# Use of human iPSCs and kidney organoids to develop a cysteamine/mTOR inhibition combination therapy to treat cystinosis

**DOI:** 10.1101/595264

**Authors:** Jennifer A. Hollywood, Aneta Przepiorski, Patrick T. Harrison, Ernst J. Wolvetang, Alan J. Davidson, Teresa M. Holm

## Abstract

Cystinosis is a lysosomal storage disease caused by mutations in *CTNS*, encoding a cystine transporter, and in its severest form is characterized by cystine accumulation, renal proximal tubule dysfunction and kidney failure. Cystinosis is treated with the cystine-depleting drug cysteamine, however this only slows progression of the disease and there is an urgent need for better treatments. Here, we have generated and characterized the first human induced pluripotent stem cell (iPSC) and kidney organoid models of cystinosis. These models exhibit elevated cystine and cysteine levels, enlarged lysosomes and a block in basal autophagy flux. Cysteamine treatment ameliorates this phenotype except for the basal autophagy flux defect. We found that treatment with Everolimus, an inhibitor of the mTOR pathway, reduces the number of large lysosomes and activates autophagy but does not rescue the cystine/cysteine loading defect. However, dual treatment of cystinotic iPSCs or kidney organoids with cysteamine and Everolimus corrects all of the observed phenotypes indicating that a combination therapy has therapeutic potential to improve the treatment of cystinosis.

## Introduction

Cystinosis is a rare autosomal recessive lysosomal storage disease caused by mutations in the *CYSTINOSIN* (*CTNS*) gene which encodes a cystine transporter.^1,2^ In the absence of CYSTINOSIN, cystine accumulates within the lysosome where it causes lysosomal dysfunction. Nephropathic cystinosis is the most severe form of cystinosis and is initially associated with the renal proximal tubule failing to reabsorb essential metabolites from the urine (Fanconi syndrome). Kidney defects present between 6-18 months of age and progress to renal failure by the end of the first decade of life.^1,3^ Other complications include derangements in non-renal tissues such as widespread cystine crystal formation (notably in the cornea), hypothyroidism, and neurological and muscular symptoms.^4–7^

The current treatment for cystinosis is life-long therapy with cysteamine, a molecule that cleaves the cystine disulphide bond to produce mixed disulphides that can escape the lysosome through alternative transporters.^8,9^ However, cysteamine only slows the progression of renal injury and kidney transplantation is inevitably required later in life.^9,10^ As a result, there remains a pressing need to develop more effective therapies for cystinosis.

While there is not yet a complete understanding of the pathogenesis of cystinosis, several lines of evidence indicate that cystine loading causes lysosomal enlargement, impaired proteolysis and delayed fusion with cargo-loaded vesicles.^11–14^ Other cellular features of cystinotic cells that are variably present in different cell types include reduced ATP and glutathione levels, mitochondrial damage, oxidative stress, increased apoptosis, and proximal tubule cell dedifferentiation.^9,15–27^ In addition, cystinotic proximal tubule cells display decreased expression of the endocytic receptors megalin and cubulin, and impaired megalin recycling.^11,12^

Defects in macroautophagy (herein referred to as autophagy) are also found in cystinotic cells. Autophagy involves the sequestration of a portion of the cytoplasm by a double-layered membrane known as an autophagosome, followed by fusion with a lysosome to form an autolysosome.^28^ This final step can be modulated by bafilomycin A1, an inhibitor of autophagolysosome acidification that disrupts autophagosome-lysosome fusion.^29^ Under, resting conditions, basal levels of autophagy are required in a ‘house-keeping’ capacity to degrade long-lived and ubiquitinated proteins, N-linked glycans, damaged organelles such as mitochondria, and to dampen certain pathways such as inflammatory, Notch and Wnt signaling.^30–32^ Under stress conditions, such as starvation, autophagy is greatly upregulated to ensure metabolically useful molecules are recycled in order to maintain cellular homeostasis. While starvation-induced autophagy is normal in cystinotic cells,^33^ basal autophagy flux is reduced in a number of cystinotic cell lines, resulting in a build-up of autophagosomes that frequently contain mitochondria.^26,33,34^ In addition, the separate pathway of chaperone-mediated autophagy, in which specific cytosolic proteins are directly delivered across the lysosomal membrane for degradation, is also defective in fibroblasts from the *Ctns*-knockout mouse.^33^

More recently, CYSTINOSIN has been implicated in modulating the mTORC1 signalling pathway, which integrates both intracellular and extracellular signals and serves as a core regulator of cellular metabolism, proliferation, survival and autophagy.^35^ mTORC1 switches between active and inactive states in response to nutrient availability.^36,37^ Inhibition of mTORC1 by a class of drugs that include Everolimus, which is used clinically as an immunosuppressant and anti-cancer agent, results in activation of autophagy.^38,39^ CYSTINOSIN physically interacts with mTORC1 binding partners, including the vacuolar ATPase proton pump and the Ragulator complex, which are necessary for mTORC1 activation by amino acids^40^. Loss of CYSTINOSIN function in conditionally immortalised human and mouse proximal tubule cell lines leads to a reduction in mTORC1 activity as well as delayed reactivation following a return to amino acid rich conditions.^40,41^

One of the challenges of the cystinosis field is a lack of good human-based cell culture models. To address this, we generated patient-specific and CRISPR/Cas9-edited cystinotic induced pluripotent stem cells (iPSCs). These cells have the advantages of being a renewable source of non-immortalised cystinotic cells and can be differentiated into numerous tissues, including kidney organoids. Our analysis of *CTNS*-iPSCs and kidney organoids revealed increased cystine and cysteine levels, enlarged lysosomes, abnormal basal autophagy flux and altered gene expression compared to healthy controls, consistent with modelling key aspects of the cystinotic phenotype. We further discovered that some of these defects can be rescued by treatment with cysteamine or Everolimus alone, but that combination therapy was the most efficacious. These results suggest that a cysteamine/Everolimus combination therapy holds tremendous therapeutic potential to improve the treatment, and health outcomes, of individuals with cystinosis.

## 2. Materials and Methods

### iPSC lines and maintenance

All work was carried out with the approval of the University of Auckland Human Participants Ethics Committee (UAHPEC 8712), Human and Disability Ethics Committee (17/NTA/204) and biosafety approval (GMO05). The CRL1502 clone C32 and the cystinosis iPSC lines were developed in Dr Wolvetang’s^42^ and Dr Davidson’s laboratory, respectively. For the patient-derived cystinosis lines (*CTNS*^−/−^), adipose-derived mesenchymal cells were derived from an individual with nephropathic cystinosis using a lentiviral doxycycline-inducible polycistronic vector containing *OCT4*, *SOX2*, *CMYC*, *KLF4* and *NANOG*. Five *CTNS*^−/−^ iPSC lines were generated, three of which (36, 108, 157) displayed a normal karyotype (determined by LabPLUS, New Zealand). These lines were confirmed to be pluripotent based on immunostaining of cell surface markers (SSEA-4, TRA-1-6-, TRA-1-81) and the formation of teratomas following transplantation of 1× 10^6^ cells under the kidney capsule of 8 wk old SCID mice (n=3 mice per line), according to established procedures^43^. All iPSC lines were cultured on LDEV-free hESC-qualified Geltrex-coated tissue culture dishes in mTeSR1 (Stemcell Technologies) medium supplemented with 1% penicillin-streptomycin, and 2.5 μg/mL Plasomcin (InvivoGen). At ~70% confluence, cells were dissociated using 1/3 Accutase in DPBS. Cells were scraped from the dish, pelleted at 800 rpm for 5 min and resuspended in mTeSR1 plus 5μM Y27632 dihydrochloride (Stemcell Technologies) for the first 24 hours to facilitate cell survival. Unless otherwise stated, all drug treatments (1 μM Cysteamine; Sigma, 100 nM Everolimus; RAD001 Selleckchem, 30 mM 3-methyladenine; Sigma, 50 mM Sucrose; Sigma) were added to cell culture medium and incubated with the cells for 24 hrs.

For starvation/refeeding experiments, cells were grown on 10 cm culture dishes until 70% confluent and incubated for 2 hrs in fresh culture medium (basal condition). For starvation, cells were washed twice in phosphate-buffered solution (PBS) and incubated in Hank’s balanced salt solution (HBSS) for 1 hr. Refeeding was performed by incubating cells in normal culture medium for the indicated time points.

### Generation of CTNS knockout lines by gene editing

Guide (g) RNA pairs targeted to introduce a 257 bp deletion in exon 8 and 9 of the *CTNS* gene were designed using RGEN and COSMID online tools^44,45^ (http://www.rgenome.net/cas-designer/; http://crispr.bme.gatech.edu/). Knockout efficiencies were first evaluated in HEK293 cells. Optimal gRNAs (gRNA_ex81.0: 5’-TCCACCCCCTGCAGTGTCATTGG-3’; gRNA_ex93.0: 5’-GCGTGAGGACAACCGCGTGCAGG-3’) were cloned into the pSPCas9(BB)-2A-GFP plasmid (Addgene 48138) and introduced into CRL1502 iPSCs by reverse transfection using *Trans*IT-LT1 (Mirus Bio). GFP positive cells were isolated 48 hrs later by flow cytometric sorting and 8,000 cells were plated on a 10 cm Geltrex-coated dish into pre-warmed mTeSR1 containing 5 μM Y27632. Medium changes were carried out daily using mTeSR1 without Y27632. Single colonies were manually picked when they reached a suitable size (~10 days post plating), clonally expanded and screened for biallelic deletions using PCR primers flanking the deleted region (F.CTNS1_primer 5’-CTCCACCCCCGCCAGTCCTC-3’; R.CTNS_1primer 5’ TCTGTGCACGGCTCTCAGC-3’). Homozygote deletions were verified by Sanger sequencing (Peters et al., 2013). Three clones, KO 15, 32 and 73 were expanded and karyotyped.

### Generation of cystinotic kidney organoids

We used an in-house developed protocol to convert iPSCs into kidney organoids^46^. Briefly, iPSCs were cultured on 10 cm Geltrex-coated dishes to ~40-50% confluency then washed twice with 1 × PBS and treated with 1mg/ml Dispase for 6 mins at 37**°**C. Dispase was removed and cells were washed 2 × PBS. Using a cell scraper, cells are manually lifted from the dish, resuspended in BPEL medium^47^ containing 8 μM CHIR99021, 3 μM Y27632 and 1 mM beta-mercaptoethanol and evenly distributed into ultra-low attachment 6 well plates (Corning). Half medium changes were carried out on day 2 with BPEL supplemented with 8 μM CHIR99021. On day 3, embryoid bodies (EB’s) were allowed to sediment in a 50 ml tube and washed 2 × PBS. EB’s were returned to the ultra-low 6 well plate and transferred to Stage II media (DMEM, 15% KnockOut Serum Replacement (KOSR; Thermo Fisher), 1% NEAA, 1% penicillin/streptomycin, 1% HEPES, 1% GlutaMAX, 0.05% PVA, 2.5 μg.ml Plasmoscin). The EBs are transferred to bioreactor flasks and grown in stage 2 media for various periods of time (up to 2 weeks). Tubule formation was observed on day 7-8. Typically, 60-80% of the EBs become kidney organoids. All drug treatments on organoids were administered on Day 10 until Day 14 when organoids were harvested for downstream analysis.

### Immunostaining

Cells were washed with TBS and fixed in 4 % paraformaldehyde (PFA)/PBS (w/v) for 10 min at RT. Following 3 washes; fixed cells (or cryosections) were blocked at room temperature (RT) for at least an hour in blocking solution (TBS containing 3 % BSA (w/v) and 5 % normal goat serum with 0.3 % Triton X100 (v/v)). Cells were incubated with primary antibody (Supplemental Table S2) in the blocking solution overnight at 4°C in a humidified chamber. Twenty-four hours later, cells were washed 3 × TBST (TBS containing 0.1% Triton X100 (v/v)) and incubated with secondary antibodies (Supplemental Table S3) at 1:500 dilution in the blocking solution for 2 hrs at RT. Cells were incubated with 10 μg/mL Hoechst 33258 for 5 mins, washed 2 × TBST and mounted with Prolong Gold (Thermo Fisher) before imaging. Cells were imaged using Zeiss LSM710 confocal microscope.

### Magic Red^TM^-Cathepsin B staining

iPSC imaging assays were set up a day before by seeding cells onto Geltrex coated 35mm Fluro dishes (WPI). Prior to staining, cells were washed 1 × DPBS. Cells were incubated for 1 hr with 26× MR-Cathepsin B in mTeSR1. Hoechst 33258 was added for the final 15 mins. Once staining was completed the dyes were washed off with 1 × DPBS and the cells were fixed with 4% PFA for 10mins. Images were taken using Zeiss LSM710 confocal microscope.

### Transmission Electron Microscopy

Samples were fixed in 2.5 % glutaraldehyde and 0.1 M phosphate buffer pH 7.4 (dissociated iPSCs or whole kidney organoids) at 4ºC, and kept in the fixative until processing. Samples were washed 3 × 0.1 M phosphate buffer for 10 min, then fixed in 1% osmium tetroxide in 0.1 M phosphate buffer for an hour at RT and washed twice in 0.1 M phosphate buffer for 5 min. The samples were then dehydrated in a graded series of ethanol washes 10 min each at RT (50%, 70%, 90% and twice 100%), following with 2 × propylene oxide wash for 10 min at RT. The samples were then infiltrated with a graded series of propylene oxide:resin mix (2:1, 1:1, 1:2) for 30 minutes each, before being imbedded in freshly made pure resin overnight. The next day the samples were placed into molds and polymerized at 60ºC for 48 hrs. All washes were performed on a rocker. Sectioned samples were imaged using Tecnai™ G² Spirit Twin transmission electron microscope.

### Transient Transfection of kidney organoids

Organoids were cultured as described above until the day of interest. Organoids were first dissociated by incubating them in 100 μl TrypLE Express (gibco) at 37°C for up 10 mins. Once dissociated cells are centrifuged at 800 g for 5 min and resuspended in stage 2 media and reverse transfected using *Trans*It-LT1.

### Western Blotting

Cells were washed 2 × ice-cold PBS and scraped on ice into 200 μl of ice-cold radioimmunoprecipitation assay buffer supplemented with protease (cOmplete mini, ROCHE) and phosphatase inhibitors (1 mM Sodium Orthoranadate, 100 mM Sodium Fluoride, 1 mM B-glycerolphosphate, 2.5 mM Sodium pyrophosphate). Protein extracts were analysed by SDS-polyacrylamide gel electrophoresis followed by blotting on nitrocellulose membranes. Primary and Secondary antibodies were used according to manufacturers’ instructions (Supplemental Table S2, S3). Blots were developed using ImageQuant^TM^ LAS 4000 image reader (Fujifilm) and analysed using image J software.

### Image analysis of lysosomes and fluorescent puncta

Magic Red^TM^ 63× magnification confocal raw images (~10 random fields per condition) were analysed using Image J analysis software. Nuclei were manually counted. To obtain a cross-sectional area of the enlarged vesicles, particle analysis was performed and the number of vesicles >10 μm^2^ were determined per field. Data were expressed as average number of enlarged vesicles per cell and statistically analysed. For the measurement of autophagic puncta, cells were transfected with the LC3-mCherry-GFP vector and imaged by confocal microscopy (10 random fields per condition containing ~1-3 cells in 3 independent experiments) and analysed using Image J. Nuclei and red and yellow puncta were manually counted and the percentage of each puncta per cell was calculated.

### Mass Spectrometry

Cells were lysed on ice with cold 50% acetonitrile and centrifuged at 16,000g for 10 min at 4°C. The supernatant was transferred to a cold Eppendorf tube and stored at −80°C until samples were ready to be processed. 2 μl of sample was added to 5 μl of a 50% solution of acetonitrile and 50 mM ammonium bicarbonate (ACN/ABC). 3 μl of sample was treated with either 3 μl 1 mM TCEP or 2 mM monobromobimane (MBrB) in 50 % ACN/ABC and incubated at RT in the dark for 20 mins. 3 μl 2 mM MBrB and 3 μl 50 mM ABC was added to the TCEP treated samples. 6 μl 25 % ACN was added to the MBrB treated set. Samples were incubated at RT in the dark for 20 mins. Following incubation, 950 μl 0.1% formic acid and 5 μl 4.292 μM CSH heavy standard was added to all samples. 10 mg HLB SPE cartridges were conditioned with 0.5 ml methanol followed by 0.5 ml 0.1% formic acid. The entire sample was loaded onto the conditioned cartridge and washed with 1 ml 0.1% formic acid. Samples were eluted into clean tubes with 0.3 ml 10% ACN in 0.1% formic acid. Samples were spun in a speedvac until volumes were reduced to 50-100 μl. Samples were either injected neat or diluted 1:3 in 0.1% formic acid and run on a QStar XL LC-MS instrument and through a LC column Zorbax SB-C18 3.5 μm 150×0.3mm (Aligent).

### RNA extraction, cDNA synthesis and quantitative PCR analysis

Cells were first washed 1 × PBS before being lysed in GENEzol for 5 mins. RNA was extracted using GENEzol^TM^ TriRNA Pure kit (Geneaid). cDNA was synthesised using qScript cDNA SuperMix (Quanta). For quantitative (q) PCR, PerfeCTa SYBR Green FastMix (Quanta) was used. qPCR was performed on a QuantStudio 6 Flex Real-Time PCR machine. Primers used are listed in Supplemental Table S1. Samples were normalised to *HPRT1* and *CREBP* expression. Gene expression was calculated using the ddCt method.^48^ Error bars represent standard deviation from technical triplicates.

### RNA-Sequencing and analysis

Total RNA from 4 samples per iPSC line were prepared using the GENEzol^TM^ TriRNA Pure kit. The quality (RIN), concentration and purity (OD260/230 and OD260/280) of the RNA was determined on Bioanalyser (RNA nano chip), Qubit and Nanodrop instruments. Library preparation and sequencing were performed commercially (New Zealand Genomics Limited, Otago). Libraries were prepared using the TruSeq standard total RNA kit with standard protocols (Illumina). 2 × 125bp paired-end sequencing was performed on an Illumina HiSeq 2500 sequencer. Reads were adapter filtered and quality trimmed using BBDuk version 37.75 (1) with a quality cut-off of phred=10 (trimq=10) and to reduce the potential mapping errors any reads less than 50bp after trimming were removed. QC filtered reads were mapped to the Human genome (GRCh38) downloaded from ENSEMBL (www.ensembl.org/Homo_sapiens/Info/Index) using HISAT2 (version 2.0.5) in stranded mapping mode (–rna-strandness RF). Read counts were generated from the alignment files using HT-Seq (version 0.6.0) under the Union mode and strand option set to “reverse”. DESeq2 (2) was used to generate differential expression calls and statistics for control versus knockout comparison based on the observed read counts for each gene. Expression changes were declared significant for q-value < 0.05. Heatmaps were generated in R using the pheatmap_1.0.8 package. GO terms enrichments were analysed using the R package goseq (version 1.22). Enrichment was tested for all differentially expressed genes with an FDR (False Discovery Rate) corrected p-value less than 0.05. The GO terms themselves were also FDR corrected at the same rate.

### Statistical analysis

Data are presented as the mean ± SEM. GraphPad PRISM software version 7 (GraphPad Software) was used for all statistical analyses. The statistical significance of the differences between two groups was calculated using an unpaired Student’s *t*-test. For between 3 or more groups one-way Anova was used. A *p* value <0.05 was considered to be statistically significant.

## Results

### Generation of *CTNS*-iPSC lines

To generate patient-specific *CTNS*-iPS cells, adipose-derived mesenchymal cells were grown from a fat sample from a nephropathic cystinosis patient undergoing renal transplantation. Exon sequencing determined that the individual was compound heterozygote for two previously described *CTNS* mutations: a 57 kb deletion and a L158P missense mutation in exon 8^2^. The mesenchymal cells were reprogrammed into iPSCs and three *CTNS*^−/−^ lines, referred to as 36, 108, 157, possessed normal karyotypes (results not shown). All three lines stained positive for the pluripotency markers alkaline phosphatase, SSEA-4, Tra-1-60 and Tra-1-81, and gave rise to teratomas containing tissues from all three germ layers (Figure 1A, B and D, data for *CTNS*^−/−^36 shown). Re-expression of endogenous *OCT4*, *NANOG*, *SOX2*, *CMYC* and *KLF4* was confirmed by qPCR (Figure 1C for representative data for *CTNS*^−/−^36). As all of these lines displayed similar phenotypes, line 36 was used for subsequent analyses (herein referred to as *CTNS*^−/−^). In addition to patient-specific lines, we also generated independent *CTNS* knockout (KO) lines by performing CRISPR/Cas9 gene editing in CRL1502 iPSCs^42^ (Briggs et al., 2013). Guide RNAs were used to introduce a 257 bp deletion in exon 8 and 9 of the *CTNS* gene, resulting in deletion of the second transmembrane domain (Figure 1E and F). Three lines with homozygote deletions (KO 15, 32 and 73) were identified by Sanger sequencing. As all three *CTNS*^KO^ lines displayed a similar phenotype (Supplemental Figure S1G, H and I). The *CTNS*^KO^ line KO73 was used for subsequent experiments.

**Figure 1.**
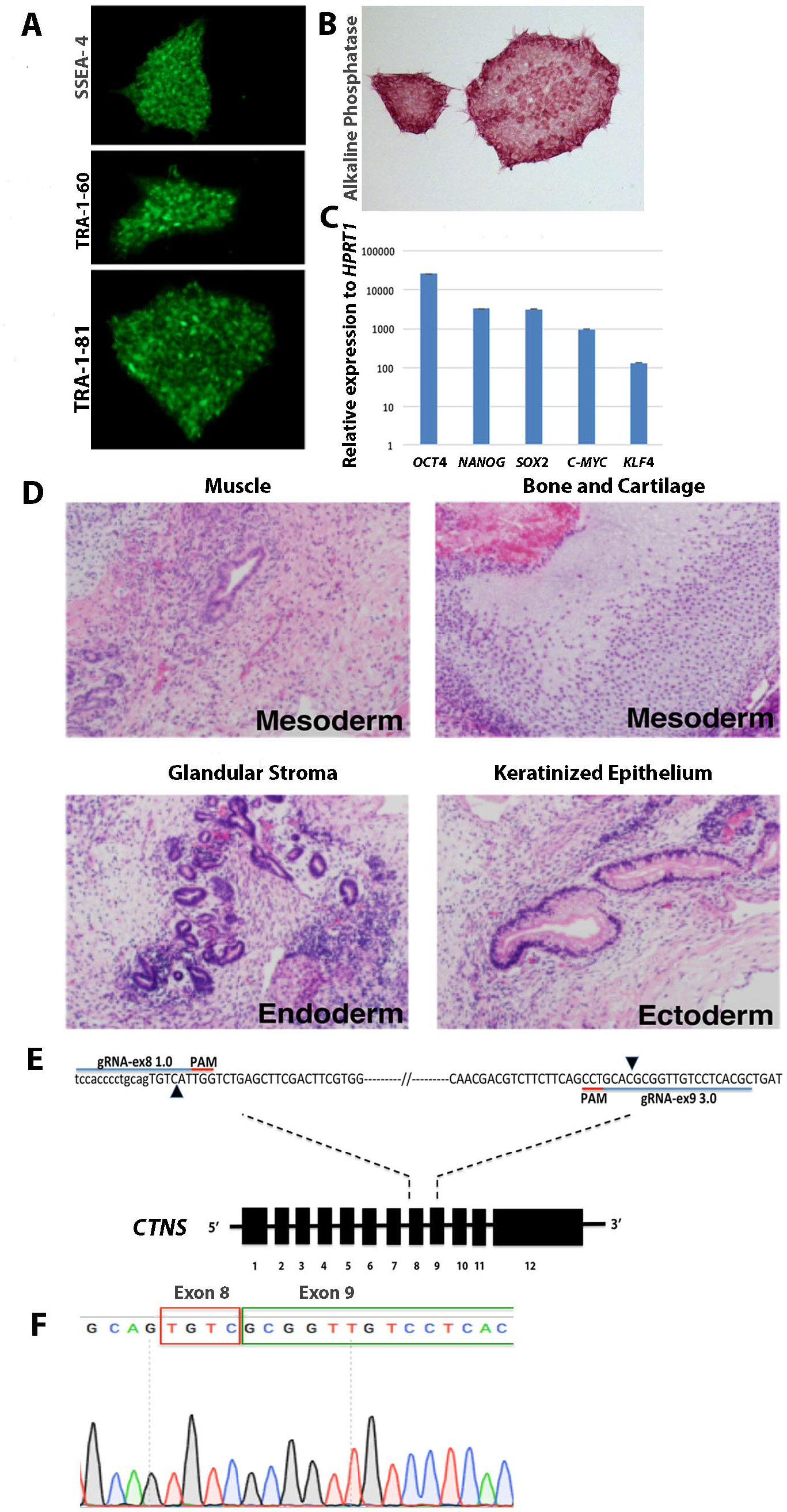
Characterization of patient-derived *CTNS*^−/−^ and *CTNS*^KO^ iPSCs. (A) *CTNS*^−/−^-iPSCs stained for stem cell surface antigens SSEA-4, TRA-1-60 and TRA-1-81. (B) *CTNS*^−/−^ iPSCs stained for alkaline phosphatase. (**C)** Quantitative PCR of endogenous genes relative to *HPRT1* expression. Plotted data are mean ± SD. (**D)** Hematoxylin and Eosin stained histological sections of tumours derived from SCID mice following injection of *CTNS*^−/−^-iPSCs under kidney capsule. All three germ layers were identified, mesoderm, endoderm and ectoderm (n=3). (**E)** Schematic overview of the CRISPR-based strategy to disrupt the *CTNS* gene in WT-iPSCs. The extent of the deletion in exon 8 and exon 9 is marked with black arrowheads. (**F)** Sanger sequencing chromatogram shows resulting sequence in *CTNS*^KO^-iPSCs.

### *CTNS*-iPSCs load cystine and cysteine

Accumulation of cystine in lysosomes in all cells of the body is a hallmark of cystinosis. Cystine was measured in cystinotic and control iPSCs by mass spectrometry, revealing 3-5 fold higher levels in *CTNS*^−/−^ and *CTNS*^KO^ iPSCs compared to the CRL1502 control cells (Figure 2A and B). Cysteine was also elevated ~6-7 fold in cystinotic iPSCs compared to controls (Figure 2A and B) in agreement with observations made by others.^19^ To assess whether cystine/cysteine levels were responsive to cysteamine treatment, we treated *CTNS*-iPSCs with 1 µM of cysteamine for 24 hrs, resulting in significant reductions in the levels of both cystine and cysteine (Figure 2A and B).

**Figure 2.**
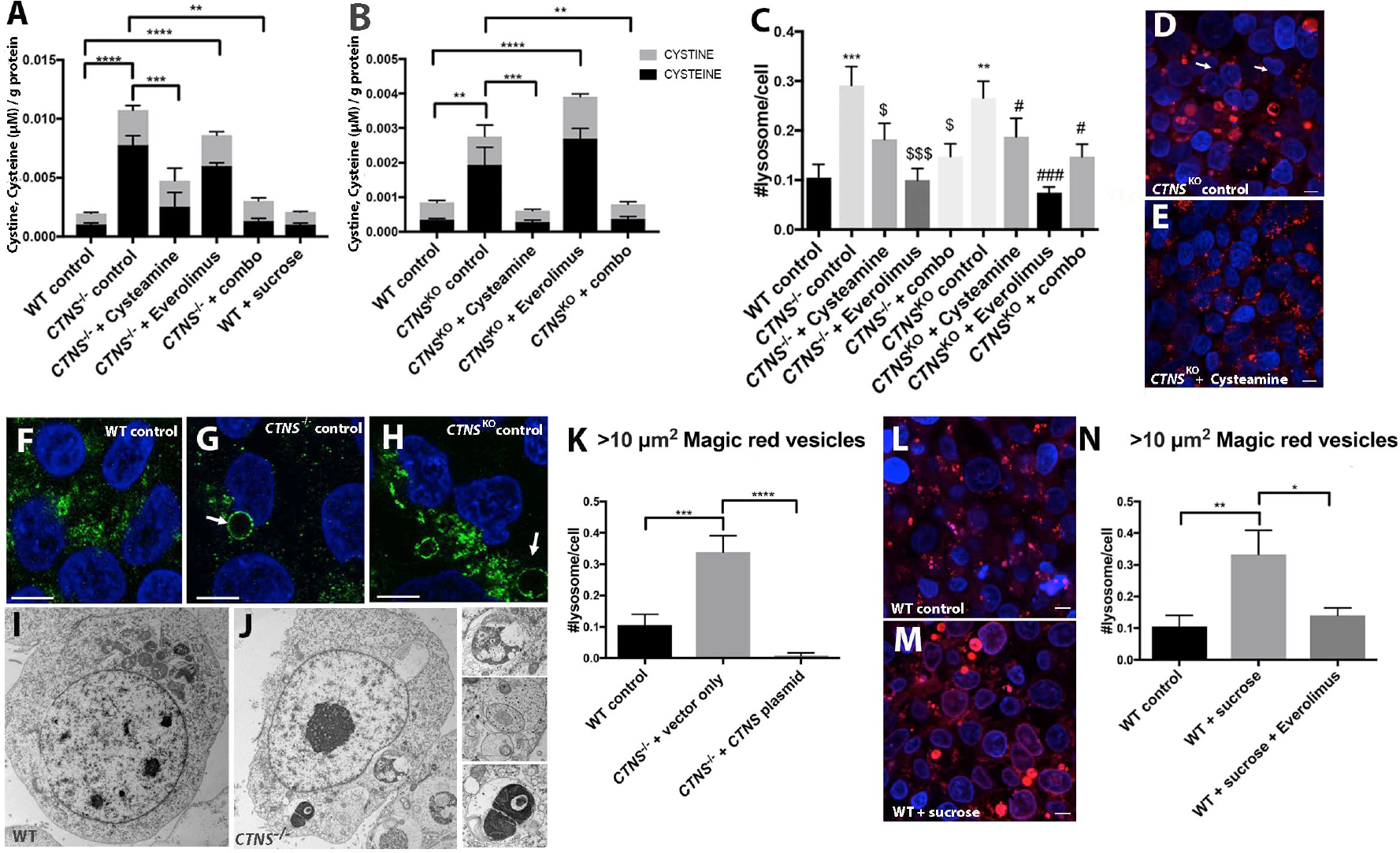
*CTNS*-iPS cells have increased cystine and cysteine levels and enlarged lysosomes. **(A, B)** Amount of cysteine (black) and cystine (grey) (µM) per gram of protein in WT and *CTNS*^−/−^-iPSCs or *CTNS*^KO^-iPSCs with various treatments. One-way analysis of variance (ANOVA) performed, **p<0.01, ***p<0.001, **** p<0.0001, data plotted as mean ± SEM, 3 independent experiments. **(C)** Graph displaying quantification of the average number of Magic Red vesicles (lysosomes) per cell over 10 µm^2^. One-way ANOVA performed, *** p=0.001, WT control vs *CTNS*^−/−^ control and ** p=0.0054, WT control vs *CTNS*^KO^ control, $ p=0.045, *CTNS*^−/−^ control vs *CTNS*^−/−^ 1 µM cysteamine and *CTNS*^−/−^combination, $$$ p<0.001, *CTNS*^−/−^ control vs CTNS^−/−^100 nM Everolimus, # p=0.047, *CTNS*^KO^ control vs *CTNS*^KO^ 1 µM Cysteamine and *CTNS*^−/−^combination, ### p=0.0004, *CTNS*^KO^ control vs *CTNS*^KO^ 100 nM Everolimus (n=600 cells from10 random fields per condition, 20 cells/field, 3 independent experiments), data plotted as mean ± SEM. **(D, E)** Representative images of fluorescent staining with Magic red in control and cysteamine treated *CTNS*^KO^-iPSCs. **(F)** Representative immunofluorescent stainings with anti-LAMP1 (green) in WT-iPSCs, and **(G, H)** *CTNS*^−/−^ and *CTNS*^KO^-iPSCs respectively. Arrows indicate enlarged vesicles. **(I, J)** Transmission electron micrograph (TEM) of WT and *CTNS*^−/−^-iPSCs showing enlarged vesicles. **(K)** Graph displaying quantification of the average number of Magic Red vesicles per cell over 10 µm^2^ in WT-iPSCs and *CTNS^−/−^*-iPSCs over-expressing vehicle or exogenous *CTNS.* One-way ANOVA performed, ***p<0.001, **** p<0.0001(n=600 cells from 10 random fields per condition, 20 cells/field, 3 independent experiments), data plotted as mean ± SEM. (**L, M)** Representative images of fluorescent staining with Magic red in WT-iPSCs and WT-iPSCs treated with 50 mM sucrose. **(N)** Average number of Magic Red vesicles per cell over 10 µm^2^ WT-iPSCs treated with 50 mM sucrose or sucrose and 100 nM Everolimus. One-way ANOVA performed, *p<0.05, **p<0.01, (n=600 cells from 10 random fields per condition, 20 cells/field, 3 independent experiments). All data are plotted mean ± SEM. *CTNS*^−/−^; patient-derived cystinotic iPS cells, *CTN*S^KO^; CRISPR generated cystinotic knock out iPS cells, Scale bars 10µm. Nuclei counter stain in panels F-H: DAPI; D, E and L, M: Hoechst.

### *CTNS*-iPSCs display enlarged lysosomes

To assess the size and distribution of lysosomes in the cystinotic iPSCs and to functionally confirm their lysosomal identity, the *CTNS*-iPSC lines were incubated with the Magic Red substrate that is degraded by cathepsin B and releases fluorescent peptides inside lysosomes and endolysosomes. Enlarged Magic Red^+^ puncta were observed more frequently in the *CTNS*-iPSCs, compared to controls (Figure 2C and D). Quantification of the enlarged lysosomes, which we defined as having an optical cross-sectional area of >10 µm^2^, showed that the average number per cell was ~2.5 fold higher in *CTNS*-iPSCs compared to controls (Figure 2C) To further confirm that these magic red structures are enlarged lysosomes we examined cystinotic iPSCs with the lysosomal marker LAMP1 by immunofluorescence and at the ultrastructural level by electron microscopy. *CTNS*^−/−^ and *CTNS^KO^* iPSCs were found to contain a mixture of small to enlarged LAMP1^+^ puncta whereas control iPSCs show qualitatively fewer enlarged LAMP1^+^ puncta (Figure 2F, G and H). Consistent with the LAMP1 and Magic Red data, we observed large degradative/storage-like bodies surrounded by a single limiting membrane in *CTNS*^−/−^ but not control iPSCs by electron microscopy (Figure 2I and J). As expected for dysfunctional lysosomes, these bodies were filled with electron-dense material, intra-luminal vesicles and undigested membranes, and therefore likely represent enlarged lysosomes and/or amphisomes (Figure 3J). To show that this phenotype is due to loss of CYSTINOSIN, we performed rescue experiments by transfecting *CTNS*-iPSCs with a plasmid encoding Cystinosin-GFP.^49^ Overexpression of this construct was capable of reducing the number of enlarged lysosomes to levels below that seen in control iPSCs (Figure 2K).

**Figure 3.**
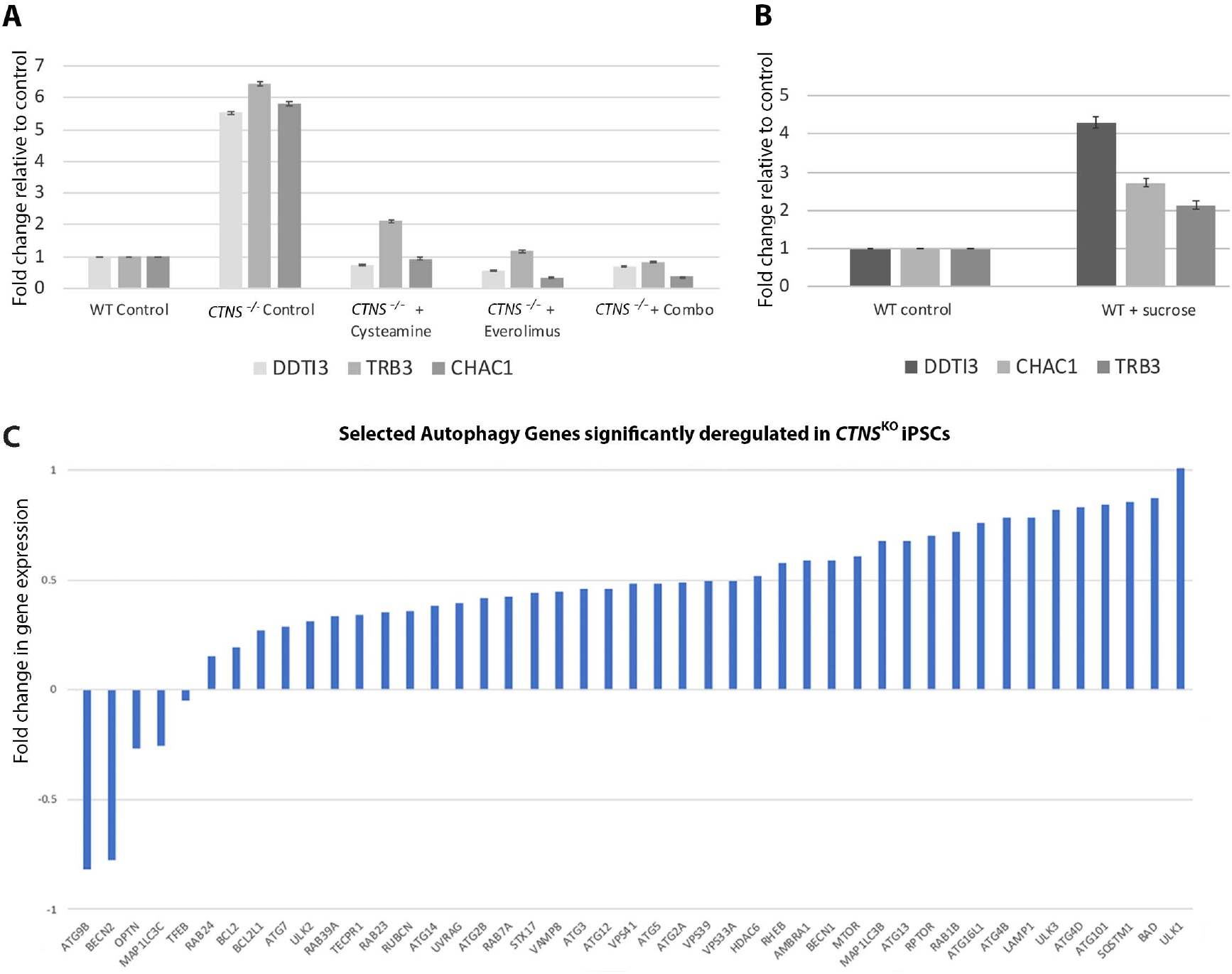
*CTNS*-iPSCs display enlarged vesicles at ultrastructural level and deregulation of genes. **(A)** Quantitative PCR of genes of interest; *DDTI3*, *TRB3* and *CHAC1*, in WT and *CTNS^−/−^*-iPSCs with various treatments normalised to *HPRT* and *CREEBP* and expressed as fold change to WT. Data plotted as mean ± SD. **(B)** Quantitative PCR analysis of target genes in WT-iPSCs treated with 50 mM sucrose, data plotted as mean ± SD. **(C)** Fold changes of selected autophagy genes deregulated in *CTNS*^KO^-iPSCs.

As the enlarged lysosome phenotype of *CTNS*-iPSCs is most likely caused by cystine accumulation, we next sought to phenocopy this using sucrose, which accumulates within the of lysosomes of normal cells^50,51^. We treated control iPSCs with 50 mM sucrose for 24 hrs and found that this led to an increase in the number of enlarged lysosomes, similar to that seen in *CTNS*-iPSCs (Figure 2L, M and N). As expeceted, sucrose had no effect on cystine loading (Figure 2A). Together, these observations are consistent with *CTNS*-iPSCs displaying an increased number of enlarged lysosomes due to cystine accumulation arising from defective CYSTINOSIN activity. In support of this, treatment of *CTNS*-iPSCs with 1 µM cysteamine for 24 hrs resulted in a reduction in the average number of enlarged lysosomes per cell although it did not completely rescue to control levels (Figure 2C and E).

### RNA-Seq analysis reveals differentially regulated genes in *CTNS*^KO^-iPSCs

To gain further insights into the phenotype of *CTNS*-iPSCs, we performed RNA-Seq to identify differentially expressed genes between *CTNS*^KO^-iPSCs and their isogenic control cells (n=4 biological repeats for each). We found a total of 12,750 differentially expressed genes with 8792 significantly up-regulated and 3958 significantly down-regulated (p < 0.05), compared to controls (Supplemental Table S4). KEGG Pathway analysis revealed several significantly enriched pathways in *CTNS*^KO^-iPSCs that include the ribosome, spliceosome, proteasome, oxidative phosphorylation, protein processing in the ER and ubiquitin mediated proteolysis (cut-off p < 1×10^−6^; Table 1). Interestingly, pathways linked to Huntington’s, Parkinson’s and Alzheimer’s disease were also enriched, perhaps reflecting common features of these disorders with lysosomal storage diseases.^52^ GO term enrichment analysis yielded a much more extensive list of gene sets that was more difficult to summarise (data not shown). However, in the ‘Biological Process’ category, we found enrichment for pathways implicated in cystinosis including autophagy, vesicle trafficking, redox homeostasis, the mTOR pathway and protein catabolism (Table 2).

**Table 1.** KEGG pathways significantly enriched in *CTNS*^KO^-iPSCs (cut-off p < 1×10^−6^)

| ENTRY | NAME | CLASS | P VALUE | NUMDEINCAT | NUMINCAT |
| --- | --- | --- | --- | --- | --- |
| **HSA03010** | Ribosome - Homo sapiens (human) | Genetic Information Processing; Translation | 2.06E-31 | 81 | 87 |
| **HSA05016** | Huntington's disease - Homo sapiens (human) | Human Diseases; Neurodegenerative diseases | 1.41E-13 | 134 | 182 |
| **HSA05012** | Parkinson's disease - Homo sapiens (human) | Human Diseases; Neurodegenerative diseases | 3.30E-12 | 93 | 129 |
| **HSA04110** | Cell cycle - Homo sapiens (human) | Cellular Processes; Cell growth and death | 1.00E-11 | 105 | 124 |
| **HSA03040** | Spliceosome - Homo sapiens (human) | Genetic Information Processing; Transcription | 1.39E-10 | 101 | 127 |
| **HSA03008** | Ribosome biogenesis in eukaryotes - Homo sapiens (human) | Genetic Information Processing; Transcription | 3.08E-10 | 66 | 76 |
| **HSA00190** | Oxidative phosphorylation - Homo sapiens (human) | Metabolism; Energy metabolism | 3.10E-10 | 90 | 131 |
| **HSA04141** | Protein processing in endoplasmic reticulum - Homo sapiens (human) | Genetic Information Processing; Folding, sorting and degradation | 8.01E-10 | 130 | 166 |
| **HSA04120** | Ubiquitin mediated proteolysis - Homo sapiens (human) | Genetic Information Processing; Folding, sorting and degradation | 1.91E-09 | 111 | 135 |
| **HSA03013** | RNA transport - Homo sapiens (human) | Genetic Information Processing; Translation | 2.05E-09 | 116 | 150 |
| **HSA05010** | Alzheimer's disease - Homo sapiens (human) | Human Diseases; Neurodegenerative diseases | 6.03E-09 | 116 | 166 |
| **HSA03050** | Proteasome - Homo sapiens (human) | Genetic Information Processing; Folding sorting and degradation | 2.18E-07 | 37 | 44 |

**Table 2.** GO Terms in ‘Biological Process’ category enriched in *CTNS*^KO^-iPSCs.

| GO TERM | TERM | P-VALUE | NUMDEINCAT | NUMINCAT |
| --- | --- | --- | --- | --- |
| GO:0016192 | vesicle-mediated transport | 1.30E-08 | 769 | 1281 |
| GO:0016236 | Macroautophagy | 2.28E-08 | 202 | 299 |
| GO:0006914 | Autophagy | 3.84E-08 | 269 | 416 |
| GO:0045454 | cell redox homeostasis | 5.43E-07 | 46 | 58 |
| GO:0000045 | autophagosome assembly | 7.60E-06 | 57 | 74 |
| GO:0045022 | early endosome to late endosome transport | 1.36E-05 | 32 | 36 |
| GO:0006900 | membrane budding | 1.58E-05 | 56 | 72 |
| GO:0000422 | mitophagy | 1.99E-05 | 132 | 200 |
| GO:0032006 | Regulation of Tor signalling | 3.06E-05 | 50 | 63 |
| GO:0006890 | retrograde vesicle-mediated transport, Golgi to ER | 4.54E-05 | 31 | 36 |
| GO:1901800 | positive regulation of proteasomal protein catabolic process | 5.15E-05 | 67 | 93 |
| GO:0032008 | positive regulation of TOR signalling | 5.96E-05 | 23 | 26 |
| GO:0032436 | positive regulation of proteasomal ubiquitin-dependent protein catabolic process | 6.88E-05 | 59 | 80 |
| GO:0006623 | protein targeting to vacuole | 7.12E-05 | 26 | 29 |
| GO:0045324 | late endosome to vacuole transport | 7.18E-05 | 1 | 14 |
| GO:0031929 | TOR signalling | 8.68E-05 | 57 | 75 |
| GO:0070534 | protein K63-linked ubiquitination | 9.68E-05 | 33 | 42 |

We next examined whether some of the differentially expressed genes would have utility as molecular biomarkers of the cystinotic phenotype. From the top 50 differentially expressed genes (Supplemental Figure S2), we focused on *DDIT3* (aka *CHOP*), which encodes a transcription factor belonging to the ‘integrated stress response’ involved in cellular adaptation to stress.^53^ In addition, we identified two downstream targets of *DDIT3*: *TRIB3*, encoding a pseudokinase that acts as a negative feedback regulator of *DDIT3*^54^ and *CHAC1*, encoding an enzyme that degrades glutathione.^55^ Using qPCR we independently confirmed that *DDIT3*, *TRIB3* and *CHAC1* were upregulated in *CTNS*-iPSCs compared to control iPSCs (Figure 3A and Supplemental Figure S1A). To assess whether the expression of this gene triad is responsive to cysteamine, we treated *CTNS*-iPSCs with 1 µM cysteamine for 24 hrs and found that they decreased to near-control levels. Incubation of control iPSCs with 50 mM sucrose for 24 hrs also resulted in an up-regulation of *DDIT3*, *TRIB3* and *CHAC1*, indicating that these genes, while not specific biomarkers of cystinotic cells *per se*, may ‘read-out’ lysosomal dysfunction caused by the accumulation of indigestible substrates (Figure 3B).

### Glutathione levels are unchanged in *CTNS*-iPSCs

Given the role of *CHAC1* in the degradation of GSH and prior reports that suggested that impaired lysosomal cystine efflux may alter GSH homeostasis,^16, 20,22,56^ we examined the levels of GSH in cystinotic iPSCs by mass spectrometry. No significant difference was seen in the total amount of GSH in *CTNS*-iPSCs compared to controls. Furthermore, no difference was observed in the ratio of oxidized (GSSG) to reduced GSH, indicating that *CTNS*-iPSCs are unlikely to be under significant oxidative stress (Supplemental Figure S1C and D).

### The mTORC1 pathway appears unaffected in *CTNS*-iPSCs

Closer scrutiny of the autophagy genes identified from the GO term analysis revealed that *CTNS*^KO^-iPSCs show a slight upregulation of *MTOR* as well as two of its downstream targets (*ULK1* and *ATG13*) compared to control cells (Figure 3C). To assess mTORC1 activity in *CTNS*-iPSCs we performed western blotting for phosphorylated S6 (pS6), a downstream target of mTORC1, under basal conditions and after starvation for 60 mins, followed by re-feeding.^57,58^ We found no difference in pS6 levels between the cystinotic iPSCs and control cells under basal conditions (Supplemental Figure S1F) and unlike prior reports,^40,41^ we did not detect a delay in the re-activation of mTORC1 at 2.5, 7, 12 or 15 mins following re-feeding (Supplemental Figure S1F and data not shown). Together these observations indicate that there are no significant defects in mTORC1 activity in *CTNS*^KO^-iPSCs under basal conditions or following starvation in terms of S6 phosphorylation.

### Basal autophagy flux is perturbed in *CTNS*-iPSCs

Of the autophagy-related genes identified in the GO term analysis, we noted that there are genes involved in early through to late processes of autophagy including autophagosome formation (*SQSTM1*, *BECLIN1*, *LC3B*), movement (*HDAC6*) and tethering and fusion (*RUBICON*, *UVRAG*, *VPS16*, *VAMP8*, *STX17*, *SNAP29*; these and other selected genes are shown in Figure 3C). In most cases, these genes are upregulated in *CTNS*^KO^-iPSCs compared to controls (Figure 3C). Notably, an increase in *SQSTM1/p62* can be indicative of a block in autophagy flux.

To explore basal autophagy levels, we first measured the levels of the autophagosome specific protein LC3B-II by western blotting. Consistent with the RNA-Seq data, we found higher levels of LC3B-II in *CTNS*^−/−^-iPSCs compared to control iPSCs, indicating either an increase in the number of autophagosomes or a decrease in autophagosome degradation (Figure 4A). To quantify autophagosome and autolysosome numbers, we transfected *CTNS*-iPSCs and control iPSCs with a plasmid encoding the mCherry-LC3B-GFP sensor that fluorescently labels autophagosomes in yellow and autolysosomes in red^59^. Under basal conditions, *CTNS*^−/−^ cells were found to have a ~2.6-fold higher levels of yellow puncta (autophagosomes) compared to control iPSCs (Figure 4B and C).

**Figure 4.**
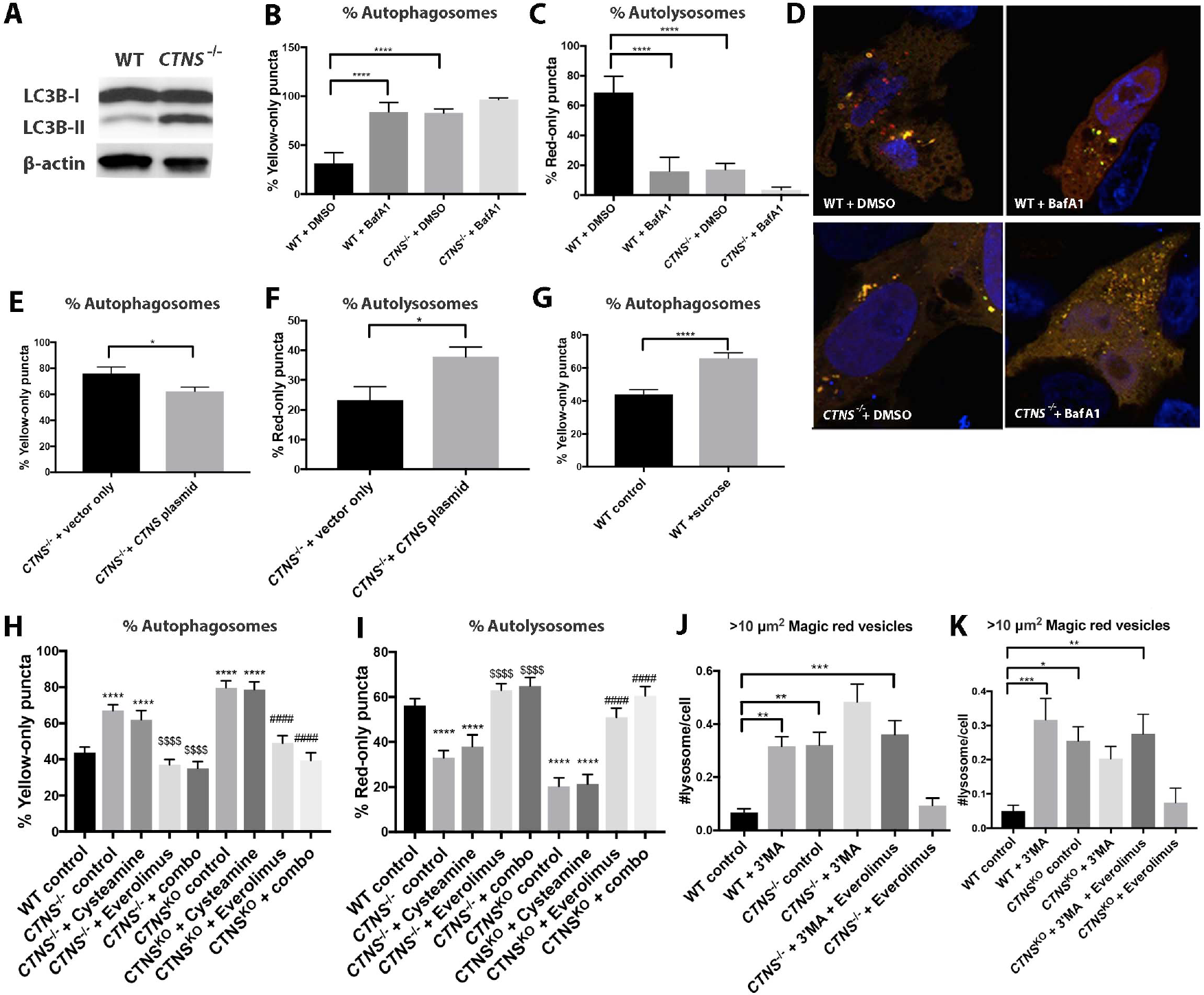
Autophagy Flux is delayed in *CTN*S-iPSCs. **(A)** Representative Western blot against autophagosome marker LC3B-II and β-actin from WT and *CTN*S-iPSCs (representative of 3 independent experiments) **(B, C)** Percentage of cells with yellow-only puncta (autophagosomes) and red-only puncta (autolysosomes) untreated or treated with Bafilomycin A1 (representative of n=30 cells, from 10 random fields per condition containing ~1-3 cells in 3 independent experiments). One-way AVOVA performed, ****p<0.0001 relative to WT, the data is mean ± SEM. **(D)** Cells transfected with tandem mCherry-LC3B-GFP plasmid showing red and yellow puncta. **(E, F)** Percentage of yellow-only and red-only puncta following exogenous expression of *CTNS* in *CTNS*^−/−^-iPSCs. Two-tailed unpaired Student *t*-test performed, *p<0.05, the data is mean ± SEM. **(G)** Sucrose treatment on WT-iPSCs to induce a cystinotic phenotype. Percentage of yellow-only puncta shown. Two-tailed unpaired Student *t*-test performed, ****p<0.0001, the data is mean ± SEM. **(H, I)** Effects of drug treatments on numbers of yellow and red puncta; *CTNS*-iPS cells treated with cysteamine alone, Everolimus alone or a combination of both for 24 hrs. One-way ANOVA performed, **** p<0.0001, WT vs *CTNS*^−/−^ & *CTNS*^KO^, $$$$ p<0.0001, *CTNS*^−/−^ vs *CTNS*^−/−^ 100 nM Everolimus & *CTNS^−^*^/−^ combination, #### p<0.0001 *CTNS*^KO^ vs *CTNS*^KO^ 100 nM Everolimus & *CTNS*^KO^ combination, data plotted as mean ± SEM (n= 30 cells from 10 random fields per condition containing ~1-3 cells in 3 independent experiments). **(J, K)** Average number of Magic Red vesicles per cell over 10 µm^2^ in WT-iPSCs and *CTNS*^−/−^ or *CTNS*^KO^-iPSCs treated with 3 mM 3’methlyadenine and Everolimus. One-way AVOVA performed *p<0.05, **p<0.01, ***p<0.001. The values are mean ± SEM, (n=600 cells from 10 random fields per condition, 20 cells/field, 3 independent experiments). *CTNS*^−/−^; patient-derived cystinotic iPS cells, *CTNS*^KO^; CRISPR generated cystinotic knockout iPS cells, BafA1; Bafilomycin A1, 3’MA; 3’methlyadenine. Nuclei counter stain in panel D: DAPI.

To assess the flux through the autophagy pathway, we treated *CTNS*-iPSCs and control iPSCs expressing the mCherry-LC3B-GFP sensor with Bafilomycin A1 (BafA1), which inhibits fusion of lysosomes with autophagosomes^60^. While BafA1 induced a 2.7-fold increase in the percentage of yellow puncta in control iPSCs compared to vehicle (DMSO) treated cells, only a slight, but non-significant increase was seen in *CTNS*-iPSCs (Figure 4B, C and D). To confirm that the autophagy defect was specific to a loss of CYSTINOSIN activity we transfected *CTNS*^−/−^-iPSCs with the Cystinosin*-*GFP plasmid, resulting in a ~1.2 fold reduction in the percentage of yellow puncta (Figure 4E and F). Taken together, these results indicate that loss of CYSTINOSIN function in iPSCs causes an accumulation of autophagosomes under basal conditions due to reduced fusion of lysosomes with autophagosomes. We speculate that this block leads to a compensatory upregulation of numerous autophagy pathway genes, as a part of a feedback response.

The basal autophagy block in *CTNS*-iPSCs may be caused by the accumulation of cystine in the lysosome. To explore this, we treated control iPSCs with 50 mM sucrose for 24 hrs and then transfected them with the mCherry-LC3B-GFP sensor. We found that the percentage of yellow puncta increased 1.5-fold in sucrose loaded cells compared to control cells, indicative of a reduction in basal autophagic flux (Figure 4G). Given this, we next tested whether treatment with cysteamine would ameliorate the basal autophagy flux defect *of CTNS*-iPSCs transfected with the mCherry-LC3B-GFP sensor. Unexpectedly, we found that treating *CTNS*-iPSCs with 1 µM cysteamine for 24 hrs did not greatly improve basal autophagy flux (Figure 4H and I). We conclude from these findings that the basal autophagy defect in cystinotic iPSCs is caused by a loss of CYSTINOSIN but this cannot be ameliorated by cysteamine treatment.

### Basal autophagy flux defects are rescued in *CTNS*-iPSCs by mTORC1 inhibition

The failure of cysteamine to restore basal autophagy flux in *CTNS*-iPSCs may provide a rationale for why cysteamine therapy is not curative and led us to speculate that activating autophagy via mTORC1 inhibition may provide additional therapeutic benefit. To test this, we treated *CTNS*-iPSCs for 24 hrs with 100 nM Everolimus and examined basal autophagy flux. We found that Everolimus restores the number of yellow puncta (autophagosomes) to control levels and correspondingly increased the number of autolysosomes, in agreement with similar results reported using *Ctns*^−/−^ mouse fibroblasts^33^ (Figure 4H and I). Importantly, dual treatment of 1 µM cysteamine and 100 nM Everolimus had similar effects as Everolimus alone, without any sign of combination toxicity (Figure 4H and I).

### Cystine levels remain high in *CTNS*-iPSCs following mTORC1 inhibition

We then assessed cystine and cysteine levels in Everolimus and combined Everolimus/cysteamine treated cells. Everolimus alone had no significant effect on cystine/cysteine levels when *CTNS*^−/−^ iPSCs were treated but caused a 1.3 fold increase in *CTNS*^KO^ iPSCs (Figure 2A and B). Combination treatment decreased cystine/cysteine to levels similar to that seen with cysteamine treatment alone (Figure 2A and B), indicating that activation of the mTORC1 pathway does not interfere with the ability of cysteamine to deplete cystine/cysteine.

### mTORC1 inhibition reduces enlarged lysosomes in *CTNS*-iPSCs via autophagy

Next we examined the effect of Everolimus treatment on the enlarged lysosome phenotype. We found that Everolimus reduces the average number of enlarged lysosomes to near-normal levels, making it more effective than cysteamine alone (Figure 2C). Combined Everolimus/cysteamine treatment yielded intermediate results with a ~2 fold reduction compared to untreated *CTNS*-iPSCs, indicating that cysteamine interferes with the ability of Everolimus to reduce the number of enlarged lysosomes (Figure 2C). In addition, Everolimus treatment reduced the number of enlarged lysosomes induced by sucrose-loading, suggesting that its effects on the lysosome are not specific to cystinotic cells (Figure 2N).

To determine if the effects of Everolimus on enlarged lysosomes was dependent on autophagy, we investigated the effects of 3-methyladenine (3-MA), an inhibitor of autophagy that acts downstream of mTORC1.^61^ First, we treated control and *CTNS*-iPSCs for 24 hours with 30 mM of 3-MA alone. We observed a ~4-5 fold increase in the average number of enlarged lysosomes per cell in control iPSCs while levels in *CTNS^−^*^/-^ and *CTNS*^KO^ iPSCs did not significantly increase further (Figure 4J and K). Treating *CTNS*-iPSCs with both 3-MA and Everolimus failed to have any significant effect on the number of enlarged lysosomes (Figure 4J and K), providing strong evidence that the effects of Everolimus are mediated via stimulation of autophagy.

### Everolimus reduces expression of *CHOP*, *TRB3*, and *CHAC1* in *CTNS*-iPSCs

We then assessed the effects of Everolimus alone and combined Everolimus/cysteamine treatment on the expression of *DDIT3*, *TRB3*, and *CHAC1*. In both *CTNS*^−/−^ and *CTNS*^KO^ iPSCs, Everolimus alone and combined treatment reduced the expression levels of the gene triad to near-control levels, in keeping with the notion that these genes are providing a read-out of lysosome dysfunction in iPSCs (Figure 3A and Supplemental Figure S1A).

### Characterisation of cystinotic kidney organoids

Having established the potential therapeutic effects of combined Everolimus/cysteamine treatment in *CTNS*-iPSCs, we next assessed whether these compounds would show efficacy on human cystinotic kidney tissue, using a kidney organoid protocol we recently developed^46^. Using this approach, we matured *CTNS*^−/−^, *CTNS*^KO^ and control cells into kidney organoids for 14 days^46^. Similar to our results obtained with undifferentiated *CTNS*-iPSCs, we found that the *CTNS*^−/−^ and *CTNS*^KO^ organoids also display cysteine/cysteine loading but no differences in the ratio of GSH/GSSG compared to isogenic control organoids (Figure 5A and Supplemental Figure S1E).

**Figure 5.**
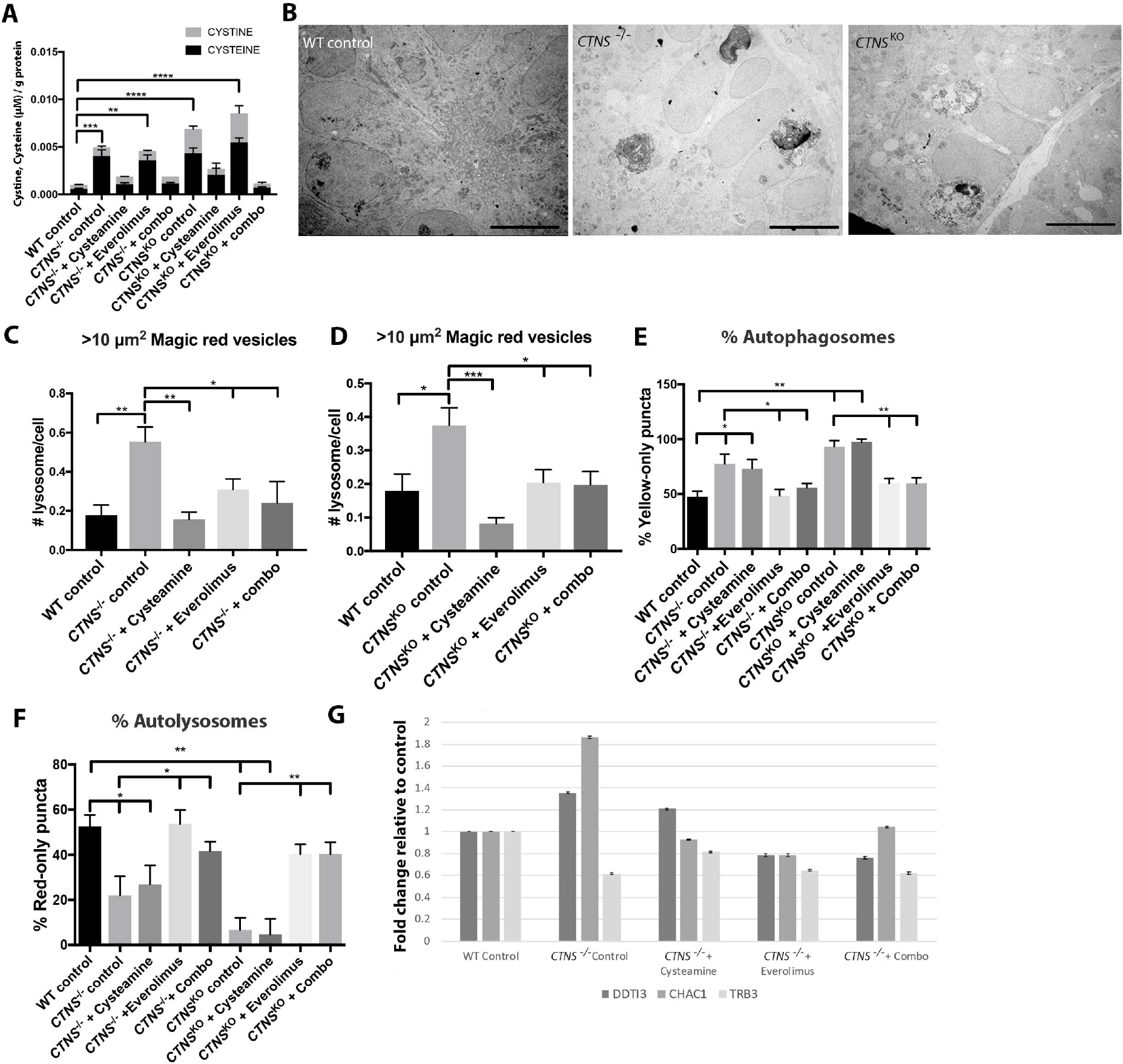
Characterization of cystinotic kidney organoids. **(A)** Amount of cysteine (black) and cystine (grey) (µM) /g of protein in WT and *CTNS*^−/−^ and *CTNS*^KO^ organoids with various treatments. One-way ANOVA performed, **p<0.01, ***p<0.001, **** p<0.0001, data plotted as mean ± SEM, n=30 organoids per experiment, 3 independent experiments. **(B)** Representative transmission electron microscope images of WT, *CTNS*^−/−^ and *CTNS*^KO^ organoids displaying enlarged multivesicular bodies. Scale bar: 5 µm. **(C)** Average number of Magic Red vesicles per cell over 10 µm^2^ in WT, *CTNS*^−/−^ and **(D)** *CTNS*^KO^ organoids. One-way AVOVA performed, *p<0.05, **p<0.01, ***p<0.001, The values are mean ± SEM (n=300 cells from 10 random fields per condition, 10 cells/field, 3 independent experiments). **(E, F)** Effects of cysteamine, Everolimus and combination treatments on autophagy flux as determined by the percentage of yellow and red puncta in day 14 *CTNS*^−/−^ and *CTNS*^KO^ organoids. One way ANOVA performed, *p<0.05, **p<0.01, all data plotted as mean ± SEM, (n= 30 cells from 10 random fields per condition containing ~1-3 cells in 3 independent experiments). **(G)** Quantitative PCR of genes of interest in *CTNS*^−/−^ organoids with various treatments expressed as fold change relative to control, data plotted as mean ± SD.

At the level of light microscopy, both the *CTNS*^−/−^ and *CTNS*^KO^ kidney organoids appear equivalent to the control organoids and we see no evidence of abnormalities such as the characteristic cystinotic ‘swan neck’ lesion (data not shown). However, at the ultrastructural level, we qualitatively observed the presence of numerous enlarged vacuoles in the tubules of cystinotic organoids, reminiscent of the degradative/storage-like bodies seen in *CTNS*-iPSCs, whereas these are rarely seen in control organoids (Figure 5B).

To quantify the enlarged lysosomes in cystinotic organoids, we dissociated the tissue at day 12 into single cells and then incubated them in Magic Red. We found that *CTNS* organoids display ~2-3 fold more enlarged lysosomes compared to control organoids (Figure 5C and D).

To determine if basal autophagy flux is affected in *CTNS* kidney organoids we transfected dissociated single cells with the mCherry-LC3B-GFP sensor plasmid. We found that cystinotic kidney organoid cells display ~1.5-2-fold more autophagosomes compared to control organoid cells, consistent with a defect in basal autophagy (Figure 5E and F).

We next assessed mRNA expression of *DDIT3*, *TRB3*, and *CHAC1* in cystinotic and control kidney organoids using qPCR. We found that *DDIT3* and *CHAC1* were increased in cystinotic kidney organoids compared to controls (Figure 5G and Supplemental Figure S1B). *TRB3* was not up-regulated, indicating that this gene may only be useful as a biomarker of lysosomal dysfunction in certain cell types.

We next examined the effects of single and combination treatment of cysteamine and Everolimus on *CTNS* kidney organoids with respect to the phenotypes of cystine/cysteine loading, basal autophagy flux, and *DDIT3*, *TRB3*, and *CHAC1* expression. To this end we treated day 13 *CTNS* and control kidney organoids with 1 µM cysteamine or 100 nM Everolimus and a combination of both drugs for 24 hrs. In keeping with our observations in *CTNS*-iPSCs, we found that cysteamine alone and when combined with Everolimus reduced cystine and cysteine levels whereas no effect was seen with Everolimus treatment alone (Figure 5A). Both cysteamine or Everolimus alone and combined treatments reduced the numbers of enlarged lysosomes (Figure 5C and D). Everolimus alone activated autophagy, as did combined treatment, but cysteamine alone had no effect (Figure 5E and F). Finally, qPCR analysis of the treated kidney organoids showed that the expression levels of *DDIT3* and *CHAC1* were normalised to near-control values in response to cysteamine alone and combined treatments, as well as Everolimus alone treatment in the case of *CTNS*^−/−^ organoids. However, Everolimus alone treatment had only modest effects on reducing *DDIT3* and *CHAC1* in *CTNS*^KO^ organoids (Figure 5G and Supplemental Figure S1B).

## Discussion

In this report we generated new human models of cystinosis in the form of cystinotic iPSCs and kidney organoids and demonstrated a phenotype characterized by cystine and cysteine loading, enlarged lysosomes, altered gene expression and defective basal autophagy. Using this unique human-based platform we tested the therapeutic effects of cysteamine and Everolimus and found that a combination therapy was able to mitigate all of the observed cystinotic phenotypes in a beneficial and effective manner. Furthermore, these models have the potential to be used as a preclinical model for testing other therapeutics as well as a novel tool to deepen our understanding of the pathogenic mechanisms at play in cystinosis.

Of the defects observed in cystinotic iPSCs and kidney organoids, the block in basal autophagy flux appears most significant as it is the only phenotype that cysteamine is unable to rescue. The finding that the number of autophagosomes in *CTNS*-iPSCs does not significantly increase in the presence of BafA1 may indicate an accumulation of autophagosomes under basal conditions due to a defect in lysosomal fusion. Although CYSTINOSIN is a H^+^ driven transporter^49^ and lysosomal pH is important for lysosomal fusion^62^ previous reports have shown that the pH of cystinotic lysosomes is normal.^12^ Thus, the underlying cause for the failure of lysosomes to fuse with autophagosomes remains unclear. There are a number of protein complexes that coordinate this fusion and a requirement for the physical movement of both autophagosomes and lysosomes. Therefore, is possible that one or more of these processes is affected in cystinotic cells.^63^ The observation that sucrose loading induces the equivalent phenotype in non-cystinotic iPSCs and that other lysosomal storage diseases show basal autophagy flux defects points to the intralysosomal accumulation of indigestible material as a generic cause of the flux defect.^64–67^ Our finding that *CTNS*-iPSCs upregulate a number of genes in the autophagy pathway suggests that compromised basal autophagy leads to the activation of transcriptional feedback mechanisms to compensate for the reduced flux.

If cystine accumulation impairs autophagosome-lysosome fusion and reduces basal autophagy flux then cysteamine treatment would have been expected to have rescued the basal autophagy phenotype of *CTNS*-iPSCs. Interpreting the failure of cysteamine to restore basal autophagy is complicated by a report showing that cysteamine treatment of HeLa cells under basal conditions affects the autophagy pathway in two places: first by acting early to induce autophagosome formation but then acting later to inhibit autolysosome maturation^68^. Thus, it is likely that while cysteamine may restore lysosomal functionality in cystinotic cells by depleting cystine, its independent inhibitory effects on autolysosome maturation induces an equivalent block in basal autophagy flux. This additional activity of cysteamine may be related to its antioxidant properties as reactive oxygen species are known to regulate autophagy via the redox-modification of several autophagy components and antioxidants can inhibit basal autophagy.^69,70^

Several mouse studies have demonstrated a critical role for basal autophagy in maintaining proximal tubule function and support the notion that continued renal dysfunction in cysteamine-treated cystinosis patients is linked to a failure to restore basal autophagy. Specifically, it has been shown that blocking autophagy in kidney cells *in vivo* and *in vitro* under nutrient replete conditions results in the accumulation of degradative vacuoles, deformed mitochondria, and p62- and ubiquitin-positive inclusion bodies.^71–73^ Similar observations have been reported for kidney biopsies and urinary cells from cystinotic patients and in primary proximal tubule cells from the *Ctns*^−/−^ mouse.^26,27,71,74^ Consistent with one of the functions of basal autophagy being the removal of damaged mitochondria, work using cystinotic mouse primary proximal tubule cells has led to a model in which defective autophagy-mediated clearance of damaged mitochondria (mitophagy) causes oxidative stress and triggers renal epithelial cell dysfunction.^27^ While we have not detected an increased level of oxidative stress or deformed mitochondria in *CTNS*-iPSCs (unpublished observations), this may be due to iPSCs relying on glycolysis rather than oxidative phosphorylation for their energy needs.^75^ By contrast, proximal tubule cells contain a large quantity of mitochondria in order to drive membrane transport processes and it is reasonable to conclude that these cells would be highly dependent on efficient basal autophagy for mitochondrial quality control. Other tissues such as hepatocytes, neurons, heart and skeletal muscle have also been found to be dependent on basal autophagy^76^ and although neural and muscle tissues are eventually affected in individuals with cystinosis, it is the proximal tubule that is affected early in the disease. One explanation for this is that the lysosomal system of proximal tubule cells is under a high degradative load due to the uptake and degradation of albumin and other plasma proteins harbouring disulphide bonds.^77,78^ Thus, the level of lysosomal cystine accumulation and the resulting basal autophagy dysfunction (assuming it is proportional to cystine load) would be expected to be pronounced in cystinotic proximal tubule cells in comparison to other tissues.

Prior studies using immortalized mouse and human proximal tubule cells found a transport-independent role for CYSTINOSIN that implicated it in the positive regulation of mTORC1.^40,41^ Despite detecting a slight upregulation of *MTOR* RNA and two of its downstream targets (*ULK1* and *ATG13*) in *CTNS*-iPSCs, we found no evidence to indicate that mTORC1 protein activity is altered. Similarly, no effect in mTORC1 activity was found in mouse cystinotic fibroblasts.^33^ At this stage, the cause for these discrepancies is not known but may be related to metabolic differences between cell lines. Alternatively, there may be confounding influences of the viral antigens used for immortalization as these can influence mTORC1 levels.^79^

We found that inhibiting mTORC1 with Everolimus was capable of overcoming the basal autophagy block, presumably via activation of the stress-induced autophagy pathway. Everolimus also reduced the frequency of enlarged lysosomes in *CTNS*-iPSCs and because this effect was abrogated by 3-MA it is likely that these structures are being cleared by autophagy. This notion is in-line with the discovery that autophagy is involved in the removal/recycling of aged and dysfunctional lysosomes (lysophagy) in order to prevent cell damage caused by the leakage of hydrolytic enzymes^80,81^. At this stage it remains unclear if the basal autophagy block seen in cystinotic cells also results in a lysophagy defect. However, it is an attractive hypothesis as large lysosomes are particularly vulnerable to rupture and some cystinotic cells have been found to be sensitive to apoptosis.^14,24,25,82,83^ Interestingly, if dysfunctional lysosomes are not removed, the total number of lysosomes remains unchanged, even if some of them are non-functional.^81^ Thus, lysosome quality control is critical for maintaining cellular degradative capabilities and therefore lysophagy defects could contribute additional stress to cystinotic proximal tubule cells.

At present, mTOR inhibitors are used in the clinic as immunosuppressants and as treatments for some cancers.^84,85^ Our work suggests that Everolimus, and related Rapamycin derivatives, may also have therapeutic potential to treat cystinosis. This notion is consistent with studies of the lysosomal disorders mucopolysaccharidosis and Niemann-Pick disease, where overcoming a block in autophagic flux improves cell viability.^64,86,87^ In the case of Niemann-Pick disease Type C, a combination therapy of low dose hydroxypropyl-β-cyclodextrin (which depletes the cholesterol accumulating in the lysosome) coupled with an autophagy stimulator has been proposed^88^. We have arrived at the same conclusion with our findings and hypothesize that dual treatment of cystinotic individuals with cysteamine and an autophagy inducer such as Everolimus will improve long-term outcomes. The next step in testing this hypothesis requires animal studies, although this will be challenging with the current *Ctns*^−/−^ mouse model as it does not fully recapitulate the progression of the human disease and shows a relatively late chronic kidney failure phenotype.^14,89^ In considering these experiments, careful attention will need to be given to finding the lowest effective dosing schedule, as mTOR inhibitors have metabolic side-effects that include dyslipidemia and impaired glucose homeostasis which may complicate their long-term use in cystinosis patients.^90^

## Author Contribution

J.A.H, P.T.H., T.M.H and A.J.D. conceptualized the study, J.A.H. and T.M.H. and A.P. designed and performed experiments. T.M.H. established the cystinotic patient lines. E.J.W. generated the CRL1502 iPSC line and reviewed manuscript. A.J.D and T.M.H co-supervised the study. J.A.H. and A.J.D. wrote the manuscript. P.T.H., T.M.H and A.J.D. acquired funding.

## Supplemental Material Table of Contents

**Supplemental Figure S1 2**

**Supplemental Figure S2 3**

**Figure Legends 4**

**Supplemental Information 5**

*Supplemental Table 1. List of primers for qPCR*

*Supplemental Table 2. List of Primary antibodies for western blot and immunohistochemistry*

*Supplemental Table 3. List of Secondary antibodies for western blot and immunohistochemistry*

**Supplemental Table S4 – List of differentially expressed genes**

## References

1. Gahl WA, Thoene JG & Schneider JA: Cystinosis. N. Engl. J. Med. 347: 111–121, 2002.

2. Kalatzis V, Nevo N, Cherqui S, Gasnier B, Antignac C: Molecular pathogenesis of cystinosis: effect of CTNS mutations on the transport activity and subcellular localization of cystinosin. Hum Mol Genet. 13: 1361–1371, 2004.

3. Emma F, Nesterova G, Langman C, Labbé A, Cherqui S, Goodyear P, et al.: Nephropathic cystinosis: an international consensus document. Nephrol Dial Transpl. 29: 87–94, 2014.

4. Scarvie KM, Ballantyne AO, Trauner DA: Visuomotor performance in children with infantile nephropathy cystinosis. Percept Motor Skill. 82: 67–75. 1996.

5. Trauner DA, Williams J, Ballantyne AO, Spilkin AM, Crowhurst J, Hesselink J: Neurological impairment in nephropathic cystinosis: motor coordination deficits. Pediatric Nephrol Berlin Ger. 25: 2061–2066, 2010.

6. Wilmer MJ, Emma F, Levtchenko EN: The pathogenesis of cystinosis: mechanisms beyond cystine accumulation. Am J Physiol-renal. 299: 905–916, 2010.

7. Ivanova E, Leo M, Matteis M, Levtchenko E: Cystinosis: clinical presentation, pathogenesis and treatment. Pediatric Endocrinol Rev Per 12 Suppl. 1: 176–84, 2014.

8. Gahl WA, Reed GF, Thoene JG, Schulam JD, Rizzo WB, Jonas AJ, et al.: Cysteamine Therapy for Children with Nephropathic Cystinosis. New Engl J Medicine. 316: 971–977, 1987.

9. Levtchenko EN, van Dael CM, de Graaf-Hess AC, Wilmer MJ, van den Heuvel LP, Monnens, L.A, et al.: Strict cysteamine dose regimen is required to prevent nocturnal cystine accumulation in cystinosis. Pediatr Nephrol. 21: 110–113, 2006.

10. Gahl WA, Balog JZ, Kleta R: Nephropathic cystinosis in adults: natural history and effects of oral cysteamine therapy. Ann Intern Med.147: 242–250, 2007.

11. Raggi C, Luciani A, Nevo N, Antignac C, Terryn S, Devuyst O: Dedifferentiation and aberrations of the endolysosomal compartment characterize the early stage of nephropathic cystinosis. Hum Mol Genet. 23: 2266–2278, 2014.

12. Ivanova EA, De Leo MG, Van Den Heuvel L, Pastore L, Dijkman H, De Matteis MA, Levtchenko E.N: Endo-lysosomal dysfunction in human proximal tubular epithelial cells deficient for lysosomal cystine transporter cystinosin. Plos One. 26: 10(3):0120998, 2015.

13. Schulman JD, Bradley KH, Seegmiller JE: Cystine: Compartmentalization within lysosomes in cystinotic leukocytes. Science. 166: 1152–1154, 1969.

14. Gaide Chevronnay HP, Janssens V, Van Der Smissen P, N’Kuli F, Nevo N, Levtchenko E, et al.: Time course of pathogenic and adaptation mechanisms in cystinotic mouse kidneys. J Am Soc Nephrol. 25: 1256–1269, 2014.

15. Coor C, Salmon RF, Quigley R, Marver D, Baum M: Role of adenosine triphosphate (ATP) and NaK ATPase in the inhibition of proximal tubule transport with intracellular cystine loading. J Clin Invest. 87: 955–61, 1991.

16. Levtchenko E, de Graaf-Hess A, Wilmer M, van den Heuvel L, Monnens L, Bloom H: Altered status of glutathione and its metabolites in cystinotic cells. Nephrol Dial Transpl. 20: 1828–1832, 2005.

17. Wilmer MJ, de Graaf-Hess A, Blom HJ, Dijkman HB, Monnens LA, van den Heuvel LP, et al.: Elevated oxidised glutathione in cystinotic proximal tubular epithelial cells. Biochem Biophys Res Commun.8: 610–4, 2005.

18. Wilmer MJ, Christensen EI, van den Heuvel LP, Monnens LA, Levtchenko EN: Urinary protein excretion pattern and renal expression of megalin and cubilin in nephropathic cystinosis. Am J Kidney Dis. 51: 893–903, 2008.

19. Wilmer MJ, Kluijtmans L, van der Velden TJ, Willems PH, Scheffer PG, Masereeuw R, et al.: Cysteamine restores glutathione redox status in cultured cystinotic proximal tubular epithelial cells. Biochimica Et Biophysica Acta. 1812: 643–651, 2011.

20. Chol M, Nevo N, Cherqui S, Antignac C, Rustin P: Glutathione precursors replenish decreased glutathione pool in cystinotic cell lines. Biochem Bioph Res Co. 324: 231–235, 2004.

21. Mannucci L, Pastore A, Rizzo C, Piemonte F, Rizzoni G, Emma F: Impaired activity of the gamma-glutamyl cycle in nephropathic cystinosis fibroblasts. Pediatr res. 59: 332–5, 2006.

22. Laube GF, Shah V, Stewart VC, Hargreaves IP, Haq MR, Heales SJ, et al.: Glutathione depletion and increased apoptosis rate in human cystinotic proximal tubular cells. Pediatr Nephrol. 21: 503–509, 2006.

23. Cherqui S, Sevin C, Hamard G, Kalatzis V, Sich M, Pequignot MO, et al.: Intralysosomal Cystine Accumulation in Mice Lacking Cystinosin, the Protein Defective in Cystinosis. Mol Cell Biol. 22: 7622–7632, 2002.

24. Park M, Helip-Wooley A, Thoene J: Lysosomal cystine storage augments apoptosis in cultured fibroblasts and renal tubular epithelial cells. J Am Soc Nephrol. 13: 878–87, 2002.

25. Park MA, Pejovic V, Kerisit KG, Junis S, Thoene JG: Increased apoptosis in cystinotic fibrolblasts and renal proximal tubule epithelial cells results from cysteinylation of protein kinase Cdelta. J Am Soc Nephrol. 17: 3167–75, 2006.

26. Sansanwal P, Yen B, Gahl WA, Ma Y, Ying l, Wong LJ, et al.: Mitochondrial autophagy promotes cellular injury in nephropathic cystinosis. J Am Soc Nephrol. 21: 272–283, 2010.

27. Festa BP, Chen Z, Berquez M, Debaix H, Tokonami N, Prange JA, et al.: Impaired autophagy bridges lysosomal storage disease and epithelial dysfunction in the kidney. Nat Commun.9: 161–177, 2018.

28. Axe EL, Walker SA, Manifava M, Chandra P, Roderick HL, Habermann A, et al.: Autophagosome formation from membrane compartments enriched in phosphatidylinositol 3-phosphate and dynamically connected to the endoplasmic reticulum. J Cell Biology 182: 685–701, 2008.

29. Mauvezin C & Neufeld TP: Bafilomycin A1 disrupts autophagic flux by inhibiting both V-ATPase-dependent acidification and Ca-P60A/SERCA-dependent autophagosome-lysosome fusion. Autophagy 11: 1437–1438, 2015.

30. Into T, Horie T, Inomata M, Gohda J, Inoue J, Murakami Y, et al.: Basal autophagy prevents autoactivation or enhancement of inflammatory signals by targeting monomeric MyD88. Sci Rep. 7: 1009, 2017.

31. Casares-Crespo L, Calatayud-Baselga I, García-Corzo L, Mira H: On the role of basal autophagy in adult neural stem cells and neurogenesis. Frontiers in cellular neuroscience, 12: 339, 2018.

32. Seino J, Wang L, Harada Y, Huang C, Ishii K, Mizushima N, Suzuki T; Basal autophagy is required for the efficient catabolism of sialyloligosaccharides. The Journal of Biological Chemistry. 288: 26898–26907, 2013.

33. Napolitano G, Johnson JL, He J, Rocca CJ, Monfregola J, Pestonjamasp K, et al.: Impairment of chaperone-mediated autophagy leads to selective lysosomal degradation defects in the lysosomal storage disease cystinosis. Embo Mol Med. 7: 158–174, 2015.

34. Lieberman AP, Puertollano R, Raben N, Slaughenhaupt S, Walkley SU, Ballabio A: Autophagy in lysosomal storage disorders. Autophagy. 8: 719–30, 2012.

35. Efeyan A, Zoncu R, Sabatini DM: Amino acids and mTORC1: from lysosomes to disease. Trends Mol Med. 18: 524–533, 2012.

36. Sancak Y, Peterson TR, Shaul YD, Lindquist RA, Thoreen CC, Bar-Peied L, et al.: The rag GTPases bind raptor and mediate amino acid signalling to mTORC1. Science. 320: 1496–1501, 2008.

37. Sancak Y, Bar-Peled L, Zoncu R, Markhard AL, Nada S & Sabatini DM: Ragulator-Rag complex targets mTORC1 to the lysosomal surface and is necessary for its activation by amino acids. Cell. 141: 290–303, 2010.

38. Gonzalez E, Andres A, Polanco N, Hernández A, Morales E, Hernadez E, et al.: Everolimus represents an advance in immunosuppression for patients who have developed cancer after renal transplantation. Transplant Proc. 41: 2332–3, 2009.

39. Gude E, Gullestad L, Andreassen AK: Everolimus immunosuppression for renal protection, reduction of allograft vasculopathy and prevention of allograft rejection in de-novo heart transplant recipients: could we have it all? Curr Opin Organ Transplant. 22: 198–206, 2017.

40. Andrzejewska Z, Nevo N, Thomas L, Chhuon C, Bailleux A, Chauvet V, et al.: Cystinosin is a component of the vacuolar H+-ATPase-Ragulator-Rag complex controlling mammalian target of rapamycin complex 1 signaling. J Am Soc Nephrol. 27: 1678–1688, 2015.

41. Ivanova EA, van den Heuvel LP, Elmonem MA, De Smedt H, Missiaen L, Pastore A, et al.: Altered mTOR signalling in nephropathic cystinosis. J Inherit Metab Dis. 39: 457–464, 2016.

42. Briggs JA, Sun J, Shepherd J, Ovchinnikov DA, Chung TL, Nayler SP, et al.: Integration-free induced pluripotent stem cells model genetic and neural developmental features of down syndrome etiology. Stem Cells. 31: 467–478, 2013.

43. Zhu F, Sun B, Wen Y, Wang Z, Reijo Pera R, Chen B: A modified method for implantation of pluripotent stem cells under the rodent kidney capsule. Stem Cells Dev. 23: 2119–25, 2014.

44. Bae S, Park J & Kim JS: Cas-OFFinder: A fast and versatile algorithm that searches for potential off-target sites of Cas9 RNA-guided endonucleases. Bioinformatics. 30: 1473–1475, 2014.

45. Cradick TJ, Qiu P, Lee CM, Fine EJ, Bao G: COSMID: A web-based tool for identifying and validating CRISPR/Cas off-target sites. Molecular therapy. Nucleic acids 3: e214, 2014.

46. Przepiorski A, Sander V, Tran T, Hollywood JA, Sorrenson B, Shih JH, et al.: A simple bioreactor-based method to generate kidney organoids from pluripotent stem cells Stem cell reports. 11: 470–484, 2018.

47. Ng ES, Davis R, Stanley EG, Elefanty AG: A protocol describing the use of recombinant protein-based, animal product-free medium (APEL) for human embryonic stem cell differentiation as spin embryoid bodies. Nat. Protoc. 3: 768–76, 2008.

48. Livak KJ & Schmittgen: T.D. Analysis of relative gene expression data using real-time quantitative PCR and 2(-Delta Delta C(T)) method. Methods. 25: 402–8, 2001.

49. Kalatzis V, Cherqui S, Antignac C, Gasnier B: Cystinosin, the protein defective in cystinosis, is a H+-driven lysosomal cystine transporter. Embo J. 20: 5940–5949, 2001.

50. Cohn ZA, Ehrenreich BA: The uptake, storage, and intracellular hydrolysis of carbohydrates by macrophages. J Exp Med. 129: 201–25, 1969.

51. DeCourcy K, Storrie B: Osmotic swelling of endocytic compartments induced by internalized sucrose is restricted to mature lysosomes in cultured mammalian cells. Exp Cell Res. 192: 52–60, 1991.

52. Beck M; The link between lysosomal storage disorders and more common diseases. Journal of Inborn Errors of Metabolism and Screening. 2016.

53. Ryoo H: Long and short (timeframe) of endoplasmic reticulum stress-induced cell death. Febs J. 283: 3718–3722, 2016.

54. Ohoka N, Yoshii S, Hattori T, Onozaki K, Hayashi H: TRB3, a novel ER stress-inducible gene, is induced via ATF4-CHOP pathway and is involved in cell death. EMBO J. 24: 1243–55, 2005.

55. Kumar A, Tikoo S, Maity S, Sengupta S, Sengupta S, Kaur A & Bachhawat AK: Mammalian proapoptotic factor ChaC1 and its homologues function as γ-glutamyl cyclotransferases acting specifically on glutathione. EMBO reports. 13: 1095–101, 2012.

56. Bellomo F, Corallini S, Pastore A, Palma A, Laurenzi C, Emma F, Taranta A: Modulation of CTNS gene expression by intracellular thiols. Free Radical Bio Med. 48: 865–872, 2010.

57. Pearson R, Dennis P, Han J, Williamson NA, Kozma SC, Wettenhall RE, et al.: The principal target of rapamycin-induced p70s6k inactivation is a novel phosphorylation site within a conserved hydrophobic domain. Embo J. 14: 5279–5287, 1995.

58. Burnett PE, Barrow RK, Cohen NA, Snyder SH, Sabantini DM: RAFT1 phosphorylation of the translational regulators p70 S6 kinase and 4E-BP1. Proc National Acad Sci. 95: 1432–1437, 1998.

59. Kimura S, Noda T, Yoshimori T: Dissection of the autophagosome maturation process by a novel reporter protein, tandem fluorescent-tagged LC3. Autophagy. 3: 452–460, 2007.

60. Yoshimori T, Yamamoto A, Moriyama Y, Futai M & Tashiro Y: Bafilomycin A1, a specific inhibitor of vacuolar-type H^+^-ATPase, inhibits acidification and protein degradation in lysosomes of cultured cells. J Biol Chem. 266: 17707–17712, 1999.

61. Seglen PO, Gordon PB: 3-Methyladenine: specific inhibitor of autophagic/lysosomal protein degradation in isolated rat hepatocytes. Proc Natl Acad Sci U S A. 79: 1889–92, 1982.

62. Kawai A, Uchiyama H, Takano S, Nakamura N, Ohkuma S: Autophagosome-lysosome fusion depends on the pH in acidic compartments in CHO cells. Autophagy 2: 154–7, 2007.

63. Nakamura S & Yoshimori T: New insights into autophagosome–lysosome fusion. J Cell Sci. jcs.196352, 2017.

64. Sarkar S, Carroll B, Buganim Y, Maetzel D, Ng AHM, Cassady JP, et al.: Impaired autophagy in the lipid-storage disorder Niemann-Pick Type C1 disease. Cell Rep. 5: 1302–1315, 2013.

65. Guo H, Zhao M, Qiu X, Deis JA, Huang H, Tang QQ, et al.: Niemann-Pick type C2 deficiency impairs autophagy-lysosomal activity, mitochondrial function, and TLR signalling in adipocytes. J. Lipid Res. 57: 1644–1658, 2016.

66. Award O, Sarkar C, Panicker LM, Miller D, Zeng X, Sgambato JA, et al.: Altered TFEB-mediated lysosomal biogenesis in Gaucher disease iPSC-derived neuronal cells. Hum. Mol. Genet. 24: 5775–5788, 2015.

67. Seranova E, Connolly KJ, Zatyka M, et al.: Dysregulation of autophagy as a common mechanism in lysosomal storage diseases. Essays Biochem. 61: 733–749, 2017.

68. Wan XM, Zheng F, Zhang L, Miao YY, Man N, Wen LP: Autophagy-mediated chemosensitization by Cysteamine in cancer cells. Int J Cancer. 129: 1087–95, 2011.

69. Pajares M, Jiménez-Moreno N, Dias IH, Debelec B, Vucetic M, Fladmark KE, et al.: Redox control of protein degradation. Redox biology. 6: 409–20, 2015.

70. Underwood BR, Imariso S, Fleming A, Rose C, Krishna G, Heard P, et al.: Antioxidants can inhibit basal autophagy and enhance neurodegeneration in models of polyglutamine disease. Hum Mol Genet. 19: 3413–3429, 2010.

71. Kimura T, Takabatake Y, Takahashi A, Kaimori JY, Matsui I, Namba T, et al.: Autophagy protects the proximal tubule from degeneration and acute ischemic injury. J Am Soc Nephrol. 22: 902–13, 2011.

72. Isaka Y, Kimura T, Takabatake Y: The protective role of autophagy against aging and acute ischemic injury in kidney proximal tubular cells. Autophagy. 7: 1085–7, 2011.

73. Komatsu M, Waguri S, Ueno T, et al.: Impairment of starvation-induced and constitutive autophagy in Atg7-deficient mice. J Cell Biol. 169: 425–34, 2005.

74. Jiang M, Wei Q, Dong G, Komatsu M, Su Y, Dong Z. Autophagy in proximal tubules protects against acute kidney injury. Kidney Int. 82: 1271–83, 2012.

75. Varum S, Rodrigues AS, Moura MB, Momcilovic O, Easley CA, Ramalho-Santos J, et al.: Energy metabolism in human pluripotent stem cells and their differentiated counterparts. PloS one. 6: e20914, 2011.

76. Levine B & Kroemer G: Autophagy in the pathogenesis of disease. Cell. 132: 27–42, 2008.

77. Liu S, Hartleben B, Kretz O, Wiech T, Igarashi T, Mizushima N, et al.: Autophagy plays a critical role in kidney tubule maintenance, aging and ischemia-reperfusion injury. Autophagy. 8: 826–37, 2012.

78. Cherqui S & Courtoy PJ: The renal Fanconi syndrome in cystinosis: pathogenic insights and therapeutic perspectives. Nature reviews. Nephrology. 13: 115–131, 2016.

79. Yu Y, Kudchodkar SB, Alwine JC: Effects of simian virus 40 large and small tumor antigens on mammalian target of rapamycin signaling: small tumor antigen mediates hypophosphorylation of eIF4E-binding protein 1 late in infection. J Virol. 79: 6882–9, 2005.

80. Papadopoulos C & Meyer H: Detection and clearance of damaged lysosomes by the endo-lysosomal damage response and lysophagy. Curr Biol. 27: R1330–R1341, 2017.

81. Hung YH, Chen LM, Yang JY, Yang WY: Spatiotemporally controlled induction of autophagy-mediated lysosome turnover. Nat Commun. 4: 2111, 2013.

82. Ono K, Kim SO, Han J: Susceptibility of lysosomes to rupture is a determinant for plasma membrane disruption in tumor necrosis factor alpha-induced cell death. Molecular and cellular biology. 23: 665–76, 2003.

83. Galarreta CI, Forbes MS, Thornhill BA, Antignac C, Gubler MC, Nevo N, et al.: The swan-neck lesion: proximal tubular adaptation to oxidative stress in nephropathic cystinosis. Am J Physiol Renal Physiol. 308: F1155–1166, 2015.

84. Thomson AW, Turnquist HR & Raimondi G: Immunoregulatory functions of mTOR inhibition. Nature reviews. Immunology. 9: 324–37, 2009.

85. Zaytseva YY, Valentino JD, Gulhati P, Evers BM: mTOR inhibitors in cancer therapy. Cancer Lett. 319: 1–7, 2012.

86. Bartolomeo R, Cinque L, De Leonibus C, Forrester A, Salzano AC, Monfregola J, et al.: mTORC1 hyperactivation arrests bone growth in lysosomal storage disorders by suppressing autophagy. The Journal of clinical investigation. 127: 3717–3729, 2017.

87. Raben N, Hill V, Shea L, Takikita S, Baum R, Mizushima N, et al.: Suppression of autophagy in skeletal muscle uncovers the accumulation of ubiquitinated proteins and their potential role in muscle damage in Pompe disease. Human molecular genetics. 17: 3897–908, 2008.

88. Maetzel D, Sarkar S, Wang H, Abi-Mosleh L, Xu P, Cheng AW, et al.: Genetic and chemical correction of cholesterol accumulation and impaired autophagy in hepatic and neural cells derived from Niemann–Pick type C patient-specific iPS cells. Stem Cell Rep. 2: 866–880, 2014.

89. Nevo N, Chol M, Bailleux A, Kalatzis V, Morisset L, Devuyst O, et al.: Renal phenotype of the cystinosis mouse model is dependent upon genetic background. Nephrol Dial Transpl. 25: 1059–1066, 2010.

90. Morviducci L, Rota F, Rizza L, Di Giacinto P, Ramponi S, Nardone MR, et al.: Everolimus is a new anti-cancer molecule: Metabolic side effects as lipid disorders and hyperglycemia. Diabetes Res. Clin. Pract. 143: 428–431, 2018.

