## Supplemental information and Figures S1,S2 for "Use of human iPSCs and kidney organoids to develop a cysteamine/mTOR inhibition combination therapy to treat cystinosis"

***Supplemental Table 1. List of primers for qPCR***

***Supplemental Table 2. List of Primary antibodies for western blot and immunohistochemistry***

***Supplemental Table 3. List of Secondary antibodies for western blot and immunohistochemistry***

**Supplemental Table S4 – List of differentially expressed genes**

### Supplementary Figure S1

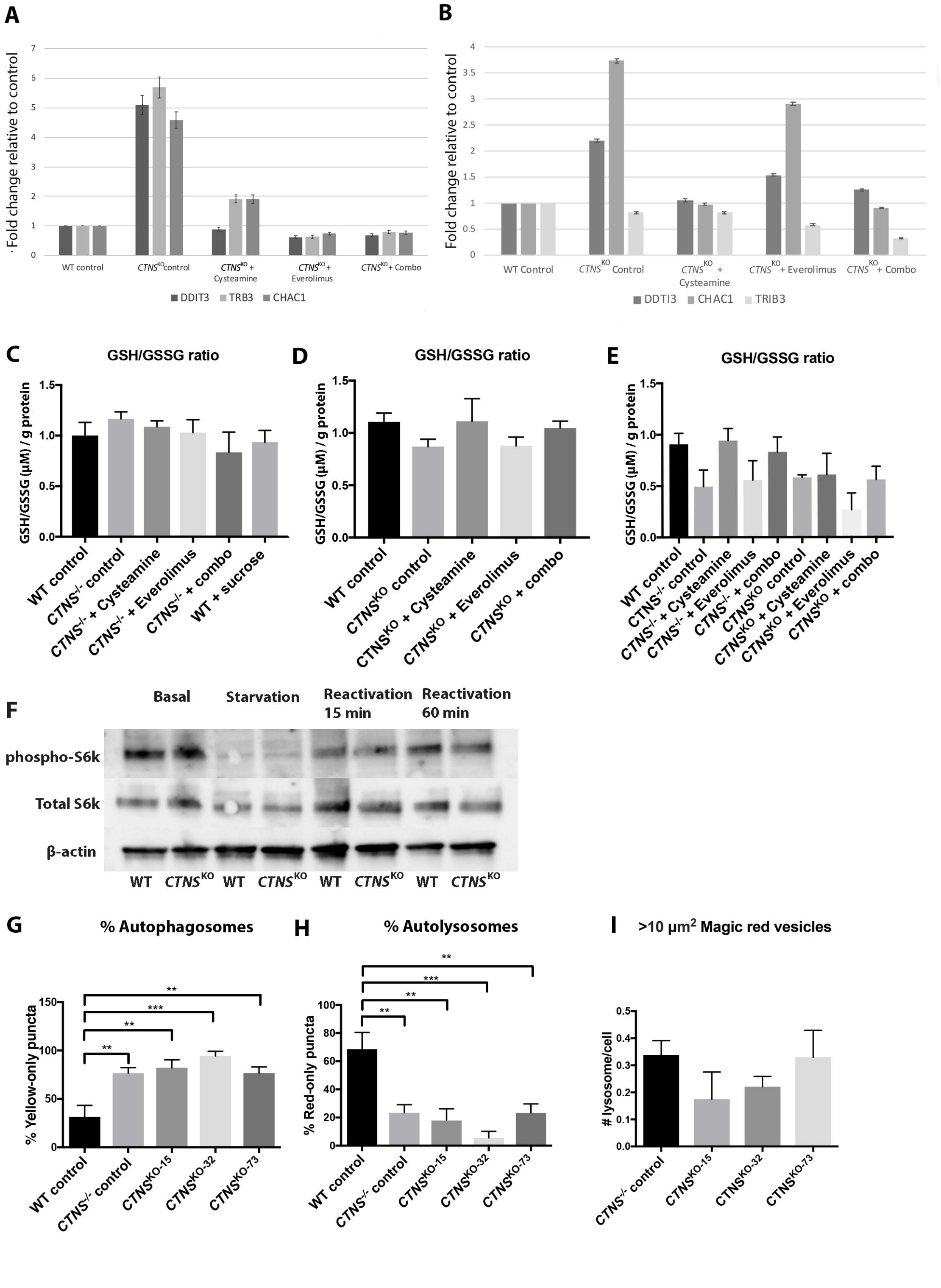

**Supplemental Figure S2**

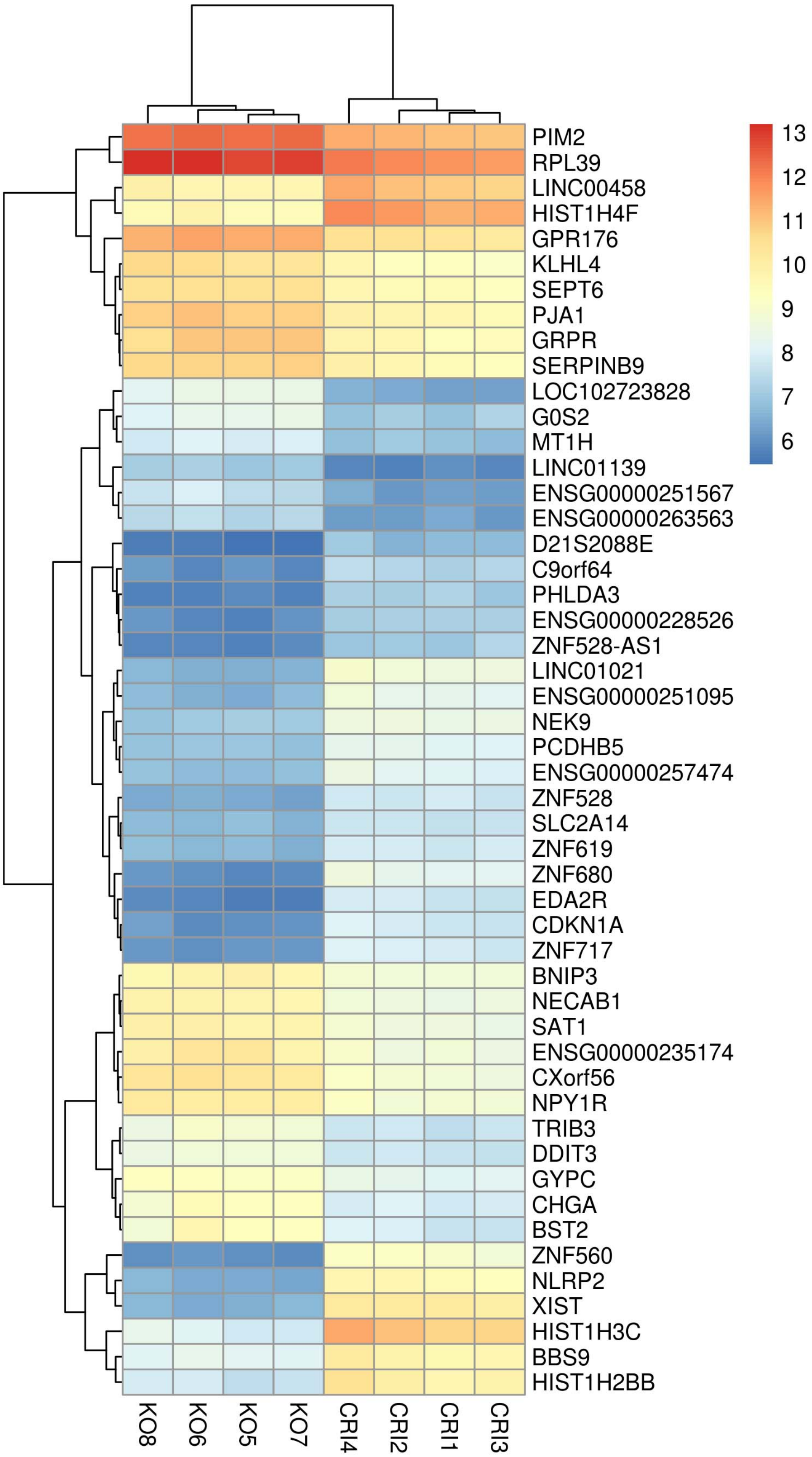

**Figure Legends:**

**Supplemental Figure S1. *CTNS*-iPS cells and organoids have unchanged GSH/GSSG ratio**

**(A)** Quantitative PCR with various treatments in $CTNS^{KO}$ -iPSCs and **(B)** $CTNS^{KO}$ organoids expressed as fold change relative to control, data plotted as mean ± SD. **(C)** Ratio of GSH/GSSG(µM) /g of protein in WT and $CTNS^{-/-}$ iPSCs with various treatments **(D)** Ratio of GSH/GSSG in WT and $CTNS^{KO}$ iPS cells with various treatments. **(E)** Ratio of GSH/GSSG/g of protein in WT and $CTNS^{-/-}$ and $CTNS^{KO}$ organoids with various treatments. One-way ANOVA performed, non-significant**,** data plotted as mean ± SEM, n=3, 3 independent experiments. **(F)** Representative Western blot against phosphorylated and total S6K under different feeding conditions in WT and $CTNS^{KO}$-iPSCs (representative of 3 independent experiments). **(G, H)** Percentage of yellow-only and red-only puncta in all three $CTNS^{KO}$ iPS cell lines compared to $CTNS^{-/-}$ and WT iPSCs. One-way ANOVA performed, **p<0.01, ***p<0.001, data is plotted as mean ± SEM, (n= 30 cells from 10 random fields per condition containing ~1-3 cells in 3 independent experiments). **(I)** Bar graph showing the average number of cells with lysosomes over 10 $\mu m^2$ in $CTNS^{-/-}$ and all three $CTNS^{KO}$-iPSCs. One-way ANOVA performed, non-significant**,** data plotted as mean ± SEM, (n=600 cells from 10 random fields per condition, 20 cells/field, 3 independent experiments).

**Supplemental Figure S2**. Top 50 differentially expressed genes in $CTNS^{KO}$-iPSCs and isogenic WT control cells. (FDR<0.05)

### Supplemental Information:

*Supplemental Table 1. List of primers for qPCR*

| Gene | *Forward primer 5'-3'* | *Reverse primer 5'-3'* |
|---|---|---|
| ***DDTI*3** | agaaccaggaaacggaaacaga | tctccttcatgcgctgcttt |
| ***CHAC*1** | gccctgtggattttcgggta | atcttgtcgctgcccctatg |
| ***TRB*3** | cccacctactgctccagatcgtgcaa | cctggacggggtacaccttgcaggtata |

*Supplemental Table 2. List of Primary antibodies for western blot and immunohistochemistry*

| Primary Antibody | Source | Product code | Dilution used |
|---|---|---|---|
| **Rabbit anti-LC3B** | Cell signalling | 3868S | 1:1000 |
| **Rabbit anti-P70 S6 kinase** | Cell signalling | 2708S | 1:1000 |
| **Rabbit anti-P-p70 S6 kinase (T389)** | Cell signalling | 9205S | 1:500 |
| **Rat anti-LAMP1** | Abcam | Ab25245 | 1:100 |
| **Mouse-anti β -actin** | Sigma | A1978 | 1:20,000 |

*Supplemental Table 3. List of Secondary antibodies for western blot and immunohistochemistry*

| Secondary Antibody | Source | Product code | Dilution |
|---|---|---|---|
| **Goat anti-Rabbit IgG-HRP** | Santa Cruz | Sc-2054 | 1:20,000 |
| **Anti-mouse IGg** | Sigma | A9044 | 1:20,000 |
| **Anti- rat Alexa Fluor 488** | Invitrogen | A-21210 | 1:500 |
