## Supplemental Table S4 for "Use of human iPSCs and kidney organoids to develop a cysteamine/mTOR inhibition combination therapy to treat cystinosis"

**Supplemental Table S4 – List of differentially expressed genes**

| | baseMean | log2FoldChange | lfcSE | stat | pvalue | padj | hgnc_symbol | entrez |
|---|---|---|---|---|---|---|---|---|
| ENSG000002 | 530.164071 | -4.519618012 | 0.1543327 | -29.284903 | 1.61E-188 | 6.55E-184 | XIST | 7503 |
| ENSG000000 | 359.043624 | -4.190680331 | 0.16845556 | -24.877067 | 1.32E-136 | 2.67E-132 | NLRP2 | 55655 |
| ENSG000001 | 239.935738 | -4.754609863 | 0.2092319 | -22.724116 | 2.59E-114 | 3.50E-110 | ZNF560 | 147741 |
| ENSG000002 | 1127.5507 | -3.21178865 | 0.17006998 | -18.885101 | 1.51E-79 | 1.53E-75 | HIST1H3C | 8352 |
| ENSG000002 | 202.862934 | -3.125251911 | 0.16913041 | -18.478356 | 3.08E-76 | 2.50E-72 | LINC01021 | 643401 |
| ENSG000001 | 136.359574 | -4.059132533 | 0.22297934 | -18.204075 | 4.79E-74 | 3.24E-70 | ZNF680 | 340252 |
| ENSG000002 | 148.153495 | 2.998121958 | 0.18699359 | 16.0332871 | 7.48E-58 | 4.33E-54 | LOC102723828 | 102723828 |
| ENSG000001 | 79.8025928 | -3.883056397 | 0.25010586 | -15.525651 | 2.33E-54 | 1.18E-50 | EDA2R | 60401 |
| ENSG000002 | 96.7985154 | -3.256302766 | 0.21194158 | -15.364153 | 2.85E-53 | 1.28E-49 | ZNF717 | 100131827 |
| ENSG000002 | 545.708404 | -2.536348455 | 0.16983839 | -14.933893 | 1.98E-50 | 8.04E-47 | HIST1H2BB | 3018 |
| ENSG000001 | 90.2534776 | -3.094859182 | 0.22300088 | -13.878237 | 8.58E-44 | 3.16E-40 | CDKN1A | 1026 |
| ENSG000001 | 190.431291 | -2.084979086 | 0.15109277 | -13.799331 | 2.57E-43 | 8.69E-40 | NEK9 | 91754 |
| ENSG000002 | 154.589929 | -2.592821198 | 0.18994414 | -13.650441 | 2.01E-42 | 6.26E-39 | NA | NA |
| ENSG000002 | 1854.09128 | -2.033737942 | 0.14977642 | -13.578492 | 5.37E-42 | 1.56E-38 | HIST1H4F | 8361 |
| ENSG000001 | 535.374351 | -1.807721828 | 0.14522789 | -12.447484 | 1.44E-35 | 3.90E-32 | BBS9 | 27241 |
| ENSG000001 | 92.0732906 | -2.237549899 | 0.18963802 | -11.799057 | 3.95E-32 | 1.00E-28 | ZNF528 | 84436 |
| ENSG000000 | 814.114002 | 1.575522045 | 0.13645534 | 11.5460637 | 7.73E-31 | 1.84E-27 | CXorf56 | 63932 |
| ENSG000002 | 142.841311 | -2.076323831 | 0.18114664 | -11.462116 | 2.04E-30 | 4.61E-27 | NA | NA |
| ENSG000001 | 44.4718355 | -2.98183992 | 0.26385973 | -11.300853 | 1.30E-29 | 2.77E-26 | PHLDA3 | 23612 |
| ENSG000001 | 142.562007 | -1.831308054 | 0.16257994 | -11.264047 | 1.97E-29 | 4.00E-26 | PCDHB5 | 26167 |
| ENSG000001 | 57.7644533 | -2.606873261 | 0.2409884 | -10.817422 | 2.85E-27 | 5.50E-24 | C9orf64 | 84267 |
| ENSG000001 | 559.533722 | 1.307742891 | 0.12445223 | 10.5079905 | 7.94E-26 | 1.46E-22 | NECAB1 | 64168 |
| ENSG000002 | 80.4177608 | 2.325417602 | 0.22540606 | 10.3165708 | 5.93E-25 | 1.05E-21 | NA | NA |
| ENSG000001 | 361.775572 | 1.549407036 | 0.15153918 | 10.2244649 | 1.54E-24 | 2.60E-21 | CHGA | 1113 |
| ENSG000002 | 41.0714723 | -2.707869228 | 0.26541335 | -10.202461 | 1.93E-24 | 3.14E-21 | ZNF528-AS1 | 102724105 |
| ENSG000001 | 105.827759 | -1.780341869 | 0.17588569 | -10.122153 | 4.41E-24 | 6.87E-21 | ZNF619 | 285267 |
| ENSG000002 | 47.642474 | -2.49788089 | 0.24707437 | -10.109834 | 5.00E-24 | 7.50E-21 | NA | NA |

| | | | | | | | | |
|---|---|---|---|---|---|---|---|---|
| ENSG000001 | 1312.41219 | 1.21159871 | 0.12019844 | 10.0799873 | 6.77E-24 | 9.81E-21 | PJA1 | 64219 |
| ENSG000001 | 3641.63111 | 1.161532164 | 0.11580132 | 10.0303877 | 1.12E-23 | 1.57E-20 | PIM2 | 11040 |
| ENSG000002 | 41.2897417 | 2.638406795 | 0.26337225 | 10.0177857 | 1.27E-23 | 1.72E-20 | LINC01139 | 339535 |
| ENSG000001 | 162.078201 | 1.723292312 | 0.17324328 | 9.94723879 | 2.59E-23 | 3.39E-20 | G0S2 | 50486 |
| ENSG000001 | 717.562925 | 1.239950532 | 0.12490622 | 9.92705159 | 3.18E-23 | 4.02E-20 | NPY1R | 4886 |
| ENSG000002 | 1440.35551 | -1.360878654 | 0.13909579 | -9.7837517 | 1.32E-22 | 1.62E-19 | LINC00458 | 100507428 |
| ENSG000002 | 65.6863537 | 2.114918251 | 0.21772275 | 9.71381369 | 2.63E-22 | 3.14E-19 | NA | NA |
| ENSG000001 | 589.791151 | 1.243973908 | 0.12864484 | 9.66983116 | 4.05E-22 | 4.69E-19 | SAT1 | 6303 |
| ENSG000002 | 27.0937181 | -2.99720099 | 0.31031127 | -9.6586922 | 4.52E-22 | 5.09E-19 | D21S2088E | 266917 |
| ENSG000001 | 1242.54867 | 1.338935565 | 0.13876869 | 9.64868626 | 4.98E-22 | 5.46E-19 | GRPR | 2925 |
| ENSG000001 | 366.353997 | 1.695749456 | 0.17826003 | 9.51278558 | 1.86E-21 | 1.98E-18 | BST2 | 684 |
| ENSG000002 | 119.771237 | 1.591136504 | 0.17036266 | 9.33970236 | 9.66E-21 | 1.00E-17 | MT1H | 4496 |
| ENSG000001 | 1006.37046 | 1.032350138 | 0.11158924 | 9.25134128 | 2.22E-20 | 2.25E-17 | Sep-06 | 23157 |
| ENSG000001 | 1193.78811 | 1.157078448 | 0.12573364 | 9.2026164 | 3.49E-20 | 3.46E-17 | SERPINB9 | 5272 |
| ENSG000001 | 388.871196 | 1.130977521 | 0.12475344 | 9.06570184 | 1.24E-19 | 1.20E-16 | GYPC | 2995 |
| ENSG000001 | 588.271523 | 1.067481701 | 0.1190635 | 8.96564994 | 3.08E-19 | 2.91E-16 | BNIP3 | 664 |
| ENSG000002 | 692.412829 | 1.419197977 | 0.15918976 | 8.91513335 | 4.87E-19 | 4.49E-16 | NA | NA |
| ENSG000001 | 991.397004 | 1.194447476 | 0.1340511 | 8.91038929 | 5.09E-19 | 4.58E-16 | KLHL4 | 56062 |
| ENSG000001 | 246.097882 | 1.232938487 | 0.1389687 | 8.87205857 | 7.18E-19 | 6.33E-16 | DDIT3 | 1649 |
| ENSG000001 | 94.5369371 | -1.563852179 | 0.17795272 | -8.7880205 | 1.52E-18 | 1.31E-15 | SLC2A14 | 144195 |
| ENSG000001 | 5963.43358 | 1.126133386 | 0.12863955 | 8.75417663 | 2.06E-18 | 1.74E-15 | RPL39 | 6170 |
| ENSG000001 | 1963.80219 | 0.998503641 | 0.11437547 | 8.73005036 | 2.55E-18 | 2.11E-15 | GPR176 | 11245 |
| ENSG000001 | 265.85223 | 1.368123887 | 0.15762388 | 8.67967414 | 3.97E-18 | 3.22E-15 | TRIB3 | 57761 |
| ENSG000001 | 1094.86483 | 1.10515429 | 0.12869307 | 8.58751962 | 8.89E-18 | 7.07E-15 | INHBE | 83729 |
| ENSG000001 | 556.927178 | 1.108756286 | 0.12956855 | 8.5572948 | 1.16E-17 | 9.01E-15 | NKAP | 79576 |
| ENSG00000C | 838.213252 | 1.001164605 | 0.11715454 | 8.54567467 | 1.28E-17 | 9.78E-15 | SMARCA2 | 6595 |
| ENSG000001 | 76.6674769 | -1.684058949 | 0.19866908 | -8.4767037 | 2.32E-17 | 1.74E-14 | TRIM4 | 89122 |
| ENSG000001 | 1900.6705 | 1.005871285 | 0.12026181 | 8.36401257 | 6.06E-17 | 4.47E-14 | TMEM64 | 169200 |
| ENSG000001 | 1686.85106 | 1.290867075 | 0.15527094 | 8.31364235 | 9.28E-17 | 6.72E-14 | DDIT4 | 54541 |

| | | | | | | | | |
|---|---|---|---|---|---|---|---|---|
| ENSG000001 | 1237.55001 | 1.100577282 | 0.13244912 | 8.30943438 | 9.62E-17 | 6.84E-14 | CDK16 | 5127 |
| ENSG000001 | 36.4881997 | -2.11536697 | 0.25573671 | -8.2716594 | 1.32E-16 | 9.24E-14 | DNAJA4 | 55466 |
| ENSG000001 | 302.115479 | 1.147418607 | 0.13912704 | 8.24727274 | 1.62E-16 | 1.10E-13 | PRRG1 | 5638 |
| ENSG000001 | 115.681934 | 1.45183308 | 0.17599886 | 8.24910516 | 1.60E-16 | 1.10E-13 | HLA-DQB1 | 3119 |
| ENSG000001 | 364.539375 | 1.149367254 | 0.13990747 | 8.21519603 | 2.12E-16 | 1.41E-13 | CXCL5 | 6374 |
| ENSG00000C | 3555.8019 | 0.964926642 | 0.11818993 | 8.16420353 | 3.24E-16 | 2.05E-13 | SARS | 6301 |
| ENSG000001 | 855.73113 | 1.081795051 | 0.13245689 | 8.1671479 | 3.16E-16 | 2.05E-13 | DMXL2 | 23312 |
| ENSG000001 | 324.17198 | 1.123994042 | 0.137649 | 8.1656537 | 3.20E-16 | 2.05E-13 | TCN2 | 6948 |
| ENSG000001 | 690.302354 | 1.144942294 | 0.14055184 | 8.14604958 | 3.76E-16 | 2.35E-13 | LCK | 3932 |
| ENSG000002 | 191.337429 | -1.236009349 | 0.15230437 | -8.1153902 | 4.84E-16 | 2.98E-13 | NA | NA |
| ENSG000002 | 49.3592911 | -1.936114294 | 0.23888028 | -8.1049566 | 5.28E-16 | 3.19E-13 | LINC00662 | 148189 |
| ENSG000001 | 1973.80308 | 0.883430497 | 0.10903366 | 8.1023652 | 5.39E-16 | 3.21E-13 | MMGT1 | 93380 |
| ENSG00000C | 1047.60864 | 1.024149066 | 0.12779523 | 8.01398508 | 1.11E-15 | 6.53E-13 | UBE2A | 7319 |
| ENSG000002 | 137.38624 | -1.220204145 | 0.15239794 | -8.0066971 | 1.18E-15 | 6.83E-13 | ZNF578 | 147660 |
| ENSG000001 | 504.942881 | 1.162594157 | 0.14558623 | 7.98560514 | 1.40E-15 | 7.99E-13 | RARRES2 | 5919 |
| ENSG000001 | 1798.28523 | 1.032779146 | 0.12993251 | 7.94858153 | 1.89E-15 | 1.06E-12 | DSEL | 92126 |
| ENSG000001 | 1920.72216 | 0.926382127 | 0.11671847 | 7.93689409 | 2.07E-15 | 1.15E-12 | KDR | 3791 |
| ENSG000002 | 427.949668 | 1.121867241 | 0.14148128 | 7.92943924 | 2.20E-15 | 1.21E-12 | LOC101927746 | 101927746 |
| ENSG000001 | 586.036287 | 1.110916567 | 0.14032668 | 7.9166453 | 2.44E-15 | 1.32E-12 | AKAP7 | 9465 |
| ENSG00000C | 9096.55668 | 1.061117976 | 0.13417301 | 7.9085798 | 2.60E-15 | 1.39E-12 | FTL | 2512 |
| ENSG000001 | 1154.1691 | 1.043367548 | 0.13214572 | 7.89558316 | 2.89E-15 | 1.52E-12 | CHAC1 | 79094 |
| ENSG00000C | 603.30517 | 0.941927352 | 0.11958727 | 7.87648524 | 3.37E-15 | 1.75E-12 | MBNL3 | 55796 |
| ENSG000002 | 2047.35049 | 1.263550648 | 0.16055594 | 7.8698468 | 3.55E-15 | 1.82E-12 | HIST1H4I | 8294 |
| ENSG000001 | 1228.20227 | 1.098225222 | 0.14018461 | 7.83413561 | 4.72E-15 | 2.36E-12 | ASS1 | 445 |
| ENSG000001 | 3320.25004 | 0.999707252 | 0.12759769 | 7.83483825 | 4.69E-15 | 2.36E-12 | PMAIP1 | 5366 |
| ENSG000001 | 836.488632 | 0.945431536 | 0.12135859 | 7.79039643 | 6.68E-15 | 3.30E-12 | TSC22D3 | 1831 |
| ENSG000001 | 1245.10573 | 0.875219998 | 0.11282032 | 7.75764484 | 8.65E-15 | 4.23E-12 | DOCK11 | 139818 |
| ENSG000001 | 5612.47312 | 0.904578908 | 0.11709897 | 7.72490887 | 1.12E-14 | 5.40E-12 | SMS | 6611 |
| ENSG000001 | 489.084238 | 1.001992705 | 0.13009768 | 7.7018491 | 1.34E-14 | 6.40E-12 | ADGRG2 | 10149 |

| | | | | | | | | |
|---|---|---|---|---|---|---|---|---|
| ENSG000001 | 469.776451 | 1.087629289 | 0.14149237 | 7.68684035 | 1.51E-14 | 7.11E-12 | TIMM17B | 10245 |
| ENSG000002 | 3765.75303 | 1.017941943 | 0.13249727 | 7.68273916 | 1.56E-14 | 7.26E-12 | PEG10 | 23089 |
| ENSG000002 | 246.420727 | 1.27744311 | 0.1669702 | 7.65072506 | 2.00E-14 | 9.21E-12 | C9orf135 | 138255 |
| ENSG000001 | 520.704252 | 1.080618838 | 0.14158812 | 7.63212929 | 2.31E-14 | 1.05E-11 | RIMS3 | 9783 |
| ENSG000001 | 43.5975188 | -1.968482983 | 0.25883055 | -7.6052963 | 2.84E-14 | 1.28E-11 | FOSL1 | 8061 |
| ENSG000001 | 1014.55548 | 0.873898498 | 0.11502888 | 7.59720975 | 3.03E-14 | 1.35E-11 | PRUNE2 | 158471 |
| ENSG000001 | 2552.14506 | 0.978264529 | 0.12918172 | 7.57277807 | 3.65E-14 | 1.61E-11 | PRPS1 | 5631 |
| ENSG000001 | 33.6080693 | -2.017046405 | 0.26702862 | -7.5536711 | 4.23E-14 | 1.85E-11 | CCL28 | 56477 |
| ENSG000001 | 268.937252 | 1.15124735 | 0.15254761 | 7.5468071 | 4.46E-14 | 1.92E-11 | ZNF248 | 57209 |
| ENSG000002 | 3332.56361 | 0.842191642 | 0.11160965 | 7.54586778 | 4.49E-14 | 1.92E-11 | FIRRE | 286467 |
| ENSG000002 | 1220.85764 | 1.185652955 | 0.15810666 | 7.49907051 | 6.43E-14 | 2.71E-11 | RN7SKP203 | 106479185 |
| ENSG000001 | 240.025688 | 1.256803052 | 0.16764702 | 7.49672191 | 6.54E-14 | 2.74E-11 | SLC35A2 | 7355 |
| ENSG000001 | 1775.60299 | 0.921326575 | 0.12313367 | 7.48232859 | 7.30E-14 | 3.02E-11 | PLP1 | 5354 |
| ENSG000001 | 457.524458 | -0.966281866 | 0.12919377 | -7.4793222 | 7.47E-14 | 3.06E-11 | TNFRSF10B | 8795 |
| ENSG000002 | 33.1601747 | -1.916602783 | 0.25701429 | -7.4571836 | 8.84E-14 | 3.58E-11 | NA | NA |
| ENSG000001 | 102.793893 | -1.313613298 | 0.1762517 | -7.453053 | 9.12E-14 | 3.66E-11 | PIWIL2 | 55124 |
| ENSG000001 | 97.7347842 | -1.324581816 | 0.17822088 | -7.4322482 | 1.07E-13 | 4.24E-11 | SPTA1 | 6708 |
| ENSG000002 | 1070.58514 | 1.022982357 | 0.13846604 | 7.38796578 | 1.49E-13 | 5.87E-11 | HLA-A | 3105 |
| ENSG00000C | 652.680438 | 1.064261426 | 0.14421803 | 7.37953082 | 1.59E-13 | 6.19E-11 | GRIPAP1 | 56850 |
| ENSG000001 | 407.129788 | 1.020454798 | 0.13874834 | 7.35471733 | 1.91E-13 | 7.39E-11 | COX8A | 1351 |
| ENSG000001 | 763.918716 | 0.870945402 | 0.11870851 | 7.33684036 | 2.19E-13 | 8.37E-11 | ESYT2 | 57488 |
| ENSG00000C | 296.85881 | 0.989129221 | 0.1350865 | 7.32219176 | 2.44E-13 | 9.25E-11 | GSTO2 | 119391 |
| ENSG000001 | 306.355706 | 1.182299996 | 0.16196546 | 7.29970455 | 2.88E-13 | 1.08E-10 | HMOX1 | 3162 |
| ENSG000001 | 86.2846503 | -1.305964436 | 0.17944977 | -7.2776045 | 3.40E-13 | 1.26E-10 | INPP5D | 3635 |
| ENSG000001 | 35.5029294 | -1.835974143 | 0.25239927 | -7.2740865 | 3.49E-13 | 1.29E-10 | TNFRSF10C | 8794 |
| ENSG000001 | 1158.97706 | 0.91738565 | 0.1261422 | 7.2726309 | 3.53E-13 | 1.29E-10 | KLHL15 | 80311 |
| ENSG000001 | 17841.2803 | 0.878192013 | 0.12097832 | 7.25908584 | 3.90E-13 | 1.41E-10 | RPS4X | 6191 |
| ENSG000002 | 9395.2005 | 0.99308295 | 0.13687936 | 7.25516948 | 4.01E-13 | 1.44E-10 | TMSB4X | 7114 |
| ENSG000001 | 1520.45912 | 1.092653597 | 0.15081418 | 7.2450321 | 4.32E-13 | 1.54E-10 | GPT2 | 84706 |

| | | | | | | | | |
|---|---|---|---|---|---|---|---|---|
| ENSG000001 | 593.323699 | 0.974398486 | 0.13456517 | 7.2410897 | 4.45E-13 | 1.57E-10 | MOSPD1 | 56180 |
| ENSG000001 | 378.503701 | 0.945518233 | 0.13061535 | 7.23895184 | 4.52E-13 | 1.58E-10 | MTM1 | 4534 |
| ENSG000001 | 397.846714 | 0.860652945 | 0.11910679 | 7.22589317 | 4.98E-13 | 1.73E-10 | ARHGEF6 | 9459 |
| ENSG000001 | 1517.7795 | 1.225635012 | 0.16966486 | 7.22385902 | 5.05E-13 | 1.74E-10 | HIST4H4 | 121504 |
| ENSG000001 | 113.069137 | 1.228223032 | 0.17012922 | 7.21935369 | 5.22E-13 | 1.78E-10 | NPY5R | 4889 |
| ENSG00000C | 233.739022 | 1.189322164 | 0.16487258 | 7.21358382 | 5.45E-13 | 1.84E-10 | NMRK2 | 27231 |
| ENSG000002 | 48.5114204 | -1.592329098 | 0.22207832 | -7.1701241 | 7.49E-13 | 2.51E-10 | NA | NA |
| ENSG000001 | 1541.34769 | 0.978783261 | 0.13657645 | 7.16655976 | 7.69E-13 | 2.56E-10 | PRDX4 | 10549 |
| ENSG000001 | 492.390464 | -1.034113359 | 0.14451227 | -7.1558864 | 8.31E-13 | 2.74E-10 | RPS27L | 51065 |
| ENSG000001 | 653.914655 | 0.845287412 | 0.11840426 | 7.13899478 | 9.40E-13 | 3.07E-10 | GPC6 | 10082 |
| ENSG000001 | 1406.63979 | 0.904805728 | 0.12715878 | 7.11555867 | 1.11E-12 | 3.59E-10 | PSMD10 | 5716 |
| ENSG000001 | 173.28433 | 1.166044032 | 0.16386887 | 7.11571412 | 1.11E-12 | 3.59E-10 | MT1G | 4495 |
| ENSG000001 | 144.710751 | 1.079871222 | 0.15181511 | 7.11306827 | 1.13E-12 | 3.62E-10 | LCP1 | 3936 |
| ENSG000002 | 268.264533 | 0.992094919 | 0.13974345 | 7.09940205 | 1.25E-12 | 3.97E-10 | NA | NA |
| ENSG000001 | 1182.43809 | 0.936472595 | 0.13199424 | 7.09479878 | 1.30E-12 | 4.07E-10 | MAGED2 | 10916 |
| ENSG000002 | 275.058911 | 1.244682744 | 0.17623239 | 7.0627356 | 1.63E-12 | 5.09E-10 | RN7SKP71 | 106479126 |
| ENSG000001 | 272.83691 | 0.925767424 | 0.13150561 | 7.03975611 | 1.93E-12 | 5.96E-10 | TNFSF11 | 8600 |
| ENSG000001 | 2000.42244 | 0.915376239 | 0.130162 | 7.0325918 | 2.03E-12 | 6.23E-10 | PRDX2 | 7001 |
| ENSG00000C | 357.102175 | 1.01878828 | 0.14492591 | 7.02971809 | 2.07E-12 | 6.31E-10 | H2AFY2 | 55506 |
| ENSG000001 | 402.406325 | 1.168180816 | 0.16626475 | 7.02602823 | 2.12E-12 | 6.43E-10 | ATF5 | 22809 |
| ENSG000001 | 685.301218 | 0.881813261 | 0.12563582 | 7.01880444 | 2.24E-12 | 6.72E-10 | S100A11 | 6282 |
| ENSG00000C | 1930.58745 | 0.806187284 | 0.11499389 | 7.01069646 | 2.37E-12 | 7.02E-10 | IFRD1 | 3475 |
| ENSG000001 | 527.604974 | 1.176529477 | 0.16781765 | 7.01076106 | 2.37E-12 | 7.02E-10 | NA | NA |
| ENSG00000C | 1388.16371 | 0.763807603 | 0.10908964 | 7.00165083 | 2.53E-12 | 7.43E-10 | CDKL5 | 6792 |
| ENSG000001 | 448.441612 | 0.901044156 | 0.12885415 | 6.9927444 | 2.70E-12 | 7.86E-10 | NA | NA |
| ENSG000001 | 46.1197803 | -1.554948221 | 0.22353687 | -6.9561151 | 3.50E-12 | 1.01E-09 | GRHL3 | 57822 |
| ENSG00000C | 985.286517 | 0.811605141 | 0.11691197 | 6.94201897 | 3.87E-12 | 1.11E-09 | NDUFB4 | 4710 |
| ENSG000001 | 3785.32216 | 0.87820901 | 0.12666851 | 6.93312813 | 4.12E-12 | 1.18E-09 | PHF6 | 84295 |
| ENSG000001 | 2771.49069 | 1.016823039 | 0.1467213 | 6.93030289 | 4.20E-12 | 1.19E-09 | SLC1A5 | 6510 |

| | | | | | | | | |
|---|---|---|---|---|---|---|---|---|
| ENSG000001 | 6414.80662 | 0.884554717 | 0.12782492 | 6.92004927 | 4.51E-12 | 1.27E-09 | HIST1H1E | 3008 |
| ENSG000001 | 669.156246 | 0.80329902 | 0.11613866 | 6.91672391 | 4.62E-12 | 1.29E-09 | MET | 4233 |
| ENSG000002 | 200.792016 | 1.098833556 | 0.15888813 | 6.91576862 | 4.65E-12 | 1.29E-09 | LINC00698 | 285401 |
| ENSG000000 | 4814.10971 | 1.005703186 | 0.14554359 | 6.90997928 | 4.85E-12 | 1.34E-09 | PHGDH | 26227 |
| ENSG000001 | 589.453512 | 0.897653437 | 0.12997495 | 6.90635703 | 4.97E-12 | 1.36E-09 | NDUFV1 | 4723 |
| ENSG000001 | 4508.35397 | 0.981346755 | 0.142278 | 6.89738912 | 5.30E-12 | 1.44E-09 | HIST1H1C | 3006 |
| ENSG000001 | 63.3465856 | 1.493856262 | 0.21666119 | 6.89489551 | 5.39E-12 | 1.46E-09 | GSTT2B | 653689 |
| ENSG000001 | 3178.31834 | 1.081906877 | 0.15699677 | 6.89126828 | 5.53E-12 | 1.49E-09 | UBA1 | 7317 |
| ENSG000001 | 452.853525 | 1.047549274 | 0.15249284 | 6.86949811 | 6.44E-12 | 1.72E-09 | ZDHHC22 | 283576 |
| ENSG000001 | 3908.02868 | 0.867703773 | 0.1263321 | 6.86843465 | 6.49E-12 | 1.72E-09 | RBBP7 | 5931 |
| ENSG000001 | 9238.03047 | 0.830560067 | 0.12099446 | 6.86444735 | 6.67E-12 | 1.75E-09 | PSAT1 | 29968 |
| ENSG000002 | 951.332523 | 0.80112696 | 0.11669954 | 6.86486839 | 6.66E-12 | 1.75E-09 | RAB12 | 201475 |
| ENSG000001 | 196.17203 | 1.069372274 | 0.15582259 | 6.8627549 | 6.75E-12 | 1.76E-09 | HRK | 8739 |
| ENSG000000 | 341.115521 | 0.866963247 | 0.12636589 | 6.86073772 | 6.85E-12 | 1.76E-09 | P3H2 | 55214 |
| ENSG000001 | 40.0102516 | -1.631650579 | 0.23779725 | -6.8615199 | 6.81E-12 | 1.76E-09 | BTG2 | 7832 |
| ENSG000001 | 266.140269 | 0.94634049 | 0.13813494 | 6.85084101 | 7.34E-12 | 1.87E-09 | TMEM115 | 11070 |
| ENSG000001 | 543.456997 | 0.963743053 | 0.14071729 | 6.84878923 | 7.45E-12 | 1.89E-09 | VPS18 | 57617 |
| ENSG000001 | 554.5939 | 0.835608564 | 0.12209132 | 6.84412749 | 7.69E-12 | 1.94E-09 | CLDN7 | 1366 |
| ENSG000002 | 22.9842265 | -2.065632469 | 0.30255176 | -6.827369 | 8.65E-12 | 2.16E-09 | NA | NA |
| ENSG000001 | 913.997664 | 1.124438384 | 0.16499502 | 6.81498381 | 9.43E-12 | 2.35E-09 | USP11 | 8237 |
| ENSG000001 | 1263.96167 | 0.988849962 | 0.14516329 | 6.81198378 | 9.63E-12 | 2.38E-09 | PRDX5 | 25824 |
| ENSG000001 | 1492.58003 | 0.936891683 | 0.13757774 | 6.80990768 | 9.77E-12 | 2.40E-09 | TALDO1 | 6888 |
| ENSG000001 | 10412.6151 | 0.883750271 | 0.12997075 | 6.79960871 | 1.05E-11 | 2.56E-09 | EIF2S3 | 1968 |
| ENSG000002 | 293.064829 | -1.039669098 | 0.15340505 | -6.777281 | 1.22E-11 | 2.97E-09 | NA | NA |
| ENSG000000 | 68.6287626 | 1.325240903 | 0.19568812 | 6.77220916 | 1.27E-11 | 3.06E-09 | NPFFR2 | 10886 |
| ENSG000000 | 476.473973 | 0.918382189 | 0.13582214 | 6.76165318 | 1.36E-11 | 3.27E-09 | PHKA2 | 5256 |
| ENSG000002 | 156.831475 | 1.079467498 | 0.15976779 | 6.7564776 | 1.41E-11 | 3.37E-09 | PNMA2 | 10687 |
| ENSG000002 | 3463.02603 | 1.024719789 | 0.15168269 | 6.75568044 | 1.42E-11 | 3.37E-09 | HIST1H3G | 8355 |
| ENSG000001 | 1958.00795 | 0.806949688 | 0.11962623 | 6.74559153 | 1.52E-11 | 3.59E-09 | BOD1 | 91272 |

| | | | | | | | | |
|---|---|---|---|---|---|---|---|---|
| ENSG00000C | 687.983778 | 1.075191083 | 0.15952856 | 6.73980319 | 1.59E-11 | 3.72E-09 | TFE3 | 7030 |
| ENSG000001 | 1513.28035 | 0.793159307 | 0.11778244 | 6.73410489 | 1.65E-11 | 3.84E-09 | ARMT1 | 79624 |
| ENSG000001 | 1000.51642 | 1.069061575 | 0.15923625 | 6.7136824 | 1.90E-11 | 4.40E-09 | PYCR1 | 5831 |
| ENSG00000C | 453.665853 | 0.987933806 | 0.14719744 | 6.71162338 | 1.92E-11 | 4.40E-09 | CDC34 | 997 |
| ENSG000001 | 1675.27635 | 0.77538546 | 0.11553292 | 6.71138142 | 1.93E-11 | 4.40E-09 | XIAP | 331 |
| ENSG000001 | 438.254104 | 1.01234129 | 0.150865 | 6.71024614 | 1.94E-11 | 4.40E-09 | ATP6V0B | 533 |
| ENSG000001 | 4962.89068 | 0.872438244 | 0.13003146 | 6.7094395 | 1.95E-11 | 4.40E-09 | STK26 | 51765 |
| ENSG000001 | 554.850611 | 0.845346258 | 0.1259837 | 6.70996508 | 1.95E-11 | 4.40E-09 | RRP7A | 27341 |
| ENSG000002 | 904.463935 | 0.794600225 | 0.11845361 | 6.70811313 | 1.97E-11 | 4.42E-09 | ZNF254 | 9534 |
| ENSG00000C | 3532.40857 | 0.808322378 | 0.12062185 | 6.70129294 | 2.07E-11 | 4.60E-09 | ACSL4 | 2182 |
| ENSG000001 | 68.4935165 | -1.330941855 | 0.19866655 | -6.6993756 | 2.09E-11 | 4.64E-09 | MDGA2 | 161357 |
| ENSG000001 | 1584.23266 | 0.858652079 | 0.12826775 | 6.69421631 | 2.17E-11 | 4.78E-09 | SQSTM1 | 8878 |
| ENSG000001 | 1096.68752 | 0.848810207 | 0.12683701 | 6.69213373 | 2.20E-11 | 4.82E-09 | PSPH | 5723 |
| ENSG000002 | 3176.79602 | 0.943500502 | 0.14100771 | 6.69112696 | 2.21E-11 | 4.83E-09 | HIST1H4D | 8360 |
| ENSG000001 | 1214.30037 | 1.026769786 | 0.15355251 | 6.68676648 | 2.28E-11 | 4.95E-09 | LONP1 | 9361 |
| ENSG000002 | 15.1141216 | -2.204590494 | 0.32975492 | -6.6855425 | 2.30E-11 | 4.96E-09 | C5orf17 | 439936 |
| ENSG000001 | 337.31179 | 1.067858683 | 0.15986386 | 6.67980051 | 2.39E-11 | 5.13E-09 | ZNF516 | 9658 |
| ENSG000001 | 3306.56765 | 0.879452374 | 0.13177548 | 6.67386965 | 2.49E-11 | 5.29E-09 | SLC3A2 | 6520 |
| ENSG000001 | 241.281636 | 0.917121503 | 0.137407 | 6.67448912 | 2.48E-11 | 5.29E-09 | MT1X | 4501 |
| ENSG000002 | 2464.10898 | 1.017890044 | 0.15271975 | 6.66508464 | 2.65E-11 | 5.59E-09 | HIST1H3A | 8350 |
| ENSG000001 | 389.487498 | 0.927794 | 0.13963325 | 6.64450636 | 3.04E-11 | 6.39E-09 | SLC43A1 | 8501 |
| ENSG00000C | 3705.1504 | 0.762734777 | 0.11481743 | 6.64302261 | 3.07E-11 | 6.42E-09 | ANOS1 | 3730 |
| ENSG000001 | 355.646565 | 1.001951674 | 0.1508818 | 6.64063982 | 3.12E-11 | 6.49E-09 | FUT1 | 2523 |
| ENSG000001 | 352.208739 | 1.021319017 | 0.1538554 | 6.63817472 | 3.18E-11 | 6.57E-09 | CLIP2 | 7461 |
| ENSG000002 | 778.478368 | 0.95995064 | 0.14463031 | 6.63727151 | 3.20E-11 | 6.58E-09 | C6orf48 | 50854 |
| ENSG00000C | 224.207621 | 0.974081448 | 0.14679488 | 6.63566342 | 3.23E-11 | 6.62E-09 | AKR7A2 | 8574 |
| ENSG000001 | 242.984303 | -0.923344888 | 0.13930275 | -6.628332 | 3.40E-11 | 6.89E-09 | AHNAK | 79026 |
| ENSG000001 | 3236.66942 | 0.859842112 | 0.12972454 | 6.62821476 | 3.40E-11 | 6.89E-09 | THY1 | 7070 |
| ENSG000002 | 4726.69607 | 0.815440189 | 0.12305153 | 6.62681885 | 3.43E-11 | 6.92E-09 | LOC441666 | 441666 |

| | | | | | | | | |
|---|---|---|---|---|---|---|---|---|
| ENSG000001 | 1217.6461 | 0.757776707 | 0.11443212 | 6.6220631 | 3.54E-11 | 7.11E-09 | STX3 | 6809 |
| ENSG000001 | 3046.2039 | 0.733777882 | 0.11085402 | 6.61931658 | 3.61E-11 | 7.21E-09 | POLA1 | 5422 |
| ENSG000002 | 2082.53697 | 0.864699118 | 0.13066933 | 6.61746017 | 3.65E-11 | 7.26E-09 | NA | NA |
| ENSG000001 | 232.361981 | -1.064699992 | 0.16104854 | -6.6110503 | 3.82E-11 | 7.55E-09 | DDB2 | 1643 |
| ENSG000001 | 317.431625 | 0.95296904 | 0.14427238 | 6.60534632 | 3.97E-11 | 7.81E-09 | SAPCD2 | 89958 |
| ENSG000001 | 1110.14467 | 0.852287745 | 0.12933002 | 6.59002268 | 4.40E-11 | 8.61E-09 | B2M | 567 |
| ENSG000001 | 401.792792 | 1.024598754 | 0.15550325 | 6.58892167 | 4.43E-11 | 8.64E-09 | GRK6 | 2870 |
| ENSG000001 | 537.971325 | 0.854446153 | 0.12978338 | 6.5836333 | 4.59E-11 | 8.88E-09 | CDS1 | 1040 |
| ENSG000001 | 494.924123 | 0.980206332 | 0.14889058 | 6.58340059 | 4.60E-11 | 8.88E-09 | APRT | 353 |
| ENSG000001 | 3695.24441 | 1.049574415 | 0.15953072 | 6.57913679 | 4.73E-11 | 9.09E-09 | SHMT2 | 6472 |
| ENSG000001 | 3252.5599 | 0.814049185 | 0.12382704 | 6.57408255 | 4.90E-11 | 9.36E-09 | WARS | 7453 |
| ENSG000001 | 2286.34957 | 0.899686257 | 0.13692271 | 6.57075969 | 5.01E-11 | 9.53E-09 | HIST1H3J | 8356 |
| ENSG000002 | 14.0205277 | -2.215406243 | 0.33750153 | -6.564137 | 5.23E-11 | 9.92E-09 | GUSBP5 | 441046 |
| ENSG000001 | 1908.74407 | 0.840479659 | 0.12807025 | 6.56264553 | 5.29E-11 | 9.97E-09 | SNRPD2 | 6633 |
| ENSG000001 | 259.712967 | 0.957700655 | 0.14608021 | 6.55599163 | 5.53E-11 | 1.04E-08 | PINK1 | 65018 |
| ENSG000001 | 347.136596 | 1.001875849 | 0.15286446 | 6.55401424 | 5.60E-11 | 1.05E-08 | NAA38 | 84316 |
| ENSG000001 | 537.934638 | 0.913063323 | 0.13978184 | 6.5320597 | 6.49E-11 | 1.21E-08 | NDUFB11 | 54539 |
| ENSG000001 | 5858.58852 | 0.924861871 | 0.14163178 | 6.53004464 | 6.58E-11 | 1.22E-08 | RPL13 | 6137 |
| ENSG000001 | 1177.4577 | 0.875102121 | 0.13402436 | 6.52942581 | 6.60E-11 | 1.22E-08 | EIF4EBP1 | 1978 |
| ENSG000001 | 1019.71239 | 0.830334485 | 0.12726404 | 6.52450224 | 6.82E-11 | 1.25E-08 | SMARCD2 | 6603 |
| ENSG000001 | 577.1565 | 0.918623953 | 0.14090134 | 6.51962536 | 7.05E-11 | 1.29E-08 | MAFG | 4097 |
| ENSG000001 | 2346.99944 | 0.71175706 | 0.10930149 | 6.51186989 | 7.42E-11 | 1.35E-08 | XBP1 | 7494 |
| ENSG00000C | 5559.77166 | 0.894294383 | 0.1374848 | 6.50467826 | 7.79E-11 | 1.41E-08 | GSTP1 | 2950 |
| ENSG000001 | 167.022867 | 1.013805774 | 0.15590846 | 6.50257053 | 7.90E-11 | 1.42E-08 | CHRNA9 | 55584 |
| ENSG000001 | 5100.03164 | 0.868174453 | 0.13356259 | 6.500132 | 8.02E-11 | 1.44E-08 | UBC | 7316 |
| ENSG000002 | 584.664382 | 0.80161189 | 0.12338279 | 6.49695042 | 8.20E-11 | 1.46E-08 | HLA-DOA | 3111 |
| ENSG000001 | 714.3121 | 0.70498233 | 0.10852153 | 6.49624377 | 8.24E-11 | 1.46E-08 | MAX | 4149 |
| ENSG000001 | 1186.64844 | 0.872780366 | 0.13437504 | 6.49510758 | 8.30E-11 | 1.47E-08 | GRB10 | 2887 |
| ENSG000001 | 721.409582 | 0.974881497 | 0.1501123 | 6.49434806 | 8.34E-11 | 1.47E-08 | RAB17 | 64284 |

| | | | | | | | | |
|---|---|---|---|---|---|---|---|---|
| ENSG00000C | 5769.47636 | 0.812554768 | 0.12513542 | 6.49340365 | 8.39E-11 | 1.47E-08 | RPL18 | 6141 |
| ENSG00000C | 783.6446 | 0.834763804 | 0.12875435 | 6.48338325 | 8.97E-11 | 1.57E-08 | LAMP2 | 3920 |
| ENSG000002 | 45.8365 | 1.455087861 | 0.22448796 | 6.48180798 | 9.06E-11 | 1.58E-08 | PCAT7 | 101928099 |
| ENSG000001 | 276.435408 | 0.90734349 | 0.14008455 | 6.47711329 | 9.35E-11 | 1.62E-08 | RITA1 | 84934 |
| ENSG000001 | 1333.9884 | 0.819907897 | 0.12666768 | 6.47290512 | 9.61E-11 | 1.66E-08 | VLDLR | 7436 |
| ENSG00000C | 620.871144 | 0.806092909 | 0.12461569 | 6.46863099 | 9.89E-11 | 1.70E-08 | ARF5 | 381 |
| ENSG00000C | 4322.28138 | 0.774168241 | 0.11970707 | 6.46718882 | 9.98E-11 | 1.71E-08 | GPM6B | 2824 |
| ENSG000002 | 204.402138 | 1.101989441 | 0.17050747 | 6.46299807 | 1.03E-10 | 1.75E-08 | RHOXF1-AS1 | 101928969 |
| ENSG000001 | 2596.40276 | 0.925988028 | 0.14328976 | 6.46234624 | 1.03E-10 | 1.75E-08 | RPLP2 | 6181 |
| ENSG000002 | 473.128144 | 1.0298962 | 0.15954429 | 6.45523716 | 1.08E-10 | 1.83E-08 | NA | NA |
| ENSG000001 | 1315.686 | 0.766457506 | 0.1187754 | 6.45299892 | 1.10E-10 | 1.85E-08 | PPM1H | 57460 |
| ENSG000001 | 2663.04487 | 0.801115982 | 0.12419446 | 6.45049674 | 1.11E-10 | 1.87E-08 | RPS15 | 6209 |
| ENSG000001 | 1369.54128 | 0.764069659 | 0.11848165 | 6.44884385 | 1.13E-10 | 1.88E-08 | WDR44 | 54521 |
| ENSG000001 | 1006.32414 | 0.798823793 | 0.1238962 | 6.44752443 | 1.14E-10 | 1.89E-08 | COX7A2 | 1347 |
| ENSG000001 | 189.604232 | 1.070288742 | 0.16605173 | 6.44551391 | 1.15E-10 | 1.91E-08 | MFSD3 | 113655 |
| ENSG000001 | 1183.62796 | 0.811738504 | 0.12596734 | 6.44403941 | 1.16E-10 | 1.92E-08 | CERK | 64781 |
| ENSG000001 | 3224.9357 | 0.751667057 | 0.11665617 | 6.44344045 | 1.17E-10 | 1.92E-08 | CUL4B | 8450 |
| ENSG000001 | 909.237375 | 0.755015377 | 0.11719265 | 6.44251478 | 1.18E-10 | 1.92E-08 | PYGL | 5836 |
| ENSG000001 | 492.727259 | 1.051483096 | 0.16350694 | 6.4308164 | 1.27E-10 | 2.07E-08 | IRAK1 | 3654 |
| ENSG000001 | 1104.55874 | 1.144245289 | 0.17809077 | 6.42506804 | 1.32E-10 | 2.14E-08 | PCK2 | 5106 |
| ENSG000001 | 1056.86241 | 0.875812302 | 0.13654996 | 6.4138598 | 1.42E-10 | 2.29E-08 | B3GNT7 | 93010 |
| ENSG000001 | 231.276419 | 0.969266387 | 0.15121086 | 6.41003156 | 1.45E-10 | 2.33E-08 | RABAC1 | 10567 |
| ENSG000001 | 122.537519 | 1.067653085 | 0.16655426 | 6.41024193 | 1.45E-10 | 2.33E-08 | TFPI2 | 7980 |
| ENSG000001 | 1043.35584 | 0.750866727 | 0.11715834 | 6.40899095 | 1.46E-10 | 2.34E-08 | ACO1 | 48 |
| ENSG000002 | 43.0218861 | 1.744849875 | 0.27242327 | 6.40492224 | 1.50E-10 | 2.39E-08 | RNU2-63P | 106480225 |
| ENSG00000C | 104.719812 | 1.177866925 | 0.18420564 | 6.39430426 | 1.61E-10 | 2.55E-08 | TRAPPC6A | 79090 |
| ENSG000001 | 10275.0248 | 0.88578422 | 0.13870371 | 6.38616093 | 1.70E-10 | 2.68E-08 | HIST1H1B | 3009 |
| ENSG000002 | 271.456069 | 0.895075048 | 0.14035514 | 6.3772161 | 1.80E-10 | 2.83E-08 | HLA-E | 3133 |
| ENSG000002 | 540.837426 | 0.917714423 | 0.14391513 | 6.37677534 | 1.81E-10 | 2.83E-08 | ARPC4 | 10093 |

| | | | | | | | | | |
|---|---|---|---|---|---|---|---|---|---|
| ENSG000002 | 1239.40088 | 0.715312927 | 0.11218792 | 6.37602452 | 1.82E-10 | 2.83E-08 | NKX1-2 | 390010 | |
| ENSG000001 | 406.733293 | 0.884919447 | 0.13886633 | 6.3724552 | 1.86E-10 | 2.89E-08 | KCTD5 | 54442 | |
| ENSG000001 | 1553.32345 | 0.89640206 | 0.14073348 | 6.36950113 | 1.90E-10 | 2.94E-08 | CDK4 | 1019 | |
| ENSG000000 | 6487.91783 | 0.760738916 | 0.11951827 | 6.36504297 | 1.95E-10 | 3.00E-08 | MTHFD2 | 10797 | |
| ENSG000002 | 1683.24545 | 0.89504456 | 0.14061457 | 6.36523352 | 1.95E-10 | 3.00E-08 | DDOST | 1650 | |
| ENSG000002 | 352.138534 | 0.914588896 | 0.14376469 | 6.36170754 | 2.00E-10 | 3.05E-08 | TMEM147-AS1 | 100506469 | |
| ENSG000001 | 44.7775406 | 1.449897823 | 0.22796508 | 6.3601752 | 2.02E-10 | 3.07E-08 | CYTL1 | 54360 | |
| ENSG000001 | 406.497101 | 0.922739418 | 0.14514342 | 6.35743214 | 2.05E-10 | 3.12E-08 | PMEL | 6490 | |
| ENSG000001 | 184.487981 | -1.173158162 | 0.18459891 | -6.3551739 | 2.08E-10 | 3.15E-08 | TRIM22 | 10346 | |
| ENSG000001 | 53.9344295 | -1.354609153 | 0.21316165 | -6.3548447 | 2.09E-10 | 3.15E-08 | CCDC125 | 202243 | |
| ENSG000001 | 60.8229654 | 1.36843863 | 0.21539341 | 6.35320557 | 2.11E-10 | 3.17E-08 | DHRS2 | 10202 | |
| ENSG000000 | 938.213122 | 0.778923804 | 0.12262446 | 6.35210771 | 2.12E-10 | 3.17E-08 | CPS1 | 1373 | |
| ENSG000001 | 99.9793379 | -1.272982627 | 0.20040495 | -6.3520517 | 2.12E-10 | 3.17E-08 | MUC16 | 94025 | |
| ENSG000001 | 5011.82185 | 0.829856203 | 0.13067865 | 6.3503578 | 2.15E-10 | 3.18E-08 | PGK1 | 5230 | |
| ENSG000001 | 1937.59685 | 0.729354913 | 0.1148461 | 6.35071529 | 2.14E-10 | 3.18E-08 | MID1IP1 | 58526 | |
| ENSG000001 | 476.512185 | 0.811287789 | 0.12776617 | 6.34978555 | 2.16E-10 | 3.18E-08 | APOO | 79135 | |
| ENSG000002 | 1398.0834 | 0.886803797 | 0.1398121 | 6.34282565 | 2.26E-10 | 3.31E-08 | RPS28 | 6234 | |
| ENSG000001 | 2780.06206 | 0.759125675 | 0.1197824 | 6.33753932 | 2.33E-10 | 3.42E-08 | ODC1 | 4953 | |
| ENSG000001 | 549.913008 | 0.79418485 | 0.12534421 | 6.33603115 | 2.36E-10 | 3.44E-08 | CHRDL1 | 91851 | |
| ENSG000001 | 1376.22712 | 0.766087243 | 0.1209255 | 6.33520024 | 2.37E-10 | 3.45E-08 | DYNC2H1 | 79659 | |
| ENSG000001 | 220.103383 | 0.990466988 | 0.15638679 | 6.3334441 | 2.40E-10 | 3.47E-08 | BRAT1 | 221927 | |
| ENSG000001 | 2271.80289 | 0.765723207 | 0.12096268 | 6.33024334 | 2.45E-10 | 3.53E-08 | APOE | 348 | |
| ENSG000001 | 588.580218 | 0.797111579 | 0.12599566 | 6.32650012 | 2.51E-10 | 3.61E-08 | HMGN4 | 10473 | |
| ENSG000002 | 3222.6607 | 1.07372446 | 0.16980736 | 6.32319139 | 2.56E-10 | 3.67E-08 | RN7SL471P | 106481051 | |
| ENSG000001 | 2981.73414 | 0.739421142 | 0.11697936 | 6.32095405 | 2.60E-10 | 3.70E-08 | MAL2 | 114569 | |
| ENSG000002 | 2441.30619 | 0.952572737 | 0.15069802 | 6.32106999 | 2.60E-10 | 3.70E-08 | NA | | NA |
| ENSG000001 | 597.720857 | 0.822942155 | 0.13022866 | 6.31920948 | 2.63E-10 | 3.70E-08 | NDUFA1 | 4694 | |
| ENSG000001 | 1732.47369 | 0.747393697 | 0.11826632 | 6.31958183 | 2.62E-10 | 3.70E-08 | UTP14A | 10813 | |
| ENSG000001 | 1487.14722 | 0.796288196 | 0.12599633 | 6.31993173 | 2.62E-10 | 3.70E-08 | APEH | 327 | |

| | | | | | | | | |
|---|---|---|---|---|---|---|---|---|
| ENSG000002 | 417.197857 | 0.95923226 | 0.15190682 | 6.3146095 | 2.71E-10 | 3.80E-08 | HIST1H4K | 8362 |
| ENSG000001 | 233.478672 | 0.844889218 | 0.13383628 | 6.31285646 | 2.74E-10 | 3.83E-08 | TMEM30B | 161291 |
| ENSG000001 | 1924.00403 | 0.846086961 | 0.13405232 | 6.31161723 | 2.76E-10 | 3.85E-08 | HSPA13 | 6782 |
| ENSG000001 | 3084.88731 | 1.004199245 | 0.15933154 | 6.30257658 | 2.93E-10 | 4.07E-08 | RPS9 | 6203 |
| ENSG000001 | 1577.07406 | 0.739473937 | 0.1173545 | 6.30119776 | 2.95E-10 | 4.09E-08 | GPAM | 57678 |
| ENSG000001 | 1577.46421 | 0.67752112 | 0.10767885 | 6.29205408 | 3.13E-10 | 4.32E-08 | PCYT1B | 9468 |
| ENSG000001 | 8017.23393 | 0.710308445 | 0.11314637 | 6.27778378 | 3.43E-10 | 4.72E-08 | GARS | 2617 |
| ENSG000001 | 8950.97494 | 0.798317708 | 0.12718564 | 6.276791 | 3.46E-10 | 4.73E-08 | PLS3 | 5358 |
| ENSG000001 | 674.090429 | 0.749599775 | 0.11943176 | 6.27638536 | 3.47E-10 | 4.73E-08 | F2RL1 | 2150 |
| ENSG000001 | 1975.30987 | 0.796278461 | 0.12692658 | 6.27353602 | 3.53E-10 | 4.80E-08 | CASP3 | 836 |
| ENSG00000C | 609.87779 | 0.825623166 | 0.13182659 | 6.26294883 | 3.78E-10 | 5.11E-08 | MRPS18A | 55168 |
| ENSG000001 | 250.554942 | 0.974825059 | 0.15565495 | 6.26273097 | 3.78E-10 | 5.11E-08 | HAPLN3 | 145864 |
| ENSG000001 | 315.021233 | 0.881747871 | 0.14089488 | 6.25819664 | 3.89E-10 | 5.25E-08 | CXorf57 | 55086 |
| ENSG000001 | 421.340151 | 0.874114742 | 0.13968909 | 6.25757341 | 3.91E-10 | 5.25E-08 | SNX17 | 9784 |
| ENSG000002 | 1869.53152 | 1.037997313 | 0.16596496 | 6.25431589 | 3.99E-10 | 5.34E-08 | HIST1H3H | 8357 |
| ENSG000001 | 234.460205 | 0.868807242 | 0.13898637 | 6.25102474 | 4.08E-10 | 5.44E-08 | GNA14 | 9630 |
| ENSG000001 | 1385.91715 | 0.924081175 | 0.14808696 | 6.24012503 | 4.37E-10 | 5.81E-08 | GPX4 | 2879 |
| ENSG000001 | 513.757745 | 0.779682107 | 0.12496567 | 6.23917034 | 4.40E-10 | 5.83E-08 | YPEL5 | 51646 |
| ENSG000002 | 9162.86066 | 0.842405859 | 0.1350728 | 6.23668013 | 4.47E-10 | 5.90E-08 | RPS18 | 6222 |
| ENSG000001 | 3830.60709 | 0.784079294 | 0.12576274 | 6.2345914 | 4.53E-10 | 5.96E-08 | RPL36 | 25873 |
| ENSG000001 | 16.5333767 | -1.984344529 | 0.31833888 | -6.2334345 | 4.56E-10 | 5.99E-08 | LRRC61 | 65999 |
| ENSG000001 | 109.548454 | -1.042237981 | 0.16729621 | -6.229896 | 4.67E-10 | 6.09E-08 | ATP10D | 57205 |
| ENSG000002 | 70.1010384 | 1.282355331 | 0.20582379 | 6.2303554 | 4.65E-10 | 6.09E-08 | NA | NA |
| ENSG000002 | 4008.77087 | 0.994286182 | 0.15963304 | 6.22857396 | 4.71E-10 | 6.12E-08 | HIST1H4E | 8367 |
| ENSG000001 | 4119.9669 | 0.802133521 | 0.12888254 | 6.22375617 | 4.85E-10 | 6.29E-08 | HIST1H2BK | 85236 |
| ENSG000001 | 209.35412 | 1.053064941 | 0.16928376 | 6.22070868 | 4.95E-10 | 6.39E-08 | LHFPL4 | 375323 |
| ENSG000001 | 1013.61136 | 0.803092798 | 0.12942662 | 6.20500478 | 5.47E-10 | 7.04E-08 | BEX2 | 84707 |
| ENSG000001 | 1717.36484 | 0.668883208 | 0.10785594 | 6.20163529 | 5.59E-10 | 7.15E-08 | TSPAN3 | 10099 |
| ENSG000001 | 371.447885 | 0.869543914 | 0.14021407 | 6.20154541 | 5.59E-10 | 7.15E-08 | NA | NA |

| | | | | | | | | |
|---|---|---|---|---|---|---|---|---|
| ENSG000001 | 495.380837 | 0.743951078 | 0.12007634 | 6.19565064 | 5.80E-10 | 7.40E-08 | PQBP1 | 10084 |
| ENSG000001 | 758.901249 | 0.743522827 | 0.12004324 | 6.19379185 | 5.87E-10 | 7.47E-08 | SLC9A6 | 10479 |
| ENSG000001 | 1079.83968 | 0.935538654 | 0.15109647 | 6.19166463 | 5.95E-10 | 7.54E-08 | FKBP8 | 23770 |
| ENSG000001 | 311.14347 | 1.062594151 | 0.17174064 | 6.18720252 | 6.12E-10 | 7.74E-08 | COL26A1 | 136227 |
| ENSG000001 | 5422.4137 | 0.75152176 | 0.12159229 | 6.18066968 | 6.38E-10 | 8.01E-08 | SLC7A11 | 23657 |
| ENSG000001 | 1456.78651 | 0.96287696 | 0.15578564 | 6.18078116 | 6.38E-10 | 8.01E-08 | RPS21 | 6227 |
| ENSG00000C | 943.809049 | 0.736694671 | 0.11921261 | 6.1796709 | 6.42E-10 | 8.04E-08 | LIMA1 | 51474 |
| ENSG000001 | 721.568764 | 0.84777095 | 0.13719928 | 6.17912113 | 6.45E-10 | 8.04E-08 | COX6C | 1345 |
| ENSG000001 | 1180.18634 | 0.744295042 | 0.12046971 | 6.17827527 | 6.48E-10 | 8.06E-08 | CENPV | 201161 |
| ENSG000001 | 4758.52879 | 0.839875034 | 0.13602372 | 6.17447486 | 6.64E-10 | 8.23E-08 | SERPINH1 | 871 |
| ENSG000001 | 1153.42495 | 0.733259766 | 0.11878888 | 6.172798 | 6.71E-10 | 8.29E-08 | TBC1D14 | 57533 |
| ENSG000001 | 455.603499 | 0.954309795 | 0.15464535 | 6.1709569 | 6.79E-10 | 8.37E-08 | NA | NA |
| ENSG000001 | 720.726217 | 0.836598514 | 0.13561122 | 6.16909509 | 6.87E-10 | 8.44E-08 | ANAPC16 | 119504 |
| ENSG000001 | 1055.61608 | 0.854755534 | 0.13858122 | 6.16790305 | 6.92E-10 | 8.48E-08 | IGFBP2 | 3485 |
| ENSG000002 | 798.190351 | 0.837190148 | 0.1357541 | 6.16696052 | 6.96E-10 | 8.50E-08 | NA | NA |
| ENSG000002 | 821.707537 | 0.972226441 | 0.157684 | 6.16566342 | 7.02E-10 | 8.55E-08 | RN7SKP255 | 106479209 |
| ENSG00000C | 190.874381 | 0.946269838 | 0.15351051 | 6.16420217 | 7.08E-10 | 8.60E-08 | BLVRB | 645 |
| ENSG00000C | 544.67501 | 0.842613481 | 0.13678675 | 6.16005181 | 7.27E-10 | 8.80E-08 | GBA2 | 57704 |
| ENSG000001 | 779.430198 | 0.895377342 | 0.14545512 | 6.155695 | 7.47E-10 | 9.02E-08 | AAMP | 14 |
| ENSG000001 | 7993.90237 | 0.786071349 | 0.12770866 | 6.15519202 | 7.50E-10 | 9.02E-08 | RPL35 | 11224 |
| ENSG000001 | 1549.52859 | 0.862167196 | 0.1400924 | 6.15427508 | 7.54E-10 | 9.05E-08 | STOML2 | 30968 |
| ENSG000001 | 2015.766 | 0.806978013 | 0.13140421 | 6.14118819 | 8.19E-10 | 9.80E-08 | CCNB1IP1 | 57820 |
| ENSG000001 | 1081.69631 | 0.856257851 | 0.13943996 | 6.14069198 | 8.22E-10 | 9.80E-08 | SNRPB | 6628 |
| ENSG000001 | 446.42773 | 0.838616529 | 0.13658443 | 6.13991329 | 8.26E-10 | 9.82E-08 | MPP2 | 4355 |
| ENSG000001 | 1490.17937 | 0.71483736 | 0.11643475 | 6.13938174 | 8.28E-10 | 9.82E-08 | WBP5 | 51186 |
| ENSG00000C | 4331.5475 | 0.943962134 | 0.15378077 | 6.13836269 | 8.34E-10 | 9.86E-08 | SLC25A5 | 292 |
| ENSG00000C | 126.551818 | -1.066534358 | 0.17376802 | -6.1376909 | 8.37E-10 | 9.87E-08 | USH2A | 7399 |
| ENSG000001 | 199.653194 | 0.869028389 | 0.14161181 | 6.13669415 | 8.43E-10 | 9.90E-08 | CLGN | 1047 |
| ENSG00000C | 648.08195 | 0.775957083 | 0.12652916 | 6.13263432 | 8.64E-10 | 1.01E-07 | HEPH | 9843 |

| | | | | | | | | |
|---|---|---|---|---|---|---|---|---|
| ENSG000001 | 845.827275 | 0.879183945 | 0.14347407 | 6.12782481 | 8.91E-10 | 1.04E-07 | HIST2H3D | 653604 |
| ENSG000001 | 408.286521 | 0.796438033 | 0.12998859 | 6.12698424 | 8.96E-10 | 1.04E-07 | NAA40 | 79829 |
| ENSG000001 | 269.642828 | 0.945009796 | 0.15426533 | 6.12587293 | 9.02E-10 | 1.05E-07 | CTNNBIP1 | 56998 |
| ENSG000001 | 659.586003 | 0.875804524 | 0.14298067 | 6.12533505 | 9.05E-10 | 1.05E-07 | RNASEH2A | 10535 |
| ENSG000001 | 1975.23259 | 0.936974848 | 0.15300502 | 6.12381763 | 9.14E-10 | 1.06E-07 | VAT1 | 10493 |
| ENSG00000C | 4470.82524 | 0.841154575 | 0.13738609 | 6.12256016 | 9.21E-10 | 1.06E-07 | RPS5 | 6193 |
| ENSG00000C | 558.306004 | 0.808494153 | 0.13209643 | 6.1204845 | 9.33E-10 | 1.07E-07 | PDK3 | 5165 |
| ENSG000001 | 213.338781 | 1.0221393 | 0.16709301 | 6.11718768 | 9.52E-10 | 1.09E-07 | TMEM129 | 92305 |
| ENSG000002 | 239.009053 | 1.024833829 | 0.16753509 | 6.11712955 | 9.53E-10 | 1.09E-07 | NA | NA |
| ENSG000001 | 1451.48014 | 0.814823567 | 0.13327954 | 6.11364308 | 9.74E-10 | 1.11E-07 | HAX1 | 10456 |
| ENSG000002 | 241.27483 | 0.886972334 | 0.14508969 | 6.11326911 | 9.76E-10 | 1.11E-07 | SEPP1 | 6414 |
| ENSG000002 | 695.562154 | 0.877697121 | 0.14355579 | 6.11397934 | 9.72E-10 | 1.11E-07 | NA | NA |
| ENSG000001 | 9975.136 | 0.699324986 | 0.11447245 | 6.10911175 | 1.00E-09 | 1.13E-07 | STAG2 | 10735 |
| ENSG000001 | 324.516359 | 0.851466508 | 0.13939587 | 6.10826219 | 1.01E-09 | 1.13E-07 | RNF185 | 91445 |
| ENSG000001 | 7726.05886 | 0.976432953 | 0.15989759 | 6.10661456 | 1.02E-09 | 1.14E-07 | TKT | 7086 |
| ENSG000001 | 242.78276 | 0.866858454 | 0.14204775 | 6.10258476 | 1.04E-09 | 1.17E-07 | ASF1B | 55723 |
| ENSG000001 | 211.036901 | 0.928619764 | 0.15218461 | 6.10192952 | 1.05E-09 | 1.17E-07 | CYSTM1 | 84418 |
| ENSG000001 | 174.69405 | 1.036393621 | 0.17000076 | 6.09640584 | 1.08E-09 | 1.21E-07 | GCAT | 23464 |
| ENSG000001 | 263.925058 | 0.893597754 | 0.14671701 | 6.09062135 | 1.12E-09 | 1.25E-07 | TMEM219 | 124446 |
| ENSG000001 | 1224.52735 | 0.850393622 | 0.13963263 | 6.09022137 | 1.13E-09 | 1.25E-07 | RUVBL2 | 10856 |
| ENSG000001 | 375.696553 | 0.847173648 | 0.13927741 | 6.082635 | 1.18E-09 | 1.31E-07 | TCEAL7 | 56849 |
| ENSG000001 | 1305.95961 | 0.78764315 | 0.12950844 | 6.08179013 | 1.19E-09 | 1.31E-07 | LAMP1 | 3916 |
| ENSG000001 | 231.900909 | 0.91164788 | 0.15012956 | 6.07240749 | 1.26E-09 | 1.38E-07 | CASP9 | 842 |
| ENSG000001 | 134.516804 | -0.950679413 | 0.15655005 | -6.0726867 | 1.26E-09 | 1.38E-07 | ENPP2 | 5168 |
| ENSG000001 | 211.275197 | 1.055028842 | 0.17371592 | 6.07329954 | 1.25E-09 | 1.38E-07 | MT1E | 4493 |
| ENSG000001 | 400.590254 | 0.809681788 | 0.13334438 | 6.07211013 | 1.26E-09 | 1.38E-07 | BRMS1 | 25855 |
| ENSG00000C | 516.328307 | 0.741857482 | 0.12219273 | 6.07120822 | 1.27E-09 | 1.38E-07 | CERS4 | 79603 |
| ENSG000001 | 611.132673 | 0.776731975 | 0.12800662 | 6.06790467 | 1.30E-09 | 1.41E-07 | SLC25A4 | 291 |
| ENSG000001 | 2495.52898 | 1.014673432 | 0.16727858 | 6.06577014 | 1.31E-09 | 1.42E-07 | HIST1H2AI | 8329 |

| | | | | | | | | |
|---|---|---|---|---|---|---|---|---|
| ENSG000001 | 2460.59693 | 0.797032082 | 0.13147285 | 6.06233229 | 1.34E-09 | 1.45E-07 | OAZ1 | 4946 |
| ENSG000001 | 161.58673 | 1.099282771 | 0.18138634 | 6.06044961 | 1.36E-09 | 1.46E-07 | PTP4A3 | 11156 |
| ENSG00000C | 208.871993 | 0.905270421 | 0.14945873 | 6.05699268 | 1.39E-09 | 1.48E-07 | NA | NA |
| ENSG000001 | 3073.33648 | 0.689377157 | 0.11380864 | 6.0573358 | 1.38E-09 | 1.48E-07 | EIF2AK4 | 440275 |
| ENSG000001 | 1289.03924 | 0.75749655 | 0.12506051 | 6.05704028 | 1.39E-09 | 1.48E-07 | NIPSNAP1 | 8508 |
| ENSG00000C | 1263.52621 | 0.800457798 | 0.13220425 | 6.05470559 | 1.41E-09 | 1.50E-07 | SLC25A39 | 51629 |
| ENSG000001 | 603.351055 | 0.850732451 | 0.14056777 | 6.05211582 | 1.43E-09 | 1.52E-07 | BAP1 | 8314 |
| ENSG000001 | 26.7816114 | -1.685460716 | 0.27857082 | -6.0503851 | 1.45E-09 | 1.53E-07 | ZNF572 | 137209 |
| ENSG000001 | 818.521959 | 0.815405767 | 0.13478365 | 6.04973804 | 1.45E-09 | 1.53E-07 | C8orf33 | 65265 |
| ENSG00000C | 227.89892 | 0.953483538 | 0.15762601 | 6.04902399 | 1.46E-09 | 1.53E-07 | RPUSD1 | 113000 |
| ENSG000001 | 485.843688 | 0.77797768 | 0.12861717 | 6.04878542 | 1.46E-09 | 1.53E-07 | YIPF2 | 78992 |
| ENSG000001 | 1056.03482 | 0.836111923 | 0.13825264 | 6.04771019 | 1.47E-09 | 1.54E-07 | TCEB2 | 6923 |
| ENSG000001 | 391.29378 | 0.966706188 | 0.15985581 | 6.04736366 | 1.47E-09 | 1.54E-07 | ZNF768 | 79724 |
| ENSG000001 | 1053.59097 | 0.776111184 | 0.12841015 | 6.04400177 | 1.50E-09 | 1.57E-07 | ATP6V1D | 51382 |
| ENSG000002 | 1060.28901 | 0.749602692 | 0.12407306 | 6.04162335 | 1.53E-09 | 1.59E-07 | THAP9-AS1 | 100499177 |
| ENSG000001 | 283.057843 | 0.932082579 | 0.15430502 | 6.04052 | 1.54E-09 | 1.59E-07 | FAHD2A | 51011 |
| ENSG000001 | 324.726067 | 0.838865377 | 0.13888306 | 6.04008436 | 1.54E-09 | 1.59E-07 | GSN | 2934 |
| ENSG000001 | 133.709821 | 0.977097432 | 0.16206563 | 6.02902305 | 1.65E-09 | 1.70E-07 | MX2 | 4600 |
| ENSG000001 | 5299.23375 | 0.688673954 | 0.11439948 | 6.01990463 | 1.75E-09 | 1.80E-07 | NGFRAP1 | 27018 |
| ENSG000001 | 1160.55505 | 0.791983752 | 0.13157368 | 6.01931744 | 1.75E-09 | 1.80E-07 | DHTKD1 | 55526 |
| ENSG000001 | 453.671514 | 0.838541081 | 0.13942415 | 6.01431736 | 1.81E-09 | 1.85E-07 | IRF3 | 3661 |
| ENSG000001 | 1094.77932 | 0.887400371 | 0.14764747 | 6.01026483 | 1.85E-09 | 1.89E-07 | FAM46B | 115572 |
| ENSG000002 | 3091.38644 | 0.810973438 | 0.1349554 | 6.0091958 | 1.86E-09 | 1.90E-07 | UBA52 | 7311 |
| ENSG000001 | 1206.50482 | 0.711116573 | 0.11835952 | 6.00810613 | 1.88E-09 | 1.91E-07 | WDR45B | 56270 |
| ENSG000002 | 702.544771 | 0.81328503 | 0.13538625 | 6.00714629 | 1.89E-09 | 1.91E-07 | NELFE | 7936 |
| ENSG00000C | 3432.71404 | 0.727642427 | 0.12121091 | 6.00311002 | 1.94E-09 | 1.96E-07 | NA | NA |
| ENSG00000C | 912.923933 | 0.690442979 | 0.11506179 | 6.00062805 | 1.97E-09 | 1.98E-07 | PITHD1 | 57095 |
| ENSG000001 | 1045.65721 | 0.86339366 | 0.1438962 | 6.00011457 | 1.97E-09 | 1.98E-07 | MAD2L2 | 10459 |
| ENSG000001 | 147.499078 | 1.027490207 | 0.17124268 | 6.00019932 | 1.97E-09 | 1.98E-07 | BRI3 | 25798 |

| | | | | | | | | |
|---|---|---|---|---|---|---|---|---|
| ENSG000001 | 1147.30068 | 0.734102078 | 0.12237997 | 5.99854781 | 1.99E-09 | 1.99E-07 | BFAR | 51283 |
| ENSG000001 | 680.017697 | 0.812735274 | 0.13550615 | 5.99777398 | 2.00E-09 | 1.99E-07 | CTH | 1491 |
| ENSG000001 | 302.812792 | 0.941411312 | 0.15696847 | 5.9974547 | 2.00E-09 | 1.99E-07 | SIVA1 | 10572 |
| ENSG000002 | 880.046037 | 0.716232485 | 0.11942198 | 5.99749295 | 2.00E-09 | 1.99E-07 | NA | NA |
| ENSG000001 | 1225.12216 | 0.785563252 | 0.13099968 | 5.99668067 | 2.01E-09 | 2.00E-07 | PSMA7 | 5688 |
| ENSG000002 | 3035.77553 | 0.947724924 | 0.15805555 | 5.99615083 | 2.02E-09 | 2.00E-07 | HIST1H2AH | 85235 |
| ENSG00000C | 238.026245 | 0.954024996 | 0.15918863 | 5.99304719 | 2.06E-09 | 2.03E-07 | RNH1 | 6050 |
| ENSG000001 | 1566.39606 | 0.837123651 | 0.13968488 | 5.99294379 | 2.06E-09 | 2.03E-07 | ATP5G2 | 517 |
| ENSG000001 | 1148.64737 | 0.803435329 | 0.13410967 | 5.99088312 | 2.09E-09 | 2.05E-07 | VEGFA | 7422 |
| ENSG000001 | 839.220819 | 0.919167407 | 0.15347985 | 5.9888475 | 2.11E-09 | 2.07E-07 | C19orf48 | 84798 |
| ENSG000002 | 599.229416 | 0.786508138 | 0.13138986 | 5.98606399 | 2.15E-09 | 2.10E-07 | MCTS1 | 28985 |
| ENSG000001 | 753.888892 | 0.875170988 | 0.14622926 | 5.98492402 | 2.16E-09 | 2.11E-07 | GPS1 | 2873 |
| ENSG000001 | 2185.27275 | 0.77625319 | 0.12973958 | 5.98316411 | 2.19E-09 | 2.13E-07 | ATP6V1G1 | 9550 |
| ENSG000001 | 467.071612 | 0.815617415 | 0.13635169 | 5.98171842 | 2.21E-09 | 2.14E-07 | NA | NA |
| ENSG000001 | 1659.95026 | 0.68232551 | 0.11411601 | 5.97922699 | 2.24E-09 | 2.17E-07 | NIPA2 | 81614 |
| ENSG000001 | 98.2082853 | 1.051951404 | 0.175985 | 5.97750614 | 2.27E-09 | 2.19E-07 | HSPB8 | 26353 |
| ENSG000001 | 3025.65821 | 0.721588471 | 0.12072694 | 5.97702948 | 2.27E-09 | 2.19E-07 | BUB3 | 9184 |
| ENSG000002 | 2843.72591 | 0.819163172 | 0.13716868 | 5.97194049 | 2.34E-09 | 2.25E-07 | HIST1H2BO | 8348 |
| ENSG000001 | 5189.54305 | 0.793915575 | 0.13295566 | 5.97128094 | 2.35E-09 | 2.26E-07 | HIST1H2BD | 3017 |
| ENSG00000C | 3311.31728 | 0.714186495 | 0.11964899 | 5.96901374 | 2.39E-09 | 2.28E-07 | EPS15 | 2060 |
| ENSG000001 | 251.47458 | 0.877603428 | 0.14703595 | 5.96863157 | 2.39E-09 | 2.28E-07 | MIIP | 60672 |
| ENSG000002 | 11.9214003 | -2.095340848 | 0.35106264 | -5.9685669 | 2.39E-09 | 2.28E-07 | NA | NA |
| ENSG000002 | 3220.48887 | 0.734165941 | 0.12308733 | 5.96459408 | 2.45E-09 | 2.33E-07 | HIST1H3I | 8354 |
| ENSG000002 | 390.806594 | 0.812061405 | 0.13619876 | 5.96232584 | 2.49E-09 | 2.36E-07 | PFDN6 | 10471 |
| ENSG000001 | 716.692553 | 0.888875391 | 0.14915124 | 5.9595574 | 2.53E-09 | 2.39E-07 | NUCB1 | 4924 |
| ENSG000002 | 1167.09777 | 0.739219765 | 0.12403977 | 5.95953818 | 2.53E-09 | 2.39E-07 | NA | NA |
| ENSG000001 | 2588.16934 | 0.92879181 | 0.1558856 | 5.95816293 | 2.55E-09 | 2.40E-07 | HIST1H4C | 8364 |
| ENSG000001 | 765.940247 | 0.733347907 | 0.1231464 | 5.95508994 | 2.60E-09 | 2.44E-07 | OS9 | 10956 |
| ENSG000001 | 555.340457 | 0.697326696 | 0.11715126 | 5.95236183 | 2.64E-09 | 2.48E-07 | STC1 | 6781 |

| | | | | | | | | |
|---|---|---|---|---|---|---|---|---|
| ENSG000001 | 481.696006 | 0.7766879 | 0.13051554 | 5.95092291 | 2.67E-09 | 2.49E-07 | CNPY3 | 10695 |
| ENSG000002 | 838.380236 | 0.804761567 | 0.13523171 | 5.9509828 | 2.67E-09 | 2.49E-07 | E2F4 | 1874 |
| ENSG000001 | 1049.49805 | 0.833275271 | 0.14012161 | 5.94680038 | 2.73E-09 | 2.54E-07 | ATP5I | 521 |
| ENSG000001 | 2704.95463 | 0.849459289 | 0.14300602 | 5.94002476 | 2.85E-09 | 2.64E-07 | RPL10 | 6134 |
| ENSG000001 | 1493.77176 | 0.682835247 | 0.11500279 | 5.93755361 | 2.89E-09 | 2.68E-07 | GTF2E1 | 2960 |
| ENSG000001 | 2028.4385 | 0.886169219 | 0.14929762 | 5.93558844 | 2.93E-09 | 2.70E-07 | STC2 | 8614 |
| ENSG000001 | 374.082605 | 0.894279242 | 0.15071082 | 5.93374286 | 2.96E-09 | 2.73E-07 | SLC35B2 | 347734 |
| ENSG000001 | 166.751722 | 0.920533651 | 0.1552202 | 5.93050158 | 3.02E-09 | 2.78E-07 | SLC1A4 | 6509 |
| ENSG000001 | 837.721753 | 0.823069129 | 0.13887019 | 5.92689573 | 3.09E-09 | 2.83E-07 | CUTA | 51596 |
| ENSG000001 | 10547.8474 | 0.784929589 | 0.13244496 | 5.92645851 | 3.10E-09 | 2.83E-07 | RPL7A | 6130 |
| ENSG000001 | 720.21775 | 0.761804962 | 0.12859305 | 5.92415336 | 3.14E-09 | 2.87E-07 | MRPL10 | 124995 |
| ENSG000001 | 3415.24225 | 0.854352154 | 0.14427788 | 5.92157395 | 3.19E-09 | 2.91E-07 | HIST1H2AC | 8334 |
| ENSG00000C | 749.904016 | 0.715614902 | 0.12103351 | 5.91253526 | 3.37E-09 | 3.06E-07 | POLDIP2 | 26073 |
| ENSG000001 | 2843.60699 | 0.862211139 | 0.14582121 | 5.91279631 | 3.36E-09 | 3.06E-07 | PSAP | 5660 |
| ENSG000001 | 449.510237 | 0.765119936 | 0.12943651 | 5.91116021 | 3.40E-09 | 3.07E-07 | FAM83F | 113828 |
| ENSG000001 | 5022.04455 | 0.736114549 | 0.12462096 | 5.9068277 | 3.49E-09 | 3.15E-07 | ALDOA | 226 |
| ENSG000001 | 2333.30707 | 0.780234741 | 0.13219215 | 5.90227731 | 3.59E-09 | 3.23E-07 | HINT1 | 3094 |
| ENSG000002 | 409.593821 | 0.825713746 | 0.13991826 | 5.90140075 | 3.60E-09 | 3.24E-07 | PISD | 23761 |
| ENSG000001 | 385.368111 | 0.774413322 | 0.1313059 | 5.89778011 | 3.68E-09 | 3.31E-07 | CLPB | 81570 |
| ENSG000001 | 520.10236 | 0.884033764 | 0.14991816 | 5.89677573 | 3.71E-09 | 3.32E-07 | AURKAIP1 | 54998 |
| ENSG000001 | 410.878536 | 0.793054942 | 0.13452324 | 5.89530061 | 3.74E-09 | 3.34E-07 | C7orf50 | 84310 |
| ENSG000001 | 97.0117608 | 1.128450669 | 0.19159574 | 5.88974807 | 3.87E-09 | 3.45E-07 | SCRN2 | 90507 |
| ENSG00000C | 587.800363 | 0.764174663 | 0.12977954 | 5.888252 | 3.90E-09 | 3.47E-07 | CD59 | 966 |
| ENSG000001 | 5740.31031 | 0.772127783 | 0.13115771 | 5.88701808 | 3.93E-09 | 3.49E-07 | TPI1 | 7167 |
| ENSG000001 | 1237.89711 | 0.74848166 | 0.12717389 | 5.88549809 | 3.97E-09 | 3.51E-07 | RAP2C | 57826 |
| ENSG000001 | 265.395931 | 0.863149555 | 0.146675 | 5.88477616 | 3.99E-09 | 3.52E-07 | SSSCA1 | 10534 |
| ENSG000001 | 918.195866 | 0.879653265 | 0.14952079 | 5.8831501 | 4.03E-09 | 3.55E-07 | ITPK1 | 3705 |
| ENSG000001 | 492.202542 | 0.833431512 | 0.14167822 | 5.88256635 | 4.04E-09 | 3.55E-07 | SF3B5 | 83443 |
| ENSG00000C | 94.4985977 | 1.030248944 | 0.17518448 | 5.88093719 | 4.08E-09 | 3.58E-07 | TAC1 | 6863 |

| | | | | | | | | |
|---|---|---|---|---|---|---|---|---|
| ENSG000002 | 319.267617 | 0.971695807 | 0.16524521 | 5.88032645 | 4.09E-09 | 3.59E-07 | FAM195B | 348262 |
| ENSG000001 | 2755.11752 | 0.760109774 | 0.12930127 | 5.87859485 | 4.14E-09 | 3.62E-07 | PHB | 5245 |
| ENSG000001 | 313.533462 | 0.948169768 | 0.16132536 | 5.87737592 | 4.17E-09 | 3.63E-07 | MGAT4B | 11282 |
| ENSG000001 | 10302.4258 | 0.642259584 | 0.10927284 | 5.87757759 | 4.16E-09 | 3.63E-07 | IARS | 3376 |
| ENSG000001 | 772.03079 | 0.86352352 | 0.14693686 | 5.87683405 | 4.18E-09 | 3.63E-07 | UROD | 7389 |
| ENSG000000 | 668.583302 | 0.781199808 | 0.13294061 | 5.87630673 | 4.20E-09 | 3.64E-07 | NDUFAF7 | 55471 |
| ENSG000001 | 328.349941 | 0.771690299 | 0.13133775 | 5.87561679 | 4.21E-09 | 3.64E-07 | ANKRD46 | 157567 |
| ENSG000001 | 829.373826 | 0.780182175 | 0.13287447 | 5.87157315 | 4.32E-09 | 3.72E-07 | RNF7 | 9616 |
| ENSG000001 | 6768.5591 | 0.776928614 | 0.13231759 | 5.87169582 | 4.31E-09 | 3.72E-07 | PRDX1 | 5052 |
| ENSG000001 | 8232.55238 | 0.846052503 | 0.14410017 | 5.87128049 | 4.32E-09 | 3.72E-07 | RPS7 | 6201 |
| ENSG000001 | 270.871151 | 0.797150223 | 0.13580664 | 5.86974389 | 4.36E-09 | 3.74E-07 | MGAT4C | 25834 |
| ENSG000001 | 340.951397 | 0.890171417 | 0.15169626 | 5.86811715 | 4.41E-09 | 3.77E-07 | PGP | 283871 |
| ENSG000001 | 649.724494 | 0.757866724 | 0.12917153 | 5.86713451 | 4.43E-09 | 3.79E-07 | MRPL44 | 65080 |
| ENSG000001 | 1545.78977 | 0.800082959 | 0.13638115 | 5.86652168 | 4.45E-09 | 3.79E-07 | EIF3K | 27335 |
| ENSG000001 | 837.855686 | 0.690196264 | 0.11767482 | 5.86528414 | 4.48E-09 | 3.81E-07 | TXNL4A | 10907 |
| ENSG000000 | 296.436135 | 0.896308979 | 0.15285542 | 5.8637695 | 4.52E-09 | 3.83E-07 | MUL1 | 79594 |
| ENSG000002 | 733.107702 | 0.759711625 | 0.1295594 | 5.86380945 | 4.52E-09 | 3.83E-07 | NA | NA |
| ENSG000001 | 321.317869 | 0.92650742 | 0.15801939 | 5.86325128 | 4.54E-09 | 3.83E-07 | RPS6KB2 | 6199 |
| ENSG000001 | 2265.65955 | -0.739237676 | 0.12609698 | -5.8624533 | 4.56E-09 | 3.84E-07 | INPP5F | 22876 |
| ENSG000001 | 5946.51572 | 0.775269317 | 0.1323245 | 5.85884919 | 4.66E-09 | 3.92E-07 | CCT7 | 10574 |
| ENSG000000 | 17173.9856 | 0.767709789 | 0.13104924 | 5.8581781 | 4.68E-09 | 3.93E-07 | ENO1 | 2023 |
| ENSG000001 | 569.341918 | 0.741491078 | 0.12660427 | 5.85676199 | 4.72E-09 | 3.95E-07 | ANXA11 | 311 |
| ENSG000001 | 340.214317 | 0.760790866 | 0.12993433 | 5.85519529 | 4.76E-09 | 3.98E-07 | PCDH11X | 27328 |
| ENSG000001 | 1331.98196 | 0.693873862 | 0.11850859 | 5.8550511 | 4.77E-09 | 3.98E-07 | ERGIC1 | 57222 |
| ENSG000001 | 1561.59024 | 0.699332266 | 0.11944957 | 5.85462349 | 4.78E-09 | 3.98E-07 | GLS | 2744 |
| ENSG000002 | 84.9727558 | -1.112000441 | 0.19000345 | -5.8525277 | 4.84E-09 | 4.02E-07 | NA | NA |
| ENSG000000 | 1384.84038 | 0.954878249 | 0.16317158 | 5.85198875 | 4.86E-09 | 4.03E-07 | PDZD4 | 57595 |
| ENSG000001 | 5085.04296 | 0.772665449 | 0.13206327 | 5.85072169 | 4.89E-09 | 4.04E-07 | RPL18A | 6142 |
| ENSG000001 | 179.542243 | 0.943378549 | 0.16125029 | 5.85039905 | 4.90E-09 | 4.04E-07 | TYSND1 | 219743 |

| | | | | | | | | |
|---|---|---|---|---|---|---|---|---|
| ENSG000001 | 228.490921 | 0.809462321 | 0.13835318 | 5.8506954 | 4.90E-09 | 4.04E-07 | COQ4 | 51117 |
| ENSG000000 | 2202.61763 | 0.774052466 | 0.13236468 | 5.84787767 | 4.98E-09 | 4.10E-07 | HADHA | 3030 |
| ENSG000001 | 3453.46406 | 0.78122442 | 0.13365699 | 5.84499479 | 5.07E-09 | 4.16E-07 | TCEA1 | 6917 |
| ENSG000001 | 11691.9817 | 0.827784476 | 0.14166376 | 5.84330443 | 5.12E-09 | 4.19E-07 | RPSA | 3921 |
| ENSG000001 | 3201.37608 | 0.750829209 | 0.12853691 | 5.8413511 | 5.18E-09 | 4.23E-07 | HIST2H2BE | 8349 |
| ENSG000001 | 919.061639 | 0.772922056 | 0.13236424 | 5.839357 | 5.24E-09 | 4.28E-07 | NA | NA |
| ENSG000001 | 1946.33166 | 0.715991443 | 0.12263505 | 5.83839167 | 5.27E-09 | 4.29E-07 | CDK6 | 1021 |
| ENSG000001 | 1679.00297 | 0.897995133 | 0.15388932 | 5.83533125 | 5.37E-09 | 4.36E-07 | CD63 | 967 |
| ENSG000001 | 283.347372 | 0.810394057 | 0.13891609 | 5.83369466 | 5.42E-09 | 4.40E-07 | COX6A1 | 1337 |
| ENSG000001 | 16486.7905 | 0.777765529 | 0.13333345 | 5.83323643 | 5.44E-09 | 4.40E-07 | RPL13A | 23521 |
| ENSG000001 | 1757.49609 | 0.689867373 | 0.11829361 | 5.83182287 | 5.48E-09 | 4.42E-07 | SYAP1 | 94056 |
| ENSG000001 | 116.782226 | -1.048223012 | 0.17974777 | -5.831633 | 5.49E-09 | 4.42E-07 | DNAH12 | 201625 |
| ENSG000001 | 8593.60384 | 0.762296532 | 0.13070801 | 5.83205679 | 5.47E-09 | 4.42E-07 | RPL37A | 6168 |
| ENSG000001 | 1260.747 | 0.790546152 | 0.13557815 | 5.83092605 | 5.51E-09 | 4.43E-07 | FAU | 2197 |
| ENSG000001 | 2086.27571 | 0.794915408 | 0.13633559 | 5.83057883 | 5.52E-09 | 4.43E-07 | NDUFA4 | 4697 |
| ENSG000001 | 206.27163 | 0.848547112 | 0.1455887 | 5.82838562 | 5.60E-09 | 4.48E-07 | HPS1 | 3257 |
| ENSG000001 | 6720.97456 | 0.852733579 | 0.14634576 | 5.82684187 | 5.65E-09 | 4.51E-07 | RPS14 | 6208 |
| ENSG000000 | 825.672865 | 0.752000356 | 0.12908311 | 5.82570675 | 5.69E-09 | 4.53E-07 | PGM1 | 5236 |
| ENSG000001 | 239.938654 | 0.950055999 | 0.16309218 | 5.82526995 | 5.70E-09 | 4.53E-07 | CHCHD2 | 51142 |
| ENSG000001 | 673.105275 | 0.659944953 | 0.11330335 | 5.82458446 | 5.73E-09 | 4.54E-07 | FAM21C | 253725 |
| ENSG000001 | 321.367309 | 0.874407191 | 0.15018661 | 5.82213825 | 5.81E-09 | 4.60E-07 | DAPK3 | 1613 |
| ENSG000000 | 480.410775 | 0.854973163 | 0.14689393 | 5.8203438 | 5.87E-09 | 4.64E-07 | OTUD5 | 55593 |
| ENSG000002 | 50.5291301 | 1.297544519 | 0.22295924 | 5.81964911 | 5.90E-09 | 4.65E-07 | NA | NA |
| ENSG000000 | 168.774003 | 0.893178926 | 0.15357277 | 5.8159979 | 6.03E-09 | 4.74E-07 | WIPI1 | 55062 |
| ENSG000001 | 549.228706 | 0.823127097 | 0.14152899 | 5.81596101 | 6.03E-09 | 4.74E-07 | NA | NA |
| ENSG000001 | 784.118507 | 0.786236573 | 0.13520984 | 5.81493614 | 6.07E-09 | 4.76E-07 | AP1M2 | 10053 |
| ENSG000001 | 322.367731 | 0.822840348 | 0.14151492 | 5.81451297 | 6.08E-09 | 4.76E-07 | COPE | 11316 |
| ENSG000001 | 645.786167 | 0.640242039 | 0.11014911 | 5.81250313 | 6.15E-09 | 4.81E-07 | PLCB1 | 23236 |
| ENSG000001 | 8902.47802 | 0.90501227 | 0.15571376 | 5.81202502 | 6.17E-09 | 4.81E-07 | RPL8 | 6132 |

| | | | | | | | | |
|---|---|---|---|---|---|---|---|---|
| ENSG000001 | 438.556692 | 0.792135342 | 0.13632356 | 5.81070026 | 6.22E-09 | 4.84E-07 | ERI3 | 79033 |
| ENSG000001 | 3265.49537 | 0.74036248 | 0.12741793 | 5.81050467 | 6.23E-09 | 4.84E-07 | FSTL1 | 11167 |
| ENSG00000C | 447.360543 | 0.834220324 | 0.14358675 | 5.8098698 | 6.25E-09 | 4.85E-07 | AP2S1 | 1175 |
| ENSG000001 | 1706.57655 | 0.819795278 | 0.14112148 | 5.80914583 | 6.28E-09 | 4.86E-07 | LSM4 | 25804 |
| ENSG000002 | 1691.56752 | 0.697586102 | 0.12011813 | 5.80750053 | 6.34E-09 | 4.90E-07 | NA | NA |
| ENSG000001 | 616.674095 | 0.812642174 | 0.13995254 | 5.80655544 | 6.38E-09 | 4.92E-07 | B4GALT2 | 8704 |
| ENSG000001 | 1073.02024 | 0.860673537 | 0.148251 | 5.80551604 | 6.42E-09 | 4.94E-07 | PLD3 | 23646 |
| ENSG000002 | 611.706692 | 0.799755336 | 0.1378806 | 5.80034688 | 6.62E-09 | 5.08E-07 | APELA | 100506013 |
| ENSG000002 | 554.321649 | 0.852500214 | 0.14705939 | 5.79697902 | 6.75E-09 | 5.18E-07 | EIF6 | 3692 |
| ENSG000001 | 427.649614 | 0.837721302 | 0.14453403 | 5.79601436 | 6.79E-09 | 5.19E-07 | NA | NA |
| ENSG000002 | 2810.50076 | 0.73559037 | 0.12691085 | 5.79611902 | 6.79E-09 | 5.19E-07 | LINC00678 | 101410541 |
| ENSG000001 | 470.020914 | 0.873473242 | 0.15076009 | 5.79379605 | 6.88E-09 | 5.25E-07 | TRMT1 | 55621 |
| ENSG000001 | 584.731879 | 0.81521843 | 0.14072844 | 5.79284762 | 6.92E-09 | 5.27E-07 | THAP4 | 51078 |
| ENSG000001 | 1669.0267 | 0.760423166 | 0.13130701 | 5.79118484 | 6.99E-09 | 5.31E-07 | ATP5G3 | 518 |
| ENSG000001 | 1044.48273 | 0.649729662 | 0.11223272 | 5.78912894 | 7.08E-09 | 5.36E-07 | QPRT | 23475 |
| ENSG000001 | 331.986996 | 0.857955212 | 0.14826821 | 5.78650819 | 7.19E-09 | 5.44E-07 | MPST | 4357 |
| ENSG000001 | 582.790577 | 0.699765492 | 0.12094989 | 5.78558177 | 7.23E-09 | 5.46E-07 | SNX30 | 401548 |
| ENSG000001 | 557.908053 | 0.832865796 | 0.14400398 | 5.78363021 | 7.31E-09 | 5.51E-07 | MYL6B | 140465 |
| ENSG000001 | 801.610261 | 0.732159053 | 0.1266754 | 5.77980454 | 7.48E-09 | 5.63E-07 | APMAP | 57136 |
| ENSG000001 | 1379.49084 | 0.77996361 | 0.13498288 | 5.77824082 | 7.55E-09 | 5.67E-07 | TRMT112 | 51504 |
| ENSG000001 | 83.2522521 | 1.097287668 | 0.18994145 | 5.77697857 | 7.61E-09 | 5.70E-07 | FUOM | 282969 |
| ENSG000002 | 146.93766 | 1.115315709 | 0.19308249 | 5.77636905 | 7.63E-09 | 5.71E-07 | SCARNA5 | 677775 |
| ENSG00000C | 643.749713 | 0.804154512 | 0.13926653 | 5.77421225 | 7.73E-09 | 5.76E-07 | CNTN1 | 1272 |
| ENSG00000C | 314.183278 | 0.845492255 | 0.14642152 | 5.77437169 | 7.72E-09 | 5.76E-07 | SLC35C2 | 51006 |
| ENSG000002 | 1809.32252 | 0.980964564 | 0.16990946 | 5.77345465 | 7.77E-09 | 5.78E-07 | HIST1H4B | 8366 |
| ENSG000002 | 152.332724 | 1.047226552 | 0.18165596 | 5.76488961 | 8.17E-09 | 6.07E-07 | PSENEN | 55851 |
| ENSG000001 | 963.726907 | 0.716842348 | 0.12438002 | 5.76332379 | 8.25E-09 | 6.11E-07 | ZNF697 | 90874 |
| ENSG000001 | 150.495342 | 1.112315705 | 0.19302056 | 5.76267985 | 8.28E-09 | 6.13E-07 | SLC52A2 | 79581 |
| ENSG000001 | 2366.30532 | 0.627284869 | 0.10886761 | 5.76190564 | 8.32E-09 | 6.14E-07 | DFFA | 1676 |

| | | | | | | | | |
|---|---|---|---|---|---|---|---|---|
| ENSG000002 | 2718.90659 | 0.784272389 | 0.13617022 | 5.75950003 | 8.44E-09 | 6.22E-07 | HIST1H2BE | 8344 |
| ENSG000001 | 2281.15464 | 0.665744388 | 0.11564293 | 5.75689668 | 8.57E-09 | 6.31E-07 | SUDS3 | 64426 |
| ENSG000001 | 361.520642 | 0.844103115 | 0.1466586 | 5.7555651 | 8.64E-09 | 6.34E-07 | ATAD3B | 83858 |
| ENSG000001 | 487.094591 | 0.68842356 | 0.11962767 | 5.75471835 | 8.68E-09 | 6.36E-07 | GZF1 | 64412 |
| ENSG000001 | 1310.6224 | 0.695733496 | 0.12093726 | 5.75284637 | 8.78E-09 | 6.42E-07 | AHSA1 | 10598 |
| ENSG000001 | 1145.64985 | 0.841053503 | 0.14621995 | 5.75197522 | 8.82E-09 | 6.44E-07 | EXOC7 | 23265 |
| ENSG000002 | 2002.1666 | 0.734888227 | 0.12781925 | 5.74943303 | 8.95E-09 | 6.53E-07 | NA | NA |
| ENSG000001 | 1344.53373 | 0.724018973 | 0.12600875 | 5.74578353 | 9.15E-09 | 6.66E-07 | GFPT2 | 9945 |
| ENSG00000C | 3217.06897 | 0.6113579 | 0.10641521 | 5.74502363 | 9.19E-09 | 6.67E-07 | DNAJA2 | 10294 |
| ENSG000001 | 299.79035 | 0.926030304 | 0.16118393 | 5.74517765 | 9.18E-09 | 6.67E-07 | FGD5 | 152273 |
| ENSG000001 | 263.048763 | 0.888236858 | 0.15464329 | 5.74377868 | 9.26E-09 | 6.69E-07 | SERPINB6 | 5269 |
| ENSG000001 | 1245.92373 | 0.606052823 | 0.10551547 | 5.74373409 | 9.26E-09 | 6.69E-07 | FGF13 | 2258 |
| ENSG000002 | 3465.57859 | 0.68087428 | 0.11855833 | 5.74294763 | 9.30E-09 | 6.71E-07 | HIST2H2BF | 440689 |
| ENSG000001 | 405.355223 | 0.74415789 | 0.12961951 | 5.74109474 | 9.41E-09 | 6.78E-07 | NRK | 203447 |
| ENSG000001 | 246.334531 | 0.944418473 | 0.16452476 | 5.74028173 | 9.45E-09 | 6.80E-07 | TIMM10 | 26519 |
| ENSG00000C | 560.733576 | 0.788550294 | 0.137419 | 5.73829163 | 9.56E-09 | 6.86E-07 | FUCA2 | 2519 |
| ENSG000001 | 407.125619 | 0.798857117 | 0.13925329 | 5.73671978 | 9.65E-09 | 6.92E-07 | EMC8 | 10328 |
| ENSG000001 | 625.445405 | 0.836466514 | 0.14583638 | 5.7356507 | 9.71E-09 | 6.95E-07 | THOP1 | 7064 |
| ENSG000001 | 728.51524 | 0.67997565 | 0.11858443 | 5.73410571 | 9.80E-09 | 7.00E-07 | NQO2 | 4835 |
| ENSG00000C | 164.576249 | 0.944610111 | 0.16475748 | 5.73333689 | 9.85E-09 | 7.02E-07 | TMED1 | 11018 |
| ENSG000001 | 327.433455 | 0.861101371 | 0.15020175 | 5.73296511 | 9.87E-09 | 7.02E-07 | ATAD3A | 55210 |
| ENSG000002 | 14948.0427 | 0.736784328 | 0.12852751 | 5.73250292 | 9.90E-09 | 7.03E-07 | GNB2L1 | 10399 |
| ENSG000001 | 1795.74101 | 0.836685592 | 0.14599368 | 5.7309714 | 9.99E-09 | 7.08E-07 | TBC1D16 | 125058 |
| ENSG000001 | 259.565066 | 0.926978823 | 0.16179738 | 5.72925741 | 1.01E-08 | 7.11E-07 | ROMO1 | 140823 |
| ENSG000001 | 389.519043 | 0.896241568 | 0.15642361 | 5.72957988 | 1.01E-08 | 7.11E-07 | KATNB1 | 10300 |
| ENSG000001 | 573.497913 | 0.86237132 | 0.15052132 | 5.72923049 | 1.01E-08 | 7.11E-07 | SSR4 | 6748 |
| ENSG000001 | 156.09355 | 0.899463998 | 0.1570041 | 5.7289206 | 1.01E-08 | 7.12E-07 | SWI5 | 375757 |
| ENSG000001 | 942.78903 | 0.742267215 | 0.12962426 | 5.72629838 | 1.03E-08 | 7.21E-07 | GPR160 | 26996 |
| ENSG000001 | 3945.34991 | 0.777792284 | 0.13589164 | 5.72362119 | 1.04E-08 | 7.30E-07 | FADS2 | 9415 |

| | | | | | | | | |
|---|---|---|---|---|---|---|---|---|
| ENSG000001 | 350.95292 | 0.788281113 | 0.13771794 | 5.72388115 | 1.04E-08 | 7.30E-07 | ICT1 | 3396 |
| ENSG000001 | 460.192462 | 0.765676244 | 0.13384166 | 5.72076155 | 1.06E-08 | 7.40E-07 | CHST10 | 9486 |
| ENSG000001 | 4611.66268 | 0.690013138 | 0.12061565 | 5.72075971 | 1.06E-08 | 7.40E-07 | DDB1 | 1642 |
| ENSG000001 | 2492.10897 | 0.692393445 | 0.12105339 | 5.7197362 | 1.07E-08 | 7.43E-07 | SOD1 | 6647 |
| ENSG00000C | 1006.98049 | 0.682655791 | 0.11938238 | 5.71822897 | 1.08E-08 | 7.49E-07 | PHLPP1 | 23239 |
| ENSG000001 | 269.326895 | 0.856252655 | 0.14981074 | 5.71556264 | 1.09E-08 | 7.58E-07 | DVL1 | 1855 |
| ENSG000001 | 634.510954 | 0.713872889 | 0.12490158 | 5.71548329 | 1.09E-08 | 7.58E-07 | SLC25A38 | 54977 |
| ENSG000001 | 4145.28607 | 0.718037823 | 0.12576981 | 5.70914295 | 1.14E-08 | 7.80E-07 | SLC39A14 | 23516 |
| ENSG000001 | 528.910901 | 0.739233107 | 0.12947629 | 5.70940915 | 1.13E-08 | 7.80E-07 | C19orf53 | 28974 |
| ENSG000001 | 5333.92279 | 0.82167668 | 0.1439247 | 5.70907337 | 1.14E-08 | 7.80E-07 | RPS19 | 6223 |
| ENSG000001 | 440.338683 | 0.858042437 | 0.1502767 | 5.70975035 | 1.13E-08 | 7.80E-07 | ZNHIT1 | 10467 |
| ENSG000001 | 5133.88083 | 0.758680086 | 0.13288488 | 5.70930338 | 1.13E-08 | 7.80E-07 | LRRC75A-AS1 | 125144 |
| ENSG000001 | 701.076507 | 0.909503071 | 0.15931316 | 5.70890108 | 1.14E-08 | 7.80E-07 | UBL5 | 59286 |
| ENSG000001 | 636.582893 | 0.754842504 | 0.13224063 | 5.7080981 | 1.14E-08 | 7.83E-07 | WASF3 | 10810 |
| ENSG000001 | 493.539519 | 0.753178073 | 0.13197428 | 5.70700628 | 1.15E-08 | 7.86E-07 | TMED4 | 222068 |
| ENSG000001 | 1562.932 | 0.728622411 | 0.12771393 | 5.70511315 | 1.16E-08 | 7.91E-07 | RTCB | 51493 |
| ENSG000001 | 9733.90716 | 0.646456646 | 0.11330065 | 5.70567454 | 1.16E-08 | 7.91E-07 | CLIC4 | 25932 |
| ENSG000002 | 149.338808 | 0.903893661 | 0.15843495 | 5.70514048 | 1.16E-08 | 7.91E-07 | NA | NA |
| ENSG000001 | 1164.3979 | 0.728042857 | 0.12770938 | 5.70077842 | 1.19E-08 | 8.10E-07 | PRICKLE1 | 144165 |
| ENSG000001 | 385.635292 | 0.723491548 | 0.12692147 | 5.70030861 | 1.20E-08 | 8.11E-07 | IL13RA1 | 3597 |
| ENSG000001 | 1800.74619 | 0.697698749 | 0.12250701 | 5.69517406 | 1.23E-08 | 8.34E-07 | PSMD3 | 5709 |
| ENSG000001 | 1060.29732 | 0.724070265 | 0.12725334 | 5.68999016 | 1.27E-08 | 8.57E-07 | PATZ1 | 23598 |
| ENSG000001 | 1142.71065 | 0.83569394 | 0.14686648 | 5.69016123 | 1.27E-08 | 8.57E-07 | LRRC47 | 57470 |
| ENSG000001 | 1056.29483 | 0.805353401 | 0.14155593 | 5.68929488 | 1.28E-08 | 8.59E-07 | PPP1R14B | 26472 |
| ENSG000001 | 591.663227 | 0.738790545 | 0.12992409 | 5.68632447 | 1.30E-08 | 8.73E-07 | DCAF11 | 80344 |
| ENSG000002 | 5556.27654 | 0.856169464 | 0.15061903 | 5.68433778 | 1.31E-08 | 8.82E-07 | RPL41 | 6171 |
| ENSG000001 | 379.914957 | 0.89831301 | 0.15805445 | 5.68356678 | 1.32E-08 | 8.84E-07 | MMP24 | 10893 |
| ENSG000001 | 3737.67041 | 0.792009945 | 0.13937773 | 5.68247112 | 1.33E-08 | 8.88E-07 | HIST2H2AC | 8338 |
| ENSG000001 | 733.455751 | 0.7431992 | 0.13080112 | 5.68190263 | 1.33E-08 | 8.90E-07 | DZIP1 | 22873 |

| | | | | | | | | |
|---|---|---|---|---|---|---|---|---|
| ENSG000001 | 1951.83328 | 0.594231962 | 0.10459774 | 5.68111672 | 1.34E-08 | 8.93E-07 | AMMECR1 | 9949 |
| ENSG000001 | 6560.25297 | 0.709346542 | 0.12497847 | 5.67575006 | 1.38E-08 | 9.18E-07 | MORF4L2 | 9643 |
| ENSG000001 | 740.641864 | 0.690004945 | 0.12157145 | 5.67571517 | 1.38E-08 | 9.18E-07 | POLR2G | 5436 |
| ENSG000001 | 1667.83926 | 0.832747208 | 0.14674299 | 5.67486859 | 1.39E-08 | 9.21E-07 | CNDP2 | 55748 |
| ENSG000001 | 460.118877 | 0.888628765 | 0.15664971 | 5.67271258 | 1.41E-08 | 9.30E-07 | CHMP1A | 5119 |
| ENSG000001 | 945.251469 | 0.787148235 | 0.13876174 | 5.67266063 | 1.41E-08 | 9.30E-07 | COX5B | 1329 |
| ENSG000001 | 313.449347 | 0.828318194 | 0.14606013 | 5.67107677 | 1.42E-08 | 9.34E-07 | MRPS26 | 64949 |
| ENSG000001 | 155.686759 | 0.880965169 | 0.15534143 | 5.67115407 | 1.42E-08 | 9.34E-07 | KLF15 | 28999 |
| ENSG000002 | 4643.73531 | 0.764073907 | 0.13472023 | 5.6715605 | 1.42E-08 | 9.34E-07 | HIST1H2AE | 3012 |
| ENSG000001 | 6046.8754 | 0.643349607 | 0.11350296 | 5.66813063 | 1.44E-08 | 9.49E-07 | MCM4 | 4173 |
| ENSG000001 | 7992.50204 | 0.738386623 | 0.13032449 | 5.66575511 | 1.46E-08 | 9.59E-07 | RPL30 | 6156 |
| ENSG000001 | 227.452937 | 1.013458774 | 0.17887015 | 5.66589112 | 1.46E-08 | 9.59E-07 | SLC6A9 | 6536 |
| ENSG000001 | 1851.58835 | 0.719588987 | 0.12702627 | 5.66488304 | 1.47E-08 | 9.62E-07 | PTTG1IP | 754 |
| ENSG000002 | 837.005365 | 0.796677887 | 0.1406541 | 5.66409273 | 1.48E-08 | 9.65E-07 | SNORD116-18 | 100033430 |
| ENSG000001 | 75.4050081 | 1.073229075 | 0.18952637 | 5.66268995 | 1.49E-08 | 9.72E-07 | HRC | 3270 |
| ENSG00000C | 839.616083 | 0.78160394 | 0.13805642 | 5.66148184 | 1.50E-08 | 9.77E-07 | ADCY2 | 108 |
| ENSG000002 | 426.487237 | 0.742809575 | 0.13124506 | 5.65971468 | 1.52E-08 | 9.85E-07 | NEU1 | 4758 |
| ENSG000001 | 1082.15173 | 0.772071872 | 0.1364689 | 5.65749326 | 1.54E-08 | 9.95E-07 | TIMM44 | 10469 |
| ENSG000001 | 332.153269 | 0.886925232 | 0.15676949 | 5.65751164 | 1.54E-08 | 9.95E-07 | NKRF | 55922 |
| ENSG00000C | 3326.35487 | 0.697272436 | 0.12328158 | 5.65593353 | 1.55E-08 | 9.99E-07 | LMAN1 | 3998 |
| ENSG000001 | 291.331237 | 0.815851946 | 0.14423655 | 5.65634687 | 1.55E-08 | 9.99E-07 | GCDH | 2639 |
| ENSG000001 | 4490.82132 | 0.768499531 | 0.13587093 | 5.65609971 | 1.55E-08 | 9.99E-07 | MCM7 | 4176 |
| ENSG000001 | 1003.42095 | 0.744617356 | 0.13165862 | 5.65566749 | 1.55E-08 | 9.99E-07 | CHEK2 | 11200 |
| ENSG00000C | 413.010728 | 0.894282664 | 0.15818788 | 5.6532946 | 1.57E-08 | 1.01E-06 | HMG20B | 10362 |
| ENSG000001 | 403.702507 | 0.746763575 | 0.13212218 | 5.65206804 | 1.59E-08 | 1.02E-06 | TRIM65 | 201292 |
| ENSG00000C | 451.167532 | 0.714191688 | 0.12643045 | 5.64888984 | 1.61E-08 | 1.03E-06 | ATP6V1H | 51606 |
| ENSG00000C | 4264.95637 | 0.710456908 | 0.12576621 | 5.64902862 | 1.61E-08 | 1.03E-06 | PEBP1 | 5037 |
| ENSG000001 | 1022.09764 | 0.677836037 | 0.1199925 | 5.64898679 | 1.61E-08 | 1.03E-06 | GMPR2 | 51292 |
| ENSG000001 | 347.639665 | 0.733905547 | 0.12991658 | 5.64905243 | 1.61E-08 | 1.03E-06 | SLC37A1 | 54020 |

| | | | | | | | | |
|---|---|---|---|---|---|---|---|---|
| ENSG000001 | 240.449361 | 0.901491448 | 0.15969347 | 5.64513665 | 1.65E-08 | 1.05E-06 | RNMTL1 | 55178 |
| ENSG000001 | 508.318329 | 0.666814819 | 0.11816739 | 5.64296838 | 1.67E-08 | 1.06E-06 | LRP11 | 84918 |
| ENSG000001 | 479.361242 | 0.741073936 | 0.13137844 | 5.64075773 | 1.69E-08 | 1.07E-06 | SCYL1 | 57410 |
| ENSG000001 | 305.807285 | 0.948839068 | 0.16820506 | 5.64096611 | 1.69E-08 | 1.07E-06 | CD151 | 977 |
| ENSG000001 | 1186.17642 | 0.677847438 | 0.12018626 | 5.63997453 | 1.70E-08 | 1.08E-06 | RNF41 | 10193 |
| ENSG000001 | 1642.78037 | 0.729648552 | 0.12940218 | 5.63861104 | 1.71E-08 | 1.08E-06 | GPX8 | 493869 |
| ENSG00000C | 203.962082 | 1.020198958 | 0.18108978 | 5.63366406 | 1.76E-08 | 1.11E-06 | ARHGAP4 | 393 |
| ENSG000001 | 1161.10408 | 0.755653777 | 0.13413557 | 5.63350758 | 1.77E-08 | 1.11E-06 | SEC13 | 6396 |
| ENSG000001 | 1291.9757 | 0.817438741 | 0.14510528 | 5.63341817 | 1.77E-08 | 1.11E-06 | TOMM7 | 54543 |
| ENSG000002 | 617.391232 | 0.732744673 | 0.13008032 | 5.63301707 | 1.77E-08 | 1.11E-06 | LINC00998 | 401397 |
| ENSG000001 | 410.166018 | 0.72216642 | 0.12824082 | 5.63133046 | 1.79E-08 | 1.12E-06 | C1GALT1C1 | 29071 |
| ENSG00000C | 1160.80032 | 0.645065788 | 0.11461279 | 5.62821827 | 1.82E-08 | 1.14E-06 | TMEM230 | 29058 |
| ENSG000001 | 229.447068 | 0.956806152 | 0.17007094 | 5.62592381 | 1.85E-08 | 1.15E-06 | MRPS12 | 6183 |
| ENSG000001 | 4074.88748 | 0.749665328 | 0.13326841 | 5.62522907 | 1.85E-08 | 1.16E-06 | EIF3B | 8662 |
| ENSG000001 | 1216.27076 | 0.711668439 | 0.12656423 | 5.62298254 | 1.88E-08 | 1.17E-06 | PHF5A | 84844 |
| ENSG000001 | 283.08006 | 0.776229601 | 0.13805277 | 5.62270227 | 1.88E-08 | 1.17E-06 | MED8 | 112950 |
| ENSG000001 | 982.520803 | 0.774756345 | 0.13780534 | 5.62210683 | 1.89E-08 | 1.17E-06 | GUCD1 | 83606 |
| ENSG000001 | 1418.7317 | 0.681420213 | 0.12120581 | 5.62200956 | 1.89E-08 | 1.17E-06 | LRRC8D | 55144 |
| ENSG000001 | 2088.00647 | 0.717034338 | 0.12759285 | 5.61970633 | 1.91E-08 | 1.18E-06 | TUFM | 7284 |
| ENSG000001 | 334.301538 | 0.844834193 | 0.15038447 | 5.61782879 | 1.93E-08 | 1.19E-06 | ATG101 | 60673 |
| ENSG000002 | 2549.2259 | 0.725035564 | 0.12906243 | 5.61771204 | 1.94E-08 | 1.19E-06 | NA | NA |
| ENSG000001 | 153.663102 | 0.998860114 | 0.17782059 | 5.61723525 | 1.94E-08 | 1.20E-06 | GPAA1 | 8733 |
| ENSG000001 | 1941.73331 | 0.66177807 | 0.11781768 | 5.61696729 | 1.94E-08 | 1.20E-06 | PES1 | 23481 |
| ENSG000001 | 3558.3914 | 0.697854794 | 0.12425209 | 5.61644295 | 1.95E-08 | 1.20E-06 | MSN | 4478 |
| ENSG000001 | 1689.48647 | 0.700053906 | 0.12466282 | 5.61557874 | 1.96E-08 | 1.20E-06 | CEBPG | 1054 |
| ENSG000001 | 737.541008 | 0.694214238 | 0.12363959 | 5.61482139 | 1.97E-08 | 1.21E-06 | MAPKAPK3 | 7867 |
| ENSG00000C | 76.3718618 | 1.115907768 | 0.19876908 | 5.61409127 | 1.98E-08 | 1.21E-06 | GUCA1A | 2978 |
| ENSG000001 | 3535.49252 | 0.641799377 | 0.11433659 | 5.61324576 | 1.99E-08 | 1.21E-06 | RPN2 | 6185 |
| ENSG000001 | 539.662463 | 0.685083164 | 0.12205936 | 5.61270507 | 1.99E-08 | 1.21E-06 | LGMN | 5641 |

| | | | | | | | | |
|---|---|---|---|---|---|---|---|---|
| ENSG000001 | 1276.01483 | 0.718236901 | 0.12796743 | 5.61265376 | 1.99E-08 | 1.21E-06 | CSRP2 | 1466 |
| ENSG00000C | 227.638055 | 0.84852427 | 0.15123699 | 5.61056034 | 2.02E-08 | 1.22E-06 | PGS1 | 9489 |
| ENSG000001 | 384.281015 | 0.746468853 | 0.13304384 | 5.61069818 | 2.02E-08 | 1.22E-06 | AMFR | 267 |
| ENSG000001 | 1013.53894 | 0.773364474 | 0.13786065 | 5.6097551 | 2.03E-08 | 1.23E-06 | IDH3B | 3420 |
| ENSG000001 | 611.757273 | 0.70268744 | 0.12525747 | 5.60994442 | 2.02E-08 | 1.23E-06 | SNX12 | 29934 |
| ENSG00000C | 23932.9665 | 0.770908154 | 0.13746158 | 5.60817195 | 2.04E-08 | 1.24E-06 | RPLP0 | 6175 |
| ENSG000001 | 3118.32764 | 0.70863744 | 0.12637512 | 5.60741239 | 2.05E-08 | 1.24E-06 | BSG | 682 |
| ENSG000001 | 556.25394 | 0.723214571 | 0.12901213 | 5.60578751 | 2.07E-08 | 1.25E-06 | TMEM164 | 84187 |
| ENSG000001 | 357.516105 | 0.749730297 | 0.13376198 | 5.60495828 | 2.08E-08 | 1.25E-06 | KCND2 | 3751 |
| ENSG000001 | 686.015773 | 0.822655347 | 0.14689885 | 5.60014827 | 2.14E-08 | 1.29E-06 | FAM64A | 54478 |
| ENSG000001 | 495.069779 | 0.834162815 | 0.14897866 | 5.59921018 | 2.15E-08 | 1.29E-06 | PPIF | 10105 |
| ENSG000002 | 2584.53045 | 0.895348966 | 0.15993224 | 5.59830205 | 2.16E-08 | 1.30E-06 | HIST1H3F | 8968 |
| ENSG000001 | 1417.31862 | 0.766688799 | 0.13699684 | 5.59639767 | 2.19E-08 | 1.31E-06 | CDC37 | 11140 |
| ENSG000001 | 288.992909 | 0.970264898 | 0.17336246 | 5.59674178 | 2.18E-08 | 1.31E-06 | TK1 | 7083 |
| ENSG000001 | 12707.005 | 0.717852306 | 0.12827105 | 5.59637057 | 2.19E-08 | 1.31E-06 | RPL12 | 6136 |
| ENSG000001 | 201.051665 | 0.909565534 | 0.1625551 | 5.59542909 | 2.20E-08 | 1.31E-06 | GPR143 | 4935 |
| ENSG000001 | 413.455218 | 0.892505686 | 0.15951112 | 5.59525675 | 2.20E-08 | 1.31E-06 | GIPC1 | 10755 |
| ENSG000001 | 542.744989 | 0.812296782 | 0.14517369 | 5.59534417 | 2.20E-08 | 1.31E-06 | NR2F6 | 2063 |
| ENSG000001 | 183.285032 | 0.795151439 | 0.14212274 | 5.59482204 | 2.21E-08 | 1.31E-06 | SELK | 58515 |
| ENSG00000C | 11709.2356 | 0.739245493 | 0.13214794 | 5.59407524 | 2.22E-08 | 1.31E-06 | RPS20 | 6224 |
| ENSG000001 | 1018.55345 | 0.748670927 | 0.13389591 | 5.59144004 | 2.25E-08 | 1.33E-06 | TBRG4 | 9238 |
| ENSG000001 | 114.829961 | 1.015747455 | 0.18171328 | 5.58983598 | 2.27E-08 | 1.34E-06 | MT2A | 4502 |
| ENSG000001 | 3698.80292 | 0.677908655 | 0.12130887 | 5.58828614 | 2.29E-08 | 1.35E-06 | TXNRD1 | 7296 |
| ENSG000001 | 1647.35961 | 0.884186578 | 0.15823368 | 5.58785338 | 2.30E-08 | 1.35E-06 | MFGE8 | 4240 |
| ENSG000001 | 216.142454 | 0.814843691 | 0.14583291 | 5.5875157 | 2.30E-08 | 1.35E-06 | THAP7 | 80764 |
| ENSG000001 | 664.409108 | 0.795408469 | 0.1423965 | 5.58587098 | 2.33E-08 | 1.36E-06 | GRWD1 | 83743 |
| ENSG000001 | 639.047913 | 1.004514152 | 0.17984931 | 5.58531003 | 2.33E-08 | 1.37E-06 | CD81 | 975 |
| ENSG000001 | 1225.08624 | 0.693401127 | 0.12420693 | 5.58262854 | 2.37E-08 | 1.39E-06 | NME4 | 4833 |
| ENSG000001 | 959.242007 | 0.675866254 | 0.12112599 | 5.57986168 | 2.41E-08 | 1.40E-06 | COX7A2L | 9167 |

| | | | | | | | | |
|---|---|---|---|---|---|---|---|---|
| ENSG000001 | 334.899598 | 0.863980624 | 0.15483172 | 5.58012673 | 2.40E-08 | 1.40E-06 | THAP11 | 57215 |
| ENSG000001 | 207.606533 | 0.857995232 | 0.15376822 | 5.57979554 | 2.41E-08 | 1.40E-06 | SYN3 | 8224 |
| ENSG000001 | 1617.9622 | 0.774964263 | 0.13894961 | 5.57730433 | 2.44E-08 | 1.42E-06 | COX4I1 | 1327 |
| ENSG00000C | 1246.83142 | 0.629412818 | 0.11286821 | 5.57652884 | 2.45E-08 | 1.43E-06 | CDC23 | 8697 |
| ENSG000001 | 9520.5462 | 0.681395187 | 0.12221067 | 5.57557859 | 2.47E-08 | 1.43E-06 | RPS8 | 6202 |
| ENSG000001 | 405.608922 | 0.697254174 | 0.12508027 | 5.57445377 | 2.48E-08 | 1.44E-06 | EMG1 | 10436 |
| ENSG000001 | 869.038839 | 0.667071182 | 0.1196842 | 5.57359422 | 2.50E-08 | 1.44E-06 | POFUT1 | 23509 |
| ENSG000002 | 244.328166 | 0.87446473 | 0.15690401 | 5.57324659 | 2.50E-08 | 1.44E-06 | C8orf88 | 100127983 |
| ENSG00000C | 113.859598 | 1.033668466 | 0.18549402 | 5.57251651 | 2.51E-08 | 1.45E-06 | ICAM3 | 3385 |
| ENSG000002 | 2439.46471 | 0.776000908 | 0.13926309 | 5.57219384 | 2.52E-08 | 1.45E-06 | RPS29 | 6235 |
| ENSG000001 | 805.760715 | 0.632743346 | 0.11356493 | 5.57164376 | 2.52E-08 | 1.45E-06 | ADAM9 | 8754 |
| ENSG000001 | 1007.71799 | 0.648216943 | 0.11636423 | 5.57058593 | 2.54E-08 | 1.46E-06 | SORD | 6652 |
| ENSG000001 | 11366.8713 | 0.712967692 | 0.12800048 | 5.57003937 | 2.55E-08 | 1.46E-06 | RPS11 | 6205 |
| ENSG000001 | 4954.62864 | 0.795156915 | 0.14295858 | 5.56214885 | 2.66E-08 | 1.52E-06 | H2AFZ | 3015 |
| ENSG000002 | 5036.94155 | 0.820721561 | 0.14755212 | 5.56224849 | 2.66E-08 | 1.52E-06 | HIST1H3B | 8358 |
| ENSG000001 | 816.990213 | 0.744549258 | 0.13388826 | 5.56097496 | 2.68E-08 | 1.53E-06 | ILK | 3611 |
| ENSG000001 | 1387.86158 | 0.811201172 | 0.14588728 | 5.56046555 | 2.69E-08 | 1.53E-06 | TOMM40 | 10452 |
| ENSG000001 | 434.321514 | 0.749205922 | 0.13477094 | 5.55910567 | 2.71E-08 | 1.54E-06 | TRAPPC2L | 51693 |
| ENSG00000C | 977.762256 | 0.743639965 | 0.13379052 | 5.55824112 | 2.73E-08 | 1.55E-06 | NA | NA |
| ENSG000001 | 1984.86276 | 0.615819834 | 0.11082708 | 5.55658267 | 2.75E-08 | 1.56E-06 | ARIH2 | 10425 |
| ENSG000001 | 1146.03755 | 0.675455448 | 0.12161648 | 5.55397966 | 2.79E-08 | 1.58E-06 | TIMM50 | 92609 |
| ENSG000001 | 268.287368 | 0.767511385 | 0.13821767 | 5.552918 | 2.81E-08 | 1.59E-06 | MOCS1 | 4337 |
| ENSG000001 | 186.507437 | 0.828278964 | 0.14915663 | 5.55308174 | 2.81E-08 | 1.59E-06 | SGPP1 | 81537 |
| ENSG00000C | 2072.68503 | 0.685926076 | 0.123538 | 5.55234885 | 2.82E-08 | 1.59E-06 | ALDH18A1 | 5832 |
| ENSG000001 | 524.889008 | 0.823887542 | 0.14838988 | 5.55218131 | 2.82E-08 | 1.59E-06 | ARMC6 | 93436 |
| ENSG000001 | 955.111826 | 0.795834095 | 0.14334176 | 5.55200448 | 2.82E-08 | 1.59E-06 | PTK7 | 5754 |
| ENSG000001 | 403.552451 | 0.68237004 | 0.12296799 | 5.54916809 | 2.87E-08 | 1.61E-06 | URM1 | 81605 |
| ENSG000001 | 1544.36952 | 0.838791205 | 0.1511812 | 5.54825081 | 2.89E-08 | 1.62E-06 | EPHA1 | 2041 |
| ENSG00000C | 1086.94129 | 0.707529404 | 0.12754906 | 5.54711587 | 2.90E-08 | 1.63E-06 | DPM1 | 8813 |

| | | | | | | | | |
|---|---|---|---|---|---|---|---|---|
| ENSG000001 | 330.084292 | 0.864422037 | 0.1558392 | 5.54688433 | 2.91E-08 | 1.63E-06 | MOGS | 7841 |
| ENSG000001 | 99.8167382 | 1.003270407 | 0.18087165 | 5.54686372 | 2.91E-08 | 1.63E-06 | NA | NA |
| ENSG000001 | 479.948038 | 0.793436513 | 0.14306879 | 5.54583932 | 2.93E-08 | 1.63E-06 | OCIAD2 | 132299 |
| ENSG000001 | 484.684283 | 0.88481416 | 0.15955478 | 5.54551973 | 2.93E-08 | 1.63E-06 | TMEM8A | 58986 |
| ENSG000001 | 561.226985 | 0.697822441 | 0.1258473 | 5.54499321 | 2.94E-08 | 1.64E-06 | SMYD5 | 10322 |
| ENSG000001 | 4767.19919 | 0.68273397 | 0.12315447 | 5.54372072 | 2.96E-08 | 1.65E-06 | MCM3 | 4172 |
| ENSG000001 | 394.609534 | 0.898410876 | 0.16208202 | 5.54293984 | 2.97E-08 | 1.65E-06 | SLC25A11 | 8402 |
| ENSG000001 | 106.584271 | 0.978428284 | 0.17662817 | 5.53948039 | 3.03E-08 | 1.68E-06 | TOR4A | 54863 |
| ENSG000001 | 314.687558 | 0.830300763 | 0.14995257 | 5.53708915 | 3.08E-08 | 1.70E-06 | C20orf27 | 54976 |
| ENSG000001 | 214.658458 | 0.846454356 | 0.15290742 | 5.53573095 | 3.10E-08 | 1.71E-06 | SCG5 | 6447 |
| ENSG000001 | 271.666382 | -0.767389655 | 0.13863597 | -5.5352853 | 3.11E-08 | 1.72E-06 | HHLA1 | 10086 |
| ENSG000001 | 1734.81442 | 0.663069148 | 0.11979766 | 5.53490898 | 3.11E-08 | 1.72E-06 | FAM171B | 165215 |
| ENSG000001 | 98.8258815 | 1.004671688 | 0.18152399 | 5.53464963 | 3.12E-08 | 1.72E-06 | SHPK | 23729 |
| ENSG000001 | 73.2076326 | 1.028331801 | 0.18590105 | 5.53160827 | 3.17E-08 | 1.75E-06 | NA | NA |
| ENSG000001 | 1325.76839 | 0.809260676 | 0.14631162 | 5.53107578 | 3.18E-08 | 1.75E-06 | MTCH1 | 23787 |
| ENSG000001 | 811.288735 | 0.649759008 | 0.11747895 | 5.53085494 | 3.19E-08 | 1.75E-06 | HIGD1A | 25994 |
| ENSG000001 | 2098.24324 | 0.757796256 | 0.13702149 | 5.53049212 | 3.19E-08 | 1.75E-06 | TUBB2B | 347733 |
| ENSG000001 | 482.931183 | 0.785219411 | 0.142004 | 5.52955822 | 3.21E-08 | 1.76E-06 | ABT1 | 29777 |
| ENSG000001 | 1885.88912 | 0.677518025 | 0.12253881 | 5.52900777 | 3.22E-08 | 1.76E-06 | PPP1CA | 5499 |
| ENSG000001 | 8864.20224 | 0.759086168 | 0.13730919 | 5.52829838 | 3.23E-08 | 1.76E-06 | RPL24 | 6152 |
| ENSG000001 | 71.6102084 | -1.198343721 | 0.21678448 | -5.5278114 | 3.24E-08 | 1.77E-06 | ARPP21 | 10777 |
| ENSG000001 | 655.741266 | 0.752111395 | 0.13609066 | 5.52654663 | 3.27E-08 | 1.78E-06 | PUDP | 8226 |
| ENSG000001 | 446.648434 | 0.721751154 | 0.13060653 | 5.52614897 | 3.27E-08 | 1.78E-06 | CALB1 | 793 |
| ENSG000001 | 575.897716 | 0.718009768 | 0.12997317 | 5.52429228 | 3.31E-08 | 1.80E-06 | METTL17 | 64745 |
| ENSG000001 | 150.812154 | 0.874274115 | 0.15827093 | 5.52390843 | 3.32E-08 | 1.80E-06 | TMEM203 | 94107 |
| ENSG000001 | 3716.10009 | 0.770258407 | 0.13945247 | 5.52344761 | 3.32E-08 | 1.80E-06 | MARS | 4141 |
| ENSG000001 | 310.894014 | 0.905098011 | 0.16395773 | 5.52031304 | 3.38E-08 | 1.82E-06 | RHBDD3 | 25807 |
| ENSG000001 | 2980.27717 | 0.783819309 | 0.1419808 | 5.52060088 | 3.38E-08 | 1.82E-06 | ITM2C | 81618 |
| ENSG000001 | 780.325746 | 0.70327407 | 0.12739697 | 5.52033585 | 3.38E-08 | 1.82E-06 | IQCB1 | 9657 |

| | | | | | | | | |
|---|---|---|---|---|---|---|---|---|
| ENSG000002 | 442.960707 | 0.660928872 | 0.1197141 | 5.52089429 | 3.37E-08 | 1.82E-06 | NA | NA |
| ENSG000001 | 2104.70981 | 0.671897318 | 0.12174189 | 5.51903145 | 3.41E-08 | 1.83E-06 | CLU | 1191 |
| ENSG000000 | 499.045456 | 0.983802796 | 0.17828097 | 5.51827149 | 3.42E-08 | 1.84E-06 | GRN | 2896 |
| ENSG000001 | 6053.14329 | 0.631293779 | 0.11439939 | 5.51833148 | 3.42E-08 | 1.84E-06 | TCP1 | 6950 |
| ENSG000000 | 122.17925 | 0.937753394 | 0.16996934 | 5.51719151 | 3.44E-08 | 1.85E-06 | KCNG1 | 3755 |
| ENSG000002 | 133.075047 | 0.929194842 | 0.16846115 | 5.51578108 | 3.47E-08 | 1.86E-06 | ZNF425 | 155054 |
| ENSG000001 | 3404.13444 | 0.740391092 | 0.13424182 | 5.51535352 | 3.48E-08 | 1.86E-06 | YARS | 8565 |
| ENSG000001 | 239.700824 | 0.88873991 | 0.16116818 | 5.51436324 | 3.50E-08 | 1.86E-06 | MPP1 | 4354 |
| ENSG000001 | 786.131517 | 0.673206381 | 0.12207865 | 5.51452997 | 3.50E-08 | 1.86E-06 | DSCR3 | 10311 |
| ENSG000001 | 2715.73463 | 0.710998163 | 0.12892798 | 5.51469261 | 3.49E-08 | 1.86E-06 | LMNB2 | 84823 |
| ENSG000001 | 162.273374 | 0.948347277 | 0.17202939 | 5.5127051 | 3.53E-08 | 1.88E-06 | CXorf40B | 541578 |
| ENSG000001 | 3564.87488 | 0.650958406 | 0.11813224 | 5.51042147 | 3.58E-08 | 1.90E-06 | ATP11C | 286410 |
| ENSG000001 | 213.863167 | 0.776298488 | 0.14089713 | 5.50968271 | 3.59E-08 | 1.91E-06 | PI4K2A | 55361 |
| ENSG000001 | 1146.62275 | 0.6249577 | 0.11349476 | 5.50648929 | 3.66E-08 | 1.94E-06 | CRTAP | 10491 |
| ENSG000000 | 147.945446 | 0.919299668 | 0.166996 | 5.50492036 | 3.69E-08 | 1.95E-06 | NAT14 | 57106 |
| ENSG000001 | 5572.79469 | 0.737636135 | 0.13399037 | 5.50514297 | 3.69E-08 | 1.95E-06 | RPL35A | 6165 |
| ENSG000001 | 704.447698 | 0.767776057 | 0.13949151 | 5.50410587 | 3.71E-08 | 1.96E-06 | TUBGCP2 | 10844 |
| ENSG000001 | 459.824209 | 0.793969937 | 0.14431098 | 5.50179837 | 3.76E-08 | 1.98E-06 | NOSIP | 51070 |
| ENSG000002 | 16.8428548 | -1.772572109 | 0.32220989 | -5.5012965 | 3.77E-08 | 1.98E-06 | NA | NA |
| ENSG000001 | 1770.95237 | 0.68115732 | 0.12384687 | 5.49999615 | 3.80E-08 | 1.99E-06 | MRS2 | 57380 |
| ENSG000001 | 255.927933 | 0.809259844 | 0.14716806 | 5.49888221 | 3.82E-08 | 2.00E-06 | MMP15 | 4324 |
| ENSG000001 | 6261.04931 | 0.746846356 | 0.13581088 | 5.49916428 | 3.82E-08 | 2.00E-06 | RPL29 | 6159 |
| ENSG000001 | 181.155185 | 0.819229825 | 0.14898135 | 5.49887511 | 3.82E-08 | 2.00E-06 | CCDC172 | 374355 |
| ENSG000002 | 1351.02036 | 0.634229466 | 0.11533042 | 5.49923851 | 3.81E-08 | 2.00E-06 | FGD5-AS1 | 100505641 |
| ENSG000001 | 582.362098 | 0.748094297 | 0.13605651 | 5.49840865 | 3.83E-08 | 2.00E-06 | MRPL17 | 63875 |
| ENSG000001 | 354.052178 | 0.861543444 | 0.15668501 | 5.49856982 | 3.83E-08 | 2.00E-06 | STK40 | 83931 |
| ENSG000001 | 645.759859 | 0.67582313 | 0.12293152 | 5.49755793 | 3.85E-08 | 2.00E-06 | TTC9 | 23508 |
| ENSG000002 | 2911.17279 | 0.723331214 | 0.13157705 | 5.4973965 | 3.85E-08 | 2.00E-06 | CLIC1 | 1192 |
| ENSG000002 | 273.61916 | 0.847432523 | 0.15424549 | 5.49405064 | 3.93E-08 | 2.04E-06 | RGL2 | 5863 |

| | | | | | | | | |
|---|---|---|---|---|---|---|---|---|
| ENSG00000C | 253.52316 | 0.839917083 | 0.15293488 | 5.49199151 | 3.97E-08 | 2.06E-06 | TSPAN9 | 10867 |
| ENSG00000C | 625.372557 | 0.762954671 | 0.13893009 | 5.49164452 | 3.98E-08 | 2.06E-06 | NA | NA |
| ENSG000001 | 814.922694 | 0.840554791 | 0.15312323 | 5.48940093 | 4.03E-08 | 2.09E-06 | CDCA5 | 113130 |
| ENSG000001 | 326.29915 | 0.864710673 | 0.15755283 | 5.48838565 | 4.06E-08 | 2.09E-06 | PPP1R12C | 54776 |
| ENSG000001 | 865.469068 | 0.846567471 | 0.15425116 | 5.48824071 | 4.06E-08 | 2.09E-06 | H2AFX | 3014 |
| ENSG000001 | 274.994209 | 0.772520426 | 0.14081917 | 5.48590369 | 4.11E-08 | 2.12E-06 | DNASE2 | 1777 |
| ENSG000001 | 848.244228 | 0.710491048 | 0.12952186 | 5.48549154 | 4.12E-08 | 2.12E-06 | RNF114 | 55905 |
| ENSG000001 | 61.6290107 | -1.115591413 | 0.20341438 | -5.484329 | 4.15E-08 | 2.13E-06 | MNS1 | 55329 |
| ENSG000001 | 462.369748 | 0.854149984 | 0.15575471 | 5.48394319 | 4.16E-08 | 2.14E-06 | DPH2 | 1802 |
| ENSG000001 | 694.815912 | 0.896802584 | 0.16356321 | 5.48291126 | 4.18E-08 | 2.14E-06 | SESN2 | 83667 |
| ENSG000001 | 7434.9459 | 0.713498429 | 0.13024681 | 5.47804896 | 4.30E-08 | 2.20E-06 | CTNNB1 | 1499 |
| ENSG000001 | 692.991814 | 0.643789615 | 0.11769062 | 5.47018607 | 4.50E-08 | 2.30E-06 | UST | 10090 |
| ENSG000001 | 2187.49569 | 0.62670397 | 0.11457894 | 5.46962631 | 4.51E-08 | 2.30E-06 | KIF5C | 3800 |
| ENSG000001 | 6463.9999 | 0.628875706 | 0.11505843 | 5.46570734 | 4.61E-08 | 2.35E-06 | TMBIM6 | 7009 |
| ENSG000001 | 2605.21062 | 0.652544233 | 0.11941447 | 5.46453251 | 4.64E-08 | 2.36E-06 | ATP6V1A | 523 |
| ENSG000001 | 254.14093 | 0.882937429 | 0.16160964 | 5.46339575 | 4.67E-08 | 2.38E-06 | ZNF581 | 51545 |
| ENSG000001 | 610.69759 | 0.741085788 | 0.1356959 | 5.46137196 | 4.72E-08 | 2.39E-06 | SLC25A1 | 6576 |
| ENSG000001 | 1188.78121 | 0.688082801 | 0.12598578 | 5.46159084 | 4.72E-08 | 2.39E-06 | SNU13 | 4809 |
| ENSG000001 | 260.8581 | 0.786673577 | 0.14403403 | 5.4617204 | 4.72E-08 | 2.39E-06 | HIGD2A | 192286 |
| ENSG000001 | 1519.15105 | 0.607564872 | 0.11126036 | 5.46074876 | 4.74E-08 | 2.40E-06 | SNRPG | 6637 |
| ENSG000002 | 79.3287667 | 1.059459858 | 0.19407406 | 5.45904941 | 4.79E-08 | 2.42E-06 | OXLD1 | 339229 |
| ENSG000001 | 419.014833 | 0.677048693 | 0.12406426 | 5.4572422 | 4.84E-08 | 2.44E-06 | PIN1 | 5300 |
| ENSG000002 | 381.730002 | 0.803479237 | 0.14727219 | 5.45574311 | 4.88E-08 | 2.46E-06 | FIS1 | 51024 |
| ENSG000001 | 6346.81332 | 0.662291466 | 0.12141618 | 5.45472175 | 4.90E-08 | 2.47E-06 | UBB | 7314 |
| ENSG000001 | 126.840908 | 0.872132895 | 0.15996099 | 5.45215997 | 4.98E-08 | 2.50E-06 | SH3BGR | 6450 |
| ENSG000002 | 1129.88785 | 0.767073342 | 0.14069203 | 5.45214496 | 4.98E-08 | 2.50E-06 | PGAM5 | 192111 |
| ENSG000001 | 582.914661 | 0.700734146 | 0.12854685 | 5.45119674 | 5.00E-08 | 2.51E-06 | UBE2L6 | 9246 |
| ENSG00000C | 1361.45743 | 0.736577854 | 0.13514873 | 5.45012793 | 5.03E-08 | 2.52E-06 | UQCRC1 | 7384 |
| ENSG000001 | 268.276615 | 0.805891254 | 0.14786546 | 5.45016562 | 5.03E-08 | 2.52E-06 | LMAN2L | 81562 |

| | | | | | | | | |
|---|---|---|---|---|---|---|---|---|
| ENSG000001 | 420.949473 | 0.737558564 | 0.13536206 | 5.44878347 | 5.07E-08 | 2.54E-06 | MINPP1 | 9562 |
| ENSG000001 | 704.165733 | 0.711513869 | 0.13061256 | 5.44751486 | 5.11E-08 | 2.55E-06 | VPS25 | 84313 |
| ENSG000001 | 612.297392 | 0.787785301 | 0.14462943 | 5.44692238 | 5.12E-08 | 2.56E-06 | TMEM147 | 10430 |
| ENSG000001 | 657.416577 | 0.893309346 | 0.16401883 | 5.44638273 | 5.14E-08 | 2.56E-06 | MAP2K2 | 5605 |
| ENSG000002 | 11128.8853 | 0.78757354 | 0.14460361 | 5.44643067 | 5.14E-08 | 2.56E-06 | POU5F1 | 5460 |
| ENSG000001 | 1381.64465 | 0.669890906 | 0.12303694 | 5.44463255 | 5.19E-08 | 2.58E-06 | FAM32A | 26017 |
| ENSG000001 | 13481.7828 | 0.724416877 | 0.13307252 | 5.44377534 | 5.22E-08 | 2.59E-06 | RPL11 | 6135 |
| ENSG000001 | 175.239803 | 0.881908804 | 0.16200218 | 5.44380811 | 5.22E-08 | 2.59E-06 | PAQR4 | 124222 |
| ENSG000002 | 3713.43971 | 0.651826836 | 0.11977616 | 5.44204158 | 5.27E-08 | 2.61E-06 | PHB2 | 11331 |
| ENSG000001 | 337.703606 | 0.785337585 | 0.14431927 | 5.44166805 | 5.28E-08 | 2.61E-06 | SPECC1L | 23384 |
| ENSG000001 | 1300.90135 | 0.695579418 | 0.12786172 | 5.44009123 | 5.33E-08 | 2.63E-06 | KIAA1191 | 57179 |
| ENSG000001 | 707.021784 | 0.61420034 | 0.11293576 | 5.4384932 | 5.37E-08 | 2.65E-06 | SLC30A5 | 64924 |
| ENSG000001 | 565.593369 | 0.736410925 | 0.13540288 | 5.4386652 | 5.37E-08 | 2.65E-06 | C9orf72 | 203228 |
| ENSG000001 | 1798.53108 | 0.709148184 | 0.13041617 | 5.43757887 | 5.40E-08 | 2.66E-06 | GNAI2 | 2771 |
| ENSG000002 | 298.552474 | 0.974879978 | 0.17928892 | 5.43748042 | 5.40E-08 | 2.66E-06 | RPSAP15 | 220885 |
| ENSG000001 | 9347.2968 | 0.619806713 | 0.11401859 | 5.43601457 | 5.45E-08 | 2.67E-06 | XPOT | 11260 |
| ENSG000002 | 2772.34805 | 0.672082966 | 0.12364697 | 5.43549873 | 5.46E-08 | 2.68E-06 | HIST1H2BF | 8343 |
| ENSG000001 | 330.699763 | 0.823382472 | 0.15154002 | 5.43343269 | 5.53E-08 | 2.71E-06 | KLHL21 | 9903 |
| ENSG000001 | 8805.30248 | 0.618580134 | 0.11388981 | 5.43139139 | 5.59E-08 | 2.73E-06 | TPM4 | 7171 |
| ENSG000001 | 457.773539 | 0.787293078 | 0.14495829 | 5.4311697 | 5.60E-08 | 2.73E-06 | ATG4B | 23192 |
| ENSG000001 | 215.807099 | 0.938548909 | 0.17280214 | 5.43134988 | 5.59E-08 | 2.73E-06 | PIM3 | 415116 |
| ENSG000002 | 1482.26838 | 0.738473439 | 0.1359851 | 5.43054653 | 5.62E-08 | 2.74E-06 | NA | NA |
| ENSG000001 | 382.100403 | 0.715599044 | 0.13178314 | 5.43012571 | 5.63E-08 | 2.74E-06 | ACTR10 | 55860 |
| ENSG00000C | 313.972267 | 0.78813582 | 0.14520946 | 5.42757895 | 5.71E-08 | 2.78E-06 | DHX32 | 55760 |
| ENSG000001 | 1937.80851 | 0.849213875 | 0.15649215 | 5.42655891 | 5.75E-08 | 2.78E-06 | HIST1H2BJ | 8970 |
| ENSG000001 | 5477.73017 | 0.574626782 | 0.1058909 | 5.42659254 | 5.74E-08 | 2.78E-06 | UCHL1 | 7345 |
| ENSG000001 | 5609.2698 | 0.654778575 | 0.12065391 | 5.42691548 | 5.73E-08 | 2.78E-06 | AP1S2 | 8905 |
| ENSG000001 | 898.842072 | 0.694280101 | 0.12795733 | 5.42587205 | 5.77E-08 | 2.79E-06 | SLC7A3 | 84889 |
| ENSG000001 | 288.441874 | 0.720481898 | 0.13281942 | 5.42452235 | 5.81E-08 | 2.81E-06 | JOSD1 | 9929 |

| | | | | | | | | |
|---|---|---|---|---|---|---|---|---|
| ENSG000001 | 106.364218 | 0.902689227 | 0.16640411 | 5.42468121 | 5.81E-08 | 2.81E-06 | FLVCR2 | 55640 |
| ENSG000002 | 956.115484 | 0.775617228 | 0.14298992 | 5.42427906 | 5.82E-08 | 2.81E-06 | NA | NA |
| ENSG000001 | 722.543073 | 0.731471184 | 0.13487663 | 5.42326119 | 5.85E-08 | 2.82E-06 | NQO1 | 1728 |
| ENSG000001 | 433.293453 | 0.865999035 | 0.1597227 | 5.42189066 | 5.90E-08 | 2.84E-06 | COLGALT1 | 79709 |
| ENSG000001 | 668.179193 | 0.617336976 | 0.11386987 | 5.42142526 | 5.91E-08 | 2.84E-06 | CFDP1 | 10428 |
| ENSG000001 | 84.7885214 | 1.036362141 | 0.19120298 | 5.42021964 | 5.95E-08 | 2.84E-06 | AURKC | 6795 |
| ENSG000001 | 114.046781 | -0.946676998 | 0.17465862 | -5.4201561 | 5.95E-08 | 2.84E-06 | TEK | 7010 |
| ENSG000001 | 173.187352 | 0.811687172 | 0.14973316 | 5.42089116 | 5.93E-08 | 2.84E-06 | LRP3 | 4037 |
| ENSG000001 | 242.831132 | 0.852145449 | 0.15721013 | 5.42042331 | 5.95E-08 | 2.84E-06 | ANAPC11 | 51529 |
| ENSG000001 | 916.064139 | 0.746294821 | 0.13768709 | 5.42022361 | 5.95E-08 | 2.84E-06 | CD2BP2 | 10421 |
| ENSG000001 | 706.285579 | 0.652647376 | 0.12043118 | 5.41925588 | 5.98E-08 | 2.86E-06 | FAM120AOS | 158293 |
| ENSG000002 | 122.184968 | 0.916859721 | 0.1692049 | 5.41863583 | 6.01E-08 | 2.86E-06 | C19orf24 | 55009 |
| ENSG000001 | 470.71709 | 0.675126314 | 0.12462746 | 5.41715554 | 6.06E-08 | 2.88E-06 | ARRDC4 | 91947 |
| ENSG000001 | 524.962971 | 0.83259284 | 0.15370107 | 5.41696206 | 6.06E-08 | 2.88E-06 | ATP5G1 | 516 |
| ENSG000002 | 274.554055 | 0.885211824 | 0.16343243 | 5.41637808 | 6.08E-08 | 2.89E-06 | NA | NA |
| ENSG000001 | 1247.47486 | 0.604199702 | 0.11156134 | 5.41585191 | 6.10E-08 | 2.89E-06 | NEMP1 | 23306 |
| ENSG000001 | 1510.15371 | 0.617087149 | 0.11395826 | 5.4150277 | 6.13E-08 | 2.90E-06 | MCCC2 | 64087 |
| ENSG000001 | 25875.9554 | 0.660258315 | 0.12193067 | 5.41503052 | 6.13E-08 | 2.90E-06 | RPL4 | 6124 |
| ENSG000001 | 372.045386 | 0.781998175 | 0.14447215 | 5.41279537 | 6.20E-08 | 2.93E-06 | FAM122B | 159090 |
| ENSG00000C | 987.533047 | 0.805738579 | 0.1488761 | 5.41214175 | 6.23E-08 | 2.94E-06 | ATP1A2 | 477 |
| ENSG000002 | 30.1962058 | -1.364939637 | 0.25221303 | -5.4118523 | 6.24E-08 | 2.94E-06 | POU5F2 | 134187 |
| ENSG000002 | 209.17544 | 0.902259677 | 0.16681197 | 5.40884264 | 6.34E-08 | 2.99E-06 | NA | NA |
| ENSG00000C | 637.113877 | 0.728810858 | 0.13480759 | 5.40630417 | 6.43E-08 | 3.03E-06 | AGBL5 | 60509 |
| ENSG000001 | 812.049622 | 0.693693543 | 0.12836275 | 5.40416566 | 6.51E-08 | 3.06E-06 | ARFGAP2 | 84364 |
| ENSG000001 | 478.601371 | 0.686723807 | 0.12707271 | 5.40417998 | 6.51E-08 | 3.06E-06 | TMEM161B | 153396 |
| ENSG000001 | 1355.07027 | 0.660463371 | 0.12228056 | 5.40121329 | 6.62E-08 | 3.10E-06 | KIF2C | 11004 |
| ENSG000001 | 27201.0852 | 0.821151822 | 0.1520833 | 5.39935563 | 6.69E-08 | 3.13E-06 | EEF2 | 1938 |
| ENSG000001 | 825.022911 | 0.703150354 | 0.13025424 | 5.39829168 | 6.73E-08 | 3.15E-06 | SPINT2 | 10653 |
| ENSG000001 | 176.406489 | 0.882601988 | 0.16351105 | 5.39781252 | 6.75E-08 | 3.15E-06 | WDR83OS | 51398 |

| | | | | | | | | |
|---|---|---|---|---|---|---|---|---|
| ENSG000002 | 579.109588 | 0.753620306 | 0.13963543 | 5.39705649 | 6.77E-08 | 3.16E-06 | NA | NA |
| ENSG000001 | 446.495652 | 0.67507867 | 0.12509694 | 5.39644451 | 6.80E-08 | 3.17E-06 | C2orf68 | 388969 |
| ENSG000001 | 948.015745 | 0.766959333 | 0.14213395 | 5.39603189 | 6.81E-08 | 3.17E-06 | CYC1 | 1537 |
| ENSG000001 | 648.116345 | 0.776190981 | 0.14387213 | 5.395006 | 6.85E-08 | 3.19E-06 | SCOC | 60592 |
| ENSG000001 | 1311.79998 | 0.810329812 | 0.15024124 | 5.39352462 | 6.91E-08 | 3.21E-06 | SEPW1 | 6415 |
| ENSG000001 | 160.604038 | 0.861805903 | 0.15985729 | 5.39109538 | 7.00E-08 | 3.25E-06 | MAGEH1 | 28986 |
| ENSG000001 | 796.876376 | 0.658076642 | 0.12207826 | 5.3906127 | 7.02E-08 | 3.25E-06 | GABARAPL1 | 23710 |
| ENSG000000 | 778.715871 | 0.841021104 | 0.15602838 | 5.39018025 | 7.04E-08 | 3.25E-06 | POLR2E | 5434 |
| ENSG000001 | 596.080649 | 0.771042973 | 0.14306245 | 5.38955538 | 7.06E-08 | 3.25E-06 | ADM | 133 |
| ENSG000001 | 1872.09688 | 0.625126498 | 0.11598824 | 5.38956778 | 7.06E-08 | 3.25E-06 | SAP18 | 10284 |
| ENSG000001 | 12040.6724 | 0.618070593 | 0.11467878 | 5.389581 | 7.06E-08 | 3.25E-06 | CALR | 811 |
| ENSG000001 | 71.2303259 | -1.046128276 | 0.19409602 | -5.3897462 | 7.06E-08 | 3.25E-06 | MYT1L | 23040 |
| ENSG000000 | 271.793107 | 0.798750508 | 0.14822083 | 5.38892229 | 7.09E-08 | 3.26E-06 | CYBA | 1535 |
| ENSG000001 | 782.235922 | 0.580770399 | 0.10778249 | 5.38835566 | 7.11E-08 | 3.27E-06 | MALT1 | 10892 |
| ENSG000001 | 433.590909 | 0.868119482 | 0.16112603 | 5.38782883 | 7.13E-08 | 3.28E-06 | CORO1B | 57175 |
| ENSG000000 | 394.447622 | 0.719056499 | 0.13347385 | 5.38724608 | 7.15E-08 | 3.28E-06 | KLHL13 | 90293 |
| ENSG000002 | 193.834861 | 0.883788997 | 0.1640985 | 5.38572274 | 7.22E-08 | 3.31E-06 | NA | NA |
| ENSG000001 | 125.235955 | -0.907059491 | 0.16845323 | -5.384637 | 7.26E-08 | 3.32E-06 | LYST | 1130 |
| ENSG000002 | 396.879186 | 0.756184813 | 0.14044604 | 5.384166 | 7.28E-08 | 3.33E-06 | NA | NA |
| ENSG000001 | 1768.19476 | 0.641223949 | 0.11912179 | 5.38292738 | 7.33E-08 | 3.35E-06 | LECT1 | 11061 |
| ENSG000001 | 19.622582 | -1.6490853 | 0.30640226 | -5.3820925 | 7.36E-08 | 3.36E-06 | SLITRK4 | 139065 |
| ENSG000001 | 5020.75691 | 0.62377988 | 0.11590763 | 5.38169829 | 7.38E-08 | 3.36E-06 | HSD17B4 | 3295 |
| ENSG000001 | 232.264778 | 0.812140173 | 0.15096875 | 5.37952521 | 7.47E-08 | 3.39E-06 | BRF2 | 55290 |
| ENSG000001 | 355.117112 | 0.805218787 | 0.14966673 | 5.38007867 | 7.45E-08 | 3.39E-06 | LGALS3BP | 3959 |
| ENSG000001 | 462.586083 | 0.70462147 | 0.13098144 | 5.37955213 | 7.47E-08 | 3.39E-06 | NEDD8 | 4738 |
| ENSG000002 | 46.7899176 | 1.237893375 | 0.2300955 | 5.37991122 | 7.45E-08 | 3.39E-06 | NA | NA |
| ENSG000000 | 793.823954 | 0.688175058 | 0.12793972 | 5.37890079 | 7.49E-08 | 3.39E-06 | LAP3 | 51056 |
| ENSG000001 | 216.495425 | 0.798011537 | 0.1483602 | 5.37887882 | 7.50E-08 | 3.39E-06 | MUS81 | 80198 |
| ENSG000001 | 973.604811 | 0.798231246 | 0.1484511 | 5.37706532 | 7.57E-08 | 3.42E-06 | STK25 | 10494 |

| | | | | | | | | |
|---|---|---|---|---|---|---|---|---|
| ENSG000002 | 1408.96084 | 0.877851339 | 0.16325564 | 5.3771578 | 7.57E-08 | 3.42E-06 | HIST1H2AB | 8335 |
| ENSG000001 | 266.110172 | 0.902409486 | 0.16788429 | 5.37518715 | 7.65E-08 | 3.45E-06 | UCK1 | 83549 |
| ENSG000001 | 1071.92047 | 0.594437725 | 0.11060296 | 5.37451894 | 7.68E-08 | 3.46E-06 | ATP6V0A2 | 23545 |
| ENSG000001 | 333.532456 | 0.755801027 | 0.1406567 | 5.3733738 | 7.73E-08 | 3.48E-06 | ZNF787 | 126208 |
| ENSG000001 | 280.623868 | 0.831141397 | 0.15468811 | 5.37301412 | 7.74E-08 | 3.48E-06 | YIF1B | 90522 |
| ENSG000002 | 125.169542 | 0.905752976 | 0.16856843 | 5.37320634 | 7.73E-08 | 3.48E-06 | LOC730101 | 730101 |
| ENSG000001 | 340.467357 | 0.791747996 | 0.14743887 | 5.37000864 | 7.87E-08 | 3.53E-06 | XRCC3 | 7517 |
| ENSG000001 | 195.013103 | 0.752440372 | 0.14015106 | 5.36878102 | 7.93E-08 | 3.55E-06 | RGS3 | 5998 |
| ENSG000001 | 806.164278 | 0.708290617 | 0.13196672 | 5.36719116 | 8.00E-08 | 3.58E-06 | AUP1 | 550 |
| ENSG000001 | 160.609426 | 0.871031975 | 0.16230788 | 5.36654145 | 8.03E-08 | 3.59E-06 | ZNF593 | 51042 |
| ENSG000001 | 3342.74998 | 0.709464785 | 0.13225704 | 5.36428744 | 8.13E-08 | 3.63E-06 | MCM5 | 4174 |
| ENSG000001 | 221.628522 | 0.818874939 | 0.1527 | 5.36263881 | 8.20E-08 | 3.66E-06 | CD99L2 | 83692 |
| ENSG00000C | 197.255732 | 0.795385571 | 0.14834006 | 5.36190674 | 8.23E-08 | 3.67E-06 | CD74 | 972 |
| ENSG00000C | 5875.38935 | 0.677754784 | 0.1263974 | 5.36209422 | 8.23E-08 | 3.67E-06 | RPL31 | 6160 |
| ENSG000001 | 530.248438 | -0.68598356 | 0.1279629 | -5.3608003 | 8.29E-08 | 3.68E-06 | ZNF300 | 91975 |
| ENSG000001 | 606.768109 | 0.705563199 | 0.13162845 | 5.36026383 | 8.31E-08 | 3.69E-06 | ECHS1 | 1892 |
| ENSG000001 | 615.216516 | 0.789634622 | 0.14732949 | 5.35965088 | 8.34E-08 | 3.70E-06 | FOXO3 | 2309 |
| ENSG000002 | 30.7004192 | -1.365099935 | 0.25469724 | -5.3596966 | 8.34E-08 | 3.70E-06 | NA | NA |
| ENSG000001 | 4944.16418 | 0.674913014 | 0.12593459 | 5.35923475 | 8.36E-08 | 3.70E-06 | CTSC | 1075 |
| ENSG000001 | 3002.90006 | 0.719336801 | 0.13423746 | 5.35868912 | 8.38E-08 | 3.71E-06 | SAE1 | 10055 |
| ENSG000001 | 2227.03957 | 0.630491419 | 0.11770243 | 5.35665603 | 8.48E-08 | 3.74E-06 | SKP2 | 6502 |
| ENSG000002 | 135.688434 | -0.902630513 | 0.16854988 | -5.3552724 | 8.54E-08 | 3.77E-06 | NA | NA |
| ENSG000001 | 9635.68827 | 0.65966894 | 0.12320344 | 5.35430615 | 8.59E-08 | 3.79E-06 | RPL37 | 6167 |
| ENSG00000C | 499.041127 | 0.64875947 | 0.12117463 | 5.35392157 | 8.61E-08 | 3.79E-06 | TUBE1 | 51175 |
| ENSG000001 | 311.281 | 0.809796309 | 0.15128049 | 5.35294595 | 8.65E-08 | 3.81E-06 | ADRBK1 | 156 |
| ENSG000001 | 453.688716 | 0.824920095 | 0.1541859 | 5.3501655 | 8.79E-08 | 3.86E-06 | FAM129B | 64855 |
| ENSG00000C | 5988.95703 | 0.777402204 | 0.14531285 | 5.34985178 | 8.80E-08 | 3.86E-06 | AARS | 16 |
| ENSG000002 | 960.6289 | -0.644290898 | 0.12043488 | -5.3497036 | 8.81E-08 | 3.86E-06 | NA | NA |
| ENSG000001 | 12519.1973 | 0.711053706 | 0.13293867 | 5.34873496 | 8.86E-08 | 3.88E-06 | RPL27A | 6157 |

| | | | | | | | | |
|---|---|---|---|---|---|---|---|---|
| ENSG000001 | 2074.01968 | 0.709838085 | 0.13275305 | 5.3470568 | 8.94E-08 | 3.91E-06 | NA | NA |
| ENSG00000C | 3200.74246 | 0.587887961 | 0.10996436 | 5.34616833 | 8.98E-08 | 3.93E-06 | CDC6 | 990 |
| ENSG000001 | 266.987666 | -0.741869793 | 0.13879523 | -5.3450669 | 9.04E-08 | 3.95E-06 | SYNE1 | 23345 |
| ENSG00000C | 2021.80574 | 0.630801143 | 0.11804041 | 5.34394249 | 9.09E-08 | 3.97E-06 | UNG | 7374 |
| ENSG000002 | 847.785421 | 0.728516869 | 0.13633096 | 5.34373737 | 9.10E-08 | 3.97E-06 | MRPL20 | 55052 |
| ENSG000001 | 3007.48148 | 0.699391106 | 0.1308909 | 5.34331359 | 9.13E-08 | 3.97E-06 | SND1 | 27044 |
| ENSG00000C | 390.387082 | 0.785341545 | 0.14699904 | 5.34249427 | 9.17E-08 | 3.97E-06 | MOCOS | 55034 |
| ENSG000001 | 548.360236 | 0.665018521 | 0.1244698 | 5.34281045 | 9.15E-08 | 3.97E-06 | TXN2 | 25828 |
| ENSG000001 | 112.565268 | 0.979701499 | 0.18337482 | 5.34261737 | 9.16E-08 | 3.97E-06 | PYCARD | 29108 |
| ENSG000001 | 1059.11833 | 0.783373627 | 0.14662715 | 5.34262326 | 9.16E-08 | 3.97E-06 | PLCXD1 | 55344 |
| ENSG000002 | 415.455887 | 0.82599496 | 0.15473255 | 5.33821071 | 9.39E-08 | 4.06E-06 | SCARNA13 | 677768 |
| ENSG00000C | 724.873083 | 0.683726591 | 0.12810685 | 5.3371586 | 9.44E-08 | 4.08E-06 | SLC25A24 | 29957 |
| ENSG000001 | 438.022489 | 0.686914673 | 0.12871947 | 5.33652516 | 9.47E-08 | 4.09E-06 | PSME2 | 5721 |
| ENSG000001 | 621.700412 | 0.653623582 | 0.1224896 | 5.33615562 | 9.49E-08 | 4.10E-06 | TMED8 | 283578 |
| ENSG000001 | 2302.74752 | 0.661294502 | 0.12394654 | 5.33532041 | 9.54E-08 | 4.11E-06 | GHITM | 27069 |
| ENSG000001 | 85.3300108 | 0.979261545 | 0.18357144 | 5.33449832 | 9.58E-08 | 4.12E-06 | GCHFR | 2644 |
| ENSG000001 | 783.750169 | 0.677745915 | 0.12708513 | 5.33300728 | 9.66E-08 | 4.15E-06 | TOM1L1 | 10040 |
| ENSG000002 | 507.352877 | 0.823498168 | 0.15442334 | 5.33273128 | 9.67E-08 | 4.16E-06 | ZBED1 | 9189 |
| ENSG00000C | 13706.4654 | 0.647486235 | 0.12143346 | 5.33202474 | 9.71E-08 | 4.17E-06 | RPL6 | 6128 |
| ENSG00000C | 1832.43257 | 0.609099639 | 0.11424788 | 5.33138666 | 9.75E-08 | 4.18E-06 | FBXW11 | 23291 |
| ENSG000001 | 820.261814 | 0.663195745 | 0.12439901 | 5.33119789 | 9.76E-08 | 4.18E-06 | NDUFS8 | 4728 |
| ENSG00000C | 3377.26835 | 0.7631061 | 0.1431881 | 5.32939594 | 9.85E-08 | 4.20E-06 | GPC4 | 2239 |
| ENSG000001 | 393.299015 | 0.764654472 | 0.14347663 | 5.32947052 | 9.85E-08 | 4.20E-06 | NINL | 22981 |
| ENSG000001 | 2876.63009 | 0.650978171 | 0.12213663 | 5.32991773 | 9.83E-08 | 4.20E-06 | PPIB | 5479 |
| ENSG000001 | 298.727694 | 0.689087645 | 0.12930823 | 5.3290315 | 9.87E-08 | 4.20E-06 | KRCC1 | 51315 |
| ENSG000001 | 673.869355 | 0.68404074 | 0.12835433 | 5.32931585 | 9.86E-08 | 4.20E-06 | C11orf95 | 65998 |
| ENSG000002 | 77.6011693 | 1.089372909 | 0.20442422 | 5.32898153 | 9.88E-08 | 4.20E-06 | SNORA12 | 677800 |
| ENSG000002 | 16.682508 | -1.683505405 | 0.31585128 | -5.3300572 | 9.82E-08 | 4.20E-06 | LOC101927151 | 101927151 |
| ENSG000001 | 483.031886 | 0.723139405 | 0.13571845 | 5.32823217 | 9.92E-08 | 4.21E-06 | UQCRQ | 27089 |

| | | | | | | | | |
|---|---|---|---|---|---|---|---|---|
| ENSG000001 | 736.004189 | 0.673238321 | 0.12637356 | 5.32736711 | 9.96E-08 | 4.23E-06 | MAF1 | 84232 |
| ENSG00000C | 1429.05761 | 0.620624068 | 0.11651695 | 5.32647032 | 1.00E-07 | 4.24E-06 | ALDH3A2 | 224 |
| ENSG000002 | 3300.48906 | 0.81988881 | 0.15396745 | 5.32507876 | 1.01E-07 | 4.27E-06 | HIST1H2BI | 8346 |
| ENSG000001 | 207.208688 | 0.853788777 | 0.16036657 | 5.32398227 | 1.02E-07 | 4.29E-06 | MPG | 4350 |
| ENSG000001 | 1968.37483 | 0.637064224 | 0.11967067 | 5.3234784 | 1.02E-07 | 4.30E-06 | TXNDC12 | 51060 |
| ENSG00000C | 2246.39906 | 0.663559421 | 0.12468216 | 5.32200779 | 1.03E-07 | 4.33E-06 | MCM2 | 4171 |
| ENSG00000C | 652.125296 | 0.895036714 | 0.16820613 | 5.32107077 | 1.03E-07 | 4.34E-06 | PRSS8 | 5652 |
| ENSG000001 | 527.678138 | 0.756559877 | 0.14218147 | 5.32108634 | 1.03E-07 | 4.34E-06 | TIMM8B | 26521 |
| ENSG000001 | 693.240828 | 0.731428103 | 0.13749432 | 5.31969692 | 1.04E-07 | 4.37E-06 | CYB5A | 1528 |
| ENSG000001 | 289.908231 | 0.728731136 | 0.13704192 | 5.31757828 | 1.05E-07 | 4.42E-06 | RASA3 | 22821 |
| ENSG000001 | 106.071568 | 0.89888607 | 0.1690864 | 5.3161347 | 1.06E-07 | 4.45E-06 | DEXI | 28955 |
| ENSG00000C | 891.673283 | 0.772189967 | 0.14528273 | 5.31508438 | 1.07E-07 | 4.47E-06 | MRTO4 | 51154 |
| ENSG000001 | 619.616784 | 0.673802955 | 0.12679036 | 5.3143073 | 1.07E-07 | 4.49E-06 | FAM101B | 359845 |
| ENSG000001 | 189.110358 | 0.835864956 | 0.15730671 | 5.31360026 | 1.07E-07 | 4.50E-06 | RAB25 | 57111 |
| ENSG000001 | 1807.45594 | 0.760618273 | 0.14317403 | 5.31254348 | 1.08E-07 | 4.52E-06 | HIST1H4H | 8365 |
| ENSG000001 | 1012.32873 | 0.705521493 | 0.13283401 | 5.31130179 | 1.09E-07 | 4.55E-06 | C11orf84 | 144097 |
| ENSG000001 | 467.471614 | 0.72517444 | 0.13653926 | 5.31110554 | 1.09E-07 | 4.55E-06 | IDH2 | 3418 |
| ENSG000001 | 526.613625 | 0.663641734 | 0.12496795 | 5.31049564 | 1.09E-07 | 4.56E-06 | ZNF518B | 85460 |
| ENSG00000C | 665.050718 | 0.681706504 | 0.12840799 | 5.30891025 | 1.10E-07 | 4.59E-06 | AAAS | 8086 |
| ENSG000001 | 740.258962 | 0.796316061 | 0.15000412 | 5.30862787 | 1.10E-07 | 4.59E-06 | FEM1A | 55527 |
| ENSG000001 | 1712.94249 | 0.661099566 | 0.12455815 | 5.30755774 | 1.11E-07 | 4.60E-06 | CYB5B | 80777 |
| ENSG000001 | 4758.01212 | 0.703028131 | 0.13244957 | 5.30789288 | 1.11E-07 | 4.60E-06 | SRSF7 | 6432 |
| ENSG000001 | 848.555429 | 0.679768346 | 0.12806919 | 5.30782104 | 1.11E-07 | 4.60E-06 | TPST2 | 8459 |
| ENSG000001 | 1078.50746 | 0.642040896 | 0.12096496 | 5.30766021 | 1.11E-07 | 4.60E-06 | FBXO21 | 23014 |
| ENSG000001 | 3103.93727 | 0.677933778 | 0.12774705 | 5.30684486 | 1.12E-07 | 4.61E-06 | AHCY | 191 |
| ENSG000002 | 56.1567804 | 1.100688953 | 0.20740918 | 5.30684779 | 1.12E-07 | 4.61E-06 | PIK3CD-AS2 | 101929074 |
| ENSG000001 | 407.540073 | 0.797433169 | 0.15027036 | 5.30665637 | 1.12E-07 | 4.61E-06 | RRP1 | 8568 |
| ENSG000001 | 4462.62352 | 0.646332357 | 0.12181089 | 5.30603098 | 1.12E-07 | 4.62E-06 | NR6A1 | 2649 |
| ENSG000001 | 505.585655 | 0.786725118 | 0.1482928 | 5.30521447 | 1.13E-07 | 4.64E-06 | DGKZ | 8525 |

| | | | | | | | | |
|---|---|---|---|---|---|---|---|---|
| ENSG000002 | 2908.4312 | 0.808790539 | 0.15250263 | 5.30345316 | 1.14E-07 | 4.68E-06 | HIST1H3E | 8353 |
| ENSG000001 | 803.346112 | 0.785135598 | 0.14808565 | 5.30190196 | 1.15E-07 | 4.71E-06 | RABL6 | 55684 |
| ENSG000002 | 99.7466206 | 0.899832278 | 0.16971587 | 5.30199244 | 1.15E-07 | 4.71E-06 | NA | NA |
| ENSG000001 | 797.885997 | 0.754948182 | 0.14243107 | 5.30044574 | 1.16E-07 | 4.74E-06 | RBM19 | 9904 |
| ENSG000002 | 2952.39262 | 0.730736011 | 0.13786352 | 5.30043052 | 1.16E-07 | 4.74E-06 | HIST1H2BH | 8345 |
| ENSG000001 | 412.372926 | 0.725313512 | 0.13686617 | 5.29943587 | 1.16E-07 | 4.75E-06 | WRNIP1 | 56897 |
| ENSG000001 | 103.520401 | 0.960178578 | 0.1812003 | 5.29898989 | 1.16E-07 | 4.75E-06 | SSC4D | 136853 |
| ENSG000001 | 1873.02399 | 0.780928657 | 0.14737116 | 5.29906033 | 1.16E-07 | 4.75E-06 | MDH2 | 4191 |
| ENSG000001 | 257.711692 | 0.722311631 | 0.13630099 | 5.29938643 | 1.16E-07 | 4.75E-06 | STRA13 | 201254 |
| ENSG000002 | 387.270876 | 0.685091443 | 0.12928275 | 5.29917139 | 1.16E-07 | 4.75E-06 | TBC1D8 | 11138 |
| ENSG000001 | 293.471184 | -0.685075221 | 0.12932078 | -5.2974875 | 1.17E-07 | 4.78E-06 | MYADM | 91663 |
| ENSG000001 | 10422.9054 | 0.703063948 | 0.13271572 | 5.29751828 | 1.17E-07 | 4.78E-06 | RPS23 | 6228 |
| ENSG00000C | 1343.8394 | 0.625621851 | 0.11811875 | 5.29654992 | 1.18E-07 | 4.80E-06 | TTC19 | 54902 |
| ENSG000001 | 297.013676 | 0.730031966 | 0.13785689 | 5.29557818 | 1.19E-07 | 4.82E-06 | STX8 | 9482 |
| ENSG000001 | 661.838413 | 0.653706442 | 0.12347844 | 5.2940939 | 1.20E-07 | 4.86E-06 | FLAD1 | 80308 |
| ENSG000001 | 1208.0051 | 0.708726946 | 0.13388458 | 5.29356653 | 1.20E-07 | 4.86E-06 | FAM199X | 139231 |
| ENSG000001 | 1681.72787 | 0.699130039 | 0.13220856 | 5.28808466 | 1.24E-07 | 5.01E-06 | UBE2E1 | 7324 |
| ENSG000001 | 211.047349 | 0.801891481 | 0.15164899 | 5.28781288 | 1.24E-07 | 5.01E-06 | TMEM205 | 374882 |
| ENSG000001 | 86.8747753 | 1.037615757 | 0.19625302 | 5.28713275 | 1.24E-07 | 5.02E-06 | PCED1A | 64773 |
| ENSG000001 | 1200.64241 | 0.59660249 | 0.11285376 | 5.28650982 | 1.25E-07 | 5.04E-06 | WDR11 | 55717 |
| ENSG000001 | 28986.4274 | 0.715415775 | 0.13533978 | 5.28607161 | 1.25E-07 | 5.04E-06 | ACTG1 | 71 |
| ENSG000001 | 2677.7 | 0.813890091 | 0.15398554 | 5.28549677 | 1.25E-07 | 5.05E-06 | KRT8 | 3856 |
| ENSG000001 | 1320.36887 | 0.627461718 | 0.11873082 | 5.28474192 | 1.26E-07 | 5.07E-06 | UBE2S | 27338 |
| ENSG000001 | 874.280361 | 0.638805022 | 0.12096014 | 5.28111987 | 1.28E-07 | 5.17E-06 | RNF11 | 26994 |
| ENSG000002 | 1764.4697 | 0.945512265 | 0.179062 | 5.28036248 | 1.29E-07 | 5.18E-06 | SNORD17 | 692086 |
| ENSG000001 | 502.06333 | 0.731419862 | 0.13854244 | 5.27939218 | 1.30E-07 | 5.20E-06 | VPS51 | 738 |
| ENSG000001 | 1047.25085 | 0.74193704 | 0.14054317 | 5.27906879 | 1.30E-07 | 5.20E-06 | ALPL | 249 |
| ENSG000001 | 1964.53255 | 0.611714427 | 0.11587232 | 5.27921107 | 1.30E-07 | 5.20E-06 | MEX3C | 51320 |
| ENSG000002 | 26.7722053 | -1.427271793 | 0.27044485 | -5.2774967 | 1.31E-07 | 5.24E-06 | ZNF571-AS1 | 100507433 |

| | | | | | | | | |
|---|---|---|---|---|---|---|---|---|
| ENSG000001 | 1244.53629 | 0.678664675 | 0.1286315 | 5.27603776 | 1.32E-07 | 5.28E-06 | PSMD13 | 5719 |
| ENSG000001 | 1950.60725 | 0.732080636 | 0.13877961 | 5.27513113 | 1.33E-07 | 5.30E-06 | TRAP1 | 10131 |
| ENSG000001 | 10383.2232 | 0.720152578 | 0.13652451 | 5.27489592 | 1.33E-07 | 5.30E-06 | RPS27A | 6233 |
| ENSG000001 | 505.431216 | 0.71719098 | 0.13596291 | 5.27490172 | 1.33E-07 | 5.30E-06 | WIBG | 84305 |
| ENSG000001 | 543.788825 | 0.743040011 | 0.14088982 | 5.27390827 | 1.34E-07 | 5.31E-06 | PYGB | 5834 |
| ENSG000001 | 1159.02244 | 0.685955132 | 0.13006001 | 5.27414323 | 1.33E-07 | 5.31E-06 | ACTR1A | 10121 |
| ENSG000001 | 382.311062 | 0.752895567 | 0.14275431 | 5.27406537 | 1.33E-07 | 5.31E-06 | GCC1 | 79571 |
| ENSG000001 | 767.572633 | 0.630123168 | 0.11952499 | 5.27189475 | 1.35E-07 | 5.36E-06 | P4HA1 | 5033 |
| ENSG000001 | 457.160073 | 0.744238048 | 0.14116601 | 5.27207674 | 1.35E-07 | 5.36E-06 | PREB | 10113 |
| ENSG000001 | 707.86813 | 0.626534069 | 0.11884692 | 5.27177367 | 1.35E-07 | 5.36E-06 | ZDHHC7 | 55625 |
| ENSG000001 | 1371.93198 | 0.730568782 | 0.13861363 | 5.27054084 | 1.36E-07 | 5.39E-06 | COX6B1 | 1340 |
| ENSG00000C | 760.191776 | 0.657454465 | 0.1247602 | 5.26974501 | 1.37E-07 | 5.40E-06 | BCL2L13 | 23786 |
| ENSG000001 | 707.805145 | 0.686881619 | 0.13037393 | 5.2685502 | 1.38E-07 | 5.43E-06 | SDCCAG3 | 10807 |
| ENSG000001 | 2053.25733 | 0.768615414 | 0.14588344 | 5.26869547 | 1.37E-07 | 5.43E-06 | HIST1H2BL | 8340 |
| ENSG000001 | 426.75311 | 0.732855389 | 0.13911971 | 5.26780427 | 1.38E-07 | 5.45E-06 | TCTN3 | 26123 |
| ENSG000001 | 343.652846 | 0.686005173 | 0.13023073 | 5.26761375 | 1.38E-07 | 5.45E-06 | ATP6V0D1 | 9114 |
| ENSG000001 | 337.366151 | 0.775593508 | 0.14726038 | 5.2668171 | 1.39E-07 | 5.46E-06 | GYS1 | 2997 |
| ENSG000001 | 5036.44362 | 0.673767585 | 0.12793901 | 5.26631841 | 1.39E-07 | 5.47E-06 | RPS13 | 6207 |
| ENSG000001 | 325.575576 | 0.838962287 | 0.15931729 | 5.26598399 | 1.39E-07 | 5.48E-06 | SLC25A33 | 84275 |
| ENSG000001 | 963.379865 | 0.719777141 | 0.13669995 | 5.26537983 | 1.40E-07 | 5.49E-06 | NA | NA |
| ENSG000001 | 980.303214 | 0.647703508 | 0.1230276 | 5.26470092 | 1.40E-07 | 5.51E-06 | SAP30BP | 29115 |
| ENSG000001 | 864.154042 | 0.667657992 | 0.12686693 | 5.26266374 | 1.42E-07 | 5.56E-06 | ERGIC3 | 51614 |
| ENSG00000C | 5123.84162 | 0.608534112 | 0.11569702 | 5.25972167 | 1.44E-07 | 5.65E-06 | KDM1A | 23028 |
| ENSG000001 | 945.544421 | 0.736605935 | 0.14007896 | 5.25850523 | 1.45E-07 | 5.68E-06 | DHX30 | 22907 |
| ENSG000001 | 401.4786 | 0.806792924 | 0.15345116 | 5.25765266 | 1.46E-07 | 5.70E-06 | JDP2 | 122953 |
| ENSG000001 | 635.803159 | 0.727688493 | 0.13840923 | 5.25751416 | 1.46E-07 | 5.70E-06 | FAM134C | 162427 |
| ENSG000001 | 158.358049 | 0.869646689 | 0.16542124 | 5.25716463 | 1.46E-07 | 5.70E-06 | C9orf16 | 79095 |
| ENSG000001 | 1184.79199 | 0.67509626 | 0.12844086 | 5.25608643 | 1.47E-07 | 5.73E-06 | MMP16 | 4325 |
| ENSG000001 | 229.265362 | 0.758416967 | 0.14431045 | 5.25545437 | 1.48E-07 | 5.75E-06 | FLCN | 201163 |

| | | | | | | | | |
|---|---|---|---|---|---|---|---|---|
| ENSG000001 | 570.978134 | 0.703782721 | 0.133921 | 5.25520824 | 1.48E-07 | 5.75E-06 | ADPGK | 83440 |
| ENSG000001 | 1205.73765 | 0.622720638 | 0.11853468 | 5.25348888 | 1.49E-07 | 5.80E-06 | ATAD1 | 84896 |
| ENSG000001 | 241.563806 | 0.75070448 | 0.14294165 | 5.25182482 | 1.51E-07 | 5.84E-06 | PTGR1 | 22949 |
| ENSG000001 | 1198.14726 | 0.732282777 | 0.13945915 | 5.2508765 | 1.51E-07 | 5.87E-06 | IDO1 | 3620 |
| ENSG000002 | 129.497321 | 0.857398569 | 0.16329742 | 5.25053346 | 1.52E-07 | 5.87E-06 | LOC101928000 | 101928000 |
| ENSG000001 | 1412.96934 | 0.588904684 | 0.11216712 | 5.25024324 | 1.52E-07 | 5.88E-06 | UBE2D2 | 7322 |
| ENSG000001 | 1121.26763 | 0.821654313 | 0.1565327 | 5.24909064 | 1.53E-07 | 5.91E-06 | NHSL1 | 57224 |
| ENSG000001 | 4113.41072 | 0.692441811 | 0.13193784 | 5.24824256 | 1.54E-07 | 5.93E-06 | KRT18 | 3875 |
| ENSG000001 | 5156.93806 | 0.689459788 | 0.13139357 | 5.2472871 | 1.54E-07 | 5.96E-06 | VCP | 7415 |
| ENSG000002 | 231.349509 | 0.78317999 | 0.1492607 | 5.24706084 | 1.55E-07 | 5.96E-06 | TSPAN4 | 7106 |
| ENSG00000C | 562.690895 | 0.710840518 | 0.13549861 | 5.24610919 | 1.55E-07 | 5.98E-06 | AKAP8L | 26993 |
| ENSG00000C | 473.045648 | 0.810195391 | 0.15446596 | 5.24513884 | 1.56E-07 | 6.01E-06 | NDUFB7 | 4713 |
| ENSG000001 | 896.271062 | 0.666273658 | 0.12704194 | 5.24451719 | 1.57E-07 | 6.02E-06 | ZFAND1 | 79752 |
| ENSG000001 | 374.989582 | 0.687930752 | 0.13120824 | 5.24304531 | 1.58E-07 | 6.07E-06 | MOB3B | 79817 |
| ENSG00000C | 2212.31159 | 0.745040411 | 0.14211234 | 5.24261585 | 1.58E-07 | 6.07E-06 | CTTN | 2017 |
| ENSG000001 | 265.049463 | 0.785382583 | 0.1498669 | 5.24053384 | 1.60E-07 | 6.14E-06 | ELK1 | 2002 |
| ENSG00000C | 459.331929 | 0.709987418 | 0.13548907 | 5.24018211 | 1.60E-07 | 6.14E-06 | ZFP64 | 55734 |
| ENSG000001 | 409.202108 | 0.835830999 | 0.15953186 | 5.23927331 | 1.61E-07 | 6.17E-06 | CPSF4 | 10898 |
| ENSG000001 | 958.744861 | 0.679003895 | 0.12961129 | 5.23877118 | 1.62E-07 | 6.18E-06 | TUSC3 | 7991 |
| ENSG000001 | 1380.27632 | 0.586709126 | 0.11200145 | 5.2384065 | 1.62E-07 | 6.18E-06 | CASC4 | 113201 |
| ENSG000001 | 52.5245594 | -1.126461503 | 0.21504255 | -5.2383191 | 1.62E-07 | 6.18E-06 | KIAA2012 | 100652824 |
| ENSG000001 | 3875.97772 | 0.675928539 | 0.12905871 | 5.23737245 | 1.63E-07 | 6.21E-06 | PRMT1 | 3276 |
| ENSG000002 | 112.847213 | 1.056043714 | 0.20166195 | 5.23670293 | 1.63E-07 | 6.22E-06 | AP5Z1 | 9907 |
| ENSG000002 | 165.325294 | 0.835542065 | 0.15956991 | 5.23621313 | 1.64E-07 | 6.23E-06 | LOC148413 | 148413 |
| ENSG000001 | 1300.62481 | 0.692619852 | 0.13230989 | 5.23483045 | 1.65E-07 | 6.28E-06 | FEN1 | 2237 |
| ENSG000001 | 487.985742 | 0.95275523 | 0.18205836 | 5.23324078 | 1.67E-07 | 6.32E-06 | AKNA | 80709 |
| ENSG000001 | 94.0382089 | -0.983836344 | 0.18799845 | -5.2332152 | 1.67E-07 | 6.32E-06 | IGFN1 | 91156 |
| ENSG000001 | 232.380289 | 0.791941819 | 0.15135756 | 5.23225812 | 1.67E-07 | 6.35E-06 | GRINA | 2907 |
| ENSG000001 | 1506.48662 | 0.636889115 | 0.12176137 | 5.23063341 | 1.69E-07 | 6.40E-06 | UNC5D | 137970 |

| | | | | | | | | |
|---|---|---|---|---|---|---|---|---|
| ENSG000001 | 532.954651 | 0.647833271 | 0.1238592 | 5.23040079 | 1.69E-07 | 6.40E-06 | MTMR14 | 64419 |
| ENSG000001 | 10017.3829 | 0.648605031 | 0.12401104 | 5.23022001 | 1.69E-07 | 6.40E-06 | RPL9 | 6133 |
| ENSG000001 | 322.747958 | 0.767422868 | 0.14674049 | 5.22979633 | 1.70E-07 | 6.41E-06 | RBM42 | 79171 |
| ENSG000001 | 3584.10085 | 0.603122434 | 0.11533161 | 5.22946332 | 1.70E-07 | 6.41E-06 | PTPN13 | 5783 |
| ENSG000001 | 503.750414 | 0.604971665 | 0.115682 | 5.22960942 | 1.70E-07 | 6.41E-06 | ZNF160 | 90338 |
| ENSG000001 | 581.77288 | 0.710504497 | 0.13589897 | 5.22818171 | 1.71E-07 | 6.44E-06 | UBAC2 | 337867 |
| ENSG000001 | 630.144673 | 0.699774709 | 0.13384985 | 5.22805747 | 1.71E-07 | 6.44E-06 | ACADSB | 36 |
| ENSG000001 | 77.7495864 | 0.960784388 | 0.18378296 | 5.22782079 | 1.72E-07 | 6.45E-06 | ALDH1A1 | 216 |
| ENSG00000C | 264.115131 | 0.839016876 | 0.16050351 | 5.2274052 | 1.72E-07 | 6.45E-06 | ARHGEF10L | 55160 |
| ENSG000001 | 283.801649 | 0.741836283 | 0.14192076 | 5.22711609 | 1.72E-07 | 6.46E-06 | PRKAB1 | 5564 |
| ENSG00000C | 709.44344 | 0.727808969 | 0.13925583 | 5.2264166 | 1.73E-07 | 6.48E-06 | NDUFAB1 | 4706 |
| ENSG000001 | 619.940956 | 0.658123569 | 0.12593966 | 5.22570535 | 1.73E-07 | 6.50E-06 | IFT57 | 55081 |
| ENSG000001 | 710.684754 | 0.699814345 | 0.13401041 | 5.22208945 | 1.77E-07 | 6.62E-06 | GTF2H3 | 2967 |
| ENSG000001 | 21.6434897 | -1.506060086 | 0.28846682 | -5.2209126 | 1.78E-07 | 6.65E-06 | OR2B11 | 127623 |
| ENSG000001 | 529.842756 | 0.728023405 | 0.13944927 | 5.22070431 | 1.78E-07 | 6.66E-06 | FIBP | 9158 |
| ENSG000001 | 622.778512 | 0.630513685 | 0.12079415 | 5.21973702 | 1.79E-07 | 6.68E-06 | CPEB4 | 80315 |
| ENSG000001 | 723.753263 | 0.67656003 | 0.12963595 | 5.2189228 | 1.80E-07 | 6.71E-06 | GSKIP | 51527 |
| ENSG00000C | 4672.46265 | 0.869259483 | 0.16658457 | 5.21812728 | 1.81E-07 | 6.73E-06 | NFE2L1 | 4779 |
| ENSG00000C | 232.50515 | 0.892831407 | 0.17115329 | 5.21655991 | 1.82E-07 | 6.78E-06 | CPXM1 | 56265 |
| ENSG00000C | 1401.69893 | 0.758144134 | 0.14537647 | 5.21504009 | 1.84E-07 | 6.83E-06 | SUGP2 | 10147 |
| ENSG000002 | 1121.61108 | 0.563937065 | 0.10814165 | 5.21479971 | 1.84E-07 | 6.83E-06 | TMX2 | 51075 |
| ENSG00000C | 100.9815 | 0.904380408 | 0.17349062 | 5.21284898 | 1.86E-07 | 6.90E-06 | COMT | 1312 |
| ENSG000001 | 1102.03235 | 0.666833522 | 0.12793184 | 5.21241234 | 1.86E-07 | 6.91E-06 | EARS2 | 124454 |
| ENSG000002 | 6.32242235 | -1.956294787 | 0.37534167 | -5.2120373 | 1.87E-07 | 6.92E-06 | LOC440446 | 440446 |
| ENSG000002 | 1049.78092 | 0.685451879 | 0.13153993 | 5.21097946 | 1.88E-07 | 6.95E-06 | HIST1H2BN | 8341 |
| ENSG000001 | 498.096921 | 0.885440878 | 0.16993414 | 5.21049438 | 1.88E-07 | 6.95E-06 | SBF1 | 6305 |
| ENSG000001 | 882.428641 | 0.585756986 | 0.11241362 | 5.21072975 | 1.88E-07 | 6.95E-06 | CEP120 | 153241 |
| ENSG000001 | 7282.89909 | 0.641075715 | 0.12303714 | 5.21042456 | 1.88E-07 | 6.95E-06 | NACA | 4666 |
| ENSG000001 | 209.891556 | 0.818889856 | 0.15723616 | 5.20802498 | 1.91E-07 | 7.04E-06 | GPR108 | 56927 |

| | | | | | | | | |
|---|---|---|---|---|---|---|---|---|
| ENSG000001 | 644.591899 | 0.759859973 | 0.14595804 | 5.20601674 | 1.93E-07 | 7.11E-06 | TBL1X | 6907 |
| ENSG00000C | 4273.4763 | 0.673863013 | 0.12944465 | 5.2058006 | 1.93E-07 | 7.11E-06 | TMED2 | 10959 |
| ENSG000001 | 9010.18909 | 0.698082345 | 0.13411724 | 5.20501577 | 1.94E-07 | 7.13E-06 | RPL10A | 4736 |
| ENSG00000C | 22275.6882 | 0.605279642 | 0.11629239 | 5.20480873 | 1.94E-07 | 7.13E-06 | BCAT1 | 586 |
| ENSG000001 | 197.791031 | 0.838055199 | 0.16107575 | 5.20286371 | 1.96E-07 | 7.20E-06 | MFSD12 | 126321 |
| ENSG000001 | 29.4757492 | -1.377575674 | 0.26488797 | -5.2005973 | 1.99E-07 | 7.28E-06 | PLG | 5340 |
| ENSG000001 | 1428.559 | 0.760269604 | 0.14621471 | 5.19967948 | 2.00E-07 | 7.31E-06 | PSMB5 | 5693 |
| ENSG000001 | 1581.2114 | 0.643708063 | 0.12379507 | 5.19978746 | 2.00E-07 | 7.31E-06 | USMG5 | 84833 |
| ENSG000001 | 574.795507 | 0.687787109 | 0.13229608 | 5.19884728 | 2.01E-07 | 7.33E-06 | SLC18B1 | 116843 |
| ENSG000001 | 367.271637 | 0.875908052 | 0.16849792 | 5.19833144 | 2.01E-07 | 7.35E-06 | GNB2 | 2783 |
| ENSG000001 | 11040.0753 | 0.647345702 | 0.12455326 | 5.19734066 | 2.02E-07 | 7.38E-06 | RPL19 | 6143 |
| ENSG00000C | 347.106441 | 0.940078181 | 0.18089516 | 5.19681237 | 2.03E-07 | 7.38E-06 | MBD3 | 53615 |
| ENSG000001 | 7643.2431 | 0.621411128 | 0.11957523 | 5.19682144 | 2.03E-07 | 7.38E-06 | ATF4 | 468 |
| ENSG000001 | 801.634574 | 0.609709346 | 0.11732549 | 5.19673375 | 2.03E-07 | 7.38E-06 | GOLGA7 | 51125 |
| ENSG000001 | 717.864256 | 0.620684534 | 0.11949447 | 5.194253 | 2.06E-07 | 7.48E-06 | ALDH6A1 | 4329 |
| ENSG00000C | 1719.56464 | 0.592465247 | 0.11407212 | 5.19377762 | 2.06E-07 | 7.48E-06 | POLQ | 10721 |
| ENSG000001 | 3605.99971 | 0.665121262 | 0.12806222 | 5.19373536 | 2.06E-07 | 7.48E-06 | DNAJB6 | 10049 |
| ENSG000001 | 762.217552 | 0.690767192 | 0.13302459 | 5.19277833 | 2.07E-07 | 7.51E-06 | NXT2 | 55916 |
| ENSG000001 | 937.722191 | 0.72187074 | 0.13901096 | 5.19290524 | 2.07E-07 | 7.51E-06 | SIGMAR1 | 10280 |
| ENSG000001 | 10147.4817 | 0.722262064 | 0.13913682 | 5.19102026 | 2.09E-07 | 7.57E-06 | RPL32 | 6161 |
| ENSG000001 | 393.366733 | 0.847460926 | 0.16328594 | 5.19004212 | 2.10E-07 | 7.61E-06 | HGS | 9146 |
| ENSG000001 | 222.813462 | 0.795163016 | 0.15321912 | 5.18971133 | 2.11E-07 | 7.61E-06 | NDUFA2 | 4695 |
| ENSG000001 | 170.155704 | 0.958650186 | 0.18474488 | 5.18904871 | 2.11E-07 | 7.63E-06 | RAB38 | 23682 |
| ENSG000002 | 5574.71559 | 0.560610324 | 0.10806534 | 5.18769755 | 2.13E-07 | 7.68E-06 | HMGN1 | 3150 |
| ENSG000001 | 657.035854 | 0.629184468 | 0.12129734 | 5.1871252 | 2.14E-07 | 7.70E-06 | JTB | 10899 |
| ENSG000001 | 556.182757 | 0.635907 | 0.12260022 | 5.186834 | 2.14E-07 | 7.70E-06 | MRPL4 | 51073 |
| ENSG000001 | 227.620886 | 0.84100953 | 0.16220891 | 5.18473065 | 2.16E-07 | 7.78E-06 | DNAJB2 | 3300 |
| ENSG000001 | 614.089485 | 0.727514311 | 0.14033204 | 5.18423511 | 2.17E-07 | 7.80E-06 | CDCA8 | 55143 |
| ENSG000001 | 443.765298 | 0.710141447 | 0.13699788 | 5.18359424 | 2.18E-07 | 7.81E-06 | P3H1 | 64175 |

| | | | | | | | | |
|---|---|---|---|---|---|---|---|---|
| ENSG000001 | 3096.90673 | 0.736391761 | 0.14205854 | 5.18372037 | 2.18E-07 | 7.81E-06 | FLNB | 2317 |
| ENSG00000C | 424.466405 | 0.757144485 | 0.14612354 | 5.1815366 | 2.20E-07 | 7.89E-06 | ISOC2 | 79763 |
| ENSG000002 | 236.607471 | 0.743977182 | 0.14358976 | 5.18126898 | 2.20E-07 | 7.89E-06 | CCHCR1 | 54535 |
| ENSG000001 | 118.186174 | -0.969279466 | 0.1870833 | -5.1810046 | 2.21E-07 | 7.89E-06 | DNAH6 | 1768 |
| ENSG000002 | 229.762458 | 0.989210965 | 0.19093274 | 5.18093945 | 2.21E-07 | 7.89E-06 | NA | NA |
| ENSG000001 | 2285.58692 | 0.62497537 | 0.12063907 | 5.1805385 | 2.21E-07 | 7.90E-06 | MID1 | 4281 |
| ENSG000001 | 542.122803 | 0.656830817 | 0.12680427 | 5.17987941 | 2.22E-07 | 7.93E-06 | LRRTM4 | 80059 |
| ENSG000001 | 255.03809 | 0.794370604 | 0.15337074 | 5.17941446 | 2.23E-07 | 7.93E-06 | ITPA | 3704 |
| ENSG000001 | 118.047264 | 0.873962637 | 0.16873559 | 5.17948018 | 2.23E-07 | 7.93E-06 | YDJC | 150223 |
| ENSG000001 | 302.035443 | 0.828813494 | 0.16003401 | 5.17898358 | 2.23E-07 | 7.94E-06 | TNFRSF8 | 943 |
| ENSG00000C | 428.473561 | 0.660932815 | 0.12762894 | 5.17854964 | 2.24E-07 | 7.95E-06 | YIPF1 | 54432 |
| ENSG000001 | 948.618635 | 0.7439594 | 0.14366416 | 5.17846221 | 2.24E-07 | 7.95E-06 | TPD52L2 | 7165 |
| ENSG00000C | 1609.30261 | 0.6355349 | 0.12276948 | 5.17665239 | 2.26E-07 | 8.01E-06 | M6PR | 4074 |
| ENSG000001 | 170.585439 | 0.760884175 | 0.14698266 | 5.17669335 | 2.26E-07 | 8.01E-06 | DHRSX | 207063 |
| ENSG000001 | 1357.19095 | 0.716422273 | 0.1384162 | 5.17585558 | 2.27E-07 | 8.04E-06 | NA | NA |
| ENSG000001 | 443.795322 | 0.716479272 | 0.13843077 | 5.17572276 | 2.27E-07 | 8.04E-06 | CENPB | 1059 |
| ENSG000001 | 194.929498 | 0.887345974 | 0.17144538 | 5.1756775 | 2.27E-07 | 8.04E-06 | TSSC4 | 10078 |
| ENSG00000C | 1993.74135 | 0.636000492 | 0.12291427 | 5.17434206 | 2.29E-07 | 8.09E-06 | CDH3 | 1001 |
| ENSG00000C | 1971.70968 | 0.600148168 | 0.11598931 | 5.17416805 | 2.29E-07 | 8.09E-06 | CTPS2 | 56474 |
| ENSG00000C | 31.2835633 | -1.359339133 | 0.26272514 | -5.1739971 | 2.29E-07 | 8.09E-06 | C6 | 729 |
| ENSG000001 | 1145.3682 | 0.628672806 | 0.12153638 | 5.17271295 | 2.31E-07 | 8.13E-06 | RNF167 | 26001 |
| ENSG000001 | 4151.37855 | 0.677285545 | 0.13093394 | 5.17272703 | 2.31E-07 | 8.13E-06 | TCEAL8 | 90843 |
| ENSG000001 | 631.971832 | 0.677109465 | 0.13090546 | 5.17250731 | 2.31E-07 | 8.13E-06 | DCAF12 | 25853 |
| ENSG000001 | 668.545693 | 0.770713315 | 0.14901366 | 5.17209852 | 2.31E-07 | 8.14E-06 | LLGL1 | 3996 |
| ENSG000001 | 3122.90188 | 0.631304218 | 0.1220845 | 5.17104312 | 2.33E-07 | 8.18E-06 | RSL24D1 | 51187 |
| ENSG000001 | 506.584638 | 0.658389704 | 0.12734679 | 5.1700534 | 2.34E-07 | 8.22E-06 | DGUOK | 1716 |
| ENSG000002 | 737.116779 | 0.729896976 | 0.14119961 | 5.16925617 | 2.35E-07 | 8.24E-06 | NA | NA |
| ENSG000001 | 3414.50994 | 0.685946272 | 0.13270603 | 5.16891548 | 2.35E-07 | 8.25E-06 | EPCAM | 4072 |
| ENSG000001 | 406.232846 | 0.690691383 | 0.13362591 | 5.16884341 | 2.36E-07 | 8.25E-06 | SIAH2 | 6478 |

| | | | | | | | | |
|---|---|---|---|---|---|---|---|---|
| ENSG000001 | 200.667438 | 0.774588755 | 0.14986339 | 5.1686322 | 2.36E-07 | 8.25E-06 | PTER | 9317 |
| ENSG000001 | 724.507486 | 0.711657826 | 0.1377227 | 5.16732398 | 2.37E-07 | 8.30E-06 | WIPI2 | 26100 |
| ENSG000001 | 114.824043 | -0.881836726 | 0.1706654 | -5.1670503 | 2.38E-07 | 8.31E-06 | LRP1B | 53353 |
| ENSG000001 | 2614.24468 | 0.669326551 | 0.12955363 | 5.16640518 | 2.39E-07 | 8.33E-06 | EEF1D | 1936 |
| ENSG000001 | 669.720245 | 0.617736314 | 0.11957368 | 5.16615605 | 2.39E-07 | 8.33E-06 | JAM3 | 83700 |
| ENSG00000C | 950.60397 | 0.779869066 | 0.15096547 | 5.16587722 | 2.39E-07 | 8.33E-06 | TIMM13 | 26517 |
| ENSG000001 | 986.544223 | 0.576366932 | 0.11157772 | 5.1656093 | 2.40E-07 | 8.33E-06 | ELP3 | 55140 |
| ENSG000002 | 4666.67075 | 0.767281201 | 0.14853534 | 5.16564758 | 2.40E-07 | 8.33E-06 | HIST1H2BG | 8339 |
| ENSG000001 | 7021.84163 | 0.678542103 | 0.13136482 | 5.16532602 | 2.40E-07 | 8.34E-06 | RPS12 | 6206 |
| ENSG000001 | 1490.49706 | 0.683338836 | 0.13229875 | 5.16511946 | 2.40E-07 | 8.34E-06 | PFDN5 | 5204 |
| ENSG000001 | 343.498869 | 0.733872568 | 0.1420854 | 5.16501034 | 2.40E-07 | 8.34E-06 | FPGS | 2356 |
| ENSG000001 | 400.846109 | 0.729554483 | 0.14125414 | 5.16483615 | 2.41E-07 | 8.34E-06 | INO80E | 283899 |
| ENSG00000C | 257.060243 | 0.743666699 | 0.14401774 | 5.16371589 | 2.42E-07 | 8.38E-06 | RPS6KA2 | 6196 |
| ENSG00000C | 1247.70855 | 0.621552562 | 0.1203732 | 5.16354625 | 2.42E-07 | 8.38E-06 | ORC1 | 4998 |
| ENSG00000C | 169.114837 | 0.88765312 | 0.17194533 | 5.16241487 | 2.44E-07 | 8.43E-06 | TMEM161A | 54929 |
| ENSG000001 | 705.527598 | 0.674148361 | 0.13059391 | 5.16217293 | 2.44E-07 | 8.43E-06 | XPNPEP1 | 7511 |
| ENSG000001 | 501.141415 | 0.699645133 | 0.13553449 | 5.16211885 | 2.44E-07 | 8.43E-06 | CMTR1 | 23070 |
| ENSG000001 | 2897.76295 | 0.697108518 | 0.13506915 | 5.16112302 | 2.45E-07 | 8.46E-06 | P4HB | 5034 |
| ENSG000002 | 791.581417 | 0.65858866 | 0.1276096 | 5.16096477 | 2.46E-07 | 8.46E-06 | NA | NA |
| ENSG00000C | 420.193378 | 0.72053865 | 0.1396268 | 5.16046105 | 2.46E-07 | 8.48E-06 | PAFAH1B3 | 5050 |
| ENSG000001 | 810.890965 | 0.793480377 | 0.15376491 | 5.16034767 | 2.46E-07 | 8.48E-06 | RNF40 | 9810 |
| ENSG000001 | 587.721647 | 0.599881312 | 0.11626986 | 5.15938787 | 2.48E-07 | 8.51E-06 | MARVELD2 | 153562 |
| ENSG000002 | 458.816562 | 0.667945913 | 0.12946763 | 5.15917309 | 2.48E-07 | 8.52E-06 | TRNP1 | 388610 |
| ENSG000002 | 506.819401 | 0.724544416 | 0.14045973 | 5.15837842 | 2.49E-07 | 8.55E-06 | NA | NA |
| ENSG000001 | 1426.16709 | 0.637772839 | 0.12367134 | 5.15699791 | 2.51E-07 | 8.60E-06 | CNIH1 | 10175 |
| ENSG000001 | 476.9899 | 0.706742731 | 0.13707707 | 5.15580559 | 2.53E-07 | 8.63E-06 | RGP1 | 9827 |
| ENSG000001 | 10343.4691 | 0.658886723 | 0.12779001 | 5.15601133 | 2.52E-07 | 8.63E-06 | USP9X | 8239 |
| ENSG000001 | 918.625121 | 0.642513376 | 0.12461847 | 5.15584374 | 2.52E-07 | 8.63E-06 | CSNK1G2 | 1455 |
| ENSG000001 | 1296.50746 | 0.765015061 | 0.14837057 | 5.15611064 | 2.52E-07 | 8.63E-06 | FLII | 2314 |

| | | | | | | | | |
|---|---|---|---|---|---|---|---|---|
| ENSG000001 | 1384.00662 | 0.649758321 | 0.12604792 | 5.15485169 | 2.54E-07 | 8.66E-06 | PSMB6 | 5694 |
| ENSG000001 | 2106.81734 | 0.624107543 | 0.12107923 | 5.15453854 | 2.54E-07 | 8.67E-06 | CUL4A | 8451 |
| ENSG000001 | 122.647969 | 0.846724402 | 0.16433003 | 5.15258485 | 2.57E-07 | 8.75E-06 | LRRTM3 | 347731 |
| ENSG000001 | 281.693283 | 0.851268416 | 0.1652197 | 5.15234201 | 2.57E-07 | 8.76E-06 | E2F1 | 1869 |
| ENSG00000C | 287.437492 | 0.858834146 | 0.1667165 | 5.15146446 | 2.58E-07 | 8.77E-06 | DVL2 | 1856 |
| ENSG000001 | 252.678455 | 0.694089022 | 0.13474258 | 5.15122266 | 2.59E-07 | 8.77E-06 | PAQR5 | 54852 |
| ENSG000001 | 20501.0369 | 0.626539738 | 0.12162933 | 5.15122242 | 2.59E-07 | 8.77E-06 | TUBB | 203068 |
| ENSG000002 | 1611.94383 | 0.621623572 | 0.12066113 | 5.15181278 | 2.58E-07 | 8.77E-06 | ANKRD28 | 23243 |
| ENSG000002 | 3512.47341 | 0.585647171 | 0.11368269 | 5.15159507 | 2.58E-07 | 8.77E-06 | NA | NA |
| ENSG000001 | 1762.01569 | 0.63933584 | 0.12414854 | 5.14976535 | 2.61E-07 | 8.84E-06 | TSPYL1 | 7259 |
| ENSG000001 | 898.176912 | 0.742102831 | 0.14412196 | 5.14913076 | 2.62E-07 | 8.86E-06 | APH1A | 51107 |
| ENSG000001 | 2245.3811 | 0.647365494 | 0.12575295 | 5.1479151 | 2.63E-07 | 8.90E-06 | UQCRC2 | 7385 |
| ENSG000001 | 158.141698 | 0.835815054 | 0.16236238 | 5.14783687 | 2.64E-07 | 8.90E-06 | MZT2B | 80097 |
| ENSG000001 | 213.776994 | 0.816429202 | 0.15860072 | 5.1477018 | 2.64E-07 | 8.90E-06 | RNASEH2C | 84153 |
| ENSG000001 | 283.045166 | 0.829227167 | 0.16113159 | 5.14627319 | 2.66E-07 | 8.96E-06 | PTDSS2 | 81490 |
| ENSG000001 | 48.840623 | 1.11895441 | 0.21754144 | 5.14363808 | 2.69E-07 | 9.08E-06 | BCHE | 590 |
| ENSG000001 | 458.305752 | 0.703030545 | 0.13673477 | 5.14156398 | 2.72E-07 | 9.18E-06 | STAT6 | 6778 |
| ENSG000001 | 167.945049 | 0.878057387 | 0.1707886 | 5.1411943 | 2.73E-07 | 9.19E-06 | ADPRHL2 | 54936 |
| ENSG000001 | 1895.14371 | 0.607219367 | 0.11811274 | 5.14101486 | 2.73E-07 | 9.19E-06 | NA | NA |
| ENSG000001 | 4513.19011 | 0.828108528 | 0.16109456 | 5.14051219 | 2.74E-07 | 9.21E-06 | HIST1H2AG | 8969 |
| ENSG000001 | 721.27324 | 0.601572881 | 0.11703339 | 5.14018141 | 2.74E-07 | 9.21E-06 | GPALPP1 | 55425 |
| ENSG000001 | 1169.97512 | 0.622651011 | 0.1211329 | 5.14023018 | 2.74E-07 | 9.21E-06 | PKIB | 5570 |
| ENSG000001 | 370.334986 | 0.622180649 | 0.12105441 | 5.13967778 | 2.75E-07 | 9.22E-06 | NCOA7 | 135112 |
| ENSG000001 | 182.218668 | -0.801771279 | 0.15599807 | -5.139623 | 2.75E-07 | 9.22E-06 | ZMAT3 | 64393 |
| ENSG000001 | 96.5327157 | 1.008273949 | 0.19619089 | 5.1392497 | 2.76E-07 | 9.23E-06 | TST | 7263 |
| ENSG000001 | 888.543002 | 0.680700108 | 0.13248328 | 5.13800775 | 2.78E-07 | 9.28E-06 | ATG13 | 9776 |
| ENSG00000C | 344.819145 | 0.837149625 | 0.16295034 | 5.13745237 | 2.78E-07 | 9.30E-06 | HYAL2 | 8692 |
| ENSG000001 | 804.407524 | 0.656502745 | 0.12779059 | 5.13733253 | 2.79E-07 | 9.30E-06 | PSRC1 | 84722 |
| ENSG000001 | 1093.16676 | 0.784758544 | 0.15278422 | 5.13638478 | 2.80E-07 | 9.34E-06 | RANGAP1 | 5905 |

| | | | | | | | | |
|---|---|---|---|---|---|---|---|---|
| ENSG000001 | 308.540458 | 0.770718516 | 0.15006286 | 5.13597122 | 2.81E-07 | 9.35E-06 | KCNK5 | 8645 |
| ENSG000001 | 673.183791 | 0.623949559 | 0.12150117 | 5.13533787 | 2.82E-07 | 9.38E-06 | GLB1 | 2720 |
| ENSG000001 | 408.298022 | 0.749886173 | 0.14603817 | 5.13486424 | 2.82E-07 | 9.39E-06 | TARBP2 | 6895 |
| ENSG000001 | 701.094499 | 0.628615313 | 0.12242218 | 5.1348156 | 2.82E-07 | 9.39E-06 | KCTD15 | 79047 |
| ENSG000001 | 2052.04455 | 0.543486962 | 0.10585015 | 5.13449425 | 2.83E-07 | 9.40E-06 | CLTA | 1211 |
| ENSG000001 | 1645.61356 | 0.605342973 | 0.11792015 | 5.13349922 | 2.84E-07 | 9.44E-06 | MKRN1 | 23608 |
| ENSG000001 | 93.3467597 | -0.979075836 | 0.19078378 | -5.1318611 | 2.87E-07 | 9.50E-06 | SOX6 | 55553 |
| ENSG000001 | 456.000745 | 0.616599474 | 0.12015353 | 5.13176319 | 2.87E-07 | 9.50E-06 | RNF146 | 81847 |
| ENSG000001 | 664.31434 | 0.630026622 | 0.12277167 | 5.13169377 | 2.87E-07 | 9.50E-06 | DTD1 | 92675 |
| ENSG000001 | 158.893978 | 1.009482712 | 0.19670497 | 5.13196339 | 2.87E-07 | 9.50E-06 | ULK1 | 8408 |
| ENSG000001 | 1169.87763 | 0.648952048 | 0.12646599 | 5.13143545 | 2.88E-07 | 9.50E-06 | CRK | 1398 |
| ENSG000002 | 2032.65714 | 0.625633041 | 0.12193466 | 5.13088747 | 2.88E-07 | 9.52E-06 | DANCR | 57291 |
| ENSG000001 | 549.763701 | 0.662528209 | 0.12914285 | 5.13019656 | 2.89E-07 | 9.55E-06 | ELOF1 | 84337 |
| ENSG000001 | 3162.09606 | 0.574446575 | 0.11199404 | 5.12926024 | 2.91E-07 | 9.59E-06 | BUB1B | 701 |
| ENSG000002 | 344.055603 | 0.764146859 | 0.14899612 | 5.12863599 | 2.92E-07 | 9.61E-06 | ARL2 | 402 |
| ENSG000001 | 3115.7755 | 0.647250209 | 0.12622378 | 5.12779948 | 2.93E-07 | 9.65E-06 | AIF1L | 83543 |
| ENSG000001 | 32.7762305 | -1.336976346 | 0.26077972 | -5.1268417 | 2.95E-07 | 9.69E-06 | SPATA18 | 132671 |
| ENSG000001 | 238.404927 | 0.804302426 | 0.15690218 | 5.12613926 | 2.96E-07 | 9.72E-06 | RGS16 | 6004 |
| ENSG000001 | 97.1096332 | -0.854300719 | 0.16667398 | -5.1255795 | 2.97E-07 | 9.72E-06 | IQCA1 | 79781 |
| ENSG000001 | 1177.58134 | 0.664717851 | 0.12968269 | 5.12572525 | 2.96E-07 | 9.72E-06 | ABRACL | 58527 |
| ENSG000001 | 662.727474 | 0.799176714 | 0.15591989 | 5.12555982 | 2.97E-07 | 9.72E-06 | PPP4C | 5531 |
| ENSG00000C | 3677.56451 | 0.639058374 | 0.12470055 | 5.12474363 | 2.98E-07 | 9.76E-06 | FKBP4 | 2288 |
| ENSG00000C | 2770.03837 | 0.630639475 | 0.12307169 | 5.1241634 | 2.99E-07 | 9.78E-06 | SLC30A9 | 10463 |
| ENSG000001 | 532.497236 | 0.683502292 | 0.13340899 | 5.12336007 | 3.00E-07 | 9.82E-06 | NARS2 | 79731 |
| ENSG000002 | 757.916407 | 0.751802334 | 0.14678447 | 5.12181118 | 3.03E-07 | 9.88E-06 | EXOSC6 | 118460 |
| ENSG000002 | 23.8900904 | -1.395677136 | 0.27249474 | -5.1218498 | 3.03E-07 | 9.88E-06 | SLC25A3P1 | 163742 |
| ENSG000002 | 312.779657 | 0.63792318 | 0.12457739 | 5.12069793 | 3.04E-07 | 9.93E-06 | MPV17L2 | 84769 |
| ENSG00000C | 17826.3672 | 0.640434583 | 0.12507343 | 5.12046864 | 3.05E-07 | 9.93E-06 | SCD | 6319 |
| ENSG000001 | 625.837369 | 0.611070148 | 0.11934315 | 5.12027818 | 3.05E-07 | 9.94E-06 | ENTPD4 | 9583 |

| | | | | | | | | |
|---|---|---|---|---|---|---|---|---|
| ENSG000002 | 976.077426 | 0.798603904 | 0.15598874 | 5.11962549 | 3.06E-07 | 9.96E-06 | NA | NA |
| ENSG000001 | 1145.48554 | 0.595355431 | 0.11630442 | 5.11894058 | 3.07E-07 | 9.98E-06 | ZNF273 | 10793 |
| ENSG000001 | 1150.68963 | 0.628977008 | 0.12286918 | 5.11907871 | 3.07E-07 | 9.98E-06 | COPS8 | 10920 |
| ENSG000001 | 620.430493 | 0.58058457 | 0.11344299 | 5.11785339 | 3.09E-07 | 1.00E-05 | IGSF1 | 3547 |
| ENSG000001 | 1447.31015 | 0.650770096 | 0.12717866 | 5.1169754 | 3.10E-07 | 1.01E-05 | DDX23 | 9416 |
| ENSG00000C | 2070.0554 | 0.615312037 | 0.12026292 | 5.11639045 | 3.11E-07 | 1.01E-05 | EIF3I | 8668 |
| ENSG000001 | 746.92448 | 0.627616916 | 0.12267273 | 5.1161892 | 3.12E-07 | 1.01E-05 | MRPL18 | 29074 |
| ENSG00000C | 273.009823 | 0.803170569 | 0.15703004 | 5.11475747 | 3.14E-07 | 1.02E-05 | DHPS | 1725 |
| ENSG000001 | 1606.54311 | 0.589877631 | 0.11532994 | 5.11469634 | 3.14E-07 | 1.02E-05 | RAB3GAP1 | 22930 |
| ENSG000001 | 2231.98247 | 0.700656068 | 0.13698307 | 5.11490981 | 3.14E-07 | 1.02E-05 | SURF4 | 6836 |
| ENSG000001 | 1462.64225 | 0.646281411 | 0.12637142 | 5.114142 | 3.15E-07 | 1.02E-05 | SHISA5 | 51246 |
| ENSG000001 | 1200.20568 | 0.697823613 | 0.13645634 | 5.11389652 | 3.16E-07 | 1.02E-05 | CARHSP1 | 23589 |
| ENSG00000C | 5285.8244 | 0.62798632 | 0.12281111 | 5.1134326 | 3.16E-07 | 1.02E-05 | MSH2 | 4436 |
| ENSG000001 | 398.282365 | 0.738539236 | 0.14443993 | 5.11312366 | 3.17E-07 | 1.02E-05 | GSTO1 | 9446 |
| ENSG000001 | 1106.92551 | 0.704713448 | 0.13783471 | 5.11274309 | 3.18E-07 | 1.02E-05 | TGIF2 | 60436 |
| ENSG000001 | 572.344785 | 0.629437267 | 0.12311338 | 5.11266337 | 3.18E-07 | 1.02E-05 | ADGRL3 | 23284 |
| ENSG000001 | 830.275822 | 0.705237091 | 0.13800944 | 5.11006416 | 3.22E-07 | 1.03E-05 | UGT8 | 7368 |
| ENSG000001 | 9124.41795 | 0.630632684 | 0.12342586 | 5.10940502 | 3.23E-07 | 1.04E-05 | EIF4A2 | 1974 |
| ENSG000001 | 262.627015 | 0.827025773 | 0.16187284 | 5.10910755 | 3.24E-07 | 1.04E-05 | FAM127A | 8933 |
| ENSG00000C | 1619.1573 | 0.630206971 | 0.12336757 | 5.10836811 | 3.25E-07 | 1.04E-05 | PDCD2 | 5134 |
| ENSG000001 | 739.598832 | 0.766362398 | 0.15005677 | 5.10714964 | 3.27E-07 | 1.05E-05 | SCAP | 22937 |
| ENSG000001 | 1183.9607 | 0.711630036 | 0.13933855 | 5.10720155 | 3.27E-07 | 1.05E-05 | CDC20 | 991 |
| ENSG00000C | 1312.56829 | 0.668344075 | 0.13086978 | 5.10693955 | 3.27E-07 | 1.05E-05 | KIF22 | 3835 |
| ENSG000001 | 1150.34867 | 0.665236956 | 0.13027876 | 5.10625783 | 3.29E-07 | 1.05E-05 | MMS19 | 64210 |
| ENSG000001 | 195.820214 | 0.870237743 | 0.17042551 | 5.10626449 | 3.29E-07 | 1.05E-05 | VWA1 | 64856 |
| ENSG000001 | 1905.92718 | 0.678663168 | 0.13292344 | 5.10566961 | 3.30E-07 | 1.05E-05 | EDF1 | 8721 |
| ENSG000001 | 1037.9945 | 0.643524676 | 0.12603772 | 5.10581002 | 3.29E-07 | 1.05E-05 | GFM2 | 84340 |
| ENSG000001 | 462.958869 | 0.664133508 | 0.1301299 | 5.10361974 | 3.33E-07 | 1.06E-05 | PLA2G12A | 81579 |
| ENSG000001 | 1240.8261 | 0.608225569 | 0.11921212 | 5.1020447 | 3.36E-07 | 1.07E-05 | ENOPH1 | 58478 |

| | | | | | | | | |
|---|---|---|---|---|---|---|---|---|
| ENSG000001 | 6158.95317 | 0.624153085 | 0.12234683 | 5.10150577 | 3.37E-07 | 1.07E-05 | CFL1 | 1072 |
| ENSG000001 | 103.779415 | -0.871961906 | 0.17092963 | -5.1012917 | 3.37E-07 | 1.07E-05 | ABCA13 | 154664 |
| ENSG000001 | 5829.38754 | 0.685909997 | 0.13446456 | 5.10104679 | 3.38E-07 | 1.07E-05 | RPL27 | 6155 |
| ENSG000001 | 268.526842 | 0.768330942 | 0.15062746 | 5.10086902 | 3.38E-07 | 1.07E-05 | NODAL | 4838 |
| ENSG000002 | 544.135634 | 0.739787764 | 0.14507722 | 5.09926899 | 3.41E-07 | 1.08E-05 | GNL1 | 2794 |
| ENSG000001 | 908.8861 | 0.605516124 | 0.11875464 | 5.09888375 | 3.42E-07 | 1.08E-05 | EMC10 | 284361 |
| ENSG000001 | 472.523297 | 0.660009641 | 0.12947682 | 5.09751184 | 3.44E-07 | 1.09E-05 | UQCC1 | 55245 |
| ENSG000001 | 190.899021 | 0.937044308 | 0.18382202 | 5.09756277 | 3.44E-07 | 1.09E-05 | PGLS | 25796 |
| ENSG00000C | 509.719802 | 0.73954522 | 0.1450863 | 5.09727822 | 3.45E-07 | 1.09E-05 | UBE2T | 29089 |
| ENSG000001 | 646.418019 | 0.691049268 | 0.13558905 | 5.09664494 | 3.46E-07 | 1.09E-05 | SPP1 | 6696 |
| ENSG000001 | 870.838614 | 0.651269393 | 0.1277985 | 5.09606463 | 3.47E-07 | 1.09E-05 | GINS2 | 51659 |
| ENSG000001 | 750.296731 | 0.626074847 | 0.12285243 | 5.09615372 | 3.47E-07 | 1.09E-05 | ADO | 84890 |
| ENSG000001 | 854.646554 | 0.579392562 | 0.11372234 | 5.0947997 | 3.49E-07 | 1.10E-05 | PPP2R2B | 5521 |
| ENSG000001 | 383.157562 | 0.751579407 | 0.14756227 | 5.09330322 | 3.52E-07 | 1.11E-05 | FOXRED1 | 55572 |
| ENSG000001 | 394.978129 | 0.746982134 | 0.14672968 | 5.09087264 | 3.56E-07 | 1.12E-05 | REXO4 | 57109 |
| ENSG000001 | 236.521061 | 0.688858475 | 0.13535121 | 5.08941508 | 3.59E-07 | 1.13E-05 | TTC31 | 64427 |
| ENSG000001 | 303.545608 | 0.867460876 | 0.17045103 | 5.08920863 | 3.60E-07 | 1.13E-05 | DUS1L | 64118 |
| ENSG000002 | 260.570694 | 0.739147996 | 0.14523598 | 5.0892898 | 3.59E-07 | 1.13E-05 | HGH1 | 51236 |
| ENSG000001 | 1785.94489 | 0.572898794 | 0.11257717 | 5.08894312 | 3.60E-07 | 1.13E-05 | PPP6C | 5537 |
| ENSG000001 | 1070.61955 | 0.745054925 | 0.14641076 | 5.08879914 | 3.60E-07 | 1.13E-05 | SMPD4 | 55627 |
| ENSG00000C | 99.4054423 | 0.912292254 | 0.17930055 | 5.08806162 | 3.62E-07 | 1.13E-05 | PIGQ | 9091 |
| ENSG000001 | 846.045181 | 0.740820843 | 0.14559597 | 5.08819601 | 3.61E-07 | 1.13E-05 | AKR1A1 | 10327 |
| ENSG000001 | 544.653288 | 0.789815125 | 0.15524141 | 5.08765764 | 3.63E-07 | 1.13E-05 | INPPL1 | 3636 |
| ENSG000001 | 365.739758 | 0.697353929 | 0.13716125 | 5.08419038 | 3.69E-07 | 1.15E-05 | DUSP3 | 1845 |
| ENSG000001 | 568.267672 | 0.766415772 | 0.15076179 | 5.08362078 | 3.70E-07 | 1.16E-05 | MRPS2 | 51116 |
| ENSG000001 | 716.659821 | 0.681823659 | 0.13420353 | 5.08051958 | 3.76E-07 | 1.17E-05 | CIAO1 | 9391 |
| ENSG000001 | 317.559369 | 0.710500852 | 0.13986397 | 5.07994194 | 3.78E-07 | 1.18E-05 | KCNMB4 | 27345 |
| ENSG000001 | 60.3015318 | -1.051186779 | 0.20697329 | -5.0788523 | 3.80E-07 | 1.18E-05 | CFAP47 | 286464 |
| ENSG000001 | 1282.08684 | 0.550399467 | 0.10838033 | 5.07840718 | 3.81E-07 | 1.18E-05 | BCAS2 | 10286 |

| | | | | | | | | |
|---|---|---|---|---|---|---|---|---|
| ENSG000001 | 2731.69265 | 0.640073987 | 0.12604871 | 5.077989 | 3.81E-07 | 1.19E-05 | ARL6IP1 | 23204 |
| ENSG000002 | 118.795899 | 0.917493706 | 0.18068159 | 5.0779591 | 3.82E-07 | 1.19E-05 | VTRNA1-3 | 56662 |
| ENSG000001 | 683.285695 | 0.714640983 | 0.14075812 | 5.07708534 | 3.83E-07 | 1.19E-05 | DNAJB1 | 3337 |
| ENSG000001 | 842.256226 | 0.726577262 | 0.14315128 | 5.0755904 | 3.86E-07 | 1.20E-05 | DNAJA3 | 9093 |
| ENSG000002 | 502.412727 | 0.651217509 | 0.12830206 | 5.07565905 | 3.86E-07 | 1.20E-05 | PDCD6 | 10016 |
| ENSG000001 | 641.164115 | 0.676605054 | 0.13331427 | 5.07526351 | 3.87E-07 | 1.20E-05 | HSPA14 | 51182 |
| ENSG000001 | 1484.24284 | 0.713997102 | 0.14076568 | 5.07223858 | 3.93E-07 | 1.22E-05 | SLC2A1 | 6513 |
| ENSG000001 | 544.510765 | 0.744419643 | 0.14678866 | 5.07137037 | 3.95E-07 | 1.22E-05 | POP7 | 10248 |
| ENSG000001 | 118.474707 | 0.870355664 | 0.17163094 | 5.07108837 | 3.96E-07 | 1.22E-05 | PUSL1 | 126789 |
| ENSG000001 | 217.285164 | 0.85496376 | 0.16859868 | 5.070999 | 3.96E-07 | 1.22E-05 | GTPBP6 | 8225 |
| ENSG000000 | 3973.30803 | 0.622987511 | 0.12286248 | 5.07060817 | 3.97E-07 | 1.22E-05 | KARS | 3735 |
| ENSG000001 | 908.429793 | 0.538843757 | 0.10627155 | 5.07044238 | 3.97E-07 | 1.22E-05 | METTL7A | 25840 |
| ENSG000001 | 64.4037489 | -1.030526216 | 0.20328137 | -5.0694572 | 3.99E-07 | 1.23E-05 | RGSL1 | 353299 |
| ENSG000001 | 1936.32365 | 0.559205071 | 0.11030563 | 5.06959679 | 3.99E-07 | 1.23E-05 | XPO5 | 57510 |
| ENSG000001 | 1911.33296 | 0.657126821 | 0.1296327 | 5.06914386 | 4.00E-07 | 1.23E-05 | EIF3G | 8666 |
| ENSG000002 | 89.91286 | 1.081433208 | 0.2133329 | 5.06922851 | 3.99E-07 | 1.23E-05 | NA | NA |
| ENSG000000 | 326.396983 | 0.76840132 | 0.15162405 | 5.06780626 | 4.02E-07 | 1.24E-05 | FAM50A | 9130 |
| ENSG000001 | 551.606365 | 0.633851543 | 0.12509169 | 5.06709547 | 4.04E-07 | 1.24E-05 | BICD1 | 636 |
| ENSG000000 | 1501.4713 | 0.682377556 | 0.13467511 | 5.06684235 | 4.04E-07 | 1.24E-05 | MRPS34 | 65993 |
| ENSG000001 | 244.880116 | 0.737252556 | 0.14553364 | 5.06585679 | 4.07E-07 | 1.25E-05 | CDIPT | 10423 |
| ENSG000001 | 643.647458 | 0.598300049 | 0.11813346 | 5.06461127 | 4.09E-07 | 1.25E-05 | JAK1 | 3716 |
| ENSG000001 | 421.202431 | 0.765776014 | 0.15121287 | 5.06422516 | 4.10E-07 | 1.25E-05 | PTGES2 | 80142 |
| ENSG000001 | 72.2384599 | -0.945901311 | 0.18678826 | -5.0640299 | 4.10E-07 | 1.25E-05 | PAPPA | 5069 |
| ENSG000001 | 728.38787 | 0.635232225 | 0.12543804 | 5.06411146 | 4.10E-07 | 1.25E-05 | MCRS1 | 10445 |
| ENSG000001 | 353.984548 | 0.778519622 | 0.15378391 | 5.06242586 | 4.14E-07 | 1.26E-05 | C12orf57 | 113246 |
| ENSG000001 | 334.651904 | 0.73559557 | 0.14532148 | 5.06185039 | 4.15E-07 | 1.27E-05 | AARS2 | 57505 |
| ENSG000001 | 434.025348 | 0.68453092 | 0.1354006 | 5.05559723 | 4.29E-07 | 1.31E-05 | ACAD8 | 27034 |
| ENSG000001 | 700.771006 | 0.670149897 | 0.13257046 | 5.05504697 | 4.30E-07 | 1.31E-05 | KCTD2 | 23510 |
| ENSG000002 | 166.885475 | 0.752733731 | 0.14891968 | 5.05462913 | 4.31E-07 | 1.31E-05 | POU5F1P3 | 642559 |

| | | | | | | | | |
|---|---|---|---|---|---|---|---|---|
| ENSG000001 | 321.978652 | 0.663783832 | 0.13133315 | 5.05419881 | 4.32E-07 | 1.31E-05 | CTSB | 1508 |
| ENSG000002 | 443.272815 | 0.732974856 | 0.14508003 | 5.05221064 | 4.37E-07 | 1.33E-05 | NDUFS3 | 4722 |
| ENSG000002 | 193.09463 | -0.862501894 | 0.17071554 | -5.0522753 | 4.37E-07 | 1.33E-05 | TSIX | 9383 |
| ENSG000001 | 984.443607 | 0.635782022 | 0.12585038 | 5.05188807 | 4.37E-07 | 1.33E-05 | ZNF3 | 7551 |
| ENSG000001 | 14554.9955 | 0.587842652 | 0.11637793 | 5.05115246 | 4.39E-07 | 1.33E-05 | TUBA1B | 10376 |
| ENSG000001 | 1794.29916 | 0.629325019 | 0.12458889 | 5.05121291 | 4.39E-07 | 1.33E-05 | MRPS16 | 51021 |
| ENSG000001 | 152.630507 | 0.762248556 | 0.15093949 | 5.05002744 | 4.42E-07 | 1.34E-05 | CUZD1 | 50624 |
| ENSG000001 | 5243.50416 | 0.594500146 | 0.11772945 | 5.04971462 | 4.42E-07 | 1.34E-05 | PDIA6 | 10130 |
| ENSG000001 | 156.223142 | 0.873143395 | 0.1729346 | 5.04898041 | 4.44E-07 | 1.34E-05 | PSMG3 | 84262 |
| ENSG000001 | 1119.35414 | 0.605429987 | 0.11991923 | 5.04864797 | 4.45E-07 | 1.34E-05 | FOCAD | 54914 |
| ENSG000002 | 392.783839 | 0.677454097 | 0.1342127 | 5.04761545 | 4.47E-07 | 1.35E-05 | NA | NA |
| ENSG000001 | 58.677824 | 1.054812966 | 0.20904133 | 5.04595423 | 4.51E-07 | 1.36E-05 | HES6 | 55502 |
| ENSG000001 | 1233.797 | 0.661059755 | 0.1310249 | 5.04529849 | 4.53E-07 | 1.37E-05 | ADRM1 | 11047 |
| ENSG000001 | 1171.25897 | 0.640279156 | 0.12694019 | 5.04394356 | 4.56E-07 | 1.37E-05 | ATP6V1B2 | 526 |
| ENSG000001 | 213.843259 | 0.817422523 | 0.16208749 | 5.04309436 | 4.58E-07 | 1.38E-05 | NUDT1 | 4521 |
| ENSG000001 | 313.60675 | 0.673741867 | 0.13360908 | 5.04263533 | 4.59E-07 | 1.38E-05 | DRG2 | 1819 |
| ENSG000001 | 2447.25747 | 0.738218524 | 0.14640179 | 5.04241474 | 4.60E-07 | 1.38E-05 | AP2M1 | 1173 |
| ENSG000002 | 32.4954986 | -1.31872883 | 0.26154868 | -5.0420015 | 4.61E-07 | 1.38E-05 | LOC100422737 | 100422737 |
| ENSG000001 | 27.4953747 | -1.320186377 | 0.26184888 | -5.0417874 | 4.61E-07 | 1.38E-05 | GABRA4 | 2557 |
| ENSG000001 | 303.231237 | 0.77462209 | 0.153697 | 5.03992991 | 4.66E-07 | 1.40E-05 | POC1A | 25886 |
| ENSG000001 | 1343.00723 | 0.55678721 | 0.11048063 | 5.03968162 | 4.66E-07 | 1.40E-05 | CHP1 | 11261 |
| ENSG000001 | 378.832569 | 0.736928126 | 0.14625078 | 5.03879788 | 4.68E-07 | 1.40E-05 | GLA | 2717 |
| ENSG000001 | 525.033848 | 0.760359068 | 0.15090075 | 5.03880238 | 4.68E-07 | 1.40E-05 | PNMAL1 | 55228 |
| ENSG000001 | 546.926043 | 0.673714739 | 0.1337095 | 5.03864526 | 4.69E-07 | 1.40E-05 | ACSS3 | 79611 |
| ENSG000001 | 1606.68629 | 0.647353092 | 0.12848175 | 5.03848277 | 4.69E-07 | 1.40E-05 | PSMC3 | 5702 |
| ENSG000001 | 503.99693 | 0.724096765 | 0.14372893 | 5.03793336 | 4.71E-07 | 1.41E-05 | RRP9 | 9136 |
| ENSG000002 | 375.649486 | 0.659381919 | 0.13088819 | 5.03774961 | 4.71E-07 | 1.41E-05 | PRR13 | 54458 |
| ENSG000001 | 789.590316 | 0.649107925 | 0.12885966 | 5.03732466 | 4.72E-07 | 1.41E-05 | NAPG | 8774 |
| ENSG000001 | 326.622543 | 0.736914726 | 0.14633181 | 5.03591598 | 4.76E-07 | 1.42E-05 | RPS19BP1 | 91582 |

| | | | | | | | | | |
|---|---|---|---|---|---|---|---|---|---|
| ENSG000001 | 1491.97516 | 0.623833337 | 0.12388674 | 5.03551349 | 4.77E-07 | 1.42E-05 | ADNP2 | | 22850 |
| ENSG000001 | 1217.20953 | 0.701288492 | 0.13926552 | 5.035622 | 4.76E-07 | 1.42E-05 | POLD2 | | 5425 |
| ENSG000002 | 430.473992 | 0.671812839 | 0.13341774 | 5.03540877 | 4.77E-07 | 1.42E-05 | NA | NA | |
| ENSG000001 | 492.031953 | 0.640282051 | 0.12716014 | 5.03524166 | 4.77E-07 | 1.42E-05 | CALCOCO2 | | 10241 |
| ENSG000001 | 679.056832 | 0.768945575 | 0.15275646 | 5.03380079 | 4.81E-07 | 1.43E-05 | POR | | 5447 |
| ENSG000001 | 707.72284 | 0.638874585 | 0.12693318 | 5.03315678 | 4.82E-07 | 1.43E-05 | RIC8A | | 60626 |
| ENSG000001 | 678.897995 | 0.582290037 | 0.11570354 | 5.03260363 | 4.84E-07 | 1.43E-05 | NGDN | | 25983 |
| ENSG000001 | 4671.47907 | 0.620368683 | 0.12330044 | 5.03135813 | 4.87E-07 | 1.44E-05 | PGD | | 5226 |
| ENSG000001 | 3385.82158 | 0.612874827 | 0.12182532 | 5.0307674 | 4.89E-07 | 1.45E-05 | COX7C | | 1350 |
| ENSG000001 | 651.061662 | 0.677747018 | 0.1347235 | 5.03065191 | 4.89E-07 | 1.45E-05 | PLA2G16 | | 11145 |
| ENSG00000C | 2227.16476 | 0.712223421 | 0.14158641 | 5.03030924 | 4.90E-07 | 1.45E-05 | OAT | | 4942 |
| ENSG00000C | 1715.65498 | 0.629301362 | 0.12510426 | 5.03021539 | 4.90E-07 | 1.45E-05 | AURKA | | 6790 |
| ENSG00000C | 323.907855 | 0.698877094 | 0.1389886 | 5.02830512 | 4.95E-07 | 1.46E-05 | SLCO1A2 | | 6579 |
| ENSG000001 | 361.211097 | 0.723069709 | 0.14380393 | 5.0281639 | 4.95E-07 | 1.46E-05 | SYT6 | | 148281 |
| ENSG00000C | 462.874985 | 0.601367148 | 0.1196203 | 5.02729996 | 4.97E-07 | 1.46E-05 | FECH | | 2235 |
| ENSG000001 | 1089.06177 | 0.54349785 | 0.10810884 | 5.02732127 | 4.97E-07 | 1.46E-05 | CNOT11 | | 55571 |
| ENSG000001 | 114.330895 | 0.888995098 | 0.17683887 | 5.02714751 | 4.98E-07 | 1.46E-05 | BLOC1S1 | | 2647 |
| ENSG000001 | 289.4577 | 0.68655583 | 0.13658628 | 5.0265358 | 4.99E-07 | 1.47E-05 | DHRS7 | | 51635 |
| ENSG00000C | 733.100286 | 0.632118481 | 0.12576918 | 5.02602059 | 5.01E-07 | 1.47E-05 | VAMP3 | | 9341 |
| ENSG000001 | 2108.92684 | 0.571482256 | 0.11372069 | 5.02531484 | 5.03E-07 | 1.48E-05 | DOCK1 | | 1793 |
| ENSG00000C | 93.9630668 | 0.870761433 | 0.17329799 | 5.02464812 | 5.04E-07 | 1.48E-05 | BAD | | 572 |
| ENSG00000C | 194.415323 | 0.7536911 | 0.14999681 | 5.02471418 | 5.04E-07 | 1.48E-05 | SNX10 | | 29887 |
| ENSG000001 | 160.650056 | 0.79170134 | 0.15755576 | 5.02489621 | 5.04E-07 | 1.48E-05 | RASSF4 | | 83937 |
| ENSG000001 | 217.719478 | 0.836169678 | 0.16640362 | 5.02494883 | 5.04E-07 | 1.48E-05 | TACR3 | | 6870 |
| ENSG000001 | 409.173888 | 0.697665735 | 0.13886553 | 5.02403812 | 5.06E-07 | 1.48E-05 | LY6E | | 4061 |
| ENSG000001 | 25.8113131 | -1.383945591 | 0.27548486 | -5.0236721 | 5.07E-07 | 1.48E-05 | NLRP4 | | 147945 |
| ENSG000001 | 732.289217 | 0.751180934 | 0.14958845 | 5.02165068 | 5.12E-07 | 1.50E-05 | GADD45GIP1 | | 90480 |
| ENSG000001 | 1890.91316 | 0.745031747 | 0.14836896 | 5.02147976 | 5.13E-07 | 1.50E-05 | PFAS | | 5198 |
| ENSG000001 | 1475.48343 | 0.67846246 | 0.13512032 | 5.02117276 | 5.14E-07 | 1.50E-05 | FOXM1 | | 2305 |

| | | | | | | | | |
|---|---|---|---|---|---|---|---|---|
| ENSG000001 | 1582.99369 | 0.695538102 | 0.13854115 | 5.02044429 | 5.16E-07 | 1.50E-05 | CKS2 | 1164 |
| ENSG000001 | 207.214514 | 0.720082398 | 0.1434921 | 5.01827221 | 5.21E-07 | 1.52E-05 | BET1L | 51272 |
| ENSG00000C | 127.136737 | 0.874816213 | 0.17434417 | 5.01775423 | 5.23E-07 | 1.52E-05 | ASNS | 440 |
| ENSG00000C | 294.830506 | 0.703765794 | 0.14027072 | 5.01719657 | 5.24E-07 | 1.53E-05 | LZTS3 | 9762 |
| ENSG000001 | 215.406325 | 0.881703133 | 0.17580028 | 5.0153683 | 5.29E-07 | 1.54E-05 | THBS4 | 7060 |
| ENSG000001 | 944.758356 | 0.760733371 | 0.15168042 | 5.01536953 | 5.29E-07 | 1.54E-05 | CTDSP2 | 10106 |
| ENSG000001 | 703.363575 | 0.633293868 | 0.12628991 | 5.0146039 | 5.31E-07 | 1.54E-05 | RCBTB1 | 55213 |
| ENSG000001 | 209.205367 | 0.792760613 | 0.15809493 | 5.01445958 | 5.32E-07 | 1.54E-05 | NDUFS7 | 374291 |
| ENSG000002 | 184.703779 | 0.72219404 | 0.14403878 | 5.01388622 | 5.33E-07 | 1.55E-05 | NA | NA |
| ENSG000002 | 733.009839 | 0.848115125 | 0.16915141 | 5.01394053 | 5.33E-07 | 1.55E-05 | HIST1H2AK | 8330 |
| ENSG000001 | 3061.48267 | 0.630291578 | 0.1257148 | 5.01366259 | 5.34E-07 | 1.55E-05 | RPL7L1 | 285855 |
| ENSG000001 | 273.576576 | 0.709429674 | 0.14150434 | 5.01348347 | 5.35E-07 | 1.55E-05 | PROCR | 10544 |
| ENSG000001 | 3835.76572 | 0.636752062 | 0.12701397 | 5.01324435 | 5.35E-07 | 1.55E-05 | EIF4H | 7458 |
| ENSG000001 | 453.801246 | 0.691461326 | 0.13793107 | 5.01309338 | 5.36E-07 | 1.55E-05 | TMEM259 | 91304 |
| ENSG000001 | 323.927088 | 0.854815569 | 0.17055074 | 5.01208953 | 5.38E-07 | 1.55E-05 | PLEKHJ1 | 55111 |
| ENSG000001 | 311.343469 | 0.790370801 | 0.15770657 | 5.01165413 | 5.40E-07 | 1.56E-05 | AGTRAP | 57085 |
| ENSG000001 | 283.550508 | 0.641140689 | 0.12793613 | 5.01141244 | 5.40E-07 | 1.56E-05 | SULF1 | 23213 |
| ENSG000002 | 1021.65314 | 0.783909788 | 0.15643643 | 5.01104365 | 5.41E-07 | 1.56E-05 | MIF | 4282 |
| ENSG000001 | 2796.43006 | 0.61614901 | 0.12296626 | 5.01071596 | 5.42E-07 | 1.56E-05 | NDUFS5 | 4725 |
| ENSG000001 | 5826.44378 | 0.706938392 | 0.14115484 | 5.00824757 | 5.49E-07 | 1.58E-05 | DHCR24 | 1718 |
| ENSG000001 | 556.869703 | 0.65046371 | 0.12989714 | 5.00752919 | 5.51E-07 | 1.58E-05 | EIF2B2 | 8892 |
| ENSG000001 | 332.678992 | 0.673601132 | 0.13453946 | 5.00671812 | 5.54E-07 | 1.59E-05 | NSG1 | 27065 |
| ENSG000001 | 159.289202 | 0.880044931 | 0.17580686 | 5.00574856 | 5.56E-07 | 1.60E-05 | MRPL54 | 116541 |
| ENSG000001 | 717.179901 | 0.702675064 | 0.14037732 | 5.00561661 | 5.57E-07 | 1.60E-05 | ASNA1 | 439 |
| ENSG000001 | 262.203863 | 0.797573173 | 0.15934268 | 5.00539579 | 5.57E-07 | 1.60E-05 | DDX49 | 54555 |
| ENSG000001 | 2878.65752 | 0.590361468 | 0.11796772 | 5.00443232 | 5.60E-07 | 1.60E-05 | TM9SF2 | 9375 |
| ENSG000001 | 440.925441 | 0.688285381 | 0.13754569 | 5.00404889 | 5.61E-07 | 1.61E-05 | PEF1 | 553115 |
| ENSG000001 | 184.716328 | 0.861780531 | 0.17222153 | 5.00390702 | 5.62E-07 | 1.61E-05 | MZT2A | 653784 |
| ENSG000001 | 2441.96655 | 0.599399742 | 0.1198093 | 5.00294837 | 5.65E-07 | 1.61E-05 | EIF3L | 51386 |

| | | | | | | | | |
|---|---|---|---|---|---|---|---|---|
| ENSG000001 | 253.898926 | 0.703589279 | 0.14063151 | 5.00306993 | 5.64E-07 | 1.61E-05 | C16orf13 | 84326 |
| ENSG000001 | 2055.18062 | 0.674701294 | 0.13486644 | 5.00273672 | 5.65E-07 | 1.61E-05 | TXN | 7295 |
| ENSG000001 | 1637.86549 | 0.633546406 | 0.12664379 | 5.00258555 | 5.66E-07 | 1.61E-05 | SSU72 | 29101 |
| ENSG000001 | 629.681684 | 0.701396859 | 0.1402075 | 5.00256311 | 5.66E-07 | 1.61E-05 | ZNF48 | 197407 |
| ENSG000002 | 43.730735 | -1.13058672 | 0.22599943 | -5.0026088 | 5.66E-07 | 1.61E-05 | CFI | 3426 |
| ENSG000001 | 517.616542 | 0.688520267 | 0.13764295 | 5.00221955 | 5.67E-07 | 1.61E-05 | TSTD2 | 158427 |
| ENSG000001 | 640.993233 | 0.637278103 | 0.12740874 | 5.00183959 | 5.68E-07 | 1.61E-05 | ZBTB8OS | 339487 |
| ENSG000001 | 2719.8275 | 0.643389119 | 0.12863754 | 5.00156595 | 5.69E-07 | 1.62E-05 | DKC1 | 1736 |
| ENSG00000C | 2168.92549 | 0.594268248 | 0.1188217 | 5.00134447 | 5.69E-07 | 1.62E-05 | PREX2 | 80243 |
| ENSG000001 | 2634.63165 | 0.527365533 | 0.10545082 | 5.00105677 | 5.70E-07 | 1.62E-05 | BRD7 | 29117 |
| ENSG000001 | 1829.42267 | 0.57251223 | 0.11449954 | 5.0001269 | 5.73E-07 | 1.62E-05 | PPIL1 | 51645 |
| ENSG000001 | 1073.91925 | 0.627752961 | 0.12556715 | 4.99934073 | 5.75E-07 | 1.63E-05 | C12orf49 | 79794 |
| ENSG000001 | 1231.19589 | 0.739542243 | 0.14794685 | 4.99870206 | 5.77E-07 | 1.63E-05 | ZC3H7B | 23264 |
| ENSG000001 | 1182.12737 | 0.748965167 | 0.14983976 | 4.99844074 | 5.78E-07 | 1.64E-05 | DDX54 | 79039 |
| ENSG00000C | 776.242291 | 0.655780243 | 0.13120071 | 4.99829803 | 5.78E-07 | 1.64E-05 | FBXL19 | 54620 |
| ENSG00000C | 6317.21229 | 0.615926196 | 0.12324641 | 4.99751832 | 5.81E-07 | 1.64E-05 | SLC2A3 | 6515 |
| ENSG00000C | 148.308766 | 0.774596662 | 0.15502119 | 4.99671477 | 5.83E-07 | 1.65E-05 | BCL7C | 9274 |
| ENSG000001 | 2236.975 | 0.6220745 | 0.1244949 | 4.99678716 | 5.83E-07 | 1.65E-05 | NMT1 | 4836 |
| ENSG000001 | 300.30817 | 0.698350316 | 0.13976672 | 4.99654219 | 5.84E-07 | 1.65E-05 | TSPAN33 | 340348 |
| ENSG000001 | 1297.66185 | 0.718890798 | 0.14389962 | 4.99577956 | 5.86E-07 | 1.65E-05 | SDCBP | 6386 |
| ENSG000001 | 6320.41608 | 0.612203639 | 0.12256008 | 4.99513094 | 5.88E-07 | 1.66E-05 | H3F3B | 3021 |
| ENSG000001 | 14818.1111 | 0.637550183 | 0.12763637 | 4.99505123 | 5.88E-07 | 1.66E-05 | PPIA | 5478 |
| ENSG000001 | 21341.6148 | 0.617269732 | 0.12360295 | 4.99397244 | 5.91E-07 | 1.66E-05 | HSPD1 | 3329 |
| ENSG000001 | 1781.52887 | 0.63788724 | 0.12773509 | 4.99382918 | 5.92E-07 | 1.66E-05 | ATP5H | 10476 |
| ENSG00000C | 582.483595 | 0.702482191 | 0.14067855 | 4.99352742 | 5.93E-07 | 1.66E-05 | BRINP1 | 1620 |
| ENSG000001 | 1068.89589 | 0.614779415 | 0.12313675 | 4.99265583 | 5.96E-07 | 1.67E-05 | MEA1 | 4201 |
| ENSG000001 | 763.915758 | 0.704952061 | 0.14119915 | 4.99260854 | 5.96E-07 | 1.67E-05 | CTBP1 | 1487 |
| ENSG00000C | 832.632958 | 0.767995286 | 0.15383127 | 4.99245232 | 5.96E-07 | 1.67E-05 | CABIN1 | 23523 |
| ENSG00000C | 60.0185895 | -1.010087838 | 0.20234487 | -4.9919123 | 5.98E-07 | 1.67E-05 | DCN | 1634 |

| | | | | | | | | | |
|---|---|---|---|---|---|---|---|---|---|
| ENSG000001 | 1042.1406 | 0.675661994 | 0.13535157 | 4.99190348 | 5.98E-07 | 1.67E-05 | ZWINT | | 11130 |
| ENSG000001 | 3541.88451 | 0.56654898 | 0.11349131 | 4.99200318 | 5.98E-07 | 1.67E-05 | HMGB2 | | 3148 |
| ENSG000001 | 547.662196 | 0.655514761 | 0.13134107 | 4.99093524 | 6.01E-07 | 1.68E-05 | SLC35B1 | | 10237 |
| ENSG000001 | 931.466185 | 0.60790377 | 0.12180149 | 4.99093864 | 6.01E-07 | 1.68E-05 | NDUFB5 | | 4711 |
| ENSG000001 | 879.071078 | 0.629847012 | 0.12620957 | 4.99048528 | 6.02E-07 | 1.68E-05 | GPAT4 | | 137964 |
| ENSG000001 | 7557.952 | 0.552988821 | 0.11081068 | 4.99039267 | 6.03E-07 | 1.68E-05 | NA | NA | |
| ENSG000001 | 897.259333 | 0.796718808 | 0.15966737 | 4.98986627 | 6.04E-07 | 1.68E-05 | SLC44A2 | | 57153 |
| ENSG000001 | 4530.81567 | 0.612790949 | 0.12281385 | 4.98959169 | 6.05E-07 | 1.69E-05 | SSRP1 | | 6749 |
| ENSG000001 | 1121.45643 | 0.611165862 | 0.12249252 | 4.98941367 | 6.06E-07 | 1.69E-05 | FOPNL | | 123811 |
| ENSG000001 | 955.654312 | 0.677076646 | 0.13570716 | 4.98924777 | 6.06E-07 | 1.69E-05 | PFKM | | 5213 |
| ENSG00000C | 261.82677 | 0.808759125 | 0.16213314 | 4.98824082 | 6.09E-07 | 1.69E-05 | NCAPH2 | | 29781 |
| ENSG000001 | 211.984191 | 0.815235539 | 0.1634253 | 4.98842932 | 6.09E-07 | 1.69E-05 | DGCR6L | | 85359 |
| ENSG000001 | 370.14898 | 0.631216674 | 0.12654318 | 4.98815262 | 6.10E-07 | 1.69E-05 | DDR2 | | 4921 |
| ENSG000001 | 913.54432 | 0.65722705 | 0.13175228 | 4.98835419 | 6.09E-07 | 1.69E-05 | ZNF138 | | 7697 |
| ENSG000001 | 459.447557 | 0.708525941 | 0.14205965 | 4.98752411 | 6.12E-07 | 1.70E-05 | UQCC2 | | 84300 |
| ENSG000001 | 874.676407 | 0.635935879 | 0.12751632 | 4.98709399 | 6.13E-07 | 1.70E-05 | TES | | 26136 |
| ENSG000001 | 1943.37196 | 0.587855136 | 0.11788455 | 4.98670214 | 6.14E-07 | 1.70E-05 | PLRG1 | | 5356 |
| ENSG000001 | 2104.38563 | 0.609075388 | 0.12214799 | 4.98637238 | 6.15E-07 | 1.70E-05 | PSMA3 | | 5684 |
| ENSG000001 | 101.928718 | -0.895839293 | 0.17966197 | -4.9862488 | 6.16E-07 | 1.70E-05 | CACNA1E | | 777 |
| ENSG000001 | 857.102186 | 0.633292714 | 0.12702809 | 4.98545413 | 6.18E-07 | 1.71E-05 | KNSTRN | | 90417 |
| ENSG000001 | 1334.37535 | 0.602146757 | 0.12078144 | 4.9854246 | 6.18E-07 | 1.71E-05 | DMKN | | 93099 |
| ENSG000001 | 427.942203 | 0.750160641 | 0.15047604 | 4.98524971 | 6.19E-07 | 1.71E-05 | PALD1 | | 27143 |
| ENSG000001 | 681.746999 | 0.884666926 | 0.17747366 | 4.98477876 | 6.20E-07 | 1.71E-05 | NUP210 | | 23225 |
| ENSG000001 | 552.640535 | 0.6485037 | 0.13009491 | 4.98485065 | 6.20E-07 | 1.71E-05 | APTX | | 54840 |
| ENSG000001 | 1306.89644 | 0.59269371 | 0.11895396 | 4.98254728 | 6.28E-07 | 1.73E-05 | RAD17 | | 5884 |
| ENSG000001 | 249.263784 | 0.771895466 | 0.1549279 | 4.98228833 | 6.28E-07 | 1.73E-05 | FOXA3 | | 3171 |
| ENSG000001 | 947.305694 | 0.727481454 | 0.14602248 | 4.98198255 | 6.29E-07 | 1.73E-05 | DAD1 | | 1603 |
| ENSG000002 | 702.869994 | 0.851814249 | 0.17098747 | 4.98173486 | 6.30E-07 | 1.73E-05 | HIST1H4A | | 8359 |
| ENSG000001 | 123.063149 | 0.813901492 | 0.16339012 | 4.98133834 | 6.31E-07 | 1.73E-05 | HTRA1 | | 5654 |

| | | | | | | | | | |
|---|---|---|---|---|---|---|---|---|---|
| ENSG000001 | 353.680526 | 0.728984151 | 0.14635327 | 4.98098985 | 6.33E-07 | 1.74E-05 | ACAD9 | | 28976 |
| ENSG000002 | 847.716633 | 0.708712516 | 0.14230179 | 4.98034873 | 6.35E-07 | 1.74E-05 | NA | NA | |
| ENSG00000C | 291.948277 | 0.704452167 | 0.14147535 | 4.97932782 | 6.38E-07 | 1.74E-05 | CLCN4 | | 1183 |
| ENSG000001 | 767.409248 | 0.569116576 | 0.11429865 | 4.97920646 | 6.38E-07 | 1.74E-05 | NA | NA | |
| ENSG000001 | 588.851138 | 0.682803403 | 0.13713057 | 4.9792209 | 6.38E-07 | 1.74E-05 | C1orf109 | | 54955 |
| ENSG000001 | 157.683984 | 0.857580408 | 0.17222091 | 4.9795372 | 6.37E-07 | 1.74E-05 | SAT2 | | 112483 |
| ENSG000001 | 240.615266 | 0.81090348 | 0.16284246 | 4.97968088 | 6.37E-07 | 1.74E-05 | CHID1 | | 66005 |
| ENSG000001 | 14716.9813 | 0.561291298 | 0.11274475 | 4.97842518 | 6.41E-07 | 1.75E-05 | HSPA9 | | 3313 |
| ENSG00000C | 4323.57692 | 0.614422847 | 0.12346026 | 4.97668521 | 6.47E-07 | 1.77E-05 | SLC25A3 | | 5250 |
| ENSG000001 | 1278.00174 | 0.611892909 | 0.12298039 | 4.9755325 | 6.51E-07 | 1.77E-05 | PDHA1 | | 5160 |
| ENSG000001 | 313.256781 | 0.728642958 | 0.1464615 | 4.97497949 | 6.53E-07 | 1.78E-05 | CACTIN | | 58509 |
| ENSG000002 | 979.188643 | 0.573626347 | 0.11530093 | 4.97503666 | 6.52E-07 | 1.78E-05 | SUPT4H1 | | 6827 |
| ENSG000001 | 35162.9509 | 0.611340817 | 0.12289103 | 4.97465784 | 6.54E-07 | 1.78E-05 | GAPDH | | 2597 |
| ENSG000001 | 277.166171 | 0.69631869 | 0.13998419 | 4.97426653 | 6.55E-07 | 1.78E-05 | NSUN5 | | 55695 |
| ENSG000001 | 1087.84204 | 0.650188763 | 0.13071421 | 4.97412452 | 6.55E-07 | 1.78E-05 | CMAS | | 55907 |
| ENSG000002 | 28.4470277 | -1.301915417 | 0.2617529 | -4.9738337 | 6.56E-07 | 1.78E-05 | PCDHA3 | | 56145 |
| ENSG000001 | 733.725599 | 0.849604036 | 0.1708421 | 4.97303685 | 6.59E-07 | 1.79E-05 | PKN1 | | 5585 |
| ENSG000001 | 179.429044 | 0.727249864 | 0.14626088 | 4.9722788 | 6.62E-07 | 1.79E-05 | TBL2 | | 26608 |
| ENSG000002 | 139.212913 | 0.813097916 | 0.16358278 | 4.97055936 | 6.68E-07 | 1.81E-05 | NA | NA | |
| ENSG000001 | 870.759746 | 0.622004577 | 0.12514102 | 4.97042923 | 6.68E-07 | 1.81E-05 | SRPRB | | 58477 |
| ENSG00000C | 372.540958 | 0.900618182 | 0.18121791 | 4.96980772 | 6.70E-07 | 1.81E-05 | PLOD1 | | 5351 |
| ENSG000001 | 1047.6235 | 0.540859884 | 0.10884117 | 4.96925807 | 6.72E-07 | 1.82E-05 | TTL | | 150465 |
| ENSG000001 | 2322.08642 | 0.616444597 | 0.12406738 | 4.96862771 | 6.74E-07 | 1.82E-05 | SLC20A1 | | 6574 |
| ENSG000001 | 20.0425989 | -1.432856433 | 0.2885109 | -4.9663859 | 6.82E-07 | 1.84E-05 | FCRL6 | | 343413 |
| ENSG000001 | 2061.30302 | 0.617628239 | 0.12437956 | 4.96567293 | 6.85E-07 | 1.85E-05 | RTN3 | | 10313 |
| ENSG000001 | 142.513929 | 0.886169477 | 0.17851589 | 4.964093 | 6.90E-07 | 1.86E-05 | CLDN4 | | 1364 |
| ENSG00000C | 224.083007 | 0.731335197 | 0.14732893 | 4.96396197 | 6.91E-07 | 1.86E-05 | NTHL1 | | 4913 |
| ENSG00000C | 764.358311 | 0.634889051 | 0.12790551 | 4.96373488 | 6.92E-07 | 1.86E-05 | AGK | | 55750 |
| ENSG000001 | 6244.28917 | 0.575399805 | 0.11592325 | 4.96362722 | 6.92E-07 | 1.86E-05 | RAD21 | | 5885 |

| | | | | | | | | |
|---|---|---|---|---|---|---|---|---|
| ENSG000001 | 267.039202 | 0.758583746 | 0.15288217 | 4.96188509 | 6.98E-07 | 1.88E-05 | SLC37A4 | 2542 |
| ENSG000001 | 238.381851 | 0.757914221 | 0.15275993 | 4.96147279 | 7.00E-07 | 1.88E-05 | AFMID | 125061 |
| ENSG000002 | 1486.93892 | 0.661630224 | 0.13336334 | 4.96111035 | 7.01E-07 | 1.88E-05 | TIMM23 | 100287932 |
| ENSG000000 | 713.199415 | 0.740813518 | 0.14933188 | 4.96085295 | 7.02E-07 | 1.88E-05 | MKNK2 | 2872 |
| ENSG000001 | 684.238175 | 0.605464512 | 0.12204721 | 4.96090414 | 7.02E-07 | 1.88E-05 | CCBL2 | 56267 |
| ENSG000001 | 1013.13149 | 0.682512996 | 0.13758514 | 4.96065923 | 7.03E-07 | 1.88E-05 | PPP2R4 | 5524 |
| ENSG000001 | 342.612436 | 0.710408023 | 0.14324562 | 4.95936995 | 7.07E-07 | 1.90E-05 | SMIM14 | 201895 |
| ENSG000001 | 957.8711 | 0.582547256 | 0.11748169 | 4.95862186 | 7.10E-07 | 1.90E-05 | EFTUD1 | 79631 |
| ENSG000001 | 43.8932407 | -1.141660361 | 0.23044272 | -4.9542045 | 7.26E-07 | 1.94E-05 | HTR4 | 3360 |
| ENSG000001 | 884.464542 | 0.589530751 | 0.11900441 | 4.95385643 | 7.28E-07 | 1.94E-05 | RNF130 | 55819 |
| ENSG000001 | 1376.64005 | 0.638361302 | 0.12886039 | 4.95389867 | 7.27E-07 | 1.94E-05 | HIST2H2AB | 317772 |
| ENSG000001 | 772.502263 | 0.763527643 | 0.15414051 | 4.95345208 | 7.29E-07 | 1.95E-05 | AP1B1 | 162 |
| ENSG000001 | 1915.59924 | 0.66690013 | 0.13463066 | 4.9535533 | 7.29E-07 | 1.95E-05 | QARS | 5859 |
| ENSG000001 | 478.689352 | 0.718046721 | 0.14497115 | 4.95303182 | 7.31E-07 | 1.95E-05 | FAM45A | 404636 |
| ENSG000001 | 194.985602 | 0.71869289 | 0.14511343 | 4.95262848 | 7.32E-07 | 1.95E-05 | JAGN1 | 84522 |
| ENSG000001 | 27.9646318 | -1.312047598 | 0.26493833 | -4.9522755 | 7.34E-07 | 1.95E-05 | ZNF439 | 90594 |
| ENSG000001 | 345.752407 | 0.746397361 | 0.1507322 | 4.95181097 | 7.35E-07 | 1.96E-05 | SETD8 | 387893 |
| ENSG000001 | 2168.17428 | 0.57651455 | 0.1164347 | 4.9513982 | 7.37E-07 | 1.96E-05 | CACHD1 | 57685 |
| ENSG000000 | 695.609776 | 0.651794177 | 0.1316667 | 4.95033415 | 7.41E-07 | 1.97E-05 | NUDCD3 | 23386 |
| ENSG000001 | 155.976075 | 0.891144199 | 0.18001448 | 4.95040279 | 7.41E-07 | 1.97E-05 | HPS6 | 79803 |
| ENSG000001 | 1014.00538 | 0.669928408 | 0.13532994 | 4.95033411 | 7.41E-07 | 1.97E-05 | LDB1 | 8861 |
| ENSG000002 | 273.4639 | 0.785498002 | 0.15867443 | 4.95037553 | 7.41E-07 | 1.97E-05 | C11orf31 | 280636 |
| ENSG000001 | 1044.74365 | 0.710930575 | 0.14364237 | 4.94930987 | 7.45E-07 | 1.98E-05 | ALYREF | 10189 |
| ENSG000002 | 257.064091 | 0.760765377 | 0.15374244 | 4.94831068 | 7.49E-07 | 1.98E-05 | TSPY26P | 128854 |
| ENSG000002 | 2209.74865 | 0.659045587 | 0.13320798 | 4.94749324 | 7.52E-07 | 1.99E-05 | HSBP1 | 3281 |
| ENSG000001 | 1551.6574 | 0.707480063 | 0.14300631 | 4.94719474 | 7.53E-07 | 1.99E-05 | NOP10 | 55505 |
| ENSG000002 | 259.86068 | 0.692457568 | 0.13998477 | 4.94666365 | 7.55E-07 | 2.00E-05 | NA | NA |
| ENSG000001 | 1016.72813 | 0.69891278 | 0.14130972 | 4.94596404 | 7.58E-07 | 2.00E-05 | LMAN2 | 10960 |
| ENSG000001 | 218.652005 | 0.737669495 | 0.14915838 | 4.94554496 | 7.59E-07 | 2.01E-05 | RBCK1 | 10616 |

| | | | | | | | | |
|---|---|---|---|---|---|---|---|---|
| ENSG000001 | 1090.91017 | 0.766626423 | 0.15504235 | 4.94462579 | 7.63E-07 | 2.01E-05 | EPHB4 | 2050 |
| ENSG000001 | 4353.69842 | 0.619121909 | 0.12521907 | 4.94431001 | 7.64E-07 | 2.02E-05 | TUBA1C | 84790 |
| ENSG000001 | 315.764096 | 0.862000676 | 0.17435715 | 4.94387914 | 7.66E-07 | 2.02E-05 | HIST1H4J | 8363 |
| ENSG000001 | 41.0777051 | 1.16096425 | 0.23483742 | 4.94369367 | 7.67E-07 | 2.02E-05 | TEX13B | 56156 |
| ENSG000002 | 420.783489 | 0.837865562 | 0.16951237 | 4.94279894 | 7.70E-07 | 2.03E-05 | SNORD116-13 | 100033425 |
| ENSG000002 | 2326.6752 | 0.606244427 | 0.122658 | 4.94255906 | 7.71E-07 | 2.03E-05 | NAP1L4 | 4676 |
| ENSG000001 | 301.009801 | 0.60350428 | 0.12211416 | 4.94213194 | 7.73E-07 | 2.03E-05 | C11orf1 | 64776 |
| ENSG000001 | 1484.98115 | 0.605486671 | 0.12251468 | 4.94215625 | 7.73E-07 | 2.03E-05 | FAM84B | 157638 |
| ENSG00000C | 22054.7579 | 0.554216714 | 0.11218736 | 4.94009939 | 7.81E-07 | 2.05E-05 | EIF4B | 1975 |
| ENSG000001 | 1860.64681 | 0.506074614 | 0.10245948 | 4.93926578 | 7.84E-07 | 2.06E-05 | MAP7D3 | 79649 |
| ENSG000001 | 479.928087 | 0.715936376 | 0.14496423 | 4.93871074 | 7.86E-07 | 2.06E-05 | MRPL52 | 122704 |
| ENSG000001 | 523.315329 | 0.67615685 | 0.13693056 | 4.93795418 | 7.89E-07 | 2.07E-05 | OSER1 | 51526 |
| ENSG00000C | 504.913958 | 0.648197699 | 0.13128064 | 4.9374966 | 7.91E-07 | 2.07E-05 | SLC9A7 | 84679 |
| ENSG00000C | 2106.34352 | 0.688061925 | 0.13935623 | 4.93743228 | 7.92E-07 | 2.07E-05 | ERP29 | 10961 |
| ENSG000001 | 568.110923 | 0.669413288 | 0.13558696 | 4.93715098 | 7.93E-07 | 2.07E-05 | COL4A5 | 1287 |
| ENSG000001 | 679.011189 | 0.744688424 | 0.15085462 | 4.93646414 | 7.96E-07 | 2.08E-05 | SAMM50 | 25813 |
| ENSG000001 | 496.519047 | 0.697744071 | 0.14135327 | 4.936172 | 7.97E-07 | 2.08E-05 | DNAJC5 | 80331 |
| ENSG000001 | 293.51791 | 0.666829266 | 0.13509862 | 4.93587031 | 7.98E-07 | 2.08E-05 | GORASP1 | 64689 |
| ENSG000001 | 654.238233 | 0.661810149 | 0.13408807 | 4.9356378 | 7.99E-07 | 2.08E-05 | STON2 | 85439 |
| ENSG000001 | 264.506832 | 0.705794006 | 0.14299065 | 4.93594515 | 7.98E-07 | 2.08E-05 | FOXD3 | 27022 |
| ENSG000002 | 122.278061 | 0.797478941 | 0.16157464 | 4.93566911 | 7.99E-07 | 2.08E-05 | CCDC160 | 347475 |
| ENSG000001 | 185.808499 | 0.750350396 | 0.15204709 | 4.93498701 | 8.02E-07 | 2.09E-05 | FASTK | 10922 |
| ENSG000002 | 293.932912 | 0.773036069 | 0.15667021 | 4.93416128 | 8.05E-07 | 2.10E-05 | MARS2 | 92935 |
| ENSG000001 | 444.516723 | 0.678717509 | 0.13756055 | 4.93395446 | 8.06E-07 | 2.10E-05 | COQ5 | 84274 |
| ENSG000001 | 1928.27817 | 0.750107626 | 0.15207337 | 4.93253774 | 8.12E-07 | 2.11E-05 | VRTN | 55237 |
| ENSG000001 | 1974.0245 | 0.558814045 | 0.11333067 | 4.93082788 | 8.19E-07 | 2.13E-05 | ELP2 | 55250 |
| ENSG000002 | 378.754958 | 0.710461303 | 0.14408571 | 4.93082431 | 8.19E-07 | 2.13E-05 | FANCG | 2189 |
| ENSG000001 | 291.261082 | 0.71054784 | 0.14412604 | 4.93004486 | 8.22E-07 | 2.13E-05 | DCAKD | 79877 |
| ENSG000001 | 1037.55997 | 0.567754505 | 0.11516891 | 4.9297548 | 8.23E-07 | 2.13E-05 | RHNO1 | 83695 |

| | | | | | | | | |
|---|---|---|---|---|---|---|---|---|
| ENSG000001 | 975.265434 | 0.63203858 | 0.12821795 | 4.92940811 | 8.25E-07 | 2.14E-05 | TMEM165 | 55858 |
| ENSG000001 | 1323.83794 | 0.630789073 | 0.12796838 | 4.92925738 | 8.25E-07 | 2.14E-05 | GNPDA1 | 10007 |
| ENSG000001 | 5255.56124 | 0.645467915 | 0.13096462 | 4.92856704 | 8.28E-07 | 2.14E-05 | IGF2BP3 | 10643 |
| ENSG000001 | 2601.30251 | 0.582856202 | 0.11832332 | 4.92596207 | 8.39E-07 | 2.17E-05 | CHD9 | 80205 |
| ENSG000001 | 313.101073 | 0.685354005 | 0.13913962 | 4.92565655 | 8.41E-07 | 2.17E-05 | NA | NA |
| ENSG000001 | 351.965688 | 0.728503486 | 0.14789674 | 4.92575769 | 8.40E-07 | 2.17E-05 | NKIRAS2 | 28511 |
| ENSG000001 | 697.607121 | 0.602542642 | 0.12233164 | 4.92548482 | 8.42E-07 | 2.17E-05 | C6orf120 | 387263 |
| ENSG000001 | 655.263132 | 0.665628187 | 0.13514779 | 4.92518741 | 8.43E-07 | 2.17E-05 | SEC61B | 10952 |
| ENSG000001 | 5190.55666 | 0.568397118 | 0.11542488 | 4.92438993 | 8.46E-07 | 2.18E-05 | ACLY | 47 |
| ENSG000001 | 626.845331 | 0.665714003 | 0.13520511 | 4.92373414 | 8.49E-07 | 2.19E-05 | PIP4K2A | 5305 |
| ENSG000001 | 978.317599 | 0.578421806 | 0.11747821 | 4.9236518 | 8.49E-07 | 2.19E-05 | TMX3 | 54495 |
| ENSG000002 | 69.3324553 | -0.984259296 | 0.19990191 | -4.9237113 | 8.49E-07 | 2.19E-05 | NA | NA |
| ENSG000001 | 1104.41613 | 0.53786519 | 0.10925598 | 4.9229819 | 8.52E-07 | 2.19E-05 | ZCCHC17 | 51538 |
| ENSG000001 | 621.610185 | 0.801696436 | 0.16287149 | 4.9222637 | 8.55E-07 | 2.20E-05 | AP2A1 | 160 |
| ENSG000001 | 1635.37149 | 0.65176127 | 0.1324284 | 4.9216126 | 8.58E-07 | 2.20E-05 | CARS | 833 |
| ENSG000001 | 851.086055 | 0.555576451 | 0.11288626 | 4.92155936 | 8.59E-07 | 2.20E-05 | NEK4 | 6787 |
| ENSG000002 | 50.6712612 | 1.123852 | 0.22835411 | 4.92153166 | 8.59E-07 | 2.20E-05 | NA | NA |
| ENSG000001 | 146.813861 | 0.763932347 | 0.15525157 | 4.92060949 | 8.63E-07 | 2.21E-05 | ZFYVE27 | 118813 |
| ENSG000001 | 1223.2451 | 0.61586045 | 0.1251695 | 4.92021164 | 8.65E-07 | 2.21E-05 | CREBZF | 58487 |
| ENSG000001 | 286.332257 | 0.698715418 | 0.14205299 | 4.91869552 | 8.71E-07 | 2.23E-05 | ECSIT | 51295 |
| ENSG000002 | 614.781644 | 0.641158697 | 0.13036195 | 4.91829619 | 8.73E-07 | 2.23E-05 | RPL13AP5 | 728658 |
| ENSG000001 | 1605.38333 | 0.609878417 | 0.12400957 | 4.91799471 | 8.74E-07 | 2.24E-05 | HN1 | 51155 |
| ENSG000002 | 1622.53598 | 0.585571436 | 0.11907089 | 4.91783883 | 8.75E-07 | 2.24E-05 | TRIM27 | 5987 |
| ENSG000001 | 2024.1988 | 0.700044649 | 0.1423541 | 4.91762893 | 8.76E-07 | 2.24E-05 | GPI | 2821 |
| ENSG000002 | 362.446551 | 0.735610867 | 0.14960688 | 4.91695878 | 8.79E-07 | 2.24E-05 | NA | NA |
| ENSG000001 | 801.213953 | 0.626308792 | 0.12740161 | 4.91601956 | 8.83E-07 | 2.25E-05 | MRPS22 | 56945 |
| ENSG00000C | 1456.58537 | 0.667234659 | 0.13573774 | 4.91561641 | 8.85E-07 | 2.26E-05 | ATP1B3 | 483 |
| ENSG00000C | 1017.5201 | 0.561208857 | 0.11417131 | 4.91549801 | 8.86E-07 | 2.26E-05 | OXCT1 | 5019 |
| ENSG000001 | 245.348806 | 0.824098955 | 0.16765771 | 4.91536579 | 8.86E-07 | 2.26E-05 | LRFN4 | 78999 |

| | | | | | | | | |
|---|---|---|---|---|---|---|---|---|
| ENSG000001 | 1685.20601 | 0.679371512 | 0.13822222 | 4.91506728 | 8.88E-07 | 2.26E-05 | LRRC59 | 55379 |
| ENSG00000C | 638.227705 | 0.778634846 | 0.1584621 | 4.91369754 | 8.94E-07 | 2.27E-05 | AKR1B1 | 231 |
| ENSG000001 | 338.982917 | 0.647280504 | 0.13173288 | 4.91358357 | 8.94E-07 | 2.27E-05 | LRCH2 | 57631 |
| ENSG000001 | 102.071739 | 0.891162414 | 0.1813673 | 4.91357808 | 8.94E-07 | 2.27E-05 | PRRT3 | 285368 |
| ENSG00000C | 398.368628 | 0.67483261 | 0.13735156 | 4.91317767 | 8.96E-07 | 2.27E-05 | HSD17B10 | 3028 |
| ENSG000001 | 197.753829 | 0.786739455 | 0.16013132 | 4.91308931 | 8.97E-07 | 2.27E-05 | BCAT2 | 587 |
| ENSG000001 | 106.922126 | 0.854530596 | 0.17395972 | 4.91223258 | 9.00E-07 | 2.28E-05 | EEF1A2 | 1917 |
| ENSG000001 | 1683.47207 | 0.641254868 | 0.13053963 | 4.91233852 | 9.00E-07 | 2.28E-05 | UBFD1 | 56061 |
| ENSG000001 | 660.230805 | 0.719601328 | 0.14649403 | 4.91215473 | 9.01E-07 | 2.28E-05 | KRT19 | 3880 |
| ENSG000001 | 1320.20218 | 0.636797516 | 0.12964588 | 4.91182206 | 9.02E-07 | 2.28E-05 | COPZ1 | 22818 |
| ENSG000001 | 529.688509 | 0.71874502 | 0.14632744 | 4.91189487 | 9.02E-07 | 2.28E-05 | MAP2K1 | 5604 |
| ENSG000001 | 2056.87183 | 0.589609877 | 0.12005913 | 4.91099554 | 9.06E-07 | 2.29E-05 | BCCIP | 56647 |
| ENSG000002 | 215.680271 | 0.78893606 | 0.16072897 | 4.90848703 | 9.18E-07 | 2.32E-05 | RPL17 | 6139 |
| ENSG000001 | 1407.1551 | 0.6178815 | 0.12590569 | 4.90749473 | 9.22E-07 | 2.33E-05 | PDPN | 10630 |
| ENSG00000C | 386.900712 | 0.610453435 | 0.12440517 | 4.90697808 | 9.25E-07 | 2.33E-05 | RFFL | 117584 |
| ENSG000001 | 157.086433 | 0.886195824 | 0.18060853 | 4.90672185 | 9.26E-07 | 2.33E-05 | LHPP | 64077 |
| ENSG000001 | 1481.80582 | 0.614426531 | 0.12522978 | 4.90639299 | 9.28E-07 | 2.34E-05 | LRRC58 | 116064 |
| ENSG000001 | 422.194215 | 0.657320014 | 0.13397912 | 4.90613757 | 9.29E-07 | 2.34E-05 | PAPSS2 | 9060 |
| ENSG000001 | 177.852467 | 0.765350904 | 0.15604628 | 4.90464063 | 9.36E-07 | 2.35E-05 | CMTM3 | 123920 |
| ENSG000001 | 96.591651 | 0.900449057 | 0.1835915 | 4.90463357 | 9.36E-07 | 2.35E-05 | MUM1L1 | 139221 |
| ENSG000001 | 123.468069 | 0.950501234 | 0.19382835 | 4.9038298 | 9.40E-07 | 2.36E-05 | CENPM | 79019 |
| ENSG000001 | 532.073759 | 0.606600182 | 0.1236941 | 4.90403497 | 9.39E-07 | 2.36E-05 | VGLL4 | 9686 |
| ENSG000001 | 70.4482612 | 0.951815933 | 0.1940926 | 4.90392706 | 9.39E-07 | 2.36E-05 | HPGD | 3248 |
| ENSG00000C | 1953.02446 | 0.563217516 | 0.11486148 | 4.90345 | 9.42E-07 | 2.36E-05 | CS | 1431 |
| ENSG00000C | 313.209305 | 0.675051301 | 0.1376684 | 4.90345846 | 9.42E-07 | 2.36E-05 | SPHK2 | 56848 |
| ENSG000001 | 15676.6097 | 0.626960133 | 0.12785881 | 4.90353477 | 9.41E-07 | 2.36E-05 | RPS3A | 6189 |
| ENSG000001 | 577.697821 | 0.623162994 | 0.12709272 | 4.90321532 | 9.43E-07 | 2.36E-05 | GNS | 2799 |
| ENSG000001 | 443.80242 | 0.721369196 | 0.14713862 | 4.90265015 | 9.46E-07 | 2.37E-05 | DCTD | 1635 |
| ENSG000001 | 547.138885 | 0.633711591 | 0.12926749 | 4.90232767 | 9.47E-07 | 2.37E-05 | RBX1 | 9978 |

| | | | | | | | | |
|---|---|---|---|---|---|---|---|---|
| ENSG000001 | 350.75502 | 0.799517493 | 0.16309486 | 4.90216255 | 9.48E-07 | 2.37E-05 | TBL3 | 10607 |
| ENSG00000C | 310.694905 | 0.649274467 | 0.13246101 | 4.90162703 | 9.50E-07 | 2.37E-05 | GINM1 | 116254 |
| ENSG000001 | 2402.34071 | 0.547057694 | 0.1116143 | 4.90132271 | 9.52E-07 | 2.37E-05 | DR1 | 1810 |
| ENSG000001 | 1155.78226 | 0.760967197 | 0.15525773 | 4.90131594 | 9.52E-07 | 2.37E-05 | MYC | 4609 |
| ENSG000001 | 224.510207 | 0.66608853 | 0.13591838 | 4.900651 | 9.55E-07 | 2.38E-05 | CPE | 1363 |
| ENSG000001 | 2877.22364 | 0.555216013 | 0.11329886 | 4.9004553 | 9.56E-07 | 2.38E-05 | ICMT | 23463 |
| ENSG000001 | 1353.72733 | 0.59259371 | 0.12094734 | 4.89960115 | 9.60E-07 | 2.39E-05 | NDUFB9 | 4715 |
| ENSG000001 | 771.921116 | 0.649697517 | 0.13260887 | 4.89935194 | 9.62E-07 | 2.39E-05 | FLOT2 | 2319 |
| ENSG000001 | 1168.30044 | 0.681269148 | 0.13906257 | 4.89901163 | 9.63E-07 | 2.39E-05 | LANCL1 | 10314 |
| ENSG000002 | 16.6942286 | 1.554763628 | 0.31735854 | 4.89907603 | 9.63E-07 | 2.39E-05 | NA | NA |
| ENSG000001 | 540.98556 | 0.682947647 | 0.13940964 | 4.89885521 | 9.64E-07 | 2.39E-05 | HDAC5 | 10014 |
| ENSG000002 | 224.026784 | 0.738312988 | 0.15072917 | 4.89827551 | 9.67E-07 | 2.40E-05 | NA | NA |
| ENSG000001 | 552.933334 | 0.668150917 | 0.13641445 | 4.89794816 | 9.68E-07 | 2.40E-05 | GM2A | 2760 |
| ENSG000001 | 207.100267 | 0.801655159 | 0.16373804 | 4.89596159 | 9.78E-07 | 2.42E-05 | NA | NA |
| ENSG000001 | 115.922594 | 0.87692339 | 0.17913207 | 4.89540135 | 9.81E-07 | 2.43E-05 | ALKBH7 | 84266 |
| ENSG000001 | 341.683951 | 0.669117391 | 0.13672018 | 4.89406473 | 9.88E-07 | 2.45E-05 | DNAJC14 | 85406 |
| ENSG000001 | 571.347362 | 0.585646583 | 0.11966792 | 4.89393126 | 9.88E-07 | 2.45E-05 | COL14A1 | 7373 |
| ENSG000001 | 1468.09165 | 0.584238687 | 0.11940137 | 4.89306513 | 9.93E-07 | 2.45E-05 | GSR | 2936 |
| ENSG000001 | 2853.99199 | 0.59926896 | 0.12249266 | 4.89228445 | 9.97E-07 | 2.46E-05 | GTPBP4 | 23560 |
| ENSG000001 | 659.756111 | 0.661572283 | 0.1352323 | 4.89211753 | 9.98E-07 | 2.46E-05 | GRHPR | 9380 |
| ENSG00000C | 6179.81189 | 0.562631816 | 0.11502488 | 4.89139251 | 1.00E-06 | 2.47E-05 | VCL | 7414 |
| ENSG000001 | 4131.57601 | 0.576212522 | 0.11780324 | 4.89131308 | 1.00E-06 | 2.47E-05 | PGAM1 | 5223 |
| ENSG00000C | 2282.41344 | 0.623284057 | 0.12744871 | 4.89046988 | 1.01E-06 | 2.48E-05 | PSMB1 | 5689 |
| ENSG000001 | 357.768515 | 0.64478668 | 0.13185437 | 4.89014268 | 1.01E-06 | 2.48E-05 | POLR1C | 9533 |
| ENSG000001 | 1711.58762 | 0.685404475 | 0.14019595 | 4.88890344 | 1.01E-06 | 2.50E-05 | PSMA5 | 5686 |
| ENSG000001 | 511.739458 | 0.67473291 | 0.13801946 | 4.88867955 | 1.02E-06 | 2.50E-05 | DTYMK | 1841 |
| ENSG000001 | 151.119753 | 0.771076919 | 0.15773146 | 4.88854234 | 1.02E-06 | 2.50E-05 | FAM69B | 138311 |
| ENSG000001 | 179.122214 | 0.729725516 | 0.14928439 | 4.8881569 | 1.02E-06 | 2.50E-05 | SLC13A3 | 64849 |
| ENSG000001 | 441.834635 | 0.666476125 | 0.13634573 | 4.88813357 | 1.02E-06 | 2.50E-05 | DOLPP1 | 57171 |

| | | | | | | | | |
|---|---|---|---|---|---|---|---|---|
| ENSG000001 | 499.17015 | 0.698579311 | 0.14292777 | 4.88763869 | 1.02E-06 | 2.51E-05 | RAP1GAP2 | 23108 |
| ENSG000001 | 6560.38407 | 0.537997817 | 0.11008718 | 4.88701615 | 1.02E-06 | 2.51E-05 | ST13 | 6767 |
| ENSG000001 | 374.039543 | 0.667495293 | 0.13658243 | 4.88712428 | 1.02E-06 | 2.51E-05 | ABCD4 | 5826 |
| ENSG000001 | 979.10142 | 0.551299818 | 0.1128336 | 4.88595415 | 1.03E-06 | 2.52E-05 | NDUFB10 | 4716 |
| ENSG000001 | 56.3775599 | 1.062766718 | 0.21751221 | 4.88600939 | 1.03E-06 | 2.52E-05 | ADRA2A | 150 |
| ENSG000001 | 1048.36977 | 0.626247519 | 0.1281973 | 4.88502903 | 1.03E-06 | 2.53E-05 | COA1 | 55744 |
| ENSG00000C | 640.206558 | 0.750933013 | 0.15375074 | 4.88409354 | 1.04E-06 | 2.54E-05 | MRPL28 | 10573 |
| ENSG000001 | 1089.19785 | 0.654965916 | 0.13411857 | 4.88348433 | 1.04E-06 | 2.55E-05 | EIF4A3 | 9775 |
| ENSG00000C | 604.932271 | 0.674124841 | 0.13805293 | 4.88308962 | 1.04E-06 | 2.55E-05 | CIAPIN1 | 57019 |
| ENSG00000C | 503.524099 | 0.724122755 | 0.14828885 | 4.88319075 | 1.04E-06 | 2.55E-05 | TECR | 9524 |
| ENSG00000C | 1592.74505 | 0.646345075 | 0.13238173 | 4.88243413 | 1.05E-06 | 2.56E-05 | BIRC5 | 332 |
| ENSG000001 | 5430.61734 | 0.579535434 | 0.11871008 | 4.88193949 | 1.05E-06 | 2.56E-05 | ETF1 | 2107 |
| ENSG000001 | 168.751065 | 0.856524353 | 0.1754769 | 4.88112324 | 1.05E-06 | 2.57E-05 | TMEM222 | 84065 |
| ENSG000002 | 43.5099507 | -1.065690826 | 0.21835542 | -4.8805329 | 1.06E-06 | 2.58E-05 | PECAM1 | 5175 |
| ENSG000001 | 346.493115 | 0.676234773 | 0.13857257 | 4.88000458 | 1.06E-06 | 2.58E-05 | SLC25A26 | 115286 |
| ENSG000001 | 148.578378 | 0.839362403 | 0.17201913 | 4.87947139 | 1.06E-06 | 2.59E-05 | ARFRP1 | 10139 |
| ENSG000001 | 51.7591256 | 1.0987345 | 0.2251886 | 4.87917469 | 1.07E-06 | 2.59E-05 | RENBP | 5973 |
| ENSG000001 | 739.960795 | 0.610333173 | 0.12509317 | 4.87902871 | 1.07E-06 | 2.59E-05 | WDR41 | 55255 |
| ENSG000001 | 169.241498 | 0.811646406 | 0.16635522 | 4.87899565 | 1.07E-06 | 2.59E-05 | SHARPIN | 81858 |
| ENSG000001 | 774.331323 | 0.821830253 | 0.16846752 | 4.87827116 | 1.07E-06 | 2.60E-05 | COL18A1 | 80781 |
| ENSG000002 | 230.43709 | 0.823048453 | 0.16871526 | 4.87832851 | 1.07E-06 | 2.60E-05 | NA | NA |
| ENSG000001 | 1438.73817 | 0.676282292 | 0.13864106 | 4.87793645 | 1.07E-06 | 2.60E-05 | GLDC | 2731 |
| ENSG000002 | 68.4774375 | 1.01826823 | 0.20875914 | 4.87771812 | 1.07E-06 | 2.60E-05 | C17orf107 | 100130311 |
| ENSG000001 | 794.502814 | 0.609439281 | 0.1249468 | 4.8775902 | 1.07E-06 | 2.60E-05 | KLF3 | 51274 |
| ENSG000001 | 443.574318 | 0.66628083 | 0.13664452 | 4.87601566 | 1.08E-06 | 2.62E-05 | CAMLG | 819 |
| ENSG000001 | 367.527031 | 0.650748146 | 0.13346694 | 4.87572542 | 1.08E-06 | 2.62E-05 | TRAPPC4 | 51399 |
| ENSG000002 | 419.834744 | 0.683065333 | 0.14010313 | 4.87544651 | 1.09E-06 | 2.62E-05 | NA | NA |
| ENSG00000C | 352.872649 | 0.702999861 | 0.14421356 | 4.87471404 | 1.09E-06 | 2.63E-05 | RRAGD | 58528 |
| ENSG000001 | 1459.81437 | 0.650072317 | 0.13335471 | 4.87476099 | 1.09E-06 | 2.63E-05 | PRPF19 | 27339 |

| | | | | | | | | |
|---|---|---|---|---|---|---|---|---|
| ENSG000001 | 501.956121 | 0.60503261 | 0.1241402 | 4.87378466 | 1.09E-06 | 2.64E-05 | LRRC41 | 10489 |
| ENSG000001 | 1855.72509 | 0.768320117 | 0.15764362 | 4.87377873 | 1.09E-06 | 2.64E-05 | BCOR | 54880 |
| ENSG00000C | 643.313563 | 0.660009336 | 0.13544367 | 4.87294329 | 1.10E-06 | 2.64E-05 | SPAG7 | 9552 |
| ENSG000001 | 1356.13236 | 0.675837863 | 0.13868109 | 4.87332372 | 1.10E-06 | 2.64E-05 | PDAP1 | 11333 |
| ENSG000001 | 772.874844 | 0.628135844 | 0.1289034 | 4.8729193 | 1.10E-06 | 2.64E-05 | DDX56 | 54606 |
| ENSG000001 | 167.295914 | 0.765063045 | 0.15699471 | 4.87317718 | 1.10E-06 | 2.64E-05 | CCDC115 | 84317 |
| ENSG000001 | 632.242929 | 0.651319258 | 0.13365624 | 4.87309294 | 1.10E-06 | 2.64E-05 | PEX13 | 5194 |
| ENSG000002 | 4145.57342 | 0.596490595 | 0.12240281 | 4.87317715 | 1.10E-06 | 2.64E-05 | VDAC1 | 7416 |
| ENSG00000C | 299.493255 | 0.652345519 | 0.13392302 | 4.87104843 | 1.11E-06 | 2.67E-05 | INPP5A | 3632 |
| ENSG000001 | 3213.73451 | 0.629815824 | 0.12930557 | 4.87075556 | 1.11E-06 | 2.67E-05 | SMARCA1 | 6594 |
| ENSG000001 | 1737.30177 | 0.57333499 | 0.11771737 | 4.87043658 | 1.11E-06 | 2.67E-05 | MAPK1 | 5594 |
| ENSG000001 | 1076.54731 | 0.625621155 | 0.12845376 | 4.8703998 | 1.11E-06 | 2.67E-05 | TMEM41B | 440026 |
| ENSG000001 | 1192.67817 | 0.647045323 | 0.13286484 | 4.86995157 | 1.12E-06 | 2.67E-05 | LRIG1 | 26018 |
| ENSG00000C | 2707.76497 | 0.606595611 | 0.1245786 | 4.86917989 | 1.12E-06 | 2.68E-05 | CDH1 | 999 |
| ENSG000001 | 803.471765 | 0.66066356 | 0.13570161 | 4.86850191 | 1.12E-06 | 2.69E-05 | CTSV | 1515 |
| ENSG000001 | 1645.18236 | 0.618198398 | 0.12700515 | 4.86750661 | 1.13E-06 | 2.70E-05 | GNPTAB | 79158 |
| ENSG000001 | 440.557671 | 0.713388879 | 0.14656136 | 4.86751006 | 1.13E-06 | 2.70E-05 | ZSCAN25 | 221785 |
| ENSG000002 | 101.693677 | 0.845196175 | 0.1736401 | 4.86751731 | 1.13E-06 | 2.70E-05 | NA | NA |
| ENSG000001 | 733.376581 | 0.712449201 | 0.14637808 | 4.86718507 | 1.13E-06 | 2.70E-05 | MED22 | 6837 |
| ENSG000001 | 8061.57232 | 0.592931374 | 0.12182515 | 4.86706879 | 1.13E-06 | 2.70E-05 | ATP5A1 | 498 |
| ENSG000001 | 298.783613 | 0.673128336 | 0.13832495 | 4.86628296 | 1.14E-06 | 2.71E-05 | DDRGK1 | 65992 |
| ENSG00000C | 822.983117 | 0.653915053 | 0.13438819 | 4.86586701 | 1.14E-06 | 2.72E-05 | ORC6 | 23594 |
| ENSG000001 | 903.167062 | 0.596347757 | 0.12257717 | 4.86508031 | 1.14E-06 | 2.72E-05 | GRID2 | 2895 |
| ENSG000001 | 1008.68151 | 0.603621069 | 0.12408692 | 4.86450201 | 1.15E-06 | 2.73E-05 | POLR2C | 5432 |
| ENSG000001 | 1015.88676 | 0.659061495 | 0.13548985 | 4.86428687 | 1.15E-06 | 2.73E-05 | ATRAID | 51374 |
| ENSG00000C | 482.599784 | 0.793625181 | 0.16316649 | 4.86389824 | 1.15E-06 | 2.74E-05 | SDF4 | 51150 |
| ENSG000001 | 2136.71362 | 0.672839895 | 0.13836369 | 4.86283559 | 1.16E-06 | 2.75E-05 | PCDH18 | 54510 |
| ENSG000002 | 142.711135 | 0.783498312 | 0.16113927 | 4.86224321 | 1.16E-06 | 2.76E-05 | C17orf51 | 339263 |
| ENSG00000C | 1328.21187 | 0.579393537 | 0.11918223 | 4.86140864 | 1.17E-06 | 2.77E-05 | TTC17 | 55761 |

| | | | | | | | | |
|---|---|---|---|---|---|---|---|---|
| ENSG000001 | 196.825342 | 0.769587273 | 0.15831793 | 4.86102402 | 1.17E-06 | 2.77E-05 | SPTSSB | 165679 |
| ENSG000001 | 1654.85081 | 0.620581768 | 0.12771231 | 4.85921665 | 1.18E-06 | 2.79E-05 | SNRPC | 6631 |
| ENSG000001 | 163.452924 | 0.772679781 | 0.15903663 | 4.85850183 | 1.18E-06 | 2.80E-05 | RCAN2 | 10231 |
| ENSG000001 | 339.943591 | 0.711261742 | 0.1464264 | 4.85746936 | 1.19E-06 | 2.81E-05 | JMJD8 | 339123 |
| ENSG000001 | 41.4394239 | -1.147409847 | 0.23629485 | -4.8558393 | 1.20E-06 | 2.84E-05 | PIK3C2G | 5288 |
| ENSG000001 | 316.861495 | 0.727145754 | 0.14976247 | 4.85532705 | 1.20E-06 | 2.84E-05 | PEX10 | 5192 |
| ENSG000001 | 1481.33694 | 0.584718221 | 0.12043957 | 4.85486799 | 1.20E-06 | 2.85E-05 | GLRX3 | 10539 |
| ENSG000001 | 51.468924 | -1.013369244 | 0.20874245 | -4.8546391 | 1.21E-06 | 2.85E-05 | UNC13C | 440279 |
| ENSG000001 | 561.500726 | 0.730523901 | 0.15048617 | 4.85442555 | 1.21E-06 | 2.85E-05 | PELO | 53918 |
| ENSG000001 | 216.617991 | 0.799152473 | 0.16468157 | 4.85271353 | 1.22E-06 | 2.87E-05 | C19orf25 | 148223 |
| ENSG000001 | 363.47642 | 0.68412762 | 0.14098503 | 4.85248402 | 1.22E-06 | 2.87E-05 | POLG | 5428 |
| ENSG000001 | 90.1139584 | 0.964313727 | 0.1987222 | 4.85257179 | 1.22E-06 | 2.87E-05 | CIB1 | 10519 |
| ENSG000001 | 1653.408 | 0.643211456 | 0.1325718 | 4.85179693 | 1.22E-06 | 2.88E-05 | ATP5L | 10632 |
| ENSG000001 | 1689.58917 | 0.54132318 | 0.11159397 | 4.85082817 | 1.23E-06 | 2.89E-05 | ANO6 | 196527 |
| ENSG00000C | 469.779194 | 0.789255177 | 0.16271453 | 4.85055145 | 1.23E-06 | 2.90E-05 | HERPUD1 | 9709 |
| ENSG000001 | 6012.02758 | 0.527167447 | 0.10868995 | 4.85019511 | 1.23E-06 | 2.90E-05 | MLEC | 9761 |
| ENSG000001 | 852.457256 | 0.791472714 | 0.16319006 | 4.85000572 | 1.23E-06 | 2.90E-05 | BCAP31 | 10134 |
| ENSG000001 | 354.23179 | 0.671231984 | 0.13840558 | 4.8497467 | 1.24E-06 | 2.90E-05 | PROSC | 11212 |
| ENSG00000C | 349.310123 | 0.902926433 | 0.18618762 | 4.84955132 | 1.24E-06 | 2.90E-05 | COASY | 80347 |
| ENSG000001 | 241.050615 | 0.709649941 | 0.14639834 | 4.8473905 | 1.25E-06 | 2.93E-05 | PGAP2 | 27315 |
| ENSG00000C | 672.27718 | 0.689184493 | 0.14218695 | 4.84703056 | 1.25E-06 | 2.94E-05 | LYPLA2 | 11313 |
| ENSG00000C | 493.831198 | 0.635177464 | 0.1310441 | 4.84705138 | 1.25E-06 | 2.94E-05 | PSME1 | 5720 |
| ENSG000001 | 635.463494 | 0.574763348 | 0.11858513 | 4.8468416 | 1.25E-06 | 2.94E-05 | SLC25A32 | 81034 |
| ENSG000001 | 2485.4694 | 0.543758834 | 0.11219356 | 4.8466136 | 1.26E-06 | 2.94E-05 | EIF1AX | 1964 |
| ENSG00000C | 825.565528 | 0.683998119 | 0.14114341 | 4.84612142 | 1.26E-06 | 2.94E-05 | RFC2 | 5982 |
| ENSG000001 | 512.404989 | 0.861234592 | 0.17770954 | 4.84630488 | 1.26E-06 | 2.94E-05 | NCLN | 56926 |
| ENSG000001 | 1015.64527 | 0.615382477 | 0.12698547 | 4.84608588 | 1.26E-06 | 2.94E-05 | OXA1L | 5018 |
| ENSG000001 | 2735.37358 | 0.537287454 | 0.11089735 | 4.8449081 | 1.27E-06 | 2.96E-05 | CSNK1A1 | 1452 |
| ENSG000001 | 361.08065 | 0.641347596 | 0.13237581 | 4.84490043 | 1.27E-06 | 2.96E-05 | IRAK1BP1 | 134728 |

| | | | | | | | | |
|---|---|---|---|---|---|---|---|---|
| ENSG000001 | 1408.13271 | 0.561417359 | 0.115913 | 4.84343742 | 1.28E-06 | 2.98E-05 | ATPAF1 | 64756 |
| ENSG000001 | 1486.14913 | 0.569636161 | 0.1176182 | 4.84309528 | 1.28E-06 | 2.98E-05 | SARAF | 51669 |
| ENSG000001 | 21.8344611 | -1.3633 | 0.28151978 | -4.8426438 | 1.28E-06 | 2.98E-05 | GDF15 | 9518 |
| ENSG000001 | 35.7739722 | 1.226714249 | 0.25333694 | 4.84222421 | 1.28E-06 | 2.99E-05 | CALCA | 796 |
| ENSG000001 | 454.553109 | 0.619078935 | 0.12784943 | 4.84225014 | 1.28E-06 | 2.99E-05 | CPT2 | 1376 |
| ENSG000001 | 282.892553 | 0.780206584 | 0.16113131 | 4.84205463 | 1.29E-06 | 2.99E-05 | STUB1 | 10273 |
| ENSG000001 | 1481.29168 | 0.516953477 | 0.10677824 | 4.84137468 | 1.29E-06 | 3.00E-05 | PARN | 5073 |
| ENSG00000C | 487.495274 | 0.705214411 | 0.1456845 | 4.84069629 | 1.29E-06 | 3.00E-05 | MED24 | 9862 |
| ENSG000002 | 100.537566 | -0.855162668 | 0.17667191 | -4.8403997 | 1.30E-06 | 3.01E-05 | NA | NA |
| ENSG000001 | 11717.1777 | 0.542176845 | 0.11201438 | 4.84024303 | 1.30E-06 | 3.01E-05 | DPPA4 | 55211 |
| ENSG000001 | 234.607427 | 0.661793322 | 0.13673363 | 4.8400185 | 1.30E-06 | 3.01E-05 | DPH7 | 92715 |
| ENSG000001 | 464.336924 | 0.696755211 | 0.143971 | 4.83955263 | 1.30E-06 | 3.02E-05 | ARHGEF40 | 55701 |
| ENSG000001 | 168.837603 | 0.857031533 | 0.17713418 | 4.83831836 | 1.31E-06 | 3.03E-05 | FAM96B | 51647 |
| ENSG000002 | 23.8695904 | -1.345736247 | 0.27814512 | -4.8382522 | 1.31E-06 | 3.03E-05 | DNM3OS | 100628315 |
| ENSG000001 | 314.28765 | 0.691377611 | 0.14292034 | 4.83750336 | 1.31E-06 | 3.04E-05 | ABHD12 | 26090 |
| ENSG000001 | 1970.53184 | 0.608480748 | 0.12578943 | 4.83729639 | 1.32E-06 | 3.04E-05 | STAU1 | 6780 |
| ENSG000001 | 981.654016 | 0.595724416 | 0.12315584 | 4.83715923 | 1.32E-06 | 3.04E-05 | SENP5 | 205564 |
| ENSG000002 | 243.079577 | 0.709833694 | 0.14676445 | 4.83655054 | 1.32E-06 | 3.05E-05 | NA | NA |
| ENSG000001 | 40.4185118 | -1.10222628 | 0.22793787 | -4.8356436 | 1.33E-06 | 3.06E-05 | GAD2 | 2572 |
| ENSG000001 | 546.195274 | 0.606685233 | 0.12548045 | 4.8348986 | 1.33E-06 | 3.07E-05 | AUNIP | 79000 |
| ENSG000001 | 1545.52831 | 0.521940772 | 0.10795513 | 4.83479372 | 1.33E-06 | 3.07E-05 | ANK2 | 287 |
| ENSG000001 | 2019.55153 | 0.584333465 | 0.12088744 | 4.83369864 | 1.34E-06 | 3.09E-05 | MRPS27 | 23107 |
| ENSG000002 | 65.6813063 | -1.057360982 | 0.21877157 | -4.8331736 | 1.34E-06 | 3.09E-05 | NA | NA |
| ENSG000001 | 63.8335711 | -0.958434862 | 0.19833272 | -4.8324597 | 1.35E-06 | 3.10E-05 | PPFIA4 | 8497 |
| ENSG000001 | 615.533494 | 0.670414896 | 0.13872947 | 4.83253399 | 1.35E-06 | 3.10E-05 | BSDC1 | 55108 |
| ENSG000001 | 14178.1818 | 0.609989977 | 0.12623426 | 4.83220627 | 1.35E-06 | 3.10E-05 | SLC16A1 | 6566 |
| ENSG000001 | 567.303479 | 0.660323447 | 0.1366769 | 4.83127309 | 1.36E-06 | 3.12E-05 | RER1 | 11079 |
| ENSG000001 | 540.964392 | 0.749852708 | 0.15521201 | 4.83115124 | 1.36E-06 | 3.12E-05 | USP19 | 10869 |
| ENSG00000C | 2898.90021 | 0.619742565 | 0.12828532 | 4.83097022 | 1.36E-06 | 3.12E-05 | KIAA0100 | 9703 |

| | | | | | | | | |
|---|---|---|---|---|---|---|---|---|
| ENSG000001 | 109.274058 | -0.811162393 | 0.16797154 | -4.8291657 | 1.37E-06 | 3.14E-05 | ZNF502 | 91392 |
| ENSG000001 | 378.645317 | 0.709198247 | 0.14689935 | 4.82778357 | 1.38E-06 | 3.16E-05 | TSPAN13 | 27075 |
| ENSG000001 | 112.765136 | -0.837925047 | 0.17358527 | -4.8271668 | 1.38E-06 | 3.17E-05 | ILDR2 | 387597 |
| ENSG000001 | 6690.15317 | 0.654025294 | 0.13549785 | 4.82683162 | 1.39E-06 | 3.18E-05 | ATP5B | 506 |
| ENSG000001 | 259.734169 | 0.645743993 | 0.13380861 | 4.82587768 | 1.39E-06 | 3.19E-05 | TMBIM1 | 64114 |
| ENSG000001 | 71.9016882 | -0.916401716 | 0.18989868 | -4.8257404 | 1.39E-06 | 3.19E-05 | CNTN2 | 6900 |
| ENSG000001 | 455.101248 | 0.920513254 | 0.19082076 | 4.82396817 | 1.41E-06 | 3.22E-05 | UBE2M | 9040 |
| ENSG000001 | 2096.42925 | 0.660626295 | 0.13694968 | 4.8238615 | 1.41E-06 | 3.22E-05 | FDPS | 2224 |
| ENSG00000C | 1306.14108 | 0.657307448 | 0.13627837 | 4.82327045 | 1.41E-06 | 3.22E-05 | CIRBP | 1153 |
| ENSG000001 | 486.975503 | 0.654948797 | 0.13581551 | 4.8223417 | 1.42E-06 | 3.24E-05 | LIMD2 | 80774 |
| ENSG000001 | 744.341631 | 0.584598385 | 0.12123589 | 4.82199101 | 1.42E-06 | 3.24E-05 | DESI1 | 27351 |
| ENSG000001 | 472.453579 | 0.603365398 | 0.12513303 | 4.8217916 | 1.42E-06 | 3.24E-05 | C7orf49 | 78996 |
| ENSG000002 | 135.776963 | 0.88315253 | 0.18315763 | 4.82181669 | 1.42E-06 | 3.24E-05 | NA | NA |
| ENSG000001 | 112.715949 | 0.89830956 | 0.18640956 | 4.81901001 | 1.44E-06 | 3.28E-05 | ADAMTS8 | 11095 |
| ENSG00000C | 1068.30628 | 0.538458818 | 0.11174148 | 4.81879102 | 1.44E-06 | 3.29E-05 | ZCCHC8 | 55596 |
| ENSG000001 | 3122.4153 | 0.595707103 | 0.12364556 | 4.817861 | 1.45E-06 | 3.30E-05 | GOLGB1 | 2804 |
| ENSG000001 | 698.121574 | 0.579777949 | 0.12037268 | 4.81652459 | 1.46E-06 | 3.32E-05 | CLMP | 79827 |
| ENSG000001 | 228.8028 | 0.884124952 | 0.1835665 | 4.81637416 | 1.46E-06 | 3.32E-05 | COL6A2 | 1292 |
| ENSG000001 | 212.873936 | 0.674096571 | 0.13996612 | 4.81614092 | 1.46E-06 | 3.32E-05 | TSPAN2 | 10100 |
| ENSG00000C | 827.633549 | 0.595584987 | 0.12371668 | 4.81410422 | 1.48E-06 | 3.36E-05 | SLC7A2 | 6542 |
| ENSG00000C | 870.102321 | 0.569626703 | 0.11834263 | 4.81336847 | 1.48E-06 | 3.36E-05 | TDP1 | 55775 |
| ENSG000001 | 921.383261 | 0.576781145 | 0.11982381 | 4.81357724 | 1.48E-06 | 3.36E-05 | FBXO7 | 25793 |
| ENSG000001 | 539.293179 | 0.664214462 | 0.13800305 | 4.8130419 | 1.49E-06 | 3.36E-05 | ZNF629 | 23361 |
| ENSG000001 | 371.509616 | 0.578048344 | 0.12009771 | 4.81315061 | 1.49E-06 | 3.36E-05 | SETD7 | 80854 |
| ENSG000001 | 1252.09249 | 0.529272265 | 0.10996322 | 4.81317554 | 1.49E-06 | 3.36E-05 | LONRF1 | 91694 |
| ENSG000001 | 1073.15074 | 0.711140563 | 0.14774218 | 4.81338884 | 1.48E-06 | 3.36E-05 | PUF60 | 22827 |
| ENSG000001 | 77.3159474 | 0.924580023 | 0.19208409 | 4.81341277 | 1.48E-06 | 3.36E-05 | BLOC1S4 | 55330 |
| ENSG000001 | 1520.90498 | 0.569297493 | 0.11829098 | 4.81268712 | 1.49E-06 | 3.36E-05 | ZNF267 | 10308 |
| ENSG000002 | 517.69012 | 0.673052954 | 0.13986494 | 4.8121636 | 1.49E-06 | 3.37E-05 | ZNHIT3 | 9326 |

| | | | | | | | | |
|---|---|---|---|---|---|---|---|---|
| ENSG000001 | 376.747546 | 0.689821455 | 0.14337671 | 4.81125182 | 1.50E-06 | 3.38E-05 | MAU2 | 23383 |
| ENSG000001 | 247.60018 | 0.629219884 | 0.1307813 | 4.81123754 | 1.50E-06 | 3.38E-05 | CHST15 | 51363 |
| ENSG00000C | 298.085913 | 0.643726427 | 0.13384003 | 4.80967051 | 1.51E-06 | 3.41E-05 | LETMD1 | 25875 |
| ENSG000001 | 1416.59757 | 0.558780338 | 0.11617586 | 4.80978005 | 1.51E-06 | 3.41E-05 | ERLIN2 | 11160 |
| ENSG000002 | 127.438603 | 0.801293935 | 0.166609 | 4.80942753 | 1.51E-06 | 3.41E-05 | NA | NA |
| ENSG000001 | 3653.86353 | 0.605141282 | 0.12584157 | 4.80875505 | 1.52E-06 | 3.42E-05 | ADH5 | 128 |
| ENSG000001 | 67.2774667 | 1.047248 | 0.2177907 | 4.80850658 | 1.52E-06 | 3.42E-05 | ISG15 | 9636 |
| ENSG000001 | 3346.2277 | 0.553628442 | 0.11514958 | 4.80790673 | 1.53E-06 | 3.43E-05 | RMND5A | 64795 |
| ENSG000001 | 385.76914 | 0.653876333 | 0.13602985 | 4.80685913 | 1.53E-06 | 3.44E-05 | DBNL | 28988 |
| ENSG000001 | 696.344143 | 0.573802562 | 0.11937486 | 4.80672879 | 1.53E-06 | 3.44E-05 | TRUB2 | 26995 |
| ENSG000001 | 188.638354 | 0.760528891 | 0.15822518 | 4.80662356 | 1.54E-06 | 3.44E-05 | MRPL41 | 64975 |
| ENSG000001 | 7173.37771 | 0.559649426 | 0.1164288 | 4.80679532 | 1.53E-06 | 3.44E-05 | HMGB1 | 3146 |
| ENSG000002 | 521.982402 | 0.65026042 | 0.13532378 | 4.80521922 | 1.55E-06 | 3.47E-05 | TSSC2 | 650368 |
| ENSG000001 | 1030.72567 | 0.651129364 | 0.13552246 | 4.80458621 | 1.55E-06 | 3.47E-05 | YKT6 | 10652 |
| ENSG000002 | 122.517723 | 0.845568329 | 0.17598991 | 4.80464083 | 1.55E-06 | 3.47E-05 | FGF13-AS1 | 100129662 |
| ENSG000001 | 39.5424134 | -1.110087929 | 0.23106291 | -4.804267 | 1.55E-06 | 3.48E-05 | WDR64 | 128025 |
| ENSG000001 | 375.661173 | 0.714430943 | 0.14873013 | 4.80353868 | 1.56E-06 | 3.49E-05 | COMMD4 | 54939 |
| ENSG00000C | 1621.49174 | 0.685314906 | 0.14268894 | 4.80285947 | 1.56E-06 | 3.50E-05 | U2AF2 | 11338 |
| ENSG00000C | 1023.74444 | 0.719780873 | 0.14989005 | 4.80205898 | 1.57E-06 | 3.51E-05 | PFKP | 5214 |
| ENSG00000C | 1387.17464 | 0.674587322 | 0.14048344 | 4.80189934 | 1.57E-06 | 3.51E-05 | ALKBH5 | 54890 |
| ENSG000001 | 739.967657 | 0.577885659 | 0.12034939 | 4.8017333 | 1.57E-06 | 3.51E-05 | ORMDL1 | 94101 |
| ENSG000001 | 149.733061 | 0.741529886 | 0.15442823 | 4.80177664 | 1.57E-06 | 3.51E-05 | NA | NA |
| ENSG000001 | 537.213527 | 0.633336148 | 0.13189088 | 4.80197081 | 1.57E-06 | 3.51E-05 | EXT2 | 2132 |
| ENSG00000C | 352.76991 | 0.706827643 | 0.1472484 | 4.80023986 | 1.58E-06 | 3.53E-05 | ERCC1 | 2067 |
| ENSG00000C | 3655.05724 | 0.606306346 | 0.12630763 | 4.80023542 | 1.58E-06 | 3.53E-05 | BZW1 | 9689 |
| ENSG000001 | 2077.63282 | 0.595782714 | 0.12411742 | 4.80015403 | 1.59E-06 | 3.53E-05 | MTPN | 136319 |
| ENSG000001 | 16943.1155 | 0.51302111 | 0.10689188 | 4.79943949 | 1.59E-06 | 3.54E-05 | CANX | 821 |
| ENSG000001 | 290.437248 | 0.642094108 | 0.13381745 | 4.79828385 | 1.60E-06 | 3.56E-05 | RALB | 5899 |
| ENSG000001 | 268.348115 | 0.706946154 | 0.14733227 | 4.79831178 | 1.60E-06 | 3.56E-05 | RPP25 | 54913 |

| | | | | | | | | |
|---|---|---|---|---|---|---|---|---|
| ENSG000001 | 3715.79283 | 0.588668745 | 0.12269956 | 4.79764349 | 1.61E-06 | 3.56E-05 | NOP56 | 10528 |
| ENSG000001 | 4962.94985 | 0.748439101 | 0.15599913 | 4.79771335 | 1.60E-06 | 3.56E-05 | TRIM28 | 10155 |
| ENSG000001 | 1388.00075 | 0.663814416 | 0.1383735 | 4.7972655 | 1.61E-06 | 3.57E-05 | PARK7 | 11315 |
| ENSG000002 | 107.597166 | 0.830907581 | 0.17321039 | 4.79710003 | 1.61E-06 | 3.57E-05 | PITPNA-AS1 | 100306951 |
| ENSG000001 | 2244.25811 | 0.639610895 | 0.1333498 | 4.79648923 | 1.61E-06 | 3.58E-05 | GOT2 | 2806 |
| ENSG000001 | 186.217649 | 0.711766641 | 0.14840521 | 4.79610272 | 1.62E-06 | 3.58E-05 | CEBPB | 1051 |
| ENSG000001 | 516.727552 | 0.624128436 | 0.13014044 | 4.79580686 | 1.62E-06 | 3.59E-05 | EMC7 | 56851 |
| ENSG000002 | 155.182973 | 0.713524385 | 0.14879933 | 4.79521234 | 1.63E-06 | 3.60E-05 | TSTD1 | 100131187 |
| ENSG000001 | 711.48236 | 0.688270598 | 0.14354056 | 4.79495554 | 1.63E-06 | 3.60E-05 | GOT1 | 2805 |
| ENSG000001 | 333.562188 | 0.756182126 | 0.15775443 | 4.79341282 | 1.64E-06 | 3.62E-05 | ELP6 | 54859 |
| ENSG000002 | 405.32398 | 0.681532937 | 0.14218204 | 4.79338259 | 1.64E-06 | 3.62E-05 | SMIM7 | 79086 |
| ENSG000002 | 123.568862 | 0.825999011 | 0.1723196 | 4.79341313 | 1.64E-06 | 3.62E-05 | NA | NA |
| ENSG000001 | 779.04197 | 0.624910954 | 0.13043198 | 4.79108706 | 1.66E-06 | 3.66E-05 | PACSIN2 | 11252 |
| ENSG000001 | 492.04482 | 0.680358843 | 0.14202025 | 4.79057643 | 1.66E-06 | 3.67E-05 | ALDH1L2 | 160428 |
| ENSG000001 | 1511.32759 | 0.587999431 | 0.12276932 | 4.78946541 | 1.67E-06 | 3.69E-05 | MAP3K7 | 6885 |
| ENSG000001 | 2285.60024 | 0.540050028 | 0.11277707 | 4.78865101 | 1.68E-06 | 3.70E-05 | TCF12 | 6938 |
| ENSG000001 | 397.954221 | 0.791930715 | 0.16537756 | 4.78862254 | 1.68E-06 | 3.70E-05 | ATP13A2 | 23400 |
| ENSG000001 | 253.506956 | 0.712619315 | 0.14883615 | 4.7879451 | 1.68E-06 | 3.70E-05 | COMMD9 | 29099 |
| ENSG000001 | 880.118378 | 0.669779127 | 0.13989102 | 4.78786371 | 1.69E-06 | 3.70E-05 | GTF3C5 | 9328 |
| ENSG000001 | 378.78615 | 0.739915439 | 0.15453382 | 4.78804849 | 1.68E-06 | 3.70E-05 | FABP7 | 2173 |
| ENSG000002 | 132.021606 | 0.779111172 | 0.16271525 | 4.78818765 | 1.68E-06 | 3.70E-05 | ZBTB48 | 3104 |
| ENSG000001 | 427.670001 | 0.7539015 | 0.15748134 | 4.78724351 | 1.69E-06 | 3.71E-05 | GYG1 | 2992 |
| ENSG000001 | 381.838561 | 0.644266958 | 0.1345851 | 4.78705997 | 1.69E-06 | 3.71E-05 | TSPAN5 | 10098 |
| ENSG000001 | 2928.55632 | 0.591786788 | 0.12363521 | 4.78655548 | 1.70E-06 | 3.72E-05 | NA | NA |
| ENSG000001 | 427.409434 | 0.69069487 | 0.14430453 | 4.78636988 | 1.70E-06 | 3.72E-05 | TOE1 | 114034 |
| ENSG00000C | 1834.78632 | 0.546543471 | 0.11419565 | 4.78602725 | 1.70E-06 | 3.73E-05 | RABEP1 | 9135 |
| ENSG000002 | 221.3615 | 0.661645625 | 0.13825111 | 4.78582491 | 1.70E-06 | 3.73E-05 | SLC26A6 | 65010 |
| ENSG000001 | 16027.4033 | 0.566686696 | 0.11841356 | 4.78565724 | 1.70E-06 | 3.73E-05 | TARS | 6897 |
| ENSG000001 | 587.083693 | 0.642193266 | 0.13424982 | 4.78356874 | 1.72E-06 | 3.77E-05 | MEGF10 | 84466 |

| | | | | | | | | |
|---|---|---|---|---|---|---|---|---|
| ENSG000001 | 1365.11795 | 0.571886751 | 0.11956533 | 4.78304846 | 1.73E-06 | 3.77E-05 | MTMR9 | 66036 |
| ENSG000002 | 479.010056 | 0.733246208 | 0.1533194 | 4.78247512 | 1.73E-06 | 3.78E-05 | NA | NA |
| ENSG000001 | 1746.11202 | 0.564296473 | 0.11800316 | 4.78204556 | 1.74E-06 | 3.79E-05 | IST1 | 9798 |
| ENSG000001 | 2629.45482 | 0.589274763 | 0.12323013 | 4.78190474 | 1.74E-06 | 3.79E-05 | SCRN1 | 9805 |
| ENSG000001 | 438.430425 | 0.591710168 | 0.1237529 | 4.78138425 | 1.74E-06 | 3.80E-05 | PLSCR1 | 5359 |
| ENSG000001 | 1333.65696 | 0.551483046 | 0.11534434 | 4.78118845 | 1.74E-06 | 3.80E-05 | PRC1 | 9055 |
| ENSG000001 | 656.052426 | 0.703408383 | 0.14713418 | 4.78072739 | 1.75E-06 | 3.81E-05 | LSS | 4047 |
| ENSG000001 | 33.4726709 | -1.166922434 | 0.24415258 | -4.7794803 | 1.76E-06 | 3.83E-05 | PPEF2 | 5470 |
| ENSG000001 | 930.358559 | 0.611013145 | 0.1278844 | 4.77785536 | 1.77E-06 | 3.86E-05 | GDAP1 | 54332 |
| ENSG000001 | 478.25318 | 0.676503473 | 0.14161036 | 4.77721727 | 1.78E-06 | 3.87E-05 | LEMD2 | 221496 |
| ENSG000001 | 327.369041 | 0.68014385 | 0.14238197 | 4.77689596 | 1.78E-06 | 3.87E-05 | ZSCAN21 | 7589 |
| ENSG000001 | 456.891711 | 0.551322094 | 0.11544207 | 4.77574663 | 1.79E-06 | 3.89E-05 | RABL3 | 285282 |
| ENSG00000C | 1382.24589 | 0.498729723 | 0.10443565 | 4.77547401 | 1.79E-06 | 3.89E-05 | LRRC16A | 55604 |
| ENSG000001 | 2708.2798 | 0.655041309 | 0.13716903 | 4.77543168 | 1.79E-06 | 3.89E-05 | UQCRH | 7388 |
| ENSG000001 | 216.626272 | 0.784252383 | 0.16423613 | 4.77515124 | 1.80E-06 | 3.90E-05 | MTG2 | 26164 |
| ENSG000001 | 1378.72396 | 0.566137619 | 0.11857485 | 4.77451697 | 1.80E-06 | 3.91E-05 | COPG1 | 22820 |
| ENSG000002 | 148.152492 | 0.800793588 | 0.16775781 | 4.77350991 | 1.81E-06 | 3.92E-05 | ARL16 | 339231 |
| ENSG00000C | 1418.82755 | 0.590143639 | 0.12365789 | 4.77238965 | 1.82E-06 | 3.94E-05 | CDK13 | 8621 |
| ENSG000001 | 6373.74528 | 0.55720118 | 0.11675899 | 4.7722338 | 1.82E-06 | 3.94E-05 | RBBP4 | 5928 |
| ENSG00000C | 3942.62989 | 0.606804912 | 0.12718451 | 4.77105978 | 1.83E-06 | 3.96E-05 | TMSB10 | 9168 |
| ENSG000001 | 992.763936 | 0.607308932 | 0.12729215 | 4.77098501 | 1.83E-06 | 3.96E-05 | UBXN1 | 51035 |
| ENSG000001 | 12051.0473 | 0.657991791 | 0.13791587 | 4.77096512 | 1.83E-06 | 3.96E-05 | RPL15 | 6138 |
| ENSG000001 | 549.820645 | 0.568868439 | 0.11923469 | 4.7709976 | 1.83E-06 | 3.96E-05 | HKR1 | 284459 |
| ENSG000001 | 346.984073 | 0.654485148 | 0.13721204 | 4.76988123 | 1.84E-06 | 3.98E-05 | CHCHD1 | 118487 |
| ENSG000001 | 84.5142738 | -0.871518641 | 0.18276049 | -4.7686382 | 1.85E-06 | 4.00E-05 | VWF | 7450 |
| ENSG000001 | 571.255922 | 0.84986542 | 0.17823047 | 4.76835089 | 1.86E-06 | 4.01E-05 | TMED9 | 54732 |
| ENSG000001 | 228.706578 | 0.633030069 | 0.13276468 | 4.76806083 | 1.86E-06 | 4.01E-05 | REPS2 | 9185 |
| ENSG00000C | 318.67828 | 0.851055674 | 0.17851107 | 4.76752324 | 1.87E-06 | 4.02E-05 | BAK1 | 578 |
| ENSG000001 | 3896.20183 | 0.521465191 | 0.10938396 | 4.76729137 | 1.87E-06 | 4.02E-05 | EIF4EBP2 | 1979 |

| | | | | | | | | |
|---|---|---|---|---|---|---|---|---|
| ENSG000001 | 2382.47357 | 0.517420641 | 0.10854868 | 4.7667152 | 1.87E-06 | 4.03E-05 | EIF3F | 8665 |
| ENSG00000C | 566.001771 | 0.609126925 | 0.12780285 | 4.7661449 | 1.88E-06 | 4.04E-05 | TOMM34 | 10953 |
| ENSG000001 | 213.240131 | 0.701195697 | 0.14713792 | 4.76556753 | 1.88E-06 | 4.05E-05 | STK16 | 8576 |
| ENSG00000C | 1005.53826 | 0.638105197 | 0.13391051 | 4.76516158 | 1.89E-06 | 4.05E-05 | LAS1L | 81887 |
| ENSG000001 | 795.174645 | 0.532477897 | 0.11174642 | 4.76505544 | 1.89E-06 | 4.05E-05 | AK3 | 50808 |
| ENSG000001 | 233.993381 | 0.72805279 | 0.15279658 | 4.76485003 | 1.89E-06 | 4.05E-05 | LRWD1 | 222229 |
| ENSG000001 | 2318.11876 | 0.618308006 | 0.12976166 | 4.76495157 | 1.89E-06 | 4.05E-05 | OAZ2 | 4947 |
| ENSG000002 | 128.143711 | -0.788644043 | 0.16558763 | -4.762699 | 1.91E-06 | 4.10E-05 | MGAM | 8972 |
| ENSG000001 | 241.298257 | 0.722678941 | 0.15186005 | 4.75884833 | 1.95E-06 | 4.17E-05 | ALKBH3 | 221120 |
| ENSG000001 | 2202.44512 | 0.593097674 | 0.12463392 | 4.75871788 | 1.95E-06 | 4.17E-05 | SLIRP | 81892 |
| ENSG000001 | 2485.51281 | 0.600384394 | 0.12616933 | 4.75856046 | 1.95E-06 | 4.17E-05 | MYL12B | 103910 |
| ENSG000001 | 237.861966 | 0.717016752 | 0.15068318 | 4.75843911 | 1.95E-06 | 4.17E-05 | MRPL34 | 64981 |
| ENSG000001 | 1358.1107 | 0.543819131 | 0.11428873 | 4.75829193 | 1.95E-06 | 4.18E-05 | YIPF6 | 286451 |
| ENSG000001 | 827.496589 | 0.590100508 | 0.12402658 | 4.75785514 | 1.96E-06 | 4.18E-05 | TFDP1 | 7027 |
| ENSG00000C | 251.636948 | 0.729226749 | 0.15332165 | 4.75618917 | 1.97E-06 | 4.21E-05 | RNF126 | 55658 |
| ENSG000001 | 1122.70959 | 0.606329884 | 0.12748036 | 4.75626108 | 1.97E-06 | 4.21E-05 | KDELR1 | 10945 |
| ENSG000001 | 247.368686 | 0.671179595 | 0.14113762 | 4.75549738 | 1.98E-06 | 4.22E-05 | TCF19 | 6941 |
| ENSG000001 | 1307.61681 | 0.59314603 | 0.12475938 | 4.75432027 | 1.99E-06 | 4.25E-05 | CCNB2 | 9133 |
| ENSG000001 | 3989.22372 | 0.560000806 | 0.11780262 | 4.75372113 | 2.00E-06 | 4.26E-05 | API5 | 8539 |
| ENSG000001 | 548.54296 | 0.614151596 | 0.12920031 | 4.7534838 | 2.00E-06 | 4.26E-05 | TOR1A | 1861 |
| ENSG000001 | 61.0449249 | -0.939133227 | 0.197643 | -4.7516645 | 2.02E-06 | 4.30E-05 | ADCY10 | 55811 |
| ENSG00000C | 858.112128 | 0.55128897 | 0.11604206 | 4.75076861 | 2.03E-06 | 4.31E-05 | CLK1 | 1195 |
| ENSG000001 | 18.509029 | -1.47571635 | 0.3106402 | -4.7505647 | 2.03E-06 | 4.32E-05 | KRT5 | 3852 |
| ENSG000001 | 463.184368 | 0.792791641 | 0.16695144 | 4.7486361 | 2.05E-06 | 4.35E-05 | ORMDL3 | 94103 |
| ENSG00000C | 2319.32403 | 0.586547162 | 0.1235433 | 4.74770501 | 2.06E-06 | 4.37E-05 | SLC4A7 | 9497 |
| ENSG000001 | 1275.12838 | 0.632312027 | 0.13320607 | 4.74687111 | 2.07E-06 | 4.39E-05 | C1orf43 | 25912 |
| ENSG000001 | 225.658937 | 0.771399674 | 0.16252344 | 4.74639035 | 2.07E-06 | 4.39E-05 | C17orf62 | 79415 |
| ENSG000002 | 967.049535 | 0.581135056 | 0.12243566 | 4.74645253 | 2.07E-06 | 4.39E-05 | DNAJC9 | 23234 |
| ENSG000002 | 68.78903 | -0.899784327 | 0.18958411 | -4.7460957 | 2.07E-06 | 4.40E-05 | NA | NA |

| | | | | | | | | |
|---|---|---|---|---|---|---|---|---|
| ENSG00000( | 704.537421 | 0.665816746 | 0.14030547 | 4.74547971 | 2.08E-06 | 4.41E-05 | TRNT1 | 51095 |
| ENSG000002 | 312.778408 | 0.648892066 | 0.13673759 | 4.7455281 | 2.08E-06 | 4.41E-05 | RASL10B | 91608 |
| ENSG000001 | 105.419568 | -0.798458016 | 0.1682764 | -4.7449199 | 2.09E-06 | 4.42E-05 | CUBN | 8029 |
| ENSG000002 | 2186.36407 | 0.610846705 | 0.12873905 | 4.74484414 | 2.09E-06 | 4.42E-05 | TSN | 7247 |
| ENSG000001 | 3585.43988 | 0.630399144 | 0.1328668 | 4.74459493 | 2.09E-06 | 4.42E-05 | HYOU1 | 10525 |
| ENSG000001 | 380.683049 | 0.694217362 | 0.14632221 | 4.74444299 | 2.09E-06 | 4.42E-05 | SCAMP5 | 192683 |
| ENSG000001 | 114.072068 | 0.88631381 | 0.18685936 | 4.74321327 | 2.10E-06 | 4.45E-05 | PPCDC | 60490 |
| ENSG000001 | 880.058428 | 0.652849254 | 0.13764513 | 4.74298827 | 2.11E-06 | 4.45E-05 | HDAC3 | 8841 |
| ENSG000001 | 722.916369 | 0.569485602 | 0.12007184 | 4.74287379 | 2.11E-06 | 4.45E-05 | ZKSCAN5 | 23660 |
| ENSG000001 | 1419.30059 | 0.666018531 | 0.14043179 | 4.74264777 | 2.11E-06 | 4.45E-05 | MAP7 | 9053 |
| ENSG000001 | 216.525524 | 0.78987691 | 0.16657171 | 4.74196317 | 2.12E-06 | 4.46E-05 | LSM10 | 84967 |
| ENSG000001 | 629.606507 | 0.568832123 | 0.11996402 | 4.74168938 | 2.12E-06 | 4.47E-05 | SAMD8 | 142891 |
| ENSG000001 | 2471.50117 | 0.5212497 | 0.10995466 | 4.74058764 | 2.13E-06 | 4.49E-05 | ARF3 | 377 |
| ENSG000001 | 375.393927 | 0.688600457 | 0.14528888 | 4.73952609 | 2.14E-06 | 4.51E-05 | NDUFB6 | 4712 |
| ENSG000001 | 444.186506 | 0.577863515 | 0.12195828 | 4.73820663 | 2.16E-06 | 4.54E-05 | MED6 | 10001 |
| ENSG000001 | 483.890013 | 0.667723585 | 0.14093951 | 4.73766084 | 2.16E-06 | 4.55E-05 | PDCL | 5082 |
| ENSG000001 | 862.533939 | 0.581732839 | 0.12283396 | 4.73592835 | 2.18E-06 | 4.58E-05 | AIMP2 | 7965 |
| ENSG000001 | 2033.53676 | 0.631199926 | 0.1332835 | 4.73576945 | 2.18E-06 | 4.58E-05 | HIP1 | 3092 |
| ENSG000001 | 1978.79156 | 0.63727105 | 0.13456831 | 4.73566944 | 2.18E-06 | 4.58E-05 | GOLM1 | 51280 |
| ENSG000001 | 863.802161 | 0.692231777 | 0.14620896 | 4.73453723 | 2.20E-06 | 4.61E-05 | TRAF7 | 84231 |
| ENSG000001 | 891.78296 | 0.603183895 | 0.12739899 | 4.73460504 | 2.19E-06 | 4.61E-05 | TMEM126B | 55863 |
| ENSG000001 | 516.294156 | 0.604409942 | 0.12767052 | 4.73413878 | 2.20E-06 | 4.61E-05 | TBRG1 | 84897 |
| ENSG000001 | 1463.12032 | 0.545223339 | 0.11517118 | 4.73402578 | 2.20E-06 | 4.61E-05 | SETD3 | 84193 |
| ENSG000001 | 1257.00055 | 0.690912668 | 0.1459513 | 4.73385757 | 2.20E-06 | 4.61E-05 | MFN2 | 9927 |
| ENSG000001 | 740.514128 | 0.673022357 | 0.14219041 | 4.73324723 | 2.21E-06 | 4.63E-05 | NHP2 | 55651 |
| ENSG000001 | 1796.46609 | 0.610071527 | 0.12889694 | 4.73301799 | 2.21E-06 | 4.63E-05 | ACP1 | 52 |
| ENSG000001 | 74.5161559 | 0.981841148 | 0.20745552 | 4.7327792 | 2.21E-06 | 4.63E-05 | RRAS | 6237 |
| ENSG000001 | 2687.28248 | 0.563022351 | 0.11896034 | 4.73285744 | 2.21E-06 | 4.63E-05 | PA2G4 | 5036 |
| ENSG000001 | 837.088541 | 0.606350132 | 0.1281443 | 4.73177588 | 2.23E-06 | 4.65E-05 | DDX41 | 51428 |

| | | | | | | | | |
|---|---|---|---|---|---|---|---|---|
| ENSG00000( | 922.507956 | 0.617949781 | 0.13060922 | 4.73128754 | 2.23E-06 | 4.66E-05 | PPP5C | 5536 |
| ENSG00000( | 1471.88875 | 0.627139458 | 0.13257429 | 4.73047585 | 2.24E-06 | 4.66E-05 | CD9 | 928 |
| ENSG000001 | 297.233859 | 0.63622654 | 0.13449731 | 4.73040357 | 2.24E-06 | 4.66E-05 | SLC25A15 | 10166 |
| ENSG000001 | 2000.62229 | 0.620825606 | 0.1312243 | 4.73102619 | 2.23E-06 | 4.66E-05 | MAZ | 4150 |
| ENSG000001 | 275.677991 | 0.737619815 | 0.15591815 | 4.73081437 | 2.24E-06 | 4.66E-05 | CIB2 | 10518 |
| ENSG000001 | 660.477931 | 0.562424793 | 0.11889039 | 4.73061627 | 2.24E-06 | 4.66E-05 | SLAIN1 | 122060 |
| ENSG000001 | 451.979634 | 0.642097989 | 0.13573696 | 4.73045811 | 2.24E-06 | 4.66E-05 | FAM103A1 | 83640 |
| ENSG000002 | 173.543458 | 0.680452098 | 0.14383935 | 4.73063944 | 2.24E-06 | 4.66E-05 | PCDHAC1 | 56135 |
| ENSG00000( | 361.722525 | 0.568748998 | 0.12023776 | 4.73020293 | 2.24E-06 | 4.66E-05 | NEDD4 | 4734 |
| ENSG000001 | 940.69504 | 0.605529514 | 0.12801848 | 4.73001657 | 2.25E-06 | 4.67E-05 | ANXA7 | 310 |
| ENSG00000( | 577.583737 | 0.618444937 | 0.13077124 | 4.72921211 | 2.25E-06 | 4.68E-05 | EXOSC7 | 23016 |
| ENSG000001 | 151.339029 | 0.765938282 | 0.16197559 | 4.72872647 | 2.26E-06 | 4.69E-05 | RHOF | 54509 |
| ENSG000001 | 949.777153 | 0.701918365 | 0.14844336 | 4.72852652 | 2.26E-06 | 4.69E-05 | UQCR10 | 29796 |
| ENSG000001 | 1901.05809 | 0.591279125 | 0.12506082 | 4.72793252 | 2.27E-06 | 4.70E-05 | PSMB2 | 5690 |
| ENSG00000( | 4077.35199 | 0.664500958 | 0.14056428 | 4.72738151 | 2.27E-06 | 4.72E-05 | CNN2 | 1265 |
| ENSG000001 | 83.1935726 | 0.955525833 | 0.20217567 | 4.72621567 | 2.29E-06 | 4.74E-05 | AMDHD2 | 51005 |
| ENSG00000( | 405.091164 | 0.797215677 | 0.16869279 | 4.72584316 | 2.29E-06 | 4.75E-05 | POLRMT | 5442 |
| ENSG000001 | 1745.39805 | 0.646039435 | 0.13671146 | 4.72556896 | 2.29E-06 | 4.75E-05 | HMGN2 | 3151 |
| ENSG000001 | 914.809904 | 0.775965008 | 0.16421235 | 4.72537541 | 2.30E-06 | 4.75E-05 | C6orf106 | 64771 |
| ENSG000001 | 1146.1005 | 0.611852685 | 0.12948546 | 4.7252616 | 2.30E-06 | 4.75E-05 | HDAC1 | 3065 |
| ENSG000001 | 658.314477 | 0.705670398 | 0.14936861 | 4.72435555 | 2.31E-06 | 4.77E-05 | C14orf2 | 9556 |
| ENSG00000( | 54.7096798 | -0.987331457 | 0.20899973 | -4.7240801 | 2.31E-06 | 4.78E-05 | ABCB5 | 340273 |
| ENSG000001 | 1996.75839 | 0.470962558 | 0.0997011 | 4.72374503 | 2.32E-06 | 4.78E-05 | RAB6A | 5870 |
| ENSG000001 | 336.180004 | 0.685693498 | 0.14520322 | 4.72230239 | 2.33E-06 | 4.81E-05 | MPLKIP | 136647 |
| ENSG000001 | 2701.01415 | 0.551407137 | 0.11677531 | 4.72194956 | 2.34E-06 | 4.82E-05 | PTCD3 | 55037 |
| ENSG00000( | 485.561638 | 0.618421226 | 0.13097918 | 4.72152311 | 2.34E-06 | 4.83E-05 | ALAS1 | 211 |
| ENSG000002 | 223.821802 | 0.642858924 | 0.13618102 | 4.72062052 | 2.35E-06 | 4.84E-05 | PCDHGA10 | 56106 |
| ENSG000001 | 235.826355 | 0.722356636 | 0.15304781 | 4.71981027 | 2.36E-06 | 4.86E-05 | PCYT2 | 5833 |
| ENSG000001 | 566.001527 | 0.591100837 | 0.12525394 | 4.71921949 | 2.37E-06 | 4.87E-05 | MGST3 | 4259 |

| | | | | | | | | |
|---|---|---|---|---|---|---|---|---|
| ENSG000001 | 607.788493 | 0.656287637 | 0.139086 | 4.7185744 | 2.38E-06 | 4.88E-05 | NA | NA |
| ENSG000001 | 84.9074949 | 0.913085515 | 0.19350544 | 4.71865561 | 2.37E-06 | 4.88E-05 | ASL | 435 |
| ENSG000001 | 1193.846 | 0.589104101 | 0.12486116 | 4.71807345 | 2.38E-06 | 4.89E-05 | GTF3C2 | 2976 |
| ENSG000002 | 1007.37387 | 0.676402294 | 0.14336633 | 4.71799972 | 2.38E-06 | 4.89E-05 | PSMB3 | 5691 |
| ENSG000002 | 873.311998 | 0.546620898 | 0.11586395 | 4.71778225 | 2.38E-06 | 4.90E-05 | AGPAT1 | 10554 |
| ENSG000000 | 845.995816 | 0.65408026 | 0.13870781 | 4.71552592 | 2.41E-06 | 4.94E-05 | CLPTM1L | 81037 |
| ENSG000001 | 55.3233182 | -0.954683493 | 0.20245465 | -4.7155424 | 2.41E-06 | 4.94E-05 | RFX4 | 5992 |
| ENSG000001 | 190.858293 | 0.868254473 | 0.18412291 | 4.71562422 | 2.41E-06 | 4.94E-05 | TRMT61A | 115708 |
| ENSG000001 | 705.018124 | 0.593045616 | 0.12580026 | 4.71418451 | 2.43E-06 | 4.97E-05 | COX18 | 285521 |
| ENSG000001 | 4324.87059 | 0.590259453 | 0.12521101 | 4.71411783 | 2.43E-06 | 4.97E-05 | RPN1 | 6184 |
| ENSG000001 | 592.83925 | 0.62339598 | 0.13226757 | 4.71314312 | 2.44E-06 | 4.99E-05 | GINS4 | 84296 |
| ENSG000001 | 213.140358 | 0.742363612 | 0.15753884 | 4.71225767 | 2.45E-06 | 5.01E-05 | PICK1 | 9463 |
| ENSG000001 | 1508.66659 | 0.618200745 | 0.13120608 | 4.7116777 | 2.46E-06 | 5.02E-05 | MTA2 | 9219 |
| ENSG000001 | 54.5859411 | 1.010918551 | 0.21457875 | 4.71117732 | 2.46E-06 | 5.03E-05 | C14orf80 | 283643 |
| ENSG000001 | 1192.79114 | 0.552378784 | 0.11726567 | 4.71049025 | 2.47E-06 | 5.05E-05 | PLXDC2 | 84898 |
| ENSG000001 | 665.146653 | 0.586815318 | 0.12458715 | 4.71007904 | 2.48E-06 | 5.06E-05 | NSMCE4A | 54780 |
| ENSG000001 | 1760.85346 | 0.501853501 | 0.10655105 | 4.70998177 | 2.48E-06 | 5.06E-05 | KIF20A | 10112 |
| ENSG000001 | 4147.43427 | 0.594449668 | 0.12621517 | 4.70981155 | 2.48E-06 | 5.06E-05 | LRRN1 | 57633 |
| ENSG000000 | 58.7098782 | -0.971262503 | 0.20625084 | -4.7091323 | 2.49E-06 | 5.07E-05 | COL19A1 | 1310 |
| ENSG000001 | 501.601929 | 0.735339323 | 0.15615467 | 4.70904482 | 2.49E-06 | 5.07E-05 | LZTS2 | 84445 |
| ENSG000001 | 268.485207 | 0.738187493 | 0.15676097 | 4.70900044 | 2.49E-06 | 5.07E-05 | SGSM2 | 9905 |
| ENSG000001 | 753.389951 | 0.580717836 | 0.12335475 | 4.70770559 | 2.51E-06 | 5.10E-05 | CENPN | 55839 |
| ENSG000000 | 1145.64749 | 0.58796955 | 0.12492044 | 4.70675214 | 2.52E-06 | 5.12E-05 | DHX8 | 1659 |
| ENSG000001 | 250.378865 | 0.90104378 | 0.19143364 | 4.70682059 | 2.52E-06 | 5.12E-05 | PUS1 | 80324 |
| ENSG000001 | 3429.29662 | 0.60176978 | 0.12784864 | 4.70689233 | 2.52E-06 | 5.12E-05 | TUBB4B | 10383 |
| ENSG000001 | 2827.52719 | 0.650923168 | 0.13829968 | 4.70661365 | 2.52E-06 | 5.12E-05 | PPM1G | 5496 |
| ENSG000001 | 699.49818 | 0.578383947 | 0.12290222 | 4.70604959 | 2.53E-06 | 5.13E-05 | PIGN | 23556 |
| ENSG000000 | 177.404227 | 0.787810441 | 0.1674128 | 4.70579569 | 2.53E-06 | 5.13E-05 | NUBP2 | 10101 |
| ENSG000001 | 1259.04487 | 0.569768401 | 0.12108779 | 4.70541591 | 2.53E-06 | 5.14E-05 | NAP1L3 | 4675 |

| | | | | | | | | |
|---|---|---|---|---|---|---|---|---|
| ENSG000001 | 1140.67149 | 0.576476079 | 0.12254254 | 4.70429363 | 2.55E-06 | 5.17E-05 | AAGAB | 79719 |
| ENSG000001 | 195.245177 | 0.809805996 | 0.1721558 | 4.70391346 | 2.55E-06 | 5.17E-05 | PLOD3 | 8985 |
| ENSG000001 | 558.257109 | 0.570812983 | 0.1213559 | 4.70362798 | 2.56E-06 | 5.18E-05 | CCDC50 | 152137 |
| ENSG000002 | 928.532135 | 0.698035016 | 0.14840758 | 4.70349986 | 2.56E-06 | 5.18E-05 | GPX1 | 2876 |
| ENSG000001 | 827.039913 | 0.735261809 | 0.15633566 | 4.70309708 | 2.56E-06 | 5.19E-05 | NA | NA |
| ENSG000001 | 1581.71778 | 0.590253011 | 0.12550636 | 4.70297305 | 2.56E-06 | 5.19E-05 | UQCRFS1 | 7386 |
| ENSG000001 | 308.415068 | 0.671556256 | 0.14281207 | 4.70237734 | 2.57E-06 | 5.20E-05 | DECR1 | 1666 |
| ENSG000001 | 62.4714699 | 1.014192581 | 0.21567601 | 4.70238936 | 2.57E-06 | 5.20E-05 | ART5 | 116969 |
| ENSG000001 | 673.260492 | 0.634417534 | 0.13492556 | 4.70198177 | 2.58E-06 | 5.20E-05 | GLRX5 | 51218 |
| ENSG000001 | 3806.80413 | 0.620199149 | 0.13192438 | 4.70117169 | 2.59E-06 | 5.22E-05 | PGRMC1 | 10857 |
| ENSG000001 | 198.435839 | 0.687387028 | 0.14623016 | 4.70072007 | 2.59E-06 | 5.23E-05 | HMHA1 | 23526 |
| ENSG000001 | 593.867438 | 0.665992001 | 0.141683 | 4.70057811 | 2.59E-06 | 5.23E-05 | LAMTOR5 | 10542 |
| ENSG000001 | 7337.53628 | 0.54202393 | 0.11531409 | 4.70041386 | 2.60E-06 | 5.23E-05 | CNBP | 7555 |
| ENSG000001 | 3544.81278 | 0.60277106 | 0.1282702 | 4.69922925 | 2.61E-06 | 5.26E-05 | SUMO2 | 6613 |
| ENSG000001 | 254.571111 | 0.764502684 | 0.16269833 | 4.69889705 | 2.62E-06 | 5.27E-05 | UPP1 | 7378 |
| ENSG000001 | 735.934564 | 0.582322268 | 0.1239468 | 4.69816305 | 2.63E-06 | 5.28E-05 | E2F5 | 1875 |
| ENSG000001 | 85.6875795 | 0.893134985 | 0.19011594 | 4.69784384 | 2.63E-06 | 5.29E-05 | LHFPL1 | 340596 |
| ENSG000001 | 1277.86725 | 0.59800501 | 0.12729663 | 4.69772852 | 2.63E-06 | 5.29E-05 | MTMR12 | 54545 |
| ENSG000001 | 5448.63128 | 0.553946494 | 0.11792495 | 4.69744935 | 2.63E-06 | 5.29E-05 | NUP205 | 23165 |
| ENSG000001 | 729.726057 | 0.562867799 | 0.11982384 | 4.69746087 | 2.63E-06 | 5.29E-05 | RAB11FIP1 | 80223 |
| ENSG000001 | 2094.01839 | 0.609509847 | 0.12975453 | 4.69740693 | 2.63E-06 | 5.29E-05 | MAD2L1 | 4085 |
| ENSG000001 | 207.364312 | 0.718301406 | 0.15299468 | 4.69494368 | 2.67E-06 | 5.35E-05 | CDC25B | 994 |
| ENSG000001 | 1421.78533 | 0.600846473 | 0.12798464 | 4.69467647 | 2.67E-06 | 5.35E-05 | SF3B6 | 51639 |
| ENSG000001 | 433.356476 | 0.578844872 | 0.12329682 | 4.69472657 | 2.67E-06 | 5.35E-05 | NDUFB3 | 4709 |
| ENSG000001 | 50.8228485 | -1.015220675 | 0.21625529 | -4.6945472 | 2.67E-06 | 5.35E-05 | EYA4 | 2070 |
| ENSG000001 | 501.481152 | 0.642906792 | 0.13695451 | 4.69430889 | 2.68E-06 | 5.36E-05 | ATP5SL | 55101 |
| ENSG000001 | 1323.61605 | 0.525872146 | 0.11202704 | 4.69415386 | 2.68E-06 | 5.36E-05 | ALG13 | 79868 |
| ENSG00000C | 310.00125 | 0.647399852 | 0.13793673 | 4.69345511 | 2.69E-06 | 5.37E-05 | UNKL | 64718 |
| ENSG000001 | 887.925169 | 0.587295319 | 0.12517195 | 4.69190825 | 2.71E-06 | 5.41E-05 | KIAA1468 | 57614 |

| | | | | | | | | |
|---|---|---|---|---|---|---|---|---|
| ENSG000001 | 235.431353 | 0.731449965 | 0.15589943 | 4.69180657 | 2.71E-06 | 5.41E-05 | SHKBP1 | 92799 |
| ENSG000001 | 271.748601 | 0.772832909 | 0.16473228 | 4.69144786 | 2.71E-06 | 5.42E-05 | ALDH2 | 217 |
| ENSG000001 | 364.81228 | 0.609661764 | 0.1299823 | 4.6903444 | 2.73E-06 | 5.45E-05 | LRRC42 | 115353 |
| ENSG000001 | 245.295445 | 0.700528545 | 0.14938111 | 4.68953916 | 2.74E-06 | 5.46E-05 | PODXL2 | 50512 |
| ENSG000001 | 26.57827 | -1.27201011 | 0.27126561 | -4.6891683 | 2.74E-06 | 5.47E-05 | TRIM67 | 440730 |
| ENSG000001 | 263.046314 | 0.665580428 | 0.14194257 | 4.68908261 | 2.74E-06 | 5.47E-05 | RPUSD2 | 27079 |
| ENSG000001 | 534.579897 | 0.682375723 | 0.14553826 | 4.68863475 | 2.75E-06 | 5.48E-05 | TXNDC17 | 84817 |
| ENSG00000C | 895.564167 | 0.515567465 | 0.11002473 | 4.68592365 | 2.79E-06 | 5.55E-05 | GDE1 | 51573 |
| ENSG000001 | 662.452914 | 0.538705291 | 0.11496813 | 4.6856925 | 2.79E-06 | 5.55E-05 | SLC16A9 | 220963 |
| ENSG00000C | 1414.76898 | 0.576595808 | 0.12306071 | 4.68545824 | 2.79E-06 | 5.56E-05 | ANAPC5 | 51433 |
| ENSG000001 | 353.198415 | 0.605556717 | 0.12928998 | 4.68370939 | 2.82E-06 | 5.60E-05 | ARL6IP6 | 151188 |
| ENSG00000C | 178.780086 | 0.688751509 | 0.14705939 | 4.6834922 | 2.82E-06 | 5.61E-05 | BCS1L | 617 |
| ENSG000001 | 253.340005 | 0.758622825 | 0.16199705 | 4.68294219 | 2.83E-06 | 5.62E-05 | TMEM120B | 144404 |
| ENSG000001 | 274.744622 | 0.615818163 | 0.13150922 | 4.68269964 | 2.83E-06 | 5.62E-05 | RNF121 | 55298 |
| ENSG000001 | 699.366746 | 0.546005882 | 0.11660393 | 4.68256859 | 2.83E-06 | 5.62E-05 | AREL1 | 9870 |
| ENSG000002 | 233.056945 | 0.785091567 | 0.16766816 | 4.68241308 | 2.84E-06 | 5.62E-05 | SNORD116-19 | 727708 |
| ENSG000001 | 573.275463 | 0.628307785 | 0.13420906 | 4.68156027 | 2.85E-06 | 5.65E-05 | RFC5 | 5985 |
| ENSG000001 | 579.315633 | 0.614920901 | 0.13137547 | 4.68063723 | 2.86E-06 | 5.67E-05 | HIBADH | 11112 |
| ENSG000001 | 7805.94457 | 0.526986782 | 0.11258967 | 4.6805964 | 2.86E-06 | 5.67E-05 | CEBPZ | 10153 |
| ENSG000002 | 331.089243 | 0.577487405 | 0.12338036 | 4.68054575 | 2.86E-06 | 5.67E-05 | FAM216A | 29902 |
| ENSG000001 | 779.234381 | 0.67291491 | 0.14377641 | 4.68028722 | 2.86E-06 | 5.67E-05 | SH3PXD2B | 285590 |
| ENSG000001 | 1310.44664 | 0.689422071 | 0.14730572 | 4.68021262 | 2.87E-06 | 5.67E-05 | RCC1 | 1104 |
| ENSG000002 | 87.5159825 | -0.941940031 | 0.20125316 | -4.680374 | 2.86E-06 | 5.67E-05 | NA | NA |
| ENSG000002 | 1109.30396 | 0.708137484 | 0.15131043 | 4.6800308 | 2.87E-06 | 5.67E-05 | NA | NA |
| ENSG000001 | 123.152654 | 0.857248163 | 0.18318021 | 4.67980768 | 2.87E-06 | 5.67E-05 | GALE | 2582 |
| ENSG000001 | 301.333001 | 0.764353418 | 0.16333603 | 4.67963754 | 2.87E-06 | 5.67E-05 | TOP3B | 8940 |
| ENSG000001 | 621.989511 | 0.583484119 | 0.1246949 | 4.67929427 | 2.88E-06 | 5.68E-05 | GBAS | 2631 |
| ENSG000002 | 363.865273 | 0.678899288 | 0.14509487 | 4.67900281 | 2.88E-06 | 5.69E-05 | COL5A2 | 1290 |
| ENSG000001 | 624.871813 | 0.604681775 | 0.12925997 | 4.67802795 | 2.90E-06 | 5.71E-05 | PRPF31 | 26121 |

| | | | | | | | | |
|---|---|---|---|---|---|---|---|---|
| ENSG000002 | 111.082676 | -0.778190739 | 0.16635195 | -4.677978 | 2.90E-06 | 5.71E-05 | ZNF350 | 59348 |
| ENSG000001 | 135.47882 | 0.870539413 | 0.1861279 | 4.67710328 | 2.91E-06 | 5.73E-05 | SNX8 | 29886 |
| ENSG000001 | 607.837876 | 0.568567129 | 0.12156645 | 4.67700674 | 2.91E-06 | 5.73E-05 | NDUFAF4 | 29078 |
| ENSG00000C | 115.415851 | 0.799944228 | 0.17106424 | 4.67627972 | 2.92E-06 | 5.75E-05 | TMPRSS11E | 28983 |
| ENSG000001 | 1820.96966 | 0.563341021 | 0.12047437 | 4.67602364 | 2.92E-06 | 5.75E-05 | PKP4 | 8502 |
| ENSG000001 | 11425.9282 | 0.514369904 | 0.10999985 | 4.67609627 | 2.92E-06 | 5.75E-05 | RBMX | 27316 |
| ENSG000001 | 1627.26038 | 0.673361875 | 0.14400495 | 4.67596329 | 2.93E-06 | 5.75E-05 | NOC2L | 26155 |
| ENSG000001 | 723.19444 | 0.647685355 | 0.13854725 | 4.67483382 | 2.94E-06 | 5.78E-05 | CD3EAP | 10849 |
| ENSG000001 | 31.5032235 | -1.227104771 | 0.26251539 | -4.6744108 | 2.95E-06 | 5.79E-05 | GRM7 | 2917 |
| ENSG000001 | 2719.0094 | 0.541143832 | 0.11578618 | 4.6736479 | 2.96E-06 | 5.80E-05 | VMA21 | 203547 |
| ENSG00000C | 678.783504 | 0.618563312 | 0.1323622 | 4.67326271 | 2.96E-06 | 5.81E-05 | IP6K2 | 51447 |
| ENSG000001 | 2321.68926 | 0.57861717 | 0.12383144 | 4.67261928 | 2.97E-06 | 5.83E-05 | GLUD1 | 2746 |
| ENSG000001 | 1502.16506 | 0.58238498 | 0.12464763 | 4.67225069 | 2.98E-06 | 5.83E-05 | NUP93 | 9688 |
| ENSG000001 | 169.268314 | 0.760771958 | 0.16282682 | 4.67227679 | 2.98E-06 | 5.83E-05 | ASB6 | 140459 |
| ENSG000001 | 403.842067 | 0.589229279 | 0.12610722 | 4.67244685 | 2.98E-06 | 5.83E-05 | PIEZO2 | 63895 |
| ENSG000001 | 869.959238 | 0.511499591 | 0.10948 | 4.67208264 | 2.98E-06 | 5.83E-05 | USP54 | 159195 |
| ENSG000001 | 429.296994 | 0.673009 | 0.14405568 | 4.67186705 | 2.98E-06 | 5.84E-05 | TMEM9 | 252839 |
| ENSG000001 | 2207.25446 | 0.529198784 | 0.11328521 | 4.67138478 | 2.99E-06 | 5.85E-05 | USP25 | 29761 |
| ENSG000002 | 2485.67132 | 0.580755108 | 0.12434411 | 4.67054776 | 3.00E-06 | 5.87E-05 | HN1L | 90861 |
| ENSG000002 | 528.611635 | 0.579468254 | 0.12407754 | 4.67021063 | 3.01E-06 | 5.87E-05 | LSM2 | 57819 |
| ENSG000001 | 2076.6525 | 0.561937427 | 0.12034457 | 4.66940392 | 3.02E-06 | 5.89E-05 | CTNNAL1 | 8727 |
| ENSG000001 | 3408.73685 | 0.609072774 | 0.13044515 | 4.66918669 | 3.02E-06 | 5.90E-05 | IMPDH2 | 3615 |
| ENSG000001 | 9709.10051 | 0.576724169 | 0.12351713 | 4.66918381 | 3.02E-06 | 5.90E-05 | XRCC6 | 2547 |
| ENSG00000C | 775.200573 | 0.653948508 | 0.14008325 | 4.66828487 | 3.04E-06 | 5.92E-05 | SMARCB1 | 6598 |
| ENSG000001 | 1463.70062 | 0.585165147 | 0.12536088 | 4.6678451 | 3.04E-06 | 5.93E-05 | MDK | 4192 |
| ENSG000001 | 2041.39453 | 0.580257899 | 0.12430985 | 4.66783532 | 3.04E-06 | 5.93E-05 | MCFD2 | 90411 |
| ENSG000001 | 740.421207 | 0.599690569 | 0.12848556 | 4.66737722 | 3.05E-06 | 5.93E-05 | PCCB | 5096 |
| ENSG000001 | 121.715083 | 0.795197631 | 0.17037208 | 4.66741746 | 3.05E-06 | 5.93E-05 | MIF4GD | 57409 |
| ENSG000001 | 1133.87795 | 0.760542615 | 0.16295455 | 4.66720707 | 3.05E-06 | 5.94E-05 | KDM5C | 8242 |

| | | | | | | | | |
|---|---|---|---|---|---|---|---|---|
| ENSG000002 | 1339.53748 | 0.54741335 | 0.11730823 | 4.66645324 | 3.06E-06 | 5.95E-05 | MRPS21 | 54460 |
| ENSG000001 | 351.7998 | 0.624092779 | 0.13374739 | 4.6662054 | 3.07E-06 | 5.96E-05 | SLC25A37 | 51312 |
| ENSG000001 | 849.539578 | 0.660031588 | 0.14146796 | 4.66559055 | 3.08E-06 | 5.97E-05 | NIP7 | 51388 |
| ENSG000001 | 2948.43236 | 0.510076448 | 0.10933263 | 4.66536348 | 3.08E-06 | 5.98E-05 | TMEM123 | 114908 |
| ENSG000001 | 512.443258 | 0.653401076 | 0.14007775 | 4.66456001 | 3.09E-06 | 6.00E-05 | FAM162A | 26355 |
| ENSG000001 | 362.804145 | 0.690724719 | 0.14810597 | 4.66371952 | 3.11E-06 | 6.02E-05 | CHPF | 79586 |
| ENSG000002 | 21.3183388 | -1.320007806 | 0.28304451 | -4.6636051 | 3.11E-06 | 6.02E-05 | ZNF726 | 730087 |
| ENSG000002 | 23.0464178 | -1.340389583 | 0.2874191 | -4.663537 | 3.11E-06 | 6.02E-05 | LOC339529 | 339529 |
| ENSG000001 | 233.173642 | 0.66675081 | 0.1429794 | 4.66326487 | 3.11E-06 | 6.02E-05 | DCUN1D2 | 55208 |
| ENSG000001 | 378.203271 | 0.687292665 | 0.14740709 | 4.66254836 | 3.12E-06 | 6.04E-05 | RAB39B | 116442 |
| ENSG000001 | 48.2556359 | -1.123074975 | 0.24089976 | -4.6620013 | 3.13E-06 | 6.06E-05 | ARMC3 | 219681 |
| ENSG000001 | 43.5086833 | -1.14030706 | 0.24460823 | -4.6617689 | 3.14E-06 | 6.06E-05 | CHL1 | 10752 |
| ENSG000001 | 75.9655355 | 0.886431004 | 0.19018738 | 4.66082986 | 3.15E-06 | 6.08E-05 | MKX | 283078 |
| ENSG000002 | 1052.16841 | 0.522692547 | 0.11215021 | 4.66064716 | 3.15E-06 | 6.09E-05 | CHMP1B | 57132 |
| ENSG000001 | 60.3470971 | -1.085656363 | 0.23295651 | -4.6603393 | 3.16E-06 | 6.09E-05 | NUP210L | 91181 |
| ENSG000001 | 1978.69372 | 0.544924292 | 0.11696201 | 4.65898536 | 3.18E-06 | 6.13E-05 | TYW3 | 127253 |
| ENSG000001 | 205.455997 | 0.816668162 | 0.17529548 | 4.65880901 | 3.18E-06 | 6.13E-05 | NAT9 | 26151 |
| ENSG000001 | 1202.24 | 0.622308552 | 0.13361817 | 4.65736472 | 3.20E-06 | 6.17E-05 | PPP3CA | 5530 |
| ENSG000000 | 1091.6958 | 0.510115158 | 0.10953433 | 4.65712574 | 3.21E-06 | 6.17E-05 | NLK | 51701 |
| ENSG000001 | 48.2881085 | -1.037239628 | 0.22271688 | -4.6572116 | 3.21E-06 | 6.17E-05 | SLC9C2 | 284525 |
| ENSG000001 | 365.232871 | 0.627687769 | 0.13480903 | 4.65612562 | 3.22E-06 | 6.20E-05 | OLFM2 | 93145 |
| ENSG000001 | 6717.92067 | 0.553857045 | 0.11897751 | 4.65514052 | 3.24E-06 | 6.23E-05 | HDLBP | 3069 |
| ENSG000001 | 757.617603 | 0.59041953 | 0.12683192 | 4.65513351 | 3.24E-06 | 6.23E-05 | HARS | 3035 |
| ENSG000001 | 2824.59173 | 0.538206458 | 0.11565347 | 4.65361283 | 3.26E-06 | 6.27E-05 | GRPEL2 | 134266 |
| ENSG000001 | 543.003792 | 0.627321692 | 0.13480768 | 4.65345672 | 3.26E-06 | 6.27E-05 | ARFIP2 | 23647 |
| ENSG000001 | 219.34594 | 0.851070223 | 0.18290828 | 4.65298899 | 3.27E-06 | 6.28E-05 | CUEDC2 | 79004 |
| ENSG000001 | 704.853373 | 0.588666634 | 0.12654368 | 4.65188508 | 3.29E-06 | 6.31E-05 | NOP2 | 4839 |
| ENSG000001 | 113.969844 | 0.750011674 | 0.16129163 | 4.65003457 | 3.32E-06 | 6.37E-05 | SYNE4 | 163183 |
| ENSG000001 | 822.133392 | 0.579994172 | 0.12475305 | 4.64913804 | 3.33E-06 | 6.39E-05 | KIAA0319L | 79932 |

| | | | | | | | | |
|---|---|---|---|---|---|---|---|---|
| ENSG000001 | 492.296656 | 0.615902 | 0.13248059 | 4.64899805 | 3.34E-06 | 6.39E-05 | CDK2 | 1017 |
| ENSG000001 | 623.128863 | 0.560657999 | 0.12060114 | 4.64886155 | 3.34E-06 | 6.39E-05 | SUPT7L | 9913 |
| ENSG000001 | 1211.19866 | 0.599988978 | 0.12907571 | 4.6483494 | 3.35E-06 | 6.40E-05 | PARP4 | 143 |
| ENSG000001 | 119853.489 | 0.655217534 | 0.14095429 | 4.64843983 | 3.34E-06 | 6.40E-05 | EEF1A1 | 1915 |
| ENSG000001 | 1619.1196 | 0.560359083 | 0.12055905 | 4.64800511 | 3.35E-06 | 6.41E-05 | RBM47 | 54502 |
| ENSG000001 | 359.297267 | 0.630188176 | 0.13559031 | 4.64773767 | 3.36E-06 | 6.42E-05 | SMYD4 | 114826 |
| ENSG000001 | 554.438806 | 0.525392352 | 0.11306374 | 4.64686875 | 3.37E-06 | 6.44E-05 | CTAGE5 | 4253 |
| ENSG000001 | 54.1431172 | 0.984458904 | 0.21189121 | 4.64605826 | 3.38E-06 | 6.46E-05 | ONECUT1 | 3175 |
| ENSG000002 | 220.390536 | 0.628914946 | 0.13537835 | 4.64560934 | 3.39E-06 | 6.47E-05 | NA | NA |
| ENSG000001 | 714.212743 | 0.613680953 | 0.13212287 | 4.64477475 | 3.40E-06 | 6.50E-05 | TM2D2 | 83877 |
| ENSG000001 | 130.309174 | 0.807771423 | 0.17395449 | 4.64357895 | 3.42E-06 | 6.53E-05 | ZNF250 | 58500 |
| ENSG000001 | 967.19644 | 0.554840156 | 0.11949273 | 4.6432965 | 3.43E-06 | 6.54E-05 | RPP30 | 10556 |
| ENSG000001 | 948.828199 | 0.531528082 | 0.1144866 | 4.64270989 | 3.44E-06 | 6.55E-05 | ERO1A | 30001 |
| ENSG00000C | 4361.05113 | 0.524820271 | 0.11304835 | 4.6424408 | 3.44E-06 | 6.56E-05 | WDFY1 | 57590 |
| ENSG000001 | 508.914323 | 0.579025822 | 0.12477291 | 4.64063716 | 3.47E-06 | 6.61E-05 | LANCL2 | 55915 |
| ENSG00000C | 363.349382 | 0.701163535 | 0.15110047 | 4.64037948 | 3.48E-06 | 6.61E-05 | EDC4 | 23644 |
| ENSG000001 | 237.757867 | 0.707144363 | 0.15238727 | 4.64044241 | 3.48E-06 | 6.61E-05 | UBL4A | 8266 |
| ENSG000001 | 386.32742 | 0.635594266 | 0.13698342 | 4.63993588 | 3.49E-06 | 6.63E-05 | COA3 | 28958 |
| ENSG000001 | 1004.85757 | 0.521444154 | 0.11238751 | 4.63969838 | 3.49E-06 | 6.63E-05 | LRIG2 | 9860 |
| ENSG00000C | 36750.0003 | 0.487588361 | 0.10509549 | 4.63947925 | 3.49E-06 | 6.63E-05 | HSP90AB1 | 3326 |
| ENSG000001 | 249.812108 | 0.690714165 | 0.14893636 | 4.63764621 | 3.52E-06 | 6.69E-05 | ZNF362 | 149076 |
| ENSG000001 | 6439.91591 | 0.553280908 | 0.11933612 | 4.63632411 | 3.55E-06 | 6.73E-05 | Sep-02 | 4735 |
| ENSG00000C | 235.176876 | 0.649824511 | 0.14017252 | 4.635891 | 3.55E-06 | 6.74E-05 | SCMH1 | 22955 |
| ENSG000001 | 717.348556 | 0.575678029 | 0.12417852 | 4.6358906 | 3.55E-06 | 6.74E-05 | RNF141 | 50862 |
| ENSG000001 | 1092.46732 | 0.633314072 | 0.13661452 | 4.63577408 | 3.56E-06 | 6.74E-05 | TDP2 | 51567 |
| ENSG00000C | 1042.20691 | 0.753749397 | 0.16260798 | 4.63537755 | 3.56E-06 | 6.75E-05 | CAPN1 | 823 |
| ENSG000001 | 1327.637 | 0.603994747 | 0.13032297 | 4.63459933 | 3.58E-06 | 6.77E-05 | CLCN3 | 1182 |
| ENSG000002 | 6801.74623 | 0.636674354 | 0.13738498 | 4.63423558 | 3.58E-06 | 6.78E-05 | HIST1H2AJ | 8331 |
| ENSG000001 | 988.27031 | 0.637121747 | 0.13748475 | 4.63412664 | 3.58E-06 | 6.78E-05 | GRB2 | 2885 |

| | | | | | | | | |
|---|---|---|---|---|---|---|---|---|
| ENSG00000C | 1367.16906 | 0.595567658 | 0.12852335 | 4.63392569 | 3.59E-06 | 6.78E-05 | RRN3 | 54700 |
| ENSG000001 | 5257.29535 | 0.551552169 | 0.11902549 | 4.63389951 | 3.59E-06 | 6.78E-05 | CKAP5 | 9793 |
| ENSG000001 | 4101.97002 | 0.544782628 | 0.11756847 | 4.63374773 | 3.59E-06 | 6.78E-05 | TUBA1A | 7846 |
| ENSG000001 | 240.892392 | 0.599938511 | 0.12948341 | 4.63332337 | 3.60E-06 | 6.79E-05 | CYP2U1 | 113612 |
| ENSG000001 | 298.031176 | 0.643497507 | 0.13891966 | 4.63215591 | 3.62E-06 | 6.82E-05 | RNF170 | 81790 |
| ENSG000001 | 479.090426 | 0.560687 | 0.12103964 | 4.63225942 | 3.62E-06 | 6.82E-05 | RPUSD3 | 285367 |
| ENSG000001 | 2049.51391 | 0.574223868 | 0.12397093 | 4.63192361 | 3.62E-06 | 6.82E-05 | ZBTB33 | 10009 |
| ENSG000001 | 52.9872242 | -1.028851057 | 0.22211532 | -4.6320581 | 3.62E-06 | 6.82E-05 | ERICH3 | 127254 |
| ENSG000001 | 107.645088 | 0.791445619 | 0.17086674 | 4.6319466 | 3.62E-06 | 6.82E-05 | EME2 | 197342 |
| ENSG000002 | 60.0669089 | -0.948620214 | 0.20478386 | -4.6322997 | 3.62E-06 | 6.82E-05 | SLC8A1-AS1 | 100128590 |
| ENSG00000C | 76.8162698 | 0.900669464 | 0.19445427 | 4.6317804 | 3.63E-06 | 6.82E-05 | SIRT6 | 51548 |
| ENSG000001 | 1500.11574 | 0.639236341 | 0.13802955 | 4.63115583 | 3.64E-06 | 6.84E-05 | NT5DC2 | 64943 |
| ENSG00000C | 1232.68928 | 0.651228927 | 0.1406247 | 4.63097127 | 3.64E-06 | 6.84E-05 | UHRF1BP1 | 54887 |
| ENSG000001 | 57.5980526 | 0.980837254 | 0.21185216 | 4.6298195 | 3.66E-06 | 6.88E-05 | ZNHIT2 | 741 |
| ENSG000002 | 775.322706 | 0.579999007 | 0.12527767 | 4.62970798 | 3.66E-06 | 6.88E-05 | SOCS7 | 30837 |
| ENSG000001 | 4642.19812 | 0.826068909 | 0.1784418 | 4.62934635 | 3.67E-06 | 6.89E-05 | RPLP1 | 6176 |
| ENSG000001 | 868.859513 | 0.537338677 | 0.11607871 | 4.62908883 | 3.67E-06 | 6.89E-05 | RNF34 | 80196 |
| ENSG000001 | 2121.21595 | 0.588439706 | 0.12712998 | 4.62864631 | 3.68E-06 | 6.90E-05 | ERH | 2079 |
| ENSG000001 | 27.5478722 | -1.234063916 | 0.26662482 | -4.628466 | 3.68E-06 | 6.91E-05 | MUC13 | 56667 |
| ENSG00000C | 281.896878 | 0.651627683 | 0.14079008 | 4.62836359 | 3.69E-06 | 6.91E-05 | SH3BP2 | 6452 |
| ENSG000001 | 770.06342 | 0.655095852 | 0.14154262 | 4.62825851 | 3.69E-06 | 6.91E-05 | TMEM14C | 51522 |
| ENSG000001 | 244.653885 | 0.665144501 | 0.1437224 | 4.62798067 | 3.69E-06 | 6.91E-05 | VPS4A | 27183 |
| ENSG00000C | 28.2854945 | -1.260393989 | 0.27235801 | -4.6277103 | 3.70E-06 | 6.92E-05 | PDE6C | 5146 |
| ENSG000001 | 3295.00628 | 0.524062132 | 0.11324684 | 4.62760911 | 3.70E-06 | 6.92E-05 | OGT | 8473 |
| ENSG000001 | 118.390467 | 0.852450629 | 0.18421528 | 4.62746965 | 3.70E-06 | 6.92E-05 | WDR24 | 84219 |
| ENSG000001 | 934.809701 | 0.724311548 | 0.15655264 | 4.6266327 | 3.72E-06 | 6.94E-05 | POLDIP3 | 84271 |
| ENSG000001 | 1223.34575 | 0.516746502 | 0.11168906 | 4.62665278 | 3.72E-06 | 6.94E-05 | FAM192A | 80011 |
| ENSG000001 | 190.777536 | 0.693543023 | 0.14990634 | 4.62650879 | 3.72E-06 | 6.94E-05 | DHRS7B | 25979 |
| ENSG000001 | 658.882194 | 0.758327254 | 0.16396887 | 4.62482445 | 3.75E-06 | 7.00E-05 | GTF2F1 | 2962 |

| | | | | | | | | |
|---|---|---|---|---|---|---|---|---|
| ENSG000001 | 2253.36202 | 0.556127766 | 0.12027801 | 4.62368596 | 3.77E-06 | 7.02E-05 | CPD | 1362 |
| ENSG000001 | 32.9436836 | -1.15104656 | 0.24893552 | -4.6238744 | 3.77E-06 | 7.02E-05 | SLC2A5 | 6518 |
| ENSG000001 | 2041.29513 | 0.528612117 | 0.11432597 | 4.62372741 | 3.77E-06 | 7.02E-05 | CEP57 | 9702 |
| ENSG000001 | 532.488965 | 0.621632945 | 0.13444513 | 4.62369243 | 3.77E-06 | 7.02E-05 | CDCA4 | 55038 |
| ENSG00000C | 551.584835 | 0.625674966 | 0.13534627 | 4.62277226 | 3.79E-06 | 7.03E-05 | CWF19L1 | 55280 |
| ENSG00000C | 1899.02896 | 0.581758888 | 0.1258334 | 4.62324701 | 3.78E-06 | 7.03E-05 | PSMD8 | 5714 |
| ENSG000001 | 795.143536 | 0.592261577 | 0.12810913 | 4.62310206 | 3.78E-06 | 7.03E-05 | PANK2 | 80025 |
| ENSG000001 | 440.864938 | 0.809112934 | 0.17502023 | 4.62296805 | 3.78E-06 | 7.03E-05 | UNC45A | 55898 |
| ENSG000001 | 1611.28299 | 0.564991223 | 0.12222157 | 4.6226803 | 3.79E-06 | 7.03E-05 | CDYL | 9425 |
| ENSG000001 | 331.745922 | 0.664598133 | 0.1437568 | 4.62307262 | 3.78E-06 | 7.03E-05 | MEX3D | 399664 |
| ENSG000001 | 5434.23742 | 0.570283563 | 0.12336253 | 4.62282637 | 3.79E-06 | 7.03E-05 | EWSR1 | 2130 |
| ENSG000002 | 149.920584 | 0.867756357 | 0.18771791 | 4.62266142 | 3.79E-06 | 7.03E-05 | NA | NA |
| ENSG000001 | 202.633876 | 0.79312299 | 0.17159751 | 4.62199583 | 3.80E-06 | 7.05E-05 | ECI1 | 1632 |
| ENSG000002 | 8.30079143 | -1.686491563 | 0.36487817 | -4.6220676 | 3.80E-06 | 7.05E-05 | BLACE | 338436 |
| ENSG000001 | 860.934238 | 0.647493283 | 0.14009472 | 4.62182494 | 3.80E-06 | 7.05E-05 | ESYT1 | 23344 |
| ENSG000001 | 683.082741 | 0.614605016 | 0.13300756 | 4.62082767 | 3.82E-06 | 7.08E-05 | CPSF3L | 54973 |
| ENSG000001 | 2064.38347 | 0.786222441 | 0.17015622 | 4.62059197 | 3.83E-06 | 7.08E-05 | CGNL1 | 84952 |
| ENSG000001 | 37.6891315 | -1.187862815 | 0.2570783 | -4.6206266 | 3.83E-06 | 7.08E-05 | PTPRT | 11122 |
| ENSG000001 | 377.612879 | 0.680340334 | 0.14725103 | 4.62027566 | 3.83E-06 | 7.09E-05 | SNX7 | 51375 |
| ENSG00000C | 1521.20986 | 0.56757263 | 0.12285019 | 4.62003873 | 3.84E-06 | 7.09E-05 | SPAG5 | 10615 |
| ENSG000001 | 745.039156 | 0.630514844 | 0.13648427 | 4.61968858 | 3.84E-06 | 7.10E-05 | C4orf3 | 401152 |
| ENSG000002 | 94.0683672 | -0.844017473 | 0.18269752 | -4.6197534 | 3.84E-06 | 7.10E-05 | LINC01206 | 100996490 |
| ENSG000001 | 295.686301 | 0.640855143 | 0.13873368 | 4.61931913 | 3.85E-06 | 7.11E-05 | MICAL1 | 64780 |
| ENSG000001 | 799.889273 | 0.536069426 | 0.11605588 | 4.61906306 | 3.85E-06 | 7.11E-05 | ARFIP1 | 27236 |
| ENSG000002 | 33.864422 | -1.184532389 | 0.25644422 | -4.6190644 | 3.85E-06 | 7.11E-05 | ZFP57 | 346171 |
| ENSG000001 | 486.348533 | 0.543752093 | 0.11772455 | 4.61885052 | 3.86E-06 | 7.12E-05 | RIOK3 | 8780 |
| ENSG000001 | 1019.74736 | 0.553476733 | 0.11985123 | 4.61803134 | 3.87E-06 | 7.14E-05 | VPS4B | 9525 |
| ENSG000002 | 47.2471912 | 1.025199298 | 0.22200312 | 4.61794992 | 3.88E-06 | 7.14E-05 | NA | NA |
| ENSG000001 | 692.965 | 0.598927603 | 0.12970374 | 4.61765859 | 3.88E-06 | 7.15E-05 | BAG2 | 9532 |

| | | | | | | | | |
|---|---|---|---|---|---|---|---|---|
| ENSG000001 | 646.893423 | 0.631399436 | 0.13673726 | 4.61761056 | 3.88E-06 | 7.15E-05 | NUP85 | 79902 |
| ENSG000001 | 1044.63637 | 0.650171117 | 0.14080617 | 4.61749017 | 3.88E-06 | 7.15E-05 | ZDHHC5 | 25921 |
| ENSG000002 | 102.96659 | 0.789154114 | 0.17095339 | 4.61619454 | 3.91E-06 | 7.19E-05 | NA | NA |
| ENSG000001 | 650.414837 | 0.598610043 | 0.12968066 | 4.61603159 | 3.91E-06 | 7.19E-05 | KANSL2 | 54934 |
| ENSG000001 | 524.497937 | 0.743637348 | 0.16110262 | 4.61592346 | 3.91E-06 | 7.19E-05 | BTBD2 | 55643 |
| ENSG000001 | 5406.5289 | 0.53914151 | 0.1168065 | 4.61568082 | 3.92E-06 | 7.19E-05 | SNHG16 | 100507246 |
| ENSG000001 | 1047.16727 | 0.586644577 | 0.12709633 | 4.61574744 | 3.92E-06 | 7.19E-05 | TPM2 | 7169 |
| ENSG000001 | 360.054317 | 0.633000424 | 0.1371488 | 4.61542813 | 3.92E-06 | 7.20E-05 | ALDOC | 230 |
| ENSG000002 | 137.409017 | 0.801776619 | 0.17372132 | 4.6153035 | 3.93E-06 | 7.20E-05 | C17orf89 | 284184 |
| ENSG000001 | 104.372243 | 0.824647982 | 0.17868207 | 4.61516916 | 3.93E-06 | 7.20E-05 | DENND2A | 27147 |
| ENSG000001 | 1080.89903 | 0.575510637 | 0.12471004 | 4.61478999 | 3.93E-06 | 7.21E-05 | NA | NA |
| ENSG000001 | 1144.40647 | 0.590345796 | 0.12793105 | 4.61456216 | 3.94E-06 | 7.21E-05 | TM9SF4 | 9777 |
| ENSG000001 | 190.100379 | -0.679794084 | 0.1473154 | -4.6145486 | 3.94E-06 | 7.21E-05 | TNFRSF10D | 8793 |
| ENSG000001 | 1275.78132 | 0.689208065 | 0.14936265 | 4.61432671 | 3.94E-06 | 7.22E-05 | NPLOC4 | 55666 |
| ENSG000001 | 1194.95437 | 0.590671086 | 0.12801611 | 4.61403712 | 3.95E-06 | 7.22E-05 | EXOSC8 | 11340 |
| ENSG000001 | 468.976317 | 0.582746656 | 0.12631512 | 4.61343544 | 3.96E-06 | 7.23E-05 | ATG4C | 84938 |
| ENSG000001 | 10622.0479 | 0.542550667 | 0.11760058 | 4.61350334 | 3.96E-06 | 7.23E-05 | LARS | 51520 |
| ENSG000001 | 6718.00257 | 0.556848712 | 0.12069938 | 4.61351749 | 3.96E-06 | 7.23E-05 | HNRNPF | 3185 |
| ENSG000001 | 796.296103 | 0.560712819 | 0.12153276 | 4.6136762 | 3.96E-06 | 7.23E-05 | SRPR | 6734 |
| ENSG000001 | 714.78153 | 0.55030603 | 0.11927618 | 4.6137126 | 3.96E-06 | 7.23E-05 | ZNF286A | 57335 |
| ENSG000001 | 704.748106 | 0.642617591 | 0.13930205 | 4.61312367 | 3.97E-06 | 7.24E-05 | SH2B3 | 10019 |
| ENSG000001 | 574.487084 | 0.55645279 | 0.1206293 | 4.61291552 | 3.97E-06 | 7.24E-05 | COX11 | 1353 |
| ENSG000002 | 192.801322 | 0.884438603 | 0.19174108 | 4.61267152 | 3.98E-06 | 7.24E-05 | NA | NA |
| ENSG000001 | 798.463973 | 0.588535487 | 0.12761612 | 4.6117644 | 3.99E-06 | 7.27E-05 | BECN1 | 8678 |
| ENSG000001 | 493.38305 | 0.665692501 | 0.14436076 | 4.61131194 | 4.00E-06 | 7.29E-05 | DVL3 | 1857 |
| ENSG000001 | 4300.92015 | 0.513537824 | 0.11138068 | 4.61065446 | 4.01E-06 | 7.31E-05 | PANK3 | 79646 |
| ENSG000001 | 2818.35321 | 0.53247229 | 0.11549458 | 4.61036608 | 4.02E-06 | 7.31E-05 | NUP98 | 4928 |
| ENSG000001 | 2838.55984 | 0.518143827 | 0.11239452 | 4.61004508 | 4.03E-06 | 7.32E-05 | FADS1 | 3992 |
| ENSG000001 | 517.168353 | 0.821262979 | 0.17818444 | 4.60906111 | 4.04E-06 | 7.35E-05 | KIF18B | 146909 |

| | | | | | | | | |
|---|---|---|---|---|---|---|---|---|
| ENSG000001 | 1906.03766 | 0.552511884 | 0.11988018 | 4.60886784 | 4.05E-06 | 7.36E-05 | ORC3 | 23595 |
| ENSG000002 | 342.860697 | 0.613099405 | 0.13305435 | 4.60788697 | 4.07E-06 | 7.39E-05 | ALG3 | 10195 |
| ENSG000001 | 595.573801 | 0.574660317 | 0.12472133 | 4.6075545 | 4.07E-06 | 7.40E-05 | CISD1 | 55847 |
| ENSG000002 | 1088.66896 | 0.559820769 | 0.12151328 | 4.60707484 | 4.08E-06 | 7.41E-05 | CYFIP1 | 23191 |
| ENSG000001 | 125.204861 | 0.862483683 | 0.1872125 | 4.60697702 | 4.09E-06 | 7.41E-05 | SPSB2 | 84727 |
| ENSG000000 | 1196.96105 | 0.61221902 | 0.13289612 | 4.60674858 | 4.09E-06 | 7.41E-05 | LARS2 | 23395 |
| ENSG000001 | 3967.84653 | 0.547923658 | 0.11893918 | 4.60675501 | 4.09E-06 | 7.41E-05 | BUB1 | 699 |
| ENSG000001 | 1217.63502 | 0.618891517 | 0.13436939 | 4.60589667 | 4.11E-06 | 7.44E-05 | RBM15 | 64783 |
| ENSG000001 | 2375.32711 | 0.575793708 | 0.12501579 | 4.60576779 | 4.11E-06 | 7.44E-05 | RUVBL1 | 8607 |
| ENSG000001 | 100.373092 | -0.791565476 | 0.17187374 | -4.6055056 | 4.11E-06 | 7.45E-05 | CCDC168 | 643677 |
| ENSG000000 | 370.951119 | 0.629875617 | 0.13678262 | 4.60493896 | 4.13E-06 | 7.46E-05 | CAPG | 822 |
| ENSG000001 | 181.074795 | 0.798059826 | 0.17331692 | 4.60462734 | 4.13E-06 | 7.47E-05 | PDDC1 | 347862 |
| ENSG000001 | 1282.85589 | 0.497080177 | 0.10796018 | 4.604292 | 4.14E-06 | 7.48E-05 | ZNF664 | 144348 |
| ENSG000001 | 529.905148 | 0.564236541 | 0.12254935 | 4.60415758 | 4.14E-06 | 7.48E-05 | CLCC1 | 23155 |
| ENSG000001 | 814.199867 | 0.618254134 | 0.13429835 | 4.60358691 | 4.15E-06 | 7.50E-05 | COX5A | 9377 |
| ENSG000001 | 747.045898 | 0.616353151 | 0.13388847 | 4.60348183 | 4.15E-06 | 7.50E-05 | DCTN2 | 10540 |
| ENSG000001 | 462.349152 | 0.562765844 | 0.12225868 | 4.60307495 | 4.16E-06 | 7.51E-05 | PPCS | 79717 |
| ENSG000001 | 1555.18526 | 0.64538656 | 0.14021339 | 4.60288833 | 4.17E-06 | 7.51E-05 | FUBP3 | 8939 |
| ENSG000001 | 539.792207 | 0.661583062 | 0.1437333 | 4.60285171 | 4.17E-06 | 7.51E-05 | SMO | 6608 |
| ENSG000001 | 554.787397 | 0.635064306 | 0.13799226 | 4.60217345 | 4.18E-06 | 7.53E-05 | L3MBTL2 | 83746 |
| ENSG000002 | 3099.32848 | 0.49920107 | 0.10849225 | 4.60126032 | 4.20E-06 | 7.56E-05 | NA | NA |
| ENSG000000 | 1480.21143 | 0.498353785 | 0.10832394 | 4.60058773 | 4.21E-06 | 7.58E-05 | SPAG9 | 9043 |
| ENSG000001 | 589.007626 | 0.669447572 | 0.14551086 | 4.60067083 | 4.21E-06 | 7.58E-05 | IMPA2 | 3613 |
| ENSG000001 | 1446.50021 | 0.681997419 | 0.14825137 | 4.60027728 | 4.22E-06 | 7.59E-05 | CCAR2 | 57805 |
| ENSG000002 | 59.7872449 | -0.922144582 | 0.20045891 | -4.6001675 | 4.22E-06 | 7.59E-05 | A2MP1 | 3 |
| ENSG000001 | 401.178845 | 0.784333578 | 0.1705168 | 4.59974362 | 4.23E-06 | 7.60E-05 | NELFB | 25920 |
| ENSG000000 | 2965.57948 | 0.691496935 | 0.15034345 | 4.59944835 | 4.24E-06 | 7.60E-05 | AGO1 | 26523 |
| ENSG000001 | 278.769618 | 0.705079199 | 0.15329907 | 4.59937028 | 4.24E-06 | 7.60E-05 | NT5C | 30833 |
| ENSG000001 | 4071.27647 | 0.534248041 | 0.11615556 | 4.59941874 | 4.24E-06 | 7.60E-05 | NLGN4X | 57502 |

| | | | | | | | | |
|---|---|---|---|---|---|---|---|---|
| ENSG000002 | 190.442037 | 0.747566019 | 0.16255836 | 4.59875472 | 4.25E-06 | 7.62E-05 | UQCC3 | 790955 |
| ENSG000001 | 311.848383 | 0.705802064 | 0.15349045 | 4.59834503 | 4.26E-06 | 7.63E-05 | CAPN15 | 6650 |
| ENSG000001 | 1083.83213 | 0.64074392 | 0.13935098 | 4.59805838 | 4.26E-06 | 7.64E-05 | RND2 | 8153 |
| ENSG000001 | 994.430912 | 0.574043795 | 0.12485161 | 4.59780868 | 4.27E-06 | 7.65E-05 | ARL8B | 55207 |
| ENSG000001 | 13317.7233 | 0.605525617 | 0.13170736 | 4.59750765 | 4.28E-06 | 7.66E-05 | LDHB | 3945 |
| ENSG000001 | 462.804573 | 0.781376815 | 0.16996949 | 4.59715914 | 4.28E-06 | 7.66E-05 | TSR2 | 90121 |
| ENSG000001 | 5271.40449 | 0.561829191 | 0.12221071 | 4.59721713 | 4.28E-06 | 7.66E-05 | AKIRIN1 | 79647 |
| ENSG000001 | 150.49315 | 0.70646704 | 0.15368027 | 4.59699237 | 4.29E-06 | 7.66E-05 | STAG3L4 | 64940 |
| ENSG000001 | 5567.21335 | 0.626121481 | 0.13621212 | 4.59666495 | 4.29E-06 | 7.67E-05 | USP22 | 23326 |
| ENSG000001 | 2269.4085 | 0.609258813 | 0.13255631 | 4.59622656 | 4.30E-06 | 7.69E-05 | MTOR | 2475 |
| ENSG000001 | 41.6992191 | -1.043937795 | 0.22713389 | -4.5961341 | 4.30E-06 | 7.69E-05 | SDR42E1 | 93517 |
| ENSG000001 | 127.892432 | 0.895412948 | 0.1948258 | 4.59596694 | 4.31E-06 | 7.69E-05 | RNU5E-1 | 26829 |
| ENSG00000C | 312.930371 | 0.674125094 | 0.14669384 | 4.59545605 | 4.32E-06 | 7.70E-05 | NDUFB2 | 4708 |
| ENSG000002 | 148.863205 | 0.749478124 | 0.16309623 | 4.59531249 | 4.32E-06 | 7.71E-05 | NA | NA |
| ENSG000001 | 1135.38674 | 0.562132252 | 0.12234314 | 4.59471812 | 4.33E-06 | 7.72E-05 | PTDSS1 | 9791 |
| ENSG000001 | 273.427928 | 0.641446444 | 0.13962786 | 4.59397187 | 4.35E-06 | 7.75E-05 | MRPL36 | 64979 |
| ENSG000001 | 1190.5376 | 0.533761893 | 0.11619994 | 4.59347809 | 4.36E-06 | 7.76E-05 | SLC30A6 | 55676 |
| ENSG000001 | 71.5141861 | -0.886998202 | 0.19310372 | -4.5933771 | 4.36E-06 | 7.76E-05 | ST18 | 9705 |
| ENSG000001 | 264.561846 | 0.667198967 | 0.14528891 | 4.59222241 | 4.39E-06 | 7.80E-05 | GSS | 2937 |
| ENSG000001 | 1042.19637 | 0.598761103 | 0.13039496 | 4.59190376 | 4.39E-06 | 7.81E-05 | CBR4 | 84869 |
| ENSG00000C | 78.3367696 | 0.900493847 | 0.19613587 | 4.59117359 | 4.41E-06 | 7.83E-05 | FAM3A | 60343 |
| ENSG00000C | 882.611421 | 0.557124598 | 0.12134604 | 4.59120541 | 4.41E-06 | 7.83E-05 | CD200 | 4345 |
| ENSG000001 | 745.127029 | 0.548309197 | 0.1194244 | 4.59126612 | 4.41E-06 | 7.83E-05 | FAM118B | 79607 |
| ENSG00000C | 175.436901 | 0.748564866 | 0.16306429 | 4.59061177 | 4.42E-06 | 7.85E-05 | EDEM2 | 55741 |
| ENSG000002 | 567.22377 | 0.714012924 | 0.15554052 | 4.59052675 | 4.42E-06 | 7.85E-05 | REPIN1 | 29803 |
| ENSG000001 | 1604.14647 | 0.50615057 | 0.110262 | 4.5904353 | 4.42E-06 | 7.85E-05 | MRPL37 | 51253 |
| ENSG000001 | 18.3459578 | -1.396494633 | 0.30425958 | -4.5898132 | 4.44E-06 | 7.86E-05 | CNTN6 | 27255 |
| ENSG000001 | 838.01664 | 0.555620599 | 0.12104972 | 4.59001983 | 4.43E-06 | 7.86E-05 | NA | NA |
| ENSG000001 | 839.683691 | 0.603900495 | 0.13157266 | 4.58986318 | 4.44E-06 | 7.86E-05 | GLOD4 | 51031 |

| | | | | | | | | | |
|---|---|---|---|---|---|---|---|---|---|
| ENSG000002 | 287.934939 | 0.731345949 | 0.15934058 | 4.58982856 | 4.44E-06 | 7.86E-05 | CARKD | | 55739 |
| ENSG000001 | 559.347418 | 0.781749675 | 0.17034472 | 4.58922156 | 4.45E-06 | 7.87E-05 | RCOR2 | | 283248 |
| ENSG000001 | 12183.1169 | 0.60322886 | 0.13146725 | 4.58843438 | 4.47E-06 | 7.90E-05 | RPL7 | | 6129 |
| ENSG000001 | 894.972287 | 0.567696523 | 0.12372771 | 4.5882732 | 4.47E-06 | 7.90E-05 | PAPSS1 | | 9061 |
| ENSG000001 | 489.094175 | 0.566317631 | 0.12344645 | 4.58755696 | 4.48E-06 | 7.93E-05 | TAF12 | | 6883 |
| ENSG000001 | 734.344446 | 0.618491688 | 0.13483302 | 4.58709356 | 4.49E-06 | 7.94E-05 | METTL3 | | 56339 |
| ENSG000001 | 804.861495 | 0.615954073 | 0.13428248 | 4.5870026 | 4.50E-06 | 7.94E-05 | MRPS23 | | 51649 |
| ENSG000002 | 710.652112 | 0.529880324 | 0.11551915 | 4.58694788 | 4.50E-06 | 7.94E-05 | PCMTD2 | | 55251 |
| ENSG000001 | 439.139821 | 0.76212309 | 0.16618734 | 4.58592738 | 4.52E-06 | 7.97E-05 | ARFGAP1 | | 55738 |
| ENSG000001 | 1159.32129 | 0.633944391 | 0.13823613 | 4.58595293 | 4.52E-06 | 7.97E-05 | DLST | | 1743 |
| ENSG000001 | 926.531301 | 0.625387749 | 0.13636841 | 4.58601637 | 4.52E-06 | 7.97E-05 | NIF3L1 | | 60491 |
| ENSG000001 | 411.740512 | 0.726119235 | 0.15840361 | 4.58398151 | 4.56E-06 | 8.04E-05 | RNF181 | | 51255 |
| ENSG000001 | 430.47126 | 0.697352414 | 0.15218839 | 4.58216571 | 4.60E-06 | 8.11E-05 | CORO2A | | 7464 |
| ENSG000001 | 970.048612 | 0.581597611 | 0.12693516 | 4.58184787 | 4.61E-06 | 8.12E-05 | TMEM248 | | 55069 |
| ENSG000001 | 743.606298 | 0.540350534 | 0.11793843 | 4.58163242 | 4.61E-06 | 8.12E-05 | GSTCD | | 79807 |
| ENSG000002 | 2119.28334 | 0.551222812 | 0.12031339 | 4.5815583 | 4.62E-06 | 8.12E-05 | TMA7 | | 51372 |
| ENSG000001 | 10072.0812 | 0.554945544 | 0.12113729 | 4.58112893 | 4.62E-06 | 8.13E-05 | RPS25 | | 6230 |
| ENSG000001 | 1840.47046 | 0.580787575 | 0.12678739 | 4.58079932 | 4.63E-06 | 8.14E-05 | CD164 | | 8763 |
| ENSG000001 | 4627.24194 | 0.551686431 | 0.12043508 | 4.58077853 | 4.63E-06 | 8.14E-05 | FEM1B | | 10116 |
| ENSG000001 | 920.787172 | 0.495405606 | 0.10815568 | 4.58048606 | 4.64E-06 | 8.14E-05 | VTI1B | | 10490 |
| ENSG000001 | 905.887493 | 0.581915705 | 0.12704156 | 4.58051435 | 4.64E-06 | 8.14E-05 | ZYG11B | | 79699 |
| ENSG000002 | 260.911881 | 0.811241779 | 0.17714059 | 4.57964937 | 4.66E-06 | 8.17E-05 | NA | NA | |
| ENSG000001 | 902.800476 | 0.604857452 | 0.13208697 | 4.57923629 | 4.67E-06 | 8.18E-05 | SLC31A1 | | 1317 |
| ENSG000001 | 879.117272 | 0.684697052 | 0.14952188 | 4.57924325 | 4.67E-06 | 8.18E-05 | SH3GL1 | | 6455 |
| ENSG000001 | 7390.19455 | 0.635177796 | 0.13872145 | 4.57880009 | 4.68E-06 | 8.20E-05 | SPTBN1 | | 6711 |
| ENSG000001 | 3546.34972 | 0.556592358 | 0.12156686 | 4.57848742 | 4.68E-06 | 8.20E-05 | CELF1 | | 10658 |
| ENSG00000C | 314.861221 | 0.559877769 | 0.12228731 | 4.57837991 | 4.69E-06 | 8.20E-05 | MKRN2 | | 23609 |
| ENSG000001 | 457.892436 | 0.706558582 | 0.1543299 | 4.57823534 | 4.69E-06 | 8.21E-05 | MGAT1 | | 4245 |
| ENSG000001 | 24.9888132 | -1.31274558 | 0.28674943 | -4.5780234 | 4.69E-06 | 8.21E-05 | NXF4 | | 55999 |

| | | | | | | | | |
|---|---|---|---|---|---|---|---|---|
| ENSG000001 | 486.600366 | 0.632395373 | 0.13814382 | 4.57780424 | 4.70E-06 | 8.22E-05 | TEAD4 | 7004 |
| ENSG000001 | 583.145429 | 0.615604858 | 0.1344993 | 4.57701164 | 4.72E-06 | 8.24E-05 | TMEM43 | 79188 |
| ENSG00000C | 1248.08133 | 0.536386654 | 0.11719377 | 4.57692116 | 4.72E-06 | 8.24E-05 | LSG1 | 55341 |
| ENSG000001 | 1083.76267 | 0.542162023 | 0.11846365 | 4.57661095 | 4.73E-06 | 8.25E-05 | BNIP3L | 665 |
| ENSG000001 | 613.022849 | 0.671133343 | 0.14664144 | 4.57669626 | 4.72E-06 | 8.25E-05 | ERAL1 | 26284 |
| ENSG000001 | 1706.25135 | 0.6190136 | 0.13526084 | 4.57644342 | 4.73E-06 | 8.25E-05 | SNRNP70 | 6625 |
| ENSG000001 | 824.094213 | 0.584391143 | 0.12771087 | 4.575892 | 4.74E-06 | 8.26E-05 | MFAP2 | 4237 |
| ENSG000001 | 478.661304 | 0.647016026 | 0.14139163 | 4.57605619 | 4.74E-06 | 8.26E-05 | STRIP1 | 85369 |
| ENSG000001 | 422.571906 | 0.779094612 | 0.17026103 | 4.57588346 | 4.74E-06 | 8.26E-05 | TBC1D9B | 23061 |
| ENSG000002 | 37.0798796 | -1.092755456 | 0.23880017 | -4.5760245 | 4.74E-06 | 8.26E-05 | NA | NA |
| ENSG00000C | 1024.92451 | 0.595383919 | 0.13012823 | 4.57536316 | 4.75E-06 | 8.28E-05 | LRRC40 | 55631 |
| ENSG000001 | 309.112606 | 0.710376344 | 0.15526899 | 4.57513331 | 4.76E-06 | 8.28E-05 | MRPL14 | 64928 |
| ENSG000001 | 648.718795 | 0.641130194 | 0.14013826 | 4.57498314 | 4.76E-06 | 8.29E-05 | LAMTOR1 | 55004 |
| ENSG00000C | 3529.72271 | 0.511373657 | 0.11179499 | 4.57420912 | 4.78E-06 | 8.31E-05 | TXLNG | 55787 |
| ENSG000001 | 3208.33645 | 0.593278381 | 0.1297192 | 4.57355888 | 4.80E-06 | 8.33E-05 | CCNB1 | 891 |
| ENSG000001 | 599.418286 | 0.597125574 | 0.13059279 | 4.57242373 | 4.82E-06 | 8.38E-05 | USP45 | 85015 |
| ENSG000001 | 574.900337 | 0.683053945 | 0.14940449 | 4.57184355 | 4.83E-06 | 8.40E-05 | PMPCA | 23203 |
| ENSG00000C | 303.877359 | 0.730106579 | 0.1597151 | 4.57130601 | 4.85E-06 | 8.41E-05 | ITM2A | 9452 |
| ENSG000001 | 246.899511 | 0.612334641 | 0.1339508 | 4.57133994 | 4.85E-06 | 8.41E-05 | LARP6 | 55323 |
| ENSG000001 | 800.229984 | 0.516611566 | 0.11301489 | 4.57118158 | 4.85E-06 | 8.41E-05 | LIN54 | 132660 |
| ENSG00000C | 1244.94945 | 0.507547464 | 0.11103477 | 4.57106789 | 4.85E-06 | 8.41E-05 | UBE3C | 9690 |
| ENSG000001 | 3974.31777 | 0.559341355 | 0.12236875 | 4.57094951 | 4.86E-06 | 8.41E-05 | YME1L1 | 10730 |
| ENSG000002 | 653.240896 | 0.630874411 | 0.13802439 | 4.57074582 | 4.86E-06 | 8.42E-05 | MRPS18B | 28973 |
| ENSG000001 | 448.657787 | 0.800652911 | 0.17518412 | 4.57035088 | 4.87E-06 | 8.43E-05 | CNP | 1267 |
| ENSG000001 | 531.265796 | 0.765864558 | 0.16757761 | 4.57020804 | 4.87E-06 | 8.43E-05 | ALPK3 | 57538 |
| ENSG000002 | 59.4651512 | 0.943102162 | 0.20636914 | 4.56997661 | 4.88E-06 | 8.44E-05 | NA | NA |
| ENSG000001 | 3970.17709 | 0.528031594 | 0.11557162 | 4.56886896 | 4.90E-06 | 8.48E-05 | CCNA2 | 890 |
| ENSG000001 | 307.458382 | 0.592430541 | 0.12966732 | 4.5688502 | 4.90E-06 | 8.48E-05 | AKTIP | 64400 |
| ENSG000001 | 1481.34764 | 0.574491747 | 0.12574361 | 4.56875488 | 4.91E-06 | 8.48E-05 | H2AFY | 9555 |

| | | | | | | | | |
|---|---|---|---|---|---|---|---|---|
| ENSG000001 | 1786.67982 | 0.54359064 | 0.11900066 | 4.5679632 | 4.92E-06 | 8.50E-05 | DARS | 1615 |
| ENSG000001 | 1797.14618 | 0.598015899 | 0.13091478 | 4.56797845 | 4.92E-06 | 8.50E-05 | CDK1 | 983 |
| ENSG000001 | 9766.89618 | 0.596741338 | 0.13063174 | 4.56811917 | 4.92E-06 | 8.50E-05 | RPS27 | 6232 |
| ENSG000001 | 310.763037 | 0.617832852 | 0.13526409 | 4.5676043 | 4.93E-06 | 8.51E-05 | TRMT61B | 55006 |
| ENSG000001 | 390.82911 | 0.619530788 | 0.13564653 | 4.56724399 | 4.94E-06 | 8.51E-05 | TMEM258 | 746 |
| ENSG000001 | 2223.06266 | 0.565329691 | 0.12377633 | 4.56734908 | 4.94E-06 | 8.51E-05 | EPB41 | 2035 |
| ENSG000001 | 174.602928 | 0.674048526 | 0.14758126 | 4.56730423 | 4.94E-06 | 8.51E-05 | SVBP | 374969 |
| ENSG00000C | 139.081926 | 0.727121849 | 0.15923573 | 4.56632341 | 4.96E-06 | 8.55E-05 | C19orf60 | 55049 |
| ENSG000001 | 799.101851 | 0.561770696 | 0.12303026 | 4.56611813 | 4.97E-06 | 8.55E-05 | SNX18 | 112574 |
| ENSG000002 | 2712.04886 | 0.591643709 | 0.12957317 | 4.56609752 | 4.97E-06 | 8.55E-05 | HIST1H2AL | 8332 |
| ENSG000001 | 3327.86504 | 0.548351767 | 0.12009586 | 4.56595058 | 4.97E-06 | 8.55E-05 | FLT1 | 2321 |
| ENSG000001 | 544.617061 | 0.623387787 | 0.1365407 | 4.56558224 | 4.98E-06 | 8.56E-05 | SMG8 | 55181 |
| ENSG000002 | 1095.47365 | 0.511544223 | 0.112044 | 4.56556569 | 4.98E-06 | 8.56E-05 | NA | NA |
| ENSG000001 | 1087.43245 | 0.56419428 | 0.12358141 | 4.56536527 | 4.99E-06 | 8.56E-05 | COMMD2 | 51122 |
| ENSG000001 | 1295.23489 | 0.757737265 | 0.16599696 | 4.56476593 | 5.00E-06 | 8.58E-05 | GPRC5B | 51704 |
| ENSG000001 | 181.26792 | 0.730912257 | 0.16016653 | 4.56345184 | 5.03E-06 | 8.63E-05 | PKMYT1 | 9088 |
| ENSG000001 | 95.2986851 | -0.878546968 | 0.19252064 | -4.563391 | 5.03E-06 | 8.63E-05 | PCDH15 | 65217 |
| ENSG000001 | 592.483985 | 0.829783496 | 0.18187725 | 4.56232714 | 5.06E-06 | 8.67E-05 | NA | NA |
| ENSG000001 | 171.741347 | 0.65591887 | 0.14377915 | 4.56198891 | 5.07E-06 | 8.68E-05 | TMEM243 | 79161 |
| ENSG000001 | 473.756461 | 0.569342096 | 0.12480105 | 4.5619976 | 5.07E-06 | 8.68E-05 | PRKACB | 5567 |
| ENSG000002 | 1010.44474 | 0.61312336 | 0.13442045 | 4.56123575 | 5.09E-06 | 8.71E-05 | TXNIP | 10628 |
| ENSG000001 | 38.9468315 | -1.045584056 | 0.22926386 | -4.5606143 | 5.10E-06 | 8.73E-05 | NA | NA |
| ENSG000002 | 312.163434 | 0.628792687 | 0.13787402 | 4.56063204 | 5.10E-06 | 8.73E-05 | SNHG15 | 285958 |
| ENSG000001 | 4335.11189 | 0.557042128 | 0.12215252 | 4.56021818 | 5.11E-06 | 8.73E-05 | UBA2 | 10054 |
| ENSG000001 | 141.638375 | 0.909437204 | 0.19942232 | 4.56035815 | 5.11E-06 | 8.73E-05 | FAM234A | 83986 |
| ENSG000002 | 86.1410679 | 0.826305475 | 0.18119871 | 4.56021727 | 5.11E-06 | 8.73E-05 | ATP2A1-AS1 | 100289092 |
| ENSG000001 | 208.47415 | 0.716900196 | 0.15724794 | 4.55904357 | 5.14E-06 | 8.78E-05 | LAMTOR2 | 28956 |
| ENSG000001 | 867.228261 | 0.578608986 | 0.12692047 | 4.5588312 | 5.14E-06 | 8.78E-05 | PERP | 64065 |
| ENSG00000C | 495.711306 | 0.646799697 | 0.14191261 | 4.55773248 | 5.17E-06 | 8.83E-05 | ACADVL | 37 |

| | | | | | | | | | |
|---|---|---|---|---|---|---|---|---|---|
| ENSG000001 | 331.834949 | 0.706509087 | 0.15502236 | 4.55746559 | 5.18E-06 | 8.83E-05 | EDEM1 | | 9695 |
| ENSG000001 | 453.722633 | 0.541707772 | 0.11889188 | 4.55630573 | 5.21E-06 | 8.88E-05 | TSEN2 | | 80746 |
| ENSG000001 | 614.242786 | 0.628201638 | 0.13787858 | 4.55619446 | 5.21E-06 | 8.88E-05 | HADHB | | 3032 |
| ENSG000001 | 825.557345 | 0.577654058 | 0.12680024 | 4.55562272 | 5.22E-06 | 8.90E-05 | FYTTD1 | | 84248 |
| ENSG000001 | 431.528825 | 0.711188611 | 0.15613358 | 4.55500096 | 5.24E-06 | 8.92E-05 | TBC1D8B | | 54885 |
| ENSG000001 | 480.049622 | 0.542362954 | 0.11908029 | 4.55459869 | 5.25E-06 | 8.94E-05 | GNAI1 | | 2770 |
| ENSG000001 | 1298.31312 | 0.515004021 | 0.1130826 | 4.55422854 | 5.26E-06 | 8.95E-05 | ZIC5 | | 85416 |
| ENSG000001 | 1682.92943 | 0.593723323 | 0.130383 | 4.5536867 | 5.27E-06 | 8.97E-05 | RNF168 | | 165918 |
| ENSG00000C | 1283.12738 | 0.553101361 | 0.12146772 | 4.55348444 | 5.28E-06 | 8.97E-05 | NDC80 | | 10403 |
| ENSG000001 | 2338.15705 | 0.619970537 | 0.13617714 | 4.55267697 | 5.30E-06 | 9.00E-05 | PDCD11 | | 22984 |
| ENSG000001 | 191.209792 | 0.776710005 | 0.17065603 | 4.55131871 | 5.33E-06 | 9.06E-05 | GRTP1 | | 79774 |
| ENSG000002 | 23.0703072 | -1.281605888 | 0.28162181 | -4.5508048 | 5.34E-06 | 9.07E-05 | NA | NA | |
| ENSG00000C | 1933.50879 | 0.654083544 | 0.14375091 | 4.55011764 | 5.36E-06 | 9.10E-05 | FSCN1 | | 6624 |
| ENSG000001 | 378.773269 | 0.633834817 | 0.13929943 | 4.55016074 | 5.36E-06 | 9.10E-05 | FAM134A | | 79137 |
| ENSG000002 | 2941.8741 | 0.573925594 | 0.12617972 | 4.54847717 | 5.40E-06 | 9.16E-05 | NA | NA | |
| ENSG00000C | 15013.5019 | 0.499687065 | 0.10986378 | 4.54824222 | 5.41E-06 | 9.17E-05 | YBX1 | | 4904 |
| ENSG000001 | 293.382755 | 0.693502966 | 0.1524743 | 4.54832698 | 5.41E-06 | 9.17E-05 | NELFA | | 7469 |
| ENSG000001 | 446.889044 | 0.622856317 | 0.13697678 | 4.54716732 | 5.44E-06 | 9.19E-05 | RASGRF2 | | 5924 |
| ENSG000001 | 268.758388 | 0.642152959 | 0.14121807 | 4.54724345 | 5.44E-06 | 9.19E-05 | UNC50 | | 25972 |
| ENSG000001 | 196.12447 | 0.686232426 | 0.15090335 | 4.54749619 | 5.43E-06 | 9.19E-05 | IFT22 | | 64792 |
| ENSG000001 | 1691.33164 | 0.635062245 | 0.13965386 | 4.54740207 | 5.43E-06 | 9.19E-05 | WDR82 | | 80335 |
| ENSG000001 | 303.935366 | 0.631883439 | 0.13896278 | 4.54714169 | 5.44E-06 | 9.19E-05 | YIF1A | | 10897 |
| ENSG000002 | 649.035414 | 0.720389525 | 0.15841752 | 4.54741083 | 5.43E-06 | 9.19E-05 | SALL3 | | 27164 |
| ENSG000001 | 401.826177 | 0.59836372 | 0.13160178 | 4.54677534 | 5.45E-06 | 9.20E-05 | NUPL2 | | 11097 |
| ENSG000001 | 540.231436 | 0.634989254 | 0.13966118 | 4.54664093 | 5.45E-06 | 9.21E-05 | TOX | | 9760 |
| ENSG00000C | 995.975221 | 0.664662291 | 0.14620005 | 4.54625211 | 5.46E-06 | 9.22E-05 | GYG2 | | 8908 |
| ENSG000001 | 453.086087 | 0.72879402 | 0.16035486 | 4.54488272 | 5.50E-06 | 9.28E-05 | CBX6 | | 23466 |
| ENSG00000C | 1235.41945 | 0.547937956 | 0.12057741 | 4.54428379 | 5.51E-06 | 9.29E-05 | CDC14B | | 8555 |
| ENSG000002 | 441.768425 | 0.689775514 | 0.15178825 | 4.5443276 | 5.51E-06 | 9.29E-05 | WBSCR16 | | 81554 |

| | | | | | | | | | |
|---|---|---|---|---|---|---|---|---|---|
| ENSG000001 | 11651.3992 | 0.579362477 | 0.12750032 | 4.54400819 | 5.52E-06 | 9.30E-05 | SLC38A2 | | 54407 |
| ENSG000001 | 54.3148186 | -0.991800198 | 0.21831185 | -4.5430433 | 5.54E-06 | 9.34E-05 | TNR | | 7143 |
| ENSG000002 | 1322.6767 | 0.571283822 | 0.1257766 | 4.54205184 | 5.57E-06 | 9.38E-05 | DHFR | | 1719 |
| ENSG000001 | 355.052407 | 0.655208607 | 0.14427063 | 4.54152448 | 5.58E-06 | 9.40E-05 | NSMF | | 26012 |
| ENSG000001 | 576.683453 | 0.598474724 | 0.13177869 | 4.54151381 | 5.59E-06 | 9.40E-05 | NDUFB1 | | 4707 |
| ENSG00000C | 217.638398 | 0.714345881 | 0.15730746 | 4.54108088 | 5.60E-06 | 9.41E-05 | EXOSC5 | | 56915 |
| ENSG00000C | 4767.11518 | 0.540262541 | 0.11898731 | 4.54050554 | 5.61E-06 | 9.43E-05 | C14orf166 | | 51637 |
| ENSG000001 | 49.85807 | -1.022188058 | 0.2251214 | -4.5406081 | 5.61E-06 | 9.43E-05 | ALDH1A2 | | 8854 |
| ENSG000001 | 386.124921 | 0.683753523 | 0.15059232 | 4.54042752 | 5.61E-06 | 9.43E-05 | CCDC137 | | 339230 |
| ENSG000002 | 279.05216 | 0.616948066 | 0.13587744 | 4.54047464 | 5.61E-06 | 9.43E-05 | SMIM13 | | 221710 |
| ENSG000001 | 77.2778759 | -0.953067482 | 0.20991744 | -4.5402015 | 5.62E-06 | 9.43E-05 | NTRK2 | | 4915 |
| ENSG000001 | 269.385091 | 0.618385002 | 0.13621982 | 4.53961118 | 5.64E-06 | 9.45E-05 | ACD | | 65057 |
| ENSG000001 | 1128.1512 | 0.660202302 | 0.14543318 | 4.53955768 | 5.64E-06 | 9.45E-05 | BCR | | 613 |
| ENSG00000C | 1465.64279 | 0.565149622 | 0.12450204 | 4.53928013 | 5.64E-06 | 9.46E-05 | SMARCD1 | | 6602 |
| ENSG000001 | 560.545906 | 0.62181664 | 0.13698451 | 4.53932067 | 5.64E-06 | 9.46E-05 | AKT1 | | 207 |
| ENSG000002 | 139.980469 | 0.807021178 | 0.17781042 | 4.53866082 | 5.66E-06 | 9.48E-05 | PSMB8 | | 5696 |
| ENSG000001 | 963.276227 | 0.656242471 | 0.14460518 | 4.53816702 | 5.67E-06 | 9.50E-05 | YTHDF1 | | 54915 |
| ENSG000002 | 648.946649 | 0.701698065 | 0.1546395 | 4.53763793 | 5.69E-06 | 9.52E-05 | WDR46 | | 9277 |
| ENSG000001 | 113.078014 | -0.834165579 | 0.18383901 | -4.5374787 | 5.69E-06 | 9.52E-05 | DNAH8 | | 1769 |
| ENSG000001 | 1417.37465 | 0.540717652 | 0.11917531 | 4.53716184 | 5.70E-06 | 9.53E-05 | NA | NA | |
| ENSG00000C | 481.448393 | 0.734715108 | 0.16198811 | 4.53561127 | 5.74E-06 | 9.60E-05 | CPSF1 | | 29894 |
| ENSG000001 | 508.590762 | 0.555694694 | 0.12252193 | 4.53547123 | 5.75E-06 | 9.60E-05 | BET1 | | 10282 |
| ENSG000001 | 1696.53079 | 0.541271147 | 0.11934728 | 4.5352616 | 5.75E-06 | 9.61E-05 | EFR3A | | 23167 |
| ENSG000001 | 390.363299 | 0.667023449 | 0.14714973 | 4.53295725 | 5.82E-06 | 9.71E-05 | ATP6V1F | | 9296 |
| ENSG000001 | 2158.7527 | 0.543456086 | 0.11990416 | 4.53242047 | 5.83E-06 | 9.73E-05 | ARPC2 | | 10109 |
| ENSG000001 | 339.488044 | 0.615295138 | 0.13575883 | 4.53226611 | 5.84E-06 | 9.73E-05 | UNC119 | | 9094 |
| ENSG000001 | 40.088618 | -1.111293139 | 0.2452063 | -4.5320742 | 5.84E-06 | 9.74E-05 | MROH9 | | 80133 |
| ENSG000002 | 2409.48311 | 0.633487441 | 0.13978257 | 4.53194876 | 5.84E-06 | 9.74E-05 | SNHG5 | | 387066 |
| ENSG000002 | 20.5128109 | -1.292582574 | 0.28521131 | -4.5320173 | 5.84E-06 | 9.74E-05 | GNG12-AS1 | | 100289178 |

| | | | | | | | | |
|---|---|---|---|---|---|---|---|---|
| ENSG000002 | 304.866137 | 0.65802047 | 0.14519941 | 4.53183971 | 5.85E-06 | 9.74E-05 | SMIM20 | 389203 |
| ENSG000001 | 35.1437107 | -1.11763217 | 0.24663122 | -4.5315925 | 5.85E-06 | 9.74E-05 | LIF | 3976 |
| ENSG000002 | 480.693928 | 0.542513766 | 0.11971857 | 4.53157567 | 5.85E-06 | 9.74E-05 | USP51 | 158880 |
| ENSG000001 | 113.448807 | 0.849259706 | 0.18742301 | 4.5312457 | 5.86E-06 | 9.75E-05 | C1orf122 | 127687 |
| ENSG000001 | 2645.00249 | 0.618388259 | 0.1364764 | 4.53110032 | 5.87E-06 | 9.76E-05 | RRP1B | 23076 |
| ENSG000001 | 380.58188 | 0.667258096 | 0.14729431 | 4.53010102 | 5.90E-06 | 9.80E-05 | CCZ1B | 221960 |
| ENSG000001 | 909.307833 | 0.51213321 | 0.11306076 | 4.52971651 | 5.91E-06 | 9.81E-05 | FAM111A | 63901 |
| ENSG000001 | 225.003032 | 0.61710214 | 0.13624293 | 4.529425 | 5.91E-06 | 9.82E-05 | MDFIC | 29969 |
| ENSG000001 | 32.3023841 | -1.151691772 | 0.25426818 | -4.5294373 | 5.91E-06 | 9.82E-05 | DMBT1P1 | 375940 |
| ENSG000001 | 1218.57528 | 0.55523399 | 0.12259358 | 4.5290625 | 5.92E-06 | 9.83E-05 | ATPIF1 | 93974 |
| ENSG000001 | 437.790065 | 0.569262236 | 0.12569445 | 4.52893697 | 5.93E-06 | 9.83E-05 | PPP2R2D | 55844 |
| ENSG000001 | 790.775728 | 0.701201055 | 0.15489584 | 4.52691978 | 5.98E-06 | 9.92E-05 | GTPBP1 | 9567 |
| ENSG000001 | 258.552871 | 0.62087062 | 0.13717059 | 4.52626624 | 6.00E-06 | 9.95E-05 | RASEF | 158158 |
| ENSG000001 | 107.325207 | 0.778911558 | 0.17209538 | 4.52604583 | 6.01E-06 | 9.96E-05 | S100A16 | 140576 |
| ENSG000001 | 256.34867 | 0.703671354 | 0.15549235 | 4.52544026 | 6.03E-06 | 9.98E-05 | DEF8 | 54849 |
| ENSG000001 | 678.496214 | 0.610198042 | 0.13484635 | 4.52513575 | 6.04E-06 | 9.99E-05 | CDKN3 | 1033 |
| ENSG000001 | 30.4464777 | -1.163586493 | 0.25714321 | -4.5250523 | 6.04E-06 | 9.99E-05 | TSPAN11 | 441631 |
| ENSG000001 | 1891.37344 | 0.496714163 | 0.10977793 | 4.52471805 | 6.05E-06 | 0.00010001 | KIF21A | 55605 |
| ENSG000001 | 561.371701 | 0.597443712 | 0.1320526 | 4.52428592 | 6.06E-06 | 0.00010018 | ZNF780A | 284323 |
| ENSG000001 | 372.209161 | 0.755104861 | 0.16693191 | 4.52343026 | 6.08E-06 | 0.00010054 | POLR2L | 5441 |
| ENSG000001 | 3045.20625 | 0.590216807 | 0.13048471 | 4.52326416 | 6.09E-06 | 0.00010058 | LYPLA1 | 10434 |
| ENSG00000C | 494.547267 | 0.577173591 | 0.12760862 | 4.52299835 | 6.10E-06 | 0.00010067 | TRIP13 | 9319 |
| ENSG000002 | 945.3715 | 0.576859564 | 0.12754212 | 4.52289452 | 6.10E-06 | 0.00010068 | MYO19 | 80179 |
| ENSG000001 | 1578.52927 | 0.594755568 | 0.13150151 | 4.522804 | 6.10E-06 | 0.00010068 | RBM3 | 5935 |
| ENSG00000C | 219.577606 | 0.742745098 | 0.1642256 | 4.52271201 | 6.11E-06 | 0.00010068 | IDH3G | 3421 |
| ENSG000001 | 1008.21754 | 0.541681921 | 0.11978113 | 4.52226425 | 6.12E-06 | 0.00010085 | EPB41L5 | 57669 |
| ENSG000001 | 6654.07635 | 0.563568123 | 0.12463159 | 4.52187215 | 6.13E-06 | 0.000101 | FAM60A | 58516 |
| ENSG000001 | 49.3777012 | -1.047772218 | 0.23174285 | -4.521271 | 6.15E-06 | 0.00010124 | CORIN | 10699 |
| ENSG000001 | 111.784674 | 0.76155473 | 0.16845743 | 4.52075475 | 6.16E-06 | 0.00010145 | FLYWCH2 | 114984 |

| | | | | | | | | | |
|---|---|---|---|---|---|---|---|---|---|
| ENSG000001 | 2025.46012 | 0.513611272 | 0.11363048 | 4.52001315 | 6.18E-06 | 0.00010177 | VDAC2 | | 7417 |
| ENSG000001 | 370.339504 | 0.645904185 | 0.1429187 | 4.51938175 | 6.20E-06 | 0.00010203 | WDR77 | | 79084 |
| ENSG000001 | 3340.65194 | 0.514226068 | 0.11378791 | 4.51916264 | 6.21E-06 | 0.00010209 | OPA1 | | 4976 |
| ENSG000001 | 561.307559 | 0.613969831 | 0.13586647 | 4.51892085 | 6.22E-06 | 0.00010213 | HACL1 | | 26061 |
| ENSG000001 | 90.8892012 | 0.851967191 | 0.18853009 | 4.51899859 | 6.21E-06 | 0.00010213 | PPP1R35 | | 221908 |
| ENSG000001 | 1214.43913 | 0.532647412 | 0.11792689 | 4.51675976 | 6.28E-06 | 0.00010313 | MTCH2 | | 23788 |
| ENSG00000C | 390.063312 | 0.594952808 | 0.13172927 | 4.51648137 | 6.29E-06 | 0.00010319 | RPL26L1 | | 51121 |
| ENSG000001 | 565.935837 | 0.549096783 | 0.1215766 | 4.51646757 | 6.29E-06 | 0.00010319 | NIPA1 | | 123606 |
| ENSG000001 | 1043.23657 | 0.674832021 | 0.14942257 | 4.51626567 | 6.29E-06 | 0.00010325 | USP5 | | 8078 |
| ENSG000001 | 356.402944 | 0.69929486 | 0.15486405 | 4.51554026 | 6.32E-06 | 0.00010356 | PACS2 | | 23241 |
| ENSG000001 | 584.593983 | 0.696354277 | 0.1542183 | 4.51538023 | 6.32E-06 | 0.0001036 | MLST8 | | 64223 |
| ENSG000002 | 393.272222 | 0.558540674 | 0.12370871 | 4.51496657 | 6.33E-06 | 0.00010376 | PPP1R11 | | 6992 |
| ENSG000001 | 145.039895 | 0.821392297 | 0.1819408 | 4.51461307 | 6.34E-06 | 0.00010389 | ULK3 | | 25989 |
| ENSG000002 | 299.850274 | 0.648695517 | 0.14369305 | 4.51445285 | 6.35E-06 | 0.00010392 | NA | NA | |
| ENSG00000C | 230.105014 | 0.815908023 | 0.18075009 | 4.5140118 | 6.36E-06 | 0.0001041 | ATP6AP1 | | 537 |
| ENSG000001 | 383.379928 | 0.613603576 | 0.13594419 | 4.51364316 | 6.37E-06 | 0.00010423 | CDC25C | | 995 |
| ENSG000002 | 238.439866 | 0.632520006 | 0.14013745 | 4.5135686 | 6.37E-06 | 0.00010423 | ZNF805 | | 390980 |
| ENSG000001 | 88.3985726 | 0.787478946 | 0.17448446 | 4.5131752 | 6.39E-06 | 0.00010438 | MAP3K8 | | 1326 |
| ENSG000001 | 16272.6132 | 0.662681786 | 0.14683757 | 4.5130262 | 6.39E-06 | 0.00010441 | RPS3 | | 6188 |
| ENSG000001 | 144.240793 | 0.747847212 | 0.16572001 | 4.51271528 | 6.40E-06 | 0.00010453 | HDDC3 | | 374659 |
| ENSG000001 | 195.938727 | -0.635144734 | 0.14075212 | -4.5125057 | 6.41E-06 | 0.00010459 | NFIB | | 4781 |
| ENSG000001 | 936.673302 | 0.555131779 | 0.12304353 | 4.51166984 | 6.43E-06 | 0.00010487 | GTF2H1 | | 2965 |
| ENSG000001 | 211.06169 | 0.778597113 | 0.17257684 | 4.51159679 | 6.43E-06 | 0.00010487 | FHOD1 | | 29109 |
| ENSG000001 | 18.5046504 | -1.353579092 | 0.30001452 | -4.5117119 | 6.43E-06 | 0.00010487 | OR3A3 | | 8392 |
| ENSG000001 | 474.632091 | 0.58026149 | 0.12861067 | 4.51176797 | 6.43E-06 | 0.00010487 | RPP14 | | 11102 |
| ENSG000001 | 289.210692 | 0.666316916 | 0.14769347 | 4.5114854 | 6.44E-06 | 0.00010488 | HSF1 | | 3297 |
| ENSG000001 | 761.873228 | 0.612963856 | 0.13589236 | 4.51065713 | 6.46E-06 | 0.00010525 | DDX39A | | 10212 |
| ENSG00000C | 2395.56051 | 0.676021126 | 0.14989428 | 4.50998627 | 6.48E-06 | 0.00010554 | GANAB | | 23193 |
| ENSG00000C | 63.25828 | -0.864673154 | 0.19174211 | -4.5095633 | 6.50E-06 | 0.00010571 | SCIN | | 85477 |

| | | | | | | | | | |
|---|---|---|---|---|---|---|---|---|---|
| ENSG000001 | 156.732136 | 0.719034952 | 0.15948426 | 4.50850097 | 6.53E-06 | 0.0001062 | FAM207A | | 85395 |
| ENSG000002 | 374.274384 | 0.57263978 | 0.12702956 | 4.50792534 | 6.55E-06 | 0.00010644 | NA | NA | |
| ENSG000001 | 200.369847 | 0.693968195 | 0.15396667 | 4.50726257 | 6.57E-06 | 0.00010669 | P3H4 | | 10609 |
| ENSG000002 | 283.807255 | 0.617562628 | 0.13701494 | 4.50726497 | 6.57E-06 | 0.00010669 | ARFGAP3 | | 26286 |
| ENSG000001 | 360.627329 | 0.568550055 | 0.12614937 | 4.50695923 | 6.58E-06 | 0.0001068 | POP4 | | 10775 |
| ENSG000001 | 24.1120901 | 1.264800277 | 0.28065362 | 4.50662375 | 6.59E-06 | 0.00010692 | KCNE5 | | 23630 |
| ENSG00000C | 276.188872 | 0.683210976 | 0.15161007 | 4.50636931 | 6.59E-06 | 0.00010701 | CTSA | | 5476 |
| ENSG000001 | 442.53292 | 0.634663764 | 0.14084272 | 4.5061879 | 6.60E-06 | 0.00010706 | FXR2 | | 9513 |
| ENSG000001 | 1176.18622 | 0.551253762 | 0.12234157 | 4.50585807 | 6.61E-06 | 0.00010718 | MRFAP1L1 | | 114932 |
| ENSG000001 | 2383.46426 | 0.549157398 | 0.12190694 | 4.50472622 | 6.65E-06 | 0.00010771 | KIAA1033 | | 23325 |
| ENSG000001 | 618.101194 | 0.553899483 | 0.12296224 | 4.50463062 | 6.65E-06 | 0.00010772 | HPS5 | | 11234 |
| ENSG000001 | 279.085112 | 0.752113235 | 0.16698877 | 4.50397501 | 6.67E-06 | 0.00010801 | SH3GLB2 | | 56904 |
| ENSG000001 | 480.616932 | 0.579276497 | 0.1286418 | 4.5030192 | 6.70E-06 | 0.00010841 | CSRNP2 | | 81566 |
| ENSG000001 | 1860.26972 | 0.577135826 | 0.12816537 | 4.5030559 | 6.70E-06 | 0.00010841 | KDM6A | | 7403 |
| ENSG000001 | 1456.65269 | 0.518157819 | 0.11507838 | 4.50265142 | 6.71E-06 | 0.00010855 | UBE4A | | 9354 |
| ENSG000001 | 1268.40195 | 0.647710945 | 0.14386096 | 4.50233996 | 6.72E-06 | 0.00010867 | DCUN1D5 | | 84259 |
| ENSG000001 | 2462.76768 | 0.605904217 | 0.1345793 | 4.50220976 | 6.73E-06 | 0.00010869 | PSMB4 | | 5692 |
| ENSG00000C | 484.454872 | 0.675523062 | 0.15006454 | 4.5015501 | 6.75E-06 | 0.00010886 | WBSCR22 | | 114049 |
| ENSG00000C | 452.643741 | 0.639801131 | 0.14212222 | 4.50176701 | 6.74E-06 | 0.00010886 | NLE1 | | 54475 |
| ENSG000001 | 502.190127 | 0.619718627 | 0.13767054 | 4.50146133 | 6.75E-06 | 0.00010886 | CHKA | | 1119 |
| ENSG000001 | 1368.02928 | 0.538805142 | 0.11968938 | 4.50169533 | 6.74E-06 | 0.00010886 | DCAF7 | | 10238 |
| ENSG000001 | 1217.08284 | 0.648139342 | 0.14398195 | 4.50153188 | 6.75E-06 | 0.00010886 | PNP | | 4860 |
| ENSG000002 | 96.294221 | 0.827075646 | 0.18374029 | 4.50132988 | 6.75E-06 | 0.00010888 | NA | NA | |
| ENSG000002 | 72.3909678 | -0.906874265 | 0.2014845 | -4.5009628 | 6.76E-06 | 0.00010903 | NA | NA | |
| ENSG000001 | 2792.72774 | 0.54800887 | 0.12176458 | 4.50056049 | 6.78E-06 | 0.00010919 | SPCS3 | | 60559 |
| ENSG000002 | 220.961642 | 0.664101154 | 0.14756821 | 4.50029954 | 6.79E-06 | 0.00010928 | NA | NA | |
| ENSG000001 | 692.406134 | 0.605508707 | 0.13455305 | 4.5001485 | 6.79E-06 | 0.00010932 | NDUFS6 | | 4726 |
| ENSG00000C | 682.260715 | 0.673981413 | 0.14978634 | 4.49961881 | 6.81E-06 | 0.00010954 | DDX11 | | 1663 |
| ENSG000002 | 125.073219 | 0.724619392 | 0.16104458 | 4.49949564 | 6.81E-06 | 0.00010956 | NA | NA | |

| | | | | | | | | | |
|---|---|---|---|---|---|---|---|---|---|
| ENSG000001 | 76.4225951 | 0.844141275 | 0.18762415 | 4.49910779 | 6.82E-06 | 0.00010972 | FBXO36 | | 130888 |
| ENSG000001 | 569.5505 | 0.592669002 | 0.13173641 | 4.49890055 | 6.83E-06 | 0.00010978 | CEP85 | | 64793 |
| ENSG000001 | 372.406935 | 0.636207619 | 0.14142704 | 4.49848635 | 6.84E-06 | 0.00010996 | LZIC | | 84328 |
| ENSG000001 | 25.0299567 | -1.225916489 | 0.27253795 | -4.4981496 | 6.85E-06 | 0.00011009 | OTOF | | 9381 |
| ENSG000001 | 1029.15358 | 0.541982305 | 0.12049943 | 4.49779959 | 6.87E-06 | 0.00011022 | EXTL2 | | 2135 |
| ENSG000001 | 372.979953 | 0.652687157 | 0.14511682 | 4.49766726 | 6.87E-06 | 0.00011025 | FAM213B | | 127281 |
| ENSG000001 | 294.432791 | 0.672507109 | 0.14953336 | 4.49737175 | 6.88E-06 | 0.00011036 | GSTZ1 | | 2954 |
| ENSG000001 | 263.038742 | 0.601895356 | 0.13384356 | 4.4970064 | 6.89E-06 | 0.0001105 | PTRHD1 | | 391356 |
| ENSG000001 | 234.771605 | 0.684451756 | 0.1522136 | 4.496653 | 6.90E-06 | 0.00011064 | DTX3 | | 196403 |
| ENSG000001 | 6849.28624 | 0.621087683 | 0.13812583 | 4.49653526 | 6.91E-06 | 0.00011066 | RPL21 | | 6144 |
| ENSG000001 | 8218.31784 | 0.486206232 | 0.10814298 | 4.49595741 | 6.93E-06 | 0.00011088 | NARS | | 4677 |
| ENSG000002 | 157.049493 | 0.880723588 | 0.19588983 | 4.49601491 | 6.92E-06 | 0.00011088 | NA | NA | |
| ENSG00000C | 890.940896 | 0.754358166 | 0.16780929 | 4.49533019 | 6.95E-06 | 0.00011116 | DNM2 | | 1785 |
| ENSG000002 | 323.71649 | 0.583616631 | 0.12983425 | 4.4950898 | 6.95E-06 | 0.00011124 | PINX1 | | 54984 |
| ENSG00000C | 283.284252 | 0.635630072 | 0.14142921 | 4.49433379 | 6.98E-06 | 0.00011155 | DERA | | 51071 |
| ENSG000001 | 1063.01584 | 0.527633132 | 0.11739781 | 4.49440336 | 6.98E-06 | 0.00011155 | KXD1 | | 79036 |
| ENSG00000C | 176.96451 | 0.686004858 | 0.15265439 | 4.49384292 | 6.99E-06 | 0.00011172 | FAM160A2 | | 84067 |
| ENSG000002 | 701.259653 | 0.568313307 | 0.12646491 | 4.49384179 | 6.99E-06 | 0.00011172 | FNIP1 | | 96459 |
| ENSG000001 | 433.873106 | 0.558668368 | 0.12432115 | 4.49375176 | 7.00E-06 | 0.00011172 | FILIP1 | | 27145 |
| ENSG000001 | 690.996494 | 0.631730165 | 0.14059937 | 4.49312225 | 7.02E-06 | 0.00011201 | TMEM261 | | 90871 |
| ENSG000001 | 173.444263 | 0.679387595 | 0.15121337 | 4.49290693 | 7.03E-06 | 0.00011208 | KLHL25 | | 64410 |
| ENSG00000C | 36.8300791 | -1.106434573 | 0.24628038 | -4.4925811 | 7.04E-06 | 0.00011216 | MYH13 | | 8735 |
| ENSG000001 | 116.268883 | 0.779351889 | 0.17347826 | 4.49250468 | 7.04E-06 | 0.00011216 | TMEM208 | | 29100 |
| ENSG000002 | 334.633345 | 0.743515796 | 0.16550177 | 4.49249441 | 7.04E-06 | 0.00011216 | C9orf69 | | 90120 |
| ENSG00000C | 1041.69278 | 0.563668904 | 0.12547608 | 4.4922418 | 7.05E-06 | 0.00011225 | MRPS35 | | 60488 |
| ENSG00000C | 392.657848 | 0.566586723 | 0.12614268 | 4.4916337 | 7.07E-06 | 0.00011253 | TXNDC16 | | 57544 |
| ENSG000001 | 167.082431 | 0.769808738 | 0.17141211 | 4.49098232 | 7.09E-06 | 0.00011283 | NA | NA | |
| ENSG000001 | 2696.89288 | 0.545829649 | 0.1215527 | 4.49047735 | 7.11E-06 | 0.00011301 | FANCD2 | | 2177 |
| ENSG000001 | 215.587161 | 0.745323307 | 0.16597755 | 4.4905067 | 7.11E-06 | 0.00011301 | PDLIM7 | | 9260 |

| | | | | | | | | |
|---|---|---|---|---|---|---|---|---|
| ENSG000002 | 655.665826 | 0.584482473 | 0.13016477 | 4.49032768 | 7.11E-06 | 0.00011304 | NUP62 | 23636 |
| ENSG000001 | 998.422868 | 0.586066122 | 0.13057114 | 4.48848118 | 7.17E-06 | 0.00011398 | LDB2 | 9079 |
| ENSG000001 | 854.641321 | 0.576549877 | 0.12847623 | 4.48759944 | 7.20E-06 | 0.00011441 | RARS2 | 57038 |
| ENSG000000 | 2241.78543 | 0.603995942 | 0.13459733 | 4.48742896 | 7.21E-06 | 0.00011446 | DAZAP1 | 26528 |
| ENSG000001 | 544.579936 | 0.580438737 | 0.12936769 | 4.48673655 | 7.23E-06 | 0.00011478 | EXOSC1 | 51013 |
| ENSG000001 | 3887.67342 | 0.576092615 | 0.12840841 | 4.4864086 | 7.24E-06 | 0.00011492 | PDIA4 | 9601 |
| ENSG000000 | 372.251233 | 0.548833409 | 0.12235036 | 4.48575231 | 7.27E-06 | 0.0001152 | MSL3 | 10943 |
| ENSG000001 | 247.423411 | 0.649403992 | 0.14477287 | 4.48567457 | 7.27E-06 | 0.0001152 | FBXO33 | 254170 |
| ENSG000001 | 191.452917 | -0.621890513 | 0.13864034 | -4.4856391 | 7.27E-06 | 0.0001152 | ZNF667-AS1 | 100128252 |
| ENSG000001 | 367.682775 | 0.580848881 | 0.12950045 | 4.48530388 | 7.28E-06 | 0.00011533 | CAMK2D | 817 |
| ENSG000000 | 316.707422 | 0.636745765 | 0.14198152 | 4.48470871 | 7.30E-06 | 0.00011561 | LMCD1 | 29995 |
| ENSG000001 | 460.463999 | 0.59298806 | 0.13223779 | 4.4842556 | 7.32E-06 | 0.00011581 | ZNF277 | 11179 |
| ENSG000001 | 1232.86144 | 0.60500604 | 0.1349326 | 4.48376493 | 7.33E-06 | 0.00011596 | FN1 | 2335 |
| ENSG000001 | 413.689612 | 0.582212461 | 0.12984976 | 4.4837393 | 7.33E-06 | 0.00011596 | CARS2 | 79587 |
| ENSG000001 | 402.808279 | 0.8453333 | 0.18852615 | 4.48390485 | 7.33E-06 | 0.00011596 | PFKL | 5211 |
| ENSG000001 | 1490.46331 | 0.555120762 | 0.1238171 | 4.48339321 | 7.35E-06 | 0.0001161 | DCTN5 | 84516 |
| ENSG000001 | 36.5851275 | -1.051820905 | 0.23460909 | -4.4832914 | 7.35E-06 | 0.00011611 | KMO | 8564 |
| ENSG000001 | 1950.83546 | 0.598919842 | 0.13360197 | 4.48286687 | 7.36E-06 | 0.00011625 | SCG3 | 29106 |
| ENSG000001 | 1282.37225 | 0.550273734 | 0.12274824 | 4.48294586 | 7.36E-06 | 0.00011625 | GTF3C3 | 9330 |
| ENSG000001 | 1308.94264 | 0.63503122 | 0.14166693 | 4.4825649 | 7.38E-06 | 0.00011637 | PLEKHB2 | 55041 |
| ENSG000001 | 295.31776 | 0.706041656 | 0.15752321 | 4.48214377 | 7.39E-06 | 0.00011655 | HMCES | 56941 |
| ENSG000000 | 2711.20125 | 0.492022762 | 0.10978127 | 4.48184611 | 7.40E-06 | 0.00011667 | KPNA6 | 23633 |
| ENSG000001 | 372.682579 | 0.593638226 | 0.13246798 | 4.48137132 | 7.42E-06 | 0.00011684 | KDSR | 2531 |
| ENSG000002 | 513.951321 | 0.750253915 | 0.16741642 | 4.48136395 | 7.42E-06 | 0.00011684 | TRIM26 | 7726 |
| ENSG000001 | 138.883513 | 0.791660113 | 0.17669462 | 4.48038608 | 7.45E-06 | 0.00011734 | CTU2 | 348180 |
| ENSG000001 | 78.1333988 | 0.865877804 | 0.1932849 | 4.47980053 | 7.47E-06 | 0.00011757 | NFKBIE | 4794 |
| ENSG000001 | 202.746361 | 0.670253767 | 0.14961676 | 4.47980395 | 7.47E-06 | 0.00011757 | GCNT1 | 2650 |
| ENSG000000 | 620.762553 | 0.56291238 | 0.12565972 | 4.47965642 | 7.48E-06 | 0.0001176 | UBR7 | 55148 |
| ENSG000001 | 351.558536 | 0.68535046 | 0.15299593 | 4.47953392 | 7.48E-06 | 0.00011762 | ESRP2 | 80004 |

| | | | | | | | | |
|---|---|---|---|---|---|---|---|---|
| ENSG000001 | 482.525024 | 0.632254971 | 0.14115141 | 4.47926768 | 7.49E-06 | 0.00011769 | RAE1 | 8480 |
| ENSG000001 | 1990.8499 | 0.575910911 | 0.12857326 | 4.47924324 | 7.49E-06 | 0.00011769 | CAPZA2 | 830 |
| ENSG00000C | 9998.19182 | 0.547928858 | 0.12233092 | 4.47907083 | 7.50E-06 | 0.00011769 | PKM | 5315 |
| ENSG000001 | 781.295499 | 0.583565739 | 0.13028946 | 4.4789944 | 7.50E-06 | 0.00011769 | ETFA | 2108 |
| ENSG000001 | 136.787548 | 0.757941807 | 0.16921574 | 4.47914484 | 7.49E-06 | 0.00011769 | NA | NA |
| ENSG00000C | 118.895064 | 0.755994996 | 0.16880393 | 4.47853897 | 7.52E-06 | 0.00011785 | SPATA20 | 64847 |
| ENSG000001 | 531.874633 | 0.62479224 | 0.13950789 | 4.4785442 | 7.52E-06 | 0.00011785 | RNASEH1 | 246243 |
| ENSG000001 | 518.383058 | 0.673227483 | 0.15032782 | 4.47839583 | 7.52E-06 | 0.00011788 | AMMECR1L | 83607 |
| ENSG00000C | 334.853961 | 0.739552089 | 0.16516995 | 4.47752191 | 7.55E-06 | 0.00011832 | ASB1 | 51665 |
| ENSG000001 | 464.672539 | 0.596610489 | 0.13325689 | 4.47714569 | 7.56E-06 | 0.00011844 | MTX2 | 10651 |
| ENSG000001 | 3239.92526 | 0.467984662 | 0.10452732 | 4.47715163 | 7.56E-06 | 0.00011844 | ANAPC1 | 64682 |
| ENSG00000C | 37.4796027 | -1.046452008 | 0.23376475 | -4.4765176 | 7.59E-06 | 0.0001187 | MYH7 | 4625 |
| ENSG000001 | 1642.83771 | 0.518574506 | 0.11584269 | 4.47654064 | 7.59E-06 | 0.0001187 | ATP6V1C1 | 528 |
| ENSG00000C | 1318.75559 | 0.549309063 | 0.12271805 | 4.47618816 | 7.60E-06 | 0.00011883 | SNRPA | 6626 |
| ENSG00000C | 1075.63126 | 0.559882669 | 0.12508573 | 4.47599149 | 7.61E-06 | 0.0001189 | PHF23 | 79142 |
| ENSG000001 | 5157.06011 | 0.540109274 | 0.12067526 | 4.47572514 | 7.62E-06 | 0.00011899 | HIST1H1A | 3024 |
| ENSG000001 | 690.713795 | 0.646075679 | 0.14435326 | 4.47565713 | 7.62E-06 | 0.00011899 | EDNRB | 1910 |
| ENSG00000C | 19.732979 | -1.317592875 | 0.29442618 | -4.4751213 | 7.64E-06 | 0.00011915 | FMO1 | 2326 |
| ENSG000001 | 339.56331 | 0.632569702 | 0.14135113 | 4.47516566 | 7.64E-06 | 0.00011915 | FADD | 8772 |
| ENSG000002 | 103.099049 | 0.794261433 | 0.1774811 | 4.4751889 | 7.63E-06 | 0.00011915 | SNORD15B | 114599 |
| ENSG000001 | 85.8921442 | 0.908015096 | 0.20291757 | 4.47479775 | 7.65E-06 | 0.00011929 | NDUFAF3 | 25915 |
| ENSG000001 | 821.697559 | 0.663655473 | 0.14835667 | 4.47337792 | 7.70E-06 | 0.00012004 | MRPL16 | 54948 |
| ENSG00000C | 477.435414 | 0.564793351 | 0.12625977 | 4.47326467 | 7.70E-06 | 0.00012005 | OSGEP | 55644 |
| ENSG00000C | 4645.46947 | 0.488878825 | 0.10929938 | 4.4728417 | 7.72E-06 | 0.00012025 | APLP2 | 334 |
| ENSG00000C | 420.309434 | 0.561587401 | 0.12558714 | 4.471695 | 7.76E-06 | 0.00012085 | KCNN2 | 3781 |
| ENSG000001 | 343.478797 | 0.632659094 | 0.14149268 | 4.47132025 | 7.77E-06 | 0.00012101 | GPR19 | 2842 |
| ENSG000001 | 329.468271 | 0.576531559 | 0.12895946 | 4.4706417 | 7.80E-06 | 0.00012121 | SCPEP1 | 59342 |
| ENSG000001 | 34.8418272 | -1.090205902 | 0.24385321 | -4.4707466 | 7.79E-06 | 0.00012121 | MR1 | 3140 |
| ENSG000001 | 193.691287 | 0.648161393 | 0.14498127 | 4.47065603 | 7.80E-06 | 0.00012121 | SAP30 | 8819 |

| | | | | | | | | |
|---|---|---|---|---|---|---|---|---|
| ENSG000002 | 85.3632033 | 0.873605673 | 0.19540186 | 4.47081543 | 7.79E-06 | 0.00012121 | SNORA22 | 677807 |
| ENSG000001 | 1722.51975 | 0.523250319 | 0.11706054 | 4.46991215 | 7.83E-06 | 0.00012158 | AFG3L2 | 10939 |
| ENSG000001 | 318.837288 | 0.619178582 | 0.13853716 | 4.46940424 | 7.84E-06 | 0.00012182 | OLFM1 | 10439 |
| ENSG000002 | 28.4018307 | -1.153527593 | 0.25812407 | -4.4688881 | 7.86E-06 | 0.00012207 | NA | NA |
| ENSG000001 | 1269.90411 | 0.646271581 | 0.14463205 | 4.46838445 | 7.88E-06 | 0.00012226 | EDIL3 | 10085 |
| ENSG000001 | 641.990523 | 0.619841793 | 0.13871688 | 4.46839489 | 7.88E-06 | 0.00012226 | ECI2 | 10455 |
| ENSG00000C | 288.538984 | 0.635437509 | 0.14221963 | 4.46800156 | 7.90E-06 | 0.00012242 | PVR | 5817 |
| ENSG000001 | 81.2558458 | -0.856413279 | 0.19167961 | -4.4679415 | 7.90E-06 | 0.00012242 | LRRK2 | 120892 |
| ENSG000001 | 1051.56768 | 0.528405287 | 0.11827772 | 4.46749634 | 7.91E-06 | 0.00012263 | REEP5 | 7905 |
| ENSG00000C | 1013.71764 | 0.544259337 | 0.12182898 | 4.46740465 | 7.92E-06 | 0.00012264 | CHMP2B | 25978 |
| ENSG00000C | 1531.99645 | 0.507037375 | 0.11350391 | 4.46713561 | 7.93E-06 | 0.00012274 | GEMIN5 | 25929 |
| ENSG00000C | 1807.59901 | 0.58872976 | 0.13179679 | 4.46695084 | 7.93E-06 | 0.0001228 | SART3 | 9733 |
| ENSG000001 | 271.033241 | 0.615138107 | 0.13772064 | 4.46656424 | 7.95E-06 | 0.00012298 | HEY2 | 23493 |
| ENSG00000C | 670.452721 | 0.516770849 | 0.11571405 | 4.46593018 | 7.97E-06 | 0.00012329 | ZNF280C | 55609 |
| ENSG000002 | 123.095015 | 0.74649979 | 0.16716756 | 4.46557815 | 7.99E-06 | 0.00012345 | FBXO17 | 115290 |
| ENSG00000C | 2803.88536 | 0.662065606 | 0.14827109 | 4.46523725 | 8.00E-06 | 0.00012355 | FGFR1 | 2260 |
| ENSG000001 | 251.412018 | 0.661217138 | 0.14808116 | 4.46523485 | 8.00E-06 | 0.00012355 | CASP6 | 839 |
| ENSG000001 | 648.463907 | 0.546927968 | 0.12249283 | 4.4649793 | 8.01E-06 | 0.00012366 | SLC37A3 | 84255 |
| ENSG000001 | 285.093526 | 0.753966658 | 0.16886631 | 4.46487318 | 8.01E-06 | 0.00012367 | LAMTOR4 | 389541 |
| ENSG000001 | 937.959954 | 0.53415287 | 0.11963948 | 4.46468729 | 8.02E-06 | 0.00012373 | OSBPL1A | 114876 |
| ENSG000001 | 316.846923 | 0.663008636 | 0.14852908 | 4.46383045 | 8.05E-06 | 0.00012418 | NDUFA8 | 4702 |
| ENSG000001 | 1455.39784 | 0.642626431 | 0.14402849 | 4.46180067 | 8.13E-06 | 0.00012531 | DUSP16 | 80824 |
| ENSG000001 | 669.06465 | 0.606647675 | 0.13596712 | 4.46172334 | 8.13E-06 | 0.00012531 | AURKB | 9212 |
| ENSG000001 | 68.9792636 | -0.892300474 | 0.20001103 | -4.4612562 | 8.15E-06 | 0.00012548 | AXDND1 | 126859 |
| ENSG000001 | 2784.47444 | 0.604769621 | 0.13556244 | 4.46118863 | 8.15E-06 | 0.00012548 | Sep-09 | 10801 |
| ENSG000001 | 346.653223 | 0.706242449 | 0.15830381 | 4.46131052 | 8.15E-06 | 0.00012548 | ATG9A | 79065 |
| ENSG00000C | 5518.13125 | 0.477185603 | 0.10698025 | 4.46050173 | 8.18E-06 | 0.00012569 | ITGA6 | 3655 |
| ENSG000001 | 581.63148 | 0.600422252 | 0.13460301 | 4.46068947 | 8.17E-06 | 0.00012569 | NUDT12 | 83594 |
| ENSG000001 | 179.458419 | 0.667896157 | 0.14973472 | 4.46052979 | 8.18E-06 | 0.00012569 | DPCD | 25911 |

| | | | | | | | | |
|---|---|---|---|---|---|---|---|---|
| ENSG000002 | 561.200972 | 0.797543478 | 0.17879771 | 4.46059117 | 8.17E-06 | 0.00012569 | NA | NA |
| ENSG000000 | 264.330396 | 0.660742159 | 0.14814576 | 4.46008147 | 8.19E-06 | 0.00012589 | CYB561 | 1534 |
| ENSG000001 | 926.899891 | 0.603479505 | 0.13536411 | 4.45819416 | 8.27E-06 | 0.00012696 | PIH1D1 | 55011 |
| ENSG000001 | 477.03639 | 0.565818213 | 0.12695391 | 4.45687886 | 8.32E-06 | 0.00012769 | TIMM22 | 29928 |
| ENSG000001 | 712.916114 | 0.535369528 | 0.12013946 | 4.45623389 | 8.34E-06 | 0.00012803 | CNOT8 | 9337 |
| ENSG000001 | 203.679028 | 0.784687465 | 0.17609543 | 4.45603543 | 8.35E-06 | 0.00012805 | RECQL4 | 9401 |
| ENSG000002 | 640.874306 | 0.613647804 | 0.1377117 | 4.45603256 | 8.35E-06 | 0.00012805 | CDKN2AIPNL | 91368 |
| ENSG000001 | 99.6702769 | 0.77418 | 0.17376622 | 4.45529624 | 8.38E-06 | 0.00012844 | TMEM37 | 140738 |
| ENSG000001 | 208.19844 | -0.664394834 | 0.14914217 | -4.4547753 | 8.40E-06 | 0.0001287 | PLK2 | 10769 |
| ENSG000001 | 2769.73329 | 0.542183221 | 0.12171352 | 4.45458502 | 8.41E-06 | 0.00012872 | SEC11A | 23478 |
| ENSG000001 | 2409.40517 | 0.552290521 | 0.12398214 | 4.45459749 | 8.41E-06 | 0.00012872 | RPRD1A | 55197 |
| ENSG000001 | 2372.90676 | 0.563080856 | 0.12642108 | 4.45401063 | 8.43E-06 | 0.00012902 | KNOP1 | 400506 |
| ENSG000001 | 945.884684 | 0.614807103 | 0.13805147 | 4.45346278 | 8.45E-06 | 0.0001293 | COA4 | 51287 |
| ENSG000000 | 328.359485 | 0.615675889 | 0.13828598 | 4.45219318 | 8.50E-06 | 0.00012998 | TSR3 | 115939 |
| ENSG000001 | 25.6739846 | 1.219688091 | 0.27395477 | 4.45215139 | 8.50E-06 | 0.00012998 | GADD45G | 10912 |
| ENSG000001 | 1095.16582 | 0.713382476 | 0.1602351 | 4.45209877 | 8.50E-06 | 0.00012998 | CBX2 | 84733 |
| ENSG000001 | 970.469775 | 0.521988969 | 0.11725066 | 4.45190643 | 8.51E-06 | 0.00012999 | KCTD20 | 222658 |
| ENSG000001 | 6330.40665 | 0.564671353 | 0.12683708 | 4.45194217 | 8.51E-06 | 0.00012999 | RPL22 | 6146 |
| ENSG000001 | 319.279264 | 0.638990927 | 0.14353516 | 4.45180776 | 8.52E-06 | 0.00013 | TOMM40L | 84134 |
| ENSG000002 | 5.53966048 | -1.677640472 | 0.37685107 | -4.4517333 | 8.52E-06 | 0.00013 | NA | NA |
| ENSG000001 | 2949.54843 | 0.53971275 | 0.12124489 | 4.45142694 | 8.53E-06 | 0.00013014 | CDC123 | 8872 |
| ENSG000002 | 2219.98932 | 0.647934771 | 0.14555958 | 4.45133704 | 8.53E-06 | 0.00013014 | RNPS1 | 10921 |
| ENSG000001 | 167.931963 | 0.784537124 | 0.1762858 | 4.45037053 | 8.57E-06 | 0.00013068 | COMMD5 | 28991 |
| ENSG000001 | 1001.82343 | 0.695192683 | 0.15622102 | 4.45005852 | 8.58E-06 | 0.00013082 | MPRIP | 23164 |
| ENSG000001 | 185.880871 | 0.711570903 | 0.1599061 | 4.44992965 | 8.59E-06 | 0.00013085 | BABAM1 | 29086 |
| ENSG000000 | 213.035871 | 0.65181622 | 0.14649342 | 4.44945745 | 8.61E-06 | 0.00013094 | LLGL2 | 3993 |
| ENSG000001 | 96.8842159 | -0.83175551 | 0.18693059 | -4.4495419 | 8.61E-06 | 0.00013094 | PTPRB | 5787 |
| ENSG000001 | 274.901581 | 0.727712547 | 0.16354251 | 4.44968444 | 8.60E-06 | 0.00013094 | MTHFR | 4524 |
| ENSG000002 | 86.1321491 | -0.850255371 | 0.19109004 | -4.4495012 | 8.61E-06 | 0.00013094 | NA | NA |

| | | | | | | | | |
|---|---|---|---|---|---|---|---|---|
| ENSG000001 | 3483.16656 | 0.547147842 | 0.12297798 | 4.44915306 | 8.62E-06 | 0.00013108 | SF3A3 | 10946 |
| ENSG000001 | 1120.41024 | 0.519306656 | 0.11672345 | 4.44903454 | 8.63E-06 | 0.0001311 | SPTLC2 | 9517 |
| ENSG000001 | 83.6635723 | 0.830625144 | 0.18671046 | 4.44873374 | 8.64E-06 | 0.00013124 | GDAP1L1 | 78997 |
| ENSG000001 | 4826.06953 | 0.528513348 | 0.11880339 | 4.44863873 | 8.64E-06 | 0.00013125 | SRSF2 | 6427 |
| ENSG000001 | 2725.7822 | 0.512032695 | 0.11510576 | 4.44836743 | 8.65E-06 | 0.00013133 | PDCD6IP | 10015 |
| ENSG000001 | 57.7339093 | -0.878347429 | 0.19745523 | -4.4483371 | 8.65E-06 | 0.00013133 | PAX8-AS1 | 654433 |
| ENSG000001 | 524.069001 | 0.599056699 | 0.13467478 | 4.44817288 | 8.66E-06 | 0.00013138 | TRAPPC10 | 7109 |
| ENSG000001 | 376.639517 | 0.605245751 | 0.13607256 | 4.44796331 | 8.67E-06 | 0.00013146 | NARF | 26502 |
| ENSG000001 | 595.216862 | 0.582172365 | 0.13091499 | 4.44694945 | 8.71E-06 | 0.00013203 | LRPAP1 | 4043 |
| ENSG000001 | 387.293329 | 0.706462569 | 0.15889951 | 4.44597066 | 8.75E-06 | 0.00013259 | PLEKHG3 | 26030 |
| ENSG00000C | 1038.32914 | 0.550012049 | 0.12371866 | 4.44566773 | 8.76E-06 | 0.00013273 | SNAP23 | 8773 |
| ENSG000001 | 86.8665244 | 0.956751429 | 0.21525242 | 4.44478826 | 8.80E-06 | 0.00013312 | GRIN2D | 2906 |
| ENSG000001 | 1905.56554 | 0.600689484 | 0.1351488 | 4.44465282 | 8.80E-06 | 0.00013312 | MRPL51 | 51258 |
| ENSG000001 | 982.40405 | 0.513883475 | 0.11561974 | 4.4445998 | 8.81E-06 | 0.00013312 | LMO4 | 8543 |
| ENSG000001 | 302.425371 | 0.644144568 | 0.14492893 | 4.44455493 | 8.81E-06 | 0.00013312 | RFT1 | 91869 |
| ENSG000001 | 35.5614671 | -1.096727271 | 0.24675677 | -4.4445682 | 8.81E-06 | 0.00013312 | PEG3 | 5178 |
| ENSG000002 | 22.6846638 | -1.297322675 | 0.29187203 | -4.4448338 | 8.80E-06 | 0.00013312 | LOC401312 | 401312 |
| ENSG000001 | 1965.1349 | 0.530493048 | 0.11939125 | 4.44331594 | 8.86E-06 | 0.00013383 | DBI | 1622 |
| ENSG000001 | 1077.60233 | 0.550998127 | 0.12401934 | 4.44284044 | 8.88E-06 | 0.00013408 | ZPR1 | 8882 |
| ENSG00000C | 789.371291 | 0.726934194 | 0.16363035 | 4.44253891 | 8.89E-06 | 0.00013422 | GOLGA3 | 2802 |
| ENSG000001 | 911.78497 | 0.61439805 | 0.13830988 | 4.44218477 | 8.91E-06 | 0.00013439 | FAM96A | 84191 |
| ENSG000001 | 297.795849 | 0.656541961 | 0.14780214 | 4.44203276 | 8.91E-06 | 0.00013444 | PSMA1 | 5682 |
| ENSG000001 | 70.9482287 | -0.875248374 | 0.19705697 | -4.4416008 | 8.93E-06 | 0.00013466 | MYH15 | 22989 |
| ENSG000001 | 921.429218 | 0.52115996 | 0.1173468 | 4.44119433 | 8.95E-06 | 0.00013481 | ISCA1 | 81689 |
| ENSG000002 | 25.5863553 | -1.213001241 | 0.2731229 | -4.4412287 | 8.94E-06 | 0.00013481 | NA | NA |
| ENSG000001 | 1215.75516 | 0.579030359 | 0.13038096 | 4.4410653 | 8.95E-06 | 0.00013484 | RHEB | 6009 |
| ENSG000001 | 21.6515724 | -1.272402886 | 0.28652413 | -4.4408228 | 8.96E-06 | 0.00013494 | NCF2 | 4688 |
| ENSG000001 | 4420.74737 | 0.527945854 | 0.11889522 | 4.44042955 | 8.98E-06 | 0.00013514 | GNL3 | 26354 |
| ENSG000001 | 1169.16555 | 0.487642358 | 0.10982144 | 4.44032018 | 8.98E-06 | 0.00013516 | SAMHD1 | 25939 |

| | | | | | | | | | |
|---|---|---|---|---|---|---|---|---|---|
| ENSG00000C | 899.252599 | 0.543967011 | 0.12252578 | 4.43961288 | 9.01E-06 | 0.0001355 | ECHDC1 | | 55862 |
| ENSG000001 | 373.491463 | 0.590009923 | 0.13289517 | 4.43966419 | 9.01E-06 | 0.0001355 | RABEPK | | 10244 |
| ENSG000001 | 3730.86083 | 0.552863435 | 0.12453459 | 4.43943668 | 9.02E-06 | 0.00013556 | GART | | 2618 |
| ENSG000001 | 876.087898 | 0.702374935 | 0.15824388 | 4.43855977 | 9.06E-06 | 0.00013602 | FKBP10 | | 60681 |
| ENSG000001 | 469.814711 | 0.590035788 | 0.13293184 | 4.43863402 | 9.05E-06 | 0.00013602 | DNAAF5 | | 54919 |
| ENSG000001 | 393.878675 | 0.55191523 | 0.12435671 | 4.43816203 | 9.07E-06 | 0.00013622 | FKBP2 | | 2286 |
| ENSG000001 | 4674.83332 | 0.573006708 | 0.12912277 | 4.43768905 | 9.09E-06 | 0.00013647 | NUP155 | | 9631 |
| ENSG000001 | 421.774252 | 0.561226224 | 0.12647489 | 4.4374519 | 9.10E-06 | 0.00013657 | STAT2 | | 6773 |
| ENSG000001 | 2351.69718 | 0.527621514 | 0.1189064 | 4.43728437 | 9.11E-06 | 0.00013662 | C6orf62 | | 81688 |
| ENSG000001 | 555.324779 | 0.809054758 | 0.1823457 | 4.4369282 | 9.13E-06 | 0.00013675 | CLUH | | 23277 |
| ENSG000001 | 1343.42912 | 0.540453502 | 0.12180717 | 4.43695956 | 9.12E-06 | 0.00013675 | NUDT4 | | 11163 |
| ENSG000001 | 5910.71134 | 0.51529604 | 0.11615189 | 4.43639811 | 9.15E-06 | 0.00013703 | RBM39 | | 9584 |
| ENSG000001 | 312.302799 | 0.589548188 | 0.13289773 | 4.43610438 | 9.16E-06 | 0.00013717 | PYURF | | 100996939 |
| ENSG000001 | 1262.09485 | 0.484879348 | 0.10933621 | 4.43475528 | 9.22E-06 | 0.00013798 | MRPS9 | | 64965 |
| ENSG000002 | 113.185343 | 0.785251454 | 0.17707322 | 4.43461446 | 9.22E-06 | 0.00013802 | FAM58A | | 92002 |
| ENSG000001 | 268.004006 | 0.758067433 | 0.17096198 | 4.43412865 | 9.24E-06 | 0.00013828 | DNPEP | | 23549 |
| ENSG000001 | 473.317915 | 0.786976275 | 0.17748777 | 4.43397475 | 9.25E-06 | 0.00013833 | IER2 | | 9592 |
| ENSG000001 | 4553.00398 | 0.704825847 | 0.15896646 | 4.43380226 | 9.26E-06 | 0.00013839 | EIF5A | | 1984 |
| ENSG000001 | 438.1929 | 0.635133991 | 0.14327205 | 4.43306292 | 9.29E-06 | 0.00013881 | VAC14 | | 55697 |
| ENSG00000C | 121.418486 | 0.747815151 | 0.16872891 | 4.4320511 | 9.33E-06 | 0.00013931 | B4GALT7 | | 11285 |
| ENSG000001 | 973.962461 | 0.711226077 | 0.16047098 | 4.43211654 | 9.33E-06 | 0.00013931 | MYBBP1A | | 10514 |
| ENSG000001 | 6009.8325 | 0.514903584 | 0.11617474 | 4.43214753 | 9.33E-06 | 0.00013931 | NOLC1 | | 9221 |
| ENSG00000C | 418.093385 | 0.620491157 | 0.14001675 | 4.43154936 | 9.36E-06 | 0.00013958 | CBFB | | 865 |
| ENSG000001 | 4765.95229 | 0.546378027 | 0.12329651 | 4.43141514 | 9.36E-06 | 0.00013962 | SF3B3 | | 23450 |
| ENSG000001 | 1407.12585 | 0.477376396 | 0.10774786 | 4.4304952 | 9.40E-06 | 0.00014016 | C19orf43 | | 79002 |
| ENSG00000C | 1110.28572 | 0.52190497 | 0.11780831 | 4.43012018 | 9.42E-06 | 0.00014029 | RBM5 | | 10181 |
| ENSG000001 | 970.027604 | 0.524264333 | 0.11834465 | 4.42997899 | 9.42E-06 | 0.00014029 | ASAH1 | | 427 |
| ENSG000001 | 2102.49052 | 0.536250666 | 0.12104852 | 4.43004725 | 9.42E-06 | 0.00014029 | PPAT | | 5471 |
| ENSG000002 | 80.5337654 | -0.883986049 | 0.19954484 | -4.430012 | 9.42E-06 | 0.00014029 | NA | NA | |

| | | | | | | | | |
|---|---|---|---|---|---|---|---|---|
| ENSG000001 | 2859.01938 | 0.530877209 | 0.1198472 | 4.42961727 | 9.44E-06 | 0.00014048 | PRKAR1A | 5573 |
| ENSG000001 | 436.785802 | 0.68487394 | 0.15461926 | 4.42942189 | 9.45E-06 | 0.00014055 | RPUSD4 | 84881 |
| ENSG000001 | 2366.12463 | 0.515229372 | 0.11633758 | 4.42874408 | 9.48E-06 | 0.00014094 | WDR36 | 134430 |
| ENSG000001 | 496.39254 | 0.60477315 | 0.13656504 | 4.42846246 | 9.49E-06 | 0.00014108 | HPS4 | 89781 |
| ENSG000001 | 605.203607 | 0.603016678 | 0.13617984 | 4.42809052 | 9.51E-06 | 0.00014127 | SPG7 | 6687 |
| ENSG000002 | 18.2518901 | -1.332973977 | 0.30104858 | -4.4277704 | 9.52E-06 | 0.00014143 | LINC00540 | 100506622 |
| ENSG000001 | 1787.53415 | 0.535654078 | 0.12100247 | 4.42680266 | 9.56E-06 | 0.00014201 | QTRTD1 | 79691 |
| ENSG000001 | 254.07241 | 0.722446898 | 0.16320755 | 4.4265532 | 9.58E-06 | 0.00014212 | MAP7D1 | 55700 |
| ENSG000002 | 67.634089 | -0.912026259 | 0.20606564 | -4.4259017 | 9.60E-06 | 0.0001425 | NA | NA |
| ENSG00000C | 137.883116 | -0.695120284 | 0.1570616 | -4.4257813 | 9.61E-06 | 0.00014253 | DNAH5 | 1767 |
| ENSG000001 | 670.385689 | 0.620375865 | 0.14018146 | 4.42552013 | 9.62E-06 | 0.00014265 | ZBTB34 | 403341 |
| ENSG000001 | 1013.03429 | 0.520167105 | 0.11754461 | 4.42527385 | 9.63E-06 | 0.00014273 | TMED5 | 50999 |
| ENSG000001 | 364.022668 | 0.635711481 | 0.14365577 | 4.42524155 | 9.63E-06 | 0.00014273 | MPI | 4351 |
| ENSG000001 | 1649.91347 | 0.620484771 | 0.14022713 | 4.42485535 | 9.65E-06 | 0.00014293 | PPP2R1A | 5518 |
| ENSG000001 | 331.757757 | 0.693270422 | 0.1566874 | 4.42454489 | 9.66E-06 | 0.00014308 | SHROOM2 | 357 |
| ENSG000001 | 5858.06119 | 0.567825103 | 0.12833877 | 4.42442374 | 9.67E-06 | 0.00014309 | HIST1H2BC | 8347 |
| ENSG000001 | 208.369957 | 0.652681142 | 0.1475194 | 4.42437511 | 9.67E-06 | 0.00014309 | TRAIP | 10293 |
| ENSG000001 | 1268.0697 | 0.602717453 | 0.13623814 | 4.42399941 | 9.69E-06 | 0.00014329 | TMEM245 | 23731 |
| ENSG000001 | 1666.99976 | 0.488183448 | 0.1103595 | 4.42357422 | 9.71E-06 | 0.00014352 | ARGLU1 | 55082 |
| ENSG000001 | 273.044975 | 0.693734998 | 0.15683798 | 4.42325886 | 9.72E-06 | 0.00014368 | PITPNM1 | 9600 |
| ENSG000001 | 72.0916158 | -0.828937396 | 0.18744113 | -4.4223879 | 9.76E-06 | 0.0001442 | ANXA1 | 301 |
| ENSG000001 | 773.866617 | 0.688414362 | 0.15569017 | 4.42169437 | 9.79E-06 | 0.00014462 | HDGFRP2 | 84717 |
| ENSG000001 | 297.259492 | 0.699108372 | 0.15813176 | 4.4210496 | 9.82E-06 | 0.000145 | C12orf10 | 60314 |
| ENSG00000C | 673.048187 | 0.575231719 | 0.13012206 | 4.42070878 | 9.84E-06 | 0.00014513 | TRIT1 | 54802 |
| ENSG00000C | 3046.61161 | 0.63151658 | 0.14285466 | 4.42069283 | 9.84E-06 | 0.00014513 | SF3B2 | 10992 |
| ENSG00000C | 283.144576 | 0.62438633 | 0.14124789 | 4.42050038 | 9.85E-06 | 0.00014514 | TTC38 | 55020 |
| ENSG000001 | 1511.25783 | 0.506846032 | 0.11465946 | 4.42044667 | 9.85E-06 | 0.00014514 | FGD6 | 55785 |
| ENSG000002 | 1468.0563 | 0.496755563 | 0.11237592 | 4.42048053 | 9.85E-06 | 0.00014514 | PDE7A | 5150 |
| ENSG00000C | 326.758122 | 0.712589895 | 0.16120804 | 4.42031251 | 9.86E-06 | 0.00014514 | POMGNT1 | 55624 |

| | | | | | | | | |
|---|---|---|---|---|---|---|---|---|
| ENSG000001 | 230.097129 | 0.760879949 | 0.17213373 | 4.42028374 | 9.86E-06 | 0.00014514 | NTMT1 | 28989 |
| ENSG000001 | 740.200181 | 0.510928799 | 0.11559719 | 4.41990682 | 9.87E-06 | 0.00014518 | PFDN1 | 5201 |
| ENSG000001 | 1046.01271 | 0.590271594 | 0.13354692 | 4.41995674 | 9.87E-06 | 0.00014518 | VASP | 7408 |
| ENSG000001 | 719.14618 | 0.534074773 | 0.12082757 | 4.42014006 | 9.86E-06 | 0.00014518 | ZNF623 | 9831 |
| ENSG000002 | 1350.6754 | 0.481407229 | 0.10891653 | 4.41996495 | 9.87E-06 | 0.00014518 | SYNJ2BP | 55333 |
| ENSG000001 | 355.571371 | 0.780700754 | 0.17664817 | 4.41952348 | 9.89E-06 | 0.00014534 | RIMS4 | 140730 |
| ENSG000001 | 1480.51456 | 0.506564841 | 0.11462266 | 4.41941291 | 9.90E-06 | 0.00014534 | GOLPH3 | 64083 |
| ENSG000001 | 317.317423 | 0.624995449 | 0.14142133 | 4.41938605 | 9.90E-06 | 0.00014534 | SNRNP25 | 79622 |
| ENSG000001 | 177.740192 | 0.731982983 | 0.16563172 | 4.41934069 | 9.90E-06 | 0.00014534 | GPATCH3 | 63906 |
| ENSG000002 | 58.7525637 | 0.937787771 | 0.21220342 | 4.41928671 | 9.90E-06 | 0.00014534 | NA | NA |
| ENSG000001 | 3771.70553 | 0.546211042 | 0.12361545 | 4.41863099 | 9.93E-06 | 0.00014573 | MGEA5 | 10724 |
| ENSG000001 | 1081.12812 | 0.551350789 | 0.12478757 | 4.41831493 | 9.95E-06 | 0.00014589 | AASDHPPT | 60496 |
| ENSG000001 | 1895.88542 | 0.548744424 | 0.1242087 | 4.41792258 | 9.97E-06 | 0.0001461 | PSMD4 | 5710 |
| ENSG000000 | 886.318648 | 0.618658968 | 0.14003828 | 4.41778466 | 9.97E-06 | 0.00014614 | ARG2 | 384 |
| ENSG000001 | 998.111844 | 0.623239005 | 0.14108013 | 4.41762416 | 9.98E-06 | 0.00014619 | RAD54L2 | 23132 |
| ENSG000001 | 52.863135 | -0.897707757 | 0.20321694 | -4.4174849 | 9.99E-06 | 0.00014624 | RGS8 | 85397 |
| ENSG000001 | 418.184331 | 0.60543353 | 0.13706102 | 4.41725535 | 1.00E-05 | 0.00014634 | KAT5 | 10524 |
| ENSG000001 | 1258.43927 | 0.521613397 | 0.11811439 | 4.4161716 | 1.00E-05 | 0.00014702 | MRPL19 | 9801 |
| ENSG000001 | 225.87254 | 0.807581563 | 0.18288282 | 4.41584163 | 1.01E-05 | 0.00014719 | INTS5 | 80789 |
| ENSG000000 | 1857.61078 | 0.591884912 | 0.13403949 | 4.41575034 | 1.01E-05 | 0.0001472 | SEC61A1 | 29927 |
| ENSG000000 | 147.717105 | 0.660143762 | 0.14952697 | 4.41488092 | 1.01E-05 | 0.00014774 | SNAPC1 | 6617 |
| ENSG000001 | 1916.94609 | 0.541624075 | 0.12268814 | 4.41464093 | 1.01E-05 | 0.00014785 | PSMC6 | 5706 |
| ENSG000001 | 452.411494 | 0.615158621 | 0.13935956 | 4.41418315 | 1.01E-05 | 0.00014811 | LIMK2 | 3985 |
| ENSG000000 | 360.095663 | 0.604913052 | 0.13705027 | 4.41380411 | 1.02E-05 | 0.00014832 | FBXO42 | 54455 |
| ENSG000002 | 116.536783 | -0.776235919 | 0.17587495 | -4.4135672 | 1.02E-05 | 0.00014843 | PKHD1L1 | 93035 |
| ENSG000001 | 1100.64177 | 0.589159387 | 0.13349557 | 4.41332539 | 1.02E-05 | 0.00014854 | ASF1A | 25842 |
| ENSG000001 | 1306.96244 | 0.573789095 | 0.13002822 | 4.41280437 | 1.02E-05 | 0.00014884 | TMEM87A | 25963 |
| ENSG000001 | 388.789524 | 0.649178111 | 0.14712111 | 4.41254218 | 1.02E-05 | 0.00014892 | PEPD | 5184 |
| ENSG000001 | 6044.90206 | 0.478371944 | 0.10841128 | 4.4125663 | 1.02E-05 | 0.00014892 | EIF2S2 | 8894 |

| | | | | | | | | | |
|---|---|---|---|---|---|---|---|---|---|
| ENSG000001 | 604.430909 | 0.633680438 | 0.14366108 | 4.41094034 | 1.03E-05 | 0.00014991 | CPVL | | 54504 |
| ENSG000001 | 267.442628 | 0.624026017 | 0.14147235 | 4.41093991 | 1.03E-05 | 0.00014991 | TUSC2 | | 11334 |
| ENSG000002 | 28.9502348 | -1.168167575 | 0.26484257 | -4.4107998 | 1.03E-05 | 0.00014996 | NA | NA | |
| ENSG000002 | 18.5225376 | -1.293208904 | 0.29322957 | -4.4102268 | 1.03E-05 | 0.0001503 | NA | NA | |
| ENSG00000C | 3477.09956 | 0.618181512 | 0.14019051 | 4.40958182 | 1.04E-05 | 0.0001507 | MYL6 | | 4637 |
| ENSG000001 | 237.684033 | 0.638979666 | 0.14491167 | 4.40944245 | 1.04E-05 | 0.00015074 | RHPN2 | | 85415 |
| ENSG000002 | 210.466437 | 0.79047744 | 0.17927358 | 4.40933592 | 1.04E-05 | 0.00015076 | DDAH2 | | 23564 |
| ENSG000001 | 479.698778 | 0.65816279 | 0.1492899 | 4.40862248 | 1.04E-05 | 0.0001512 | IGDCC3 | | 9543 |
| ENSG000001 | 649.367669 | 0.528755953 | 0.11994624 | 4.40827436 | 1.04E-05 | 0.00015139 | SCGB3A2 | | 117156 |
| ENSG000001 | 258.713898 | 0.606651735 | 0.13761962 | 4.40817779 | 1.04E-05 | 0.0001514 | DCLRE1B | | 64858 |
| ENSG000001 | 1251.76451 | 0.58352851 | 0.13237768 | 4.4080583 | 1.04E-05 | 0.00015143 | FAM168B | | 130074 |
| ENSG00000C | 1187.00969 | 0.581186354 | 0.13185885 | 4.40764004 | 1.05E-05 | 0.00015162 | TXLNA | | 200081 |
| ENSG000001 | 288.718515 | 0.683616976 | 0.15509592 | 4.40770444 | 1.04E-05 | 0.00015162 | NSDHL | | 50814 |
| ENSG000001 | 839.292639 | 0.662328864 | 0.15027283 | 4.40750916 | 1.05E-05 | 0.00015165 | NOMO1 | | 23420 |
| ENSG000001 | 265.895893 | 0.614877629 | 0.13952432 | 4.40695656 | 1.05E-05 | 0.00015199 | PCBD1 | | 5092 |
| ENSG000001 | 667.277893 | 0.737999309 | 0.16748553 | 4.40634679 | 1.05E-05 | 0.00015236 | DHCR7 | | 1717 |
| ENSG000002 | 20.1642491 | -1.261092733 | 0.28622621 | -4.4059303 | 1.05E-05 | 0.0001526 | NA | NA | |
| ENSG000001 | 1380.91573 | 0.516219583 | 0.11717621 | 4.40549814 | 1.06E-05 | 0.00015285 | POLR1D | | 51082 |
| ENSG00000C | 4166.40139 | 0.505620078 | 0.11477736 | 4.40522496 | 1.06E-05 | 0.00015299 | DNAJC10 | | 54431 |
| ENSG000001 | 1681.38293 | 0.524697478 | 0.1191233 | 4.40465883 | 1.06E-05 | 0.00015333 | UNC13B | | 10497 |
| ENSG000001 | 236.166471 | 0.694123147 | 0.15766694 | 4.40246474 | 1.07E-05 | 0.00015484 | STX10 | | 8677 |
| ENSG00000C | 2148.46189 | 0.57565426 | 0.13076743 | 4.40212264 | 1.07E-05 | 0.00015503 | DSP | | 1832 |
| ENSG000001 | 5706.89406 | 0.554793865 | 0.12605058 | 4.401359 | 1.08E-05 | 0.00015527 | EEF1B2 | | 1933 |
| ENSG000001 | 654.051789 | 0.608405569 | 0.13823208 | 4.40133409 | 1.08E-05 | 0.00015527 | RAB11FIP4 | | 84440 |
| ENSG000001 | 17507.4406 | 0.541984158 | 0.12314122 | 4.40132201 | 1.08E-05 | 0.00015527 | RPS6 | | 6194 |
| ENSG000001 | 23104.7392 | 0.518254111 | 0.11774299 | 4.40157076 | 1.07E-05 | 0.00015527 | RPS24 | | 6229 |
| ENSG000001 | 1463.9676 | 0.50802726 | 0.11542622 | 4.40131599 | 1.08E-05 | 0.00015527 | CENPU | | 79682 |
| ENSG000001 | 781.95023 | 0.535819207 | 0.12173025 | 4.40169311 | 1.07E-05 | 0.00015527 | EIF2AK3 | | 9451 |
| ENSG000001 | 170.256043 | 0.688632102 | 0.15646451 | 4.40120323 | 1.08E-05 | 0.0001553 | CLTB | | 1212 |

| | | | | | | | | | |
|---|---|---|---|---|---|---|---|---|---|
| ENSG000001 | 577.387431 | 0.570865366 | 0.12972317 | 4.40064288 | 1.08E-05 | 0.00015564 | IMPA1 | | 3612 |
| ENSG000001 | 2783.33947 | 0.516165356 | 0.11729512 | 4.40056957 | 1.08E-05 | 0.00015564 | WWC2 | | 80014 |
| ENSG000001 | 1199.58673 | 0.551623344 | 0.12536464 | 4.40015085 | 1.08E-05 | 0.00015589 | ARL1 | | 400 |
| ENSG00000C | 495.543491 | 0.73211932 | 0.16639246 | 4.39995494 | 1.08E-05 | 0.00015593 | POLD1 | | 5424 |
| ENSG000001 | 840.291586 | 0.593022998 | 0.13478463 | 4.39978218 | 1.08E-05 | 0.00015593 | RHOQ | | 23433 |
| ENSG000001 | 241.624324 | 0.735487093 | 0.16716012 | 4.39989581 | 1.08E-05 | 0.00015593 | TM7SF2 | | 7108 |
| ENSG000002 | 58.4693606 | -0.8691446 | 0.19754295 | -4.3997753 | 1.08E-05 | 0.00015593 | NA | NA | |
| ENSG000001 | 567.480641 | 0.525734929 | 0.11949798 | 4.39952981 | 1.08E-05 | 0.00015603 | NA | NA | |
| ENSG000001 | 2234.73298 | 0.52604152 | 0.11956867 | 4.39949303 | 1.09E-05 | 0.00015603 | GRSF1 | | 2926 |
| ENSG000002 | 7477.14425 | 0.595028142 | 0.13525968 | 4.39915379 | 1.09E-05 | 0.00015622 | SNORD3A | | 780851 |
| ENSG000001 | 1769.17198 | 0.588196892 | 0.13371077 | 4.39902397 | 1.09E-05 | 0.00015625 | NAT10 | | 55226 |
| ENSG000001 | 30.1876072 | -1.141307628 | 0.25945037 | -4.3989439 | 1.09E-05 | 0.00015626 | SLC5A1 | | 6523 |
| ENSG000001 | 146.110999 | 0.677241557 | 0.15396355 | 4.39871371 | 1.09E-05 | 0.00015637 | ZNF792 | | 126375 |
| ENSG00000C | 492.590415 | 0.687465007 | 0.156303 | 4.39828423 | 1.09E-05 | 0.00015662 | CECR5 | | 27440 |
| ENSG000001 | 202.325349 | 0.800185234 | 0.18194556 | 4.39793768 | 1.09E-05 | 0.00015681 | CTSD | | 1509 |
| ENSG000001 | 84.7799728 | 0.836552397 | 0.19021881 | 4.3978427 | 1.09E-05 | 0.00015683 | GALK1 | | 2584 |
| ENSG000001 | 116.747491 | 0.770372841 | 0.17517927 | 4.39762571 | 1.09E-05 | 0.00015686 | MRAP2 | | 112609 |
| ENSG000001 | 333.800286 | 0.698448062 | 0.1588233 | 4.39764241 | 1.09E-05 | 0.00015686 | GUK1 | | 2987 |
| ENSG000001 | 153.819942 | 0.655197627 | 0.14899098 | 4.39756564 | 1.09E-05 | 0.00015686 | WDYHV1 | | 55093 |
| ENSG000002 | 518.29895 | 0.719813301 | 0.16373103 | 4.39631582 | 1.10E-05 | 0.00015771 | BOP1 | | 23246 |
| ENSG000002 | 174.136745 | 0.644704105 | 0.14669559 | 4.39484314 | 1.11E-05 | 0.00015873 | NA | NA | |
| ENSG000001 | 18274.2788 | 0.574300778 | 0.13070276 | 4.39394518 | 1.11E-05 | 0.0001593 | PODXL | | 5420 |
| ENSG000001 | 340.110166 | 0.705901181 | 0.16065469 | 4.39390331 | 1.11E-05 | 0.0001593 | FAM219B | | 57184 |
| ENSG000001 | 6366.42123 | 0.54590009 | 0.12425405 | 4.39341895 | 1.12E-05 | 0.0001596 | RPL14 | | 9045 |
| ENSG00000C | 403.241602 | 0.613220127 | 0.13958161 | 4.393273 | 1.12E-05 | 0.00015962 | PIR | | 8544 |
| ENSG000001 | 340.044654 | 0.597831054 | 0.13607958 | 4.39324603 | 1.12E-05 | 0.00015962 | GSG2 | | 83903 |
| ENSG000001 | 3106.16217 | 0.525470581 | 0.11961159 | 4.39314087 | 1.12E-05 | 0.00015964 | YES1 | | 7525 |
| ENSG000001 | 309.21182 | 0.577553803 | 0.1315162 | 4.39150312 | 1.13E-05 | 0.00016075 | PAN2 | | 9924 |
| ENSG000001 | 595.242125 | 0.579336293 | 0.13192493 | 4.39140885 | 1.13E-05 | 0.00016075 | C11orf73 | | 51501 |

| | | | | | | | | |
|---|---|---|---|---|---|---|---|---|
| ENSG000001 | 585.318366 | 0.607464791 | 0.13833063 | 4.39139755 | 1.13E-05 | 0.00016075 | SPINT1 | 6692 |
| ENSG000001 | 2246.35317 | 0.475743837 | 0.10833749 | 4.39131327 | 1.13E-05 | 0.00016076 | CCNT1 | 904 |
| ENSG000001 | 6115.65466 | 0.512792812 | 0.11678796 | 4.39080201 | 1.13E-05 | 0.00016108 | CALU | 813 |
| ENSG000001 | 1428.9622 | 0.55110357 | 0.12552713 | 4.39031438 | 1.13E-05 | 0.00016134 | ALDH1B1 | 219 |
| ENSG000001 | 6646.07764 | 0.542829244 | 0.12364277 | 4.39030326 | 1.13E-05 | 0.00016134 | CCT8 | 10694 |
| ENSG000001 | 299.084708 | 0.663560715 | 0.15115296 | 4.38999488 | 1.13E-05 | 0.00016151 | ATCAY | 85300 |
| ENSG000001 | 427.064803 | 0.589699409 | 0.13435884 | 4.38898869 | 1.14E-05 | 0.00016216 | POLR2I | 5438 |
| ENSG000002 | 11.9989717 | -1.453567547 | 0.33118623 | -4.3889734 | 1.14E-05 | 0.00016216 | NA | NA |
| ENSG00000C | 1754.13186 | 0.580318981 | 0.1322288 | 4.38874859 | 1.14E-05 | 0.00016221 | MAP4 | 4134 |
| ENSG000002 | 38.2402568 | -1.052854703 | 0.23989811 | -4.3887577 | 1.14E-05 | 0.00016221 | LINC00624 | 100289211 |
| ENSG000001 | 50.1700195 | -0.94291918 | 0.21485949 | -4.3885387 | 1.14E-05 | 0.0001623 | VWA3B | 200403 |
| ENSG000001 | 617.81349 | 0.54880966 | 0.12505691 | 4.3884792 | 1.14E-05 | 0.0001623 | SLC33A1 | 9197 |
| ENSG000001 | 299.67219 | 0.644741875 | 0.14692242 | 4.38831516 | 1.14E-05 | 0.00016236 | TRMU | 55687 |
| ENSG000001 | 75.9005203 | 0.855306945 | 0.19491512 | 4.38809949 | 1.14E-05 | 0.00016247 | RHOD | 29984 |
| ENSG000001 | 2825.66055 | 0.540619613 | 0.12320576 | 4.38794118 | 1.14E-05 | 0.00016253 | MAPRE1 | 22919 |
| ENSG000001 | 949.856757 | 0.522463343 | 0.11909551 | 4.38692731 | 1.15E-05 | 0.00016312 | CBR1 | 873 |
| ENSG000001 | 2764.91504 | 0.649854682 | 0.14813444 | 4.38692493 | 1.15E-05 | 0.00016312 | PSMD2 | 5708 |
| ENSG000002 | 363.856574 | 0.608219573 | 0.13864351 | 4.38693136 | 1.15E-05 | 0.00016312 | TMEM189 | 387521 |
| ENSG00000C | 1494.25901 | 0.552178952 | 0.12587735 | 4.38664262 | 1.15E-05 | 0.00016327 | SCAMP1 | 9522 |
| ENSG000001 | 613.76779 | 0.759636557 | 0.17321182 | 4.38559304 | 1.16E-05 | 0.000164 | CMTM4 | 146223 |
| ENSG000001 | 346.391239 | 0.563560625 | 0.12852249 | 4.38491823 | 1.16E-05 | 0.00016446 | ZFP69B | 65243 |
| ENSG000001 | 1654.34986 | 0.544866952 | 0.12426182 | 4.38483008 | 1.16E-05 | 0.00016447 | DYNC1LI2 | 1783 |
| ENSG000001 | 154.148698 | 0.75301872 | 0.17176509 | 4.38400323 | 1.17E-05 | 0.00016503 | OGDHL | 55753 |
| ENSG000001 | 95.2007091 | 0.787764258 | 0.17969566 | 4.38388034 | 1.17E-05 | 0.00016507 | MVB12A | 93343 |
| ENSG000001 | 4739.15839 | 0.567170074 | 0.12939002 | 4.38341426 | 1.17E-05 | 0.00016537 | PTPN11 | 5781 |
| ENSG000001 | 241.707425 | 0.637802962 | 0.14552933 | 4.38264195 | 1.17E-05 | 0.00016584 | CCDC71 | 64925 |
| ENSG000002 | 1504.61359 | 0.624857313 | 0.14257557 | 4.38263949 | 1.17E-05 | 0.00016584 | RBM15B | 29890 |
| ENSG00000C | 46.7234883 | -1.001370635 | 0.22849408 | -4.3824795 | 1.17E-05 | 0.0001659 | ACSM2B | 348158 |
| ENSG00000C | 2055.25528 | 0.585554139 | 0.13363578 | 4.3817169 | 1.18E-05 | 0.00016643 | CMTM6 | 54918 |

| | | | | | | | | |
|---|---|---|---|---|---|---|---|---|
| ENSG000001 | 584.53854 | 0.606188456 | 0.138386 | 4.3804175 | 1.18E-05 | 0.00016731 | GATAD1 | 57798 |
| ENSG000001 | 69.9810053 | 0.888499006 | 0.20283262 | 4.38045412 | 1.18E-05 | 0.00016731 | APOBEC3B | 9582 |
| ENSG000001 | 2324.32554 | 0.556166893 | 0.12697828 | 4.38001582 | 1.19E-05 | 0.00016756 | ARF4 | 378 |
| ENSG000001 | 1773.82832 | 0.505095281 | 0.11532092 | 4.37991015 | 1.19E-05 | 0.00016758 | DLAT | 1737 |
| ENSG000001 | 1262.18466 | 0.501932583 | 0.11460923 | 4.37951268 | 1.19E-05 | 0.00016783 | CDCA2 | 157313 |
| ENSG000002 | 747.595034 | 0.551261194 | 0.12588096 | 4.37922602 | 1.19E-05 | 0.00016799 | ZNF512 | 84450 |
| ENSG000001 | 111.056107 | 0.828138764 | 0.18913231 | 4.37862126 | 1.19E-05 | 0.0001684 | NA | NA |
| ENSG000001 | 186.28179 | 0.65866117 | 0.15043404 | 4.37840514 | 1.20E-05 | 0.00016851 | STX5 | 6811 |
| ENSG000001 | 665.586112 | 0.729041489 | 0.16656779 | 4.37684547 | 1.20E-05 | 0.00016966 | BHLHB9 | 80823 |
| ENSG000002 | 526.302302 | 0.622415926 | 0.14220972 | 4.3767468 | 1.20E-05 | 0.00016967 | GDI1 | 2664 |
| ENSG000000 | 77.5559242 | 0.921168692 | 0.21050057 | 4.37608638 | 1.21E-05 | 0.00017013 | SELO | 83642 |
| ENSG000001 | 39.3744358 | -1.044965543 | 0.23882534 | -4.3754383 | 1.21E-05 | 0.00017058 | MYH1 | 4619 |
| ENSG000001 | 1520.9885 | 0.675307045 | 0.15436129 | 4.37484715 | 1.22E-05 | 0.00017098 | CDC16 | 8881 |
| ENSG000001 | 904.045357 | 0.533110088 | 0.12186039 | 4.374761 | 1.22E-05 | 0.00017099 | ZNF12 | 7559 |
| ENSG000001 | 69.7554195 | -0.91906916 | 0.21008847 | -4.3746769 | 1.22E-05 | 0.000171 | STAB2 | 55576 |
| ENSG000001 | 76.5975075 | 0.949085476 | 0.21695625 | 4.37454768 | 1.22E-05 | 0.00017104 | DOHH | 83475 |
| ENSG000001 | 190.170121 | 0.669635846 | 0.15311679 | 4.37336652 | 1.22E-05 | 0.00017191 | LRRC14 | 9684 |
| ENSG000001 | 711.917199 | 0.541849656 | 0.12392785 | 4.37229932 | 1.23E-05 | 0.0001726 | PAPD4 | 167153 |
| ENSG000001 | 628.213365 | 0.555116384 | 0.12696307 | 4.37226644 | 1.23E-05 | 0.0001726 | ORC5 | 5001 |
| ENSG000001 | 215.809194 | 0.667198957 | 0.15259579 | 4.37232874 | 1.23E-05 | 0.0001726 | ZFYVE1 | 53349 |
| ENSG000001 | 40.3163437 | -0.994687569 | 0.22750779 | -4.3721033 | 1.23E-05 | 0.00017266 | FAM153A | 285596 |
| ENSG000001 | 10777.028 | 0.546933823 | 0.12513629 | 4.37070513 | 1.24E-05 | 0.00017371 | CCT5 | 22948 |
| ENSG000002 | 380.386115 | 0.622882105 | 0.14251513 | 4.37063854 | 1.24E-05 | 0.00017371 | MZT1 | 440145 |
| ENSG000001 | 620.798463 | 0.603348301 | 0.13805698 | 4.37028475 | 1.24E-05 | 0.00017387 | HRSP12 | 10247 |
| ENSG000002 | 39.5091222 | -1.053745201 | 0.2411157 | -4.3702887 | 1.24E-05 | 0.00017387 | COL15A1 | 1306 |
| ENSG000001 | 478.329364 | 0.671028424 | 0.15354912 | 4.37012214 | 1.24E-05 | 0.00017394 | RAB35 | 11021 |
| ENSG000001 | 850.463855 | 0.567248963 | 0.12981425 | 4.36969709 | 1.24E-05 | 0.00017422 | ETS1 | 2113 |
| ENSG000002 | 52.0345803 | -0.915976017 | 0.20962378 | -4.3696189 | 1.24E-05 | 0.00017422 | LOC285626 | 285626 |
| ENSG000001 | 469.270794 | 0.560738087 | 0.12834192 | 4.36909523 | 1.25E-05 | 0.00017458 | TMEM18 | 129787 |

| | | | | | | | | |
|---|---|---|---|---|---|---|---|---|
| ENSG000001 | 322.49876 | 0.690698634 | 0.15809514 | 4.36887956 | 1.25E-05 | 0.00017469 | SHMT1 | 6470 |
| ENSG000002 | 281.851681 | 0.683630027 | 0.15650614 | 4.36807159 | 1.25E-05 | 0.00017528 | TP73-AS1 | 57212 |
| ENSG00000C | 929.615293 | 0.593836878 | 0.1359643 | 4.36759411 | 1.26E-05 | 0.0001756 | PHF20 | 51230 |
| ENSG000001 | 105.264866 | -0.737565787 | 0.16887972 | -4.367403 | 1.26E-05 | 0.00017569 | CACNA1D | 776 |
| ENSG00000C | 4976.60616 | 0.533590818 | 0.12220281 | 4.36643661 | 1.26E-05 | 0.00017641 | STARD7 | 56910 |
| ENSG000001 | 964.985978 | 0.525278176 | 0.12031974 | 4.3656856 | 1.27E-05 | 0.00017696 | ATP6V0E1 | 8992 |
| ENSG000001 | 1332.35205 | 0.525891106 | 0.1204628 | 4.36558926 | 1.27E-05 | 0.00017697 | MRPS15 | 64960 |
| ENSG000001 | 336.214504 | 0.793559639 | 0.18179338 | 4.36517341 | 1.27E-05 | 0.00017725 | ASMTL | 8623 |
| ENSG00000C | 380.877291 | 0.574134021 | 0.13157111 | 4.36367856 | 1.28E-05 | 0.0001784 | PIGS | 94005 |
| ENSG000001 | 459.20504 | 0.650002647 | 0.14901576 | 4.3619726 | 1.29E-05 | 0.00017974 | FBLIM1 | 54751 |
| ENSG000001 | 289.800675 | 0.81368985 | 0.18657012 | 4.36130859 | 1.29E-05 | 0.00018022 | LSM7 | 51690 |
| ENSG000001 | 6495.87239 | 0.558599911 | 0.12808317 | 4.36122809 | 1.29E-05 | 0.00018023 | CCT4 | 10575 |
| ENSG000001 | 523.095881 | 0.5802577 | 0.1330558 | 4.36101016 | 1.29E-05 | 0.00018023 | METTL21A | 151194 |
| ENSG000001 | 364.875182 | 0.634608142 | 0.1455169 | 4.3610613 | 1.29E-05 | 0.00018023 | TMEM11 | 8834 |
| ENSG000002 | 250.973531 | 0.579969247 | 0.13299012 | 4.36099499 | 1.29E-05 | 0.00018023 | HLA-DPB1 | 3115 |
| ENSG000001 | 2865.37699 | 0.570117689 | 0.13074077 | 4.36067268 | 1.30E-05 | 0.00018044 | YTHDF2 | 51441 |
| ENSG000001 | 2276.68321 | 0.488411441 | 0.11204195 | 4.35918364 | 1.31E-05 | 0.00018161 | RQCD1 | 9125 |
| ENSG000002 | 143.692671 | 0.813733662 | 0.18667761 | 4.35903194 | 1.31E-05 | 0.00018167 | NA | NA |
| ENSG000001 | 118.14109 | 0.751437141 | 0.17239208 | 4.35888444 | 1.31E-05 | 0.00018173 | BOK | 666 |
| ENSG000001 | 800.372139 | 0.493891548 | 0.11331767 | 4.35846889 | 1.31E-05 | 0.00018195 | BEX1 | 55859 |
| ENSG000001 | 269.805091 | 0.607119532 | 0.13929523 | 4.35850928 | 1.31E-05 | 0.00018195 | SLC25A25 | 114789 |
| ENSG000001 | 1222.9328 | 0.514124428 | 0.1179782 | 4.35779172 | 1.31E-05 | 0.00018245 | EXOC2 | 55770 |
| ENSG000001 | 1460.29672 | 0.506980213 | 0.11634221 | 4.35766361 | 1.31E-05 | 0.0001825 | NUDT5 | 11164 |
| ENSG000001 | 344.605947 | 0.659796739 | 0.15143094 | 4.35708003 | 1.32E-05 | 0.00018292 | LPIN2 | 9663 |
| ENSG000001 | 223.322321 | 0.714106878 | 0.16391852 | 4.35647459 | 1.32E-05 | 0.00018326 | PPM1F | 9647 |
| ENSG000001 | 1131.76259 | 0.741460205 | 0.17019692 | 4.3564843 | 1.32E-05 | 0.00018326 | PLCG1 | 5335 |
| ENSG000001 | 815.729718 | 0.478265229 | 0.10978309 | 4.3564562 | 1.32E-05 | 0.00018326 | SBDS | 51119 |
| ENSG000001 | 1628.22435 | 0.540943369 | 0.12417777 | 4.35620128 | 1.32E-05 | 0.00018334 | FANCI | 55215 |
| ENSG000001 | 198.256078 | 0.722175242 | 0.16577986 | 4.35623014 | 1.32E-05 | 0.00018334 | SSNA1 | 8636 |

| | | | | | | | | | |
|---|---|---|---|---|---|---|---|---|---|
| ENSG000001 | 1098.18271 | 0.531856755 | 0.12210066 | 4.35588772 | 1.33E-05 | 0.00018354 | DHX40 | | 79665 |
| ENSG000001 | 242.727261 | 0.624431257 | 0.14338552 | 4.35491154 | 1.33E-05 | 0.0001843 | SLC17A5 | | 26503 |
| ENSG000001 | 791.136653 | 0.618701325 | 0.14207577 | 4.35472794 | 1.33E-05 | 0.00018439 | FARSA | | 2193 |
| ENSG000001 | 19.9983929 | -1.293300759 | 0.29702357 | -4.3542024 | 1.34E-05 | 0.00018477 | SIRPB2 | | 284759 |
| ENSG000001 | 37.0823799 | -1.008795314 | 0.23169945 | -4.3538961 | 1.34E-05 | 0.0001849 | HKDC1 | | 80201 |
| ENSG000002 | 23.4222121 | -1.181781507 | 0.27142834 | -4.3539356 | 1.34E-05 | 0.0001849 | NA | NA | |
| ENSG000001 | 1523.01574 | 0.512994495 | 0.11784979 | 4.35295231 | 1.34E-05 | 0.00018562 | CNIH4 | | 29097 |
| ENSG000001 | 193.888745 | 0.654413755 | 0.15034221 | 4.35282773 | 1.34E-05 | 0.00018562 | NETO1 | | 81832 |
| ENSG000002 | 53.1780631 | -0.940401251 | 0.21604305 | -4.352842 | 1.34E-05 | 0.00018562 | NOX5 | | 79400 |
| ENSG000001 | 7662.91552 | 0.566838891 | 0.13023048 | 4.35258226 | 1.35E-05 | 0.0001857 | DPYSL3 | | 1809 |
| ENSG000001 | 1202.5875 | 0.60866484 | 0.13983977 | 4.35258757 | 1.35E-05 | 0.0001857 | THYN1 | | 29087 |
| ENSG000001 | 79.6651681 | -0.81170177 | 0.18649364 | -4.3524367 | 1.35E-05 | 0.00018576 | ST8SIA1 | | 6489 |
| ENSG000001 | 224.921506 | 0.679032259 | 0.15602536 | 4.35206335 | 1.35E-05 | 0.00018601 | NPTX2 | | 4885 |
| ENSG000001 | 129.494153 | 0.745767987 | 0.17136849 | 4.35183836 | 1.35E-05 | 0.00018609 | SLC17A9 | | 63910 |
| ENSG000001 | 2037.8258 | 0.48473862 | 0.11138936 | 4.35174973 | 1.35E-05 | 0.00018609 | IDE | | 3416 |
| ENSG000001 | 1264.5957 | 0.655112475 | 0.15053828 | 4.35180003 | 1.35E-05 | 0.00018609 | SREBF2 | | 6721 |
| ENSG000001 | 608.316348 | 0.517571353 | 0.118948 | 4.35124062 | 1.35E-05 | 0.00018627 | TNNT1 | | 7138 |
| ENSG000001 | 2057.85821 | 0.584391459 | 0.13429882 | 4.35142673 | 1.35E-05 | 0.00018627 | ARHGAP11A | | 9824 |
| ENSG000002 | 283.006551 | 0.60472062 | 0.13897628 | 4.35125068 | 1.35E-05 | 0.00018627 | CENPW | | 387103 |
| ENSG000002 | 9.93376151 | -1.5174045 | 0.34872056 | -4.351348 | 1.35E-05 | 0.00018627 | NA | NA | |
| ENSG000001 | 257.452816 | 0.649742002 | 0.14933974 | 4.35076425 | 1.36E-05 | 0.00018661 | COPRS | | 55352 |
| ENSG000001 | 126.109052 | 0.79071295 | 0.18175479 | 4.35043802 | 1.36E-05 | 0.00018683 | NUDT16L1 | | 84309 |
| ENSG000001 | 721.320516 | 0.540887109 | 0.12433692 | 4.35017285 | 1.36E-05 | 0.00018699 | SH3BGRL2 | | 83699 |
| ENSG00000C | 41.0113098 | -1.002241373 | 0.23042731 | -4.3494905 | 1.36E-05 | 0.00018751 | FAP | | 2191 |
| ENSG000001 | 1479.73489 | 0.54000367 | 0.12415877 | 4.34929951 | 1.37E-05 | 0.00018761 | NA | NA | |
| ENSG000001 | 2393.43603 | 0.533432972 | 0.12266191 | 4.34880717 | 1.37E-05 | 0.00018797 | SERP1 | | 27230 |
| ENSG000001 | 4.88065731 | -1.644896019 | 0.378274 | -4.3484248 | 1.37E-05 | 0.00018823 | CRYGD | | 1421 |
| ENSG000001 | 1300.75988 | 0.500656698 | 0.11514392 | 4.34809482 | 1.37E-05 | 0.00018839 | DCTN4 | | 51164 |
| ENSG000001 | 422.042776 | 0.679530775 | 0.15628183 | 4.34811106 | 1.37E-05 | 0.00018839 | CCDC8 | | 83987 |

| | | | | | | | | | |
|---|---|---|---|---|---|---|---|---|---|
| ENSG000001 | 568.998035 | 0.548078582 | 0.12606006 | 4.34775762 | 1.38E-05 | 0.00018855 | LAMTOR3 | | 8649 |
| ENSG000001 | 997.502819 | 0.53564837 | 0.12320067 | 4.34777174 | 1.38E-05 | 0.00018855 | EMC1 | | 23065 |
| ENSG00000C | 1933.60734 | 0.455168945 | 0.10470241 | 4.34726344 | 1.38E-05 | 0.00018891 | EIF4G3 | | 8672 |
| ENSG000001 | 149.778886 | 0.746086947 | 0.17164065 | 4.34679629 | 1.38E-05 | 0.00018925 | SLC2A8 | | 29988 |
| ENSG000001 | 209.984397 | 0.690803431 | 0.15897643 | 4.34531993 | 1.39E-05 | 0.00019046 | ITPRIP | | 85450 |
| ENSG000001 | 741.120009 | 0.552993774 | 0.1272656 | 4.3451944 | 1.39E-05 | 0.00019051 | FAM124A | | 220108 |
| ENSG00000C | 986.251758 | 0.601637233 | 0.13848678 | 4.34436596 | 1.40E-05 | 0.00019103 | TYRO3 | | 7301 |
| ENSG000001 | 4651.4986 | 0.521597916 | 0.12006292 | 4.3443714 | 1.40E-05 | 0.00019103 | C1QBP | | 708 |
| ENSG000002 | 36.4447832 | -1.039767689 | 0.239332 | -4.3444574 | 1.40E-05 | 0.00019103 | NA | NA | |
| ENSG000001 | 1285.69114 | 0.641571212 | 0.14770499 | 4.34359862 | 1.40E-05 | 0.00019164 | NCAPG2 | | 54892 |
| ENSG00000C | 143.774402 | 0.691753707 | 0.15928516 | 4.3428634 | 1.41E-05 | 0.00019218 | ST3GAL1 | | 6482 |
| ENSG000001 | 533.366605 | 0.634534346 | 0.14611347 | 4.34275058 | 1.41E-05 | 0.00019218 | STK11 | | 6794 |
| ENSG000002 | 753.811898 | 0.551729767 | 0.12704553 | 4.34277201 | 1.41E-05 | 0.00019218 | MRPL45 | | 84311 |
| ENSG000001 | 233.278429 | 0.701619517 | 0.16159414 | 4.34186233 | 1.41E-05 | 0.0001929 | TMC6 | | 11322 |
| ENSG00000C | 494.038755 | 0.546753325 | 0.12592941 | 4.34174435 | 1.41E-05 | 0.00019293 | LTBP1 | | 4052 |
| ENSG000001 | 2941.35842 | 0.50176154 | 0.11556855 | 4.34167883 | 1.41E-05 | 0.00019293 | SQLE | | 6713 |
| ENSG000001 | 53.3862652 | 0.93051337 | 0.2143254 | 4.34159172 | 1.41E-05 | 0.00019294 | ARRDC1 | | 92714 |
| ENSG000001 | 137.357517 | 0.702381235 | 0.16179006 | 4.34131268 | 1.42E-05 | 0.00019312 | C7orf26 | | 79034 |
| ENSG000001 | 277.554235 | 0.586692338 | 0.13514689 | 4.34114569 | 1.42E-05 | 0.00019314 | GNAL | | 2774 |
| ENSG000001 | 80.6428841 | 0.898674193 | 0.20701317 | 4.341145 | 1.42E-05 | 0.00019314 | RPP25L | | 138716 |
| ENSG000001 | 235.355635 | 0.760542415 | 0.17521048 | 4.34073594 | 1.42E-05 | 0.00019343 | MAPK7 | | 5598 |
| ENSG000001 | 541.803214 | 0.585950316 | 0.13500169 | 4.34031846 | 1.42E-05 | 0.00019374 | NGRN | | 51335 |
| ENSG000001 | 573.475568 | 0.544918407 | 0.1255512 | 4.34020869 | 1.42E-05 | 0.00019377 | FEZ2 | | 9637 |
| ENSG000001 | 457.860397 | 0.583534444 | 0.13445267 | 4.34007318 | 1.42E-05 | 0.00019382 | POC1B | | 282809 |
| ENSG00000C | 2068.3503 | 0.496305482 | 0.11436135 | 4.33980094 | 1.43E-05 | 0.000194 | TCEB3 | | 6924 |
| ENSG00000C | 137.937241 | 0.737604771 | 0.16996707 | 4.33969232 | 1.43E-05 | 0.00019403 | SLC22A17 | | 51310 |
| ENSG000001 | 1276.80653 | 0.525025712 | 0.12099388 | 4.3392751 | 1.43E-05 | 0.00019433 | FAF2 | | 23197 |
| ENSG000001 | 737.48409 | 0.624758337 | 0.14399101 | 4.3388704 | 1.43E-05 | 0.00019462 | SUN1 | | 23353 |
| ENSG000001 | 937.498268 | 0.548033968 | 0.12631212 | 4.33872829 | 1.43E-05 | 0.00019468 | PRPF4 | | 9128 |

| | | | | | | | | | |
|---|---|---|---|---|---|---|---|---|---|
| ENSG000001 | 1808.00001 | 0.576702485 | 0.13292922 | 4.33841779 | 1.44E-05 | 0.00019489 | TP53 | | 7157 |
| ENSG000001 | 186.867623 | 0.686194272 | 0.15820947 | 4.33725148 | 1.44E-05 | 0.00019587 | TMEM223 | | 79064 |
| ENSG000001 | 239.346433 | 0.590059727 | 0.13605791 | 4.33682793 | 1.45E-05 | 0.00019618 | ZSCAN32 | | 54925 |
| ENSG000001 | 240.299734 | 0.590117667 | 0.13607452 | 4.33672429 | 1.45E-05 | 0.0001962 | EME1 | | 146956 |
| ENSG000001 | 107.294283 | 0.717178368 | 0.16539744 | 4.33609107 | 1.45E-05 | 0.0001967 | MOB2 | | 81532 |
| ENSG000001 | 216.434667 | 0.792312696 | 0.18273585 | 4.33583599 | 1.45E-05 | 0.00019675 | ZER1 | | 10444 |
| ENSG000001 | 498.744553 | 0.689711019 | 0.15907038 | 4.33588593 | 1.45E-05 | 0.00019675 | C9orf114 | | 51490 |
| ENSG000002 | 427.350694 | 0.66874695 | 0.15423753 | 4.33582495 | 1.45E-05 | 0.00019675 | NA | NA | |
| ENSG000001 | 308.154443 | 0.760649124 | 0.17544862 | 4.33545235 | 1.45E-05 | 0.00019701 | PCNXL3 | | 399909 |
| ENSG000001 | 1063.67838 | 0.547483015 | 0.12631103 | 4.33440377 | 1.46E-05 | 0.00019789 | UBE2G2 | | 7327 |
| ENSG000001 | 570.195368 | 0.539941174 | 0.124583 | 4.33398749 | 1.46E-05 | 0.00019818 | SRD5A1 | | 6715 |
| ENSG000002 | 270.274778 | 0.647109663 | 0.14931222 | 4.33393629 | 1.46E-05 | 0.00019818 | NA | NA | |
| ENSG000002 | 22.0534561 | -1.24392274 | 0.28702978 | -4.3337758 | 1.47E-05 | 0.00019826 | NA | NA | |
| ENSG000001 | 581.32205 | 0.576282374 | 0.13297972 | 4.33361097 | 1.47E-05 | 0.0001983 | UBAC1 | | 10422 |
| ENSG000001 | 84.3184686 | -0.836715567 | 0.19307738 | -4.3335764 | 1.47E-05 | 0.0001983 | CFAP54 | | 144535 |
| ENSG000001 | 1087.73952 | 0.662119424 | 0.15279093 | 4.33349949 | 1.47E-05 | 0.00019831 | MCM3AP | | 8888 |
| ENSG000002 | 1082.34904 | 0.61942752 | 0.14295646 | 4.33298037 | 1.47E-05 | 0.00019871 | NA | NA | |
| ENSG000001 | 788.935048 | 0.566122307 | 0.13066097 | 4.33275771 | 1.47E-05 | 0.00019884 | JAG1 | | 182 |
| ENSG000001 | 99.6269205 | 0.7916268 | 0.18271403 | 4.33260007 | 1.47E-05 | 0.00019892 | NME3 | | 4832 |
| ENSG000001 | 259.845785 | 0.58490866 | 0.13500788 | 4.33240386 | 1.47E-05 | 0.00019903 | PKIA | | 5569 |
| ENSG000001 | 295.301689 | 0.693662845 | 0.16012074 | 4.3321237 | 1.48E-05 | 0.00019922 | ENTPD6 | | 955 |
| ENSG000001 | 15.3259351 | -1.345177793 | 0.31052379 | -4.3319637 | 1.48E-05 | 0.0001993 | NA | NA | |
| ENSG00000C | 1422.88232 | 0.512929855 | 0.11841342 | 4.33168689 | 1.48E-05 | 0.00019936 | SNRNP40 | | 9410 |
| ENSG000001 | 187.43351 | 0.786529429 | 0.18157625 | 4.33167565 | 1.48E-05 | 0.00019936 | ILVBL | | 10994 |
| ENSG000001 | 1447.94348 | 0.495037912 | 0.11427991 | 4.33180165 | 1.48E-05 | 0.00019936 | METAP1 | | 23173 |
| ENSG000001 | 3433.18265 | 0.498918984 | 0.11518154 | 4.33158803 | 1.48E-05 | 0.00019937 | TSR1 | | 55720 |
| ENSG000001 | 3446.18238 | 0.576291301 | 0.13304666 | 4.33149753 | 1.48E-05 | 0.00019939 | SSR3 | | 6747 |
| ENSG000001 | 669.281702 | 0.521430663 | 0.12038381 | 4.33140173 | 1.48E-05 | 0.00019941 | ZBTB2 | | 57621 |
| ENSG000001 | 989.451499 | 0.561444429 | 0.12966977 | 4.32980212 | 1.49E-05 | 0.00020073 | DDX50 | | 79009 |

| | | | | | | | | |
|---|---|---|---|---|---|---|---|---|
| ENSG000001 | 354.515601 | 0.531768142 | 0.12281384 | 4.32987138 | 1.49E-05 | 0.00020073 | LIN7A | 8825 |
| ENSG000001 | 394.612449 | 0.598908659 | 0.13832637 | 4.3296782 | 1.49E-05 | 0.00020078 | SLC16A2 | 6567 |
| ENSG000001 | 1381.19406 | 0.500559992 | 0.11563128 | 4.32893241 | 1.50E-05 | 0.00020139 | BUD31 | 8896 |
| ENSG000001 | 743.2009 | 0.57591535 | 0.13304898 | 4.32859657 | 1.50E-05 | 0.00020163 | GALNT3 | 2591 |
| ENSG000001 | 1799.86917 | 0.488279798 | 0.11281045 | 4.3283205 | 1.50E-05 | 0.00020182 | ENPP1 | 5167 |
| ENSG000001 | 67.9404207 | 0.860943911 | 0.19891839 | 4.32812625 | 1.50E-05 | 0.00020186 | SPSB3 | 90864 |
| ENSG000001 | 104.55259 | 0.76783052 | 0.17740227 | 4.3281889 | 1.50E-05 | 0.00020186 | TMEM150A | 129303 |
| ENSG000001 | 467.451837 | 0.548946978 | 0.12684342 | 4.32775289 | 1.51E-05 | 0.00020214 | SNRK | 54861 |
| ENSG00000C | 480.117167 | 0.639632949 | 0.14782428 | 4.32698158 | 1.51E-05 | 0.00020278 | ANKS1A | 23294 |
| ENSG000001 | 1601.08275 | 0.605315727 | 0.13990045 | 4.32676047 | 1.51E-05 | 0.00020291 | RPL22L1 | 200916 |
| ENSG000001 | 317.204094 | 0.59234328 | 0.13690583 | 4.3266476 | 1.51E-05 | 0.00020295 | ZNF543 | 125919 |
| ENSG000001 | 202.172878 | 0.636347273 | 0.14708764 | 4.32631368 | 1.52E-05 | 0.00020319 | MOAP1 | 64112 |
| ENSG00000C | 61.3638589 | -0.87843962 | 0.20305476 | -4.3261217 | 1.52E-05 | 0.0002033 | CFAP61 | 26074 |
| ENSG000001 | 1044.99531 | 0.502436758 | 0.11614342 | 4.32600274 | 1.52E-05 | 0.00020334 | ALG8 | 79053 |
| ENSG00000C | 2129.33339 | 0.542569968 | 0.12542873 | 4.32572305 | 1.52E-05 | 0.00020353 | VPS35 | 55737 |
| ENSG000001 | 762.485428 | 0.502459538 | 0.11615924 | 4.32560964 | 1.52E-05 | 0.00020357 | CEP70 | 80321 |
| ENSG00000C | 4240.42153 | 0.50687866 | 0.11719007 | 4.3252697 | 1.52E-05 | 0.00020375 | CTNNA1 | 1495 |
| ENSG00000C | 1975.60968 | 0.53043985 | 0.12263604 | 4.32531778 | 1.52E-05 | 0.00020375 | ZMPSTE24 | 10269 |
| ENSG00000C | 329.677153 | 0.727934482 | 0.1683451 | 4.32406108 | 1.53E-05 | 0.0002048 | NCDN | 23154 |
| ENSG000001 | 302.738133 | 0.655119636 | 0.15152482 | 4.32351381 | 1.54E-05 | 0.00020525 | POLR3H | 171568 |
| ENSG000001 | 25.5254147 | -1.157283087 | 0.26768161 | -4.3233567 | 1.54E-05 | 0.00020532 | ADAM2 | 2515 |
| ENSG000001 | 16172.5237 | 0.539127927 | 0.12470807 | 4.32311965 | 1.54E-05 | 0.00020548 | RPL23 | 9349 |
| ENSG000001 | 1217.04492 | 0.602305312 | 0.13934172 | 4.32250537 | 1.54E-05 | 0.00020598 | COTL1 | 23406 |
| ENSG000001 | 245.200791 | 0.633671197 | 0.1466068 | 4.32224979 | 1.54E-05 | 0.00020615 | TIMP1 | 7076 |
| ENSG000001 | 2570.02656 | 0.550894767 | 0.12746985 | 4.32176528 | 1.55E-05 | 0.00020651 | RCN2 | 5955 |
| ENSG000001 | 793.156025 | 0.61259236 | 0.14174731 | 4.32172116 | 1.55E-05 | 0.00020651 | CENPO | 79172 |
| ENSG000001 | 130.075835 | 0.71440664 | 0.16533135 | 4.32105972 | 1.55E-05 | 0.000207 | DNPH1 | 10591 |
| ENSG000001 | 31.9767139 | -1.135197893 | 0.26270983 | -4.3211093 | 1.55E-05 | 0.000207 | LY9 | 4063 |
| ENSG000001 | 111.435532 | 0.681275371 | 0.15767372 | 4.32079216 | 1.55E-05 | 0.00020718 | STARD13 | 90627 |

| | | | | | | | | |
|---|---|---|---|---|---|---|---|---|
| ENSG000002 | 1027.05806 | 0.597668822 | 0.13833311 | 4.32050443 | 1.56E-05 | 0.00020738 | SACM1L | 22908 |
| ENSG000001 | 594.729192 | 0.612679891 | 0.14181194 | 4.32036888 | 1.56E-05 | 0.00020744 | ARL6IP5 | 10550 |
| ENSG000001 | 99.3954917 | 0.743044955 | 0.17199304 | 4.32020353 | 1.56E-05 | 0.00020753 | STK32C | 282974 |
| ENSG000001 | 3294.34133 | 0.513697029 | 0.11891608 | 4.31982817 | 1.56E-05 | 0.00020781 | CDCA7L | 55536 |
| ENSG000001 | 718.328566 | 0.575976528 | 0.13335634 | 4.31907861 | 1.57E-05 | 0.00020845 | KBTBD2 | 25948 |
| ENSG00000C | 47.4406556 | -1.01219584 | 0.2343672 | -4.3188459 | 1.57E-05 | 0.00020853 | ESR1 | 2099 |
| ENSG000001 | 1127.29263 | 0.514732502 | 0.1191828 | 4.31884888 | 1.57E-05 | 0.00020853 | DUT | 1854 |
| ENSG000001 | 1405.78621 | 0.6130967 | 0.14198903 | 4.31791593 | 1.58E-05 | 0.00020935 | OSTC | 58505 |
| ENSG000001 | 7938.11555 | 0.548728724 | 0.12709723 | 4.31739335 | 1.58E-05 | 0.00020975 | IDH1 | 3417 |
| ENSG000001 | 970.33607 | 0.547491816 | 0.12681217 | 4.31734448 | 1.58E-05 | 0.00020975 | MRPL49 | 740 |
| ENSG000001 | 184.538681 | 0.707217048 | 0.16381238 | 4.3172382 | 1.58E-05 | 0.00020978 | SERTAD3 | 29946 |
| ENSG000001 | 1905.90904 | 0.518900841 | 0.12020546 | 4.31678246 | 1.58E-05 | 0.00021015 | SCO1 | 6341 |
| ENSG000001 | 1121.03527 | 0.647162195 | 0.14992475 | 4.31658009 | 1.58E-05 | 0.00021022 | OGDH | 4967 |
| ENSG000002 | 233.486745 | -0.645095708 | 0.14944676 | -4.3165587 | 1.58E-05 | 0.00021022 | LINC-ROR | 100885779 |
| ENSG000001 | 286.554695 | 0.711168051 | 0.16478182 | 4.31581623 | 1.59E-05 | 0.00021075 | FGFR4 | 2264 |
| ENSG000002 | 18.2714494 | -1.290814347 | 0.2990911 | -4.3157899 | 1.59E-05 | 0.00021075 | NA | NA |
| ENSG000002 | 45.9274823 | 0.977959481 | 0.22659715 | 4.31585082 | 1.59E-05 | 0.00021075 | NA | NA |
| ENSG000001 | 376.618273 | 0.631037483 | 0.14624147 | 4.31503793 | 1.60E-05 | 0.0002114 | RIPK2 | 8767 |
| ENSG000001 | 78.1999279 | 0.796376238 | 0.18457683 | 4.31460565 | 1.60E-05 | 0.00021174 | MT1F | 4494 |
| ENSG00000C | 274.235019 | 0.660614036 | 0.1531188 | 4.31438866 | 1.60E-05 | 0.00021176 | METTL22 | 79091 |
| ENSG000001 | 165.650114 | 0.658812813 | 0.15270188 | 4.31437252 | 1.60E-05 | 0.00021176 | TMEM125 | 128218 |
| ENSG000001 | 346.970229 | 0.68187785 | 0.15804522 | 4.31444774 | 1.60E-05 | 0.00021176 | MYO1C | 4641 |
| ENSG000001 | 401.606827 | 0.799256277 | 0.18528886 | 4.3135689 | 1.61E-05 | 0.00021246 | CDK9 | 1025 |
| ENSG000001 | 431.979519 | 0.714186715 | 0.16558124 | 4.31321045 | 1.61E-05 | 0.00021274 | ABR | 29 |
| ENSG000001 | 340.070864 | 0.669023453 | 0.15511837 | 4.31298671 | 1.61E-05 | 0.00021288 | SLC35E1 | 79939 |
| ENSG000001 | 1706.51667 | 0.594296281 | 0.13780943 | 4.31245008 | 1.61E-05 | 0.00021333 | CDC25A | 993 |
| ENSG000002 | 196.979868 | 0.675312539 | 0.15660869 | 4.31210119 | 1.62E-05 | 0.0002136 | NA | NA |
| ENSG000001 | 108.025454 | 0.773503407 | 0.17938313 | 4.31201869 | 1.62E-05 | 0.00021361 | CHMP6 | 79643 |
| ENSG000001 | 854.010512 | 0.571720123 | 0.13260213 | 4.31154542 | 1.62E-05 | 0.000214 | PNO1 | 56902 |

| | | | | | | | | |
|---|---|---|---|---|---|---|---|---|
| ENSG000001 | 61.7365987 | 0.885295549 | 0.20534459 | 4.31126794 | 1.62E-05 | 0.0002142 | BHLHE40 | 8553 |
| ENSG000001 | 1816.88465 | 0.594974515 | 0.13801603 | 4.31090879 | 1.63E-05 | 0.00021441 | ARF1 | 375 |
| ENSG000001 | 15.860013 | 1.353442542 | 0.31395546 | 4.31093809 | 1.63E-05 | 0.00021441 | PF4 | 5196 |
| ENSG00000C | 56.2912312 | -0.876411609 | 0.20330973 | -4.3107213 | 1.63E-05 | 0.00021452 | CFH | 3075 |
| ENSG000001 | 546.623762 | 0.510054526 | 0.11833653 | 4.3102034 | 1.63E-05 | 0.00021495 | CLCN5 | 1184 |
| ENSG000001 | 209.419093 | 0.653623797 | 0.15166027 | 4.30978909 | 1.63E-05 | 0.00021521 | PHYH | 5264 |
| ENSG000001 | 2200.2084 | 0.513607903 | 0.11917194 | 4.30980552 | 1.63E-05 | 0.00021521 | DENR | 8562 |
| ENSG000001 | 59.8662585 | -0.923970705 | 0.21441018 | -4.3093603 | 1.64E-05 | 0.00021556 | KLRD1 | 3824 |
| ENSG000001 | 702.214774 | 0.636957561 | 0.14782872 | 4.30875392 | 1.64E-05 | 0.00021608 | NPM3 | 10360 |
| ENSG000001 | 1435.11174 | 0.461687103 | 0.10715417 | 4.30862453 | 1.64E-05 | 0.00021614 | EMB | 133418 |
| ENSG000001 | 13.1917883 | -1.414548653 | 0.3283155 | -4.308504 | 1.64E-05 | 0.00021619 | OR10K2 | 391107 |
| ENSG00000C | 2680.75419 | 0.507477171 | 0.11778991 | 4.3083248 | 1.64E-05 | 0.00021629 | PTPN4 | 5775 |
| ENSG000001 | 1491.75543 | 0.489165204 | 0.11355081 | 4.3078971 | 1.65E-05 | 0.00021664 | PPID | 5481 |
| ENSG000001 | 4390.16697 | 0.523310814 | 0.12149203 | 4.30736727 | 1.65E-05 | 0.00021709 | CYCS | 54205 |
| ENSG00000C | 116.528986 | 0.685113057 | 0.15908882 | 4.3064814 | 1.66E-05 | 0.00021772 | RHBDF1 | 64285 |
| ENSG000001 | 12392.4762 | 0.498214714 | 0.11568601 | 4.30661144 | 1.66E-05 | 0.00021772 | SLC38A1 | 81539 |
| ENSG000001 | 2012.19094 | 0.57299513 | 0.1330553 | 4.30644348 | 1.66E-05 | 0.00021772 | PRDX3 | 10935 |
| ENSG000001 | 550.478438 | 0.557066391 | 0.12935643 | 4.30644545 | 1.66E-05 | 0.00021772 | MRPS11 | 64963 |
| ENSG000001 | 1600.06812 | 0.605736787 | 0.14066241 | 4.30631592 | 1.66E-05 | 0.00021777 | LRRC8A | 56262 |
| ENSG00000C | 268.520995 | 0.684589821 | 0.1589898 | 4.30587262 | 1.66E-05 | 0.00021789 | GTF2IRD1 | 9569 |
| ENSG000001 | 1757.20124 | 0.56073643 | 0.13022116 | 4.30603176 | 1.66E-05 | 0.00021789 | TJP2 | 9414 |
| ENSG000001 | 309.101366 | 0.712275403 | 0.16540985 | 4.30612446 | 1.66E-05 | 0.00021789 | ZNF282 | 8427 |
| ENSG000001 | 299.807593 | 0.601222931 | 0.13962962 | 4.30584106 | 1.66E-05 | 0.00021789 | NDNL2 | 56160 |
| ENSG000001 | 298.240463 | 0.739222838 | 0.17167485 | 4.3059471 | 1.66E-05 | 0.00021789 | PPP1R26 | 9858 |
| ENSG000001 | 206.620507 | 0.6582133 | 0.15289186 | 4.30509042 | 1.67E-05 | 0.00021856 | ACTR1B | 10120 |
| ENSG000001 | 3094.71772 | 0.482283311 | 0.11203761 | 4.30465549 | 1.67E-05 | 0.00021891 | DNAJC8 | 22826 |
| ENSG000001 | 1027.89092 | 0.563057421 | 0.13080754 | 4.30447229 | 1.67E-05 | 0.00021903 | STAT3 | 6774 |
| ENSG000001 | 757.503867 | 0.676908861 | 0.15726563 | 4.30423894 | 1.68E-05 | 0.00021919 | JUP | 3728 |
| ENSG000002 | 70.5376856 | 0.841942387 | 0.19562112 | 4.30394411 | 1.68E-05 | 0.00021934 | NA | NA |

| | | | | | | | | |
|---|---|---|---|---|---|---|---|---|
| ENSG000002 | 2115.41415 | 0.496155556 | 0.11527911 | 4.30395032 | 1.68E-05 | 0.00021934 | NA | NA |
| ENSG000001 | 9673.6401 | 0.550037838 | 0.12782 | 4.30322196 | 1.68E-05 | 0.00021992 | SKIL | 6498 |
| ENSG000001 | 387.562141 | 0.607895092 | 0.14126552 | 4.30320936 | 1.68E-05 | 0.00021992 | NMRAL1 | 57407 |
| ENSG000001 | 677.840724 | 0.541325359 | 0.12580682 | 4.30283015 | 1.69E-05 | 0.00022023 | SEC11C | 90701 |
| ENSG000000 | 38.4329005 | -1.032401534 | 0.23997842 | -4.3020599 | 1.69E-05 | 0.00022089 | FYB | 2533 |
| ENSG000002 | 15831.5179 | 0.526613433 | 0.12241059 | 4.30202527 | 1.69E-05 | 0.00022089 | NA | NA |
| ENSG000002 | 300.19496 | 0.639749259 | 0.14871293 | 4.30190755 | 1.69E-05 | 0.00022094 | NA | NA |
| ENSG000001 | 651.564731 | 0.603204427 | 0.14022235 | 4.30177104 | 1.69E-05 | 0.000221 | CDKN1B | 1027 |
| ENSG000001 | 59.2175326 | 0.886352166 | 0.20606473 | 4.30132875 | 1.70E-05 | 0.00022137 | TRIM54 | 57159 |
| ENSG000000 | 448.64453 | 0.627292745 | 0.14584994 | 4.30094611 | 1.70E-05 | 0.00022164 | MYLIP | 29116 |
| ENSG000001 | 2009.15845 | 0.492524287 | 0.1145161 | 4.30091755 | 1.70E-05 | 0.00022164 | ZNF107 | 51427 |
| ENSG000001 | 766.926706 | 0.509027548 | 0.11836673 | 4.30042757 | 1.70E-05 | 0.00022206 | RHOT1 | 55288 |
| ENSG000001 | 1220.41189 | 0.539727191 | 0.12551811 | 4.2999947 | 1.71E-05 | 0.00022242 | RFC3 | 5983 |
| ENSG000001 | 435.003529 | 0.675847171 | 0.15723537 | 4.29831521 | 1.72E-05 | 0.00022404 | POLR2H | 5437 |
| ENSG000001 | 2569.57102 | 0.514157235 | 0.11963363 | 4.29776513 | 1.73E-05 | 0.00022453 | UTRN | 7402 |
| ENSG000001 | 854.584572 | 0.559867831 | 0.13027697 | 4.29751974 | 1.73E-05 | 0.0002247 | PHKB | 5257 |
| ENSG000000 | 679.341902 | 0.684821936 | 0.15935908 | 4.29735115 | 1.73E-05 | 0.0002248 | DGCR2 | 9993 |
| ENSG000001 | 1193.87403 | 0.520729431 | 0.12118523 | 4.29697105 | 1.73E-05 | 0.00022511 | FREM2 | 341640 |
| ENSG000001 | 306.759263 | 0.642744896 | 0.14959773 | 4.29648836 | 1.74E-05 | 0.0002255 | C16orf58 | 64755 |
| ENSG000001 | 1168.41952 | 0.515743187 | 0.12003935 | 4.29645094 | 1.74E-05 | 0.0002255 | RAB8A | 4218 |
| ENSG000000 | 83.9416375 | 0.774830728 | 0.18034792 | 4.29631089 | 1.74E-05 | 0.00022554 | TMEM38A | 79041 |
| ENSG000001 | 229.067438 | 0.742408151 | 0.17280305 | 4.29626768 | 1.74E-05 | 0.00022554 | TAF1C | 9013 |
| ENSG000001 | 7029.92222 | 0.519791178 | 0.12099592 | 4.29593978 | 1.74E-05 | 0.0002258 | SRSF1 | 6426 |
| ENSG000001 | 41.3395595 | -1.038885376 | 0.24184752 | -4.2956213 | 1.74E-05 | 0.00022605 | PDE11A | 50940 |
| ENSG000001 | 298.413967 | 0.631527485 | 0.14705195 | 4.29458756 | 1.75E-05 | 0.00022704 | ZNF574 | 64763 |
| ENSG000001 | 2133.91454 | 0.458778477 | 0.10683919 | 4.29410298 | 1.75E-05 | 0.00022746 | OCIAD1 | 54940 |
| ENSG000001 | 609.335518 | 0.567130057 | 0.13207848 | 4.29388689 | 1.76E-05 | 0.00022761 | GPC3 | 2719 |
| ENSG000002 | 226.265562 | 0.693437197 | 0.16150641 | 4.29355832 | 1.76E-05 | 0.00022781 | SIPA1 | 6494 |
| ENSG000002 | 550.95542 | 0.651552943 | 0.15175173 | 4.29354544 | 1.76E-05 | 0.00022781 | NA | NA |

| | | | | | | | | |
|---|---|---|---|---|---|---|---|---|
| ENSG000002 | 74.1597521 | 0.932985846 | 0.21730428 | 4.29345366 | 1.76E-05 | 0.00022784 | LOC100506691 | 100506691 |
| ENSG000001 | 952.047342 | 0.637380231 | 0.14846484 | 4.29313932 | 1.76E-05 | 0.00022794 | NFE2L2 | 4780 |
| ENSG000001 | 1004.45335 | 0.580239924 | 0.13515673 | 4.29308937 | 1.76E-05 | 0.00022794 | STK35 | 140901 |
| ENSG000001 | 856.088706 | 0.510602387 | 0.11893164 | 4.29324266 | 1.76E-05 | 0.00022794 | KIAA1324L | 222223 |
| ENSG000002 | 35.3630535 | 1.05035405 | 0.24466276 | 4.29306876 | 1.76E-05 | 0.00022794 | NA | NA |
| ENSG000001 | 2934.006 | 0.511712111 | 0.11920459 | 4.2927216 | 1.76E-05 | 0.00022817 | STT3A | 3703 |
| ENSG000001 | 216.549371 | 0.639259172 | 0.14891777 | 4.29269904 | 1.77E-05 | 0.00022817 | ATP6V0E2 | 155066 |
| ENSG00000C | 460.997584 | 0.52582269 | 0.12249614 | 4.29256548 | 1.77E-05 | 0.00022824 | EPB41L3 | 23136 |
| ENSG00000C | 906.024621 | 0.605070161 | 0.14098842 | 4.29163023 | 1.77E-05 | 0.00022913 | MAVS | 57506 |
| ENSG000001 | 534.943021 | 0.587101499 | 0.13681782 | 4.29111874 | 1.78E-05 | 0.00022958 | VOPP1 | 81552 |
| ENSG000001 | 747.146992 | 0.558381085 | 0.13013855 | 4.29066621 | 1.78E-05 | 0.00022995 | DBT | 1629 |
| ENSG000001 | 2793.79105 | 0.474581039 | 0.11060891 | 4.29062226 | 1.78E-05 | 0.00022995 | STRBP | 55342 |
| ENSG000001 | 842.320937 | 0.586352064 | 0.13667882 | 4.28999939 | 1.79E-05 | 0.00023045 | MGAT5 | 4249 |
| ENSG000001 | 1626.49287 | 0.484260295 | 0.11287962 | 4.29005943 | 1.79E-05 | 0.00023045 | DZIP3 | 9666 |
| ENSG00000C | 494.638599 | 0.567802462 | 0.13237587 | 4.28931984 | 1.79E-05 | 0.00023101 | MFAP3 | 4238 |
| ENSG000002 | 540.336576 | 0.687174335 | 0.1602042 | 4.28936529 | 1.79E-05 | 0.00023101 | MIAT | 440823 |
| ENSG00000C | 541.806871 | 0.5765977 | 0.13442893 | 4.28923828 | 1.79E-05 | 0.00023102 | TRO | 7216 |
| ENSG000001 | 1523.62343 | 0.716144175 | 0.16698607 | 4.28864628 | 1.80E-05 | 0.00023157 | TTLL12 | 23170 |
| ENSG00000C | 975.718448 | 0.562182292 | 0.13111756 | 4.28762005 | 1.81E-05 | 0.00023251 | STRN4 | 29888 |
| ENSG000001 | 1849.67408 | 0.519987102 | 0.12127886 | 4.28753284 | 1.81E-05 | 0.00023251 | METTL9 | 51108 |
| ENSG000002 | 141.400167 | 0.672488457 | 0.15684591 | 4.28757397 | 1.81E-05 | 0.00023251 | NA | NA |
| ENSG000001 | 1014.87245 | 0.517263553 | 0.12066092 | 4.28691859 | 1.81E-05 | 0.00023308 | EAF1 | 85403 |
| ENSG000001 | 323.156645 | 0.743910262 | 0.17355214 | 4.28637905 | 1.82E-05 | 0.00023357 | GGA1 | 26088 |
| ENSG000001 | 17783.9529 | 0.577018765 | 0.13462233 | 4.28620392 | 1.82E-05 | 0.0002336 | TPT1 | 7178 |
| ENSG000002 | 69.6078761 | 0.853263053 | 0.19907102 | 4.28622438 | 1.82E-05 | 0.0002336 | MRM1 | 79922 |
| ENSG000001 | 345.285079 | 0.550558307 | 0.12845245 | 4.28608641 | 1.82E-05 | 0.00023365 | NIT2 | 56954 |
| ENSG000001 | 70.7293804 | 0.905800333 | 0.2113654 | 4.28547122 | 1.82E-05 | 0.00023423 | LGALS1 | 3956 |
| ENSG00000C | 2351.63508 | 0.567608388 | 0.13246215 | 4.28506088 | 1.83E-05 | 0.00023459 | TFRC | 7037 |
| ENSG000001 | 54.7719689 | 1.005854607 | 0.23475849 | 4.28463568 | 1.83E-05 | 0.00023496 | RNU5A-1 | 26831 |

| | | | | | | | | |
|---|---|---|---|---|---|---|---|---|
| ENSG000001 | 901.531383 | 0.574070691 | 0.13398881 | 4.28446729 | 1.83E-05 | 0.00023504 | CCNF | 899 |
| ENSG000002 | 145.336856 | 0.647180231 | 0.15105429 | 4.28442153 | 1.83E-05 | 0.00023504 | TIMM23B | 100652748 |
| ENSG000000 | 2119.55918 | 0.502742522 | 0.11735349 | 4.28400138 | 1.84E-05 | 0.00023541 | CDC7 | 8317 |
| ENSG000001 | 160.188207 | 0.633409641 | 0.14789087 | 4.28295306 | 1.84E-05 | 0.00023637 | PEX5L | 51555 |
| ENSG000001 | 458.498761 | 0.671427779 | 0.15676562 | 4.28300393 | 1.84E-05 | 0.00023637 | OBSL1 | 23363 |
| ENSG000001 | 913.160513 | 0.57966943 | 0.13537949 | 4.2818114 | 1.85E-05 | 0.00023751 | PCID2 | 55795 |
| ENSG000001 | 265.511451 | 0.566599663 | 0.13234156 | 4.28134332 | 1.86E-05 | 0.00023794 | TMEM192 | 201931 |
| ENSG000001 | 288.54323 | 0.654951616 | 0.15298868 | 4.28104632 | 1.86E-05 | 0.0002381 | PRKRIP1 | 79706 |
| ENSG000001 | 614.519704 | 0.574339989 | 0.13415865 | 4.28105088 | 1.86E-05 | 0.0002381 | PPA2 | 27068 |
| ENSG000001 | 419.669414 | 0.589795623 | 0.13777447 | 4.28087758 | 1.86E-05 | 0.00023821 | UBIAD1 | 29914 |
| ENSG000001 | 535.193425 | 0.602449595 | 0.14073735 | 4.28066614 | 1.86E-05 | 0.00023836 | NT5C3B | 115024 |
| ENSG000000 | 518.235028 | 0.604080873 | 0.14112792 | 4.2803783 | 1.87E-05 | 0.00023852 | USP36 | 57602 |
| ENSG000001 | 3381.73322 | 0.525005331 | 0.12265387 | 4.28038119 | 1.87E-05 | 0.00023852 | BMS1 | 9790 |
| ENSG000000 | 1199.34354 | 0.518707839 | 0.12118492 | 4.28030012 | 1.87E-05 | 0.00023853 | PSMC4 | 5704 |
| ENSG000001 | 355.693691 | 0.564054886 | 0.1317923 | 4.27987734 | 1.87E-05 | 0.00023891 | COQ7 | 10229 |
| ENSG000001 | 116.018746 | -0.6700057 | 0.15656265 | -4.2794734 | 1.87E-05 | 0.00023927 | GRIA1 | 2890 |
| ENSG000001 | 936.095282 | 0.673539408 | 0.15740073 | 4.27913778 | 1.88E-05 | 0.00023955 | MAGED1 | 9500 |
| ENSG000001 | 168.777834 | 0.673510958 | 0.15739721 | 4.27905274 | 1.88E-05 | 0.00023957 | ACOXL | 55289 |
| ENSG000001 | 264.55084 | 0.631152895 | 0.14750703 | 4.2787987 | 1.88E-05 | 0.00023976 | PMEPA1 | 56937 |
| ENSG000001 | 363.572378 | 0.578964683 | 0.13532255 | 4.27840507 | 1.88E-05 | 0.00024011 | DSN1 | 79980 |
| ENSG000001 | 435.949818 | 0.538396584 | 0.12585239 | 4.27800054 | 1.89E-05 | 0.00024047 | RBMX2 | 51634 |
| ENSG000001 | 279.232465 | 0.624695063 | 0.14606944 | 4.27669914 | 1.90E-05 | 0.00024181 | FBXL12 | 54850 |
| ENSG000002 | 378.448253 | 0.72684562 | 0.1699727 | 4.27624913 | 1.90E-05 | 0.00024222 | IFRD2 | 7866 |
| ENSG000000 | 14163.9873 | 0.518295695 | 0.12120583 | 4.27616151 | 1.90E-05 | 0.00024224 | SEPHS1 | 22929 |
| ENSG000001 | 97.860436 | -0.781808327 | 0.18283822 | -4.2759567 | 1.90E-05 | 0.00024235 | CSMD3 | 114788 |
| ENSG000002 | 126.679518 | 0.84353882 | 0.19727673 | 4.27591653 | 1.90E-05 | 0.00024235 | TWF2 | 11344 |
| ENSG000002 | 50.765454 | -0.91795354 | 0.21468514 | -4.2758132 | 1.90E-05 | 0.00024239 | TTC41P | 253724 |
| ENSG000001 | 366.939394 | 0.575058971 | 0.13449639 | 4.27564607 | 1.91E-05 | 0.0002425 | KLF10 | 7071 |
| ENSG000001 | 1473.54525 | 0.719002886 | 0.16817818 | 4.2752448 | 1.91E-05 | 0.00024286 | PRKCSH | 5589 |

| | | | | | | | | |
|---|---|---|---|---|---|---|---|---|
| ENSG000001 | 1673.86441 | 0.527056261 | 0.12329446 | 4.27477649 | 1.91E-05 | 0.00024329 | NUP43 | 348995 |
| ENSG000001 | 184.439689 | 0.692019606 | 0.16189283 | 4.2745537 | 1.92E-05 | 0.00024346 | ACSF3 | 197322 |
| ENSG00000C | 1202.73707 | 0.523274374 | 0.12242295 | 4.2743161 | 1.92E-05 | 0.00024364 | MAPKAPK5 | 8550 |
| ENSG000001 | 688.05922 | 0.549397727 | 0.12854731 | 4.27389503 | 1.92E-05 | 0.00024403 | ARL14EP | 120534 |
| ENSG000001 | 1828.03878 | 0.511562758 | 0.11969904 | 4.27374137 | 1.92E-05 | 0.00024412 | C16orf72 | 29035 |
| ENSG000001 | 734.43151 | 0.505978395 | 0.11840469 | 4.27329674 | 1.93E-05 | 0.00024445 | LHFPL2 | 10184 |
| ENSG000002 | 313.97101 | 0.536908736 | 0.12564175 | 4.27333047 | 1.93E-05 | 0.00024445 | NA | NA |
| ENSG000001 | 1521.46339 | 0.525180831 | 0.12291317 | 4.2727792 | 1.93E-05 | 0.00024494 | RAB5A | 5868 |
| ENSG00000C | 5569.96056 | 0.504089211 | 0.11798103 | 4.2726295 | 1.93E-05 | 0.00024503 | HMGB3 | 3149 |
| ENSG000002 | 29.9582579 | -1.053908897 | 0.24670332 | -4.2719689 | 1.94E-05 | 0.00024568 | NA | NA |
| ENSG00000C | 4647.42974 | 0.525621161 | 0.12304199 | 4.27188438 | 1.94E-05 | 0.0002457 | TM9SF3 | 56889 |
| ENSG000002 | 298.516333 | 0.585190543 | 0.13701611 | 4.27096171 | 1.95E-05 | 0.00024664 | LBH | 81606 |
| ENSG000001 | 413.81329 | 0.550730649 | 0.12897904 | 4.26992373 | 1.96E-05 | 0.00024771 | SLC7A6OS | 84138 |
| ENSG000001 | 728.652837 | 0.660416404 | 0.15467685 | 4.26965247 | 1.96E-05 | 0.00024794 | OCRL | 4952 |
| ENSG000002 | 44.5394457 | 0.959102705 | 0.22464624 | 4.26939127 | 1.96E-05 | 0.00024807 | NA | NA |
| ENSG000002 | 90.7974053 | 0.765730496 | 0.17935114 | 4.26944874 | 1.96E-05 | 0.00024807 | HEIH | 100859930 |
| ENSG00000C | 532.350483 | 0.608974442 | 0.14264167 | 4.26926049 | 1.96E-05 | 0.00024814 | KIF26A | 26153 |
| ENSG000001 | 126.913229 | 0.785045904 | 0.18390195 | 4.26882867 | 1.97E-05 | 0.00024855 | POMGNT2 | 84892 |
| ENSG000001 | 199.928892 | 0.664454628 | 0.15566936 | 4.26837128 | 1.97E-05 | 0.00024884 | PAFAH2 | 5051 |
| ENSG000001 | 3.86194651 | -1.607684516 | 0.37665778 | -4.2682897 | 1.97E-05 | 0.00024884 | OTOP3 | 347741 |
| ENSG000001 | 817.033925 | 0.623200358 | 0.14600541 | 4.26833752 | 1.97E-05 | 0.00024884 | NOP9 | 161424 |
| ENSG000002 | 220.577253 | 0.639601818 | 0.14984672 | 4.26837391 | 1.97E-05 | 0.00024884 | LINC00888 | 100505687 |
| ENSG00000C | 73.2789279 | 0.823666963 | 0.19305471 | 4.26649499 | 1.99E-05 | 0.00025077 | ICAM1 | 3383 |
| ENSG000001 | 365.639144 | 0.626538013 | 0.14685854 | 4.26626892 | 1.99E-05 | 0.00025094 | SFT2D1 | 113402 |
| ENSG000001 | 200.29509 | 0.634210491 | 0.14866107 | 4.26615037 | 1.99E-05 | 0.000251 | SHISA3 | 152573 |
| ENSG000001 | 283.162265 | 0.575274525 | 0.13485015 | 4.26602814 | 1.99E-05 | 0.00025106 | NUBP1 | 4682 |
| ENSG00000C | 2107.71968 | 0.519584126 | 0.1218146 | 4.2653684 | 2.00E-05 | 0.00025172 | NSUN2 | 54888 |
| ENSG000002 | 165.207082 | 0.693976666 | 0.1627191 | 4.26487537 | 2.00E-05 | 0.0002522 | LINC00339 | 29092 |
| ENSG000002 | 1247.51918 | 0.716587794 | 0.16802717 | 4.26471377 | 2.00E-05 | 0.0002523 | BAG6 | 7917 |

| | | | | | | | | |
|---|---|---|---|---|---|---|---|---|
| ENSG000001 | 877.806917 | 0.618067124 | 0.14493174 | 4.26453936 | 2.00E-05 | 0.00025234 | ANXA6 | 309 |
| ENSG000002 | 114.317783 | -0.766620141 | 0.17976562 | -4.2645536 | 2.00E-05 | 0.00025234 | NA | NA |
| ENSG000001 | 394.444024 | 0.538627987 | 0.12630841 | 4.26438726 | 2.00E-05 | 0.00025244 | G6PC3 | 92579 |
| ENSG000001 | 543.372307 | 0.71877895 | 0.16855965 | 4.26424079 | 2.01E-05 | 0.00025253 | SEMA4C | 54910 |
| ENSG000001 | 990.46063 | 0.648872113 | 0.15217113 | 4.26409483 | 2.01E-05 | 0.00025253 | RALY | 22913 |
| ENSG000001 | 297.163951 | 0.686547753 | 0.16100664 | 4.26409588 | 2.01E-05 | 0.00025253 | MXRA7 | 439921 |
| ENSG000001 | 150.741874 | -0.639124561 | 0.14990142 | -4.2636326 | 2.01E-05 | 0.00025298 | TNFRSF10A | 8797 |
| ENSG000001 | 732.429532 | 0.531508341 | 0.12468953 | 4.26265407 | 2.02E-05 | 0.00025401 | TUBG1 | 7283 |
| ENSG00000C | 672.486856 | 0.546622178 | 0.12823773 | 4.26256891 | 2.02E-05 | 0.00025403 | MED29 | 55588 |
| ENSG000001 | 275.994499 | 0.574186787 | 0.13470956 | 4.2624057 | 2.02E-05 | 0.00025414 | RNF144A | 9781 |
| ENSG000001 | 484.320253 | 0.642331826 | 0.15070766 | 4.26210464 | 2.03E-05 | 0.0002544 | USF2 | 7392 |
| ENSG000001 | 849.208473 | 0.629783107 | 0.14776804 | 4.26197099 | 2.03E-05 | 0.00025447 | PSMF1 | 9491 |
| ENSG000001 | 525.980904 | 0.546481456 | 0.12823554 | 4.2615443 | 2.03E-05 | 0.0002548 | MICU1 | 10367 |
| ENSG000002 | 472.280075 | 0.548261143 | 0.12865125 | 4.26160768 | 2.03E-05 | 0.0002548 | NA | NA |
| ENSG00000C | 73.8968956 | -0.878260919 | 0.20609653 | -4.2614057 | 2.03E-05 | 0.00025481 | LRRC7 | 57554 |
| ENSG000001 | 1131.41366 | 0.556368912 | 0.13056015 | 4.26139922 | 2.03E-05 | 0.00025481 | SLC5A6 | 8884 |
| ENSG000001 | 13027.7604 | 0.497394365 | 0.11672785 | 4.26114547 | 2.03E-05 | 0.00025502 | DDX21 | 9188 |
| ENSG000001 | 1243.86236 | 0.522605716 | 0.12265463 | 4.26079075 | 2.04E-05 | 0.00025527 | HP1BP3 | 50809 |
| ENSG000001 | 466.964411 | 0.526847106 | 0.12365001 | 4.26079313 | 2.04E-05 | 0.00025527 | SMIM12 | 113444 |
| ENSG000001 | 2213.12753 | 0.510159173 | 0.11974076 | 4.26053067 | 2.04E-05 | 0.00025549 | SMU1 | 55234 |
| ENSG00000C | 85.8887795 | -0.804227091 | 0.18877881 | -4.2601555 | 2.04E-05 | 0.00025576 | PAX6 | 5080 |
| ENSG000001 | 156.960544 | 0.796969245 | 0.18707331 | 4.26019759 | 2.04E-05 | 0.00025576 | C16orf59 | 80178 |
| ENSG000002 | 765.76684 | 0.532947213 | 0.12512956 | 4.2591631 | 2.05E-05 | 0.00025682 | NOL7 | 51406 |
| ENSG00000C | 832.436172 | 0.503855628 | 0.11830285 | 4.25903214 | 2.05E-05 | 0.00025689 | GOLGA5 | 9950 |
| ENSG000001 | 1666.34895 | 0.597036144 | 0.14019026 | 4.2587563 | 2.06E-05 | 0.00025697 | POMP | 51371 |
| ENSG000001 | 405.950357 | 0.636162973 | 0.14937282 | 4.25889387 | 2.05E-05 | 0.00025697 | C9orf91 | 203197 |
| ENSG000001 | 2061.58874 | 0.566062601 | 0.13291527 | 4.25882288 | 2.06E-05 | 0.00025697 | RYK | 6259 |
| ENSG000001 | 789.665193 | 0.560437884 | 0.13164034 | 4.25734137 | 2.07E-05 | 0.00025852 | TGOLN2 | 10618 |
| ENSG00000C | 250.587499 | 0.567860561 | 0.13340596 | 4.25663565 | 2.08E-05 | 0.00025925 | STK10 | 6793 |

| | | | | | | | | |
|---|---|---|---|---|---|---|---|---|
| ENSG000001 | 1121.55143 | 0.578481569 | 0.13591786 | 4.25611149 | 2.08E-05 | 0.00025974 | GGA2 | 23062 |
| ENSG000001 | 153.097094 | 0.747605357 | 0.17565587 | 4.25607952 | 2.08E-05 | 0.00025974 | FZR1 | 51343 |
| ENSG00000C | 419.442215 | 0.529385816 | 0.1243874 | 4.2559442 | 2.08E-05 | 0.00025982 | MRPS33 | 51650 |
| ENSG000001 | 601.24949 | 0.506199036 | 0.11896449 | 4.25504329 | 2.09E-05 | 0.00026078 | ZFAS1 | 441951 |
| ENSG000002 | 231.744639 | 0.743366646 | 0.1747424 | 4.25407136 | 2.10E-05 | 0.00026184 | PLEKHO2 | 80301 |
| ENSG00000C | 294.187674 | 0.655720203 | 0.15415396 | 4.25367073 | 2.10E-05 | 0.00026223 | ME1 | 4199 |
| ENSG000001 | 356.507367 | 0.576265403 | 0.13547948 | 4.2535255 | 2.10E-05 | 0.00026232 | KLHL36 | 79786 |
| ENSG00000C | 9989.11689 | 0.509017879 | 0.11968039 | 4.25314343 | 2.11E-05 | 0.0002626 | SUPT16H | 11198 |
| ENSG000001 | 690.820626 | 0.506525078 | 0.11909316 | 4.25318358 | 2.11E-05 | 0.0002626 | TAF4B | 6875 |
| ENSG000001 | 843.237522 | 0.520068228 | 0.12229549 | 4.25255437 | 2.11E-05 | 0.00026322 | SNX9 | 51429 |
| ENSG000001 | 46.8525478 | -0.957168027 | 0.22511819 | -4.2518467 | 2.12E-05 | 0.00026397 | CD163 | 9332 |
| ENSG000001 | 29.3944622 | -1.122498277 | 0.2640133 | -4.2516733 | 2.12E-05 | 0.00026409 | GBP5 | 115362 |
| ENSG000001 | 894.929075 | 0.531614585 | 0.12503971 | 4.25156598 | 2.12E-05 | 0.00026414 | RPF1 | 80135 |
| ENSG000001 | 3480.61296 | 0.520218433 | 0.12237013 | 4.25118797 | 2.13E-05 | 0.00026449 | SNRPD3 | 6634 |
| ENSG000001 | 641.302728 | 0.547909778 | 0.12888576 | 4.25112721 | 2.13E-05 | 0.00026449 | ZBTB14 | 7541 |
| ENSG000001 | 601.043794 | 0.528000087 | 0.12420643 | 4.25098828 | 2.13E-05 | 0.00026458 | NGLY1 | 55768 |
| ENSG000002 | 7.66755373 | -1.540921903 | 0.36249473 | -4.2508809 | 2.13E-05 | 0.00026462 | NA | NA |
| ENSG000001 | 1140.93674 | 0.543351075 | 0.12784113 | 4.25020552 | 2.14E-05 | 0.00026534 | MRPL30 | 51263 |
| ENSG000001 | 1409.46314 | 0.503359707 | 0.11844756 | 4.24964176 | 2.14E-05 | 0.00026593 | PNPT1 | 87178 |
| ENSG000001 | 479.273673 | 0.598803899 | 0.14092609 | 4.24906338 | 2.15E-05 | 0.00026653 | CNOT6L | 246175 |
| ENSG000001 | 469.961787 | 0.602317875 | 0.1417669 | 4.24864953 | 2.15E-05 | 0.00026694 | HMGXB3 | 22993 |
| ENSG000001 | 327.383754 | 0.703506436 | 0.16558696 | 4.24856173 | 2.15E-05 | 0.00026694 | ARAP1 | 116985 |
| ENSG000002 | 2135.52747 | 0.696695264 | 0.16398571 | 4.24851216 | 2.15E-05 | 0.00026694 | NA | NA |
| ENSG000001 | 425.870696 | 0.546070785 | 0.12853697 | 4.24835586 | 2.15E-05 | 0.00026705 | APH1B | 83464 |
| ENSG000001 | 1317.83545 | 0.535730021 | 0.12612387 | 4.24764955 | 2.16E-05 | 0.00026774 | TDG | 6996 |
| ENSG000001 | 1071.69217 | 0.500919724 | 0.11792901 | 4.24763785 | 2.16E-05 | 0.00026774 | ZNF655 | 79027 |
| ENSG000001 | 2808.263 | 0.516926623 | 0.12175146 | 4.24575301 | 2.18E-05 | 0.00026984 | LTA4H | 4048 |
| ENSG000001 | 100.350236 | 0.71388358 | 0.1681407 | 4.24575115 | 2.18E-05 | 0.00026984 | AKIP1 | 56672 |
| ENSG000001 | 559.315627 | 0.624055879 | 0.14701708 | 4.24478489 | 2.19E-05 | 0.00027092 | ETS2 | 2114 |

| | | | | | | | | |
|---|---|---|---|---|---|---|---|---|
| ENSG000001 | 128.630261 | 0.6865984 | 0.16176259 | 4.24448205 | 2.19E-05 | 0.00027114 | APOC1 | 341 |
| ENSG000001 | 115.287499 | 0.712006559 | 0.16774949 | 4.24446343 | 2.19E-05 | 0.00027114 | P4HTM | 54681 |
| ENSG000001 | 464.412722 | 0.603330427 | 0.14214865 | 4.24436264 | 2.19E-05 | 0.00027118 | TFIP11 | 24144 |
| ENSG00000C | 169.907199 | 0.693261253 | 0.16334155 | 4.24424302 | 2.19E-05 | 0.00027125 | GAL | 51083 |
| ENSG000001 | 498.993372 | 0.565092917 | 0.13316477 | 4.2435617 | 2.20E-05 | 0.00027199 | CDKN2AIP | 55602 |
| ENSG000001 | 65.3372191 | -0.879695656 | 0.20732835 | -4.2430072 | 2.21E-05 | 0.00027258 | SCN3A | 6328 |
| ENSG000002 | 43.147453 | -0.92174773 | 0.217296 | -4.2418993 | 2.22E-05 | 0.00027384 | NA | NA |
| ENSG00000C | 100.869625 | 0.803679849 | 0.18948672 | 4.24135178 | 2.22E-05 | 0.00027426 | PQLC2 | 54896 |
| ENSG00000C | 153.923687 | 0.781071682 | 0.18415075 | 4.24147975 | 2.22E-05 | 0.00027426 | LZTR1 | 8216 |
| ENSG000001 | 37.0648377 | 0.996191523 | 0.23487529 | 4.24136358 | 2.22E-05 | 0.00027426 | FAM71E1 | 112703 |
| ENSG000001 | 44.621691 | -1.007174825 | 0.23747989 | -4.2410952 | 2.22E-05 | 0.00027449 | EMP1 | 2012 |
| ENSG000002 | 50.1272817 | -0.904647036 | 0.21332907 | -4.2406176 | 2.23E-05 | 0.00027499 | NA | NA |
| ENSG000001 | 1770.92805 | 0.488866149 | 0.11528957 | 4.24033274 | 2.23E-05 | 0.00027526 | ASPH | 444 |
| ENSG000001 | 933.648163 | 0.551160759 | 0.12998432 | 4.24020963 | 2.23E-05 | 0.00027533 | ANKLE2 | 23141 |
| ENSG000001 | 688.44222 | 0.523114997 | 0.12340629 | 4.23896538 | 2.25E-05 | 0.00027677 | TMEM14B | 81853 |
| ENSG000001 | 439.609912 | 0.541144731 | 0.12766309 | 4.23885038 | 2.25E-05 | 0.00027683 | ARMC9 | 80210 |
| ENSG000001 | 25.0185324 | -1.153308479 | 0.27213515 | -4.2379989 | 2.26E-05 | 0.0002778 | LOC100129940 | 100129940 |
| ENSG000002 | 3955.16371 | 0.473978472 | 0.11185757 | 4.23733911 | 2.26E-05 | 0.00027853 | ESRG | 790952 |
| ENSG00000C | 69.5787211 | 0.833963064 | 0.19685971 | 4.23633185 | 2.27E-05 | 0.0002797 | CYP51A1 | 1595 |
| ENSG000001 | 594.116586 | 0.563840221 | 0.13310084 | 4.23618822 | 2.27E-05 | 0.00027979 | NDUFAF2 | 91942 |
| ENSG000001 | 334.822596 | 0.540694284 | 0.12763981 | 4.23609427 | 2.27E-05 | 0.00027982 | ELOVL4 | 6785 |
| ENSG000001 | 156.445569 | 0.651786095 | 0.15387498 | 4.23581603 | 2.28E-05 | 0.00028008 | WDTC1 | 23038 |
| ENSG00000C | 312.462494 | 0.755944955 | 0.17848952 | 4.23523437 | 2.28E-05 | 0.00028073 | CTDP1 | 9150 |
| ENSG000001 | 40.9712756 | -0.956039102 | 0.22575645 | -4.2348252 | 2.29E-05 | 0.00028113 | C1orf101 | 257044 |
| ENSG000001 | 5168.79014 | 0.458418495 | 0.10825107 | 4.23477117 | 2.29E-05 | 0.00028113 | NA | NA |
| ENSG000001 | 267.07865 | 0.630970156 | 0.14900097 | 4.23467152 | 2.29E-05 | 0.00028117 | DCPS | 28960 |
| ENSG000001 | 3079.32785 | 0.503528252 | 0.11891108 | 4.23449413 | 2.29E-05 | 0.00028131 | GALNT1 | 2589 |
| ENSG000001 | 613.159605 | 0.607157406 | 0.14341922 | 4.23344527 | 2.30E-05 | 0.00028254 | IGSF3 | 3321 |
| ENSG00000C | 1584.2619 | 0.651397188 | 0.15388989 | 4.23287826 | 2.31E-05 | 0.00028317 | ITGB5 | 3693 |

| | | | | | | | | |
|---|---|---|---|---|---|---|---|---|
| ENSG000001 | 789.035897 | 0.507744111 | 0.11996253 | 4.23252236 | 2.31E-05 | 0.00028353 | FAM91A1 | 157769 |
| ENSG000001 | 835.447791 | 0.528981434 | 0.12498257 | 4.23244167 | 2.31E-05 | 0.00028355 | MRPL13 | 28998 |
| ENSG000001 | 1599.28308 | 0.508652036 | 0.1201905 | 4.23204874 | 2.32E-05 | 0.00028381 | RNMT | 8731 |
| ENSG000001 | 21.1910135 | -1.202463101 | 0.28413102 | -4.2320725 | 2.32E-05 | 0.00028381 | CYSLTR2 | 57105 |
| ENSG000002 | 40.2423097 | -0.992104743 | 0.23442774 | -4.232028 | 2.32E-05 | 0.00028381 | C6orf10 | 10665 |
| ENSG000001 | 2727.32275 | 0.516105852 | 0.12196422 | 4.23161705 | 2.32E-05 | 0.00028425 | CBX1 | 10951 |
| ENSG000001 | 495.661819 | 0.476353855 | 0.11258115 | 4.2312045 | 2.32E-05 | 0.00028468 | ERAP1 | 51752 |
| ENSG000001 | 2719.14252 | 0.750927833 | 0.17747644 | 4.23114085 | 2.33E-05 | 0.00028468 | SLC25A6 | 293 |
| ENSG00000C | 18.8151176 | -1.269144183 | 0.29996845 | -4.2309256 | 2.33E-05 | 0.00028486 | PCDHA6 | 56142 |
| ENSG00000C | 1933.88789 | 0.519407835 | 0.12276882 | 4.23077966 | 2.33E-05 | 0.00028496 | SLC25A13 | 10165 |
| ENSG000001 | 88.0944364 | -0.831365405 | 0.19653415 | -4.230132 | 2.34E-05 | 0.00028561 | KYNU | 8942 |
| ENSG000001 | 1113.53184 | 0.498734936 | 0.1178988 | 4.23019509 | 2.33E-05 | 0.00028561 | SFXN1 | 94081 |
| ENSG000001 | 195.05707 | 0.627288084 | 0.14829923 | 4.22988082 | 2.34E-05 | 0.00028584 | SENP3 | 26168 |
| ENSG000001 | 182.892585 | 0.675650042 | 0.1597379 | 4.22974151 | 2.34E-05 | 0.00028593 | E4F1 | 1877 |
| ENSG000001 | 170.450688 | 0.716674852 | 0.16944448 | 4.22955571 | 2.34E-05 | 0.00028608 | SLC25A22 | 79751 |
| ENSG000001 | 537.000901 | 0.62375103 | 0.14747824 | 4.22944437 | 2.34E-05 | 0.00028614 | COMMD10 | 51397 |
| ENSG000001 | 88.9150317 | -0.758116456 | 0.17925504 | -4.2292616 | 2.34E-05 | 0.00028629 | DOCK8 | 81704 |
| ENSG00000C | 662.8102 | 0.473206047 | 0.11190775 | 4.22853672 | 2.35E-05 | 0.00028712 | STX7 | 8417 |
| ENSG000001 | 1296.42274 | 0.512264193 | 0.12114862 | 4.22839485 | 2.35E-05 | 0.00028722 | PPFIA1 | 8500 |
| ENSG000001 | 163.283189 | -0.610340626 | 0.14438369 | -4.227213 | 2.37E-05 | 0.00028864 | SCN2A | 6326 |
| ENSG00000C | 3822.96085 | 0.462421864 | 0.10939967 | 4.22690345 | 2.37E-05 | 0.00028878 | HECTD1 | 25831 |
| ENSG000001 | 104.921326 | 0.744691535 | 0.17617776 | 4.22693274 | 2.37E-05 | 0.00028878 | SNCG | 6623 |
| ENSG000001 | 1254.393 | 0.553374755 | 0.13091423 | 4.22700236 | 2.37E-05 | 0.00028878 | MRPL11 | 65003 |
| ENSG000001 | 752.149634 | 0.531651244 | 0.1257838 | 4.22670688 | 2.37E-05 | 0.00028891 | FAM35A | 54537 |
| ENSG000001 | 4933.41023 | 0.504746075 | 0.11941946 | 4.22666532 | 2.37E-05 | 0.00028891 | GABRB3 | 2562 |
| ENSG000001 | 2125.95405 | 0.52406584 | 0.12399883 | 4.22637739 | 2.37E-05 | 0.0002892 | EIF2S1 | 1965 |
| ENSG000001 | 669.088114 | 0.708098097 | 0.16756673 | 4.22576781 | 2.38E-05 | 0.00028989 | TTYH3 | 80727 |
| ENSG000001 | 2000.95752 | 0.534231777 | 0.12642705 | 4.22561297 | 2.38E-05 | 0.00029 | PSMD14 | 10213 |
| ENSG000001 | 254.48381 | 0.705336774 | 0.16692259 | 4.22553206 | 2.38E-05 | 0.00029002 | MAPK3 | 5595 |

| | | | | | | | | |
|---|---|---|---|---|---|---|---|---|
| ENSG00000C | 342.377995 | 0.675240997 | 0.15980461 | 4.22541625 | 2.38E-05 | 0.00029008 | JPH4 | 84502 |
| ENSG000001 | 1368.51777 | 0.571679834 | 0.13530156 | 4.22522711 | 2.39E-05 | 0.00029024 | SSR2 | 6746 |
| ENSG000001 | 47.2206642 | 0.911738408 | 0.21580528 | 4.22481979 | 2.39E-05 | 0.00029068 | C9orf89 | 84270 |
| ENSG00000C | 286.491971 | 0.71371529 | 0.16894954 | 4.22442865 | 2.40E-05 | 0.0002911 | ATP5D | 513 |
| ENSG000001 | 443.489526 | 0.618380189 | 0.14638952 | 4.22421079 | 2.40E-05 | 0.00029126 | MLX | 6945 |
| ENSG000001 | 124.349041 | 0.733379062 | 0.17361789 | 4.22409836 | 2.40E-05 | 0.00029126 | ZNF205 | 7755 |
| ENSG000001 | 98.6607503 | 0.834480104 | 0.19755137 | 4.22411704 | 2.40E-05 | 0.00029126 | ATG4D | 84971 |
| ENSG000001 | 510.082843 | 0.548555798 | 0.12986557 | 4.22402782 | 2.40E-05 | 0.00029127 | CCNE1 | 898 |
| ENSG000001 | 577.644078 | 0.571959151 | 0.13543913 | 4.22299772 | 2.41E-05 | 0.00029251 | KLHL24 | 54800 |
| ENSG000001 | 1511.53655 | 0.558264817 | 0.13220108 | 4.22284603 | 2.41E-05 | 0.00029262 | VRK1 | 7443 |
| ENSG000001 | 1095.12277 | 0.495661497 | 0.1173835 | 4.22258251 | 2.42E-05 | 0.00029288 | CNOT10 | 25904 |
| ENSG000001 | 450.435042 | 0.595691168 | 0.14107551 | 4.22249863 | 2.42E-05 | 0.0002929 | C1orf52 | 148423 |
| ENSG000002 | 55.3882696 | -0.927296817 | 0.21961218 | -4.2224289 | 2.42E-05 | 0.0002929 | LINC01588 | 283551 |
| ENSG000001 | 190.146023 | 0.625230744 | 0.14808462 | 4.22211815 | 2.42E-05 | 0.00029322 | FARS2 | 10667 |
| ENSG000001 | 571.798802 | 0.566040179 | 0.1340705 | 4.22195914 | 2.42E-05 | 0.00029334 | CHMP2A | 27243 |
| ENSG00000C | 718.724239 | 0.476851136 | 0.11295321 | 4.22166971 | 2.42E-05 | 0.00029352 | MRPS10 | 55173 |
| ENSG00000C | 227.63689 | 0.664883573 | 0.15749506 | 4.22161544 | 2.43E-05 | 0.00029352 | HOOK2 | 29911 |
| ENSG000001 | 140.76168 | 0.721908509 | 0.17100269 | 4.22162081 | 2.43E-05 | 0.00029352 | KCTD17 | 79734 |
| ENSG000001 | 1016.76618 | 0.54659504 | 0.12948395 | 4.22133447 | 2.43E-05 | 0.0002938 | SLC39A7 | 7922 |
| ENSG000001 | 7424.21263 | 0.582866912 | 0.13810019 | 4.22060911 | 2.44E-05 | 0.00029466 | CHD4 | 1108 |
| ENSG000001 | 3051.94456 | 0.524508352 | 0.12428204 | 4.2203069 | 2.44E-05 | 0.00029497 | WDHD1 | 11169 |
| ENSG000001 | 213.264978 | 0.729451888 | 0.17285587 | 4.22000074 | 2.44E-05 | 0.00029528 | NAGA | 4668 |
| ENSG000001 | 230.293925 | 0.65380728 | 0.15494744 | 4.21954241 | 2.45E-05 | 0.00029579 | PIGO | 84720 |
| ENSG000001 | 1428.15634 | 0.557410914 | 0.13210984 | 4.21929894 | 2.45E-05 | 0.00029602 | CKS1B | 1163 |
| ENSG000001 | 2663.41552 | 0.533624188 | 0.12649279 | 4.21861331 | 2.46E-05 | 0.00029684 | CCNC | 892 |
| ENSG000001 | 750.27255 | 0.482977298 | 0.11449224 | 4.21842816 | 2.46E-05 | 0.00029699 | VPS36 | 51028 |
| ENSG00000C | 2657.22247 | 0.573298503 | 0.13591202 | 4.21815889 | 2.46E-05 | 0.00029699 | MDH1 | 4190 |
| ENSG00000C | 293.713154 | 0.648908501 | 0.15383542 | 4.21819949 | 2.46E-05 | 0.00029699 | GEMIN2 | 8487 |
| ENSG000001 | 1748.84561 | 0.502351627 | 0.11908983 | 4.21825793 | 2.46E-05 | 0.00029699 | ADRBK2 | 157 |

| | | | | | | | | |
|---|---|---|---|---|---|---|---|---|
| ENSG000001 | 203.468992 | 0.634724969 | 0.1504723 | 4.21821795 | 2.46E-05 | 0.00029699 | HRASLS5 | 117245 |
| ENSG000001 | 1097.59381 | 0.550080471 | 0.130449 | 4.2168239 | 2.48E-05 | 0.00029858 | RPA2 | 6118 |
| ENSG000001 | 2465.06549 | 0.586819983 | 0.13916015 | 4.21686793 | 2.48E-05 | 0.00029858 | SLC39A1 | 27173 |
| ENSG00000C | 802.514576 | 0.519741983 | 0.12328144 | 4.21589816 | 2.49E-05 | 0.0002997 | SRI | 6717 |
| ENSG000001 | 356.166772 | 0.814716452 | 0.19325119 | 4.21584179 | 2.49E-05 | 0.0002997 | LRP5 | 4041 |
| ENSG00000C | 2788.91644 | 0.436807121 | 0.10361911 | 4.21550749 | 2.49E-05 | 0.00030006 | RDH11 | 51109 |
| ENSG00000C | 339.519858 | 0.550978226 | 0.13071207 | 4.21520554 | 2.50E-05 | 0.00030037 | BCKDHB | 594 |
| ENSG000001 | 86.7534893 | 0.761749942 | 0.18073494 | 4.2147353 | 2.50E-05 | 0.00030091 | CCDC28B | 79140 |
| ENSG000001 | 20817.3612 | 0.686841171 | 0.16298154 | 4.2142268 | 2.51E-05 | 0.0003015 | RPS2 | 6187 |
| ENSG000001 | 1687.98392 | 0.515915318 | 0.12246925 | 4.21261123 | 2.52E-05 | 0.00030357 | UBE4B | 10277 |
| ENSG000001 | 2560.18376 | 0.503209389 | 0.11945819 | 4.21243114 | 2.53E-05 | 0.00030372 | IQGAP2 | 10788 |
| ENSG000001 | 4427.94615 | 0.462798195 | 0.10986645 | 4.2123705 | 2.53E-05 | 0.00030372 | KIF5B | 3799 |
| ENSG000001 | 228.934596 | 0.677255417 | 0.16078902 | 4.21207494 | 2.53E-05 | 0.00030403 | CBY1 | 25776 |
| ENSG000001 | 134.669715 | 0.676923555 | 0.16073764 | 4.21135689 | 2.54E-05 | 0.00030481 | SHF | 90525 |
| ENSG000001 | 46.9414016 | -0.90914649 | 0.21587723 | -4.2114052 | 2.54E-05 | 0.00030481 | CD34 | 947 |
| ENSG000001 | 980.212523 | 0.605207378 | 0.14371135 | 4.21127062 | 2.54E-05 | 0.00030484 | MRPS7 | 51081 |
| ENSG00000C | 469.614489 | 0.587004788 | 0.13939439 | 4.21110757 | 2.54E-05 | 0.00030497 | ALG1 | 56052 |
| ENSG000001 | 1079.99397 | 0.722844519 | 0.17166088 | 4.21088665 | 2.54E-05 | 0.00030518 | SUPT5H | 6829 |
| ENSG000002 | 20.3115486 | -1.248924331 | 0.2966299 | -4.2103791 | 2.55E-05 | 0.00030577 | LOC283038 | 283038 |
| ENSG000001 | 841.930302 | 0.524571758 | 0.12459881 | 4.21008644 | 2.55E-05 | 0.00030599 | UBE2W | 55284 |
| ENSG000001 | 819.521869 | 0.504681931 | 0.11987383 | 4.21010925 | 2.55E-05 | 0.00030599 | TC2N | 123036 |
| ENSG000001 | 3741.0417 | 0.527217671 | 0.12523863 | 4.20970486 | 2.56E-05 | 0.00030641 | PCNA | 5111 |
| ENSG000002 | 26193.1735 | 0.467529434 | 0.11106814 | 4.20939271 | 2.56E-05 | 0.00030675 | CD24 | 100133941 |
| ENSG00000C | 2324.14901 | 0.623066924 | 0.14805431 | 4.20836723 | 2.57E-05 | 0.00030805 | GCN1 | 10985 |
| ENSG000001 | 479.172264 | 0.594910051 | 0.14136609 | 4.20829388 | 2.57E-05 | 0.00030806 | NRBF2 | 29982 |
| ENSG00000C | 376.011623 | 0.754339397 | 0.17927497 | 4.207723 | 2.58E-05 | 0.00030866 | CELSR3 | 1951 |
| ENSG000001 | 175.466082 | 0.594938277 | 0.14139026 | 4.20777426 | 2.58E-05 | 0.00030866 | TAF6L | 10629 |
| ENSG00000C | 1154.75151 | 0.502886859 | 0.11953168 | 4.20714303 | 2.59E-05 | 0.0003092 | ANKRD10 | 55608 |
| ENSG000001 | 627.658871 | 0.588700696 | 0.13992934 | 4.20712834 | 2.59E-05 | 0.0003092 | NPC2 | 10577 |

| | | | | | | | | |
|---|---|---|---|---|---|---|---|---|
| ENSG000001 | 566.550785 | 0.506660187 | 0.12042625 | 4.20722366 | 2.59E-05 | 0.0003092 | CCDC77 | 84318 |
| ENSG00000( | 17990.4573 | 0.527252361 | 0.12533513 | 4.20674051 | 2.59E-05 | 0.00030964 | CSDE1 | 7812 |
| ENSG00000( | 1939.96098 | 0.502256334 | 0.11940691 | 4.20625871 | 2.60E-05 | 0.00031021 | WDR3 | 10885 |
| ENSG000001 | 480.651991 | 0.595114881 | 0.14152427 | 4.20503772 | 2.61E-05 | 0.00031179 | P2RY1 | 5028 |
| ENSG000001 | 737.525965 | 0.546154954 | 0.12988678 | 4.20485408 | 2.61E-05 | 0.00031195 | SEC24C | 9632 |
| ENSG00000( | 727.477914 | 0.670248222 | 0.15940795 | 4.20460975 | 2.62E-05 | 0.00031211 | KEAP1 | 9817 |
| ENSG000001 | 719.517134 | 0.513906722 | 0.12222353 | 4.20464636 | 2.61E-05 | 0.00031211 | LDAH | 60526 |
| ENSG00000( | 2657.90614 | 0.487345079 | 0.11592345 | 4.20402492 | 2.62E-05 | 0.00031277 | NDUFS1 | 4719 |
| ENSG000001 | 37.6566778 | 0.985059552 | 0.23431517 | 4.2039939 | 2.62E-05 | 0.00031277 | TSPO | 706 |
| ENSG00000( | 451.095794 | 0.607143212 | 0.14443094 | 4.20369206 | 2.63E-05 | 0.00031296 | EPN1 | 29924 |
| ENSG00000( | 1684.74457 | 0.606422828 | 0.14426304 | 4.20359109 | 2.63E-05 | 0.00031296 | FKBP1A | 2280 |
| ENSG000001 | 2494.04957 | 0.500614225 | 0.11909034 | 4.20365101 | 2.63E-05 | 0.00031296 | ZNF451 | 26036 |
| ENSG000001 | 199.872246 | 0.641387909 | 0.15258046 | 4.20360457 | 2.63E-05 | 0.00031296 | MED9 | 55090 |
| ENSG000001 | 512.644296 | 0.655780491 | 0.15600908 | 4.20347656 | 2.63E-05 | 0.00031303 | PGRMC2 | 10424 |
| ENSG00000( | 672.566912 | 0.496099502 | 0.11802521 | 4.20333488 | 2.63E-05 | 0.00031313 | PPAP2A | 8611 |
| ENSG000001 | 326.754649 | 0.576414693 | 0.13714103 | 4.20307967 | 2.63E-05 | 0.0003134 | ARMCX5 | 64860 |
| ENSG000001 | 524.847994 | 0.58150985 | 0.13837993 | 4.20227012 | 2.64E-05 | 0.00031443 | COG4 | 25839 |
| ENSG00000( | 465.532635 | 0.68893288 | 0.16396142 | 4.20179861 | 2.65E-05 | 0.00031488 | XAB2 | 56949 |
| ENSG000001 | 3488.29012 | 0.467128031 | 0.1111748 | 4.20174395 | 2.65E-05 | 0.00031488 | PRKCI | 5584 |
| ENSG000001 | 510.528992 | 0.600934912 | 0.14301887 | 4.20178747 | 2.65E-05 | 0.00031488 | SNHG17 | 388796 |
| ENSG000001 | 2210.00203 | 0.502171729 | 0.11952312 | 4.20146113 | 2.65E-05 | 0.00031518 | RLIM | 51132 |
| ENSG00000( | 270.938453 | 0.631024185 | 0.15021167 | 4.20089997 | 2.66E-05 | 0.0003158 | MAD1L1 | 8379 |
| ENSG000001 | 870.360296 | 0.532884247 | 0.12685042 | 4.20088673 | 2.66E-05 | 0.0003158 | NPTN | 27020 |
| ENSG00000( | 820.607391 | 0.450485407 | 0.10724341 | 4.20058821 | 2.66E-05 | 0.00031612 | BRAP | 8315 |
| ENSG000001 | 26.3375547 | -1.131916176 | 0.26949354 | -4.2001607 | 2.67E-05 | 0.00031663 | BRINP3 | 339479 |
| ENSG000001 | 571.417191 | 0.612716613 | 0.14589866 | 4.19960404 | 2.67E-05 | 0.00031728 | N4BP3 | 23138 |
| ENSG000001 | 1341.3818 | 0.513805798 | 0.12234932 | 4.1994987 | 2.68E-05 | 0.00031728 | NBAS | 51594 |
| ENSG000002 | 31.1160278 | -1.107335191 | 0.26368136 | -4.1995202 | 2.67E-05 | 0.00031728 | PKD1L3 | 342372 |
| ENSG000001 | 635.177352 | 0.541298267 | 0.12891146 | 4.19899262 | 2.68E-05 | 0.00031789 | VWA9 | 81556 |

| | | | | | | | | | |
|---|---|---|---|---|---|---|---|---|---|
| ENSG00000C | 2114.8443 | 0.697886964 | 0.16621018 | 4.19882191 | 2.68E-05 | 0.00031804 | DYNLL1 | | 8655 |
| ENSG00000C | 1260.31575 | 0.497845026 | 0.11857152 | 4.1986898 | 2.68E-05 | 0.00031813 | GCLC | | 2729 |
| ENSG000001 | 422.375444 | 0.638975287 | 0.1522306 | 4.19741688 | 2.70E-05 | 0.00031983 | KHNYN | | 23351 |
| ENSG00000C | 570.79028 | 0.609072891 | 0.14512751 | 4.196812 | 2.71E-05 | 0.00032032 | STARD3NL | | 83930 |
| ENSG000001 | 754.803639 | 0.606724718 | 0.14456826 | 4.19680437 | 2.71E-05 | 0.00032032 | LIG1 | | 3978 |
| ENSG000001 | 438.32087 | 0.575795772 | 0.13719264 | 4.19698725 | 2.70E-05 | 0.00032032 | WDR55 | | 54853 |
| ENSG000002 | 8.66326674 | -1.513190425 | 0.36055409 | -4.1968472 | 2.71E-05 | 0.00032032 | NA | NA | |
| ENSG000002 | 98.628551 | -0.742937445 | 0.17703153 | -4.1966391 | 2.71E-05 | 0.00032042 | LINC00598 | | 646982 |
| ENSG000002 | 316.343032 | 0.651635137 | 0.15527678 | 4.19660377 | 2.71E-05 | 0.00032042 | NA | NA | |
| ENSG000001 | 543.763684 | 0.500746185 | 0.11932764 | 4.19639721 | 2.71E-05 | 0.00032062 | DIS3L | | 115752 |
| ENSG000001 | 270.144406 | 0.568824265 | 0.13555948 | 4.19612295 | 2.72E-05 | 0.00032091 | GSTA4 | | 2941 |
| ENSG000001 | 485.221648 | -0.580120088 | 0.13825647 | -4.1959707 | 2.72E-05 | 0.00032104 | DNAJC15 | | 29103 |
| ENSG000001 | 3951.72014 | 0.491507644 | 0.11714208 | 4.19582485 | 2.72E-05 | 0.00032115 | PTAR1 | | 375743 |
| ENSG000001 | 229.959007 | 0.678058872 | 0.1616577 | 4.19441114 | 2.74E-05 | 0.00032306 | TMEM55B | | 90809 |
| ENSG000001 | 248.059376 | 0.581745727 | 0.13871145 | 4.19392723 | 2.74E-05 | 0.00032366 | UBXN8 | | 7993 |
| ENSG000001 | 2918.22472 | 0.492590531 | 0.11746513 | 4.1935044 | 2.75E-05 | 0.00032417 | PAK1 | | 5058 |
| ENSG000001 | 1282.70423 | 0.490193958 | 0.11689645 | 4.1934033 | 2.75E-05 | 0.00032422 | TTF2 | | 8458 |
| ENSG000001 | 790.689208 | 0.532323677 | 0.12695519 | 4.19300435 | 2.75E-05 | 0.0003247 | SNX19 | | 399979 |
| ENSG000002 | 23.4864368 | -1.204371254 | 0.28726125 | -4.1925991 | 2.76E-05 | 0.00032518 | NA | NA | |
| ENSG000001 | 479.200922 | 0.730416088 | 0.17422344 | 4.19241009 | 2.76E-05 | 0.00032536 | DHX37 | | 57647 |
| ENSG000001 | 3194.80671 | 0.498954192 | 0.11901629 | 4.19231839 | 2.76E-05 | 0.00032536 | EIF3H | | 8667 |
| ENSG000001 | 582.752263 | 0.518328127 | 0.1236388 | 4.19227729 | 2.76E-05 | 0.00032536 | RPL39L | | 116832 |
| ENSG000002 | 667.780843 | 0.547122756 | 0.13051662 | 4.1919777 | 2.77E-05 | 0.0003257 | TUNAR | | 100507043 |
| ENSG000001 | 1191.96151 | 0.463049434 | 0.11047369 | 4.19149073 | 2.77E-05 | 0.0003263 | NOL10 | | 79954 |
| ENSG00000C | 113.063236 | 0.716319902 | 0.17091219 | 4.19115752 | 2.78E-05 | 0.00032669 | SMOX | | 54498 |
| ENSG000001 | 8002.19704 | 0.645134136 | 0.1539524 | 4.19047808 | 2.78E-05 | 0.00032757 | MDN1 | | 23195 |
| ENSG000001 | 638.046905 | 0.542862978 | 0.12954873 | 4.19041528 | 2.78E-05 | 0.00032757 | TRMT5 | | 57570 |
| ENSG000001 | 2636.79272 | 0.523609461 | 0.12495787 | 4.19028793 | 2.79E-05 | 0.00032765 | USP10 | | 9100 |
| ENSG00000C | 3690.81967 | 0.494937884 | 0.11813203 | 4.18970104 | 2.79E-05 | 0.00032841 | POLR2B | | 5431 |

| | | | | | | | | |
|---|---|---|---|---|---|---|---|---|
| ENSG000001 | 376.236153 | 0.53653781 | 0.12806457 | 4.18958808 | 2.79E-05 | 0.00032848 | MND1 | 84057 |
| ENSG000001 | 604.201389 | 0.523702586 | 0.12500598 | 4.18942018 | 2.80E-05 | 0.00032862 | METTL2B | 55798 |
| ENSG000001 | 288.431441 | 0.61316206 | 0.14636224 | 4.18934605 | 2.80E-05 | 0.00032864 | ZFYVE21 | 79038 |
| ENSG000001 | 176.869315 | 0.599583639 | 0.14314289 | 4.18870717 | 2.81E-05 | 0.00032947 | THRB | 7068 |
| ENSG000001 | 1508.98066 | 0.506266432 | 0.12087006 | 4.18851817 | 2.81E-05 | 0.00032965 | HJURP | 55355 |
| ENSG000001 | 11.351515 | -1.413488863 | 0.33751134 | -4.1879745 | 2.81E-05 | 0.00033034 | CTSF | 8722 |
| ENSG00000C | 156.846707 | 0.760851733 | 0.18168104 | 4.18784336 | 2.82E-05 | 0.00033044 | ITPKC | 80271 |
| ENSG000001 | 1895.75087 | 0.466510888 | 0.11139861 | 4.18776223 | 2.82E-05 | 0.00033046 | MASTL | 84930 |
| ENSG000001 | 759.607891 | 0.586059866 | 0.13995579 | 4.18746433 | 2.82E-05 | 0.0003308 | FAM213A | 84293 |
| ENSG00000C | 1820.24537 | 0.500904881 | 0.11962233 | 4.1873861 | 2.82E-05 | 0.00033082 | AK2 | 204 |
| ENSG000001 | 445.683776 | 0.514778003 | 0.12295789 | 4.18662044 | 2.83E-05 | 0.00033184 | PIGA | 5277 |
| ENSG000001 | 231.215291 | 0.633370419 | 0.1512919 | 4.18641335 | 2.83E-05 | 0.00033204 | TNIP2 | 79155 |
| ENSG000001 | 461.078694 | 0.779700906 | 0.18624851 | 4.18634698 | 2.83E-05 | 0.00033204 | C10orf2 | 56652 |
| ENSG000001 | 790.920254 | 0.52105997 | 0.12447211 | 4.1861585 | 2.84E-05 | 0.00033222 | KLHL9 | 55958 |
| ENSG00000C | 1633.18076 | 0.465501029 | 0.11120884 | 4.18582761 | 2.84E-05 | 0.00033252 | PUS7 | 54517 |
| ENSG000001 | 487.761734 | 0.575996266 | 0.13760468 | 4.18587712 | 2.84E-05 | 0.00033252 | BICD2 | 23299 |
| ENSG00000C | 3049.40208 | 0.495620671 | 0.11841223 | 4.18555294 | 2.84E-05 | 0.00033282 | DDX24 | 57062 |
| ENSG000001 | 147.938427 | 0.699869753 | 0.1672199 | 4.18532583 | 2.85E-05 | 0.00033306 | REEP6 | 92840 |
| ENSG000001 | 2385.20538 | 0.509830398 | 0.12181607 | 4.18524763 | 2.85E-05 | 0.00033308 | LIN7C | 55327 |
| ENSG000001 | 203.417945 | 0.565496823 | 0.13512406 | 4.18501941 | 2.85E-05 | 0.00033332 | IMMP1L | 196294 |
| ENSG000001 | 1375.69019 | 0.553212572 | 0.13221513 | 4.18418501 | 2.86E-05 | 0.00033437 | METTL16 | 79066 |
| ENSG000002 | 1002.69517 | 0.483758781 | 0.11561629 | 4.18417506 | 2.86E-05 | 0.00033437 | NA | NA |
| ENSG000001 | 146.736192 | 0.677906601 | 0.16202671 | 4.18391889 | 2.87E-05 | 0.00033465 | ZNF747 | 65988 |
| ENSG000001 | 611.719372 | 0.494165389 | 0.11812041 | 4.18357313 | 2.87E-05 | 0.00033506 | KDM4C | 23081 |
| ENSG000001 | 94.8787706 | 0.73348879 | 0.17533304 | 4.18340325 | 2.87E-05 | 0.00033521 | PLEKHG4 | 25894 |
| ENSG00000C | 317.778118 | 0.551250561 | 0.13180909 | 4.18218933 | 2.89E-05 | 0.00033682 | CD99 | 4267 |
| ENSG000001 | 671.054303 | 0.501283622 | 0.11986145 | 4.18219214 | 2.89E-05 | 0.00033682 | KANK1 | 23189 |
| ENSG00000C | 3733.13492 | 0.53750011 | 0.12854168 | 4.18152396 | 2.90E-05 | 0.00033771 | ZNF532 | 55205 |
| ENSG000002 | 39.7666809 | 0.977357195 | 0.2337534 | 4.18114643 | 2.90E-05 | 0.00033817 | NA | NA |

| | | | | | | | | |
|---|---|---|---|---|---|---|---|---|
| ENSG000001 | 787.626607 | 0.5783208 | 0.13833563 | 4.18056278 | 2.91E-05 | 0.00033894 | PLOD2 | 5352 |
| ENSG000001 | 1019.82923 | 0.567702553 | 0.13580451 | 4.18029233 | 2.91E-05 | 0.00033925 | BZW2 | 28969 |
| ENSG000001 | 2878.8153 | 0.484988134 | 0.11602005 | 4.1802097 | 2.91E-05 | 0.00033927 | MYO1B | 4430 |
| ENSG00000C | 1179.88622 | 0.542600187 | 0.12980544 | 4.18010367 | 2.91E-05 | 0.00033933 | TRIO | 7204 |
| ENSG000001 | 4062.08358 | 0.512472649 | 0.12260339 | 4.17992242 | 2.92E-05 | 0.00033951 | CPSF6 | 11052 |
| ENSG00000C | 201.063529 | 0.776110677 | 0.18568243 | 4.17977451 | 2.92E-05 | 0.00033963 | PDRG1 | 81572 |
| ENSG000001 | 306.927559 | 0.547986113 | 0.1311111 | 4.17955548 | 2.92E-05 | 0.00033986 | HINT3 | 135114 |
| ENSG000002 | 21.50741 | -1.215154031 | 0.29075583 | -4.1792938 | 2.92E-05 | 0.00034015 | LOC100506497 | 100506497 |
| ENSG000001 | 2980.38533 | 0.533398759 | 0.12763332 | 4.17914969 | 2.93E-05 | 0.00034027 | COPB2 | 9276 |
| ENSG00000C | 3207.44421 | 0.504774258 | 0.12078812 | 4.17900585 | 2.93E-05 | 0.00034039 | SAR1A | 56681 |
| ENSG000001 | 2372.55991 | 0.617704163 | 0.14782527 | 4.17861014 | 2.93E-05 | 0.00034088 | SPTAN1 | 6709 |
| ENSG00000C | 525.149109 | 0.526463226 | 0.12599922 | 4.17830534 | 2.94E-05 | 0.00034123 | KIAA0141 | 9812 |
| ENSG000001 | 852.307122 | 0.519065882 | 0.12423049 | 4.17824869 | 2.94E-05 | 0.00034123 | SRPK2 | 6733 |
| ENSG000001 | 890.657455 | 0.513107413 | 0.12281538 | 4.17787602 | 2.94E-05 | 0.00034169 | VPS8 | 23355 |
| ENSG000001 | 941.683155 | 0.489140623 | 0.11708638 | 4.17760474 | 2.95E-05 | 0.000342 | SUV39H2 | 79723 |
| ENSG000001 | 436.610595 | 0.748719845 | 0.17923377 | 4.1773369 | 2.95E-05 | 0.00034221 | MEGF8 | 1954 |
| ENSG000001 | 1179.11961 | 0.509032277 | 0.12185508 | 4.17735804 | 2.95E-05 | 0.00034221 | TMA16 | 55319 |
| ENSG000001 | 1046.97725 | 0.487926702 | 0.11681384 | 4.17695979 | 2.95E-05 | 0.00034268 | SPG11 | 80208 |
| ENSG000001 | 281.21265 | 0.829608247 | 0.19866414 | 4.17593363 | 2.97E-05 | 0.00034413 | BYSL | 705 |
| ENSG00000C | 8699.90406 | 0.578099965 | 0.13847616 | 4.17472565 | 2.98E-05 | 0.00034576 | CBX5 | 23468 |
| ENSG000001 | 337.881081 | 0.731780629 | 0.17528587 | 4.17478397 | 2.98E-05 | 0.00034576 | G6PD | 2539 |
| ENSG000001 | 118.330195 | 0.810807392 | 0.19422197 | 4.17464312 | 2.98E-05 | 0.00034579 | MED10 | 84246 |
| ENSG000001 | 1162.64473 | 0.533903737 | 0.12790299 | 4.17428671 | 2.99E-05 | 0.00034617 | ZSCAN10 | 84891 |
| ENSG000001 | 2045.11484 | 0.630509812 | 0.15104806 | 4.17423305 | 2.99E-05 | 0.00034617 | PPM1B | 5495 |
| ENSG000001 | 706.567732 | 0.510320491 | 0.12225606 | 4.17419384 | 2.99E-05 | 0.00034617 | PIGK | 10026 |
| ENSG000001 | 12083.8474 | 0.73080507 | 0.175082 | 4.17407316 | 2.99E-05 | 0.00034626 | RPL3 | 6122 |
| ENSG000001 | 56.6197861 | 0.875950243 | 0.20991187 | 4.17294278 | 3.01E-05 | 0.00034788 | CRB3 | 92359 |
| ENSG000001 | 2390.29203 | 0.507701524 | 0.12166928 | 4.17279977 | 3.01E-05 | 0.000348 | RAB11A | 8766 |
| ENSG000002 | 229.621578 | 0.680825747 | 0.16317827 | 4.17228198 | 3.02E-05 | 0.00034869 | NA | NA |

| | | | | | | | | |
|---|---|---|---|---|---|---|---|---|
| ENSG00000( | 2781.76485 | 0.509716861 | 0.12217417 | 4.1720509 | 3.02E-05 | 0.00034895 | SNX5 | 27131 |
| ENSG000001 | 4920.5542 | 0.576452519 | 0.13818938 | 4.17146763 | 3.03E-05 | 0.00034974 | YWHAB | 7529 |
| ENSG000001 | 225.654375 | 0.675533366 | 0.16194603 | 4.17134868 | 3.03E-05 | 0.00034983 | DGCR14 | 8220 |
| ENSG000001 | 18546.0915 | 0.443417954 | 0.10630688 | 4.17111262 | 3.03E-05 | 0.00035002 | EIF4G2 | 1982 |
| ENSG000002 | 73.2424582 | 0.79303328 | 0.19012598 | 4.17109362 | 3.03E-05 | 0.00035002 | NSUN5P1 | 155400 |
| ENSG000001 | 360.791251 | 0.520005483 | 0.12467773 | 4.17079684 | 3.04E-05 | 0.00035038 | UGT3A2 | 167127 |
| ENSG00000( | 1367.63308 | 0.521538044 | 0.12505316 | 4.17053069 | 3.04E-05 | 0.00035069 | SPTLC1 | 10558 |
| ENSG000001 | 1122.39249 | 0.515286872 | 0.1235636 | 4.17021569 | 3.04E-05 | 0.00035107 | XPO6 | 23214 |
| ENSG00000( | 701.850321 | 0.593428106 | 0.14230888 | 4.1700005 | 3.05E-05 | 0.0003513 | RANBP3 | 8498 |
| ENSG000001 | 2219.68855 | 0.518183206 | 0.12426947 | 4.1698351 | 3.05E-05 | 0.00035146 | RABGGTB | 5876 |
| ENSG000001 | 1025.43542 | 0.486399213 | 0.11664918 | 4.16976116 | 3.05E-05 | 0.00035147 | ZMYM1 | 79830 |
| ENSG00000( | 144.228126 | 0.754386247 | 0.18092481 | 4.16961199 | 3.05E-05 | 0.0003516 | MYDGF | 56005 |
| ENSG000001 | 535.703574 | 0.53774567 | 0.12897549 | 4.16936326 | 3.05E-05 | 0.00035189 | SUCLG1 | 8802 |
| ENSG00000( | 368.235613 | 0.671582202 | 0.16108618 | 4.16908645 | 3.06E-05 | 0.00035221 | TSPAN17 | 26262 |
| ENSG000001 | 964.880172 | 0.534410116 | 0.12820748 | 4.16832255 | 3.07E-05 | 0.0003533 | THOC5 | 8563 |
| ENSG000001 | 5062.34688 | 0.52227368 | 0.12529981 | 4.16819202 | 3.07E-05 | 0.0003534 | DCP2 | 167227 |
| ENSG000001 | 1481.22843 | 0.583628292 | 0.1400396 | 4.16759475 | 3.08E-05 | 0.00035422 | RND3 | 390 |
| ENSG00000( | 1752.05631 | 0.698104712 | 0.16752436 | 4.16718336 | 3.08E-05 | 0.00035466 | AP3D1 | 8943 |
| ENSG000001 | 273.878731 | 0.56888729 | 0.13651526 | 4.16720649 | 3.08E-05 | 0.00035466 | ZNF274 | 10782 |
| ENSG000002 | 887.133758 | 0.631066085 | 0.15145981 | 4.16655796 | 3.09E-05 | 0.00035554 | PIP4K2B | 8396 |
| ENSG000001 | 489.91577 | 0.522719536 | 0.1254669 | 4.1661947 | 3.10E-05 | 0.000356 | PLAG1 | 5324 |
| ENSG000001 | 8082.55768 | 0.491504585 | 0.11797838 | 4.16605653 | 3.10E-05 | 0.00035612 | KPNA2 | 3838 |
| ENSG000002 | 271.080532 | 0.713063931 | 0.17117657 | 4.16566317 | 3.10E-05 | 0.00035663 | NCKIPSD | 51517 |
| ENSG000001 | 962.866531 | 0.500949564 | 0.12026238 | 4.16547177 | 3.11E-05 | 0.00035683 | MRPS5 | 64969 |
| ENSG000001 | 187.138775 | 0.654901858 | 0.15722579 | 4.16535902 | 3.11E-05 | 0.00035688 | MAN2B1 | 4125 |
| ENSG000002 | 1058.23649 | 0.532255852 | 0.1277831 | 4.16530703 | 3.11E-05 | 0.00035688 | DAXX | 1616 |
| ENSG000001 | 217.18585 | 0.618701828 | 0.14855091 | 4.16491451 | 3.11E-05 | 0.00035728 | CDO1 | 1036 |
| ENSG000001 | 851.481031 | 0.499869223 | 0.12001817 | 4.16494612 | 3.11E-05 | 0.00035728 | SOCS6 | 9306 |
| ENSG000002 | 722.253197 | 0.534846686 | 0.1284189 | 4.16485951 | 3.12E-05 | 0.00035728 | ZNF542P | 147947 |

| | | | | | | | | |
|---|---|---|---|---|---|---|---|---|
| ENSG000001 | 526.318887 | 0.524614637 | 0.12598604 | 4.16406954 | 3.13E-05 | 0.00035832 | LIPA | 3988 |
| ENSG000001 | 95.6091229 | 0.760079951 | 0.18253034 | 4.16412931 | 3.13E-05 | 0.00035832 | SDHAF4 | 135154 |
| ENSG000001 | 2031.92814 | 0.523962898 | 0.12583363 | 4.16393386 | 3.13E-05 | 0.00035843 | ACAT2 | 39 |
| ENSG00000C | 710.64607 | 0.461901118 | 0.11093123 | 4.16385094 | 3.13E-05 | 0.00035846 | MCUR1 | 63933 |
| ENSG000002 | 206.187464 | 0.546601651 | 0.13127914 | 4.16365954 | 3.13E-05 | 0.00035866 | LIN52 | 91750 |
| ENSG000001 | 716.589359 | 0.624333593 | 0.14995313 | 4.16352489 | 3.13E-05 | 0.00035877 | VHL | 7428 |
| ENSG000001 | 189.862669 | 0.673913635 | 0.16186709 | 4.16337655 | 3.14E-05 | 0.0003588 | BCL7A | 605 |
| ENSG000001 | 761.348993 | 0.628024613 | 0.15084349 | 4.16341863 | 3.14E-05 | 0.0003588 | PITPNA | 5306 |
| ENSG000001 | 2297.05374 | 0.572526332 | 0.13755752 | 4.16208677 | 3.15E-05 | 0.00036073 | NACC1 | 112939 |
| ENSG000001 | 197.108058 | 0.719651357 | 0.17296646 | 4.16064115 | 3.17E-05 | 0.00036292 | CD320 | 51293 |
| ENSG000001 | 2364.82889 | 0.497452191 | 0.11957075 | 4.16031656 | 3.18E-05 | 0.00036333 | TMEM33 | 55161 |
| ENSG000001 | 1285.62074 | 0.447461146 | 0.10756065 | 4.16008237 | 3.18E-05 | 0.0003636 | UTP11L | 51118 |
| ENSG00000C | 1608.99263 | 0.498321803 | 0.11978886 | 4.16000105 | 3.18E-05 | 0.00036363 | PROM1 | 8842 |
| ENSG000001 | 9588.53219 | 0.520303869 | 0.12508386 | 4.15964038 | 3.19E-05 | 0.00036409 | HSP90B1 | 7184 |
| ENSG000001 | 55.8968448 | -0.895443022 | 0.2152723 | -4.159583 | 3.19E-05 | 0.00036409 | SH3TC2 | 79628 |
| ENSG000001 | 181.808171 | 0.629911085 | 0.15143902 | 4.15950331 | 3.19E-05 | 0.00036411 | NA | NA |
| ENSG000001 | 1560.23057 | 0.473192666 | 0.11376835 | 4.15926444 | 3.19E-05 | 0.00036439 | TTC1 | 7265 |
| ENSG000001 | 5724.26245 | 0.507482665 | 0.12201499 | 4.159183 | 3.19E-05 | 0.00036442 | EIF3M | 10480 |
| ENSG000001 | 1449.5698 | 0.573343389 | 0.1378622 | 4.15881516 | 3.20E-05 | 0.00036475 | PTPRS | 5802 |
| ENSG000001 | 531.931887 | 0.665814588 | 0.16009836 | 4.15878458 | 3.20E-05 | 0.00036475 | QSOX1 | 5768 |
| ENSG000001 | 628.487654 | 0.531475555 | 0.127792 | 4.15891117 | 3.20E-05 | 0.00036475 | PTGFRN | 5738 |
| ENSG000002 | 7.16297693 | -1.543973321 | 0.37132629 | -4.1579963 | 3.21E-05 | 0.00036591 | LOC101927843 | 101927843 |
| ENSG000001 | 32.7486446 | -1.00167759 | 0.240964 | -4.1569596 | 3.23E-05 | 0.00036726 | TFEC | 22797 |
| ENSG000001 | 410.501562 | 0.56325784 | 0.13549705 | 4.15697484 | 3.22E-05 | 0.00036726 | KIAA0355 | 9710 |
| ENSG000001 | 284.724939 | 0.56608582 | 0.13617631 | 4.1570067 | 3.22E-05 | 0.00036726 | MRPL48 | 51642 |
| ENSG000001 | 66.8168835 | 0.793829338 | 0.19099261 | 4.15633542 | 3.23E-05 | 0.00036816 | LOC100505501 | 100505501 |
| ENSG000001 | 550.085154 | 0.585417616 | 0.14085616 | 4.15613773 | 3.24E-05 | 0.00036838 | COPS5 | 10987 |
| ENSG00000C | 3123.82005 | 0.479016677 | 0.1152778 | 4.15532452 | 3.25E-05 | 0.00036948 | TNPO3 | 23534 |
| ENSG000001 | 25.3692162 | -1.102688317 | 0.26536725 | -4.1553293 | 3.25E-05 | 0.00036948 | ACTA2 | 59 |

| | | | | | | | | |
|---|---|---|---|---|---|---|---|---|
| ENSG000001 | 1432.65513 | 0.501008938 | 0.12058815 | 4.15471105 | 3.26E-05 | 0.00037037 | PSMD6 | 9861 |
| ENSG000001 | 539.974939 | 0.582053433 | 0.14010155 | 4.15451115 | 3.26E-05 | 0.00037059 | DPY19L4 | 286148 |
| ENSG000001 | 497.756188 | 0.582891894 | 0.14031535 | 4.15415621 | 3.26E-05 | 0.00037106 | FMNL3 | 91010 |
| ENSG000002 | 301.009121 | 0.556791109 | 0.13404269 | 4.15383431 | 3.27E-05 | 0.00037148 | UQCRHL | 440567 |
| ENSG000001 | 2131.43133 | 0.516540329 | 0.12436611 | 4.15338498 | 3.28E-05 | 0.00037211 | TGIF1 | 7050 |
| ENSG000001 | 529.199341 | 0.528189786 | 0.12717765 | 4.15316512 | 3.28E-05 | 0.00037236 | TPRKB | 51002 |
| ENSG000001 | 790.028622 | 0.503275403 | 0.12118744 | 4.15286768 | 3.28E-05 | 0.00037274 | SLC38A9 | 153129 |
| ENSG000001 | 210.257307 | 0.703731104 | 0.16946204 | 4.15273583 | 3.29E-05 | 0.00037285 | YRDC | 79693 |
| ENSG000001 | 667.835787 | 0.575043431 | 0.13849338 | 4.15213671 | 3.29E-05 | 0.00037372 | SRF | 6722 |
| ENSG000001 | 2974.21407 | 0.575355159 | 0.13858921 | 4.15151484 | 3.30E-05 | 0.00037464 | CHD7 | 55636 |
| ENSG00000C | 238.078487 | 0.593663563 | 0.1430178 | 4.15097682 | 3.31E-05 | 0.00037541 | TSSC1 | 7260 |
| ENSG000001 | 2577.35632 | 0.509413936 | 0.12272705 | 4.15078785 | 3.31E-05 | 0.00037562 | COPA | 1314 |
| ENSG000001 | 490.085088 | 0.650701573 | 0.15678613 | 4.1502496 | 3.32E-05 | 0.0003764 | PABPN1 | 8106 |
| ENSG000001 | 529.246463 | 0.590052278 | 0.14221634 | 4.14897667 | 3.34E-05 | 0.00037829 | NXPE3 | 91775 |
| ENSG000001 | 22.9006456 | -1.171187238 | 0.2822831 | -4.1489811 | 3.34E-05 | 0.00037829 | OR51D1 | 390038 |
| ENSG000001 | 1485.67489 | 0.533730803 | 0.12864511 | 4.14886205 | 3.34E-05 | 0.00037837 | RPE | 6120 |
| ENSG000002 | 10.1710833 | -1.431137301 | 0.3449615 | -4.1486871 | 3.34E-05 | 0.00037855 | LINC00661 | 126536 |
| ENSG000001 | 1062.42883 | 0.503174601 | 0.12128758 | 4.1486079 | 3.35E-05 | 0.00037858 | THUMPD3 | 25917 |
| ENSG000002 | 70.4290132 | 0.836533877 | 0.20165806 | 4.14827895 | 3.35E-05 | 0.00037891 | NA | NA |
| ENSG000002 | 368.297112 | 0.676756112 | 0.16313993 | 4.14831687 | 3.35E-05 | 0.00037891 | TAPBP | 6892 |
| ENSG00000C | 2302.29833 | 0.560243492 | 0.13506137 | 4.14806597 | 3.35E-05 | 0.00037905 | CRKL | 1399 |
| ENSG000001 | 578.035375 | 0.511876346 | 0.1234004 | 4.148093 | 3.35E-05 | 0.00037905 | PRPSAP2 | 5636 |
| ENSG00000C | 271.261792 | 0.806225911 | 0.1943885 | 4.14749802 | 3.36E-05 | 0.00037989 | WDR18 | 57418 |
| ENSG00000C | 14040.1989 | 0.476933077 | 0.11502255 | 4.14643114 | 3.38E-05 | 0.00038148 | IPO5 | 3843 |
| ENSG000001 | 1047.31501 | 0.53693338 | 0.12949349 | 4.14641222 | 3.38E-05 | 0.00038148 | TIMMDC1 | 51300 |
| ENSG000001 | 1858.38421 | 0.490156234 | 0.11821473 | 4.14632119 | 3.38E-05 | 0.00038152 | CHEK1 | 1111 |
| ENSG000001 | 76.7159026 | -0.777947148 | 0.18763853 | -4.1459883 | 3.38E-05 | 0.00038197 | SLIT3 | 6586 |
| ENSG000002 | 36.1833325 | 1.056463575 | 0.25484044 | 4.14558836 | 3.39E-05 | 0.00038253 | MIR621 | 693206 |
| ENSG000001 | 706.902198 | 0.569680287 | 0.13742554 | 4.14537416 | 3.39E-05 | 0.00038279 | TTLL4 | 9654 |

| | | | | | | | | |
|---|---|---|---|---|---|---|---|---|
| ENSG000001 | 288.498394 | 0.585239933 | 0.14118365 | 4.14523865 | 3.39E-05 | 0.00038291 | COA5 | 493753 |
| ENSG000001 | 801.398994 | 0.53722089 | 0.12960684 | 4.14500406 | 3.40E-05 | 0.00038315 | WDR74 | 54663 |
| ENSG000001 | 162.916294 | 0.690920859 | 0.16668918 | 4.14496531 | 3.40E-05 | 0.00038315 | CYHR1 | 50626 |
| ENSG00000C | 1304.5139 | 0.503393724 | 0.12146282 | 4.1444263 | 3.41E-05 | 0.00038394 | CLASP1 | 23332 |
| ENSG000001 | 768.415186 | 0.568261161 | 0.13712495 | 4.14411208 | 3.41E-05 | 0.00038436 | SLC39A8 | 64116 |
| ENSG00000C | 2334.44259 | 0.52288328 | 0.12618729 | 4.14370809 | 3.42E-05 | 0.00038466 | ZNF195 | 7748 |
| ENSG000001 | 8.4453176 | -1.515091256 | 0.36563893 | -4.1436814 | 3.42E-05 | 0.00038466 | PERM1 | 84808 |
| ENSG000001 | 1079.89064 | 0.52828542 | 0.12748673 | 4.14384622 | 3.42E-05 | 0.00038466 | NBR1 | 4077 |
| ENSG000001 | 741.824696 | 0.547309979 | 0.13207976 | 4.14378385 | 3.42E-05 | 0.00038466 | FAN1 | 22909 |
| ENSG000001 | 1794.21079 | 0.527465538 | 0.12731203 | 4.14309265 | 3.43E-05 | 0.00038554 | TOMM22 | 56993 |
| ENSG000002 | 45.4567327 | 1.098868046 | 0.26524451 | 4.14284933 | 3.43E-05 | 0.00038584 | P2RY11 | 5032 |
| ENSG000001 | 28.7368274 | -1.087558938 | 0.26252733 | -4.1426503 | 3.43E-05 | 0.00038607 | NCAM2 | 4685 |
| ENSG000001 | 933.585836 | 0.530188966 | 0.12799418 | 4.14228979 | 3.44E-05 | 0.00038657 | RAB15 | 376267 |
| ENSG00000C | 1286.64726 | 0.483571424 | 0.11676223 | 4.1415054 | 3.45E-05 | 0.00038776 | ALS2 | 57679 |
| ENSG000002 | 40.3270808 | -0.927412895 | 0.22393372 | -4.1414616 | 3.45E-05 | 0.00038776 | LINC01435 | 103695365 |
| ENSG000001 | 175.202345 | 0.596430702 | 0.1440441 | 4.1406119 | 3.46E-05 | 0.00038909 | SLC25A16 | 8034 |
| ENSG00000C | 72.7853753 | -0.817762344 | 0.19751085 | -4.1403413 | 3.47E-05 | 0.00038925 | TG | 7038 |
| ENSG000002 | 759.869851 | 0.487960297 | 0.11785549 | 4.14032716 | 3.47E-05 | 0.00038925 | SNHG8 | 100093630 |
| ENSG000002 | 45.9620967 | -0.926851484 | 0.2238527 | -4.1404525 | 3.47E-05 | 0.00038925 | NA | NA |
| ENSG000001 | 876.212112 | 0.711442763 | 0.17183572 | 4.14024953 | 3.47E-05 | 0.00038927 | GIGYF1 | 64599 |
| ENSG000001 | 21.8487483 | -1.166919865 | 0.28186785 | -4.1399537 | 3.47E-05 | 0.00038967 | GALNT15 | 117248 |
| ENSG000001 | 861.331236 | 0.474427411 | 0.11460539 | 4.13966057 | 3.48E-05 | 0.00039006 | RNF6 | 6049 |
| ENSG00000C | 1165.35286 | 0.485410689 | 0.11727944 | 4.13892384 | 3.49E-05 | 0.0003912 | PMS1 | 5378 |
| ENSG00000C | 325.327177 | 0.729860473 | 0.17635073 | 4.13868692 | 3.49E-05 | 0.00039145 | GTSE1 | 51512 |
| ENSG000001 | 1046.68243 | 0.498948265 | 0.12055809 | 4.1386544 | 3.49E-05 | 0.00039145 | VBP1 | 7411 |
| ENSG000001 | 5468.34695 | 0.4437937 | 0.1072392 | 4.13835326 | 3.50E-05 | 0.00039185 | RSL1D1 | 26156 |
| ENSG000001 | 1741.05827 | 0.487501923 | 0.11781411 | 4.13789098 | 3.51E-05 | 0.00039243 | SSR1 | 6745 |
| ENSG000001 | 425.506105 | 0.575169641 | 0.13899898 | 4.13794132 | 3.50E-05 | 0.00039243 | PRKAR1B | 5575 |
| ENSG000001 | 394.507789 | 0.570465761 | 0.13787468 | 4.13756731 | 3.51E-05 | 0.00039287 | CDK5RAP1 | 51654 |

| | | | | | | | | |
|---|---|---|---|---|---|---|---|---|
| ENSG000002 | 51.9885856 | -0.85648195 | 0.20700909 | -4.1374122 | 3.51E-05 | 0.00039303 | LOC101927661 | 101927661 |
| ENSG000001 | 1370.4485 | 0.51621584 | 0.12477087 | 4.13731041 | 3.51E-05 | 0.00039309 | PPP1R2 | 5504 |
| ENSG000000 | 1296.92649 | 0.602470498 | 0.14562789 | 4.1370544 | 3.52E-05 | 0.00039342 | MAT2B | 27430 |
| ENSG000001 | 662.5917 | 0.479390501 | 0.1158793 | 4.13698132 | 3.52E-05 | 0.00039344 | C9orf78 | 51759 |
| ENSG000001 | 1111.53745 | 0.430728679 | 0.10414048 | 4.13603527 | 3.53E-05 | 0.00039469 | STAMBP | 10617 |
| ENSG000001 | 1510.44308 | 0.561293793 | 0.13570814 | 4.13603641 | 3.53E-05 | 0.00039469 | KLHDC3 | 116138 |
| ENSG000001 | 90.297634 | -0.715681995 | 0.1730372 | -4.1360007 | 3.53E-05 | 0.00039469 | NEK10 | 152110 |
| ENSG000001 | 578.675868 | 0.572232241 | 0.13834808 | 4.13617774 | 3.53E-05 | 0.00039469 | NUDCD2 | 134492 |
| ENSG000001 | 34.7865737 | -1.004442433 | 0.2428626 | -4.1358465 | 3.54E-05 | 0.00039485 | GRIA2 | 2891 |
| ENSG000000 | 437.202462 | 0.567217701 | 0.13716861 | 4.13518593 | 3.55E-05 | 0.00039587 | NCK2 | 8440 |
| ENSG000001 | 300.100607 | 0.550950129 | 0.13324611 | 4.13483085 | 3.55E-05 | 0.00039638 | HINFP | 25988 |
| ENSG000001 | 330.716972 | 0.567428567 | 0.13723648 | 4.13467749 | 3.55E-05 | 0.00039644 | IFT52 | 51098 |
| ENSG000001 | 273.209716 | 0.570445421 | 0.13796643 | 4.13466837 | 3.55E-05 | 0.00039644 | C14orf159 | 80017 |
| ENSG000001 | 1533.45087 | 0.51452194 | 0.12445723 | 4.13412647 | 3.56E-05 | 0.00039727 | TGFBR1 | 7046 |
| ENSG000000 | 245.168178 | 0.681301111 | 0.164804 | 4.1340084 | 3.56E-05 | 0.00039736 | PNKP | 11284 |
| ENSG000000 | 287.139583 | 0.602522472 | 0.14575025 | 4.13393779 | 3.57E-05 | 0.00039738 | ANGEL1 | 23357 |
| ENSG000001 | 672.919773 | 0.716344783 | 0.17330657 | 4.13339658 | 3.57E-05 | 0.0003982 | PLXNA1 | 5361 |
| ENSG000002 | 37.1775378 | 0.982466082 | 0.23772237 | 4.13282982 | 3.58E-05 | 0.00039908 | HIST2H2AA3 | 8337 |
| ENSG000001 | 521.248259 | 0.698676442 | 0.16906627 | 4.13255955 | 3.59E-05 | 0.00039944 | MTA1 | 9112 |
| ENSG000001 | 458.015766 | 0.58683588 | 0.14201791 | 4.13212577 | 3.59E-05 | 0.00040008 | HIC2 | 23119 |
| ENSG000001 | 1430.10697 | 0.443821589 | 0.10741376 | 4.13188777 | 3.60E-05 | 0.00040039 | UTP15 | 84135 |
| ENSG000000 | 1303.30057 | 0.469751609 | 0.11369798 | 4.13157387 | 3.60E-05 | 0.00040082 | VEZT | 55591 |
| ENSG000001 | 1239.16131 | 0.52933683 | 0.12813003 | 4.13124717 | 3.61E-05 | 0.00040128 | ZNF460 | 10794 |
| ENSG000001 | 644.102784 | 0.649352722 | 0.15719132 | 4.13097061 | 3.61E-05 | 0.00040166 | IRGQ | 126298 |
| ENSG000001 | 999.922947 | 0.494163732 | 0.11964204 | 4.130352 | 3.62E-05 | 0.00040241 | IFITM3 | 10410 |
| ENSG000001 | 105.671951 | -0.716389088 | 0.17344427 | -4.1303705 | 3.62E-05 | 0.00040241 | CCDC30 | 728621 |
| ENSG000001 | 469.347933 | 0.561144566 | 0.13585671 | 4.13041483 | 3.62E-05 | 0.00040241 | SPIN4 | 139886 |
| ENSG000001 | 1895.21782 | 0.43336904 | 0.1049396 | 4.12969957 | 3.63E-05 | 0.00040344 | ROR1 | 4919 |
| ENSG000001 | 363.44941 | 0.618798441 | 0.14985355 | 4.12935443 | 3.64E-05 | 0.0004039 | SUMF2 | 25870 |

| | | | | | | | | |
|---|---|---|---|---|---|---|---|---|
| ENSG000001 | 47.8939849 | -0.913196522 | 0.22115264 | -4.129259 | 3.64E-05 | 0.0004039 | ATP13A5 | 344905 |
| ENSG000002 | 33.7356595 | -1.040649883 | 0.25201905 | -4.129251 | 3.64E-05 | 0.0004039 | CAPN8 | 388743 |
| ENSG000001 | 280.044946 | 0.598707182 | 0.14500648 | 4.12883056 | 3.65E-05 | 0.00040444 | MMAB | 326625 |
| ENSG000002 | 1068.95646 | 0.465005977 | 0.11262449 | 4.12881764 | 3.65E-05 | 0.00040444 | UBXN2B | 137886 |
| ENSG000000 | 462.856552 | 0.560895235 | 0.13586097 | 4.12845001 | 3.65E-05 | 0.00040497 | RTFDC1 | 51507 |
| ENSG000001 | 3817.56778 | 0.469541064 | 0.11374511 | 4.12801085 | 3.66E-05 | 0.00040564 | NUP153 | 9972 |
| ENSG000002 | 80.2473867 | -0.793955227 | 0.19233976 | -4.1278789 | 3.66E-05 | 0.00040576 | ABCC13 | 150000 |
| ENSG000001 | 1094.55159 | 0.496387584 | 0.12025834 | 4.12767695 | 3.66E-05 | 0.000406 | TMEM50A | 23585 |
| ENSG000001 | 65.6947282 | -0.818500865 | 0.19835681 | -4.1264067 | 3.68E-05 | 0.00040814 | ATP8B4 | 79895 |
| ENSG000001 | 390.054776 | 0.493479274 | 0.1196238 | 4.12525998 | 3.70E-05 | 0.00041007 | NAP1L2 | 4674 |
| ENSG000002 | 26.6684359 | -1.067884071 | 0.25888127 | -4.1249955 | 3.71E-05 | 0.00041043 | ZNF736 | 728927 |
| ENSG000001 | 390.445413 | 0.540509449 | 0.13103932 | 4.12478815 | 3.71E-05 | 0.00041069 | TRAPPC13 | 80006 |
| ENSG000001 | 518.432059 | 0.517358224 | 0.12543182 | 4.1246171 | 3.71E-05 | 0.00041088 | SNX4 | 8723 |
| ENSG000001 | 437.566544 | 0.553051981 | 0.13410921 | 4.12389265 | 3.73E-05 | 0.00041206 | ZDHHC9 | 51114 |
| ENSG000001 | 186.147389 | 0.670260924 | 0.1625529 | 4.12334027 | 3.73E-05 | 0.00041291 | DUS3L | 56931 |
| ENSG000001 | 17.6087081 | -1.262553393 | 0.30620039 | -4.1232913 | 3.73E-05 | 0.00041291 | KIAA1211L | 343990 |
| ENSG000001 | 1583.55716 | 0.640200182 | 0.15527803 | 4.12292827 | 3.74E-05 | 0.00041345 | PXDN | 7837 |
| ENSG000000 | 1823.38184 | 0.690784295 | 0.16755554 | 4.12271828 | 3.74E-05 | 0.00041372 | KDM4A | 9682 |
| ENSG000001 | 238.239022 | 0.563530971 | 0.13670211 | 4.12232818 | 3.75E-05 | 0.00041431 | NSUN7 | 79730 |
| ENSG000002 | 37.0664904 | -1.036536839 | 0.25147076 | -4.121898 | 3.76E-05 | 0.00041497 | NA | NA |
| ENSG000000 | 2250.43394 | 0.476319895 | 0.11556525 | 4.12165331 | 3.76E-05 | 0.0004153 | DEPDC1 | 55635 |
| ENSG000001 | 222.624562 | 0.604774982 | 0.1467347 | 4.12155405 | 3.76E-05 | 0.00041536 | STEAP3 | 55240 |
| ENSG000001 | 6421.94174 | 0.491330615 | 0.11921205 | 4.12148436 | 3.76E-05 | 0.00041537 | PPP1CC | 5501 |
| ENSG000002 | 150.168847 | 0.619484434 | 0.15032418 | 4.12099001 | 3.77E-05 | 0.00041615 | S1PR2 | 9294 |
| ENSG000000 | 453.350687 | 0.60365682 | 0.14649552 | 4.12065044 | 3.78E-05 | 0.00041665 | WDR62 | 284403 |
| ENSG000000 | 51.1438399 | -0.924162075 | 0.22428986 | -4.1203916 | 3.78E-05 | 0.00041701 | BCAS1 | 8537 |
| ENSG000000 | 7459.07139 | 0.518356524 | 0.12581123 | 4.12011341 | 3.79E-05 | 0.00041706 | THRAP3 | 9967 |
| ENSG000000 | 1691.41421 | 0.46527493 | 0.11292668 | 4.12015067 | 3.79E-05 | 0.00041706 | KDM5A | 5927 |
| ENSG000001 | 2011.42929 | 0.573679042 | 0.13923639 | 4.12018042 | 3.79E-05 | 0.00041706 | SHC1 | 6464 |

| | | | | | | | | | |
|---|---|---|---|---|---|---|---|---|---|
| ENSG000001 | 1753.21748 | 0.516129815 | 0.1252664 | 4.12025751 | 3.78E-05 | 0.00041706 | PDZD8 | | 118987 |
| ENSG00000C | 462.317947 | 0.51652484 | 0.12538626 | 4.11946927 | 3.80E-05 | 0.00041811 | ANKRD13A | | 88455 |
| ENSG000001 | 102.938126 | 0.727450239 | 0.17659391 | 4.11933928 | 3.80E-05 | 0.00041823 | PEX16 | | 9409 |
| ENSG000001 | 1150.16145 | 0.716827368 | 0.17404312 | 4.11867686 | 3.81E-05 | 0.00041932 | SRSF9 | | 8683 |
| ENSG000001 | 30.949467 | -1.064113387 | 0.25838119 | -4.1183857 | 3.82E-05 | 0.00041974 | TPH1 | | 7166 |
| ENSG00000C | 3042.29024 | 0.482636137 | 0.1171957 | 4.11820677 | 3.82E-05 | 0.00041984 | NUP50 | | 10762 |
| ENSG000001 | 46.6482649 | -0.942791128 | 0.2289293 | -4.1182632 | 3.82E-05 | 0.00041984 | MDGA1 | | 266727 |
| ENSG00000C | 140.024056 | 0.706194512 | 0.17149155 | 4.1179551 | 3.82E-05 | 0.00042018 | PCBP4 | | 57060 |
| ENSG000001 | 63.1822147 | -0.858467322 | 0.2084905 | -4.117537 | 3.83E-05 | 0.00042083 | NCKAP1L | | 3071 |
| ENSG000002 | 306.824105 | 0.659639146 | 0.16020521 | 4.11746371 | 3.83E-05 | 0.00042085 | NA | NA | |
| ENSG00000C | 183.654305 | 0.727422682 | 0.17667453 | 4.11730369 | 3.83E-05 | 0.00042103 | JMJD4 | | 65094 |
| ENSG000001 | 52.1203283 | -0.858550788 | 0.20854826 | -4.1167967 | 3.84E-05 | 0.00042184 | ATP13A4 | | 84239 |
| ENSG000001 | 1034.47733 | 0.479166339 | 0.11639715 | 4.11664994 | 3.84E-05 | 0.000422 | MELK | | 9833 |
| ENSG000001 | 475.381478 | 0.630165447 | 0.15314934 | 4.11471224 | 3.88E-05 | 0.00042533 | NA | NA | |
| ENSG000002 | 23.0528318 | -1.136192136 | 0.27612826 | -4.114726 | 3.88E-05 | 0.00042533 | NA | NA | |
| ENSG000001 | 682.813065 | 0.477657098 | 0.1160972 | 4.11428615 | 3.88E-05 | 0.000426 | SAAL1 | | 113174 |
| ENSG000001 | 230.551694 | 0.655611161 | 0.15935298 | 4.11420709 | 3.89E-05 | 0.00042603 | ADCK4 | | 79934 |
| ENSG000001 | 52.9319624 | -0.851570452 | 0.20702376 | -4.1133948 | 3.90E-05 | 0.0004273 | CFAP221 | | 200373 |
| ENSG000002 | 1041.59574 | 0.474125145 | 0.11526236 | 4.11344293 | 3.90E-05 | 0.0004273 | JPX | | 554203 |
| ENSG000002 | 5.52872863 | -1.549513393 | 0.37673982 | -4.1129536 | 3.91E-05 | 0.000428 | NA | NA | |
| ENSG000001 | 415.786853 | 0.550890819 | 0.13395109 | 4.11262669 | 3.91E-05 | 0.00042849 | PRMT7 | | 54496 |
| ENSG000001 | 62.857671 | 0.88015444 | 0.21403858 | 4.11212995 | 3.92E-05 | 0.0004293 | ACP5 | | 54 |
| ENSG000001 | 473.568829 | 0.640262741 | 0.15571299 | 4.11181331 | 3.93E-05 | 0.00042977 | EHD4 | | 30844 |
| ENSG00000C | 110.143198 | 0.83729006 | 0.20363536 | 4.11171261 | 3.93E-05 | 0.00042985 | QPCTL | | 54814 |
| ENSG000001 | 870.187358 | 0.626932479 | 0.15249484 | 4.11117163 | 3.94E-05 | 0.00043074 | KDELR2 | | 11014 |
| ENSG000001 | 668.841509 | 0.535889471 | 0.13035343 | 4.11104999 | 3.94E-05 | 0.00043085 | MAPK14 | | 1432 |
| ENSG000001 | 725.742409 | 0.653858432 | 0.15905314 | 4.11094324 | 3.94E-05 | 0.00043093 | ACO2 | | 50 |
| ENSG000002 | 67437.48 | 1.073428853 | 0.261135 | 4.11062808 | 3.95E-05 | 0.0004314 | RN7SL2 | | 378706 |
| ENSG000001 | 1453.88748 | 0.480817873 | 0.11698772 | 4.10998599 | 3.96E-05 | 0.00043237 | ZNF426 | | 79088 |

| | | | | | | | | |
|---|---|---|---|---|---|---|---|---|
| ENSG000001 | 2390.63889 | 0.700051205 | 0.17032871 | 4.11000126 | 3.96E-05 | 0.00043237 | MRFAP1 | 93621 |
| ENSG000001 | 292.219032 | 0.65137436 | 0.15848892 | 4.10990481 | 3.96E-05 | 0.00043241 | MYEOV2 | 150678 |
| ENSG000001 | 1069.14984 | 0.496196302 | 0.12077294 | 4.10850571 | 3.98E-05 | 0.00043492 | BLOC1S6 | 26258 |
| ENSG00000C | 258.4669 | 0.543871486 | 0.13239356 | 4.10799035 | 3.99E-05 | 0.00043577 | ZNF200 | 7752 |
| ENSG000002 | 29.1686448 | -1.037036688 | 0.2524744 | -4.1074924 | 4.00E-05 | 0.0004366 | NA | NA |
| ENSG000002 | 88.8384491 | 0.776084948 | 0.18894884 | 4.10738138 | 4.00E-05 | 0.00043669 | SDHAF1 | 644096 |
| ENSG000001 | 41.3987351 | -0.942082807 | 0.2293737 | -4.1071964 | 4.00E-05 | 0.00043692 | KLHL6 | 89857 |
| ENSG000001 | 3207.0867 | 0.481743857 | 0.11730643 | 4.10671324 | 4.01E-05 | 0.0004376 | NOL11 | 25926 |
| ENSG000001 | 789.38279 | 0.595874218 | 0.14509734 | 4.10672051 | 4.01E-05 | 0.0004376 | RAC3 | 5881 |
| ENSG000001 | 316.794914 | 0.550729853 | 0.13411798 | 4.10630894 | 4.02E-05 | 0.00043825 | TSFM | 10102 |
| ENSG000001 | 1620.07771 | 0.591819921 | 0.14412859 | 4.10619372 | 4.02E-05 | 0.00043835 | CDS2 | 8760 |
| ENSG000001 | 369.123814 | 0.559597507 | 0.13628687 | 4.10602662 | 4.03E-05 | 0.00043855 | NDUFC2 | 4718 |
| ENSG000001 | 562.189394 | 0.496656871 | 0.12096843 | 4.10567338 | 4.03E-05 | 0.0004391 | JAKMIP2 | 9832 |
| ENSG000001 | 6616.79627 | 0.668384675 | 0.16280328 | 4.10547424 | 4.03E-05 | 0.00043936 | EIF4G1 | 1981 |
| ENSG000001 | 462.246924 | 0.551713312 | 0.13438752 | 4.10539106 | 4.04E-05 | 0.0004394 | ANAPC13 | 25847 |
| ENSG000002 | 557.764865 | 0.566505485 | 0.1380256 | 4.10435093 | 4.05E-05 | 0.00044126 | CRCP | 27297 |
| ENSG000001 | 63.8589193 | 0.842467903 | 0.20527259 | 4.10414226 | 4.06E-05 | 0.00044154 | HPN | 3249 |
| ENSG000001 | 176.496403 | 0.689928568 | 0.16812797 | 4.10359192 | 4.07E-05 | 0.00044248 | SH2B1 | 25970 |
| ENSG000001 | 474.570145 | 0.579031586 | 0.1411079 | 4.1034667 | 4.07E-05 | 0.00044251 | KIAA0930 | 23313 |
| ENSG000001 | 511.276077 | 0.525672318 | 0.1281049 | 4.10345195 | 4.07E-05 | 0.00044251 | DPY19L1 | 23333 |
| ENSG000001 | 3729.39026 | 0.514976317 | 0.12550531 | 4.10322348 | 4.07E-05 | 0.00044283 | C11orf58 | 10944 |
| ENSG000001 | 30.902121 | 1.07287756 | 0.26151417 | 4.10255993 | 4.09E-05 | 0.00044398 | PPAN | 56342 |
| ENSG000001 | 574.078844 | 0.499998382 | 0.1218794 | 4.10240278 | 4.09E-05 | 0.00044416 | MIS12 | 79003 |
| ENSG000001 | 187.009758 | 0.627969769 | 0.15307838 | 4.10227594 | 4.09E-05 | 0.00044429 | RAD51D | 5892 |
| ENSG00000C | 1013.55782 | 0.483115167 | 0.11777176 | 4.10213067 | 4.09E-05 | 0.00044445 | VPS41 | 27072 |
| ENSG000001 | 256.137739 | 0.66383358 | 0.161833 | 4.10196668 | 4.10E-05 | 0.00044464 | FBXO46 | 23403 |
| ENSG000001 | 289.70849 | 0.571231614 | 0.13926238 | 4.10183724 | 4.10E-05 | 0.00044477 | COQ10B | 80219 |
| ENSG000001 | 694.596815 | 0.532388103 | 0.12979827 | 4.10165797 | 4.10E-05 | 0.000445 | CXCL12 | 6387 |
| ENSG000001 | 138.436572 | 0.766077368 | 0.18677972 | 4.10150178 | 4.10E-05 | 0.00044518 | SIX5 | 147912 |

| | | | | | | | | |
|---|---|---|---|---|---|---|---|---|
| ENSG000001 | 2499.28035 | 0.551782872 | 0.13456262 | 4.10056569 | 4.12E-05 | 0.00044686 | CTBP2 | 1488 |
| ENSG00000C | 820.162058 | 0.49670745 | 0.12113806 | 4.10034192 | 4.13E-05 | 0.00044706 | NSMAF | 8439 |
| ENSG000001 | 1281.29452 | 0.454939572 | 0.11095143 | 4.10034877 | 4.13E-05 | 0.00044706 | RFC4 | 5984 |
| ENSG000001 | 1418.38462 | 0.496776376 | 0.12115867 | 4.10021331 | 4.13E-05 | 0.00044719 | ANK3 | 288 |
| ENSG00000C | 23.9227061 | -1.142194904 | 0.27857477 | -4.1001376 | 4.13E-05 | 0.00044721 | NOX1 | 27035 |
| ENSG000001 | 4121.25671 | 0.500228898 | 0.12202444 | 4.0994156 | 4.14E-05 | 0.00044838 | CAP1 | 10487 |
| ENSG000001 | 378.062552 | 0.531212308 | 0.12958256 | 4.09941221 | 4.14E-05 | 0.00044838 | C12orf43 | 64897 |
| ENSG000001 | 374.066133 | 0.504057443 | 0.12296249 | 4.09927806 | 4.14E-05 | 0.00044852 | ANAPC15 | 25906 |
| ENSG000001 | 292.956571 | 0.735647518 | 0.17952199 | 4.09781287 | 4.17E-05 | 0.00045125 | PLIN3 | 10226 |
| ENSG00000C | 201.893617 | 0.548348131 | 0.1338376 | 4.09711581 | 4.18E-05 | 0.00045249 | TMEM206 | 55248 |
| ENSG000001 | 761.256207 | 0.49077753 | 0.11979207 | 4.0969118 | 4.19E-05 | 0.00045276 | TSC1 | 7248 |
| ENSG00000C | 2455.45917 | 0.468748302 | 0.11441892 | 4.09677265 | 4.19E-05 | 0.00045279 | THUMPD1 | 55623 |
| ENSG000001 | 695.667794 | 0.566726205 | 0.13833275 | 4.09683311 | 4.19E-05 | 0.00045279 | ZKSCAN1 | 7586 |
| ENSG000001 | 2134.60263 | 0.608780993 | 0.14863459 | 4.09582328 | 4.21E-05 | 0.00045453 | EFTUD2 | 9343 |
| ENSG000001 | 405.693023 | 0.544580901 | 0.13297733 | 4.09529125 | 4.22E-05 | 0.00045546 | KAT8 | 84148 |
| ENSG000001 | 249.703829 | 0.643693816 | 0.15718222 | 4.09520746 | 4.22E-05 | 0.0004555 | GLB1L3 | 112937 |
| ENSG000001 | 2555.94888 | 0.664154342 | 0.16219599 | 4.09476418 | 4.23E-05 | 0.00045606 | HNRNPL | 3191 |
| ENSG000001 | 170.528754 | 0.628250583 | 0.15342455 | 4.0948504 | 4.22E-05 | 0.00045606 | MORC4 | 79710 |
| ENSG000002 | 1576.41828 | 0.547820684 | 0.13378658 | 4.09473576 | 4.23E-05 | 0.00045606 | ATXN7L3B | 552889 |
| ENSG00000C | 188.269815 | 0.750471937 | 0.1832815 | 4.09464088 | 4.23E-05 | 0.00045613 | CHPF2 | 54480 |
| ENSG000001 | 814.679725 | 0.680125689 | 0.16610817 | 4.09447453 | 4.23E-05 | 0.00045634 | DHX38 | 9785 |
| ENSG000001 | 1421.97901 | 0.443257177 | 0.10826525 | 4.09417765 | 4.24E-05 | 0.0004568 | YEATS2 | 55689 |
| ENSG000001 | 4400.20802 | 0.494822087 | 0.12087194 | 4.09377152 | 4.24E-05 | 0.00045748 | CNN3 | 1266 |
| ENSG000001 | 293.672148 | 0.691150886 | 0.16883341 | 4.09368546 | 4.25E-05 | 0.00045753 | H1FX | 8971 |
| ENSG000001 | 412.8215 | 0.527886384 | 0.12895531 | 4.09356053 | 4.25E-05 | 0.00045753 | ENOX2 | 10495 |
| ENSG000001 | 129.861546 | 0.676290136 | 0.16520636 | 4.09360828 | 4.25E-05 | 0.00045753 | ZNF511 | 118472 |
| ENSG000001 | 440.432344 | 0.686061947 | 0.16761246 | 4.09314397 | 4.26E-05 | 0.00045823 | ZNF512B | 57473 |
| ENSG000001 | 730.719343 | 0.507478687 | 0.12400784 | 4.09231135 | 4.27E-05 | 0.00045976 | KLHDC2 | 23588 |
| ENSG000001 | 1491.50848 | 0.678062789 | 0.16570512 | 4.09198457 | 4.28E-05 | 0.00046016 | RRBP1 | 6238 |

| | | | | | | | | |
|---|---|---|---|---|---|---|---|---|
| ENSG000001 | 508.014585 | 0.552979269 | 0.13513535 | 4.09204002 | 4.28E-05 | 0.00046016 | SNF8 | 11267 |
| ENSG000001 | 56.2838015 | 0.84093201 | 0.20552813 | 4.09156656 | 4.28E-05 | 0.00046087 | SPATA24 | 202051 |
| ENSG000001 | 1169.37298 | 0.483927253 | 0.11830304 | 4.09057324 | 4.30E-05 | 0.00046273 | MACC1 | 346389 |
| ENSG00000C | 160.262978 | 0.671093168 | 0.16406813 | 4.09033229 | 4.31E-05 | 0.00046309 | RHBDD2 | 57414 |
| ENSG000001 | 183.128041 | 0.630071375 | 0.15405234 | 4.0899825 | 4.31E-05 | 0.00046361 | NDRG4 | 65009 |
| ENSG000001 | 123.364977 | 0.711284026 | 0.17391086 | 4.0899344 | 4.31E-05 | 0.00046361 | THEMIS2 | 9473 |
| ENSG000001 | 2096.80369 | 0.47766851 | 0.1167927 | 4.08988313 | 4.32E-05 | 0.00046361 | ZNF714 | 148206 |
| ENSG000001 | 798.699433 | 0.517142203 | 0.12646895 | 4.08908423 | 4.33E-05 | 0.00046509 | PANX1 | 24145 |
| ENSG000001 | 369.832268 | 0.702250093 | 0.17175177 | 4.08875036 | 4.34E-05 | 0.00046556 | TSC2 | 7249 |
| ENSG000001 | 823.106496 | 0.524288898 | 0.12822786 | 4.08872851 | 4.34E-05 | 0.00046556 | E2F3 | 1871 |
| ENSG000001 | 354.911012 | 0.597668665 | 0.14618577 | 4.08841877 | 4.34E-05 | 0.00046606 | ST3GAL2 | 6483 |
| ENSG000001 | 61.326915 | -0.860010109 | 0.21038269 | -4.0878368 | 4.35E-05 | 0.0004671 | A2M | 2 |
| ENSG000001 | 688.081389 | 0.448461177 | 0.10971083 | 4.08766549 | 4.36E-05 | 0.00046732 | RAVER2 | 55225 |
| ENSG000001 | 2945.81913 | 0.513732467 | 0.12568421 | 4.08748599 | 4.36E-05 | 0.00046752 | ESCO1 | 114799 |
| ENSG000002 | 42.8463998 | 0.950561152 | 0.23255622 | 4.0874467 | 4.36E-05 | 0.00046752 | NA | NA |
| ENSG00000C | 715.458877 | 0.606114918 | 0.14832482 | 4.0864025 | 4.38E-05 | 0.00046876 | ATXN7L3 | 56970 |
| ENSG00000C | 127.386698 | 0.685765093 | 0.16781143 | 4.086522 | 4.38E-05 | 0.00046876 | GNPTG | 84572 |
| ENSG000001 | 1762.51033 | 0.51085622 | 0.12500437 | 4.08670682 | 4.38E-05 | 0.00046876 | SCYL2 | 55681 |
| ENSG000001 | 1534.87933 | 0.43531657 | 0.10652416 | 4.08655249 | 4.38E-05 | 0.00046876 | TMOD3 | 29766 |
| ENSG000001 | 189.300318 | -0.578142762 | 0.14146732 | -4.0867583 | 4.37E-05 | 0.00046876 | PCDH10 | 57575 |
| ENSG000001 | 28.7112481 | -1.05230059 | 0.25750734 | -4.0864878 | 4.38E-05 | 0.00046876 | C12orf50 | 160419 |
| ENSG000001 | 271.74957 | 0.665613919 | 0.16288321 | 4.08644899 | 4.38E-05 | 0.00046876 | MAP3K11 | 4296 |
| ENSG000001 | 1052.32027 | 0.491020449 | 0.12017125 | 4.08600592 | 4.39E-05 | 0.00046944 | TTI1 | 9675 |
| ENSG00000C | 766.389254 | 0.497005121 | 0.1216409 | 4.0858389 | 4.39E-05 | 0.00046965 | RNF13 | 11342 |
| ENSG000002 | 321.92268 | 0.650024787 | 0.15911412 | 4.08527415 | 4.40E-05 | 0.00047067 | SKIV2L | 6499 |
| ENSG000001 | 641.837476 | 0.638841664 | 0.15639568 | 4.08477815 | 4.41E-05 | 0.00047155 | ZNF646 | 9726 |
| ENSG00000C | 38.5045456 | -0.92786179 | 0.22715692 | -4.0846733 | 4.41E-05 | 0.00047164 | PHACTR3 | 116154 |
| ENSG000001 | 139.283162 | 0.617615987 | 0.15122921 | 4.08397274 | 4.43E-05 | 0.00047294 | ATOX1 | 475 |
| ENSG000001 | 1327.54221 | 0.552396583 | 0.13526718 | 4.08374439 | 4.43E-05 | 0.00047328 | CAB39 | 51719 |

| | | | | | | | | |
|---|---|---|---|---|---|---|---|---|
| ENSG000001 | 627.843961 | 0.527550334 | 0.1292029 | 4.08311537 | 4.44E-05 | 0.00047444 | GNL3L | 54552 |
| ENSG000002 | 27.8696086 | -1.049741051 | 0.25713967 | -4.0823769 | 4.46E-05 | 0.00047583 | NA | NA |
| ENSG000001 | 359.472859 | 0.520299681 | 0.12745325 | 4.08227856 | 4.46E-05 | 0.0004759 | LCMT2 | 9836 |
| ENSG000001 | 15.3800325 | -1.30456657 | 0.31958035 | -4.0821238 | 4.46E-05 | 0.00047609 | AADACL2 | 344752 |
| ENSG00000C | 1077.68957 | 0.532264909 | 0.13039858 | 4.08183055 | 4.47E-05 | 0.00047657 | CRMP1 | 1400 |
| ENSG000002 | 479.031848 | 0.623223765 | 0.15269328 | 4.08154034 | 4.47E-05 | 0.00047704 | LCMT1 | 51451 |
| ENSG000001 | 208.163346 | 0.565719402 | 0.13861263 | 4.08129753 | 4.48E-05 | 0.00047741 | ZKSCAN3 | 80317 |
| ENSG000001 | 7523.00317 | 0.447503717 | 0.10965636 | 4.08096469 | 4.48E-05 | 0.00047785 | NUDT21 | 11051 |
| ENSG000001 | 501.793262 | 0.561169436 | 0.13750868 | 4.08097453 | 4.48E-05 | 0.00047785 | NHS | 4810 |
| ENSG000001 | 467.578382 | 0.557152976 | 0.13653467 | 4.08067028 | 4.49E-05 | 0.00047833 | CHAF1B | 8208 |
| ENSG00000C | 584.937959 | 0.46791354 | 0.11466888 | 4.0805627 | 4.49E-05 | 0.00047842 | EXOC1 | 55763 |
| ENSG00000C | 31.0452733 | -1.016278954 | 0.24907312 | -4.0802435 | 4.50E-05 | 0.00047895 | PKP1 | 5317 |
| ENSG000001 | 669.163251 | 0.564990364 | 0.13847834 | 4.07999098 | 4.50E-05 | 0.00047935 | DCLRE1A | 9937 |
| ENSG000001 | 516.822971 | 0.503292111 | 0.12336045 | 4.07984969 | 4.51E-05 | 0.00047951 | THAP9 | 79725 |
| ENSG00000C | 1260.57436 | 0.489633716 | 0.12002007 | 4.07959877 | 4.51E-05 | 0.0004799 | SEL1L3 | 23231 |
| ENSG000001 | 103.015001 | 0.730795187 | 0.17915245 | 4.07918052 | 4.52E-05 | 0.00048052 | FAM171A2 | 284069 |
| ENSG000001 | 954.884002 | 0.510672174 | 0.12518912 | 4.0792057 | 4.52E-05 | 0.00048052 | NELL2 | 4753 |
| ENSG000001 | 902.552754 | 0.562735336 | 0.13796502 | 4.07882619 | 4.53E-05 | 0.00048087 | TRPC4AP | 26133 |
| ENSG000001 | 110.085971 | 0.718827316 | 0.17623092 | 4.07889435 | 4.53E-05 | 0.00048087 | KLRG2 | 346689 |
| ENSG000002 | 40.7787666 | 0.947838784 | 0.23237952 | 4.0788396 | 4.53E-05 | 0.00048087 | BBOX1-AS1 | 103695435 |
| ENSG000002 | 85.9678634 | 0.724811218 | 0.17773606 | 4.07802015 | 4.54E-05 | 0.00048241 | LINC00890 | 401613 |
| ENSG00000C | 364.021799 | 0.552039454 | 0.13537404 | 4.07788284 | 4.54E-05 | 0.00048257 | MRPL43 | 84545 |
| ENSG000001 | 267.944411 | -0.561063669 | 0.137621 | -4.0768754 | 4.56E-05 | 0.00048446 | ZNF667 | 63934 |
| ENSG000002 | 133.16899 | 0.743981464 | 0.18248914 | 4.07685329 | 4.56E-05 | 0.00048446 | NA | NA |
| ENSG000001 | 1672.44439 | 0.559123547 | 0.13714839 | 4.07677796 | 4.57E-05 | 0.00048449 | KIF1C | 10749 |
| ENSG00000C | 3815.53058 | 0.488822588 | 0.11991177 | 4.07651877 | 4.57E-05 | 0.00048468 | CDV3 | 55573 |
| ENSG000001 | 441.24809 | 0.548915099 | 0.13465616 | 4.07642022 | 4.57E-05 | 0.00048468 | TAF8 | 129685 |
| ENSG000001 | 301.751039 | 0.635002854 | 0.15577611 | 4.07638141 | 4.57E-05 | 0.00048468 | GOLGA2 | 2801 |
| ENSG000001 | 2160.87011 | 0.478881035 | 0.11747245 | 4.07653916 | 4.57E-05 | 0.00048468 | PTEN | 5728 |

| | | | | | | | | |
|---|---|---|---|---|---|---|---|---|
| ENSG000002 | 210.633925 | 0.567545113 | 0.13922707 | 4.07639928 | 4.57E-05 | 0.00048468 | CCDC71L | 168455 |
| ENSG000001 | 1226.39236 | 0.467031483 | 0.11457515 | 4.07620213 | 4.58E-05 | 0.00048493 | LEPROTL1 | 23484 |
| ENSG000001 | 580.858383 | 0.464234276 | 0.11389346 | 4.07603994 | 4.58E-05 | 0.00048514 | ALG2 | 85365 |
| ENSG000001 | 626.395291 | 0.587813066 | 0.14421952 | 4.07582194 | 4.59E-05 | 0.00048524 | AMBRA1 | 55626 |
| ENSG000001 | 510.751779 | 0.580302936 | 0.1423773 | 4.07581066 | 4.59E-05 | 0.00048524 | TOP3A | 7156 |
| ENSG000002 | 97.5447317 | 0.710462635 | 0.17431039 | 4.07584787 | 4.58E-05 | 0.00048524 | NA | NA |
| ENSG000001 | 714.191935 | 0.511270268 | 0.12544222 | 4.07574323 | 4.59E-05 | 0.00048525 | QRSL1 | 55278 |
| ENSG000000 | 3276.86819 | 0.484618708 | 0.11890825 | 4.07556846 | 4.59E-05 | 0.00048537 | PSMA4 | 5685 |
| ENSG000001 | 3128.50964 | 0.487336538 | 0.11957486 | 4.07557687 | 4.59E-05 | 0.00048537 | YWHAH | 7533 |
| ENSG000000 | 684.430164 | 0.465312323 | 0.11417972 | 4.07526238 | 4.60E-05 | 0.00048588 | FHL1 | 2273 |
| ENSG000001 | 17.0307182 | -1.24930632 | 0.30658503 | -4.0749097 | 4.60E-05 | 0.00048649 | DCST1 | 149095 |
| ENSG000002 | 24.2913973 | -1.105827392 | 0.27139278 | -4.0746382 | 4.61E-05 | 0.00048693 | MTND6P4 | 106478943 |
| ENSG000001 | 206.958992 | 0.774123337 | 0.1900114 | 4.07408892 | 4.62E-05 | 0.00048795 | DDX28 | 55794 |
| ENSG000001 | 785.265679 | 0.593390181 | 0.14567426 | 4.07340448 | 4.63E-05 | 0.00048926 | RNF187 | 149603 |
| ENSG000000 | 778.45434 | 0.565057338 | 0.13872311 | 4.07327479 | 4.64E-05 | 0.00048941 | CFAP20 | 29105 |
| ENSG000002 | 710.041988 | 0.483356912 | 0.11866797 | 4.0731876 | 4.64E-05 | 0.00048946 | INPP5B | 3633 |
| ENSG000000 | 632.656908 | 0.567531349 | 0.13933791 | 4.07305786 | 4.64E-05 | 0.00048961 | NUP37 | 79023 |
| ENSG000001 | 203.531002 | 0.586156013 | 0.14391589 | 4.0729067 | 4.64E-05 | 0.0004898 | EXOSC4 | 54512 |
| ENSG000000 | 44.5719058 | -0.901245917 | 0.22128502 | -4.0727833 | 4.65E-05 | 0.00048993 | TRAF3IP3 | 80342 |
| ENSG000001 | 967.900574 | 0.534186853 | 0.13116341 | 4.07268191 | 4.65E-05 | 0.00049002 | DPY30 | 84661 |
| ENSG000001 | 68.8779083 | 0.779875691 | 0.19150766 | 4.07229512 | 4.66E-05 | 0.0004907 | PTPMT1 | 114971 |
| ENSG000001 | 6563.00895 | 0.576988088 | 0.14169527 | 4.0720349 | 4.66E-05 | 0.00049112 | NA | NA |
| ENSG000001 | 2043.47819 | 0.480114521 | 0.11790953 | 4.07188906 | 4.66E-05 | 0.0004913 | DIAPH2 | 1730 |
| ENSG000001 | 764.033698 | 0.554058028 | 0.13607174 | 4.07180813 | 4.66E-05 | 0.00049135 | SDHB | 6390 |
| ENSG000000 | 1196.99439 | 0.489254141 | 0.12018115 | 4.07097233 | 4.68E-05 | 0.00049273 | AP5M1 | 55745 |
| ENSG000000 | 67.963602 | -0.801496045 | 0.19687811 | -4.0710268 | 4.68E-05 | 0.00049273 | ACTN2 | 88 |
| ENSG000001 | 1332.81762 | 0.499000719 | 0.12257241 | 4.0710689 | 4.68E-05 | 0.00049273 | RAF1 | 5894 |
| ENSG000001 | 628.120383 | 0.590611268 | 0.14509108 | 4.07062426 | 4.69E-05 | 0.00049334 | OTUB1 | 55611 |
| ENSG000000 | 1507.08373 | 0.489821838 | 0.12034674 | 4.0700881 | 4.70E-05 | 0.00049422 | MTIF2 | 4528 |

| | | | | | | | | |
|---|---|---|---|---|---|---|---|---|
| ENSG000001 | 468.237645 | 0.716279215 | 0.17598606 | 4.07009055 | 4.70E-05 | 0.00049422 | CDT1 | 81620 |
| ENSG000001 | 2079.32156 | 0.480297437 | 0.11801532 | 4.06978885 | 4.71E-05 | 0.0004946 | NCBP1 | 4686 |
| ENSG000001 | 64.7087778 | -0.861906072 | 0.21177843 | -4.0698483 | 4.70E-05 | 0.0004946 | KCNB1 | 3745 |
| ENSG000001 | 27.2907798 | -1.078955566 | 0.26512759 | -4.0695711 | 4.71E-05 | 0.00049493 | DPP6 | 1804 |
| ENSG000001 | 269.681999 | 0.52950454 | 0.13013882 | 4.06876689 | 4.73E-05 | 0.00049651 | MYO5B | 4645 |
| ENSG000001 | 205.017314 | 0.743258914 | 0.18268026 | 4.06863286 | 4.73E-05 | 0.00049667 | HCN4 | 10021 |
| ENSG000001 | 28.0396304 | -1.052389793 | 0.25870815 | -4.0678649 | 4.74E-05 | 0.00049818 | ANKRD7 | 56311 |
| ENSG00000C | 678.245048 | 0.531375372 | 0.13063603 | 4.06760201 | 4.75E-05 | 0.00049842 | MAPK9 | 5601 |
| ENSG000001 | 482.811967 | 0.516388111 | 0.12694993 | 4.06765176 | 4.75E-05 | 0.00049842 | CTSL | 1514 |
| ENSG000001 | 627.344126 | 0.642745533 | 0.15801939 | 4.06751042 | 4.75E-05 | 0.00049842 | NOL6 | 65083 |
| ENSG000001 | 1082.6126 | 0.54007145 | 0.13277609 | 4.06753528 | 4.75E-05 | 0.00049842 | DRG1 | 4733 |
| ENSG000001 | 2048.81409 | 0.55681584 | 0.13690394 | 4.06720087 | 4.76E-05 | 0.00049883 | PPT1 | 5538 |
| ENSG000001 | 4919.66915 | 0.697560077 | 0.17150756 | 4.0672263 | 4.76E-05 | 0.00049883 | DYNC1H1 | 1778 |
| ENSG000001 | 774.833476 | 0.462575593 | 0.113735 | 4.06713506 | 4.76E-05 | 0.00049884 | INTU | 27152 |
| ENSG000001 | 404.450888 | 0.600911451 | 0.14775084 | 4.06705942 | 4.76E-05 | 0.00049887 | MAGEF1 | 64110 |
| ENSG000001 | 205.442491 | 0.613833673 | 0.15093308 | 4.06692595 | 4.76E-05 | 0.0004989 | ABHD17A | 81926 |
| ENSG000001 | 902.659151 | 0.498646153 | 0.12260975 | 4.06693705 | 4.76E-05 | 0.0004989 | TMEM39A | 55254 |
| ENSG000001 | 185.672369 | 0.66882257 | 0.16446317 | 4.06670111 | 4.77E-05 | 0.00049925 | TMEM216 | 51259 |
| ENSG000001 | 106.728105 | -0.694984801 | 0.17091791 | -4.0661906 | 4.78E-05 | 0.00050022 | ZBTB20 | 26137 |
| ENSG000001 | 127.113342 | -0.758175036 | 0.18647052 | -4.0659243 | 4.78E-05 | 0.00050053 | DNAH7 | 56171 |
| ENSG000001 | 941.126786 | 0.492774262 | 0.12119565 | 4.06594012 | 4.78E-05 | 0.00050053 | PRIM2 | 5558 |
| ENSG000001 | 218.040867 | 0.555270472 | 0.13657594 | 4.06565365 | 4.79E-05 | 0.00050085 | ZNF691 | 51058 |
| ENSG000001 | 36.1591586 | -0.98081875 | 0.24124497 | -4.0656548 | 4.79E-05 | 0.00050085 | IL31RA | 133396 |
| ENSG000001 | 1839.28567 | 0.511874695 | 0.12591876 | 4.06511859 | 4.80E-05 | 0.00050188 | GALNT7 | 51809 |
| ENSG000001 | 2463.703 | 0.443141974 | 0.10903617 | 4.06417405 | 4.82E-05 | 0.00050378 | UBE2G1 | 7326 |
| ENSG000001 | 623.433477 | 0.585534031 | 0.14408493 | 4.06381187 | 4.83E-05 | 0.00050443 | HTATIP2 | 10553 |
| ENSG000001 | 259.299686 | 0.597027517 | 0.14694042 | 4.06305845 | 4.84E-05 | 0.00050594 | DCXR | 51181 |
| ENSG000001 | 110.008542 | 0.751341928 | 0.18492338 | 4.06299037 | 4.84E-05 | 0.00050595 | ZNF358 | 140467 |
| ENSG000001 | 571.023364 | 0.561881419 | 0.13829558 | 4.06290219 | 4.85E-05 | 0.00050601 | YIPF3 | 25844 |

| | | | | | | | | |
|---|---|---|---|---|---|---|---|---|
| ENSG000001 | 16161.7623 | 0.515698736 | 0.12695011 | 4.06221584 | 4.86E-05 | 0.00050737 | SERBP1 | 26135 |
| ENSG000001 | 160.172504 | 0.678518167 | 0.16704612 | 4.06186121 | 4.87E-05 | 0.00050801 | ESRRA | 2101 |
| ENSG000001 | 362.647363 | 0.494118234 | 0.12165243 | 4.06172097 | 4.87E-05 | 0.00050819 | SIRT5 | 23408 |
| ENSG000001 | 866.330677 | 0.536145197 | 0.13201063 | 4.06137884 | 4.88E-05 | 0.0005088 | ATP6V1E1 | 529 |
| ENSG00000C | 676.762999 | 0.541545671 | 0.13334656 | 4.06118964 | 4.88E-05 | 0.00050909 | FAM21A | 387680 |
| ENSG000001 | 283.15534 | 0.620194042 | 0.15272685 | 4.06080565 | 4.89E-05 | 0.00050979 | DOLK | 22845 |
| ENSG000001 | 30.1252318 | -0.996016388 | 0.2453079 | -4.0602703 | 4.90E-05 | 0.00051083 | EFCAB3 | 146779 |
| ENSG00000C | 127.846329 | 0.685062792 | 0.16874893 | 4.05965707 | 4.91E-05 | 0.00051204 | XK | 7504 |
| ENSG000002 | 3.51202847 | -1.52476207 | 0.37563483 | -4.0591605 | 4.92E-05 | 0.000513 | NA | NA |
| ENSG000001 | 237.972551 | 0.661687512 | 0.16303278 | 4.05861645 | 4.94E-05 | 0.00051407 | DPP9 | 91039 |
| ENSG000001 | 644.842185 | 0.536606129 | 0.13221885 | 4.05846912 | 4.94E-05 | 0.00051426 | PAIP2B | 400961 |
| ENSG000001 | 11396.7025 | 0.450638246 | 0.11104275 | 4.05824094 | 4.94E-05 | 0.00051463 | DDX17 | 10521 |
| ENSG000001 | 336.054899 | 0.711131918 | 0.17524626 | 4.05790062 | 4.95E-05 | 0.00051525 | PCSK9 | 255738 |
| ENSG000002 | 328.605252 | 0.544468662 | 0.13417716 | 4.05783424 | 4.95E-05 | 0.00051526 | TMEM199 | 147007 |
| ENSG00000C | 1666.52179 | 0.487210307 | 0.12007155 | 4.05766638 | 4.96E-05 | 0.00051537 | FAM136A | 84908 |
| ENSG000001 | 773.195522 | 0.56264641 | 0.13866088 | 4.05771552 | 4.96E-05 | 0.00051537 | PCMT1 | 5110 |
| ENSG000001 | 766.819293 | 0.536235674 | 0.13216635 | 4.05727852 | 4.96E-05 | 0.00051609 | TWF1 | 5756 |
| ENSG000001 | 482.331672 | 0.541760241 | 0.13353278 | 4.05713291 | 4.97E-05 | 0.00051628 | TMED3 | 23423 |
| ENSG000001 | 48.5863855 | -0.879114107 | 0.21669759 | -4.0568708 | 4.97E-05 | 0.00051673 | SOX15 | 6665 |
| ENSG000002 | 3079.22999 | 0.527607029 | 0.13007822 | 4.05607495 | 4.99E-05 | 0.00051834 | BRD2 | 6046 |
| ENSG000002 | 21.0107666 | -1.158912137 | 0.28572631 | -4.0560218 | 4.99E-05 | 0.00051834 | NA | NA |
| ENSG000002 | 36.7187241 | 1.000405443 | 0.24666819 | 4.05567267 | 5.00E-05 | 0.00051898 | NA | NA |
| ENSG000001 | 1990.5787 | 0.482908131 | 0.11907585 | 4.05546673 | 5.00E-05 | 0.00051931 | MTDH | 92140 |
| ENSG000001 | 1052.33879 | 0.510884869 | 0.12597986 | 4.05528992 | 5.01E-05 | 0.0005195 | FASTKD1 | 79675 |
| ENSG000001 | 645.514319 | 0.608898438 | 0.15015239 | 4.05520318 | 5.01E-05 | 0.0005195 | ARHGDIA | 396 |
| ENSG000001 | 701.603822 | 0.501954506 | 0.12377939 | 4.05523478 | 5.01E-05 | 0.0005195 | ACOX1 | 51 |
| ENSG00000C | 594.356498 | 0.527574447 | 0.13012087 | 4.05449529 | 5.02E-05 | 0.00052077 | GGCT | 79017 |
| ENSG000001 | 341.43698 | 0.538547534 | 0.13282596 | 4.0545351 | 5.02E-05 | 0.00052077 | SLC25A17 | 10478 |
| ENSG000001 | 19.2289396 | -1.176632591 | 0.29020771 | -4.0544498 | 5.03E-05 | 0.00052077 | MS4A1 | 931 |

| | | | | | | | | |
|---|---|---|---|---|---|---|---|---|
| ENSG00000C | 67.8882334 | -0.813169242 | 0.2005949 | -4.0537882 | 5.04E-05 | 0.00052212 | SP140 | 11262 |
| ENSG000002 | 4039.51092 | 0.516005533 | 0.12730509 | 4.05329841 | 5.05E-05 | 0.00052308 | HIST1H2BM | 8342 |
| ENSG000001 | 699.111503 | 0.508069155 | 0.12535528 | 4.05303343 | 5.06E-05 | 0.00052354 | POT1 | 25913 |
| ENSG000001 | 29.3654476 | -1.022048062 | 0.25217707 | -4.0528984 | 5.06E-05 | 0.0005237 | NPFFR1 | 64106 |
| ENSG000001 | 680.129006 | 0.518240382 | 0.12787381 | 4.05274832 | 5.06E-05 | 0.00052391 | RPAIN | 84268 |
| ENSG000001 | 700.564365 | 0.523208639 | 0.12910377 | 4.05262098 | 5.06E-05 | 0.00052406 | OXNAD1 | 92106 |
| ENSG000001 | 467.943386 | 0.545970659 | 0.13474779 | 4.05179677 | 5.08E-05 | 0.00052577 | SMIM19 | 114926 |
| ENSG000001 | 442.757571 | 0.561144167 | 0.13850647 | 4.05139308 | 5.09E-05 | 0.00052655 | ABHD4 | 63874 |
| ENSG000002 | 192.222418 | 0.670082904 | 0.16539965 | 4.05129577 | 5.09E-05 | 0.00052663 | NA | NA |
| ENSG000001 | 378.502731 | 0.576920492 | 0.14240619 | 4.05123185 | 5.09E-05 | 0.00052664 | MUTYH | 4595 |
| ENSG000001 | 783.751937 | 0.515217998 | 0.12718494 | 4.05093546 | 5.10E-05 | 0.00052718 | RAD23A | 5886 |
| ENSG000001 | 173.689184 | 0.788322975 | 0.19461546 | 4.05066976 | 5.11E-05 | 0.00052764 | CLCN7 | 1186 |
| ENSG000001 | 594.117768 | 0.601422254 | 0.14850799 | 4.04976367 | 5.13E-05 | 0.00052955 | TADA3 | 10474 |
| ENSG000001 | 53.0397093 | -0.90925235 | 0.22453253 | -4.049535 | 5.13E-05 | 0.00052994 | CRB1 | 23418 |
| ENSG000001 | 184.174717 | 0.594506432 | 0.14683061 | 4.04892699 | 5.15E-05 | 0.00053118 | TRIM8 | 81603 |
| ENSG000001 | 1439.51788 | 0.504383151 | 0.12457972 | 4.04867799 | 5.15E-05 | 0.00053161 | UBE2C | 11065 |
| ENSG000001 | 239.191409 | 0.668571602 | 0.16514288 | 4.04844328 | 5.16E-05 | 0.00053201 | STYXL1 | 51657 |
| ENSG000001 | 132.271152 | 0.743904829 | 0.18376188 | 4.04819982 | 5.16E-05 | 0.00053242 | ARHGEF16 | 27237 |
| ENSG000001 | 2169.23273 | 0.480105781 | 0.11861067 | 4.04774537 | 5.17E-05 | 0.00053319 | BAG4 | 9530 |
| ENSG000001 | 210.657891 | 0.622011615 | 0.15366799 | 4.04776316 | 5.17E-05 | 0.00053319 | TBC1D24 | 57465 |
| ENSG000001 | 341.46496 | 0.67307016 | 0.16628988 | 4.04757148 | 5.18E-05 | 0.00053345 | ZMYND19 | 116225 |
| ENSG00000C | 187.61617 | 0.600476092 | 0.14835834 | 4.04747122 | 5.18E-05 | 0.00053354 | ZNF275 | 10838 |
| ENSG000001 | 659.243047 | 0.470268137 | 0.11619621 | 4.04719006 | 5.18E-05 | 0.00053396 | RAB3IP | 117177 |
| ENSG000001 | 1625.08062 | 0.537912225 | 0.13291068 | 4.04717067 | 5.18E-05 | 0.00053396 | COX7B | 1349 |
| ENSG000002 | 150.951933 | 0.593980697 | 0.14677789 | 4.04679945 | 5.19E-05 | 0.00053467 | NA | NA |
| ENSG00000C | 679.504131 | 0.53268498 | 0.13166256 | 4.04583475 | 5.21E-05 | 0.00053674 | COX15 | 1355 |
| ENSG000001 | 208.669651 | 0.669442194 | 0.16547927 | 4.04547468 | 5.22E-05 | 0.00053729 | ZNF598 | 90850 |
| ENSG000001 | 91.4295369 | 0.757429528 | 0.18722618 | 4.04553224 | 5.22E-05 | 0.00053729 | COX14 | 84987 |
| ENSG00000C | 124.761775 | -0.660777282 | 0.16335314 | -4.0450847 | 5.23E-05 | 0.00053805 | LAMA3 | 3909 |

| | | | | | | | | |
|---|---|---|---|---|---|---|---|---|
| ENSG000001 | 2136.75659 | 0.438533261 | 0.10841433 | 4.04497493 | 5.23E-05 | 0.00053816 | MAPK1IP1L | 93487 |
| ENSG000001 | 1416.00946 | 0.480051207 | 0.11868135 | 4.04487474 | 5.24E-05 | 0.00053826 | SRP54 | 6729 |
| ENSG000001 | 1236.56936 | 0.465248053 | 0.11502597 | 4.04472193 | 5.24E-05 | 0.00053847 | MYO1E | 4643 |
| ENSG000002 | 469.570157 | 0.53907746 | 0.13329036 | 4.0443844 | 5.25E-05 | 0.00053911 | DPP3 | 10072 |
| ENSG000001 | 1006.82168 | 0.566156031 | 0.13999539 | 4.04410487 | 5.25E-05 | 0.00053962 | MTHFD1L | 25902 |
| ENSG000001 | 1159.48304 | 0.469687158 | 0.11614791 | 4.04387098 | 5.26E-05 | 0.00054002 | HPS3 | 84343 |
| ENSG000001 | 2994.44327 | 0.49548933 | 0.12253684 | 4.04359479 | 5.26E-05 | 0.00054052 | RARS | 5917 |
| ENSG000001 | 669.097956 | 0.709469258 | 0.17546419 | 4.04338493 | 5.27E-05 | 0.00054087 | TBCD | 6904 |
| ENSG000001 | 654.770352 | 0.558294611 | 0.13808154 | 4.04322407 | 5.27E-05 | 0.0005411 | SCP2 | 6342 |
| ENSG000001 | 355.855955 | 0.557929313 | 0.13799999 | 4.04296634 | 5.28E-05 | 0.00054156 | CCDC127 | 133957 |
| ENSG000001 | 3740.07729 | 0.478481509 | 0.11835853 | 4.04264502 | 5.29E-05 | 0.00054203 | EIF3D | 8664 |
| ENSG000001 | 737.435644 | 0.538882323 | 0.133298 | 4.04268872 | 5.28E-05 | 0.00054203 | CMPK1 | 51727 |
| ENSG000001 | 1109.66998 | 0.471475507 | 0.11663064 | 4.04246701 | 5.29E-05 | 0.00054217 | RRP36 | 88745 |
| ENSG000002 | 21.4465712 | -1.17444623 | 0.29052285 | -4.0425262 | 5.29E-05 | 0.00054217 | CEACAM22P | 388550 |
| ENSG000001 | 13956.0455 | 0.470186176 | 0.11631428 | 4.0423771 | 5.29E-05 | 0.00054224 | PAICS | 10606 |
| ENSG000001 | 572.45683 | 0.456582846 | 0.11295341 | 4.04222267 | 5.29E-05 | 0.00054246 | SKA1 | 220134 |
| ENSG000001 | 797.521868 | 0.512192499 | 0.1267132 | 4.04213999 | 5.30E-05 | 0.00054251 | CLDN12 | 9069 |
| ENSG000001 | 1038.07978 | 0.542260443 | 0.13418648 | 4.04109586 | 5.32E-05 | 0.0005448 | UMPS | 7372 |
| ENSG000001 | 630.137942 | 0.532783068 | 0.13186136 | 4.04047909 | 5.33E-05 | 0.00054596 | ZNF334 | 55713 |
| ENSG000002 | 27.5901259 | -1.086796805 | 0.26897362 | -4.0405331 | 5.33E-05 | 0.00054596 | NA | NA |
| ENSG000001 | 991.262871 | 0.560281191 | 0.13866998 | 4.04039276 | 5.34E-05 | 0.00054602 | ELMOD2 | 255520 |
| ENSG000001 | 94.6233097 | 0.812566815 | 0.20111424 | 4.04032467 | 5.34E-05 | 0.00054604 | TATDN2 | 9797 |
| ENSG000001 | 426.902249 | 0.658802958 | 0.16306324 | 4.04016845 | 5.34E-05 | 0.00054627 | IGSF8 | 93185 |
| ENSG000001 | 336.085234 | 0.570591265 | 0.14124964 | 4.03959434 | 5.35E-05 | 0.00054747 | GLTP | 51228 |
| ENSG000001 | 680.159084 | 0.485382603 | 0.12015817 | 4.03953054 | 5.36E-05 | 0.00054748 | ASCC1 | 51008 |
| ENSG000001 | 490.303896 | 0.579443427 | 0.1434674 | 4.0388509 | 5.37E-05 | 0.00054893 | BRPF1 | 7862 |
| ENSG000001 | 363.043299 | 0.567761108 | 0.14060464 | 4.03799693 | 5.39E-05 | 0.00055079 | ACBD7 | 414149 |
| ENSG000001 | 707.694021 | 0.534185394 | 0.13229373 | 4.037874 | 5.39E-05 | 0.00055082 | ZNF761 | 388561 |
| ENSG000001 | 280.348324 | 0.523802738 | 0.12972265 | 4.03786639 | 5.39E-05 | 0.00055082 | AVEN | 57099 |

| | | | | | | | | |
|---|---|---|---|---|---|---|---|---|
| ENSG00000C | 335.411033 | 0.546749952 | 0.13540767 | 4.03780637 | 5.40E-05 | 0.00055082 | GATB | 5188 |
| ENSG000001 | 1851.84091 | 0.515602935 | 0.12771968 | 4.03698887 | 5.41E-05 | 0.0005526 | SKP1 | 6500 |
| ENSG000001 | 948.152522 | 0.685829261 | 0.16990379 | 4.03657416 | 5.42E-05 | 0.00055344 | SGTA | 6449 |
| ENSG00000C | 4434.77023 | 0.467305152 | 0.11577039 | 4.03648252 | 5.43E-05 | 0.00055352 | RAI14 | 26064 |
| ENSG000002 | 337.226757 | 0.581604644 | 0.14410641 | 4.03593879 | 5.44E-05 | 0.00055466 | NA | NA |
| ENSG000001 | 33.8920475 | -1.009643746 | 0.25018205 | -4.0356362 | 5.45E-05 | 0.00055524 | KCNJ15 | 3772 |
| ENSG000001 | 1208.06822 | 0.490320013 | 0.12152071 | 4.03486782 | 5.46E-05 | 0.00055678 | TSC22D1 | 8848 |
| ENSG000001 | 288.307527 | 0.635128302 | 0.1574081 | 4.0349149 | 5.46E-05 | 0.00055678 | R3HDM4 | 91300 |
| ENSG000001 | 40.9245867 | -0.923119846 | 0.22880549 | -4.0345179 | 5.47E-05 | 0.00055747 | ABCC11 | 85320 |
| ENSG000001 | 222.081896 | 0.620590784 | 0.15382253 | 4.03445973 | 5.47E-05 | 0.00055747 | STARD3 | 10948 |
| ENSG000001 | 8314.03343 | 0.475567447 | 0.11789035 | 4.03398116 | 5.48E-05 | 0.00055846 | BTF3 | 689 |
| ENSG000001 | 122.788401 | 0.636378117 | 0.15776152 | 4.03379806 | 5.49E-05 | 0.00055876 | PDGFD | 80310 |
| ENSG000001 | 409.954983 | 0.64283274 | 0.15940887 | 4.03260339 | 5.52E-05 | 0.00056147 | DHX34 | 9704 |
| ENSG000001 | 427.475845 | 0.639147754 | 0.15850483 | 4.03235499 | 5.52E-05 | 0.00056192 | CXXC1 | 30827 |
| ENSG00000C | 657.65241 | 0.554785237 | 0.13759381 | 4.03205089 | 5.53E-05 | 0.00056251 | UFD1L | 7353 |
| ENSG000001 | 36.5203177 | -0.983096377 | 0.24384527 | -4.0316401 | 5.54E-05 | 0.00056335 | EBF1 | 1879 |
| ENSG000001 | 20.7313174 | -1.140138266 | 0.28281827 | -4.0313458 | 5.55E-05 | 0.00056391 | PLCH2 | 9651 |
| ENSG000001 | 1428.11584 | 0.491723332 | 0.12198209 | 4.03111097 | 5.55E-05 | 0.00056434 | UTP6 | 55813 |
| ENSG000001 | 49.0180034 | -0.851606553 | 0.21131832 | -4.0299704 | 5.58E-05 | 0.00056694 | CCDC129 | 223075 |
| ENSG000001 | 940.874018 | 0.459685285 | 0.11409215 | 4.02907024 | 5.60E-05 | 0.00056897 | ATG3 | 64422 |
| ENSG000001 | 54.184007 | -0.837492245 | 0.20788268 | -4.0286774 | 5.61E-05 | 0.00056977 | GLIS3 | 169792 |
| ENSG000001 | 317.823849 | 0.529982258 | 0.13155421 | 4.02862249 | 5.61E-05 | 0.00056977 | CCZ1 | 51622 |
| ENSG000001 | 1420.03809 | 0.461959747 | 0.11467159 | 4.02854562 | 5.61E-05 | 0.00056982 | ATL2 | 64225 |
| ENSG000001 | 273.289034 | 0.608033751 | 0.15093647 | 4.02840849 | 5.62E-05 | 0.00057001 | NA | NA |
| ENSG000001 | 685.343049 | 0.532784973 | 0.13226757 | 4.02808474 | 5.62E-05 | 0.00057058 | FLOT1 | 10211 |
| ENSG000001 | 2620.8837 | 0.525251363 | 0.1303983 | 4.02805384 | 5.62E-05 | 0.00057058 | AK4 | 205 |
| ENSG000001 | 2153.73608 | 0.476840966 | 0.1183853 | 4.02787296 | 5.63E-05 | 0.00057088 | SSBP1 | 6742 |
| ENSG000001 | 1122.41902 | 0.468671913 | 0.11636731 | 4.02752218 | 5.64E-05 | 0.00057158 | CRABP1 | 1381 |
| ENSG000001 | 856.472416 | 0.480793024 | 0.1193786 | 4.02746394 | 5.64E-05 | 0.00057158 | TRAPPC6B | 122553 |

| | | | | | | | | |
|---|---|---|---|---|---|---|---|---|
| ENSG000001 | 312.641147 | 0.579340362 | 0.1438502 | 4.02738667 | 5.64E-05 | 0.00057163 | PPP1R16B | 26051 |
| ENSG000001 | 37.9786659 | -0.952568482 | 0.23652806 | -4.0272959 | 5.64E-05 | 0.00057171 | DSCAM | 1826 |
| ENSG000001 | 1328.85923 | 0.690837265 | 0.17155505 | 4.02691307 | 5.65E-05 | 0.00057249 | NUMA1 | 4926 |
| ENSG000001 | 150.100822 | 0.706790195 | 0.17553489 | 4.02649402 | 5.66E-05 | 0.00057337 | CCS | 9973 |
| ENSG000001 | 6328.63675 | 0.470889888 | 0.11695133 | 4.02637476 | 5.66E-05 | 0.00057352 | YWHAQ | 10971 |
| ENSG000001 | 462.210204 | 0.696145902 | 0.17291606 | 4.02591807 | 5.68E-05 | 0.00057435 | PEX5 | 5830 |
| ENSG000001 | 89.8521261 | -0.7479192 | 0.18577469 | -4.0259478 | 5.67E-05 | 0.00057435 | NA | NA |
| ENSG000001 | 435.888246 | 0.490313414 | 0.12179551 | 4.02570996 | 5.68E-05 | 0.00057471 | SMPDL3B | 27293 |
| ENSG000001 | 22.8631943 | -1.143351225 | 0.28404661 | -4.025224 | 5.69E-05 | 0.00057561 | GBP3 | 2635 |
| ENSG000001 | 3342.00532 | 0.514900945 | 0.12791814 | 4.02523785 | 5.69E-05 | 0.00057561 | UBQLN1 | 29979 |
| ENSG00000C | 703.855235 | 0.510995649 | 0.12695749 | 4.02493512 | 5.70E-05 | 0.00057618 | DOPEY1 | 23033 |
| ENSG000001 | 648.443925 | 0.526823425 | 0.13090166 | 4.0245742 | 5.71E-05 | 0.0005765 | YARS2 | 51067 |
| ENSG000001 | 1359.97504 | 0.533738892 | 0.13261511 | 4.02472163 | 5.70E-05 | 0.0005765 | TRUB1 | 142940 |
| ENSG000001 | 12564.4717 | 0.442737601 | 0.11000868 | 4.02456977 | 5.71E-05 | 0.0005765 | HSPA4 | 3308 |
| ENSG000002 | 31.5849271 | -1.023132991 | 0.25421568 | -4.0246652 | 5.71E-05 | 0.0005765 | NAMA | 100996569 |
| ENSG000002 | 29.1223153 | -1.075183644 | 0.26716848 | -4.0243656 | 5.71E-05 | 0.00057685 | LINC00964 | 157381 |
| ENSG000001 | 186.193299 | 0.599594685 | 0.14899843 | 4.02416788 | 5.72E-05 | 0.00057719 | PLEKHG5 | 57449 |
| ENSG000001 | 502.012616 | 0.497167507 | 0.12355432 | 4.023878 | 5.72E-05 | 0.00057776 | PITRM1 | 10531 |
| ENSG000001 | 868.858497 | 0.57955517 | 0.14403428 | 4.02373089 | 5.73E-05 | 0.00057798 | NF2 | 4771 |
| ENSG000001 | 727.241344 | 0.543991727 | 0.13520265 | 4.02352873 | 5.73E-05 | 0.00057833 | ARL4A | 10124 |
| ENSG00000C | 571.526544 | 0.564331481 | 0.14026665 | 4.0232763 | 5.74E-05 | 0.00057881 | METTL2A | 339175 |
| ENSG000001 | 230.321981 | 0.665756434 | 0.16548653 | 4.02302499 | 5.75E-05 | 0.00057924 | CCDC51 | 79714 |
| ENSG000001 | 1220.63193 | 0.490087386 | 0.12182364 | 4.02292527 | 5.75E-05 | 0.00057924 | TMEM170A | 124491 |
| ENSG000002 | 34.864642 | -1.061214347 | 0.26378872 | -4.022971 | 5.75E-05 | 0.00057924 | LINC00971 | 440970 |
| ENSG000001 | 2895.31518 | 0.543601038 | 0.13513048 | 4.02278618 | 5.75E-05 | 0.00057932 | MYL12A | 10627 |
| ENSG000001 | 1099.15017 | 0.543227763 | 0.13503805 | 4.02277548 | 5.75E-05 | 0.00057932 | CISD2 | 493856 |
| ENSG000001 | 235.843319 | 0.560079062 | 0.13923429 | 4.02256549 | 5.76E-05 | 0.00057969 | FUT11 | 170384 |
| ENSG000001 | 523.075184 | 0.514489592 | 0.12790641 | 4.02239097 | 5.76E-05 | 0.00057998 | TUBA4A | 7277 |
| ENSG00000C | 1609.37134 | 0.459366073 | 0.11421411 | 4.02197321 | 5.77E-05 | 0.00058086 | GPATCH2L | 55668 |

| | | | | | | | | |
|---|---|---|---|---|---|---|---|---|
| ENSG000001 | 2107.25205 | 0.491142487 | 0.1221642 | 4.0203472 | 5.81E-05 | 0.00058474 | AGTPBP1 | 23287 |
| ENSG000001 | 1169.05161 | 0.481152459 | 0.11968765 | 4.02006784 | 5.82E-05 | 0.00058529 | SOCS4 | 122809 |
| ENSG000001 | 543.528517 | 0.732213431 | 0.18214605 | 4.01992491 | 5.82E-05 | 0.00058544 | SRM | 6723 |
| ENSG000001 | 989.421976 | 0.501230452 | 0.12468751 | 4.019893 | 5.82E-05 | 0.00058544 | RACGAP1 | 29127 |
| ENSG000002 | 89.8500073 | -0.790999476 | 0.19678477 | -4.0196175 | 5.83E-05 | 0.00058598 | LINC01212 | 101927152 |
| ENSG00000C | 2572.28317 | 0.479062941 | 0.11918487 | 4.01949456 | 5.83E-05 | 0.00058614 | BTAF1 | 9044 |
| ENSG000001 | 3621.80163 | 0.467917962 | 0.11641583 | 4.01936725 | 5.84E-05 | 0.00058631 | CNOT7 | 29883 |
| ENSG000001 | 592.330072 | 0.555356043 | 0.13817625 | 4.01918601 | 5.84E-05 | 0.00058651 | RPA3 | 6119 |
| ENSG000001 | 402.029018 | 0.652064652 | 0.16223863 | 4.01917012 | 5.84E-05 | 0.00058651 | TMEM184B | 25829 |
| ENSG00000C | 1220.96602 | 0.484770849 | 0.12062362 | 4.01887162 | 5.85E-05 | 0.00058711 | NEDD4L | 23327 |
| ENSG000001 | 222.541962 | 0.671676947 | 0.16715599 | 4.01826437 | 5.86E-05 | 0.00058848 | SCAMP2 | 10066 |
| ENSG000001 | 267.328197 | 0.597654867 | 0.14875714 | 4.01765511 | 5.88E-05 | 0.00058985 | RMDN3 | 55177 |
| ENSG000001 | 551.911124 | 0.594087222 | 0.14791157 | 4.01650274 | 5.91E-05 | 0.00059256 | CA14 | 23632 |
| ENSG000001 | 234.549687 | 0.6132559 | 0.15268567 | 4.0164601 | 5.91E-05 | 0.00059256 | CCDC97 | 90324 |
| ENSG000001 | 539.844948 | 0.502515037 | 0.12511657 | 4.01637469 | 5.91E-05 | 0.00059262 | NUDT11 | 55190 |
| ENSG000001 | 464.340445 | 0.463432707 | 0.11538926 | 4.0162551 | 5.91E-05 | 0.00059266 | MOSPD2 | 158747 |
| ENSG000002 | 26.0541091 | -1.116455771 | 0.27798502 | -4.0162443 | 5.91E-05 | 0.00059266 | LOC253573 | 253573 |
| ENSG00000C | 491.722985 | 0.627257414 | 0.15618568 | 4.01610069 | 5.92E-05 | 0.00059287 | TAF11 | 6882 |
| ENSG000001 | 146.018198 | 0.697163217 | 0.17359878 | 4.01594533 | 5.92E-05 | 0.00059312 | ADM2 | 79924 |
| ENSG000001 | 187.639736 | 0.767687371 | 0.19116538 | 4.01582846 | 5.92E-05 | 0.00059327 | BCL2L12 | 83596 |
| ENSG00000C | 737.897143 | 0.556511141 | 0.13858679 | 4.01561451 | 5.93E-05 | 0.00059344 | TRIB2 | 28951 |
| ENSG000001 | 237.005853 | 0.581251518 | 0.14474528 | 4.01568534 | 5.93E-05 | 0.00059344 | PPP6R2 | 9701 |
| ENSG000002 | 61.0142151 | -0.85563693 | 0.21307896 | -4.0155862 | 5.93E-05 | 0.00059344 | NA | NA |
| ENSG000001 | 829.974969 | 0.587404016 | 0.14629689 | 4.01515031 | 5.94E-05 | 0.00059439 | OCLN | 100506658 |
| ENSG000001 | 1647.71024 | 0.540115216 | 0.1345422 | 4.01446685 | 5.96E-05 | 0.00059557 | RBM17 | 84991 |
| ENSG000001 | 2262.05528 | 0.474108884 | 0.11810067 | 4.01444706 | 5.96E-05 | 0.00059557 | TBCA | 6902 |
| ENSG000001 | 838.360299 | 0.46709953 | 0.11635401 | 4.01446865 | 5.96E-05 | 0.00059557 | IMP3 | 55272 |
| ENSG000002 | 676.791954 | 0.566614818 | 0.14113998 | 4.01455919 | 5.96E-05 | 0.00059557 | NA | NA |
| ENSG000001 | 470.978308 | 0.51642651 | 0.12865057 | 4.01417987 | 5.97E-05 | 0.0005961 | WBP4 | 11193 |

| | | | | | | | | |
|---|---|---|---|---|---|---|---|---|
| ENSG000001 | 310.496776 | 0.53766125 | 0.13394549 | 4.01403031 | 5.97E-05 | 0.00059633 | TCEAL3 | 85012 |
| ENSG000001 | 2301.66719 | 0.486928594 | 0.12131156 | 4.01386802 | 5.97E-05 | 0.0005966 | PTK2 | 5747 |
| ENSG000001 | 617.896399 | 0.515809065 | 0.12851013 | 4.01376187 | 5.98E-05 | 0.00059672 | RGMB | 285704 |
| ENSG000002 | 836.77497 | 0.546659528 | 0.13620145 | 4.01361017 | 5.98E-05 | 0.00059695 | FOXD3-AS1 | 100996301 |
| ENSG000001 | 1207.10506 | 0.480311612 | 0.11968899 | 4.01299729 | 6.00E-05 | 0.00059836 | PRIM1 | 5557 |
| ENSG000000 | 3365.65779 | 0.438954895 | 0.10939203 | 4.01267726 | 6.00E-05 | 0.00059888 | ZC3H15 | 55854 |
| ENSG000001 | 2938.18663 | 0.480729639 | 0.11980218 | 4.01269536 | 6.00E-05 | 0.00059888 | DCAF13 | 25879 |
| ENSG000000 | 1831.64874 | 0.504381393 | 0.12571096 | 4.01223092 | 6.01E-05 | 0.00059986 | USP33 | 23032 |
| ENSG000001 | 1729.08919 | 0.45580746 | 0.11360692 | 4.01214535 | 6.02E-05 | 0.00059993 | ZNF33B | 7582 |
| ENSG000001 | 63.1869617 | -0.827843471 | 0.20634679 | -4.0119037 | 6.02E-05 | 0.0006004 | KCNN3 | 3782 |
| ENSG000001 | 847.975672 | 0.49300225 | 0.12290389 | 4.01128274 | 6.04E-05 | 0.00060183 | CWC15 | 51503 |
| ENSG000001 | 2196.76895 | 0.488105105 | 0.12169753 | 4.01080545 | 6.05E-05 | 0.0006029 | PSMC2 | 5701 |
| ENSG000000 | 239.124262 | 0.575502353 | 0.14349055 | 4.01073351 | 6.05E-05 | 0.00060294 | METTL1 | 4234 |
| ENSG000001 | 286.888348 | 0.510559708 | 0.12730684 | 4.01046555 | 6.06E-05 | 0.00060333 | EPHA4 | 2043 |
| ENSG000002 | 1403.11599 | 0.560750738 | 0.1398201 | 4.01051581 | 6.06E-05 | 0.00060333 | DCTN1 | 1639 |
| ENSG000001 | 1406.03602 | 0.487238037 | 0.12149443 | 4.01037332 | 6.06E-05 | 0.00060342 | BNIP2 | 663 |
| ENSG000001 | 862.685065 | 0.482990558 | 0.12043765 | 4.01029557 | 6.06E-05 | 0.00060347 | CAAP1 | 79886 |
| ENSG000001 | 274.028788 | 0.588456726 | 0.14674147 | 4.01015977 | 6.07E-05 | 0.00060352 | WDR34 | 89891 |
| ENSG000002 | 466.148225 | 0.481280118 | 0.1200146 | 4.01017975 | 6.07E-05 | 0.00060352 | NA | NA |
| ENSG000001 | 1541.4358 | 0.588328898 | 0.14672603 | 4.00971039 | 6.08E-05 | 0.00060437 | PSMB7 | 5695 |
| ENSG000002 | 2.91754726 | -1.481038144 | 0.36935967 | -4.0097451 | 6.08E-05 | 0.00060437 | NA | NA |
| ENSG000001 | 9903.34946 | 0.523396943 | 0.13053663 | 4.00957901 | 6.08E-05 | 0.00060456 | RPL23A | 6147 |
| ENSG000001 | 681.552353 | 0.506517636 | 0.12633096 | 4.00944971 | 6.09E-05 | 0.00060474 | CRISPLD1 | 83690 |
| ENSG000001 | 161.426702 | 0.750949617 | 0.18731711 | 4.00897513 | 6.10E-05 | 0.00060581 | IFT140 | 9742 |
| ENSG000001 | 1021.28688 | 0.525460566 | 0.13108087 | 4.00867474 | 6.11E-05 | 0.00060643 | ATP8B2 | 57198 |
| ENSG000001 | 862.829677 | 0.466977811 | 0.11649633 | 4.00851966 | 6.11E-05 | 0.00060668 | ARMC10 | 83787 |
| ENSG000001 | 15.1899761 | -1.26751197 | 0.31621263 | -4.0084166 | 6.11E-05 | 0.0006068 | NA | NA |
| ENSG000001 | 503.208681 | 0.578288573 | 0.14427713 | 4.00817901 | 6.12E-05 | 0.00060726 | FTSJ2 | 29960 |
| ENSG000001 | 4784.43593 | 0.539793699 | 0.13469051 | 4.00765939 | 6.13E-05 | 0.00060819 | GLO1 | 2739 |

| | | | | | | | | |
|---|---|---|---|---|---|---|---|---|
| ENSG000002 | 63.9853259 | 0.80535347 | 0.20094867 | 4.00775712 | 6.13E-05 | 0.00060819 | NA | NA |
| ENSG000002 | 54.2783606 | 0.886141209 | 0.22111282 | 4.00764289 | 6.13E-05 | 0.00060819 | THTPA | 79178 |
| ENSG000001 | 454.697927 | 0.507274927 | 0.12658108 | 4.00750987 | 6.14E-05 | 0.00060824 | MTRF1L | 54516 |
| ENSG000001 | 294.125602 | 0.512474437 | 0.12787741 | 4.00754463 | 6.14E-05 | 0.00060824 | NDN | 4692 |
| ENSG000001 | 283.752318 | 0.574623132 | 0.1433893 | 4.00743372 | 6.14E-05 | 0.00060829 | ZNF211 | 10520 |
| ENSG000001 | 443.684007 | 0.536887334 | 0.13397623 | 4.00733265 | 6.14E-05 | 0.0006084 | TCF25 | 22980 |
| ENSG000001 | 28.2507278 | -1.021681024 | 0.25497145 | -4.0070409 | 6.15E-05 | 0.000609 | AOAH | 313 |
| ENSG000002 | 20.9803627 | -1.154904032 | 0.28822846 | -4.006905 | 6.15E-05 | 0.0006092 | FLJ16779 | 100192386 |
| ENSG000001 | 327.360469 | 0.63944085 | 0.15960649 | 4.00635864 | 6.17E-05 | 0.00061002 | MRPL2 | 51069 |
| ENSG000001 | 193.942433 | 0.60822849 | 0.15181186 | 4.00646239 | 6.16E-05 | 0.00061002 | WDR45 | 11152 |
| ENSG000001 | 10.3253674 | -1.383214307 | 0.34525374 | -4.0063702 | 6.17E-05 | 0.00061002 | MYL4 | 4635 |
| ENSG000002 | 111.768143 | 0.735148269 | 0.183491 | 4.00645413 | 6.16E-05 | 0.00061002 | NA | NA |
| ENSG000001 | 5493.0122 | 0.489867593 | 0.1222771 | 4.00620863 | 6.17E-05 | 0.00061025 | PSMD1 | 5707 |
| ENSG00000C | 748.436839 | 0.473039392 | 0.11808135 | 4.00604656 | 6.17E-05 | 0.00061052 | NFATC3 | 4775 |
| ENSG000001 | 2337.67009 | 0.484790367 | 0.12102022 | 4.00586261 | 6.18E-05 | 0.00061085 | SENP2 | 59343 |
| ENSG000001 | 1491.34306 | 0.552823702 | 0.1380231 | 4.00529849 | 6.19E-05 | 0.00061216 | GNPNAT1 | 64841 |
| ENSG000001 | 509.058635 | 0.537454537 | 0.13418811 | 4.00523206 | 6.20E-05 | 0.00061218 | RUFY1 | 80230 |
| ENSG000001 | 224.121154 | 0.572240045 | 0.14287953 | 4.00505273 | 6.20E-05 | 0.0006125 | CC2D1B | 200014 |
| ENSG000001 | 550.201504 | 0.504707609 | 0.12602978 | 4.00466953 | 6.21E-05 | 0.00061334 | USPL1 | 10208 |
| ENSG000001 | 2707.09966 | 0.51953518 | 0.12973611 | 4.00455354 | 6.21E-05 | 0.00061335 | TMX1 | 81542 |
| ENSG000001 | 428.223555 | 0.486758895 | 0.12155069 | 4.00457526 | 6.21E-05 | 0.00061335 | WDR19 | 57728 |
| ENSG00000C | 470.851147 | 0.508966348 | 0.12710705 | 4.00423376 | 6.22E-05 | 0.00061403 | FAR2 | 55711 |
| ENSG00000C | 2100.19814 | 0.482652118 | 0.12054915 | 4.00377872 | 6.23E-05 | 0.00061491 | USP28 | 57646 |
| ENSG000001 | 458.977127 | 0.660897302 | 0.16506712 | 4.00380956 | 6.23E-05 | 0.00061491 | SIN3B | 23309 |
| ENSG00000C | 2812.31839 | 0.6449739 | 0.16110608 | 4.00341128 | 6.24E-05 | 0.00061542 | GNB1 | 2782 |
| ENSG000001 | 21.9465872 | -1.097769317 | 0.27420397 | -4.0034771 | 6.24E-05 | 0.00061542 | MS4A4A | 51338 |
| ENSG000001 | 766.555768 | 0.51074593 | 0.12757634 | 4.00345331 | 6.24E-05 | 0.00061542 | WSB2 | 55884 |
| ENSG00000C | 2319.20106 | 0.451698522 | 0.11284363 | 4.00287126 | 6.26E-05 | 0.00061652 | WBP11 | 51729 |
| ENSG000001 | 1039.45744 | 0.56210268 | 0.14042452 | 4.00288131 | 6.26E-05 | 0.00061652 | STOX2 | 56977 |

| | | | | | | | | |
|---|---|---|---|---|---|---|---|---|
| ENSG000001 | 171.829345 | 0.619054062 | 0.15466115 | 4.00264749 | 6.26E-05 | 0.00061696 | SPSB4 | 92369 |
| ENSG000002 | 4294.54902 | 0.384181653 | 0.09598579 | 4.00248453 | 6.27E-05 | 0.00061723 | MARCKS | 4082 |
| ENSG000001 | 465.458475 | 0.513876331 | 0.12840026 | 4.00214388 | 6.28E-05 | 0.00061797 | MMD | 23531 |
| ENSG000002 | 25.6242201 | -1.080562611 | 0.27001431 | -4.0018716 | 6.28E-05 | 0.00061853 | NA | NA |
| ENSG00000C | 80.1908664 | 0.719547373 | 0.17983693 | 4.00111015 | 6.30E-05 | 0.00062038 | MID2 | 11043 |
| ENSG000001 | 6489.08339 | 0.450526017 | 0.11261401 | 4.00062126 | 6.32E-05 | 0.00062151 | POLR3G | 10622 |
| ENSG000001 | 469.027531 | 0.616675634 | 0.15415094 | 4.00046636 | 6.32E-05 | 0.00062177 | MRPL27 | 51264 |
| ENSG000001 | 579.549302 | 0.559378465 | 0.13983251 | 4.0003463 | 6.32E-05 | 0.00062193 | LSM1 | 27257 |
| ENSG000002 | 244.30528 | 0.652127271 | 0.16303484 | 3.99992594 | 6.34E-05 | 0.00062279 | SNORD116-7 | 100033419 |
| ENSG000002 | 34.0375419 | -0.951175576 | 0.23779955 | -3.9999048 | 6.34E-05 | 0.00062279 | NA | NA |
| ENSG00000C | 93.2586888 | 0.726646837 | 0.18167357 | 3.99973878 | 6.34E-05 | 0.00062308 | SPPL2B | 56928 |
| ENSG000001 | 757.194418 | 0.538130295 | 0.13455396 | 3.9993642 | 6.35E-05 | 0.00062391 | LSM5 | 23658 |
| ENSG000001 | 783.727023 | 0.493926728 | 0.12350867 | 3.99912604 | 6.36E-05 | 0.00062439 | RAD1 | 5810 |
| ENSG000001 | 771.437877 | 0.549910109 | 0.13754085 | 3.99815833 | 6.38E-05 | 0.0006268 | ZBTB3 | 79842 |
| ENSG000001 | 249.438665 | 0.774297155 | 0.19367256 | 3.99797029 | 6.39E-05 | 0.00062714 | EMC6 | 83460 |
| ENSG000001 | 399.460068 | 0.574998973 | 0.14383154 | 3.99772511 | 6.40E-05 | 0.00062764 | NDUFS4 | 4724 |
| ENSG000001 | 931.858145 | 0.430526178 | 0.10769528 | 3.99763281 | 6.40E-05 | 0.00062773 | NDUFA10 | 4705 |
| ENSG00000C | 50.3766103 | -0.886295083 | 0.22170863 | -3.9975669 | 6.40E-05 | 0.00062775 | LAMB4 | 22798 |
| ENSG000001 | 297.006352 | 0.563135866 | 0.14087184 | 3.99750473 | 6.40E-05 | 0.00062777 | HARS2 | 23438 |
| ENSG000001 | 646.268292 | 0.578199089 | 0.14465518 | 3.99708524 | 6.41E-05 | 0.00062873 | IRF2BPL | 64207 |
| ENSG00000C | 1495.7383 | 0.513496607 | 0.12849409 | 3.9962664 | 6.43E-05 | 0.00063075 | SLC39A9 | 55334 |
| ENSG000001 | 4075.44472 | 0.427310605 | 0.10695628 | 3.99518945 | 6.46E-05 | 0.00063332 | CSNK2A1 | 1457 |
| ENSG000001 | 73.4915905 | -0.758992889 | 0.18997533 | -3.9952181 | 6.46E-05 | 0.00063332 | LRMP | 4033 |
| ENSG000001 | 275.680785 | 0.688541832 | 0.17234716 | 3.99508662 | 6.47E-05 | 0.00063334 | PIEZO1 | 9780 |
| ENSG000001 | 710.638382 | 0.503990804 | 0.12615471 | 3.99502175 | 6.47E-05 | 0.00063334 | ANKRD13C | 81573 |
| ENSG000002 | 78.8707311 | 0.759932126 | 0.19022028 | 3.99501111 | 6.47E-05 | 0.00063334 | SNORA74A | 26821 |
| ENSG000001 | 160.422138 | 0.602885769 | 0.1509128 | 3.99492802 | 6.47E-05 | 0.00063341 | MECR | 51102 |
| ENSG000001 | 424.797162 | 0.582931802 | 0.14595389 | 3.99394507 | 6.50E-05 | 0.00063589 | KDELC2 | 143888 |
| ENSG000001 | 6882.18337 | 0.595388338 | 0.14908902 | 3.99350897 | 6.51E-05 | 0.00063691 | MYH10 | 4628 |

| | | | | | | | | | |
|---|---|---|---|---|---|---|---|---|---|
| ENSG000001 | 364.960478 | 0.495800101 | 0.12415698 | 3.99333254 | 6.52E-05 | 0.00063713 | AHCYL2 | | 23382 |
| ENSG000002 | 334.777319 | 0.593980807 | 0.14874395 | 3.99331075 | 6.52E-05 | 0.00063713 | LINC00667 | | 339290 |
| ENSG000001 | 1038.9255 | 0.460392235 | 0.11529693 | 3.99310072 | 6.52E-05 | 0.00063753 | ATG12 | | 9140 |
| ENSG000001 | 2205.16731 | 0.608254668 | 0.15232839 | 3.99304861 | 6.52E-05 | 0.00063753 | GATAD2A | | 54815 |
| ENSG000001 | 135.672292 | 0.703002421 | 0.17605975 | 3.99297631 | 6.52E-05 | 0.00063757 | RRP7BP | | 91695 |
| ENSG000001 | 1281.48567 | 0.481379735 | 0.12056449 | 3.99271583 | 6.53E-05 | 0.00063812 | SLC29A1 | | 2030 |
| ENSG000001 | 260.514935 | 0.630490505 | 0.15792326 | 3.99238525 | 6.54E-05 | 0.00063885 | RUNDC1 | | 146923 |
| ENSG000001 | 420.53848 | 0.533246817 | 0.13357344 | 3.99216215 | 6.55E-05 | 0.00063922 | POLR1E | | 64425 |
| ENSG000001 | 903.794051 | 0.684627612 | 0.1714941 | 3.99213496 | 6.55E-05 | 0.00063922 | SMCR8 | | 140775 |
| ENSG000001 | 2549.10536 | 0.504847772 | 0.12648138 | 3.99147894 | 6.57E-05 | 0.00064084 | TTC37 | | 9652 |
| ENSG000002 | 15.0210586 | -1.254545099 | 0.31431151 | -3.9914068 | 6.57E-05 | 0.00064088 | NA | NA | |
| ENSG000001 | 340.009242 | 0.602635178 | 0.15099329 | 3.99113868 | 6.58E-05 | 0.00064145 | CCDC94 | | 55702 |
| ENSG00000C | 25.5004198 | -1.054648959 | 0.26425381 | -3.9910454 | 6.58E-05 | 0.00064155 | AFM | | 173 |
| ENSG00000C | 102.199244 | -0.713936929 | 0.1788988 | -3.9907307 | 6.59E-05 | 0.00064194 | DNAH9 | | 1770 |
| ENSG00000C | 2667.85719 | 0.456021622 | 0.11426979 | 3.99074538 | 6.59E-05 | 0.00064194 | CLNS1A | | 1207 |
| ENSG000001 | 1026.95603 | 0.501278716 | 0.12560973 | 3.99076327 | 6.59E-05 | 0.00064194 | CPSF7 | | 79869 |
| ENSG000001 | 45.7193973 | -0.891373601 | 0.22336719 | -3.9906202 | 6.59E-05 | 0.00064208 | DUOX1 | | 53905 |
| ENSG000001 | 729.090543 | 0.521147309 | 0.13062014 | 3.98979301 | 6.61E-05 | 0.00064417 | MTMR4 | | 9110 |
| ENSG000002 | 19.4321722 | -1.18299911 | 0.29652498 | -3.9895428 | 6.62E-05 | 0.00064469 | NA | NA | |
| ENSG000001 | 1837.56977 | 0.50755546 | 0.12724067 | 3.98894057 | 6.64E-05 | 0.00064618 | TIA1 | | 7072 |
| ENSG000001 | 662.674183 | 0.467053934 | 0.11709975 | 3.9885134 | 6.65E-05 | 0.00064624 | EAPP | | 55837 |
| ENSG000001 | 53.997066 | 0.857392046 | 0.21496807 | 3.98846236 | 6.65E-05 | 0.00064624 | RAB19 | | 401409 |
| ENSG000001 | 593.677504 | 0.563662482 | 0.14130999 | 3.98883669 | 6.64E-05 | 0.00064624 | DPH3 | | 285381 |
| ENSG000001 | 99.9771724 | 0.706300289 | 0.1770714 | 3.98878804 | 6.64E-05 | 0.00064624 | SYVN1 | | 84447 |
| ENSG000001 | 1190.79473 | 0.563331833 | 0.14123369 | 3.98865056 | 6.65E-05 | 0.00064624 | TLE1 | | 7088 |
| ENSG000001 | 993.598114 | 0.513824565 | 0.12882292 | 3.98861136 | 6.65E-05 | 0.00064624 | ZNF544 | | 27300 |
| ENSG000001 | 238.542349 | 0.60867225 | 0.15260513 | 3.98854384 | 6.65E-05 | 0.00064624 | C1orf174 | | 339448 |
| ENSG000002 | 119.547749 | 0.693700489 | 0.17392332 | 3.98854217 | 6.65E-05 | 0.00064624 | ORAI1 | | 84876 |
| ENSG000001 | 1748.30942 | 0.440251605 | 0.11038455 | 3.98834457 | 6.65E-05 | 0.00064641 | USP47 | | 55031 |

| | | | | | | | | |
|---|---|---|---|---|---|---|---|---|
| ENSG00000C | 1045.17019 | 0.472081822 | 0.11837716 | 3.98794701 | 6.66E-05 | 0.00064734 | MARK3 | 4140 |
| ENSG000001 | 282.241665 | 0.5979654 | 0.14995211 | 3.98770926 | 6.67E-05 | 0.00064767 | BCKDK | 10295 |
| ENSG000001 | 570.713924 | 0.516781181 | 0.12959225 | 3.98774753 | 6.67E-05 | 0.00064767 | ZNF304 | 57343 |
| ENSG000001 | 286.821019 | 0.590742364 | 0.14816374 | 3.98709127 | 6.69E-05 | 0.00064905 | RTKN | 6242 |
| ENSG000001 | 589.44206 | 0.49066006 | 0.12306158 | 3.98711006 | 6.69E-05 | 0.00064905 | SGMS1 | 259230 |
| ENSG000001 | 45.417617 | -0.899611707 | 0.22564084 | -3.9869188 | 6.69E-05 | 0.00064937 | ACSM2A | 123876 |
| ENSG000001 | 713.737812 | 0.462247637 | 0.11594585 | 3.98675438 | 6.70E-05 | 0.00064966 | MICU2 | 221154 |
| ENSG00000C | 2150.78187 | 0.526901778 | 0.13217701 | 3.98633438 | 6.71E-05 | 0.00065066 | NCAPD2 | 9918 |
| ENSG000001 | 990.13693 | 0.518418627 | 0.1300509 | 3.98627478 | 6.71E-05 | 0.00065067 | OSBPL10 | 114884 |
| ENSG000001 | 15.452227 | 1.316786459 | 0.33034184 | 3.98613283 | 6.72E-05 | 0.0006509 | FAM86B2 | 653333 |
| ENSG000001 | 873.253953 | 0.548891582 | 0.13770519 | 3.98599048 | 6.72E-05 | 0.00065113 | PRMT6 | 55170 |
| ENSG000001 | 344.302882 | 0.549647355 | 0.13789938 | 3.98585794 | 6.72E-05 | 0.00065134 | SNIP1 | 79753 |
| ENSG000001 | 247.537023 | 0.64073142 | 0.16077492 | 3.98526973 | 6.74E-05 | 0.0006528 | TROAP | 10024 |
| ENSG000001 | 774.295349 | 0.441176739 | 0.11071232 | 3.98489285 | 6.75E-05 | 0.00065368 | THAP5 | 168451 |
| ENSG000001 | 518.950174 | 0.511025868 | 0.12824574 | 3.98473949 | 6.76E-05 | 0.00065395 | DYNLT1 | 6993 |
| ENSG00000C | 1023.83034 | 0.68234636 | 0.17124499 | 3.98462076 | 6.76E-05 | 0.00065396 | SEC63 | 11231 |
| ENSG00000C | 996.455449 | 0.522620528 | 0.13115905 | 3.98463195 | 6.76E-05 | 0.00065396 | KIF4A | 24137 |
| ENSG000001 | 76.5068225 | 0.742369176 | 0.18635185 | 3.98369623 | 6.79E-05 | 0.00065636 | CHMP4C | 92421 |
| ENSG000001 | 41.5759142 | -0.898581544 | 0.22557188 | -3.9835707 | 6.79E-05 | 0.00065655 | CNGB3 | 54714 |
| ENSG000001 | 99.9635444 | -0.805684941 | 0.20227592 | -3.9830987 | 6.80E-05 | 0.0006577 | SPEF2 | 79925 |
| ENSG00000C | 2079.10736 | 0.447402433 | 0.11235948 | 3.98188405 | 6.84E-05 | 0.00066091 | MTAP | 4507 |
| ENSG000001 | 3021.96269 | 0.46810815 | 0.11757374 | 3.98140064 | 6.85E-05 | 0.00066194 | Sep-11 | 55752 |
| ENSG000002 | 914.078577 | 0.530460168 | 0.13323411 | 3.98141419 | 6.85E-05 | 0.00066194 | BRK1 | 55845 |
| ENSG000001 | 2018.01098 | 0.589393272 | 0.14805038 | 3.98103171 | 6.86E-05 | 0.00066266 | AGO2 | 27161 |
| ENSG000001 | 586.939258 | 0.547005896 | 0.13740319 | 3.98102749 | 6.86E-05 | 0.00066266 | APOA1BP | 128240 |
| ENSG00000C | 125.705795 | 0.646638771 | 0.16244859 | 3.98057496 | 6.87E-05 | 0.00066377 | MCOLN3 | 55283 |
| ENSG000001 | 376.652388 | 0.648091513 | 0.16283039 | 3.98016314 | 6.89E-05 | 0.00066476 | NTS | 4922 |
| ENSG000002 | 186.810949 | 0.644593707 | 0.16195519 | 3.9800743 | 6.89E-05 | 0.00066485 | ZDHHC18 | 84243 |
| ENSG000001 | 858.353956 | 0.530897942 | 0.13339428 | 3.97991532 | 6.89E-05 | 0.00066514 | SLC35A5 | 55032 |

| | | | | | | | | |
|---|---|---|---|---|---|---|---|---|
| ENSG000001 | 2192.91555 | 0.453098992 | 0.11384806 | 3.97985704 | 6.90E-05 | 0.00066514 | ADD3 | 120 |
| ENSG000001 | 230.156296 | 0.624121307 | 0.15682306 | 3.97978007 | 6.90E-05 | 0.0006652 | CREB3 | 10488 |
| ENSG000001 | 968.612442 | 0.496265594 | 0.12469964 | 3.97968738 | 6.90E-05 | 0.0006653 | ARPC3 | 10094 |
| ENSG000001 | 403.916634 | 0.666424499 | 0.16746085 | 3.97958394 | 6.90E-05 | 0.00066543 | SLC4A2 | 6522 |
| ENSG000002 | 942.559014 | 0.591633626 | 0.14867005 | 3.97950793 | 6.91E-05 | 0.00066549 | AATF | 26574 |
| ENSG000001 | 154.530245 | 0.688933739 | 0.17312535 | 3.97939259 | 6.91E-05 | 0.00066549 | PCOLCE | 5118 |
| ENSG000001 | 341.313245 | 0.541180691 | 0.13599534 | 3.97940618 | 6.91E-05 | 0.00066549 | NHLRC3 | 387921 |
| ENSG000001 | 947.531973 | 0.462456591 | 0.11621613 | 3.97928074 | 6.91E-05 | 0.00066565 | SLU7 | 10569 |
| ENSG000001 | 15.6000164 | -1.233774383 | 0.31006539 | -3.9790781 | 6.92E-05 | 0.00066606 | NA | NA |
| ENSG000001 | 1354.024 | 0.461677326 | 0.11602983 | 3.97895389 | 6.92E-05 | 0.00066625 | MESDC2 | 23184 |
| ENSG000002 | 57.4030029 | 0.795311767 | 0.19990258 | 3.97849672 | 6.94E-05 | 0.00066737 | NA | NA |
| ENSG000001 | 1221.83291 | 0.438987468 | 0.11034887 | 3.97817803 | 6.94E-05 | 0.00066811 | REPS1 | 85021 |
| ENSG000001 | 1110.97249 | 0.463547872 | 0.11655138 | 3.97719771 | 6.97E-05 | 0.00067071 | DAP3 | 7818 |
| ENSG000001 | 1014.82758 | 0.461730002 | 0.11611054 | 3.97664153 | 6.99E-05 | 0.00067212 | LRP8 | 7804 |
| ENSG000001 | 1735.29039 | 0.477433782 | 0.12006943 | 3.97631427 | 7.00E-05 | 0.00067289 | KCMF1 | 56888 |
| ENSG00000C | 1717.63798 | 0.528449183 | 0.13291729 | 3.97577444 | 7.02E-05 | 0.00067417 | SIRT1 | 23411 |
| ENSG000001 | 2833.47681 | 0.490507283 | 0.12337659 | 3.97569184 | 7.02E-05 | 0.00067417 | SRSF5 | 6430 |
| ENSG000001 | 1174.92919 | 0.470329498 | 0.1182998 | 3.9757422 | 7.02E-05 | 0.00067417 | PRPF39 | 55015 |
| ENSG000001 | 1076.10623 | 0.542494838 | 0.136458 | 3.97554429 | 7.02E-05 | 0.00067443 | ACADM | 34 |
| ENSG00000C | 1214.18913 | 0.449913386 | 0.11317193 | 3.97548551 | 7.02E-05 | 0.00067443 | RNF4 | 6047 |
| ENSG00000C | 5814.49584 | 0.468761394 | 0.1179218 | 3.97518864 | 7.03E-05 | 0.00067487 | MAPK6 | 5597 |
| ENSG000001 | 177.278525 | 0.69486315 | 0.17479857 | 3.97522218 | 7.03E-05 | 0.00067487 | CHTF18 | 63922 |
| ENSG000001 | 902.996641 | 0.47380161 | 0.1191905 | 3.97516242 | 7.03E-05 | 0.00067487 | ANAPC7 | 51434 |
| ENSG000001 | 172.252709 | 0.669202785 | 0.16835537 | 3.97494181 | 7.04E-05 | 0.00067502 | TUBGCP6 | 85378 |
| ENSG000001 | 404.12842 | 0.545207469 | 0.13715923 | 3.97499665 | 7.04E-05 | 0.00067502 | SNTB1 | 6641 |
| ENSG000002 | 10.2120882 | -1.374163654 | 0.34570432 | -3.9749682 | 7.04E-05 | 0.00067502 | LINC01480 | 101927931 |
| ENSG000002 | 2423.00061 | 0.476072525 | 0.11979485 | 3.97406495 | 7.07E-05 | 0.00067735 | EID1 | 23741 |
| ENSG000001 | 253.027368 | 0.610085324 | 0.15352256 | 3.9739131 | 7.07E-05 | 0.00067762 | CDR2L | 30850 |
| ENSG000001 | 85.3500638 | 0.720903885 | 0.18144112 | 3.97321112 | 7.09E-05 | 0.00067946 | ZNF251 | 90987 |

| | | | | | | | | | |
|---|---|---|---|---|---|---|---|---|---|
| ENSG000000 | 1102.81661 | 0.511526075 | 0.12874607 | 3.97313922 | 7.09E-05 | 0.00067951 | RAB21 | | 23011 |
| ENSG000001 | 393.423421 | 0.565401885 | 0.1423108 | 3.9730077 | 7.10E-05 | 0.00067972 | DNAJB12 | | 54788 |
| ENSG000001 | 80.1004265 | 0.723132394 | 0.18202225 | 3.97276929 | 7.10E-05 | 0.00068024 | MANEAL | | 149175 |
| ENSG000000 | 2173.27998 | 0.494746668 | 0.12453742 | 3.97267488 | 7.11E-05 | 0.00068035 | TP53BP1 | | 7158 |
| ENSG000000 | 2919.12496 | 0.487347223 | 0.12268354 | 3.97239305 | 7.12E-05 | 0.00068085 | GPBP1 | | 65056 |
| ENSG000001 | 484.015331 | 0.54555544 | 0.13733691 | 3.97238775 | 7.12E-05 | 0.00068085 | LPAR3 | | 23566 |
| ENSG000001 | 221.189961 | 0.633884422 | 0.15957599 | 3.97230457 | 7.12E-05 | 0.00068093 | SYNGR2 | | 9144 |
| ENSG000000 | 114.923034 | 0.674759973 | 0.16987161 | 3.97217625 | 7.12E-05 | 0.00068113 | TRAF1 | | 7185 |
| ENSG000001 | 2367.02846 | 0.446470401 | 0.11240493 | 3.9719824 | 7.13E-05 | 0.00068137 | CPSF3 | | 51692 |
| ENSG000001 | 442.834781 | 0.485803251 | 0.1223073 | 3.97198899 | 7.13E-05 | 0.00068137 | C4orf29 | | 80167 |
| ENSG000001 | 178.181521 | 0.563515238 | 0.14188664 | 3.97158771 | 7.14E-05 | 0.00068233 | NMT2 | | 9397 |
| ENSG000001 | 157.420779 | 0.643056301 | 0.16192131 | 3.97141232 | 7.14E-05 | 0.00068268 | FRMD8 | | 83786 |
| ENSG000002 | 29.5477756 | -1.03928985 | 0.26170776 | -3.9711847 | 7.15E-05 | 0.00068317 | NA | NA | |
| ENSG000001 | 576.321361 | 0.546348036 | 0.13759878 | 3.97058775 | 7.17E-05 | 0.00068472 | TBCB | | 1155 |
| ENSG000001 | 103.780062 | 0.726310535 | 0.18293463 | 3.9703282 | 7.18E-05 | 0.00068531 | FBXL14 | | 144699 |
| ENSG000001 | 562.538944 | 0.486096016 | 0.12243682 | 3.97017828 | 7.18E-05 | 0.00068558 | NR2C1 | | 7181 |
| ENSG000001 | 42.8389812 | 0.954768794 | 0.24052879 | 3.96945744 | 7.20E-05 | 0.00068749 | UAP1L1 | | 91373 |
| ENSG000001 | 183.239869 | 0.552049715 | 0.1390806 | 3.96927894 | 7.21E-05 | 0.00068784 | ZNF75D | | 7626 |
| ENSG000001 | 503.214956 | 0.468255585 | 0.11799494 | 3.96843781 | 7.23E-05 | 0.00069011 | ZNF720 | | 124411 |
| ENSG000002 | 25.39877 | -1.171026233 | 0.29511457 | -3.9680394 | 7.25E-05 | 0.00069111 | NA | NA | |
| ENSG000001 | 2060.28166 | 0.441996459 | 0.11139828 | 3.96771348 | 7.26E-05 | 0.00069189 | PAIP1 | | 10605 |
| ENSG000002 | 310.778517 | 0.556385683 | 0.14023114 | 3.96763282 | 7.26E-05 | 0.00069196 | NA | NA | |
| ENSG000001 | 210.36634 | 0.621388804 | 0.156648 | 3.96678411 | 7.28E-05 | 0.00069426 | GLI1 | | 2735 |
| ENSG000001 | 869.305333 | 0.44800782 | 0.11298938 | 3.96504353 | 7.34E-05 | 0.00069919 | LRCH3 | | 84859 |
| ENSG000001 | 82.7145863 | 0.772092905 | 0.19474115 | 3.96471377 | 7.35E-05 | 0.00069999 | CHST8 | | 64377 |
| ENSG000001 | 409.684114 | 0.548885926 | 0.13845359 | 3.96440383 | 7.36E-05 | 0.00070064 | TTPAL | | 79183 |
| ENSG000001 | 68.6012844 | 0.82138592 | 0.20719149 | 3.96438052 | 7.36E-05 | 0.00070064 | ENDOG | | 2021 |
| ENSG000000 | 464.323413 | 0.506332875 | 0.12772308 | 3.96430226 | 7.36E-05 | 0.0007007 | MTFR1 | | 9650 |
| ENSG000001 | 212.469595 | 0.65109633 | 0.16424742 | 3.96411906 | 7.37E-05 | 0.00070075 | NR1H2 | | 7376 |

| | | | | | | | | |
|---|---|---|---|---|---|---|---|---|
| ENSG000001 | 326.176671 | 0.616124761 | 0.1554253 | 3.96412149 | 7.37E-05 | 0.00070075 | DAB2IP | 153090 |
| ENSG000001 | 1883.07626 | 0.458461848 | 0.11564979 | 3.96422555 | 7.36E-05 | 0.00070075 | ZNF43 | 7594 |
| ENSG000001 | 646.916823 | 0.50515744 | 0.12745192 | 3.96351376 | 7.39E-05 | 0.00070236 | VAMP7 | 6845 |
| ENSG000001 | 73.1379996 | -0.811593518 | 0.20477085 | -3.9634231 | 7.39E-05 | 0.00070247 | EYS | 346007 |
| ENSG000001 | 346.62791 | 0.503755634 | 0.12711912 | 3.96286296 | 7.41E-05 | 0.00070395 | IFIT5 | 24138 |
| ENSG000001 | 284.440352 | 0.573353141 | 0.14469849 | 3.9623988 | 7.42E-05 | 0.00070516 | DEAF1 | 10522 |
| ENSG000001 | 1204.92528 | 0.520413882 | 0.13134989 | 3.96204285 | 7.43E-05 | 0.00070604 | CAMSAP1 | 157922 |
| ENSG000001 | 459.306969 | 0.506119506 | 0.12774506 | 3.96194978 | 7.43E-05 | 0.00070615 | CYB5R2 | 51700 |
| ENSG000001 | 62.4584301 | -0.792783049 | 0.20011982 | -3.9615419 | 7.45E-05 | 0.0007072 | MYH11 | 4629 |
| ENSG000001 | 270.033569 | 0.580464831 | 0.14653101 | 3.9613787 | 7.45E-05 | 0.00070739 | ZNF394 | 84124 |
| ENSG000001 | 143.25954 | 0.603613294 | 0.15237511 | 3.96136414 | 7.45E-05 | 0.00070739 | TMEM107 | 84314 |
| ENSG000001 | 207.337714 | 0.661556946 | 0.16700505 | 3.96129906 | 7.45E-05 | 0.00070742 | ADCY3 | 109 |
| ENSG000001 | 276.673149 | 0.58284691 | 0.14715311 | 3.96081952 | 7.47E-05 | 0.00070867 | TMSB15A | 11013 |
| ENSG000001 | 7347.68795 | 0.491561032 | 0.12413471 | 3.95990006 | 7.50E-05 | 0.00071124 | CCT2 | 10576 |
| ENSG000001 | 30.4107293 | -1.011645455 | 0.2555183 | -3.9591899 | 7.52E-05 | 0.00071319 | GABRR3 | 200959 |
| ENSG00000C | 1080.53688 | 0.60330717 | 0.15239916 | 3.95873018 | 7.53E-05 | 0.00071423 | SZRD1 | 26099 |
| ENSG00000Z | 16.8785649 | -1.222845086 | 0.30889738 | -3.9587422 | 7.53E-05 | 0.00071423 | NA | NA |
| ENSG00000C | 278.174756 | -0.54451591 | 0.13755277 | -3.9585963 | 7.54E-05 | 0.00071439 | SESN1 | 27244 |
| ENSG000001 | 41.8990732 | -0.889032376 | 0.22459389 | -3.9583997 | 7.55E-05 | 0.00071439 | SLC9A2 | 6549 |
| ENSG000001 | 3263.65715 | 0.617581667 | 0.15601614 | 3.9584474 | 7.54E-05 | 0.00071439 | SNRNP200 | 23020 |
| ENSG000001 | 808.138079 | 0.520888751 | 0.13159039 | 3.95841024 | 7.55E-05 | 0.00071439 | TCEB1 | 6921 |
| ENSG000001 | 419.236024 | 0.514010606 | 0.12985271 | 3.95841273 | 7.54E-05 | 0.00071439 | WDR73 | 84942 |
| ENSG000001 | 333.472334 | 0.598798672 | 0.15130462 | 3.9575704 | 7.57E-05 | 0.0007167 | PTGIS | 5740 |
| ENSG000001 | 116.411884 | 0.680197676 | 0.17188542 | 3.95727394 | 7.58E-05 | 0.00071743 | NRM | 11270 |
| ENSG00000C | 596.72625 | 0.680512871 | 0.17196956 | 3.95717045 | 7.58E-05 | 0.00071757 | NISCH | 11188 |
| ENSG000001 | 189.878696 | 0.809214301 | 0.20449657 | 3.95710447 | 7.59E-05 | 0.0007176 | L1CAM | 3897 |
| ENSG000001 | 2240.69002 | 0.476274329 | 0.12037942 | 3.95644319 | 7.61E-05 | 0.00071942 | NUP58 | 9818 |
| ENSG000001 | 1151.84614 | 0.446941679 | 0.11297402 | 3.95614551 | 7.62E-05 | 0.00072015 | CENPH | 64946 |
| ENSG00000C | 1445.88764 | 0.470625493 | 0.11896642 | 3.95595225 | 7.62E-05 | 0.0007204 | BRCA1 | 672 |

| | | | | | | | | |
|---|---|---|---|---|---|---|---|---|
| ENSG000001 | 508.703636 | 0.548358854 | 0.13861454 | 3.95599794 | 7.62E-05 | 0.0007204 | HSPE1 | 3336 |
| ENSG000001 | 30.1365134 | -1.0704317 | 0.2705962 | -3.9558268 | 7.63E-05 | 0.00072061 | MUC17 | 140453 |
| ENSG000001 | 2300.19333 | 0.608444549 | 0.15382304 | 3.95548394 | 7.64E-05 | 0.00072147 | UBTF | 7343 |
| ENSG000001 | 1260.62911 | 0.504449406 | 0.1275375 | 3.95530252 | 7.64E-05 | 0.00072185 | FBXO30 | 84085 |
| ENSG000001 | 422.889098 | 0.56409399 | 0.14262393 | 3.95511461 | 7.65E-05 | 0.00072225 | CEP44 | 80817 |
| ENSG000001 | 1832.48027 | 0.460013641 | 0.11631357 | 3.95494389 | 7.66E-05 | 0.0007226 | MAPKAP1 | 79109 |
| ENSG00000C | 1997.01536 | 0.474333559 | 0.11993604 | 3.95488746 | 7.66E-05 | 0.0007226 | RC3H2 | 54542 |
| ENSG000001 | 2015.26223 | 0.55734086 | 0.14093416 | 3.95461876 | 7.67E-05 | 0.00072308 | PTTG1 | 9232 |
| ENSG000001 | 113.676639 | 0.711936993 | 0.18002523 | 3.9546512 | 7.66E-05 | 0.00072308 | SMIM4 | 440957 |
| ENSG000001 | 2083.52032 | 0.519065686 | 0.13125746 | 3.95456132 | 7.67E-05 | 0.00072308 | ZNF398 | 57541 |
| ENSG000001 | 599.857731 | 0.514897803 | 0.13021629 | 3.95417352 | 7.68E-05 | 0.00072409 | TPMT | 7172 |
| ENSG000001 | 800.196581 | 0.489652947 | 0.1238403 | 3.95390644 | 7.69E-05 | 0.00072473 | LCORL | 254251 |
| ENSG000002 | 502.575363 | 0.572576672 | 0.1448218 | 3.95366359 | 7.70E-05 | 0.0007253 | NA | NA |
| ENSG00000C | 975.616396 | 0.458689576 | 0.11602251 | 3.95345337 | 7.70E-05 | 0.00072577 | YTHDC2 | 64848 |
| ENSG00000C | 1113.3608 | 0.56175126 | 0.14210609 | 3.95304155 | 7.72E-05 | 0.00072685 | ELAC2 | 60528 |
| ENSG000001 | 573.424325 | 0.482940059 | 0.12219421 | 3.95223354 | 7.74E-05 | 0.00072914 | ANKEF1 | 63926 |
| ENSG000002 | 40.3613358 | -0.904208897 | 0.22879794 | -3.9519975 | 7.75E-05 | 0.00072969 | NA | NA |
| ENSG000001 | 4104.37335 | 0.502047844 | 0.1270411 | 3.95185364 | 7.75E-05 | 0.00072996 | AKAP12 | 9590 |
| ENSG000001 | 604.570368 | 0.488995788 | 0.12374117 | 3.95176298 | 7.76E-05 | 0.00073006 | ZNF232 | 7775 |
| ENSG00000C | 646.8533 | 0.4868237 | 0.12320095 | 3.95146074 | 7.77E-05 | 0.00073065 | MNAT1 | 4331 |
| ENSG000001 | 613.273515 | 0.52190867 | 0.13207844 | 3.95150527 | 7.77E-05 | 0.00073065 | CREBL2 | 1389 |
| ENSG000001 | 2824.55003 | 0.584304006 | 0.14788328 | 3.95111621 | 7.78E-05 | 0.00073136 | LPCAT1 | 79888 |
| ENSG000001 | 906.146879 | 0.538362717 | 0.13625524 | 3.95113416 | 7.78E-05 | 0.00073136 | PGAP1 | 80055 |
| ENSG000001 | 584.845425 | 0.697877389 | 0.176636 | 3.95093529 | 7.78E-05 | 0.00073175 | EIF1B | 10289 |
| ENSG000001 | 1309.5171 | 0.546501656 | 0.13832663 | 3.95080571 | 7.79E-05 | 0.00073197 | ACTL6A | 86 |
| ENSG00000C | 1580.17197 | 0.519080029 | 0.13139489 | 3.95053447 | 7.80E-05 | 0.00073263 | FGFR2 | 2263 |
| ENSG000002 | 1155.03367 | 0.651067184 | 0.16480742 | 3.95047254 | 7.80E-05 | 0.00073265 | EHMT2 | 10919 |
| ENSG000001 | 3043.64526 | 0.493445328 | 0.12491015 | 3.95040208 | 7.80E-05 | 0.0007327 | ATP5F1 | 515 |
| ENSG000001 | 34.2150272 | -1.091932744 | 0.27643563 | -3.9500434 | 7.81E-05 | 0.00073363 | LHX9 | 56956 |

| | | | | | | | | |
|---|---|---|---|---|---|---|---|---|
| ENSG000000 | 547.095053 | 0.53686727 | 0.13591899 | 3.94990612 | 7.82E-05 | 0.00073388 | NANS | 54187 |
| ENSG000001 | 880.711576 | 0.497073675 | 0.12585369 | 3.94961527 | 7.83E-05 | 0.0007346 | C17orf80 | 55028 |
| ENSG000001 | 571.227465 | 0.533277384 | 0.13502339 | 3.94951869 | 7.83E-05 | 0.00073473 | ERLIN1 | 10613 |
| ENSG000001 | 447.486749 | 0.578165326 | 0.14639196 | 3.94943371 | 7.83E-05 | 0.00073482 | UBL7 | 84993 |
| ENSG000001 | 9751.7316 | 0.471198241 | 0.11931455 | 3.9492103 | 7.84E-05 | 0.00073533 | SRSF3 | 6428 |
| ENSG000000 | 2658.84772 | 0.517717723 | 0.13109597 | 3.94915063 | 7.84E-05 | 0.00073535 | AFF4 | 27125 |
| ENSG000002 | 33.9960139 | 0.93933311 | 0.23786488 | 3.94901983 | 7.85E-05 | 0.00073558 | NA | NA |
| ENSG000001 | 611.363365 | 0.521922536 | 0.13216938 | 3.94889156 | 7.85E-05 | 0.0007358 | TMCO3 | 55002 |
| ENSG000001 | 421.59896 | 0.504033439 | 0.12764219 | 3.94879957 | 7.85E-05 | 0.00073592 | NADK2 | 133686 |
| ENSG000001 | 2837.9236 | 0.459916086 | 0.11647363 | 3.94867137 | 7.86E-05 | 0.00073614 | RRAS2 | 22800 |
| ENSG000001 | 75.198619 | 0.717501677 | 0.18172505 | 3.94828163 | 7.87E-05 | 0.00073717 | TIGD5 | 84948 |
| ENSG000001 | 1124.53403 | 0.473199127 | 0.11985436 | 3.94811764 | 7.88E-05 | 0.0007374 | LEMD3 | 23592 |
| ENSG000001 | 906.190537 | 0.487698018 | 0.12352738 | 3.9480965 | 7.88E-05 | 0.0007374 | DENND5A | 23258 |
| ENSG000001 | 537.007802 | 0.542077456 | 0.13732558 | 3.94738869 | 7.90E-05 | 0.00073924 | RPS6KA1 | 6195 |
| ENSG000001 | 446.048943 | 0.567541894 | 0.1437763 | 3.94739544 | 7.90E-05 | 0.00073924 | KBTBD6 | 89890 |
| ENSG000001 | 398.929863 | 0.513285583 | 0.13004209 | 3.94707257 | 7.91E-05 | 0.00074004 | KLHDC10 | 23008 |
| ENSG000001 | 698.364699 | 0.502615895 | 0.12734261 | 3.94695772 | 7.92E-05 | 0.00074023 | FBXO45 | 200933 |
| ENSG000001 | 295.829624 | 0.55001405 | 0.13936177 | 3.94666366 | 7.92E-05 | 0.00074097 | ISCA2 | 122961 |
| ENSG000001 | 63.6812067 | -0.76579671 | 0.19404276 | -3.9465359 | 7.93E-05 | 0.00074119 | RP1 | 6101 |
| ENSG000001 | 117.421068 | 0.729839317 | 0.18494318 | 3.94628948 | 7.94E-05 | 0.00074178 | NUDT22 | 84304 |
| ENSG000001 | 88.6691493 | 0.743256116 | 0.18836328 | 3.94586514 | 7.95E-05 | 0.00074293 | ZNF823 | 55552 |
| ENSG000001 | 5865.68101 | 0.426445652 | 0.10807747 | 3.94574043 | 7.96E-05 | 0.00074314 | PPP6R3 | 55291 |
| ENSG000000 | 826.764172 | 0.559311456 | 0.1417581 | 3.94553439 | 7.96E-05 | 0.00074323 | RRP12 | 23223 |
| ENSG000001 | 488.064933 | 0.498226053 | 0.12627483 | 3.94556886 | 7.96E-05 | 0.00074323 | ATXN7L1 | 222255 |
| ENSG000001 | 1426.28802 | 0.43087273 | 0.1092057 | 3.94551511 | 7.96E-05 | 0.00074323 | ATAD2 | 29028 |
| ENSG000001 | 240.693407 | 0.55604189 | 0.14093092 | 3.94549255 | 7.96E-05 | 0.00074323 | ZNF816 | 125893 |
| ENSG000001 | 606.682043 | 0.523561881 | 0.13271525 | 3.94500157 | 7.98E-05 | 0.00074458 | PRMT3 | 10196 |
| ENSG000001 | 1482.58294 | 0.470193729 | 0.11923528 | 3.94341101 | 8.03E-05 | 0.00074937 | IK | 3550 |
| ENSG000001 | 518.869671 | 0.508126492 | 0.12887977 | 3.94263977 | 8.06E-05 | 0.00075161 | IFITM1 | 8519 |

| | | | | | | | | | |
|---|---|---|---|---|---|---|---|---|---|
| ENSG000001 | 112.13249 | 0.775034736 | 0.19659538 | 3.94228361 | 8.07E-05 | 0.00075255 | EMC9 | | 51016 |
| ENSG000001 | 485.062179 | 0.493293417 | 0.1251381 | 3.94199231 | 8.08E-05 | 0.00075312 | EXOSC3 | | 51010 |
| ENSG000001 | 508.963914 | 0.548992426 | 0.13926648 | 3.94202831 | 8.08E-05 | 0.00075312 | ZNF45 | | 7596 |
| ENSG000001 | 571.203393 | 0.649550598 | 0.16481666 | 3.94104934 | 8.11E-05 | 0.00075591 | ZNF202 | | 7753 |
| ENSG000001 | 77.6320103 | 0.74405624 | 0.18881454 | 3.94067237 | 8.13E-05 | 0.00075693 | RAD9A | | 5883 |
| ENSG000001 | 104.54042 | 0.693037004 | 0.17588242 | 3.94034271 | 8.14E-05 | 0.0007578 | ADRA2C | | 152 |
| ENSG000001 | 924.332764 | 0.487330455 | 0.12368176 | 3.94019661 | 8.14E-05 | 0.00075791 | EBPL | | 84650 |
| ENSG000002 | 21.9874172 | -1.08814374 | 0.27616185 | -3.9402391 | 8.14E-05 | 0.00075791 | ANKRD20A11P | | 391267 |
| ENSG000001 | 387.060531 | 0.54673682 | 0.13876101 | 3.94013276 | 8.14E-05 | 0.00075794 | MUT | | 4594 |
| ENSG000001 | 996.029676 | 0.517726036 | 0.13140297 | 3.93998732 | 8.15E-05 | 0.00075822 | SGPL1 | | 8879 |
| ENSG000000 | 1030.225 | 0.622335492 | 0.15795672 | 3.93991156 | 8.15E-05 | 0.00075829 | RNF10 | | 9921 |
| ENSG000002 | 18.9984637 | -1.172205582 | 0.29753079 | -3.9397791 | 8.16E-05 | 0.00075853 | NA | NA | |
| ENSG000001 | 1122.36797 | 0.599649682 | 0.15220911 | 3.93964372 | 8.16E-05 | 0.00075879 | CDC42BPB | | 9578 |
| ENSG000001 | 161.154165 | 0.681637468 | 0.17302964 | 3.93942609 | 8.17E-05 | 0.0007593 | MAN2C1 | | 4123 |
| ENSG000001 | 1153.28128 | 0.501577803 | 0.12732477 | 3.9393575 | 8.17E-05 | 0.00075934 | SMIM15 | | 643155 |
| ENSG000001 | 440.97547 | 0.486074247 | 0.1233956 | 3.93915384 | 8.18E-05 | 0.00075982 | ATRNL1 | | 26033 |
| ENSG000001 | 1408.56863 | 0.551336741 | 0.13997079 | 3.9389413 | 8.18E-05 | 0.00076031 | ANKRD50 | | 57182 |
| ENSG000002 | 5.23266594 | -1.485878531 | 0.37725564 | -3.9386516 | 8.19E-05 | 0.00076106 | NA | NA | |
| ENSG000001 | 7300.69918 | 0.432735669 | 0.10988478 | 3.93808546 | 8.21E-05 | 0.00076268 | RPL38 | | 6169 |
| ENSG000000 | 1008.27537 | 0.633721831 | 0.1609292 | 3.93789205 | 8.22E-05 | 0.00076312 | ELAVL1 | | 1994 |
| ENSG000001 | 1498.06056 | 0.465689396 | 0.11826184 | 3.93778235 | 8.22E-05 | 0.0007633 | SEC23A | | 10484 |
| ENSG000001 | 1686.6562 | 0.478912026 | 0.12162363 | 3.93765621 | 8.23E-05 | 0.00076352 | TMEM47 | | 83604 |
| ENSG000001 | 1255.75767 | 0.519767705 | 0.13200338 | 3.93753343 | 8.23E-05 | 0.00076374 | ETV4 | | 2118 |
| ENSG000002 | 135.969242 | 0.687723952 | 0.17466239 | 3.93744731 | 8.24E-05 | 0.00076384 | DHRS11 | | 79154 |
| ENSG000000 | 282.349937 | 0.537301593 | 0.13647085 | 3.93711632 | 8.25E-05 | 0.00076472 | PPIE | | 10450 |
| ENSG000001 | 1217.14644 | 0.458090434 | 0.11636182 | 3.936776 | 8.26E-05 | 0.00076563 | SERINC1 | | 57515 |
| ENSG000001 | 384.010561 | 0.706893073 | 0.17956819 | 3.93662759 | 8.26E-05 | 0.00076592 | SLC35E2B | | 728661 |
| ENSG000000 | 686.300691 | 0.484706761 | 0.12314607 | 3.93603098 | 8.28E-05 | 0.00076765 | ERP44 | | 23071 |
| ENSG000001 | 561.748597 | 0.499791882 | 0.12698823 | 3.93573378 | 8.29E-05 | 0.00076843 | TMEM181 | | 57583 |

| | | | | | | | | | |
|---|---|---|---|---|---|---|---|---|---|
| ENSG000001 | 737.919881 | 0.502963099 | 0.12780352 | 3.93544001 | 8.30E-05 | 0.00076902 | COPG2 | | 26958 |
| ENSG000001 | 2311.86174 | 0.451943149 | 0.1148381 | 3.93548098 | 8.30E-05 | 0.00076902 | UBE2L3 | | 7332 |
| ENSG00000C | 1081.57922 | 0.457310649 | 0.11620518 | 3.93537243 | 8.31E-05 | 0.00076906 | RBM41 | | 55285 |
| ENSG000001 | 1218.99202 | 0.523415804 | 0.13301893 | 3.93489715 | 8.32E-05 | 0.00077041 | SEC23B | | 10483 |
| ENSG000001 | 153.033539 | 0.659716682 | 0.16767751 | 3.93443758 | 8.34E-05 | 0.00077171 | NXT1 | | 29107 |
| ENSG000001 | 1608.35829 | 0.487171901 | 0.12383879 | 3.93392003 | 8.36E-05 | 0.00077319 | NDUFA5 | | 4698 |
| ENSG000001 | 756.551005 | 0.697784479 | 0.17739338 | 3.93354291 | 8.37E-05 | 0.00077423 | SEC16A | | 9919 |
| ENSG000001 | 19670.3126 | 0.454904353 | 0.11565787 | 3.93318962 | 8.38E-05 | 0.0007752 | RPL5 | | 6125 |
| ENSG000001 | 2344.38877 | 0.446904417 | 0.1136266 | 3.93309676 | 8.39E-05 | 0.00077532 | CCNI | | 10983 |
| ENSG000001 | 570.793399 | 0.729169661 | 0.18541903 | 3.93255036 | 8.40E-05 | 0.00077655 | SYMPK | | 8189 |
| ENSG000001 | 420.497668 | 0.539975078 | 0.13730559 | 3.9326519 | 8.40E-05 | 0.00077655 | ANKZF1 | | 55139 |
| ENSG000001 | 165.259181 | 0.598969026 | 0.15230901 | 3.93259098 | 8.40E-05 | 0.00077655 | DUSP7 | | 1849 |
| ENSG00000C | 674.830555 | 0.503240678 | 0.12798373 | 3.93206774 | 8.42E-05 | 0.00077749 | LUC7L | | 55692 |
| ENSG00000C | 344.202123 | 0.55310888 | 0.1406629 | 3.93215899 | 8.42E-05 | 0.00077749 | ADAT1 | | 23536 |
| ENSG000001 | 21.8232474 | 1.1043864 | 0.28086839 | 3.93204234 | 8.42E-05 | 0.00077749 | EVA1B | | 55194 |
| ENSG000001 | 542.966204 | 0.516652574 | 0.13139401 | 3.93208631 | 8.42E-05 | 0.00077749 | PRKAA2 | | 5563 |
| ENSG000001 | 1195.22769 | 0.460129735 | 0.11704468 | 3.93123143 | 8.45E-05 | 0.00077994 | TRIB1 | | 10221 |
| ENSG000001 | 357.419241 | 0.498707758 | 0.12688113 | 3.93051172 | 8.48E-05 | 0.0007821 | LMBRD2 | | 92255 |
| ENSG00000C | 197.408179 | 0.641633168 | 0.16327706 | 3.9297202 | 8.50E-05 | 0.0007845 | NA | NA | |
| ENSG00000C | 4140.5626 | 0.433371512 | 0.11028236 | 3.92965391 | 8.51E-05 | 0.00078454 | RANBP1 | | 5902 |
| ENSG000001 | 34.3895332 | -0.986529558 | 0.25106284 | -3.9294129 | 8.52E-05 | 0.00078514 | POU2AF1 | | 5450 |
| ENSG000001 | 266.240205 | 0.516308484 | 0.13140221 | 3.92922237 | 8.52E-05 | 0.00078559 | SDF2 | | 6388 |
| ENSG000001 | 836.09413 | 0.502802481 | 0.1279736 | 3.92895459 | 8.53E-05 | 0.00078628 | N4BP1 | | 9683 |
| ENSG00000C | 278.042557 | 0.623340483 | 0.15865925 | 3.92880005 | 8.54E-05 | 0.00078661 | RAD51 | | 5888 |
| ENSG000001 | 313.044831 | 0.50411373 | 0.12831435 | 3.92873998 | 8.54E-05 | 0.00078663 | ACSS2 | | 55902 |
| ENSG000001 | 420.114728 | 0.70412076 | 0.17924407 | 3.92827917 | 8.56E-05 | 0.00078796 | RPTOR | | 57521 |
| ENSG000001 | 595.62493 | 0.507883082 | 0.12930109 | 3.92791033 | 8.57E-05 | 0.00078899 | ANO10 | | 55129 |
| ENSG000001 | 614.25207 | 0.478263879 | 0.12176223 | 3.92785072 | 8.57E-05 | 0.000789 | PTPN9 | | 5780 |
| ENSG000001 | 820.054964 | 0.494217634 | 0.12584601 | 3.92716179 | 8.60E-05 | 0.00079109 | ZNF711 | | 7552 |

| | | | | | | | | |
|---|---|---|---|---|---|---|---|---|
| ENSG000001 | 1710.6247 | 0.52743507 | 0.13431376 | 3.92688771 | 8.61E-05 | 0.00079181 | DAPK1 | 1612 |
| ENSG000001 | 1089.4182 | 0.473855177 | 0.12068035 | 3.92653132 | 8.62E-05 | 0.00079244 | PRCP | 5547 |
| ENSG000001 | 1274.47934 | 0.435929439 | 0.11102004 | 3.92658327 | 8.62E-05 | 0.00079244 | ZDHHC3 | 51304 |
| ENSG000001 | 42.968812 | -0.896595324 | 0.22833994 | -3.9265812 | 8.62E-05 | 0.00079244 | ITGBL1 | 9358 |
| ENSG000001 | 160.73846 | 0.625991252 | 0.15943978 | 3.92619241 | 8.63E-05 | 0.00079338 | TPST1 | 8460 |
| ENSG000001 | 791.582259 | 0.542688231 | 0.13823097 | 3.92595253 | 8.64E-05 | 0.00079399 | MPHOSPH8 | 54737 |
| ENSG000001 | 2467.01571 | 0.398907802 | 0.1016335 | 3.92496391 | 8.67E-05 | 0.00079708 | ATMIN | 23300 |
| ENSG000001 | 39.3080053 | -0.941555479 | 0.23993076 | -3.9242801 | 8.70E-05 | 0.00079916 | COL3A1 | 1281 |
| ENSG00000C | 3328.80004 | 0.473669728 | 0.12070617 | 3.92415516 | 8.70E-05 | 0.0007994 | DDX18 | 8886 |
| ENSG000001 | 33.5787898 | -0.97632114 | 0.24880779 | -3.9239975 | 8.71E-05 | 0.00079956 | DUOX2 | 50506 |
| ENSG000002 | 84.2229542 | 0.730955111 | 0.18627711 | 3.92402012 | 8.71E-05 | 0.00079956 | LOC105375014 | 105375014 |
| ENSG000001 | 51.4258563 | 0.902430586 | 0.22999643 | 3.9236721 | 8.72E-05 | 0.00080046 | ENHO | 375704 |
| ENSG000001 | 165.948409 | 0.61077601 | 0.15566702 | 3.92360567 | 8.72E-05 | 0.0008005 | SNX21 | 90203 |
| ENSG00000C | 1030.2277 | 0.517873267 | 0.13200924 | 3.92300776 | 8.75E-05 | 0.00080231 | TSPAN6 | 7105 |
| ENSG00000C | 441.191331 | 0.484104004 | 0.12341365 | 3.92261325 | 8.76E-05 | 0.00080344 | LYRM2 | 57226 |
| ENSG000002 | 24.4889267 | -1.035821951 | 0.26407224 | -3.9224946 | 8.76E-05 | 0.00080365 | LRRC37A11P | 342666 |
| ENSG000002 | 598.658992 | 0.528805241 | 0.13481949 | 3.92232047 | 8.77E-05 | 0.00080405 | DHX16 | 8449 |
| ENSG000001 | 281.02186 | 0.512522016 | 0.13068271 | 3.92188071 | 8.79E-05 | 0.00080534 | SETDB2 | 83852 |
| ENSG000001 | 914.590129 | 0.493945363 | 0.1259478 | 3.92182591 | 8.79E-05 | 0.00080534 | SUPV3L1 | 6832 |
| ENSG000001 | 252.691022 | 0.614823919 | 0.15678345 | 3.92148486 | 8.80E-05 | 0.0008063 | OIP5 | 11339 |
| ENSG000002 | 43.9352061 | 0.928844811 | 0.23687171 | 3.92129897 | 8.81E-05 | 0.00080674 | SCARNA6 | 677772 |
| ENSG000001 | 204.369821 | 0.635560971 | 0.16208409 | 3.92118055 | 8.81E-05 | 0.00080696 | VARS2 | 57176 |
| ENSG000001 | 2024.41641 | 0.444645195 | 0.1134055 | 3.92084329 | 8.82E-05 | 0.0008079 | URI1 | 8725 |
| ENSG000001 | 1306.61284 | 0.525912659 | 0.13414287 | 3.92054132 | 8.84E-05 | 0.00080863 | KLHL7 | 55975 |
| ENSG000001 | 390.816224 | 0.4969817 | 0.12676434 | 3.9205165 | 8.84E-05 | 0.00080863 | MANEA | 79694 |
| ENSG00000C | 581.017033 | 0.544942358 | 0.13900059 | 3.92043196 | 8.84E-05 | 0.00080874 | RCOR1 | 23186 |
| ENSG000001 | 51.7620139 | -0.831608327 | 0.21215366 | -3.9198397 | 8.86E-05 | 0.00081054 | SSPO | 23145 |
| ENSG00000C | 635.905463 | 0.568704755 | 0.1450883 | 3.91971488 | 8.87E-05 | 0.00081078 | SDHA | 6389 |
| ENSG00000C | 3892.35037 | 0.411969593 | 0.10511252 | 3.91931976 | 8.88E-05 | 0.00081193 | RFC1 | 5981 |

| | | | | | | | | |
|---|---|---|---|---|---|---|---|---|
| ENSG000001 | 16.6007146 | -1.183455284 | 0.30197465 | -3.9190551 | 8.89E-05 | 0.00081264 | OR2G6 | 391211 |
| ENSG000001 | 3955.12033 | 0.480214564 | 0.12254083 | 3.91881257 | 8.90E-05 | 0.00081327 | ARL5B | 221079 |
| ENSG000001 | 24.6413126 | -1.047101427 | 0.26723796 | -3.9182362 | 8.92E-05 | 0.00081503 | HOXB3 | 3213 |
| ENSG000001 | 459.734666 | 0.569409087 | 0.14533725 | 3.91784674 | 8.93E-05 | 0.00081598 | TAGLN | 6876 |
| ENSG000001 | 135.22696 | 0.6883074 | 0.17568417 | 3.91786801 | 8.93E-05 | 0.00081598 | ALKBH4 | 54784 |
| ENSG000001 | 38.2779928 | 0.95911676 | 0.24482534 | 3.91755503 | 8.95E-05 | 0.0008166 | PILRB | 29990 |
| ENSG000001 | 559.553125 | 0.558560772 | 0.1425772 | 3.91760246 | 8.94E-05 | 0.0008166 | WDR61 | 80349 |
| ENSG000001 | 380.519177 | 0.519322826 | 0.13256547 | 3.91748183 | 8.95E-05 | 0.00081667 | PNPLA8 | 50640 |
| ENSG000001 | 1782.85065 | 0.51972796 | 0.13267678 | 3.91724885 | 8.96E-05 | 0.00081727 | NCBP2 | 22916 |
| ENSG00000C | 286.401468 | 0.576715472 | 0.1472386 | 3.91687688 | 8.97E-05 | 0.00081813 | CPOX | 1371 |
| ENSG000001 | 471.731141 | 0.50019705 | 0.12770448 | 3.91683245 | 8.97E-05 | 0.00081813 | MRRF | 92399 |
| ENSG000001 | 46.2010442 | -0.913566206 | 0.23323693 | -3.9169021 | 8.97E-05 | 0.00081813 | AGBL4 | 84871 |
| ENSG000001 | 234.326012 | 0.544056631 | 0.13891105 | 3.91658278 | 8.98E-05 | 0.00081861 | FDX1 | 2230 |
| ENSG000001 | 134.267951 | 0.738841415 | 0.18864438 | 3.91658316 | 8.98E-05 | 0.00081861 | FBXL18 | 80028 |
| ENSG00000C | 4692.71463 | 0.456603272 | 0.11658613 | 3.91644594 | 8.99E-05 | 0.00081889 | XRN2 | 22803 |
| ENSG000001 | 13.0750645 | -1.262760521 | 0.32245992 | -3.9160232 | 9.00E-05 | 0.00081996 | PRSS54 | 221191 |
| ENSG000001 | 1467.80551 | 0.417660181 | 0.10665386 | 3.91603429 | 9.00E-05 | 0.00081996 | SECISBP2 | 79048 |
| ENSG000001 | 87.2750125 | 0.855746403 | 0.21854718 | 3.91561396 | 9.02E-05 | 0.00082117 | NA | NA |
| ENSG000001 | 361.703906 | 0.602283607 | 0.15383214 | 3.91519997 | 9.03E-05 | 0.00082239 | PTOV1 | 53635 |
| ENSG000001 | 1363.92509 | 0.484739129 | 0.12381368 | 3.91506924 | 9.04E-05 | 0.00082266 | CSTF1 | 1477 |
| ENSG000001 | 256.992611 | 0.559994311 | 0.14305377 | 3.91457224 | 9.06E-05 | 0.00082417 | DGKE | 8526 |
| ENSG000001 | 835.891098 | 0.492804069 | 0.12589374 | 3.91444449 | 9.06E-05 | 0.00082442 | THOC7 | 80145 |
| ENSG00000C | 292.926483 | 0.61998101 | 0.15838567 | 3.91437576 | 9.06E-05 | 0.00082447 | NADK | 65220 |
| ENSG00000C | 1055.46803 | 0.467369477 | 0.11940529 | 3.91414394 | 9.07E-05 | 0.00082489 | ZRANB1 | 54764 |
| ENSG000001 | 24.8040468 | -1.053548039 | 0.2691635 | -3.9141564 | 9.07E-05 | 0.00082489 | SCML4 | 256380 |
| ENSG00000C | 1493.58324 | 0.620945974 | 0.1586624 | 3.9136304 | 9.09E-05 | 0.00082646 | TCF3 | 6929 |
| ENSG000002 | 8.26520845 | -1.401460035 | 0.35814236 | -3.9131367 | 9.11E-05 | 0.00082797 | NA | NA |
| ENSG00000C | 270.977767 | 0.523993247 | 0.13391128 | 3.91298806 | 9.12E-05 | 0.00082829 | HSPB11 | 51668 |
| ENSG000001 | 2053.76797 | 0.443456196 | 0.11333384 | 3.91283131 | 9.12E-05 | 0.00082865 | CFAP97 | 57587 |

| | | | | | | | | |
|---|---|---|---|---|---|---|---|---|
| ENSG000001 | 25.1531662 | -1.125030798 | 0.2875304 | -3.9127369 | 9.13E-05 | 0.00082878 | KCNT1 | 57582 |
| ENSG000001 | 20.7863549 | -1.107083725 | 0.28295062 | -3.9126393 | 9.13E-05 | 0.00082892 | TMPRSS11D | 9407 |
| ENSG000002 | 43.4041016 | 0.876326852 | 0.22397624 | 3.91258853 | 9.13E-05 | 0.00082892 | MIR503HG | 84848 |
| ENSG000001 | 584.013412 | 0.475842737 | 0.12162115 | 3.91249978 | 9.13E-05 | 0.00082904 | MEAF6 | 64769 |
| ENSG000001 | 351.297544 | 0.502584062 | 0.12846127 | 3.9123393 | 9.14E-05 | 0.00082941 | SETD4 | 54093 |
| ENSG000001 | 110.29972 | 0.78202137 | 0.19988881 | 3.91228196 | 9.14E-05 | 0.00082942 | PNPLA2 | 57104 |
| ENSG000002 | 16.0683181 | -1.199998055 | 0.30676688 | -3.9117589 | 9.16E-05 | 0.00083103 | TECRL | 253017 |
| ENSG000001 | 468.495183 | 0.486684488 | 0.12442971 | 3.91132056 | 9.18E-05 | 0.00083217 | TMEM55A | 55529 |
| ENSG000001 | 492.017746 | 0.465616036 | 0.1190431 | 3.91132323 | 9.18E-05 | 0.00083217 | ZNF620 | 253639 |
| ENSG00000C | 5.5380331 | -1.46852245 | 0.37548307 | -3.9110217 | 9.19E-05 | 0.00083283 | KRT31 | 3881 |
| ENSG000001 | 10.8136374 | -1.328742288 | 0.33974026 | -3.9110534 | 9.19E-05 | 0.00083283 | TMEM212 | 389177 |
| ENSG00000C | 706.647222 | 0.646468263 | 0.16530799 | 3.9106897 | 9.20E-05 | 0.00083379 | ADIPOR2 | 79602 |
| ENSG000001 | 67.2493036 | -0.781019452 | 0.19972553 | -3.9104638 | 9.21E-05 | 0.00083438 | NEK5 | 341676 |
| ENSG00000C | 104.457928 | 0.673330319 | 0.17220808 | 3.90998102 | 9.23E-05 | 0.00083586 | SLC27A5 | 10998 |
| ENSG00000C | 1573.57952 | 0.496427687 | 0.12696754 | 3.90987863 | 9.23E-05 | 0.00083603 | VTA1 | 51534 |
| ENSG00000C | 1436.12202 | 0.498112766 | 0.12740613 | 3.90964529 | 9.24E-05 | 0.00083665 | ATP2C1 | 27032 |
| ENSG000001 | 238.602241 | 0.59728242 | 0.15279382 | 3.90907434 | 9.27E-05 | 0.00083844 | TSKU | 25987 |
| ENSG000001 | 1120.3194 | 0.502737019 | 0.12861458 | 3.90886479 | 9.27E-05 | 0.0008388 | CLPX | 10845 |
| ENSG000002 | 100.376526 | 0.758711382 | 0.19409872 | 3.90889435 | 9.27E-05 | 0.0008388 | NFKBIL1 | 4795 |
| ENSG000001 | 152.246117 | 0.57554448 | 0.14724393 | 3.90878249 | 9.28E-05 | 0.0008389 | ZNF18 | 7566 |
| ENSG00000C | 546.770064 | 0.727635301 | 0.18618983 | 3.90802917 | 9.31E-05 | 0.00084133 | PHRF1 | 57661 |
| ENSG000001 | 709.075112 | 0.50435132 | 0.12905938 | 3.90790139 | 9.31E-05 | 0.00084142 | AP1AR | 55435 |
| ENSG000002 | 29.6637313 | -0.97910886 | 0.25054645 | -3.9078936 | 9.31E-05 | 0.00084142 | LOC400794 | 400794 |
| ENSG000001 | 1446.51981 | 0.473778106 | 0.12124627 | 3.90756837 | 9.32E-05 | 0.00084237 | CDC42SE1 | 56882 |
| ENSG000001 | 1036.31408 | 0.43907159 | 0.11236878 | 3.90741615 | 9.33E-05 | 0.00084271 | MRPL57 | 78988 |
| ENSG000002 | 206.945651 | 0.576496679 | 0.14754933 | 3.90714528 | 9.34E-05 | 0.00084347 | CNEP1R1 | 255919 |
| ENSG00000C | 265.270645 | 0.63477002 | 0.16246866 | 3.9070306 | 9.34E-05 | 0.00084368 | GNA11 | 2767 |
| ENSG000001 | 1089.07114 | 0.476506588 | 0.121967 | 3.90684855 | 9.35E-05 | 0.00084413 | PPM1A | 5494 |
| ENSG000001 | 400.83096 | 0.598072396 | 0.15309577 | 3.90652457 | 9.36E-05 | 0.00084489 | EMC3 | 55831 |

| | | | | | | | | | |
|---|---|---|---|---|---|---|---|---|---|
| ENSG000001 | 1433.66647 | 0.516070047 | 0.13210356 | 3.90655682 | 9.36E-05 | 0.00084489 | RAP2A | | 5911 |
| ENSG00000C | 450.530194 | 0.442981918 | 0.11341798 | 3.90574672 | 9.39E-05 | 0.00084742 | HDAC9 | | 9734 |
| ENSG000001 | 173.090877 | 0.560426984 | 0.14350479 | 3.9052843 | 9.41E-05 | 0.00084885 | GALM | | 130589 |
| ENSG000002 | 234.054644 | 0.581109044 | 0.14880586 | 3.90514899 | 9.42E-05 | 0.00084914 | ZSWIM8 | | 23053 |
| ENSG000001 | 13480.4513 | 0.505875348 | 0.12956228 | 3.90449552 | 9.44E-05 | 0.00085115 | HMGA1 | | 3159 |
| ENSG000002 | 24.5534036 | -1.10023569 | 0.2817888 | -3.9044692 | 9.44E-05 | 0.00085115 | NA | NA | |
| ENSG000001 | 1715.18366 | 0.519045271 | 0.13295269 | 3.90398475 | 9.46E-05 | 0.00085267 | TCF20 | | 6942 |
| ENSG000001 | 1307.47313 | 0.607462052 | 0.15560388 | 3.90390035 | 9.47E-05 | 0.00085278 | TLN1 | | 7094 |
| ENSG000001 | 1296.75252 | 0.537756266 | 0.13776118 | 3.90353996 | 9.48E-05 | 0.00085386 | FTSJ3 | | 117246 |
| ENSG000001 | 115.946481 | 0.65457429 | 0.16770281 | 3.90318033 | 9.49E-05 | 0.00085494 | GSDMD | | 79792 |
| ENSG000002 | 213.224205 | -0.66111303 | 0.16939994 | -3.9026758 | 9.51E-05 | 0.00085642 | RNU1-1 | | 26871 |
| ENSG000002 | 624.499899 | 0.470738003 | 0.12061996 | 3.90265431 | 9.51E-05 | 0.00085642 | LYRM4 | | 57128 |
| ENSG000001 | 2684.29106 | 0.693699501 | 0.17776919 | 3.90224813 | 9.53E-05 | 0.00085767 | SLC7A5 | | 8140 |
| ENSG000001 | 546.556239 | 0.456076489 | 0.1168803 | 3.90208172 | 9.54E-05 | 0.00085807 | SLC35F5 | | 80255 |
| ENSG000001 | 54.5240416 | -0.79175519 | 0.20291062 | -3.9019899 | 9.54E-05 | 0.0008582 | ATP1A4 | | 480 |
| ENSG000001 | 112.603919 | 0.710389495 | 0.18207556 | 3.90161927 | 9.56E-05 | 0.00085933 | MAPK12 | | 6300 |
| ENSG000001 | 664.133781 | 0.522834041 | 0.13402002 | 3.9011636 | 9.57E-05 | 0.00086075 | UBE2Q2 | | 92912 |
| ENSG000001 | 529.630671 | 0.730506851 | 0.18727446 | 3.90072873 | 9.59E-05 | 0.00086211 | INTS1 | | 26173 |
| ENSG000001 | 2777.47595 | 0.47257108 | 0.12115334 | 3.90060297 | 9.60E-05 | 0.00086237 | NA | NA | |
| ENSG00000C | 1494.83518 | 0.615739808 | 0.15787623 | 3.90014257 | 9.61E-05 | 0.00086364 | LAPTM4A | | 9741 |
| ENSG000001 | 451.812406 | 0.600228707 | 0.15389929 | 3.90013966 | 9.61E-05 | 0.00086364 | ACOT9 | | 23597 |
| ENSG00000C | 565.222638 | 0.464183832 | 0.11902728 | 3.89981049 | 9.63E-05 | 0.00086461 | RBL1 | | 5933 |
| ENSG000001 | 7815.91783 | 0.483977895 | 0.12410453 | 3.89976002 | 9.63E-05 | 0.00086461 | AMD1 | | 262 |
| ENSG000001 | 277.255639 | 0.661960715 | 0.16975221 | 3.89957056 | 9.64E-05 | 0.0008651 | FURIN | | 5045 |
| ENSG000001 | 419.166253 | 0.601365895 | 0.15422394 | 3.89930306 | 9.65E-05 | 0.00086586 | RPAP1 | | 26015 |
| ENSG000001 | 185.393286 | 0.756776913 | 0.19414708 | 3.89795669 | 9.70E-05 | 0.00087049 | MSI1 | | 4440 |
| ENSG00000C | 143.98945 | 0.639890394 | 0.16418447 | 3.89738679 | 9.72E-05 | 0.00087203 | PNPLA4 | | 8228 |
| ENSG00000C | 66.2053874 | -0.732193342 | 0.18786364 | -3.8974724 | 9.72E-05 | 0.00087203 | COL5A3 | | 50509 |
| ENSG000001 | 157.182614 | 0.630478661 | 0.16177031 | 3.89736941 | 9.72E-05 | 0.00087203 | LRRC45 | | 201255 |

| | | | | | | | | |
|---|---|---|---|---|---|---|---|---|
| ENSG000001 | 6614.79058 | 0.465682425 | 0.11951084 | 3.89657054 | 9.76E-05 | 0.00087471 | TRA2B | 6434 |
| ENSG000002 | 118.896424 | 0.63658885 | 0.16338705 | 3.89620127 | 9.77E-05 | 0.00087585 | LINC01090 | 104355152 |
| ENSG000001 | 622.584782 | 0.587836569 | 0.1509055 | 3.89539534 | 9.80E-05 | 0.00087838 | ING5 | 84289 |
| ENSG000001 | 156.046829 | 0.59594491 | 0.15298656 | 3.89540687 | 9.80E-05 | 0.00087838 | WDR53 | 348793 |
| ENSG000002 | 174.258263 | 0.733369949 | 0.18827955 | 3.89511201 | 9.82E-05 | 0.00087922 | NEURL4 | 84461 |
| ENSG000001 | 2796.83805 | 0.458392827 | 0.1177017 | 3.89453035 | 9.84E-05 | 0.00088114 | ZDHHC20 | 253832 |
| ENSG000001 | 941.704498 | 0.449277442 | 0.11536666 | 3.89434381 | 9.85E-05 | 0.00088162 | FAF1 | 11124 |
| ENSG000001 | 75.0478546 | 0.856273343 | 0.21988731 | 3.89414628 | 9.85E-05 | 0.00088214 | SLC25A28 | 81894 |
| ENSG000001 | 3447.00607 | 0.597859824 | 0.15354313 | 3.8937583 | 9.87E-05 | 0.00088317 | H3F3A | 3020 |
| ENSG000001 | 237.524246 | 0.656532175 | 0.16861105 | 3.89376726 | 9.87E-05 | 0.00088317 | SEZ6L2 | 26470 |
| ENSG00000C | 37008.3985 | 0.573106409 | 0.14721558 | 3.89297387 | 9.90E-05 | 0.00088576 | ACTB | 60 |
| ENSG000001 | 32.2231121 | -0.97616756 | 0.25075324 | -3.8929409 | 9.90E-05 | 0.00088576 | PVRL4 | 81607 |
| ENSG000001 | 10992.1991 | 0.514895609 | 0.13227321 | 3.89266744 | 9.91E-05 | 0.00088636 | YWHAZ | 7534 |
| ENSG000001 | 583.435638 | 0.473133414 | 0.1215446 | 3.89267327 | 9.91E-05 | 0.00088636 | SNAPC3 | 6619 |
| ENSG000001 | 433.763205 | 0.496762023 | 0.12762867 | 3.89224474 | 9.93E-05 | 0.00088772 | FBXO18 | 84893 |
| ENSG000001 | 1439.32255 | 0.43388993 | 0.11147909 | 3.89211926 | 9.94E-05 | 0.00088778 | SUPT20H | 55578 |
| ENSG000002 | 808.408554 | 0.562226609 | 0.14445181 | 3.89213967 | 9.94E-05 | 0.00088778 | HLA-C | 3107 |
| ENSG000001 | 579.602037 | 0.528005043 | 0.13566766 | 3.89190068 | 9.95E-05 | 0.00088839 | RBSN | 64145 |
| ENSG000001 | 209.910257 | 0.548871452 | 0.14103798 | 3.89165716 | 9.96E-05 | 0.00088908 | LOXL3 | 84695 |
| ENSG000001 | 273.784786 | 0.626756559 | 0.16105467 | 3.8915765 | 9.96E-05 | 0.00088918 | TJAP1 | 93643 |
| ENSG000001 | 1611.26405 | 0.522891364 | 0.13437519 | 3.89127911 | 9.97E-05 | 0.00089008 | RRM2 | 6241 |
| ENSG000001 | 272.221849 | 0.533741185 | 0.13716841 | 3.89113791 | 9.98E-05 | 0.0008904 | EFEMP1 | 2202 |
| ENSG000001 | 710.349273 | 0.466911742 | 0.11999538 | 3.8910811 | 9.98E-05 | 0.00089041 | DPH5 | 51611 |
| ENSG000001 | 324.159803 | 0.622333977 | 0.15994775 | 3.89085796 | 9.99E-05 | 0.00089104 | PPAP2C | 8612 |
| ENSG00000C | 742.107548 | 0.515998958 | 0.13264 | 3.89022126 | 0.00010015 | 0.00089318 | SPG21 | 51324 |
| ENSG000001 | 218.817379 | 0.708866155 | 0.1822232 | 3.89009829 | 0.0001002 | 0.00089344 | POFUT2 | 23275 |
| ENSG00000C | 818.114223 | 0.427027678 | 0.10977736 | 3.88994314 | 0.00010027 | 0.00089381 | NUB1 | 51667 |
| ENSG000001 | 1167.38689 | 0.47249242 | 0.12146804 | 3.88984973 | 0.00010031 | 0.00089396 | PRKAR2A | 5576 |
| ENSG000001 | 629.788385 | 0.503898382 | 0.12954801 | 3.88966505 | 0.00010038 | 0.00089441 | IDH3A | 3419 |

| | | | | | | | | | |
|---|---|---|---|---|---|---|---|---|---|
| ENSG000001 | 1533.15436 | 0.471762515 | 0.12128753 | 3.88962095 | 0.0001004 | 0.00089441 | Sep-15 | | 9403 |
| ENSG000001 | 13.6237118 | -1.243723026 | 0.31977712 | -3.8893434 | 0.00010052 | 0.00089493 | NA | NA | |
| ENSG000002 | 521.674554 | 0.58487578 | 0.15037999 | 3.88931915 | 0.00010053 | 0.00089493 | NA | NA | |
| ENSG000002 | 59.3089799 | 0.830515904 | 0.21353313 | 3.88940073 | 0.00010049 | 0.00089493 | CISD3 | | 284106 |
| ENSG00000C | 401.015592 | 0.501798303 | 0.12902194 | 3.88924774 | 0.00010056 | 0.000895 | EXD2 | | 55218 |
| ENSG000001 | 8.7606359 | -1.393386324 | 0.35827292 | -3.8891757 | 0.00010059 | 0.00089507 | TBX22 | | 50945 |
| ENSG000002 | 264.412803 | 0.540131362 | 0.13889116 | 3.88888211 | 0.00010071 | 0.00089596 | TMEM242 | | 729515 |
| ENSG000001 | 60.4465026 | -0.777439795 | 0.19992194 | -3.8887167 | 0.00010078 | 0.00089637 | TNNT2 | | 7139 |
| ENSG000001 | 9.18458375 | 1.389346397 | 0.35729422 | 3.88852192 | 0.00010086 | 0.00089689 | PYDC1 | | 260434 |
| ENSG000001 | 461.416924 | 0.576262124 | 0.14820679 | 3.88823015 | 0.00010098 | 0.00089778 | ESPL1 | | 9700 |
| ENSG000001 | 1020.26036 | 0.537710104 | 0.1382994 | 3.8880148 | 0.00010107 | 0.00089838 | ESD | | 2098 |
| ENSG000001 | 958.592127 | 0.515964701 | 0.13271405 | 3.88779259 | 0.00010116 | 0.000899 | FAT4 | | 79633 |
| ENSG000001 | 4044.06158 | 0.518375432 | 0.1333438 | 3.88751056 | 0.00010128 | 0.00089985 | HK1 | | 3098 |
| ENSG000001 | 1155.57955 | 0.471324073 | 0.12124681 | 3.88731101 | 0.00010136 | 0.00090039 | DNAJB14 | | 79982 |
| ENSG00000C | 421.080613 | 0.510761748 | 0.13139924 | 3.88709828 | 0.00010145 | 0.00090098 | L2HGDH | | 79944 |
| ENSG000001 | 1027.79747 | 0.500870118 | 0.12885797 | 3.8869937 | 0.00010149 | 0.00090117 | TULP4 | | 56995 |
| ENSG000001 | 25.2571752 | -1.034766131 | 0.26622402 | -3.8868248 | 0.00010156 | 0.0009016 | ADIPOQ | | 9370 |
| ENSG000001 | 1754.35624 | 0.563490413 | 0.14497836 | 3.88672078 | 0.00010161 | 0.00090179 | PRELID1 | | 27166 |
| ENSG00000C | 1186.22014 | 0.496301809 | 0.12771006 | 3.88616052 | 0.00010184 | 0.00090368 | ST6GAL1 | | 6480 |
| ENSG000001 | 10685.6011 | 0.492927057 | 0.12685305 | 3.88581163 | 0.00010199 | 0.00090467 | CLTC | | 1213 |
| ENSG000001 | 369.871945 | 0.577260466 | 0.14855686 | 3.88578806 | 0.000102 | 0.00090467 | GREB1 | | 9687 |
| ENSG000001 | 21.1169127 | -1.100607553 | 0.28328024 | -3.8852253 | 0.00010224 | 0.00090657 | NA | NA | |
| ENSG000001 | 1196.9248 | 0.508607409 | 0.13091572 | 3.88499884 | 0.00010233 | 0.00090721 | PRPF6 | | 24148 |
| ENSG000001 | 1131.86993 | 0.504641502 | 0.12989917 | 3.88487088 | 0.00010238 | 0.00090749 | RAB34 | | 83871 |
| ENSG000001 | 143.408892 | -0.60319051 | 0.15527683 | -3.8846138 | 0.00010249 | 0.00090786 | COL12A1 | | 1303 |
| ENSG000001 | 977.991119 | 0.539314652 | 0.13883268 | 3.88463761 | 0.00010248 | 0.00090786 | TMEM59 | | 9528 |
| ENSG000002 | 42.4342795 | -0.879051959 | 0.22628967 | -3.8846314 | 0.00010249 | 0.00090786 | ADGRG1 | | 9289 |
| ENSG00000C | 431.930781 | 0.533477637 | 0.13736481 | 3.88365574 | 0.0001029 | 0.00091125 | GCLM | | 2730 |
| ENSG000001 | 139.129036 | -0.604432644 | 0.15564414 | -3.883427 | 0.00010299 | 0.00091191 | UNC80 | | 285175 |

| | | | | | | | | |
|---|---|---|---|---|---|---|---|---|
| ENSG00000C | 130.17104 | 0.601840361 | 0.15499112 | 3.88306363 | 0.00010315 | 0.00091307 | SLC25A43 | 203427 |
| ENSG000001 | 651.072739 | 0.490241221 | 0.12625344 | 3.88299305 | 0.00010318 | 0.00091314 | TWSG1 | 57045 |
| ENSG000002 | 15.7577573 | -1.183153358 | 0.30470884 | -3.882898 | 0.00010322 | 0.00091329 | LOC101929717 | 101929717 |
| ENSG000001 | 665.207391 | 0.531656035 | 0.13693813 | 3.88245438 | 0.00010341 | 0.00091463 | MED4 | 29079 |
| ENSG000001 | 425.876923 | 0.512650283 | 0.1320434 | 3.88243778 | 0.00010341 | 0.00091463 | LRR1 | 122769 |
| ENSG000001 | 8999.77376 | 0.490310874 | 0.12629452 | 3.88228142 | 0.00010348 | 0.00091502 | LRPPRC | 10128 |
| ENSG000001 | 92.308368 | 0.703299824 | 0.18116114 | 3.88217816 | 0.00010352 | 0.0009152 | TP53I13 | 90313 |
| ENSG000001 | 494.503034 | 0.486083838 | 0.12521389 | 3.88202819 | 0.00010359 | 0.00091557 | ZNF330 | 27309 |
| ENSG000001 | 45.1233527 | -0.900879883 | 0.23207856 | -3.8817884 | 0.00010369 | 0.00091627 | STK31 | 56164 |
| ENSG00000C | 903.310936 | 0.579970321 | 0.14941817 | 3.88152466 | 0.0001038 | 0.00091687 | EXTL3 | 2137 |
| ENSG000002 | 3.24830003 | -1.452447966 | 0.37419119 | -3.8815664 | 0.00010379 | 0.00091687 | NA | NA |
| ENSG000001 | 1762.67933 | 0.442262795 | 0.11394277 | 3.88144677 | 0.00010384 | 0.00091696 | PRKRIR | 5612 |
| ENSG000002 | 65.097684 | 0.741116482 | 0.19095396 | 3.88112652 | 0.00010397 | 0.00091797 | NA | NA |
| ENSG000001 | 527.012954 | 0.561421687 | 0.14466202 | 3.8809197 | 0.00010406 | 0.00091855 | C14orf169 | 79697 |
| ENSG000001 | 148.734333 | 0.742512084 | 0.19133283 | 3.88073544 | 0.00010414 | 0.00091905 | FAM73B | 84895 |
| ENSG000002 | 138.137022 | 0.605281091 | 0.15597839 | 3.88054453 | 0.00010422 | 0.00091957 | NA | NA |
| ENSG000001 | 18.9183032 | -1.121464439 | 0.28900504 | -3.8804322 | 0.00010427 | 0.00091979 | OR8D4 | 338662 |
| ENSG000001 | 1505.81556 | 0.491214889 | 0.12660711 | 3.87983638 | 0.00010453 | 0.00092185 | HSD17B12 | 51144 |
| ENSG000001 | 303.068288 | 0.750529014 | 0.19354737 | 3.87775361 | 0.00010543 | 0.00092957 | MYH14 | 79784 |
| ENSG000001 | 3232.24122 | 0.437933101 | 0.11295155 | 3.87717638 | 0.00010568 | 0.00093158 | ACTR3 | 10096 |
| ENSG000002 | 184.20246 | 0.640921916 | 0.16531282 | 3.87702485 | 0.00010574 | 0.00093196 | C6orf47 | 57827 |
| ENSG000001 | 536.888797 | 0.562451045 | 0.14507605 | 3.87693931 | 0.00010578 | 0.00093208 | STAT5B | 6777 |
| ENSG000002 | 24.6600816 | -1.059603103 | 0.27332573 | -3.8767046 | 0.00010588 | 0.00093278 | LINC01344 | 400799 |
| ENSG000001 | 614.848366 | 0.51874212 | 0.13382175 | 3.87636627 | 0.00010603 | 0.00093367 | PPP2R5D | 5528 |
| ENSG000001 | 138.283755 | 0.615825109 | 0.15886465 | 3.87641375 | 0.00010601 | 0.00093367 | TMEM60 | 85025 |
| ENSG000001 | 1300.62847 | 0.481018888 | 0.12409691 | 3.87615526 | 0.00010612 | 0.00093428 | KIAA0196 | 9897 |
| ENSG000001 | 1477.32205 | 0.451976421 | 0.11661909 | 3.87566416 | 0.00010633 | 0.00093596 | MAGT1 | 84061 |
| ENSG00000C | 624.153751 | 0.583032231 | 0.15046052 | 3.87498487 | 0.00010663 | 0.00093837 | FAM107B | 83641 |
| ENSG00000C | 3146.0789 | 0.653964655 | 0.1687746 | 3.8747813 | 0.00010672 | 0.00093895 | ADD2 | 119 |

| | | | | | | | | |
|---|---|---|---|---|---|---|---|---|
| ENSG00000( | 1167.10898 | 0.556684255 | 0.14367642 | 3.87456948 | 0.00010681 | 0.00093956 | DNAJC11 | 55735 |
| ENSG00000( | 1713.52862 | 0.483364325 | 0.124765 | 3.87419819 | 0.00010698 | 0.00094059 | LASP1 | 3927 |
| ENSG000001 | 382.588846 | 0.513754674 | 0.13260895 | 3.87420811 | 0.00010697 | 0.00094059 | PMS2 | 5395 |
| ENSG00000( | 99.1217245 | 0.702473482 | 0.18132507 | 3.87411115 | 0.00010701 | 0.00094072 | NA | NA |
| ENSG000001 | 3301.87346 | 0.437659857 | 0.11297344 | 3.87400666 | 0.00010706 | 0.00094072 | PDS5A | 23244 |
| ENSG000002 | 27.5321847 | -1.000323535 | 0.25821355 | -3.8740164 | 0.00010706 | 0.00094072 | MIR663AHG | 284801 |
| ENSG000001 | 1446.80126 | 0.524715592 | 0.13544961 | 3.87388052 | 0.00010712 | 0.000941 | NRBP1 | 29959 |
| ENSG000001 | 670.752033 | 0.579176367 | 0.14951772 | 3.87363039 | 0.00010723 | 0.00094154 | PVRL2 | 5819 |
| ENSG000001 | 460.226538 | 0.493034482 | 0.12727856 | 3.87366475 | 0.00010721 | 0.00094154 | UTP23 | 84294 |
| ENSG000001 | 173.360831 | 0.630042179 | 0.16265097 | 3.8735838 | 0.00010725 | 0.00094154 | ORAI2 | 80228 |
| ENSG000001 | 425.311868 | 0.526789223 | 0.13600015 | 3.87344596 | 0.00010731 | 0.00094186 | KLHL18 | 23276 |
| ENSG000001 | 284.200817 | 0.625085959 | 0.16138837 | 3.87317858 | 0.00010743 | 0.00094269 | MAN1B1 | 11253 |
| ENSG00000( | 753.350043 | 0.487828585 | 0.12596298 | 3.8727932 | 0.0001076 | 0.00094398 | STRADB | 55437 |
| ENSG000001 | 265.151529 | 0.713480047 | 0.18424372 | 3.87247964 | 0.00010773 | 0.00094499 | DCAF15 | 90379 |
| ENSG00000( | 1052.05668 | 0.458632013 | 0.11844024 | 3.87226518 | 0.00010783 | 0.00094562 | SRBD1 | 55133 |
| ENSG000001 | 68.2256275 | -0.719955432 | 0.18593032 | -3.8721788 | 0.00010787 | 0.00094575 | XIRP2 | 129446 |
| ENSG00000( | 642.991159 | 0.474890561 | 0.12265823 | 3.87165684 | 0.0001081 | 0.00094758 | IGBP1 | 3476 |
| ENSG000001 | 37.9188774 | 0.96905335 | 0.25033809 | 3.8709784 | 0.0001084 | 0.00095001 | GHDC | 84514 |
| ENSG000001 | 1416.35925 | 0.481013196 | 0.12427142 | 3.87066641 | 0.00010854 | 0.00095102 | SPRED1 | 161742 |
| ENSG000001 | 1998.1241 | 0.460547407 | 0.11898697 | 3.87057003 | 0.00010858 | 0.00095119 | ACAA2 | 10449 |
| ENSG000001 | 5007.04207 | 0.417400262 | 0.10784103 | 3.87051452 | 0.00010861 | 0.0009512 | EIF5 | 1983 |
| ENSG000001 | 478.406733 | 0.583944745 | 0.15087553 | 3.8703741 | 0.00010867 | 0.00095155 | PPIH | 10465 |
| ENSG000002 | 3046.37451 | 0.472201017 | 0.12200998 | 3.87018342 | 0.00010875 | 0.00095209 | SNHG1 | 23642 |
| ENSG00000( | 376.775624 | 0.506417105 | 0.13088885 | 3.86906229 | 0.00010925 | 0.00095627 | TFB1M | 51106 |
| ENSG000001 | 212.937201 | -0.606906408 | 0.15687589 | -3.8687042 | 0.00010942 | 0.00095747 | TP53INP1 | 94241 |
| ENSG00000( | 1693.06408 | 0.616624295 | 0.15939068 | 3.86863464 | 0.00010945 | 0.00095753 | PFN2 | 5217 |
| ENSG000001 | 729.221884 | 0.475266138 | 0.122857 | 3.86844994 | 0.00010953 | 0.00095805 | HSDL1 | 83693 |
| ENSG000001 | 3268.8481 | 0.506138133 | 0.13084199 | 3.86831575 | 0.00010959 | 0.00095837 | PHC1 | 1911 |
| ENSG000001 | 52.9019078 | -0.85027649 | 0.21983056 | -3.8678721 | 0.00010979 | 0.00095991 | PLCL1 | 5334 |

| | | | | | | | | | |
|---|---|---|---|---|---|---|---|---|---|
| ENSG000001 | 291.527237 | 0.489315807 | 0.12651075 | 3.86778052 | 0.00010983 | 0.00096006 | INTS9 | | 55756 |
| ENSG000001 | 471.090853 | 0.56651046 | 0.14647142 | 3.86772026 | 0.00010986 | 0.00096009 | MAPKAPK2 | | 9261 |
| ENSG000001 | 508.996915 | 0.551853623 | 0.14269884 | 3.86726063 | 0.00011006 | 0.0009617 | BAHD1 | | 22893 |
| ENSG000001 | 1213.52924 | 0.518025663 | 0.1339556 | 3.86714455 | 0.00011012 | 0.00096195 | H2AFV | | 94239 |
| ENSG000001 | 1121.94255 | 0.559317033 | 0.14463711 | 3.86703681 | 0.00011017 | 0.00096196 | SURF6 | | 6838 |
| ENSG000001 | 813.455651 | 0.452021956 | 0.11689047 | 3.86705571 | 0.00011016 | 0.00096196 | TTC14 | | 151613 |
| ENSG000001 | 21.1256871 | -1.082907143 | 0.28005366 | -3.8667845 | 0.00011028 | 0.00096275 | ZNF596 | | 169270 |
| ENSG000001 | 837.425395 | 0.562408831 | 0.14545735 | 3.86648603 | 0.00011041 | 0.00096372 | ZSCAN29 | | 146050 |
| ENSG000001 | 335.985094 | 0.561447236 | 0.14522769 | 3.86597924 | 0.00011064 | 0.00096551 | TBCC | | 6903 |
| ENSG000001 | 1256.02329 | 0.481339433 | 0.12452098 | 3.86552893 | 0.00011085 | 0.00096709 | CRNKL1 | | 51340 |
| ENSG00000C | 3337.23641 | 0.461048602 | 0.11929466 | 3.8647884 | 0.00011119 | 0.00096961 | NDC1 | | 55706 |
| ENSG000001 | 2319.92999 | 0.499247635 | 0.1291768 | 3.86483975 | 0.00011116 | 0.00096961 | AGPAT5 | | 55326 |
| ENSG000001 | 99.5134445 | 0.6865037 | 0.17763943 | 3.86459067 | 0.00011128 | 0.00097019 | ROBO3 | | 64221 |
| ENSG000002 | 24.5131636 | -1.050438417 | 0.27182212 | -3.8644332 | 0.00011135 | 0.0009706 | NA | NA | |
| ENSG000001 | 45.3737805 | -0.881052838 | 0.22802289 | -3.8638789 | 0.0001116 | 0.0009726 | LVRN | | 206338 |
| ENSG000001 | 23.9465926 | -1.061182782 | 0.2746563 | -3.8636753 | 0.00011169 | 0.0009732 | PPARG | | 5468 |
| ENSG000001 | 813.447863 | 0.503134966 | 0.13023165 | 3.86338483 | 0.00011183 | 0.00097415 | HDDC2 | | 51020 |
| ENSG000001 | 106.175245 | 0.714787126 | 0.18503693 | 3.86294314 | 0.00011203 | 0.00097564 | MCAT | | 27349 |
| ENSG000001 | 1593.25486 | 0.446449335 | 0.11557338 | 3.86290784 | 0.00011205 | 0.00097564 | NFYB | | 4801 |
| ENSG000001 | 50.3117785 | 0.81660495 | 0.21142195 | 3.86244163 | 0.00011226 | 0.00097729 | ACOT2 | | 10965 |
| ENSG000001 | 75.8890964 | -0.780803155 | 0.20216454 | -3.8622161 | 0.00011236 | 0.00097799 | IAH1 | | 285148 |
| ENSG000001 | 472.197106 | 0.572186392 | 0.14815515 | 3.86207558 | 0.00011243 | 0.00097834 | EFNB2 | | 1948 |
| ENSG000001 | 2701.43178 | 0.436933689 | 0.11314539 | 3.86170133 | 0.0001126 | 0.00097963 | USP1 | | 7398 |
| ENSG000001 | 632.223884 | 0.616158822 | 0.15956651 | 3.86145462 | 0.00011271 | 0.00098041 | LETM1 | | 3954 |
| ENSG00000C | 30.5712629 | -0.964791916 | 0.24985834 | -3.8613557 | 0.00011276 | 0.0009806 | LTF | | 4057 |
| ENSG000001 | 1506.6651 | 0.425345119 | 0.11015758 | 3.86124226 | 0.00011281 | 0.00098084 | ARHGAP21 | | 57584 |
| ENSG000001 | 105.967783 | -0.630590208 | 0.16333073 | -3.860818 | 0.00011301 | 0.00098233 | DOCK10 | | 55619 |
| ENSG000002 | 395.263961 | 0.573905683 | 0.14865103 | 3.86075811 | 0.00011304 | 0.00098236 | NA | NA | |
| ENSG000001 | 968.442021 | 0.489189229 | 0.12672425 | 3.86026521 | 0.00011326 | 0.00098414 | CDK7 | | 1022 |

| | | | | | | | | | |
|---|---|---|---|---|---|---|---|---|---|
| ENSG000001 | 107.929524 | 0.68751807 | 0.17810798 | 3.86011951 | 0.00011333 | 0.00098451 | XKR8 | | 55113 |
| ENSG000001 | 302.85185 | 0.524237603 | 0.13581446 | 3.8599544 | 0.00011341 | 0.00098497 | TMEM87B | | 84910 |
| ENSG000001 | 42.538052 | -0.936833405 | 0.2427205 | -3.8597209 | 0.00011352 | 0.0009857 | GADL1 | | 339896 |
| ENSG000001 | 297.228937 | 0.589474304 | 0.15274565 | 3.8591888 | 0.00011376 | 0.00098764 | PHF19 | | 26147 |
| ENSG000001 | 277.33217 | 0.568382321 | 0.14729593 | 3.85877822 | 0.00011396 | 0.00098887 | MAPK8IP1 | | 9479 |
| ENSG000001 | 251.693994 | 0.538175672 | 0.13946802 | 3.85877469 | 0.00011396 | 0.00098887 | DNAJC1 | | 64215 |
| ENSG000001 | 719.916979 | 0.61897186 | 0.16040835 | 3.85872605 | 0.00011398 | 0.00098887 | ADCY1 | | 107 |
| ENSG000001 | 849.807493 | 0.512414858 | 0.13280771 | 3.85832169 | 0.00011417 | 0.0009903 | AES | | 166 |
| ENSG000001 | 44.8344363 | -0.837935439 | 0.21719943 | -3.8579081 | 0.00011436 | 0.00099176 | ATP8B5P | | 158381 |
| ENSG000001 | 363.034361 | 0.504890292 | 0.13087832 | 3.85770764 | 0.00011446 | 0.00099236 | GINS3 | | 64785 |
| ENSG000001 | 361.523178 | 0.544437022 | 0.14113483 | 3.85756677 | 0.00011452 | 0.00099272 | NECAP2 | | 55707 |
| ENSG000002 | 23.6943206 | -1.069203762 | 0.27719623 | -3.8572089 | 0.00011469 | 0.00099396 | NA | NA | |
| ENSG000001 | 454.451985 | 0.501243265 | 0.12995762 | 3.85697484 | 0.0001148 | 0.0009947 | ZNF320 | | 162967 |
| ENSG000001 | 70.4115815 | 0.779110032 | 0.20200776 | 3.85683229 | 0.00011487 | 0.00099507 | SLC10A3 | | 8273 |
| ENSG00000C | 496.698253 | 0.463215695 | 0.12010823 | 3.85665243 | 0.00011495 | 0.00099559 | NUFIP1 | | 26747 |
| ENSG000001 | 22.8591143 | -1.075568953 | 0.27890443 | -3.8564068 | 0.00011507 | 0.00099638 | SLC39A12 | | 221074 |
| ENSG00000C | 140.47658 | 0.591506266 | 0.15340137 | 3.85593852 | 0.00011529 | 0.00099807 | E2F2 | | 1870 |
| ENSG000001 | 42.4759708 | -0.866779795 | 0.22479876 | -3.8558033 | 0.00011535 | 0.00099841 | COL22A1 | | 169044 |
| ENSG000001 | 476.28056 | 0.526046424 | 0.13643398 | 3.85568477 | 0.00011541 | 0.00099868 | NEU3 | | 10825 |
| ENSG000001 | 477.832305 | 0.517505204 | 0.13422916 | 3.85538583 | 0.00011555 | 0.00099969 | YIPF4 | | 84272 |
| ENSG000001 | 364.156973 | 0.488739108 | 0.12677193 | 3.85526296 | 0.00011561 | 0.00099998 | TMEM41A | | 90407 |
| ENSG000001 | 159.480797 | 0.592519402 | 0.15369735 | 3.85510498 | 0.00011568 | 0.00100041 | NA | NA | |
| ENSG000001 | 658.123773 | 0.544830403 | 0.14133442 | 3.8549024 | 0.00011578 | 0.00100103 | LDOC1L | | 84247 |
| ENSG000001 | 216.804713 | 0.684422853 | 0.17756069 | 3.85458543 | 0.00011593 | 0.00100211 | MLLT1 | | 4298 |
| ENSG000001 | 295.868434 | 0.689982936 | 0.17902077 | 3.85420595 | 0.00011611 | 0.00100324 | TSPYL2 | | 64061 |
| ENSG000002 | 12.2574861 | -1.264892309 | 0.32818212 | -3.8542389 | 0.00011609 | 0.00100324 | FMO9P | | 116123 |
| ENSG000001 | 2727.37557 | 0.459442878 | 0.11920996 | 3.85406464 | 0.00011617 | 0.00100361 | RFWD3 | | 55159 |
| ENSG000001 | 2624.88954 | 0.499483398 | 0.12961165 | 3.85369213 | 0.00011635 | 0.00100492 | ZFAND5 | | 7763 |
| ENSG00000C | 326.682548 | 0.56265564 | 0.14600983 | 3.85354635 | 0.00011642 | 0.00100531 | ZXDC | | 79364 |

| | | | | | | | | |
|---|---|---|---|---|---|---|---|---|
| ENSG000001 | 2895.09198 | 0.487522772 | 0.12652356 | 3.85321724 | 0.00011658 | 0.00100625 | MLLT4 | 4301 |
| ENSG000001 | 1800.00745 | 0.504038991 | 0.13081009 | 3.85321187 | 0.00011658 | 0.00100625 | NUP54 | 53371 |
| ENSG000001 | 184.312293 | 0.59080428 | 0.15333193 | 3.85310659 | 0.00011663 | 0.00100626 | STX12 | 23673 |
| ENSG000001 | 13.0402412 | -1.251166316 | 0.32471222 | -3.8531544 | 0.00011661 | 0.00100626 | KLK11 | 11012 |
| ENSG000001 | 411.175427 | 0.520793079 | 0.13517455 | 3.85274509 | 0.0001168 | 0.00100753 | KIAA1919 | 91749 |
| ENSG000001 | 46.7847331 | -0.881487068 | 0.22881475 | -3.8524049 | 0.00011696 | 0.00100847 | LGI1 | 9211 |
| ENSG000001 | 353.684358 | 0.567500595 | 0.14731244 | 3.85236034 | 0.00011698 | 0.00100847 | NDUFC1 | 4717 |
| ENSG000002 | 340.539754 | 0.486640923 | 0.12632029 | 3.8524366 | 0.00011695 | 0.00100847 | NA | NA |
| ENSG00000C | 573.310734 | 0.524334163 | 0.13611522 | 3.85213458 | 0.00011709 | 0.00100897 | UBE2D1 | 7321 |
| ENSG000001 | 2023.64069 | 0.568958024 | 0.14769804 | 3.85217039 | 0.00011708 | 0.00100897 | KDM3B | 51780 |
| ENSG000002 | 43.4068292 | -0.884543415 | 0.22966743 | -3.8514099 | 0.00011744 | 0.00101175 | FER1L6 | 654463 |
| ENSG00000C | 202.003168 | 0.560148545 | 0.14545043 | 3.85113016 | 0.00011757 | 0.00101263 | CUL7 | 9820 |
| ENSG000001 | 21.3469102 | -1.093231319 | 0.28387564 | -3.8510923 | 0.00011759 | 0.00101263 | VWC2L | 402117 |
| ENSG000001 | 189.678611 | 0.600051929 | 0.15583003 | 3.85068232 | 0.00011779 | 0.0010139 | C9orf40 | 55071 |
| ENSG000001 | 1333.03054 | 0.488062117 | 0.12674619 | 3.85070434 | 0.00011778 | 0.0010139 | ADGRA3 | 166647 |
| ENSG00000C | 1351.41144 | 0.503487736 | 0.13079639 | 3.849401 | 0.00011841 | 0.001019 | SEH1L | 81929 |
| ENSG000001 | 915.419615 | 0.467697652 | 0.1215266 | 3.84852094 | 0.00011883 | 0.00102245 | BRD8 | 10902 |
| ENSG000001 | 56.1440098 | -0.870903804 | 0.22633572 | -3.8478407 | 0.00011916 | 0.00102507 | CD36 | 948 |
| ENSG000001 | 981.889576 | 0.483654899 | 0.12570002 | 3.84769143 | 0.00011924 | 0.00102548 | B3GNT2 | 10678 |
| ENSG00000C | 601.358247 | 0.449156034 | 0.1167428 | 3.84739802 | 0.00011938 | 0.00102649 | RANBP9 | 10048 |
| ENSG000001 | 5164.56758 | 0.459937808 | 0.1195562 | 3.84704268 | 0.00011955 | 0.00102776 | RAD23B | 5887 |
| ENSG00000C | 5339.2124 | 0.433038949 | 0.11257431 | 3.84669407 | 0.00011972 | 0.00102896 | TPX2 | 22974 |
| ENSG000002 | 37.6950515 | 0.914078363 | 0.23762954 | 3.84665285 | 0.00011974 | 0.00102896 | LINC01315 | 102723775 |
| ENSG00000C | 280.845372 | 0.492509145 | 0.12804514 | 3.8463711 | 0.00011988 | 0.00102993 | OSBPL6 | 114880 |
| ENSG000001 | 2800.84518 | 0.420427112 | 0.10931265 | 3.84609735 | 0.00012001 | 0.00103086 | MRPL42 | 28977 |
| ENSG000002 | 80.1534514 | 0.703329862 | 0.18287585 | 3.84594178 | 0.00012009 | 0.0010313 | SNORD13 | 692084 |
| ENSG000001 | 16.4855017 | -1.169106229 | 0.30402855 | -3.845383 | 0.00012036 | 0.00103343 | KRT3 | 3850 |
| ENSG00000C | 220.255953 | 0.612080198 | 0.15918538 | 3.84507798 | 0.00012051 | 0.0010345 | XYLT2 | 64132 |
| ENSG000001 | 1745.89367 | 0.509650096 | 0.13256264 | 3.8445982 | 0.00012075 | 0.0010363 | SLC25A36 | 55186 |

| | | | | | | | | |
|---|---|---|---|---|---|---|---|---|
| ENSG000002 | 75.7606508 | 0.709558587 | 0.18457462 | 3.8442913 | 0.0001209 | 0.00103738 | NA | NA |
| ENSG000001 | 1970.3469 | 0.46969564 | 0.12218456 | 3.84414901 | 0.00012097 | 0.00103776 | FABP5 | 2171 |
| ENSG000001 | 3545.60605 | 0.446515356 | 0.11615618 | 3.84409467 | 0.000121 | 0.00103778 | PSMD7 | 5713 |
| ENSG000001 | 1039.56848 | 0.488458745 | 0.12708385 | 3.84359411 | 0.00012125 | 0.00103968 | FAM63B | 54629 |
| ENSG000001 | 132.265254 | -0.666870162 | 0.17352597 | -3.8430568 | 0.00012151 | 0.00104173 | ABHD12B | 145447 |
| ENSG000001 | 224.378541 | 0.671410102 | 0.17471729 | 3.84283723 | 0.00012162 | 0.00104223 | KIAA0195 | 9772 |
| ENSG000001 | 680.305715 | 0.506066782 | 0.13169046 | 3.84285079 | 0.00012161 | 0.00104223 | UBE2J1 | 51465 |
| ENSG000001 | 1248.67982 | 0.474918318 | 0.12359039 | 3.84267989 | 0.0001217 | 0.00104268 | USP39 | 10713 |
| ENSG000002 | 3214.6671 | 0.565578349 | 0.14719696 | 3.84232365 | 0.00012187 | 0.00104397 | ACACA | 31 |
| ENSG000001 | 137.090414 | 0.759049918 | 0.19757271 | 3.84187636 | 0.0001221 | 0.00104565 | MFSD10 | 10227 |
| ENSG000001 | 106.139975 | 0.713655947 | 0.18576135 | 3.84178917 | 0.00012214 | 0.0010458 | ZNF500 | 26048 |
| ENSG00000C | 3062.61535 | 0.452705465 | 0.11784653 | 3.84148318 | 0.00012229 | 0.00104689 | CDC5L | 988 |
| ENSG000001 | 332.574177 | 0.580216764 | 0.15104316 | 3.84139702 | 0.00012234 | 0.00104703 | ITGA5 | 3678 |
| ENSG000001 | 610.029354 | 0.510091186 | 0.13279081 | 3.84131401 | 0.00012238 | 0.00104717 | COMMD6 | 170622 |
| ENSG000002 | 21.5763696 | -1.083988915 | 0.28220165 | -3.8411857 | 0.00012244 | 0.00104749 | FLJ12825 | 440101 |
| ENSG000001 | 99.2920509 | 0.67877585 | 0.1767205 | 3.84095694 | 0.00012256 | 0.00104806 | TSEN34 | 79042 |
| ENSG000001 | 153.575498 | 0.627998175 | 0.16350071 | 3.84095079 | 0.00012256 | 0.00104806 | CCBL1 | 883 |
| ENSG000001 | 148.049385 | 0.685919575 | 0.17858629 | 3.84083002 | 0.00012262 | 0.00104835 | CC2D1A | 54862 |
| ENSG00000C | 480.413818 | 0.496689111 | 0.12932482 | 3.84063243 | 0.00012272 | 0.00104897 | OSTM1 | 28962 |
| ENSG000001 | 119.129748 | 0.644548303 | 0.16782836 | 3.84052086 | 0.00012277 | 0.00104923 | PIK3CD | 5293 |
| ENSG000001 | 50.6684576 | -0.82356941 | 0.21444976 | -3.840384 | 0.00012284 | 0.00104959 | TSHZ2 | 128553 |
| ENSG000001 | 2123.01159 | 0.417710282 | 0.10877079 | 3.84027997 | 0.00012289 | 0.00104982 | CACUL1 | 143384 |
| ENSG00000C | 5810.97153 | 0.46559076 | 0.1212416 | 3.84018977 | 0.00012294 | 0.00104998 | TOP2B | 7155 |
| ENSG000002 | 61.3136057 | 0.911889809 | 0.2375001 | 3.83953447 | 0.00012327 | 0.00105257 | TMEM256 | 254863 |
| ENSG000001 | 206.585516 | 0.566931842 | 0.14767248 | 3.83911649 | 0.00012348 | 0.00105414 | CDK10 | 8558 |
| ENSG000001 | 815.500713 | 0.562649794 | 0.14656517 | 3.83890519 | 0.00012358 | 0.00105482 | WDR5 | 11091 |
| ENSG00000C | 1140.20884 | 0.515492019 | 0.13428586 | 3.83876629 | 0.00012365 | 0.0010552 | DYNC1I2 | 1781 |
| ENSG00000C | 28.2265034 | -1.012581256 | 0.26378393 | -3.8386768 | 0.0001237 | 0.00105536 | HAL | 3034 |
| ENSG00000C | 104.863709 | 0.726584674 | 0.18928931 | 3.83848766 | 0.00012379 | 0.00105595 | LZTS1 | 11178 |

| | | | | | | | | |
|---|---|---|---|---|---|---|---|---|
| ENSG000001 | 474.200204 | 0.498597012 | 0.12990483 | 3.83817146 | 0.00012395 | 0.00105709 | GLE1 | 2733 |
| ENSG000001 | 122.658432 | 0.645284866 | 0.16816463 | 3.83722117 | 0.00012443 | 0.00106096 | RAB40C | 57799 |
| ENSG000001 | 382.981689 | 0.556154473 | 0.1449407 | 3.83711725 | 0.00012449 | 0.00106119 | CEP104 | 9731 |
| ENSG000002 | 13.605649 | -1.278285854 | 0.33314843 | -3.8369859 | 0.00012455 | 0.00106153 | LOC101926897 | 101926897 |
| ENSG000001 | 1192.69146 | 0.426800011 | 0.11124478 | 3.83658452 | 0.00012476 | 0.00106282 | MFF | 56947 |
| ENSG000002 | 65.1471807 | 0.73309187 | 0.19107924 | 3.83658573 | 0.00012476 | 0.00106282 | STK19 | 8859 |
| ENSG00000C | 525.938817 | 0.563519496 | 0.14689474 | 3.83621279 | 0.00012495 | 0.00106421 | TMEM98 | 26022 |
| ENSG000002 | 9739.7751 | 0.553305955 | 0.14424186 | 3.83595967 | 0.00012507 | 0.00106508 | PRKDC | 5591 |
| ENSG000002 | 24.5969351 | -1.057199185 | 0.27562891 | -3.8355889 | 0.00012526 | 0.00106646 | TCEAL5 | 340543 |
| ENSG000001 | 376.216829 | 0.502688907 | 0.13108517 | 3.83482675 | 0.00012565 | 0.00106955 | TRIM35 | 23087 |
| ENSG000001 | 44.7297677 | -0.868568388 | 0.22651145 | -3.8345452 | 0.0001258 | 0.00107055 | BMPER | 168667 |
| ENSG000002 | 11.8247715 | -1.316392664 | 0.34333814 | -3.8340997 | 0.00012603 | 0.00107227 | NA | NA |
| ENSG000001 | 7998.55994 | 0.473689516 | 0.12355306 | 3.83389559 | 0.00012613 | 0.00107294 | CCT6A | 908 |
| ENSG000001 | 559.086094 | 0.462925468 | 0.12075851 | 3.8334811 | 0.00012634 | 0.00107452 | UFSP2 | 55325 |
| ENSG000001 | 1368.56374 | 0.445529608 | 0.11623047 | 3.83315672 | 0.00012651 | 0.00107571 | TSEN15 | 116461 |
| ENSG00000C | 1472.36218 | 0.467434278 | 0.12196091 | 3.83265654 | 0.00012677 | 0.00107768 | PALB2 | 79728 |
| ENSG000001 | 421.46228 | 0.503393575 | 0.13134603 | 3.83257557 | 0.00012681 | 0.00107781 | UBE3B | 89910 |
| ENSG000001 | 189.346484 | 0.57343687 | 0.14963492 | 3.83223962 | 0.00012698 | 0.00107905 | RCE1 | 9986 |
| ENSG000001 | 98.1005626 | 0.693532476 | 0.18098549 | 3.83197837 | 0.00012712 | 0.00107975 | RELL2 | 285613 |
| ENSG000001 | 20.854351 | -1.104029634 | 0.28810636 | -3.8320211 | 0.00012709 | 0.00107975 | KLHL33 | 123103 |
| ENSG000001 | 1075.3005 | 0.503860629 | 0.13149987 | 3.8316436 | 0.00012729 | 0.00108099 | CHD1L | 9557 |
| ENSG000001 | 19.7387086 | -1.132537368 | 0.29564318 | -3.8307577 | 0.00012775 | 0.00108467 | FSCN3 | 29999 |
| ENSG000001 | 1360.41506 | 0.465823743 | 0.12161682 | 3.83025759 | 0.00012801 | 0.00108664 | CLASP2 | 23122 |
| ENSG000001 | 43.9684875 | -0.864355557 | 0.22567299 | -3.8301241 | 0.00012808 | 0.00108701 | C7 | 730 |
| ENSG000001 | 2932.54013 | 0.483099897 | 0.12614181 | 3.8298159 | 0.00012824 | 0.00108814 | ATP5C1 | 509 |
| ENSG000001 | 1222.71929 | 0.433793482 | 0.11327364 | 3.82960663 | 0.00012835 | 0.00108884 | PRPF3 | 9129 |
| ENSG000001 | 55.9087042 | 0.933794195 | 0.24385282 | 3.82933529 | 0.00012849 | 0.00108981 | TMEM134 | 80194 |
| ENSG000001 | 198.644272 | 0.638841267 | 0.16683192 | 3.82925071 | 0.00012853 | 0.00108996 | DNM1 | 1759 |
| ENSG000001 | 174.748192 | 0.650047091 | 0.16977643 | 3.82884173 | 0.00012875 | 0.00109131 | FAAP100 | 80233 |

| | | | | | | | | |
|---|---|---|---|---|---|---|---|---|
| ENSG000002 | 4.30144583 | -1.448401788 | 0.37828563 | -3.8288575 | 0.00012874 | 0.00109131 | NA | NA |
| ENSG000001 | 405.562654 | 0.462820197 | 0.12088336 | 3.82865087 | 0.00012885 | 0.00109193 | BBS7 | 55212 |
| ENSG000001 | 472.681803 | 0.536653177 | 0.140217 | 3.82730479 | 0.00012955 | 0.00109769 | FZD6 | 8323 |
| ENSG000001 | 2610.73986 | 0.498220159 | 0.1301778 | 3.82722822 | 0.00012959 | 0.0010978 | GULP1 | 51454 |
| ENSG000001 | 2085.78178 | 0.494063893 | 0.12911258 | 3.82661309 | 0.00012992 | 0.00110032 | COPB1 | 1315 |
| ENSG000002 | 91.7315131 | 0.738318554 | 0.19295518 | 3.82637329 | 0.00013005 | 0.00110116 | RN7SKP80 | 106479130 |
| ENSG000001 | 1309.78656 | 0.643539739 | 0.16818825 | 3.82630617 | 0.00013008 | 0.00110123 | USP14 | 9097 |
| ENSG00000C | 1266.54739 | 0.465044464 | 0.121566 | 3.82544829 | 0.00013053 | 0.00110484 | SPDL1 | 54908 |
| ENSG000001 | 166.440802 | 0.548895537 | 0.14348804 | 3.82537492 | 0.00013057 | 0.00110494 | TMEM179B | 374395 |
| ENSG000001 | 3194.83337 | 0.463902556 | 0.12127205 | 3.82530483 | 0.00013061 | 0.00110502 | CHCHD3 | 54927 |
| ENSG000001 | 244.420386 | 0.727674078 | 0.1902384 | 3.82506418 | 0.00013074 | 0.00110587 | MED16 | 10025 |
| ENSG00000C | 9750.87701 | 0.461515236 | 0.1206829 | 3.82419752 | 0.0001312 | 0.00110922 | TNPO1 | 3842 |
| ENSG000001 | 489.373275 | 0.485888182 | 0.12705752 | 3.82415932 | 0.00013122 | 0.00110922 | PDE3B | 5140 |
| ENSG000001 | 729.139762 | 0.534884456 | 0.13986715 | 3.82423222 | 0.00013118 | 0.00110922 | DRAXIN | 374946 |
| ENSG000001 | 106.251311 | 0.67723972 | 0.17709719 | 3.82411323 | 0.00013124 | 0.00110922 | GBA | 2629 |
| ENSG000002 | 179.357414 | 0.561690598 | 0.14688342 | 3.82405723 | 0.00013127 | 0.00110925 | GPX3 | 2878 |
| ENSG00000C | 2118.88583 | 0.579028885 | 0.15141985 | 3.82399592 | 0.00013131 | 0.00110929 | YTHDC1 | 91746 |
| ENSG00000C | 1659.87882 | 0.398343423 | 0.10417954 | 3.82362443 | 0.0001315 | 0.00111073 | OGFOD1 | 55239 |
| ENSG000001 | 2945.35617 | 0.478680512 | 0.12519247 | 3.82355662 | 0.00013154 | 0.00111081 | MIB1 | 57534 |
| ENSG000001 | 38.6974938 | -0.903038158 | 0.23618779 | -3.8233905 | 0.00013163 | 0.00111132 | CNTN4 | 152330 |
| ENSG000001 | 3196.67244 | 0.463862417 | 0.12132708 | 3.82323884 | 0.00013171 | 0.00111178 | TMED10 | 10972 |
| ENSG00000C | 2345.08235 | 0.591706618 | 0.15477845 | 3.8229263 | 0.00013188 | 0.00111274 | TCOF1 | 6949 |
| ENSG000001 | 3581.88898 | 0.530636787 | 0.13880397 | 3.82292226 | 0.00013188 | 0.00111274 | SFRP2 | 6423 |
| ENSG000001 | 1175.53315 | 0.509651361 | 0.13335105 | 3.82187743 | 0.00013244 | 0.001117 | COA7 | 65260 |
| ENSG000001 | 531.617006 | 0.490219224 | 0.12826573 | 3.82190333 | 0.00013243 | 0.001117 | POLR3D | 661 |
| ENSG000002 | 33.8654032 | 0.936585335 | 0.24508219 | 3.82151522 | 0.00013263 | 0.00111841 | NA | NA |
| ENSG000001 | 837.354582 | 0.498793972 | 0.13053399 | 3.82118076 | 0.00013281 | 0.0011197 | ZNF317 | 57693 |
| ENSG00000C | 10414.0882 | 0.485147162 | 0.12697419 | 3.820833 | 0.000133 | 0.00112081 | GDI2 | 2665 |
| ENSG000001 | 1003.98819 | 0.514509407 | 0.1346588 | 3.8208375 | 0.000133 | 0.00112081 | MRPL47 | 57129 |

| | | | | | | | | |
|---|---|---|---|---|---|---|---|---|
| ENSG000001 | 107.285831 | 0.682905769 | 0.17873707 | 3.82072823 | 0.00013306 | 0.00112106 | IGDCC4 | 57722 |
| ENSG000001 | 1813.06817 | 0.478736474 | 0.12530633 | 3.82052907 | 0.00013317 | 0.00112173 | IMMT | 10989 |
| ENSG000001 | 30.1038207 | -1.011459416 | 0.26476661 | -3.8201925 | 0.00013335 | 0.00112303 | C1QTNF7 | 114905 |
| ENSG000002 | 24.6796056 | -1.036828323 | 0.27141891 | -3.8200298 | 0.00013344 | 0.00112354 | NA | NA |
| ENSG000000 | 538.0246 | 0.547508205 | 0.14332929 | 3.81993234 | 0.00013349 | 0.00112375 | ASAP3 | 55616 |
| ENSG000000 | 17.9885232 | -1.127592683 | 0.29521491 | -3.8195655 | 0.00013369 | 0.00112518 | CHRD | 8646 |
| ENSG000001 | 92.2146764 | -0.70331261 | 0.18414083 | -3.8194278 | 0.00013376 | 0.00112558 | FAM19A2 | 338811 |
| ENSG000002 | 17.2215463 | -1.155004672 | 0.30244199 | -3.8189297 | 0.00013403 | 0.00112762 | NA | NA |
| ENSG000000 | 381.318126 | 0.514841542 | 0.13482535 | 3.81858118 | 0.00013422 | 0.00112898 | SH3YL1 | 26751 |
| ENSG000000 | 2499.14234 | 0.432508436 | 0.11326609 | 3.81851644 | 0.00013426 | 0.00112904 | KIF2A | 3796 |
| ENSG000001 | 1600.8975 | 0.434079562 | 0.11368194 | 3.81836886 | 0.00013434 | 0.00112948 | FARSB | 10056 |
| ENSG000001 | 55.6187354 | -0.78846185 | 0.20649891 | -3.8182373 | 0.00013441 | 0.00112985 | SLC28A2 | 9153 |
| ENSG000002 | 203.715974 | 0.711765464 | 0.18643694 | 3.81772756 | 0.00013469 | 0.00113195 | SNORD3B-1 | 26851 |
| ENSG000000 | 370.238279 | 0.553838148 | 0.14508287 | 3.81739179 | 0.00013487 | 0.0011332 | SS18L2 | 51188 |
| ENSG000001 | 86.1404157 | 0.725862151 | 0.19014799 | 3.81735375 | 0.00013489 | 0.0011332 | SLC39A13 | 91252 |
| ENSG000001 | 81.073902 | -0.688843728 | 0.18047556 | -3.8168255 | 0.00013518 | 0.00113539 | GSAP | 54103 |
| ENSG000001 | 236.381336 | 0.684036908 | 0.17922329 | 3.81667412 | 0.00013526 | 0.00113585 | ADAM15 | 8751 |
| ENSG000001 | 546.119557 | 0.564183605 | 0.14783392 | 3.81633391 | 0.00013545 | 0.00113718 | PLAGL2 | 5326 |
| ENSG000002 | 29.7648416 | 0.957351816 | 0.25087535 | 3.81604575 | 0.00013561 | 0.00113828 | NA | NA |
| ENSG000002 | 20.9882085 | -1.07311795 | 0.28121626 | -3.8159882 | 0.00013564 | 0.00113831 | DCDC2C | 728597 |
| ENSG000001 | 876.996781 | 0.522111554 | 0.13683044 | 3.81575582 | 0.00013577 | 0.00113914 | C14orf119 | 55017 |
| ENSG000001 | 722.069102 | 0.501924254 | 0.1315515 | 3.81542028 | 0.00013595 | 0.00114017 | ZDHHC2 | 51201 |
| ENSG000001 | 25.9353728 | -1.039450184 | 0.27243282 | -3.8154367 | 0.00013594 | 0.00114017 | IL1RL1 | 9173 |
| ENSG000001 | 5750.98314 | 0.425557679 | 0.11153744 | 3.81537956 | 0.00013597 | 0.00114017 | NAA50 | 80218 |
| ENSG000000 | 2234.2848 | 0.598600991 | 0.15690451 | 3.81506544 | 0.00013615 | 0.00114139 | GTF3C1 | 2975 |
| ENSG000002 | 2090.2083 | 0.611924077 | 0.16041976 | 3.81451799 | 0.00013645 | 0.00114368 | ABCF1 | 23 |
| ENSG000001 | 278.219144 | 0.537427476 | 0.14089385 | 3.81441396 | 0.00013651 | 0.00114393 | OPA3 | 80207 |
| ENSG000001 | 1908.33232 | 0.416766987 | 0.10927039 | 3.81408888 | 0.00013669 | 0.0011452 | STK24 | 8428 |
| ENSG000000 | 361.651244 | 0.503224704 | 0.1319488 | 3.81378752 | 0.00013685 | 0.00114636 | FNIP2 | 57600 |

| | | | | | | | | |
|---|---|---|---|---|---|---|---|---|
| ENSG000001 | 970.385026 | 0.464554603 | 0.12181412 | 3.81363502 | 0.00013694 | 0.00114683 | SIX4 | 51804 |
| ENSG000001 | 364.530436 | 0.612198493 | 0.16053771 | 3.81342494 | 0.00013705 | 0.00114757 | SH3KBP1 | 30011 |
| ENSG000001 | 1016.5559 | 0.483270123 | 0.12673383 | 3.8132685 | 0.00013714 | 0.00114792 | DNMBP | 23268 |
| ENSG000001 | 166.353723 | 0.580775127 | 0.1523046 | 3.81324748 | 0.00013715 | 0.00114792 | CXorf38 | 159013 |
| ENSG000002 | 9.21545491 | -1.339716175 | 0.35140759 | -3.8124281 | 0.00013761 | 0.0011515 | NA | NA |
| ENSG000002 | 51.5940233 | 0.843744919 | 0.22132873 | 3.81217987 | 0.00013775 | 0.00115242 | TMEM141 | 85014 |
| ENSG000001 | 21.9847019 | -1.084584243 | 0.28459247 | -3.8110082 | 0.0001384 | 0.00115765 | LAX1 | 54900 |
| ENSG000001 | 1786.22142 | 0.461794452 | 0.12117942 | 3.81083245 | 0.0001385 | 0.00115824 | HNRNPH2 | 3188 |
| ENSG000001 | 582.435747 | 0.480848126 | 0.12619895 | 3.81023883 | 0.00013883 | 0.00116078 | C14orf1 | 11161 |
| ENSG000002 | 358.587662 | 0.5069832 | 0.13306529 | 3.81003333 | 0.00013895 | 0.00116151 | MYCBP | 26292 |
| ENSG000001 | 1994.54943 | 0.508651723 | 0.13351113 | 3.80980753 | 0.00013908 | 0.00116233 | MMADHC | 27249 |
| ENSG000001 | 542.543728 | 0.466651164 | 0.12249049 | 3.80969292 | 0.00013914 | 0.00116263 | MINA | 84864 |
| ENSG000001 | 252.455797 | 0.596646721 | 0.15662802 | 3.80932291 | 0.00013935 | 0.00116413 | TACO1 | 51204 |
| ENSG00000C | 1263.33267 | 0.622926123 | 0.16353027 | 3.80924036 | 0.00013939 | 0.00116428 | ABL1 | 25 |
| ENSG00000C | 43.6382408 | -0.84455511 | 0.22173533 | -3.8088433 | 0.00013962 | 0.00116519 | MSR1 | 4481 |
| ENSG00000C | 361.697195 | 0.584871475 | 0.15355463 | 3.80888211 | 0.0001396 | 0.00116519 | PLEKHG2 | 64857 |
| ENSG000001 | 326.252248 | 0.519685801 | 0.13643779 | 3.80895793 | 0.00013955 | 0.00116519 | RHBDD1 | 84236 |
| ENSG000001 | 420.402437 | 0.604592001 | 0.15873212 | 3.80888246 | 0.0001396 | 0.00116519 | SDC3 | 9672 |
| ENSG00000C | 460.892106 | 0.492078613 | 0.12924653 | 3.80728672 | 0.0001405 | 0.00117189 | ISOC1 | 51015 |
| ENSG00000C | 1264.76951 | 0.594764619 | 0.15621397 | 3.80737144 | 0.00014045 | 0.00117189 | NUDC | 10726 |
| ENSG000001 | 201.502238 | 0.633101974 | 0.16628975 | 3.80722192 | 0.00014054 | 0.00117189 | FKRP | 79147 |
| ENSG000001 | 6973.36437 | 0.427512799 | 0.11228931 | 3.80724384 | 0.00014052 | 0.00117189 | LIN28B | 389421 |
| ENSG000001 | 464.971369 | 0.488275464 | 0.12826746 | 3.80669793 | 0.00014083 | 0.00117413 | ERI2 | 112479 |
| ENSG000001 | 195.820046 | 0.607748955 | 0.15967237 | 3.80622491 | 0.0001411 | 0.00117614 | SMUG1 | 23583 |
| ENSG000001 | 2592.25392 | 0.412691248 | 0.10843423 | 3.805913 | 0.00014128 | 0.00117738 | FAM208B | 54906 |
| ENSG000001 | 84.7747414 | 0.72092137 | 0.18942604 | 3.8058198 | 0.00014134 | 0.00117758 | RNF135 | 84282 |
| ENSG00000C | 1742.91809 | 0.486500896 | 0.12784432 | 3.80541658 | 0.00014157 | 0.00117926 | ZFX | 7543 |
| ENSG000001 | 722.021061 | 0.532563167 | 0.13995069 | 3.8053628 | 0.0001416 | 0.00117927 | GNG5 | 2787 |
| ENSG00000C | 64.386805 | -0.751534015 | 0.19750941 | -3.8050543 | 0.00014177 | 0.0011805 | ABCC9 | 10060 |

| | | | | | | | | |
|---|---|---|---|---|---|---|---|---|
| ENSG000001 | 1050.53479 | 0.410530678 | 0.10789422 | 3.80493659 | 0.00014184 | 0.00118082 | PDLIM5 | 10611 |
| ENSG000001 | 966.569931 | 0.531543526 | 0.13970611 | 3.80472627 | 0.00014196 | 0.00118158 | RAB2A | 5862 |
| ENSG000001 | 720.718488 | 0.488960217 | 0.12854024 | 3.80394666 | 0.00014241 | 0.00118506 | CSRP2BP | 57325 |
| ENSG000001 | 338.021077 | 0.556567457 | 0.1463265 | 3.80359987 | 0.00014261 | 0.00118648 | PIAS4 | 51588 |
| ENSG00000C | 2827.48614 | 0.507354944 | 0.13339533 | 3.80339355 | 0.00014273 | 0.00118723 | ELOVL5 | 60481 |
| ENSG000001 | 116.272148 | -0.623936548 | 0.16405369 | -3.803246 | 0.00014281 | 0.00118769 | DNAH10 | 196385 |
| ENSG00000C | 657.109168 | 0.443679594 | 0.11666143 | 3.80313853 | 0.00014287 | 0.00118796 | MEF2A | 4205 |
| ENSG00000C | 1110.98504 | 0.491483207 | 0.12924462 | 3.80273635 | 0.00014311 | 0.00118965 | ABCF2 | 10061 |
| ENSG000002 | 26.392425 | -1.006876105 | 0.26478199 | -3.8026608 | 0.00014315 | 0.00118977 | NA | NA |
| ENSG000001 | 30.7813128 | -0.941412091 | 0.24763005 | -3.8016875 | 0.00014371 | 0.00119421 | TMPRSS4 | 56649 |
| ENSG000001 | 604.656105 | 0.479939891 | 0.12624841 | 3.8015521 | 0.00014379 | 0.00119462 | ADI1 | 55256 |
| ENSG000001 | 118.347359 | 0.606404479 | 0.15952205 | 3.80138352 | 0.00014389 | 0.00119519 | FAM219A | 203259 |
| ENSG000002 | 64.5465968 | 0.740585468 | 0.1948302 | 3.80118424 | 0.00014401 | 0.0011959 | NA | NA |
| ENSG000001 | 208.171172 | 0.700009064 | 0.18417392 | 3.8008044 | 0.00014423 | 0.00119749 | FBXW4 | 6468 |
| ENSG00000C | 48.7875398 | -0.837764921 | 0.22043185 | -3.8005621 | 0.00014437 | 0.00119842 | RPH3A | 22895 |
| ENSG000001 | 108.10079 | 0.646664277 | 0.17015879 | 3.80035766 | 0.00014449 | 0.00119916 | LAGE3 | 8270 |
| ENSG000001 | 523.650882 | 0.502136874 | 0.13213531 | 3.8001718 | 0.0001446 | 0.00119982 | RAP1GDS1 | 5910 |
| ENSG000001 | 446.780542 | 0.487212087 | 0.12821561 | 3.79994366 | 0.00014473 | 0.00120068 | GTF2E2 | 2961 |
| ENSG000001 | 1334.11462 | 0.451170439 | 0.11873561 | 3.79979048 | 0.00014482 | 0.00120117 | RLF | 6018 |
| ENSG000001 | 152.305643 | -0.611440599 | 0.16092346 | -3.7995741 | 0.00014494 | 0.00120198 | ZEB2 | 9839 |
| ENSG000001 | 57.9753701 | -0.766779206 | 0.20181386 | -3.7994377 | 0.00014502 | 0.00120239 | AMICA1 | 120425 |
| ENSG000001 | 979.712243 | 0.491176101 | 0.12928439 | 3.79919106 | 0.00014517 | 0.00120334 | IMP4 | 92856 |
| ENSG000001 | 3630.5793 | 0.478915783 | 0.12606357 | 3.79900228 | 0.00014528 | 0.00120401 | PAWR | 5074 |
| ENSG000001 | 278.694814 | 0.578033456 | 0.1521871 | 3.79817639 | 0.00014576 | 0.00120778 | FHL2 | 2274 |
| ENSG000001 | 186.201867 | 0.574623319 | 0.15129714 | 3.79797862 | 0.00014588 | 0.0012085 | TBC1D13 | 54662 |
| ENSG00000C | 120.25412 | 0.656202721 | 0.17278808 | 3.79773143 | 0.00014603 | 0.00120946 | PLEKHH3 | 79990 |
| ENSG000001 | 34.9581608 | 0.982190581 | 0.25863492 | 3.79759465 | 0.00014611 | 0.00120988 | SPATA2L | 124044 |
| ENSG000001 | 218.596363 | 0.576995913 | 0.15194792 | 3.7973269 | 0.00014626 | 0.00121094 | NRIP3 | 56675 |
| ENSG000001 | 365.71233 | 0.475741046 | 0.1252882 | 3.79717375 | 0.00014636 | 0.00121144 | TCAIM | 285343 |

| | | | | | | | | | |
|---|---|---|---|---|---|---|---|---|---|
| ENSG000001 | 212.498218 | 0.578116689 | 0.15226986 | 3.79665866 | 0.00014666 | 0.00121371 | METTL4 | | 64863 |
| ENSG00000C | 110.810429 | 0.634400086 | 0.16709823 | 3.79656982 | 0.00014671 | 0.0012139 | RRAGB | | 10325 |
| ENSG000001 | 366.45341 | 0.545497325 | 0.14370246 | 3.79601943 | 0.00014704 | 0.00121635 | SEMA6B | | 10501 |
| ENSG000001 | 11930.5691 | 0.489479032 | 0.12895601 | 3.79570554 | 0.00014722 | 0.00121764 | RAN | | 5901 |
| ENSG000001 | 199.269345 | 0.657436298 | 0.17320774 | 3.7956519 | 0.00014726 | 0.00121765 | DDX51 | | 317781 |
| ENSG000001 | 393.03122 | 0.461695032 | 0.12164853 | 3.79531931 | 0.00014745 | 0.00121904 | GIT2 | | 9815 |
| ENSG000002 | 11.1980853 | -1.28042303 | 0.33741212 | -3.7948341 | 0.00014774 | 0.00122118 | PPIAL4G | | 644591 |
| ENSG000001 | 1333.70166 | 0.51658915 | 0.13613965 | 3.79455323 | 0.00014791 | 0.00122231 | CCNJ | | 54619 |
| ENSG000001 | 1727.81879 | 0.484894951 | 0.12779931 | 3.79419067 | 0.00014813 | 0.00122385 | SNRPB2 | | 6629 |
| ENSG000001 | 1509.4869 | 0.519691114 | 0.13698246 | 3.79385155 | 0.00014833 | 0.00122502 | LDLR | | 3949 |
| ENSG000002 | 472.700039 | 0.496013516 | 0.13074015 | 3.79388822 | 0.00014831 | 0.00122502 | GATC | | 283459 |
| ENSG000001 | 2050.32919 | 0.444771911 | 0.11723757 | 3.79376619 | 0.00014838 | 0.00122508 | CTDSPL2 | | 51496 |
| ENSG000002 | 29.0878999 | -0.990360294 | 0.26105129 | -3.7937384 | 0.0001484 | 0.00122508 | NA | NA | |
| ENSG000002 | 27.7610224 | 1.012121407 | 0.26682585 | 3.793191 | 0.00014872 | 0.00122754 | NA | NA | |
| ENSG000001 | 39.7811435 | -0.925852809 | 0.24409843 | -3.7929487 | 0.00014887 | 0.00122849 | COMMD8 | | 54951 |
| ENSG00000C | 4138.03518 | 0.492260744 | 0.12978613 | 3.79286104 | 0.00014892 | 0.00122867 | STRAP | | 11171 |
| ENSG000001 | 46.9377745 | -0.826916898 | 0.21803104 | -3.7926567 | 0.00014904 | 0.00122918 | FRMPD2 | | 143162 |
| ENSG000002 | 172.476112 | 0.560226218 | 0.14771192 | 3.79269482 | 0.00014902 | 0.00122918 | NA | NA | |
| ENSG000001 | 20.2870409 | -1.090638018 | 0.28758555 | -3.792395 | 0.0001492 | 0.00123023 | ACSM6 | | 142827 |
| ENSG000001 | 422.751911 | 0.494171247 | 0.13031003 | 3.79227336 | 0.00014927 | 0.00123058 | TMTC4 | | 84899 |
| ENSG00000C | 35.490937 | -0.889094192 | 0.23447617 | -3.7918318 | 0.00014954 | 0.00123252 | IYD | | 389434 |
| ENSG000001 | 166.068492 | 0.581595662 | 0.15339856 | 3.79140236 | 0.0001498 | 0.0012344 | PNPO | | 55163 |
| ENSG000001 | 1408.28708 | 0.415174158 | 0.10951043 | 3.79118365 | 0.00014993 | 0.00123516 | ZBTB1 | | 22890 |
| ENSG000001 | 1738.56571 | 0.480689078 | 0.12679244 | 3.79114938 | 0.00014995 | 0.00123516 | KIAA1143 | | 57456 |
| ENSG000001 | 977.712122 | 0.406240292 | 0.10715647 | 3.79109452 | 0.00014998 | 0.00123518 | NDFIP1 | | 80762 |
| ENSG000001 | 494.786083 | 0.708409355 | 0.18687737 | 3.79077116 | 0.00015018 | 0.00123654 | PLXNB1 | | 5364 |
| ENSG000001 | 194.240593 | 0.608589972 | 0.16055111 | 3.79063065 | 0.00015027 | 0.00123699 | C5orf45 | | 51149 |
| ENSG000002 | 11.1897417 | -1.30242965 | 0.34360043 | -3.7905356 | 0.00015032 | 0.00123721 | NA | NA | |
| ENSG000001 | 1140.14309 | 0.552497169 | 0.14576396 | 3.79035498 | 0.00015043 | 0.00123786 | RBM10 | | 8241 |

| | | | | | | | | |
|---|---|---|---|---|---|---|---|---|
| ENSG000001 | 2175.53533 | 0.413017569 | 0.10897156 | 3.79014091 | 0.00015056 | 0.00123844 | DMTF1 | 9988 |
| ENSG000001 | 380.064972 | 0.490385182 | 0.12938454 | 3.79013726 | 0.00015056 | 0.00123844 | ZNF607 | 84775 |
| ENSG000001 | 645.245865 | 0.520299677 | 0.13731429 | 3.7891152 | 0.00015118 | 0.0012433 | RTCA | 8634 |
| ENSG00000C | 19.0096014 | -1.114984102 | 0.29427116 | -3.7889683 | 0.00015127 | 0.00124353 | SLAMF7 | 57823 |
| ENSG000002 | 710.292722 | 0.471404045 | 0.12441325 | 3.78901797 | 0.00015124 | 0.00124353 | ARHGAP19 | 84986 |
| ENSG00000C | 115.160717 | 0.690092584 | 0.18214476 | 3.78870409 | 0.00015144 | 0.0012446 | SYT7 | 9066 |
| ENSG000001 | 146.842283 | 0.641676083 | 0.16939542 | 3.78803679 | 0.00015184 | 0.00124769 | C21orf58 | 54058 |
| ENSG000001 | 55.8487281 | 0.856314913 | 0.22606351 | 3.78793959 | 0.0001519 | 0.00124793 | THAP8 | 199745 |
| ENSG000001 | 115.25881 | 0.717545019 | 0.18943441 | 3.78782823 | 0.00015197 | 0.00124823 | NDOR1 | 27158 |
| ENSG000002 | 253.47108 | 0.507248759 | 0.13391768 | 3.78776547 | 0.00015201 | 0.0012483 | SYS1 | 90196 |
| ENSG000002 | 20.510525 | -1.100503601 | 0.29055209 | -3.7876292 | 0.00015209 | 0.00124873 | C1orf141 | 400757 |
| ENSG000001 | 2233.64902 | 0.525158239 | 0.13865277 | 3.78757841 | 0.00015212 | 0.00124873 | UBXN4 | 23190 |
| ENSG000001 | 2341.7672 | 0.421119147 | 0.11118869 | 3.78742801 | 0.00015221 | 0.00124923 | KIF23 | 9493 |
| ENSG000002 | 352.636078 | 0.544417229 | 0.14375578 | 3.78709807 | 0.00015242 | 0.00125064 | NA | NA |
| ENSG000001 | 2992.47374 | 0.517307915 | 0.13662146 | 3.7864324 | 0.00015283 | 0.00125374 | GMFB | 2764 |
| ENSG00000C | 579.370803 | 0.454364578 | 0.1200104 | 3.78604347 | 0.00015306 | 0.00125519 | HEATR6 | 63897 |
| ENSG000001 | 492.456493 | 0.496573739 | 0.13115752 | 3.7860868 | 0.00015304 | 0.00125519 | ZNF215 | 7762 |
| ENSG000001 | 1180.45756 | 0.468950335 | 0.12386483 | 3.78598463 | 0.0001531 | 0.00125524 | EXOSC2 | 23404 |
| ENSG000001 | 1327.28345 | 0.501211421 | 0.13239953 | 3.78559808 | 0.00015334 | 0.00125694 | USP46 | 64854 |
| ENSG00000C | 1121.77291 | 0.426175799 | 0.11258107 | 3.78550146 | 0.0001534 | 0.00125717 | FAM135A | 57579 |
| ENSG000001 | 644.198229 | 0.480612578 | 0.12696704 | 3.78533353 | 0.0001535 | 0.00125777 | OLFML3 | 56944 |
| ENSG000001 | 257.485309 | 0.684718748 | 0.1808994 | 3.78508017 | 0.00015366 | 0.00125879 | ZHX2 | 22882 |
| ENSG00000C | 1945.78948 | 0.662500758 | 0.1750415 | 3.78482107 | 0.00015382 | 0.00125985 | KHSRP | 8570 |
| ENSG000001 | 814.628226 | 0.528584185 | 0.13966119 | 3.78476075 | 0.00015386 | 0.0012599 | MRPL15 | 29088 |
| ENSG000001 | 328.660128 | 0.536233259 | 0.14171781 | 3.78380982 | 0.00015445 | 0.00126447 | VIMP | 55829 |
| ENSG00000C | 3217.46393 | 0.566715804 | 0.14979377 | 3.78330688 | 0.00015476 | 0.00126678 | MCM6 | 4175 |
| ENSG00000C | 558.008674 | 0.467774748 | 0.12365702 | 3.78284025 | 0.00015505 | 0.0012689 | RUFY3 | 22902 |
| ENSG000001 | 11.1389277 | -1.295264714 | 0.34242482 | -3.7826251 | 0.00015518 | 0.00126974 | SYCE1 | 93426 |
| ENSG000001 | 140.633346 | 0.575264475 | 0.15209994 | 3.782148 | 0.00015548 | 0.00127192 | MX1 | 4599 |

| | | | | | | | | | |
|---|---|---|---|---|---|---|---|---|---|
| ENSG000001 | 1224.24041 | 0.470901544 | 0.1245123 | 3.78196794 | 0.00015559 | 0.00127233 | NPAT | | 4863 |
| ENSG000001 | 211.386547 | 0.609975769 | 0.16128491 | 3.78197673 | 0.00015559 | 0.00127233 | DHRS13 | | 147015 |
| ENSG000001 | 576.424822 | 0.534341552 | 0.14130759 | 3.78140721 | 0.00015594 | 0.00127443 | SUGP1 | | 57794 |
| ENSG000001 | 572.297827 | 0.570284877 | 0.15081198 | 3.7814295 | 0.00015593 | 0.00127443 | TRIP10 | | 9322 |
| ENSG000001 | 24.3032467 | 1.085785815 | 0.28713702 | 3.78142049 | 0.00015594 | 0.00127443 | SNORD104 | | 692227 |
| ENSG000001 | 697.619649 | 0.427366677 | 0.11303862 | 3.78071391 | 0.00015638 | 0.00127747 | TATDN1 | | 83940 |
| ENSG000001 | 196.324158 | 0.53035152 | 0.14027706 | 3.78074299 | 0.00015636 | 0.00127747 | FAM221A | | 340277 |
| ENSG000002 | 142.93255 | 0.564259861 | 0.14925707 | 3.78045654 | 0.00015654 | 0.00127853 | NA | NA | |
| ENSG00000C | 620.320509 | 0.485201628 | 0.12835016 | 3.78029619 | 0.00015664 | 0.0012791 | HEXB | | 3074 |
| ENSG00000C | 2383.31749 | 0.463310228 | 0.12256299 | 3.78018059 | 0.00015671 | 0.00127917 | TRPM7 | | 54822 |
| ENSG000001 | 2178.40788 | 0.487601115 | 0.12898754 | 3.78021879 | 0.00015669 | 0.00127917 | ZNF589 | | 51385 |
| ENSG00000C | 1004.9893 | 0.457320838 | 0.12099009 | 3.77982059 | 0.00015694 | 0.00128065 | PSEN1 | | 5663 |
| ENSG000001 | 125.811413 | 0.652412759 | 0.1726054 | 3.77979338 | 0.00015696 | 0.00128065 | TPRN | | 286262 |
| ENSG000002 | 114.292684 | 0.649559021 | 0.17185336 | 3.77972831 | 0.000157 | 0.00128073 | NA | NA | |
| ENSG000001 | 728.37832 | 0.432034356 | 0.1143133 | 3.77938844 | 0.00015721 | 0.00128222 | CD55 | | 1604 |
| ENSG000001 | 460.578719 | 0.489874234 | 0.12962303 | 3.77922236 | 0.00015732 | 0.00128281 | CTDSPL | | 10217 |
| ENSG000001 | 59.7439763 | -0.75167852 | 0.19890792 | -3.7790277 | 0.00015744 | 0.00128356 | TPRG1 | | 285386 |
| ENSG000001 | 167.847439 | 0.604213887 | 0.15989467 | 3.77882443 | 0.00015757 | 0.00128435 | PARS2 | | 25973 |
| ENSG000001 | 225.896555 | 0.599334598 | 0.15862294 | 3.77836009 | 0.00015786 | 0.0012863 | DBF4B | | 80174 |
| ENSG000002 | 25.2919612 | -1.04863594 | 0.27753829 | -3.7783469 | 0.00015787 | 0.0012863 | NA | NA | |
| ENSG000001 | 121.450152 | 0.615324554 | 0.16286964 | 3.77801877 | 0.00015808 | 0.00128773 | IL27RA | | 9466 |
| ENSG000001 | 477.01986 | 0.48769418 | 0.12909012 | 3.77793566 | 0.00015813 | 0.00128791 | SPOP | | 8405 |
| ENSG000002 | 24.2472097 | -1.045729311 | 0.27683252 | -3.77748 | 0.00015842 | 0.00129 | C1orf186 | | 440712 |
| ENSG000001 | 56.9806665 | -0.781108573 | 0.20679862 | -3.7771459 | 0.00015864 | 0.00129125 | COL4A3 | | 1285 |
| ENSG000002 | 139.984527 | 0.65536186 | 0.17350747 | 3.77713908 | 0.00015864 | 0.00129125 | NA | NA | |
| ENSG000001 | 400.033251 | 0.480670885 | 0.12726829 | 3.77683162 | 0.00015884 | 0.00129233 | CUTC | | 51076 |
| ENSG000001 | 779.119445 | 0.571383944 | 0.15128488 | 3.77687416 | 0.00015881 | 0.00129233 | FOXK2 | | 3607 |
| ENSG000001 | 515.823933 | 0.678407447 | 0.1796376 | 3.77653361 | 0.00015903 | 0.00129361 | MAP1LC3B | | 81631 |
| ENSG000001 | 890.037198 | 0.525382918 | 0.13914657 | 3.77575175 | 0.00015953 | 0.00129742 | ZSCAN2 | | 54993 |

| | | | | | | | | |
|---|---|---|---|---|---|---|---|---|
| ENSG000001 | 25.7838542 | -1.001077509 | 0.26521143 | -3.7746393 | 0.00016024 | 0.00130296 | TMPRSS15 | 5651 |
| ENSG000001 | 2249.87126 | -0.440246945 | 0.11663505 | -3.774568 | 0.00016029 | 0.00130307 | TTN | 7273 |
| ENSG000002 | 25.8816949 | -1.041352731 | 0.27593876 | -3.7738545 | 0.00016074 | 0.00130654 | REXO1L1P | 254958 |
| ENSG000001 | 833.842781 | 0.439839762 | 0.1165528 | 3.77373814 | 0.00016082 | 0.00130689 | NDUFV3 | 4731 |
| ENSG000001 | 28.1777162 | -1.003493028 | 0.26592384 | -3.7736107 | 0.0001609 | 0.0013073 | TDRD6 | 221400 |
| ENSG000001 | 651.807431 | 0.438860716 | 0.11630014 | 3.77351839 | 0.00016096 | 0.00130752 | ZKSCAN2 | 342357 |
| ENSG00000C | 1092.58791 | 0.458179418 | 0.12142239 | 3.77343439 | 0.00016102 | 0.0013077 | UTP18 | 51096 |
| ENSG000001 | 431.244738 | 0.516892572 | 0.13698536 | 3.7733417 | 0.00016108 | 0.00130792 | TRAFD1 | 10906 |
| ENSG000001 | 406.320037 | 0.566110084 | 0.15004913 | 3.77283152 | 0.00016141 | 0.00131034 | AXIN1 | 8312 |
| ENSG000001 | 33.1102177 | -0.915691768 | 0.24271318 | -3.772732 | 0.00016147 | 0.0013106 | P4HA3 | 283208 |
| ENSG000001 | 599.056375 | 0.513707191 | 0.13617225 | 3.77248088 | 0.00016163 | 0.0013114 | PIK3R1 | 5295 |
| ENSG000002 | 8.08141867 | -1.356465083 | 0.35956451 | -3.7725221 | 0.00016161 | 0.0013114 | SDR16C6P | 442388 |
| ENSG00000C | 54.3171375 | -0.857279597 | 0.22725874 | -3.7722623 | 0.00016177 | 0.00131228 | NA | NA |
| ENSG000002 | 30.3792728 | -0.968852877 | 0.25687692 | -3.7716619 | 0.00016216 | 0.00131518 | NA | NA |
| ENSG000001 | 23.0557275 | -1.031549169 | 0.27351345 | -3.7714751 | 0.00016229 | 0.0013159 | NA | NA |
| ENSG000001 | 4234.17908 | 0.465981758 | 0.12358588 | 3.77050967 | 0.00016291 | 0.00132048 | GSPT1 | 2935 |
| ENSG000001 | 194.365958 | 0.532476285 | 0.14122003 | 3.77054361 | 0.00016289 | 0.00132048 | BCAS4 | 55653 |
| ENSG000001 | 35.278978 | -0.903600035 | 0.23965747 | -3.7703812 | 0.000163 | 0.00132089 | NA | NA |
| ENSG000001 | 1185.97357 | 0.47829523 | 0.12686227 | 3.77019279 | 0.00016312 | 0.00132163 | SSX2IP | 117178 |
| ENSG00000C | 102.925618 | 0.64266697 | 0.1704729 | 3.76990684 | 0.00016331 | 0.00132288 | TESK2 | 10420 |
| ENSG000001 | 815.917139 | 0.520359415 | 0.13803525 | 3.76975742 | 0.00016341 | 0.00132302 | TBC1D4 | 9882 |
| ENSG000001 | 978.676475 | 0.460240025 | 0.12208675 | 3.76977866 | 0.00016339 | 0.00132302 | LARP7 | 51574 |
| ENSG000001 | 467.436999 | 0.600397611 | 0.15926807 | 3.76973004 | 0.00016342 | 0.00132302 | RRS1 | 23212 |
| ENSG000001 | 609.860436 | 0.590368911 | 0.15665876 | 3.7685024 | 0.00016423 | 0.00132928 | DCAF5 | 8816 |
| ENSG000001 | 6131.43677 | 0.523171217 | 0.13883131 | 3.76839491 | 0.0001643 | 0.00132932 | APP | 351 |
| ENSG000001 | 192.725116 | 0.569533236 | 0.15113265 | 3.76843287 | 0.00016428 | 0.00132932 | CCDC69 | 26112 |
| ENSG000001 | 392.411462 | 0.477811686 | 0.1267992 | 3.76825461 | 0.00016439 | 0.0013298 | ARMCX1 | 51309 |
| ENSG000001 | 875.712335 | 0.487311234 | 0.12932757 | 3.76803839 | 0.00016454 | 0.00133018 | SCARB2 | 950 |
| ENSG000001 | 17507.4806 | 0.430000227 | 0.11411791 | 3.76803465 | 0.00016454 | 0.00133018 | NAP1L1 | 4673 |

| | | | | | | | | |
|---|---|---|---|---|---|---|---|---|
| ENSG000002 | 2786.37249 | 0.493047152 | 0.13084831 | 3.76808181 | 0.00016451 | 0.00133018 | NCOA4 | 8031 |
| ENSG000002 | 3259.67935 | 0.446128877 | 0.11840631 | 3.76777962 | 0.00016471 | 0.00133127 | SCARNA7 | 677767 |
| ENSG000001 | 475.843418 | 0.468638142 | 0.12438464 | 3.76765286 | 0.00016479 | 0.00133168 | CYYR1 | 116159 |
| ENSG000001 | 1686.35401 | 0.507469143 | 0.13473087 | 3.76653942 | 0.00016553 | 0.00133737 | GLYR1 | 84656 |
| ENSG00000C | 152.035772 | 0.644472151 | 0.17110944 | 3.76643257 | 0.0001656 | 0.00133767 | TMEM101 | 84336 |
| ENSG00000C | 170.618055 | 0.610416203 | 0.16207103 | 3.76634987 | 0.00016565 | 0.00133785 | SBNO2 | 22904 |
| ENSG000001 | 225.999897 | 0.5628001 | 0.14944586 | 3.76591303 | 0.00016594 | 0.00133966 | C14orf37 | 145407 |
| ENSG000002 | 270.629044 | 0.621099901 | 0.16492677 | 3.76591311 | 0.00016594 | 0.00133966 | NA | NA |
| ENSG000002 | 26.5394028 | -1.020480311 | 0.2710109 | -3.7654586 | 0.00016624 | 0.00134183 | NA | NA |
| ENSG000001 | 224.066054 | 0.5497194 | 0.14599767 | 3.76526143 | 0.00016637 | 0.00134262 | GPN2 | 54707 |
| ENSG000001 | 584.498626 | 0.634044628 | 0.16840596 | 3.76497728 | 0.00016656 | 0.00134388 | IVD | 3712 |
| ENSG000001 | 377.533075 | 0.621389905 | 0.16505568 | 3.76472891 | 0.00016673 | 0.00134495 | CYB5R3 | 1727 |
| ENSG000001 | 142.714332 | 0.60249326 | 0.1600488 | 3.76443469 | 0.00016693 | 0.00134607 | ING4 | 51147 |
| ENSG000001 | 954.924439 | 0.414571974 | 0.11012966 | 3.76439876 | 0.00016695 | 0.00134607 | BTBD3 | 22903 |
| ENSG000001 | 144.773281 | 0.557385883 | 0.14806876 | 3.76437187 | 0.00016697 | 0.00134607 | SCARA3 | 51435 |
| ENSG000001 | 114.69345 | -0.755879374 | 0.20080839 | -3.7641823 | 0.00016709 | 0.00134656 | DNAH2 | 146754 |
| ENSG000001 | 718.083025 | 0.433723198 | 0.11522269 | 3.76421705 | 0.00016707 | 0.00134656 | PTPN1 | 5770 |
| ENSG000002 | 458.680444 | 0.560851142 | 0.1490185 | 3.76363423 | 0.00016746 | 0.00134915 | HLA-DPB2 | 3116 |
| ENSG000002 | 18.2543955 | -1.093175977 | 0.29046006 | -3.7636017 | 0.00016748 | 0.00134915 | AKR1B15 | 441282 |
| ENSG000001 | 544.254885 | 0.514030847 | 0.13658196 | 3.76353402 | 0.00016753 | 0.00134925 | PDK1 | 5163 |
| ENSG000001 | 760.752084 | 0.458860207 | 0.12193771 | 3.76307056 | 0.00016784 | 0.00135148 | DPF2 | 5977 |
| ENSG00000C | 197.31121 | 0.517517113 | 0.13754089 | 3.76264176 | 0.00016813 | 0.00135273 | CAPN6 | 827 |
| ENSG00000C | 138.975848 | 0.61937819 | 0.16460913 | 3.76272082 | 0.00016807 | 0.00135273 | RAB11FIP3 | 9727 |
| ENSG000001 | 15.7935979 | -1.165004969 | 0.3096212 | -3.7626783 | 0.0001681 | 0.00135273 | FRMD1 | 79981 |
| ENSG000001 | 1786.29359 | 0.461433576 | 0.12263104 | 3.76277954 | 0.00016804 | 0.00135273 | BLM | 641 |
| ENSG000001 | 1283.11264 | 0.479892796 | 0.12754789 | 3.76245172 | 0.00016826 | 0.00135349 | MYCN | 4613 |
| ENSG000001 | 5463.7748 | 0.482674955 | 0.12829352 | 3.76227083 | 0.00016838 | 0.0013542 | CXADR | 1525 |
| ENSG00000C | 202.080598 | 0.531752553 | 0.14134463 | 3.76209935 | 0.00016849 | 0.00135463 | TINF2 | 26277 |
| ENSG000001 | 179.251899 | 0.579021282 | 0.15390941 | 3.7620915 | 0.0001685 | 0.00135463 | ANKRD9 | 122416 |

| | | | | | | | | |
|---|---|---|---|---|---|---|---|---|
| ENSG000001 | 440.896161 | 0.491543735 | 0.13066936 | 3.76173666 | 0.00016874 | 0.00135629 | ZNF689 | 115509 |
| ENSG000001 | 221.847567 | 0.544846678 | 0.14485377 | 3.76135667 | 0.00016899 | 0.00135789 | TEX261 | 113419 |
| ENSG000001 | 1025.02897 | 0.521028894 | 0.13852207 | 3.76134221 | 0.000169 | 0.00135789 | ZBTB39 | 9880 |
| ENSG000001 | 13.2994334 | -1.225696043 | 0.32587975 | -3.7611912 | 0.00016911 | 0.00135844 | TNFSF14 | 8740 |
| ENSG00000C | 798.918442 | 0.645448774 | 0.17161071 | 3.76112183 | 0.00016915 | 0.00135855 | ADGRL1 | 22859 |
| ENSG000001 | 430.218968 | 0.454993314 | 0.12097685 | 3.76099491 | 0.00016924 | 0.00135897 | MTERF1 | 7978 |
| ENSG000001 | 2867.05752 | 0.494213997 | 0.13140719 | 3.76093585 | 0.00016928 | 0.00135902 | FAM169A | 26049 |
| ENSG000001 | 3476.26858 | 0.437129485 | 0.11623459 | 3.76075206 | 0.0001694 | 0.00135975 | ICE1 | 23379 |
| ENSG000001 | 9630.07967 | 0.387502418 | 0.10304107 | 3.76065977 | 0.00016947 | 0.00135999 | TPM3 | 7170 |
| ENSG00000C | 203.9178 | 0.543201946 | 0.14444553 | 3.76060064 | 0.00016951 | 0.00136004 | EHD3 | 30845 |
| ENSG000001 | 26.0770219 | -1.013374032 | 0.26948078 | -3.7604686 | 0.0001696 | 0.00136025 | KCNJ9 | 3765 |
| ENSG000002 | 21.1787697 | -1.056921401 | 0.28106518 | -3.7604138 | 0.00016963 | 0.00136025 | NA | NA |
| ENSG000002 | 68.5983391 | 0.726693756 | 0.19324773 | 3.76042584 | 0.00016962 | 0.00136025 | NA | NA |
| ENSG00000C | 310.857372 | 0.491871863 | 0.13080462 | 3.76035542 | 0.00016967 | 0.0013603 | DCX | 1641 |
| ENSG000001 | 3071.6269 | 0.465877565 | 0.12389675 | 3.76020806 | 0.00016977 | 0.00136083 | ECT2 | 1894 |
| ENSG000001 | 80.5927153 | -0.701748616 | 0.18663263 | -3.7600533 | 0.00016988 | 0.0013614 | ZNF471 | 57573 |
| ENSG00000C | 43.5248901 | -0.834291762 | 0.22189834 | -3.7597928 | 0.00017005 | 0.00136255 | ADGRA2 | 25960 |
| ENSG00000C | 1379.17791 | 0.471002476 | 0.12527531 | 3.759739 | 0.00017009 | 0.00136257 | IGF2BP2 | 10644 |
| ENSG000001 | 627.137968 | 0.465886229 | 0.12392037 | 3.75956119 | 0.00017021 | 0.00136327 | ANKRD27 | 84079 |
| ENSG000002 | 2055.07413 | 0.458999839 | 0.12210137 | 3.75917027 | 0.00017048 | 0.00136514 | PPP1R10 | 5514 |
| ENSG00000C | 731.497098 | 0.454545734 | 0.12092199 | 3.75899991 | 0.00017059 | 0.0013658 | FXYD5 | 53827 |
| ENSG000001 | 889.475547 | 0.406299063 | 0.10809484 | 3.75872781 | 0.00017078 | 0.00136701 | BRIP1 | 83990 |
| ENSG000001 | 82.0166974 | -0.752631985 | 0.20024402 | -3.7585741 | 0.00017088 | 0.00136758 | PDE1C | 5137 |
| ENSG000002 | 31.3737372 | -0.936957606 | 0.24928959 | -3.7585108 | 0.00017093 | 0.00136766 | GDNF-AS1 | 100861519 |
| ENSG000001 | 640.167643 | 0.604643877 | 0.1608816 | 3.75831591 | 0.00017106 | 0.00136842 | KCTD10 | 83892 |
| ENSG000001 | 803.528588 | 0.482868582 | 0.1284815 | 3.75827328 | 0.00017109 | 0.00136842 | RAD51AP1 | 10635 |
| ENSG000002 | 62.8762413 | 0.733064318 | 0.19510003 | 3.75737684 | 0.0001717 | 0.00137305 | C8orf82 | 414919 |
| ENSG000001 | 90.7299727 | -0.676779522 | 0.18012671 | -3.7572413 | 0.0001718 | 0.00137353 | AKAP6 | 9472 |
| ENSG00000C | 45.4850018 | -0.858401207 | 0.22847295 | -3.757124 | 0.00017188 | 0.0013739 | SMC1B | 27127 |

| | | | | | | | | |
|---|---|---|---|---|---|---|---|---|
| ENSG000002 | 62.974909 | 0.809586643 | 0.21550233 | 3.75674194 | 0.00017214 | 0.00137573 | MIR3609 | 100500819 |
| ENSG000002 | 13.0585959 | -1.216002234 | 0.32369254 | -3.7566582 | 0.0001722 | 0.00137592 | LINC01350 | 101929093 |
| ENSG000001 | 2390.30762 | 0.423001283 | 0.11260623 | 3.75646427 | 0.00017233 | 0.00137671 | NUTF2 | 10204 |
| ENSG000001 | 181.211571 | 0.551565277 | 0.1468363 | 3.75632769 | 0.00017242 | 0.00137719 | SLC25A14 | 9016 |
| ENSG000001 | 44.5815161 | -0.834154549 | 0.22207282 | -3.7562208 | 0.0001725 | 0.00137751 | F5 | 2153 |
| ENSG000002 | 15.582839 | -1.171005829 | 0.31175973 | -3.7561164 | 0.00017257 | 0.00137781 | LINC01183 | 728586 |
| ENSG000001 | 366.044551 | 0.464415991 | 0.12364929 | 3.75591313 | 0.00017271 | 0.00137842 | C18orf21 | 83608 |
| ENSG000001 | 147.024164 | 0.629434757 | 0.16758525 | 3.75590783 | 0.00017271 | 0.00137842 | ALDH16A1 | 126133 |
| ENSG000001 | 382.561292 | 0.583832959 | 0.15544851 | 3.7557964 | 0.00017279 | 0.00137876 | LAMB2 | 3913 |
| ENSG000002 | 98.3613714 | 0.637888912 | 0.16985247 | 3.75554679 | 0.00017296 | 0.00137986 | NA | NA |
| ENSG00000C | 2236.38554 | 0.403597401 | 0.10746926 | 3.75546847 | 0.00017302 | 0.00138002 | DBF4 | 10926 |
| ENSG000001 | 2068.88687 | 0.468123101 | 0.12465606 | 3.75531766 | 0.00017312 | 0.00138058 | ZIC3 | 7547 |
| ENSG000001 | 50.0777071 | -0.823320962 | 0.21924809 | -3.7552024 | 0.0001732 | 0.00138095 | NPAS3 | 64067 |
| ENSG000001 | 494.806537 | 0.533065468 | 0.14195671 | 3.75512699 | 0.00017325 | 0.00138109 | CCDC93 | 54520 |
| ENSG000001 | 385.220746 | 0.519845389 | 0.13844877 | 3.75478526 | 0.00017349 | 0.00138259 | CINP | 51550 |
| ENSG000001 | 1295.61795 | 0.46177032 | 0.12298274 | 3.75475719 | 0.00017351 | 0.00138259 | EBNA1BP2 | 10969 |
| ENSG000001 | 19.9882846 | -1.078591051 | 0.28726638 | -3.7546721 | 0.00017357 | 0.00138279 | CD244 | 51744 |
| ENSG00000C | 714.600653 | 0.497742934 | 0.13256879 | 3.75460106 | 0.00017362 | 0.00138291 | VIM | 7431 |
| ENSG000001 | 1656.30919 | 0.511562143 | 0.13627221 | 3.75397251 | 0.00017405 | 0.00138611 | GID8 | 54994 |
| ENSG000002 | 31.9890145 | -0.954018079 | 0.25414578 | -3.7538223 | 0.00017416 | 0.00138667 | LOC643711 | 643711 |
| ENSG000001 | 2547.77605 | 0.494227991 | 0.13166943 | 3.75355165 | 0.00017435 | 0.00138762 | GTF2A1 | 2957 |
| ENSG000001 | 1209.03105 | 0.446885832 | 0.11905664 | 3.75355649 | 0.00017434 | 0.00138762 | BAG5 | 9529 |
| ENSG000001 | 1551.23746 | 0.419430968 | 0.11175966 | 3.75297276 | 0.00017475 | 0.00139056 | BBX | 56987 |
| ENSG000001 | 839.064302 | 0.440297684 | 0.11733855 | 3.75237035 | 0.00017517 | 0.00139363 | MTPAP | 55149 |
| ENSG000001 | 454.942841 | 0.508138544 | 0.13544276 | 3.75168475 | 0.00017565 | 0.0013969 | POLR3F | 10621 |
| ENSG000002 | 119.96544 | 0.657678107 | 0.1753015 | 3.75169698 | 0.00017564 | 0.0013969 | SMG1P7 | 100506060 |
| ENSG000001 | 335.989194 | 0.472648085 | 0.12598636 | 3.75158153 | 0.00017572 | 0.0013972 | VPS26B | 112936 |
| ENSG00000C | 2031.44172 | 0.539183524 | 0.14373142 | 3.75132678 | 0.0001759 | 0.00139835 | LIG3 | 3980 |
| ENSG000001 | 1385.43274 | 0.425868776 | 0.11353404 | 3.75102274 | 0.00017611 | 0.00139977 | CCDC117 | 150275 |

| | | | | | | | | |
|---|---|---|---|---|---|---|---|---|
| ENSG000002 | 767.90828 | 0.527486377 | 0.14063715 | 3.7506902 | 0.00017635 | 0.00140135 | ATXN1L | 342371 |
| ENSG000001 | 6576.71021 | 0.419655813 | 0.11191151 | 3.7498896 | 0.00017691 | 0.00140556 | PDIA3 | 2923 |
| ENSG000001 | 932.105113 | 0.442376779 | 0.11797235 | 3.74983444 | 0.00017695 | 0.00140559 | MRPL39 | 54148 |
| ENSG000001 | 10319.5324 | 0.501927394 | 0.13386067 | 3.74962571 | 0.0001771 | 0.00140614 | DDX5 | 1655 |
| ENSG000001 | 1182.46198 | 0.486637644 | 0.1297822 | 3.74964852 | 0.00017708 | 0.00140614 | ECD | 11319 |
| ENSG000002 | 351.876863 | 0.598398606 | 0.15959042 | 3.74958985 | 0.00017712 | 0.00140614 | RNU4-1 | 26835 |
| ENSG000001 | 767.330409 | 0.461363926 | 0.12305656 | 3.74920225 | 0.0001774 | 0.00140803 | TBC1D5 | 9779 |
| ENSG000002 | 207.496565 | 0.552037692 | 0.14724324 | 3.74915471 | 0.00017743 | 0.00140803 | NA | NA |
| ENSG000001 | 1040.2691 | 0.415106325 | 0.11072406 | 3.74901659 | 0.00017753 | 0.00140825 | NA | NA |
| ENSG000001 | 16.484366 | -1.131677135 | 0.3018569 | -3.7490517 | 0.0001775 | 0.00140825 | C4BPB | 725 |
| ENSG000001 | 539.32079 | 0.468605534 | 0.1250083 | 3.7485953 | 0.00017783 | 0.00141034 | KANSL3 | 55683 |
| ENSG000002 | 9.00410272 | -1.318262616 | 0.35169457 | -3.7483167 | 0.00017803 | 0.00141164 | NA | NA |
| ENSG000001 | 214.164989 | 0.573189602 | 0.15293726 | 3.74787425 | 0.00017834 | 0.00141385 | RNF149 | 284996 |
| ENSG00000C | 430.367548 | 0.431416757 | 0.11511239 | 3.74778717 | 0.0001784 | 0.001414 | DDHD2 | 23259 |
| ENSG000002 | 211.609822 | 0.60624413 | 0.16176215 | 3.74775014 | 0.00017843 | 0.001414 | INF2 | 64423 |
| ENSG000001 | 172.217028 | 0.611303161 | 0.16312699 | 3.74740653 | 0.00017867 | 0.00141566 | IFITM2 | 10581 |
| ENSG00000C | 115.965782 | 0.606821896 | 0.16194135 | 3.74717071 | 0.00017884 | 0.00141671 | DNASE1L1 | 1774 |
| ENSG00000C | 101.600387 | 0.651924464 | 0.17401132 | 3.74644853 | 0.00017936 | 0.00142052 | HOMER3 | 9454 |
| ENSG000001 | 375.962749 | 0.643366587 | 0.17174253 | 3.746111 | 0.0001796 | 0.00142215 | FAM83G | 644815 |
| ENSG000001 | 1410.41038 | 0.452805246 | 0.12087676 | 3.74600755 | 0.00017967 | 0.00142246 | VSIG10 | 54621 |
| ENSG000001 | 2272.92065 | 0.447989004 | 0.11959507 | 3.74588192 | 0.00017976 | 0.00142278 | NUP107 | 57122 |
| ENSG000001 | 516.980194 | 0.554160153 | 0.1479396 | 3.74585417 | 0.00017978 | 0.00142278 | UBAP1 | 51271 |
| ENSG000001 | 315.102676 | 0.524821999 | 0.14011181 | 3.74573713 | 0.00017986 | 0.00142301 | PTN | 5764 |
| ENSG000001 | 135.529541 | -0.610323559 | 0.16293911 | -3.7457156 | 0.00017988 | 0.00142301 | PKHD1 | 5314 |
| ENSG000002 | 49.7437309 | -0.797516693 | 0.21291933 | -3.7456283 | 0.00017994 | 0.00142322 | C2orf16 | 84226 |
| ENSG000001 | 315.021599 | 0.530623906 | 0.14167419 | 3.74538151 | 0.00018012 | 0.00142434 | NABP2 | 79035 |
| ENSG00000C | 682.460716 | 0.529920178 | 0.1414999 | 3.74502162 | 0.00018038 | 0.00142611 | UBA5 | 79876 |
| ENSG00000C | 1907.60274 | 0.435016356 | 0.11617766 | 3.74440623 | 0.00018082 | 0.00142933 | UBA6 | 55236 |
| ENSG000001 | 185.847612 | 0.583057176 | 0.15571844 | 3.74430394 | 0.00018089 | 0.00142963 | CANT1 | 124583 |

| | | | | | | | | | |
|---|---|---|---|---|---|---|---|---|---|
| ENSG000000 | 390.459559 | 0.464130477 | 0.12396109 | 3.74416267 | 0.000181 | 0.00143016 | TMEM260 | | 54916 |
| ENSG000000 | 2097.57386 | 0.50901995 | 0.1359535 | 3.74407387 | 0.00018106 | 0.00143039 | PAFAH1B1 | | 5048 |
| ENSG000001 | 1228.23975 | 0.452757222 | 0.1209524 | 3.74326764 | 0.00018164 | 0.00143471 | CCDC25 | | 55246 |
| ENSG000001 | 23.8817713 | -1.05256232 | 0.28121825 | -3.7428663 | 0.00018193 | 0.00143672 | NA | NA | |
| ENSG000001 | 79.4403503 | -0.698081092 | 0.18653623 | -3.7423351 | 0.00018232 | 0.00143948 | ESRRG | | 2104 |
| ENSG000000 | 50.5346937 | -0.816168422 | 0.21812077 | -3.7418188 | 0.00018269 | 0.00144188 | HGF | | 3082 |
| ENSG000001 | 5700.50366 | 0.458374114 | 0.12249974 | 3.74183757 | 0.00018268 | 0.00144188 | CAND1 | | 55832 |
| ENSG000001 | 13.0082601 | -1.209839036 | 0.32335245 | -3.741549 | 0.00018289 | 0.00144287 | LINC00303 | | 284573 |
| ENSG000001 | 431.638895 | 0.500166597 | 0.13367888 | 3.74155299 | 0.00018289 | 0.00144287 | ZDHHC13 | | 54503 |
| ENSG000000 | 642.012016 | 0.415304476 | 0.11100963 | 3.74115721 | 0.00018317 | 0.00144484 | HEBP2 | | 23593 |
| ENSG000000 | 660.986642 | 0.492593286 | 0.13167659 | 3.7409328 | 0.00018334 | 0.00144557 | JKAMP | | 51528 |
| ENSG000002 | 17.8509368 | -1.111037252 | 0.29699251 | -3.7409605 | 0.00018332 | 0.00144557 | SRD5A2 | | 6716 |
| ENSG000001 | 2087.41436 | 0.443000995 | 0.11843114 | 3.74057866 | 0.0001836 | 0.00144704 | USP24 | | 23358 |
| ENSG000001 | 800.914101 | 0.427253404 | 0.11422078 | 3.74059248 | 0.00018359 | 0.00144704 | ZFP91 | | 80829 |
| ENSG000001 | 19.983654 | -1.077060033 | 0.28794705 | -3.7404795 | 0.00018367 | 0.00144733 | PM20D1 | | 148811 |
| ENSG000001 | 1491.71525 | 0.454428647 | 0.12152284 | 3.73945056 | 0.00018442 | 0.00145299 | HAT1 | | 8520 |
| ENSG000001 | 2984.55401 | 0.45502172 | 0.12168746 | 3.7392656 | 0.00018456 | 0.00145378 | TFAM | | 7019 |
| ENSG000002 | 15.2710592 | -1.157633715 | 0.30960218 | -3.7391007 | 0.00018468 | 0.00145445 | NA | NA | |
| ENSG000002 | 236.874664 | 0.512553966 | 0.13708706 | 3.73889393 | 0.00018483 | 0.00145536 | NA | NA | |
| ENSG000000 | 1403.52938 | 0.53284113 | 0.1425184 | 3.73875327 | 0.00018494 | 0.00145589 | ADD1 | | 118 |
| ENSG000001 | 779.780695 | 0.461998509 | 0.12357606 | 3.73857604 | 0.00018507 | 0.0014564 | PCYT1A | | 5130 |
| ENSG000001 | 400.082677 | 0.465487736 | 0.12450965 | 3.73856751 | 0.00018507 | 0.0014564 | CDC26 | | 246184 |
| ENSG000001 | 1070.12732 | 0.478131471 | 0.12789587 | 3.73844326 | 0.00018516 | 0.00145684 | INO80 | | 54617 |
| ENSG000001 | 214.759835 | 0.531886963 | 0.1422811 | 3.73828271 | 0.00018528 | 0.00145749 | FAHD1 | | 81889 |
| ENSG000001 | 778.862421 | 0.455338131 | 0.1218187 | 3.7378344 | 0.00018561 | 0.00145981 | SUCLG2 | | 8801 |
| ENSG000001 | 50.5194488 | -0.780203824 | 0.20877766 | -3.7370082 | 0.00018622 | 0.00146432 | RXFP1 | | 59350 |
| ENSG000002 | 17.0370315 | -1.114789226 | 0.29832866 | -3.7367822 | 0.00018639 | 0.00146536 | NA | NA | |
| ENSG000001 | 725.245113 | 0.444534042 | 0.11896931 | 3.73654399 | 0.00018657 | 0.00146646 | PKD2 | | 5311 |
| ENSG000001 | 3447.62185 | 0.511236209 | 0.13682276 | 3.73648513 | 0.00018661 | 0.00146652 | MTHFD1 | | 4522 |

| | | | | | | | | |
|---|---|---|---|---|---|---|---|---|
| ENSG000001 | 6110.08841 | 0.475730258 | 0.12733284 | 3.73611591 | 0.00018688 | 0.00146839 | RPL34 | 6164 |
| ENSG000002 | 10.1169575 | -1.288585929 | 0.34491859 | -3.7359133 | 0.00018704 | 0.00146929 | NA | NA |
| ENSG000001 | 152.353586 | 0.561608918 | 0.15034802 | 3.73539276 | 0.00018742 | 0.00147204 | SLC25A21 | 89874 |
| ENSG000001 | 168.956529 | 0.599797165 | 0.16058291 | 3.73512455 | 0.00018762 | 0.00147333 | FAM53B | 9679 |
| ENSG000001 | 427.447552 | 0.49990312 | 0.13384111 | 3.73504915 | 0.00018768 | 0.00147349 | DPAGT1 | 1798 |
| ENSG000001 | 303.0823 | 0.526659265 | 0.14100949 | 3.73492068 | 0.00018777 | 0.00147395 | SLC25A29 | 123096 |
| ENSG000001 | 824.181873 | 0.594988802 | 0.15934714 | 3.73391571 | 0.00018853 | 0.00147956 | LSR | 51599 |
| ENSG000001 | 1751.94604 | 0.42927869 | 0.11498263 | 3.73342192 | 0.0001889 | 0.00148218 | UBE2V2 | 7336 |
| ENSG000001 | 147.938147 | 0.576885759 | 0.15452572 | 3.73326695 | 0.00018901 | 0.00148281 | ATG4A | 115201 |
| ENSG00000C | 986.416167 | 0.459243985 | 0.12303023 | 3.73277344 | 0.00018938 | 0.00148543 | ZMYND11 | 10771 |
| ENSG000002 | 185.740058 | 0.569075538 | 0.15246263 | 3.73255762 | 0.00018955 | 0.00148642 | AP1G2 | 8906 |
| ENSG000001 | 107.727446 | 0.660543045 | 0.1769729 | 3.73245307 | 0.00018962 | 0.00148674 | GALNS | 2588 |
| ENSG000001 | 2337.5568 | 0.435031033 | 0.11655667 | 3.73235637 | 0.0001897 | 0.00148703 | SS18 | 6760 |
| ENSG000001 | 867.711808 | 0.491839774 | 0.1318202 | 3.73114115 | 0.00019061 | 0.00149364 | FNTA | 2339 |
| ENSG000001 | 566.338086 | 0.502121689 | 0.13457566 | 3.73114801 | 0.00019061 | 0.00149364 | CXXC5 | 51523 |
| ENSG000001 | 678.970167 | 0.475105528 | 0.12735157 | 3.73066105 | 0.00019098 | 0.0014962 | SLC9A3R1 | 9368 |
| ENSG00000C | 4145.68458 | 0.460626987 | 0.12347635 | 3.73048743 | 0.00019111 | 0.00149637 | CDC42 | 998 |
| ENSG000001 | 805.55234 | 0.479433077 | 0.12851749 | 3.73048888 | 0.00019111 | 0.00149637 | AKAP1 | 8165 |
| ENSG000001 | 106.764226 | -0.653401009 | 0.1751506 | -3.7305097 | 0.00019109 | 0.00149637 | GREM1 | 26585 |
| ENSG000001 | 635.577398 | 0.454161213 | 0.1217542 | 3.73014816 | 0.00019137 | 0.00149809 | OSGIN2 | 734 |
| ENSG000001 | 275.344836 | 0.650154035 | 0.17434904 | 3.72903711 | 0.00019221 | 0.00150442 | BIN1 | 274 |
| ENSG000001 | 65.3410283 | -0.792695944 | 0.21259426 | -3.7286798 | 0.00019249 | 0.00150569 | MYOM1 | 8736 |
| ENSG000001 | 993.077602 | 0.468609422 | 0.12567673 | 3.72868872 | 0.00019248 | 0.00150569 | PPHLN1 | 51535 |
| ENSG000001 | 641.3002 | 0.485471623 | 0.13019874 | 3.72869667 | 0.00019247 | 0.00150569 | EIF2B5 | 8893 |
| ENSG000001 | 494.424693 | 0.548377686 | 0.1470738 | 3.72858854 | 0.00019256 | 0.00150594 | MON1B | 22879 |
| ENSG000001 | 1941.19424 | 0.446322073 | 0.11970701 | 3.72845387 | 0.00019266 | 0.00150646 | NBN | 4683 |
| ENSG000001 | 102.341554 | 0.771464257 | 0.20693264 | 3.72809365 | 0.00019293 | 0.0015081 | ATHL1 | 80162 |
| ENSG000001 | 241.926963 | 0.579726711 | 0.15550265 | 3.7280824 | 0.00019294 | 0.0015081 | JPH3 | 57338 |
| ENSG000001 | 2175.24906 | 0.45317357 | 0.1215589 | 3.72801643 | 0.00019299 | 0.0015082 | WTAP | 9589 |

| | | | | | | | | |
|---|---|---|---|---|---|---|---|---|
| ENSG000001 | 576.142264 | 0.440311438 | 0.1181118 | 3.72792095 | 0.00019307 | 0.00150848 | MTIF3 | 219402 |
| ENSG00000C | 490.40845 | 0.43221943 | 0.11594723 | 3.7277253 | 0.00019322 | 0.00150913 | TIPIN | 54962 |
| ENSG00000C | 58.878116 | -0.788916099 | 0.21163645 | -3.7276948 | 0.00019324 | 0.00150913 | PTPRC | 5788 |
| ENSG000001 | 4269.06124 | 0.409583696 | 0.10988012 | 3.72755043 | 0.00019335 | 0.00150913 | KAT7 | 11143 |
| ENSG000001 | 609.532652 | 0.475706699 | 0.12761847 | 3.72756936 | 0.00019334 | 0.00150913 | ZNF219 | 51222 |
| ENSG000001 | 4426.11267 | 0.442069269 | 0.11859541 | 3.72754101 | 0.00019336 | 0.00150913 | SRSF10 | 10772 |
| ENSG000001 | 40.3651473 | -0.863852119 | 0.23174978 | -3.7275207 | 0.00019337 | 0.00150913 | AADACL3 | 126767 |
| ENSG000001 | 28.48449 | -1.009604893 | 0.27086021 | -3.7274021 | 0.00019346 | 0.00150955 | CCDC175 | 729665 |
| ENSG000001 | 4285.33949 | 0.466534303 | 0.125168 | 3.72726503 | 0.00019357 | 0.00151008 | USP7 | 7874 |
| ENSG000001 | 2167.30208 | 0.496014373 | 0.13307957 | 3.72720149 | 0.00019362 | 0.00151017 | COPS2 | 9318 |
| ENSG000001 | 662.118884 | 0.576857549 | 0.15479053 | 3.726698 | 0.000194 | 0.0015129 | ERCC2 | 2068 |
| ENSG000001 | 1462.69655 | 0.437142874 | 0.11730436 | 3.72656982 | 0.0001941 | 0.00151338 | LMBR1 | 64327 |
| ENSG000001 | 59.5966726 | 0.745441607 | 0.20010269 | 3.72529521 | 0.00019509 | 0.00152075 | HS3ST3A1 | 9955 |
| ENSG000002 | 21.4419202 | -1.079517211 | 0.28979339 | -3.7251271 | 0.00019522 | 0.00152148 | NA | NA |
| ENSG000001 | 1251.87973 | 0.494021943 | 0.13262914 | 3.72483722 | 0.00019544 | 0.00152264 | VCPIP1 | 80124 |
| ENSG000001 | 26.8846782 | -0.958378674 | 0.25729164 | -3.724873 | 0.00019541 | 0.00152264 | C5orf56 | 441108 |
| ENSG000002 | 35.4550588 | -0.878599341 | 0.23589223 | -3.7245794 | 0.00019564 | 0.0015239 | LINC00607 | 646324 |
| ENSG00000C | 1565.61586 | 0.470629584 | 0.12638485 | 3.72378158 | 0.00019626 | 0.00152814 | IBTK | 25998 |
| ENSG000002 | 1408.18894 | 0.419399855 | 0.11262597 | 3.72382916 | 0.00019622 | 0.00152814 | TENM3 | 55714 |
| ENSG000001 | 1338.88743 | 0.472634653 | 0.12693084 | 3.72356045 | 0.00019643 | 0.00152919 | ITGAV | 3685 |
| ENSG000001 | 554.83819 | 0.447967864 | 0.1203166 | 3.7232425 | 0.00019668 | 0.00153082 | SLC25A23 | 79085 |
| ENSG000001 | 88.5134433 | 0.677311705 | 0.18192166 | 3.72309536 | 0.0001968 | 0.00153142 | TRPT1 | 83707 |
| ENSG000002 | 37.7704847 | -0.860713393 | 0.23118969 | -3.7229747 | 0.00019689 | 0.00153186 | KCNIP4-IT1 | 359822 |
| ENSG000002 | 207.109703 | 0.546771928 | 0.14687966 | 3.72258435 | 0.00019719 | 0.00153394 | FADS3 | 3995 |
| ENSG000001 | 136.454359 | 0.711135016 | 0.19104059 | 3.72242886 | 0.00019732 | 0.00153459 | SURF1 | 6834 |
| ENSG000002 | 42.0958796 | 0.865125074 | 0.23242771 | 3.72212536 | 0.00019755 | 0.00153614 | MYL5 | 4636 |
| ENSG000001 | 627.717004 | 0.53402007 | 0.14348902 | 3.72167893 | 0.0001979 | 0.00153856 | CRYZ | 1429 |
| ENSG000001 | 25.3922761 | -1.026864972 | 0.27592822 | -3.7214931 | 0.00019805 | 0.0015394 | LMX1A | 4009 |
| ENSG000001 | 1300.3522 | 0.552553476 | 0.14848026 | 3.7213934 | 0.00019813 | 0.00153971 | EHMT1 | 79813 |

| | | | | | | | | |
|---|---|---|---|---|---|---|---|---|
| ENSG00000( | 3499.49792 | 0.597257009 | 0.16050745 | 3.72105476 | 0.00019839 | 0.00154149 | AP2B1 | 163 |
| ENSG000001 | 1154.34061 | 0.541907848 | 0.14563701 | 3.72094881 | 0.00019848 | 0.00154184 | POLR3A | 11128 |
| ENSG000002 | 50.7085445 | -0.805374229 | 0.216449 | -3.7208498 | 0.00019855 | 0.00154215 | MBNL1-AS1 | 401093 |
| ENSG00000( | 220.208975 | 0.681653505 | 0.18321418 | 3.72052807 | 0.00019881 | 0.00154331 | NA | NA |
| ENSG000001 | 295.247622 | 0.515352807 | 0.1385157 | 3.72053712 | 0.0001988 | 0.00154331 | CCDC113 | 29070 |
| ENSG000001 | 452.478474 | 0.593250626 | 0.15945391 | 3.72051478 | 0.00019882 | 0.00154331 | TAF6 | 6878 |
| ENSG000002 | 171.328379 | 0.579765885 | 0.15583239 | 3.7204452 | 0.00019887 | 0.00154344 | NA | NA |
| ENSG000001 | 399.485015 | 0.565513503 | 0.15200795 | 3.72028891 | 0.000199 | 0.00154351 | GCA | 25801 |
| ENSG000001 | 2390.79061 | 0.430151923 | 0.1156203 | 3.72038401 | 0.00019892 | 0.00154351 | SDAD1 | 55153 |
| ENSG000002 | 1080.87135 | 0.42380179 | 0.11391625 | 3.72029281 | 0.00019899 | 0.00154351 | SLC35B4 | 84912 |
| ENSG000001 | 6432.43153 | 0.392218721 | 0.10543432 | 3.72002878 | 0.0001992 | 0.0015448 | KTN1 | 3895 |
| ENSG000001 | 263.355372 | 0.688524914 | 0.18513442 | 3.71905396 | 0.00019997 | 0.00155048 | MAP1S | 55201 |
| ENSG000001 | 46.8719998 | 0.814380832 | 0.21901506 | 3.71837821 | 0.00020051 | 0.00155433 | NA | NA |
| ENSG000001 | 750.365203 | 0.458442094 | 0.12329589 | 3.71822685 | 0.00020063 | 0.00155497 | VPS37A | 137492 |
| ENSG00000( | 1349.47262 | 0.480557458 | 0.12924973 | 3.71805389 | 0.00020076 | 0.00155559 | PGM3 | 5238 |
| ENSG000001 | 5.57536385 | -1.403105427 | 0.37737884 | -3.7180289 | 0.00020078 | 0.00155559 | AVPR1B | 553 |
| ENSG00000( | 1587.1747 | 0.453286343 | 0.12191918 | 3.71792468 | 0.00020087 | 0.00155594 | CPNE3 | 8895 |
| ENSG000001 | 5103.06765 | 0.42096914 | 0.11324433 | 3.71735292 | 0.00020132 | 0.00155901 | STIP1 | 10963 |
| ENSG000002 | 897.319652 | 0.451511696 | 0.12146132 | 3.71732912 | 0.00020134 | 0.00155901 | NA | NA |
| ENSG000001 | 331.85582 | 0.559500919 | 0.15052248 | 3.71705884 | 0.00020156 | 0.00156038 | GTF2B | 2959 |
| ENSG000001 | 502.311956 | 0.486981949 | 0.13102587 | 3.71668536 | 0.00020185 | 0.00156239 | TP53RK | 112858 |
| ENSG000001 | 15.4747564 | -1.142989639 | 0.30754065 | -3.7165482 | 0.00020196 | 0.00156264 | FCRLA | 84824 |
| ENSG000002 | 176.391373 | 0.65268449 | 0.17561402 | 3.71658538 | 0.00020193 | 0.00156264 | SLC35F6 | 54978 |
| ENSG000001 | 19115.2414 | 0.43830392 | 0.11795013 | 3.71601046 | 0.00020239 | 0.0015655 | GJA1 | 2697 |
| ENSG000001 | 1372.33051 | 0.642254869 | 0.17283547 | 3.71598996 | 0.00020241 | 0.0015655 | SOX11 | 6664 |
| ENSG00000( | 14.1758022 | -1.193812707 | 0.3212725 | -3.7158882 | 0.00020249 | 0.00156576 | NRIP2 | 83714 |
| ENSG000001 | 276.869056 | 0.676762808 | 0.18212858 | 3.71585172 | 0.00020252 | 0.00156576 | RHOT2 | 89941 |
| ENSG000001 | 49.5737601 | -0.806211128 | 0.21697109 | -3.7157536 | 0.0002026 | 0.00156607 | ITGA10 | 8515 |
| ENSG000001 | 587.293189 | 0.487031284 | 0.13109665 | 3.71505507 | 0.00020316 | 0.0015701 | RNF139 | 11236 |

| | | | | | | | | |
|---|---|---|---|---|---|---|---|---|
| ENSG000001 | 105.539694 | 0.631704646 | 0.17004654 | 3.71489277 | 0.00020329 | 0.00157081 | KDM8 | 79831 |
| ENSG000001 | 752.681036 | 0.497972267 | 0.13405031 | 3.71481615 | 0.00020335 | 0.00157099 | SPCS1 | 28972 |
| ENSG000001 | 433.475148 | 0.520425421 | 0.14010845 | 3.71444708 | 0.00020365 | 0.00157298 | NFATC2IP | 84901 |
| ENSG000001 | 627.122239 | 0.441836132 | 0.11897084 | 3.71381867 | 0.00020416 | 0.00157659 | REEP3 | 221035 |
| ENSG00000C | 2401.87266 | 0.430168847 | 0.11583387 | 3.7136706 | 0.00020427 | 0.00157722 | GNAI3 | 2773 |
| ENSG000001 | 271.449452 | 0.500871464 | 0.13490144 | 3.71286975 | 0.00020492 | 0.00158173 | VWA5A | 4013 |
| ENSG000001 | 267.504594 | 0.51792528 | 0.1394971 | 3.71280327 | 0.00020498 | 0.00158173 | VPS11 | 55823 |
| ENSG000002 | 11.6728684 | -1.233084616 | 0.33211565 | -3.7128169 | 0.00020497 | 0.00158173 | NA | NA |
| ENSG000001 | 7936.6693 | 0.414322137 | 0.11159749 | 3.7126476 | 0.0002051 | 0.0015824 | UBE2D3 | 7323 |
| ENSG00000C | 510.965268 | 0.436713333 | 0.11763287 | 3.71251101 | 0.00020521 | 0.00158278 | IFNGR1 | 3459 |
| ENSG000002 | 557.603036 | 0.569696004 | 0.15345381 | 3.71249171 | 0.00020523 | 0.00158278 | ATP5O | 539 |
| ENSG000001 | 10.5977504 | -1.25787293 | 0.33884393 | -3.7122487 | 0.00020543 | 0.00158399 | TCHHL1 | 126637 |
| ENSG00000C | 432.048436 | 0.490048815 | 0.13201985 | 3.71193273 | 0.00020568 | 0.00158537 | AGPAT4 | 56895 |
| ENSG000001 | 69.2247292 | -0.757248025 | 0.20400147 | -3.7119734 | 0.00020565 | 0.00158537 | GREB1L | 80000 |
| ENSG000001 | 1776.95579 | 0.419824682 | 0.11310369 | 3.71185654 | 0.00020574 | 0.00158555 | ZMAT2 | 153527 |
| ENSG000001 | 1006.22453 | 0.439525937 | 0.11841527 | 3.71173374 | 0.00020584 | 0.00158602 | FCF1 | 51077 |
| ENSG000001 | 624.605041 | 0.562602519 | 0.15159143 | 3.71130816 | 0.00020619 | 0.00158808 | UBN1 | 29855 |
| ENSG000001 | 474.953854 | 0.433194164 | 0.11672158 | 3.71134596 | 0.00020616 | 0.00158808 | WDR20 | 91833 |
| ENSG000001 | 2496.14958 | 0.490044406 | 0.13204364 | 3.71123071 | 0.00020625 | 0.00158827 | PPP2CA | 5515 |
| ENSG000001 | 995.105126 | 0.553915968 | 0.14926602 | 3.71093137 | 0.0002065 | 0.00158984 | LIMK1 | 3984 |
| ENSG000002 | 230.485193 | 0.486629357 | 0.13114399 | 3.71064939 | 0.00020673 | 0.00159131 | TMEM200C | 645369 |
| ENSG000001 | 433.83833 | 0.569066065 | 0.15337651 | 3.71025566 | 0.00020705 | 0.00159349 | MBD1 | 4152 |
| ENSG000002 | 112.400786 | 0.668823457 | 0.18027719 | 3.70997263 | 0.00020728 | 0.00159497 | SNORA53 | 677832 |
| ENSG00000C | 1214.84422 | 0.583703586 | 0.15735174 | 3.70954639 | 0.00020763 | 0.00159735 | POLR1A | 25885 |
| ENSG00000C | 142.92711 | 0.637290784 | 0.17180405 | 3.7094048 | 0.00020775 | 0.00159794 | MATK | 4145 |
| ENSG000001 | 1289.73879 | 0.400719667 | 0.10803489 | 3.70916916 | 0.00020794 | 0.00159913 | EFCAB14 | 9813 |
| ENSG000001 | 11.5768031 | -1.254414639 | 0.33822033 | -3.7088682 | 0.00020819 | 0.00160072 | TGM3 | 7053 |
| ENSG000001 | 486.030499 | 0.49561374 | 0.1336352 | 3.70870655 | 0.00020832 | 0.00160144 | DHDDS | 79947 |
| ENSG00000C | 1207.17345 | 0.423283862 | 0.11413716 | 3.70855449 | 0.00020845 | 0.00160197 | SMARCE1 | 6605 |

| | | | | | | | | |
|---|---|---|---|---|---|---|---|---|
| ENSG000002 | 306.384358 | 0.512966251 | 0.13832075 | 3.70852713 | 0.00020847 | 0.00160197 | NA | NA |
| ENSG000001 | 1186.71697 | 0.484771364 | 0.13073212 | 3.70812758 | 0.0002088 | 0.0016042 | ZNF22 | 7570 |
| ENSG000001 | 71.8271421 | 0.711977326 | 0.19201493 | 3.70792682 | 0.00020896 | 0.00160516 | ARHGAP9 | 64333 |
| ENSG00000C | 541.309273 | 0.481969051 | 0.12998638 | 3.70784262 | 0.00020903 | 0.00160539 | ERLEC1 | 27248 |
| ENSG000001 | 1400.07077 | 0.456215798 | 0.12304565 | 3.70769531 | 0.00020915 | 0.00160602 | PWP1 | 11137 |
| ENSG000002 | 53.5926685 | 0.822590042 | 0.22186318 | 3.70764556 | 0.0002092 | 0.00160603 | NA | NA |
| ENSG000002 | 29.893544 | -0.914107048 | 0.2465627 | -3.707402 | 0.0002094 | 0.00160727 | PGM5P2 | 595135 |
| ENSG000001 | 216.651888 | 0.697978055 | 0.18827932 | 3.70714142 | 0.00020961 | 0.00160814 | MTSS1L | 92154 |
| ENSG000001 | 25.718294 | -0.98407047 | 0.26545408 | -3.7071212 | 0.00020963 | 0.00160814 | SPOCK3 | 50859 |
| ENSG000002 | 50.0940809 | -0.807751051 | 0.21788979 | -3.7071543 | 0.0002096 | 0.00160814 | NA | NA |
| ENSG000002 | 263.899876 | 0.70112876 | 0.18913536 | 3.70702114 | 0.00020971 | 0.00160848 | JRK | 8629 |
| ENSG000001 | 1135.71293 | 0.438987257 | 0.11842242 | 3.70696075 | 0.00020976 | 0.00160855 | ZFYVE9 | 9372 |
| ENSG00000C | 3964.4609 | 0.494977826 | 0.13353107 | 3.70683637 | 0.00020986 | 0.00160904 | RHOA | 387 |
| ENSG000001 | 389.278249 | 0.588226074 | 0.15869427 | 3.70666244 | 0.00021001 | 0.00160954 | MYL9 | 10398 |
| ENSG000001 | 65.1491804 | -0.71169468 | 0.19200362 | -3.7066732 | 0.00021 | 0.00160954 | PCDHB12 | 56124 |
| ENSG000002 | 1334.57417 | 0.428518041 | 0.11561947 | 3.70627928 | 0.00021033 | 0.00161167 | PARG | 8505 |
| ENSG00000C | 816.873694 | 0.422542466 | 0.11402658 | 3.70564878 | 0.00021085 | 0.00161507 | SEL1L | 6400 |
| ENSG00000C | 1422.12802 | 0.633295921 | 0.17089854 | 3.70568367 | 0.00021082 | 0.00161507 | SRRT | 51593 |
| ENSG000001 | 1144.56892 | 0.463621353 | 0.12513404 | 3.70499788 | 0.00021139 | 0.00161892 | MOB4 | 25843 |
| ENSG000001 | 181.258636 | 0.548963666 | 0.14819398 | 3.70435868 | 0.00021193 | 0.00162239 | SPATC1L | 84221 |
| ENSG000001 | 1787.85406 | 0.453043853 | 0.12230015 | 3.70436066 | 0.00021192 | 0.00162239 | PJA2 | 9867 |
| ENSG000001 | 990.964864 | 0.428134982 | 0.11558653 | 3.70402136 | 0.00021221 | 0.00162425 | ARHGAP42 | 143872 |
| ENSG000001 | 461.569877 | 0.529762633 | 0.14303752 | 3.70366204 | 0.00021251 | 0.00162624 | RMI2 | 116028 |
| ENSG000001 | 466.05979 | 0.452377797 | 0.12216047 | 3.70314394 | 0.00021294 | 0.00162926 | NEK3 | 4752 |
| ENSG000001 | 3066.3532 | 0.468947736 | 0.12663909 | 3.70302525 | 0.00021304 | 0.00162972 | HNRNPUL1 | 11100 |
| ENSG000001 | 6567.42702 | 0.370711741 | 0.10011458 | 3.70287457 | 0.00021317 | 0.00162977 | PHF3 | 23469 |
| ENSG000001 | 1148.88359 | 0.446265761 | 0.12051592 | 3.70296104 | 0.0002131 | 0.00162977 | SNX6 | 58533 |
| ENSG000001 | 609.347037 | 0.537723948 | 0.145218 | 3.70287383 | 0.00021317 | 0.00162977 | BORA | 79866 |
| ENSG000001 | 534.144808 | 0.535126856 | 0.14453577 | 3.70238353 | 0.00021358 | 0.00163261 | COPS7B | 64708 |

| | | | | | | | | |
|---|---|---|---|---|---|---|---|---|
| ENSG000002 | 3491.08194 | 0.447311876 | 0.12082127 | 3.70226114 | 0.00021369 | 0.00163309 | NRAS | 4893 |
| ENSG000001 | 428.710209 | 0.540534126 | 0.14602194 | 3.7017322 | 0.00021413 | 0.00163589 | ZNF7 | 7553 |
| ENSG000002 | 84.4765675 | 0.65470503 | 0.1768644 | 3.7017344 | 0.00021413 | 0.00163589 | PXN-AS1 | 100506649 |
| ENSG000001 | 111.344549 | 0.631268434 | 0.17055617 | 3.70123482 | 0.00021455 | 0.00163879 | SIGIRR | 59307 |
| ENSG000001 | 20.7109234 | -1.081261203 | 0.29214362 | -3.701129 | 0.00021464 | 0.00163916 | HHLA2 | 11148 |
| ENSG000001 | 821.662603 | 0.416850519 | 0.11264692 | 3.70050538 | 0.00021517 | 0.00164289 | RASSF3 | 283349 |
| ENSG000001 | 491.575767 | 0.454343255 | 0.12279365 | 3.70005508 | 0.00021555 | 0.0016455 | INIP | 58493 |
| ENSG000001 | 2088.37076 | 0.511693014 | 0.13829853 | 3.69991658 | 0.00021567 | 0.00164608 | MED28 | 80306 |
| ENSG000001 | 10910.7031 | 0.438078094 | 0.11840386 | 3.69986328 | 0.00021572 | 0.00164612 | SF3B1 | 23451 |
| ENSG00000C | 385.807348 | 0.485764548 | 0.13130016 | 3.69964919 | 0.0002159 | 0.0016472 | XRCC1 | 7515 |
| ENSG000001 | 177.513455 | 0.570616423 | 0.15424443 | 3.69942973 | 0.00021608 | 0.00164769 | SNX16 | 64089 |
| ENSG000002 | 533.434609 | 0.58148322 | 0.15717894 | 3.69949835 | 0.00021603 | 0.00164769 | CCNL2 | 81669 |
| ENSG000002 | 28.1449224 | -0.967883831 | 0.26162728 | -3.699476 | 0.00021605 | 0.00164769 | LOC285692 | 285692 |
| ENSG000001 | 422.264141 | 0.506068491 | 0.1368069 | 3.69914438 | 0.00021633 | 0.00164924 | SLCO4C1 | 353189 |
| ENSG00000C | 447.661174 | 0.577285591 | 0.15608373 | 3.69856347 | 0.00021682 | 0.0016524 | MARK2 | 2011 |
| ENSG000002 | 16.6448161 | -1.172547858 | 0.31702745 | -3.6985689 | 0.00021682 | 0.0016524 | NA | NA |
| ENSG000001 | 1970.90397 | 0.448237627 | 0.12119671 | 3.69843054 | 0.00021694 | 0.00165295 | TRIM37 | 4591 |
| ENSG000002 | 1074.02437 | 0.465136322 | 0.12576884 | 3.69834303 | 0.00021701 | 0.00165321 | ZNF788 | 388507 |
| ENSG000001 | 3316.625 | 0.451150111 | 0.12199081 | 3.69823042 | 0.00021711 | 0.00165363 | UBE2N | 7334 |
| ENSG000001 | 143.314064 | 0.616139502 | 0.16661907 | 3.69789309 | 0.0002174 | 0.00165552 | CHCHD6 | 84303 |
| ENSG000001 | 1149.45666 | 0.443061126 | 0.11984656 | 3.69690304 | 0.00021825 | 0.00166167 | ZNF252P | 286101 |
| ENSG000001 | 392.434221 | 0.444513212 | 0.12025249 | 3.69649897 | 0.00021859 | 0.00166401 | STX17 | 55014 |
| ENSG00000C | 3935.54217 | 0.433140987 | 0.11717894 | 3.69640651 | 0.00021867 | 0.0016643 | DNAJA1 | 3301 |
| ENSG000001 | 213.75547 | 0.518862749 | 0.14037952 | 3.69614284 | 0.0002189 | 0.0016654 | CNPPD1 | 27013 |
| ENSG000001 | 2366.14597 | 0.443823661 | 0.12007695 | 3.69616035 | 0.00021888 | 0.0016654 | PUM1 | 9698 |
| ENSG00000C | 281.511521 | 0.641112385 | 0.17345888 | 3.69604813 | 0.00021898 | 0.00166571 | SCARB1 | 949 |
| ENSG000002 | 17.4629955 | -1.108429278 | 0.2999702 | -3.6951313 | 0.00021977 | 0.00167142 | LINC01136 | 730227 |
| ENSG000001 | 4004.08352 | 0.410096843 | 0.11099088 | 3.69486973 | 0.00022 | 0.00167252 | EML4 | 27436 |
| ENSG000002 | 50.3375466 | 0.777852183 | 0.21052215 | 3.69487102 | 0.00022 | 0.00167252 | LOC729970 | 729970 |

| | | | | | | | | |
|---|---|---|---|---|---|---|---|---|
| ENSG000001 | 374.308941 | 0.453917697 | 0.1228549 | 3.69474629 | 0.00022011 | 0.00167302 | ZNF557 | 79230 |
| ENSG000002 | 4.3684223 | -1.398550576 | 0.37854416 | -3.6945507 | 0.00022028 | 0.00167399 | LOC102723362 | 102723362 |
| ENSG000000 | 26768.9121 | 0.412897019 | 0.11176402 | 3.69436458 | 0.00022044 | 0.0016749 | HNRNPC | 3183 |
| ENSG000001 | 279.840234 | 0.533174214 | 0.14433537 | 3.69399557 | 0.00022076 | 0.00167702 | RTN4IP1 | 84816 |
| ENSG000000 | 1972.63456 | 0.526407131 | 0.14250626 | 3.69392279 | 0.00022082 | 0.00167715 | GLG1 | 2734 |
| ENSG000001 | 1081.53636 | 0.472369718 | 0.12787897 | 3.69388104 | 0.00022086 | 0.00167715 | PPP1R8 | 5511 |
| ENSG000002 | 17.2945386 | -1.136204137 | 0.30761396 | -3.693604 | 0.0002211 | 0.00167866 | NA | NA |
| ENSG000001 | 198.440662 | 0.559556645 | 0.15150611 | 3.69329424 | 0.00022137 | 0.0016804 | COMMD1 | 150684 |
| ENSG000001 | 130.796166 | 0.637653107 | 0.17265676 | 3.69318359 | 0.00022146 | 0.00168081 | NA | NA |
| ENSG000001 | 909.16909 | 0.408215847 | 0.11054377 | 3.69279842 | 0.0002218 | 0.00168305 | PSMG1 | 8624 |
| ENSG000002 | 56.9457254 | 0.782144782 | 0.21183359 | 3.69226038 | 0.00022227 | 0.0016863 | LOC101926975 | 101926975 |
| ENSG000002 | 160.89544 | 0.647376857 | 0.17534057 | 3.69211105 | 0.0002224 | 0.00168697 | NA | NA |
| ENSG000001 | 265.752437 | 0.556821644 | 0.1508255 | 3.69182692 | 0.00022265 | 0.00168854 | PDZD11 | 51248 |
| ENSG000001 | 41.2129857 | -0.897711196 | 0.24316541 | -3.6917717 | 0.0002227 | 0.00168859 | CATSPERB | 79820 |
| ENSG000000 | 72.9689194 | -0.760094101 | 0.20589687 | -3.6916253 | 0.00022283 | 0.00168925 | ALPK1 | 80216 |
| ENSG000001 | 29.6559365 | -0.950416673 | 0.25746837 | -3.6913919 | 0.00022303 | 0.00169049 | NME9 | 347736 |
| ENSG000000 | 340.011496 | 0.512187161 | 0.13875983 | 3.69117755 | 0.00022322 | 0.00169096 | PTPRU | 10076 |
| ENSG000001 | 464.246119 | 0.424625655 | 0.11503724 | 3.69120175 | 0.0002232 | 0.00169096 | RBL2 | 5934 |
| ENSG000001 | 1583.95191 | 0.441220401 | 0.11953323 | 3.69119435 | 0.0002232 | 0.00169096 | MIOS | 54468 |
| ENSG000000 | 4186.70667 | 0.484302802 | 0.13121047 | 3.69103788 | 0.00022334 | 0.00169158 | FDFT1 | 2222 |
| ENSG000000 | 1292.22846 | 0.429988192 | 0.11650812 | 3.6906284 | 0.0002237 | 0.00169398 | ITCH | 83737 |
| ENSG000001 | 246.199891 | 0.544927922 | 0.14765525 | 3.69054219 | 0.00022378 | 0.00169424 | ST6GALNAC6 | 30815 |
| ENSG000001 | 207.909191 | 0.565765054 | 0.15331757 | 3.69015143 | 0.00022412 | 0.00169621 | NFKBIB | 4793 |
| ENSG000001 | 55.1202939 | -0.777016744 | 0.2105644 | -3.690162 | 0.00022411 | 0.00169621 | C9orf84 | 158401 |
| ENSG000001 | 6820.34651 | 0.449467329 | 0.12181002 | 3.68990451 | 0.00022434 | 0.00169723 | LAPTM4B | 55353 |
| ENSG000002 | 34.454415 | -0.880396474 | 0.23859427 | -3.6899313 | 0.00022431 | 0.00169723 | TARID | 100507308 |
| ENSG000000 | 1316.0883 | 0.494265801 | 0.13395575 | 3.68976913 | 0.00022446 | 0.00169781 | NUP188 | 23511 |
| ENSG000001 | 813.248483 | 0.443026198 | 0.12007939 | 3.68944402 | 0.00022474 | 0.00169967 | TBC1D31 | 93594 |
| ENSG000001 | 687.043505 | 0.490100744 | 0.13284339 | 3.68931222 | 0.00022486 | 0.00170023 | TMEM237 | 65062 |

| | | | | | | | | | |
|---|---|---|---|---|---|---|---|---|---|
| ENSG000001 | 483.173572 | 0.454326985 | 0.12315296 | 3.68912749 | 0.00022502 | 0.00170115 | FAM179B | | 23116 |
| ENSG000001 | 1072.58448 | 0.566898462 | 0.15367929 | 3.68884102 | 0.00022528 | 0.00170275 | POLE | | 5426 |
| ENSG000001 | 114.012624 | 0.748430489 | 0.20291316 | 3.68842758 | 0.00022564 | 0.0017052 | LMF2 | | 91289 |
| ENSG000002 | 19.3219328 | -1.095475437 | 0.29701229 | -3.6883169 | 0.00022574 | 0.00170562 | NA | NA | |
| ENSG00000C | 763.822822 | 0.496964236 | 0.13477367 | 3.68739846 | 0.00022656 | 0.00171027 | MYNN | | 55892 |
| ENSG000001 | 1177.04389 | 0.483873076 | 0.13122082 | 3.68747187 | 0.00022649 | 0.00171027 | WSB1 | | 26118 |
| ENSG000002 | 11.460095 | -1.218381075 | 0.33040684 | -3.6875177 | 0.00022645 | 0.00171027 | LRIT2 | | 340745 |
| ENSG000002 | 209.335688 | 0.578867829 | 0.15698593 | 3.68738679 | 0.00022657 | 0.00171027 | NA | NA | |
| ENSG000002 | 42.069549 | -0.893849227 | 0.24240289 | -3.6874529 | 0.00022651 | 0.00171027 | NA | NA | |
| ENSG00000C | 553.864308 | 0.497149798 | 0.13485757 | 3.68648048 | 0.00022738 | 0.00171542 | POLA2 | | 23649 |
| ENSG00000C | 1544.83774 | 0.43755737 | 0.1186922 | 3.68648785 | 0.00022737 | 0.00171542 | PKN2 | | 5586 |
| ENSG000002 | 724.647454 | 0.483873311 | 0.13125575 | 3.68649218 | 0.00022737 | 0.00171542 | ZDBF2 | | 57683 |
| ENSG000001 | 1793.16064 | 0.497998219 | 0.13510636 | 3.68597165 | 0.00022783 | 0.00171853 | PCDH1 | | 5097 |
| ENSG000001 | 563.235429 | 0.669068054 | 0.18152171 | 3.68588445 | 0.00022791 | 0.0017188 | CCDC86 | | 79080 |
| ENSG000001 | 308.060079 | -0.490798735 | 0.13315924 | -3.6858031 | 0.00022798 | 0.00171903 | ZBTB6 | | 10773 |
| ENSG000001 | 23.3818065 | -1.020268887 | 0.27682591 | -3.6855976 | 0.00022817 | 0.00171995 | HORMAD1 | | 84072 |
| ENSG000002 | 15.0007676 | -1.133009813 | 0.30741766 | -3.6855717 | 0.00022819 | 0.00171995 | LOC100130331 | | 100130331 |
| ENSG00000C | 1324.57664 | 0.46544424 | 0.12629027 | 3.68551152 | 0.00022824 | 0.00172004 | BTBD1 | | 53339 |
| ENSG000001 | 570.214304 | 0.46592665 | 0.12643457 | 3.68512079 | 0.00022859 | 0.00172204 | TTC39B | | 158219 |
| ENSG000001 | 99.6013793 | 0.687547539 | 0.18657352 | 3.68512924 | 0.00022859 | 0.00172204 | BEX5 | | 340542 |
| ENSG000001 | 877.859139 | 0.467939214 | 0.12698396 | 3.6850261 | 0.00022868 | 0.00172205 | CCNY | | 219771 |
| ENSG000001 | 93.0275881 | 0.691477241 | 0.18764524 | 3.68502406 | 0.00022868 | 0.00172205 | FBXO27 | | 126433 |
| ENSG00000C | 160.425241 | 0.583051664 | 0.15823897 | 3.68462748 | 0.00022904 | 0.00172409 | GPKOW | | 27238 |
| ENSG000001 | 5430.86654 | 0.397531426 | 0.10788816 | 3.68466237 | 0.00022901 | 0.00172409 | PAFAH1B2 | | 5049 |
| ENSG000001 | 481.916127 | 0.482430712 | 0.13094015 | 3.68436046 | 0.00022928 | 0.00172558 | COG3 | | 83548 |
| ENSG000001 | 4033.93654 | 0.427033808 | 0.1159101 | 3.68418107 | 0.00022944 | 0.00172616 | MRPL3 | | 11222 |
| ENSG000001 | 81.2478451 | 0.690797864 | 0.18750338 | 3.68418879 | 0.00022943 | 0.00172616 | ANKRD34B | | 340120 |
| ENSG000001 | 120.027235 | 0.633652371 | 0.1720034 | 3.68395256 | 0.00022964 | 0.00172739 | MIEN1 | | 84299 |
| ENSG000002 | 859.57029 | 0.519195281 | 0.14093822 | 3.68385026 | 0.00022974 | 0.00172776 | C4orf46 | | 201725 |

| | | | | | | | | | |
|---|---|---|---|---|---|---|---|---|---|
| ENSG000001 | 1764.72307 | 0.631950767 | 0.17155761 | 3.68360681 | 0.00022996 | 0.00172909 | MYBL2 | | 4605 |
| ENSG000002 | 886.003858 | 0.520592841 | 0.14134295 | 3.68318919 | 0.00023033 | 0.00173161 | PEX26 | | 55670 |
| ENSG000001 | 10735.4288 | 0.472586788 | 0.12831123 | 3.68312895 | 0.00023039 | 0.00173169 | TOP2A | | 7153 |
| ENSG000002 | 204.299599 | 0.554061285 | 0.15044145 | 3.68290312 | 0.00023059 | 0.00173291 | LOC101929709 | | 101929709 |
| ENSG000001 | 193.999379 | 0.614744983 | 0.16694129 | 3.68240235 | 0.00023105 | 0.00173599 | CRY2 | | 1408 |
| ENSG000002 | 5.32233619 | -1.388216927 | 0.37700219 | -3.6822516 | 0.00023118 | 0.0017367 | NA | NA | |
| ENSG000001 | 417.000786 | 0.483548961 | 0.13132638 | 3.68203992 | 0.00023138 | 0.00173782 | TRPC1 | | 7220 |
| ENSG000001 | 36.5043144 | -0.902270142 | 0.24505762 | -3.6818694 | 0.00023153 | 0.00173866 | OR2C3 | | 81472 |
| ENSG000001 | 51.0475487 | 0.769408133 | 0.20898543 | 3.68163529 | 0.00023174 | 0.00173991 | FBXL15 | | 79176 |
| ENSG000001 | 866.885115 | 0.446243207 | 0.12120931 | 3.68159178 | 0.00023178 | 0.00173991 | BEND4 | | 389206 |
| ENSG000001 | 251.51069 | 0.487715243 | 0.13249528 | 3.68100091 | 0.00023232 | 0.00174363 | APEX2 | | 27301 |
| ENSG000001 | 2193.76974 | 0.489845006 | 0.13312284 | 3.67964654 | 0.00023356 | 0.00175259 | DSTN | | 11034 |
| ENSG000002 | 131.865505 | 0.570698782 | 0.15511125 | 3.67928682 | 0.00023389 | 0.00175474 | ILF3-AS1 | | 147727 |
| ENSG000001 | 29.3183669 | -0.932256991 | 0.25338616 | -3.6791947 | 0.00023397 | 0.00175505 | CNTN3 | | 5067 |
| ENSG000002 | 71.7095412 | -0.707300606 | 0.19226451 | -3.6787891 | 0.00023434 | 0.00175751 | NA | NA | |
| ENSG000001 | 762.757749 | 0.480259969 | 0.13055023 | 3.67873702 | 0.00023439 | 0.00175755 | TXNDC9 | | 10190 |
| ENSG000001 | 60.7831201 | 0.718762436 | 0.19541404 | 3.67815151 | 0.00023493 | 0.00176126 | CRIP3 | | 401262 |
| ENSG000001 | 412.009606 | 0.69963337 | 0.19021582 | 3.67810304 | 0.00023498 | 0.00176127 | SEPN1 | | 57190 |
| ENSG000001 | 132.156822 | 0.724323052 | 0.19693281 | 3.67802119 | 0.00023505 | 0.00176151 | GAA | | 2548 |
| ENSG00000C | 50.4008701 | -0.805549277 | 0.21902623 | -3.6778667 | 0.00023519 | 0.00176225 | ROS1 | | 6098 |
| ENSG000002 | 8923.69654 | 0.446077873 | 0.12129343 | 3.67767558 | 0.00023537 | 0.00176325 | DDX3X | | 1654 |
| ENSG000001 | 1071.58811 | 0.515026869 | 0.14004353 | 3.67761995 | 0.00023542 | 0.0017633 | GOLT1B | | 51026 |
| ENSG000002 | 49.7620211 | -0.809470035 | 0.22011473 | -3.6774914 | 0.00023554 | 0.00176387 | NA | NA | |
| ENSG000002 | 34.1780095 | -0.908051272 | 0.24695041 | -3.6770592 | 0.00023594 | 0.00176653 | NA | NA | |
| ENSG000001 | 1525.2985 | 0.430149653 | 0.11698551 | 3.67694823 | 0.00023604 | 0.00176697 | ZNF675 | | 171392 |
| ENSG00000C | 34.3824084 | -0.891215069 | 0.24239502 | -3.6767053 | 0.00023627 | 0.00176816 | CACNA1S | | 779 |
| ENSG000001 | 3768.66527 | 0.44980533 | 0.12234 | 3.67668258 | 0.00023629 | 0.00176816 | ZNF24 | | 7572 |
| ENSG000001 | 214.402987 | -0.49465884 | 0.1345433 | -3.6765771 | 0.00023638 | 0.00176857 | KCNMA1 | | 3778 |
| ENSG000001 | 822.643691 | 0.448699394 | 0.12204759 | 3.67642982 | 0.00023652 | 0.00176926 | PDE5A | | 8654 |

| | | | | | | | | |
|---|---|---|---|---|---|---|---|---|
| ENSG000001 | 72.8603894 | 0.754795036 | 0.20530922 | 3.67638159 | 0.00023657 | 0.00176927 | C16orf91 | 283951 |
| ENSG000002 | 16.5559939 | -1.129999102 | 0.30737657 | -3.6762695 | 0.00023667 | 0.00176972 | NA | NA |
| ENSG000001 | 285.195606 | 0.530388525 | 0.14428236 | 3.67604548 | 0.00023688 | 0.00177095 | ZNF266 | 10781 |
| ENSG000002 | 8.99058159 | -1.321403189 | 0.35949525 | -3.6757181 | 0.00023718 | 0.00177276 | LOC440704 | 440704 |
| ENSG000002 | 5.87658924 | -1.367669481 | 0.3720851 | -3.67569 | 0.00023721 | 0.00177276 | LOC101929412 | 101929412 |
| ENSG000002 | 32.3955287 | -0.921496202 | 0.25073524 | -3.6751762 | 0.00023769 | 0.001776 | OR10A6 | 390093 |
| ENSG000001 | 20.2862161 | -1.041710986 | 0.2834584 | -3.6750048 | 0.00023784 | 0.00177687 | CLEC9A | 283420 |
| ENSG000001 | 160.154506 | 0.591924253 | 0.16109424 | 3.67439727 | 0.00023841 | 0.00178077 | CTDNEP1 | 23399 |
| ENSG000001 | 278.656518 | 0.597685156 | 0.16268067 | 3.67397764 | 0.0002388 | 0.00178307 | MBOAT7 | 79143 |
| ENSG000002 | 239.05338 | 0.565886593 | 0.15402573 | 3.6739745 | 0.00023881 | 0.00178307 | NA | NA |
| ENSG000001 | 294.184667 | 0.502626033 | 0.13681099 | 3.67387177 | 0.0002389 | 0.00178346 | GATM | 2628 |
| ENSG000001 | 137.623135 | 0.706494036 | 0.19232154 | 3.67350446 | 0.00023925 | 0.0017857 | CEP72 | 55722 |
| ENSG000002 | 3.50671402 | -1.382221391 | 0.37629493 | -3.6732395 | 0.00023949 | 0.00178689 | NA | NA |
| ENSG000002 | 779.345761 | 0.457346169 | 0.12450704 | 3.67325556 | 0.00023948 | 0.00178689 | EBLN3 | 100506710 |
| ENSG000001 | 27.5505399 | -0.965418419 | 0.26283639 | -3.6730775 | 0.00023965 | 0.0017877 | SLC6A20 | 54716 |
| ENSG000001 | 64.0924658 | -0.723761445 | 0.19705388 | -3.6729114 | 0.0002398 | 0.00178853 | NA | NA |
| ENSG000001 | 120.319703 | 0.606523851 | 0.1651618 | 3.672301 | 0.00024038 | 0.00179248 | PCDH19 | 57526 |
| ENSG000002 | 713.672488 | 0.57237237 | 0.15586591 | 3.67221013 | 0.00024046 | 0.00179279 | RBM14 | 10432 |
| ENSG00000C | 78.6689296 | -0.666189198 | 0.18141851 | -3.6721126 | 0.00024055 | 0.00179286 | CFTR | 1080 |
| ENSG000001 | 501.329178 | 0.497637018 | 0.13551842 | 3.67209861 | 0.00024057 | 0.00179286 | PTRH2 | 51651 |
| ENSG000002 | 40.8892013 | -0.836926305 | 0.2279175 | -3.6720582 | 0.0002406 | 0.00179286 | LOC101927829 | 101927829 |
| ENSG000001 | 22.6636911 | -1.028884181 | 0.28019954 | -3.6719696 | 0.00024069 | 0.00179316 | OR10Z1 | 128368 |
| ENSG000002 | 16.4151616 | -1.116135643 | 0.30397114 | -3.6718474 | 0.0002408 | 0.00179369 | HOTAIR | 100124700 |
| ENSG000001 | 394.301565 | 0.644516648 | 0.17553239 | 3.6717819 | 0.00024087 | 0.00179382 | VAV2 | 7410 |
| ENSG000001 | 328.10609 | 0.494936449 | 0.13480603 | 3.67147126 | 0.00024116 | 0.00179567 | ENOSF1 | 55556 |
| ENSG000001 | 609.112736 | 0.451177746 | 0.12289533 | 3.67123594 | 0.00024138 | 0.00179699 | ZNF117 | 51351 |
| ENSG000001 | 174.253717 | 0.528433659 | 0.14394343 | 3.67112048 | 0.00024149 | 0.00179714 | TNFAIP1 | 7126 |
| ENSG000001 | 284.754085 | 0.513725642 | 0.13993611 | 3.67114427 | 0.00024147 | 0.00179714 | TLCD1 | 116238 |
| ENSG000001 | 9.41287491 | -1.2919517 | 0.35195029 | -3.6708358 | 0.00024176 | 0.00179882 | RAB41 | 347517 |

| | | | | | | | | |
|---|---|---|---|---|---|---|---|---|
| ENSG000001 | 33.0428477 | -0.93424782 | 0.25451199 | -3.6707419 | 0.00024185 | 0.00179882 | FAM135B | 51059 |
| ENSG000001 | 460.556466 | 0.476860524 | 0.12990765 | 3.6707654 | 0.00024183 | 0.00179882 | AASDH | 132949 |
| ENSG000001 | 746.545173 | 0.497437295 | 0.13553918 | 3.67006287 | 0.00024249 | 0.00180327 | COPS4 | 51138 |
| ENSG00000C | 778.70395 | 0.430510126 | 0.11730857 | 3.66989497 | 0.00024265 | 0.00180413 | CLDND1 | 56650 |
| ENSG00000C | 804.381376 | 0.443848289 | 0.12097056 | 3.66906031 | 0.00024344 | 0.00180969 | ZFYVE26 | 23503 |
| ENSG00000C | 253.183882 | 0.587406094 | 0.16011455 | 3.66866158 | 0.00024382 | 0.00181219 | APBA2 | 321 |
| ENSG000001 | 509.329372 | 0.441019406 | 0.12021418 | 3.66861384 | 0.00024387 | 0.00181219 | GRPEL1 | 80273 |
| ENSG000002 | 8.57052934 | -1.303407013 | 0.35531087 | -3.6683567 | 0.00024411 | 0.00181368 | NA | NA |
| ENSG000002 | 4.32066721 | -1.388563958 | 0.37855449 | -3.6680689 | 0.00024439 | 0.00181539 | NA | NA |
| ENSG000001 | 360.391594 | 0.433996497 | 0.11832597 | 3.66780412 | 0.00024464 | 0.00181694 | SNX25 | 83891 |
| ENSG000001 | 234.9225 | 0.633449067 | 0.17271152 | 3.66767109 | 0.00024477 | 0.00181722 | DPM2 | 8818 |
| ENSG000001 | 2932.89628 | 0.433345656 | 0.1181528 | 3.66767163 | 0.00024477 | 0.00181722 | HAUS6 | 54801 |
| ENSG000001 | 63.7679476 | 0.788586075 | 0.21506975 | 3.6666526 | 0.00024575 | 0.00182414 | NA | NA |
| ENSG000001 | 517.040263 | 0.52319471 | 0.14269237 | 3.66659198 | 0.0002458 | 0.00182423 | UBL3 | 5412 |
| ENSG000001 | 685.045997 | 0.484596514 | 0.13218915 | 3.66593268 | 0.00024644 | 0.00182861 | MIS18A | 54069 |
| ENSG000001 | 450.081333 | 0.448144781 | 0.12225679 | 3.66560237 | 0.00024676 | 0.00183063 | ACTR8 | 93973 |
| ENSG00000C | 5466.08227 | 0.566868918 | 0.15466878 | 3.66505076 | 0.00024729 | 0.00183391 | HUWE1 | 10075 |
| ENSG000001 | 817.806466 | 0.506364065 | 0.13815848 | 3.66509567 | 0.00024725 | 0.00183391 | CKAP4 | 10970 |
| ENSG000001 | 1049.01358 | 0.471418025 | 0.12863246 | 3.66484506 | 0.00024749 | 0.00183505 | GTF3A | 2971 |
| ENSG000001 | 549.51224 | 0.468144989 | 0.12774506 | 3.66468169 | 0.00024765 | 0.00183589 | BBS4 | 585 |
| ENSG000001 | 544.368186 | 0.574725977 | 0.15683279 | 3.66457788 | 0.00024775 | 0.0018363 | NDFIP2 | 54602 |
| ENSG00000C | 1316.34982 | 0.484676507 | 0.13226258 | 3.66450201 | 0.00024782 | 0.00183651 | VMP1 | 81671 |
| ENSG000001 | 1529.32397 | 0.495309226 | 0.13517305 | 3.66426018 | 0.00024805 | 0.00183791 | WDR33 | 55339 |
| ENSG000001 | 273.978188 | 0.540131576 | 0.14743801 | 3.66344853 | 0.00024884 | 0.00184341 | GFER | 2671 |
| ENSG000002 | 702.012339 | 0.439634056 | 0.1200072 | 3.66339737 | 0.00024889 | 0.00184344 | ZNF271P | 10778 |
| ENSG000001 | 209.556864 | 0.713956685 | 0.194909 | 3.66302577 | 0.00024925 | 0.00184578 | C15orf39 | 56905 |
| ENSG00000C | 680.295983 | 0.438395466 | 0.1196828 | 3.66297802 | 0.0002493 | 0.00184578 | KRIT1 | 889 |
| ENSG000001 | 228.557978 | 0.603955013 | 0.16490391 | 3.66246636 | 0.0002498 | 0.00184914 | SUV39H1 | 6839 |
| ENSG000001 | 77.3742729 | 0.733932301 | 0.20040884 | 3.66217533 | 0.00025008 | 0.0018509 | UBTD1 | 80019 |

| | | | | | | | | |
|---|---|---|---|---|---|---|---|---|
| ENSG00000C | 575.692811 | 0.486164807 | 0.13275852 | 3.66202336 | 0.00025023 | 0.00185166 | DERL2 | 51009 |
| ENSG000001 | 244.286335 | 0.494967936 | 0.13517466 | 3.66169179 | 0.00025056 | 0.00185372 | ZMYM5 | 9205 |
| ENSG000001 | 7.48440676 | -1.330186112 | 0.3632827 | -3.661573 | 0.00025067 | 0.00185425 | FGFBP1 | 9982 |
| ENSG00000C | 100.009258 | 0.658051523 | 0.17974494 | 3.66102954 | 0.0002512 | 0.00185785 | APBA3 | 9546 |
| ENSG000001 | 2635.4503 | 0.422255948 | 0.11534006 | 3.66096533 | 0.00025127 | 0.00185797 | TOMM70A | 9868 |
| ENSG00000C | 668.274948 | 0.448053021 | 0.12239725 | 3.660646 | 0.00025158 | 0.00185859 | TTC27 | 55622 |
| ENSG00000C | 130.692549 | 0.571558635 | 0.15613146 | 3.66075244 | 0.00025148 | 0.00185859 | GRAMD4 | 23151 |
| ENSG000001 | 1677.78797 | 0.408990719 | 0.111721 | 3.66082206 | 0.00025141 | 0.00185859 | NUP88 | 4927 |
| ENSG000001 | 17.8958941 | -1.08024787 | 0.29509529 | -3.6606747 | 0.00025155 | 0.00185859 | ENPP6 | 133121 |
| ENSG000002 | 24.5619434 | -1.001319654 | 0.2735316 | -3.6607092 | 0.00025152 | 0.00185859 | NA | NA |
| ENSG000001 | 1098.66251 | 0.429365731 | 0.11729438 | 3.66058221 | 0.00025164 | 0.00185872 | DCAF16 | 54876 |
| ENSG000001 | 184.724645 | -0.515694463 | 0.14088641 | -3.6603562 | 0.00025186 | 0.00186002 | RYR3 | 6263 |
| ENSG000001 | 1828.17417 | -0.491868515 | 0.13439205 | -3.6599525 | 0.00025226 | 0.00186261 | ZNF649 | 65251 |
| ENSG000001 | 3574.98148 | 0.388823234 | 0.10625253 | 3.65942554 | 0.00025278 | 0.00186611 | HDGFRP3 | 50810 |
| ENSG000001 | 14.5138304 | -1.174765798 | 0.32103691 | -3.6592857 | 0.00025292 | 0.00186679 | SLC17A2 | 10246 |
| ENSG000001 | 66.1730927 | -0.681627305 | 0.18629688 | -3.658823 | 0.00025338 | 0.00186982 | SLC22A2 | 6582 |
| ENSG000001 | 2069.97957 | 0.441134721 | 0.12057343 | 3.65863941 | 0.00025356 | 0.00187082 | BRIX1 | 55299 |
| ENSG000001 | 16.148941 | -1.128205213 | 0.30842962 | -3.6579016 | 0.00025429 | 0.00187587 | NLRP5 | 126206 |
| ENSG000001 | 198.969642 | 0.73059343 | 0.19975625 | 3.65742458 | 0.00025476 | 0.00187902 | DAGLB | 221955 |
| ENSG000001 | 17.3599998 | -1.104918928 | 0.3021303 | -3.6570941 | 0.00025509 | 0.0018811 | C1orf110 | 339512 |
| ENSG00000C | 169.611111 | 0.623070041 | 0.17041084 | 3.65628183 | 0.0002559 | 0.00188673 | SCN4A | 6329 |
| ENSG000001 | 69.4889757 | -0.700743484 | 0.19166182 | -3.6561453 | 0.00025604 | 0.00188739 | SOX5 | 6660 |
| ENSG000001 | 208.319127 | 0.67867285 | 0.18562877 | 3.65607583 | 0.00025611 | 0.00188756 | TRAF2 | 7186 |
| ENSG000001 | 25.56316 | -0.960694649 | 0.26277557 | -3.6559511 | 0.00025623 | 0.00188813 | SAGE1 | 55511 |
| ENSG00000C | 2534.47602 | 0.407261864 | 0.11140403 | 3.6557192 | 0.00025646 | 0.00188915 | ETV1 | 2115 |
| ENSG000001 | 1377.41303 | 0.480460093 | 0.1314256 | 3.65575717 | 0.00025642 | 0.00188915 | CGGBP1 | 8545 |
| ENSG000002 | 10.7066852 | -1.250463808 | 0.34206767 | -3.6556036 | 0.00025658 | 0.00188966 | NA | NA |
| ENSG000001 | 301.893799 | 0.559512935 | 0.15306548 | 3.65538279 | 0.0002568 | 0.00189095 | SURF2 | 6835 |
| ENSG00000C | 580.908285 | 0.495928657 | 0.13569451 | 3.65474368 | 0.00025744 | 0.00189532 | CHMP5 | 51510 |

| | | | | | | | | | |
|---|---|---|---|---|---|---|---|---|---|
| ENSG000001 | 32.3093565 | -0.894300975 | 0.24472901 | -3.6542499 | 0.00025793 | 0.00189862 | NWD1 | | 284434 |
| ENSG00000C | 446.944441 | 0.460590514 | 0.12604457 | 3.65418778 | 0.000258 | 0.00189874 | TRAPPC3 | | 27095 |
| ENSG000001 | 879.736519 | 0.457366544 | 0.12518005 | 3.65366952 | 0.00025852 | 0.00190223 | PTPRA | | 5786 |
| ENSG000001 | 7296.84007 | 0.451326373 | 0.123532 | 3.65351785 | 0.00025867 | 0.00190301 | ITGB1 | | 3688 |
| ENSG000001 | 311.309409 | 0.52242733 | 0.14302833 | 3.65261438 | 0.00025958 | 0.00190903 | C5orf30 | | 90355 |
| ENSG000002 | 48.4867018 | 0.806826873 | 0.22088817 | 3.65264863 | 0.00025955 | 0.00190903 | NA | NA | |
| ENSG000001 | 645.803137 | 0.46190513 | 0.12646507 | 3.65243247 | 0.00025977 | 0.00191004 | FAM104A | | 84923 |
| ENSG000001 | 227.69811 | 0.5082854 | 0.13917057 | 3.65224787 | 0.00025995 | 0.00191072 | MRPS36 | | 92259 |
| ENSG000002 | 26.1985581 | -0.952270992 | 0.26073257 | -3.6522902 | 0.00025991 | 0.00191072 | NA | NA | |
| ENSG000001 | 1935.23389 | 0.457459131 | 0.1252598 | 3.65208262 | 0.00026012 | 0.0019116 | SC5D | | 6309 |
| ENSG000002 | 19.5072883 | -1.096388511 | 0.30023697 | -3.6517438 | 0.00026047 | 0.00191378 | MUC5AC | | 4586 |
| ENSG000001 | 52.152729 | -0.758868937 | 0.20781398 | -3.6516741 | 0.00026054 | 0.00191395 | MEIS1 | | 4211 |
| ENSG000001 | 1263.85764 | 0.458456815 | 0.12555133 | 3.65154879 | 0.00026066 | 0.00191454 | LYAR | | 55646 |
| ENSG000001 | 383.699488 | 0.439714366 | 0.12043555 | 3.65103476 | 0.00026119 | 0.00191803 | ZRANB3 | | 84083 |
| ENSG000002 | 84.0402871 | 0.708342158 | 0.19401396 | 3.65098554 | 0.00026124 | 0.00191805 | SMIM3 | | 85027 |
| ENSG000001 | 47.3903302 | -0.804435844 | 0.22034599 | -3.650785 | 0.00026144 | 0.0019192 | DRP2 | | 1821 |
| ENSG000001 | 103.627778 | 0.706640328 | 0.19357738 | 3.65042818 | 0.0002618 | 0.00192152 | ARMC5 | | 79798 |
| ENSG000001 | 175.953708 | 0.597920081 | 0.16380206 | 3.65025989 | 0.00026198 | 0.00192243 | NRSN2 | | 80023 |
| ENSG00000C | 935.825134 | 0.488562091 | 0.13385112 | 3.65004115 | 0.0002622 | 0.00192337 | TOX4 | | 9878 |
| ENSG000001 | 857.894785 | 0.546288598 | 0.14966611 | 3.65004881 | 0.00026219 | 0.00192337 | POLR2D | | 5433 |
| ENSG000001 | 796.091238 | 0.500757969 | 0.13721123 | 3.64954059 | 0.00026271 | 0.00192678 | EDRF1 | | 26098 |
| ENSG00000C | 143.242795 | 0.569534856 | 0.15606284 | 3.64939449 | 0.00026286 | 0.00192752 | ARSD | | 414 |
| ENSG000001 | 506.195728 | 0.435710378 | 0.11940178 | 3.64911122 | 0.00026315 | 0.0019293 | FAM13A | | 10144 |
| ENSG000001 | 575.227558 | 0.454138527 | 0.12446498 | 3.64872532 | 0.00026354 | 0.00193171 | DDX55 | | 57696 |
| ENSG000002 | 6177.88441 | 0.477681127 | 0.13091826 | 3.64869757 | 0.00026357 | 0.00193171 | LINC00657 | | 647979 |
| ENSG000001 | 153.284995 | 0.617788842 | 0.16932069 | 3.64863183 | 0.00026364 | 0.00193186 | TMPRSS2 | | 7113 |
| ENSG000001 | 1039.40155 | 0.407132326 | 0.11159016 | 3.64846094 | 0.00026382 | 0.00193279 | PEX19 | | 5824 |
| ENSG000001 | 892.597656 | 0.475987442 | 0.13046905 | 3.6482786 | 0.000264 | 0.00193381 | C8orf59 | | 401466 |
| ENSG000001 | 13.8779858 | -1.187078581 | 0.32538752 | -3.6481995 | 0.00026408 | 0.00193406 | F9 | | 2158 |

| | | | | | | | | |
|---|---|---|---|---|---|---|---|---|
| ENSG000001 | 8.96446774 | -1.313910253 | 0.36016768 | -3.6480515 | 0.00026424 | 0.00193483 | DPT | 1805 |
| ENSG000001 | 923.746607 | 0.513662477 | 0.14082916 | 3.64741559 | 0.00026489 | 0.00193927 | MFHAS1 | 9258 |
| ENSG000001 | 3983.3415 | 0.547716229 | 0.1501811 | 3.64703834 | 0.00026528 | 0.00194177 | ATP1A1 | 476 |
| ENSG000001 | 967.751525 | 0.557599245 | 0.15290744 | 3.64664553 | 0.00026569 | 0.00194438 | URB1 | 9875 |
| ENSG000001 | 50.8116404 | 0.866967603 | 0.23779127 | 3.64591859 | 0.00026644 | 0.00194954 | FBXW9 | 84261 |
| ENSG000001 | 1326.11905 | 0.487901314 | 0.13383155 | 3.64563753 | 0.00026673 | 0.00195132 | FNDC3A | 22862 |
| ENSG00000C | 4193.72604 | 0.438564686 | 0.12031836 | 3.64503548 | 0.00026735 | 0.00195379 | SRPK1 | 6732 |
| ENSG000001 | 3575.29257 | 0.397456586 | 0.10904017 | 3.64504722 | 0.00026734 | 0.00195379 | ESRP1 | 54845 |
| ENSG000001 | 700.63308 | 0.471687242 | 0.12940546 | 3.64503343 | 0.00026736 | 0.00195379 | AKAP8 | 10270 |
| ENSG000001 | 373.42831 | 0.478695853 | 0.13132764 | 3.6450504 | 0.00026734 | 0.00195379 | FAM188A | 80013 |
| ENSG000001 | 25.0185681 | -1.032533765 | 0.28326961 | -3.6450566 | 0.00026733 | 0.00195379 | ADAM18 | 8749 |
| ENSG000001 | 737.455738 | 0.437331651 | 0.11997286 | 3.64525476 | 0.00026713 | 0.00195379 | PRKRA | 8575 |
| ENSG00000C | 464.874803 | 0.497754699 | 0.13657033 | 3.64467661 | 0.00026773 | 0.00195615 | DNAJB11 | 51726 |
| ENSG000001 | 1381.82353 | 0.447679523 | 0.12284165 | 3.64436254 | 0.00026806 | 0.00195819 | NOL8 | 55035 |
| ENSG000001 | 664.80499 | 0.48483091 | 0.13303932 | 3.6442679 | 0.00026815 | 0.00195856 | ADAMTS19 | 171019 |
| ENSG00000C | 395.612341 | 0.49055961 | 0.13461991 | 3.64403468 | 0.0002684 | 0.00195963 | FUT8 | 2530 |
| ENSG000001 | 414.189194 | 0.43882886 | 0.12042391 | 3.64403416 | 0.0002684 | 0.00195963 | FKBP15 | 23307 |
| ENSG000001 | 1669.17194 | 0.406090143 | 0.11144763 | 3.64377545 | 0.00026867 | 0.00196085 | SNW1 | 22938 |
| ENSG000001 | 62.1830725 | 0.759382272 | 0.2084058 | 3.64376738 | 0.00026868 | 0.00196085 | DNAJC4 | 3338 |
| ENSG000001 | 232.695446 | 0.59184006 | 0.16243027 | 3.64365613 | 0.00026879 | 0.00196085 | ARHGEF19 | 128272 |
| ENSG000001 | 43.3965013 | -0.876234143 | 0.24048301 | -3.6436426 | 0.00026881 | 0.00196085 | PDE7B | 27115 |
| ENSG000002 | 3684.95558 | 0.469387727 | 0.12882185 | 3.64369664 | 0.00026875 | 0.00196085 | PPP1CB | 5500 |
| ENSG000001 | 114.771876 | 0.627613241 | 0.17225509 | 3.64351053 | 0.00026894 | 0.00196115 | HTR3A | 3359 |
| ENSG000001 | 327.533211 | 0.4950056 | 0.13585918 | 3.64351973 | 0.00026894 | 0.00196115 | ASB7 | 140460 |
| ENSG000001 | 868.793405 | 0.426616012 | 0.11710672 | 3.6429679 | 0.00026951 | 0.00196472 | C5orf15 | 56951 |
| ENSG000001 | 188.78456 | 0.551957804 | 0.15151396 | 3.64295023 | 0.00026953 | 0.00196472 | PKIG | 11142 |
| ENSG00000C | 45.4477463 | -0.802965928 | 0.22045159 | -3.6423685 | 0.00027014 | 0.00196882 | ABCB11 | 8647 |
| ENSG000001 | 1166.03244 | 0.438074524 | 0.12027487 | 3.64227802 | 0.00027024 | 0.00196915 | PARP8 | 79668 |
| ENSG00000C | 1507.76648 | 0.378870551 | 0.10403169 | 3.64187642 | 0.00027066 | 0.00197187 | TXNL1 | 9352 |

| | | | | | | | | |
|---|---|---|---|---|---|---|---|---|
| ENSG000001 | 1996.47012 | 0.525617116 | 0.14433124 | 3.64174178 | 0.0002708 | 0.00197255 | SERINC5 | 256987 |
| ENSG000001 | 331.682892 | 0.49260209 | 0.13529296 | 3.64100326 | 0.00027158 | 0.00197786 | PRKCB | 5579 |
| ENSG00000C | 152.467523 | 0.556830664 | 0.15294687 | 3.64068037 | 0.00027192 | 0.00197975 | LPAR2 | 9170 |
| ENSG000001 | 288.155703 | 0.524916258 | 0.14418143 | 3.64066488 | 0.00027193 | 0.00197975 | DEDD2 | 162989 |
| ENSG000001 | 801.984413 | 0.481349129 | 0.13221753 | 3.64058487 | 0.00027202 | 0.00198001 | UBE2R2 | 54926 |
| ENSG000001 | 2768.74305 | 0.412840288 | 0.11341044 | 3.64023175 | 0.00027239 | 0.00198238 | MOB1A | 55233 |
| ENSG000001 | 195.560974 | 0.545008238 | 0.14972919 | 3.63995992 | 0.00027268 | 0.00198411 | RNF216P1 | 441191 |
| ENSG000001 | 806.567982 | 0.44972113 | 0.12355878 | 3.63973432 | 0.00027292 | 0.0019855 | ATP6AP2 | 10159 |
| ENSG000002 | 14.7474382 | -1.137547485 | 0.31257066 | -3.6393291 | 0.00027335 | 0.00198827 | NA | NA |
| ENSG00000C | 1177.25236 | 0.428939316 | 0.11786826 | 3.63914187 | 0.00027355 | 0.00198936 | ZNF302 | 55900 |
| ENSG000001 | 30.9926925 | -0.935471939 | 0.25708285 | -3.6387956 | 0.00027392 | 0.00199167 | ALS2CR12 | 130540 |
| ENSG000002 | 991.445553 | 0.553185904 | 0.15205609 | 3.63803835 | 0.00027472 | 0.00199718 | DDR1 | 780 |
| ENSG00000C | 57.8752524 | -0.75882415 | 0.20859316 | -3.6378189 | 0.00027496 | 0.00199852 | MYO15A | 51168 |
| ENSG000001 | 1056.95445 | 0.452172882 | 0.12430009 | 3.63775192 | 0.00027503 | 0.00199869 | DSCC1 | 79075 |
| ENSG000001 | 1501.21154 | 0.422792964 | 0.11623572 | 3.63737541 | 0.00027543 | 0.00200089 | ACAP2 | 23527 |
| ENSG000001 | 346.993485 | 0.496324252 | 0.13645017 | 3.637403 | 0.0002754 | 0.00200089 | VPS33A | 65082 |
| ENSG000001 | 748.878791 | 0.452780997 | 0.12448241 | 3.63730907 | 0.0002755 | 0.00200105 | STAM | 8027 |
| ENSG000001 | 508.384096 | 0.603248505 | 0.1658616 | 3.63705946 | 0.00027577 | 0.00200263 | RAPGEF1 | 2889 |
| ENSG00000C | 451.351564 | 0.531081405 | 0.14604243 | 3.63648713 | 0.00027638 | 0.00200672 | ACTR6 | 64431 |
| ENSG000001 | 664.157644 | 0.453501792 | 0.12471086 | 3.63642584 | 0.00027645 | 0.00200684 | HAUS2 | 55142 |
| ENSG000002 | 7.87219434 | -1.313947093 | 0.36134153 | -3.6363025 | 0.00027658 | 0.00200744 | NA | NA |
| ENSG000001 | 90.1335008 | 0.715890081 | 0.1969109 | 3.63560406 | 0.00027733 | 0.00201253 | CCDC64B | 146439 |
| ENSG000002 | 467.173449 | 0.490892188 | 0.13504395 | 3.63505502 | 0.00027792 | 0.00201646 | ZNF844 | 284391 |
| ENSG00000C | 773.251788 | 0.476748537 | 0.13115592 | 3.63497547 | 0.00027801 | 0.00201672 | FYN | 2534 |
| ENSG000002 | 5.73194045 | -1.366217259 | 0.3758699 | -3.6348142 | 0.00027818 | 0.00201762 | NA | NA |
| ENSG000002 | 16.0736962 | -1.09366442 | 0.30091151 | -3.6345052 | 0.00027851 | 0.00201968 | LOC101928535 | 101928535 |
| ENSG000001 | 2690.96947 | 0.464182096 | 0.12771844 | 3.63441729 | 0.00027861 | 0.00202001 | MTF2 | 22823 |
| ENSG000001 | 1968.22589 | 0.603069761 | 0.1659372 | 3.63432535 | 0.00027871 | 0.00202037 | SMARCA4 | 6597 |
| ENSG000001 | 1908.37377 | 0.496158955 | 0.13653677 | 3.6338852 | 0.00027919 | 0.00202346 | FRAS1 | 80144 |

| | | | | | | | | |
|---|---|---|---|---|---|---|---|---|
| ENSG000001 | 57.1999357 | -0.738310817 | 0.20320841 | -3.633269 | 0.00027985 | 0.00202793 | LRFN5 | 145581 |
| ENSG000001 | 29.6011336 | -0.966354493 | 0.26598626 | -3.6330993 | 0.00028004 | 0.00202879 | SLC17A4 | 10050 |
| ENSG000002 | 15.5139149 | -1.128941416 | 0.31074056 | -3.6330674 | 0.00028007 | 0.00202879 | LOC100506444 | 100506444 |
| ENSG000001 | 167.496454 | 0.652302057 | 0.17955836 | 3.63281359 | 0.00028035 | 0.00203043 | FN3KRP | 79672 |
| ENSG000001 | 528.793682 | 0.475421616 | 0.13087157 | 3.63273403 | 0.00028043 | 0.00203069 | ZBTB43 | 23099 |
| ENSG000002 | 29.4697006 | -0.952270346 | 0.26219085 | -3.631974 | 0.00028126 | 0.00203632 | LOC643733 | 643733 |
| ENSG000001 | 434.329325 | 0.543284517 | 0.14960768 | 3.63139461 | 0.00028189 | 0.00204053 | MUM1 | 84939 |
| ENSG000001 | 391.294599 | 0.484405496 | 0.1334129 | 3.63087463 | 0.00028246 | 0.00204429 | FAM8A1 | 51439 |
| ENSG000001 | 234.83606 | 0.542625709 | 0.14945701 | 3.63064741 | 0.00028271 | 0.00204572 | HDHD2 | 84064 |
| ENSG000001 | 337.525721 | 0.583417978 | 0.16070189 | 3.63043632 | 0.00028294 | 0.00204703 | ENO2 | 2026 |
| ENSG000001 | 52464.166 | 0.451417215 | 0.12438185 | 3.62928515 | 0.00028421 | 0.00205581 | NPM1 | 4869 |
| ENSG000001 | 505.316869 | 0.457888081 | 0.12617795 | 3.62890725 | 0.00028462 | 0.00205846 | ZNF189 | 7743 |
| ENSG000001 | 390.720418 | 0.593283699 | 0.16349131 | 3.62883932 | 0.0002847 | 0.00205863 | ATF3 | 467 |
| ENSG00000C | 47.5835384 | -0.794845934 | 0.21905555 | -3.6285131 | 0.00028506 | 0.0020604 | PGR | 5241 |
| ENSG000001 | 76.4938913 | 0.723340269 | 0.19935083 | 3.62847892 | 0.0002851 | 0.0020604 | IL20RB | 53833 |
| ENSG000002 | 741.796309 | 0.445592133 | 0.122804 | 3.62848229 | 0.00028509 | 0.0020604 | RNF5 | 6048 |
| ENSG000002 | 12.8309708 | -1.179265577 | 0.32500873 | -3.628412 | 0.00028517 | 0.00206057 | PADI6 | 353238 |
| ENSG000002 | 147.618035 | 0.646236982 | 0.17811542 | 3.62819221 | 0.00028541 | 0.00206196 | NA | NA |
| ENSG000001 | 166.348867 | 0.58720744 | 0.16185894 | 3.62789626 | 0.00028574 | 0.00206396 | RANGRF | 29098 |
| ENSG000001 | 494.273889 | 0.453381645 | 0.1249796 | 3.62764525 | 0.00028602 | 0.00206556 | PRDM4 | 11108 |
| ENSG000001 | 20.0776557 | -1.028293404 | 0.28346481 | -3.6275875 | 0.00028608 | 0.00206556 | CYP4F12 | 66002 |
| ENSG000001 | 53.8333659 | 0.743292481 | 0.20490163 | 3.62755769 | 0.00028611 | 0.00206556 | C17orf97 | 400566 |
| ENSG000001 | 2312.08734 | 0.414537435 | 0.11427821 | 3.62744064 | 0.00028624 | 0.00206576 | RIMKLB | 57494 |
| ENSG000002 | 78.6700073 | 0.69248389 | 0.19090141 | 3.62744248 | 0.00028624 | 0.00206576 | SNORD116-14 | 100033426 |
| ENSG000001 | 227.010697 | 0.516433354 | 0.14237575 | 3.62725654 | 0.00028645 | 0.00206686 | FAM220A | 84792 |
| ENSG00000C | 834.062653 | 0.484603032 | 0.13361451 | 3.62687422 | 0.00028687 | 0.00206919 | ATG5 | 9474 |
| ENSG000001 | 611.688685 | 0.481237711 | 0.13268527 | 3.6269113 | 0.00028683 | 0.00206919 | MANF | 7873 |
| ENSG000001 | 38.1260514 | -0.885080579 | 0.24405982 | -3.6264904 | 0.0002873 | 0.00207176 | SAG | 6295 |
| ENSG000001 | 4.33381171 | -1.372954347 | 0.37859339 | -3.6264615 | 0.00028733 | 0.00207176 | C9orf66 | 157983 |

| | | | | | | | | |
|---|---|---|---|---|---|---|---|---|
| ENSG00000C | 1054.38323 | 0.410653244 | 0.11324614 | 3.62620078 | 0.00028762 | 0.00207348 | MAP3K1 | 4214 |
| ENSG00000C | 336.238749 | 0.4959148 | 0.13676317 | 3.62608455 | 0.00028775 | 0.00207405 | SLC38A5 | 92745 |
| ENSG000002 | 267.95185 | 0.497363181 | 0.13716734 | 3.62595934 | 0.00028789 | 0.00207469 | NA | NA |
| ENSG000001 | 24.460836 | -0.976918807 | 0.2694285 | -3.6258927 | 0.00028796 | 0.00207485 | SELP | 6403 |
| ENSG000001 | 684.550048 | 0.437620921 | 0.12069619 | 3.62580572 | 0.00028806 | 0.00207518 | SLC35F2 | 54733 |
| ENSG000002 | 36.7563437 | -0.854864723 | 0.23578578 | -3.6255991 | 0.00028829 | 0.00207647 | DIO2 | 1734 |
| ENSG000001 | 875.497263 | 0.498060868 | 0.13737684 | 3.62550822 | 0.00028839 | 0.00207683 | CIT | 11113 |
| ENSG000002 | 24.1194819 | -0.980525865 | 0.27045582 | -3.6254567 | 0.00028845 | 0.00207688 | NA | NA |
| ENSG000001 | 2340.09439 | 0.478988446 | 0.1321543 | 3.62446364 | 0.00028956 | 0.00208451 | SETD5 | 55209 |
| ENSG000001 | 1116.13456 | 0.457733399 | 0.12629884 | 3.62420908 | 0.00028985 | 0.00208619 | NDUFA6 | 4700 |
| ENSG00000C | 148.111629 | 0.663433036 | 0.1830681 | 3.62396847 | 0.00029012 | 0.00208776 | CALCRL | 10203 |
| ENSG000001 | 49.6636203 | 0.752091541 | 0.20753724 | 3.62388726 | 0.00029021 | 0.00208805 | DUSP2 | 1844 |
| ENSG00000C | 1221.52871 | 0.423390085 | 0.11686677 | 3.62284415 | 0.00029138 | 0.00209612 | ERGIC2 | 51290 |
| ENSG000001 | 80.7349143 | 0.708604202 | 0.19560846 | 3.6225641 | 0.0002917 | 0.00209802 | ZNF428 | 126299 |
| ENSG00000C | 493.884037 | 0.647042695 | 0.17862141 | 3.6224251 | 0.00029185 | 0.00209877 | ACOT7 | 11332 |
| ENSG000002 | 162.100099 | 0.60961074 | 0.16830166 | 3.62213155 | 0.00029219 | 0.00210078 | NA | NA |
| ENSG000001 | 199.539736 | 0.609914179 | 0.16840389 | 3.62173446 | 0.00029263 | 0.00210364 | PREX1 | 57580 |
| ENSG00000C | 1570.73449 | 0.449833671 | 0.12421826 | 3.62131687 | 0.00029311 | 0.00210666 | SLC7A8 | 23428 |
| ENSG000001 | 151.882871 | 0.571315795 | 0.15778271 | 3.62090252 | 0.00029358 | 0.00210967 | RELL1 | 768211 |
| ENSG000001 | 29.0270831 | -0.94879909 | 0.26207488 | -3.6203359 | 0.00029422 | 0.00211392 | CHIT1 | 1118 |
| ENSG00000C | 858.612225 | 0.421195064 | 0.11637412 | 3.619319 | 0.00029538 | 0.00212111 | ATG2B | 55102 |
| ENSG000001 | 586.227127 | 0.406537931 | 0.11232435 | 3.61932161 | 0.00029538 | 0.00212111 | FKTN | 2218 |
| ENSG000001 | 41.3668232 | -0.802434056 | 0.22170537 | -3.6193713 | 0.00029532 | 0.00212111 | FNDC1 | 84624 |
| ENSG000001 | 1454.92495 | 0.472339152 | 0.13052624 | 3.61872953 | 0.00029605 | 0.00212557 | WDR48 | 57599 |
| ENSG000001 | 21.0469909 | -1.033493995 | 0.28560588 | -3.618602 | 0.0002962 | 0.00212624 | CYP4F8 | 11283 |
| ENSG000002 | 210.186903 | 0.577269565 | 0.15953205 | 3.61851778 | 0.0002963 | 0.00212656 | LINC00664 | 400680 |
| ENSG000001 | 1638.94211 | 0.442335926 | 0.12225629 | 3.61810354 | 0.00029677 | 0.00212959 | PHAX | 51808 |
| ENSG000001 | 19.1077257 | -1.092167725 | 0.30189871 | -3.6176627 | 0.00029728 | 0.00213284 | DCAF4L2 | 138009 |
| ENSG000001 | 51.3734334 | -0.761365812 | 0.21047505 | -3.6173684 | 0.00029761 | 0.00213467 | ABCA10 | 10349 |

| | | | | | | | | | |
|---|---|---|---|---|---|---|---|---|---|
| ENSG000002 | 44.2853526 | -0.942097663 | 0.26043869 | -3.6173491 | 0.00029764 | 0.00213467 | ABCA17P | | 650655 |
| ENSG000001 | 4167.60439 | 0.574013589 | 0.15869003 | 3.61720005 | 0.00029781 | 0.00213514 | MYO10 | | 4651 |
| ENSG000001 | 3191.28448 | 0.432126744 | 0.11946299 | 3.61724352 | 0.00029776 | 0.00213514 | PTP4A2 | | 8073 |
| ENSG000000 | 461.912804 | 0.497261026 | 0.13747479 | 3.61710696 | 0.00029791 | 0.00213516 | MAP3K9 | | 4293 |
| ENSG000001 | 767.286758 | 0.450813197 | 0.12463289 | 3.61712867 | 0.00029789 | 0.00213516 | ZBTB21 | | 49854 |
| ENSG000001 | 104.099228 | 0.653421921 | 0.18065294 | 3.61700139 | 0.00029804 | 0.00213565 | SLC29A3 | | 55315 |
| ENSG000001 | 43.5428194 | -0.805531878 | 0.2227252 | -3.6167073 | 0.00029837 | 0.0021377 | DPYD | | 1806 |
| ENSG000002 | 66.5142411 | 0.779401579 | 0.2155351 | 3.61612373 | 0.00029905 | 0.00214214 | NA | NA | |
| ENSG000001 | 1003.19784 | 0.646093913 | 0.17867445 | 3.61603983 | 0.00029914 | 0.00214246 | HM13 | | 81502 |
| ENSG000001 | 70.7282159 | 0.759769655 | 0.21012094 | 3.61586834 | 0.00029934 | 0.0021435 | DCAF12L1 | | 139170 |
| ENSG000001 | 2120.86048 | 0.418129036 | 0.11565536 | 3.61530176 | 0.0003 | 0.00214762 | TRIM2 | | 23321 |
| ENSG000002 | 60.6415147 | 0.753562214 | 0.20843813 | 3.61528 | 0.00030002 | 0.00214762 | NA | NA | |
| ENSG000001 | 317.360719 | 0.590877284 | 0.16347433 | 3.61449574 | 0.00030093 | 0.00215375 | THOC6 | | 79228 |
| ENSG000001 | 204.42116 | 0.543954519 | 0.15050757 | 3.6141339 | 0.00030135 | 0.00215638 | STRIP2 | | 57464 |
| ENSG000001 | 426.956824 | 0.600947052 | 0.16628206 | 3.61402223 | 0.00030148 | 0.00215692 | TRIM25 | | 7706 |
| ENSG000001 | 19.4618123 | -1.036969903 | 0.28694939 | -3.6137728 | 0.00030177 | 0.00215862 | TREM1 | | 54210 |
| ENSG000001 | 4641.04417 | 0.438948658 | 0.12147782 | 3.61340593 | 0.0003022 | 0.00216129 | SUMO1 | | 7341 |
| ENSG000001 | 46.9085071 | 0.795350659 | 0.22011383 | 3.61336061 | 0.00030225 | 0.00216129 | SELV | | 348303 |
| ENSG000001 | 455.361952 | 0.470856416 | 0.13031487 | 3.61322082 | 0.00030242 | 0.00216208 | PPP1R7 | | 5510 |
| ENSG000001 | 167.120846 | 0.675611547 | 0.18699794 | 3.61293584 | 0.00030275 | 0.00216408 | MORN4 | | 118812 |
| ENSG000001 | 61.6157469 | 0.823612891 | 0.22799404 | 3.61243163 | 0.00030334 | 0.00216791 | C10orf35 | | 219738 |
| ENSG000001 | 34.5826896 | -0.926406653 | 0.25646789 | -3.6121741 | 0.00030364 | 0.0021693 | FCRL3 | | 115352 |
| ENSG000001 | 4009.02121 | 0.401805822 | 0.11123659 | 3.61217319 | 0.00030364 | 0.0021693 | CPSF2 | | 53981 |
| ENSG000001 | 23.0332842 | -1.019728675 | 0.28231672 | -3.6120024 | 0.00030384 | 0.00217035 | FLG2 | | 388698 |
| ENSG000001 | 585.19458 | 0.470158541 | 0.13018332 | 3.61151133 | 0.00030442 | 0.00217408 | PCGF3 | | 10336 |
| ENSG000001 | 1804.93355 | 0.415887044 | 0.11515833 | 3.61143681 | 0.00030451 | 0.00217433 | DNAJC13 | | 23317 |
| ENSG000001 | 1033.10089 | 0.455735963 | 0.12620784 | 3.61099556 | 0.00030502 | 0.00217764 | LSM8 | | 51691 |
| ENSG000002 | 15.8071581 | -1.13999038 | 0.31572726 | -3.6106809 | 0.00030539 | 0.00217991 | NA | NA | |
| ENSG000000 | 1329.14633 | 0.431871397 | 0.11961275 | 3.61057983 | 0.00030551 | 0.00218016 | DDX20 | | 11218 |

| | | | | | | | | |
|---|---|---|---|---|---|---|---|---|
| ENSG000001 | 43.3164612 | -0.883642477 | 0.24473838 | -3.6105595 | 0.00030554 | 0.00218016 | GRM5 | 2915 |
| ENSG000001 | 18.9858629 | -1.069527057 | 0.29623746 | -3.6103707 | 0.00030576 | 0.00218116 | IGSF6 | 10261 |
| ENSG000001 | 115.038307 | -0.687719503 | 0.19048559 | -3.6103493 | 0.00030578 | 0.00218116 | OR52A1 | 23538 |
| ENSG000001 | 816.114497 | 0.493099422 | 0.13660309 | 3.60972372 | 0.00030652 | 0.00218604 | CDK2AP1 | 8099 |
| ENSG000001 | 853.696074 | 0.489778951 | 0.13570859 | 3.60904893 | 0.00030732 | 0.00219131 | MED20 | 9477 |
| ENSG000001 | 262.997486 | 0.528859176 | 0.14653867 | 3.6090075 | 0.00030737 | 0.00219131 | SDHAF3 | 57001 |
| ENSG000002 | 240.950442 | 0.495469631 | 0.13733548 | 3.60773231 | 0.00030888 | 0.00220172 | BBIP1 | 92482 |
| ENSG000001 | 601.999355 | 0.470200602 | 0.13034168 | 3.60744617 | 0.00030923 | 0.00220376 | ZNF227 | 7770 |
| ENSG000001 | 125.827972 | 0.55103245 | 0.15275257 | 3.60735298 | 0.00030934 | 0.00220417 | WDR25 | 79446 |
| ENSG00000C | 465.492428 | 0.48858674 | 0.13546145 | 3.60683239 | 0.00030996 | 0.0022081 | SEC22C | 9117 |
| ENSG000001 | 694.963503 | 0.551241999 | 0.15283546 | 3.60676764 | 0.00031004 | 0.0022081 | TACC2 | 10579 |
| ENSG000001 | 1023.84771 | 0.403340018 | 0.11182911 | 3.60675328 | 0.00031005 | 0.0022081 | METTL14 | 57721 |
| ENSG000001 | 96.5715741 | 0.627945033 | 0.17411175 | 3.60656315 | 0.00031028 | 0.00220933 | HIP1R | 9026 |
| ENSG000001 | 688.788577 | 0.492205236 | 0.13647806 | 3.60647875 | 0.00031038 | 0.00220966 | LARGE | 9215 |
| ENSG000001 | 19.7776505 | -1.10421317 | 0.30619247 | -3.6062714 | 0.00031063 | 0.00221104 | PLA2G4F | 255189 |
| ENSG000001 | 179.712917 | 0.558285496 | 0.1548188 | 3.6060576 | 0.00031088 | 0.00221243 | AGAP3 | 116988 |
| ENSG000001 | 262.915825 | 0.557265316 | 0.15453763 | 3.60601706 | 0.00031093 | 0.00221243 | NA | NA |
| ENSG000002 | 557.757369 | 0.430467691 | 0.11937741 | 3.60593936 | 0.00031103 | 0.0022127 | PPIL3 | 53938 |
| ENSG000002 | 15.4920194 | -1.127769229 | 0.31277399 | -3.6057002 | 0.00031131 | 0.00221435 | NA | NA |
| ENSG000002 | 54.5516471 | 0.758853417 | 0.21048143 | 3.60532246 | 0.00031177 | 0.00221719 | NA | NA |
| ENSG000001 | 916.550402 | 0.419887583 | 0.1164705 | 3.60509821 | 0.00031203 | 0.00221871 | PDHX | 8050 |
| ENSG000001 | 70.5466661 | -0.665087578 | 0.18449689 | -3.6048715 | 0.00031231 | 0.00222026 | MYO18B | 84700 |
| ENSG000001 | 3132.85091 | 0.390316763 | 0.10827956 | 3.60471334 | 0.0003125 | 0.00222123 | TRIM33 | 51592 |
| ENSG000001 | 147.348278 | 0.567916559 | 0.15755378 | 3.60458862 | 0.00031265 | 0.0022219 | INPP5K | 51763 |
| ENSG000001 | 250.41845 | 0.552290305 | 0.15322295 | 3.60448825 | 0.00031277 | 0.00222237 | DTX4 | 23220 |
| ENSG00000C | 11.7742183 | -1.197605023 | 0.33229234 | -3.6040705 | 0.00031327 | 0.00222556 | FETUB | 26998 |
| ENSG000001 | 2591.03858 | 0.461530437 | 0.12808131 | 3.60341764 | 0.00031406 | 0.00223038 | EIF2A | 83939 |
| ENSG000002 | 33.4546768 | -0.882505233 | 0.24490576 | -3.6034482 | 0.00031402 | 0.00223038 | NA | NA |
| ENSG00000C | 1998.31978 | 0.453503399 | 0.12587104 | 3.60292076 | 0.00031466 | 0.00223426 | DLD | 1738 |

| | | | | | | | | |
|---|---|---|---|---|---|---|---|---|
| ENSG000001 | 62.6149141 | -0.783300694 | 0.21741081 | -3.6028599 | 0.00031474 | 0.00223439 | PZP | 5858 |
| ENSG000001 | 414.131156 | 0.481230419 | 0.13359179 | 3.60224536 | 0.00031548 | 0.00223929 | TSPYL4 | 23270 |
| ENSG000001 | 988.731229 | 0.575664504 | 0.15982607 | 3.60181847 | 0.000316 | 0.00224258 | PLK1 | 5347 |
| ENSG00000C | 37.9286099 | 0.876987905 | 0.243517 | 3.60134165 | 0.00031658 | 0.00224591 | HSD17B14 | 51171 |
| ENSG000001 | 1177.71988 | 0.432324866 | 0.12004453 | 3.60137078 | 0.00031654 | 0.00224591 | DUSP6 | 1848 |
| ENSG000001 | 39.7926739 | -0.83486819 | 0.23186742 | -3.6006275 | 0.00031745 | 0.00225169 | MYOM3 | 127294 |
| ENSG00000C | 640.797682 | 0.445303173 | 0.12367649 | 3.60054831 | 0.00031755 | 0.00225181 | NOA1 | 84273 |
| ENSG000001 | 14877.3719 | 0.431194291 | 0.11975879 | 3.60052315 | 0.00031758 | 0.00225181 | G3BP1 | 10146 |
| ENSG000001 | 1614.4599 | 0.443904843 | 0.12329492 | 3.60034995 | 0.00031779 | 0.00225291 | EIF4E2 | 9470 |
| ENSG00000C | 387.844365 | 0.466570972 | 0.1296002 | 3.60007898 | 0.00031812 | 0.00225432 | ZNF586 | 54807 |
| ENSG000001 | 289.368979 | 0.626304794 | 0.17396912 | 3.60009179 | 0.0003181 | 0.00225432 | NCKAP5L | 57701 |
| ENSG000001 | 82.2269534 | 0.663654146 | 0.18434573 | 3.60005164 | 0.00031815 | 0.00225432 | TMEM99 | 147184 |
| ENSG000001 | 742.807601 | 0.445316031 | 0.12373867 | 3.59884277 | 0.00031964 | 0.00226443 | ELF2 | 1998 |
| ENSG000001 | 53.0833016 | -0.764520302 | 0.21253403 | -3.5971665 | 0.0003217 | 0.00227867 | FAM47E | 100129583 |
| ENSG00000C | 27.3568157 | -0.968555364 | 0.26926471 | -3.5970379 | 0.00032186 | 0.00227939 | CLUL1 | 27098 |
| ENSG000001 | 390.274928 | 0.584320243 | 0.16245799 | 3.59674679 | 0.00032222 | 0.00228075 | GYLTL1B | 120071 |
| ENSG000001 | 155.693936 | 0.591714456 | 0.16451028 | 3.59682353 | 0.00032213 | 0.00228075 | APBB1 | 322 |
| ENSG000001 | 12.828593 | -1.173144516 | 0.32616701 | -3.5967602 | 0.00032221 | 0.00228075 | OR13G1 | 441933 |
| ENSG00000C | 1611.72247 | 0.41625213 | 0.11573878 | 3.59647935 | 0.00032255 | 0.0022827 | MCM10 | 55388 |
| ENSG000001 | 612.302538 | 0.441037215 | 0.12263266 | 3.59640901 | 0.00032264 | 0.00228292 | B4GALT4 | 8702 |
| ENSG000001 | 220.430166 | 0.57951829 | 0.16116013 | 3.59591595 | 0.00032325 | 0.00228685 | NOC4L | 79050 |
| ENSG000001 | 1840.29641 | 0.515603364 | 0.14340057 | 3.59554623 | 0.00032371 | 0.0022897 | MYEF2 | 50804 |
| ENSG00000C | 533.536798 | 0.540465293 | 0.15032217 | 3.59537968 | 0.00032392 | 0.00229013 | FTSJ1 | 24140 |
| ENSG000001 | 1755.3747 | 0.469897593 | 0.13069554 | 3.59536053 | 0.00032394 | 0.00229013 | B4GALT5 | 9334 |
| ENSG000001 | 3231.37596 | 0.448790456 | 0.12482384 | 3.59539048 | 0.00032391 | 0.00229013 | PPP4R2 | 151987 |
| ENSG000001 | 128.067344 | 0.583424315 | 0.16228795 | 3.59499468 | 0.0003244 | 0.00229295 | ANKRD54 | 129138 |
| ENSG000001 | 1429.08868 | 0.403683525 | 0.11229966 | 3.59469936 | 0.00032477 | 0.00229516 | MCM8 | 84515 |
| ENSG00000C | 86.313554 | 0.773801697 | 0.2152718 | 3.59453345 | 0.00032497 | 0.00229622 | MMP25 | 64386 |
| ENSG00000C | 3736.20272 | 0.552654683 | 0.15377018 | 3.59403031 | 0.0003256 | 0.00230026 | PTBP1 | 5725 |

| | | | | | | | | |
|---|---|---|---|---|---|---|---|---|
| ENSG000002 | 377.054592 | 0.501650755 | 0.13958221 | 3.59394469 | 0.00032571 | 0.00230061 | SRA1 | 10011 |
| ENSG000001 | 1384.4301 | 0.447955957 | 0.12464475 | 3.59386149 | 0.00032581 | 0.00230076 | PLEKHA8 | 84725 |
| ENSG000002 | 32.0070004 | -0.91818978 | 0.2554901 | -3.593837 | 0.00032584 | 0.00230076 | ANKUB1 | 389161 |
| ENSG000001 | 1231.96219 | 0.586376048 | 0.16316675 | 3.59372257 | 0.00032599 | 0.00230097 | PLCB3 | 5331 |
| ENSG000001 | 1557.36716 | 0.403140388 | 0.11217904 | 3.59372288 | 0.00032599 | 0.00230097 | ARIH1 | 25820 |
| ENSG000001 | 575.507428 | 0.444451284 | 0.12368004 | 3.59355714 | 0.00032619 | 0.00230204 | CHCHD7 | 79145 |
| ENSG00000C | 341.171992 | 0.448699946 | 0.12486737 | 3.59341235 | 0.00032638 | 0.0023028 | PDCD7 | 10081 |
| ENSG000001 | 92.2693955 | -0.664456342 | 0.18491125 | -3.5933797 | 0.00032642 | 0.0023028 | MIR4435-2HG | 541471 |
| ENSG00000C | 18863.4167 | 0.444387585 | 0.12368814 | 3.59280683 | 0.00032714 | 0.00230747 | XRCC5 | 7520 |
| ENSG000001 | 91.3938596 | 0.637643628 | 0.17748848 | 3.59259162 | 0.00032741 | 0.00230898 | PQLC1 | 80148 |
| ENSG000001 | 1303.2125 | 0.447118101 | 0.12448357 | 3.59178403 | 0.00032842 | 0.00231574 | ARFGEF2 | 10564 |
| ENSG00000C | 2505.235 | 0.583414757 | 0.1624417 | 3.59153314 | 0.00032874 | 0.00231717 | SMC1A | 8243 |
| ENSG000001 | 185.46636 | 0.578420767 | 0.16105009 | 3.59155825 | 0.00032871 | 0.00231717 | AMPD2 | 271 |
| ENSG000001 | 14.3027149 | -1.129881076 | 0.31461174 | -3.5913507 | 0.00032897 | 0.00231802 | PSG6 | 5675 |
| ENSG000002 | 1242.83806 | 0.415519548 | 0.11570022 | 3.59134627 | 0.00032897 | 0.00231802 | C5orf51 | 285636 |
| ENSG000001 | 262.227472 | 0.465890864 | 0.12974538 | 3.59080902 | 0.00032965 | 0.00232241 | APPL2 | 55198 |
| ENSG000001 | 886.437536 | 0.536912361 | 0.14952899 | 3.59069075 | 0.0003298 | 0.00232306 | BLMH | 642 |
| ENSG000001 | 77.3413154 | 0.750237941 | 0.20894666 | 3.59057165 | 0.00032995 | 0.00232371 | PYCRL | 65263 |
| ENSG00000C | 26.3270282 | -0.938548851 | 0.26142647 | -3.5901065 | 0.00033054 | 0.00232728 | CHI3L2 | 1117 |
| ENSG000001 | 149.75674 | 0.526867994 | 0.14675657 | 3.59008112 | 0.00033058 | 0.00232728 | CHST4 | 10164 |
| ENSG000001 | 53.5093971 | -0.763984671 | 0.21282694 | -3.5896991 | 0.00033106 | 0.00233029 | TRPA1 | 8989 |
| ENSG000001 | 166.30673 | 0.562719148 | 0.15676452 | 3.58958227 | 0.00033121 | 0.00233093 | OGFOD3 | 79701 |
| ENSG000001 | 21.1304639 | -1.021013754 | 0.28444637 | -3.5894772 | 0.00033134 | 0.00233147 | AIFM3 | 150209 |
| ENSG000001 | 378.679646 | 0.473042982 | 0.13179553 | 3.58921861 | 0.00033167 | 0.00233338 | ACBD5 | 91452 |
| ENSG000002 | 138.179325 | 0.637753632 | 0.17769627 | 3.58900975 | 0.00033194 | 0.00233484 | CENPBD1P1 | 65996 |
| ENSG000002 | 14.9459844 | -1.114504407 | 0.31055647 | -3.5887335 | 0.00033229 | 0.00233691 | NA | NA |
| ENSG000001 | 63.8044095 | -0.80134569 | 0.2233231 | -3.5882794 | 0.00033287 | 0.00234017 | COL6A5 | 256076 |
| ENSG000002 | 2.38380103 | -1.298383366 | 0.36183709 | -3.5883092 | 0.00033283 | 0.00234017 | NA | NA |
| ENSG000001 | 197.727876 | 0.600771575 | 0.16743715 | 3.58804221 | 0.00033317 | 0.00234189 | IRF2BP1 | 26145 |

| | | | | | | | | | |
|---|---|---|---|---|---|---|---|---|---|
| ENSG000002 | 80.6344946 | 0.715003557 | 0.19930183 | 3.58754128 | 0.00033381 | 0.00234599 | ZBTB22 | | 9278 |
| ENSG000001 | 31.2421676 | -0.918666426 | 0.25608437 | -3.5873584 | 0.00033404 | 0.00234723 | TFAP2A | | 7020 |
| ENSG000001 | 355.956619 | 0.497648589 | 0.13873758 | 3.58697752 | 0.00033453 | 0.00235025 | PMF1 | | 11243 |
| ENSG000002 | 22.1678848 | -1.014209773 | 0.28276274 | -3.5867872 | 0.00033478 | 0.00235156 | LY86-AS1 | | 285780 |
| ENSG000001 | 1937.86879 | 0.41447687 | 0.11555925 | 3.5867043 | 0.00033488 | 0.0023519 | VAPA | | 9218 |
| ENSG00000C | 516.638205 | 0.502545815 | 0.14011575 | 3.58664751 | 0.00033496 | 0.002352 | PHLPP2 | | 23035 |
| ENSG000001 | 1065.10616 | 0.493302902 | 0.1375475 | 3.58641847 | 0.00033525 | 0.00235366 | CASK | | 8573 |
| ENSG000001 | 716.293493 | 0.424444945 | 0.11835364 | 3.58624339 | 0.00033548 | 0.00235443 | PARD6B | | 84612 |
| ENSG000001 | 230.850218 | 0.542858554 | 0.15137181 | 3.58625921 | 0.00033546 | 0.00235443 | UQCR11 | | 10975 |
| ENSG000001 | 320.259776 | 0.473716898 | 0.13210154 | 3.58600588 | 0.00033578 | 0.00235576 | TMEM127 | | 55654 |
| ENSG000001 | 36.1324487 | -0.846076658 | 0.2359369 | -3.5860294 | 0.00033575 | 0.00235576 | C1orf168 | | 199920 |
| ENSG000002 | 48.0781244 | 0.821559023 | 0.22910479 | 3.58595313 | 0.00033585 | 0.00235582 | HIST2H3C | | 126961 |
| ENSG000001 | 14.2480326 | -1.131087177 | 0.31542614 | -3.5859019 | 0.00033592 | 0.00235588 | SLC22A14 | | 9389 |
| ENSG000001 | 829.743579 | 0.45665028 | 0.12734856 | 3.58582982 | 0.00033601 | 0.00235612 | HAUS1 | | 115106 |
| ENSG000001 | 23.2531382 | -1.021657161 | 0.28492487 | -3.5857072 | 0.00033617 | 0.00235682 | CCR2 | | 729230 |
| ENSG000001 | 372.710863 | 0.463674266 | 0.12931497 | 3.58561937 | 0.00033628 | 0.00235721 | NA | NA | |
| ENSG000002 | 12.531651 | -1.178519631 | 0.32871807 | -3.5851988 | 0.00033682 | 0.0023606 | NA | NA | |
| ENSG000001 | 391.107367 | 0.56696258 | 0.1581559 | 3.58483354 | 0.00033729 | 0.0023635 | ZNF74 | | 7625 |
| ENSG000001 | 2003.83673 | 0.413957055 | 0.11548302 | 3.58457074 | 0.00033763 | 0.00236547 | ARHGAP12 | | 94134 |
| ENSG000001 | 1188.38941 | 0.419011512 | 0.1169034 | 3.58425416 | 0.00033804 | 0.00236793 | DCAF17 | | 80067 |
| ENSG000001 | 161.891189 | 0.602139059 | 0.16801492 | 3.58384285 | 0.00033858 | 0.00237126 | CCND3 | | 896 |
| ENSG000001 | 993.561784 | 0.444909171 | 0.1241565 | 3.58345441 | 0.00033908 | 0.00237438 | ZNF100 | | 163227 |
| ENSG000001 | 272.7488 | 0.506038212 | 0.14123005 | 3.58307739 | 0.00033957 | 0.0023774 | PPM1K | | 152926 |
| ENSG000001 | 301.22701 | 0.524334001 | 0.14633956 | 3.58299563 | 0.00033968 | 0.00237773 | CHPT1 | | 56994 |
| ENSG000002 | 14.6096106 | -1.119898537 | 0.31257447 | -3.5828216 | 0.0003399 | 0.00237891 | HNF1B | | 6928 |
| ENSG00000C | 104.901582 | 0.665199506 | 0.18567797 | 3.58254402 | 0.00034026 | 0.00238103 | ZDHHC8 | | 29801 |
| ENSG000001 | 1065.3975 | 0.446259658 | 0.12456938 | 3.58241866 | 0.00034043 | 0.00238176 | NUP35 | | 129401 |
| ENSG000001 | 34.7409154 | -0.900477771 | 0.25137809 | -3.5821648 | 0.00034076 | 0.00238366 | PLCXD3 | | 345557 |
| ENSG00000C | 450.866813 | 0.584015716 | 0.16304037 | 3.58203134 | 0.00034093 | 0.00238447 | TNFRSF1A | | 7132 |

| | | | | | | | | |
|---|---|---|---|---|---|---|---|---|
| ENSG000001 | 332.813102 | 0.499659373 | 0.13949805 | 3.58183758 | 0.00034119 | 0.00238542 | ZNF175 | 7728 |
| ENSG000001 | 244.100474 | 0.564189909 | 0.15751387 | 3.58184262 | 0.00034118 | 0.00238542 | RXRA | 6256 |
| ENSG000001 | 253.439347 | 0.533843144 | 0.14905122 | 3.58160877 | 0.00034148 | 0.00238705 | KBTBD4 | 55709 |
| ENSG000002 | 1689.52072 | 0.410108494 | 0.11450525 | 3.58156937 | 0.00034154 | 0.00238705 | DDX52 | 11056 |
| ENSG000001 | 136.364598 | 0.563489615 | 0.15733569 | 3.58144808 | 0.0003417 | 0.00238774 | CENPA | 1058 |
| ENSG000002 | 33.7273955 | -0.990802927 | 0.27666015 | -3.5812998 | 0.00034189 | 0.00238869 | SLFN12L | 100506736 |
| ENSG000001 | 182.376825 | 0.723071911 | 0.20190971 | 3.58116456 | 0.00034207 | 0.00238916 | DAP | 1611 |
| ENSG000002 | 14.1525116 | -1.165879878 | 0.32555943 | -3.5811584 | 0.00034207 | 0.00238916 | LOC100996624 | 100996624 |
| ENSG000001 | 633.487791 | 0.536727791 | 0.14988482 | 3.5809349 | 0.00034237 | 0.00239079 | CTNNBL1 | 56259 |
| ENSG000001 | 98.5867393 | 0.608805559 | 0.17002222 | 3.58074119 | 0.00034262 | 0.00239174 | ADCK2 | 90956 |
| ENSG000001 | 3192.35102 | 0.436885618 | 0.12200976 | 3.58074316 | 0.00034262 | 0.00239174 | SRP14 | 6727 |
| ENSG000001 | 1583.0251 | 0.464798391 | 0.12981267 | 3.58053169 | 0.0003429 | 0.00239284 | PIP5K1A | 8394 |
| ENSG000001 | 1157.59492 | 0.469194074 | 0.13104013 | 3.58053732 | 0.00034289 | 0.00239284 | ZNF397 | 84307 |
| ENSG000001 | 31.1227261 | -0.895861396 | 0.25021369 | -3.5803852 | 0.00034309 | 0.00239377 | SLC40A1 | 30061 |
| ENSG000001 | 1889.09641 | 0.47063601 | 0.13145213 | 3.58028446 | 0.00034322 | 0.00239428 | YY1 | 7528 |
| ENSG000001 | 8.99140088 | -1.254120113 | 0.35037469 | -3.5793685 | 0.00034443 | 0.00240227 | KRT13 | 3860 |
| ENSG000001 | 111.606085 | 0.617468117 | 0.17251637 | 3.57918569 | 0.00034467 | 0.00240354 | TMUB2 | 79089 |
| ENSG000001 | 2109.25284 | 0.416341439 | 0.11632963 | 3.57898006 | 0.00034494 | 0.00240502 | ERBB2IP | 55914 |
| ENSG000002 | 64.5047032 | -0.776657691 | 0.21702244 | -3.5786976 | 0.00034531 | 0.0024072 | MGAM2 | 93432 |
| ENSG00000C | 195.32938 | 0.617791779 | 0.17264089 | 3.57847893 | 0.0003456 | 0.00240854 | MRI1 | 84245 |
| ENSG000001 | 6236.91193 | 0.533066469 | 0.14896524 | 3.57846209 | 0.00034562 | 0.00240854 | SMARCC1 | 6599 |
| ENSG000001 | 260.620594 | 0.555397773 | 0.15524373 | 3.57758577 | 0.00034678 | 0.00241622 | HCFC1R1 | 54985 |
| ENSG000002 | 114.570153 | 0.578087748 | 0.16159491 | 3.5773883 | 0.00034704 | 0.00241763 | NA | NA |
| ENSG000001 | 103.932031 | 0.737587801 | 0.20619638 | 3.57711329 | 0.00034741 | 0.00241975 | HINT2 | 84681 |
| ENSG000002 | 2543.30678 | 0.408049363 | 0.1140817 | 3.57681703 | 0.0003478 | 0.00242208 | LGR4 | 55366 |
| ENSG000001 | 622.579142 | 0.520434627 | 0.14552957 | 3.57614349 | 0.0003487 | 0.00242791 | SCNN1A | 6337 |
| ENSG000001 | 326.612596 | 0.49345538 | 0.13799804 | 3.57581447 | 0.00034914 | 0.00243055 | ZNF749 | 388567 |
| ENSG000002 | 40.5274002 | -0.85585858 | 0.23935872 | -3.5756316 | 0.00034938 | 0.00243183 | NA | NA |
| ENSG000002 | 649.193398 | 0.441368586 | 0.12346442 | 3.57486455 | 0.00035041 | 0.00243855 | FAM133B | 257415 |

| | | | | | | | | |
|---|---|---|---|---|---|---|---|---|
| ENSG000002 | 34.4068639 | -0.86590036 | 0.24223748 | -3.5745929 | 0.00035077 | 0.00244067 | LINC01606 | 100507651 |
| ENSG000001 | 689.772251 | 0.473187853 | 0.13239236 | 3.57413273 | 0.00035139 | 0.00244454 | DRAP1 | 10589 |
| ENSG000001 | 271.250433 | 0.4531388 | 0.12679283 | 3.57385183 | 0.00035177 | 0.00244675 | MBLAC2 | 153364 |
| ENSG000001 | 239.09826 | 0.490431896 | 0.13723758 | 3.57359776 | 0.00035211 | 0.00244871 | MTFMT | 123263 |
| ENSG000001 | 111.183163 | -0.568589199 | 0.15911622 | -3.5734208 | 0.00035235 | 0.00244994 | CYP2R1 | 120227 |
| ENSG000000 | 5094.09375 | 0.397873225 | 0.11134801 | 3.57324059 | 0.00035259 | 0.00245048 | DPYSL2 | 1808 |
| ENSG000001 | 20.840467 | -0.997634165 | 0.27919683 | -3.5732288 | 0.00035261 | 0.00245048 | CDHR4 | 389118 |
| ENSG000002 | 22.6637503 | -0.972316893 | 0.27210668 | -3.573293 | 0.00035252 | 0.00245048 | NA | NA |
| ENSG000001 | 178.805168 | 0.533121219 | 0.14920412 | 3.57309985 | 0.00035278 | 0.00245127 | KLC4 | 89953 |
| ENSG000001 | 38.2678555 | -0.808389822 | 0.22626011 | -3.5728341 | 0.00035314 | 0.0024532 | PCDHB1 | 29930 |
| ENSG000002 | 11.1197624 | -1.22620642 | 0.34320568 | -3.5728034 | 0.00035318 | 0.0024532 | NA | NA |
| ENSG000000 | 1371.16526 | 0.44062687 | 0.12333986 | 3.57246135 | 0.00035364 | 0.00245599 | AKAP11 | 11215 |
| ENSG000001 | 961.496654 | 0.470718833 | 0.13179091 | 3.57170944 | 0.00035466 | 0.00246263 | LAMA1 | 284217 |
| ENSG000001 | 9874.54933 | 0.38865753 | 0.10881882 | 3.57160227 | 0.0003548 | 0.00246322 | PFN1 | 5216 |
| ENSG000001 | 262.816875 | 0.496708409 | 0.1390765 | 3.57147625 | 0.00035497 | 0.00246356 | MOCS3 | 27304 |
| ENSG000002 | 296.973303 | 0.504725425 | 0.14131963 | 3.57151683 | 0.00035492 | 0.00246356 | NA | NA |
| ENSG000000 | 2491.87965 | 0.427132397 | 0.11961785 | 3.57080832 | 0.00035588 | 0.00246943 | ESF1 | 51575 |
| ENSG000001 | 1706.36112 | 0.439737695 | 0.12315048 | 3.57073479 | 0.00035598 | 0.0024697 | POLE3 | 54107 |
| ENSG000001 | 229.548669 | 0.477718937 | 0.13381079 | 3.5701077 | 0.00035683 | 0.0024752 | NA | NA |
| ENSG000001 | 767.517564 | 0.445778002 | 0.12488288 | 3.56956865 | 0.00035757 | 0.00247987 | DNAJC16 | 23341 |
| ENSG000001 | 884.680319 | 0.476661334 | 0.13353715 | 3.56950358 | 0.00035766 | 0.00248006 | ZNF496 | 84838 |
| ENSG000000 | 2642.58139 | 0.422318618 | 0.11831645 | 3.56939903 | 0.0003578 | 0.00248063 | ZZZ3 | 26009 |
| ENSG000000 | 144.103896 | 0.5942788 | 0.16649507 | 3.56934765 | 0.00035787 | 0.00248069 | CYTH3 | 9265 |
| ENSG000001 | 271.895396 | 0.478954893 | 0.1341909 | 3.56920559 | 0.00035807 | 0.00248161 | TMEM70 | 54968 |
| ENSG000001 | 323.590802 | 0.46667351 | 0.13075345 | 3.56911058 | 0.0003582 | 0.00248209 | CHCHD4 | 131474 |
| ENSG000001 | 1478.54695 | 0.442773274 | 0.12406 | 3.56902528 | 0.00035831 | 0.00248247 | MTX3 | 345778 |
| ENSG000002 | 183.303388 | 0.61845516 | 0.17329029 | 3.56889689 | 0.00035849 | 0.00248299 | ATF6B | 1388 |
| ENSG000002 | 49.1433766 | 0.792259424 | 0.22199099 | 3.56888095 | 0.00035851 | 0.00248299 | NA | NA |
| ENSG000002 | 15.8810526 | -1.094960146 | 0.30681583 | -3.5687864 | 0.00035864 | 0.00248346 | LOC101929517 | 101929517 |

| | | | | | | | | |
|---|---|---|---|---|---|---|---|---|
| ENSG00000C | 793.899163 | 0.463762172 | 0.12995993 | 3.56850129 | 0.00035903 | 0.00248574 | DPP8 | 54878 |
| ENSG000002 | 332.820776 | 0.448597014 | 0.12571569 | 3.5683454 | 0.00035924 | 0.0024868 | EMP2 | 2013 |
| ENSG000001 | 1493.34749 | 0.590422872 | 0.16547824 | 3.56797898 | 0.00035975 | 0.00248985 | AHCYL1 | 10768 |
| ENSG000001 | 2256.79756 | 0.449681654 | 0.12603995 | 3.56777092 | 0.00036003 | 0.0024914 | NAA25 | 80018 |
| ENSG000001 | 603.810864 | 0.469578 | 0.13162349 | 3.56758502 | 0.00036029 | 0.00249274 | SPTY2D1 | 144108 |
| ENSG000001 | 790.807847 | 0.433537165 | 0.12153379 | 3.56721493 | 0.0003608 | 0.00249584 | CCNH | 902 |
| ENSG000002 | 21.9672378 | -0.976453137 | 0.27374033 | -3.5670781 | 0.00036098 | 0.00249672 | LOC101927580 | 101927580 |
| ENSG00000C | 2198.41444 | 0.363727621 | 0.1019735 | 3.56688379 | 0.00036125 | 0.00249777 | ARCN1 | 372 |
| ENSG000001 | 1116.60629 | 0.504774618 | 0.14151721 | 3.56687799 | 0.00036126 | 0.00249777 | OXSR1 | 9943 |
| ENSG000001 | 1742.10601 | 0.464738043 | 0.13029881 | 3.56670971 | 0.00036149 | 0.00249895 | KIAA0232 | 9778 |
| ENSG00000C | 3885.83986 | 0.370093224 | 0.10378788 | 3.56586156 | 0.00036266 | 0.00250662 | FERMT2 | 10979 |
| ENSG000001 | 21.6425338 | -0.984849532 | 0.2762274 | -3.5653578 | 0.00036336 | 0.00251089 | TAF7L | 54457 |
| ENSG000002 | 297.022816 | 0.447610547 | 0.12554549 | 3.56532552 | 0.0003634 | 0.00251089 | KIAA0040 | 9674 |
| ENSG000001 | 1012.73474 | 0.459859488 | 0.12898472 | 3.56522442 | 0.00036354 | 0.00251143 | FBXO38 | 81545 |
| ENSG000001 | 10.6993099 | -1.227631733 | 0.34434086 | -3.5651642 | 0.00036363 | 0.00251158 | DPEP3 | 64180 |
| ENSG000001 | 320.777623 | 0.492354293 | 0.13811025 | 3.56493676 | 0.00036394 | 0.00251333 | DBR1 | 51163 |
| ENSG000001 | 218.766299 | 0.569166856 | 0.15968924 | 3.56421534 | 0.00036495 | 0.00251983 | NRF1 | 4899 |
| ENSG000001 | 1044.17635 | 0.597318961 | 0.16760455 | 3.56385891 | 0.00036544 | 0.00252239 | SRSF4 | 6429 |
| ENSG000001 | 1349.69082 | 0.418280849 | 0.11736698 | 3.56387171 | 0.00036542 | 0.00252239 | GUF1 | 60558 |
| ENSG000001 | 56.6862262 | -0.749168444 | 0.21021848 | -3.5637611 | 0.00036558 | 0.00252247 | MROH2B | 133558 |
| ENSG000001 | 134.458632 | 0.582771987 | 0.16352528 | 3.56380362 | 0.00036552 | 0.00252247 | ZNF395 | 55893 |
| ENSG000001 | 335.311975 | 0.464768702 | 0.13043576 | 3.56319994 | 0.00036636 | 0.00252745 | TMEM67 | 91147 |
| ENSG00000C | 3241.22764 | 0.556063461 | 0.15606188 | 3.56309599 | 0.00036651 | 0.00252802 | WDR1 | 9948 |
| ENSG000001 | 510.303088 | 0.439031332 | 0.12322167 | 3.56293932 | 0.00036673 | 0.0025291 | CETN2 | 1069 |
| ENSG000002 | 16.7612655 | -1.078399424 | 0.3027055 | -3.5625366 | 0.00036729 | 0.00253255 | NA | NA |
| ENSG000001 | 340.127351 | 0.479040973 | 0.1344766 | 3.56226278 | 0.00036767 | 0.00253476 | CEP76 | 79959 |
| ENSG000001 | 28.37814 | -0.959399002 | 0.26934382 | -3.5619864 | 0.00036806 | 0.002537 | OR7C1 | 26664 |
| ENSG000001 | 75.8202534 | 0.711953422 | 0.19988775 | 3.56176614 | 0.00036837 | 0.0025387 | SLC34A2 | 10568 |
| ENSG000001 | 2052.09573 | 0.420693476 | 0.11814901 | 3.56070246 | 0.00036986 | 0.00254857 | TAX1BP1 | 8887 |

| | | | | | | | | |
|---|---|---|---|---|---|---|---|---|
| ENSG00000C | 889.404914 | 0.426925227 | 0.11991125 | 3.56034344 | 0.00037037 | 0.00255102 | SLC6A15 | 55117 |
| ENSG000001 | 367.843719 | 0.487764766 | 0.1370021 | 3.56027214 | 0.00037047 | 0.00255102 | BAG1 | 573 |
| ENSG000001 | 150.335404 | 0.573124283 | 0.16097718 | 3.56028273 | 0.00037046 | 0.00255102 | AIP | 9049 |
| ENSG000002 | 29.5263201 | -0.917482092 | 0.25769841 | -3.560294 | 0.00037044 | 0.00255102 | LOC157273 | 157273 |
| ENSG000001 | 19.6224562 | -1.072786008 | 0.30132614 | -3.5602156 | 0.00037055 | 0.00255114 | MYOZ2 | 51778 |
| ENSG000001 | 159.655269 | 0.543982415 | 0.15281435 | 3.55976002 | 0.00037119 | 0.00255513 | YBEY | 54059 |
| ENSG000001 | 661.725858 | 0.562788593 | 0.1581047 | 3.55959429 | 0.00037143 | 0.00255631 | GPR107 | 57720 |
| ENSG000002 | 127.365692 | 0.612971329 | 0.17220517 | 3.55954089 | 0.0003715 | 0.0025564 | EGLN2 | 112398 |
| ENSG000001 | 104.043526 | 0.622047641 | 0.17476658 | 3.55930542 | 0.00037184 | 0.00255826 | USP20 | 10868 |
| ENSG000001 | 40.2417214 | -0.81976218 | 0.23033199 | -3.5590462 | 0.0003722 | 0.00256035 | ZFPM2 | 23414 |
| ENSG00000C | 198.303278 | 0.591599962 | 0.16622844 | 3.55895754 | 0.00037233 | 0.00256078 | MAP2K7 | 5609 |
| ENSG00000C | 826.097871 | 0.413721899 | 0.11625387 | 3.55877964 | 0.00037258 | 0.00256199 | FAM76B | 143684 |
| ENSG000002 | 201.232354 | 0.523726533 | 0.14716613 | 3.5587436 | 0.00037263 | 0.00256199 | RNASEH1-AS1 | 100506054 |
| ENSG000001 | 631.834862 | 0.458227635 | 0.12878096 | 3.55819401 | 0.00037341 | 0.00256692 | PHTF1 | 10745 |
| ENSG000001 | 45.5661506 | -0.772611972 | 0.21713887 | -3.5581467 | 0.00037348 | 0.00256695 | SCN11A | 11280 |
| ENSG00000C | 681.431843 | 0.410957303 | 0.11550072 | 3.55804957 | 0.00037362 | 0.00256715 | TSG101 | 7251 |
| ENSG000001 | 118.71381 | -0.61204701 | 0.17202031 | -3.5579926 | 0.0003737 | 0.00256715 | POLH | 5429 |
| ENSG000002 | 36.3064863 | -0.871333171 | 0.24489361 | -3.558007 | 0.00037368 | 0.00256715 | AMY2B | 280 |
| ENSG000002 | 11.865008 | -1.191977649 | 0.33506491 | -3.5574529 | 0.00037447 | 0.002572 | LOC100507156 | 100507156 |
| ENSG000001 | 1065.71448 | 0.389352263 | 0.1094521 | 3.55728444 | 0.00037471 | 0.00257321 | UACA | 55075 |
| ENSG000001 | 196.218922 | 0.64179226 | 0.1804269 | 3.55707629 | 0.00037501 | 0.00257481 | KLC2 | 64837 |
| ENSG000001 | 865.393808 | 0.430362969 | 0.12099436 | 3.55688444 | 0.00037528 | 0.00257602 | SH3BGRL | 6451 |
| ENSG000001 | 1095.78469 | 0.468888344 | 0.131827 | 3.55684616 | 0.00037533 | 0.00257602 | POLR2K | 5440 |
| ENSG000001 | 24.8604817 | 1.020241224 | 0.28684087 | 3.55681957 | 0.00037537 | 0.00257602 | SLC25A10 | 1468 |
| ENSG000001 | 326.64689 | 0.496510989 | 0.13963782 | 3.5557058 | 0.00037697 | 0.00258652 | ZNF322 | 79692 |
| ENSG000001 | 534.328533 | 0.464976293 | 0.13077181 | 3.55563088 | 0.00037707 | 0.00258682 | MTMR6 | 9107 |
| ENSG000002 | 21.158064 | -1.006553817 | 0.28312669 | -3.5551358 | 0.00037778 | 0.00259126 | NA | NA |
| ENSG000001 | 850.841407 | 0.427835547 | 0.1203474 | 3.55500441 | 0.00037797 | 0.00259211 | CDC42SE2 | 56990 |
| ENSG000001 | 1867.11839 | 0.462988445 | 0.13026132 | 3.55430492 | 0.00037898 | 0.00259858 | COPS3 | 8533 |

| | | | | | | | | |
|---|---|---|---|---|---|---|---|---|
| ENSG000001 | 157.341352 | 0.590555798 | 0.16616325 | 3.55406994 | 0.00037932 | 0.00260046 | KLF16 | 83855 |
| ENSG000001 | 260.654309 | 0.519676908 | 0.14622716 | 3.55390131 | 0.00037956 | 0.00260169 | ZNF32 | 7580 |
| ENSG000002 | 44.8669218 | -0.837098615 | 0.23556264 | -3.5536137 | 0.00037998 | 0.0026041 | NA | NA |
| ENSG000001 | 2048.09971 | 0.419443468 | 0.11803575 | 3.553529 | 0.0003801 | 0.0026045 | PSPC1 | 55269 |
| ENSG000001 | 284.643898 | 0.554702582 | 0.15613155 | 3.55278985 | 0.00038117 | 0.00261138 | CCDC12 | 151903 |
| ENSG000001 | 3324.85032 | 0.595719494 | 0.16768077 | 3.55270015 | 0.0003813 | 0.00261148 | ACIN1 | 22985 |
| ENSG000001 | 329.684179 | 0.467567443 | 0.13160936 | 3.5526913 | 0.00038131 | 0.00261148 | FBXW8 | 26259 |
| ENSG000001 | 1077.12002 | 0.517677183 | 0.14572371 | 3.55245676 | 0.00038165 | 0.00261337 | NCOA5 | 57727 |
| ENSG000001 | 104.103377 | -0.609953955 | 0.17170723 | -3.5522905 | 0.00038189 | 0.00261458 | DNAH3 | 55567 |
| ENSG000001 | 142.741445 | 0.669937203 | 0.18860533 | 3.55205863 | 0.00038223 | 0.00261644 | CCDC134 | 79879 |
| ENSG000001 | 192.152085 | 0.52210282 | 0.14700435 | 3.55161472 | 0.00038288 | 0.00261944 | TUFT1 | 7286 |
| ENSG000001 | 398.871823 | 0.503490664 | 0.14176529 | 3.55157919 | 0.00038293 | 0.00261944 | SH3RF1 | 57630 |
| ENSG000001 | 655.849988 | 0.547358623 | 0.15411264 | 3.55167907 | 0.00038278 | 0.00261944 | MMP14 | 4323 |
| ENSG000001 | 369.785233 | 0.481661177 | 0.13561554 | 3.55166642 | 0.0003828 | 0.00261944 | ANO5 | 203859 |
| ENSG000002 | 21.9152404 | -0.972266396 | 0.27379946 | -3.5510165 | 0.00038375 | 0.00262461 | LOC101929450 | 101929450 |
| ENSG000001 | 382.465799 | 0.55483783 | 0.15625004 | 3.55096119 | 0.00038383 | 0.00262472 | TGFBRAP1 | 9392 |
| ENSG000001 | 21.0398618 | -1.026484211 | 0.28907966 | -3.5508697 | 0.00038396 | 0.00262519 | GREM2 | 64388 |
| ENSG000001 | 269.075617 | 0.564047934 | 0.15886446 | 3.55049793 | 0.0003845 | 0.00262845 | WBP2 | 23558 |
| ENSG000001 | 148.123703 | 0.560229859 | 0.15779872 | 3.55028132 | 0.00038482 | 0.00263017 | PEMT | 10400 |
| ENSG000001 | 49.4358881 | 0.766173266 | 0.21582155 | 3.55003132 | 0.00038519 | 0.00263208 | LOXL1 | 4016 |
| ENSG000002 | 10.5054111 | -1.217341153 | 0.34291283 | -3.5500018 | 0.00038523 | 0.00263208 | KIR2DL3 | 3804 |
| ENSG000001 | 476.172532 | 0.453116829 | 0.12764103 | 3.5499309 | 0.00038533 | 0.00263235 | SLC50A1 | 55974 |
| ENSG000002 | 134.797153 | 0.547635825 | 0.15427341 | 3.54977457 | 0.00038556 | 0.00263347 | NUDT16P1 | 152195 |
| ENSG000001 | 6711.30378 | 0.440061942 | 0.12397155 | 3.54970117 | 0.00038567 | 0.00263376 | ZNF770 | 54989 |
| ENSG000001 | 20.0458031 | -1.146936909 | 0.32312991 | -3.5494607 | 0.00038602 | 0.00263572 | NA | NA |
| ENSG000001 | 608.19728 | 0.466626594 | 0.1314669 | 3.54938473 | 0.00038613 | 0.00263604 | BUD13 | 84811 |
| ENSG000002 | 8.07378058 | -1.275314173 | 0.35932101 | -3.5492335 | 0.00038635 | 0.00263711 | C8orf49 | 606553 |
| ENSG000001 | 30.5265382 | -0.909189131 | 0.25617818 | -3.5490499 | 0.00038662 | 0.0026385 | EMCN | 51705 |
| ENSG000001 | 337.51168 | 0.544843822 | 0.15352886 | 3.54880402 | 0.00038698 | 0.00264039 | FOXH1 | 8928 |

| | | | | | | | | |
|---|---|---|---|---|---|---|---|---|
| ENSG000001 | 1191.1753 | 0.430738717 | 0.1213768 | 3.54877293 | 0.00038703 | 0.00264039 | CCDC43 | 124808 |
| ENSG00000C | 507.935601 | 0.468624053 | 0.13205618 | 3.54867194 | 0.00038718 | 0.00264051 | CADPS2 | 93664 |
| ENSG000001 | 111.62869 | 0.654299082 | 0.18437783 | 3.54868636 | 0.00038716 | 0.00264051 | KNDC1 | 85442 |
| ENSG000002 | 50.1195119 | -0.835267049 | 0.23538401 | -3.5485292 | 0.00038739 | 0.0026415 | LOC100287225 | 100287225 |
| ENSG000001 | 536.780913 | 0.648691712 | 0.18281327 | 3.54838414 | 0.0003876 | 0.00264251 | ABCC1 | 4363 |
| ENSG000002 | 14.787155 | -1.102371175 | 0.31067427 | -3.5483182 | 0.0003877 | 0.00264273 | MTND6P2 | 106478941 |
| ENSG000001 | 29.4473868 | -0.91358993 | 0.25748048 | -3.5481911 | 0.00038789 | 0.00264356 | IL10RA | 3587 |
| ENSG000002 | 10.2459984 | -1.206174669 | 0.33996173 | -3.5479719 | 0.00038821 | 0.00264532 | NA | NA |
| ENSG000001 | 648.800484 | 0.467942589 | 0.13190581 | 3.54755093 | 0.00038883 | 0.0026491 | TNFRSF21 | 27242 |
| ENSG000002 | 52.6660928 | -0.730577461 | 0.205948 | -3.5473879 | 0.00038907 | 0.0026503 | COL11A2 | 1302 |
| ENSG00000C | 43.1558059 | -0.781935536 | 0.2204632 | -3.5467848 | 0.00038996 | 0.00265548 | HHAT | 55733 |
| ENSG000001 | 1568.35058 | 0.475158254 | 0.13396707 | 3.54682882 | 0.0003899 | 0.00265548 | CUL5 | 8065 |
| ENSG000001 | 102.778236 | 0.628231801 | 0.17713222 | 3.54668277 | 0.00039011 | 0.00265606 | NECAB3 | 63941 |
| ENSG00000C | 888.624301 | 0.529764428 | 0.14938611 | 3.54627636 | 0.00039072 | 0.00265972 | USP13 | 8975 |
| ENSG000001 | 395.468628 | 0.483521233 | 0.13635311 | 3.54609619 | 0.00039098 | 0.00266109 | C12orf29 | 91298 |
| ENSG000001 | 81.8276603 | 0.702471925 | 0.19811579 | 3.54576438 | 0.00039148 | 0.00266399 | HID1 | 283987 |
| ENSG000002 | 18.2020683 | -1.038976367 | 0.29302414 | -3.5457023 | 0.00039157 | 0.00266418 | NA | NA |
| ENSG000001 | 639.142133 | 0.414918095 | 0.11702278 | 3.54561809 | 0.00039169 | 0.00266458 | RBM23 | 55147 |
| ENSG000001 | 35.0830377 | -0.919447736 | 0.25933493 | -3.5454064 | 0.00039201 | 0.00266627 | CSGALNACT1 | 55790 |
| ENSG000001 | 3067.20141 | 0.531749223 | 0.1499858 | 3.54533041 | 0.00039212 | 0.0026666 | DBN1 | 1627 |
| ENSG000001 | 426.812923 | 0.454984302 | 0.12833887 | 3.54517922 | 0.00039235 | 0.00266768 | ZNF594 | 84622 |
| ENSG000001 | 24.2986155 | -0.995024015 | 0.28068309 | -3.5450088 | 0.0003926 | 0.00266896 | ADGRF4 | 221393 |
| ENSG000001 | 918.870783 | 0.434746402 | 0.12264908 | 3.54463645 | 0.00039315 | 0.00267139 | C2orf69 | 205327 |
| ENSG000001 | 113.689942 | 0.573878196 | 0.16190016 | 3.54464246 | 0.00039315 | 0.00267139 | MFSD5 | 84975 |
| ENSG000001 | 103.62771 | 0.620175169 | 0.17496006 | 3.54466708 | 0.00039311 | 0.00267139 | ABCB8 | 11194 |
| ENSG000001 | 1535.62095 | 0.617856279 | 0.1743129 | 3.54452422 | 0.00039332 | 0.00267207 | HERC2 | 8924 |
| ENSG000001 | 2513.61725 | 0.451966851 | 0.12751552 | 3.54440663 | 0.0003935 | 0.00267282 | USP32 | 84669 |
| ENSG000001 | 169.489211 | 0.567542476 | 0.16013342 | 3.54418508 | 0.00039383 | 0.00267462 | CCM2 | 83605 |
| ENSG000001 | 7225.29018 | 0.506285736 | 0.14285659 | 3.54401396 | 0.00039408 | 0.00267591 | CNOT1 | 23019 |

| | | | | | | | | |
|---|---|---|---|---|---|---|---|---|
| ENSG00000C | 5714.69818 | 0.51352936 | 0.14490943 | 3.54379528 | 0.00039441 | 0.00267768 | EZR | 7430 |
| ENSG00000C | 22.3559071 | -0.975623259 | 0.27533737 | -3.543374 | 0.00039504 | 0.00268151 | NLRC4 | 58484 |
| ENSG000001 | 936.634717 | 0.408877363 | 0.11539614 | 3.54324985 | 0.00039523 | 0.00268233 | SPATA5 | 166378 |
| ENSG000001 | 807.466486 | 0.596280856 | 0.16828861 | 3.54320393 | 0.0003953 | 0.00268234 | MEPCE | 56257 |
| ENSG000001 | 1947.64231 | 0.430077294 | 0.1213922 | 3.54287424 | 0.00039579 | 0.00268525 | ROBO1 | 6091 |
| ENSG000001 | 84.095548 | 0.66234247 | 0.1869664 | 3.54257489 | 0.00039624 | 0.00268785 | CTBS | 1486 |
| ENSG000001 | 124.094401 | 0.640924587 | 0.18094388 | 3.54211816 | 0.00039693 | 0.00269206 | SELENBP1 | 8991 |
| ENSG000001 | 677.898845 | 0.409999422 | 0.11575127 | 3.54207288 | 0.000397 | 0.00269207 | RAB5B | 5869 |
| ENSG000001 | 1768.84193 | 0.418243609 | 0.11810317 | 3.54134119 | 0.0003981 | 0.0026991 | HOMER1 | 9456 |
| ENSG000001 | 179.811395 | 0.533143852 | 0.1505505 | 3.54129569 | 0.00039817 | 0.00269911 | POMT1 | 10585 |
| ENSG000001 | 105.659449 | 0.633962073 | 0.17902715 | 3.54115053 | 0.00039839 | 0.00270014 | NAA10 | 8260 |
| ENSG000001 | 225.248156 | 0.479680633 | 0.13547023 | 3.54085643 | 0.00039883 | 0.0027027 | PLBD1 | 79887 |
| ENSG000001 | 681.02399 | 0.510571263 | 0.14424234 | 3.53967684 | 0.00040062 | 0.00271436 | FAM83D | 81610 |
| ENSG000001 | 947.543261 | 0.626832885 | 0.17709221 | 3.53958474 | 0.00040076 | 0.00271485 | NOB1 | 28987 |
| ENSG000001 | 193.279392 | 0.555590348 | 0.1569713 | 3.53943912 | 0.00040098 | 0.0027159 | GTPBP3 | 84705 |
| ENSG000001 | 28.9081291 | -0.917770344 | 0.25932164 | -3.5391198 | 0.00040146 | 0.00271873 | WDR93 | 56964 |
| ENSG00000C | 2558.5881 | 0.516914573 | 0.14607357 | 3.53872762 | 0.00040206 | 0.00272232 | DDX1 | 1653 |
| ENSG000001 | 640.224625 | 0.417881333 | 0.1180929 | 3.53858146 | 0.00040228 | 0.00272337 | KIAA0907 | 22889 |
| ENSG000001 | 51.498672 | -0.749765826 | 0.21189054 | -3.5384583 | 0.00040247 | 0.00272342 | PRRX1 | 5396 |
| ENSG000001 | 19.4993918 | -1.046919272 | 0.29586726 | -3.5384762 | 0.00040244 | 0.00272342 | ARR3 | 407 |
| ENSG000001 | 1895.98075 | 0.476129452 | 0.13455898 | 3.53844438 | 0.00040249 | 0.00272342 | GTF3C4 | 9329 |
| ENSG000001 | 64.3862169 | 0.732394327 | 0.20700746 | 3.53800929 | 0.00040316 | 0.00272746 | PARD6A | 50855 |
| ENSG000001 | 1469.32272 | 0.427141372 | 0.12075234 | 3.53733425 | 0.00040419 | 0.00273372 | MRPL32 | 64983 |
| ENSG000001 | 632.595847 | 0.432403968 | 0.12224137 | 3.53729649 | 0.00040425 | 0.00273372 | PRKAG1 | 5571 |
| ENSG000002 | 41.1838003 | 0.835879486 | 0.23630629 | 3.53727146 | 0.00040428 | 0.00273372 | NA | NA |
| ENSG00000C | 78.7855222 | -0.657495568 | 0.18589457 | -3.5369272 | 0.00040481 | 0.00273683 | ANKRD44 | 91526 |
| ENSG000001 | 1134.51687 | 0.43169261 | 0.12205816 | 3.53677793 | 0.00040504 | 0.00273792 | PMPCB | 9512 |
| ENSG000001 | 150.738636 | 0.60366541 | 0.17068826 | 3.53665445 | 0.00040523 | 0.00273874 | HTRA2 | 27429 |
| ENSG000001 | 579.42722 | 0.430350933 | 0.12169142 | 3.5364116 | 0.0004056 | 0.00274023 | POLI | 11201 |

| | | | | | | | | |
|---|---|---|---|---|---|---|---|---|
| ENSG000001 | 1065.76256 | 0.443305622 | 0.12535581 | 3.53637884 | 0.00040565 | 0.00274023 | PEX1 | 5189 |
| ENSG000001 | 876.29695 | 0.434910006 | 0.12297909 | 3.53645486 | 0.00040554 | 0.00274023 | TMEM209 | 84928 |
| ENSG000001 | 1747.48999 | 0.477859088 | 0.13513219 | 3.53623432 | 0.00040587 | 0.00274128 | HTATSF1 | 27336 |
| ENSG000002 | 570.534693 | 0.502845015 | 0.14219976 | 3.53618745 | 0.00040595 | 0.00274131 | CPNE1 | 8904 |
| ENSG000001 | 143.564645 | 0.568201379 | 0.16068888 | 3.53603419 | 0.00040618 | 0.00274165 | SLC35B3 | 51000 |
| ENSG000001 | 415.444432 | 0.609915392 | 0.17248631 | 3.53602211 | 0.0004062 | 0.00274165 | FAM171A1 | 221061 |
| ENSG000001 | 35.7646521 | 0.888244084 | 0.2511943 | 3.53608378 | 0.00040611 | 0.00274165 | NA | NA |
| ENSG000001 | 135.123693 | 0.589400228 | 0.16669051 | 3.53589556 | 0.0004064 | 0.00274251 | SH3BP5 | 9467 |
| ENSG000001 | 226.764401 | 0.491378845 | 0.13900327 | 3.53501653 | 0.00040775 | 0.00275119 | DUS2 | 54920 |
| ENSG000001 | 585.935717 | 0.458704587 | 0.12976485 | 3.53489091 | 0.00040794 | 0.00275204 | MRPS18C | 51023 |
| ENSG000002 | 1720.59651 | 0.41888756 | 0.11850885 | 3.53465207 | 0.00040831 | 0.00275407 | N4BP2L2 | 10443 |
| ENSG000001 | 1768.40359 | 0.425529781 | 0.12039358 | 3.53448893 | 0.00040856 | 0.0027544 | FASTKD2 | 22868 |
| ENSG000001 | 815.625611 | 0.421518073 | 0.11925584 | 3.5345697 | 0.00040844 | 0.0027544 | CAST | 831 |
| ENSG000001 | 676.137083 | 0.445099541 | 0.12592962 | 3.53451033 | 0.00040853 | 0.0027544 | C2orf44 | 80304 |
| ENSG000001 | 1127.31244 | 0.469280551 | 0.13278596 | 3.53411274 | 0.00040915 | 0.0027574 | ETNK1 | 55500 |
| ENSG000002 | 201.74857 | 0.608951194 | 0.17230616 | 3.53412315 | 0.00040913 | 0.0027574 | NA | NA |
| ENSG000001 | 1551.76071 | 0.526356657 | 0.14893821 | 3.53406066 | 0.00040923 | 0.00275749 | WHSC1 | 7468 |
| ENSG00000C | 1592.45679 | 0.57722154 | 0.163337 | 3.53393018 | 0.00040943 | 0.00275839 | EPB41L2 | 2037 |
| ENSG00000C | 2068.01117 | 0.414928603 | 0.11741592 | 3.53383609 | 0.00040957 | 0.0027589 | DNM1L | 10059 |
| ENSG00000C | 342.817278 | 0.461300092 | 0.13053964 | 3.53379329 | 0.00040964 | 0.0027589 | ZC3HC1 | 51530 |
| ENSG000001 | 415.972011 | 0.52778424 | 0.14935804 | 3.53368492 | 0.00040981 | 0.00275937 | CERS5 | 91012 |
| ENSG000001 | 160.683468 | 0.548343909 | 0.1551773 | 3.53366069 | 0.00040985 | 0.00275937 | ANGPT1 | 284 |
| ENSG000001 | 4275.32663 | 0.452152872 | 0.12797014 | 3.53326869 | 0.00041046 | 0.00276209 | CBX3 | 11335 |
| ENSG000001 | 741.314547 | 0.44331294 | 0.12546756 | 3.53328738 | 0.00041043 | 0.00276209 | ZNF562 | 54811 |
| ENSG000002 | 58.0121894 | -0.743811928 | 0.21051345 | -3.5333226 | 0.00041037 | 0.00276209 | MAMDC2-AS1 | 100507244 |
| ENSG000001 | 572.383671 | 0.447333234 | 0.1266136 | 3.53305846 | 0.00041078 | 0.00276383 | PCMTD1 | 115294 |
| ENSG000001 | 222.39879 | 0.630915103 | 0.17859087 | 3.53273998 | 0.00041128 | 0.0027667 | SH3PXD2A | 9644 |
| ENSG000001 | 509.932417 | 0.397708037 | 0.11258607 | 3.53247987 | 0.00041168 | 0.00276897 | YEATS4 | 8089 |
| ENSG000001 | 260.318774 | 0.535548492 | 0.15160948 | 3.53242097 | 0.00041177 | 0.00276912 | PLD2 | 5338 |

| | | | | | | | | |
|---|---|---|---|---|---|---|---|---|
| ENSG000002 | 33.4788383 | 0.935061011 | 0.26472681 | 3.53217341 | 0.00041216 | 0.00277126 | NA | NA |
| ENSG000002 | 3.88545111 | -1.333741438 | 0.37762036 | -3.5319637 | 0.00041249 | 0.002773 | NA | NA |
| ENSG000001 | 177.508015 | 0.640708571 | 0.18140604 | 3.53190324 | 0.00041258 | 0.00277317 | SH3BP1 | 23616 |
| ENSG000001 | 380.594888 | 0.490243234 | 0.13880745 | 3.53182219 | 0.00041271 | 0.00277356 | SLC35A4 | 113829 |
| ENSG000001 | 2282.74009 | 0.37748512 | 0.10688652 | 3.53164373 | 0.00041299 | 0.00277497 | BAZ2B | 29994 |
| ENSG000002 | 44.8926092 | 0.805465441 | 0.22809103 | 3.53133333 | 0.00041347 | 0.00277777 | NA | NA |
| ENSG00000C | 1032.78244 | 0.404173594 | 0.11445753 | 3.53121014 | 0.00041366 | 0.00277815 | SPAST | 6683 |
| ENSG000001 | 378.734089 | 0.422582426 | 0.11967059 | 3.53121372 | 0.00041366 | 0.00277815 | ARL13B | 200894 |
| ENSG00000C | 13.3386764 | -1.168831306 | 0.3310057 | -3.5311516 | 0.00041375 | 0.0027783 | TBX21 | 30009 |
| ENSG000001 | 2188.15488 | 0.448042215 | 0.12688764 | 3.53101535 | 0.00041397 | 0.00277835 | XPO7 | 23039 |
| ENSG000001 | 474.446398 | 0.554127372 | 0.15692992 | 3.53104984 | 0.00041391 | 0.00277835 | GAK | 2580 |
| ENSG000002 | 33.9734994 | -0.954460021 | 0.27030166 | -3.531092 | 0.00041385 | 0.00277835 | LOC101927009 | 101927009 |
| ENSG000001 | 685.30673 | 0.524916123 | 0.14866582 | 3.530846 | 0.00041423 | 0.00277967 | NXN | 64359 |
| ENSG00000C | 68.0277348 | -0.698649375 | 0.19787645 | -3.5307353 | 0.00041441 | 0.00278038 | FAS | 355 |
| ENSG000001 | 688.969366 | 0.594135606 | 0.1682867 | 3.53049654 | 0.00041478 | 0.00278243 | CEP250 | 11190 |
| ENSG000001 | 1618.82643 | 0.549418674 | 0.15562944 | 3.53030035 | 0.00041509 | 0.00278403 | ARID3B | 10620 |
| ENSG000001 | 457.56811 | 0.464085999 | 0.13146354 | 3.53014976 | 0.00041532 | 0.00278515 | FAM160B1 | 57700 |
| ENSG000001 | 89.5732528 | 0.671901824 | 0.19033799 | 3.53004576 | 0.00041549 | 0.00278579 | TPRA1 | 131601 |
| ENSG000001 | 3225.29452 | 0.420074009 | 0.11902166 | 3.52939122 | 0.00041652 | 0.00279223 | ADGRL2 | 23266 |
| ENSG00000C | 1063.19469 | 0.434413482 | 0.12309209 | 3.52917473 | 0.00041686 | 0.00279405 | STAU2 | 27067 |
| ENSG000001 | 114.311679 | 0.559592394 | 0.15856548 | 3.5290934 | 0.00041699 | 0.00279445 | HRAS | 3265 |
| ENSG000001 | 1352.92536 | 0.47651783 | 0.13503314 | 3.52889531 | 0.0004173 | 0.00279608 | ARMC1 | 55156 |
| ENSG000001 | 28.870099 | -0.884404932 | 0.25063077 | -3.5287165 | 0.00041758 | 0.0027975 | ABLIM2 | 84448 |
| ENSG000002 | 9.23767457 | -1.227838141 | 0.34800589 | -3.5282108 | 0.00041838 | 0.00280239 | DACT3-AS1 | 100506068 |
| ENSG000001 | 2160.91956 | 0.388333254 | 0.11006776 | 3.52812906 | 0.00041851 | 0.00280279 | OSBPL9 | 114883 |
| ENSG000001 | 128.95792 | 0.567360615 | 0.16082196 | 3.52788031 | 0.0004189 | 0.00280497 | KRBOX4 | 55634 |
| ENSG000001 | 269.318911 | 0.501098209 | 0.14205571 | 3.5274768 | 0.00041954 | 0.00280878 | R3HCC1 | 203069 |
| ENSG000001 | 412.864652 | 0.513175305 | 0.1454844 | 3.52735615 | 0.00041973 | 0.0028096 | BRI3BP | 140707 |
| ENSG00000C | 734.63835 | 0.42517989 | 0.12054172 | 3.52724265 | 0.00041991 | 0.00281034 | CRLS1 | 54675 |

| | | | | | | | | |
|---|---|---|---|---|---|---|---|---|
| ENSG00000( | 508.36807 | 0.45460172 | 0.12888861 | 3.52708993 | 0.00042015 | 0.00281103 | ZNF264 | 9422 |
| ENSG000002 | 888.869157 | 0.628397642 | 0.17816269 | 3.52710002 | 0.00042014 | 0.00281103 | PBX2 | 5089 |
| ENSG000001 | 36.0779087 | -0.823520103 | 0.23350246 | -3.5268155 | 0.00042059 | 0.00281348 | SLC26A5 | 375611 |
| ENSG000001 | 636.561445 | 0.422139189 | 0.11971482 | 3.52620669 | 0.00042156 | 0.00281949 | PDGFA | 5154 |
| ENSG000001 | 1157.62014 | 0.428245205 | 0.12145953 | 3.52582638 | 0.00042216 | 0.00282308 | DYNC1LI1 | 51143 |
| ENSG000001 | 1621.53504 | 0.581380367 | 0.16490949 | 3.52545126 | 0.00042276 | 0.00282662 | ARHGAP35 | 2909 |
| ENSG000001 | 298.549198 | 0.528317969 | 0.14987052 | 3.52516268 | 0.00042322 | 0.00282876 | TBC1D2B | 23102 |
| ENSG000001 | 157.212306 | 0.582231684 | 0.16516381 | 3.52517701 | 0.0004232 | 0.00282876 | ZNF408 | 79797 |
| ENSG000001 | 375.577447 | 0.46477433 | 0.13184842 | 3.52506549 | 0.00042338 | 0.00282934 | ZNF226 | 7769 |
| ENSG000001 | 899.838009 | 0.545325543 | 0.15470724 | 3.52488694 | 0.00042366 | 0.00283078 | UNC119B | 84747 |
| ENSG000001 | 664.48079 | 0.484430499 | 0.13743605 | 3.52477029 | 0.00042385 | 0.00283156 | NR2C2 | 7182 |
| ENSG000001 | 123.237604 | 0.647487315 | 0.18371069 | 3.52449459 | 0.00042429 | 0.00283401 | B9D1 | 27077 |
| ENSG000001 | 28.6899916 | 0.911008107 | 0.25848211 | 3.52445327 | 0.00042436 | 0.00283401 | INSM1 | 3642 |
| ENSG000001 | 744.643169 | 0.423973814 | 0.12029877 | 3.52434038 | 0.00042454 | 0.00283475 | CHN1 | 1123 |
| ENSG000001 | 294.795018 | 0.463042785 | 0.13139736 | 3.52398856 | 0.0004251 | 0.00283805 | DAZAP2 | 9802 |
| ENSG000002 | 19.3782303 | -1.042802483 | 0.29595703 | -3.5234929 | 0.0004259 | 0.0028429 | NAV2-AS5 | 100874012 |
| ENSG000001 | 2885.42622 | 0.412725893 | 0.11714067 | 3.52333549 | 0.00042615 | 0.00284412 | CTNND1 | 1500 |
| ENSG000001 | 1242.02599 | 0.590332632 | 0.16755807 | 3.52315241 | 0.00042645 | 0.0028455 | TIMELESS | 8914 |
| ENSG000001 | 957.838731 | 0.473543713 | 0.13441033 | 3.52312005 | 0.0004265 | 0.0028455 | PARPBP | 55010 |
| ENSG000001 | 893.37179 | 0.447947271 | 0.1271613 | 3.52266987 | 0.00042722 | 0.00284986 | HNRNPLL | 92906 |
| ENSG000002 | 20.4604552 | 1.040168742 | 0.2953029 | 3.52237898 | 0.00042769 | 0.00285252 | PAGR1 | 79447 |
| ENSG000001 | 55.945906 | -0.777543399 | 0.22075225 | -3.5222444 | 0.00042791 | 0.00285303 | ABCA8 | 10351 |
| ENSG000001 | 26.487402 | -0.906867705 | 0.25746695 | -3.5222684 | 0.00042787 | 0.00285303 | RCSD1 | 92241 |
| ENSG000001 | 296.568677 | 0.480646462 | 0.13646277 | 3.52218025 | 0.00042801 | 0.00285326 | PPAPDC1B | 84513 |
| ENSG000001 | 426.011004 | 0.550498283 | 0.15630628 | 3.5219204 | 0.00042843 | 0.00285558 | ZNF740 | 283337 |
| ENSG000001 | 39.7545567 | -0.78806159 | 0.22378001 | -3.5215906 | 0.00042897 | 0.00285867 | NA | NA |
| ENSG00000( | 1546.89091 | 0.452733306 | 0.12856118 | 3.52153972 | 0.00042905 | 0.00285875 | PICALM | 8301 |
| ENSG000001 | 1112.53154 | 0.394473147 | 0.1120594 | 3.52021455 | 0.0004312 | 0.0028726 | RNF38 | 152006 |
| ENSG000001 | 3368.25584 | 0.42661342 | 0.12120087 | 3.51988758 | 0.00043173 | 0.00287567 | SBNO1 | 55206 |

| | | | | | | | | |
|---|---|---|---|---|---|---|---|---|
| ENSG000001 | 52.6938945 | -0.739844451 | 0.21019448 | -3.5198091 | 0.00043186 | 0.00287605 | ESR2 | 2100 |
| ENSG000001 | 2255.57514 | 0.362585928 | 0.1030153 | 3.51972882 | 0.00043199 | 0.00287645 | RPF2 | 84154 |
| ENSG000001 | 632.733937 | 0.457597678 | 0.13001845 | 3.51948257 | 0.00043239 | 0.0028777 | MSANTD4 | 84437 |
| ENSG000001 | 611.581825 | 0.496716011 | 0.14113118 | 3.51953427 | 0.00043231 | 0.0028777 | INSR | 3643 |
| ENSG000001 | 7.94016009 | -1.26308731 | 0.3588839 | -3.5194872 | 0.00043238 | 0.0028777 | NA | NA |
| ENSG000001 | 444.306823 | 0.437512436 | 0.1243155 | 3.51937146 | 0.00043257 | 0.00287843 | CENPK | 64105 |
| ENSG000001 | 693.984547 | 0.43808889 | 0.12450733 | 3.5185791 | 0.00043386 | 0.00288657 | DYM | 54808 |
| ENSG000001 | 6404.4883 | 0.376114033 | 0.10689698 | 3.51847191 | 0.00043404 | 0.00288726 | TMPO | 7112 |
| ENSG000001 | 790.750117 | 0.424638935 | 0.12069204 | 3.51836744 | 0.00043421 | 0.00288792 | PAQR3 | 152559 |
| ENSG000001 | 1429.59952 | 0.410961731 | 0.11682536 | 3.51774422 | 0.00043523 | 0.00289424 | TNKS | 8658 |
| ENSG00000C | 1616.93807 | 0.453071493 | 0.12880785 | 3.51742151 | 0.00043576 | 0.00289729 | ZFAND6 | 54469 |
| ENSG000001 | 384.309299 | 0.531692944 | 0.15116293 | 3.51735004 | 0.00043588 | 0.00289759 | GIT1 | 28964 |
| ENSG000001 | 11.7854138 | -1.155990485 | 0.32866363 | -3.5172449 | 0.00043605 | 0.00289779 | CALHM2 | 51063 |
| ENSG000001 | 32.8738414 | -0.844645441 | 0.24014113 | -3.5172877 | 0.00043598 | 0.00289779 | GABRR1 | 2569 |
| ENSG000001 | 217.384832 | 0.601353906 | 0.17098312 | 3.51703672 | 0.00043639 | 0.00289959 | POLR3K | 51728 |
| ENSG000001 | 13.6988855 | -1.111027558 | 0.31597794 | -3.5161555 | 0.00043784 | 0.00290836 | GABRR2 | 2570 |
| ENSG000001 | 614.62974 | 0.442915936 | 0.12596624 | 3.51614799 | 0.00043786 | 0.00290836 | ZNF813 | 126017 |
| ENSG000001 | 188.862868 | 0.716738754 | 0.20384612 | 3.51607743 | 0.00043797 | 0.00290866 | PPDPF | 79144 |
| ENSG000001 | 562.844964 | 0.520100162 | 0.14792768 | 3.51590826 | 0.00043825 | 0.00291003 | SAMD4B | 55095 |
| ENSG00000C | 50.0733872 | 0.793804458 | 0.22578674 | 3.51572671 | 0.00043855 | 0.00291107 | TNMD | 64102 |
| ENSG000001 | 197.096537 | 0.57580594 | 0.16377901 | 3.51574928 | 0.00043851 | 0.00291107 | FABP3 | 2170 |
| ENSG000002 | 36.4707392 | -0.829767451 | 0.23603915 | -3.5153806 | 0.00043912 | 0.00291439 | CTD-2201I18.1 | 101929215 |
| ENSG000001 | 733.24204 | 0.432570453 | 0.12305509 | 3.51525842 | 0.00043933 | 0.00291525 | MAGOH | 4116 |
| ENSG00000C | 1424.35018 | 0.401619687 | 0.11426467 | 3.51481956 | 0.00044005 | 0.0029196 | ATP2B1 | 490 |
| ENSG000001 | 4580.67673 | 0.561799074 | 0.15984357 | 3.51468051 | 0.00044028 | 0.00292065 | SALL4 | 57167 |
| ENSG000001 | 18.5128402 | -1.034220193 | 0.29427903 | -3.5144203 | 0.00044071 | 0.00292303 | PDZD3 | 79849 |
| ENSG000002 | 15.9366295 | -1.074910144 | 0.30588157 | -3.5141383 | 0.00044118 | 0.00292566 | NFIA-AS2 | 100996570 |
| ENSG000001 | 1008.61825 | 0.369274176 | 0.10510413 | 3.51341245 | 0.00044239 | 0.00293319 | SECISBP2L | 9728 |
| ENSG000001 | 6588.79322 | 0.40232083 | 0.1145249 | 3.51295493 | 0.00044315 | 0.00293776 | ARPP19 | 10776 |

| | | | | | | | | | |
|---|---|---|---|---|---|---|---|---|---|
| ENSG000001 | 1158.07388 | 0.490463443 | 0.13962009 | 3.51284292 | 0.00044334 | 0.00293852 | CAPNS1 | | 826 |
| ENSG000001 | 1391.82269 | 0.406957345 | 0.11585443 | 3.51266106 | 0.00044364 | 0.00294005 | IMPACT | | 55364 |
| ENSG00000C | 13.315165 | -1.13607584 | 0.32343095 | -3.5125762 | 0.00044378 | 0.00294019 | CD5L | | 922 |
| ENSG000002 | 5.05008494 | -1.325587892 | 0.37738493 | -3.5125618 | 0.00044381 | 0.00294019 | NA | NA | |
| ENSG000001 | 18.633937 | -1.021801181 | 0.29093437 | -3.5121363 | 0.00044452 | 0.00294442 | CASP1 | | 834 |
| ENSG00000C | 28.67321 | -0.967902338 | 0.27559623 | -3.5120304 | 0.0004447 | 0.00294463 | CD84 | | 8832 |
| ENSG000001 | 66.1012495 | 0.715984039 | 0.2038638 | 3.51207054 | 0.00044463 | 0.00294463 | PPP1R16A | | 84988 |
| ENSG000001 | 593.949542 | -0.416938962 | 0.11872285 | -3.5118679 | 0.00044497 | 0.00294522 | EMC4 | | 51234 |
| ENSG000001 | 817.041267 | 0.434610024 | 0.12375336 | 3.51190486 | 0.00044491 | 0.00294522 | FAM102B | | 284611 |
| ENSG000001 | 29.1212576 | -0.916137566 | 0.26087057 | -3.5118471 | 0.000445 | 0.00294522 | DNAJB7 | | 150353 |
| ENSG000001 | 150.085328 | 0.556550544 | 0.15850238 | 3.51130707 | 0.00044591 | 0.00295073 | OBFC1 | | 79991 |
| ENSG000001 | 1151.37561 | 0.453224626 | 0.12907771 | 3.51125393 | 0.000446 | 0.00295084 | KIF18A | | 81930 |
| ENSG00000C | 792.06408 | 0.550195614 | 0.15671876 | 3.51071951 | 0.0004469 | 0.00295581 | NSFL1C | | 55968 |
| ENSG000001 | 9.01509786 | -1.237930975 | 0.35261404 | -3.5107252 | 0.00044689 | 0.00295581 | FAM71A | | 149647 |
| ENSG000001 | 506.063937 | 0.515219416 | 0.14676114 | 3.51059842 | 0.0004471 | 0.0029562 | COMMD7 | | 149951 |
| ENSG000002 | 1132.43435 | 0.423280673 | 0.12057226 | 3.51059758 | 0.0004471 | 0.0029562 | ZNF551 | | 90233 |
| ENSG000001 | 4893.58597 | 0.545722618 | 0.15548504 | 3.50980794 | 0.00044843 | 0.00296451 | MAT2A | | 4144 |
| ENSG000001 | 103.311521 | 0.609526668 | 0.17368192 | 3.50944221 | 0.00044905 | 0.00296793 | COQ10A | | 93058 |
| ENSG000001 | 41.861481 | -0.789289906 | 0.22490921 | -3.5093712 | 0.00044917 | 0.00296793 | MYPN | | 84665 |
| ENSG000001 | 333.541924 | 0.589575319 | 0.16799969 | 3.50938346 | 0.00044915 | 0.00296793 | ABCF3 | | 55324 |
| ENSG000001 | 3338.62507 | 0.445086873 | 0.12684589 | 3.50887889 | 0.00045 | 0.00297295 | BMPR1A | | 657 |
| ENSG000001 | 967.184628 | 0.434387148 | 0.12380026 | 3.50877423 | 0.00045018 | 0.00297316 | ACOT13 | | 55856 |
| ENSG000001 | 19.6109549 | -1.010152739 | 0.28789342 | -3.5087733 | 0.00045018 | 0.00297316 | MOGAT2 | | 80168 |
| ENSG000001 | 77.7600541 | 0.718867097 | 0.20488863 | 3.50857491 | 0.00045051 | 0.00297478 | CCDC74A | | 90557 |
| ENSG000001 | 163.804957 | 0.699727766 | 0.19943548 | 3.50854199 | 0.00045057 | 0.00297478 | UBXN6 | | 80700 |
| ENSG000001 | 569.423234 | 0.440099615 | 0.12544317 | 3.50835857 | 0.00045088 | 0.00297634 | TYW1 | | 55253 |
| ENSG000001 | 245.392387 | 0.49658152 | 0.14156385 | 3.50782721 | 0.00045178 | 0.00298168 | IFT46 | | 56912 |
| ENSG000001 | 51.9969566 | 0.777279351 | 0.22158626 | 3.50779572 | 0.00045184 | 0.00298168 | PIP5KL1 | | 138429 |
| ENSG000001 | 16.8188669 | -1.049178574 | 0.2991241 | -3.5075027 | 0.00045233 | 0.00298448 | MYBPC3 | | 4607 |

| | | | | | | | | |
|---|---|---|---|---|---|---|---|---|
| ENSG000001 | 486.84099 | 0.482508674 | 0.13757974 | 3.50712014 | 0.00045298 | 0.00298828 | FBN2 | 2201 |
| ENSG000002 | 64.3007082 | 0.694396282 | 0.19801255 | 3.50682964 | 0.00045348 | 0.00299106 | NA | NA |
| ENSG000001 | 619.184596 | 0.439138059 | 0.12523489 | 3.50651533 | 0.00045402 | 0.00299362 | ICK | 22858 |
| ENSG000001 | 3959.97447 | 0.381196174 | 0.10871022 | 3.50653487 | 0.00045398 | 0.00299362 | ANXA5 | 308 |
| ENSG000001 | 197.592649 | 0.501044194 | 0.14290883 | 3.50604091 | 0.00045483 | 0.0029981 | CRIPT | 9419 |
| ENSG000002 | 3.22869095 | -1.316196205 | 0.37540923 | -3.5060305 | 0.00045484 | 0.0029981 | VN1R4 | 317703 |
| ENSG000001 | 1988.88355 | 0.446589168 | 0.12738173 | 3.50591238 | 0.00045504 | 0.00299895 | NIFK | 84365 |
| ENSG000001 | 187.317598 | 0.516007928 | 0.1471966 | 3.50556968 | 0.00045563 | 0.00300232 | ACOT8 | 10005 |
| ENSG00000C | 608.838735 | 0.519022089 | 0.14805847 | 3.50552109 | 0.00045571 | 0.00300238 | KIF3C | 3797 |
| ENSG000001 | 21.8952071 | -0.985451433 | 0.28114159 | -3.5051784 | 0.0004563 | 0.00300576 | KCNJ16 | 3773 |
| ENSG000001 | 195.850305 | 0.516046879 | 0.14723277 | 3.50497304 | 0.00045665 | 0.00300742 | IGFBP3 | 3486 |
| ENSG000001 | 22.1240047 | -0.975250786 | 0.27824994 | -3.5049451 | 0.0004567 | 0.00300742 | RAB40A | 142684 |
| ENSG000001 | 293.89048 | 0.584450839 | 0.16679902 | 3.50392248 | 0.00045846 | 0.0030185 | TONSL | 4796 |
| ENSG00000C | 433.589449 | 0.451036615 | 0.12873316 | 3.5036554 | 0.00045892 | 0.00302022 | CHFR | 55743 |
| ENSG000001 | 3907.00832 | 0.436226824 | 0.12450607 | 3.50365917 | 0.00045891 | 0.00302022 | PNISR | 25957 |
| ENSG000001 | 39.6516945 | -0.893334244 | 0.2549731 | -3.5036412 | 0.00045894 | 0.00302022 | OR4K2 | 390431 |
| ENSG00000C | 846.914276 | 0.416854668 | 0.11898435 | 3.50344109 | 0.00045929 | 0.00302189 | ZBTB11 | 27107 |
| ENSG000001 | 2281.2309 | 0.459219806 | 0.13107802 | 3.50340811 | 0.00045935 | 0.00302189 | HMGN3 | 9324 |
| ENSG000001 | 46.3826108 | -0.771707399 | 0.22029323 | -3.5030918 | 0.00045989 | 0.00302499 | IL16 | 3603 |
| ENSG00000C | 665.277528 | 0.415167926 | 0.11851892 | 3.5029676 | 0.00046011 | 0.00302591 | SLC12A2 | 6558 |
| ENSG000002 | 11.8881052 | -1.157374759 | 0.33040435 | -3.5029042 | 0.00046022 | 0.00302614 | NA | NA |
| ENSG000001 | 2015.07275 | 0.419884031 | 0.11988927 | 3.50226518 | 0.00046132 | 0.00303291 | PPP2R1B | 5519 |
| ENSG000001 | 462.608281 | 0.426346581 | 0.12174136 | 3.5020686 | 0.00046166 | 0.00303466 | ITFG1 | 81533 |
| ENSG000001 | 6641.70737 | 0.4353364 | 0.12433372 | 3.5013542 | 0.0004629 | 0.00304231 | ZFP42 | 132625 |
| ENSG000001 | 88.3060266 | 0.659633705 | 0.18843319 | 3.5006238 | 0.00046417 | 0.00305017 | MFAP4 | 4239 |
| ENSG000001 | 22.8440983 | -0.97968825 | 0.27990428 | -3.5000832 | 0.00046511 | 0.00305586 | PRG4 | 10216 |
| ENSG000001 | 728.855954 | 0.417917876 | 0.11940596 | 3.49997493 | 0.0004653 | 0.00305661 | ZCCHC9 | 84240 |
| ENSG000001 | 142.612633 | 0.657320427 | 0.18780961 | 3.49992969 | 0.00046538 | 0.00305663 | HIRA | 7290 |
| ENSG000001 | 24.2295656 | -0.999110808 | 0.28548428 | -3.4997051 | 0.00046577 | 0.0030575 | ACSBG2 | 81616 |

| | | | | | | | | |
|---|---|---|---|---|---|---|---|---|
| ENSG000001 | 748.209861 | 0.399789536 | 0.11423597 | 3.49968177 | 0.00046581 | 0.0030575 | SLBP | 7884 |
| ENSG000001 | 492.930523 | 0.884925019 | 0.25285445 | 3.49974076 | 0.00046571 | 0.0030575 | HPRT1 | 3251 |
| ENSG000001 | 545.532437 | 0.603359897 | 0.17240014 | 3.49976453 | 0.00046567 | 0.0030575 | SNN | 8303 |
| ENSG000001 | 76.3462674 | 0.663478467 | 0.18961115 | 3.49915315 | 0.00046674 | 0.00306298 | DNAL4 | 10126 |
| ENSG000001 | 3804.87058 | 0.397192739 | 0.11351226 | 3.49911766 | 0.0004668 | 0.00306298 | SRP72 | 6731 |
| ENSG000001 | 219.450047 | 0.506960346 | 0.14489725 | 3.49875745 | 0.00046743 | 0.00306663 | GRB14 | 2888 |
| ENSG000001 | 447.958547 | 0.469248951 | 0.13412363 | 3.49862994 | 0.00046766 | 0.0030671 | PRNP | 5621 |
| ENSG000002 | 14.2709448 | -1.112249294 | 0.31790993 | -3.4986303 | 0.00046765 | 0.0030671 | NA | NA |
| ENSG000001 | 348.48086 | 0.436495749 | 0.12476385 | 3.49857543 | 0.00046775 | 0.00306723 | ARSB | 411 |
| ENSG000001 | 26.6748694 | -0.940110549 | 0.26872328 | -3.4984336 | 0.000468 | 0.00306837 | TSHR | 7253 |
| ENSG00000C | 22.5154529 | -0.965190033 | 0.27591145 | -3.4981876 | 0.00046843 | 0.00307026 | P2RX7 | 5027 |
| ENSG000001 | 153.806681 | 0.532413738 | 0.15219724 | 3.4981826 | 0.00046844 | 0.00307026 | NCEH1 | 57552 |
| ENSG000001 | 3491.57912 | 0.418520681 | 0.11964542 | 3.49800852 | 0.00046875 | 0.00307153 | MEST | 4232 |
| ENSG000002 | 1087.44046 | 0.385844778 | 0.11030481 | 3.49798677 | 0.00046878 | 0.00307153 | OTUD6B-AS1 | 100506365 |
| ENSG000001 | 163.479847 | 0.547518518 | 0.15654262 | 3.49756838 | 0.00046952 | 0.00307585 | MTX1 | 4580 |
| ENSG000001 | 24.5314493 | -0.930161774 | 0.26599307 | -3.4969398 | 0.00047063 | 0.00308226 | SAMSN1 | 64092 |
| ENSG000001 | 11.3453288 | -1.175669547 | 0.33620081 | -3.4969266 | 0.00047065 | 0.00308226 | LRRC71 | 149499 |
| ENSG000001 | 317.166674 | 0.530114165 | 0.15159796 | 3.49684239 | 0.0004708 | 0.00308274 | GGH | 8836 |
| ENSG000001 | 21.3606055 | -0.975105526 | 0.27886272 | -3.4967224 | 0.00047101 | 0.00308313 | SLC17A3 | 10786 |
| ENSG000001 | 41.2525806 | -0.878261338 | 0.25116496 | -3.496751 | 0.00047096 | 0.00308313 | CSMD1 | 64478 |
| ENSG000001 | 94.5500895 | 0.67804438 | 0.19391736 | 3.49656353 | 0.00047129 | 0.00308447 | EVI5L | 115704 |
| ENSG000001 | 394.498447 | 0.500313548 | 0.14309767 | 3.49630815 | 0.00047174 | 0.00308692 | GPN3 | 51184 |
| ENSG000001 | 879.736211 | 0.470026833 | 0.13445113 | 3.4958934 | 0.00047248 | 0.00309089 | SREK1IP1 | 285672 |
| ENSG000001 | 986.581468 | 0.441861984 | 0.12639507 | 3.49587985 | 0.0004725 | 0.00309089 | RNF220 | 55182 |
| ENSG000002 | 13.0921975 | -1.121518282 | 0.32081709 | -3.4958184 | 0.00047261 | 0.0030911 | NA | NA |
| ENSG000001 | 402.096073 | 0.624210979 | 0.17856858 | 3.49563723 | 0.00047293 | 0.0030927 | EDC3 | 80153 |
| ENSG000001 | 388.15514 | 0.458185202 | 0.13109401 | 3.49508877 | 0.0004739 | 0.00309856 | RMND1 | 55005 |
| ENSG000001 | 1071.31053 | 0.428380926 | 0.12257149 | 3.49494762 | 0.00047415 | 0.00309917 | CCDC58 | 131076 |
| ENSG000001 | 33.405571 | 0.864671218 | 0.24740888 | 3.49490773 | 0.00047423 | 0.00309917 | CXCL3 | 2921 |

| | | | | | | | | |
|---|---|---|---|---|---|---|---|---|
| ENSG000001 | 176.465662 | 0.62303893 | 0.17826821 | 3.49495247 | 0.00047415 | 0.00309917 | MRPL40 | 64976 |
| ENSG000001 | 762.730004 | 0.595878262 | 0.17050931 | 3.49469636 | 0.0004746 | 0.0031007 | ATXN2L | 11273 |
| ENSG000002 | 193.752026 | 0.491853019 | 0.14074297 | 3.49468968 | 0.00047461 | 0.0031007 | FAM86DP | 692099 |
| ENSG00000C | 1068.18317 | 0.404080107 | 0.11563908 | 3.4943215 | 0.00047527 | 0.00310398 | FH | 2271 |
| ENSG000001 | 145.315409 | 0.681655087 | 0.19507359 | 3.49434843 | 0.00047522 | 0.00310398 | EGFL7 | 51162 |
| ENSG000001 | 83.0415544 | 0.678344509 | 0.19414557 | 3.49399941 | 0.00047584 | 0.00310723 | DPM3 | 54344 |
| ENSG000001 | 16.1751739 | -1.057513089 | 0.3026707 | -3.4939394 | 0.00047595 | 0.00310742 | ADH7 | 131 |
| ENSG000001 | 479.536894 | 0.611533125 | 0.17504857 | 3.49350544 | 0.00047672 | 0.00311198 | CLPTM1 | 1209 |
| ENSG000001 | 321.145393 | 0.481422452 | 0.13780886 | 3.4934072 | 0.0004769 | 0.00311262 | LYRM1 | 57149 |
| ENSG000001 | 605.429935 | 0.457568861 | 0.13098393 | 3.49332052 | 0.00047705 | 0.00311313 | KATNBL1 | 79768 |
| ENSG000002 | 316.103981 | 0.479561985 | 0.1372841 | 3.49320846 | 0.00047725 | 0.00311394 | NA | NA |
| ENSG000001 | 22.1900042 | -0.975538914 | 0.27927092 | -3.4931632 | 0.00047733 | 0.00311396 | CD160 | 11126 |
| ENSG00000C | 1516.49984 | 0.383030409 | 0.10966605 | 3.49269804 | 0.00047817 | 0.00311889 | MAP3K4 | 4216 |
| ENSG000001 | 12.6666691 | -1.17091101 | 0.33525047 | -3.4926454 | 0.00047826 | 0.003119 | ALOX5AP | 241 |
| ENSG000001 | 3739.06033 | 0.399420132 | 0.11436294 | 3.49256626 | 0.0004784 | 0.00311943 | GMPS | 8833 |
| ENSG000002 | 15.3128196 | -1.117215324 | 0.31989435 | -3.492451 | 0.00047861 | 0.00312027 | RAB7B | 338382 |
| ENSG000001 | 966.920285 | 0.480149299 | 0.13749815 | 3.49204182 | 0.00047934 | 0.00312455 | MBTPS1 | 8720 |
| ENSG000001 | 1220.48435 | 0.417648896 | 0.11960823 | 3.49180736 | 0.00047976 | 0.00312679 | ATR | 545 |
| ENSG000001 | 1008.5659 | 0.46591932 | 0.13344157 | 3.49156064 | 0.00048021 | 0.00312918 | NUDT15 | 55270 |
| ENSG000001 | 149.164851 | 0.532435006 | 0.15250119 | 3.49134973 | 0.00048059 | 0.00313115 | KBTBD11 | 9920 |
| ENSG000001 | 2699.25891 | 0.374651497 | 0.107324 | 3.49084561 | 0.00048149 | 0.00313656 | OLA1 | 29789 |
| ENSG000001 | 411.3566 | 0.428111041 | 0.12264483 | 3.49065718 | 0.00048183 | 0.00313801 | SPIRE1 | 56907 |
| ENSG000001 | 410.657379 | 0.513455823 | 0.14709519 | 3.49063639 | 0.00048187 | 0.00313801 | PI4KB | 5298 |
| ENSG000001 | 847.690902 | 0.464377556 | 0.13303929 | 3.49052947 | 0.00048206 | 0.00313876 | RFX5 | 5993 |
| ENSG000001 | 505.965877 | 0.588757584 | 0.16868923 | 3.49019062 | 0.00048268 | 0.00314224 | NUDT9 | 53343 |
| ENSG00000C | 2766.63382 | 0.42749869 | 0.12250289 | 3.48970281 | 0.00048356 | 0.00314747 | EXOC5 | 10640 |
| ENSG000001 | 479.044747 | 0.525930652 | 0.15071807 | 3.48949954 | 0.00048393 | 0.00314924 | RASL11B | 65997 |
| ENSG000001 | 143.562484 | 0.64564607 | 0.18502711 | 3.48946738 | 0.00048398 | 0.00314924 | ZNF212 | 7988 |
| ENSG000001 | 157.249169 | 0.596447609 | 0.17093223 | 3.48938059 | 0.00048414 | 0.00314975 | ZNF786 | 136051 |

| | | | | | | | | |
|---|---|---|---|---|---|---|---|---|
| ENSG000001 | 1430.88837 | 0.499300102 | 0.14310093 | 3.48914654 | 0.00048457 | 0.00315201 | FBXW2 | 26190 |
| ENSG00000C | 166.430904 | 0.615085932 | 0.1762896 | 3.48906525 | 0.00048471 | 0.00315246 | TRMT2A | 27037 |
| ENSG000002 | 15.5544904 | -1.093404022 | 0.31340356 | -3.4888054 | 0.00048518 | 0.00315502 | NA | NA |
| ENSG00000C | 471.742321 | 0.471377591 | 0.13511733 | 3.48865393 | 0.00048546 | 0.0031558 | POLB | 5423 |
| ENSG000001 | 14.5879 | -1.133274603 | 0.32484415 | -3.4886717 | 0.00048543 | 0.0031558 | IL1B | 3553 |
| ENSG000001 | 11102.2024 | 0.416273017 | 0.11932881 | 3.48845349 | 0.00048582 | 0.00315766 | JMJD1C | 221037 |
| ENSG00000C | 504.561245 | 0.436181295 | 0.12504875 | 3.48809009 | 0.00048648 | 0.00316145 | ATRN | 8455 |
| ENSG000002 | 8.47789213 | -1.233100453 | 0.35354189 | -3.4878482 | 0.00048692 | 0.0031638 | NA | NA |
| ENSG000001 | 76.2530269 | 0.661865493 | 0.18976799 | 3.48776158 | 0.00048708 | 0.00316394 | DHRS4L2 | 317749 |
| ENSG000001 | 331.186041 | 0.516617563 | 0.14812341 | 3.48775104 | 0.0004871 | 0.00316394 | TMEM63A | 9725 |
| ENSG000001 | 46.9647033 | -0.803193707 | 0.23029579 | -3.4876612 | 0.00048727 | 0.00316449 | NRAP | 4892 |
| ENSG00000C | 132.020318 | 0.552846784 | 0.15852595 | 3.48742143 | 0.0004877 | 0.00316632 | ZNF112 | 7771 |
| ENSG000001 | 6391.01612 | 0.404028393 | 0.115852 | 3.48745287 | 0.00048764 | 0.00316632 | ZNF146 | 7705 |
| ENSG000002 | 38.2403928 | 0.788076127 | 0.22598357 | 3.48731607 | 0.00048789 | 0.00316706 | NA | NA |
| ENSG000001 | 46.304789 | -0.841841562 | 0.24142656 | -3.4869468 | 0.00048857 | 0.00317093 | MYH8 | 4626 |
| ENSG000001 | 61.5059919 | -0.692777844 | 0.19868439 | -3.4868257 | 0.00048879 | 0.00317185 | RASGRP3 | 25780 |
| ENSG000001 | 269.615097 | 0.473527041 | 0.13580895 | 3.48671461 | 0.00048899 | 0.00317266 | PTCHD1 | 139411 |
| ENSG000001 | 457.975031 | 0.472163835 | 0.13542012 | 3.4866594 | 0.00048909 | 0.00317281 | GLCE | 26035 |
| ENSG000002 | 333.693228 | 0.505720289 | 0.14510396 | 3.48522741 | 0.00049172 | 0.00318933 | LEPROT | 54741 |
| ENSG000001 | 1589.19946 | 0.361134867 | 0.10362337 | 3.48507155 | 0.00049201 | 0.00319068 | SMC5 | 23137 |
| ENSG000001 | 546.144112 | 0.493912608 | 0.14172988 | 3.48488687 | 0.00049235 | 0.00319214 | HS2ST1 | 9653 |
| ENSG000001 | 2788.97023 | 0.460387454 | 0.13211061 | 3.48486359 | 0.00049239 | 0.00319214 | KNTC1 | 9735 |
| ENSG00000C | 115.438803 | 0.651155385 | 0.18687633 | 3.48441877 | 0.00049321 | 0.00319694 | NFKB2 | 4791 |
| ENSG000001 | 133.498344 | 0.545042072 | 0.15642737 | 3.48431407 | 0.0004934 | 0.00319768 | SLC2A10 | 81031 |
| ENSG000001 | 846.89654 | 0.450424333 | 0.12929268 | 3.48375736 | 0.00049443 | 0.00320382 | C3orf17 | 25871 |
| ENSG000001 | 3.25800945 | -1.305907868 | 0.37487145 | -3.4836152 | 0.00049469 | 0.00320501 | CST4 | 1472 |
| ENSG000001 | 178.856841 | 0.539108638 | 0.15476532 | 3.48339432 | 0.0004951 | 0.00320715 | TOR1B | 27348 |
| ENSG000001 | 3860.05772 | 0.441496913 | 0.12674553 | 3.48333307 | 0.00049521 | 0.00320737 | LRBA | 987 |
| ENSG000001 | 8665.6598 | 0.393031765 | 0.11285786 | 3.4825379 | 0.00049668 | 0.0032164 | SMARCA5 | 8467 |

| | | | | | | | | |
|---|---|---|---|---|---|---|---|---|
| ENSG000001 | 90.7460188 | 0.618688535 | 0.17767035 | 3.48222728 | 0.00049726 | 0.00321859 | NAGPA | 51172 |
| ENSG000001 | 861.427094 | 0.497205458 | 0.14278042 | 3.48230832 | 0.00049711 | 0.00321859 | SWAP70 | 23075 |
| ENSG000001 | 106.598013 | 0.642571957 | 0.18452832 | 3.48224033 | 0.00049724 | 0.00321859 | PPIP5K1 | 9677 |
| ENSG000001 | 259.28668 | 0.661237599 | 0.189903 | 3.48197545 | 0.00049773 | 0.00322066 | FGD1 | 2245 |
| ENSG000001 | 173.59752 | 0.506353973 | 0.14542198 | 3.48196304 | 0.00049775 | 0.00322066 | DYNC2LI1 | 51626 |
| ENSG000001 | 10244.604 | 0.491563716 | 0.14117578 | 3.48192666 | 0.00049782 | 0.00322066 | RPS17 | 6218 |
| ENSG000001 | 292.592044 | 0.491231958 | 0.14108381 | 3.48184506 | 0.00049797 | 0.00322113 | WBSCR17 | 64409 |
| ENSG00000C | 376.035816 | 0.519301914 | 0.14915805 | 3.48155473 | 0.00049851 | 0.00322206 | SLC1A3 | 6507 |
| ENSG000001 | 318.605467 | 0.505929053 | 0.14532094 | 3.48146014 | 0.00049869 | 0.00322206 | SFXN2 | 118980 |
| ENSG000001 | 37.649449 | -0.848141859 | 0.24361907 | -3.4814264 | 0.00049875 | 0.00322206 | OR2AT4 | 341152 |
| ENSG000001 | 339.955548 | 0.520109568 | 0.14939535 | 3.48143074 | 0.00049874 | 0.00322206 | FANCA | 2175 |
| ENSG000002 | 31.845069 | -0.884925377 | 0.25416569 | -3.481687 | 0.00049827 | 0.00322206 | LHFPL3-AS1 | 645591 |
| ENSG000002 | 196.593395 | 0.64956679 | 0.18657805 | 3.4814748 | 0.00049866 | 0.00322206 | SNHG7 | 84973 |
| ENSG000002 | 17.9778747 | -1.038108699 | 0.29817307 | -3.4815643 | 0.00049849 | 0.00322206 | MUC22 | 100507679 |
| ENSG000002 | 7.31473581 | -1.276386278 | 0.36661112 | -3.4815809 | 0.00049846 | 0.00322206 | NA | NA |
| ENSG000001 | 819.944249 | 0.434490018 | 0.12480748 | 3.4812818 | 0.00049902 | 0.00322328 | REV1 | 51455 |
| ENSG00000C | 285.576945 | 0.505225331 | 0.14513454 | 3.4810827 | 0.00049939 | 0.00322517 | PHKA1 | 5255 |
| ENSG000001 | 145.041408 | 0.529814795 | 0.1522157 | 3.48068426 | 0.00050013 | 0.00322945 | FAM227B | 196951 |
| ENSG000002 | 40.6763297 | 0.845279733 | 0.24286787 | 3.48040994 | 0.00050065 | 0.00323225 | CD177 | 57126 |
| ENSG000001 | 23.5203512 | -0.955855842 | 0.27465654 | -3.480186 | 0.00050107 | 0.00323443 | CHRM5 | 1133 |
| ENSG000001 | 396.252915 | 0.51514256 | 0.14803083 | 3.47996795 | 0.00050147 | 0.00323604 | NA | NA |
| ENSG000001 | 78.2264416 | -0.675044117 | 0.1939789 | -3.4799873 | 0.00050144 | 0.00323604 | RELN | 5649 |
| ENSG00000C | 576.086773 | 0.573906659 | 0.16492326 | 3.47984051 | 0.00050171 | 0.00323673 | MYO9B | 4650 |
| ENSG000001 | 7321.91642 | 0.418144845 | 0.1201634 | 3.47980213 | 0.00050178 | 0.00323673 | ILF2 | 3608 |
| ENSG000002 | 690.033733 | 0.490929471 | 0.14108048 | 3.47978306 | 0.00050182 | 0.00323673 | UTP14C | 9724 |
| ENSG000001 | 688.493076 | 0.411450378 | 0.11825639 | 3.47930773 | 0.00050271 | 0.00324196 | CDKAL1 | 54901 |
| ENSG000002 | 109.743203 | 0.606674945 | 0.17441358 | 3.47836999 | 0.00050447 | 0.0032528 | APOPT1 | 84334 |
| ENSG000001 | 2522.52583 | 0.453906509 | 0.13050901 | 3.4779706 | 0.00050523 | 0.00325714 | PSMD11 | 5717 |
| ENSG000002 | 10.1700804 | -1.211825192 | 0.34843915 | -3.4778675 | 0.00050542 | 0.00325735 | NA | NA |

| | | | | | | | | |
|---|---|---|---|---|---|---|---|---|
| ENSG000002 | 434.268197 | 0.425828208 | 0.12243814 | 3.47790503 | 0.00050535 | 0.00325735 | TUBGCP5 | 114791 |
| ENSG000001 | 26.1824687 | -0.918810397 | 0.26420327 | -3.4776648 | 0.0005058 | 0.0032593 | FBXW12 | 285231 |
| ENSG000001 | 6978.22075 | 0.455448706 | 0.13097223 | 3.47744484 | 0.00050622 | 0.00326146 | CCT3 | 7203 |
| ENSG000001 | 1772.23194 | 0.390417142 | 0.11227964 | 3.47718559 | 0.00050671 | 0.00326409 | FAM49B | 51571 |
| ENSG00000C | 3829.38911 | 0.404781716 | 0.11642405 | 3.47678794 | 0.00050746 | 0.0032679 | SKIV2L2 | 23517 |
| ENSG000002 | 333.078991 | 0.483714187 | 0.1391265 | 3.47679412 | 0.00050745 | 0.0032679 | HLA-B | 3106 |
| ENSG000001 | 15.8555053 | -1.058673714 | 0.30450202 | -3.4767379 | 0.00050755 | 0.00326799 | LRRC46 | 90506 |
| ENSG000001 | 183.458663 | 0.557847949 | 0.16046225 | 3.47650583 | 0.00050799 | 0.0032703 | TTC9C | 283237 |
| ENSG000001 | 230.591765 | 0.658579862 | 0.18945738 | 3.47613732 | 0.00050869 | 0.00327428 | UBE2B | 7320 |
| ENSG000001 | 683.05126 | 0.48220926 | 0.13872728 | 3.47595117 | 0.00050904 | 0.00327542 | SPRY2 | 10253 |
| ENSG000001 | 1247.76804 | 0.440551227 | 0.1267439 | 3.47591655 | 0.00050911 | 0.00327542 | FBXO11 | 80204 |
| ENSG000001 | 179.8289 | 0.68420866 | 0.19683793 | 3.47600011 | 0.00050895 | 0.00327542 | FAAP20 | 199990 |
| ENSG000001 | 10153.672 | 0.414238218 | 0.11917916 | 3.47576042 | 0.00050941 | 0.00327628 | G3BP2 | 9908 |
| ENSG000001 | 64.1659589 | -0.66259298 | 0.19063201 | -3.4757697 | 0.00050939 | 0.00327628 | DNAH17 | 8632 |
| ENSG000002 | 567.645549 | 0.480494971 | 0.13827041 | 3.47503821 | 0.00051078 | 0.0032846 | TRIQK | 286144 |
| ENSG00000C | 187.373924 | 0.561544038 | 0.16161626 | 3.47455162 | 0.00051171 | 0.00329004 | ZNRD1 | 30834 |
| ENSG000001 | 2618.92115 | 0.413721394 | 0.11908244 | 3.4742434 | 0.0005123 | 0.0032933 | DMXL1 | 1657 |
| ENSG000001 | 836.953412 | 0.470633205 | 0.1354828 | 3.47374873 | 0.00051324 | 0.00329885 | CKAP2L | 150468 |
| ENSG000001 | 1828.06692 | 0.415020365 | 0.11948224 | 3.47348997 | 0.00051374 | 0.00330099 | WDR12 | 55759 |
| ENSG000002 | 1328.78394 | 0.562945247 | 0.16206827 | 3.47350687 | 0.0005137 | 0.00330099 | DYNLL2 | 140735 |
| ENSG000002 | 18.113496 | -1.047396454 | 0.30155432 | -3.4733259 | 0.00051405 | 0.00330222 | CHRM3-AS2 | 100506915 |
| ENSG000002 | 341.189575 | 0.540080126 | 0.1554946 | 3.47330479 | 0.00051409 | 0.00330222 | MRPL33 | 9553 |
| ENSG000001 | 184.400659 | 0.509123847 | 0.14658744 | 3.47317504 | 0.00051434 | 0.00330329 | SFXN4 | 119559 |
| ENSG000001 | 54.3216982 | -0.78588281 | 0.22628187 | -3.473026 | 0.00051463 | 0.00330461 | RNF17 | 56163 |
| ENSG00000C | 307.97395 | 0.632453499 | 0.1821677 | 3.47182023 | 0.00051694 | 0.00331896 | TFAP4 | 7023 |
| ENSG000002 | 16.9742134 | -1.051586541 | 0.30289873 | -3.471743 | 0.00051709 | 0.00331939 | LOC101927849 | 101927849 |
| ENSG000002 | 277.703119 | 0.568600405 | 0.16379964 | 3.47131662 | 0.00051791 | 0.00332414 | NA | NA |
| ENSG000002 | 2.69423839 | -1.283674051 | 0.36987197 | -3.4705902 | 0.00051932 | 0.00333261 | NA | NA |
| ENSG00000C | 751.462451 | 0.471032658 | 0.13573177 | 3.47031991 | 0.00051984 | 0.00333463 | METTL13 | 51603 |

| | | | | | | | | |
|---|---|---|---|---|---|---|---|---|
| ENSG00000C | 3511.09737 | 0.419793817 | 0.12096759 | 3.47029993 | 0.00051988 | 0.00333463 | NFE2L3 | 9603 |
| ENSG000001 | 2861.56572 | 0.40273994 | 0.11605193 | 3.47034259 | 0.00051979 | 0.00333463 | PNRC2 | 55629 |
| ENSG00000C | 210.248131 | 0.534058839 | 0.15390065 | 3.47015318 | 0.00052016 | 0.0033354 | PI4K2B | 55300 |
| ENSG000001 | 850.878797 | 0.472083743 | 0.13604067 | 3.47016638 | 0.00052014 | 0.0033354 | TMEM135 | 65084 |
| ENSG000001 | 919.916398 | 0.448887671 | 0.12936972 | 3.46980478 | 0.00052084 | 0.0033392 | AVL9 | 23080 |
| ENSG00000C | 1379.64646 | 0.492299248 | 0.14190065 | 3.4693234 | 0.00052177 | 0.00334413 | VDAC3 | 7419 |
| ENSG000001 | 427.014157 | 0.45315607 | 0.13061708 | 3.46934763 | 0.00052172 | 0.00334413 | COQ2 | 27235 |
| ENSG000001 | 2617.47062 | 0.415944257 | 0.1198965 | 3.46919431 | 0.00052202 | 0.00334503 | MAN2A1 | 4124 |
| ENSG000001 | 125.785277 | 0.632596986 | 0.1823484 | 3.46916665 | 0.00052208 | 0.00334503 | ZNF785 | 146540 |
| ENSG00000C | 5441.41012 | 0.370517173 | 0.1068088 | 3.46897613 | 0.00052245 | 0.00334628 | PCM1 | 5108 |
| ENSG000001 | 290.827619 | 0.482107959 | 0.13897846 | 3.46894023 | 0.00052252 | 0.00334628 | TIGD2 | 166815 |
| ENSG000002 | 18.0312474 | -1.047446898 | 0.30195023 | -3.4689389 | 0.00052252 | 0.00334628 | NA | NA |
| ENSG000001 | 119.99334 | 0.555137926 | 0.16003777 | 3.46879308 | 0.0005228 | 0.00334757 | PSMA6 | 5687 |
| ENSG000001 | 228.193569 | 0.54861695 | 0.15816556 | 3.46862464 | 0.00052313 | 0.00334914 | ACVR1 | 90 |
| ENSG000001 | 2527.17415 | 0.385258933 | 0.11108204 | 3.4682378 | 0.00052388 | 0.00335278 | SPECC1 | 92521 |
| ENSG000002 | 57.5050663 | -0.708131415 | 0.20417807 | -3.468205 | 0.00052395 | 0.00335278 | MEF2C-AS1 | 101929423 |
| ENSG000002 | 5.78599288 | -1.300198249 | 0.37488289 | -3.4682785 | 0.0005238 | 0.00335278 | NA | NA |
| ENSG000001 | 308.544973 | 0.498132597 | 0.14364095 | 3.46790109 | 0.00052454 | 0.00335605 | CHRNA5 | 1138 |
| ENSG000001 | 260.119862 | 0.532990035 | 0.15370386 | 3.46764245 | 0.00052505 | 0.00335875 | GMDS | 2762 |
| ENSG00000C | 4062.20922 | 0.421473276 | 0.12154691 | 3.46757701 | 0.00052517 | 0.00335883 | NCKAP1 | 10787 |
| ENSG000002 | 1703.79263 | 0.431137412 | 0.12433483 | 3.46755138 | 0.00052522 | 0.00335883 | PPP4R3B | 57223 |
| ENSG000002 | 12.8834001 | -1.155613285 | 0.33329663 | -3.4672216 | 0.00052587 | 0.00336242 | LOC100506159 | 100506159 |
| ENSG000002 | 45.9670763 | -0.769360199 | 0.22193028 | -3.4666752 | 0.00052694 | 0.00336873 | MYH4 | 4622 |
| ENSG000001 | 2848.80529 | 0.385492521 | 0.11120213 | 3.46659281 | 0.0005271 | 0.00336923 | CDK12 | 51755 |
| ENSG000002 | 13.1801832 | -1.128569153 | 0.32556201 | -3.4665259 | 0.00052723 | 0.00336954 | NA | NA |
| ENSG000001 | 424.391891 | 0.451680364 | 0.13030691 | 3.46628094 | 0.00052771 | 0.00337208 | TACC1 | 6867 |
| ENSG000001 | 209.354817 | 0.528290804 | 0.15241123 | 3.46621971 | 0.00052783 | 0.00337232 | CCNDBP1 | 23582 |
| ENSG000001 | 358.035792 | 0.517144183 | 0.14919893 | 3.46613859 | 0.00052799 | 0.0033728 | PYCR2 | 29920 |
| ENSG00000C | 244.964952 | 0.519491059 | 0.14988026 | 3.46604062 | 0.00052818 | 0.0033735 | HDAC6 | 10013 |

| | | | | | | | | |
|---|---|---|---|---|---|---|---|---|
| ENSG000001 | 576.165288 | 0.442619615 | 0.12771418 | 3.4657045 | 0.00052884 | 0.00337719 | ZBTB40 | 9923 |
| ENSG000001 | 110.934709 | 0.621171572 | 0.17924217 | 3.46554371 | 0.00052916 | 0.00337815 | PPIC | 5480 |
| ENSG000002 | 7.32107675 | -1.269075314 | 0.36619727 | -3.465551 | 0.00052915 | 0.00337815 | NA | NA |
| ENSG000001 | 619.685183 | 0.45089238 | 0.13011164 | 3.46542684 | 0.00052939 | 0.00337842 | INTS10 | 55174 |
| ENSG000001 | 102.810873 | 0.705953599 | 0.20371521 | 3.46539463 | 0.00052945 | 0.00337842 | NR4A1 | 3164 |
| ENSG000001 | 129.003168 | 0.566054432 | 0.16334307 | 3.46543285 | 0.00052938 | 0.00337842 | NA | NA |
| ENSG000002 | 24.5145379 | -0.925069259 | 0.26696348 | -3.4651528 | 0.00052993 | 0.00338093 | KIAA1210 | 57481 |
| ENSG000001 | 498.466905 | 0.455737466 | 0.13154366 | 3.46453378 | 0.00053115 | 0.00338819 | ZNF622 | 90441 |
| ENSG000001 | 87.1847548 | 0.612036185 | 0.17666253 | 3.46443679 | 0.00053134 | 0.00338878 | TMEM234 | 56063 |
| ENSG000002 | 7.88141896 | -1.247643604 | 0.36013245 | -3.464402 | 0.00053141 | 0.00338878 | NA | NA |
| ENSG00000C | 878.895955 | 0.426657959 | 0.12315712 | 3.46433866 | 0.00053154 | 0.00338905 | REXO2 | 25996 |
| ENSG000001 | 37.3018533 | -0.864101832 | 0.24945078 | -3.4640173 | 0.00053217 | 0.00339203 | ITK | 3702 |
| ENSG000001 | 31.5323498 | -0.849941958 | 0.24536056 | -3.4640529 | 0.0005321 | 0.00339203 | GBP2 | 2634 |
| ENSG000002 | 10.6595097 | -1.183435528 | 0.34164177 | -3.4639661 | 0.00053227 | 0.00339215 | LOC105378057 | 105378057 |
| ENSG000002 | 4.48898089 | -1.310570773 | 0.37837258 | -3.4637044 | 0.00053279 | 0.00339491 | NA | NA |
| ENSG00000C | 654.033343 | 0.421918535 | 0.12181872 | 3.46349511 | 0.00053321 | 0.00339702 | SYT1 | 6857 |
| ENSG000002 | 736.798483 | 0.514297583 | 0.14849626 | 3.46337064 | 0.00053345 | 0.00339806 | GGNBP2 | 79893 |
| ENSG000001 | 1281.66006 | 0.394197542 | 0.11382894 | 3.46306955 | 0.00053405 | 0.00340133 | PCNXL4 | 64430 |
| ENSG000001 | 46.4774773 | -0.75424129 | 0.21780543 | -3.4629132 | 0.00053436 | 0.00340277 | PLB1 | 151056 |
| ENSG000001 | 1668.82811 | 0.387347997 | 0.11185845 | 3.4628408 | 0.0005345 | 0.00340316 | CEP295 | 85459 |
| ENSG000001 | 2186.43179 | 0.416857156 | 0.1203906 | 3.46253897 | 0.0005351 | 0.00340644 | CREB1 | 1385 |
| ENSG000002 | 62.8901765 | 0.71847953 | 0.20751174 | 3.46235599 | 0.00053547 | 0.00340822 | SNORA75 | 654321 |
| ENSG000001 | 1217.45793 | 0.572455644 | 0.16536235 | 3.46182574 | 0.00053652 | 0.00341441 | BEND3 | 57673 |
| ENSG000001 | 2352.33419 | 0.423379439 | 0.12230352 | 3.46171087 | 0.00053675 | 0.00341533 | DNMT1 | 1786 |
| ENSG000001 | 47.6082106 | -0.771471267 | 0.22289248 | -3.4611812 | 0.00053781 | 0.00342152 | SGCD | 6444 |
| ENSG000002 | 24.8527462 | -0.905289849 | 0.26156006 | -3.4611166 | 0.00053794 | 0.0034218 | NA | NA |
| ENSG000001 | 25.9564945 | -0.958898324 | 0.27705347 | -3.4610588 | 0.00053806 | 0.003422 | PIK3CG | 5294 |
| ENSG000001 | 538.481224 | 0.388452265 | 0.11224301 | 3.46081463 | 0.00053854 | 0.00342457 | KIN | 22944 |
| ENSG000002 | 601.456668 | 0.517208827 | 0.14945367 | 3.46066331 | 0.00053885 | 0.00342596 | DNAJC19 | 131118 |

| | | | | | | | | |
|---|---|---|---|---|---|---|---|---|
| ENSG000001 | 610.979785 | 0.478977383 | 0.13841397 | 3.46046986 | 0.00053923 | 0.00342788 | EIF2B1 | 1967 |
| ENSG000001 | 469.667309 | 0.484275862 | 0.13995273 | 3.46028159 | 0.00053961 | 0.00342974 | GBF1 | 8729 |
| ENSG000002 | 17.6774657 | -1.026188391 | 0.29659545 | -3.4598925 | 0.00054039 | 0.00343416 | NA | NA |
| ENSG000000 | 1904.21987 | 0.396383652 | 0.11457162 | 3.45970204 | 0.00054077 | 0.00343605 | SIKE1 | 80143 |
| ENSG000001 | 460.413478 | 0.466743271 | 0.13491309 | 3.45958473 | 0.00054101 | 0.00343648 | RIPK1 | 8737 |
| ENSG000002 | 10.2961984 | -1.190210539 | 0.34402926 | -3.4596201 | 0.00054094 | 0.00343648 | NA | NA |
| ENSG000001 | 982.874395 | 0.432473723 | 0.12501033 | 3.45950397 | 0.00054117 | 0.00343697 | EXOC4 | 60412 |
| ENSG000001 | 1854.07364 | 0.564257538 | 0.16310791 | 3.45941239 | 0.00054136 | 0.0034376 | SUPT6H | 6830 |
| ENSG000001 | 1132.46361 | 0.39029717 | 0.11282677 | 3.45926046 | 0.00054166 | 0.0034388 | S100PBP | 64766 |
| ENSG000001 | 38.1247464 | -0.82014952 | 0.23708991 | -3.4592342 | 0.00054171 | 0.0034388 | TDRD5 | 163589 |
| ENSG000001 | 1294.07259 | 0.483654855 | 0.1398282 | 3.45892223 | 0.00054234 | 0.00344224 | PRPF38A | 84950 |
| ENSG000000 | 1718.40049 | 0.407640476 | 0.11786218 | 3.45862 | 0.00054295 | 0.00344497 | RALBP1 | 10928 |
| ENSG000001 | 604.798777 | 0.527259779 | 0.15245029 | 3.45856865 | 0.00054305 | 0.00344497 | PPP2CB | 5516 |
| ENSG000001 | 330.009701 | 0.472867847 | 0.1367247 | 3.45854011 | 0.00054311 | 0.00344497 | ANAPC10 | 10393 |
| ENSG000001 | 140.301741 | 0.521378278 | 0.15074696 | 3.45863202 | 0.00054293 | 0.00344497 | RCCD1 | 91433 |
| ENSG000001 | 263.698058 | 0.505222731 | 0.14609228 | 3.45824385 | 0.00054371 | 0.00344822 | ETNK2 | 55224 |
| ENSG000001 | 2617.33401 | 0.404157126 | 0.11687158 | 3.45813001 | 0.00054394 | 0.00344914 | CHORDC1 | 26973 |
| ENSG000000 | 264.115324 | 0.493495952 | 0.14272909 | 3.45757083 | 0.00054507 | 0.00345577 | NIPAL3 | 57185 |
| ENSG000002 | 10327.6569 | 0.424354764 | 0.12274922 | 3.45708732 | 0.00054605 | 0.00346143 | IPO7 | 10527 |
| ENSG000001 | 360.99683 | 0.497630869 | 0.14394865 | 3.45700258 | 0.00054622 | 0.00346198 | GGCX | 2677 |
| ENSG000001 | 1999.59412 | 0.389857856 | 0.11277521 | 3.45694631 | 0.00054633 | 0.00346216 | CNOT2 | 4848 |
| ENSG000000 | 11.9099867 | -1.150555783 | 0.33283993 | -3.4567841 | 0.00054666 | 0.00346316 | CEACAM6 | 4680 |
| ENSG000001 | 1495.16608 | 0.38385239 | 0.11104221 | 3.45681513 | 0.0005466 | 0.00346316 | WEE1 | 7465 |
| ENSG000000 | 167.981124 | 0.558321587 | 0.16153755 | 3.45629603 | 0.00054765 | 0.0034689 | WDR54 | 84058 |
| ENSG000002 | 87.2832006 | -0.624698643 | 0.1807466 | -3.4562123 | 0.00054782 | 0.00346893 | CCDC162P | 221262 |
| ENSG000002 | 24.4341781 | -0.927236688 | 0.2682814 | -3.4562094 | 0.00054783 | 0.00346893 | OR52I2 | 143502 |
| ENSG000001 | 313.692597 | 0.467328774 | 0.13521784 | 3.45611761 | 0.00054802 | 0.00346957 | DNAJC18 | 202052 |
| ENSG000000 | 89.0782068 | 0.624573491 | 0.18073645 | 3.45571398 | 0.00054884 | 0.00347369 | RNF215 | 200312 |
| ENSG000001 | 167.458368 | 0.579689161 | 0.1677468 | 3.45573906 | 0.00054879 | 0.00347369 | HSD17B11 | 51170 |

| | | | | | | | | | |
|---|---|---|---|---|---|---|---|---|---|
| ENSG000001 | 1053.91108 | 0.425292222 | 0.12307103 | 3.45566465 | 0.00054894 | 0.00347378 | DNAJC3 | | 5611 |
| ENSG000001 | 245.606967 | 0.496766115 | 0.14375601 | 3.45561974 | 0.00054903 | 0.00347382 | HECTD3 | | 79654 |
| ENSG000001 | 1094.14248 | 0.411557261 | 0.1191028 | 3.4554793 | 0.00054932 | 0.00347476 | CCDC18 | | 343099 |
| ENSG000001 | 1897.20222 | 0.432748618 | 0.12523609 | 3.4554625 | 0.00054935 | 0.00347476 | QRICH1 | | 54870 |
| ENSG000001 | 80.7636598 | 0.635334749 | 0.18386993 | 3.45534878 | 0.00054958 | 0.00347568 | THEM6 | | 51337 |
| ENSG000002 | 39.7020266 | 0.782464685 | 0.22645741 | 3.45523989 | 0.0005498 | 0.00347654 | NA | NA | |
| ENSG000002 | 5.66135365 | -1.292885237 | 0.37419076 | -3.4551501 | 0.00054999 | 0.00347716 | NA | NA | |
| ENSG000001 | 2051.95399 | 0.361199834 | 0.10454979 | 3.45481167 | 0.00055068 | 0.00348098 | PSME3 | | 10197 |
| ENSG00000C | 11.734941 | -1.160287324 | 0.33586402 | -3.4546342 | 0.00055104 | 0.00348273 | OR1I1 | | 126370 |
| ENSG00000C | 1169.2825 | 0.496919526 | 0.14385744 | 3.45424969 | 0.00055183 | 0.00348662 | RECQL | | 5965 |
| ENSG000001 | 897.295953 | 0.465182339 | 0.13466955 | 3.45425047 | 0.00055182 | 0.00348662 | NR1D2 | | 9975 |
| ENSG000001 | 77.4754604 | -0.651768856 | 0.18869039 | -3.4541709 | 0.00055199 | 0.00348709 | FRK | | 2444 |
| ENSG000001 | 2302.48629 | 0.389130209 | 0.11266535 | 3.45385891 | 0.00055263 | 0.00349058 | NAMPT | | 10135 |
| ENSG000001 | 92.0974152 | 0.645146472 | 0.18679309 | 3.45380269 | 0.00055274 | 0.00349077 | C19orf12 | | 83636 |
| ENSG000001 | 40.5997626 | -0.845594196 | 0.24483854 | -3.453681 | 0.00055299 | 0.0034918 | ADGB | | 79747 |
| ENSG000001 | 1949.53665 | 0.385521926 | 0.11162809 | 3.45362837 | 0.0005531 | 0.00349194 | ZNF91 | | 7644 |
| ENSG000001 | 1475.02186 | 0.448028268 | 0.1297388 | 3.4533097 | 0.00055375 | 0.00349518 | MPP6 | | 51678 |
| ENSG000002 | 105.342197 | 0.647405923 | 0.1874749 | 3.45329384 | 0.00055379 | 0.00349518 | RGL3 | | 57139 |
| ENSG000001 | 324.777667 | 0.465566268 | 0.13484352 | 3.4526411 | 0.00055513 | 0.00350311 | SGCE | | 8910 |
| ENSG000001 | 4374.64019 | 0.439447181 | 0.12729476 | 3.45220164 | 0.00055603 | 0.00350827 | DLGAP5 | | 9787 |
| ENSG000001 | 33.9378359 | -0.829685749 | 0.24033828 | -3.4521581 | 0.00055612 | 0.00350829 | SELL | | 6402 |
| ENSG000001 | 563.743019 | 0.430093344 | 0.12459447 | 3.45194558 | 0.00055656 | 0.00351051 | ZNF197 | | 10168 |
| ENSG000001 | 33.3959258 | -0.864857934 | 0.25054913 | -3.4518497 | 0.00055676 | 0.00351121 | TYRP1 | | 7306 |
| ENSG00000C | 263.683042 | 0.512631289 | 0.1485117 | 3.45179048 | 0.00055688 | 0.00351144 | TBC1D22B | | 55633 |
| ENSG000001 | 3244.94307 | 0.397885333 | 0.11527902 | 3.45149827 | 0.00055748 | 0.00351469 | MAK16 | | 84549 |
| ENSG000002 | 51.7379542 | 0.742960426 | 0.2152933 | 3.45092214 | 0.00055867 | 0.00352166 | ADRA2B | | 151 |
| ENSG000001 | 12485.5498 | 0.351316291 | 0.10180492 | 3.45087724 | 0.00055877 | 0.0035217 | NONO | | 4841 |
| ENSG000001 | 201.782362 | 0.529095086 | 0.15332433 | 3.45082268 | 0.00055888 | 0.00352186 | TSPAN31 | | 6302 |
| ENSG000001 | 54.2328879 | 0.729941055 | 0.21155744 | 3.4503209 | 0.00055992 | 0.00352786 | ENKD1 | | 84080 |

| | | | | | | | | |
|---|---|---|---|---|---|---|---|---|
| ENSG000001 | 430.551719 | 0.48344947 | 0.14012158 | 3.45021434 | 0.00056014 | 0.00352871 | TRIM23 | 373 |
| ENSG000001 | 5163.18702 | 0.395204135 | 0.11455379 | 3.44994389 | 0.0005607 | 0.0035317 | ZNF644 | 84146 |
| ENSG000001 | 188.349068 | 0.493746621 | 0.14312135 | 3.44984595 | 0.00056091 | 0.00353243 | GBE1 | 2632 |
| ENSG000002 | 9.472007 | -1.197338214 | 0.34708513 | -3.4496961 | 0.00056122 | 0.00353337 | C1orf234 | 729059 |
| ENSG000002 | 85.6041332 | 0.774463085 | 0.22450222 | 3.44969003 | 0.00056123 | 0.00353337 | NA | NA |
| ENSG000001 | 12.350597 | -1.225916667 | 0.3554072 | -3.4493299 | 0.00056198 | 0.00353713 | GRIN3B | 116444 |
| ENSG000002 | 21.1742393 | -0.982388358 | 0.28480647 | -3.449319 | 0.000562 | 0.00353713 | LOC101928227 | 101928227 |
| ENSG000001 | 29.5969602 | -0.923465966 | 0.26775393 | -3.4489352 | 0.0005628 | 0.00354087 | PRDM7 | 11105 |
| ENSG000001 | 2337.76284 | 0.380169687 | 0.11022898 | 3.44890857 | 0.00056286 | 0.00354087 | AP1G1 | 164 |
| ENSG000002 | 13.7419105 | -1.101140341 | 0.3192658 | -3.4489768 | 0.00056271 | 0.00354087 | NA | NA |
| ENSG000001 | 605.678342 | 0.532164767 | 0.1543237 | 3.44836711 | 0.00056399 | 0.00354742 | LSM14B | 149986 |
| ENSG000001 | 658.217805 | 0.450232327 | 0.13057819 | 3.44799035 | 0.00056477 | 0.00355182 | ZCCHC10 | 54819 |
| ENSG000001 | 2703.00678 | 0.439982289 | 0.12762362 | 3.44749884 | 0.0005658 | 0.00355774 | ZBTB44 | 29068 |
| ENSG000001 | 118.193619 | 0.575114674 | 0.16682627 | 3.44738682 | 0.00056604 | 0.00355866 | FAM114A1 | 92689 |
| ENSG000001 | 788.997024 | 0.425552792 | 0.12344804 | 3.44722195 | 0.00056638 | 0.00356029 | SHROOM3 | 57619 |
| ENSG000002 | 6.35532341 | -1.274850671 | 0.36983414 | -3.4470876 | 0.00056666 | 0.0035615 | NA | NA |
| ENSG00000C | 19.9487409 | -1.029152596 | 0.29857785 | -3.4468484 | 0.00056717 | 0.00356411 | SIGLEC1 | 6614 |
| ENSG000001 | 414.131929 | 0.519207886 | 0.15063971 | 3.4466868 | 0.00056751 | 0.00356569 | SRRD | 402055 |
| ENSG000002 | 12.8746308 | -1.151319303 | 0.33404375 | -3.4466123 | 0.00056766 | 0.00356612 | NA | NA |
| ENSG000001 | 1345.09576 | 0.468961636 | 0.13607499 | 3.44634698 | 0.00056822 | 0.00356907 | POLR1B | 84172 |
| ENSG000001 | 929.123007 | 0.432150451 | 0.12541738 | 3.44569823 | 0.00056959 | 0.00357667 | UHRF2 | 115426 |
| ENSG000001 | 523.223875 | 0.54840195 | 0.159156 | 3.44568829 | 0.00056961 | 0.00357667 | GEMIN4 | 50628 |
| ENSG00000C | 128.468063 | 0.657184994 | 0.19073483 | 3.4455427 | 0.00056991 | 0.00357742 | MAP2K3 | 5606 |
| ENSG000001 | 420.913622 | 0.551125985 | 0.15995114 | 3.44558958 | 0.00056982 | 0.00357742 | ATP13A1 | 57130 |
| ENSG000001 | 71.6453358 | -0.690243417 | 0.20033151 | -3.445506 | 0.00056999 | 0.00357742 | KCTD16 | 57528 |
| ENSG000001 | 6741.64144 | 0.386200171 | 0.11209395 | 3.44532589 | 0.00057037 | 0.00357925 | UBR5 | 51366 |
| ENSG000001 | 2754.62558 | 0.54138956 | 0.15715538 | 3.44493173 | 0.0005712 | 0.00358392 | MARCKSL1 | 65108 |
| ENSG000001 | 135.774006 | 0.643700769 | 0.1868746 | 3.44456005 | 0.00057199 | 0.0035883 | SLC29A2 | 3177 |
| ENSG000001 | 144.007046 | 0.569878075 | 0.16547098 | 3.44397587 | 0.00057323 | 0.0035955 | ZNF57 | 126295 |

| | | | | | | | | |
|---|---|---|---|---|---|---|---|---|
| ENSG00000( | 181.448116 | 0.553454482 | 0.16070617 | 3.44389059 | 0.00057341 | 0.00359608 | ROGDI | 79641 |
| ENSG00000( | 93.7543067 | -0.681081757 | 0.19777557 | -3.4437103 | 0.00057379 | 0.00359681 | MEF2C | 4208 |
| ENSG000001 | 264.043087 | 0.650020092 | 0.1887524 | 3.44377124 | 0.00057366 | 0.00359681 | LAMA5 | 3911 |
| ENSG000002 | 11.8899739 | -1.148567908 | 0.33352496 | -3.443724 | 0.00057376 | 0.00359681 | AQP1 | 358 |
| ENSG000001 | 19.1140134 | -0.997552305 | 0.28968083 | -3.4436256 | 0.00057397 | 0.00359683 | CHRNA2 | 1135 |
| ENSG000001 | 40.6964726 | -0.805477662 | 0.23390243 | -3.4436481 | 0.00057392 | 0.00359683 | DSG3 | 1830 |
| ENSG00000( | 280.819118 | 0.619003657 | 0.17977014 | 3.44330629 | 0.00057465 | 0.00360025 | CEP170B | 283638 |
| ENSG000001 | 49.0688705 | -0.780905182 | 0.22679079 | -3.4432843 | 0.00057469 | 0.00360025 | NA | NA |
| ENSG000001 | 483.318945 | 0.46339413 | 0.13458623 | 3.44310202 | 0.00057508 | 0.00360213 | ZNF236 | 7776 |
| ENSG000001 | 188.959834 | 0.496957209 | 0.14434068 | 3.44294632 | 0.00057541 | 0.00360364 | TMEM9B | 56674 |
| ENSG000001 | 594.5693 | 0.442172108 | 0.12843444 | 3.44278448 | 0.00057576 | 0.00360524 | USP38 | 84640 |
| ENSG000001 | 587.254504 | 0.558006697 | 0.16209175 | 3.44253608 | 0.00057629 | 0.003608 | BRD1 | 23774 |
| ENSG000001 | 1153.82705 | 0.426307382 | 0.1238403 | 3.44239627 | 0.00057658 | 0.00360931 | MRPL35 | 51318 |
| ENSG000002 | 12.3434255 | -1.145653664 | 0.33283565 | -3.4421002 | 0.00057722 | 0.0036127 | NA | NA |
| ENSG000001 | 215.669833 | 0.489044391 | 0.14212961 | 3.44083407 | 0.00057992 | 0.00362909 | PGBD1 | 84547 |
| ENSG000001 | 648.349089 | 0.495946193 | 0.14413919 | 3.44074489 | 0.00058012 | 0.00362973 | VPS29 | 51699 |
| ENSG00000( | 213.854159 | 0.559579067 | 0.16264559 | 3.44048101 | 0.00058068 | 0.00363271 | TMEM132A | 54972 |
| ENSG000001 | 419.962146 | 0.452007477 | 0.13138139 | 3.44042249 | 0.00058081 | 0.00363293 | MANBA | 4126 |
| ENSG000001 | 539.34175 | 0.501582655 | 0.14579331 | 3.44036809 | 0.00058092 | 0.0036331 | ZNF627 | 199692 |
| ENSG000001 | 599.031481 | 0.41389588 | 0.12031519 | 3.44009658 | 0.00058151 | 0.00363619 | ATF1 | 466 |
| ENSG000001 | 476.797475 | 0.525003686 | 0.15263627 | 3.43957365 | 0.00058263 | 0.00364266 | ZNF587 | 84914 |
| ENSG000001 | 46.8181049 | -0.807755345 | 0.23485914 | -3.4393183 | 0.00058318 | 0.00364553 | NOL4 | 8715 |
| ENSG000001 | 233.247009 | 0.466354332 | 0.13561051 | 3.43892469 | 0.00058403 | 0.00365008 | TMEM136 | 219902 |
| ENSG000002 | 191.822378 | 0.495707798 | 0.14414729 | 3.43889779 | 0.00058409 | 0.00365008 | NOP14-AS1 | 317648 |
| ENSG000002 | 20.1906817 | -0.971244144 | 0.28243901 | -3.4387748 | 0.00058435 | 0.00365117 | LOC100506178 | 100506178 |
| ENSG000001 | 156.784561 | 0.555866372 | 0.1616616 | 3.4384564 | 0.00058504 | 0.0036549 | FBXO44 | 93611 |
| ENSG000001 | 607.441956 | 0.471166207 | 0.13705913 | 3.4376858 | 0.00058671 | 0.00366454 | EIF2B4 | 8890 |
| ENSG000001 | 14.5263951 | -1.072185402 | 0.31189397 | -3.43766 | 0.00058676 | 0.00366454 | C6orf201 | 404220 |
| ENSG000001 | 37617.7906 | 0.35476371 | 0.10322142 | 3.43691962 | 0.00058837 | 0.00367401 | SET | 6418 |

| | | | | | | | | |
|---|---|---|---|---|---|---|---|---|
| ENSG000001 | 572.421285 | 0.447862211 | 0.13031656 | 3.43672533 | 0.00058879 | 0.00367608 | IPMK | 253430 |
| ENSG000001 | 715.783073 | 0.434519741 | 0.12643603 | 3.43667643 | 0.0005889 | 0.00367617 | KBTBD7 | 84078 |
| ENSG000001 | 88.3326015 | -0.590595218 | 0.17185511 | -3.436588 | 0.00058909 | 0.00367681 | CACNA1C | 775 |
| ENSG000001 | 1887.16257 | 0.403852116 | 0.11751696 | 3.43654321 | 0.00058919 | 0.00367685 | TOPBP1 | 11073 |
| ENSG000001 | 457.330667 | 0.500943089 | 0.1457811 | 3.4362692 | 0.00058978 | 0.00368001 | GGA3 | 23163 |
| ENSG00000C | 238.304286 | 0.563221205 | 0.16391914 | 3.43596969 | 0.00059044 | 0.00368351 | RFXANK | 8625 |
| ENSG000001 | 13.2271996 | -1.095755912 | 0.31891217 | -3.4359175 | 0.00059055 | 0.00368365 | KPNA7 | 402569 |
| ENSG000001 | 886.108936 | 0.480878396 | 0.13996604 | 3.4356792 | 0.00059107 | 0.00368633 | HIATL1 | 84641 |
| ENSG000002 | 151.010714 | 0.725274322 | 0.21112145 | 3.43534174 | 0.00059181 | 0.00369035 | NA | NA |
| ENSG000001 | 43.3432256 | 0.749748267 | 0.21827305 | 3.43490997 | 0.00059275 | 0.00369567 | COMTD1 | 118881 |
| ENSG00000C | 338.317491 | 0.50399504 | 0.14673044 | 3.43483635 | 0.00059291 | 0.00369611 | NA | NA |
| ENSG00000C | 814.465255 | 0.513712475 | 0.14956146 | 3.43479189 | 0.00059301 | 0.00369614 | MAEA | 10296 |
| ENSG000001 | 419.617423 | 0.422356862 | 0.12297715 | 3.4344336 | 0.00059379 | 0.00370046 | NNT | 23530 |
| ENSG000001 | 1465.37222 | 0.438509086 | 0.12768842 | 3.43421188 | 0.00059428 | 0.00370292 | Sep-10 | 151011 |
| ENSG000001 | 544.71411 | 0.524592896 | 0.1527782 | 3.4336895 | 0.00059543 | 0.0037095 | SPC24 | 147841 |
| ENSG000001 | 947.562901 | 0.441668497 | 0.12865769 | 3.43289633 | 0.00059717 | 0.00371962 | TTC21B | 79809 |
| ENSG000001 | 395.769659 | 0.46937826 | 0.13673067 | 3.43286736 | 0.00059723 | 0.00371962 | TMEM246 | 84302 |
| ENSG000001 | 1302.85556 | 0.432113694 | 0.12588018 | 3.43273826 | 0.00059752 | 0.00372082 | RBAK | 57786 |
| ENSG000001 | 963.037632 | 0.455647324 | 0.13276445 | 3.4319979 | 0.00059915 | 0.00373042 | ZNF101 | 94039 |
| ENSG000001 | 750.209859 | 0.375616923 | 0.10946299 | 3.43145124 | 0.00060036 | 0.00373737 | RNASEH2B | 79621 |
| ENSG000001 | 129.30594 | 0.664463153 | 0.19366652 | 3.43096554 | 0.00060144 | 0.00374344 | VPS37B | 79720 |
| ENSG000001 | 54.0497622 | -0.701819724 | 0.20455681 | -3.4309282 | 0.00060152 | 0.00374344 | SYNE3 | 161176 |
| ENSG000001 | 54.2769425 | -0.732195826 | 0.21341596 | -3.4308391 | 0.00060172 | 0.00374409 | MOBP | 4336 |
| ENSG000001 | 58.656316 | 0.752351839 | 0.21932764 | 3.4302646 | 0.00060299 | 0.00375145 | CCDC22 | 28952 |
| ENSG00000C | 39.0679306 | -0.803510633 | 0.23426732 | -3.4298879 | 0.00060383 | 0.00375609 | DDX43 | 55510 |
| ENSG000001 | 1397.65295 | 0.365996789 | 0.10671708 | 3.429599 | 0.00060447 | 0.00375951 | CENPJ | 55835 |
| ENSG000001 | 925.460387 | 0.500865757 | 0.14605088 | 3.42939215 | 0.00060493 | 0.00376065 | TICRR | 90381 |
| ENSG000001 | 2428.36612 | 0.408393854 | 0.11908539 | 3.42942025 | 0.00060487 | 0.00376065 | NAE1 | 8883 |
| ENSG000001 | 10.580224 | -1.175530665 | 0.34277847 | -3.429418 | 0.00060488 | 0.00376065 | NA | NA |

| | | | | | | | | |
|---|---|---|---|---|---|---|---|---|
| ENSG000001 | 229.67727 | 0.447473448 | 0.13048478 | 3.42931522 | 0.00060511 | 0.00376114 | GOLGA1 | 2800 |
| ENSG000001 | 138.254894 | 0.528416309 | 0.15410057 | 3.42903535 | 0.00060573 | 0.0037642 | TTC30B | 150737 |
| ENSG000002 | 9.5838369 | -1.199297202 | 0.34975013 | -3.4290114 | 0.00060578 | 0.0037642 | NA | NA |
| ENSG000001 | 471.15829 | 0.44403354 | 0.12953668 | 3.42785955 | 0.00060836 | 0.00377962 | R3HCC1L | 27291 |
| ENSG00000C | 11702.4977 | 0.3832856 | 0.11181743 | 3.42778059 | 0.00060854 | 0.00378015 | BCLAF1 | 9774 |
| ENSG000002 | 12.3440194 | -1.120101149 | 0.3267814 | -3.4276772 | 0.00060877 | 0.00378101 | SPINK13 | 153218 |
| ENSG000001 | 32.0966538 | -0.891428624 | 0.26008805 | -3.4274109 | 0.00060937 | 0.00378414 | CD33 | 945 |
| ENSG000001 | 29.6356644 | -0.904415751 | 0.26389373 | -3.4271968 | 0.00060985 | 0.00378654 | OCA2 | 4948 |
| ENSG00000C | 20.1311514 | -0.984242013 | 0.28719302 | -3.4271098 | 0.00061004 | 0.0037866 | UPP2 | 151531 |
| ENSG000001 | 21.801923 | -0.955901434 | 0.27892058 | -3.4271456 | 0.00060996 | 0.0037866 | PYHIN1 | 149628 |
| ENSG000001 | 215.570163 | 0.488329679 | 0.14250987 | 3.42663756 | 0.0006111 | 0.00379261 | RBFA | 79863 |
| ENSG000002 | 27.1871599 | -0.875752908 | 0.25559055 | -3.42639 | 0.00061166 | 0.00379548 | PTCHD4 | 442213 |
| ENSG000001 | 83.1521086 | 0.630010282 | 0.18387288 | 3.42633606 | 0.00061178 | 0.00379566 | TMCO4 | 255104 |
| ENSG00000C | 392.310271 | 0.412959118 | 0.12056073 | 3.42532033 | 0.00061407 | 0.00380929 | RNF14 | 9604 |
| ENSG000001 | 110.192792 | 0.644658106 | 0.18822173 | 3.4249929 | 0.00061482 | 0.0038133 | NARFL | 64428 |
| ENSG000001 | 38.7981064 | -0.794693453 | 0.23207027 | -3.4243656 | 0.00061624 | 0.00382126 | ACE2 | 59272 |
| ENSG000001 | 188.380693 | -0.542288538 | 0.15836277 | -3.4243435 | 0.00061629 | 0.00382126 | CABYR | 26256 |
| ENSG00000C | 426.934747 | 0.434601465 | 0.12692042 | 3.42420431 | 0.0006166 | 0.00382263 | APPBP2 | 10513 |
| ENSG000001 | 175.441461 | 0.539774238 | 0.15764312 | 3.42402671 | 0.00061701 | 0.00382455 | DCAF4 | 26094 |
| ENSG00000C | 845.543246 | 0.512762542 | 0.14976036 | 3.42388692 | 0.00061732 | 0.00382577 | NDST1 | 3340 |
| ENSG000001 | 14.192078 | -1.073820399 | 0.31362886 | -3.423857 | 0.00061739 | 0.00382577 | NA | NA |
| ENSG000002 | 216.416189 | 0.46095553 | 0.13463652 | 3.42370359 | 0.00061774 | 0.00382734 | JAKMIP2-AS1 | 153469 |
| ENSG000001 | 519.036877 | 0.482977572 | 0.14107171 | 3.42363177 | 0.0006179 | 0.00382777 | ZNF331 | 55422 |
| ENSG000002 | 308.510824 | 0.551069423 | 0.16096745 | 3.4234835 | 0.00061824 | 0.00382927 | NSUN6 | 221078 |
| ENSG000001 | 167.787419 | 0.58238798 | 0.17013537 | 3.42308585 | 0.00061915 | 0.00383429 | TCAF1 | 9747 |
| ENSG000001 | 223.129611 | 0.539481583 | 0.15763303 | 3.42238932 | 0.00062073 | 0.00384354 | FAM57A | 79850 |
| ENSG000001 | 1155.26413 | 0.404243958 | 0.11812994 | 3.42202793 | 0.00062156 | 0.00384807 | APBB2 | 323 |
| ENSG000002 | 1276.30573 | 0.449040671 | 0.13124573 | 3.42137365 | 0.00062306 | 0.00385675 | OST4 | 100128731 |
| ENSG000002 | 80.0097039 | 0.612816152 | 0.17911787 | 3.42130097 | 0.00062322 | 0.00385719 | NA | NA |

| | | | | | | | | | |
|---|---|---|---|---|---|---|---|---|---|
| ENSG000001 | 618.468133 | 0.427125197 | 0.1248512 | 3.42107408 | 0.00062374 | 0.00385982 | GPN1 | | 11321 |
| ENSG00000C | 1394.46237 | 0.441618671 | 0.12909097 | 3.42098808 | 0.00062394 | 0.00385994 | RBM6 | | 10180 |
| ENSG000001 | 379.850314 | 0.445666846 | 0.13027451 | 3.42098271 | 0.00062395 | 0.00385994 | FIGNL1 | | 63979 |
| ENSG000001 | 2279.44591 | 0.378954846 | 0.11079396 | 3.42035664 | 0.00062539 | 0.00386707 | GNL2 | | 29889 |
| ENSG000002 | 1224.80618 | 0.453826667 | 0.13268258 | 3.42039366 | 0.00062531 | 0.00386707 | TDGF1P3 | | 6998 |
| ENSG000002 | 33.6330312 | 0.819472479 | 0.23958168 | 3.42043041 | 0.00062522 | 0.00386707 | LOC100506098 | | 100506098 |
| ENSG000002 | 21.0103083 | -0.986721636 | 0.28852553 | -3.4198764 | 0.0006265 | 0.00387331 | NA | NA | |
| ENSG000001 | 2117.78363 | 0.436681933 | 0.12771082 | 3.41930256 | 0.00062782 | 0.0038809 | PPIP5K2 | | 23262 |
| ENSG00000C | 1056.93422 | 0.450388017 | 0.13172777 | 3.41908171 | 0.00062833 | 0.00388322 | PDPR | | 55066 |
| ENSG000001 | 306.651022 | 0.479444599 | 0.14022717 | 3.41905649 | 0.00062839 | 0.00388322 | NA | NA | |
| ENSG000001 | 949.189333 | 0.412625415 | 0.12069339 | 3.41879054 | 0.000629 | 0.00388643 | NCOA3 | | 8202 |
| ENSG000001 | 238.794688 | 0.517521861 | 0.15138116 | 3.41866763 | 0.00062929 | 0.00388759 | CLN6 | | 54982 |
| ENSG000001 | 1256.78429 | 0.567768659 | 0.16608967 | 3.41844647 | 0.0006298 | 0.00389016 | UGDH | | 7358 |
| ENSG000001 | 8.74072903 | -1.202058485 | 0.35166013 | -3.4182394 | 0.00063028 | 0.00389253 | SLC22A13 | | 9390 |
| ENSG000001 | 1481.45114 | 0.402395818 | 0.11772256 | 3.41817071 | 0.00063044 | 0.00389292 | ZNF131 | | 7690 |
| ENSG000001 | 2672.79572 | 0.371250009 | 0.10862921 | 3.41758928 | 0.00063178 | 0.00390065 | PCF11 | | 51585 |
| ENSG000001 | 590.548407 | 0.459095863 | 0.13434054 | 3.41740384 | 0.00063221 | 0.00390222 | FGFR1OP2 | | 26127 |
| ENSG000002 | 17.4389604 | -1.0678237 | 0.31246701 | -3.4173966 | 0.00063223 | 0.00390222 | NA | NA | |
| ENSG000001 | 393.596188 | 0.454472725 | 0.13299478 | 3.41722228 | 0.00063264 | 0.00390413 | BCL7B | | 9275 |
| ENSG000001 | 23.3372051 | -0.918370705 | 0.26875787 | -3.4170932 | 0.00063294 | 0.00390479 | CYTH4 | | 27128 |
| ENSG000001 | 495.590654 | 0.591129095 | 0.17299045 | 3.41711983 | 0.00063287 | 0.00390479 | CHMP4B | | 128866 |
| ENSG000001 | 7460.88669 | 0.410509232 | 0.12015303 | 3.4165532 | 0.00063419 | 0.00391195 | HNRNPDL | | 9987 |
| ENSG000001 | 202.33858 | 0.542160526 | 0.15871116 | 3.41602014 | 0.00063544 | 0.00391869 | PLA2G3 | | 50487 |
| ENSG000001 | 877.248909 | 0.419124395 | 0.12269592 | 3.41596027 | 0.00063558 | 0.00391869 | NCAPH | | 23397 |
| ENSG000001 | 34.7697689 | -0.893459897 | 0.2615538 | -3.4159699 | 0.00063555 | 0.00391869 | C3P1 | | 388503 |
| ENSG000001 | 1014.36885 | 0.453484605 | 0.13275771 | 3.41588144 | 0.00063576 | 0.00391923 | ALDH9A1 | | 223 |
| ENSG00000C | 2701.13831 | 0.385810132 | 0.11295313 | 3.41566559 | 0.00063626 | 0.00392174 | RAB10 | | 10890 |
| ENSG00000C | 261.088634 | 0.470066762 | 0.13763375 | 3.41534521 | 0.00063701 | 0.00392576 | SNX24 | | 28966 |
| ENSG000002 | 538.099812 | 0.474533399 | 0.13895927 | 3.41491009 | 0.00063803 | 0.00393144 | NA | NA | |

| | | | | | | | | |
|---|---|---|---|---|---|---|---|---|
| ENSG00000C | 474.999816 | 0.586631089 | 0.17183482 | 3.41392446 | 0.00064034 | 0.0039441 | ITPR3 | 3710 |
| ENSG000001 | 11629.7831 | 0.417489289 | 0.12229065 | 3.41391002 | 0.00064038 | 0.0039441 | HNRNPK | 3190 |
| ENSG000002 | 33.235662 | -0.900293361 | 0.26371272 | -3.4139171 | 0.00064036 | 0.0039441 | CDKN2B-AS1 | 100048912 |
| ENSG000001 | 316.720499 | 0.490221178 | 0.14359702 | 3.41386746 | 0.00064048 | 0.00394411 | RPS6KA3 | 6197 |
| ENSG000001 | 497.544559 | 0.470747117 | 0.13792391 | 3.41309297 | 0.0006423 | 0.00395474 | TXNDC15 | 79770 |
| ENSG00000C | 13.3581856 | -1.121462454 | 0.32862396 | -3.412601 | 0.00064346 | 0.00396128 | PAX2 | 5076 |
| ENSG000001 | 32.3053467 | -0.873064811 | 0.25584842 | -3.41243 | 0.00064386 | 0.00396317 | FAM9C | 171484 |
| ENSG000002 | 3.2053025 | -1.275924854 | 0.37395241 | -3.4119979 | 0.00064489 | 0.00396885 | RN7SL116P | 106480488 |
| ENSG000001 | 243.642378 | 0.483430197 | 0.1417206 | 3.41114983 | 0.0006469 | 0.00398061 | CSRNP3 | 80034 |
| ENSG000001 | 66.8267363 | 0.667617078 | 0.19571878 | 3.41110384 | 0.000647 | 0.00398068 | PEX6 | 5190 |
| ENSG000001 | 181.580065 | 0.518934575 | 0.15213881 | 3.41092832 | 0.00064742 | 0.00398264 | ZNF346 | 23567 |
| ENSG000001 | 9.37932565 | -1.182685296 | 0.34674467 | -3.4108248 | 0.00064767 | 0.00398355 | GZMK | 3003 |
| ENSG000001 | 24.1336314 | -0.934415243 | 0.27396449 | -3.4107166 | 0.00064792 | 0.00398453 | SHCBP1L | 81626 |
| ENSG000001 | 927.682695 | 0.518297666 | 0.15196572 | 3.41062221 | 0.00064815 | 0.0039847 | CDK5RAP2 | 55755 |
| ENSG000002 | 141.250098 | 0.556600861 | 0.16319493 | 3.41065037 | 0.00064808 | 0.0039847 | LINC00493 | 388789 |
| ENSG000001 | 11.2225394 | -1.149070754 | 0.33694007 | -3.410312 | 0.00064889 | 0.00398863 | OR2V1 | 26693 |
| ENSG000001 | 670.806098 | 0.51646016 | 0.15144583 | 3.41019739 | 0.00064916 | 0.00398909 | SAV1 | 60485 |
| ENSG000002 | 6.43298333 | -1.26427354 | 0.37072878 | -3.4102384 | 0.00064906 | 0.00398909 | NA | NA |
| ENSG000001 | 60.2796314 | -0.663986511 | 0.19472252 | -3.4099112 | 0.00064984 | 0.00399268 | VWCE | 220001 |
| ENSG000001 | 1960.06413 | 0.400564508 | 0.1174783 | 3.4096893 | 0.00065037 | 0.00399532 | ARHGEF12 | 23365 |
| ENSG000001 | 6683.36757 | 0.379425205 | 0.1112831 | 3.40954927 | 0.0006507 | 0.00399677 | HDGF | 3068 |
| ENSG000001 | 304.378494 | 0.509751409 | 0.14951159 | 3.40944421 | 0.00065095 | 0.0039977 | ZNF417 | 147687 |
| ENSG000002 | 25.0872172 | -0.923907315 | 0.27100951 | -3.4091325 | 0.0006517 | 0.00400167 | NA | NA |
| ENSG000001 | 595.206009 | 0.478189436 | 0.14028561 | 3.40868477 | 0.00065277 | 0.00400703 | ANKS6 | 203286 |
| ENSG000001 | 2585.22363 | 0.585473589 | 0.1717574 | 3.40872401 | 0.00065267 | 0.00400703 | DDX42 | 11325 |
| ENSG000001 | 295.657844 | 0.466269379 | 0.13679581 | 3.40850628 | 0.0006532 | 0.00400904 | DENND1A | 57706 |
| ENSG000001 | 3084.67192 | 0.377502977 | 0.11075584 | 3.40842512 | 0.00065339 | 0.00400963 | ZC3H14 | 79882 |
| ENSG000002 | 14.333453 | -1.058599477 | 0.31062075 | -3.4080128 | 0.00065438 | 0.00401508 | NA | NA |
| ENSG000002 | 331.560028 | 0.471889289 | 0.13847431 | 3.40777499 | 0.00065495 | 0.00401798 | NA | NA |

| | | | | | | | | |
|---|---|---|---|---|---|---|---|---|
| ENSG000001 | 156.455265 | 0.562021908 | 0.1649303 | 3.40763284 | 0.00065529 | 0.00401946 | MAP4K2 | 5871 |
| ENSG000001 | 23.0800971 | -0.964085604 | 0.28293622 | -3.4074309 | 0.00065578 | 0.00402183 | SLC26A9 | 115019 |
| ENSG00000C | 410.657468 | 0.463077575 | 0.13593174 | 3.4066921 | 0.00065755 | 0.00403212 | AP3M2 | 10947 |
| ENSG000001 | 22.1668988 | -0.949283041 | 0.27865801 | -3.4066239 | 0.00065772 | 0.00403252 | APOL1 | 8542 |
| ENSG000002 | 27.738624 | -0.972523622 | 0.28548616 | -3.4065527 | 0.00065789 | 0.00403296 | NR2F2-AS1 | 644192 |
| ENSG000001 | 54.1806508 | -0.693784477 | 0.20367052 | -3.4064059 | 0.00065824 | 0.00403452 | ADGRF1 | 266977 |
| ENSG000002 | 13.8971944 | -1.081514764 | 0.317533 | -3.4059917 | 0.00065924 | 0.00403958 | C10orf128 | 170371 |
| ENSG000002 | 68.947181 | 0.673462413 | 0.19773175 | 3.40593965 | 0.00065937 | 0.00403958 | NA | NA |
| ENSG000002 | 32.6720737 | 0.813508945 | 0.23884813 | 3.40596739 | 0.0006593 | 0.00403958 | NA | NA |
| ENSG000001 | 509.456134 | 0.469062086 | 0.13774437 | 3.40530855 | 0.00066089 | 0.00404832 | C17orf75 | 64149 |
| ENSG000001 | 883.301321 | 0.400072197 | 0.11749025 | 3.40515228 | 0.00066127 | 0.00404942 | NIN | 51199 |
| ENSG000002 | 12.7135068 | -1.109177768 | 0.32573357 | -3.4051687 | 0.00066123 | 0.00404942 | NA | NA |
| ENSG000001 | 537.959109 | 0.408347267 | 0.11992339 | 3.40506787 | 0.00066148 | 0.00405006 | STX18 | 53407 |
| ENSG000001 | 20.2603867 | -0.99096975 | 0.29104163 | -3.4049072 | 0.00066187 | 0.00405122 | PAX8 | 7849 |
| ENSG000001 | 2743.85731 | 0.421746581 | 0.12386293 | 3.40494607 | 0.00066177 | 0.00405122 | SETD2 | 29072 |
| ENSG000001 | 1178.82246 | 0.401994496 | 0.11807058 | 3.40469649 | 0.00066238 | 0.00405373 | PROSER1 | 80209 |
| ENSG000001 | 297.768669 | 0.505324829 | 0.14843212 | 3.40441686 | 0.00066305 | 0.00405727 | TBC1D20 | 128637 |
| ENSG000002 | 28.3714986 | -0.87877545 | 0.25815303 | -3.4040874 | 0.00066386 | 0.00406156 | LINC01568 | 100506172 |
| ENSG000001 | 1281.85982 | 0.429103285 | 0.12606931 | 3.40370926 | 0.00066477 | 0.00406657 | STARD4 | 134429 |
| ENSG000001 | 27.1513826 | -0.874045908 | 0.25680171 | -3.4035828 | 0.00066508 | 0.00406784 | C12orf42 | 374470 |
| ENSG00000C | 1533.15874 | 0.414851405 | 0.12192368 | 3.40254984 | 0.0006676 | 0.00408263 | IDI1 | 3422 |
| ENSG000001 | 646.361383 | 0.520983476 | 0.15312409 | 3.40236137 | 0.00066806 | 0.00408483 | NUDT16 | 131870 |
| ENSG000001 | 65.4408064 | 0.699984427 | 0.20573935 | 3.40228747 | 0.00066824 | 0.00408532 | CBLC | 23624 |
| ENSG000001 | 8.09956694 | -1.219445715 | 0.35844642 | -3.4020307 | 0.00066887 | 0.00408793 | HOXD10 | 3236 |
| ENSG000002 | 25.6981362 | -0.894249908 | 0.26285619 | -3.4020501 | 0.00066882 | 0.00408793 | NPHP3-AS1 | 348808 |
| ENSG000002 | 627.25676 | 0.482791233 | 0.14192009 | 3.4018527 | 0.00066931 | 0.00408997 | VPS52 | 6293 |
| ENSG000002 | 30.1869934 | -0.852569312 | 0.25064227 | -3.4015385 | 0.00067008 | 0.00409406 | NA | NA |
| ENSG000001 | 54.1671381 | 0.707606539 | 0.2080562 | 3.40103562 | 0.00067131 | 0.00410098 | NAPRT | 93100 |
| ENSG00000C | 256.373558 | 0.507759558 | 0.14931429 | 3.40060926 | 0.00067236 | 0.00410677 | FUNDC1 | 139341 |

| | | | | | | | | |
|---|---|---|---|---|---|---|---|---|
| ENSG000001 | 453.368995 | 0.459276491 | 0.13505939 | 3.40055221 | 0.0006725 | 0.00410679 | FANCL | 55120 |
| ENSG000001 | 654.266032 | 0.383645875 | 0.1128196 | 3.400525 | 0.00067257 | 0.00410679 | ZNF776 | 284309 |
| ENSG000001 | 2282.81095 | 0.541267676 | 0.15918197 | 3.40030759 | 0.0006731 | 0.00410944 | FZD7 | 8324 |
| ENSG000002 | 44.6030513 | -0.785809392 | 0.23112872 | -3.3998777 | 0.00067416 | 0.00411529 | LINC01330 | 646168 |
| ENSG000001 | 25.1935636 | -0.917198941 | 0.26978328 | -3.399762 | 0.00067445 | 0.00411641 | VWC2 | 375567 |
| ENSG000001 | 428.979654 | 0.478000254 | 0.14062651 | 3.3990764 | 0.00067614 | 0.00412612 | PASK | 23178 |
| ENSG000002 | 538.054458 | 0.425121755 | 0.12508442 | 3.39867861 | 0.00067712 | 0.00413151 | RMST | 196475 |
| ENSG000001 | 62.8232307 | -0.683209973 | 0.20102978 | -3.3985511 | 0.00067744 | 0.00413281 | MYBPC1 | 4604 |
| ENSG000001 | 497.475941 | 0.415152597 | 0.1221967 | 3.39741246 | 0.00068026 | 0.00414942 | ADAMTS12 | 81792 |
| ENSG00000C | 1398.85947 | 0.402798422 | 0.11857169 | 3.39708769 | 0.00068107 | 0.00415373 | SNX13 | 23161 |
| ENSG000001 | 1760.31763 | 0.412747127 | 0.12150317 | 3.39700713 | 0.00068127 | 0.00415432 | WASL | 8976 |
| ENSG000001 | 10060.8346 | 0.43132451 | 0.12697913 | 3.39681426 | 0.00068175 | 0.00415663 | KPNB1 | 3837 |
| ENSG000001 | 18.6844161 | -1.007425871 | 0.29659228 | -3.3966692 | 0.00068211 | 0.00415821 | UBE2U | 148581 |
| ENSG000001 | 373.370478 | 0.580129396 | 0.17080523 | 3.39643815 | 0.00068269 | 0.00416109 | AKAP17A | 8227 |
| ENSG000001 | 12598.5425 | 0.628928515 | 0.1852114 | 3.39573327 | 0.00068445 | 0.0041712 | HNRNPA2B1 | 3181 |
| ENSG000001 | 811.837167 | 0.384891661 | 0.11336629 | 3.39511553 | 0.000686 | 0.00417999 | ATG14 | 22863 |
| ENSG000001 | 383.000546 | 0.485561174 | 0.14302153 | 3.39502163 | 0.00068623 | 0.0041808 | POMK | 84197 |
| ENSG000001 | 2327.06754 | 0.383576912 | 0.11300073 | 3.39446398 | 0.00068763 | 0.00418869 | GPBP1L1 | 60313 |
| ENSG000002 | 7.06659927 | -1.245136642 | 0.36683322 | -3.3942854 | 0.00068808 | 0.00419079 | TSSK1B | 83942 |
| ENSG000001 | 374.881161 | 0.47093201 | 0.13874699 | 3.39417825 | 0.00068835 | 0.00419181 | CFL2 | 1073 |
| ENSG000002 | 24.5481956 | -0.906396495 | 0.26705369 | -3.3940609 | 0.00068864 | 0.00419297 | ANKRD66 | 100287718 |
| ENSG00000C | 706.557156 | 0.430976971 | 0.12699593 | 3.39362815 | 0.00068973 | 0.00419834 | GOPC | 57120 |
| ENSG000001 | 282.219199 | 0.55839009 | 0.16454033 | 3.39363667 | 0.00068971 | 0.00419834 | RRAGA | 10670 |
| ENSG000001 | 2741.96173 | 0.461878196 | 0.13610588 | 3.39352123 | 0.00069 | 0.00419935 | GFPT1 | 2673 |
| ENSG000002 | 699.311312 | 0.451581824 | 0.13309537 | 3.39291906 | 0.00069152 | 0.00420796 | CUX1 | 1523 |
| ENSG000001 | 22.842539 | 0.928423661 | 0.27364937 | 3.39274911 | 0.00069195 | 0.00420994 | NEURL2 | 140825 |
| ENSG000001 | 1212.4476 | 0.433823084 | 0.12788394 | 3.39231872 | 0.00069304 | 0.00421593 | AP3M1 | 26985 |
| ENSG000002 | 7.63287801 | -1.218381624 | 0.35919614 | -3.3919675 | 0.00069393 | 0.0042207 | NA | NA |
| ENSG000001 | 549.815893 | 0.865075565 | 0.25504834 | 3.39181024 | 0.00069433 | 0.00422249 | COL6A1 | 1291 |

| | | | | | | | | |
|---|---|---|---|---|---|---|---|---|
| ENSG00000C | 2433.27883 | 0.470193848 | 0.13865482 | 3.39111079 | 0.0006961 | 0.00423232 | KIF1B | 23095 |
| ENSG000001 | 516.630584 | 0.462936006 | 0.13651702 | 3.39104974 | 0.00069625 | 0.00423232 | NELFCD | 51497 |
| ENSG000001 | 79.5808799 | 0.651392469 | 0.19209016 | 3.39107663 | 0.00069619 | 0.00423232 | TMEM251 | 26175 |
| ENSG000001 | 1474.20865 | 0.498896159 | 0.14712612 | 3.39094203 | 0.00069653 | 0.00423335 | RBM12B | 389677 |
| ENSG000001 | 445.025146 | 0.490123934 | 0.14454423 | 3.39082313 | 0.00069683 | 0.00423455 | SMURF1 | 57154 |
| ENSG000002 | 18.7805072 | -0.989336281 | 0.29177798 | -3.3907161 | 0.0006971 | 0.00423557 | NA | NA |
| ENSG000001 | 358.490519 | 0.503757769 | 0.14857336 | 3.39063329 | 0.00069731 | 0.00423622 | WBP1L | 54838 |
| ENSG000001 | 796.673661 | 0.402720731 | 0.11877743 | 3.39054929 | 0.00069753 | 0.00423688 | ARHGEF9 | 23229 |
| ENSG000001 | 1675.51069 | 0.462438508 | 0.1363996 | 3.39032147 | 0.00069811 | 0.0042385 | GLTSCR2 | 29997 |
| ENSG000001 | 1278.28774 | 0.385388443 | 0.11367102 | 3.39038425 | 0.00069795 | 0.0042385 | FBXO5 | 26271 |
| ENSG000001 | 472.024312 | 0.494705201 | 0.14591646 | 3.39033169 | 0.00069808 | 0.0042385 | VPS39 | 23339 |
| ENSG000001 | 3004.44147 | 0.408955571 | 0.12062831 | 3.39021218 | 0.00069839 | 0.00423892 | HBS1L | 10767 |
| ENSG000001 | 2548.84047 | 0.422503496 | 0.12462446 | 3.39021326 | 0.00069838 | 0.00423892 | PPA1 | 5464 |
| ENSG000001 | 227.44945 | 0.583249763 | 0.17205815 | 3.38984103 | 0.00069933 | 0.00424403 | FAM193B | 54540 |
| ENSG000001 | 37.994752 | -0.798176118 | 0.2354652 | -3.3897838 | 0.00069948 | 0.00424428 | TXLNB | 167838 |
| ENSG000001 | 10285.4143 | 0.419700567 | 0.12381779 | 3.38966289 | 0.00069979 | 0.00424434 | UGP2 | 7360 |
| ENSG000001 | 970.547113 | 0.434223873 | 0.12810261 | 3.38965661 | 0.0006998 | 0.00424434 | GTF2F2 | 2963 |
| ENSG000001 | 2418.88234 | 0.397788675 | 0.11735136 | 3.38972354 | 0.00069963 | 0.00424434 | TOPORS | 10210 |
| ENSG00000C | 1919.51462 | 0.40485673 | 0.11944811 | 3.38939419 | 0.00070047 | 0.00424606 | AGPS | 8540 |
| ENSG000001 | 1237.68848 | 0.468415931 | 0.13819685 | 3.3894833 | 0.00070024 | 0.00424606 | ASH2L | 9070 |
| ENSG000001 | 550.759688 | 0.475391818 | 0.14025915 | 3.38938186 | 0.0007005 | 0.00424606 | PHF8 | 23133 |
| ENSG000002 | 4.52438021 | -1.283142553 | 0.37857029 | -3.3894433 | 0.00070035 | 0.00424606 | LOC105374205 | 105374205 |
| ENSG000001 | 31.125258 | -0.833994831 | 0.24606859 | -3.389278 | 0.00070077 | 0.00424703 | MS4A2 | 2206 |
| ENSG000002 | 30.3860451 | -0.857292902 | 0.25295552 | -3.3891054 | 0.00070121 | 0.00424907 | NA | NA |
| ENSG000002 | 8.76841792 | -1.20219151 | 0.35475245 | -3.388818 | 0.00070195 | 0.00425289 | NA | NA |
| ENSG000001 | 1243.73312 | 0.432053058 | 0.12749872 | 3.38868538 | 0.00070229 | 0.00425431 | GORASP2 | 26003 |
| ENSG00000C | 282.568097 | 0.538421585 | 0.15890701 | 3.38828084 | 0.00070332 | 0.00425805 | RAD54L | 8438 |
| ENSG000001 | 632.528375 | 0.442780424 | 0.13067808 | 3.38832967 | 0.0007032 | 0.00425805 | FERMT1 | 55612 |
| ENSG000001 | 1358.57194 | 0.432025797 | 0.12750523 | 3.38829865 | 0.00070328 | 0.00425805 | TIAL1 | 7073 |

| | | | | | | | | |
|---|---|---|---|---|---|---|---|---|
| ENSG000001 | 173.555075 | 0.652302132 | 0.19251291 | 3.38835531 | 0.00070313 | 0.00425805 | ZNF526 | 116115 |
| ENSG000001 | 487.61085 | 0.411970773 | 0.12159433 | 3.38807539 | 0.00070385 | 0.0042606 | SFMBT1 | 51460 |
| ENSG000001 | 28.468851 | -0.858288615 | 0.25333049 | -3.3880195 | 0.00070399 | 0.00426084 | EPB42 | 2038 |
| ENSG000002 | 35.1171866 | 0.804022411 | 0.23731777 | 3.38795706 | 0.00070415 | 0.00426117 | NA | NA |
| ENSG000001 | 67.0143912 | 0.676194472 | 0.19960883 | 3.38759797 | 0.00070508 | 0.00426612 | CCM2L | 140706 |
| ENSG000002 | 6.39341621 | -1.256303649 | 0.37086677 | -3.3874797 | 0.00070538 | 0.00426732 | NA | NA |
| ENSG000002 | 12.969142 | -1.096929145 | 0.32383996 | -3.387257 | 0.00070595 | 0.00427015 | NA | NA |
| ENSG000001 | 534.971626 | 0.43815828 | 0.12935737 | 3.38719222 | 0.00070612 | 0.00427052 | TTF1 | 7270 |
| ENSG000001 | 323.059884 | 0.555541707 | 0.16402782 | 3.38687495 | 0.00070694 | 0.00427482 | AXIN2 | 8313 |
| ENSG000001 | 1309.17521 | 0.450848348 | 0.13314395 | 3.3861722 | 0.00070875 | 0.00428515 | ARRDC3 | 57561 |
| ENSG000001 | 1675.9002 | 0.400548149 | 0.11829866 | 3.38590612 | 0.00070944 | 0.00428866 | ZGRF1 | 55345 |
| ENSG000001 | 9390.33216 | 0.399418125 | 0.11798455 | 3.38534258 | 0.00071089 | 0.00429684 | DHX9 | 1660 |
| ENSG000001 | 20.8118127 | -0.963621847 | 0.28466626 | -3.3850933 | 0.00071154 | 0.00429946 | ASB18 | 401036 |
| ENSG000002 | 71.1938714 | 0.625073041 | 0.1846532 | 3.3851189 | 0.00071147 | 0.00429946 | LOC101928254 | 101928254 |
| ENSG000001 | 121.214776 | 0.522737539 | 0.1544525 | 3.38445511 | 0.0007132 | 0.00430883 | C11orf71 | 54494 |
| ENSG000002 | 23.0516775 | -0.9215335 | 0.27232557 | -3.3839404 | 0.00071454 | 0.00431627 | ADAMTS9-AS2 | 100507098 |
| ENSG000002 | 71.7196608 | -0.66704445 | 0.19718973 | -3.3827545 | 0.00071763 | 0.0043343 | NA | NA |
| ENSG00000C | 244.140315 | 0.445927411 | 0.13183132 | 3.38256055 | 0.00071813 | 0.0043357 | ABHD5 | 51099 |
| ENSG000001 | 665.766041 | 0.47123229 | 0.139313 | 3.38254361 | 0.00071818 | 0.0043357 | ZSCAN12 | 9753 |
| ENSG000002 | 11.860435 | -1.107532967 | 0.32742364 | -3.3825687 | 0.00071811 | 0.0043357 | CLEC12B | 387837 |
| ENSG000001 | 887.248354 | 0.406925114 | 0.12030579 | 3.38242346 | 0.00071849 | 0.00433695 | LCLAT1 | 253558 |
| ENSG000001 | 145.358414 | 0.516490844 | 0.1527078 | 3.38221643 | 0.00071903 | 0.00433957 | TOM1 | 10043 |
| ENSG000001 | 207.242115 | 0.503021936 | 0.14872859 | 3.38214684 | 0.00071922 | 0.00434002 | IMPDH1 | 3614 |
| ENSG000001 | 8202.14156 | 0.82719911 | 0.24459507 | 3.3819124 | 0.00071983 | 0.00434308 | FLNA | 2316 |
| ENSG000001 | 532.881952 | 0.563548795 | 0.16665135 | 3.38160361 | 0.00072064 | 0.00434732 | ZNF592 | 9640 |
| ENSG00000C | 4730.62565 | 0.402965868 | 0.11916639 | 3.38153957 | 0.00072081 | 0.00434769 | GNAS | 2778 |
| ENSG000002 | 6.64869712 | -1.244466503 | 0.36805021 | -3.3812411 | 0.00072159 | 0.00435177 | NA | NA |
| ENSG00000C | 2411.34216 | 0.381439583 | 0.11281575 | 3.38108464 | 0.000722 | 0.00435359 | SLC44A1 | 23446 |
| ENSG000001 | 305.562187 | 0.545351098 | 0.16129664 | 3.38104446 | 0.00072211 | 0.00435359 | IP6K1 | 9807 |

| | | | | | | | | | |
|---|---|---|---|---|---|---|---|---|---|
| ENSG000001 | 535.82916 | 0.480989125 | 0.14226361 | 3.38097081 | 0.0007223 | 0.00435411 | ABHD10 | | 55347 |
| ENSG00000C | 1986.39394 | 0.398007568 | 0.11772972 | 3.38068891 | 0.00072304 | 0.00435793 | CCNT2 | | 905 |
| ENSG000002 | 69.6485119 | -0.641108039 | 0.18965184 | -3.3804472 | 0.00072368 | 0.00436111 | XKR4 | | 114786 |
| ENSG000002 | 20.2754324 | -0.970268975 | 0.28704941 | -3.3801462 | 0.00072447 | 0.00436524 | NA | NA | |
| ENSG000001 | 206.356732 | 0.560515975 | 0.16582933 | 3.3800775 | 0.00072465 | 0.00436569 | BDH2 | | 56898 |
| ENSG000001 | 81.1933955 | 0.653596585 | 0.19338765 | 3.37972241 | 0.00072559 | 0.00437068 | ST6GALNAC4 | | 27090 |
| ENSG000001 | 278.18301 | 0.526902639 | 0.15590429 | 3.37965457 | 0.00072577 | 0.00437111 | MVD | | 4597 |
| ENSG000001 | 1706.45372 | 0.449147372 | 0.13291041 | 3.37932431 | 0.00072664 | 0.00437572 | Mar-06 | | 10299 |
| ENSG000001 | 2877.61546 | 0.372554609 | 0.11026004 | 3.37887252 | 0.00072784 | 0.00438226 | INTS6 | | 26512 |
| ENSG000001 | 1295.88247 | 0.478322856 | 0.14157756 | 3.37852178 | 0.00072877 | 0.00438721 | STAT1 | | 6772 |
| ENSG000001 | 87.6776938 | -0.594952612 | 0.17610302 | -3.378435 | 0.000729 | 0.00438729 | PAPPA2 | | 60676 |
| ENSG000001 | 31.8186033 | -0.818109955 | 0.24215372 | -3.3784736 | 0.00072889 | 0.00438729 | CPB1 | | 1360 |
| ENSG000001 | 271.515916 | 0.479377266 | 0.14191303 | 3.37796505 | 0.00073024 | 0.00439414 | POP5 | | 51367 |
| ENSG000001 | 218.891802 | 0.510033942 | 0.15099388 | 3.37784505 | 0.00073056 | 0.00439541 | CDK18 | | 5129 |
| ENSG000002 | 4.20886834 | -1.277276354 | 0.37815143 | -3.3776848 | 0.00073099 | 0.00439731 | NA | NA | |
| ENSG000001 | 68.2969496 | 0.693661553 | 0.20536852 | 3.37764301 | 0.0007311 | 0.00439733 | TNNI3 | | 7137 |
| ENSG000001 | 159.399703 | 0.502478969 | 0.14877639 | 3.37741071 | 0.00073172 | 0.00440039 | CTSH | | 1512 |
| ENSG00000C | 53.6175902 | -0.744144464 | 0.22041858 | -3.3760515 | 0.00073534 | 0.00442154 | SLC26A4 | | 5172 |
| ENSG00000C | 19.4242655 | -0.980071372 | 0.29030812 | -3.3759695 | 0.00073556 | 0.0044222 | ZPBP | | 11055 |
| ENSG000002 | 49.5471593 | -0.762065363 | 0.22573722 | -3.375896 | 0.00073576 | 0.00442272 | NA | NA | |
| ENSG000001 | 46.6091297 | -0.740864966 | 0.21948606 | -3.3754534 | 0.00073694 | 0.00442919 | CPED1 | | 79974 |
| ENSG000001 | 192.354153 | 0.558029661 | 0.16534209 | 3.37500057 | 0.00073816 | 0.00443582 | TKFC | | 26007 |
| ENSG00000C | 537.770309 | 0.566402867 | 0.16782576 | 3.37494594 | 0.0007383 | 0.00443605 | SREBF1 | | 6720 |
| ENSG000001 | 1888.59416 | 0.40095015 | 0.11880388 | 3.37489096 | 0.00073845 | 0.00443628 | C1GALT1 | | 56913 |
| ENSG000001 | 20.8444855 | -0.994256552 | 0.2946214 | -3.3746922 | 0.00073898 | 0.00443882 | FAM81B | | 153643 |
| ENSG000001 | 5245.11152 | 0.363628873 | 0.10775471 | 3.37459834 | 0.00073923 | 0.00443968 | WDR43 | | 23160 |
| ENSG00000C | 966.905919 | 0.382578969 | 0.11338357 | 3.3742012 | 0.0007403 | 0.00444543 | PRR11 | | 55771 |
| ENSG000001 | 179.49254 | 0.52315832 | 0.15504887 | 3.37415117 | 0.00074044 | 0.00444558 | ZNF555 | | 148254 |
| ENSG000001 | 357.176399 | 0.502230702 | 0.14886374 | 3.37376105 | 0.00074149 | 0.00445123 | TVP23B | | 51030 |

| | | | | | | | | |
|---|---|---|---|---|---|---|---|---|
| ENSG000001 | 3170.69539 | 0.369736162 | 0.10959659 | 3.37361015 | 0.00074189 | 0.00445301 | SP1 | 6667 |
| ENSG000001 | 1538.83355 | 0.582357926 | 0.17263261 | 3.37339474 | 0.00074247 | 0.00445518 | SF1 | 7536 |
| ENSG000001 | 91.2159257 | -0.623983102 | 0.18496971 | -3.373434 | 0.00074237 | 0.00445518 | FAM179A | 165186 |
| ENSG000002 | 2.57352995 | -1.243091677 | 0.36853566 | -3.3730567 | 0.00074339 | 0.00445999 | NA | NA |
| ENSG000002 | 204.662607 | 0.533526062 | 0.15818332 | 3.37283383 | 0.00074399 | 0.00446294 | HSPA1B | 3304 |
| ENSG000000 | 4357.61515 | 0.358422613 | 0.10626917 | 3.37278073 | 0.00074413 | 0.00446314 | ZNF207 | 7756 |
| ENSG000001 | 82.9826912 | 0.632005891 | 0.18739165 | 3.37264704 | 0.00074449 | 0.00446333 | NA | NA |
| ENSG000001 | 27.2238377 | -0.872066558 | 0.25856607 | -3.372703 | 0.00074434 | 0.00446333 | ANKRD30B | 374860 |
| ENSG000002 | 191.688609 | 0.52012813 | 0.15421868 | 3.37266619 | 0.00074444 | 0.00446333 | PCDHAC2 | 56134 |
| ENSG000000 | 53.4803316 | -0.680333931 | 0.20174171 | -3.3723018 | 0.00074543 | 0.00446827 | MECOM | 2122 |
| ENSG000000 | 3634.65198 | 0.420316646 | 0.12469158 | 3.37085022 | 0.00074937 | 0.00448988 | REST | 5978 |
| ENSG000001 | 34.8601967 | -0.837873772 | 0.24856414 | -3.3708554 | 0.00074935 | 0.00448988 | SLC24A4 | 123041 |
| ENSG000001 | 76.1526377 | 0.654093609 | 0.19403979 | 3.37092519 | 0.00074916 | 0.00448988 | SPATA33 | 124045 |
| ENSG000001 | 73.0770525 | -0.656384149 | 0.19474623 | -3.3704589 | 0.00075043 | 0.0044956 | ADAMTSL3 | 57188 |
| ENSG000001 | 186.508472 | 0.523720131 | 0.15538797 | 3.37040332 | 0.00075058 | 0.00449584 | SCAND1 | 51282 |
| ENSG000001 | 5606.12631 | 0.502160442 | 0.1490197 | 3.3697587 | 0.00075234 | 0.00450571 | MAP1B | 4131 |
| ENSG000000 | 31.15671 | -0.891159096 | 0.26447589 | -3.3695287 | 0.00075297 | 0.0045081 | BLNK | 29760 |
| ENSG000001 | 909.69529 | 0.435689146 | 0.12930408 | 3.36949251 | 0.00075307 | 0.0045081 | GOSR1 | 9527 |
| ENSG000001 | 399.700786 | 0.428220775 | 0.1270877 | 3.36949041 | 0.00075307 | 0.0045081 | CLK4 | 57396 |
| ENSG000002 | 640.528135 | 0.420305906 | 0.12476621 | 3.36874793 | 0.0007551 | 0.00451959 | NA | NA |
| ENSG000001 | 217.327631 | 0.544942093 | 0.16177753 | 3.36846587 | 0.00075588 | 0.00452355 | DDA1 | 79016 |
| ENSG000001 | 350.720867 | 0.427666197 | 0.12696822 | 3.36829312 | 0.00075635 | 0.00452438 | PIN4 | 5303 |
| ENSG000001 | 296.115852 | 0.518315727 | 0.15387901 | 3.36833283 | 0.00075624 | 0.00452438 | AAR2 | 25980 |
| ENSG000001 | 226.130604 | 0.478442206 | 0.14204171 | 3.36832197 | 0.00075627 | 0.00452438 | DNAJB4 | 11080 |
| ENSG000000 | 210.658813 | 0.563461644 | 0.16728649 | 3.36824349 | 0.00075649 | 0.00452453 | MARK4 | 57787 |
| ENSG000001 | 611.879506 | 0.422418222 | 0.12542068 | 3.36801083 | 0.00075713 | 0.00452768 | RCHY1 | 25898 |
| ENSG000001 | 2291.27809 | 0.376591409 | 0.11183132 | 3.36749486 | 0.00075854 | 0.0045355 | APC | 324 |
| ENSG000002 | 26.7042534 | -0.915329986 | 0.27182106 | -3.3673991 | 0.00075881 | 0.0045364 | LINC01427 | 101929663 |
| ENSG000001 | 15.5817262 | -1.058863134 | 0.31448736 | -3.3669498 | 0.00076005 | 0.00454313 | KLRC1 | 3821 |

| | | | | | | | | |
|---|---|---|---|---|---|---|---|---|
| ENSG000001 | 145.438443 | 0.557202363 | 0.16550593 | 3.36666103 | 0.00076084 | 0.00454722 | NPRL2 | 10641 |
| ENSG000002 | 11.064599 | -1.140891549 | 0.33889178 | -3.3665365 | 0.00076118 | 0.0045486 | NA | NA |
| ENSG000002 | 9.26414403 | -1.191271437 | 0.35387283 | -3.3663829 | 0.00076161 | 0.00455047 | KIR3DL3 | 115653 |
| ENSG000001 | 2437.12089 | 0.418580442 | 0.1243454 | 3.36627211 | 0.00076192 | 0.00455161 | NUSAP1 | 51203 |
| ENSG000002 | 19.6020743 | -0.969389004 | 0.28797447 | -3.3662324 | 0.00076202 | 0.00455161 | LOC100996635 | 100996635 |
| ENSG000001 | 1743.7833 | 0.389456592 | 0.11571061 | 3.36578124 | 0.00076327 | 0.00455839 | INADL | 10207 |
| ENSG000002 | 27.8550861 | -0.890555233 | 0.26460275 | -3.3656311 | 0.00076369 | 0.0045602 | NA | NA |
| ENSG000001 | 896.654149 | 0.38222399 | 0.11357143 | 3.36549414 | 0.00076407 | 0.00456179 | MAPRE2 | 10982 |
| ENSG000002 | 26.8593811 | -0.94176433 | 0.27984629 | -3.3652915 | 0.00076463 | 0.00456447 | NA | NA |
| ENSG000002 | 11.1848289 | 1.128171638 | 0.33525519 | 3.36511307 | 0.00076512 | 0.00456675 | KRT222 | 125113 |
| ENSG000001 | 178.188973 | 0.52192695 | 0.15511248 | 3.36482897 | 0.00076591 | 0.00457078 | FAM104B | 90736 |
| ENSG000001 | 8.5896239 | -1.185324621 | 0.35227535 | -3.3647674 | 0.00076608 | 0.00457113 | SHANK2-AS3 | 220070 |
| ENSG000001 | 316.295848 | 0.520116766 | 0.15458375 | 3.36462773 | 0.00076647 | 0.00457277 | NA | NA |
| ENSG000001 | 203.662753 | 0.483405709 | 0.14369403 | 3.3641321 | 0.00076785 | 0.00458032 | ARHGEF39 | 84904 |
| ENSG000002 | 201.603224 | 0.507834263 | 0.15096351 | 3.36395372 | 0.00076834 | 0.0045826 | NA | NA |
| ENSG00000C | 966.529609 | 0.436687226 | 0.12982285 | 3.36371634 | 0.00076901 | 0.0045852 | RBM7 | 10179 |
| ENSG000001 | 11.4213029 | -1.128311817 | 0.33543375 | -3.3637397 | 0.00076894 | 0.0045852 | BPIFB6 | 128859 |
| ENSG000001 | 237.012196 | 0.5021283 | 0.14928728 | 3.36350358 | 0.0007696 | 0.00458806 | ALKBH1 | 8846 |
| ENSG00000C | 322.405311 | 0.521908116 | 0.15517149 | 3.36342777 | 0.00076981 | 0.00458865 | CRY1 | 1407 |
| ENSG000001 | 195.733461 | 0.498980802 | 0.1483567 | 3.36338573 | 0.00076993 | 0.00458867 | DUSP23 | 54935 |
| ENSG000001 | 22.0995815 | -0.974456682 | 0.28979157 | -3.3626123 | 0.00077209 | 0.00460087 | SEC14L3 | 266629 |
| ENSG000001 | 11.4627067 | -1.107873438 | 0.32947361 | -3.3625559 | 0.00077224 | 0.00460113 | ADH1A | 124 |
| ENSG000001 | 319.972714 | 0.459283205 | 0.1365936 | 3.36240658 | 0.00077266 | 0.00460294 | SUOX | 6821 |
| ENSG000001 | 110.053384 | 0.61449946 | 0.18277121 | 3.36212396 | 0.00077345 | 0.00460698 | KDF1 | 126695 |
| ENSG000002 | 7.91974956 | -1.203169744 | 0.35788294 | -3.3619086 | 0.00077406 | 0.0046099 | NA | NA |
| ENSG000001 | 62.0487093 | 0.807861381 | 0.24030709 | 3.36178754 | 0.0007744 | 0.00461057 | IFI27L2 | 83982 |
| ENSG000001 | 6.32749915 | -1.247989573 | 0.37122647 | -3.3618011 | 0.00077436 | 0.00461057 | PLGLB2 | 5342 |
| ENSG000001 | 6.17139029 | -1.244554224 | 0.37027466 | -3.361165 | 0.00077614 | 0.00462029 | TLX1 | 3195 |
| ENSG00000C | 343.109677 | 0.447515967 | 0.13315089 | 3.36096876 | 0.0007767 | 0.0046229 | COQ9 | 57017 |

| | | | | | | | | |
|---|---|---|---|---|---|---|---|---|
| ENSG000001 | 140.524373 | 0.591212167 | 0.17591154 | 3.36084926 | 0.00077703 | 0.00462422 | SDR39U1 | 56948 |
| ENSG000001 | 1648.51936 | 0.45008423 | 0.13393465 | 3.36047636 | 0.00077808 | 0.00462979 | FBXL5 | 26234 |
| ENSG00000C | 1102.39067 | 0.473621952 | 0.14094686 | 3.36028731 | 0.00077861 | 0.00463092 | KAT6A | 7994 |
| ENSG000001 | 56.1614385 | -0.711635884 | 0.21177445 | -3.3603481 | 0.00077844 | 0.00463092 | ENPEP | 2028 |
| ENSG000001 | 509.208597 | 0.448537404 | 0.1334804 | 3.36032406 | 0.00077851 | 0.00463092 | PRMT2 | 3275 |
| ENSG00000C | 3263.67974 | 0.392394105 | 0.116779 | 3.36014262 | 0.00077902 | 0.00463266 | WAPL | 23063 |
| ENSG000001 | 729.270775 | 0.393184569 | 0.11701614 | 3.36008832 | 0.00077918 | 0.0046329 | GDAP2 | 54834 |
| ENSG000001 | 122.282218 | 0.577830452 | 0.17198071 | 3.35985613 | 0.00077983 | 0.00463611 | FBXO41 | 150726 |
| ENSG000002 | 21.1578275 | -0.971675763 | 0.28920855 | -3.3597754 | 0.00078006 | 0.00463679 | NA | NA |
| ENSG000001 | 2834.07514 | 0.563323643 | 0.16768 | 3.35951608 | 0.00078079 | 0.00464046 | ANP32A | 8125 |
| ENSG000001 | 41.781192 | -0.751716124 | 0.22377153 | -3.3593019 | 0.0007814 | 0.00464338 | PMFBP1 | 83449 |
| ENSG00000C | 44.9722194 | -0.739320304 | 0.22008507 | -3.3592478 | 0.00078155 | 0.00464361 | SIDT1 | 54847 |
| ENSG000001 | 1101.82725 | 0.400185857 | 0.11913863 | 3.35899323 | 0.00078227 | 0.00464721 | TMEM106B | 54664 |
| ENSG000001 | 486.861372 | 0.421079129 | 0.12536409 | 3.35884959 | 0.00078268 | 0.00464894 | UBXN2A | 165324 |
| ENSG000001 | 75.2924008 | 0.654150497 | 0.19475917 | 3.35876615 | 0.00078291 | 0.00464966 | AMIGO2 | 347902 |
| ENSG000001 | 27.2768826 | -0.847153888 | 0.25227037 | -3.3581189 | 0.00078475 | 0.00465988 | GPR26 | 2849 |
| ENSG000001 | 307.492428 | 0.492987073 | 0.14680802 | 3.35803905 | 0.00078498 | 0.00466055 | DMAP1 | 55929 |
| ENSG000002 | 10.1289134 | -1.157869334 | 0.34483964 | -3.3577037 | 0.00078593 | 0.00466552 | LOC286370 | 286370 |
| ENSG000001 | 140.023355 | 0.5130425 | 0.15280702 | 3.35745367 | 0.00078664 | 0.00466906 | NA | NA |
| ENSG000002 | 8.5894886 | -1.181503837 | 0.35190986 | -3.3574048 | 0.00078678 | 0.0046692 | NA | NA |
| ENSG000001 | 34.2139112 | -0.858265129 | 0.25564307 | -3.357279 | 0.00078714 | 0.00467064 | CIITA | 4261 |
| ENSG000001 | 1073.48218 | 0.396220328 | 0.11802658 | 3.35704313 | 0.00078781 | 0.00467394 | CEP78 | 84131 |
| ENSG000001 | 35944.9862 | 0.385102501 | 0.11473551 | 3.35643685 | 0.00078954 | 0.00468314 | LIN28A | 79727 |
| ENSG000001 | 31.1182732 | -0.818643476 | 0.24390383 | -3.3564191 | 0.00078959 | 0.00468314 | ATG9B | 285973 |
| ENSG000001 | 668.102173 | 0.561391977 | 0.16726515 | 3.35629968 | 0.00078993 | 0.00468447 | MAML1 | 9794 |
| ENSG000002 | 3.13963738 | -1.254899126 | 0.37390225 | -3.3562225 | 0.00079015 | 0.0046851 | LOC105375759 | 105375759 |
| ENSG000002 | 37.9733419 | -0.815010501 | 0.24284027 | -3.3561587 | 0.00079033 | 0.00468549 | NA | NA |
| ENSG00000C | 272.182332 | 0.435731787 | 0.12984396 | 3.35581093 | 0.00079133 | 0.00469003 | TMEM38B | 55151 |
| ENSG000001 | 695.842638 | 0.40449903 | 0.12053693 | 3.35581005 | 0.00079133 | 0.00469003 | ZNF519 | 162655 |

| | | | | | | | | |
|---|---|---|---|---|---|---|---|---|
| ENSG000001 | 150.785482 | 0.504801917 | 0.15042816 | 3.3557675 | 0.00079145 | 0.00469007 | BRF1 | 2972 |
| ENSG000001 | 3071.34872 | 0.391598214 | 0.11670278 | 3.35551736 | 0.00079217 | 0.00469362 | ATP13A3 | 79572 |
| ENSG000001 | 86.836743 | -0.654439963 | 0.19505262 | -3.3551969 | 0.00079309 | 0.00469838 | NA | NA |
| ENSG000001 | 161.771153 | 0.601518867 | 0.17928462 | 3.35510574 | 0.00079335 | 0.00469924 | PBK | 55872 |
| ENSG000001 | 2723.71831 | 0.379670131 | 0.11316709 | 3.35495191 | 0.00079379 | 0.00470117 | CCDC6 | 8030 |
| ENSG000001 | 27.2156981 | -0.860108258 | 0.2563822 | -3.3547893 | 0.00079425 | 0.00470324 | SNAP25 | 6616 |
| ENSG000001 | 401.196767 | 0.434513635 | 0.12952474 | 3.35467683 | 0.00079458 | 0.00470378 | GLMP | 112770 |
| ENSG000002 | 7.4373757 | -1.223187948 | 0.36461969 | -3.3546953 | 0.00079452 | 0.00470378 | NA | NA |
| ENSG000001 | 691.708542 | 0.440731027 | 0.13138656 | 3.35446059 | 0.0007952 | 0.00470677 | METTL5 | 29081 |
| ENSG000001 | 464.795873 | 0.543210683 | 0.16194082 | 3.35437782 | 0.00079544 | 0.00470749 | HADH | 3033 |
| ENSG000001 | 1455.89426 | 0.447375439 | 0.13337883 | 3.35417135 | 0.00079603 | 0.00471032 | TWISTNB | 221830 |
| ENSG000001 | 7541.19507 | 0.392588758 | 0.11704842 | 3.35407136 | 0.00079632 | 0.00471134 | DHX15 | 1665 |
| ENSG000001 | 354.262588 | 0.427543596 | 0.12747424 | 3.35396074 | 0.00079664 | 0.00471185 | PIP4K2C | 79837 |
| ENSG000001 | 39.1809343 | 0.766428429 | 0.22851293 | 3.3539827 | 0.00079657 | 0.00471185 | SLC25A42 | 284439 |
| ENSG000001 | 217.49886 | 0.507924414 | 0.15144275 | 3.35390378 | 0.0007968 | 0.00471213 | CRYBB2P1 | 1416 |
| ENSG000001 | 1633.78288 | 0.373555245 | 0.111382 | 3.35382069 | 0.00079704 | 0.00471286 | ANKRD40 | 91369 |
| ENSG000002 | 1271.28205 | 0.412019114 | 0.12286764 | 3.35335737 | 0.00079838 | 0.00472006 | SNHG3 | 8420 |
| ENSG000000 | 46.045013 | -0.747092705 | 0.22279948 | -3.3532067 | 0.00079881 | 0.0047217 | BIRC3 | 330 |
| ENSG000002 | 23.4628776 | -0.972539213 | 0.29003481 | -3.3531811 | 0.00079888 | 0.0047217 | TEX26-AS1 | 100507064 |
| ENSG000001 | 336.465393 | 0.451004826 | 0.13450649 | 3.35303383 | 0.00079931 | 0.00472352 | MAP3K3 | 4215 |
| ENSG000000 | 504.522119 | 0.524256705 | 0.15635815 | 3.35292218 | 0.00079963 | 0.00472474 | GRAMD1A | 57655 |
| ENSG000000 | 28.9092969 | -0.847565299 | 0.25279029 | -3.3528396 | 0.00079987 | 0.00472546 | SLC12A3 | 6559 |
| ENSG000000 | 1484.31819 | 0.52160775 | 0.1555803 | 3.3526593 | 0.00080039 | 0.0047272 | IPO11 | 51194 |
| ENSG000001 | 5507.54437 | 0.435736882 | 0.12996763 | 3.3526569 | 0.0008004 | 0.0047272 | ADAR | 103 |
| ENSG000000 | 1569.63993 | 0.421798234 | 0.12581954 | 3.35240641 | 0.00080112 | 0.00473079 | PPP2R5C | 5527 |
| ENSG000000 | 912.63111 | 0.373374803 | 0.11138306 | 3.35216859 | 0.00080181 | 0.00473348 | POLD3 | 10714 |
| ENSG000002 | 39.844986 | -0.771145838 | 0.23004308 | -3.3521802 | 0.00080178 | 0.00473348 | NA | NA |
| ENSG000001 | 1064.93985 | 0.468276606 | 0.13969861 | 3.35204924 | 0.00080216 | 0.00473483 | PCBP2 | 5094 |
| ENSG000001 | 1475.11454 | 0.401124149 | 0.11966777 | 3.35198138 | 0.00080235 | 0.0047353 | KIDINS220 | 57498 |

| | | | | | | | | | |
|---|---|---|---|---|---|---|---|---|---|
| ENSG000002 | 179.379689 | 0.520243899 | 0.15520945 | 3.35188297 | 0.00080264 | 0.0047363 | NA | NA | |
| ENSG000001 | 1151.74463 | 0.370690157 | 0.11060481 | 3.35148301 | 0.0008038 | 0.00474245 | MSI2 | | 124540 |
| ENSG000001 | 813.48109 | 0.439222398 | 0.13106827 | 3.35109628 | 0.00080492 | 0.0047477 | ATE1 | | 11101 |
| ENSG000001 | 38.1617601 | -0.847007123 | 0.25275242 | -3.3511336 | 0.00080481 | 0.0047477 | C10orf11 | | 83938 |
| ENSG000002 | 12.3434739 | -1.104063677 | 0.32961601 | -3.349545 | 0.00080944 | 0.00477367 | NA | NA | |
| ENSG000001 | 263.494642 | 0.581554402 | 0.17363738 | 3.34924661 | 0.00081032 | 0.00477764 | PYGO2 | | 90780 |
| ENSG000001 | 807.939314 | 0.413456575 | 0.1234481 | 3.34923404 | 0.00081035 | 0.00477764 | SLC25A46 | | 91137 |
| ENSG000002 | 250.066796 | 0.552794221 | 0.16505552 | 3.34914116 | 0.00081062 | 0.00477855 | NA | NA | |
| ENSG000001 | 104.06906 | 0.630453702 | 0.1882509 | 3.34900762 | 0.00081102 | 0.00478016 | CHST14 | | 113189 |
| ENSG000002 | 85.7247042 | -0.604173776 | 0.18041293 | -3.3488385 | 0.00081151 | 0.00478238 | NA | NA | |
| ENSG000001 | 1250.74659 | 0.386577424 | 0.11543986 | 3.34873422 | 0.00081182 | 0.00478349 | FOXJ3 | | 22887 |
| ENSG000002 | 179.407013 | 0.496041621 | 0.14813743 | 3.3485231 | 0.00081244 | 0.00478644 | NRAV | | 100506668 |
| ENSG00000C | 2842.89369 | 0.452239015 | 0.13505867 | 3.3484634 | 0.00081261 | 0.00478677 | TRAM1 | | 23471 |
| ENSG00000C | 19.8807698 | -0.982936316 | 0.29355742 | -3.3483614 | 0.00081291 | 0.00478784 | RGS1 | | 5996 |
| ENSG000001 | 102.451236 | 0.566556146 | 0.16922161 | 3.34801289 | 0.00081393 | 0.00479317 | SEMG1 | | 6406 |
| ENSG000001 | 311.490969 | 0.443027756 | 0.1323293 | 3.34791887 | 0.00081421 | 0.0047941 | CYP20A1 | | 57404 |
| ENSG000001 | 725.491529 | 0.532204802 | 0.15897391 | 3.34774936 | 0.00081471 | 0.00479634 | ARID3A | | 1820 |
| ENSG000001 | 21.9676398 | -0.934460957 | 0.27915413 | -3.3474732 | 0.00081552 | 0.00480042 | CNTD1 | | 124817 |
| ENSG000001 | 122.261138 | 0.61721416 | 0.18438791 | 3.34736784 | 0.00081583 | 0.00480155 | NIPSNAP3A | | 25934 |
| ENSG000001 | 1391.88896 | 0.374276085 | 0.11182032 | 3.34712041 | 0.00081656 | 0.00480514 | NOC3L | | 64318 |
| ENSG00000C | 594.788643 | 0.463334193 | 0.13843492 | 3.34694594 | 0.00081707 | 0.00480747 | ALG6 | | 29929 |
| ENSG000002 | 511.27433 | 0.492659879 | 0.14719964 | 3.34688234 | 0.00081726 | 0.00480787 | NA | NA | |
| ENSG000001 | 1599.58142 | 0.4412027 | 0.13182771 | 3.346813 | 0.00081746 | 0.00480838 | LTN1 | | 26046 |
| ENSG000001 | 80.5431643 | 0.63270987 | 0.18905444 | 3.34670731 | 0.00081778 | 0.00480951 | CACFD1 | | 11094 |
| ENSG000001 | 30.1732444 | -0.835675614 | 0.24973008 | -3.3463155 | 0.00081893 | 0.00481422 | CREG2 | | 200407 |
| ENSG000001 | 2811.52748 | 0.451980857 | 0.1350673 | 3.3463381 | 0.00081886 | 0.00481422 | YTHDF3 | | 253943 |
| ENSG000001 | 3301.21413 | 0.391310564 | 0.11693635 | 3.34635515 | 0.00081881 | 0.00481422 | KPNA4 | | 3840 |
| ENSG000001 | 95.4206848 | 0.665248441 | 0.19880448 | 3.34624465 | 0.00081914 | 0.00481431 | RNF122 | | 79845 |
| ENSG000002 | 260.793273 | 0.627449957 | 0.18750951 | 3.34623002 | 0.00081918 | 0.00481431 | PCGF2 | | 7703 |

| | | | | | | | | |
|---|---|---|---|---|---|---|---|---|
| ENSG000001 | 51.6636394 | -0.723576 | 0.21624573 | -3.3460823 | 0.00081962 | 0.00481618 | CC2D2B | 387707 |
| ENSG00000C | 25517.3057 | 0.42341721 | 0.12655591 | 3.3456929 | 0.00082077 | 0.00482155 | DNMT3B | 1789 |
| ENSG000001 | 218.683575 | 0.479743921 | 0.14339125 | 3.34569875 | 0.00082076 | 0.00482155 | DOK4 | 55715 |
| ENSG000001 | 130.132195 | 0.554764215 | 0.16581796 | 3.3456219 | 0.00082098 | 0.00482208 | PHF1 | 5252 |
| ENSG00000C | 288.205676 | 0.503435212 | 0.1504916 | 3.3452711 | 0.00082202 | 0.004824 | MIPEP | 4285 |
| ENSG000001 | 355.206167 | 0.450778206 | 0.1347486 | 3.34532744 | 0.00082185 | 0.004824 | ZC4H2 | 55906 |
| ENSG000001 | 3685.45499 | 0.408502407 | 0.12211018 | 3.34535906 | 0.00082176 | 0.004824 | STT3B | 201595 |
| ENSG000001 | 354.957096 | 0.473655812 | 0.14158892 | 3.34528872 | 0.00082197 | 0.004824 | DNAJC24 | 120526 |
| ENSG000001 | 1245.85658 | 0.402257963 | 0.12024253 | 3.34538839 | 0.00082167 | 0.004824 | PAK2 | 5062 |
| ENSG000002 | 23.0493934 | -0.955263203 | 0.28555472 | -3.3452895 | 0.00082197 | 0.004824 | DNAJC27-AS1 | 729723 |
| ENSG000001 | 2798.73162 | 0.358712098 | 0.10723763 | 3.34502065 | 0.00082276 | 0.00482765 | ALMS1 | 7840 |
| ENSG00000C | 1179.15175 | 0.407160719 | 0.12173065 | 3.34476737 | 0.00082352 | 0.00483137 | MAP4K5 | 11183 |
| ENSG000001 | 31.6234698 | -0.855765641 | 0.25586097 | -3.344651 | 0.00082386 | 0.00483269 | MAEL | 84944 |
| ENSG000002 | 272.724344 | 0.447806065 | 0.13389087 | 3.34456025 | 0.00082413 | 0.00483288 | CHURC1 | 91612 |
| ENSG000002 | 6.94864534 | -1.225693105 | 0.36646977 | -3.3445954 | 0.00082403 | 0.00483288 | NA | NA |
| ENSG000001 | 106.614977 | -0.554449412 | 0.16578334 | -3.3444217 | 0.00082454 | 0.00483459 | CREB5 | 9586 |
| ENSG000001 | 281.330371 | 0.54785963 | 0.16381662 | 3.34434712 | 0.00082476 | 0.00483519 | ZBTB7B | 51043 |
| ENSG000002 | 55.4014933 | 0.763288795 | 0.22824157 | 3.34421467 | 0.00082516 | 0.0048368 | SNORD46 | 94161 |
| ENSG000002 | 55.5028239 | 0.739239929 | 0.22105316 | 3.3441727 | 0.00082528 | 0.00483684 | LOC105376805 | 105376805 |
| ENSG000001 | 1199.9629 | 0.474712107 | 0.14195465 | 3.34411109 | 0.00082547 | 0.00483721 | LARP4B | 23185 |
| ENSG000001 | 1592.89732 | 0.432908637 | 0.12945636 | 3.34405087 | 0.00082565 | 0.00483756 | ZWILCH | 55055 |
| ENSG000001 | 11.8615311 | -1.100681895 | 0.32915055 | -3.3440075 | 0.00082578 | 0.00483762 | TREML1 | 340205 |
| ENSG000001 | 277.825621 | 0.537752669 | 0.16082491 | 3.34371508 | 0.00082665 | 0.00484202 | ARPC5L | 81873 |
| ENSG000001 | 237.328203 | 0.455081751 | 0.13611897 | 3.34326471 | 0.00082799 | 0.00484919 | ARL2BP | 23568 |
| ENSG000001 | 240.768561 | 0.454798354 | 0.13603892 | 3.34314882 | 0.00082833 | 0.00485035 | PDSS2 | 57107 |
| ENSG000002 | 78.5982713 | 0.614472384 | 0.18380218 | 3.343118 | 0.00082843 | 0.00485035 | UBE2V1 | 7335 |
| ENSG000002 | 35715.2803 | 0.350968901 | 0.10499005 | 3.3428778 | 0.00082914 | 0.00485385 | L1TD1 | 54596 |
| ENSG000002 | 8.58240393 | -1.199578854 | 0.35885644 | -3.342782 | 0.00082943 | 0.00485482 | NA | NA |
| ENSG000002 | 111.681561 | 0.528596013 | 0.15813548 | 3.34267818 | 0.00082974 | 0.00485594 | FNTB | 2342 |

| | | | | | | | | | |
|---|---|---|---|---|---|---|---|---|---|
| ENSG000001 | 31.4737623 | -0.881673814 | 0.26377435 | -3.3425305 | 0.00083018 | 0.00485782 | PLCZ1 | | 89869 |
| ENSG000001 | 1119.54089 | 0.490874178 | 0.14686383 | 3.34237614 | 0.00083064 | 0.00485983 | CTR9 | | 9646 |
| ENSG000001 | 150.97732 | 0.590850952 | 0.17679121 | 3.34208338 | 0.00083152 | 0.00486425 | ZBTB17 | | 7709 |
| ENSG000001 | 272.264505 | 0.470328966 | 0.14075124 | 3.34156188 | 0.00083308 | 0.0048727 | TUBD1 | | 51174 |
| ENSG000001 | 71.4340432 | 0.640354137 | 0.19164777 | 3.3413076 | 0.00083385 | 0.00487646 | NMU | | 10874 |
| ENSG000001 | 1393.33648 | 0.383932107 | 0.11491011 | 3.3411516 | 0.00083432 | 0.0048785 | RBMS1 | | 5937 |
| ENSG000001 | 2447.48121 | 0.429496492 | 0.12855224 | 3.34102685 | 0.00083469 | 0.00487998 | | Sep-07 | 989 |
| ENSG000001 | 113.675046 | 0.525147406 | 0.15718515 | 3.34094789 | 0.00083493 | 0.00488067 | SLC35D2 | | 11046 |
| ENSG000001 | 478.412792 | 0.443880292 | 0.13286601 | 3.34081144 | 0.00083534 | 0.00488236 | ABHD13 | | 84945 |
| ENSG00000C | 2921.71046 | 0.555140958 | 0.1661753 | 3.34069482 | 0.00083569 | 0.00488371 | UBE2K | | 3093 |
| ENSG000001 | 393.737867 | 0.437545146 | 0.13098549 | 3.34040924 | 0.00083655 | 0.00488803 | COG6 | | 57511 |
| ENSG000001 | 493.214893 | 0.440464269 | 0.13186328 | 3.34031029 | 0.00083685 | 0.00488846 | RNF125 | | 54941 |
| ENSG000001 | 1190.77888 | 0.42546738 | 0.12737382 | 3.34030476 | 0.00083686 | 0.00488846 | GFM1 | | 85476 |
| ENSG000001 | 23.4013968 | 0.921028669 | 0.27575443 | 3.34003213 | 0.00083769 | 0.00489256 | NA | NA | |
| ENSG000001 | 87.7496792 | 0.687032089 | 0.20570008 | 3.33997004 | 0.00083787 | 0.00489295 | GGT7 | | 2686 |
| ENSG000001 | 843.476378 | 0.374902128 | 0.11225564 | 3.33971738 | 0.00083864 | 0.0048967 | PHLDB2 | | 90102 |
| ENSG000001 | 98.1135656 | 0.642456218 | 0.19238102 | 3.33949896 | 0.0008393 | 0.00489984 | RND1 | | 27289 |
| ENSG000001 | 556.153149 | 0.428559173 | 0.12833503 | 3.33937793 | 0.00083966 | 0.00490127 | SIAH1 | | 6477 |
| ENSG000001 | 14.2303398 | -1.042367985 | 0.31215199 | -3.3392963 | 0.00083991 | 0.00490167 | CXorf36 | | 79742 |
| ENSG000001 | 376.216912 | 0.441218359 | 0.13213002 | 3.3392741 | 0.00083998 | 0.00490167 | MALSU1 | | 115416 |
| ENSG000001 | 825.787069 | 0.509072241 | 0.15245172 | 3.33923581 | 0.00084009 | 0.00490167 | TET3 | | 200424 |
| ENSG000001 | 176.718287 | 0.496549904 | 0.14870458 | 3.33917012 | 0.00084029 | 0.00490212 | AP3S2 | | 10239 |
| ENSG000001 | 23.8586714 | -0.924722775 | 0.27693584 | -3.3391228 | 0.00084043 | 0.00490225 | DSCAML1 | | 57453 |
| ENSG000001 | 517.881449 | 0.507723153 | 0.15205481 | 3.33907994 | 0.00084056 | 0.0049023 | ERF | | 2077 |
| ENSG000001 | 83.2481188 | -0.66023273 | 0.19774585 | -3.3387944 | 0.00084143 | 0.00490659 | SVEP1 | | 79987 |
| ENSG000001 | 307.969072 | 0.427177165 | 0.12794498 | 3.33875684 | 0.00084154 | 0.00490659 | SVIP | | 258010 |
| ENSG000001 | 143.83741 | 0.533665187 | 0.15984215 | 3.33870126 | 0.00084171 | 0.00490687 | TYK2 | | 7297 |
| ENSG000001 | 227.360771 | 0.465718208 | 0.13950024 | 3.33847601 | 0.00084239 | 0.00491015 | VRK3 | | 51231 |
| ENSG000001 | 56.0330266 | -0.678381377 | 0.20321237 | -3.3382878 | 0.00084296 | 0.00491277 | ATP10B | | 23120 |

| | | | | | | | | | |
|---|---|---|---|---|---|---|---|---|---|
| ENSG000001 | 652.841026 | 0.432724393 | 0.12963626 | 3.33798887 | 0.00084387 | 0.00491735 | ATL3 | | 25923 |
| ENSG000001 | 153.594328 | 0.53853998 | 0.16134085 | 3.33790218 | 0.00084413 | 0.00491818 | EHD1 | | 10938 |
| ENSG00000C | 346.500303 | 0.527596139 | 0.15807101 | 3.33771609 | 0.0008447 | 0.00492076 | TNFRSF12A | | 51330 |
| ENSG000001 | 1810.96763 | 0.41000427 | 0.12284706 | 3.33751802 | 0.0008453 | 0.00492356 | USP37 | | 57695 |
| ENSG00000C | 760.3294 | 0.516319663 | 0.15471043 | 3.33732936 | 0.00084588 | 0.0049262 | CEP68 | | 23177 |
| ENSG000001 | 19.3235107 | -0.958751449 | 0.28731815 | -3.3368983 | 0.00084719 | 0.00493314 | RUFY4 | | 285180 |
| ENSG000001 | 810.134925 | 0.395300911 | 0.11846542 | 3.3368464 | 0.00084735 | 0.00493332 | ZNF639 | | 51193 |
| ENSG000001 | 437.081618 | 0.564403192 | 0.16914462 | 3.3368085 | 0.00084746 | 0.00493332 | MINK1 | | 50488 |
| ENSG000001 | 122.167991 | 0.658530835 | 0.1973636 | 3.33663775 | 0.00084798 | 0.00493564 | NA | NA | |
| ENSG000001 | 38.9125724 | -0.79285631 | 0.23766875 | -3.3359721 | 0.00085002 | 0.00494577 | GRAP2 | | 9402 |
| ENSG000001 | 251.9727 | 0.482196948 | 0.14454571 | 3.33594795 | 0.00085009 | 0.00494577 | CLPP | | 8192 |
| ENSG000002 | 11.1338326 | -1.10957369 | 0.33260482 | -3.3360121 | 0.00084989 | 0.00494577 | NA | NA | |
| ENSG000001 | 323.791592 | 0.452127019 | 0.13553666 | 3.33582819 | 0.00085046 | 0.00494719 | GMPPA | | 29926 |
| ENSG000002 | 17.5767047 | -1.001748346 | 0.30033227 | -3.3354669 | 0.00085156 | 0.00495292 | NA | NA | |
| ENSG000001 | 17.211491 | -1.007412236 | 0.30203796 | -3.3353828 | 0.00085182 | 0.0049537 | MEP1A | | 4224 |
| ENSG00000C | 236.065971 | 0.491889334 | 0.14747904 | 3.33531679 | 0.00085202 | 0.00495417 | SLC45A4 | | 57210 |
| ENSG000001 | 1297.68155 | 0.447045163 | 0.13404823 | 3.33495755 | 0.00085312 | 0.00495986 | FRAT2 | | 23401 |
| ENSG000002 | 42.2902007 | 0.751444496 | 0.22534116 | 3.33469707 | 0.00085392 | 0.0049638 | NA | NA | |
| ENSG000001 | 38.2231653 | -0.807947421 | 0.24231442 | -3.3342937 | 0.00085516 | 0.00497029 | CNTNAP4 | | 85445 |
| ENSG000001 | 1076.74729 | 0.517637631 | 0.15526053 | 3.33399381 | 0.00085609 | 0.00497494 | CD276 | | 80381 |
| ENSG000001 | 380.907651 | 0.500280545 | 0.15006584 | 3.33374026 | 0.00085687 | 0.00497876 | TCEAL2 | | 140597 |
| ENSG000001 | 384.946029 | 0.458943225 | 0.13768281 | 3.33333709 | 0.00085811 | 0.00498527 | BLCAP | | 10904 |
| ENSG000001 | 6442.34904 | 0.391717426 | 0.11754317 | 3.33254096 | 0.00086057 | 0.00499884 | TMEM167A | | 153339 |
| ENSG000001 | 808.4241 | 0.469682445 | 0.14094175 | 3.33245778 | 0.00086083 | 0.00499962 | AKT2 | | 208 |
| ENSG000001 | 1691.15827 | 0.474103104 | 0.14228019 | 3.33217933 | 0.00086169 | 0.00500391 | CERS2 | | 29956 |
| ENSG000001 | 1307.74255 | 0.406108271 | 0.12187847 | 3.33207554 | 0.00086201 | 0.00500506 | OTUD6B | | 51633 |
| ENSG000002 | 67.1239848 | 0.729400484 | 0.21891391 | 3.33190555 | 0.00086254 | 0.0050074 | NA | NA | |
| ENSG000001 | 14.7235564 | -1.045161981 | 0.31368734 | -3.331859 | 0.00086268 | 0.00500752 | CLEC4M | | 10332 |
| ENSG000002 | 12.1356979 | -1.092542225 | 0.32791241 | -3.3318112 | 0.00086283 | 0.00500766 | NA | NA | |

| | | | | | | | | |
|---|---|---|---|---|---|---|---|---|
| ENSG000001 | 52.1458231 | -0.724197085 | 0.21739408 | -3.3312641 | 0.00086453 | 0.0050168 | OTOG | 340990 |
| ENSG000001 | 66.4024423 | 0.645274595 | 0.19370688 | 3.33119092 | 0.00086475 | 0.0050174 | GPR173 | 54328 |
| ENSG00000C | 321.273801 | 0.448142861 | 0.13454703 | 3.33075242 | 0.00086612 | 0.00502358 | PRTFDC1 | 56952 |
| ENSG000001 | 176.166092 | 0.55894552 | 0.16781367 | 3.33075082 | 0.00086612 | 0.00502358 | ATP1A3 | 478 |
| ENSG000002 | 10.5219113 | -1.151100692 | 0.34560028 | -3.3307285 | 0.00086619 | 0.00502358 | NA | NA |
| ENSG000001 | 244.957568 | 0.443527135 | 0.13316962 | 3.33054284 | 0.00086677 | 0.00502621 | SCRN3 | 79634 |
| ENSG000001 | 14.5210053 | -1.059828489 | 0.31821936 | -3.3304965 | 0.00086691 | 0.00502633 | ADGRG3 | 222487 |
| ENSG000001 | 36.8801326 | -0.791675078 | 0.23773332 | -3.3300973 | 0.00086816 | 0.00503283 | SEL1L2 | 80343 |
| ENSG00000C | 1662.11594 | 0.414194824 | 0.12438525 | 3.32993518 | 0.00086866 | 0.00503504 | TEAD2 | 8463 |
| ENSG000001 | 279.195762 | 0.459902841 | 0.13811848 | 3.32977048 | 0.00086918 | 0.0050373 | SEMA5A | 9037 |
| ENSG000001 | 928.247448 | 0.388467478 | 0.11667588 | 3.32945844 | 0.00087015 | 0.00504222 | ATF6 | 22926 |
| ENSG000002 | 5.76475703 | -1.25434251 | 0.37676107 | -3.3292785 | 0.00087071 | 0.00504476 | NA | NA |
| ENSG000001 | 266.389283 | 0.549787865 | 0.16514373 | 3.32914761 | 0.00087112 | 0.00504641 | FBXW5 | 54461 |
| ENSG000001 | 8.7015946 | -1.179647435 | 0.35437248 | -3.3288347 | 0.0008721 | 0.00505117 | TNFSF18 | 8995 |
| ENSG000001 | 119.103014 | 0.641716647 | 0.19277684 | 3.32880564 | 0.00087219 | 0.00505117 | BCL2L2 | 599 |
| ENSG000001 | 2684.19088 | 0.394822582 | 0.11861449 | 3.32862019 | 0.00087277 | 0.00505381 | RBBP8 | 5932 |
| ENSG000001 | 1003.87426 | 0.457122325 | 0.13735059 | 3.32814239 | 0.00087427 | 0.00506104 | STK4 | 6789 |
| ENSG000001 | 198.403986 | 0.598708933 | 0.17989247 | 3.32814889 | 0.00087425 | 0.00506104 | TADA2B | 93624 |
| ENSG00000C | 1855.42664 | 0.429857112 | 0.12916656 | 3.32792865 | 0.00087494 | 0.0050642 | CUL1 | 8454 |
| ENSG000001 | 111.752883 | 0.545005073 | 0.16377406 | 3.32778639 | 0.00087539 | 0.00506607 | BIN3 | 55909 |
| ENSG000002 | 39.2686406 | 0.767099101 | 0.23053582 | 3.32746162 | 0.00087641 | 0.00507125 | NA | NA |
| ENSG000001 | 1966.72403 | 0.403524052 | 0.12128203 | 3.32715457 | 0.00087738 | 0.00507539 | AP3B1 | 8546 |
| ENSG000001 | 533.603499 | 0.446899723 | 0.13431743 | 3.32719074 | 0.00087726 | 0.00507539 | TMEM69 | 51249 |
| ENSG000002 | 36.5353939 | -0.803338486 | 0.24145892 | -3.3270193 | 0.0008778 | 0.00507713 | NA | NA |
| ENSG000001 | 307.688757 | 0.424801761 | 0.12769236 | 3.32675942 | 0.00087862 | 0.00508115 | FBXO34 | 55030 |
| ENSG000002 | 8.78480264 | -1.183348588 | 0.35576255 | -3.3262315 | 0.00088029 | 0.00509005 | LOC105376072 | 105376072 |
| ENSG000001 | 101.840438 | 0.558389562 | 0.16787731 | 3.32617658 | 0.00088046 | 0.00509033 | RILPL1 | 353116 |
| ENSG00000C | 22.2406653 | -0.912183953 | 0.27425694 | -3.3260196 | 0.00088096 | 0.00509247 | IGF1 | 3479 |
| ENSG000002 | 232.447719 | 0.46014165 | 0.13835033 | 3.32591652 | 0.00088128 | 0.00509363 | NA | NA |

| | | | | | | | | |
|---|---|---|---|---|---|---|---|---|
| ENSG000002 | 65.0993863 | 0.661375496 | 0.19887995 | 3.3255011 | 0.0008826 | 0.0051005 | NA | NA |
| ENSG000002 | 42.9657534 | -0.758116337 | 0.22797501 | -3.3254362 | 0.0008828 | 0.00510096 | NA | NA |
| ENSG000002 | 12.1715797 | -1.117088902 | 0.33594967 | -3.3251675 | 0.00088365 | 0.00510464 | LINC01360 | 101927295 |
| ENSG000002 | 34.7801856 | -0.809433749 | 0.24342731 | -3.3251558 | 0.00088369 | 0.00510464 | NA | NA |
| ENSG00000C | 2167.99057 | 0.387033603 | 0.1164004 | 3.32501959 | 0.00088412 | 0.00510641 | SLK | 9748 |
| ENSG000001 | 24.6028455 | -0.929402353 | 0.27958331 | -3.3242412 | 0.00088659 | 0.00511995 | PLA2G2F | 64600 |
| ENSG00000C | 202.827912 | 0.478869661 | 0.14406446 | 3.32399569 | 0.00088738 | 0.00512155 | MKS1 | 54903 |
| ENSG00000C | 314.803471 | 0.434016539 | 0.13056955 | 3.32402575 | 0.00088728 | 0.00512155 | MSANTD3 | 91283 |
| ENSG000001 | 152.651559 | 0.511467152 | 0.15386794 | 3.32406574 | 0.00088715 | 0.00512155 | ZDHHC12 | 84885 |
| ENSG000002 | 344.011283 | 0.56284236 | 0.16932705 | 3.32399564 | 0.00088738 | 0.00512155 | AKT1S1 | 84335 |
| ENSG000001 | 30.0314873 | -0.829860751 | 0.24966167 | -3.3239414 | 0.00088755 | 0.00512181 | MYO1A | 4640 |
| ENSG000001 | 203.60791 | 0.497254392 | 0.14960961 | 3.32367941 | 0.00088838 | 0.0051259 | BMP7 | 655 |
| ENSG000001 | 2490.17375 | 0.453245533 | 0.13638812 | 3.32320393 | 0.0008899 | 0.00513391 | RRM1 | 6240 |
| ENSG00000C | 1110.81155 | 0.438086768 | 0.13182997 | 3.32311967 | 0.00089017 | 0.00513473 | IKBKAP | 8518 |
| ENSG000001 | 7093.72565 | 0.439179844 | 0.13216273 | 3.32302347 | 0.00089047 | 0.00513577 | DDX6 | 1656 |
| ENSG000001 | 3249.38914 | 0.853995463 | 0.25700065 | 3.32293112 | 0.00089077 | 0.00513674 | KIF1A | 547 |
| ENSG000001 | 70.0684258 | -0.644479787 | 0.19395423 | -3.3228446 | 0.00089105 | 0.0051376 | DLGAP2 | 9228 |
| ENSG000001 | 2760.50853 | 0.413584342 | 0.12448499 | 3.3223632 | 0.00089258 | 0.00514575 | DNAJC7 | 7266 |
| ENSG000001 | 459.70849 | 0.404972143 | 0.12189841 | 3.32221011 | 0.00089307 | 0.00514784 | ZNF329 | 79673 |
| ENSG000001 | 446.959663 | 0.469614956 | 0.14137386 | 3.32179471 | 0.0008944 | 0.00515478 | LGALSL | 29094 |
| ENSG000001 | 156.516603 | 0.540995303 | 0.16289045 | 3.32122166 | 0.00089624 | 0.0051639 | MRPL55 | 128308 |
| ENSG000002 | 464.599798 | 0.423410124 | 0.12748569 | 3.32123657 | 0.0008962 | 0.0051639 | NA | NA |
| ENSG00000C | 747.746806 | 0.383983014 | 0.11563789 | 3.32056405 | 0.00089836 | 0.00517535 | TRMT6 | 51605 |
| ENSG000001 | 842.078964 | 0.610716202 | 0.18395436 | 3.31993328 | 0.00090039 | 0.00518632 | CARM1 | 10498 |
| ENSG000001 | 810.167615 | 0.423461205 | 0.12755864 | 3.31973753 | 0.00090102 | 0.00518922 | AIMP1 | 9255 |
| ENSG000002 | 86.3349559 | -0.634742344 | 0.19121346 | -3.3195484 | 0.00090163 | 0.005192 | ARHGEF28 | 64283 |
| ENSG000001 | 26.6772586 | -0.882809351 | 0.26594958 | -3.3194613 | 0.00090191 | 0.00519288 | NHLH2 | 4808 |
| ENSG000002 | 160.607188 | 0.523588429 | 0.15773519 | 3.31941411 | 0.00090207 | 0.00519302 | NA | NA |
| ENSG000002 | 19.9644983 | -0.945325265 | 0.28483098 | -3.318899 | 0.00090373 | 0.00520187 | NA | NA |

| | | | | | | | | |
|---|---|---|---|---|---|---|---|---|
| ENSG000002 | 36.1407239 | 0.801360554 | 0.24146176 | 3.31878868 | 0.00090409 | 0.00520319 | NA | NA |
| ENSG000001 | 8014.53939 | 0.368338329 | 0.11099877 | 3.3184 | 0.00090535 | 0.0052097 | DEK | 7913 |
| ENSG000002 | 36.3143169 | -0.76429183 | 0.23032961 | -3.3182526 | 0.00090583 | 0.00521171 | NA | NA |
| ENSG000001 | 344.176681 | 0.600877289 | 0.18108507 | 3.31820443 | 0.00090598 | 0.00521187 | NA | NA |
| ENSG000001 | 314.38132 | 0.419235171 | 0.12634618 | 3.31814677 | 0.00090617 | 0.0052122 | RAB33B | 83452 |
| ENSG000001 | 134.078647 | 0.506287899 | 0.15258365 | 3.31810062 | 0.00090632 | 0.00521233 | ZNF214 | 7761 |
| ENSG000002 | 7.62008241 | -1.232971186 | 0.37160256 | -3.3179836 | 0.0009067 | 0.00521377 | LINC00993 | 101929520 |
| ENSG000001 | 141.076729 | 0.554179875 | 0.16704316 | 3.31758497 | 0.00090799 | 0.00522048 | TFPT | 29844 |
| ENSG000002 | 31.2718988 | -0.806205739 | 0.24301989 | -3.3174476 | 0.00090844 | 0.0052223 | NA | NA |
| ENSG00000C | 104.533802 | 0.623780328 | 0.18803844 | 3.31730215 | 0.00090891 | 0.00522428 | LAMC3 | 10319 |
| ENSG000001 | 5.94016791 | -1.238962161 | 0.3734989 | -3.3171775 | 0.00090932 | 0.00522587 | ZNF750 | 79755 |
| ENSG000002 | 25.9785285 | -0.897389804 | 0.27058031 | -3.3165377 | 0.0009114 | 0.00523711 | HAND2-AS1 | 79804 |
| ENSG000001 | 1973.39871 | 0.554301471 | 0.16714274 | 3.31633591 | 0.00091206 | 0.00524015 | ACTN4 | 81 |
| ENSG000001 | 1216.94882 | 0.426646285 | 0.12865638 | 3.31616888 | 0.00091261 | 0.00524114 | MFAP1 | 4236 |
| ENSG000001 | 791.805091 | 0.392047637 | 0.11822322 | 3.31616432 | 0.00091262 | 0.00524114 | B3GALT1 | 8708 |
| ENSG000002 | 18.5173342 | -0.983690393 | 0.2966344 | -3.316171 | 0.0009126 | 0.00524114 | LOC101928446 | 101928446 |
| ENSG000001 | 1238.96513 | 0.398020104 | 0.12002709 | 3.3160856 | 0.00091288 | 0.00524188 | PAK1IP1 | 55003 |
| ENSG00000C | 1077.1405 | 0.415803525 | 0.12540167 | 3.31577347 | 0.0009139 | 0.00524699 | SENP1 | 29843 |
| ENSG000001 | 26.0324392 | -0.879977614 | 0.26540744 | -3.3155726 | 0.00091456 | 0.00525002 | FAM177B | 400823 |
| ENSG000002 | 30.3800187 | -0.82357901 | 0.24842076 | -3.3152584 | 0.00091558 | 0.00525518 | LOC642366 | 642366 |
| ENSG000001 | 78.0786534 | -0.607491602 | 0.18325927 | -3.3149296 | 0.00091666 | 0.00525869 | KIF6 | 221458 |
| ENSG000001 | 52.1780097 | -0.71114087 | 0.21452535 | -3.3149503 | 0.00091659 | 0.00525869 | GTSF1 | 121355 |
| ENSG000001 | 282.591455 | 0.455779398 | 0.1374936 | 3.31491344 | 0.00091672 | 0.00525869 | C5orf34 | 375444 |
| ENSG000001 | 1136.4871 | 0.398271464 | 0.12014131 | 3.31502512 | 0.00091635 | 0.00525869 | PIK3R4 | 30849 |
| ENSG000001 | 4868.43348 | 0.397657744 | 0.11997362 | 3.31454311 | 0.00091793 | 0.00526492 | BIRC6 | 57448 |
| ENSG000001 | 1402.9999 | 0.415735674 | 0.12543328 | 3.31439682 | 0.00091841 | 0.00526643 | PIKFYVE | 200576 |
| ENSG000001 | 461.238044 | 0.546672349 | 0.16493936 | 3.31438382 | 0.00091845 | 0.00526643 | CCNYL1 | 151195 |
| ENSG00000C | 4148.76943 | 0.406050906 | 0.12251429 | 3.31431475 | 0.00091868 | 0.00526699 | HACD3 | 51495 |
| ENSG000001 | 300.33879 | 0.550685106 | 0.16617593 | 3.3138681 | 0.00092015 | 0.00527466 | FANCF | 2188 |

| | | | | | | | | |
|---|---|---|---|---|---|---|---|---|
| ENSG000002 | 165.408558 | 0.507927435 | 0.15328807 | 3.31354837 | 0.0009212 | 0.00527986 | NUTM2A-AS1 | 728190 |
| ENSG000002 | 11.2370206 | -1.136481704 | 0.34298387 | -3.3135136 | 0.00092132 | 0.00527986 | NA | NA |
| ENSG000002 | 17.7871551 | -0.98675359 | 0.29780282 | -3.3134461 | 0.00092154 | 0.00528039 | DACH1 | 1602 |
| ENSG000001 | 1645.40822 | 0.47511085 | 0.14339314 | 3.31334433 | 0.00092187 | 0.00528156 | PRMT5 | 10419 |
| ENSG000002 | 4.71744275 | -1.252260073 | 0.37795963 | -3.3132112 | 0.00092231 | 0.00528333 | NA | NA |
| ENSG000001 | 1209.18034 | 0.370960459 | 0.11197313 | 3.31294186 | 0.0009232 | 0.00528768 | ATF2 | 1386 |
| ENSG000001 | 774.66531 | 0.376628505 | 0.11368698 | 3.3128551 | 0.00092349 | 0.00528857 | PRKAA1 | 5562 |
| ENSG000002 | 37.3961418 | -0.806367553 | 0.24341206 | -3.3127675 | 0.00092378 | 0.00528948 | LOC100506258 | 100506258 |
| ENSG000001 | 4751.16805 | 0.409785062 | 0.12370789 | 3.31252161 | 0.00092459 | 0.00529189 | SUB1 | 10923 |
| ENSG000001 | 236.291838 | 0.451560996 | 0.13631859 | 3.31254156 | 0.00092452 | 0.00529189 | SLFN12 | 55106 |
| ENSG000002 | 212.618583 | 0.526602782 | 0.15897277 | 3.31253439 | 0.00092455 | 0.00529189 | LOC102724159 | 102724159 |
| ENSG000001 | 94.8281367 | 0.638264192 | 0.19268603 | 3.31245707 | 0.0009248 | 0.00529236 | BTBD6 | 90135 |
| ENSG000001 | 562.509652 | 0.436461523 | 0.13177276 | 3.31222873 | 0.00092556 | 0.00529594 | AIG1 | 51390 |
| ENSG000001 | 12.1548274 | -1.088261348 | 0.32858108 | -3.3120025 | 0.00092631 | 0.00529948 | FAM167A-AS1 | 83656 |
| ENSG000001 | 1167.10553 | 0.439406373 | 0.1326747 | 3.31190789 | 0.00092662 | 0.00530052 | ORC4 | 5000 |
| ENSG000000 | 86.0053921 | 0.621224312 | 0.18760698 | 3.31130697 | 0.00092861 | 0.00531117 | IL17RB | 55540 |
| ENSG000001 | 348.898251 | 0.446650182 | 0.13489822 | 3.31101604 | 0.00092958 | 0.00531471 | CAMK2G | 818 |
| ENSG000002 | 21.1926251 | 0.993870574 | 0.30016565 | 3.31107367 | 0.00092939 | 0.00531471 | TSPEAR-AS1 | 54082 |
| ENSG000002 | 816.943912 | 0.411728725 | 0.1243517 | 3.31100209 | 0.00092963 | 0.00531471 | NA | NA |
| ENSG000001 | 196.860482 | 0.491960079 | 0.14858811 | 3.31089805 | 0.00092997 | 0.00531594 | MICU3 | 286097 |
| ENSG000001 | 1009.60168 | 0.404334025 | 0.12213476 | 3.31055661 | 0.00093111 | 0.00532168 | SRFBP1 | 153443 |
| ENSG000001 | 888.234763 | 0.496595381 | 0.15002878 | 3.31000078 | 0.00093296 | 0.0053315 | WWP1 | 11059 |
| ENSG000001 | 165.441689 | 0.509115641 | 0.1538201 | 3.30981227 | 0.00093359 | 0.00533434 | TRMT44 | 152992 |
| ENSG000001 | 133.763171 | 0.570083946 | 0.17224438 | 3.30973911 | 0.00093383 | 0.00533499 | FHL3 | 2275 |
| ENSG000000 | 94.9681273 | 0.628067391 | 0.18976808 | 3.3096577 | 0.0009341 | 0.00533579 | Mar-02 | 51257 |
| ENSG000001 | 1364.62278 | 0.387267733 | 0.11701402 | 3.3095841 | 0.00093435 | 0.00533644 | STIL | 6491 |
| ENSG000001 | 1481.69067 | 0.415808247 | 0.12564205 | 3.30946711 | 0.00093474 | 0.00533792 | SEC23IP | 11196 |
| ENSG000001 | 116.927166 | 0.572695009 | 0.17306166 | 3.30919624 | 0.00093564 | 0.00534233 | CLUHP3 | 100132341 |
| ENSG000001 | 27.4650269 | 0.85004339 | 0.25689063 | 3.30897001 | 0.0009364 | 0.0053459 | NA | NA |

| | | | | | | | | |
|---|---|---|---|---|---|---|---|---|
| ENSG000002 | 7.36515505 | -1.209393185 | 0.36553312 | -3.3085735 | 0.00093773 | 0.00535272 | NA | NA |
| ENSG000001 | 78.0359333 | 0.594855849 | 0.17980283 | 3.30837861 | 0.00093838 | 0.00535569 | ULBP1 | 80329 |
| ENSG000000 | 30.3131771 | -0.853066816 | 0.25788549 | -3.3079287 | 0.00093989 | 0.00536355 | SLC18A1 | 6570 |
| ENSG000001 | 378.072333 | 0.482646039 | 0.14591182 | 3.30779252 | 0.00094034 | 0.0053654 | C11orf24 | 53838 |
| ENSG000001 | 762.420532 | 0.389016698 | 0.11760895 | 3.30771346 | 0.00094061 | 0.00536616 | PIGT | 51604 |
| ENSG000000 | 26.5664408 | -0.910928045 | 0.27542021 | -3.3074118 | 0.00094162 | 0.00537119 | CNGB1 | 1258 |
| ENSG000001 | 1675.26122 | 0.462808972 | 0.13995562 | 3.30682668 | 0.00094359 | 0.00538166 | FAR1 | 84188 |
| ENSG000001 | 256.557298 | 0.470103344 | 0.14216901 | 3.3066513 | 0.00094418 | 0.00538374 | NA | NA |
| ENSG000002 | 13.9901738 | -1.069967956 | 0.32358165 | -3.3066398 | 0.00094422 | 0.00538374 | NA | NA |
| ENSG000002 | 373.426245 | 0.477528259 | 0.14441993 | 3.30652593 | 0.00094461 | 0.00538517 | FASTKD5 | 60493 |
| ENSG000002 | 30.0482663 | -0.890298832 | 0.26928279 | -3.3061854 | 0.00094575 | 0.00539096 | NA | NA |
| ENSG000002 | 8.56636073 | -1.169314313 | 0.35369815 | -3.3059667 | 0.00094649 | 0.00539441 | NA | NA |
| ENSG000001 | 195.782619 | 0.522002901 | 0.15790473 | 3.30580923 | 0.00094703 | 0.00539669 | MTL5 | 9633 |
| ENSG000002 | 20.9053647 | -0.934144049 | 0.28259952 | -3.3055401 | 0.00094794 | 0.00540111 | NA | NA |
| ENSG000002 | 14.680618 | -1.038797394 | 0.31426394 | -3.3054934 | 0.00094809 | 0.00540125 | NA | NA |
| ENSG000001 | 1387.20599 | 0.403407714 | 0.12204591 | 3.30537667 | 0.00094849 | 0.00540275 | ETAA1 | 54465 |
| ENSG000001 | 5.86226428 | -1.234935349 | 0.37361859 | -3.305337 | 0.00094862 | 0.00540275 | EVI2B | 2124 |
| ENSG000001 | 291.271559 | 0.468012271 | 0.14160064 | 3.30515653 | 0.00094923 | 0.00540547 | LACTB2 | 51110 |
| ENSG000002 | 803.237388 | 0.490791071 | 0.14850089 | 3.30497048 | 0.00094986 | 0.0054083 | ZCCHC3 | 85364 |
| ENSG000002 | 11.2795011 | -1.113603211 | 0.33695637 | -3.3048885 | 0.00095014 | 0.00540913 | NA | NA |
| ENSG000001 | 8306.21749 | 0.405342926 | 0.12265616 | 3.30470896 | 0.00095075 | 0.00541183 | KIAA1551 | 55196 |
| ENSG000002 | 789.571343 | 0.430303101 | 0.13024814 | 3.30371783 | 0.00095412 | 0.00543024 | NA | NA |
| ENSG000001 | 934.818809 | 0.406227753 | 0.12296349 | 3.30364536 | 0.00095436 | 0.00543088 | ZNF681 | 148213 |
| ENSG000002 | 481.9914 | 0.414425911 | 0.12545516 | 3.30337871 | 0.00095527 | 0.00543528 | CHIC1 | 53344 |
| ENSG000001 | 1887.90507 | 0.414238761 | 0.12540547 | 3.30319538 | 0.0009559 | 0.00543771 | ABI2 | 10152 |
| ENSG000001 | 688.603997 | 0.380485738 | 0.11518789 | 3.30317477 | 0.00095597 | 0.00543771 | CCDC138 | 165055 |
| ENSG000001 | 2946.31016 | 0.396042718 | 0.11990063 | 3.30309129 | 0.00095625 | 0.00543857 | KRR1 | 11103 |
| ENSG000001 | 14.9747045 | -1.011339925 | 0.3062036 | -3.3028348 | 0.00095713 | 0.00544278 | TINAG | 27283 |
| ENSG000001 | 20.6356959 | -0.934716917 | 0.28308228 | -3.3019266 | 0.00096023 | 0.00545967 | PLEK | 5341 |

| | | | | | | | | |
|---|---|---|---|---|---|---|---|---|
| ENSG00000C | 196.576179 | 0.497804162 | 0.15076626 | 3.30182742 | 0.00096057 | 0.00546083 | WDR37 | 22884 |
| ENSG000002 | 14.6072631 | -1.033413257 | 0.31299682 | -3.3016733 | 0.0009611 | 0.00546307 | CECR7 | 100130418 |
| ENSG000001 | 3181.98979 | 0.412200507 | 0.12486384 | 3.3012 | 0.00096272 | 0.00547153 | TMTC3 | 160418 |
| ENSG000001 | 21.0608972 | -0.922053152 | 0.27931522 | -3.3011203 | 0.000963 | 0.00547231 | PLA2G5 | 5322 |
| ENSG000002 | 20.8566846 | 0.918740176 | 0.27831961 | 3.3010257 | 0.00096332 | 0.00547339 | NA | NA |
| ENSG00000C | 27.6490945 | -0.887867812 | 0.26900265 | -3.3005913 | 0.00096481 | 0.00548111 | PRDM1 | 639 |
| ENSG000001 | 110.915323 | 0.528222078 | 0.1600925 | 3.29948054 | 0.00096864 | 0.00550207 | IFT20 | 90410 |
| ENSG000001 | 348.524799 | 0.401872212 | 0.12183418 | 3.29851769 | 0.00097197 | 0.00551972 | PKNOX1 | 5316 |
| ENSG000001 | 538.756843 | 0.464490334 | 0.14081853 | 3.29850303 | 0.00097202 | 0.00551972 | UBE2E3 | 10477 |
| ENSG000002 | 57.319211 | -0.671685997 | 0.20365557 | -3.2981469 | 0.00097325 | 0.00552595 | NA | NA |
| ENSG000001 | 277.480983 | 0.450357062 | 0.13655442 | 3.29800417 | 0.00097375 | 0.00552799 | FAM206A | 54942 |
| ENSG000001 | 4631.58384 | 0.373560845 | 0.11327271 | 3.29788926 | 0.00097415 | 0.0055287 | NLN | 57486 |
| ENSG000001 | 292.442656 | 0.469965648 | 0.14250434 | 3.29790414 | 0.00097409 | 0.0055287 | PCGF6 | 84108 |
| ENSG000001 | 5990.27453 | 0.412911781 | 0.1252102 | 3.29774863 | 0.00097463 | 0.0055307 | SYNCRIP | 10492 |
| ENSG000001 | 115.785338 | 0.56423232 | 0.17109883 | 3.29769833 | 0.00097481 | 0.00553091 | SERINC2 | 347735 |
| ENSG000001 | 173.138822 | 0.53200764 | 0.16133726 | 3.29748784 | 0.00097554 | 0.00553351 | CES2 | 8824 |
| ENSG000001 | 4.20579844 | -1.248184738 | 0.37852411 | -3.2975039 | 0.00097548 | 0.00553351 | NA | NA |
| ENSG000001 | 391.119578 | 0.444670302 | 0.13485932 | 3.29729005 | 0.00097623 | 0.00553546 | TMX4 | 56255 |
| ENSG000001 | 16.8740704 | -0.984022644 | 0.29843545 | -3.2972713 | 0.00097629 | 0.00553546 | DSPP | 1834 |
| ENSG000001 | 23.6475192 | -0.922364457 | 0.27973185 | -3.2973165 | 0.00097613 | 0.00553546 | FAM71D | 161142 |
| ENSG000001 | 23.1664535 | -0.919795058 | 0.27898065 | -3.2969851 | 0.00097729 | 0.00554033 | SLC4A9 | 83697 |
| ENSG000002 | 8.09251528 | -1.168073047 | 0.35430747 | -3.2967779 | 0.00097801 | 0.00554364 | NA | NA |
| ENSG000001 | 252.724679 | 0.512182925 | 0.15536586 | 3.29662465 | 0.00097854 | 0.00554589 | PACSIN3 | 29763 |
| ENSG000001 | 131.280321 | 0.612759314 | 0.18588807 | 3.29638864 | 0.00097936 | 0.00554851 | RAVER1 | 125950 |
| ENSG000001 | 67.2500989 | 0.625240639 | 0.18967527 | 3.29637414 | 0.00097941 | 0.00554851 | MED30 | 90390 |
| ENSG000002 | 47.2009691 | 0.731017427 | 0.22176258 | 3.29639662 | 0.00097934 | 0.00554851 | NA | NA |
| ENSG000001 | 17.4107586 | -0.993944533 | 0.30157585 | -3.2958359 | 0.00098129 | 0.00555838 | NDST4 | 64579 |
| ENSG000001 | 312.085282 | 0.541811771 | 0.16439848 | 3.29572247 | 0.00098169 | 0.00555984 | NA | NA |
| ENSG000001 | 45.0440316 | -0.770619312 | 0.23382694 | -3.2956823 | 0.00098183 | 0.00555986 | ADGRG4 | 139378 |

| | | | | | | | | | |
|---|---|---|---|---|---|---|---|---|---|
| ENSG000002 | 17.0794885 | 1.030832519 | 0.31279619 | 3.29554052 | 0.00098233 | 0.00556189 | NA | NA | |
| ENSG000001 | 1047.69869 | 0.375972447 | 0.11409512 | 3.29525443 | 0.00098333 | 0.00556678 | PPWD1 | | 23398 |
| ENSG000001 | 3793.33692 | 0.412730856 | 0.12525237 | 3.29519397 | 0.00098354 | 0.0055672 | CHD8 | | 57680 |
| ENSG000001 | 1401.28176 | 0.378582277 | 0.11490266 | 3.29480865 | 0.00098489 | 0.00557357 | TMEM184C | | 55751 |
| ENSG000002 | 2.95713926 | -1.231125348 | 0.37365776 | -3.294794 | 0.00098494 | 0.00557357 | NA | NA | |
| ENSG000001 | 34.4483415 | -0.815844156 | 0.2476401 | -3.2944752 | 0.00098606 | 0.00557912 | DDX4 | | 54514 |
| ENSG000001 | 18.4033828 | -0.998941659 | 0.30324836 | -3.294137 | 0.00098724 | 0.00558506 | ENTHD1 | | 150350 |
| ENSG000001 | 475.341817 | 0.406867209 | 0.12351739 | 3.29400743 | 0.0009877 | 0.00558686 | SAR1B | | 51128 |
| ENSG000001 | 89.3173307 | 0.60480139 | 0.1836321 | 3.29354932 | 0.00098931 | 0.00559519 | SLC39A3 | | 29985 |
| ENSG000002 | 1906.47861 | 0.440577022 | 0.13377836 | 3.29333542 | 0.00099006 | 0.00559867 | STK38L | | 23012 |
| ENSG000001 | 517.61153 | 0.541533856 | 0.16444525 | 3.29309514 | 0.00099091 | 0.00560267 | HSPB1 | | 3315 |
| ENSG000002 | 3.81433288 | -1.245817619 | 0.37832895 | -3.2929482 | 0.00099143 | 0.00560482 | NA | NA | |
| ENSG000001 | 853.133904 | 0.561918132 | 0.17064549 | 3.29289772 | 0.00099161 | 0.00560504 | ZNF445 | | 353274 |
| ENSG000002 | 42.5383779 | -0.744628313 | 0.22616951 | -3.2923462 | 0.00099355 | 0.00561527 | NA | NA | |
| ENSG000001 | 466.744748 | 0.409889957 | 0.1245065 | 3.29211686 | 0.00099436 | 0.00561906 | TRPS1 | | 7227 |
| ENSG000001 | 4358.16342 | 0.546518011 | 0.1660218 | 3.2918448 | 0.00099533 | 0.00562372 | YWHAG | | 7532 |
| ENSG000001 | 295.031467 | 0.431690004 | 0.13115236 | 3.29151529 | 0.00099649 | 0.00562953 | XRCC4 | | 7518 |
| ENSG000002 | 321.979496 | 0.497655792 | 0.15120934 | 3.29117108 | 0.00099771 | 0.00563564 | CWC25 | | 54883 |
| ENSG000002 | 4.38776633 | -1.245969173 | 0.37859226 | -3.2910582 | 0.00099811 | 0.00563711 | NA | NA | |
| ENSG000002 | 3.86388357 | -1.244935951 | 0.3783121 | -3.2907643 | 0.00099916 | 0.00564222 | NA | NA | |
| ENSG000001 | 253.011561 | 0.486724866 | 0.14791172 | 3.29064427 | 0.00099958 | 0.00564384 | NA | NA | |
| ENSG000001 | 86.6292455 | 0.592672996 | 0.18011618 | 3.29050401 | 0.00100008 | 0.00564457 | ECHDC3 | | 79746 |
| ENSG000001 | 278.543051 | 0.493723501 | 0.15004295 | 3.29054785 | 0.00099992 | 0.00564457 | SPC25 | | 57405 |
| ENSG000001 | 19.4275361 | -0.976117787 | 0.29664966 | -3.2904734 | 0.00100019 | 0.00564457 | ACMSD | | 130013 |
| ENSG000001 | 213.76584 | 0.452815349 | 0.13761497 | 3.29045132 | 0.00100027 | 0.00564457 | SLFN5 | | 162394 |
| ENSG000001 | 264.101487 | 0.448442946 | 0.13628965 | 3.29036692 | 0.00100057 | 0.00564548 | ARL15 | | 54622 |
| ENSG000001 | 139.398814 | 0.562334951 | 0.17090573 | 3.2903224 | 0.00100073 | 0.00564558 | DALRD3 | | 55152 |
| ENSG000001 | 680.951664 | 0.408485955 | 0.12419515 | 3.28906528 | 0.00100521 | 0.00567007 | SLC19A2 | | 10560 |
| ENSG00000C | 3069.2416 | 0.500603646 | 0.15220768 | 3.28895128 | 0.00100561 | 0.00567158 | NRD1 | | 4898 |

| | | | | | | | | |
|---|---|---|---|---|---|---|---|---|
| ENSG000001 | 2179.86831 | 0.456581742 | 0.1388283 | 3.28882336 | 0.00100607 | 0.00567337 | TNKS2 | 80351 |
| ENSG000001 | 59.3258483 | 0.708498984 | 0.21543398 | 3.28870581 | 0.00100649 | 0.00567495 | NA | NA |
| ENSG000001 | 152.829481 | 0.498650364 | 0.15165781 | 3.28799656 | 0.00100903 | 0.00568848 | FAH | 2184 |
| ENSG000001 | 29.2696122 | -0.895513096 | 0.27237975 | -3.2877374 | 0.00100996 | 0.00569292 | CD1C | 911 |
| ENSG000001 | 187.920916 | 0.556792431 | 0.1693594 | 3.28763809 | 0.00101032 | 0.00569414 | C17orf58 | 284018 |
| ENSG000002 | 11.6013591 | -1.1074123 | 0.33684825 | -3.2875703 | 0.00101056 | 0.00569472 | NA | NA |
| ENSG000001 | 3112.11193 | 0.442565343 | 0.13463878 | 3.2870571 | 0.0010124 | 0.00570432 | TMEM2 | 23670 |
| ENSG000001 | 174.625327 | 0.526175729 | 0.16008224 | 3.28690876 | 0.00101294 | 0.00570653 | NAPA | 8775 |
| ENSG000001 | 117.543242 | 0.595257768 | 0.18110556 | 3.28680001 | 0.00101333 | 0.00570782 | NLRX1 | 79671 |
| ENSG000002 | 27.7984342 | -0.830048122 | 0.25254245 | -3.2867668 | 0.00101345 | 0.00570782 | FLG-AS1 | 339400 |
| ENSG00000C | 510.333095 | 0.444695036 | 0.13530552 | 3.28659942 | 0.00101405 | 0.00571042 | ATG16L1 | 55054 |
| ENSG000001 | 513.152653 | 0.477233339 | 0.14523263 | 3.28599251 | 0.00101624 | 0.00572194 | KLHL5 | 51088 |
| ENSG00000C | 634.922972 | 0.431506575 | 0.13132366 | 3.28582499 | 0.00101684 | 0.00572455 | CDK17 | 5128 |
| ENSG000002 | 99.1580371 | 0.683665626 | 0.20807877 | 3.28560968 | 0.00101762 | 0.00572813 | C1orf233 | 643988 |
| ENSG000001 | 293.00433 | 0.47592005 | 0.14486563 | 3.28525172 | 0.00101891 | 0.00573374 | NAPB | 63908 |
| ENSG000001 | 51.1566141 | -0.677235963 | 0.2061465 | -3.2852168 | 0.00101904 | 0.00573374 | ZEB1 | 6935 |
| ENSG000002 | 6.26633357 | -1.223164751 | 0.3723222 | -3.2852318 | 0.00101898 | 0.00573374 | NA | NA |
| ENSG000001 | 906.814054 | 0.391373432 | 0.1191333 | 3.28517246 | 0.0010192 | 0.00573385 | KIF15 | 56992 |
| ENSG00000C | 417.997618 | 0.563371454 | 0.17149887 | 3.28498644 | 0.00101987 | 0.00573525 | MAST2 | 23139 |
| ENSG000001 | 1048.97385 | 0.55898371 | 0.17016119 | 3.28502467 | 0.00101973 | 0.00573525 | OSBP | 5007 |
| ENSG000002 | 27.6818825 | -0.873605922 | 0.26593787 | -3.2850001 | 0.00101982 | 0.00573525 | OR2M3 | 127062 |
| ENSG000001 | 1247.12611 | 0.395052434 | 0.12027334 | 3.28462193 | 0.00102119 | 0.00574187 | PPP1R9A | 55607 |
| ENSG000002 | 319.819545 | 0.560923559 | 0.17078235 | 3.28443516 | 0.00102187 | 0.00574488 | MIR600HG | 81571 |
| ENSG000001 | 3522.11425 | 0.346536694 | 0.10551308 | 3.28430075 | 0.00102236 | 0.00574683 | CLDN6 | 9074 |
| ENSG000001 | 455.898823 | 0.436161752 | 0.13280426 | 3.2842452 | 0.00102256 | 0.00574717 | PLTP | 5360 |
| ENSG00000C | 236.14724 | 0.507417566 | 0.15450574 | 3.284134 | 0.00102296 | 0.00574784 | B4GALT1 | 2683 |
| ENSG000002 | 19.7171233 | -0.945435947 | 0.28787973 | -3.2841352 | 0.00102296 | 0.00574784 | DAB1-AS1 | 101926890 |
| ENSG000001 | 530.058676 | 0.426879707 | 0.12999595 | 3.28379243 | 0.0010242 | 0.00575162 | SCML2 | 10389 |
| ENSG000001 | 242.497165 | 0.470376007 | 0.14323999 | 3.28383164 | 0.00102406 | 0.00575162 | GRM4 | 2914 |

| | | | | | | | | |
|---|---|---|---|---|---|---|---|---|
| ENSG000001 | 6.11490433 | -1.219192161 | 0.37127374 | -3.2838093 | 0.00102414 | 0.00575162 | DMRTB1 | 63948 |
| ENSG000001 | 128.460385 | 0.580085169 | 0.17664638 | 3.283878 | 0.00102389 | 0.00575162 | RDH13 | 112724 |
| ENSG00000C | 112.386384 | 0.576443697 | 0.17556672 | 3.28333121 | 0.00102588 | 0.00576024 | MPC1 | 51660 |
| ENSG000001 | 6015.04594 | 0.424235405 | 0.12921585 | 3.28315311 | 0.00102653 | 0.00576309 | FUBP1 | 8880 |
| ENSG000001 | 238.360306 | 0.474000754 | 0.14441201 | 3.28228062 | 0.00102971 | 0.00578015 | ANKMY2 | 57037 |
| ENSG00000C | 346.804162 | 0.4179663 | 0.12734396 | 3.28218399 | 0.00103006 | 0.00578133 | CYB5R4 | 51167 |
| ENSG000002 | 2.63319389 | -1.199897079 | 0.36561476 | -3.2818617 | 0.00103124 | 0.00578634 | NA | NA |
| ENSG000002 | 8.72239737 | -1.193004926 | 0.36351273 | -3.2818794 | 0.00103118 | 0.00578634 | NA | NA |
| ENSG00000C | 231.23913 | 0.556087842 | 0.16949481 | 3.28085473 | 0.00103493 | 0.00580519 | REXO1 | 57455 |
| ENSG000002 | 8.25857132 | -1.173838814 | 0.35778743 | -3.2808274 | 0.00103503 | 0.00580519 | NA | NA |
| ENSG000002 | 21.0319789 | -0.918826883 | 0.28005498 | -3.2808804 | 0.00103484 | 0.00580519 | LINC00836 | 101929052 |
| ENSG000001 | 2554.99283 | 0.391749083 | 0.11940824 | 3.28075415 | 0.0010353 | 0.0058059 | RTN4 | 57142 |
| ENSG000002 | 6.30163827 | -1.222271221 | 0.37259205 | -3.2804544 | 0.0010364 | 0.00581126 | LINC00626 | 79100 |
| ENSG000002 | 89.7690751 | 0.613259942 | 0.18694611 | 3.28041037 | 0.00103656 | 0.00581137 | SLC48A1 | 55652 |
| ENSG000001 | 202.269206 | 0.495107529 | 0.15093708 | 3.28022464 | 0.00103724 | 0.00581359 | PDIK1L | 149420 |
| ENSG000001 | 1352.04844 | 0.378879409 | 0.11550299 | 3.28025637 | 0.00103713 | 0.00581359 | MED14 | 9282 |
| ENSG000001 | 170.355485 | 0.513003412 | 0.15640939 | 3.27987599 | 0.00103853 | 0.00581997 | CD83 | 9308 |
| ENSG000002 | 14.8833064 | -1.026297508 | 0.31295013 | -3.2794283 | 0.00104018 | 0.00582841 | NA | NA |
| ENSG000001 | 666.583154 | 0.417205361 | 0.12722279 | 3.27932893 | 0.00104054 | 0.00582966 | PSMG2 | 56984 |
| ENSG000002 | 4.76806979 | -1.240390661 | 0.37825333 | -3.279259 | 0.0010408 | 0.00583029 | NA | NA |
| ENSG000001 | 1979.93603 | 0.432500779 | 0.13190069 | 3.27898807 | 0.0010418 | 0.00583509 | SMG5 | 23381 |
| ENSG000001 | 879.838339 | 0.580989395 | 0.17719716 | 3.27877382 | 0.00104259 | 0.00583871 | ANKFY1 | 51479 |
| ENSG00000C | 150.930778 | 0.534914709 | 0.16315711 | 3.27852526 | 0.00104351 | 0.00584305 | EFR3B | 22979 |
| ENSG000001 | 317.180872 | 0.411445251 | 0.12549966 | 3.27845712 | 0.00104376 | 0.00584366 | RHOBTB3 | 22836 |
| ENSG000001 | 1372.38622 | 0.425573727 | 0.12982271 | 3.27811456 | 0.00104503 | 0.00584914 | MRPL50 | 54534 |
| ENSG000001 | 33.1889902 | -0.847061034 | 0.25839849 | -3.2781191 | 0.00104501 | 0.00584914 | CDH12 | 1010 |
| ENSG000001 | 11.49122 | -1.116202684 | 0.34051934 | -3.2779421 | 0.00104567 | 0.00585191 | OR2L3 | 391192 |
| ENSG000002 | 29.1836244 | 0.824543492 | 0.25157724 | 3.27749642 | 0.00104732 | 0.00586034 | MAFG-AS1 | 92659 |
| ENSG000001 | 175.791594 | 0.486401555 | 0.14842094 | 3.27717611 | 0.00104851 | 0.00586618 | ZKSCAN4 | 387032 |

| | | | | | | | | |
|---|---|---|---|---|---|---|---|---|
| ENSG000001 | 269.682049 | 0.437022964 | 0.13336141 | 3.2769821 | 0.00104923 | 0.00586941 | C6orf89 | 221477 |
| ENSG000001 | 1658.81762 | 0.434119862 | 0.13248479 | 3.27675251 | 0.00105008 | 0.00587337 | NOL9 | 79707 |
| ENSG000002 | 31.2572553 | 0.848176738 | 0.25886746 | 3.27649035 | 0.00105106 | 0.00587802 | MRPL12 | 6182 |
| ENSG000002 | 2608.07123 | 0.487115485 | 0.14867462 | 3.27638624 | 0.00105145 | 0.00587937 | LSM14A | 26065 |
| ENSG000001 | 3905.591 | 0.376513618 | 0.11492076 | 3.27628888 | 0.00105181 | 0.00588059 | DICER1 | 23405 |
| ENSG000001 | 861.426332 | 0.345146312 | 0.10535103 | 3.27615496 | 0.00105231 | 0.00588257 | ZCCHC7 | 84186 |
| ENSG000001 | 1277.11321 | 0.488170008 | 0.14900955 | 3.27609891 | 0.00105252 | 0.00588292 | RNF213 | 57674 |
| ENSG00000C | 24.5579307 | -0.878966869 | 0.26830319 | -3.2760209 | 0.00105281 | 0.00588374 | IL5RA | 3568 |
| ENSG000001 | 223.134267 | 0.53393779 | 0.16300589 | 3.2755736 | 0.00105448 | 0.00589225 | ASPSCR1 | 79058 |
| ENSG000001 | 2470.27469 | 0.434222354 | 0.13257591 | 3.27527353 | 0.0010556 | 0.005897 | ADAM10 | 102 |
| ENSG000001 | 150.842641 | 0.498377315 | 0.15216382 | 3.27526816 | 0.00105562 | 0.005897 | MRAS | 22808 |
| ENSG000001 | 1401.19171 | 0.393082987 | 0.12002382 | 3.27504155 | 0.00105646 | 0.00590092 | RAB8B | 51762 |
| ENSG000001 | 28.559025 | -0.866842866 | 0.26471319 | -3.2746493 | 0.00105793 | 0.00590749 | PXDNL | 137902 |
| ENSG000001 | 160.094123 | 0.525335317 | 0.16042489 | 3.2746496 | 0.00105793 | 0.00590749 | GMPPB | 29925 |
| ENSG000001 | 1412.40243 | 0.436459168 | 0.13329562 | 3.27436992 | 0.00105898 | 0.00591252 | ASXL2 | 55252 |
| ENSG000001 | 225.328061 | 0.467256779 | 0.14271769 | 3.27399336 | 0.00106039 | 0.00591959 | CDCA3 | 83461 |
| ENSG000002 | 943.789958 | 0.422184137 | 0.1289583 | 3.27380358 | 0.0010611 | 0.00592275 | ETV5 | 2119 |
| ENSG000001 | 16.6177971 | -1.003951831 | 0.30666814 | -3.2737403 | 0.00106134 | 0.00592326 | LINC00158 | 54072 |
| ENSG000001 | 320.863853 | 0.494229344 | 0.1509974 | 3.27309843 | 0.00106375 | 0.00593592 | FAM210A | 125228 |
| ENSG00000C | 3153.78537 | 0.402609145 | 0.12301233 | 3.27291712 | 0.00106444 | 0.00593832 | PSME4 | 23198 |
| ENSG000001 | 2031.37688 | 0.37489349 | 0.11454452 | 3.27290641 | 0.00106448 | 0.00593832 | GINS1 | 9837 |
| ENSG000001 | 9.93767309 | -1.158531925 | 0.35399088 | -3.2727734 | 0.00106498 | 0.00594029 | SOAT2 | 8435 |
| ENSG000001 | 354.251064 | 0.435121068 | 0.1329667 | 3.27240619 | 0.00106636 | 0.00594556 | NFKBIA | 4792 |
| ENSG000001 | 238.499341 | 0.441676971 | 0.13496951 | 3.27242037 | 0.00106631 | 0.00594556 | FAM73A | 374986 |
| ENSG000002 | 13.6419304 | -1.044252111 | 0.31910466 | -3.272444 | 0.00106622 | 0.00594556 | C3orf67-AS1 | 101929238 |
| ENSG000002 | 42.8728594 | -0.755340771 | 0.23082486 | -3.2723545 | 0.00106656 | 0.00594583 | NA | NA |
| ENSG000001 | 494.902445 | 0.45209305 | 0.13815924 | 3.27226073 | 0.00106691 | 0.00594699 | KCTD9 | 54793 |
| ENSG000001 | 215.439147 | 0.490210046 | 0.14981001 | 3.27221145 | 0.0010671 | 0.00594721 | POLL | 27343 |
| ENSG00000C | 296.368482 | -0.503779383 | 0.15396551 | -3.2720274 | 0.00106779 | 0.00595026 | BAX | 581 |

| | | | | | | | | |
|---|---|---|---|---|---|---|---|---|
| ENSG00000C | 2151.83005 | 0.394615845 | 0.12060988 | 3.27183694 | 0.00106851 | 0.0059528 | CDC27 | 996 |
| ENSG000001 | 3043.26353 | 0.373803736 | 0.11424916 | 3.27182908 | 0.00106854 | 0.0059528 | ARID2 | 196528 |
| ENSG000001 | 429.756407 | 0.402747397 | 0.12311643 | 3.27127247 | 0.00107065 | 0.00596371 | APIP | 51074 |
| ENSG000002 | 8.13813791 | -1.184563976 | 0.36213792 | -3.2710299 | 0.00107157 | 0.00596801 | LOC101928618 | 101928618 |
| ENSG000001 | 306.224009 | 0.455256586 | 0.13918331 | 3.27091353 | 0.00107201 | 0.00596964 | POC5 | 134359 |
| ENSG00000C | 114.834266 | 0.563876013 | 0.17239884 | 3.2707645 | 0.00107257 | 0.00597197 | HAGH | 3029 |
| ENSG000001 | 116.381667 | 0.634865738 | 0.19411889 | 3.2704996 | 0.00107358 | 0.00597675 | RARA | 5914 |
| ENSG00000C | 535.19765 | 0.453369549 | 0.13863696 | 3.27019242 | 0.00107474 | 0.0059816 | NFYC | 4802 |
| ENSG000001 | 716.583974 | 0.434589195 | 0.13289322 | 3.2702134 | 0.00107466 | 0.0059816 | LIMD1 | 8994 |
| ENSG000002 | 37.5927631 | -0.795053322 | 0.24312619 | -3.2701263 | 0.00107499 | 0.00598218 | LOC101927766 | 101927766 |
| ENSG000001 | 388.533129 | 0.46281349 | 0.14156998 | 3.26915004 | 0.00107871 | 0.00600203 | OTUD3 | 23252 |
| ENSG000001 | 335.732014 | 0.414010068 | 0.12664668 | 3.26901633 | 0.00107922 | 0.00600404 | PIGG | 54872 |
| ENSG00000C | 859.37906 | 0.350527135 | 0.10723327 | 3.26882826 | 0.00107994 | 0.00600639 | RBM22 | 55696 |
| ENSG000001 | 51.8460519 | -0.726915842 | 0.22237622 | -3.268856 | 0.00107983 | 0.00600639 | NA | NA |
| ENSG000002 | 226.146023 | 0.505700631 | 0.15472968 | 3.26828469 | 0.00108201 | 0.00601711 | NA | NA |
| ENSG00000C | 1218.82623 | 0.382482538 | 0.11703116 | 3.26821116 | 0.0010823 | 0.00601785 | GSK3B | 2932 |
| ENSG000002 | 7.24201436 | -1.191331781 | 0.36453076 | -3.2681241 | 0.00108263 | 0.00601888 | NA | NA |
| ENSG000001 | 441.715555 | 0.523248977 | 0.16012484 | 3.26775648 | 0.00108404 | 0.00602587 | SEMA4D | 10507 |
| ENSG000001 | 275.476531 | 0.518422453 | 0.15867714 | 3.26715265 | 0.00108635 | 0.00603791 | EXOC3 | 11336 |
| ENSG000002 | 32.077232 | -0.808859599 | 0.24757724 | -3.2671 | 0.00108655 | 0.00603821 | NA | NA |
| ENSG00000C | 750.77513 | 0.410553411 | 0.12567641 | 3.26674998 | 0.0010879 | 0.00604478 | GRHL2 | 79977 |
| ENSG000002 | 18.3076632 | 0.957117597 | 0.29299091 | 3.26671434 | 0.00108803 | 0.00604478 | LOC101929054 | 101929054 |
| ENSG000001 | 29.0283446 | -0.82610352 | 0.25291109 | -3.2663792 | 0.00108932 | 0.00605111 | RAD9B | 144715 |
| ENSG000001 | 1474.18334 | 0.365334906 | 0.11184857 | 3.26633506 | 0.00108949 | 0.00605123 | TLK2 | 11011 |
| ENSG00000C | 7.43497885 | -1.179069195 | 0.3609962 | -3.266154 | 0.00109019 | 0.00605427 | DMRT3 | 58524 |
| ENSG000002 | 42.6479707 | 0.749516638 | 0.22948544 | 3.26607487 | 0.00109049 | 0.00605513 | SCARNA21 | 677763 |
| ENSG000002 | 680.394937 | 0.504273654 | 0.15439938 | 3.26603415 | 0.00109065 | 0.00605517 | KIFC1 | 3833 |
| ENSG000002 | 11.1113403 | -1.093484973 | 0.33481543 | -3.2659336 | 0.00109104 | 0.00605649 | OR4E2 | 26686 |
| ENSG000001 | 129674.372 | 0.35187579 | 0.10774854 | 3.26571273 | 0.00109189 | 0.00606039 | COX3 | 4514 |

| | | | | | | | | |
|---|---|---|---|---|---|---|---|---|
| ENSG000001 | 35.864992 | -0.782999891 | 0.23976708 | -3.2656689 | 0.00109206 | 0.0060605 | TMCO5A | 145942 |
| ENSG000001 | 29.2906745 | -0.808195746 | 0.24749278 | -3.2655327 | 0.00109258 | 0.00606259 | ZNF831 | 128611 |
| ENSG000001 | 21.9120879 | -0.939660056 | 0.28776771 | -3.2653422 | 0.00109332 | 0.00606583 | MATN1 | 4146 |
| ENSG000001 | 733.324504 | 0.529540792 | 0.16217258 | 3.26529176 | 0.00109351 | 0.00606608 | WASF2 | 10163 |
| ENSG000001 | 1188.57856 | 0.372429525 | 0.11406684 | 3.26501137 | 0.0010946 | 0.00607043 | HIPK3 | 10114 |
| ENSG000001 | 474.315192 | 0.436545146 | 0.13370344 | 3.26502543 | 0.00109454 | 0.00607043 | GTPBP8 | 29083 |
| ENSG000001 | 1143.10726 | 0.367131785 | 0.1124477 | 3.26491149 | 0.00109498 | 0.00607172 | RAPGEF2 | 9693 |
| ENSG000001 | 20.4418144 | -0.923728301 | 0.28292927 | -3.2648736 | 0.00109513 | 0.00607172 | SH2D1B | 117157 |
| ENSG000002 | 43.2646376 | -0.773037305 | 0.23678578 | -3.2647117 | 0.00109576 | 0.00607437 | NA | NA |
| ENSG000001 | 581.926488 | 0.519392098 | 0.15910901 | 3.2643789 | 0.00109704 | 0.00608067 | ZNF687 | 57592 |
| ENSG000001 | 1269.28375 | 0.374391464 | 0.11469601 | 3.2642064 | 0.00109771 | 0.00608355 | CCDC59 | 29080 |
| ENSG000001 | 81.9492181 | 0.63242061 | 0.19376044 | 3.26393057 | 0.00109878 | 0.00608864 | DOK1 | 1796 |
| ENSG000001 | 648.471406 | 0.40429455 | 0.12387377 | 3.26376242 | 0.00109943 | 0.00609059 | ZDHHC16 | 84287 |
| ENSG000002 | 641.417143 | 0.434146245 | 0.13302011 | 3.26376402 | 0.00109943 | 0.00609059 | NA | NA |
| ENSG00000C | 676.11506 | 0.460656028 | 0.14115496 | 3.26347751 | 0.00110054 | 0.00609588 | SYPL1 | 6856 |
| ENSG000001 | 101.951132 | 0.548501695 | 0.16807526 | 3.2634291 | 0.00110073 | 0.00609609 | TTLL1 | 25809 |
| ENSG000001 | 37.9589947 | -0.755150578 | 0.23143298 | -3.2629342 | 0.00110265 | 0.00610508 | WDR87 | 83889 |
| ENSG000002 | 683.703429 | 0.445098286 | 0.13640901 | 3.26296828 | 0.00110252 | 0.00610508 | NA | NA |
| ENSG000002 | 29.8443701 | -0.843714601 | 0.25860887 | -3.2625122 | 0.00110429 | 0.00611334 | NA | NA |
| ENSG000002 | 23.5228157 | -0.895319449 | 0.27443225 | -3.2624425 | 0.00110457 | 0.00611401 | NA | NA |
| ENSG000001 | 184.527299 | 0.58755591 | 0.18010066 | 3.26237519 | 0.00110483 | 0.00611463 | REEP2 | 51308 |
| ENSG000001 | 893.03403 | 0.408330236 | 0.1251698 | 3.2622104 | 0.00110547 | 0.00611735 | MTRR | 4552 |
| ENSG000001 | 470.102796 | 0.413414928 | 0.12673207 | 3.26211782 | 0.00110583 | 0.00611851 | ABHD2 | 11057 |
| ENSG000001 | 1104.58641 | 0.373473249 | 0.11450786 | 3.26155119 | 0.00110804 | 0.00612992 | OTX2 | 5015 |
| ENSG000001 | 723.108668 | 0.441761961 | 0.135465 | 3.26107817 | 0.00110989 | 0.00613932 | LIFR | 3977 |
| ENSG000001 | 312.714826 | 0.491654234 | 0.15077031 | 3.2609486 | 0.0011104 | 0.00614128 | CLN8 | 2055 |
| ENSG000002 | 10.9835574 | -1.089411208 | 0.33409047 | -3.260827 | 0.00111088 | 0.00614308 | LINC00571 | 100874188 |
| ENSG000001 | 1109.66783 | 0.39432761 | 0.12093437 | 3.26067447 | 0.00111148 | 0.00614555 | C11orf57 | 55216 |
| ENSG00000C | 45.8030311 | -0.81367252 | 0.24955609 | -3.2604796 | 0.00111224 | 0.00614894 | ITGA8 | 8516 |

| | | | | | | | | |
|---|---|---|---|---|---|---|---|---|
| ENSG000001 | 3250.1659 | 0.428351786 | 0.13137923 | 3.26042251 | 0.00111246 | 0.00614933 | FAM126A | 84668 |
| ENSG000001 | 131.471017 | 0.512873355 | 0.15732157 | 3.26003193 | 0.001114 | 0.00615697 | TEX264 | 51368 |
| ENSG000001 | 255.035649 | 0.575124848 | 0.17643502 | 3.25969773 | 0.00111531 | 0.00616339 | KDM4B | 23030 |
| ENSG000001 | 61.5849484 | 0.737589931 | 0.22628308 | 3.25958938 | 0.00111574 | 0.00616441 | NAGLU | 4669 |
| ENSG000001 | 663.920864 | 0.457728611 | 0.14042732 | 3.25954092 | 0.00111593 | 0.00616441 | B4GALT6 | 9331 |
| ENSG000001 | 3057.17721 | 0.376405646 | 0.11547834 | 3.25953459 | 0.00111595 | 0.00616441 | TPP2 | 7174 |
| ENSG000002 | 8.83818734 | -1.147181653 | 0.3519764 | -3.2592573 | 0.00111704 | 0.0061696 | UBOX5-AS1 | 100134015 |
| ENSG000002 | 4.66995725 | 1.233285309 | 0.37840542 | 3.25916395 | 0.00111741 | 0.00617079 | LIME1 | 54923 |
| ENSG000001 | 28.3940544 | 0.829978313 | 0.25467855 | 3.25892502 | 0.00111835 | 0.00617515 | FAM173A | 65990 |
| ENSG000000 | 382.655523 | 0.457878043 | 0.14050553 | 3.25879029 | 0.00111888 | 0.00617724 | MTMR1 | 8776 |
| ENSG000001 | 44.5746224 | 0.724140711 | 0.22221901 | 3.25868031 | 0.00111932 | 0.00617824 | EPOR | 2057 |
| ENSG000002 | 8.10588847 | -1.159568308 | 0.35584127 | -3.2586673 | 0.00111937 | 0.00617824 | HCG9 | 10255 |
| ENSG000001 | 634.835519 | 0.411786529 | 0.12638117 | 3.25829018 | 0.00112086 | 0.00618561 | MBIP | 51562 |
| ENSG000001 | 463.786119 | 0.463096929 | 0.14213972 | 3.25804018 | 0.00112185 | 0.00619022 | STMN3 | 50861 |
| ENSG000001 | 197.747749 | 0.467462465 | 0.14348172 | 3.25799324 | 0.00112203 | 0.0061904 | TMEM138 | 51524 |
| ENSG000002 | 2.98593603 | -1.214114441 | 0.37269668 | -3.2576476 | 0.0011234 | 0.00619711 | NA | NA |
| ENSG000001 | 15.9570176 | -1.011451015 | 0.31049553 | -3.2575381 | 0.00112383 | 0.00619865 | HAO2 | 51179 |
| ENSG000001 | 1761.83612 | 0.455625071 | 0.13987536 | 3.25736482 | 0.00112452 | 0.0062016 | NUDCD1 | 84955 |
| ENSG000002 | 34.8815996 | -0.758031454 | 0.23271998 | -3.2572684 | 0.0011249 | 0.00620286 | OR5H14 | 403273 |
| ENSG000001 | 98.200005 | 0.580458622 | 0.17822161 | 3.25694852 | 0.00112617 | 0.00620901 | ASB13 | 79754 |
| ENSG000000 | 2106.15322 | 0.400512821 | 0.12297453 | 3.25687628 | 0.00112646 | 0.00620942 | MBTD1 | 54799 |
| ENSG000001 | 866.594893 | 0.559424428 | 0.17176842 | 3.25685256 | 0.00112655 | 0.00620942 | LMTK2 | 22853 |
| ENSG000001 | 1481.6219 | 0.39282245 | 0.12061615 | 3.25679815 | 0.00112677 | 0.00620977 | ATAD5 | 79915 |
| ENSG000002 | 858.895534 | 0.450313326 | 0.13828678 | 3.2563729 | 0.00112845 | 0.00621823 | SDHD | 6392 |
| ENSG000001 | 29.3344102 | -0.818646511 | 0.25140853 | -3.25624 | 0.00112898 | 0.0062203 | HOPX | 84525 |
| ENSG000001 | 987.729096 | 0.397602829 | 0.1221088 | 3.25613585 | 0.0011294 | 0.00622118 | FMNL2 | 114793 |
| ENSG000001 | 413.079371 | 0.42078787 | 0.12922974 | 3.25612251 | 0.00112945 | 0.00622118 | Sep-08 | 23176 |
| ENSG000001 | 1738.36434 | 0.394665463 | 0.12122058 | 3.25576291 | 0.00113088 | 0.00622804 | CHD6 | 84181 |
| ENSG000002 | 236.599048 | 0.43586776 | 0.13387702 | 3.25573234 | 0.001131 | 0.00622804 | ASAH2B | 653308 |

| | | | | | | | | |
|---|---|---|---|---|---|---|---|---|
| ENSG000001 | 44.9131447 | -0.714253804 | 0.21941911 | -3.2552033 | 0.00113311 | 0.00623881 | CYP19A1 | 1588 |
| ENSG000001 | 165.968639 | 0.584641877 | 0.1796098 | 3.25506672 | 0.00113366 | 0.00624096 | KIF7 | 374654 |
| ENSG000002 | 28.390451 | -0.81031385 | 0.24894683 | -3.2549675 | 0.00113405 | 0.0062423 | NA | NA |
| ENSG000002 | 7.39069804 | -1.192865926 | 0.36655151 | -3.2542928 | 0.00113675 | 0.00625629 | NA | NA |
| ENSG000002 | 8.1907864 | -1.160019607 | 0.35652799 | -3.2536565 | 0.0011393 | 0.00626947 | NA | NA |
| ENSG000001 | 43.2209084 | -0.707798603 | 0.21754435 | -3.253583 | 0.00113959 | 0.00627024 | TRPM8 | 79054 |
| ENSG000001 | 9916.72285 | 0.420424802 | 0.12924859 | 3.2528385 | 0.00114258 | 0.00628584 | CAPRIN1 | 4076 |
| ENSG000001 | 182.065068 | -0.473021007 | 0.14542561 | -3.2526665 | 0.00114328 | 0.00628879 | FAM172A | 83989 |
| ENSG000002 | 29.7686615 | -0.859448467 | 0.26423993 | -3.2525306 | 0.00114382 | 0.00629095 | CFAP45 | 25790 |
| ENSG000001 | 631.558355 | 0.385853192 | 0.11863463 | 3.25244984 | 0.00114415 | 0.00629188 | MRPL21 | 219927 |
| ENSG000001 | 50.3425054 | 0.741210657 | 0.22789956 | 3.25235673 | 0.00114452 | 0.00629309 | NDUFA11 | 126328 |
| ENSG000002 | 23.146134 | -0.893073926 | 0.27462563 | -3.2519686 | 0.00114609 | 0.00630084 | PRICKLE2-AS1 | 100652759 |
| ENSG000001 | 328.966961 | 0.48725995 | 0.14983953 | 3.25187852 | 0.00114645 | 0.00630198 | KSR1 | 8844 |
| ENSG000001 | 368.862449 | 0.47802121 | 0.14700304 | 3.25177778 | 0.00114686 | 0.00630336 | FST | 10468 |
| ENSG000002 | 2.46540596 | -1.19158244 | 0.36644502 | -3.251736 | 0.00114703 | 0.00630343 | NA | NA |
| ENSG000001 | 278.182013 | 0.523455915 | 0.16098289 | 3.25162454 | 0.00114747 | 0.00630505 | TECPR2 | 9895 |
| ENSG000002 | 138.739884 | 0.529448427 | 0.16283115 | 3.25151808 | 0.0011479 | 0.00630656 | TPM3P9 | 147804 |
| ENSG000001 | 16.8299489 | -1.000270183 | 0.30763961 | -3.251435 | 0.00114824 | 0.00630755 | CYSLTR1 | 10800 |
| ENSG000001 | 21.5003455 | -0.900203445 | 0.27687476 | -3.2513019 | 0.00114878 | 0.00630964 | UPB1 | 51733 |
| ENSG000001 | 140.435469 | 0.543240042 | 0.16708929 | 3.25119607 | 0.00114921 | 0.0063109 | CHIC2 | 26511 |
| ENSG000002 | 8.03629475 | -1.17682995 | 0.36197139 | -3.2511684 | 0.00114932 | 0.0063109 | NA | NA |
| ENSG000001 | 795.615584 | 0.410763611 | 0.12635027 | 3.25099108 | 0.00115003 | 0.00631398 | C16orf62 | 57020 |
| ENSG000001 | 180.160256 | 0.559999451 | 0.17226729 | 3.250759 | 0.00115097 | 0.00631743 | SLC38A10 | 124565 |
| ENSG000001 | 1611.93285 | 0.363637707 | 0.11186167 | 3.2507802 | 0.00115089 | 0.00631743 | STRN3 | 29966 |
| ENSG000001 | 232.651284 | 0.517114208 | 0.15908101 | 3.25063444 | 0.00115148 | 0.00631934 | RECQL5 | 9400 |
| ENSG000001 | 804.131973 | 0.381591273 | 0.11739192 | 3.25057535 | 0.00115172 | 0.0063198 | TMF1 | 7110 |
| ENSG000001 | 75.9475854 | 0.599659198 | 0.18449451 | 3.25028212 | 0.00115291 | 0.00632546 | BMP2 | 650 |
| ENSG000002 | 10.5004574 | -1.11003379 | 0.3415802 | -3.2497018 | 0.00115526 | 0.00633667 | NA | NA |
| ENSG000002 | 12.8029352 | -1.041351768 | 0.32044229 | -3.2497326 | 0.00115514 | 0.00633667 | NA | NA |

| | | | | | | | | | |
|---|---|---|---|---|---|---|---|---|---|
| ENSG00000C | 691.880288 | 0.538212045 | 0.16562105 | 3.24965962 | 0.00115543 | 0.00633675 | PREP | | 5550 |
| ENSG00000C | 2766.48118 | 0.44115282 | 0.13575532 | 3.24961707 | 0.00115561 | 0.00633684 | MGST1 | | 4257 |
| ENSG000001 | 459.411431 | 0.481224785 | 0.1480931 | 3.24947478 | 0.00115618 | 0.00633915 | MRPS25 | | 64432 |
| ENSG000001 | 424.062505 | 0.391554711 | 0.1205075 | 3.24921454 | 0.00115724 | 0.0063441 | CENPI | | 2491 |
| ENSG000001 | 493.556962 | 0.48616747 | 0.14963758 | 3.24896638 | 0.00115825 | 0.00634792 | NA | NA | |
| ENSG000002 | 45.3049472 | 0.746363531 | 0.22972183 | 3.24898829 | 0.00115816 | 0.00634792 | NT5M | | 56953 |
| ENSG000001 | 115.014178 | 0.56998217 | 0.17545059 | 3.24867636 | 0.00115943 | 0.00635354 | TMLHE | | 55217 |
| ENSG000002 | 120.184648 | 0.51838939 | 0.15957482 | 3.2485664 | 0.00115988 | 0.00635513 | LOC101928464 | | 101928464 |
| ENSG000001 | 36.9892126 | -0.770669865 | 0.23725327 | -3.2483003 | 0.00116097 | 0.00636022 | PLAGL1 | | 5325 |
| ENSG000001 | 549.374004 | 0.461015146 | 0.14192673 | 3.24826165 | 0.00116112 | 0.00636023 | RINT1 | | 60561 |
| ENSG000001 | 28.5580972 | -0.846894776 | 0.26074505 | -3.2479803 | 0.00116227 | 0.00636566 | DMBT1 | | 1755 |
| ENSG000002 | 12.1774289 | -1.082277969 | 0.33326226 | -3.2475264 | 0.00116413 | 0.00637496 | ASTN2-AS1 | | 100128505 |
| ENSG000001 | 131.37336 | 0.554855629 | 0.17086667 | 3.2473017 | 0.00116505 | 0.00637871 | CAPN10 | | 11132 |
| ENSG000001 | 31.5088745 | -0.816475868 | 0.25143546 | -3.2472583 | 0.00116523 | 0.00637871 | ITIH2 | | 3698 |
| ENSG000002 | 16.5737447 | -1.006155186 | 0.30984899 | -3.2472437 | 0.00116529 | 0.00637871 | NA | NA | |
| ENSG00000C | 9708.98749 | 0.364204561 | 0.11216365 | 3.24708194 | 0.00116595 | 0.00638148 | XPO1 | | 7514 |
| ENSG00000C | 254.630996 | 0.426245066 | 0.13127186 | 3.2470406 | 0.00116612 | 0.00638154 | FAM76A | | 199870 |
| ENSG000002 | 2.49699558 | -1.188886657 | 0.36617329 | -3.246787 | 0.00116716 | 0.00638637 | NA | NA | |
| ENSG000001 | 1372.01598 | 0.401954616 | 0.12380851 | 3.24658319 | 0.00116799 | 0.00639008 | UBA3 | | 9039 |
| ENSG000001 | 1278.85017 | 0.369795258 | 0.11391235 | 3.2463139 | 0.0011691 | 0.00639527 | KIAA1524 | | 57650 |
| ENSG000001 | 150.653057 | 0.640880055 | 0.19742243 | 3.24623733 | 0.00116941 | 0.00639612 | PKN3 | | 29941 |
| ENSG000001 | 12.2993508 | -1.062874138 | 0.32742508 | -3.2461598 | 0.00116973 | 0.006397 | POTEE | | 445582 |
| ENSG000001 | 10.7271451 | -1.122176362 | 0.34571081 | -3.2459973 | 0.0011704 | 0.00639828 | HOXC12 | | 3228 |
| ENSG000001 | 118.590699 | 0.534783338 | 0.1647521 | 3.24598794 | 0.00117044 | 0.00639828 | ARHGEF4 | | 50649 |
| ENSG000001 | 554.610461 | 0.568009692 | 0.17498588 | 3.24603158 | 0.00117026 | 0.00639828 | ZYX | | 7791 |
| ENSG000001 | 376.138403 | 0.401818209 | 0.12379742 | 3.24577206 | 0.00117133 | 0.00640227 | ZNF830 | | 91603 |
| ENSG000002 | 48.4667917 | 0.728373823 | 0.22441453 | 3.24566251 | 0.00117178 | 0.00640387 | SNORA28 | | 677811 |
| ENSG000002 | 16.1276242 | -0.977239717 | 0.30109604 | -3.245608 | 0.001172 | 0.00640423 | NA | NA | |
| ENSG000002 | 159.81344 | 0.500590668 | 0.154251 | 3.2452994 | 0.00117327 | 0.00641031 | SCAMP4 | | 113178 |

| | | | | | | | | | |
|---|---|---|---|---|---|---|---|---|---|
| ENSG000001 | 3013.61054 | 0.402444955 | 0.12401116 | 3.24523172 | 0.00117355 | 0.00641097 | TTK | | 7272 |
| ENSG000001 | 531.402416 | 0.389537205 | 0.12004188 | 3.24501084 | 0.00117446 | 0.00641422 | INTS4 | | 92105 |
| ENSG000001 | 353.021826 | 0.42935413 | 0.13231088 | 3.24504018 | 0.00117434 | 0.00641422 | GTPBP2 | | 54676 |
| ENSG000001 | 656.133155 | 0.387236734 | 0.119335 | 3.24495527 | 0.00117469 | 0.00641461 | NAB1 | | 4664 |
| ENSG000002 | 4.52579406 | -1.226464514 | 0.37799029 | -3.2446985 | 0.00117575 | 0.00641953 | NA | NA | |
| ENSG000002 | 39.9606654 | -0.774056886 | 0.23856421 | -3.244648 | 0.00117596 | 0.00641981 | NA | NA | |
| ENSG000001 | 227.302449 | 0.458367524 | 0.14127688 | 3.24446228 | 0.00117673 | 0.00642313 | SUPT3H | | 8464 |
| ENSG000001 | 153.620404 | 0.673701388 | 0.20766923 | 3.24410794 | 0.00117819 | 0.00643026 | SLC27A4 | | 10999 |
| ENSG000002 | 135.66819 | 0.55632985 | 0.17150594 | 3.24379343 | 0.00117949 | 0.0064365 | POU5F1P4 | | 645682 |
| ENSG000001 | 18.6892942 | -0.999229566 | 0.30807845 | -3.2434257 | 0.00118102 | 0.00644394 | NA | NA | |
| ENSG000002 | 19.5916426 | -0.922349643 | 0.28438579 | -3.2433042 | 0.00118152 | 0.00644582 | CCDC187 | | 399693 |
| ENSG000001 | 817.847246 | 0.386607729 | 0.11921317 | 3.24299511 | 0.0011828 | 0.00645195 | SLC30A7 | | 148867 |
| ENSG000001 | 591.298548 | 0.425088736 | 0.13108545 | 3.24283686 | 0.00118346 | 0.00645467 | YIPF5 | | 81555 |
| ENSG000001 | 126.798142 | 0.592304584 | 0.18266626 | 3.2425505 | 0.00118465 | 0.00645967 | LOC150776 | | 150776 |
| ENSG000002 | 9.6817821 | -1.134834279 | 0.34998319 | -3.2425394 | 0.0011847 | 0.00645967 | NA | NA | |
| ENSG00000C | 23.4365771 | -0.875168747 | 0.26990939 | -3.2424539 | 0.00118505 | 0.00646074 | SLC4A1 | | 6521 |
| ENSG000001 | 428.185229 | 0.500034298 | 0.1542238 | 3.24226405 | 0.00118584 | 0.00646277 | QSOX2 | | 169714 |
| ENSG000001 | 451.775139 | 0.401585898 | 0.12386027 | 3.24224951 | 0.0011859 | 0.00646277 | GLMN | | 11146 |
| ENSG000002 | 16.4020395 | -0.971883035 | 0.29975376 | -3.2422713 | 0.00118581 | 0.00646277 | MORC2-AS1 | | 150291 |
| ENSG000002 | 3.90249177 | -1.225316567 | 0.37795569 | -3.2419583 | 0.00118711 | 0.00646851 | NA | NA | |
| ENSG000001 | 96.9848623 | 0.561993685 | 0.17336334 | 3.24171003 | 0.00118815 | 0.00647328 | PRR15 | | 222171 |
| ENSG00000C | 39.1807589 | -0.76449301 | 0.23583985 | -3.2415768 | 0.0011887 | 0.00647543 | ITGAL | | 3683 |
| ENSG000002 | 6.06811457 | -1.204603221 | 0.37162419 | -3.2414554 | 0.00118921 | 0.00647558 | FAM205A | | 259308 |
| ENSG000002 | 31.9847502 | 0.825386358 | 0.25463035 | 3.24150822 | 0.00118899 | 0.00647558 | NA | NA | |
| ENSG000002 | 8.09237264 | -1.17638768 | 0.36291561 | -3.241491 | 0.00118906 | 0.00647558 | NA | NA | |
| ENSG000001 | 18.0667475 | -0.954733726 | 0.29455173 | -3.2413109 | 0.00118981 | 0.006478 | NA | NA | |
| ENSG000002 | 25.1895308 | -0.870028408 | 0.26842742 | -3.2412055 | 0.00119025 | 0.00647952 | C10orf55 | | 414236 |
| ENSG000002 | 11.6219341 | -1.092851518 | 0.33718268 | -3.2411259 | 0.00119059 | 0.00648046 | NA | NA | |
| ENSG000001 | 435.664034 | 0.466659036 | 0.14398361 | 3.24105654 | 0.00119088 | 0.00648117 | WWP2 | | 11060 |

| | | | | | | | | |
|---|---|---|---|---|---|---|---|---|
| ENSG000001 | 402.207406 | 0.420303232 | 0.12968592 | 3.24093191 | 0.0011914 | 0.00648226 | KLHL11 | 55175 |
| ENSG000002 | 15.7983654 | -1.000077237 | 0.30857659 | -3.2409369 | 0.00119138 | 0.00648226 | NA | NA |
| ENSG000001 | 112.388074 | 0.548638381 | 0.16929042 | 3.24081173 | 0.0011919 | 0.00648413 | CACNG7 | 59284 |
| ENSG000001 | 9.90951677 | -1.102500133 | 0.34022285 | -3.2405235 | 0.0011931 | 0.00648982 | CD207 | 50489 |
| ENSG000001 | 69.3829938 | -0.611249943 | 0.18863348 | -3.2404106 | 0.00119358 | 0.00649152 | ABCA12 | 26154 |
| ENSG000002 | 8.45961366 | -1.14927452 | 0.35475681 | -3.2396123 | 0.00119692 | 0.00650884 | NA | NA |
| ENSG000001 | 90.2020866 | 0.569383526 | 0.17576199 | 3.23951464 | 0.00119733 | 0.0065102 | ZFAND2A | 90637 |
| ENSG000002 | 78.4941833 | 0.602256819 | 0.18591758 | 3.23937535 | 0.00119792 | 0.00651251 | NA | NA |
| ENSG000002 | 17.0235558 | -1.002519339 | 0.30948569 | -3.2393076 | 0.0011982 | 0.00651318 | LOC100132249 | 100132249 |
| ENSG000002 | 5.17329476 | -1.222743823 | 0.3774913 | -3.2391311 | 0.00119894 | 0.00651634 | NA | NA |
| ENSG000001 | 438.096901 | 0.401753377 | 0.12403287 | 3.23908798 | 0.00119913 | 0.00651645 | DDX26B | 203522 |
| ENSG00000C | 190.093909 | 0.47080185 | 0.14535943 | 3.23888078 | 0.0012 | 0.00652031 | PPP1R15A | 23645 |
| ENSG000001 | 19.9627774 | -0.931625727 | 0.28766705 | -3.2385555 | 0.00120137 | 0.00652688 | MYOT | 9499 |
| ENSG000001 | 22.6127375 | -0.935791562 | 0.2889783 | -3.2382763 | 0.00120254 | 0.00653239 | CSF3R | 1441 |
| ENSG000002 | 11.4799367 | -1.088816638 | 0.33625051 | -3.2381115 | 0.00120324 | 0.00653529 | PSG1 | 5669 |
| ENSG000002 | 56.0365821 | 0.683588538 | 0.21111098 | 3.23805292 | 0.00120348 | 0.00653576 | NA | NA |
| ENSG000001 | 478.741936 | 0.466961921 | 0.14421922 | 3.23786197 | 0.00120429 | 0.00653926 | DHX35 | 60625 |
| ENSG000001 | 139.1914 | 0.482785879 | 0.14913075 | 3.23733296 | 0.00120653 | 0.00655052 | MFSD2A | 84879 |
| ENSG000001 | 355.435935 | 0.535593227 | 0.16544747 | 3.23724034 | 0.00120692 | 0.00655177 | BRCC3 | 79184 |
| ENSG000002 | 11.0127268 | -1.085566213 | 0.33536148 | -3.2370033 | 0.00120792 | 0.00655633 | PLCE1-AS1 | 100128054 |
| ENSG000001 | 26.5298554 | -0.86949957 | 0.26862117 | -3.2368989 | 0.00120836 | 0.00655785 | C22orf34 | 348645 |
| ENSG00000C | 297.610883 | 0.454805027 | 0.14051231 | 3.23676298 | 0.00120894 | 0.00655962 | MCCC1 | 56922 |
| ENSG000002 | 21.978474 | -0.959759431 | 0.29651989 | -3.2367456 | 0.00120901 | 0.00655962 | LINC01133 | 100505633 |
| ENSG000001 | 711.588175 | 0.386645112 | 0.11946286 | 3.23652983 | 0.00120993 | 0.00656283 | EMC2 | 9694 |
| ENSG000001 | 4641.89821 | 0.375370809 | 0.11597841 | 3.23655775 | 0.00120981 | 0.00656283 | BDP1 | 55814 |
| ENSG000001 | 910.528961 | 0.386661361 | 0.1194714 | 3.23643439 | 0.00121033 | 0.00656414 | RAB31 | 11031 |
| ENSG000001 | 6146.72606 | 0.399571994 | 0.12346306 | 3.23636873 | 0.00121061 | 0.00656478 | ZMYM2 | 7750 |
| ENSG000001 | 131.451171 | 0.507097704 | 0.15669087 | 3.23629398 | 0.00121093 | 0.00656562 | KIAA0101 | 9768 |
| ENSG000001 | 134.130269 | -0.529832992 | 0.16373039 | -3.2360089 | 0.00121214 | 0.0065713 | DNM3 | 26052 |

| | | | | | | | | | |
|---|---|---|---|---|---|---|---|---|---|
| ENSG000001 | 999.477186 | 0.411394524 | 0.12713203 | 3.23596293 | 0.00121233 | 0.00657148 | UBE2Q1 | | 55585 |
| ENSG000002 | 8.08952189 | -1.152834747 | 0.3562713 | -3.2358338 | 0.00121288 | 0.00657357 | NA | NA | |
| ENSG000001 | 200.570865 | 0.550711863 | 0.17020125 | 3.23565111 | 0.00121366 | 0.0065769 | LRRC4 | | 64101 |
| ENSG000001 | 35.4581074 | -0.780296423 | 0.24119641 | -3.2351079 | 0.00121597 | 0.00658855 | PLXDC1 | | 57125 |
| ENSG000001 | 283.23967 | 0.46341322 | 0.14326302 | 3.23470221 | 0.0012177 | 0.00659704 | ZFP30 | | 22835 |
| ENSG000001 | 411.584406 | 0.407122064 | 0.12586583 | 3.23457173 | 0.00121825 | 0.00659917 | PUS3 | | 83480 |
| ENSG000001 | 265.735842 | 0.47686054 | 0.14743496 | 3.23437911 | 0.00121908 | 0.00660186 | TNIP1 | | 10318 |
| ENSG000002 | 897.116623 | 0.3833375 | 0.11851935 | 3.23438757 | 0.00121904 | 0.00660186 | NA | NA | |
| ENSG000001 | 178.910641 | 0.56180236 | 0.17370977 | 3.23414371 | 0.00122008 | 0.00660642 | CDC42EP1 | | 11135 |
| ENSG000002 | 553.042968 | 0.390440987 | 0.12073288 | 3.23392427 | 0.00122102 | 0.00661062 | ZNF8 | | 7554 |
| ENSG00000C | 448.089207 | 0.428198971 | 0.1324115 | 3.23385031 | 0.00122133 | 0.00661145 | HIVEP2 | | 3097 |
| ENSG000002 | 21.0738635 | -0.932622224 | 0.28841559 | -3.2336054 | 0.00122238 | 0.00661623 | RAB44 | | 401258 |
| ENSG000001 | 1695.46989 | 0.418530376 | 0.12945508 | 3.23301632 | 0.00122491 | 0.00662901 | PRDM14 | | 63978 |
| ENSG000001 | 879.40925 | 0.414316832 | 0.12815475 | 3.23294178 | 0.00122523 | 0.00662985 | SDC4 | | 6385 |
| ENSG000001 | 42.9670988 | -0.710798274 | 0.21988481 | -3.2325938 | 0.00122672 | 0.00663705 | PARP15 | | 165631 |
| ENSG000001 | 1641.69356 | 0.39210168 | 0.12130019 | 3.23249023 | 0.00122716 | 0.00663857 | WHSC1L1 | | 54904 |
| ENSG000001 | 53.290337 | -0.746529894 | 0.23095557 | -3.2323528 | 0.00122775 | 0.00664087 | ABCA4 | | 24 |
| ENSG00000C | 42.9395908 | -0.788151834 | 0.24383585 | -3.232305 | 0.00122796 | 0.0066411 | HSD17B2 | | 3294 |
| ENSG000001 | 222.227015 | 0.438472288 | 0.13567299 | 3.2318318 | 0.00122999 | 0.00665122 | TTC23 | | 64927 |
| ENSG000001 | 3576.91564 | 0.39714824 | 0.12289289 | 3.23166177 | 0.00123073 | 0.00665429 | FAT3 | | 120114 |
| ENSG000001 | 298.289672 | 0.430015214 | 0.13307082 | 3.23147652 | 0.00123152 | 0.00665772 | TRMT12 | | 55039 |
| ENSG000001 | 1212.17074 | 0.379344773 | 0.11739245 | 3.23142385 | 0.00123175 | 0.00665806 | USP12 | | 219333 |
| ENSG000001 | 138.439132 | 0.519455232 | 0.16077369 | 3.23097169 | 0.0012337 | 0.00666771 | MBOAT1 | | 154141 |
| ENSG000001 | 1234.90713 | 0.454169142 | 0.14057582 | 3.23077715 | 0.00123454 | 0.00667136 | CHSY1 | | 22856 |
| ENSG00000C | 1853.98125 | 0.43114665 | 0.13345958 | 3.23054113 | 0.00123556 | 0.00667509 | CUL3 | | 8452 |
| ENSG000001 | 700.781385 | 0.406223462 | 0.12574427 | 3.23055242 | 0.00123551 | 0.00667509 | STXBP3 | | 6814 |
| ENSG000001 | 12.1852072 | -1.050807893 | 0.32527942 | -3.2304776 | 0.00123584 | 0.00667569 | NA | NA | |
| ENSG000001 | 25.3501907 | -0.863784292 | 0.26740552 | -3.230241 | 0.00123686 | 0.00668032 | ABCG4 | | 64137 |
| ENSG000002 | 7.95311064 | -1.168731027 | 0.36182324 | -3.230116 | 0.0012374 | 0.00668135 | NA | NA | |

| | | | | | | | | |
|---|---|---|---|---|---|---|---|---|
| ENSG000002 | 15.0171157 | -1.041519269 | 0.32243979 | -3.2301202 | 0.00123738 | 0.00668135 | LRRD1 | 401387 |
| ENSG000002 | 48.1115009 | -0.754133136 | 0.23347177 | -3.2300827 | 0.00123754 | 0.00668135 | NBPF25P | 101929780 |
| ENSG000002 | 11.987173 | -1.065284487 | 0.32982369 | -3.2298604 | 0.00123851 | 0.00668566 | NA | NA |
| ENSG000001 | 545.784934 | 0.430833398 | 0.13340297 | 3.22956374 | 0.00123979 | 0.00669171 | JADE3 | 9767 |
| ENSG000001 | 423.828209 | 0.430620594 | 0.13333981 | 3.22949751 | 0.00124008 | 0.00669237 | ZNF778 | 197320 |
| ENSG000001 | 324.907966 | 0.554824949 | 0.1718112 | 3.22927105 | 0.00124106 | 0.00669678 | TSTA3 | 7264 |
| ENSG000001 | 59.2503699 | 0.644037184 | 0.19944391 | 3.22916443 | 0.00124152 | 0.00669839 | SLC35C1 | 55343 |
| ENSG000001 | 934.013719 | 0.41159176 | 0.12746732 | 3.22899823 | 0.00124225 | 0.00670139 | RANBP6 | 26953 |
| ENSG000001 | 87.5285284 | -0.557071056 | 0.17254021 | -3.2286448 | 0.00124378 | 0.00670789 | DENND3 | 22898 |
| ENSG000001 | 62.4296358 | 0.74091982 | 0.22948128 | 3.22867217 | 0.00124366 | 0.00670789 | AVPI1 | 60370 |
| ENSG000002 | 35.2921502 | -0.793207371 | 0.24568482 | -3.2285567 | 0.00124417 | 0.00670907 | NA | NA |
| ENSG00000C | 120.09869 | 0.542924006 | 0.16817306 | 3.22836487 | 0.001245 | 0.00671089 | EPDR1 | 54749 |
| ENSG000001 | 48.9684044 | 0.71079726 | 0.22017195 | 3.22837338 | 0.00124496 | 0.00671089 | CGREF1 | 10669 |
| ENSG000001 | 44.0438927 | -0.753958851 | 0.233538 | -3.2284204 | 0.00124476 | 0.00671089 | ADAM12 | 8038 |
| ENSG000002 | 2.46859001 | -1.179648341 | 0.36543005 | -3.2281098 | 0.00124611 | 0.00671598 | NA | NA |
| ENSG000002 | 17.3386149 | -0.957761618 | 0.29671694 | -3.227863 | 0.00124719 | 0.00672089 | LOC653786 | 653786 |
| ENSG000001 | 41.8622502 | -0.716886229 | 0.2221193 | -3.2274828 | 0.00124885 | 0.00672893 | NELL1 | 4745 |
| ENSG000001 | 294.586651 | 0.490358911 | 0.15194092 | 3.22729978 | 0.00124964 | 0.00673234 | PAAF1 | 80227 |
| ENSG000001 | 8.32007522 | -1.141210684 | 0.35364803 | -3.2269674 | 0.0012511 | 0.00673927 | CCR6 | 1235 |
| ENSG000001 | 339.50713 | 0.472853784 | 0.1465426 | 3.2267325 | 0.00125212 | 0.00674377 | CCNJL | 79616 |
| ENSG000002 | 7.21728211 | -1.181765161 | 0.36624569 | -3.2267005 | 0.00125226 | 0.00674377 | NA | NA |
| ENSG000001 | 15.3568227 | -1.060688445 | 0.32875331 | -3.2263963 | 0.0012536 | 0.00675005 | KRT4 | 3851 |
| ENSG000001 | 22.9993661 | -0.880324165 | 0.272877 | -3.2260842 | 0.00125496 | 0.00675651 | CYP2C9 | 1559 |
| ENSG000002 | 20.6861003 | -0.928107696 | 0.28771937 | -3.2257393 | 0.00125648 | 0.00676376 | ACSM4 | 341392 |
| ENSG000001 | 5211.73373 | 0.392317592 | 0.1216282 | 3.22554803 | 0.00125732 | 0.00676649 | CALM2 | 805 |
| ENSG000001 | 41.7541013 | -0.847214514 | 0.26265668 | -3.2255587 | 0.00125727 | 0.00676649 | ENPP3 | 5169 |
| ENSG00000C | 29.1931978 | -0.825038887 | 0.25579971 | -3.2253315 | 0.00125827 | 0.00677071 | FCGR2B | 2213 |
| ENSG00000C | 302.238101 | 0.453880424 | 0.14073261 | 3.22512622 | 0.00125917 | 0.00677467 | KITLG | 4254 |
| ENSG000001 | 111.591324 | 0.641522969 | 0.19892456 | 3.22495609 | 0.00125992 | 0.00677702 | C20orf24 | 55969 |

| | | | | | | | | |
|---|---|---|---|---|---|---|---|---|
| ENSG000002 | 3.35424182 | -1.214624474 | 0.3766335 | -3.2249507 | 0.00125994 | 0.00677702 | NA | NA |
| ENSG000002 | 20.7617383 | -0.91847509 | 0.28481299 | -3.2248357 | 0.00126045 | 0.00677885 | NA | NA |
| ENSG000001 | 989.255063 | 0.377312801 | 0.11700478 | 3.22476398 | 0.00126077 | 0.00677965 | MCU | 90550 |
| ENSG000002 | 37.4789789 | -0.770235128 | 0.23889366 | -3.2241756 | 0.00126336 | 0.00679225 | EMX2OS | 196047 |
| ENSG000002 | 18.1469152 | -1.015841375 | 0.315072 | -3.2241563 | 0.00126344 | 0.00679225 | LOC101927132 | 101927132 |
| ENSG000002 | 30.6051714 | -0.832810448 | 0.25831772 | -3.2239772 | 0.00126423 | 0.0067956 | NA | NA |
| ENSG000001 | 26.1144291 | -0.856348175 | 0.26562253 | -3.223929 | 0.00126445 | 0.00679584 | PLEKHS1 | 79949 |
| ENSG000001 | 6.97255295 | -1.18971733 | 0.36905066 | -3.2237236 | 0.00126535 | 0.00679981 | LRRC10 | 376132 |
| ENSG000002 | 153.517549 | 0.590240304 | 0.18309658 | 3.22365552 | 0.00126566 | 0.00680053 | QTRT1 | 81890 |
| ENSG00000C | 16.9083378 | -0.958294306 | 0.29729835 | -3.2233421 | 0.00126704 | 0.00680617 | ISL1 | 3670 |
| ENSG000002 | 10.5721376 | -1.106784469 | 0.34336336 | -3.2233622 | 0.00126695 | 0.00680617 | LOC101928782 | 101928782 |
| ENSG000001 | 2448.18972 | 0.38811375 | 0.12041341 | 3.22317712 | 0.00126777 | 0.00680919 | CCDC14 | 64770 |
| ENSG000001 | 260.010672 | 0.472477122 | 0.14659109 | 3.22309575 | 0.00126813 | 0.00681022 | BTN2A1 | 11120 |
| ENSG000001 | 661.073247 | 0.544287086 | 0.16887908 | 3.22293974 | 0.00126882 | 0.00681303 | PCNT | 5116 |
| ENSG00000C | 539.148696 | 0.430548772 | 0.1335963 | 3.22275964 | 0.00126962 | 0.00681641 | BRD9 | 65980 |
| ENSG000002 | 13.9375223 | -1.007337453 | 0.31257933 | -3.2226617 | 0.00127005 | 0.00681784 | NA | NA |
| ENSG000001 | 220.5682 | 0.470341943 | 0.14598575 | 3.22183463 | 0.00127373 | 0.00683665 | SNUPN | 10073 |
| ENSG000001 | 477.335979 | 0.429363847 | 0.13327735 | 3.22158152 | 0.00127485 | 0.00684178 | CAMTA1 | 23261 |
| ENSG000001 | 698.32874 | 0.457750926 | 0.14209965 | 3.22133742 | 0.00127594 | 0.0068458 | SMARCC2 | 6601 |
| ENSG000001 | 1673.58822 | 0.386851504 | 0.12008989 | 3.22134947 | 0.00127588 | 0.0068458 | PDHB | 5162 |
| ENSG000001 | 2118.27573 | 0.394059765 | 0.12233529 | 3.2211456 | 0.00127679 | 0.00684857 | FZD3 | 7976 |
| ENSG000001 | 164.796967 | 0.5276339 | 0.16380132 | 3.22118231 | 0.00127663 | 0.00684857 | TMEM177 | 80775 |
| ENSG000002 | 17.3556997 | -0.960340826 | 0.29817433 | -3.2207361 | 0.00127862 | 0.00685746 | FAM181A-AS1 | 283592 |
| ENSG000001 | 143.009099 | 0.657703157 | 0.2042147 | 3.22064554 | 0.00127902 | 0.00685872 | CSK | 1445 |
| ENSG000001 | 115.291483 | 0.551634553 | 0.17128521 | 3.22056154 | 0.0012794 | 0.00685982 | TAP1 | 6890 |
| ENSG000002 | 19.3671985 | -0.94563325 | 0.29362944 | -3.2204988 | 0.00127968 | 0.00686042 | NA | NA |
| ENSG000001 | 56.6059779 | -0.713144363 | 0.22146073 | -3.2201843 | 0.00128108 | 0.00686704 | WDFY4 | 57705 |
| ENSG000001 | 1194.38861 | 0.410652506 | 0.12754028 | 3.2197867 | 0.00128286 | 0.00687566 | WRN | 7486 |
| ENSG000001 | 267.44662 | 0.464348946 | 0.14423096 | 3.21948178 | 0.00128423 | 0.00688207 | CENPT | 80152 |

| | | | | | | | | | |
|---|---|---|---|---|---|---|---|---|---|
| ENSG000001 | 24.5792586 | -0.873497089 | 0.27133047 | -3.2193107 | 0.00128499 | 0.00688381 | OPRM1 | | 4988 |
| ENSG000001 | 716.241524 | 0.443589781 | 0.13778883 | 3.21934488 | 0.00128484 | 0.00688381 | USF1 | | 7391 |
| ENSG000002 | 4.23466048 | -1.218782199 | 0.37858662 | -3.2192955 | 0.00128506 | 0.00688381 | NA | NA | |
| ENSG000001 | 608.922772 | 0.379343119 | 0.1178383 | 3.2191836 | 0.00128556 | 0.00688559 | LTV1 | | 84946 |
| ENSG000002 | 58.0817542 | -0.640093802 | 0.19885031 | -3.2189731 | 0.00128651 | 0.00688974 | HLA-L | | 3139 |
| ENSG000001 | 673.366806 | 0.386358759 | 0.12002777 | 3.21891152 | 0.00128678 | 0.00689031 | LSM6 | | 11157 |
| ENSG000001 | 179.70602 | 0.456876922 | 0.14195865 | 3.21838038 | 0.00128917 | 0.00690217 | NA | NA | |
| ENSG000001 | 225.851936 | 0.613938734 | 0.190809 | 3.21755641 | 0.00129288 | 0.00692111 | FBXO2 | | 26232 |
| ENSG000002 | 8.73600777 | -1.130558818 | 0.3513922 | -3.2173703 | 0.00129372 | 0.00692469 | NA | NA | |
| ENSG000002 | 1834.13607 | 0.380268858 | 0.11819397 | 3.21732876 | 0.0012939 | 0.00692478 | GAN | | 8139 |
| ENSG000001 | 42.7323584 | -0.749107536 | 0.23284783 | -3.2171549 | 0.00129469 | 0.00692806 | APOL3 | | 80833 |
| ENSG000001 | 4115.27985 | 0.524839964 | 0.16314223 | 3.21707 | 0.00129507 | 0.0069292 | UBR4 | | 23352 |
| ENSG000001 | 2904.33793 | 0.380215861 | 0.11818845 | 3.21703048 | 0.00129525 | 0.00692924 | RPS15A | | 6210 |
| ENSG000001 | 2517.65916 | 0.380434379 | 0.11825939 | 3.21694873 | 0.00129562 | 0.00692928 | SMCHD1 | | 23347 |
| ENSG000001 | 1163.34432 | 0.477921872 | 0.14856527 | 3.2169152 | 0.00129577 | 0.00692928 | DDX27 | | 55661 |
| ENSG000001 | 1266.84373 | 0.432987682 | 0.134596 | 3.21694308 | 0.00129564 | 0.00692928 | CDCA7 | | 83879 |
| ENSG000002 | 5.64488232 | -1.200699759 | 0.37328172 | -3.2166048 | 0.00129717 | 0.00693587 | NA | NA | |
| ENSG00000C | 150.189649 | 0.560016186 | 0.17410742 | 3.2164981 | 0.00129765 | 0.00693753 | ZNF839 | | 55778 |
| ENSG000001 | 947.530671 | 0.362899521 | 0.11282885 | 3.21637164 | 0.00129823 | 0.00693967 | KIAA0586 | | 9786 |
| ENSG000001 | 116.161806 | 0.529906596 | 0.16476947 | 3.21604836 | 0.00129969 | 0.00694658 | PGAP3 | | 93210 |
| ENSG000002 | 17.4805449 | -0.95852781 | 0.29804908 | -3.2160066 | 0.00129988 | 0.00694668 | NA | NA | |
| ENSG00000C | 30.1208994 | -0.86806666 | 0.26996046 | -3.2155326 | 0.00130203 | 0.00695716 | ADAM7 | | 8756 |
| ENSG000001 | 699.881255 | 0.422418076 | 0.13136941 | 3.21549796 | 0.00130218 | 0.00695716 | DNAAF2 | | 55172 |
| ENSG00000C | 212.862906 | 0.52604069 | 0.16362105 | 3.21499405 | 0.00130447 | 0.00696847 | ARAF | | 369 |
| ENSG000001 | 141.954167 | 0.656282175 | 0.20415006 | 3.21470474 | 0.00130579 | 0.00697458 | KAT2A | | 2648 |
| ENSG000001 | 1438.58801 | 0.374501219 | 0.11649837 | 3.21464761 | 0.00130605 | 0.00697505 | MPZL1 | | 9019 |
| ENSG000001 | 920.65609 | 0.429507832 | 0.13361402 | 3.21454154 | 0.00130653 | 0.00697671 | CCDC90B | | 60492 |
| ENSG000001 | 363.224868 | 0.471024278 | 0.14654261 | 3.21424802 | 0.00130787 | 0.00698292 | PPIL2 | | 23759 |
| ENSG000001 | 7.14177854 | -1.17304965 | 0.36496433 | -3.2141487 | 0.00130832 | 0.0069835 | SERPINA13P | | 388007 |

| | | | | | | | | |
|---|---|---|---|---|---|---|---|---|
| ENSG000002 | 22.4020562 | -0.938662767 | 0.29203903 | -3.2141689 | 0.00130823 | 0.0069835 | FAM153C | 653316 |
| ENSG000001 | 11.1085744 | -1.105469287 | 0.34395311 | -3.2140116 | 0.00130894 | 0.00698592 | MPEG1 | 219972 |
| ENSG000001 | 744.157092 | 0.413352518 | 0.12861245 | 3.21393859 | 0.00130928 | 0.00698677 | KBTBD8 | 84541 |
| ENSG000001 | 461.801108 | 0.485370845 | 0.15102669 | 3.21380845 | 0.00130987 | 0.00698902 | IRS1 | 3667 |
| ENSG000001 | 66.8881521 | 0.705925306 | 0.21966531 | 3.21364042 | 0.00131064 | 0.00699219 | C9orf142 | 286257 |
| ENSG000002 | 12.809159 | -1.05983583 | 0.32981571 | -3.2134182 | 0.00131165 | 0.00699668 | NA | NA |
| ENSG000001 | 65.6669856 | 0.651339862 | 0.20271124 | 3.21314138 | 0.00131292 | 0.00700251 | HDAC11 | 79885 |
| ENSG000002 | 10.4359684 | -1.103126697 | 0.34336776 | -3.2126683 | 0.00131508 | 0.00701313 | LINC00970 | 101978719 |
| ENSG000002 | 42.1169285 | -0.725200541 | 0.22577482 | -3.2120524 | 0.0013179 | 0.00702726 | NA | NA |
| ENSG000002 | 52.1178802 | -0.67836458 | 0.21119615 | -3.2120121 | 0.00131809 | 0.00702732 | ARHGEF33 | 100271715 |
| ENSG000001 | 2007.93542 | 0.379797148 | 0.11824427 | 3.21197095 | 0.00131828 | 0.0070274 | PSMC1 | 5700 |
| ENSG000002 | 816.832521 | 0.441043593 | 0.13731536 | 3.21190287 | 0.00131859 | 0.00702814 | TRIM13 | 10206 |
| ENSG000001 | 146.750298 | 0.496356307 | 0.15454174 | 3.2117945 | 0.00131909 | 0.00702987 | FBXL7 | 23194 |
| ENSG000001 | 24.1429591 | -0.867262351 | 0.27003328 | -3.2116869 | 0.00131958 | 0.00703158 | SSC5D | 284297 |
| ENSG00000C | 64.8287179 | -0.656717506 | 0.20452193 | -3.2109881 | 0.00132279 | 0.00704778 | FAT2 | 2196 |
| ENSG000001 | 12.7817724 | -1.056028619 | 0.32888576 | -3.2109283 | 0.00132307 | 0.00704832 | HSD11B1 | 3290 |
| ENSG000001 | 180.168416 | 0.578934462 | 0.18031678 | 3.21065218 | 0.00132434 | 0.00705417 | PIP5K1C | 23396 |
| ENSG000001 | 891.916779 | 0.406695782 | 0.12667746 | 3.21048267 | 0.00132512 | 0.0070574 | MRPL9 | 65005 |
| ENSG000002 | 688.398754 | 0.41397507 | 0.12894918 | 3.21037374 | 0.00132562 | 0.00705915 | TRIM59 | 286827 |
| ENSG000001 | 248.953328 | 0.483789669 | 0.15069891 | 3.21030626 | 0.00132594 | 0.00705988 | MED27 | 9442 |
| ENSG000001 | 1668.10065 | 0.405846388 | 0.12642222 | 3.21024563 | 0.00132622 | 0.00706044 | RNF138 | 51444 |
| ENSG000001 | 359.476742 | 0.456096446 | 0.14208075 | 3.21012124 | 0.00132679 | 0.00706257 | DONSON | 29980 |
| ENSG000001 | 79.2636975 | -0.578772142 | 0.18030883 | -3.2098934 | 0.00132784 | 0.00706724 | PTPRO | 5800 |
| ENSG000001 | 277.280463 | 0.447945482 | 0.13955551 | 3.20980157 | 0.00132827 | 0.00706857 | RPP38 | 10557 |
| ENSG000001 | 368.631015 | 0.382025524 | 0.11903452 | 3.20936758 | 0.00133027 | 0.00707794 | SLC35A3 | 23443 |
| ENSG000001 | 911.358562 | 0.351739196 | 0.10959843 | 3.20934527 | 0.00133038 | 0.00707794 | KIF1BP | 26128 |
| ENSG00000C | 391.525674 | 0.385588297 | 0.12015808 | 3.20900847 | 0.00133194 | 0.00708345 | NKAIN1 | 79570 |
| ENSG000001 | 1109.76395 | 0.38135941 | 0.11883823 | 3.20906333 | 0.00133168 | 0.00708345 | SLC11A2 | 4891 |
| ENSG000001 | 5.00667669 | -1.210703194 | 0.37728247 | -3.20901 | 0.00133193 | 0.00708345 | NA | NA |

| | | | | | | | | |
|---|---|---|---|---|---|---|---|---|
| ENSG00000( | 2721.39814 | 0.440330947 | 0.13722541 | 3.20881505 | 0.00133283 | 0.00708729 | HLTF | 6596 |
| ENSG000002 | 18.1406495 | -0.932977055 | 0.2907775 | -3.20856 | 0.00133401 | 0.00709265 | NA | NA |
| ENSG000001 | 72.2026018 | 0.59909743 | 0.1867376 | 3.20823144 | 0.00133554 | 0.00709983 | SP8 | 221833 |
| ENSG000001 | 1458.74549 | 0.428110378 | 0.13344983 | 3.20802493 | 0.0013365 | 0.00710399 | WDR35 | 57539 |
| ENSG000001 | 5345.52537 | 0.370644322 | 0.11554805 | 3.2077074 | 0.00133798 | 0.00711091 | CENPE | 1062 |
| ENSG000001 | 256.021094 | 0.497905578 | 0.15522729 | 3.20759058 | 0.00133852 | 0.00711287 | BDH1 | 622 |
| ENSG00000( | 1246.94864 | 0.453375684 | 0.1413603 | 3.20723491 | 0.00134018 | 0.00712073 | DEPDC1B | 55789 |
| ENSG000001 | 53.542543 | -0.669963058 | 0.20889673 | -3.2071496 | 0.00134057 | 0.00712191 | ABCA9 | 10350 |
| ENSG000001 | 739.744047 | 0.385423929 | 0.12018535 | 3.20691279 | 0.00134168 | 0.00712684 | ADAM17 | 6868 |
| ENSG000001 | 117.456822 | 0.708857021 | 0.22104661 | 3.20682151 | 0.0013421 | 0.00712817 | ANAPC2 | 29882 |
| ENSG000001 | 294.415868 | 0.433603839 | 0.13522229 | 3.20660025 | 0.00134313 | 0.00713272 | XXYLT1 | 152002 |
| ENSG000002 | 14.2363515 | -1.018782607 | 0.31774532 | -3.2062867 | 0.0013446 | 0.00713956 | NA | NA |
| ENSG000002 | 5.36720966 | -1.206547657 | 0.37632392 | -3.2061413 | 0.00134528 | 0.00714224 | NA | NA |
| ENSG000001 | 29.7237814 | -0.805768438 | 0.25132665 | -3.2060604 | 0.00134566 | 0.00714306 | C10orf67 | 256815 |
| ENSG000002 | 11.0796122 | -1.142734066 | 0.35643242 | -3.2060329 | 0.00134579 | 0.00714306 | NA | NA |
| ENSG000002 | 6.00448394 | -1.191533047 | 0.37167514 | -3.2058454 | 0.00134666 | 0.00714678 | NA | NA |
| ENSG000001 | 544.242089 | 0.400215413 | 0.12484414 | 3.20572042 | 0.00134725 | 0.00714708 | TMTC1 | 83857 |
| ENSG000001 | 17.693054 | -0.952514608 | 0.29712917 | -3.2057257 | 0.00134722 | 0.00714708 | GABRA6 | 2559 |
| ENSG000002 | 5.74116805 | -1.201562486 | 0.37481027 | -3.2057886 | 0.00134693 | 0.00714708 | NA | NA |
| ENSG000001 | 236.383094 | 0.43024611 | 0.13422234 | 3.20547305 | 0.00134841 | 0.00715229 | FAM83B | 222584 |
| ENSG000001 | 516.702992 | 0.457024274 | 0.14258106 | 3.20536465 | 0.00134891 | 0.00715405 | TUBGCP3 | 10426 |
| ENSG000001 | 118.452831 | 0.559566105 | 0.17458355 | 3.20514789 | 0.00134993 | 0.00715756 | BRSK2 | 9024 |
| ENSG000001 | 324.820096 | 0.524495101 | 0.16363967 | 3.20518314 | 0.00134977 | 0.00715756 | TOR3A | 64222 |
| ENSG00000( | 213.512842 | 0.425521438 | 0.13277511 | 3.20482825 | 0.00135143 | 0.00716458 | TRAF3IP2 | 10758 |
| ENSG000002 | 13.0012271 | -1.046082051 | 0.32641552 | -3.2047559 | 0.00135177 | 0.00716544 | NA | NA |
| ENSG000002 | 17.4645999 | -0.963470408 | 0.30064526 | -3.2046752 | 0.00135215 | 0.00716652 | PKIA-AS1 | 101927003 |
| ENSG000002 | 55.6893655 | 0.654465607 | 0.20422823 | 3.20457953 | 0.0013526 | 0.00716796 | LOC102724362 | 102724362 |
| ENSG000002 | 36.0808975 | 0.799835202 | 0.24959858 | 3.20448618 | 0.00135304 | 0.00716935 | NA | NA |
| ENSG000002 | 7.31823711 | -1.158326348 | 0.36148658 | -3.2043412 | 0.00135372 | 0.00717202 | NA | NA |

| | | | | | | | | |
|---|---|---|---|---|---|---|---|---|
| ENSG00000C | 1949.5694 | 0.43346746 | 0.135286 | 3.20408219 | 0.00135494 | 0.00717655 | MTMR2 | 8898 |
| ENSG000001 | 2056.20802 | 0.43272983 | 0.13505862 | 3.2040149 | 0.00135525 | 0.00717655 | DIDO1 | 11083 |
| ENSG000001 | 275.346151 | 0.407269023 | 0.12711232 | 3.20400914 | 0.00135528 | 0.00717655 | LIPG | 9388 |
| ENSG000002 | 45.4803204 | 0.748221716 | 0.23352443 | 3.2040404 | 0.00135513 | 0.00717655 | NA | NA |
| ENSG000001 | 675.421272 | 0.424124034 | 0.13238288 | 3.2037681 | 0.00135642 | 0.00718069 | MRPS31 | 10240 |
| ENSG000001 | 1098.22732 | 0.383301389 | 0.11964034 | 3.20378043 | 0.00135636 | 0.00718069 | TRMT10C | 54931 |
| ENSG000002 | 31.6797048 | -0.797870979 | 0.24904557 | -3.2037148 | 0.00135667 | 0.00718108 | CBR3-AS1 | 100506428 |
| ENSG000001 | 274.63944 | 0.453402634 | 0.1415298 | 3.20358421 | 0.00135728 | 0.0071834 | N6AMT1 | 29104 |
| ENSG000001 | 1332.72758 | 0.407379778 | 0.1271658 | 3.20353259 | 0.00135753 | 0.00718375 | CSTF2T | 23283 |
| ENSG000001 | 199.284125 | 0.598118001 | 0.18671275 | 3.20341268 | 0.00135809 | 0.0071858 | FOXK1 | 221937 |
| ENSG00000C | 4337.24128 | 0.332406901 | 0.10378759 | 3.20276145 | 0.00136117 | 0.00720113 | ZFR | 51663 |
| ENSG000001 | 636.909949 | 0.40043213 | 0.12504256 | 3.20236664 | 0.00136303 | 0.00721007 | FANCM | 57697 |
| ENSG000001 | 85.7585132 | 0.628356911 | 0.19624017 | 3.20197897 | 0.00136487 | 0.00721884 | AGPAT2 | 10555 |
| ENSG000002 | 11.5004438 | -1.054679738 | 0.32939717 | -3.2018482 | 0.00136549 | 0.00722117 | NA | NA |
| ENSG000001 | 5272.42436 | 0.36132836 | 0.11286148 | 3.2015206 | 0.00136704 | 0.00722845 | STMN1 | 3925 |
| ENSG000001 | 1200.37695 | 0.332280834 | 0.10380513 | 3.20100582 | 0.00136949 | 0.00724043 | TIMM10B | 26515 |
| ENSG000001 | 2958.22107 | 0.398960721 | 0.12464181 | 3.20085778 | 0.00137019 | 0.00724304 | ZNF106 | 64397 |
| ENSG000001 | 343.901976 | 0.436881609 | 0.13649184 | 3.20078923 | 0.00137052 | 0.00724304 | CEP89 | 84902 |
| ENSG000001 | 254.126 | 0.478043723 | 0.14935041 | 3.20081966 | 0.00137037 | 0.00724304 | PIF1 | 80119 |
| ENSG000001 | 29.3491282 | -0.800704157 | 0.25019764 | -3.2002866 | 0.00137291 | 0.00725474 | NA | NA |
| ENSG00000C | 47.9456788 | -0.673744892 | 0.21054525 | -3.2000004 | 0.00137427 | 0.0072603 | FGF4 | 2249 |
| ENSG00000C | 2799.85722 | 0.45094682 | 0.14092129 | 3.19999063 | 0.00137432 | 0.0072603 | PABPC4 | 8761 |
| ENSG000001 | 8391.4106 | 0.349485151 | 0.10922172 | 3.1997769 | 0.00137534 | 0.00726474 | U2SURP | 23350 |
| ENSG000002 | 757.871223 | 0.378661785 | 0.11836667 | 3.19905761 | 0.00137878 | 0.00728194 | ZNF718 | 255403 |
| ENSG000002 | 6.36832712 | -1.181178956 | 0.36925114 | -3.1988499 | 0.00137977 | 0.00728624 | NA | NA |
| ENSG000001 | 774.654011 | 0.395874188 | 0.12377007 | 3.19846467 | 0.00138161 | 0.00729503 | CHM | 1121 |
| ENSG000002 | 19.2288312 | -0.968097487 | 0.30268088 | -3.1984098 | 0.00138188 | 0.00729547 | ADGRE4P | 326342 |
| ENSG000001 | 146.153751 | 0.686379706 | 0.21465613 | 3.19757787 | 0.00138587 | 0.0073156 | TPRG1L | 127262 |
| ENSG00000C | 1240.56214 | 0.526478225 | 0.16465459 | 3.19747073 | 0.00138638 | 0.00731736 | DHX33 | 56919 |

| | | | | | | | | |
|---|---|---|---|---|---|---|---|---|
| ENSG000002 | 207.482222 | 0.430658028 | 0.13469458 | 3.19729288 | 0.00138724 | 0.00732092 | NA | NA |
| ENSG000000 | 1354.51381 | 0.405421691 | 0.12681582 | 3.19693296 | 0.00138897 | 0.00732911 | THOC1 | 9984 |
| ENSG000002 | 20.3512474 | -0.917992817 | 0.2871985 | -3.1963705 | 0.00139168 | 0.00734211 | NA | NA |
| ENSG000002 | 26.790768 | -0.831441563 | 0.26012245 | -3.1963468 | 0.0013918 | 0.00734211 | NA | NA |
| ENSG000002 | 14.7058385 | -1.000489407 | 0.31302645 | -3.1961817 | 0.00139259 | 0.00734535 | LOC101929116 | 101929116 |
| ENSG000001 | 2837.33214 | 0.3878006 | 0.12133519 | 3.19611 | 0.00139294 | 0.00734579 | BTF3L4 | 91408 |
| ENSG000002 | 17.8708622 | -0.982401961 | 0.30737625 | -3.1960894 | 0.00139304 | 0.00734579 | LOC100506474 | 100506474 |
| ENSG000001 | 150.782708 | 0.500010645 | 0.15645089 | 3.19595907 | 0.00139367 | 0.00734816 | PLGRKT | 55848 |
| ENSG000001 | 1222.68385 | 0.46611608 | 0.14585593 | 3.19572944 | 0.00139478 | 0.00735305 | SLC6A6 | 6533 |
| ENSG000002 | 12.3921396 | -1.034469691 | 0.32372301 | -3.1955395 | 0.0013957 | 0.00735693 | LYPD4 | 147719 |
| ENSG000002 | 10.8529754 | -1.089838758 | 0.34106081 | -3.1954382 | 0.00139619 | 0.00735856 | LOC105373463 | 105373463 |
| ENSG000001 | 915.068597 | 0.500233911 | 0.15656839 | 3.19498654 | 0.00139837 | 0.00736913 | PPRC1 | 23082 |
| ENSG000002 | 200.021423 | 0.462925741 | 0.14490222 | 3.19474559 | 0.00139954 | 0.00737433 | C22orf39 | 128977 |
| ENSG000000 | 39.3183786 | 0.724652012 | 0.22683344 | 3.19464365 | 0.00140004 | 0.00737453 | SMIM24 | 284422 |
| ENSG000001 | 92.3199478 | -0.564898425 | 0.17682776 | -3.1946253 | 0.00140013 | 0.00737453 | KSR2 | 283455 |
| ENSG000002 | 33.1013408 | -0.803766213 | 0.25159404 | -3.194695 | 0.00139979 | 0.00737453 | LOC101929719 | 101929719 |
| ENSG000001 | 276.234846 | 0.435571148 | 0.13635078 | 3.19448948 | 0.00140078 | 0.00737704 | LMBRD1 | 55788 |
| ENSG000000 | 40.3056272 | -0.735572641 | 0.2302711 | -3.1943768 | 0.00140133 | 0.00737896 | TF | 7018 |
| ENSG000002 | 20.5585586 | -0.920679061 | 0.28822228 | -3.1943369 | 0.00140152 | 0.00737902 | SERPINB11 | 89778 |
| ENSG000001 | 153.032858 | 0.575765388 | 0.18025894 | 3.19410163 | 0.00140267 | 0.00738408 | GEMIN7 | 79760 |
| ENSG000001 | 962.536929 | 0.397069404 | 0.12432524 | 3.19379548 | 0.00140416 | 0.00739 | VPS72 | 6944 |
| ENSG000002 | 15.3082351 | -0.967476704 | 0.30292314 | -3.1938026 | 0.00140412 | 0.00739 | NA | NA |
| ENSG000001 | 9.65847432 | -1.099160517 | 0.34417191 | -3.1936381 | 0.00140492 | 0.00739307 | MYBPH | 4608 |
| ENSG000001 | 17.4043868 | -0.982635892 | 0.30769158 | -3.1935742 | 0.00140523 | 0.00739374 | SLAMF1 | 6504 |
| ENSG000001 | 19.1720848 | -0.944365292 | 0.29572185 | -3.1934241 | 0.00140596 | 0.00739663 | SH2D1A | 4068 |
| ENSG000000 | 287.882987 | 0.443772023 | 0.13897831 | 3.19310265 | 0.00140753 | 0.00740391 | PRKCQ | 5588 |
| ENSG000000 | 1261.16144 | 0.405467142 | 0.12700272 | 3.19258627 | 0.00141005 | 0.0074162 | SCFD1 | 23256 |
| ENSG000001 | 319.580603 | 0.440808433 | 0.13807641 | 3.19249628 | 0.00141049 | 0.00741658 | TTI2 | 80185 |
| ENSG000002 | 769.977374 | 0.419393655 | 0.131368 | 3.19250997 | 0.00141042 | 0.00741658 | HACD2 | 201562 |

| | | | | | | | | |
|---|---|---|---|---|---|---|---|---|
| ENSG000001 | 11.5189582 | -1.06034176 | 0.33214452 | -3.1924108 | 0.0014109 | 0.00741781 | IL19 | 29949 |
| ENSG000002 | 10.0847275 | -1.099349337 | 0.34439383 | -3.1921284 | 0.00141229 | 0.00742411 | DDO | 8528 |
| ENSG000001 | 2912.81269 | 0.442409822 | 0.13860442 | 3.1918882 | 0.00141346 | 0.00742932 | RAC1 | 5879 |
| ENSG000001 | 60.6913179 | 0.65761472 | 0.20606786 | 3.19125321 | 0.00141657 | 0.00744181 | HAGHL | 84264 |
| ENSG000001 | 939.667755 | 0.393953068 | 0.12344598 | 3.19129935 | 0.00141634 | 0.00744181 | TRAPPC11 | 60684 |
| ENSG000001 | 14.6042038 | -1.022564437 | 0.3204234 | -3.1912914 | 0.00141638 | 0.00744181 | ZBED6CL | 113763 |
| ENSG000002 | 165.348348 | 0.505284326 | 0.15833375 | 3.19126104 | 0.00141653 | 0.00744181 | NA | NA |
| ENSG000001 | 27.349071 | -0.82157405 | 0.25747435 | -3.1908967 | 0.00141832 | 0.00745004 | KLHL32 | 114792 |
| ENSG000001 | 63.2148838 | -0.677823727 | 0.2124446 | -3.1905905 | 0.00141982 | 0.00745697 | WNT2B | 7482 |
| ENSG000002 | 44.2094192 | -0.752075287 | 0.23572953 | -3.1904161 | 0.00142068 | 0.00746051 | LOC100128317 | 100128317 |
| ENSG00000C | 54.3023258 | -0.666030723 | 0.20877357 | -3.1902062 | 0.00142171 | 0.00746496 | DGKG | 1608 |
| ENSG000002 | 35.6305906 | 0.772049474 | 0.24201671 | 3.19006675 | 0.0014224 | 0.0074676 | GLI4 | 2738 |
| ENSG000002 | 2459.47628 | 0.435687605 | 0.13659255 | 3.18968787 | 0.00142427 | 0.00747642 | HIST1H4L | 8368 |
| ENSG000001 | 1430.28231 | 0.434774753 | 0.1363093 | 3.18961906 | 0.0014246 | 0.00747724 | RNF20 | 56254 |
| ENSG000002 | 7.02792506 | -1.162525384 | 0.36450571 | -3.1893201 | 0.00142608 | 0.00748401 | LINC01036 | 104169671 |
| ENSG000001 | 1331.28598 | 0.561495345 | 0.17608075 | 3.18885139 | 0.00142839 | 0.00749518 | ZC3H4 | 23211 |
| ENSG00000C | 26.7246369 | -0.828633117 | 0.25987301 | -3.1886079 | 0.0014296 | 0.00750053 | CHRDL2 | 25884 |
| ENSG000002 | 67.1737623 | 0.619255473 | 0.19421589 | 3.18849033 | 0.00143018 | 0.00750164 | NA | NA |
| ENSG000002 | 3.71573545 | -1.205586203 | 0.37810531 | -3.1884932 | 0.00143016 | 0.00750164 | NA | NA |
| ENSG000001 | 314.578394 | 0.437251707 | 0.13713902 | 3.1883829 | 0.00143071 | 0.00750255 | KLHL28 | 54813 |
| ENSG000002 | 6.61164347 | -1.168684609 | 0.36654493 | -3.1883802 | 0.00143072 | 0.00750255 | NA | NA |
| ENSG000002 | 466.373906 | 0.449357071 | 0.14095945 | 3.18784639 | 0.00143337 | 0.00751544 | GPANK1 | 7918 |
| ENSG000002 | 4.51420158 | 1.202905457 | 0.37736761 | 3.18762243 | 0.00143448 | 0.0075203 | NA | NA |
| ENSG000001 | 224.077907 | 0.460026007 | 0.14431924 | 3.18755844 | 0.00143479 | 0.00752041 | IPO13 | 9670 |
| ENSG000001 | 320.128488 | 0.392756555 | 0.12321607 | 3.18754321 | 0.00143487 | 0.00752041 | ZNF841 | 284371 |
| ENSG00000C | 583.981828 | 0.496149238 | 0.15566187 | 3.18735247 | 0.00143582 | 0.0075244 | BCAR1 | 9564 |
| ENSG000001 | 3.79232875 | -1.203966922 | 0.3777372 | -3.1873136 | 0.00143601 | 0.00752444 | TRIM48 | 79097 |
| ENSG000001 | 8.73258387 | -1.128507669 | 0.35408927 | -3.1870711 | 0.00143721 | 0.00752881 | CELA1 | 1990 |
| ENSG000001 | 24.4504664 | -0.916537344 | 0.28757763 | -3.1870954 | 0.00143709 | 0.00752881 | GALNT9 | 50614 |

| | | | | | | | | |
|---|---|---|---|---|---|---|---|---|
| ENSG000002 | 410.668375 | 0.466023145 | 0.14622684 | 3.18698764 | 0.00143763 | 0.00753001 | NA | NA |
| ENSG000001 | 649.270477 | 0.384148339 | 0.1205567 | 3.18645356 | 0.00144029 | 0.00754295 | POLR3C | 10623 |
| ENSG000001 | 1367.74636 | 0.407994291 | 0.128058 | 3.18601181 | 0.00144249 | 0.0075535 | PTPRK | 5796 |
| ENSG000001 | 2549.63218 | 0.383208359 | 0.12028208 | 3.18591388 | 0.00144298 | 0.00755508 | XPO4 | 64328 |
| ENSG000001 | 186.467038 | 0.476889627 | 0.14971896 | 3.18523204 | 0.00144638 | 0.00757193 | PIGX | 54965 |
| ENSG000001 | 56.8101711 | -0.674245258 | 0.2117181 | -3.1846369 | 0.00144936 | 0.00758458 | MEIS2 | 4212 |
| ENSG000001 | 39.2622253 | -0.73453942 | 0.23064655 | -3.1846972 | 0.00144906 | 0.00758458 | EGFLAM | 133584 |
| ENSG000001 | 179.785229 | 0.478212445 | 0.15016082 | 3.18466865 | 0.0014492 | 0.00758458 | FCHSD1 | 89848 |
| ENSG000002 | 112.937325 | 0.535482699 | 0.16815296 | 3.18449755 | 0.00145006 | 0.00758725 | NA | NA |
| ENSG00000C | 334.413342 | 0.492457977 | 0.15466014 | 3.18412988 | 0.0014519 | 0.00759591 | GABARAPL2 | 11345 |
| ENSG000001 | 101.280778 | 0.553678827 | 0.17390735 | 3.18375751 | 0.00145377 | 0.00760471 | TMEM104 | 54868 |
| ENSG000001 | 2466.33287 | 0.375602989 | 0.11798845 | 3.18338777 | 0.00145563 | 0.00761345 | UGGT1 | 56886 |
| ENSG000001 | 4999.13986 | 0.416498842 | 0.13083833 | 3.18330901 | 0.00145602 | 0.00761454 | SRSF6 | 6431 |
| ENSG000001 | 479.186952 | 0.465750169 | 0.14631754 | 3.1831465 | 0.00145684 | 0.00761783 | NPC1 | 4864 |
| ENSG000001 | 8.23215116 | -1.123400519 | 0.35293421 | -3.183031 | 0.00145742 | 0.00761989 | OR6F1 | 343169 |
| ENSG000001 | 58.8851081 | -0.634104094 | 0.19923279 | -3.1827295 | 0.00145894 | 0.00762684 | CLSTN2 | 64084 |
| ENSG000001 | 267.293225 | 0.46074479 | 0.14477953 | 3.18238908 | 0.00146065 | 0.00763483 | C2orf47 | 79568 |
| ENSG000002 | 64.640161 | 0.635816471 | 0.19980087 | 3.18225082 | 0.00146135 | 0.00763749 | NA | NA |
| ENSG00000C | 413.464284 | 0.38249789 | 0.12021018 | 3.18190919 | 0.00146308 | 0.00764541 | COX10 | 1352 |
| ENSG000001 | 10.002071 | -1.095510577 | 0.34429708 | -3.1818759 | 0.00146325 | 0.00764541 | C1QB | 713 |
| ENSG000001 | 136.845642 | 0.519428249 | 0.16325012 | 3.18179401 | 0.00146366 | 0.00764564 | HABP4 | 22927 |
| ENSG000001 | 86.5400124 | 0.616050447 | 0.19361741 | 3.18179261 | 0.00146367 | 0.00764564 | SMPD2 | 6610 |
| ENSG000002 | 11.5140713 | -1.083759762 | 0.34064868 | -3.1814588 | 0.00146535 | 0.00765347 | ADAM21P1 | 145241 |
| ENSG000002 | 937.599166 | 0.390482524 | 0.12275527 | 3.18098382 | 0.00146776 | 0.00766504 | ZBTB10 | 65986 |
| ENSG000001 | 5.17661779 | -1.193980568 | 0.37536636 | -3.1808406 | 0.00146848 | 0.00766785 | OR2T2 | 401992 |
| ENSG000002 | 40.6427357 | 0.871906666 | 0.27416233 | 3.18025701 | 0.00147144 | 0.00768232 | NA | NA |
| ENSG000001 | 207.131679 | 0.441628311 | 0.1388686 | 3.18018836 | 0.00147179 | 0.00768315 | ANKRD6 | 22881 |
| ENSG000001 | 123.236614 | 0.612455282 | 0.19258687 | 3.18015074 | 0.00147198 | 0.00768316 | RLTPR | 146206 |
| ENSG000001 | 569.167799 | 0.397615215 | 0.12503851 | 3.17994198 | 0.00147305 | 0.0076877 | NFU1 | 27247 |

| | | | | | | | | |
|---|---|---|---|---|---|---|---|---|
| ENSG000002 | 30.8661459 | -0.781149187 | 0.2456548 | -3.1798653 | 0.00147344 | 0.00768875 | NA | NA |
| ENSG000002 | 106.206306 | 0.545320735 | 0.17149931 | 3.17972554 | 0.00147415 | 0.00769147 | ARRDC1-AS1 | 85026 |
| ENSG000001 | 110.435009 | 0.515262442 | 0.16208097 | 3.17904347 | 0.00147762 | 0.00770672 | ZNF576 | 79177 |
| ENSG000001 | 239.869815 | 0.413984077 | 0.13022011 | 3.17911017 | 0.00147728 | 0.00770672 | GID4 | 79018 |
| ENSG000002 | 102.292282 | 0.580280013 | 0.18253312 | 3.17903964 | 0.00147764 | 0.00770672 | TMEM231 | 79583 |
| ENSG000002 | 3.68821612 | -1.201522776 | 0.37802179 | -3.1784484 | 0.00148066 | 0.00772146 | NA | NA |
| ENSG000001 | 87.2128696 | 0.581431559 | 0.18293562 | 3.17833974 | 0.00148121 | 0.00772336 | PCGF1 | 84759 |
| ENSG000001 | 430.739298 | 0.540786434 | 0.17017099 | 3.17790028 | 0.00148346 | 0.00773209 | KRI1 | 65095 |
| ENSG000001 | 375.800876 | 0.393932883 | 0.12395985 | 3.17790694 | 0.00148342 | 0.00773209 | MYO1D | 4642 |
| ENSG000002 | 232.05691 | 0.418967701 | 0.13183702 | 3.17792141 | 0.00148335 | 0.00773209 | SBDSP1 | 155370 |
| ENSG000002 | 5.82111276 | -1.17807038 | 0.37071214 | -3.1778575 | 0.00148368 | 0.00773224 | NA | NA |
| ENSG000001 | 260.914071 | 0.423910124 | 0.1334185 | 3.17729638 | 0.00148655 | 0.00774439 | MREG | 55686 |
| ENSG000001 | 635.690208 | 0.409619262 | 0.1289208 | 3.17729388 | 0.00148656 | 0.00774439 | BTBD10 | 84280 |
| ENSG000002 | 8.08344161 | -1.137857345 | 0.35812191 | -3.1772905 | 0.00148658 | 0.00774439 | NA | NA |
| ENSG000001 | 33.0530333 | -0.825813187 | 0.25991623 | -3.1772283 | 0.0014869 | 0.00774506 | CD96 | 10225 |
| ENSG000001 | 638.184481 | 0.537332628 | 0.16912225 | 3.17718463 | 0.00148712 | 0.00774523 | ERCC3 | 2071 |
| ENSG000001 | 881.173822 | 0.440181349 | 0.13855533 | 3.17693547 | 0.0014884 | 0.00775089 | RSPRY1 | 89970 |
| ENSG000001 | 845.127813 | 0.830256078 | 0.2613437 | 3.17687433 | 0.00148872 | 0.00775153 | AIFM1 | 9131 |
| ENSG000002 | 21.2742627 | -0.92899771 | 0.29245656 | -3.1765323 | 0.00149047 | 0.00775968 | LOC102724776 | 102724776 |
| ENSG000002 | 9.61544982 | -1.094335819 | 0.34451603 | -3.1764438 | 0.00149093 | 0.00776105 | NA | NA |
| ENSG000001 | 916.080639 | 0.471629773 | 0.14848055 | 3.17637407 | 0.00149129 | 0.00776192 | PURB | 5814 |
| ENSG000002 | 2.52605349 | -1.156328563 | 0.364047 | -3.1763167 | 0.00149158 | 0.00776246 | NA | NA |
| ENSG000002 | 62.2981678 | -0.653483525 | 0.20575581 | -3.176015 | 0.00149313 | 0.00776954 | COL6A6 | 131873 |
| ENSG000001 | 43.6213491 | -0.703209866 | 0.22142805 | -3.175794 | 0.00149427 | 0.00777372 | ANO3 | 63982 |
| ENSG000001 | 501.128466 | 0.399241443 | 0.12571428 | 3.17578443 | 0.00149432 | 0.00777372 | SLC20A2 | 6575 |
| ENSG000001 | 403.452769 | 0.498554541 | 0.15700541 | 3.1753972 | 0.00149632 | 0.0077831 | PDPK1 | 5170 |
| ENSG000001 | 12.2543683 | -1.050854889 | 0.33094489 | -3.1753169 | 0.00149673 | 0.00778426 | CLEC2B | 9976 |
| ENSG000001 | 45.5478168 | -0.687283441 | 0.21646217 | -3.1750742 | 0.00149798 | 0.00778977 | CLIC5 | 53405 |
| ENSG000001 | 2077.55801 | 0.362610668 | 0.11421232 | 3.17488212 | 0.00149897 | 0.00779393 | ITPR2 | 3709 |

| | | | | | | | | | |
|---|---|---|---|---|---|---|---|---|---|
| ENSG000002 | 11.5724018 | -1.061986937 | 0.33450053 | -3.1748438 | 0.00149917 | 0.00779396 | TEDDM1 | | 127670 |
| ENSG000001 | 1716.72961 | 0.379172979 | 0.11943388 | 3.17475213 | 0.00149965 | 0.00779543 | ORC2 | | 4999 |
| ENSG000002 | 10.7854808 | -1.064015681 | 0.33516054 | -3.1746449 | 0.0015002 | 0.00779731 | LOC340113 | | 340113 |
| ENSG000001 | 759.038505 | 0.407778213 | 0.12845774 | 3.17441524 | 0.00150139 | 0.00780248 | EBP | | 10682 |
| ENSG000001 | 463.362431 | 0.372056126 | 0.11721721 | 3.17407435 | 0.00150315 | 0.00780966 | PBX3 | | 5090 |
| ENSG000001 | 9.14717036 | -1.097517991 | 0.3457758 | -3.174074 | 0.00150315 | 0.00780966 | FLJ40288 | | 286023 |
| ENSG000001 | 14.8688385 | -0.987090896 | 0.31099204 | -3.174007 | 0.0015035 | 0.00781046 | OR10K1 | | 391109 |
| ENSG000001 | 449.46822 | 0.363154483 | 0.11442829 | 3.17364257 | 0.00150539 | 0.00781906 | VPS45 | | 11311 |
| ENSG000001 | 496.242576 | 0.587543357 | 0.18513605 | 3.17357617 | 0.00150573 | 0.00781906 | RNF44 | | 22838 |
| ENSG000002 | 6.5232655 | -1.179521001 | 0.37166723 | -3.1735943 | 0.00150564 | 0.00781906 | MCHR2-AS1 | | 728012 |
| ENSG000001 | 331.67053 | 0.460686668 | 0.14517375 | 3.17334687 | 0.00150692 | 0.00782323 | RAP1A | | 5906 |
| ENSG000001 | 280.090086 | -0.442693027 | 0.13950328 | -3.1733521 | 0.0015069 | 0.00782323 | ZNF682 | | 91120 |
| ENSG000002 | 8.82303974 | -1.102546701 | 0.34747257 | -3.1730467 | 0.00150848 | 0.00783032 | NA | NA | |
| ENSG00000C | 186.401217 | 0.526280525 | 0.16590252 | 3.17222745 | 0.00151274 | 0.00785144 | PDIA5 | | 10954 |
| ENSG000001 | 108.227618 | -0.563083695 | 0.17752301 | -3.1718912 | 0.0015145 | 0.00785953 | HYDIN | | 54768 |
| ENSG00000C | 599.639592 | 0.542482778 | 0.17103246 | 3.17181177 | 0.00151491 | 0.00786068 | ATP11A | | 23250 |
| ENSG000001 | 156.083601 | 0.49691708 | 0.15667054 | 3.17173267 | 0.00151532 | 0.00786181 | MXI1 | | 4601 |
| ENSG00000C | 280.00679 | 0.452695392 | 0.14273859 | 3.17149966 | 0.00151654 | 0.00786711 | GLT8D1 | | 55830 |
| ENSG000001 | 378.188098 | 0.478237778 | 0.15079682 | 3.17140491 | 0.00151704 | 0.00786867 | MLF1 | | 4291 |
| ENSG000001 | 1421.73057 | 0.434499171 | 0.13702161 | 3.17102656 | 0.00151901 | 0.00787792 | CWC27 | | 10283 |
| ENSG000001 | 488.622509 | 0.382596686 | 0.12065809 | 3.1709162 | 0.00151959 | 0.0078789 | SNX14 | | 57231 |
| ENSG000002 | 10.7006555 | -1.078806413 | 0.34021628 | -3.170943 | 0.00151945 | 0.0078789 | NA | NA | |
| ENSG000002 | 2.50974311 | -1.152666235 | 0.36354865 | -3.170597 | 0.00152126 | 0.00788655 | NA | NA | |
| ENSG00000C | 61.6354039 | 0.635350073 | 0.20039538 | 3.17048266 | 0.00152186 | 0.00788865 | EFNB1 | | 1947 |
| ENSG000001 | 2561.67813 | 0.386079576 | 0.12179041 | 3.17003262 | 0.00152422 | 0.00789987 | ENSA | | 2029 |
| ENSG000001 | 16.6493797 | -0.970805074 | 0.30628233 | -3.1696412 | 0.00152627 | 0.00790951 | TMEM156 | | 80008 |
| ENSG000001 | 85.3829422 | 0.609199131 | 0.19220319 | 3.16955791 | 0.00152671 | 0.00791076 | CDC42EP2 | | 10435 |
| ENSG00000C | 70.4123411 | 0.605257698 | 0.19098786 | 3.16908982 | 0.00152917 | 0.0079225 | PPP1R3F | | 89801 |
| ENSG000001 | 15.9660471 | -0.956016014 | 0.30169384 | -3.1688284 | 0.00153055 | 0.00792862 | OR10J1 | | 26476 |

| | | | | | | | | | |
|---|---|---|---|---|---|---|---|---|---|
| ENSG000001 | 571.852744 | 0.423934451 | 0.13380381 | 3.16832878 | 0.00153318 | 0.00794124 | FAM204A | | 63877 |
| ENSG000002 | 1574.48005 | 0.372866293 | 0.11768884 | 3.16823837 | 0.00153366 | 0.0079427 | NME1 | | 4830 |
| ENSG000002 | 7.95212341 | -1.132856076 | 0.35758806 | -3.1680479 | 0.00153466 | 0.00794689 | NA | NA | |
| ENSG000002 | 15.8533786 | -0.972471427 | 0.30700102 | -3.1676488 | 0.00153677 | 0.00795679 | NA | NA | |
| ENSG000002 | 4.92109205 | -1.195609233 | 0.37747071 | -3.1674225 | 0.00153797 | 0.00796197 | NA | NA | |
| ENSG000002 | 258.392626 | 0.435184861 | 0.13740972 | 3.16706023 | 0.00153988 | 0.00797088 | ZNF674 | | 641339 |
| ENSG000001 | 141.326925 | 0.575963243 | 0.18186324 | 3.16701297 | 0.00154013 | 0.00797115 | SIRT3 | | 23410 |
| ENSG000001 | 416.511041 | 0.443813309 | 0.14014288 | 3.16686301 | 0.00154093 | 0.00797425 | UROS | | 7390 |
| ENSG000001 | 308.11827 | 0.438422744 | 0.13844824 | 3.16669073 | 0.00154184 | 0.00797796 | SNX33 | | 257364 |
| ENSG000001 | 886.361259 | 0.339053947 | 0.10708489 | 3.1662166 | 0.00154436 | 0.00798995 | RALGAPA1 | | 253959 |
| ENSG000002 | 7.58936138 | -1.1551017 | 0.36487264 | -3.1657668 | 0.00154675 | 0.0080013 | NA | NA | |
| ENSG000001 | 17.9651392 | -0.957951804 | 0.30263292 | -3.1653919 | 0.00154874 | 0.00801059 | OR5A1 | | 219982 |
| ENSG00000C | 1550.55557 | 0.365586426 | 0.11549674 | 3.16533977 | 0.00154902 | 0.00801101 | MYCBP2 | | 23077 |
| ENSG000002 | 44.0477271 | -0.8028735 | 0.25366808 | -3.1650553 | 0.00155053 | 0.00801782 | DPF3 | | 8110 |
| ENSG000001 | 1217.79225 | 0.400286291 | 0.12648142 | 3.16478343 | 0.00155198 | 0.00802429 | SOCS2 | | 8835 |
| ENSG000001 | 253.275597 | 0.47178321 | 0.14908476 | 3.1645302 | 0.00155334 | 0.00803025 | ZBTB46 | | 140685 |
| ENSG000001 | 77.6738284 | 0.58639529 | 0.18530965 | 3.16440779 | 0.00155399 | 0.00803261 | CHCHD5 | | 84269 |
| ENSG000001 | 71.2578431 | 0.613792564 | 0.19397385 | 3.1643058 | 0.00155453 | 0.0080344 | TUSC1 | | 286319 |
| ENSG000001 | 8.34570251 | -1.148069051 | 0.36285943 | -3.1639499 | 0.00155644 | 0.00804218 | NA | NA | |
| ENSG000002 | 6.05709252 | -1.17962058 | 0.37282893 | -3.1639727 | 0.00155631 | 0.00804218 | NA | NA | |
| ENSG000001 | 137.44463 | 0.473675408 | 0.14971995 | 3.16374281 | 0.00155754 | 0.00804688 | BTD | | 686 |
| ENSG000001 | 19.2450273 | -0.966245317 | 0.3054306 | -3.1635511 | 0.00155857 | 0.00805115 | GCSAML | | 148823 |
| ENSG000001 | 52.9563308 | -0.709100092 | 0.22416151 | -3.1633446 | 0.00155968 | 0.00805584 | TDRD9 | | 122402 |
| ENSG000001 | 123.877091 | 0.531916269 | 0.16815408 | 3.16326714 | 0.00156009 | 0.00805636 | PHKG2 | | 5261 |
| ENSG000001 | 23.267456 | -0.876518148 | 0.27709402 | -3.1632518 | 0.00156017 | 0.00805636 | LRRC48 | | 83450 |
| ENSG000001 | 744.151932 | 0.362170404 | 0.11449862 | 3.16309854 | 0.00156099 | 0.00805958 | EYA3 | | 2140 |
| ENSG000001 | 25.8077976 | -0.822696439 | 0.26009799 | -3.163025 | 0.00156139 | 0.00806058 | CATSPERD | | 257062 |
| ENSG000001 | 138.594539 | 0.478533404 | 0.15130671 | 3.16267146 | 0.00156329 | 0.00806935 | MXD4 | | 10608 |
| ENSG000001 | 236.799388 | 0.44698042 | 0.14135512 | 3.16210981 | 0.0015663 | 0.00808287 | GIN1 | | 54826 |

| | | | | | | | | |
|---|---|---|---|---|---|---|---|---|
| ENSG000002 | 7.39480272 | -1.148701376 | 0.36326759 | -3.1621356 | 0.00156617 | 0.00808287 | NA | NA |
| ENSG000001 | 12.1627304 | -1.044533026 | 0.33034591 | -3.1619372 | 0.00156723 | 0.00808664 | PRAM1 | 84106 |
| ENSG000001 | 20.1437325 | -0.935968061 | 0.29604843 | -3.161537 | 0.00156939 | 0.00809673 | ENDOU | 8909 |
| ENSG000001 | 25.1106616 | -0.839537351 | 0.26556403 | -3.1613368 | 0.00157047 | 0.00810126 | KLK15 | 55554 |
| ENSG000001 | 1150.86786 | 0.396258487 | 0.12535751 | 3.16102723 | 0.00157214 | 0.00810885 | ARMC8 | 25852 |
| ENSG000001 | 451.23525 | 0.444683404 | 0.14068185 | 3.16091524 | 0.00157274 | 0.00811094 | THAP1 | 55145 |
| ENSG000001 | 399.804716 | 0.423290414 | 0.13394196 | 3.16025262 | 0.00157632 | 0.00812837 | MITD1 | 129531 |
| ENSG000001 | 220.09676 | 0.464886655 | 0.14710905 | 3.16014988 | 0.00157688 | 0.0081302 | B3GALT6 | 126792 |
| ENSG00000C | 147.910954 | 0.552035564 | 0.17470762 | 3.15976805 | 0.00157895 | 0.00813568 | PRKACA | 5566 |
| ENSG00000C | 1424.06269 | 0.344825541 | 0.10913102 | 3.15973897 | 0.00157911 | 0.00813568 | PIAS2 | 9063 |
| ENSG000001 | 383.49253 | 0.388078843 | 0.12281379 | 3.1598963 | 0.00157825 | 0.00813568 | FIG4 | 9896 |
| ENSG000001 | 8299.47701 | 0.386888263 | 0.122442 | 3.15976769 | 0.00157895 | 0.00813568 | LDHA | 3939 |
| ENSG000001 | 312.077162 | 0.418879636 | 0.13256737 | 3.15974926 | 0.00157905 | 0.00813568 | HLCS | 3141 |
| ENSG000001 | 2605.16566 | 0.381902035 | 0.12086535 | 3.15973144 | 0.00157915 | 0.00813568 | ZNF93 | 81931 |
| ENSG000001 | 603.165454 | 0.398964439 | 0.12626768 | 3.15967201 | 0.00157947 | 0.00813631 | ATP8A1 | 10396 |
| ENSG000001 | 1022.43813 | 0.397016972 | 0.1256602 | 3.15944876 | 0.00158068 | 0.00814151 | INSIG1 | 3638 |
| ENSG000001 | 469.02059 | 0.408537672 | 0.12933495 | 3.1587569 | 0.00158444 | 0.00815983 | PIGU | 128869 |
| ENSG000001 | 23.4770953 | -0.886477227 | 0.28064777 | -3.1586826 | 0.00158484 | 0.00816087 | GPA33 | 10223 |
| ENSG000001 | 22.3865343 | -0.920331528 | 0.29140538 | -3.1582517 | 0.00158718 | 0.00817191 | SIGLECL1 | 284369 |
| ENSG000002 | 42.3823075 | 0.694269407 | 0.21984492 | 3.15799617 | 0.00158858 | 0.0081776 | NA | NA |
| ENSG000002 | 10.528483 | -1.091586045 | 0.34566019 | -3.1579745 | 0.00158869 | 0.0081776 | NA | NA |
| ENSG000002 | 6.85180822 | -1.148793031 | 0.36380576 | -3.1577099 | 0.00159014 | 0.00818399 | LINC01098 | 285501 |
| ENSG000001 | 23.4451932 | -0.868761545 | 0.27522628 | -3.1565356 | 0.00159655 | 0.00821597 | TBX3 | 6926 |
| ENSG000001 | 27.3246387 | -0.849742021 | 0.26923893 | -3.1560889 | 0.001599 | 0.00822752 | SHANK1 | 50944 |
| ENSG000002 | 3.42826253 | -1.185095118 | 0.37554065 | -3.1557039 | 0.00160111 | 0.00823734 | LOC100288966 | 100288966 |
| ENSG000001 | 2383.62944 | 0.372722584 | 0.11812004 | 3.15545586 | 0.00160248 | 0.00824226 | CNTNAP2 | 26047 |
| ENSG000001 | 775.485791 | 0.376184134 | 0.11921589 | 3.15548651 | 0.00160231 | 0.00824226 | ZNF765 | 91661 |
| ENSG000001 | 902.000293 | 0.383681436 | 0.12160466 | 3.15515411 | 0.00160413 | 0.00824974 | BIRC2 | 329 |
| ENSG000002 | 21.2956093 | -0.904032301 | 0.28653913 | -3.1550047 | 0.00160496 | 0.00825292 | LOC340512 | 340512 |

| | | | | | | | | | |
|---|---|---|---|---|---|---|---|---|---|
| ENSG000001 | 20.7362855 | -0.886797054 | 0.28108866 | -3.1548659 | 0.00160572 | 0.0082558 | SCN2B | | 6327 |
| ENSG00000C | 193.515566 | 0.50520116 | 0.16014911 | 3.15456729 | 0.00160736 | 0.00826164 | TBC1D22A | | 25771 |
| ENSG000001 | 1124.40585 | 0.496400404 | 0.15735883 | 3.15457618 | 0.00160731 | 0.00826164 | DIAPH1 | | 1729 |
| ENSG000001 | 460.423554 | 0.393555761 | 0.12475818 | 3.15454868 | 0.00160747 | 0.00826164 | NA | NA | |
| ENSG000001 | 885.179692 | 0.405239275 | 0.12846524 | 3.15446623 | 0.00160792 | 0.00826293 | SEC61G | | 23480 |
| ENSG000001 | 623.34992 | 0.425712819 | 0.13497307 | 3.15405748 | 0.00161017 | 0.00827346 | FUNDC2 | | 65991 |
| ENSG000001 | 29.4534007 | -0.799189283 | 0.25339516 | -3.1539248 | 0.00161091 | 0.00827617 | ASPA | | 443 |
| ENSG000001 | 288.821315 | 0.410395245 | 0.13012484 | 3.15385788 | 0.00161128 | 0.00827702 | ANKRD49 | | 54851 |
| ENSG000001 | 128.348067 | 0.543620098 | 0.17237113 | 3.15377704 | 0.00161172 | 0.00827722 | SRGAP2C | | 653464 |
| ENSG000002 | 88.6368153 | 0.559367027 | 0.17736229 | 3.1538104 | 0.00161154 | 0.00827722 | PLEKHM1 | | 9842 |
| ENSG000001 | 189.778555 | 0.447697706 | 0.14196002 | 3.15368873 | 0.00161221 | 0.00827868 | RNFT2 | | 84900 |
| ENSG00000C | 656.186781 | 0.447950152 | 0.14206838 | 3.15306012 | 0.00161569 | 0.00829272 | SFSWAP | | 6433 |
| ENSG00000C | 22.4083482 | -0.893167019 | 0.28326644 | -3.1530986 | 0.00161547 | 0.00829272 | FMO2 | | 2327 |
| ENSG000001 | 1511.95248 | 0.47759443 | 0.15147006 | 3.15306159 | 0.00161568 | 0.00829272 | TMEM30A | | 55754 |
| ENSG000001 | 15.0386899 | -0.989832461 | 0.31392893 | -3.1530463 | 0.00161576 | 0.00829272 | CDKN2B | | 1030 |
| ENSG000001 | 138.17852 | 0.476251849 | 0.15107096 | 3.15250431 | 0.00161876 | 0.00830602 | SLC41A3 | | 54946 |
| ENSG000001 | 201.057239 | 0.495049124 | 0.15703186 | 3.15253941 | 0.00161857 | 0.00830602 | CDK2AP2 | | 10263 |
| ENSG000001 | 267.013093 | 0.425590955 | 0.13501693 | 3.1521304 | 0.00162084 | 0.00831562 | SUMF1 | | 285362 |
| ENSG000001 | 1934.32483 | 0.379046788 | 0.12025326 | 3.15207073 | 0.00162117 | 0.00831626 | NCAPG | | 64151 |
| ENSG000001 | 2979.14352 | 0.446400963 | 0.14162391 | 3.15201699 | 0.00162147 | 0.00831674 | LARP4 | | 113251 |
| ENSG00000C | 541.984688 | 0.464791269 | 0.14748397 | 3.15146971 | 0.00162451 | 0.00833024 | ERMP1 | | 79956 |
| ENSG000002 | 2.8263588 | -1.172675609 | 0.37210423 | -3.1514708 | 0.0016245 | 0.00833024 | NA | NA | |
| ENSG00000C | 439.750011 | 0.546004839 | 0.17326308 | 3.15130515 | 0.00162543 | 0.00833282 | TNRC6C | | 57690 |
| ENSG000001 | 8230.00006 | 0.475126884 | 0.1507708 | 3.15131907 | 0.00162535 | 0.00833282 | RCC2 | | 55920 |
| ENSG000001 | 319.622093 | 0.386629573 | 0.12269171 | 3.15122811 | 0.00162585 | 0.00833397 | TRAF6 | | 7189 |
| ENSG00000C | 15.7350363 | -0.967154214 | 0.30692542 | -3.1511049 | 0.00162654 | 0.00833643 | FMO3 | | 2328 |
| ENSG000001 | 1157.42436 | 0.371025244 | 0.11776272 | 3.15061708 | 0.00162926 | 0.00834931 | MAPK8 | | 5599 |
| ENSG000001 | 1261.7209 | 0.479909289 | 0.15233073 | 3.15044308 | 0.00163023 | 0.00835323 | FAM98A | | 25940 |
| ENSG000001 | 104.590369 | 0.532609608 | 0.1690634 | 3.15035431 | 0.00163073 | 0.00835471 | CLYBL | | 171425 |

| | | | | | | | | |
|---|---|---|---|---|---|---|---|---|
| ENSG000001 | 837.978339 | 0.412019992 | 0.13080116 | 3.14997203 | 0.00163286 | 0.0083646 | C21orf91 | 54149 |
| ENSG000001 | 387.003992 | 0.438964731 | 0.13939041 | 3.14917456 | 0.00163732 | 0.0083864 | CLDN10 | 9071 |
| ENSG000001 | 185.391693 | 0.576389633 | 0.1830337 | 3.14909028 | 0.0016378 | 0.00838776 | BMF | 90427 |
| ENSG000001 | 16.574334 | -0.964725978 | 0.3063612 | -3.1489822 | 0.0016384 | 0.0083898 | GLYAT | 10249 |
| ENSG000001 | 390.225962 | 0.373154722 | 0.11850242 | 3.14892078 | 0.00163875 | 0.0083905 | TAF5 | 6877 |
| ENSG000001 | 30.4731775 | -0.817297911 | 0.25959519 | -3.1483554 | 0.00164192 | 0.00840569 | B4GALNT2 | 124872 |
| ENSG000002 | 48.9068464 | -0.695626812 | 0.22096191 | -3.1481752 | 0.00164293 | 0.00840981 | LOC101928035 | 101928035 |
| ENSG000001 | 98.481831 | 0.641412533 | 0.20376576 | 3.14779354 | 0.00164508 | 0.00841974 | RTN2 | 6253 |
| ENSG000001 | 2583.40945 | 0.430435465 | 0.13674814 | 3.14765132 | 0.00164588 | 0.00842277 | ELOVL6 | 79071 |
| ENSG000001 | 155.719399 | 0.491446468 | 0.15613456 | 3.14758281 | 0.00164626 | 0.00842368 | FGFRL1 | 53834 |
| ENSG000001 | 54.7011403 | -0.738971543 | 0.23480492 | -3.1471723 | 0.00164858 | 0.00843445 | PGM5 | 5239 |
| ENSG000001 | 146.199725 | 0.494656964 | 0.15719251 | 3.14682282 | 0.00165055 | 0.00844348 | SERPINI1 | 5274 |
| ENSG00000C | 81.5958646 | -0.569331658 | 0.18092557 | -3.1467729 | 0.00165083 | 0.00844385 | MOK | 5891 |
| ENSG000001 | 65.4478037 | 0.617454738 | 0.19623892 | 3.14644378 | 0.00165269 | 0.00845229 | ADSSL1 | 122622 |
| ENSG00000C | 1407.55463 | 0.374036847 | 0.11890316 | 3.14572671 | 0.00165675 | 0.00847198 | RPAP3 | 79657 |
| ENSG000001 | 2304.44416 | 0.34540367 | 0.10980459 | 3.1456214 | 0.00165734 | 0.00847396 | NSA2 | 10412 |
| ENSG000002 | 26.9651692 | -0.851289706 | 0.27064926 | -3.1453612 | 0.00165882 | 0.00848043 | LOC284788 | 284788 |
| ENSG000001 | 3301.7821 | 0.344848203 | 0.10964108 | 3.14524642 | 0.00165947 | 0.00848269 | UTP20 | 27340 |
| ENSG000001 | 59.5524697 | 0.703402298 | 0.2236477 | 3.14513543 | 0.0016601 | 0.00848484 | MGMT | 4255 |
| ENSG000001 | 48.2718479 | -0.690420228 | 0.21953197 | -3.1449643 | 0.00166107 | 0.00848632 | TNN | 63923 |
| ENSG000001 | 17.9475378 | -0.950772711 | 0.30232351 | -3.1448851 | 0.00166152 | 0.00848632 | SLC17A1 | 6568 |
| ENSG000001 | 22.5186829 | -0.851206203 | 0.27066157 | -3.1449097 | 0.00166138 | 0.00848632 | CAPN9 | 10753 |
| ENSG000002 | 26.6038335 | -0.815870662 | 0.25941981 | -3.1449821 | 0.00166097 | 0.00848632 | NA | NA |
| ENSG000002 | 21.817985 | -0.889179079 | 0.2827394 | -3.1448715 | 0.0016616 | 0.00848632 | NA | NA |
| ENSG000002 | 15.9622804 | -0.955383817 | 0.30379185 | -3.1448632 | 0.00166165 | 0.00848632 | NA | NA |
| ENSG000001 | 103.692822 | 0.594080134 | 0.18891984 | 3.14461487 | 0.00166306 | 0.00849246 | TCEA2 | 6919 |
| ENSG000002 | 5.29421376 | -1.184474697 | 0.37667975 | -3.1445138 | 0.00166363 | 0.00849398 | NA | NA |
| ENSG000002 | 9.64518435 | -1.089611853 | 0.3465148 | -3.1444887 | 0.00166377 | 0.00849398 | MTND6P20 | 106478952 |
| ENSG000001 | 295.898463 | 0.482017916 | 0.15330835 | 3.14410736 | 0.00166594 | 0.00850399 | PDE6D | 5147 |

| | | | | | | | | | |
|---|---|---|---|---|---|---|---|---|---|
| ENSG000001 | 756.014481 | 0.382541904 | 0.12167762 | 3.14389713 | 0.00166714 | 0.00850903 | CASD1 | | 64921 |
| ENSG000001 | 25679.1111 | 0.394314194 | 0.12544356 | 3.1433594 | 0.00167021 | 0.0085236 | NA | NA | |
| ENSG000001 | 216.580481 | 0.462872802 | 0.14725627 | 3.14331463 | 0.00167046 | 0.00852383 | RCBTB2 | | 1102 |
| ENSG000001 | 901.209718 | 0.427385416 | 0.13596948 | 3.14324521 | 0.00167086 | 0.00852478 | PCYOX1 | | 51449 |
| ENSG000002 | 7.19564931 | -1.137567473 | 0.36193554 | -3.1430112 | 0.00167219 | 0.00853053 | NA | NA | |
| ENSG000001 | 14.2074304 | -1.013103319 | 0.32237163 | -3.1426566 | 0.00167422 | 0.00853979 | MEFV | | 4210 |
| ENSG000000 | 3496.29653 | 0.3925048 | 0.12491 | 3.14230084 | 0.00167626 | 0.0085491 | OSBPL8 | | 114882 |
| ENSG000002 | 247.876931 | 0.503971449 | 0.16038515 | 3.14225759 | 0.0016765 | 0.00854929 | HAUS5 | | 23354 |
| ENSG000002 | 8.37498216 | -1.122324799 | 0.3571765 | -3.1422135 | 0.00167676 | 0.0085495 | NA | NA | |
| ENSG000001 | 361.177012 | 0.404790238 | 0.12883041 | 3.14203956 | 0.00167775 | 0.00855351 | ODF2 | | 4957 |
| ENSG000001 | 373.160742 | 0.421437742 | 0.13414102 | 3.14175144 | 0.00167941 | 0.00855945 | TTC5 | | 91875 |
| ENSG000001 | 520.842391 | 0.354241213 | 0.1127537 | 3.14172579 | 0.00167955 | 0.00855945 | NA | NA | |
| ENSG000002 | 3.69652884 | -1.186760302 | 0.37774129 | -3.1417277 | 0.00167954 | 0.00855945 | NA | NA | |
| ENSG000001 | 818.816716 | 0.41606016 | 0.13243485 | 3.14162143 | 0.00168015 | 0.00856142 | NA | NA | |
| ENSG000001 | 246.845347 | 0.477243166 | 0.15193332 | 3.14113571 | 0.00168294 | 0.00857456 | FGF12 | | 2257 |
| ENSG000001 | 144.429956 | 0.488213522 | 0.15544592 | 3.14072913 | 0.00168528 | 0.00858539 | ALG12 | | 79087 |
| ENSG000002 | 63.2191724 | -0.622181946 | 0.19813967 | -3.140118 | 0.0016888 | 0.00860224 | LOC101927636 | | 101927636 |
| ENSG000001 | 1239.87944 | 0.360815943 | 0.11490963 | 3.13999755 | 0.00168949 | 0.0086047 | SNX3 | | 8724 |
| ENSG000001 | 12.4990837 | -1.007382094 | 0.32085321 | -3.1396977 | 0.00169122 | 0.00861243 | TTC24 | | 164118 |
| ENSG000001 | 16.5205687 | -0.958393034 | 0.30526312 | -3.1395638 | 0.001692 | 0.00861528 | FGA | | 2243 |
| ENSG000001 | 59.6138864 | 0.628689137 | 0.20025761 | 3.13940202 | 0.00169293 | 0.00861896 | HNMT | | 3176 |
| ENSG000001 | 7.01518095 | -1.15129255 | 0.36675366 | -3.139144 | 0.00169442 | 0.00862547 | FCGR1A | | 2209 |
| ENSG000001 | 17.4368876 | -0.946976404 | 0.30167289 | -3.1390835 | 0.00169477 | 0.00862617 | MS4A3 | | 932 |
| ENSG000001 | 213.301225 | 0.491412869 | 0.15656088 | 3.13879728 | 0.00169643 | 0.00863351 | NA | NA | |
| ENSG000001 | 97.3599363 | 0.543913504 | 0.17329202 | 3.13871058 | 0.00169693 | 0.00863498 | GAL3ST3 | | 89792 |
| ENSG000001 | 12.2817919 | -1.018529245 | 0.32451972 | -3.1385743 | 0.00169772 | 0.00863792 | RBPJL | | 11317 |
| ENSG000001 | 71.2119633 | -0.606434071 | 0.19323722 | -3.1382881 | 0.00169938 | 0.00864458 | WDR78 | | 79819 |
| ENSG000002 | 7.02157843 | -1.148484064 | 0.36596031 | -3.1382749 | 0.00169945 | 0.00864458 | NA | NA | |
| ENSG000001 | 289.998039 | 0.564137877 | 0.17979856 | 3.13761065 | 0.00170331 | 0.0086631 | SF3A2 | | 8175 |

| | | | | | | | | |
|---|---|---|---|---|---|---|---|---|
| ENSG000001 | 4611.05938 | 0.418940568 | 0.13355507 | 3.1368377 | 0.00170781 | 0.00868488 | APEX1 | 328 |
| ENSG000001 | 379.234177 | 0.495487062 | 0.15796067 | 3.13677492 | 0.00170817 | 0.00868565 | KIAA1211 | 57482 |
| ENSG000002 | 13.1601486 | -1.02415559 | 0.32658942 | -3.1359117 | 0.00171321 | 0.00871017 | LOC100505795 | 100505795 |
| ENSG000001 | 115.715017 | 0.572444933 | 0.18256518 | 3.1355647 | 0.00171524 | 0.00871938 | NA | NA |
| ENSG000002 | 10.4871483 | -1.059192805 | 0.3378151 | -3.1354217 | 0.00171607 | 0.00872254 | LOC101928748 | 101928748 |
| ENSG000001 | 686.801622 | 0.409581215 | 0.13064142 | 3.13515576 | 0.00171763 | 0.00872936 | MB21D2 | 151963 |
| ENSG000002 | 47.6908256 | -0.690498699 | 0.22026654 | -3.1348325 | 0.00171952 | 0.00873789 | LOC100631378 | 100631378 |
| ENSG000001 | 31.4270692 | -0.784921869 | 0.25039144 | -3.1347791 | 0.00171984 | 0.00873838 | CASP4 | 837 |
| ENSG000002 | 12.498892 | -1.036295791 | 0.33061824 | -3.1344181 | 0.00172195 | 0.00874805 | NA | NA |
| ENSG000001 | 1892.42753 | 0.387509008 | 0.12363705 | 3.13424672 | 0.00172296 | 0.00875206 | CASP8AP2 | 9994 |
| ENSG000001 | 13.1367273 | -1.037399702 | 0.33100064 | -3.134132 | 0.00172363 | 0.00875439 | PRSS55 | 203074 |
| ENSG000000 | 1893.24251 | 0.49336237 | 0.15743831 | 3.13368687 | 0.00172625 | 0.00876647 | CSNK2A2 | 1459 |
| ENSG000002 | 35.2860694 | -0.847760516 | 0.27053419 | -3.1336539 | 0.00172644 | 0.00876647 | NA | NA |
| ENSG000000 | 61.3904273 | 0.614986334 | 0.19626576 | 3.13343677 | 0.00172772 | 0.00877186 | DEF6 | 50619 |
| ENSG000001 | 20.5588602 | -0.894756868 | 0.28555811 | -3.1333618 | 0.00172816 | 0.008773 | GCOM1 | 145781 |
| ENSG000001 | 1733.25223 | 0.358295834 | 0.11435902 | 3.13307896 | 0.00172983 | 0.00878036 | GNAQ | 2776 |
| ENSG000002 | 53.427023 | 0.635801251 | 0.20295999 | 3.13264326 | 0.0017324 | 0.0087923 | FOXL2NB | 401089 |
| ENSG000001 | 202.199608 | 0.473467952 | 0.15114445 | 3.13255258 | 0.00173293 | 0.00879392 | PPP1R21 | 129285 |
| ENSG000002 | 8.3651779 | -1.118336526 | 0.35704188 | -3.1322278 | 0.00173485 | 0.00880255 | NA | NA |
| ENSG000001 | 69.7558237 | 0.602823784 | 0.19247144 | 3.13201688 | 0.0017361 | 0.00880778 | ARHGAP39 | 80728 |
| ENSG000001 | 591.874483 | 0.45147152 | 0.14417144 | 3.13149069 | 0.00173921 | 0.00882247 | ZNF384 | 171017 |
| ENSG000001 | 24.842034 | -0.846772965 | 0.27044449 | -3.1310417 | 0.00174187 | 0.00883486 | PEAR1 | 375033 |
| ENSG000001 | 30.0331182 | -0.799059228 | 0.25522154 | -3.1308456 | 0.00174304 | 0.00883966 | SRPX2 | 27286 |
| ENSG000001 | 372.770443 | 0.443835745 | 0.14176871 | 3.1307031 | 0.00174388 | 0.00884284 | HMOX2 | 3163 |
| ENSG000002 | 26.1934744 | -0.824640384 | 0.26342298 | -3.13048 | 0.00174521 | 0.00884846 | LINC00506 | 100846978 |
| ENSG000001 | 2513.28161 | 0.327402129 | 0.10459676 | 3.13013647 | 0.00174725 | 0.0088572 | HMGA2 | 8091 |
| ENSG000002 | 25.8802771 | -0.82369158 | 0.26315046 | -3.1301164 | 0.00174737 | 0.0088572 | KCNU1 | 157855 |
| ENSG000001 | 709.537487 | 0.420349547 | 0.1343051 | 3.12981075 | 0.00174919 | 0.00886531 | AEBP2 | 121536 |
| ENSG000002 | 3.62101187 | -1.183124889 | 0.37803659 | -3.1296571 | 0.0017501 | 0.00886774 | NA | NA |

| | | | | | | | | |
|---|---|---|---|---|---|---|---|---|
| ENSG000002 | 44.5175462 | -0.715349049 | 0.22856842 | -3.1296932 | 0.00174989 | 0.00886774 | OVCH1-AS1 | 101055625 |
| ENSG000002 | 20.3745582 | -0.909561514 | 0.29063354 | -3.1295821 | 0.00175055 | 0.00886889 | NA | NA |
| ENSG000002 | 400.617072 | 0.600282357 | 0.19181805 | 3.12943631 | 0.00175142 | 0.00887219 | SGK223 | 157285 |
| ENSG000001 | 38.8595788 | -0.737848937 | 0.23579199 | -3.1292367 | 0.00175261 | 0.00887711 | OAS3 | 4940 |
| ENSG000001 | 198.332578 | 0.439473259 | 0.14044453 | 3.12915898 | 0.00175307 | 0.00887835 | ETFDH | 2110 |
| ENSG000000 | 318.095396 | 0.414246482 | 0.1323926 | 3.12892463 | 0.00175447 | 0.008881 | MRPL22 | 29093 |
| ENSG000001 | 609.864695 | 0.417485403 | 0.13342879 | 3.12890047 | 0.00175462 | 0.008881 | MIEF1 | 54471 |
| ENSG000001 | 413.171865 | 0.420325256 | 0.13433696 | 3.12888765 | 0.00175469 | 0.008881 | SNRPN | 6638 |
| ENSG000001 | 281.32723 | 0.465074176 | 0.14863808 | 3.12890333 | 0.0017546 | 0.008881 | UBE2J2 | 118424 |
| ENSG000002 | 38.4809776 | 0.81781041 | 0.26137387 | 3.12889123 | 0.00175467 | 0.008881 | LOC401585 | 401585 |
| ENSG000002 | 89.7980145 | 0.56544819 | 0.18072088 | 3.1288481 | 0.00175493 | 0.00888109 | DXO | 1797 |
| ENSG000001 | 515.194376 | 0.372072697 | 0.1189201 | 3.12876208 | 0.00175544 | 0.00888258 | FBXL4 | 26235 |
| ENSG000002 | 726.434139 | 0.3839774 | 0.1227308 | 3.12861474 | 0.00175632 | 0.00888593 | MLLT11 | 10962 |
| ENSG000001 | 1200.00252 | 0.377982231 | 0.12082714 | 3.12828923 | 0.00175827 | 0.00889466 | TPGS2 | 25941 |
| ENSG000001 | 327.368632 | 0.411339591 | 0.13149262 | 3.12823319 | 0.00175861 | 0.00889525 | OMA1 | 115209 |
| ENSG000001 | 808.480905 | 0.383525684 | 0.12260313 | 3.12818837 | 0.00175887 | 0.0088955 | DERL1 | 79139 |
| ENSG000001 | 278.317256 | 0.463640083 | 0.14822039 | 3.12804523 | 0.00175973 | 0.00889872 | IER3IP1 | 51124 |
| ENSG000001 | 268.211737 | 0.418794926 | 0.13390718 | 3.12750163 | 0.00176299 | 0.00891408 | PSMG4 | 389362 |
| ENSG000001 | 360.83884 | 0.421036788 | 0.13463002 | 3.12736185 | 0.00176383 | 0.00891721 | E2F8 | 79733 |
| ENSG000001 | 103.793569 | 0.607459858 | 0.19425292 | 3.12715954 | 0.00176504 | 0.00892125 | HS1BP3 | 64342 |
| ENSG000002 | 12.1630904 | -1.035202782 | 0.33103658 | -3.1271553 | 0.00176507 | 0.00892125 | NA | NA |
| ENSG000002 | 13.1617308 | -1.021171615 | 0.32659365 | -3.1267344 | 0.0017676 | 0.00893292 | NA | NA |
| ENSG000001 | 1765.57185 | 0.325075116 | 0.10397718 | 3.12640825 | 0.00176956 | 0.00894172 | PREPL | 9581 |
| ENSG000001 | 8.95716647 | -1.100545266 | 0.3520455 | -3.1261449 | 0.00177114 | 0.00894862 | SFTPA1 | 653509 |
| ENSG000002 | 6.29818623 | -1.156614715 | 0.36999225 | -3.1260512 | 0.00177171 | 0.00895035 | NA | NA |
| ENSG000002 | 8.55336956 | -1.123471106 | 0.35940206 | -3.1259451 | 0.00177235 | 0.00895247 | LOC105376603 | 105376603 |
| ENSG000000 | 1895.71581 | 0.478587354 | 0.15311742 | 3.12562309 | 0.00177429 | 0.00896116 | PCNP | 57092 |
| ENSG000001 | 489.700716 | 0.497186284 | 0.15910118 | 3.12496925 | 0.00177824 | 0.00897998 | FAM83H | 286077 |
| ENSG000001 | 50.1310579 | -0.725119269 | 0.23206028 | -3.1247022 | 0.00177985 | 0.00898702 | SV2C | 22987 |

| | | | | | | | | |
|---|---|---|---|---|---|---|---|---|
| ENSG000001 | 216.597736 | 0.528164559 | 0.16903582 | 3.12457172 | 0.00178064 | 0.00898897 | PRKD2 | 25865 |
| ENSG000002 | 530.055916 | 0.41186658 | 0.13181565 | 3.12456509 | 0.00178068 | 0.00898897 | NA | NA |
| ENSG000002 | 10.6730912 | -1.045192662 | 0.33455254 | -3.1241511 | 0.00178319 | 0.00900051 | NA | NA |
| ENSG000001 | 911.769835 | 0.343637151 | 0.11001666 | 3.12350113 | 0.00178713 | 0.00901929 | CCDC66 | 285331 |
| ENSG000001 | 26.7694681 | -0.802736827 | 0.25701684 | -3.123285 | 0.00178845 | 0.00902479 | PRDM9 | 56979 |
| ENSG00000C | 16267.5344 | 0.369380204 | 0.11827227 | 3.12313459 | 0.00178936 | 0.00902828 | VCAN | 1462 |
| ENSG000001 | 28.5253475 | -0.787756222 | 0.25224508 | -3.1229795 | 0.0017903 | 0.00903192 | KCNC1 | 3746 |
| ENSG000002 | 6.37053298 | -1.158727246 | 0.37104122 | -3.122907 | 0.00179074 | 0.00903302 | NA | NA |
| ENSG000002 | 45.4616802 | -0.732053818 | 0.23442577 | -3.1227532 | 0.00179168 | 0.00903662 | NA | NA |
| ENSG000001 | 3.21696186 | -1.173281783 | 0.37578849 | -3.1221866 | 0.00179513 | 0.0090529 | PTGDR2 | 11251 |
| ENSG000001 | 347.848223 | 0.396389952 | 0.12696492 | 3.12204305 | 0.00179601 | 0.00905619 | CBWD2 | 150472 |
| ENSG000002 | 13.8069045 | -1.040298686 | 0.3332261 | -3.1219004 | 0.00179688 | 0.00905945 | NA | NA |
| ENSG000002 | 5.61168571 | -1.17909816 | 0.37769421 | -3.1218328 | 0.00179729 | 0.00906041 | NA | NA |
| ENSG000002 | 3.30119409 | -1.17093708 | 0.37514002 | -3.1213334 | 0.00180034 | 0.00907466 | TRIM64C | 646754 |
| ENSG00000C | 526.726959 | 0.410851374 | 0.13163114 | 3.12123237 | 0.00180096 | 0.00907664 | TRMT11 | 60487 |
| ENSG000002 | 15.0814972 | -0.964581164 | 0.30906443 | -3.1209712 | 0.00180256 | 0.00908357 | NA | NA |
| ENSG000001 | 31.0738397 | -0.81134252 | 0.25998324 | -3.1207494 | 0.00180391 | 0.00908873 | ASB15 | 142685 |
| ENSG000001 | 282.944024 | 0.387511955 | 0.12417347 | 3.1207306 | 0.00180403 | 0.00908873 | TMEM68 | 137695 |
| ENSG000001 | 67.1730901 | 0.61288645 | 0.19642662 | 3.12018026 | 0.0018074 | 0.0091046 | CLDN19 | 149461 |
| ENSG000002 | 3.20746309 | -1.171941203 | 0.37563587 | -3.1198863 | 0.00180921 | 0.00911256 | RN7SL465P | 106479392 |
| ENSG000001 | 708.721959 | 0.482806615 | 0.15475418 | 3.11982926 | 0.00180956 | 0.00911319 | TFCP2 | 7024 |
| ENSG000001 | 829.061047 | 0.400306501 | 0.12833285 | 3.11928316 | 0.00181292 | 0.00912897 | SDE2 | 163859 |
| ENSG000001 | 427.874138 | 0.454022164 | 0.14556291 | 3.11907874 | 0.00181417 | 0.00913417 | SIK2 | 23235 |
| ENSG000002 | 9.43027505 | -1.099799365 | 0.35262685 | -3.1188758 | 0.00181542 | 0.00913933 | FMR1-AS1 | 100126270 |
| ENSG000001 | 24.4893476 | -0.846207823 | 0.27133496 | -3.1186834 | 0.00181661 | 0.00914416 | CRX | 1406 |
| ENSG000001 | 191.975797 | 0.518263869 | 0.16619218 | 3.11846122 | 0.00181798 | 0.00914902 | SPATA2 | 9825 |
| ENSG000002 | 20.9144231 | 0.914188961 | 0.29315458 | 3.11845366 | 0.00181803 | 0.00914902 | NA | NA |
| ENSG00000C | 744.083884 | 0.451847034 | 0.14489631 | 3.11841645 | 0.00181826 | 0.00914904 | TRAF4 | 9618 |
| ENSG000001 | 317.250707 | 0.458672289 | 0.14709145 | 3.11827977 | 0.0018191 | 0.00915102 | UNK | 85451 |

| | | | | | | | | | |
|---|---|---|---|---|---|---|---|---|---|
| ENSG000002 | 16.9391489 | -0.948596486 | 0.304205 | -3.1182804 | 0.0018191 | 0.00915102 | LOC102723854 | | 102723854 |
| ENSG000001 | 116.640035 | 0.577993874 | 0.18536148 | 3.11819846 | 0.0018196 | 0.00915241 | SLC19A1 | | 6573 |
| ENSG000002 | 84.0640538 | 0.563879447 | 0.18084636 | 3.11800272 | 0.00182081 | 0.00915735 | STARD10 | | 10809 |
| ENSG000000 | 3333.39502 | 0.375020495 | 0.12027833 | 3.11793904 | 0.0018212 | 0.00915819 | HMMR | | 3161 |
| ENSG000000 | 419.940949 | 0.472952947 | 0.15170585 | 3.11756559 | 0.00182351 | 0.00916867 | NOP16 | | 51491 |
| ENSG000001 | 206.792925 | 0.504768471 | 0.16192423 | 3.11731276 | 0.00182508 | 0.00917426 | FAM117A | | 81558 |
| ENSG000001 | 27.5504092 | -0.832100708 | 0.26692729 | -3.1173309 | 0.00182497 | 0.00917426 | ENAM | | 10117 |
| ENSG000000 | 111.915019 | -0.544297327 | 0.17460997 | -3.1172179 | 0.00182567 | 0.00917552 | LMO3 | | 55885 |
| ENSG000001 | 29.9330279 | -0.789647078 | 0.25331941 | -3.1171993 | 0.00182578 | 0.00917552 | LRRTM2 | | 26045 |
| ENSG000001 | 51.5697465 | 0.687244096 | 0.22047293 | 3.11713591 | 0.00182617 | 0.00917635 | WBSCR27 | | 155368 |
| ENSG000001 | 60.6830921 | -0.613276306 | 0.19678988 | -3.1164016 | 0.00183073 | 0.0091981 | NLRC5 | | 84166 |
| ENSG000001 | 441.469915 | 0.553427307 | 0.17759785 | 3.11618242 | 0.00183209 | 0.0092038 | MYO18A | | 399687 |
| ENSG000002 | 7.13673679 | -1.129639644 | 0.36253996 | -3.1159038 | 0.00183382 | 0.00921136 | CCL18 | | 6362 |
| ENSG000001 | 969.490972 | 0.353514763 | 0.11346057 | 3.1157498 | 0.00183478 | 0.00921443 | SOS1 | | 6654 |
| ENSG000002 | 38.1042354 | -0.797051544 | 0.25581515 | -3.1157323 | 0.00183489 | 0.00921443 | MEG3 | | 55384 |
| ENSG000001 | 3057.91963 | 0.452479971 | 0.14522724 | 3.11566865 | 0.00183528 | 0.00921528 | ZNF721 | | 170960 |
| ENSG000001 | 643.70763 | 0.372572239 | 0.11958503 | 3.11554241 | 0.00183607 | 0.00921694 | ZBTB38 | | 253461 |
| ENSG000002 | 27.4242808 | -0.842992843 | 0.27057657 | -3.1155427 | 0.00183607 | 0.00921694 | NA | NA | |
| ENSG000001 | 258.885555 | 0.489862925 | 0.15725824 | 3.11502232 | 0.00183931 | 0.00923207 | MDFI | | 4188 |
| ENSG000001 | 1921.66235 | 0.385874436 | 0.12387698 | 3.11498103 | 0.00183957 | 0.00923222 | PAXBP1 | | 94104 |
| ENSG000001 | 56.8821273 | 0.663103578 | 0.21288805 | 3.11479943 | 0.0018407 | 0.00923676 | RELB | | 5971 |
| ENSG000001 | 217.333211 | 0.473194731 | 0.1519348 | 3.11445911 | 0.00184282 | 0.00924627 | TPBG | | 7162 |
| ENSG000001 | 144.260766 | 0.506805642 | 0.16277642 | 3.11350771 | 0.00184878 | 0.00927287 | TCHP | | 84260 |
| ENSG000001 | 37.5855993 | -0.787633917 | 0.25296967 | -3.1135508 | 0.00184851 | 0.00927287 | KIF26B | | 55083 |
| ENSG000002 | 10.9643612 | -1.061777859 | 0.34102365 | -3.1135021 | 0.00184881 | 0.00927287 | LOC283440 | | 283440 |
| ENSG000001 | 953.208976 | 0.34882757 | 0.11204677 | 3.1132319 | 0.00185051 | 0.00928022 | UHRF1BP1L | | 23074 |
| ENSG000002 | 722.004969 | 0.436133863 | 0.14009214 | 3.11319287 | 0.00185075 | 0.0092803 | NA | NA | |
| ENSG000002 | 7.54444802 | -1.121137317 | 0.36013476 | -3.113105 | 0.0018513 | 0.00928192 | NA | NA | |
| ENSG000001 | 328.706934 | 0.459741569 | 0.1477074 | 3.11251546 | 0.001855 | 0.00929932 | ARL6 | | 84100 |

| | | | | | | | | | |
|---|---|---|---|---|---|---|---|---|---|
| ENSG000001 | 484.109278 | 0.467555134 | 0.15022321 | 3.11240277 | 0.00185571 | 0.00929943 | SLC22A23 | | 63027 |
| ENSG000002 | 9.44823203 | -1.072566104 | 0.34460875 | -3.1124169 | 0.00185562 | 0.00929943 | LHFPL3-AS2 | | 723809 |
| ENSG000002 | 7.30528363 | -1.132976023 | 0.36401307 | -3.1124597 | 0.00185535 | 0.00929943 | LOC646268 | | 646268 |
| ENSG000000 | 567.326348 | 0.438399357 | 0.1408766 | 3.11193882 | 0.00185863 | 0.0093129 | TRIP6 | | 7205 |
| ENSG000001 | 559.355877 | 0.393798915 | 0.12654692 | 3.11188061 | 0.001859 | 0.00931359 | MGAT2 | | 4247 |
| ENSG000001 | 7.18669241 | -1.127481719 | 0.36233121 | -3.1117433 | 0.00185986 | 0.00931677 | NA | NA | |
| ENSG000000 | 1225.24184 | 0.386433278 | 0.12419143 | 3.11159367 | 0.0018608 | 0.00931934 | EIF2AK1 | | 27102 |
| ENSG000002 | 371.582932 | 0.422834249 | 0.13589014 | 3.11158881 | 0.00186083 | 0.00931934 | ZNF486 | | 90649 |
| ENSG000001 | 991.993764 | 0.843818865 | 0.27119442 | 3.11149049 | 0.00186145 | 0.00932014 | H1F0 | | 3005 |
| ENSG000002 | 8.47353717 | -1.095203651 | 0.35198293 | -3.1115249 | 0.00186124 | 0.00932014 | NA | NA | |
| ENSG000001 | 2274.33491 | 0.329105644 | 0.10577558 | 3.11135741 | 0.00186229 | 0.0093232 | USP8 | | 9101 |
| ENSG000001 | 211.802031 | 0.441065539 | 0.14176369 | 3.11127295 | 0.00186283 | 0.00932471 | ZNF354B | | 117608 |
| ENSG000001 | 1595.90263 | 0.387281399 | 0.12448193 | 3.11114561 | 0.00186363 | 0.00932758 | PFDN4 | | 5203 |
| ENSG000001 | 129.648401 | 0.536216419 | 0.17236122 | 3.11100379 | 0.00186453 | 0.00933091 | C14orf142 | | 84520 |
| ENSG000002 | 29.0976929 | -0.769400615 | 0.24732631 | -3.1108725 | 0.00186535 | 0.00933391 | NA | NA | |
| ENSG000001 | 295.042455 | 0.449594963 | 0.14454771 | 3.11035679 | 0.00186861 | 0.00934907 | NA | NA | |
| ENSG000002 | 51.3838664 | 0.638598067 | 0.20533344 | 3.11005385 | 0.00187053 | 0.00935751 | NA | NA | |
| ENSG000001 | 396.401682 | 0.486981153 | 0.15660266 | 3.1096608 | 0.00187302 | 0.00936881 | ARHGAP1 | | 392 |
| ENSG000001 | 36.0671716 | 0.730501892 | 0.23494816 | 3.1092046 | 0.00187592 | 0.00938213 | CDX4 | | 1046 |
| ENSG000002 | 10.4749536 | -1.055362605 | 0.33947079 | -3.1088466 | 0.00187819 | 0.00939235 | AOC4P | | 90586 |
| ENSG000001 | 59.6584347 | 0.647971276 | 0.20847422 | 3.10816023 | 0.00188256 | 0.00941303 | SOX3 | | 6658 |
| ENSG000001 | 26.5813963 | -0.806791643 | 0.25957752 | -3.1080952 | 0.00188297 | 0.00941394 | CA10 | | 56934 |
| ENSG000002 | 19.5529342 | -0.898303597 | 0.28902429 | -3.1080558 | 0.00188323 | 0.00941404 | PRCD | | 768206 |
| ENSG000001 | 831.766033 | 0.371924159 | 0.11967859 | 3.10769177 | 0.00188555 | 0.00942448 | HEATR3 | | 55027 |
| ENSG000001 | 7.73673666 | -1.12928749 | 0.36341426 | -3.1074386 | 0.00188716 | 0.00943139 | CYLC2 | | 1539 |
| ENSG000002 | 94.1769664 | 0.55303665 | 0.17798895 | 3.10714042 | 0.00188907 | 0.00943975 | TMEM200B | | 399474 |
| ENSG000002 | 3.73882409 | -1.172560427 | 0.37744105 | -3.1066055 | 0.00189249 | 0.00945568 | NA | NA | |
| ENSG000001 | 29.9343234 | -0.79008624 | 0.25434049 | -3.1064116 | 0.00189373 | 0.00946071 | IL7R | | 3575 |
| ENSG000001 | 76.2558079 | 0.565463892 | 0.18204046 | 3.10625385 | 0.00189474 | 0.0094646 | GXYLT2 | | 727936 |

| | | | | | | | | |
|---|---|---|---|---|---|---|---|---|
| ENSG00000C | 412.660612 | 0.43783127 | 0.14097611 | 3.10571263 | 0.00189821 | 0.00947843 | TRAM2 | 9697 |
| ENSG00000C | 26.2051282 | -0.831959898 | 0.26788007 | -3.1057178 | 0.00189818 | 0.00947843 | CAMK2A | 815 |
| ENSG00000C | 659.596405 | 0.37255595 | 0.11995748 | 3.1057334 | 0.00189808 | 0.00947843 | WDR70 | 55100 |
| ENSG000002 | 1984.718 | 0.359577564 | 0.11578247 | 3.10563052 | 0.00189874 | 0.0094799 | ZNF253 | 56242 |
| ENSG000001 | 474.305929 | 0.390708535 | 0.12581326 | 3.10546397 | 0.00189981 | 0.00948407 | GOSR2 | 9570 |
| ENSG000002 | 115.481448 | 0.547909415 | 0.17644068 | 3.10534625 | 0.00190056 | 0.00948668 | VLDLR-AS1 | 401491 |
| ENSG00000C | 887.46775 | 0.35607801 | 0.11467776 | 3.10503111 | 0.00190259 | 0.00949445 | C5orf22 | 55322 |
| ENSG000002 | 15.174983 | -0.946561953 | 0.30484474 | -3.1050625 | 0.00190239 | 0.00949445 | NA | NA |
| ENSG000002 | 7.67955834 | -1.124555237 | 0.36220175 | -3.1047759 | 0.00190423 | 0.00950148 | NA | NA |
| ENSG000001 | 1356.28288 | 0.47470246 | 0.15295669 | 3.10350902 | 0.0019124 | 0.00954108 | ZNF609 | 23060 |
| ENSG000001 | 28.7647539 | -0.800401762 | 0.25791129 | -3.1033994 | 0.00191311 | 0.00954344 | ERICH6B | 220081 |
| ENSG000001 | 386.524176 | 0.468856512 | 0.15108688 | 3.10322459 | 0.00191424 | 0.00954711 | AJUBA | 84962 |
| ENSG000001 | 154.815903 | 0.517910848 | 0.16689504 | 3.10321296 | 0.00191432 | 0.00954711 | VPS37C | 55048 |
| ENSG000001 | 14.944715 | -0.977529938 | 0.31502779 | -3.1029959 | 0.00191572 | 0.00955081 | HOXA13 | 3209 |
| ENSG000001 | 8.6199075 | -1.101057547 | 0.35483763 | -3.1029898 | 0.00191576 | 0.00955081 | TSPAN16 | 26526 |
| ENSG000001 | 656.297648 | 0.432828284 | 0.13948754 | 3.10298887 | 0.00191577 | 0.00955081 | KLKB1 | 3818 |
| ENSG00000C | 2581.8634 | 0.350865093 | 0.11307988 | 3.10280751 | 0.00191694 | 0.00955549 | KIAA0020 | 9933 |
| ENSG000002 | 24.6816201 | -0.838922997 | 0.27039159 | -3.1026223 | 0.00191814 | 0.0095603 | ADAMTS9-AS1 | 101929335 |
| ENSG000002 | 16.9183863 | -0.934523217 | 0.30121168 | -3.1025464 | 0.00191863 | 0.00956158 | NA | NA |
| ENSG000001 | 327.552897 | 0.540431984 | 0.17419537 | 3.10244746 | 0.00191928 | 0.0095636 | SBK1 | 388228 |
| ENSG000001 | 75.0277036 | -0.582233302 | 0.1876992 | -3.1019487 | 0.00192251 | 0.00957855 | LDLRAD4 | 753 |
| ENSG000001 | 195.746465 | 0.466719813 | 0.15047281 | 3.10168877 | 0.0019242 | 0.00958579 | C17orf53 | 78995 |
| ENSG000001 | 19.7365558 | -0.889831584 | 0.28689178 | -3.101628 | 0.0019246 | 0.00958658 | ITLN1 | 55600 |
| ENSG000001 | 61.9613105 | -0.631550656 | 0.2036235 | -3.1015607 | 0.00192503 | 0.00958758 | CD163L1 | 283316 |
| ENSG00000C | 69.8141268 | -0.643799843 | 0.20761461 | -3.1009371 | 0.00192909 | 0.00960388 | MYO3B | 140469 |
| ENSG000001 | 24.3523071 | 0.839756868 | 0.27080424 | 3.10097383 | 0.00192885 | 0.00960388 | NA | NA |
| ENSG000002 | 16.9559104 | -0.938950993 | 0.30279831 | -3.1009123 | 0.00192925 | 0.00960388 | NA | NA |
| ENSG000002 | 644.727025 | 0.423015425 | 0.13641627 | 3.10091618 | 0.00192923 | 0.00960388 | SEC22B | 9554 |
| ENSG000001 | 10.7213623 | -1.044984119 | 0.33700214 | -3.1008234 | 0.00192983 | 0.0096044 | FAM46D | 169966 |

| | | | | | | | | |
|---|---|---|---|---|---|---|---|---|
| ENSG000002 | 770.499365 | 0.406894416 | 0.1312202 | 3.10085201 | 0.00192965 | 0.0096044 | NA | NA |
| ENSG000001 | 647.853103 | 0.419160137 | 0.1351857 | 3.10062488 | 0.00193113 | 0.00960966 | CSTF3 | 1479 |
| ENSG000001 | 2433.45981 | 0.337868918 | 0.1089757 | 3.10040599 | 0.00193256 | 0.00961261 | RSRC2 | 65117 |
| ENSG000001 | 2127.49292 | 0.464090218 | 0.14968339 | 3.10047904 | 0.00193208 | 0.00961261 | IREB2 | 3658 |
| ENSG000001 | 15.9028798 | -0.965956919 | 0.31155651 | -3.1004228 | 0.00193245 | 0.00961261 | FLG | 2312 |
| ENSG000001 | 15.4757103 | -0.970334029 | 0.31297434 | -3.1003629 | 0.00193284 | 0.00961261 | SPON2 | 10417 |
| ENSG000002 | 2.51609927 | -1.120104924 | 0.3612831 | -3.1003524 | 0.00193291 | 0.00961261 | MYLK-AS2 | 100873940 |
| ENSG000002 | 6.69503732 | -1.148677148 | 0.37056418 | -3.0998062 | 0.00193647 | 0.00962917 | NA | NA |
| ENSG00000C | 2258.68467 | 0.506254585 | 0.16333844 | 3.099421 | 0.00193899 | 0.00964052 | NUP160 | 23279 |
| ENSG000001 | 79.7329161 | -0.612035079 | 0.1975131 | -3.0987062 | 0.00194368 | 0.00966262 | SPAG17 | 200162 |
| ENSG000002 | 160.968483 | 0.527010852 | 0.1700852 | 3.09851095 | 0.00194496 | 0.00966781 | TMEM185B | 79134 |
| ENSG000001 | 1078.40099 | 0.331551676 | 0.10701949 | 3.09804937 | 0.00194799 | 0.00968169 | DOCK7 | 85440 |
| ENSG000001 | 1459.55958 | 0.353559917 | 0.11413366 | 3.09777085 | 0.00194982 | 0.00968961 | DDX10 | 1662 |
| ENSG000001 | 1067.69558 | 0.431183424 | 0.1391954 | 3.0976844 | 0.00195039 | 0.00969125 | IPO8 | 10526 |
| ENSG000002 | 39.7245075 | -0.711099748 | 0.22956469 | -3.0976008 | 0.00195094 | 0.00969279 | NA | NA |
| ENSG000001 | 417.078918 | 0.558635449 | 0.18035162 | 3.0974796 | 0.00195174 | 0.00969557 | CSTB | 1476 |
| ENSG000001 | 18.4644358 | -0.900244227 | 0.29064635 | -3.097387 | 0.00195235 | 0.00969741 | ALDH8A1 | 64577 |
| ENSG000001 | 328.783265 | 0.402958994 | 0.13009805 | 3.09734834 | 0.0019526 | 0.00969749 | Mar-08 | 220972 |
| ENSG000001 | 162.429907 | 0.50635988 | 0.16349549 | 3.09708769 | 0.00195432 | 0.00970246 | FSD1 | 79187 |
| ENSG000001 | 1070.85256 | 0.336622291 | 0.10868773 | 3.0971507 | 0.00195391 | 0.00970246 | PPIL4 | 85313 |
| ENSG000001 | 8082.57961 | 0.390975716 | 0.12623914 | 3.09710378 | 0.00195421 | 0.00970246 | SLC39A10 | 57181 |
| ENSG000001 | 17.8792203 | -0.911511212 | 0.29432682 | -3.0969356 | 0.00195532 | 0.00970617 | ITIH6 | 347365 |
| ENSG000001 | 159.798948 | 0.454402686 | 0.14672817 | 3.09690156 | 0.00195555 | 0.00970617 | B4GAT1 | 11041 |
| ENSG000002 | 19.006342 | -0.905381777 | 0.29236681 | -3.0967325 | 0.00195666 | 0.00971052 | NA | NA |
| ENSG000001 | 16.4965275 | -0.93259634 | 0.30116333 | -3.0966464 | 0.00195723 | 0.00971215 | OR8A1 | 390275 |
| ENSG000002 | 18.3977249 | -0.93570114 | 0.30217023 | -3.0966026 | 0.00195752 | 0.0097124 | NA | NA |
| ENSG000002 | 8.67756976 | -1.098595097 | 0.35478487 | -3.0965106 | 0.00195813 | 0.00971423 | NA | NA |
| ENSG000001 | 775.163077 | 0.414007968 | 0.13370558 | 3.09641503 | 0.00195876 | 0.00971498 | FMR1 | 2332 |
| ENSG000001 | 31.0094798 | -0.804033944 | 0.25966347 | -3.0964461 | 0.00195855 | 0.00971498 | C12orf40 | 283461 |

| | | | | | | | | | |
|---|---|---|---|---|---|---|---|---|---|
| ENSG000002 | 6.86397564 | -1.131208535 | 0.3653712 | -3.0960528 | 0.00196115 | 0.00972567 | NA | NA | |
| ENSG000002 | 6.31825336 | -1.149237839 | 0.37121945 | -3.0958449 | 0.00196253 | 0.0097313 | NA | NA | |
| ENSG000001 | 170.840782 | 0.48590065 | 0.15695708 | 3.095755 | 0.00196312 | 0.00973306 | HMGCL | | 3155 |
| ENSG000000 | 47.9234017 | -0.646007049 | 0.20868084 | -3.0956701 | 0.00196369 | 0.00973465 | SEC31B | | 25956 |
| ENSG000000 | 1066.53177 | 0.476444486 | 0.15392525 | 3.09529779 | 0.00196615 | 0.00974569 | ERBB3 | | 2065 |
| ENSG000002 | 239.599585 | 0.445598355 | 0.1439696 | 3.09508641 | 0.00196756 | 0.00975145 | HEXA | | 3073 |
| ENSG000002 | 5.82493746 | 1.162889456 | 0.37573014 | 3.09501247 | 0.00196805 | 0.00975269 | CHKB-AS1 | | 100144603 |
| ENSG000001 | 26.2968427 | -0.794588505 | 0.25674742 | -3.0948256 | 0.00196929 | 0.0097569 | CCDC92 | | 80212 |
| ENSG000001 | 1415.03817 | 0.428910652 | 0.13859022 | 3.09481185 | 0.00196938 | 0.0097569 | ZNF608 | | 57507 |
| ENSG000001 | 27.5234223 | -0.792266735 | 0.2560317 | -3.0944087 | 0.00197206 | 0.00976898 | SNTN | | 132203 |
| ENSG000001 | 241.929308 | 0.472006768 | 0.15254368 | 3.09424016 | 0.00197318 | 0.00977334 | TRIM14 | | 9830 |
| ENSG000002 | 20.23388 | -0.897841814 | 0.29021096 | -3.0937557 | 0.0019764 | 0.00978811 | NA | NA | |
| ENSG000001 | 477.727376 | 0.46675279 | 0.15087201 | 3.09370044 | 0.00197677 | 0.00978874 | ISYNA1 | | 51477 |
| ENSG000002 | 13.9456369 | -1.005280088 | 0.3249808 | -3.0933522 | 0.00197909 | 0.00979904 | PNLIPRP3 | | 119548 |
| ENSG000002 | 170.466676 | 0.510909524 | 0.16516628 | 3.09330401 | 0.00197941 | 0.00979943 | NA | NA | |
| ENSG000001 | 9.85537304 | -1.066798091 | 0.34490995 | -3.0929757 | 0.0019816 | 0.00980789 | PLA2G2C | | 391013 |
| ENSG000002 | 20.2010456 | -0.877876187 | 0.28382651 | -3.0930028 | 0.00198142 | 0.00980789 | NA | NA | |
| ENSG000001 | 66.908835 | 0.642298232 | 0.20767077 | 3.09286771 | 0.00198232 | 0.00981026 | HEXIM2 | | 124790 |
| ENSG000002 | 54.6670386 | -0.630293552 | 0.20384953 | -3.0919549 | 0.00198843 | 0.00983902 | COL28A1 | | 340267 |
| ENSG000002 | 57.6175332 | -0.626894055 | 0.20275193 | -3.0919264 | 0.00198862 | 0.00983902 | NA | NA | |
| ENSG000002 | 6.6956326 | -1.142656769 | 0.36957225 | -3.091836 | 0.00198923 | 0.00984081 | NA | NA | |
| ENSG000002 | 12.090034 | -1.024026056 | 0.33121061 | -3.091767 | 0.00198969 | 0.0098419 | BVES-AS1 | | 154442 |
| ENSG000001 | 255.47617 | 0.51249017 | 0.16577926 | 3.09140094 | 0.00199214 | 0.00985284 | TBC1D7 | | 51256 |
| ENSG000001 | 187.599359 | 0.451015614 | 0.14589941 | 3.09127782 | 0.00199297 | 0.00985328 | MFSD9 | | 84804 |
| ENSG000001 | 1166.88846 | 0.394786104 | 0.12770908 | 3.09129236 | 0.00199287 | 0.00985328 | BASP1 | | 10409 |
| ENSG000002 | 356.514786 | 0.379440014 | 0.12274674 | 3.09124304 | 0.0019932 | 0.00985328 | RUFY2 | | 55680 |
| ENSG000002 | 21.7798215 | -0.882574007 | 0.28550449 | -3.0912789 | 0.00199296 | 0.00985328 | LOC102724663 | | 102724663 |
| ENSG000001 | 169.194111 | 0.47453305 | 0.15352422 | 3.09093277 | 0.00199529 | 0.00986238 | PDCD2L | | 84306 |
| ENSG000001 | 439.419836 | 0.444672765 | 0.14388286 | 3.09051934 | 0.00199807 | 0.00987491 | PDCL3 | | 79031 |

| | | | | | | | | |
|---|---|---|---|---|---|---|---|---|
| ENSG000002 | 42.3085014 | -0.685935691 | 0.22196845 | -3.0902396 | 0.00199995 | 0.00988301 | NA | NA |
| ENSG000002 | 3.59687357 | -1.167482616 | 0.37780559 | -3.0901677 | 0.00200044 | 0.0098842 | NA | NA |
| ENSG000002 | 142.537872 | 0.524981263 | 0.1698962 | 3.09001175 | 0.00200149 | 0.00988698 | NA | NA |
| ENSG000002 | 1841.80682 | 0.394076884 | 0.12753239 | 3.09001416 | 0.00200147 | 0.00988698 | ZNF587B | 100293516 |
| ENSG000001 | 30.6736341 | -0.763020538 | 0.24693426 | -3.0899743 | 0.00200174 | 0.00988702 | CCR5 | 1234 |
| ENSG000001 | 5324.69757 | 0.36787262 | 0.11906964 | 3.08955847 | 0.00200454 | 0.00989967 | FXR1 | 8087 |
| ENSG000001 | 2625.3229 | 0.400943067 | 0.12978244 | 3.0893476 | 0.00200597 | 0.00990549 | ZNF92 | 168374 |
| ENSG000001 | 144.492685 | 0.524164007 | 0.16967345 | 3.0892518 | 0.00200661 | 0.00990695 | RDH10 | 157506 |
| ENSG000001 | 2839.08747 | 0.391981142 | 0.12688629 | 3.08923163 | 0.00200675 | 0.00990695 | NCOR1 | 9611 |
| ENSG000001 | 421.997406 | 0.445628838 | 0.14426502 | 3.08895978 | 0.00200859 | 0.00991481 | GABRA5 | 2558 |
| ENSG00000C | 1109.46291 | 0.385673104 | 0.12485694 | 3.08892002 | 0.00200886 | 0.00991493 | ANKIB1 | 54467 |
| ENSG000001 | 109.311193 | 0.554815818 | 0.17962861 | 3.08868293 | 0.00201046 | 0.00992164 | BRSK1 | 84446 |
| ENSG000001 | 375.664355 | 0.415824119 | 0.13463229 | 3.08859126 | 0.00201108 | 0.00992349 | TIMM9 | 26520 |
| ENSG000002 | 9.72814922 | -1.080854583 | 0.34997462 | -3.0883799 | 0.00201251 | 0.00992935 | OR5M1 | 390168 |
| ENSG000002 | 4.71013683 | -1.165769854 | 0.37748256 | -3.0882747 | 0.00201322 | 0.00993165 | NA | NA |
| ENSG000001 | 121.529825 | 0.539843265 | 0.1748096 | 3.08817861 | 0.00201387 | 0.00993366 | ZSCAN22 | 342945 |
| ENSG000001 | 24.8584993 | -0.822496167 | 0.26635059 | -3.0880208 | 0.00201494 | 0.00993652 | KCNA7 | 3743 |
| ENSG000001 | 195.373874 | 0.518487473 | 0.16790276 | 3.08802225 | 0.00201493 | 0.00993652 | TLDC1 | 57707 |
| ENSG000002 | 195.510216 | 0.427169759 | 0.13833693 | 3.08789384 | 0.0020158 | 0.00993955 | SEPT7P2 | 641977 |
| ENSG00000C | 39.6286952 | 0.730817208 | 0.23669057 | 3.08764822 | 0.00201747 | 0.00994535 | RASSF7 | 8045 |
| ENSG000001 | 143.44248 | 0.513054972 | 0.16616188 | 3.0876816 | 0.00201724 | 0.00994535 | KLF11 | 8462 |
| ENSG000001 | 35.5901186 | -0.749083376 | 0.24261813 | -3.0874996 | 0.00201848 | 0.00994821 | ZIC4 | 84107 |
| ENSG000002 | 7.44780227 | -1.129438178 | 0.36581105 | -3.0874906 | 0.00201854 | 0.00994821 | NA | NA |
| ENSG000001 | 7.71265408 | -1.120282741 | 0.3628819 | -3.0871827 | 0.00202063 | 0.00995731 | KRT71 | 112802 |
| ENSG000001 | 108.534053 | 0.588474547 | 0.19062167 | 3.08713345 | 0.00202097 | 0.00995775 | CTDSP1 | 58190 |
| ENSG000001 | 479.77632 | 0.350292002 | 0.11347257 | 3.08701921 | 0.00202175 | 0.00996037 | MED23 | 9439 |
| ENSG000002 | 191.227161 | 0.470461518 | 0.15241158 | 3.08678338 | 0.00202335 | 0.00996706 | POU5F1B | 5462 |
| ENSG000001 | 613.895194 | 0.393928083 | 0.12763718 | 3.08631134 | 0.00202657 | 0.00998082 | SLC4A1AP | 22950 |
| ENSG000001 | 6.09186987 | -1.139659227 | 0.36926378 | -3.0863011 | 0.00202663 | 0.00998082 | POU3F3 | 5455 |

| | | | | | | | | |
|---|---|---|---|---|---|---|---|---|
| ENSG000002 | 11.2477083 | -1.0430328 | 0.33798689 | -3.0860155 | 0.00202858 | 0.0099892 | NA | NA |
| ENSG000002 | 1410.32611 | 0.449444545 | 0.14564769 | 3.08583365 | 0.00202982 | 0.0099941 | PPP2R2A | 5520 |
| ENSG000002 | 230.913761 | 0.464265641 | 0.15045267 | 3.08579204 | 0.00203011 | 0.00999428 | MAPKAPK5-AS1 | 51275 |
| ENSG000002 | 613.697197 | 0.527446698 | 0.17094185 | 3.08553294 | 0.00203188 | 0.01000178 | UHRF1 | 29128 |
| ENSG00000C | 1793.2154 | 0.481513691 | 0.1560637 | 3.08536641 | 0.00203302 | 0.01000388 | SF3A1 | 10291 |
| ENSG000001 | 1187.36926 | 0.438263183 | 0.14204388 | 3.08540691 | 0.00203274 | 0.01000388 | HOOK1 | 51361 |
| ENSG000002 | 21.1361916 | -0.900372014 | 0.29182052 | -3.0853622 | 0.00203304 | 0.01000388 | TEX35 | 84066 |
| ENSG000001 | 23.8495457 | -0.824679093 | 0.26730486 | -3.0851631 | 0.00203441 | 0.01000936 | CHRM2 | 1129 |
| ENSG000002 | 9.22124089 | -1.098604521 | 0.35612419 | -3.0848916 | 0.00203626 | 0.01001729 | SH3PXD2A-AS1 | 100505839 |
| ENSG000001 | 101.367086 | 0.556709912 | 0.18048659 | 3.08449464 | 0.00203898 | 0.01002945 | B3GAT3 | 26229 |
| ENSG00000C | 1167.05863 | 0.338201673 | 0.10965036 | 3.08436438 | 0.00203988 | 0.01003141 | TAF2 | 6873 |
| ENSG000001 | 2758.08526 | 0.419425176 | 0.13598275 | 3.08439984 | 0.00203963 | 0.01003141 | SMG1 | 23049 |
| ENSG00000C | 1054.37135 | 0.410435016 | 0.13308125 | 3.0840934 | 0.00204173 | 0.01003568 | VPS13D | 55187 |
| ENSG000001 | 23.6491614 | -0.892463601 | 0.28937317 | -3.084127 | 0.0020415 | 0.01003568 | GABRA2 | 2555 |
| ENSG000002 | 32.9875901 | 0.758637767 | 0.24597864 | 3.08416116 | 0.00204127 | 0.01003568 | NA | NA |
| ENSG000002 | 13.4865369 | -0.982160049 | 0.31845386 | -3.0841518 | 0.00204133 | 0.01003568 | NA | NA |
| ENSG000001 | 26.8412573 | -0.78704763 | 0.2551991 | -3.0840534 | 0.00204201 | 0.01003582 | GPR61 | 83873 |
| ENSG000001 | 18.3841874 | -0.894713911 | 0.29013798 | -3.0837532 | 0.00204407 | 0.01004473 | CLEC1A | 51267 |
| ENSG000001 | 6539.94973 | 0.391927951 | 0.12709727 | 3.08368501 | 0.00204454 | 0.01004582 | SRP9 | 6726 |
| ENSG000001 | 670.242021 | 0.496569687 | 0.16104315 | 3.0834573 | 0.0020461 | 0.01005229 | RREB1 | 6239 |
| ENSG000002 | 14.7648001 | -0.943937283 | 0.30615947 | -3.0831556 | 0.00204818 | 0.01006127 | NA | NA |
| ENSG00000C | 145.104489 | 0.488513589 | 0.15845694 | 3.08294211 | 0.00204965 | 0.01006727 | HAUS4 | 54930 |
| ENSG000001 | 176.140049 | 0.488226898 | 0.15836802 | 3.08286299 | 0.0020502 | 0.01006873 | RNF166 | 115992 |
| ENSG000001 | 610.01262 | 0.383760248 | 0.12449 | 3.08265922 | 0.0020516 | 0.0100744 | ZNF791 | 163049 |
| ENSG000001 | 25.9753604 | -0.792097649 | 0.25697293 | -3.0824167 | 0.00205327 | 0.01007693 | ITGAX | 3687 |
| ENSG000001 | 117.037181 | 0.554014386 | 0.17973449 | 3.08240443 | 0.00205336 | 0.01007693 | CENPBD1 | 92806 |
| ENSG000001 | 247.745627 | 0.422990866 | 0.13722707 | 3.08241564 | 0.00205328 | 0.01007693 | TRNAU1AP | 54952 |
| ENSG000002 | 51.0184563 | 0.687697246 | 0.2231039 | 3.08240794 | 0.00205333 | 0.01007693 | NA | NA |
| ENSG000002 | 10.6205548 | -1.037321802 | 0.33652147 | -3.0824832 | 0.00205281 | 0.01007693 | NA | NA |

| | | | | | | | | |
|---|---|---|---|---|---|---|---|---|
| ENSG000001 | 864.248446 | 0.421756425 | 0.13683276 | 3.08227676 | 0.00205424 | 0.01008003 | RASA1 | 5921 |
| ENSG000002 | 11.8100057 | -1.007489584 | 0.32686947 | -3.0822382 | 0.0020545 | 0.01008012 | NA | NA |
| ENSG000001 | 1077.89434 | 0.408746343 | 0.13264768 | 3.08144354 | 0.00206 | 0.01010486 | GMNN | 51053 |
| ENSG000001 | 481.693739 | 0.550142279 | 0.17853644 | 3.08140045 | 0.00206029 | 0.01010486 | FAM98B | 283742 |
| ENSG000002 | 53.2765651 | -0.678463415 | 0.22017853 | -3.081424 | 0.00206013 | 0.01010486 | NA | NA |
| ENSG000002 | 989.898527 | 0.394186879 | 0.12793037 | 3.08126123 | 0.00206126 | 0.01010621 | ZNF845 | 91664 |
| ENSG000002 | 15.0288541 | -0.976463758 | 0.31690031 | -3.0812963 | 0.00206101 | 0.01010621 | NA | NA |
| ENSG000002 | 77.4886753 | 0.571062651 | 0.18533457 | 3.08125269 | 0.00206132 | 0.01010621 | NA | NA |
| ENSG000002 | 115.84057 | 0.506811895 | 0.16449536 | 3.08101026 | 0.002063 | 0.01011293 | NA | NA |
| ENSG000002 | 26.2748606 | -0.805931841 | 0.26158272 | -3.0809827 | 0.00206319 | 0.01011293 | NA | NA |
| ENSG000002 | 36.7917797 | 0.73339144 | 0.23804308 | 3.08091891 | 0.00206363 | 0.01011387 | NA | NA |
| ENSG000001 | 253.136525 | 0.413025563 | 0.13406319 | 3.08082759 | 0.00206426 | 0.01011453 | SLC16A7 | 9194 |
| ENSG000001 | 367.204104 | 0.442478391 | 0.14362273 | 3.08083829 | 0.00206419 | 0.01011453 | PEX2 | 5828 |
| ENSG000001 | 93.5269303 | 0.572924977 | 0.18597286 | 3.08069128 | 0.00206521 | 0.01011794 | EPS8L1 | 54869 |
| ENSG000001 | 16.7792353 | -0.925786054 | 0.30052867 | -3.0805249 | 0.00206636 | 0.01012195 | ADORA2A-AS1 | 646023 |
| ENSG000002 | 9.65004823 | -1.078594596 | 0.35013605 | -3.0805014 | 0.00206652 | 0.01012195 | NA | NA |
| ENSG000001 | 10.329121 | -1.066222071 | 0.34613217 | -3.08039 | 0.0020673 | 0.01012451 | HGFAC | 3083 |
| ENSG000001 | 3050.135 | 0.374996472 | 0.1217454 | 3.08016962 | 0.00206883 | 0.01013078 | SOX4 | 6659 |
| ENSG000001 | 215.903186 | 0.445635427 | 0.14475154 | 3.07862305 | 0.0020796 | 0.0101799 | IKBKB | 3551 |
| ENSG000001 | 380.139454 | 0.401296965 | 0.13034853 | 3.07864583 | 0.00207944 | 0.0101799 | SRD5A3 | 79644 |
| ENSG000002 | 21.8201029 | -0.850038119 | 0.27611002 | -3.0786211 | 0.00207961 | 0.0101799 | NA | NA |
| ENSG000001 | 355.643542 | 0.398120932 | 0.12932119 | 3.07854364 | 0.00208015 | 0.01018131 | LZTFL1 | 54585 |
| ENSG000002 | 18.0243904 | -0.891370301 | 0.28959467 | -3.0779928 | 0.002084 | 0.01019892 | LOC101929577 | 101929577 |
| ENSG000001 | 1416.41518 | 0.372489575 | 0.12103878 | 3.07743991 | 0.00208787 | 0.01021663 | CSNK1G3 | 1456 |
| ENSG000002 | 221.670183 | 0.429702561 | 0.13963515 | 3.07732367 | 0.00208868 | 0.01021938 | GANC | 2595 |
| ENSG000002 | 9.09466732 | -1.092233801 | 0.35495001 | -3.0771483 | 0.00208991 | 0.01022416 | NA | NA |
| ENSG000001 | 106.618721 | 0.505856741 | 0.16440491 | 3.07689567 | 0.00209168 | 0.01023159 | PRIMPOL | 201973 |
| ENSG000001 | 362.489873 | 0.421162454 | 0.13691174 | 3.07616027 | 0.00209685 | 0.01025563 | MPC2 | 25874 |
| ENSG000002 | 14.5803611 | -0.952705391 | 0.3097197 | -3.0760245 | 0.00209781 | 0.01025906 | NA | NA |

| | | | | | | | | |
|---|---|---|---|---|---|---|---|---|
| ENSG000002 | 26.7547226 | -0.817644607 | 0.26584325 | -3.0756643 | 0.00210034 | 0.01027022 | NA | NA |
| ENSG000001 | 63.5458105 | -0.641250063 | 0.20849673 | -3.0755882 | 0.00210088 | 0.01027161 | KCNJ6 | 3763 |
| ENSG000001 | 1636.42345 | 0.376031584 | 0.12228395 | 3.07506909 | 0.00210454 | 0.01028826 | CCNL1 | 57018 |
| ENSG000001 | 172.547542 | 0.467601799 | 0.15206713 | 3.07496963 | 0.00210524 | 0.01029045 | SNAPIN | 23557 |
| ENSG000001 | 54.2139694 | 0.664042759 | 0.21597436 | 3.07463705 | 0.00210759 | 0.01030069 | ABHD8 | 79575 |
| ENSG00000C | 243.146995 | 0.438829545 | 0.14273032 | 3.07453635 | 0.0021083 | 0.01030226 | PEX3 | 8504 |
| ENSG000001 | 394.756455 | 0.382388979 | 0.12437501 | 3.07448391 | 0.00210867 | 0.01030226 | LYRM7 | 90624 |
| ENSG000002 | 17.3873843 | -0.918413815 | 0.29871976 | -3.0744997 | 0.00210856 | 0.01030226 | ZNF630 | 57232 |
| ENSG00000C | 1546.00723 | 0.374671627 | 0.1218773 | 3.0741706 | 0.00211089 | 0.0103106 | RAB18 | 22931 |
| ENSG000001 | 339.08281 | 0.399061888 | 0.12981052 | 3.07418761 | 0.00211077 | 0.0103106 | GAP43 | 2596 |
| ENSG000001 | 9.54031234 | -1.069637905 | 0.34799911 | -3.07368 | 0.00211436 | 0.01032632 | ARGFX | 503582 |
| ENSG000001 | 911.997893 | 0.347384365 | 0.11303025 | 3.0733751 | 0.00211652 | 0.01033563 | TBK1 | 29110 |
| ENSG00000C | 256.963865 | 0.387147169 | 0.12598529 | 3.07295529 | 0.0021195 | 0.01034877 | ST7L | 54879 |
| ENSG000001 | 2002.41088 | 0.404975746 | 0.1317884 | 3.07292417 | 0.00211972 | 0.01034877 | GNA13 | 10672 |
| ENSG000001 | 23.3564392 | -0.853854903 | 0.27787632 | -3.0727875 | 0.00212069 | 0.01034977 | DNAAF1 | 123872 |
| ENSG000001 | 49.5990529 | -0.631048692 | 0.20536487 | -3.0728172 | 0.00212048 | 0.01034977 | CATSPER2 | 117155 |
| ENSG000001 | 133.465018 | 0.47610056 | 0.15494021 | 3.0728018 | 0.00212059 | 0.01034977 | ANKS3 | 124401 |
| ENSG000001 | 415.40314 | 0.415937496 | 0.13536611 | 3.07268561 | 0.00212142 | 0.01035206 | TMCC1 | 23023 |
| ENSG000001 | 68.4882378 | 0.570956363 | 0.18584453 | 3.07222587 | 0.00212469 | 0.01036553 | ZFR2 | 23217 |
| ENSG000001 | 90.3733462 | 0.547455349 | 0.17819377 | 3.07224739 | 0.00212454 | 0.01036553 | CHST7 | 56548 |
| ENSG000002 | 24.3038117 | -0.826497029 | 0.26903021 | -3.0721346 | 0.00212534 | 0.01036745 | NA | NA |
| ENSG000001 | 810.148117 | 0.393476725 | 0.1280865 | 3.07196091 | 0.00212658 | 0.01037219 | NAA35 | 60560 |
| ENSG000001 | 988.770005 | 0.423872315 | 0.13798258 | 3.07192625 | 0.00212682 | 0.01037219 | UBE2H | 7328 |
| ENSG000002 | 4.57525537 | -1.162995479 | 0.37859288 | -3.0718895 | 0.00212708 | 0.01037222 | NA | NA |
| ENSG000002 | 12.5573724 | -1.018902159 | 0.33172329 | -3.0715425 | 0.00212956 | 0.01038304 | NA | NA |
| ENSG000001 | 8.93124785 | -1.087392153 | 0.35403687 | -3.0714094 | 0.00213051 | 0.01038642 | HOXB6 | 3216 |
| ENSG000001 | 31.444809 | -0.824970642 | 0.26861331 | -3.0712202 | 0.00213186 | 0.01039175 | SLC6A5 | 9152 |
| ENSG000001 | 482.756724 | 0.780842391 | 0.25425562 | 3.07109193 | 0.00213277 | 0.01039497 | IGFBP4 | 3487 |
| ENSG000001 | 58.7811196 | -0.657479925 | 0.21413337 | -3.0704226 | 0.00213756 | 0.01041705 | CD226 | 10666 |

| | | | | | | | | |
|---|---|---|---|---|---|---|---|---|
| ENSG00000C | 29.154992 | -0.794845431 | 0.25889612 | -3.0701326 | 0.00213964 | 0.01042591 | CDH17 | 1015 |
| ENSG000001 | 323.549483 | 0.392121099 | 0.12774458 | 3.06957122 | 0.00214366 | 0.01044427 | BTG1 | 694 |
| ENSG000001 | 29.0712756 | -0.762231946 | 0.24833403 | -3.0693817 | 0.00214502 | 0.01044964 | GDF7 | 151449 |
| ENSG000001 | 6.77637158 | -1.123691454 | 0.36611483 | -3.0692323 | 0.0021461 | 0.01045362 | DEFB129 | 140881 |
| ENSG000001 | 468.869144 | 0.38635678 | 0.12588711 | 3.06907342 | 0.00214724 | 0.01045792 | RIC8B | 55188 |
| ENSG00000C | 1133.40757 | 0.35057233 | 0.11423131 | 3.06896894 | 0.00214799 | 0.01046032 | RBM27 | 54439 |
| ENSG000002 | 5.82902087 | -1.144119739 | 0.37281081 | -3.0689017 | 0.00214847 | 0.01046142 | NA | NA |
| ENSG000002 | 39.1418833 | -0.756569034 | 0.24653266 | -3.068839 | 0.00214892 | 0.01046236 | MIR99AHG | 388815 |
| ENSG000002 | 25.3433234 | -0.81426682 | 0.26534114 | -3.0687545 | 0.00214953 | 0.01046406 | NA | NA |
| ENSG00000C | 1501.27428 | 0.366312822 | 0.11937714 | 3.06853401 | 0.00215112 | 0.01047053 | MLLT10 | 8028 |
| ENSG000001 | 163.709619 | 0.52400206 | 0.17077148 | 3.06844014 | 0.00215179 | 0.01047256 | FBXO31 | 79791 |
| ENSG000001 | 239.269278 | 0.45618013 | 0.14867248 | 3.06835623 | 0.0021524 | 0.01047299 | TAMM41 | 132001 |
| ENSG000002 | 7.92494039 | -1.105029038 | 0.36013598 | -3.0683661 | 0.00215233 | 0.01047299 | NA | NA |
| ENSG000001 | 149.599377 | 0.482403478 | 0.15723033 | 3.0681325 | 0.00215401 | 0.01047905 | KCTD18 | 130535 |
| ENSG000002 | 4.37562175 | -1.160970612 | 0.37839907 | -3.0681117 | 0.00215416 | 0.01047905 | NA | NA |
| ENSG000001 | 798.853819 | 0.393539567 | 0.12826985 | 3.06805973 | 0.00215454 | 0.01047962 | PGM2 | 55276 |
| ENSG00000C | 475.154442 | 0.377627006 | 0.1230943 | 3.06778641 | 0.00215651 | 0.01048691 | C12orf4 | 57102 |
| ENSG000002 | 11.337649 | -1.023000295 | 0.33346988 | -3.0677442 | 0.00215681 | 0.01048691 | LOC101927411 | 101927411 |
| ENSG000002 | 243.966464 | 0.426694251 | 0.1390905 | 3.0677455 | 0.0021568 | 0.01048691 | ZNF224 | 7767 |
| ENSG000001 | 10.5113211 | -1.028233987 | 0.33519608 | -3.0675597 | 0.00215814 | 0.01049213 | PDC | 5132 |
| ENSG000002 | 5.30803031 | -1.157262835 | 0.37729185 | -3.0672882 | 0.00216011 | 0.01050041 | NA | NA |
| ENSG000001 | 581.053681 | 0.42958967 | 0.14005841 | 3.06721786 | 0.00216061 | 0.01050113 | DRAM2 | 128338 |
| ENSG000001 | 1635.46362 | 0.481772928 | 0.15707275 | 3.06719607 | 0.00216077 | 0.01050113 | CHD2 | 1106 |
| ENSG000001 | 182.430753 | 0.49992935 | 0.16299562 | 3.06713355 | 0.00216122 | 0.01050207 | ZNF777 | 27153 |
| ENSG00000C | 1066.67489 | 0.45754219 | 0.14919547 | 3.0667297 | 0.00216414 | 0.01050871 | NOP14 | 8602 |
| ENSG000001 | 297.94901 | 0.421246892 | 0.13735977 | 3.06674144 | 0.00216406 | 0.01050871 | NEIL3 | 55247 |
| ENSG000001 | 59.7049697 | 0.630802916 | 0.20569041 | 3.06675907 | 0.00216393 | 0.01050871 | LDHD | 197257 |
| ENSG000002 | 11.3943704 | -1.030598163 | 0.33605275 | -3.066775 | 0.00216382 | 0.01050871 | NA | NA |
| ENSG000002 | 5.68042804 | -1.140771149 | 0.37198162 | -3.0667407 | 0.00216407 | 0.01050871 | NA | NA |

| | | | | | | | | |
|---|---|---|---|---|---|---|---|---|
| ENSG000002 | 20.4139552 | -0.866926592 | 0.28267972 | -3.0668157 | 0.00216352 | 0.01050871 | NA | NA |
| ENSG000001 | 28.9078705 | -0.784389383 | 0.25578829 | -3.0665571 | 0.00216539 | 0.01051352 | GYS2 | 2998 |
| ENSG000001 | 709.156648 | 0.441382556 | 0.14393645 | 3.06650986 | 0.00216574 | 0.01051392 | GSPT2 | 23708 |
| ENSG000001 | 2139.72331 | 0.402037991 | 0.13110969 | 3.06642476 | 0.00216635 | 0.0105144 | PBRM1 | 55193 |
| ENSG000001 | 319.878415 | 0.411779278 | 0.13428614 | 3.06643166 | 0.0021663 | 0.0105144 | STXBP5 | 134957 |
| ENSG000001 | 2343.1919 | 0.395287938 | 0.128918 | 3.06619654 | 0.00216801 | 0.01052117 | TNRC6B | 23112 |
| ENSG000000 | 29.7083088 | -0.760485789 | 0.24803636 | -3.0660255 | 0.00216925 | 0.01052593 | SLC13A1 | 6561 |
| ENSG000002 | 349.929305 | 0.376311672 | 0.12275114 | 3.06564713 | 0.00217199 | 0.010538 | RPARP-AS1 | 100505761 |
| ENSG000001 | 1613.9731 | 0.355290693 | 0.11590652 | 3.06532114 | 0.00217436 | 0.01054823 | TRAPPC8 | 22878 |
| ENSG000002 | 38.4299194 | 0.7347365 | 0.23970823 | 3.06512837 | 0.00217577 | 0.01055341 | LINC01089 | 338799 |
| ENSG000002 | 64.9417098 | 0.601511912 | 0.19624527 | 3.06510269 | 0.00217595 | 0.01055341 | LOC101928979 | 101928979 |
| ENSG000001 | 714.56386 | 0.415463044 | 0.1355531 | 3.06494689 | 0.00217709 | 0.01055512 | LIMS1 | 3987 |
| ENSG000001 | 288.394416 | 0.475617893 | 0.15517958 | 3.0649515 | 0.00217705 | 0.01055512 | ZNRF1 | 84937 |
| ENSG000002 | 51.6068238 | 0.665605299 | 0.21716249 | 3.0650104 | 0.00217662 | 0.01055512 | NA | NA |
| ENSG000000 | 449.779944 | 0.444036159 | 0.14488615 | 3.06472465 | 0.0021787 | 0.01056072 | EIF2B3 | 8891 |
| ENSG000002 | 121.689929 | 0.545495149 | 0.17799202 | 3.06471688 | 0.00217876 | 0.01056072 | ZBED8 | 63920 |
| ENSG000001 | 54.5507556 | -0.734435741 | 0.23965153 | -3.0645986 | 0.00217962 | 0.01056363 | CNTN5 | 53942 |
| ENSG000001 | 15.5167031 | -0.959915923 | 0.31323598 | -3.0645136 | 0.00218024 | 0.01056452 | C2orf81 | 388963 |
| ENSG000001 | 269.487362 | 0.436205584 | 0.14234142 | 3.06450203 | 0.00218033 | 0.01056452 | SETMAR | 6419 |
| ENSG000002 | 4.66676056 | -1.156816685 | 0.3775067 | -3.0643607 | 0.00218136 | 0.01056825 | RNU6-837P | 106479936 |
| ENSG000002 | 562.071764 | 0.384397794 | 0.12545392 | 3.06405573 | 0.00218358 | 0.01057776 | TMEM57 | 55219 |
| ENSG000001 | 96.5045038 | 0.578240201 | 0.18872664 | 3.06390338 | 0.00218469 | 0.01058189 | SOX9 | 6662 |
| ENSG000002 | 54.3789372 | -0.65014655 | 0.2122163 | -3.0636033 | 0.00218689 | 0.01059124 | CFAP57 | 149465 |
| ENSG000001 | 77.9692842 | -0.552112219 | 0.18022122 | -3.063525 | 0.00218746 | 0.01059275 | DCHS2 | 54798 |
| ENSG000001 | 636.551606 | 0.41428232 | 0.13523281 | 3.06347492 | 0.00218783 | 0.01059325 | AIM1 | 202 |
| ENSG000002 | 36.3249725 | -0.725527796 | 0.23689763 | -3.0626216 | 0.00219407 | 0.01062224 | SLC7A11-AS1 | 641364 |
| ENSG000001 | 5005.16145 | 0.339317073 | 0.1107982 | 3.0624781 | 0.00219513 | 0.01062606 | FNBP1L | 54874 |
| ENSG000002 | 102.48474 | 0.525344795 | 0.17154491 | 3.06243299 | 0.00219546 | 0.0106264 | CNPY2 | 10330 |
| ENSG000001 | 198.524154 | 0.511975063 | 0.16719257 | 3.06218782 | 0.00219726 | 0.01063311 | GMEB2 | 26205 |

| | | | | | | | | |
|---|---|---|---|---|---|---|---|---|
| ENSG000001 | 217.012815 | 0.510075526 | 0.16657308 | 3.06217254 | 0.00219737 | 0.01063311 | PML | 5371 |
| ENSG00000C | 2649.05598 | 0.349858073 | 0.11426155 | 3.06190566 | 0.00219933 | 0.01064132 | DIS3 | 22894 |
| ENSG000001 | 183.754056 | 0.445604289 | 0.14554994 | 3.06152163 | 0.00220215 | 0.01065371 | GLRX | 2745 |
| ENSG000002 | 16.0299753 | -0.929014176 | 0.30347011 | -3.0613037 | 0.00220375 | 0.0106602 | NA | NA |
| ENSG000002 | 7.61365272 | -1.11197131 | 0.36324226 | -3.0612388 | 0.00220423 | 0.01066124 | GLYATL3 | 389396 |
| ENSG000001 | 65.9726745 | 0.631223153 | 0.20621305 | 3.06102429 | 0.00220581 | 0.01066758 | RRAGC | 64121 |
| ENSG000001 | 1434.42119 | 0.332098305 | 0.10849378 | 3.06098936 | 0.00220607 | 0.01066758 | EXOSC10 | 5394 |
| ENSG000002 | 32.7201619 | 0.771121003 | 0.25194089 | 3.0607219 | 0.00220804 | 0.01067584 | NA | NA |
| ENSG000002 | 3.44998383 | -1.157026516 | 0.378035 | -3.0606333 | 0.00220869 | 0.01067773 | NA | NA |
| ENSG000001 | 1254.51764 | 0.852579597 | 0.27859231 | 3.06031271 | 0.00221106 | 0.01068789 | FBL | 2091 |
| ENSG000001 | 396.450679 | 0.507097059 | 0.16573175 | 3.05974604 | 0.00221525 | 0.01070685 | KIAA1671 | 85379 |
| ENSG000001 | 26.1744265 | -0.823023072 | 0.26901245 | -3.0594238 | 0.00221763 | 0.0107171 | MEP1B | 4225 |
| ENSG000001 | 55.1049987 | 0.65138578 | 0.2129368 | 3.0590568 | 0.00222035 | 0.01072896 | C15orf27 | 123591 |
| ENSG000001 | 24.2759972 | -0.840127527 | 0.27465398 | -3.058858 | 0.00222182 | 0.0107348 | FGL1 | 2267 |
| ENSG00000C | 132.464323 | 0.5734435 | 0.18748738 | 3.05857121 | 0.00222395 | 0.0107438 | ZNF324 | 25799 |
| ENSG000001 | 68.9380405 | 0.597685389 | 0.19542925 | 3.05832112 | 0.00222581 | 0.01075149 | MAP3K10 | 4294 |
| ENSG00000C | 226.822769 | 0.424161953 | 0.13869427 | 3.05825147 | 0.00222633 | 0.01075271 | PTPN21 | 11099 |
| ENSG000001 | 2739.68722 | 0.421856528 | 0.13794223 | 3.0582115 | 0.00222662 | 0.01075287 | SRSF11 | 9295 |
| ENSG000002 | 36.2468605 | -0.748043213 | 0.24470473 | -3.0569218 | 0.00223623 | 0.01079795 | CASC11 | 100270680 |
| ENSG000001 | 364.551446 | 0.425802002 | 0.13929273 | 3.05688609 | 0.00223649 | 0.01079795 | SLFN13 | 146857 |
| ENSG000001 | 1605.1238 | 0.369869205 | 0.12099993 | 3.0567721 | 0.00223734 | 0.01080077 | FAM208A | 23272 |
| ENSG000001 | 2501.92423 | 0.342523234 | 0.11206493 | 3.05647123 | 0.00223959 | 0.01081033 | CASC5 | 57082 |
| ENSG000001 | 15.217662 | -0.948268462 | 0.31026258 | -3.0563417 | 0.00224056 | 0.01081264 | HTR3E | 285242 |
| ENSG000002 | 7.16141031 | -1.115430936 | 0.36495693 | -3.0563358 | 0.0022406 | 0.01081264 | BACH1-AS1 | 106478962 |
| ENSG000001 | 19.1476335 | -0.87654559 | 0.28681603 | -3.0561248 | 0.00224218 | 0.01081897 | PLA2G4D | 283748 |
| ENSG000001 | 1199.45632 | 0.435081488 | 0.1423679 | 3.05603649 | 0.00224284 | 0.01082087 | KDM2A | 22992 |
| ENSG000002 | 49.2304439 | 0.677002057 | 0.22153623 | 3.05594287 | 0.00224354 | 0.01082296 | VPS9D1-AS1 | 100128881 |
| ENSG000002 | 7.78215264 | -1.098255279 | 0.35940704 | -3.0557422 | 0.00224504 | 0.01082892 | NA | NA |
| ENSG000001 | 517.939829 | 0.407165478 | 0.13325799 | 3.05546764 | 0.0022471 | 0.01083755 | LNX2 | 222484 |

| | | | | | | | | |
|---|---|---|---|---|---|---|---|---|
| ENSG000001 | 1044.72102 | 0.493330215 | 0.16146064 | 3.05542088 | 0.00224745 | 0.01083795 | UNC13A | 23025 |
| ENSG000002 | 9.14366343 | 1.064435677 | 0.34840751 | 3.05514563 | 0.00224951 | 0.01084661 | NA | NA |
| ENSG000002 | 13.2127956 | -0.988763665 | 0.32365302 | -3.0550114 | 0.00225052 | 0.01085017 | NA | NA |
| ENSG000002 | 38.4032519 | 0.738018065 | 0.24158168 | 3.05494217 | 0.00225104 | 0.01085139 | NA | NA |
| ENSG000000 | 394.62391 | 0.396805519 | 0.12989481 | 3.05482201 | 0.00225194 | 0.01085152 | ERCC8 | 1161 |
| ENSG000001 | 65.084329 | -0.714682733 | 0.23395418 | -3.0547979 | 0.00225212 | 0.01085152 | ABCA6 | 23460 |
| ENSG000002 | 2.74131026 | -1.132159526 | 0.37061708 | -3.0547959 | 0.00225214 | 0.01085152 | DDX43P3 | 106481709 |
| ENSG000002 | 134.213014 | 0.473471061 | 0.15498803 | 3.05488789 | 0.00225145 | 0.01085152 | LINC01184 | 644873 |
| ENSG000001 | 1192.9166 | 0.461609865 | 0.15111753 | 3.05464134 | 0.0022533 | 0.01085583 | VPRBP | 9730 |
| ENSG000001 | 7.78544694 | -1.096536318 | 0.35899144 | -3.0544915 | 0.00225442 | 0.01085996 | NA | NA |
| ENSG000001 | 137.01236 | 0.488107774 | 0.15982536 | 3.054007 | 0.00225807 | 0.0108746 | BACE2 | 25825 |
| ENSG000002 | 721.978136 | 0.416233824 | 0.13629132 | 3.05400086 | 0.00225811 | 0.0108746 | PPP3R1 | 5534 |
| ENSG000002 | 23.6521737 | -0.841326044 | 0.27548507 | -3.0539806 | 0.00225827 | 0.0108746 | NA | NA |
| ENSG000001 | 18.7466542 | -0.910915211 | 0.29830928 | -3.0535932 | 0.00226118 | 0.01088735 | DNAJC5B | 85479 |
| ENSG000001 | 17.9292453 | -0.9443981 | 0.30932659 | -3.0530777 | 0.00226507 | 0.01090368 | AICDA | 57379 |
| ENSG000002 | 18.4443538 | -0.950317564 | 0.31126598 | -3.0530723 | 0.00226511 | 0.01090368 | LOC101927697 | 101927697 |
| ENSG000001 | 825.860813 | 0.420278767 | 0.13767 | 3.05279852 | 0.00226718 | 0.01091104 | OARD1 | 221443 |
| ENSG000002 | 8.28974841 | -1.073220548 | 0.3515509 | -3.052817 | 0.00226704 | 0.01091104 | NA | NA |
| ENSG000001 | 1252.61385 | 0.345803068 | 0.113281 | 3.05261315 | 0.00226858 | 0.0109152 | IMPAD1 | 54928 |
| ENSG000001 | 2989.15539 | 0.340215606 | 0.11145007 | 3.05262794 | 0.00226847 | 0.0109152 | MED13 | 9969 |
| ENSG000001 | 153.573081 | 0.480958858 | 0.1575627 | 3.05249188 | 0.0022695 | 0.01091831 | FAM72B | 653820 |
| ENSG000001 | 1831.06453 | 0.363041819 | 0.11894298 | 3.05223413 | 0.00227145 | 0.01092639 | SORL1 | 6653 |
| ENSG000001 | 243.79966 | 0.475544298 | 0.15580455 | 3.05218484 | 0.00227182 | 0.01092689 | SLC38A7 | 55238 |
| ENSG000002 | 6.41475421 | -1.125211021 | 0.36868446 | -3.0519621 | 0.00227351 | 0.01093371 | LOC101928098 | 101928098 |
| ENSG000002 | 29.1376812 | -0.765011402 | 0.25068328 | -3.0517049 | 0.00227546 | 0.01094178 | NA | NA |
| ENSG000002 | 31.8986919 | 0.766359205 | 0.25114014 | 3.05152018 | 0.00227686 | 0.01094722 | NA | NA |
| ENSG000001 | 225.408872 | 0.456607082 | 0.14965562 | 3.05105208 | 0.00228041 | 0.010963 | SPRED2 | 200734 |
| ENSG000002 | 4.84561597 | -1.153797572 | 0.37819394 | -3.0508093 | 0.00228226 | 0.01097057 | NA | NA |
| ENSG000001 | 1130.68346 | 0.3321521 | 0.10888004 | 3.05062439 | 0.00228366 | 0.01097472 | PWWP2A | 114825 |

| | | | | | | | | |
|---|---|---|---|---|---|---|---|---|
| ENSG000002 | 21.1452732 | -0.843181385 | 0.27639617 | -3.0506262 | 0.00228365 | 0.01097472 | NA | NA |
| ENSG000001 | 171.244242 | 0.465819616 | 0.15270265 | 3.05050121 | 0.0022846 | 0.01097793 | FANCE | 2178 |
| ENSG000001 | 87.0539598 | 0.552833994 | 0.18123631 | 3.05034902 | 0.00228576 | 0.01098219 | STAP2 | 55620 |
| ENSG000001 | 125.91729 | 0.526523729 | 0.17261884 | 3.05021009 | 0.00228681 | 0.01098597 | PTPN6 | 5777 |
| ENSG000001 | 1147.53744 | 0.374289303 | 0.12271295 | 3.05012057 | 0.0022875 | 0.01098794 | MSL1 | 339287 |
| ENSG000001 | 435.688279 | 0.387638576 | 0.12709607 | 3.04996511 | 0.00228868 | 0.01099217 | RAPGEF5 | 9771 |
| ENSG000001 | 152.16019 | 0.490944653 | 0.16096894 | 3.04993404 | 0.00228892 | 0.01099217 | TSEN54 | 283989 |
| ENSG000001 | 131.324614 | 0.486481456 | 0.15951182 | 3.04981446 | 0.00228983 | 0.01099524 | FLVCR1-AS1 | 642946 |
| ENSG000001 | 2844.0061 | 0.317495588 | 0.10410909 | 3.04964326 | 0.00229113 | 0.01100021 | RDX | 5962 |
| ENSG000001 | 37.5000053 | 0.707173681 | 0.23189735 | 3.04951174 | 0.00229214 | 0.01100112 | CNFN | 84518 |
| ENSG000001 | 323.067862 | 0.478718036 | 0.15698163 | 3.04951623 | 0.0022921 | 0.01100112 | PAPD7 | 11044 |
| ENSG000002 | 6.77959318 | -1.113791221 | 0.3652279 | -3.0495787 | 0.00229163 | 0.01100112 | DDC-AS1 | 100129427 |
| ENSG000001 | 28.3737486 | -0.843690024 | 0.27670927 | -3.0490124 | 0.00229595 | 0.01101811 | CDH9 | 1007 |
| ENSG000001 | 25.2121843 | -0.802508681 | 0.26320886 | -3.0489425 | 0.00229648 | 0.01101865 | SLC2A9 | 56606 |
| ENSG000001 | 467.192556 | 0.46670185 | 0.15307087 | 3.0489267 | 0.00229661 | 0.01101865 | TOP1MT | 116447 |
| ENSG000001 | 371.149996 | 0.4984583 | 0.16349348 | 3.0487962 | 0.0022976 | 0.01102213 | CAMKK2 | 10645 |
| ENSG000002 | 14.5180697 | -0.948981819 | 0.31127024 | -3.0487393 | 0.00229804 | 0.01102292 | NA | NA |
| ENSG000001 | 13.2616646 | -0.986294581 | 0.32351494 | -3.0486833 | 0.00229847 | 0.01102367 | PMP2 | 5375 |
| ENSG000000 | 556.293262 | 0.385693305 | 0.12651849 | 3.04851327 | 0.00229977 | 0.011026 | ZDHHC6 | 64429 |
| ENSG000001 | 1428.1078 | 0.377718617 | 0.1239024 | 3.04851731 | 0.00229974 | 0.011026 | DYRK2 | 8445 |
| ENSG000001 | 105.923315 | 0.485918185 | 0.15939211 | 3.04857121 | 0.00229932 | 0.011026 | METTL23 | 124512 |
| ENSG000001 | 2401.03119 | 0.341153967 | 0.11191363 | 3.04836841 | 0.00230088 | 0.01102827 | TEX10 | 54881 |
| ENSG000001 | 96.2327962 | 0.549276663 | 0.18018849 | 3.04834486 | 0.00230106 | 0.01102827 | VPS28 | 51160 |
| ENSG000001 | 156.986177 | 0.459562868 | 0.15075652 | 3.048378 | 0.0023008 | 0.01102827 | SGF29 | 112869 |
| ENSG000001 | 463.482936 | 0.351607962 | 0.11534767 | 3.04824508 | 0.00230182 | 0.0110299 | IFT81 | 28981 |
| ENSG000001 | 12.8280668 | -1.009633562 | 0.33121968 | -3.0482294 | 0.00230194 | 0.0110299 | CA1 | 759 |
| ENSG000001 | 1327.19222 | 0.389087698 | 0.12766838 | 3.04764343 | 0.00230643 | 0.01104881 | RBM28 | 55131 |
| ENSG000002 | 13.8883051 | -0.986624351 | 0.32373159 | -3.0476616 | 0.00230629 | 0.01104881 | NA | NA |
| ENSG000000 | 43.532935 | -0.651525109 | 0.21378291 | -3.0476015 | 0.00230676 | 0.01104905 | MYO16 | 23026 |

| | | | | | | | | |
|---|---|---|---|---|---|---|---|---|
| ENSG000002 | 13.3202173 | -1.011477195 | 0.3319085 | -3.0474579 | 0.00230786 | 0.01105303 | NA | NA |
| ENSG000001 | 283.857349 | 0.425413464 | 0.13961506 | 3.04704568 | 0.00231103 | 0.01106689 | PCTP | 58488 |
| ENSG000001 | 158.683875 | 0.4973517 | 0.16324261 | 3.04670263 | 0.00231366 | 0.01107822 | TEFM | 79736 |
| ENSG000001 | 119.839983 | 0.538932476 | 0.17690375 | 3.04647287 | 0.00231543 | 0.01108538 | CA5BP1 | 340591 |
| ENSG000002 | 24.0572338 | -0.834413411 | 0.27391166 | -3.0462866 | 0.00231687 | 0.01109094 | FLJ41278 | 400046 |
| ENSG000001 | 610.461573 | 0.484504304 | 0.15906448 | 3.04596162 | 0.00231937 | 0.01110031 | FHDC1 | 85462 |
| ENSG000002 | 17.5928845 | 0.909176845 | 0.29848324 | 3.04598963 | 0.00231916 | 0.01110031 | LOC400684 | 400684 |
| ENSG000001 | 174.952775 | 0.473284199 | 0.15538727 | 3.04583638 | 0.00232034 | 0.01110363 | VPS33B | 26276 |
| ENSG000001 | 462.420587 | 0.439243224 | 0.14424519 | 3.04511514 | 0.00232591 | 0.01112898 | MAP2K6 | 5608 |
| ENSG000001 | 3993.32811 | 0.32474882 | 0.10664715 | 3.04507728 | 0.00232621 | 0.01112906 | METAP2 | 10988 |
| ENSG000002 | 197.457258 | 0.940555116 | 0.30892103 | 3.04464583 | 0.00232954 | 0.01114373 | NA | NA |
| ENSG00000C | 33.3185202 | -0.771821394 | 0.25350762 | -3.0445688 | 0.00233014 | 0.01114527 | SI | 6476 |
| ENSG000001 | 470.242057 | 0.504539725 | 0.16572298 | 3.04447652 | 0.00233086 | 0.01114737 | CAMKV | 79012 |
| ENSG000001 | 24.0275985 | -0.828337542 | 0.27210234 | -3.0442132 | 0.0023329 | 0.01115575 | NLRP3 | 114548 |
| ENSG000002 | 81.5140665 | 0.626367834 | 0.20575917 | 3.04417945 | 0.00233316 | 0.01115575 | NA | NA |
| ENSG00000C | 98.0002062 | -0.52022697 | 0.17090929 | -3.0438776 | 0.0023355 | 0.01116563 | NRP1 | 8829 |
| ENSG000001 | 402.753001 | 0.388107028 | 0.12751034 | 3.04372985 | 0.00233665 | 0.0111698 | ARRB2 | 409 |
| ENSG00000C | 129.149999 | 0.552983798 | 0.18169092 | 3.04354126 | 0.00233811 | 0.01117549 | MRC2 | 9902 |
| ENSG000002 | 8.12604525 | -1.090294589 | 0.35823868 | -3.0434865 | 0.00233854 | 0.01117621 | NA | NA |
| ENSG000001 | 137.159094 | 0.510649692 | 0.16781323 | 3.04296436 | 0.0023426 | 0.0111943 | ASRGL1 | 80150 |
| ENSG000002 | 7.12699724 | -1.108498219 | 0.36429967 | -3.0428197 | 0.00234373 | 0.01119836 | CYP2AC1P | 106478960 |
| ENSG000002 | 35.6844883 | 0.751621931 | 0.24704375 | 3.04246485 | 0.00234649 | 0.01121025 | OSER1-AS1 | 100505783 |
| ENSG000002 | 13.3269302 | -0.989748853 | 0.32534014 | -3.0421971 | 0.00234858 | 0.0112189 | NA | NA |
| ENSG000002 | 22.9282516 | -0.8571897 | 0.28177133 | -3.0421466 | 0.00234897 | 0.01121946 | NA | NA |
| ENSG000001 | 1298.38572 | 0.365744191 | 0.12024567 | 3.04164129 | 0.00235292 | 0.01123699 | KIAA1586 | 57691 |
| ENSG00000C | 646.827366 | 0.392852388 | 0.12916756 | 3.04141689 | 0.00235468 | 0.01124405 | PPP2R5A | 5525 |
| ENSG000001 | 22.5431598 | -0.841276627 | 0.27661714 | -3.0413033 | 0.00235556 | 0.01124697 | CYP4F3 | 4051 |
| ENSG000002 | 8.86273967 | -1.071736665 | 0.35242404 | -3.0410431 | 0.0023576 | 0.01125537 | NA | NA |
| ENSG000002 | 11.4814604 | -1.000652566 | 0.32906177 | -3.0409262 | 0.00235852 | 0.01125841 | NA | NA |

| | | | | | | | | |
|---|---|---|---|---|---|---|---|---|
| ENSG000001 | 445.259437 | 0.432737114 | 0.14231769 | 3.04064181 | 0.00236074 | 0.01126772 | BOLA3 | 388962 |
| ENSG000001 | 333.355264 | 0.452476629 | 0.14881292 | 3.04057356 | 0.00236128 | 0.01126895 | ALG10 | 84920 |
| ENSG00000C | 15.4264588 | -0.93995832 | 0.30915662 | -3.0403953 | 0.00236268 | 0.01127297 | TGM1 | 7051 |
| ENSG000002 | 3.65485346 | -1.14583485 | 0.37686828 | -3.040412 | 0.00236255 | 0.01127297 | NA | NA |
| ENSG00000C | 3806.43999 | 0.342874481 | 0.11278709 | 3.04001531 | 0.00236566 | 0.01128587 | AQR | 9716 |
| ENSG000001 | 29.7396667 | -0.75249601 | 0.24754731 | -3.0398068 | 0.0023673 | 0.01129236 | MYH6 | 4624 |
| ENSG00000C | 321.343658 | 0.409870307 | 0.13484266 | 3.03961898 | 0.00236878 | 0.01129542 | CDC45 | 8318 |
| ENSG000001 | 337.425864 | 0.404483129 | 0.13306937 | 3.03964107 | 0.0023686 | 0.01129542 | JAZF1 | 221895 |
| ENSG000001 | 48.4566598 | 0.64601163 | 0.21252621 | 3.03967988 | 0.0023683 | 0.01129542 | HS6ST3 | 266722 |
| ENSG000002 | 5.25919254 | -1.146960114 | 0.37737588 | -3.0393042 | 0.00237125 | 0.0113059 | SEPT14P1 | 106480340 |
| ENSG000001 | 975.902876 | 0.403963522 | 0.13295129 | 3.03843251 | 0.00237812 | 0.01133733 | UGCG | 7357 |
| ENSG000001 | 22.6248845 | -0.829211706 | 0.27294099 | -3.0380622 | 0.00238105 | 0.0113486 | CES5A | 221223 |
| ENSG000001 | 18.691015 | -0.922689862 | 0.30370793 | -3.0380829 | 0.00238088 | 0.0113486 | NA | NA |
| ENSG000002 | 3.82764348 | -1.149958751 | 0.37858486 | -3.0375191 | 0.00238534 | 0.01136773 | NA | NA |
| ENSG00000C | 124.097834 | 0.560595887 | 0.18457879 | 3.03716315 | 0.00238816 | 0.01137983 | IGSF9B | 22997 |
| ENSG000002 | 224.592134 | 0.426438951 | 0.14041497 | 3.03699073 | 0.00238953 | 0.011385 | TMEM170B | 100113407 |
| ENSG000001 | 1088.76704 | 0.449209688 | 0.14793338 | 3.03656744 | 0.00239289 | 0.01139832 | KPNA1 | 3836 |
| ENSG000002 | 3.14000348 | -1.139628622 | 0.37530129 | -3.03657 | 0.00239287 | 0.01139832 | NA | NA |
| ENSG000001 | 30.7108497 | -0.769114699 | 0.2532951 | -3.0364374 | 0.00239392 | 0.0114019 | STKLD1 | 169436 |
| ENSG000001 | 332.306826 | 0.405335876 | 0.13349509 | 3.03633544 | 0.00239473 | 0.01140442 | GK5 | 256356 |
| ENSG000001 | 24.4756815 | -0.80242122 | 0.26428041 | -3.0362493 | 0.00239541 | 0.01140634 | RUNX2 | 860 |
| ENSG000002 | 2702.00123 | 0.382932412 | 0.12613894 | 3.03579859 | 0.002399 | 0.01142206 | NA | NA |
| ENSG000001 | 488.318229 | 0.544757338 | 0.17945089 | 3.03569045 | 0.00239986 | 0.01142252 | VPS26A | 9559 |
| ENSG000001 | 17.2966319 | -0.930986044 | 0.30667791 | -3.0357128 | 0.00239968 | 0.01142252 | C1orf158 | 93190 |
| ENSG000002 | 26.7496902 | -0.794141158 | 0.26160238 | -3.0356801 | 0.00239994 | 0.01142252 | SRD5A3-AS1 | 100506462 |
| ENSG000002 | 9.38963217 | -1.054814808 | 0.34750219 | -3.0354192 | 0.00240202 | 0.01143107 | NA | NA |
| ENSG000002 | 76.8009409 | 0.564037599 | 0.18583647 | 3.03512872 | 0.00240433 | 0.01144074 | RPL21P28 | 100131205 |
| ENSG000001 | 559.193175 | 0.365803665 | 0.12052669 | 3.03504277 | 0.00240502 | 0.01144266 | RSRC1 | 51319 |
| ENSG000002 | 94.7067258 | 0.627543547 | 0.20678686 | 3.03473608 | 0.00240746 | 0.01145296 | GPC2 | 221914 |

| | | | | | | | | |
|---|---|---|---|---|---|---|---|---|
| ENSG000002 | 9.97783304 | -1.040066483 | 0.34272592 | -3.0346887 | 0.00240784 | 0.01145341 | NA | NA |
| ENSG000002 | 5.07798615 | -1.143006827 | 0.37665666 | -3.034612 | 0.00240845 | 0.01145498 | NA | NA |
| ENSG000001 | 200.962358 | 0.429207724 | 0.14143985 | 3.03456004 | 0.00240887 | 0.01145561 | SEMA3D | 223117 |
| ENSG000000 | 737.393731 | 0.422998563 | 0.13939681 | 3.03449242 | 0.00240941 | 0.01145684 | NFX1 | 4799 |
| ENSG000002 | 15.7427565 | -0.925416442 | 0.30497213 | -3.0344296 | 0.00240991 | 0.01145788 | NA | NA |
| ENSG000001 | 6.40223275 | -1.132523871 | 0.37325682 | -3.0341679 | 0.002412 | 0.01146648 | TBPL2 | 387332 |
| ENSG000001 | 1650.518 | 0.386363226 | 0.12734095 | 3.0340847 | 0.00241267 | 0.0114683 | ASNSD1 | 54529 |
| ENSG000001 | 2449.84453 | 0.384149384 | 0.12662046 | 3.03386506 | 0.00241442 | 0.01147486 | TCF4 | 6925 |
| ENSG000002 | 4.61088705 | -1.144733837 | 0.37732157 | -3.0338415 | 0.00241461 | 0.01147486 | NA | NA |
| ENSG000001 | 336.055333 | 0.406690337 | 0.13405342 | 3.03379313 | 0.002415 | 0.01147535 | SENP7 | 57337 |
| ENSG000001 | 269.019551 | 0.408342104 | 0.13460464 | 3.03364065 | 0.00241622 | 0.01147981 | PALLD | 23022 |
| ENSG000002 | 8.8763156 | -1.072555824 | 0.35356833 | -3.0335178 | 0.0024172 | 0.01148314 | LINC01508 | 101927873 |
| ENSG000001 | 300.020099 | 0.410979852 | 0.13548454 | 3.03340768 | 0.00241809 | 0.01148487 | STX2 | 2054 |
| ENSG000001 | 822.502009 | 0.393129316 | 0.12960016 | 3.03340147 | 0.00241814 | 0.01148487 | MFN1 | 55669 |
| ENSG000001 | 21.5383117 | -0.844273435 | 0.27834845 | -3.033153 | 0.00242013 | 0.01149299 | FBXO24 | 26261 |
| ENSG000001 | 483.128269 | 0.52876356 | 0.17433981 | 3.03294788 | 0.00242177 | 0.01149946 | DGCR8 | 54487 |
| ENSG000001 | 2469.15323 | 0.348635587 | 0.11495103 | 3.03290533 | 0.00242212 | 0.01149973 | PHF14 | 9678 |
| ENSG000002 | 7.73270563 | -1.087760177 | 0.3587047 | -3.032467 | 0.00242564 | 0.0115151 | NA | NA |
| ENSG000000 | 1075.73517 | 0.342691783 | 0.11300898 | 3.0324296 | 0.00242594 | 0.01151518 | MAP2K4 | 6416 |
| ENSG000001 | 172.384939 | 0.54951309 | 0.18121697 | 3.03234889 | 0.00242659 | 0.01151691 | NAB2 | 4665 |
| ENSG000001 | 448.299015 | 0.552008141 | 0.18204847 | 3.0322043 | 0.00242775 | 0.01152108 | RNF26 | 79102 |
| ENSG000001 | 185.178117 | 0.485025785 | 0.15997305 | 3.0319218 | 0.00243002 | 0.01153052 | PPP1R37 | 284352 |
| ENSG000002 | 9.61287927 | -1.040107131 | 0.34306375 | -3.0318188 | 0.00243085 | 0.0115331 | FGD5P1 | 100132526 |
| ENSG000002 | 4.54678878 | -1.143679349 | 0.37727392 | -3.0314297 | 0.00243399 | 0.01154663 | NA | NA |
| ENSG000001 | 476.406293 | 0.509030275 | 0.16793209 | 3.03116732 | 0.0024361 | 0.01155261 | SPTBN2 | 6712 |
| ENSG000002 | 5.78534883 | 1.128885471 | 0.37242415 | 3.03118226 | 0.00243598 | 0.01155261 | MIR130B | 406920 |
| ENSG000002 | 17.7402923 | -0.914425155 | 0.30166734 | -3.0312369 | 0.00243554 | 0.01155261 | LOC100507391 | 100507391 |
| ENSG000002 | 2.22037525 | -1.098326072 | 0.36234998 | -3.0311195 | 0.00243649 | 0.01155309 | NA | NA |
| ENSG000000 | 13.8095334 | -0.973733599 | 0.32127459 | -3.0308453 | 0.0024387 | 0.01156039 | ATP2B3 | 492 |

| | | | | | | | | | |
|---|---|---|---|---|---|---|---|---|---|
| ENSG000001 | 27.6503635 | -0.771411724 | 0.25452222 | -3.0308228 | 0.00243888 | 0.01156039 | TESPA1 | | 9840 |
| ENSG000002 | 6.67358478 | -1.108510358 | 0.36573889 | -3.0308791 | 0.00243843 | 0.01156039 | NA | NA | |
| ENSG000001 | 68.9890805 | 0.58856854 | 0.19422389 | 3.03036115 | 0.00244261 | 0.01157673 | SLC26A11 | | 284129 |
| ENSG00000C | 29.0721488 | -0.772355576 | 0.25489459 | -3.030098 | 0.00244474 | 0.01158546 | NXPE1 | | 120400 |
| ENSG000001 | 2284.52504 | 0.419078607 | 0.13830981 | 3.02999916 | 0.00244554 | 0.0115879 | ADNP | | 23394 |
| ENSG000001 | 19.8985476 | -0.90249389 | 0.29787045 | -3.0298201 | 0.00244699 | 0.01159342 | KRT1 | | 3848 |
| ENSG000001 | 127.11701 | 0.609768188 | 0.201258 | 3.02978366 | 0.00244729 | 0.01159346 | NPRL3 | | 8131 |
| ENSG000001 | 392.619808 | 0.370902632 | 0.1224442 | 3.02915646 | 0.00245238 | 0.0116162 | ZNF75A | | 7627 |
| ENSG000002 | 356.395299 | 0.443272866 | 0.14633839 | 3.02909481 | 0.00245288 | 0.01161722 | NA | NA | |
| ENSG000002 | 14.2475091 | -0.977636622 | 0.32277965 | -3.028805 | 0.00245523 | 0.01162457 | NA | NA | |
| ENSG000002 | 31.7285679 | -0.80256755 | 0.264982 | -3.0287625 | 0.00245558 | 0.01162457 | CPEB2-AS1 | | 441009 |
| ENSG000002 | 4.04503761 | -1.146167052 | 0.37842396 | -3.0287909 | 0.00245535 | 0.01162457 | NA | NA | |
| ENSG000002 | 13.4114557 | -1.019287238 | 0.33652506 | -3.0288599 | 0.00245479 | 0.01162457 | NA | NA | |
| ENSG000001 | 280.185077 | 0.433506518 | 0.14315047 | 3.02832767 | 0.00245911 | 0.01163957 | TRIOBP | | 11078 |
| ENSG000001 | 163.090458 | 0.4748098 | 0.15679075 | 3.02830241 | 0.00245932 | 0.01163957 | B4GALNT3 | | 283358 |
| ENSG00000C | 142.431953 | 0.516812192 | 0.17068308 | 3.02790518 | 0.00246255 | 0.01165009 | ZNF76 | | 7629 |
| ENSG000001 | 901.626585 | 0.368325539 | 0.12164145 | 3.02796066 | 0.0024621 | 0.01165009 | SETDB1 | | 9869 |
| ENSG000001 | 856.287743 | 0.399678584 | 0.13199912 | 3.02788831 | 0.00246269 | 0.01165009 | CENPC | | 1060 |
| ENSG000002 | 7.63054789 | -1.106268263 | 0.36535212 | -3.0279509 | 0.00246218 | 0.01165009 | NA | NA | |
| ENSG000001 | 95.5333181 | 0.510428845 | 0.16857863 | 3.02783834 | 0.0024631 | 0.01165066 | CA2 | | 760 |
| ENSG00000C | 37.8240481 | -0.693312398 | 0.22898638 | -3.0277451 | 0.00246386 | 0.01165153 | MPP4 | | 58538 |
| ENSG000001 | 362.922836 | 0.494339923 | 0.1632682 | 3.02777844 | 0.00246359 | 0.01165153 | FAM120B | | 84498 |
| ENSG000001 | 44.1098734 | 0.765657022 | 0.25289442 | 3.02757576 | 0.00246524 | 0.01165535 | SMDT1 | | 91689 |
| ENSG000002 | 60.6202901 | 0.661625116 | 0.21853191 | 3.02759039 | 0.00246512 | 0.01165535 | NA | NA | |
| ENSG000001 | 47.0730188 | -0.643038741 | 0.21242691 | -3.0271059 | 0.00246907 | 0.01167212 | CLIP4 | | 79745 |
| ENSG000001 | 24.5261067 | -0.83445723 | 0.27567236 | -3.0269891 | 0.00247003 | 0.01167391 | NA | NA | |
| ENSG000002 | 6.54296948 | -1.120106766 | 0.37003562 | -3.0270242 | 0.00246974 | 0.01167391 | NA | NA | |
| ENSG000001 | 61.3520648 | -0.608417306 | 0.20102319 | -3.0266026 | 0.00247319 | 0.01168748 | EFCAB5 | | 374786 |
| ENSG000001 | 823.720824 | 0.386908011 | 0.12783772 | 3.02655588 | 0.00247357 | 0.01168793 | NDUFA12 | | 55967 |

| | | | | | | | | | |
|---|---|---|---|---|---|---|---|---|---|
| ENSG000001 | 306.27075 | 0.494559352 | 0.16342074 | 3.02629494 | 0.00247571 | 0.01169665 | ASCC2 | | 84164 |
| ENSG00000C | 727.070238 | 0.367720611 | 0.12151415 | 3.02615456 | 0.00247686 | 0.01169936 | POLR3B | | 55703 |
| ENSG000001 | 432.679585 | 0.369561645 | 0.12212208 | 3.02616572 | 0.00247677 | 0.01169936 | GEMIN6 | | 79833 |
| ENSG000002 | 11.6071543 | -1.026498086 | 0.33926085 | -3.0256898 | 0.00248067 | 0.01171599 | NA | NA | |
| ENSG000002 | 11.3143431 | -1.054858075 | 0.34867284 | -3.025352 | 0.00248344 | 0.01172772 | NA | NA | |
| ENSG000001 | 2053.32842 | 0.375616574 | 0.1241953 | 3.02440243 | 0.00249125 | 0.01176323 | VPS13A | | 23230 |
| ENSG000001 | 12.6752611 | -0.974868453 | 0.32233835 | -3.0243639 | 0.00249157 | 0.01176336 | OR2T6 | | 254879 |
| ENSG000001 | 61.500676 | 0.717859026 | 0.2374165 | 3.02362731 | 0.00249764 | 0.01178929 | ABCD1 | | 215 |
| ENSG000001 | 355.050476 | 0.367334731 | 0.12148779 | 3.02363502 | 0.00249758 | 0.01178929 | MECP2 | | 4204 |
| ENSG000001 | 890.621747 | 0.40523203 | 0.1340281 | 3.02348556 | 0.00249881 | 0.01179344 | SUCLA2 | | 8803 |
| ENSG00000C | 183.275106 | 0.457527112 | 0.15132874 | 3.02339877 | 0.00249953 | 0.01179545 | YBX2 | | 51087 |
| ENSG000001 | 44.4614475 | -0.702038262 | 0.23224538 | -3.0228298 | 0.00250423 | 0.01181627 | ITGA11 | | 22801 |
| ENSG000001 | 35.7789592 | -0.757082086 | 0.25046056 | -3.0227597 | 0.00250481 | 0.01181693 | FAM9B | | 171483 |
| ENSG000001 | 522.166918 | 0.447717591 | 0.14811635 | 3.02274261 | 0.00250495 | 0.01181693 | AP2A2 | | 161 |
| ENSG00000C | 20.1158526 | -0.912895562 | 0.30202526 | -3.0225802 | 0.0025063 | 0.0118219 | EFCAB1 | | 79645 |
| ENSG000001 | 836.174131 | 0.38237843 | 0.12651908 | 3.02229861 | 0.00250863 | 0.01183152 | RSU1 | | 6251 |
| ENSG000002 | 154.347432 | 0.494298143 | 0.16355315 | 3.02224769 | 0.00250905 | 0.01183214 | NA | NA | |
| ENSG000001 | 585.004967 | 0.347732249 | 0.1150657 | 3.0220323 | 0.00251084 | 0.01183505 | FRG1 | | 2483 |
| ENSG000002 | 84.7123579 | 0.590627824 | 0.19543997 | 3.0220421 | 0.00251076 | 0.01183505 | SNORA33 | | 594839 |
| ENSG000002 | 25.6383888 | -0.791913393 | 0.26204144 | -3.0220923 | 0.00251034 | 0.01183505 | LINC00954 | | 400946 |
| ENSG000002 | 5.36626584 | -1.132068475 | 0.37459578 | -3.0221068 | 0.00251022 | 0.01183505 | NA | NA | |
| ENSG00000C | 803.756702 | 0.373521707 | 0.12360743 | 3.02183865 | 0.00251244 | 0.01184125 | ZNF510 | | 22869 |
| ENSG00000C | 11.6134038 | -1.009381143 | 0.33404208 | -3.0217185 | 0.00251344 | 0.01184421 | HHATL | | 57467 |
| ENSG000002 | 8.70959917 | -1.06281506 | 0.35172839 | -3.0216926 | 0.00251366 | 0.01184421 | NA | NA | |
| ENSG000001 | 2985.45384 | 0.358387072 | 0.11860981 | 3.02156362 | 0.00251473 | 0.01184788 | CKAP2 | | 26586 |
| ENSG00000C | 1274.46214 | 0.384427363 | 0.12723036 | 3.02150658 | 0.0025152 | 0.01184816 | MPHOSPH9 | | 10198 |
| ENSG000002 | 25.8874677 | -0.830377134 | 0.27482407 | -3.0214862 | 0.00251537 | 0.01184816 | NA | NA | |
| ENSG000001 | 12.6833314 | -0.972505378 | 0.32186922 | -3.0214302 | 0.00251584 | 0.01184898 | NA | NA | |
| ENSG000002 | 3.63626442 | -1.139622261 | 0.37719574 | -3.021302 | 0.0025169 | 0.01185262 | NA | NA | |

| | | | | | | | | |
|---|---|---|---|---|---|---|---|---|
| ENSG000001 | 1050.87598 | 0.376790183 | 0.12473128 | 3.02081556 | 0.00252095 | 0.0118703 | CARNMT1 | 138199 |
| ENSG000002 | 11.6974062 | -1.016101115 | 0.33639248 | -3.0205822 | 0.00252289 | 0.01187807 | NA | NA |
| ENSG000002 | 2.75279455 | -1.115128498 | 0.36918739 | -3.0204946 | 0.00252362 | 0.01188013 | NA | NA |
| ENSG000001 | 24.7340164 | -0.824064895 | 0.27287272 | -3.0199607 | 0.00252807 | 0.0118997 | LINC00238 | 440184 |
| ENSG000002 | 8.65090922 | -1.062773301 | 0.35192866 | -3.0198543 | 0.00252896 | 0.0119025 | LOC101927501 | 101927501 |
| ENSG000001 | 17.5330669 | -0.89818755 | 0.29745054 | -3.0196198 | 0.00253092 | 0.01190836 | VSIG1 | 340547 |
| ENSG000001 | 14.8656687 | -0.945334917 | 0.31305981 | -3.0196623 | 0.00253057 | 0.01190836 | CHRNB3 | 1142 |
| ENSG000002 | 6.2983968 | -1.111317356 | 0.36803464 | -3.0195998 | 0.00253109 | 0.01190836 | NA | NA |
| ENSG000001 | 18.7081949 | -0.876906918 | 0.29043407 | -3.0192977 | 0.00253361 | 0.01191886 | BBOX1 | 8424 |
| ENSG000001 | 37.8965712 | -0.69505776 | 0.23021331 | -3.0191901 | 0.00253451 | 0.01192171 | ADAMTS18 | 170692 |
| ENSG000002 | 8.54213691 | -1.075169309 | 0.3561295 | -3.0190403 | 0.00253577 | 0.01192623 | NA | NA |
| ENSG000001 | 22.2177075 | -0.840616447 | 0.27849776 | -3.0183957 | 0.00254117 | 0.01194674 | PKDREJ | 10343 |
| ENSG000001 | 32.730515 | -0.730804179 | 0.24211293 | -3.0184434 | 0.00254077 | 0.01194674 | ALB | 213 |
| ENSG000001 | 24.1280116 | -0.832491121 | 0.27580772 | -3.018375 | 0.00254134 | 0.01194674 | KBTBD12 | 166348 |
| ENSG000002 | 160.669533 | 0.436957538 | 0.14476732 | 3.01834385 | 0.0025416 | 0.01194674 | LNP1 | 348801 |
| ENSG000002 | 31.2956037 | -0.743695247 | 0.24638336 | -3.0184476 | 0.00254073 | 0.01194674 | LOC101926964 | 101926964 |
| ENSG000001 | 1358.59262 | 0.380261774 | 0.12600568 | 3.01781445 | 0.00254605 | 0.01196625 | ZNF84 | 7637 |
| ENSG000001 | 6154.03349 | 0.350094818 | 0.11601488 | 3.01767167 | 0.00254725 | 0.01196854 | HNRNPR | 10236 |
| ENSG000001 | 2396.52403 | 0.368437727 | 0.12209422 | 3.01765092 | 0.00254742 | 0.01196854 | KIAA1109 | 84162 |
| ENSG000001 | 45.9368881 | -0.687688161 | 0.22788829 | -3.0176546 | 0.00254739 | 0.01196854 | TMC1 | 117531 |
| ENSG000001 | 2949.71884 | 0.380907199 | 0.12622926 | 3.01758252 | 0.002548 | 0.01196903 | PTPRG | 5793 |
| ENSG000001 | 4534.02874 | 0.331058924 | 0.1097105 | 3.01756824 | 0.00254812 | 0.01196903 | MGA | 23269 |
| ENSG000002 | 11.0295423 | -0.999044623 | 0.3310843 | -3.0174932 | 0.00254875 | 0.01197061 | NA | NA |
| ENSG000001 | 72.6550861 | 0.661319056 | 0.21916478 | 3.01745133 | 0.0025491 | 0.01197088 | RHOG | 391 |
| ENSG000001 | 199.18649 | 0.452325497 | 0.14991268 | 3.01725982 | 0.00255071 | 0.01197706 | RNF123 | 63891 |
| ENSG000001 | 136.29656 | 0.559838525 | 0.18556118 | 3.01700236 | 0.00255288 | 0.01198585 | NENF | 29937 |
| ENSG00000C | 690.458158 | 0.395749006 | 0.13118055 | 3.01682695 | 0.00255436 | 0.0119914 | RNF216 | 54476 |
| ENSG000001 | 525.047263 | 0.557276162 | 0.18474186 | 3.01651262 | 0.00255701 | 0.01200106 | SOX12 | 6666 |
| ENSG000001 | 546.956185 | 0.406973844 | 0.13491502 | 3.01651983 | 0.00255694 | 0.01200106 | ZFP82 | 284406 |

| | | | | | | | | | |
|---|---|---|---|---|---|---|---|---|---|
| ENSG000001 | 320.028753 | 0.435035128 | 0.14422028 | 3.01646303 | 0.00255742 | 0.01200163 | FANCB | | 2187 |
| ENSG000001 | 344.427874 | 0.468846588 | 0.1554317 | 3.01641555 | 0.00255782 | 0.01200212 | AMER1 | | 139285 |
| ENSG000002 | 17.507821 | 0.905192683 | 0.30010824 | 3.01622074 | 0.00255947 | 0.01200845 | SNORD12C | | 26765 |
| ENSG000002 | 20.1354311 | -0.904731778 | 0.29996634 | -3.016111 | 0.0025604 | 0.0120114 | NA | NA | |
| ENSG000001 | 356.831537 | 0.379816212 | 0.12593342 | 3.01600802 | 0.00256126 | 0.0120141 | BLOC1S5 | | 63915 |
| ENSG000001 | 14.4963675 | -0.964265189 | 0.31971991 | -3.0159686 | 0.0025616 | 0.01201427 | CFHR5 | | 81494 |
| ENSG000001 | 641.024394 | 0.342561362 | 0.11358624 | 3.01587026 | 0.00256243 | 0.01201677 | NDEL1 | | 81565 |
| ENSG000001 | 15.2843269 | -0.918922169 | 0.30473284 | -3.0155009 | 0.00256555 | 0.01203003 | SH2D4B | | 387694 |
| ENSG000001 | 47.3400849 | -0.652452875 | 0.21639161 | -3.0151487 | 0.00256853 | 0.01204122 | GHR | | 2690 |
| ENSG000001 | 1074.79572 | 0.366152115 | 0.12143694 | 3.0151625 | 0.00256842 | 0.01204122 | TUBB2A | | 7280 |
| ENSG000001 | 13143.8375 | 0.379432934 | 0.12584745 | 3.01502291 | 0.0025696 | 0.01204483 | EIF3E | | 3646 |
| ENSG000001 | 3911.48595 | 0.385510743 | 0.12786898 | 3.0148886 | 0.00257074 | 0.01204877 | TRIP12 | | 9320 |
| ENSG000001 | 101.949761 | 0.546869555 | 0.18139274 | 3.01483713 | 0.00257117 | 0.01204942 | WDR83 | | 84292 |
| ENSG000002 | 152.044809 | 0.531156024 | 0.17618543 | 3.01475572 | 0.00257186 | 0.01205126 | GNAS-AS1 | | 149775 |
| ENSG000001 | 248.877141 | 0.440223709 | 0.14602507 | 3.01471323 | 0.00257222 | 0.01205156 | NA | NA | |
| ENSG000002 | 2.76849038 | -1.111345023 | 0.3686727 | -3.0144489 | 0.00257447 | 0.01206067 | OR13C2 | | 392376 |
| ENSG000001 | 3828.17337 | 0.354644594 | 0.11765679 | 3.01422965 | 0.00257633 | 0.01206521 | SMARCAD1 | | 56916 |
| ENSG000001 | 521.547886 | 0.386018674 | 0.12806272 | 3.01429392 | 0.00257578 | 0.01206521 | ATXN7 | | 6314 |
| ENSG000002 | 23.3640766 | -0.807792389 | 0.26799128 | -3.0142487 | 0.00257617 | 0.01206521 | ASB14 | | 142686 |
| ENSG00000C | 27.1699395 | -0.793702131 | 0.26333094 | -3.0140861 | 0.00257755 | 0.01206953 | MUSK | | 4593 |
| ENSG000002 | 85.5893899 | 0.517383584 | 0.17166559 | 3.01390394 | 0.00257909 | 0.01207538 | NA | NA | |
| ENSG000001 | 1566.69502 | 0.321264545 | 0.10659964 | 3.013749 | 0.00258041 | 0.01208015 | SLC39A6 | | 25800 |
| ENSG00000C | 1569.92256 | 0.345534835 | 0.11465588 | 3.01366872 | 0.00258109 | 0.01208195 | ME2 | | 4200 |
| ENSG000002 | 75.8376388 | 0.603524484 | 0.20026691 | 3.01360061 | 0.00258167 | 0.01208327 | PPT2 | | 9374 |
| ENSG000001 | 589.220643 | 0.377536749 | 0.12528948 | 3.01331568 | 0.0025841 | 0.01209323 | MTO1 | | 25821 |
| ENSG000001 | 2018.49185 | 0.36616982 | 0.12152005 | 3.01324618 | 0.00258469 | 0.0120946 | PLAA | | 9373 |
| ENSG000001 | 22.1890414 | -0.820452956 | 0.27229106 | -3.0131469 | 0.00258554 | 0.01209716 | SLC22A10 | | 387775 |
| ENSG000001 | 1141.54368 | 0.323438279 | 0.10734803 | 3.01298759 | 0.00258689 | 0.01210212 | PHF20L1 | | 51105 |
| ENSG000001 | 25.7213745 | -0.808826362 | 0.268477 | -3.0126467 | 0.0025898 | 0.01211298 | TRIM58 | | 25893 |

| | | | | | | | | |
|---|---|---|---|---|---|---|---|---|
| ENSG000001 | 137.639041 | 0.489418579 | 0.16245475 | 3.01264548 | 0.00258981 | 0.01211298 | ALG10B | 144245 |
| ENSG000002 | 6.36254659 | -1.110181764 | 0.36851766 | -3.0125606 | 0.00259054 | 0.01211497 | NA | NA |
| ENSG000001 | 4032.78068 | 0.440747035 | 0.14631274 | 3.01236261 | 0.00259223 | 0.01212147 | HMGCR | 3156 |
| ENSG000001 | 7.46211193 | -1.094276186 | 0.36329917 | -3.0120525 | 0.00259488 | 0.01213246 | WISP1 | 8840 |
| ENSG000001 | 190.626454 | 0.441110953 | 0.14645539 | 3.01191332 | 0.00259607 | 0.01213663 | ATG10 | 83734 |
| ENSG000002 | 41.9434892 | -0.721131927 | 0.23943693 | -3.0117824 | 0.00259719 | 0.01214046 | MSC-AS1 | 100132891 |
| ENSG000001 | 17.7965586 | -0.896601192 | 0.29770995 | -3.0116602 | 0.00259823 | 0.01214255 | STOML3 | 161003 |
| ENSG000001 | 68.0171026 | -0.570259127 | 0.18934843 | -3.0116918 | 0.00259796 | 0.01214255 | PPFIA2 | 8499 |
| ENSG000001 | 177.296034 | 0.479483417 | 0.15921833 | 3.0114838 | 0.00259974 | 0.01214821 | ORMDL2 | 29095 |
| ENSG000002 | 24.7596119 | -0.847026794 | 0.28127056 | -3.0114307 | 0.0026002 | 0.01214893 | NA | NA |
| ENSG000001 | 164.059814 | -0.432321184 | 0.14356588 | -3.0113087 | 0.00260124 | 0.01215102 | MFSD6 | 54842 |
| ENSG000001 | 790.540488 | 0.42457905 | 0.14099484 | 3.01130911 | 0.00260124 | 0.01215102 | GNG12 | 55970 |
| ENSG00000C | 1649.85201 | 0.340436818 | 0.1130555 | 3.01123628 | 0.00260186 | 0.01215252 | MAP4K3 | 8491 |
| ENSG000002 | 14.5176086 | -0.941437616 | 0.31268811 | -3.010788 | 0.00260571 | 0.01216907 | NA | NA |
| ENSG000001 | 24.8982296 | -0.801021397 | 0.26608319 | -3.0104172 | 0.00260889 | 0.01218254 | SNX31 | 169166 |
| ENSG000001 | 266.004816 | 0.533659927 | 0.17730327 | 3.00987071 | 0.00261359 | 0.01220307 | MLXIP | 22877 |
| ENSG000001 | 1274.31071 | 0.388471701 | 0.1290697 | 3.00978236 | 0.00261435 | 0.01220521 | INTS3 | 65123 |
| ENSG000001 | 3712.05743 | 0.378210605 | 0.12566554 | 3.00966044 | 0.0026154 | 0.0122073 | DESI2 | 51029 |
| ENSG000001 | 279.390618 | 0.398588349 | 0.1324356 | 3.00967679 | 0.00261526 | 0.0122073 | SLC2A13 | 114134 |
| ENSG00000C | 891.9307 | 0.35500197 | 0.11795714 | 3.00958431 | 0.00261605 | 0.01220896 | ZW10 | 9183 |
| ENSG000001 | 18.1581489 | -0.868808037 | 0.28868772 | -3.0095082 | 0.00261671 | 0.01221061 | KCNE4 | 23704 |
| ENSG000001 | 35.8776217 | -0.69372151 | 0.23052092 | -3.0093647 | 0.00261795 | 0.01221469 | ABCC3 | 8714 |
| ENSG000001 | 31.8835895 | -0.784645513 | 0.26073702 | -3.0093369 | 0.00261819 | 0.01221469 | TMC2 | 117532 |
| ENSG00000C | 12.7386104 | -0.979804024 | 0.32559987 | -3.0092273 | 0.00261913 | 0.01221657 | NA | NA |
| ENSG000001 | 110.840135 | 0.621267951 | 0.20645479 | 3.00922027 | 0.00261919 | 0.01221657 | ABCA3 | 21 |
| ENSG000001 | 21.53516 | -0.858300655 | 0.28523467 | -3.0091035 | 0.0026202 | 0.01221986 | DTHD1 | 401124 |
| ENSG000002 | 28.713942 | -0.788475216 | 0.26203913 | -3.008998 | 0.00262111 | 0.0122227 | OR2A5 | 393046 |
| ENSG000002 | 15.9438732 | -0.91358386 | 0.30366616 | -3.0085139 | 0.00262529 | 0.01224078 | NA | NA |
| ENSG000002 | 6.88812047 | -1.109892655 | 0.36893046 | -3.0084061 | 0.00262622 | 0.01224371 | NA | NA |

| | | | | | | | | |
|---|---|---|---|---|---|---|---|---|
| ENSG00000( | 185.547582 | 0.449339743 | 0.14937679 | 3.00809614 | 0.0026289 | 0.01225198 | DZANK1 | 55184 |
| ENSG000001 | 113.127132 | 0.504812963 | 0.16781729 | 3.00811061 | 0.00262877 | 0.01225198 | BORCS5 | 118426 |
| ENSG00000Z | 8.10218879 | -1.077503839 | 0.35819642 | -3.0081369 | 0.00262855 | 0.01225198 | NA | NA |
| ENSG000001 | 23.0482785 | -0.810754717 | 0.26955209 | -3.007785 | 0.00263159 | 0.01226313 | TULP2 | 7288 |
| ENSG000001 | 969.230323 | 0.40574621 | 0.13490517 | 3.00764029 | 0.00263285 | 0.01226755 | RNGTT | 8732 |
| ENSG000001 | 178.106203 | 0.437711333 | 0.14554769 | 3.00733963 | 0.00263545 | 0.01227828 | IFT27 | 11020 |
| ENSG000001 | 19.0824914 | -0.891627494 | 0.29650125 | -3.0071627 | 0.00263699 | 0.01228402 | NPAP1 | 23742 |
| ENSG000001 | 144.947138 | 0.495700281 | 0.1648471 | 3.00703066 | 0.00263813 | 0.01228795 | LINC01405 | 100131138 |
| ENSG00000Z | 4.98493233 | -1.132308093 | 0.37656117 | -3.0069698 | 0.00263866 | 0.01228899 | NA | NA |
| ENSG000001 | 13.8411421 | -0.989853895 | 0.3291947 | -3.006895 | 0.00263931 | 0.01229061 | NA | NA |
| ENSG00000Z | 10.1288777 | -1.021408156 | 0.33969552 | -3.0068343 | 0.00263984 | 0.01229165 | NA | NA |
| ENSG000001 | 78.5519976 | -0.553574985 | 0.18412334 | -3.0065443 | 0.00264236 | 0.01230143 | BMP4 | 652 |
| ENSG000001 | 29.7317904 | -0.750578951 | 0.24965018 | -3.0065227 | 0.00264254 | 0.01230143 | OR5A2 | 219981 |
| ENSG000001 | 1123.58439 | 0.344693747 | 0.11466221 | 3.00616687 | 0.00264564 | 0.01231442 | ARPC5 | 10092 |
| ENSG00000Z | 23.8773953 | -0.832044966 | 0.27679625 | -3.0059835 | 0.00264723 | 0.01232044 | CAPN14 | 440854 |
| ENSG000001 | 866.88154 | 0.359673371 | 0.11966232 | 3.00573628 | 0.00264939 | 0.01232904 | FTO | 79068 |
| ENSG000001 | 32.8897286 | -0.792070067 | 0.26352676 | -3.0056532 | 0.00265011 | 0.012331 | NOSTRIN | 115677 |
| ENSG000001 | 112.251887 | -0.507122563 | 0.16873255 | -3.0054815 | 0.00265161 | 0.01233654 | C1orf61 | 10485 |
| ENSG000001 | 6.74484152 | -1.100326318 | 0.36611467 | -3.0054145 | 0.00265219 | 0.01233785 | OR2AK2 | 391191 |
| ENSG00000( | 17.6408406 | -0.881524395 | 0.29333625 | -3.0051669 | 0.00265435 | 0.01234648 | SELE | 6401 |
| ENSG000001 | 712.503356 | 0.391780418 | 0.13037114 | 3.00511623 | 0.00265479 | 0.01234712 | GPATCH8 | 23131 |
| ENSG000001 | 380.029418 | 0.393409809 | 0.13093813 | 3.00454732 | 0.00265976 | 0.01236882 | TARSL2 | 123283 |
| ENSG000001 | 35.1732943 | -0.73756397 | 0.24548946 | -3.0044629 | 0.0026605 | 0.01236941 | PIWIL4 | 143689 |
| ENSG000001 | 418.073915 | 0.388134265 | 0.1291847 | 3.00449091 | 0.00266026 | 0.01236941 | ZNF429 | 353088 |
| ENSG000001 | 1423.02579 | 0.392562523 | 0.13066446 | 3.00435585 | 0.00266144 | 0.01237093 | ITM2B | 9445 |
| ENSG000001 | 10899.9771 | 0.353038935 | 0.11750793 | 3.0043838 | 0.00266119 | 0.01237093 | RANBP2 | 5903 |
| ENSG00000Z | 5.64808912 | -1.12044021 | 0.37296711 | -3.004126 | 0.00266345 | 0.01237886 | LOC146795 | 146795 |
| ENSG000001 | 134.315497 | 0.490898242 | 0.16341663 | 3.00396745 | 0.00266484 | 0.01238268 | LRRC20 | 55222 |
| ENSG00000Z | 28.5033011 | 0.792593894 | 0.26384947 | 3.00396239 | 0.00266488 | 0.01238268 | CEBPB-AS1 | 101927559 |

| | | | | | | | | | |
|---|---|---|---|---|---|---|---|---|---|
| ENSG000002 | 24.6675807 | 0.871783016 | 0.29023702 | 3.00369342 | 0.00266724 | 0.01239221 | SNORA5C | | 677796 |
| ENSG000001 | 81.4453773 | 0.591798526 | 0.1970504 | 3.0032851 | 0.00267082 | 0.01240743 | DNAJC30 | | 84277 |
| ENSG000001 | 334.390874 | 0.506074285 | 0.16851047 | 3.0032216 | 0.00267138 | 0.0124086 | ADK | | 132 |
| ENSG000002 | 19.2859843 | -0.857775604 | 0.28562326 | -3.0031714 | 0.00267182 | 0.01240922 | NA | NA | |
| ENSG000001 | 1439.75863 | 0.37362241 | 0.12441274 | 3.00308796 | 0.00267255 | 0.01241121 | DROSHA | | 29102 |
| ENSG000001 | 961.247872 | 0.352860932 | 0.11751119 | 3.00278589 | 0.00267521 | 0.01242069 | USP6NL | | 9712 |
| ENSG000002 | 7.74744216 | -1.101259047 | 0.36674429 | -3.0027981 | 0.0026751 | 0.01242069 | CSN1S2AP | | 286828 |
| ENSG000001 | 145.50441 | 0.490449477 | 0.16335161 | 3.00241591 | 0.00267846 | 0.01243437 | PIGL | | 9487 |
| ENSG000001 | 2595.76521 | 0.348687819 | 0.11614719 | 3.00212012 | 0.00268106 | 0.01244503 | MCMBP | | 79892 |
| ENSG000001 | 10.3127302 | -1.023060844 | 0.34078923 | -3.002034 | 0.00268182 | 0.01244713 | OR4F15 | | 390649 |
| ENSG000002 | 8.13152235 | -1.095145356 | 0.36480958 | -3.0019644 | 0.00268244 | 0.01244855 | NA | NA | |
| ENSG000001 | 14.3608088 | -0.999503009 | 0.33302433 | -3.0012913 | 0.00268837 | 0.01247325 | NEUROD2 | | 4761 |
| ENSG000001 | 182.793118 | 0.444984484 | 0.14826363 | 3.00130577 | 0.00268824 | 0.01247325 | SPATA5L1 | | 79029 |
| ENSG000001 | 126.765928 | 0.523256037 | 0.17435361 | 3.00111959 | 0.00268989 | 0.01247886 | C10orf88 | | 80007 |
| ENSG000001 | 3855.77634 | 0.366782499 | 0.12221794 | 3.00105275 | 0.00269048 | 0.01248017 | USP34 | | 9736 |
| ENSG000002 | 9.34119757 | -1.043382778 | 0.34771811 | -3.000657 | 0.00269398 | 0.01249497 | NA | NA | |
| ENSG000001 | 24.7208204 | -0.789575281 | 0.26314648 | -3.0005162 | 0.00269522 | 0.01249789 | POU4F1 | | 5457 |
| ENSG000001 | 238.341357 | 0.41857913 | 0.13950096 | 3.00054649 | 0.00269496 | 0.01249789 | DDX19A | | 55308 |
| ENSG000002 | 30.6136668 | -0.782768958 | 0.26090606 | -3.0001947 | 0.00269807 | 0.01250966 | NA | NA | |
| ENSG000001 | 3354.22302 | 0.34872876 | 0.11624023 | 3.00006946 | 0.00269918 | 0.01251337 | CCDC88A | | 55704 |
| ENSG000001 | 55.572826 | 0.632629635 | 0.21087608 | 3.00000656 | 0.00269974 | 0.01251453 | CCDC106 | | 29903 |
| ENSG000001 | 605.228446 | -0.36760633 | 0.12253924 | -2.999907 | 0.00270062 | 0.01251719 | RYR2 | | 6262 |
| ENSG000002 | 2.21869176 | -1.082302752 | 0.36083146 | -2.9994689 | 0.00270451 | 0.01253377 | NA | NA | |
| ENSG000002 | 3.13106565 | -1.124426106 | 0.37491098 | -2.9991816 | 0.00270706 | 0.01254416 | NA | NA | |
| ENSG000001 | 1055.7958 | 0.359351038 | 0.11982717 | 2.99891116 | 0.00270946 | 0.01255387 | ZNF493 | | 284443 |
| ENSG000001 | 739.857409 | 0.346334316 | 0.11549996 | 2.99856658 | 0.00271253 | 0.01256663 | SPIDR | | 23514 |
| ENSG000001 | 181.092781 | 0.587827064 | 0.196058 | 2.99823041 | 0.00271552 | 0.01257907 | PCIF1 | | 63935 |
| ENSG000001 | 264.313607 | 0.450717618 | 0.15033746 | 2.9980394 | 0.00271723 | 0.01258552 | KLF13 | | 51621 |
| ENSG000001 | 207.953737 | 0.479880092 | 0.16007888 | 2.99777275 | 0.0027196 | 0.01259366 | KLC1 | | 3831 |

| | | | | | | | | | |
|---|---|---|---|---|---|---|---|---|---|
| ENSG000002 | 19.4449041 | -0.875734431 | 0.29212687 | -2.997788 | 0.00271947 | 0.01259366 | MACROD2-IT1 | | 140848 |
| ENSG000002 | 4208.08945 | 0.355657159 | 0.11864606 | 2.99763144 | 0.00272087 | 0.01259806 | GAS5 | | 60674 |
| ENSG000001 | 1420.79171 | 0.388971273 | 0.12976136 | 2.99758928 | 0.00272124 | 0.01259836 | RAB3B | | 5865 |
| ENSG000002 | 56.0300698 | -0.622091753 | 0.20753675 | -2.9975016 | 0.00272202 | 0.01260055 | LINC01122 | | 400955 |
| ENSG000001 | 378.093196 | 0.381031269 | 0.12712021 | 2.99740899 | 0.00272285 | 0.01260294 | NETO2 | | 81831 |
| ENSG000002 | 10.0014399 | -1.031487559 | 0.34414648 | -2.997234 | 0.00272441 | 0.01260874 | NA | NA | |
| ENSG000000 | 16.3518535 | -0.914605266 | 0.30516776 | -2.9970573 | 0.00272599 | 0.01261461 | STAP1 | | 26228 |
| ENSG000002 | 4.99540461 | -1.128471096 | 0.37653784 | -2.996966 | 0.00272681 | 0.01261695 | NA | NA | |
| ENSG000002 | 4.4876073 | -1.13457331 | 0.37857852 | -2.9969299 | 0.00272713 | 0.012617 | NA | NA | |
| ENSG000002 | 39.3625398 | -0.721538962 | 0.24080376 | -2.9963775 | 0.00273208 | 0.01263844 | LINC00635 | | 151658 |
| ENSG000002 | 11.5053359 | -1.039579499 | 0.34699141 | -2.9959805 | 0.00273564 | 0.01265346 | NA | NA | |
| ENSG000000 | 147.170353 | 0.501668612 | 0.16744928 | 2.9959436 | 0.00273597 | 0.01265355 | LTBP4 | | 8425 |
| ENSG000001 | 26.8428891 | -0.798169563 | 0.26644334 | -2.9956446 | 0.00273865 | 0.01266452 | FCRL4 | | 83417 |
| ENSG000001 | 398.66599 | 0.538943493 | 0.1799222 | 2.99542526 | 0.00274062 | 0.01267074 | RBM38 | | 55544 |
| ENSG000001 | 1907.0527 | 0.366133667 | 0.12223077 | 2.99542955 | 0.00274059 | 0.01267074 | MIER3 | | 166968 |
| ENSG000002 | 1249.67295 | 0.347040811 | 0.11587217 | 2.9950315 | 0.00274416 | 0.01268566 | ZBED5 | | 58486 |
| ENSG000001 | 338.60881 | 0.537970407 | 0.17963344 | 2.99482323 | 0.00274604 | 0.01269143 | ERICH1 | | 157697 |
| ENSG000001 | 69.7596937 | 0.586108204 | 0.1957069 | 2.99482646 | 0.00274601 | 0.01269143 | ABHD14B | | 84836 |
| ENSG000001 | 227.820154 | 0.394235905 | 0.13164599 | 2.99466701 | 0.00274745 | 0.01269649 | UVRAG | | 7405 |
| ENSG000001 | 50.7690621 | -0.654801454 | 0.21868791 | -2.994228 | 0.0027514 | 0.01271333 | ANKFN1 | | 162282 |
| ENSG000001 | 582.949553 | 0.344727715 | 0.11513385 | 2.99414733 | 0.00275213 | 0.01271524 | TBCK | | 93627 |
| ENSG000001 | 573.59067 | 0.400855518 | 0.13389338 | 2.99384122 | 0.00275489 | 0.01272655 | XPNPEP3 | | 63929 |
| ENSG000002 | 3.79542876 | -1.133282252 | 0.37854781 | -2.9937625 | 0.0027556 | 0.01272838 | NA | NA | |
| ENSG000000 | 24.7940454 | -0.888048311 | 0.29664066 | -2.9936837 | 0.00275632 | 0.01273022 | SLC22A16 | | 85413 |
| ENSG000002 | 16.3174726 | -0.933819211 | 0.3119442 | -2.9935457 | 0.00275756 | 0.01273453 | NA | NA | |
| ENSG000001 | 6.71181582 | -1.098002104 | 0.36683838 | -2.9931495 | 0.00276114 | 0.01274962 | KLHL30 | | 377007 |
| ENSG000002 | 25.8993599 | -0.780930015 | 0.2609246 | -2.9929337 | 0.0027631 | 0.01275719 | NA | NA | |
| ENSG000001 | 821.539569 | 0.332264492 | 0.11102092 | 2.99280973 | 0.00276422 | 0.01276092 | CEP290 | | 80184 |
| ENSG000001 | 950.956062 | 0.373485598 | 0.12479959 | 2.99268288 | 0.00276537 | 0.01276477 | ZNF561 | | 93134 |

| | | | | | | | | |
|---|---|---|---|---|---|---|---|---|
| ENSG000002 | 18.2999671 | -0.876598841 | 0.29296754 | -2.9921364 | 0.00277032 | 0.01278619 | NA | NA |
| ENSG000001 | 956.270598 | 0.34818017 | 0.11637187 | 2.9919615 | 0.00277191 | 0.01279097 | ZNF766 | 90321 |
| ENSG000002 | 220.554651 | 0.583297514 | 0.19495546 | 2.99195263 | 0.00277199 | 0.01279097 | C1orf226 | 400793 |
| ENSG000002 | 59.2334048 | 0.610750283 | 0.20414791 | 2.99170486 | 0.00277424 | 0.0127999 | NA | NA |
| ENSG000000 | 278.839533 | 0.435122404 | 0.14545431 | 2.99147134 | 0.00277637 | 0.01280824 | PRKCZ | 5590 |
| ENSG000001 | 41.7508601 | -0.706033579 | 0.23603754 | -2.9911919 | 0.00277891 | 0.01281851 | SLC26A8 | 116369 |
| ENSG000002 | 16.6108078 | -0.926215377 | 0.30966417 | -2.9910318 | 0.00278037 | 0.01282377 | SLC39A12-AS1 | 100129213 |
| ENSG000002 | 22.7708264 | -0.827093518 | 0.27654539 | -2.9908056 | 0.00278243 | 0.01283181 | NA | NA |
| ENSG000002 | 17.235256 | -0.898419071 | 0.30040146 | -2.9907281 | 0.00278313 | 0.01283361 | LARGE-AS1 | 100506195 |
| ENSG000001 | 21.1262182 | -0.821317979 | 0.27465069 | -2.9904093 | 0.00278604 | 0.01284555 | KIRREL3 | 84623 |
| ENSG000001 | 7.05661637 | -1.089750493 | 0.36442691 | -2.9903129 | 0.00278692 | 0.01284815 | NA | NA |
| ENSG000001 | 527.034297 | 0.381871382 | 0.12773721 | 2.98950776 | 0.00279427 | 0.01288059 | CNKSR2 | 22866 |
| ENSG000001 | 368.802444 | 0.465406129 | 0.15568486 | 2.98941161 | 0.00279515 | 0.01288318 | RNPEP | 6051 |
| ENSG000001 | 42.6590408 | -0.674738704 | 0.22571779 | -2.9893022 | 0.00279615 | 0.01288633 | IL1R1 | 3554 |
| ENSG000001 | 288.604585 | 0.413645023 | 0.13840433 | 2.98867104 | 0.00280194 | 0.01291151 | ZCCHC2 | 54877 |
| ENSG000002 | 4.26179605 | -1.131119277 | 0.37847708 | -2.9886071 | 0.00280252 | 0.01291275 | NA | NA |
| ENSG000000 | 1829.62288 | 0.492920849 | 0.16495545 | 2.98820583 | 0.0028062 | 0.01292825 | ACTN1 | 87 |
| ENSG000001 | 747.859586 | 0.446433617 | 0.1494091 | 2.98799486 | 0.00280814 | 0.01293423 | ARFGEF3 | 57221 |
| ENSG000002 | 64.6149652 | 0.630875599 | 0.21113532 | 2.98801543 | 0.00280795 | 0.01293423 | RPSAP9 | 653162 |
| ENSG000000 | 37.0095477 | -0.780092823 | 0.26109432 | -2.9877817 | 0.0028101 | 0.01294179 | PRDM6 | 93166 |
| ENSG000001 | 14.6568159 | -0.928325559 | 0.31072295 | -2.9876312 | 0.00281149 | 0.01294522 | GCM1 | 8521 |
| ENSG000002 | 3.04614343 | -1.120052059 | 0.37489222 | -2.9876642 | 0.00281118 | 0.01294522 | NA | NA |
| ENSG000002 | 4.93170386 | -1.124449164 | 0.37637981 | -2.9875384 | 0.00281234 | 0.01294768 | NA | NA |
| ENSG000001 | 923.81247 | 0.382747513 | 0.12812244 | 2.98735738 | 0.00281401 | 0.01295241 | KIF3B | 9371 |
| ENSG000002 | 22.7826904 | -0.835168812 | 0.27956588 | -2.9873774 | 0.00281382 | 0.01295241 | NA | NA |
| ENSG000001 | 278.310181 | 0.577559097 | 0.19335209 | 2.98708483 | 0.00281652 | 0.01296249 | CARD11 | 84433 |
| ENSG000002 | 134.172308 | 0.476864249 | 0.15965941 | 2.98675944 | 0.00281951 | 0.01297482 | CYB5RL | 606495 |
| ENSG000002 | 16.8943491 | -0.904565015 | 0.30289117 | -2.9864357 | 0.0028225 | 0.01298709 | NA | NA |
| ENSG000002 | 14.9106929 | -0.936288126 | 0.31354177 | -2.9861671 | 0.00282498 | 0.01299703 | LINC00861 | 100130231 |

| | | | | | | | | | |
|---|---|---|---|---|---|---|---|---|---|
| ENSG000001 | 29.2876625 | -0.770014138 | 0.25787435 | -2.9860052 | 0.00282648 | 0.01300244 | ATP1B4 | | 23439 |
| ENSG000001 | 653.49678 | 0.452840073 | 0.15168811 | 2.98533658 | 0.00283266 | 0.01302942 | COL4A2 | | 1284 |
| ENSG000001 | 13.234704 | -0.956080222 | 0.32026643 | -2.9852652 | 0.00283333 | 0.01302951 | CRISP1 | | 167 |
| ENSG000001 | 19.8657931 | -0.862535048 | 0.28892853 | -2.9852886 | 0.00283311 | 0.01302951 | TMEM52B | | 120939 |
| ENSG000001 | 1151.45158 | 0.380103802 | 0.12734287 | 2.98488481 | 0.00283685 | 0.01304424 | DNA2 | | 1763 |
| ENSG000001 | 349.779153 | 0.390099169 | 0.13069637 | 2.98477421 | 0.00283788 | 0.01304748 | ERCC4 | | 2072 |
| ENSG000001 | 6620.93938 | 0.398449219 | 0.13350008 | 2.9846366 | 0.00283915 | 0.01305187 | MORF4L1 | | 10933 |
| ENSG000002 | 25.6710583 | 0.794285492 | 0.26614546 | 2.98440364 | 0.00284132 | 0.01306033 | NA | NA | |
| ENSG00000C | 57.3133732 | -0.591882819 | 0.19832928 | -2.9843441 | 0.00284187 | 0.01306139 | RPGRIP1 | | 57096 |
| ENSG000001 | 21.7396077 | -0.835848503 | 0.28010437 | -2.984061 | 0.0028445 | 0.013072 | PRDM16 | | 63976 |
| ENSG000001 | 321.237843 | 0.465871905 | 0.1561504 | 2.9834819 | 0.00284989 | 0.01309528 | SPPL3 | | 121665 |
| ENSG00000C | 304.027592 | 0.419925505 | 0.14075603 | 2.98335703 | 0.00285105 | 0.01309802 | STK17B | | 9262 |
| ENSG000001 | 37.6445879 | -0.788080726 | 0.2641598 | -2.9833485 | 0.00285113 | 0.01309802 | MAP3K19 | | 80122 |
| ENSG00000C | 198.122258 | 0.454155862 | 0.15223934 | 2.98317015 | 0.00285279 | 0.01310322 | ZNF582 | | 147948 |
| ENSG00000C | 21.4327823 | -0.854268869 | 0.28636397 | -2.9831576 | 0.00285291 | 0.01310322 | RARB | | 5915 |
| ENSG000002 | 4.49509014 | -1.128832121 | 0.37845439 | -2.9827429 | 0.00285678 | 0.01311951 | NA | NA | |
| ENSG000001 | 203.463026 | 0.42263442 | 0.14169792 | 2.98264387 | 0.0028577 | 0.01312226 | LPCAT4 | | 254531 |
| ENSG000001 | 41.2383256 | -0.698708331 | 0.23427075 | -2.9824822 | 0.00285921 | 0.01312771 | ARHGAP24 | | 83478 |
| ENSG000001 | 498.084216 | 0.376532154 | 0.12625545 | 2.98230414 | 0.00286088 | 0.01313386 | ZUFSP | | 221302 |
| ENSG000001 | 878.47222 | 0.376552377 | 0.12627785 | 2.98193527 | 0.00286433 | 0.01314821 | RNF145 | | 153830 |
| ENSG000001 | 208.080287 | 0.48865649 | 0.16388006 | 2.98179346 | 0.00286565 | 0.01315022 | TMEM109 | | 79073 |
| ENSG000001 | 144.247601 | 0.577463724 | 0.19366381 | 2.98178434 | 0.00286574 | 0.01315022 | FAHD2B | | 151313 |
| ENSG000002 | 38.9004099 | -0.734518472 | 0.24633241 | -2.9818182 | 0.00286542 | 0.01315022 | NA | NA | |
| ENSG000002 | 59.1039277 | 0.633105728 | 0.21233264 | 2.98166941 | 0.00286681 | 0.01315219 | NA | NA | |
| ENSG000002 | 9.47677447 | -1.022827641 | 0.34303596 | -2.9816922 | 0.0028666 | 0.01315219 | NA | NA | |
| ENSG000002 | 9.45216621 | -1.081726827 | 0.36282344 | -2.9814138 | 0.00286921 | 0.01316168 | NA | NA | |
| ENSG000002 | 13.5798609 | -0.954005242 | 0.31998905 | -2.9813684 | 0.00286963 | 0.01316214 | NA | NA | |
| ENSG000002 | 20.6148172 | 0.838076059 | 0.28115563 | 2.98082615 | 0.00287472 | 0.01318398 | LOC102724788 | | 102724788 |
| ENSG000001 | 392.782126 | 0.418884544 | 0.14053551 | 2.98063134 | 0.00287655 | 0.01319088 | RCL1 | | 10171 |

| | | | | | | | | |
|---|---|---|---|---|---|---|---|---|
| ENSG000001 | 20.0671471 | -0.877365765 | 0.29436139 | -2.9805735 | 0.00287709 | 0.01319188 | PPP1R3A | 5506 |
| ENSG000002 | 144.387889 | 0.505391416 | 0.16958205 | 2.98021762 | 0.00288044 | 0.01320572 | NA | NA |
| ENSG000002 | 112.216495 | 0.515493837 | 0.17297672 | 2.98013421 | 0.00288122 | 0.01320782 | NA | NA |
| ENSG000001 | 446.5407 | 0.405562117 | 0.13609113 | 2.98007755 | 0.00288175 | 0.01320877 | PPAP2B | 8613 |
| ENSG000001 | 8.21867366 | -1.091759104 | 0.36637551 | -2.9798911 | 0.00288351 | 0.01321532 | OR52N5 | 390075 |
| ENSG000001 | 36.0483571 | -0.684281141 | 0.22963734 | -2.979834 | 0.00288405 | 0.01321629 | ATP2B2 | 491 |
| ENSG000002 | 8.34713911 | -1.06045963 | 0.35588916 | -2.9797469 | 0.00288487 | 0.01321856 | NA | NA |
| ENSG000001 | 418.709863 | 0.487598493 | 0.16364405 | 2.97962866 | 0.00288598 | 0.01322207 | EMD | 2010 |
| ENSG000001 | 9.40309812 | -1.021491801 | 0.34282893 | -2.9795963 | 0.00288629 | 0.01322207 | TRY2P | 207147 |
| ENSG000002 | 18.0731093 | -0.896457214 | 0.30088013 | -2.9794497 | 0.00288767 | 0.0132269 | C17orf105 | 284067 |
| ENSG000001 | 11.0113286 | -0.992316832 | 0.33306311 | -2.9793658 | 0.00288846 | 0.01322903 | KIAA0087 | 9808 |
| ENSG00000C | 24.0975656 | -0.79344406 | 0.2663479 | -2.9789762 | 0.00289213 | 0.01324436 | NPC1L1 | 29881 |
| ENSG000001 | 240.55132 | 0.49383215 | 0.16579324 | 2.9786024 | 0.00289566 | 0.01325903 | HSPBP1 | 23640 |
| ENSG000001 | 345.032397 | 0.381530826 | 0.12809552 | 2.97848689 | 0.00289675 | 0.01326254 | VTI1A | 143187 |
| ENSG000001 | 1839.49558 | 0.356323141 | 0.11967721 | 2.97736835 | 0.00290734 | 0.01330829 | PUS7L | 83448 |
| ENSG000001 | 1227.80294 | 0.458761723 | 0.15408688 | 2.97729249 | 0.00290806 | 0.01330829 | DDI2 | 84301 |
| ENSG000001 | 190.877314 | 0.434828894 | 0.14604784 | 2.97730453 | 0.00290795 | 0.01330829 | TMEM116 | 89894 |
| ENSG000002 | 16.8583673 | -0.933822271 | 0.31365176 | -2.9772582 | 0.00290839 | 0.01330829 | OR1C1 | 26188 |
| ENSG000002 | 7.60610913 | -1.069863846 | 0.35933569 | -2.9773381 | 0.00290763 | 0.01330829 | LOC100506289 | 100506289 |
| ENSG000002 | 7.78365843 | -1.07988265 | 0.36272321 | -2.9771534 | 0.00290938 | 0.01331134 | NA | NA |
| ENSG000002 | 18.6209309 | -0.858421888 | 0.28834203 | -2.977096 | 0.00290993 | 0.01331233 | LOC100507073 | 100507073 |
| ENSG000001 | 70.2382785 | -0.621017326 | 0.20860131 | -2.9770538 | 0.00291033 | 0.01331267 | CFAP70 | 118491 |
| ENSG000002 | 32.4132082 | 0.728549382 | 0.24472937 | 2.97695933 | 0.00291123 | 0.01331526 | LTB4R2 | 56413 |
| ENSG000001 | 26.6076419 | -0.831750965 | 0.27941224 | -2.9767879 | 0.00291285 | 0.01332121 | SPATA9 | 83890 |
| ENSG000001 | 1670.82848 | 0.364453586 | 0.12243425 | 2.97672902 | 0.00291341 | 0.01332227 | FOXO1 | 2308 |
| ENSG00000C | 27.0928566 | -0.805631994 | 0.27065861 | -2.9765614 | 0.00291501 | 0.01332805 | ASB4 | 51666 |
| ENSG000001 | 195.013105 | 0.417249507 | 0.1401861 | 2.97639715 | 0.00291657 | 0.01333218 | ARMCX6 | 54470 |
| ENSG000002 | 9.4620638 | -1.019216065 | 0.342429 | -2.9764303 | 0.00291625 | 0.01333218 | NA | NA |
| ENSG000001 | 499.741926 | 0.344853725 | 0.1158734 | 2.97612502 | 0.00291916 | 0.01334251 | C16orf87 | 388272 |

| | | | | | | | | |
|---|---|---|---|---|---|---|---|---|
| ENSG000002 | 9.32461307 | -1.039346583 | 0.34924264 | -2.9760014 | 0.00292034 | 0.01334639 | NA | NA |
| ENSG000001 | 162.592585 | 0.47779408 | 0.16056527 | 2.97570011 | 0.00292321 | 0.0133565 | RNF113A | 7737 |
| ENSG000001 | 1141.88879 | 0.343842037 | 0.11554996 | 2.97570007 | 0.00292321 | 0.0133565 | PTPN2 | 5771 |
| ENSG000002 | 73.1025255 | 0.580531052 | 0.19510051 | 2.97554866 | 0.00292465 | 0.01336159 | LOC105369779 | 105369779 |
| ENSG000002 | 20.5351651 | -0.841510491 | 0.28281767 | -2.9754523 | 0.00292557 | 0.01336428 | NA | NA |
| ENSG000001 | 178.981932 | 0.434770036 | 0.14612473 | 2.9753351 | 0.00292669 | 0.01336638 | RAMP2 | 10266 |
| ENSG000002 | 26.1452013 | -0.782300491 | 0.26292585 | -2.9753655 | 0.0029264 | 0.01336638 | NA | NA |
| ENSG000001 | 21.9264738 | -0.825304503 | 0.2773938 | -2.9752089 | 0.00292789 | 0.01337037 | PROX2 | 283571 |
| ENSG000002 | 11.3034062 | -1.004646423 | 0.33769415 | -2.9750187 | 0.00292971 | 0.01337716 | MUC21 | 394263 |
| ENSG000001 | 102.174363 | -0.517065307 | 0.17380962 | -2.9748946 | 0.00293089 | 0.01338106 | PLA2R1 | 22925 |
| ENSG000002 | 40.7098455 | -0.707400763 | 0.23781224 | -2.9746188 | 0.00293353 | 0.01339159 | LINC01291 | 102724515 |
| ENSG000001 | 17806.6373 | 0.320349857 | 0.1077003 | 2.97445646 | 0.00293508 | 0.01339717 | PSIP1 | 11168 |
| ENSG000002 | 969.420587 | 0.423380035 | 0.14235011 | 2.97421635 | 0.00293738 | 0.01340615 | KIAA1147 | 57189 |
| ENSG000001 | 48.1185759 | -0.635992722 | 0.2138439 | -2.9740981 | 0.00293851 | 0.01340981 | TECTA | 7007 |
| ENSG000001 | 12.9198022 | -0.956749999 | 0.32170584 | -2.9739901 | 0.00293955 | 0.01341302 | NIPAL4 | 348938 |
| ENSG000002 | 34.004089 | -0.764255797 | 0.25698951 | -2.9738793 | 0.00294061 | 0.01341635 | C2orf91 | 400950 |
| ENSG000002 | 9.45693458 | -1.020700128 | 0.34324354 | -2.9736907 | 0.00294241 | 0.01342309 | NA | NA |
| ENSG000001 | 1825.81912 | 0.355504264 | 0.11955345 | 2.97360093 | 0.00294328 | 0.01342551 | PPP4R1 | 9989 |
| ENSG000001 | 15.599222 | -0.911009374 | 0.30637107 | -2.9735489 | 0.00294377 | 0.01342628 | TNFSF13B | 10673 |
| ENSG000001 | 4330.61795 | 0.37534075 | 0.1262469 | 2.97306903 | 0.00294838 | 0.01344565 | TBL1XR1 | 79718 |
| ENSG000002 | 8.36003732 | -1.05716571 | 0.35558441 | -2.9730373 | 0.00294869 | 0.01344565 | LINC00494 | 284749 |
| ENSG00000C | 389.800177 | 0.486931356 | 0.16378815 | 2.97293395 | 0.00294968 | 0.01344867 | WWTR1 | 25937 |
| ENSG00000C | 44.9681406 | -0.673776308 | 0.22666768 | -2.9725293 | 0.00295357 | 0.01346338 | MYOM2 | 9172 |
| ENSG000002 | 21.5054526 | -0.834704904 | 0.28080504 | -2.9725425 | 0.00295344 | 0.01346338 | NA | NA |
| ENSG000001 | 92.3477731 | 0.550193669 | 0.18511755 | 2.97213135 | 0.0029574 | 0.01347781 | KTI12 | 112970 |
| ENSG000002 | 5.84678338 | -1.122610131 | 0.37771027 | -2.9721462 | 0.00295726 | 0.01347781 | NA | NA |
| ENSG000001 | 122.524314 | 0.582034469 | 0.19583766 | 2.9720253 | 0.00295842 | 0.01347797 | STX4 | 6810 |
| ENSG000001 | 7.54945031 | -1.088354211 | 0.36619965 | -2.9720242 | 0.00295843 | 0.01347797 | NA | NA |
| ENSG000002 | 11.3391989 | -0.98564452 | 0.33163444 | -2.9720813 | 0.00295788 | 0.01347797 | NA | NA |

| | | | | | | | | |
|---|---|---|---|---|---|---|---|---|
| ENSG000001 | 362.513148 | 0.425185911 | 0.14307703 | 2.97172726 | 0.0029613 | 0.0134895 | SH3BP5L | 80851 |
| ENSG000001 | 178.977204 | -0.466067669 | 0.15685045 | -2.9714143 | 0.00296432 | 0.01350174 | RPS6KA5 | 9252 |
| ENSG000001 | 23.8428938 | -0.795038663 | 0.26757032 | -2.9713261 | 0.00296517 | 0.0135041 | RBM46 | 166863 |
| ENSG000001 | 14.0136682 | -0.952295025 | 0.32050373 | -2.9712447 | 0.00296595 | 0.01350616 | VWA5B2 | 90113 |
| ENSG000001 | 994.887312 | 0.356546025 | 0.1200008 | 2.971197 | 0.00296641 | 0.01350674 | STAG1 | 10274 |
| ENSG000001 | 76.5763092 | 0.661440266 | 0.22262097 | 2.97114987 | 0.00296687 | 0.0135073 | ZCCHC24 | 219654 |
| ENSG000001 | 187.770215 | -0.420530012 | 0.14154702 | -2.9709563 | 0.00296874 | 0.0135143 | CACNA1A | 773 |
| ENSG000002 | 17.7147719 | -0.863507456 | 0.29065505 | -2.9709013 | 0.00296927 | 0.0135152 | NA | NA |
| ENSG000001 | 268.107275 | 0.424776492 | 0.14298297 | 2.97081878 | 0.00297007 | 0.01351732 | DTNB | 1838 |
| ENSG000002 | 15.4284981 | -0.917942226 | 0.30900623 | -2.970627 | 0.00297193 | 0.01352424 | LINC01163 | 101927381 |
| ENSG000001 | 220.976205 | 0.433618937 | 0.14597304 | 2.9705412 | 0.00297276 | 0.01352499 | RBM48 | 84060 |
| ENSG000002 | 385.572147 | 0.430966254 | 0.14507937 | 2.97055505 | 0.00297262 | 0.01352499 | NA | NA |
| ENSG000001 | 341.316778 | 0.409035669 | 0.13770137 | 2.97045463 | 0.00297359 | 0.01352728 | BPHL | 670 |
| ENSG000002 | 27.746096 | -0.774875117 | 0.26090221 | -2.969983 | 0.00297816 | 0.01354655 | NA | NA |
| ENSG000001 | 1051.0152 | 0.352963591 | 0.11885209 | 2.96977185 | 0.00298021 | 0.01355434 | ZBTB24 | 9841 |
| ENSG000001 | 609.864103 | 0.413001082 | 0.13906992 | 2.96973694 | 0.00298055 | 0.01355436 | TRAK1 | 22906 |
| ENSG000002 | 18.925345 | -0.851097648 | 0.28662537 | -2.9693731 | 0.00298408 | 0.0135689 | FGF14-IT1 | 283480 |
| ENSG000002 | 25.9965 | -0.76579653 | 0.25790473 | -2.9693001 | 0.00298479 | 0.0135706 | NA | NA |
| ENSG000000 | 401.691161 | 0.409715893 | 0.13799513 | 2.96906043 | 0.00298712 | 0.01357967 | ELMO2 | 63916 |
| ENSG000002 | 6.02248726 | -1.104436796 | 0.3719952 | -2.9689544 | 0.00298815 | 0.01358283 | NA | NA |
| ENSG000000 | 467.795007 | 0.406864111 | 0.13704792 | 2.96877271 | 0.00298992 | 0.01358934 | INPP4A | 3631 |
| ENSG000001 | 732.956724 | 0.784312433 | 0.26420932 | 2.96852675 | 0.00299231 | 0.0135987 | CHTF8 | 54921 |
| ENSG000001 | 198.494095 | 0.525209908 | 0.1769384 | 2.96832062 | 0.00299432 | 0.0136063 | SRC | 6714 |
| ENSG000000 | 2143.13311 | 0.364962528 | 0.12295879 | 2.96816954 | 0.00299579 | 0.01361146 | SEC62 | 7095 |
| ENSG000002 | 12.8425375 | -0.951673089 | 0.32065516 | -2.967902 | 0.0029984 | 0.01362179 | NA | NA |
| ENSG000001 | 186.988084 | 0.454217517 | 0.15305616 | 2.96765256 | 0.00300083 | 0.01363132 | TTC33 | 23548 |
| ENSG000002 | 9.27475953 | -1.054383598 | 0.35530703 | -2.9675281 | 0.00300205 | 0.01363531 | NA | NA |
| ENSG000001 | 6505.55483 | 0.354777946 | 0.11955653 | 2.96744929 | 0.00300282 | 0.01363728 | ABCE1 | 6059 |
| ENSG000001 | 2023.30886 | 0.322574834 | 0.10870806 | 2.96734988 | 0.00300379 | 0.01363985 | NA | NA |

| | | | | | | | | | |
|---|---|---|---|---|---|---|---|---|---|
| ENSG000001 | 308.205085 | 0.427490687 | 0.14406613 | 2.96732253 | 0.00300406 | 0.01363985 | METTL15 | | 196074 |
| ENSG000001 | 456.884243 | 0.394125535 | 0.13283042 | 2.967133 | 0.00300591 | 0.01364673 | SHTN1 | | 57698 |
| ENSG000002 | 957.169603 | 0.393115778 | 0.13249755 | 2.96696636 | 0.00300754 | 0.0136526 | MIR17HG | | 407975 |
| ENSG000002 | 4.70138224 | -1.122235814 | 0.37830177 | -2.9665095 | 0.00301201 | 0.01367137 | NA | NA | |
| ENSG000001 | 215.071551 | 0.450165499 | 0.15179619 | 2.9655915 | 0.00302101 | 0.01371071 | EPHB2 | | 2048 |
| ENSG000002 | 5.78671664 | -1.109583594 | 0.37418313 | -2.9653491 | 0.0030234 | 0.01371998 | NA | NA | |
| ENSG000001 | 5.59383405 | -1.105020467 | 0.37265897 | -2.9652324 | 0.00302454 | 0.01372365 | OR8H1 | | 219469 |
| ENSG000001 | 26.4069059 | -0.820478081 | 0.27670304 | -2.9651936 | 0.00302493 | 0.01372385 | LGSN | | 51557 |
| ENSG000001 | 5.18165555 | -1.110072279 | 0.37438272 | -2.9650735 | 0.00302611 | 0.01372767 | RAX2 | | 84839 |
| ENSG000001 | 875.397141 | 0.364932715 | 0.1230929 | 2.96469357 | 0.00302985 | 0.0137431 | ZDHHC17 | | 23390 |
| ENSG00000C | 31.0309561 | -0.774884266 | 0.26137606 | -2.9646337 | 0.00303044 | 0.01374423 | CLCA4 | | 22802 |
| ENSG000001 | 883.710133 | 0.383991822 | 0.12953167 | 2.96446282 | 0.00303212 | 0.01374726 | PLIN2 | | 123 |
| ENSG000001 | 13.6699373 | -0.930570063 | 0.31390762 | -2.9644711 | 0.00303204 | 0.01374726 | CD1A | | 909 |
| ENSG000001 | 20.6972058 | -0.850042004 | 0.28673876 | -2.9645172 | 0.00303158 | 0.01374726 | HK3 | | 3101 |
| ENSG000001 | 75.1264482 | 0.565935662 | 0.19091795 | 2.96428735 | 0.00303385 | 0.01375356 | SLC2A6 | | 11182 |
| ENSG000001 | 538.6816 | 0.39910999 | 0.13464105 | 2.96425182 | 0.0030342 | 0.01375361 | ABCB7 | | 22 |
| ENSG000001 | 102.096349 | 0.569601607 | 0.19219205 | 2.96371058 | 0.00303954 | 0.01377628 | TSPAN18 | | 90139 |
| ENSG000001 | 40.9719633 | -0.654391214 | 0.22082417 | -2.963404 | 0.00304257 | 0.01378847 | CCDC148 | | 130940 |
| ENSG00000C | 539.824725 | 0.388652308 | 0.13115299 | 2.96335066 | 0.0030431 | 0.01378932 | GMCL1 | | 64395 |
| ENSG000001 | 588.897918 | 0.367039945 | 0.12388439 | 2.96276196 | 0.00304892 | 0.01381417 | ZNF780B | | 163131 |
| ENSG000001 | 429.130019 | 0.399302366 | 0.13478783 | 2.96245122 | 0.003052 | 0.01382631 | ZNF180 | | 7733 |
| ENSG000002 | 2.22390476 | -1.065113521 | 0.35954134 | -2.9624229 | 0.00305228 | 0.01382631 | NA | NA | |
| ENSG000001 | 943.254169 | 0.346763895 | 0.11706096 | 2.96225062 | 0.00305399 | 0.01383169 | USP53 | | 54532 |
| ENSG000001 | 936.962384 | 0.460987507 | 0.15562155 | 2.96223429 | 0.00305415 | 0.01383169 | NAV2 | | 89797 |
| ENSG000001 | 4.15278444 | -1.119119503 | 0.37781313 | -2.962098 | 0.00305551 | 0.01383627 | PRTN3 | | 5657 |
| ENSG000001 | 929.373202 | 0.505620177 | 0.17070282 | 2.96199069 | 0.00305657 | 0.01383955 | TOB2 | | 10766 |
| ENSG000001 | 12.3206378 | -0.986157108 | 0.332965 | -2.9617441 | 0.00305902 | 0.01384909 | ASB11 | | 140456 |
| ENSG000001 | 223.561275 | 0.418422312 | 0.14128079 | 2.96163629 | 0.00306009 | 0.01385085 | LLPH | | 84298 |
| ENSG000001 | 231.029234 | 0.405874518 | 0.13704319 | 2.96165415 | 0.00305991 | 0.01385085 | APOOL | | 139322 |

| | | | | | | | | | |
|---|---|---|---|---|---|---|---|---|---|
| ENSG000001 | 52.1060128 | -0.650655816 | 0.21970438 | -2.9615059 | 0.00306139 | 0.01385517 | TRPV6 | | 55503 |
| ENSG000002 | 2.31242276 | -1.062298637 | 0.35871408 | -2.9614077 | 0.00306236 | 0.01385804 | NA | NA | |
| ENSG000001 | 579.897028 | 0.346770876 | 0.11710926 | 2.96108853 | 0.00306554 | 0.01386931 | CEP135 | | 9662 |
| ENSG000002 | 6.3330928 | -1.092587218 | 0.36897891 | -2.9611102 | 0.00306532 | 0.01386931 | LOC101927948 | | 101927948 |
| ENSG000002 | 4.84057485 | -1.113650665 | 0.37610102 | -2.9610413 | 0.00306601 | 0.01386989 | RNU6-108P | | 106481875 |
| ENSG000001 | 385.765411 | 0.404316014 | 0.13655086 | 2.96091874 | 0.00306723 | 0.01387386 | GPCPD1 | | 56261 |
| ENSG000001 | 565.359153 | 0.401158512 | 0.13550009 | 2.96057748 | 0.00307063 | 0.0138815 | COG5 | | 10466 |
| ENSG000001 | 192.24327 | 0.572362458 | 0.19332083 | 2.96068696 | 0.00306954 | 0.0138815 | KCNS3 | | 3790 |
| ENSG000001 | 21.0897177 | -0.829747384 | 0.28026001 | -2.9606343 | 0.00307006 | 0.0138815 | NLRP11 | | 204801 |
| ENSG000001 | 34.9523075 | -0.768025716 | 0.25941619 | -2.9605928 | 0.00307048 | 0.0138815 | DPPA3 | | 359787 |
| ENSG000002 | 62.8754237 | 0.586668445 | 0.19815837 | 2.96060393 | 0.00307037 | 0.0138815 | GALT | | 2592 |
| ENSG000001 | 4636.35024 | 0.314121699 | 0.10610874 | 2.96037542 | 0.00307264 | 0.01388906 | PRPF4B | | 8899 |
| ENSG000001 | 64.8738359 | 0.644860105 | 0.21784142 | 2.9602272 | 0.00307412 | 0.0138942 | ARMC7 | | 79637 |
| ENSG000001 | 17.7658623 | -0.860214055 | 0.29066349 | -2.9594844 | 0.00308154 | 0.01392618 | GLIPR1L1 | | 256710 |
| ENSG000001 | 30.3943721 | -0.742918116 | 0.25104029 | -2.9593581 | 0.00308281 | 0.01393034 | ANKRD62 | | 342850 |
| ENSG000001 | 349.826018 | 0.388944406 | 0.13143322 | 2.95925505 | 0.00308384 | 0.01393345 | RNF25 | | 64320 |
| ENSG000002 | 20.9090294 | -0.857325331 | 0.28972865 | -2.959063 | 0.00308576 | 0.01393917 | LOC101927136 | | 101927136 |
| ENSG000002 | 13.0208015 | -0.955838846 | 0.32302112 | -2.9590599 | 0.00308579 | 0.01393917 | NA | NA | |
| ENSG000001 | 1657.23859 | 0.355769699 | 0.1202336 | 2.95898731 | 0.00308652 | 0.0139409 | PLK4 | | 10733 |
| ENSG000002 | 22.1411681 | -0.813562687 | 0.2749532 | -2.9589133 | 0.00308726 | 0.01394269 | LOC100506314 | | 100506314 |
| ENSG000001 | 1894.55082 | 0.396702851 | 0.13407507 | 2.95881138 | 0.00308828 | 0.01394575 | MAGI1 | | 9223 |
| ENSG000002 | 10.1748473 | -1.009432575 | 0.34116558 | -2.9587762 | 0.00308863 | 0.01394579 | NA | NA | |
| ENSG000001 | 63.3185507 | -0.612793068 | 0.20713306 | -2.9584512 | 0.00309189 | 0.01395584 | NPNT | | 255743 |
| ENSG000001 | 9.39136298 | -1.01483455 | 0.3430232 | -2.9585012 | 0.00309139 | 0.01395584 | GPR149 | | 344758 |
| ENSG000002 | 2.21343247 | -1.063280795 | 0.35940408 | -2.958455 | 0.00309185 | 0.01395584 | NA | NA | |
| ENSG000001 | 22.2458377 | -0.805983045 | 0.27244326 | -2.9583519 | 0.00309289 | 0.01395879 | CLN3 | | 1201 |
| ENSG000001 | 48.5010057 | -0.643335727 | 0.21749669 | -2.9579104 | 0.00309732 | 0.01397724 | BTN3A3 | | 10384 |
| ENSG000001 | 43.5686088 | 0.67705215 | 0.22890248 | 2.95781916 | 0.00309824 | 0.01397982 | PNPLA5 | | 150379 |
| ENSG000002 | 4.64041096 | -1.118469433 | 0.37817592 | -2.9575374 | 0.00310107 | 0.01399105 | NA | NA | |

| | | | | | | | | | |
|---|---|---|---|---|---|---|---|---|---|
| ENSG000001 | 445.072521 | 0.374213314 | 0.12653093 | 2.95748484 | 0.0031016 | 0.01399188 | SPOPL | | 339745 |
| ENSG000001 | 419.268544 | 0.491166954 | 0.16610961 | 2.95688463 | 0.00310764 | 0.01401758 | ARAP3 | | 64411 |
| ENSG000001 | 18.4786757 | -0.868391375 | 0.2937242 | -2.9564856 | 0.00311167 | 0.01403417 | SYNC | | 81493 |
| ENSG000002 | 3.69236503 | -1.112374192 | 0.37625747 | -2.9564176 | 0.00311235 | 0.01403571 | NA | NA | |
| ENSG000001 | 82.5302533 | 0.51634351 | 0.17465907 | 2.95629366 | 0.0031136 | 0.01403979 | NCR3LG1 | | 374383 |
| ENSG000002 | 15.3674309 | -0.912075427 | 0.30853064 | -2.9561907 | 0.00311464 | 0.01404291 | NA | NA | |
| ENSG000002 | 16.137836 | -0.931440382 | 0.31514041 | -2.9556361 | 0.00312025 | 0.01406662 | NA | NA | |
| ENSG000001 | 91.0865321 | -0.497862387 | 0.16845288 | -2.9554994 | 0.00312163 | 0.01407129 | ABI3BP | | 25890 |
| ENSG000001 | 1791.44542 | 0.439408503 | 0.14868424 | 2.95531326 | 0.00312352 | 0.01407822 | DHX36 | | 170506 |
| ENSG000001 | 53.9694665 | -0.594993616 | 0.2013327 | -2.9552756 | 0.0031239 | 0.01407838 | SCUBE2 | | 57758 |
| ENSG000002 | 11.0795318 | -1.014525358 | 0.34330077 | -2.9552085 | 0.00312458 | 0.01407987 | NA | NA | |
| ENSG000001 | 2896.08479 | 0.362091254 | 0.12253564 | 2.95498724 | 0.00312682 | 0.01408813 | QKI | | 9444 |
| ENSG000002 | 3.02823443 | -1.105788397 | 0.37421446 | -2.9549591 | 0.00312711 | 0.01408813 | CDRT15 | | 146822 |
| ENSG000001 | 2170.21735 | 0.344052812 | 0.11643471 | 2.95489906 | 0.00312771 | 0.0140893 | PTBP2 | | 58155 |
| ENSG000001 | 43.7118909 | 0.65026128 | 0.2200782 | 2.9546829 | 0.00312991 | 0.01409761 | TRIM62 | | 55223 |
| ENSG000001 | 71.1110305 | 0.562479847 | 0.19037784 | 2.95454474 | 0.00313131 | 0.01410236 | PID1 | | 55022 |
| ENSG000001 | 491.96354 | 0.476371319 | 0.16124891 | 2.95426072 | 0.00313419 | 0.01411378 | ARHGAP17 | | 55114 |
| ENSG000001 | 31.5918347 | -0.735509876 | 0.24901829 | -2.953638 | 0.00314052 | 0.01413915 | ACSL5 | | 51703 |
| ENSG000002 | 8.10608681 | -1.061468541 | 0.35937619 | -2.9536418 | 0.00314048 | 0.01413915 | NA | NA | |
| ENSG000001 | 6.83892983 | -1.093020262 | 0.37007704 | -2.9534938 | 0.00314199 | 0.01414418 | SIX6 | | 4990 |
| ENSG000001 | 9.26269312 | -1.032786425 | 0.34969504 | -2.9533917 | 0.00314303 | 0.01414572 | IL13 | | 3596 |
| ENSG000001 | 40.894946 | -0.662694516 | 0.22438247 | -2.9534148 | 0.00314279 | 0.01414572 | ITGAM | | 3684 |
| ENSG000001 | 59.3202306 | 0.616221075 | 0.20865608 | 2.95328598 | 0.00314411 | 0.014149 | SELM | | 140606 |
| ENSG000002 | 7.91855993 | -1.054746449 | 0.35725925 | -2.9523279 | 0.00315388 | 0.0141914 | NA | NA | |
| ENSG000001 | 841.938924 | 0.362864247 | 0.12291225 | 2.95222195 | 0.00315496 | 0.01419312 | NT5DC1 | | 221294 |
| ENSG000002 | 12.0397652 | -0.969452073 | 0.32837953 | -2.9522305 | 0.00315487 | 0.01419312 | NA | NA | |
| ENSG000001 | 360.701406 | 0.539806778 | 0.18285278 | 2.95213872 | 0.00315581 | 0.01419537 | DCTPP1 | | 79077 |
| ENSG000001 | 2829.12281 | 0.343582808 | 0.11638717 | 2.95206776 | 0.00315654 | 0.01419549 | WDR75 | | 84128 |
| ENSG000001 | 11.819875 | -0.963914945 | 0.32652009 | -2.9520847 | 0.00315636 | 0.01419549 | NA | NA | |

| | | | | | | | | |
|---|---|---|---|---|---|---|---|---|
| ENSG000001 | 15.4695673 | -0.902827616 | 0.30590171 | -2.951365 | 0.00316373 | 0.01422625 | ATOH8 | 84913 |
| ENSG000002 | 110.891955 | 0.500650365 | 0.16964455 | 2.95117271 | 0.0031657 | 0.01423353 | ZNF561-AS1 | 284385 |
| ENSG000001 | 75.0857894 | -0.596604548 | 0.2021638 | -2.9510949 | 0.0031665 | 0.01423554 | ADAM32 | 203102 |
| ENSG000002 | 3.02755725 | -1.104502305 | 0.37431337 | -2.9507424 | 0.00317011 | 0.01425022 | NA | NA |
| ENSG000002 | 4.02670891 | -1.115256299 | 0.37797571 | -2.9506031 | 0.00317154 | 0.01425507 | HLTF-AS1 | 100873945 |
| ENSG000002 | 9.52449224 | -1.020171352 | 0.34576761 | -2.9504538 | 0.00317307 | 0.01426037 | OR2T11 | 127077 |
| ENSG000001 | 845.569122 | 0.388680052 | 0.13173792 | 2.95040369 | 0.00317359 | 0.0142604 | NAA30 | 122830 |
| ENSG000001 | 200.916678 | 0.506505343 | 0.17167433 | 2.95038487 | 0.00317378 | 0.0142604 | SS18L1 | 26039 |
| ENSG000002 | 49.7602633 | 0.657854449 | 0.2229771 | 2.95032286 | 0.00317442 | 0.01426168 | NA | NA |
| ENSG000002 | 17.6319636 | -0.916772756 | 0.31074158 | -2.9502738 | 0.00317492 | 0.01426236 | LINC01314 | 100996492 |
| ENSG000002 | 33.6527826 | 0.729124142 | 0.24716517 | 2.94994697 | 0.00317828 | 0.01427588 | NA | NA |
| ENSG000001 | 805.471314 | 0.369349927 | 0.12521115 | 2.94981667 | 0.00317963 | 0.01428032 | ZNF28 | 7576 |
| ENSG000001 | 156.895646 | 0.436076969 | 0.14783724 | 2.94970985 | 0.00318072 | 0.01428367 | PRKG2 | 5593 |
| ENSG000002 | 219.44932 | -0.402413982 | 0.13643146 | -2.9495688 | 0.00318218 | 0.01428861 | GRIN2B | 2904 |
| ENSG000002 | 5.20314557 | -1.105421952 | 0.37478612 | -2.9494741 | 0.00318315 | 0.01429141 | NA | NA |
| ENSG000002 | 35.9649552 | -0.702098039 | 0.23806082 | -2.9492381 | 0.00318558 | 0.01429916 | NA | NA |
| ENSG000002 | 11.5657075 | -0.9679405 | 0.32819923 | -2.9492467 | 0.0031855 | 0.01429916 | NA | NA |
| ENSG00000C | 486.22308 | 0.441861648 | 0.1498303 | 2.94908066 | 0.00318721 | 0.01430487 | SMG6 | 23293 |
| ENSG000001 | 606.510857 | 0.350329339 | 0.11880523 | 2.94877035 | 0.00319041 | 0.01431765 | STK39 | 27347 |
| ENSG000001 | 10.315521 | -0.990155657 | 0.33579761 | -2.948668 | 0.00319147 | 0.01432081 | NMUR1 | 10316 |
| ENSG000001 | 79.5793385 | 0.560757344 | 0.19017611 | 2.94862133 | 0.00319195 | 0.01432139 | RABEP2 | 79874 |
| ENSG000001 | 619.258699 | 0.382384393 | 0.12969069 | 2.94843358 | 0.00319389 | 0.01432692 | ALKBH8 | 91801 |
| ENSG000001 | 230.912167 | 0.448977394 | 0.15227596 | 2.9484457 | 0.00319376 | 0.01432692 | RPIA | 22934 |
| ENSG000001 | 27.9616873 | -0.775627868 | 0.26308163 | -2.9482403 | 0.00319589 | 0.01433429 | GRIN3A | 116443 |
| ENSG000002 | 11.8798878 | -0.981120649 | 0.33278836 | -2.948182 | 0.00319649 | 0.01433541 | NA | NA |
| ENSG000001 | 13.4029309 | -0.938897915 | 0.31850131 | -2.947862 | 0.0031998 | 0.01434867 | GPR12 | 2835 |
| ENSG000001 | 492.175115 | 0.371873603 | 0.12615511 | 2.94774907 | 0.00320097 | 0.01435233 | MIPOL1 | 145282 |
| ENSG000002 | 10.2061112 | -1.028880335 | 0.34914972 | -2.946817 | 0.00321063 | 0.01439406 | NA | NA |
| ENSG00000C | 1049.0175 | 0.35299915 | 0.11979519 | 2.94668875 | 0.00321196 | 0.01439844 | DIP2B | 57609 |

| | | | | | | | | |
|---|---|---|---|---|---|---|---|---|
| ENSG000002 | 15.7480305 | -0.891110664 | 0.3024206 | -2.9465938 | 0.00321295 | 0.01440127 | HOGA1 | 112817 |
| ENSG000001 | 386.792867 | 0.376304319 | 0.1277162 | 2.94641027 | 0.00321486 | 0.01440664 | ZNF473 | 25888 |
| ENSG000001 | 66.790307 | -0.576344418 | 0.19560779 | -2.9464288 | 0.00321466 | 0.01440664 | WDR49 | 151790 |
| ENSG000001 | 74.6566906 | -0.547944542 | 0.18598216 | -2.946221 | 0.00321682 | 0.01441068 | ADAMTS9 | 56999 |
| ENSG000001 | 487.527799 | 0.373585072 | 0.12680042 | 2.94624478 | 0.00321658 | 0.01441068 | ALCAM | 214 |
| ENSG000002 | 2067.06842 | 0.348253294 | 0.11820109 | 2.94627821 | 0.00321623 | 0.01441068 | SCAF8 | 22828 |
| ENSG000001 | 18.5032867 | -0.868784479 | 0.29490468 | -2.945984 | 0.00321929 | 0.01442013 | ZNF556 | 80032 |
| ENSG000002 | 81.3821097 | 0.615235331 | 0.20884165 | 2.94594179 | 0.00321973 | 0.01442051 | NA | NA |
| ENSG000001 | 3971.93934 | 0.420846785 | 0.14286889 | 2.94568525 | 0.0032224 | 0.01443088 | HNRNPAB | 3182 |
| ENSG000001 | 224.966959 | 0.473266384 | 0.16068154 | 2.94536868 | 0.0032257 | 0.01444406 | URGCP | 55665 |
| ENSG000001 | 915.023661 | 0.363517316 | 0.12342311 | 2.9452937 | 0.00322648 | 0.01444596 | ABCD3 | 5825 |
| ENSG000001 | 1277.78676 | 0.348547392 | 0.11835917 | 2.94482802 | 0.00323134 | 0.01446612 | ENY2 | 56943 |
| ENSG000001 | 95.7433459 | 0.527789354 | 0.17924008 | 2.94459442 | 0.00323378 | 0.01447545 | ACSF2 | 80221 |
| ENSG00000C | 692.664794 | 0.468333801 | 0.15905912 | 2.94440082 | 0.00323581 | 0.01448291 | CCDC88C | 440193 |
| ENSG000001 | 53.2217828 | 0.617684728 | 0.20978751 | 2.94433512 | 0.00323649 | 0.01448439 | VASN | 114990 |
| ENSG000001 | 13.5527596 | -0.943649066 | 0.32062839 | -2.9431239 | 0.00324918 | 0.01453958 | OR8G1 | 26494 |
| ENSG000001 | 64.2418765 | 0.586976593 | 0.199443 | 2.94307942 | 0.00324965 | 0.01454006 | HYI | 81888 |
| ENSG000001 | 1011.74486 | 0.372089881 | 0.12643222 | 2.94299882 | 0.0032505 | 0.01454224 | BEX4 | 56271 |
| ENSG000002 | 516.500234 | 0.382835724 | 0.13009817 | 2.94266794 | 0.00325397 | 0.01455619 | TADA2A | 6871 |
| ENSG000002 | 25.8209238 | -0.79554961 | 0.27038104 | -2.9423276 | 0.00325755 | 0.01457059 | NA | NA |
| ENSG000002 | 97.0223075 | -0.532526732 | 0.18101656 | -2.9418675 | 0.00326239 | 0.01459064 | EML6 | 400954 |
| ENSG000001 | 276.05469 | 0.453788821 | 0.1542563 | 2.94178478 | 0.00326327 | 0.01459293 | NA | NA |
| ENSG000001 | 38.3974352 | -0.66086652 | 0.22466716 | -2.9415359 | 0.00326589 | 0.01460306 | KCNJ10 | 3766 |
| ENSG000001 | 16.3525342 | -0.905455943 | 0.30782829 | -2.9414319 | 0.00326699 | 0.01460474 | MMP8 | 4317 |
| ENSG000001 | 83.5252186 | 0.549325523 | 0.18675398 | 2.94143942 | 0.00326691 | 0.01460474 | WDR90 | 197335 |
| ENSG000002 | 304.309204 | 0.392848787 | 0.13356199 | 2.94132165 | 0.00326815 | 0.01460833 | TMEM167B | 56900 |
| ENSG000002 | 7.2152024 | -1.062790019 | 0.36133671 | -2.9412733 | 0.00326866 | 0.014609 | NA | NA |
| ENSG000001 | 420.874923 | 0.421118367 | 0.14319231 | 2.94092863 | 0.0032723 | 0.01462365 | LIAS | 11019 |
| ENSG000001 | 5.28635964 | -1.105442948 | 0.37594596 | -2.9404304 | 0.00327757 | 0.01464557 | IFNE | 338376 |

| | | | | | | | | |
|---|---|---|---|---|---|---|---|---|
| ENSG000001 | 301.616625 | 0.40803755 | 0.13877342 | 2.94031483 | 0.00327879 | 0.01464781 | KHDRBS3 | 10656 |
| ENSG000001 | 723.906932 | 0.535765686 | 0.1822119 | 2.94034408 | 0.00327848 | 0.01464781 | CLSTN1 | 22883 |
| ENSG000001 | 1408.39783 | 0.41417753 | 0.14086452 | 2.94025454 | 0.00327943 | 0.01464905 | HIPK1 | 204851 |
| ENSG00000C | 1820.81776 | 0.343648191 | 0.11688181 | 2.94013418 | 0.0032807 | 0.01465226 | ASUN | 55726 |
| ENSG000002 | 29.6161164 | -0.778701399 | 0.26485376 | -2.9401183 | 0.00328087 | 0.01465226 | NA | NA |
| ENSG000001 | 238.496091 | 0.41371249 | 0.14071693 | 2.94003361 | 0.00328177 | 0.01465465 | LOC728554 | 728554 |
| ENSG000001 | 627.4843 | 0.370755068 | 0.12612497 | 2.93958508 | 0.00328652 | 0.01467374 | GTF3C6 | 112495 |
| ENSG000002 | 7.24129196 | -1.062936758 | 0.36159698 | -2.9395621 | 0.00328676 | 0.01467374 | NA | NA |
| ENSG000001 | 874.051427 | 0.455594452 | 0.15501789 | 2.93897989 | 0.00329294 | 0.01469971 | MDC1 | 9656 |
| ENSG000001 | 28.0705979 | -0.742944758 | 0.25281608 | -2.9386769 | 0.00329617 | 0.01471247 | SH2D6 | 284948 |
| ENSG000002 | 5.01566108 | -1.108148923 | 0.37710783 | -2.9385466 | 0.00329755 | 0.01471703 | ZIM2 | 23619 |
| ENSG000002 | 4.59216916 | -1.111272598 | 0.37822638 | -2.938115 | 0.00330215 | 0.01473275 | RNU1-93P | 106480180 |
| ENSG000002 | 17.9965074 | -0.855254821 | 0.29108979 | -2.9381134 | 0.00330216 | 0.01473275 | LOC101927156 | 101927156 |
| ENSG000002 | 4.7960038 | -1.105483777 | 0.37625621 | -2.9381144 | 0.00330215 | 0.01473275 | NA | NA |
| ENSG000002 | 21.9558464 | -0.819208254 | 0.2788326 | -2.9379931 | 0.00330344 | 0.01473685 | NA | NA |
| ENSG000001 | 12.73201 | -0.963792906 | 0.32810379 | -2.9374635 | 0.00330909 | 0.01476042 | RGS18 | 64407 |
| ENSG000001 | 196.292403 | 0.43876225 | 0.14937574 | 2.93730596 | 0.00331077 | 0.01476629 | ITGB1BP1 | 9270 |
| ENSG000002 | 4.92419526 | -1.10767151 | 0.37712159 | -2.9371734 | 0.00331219 | 0.01477098 | NA | NA |
| ENSG000001 | 106.599202 | 0.488563272 | 0.16634211 | 2.93709915 | 0.00331298 | 0.01477135 | SAYSD1 | 55776 |
| ENSG000002 | 8.80653485 | -1.027175844 | 0.34972861 | -2.9370656 | 0.00331334 | 0.01477135 | LINC01031 | 101929184 |
| ENSG000002 | 13.1151672 | -0.958894991 | 0.32648085 | -2.9370635 | 0.00331336 | 0.01477135 | NA | NA |
| ENSG000001 | 181.634567 | 0.446971571 | 0.15218611 | 2.93700639 | 0.00331397 | 0.01477244 | COL6A3 | 1293 |
| ENSG000001 | 165.172211 | 0.511789208 | 0.17426544 | 2.93683705 | 0.00331578 | 0.01477889 | ACP2 | 53 |
| ENSG00000C | 663.043846 | 0.395091194 | 0.13454165 | 2.9365716 | 0.00331862 | 0.01478992 | KDM7A | 80853 |
| ENSG000001 | 437.599324 | 0.470799935 | 0.16032955 | 2.9364514 | 0.00331991 | 0.01479402 | SIPA1L3 | 23094 |
| ENSG000001 | 1428.68413 | 0.343621719 | 0.1170382 | 2.93597925 | 0.00332497 | 0.01481493 | PPP2R5E | 5529 |
| ENSG000001 | 29.2785033 | -0.735516992 | 0.25053183 | -2.9358225 | 0.00332665 | 0.01482079 | ME3 | 10873 |
| ENSG000002 | 2.96117921 | -1.100272562 | 0.37479083 | -2.9356977 | 0.00332799 | 0.01482513 | NA | NA |
| ENSG000002 | 24.7083215 | 0.818884146 | 0.27895066 | 2.93558778 | 0.00332916 | 0.01482875 | NA | NA |

| | | | | | | | | |
|---|---|---|---|---|---|---|---|---|
| ENSG000001 | 1279.87424 | 0.34946905 | 0.11905626 | 2.93532685 | 0.00333197 | 0.0148396 | USP16 | 10600 |
| ENSG000001 | 297.334363 | 0.463038955 | 0.15775971 | 2.93509003 | 0.00333451 | 0.0148493 | RYR1 | 6261 |
| ENSG000001 | 83.2579369 | 0.556260738 | 0.18953505 | 2.93486998 | 0.00333687 | 0.0148582 | C7orf43 | 55262 |
| ENSG000001 | 245.746857 | 0.397658564 | 0.13549715 | 2.93481135 | 0.00333751 | 0.01485937 | SCAMP3 | 10067 |
| ENSG00000C | 34.2718581 | -0.698763813 | 0.23813202 | -2.9343547 | 0.00334242 | 0.01487962 | CEACAM1 | 634 |
| ENSG000001 | 82.3882915 | 0.574046385 | 0.19568078 | 2.93358601 | 0.00335071 | 0.01491488 | NUDT14 | 256281 |
| ENSG00000C | 5672.10853 | 0.39938346 | 0.13617079 | 2.93295981 | 0.00335747 | 0.01494336 | JARID2 | 3720 |
| ENSG000001 | 13.8490659 | -0.915094837 | 0.3120312 | -2.932703 | 0.00336025 | 0.01495408 | DSCR8 | 84677 |
| ENSG000002 | 15.8156295 | -0.918794741 | 0.31330005 | -2.9326352 | 0.00336099 | 0.0149557 | NA | NA |
| ENSG000001 | 18.0590217 | -0.899993541 | 0.30690115 | -2.9325192 | 0.00336224 | 0.01495801 | SLAMF6 | 114836 |
| ENSG000001 | 895.966219 | 0.358674193 | 0.12230874 | 2.93253122 | 0.00336211 | 0.01495801 | NA | NA |
| ENSG000002 | 204.714587 | 0.509576966 | 0.17378237 | 2.93227086 | 0.00336493 | 0.01496833 | COG8 | 84342 |
| ENSG000001 | 10.9746296 | -0.976394998 | 0.33299879 | -2.9321277 | 0.00336648 | 0.01497359 | USP29 | 57663 |
| ENSG000001 | 22.0705773 | 0.917780292 | 0.3130221 | 2.93199842 | 0.00336788 | 0.01497818 | PRODH | 5625 |
| ENSG000001 | 465.252539 | 0.394257623 | 0.1344706 | 2.93192425 | 0.00336869 | 0.01498012 | TBP | 6908 |
| ENSG000002 | 10.5002494 | -1.018665102 | 0.3474527 | -2.9318094 | 0.00336993 | 0.01498402 | TMEM246-AS1 | 101928470 |
| ENSG000001 | 21.583534 | -0.804806015 | 0.2745131 | -2.9317581 | 0.00337049 | 0.01498485 | ROBO4 | 54538 |
| ENSG000002 | 5.65016436 | -1.097179565 | 0.37424964 | -2.9316783 | 0.00337136 | 0.01498706 | NA | NA |
| ENSG000002 | 11.3848173 | -1.022530589 | 0.34879935 | -2.9315725 | 0.00337251 | 0.01499052 | NA | NA |
| ENSG000001 | 213.428051 | 0.483244881 | 0.16484903 | 2.93143903 | 0.00337396 | 0.01499301 | CGRRF1 | 10668 |
| ENSG000001 | 205.265171 | 0.422787753 | 0.14422633 | 2.93141867 | 0.00337418 | 0.01499301 | MMACHC | 25974 |
| ENSG000002 | 8.77294759 | -1.042482446 | 0.35562339 | -2.9314226 | 0.00337413 | 0.01499301 | LINC00184 | 100302691 |
| ENSG000001 | 194.794422 | 0.423043904 | 0.14433122 | 2.93106304 | 0.00337804 | 0.01500855 | FAM174B | 400451 |
| ENSG000001 | 9.71816102 | -1.000648892 | 0.34140203 | -2.9309986 | 0.00337874 | 0.01501001 | TEX101 | 83639 |
| ENSG000002 | 26.269875 | -0.767971545 | 0.26202053 | -2.9309595 | 0.00337917 | 0.01501026 | NA | NA |
| ENSG000001 | 177.650647 | 0.46761264 | 0.15954827 | 2.93085364 | 0.00338032 | 0.01501373 | PCYOX1L | 78991 |
| ENSG00000C | 398.76208 | 0.421636839 | 0.14387856 | 2.93050497 | 0.00338412 | 0.01502782 | SYNJ2 | 8871 |
| ENSG000002 | 8.63005678 | -1.03540776 | 0.35332189 | -2.9304943 | 0.00338423 | 0.01502782 | NA | NA |
| ENSG000002 | 27.471488 | -0.771041312 | 0.26311741 | -2.9304078 | 0.00338517 | 0.01503035 | NA | NA |

| | | | | | | | | |
|---|---|---|---|---|---|---|---|---|
| ENSG000002 | 80.0174231 | 0.583915981 | 0.19926669 | 2.93032408 | 0.00338609 | 0.01503276 | C9orf129 | 445577 |
| ENSG000001 | 241.023414 | 0.401825534 | 0.13713724 | 2.93009784 | 0.00338855 | 0.01504041 | EZH1 | 2145 |
| ENSG000001 | 195.734675 | 0.420630046 | 0.14355437 | 2.93010973 | 0.00338842 | 0.01504041 | COQ3 | 51805 |
| ENSG000002 | 4.42107695 | -1.106298178 | 0.37757361 | -2.9300199 | 0.0033894 | 0.01504254 | NA | NA |
| ENSG000001 | 68.1386798 | -0.575125762 | 0.19629333 | -2.9299302 | 0.00339038 | 0.01504523 | PLCE1 | 51196 |
| ENSG000001 | 22.5683036 | 0.813738416 | 0.27773778 | 2.92988016 | 0.00339093 | 0.01504601 | P2RX1 | 5023 |
| ENSG000001 | 102.261226 | 0.549822704 | 0.18766312 | 2.92983893 | 0.00339138 | 0.01504636 | KRTCAP3 | 200634 |
| ENSG000002 | 16.2008195 | -0.920875242 | 0.31432873 | -2.9296566 | 0.00339337 | 0.01505354 | GOLGA6L6 | 727832 |
| ENSG000001 | 1060.94174 | 0.40260533 | 0.13743137 | 2.92950104 | 0.00339507 | 0.01505779 | PCNXL2 | 80003 |
| ENSG000002 | 5.63013967 | -1.095356159 | 0.37390155 | -2.929531 | 0.00339474 | 0.01505779 | NA | NA |
| ENSG000001 | 612.368455 | 0.379379413 | 0.12951224 | 2.92929386 | 0.00339733 | 0.01506618 | ITGB3BP | 23421 |
| ENSG000001 | 218.808585 | 0.480233805 | 0.16394825 | 2.92917929 | 0.00339858 | 0.01507009 | WRAP53 | 55135 |
| ENSG00000C | 37.4645106 | -0.674880779 | 0.23040893 | -2.9290566 | 0.00339993 | 0.01507302 | TBXAS1 | 6916 |
| ENSG000002 | 21.3083442 | -0.818583265 | 0.27947048 | -2.9290509 | 0.00339999 | 0.01507302 | LINC01483 | 101928122 |
| ENSG000001 | 1039.77064 | 0.409031291 | 0.13965252 | 2.92892166 | 0.0034014 | 0.01507764 | ERCC6L | 54821 |
| ENSG000002 | 24.7607125 | -0.785816328 | 0.26830562 | -2.9288105 | 0.00340262 | 0.01508138 | PNLIPRP2 | 5408 |
| ENSG000001 | 40.4750099 | -0.663015809 | 0.22638437 | -2.9287173 | 0.00340364 | 0.01508426 | GAD1 | 2571 |
| ENSG000001 | 16.9972698 | -0.918587282 | 0.31367473 | -2.9284708 | 0.00340634 | 0.01509457 | C5AR2 | 27202 |
| ENSG000002 | 9.24410831 | -1.045606842 | 0.35707366 | -2.9282665 | 0.00340858 | 0.01510284 | NA | NA |
| ENSG000001 | 16.2579374 | -0.920339852 | 0.31431952 | -2.9280391 | 0.00341107 | 0.01511224 | IL18RAP | 8807 |
| ENSG000002 | 5.84058208 | -1.084450425 | 0.37038126 | -2.9279301 | 0.00341227 | 0.0151152 | NA | NA |
| ENSG000002 | 16.4905019 | -0.908829248 | 0.310402 | -2.9279104 | 0.00341248 | 0.0151152 | YWHAEP7 | 284100 |
| ENSG000002 | 6.41182405 | -1.084891464 | 0.37054537 | -2.9278236 | 0.00341344 | 0.01511777 | NA | NA |
| ENSG000002 | 78.3839048 | 0.551635598 | 0.18841384 | 2.92778701 | 0.00341384 | 0.01511789 | FAM86EP | 348926 |
| ENSG000002 | 24.8725469 | -0.790973126 | 0.27017886 | -2.9275908 | 0.00341599 | 0.01512579 | ZNF705G | 100131980 |
| ENSG000001 | 3.413209 | -1.103398094 | 0.37691858 | -2.9274176 | 0.0034179 | 0.01513256 | NA | NA |
| ENSG000002 | 3.34439154 | -1.104774631 | 0.37741367 | -2.9272247 | 0.00342002 | 0.01514029 | HNRNPDP2 | 106481661 |
| ENSG000001 | 18.1635247 | -0.86137592 | 0.29430517 | -2.926812 | 0.00342456 | 0.01515874 | VTCN1 | 79679 |
| ENSG000002 | 83.2460124 | 0.591250818 | 0.20203468 | 2.92648179 | 0.0034282 | 0.01517319 | NATD1 | 256302 |

| | | | | | | | | | |
|---|---|---|---|---|---|---|---|---|---|
| ENSG000002 | 5.72585739 | -1.098288673 | 0.37534498 | -2.926078 | 0.00343265 | 0.01518958 | NA | NA | |
| ENSG000002 | 6.57236225 | -1.075669331 | 0.3676111 | -2.9261067 | 0.00343233 | 0.01518958 | NA | NA | |
| ENSG000002 | 5.43972671 | -1.101235689 | 0.37638801 | -2.9257991 | 0.00343573 | 0.01520155 | NA | NA | |
| ENSG000001 | 44.5696811 | -0.682521747 | 0.23329803 | -2.9255358 | 0.00343864 | 0.01521276 | SLC26A7 | | 115111 |
| ENSG000001 | 9.63278673 | -1.022799904 | 0.34962804 | -2.9253944 | 0.0034402 | 0.01521801 | OR2L5 | | 81466 |
| ENSG000001 | 25.9859547 | -0.79688799 | 0.27243757 | -2.9250297 | 0.00344423 | 0.0152342 | GPD1 | | 2819 |
| ENSG000002 | 132.676784 | 0.457751116 | 0.15649956 | 2.92493555 | 0.00344528 | 0.01523714 | HOMEZ | | 57594 |
| ENSG000001 | 263.936272 | 0.393085354 | 0.13439389 | 2.92487515 | 0.00344594 | 0.01523844 | NUBPL | | 80224 |
| ENSG000002 | 13.7810319 | -0.953809944 | 0.32614267 | -2.9245175 | 0.00344991 | 0.0152543 | NA | NA | |
| ENSG000001 | 1326.63374 | 0.35107969 | 0.12005259 | 2.92438256 | 0.0034514 | 0.01525925 | FBXO28 | | 23219 |
| ENSG000001 | 20.2129814 | -0.819320899 | 0.28017397 | -2.9243291 | 0.003452 | 0.01526021 | UTS2B | | 257313 |
| ENSG000001 | 752.449856 | 0.347508458 | 0.11884642 | 2.92401284 | 0.0034555 | 0.01527405 | RSBN1L | | 222194 |
| ENSG000002 | 22.7903002 | -0.875990063 | 0.29959595 | -2.9239049 | 0.0034567 | 0.01527602 | FAM221B | | 392307 |
| ENSG000002 | 6.55941472 | -1.072772765 | 0.3668944 | -2.9239279 | 0.00345645 | 0.01527602 | LINC01019 | | 285577 |
| ENSG000001 | 2179.57904 | 0.44636162 | 0.15266847 | 2.92373156 | 0.00345863 | 0.01528286 | PSMD12 | | 5718 |
| ENSG000001 | 198.060718 | 0.556756596 | 0.19043233 | 2.92364541 | 0.00345959 | 0.01528397 | GAR1 | | 54433 |
| ENSG000001 | 17.386146 | -0.875408731 | 0.299425 | -2.9236328 | 0.00345973 | 0.01528397 | FBLN7 | | 129804 |
| ENSG000001 | 1069.5068 | 0.35097867 | 0.12005126 | 2.92357333 | 0.00346039 | 0.01528397 | DHX57 | | 90957 |
| ENSG000002 | 4.90349817 | -1.101925646 | 0.37690958 | -2.9235809 | 0.0034603 | 0.01528397 | NA | NA | |
| ENSG00000C | 27.7247167 | -0.753835365 | 0.25787608 | -2.9232466 | 0.00346402 | 0.01529836 | CASR | | 846 |
| ENSG000001 | 15.8362039 | -0.874482811 | 0.29917607 | -2.9229704 | 0.00346709 | 0.0153086 | CPA4 | | 51200 |
| ENSG000001 | 391.646685 | 0.389076747 | 0.13310973 | 2.9229776 | 0.00346701 | 0.0153086 | ZNF577 | | 84765 |
| ENSG000001 | 329.688002 | 0.386947489 | 0.13238318 | 2.92293541 | 0.00346748 | 0.01530865 | FUT10 | | 84750 |
| ENSG000001 | 11.0086532 | -0.974321044 | 0.33334889 | -2.9228267 | 0.0034687 | 0.01531233 | ZNF804B | | 219578 |
| ENSG000001 | 235.404287 | 0.382005214 | 0.13071322 | 2.92246801 | 0.00347269 | 0.01532831 | BRE | | 9577 |
| ENSG000001 | 78.2885474 | 0.554076048 | 0.18962 | 2.92203382 | 0.00347754 | 0.01534802 | TAZ | | 6901 |
| ENSG00000C | 1068.7707 | 0.338474168 | 0.11584022 | 2.92190556 | 0.00347897 | 0.01535267 | ZCCHC6 | | 79670 |
| ENSG000001 | 1164.98579 | 0.400978623 | 0.1372368 | 2.92180095 | 0.00348014 | 0.01535616 | NMD3 | | 51068 |
| ENSG000001 | 11.4426336 | -0.974968527 | 0.33370111 | -2.921682 | 0.00348147 | 0.01536035 | ZP4 | | 57829 |

| | | | | | | | | |
|---|---|---|---|---|---|---|---|---|
| ENSG000001 | 12.9848557 | -0.932421334 | 0.31914906 | -2.9215857 | 0.00348254 | 0.01536343 | SLC39A2 | 29986 |
| ENSG000002 | 4.45266657 | -1.103500849 | 0.37772052 | -2.9214744 | 0.00348379 | 0.01536725 | NA | NA |
| ENSG000001 | 94.5704533 | 0.566705287 | 0.19399477 | 2.92124002 | 0.00348641 | 0.01537714 | FZD2 | 2535 |
| ENSG000001 | 17.7833634 | 0.880278242 | 0.30134692 | 2.9211456 | 0.00348747 | 0.01538013 | ARHGEF18 | 23370 |
| ENSG000001 | 182.608347 | 0.509254243 | 0.17433663 | 2.92109727 | 0.00348801 | 0.01538085 | FKBP11 | 51303 |
| ENSG000001 | 1409.83232 | 0.382467616 | 0.13093546 | 2.92103926 | 0.00348866 | 0.01538204 | KPNA3 | 3839 |
| ENSG000001 | 19.4350078 | -0.847160698 | 0.29003451 | -2.9208962 | 0.00349026 | 0.01538743 | HHIPL2 | 79802 |
| ENSG000001 | 3047.4497 | 0.39862492 | 0.13647982 | 2.92076093 | 0.00349178 | 0.01539244 | DDX46 | 9879 |
| ENSG000001 | 505.075638 | 0.410590445 | 0.14058096 | 2.9206689 | 0.00349281 | 0.01539391 | HS6ST1 | 9394 |
| ENSG000001 | 17.796945 | -0.860722389 | 0.29470098 | -2.9206634 | 0.00349287 | 0.01539391 | OR6C4 | 341418 |
| ENSG000002 | 2469.10662 | 0.398452833 | 0.1364292 | 2.92058331 | 0.00349377 | 0.01539619 | NA | NA |
| ENSG000001 | 78.521099 | -0.555416532 | 0.19017591 | -2.9205409 | 0.00349424 | 0.01539654 | PTPRQ | 374462 |
| ENSG000001 | 29.2080331 | -0.753650669 | 0.2580546 | -2.9205086 | 0.00349461 | 0.01539654 | NUGGC | 389643 |
| ENSG000002 | 3.00793706 | -1.09311362 | 0.37433319 | -2.9201622 | 0.00349849 | 0.01541199 | NA | NA |
| ENSG000001 | 235.119647 | 0.421884726 | 0.1444786 | 2.92004993 | 0.00349975 | 0.01541419 | GALK2 | 2585 |
| ENSG000002 | 5.97440653 | -1.088576587 | 0.37279044 | -2.9200764 | 0.00349946 | 0.01541419 | NA | NA |
| ENSG000001 | 1539.52366 | 0.449659737 | 0.15400423 | 2.91978815 | 0.00350269 | 0.01542547 | CCDC47 | 57003 |
| ENSG00000C | 165.655967 | -0.496060464 | 0.1699119 | -2.9195157 | 0.00350576 | 0.01543729 | LRP2 | 4036 |
| ENSG000001 | 480.623125 | 0.377340818 | 0.12925009 | 2.91946274 | 0.00350635 | 0.01543823 | KIAA0753 | 9851 |
| ENSG000001 | 266.321784 | 0.414934267 | 0.14214419 | 2.91910812 | 0.00351034 | 0.01545413 | HUS1 | 3364 |
| ENSG000001 | 5541.6513 | 0.294518537 | 0.10089561 | 2.91904227 | 0.00351109 | 0.01545572 | HIF1A | 3091 |
| ENSG000001 | 450.972951 | 0.351965077 | 0.1205823 | 2.91887837 | 0.00351293 | 0.01546217 | CIR1 | 9541 |
| ENSG000001 | 323.071901 | 0.399748878 | 0.13695958 | 2.91873624 | 0.00351453 | 0.01546592 | STK36 | 27148 |
| ENSG000002 | 39.1938573 | -0.683391737 | 0.2341397 | -2.918735 | 0.00351455 | 0.01546592 | RNF219-AS1 | 100874222 |
| ENSG000002 | 32442.8215 | 0.843980055 | 0.28916583 | 2.91867141 | 0.00351527 | 0.0154674 | NA | NA |
| ENSG00000C | 1284.64771 | 0.349933818 | 0.1199006 | 2.91853255 | 0.00351683 | 0.01547261 | TBC1D1 | 23216 |
| ENSG000001 | 30.4025784 | -0.75114174 | 0.25738691 | -2.918337 | 0.00351904 | 0.01548064 | FAM186A | 121006 |
| ENSG000001 | 138.504339 | 0.524792537 | 0.17984821 | 2.91797484 | 0.00352313 | 0.01549695 | NDUFB8 | 4714 |
| ENSG000002 | 5.57567711 | -1.091132416 | 0.37394567 | -2.9178902 | 0.00352408 | 0.01549947 | NA | NA |

| | | | | | | | | |
|---|---|---|---|---|---|---|---|---|
| ENSG000001 | 348.746417 | 0.375686417 | 0.12875901 | 2.91774851 | 0.00352569 | 0.01550147 | SSFA2 | 6744 |
| ENSG000001 | 12.5936442 | -0.97972854 | 0.3357806 | -2.917764 | 0.00352551 | 0.01550147 | BTNL3 | 10917 |
| ENSG000001 | 10.4837144 | -0.994578947 | 0.34086671 | -2.9177943 | 0.00352517 | 0.01550147 | NUP210P1 | 255330 |
| ENSG000002 | 15.5658707 | -0.902815299 | 0.30944939 | -2.9174893 | 0.00352862 | 0.01551268 | NA | NA |
| ENSG000001 | 982.969766 | 0.354295294 | 0.12144213 | 2.91740013 | 0.00352963 | 0.01551375 | INTS2 | 57508 |
| ENSG000002 | 11.1294735 | -0.979314261 | 0.33567733 | -2.9174274 | 0.00352932 | 0.01551375 | KRBOX1 | 100506243 |
| ENSG000001 | 1641.22852 | 0.311113556 | 0.10665363 | 2.91704612 | 0.00353364 | 0.01552969 | RAP1B | 5908 |
| ENSG000000 | 32.2649342 | 0.730879798 | 0.25059948 | 2.91652557 | 0.00353954 | 0.01555327 | TESC | 54997 |
| ENSG000001 | 64.1491184 | 0.557950335 | 0.19130784 | 2.91650529 | 0.00353977 | 0.01555327 | RAB37 | 326624 |
| ENSG000001 | 464.478117 | 0.38745336 | 0.13285766 | 2.91630431 | 0.00354205 | 0.0155616 | HSDL2 | 84263 |
| ENSG000001 | 924.977387 | 0.345123103 | 0.11834638 | 2.9162117 | 0.0035431 | 0.01556454 | TIPRL | 261726 |
| ENSG000001 | 11.8421594 | -0.951339704 | 0.32625928 | -2.9159008 | 0.00354663 | 0.01557837 | NLRP8 | 126205 |
| ENSG000001 | 99.1766995 | 0.501902956 | 0.17214941 | 2.9155078 | 0.0035511 | 0.01559631 | PAQR7 | 164091 |
| ENSG000001 | 46.2407755 | -0.665540362 | 0.22828091 | -2.9154447 | 0.00355182 | 0.01559778 | MEGF11 | 84465 |
| ENSG000001 | 78.3666413 | 0.524999034 | 0.18008167 | 2.9153385 | 0.00355303 | 0.0156014 | SLC5A12 | 159963 |
| ENSG000001 | 78.1769764 | -0.522312885 | 0.1791678 | -2.9152162 | 0.00355442 | 0.01560582 | GRIP2 | 80852 |
| ENSG000001 | 53.612031 | -0.626535944 | 0.21493366 | -2.9150201 | 0.00355666 | 0.01561394 | GDA | 9615 |
| ENSG000001 | 962.431257 | 0.359239832 | 0.12323997 | 2.91496203 | 0.00355732 | 0.01561417 | PELI1 | 57162 |
| ENSG000002 | 30.5291982 | -0.715819131 | 0.24556841 | -2.9149479 | 0.00355748 | 0.01561417 | NA | NA |
| ENSG000002 | 2.56199673 | -1.074505789 | 0.36862626 | -2.9148922 | 0.00355811 | 0.01561527 | NA | NA |
| ENSG000001 | 22.8913502 | 0.782501971 | 0.26848933 | 2.91446206 | 0.00356302 | 0.01563511 | LURAP1 | 541468 |
| ENSG000001 | 14.8775483 | -0.901378021 | 0.30928689 | -2.9143751 | 0.00356401 | 0.01563777 | TFAP2E | 339488 |
| ENSG000001 | 54.4683939 | -0.603189568 | 0.20698692 | -2.9141434 | 0.00356666 | 0.01564769 | USP6 | 9098 |
| ENSG000002 | 2013.5211 | 0.342685713 | 0.11759977 | 2.91399988 | 0.0035683 | 0.0156532 | ITSN1 | 6453 |
| ENSG000001 | 633.135961 | 0.374381999 | 0.1284827 | 2.91387085 | 0.00356978 | 0.01565628 | HERC4 | 26091 |
| ENSG000002 | 197.609922 | 0.409904403 | 0.14067189 | 2.91390418 | 0.00356939 | 0.01565628 | NFS1 | 9054 |
| ENSG000002 | 15.9647827 | -0.910847755 | 0.31265604 | -2.9132581 | 0.00357679 | 0.01568364 | NA | NA |
| ENSG000002 | 5.44669637 | -1.08552795 | 0.37261326 | -2.9132832 | 0.0035765 | 0.01568364 | NA | NA |
| ENSG000002 | 2.91641229 | -1.091109164 | 0.37458457 | -2.9128513 | 0.00358145 | 0.01570239 | NA | NA |

| | | | | | | | | |
|---|---|---|---|---|---|---|---|---|
| ENSG000001 | 102.766973 | 0.500115466 | 0.17169791 | 2.91276386 | 0.00358245 | 0.01570509 | PINK1-AS | 100861548 |
| ENSG000001 | 8.32877006 | -1.039757852 | 0.35698432 | -2.9126149 | 0.00358416 | 0.01571088 | MORN5 | 254956 |
| ENSG00000C | 10.2486231 | -1.006632049 | 0.34565802 | -2.9122195 | 0.0035887 | 0.01572569 | TRPM5 | 29850 |
| ENSG000001 | 18.8834608 | -0.859822703 | 0.295246 | -2.9122247 | 0.00358864 | 0.01572569 | SMCO2 | 341346 |
| ENSG000002 | 8.76848835 | -1.017796409 | 0.34948619 | -2.912265 | 0.00358818 | 0.01572569 | ANKRD44-IT1 | 101927547 |
| ENSG000002 | 8.7689663 | -1.038940653 | 0.35680992 | -2.9117482 | 0.00359412 | 0.01574603 | NA | NA |
| ENSG000002 | 20.3440296 | -0.854160639 | 0.29334644 | -2.9117811 | 0.00359374 | 0.01574603 | NA | NA |
| ENSG00000C | 135.400276 | 0.478920331 | 0.16448396 | 2.91165374 | 0.00359521 | 0.01574909 | ELOVL1 | 64834 |
| ENSG00000C | 529.802264 | 0.381738047 | 0.13110912 | 2.91160551 | 0.00359576 | 0.01574982 | TBPL1 | 9519 |
| ENSG000002 | 17.8652872 | -0.879910316 | 0.30222529 | -2.9114384 | 0.00359769 | 0.01575655 | NA | NA |
| ENSG000001 | 3348.05411 | 0.346640136 | 0.1190677 | 2.91128609 | 0.00359944 | 0.01576253 | CD2AP | 23607 |
| ENSG000001 | 101.465182 | 0.497568684 | 0.17092347 | 2.91106127 | 0.00360203 | 0.01577217 | ASPHD2 | 57168 |
| ENSG000002 | 8.4402504 | -1.044489537 | 0.35885195 | -2.9106419 | 0.00360687 | 0.01579165 | NA | NA |
| ENSG000001 | 207.768632 | 0.483648728 | 0.16617967 | 2.91039641 | 0.00360971 | 0.01580235 | ZNF133 | 7692 |
| ENSG000002 | 17.6526073 | -0.887616773 | 0.30504842 | -2.909757 | 0.0036171 | 0.015833 | LINC01239 | 441389 |
| ENSG000001 | 353.272616 | 0.396690042 | 0.13634064 | 2.909551 | 0.00361948 | 0.01584173 | RBM18 | 92400 |
| ENSG000001 | 829.063828 | 0.336650471 | 0.11570823 | 2.90947738 | 0.00362034 | 0.01584376 | PAXIP1 | 22976 |
| ENSG000001 | 1720.7361 | 0.33106828 | 0.11380515 | 2.90907995 | 0.00362494 | 0.0158622 | TRIP11 | 9321 |
| ENSG000002 | 9.46082594 | -1.011384669 | 0.34768276 | -2.9089296 | 0.00362668 | 0.01586812 | NA | NA |
| ENSG000002 | 7.64945247 | -1.050438263 | 0.36115382 | -2.908562 | 0.00363095 | 0.01588507 | NA | NA |
| ENSG000001 | 58.0079984 | -0.614086547 | 0.21118958 | -2.9077502 | 0.00364039 | 0.01592464 | GABBR2 | 9568 |
| ENSG000002 | 590.63455 | 0.484352789 | 0.16657839 | 2.90765677 | 0.00364148 | 0.01592768 | NA | NA |
| ENSG000001 | 37.764066 | 0.67346354 | 0.23162224 | 2.90759442 | 0.0036422 | 0.01592914 | LRRC75B | 388886 |
| ENSG000001 | 357.354301 | 0.434572871 | 0.14946718 | 2.90748018 | 0.00364353 | 0.01593324 | OSBPL2 | 9885 |
| ENSG000001 | 9.21371004 | -1.035400559 | 0.35612496 | -2.907408 | 0.00364438 | 0.0159352 | ALDH3B2 | 222 |
| ENSG000002 | 19.865887 | -0.852308505 | 0.29315751 | -2.9073398 | 0.00364517 | 0.01593696 | NA | NA |
| ENSG000001 | 120.328144 | 0.939074204 | 0.32300509 | 2.9073047 | 0.00364558 | 0.01593703 | NA | NA |
| ENSG000002 | 2.92582674 | -1.088599488 | 0.37446019 | -2.9071167 | 0.00364777 | 0.01594489 | NA | NA |
| ENSG000001 | 228.091221 | 0.408889356 | 0.14066617 | 2.90680663 | 0.00365139 | 0.01595898 | BPGM | 669 |

| | | | | | | | | |
|---|---|---|---|---|---|---|---|---|
| ENSG00000C | 931.044734 | 0.418009416 | 0.1438124 | 2.90662977 | 0.00365345 | 0.01596629 | NA | NA |
| ENSG000001 | 77.7248817 | 0.550655784 | 0.18947338 | 2.90624357 | 0.00365797 | 0.01598429 | LIPT2 | 387787 |
| ENSG000002 | 9.92351876 | -0.986017215 | 0.33929926 | -2.90604 | 0.00366035 | 0.01599296 | LOC105378753 | 105378753 |
| ENSG000002 | 13.4052257 | 0.929663149 | 0.31991131 | 2.90600274 | 0.00366078 | 0.01599315 | NA | NA |
| ENSG000001 | 40.1528614 | -0.674090442 | 0.23197038 | -2.9059332 | 0.0036616 | 0.01599498 | TIGIT | 201633 |
| ENSG000001 | 1811.4759 | 0.419288848 | 0.14428991 | 2.90587774 | 0.00366225 | 0.01599609 | PTPN12 | 5782 |
| ENSG000001 | 1327.25109 | 0.715222432 | 0.24615141 | 2.9056199 | 0.00366526 | 0.01600755 | CKB | 1152 |
| ENSG000002 | 10.5911825 | -0.980196402 | 0.33735931 | -2.9054968 | 0.00366671 | 0.01601212 | LOC101928731 | 101928731 |
| ENSG000002 | 4.6524892 | -1.099226893 | 0.37834624 | -2.9053464 | 0.00366847 | 0.01601809 | NA | NA |
| ENSG00000C | 3159.44807 | 0.334384216 | 0.11509516 | 2.90528469 | 0.00366919 | 0.01601909 | DNTTIP2 | 30836 |
| ENSG000001 | 46.0909119 | 0.61545566 | 0.21184187 | 2.9052597 | 0.00366948 | 0.01601909 | GLYCTK | 132158 |
| ENSG000001 | 24.739897 | -0.803810486 | 0.27671929 | -2.9047866 | 0.00367504 | 0.01603986 | SYCP3 | 50511 |
| ENSG000002 | 6.60079649 | -1.066721982 | 0.36722661 | -2.9048058 | 0.00367481 | 0.01603986 | LOC101928661 | 101928661 |
| ENSG000002 | 24.9370329 | -0.804973681 | 0.27712424 | -2.9047393 | 0.00367559 | 0.01604056 | NA | NA |
| ENSG00000C | 12.917812 | -0.964363049 | 0.33202963 | -2.9044488 | 0.003679 | 0.01605373 | C2orf83 | 56918 |
| ENSG000001 | 2678.03069 | 0.358733034 | 0.12351565 | 2.9043528 | 0.00368013 | 0.01605692 | EIF4E | 1977 |
| ENSG000001 | 2944.29283 | 0.301810433 | 0.10393226 | 2.90391497 | 0.00368528 | 0.01607766 | DYRK1A | 1859 |
| ENSG000002 | 7.18540727 | -1.052393179 | 0.36244679 | -2.9035798 | 0.00368923 | 0.01609315 | NA | NA |
| ENSG000001 | 22.1293382 | -0.788060128 | 0.27141426 | -2.9035325 | 0.00368979 | 0.01609385 | NR4A2 | 4929 |
| ENSG000002 | 30.2696318 | -0.726681334 | 0.25029749 | -2.9032705 | 0.00369288 | 0.01610559 | NA | NA |
| ENSG000001 | 842.454646 | 0.472921583 | 0.16289826 | 2.90317153 | 0.00369404 | 0.01610771 | ATP5J | 522 |
| ENSG000002 | 18.9461282 | -0.84370747 | 0.29061674 | -2.9031619 | 0.00369416 | 0.01610771 | NA | NA |
| ENSG000002 | 5.20360293 | -1.089492304 | 0.37528535 | -2.9031037 | 0.00369484 | 0.01610897 | NA | NA |
| ENSG000002 | 4.65665974 | -1.098322386 | 0.37835732 | -2.9028707 | 0.00369759 | 0.01611923 | NA | NA |
| ENSG000002 | 19.1533157 | -0.835710924 | 0.28791038 | -2.9026773 | 0.00369988 | 0.01612745 | LOC101928877 | 101928877 |
| ENSG00000C | 126.357717 | 0.501825088 | 0.17289805 | 2.90243351 | 0.00370276 | 0.01613828 | TMEM159 | 57146 |
| ENSG000001 | 178.167372 | 0.428535005 | 0.1476662 | 2.90205203 | 0.00370727 | 0.01615594 | TTC12 | 54970 |
| ENSG000001 | 442.873641 | 0.364103201 | 0.12546529 | 2.90202347 | 0.00370761 | 0.01615594 | NFRKB | 4798 |
| ENSG000001 | 638.08276 | 0.347923364 | 0.11989699 | 2.90185236 | 0.00370963 | 0.0161613 | OTULIN | 90268 |

| | | | | | | | | |
|---|---|---|---|---|---|---|---|---|
| ENSG000002 | 6.45080091 | -1.061867411 | 0.36592615 | -2.9018626 | 0.00370951 | 0.0161613 | NA | NA |
| ENSG00000C | 36.3207373 | -0.692367618 | 0.23865648 | -2.9011055 | 0.00371849 | 0.01619465 | NA | NA |
| ENSG000001 | 771.98004 | 0.353966817 | 0.12201044 | 2.90111921 | 0.00371832 | 0.01619465 | RAB23 | 51715 |
| ENSG000001 | 234.448252 | 0.435022642 | 0.14994944 | 2.90112883 | 0.00371821 | 0.01619465 | ZNF30 | 90075 |
| ENSG000001 | 216.75885 | 0.439429322 | 0.1514773 | 2.90095825 | 0.00372023 | 0.01620052 | DHODH | 1723 |
| ENSG000001 | 34.9518674 | -0.748758994 | 0.25813406 | -2.9006594 | 0.00372378 | 0.01621423 | ADH1B | 125 |
| ENSG000002 | 29.7215866 | 0.741450448 | 0.25564145 | 2.90035299 | 0.00372743 | 0.01622835 | LOC101929112 | 101929112 |
| ENSG000002 | 6.57618515 | -1.068960429 | 0.36856764 | -2.90031 | 0.00372794 | 0.01622883 | NA | NA |
| ENSG000001 | 6.14565687 | -1.070645641 | 0.3691628 | -2.9001992 | 0.00372926 | 0.01623282 | SIGLEC7 | 27036 |
| ENSG000002 | 98.8056925 | -0.492248201 | 0.16973105 | -2.9001658 | 0.00372965 | 0.01623282 | CFAP44 | 55779 |
| ENSG000001 | 278.519523 | 0.460248841 | 0.15870407 | 2.90004443 | 0.0037311 | 0.01623736 | SMG9 | 56006 |
| ENSG000001 | 86.1876482 | -0.533735526 | 0.18405101 | -2.8999326 | 0.00373243 | 0.01624141 | NAALADL2 | 254827 |
| ENSG000002 | 6.95652085 | -1.062362014 | 0.36638785 | -2.8995558 | 0.00373692 | 0.0162592 | NA | NA |
| ENSG000001 | 1220.83187 | 0.370715085 | 0.12785583 | 2.89947736 | 0.00373785 | 0.01626153 | ZMYM3 | 9203 |
| ENSG000002 | 6.00319989 | -1.080569218 | 0.37269828 | -2.8993136 | 0.00373981 | 0.01626828 | NA | NA |
| ENSG000001 | 4699.04099 | 0.350778331 | 0.1210373 | 2.89810104 | 0.0037543 | 0.01632956 | MCL1 | 4170 |
| ENSG000001 | 6.19172349 | -1.06983158 | 0.36916142 | -2.8980048 | 0.00375545 | 0.01633281 | CLDN20 | 49861 |
| ENSG000002 | 9.93756252 | -1.009244096 | 0.34828468 | -2.8977562 | 0.00375843 | 0.01634402 | NA | NA |
| ENSG000002 | 211.864284 | 0.863547522 | 0.29805272 | 2.89729792 | 0.00376392 | 0.01636616 | NA | NA |
| ENSG000001 | 1283.61596 | 0.376578737 | 0.12998045 | 2.89719516 | 0.00376515 | 0.01636976 | PGGT1B | 5229 |
| ENSG000001 | 23.3378687 | -0.780736643 | 0.26950166 | -2.8969641 | 0.00376793 | 0.01638007 | SLC30A8 | 169026 |
| ENSG000002 | 3.40348134 | -1.090311695 | 0.37643929 | -2.8963812 | 0.00377494 | 0.01640877 | NA | NA |
| ENSG000001 | 6148.71769 | 0.335249836 | 0.1157535 | 2.89623929 | 0.00377664 | 0.01641311 | HMGCS1 | 3157 |
| ENSG000001 | 384.744271 | 0.40317458 | 0.13920778 | 2.89620724 | 0.00377703 | 0.01641311 | SSBP2 | 23635 |
| ENSG000001 | 218.535021 | 0.445182335 | 0.15371271 | 2.89619726 | 0.00377715 | 0.01641311 | CCNE2 | 9134 |
| ENSG000002 | 14.1216869 | -0.959559093 | 0.33133015 | -2.8960814 | 0.00377854 | 0.01641741 | OR2H2 | 7932 |
| ENSG000001 | 415.149623 | 0.424941464 | 0.14673701 | 2.89593918 | 0.00378026 | 0.0164231 | ZNF154 | 7710 |
| ENSG00000C | 445.904971 | 0.356529435 | 0.12312696 | 2.89562449 | 0.00378405 | 0.01643781 | ACER3 | 55331 |
| ENSG000002 | 200.791959 | 0.485951459 | 0.16783418 | 2.89542599 | 0.00378644 | 0.01644645 | NA | NA |

| | | | | | | | | |
|---|---|---|---|---|---|---|---|---|
| ENSG00000C | 2628.25774 | 0.394740283 | 0.13634462 | 2.89516584 | 0.00378958 | 0.01645832 | YBX3 | 8531 |
| ENSG000002 | 3.68027299 | -1.095612628 | 0.37844323 | -2.8950515 | 0.00379096 | 0.01646256 | NA | NA |
| ENSG000002 | 101.450937 | 0.486370639 | 0.16801039 | 2.89488433 | 0.00379298 | 0.01646956 | FAM19A5 | 25817 |
| ENSG000001 | 155.552123 | 0.452188791 | 0.15620507 | 2.89484066 | 0.00379351 | 0.01647009 | FAM86C1 | 55199 |
| ENSG000001 | 1481.05335 | 0.50163879 | 0.17329475 | 2.8947143 | 0.00379504 | 0.0164735 | HTT | 3064 |
| ENSG000002 | 53.7661286 | 0.590885798 | 0.20412619 | 2.89470839 | 0.00379511 | 0.0164735 | TOPORS-AS1 | 100129250 |
| ENSG000001 | 202.679916 | 0.391246303 | 0.13516542 | 2.89457387 | 0.00379674 | 0.01647864 | ZNF41 | 7592 |
| ENSG000002 | 6.77197301 | -1.070474078 | 0.36982486 | -2.8945433 | 0.0037971 | 0.01647864 | LOC105373006 | 105373006 |
| ENSG000001 | 908.490153 | 0.348653296 | 0.12045667 | 2.89442924 | 0.00379849 | 0.01648286 | ZNF652 | 22834 |
| ENSG000002 | 214.964227 | 0.449787426 | 0.15540311 | 2.89432708 | 0.00379972 | 0.01648646 | NA | NA |
| ENSG000002 | 10.4428322 | -0.993259882 | 0.34318509 | -2.8942396 | 0.00380078 | 0.01648929 | LOC100505782 | 100505782 |
| ENSG000001 | 105.68061 | 0.505167085 | 0.17455001 | 2.89411085 | 0.00380234 | 0.01649429 | DFFB | 1677 |
| ENSG000001 | 61.9701985 | -0.56855166 | 0.19648579 | -2.893602 | 0.00380851 | 0.01651928 | SGIP1 | 84251 |
| ENSG000001 | 158.110375 | 0.421479183 | 0.14567044 | 2.89337491 | 0.00381126 | 0.0165293 | BEND6 | 221336 |
| ENSG000002 | 15.8513184 | -0.895680277 | 0.30956574 | -2.8933443 | 0.00381163 | 0.0165293 | NA | NA |
| ENSG00000C | 1112.6566 | 0.377584503 | 0.13053168 | 2.89266566 | 0.00381988 | 0.01656151 | MPP5 | 64398 |
| ENSG000001 | 570.707642 | 0.38905889 | 0.13449792 | 2.89267591 | 0.00381975 | 0.01656151 | RASA2 | 5922 |
| ENSG000001 | 406.32109 | 0.354414927 | 0.12252813 | 2.89251892 | 0.00382166 | 0.01656748 | RAB9A | 9367 |
| ENSG000002 | 5.01414269 | -1.093455296 | 0.37804328 | -2.8924077 | 0.00382301 | 0.01657157 | NA | NA |
| ENSG000001 | 4.58432914 | -1.093870583 | 0.37819624 | -2.892336 | 0.00382389 | 0.01657359 | C11orf53 | 341032 |
| ENSG000002 | 3.36642291 | -1.090312243 | 0.37697136 | -2.8922947 | 0.00382439 | 0.01657399 | NA | NA |
| ENSG000001 | 4.80319975 | -1.089709338 | 0.37677768 | -2.8921813 | 0.00382577 | 0.01657821 | TH | 7054 |
| ENSG000002 | 11.866257 | -0.950070572 | 0.32850085 | -2.8921404 | 0.00382627 | 0.01657859 | NA | NA |
| ENSG000001 | 6789.78624 | 0.312997389 | 0.10822958 | 2.89197627 | 0.00382827 | 0.01658548 | NUFIP2 | 57532 |
| ENSG000001 | 325.370642 | 0.413496253 | 0.14300808 | 2.89141882 | 0.00383507 | 0.01661316 | NRDE2 | 55051 |
| ENSG000001 | 155.625625 | -0.446332421 | 0.15437268 | -2.8912656 | 0.00383694 | 0.01661949 | PCDH9 | 5101 |
| ENSG000002 | 13.5621615 | -0.91030151 | 0.31488176 | -2.8909312 | 0.00384102 | 0.0166354 | LOC101929418 | 101929418 |
| ENSG000002 | 18.67463 | -0.886240597 | 0.3065893 | -2.8906442 | 0.00384453 | 0.01664882 | PTPRVP | 148713 |
| ENSG00000C | 247.188812 | 0.434482837 | 0.1503226 | 2.89033601 | 0.0038483 | 0.01666338 | MMP2 | 4313 |

| | | | | | | | | |
|---|---|---|---|---|---|---|---|---|
| ENSG000001 | 1131.18738 | 0.472971113 | 0.16364589 | 2.89021079 | 0.00384984 | 0.01666824 | TRIM44 | 54765 |
| ENSG000000 | 94.619554 | 0.532948439 | 0.18444923 | 2.88940445 | 0.00385972 | 0.01670923 | MGST2 | 4258 |
| ENSG000001 | 839.771048 | 0.356658009 | 0.12343792 | 2.88937143 | 0.00386013 | 0.01670923 | CLIP1 | 6249 |
| ENSG000001 | 18.478612 | -0.855081246 | 0.29595429 | -2.8892342 | 0.00386181 | 0.01671474 | BCO1 | 53630 |
| ENSG000001 | 3910.67906 | 0.349689232 | 0.12103738 | 2.88910116 | 0.00386345 | 0.01672003 | SP3 | 6670 |
| ENSG000001 | 1704.50463 | 0.730515526 | 0.25286339 | 2.88897309 | 0.00386502 | 0.01672149 | EIF1 | 10209 |
| ENSG000002 | 19.2128376 | -0.834840922 | 0.28897446 | -2.8889782 | 0.00386496 | 0.01672149 | ERICH3-AS1 | 101927320 |
| ENSG000002 | 3.28980523 | -1.089323038 | 0.37706018 | -2.8889899 | 0.00386481 | 0.01672149 | NA | NA |
| ENSG000001 | 8.58612622 | -1.024995069 | 0.35485562 | -2.8884848 | 0.00387103 | 0.01674568 | NA | NA |
| ENSG000001 | 433.624907 | 0.411899765 | 0.14260495 | 2.8883973 | 0.0038721 | 0.01674677 | DDX39B | 7919 |
| ENSG000002 | 5.38567595 | -1.088332041 | 0.3767915 | -2.8884198 | 0.00387183 | 0.01674677 | NA | NA |
| ENSG000001 | 137.074 | 0.483009564 | 0.16722672 | 2.88835153 | 0.00387267 | 0.01674742 | VCPKMT | 79609 |
| ENSG000001 | 1014.9963 | 0.410122557 | 0.14199998 | 2.88818749 | 0.00387469 | 0.01675437 | NCAPD3 | 23310 |
| ENSG000001 | 323.98465 | 0.430078516 | 0.14891258 | 2.88812754 | 0.00387543 | 0.01675578 | FRA10AC1 | 118924 |
| ENSG000001 | 59.3811006 | -0.584949599 | 0.20255129 | -2.8879086 | 0.00387813 | 0.01676566 | FAM13C | 220965 |
| ENSG000001 | 139.318133 | 0.445953 | 0.15442381 | 2.88785134 | 0.00387883 | 0.01676692 | KLF8 | 11279 |
| ENSG000000 | 29.8965148 | -0.715814348 | 0.24788055 | -2.8877391 | 0.00388022 | 0.01676754 | GCKR | 2646 |
| ENSG000001 | 163.600956 | 0.425632601 | 0.14739076 | 2.88778348 | 0.00387967 | 0.01676754 | IRS4 | 8471 |
| ENSG000001 | 1654.95139 | 0.469417886 | 0.16255365 | 2.88777199 | 0.00387981 | 0.01676754 | TRA2A | 29896 |
| ENSG000001 | 1157.02716 | 0.33307358 | 0.11534812 | 2.88755095 | 0.00388254 | 0.01677579 | UBLCP1 | 134510 |
| ENSG000001 | 73.2230375 | 0.556968146 | 0.1928908 | 2.88747911 | 0.00388342 | 0.01677783 | FAM86C2P | 645332 |
| ENSG000001 | 10.9591376 | -0.974058654 | 0.33735029 | -2.8873805 | 0.00388464 | 0.0167813 | DNAJB13 | 374407 |
| ENSG000001 | 20.1067191 | -0.845533012 | 0.29285908 | -2.8871668 | 0.00388728 | 0.01679092 | PRR26 | 414235 |
| ENSG000001 | 734.525161 | 0.361334839 | 0.12516213 | 2.88693428 | 0.00389015 | 0.01680154 | TDRP | 157695 |
| ENSG000001 | 9.62756821 | -1.013822449 | 0.35119591 | -2.8867718 | 0.00389216 | 0.01680843 | TCERG1L | 256536 |
| ENSG000002 | 10.7610386 | -0.959858989 | 0.33250808 | -2.8867238 | 0.00389276 | 0.0168092 | MIR8061 | 102466251 |
| ENSG000001 | 8.15196412 | -1.025421107 | 0.35527328 | -2.8862883 | 0.00389815 | 0.01683069 | NA | NA |
| ENSG000001 | 12.9651436 | -0.92205721 | 0.31948099 | -2.8861098 | 0.00390036 | 0.01683845 | UGT2B17 | 7367 |
| ENSG000002 | 21.6254153 | 0.798054291 | 0.27651912 | 2.88607278 | 0.00390082 | 0.01683863 | SNORD116-25 | 100033436 |

| | | | | | | | | |
|---|---|---|---|---|---|---|---|---|
| ENSG000001 | 431.919013 | 0.461835399 | 0.16003063 | 2.88591883 | 0.00390273 | 0.01684508 | PHF10 | 55274 |
| ENSG000001 | 29.2003032 | -0.768089684 | 0.26615865 | -2.8858341 | 0.00390378 | 0.01684603 | TRIM29 | 23650 |
| ENSG000001 | 102.791626 | 0.537306372 | 0.18618618 | 2.88585536 | 0.00390352 | 0.01684603 | GPSM1 | 26086 |
| ENSG000002 | 41.1131051 | -0.670661007 | 0.23240633 | -2.8857261 | 0.00390512 | 0.01685002 | NA | NA |
| ENSG000002 | 40.1132265 | 0.694125807 | 0.2405704 | 2.88533341 | 0.00390999 | 0.01686926 | SNORD4A | 26773 |
| ENSG000001 | 50.8063369 | -0.627215855 | 0.2173971 | -2.8851161 | 0.00391269 | 0.01687911 | GALNTL6 | 442117 |
| ENSG000002 | 115.307967 | 0.510780107 | 0.17704441 | 2.88503945 | 0.00391365 | 0.01688142 | NA | NA |
| ENSG000002 | 25.4119483 | -0.765338495 | 0.26528246 | -2.8849947 | 0.0039142 | 0.01688203 | LOC101927780 | 101927780 |
| ENSG000001 | 1674.13192 | 0.319803882 | 0.11085684 | 2.88483678 | 0.00391617 | 0.0168887 | PRRC1 | 133619 |
| ENSG000001 | 74.5935071 | 0.653390473 | 0.22652451 | 2.88441398 | 0.00392143 | 0.0169096 | SLC16A5 | 9121 |
| ENSG000002 | 4.05744283 | -1.09187776 | 0.3785783 | -2.8841531 | 0.00392468 | 0.01692181 | NA | NA |
| ENSG000001 | 4.20662156 | -1.091500645 | 0.37855789 | -2.8833124 | 0.00393517 | 0.01696524 | OR2T10 | 127069 |
| ENSG000001 | 84.6384076 | 0.528878536 | 0.18343486 | 2.8831954 | 0.00393663 | 0.01696974 | CYB5D2 | 124936 |
| ENSG000002 | 6.20866451 | -1.06695802 | 0.37008546 | -2.8830045 | 0.00393902 | 0.01697823 | NA | NA |
| ENSG000001 | 429.713824 | 0.419946704 | 0.14566567 | 2.88294895 | 0.00393971 | 0.01697941 | MADD | 8567 |
| ENSG000001 | 4.18112127 | -1.091320817 | 0.37859338 | -2.8825671 | 0.00394449 | 0.0169982 | HOXD11 | 3237 |
| ENSG00000C | 1323.76462 | 0.510295954 | 0.17703241 | 2.88250016 | 0.00394533 | 0.01700001 | RALA | 5898 |
| ENSG000002 | 3.36849815 | -1.086529116 | 0.37697052 | -2.8822655 | 0.00394827 | 0.01701086 | NA | NA |
| ENSG000001 | 113.312709 | 0.527571121 | 0.18304998 | 2.88211519 | 0.00395015 | 0.01701717 | PFKFB4 | 5210 |
| ENSG000002 | 131.052252 | 0.524666043 | 0.18205944 | 2.88183922 | 0.00395361 | 0.01703027 | NA | NA |
| ENSG000002 | 15.5590569 | -0.88060299 | 0.30557489 | -2.8817911 | 0.00395422 | 0.01703107 | NA | NA |
| ENSG00000C | 7645.51507 | 0.342726887 | 0.11893562 | 2.88161685 | 0.00395641 | 0.01703687 | AASS | 10157 |
| ENSG000001 | 21.6931519 | -0.886953736 | 0.3077961 | -2.8816276 | 0.00395627 | 0.01703687 | OR51L1 | 119682 |
| ENSG000001 | 323.46373 | 0.509452683 | 0.17682288 | 2.88114692 | 0.00396231 | 0.01706048 | C6orf132 | 647024 |
| ENSG000002 | 20.9893158 | -0.799987195 | 0.27769581 | -2.880804 | 0.00396662 | 0.01707724 | NA | NA |
| ENSG00000C | 43.4532936 | -0.639026714 | 0.22183674 | -2.8806171 | 0.00396898 | 0.01708555 | CD38 | 952 |
| ENSG000001 | 262.709486 | 0.417052867 | 0.14478753 | 2.8804474 | 0.00397111 | 0.01709112 | F2R | 2149 |
| ENSG000002 | 5.07356402 | -1.079844009 | 0.3748873 | -2.8804497 | 0.00397108 | 0.01709112 | PRAMEF15 | 653619 |
| ENSG000001 | 170.242039 | 0.461405638 | 0.16018808 | 2.88039937 | 0.00397172 | 0.01709191 | CHAC2 | 494143 |

| | | | | | | | | |
|---|---|---|---|---|---|---|---|---|
| ENSG000002 | 10.6954219 | -0.978085969 | 0.33957819 | -2.8802968 | 0.00397301 | 0.01709566 | NA | NA |
| ENSG000001 | 319.600307 | 0.41648762 | 0.14461412 | 2.87999285 | 0.00397684 | 0.01711034 | LDLRAD3 | 143458 |
| ENSG000001 | 12.4647865 | -0.935962944 | 0.32504646 | -2.8794743 | 0.00398339 | 0.01713668 | G6PC2 | 57818 |
| ENSG000002 | 22.3452446 | -0.835612792 | 0.29021154 | -2.8793231 | 0.0039853 | 0.01714308 | LOC101927379 | 101927379 |
| ENSG000001 | 657.66804 | 0.378458044 | 0.13144192 | 2.87927974 | 0.00398585 | 0.01714362 | PGM2L1 | 283209 |
| ENSG000002 | 12.942631 | 0.919924487 | 0.31953359 | 2.87896019 | 0.00398989 | 0.01715918 | NA | NA |
| ENSG00000C | 26.0453631 | -0.768181029 | 0.2668312 | -2.8789026 | 0.00399062 | 0.0171605 | SCTR | 6344 |
| ENSG000001 | 24.3352889 | -0.782820289 | 0.27193959 | -2.8786551 | 0.00399375 | 0.01717032 | KCNH1 | 3756 |
| ENSG000001 | 24.8785271 | -0.808304313 | 0.28079034 | -2.8786756 | 0.00399349 | 0.01717032 | NA | NA |
| ENSG000001 | 15.4341722 | -0.893214469 | 0.31031626 | -2.8784004 | 0.00399697 | 0.01718237 | CYP7A1 | 1581 |
| ENSG000002 | 173.806912 | 0.444193981 | 0.15432202 | 2.8783578 | 0.00399751 | 0.01718287 | NA | NA |
| ENSG000001 | 52.8133904 | -0.615387394 | 0.21382159 | -2.8780415 | 0.00400153 | 0.01719829 | PLD5 | 200150 |
| ENSG000002 | 5.14298209 | -1.08122566 | 0.37568957 | -2.8779762 | 0.00400235 | 0.01720002 | NA | NA |
| ENSG000002 | 4.13216187 | -1.089501648 | 0.37859192 | -2.8777731 | 0.00400493 | 0.01720928 | NA | NA |
| ENSG000001 | 14.5427619 | -0.923498447 | 0.32092478 | -2.8776165 | 0.00400692 | 0.017216 | RHCE | 6006 |
| ENSG000001 | 1692.03343 | 0.315416795 | 0.10962179 | 2.87731832 | 0.00401071 | 0.01723045 | NEMF | 9147 |
| ENSG000001 | 10.0498791 | -0.985645657 | 0.34256166 | -2.8772795 | 0.0040112 | 0.01723074 | ASPG | 374569 |
| ENSG000001 | 2485.34261 | 0.375688761 | 0.13058091 | 2.87705733 | 0.00401403 | 0.01724105 | EP300 | 2033 |
| ENSG000002 | 345.86089 | 0.362529404 | 0.12601727 | 2.87682324 | 0.00401701 | 0.01725202 | PPME1 | 51400 |
| ENSG000001 | 155.929313 | 0.469814681 | 0.16332757 | 2.87651785 | 0.00402089 | 0.01726141 | WFS1 | 7466 |
| ENSG000001 | 74.1945821 | 0.531502318 | 0.18477043 | 2.87655504 | 0.00402042 | 0.01726141 | RILP | 83547 |
| ENSG000001 | 114.483311 | 0.494467302 | 0.17189594 | 2.87655021 | 0.00402048 | 0.01726141 | ZBTB49 | 166793 |
| ENSG000002 | 4.88584563 | -1.08508932 | 0.37721695 | -2.8765656 | 0.00402029 | 0.01726141 | NA | NA |
| ENSG000002 | 15.9285507 | -0.870047588 | 0.30247568 | -2.8764217 | 0.00402212 | 0.01726485 | NA | NA |
| ENSG000002 | 201.697026 | 0.421864564 | 0.14667106 | 2.8762631 | 0.00402414 | 0.01727169 | DUSP14 | 11072 |
| ENSG000002 | 15.0864713 | -0.887909187 | 0.30874796 | -2.8758382 | 0.00402956 | 0.01729313 | NA | NA |
| ENSG000001 | 28.7073333 | -0.733963666 | 0.25523412 | -2.8756487 | 0.00403198 | 0.01730168 | ZBTB16 | 7704 |
| ENSG000001 | 6.23545982 | -1.065175794 | 0.37043065 | -2.8755066 | 0.0040338 | 0.01730732 | OR4B1 | 119765 |
| ENSG000002 | 16.2718445 | -0.890790799 | 0.30978865 | -2.8754791 | 0.00403415 | 0.01730732 | NA | NA |

| | | | | | | | | |
|---|---|---|---|---|---|---|---|---|
| ENSG00000C | 884.610968 | 0.333731689 | 0.11606564 | 2.87537037 | 0.00403554 | 0.01730891 | PIK3C3 | 5289 |
| ENSG000001 | 132.80455 | -0.450223299 | 0.15658036 | -2.8753498 | 0.0040358 | 0.01730891 | PCDH7 | 5099 |
| ENSG000002 | 10.5219964 | -0.966929953 | 0.33627516 | -2.875413 | 0.00403499 | 0.01730891 | NA | NA |
| ENSG000001 | 1050.28495 | 0.382789436 | 0.13313041 | 2.87529683 | 0.00403648 | 0.01730999 | C3orf38 | 285237 |
| ENSG000002 | 7.21347317 | -1.04606958 | 0.36382161 | -2.8752266 | 0.00403738 | 0.017312 | NA | NA |
| ENSG000001 | 44.833365 | -0.615812237 | 0.21418691 | -2.8751161 | 0.00403879 | 0.01731441 | SUSD6 | 9766 |
| ENSG000001 | 293.074773 | 0.393862329 | 0.13698925 | 2.8751331 | 0.00403857 | 0.01731441 | TRAPPC12 | 51112 |
| ENSG000001 | 714.460504 | 0.36039919 | 0.12535611 | 2.8750029 | 0.00404024 | 0.01731695 | MRPL1 | 65008 |
| ENSG000001 | 232.005014 | 0.407513787 | 0.14174348 | 2.87500897 | 0.00404016 | 0.01731695 | ZBTB26 | 57684 |
| ENSG00000C | 132.326885 | 0.556888042 | 0.19370675 | 2.87490266 | 0.00404152 | 0.01732062 | ARVCF | 421 |
| ENSG000002 | 21.9754421 | -0.775890107 | 0.2698992 | -2.8747403 | 0.0040436 | 0.0173277 | NA | NA |
| ENSG00000C | 3744.57022 | 0.319081155 | 0.11100234 | 2.8745445 | 0.00404611 | 0.01733657 | PPP1R12A | 4659 |
| ENSG000002 | 192.569115 | 0.402363315 | 0.13997622 | 2.87451186 | 0.00404653 | 0.01733657 | ZSCAN31 | 64288 |
| ENSG000001 | 14.5298005 | -0.895783861 | 0.31165377 | -2.8742917 | 0.00404935 | 0.01734538 | CX3CR1 | 1524 |
| ENSG000002 | 15.8948465 | -0.894320663 | 0.31114547 | -2.8742847 | 0.00404944 | 0.01734538 | NA | NA |
| ENSG000001 | 117.35371 | 0.481536765 | 0.16753823 | 2.87419037 | 0.00405065 | 0.01734873 | RHOB | 388 |
| ENSG000001 | 330.028061 | 0.368344983 | 0.12816127 | 2.87407405 | 0.00405214 | 0.01735329 | WARS2 | 10352 |
| ENSG000002 | 517.01797 | 0.380519581 | 0.1323996 | 2.87402359 | 0.00405279 | 0.01735423 | TMEM178B | 100507421 |
| ENSG000002 | 29.981432 | -0.84731478 | 0.29488625 | -2.8733614 | 0.00406129 | 0.01738881 | NA | NA |
| ENSG000002 | 12.5558809 | -0.968763391 | 0.33716511 | -2.8732611 | 0.00406258 | 0.0173925 | LOC440434 | 440434 |
| ENSG000001 | 210.730348 | 0.424757696 | 0.14785287 | 2.87284047 | 0.00406799 | 0.01741383 | TM2D1 | 83941 |
| ENSG000002 | 159.399692 | 0.455387409 | 0.15853106 | 2.87254375 | 0.00407182 | 0.01742836 | NA | NA |
| ENSG000001 | 60.1726816 | 0.573138301 | 0.19952975 | 2.87244529 | 0.00407309 | 0.01743195 | CCDC102A | 92922 |
| ENSG000001 | 5.81157101 | -1.06721778 | 0.37156132 | -2.8722521 | 0.00407558 | 0.01744077 | LDHAL6B | 92483 |
| ENSG000002 | 2.52940932 | -1.055111032 | 0.36735999 | -2.8721447 | 0.00407696 | 0.01744486 | NA | NA |
| ENSG000001 | 5056.4093 | 0.439874204 | 0.15315446 | 2.87209533 | 0.0040776 | 0.01744574 | IGF2BP1 | 10642 |
| ENSG000002 | 6.5463057 | -1.059855476 | 0.36903047 | -2.8719999 | 0.00407883 | 0.01744917 | LINC00479 | 150135 |
| ENSG000001 | 33.4553339 | -0.695326526 | 0.24212743 | -2.8717379 | 0.00408221 | 0.0174618 | GRM1 | 2911 |
| ENSG000001 | 188.938303 | 0.424261463 | 0.14773949 | 2.87168619 | 0.00408288 | 0.01746281 | RUSC1 | 23623 |

| | | | | | | | | |
|---|---|---|---|---|---|---|---|---|
| ENSG000002 | 6.07794931 | -1.057588936 | 0.36832155 | -2.8713741 | 0.00408692 | 0.01747823 | NA | NA |
| ENSG000002 | 3.70559233 | -1.08619189 | 0.37831789 | -2.8711089 | 0.00409035 | 0.01749105 | NA | NA |
| ENSG000002 | 12.7517568 | -0.934396182 | 0.32545331 | -2.8710606 | 0.00409097 | 0.01749188 | NA | NA |
| ENSG000001 | 169.898699 | 0.415749227 | 0.14481614 | 2.87087622 | 0.00409336 | 0.01750024 | ZNF235 | 9310 |
| ENSG000001 | 43.2547154 | -0.693991002 | 0.24174845 | -2.8707154 | 0.00409544 | 0.0175073 | RGS22 | 26166 |
| ENSG000001 | 24.0977183 | -0.814838445 | 0.28389815 | -2.8701787 | 0.0041024 | 0.01753519 | CARD14 | 79092 |
| ENSG00000C | 1704.63098 | 0.33561723 | 0.11694152 | 2.86995784 | 0.00410527 | 0.0175456 | IARS2 | 55699 |
| ENSG000001 | 18.8717712 | -0.833574843 | 0.29049432 | -2.8695048 | 0.00411115 | 0.0175689 | LIX1 | 167410 |
| ENSG000001 | 27.5705978 | -0.780366294 | 0.27195716 | -2.8694456 | 0.00411192 | 0.01756928 | GFI1B | 8328 |
| ENSG000001 | 290.781075 | 0.4213937 | 0.14685618 | 2.86943125 | 0.00411211 | 0.01756928 | COL4A6 | 1288 |
| ENSG000001 | 3.75841606 | -1.085054465 | 0.37817328 | -2.8691992 | 0.00411513 | 0.01758032 | CCL7 | 6354 |
| ENSG000001 | 296.042091 | 0.385098652 | 0.13422206 | 2.86911603 | 0.00411621 | 0.0175831 | ZNF746 | 155061 |
| ENSG000001 | 251.472457 | 0.430792025 | 0.15015616 | 2.86896015 | 0.00411824 | 0.01758621 | DDX19B | 11269 |
| ENSG000001 | 330.379805 | 0.420221202 | 0.14647037 | 2.86898443 | 0.00411792 | 0.01758621 | FAM92A1 | 137392 |
| ENSG000002 | 20.3364459 | -0.867992401 | 0.30254404 | -2.8689787 | 0.004118 | 0.01758621 | HNRNPA3P15 | 106480234 |
| ENSG000001 | 29.4112521 | -0.738168999 | 0.25733647 | -2.8684974 | 0.00412427 | 0.01760928 | PARK2 | 5071 |
| ENSG000001 | 6.29299644 | -1.065579242 | 0.37147887 | -2.8684788 | 0.00412451 | 0.01760928 | NA | NA |
| ENSG000001 | 67.4917903 | 0.598111539 | 0.20852646 | 2.86827652 | 0.00412715 | 0.01761869 | OPLAH | 26873 |
| ENSG000001 | 2539.3418 | 0.306460776 | 0.10687441 | 2.86748517 | 0.00413748 | 0.01765603 | FGD4 | 121512 |
| ENSG000001 | 186.395669 | 0.498514213 | 0.17384872 | 2.86751732 | 0.00413706 | 0.01765603 | RELA | 5970 |
| ENSG000002 | 12.4512951 | -0.93682538 | 0.3267076 | -2.8674735 | 0.00413763 | 0.01765603 | FCGR3A | 2214 |
| ENSG000002 | 3.74140182 | -1.084515536 | 0.37820239 | -2.8675533 | 0.00413659 | 0.01765603 | NA | NA |
| ENSG000002 | 18.3163125 | -0.842732519 | 0.29390795 | -2.8673349 | 0.00413945 | 0.0176619 | LOC101929532 | 101929532 |
| ENSG000001 | 37.3038076 | -0.65371327 | 0.22800229 | -2.8671347 | 0.00414207 | 0.01766763 | PDE1A | 5136 |
| ENSG000001 | 14.3270265 | -0.908585245 | 0.31689987 | -2.8671051 | 0.00414245 | 0.01766763 | ACSM5 | 54988 |
| ENSG000002 | 20.2839835 | -0.799953874 | 0.27900842 | -2.8671316 | 0.00414211 | 0.01766763 | NA | NA |
| ENSG000002 | 4.84151261 | -1.082157464 | 0.37743985 | -2.8670991 | 0.00414253 | 0.01766763 | NA | NA |
| ENSG000001 | 101.265908 | 0.48301584 | 0.16849579 | 2.86663444 | 0.00414862 | 0.01768987 | POSTN | 10631 |
| ENSG000002 | 21.4123506 | 0.844810569 | 0.29470466 | 2.86663454 | 0.00414862 | 0.01768987 | NA | NA |

| | | | | | | | | |
|---|---|---|---|---|---|---|---|---|
| ENSG000001 | 339.784028 | 0.408254248 | 0.14242417 | 2.86646742 | 0.00415081 | 0.01769548 | IFNGR2 | 3460 |
| ENSG000002 | 36.1699843 | -0.665531996 | 0.23217613 | -2.8664962 | 0.00415043 | 0.01769548 | LINC01091 | 285419 |
| ENSG000001 | 676.587918 | 0.439861233 | 0.15346322 | 2.86623218 | 0.00415389 | 0.01770678 | GDF11 | 10220 |
| ENSG000001 | 25.2804503 | -0.747055374 | 0.26064814 | -2.8661451 | 0.00415504 | 0.01770979 | KRT9 | 3857 |
| ENSG000002 | 17.7540169 | -0.855652324 | 0.29854605 | -2.8660648 | 0.00415609 | 0.01771242 | TBC1D29 | 26083 |
| ENSG000002 | 236.351936 | 0.495229527 | 0.17280769 | 2.86578402 | 0.00415978 | 0.01772627 | NYNRIN | 57523 |
| ENSG000002 | 22.7233837 | -0.80004756 | 0.2791928 | -2.8655738 | 0.00416254 | 0.01773619 | LINC01132 | 100506810 |
| ENSG000002 | 148.303217 | 0.484177242 | 0.16896708 | 2.86551234 | 0.00416335 | 0.01773776 | C12orf73 | 728568 |
| ENSG000002 | 13.0525066 | -0.921885266 | 0.32177035 | -2.8650411 | 0.00416955 | 0.01776231 | NA | NA |
| ENSG000002 | 21.4003882 | -0.80458145 | 0.28083951 | -2.8649155 | 0.00417121 | 0.01776749 | PCA3 | 50652 |
| ENSG000002 | 6.48883864 | -1.05477464 | 0.36819128 | -2.8647464 | 0.00417343 | 0.01777512 | NA | NA |
| ENSG000001 | 14057.7844 | 0.380257593 | 0.13274808 | 2.86450535 | 0.00417661 | 0.01778575 | YWHAE | 7531 |
| ENSG000001 | 24.4944427 | -0.786450004 | 0.27455145 | -2.8644905 | 0.00417681 | 0.01778575 | TMPRSS11A | 339967 |
| ENSG000001 | 32.7851 | -0.729773935 | 0.2547712 | -2.8644287 | 0.00417762 | 0.01778735 | EPPIN | 57119 |
| ENSG000001 | 73.3550452 | 0.598927189 | 0.20909883 | 2.86432585 | 0.00417898 | 0.01779126 | ENGASE | 64772 |
| ENSG000001 | 180.543994 | 0.456447566 | 0.15936732 | 2.8641228 | 0.00418166 | 0.0178008 | ZNF16 | 7564 |
| ENSG000002 | 9.58413785 | -1.00116893 | 0.34959542 | -2.8637931 | 0.00418601 | 0.01781747 | NA | NA |
| ENSG000001 | 17.2775658 | 0.854497097 | 0.29839707 | 2.86362425 | 0.00418825 | 0.0178251 | CCDC184 | 387856 |
| ENSG000001 | 110.553889 | 0.475804885 | 0.16617392 | 2.86329462 | 0.00419261 | 0.01784178 | AGMAT | 79814 |
| ENSG000002 | 4.46609924 | -1.082866292 | 0.37822076 | -2.8630535 | 0.0041958 | 0.01785349 | NA | NA |
| ENSG000002 | 12.3354994 | -0.926007572 | 0.32344733 | -2.8629315 | 0.00419741 | 0.01785849 | NA | NA |
| ENSG000001 | 19.5467164 | -0.83649601 | 0.29220501 | -2.8627025 | 0.00420045 | 0.01786785 | MMP3 | 4314 |
| ENSG000002 | 18.2320953 | -0.926448181 | 0.32362753 | -2.862699 | 0.00420049 | 0.01786785 | MTND6P19 | 106481788 |
| ENSG000001 | 5.92037905 | -1.068781172 | 0.37336844 | -2.8625375 | 0.00420263 | 0.01787508 | SRMS | 6725 |
| ENSG00000C | 611.786094 | 0.374762166 | 0.13092677 | 2.86237991 | 0.00420473 | 0.01788045 | UIMC1 | 51720 |
| ENSG000001 | 112.63431 | 0.455533537 | 0.15914895 | 2.8623095 | 0.00420566 | 0.01788045 | GLB1L2 | 89944 |
| ENSG000002 | 4.14886557 | -1.083612796 | 0.37857795 | -2.8623241 | 0.00420547 | 0.01788045 | NA | NA |
| ENSG000002 | 11.7314525 | -0.960502548 | 0.33556797 | -2.8623189 | 0.00420554 | 0.01788045 | NA | NA |
| ENSG000001 | 1172.81142 | 0.403393665 | 0.14094044 | 2.8621569 | 0.00420769 | 0.01788718 | UCK2 | 7371 |

| | | | | | | | | |
|---|---|---|---|---|---|---|---|---|
| ENSG000001 | 466.609094 | 0.410315478 | 0.14336188 | 2.86209601 | 0.00420849 | 0.01788875 | TSPAN14 | 81619 |
| ENSG000002 | 9.71933446 | -0.995384925 | 0.34779677 | -2.8619729 | 0.00421013 | 0.01789382 | NA | NA |
| ENSG000001 | 13.8144144 | -0.943783556 | 0.32977427 | -2.8619078 | 0.00421099 | 0.01789562 | TRIM50 | 135892 |
| ENSG000001 | 1154.38137 | 0.373739042 | 0.13059538 | 2.861809 | 0.00421231 | 0.01789932 | TRIML2 | 205860 |
| ENSG000001 | 31.9261771 | -0.776504008 | 0.27136588 | -2.8614652 | 0.00421688 | 0.01791687 | MS4A6A | 64231 |
| ENSG000002 | 10.037725 | -0.989856989 | 0.34593846 | -2.8613673 | 0.00421818 | 0.01792053 | NA | NA |
| ENSG000002 | 10.261168 | -0.972093639 | 0.33973756 | -2.8613075 | 0.00421898 | 0.01792203 | NA | NA |
| ENSG000001 | 59.5983218 | -0.583183302 | 0.20382152 | -2.861245 | 0.00421981 | 0.01792369 | MAPT | 4137 |
| ENSG000001 | 14071.7478 | 0.326724888 | 0.11419694 | 2.8610651 | 0.0042222 | 0.01793198 | TERF1 | 7013 |
| ENSG000002 | 2.56359591 | -1.050349072 | 0.36712566 | -2.861007 | 0.00422298 | 0.01793339 | NA | NA |
| ENSG000002 | 10.9113427 | -0.958147751 | 0.3349515 | -2.8605567 | 0.00422898 | 0.017957 | CHMP1B2P | 101060146 |
| ENSG000002 | 16.5192998 | -0.888807267 | 0.31071893 | -2.8604863 | 0.00422992 | 0.0179591 | ZNF503-AS1 | 253264 |
| ENSG000001 | 235.361616 | 0.535671267 | 0.18728008 | 2.86026827 | 0.00423283 | 0.01796958 | TERF2 | 7014 |
| ENSG000001 | 15.9943601 | -0.878060101 | 0.30699154 | -2.8602095 | 0.00423361 | 0.01797103 | TPRXL | 348825 |
| ENSG00000C | 245.584756 | 0.466160148 | 0.16298581 | 2.86012723 | 0.00423471 | 0.01797103 | PNPLA6 | 10908 |
| ENSG000001 | 27.1720304 | -0.789234092 | 0.27594538 | -2.8601098 | 0.00423494 | 0.01797103 | HMP19 | 51617 |
| ENSG000001 | 325.289912 | 0.506414156 | 0.17705865 | 2.86014927 | 0.00423442 | 0.01797103 | PNMA1 | 9240 |
| ENSG000001 | 320.796305 | 0.392843345 | 0.13735978 | 2.85995914 | 0.00423696 | 0.01797769 | ZNRF3 | 84133 |
| ENSG000001 | 1852.36522 | 0.469779967 | 0.16427518 | 2.85971362 | 0.00424024 | 0.01798785 | ARID1A | 8289 |
| ENSG000001 | 151.846361 | 0.452412093 | 0.15820076 | 2.85973396 | 0.00423997 | 0.01798785 | PANK4 | 55229 |
| ENSG00000C | 548.053229 | 0.491126115 | 0.1717451 | 2.85962229 | 0.00424146 | 0.01799115 | HIPK2 | 28996 |
| ENSG000001 | 35.8276186 | -0.655506937 | 0.22923316 | -2.8595642 | 0.00424224 | 0.01799136 | LGR6 | 59352 |
| ENSG000002 | 4.88795333 | -1.079006093 | 0.37733394 | -2.8595522 | 0.0042424 | 0.01799136 | NA | NA |
| ENSG000001 | 563.633069 | 0.373163281 | 0.13051442 | 2.8591728 | 0.00424747 | 0.01801101 | PIM1 | 5292 |
| ENSG000001 | 7.21400818 | -1.043046618 | 0.36482237 | -2.8590533 | 0.00424907 | 0.01801591 | LYZL1 | 84569 |
| ENSG000002 | 7.5835914 | -1.037339439 | 0.36284535 | -2.8589024 | 0.0042511 | 0.0180226 | PLGLA | 285189 |
| ENSG000001 | 1216.4323 | 0.357747057 | 0.12513617 | 2.85886214 | 0.00425164 | 0.018023 | ANGEL2 | 90806 |
| ENSG000001 | 185.553321 | 0.452185246 | 0.15817151 | 2.85882871 | 0.00425208 | 0.01802302 | TMEM128 | 85013 |
| ENSG000002 | 22.9772258 | -0.779711583 | 0.27274924 | -2.8587122 | 0.00425365 | 0.01802776 | NA | NA |

| | | | | | | | | |
|---|---|---|---|---|---|---|---|---|
| ENSG000001 | 496.661129 | 0.38051619 | 0.13311243 | 2.85860758 | 0.00425505 | 0.01803182 | NXF1 | 10482 |
| ENSG000001 | 8.40766723 | -1.009190513 | 0.35305392 | -2.85846 | 0.00425703 | 0.01803643 | LINC01551 | 387978 |
| ENSG000002 | 8.18647289 | -1.021945005 | 0.35751467 | -2.8584702 | 0.00425689 | 0.01803643 | NA | NA |
| ENSG000001 | 50.728234 | -0.597543032 | 0.20904752 | -2.8584077 | 0.00425773 | 0.01803752 | C12orf56 | 115749 |
| ENSG000001 | 5575.41741 | 0.684347028 | 0.23943187 | 2.85821194 | 0.00426036 | 0.01804677 | RPS16 | 6217 |
| ENSG000002 | 4.7404844 | -1.077969365 | 0.37716966 | -2.858049 | 0.00426255 | 0.01805227 | ABCC5-AS1 | 100873982 |
| ENSG000002 | 10.0821925 | -0.990460924 | 0.34655001 | -2.8580606 | 0.00426239 | 0.01805227 | LOC100507477 | 100507477 |
| ENSG000001 | 940.408316 | 0.377362589 | 0.1320507 | 2.85770978 | 0.0042671 | 0.01806969 | ABHD17B | 51104 |
| ENSG000001 | 948.701112 | 0.349783643 | 0.12240507 | 2.85759109 | 0.0042687 | 0.01807456 | ISCU | 23479 |
| ENSG000001 | 186.377052 | 0.479893071 | 0.16796548 | 2.85709339 | 0.0042754 | 0.01810104 | TBC1D17 | 79735 |
| ENSG00000C | 11.4478764 | -0.942736988 | 0.33000806 | -2.856709 | 0.00428058 | 0.01811831 | CYP26B1 | 56603 |
| ENSG000001 | 29.8032441 | -0.751883757 | 0.26320091 | -2.8566913 | 0.00428082 | 0.01811831 | TAT | 6898 |
| ENSG000002 | 11.5156101 | -0.956688088 | 0.33489303 | -2.8566975 | 0.00428074 | 0.01811831 | NA | NA |
| ENSG000001 | 2795.59312 | 0.327261234 | 0.11457433 | 2.85632257 | 0.00428579 | 0.01813747 | TMEM97 | 27346 |
| ENSG000001 | 361.362781 | 0.47697118 | 0.16700682 | 2.85599815 | 0.00429018 | 0.01815411 | ZNF142 | 7701 |
| ENSG000002 | 97.0226342 | 0.606179537 | 0.21225033 | 2.85596514 | 0.00429062 | 0.01815411 | NA | NA |
| ENSG000001 | 9.46365468 | 0.986079631 | 0.34534471 | 2.8553489 | 0.00429896 | 0.01818748 | MIR335 | 442904 |
| ENSG000001 | 17.2278214 | -0.892365396 | 0.3125416 | -2.8551892 | 0.00430112 | 0.01819473 | DNTT | 1791 |
| ENSG000001 | 3.78603235 | -1.079475157 | 0.378094 | -2.8550444 | 0.00430308 | 0.01820113 | NA | NA |
| ENSG000002 | 21.2548748 | -0.838046486 | 0.29354571 | -2.8549096 | 0.00430491 | 0.01820696 | LOC339874 | 339874 |
| ENSG000002 | 819.294666 | 0.341155657 | 0.11950532 | 2.8547319 | 0.00430732 | 0.01821525 | POLR2M | 81488 |
| ENSG000002 | 516.398642 | 0.347848479 | 0.12185403 | 2.85463256 | 0.00430866 | 0.01821905 | SMG1P1 | 641298 |
| ENSG000001 | 43.1091404 | -0.64911423 | 0.22740079 | -2.8544941 | 0.00431054 | 0.01822509 | UNC5C | 8633 |
| ENSG000001 | 52.841377 | -0.601128222 | 0.21062838 | -2.8539754 | 0.00431759 | 0.01825297 | ELAVL4 | 1996 |
| ENSG000001 | 430.37848 | 0.342647237 | 0.12006269 | 2.85390272 | 0.00431858 | 0.01825524 | POGLUT1 | 56983 |
| ENSG000002 | 3.70588572 | -1.079401259 | 0.37823266 | -2.8538023 | 0.00431994 | 0.01825911 | NA | NA |
| ENSG000001 | 1448.8431 | 0.390866109 | 0.13696555 | 2.85375489 | 0.00432059 | 0.01825993 | SPRY4 | 81848 |
| ENSG000001 | 586.173366 | 0.377787834 | 0.13238749 | 2.85365213 | 0.00432198 | 0.01826356 | RNF8 | 9025 |
| ENSG000002 | 629.47078 | 0.356439949 | 0.1249092 | 2.85359253 | 0.00432279 | 0.01826356 | CHUK | 1147 |

| | | | | | | | | |
|---|---|---|---|---|---|---|---|---|
| ENSG000002 | 13.4629896 | -0.902743778 | 0.31634983 | -2.853625 | 0.00432235 | 0.01826356 | NA | NA |
| ENSG000002 | 6.32196763 | -1.061746816 | 0.37213035 | -2.8531583 | 0.0043287 | 0.01828663 | MIR1183 | 100302122 |
| ENSG000001 | 337.271997 | 0.392359012 | 0.13752145 | 2.85307508 | 0.00432984 | 0.01828951 | ALG5 | 29880 |
| ENSG000001 | 1685.47161 | 0.390276145 | 0.13680039 | 2.85288765 | 0.00433239 | 0.01829649 | GJC1 | 10052 |
| ENSG000002 | 5.13597935 | -1.070942786 | 0.37538899 | -2.8528881 | 0.00433239 | 0.01829649 | NA | NA |
| ENSG000001 | 253.392303 | 0.373402963 | 0.13094548 | 2.8515911 | 0.0043501 | 0.01836912 | SPAG16 | 79582 |
| ENSG000002 | 7.14833165 | -1.038366766 | 0.36413962 | -2.8515621 | 0.0043505 | 0.01836912 | NA | NA |
| ENSG000001 | 497.216342 | 0.781845485 | 0.27421589 | 2.85120415 | 0.0043554 | 0.0183879 | SART1 | 9092 |
| ENSG000001 | 13.4192186 | -0.939504439 | 0.32951966 | -2.8511332 | 0.00435637 | 0.01838936 | NA | NA |
| ENSG000002 | 2.45620857 | -1.049943049 | 0.36825729 | -2.8511127 | 0.00435665 | 0.01838936 | NA | NA |
| ENSG000002 | 4.04830858 | -1.079389248 | 0.37859001 | -2.8510769 | 0.00435714 | 0.01838952 | NA | NA |
| ENSG000002 | 296.426806 | 0.441674044 | 0.15491986 | 2.85098398 | 0.00435842 | 0.01839298 | RNU4-2 | 26834 |
| ENSG000002 | 5.79706777 | -1.06149438 | 0.37235365 | -2.8507693 | 0.00436136 | 0.01840348 | LOC200772 | 200772 |
| ENSG00000C | 3630.07635 | 0.320627153 | 0.11247326 | 2.85069666 | 0.00436236 | 0.01840578 | ARFGEF1 | 10565 |
| ENSG000002 | 3.68900175 | -1.078175284 | 0.3782192 | -2.8506624 | 0.00436283 | 0.01840584 | NA | NA |
| ENSG000001 | 49.8454116 | -0.601455225 | 0.21099265 | -2.850598 | 0.00436371 | 0.01840766 | RFX1 | 5989 |
| ENSG000001 | 1188.97293 | 0.317796153 | 0.11149066 | 2.85042846 | 0.00436604 | 0.01841556 | AP3S1 | 1176 |
| ENSG000001 | 298.644812 | 0.406435707 | 0.1425921 | 2.85033819 | 0.00436728 | 0.01841887 | SPATS2 | 65244 |
| ENSG000002 | 22.1933244 | -0.771240726 | 0.27060785 | -2.8500308 | 0.0043715 | 0.01843476 | LINC00536 | 100859921 |
| ENSG000002 | 2.50548008 | -1.046351022 | 0.36717257 | -2.8497527 | 0.00437532 | 0.01844876 | RN7SKP177 | 106479174 |
| ENSG000002 | 13.3493907 | -0.943579184 | 0.33111258 | -2.8497232 | 0.00437573 | 0.01844876 | OSBPL10-AS1 | 100874206 |
| ENSG000002 | 2.47297542 | -1.048893163 | 0.36810472 | -2.8494423 | 0.00437959 | 0.01846314 | TPRG1-AS2 | 100874027 |
| ENSG000001 | 28.173777 | 0.730781688 | 0.25646948 | 2.84939048 | 0.00438031 | 0.01846423 | TMEM102 | 284114 |
| ENSG00000C | 3117.11793 | 0.391468749 | 0.13739576 | 2.84920537 | 0.00438286 | 0.01847306 | TNRC6A | 27327 |
| ENSG000001 | 306.234337 | 0.357568407 | 0.12551098 | 2.84890139 | 0.00438705 | 0.01848879 | VAV3 | 10451 |
| ENSG000002 | 4.44045929 | -1.077802237 | 0.37833559 | -2.8487995 | 0.00438845 | 0.01849279 | NA | NA |
| ENSG000001 | 350.459534 | 0.358483871 | 0.12585119 | 2.8484742 | 0.00439294 | 0.01850786 | GPR63 | 81491 |
| ENSG000002 | 7.58055943 | -1.031273997 | 0.36204215 | -2.8484915 | 0.0043927 | 0.01850786 | NA | NA |
| ENSG000002 | 14.2532922 | -0.880632251 | 0.30916381 | -2.8484325 | 0.00439352 | 0.01850836 | LINC00893 | 100131434 |

| | | | | | | | | |
|---|---|---|---|---|---|---|---|---|
| ENSG000001 | 146.772425 | 0.590948414 | 0.20747307 | 2.84831378 | 0.00439516 | 0.01851335 | CNNM3 | 26505 |
| ENSG000001 | 103.847926 | 0.5297287 | 0.18600224 | 2.84796949 | 0.00439991 | 0.01853146 | PLEKHA8P1 | 51054 |
| ENSG000001 | 62.8429467 | -0.560261603 | 0.19673044 | -2.8478643 | 0.00440137 | 0.01853566 | RAG1 | 5896 |
| ENSG000002 | 3.74470299 | -1.076823531 | 0.37813448 | -2.8477264 | 0.00440328 | 0.01854177 | NA | NA |
| ENSG000002 | 6.91223907 | -1.045461659 | 0.36715031 | -2.8475032 | 0.00440637 | 0.01855285 | LINC01266 | 101927215 |
| ENSG000001 | 155.244281 | 0.516961838 | 0.18156784 | 2.84721041 | 0.00441042 | 0.01856652 | DMWD | 1762 |
| ENSG000002 | 4.06512282 | -1.07792219 | 0.37858991 | -2.8472026 | 0.00441053 | 0.01856652 | NA | NA |
| ENSG000001 | 229.470657 | 0.381950306 | 0.13416237 | 2.84692577 | 0.00441437 | 0.01857882 | THAP6 | 152815 |
| ENSG000002 | 17.7083795 | 0.835629261 | 0.2935188 | 2.84693608 | 0.00441422 | 0.01857882 | SCARNA22 | 677770 |
| ENSG000001 | 44.4828732 | -0.618447863 | 0.21723807 | -2.8468669 | 0.00441518 | 0.01858033 | MAS1 | 4142 |
| ENSG000001 | 278.117872 | 0.373307627 | 0.13114254 | 2.84657925 | 0.00441917 | 0.01859472 | RRP8 | 23378 |
| ENSG000001 | 832.318162 | 0.377274977 | 0.13253743 | 2.84655424 | 0.00441952 | 0.01859472 | MYO5A | 4644 |
| ENSG000001 | 11.631258 | -0.970328476 | 0.34088891 | -2.8464653 | 0.00442075 | 0.01859799 | FAM71C | 196472 |
| ENSG00000C | 173.183107 | 0.464628434 | 0.16324106 | 2.84627184 | 0.00442344 | 0.01860736 | GAB2 | 9846 |
| ENSG000002 | 152.205475 | 0.436999183 | 0.15356025 | 2.84578317 | 0.00443024 | 0.01863401 | NA | NA |
| ENSG000001 | 120.714861 | 0.509054988 | 0.17890328 | 2.84542003 | 0.00443529 | 0.01865333 | EFS | 10278 |
| ENSG000002 | 7.42783106 | -1.02683724 | 0.36093102 | -2.8449681 | 0.00444159 | 0.01867788 | PRODH2 | 58510 |
| ENSG000001 | 937.586992 | 0.341764717 | 0.12013583 | 2.84481924 | 0.00444366 | 0.01868467 | NRIP1 | 8204 |
| ENSG000001 | 420.904765 | 0.451827832 | 0.15882902 | 2.8447436 | 0.00444472 | 0.01868717 | PPP3CB | 5532 |
| ENSG000002 | 9.00861487 | -0.994042919 | 0.34948607 | -2.8442991 | 0.00445092 | 0.01871133 | NA | NA |
| ENSG000002 | 11.3872787 | -0.959288053 | 0.3372821 | -2.8441712 | 0.00445271 | 0.0187169 | NA | NA |
| ENSG000002 | 11.6430983 | -0.928734616 | 0.32655617 | -2.8440272 | 0.00445472 | 0.01872342 | NA | NA |
| ENSG00000C | 79.5425664 | 0.551139791 | 0.1937979 | 2.84388944 | 0.00445665 | 0.01872957 | RWDD2A | 112611 |
| ENSG000001 | 151.049029 | 0.51039096 | 0.17948186 | 2.84369106 | 0.00445943 | 0.0187393 | TMEM180 | 79847 |
| ENSG000001 | 8209.10563 | 0.31151666 | 0.10954962 | 2.84361236 | 0.00446053 | 0.01874198 | MSH6 | 2956 |
| ENSG000001 | 943.830413 | 0.47268393 | 0.16626433 | 2.84296655 | 0.00446958 | 0.01877591 | ATXN10 | 25814 |
| ENSG000001 | 8.67876795 | -0.997775367 | 0.35096028 | -2.8429866 | 0.00446929 | 0.01877591 | OR4Q3 | 441669 |
| ENSG000002 | 66.4350686 | 0.620391268 | 0.21822197 | 2.84293683 | 0.00446999 | 0.01877591 | SNORD116-17 | 100033429 |
| ENSG000002 | 100.474518 | 0.487855184 | 0.17160754 | 2.84285409 | 0.00447115 | 0.01877884 | NA | NA |

| | | | | | | | | |
|---|---|---|---|---|---|---|---|---|
| ENSG000001 | 334.537381 | 0.514167551 | 0.18087049 | 2.84273875 | 0.00447277 | 0.0187837 | PHF13 | 148479 |
| ENSG000002 | 19.371089 | -0.836821403 | 0.29439143 | -2.8425467 | 0.00447547 | 0.01879307 | LINC01470 | 101927134 |
| ENSG000002 | 74.3872195 | 0.562755068 | 0.19798233 | 2.84245094 | 0.00447681 | 0.01879391 | NA | NA |
| ENSG000002 | 8.66862292 | -0.997553583 | 0.35095056 | -2.8424334 | 0.00447706 | 0.01879391 | LOC101928907 | 101928907 |
| ENSG000002 | 169.462513 | 0.455838339 | 0.16036822 | 2.84244804 | 0.00447685 | 0.01879391 | PKI55 | 150967 |
| ENSG000001 | 754.909969 | 0.34662874 | 0.12195769 | 2.84220495 | 0.00448027 | 0.01880544 | ZFP62 | 643836 |
| ENSG00000C | 819.838502 | 0.335666567 | 0.11810444 | 2.84211631 | 0.00448151 | 0.01880872 | ZFYVE16 | 9765 |
| ENSG000001 | 50.1982679 | -0.599388191 | 0.21089919 | -2.8420601 | 0.0044823 | 0.01881009 | GFAP | 2670 |
| ENSG000001 | 34.0701604 | -0.7093485 | 0.2495984 | -2.8419594 | 0.00448372 | 0.01881409 | COL8A1 | 1295 |
| ENSG000001 | 287.945563 | 0.474855505 | 0.16711702 | 2.84145505 | 0.00449082 | 0.01884192 | CAPN2 | 824 |
| ENSG00000C | 72.0613721 | -0.549794412 | 0.19349912 | -2.8413277 | 0.00449261 | 0.0188475 | ADAM28 | 10863 |
| ENSG000001 | 1983.08093 | 0.344656024 | 0.12130347 | 2.84127086 | 0.00449341 | 0.01884891 | TFDP2 | 7029 |
| ENSG00000C | 135.925933 | 0.45291288 | 0.15941248 | 2.84113824 | 0.00449528 | 0.01885374 | AGA | 175 |
| ENSG000002 | 14.3531256 | -0.906467101 | 0.31905237 | -2.8411232 | 0.00449549 | 0.01885374 | NA | NA |
| ENSG000002 | 18.1686341 | -0.850388047 | 0.29932211 | -2.8410465 | 0.00449658 | 0.01885632 | NA | NA |
| ENSG000002 | 11.9091307 | -0.944729965 | 0.33255509 | -2.8408224 | 0.00449974 | 0.01886762 | NA | NA |
| ENSG000001 | 92.5329282 | 0.576467508 | 0.20293033 | 2.84071639 | 0.00450123 | 0.01887195 | TSNARE1 | 203062 |
| ENSG000002 | 18.6214084 | -0.816811571 | 0.28756268 | -2.8404645 | 0.00450479 | 0.01888491 | LINC00616 | 641365 |
| ENSG000002 | 3.72827658 | -1.07408974 | 0.37815815 | -2.8403189 | 0.00450685 | 0.01889157 | NA | NA |
| ENSG00000C | 495.881236 | 0.348222984 | 0.1226207 | 2.83983858 | 0.00451364 | 0.01891418 | ZAK | 51776 |
| ENSG000001 | 55.970862 | -0.57493414 | 0.20245096 | -2.8398686 | 0.00451321 | 0.01891418 | GPR155 | 151556 |
| ENSG000001 | 210.084695 | 0.54866382 | 0.19320165 | 2.83985053 | 0.00451347 | 0.01891418 | GUSB | 2990 |
| ENSG000001 | 1336.02658 | 0.778138508 | 0.274069 | 2.83920659 | 0.00452259 | 0.01894972 | TNKS1BP1 | 85456 |
| ENSG000002 | 7.09001927 | -1.028974945 | 0.36245926 | -2.8388706 | 0.00452735 | 0.01896773 | NA | NA |
| ENSG000002 | 2.59014117 | -1.039768382 | 0.36628443 | -2.8386912 | 0.0045299 | 0.01897643 | NA | NA |
| ENSG000001 | 978.759043 | 0.334272823 | 0.1177611 | 2.83856732 | 0.00453166 | 0.018981 | BARD1 | 580 |
| ENSG000002 | 7.85527099 | -1.025680117 | 0.36133964 | -2.8385485 | 0.00453192 | 0.018981 | DLGAP2-AS1 | 100507435 |
| ENSG000002 | 4.75211359 | -1.070302368 | 0.37712473 | -2.8380594 | 0.00453887 | 0.01900815 | NA | NA |
| ENSG00000C | 512.506114 | 0.413898211 | 0.14584753 | 2.83788298 | 0.00454138 | 0.01901669 | ACAT1 | 38 |

| | | | | | | | | |
|---|---|---|---|---|---|---|---|---|
| ENSG000001 | 34.2183396 | -0.677711647 | 0.23883005 | -2.8376313 | 0.00454496 | 0.01902972 | IFI44L | 10964 |
| ENSG000001 | 559.603963 | 0.392952918 | 0.13849255 | 2.83735777 | 0.00454886 | 0.01904176 | ZNF484 | 83744 |
| ENSG000001 | 106.867104 | 0.524235047 | 0.18476394 | 2.83732342 | 0.00454935 | 0.01904176 | WDR81 | 124997 |
| ENSG000001 | 8.90863051 | -1.003751405 | 0.35376465 | -2.8373423 | 0.00454908 | 0.01904176 | OR14A16 | 284532 |
| ENSG000002 | 27.4449901 | -0.761757231 | 0.26847984 | -2.8372976 | 0.00454972 | 0.01904176 | LINC01498 | 102723562 |
| ENSG000001 | 497.716863 | 0.398610606 | 0.14049249 | 2.83723788 | 0.00455057 | 0.01904336 | WWC1 | 23286 |
| ENSG000001 | 8.0321509 | -1.010855237 | 0.356299 | -2.8370982 | 0.00455256 | 0.01904973 | CPSF4L | 642843 |
| ENSG000001 | 158.952925 | 0.460088688 | 0.16217067 | 2.83706473 | 0.00455304 | 0.01904976 | TOX3 | 27324 |
| ENSG000002 | 21.3802444 | -0.812683462 | 0.28648612 | -2.8367289 | 0.00455783 | 0.01906784 | ST7-AS2 | 93654 |
| ENSG000001 | 697.813265 | 0.420206974 | 0.14813411 | 2.83666578 | 0.00455873 | 0.01906964 | COL1A2 | 1278 |
| ENSG000002 | 10.3719684 | -0.974120474 | 0.34343052 | -2.8364412 | 0.00456194 | 0.01908109 | NA | NA |
| ENSG000001 | 724.45834 | 0.372851889 | 0.13145637 | 2.83631661 | 0.00456372 | 0.01908656 | ANTXR1 | 84168 |
| ENSG000002 | 85.9021377 | 0.503242131 | 0.1774372 | 2.83617044 | 0.00456581 | 0.01909333 | FAM127C | 441518 |
| ENSG000001 | 277.973464 | 0.402197708 | 0.14181968 | 2.83597951 | 0.00456854 | 0.01910081 | ZHX1 | 11244 |
| ENSG000001 | 496.649546 | 0.383497293 | 0.13522596 | 2.83597385 | 0.00456862 | 0.01910081 | PRKG1 | 5592 |
| ENSG000002 | 13.0973599 | -0.922392535 | 0.32525033 | -2.8359465 | 0.00456901 | 0.01910081 | NA | NA |
| ENSG000002 | 3.32144692 | -1.067087619 | 0.37628409 | -2.8358563 | 0.0045703 | 0.01910424 | NA | NA |
| ENSG000002 | 6.36929609 | -1.043323671 | 0.36791149 | -2.8358007 | 0.0045711 | 0.0191056 | FAM83A-AS1 | 100131726 |
| ENSG000001 | 133.626399 | 0.469205693 | 0.16546517 | 2.83567651 | 0.00457287 | 0.01910909 | ZNF668 | 79759 |
| ENSG000002 | 7.0299797 | -1.044379291 | 0.36829739 | -2.8356956 | 0.0045726 | 0.01910909 | NA | NA |
| ENSG000001 | 295.519175 | 0.412614513 | 0.14551038 | 2.83563628 | 0.00457345 | 0.01910953 | WRB | 7485 |
| ENSG000002 | 5.87393373 | -1.057023785 | 0.372795 | -2.8354022 | 0.0045768 | 0.01911762 | AZGP1P1 | 646282 |
| ENSG000002 | 119.16873 | -0.449044522 | 0.1583673 | -2.8354624 | 0.00457594 | 0.01911762 | NA | NA |
| ENSG000002 | 19.5289629 | -0.863504732 | 0.30454135 | -2.8354269 | 0.00457645 | 0.01911762 | DKFZp434J0226 | 93429 |
| ENSG000001 | 1147.00803 | 0.459001184 | 0.16191669 | 2.83479851 | 0.00458546 | 0.01915182 | SORT1 | 6272 |
| ENSG000001 | 12.2023572 | -0.935079547 | 0.32986378 | -2.8347445 | 0.00458623 | 0.01915308 | SAA2 | 6289 |
| ENSG000001 | 21.201383 | -0.829604159 | 0.29267237 | -2.8345831 | 0.00458855 | 0.01915684 | TPTE2 | 93492 |
| ENSG000002 | 39.5753205 | -0.686422063 | 0.24215897 | -2.8345928 | 0.00458841 | 0.01915684 | WEE2 | 494551 |
| ENSG000002 | 10.9424967 | -0.960588927 | 0.33887696 | -2.8346245 | 0.00458796 | 0.01915684 | LOC105374250 | 105374250 |

| | | | | | | | | | |
|---|---|---|---|---|---|---|---|---|---|
| ENSG000001 | 766.928829 | 0.396208729 | 0.13978445 | 2.83442628 | 0.0045908 | 0.01916427 | MGME1 | | 92667 |
| ENSG000002 | 7.71912529 | -1.014267342 | 0.35784668 | -2.8343629 | 0.00459172 | 0.0191661 | NA | NA | |
| ENSG000001 | 16.6365085 | -0.842770073 | 0.29736155 | -2.8341596 | 0.00459464 | 0.01917633 | SIX1 | | 6495 |
| ENSG000001 | 24.0326179 | -0.816994055 | 0.28827384 | -2.8340902 | 0.00459564 | 0.01917851 | SKAP1 | | 8631 |
| ENSG000001 | 78.3951417 | 0.547019516 | 0.19302513 | 2.83392908 | 0.00459795 | 0.01918621 | TMEM42 | | 131616 |
| ENSG000001 | 652.95208 | 0.384520527 | 0.13569535 | 2.8337045 | 0.00460119 | 0.01919772 | OSBPL11 | | 114885 |
| ENSG000001 | 414.638625 | 0.328013343 | 0.11575779 | 2.83361791 | 0.00460243 | 0.01920069 | ABCC4 | | 10257 |
| ENSG000001 | 110.487008 | -0.458719965 | 0.16188653 | -2.8335894 | 0.00460284 | 0.01920069 | ZNF836 | | 162962 |
| ENSG000002 | 8.88395292 | -0.984339406 | 0.34742039 | -2.8332805 | 0.00460729 | 0.01921648 | NA | NA | |
| ENSG000002 | 30.6464244 | -0.717413321 | 0.25321117 | -2.833261 | 0.00460758 | 0.01921648 | OR2AG1 | | 144125 |
| ENSG000002 | 8.33188106 | -1.001579115 | 0.3535224 | -2.8331419 | 0.00460929 | 0.01922166 | NA | NA | |
| ENSG000001 | 644.941393 | 0.364113338 | 0.12852232 | 2.83307473 | 0.00461026 | 0.01922372 | SV2A | | 9900 |
| ENSG000002 | 3.30129437 | -1.066453004 | 0.37644149 | -2.8329848 | 0.00461156 | 0.01922715 | NA | NA | |
| ENSG000001 | 36.8656076 | 0.662290117 | 0.23379666 | 2.83276128 | 0.00461478 | 0.01923862 | CYP27A1 | | 1593 |
| ENSG000001 | 53.8517717 | 0.588473957 | 0.20776403 | 2.83241499 | 0.00461978 | 0.01925749 | FRG1BP | | 284802 |
| ENSG000001 | 981.545446 | 0.360267274 | 0.12719793 | 2.83233604 | 0.00462093 | 0.01926027 | AMZ2 | | 51321 |
| ENSG000001 | 156.686174 | 0.465335959 | 0.16431861 | 2.83191262 | 0.00462705 | 0.01928381 | NFATC4 | | 4776 |
| ENSG000001 | 2319.20558 | 0.341571907 | 0.1206242 | 2.83170287 | 0.00463009 | 0.01929448 | SENP6 | | 26054 |
| ENSG000002 | 4.86337044 | -1.069234266 | 0.37760503 | -2.8316208 | 0.00463127 | 0.01929745 | NA | NA | |
| ENSG000001 | 1265.07273 | 0.308885659 | 0.10908884 | 2.83150568 | 0.00463294 | 0.01930241 | HMG20A | | 10363 |
| ENSG000001 | 13.6196779 | -0.908067448 | 0.32072342 | -2.8313101 | 0.00463578 | 0.01931224 | AIPL1 | | 23746 |
| ENSG000001 | 166.676313 | 0.406271095 | 0.1434973 | 2.83121073 | 0.00463722 | 0.01931625 | TMEM45A | | 55076 |
| ENSG000001 | 255.106782 | 0.464661974 | 0.16414744 | 2.83075973 | 0.00464376 | 0.01934152 | TIMM8A | | 1678 |
| ENSG000002 | 31.9263104 | 0.693810031 | 0.2451603 | 2.83002604 | 0.00465442 | 0.01938394 | SNORA71A | | 26777 |
| ENSG000001 | 70.9511169 | 0.565558831 | 0.19986356 | 2.82972461 | 0.00465881 | 0.01940022 | ZFAND2B | | 130617 |
| ENSG000001 | 361.816603 | 0.329291672 | 0.11637535 | 2.82956541 | 0.00466113 | 0.01940788 | CFAP36 | | 112942 |
| ENSG000002 | 6.38612645 | -1.055282627 | 0.37297081 | -2.8293973 | 0.00466358 | 0.01941609 | OR6C3 | | 254786 |
| ENSG000001 | 22.9311371 | -0.767459654 | 0.27127409 | -2.8290931 | 0.00466801 | 0.01943224 | BIN2 | | 51411 |
| ENSG000001 | 59.5442213 | 0.651764452 | 0.23038153 | 2.82906555 | 0.00466841 | 0.01943224 | C11orf68 | | 83638 |

| | | | | | | | | |
|---|---|---|---|---|---|---|---|---|
| ENSG00000C | 145.872314 | 0.543731411 | 0.19222748 | 2.82858314 | 0.00467546 | 0.01945955 | TOLLIP | 54472 |
| ENSG000002 | 14.7315852 | -0.880689364 | 0.3113722 | -2.8284136 | 0.00467793 | 0.01946786 | NA | NA |
| ENSG000001 | 13.7204107 | -0.907499183 | 0.32087839 | -2.8281717 | 0.00468147 | 0.01948058 | OR7D4 | 125958 |
| ENSG000001 | 525.640417 | 0.368439933 | 0.13027987 | 2.82806485 | 0.00468303 | 0.01948508 | FAM175B | 23172 |
| ENSG000002 | 10.3935311 | -0.94922372 | 0.33565487 | -2.8279754 | 0.00468434 | 0.01948853 | NA | NA |
| ENSG000001 | 520.727236 | 0.50000471 | 0.17681049 | 2.82791308 | 0.00468525 | 0.01949032 | RMND5B | 64777 |
| ENSG000001 | 527.483862 | 0.40118976 | 0.14186954 | 2.82787811 | 0.00468576 | 0.01949045 | RALGAPA2 | 57186 |
| ENSG000002 | 163.885324 | 0.445672766 | 0.15761104 | 2.82767471 | 0.00468874 | 0.01950084 | NA | NA |
| ENSG000001 | 2078.75339 | 0.329782741 | 0.11663164 | 2.82755815 | 0.00469045 | 0.01950489 | IWS1 | 55677 |
| ENSG000002 | 5.09254937 | -1.063050888 | 0.37596283 | -2.8275425 | 0.00469068 | 0.01950489 | NA | NA |
| ENSG000002 | 20.8659666 | -0.785615626 | 0.27786161 | -2.8273629 | 0.00469331 | 0.01951383 | NA | NA |
| ENSG000001 | 3690.86375 | 0.341754624 | 0.12090338 | 2.8266754 | 0.0047034 | 0.01955377 | ANKRD17 | 26057 |
| ENSG000001 | 246.366904 | 0.560582689 | 0.19833127 | 2.8264968 | 0.00470602 | 0.01956267 | DNTTIP1 | 116092 |
| ENSG000001 | 5.70785962 | -1.061835932 | 0.37571466 | -2.8261765 | 0.00471073 | 0.01958024 | MAGEB16 | 139604 |
| ENSG000001 | 99.8063435 | 0.560021202 | 0.19816537 | 2.82602967 | 0.00471289 | 0.01958721 | GPR137 | 56834 |
| ENSG000001 | 19.8311226 | -0.80286081 | 0.28410265 | -2.8259533 | 0.00471401 | 0.01958987 | PDILT | 204474 |
| ENSG000001 | 4.32737079 | -1.068250839 | 0.37802572 | -2.8258681 | 0.00471527 | 0.01959307 | PRAMEF2 | 65122 |
| ENSG000002 | 13.7261953 | -0.933184284 | 0.33025002 | -2.8256904 | 0.00471788 | 0.01960194 | NA | NA |
| ENSG000002 | 6.82747617 | -1.036146764 | 0.36669721 | -2.8256194 | 0.00471893 | 0.01960427 | NA | NA |
| ENSG000001 | 32.8699274 | -0.720468953 | 0.25501079 | -2.8252489 | 0.00472439 | 0.01962495 | CYP2E1 | 1571 |
| ENSG000001 | 135.714794 | 0.526555981 | 0.18638515 | 2.82509627 | 0.00472664 | 0.01963229 | TSHZ3 | 57616 |
| ENSG000002 | 9.67750883 | -0.978440294 | 0.34635965 | -2.8249258 | 0.00472916 | 0.01964073 | NA | NA |
| ENSG000001 | 7.32628945 | -1.015810063 | 0.35959888 | -2.8248421 | 0.00473039 | 0.01964384 | SIGLEC8 | 27181 |
| ENSG000001 | 2315.89595 | 0.386106711 | 0.13668464 | 2.82479947 | 0.00473102 | 0.01964445 | SLC30A1 | 7779 |
| ENSG000002 | 26.4712085 | -0.746081377 | 0.26412524 | -2.8247257 | 0.00473211 | 0.01964695 | ITGB3 | 3690 |
| ENSG000002 | 11.8893082 | -0.960692376 | 0.34011311 | -2.8246261 | 0.00473358 | 0.01964904 | LOC100507562 | 100507562 |
| ENSG000002 | 23.0528597 | -0.758426221 | 0.26850194 | -2.8246582 | 0.00473311 | 0.01964904 | NA | NA |
| ENSG000001 | 44.031765 | 0.660673376 | 0.23392423 | 2.824305 | 0.00473833 | 0.01966672 | EMILIN3 | 90187 |
| ENSG00000C | 41.7463928 | -0.620331949 | 0.21965636 | -2.8241019 | 0.00474133 | 0.01967249 | CATSPERG | 57828 |

| | | | | | | | | |
|---|---|---|---|---|---|---|---|---|
| ENSG000001 | 23.2412338 | -0.776938051 | 0.27510597 | -2.824141 | 0.00474075 | 0.01967249 | ITIH3 | 3699 |
| ENSG000001 | 381.800677 | 0.361523848 | 0.12801199 | 2.82414047 | 0.00474076 | 0.01967249 | ZNF114 | 163071 |
| ENSG000002 | 22.6989898 | -0.770653433 | 0.27288655 | -2.8240799 | 0.00474166 | 0.01967249 | NA | NA |
| ENSG000001 | 25.8004272 | 0.754806156 | 0.26728506 | 2.82397439 | 0.00474322 | 0.01967695 | AOC2 | 314 |
| ENSG000001 | 2351.5396 | 0.317997665 | 0.11261289 | 2.82381234 | 0.00474562 | 0.01968308 | ZNF33A | 7581 |
| ENSG000002 | 25.2366796 | 0.748146381 | 0.26494227 | 2.82380905 | 0.00474566 | 0.01968308 | NA | NA |
| ENSG000001 | 191.669877 | 0.557628392 | 0.19751745 | 2.82318543 | 0.00475491 | 0.01971939 | LOXL2 | 4017 |
| ENSG000001 | 108.657596 | 0.490582425 | 0.17377377 | 2.82310976 | 0.00475603 | 0.01972001 | PDGFRA | 5156 |
| ENSG000001 | 135.376849 | 0.604221713 | 0.2140252 | 2.82313356 | 0.00475568 | 0.01972001 | NAT8L | 339983 |
| ENSG000001 | 9.02851185 | -0.990382484 | 0.35082188 | -2.8230351 | 0.00475714 | 0.01972259 | SPIC | 121599 |
| ENSG000001 | 349.241688 | 0.400082329 | 0.14172648 | 2.8229186 | 0.00475887 | 0.01972774 | WDR27 | 253769 |
| ENSG000001 | 185.609428 | -0.416711866 | 0.147629 | -2.8226965 | 0.00476216 | 0.01973939 | IKBIP | 121457 |
| ENSG000001 | 635.793408 | 0.435536445 | 0.15429996 | 2.82266082 | 0.00476269 | 0.01973957 | ZBED4 | 9889 |
| ENSG000001 | 1790.93077 | 0.356939225 | 0.12646218 | 2.82249783 | 0.00476511 | 0.01974557 | ZNF141 | 7700 |
| ENSG000001 | 321.858937 | 0.40340756 | 0.14292561 | 2.82250013 | 0.00476508 | 0.01974557 | ZNF418 | 147686 |
| ENSG000002 | 12.2000477 | -0.931515852 | 0.33005036 | -2.8223446 | 0.00476739 | 0.01975148 | NA | NA |
| ENSG000002 | 9.33685124 | -0.983047559 | 0.34830985 | -2.8223364 | 0.00476751 | 0.01975148 | FGF10-AS1 | 101927075 |
| ENSG000001 | 88.6186198 | 0.546918435 | 0.19379612 | 2.82213311 | 0.00477054 | 0.01976198 | SSH3 | 54961 |
| ENSG000001 | 371.0557 | 0.39187515 | 0.13886401 | 2.82200659 | 0.00477242 | 0.01976776 | FAM184A | 79632 |
| ENSG000001 | 1177.63244 | 0.364368414 | 0.12912506 | 2.82182568 | 0.00477511 | 0.0197769 | EXOC6B | 23233 |
| ENSG000002 | 12.4163093 | -0.921859931 | 0.32673467 | -2.8214328 | 0.00478096 | 0.01979911 | NA | NA |
| ENSG000001 | 15.3391117 | -0.869412709 | 0.30815547 | -2.8213444 | 0.00478228 | 0.01980255 | MUC6 | 4588 |
| ENSG000001 | 260.295048 | 0.384225091 | 0.13619098 | 2.82122268 | 0.0047841 | 0.01980804 | TCFL5 | 10732 |
| ENSG000002 | 2.41763925 | -1.037199015 | 0.36764838 | -2.8211712 | 0.00478487 | 0.0198092 | NA | NA |
| ENSG000001 | 1572.3013 | 0.312836756 | 0.11089085 | 2.82112325 | 0.00478558 | 0.01981014 | SLMAP | 7871 |
| ENSG000002 | 3.67600628 | -1.066379611 | 0.37801424 | -2.8210038 | 0.00478736 | 0.01981549 | NA | NA |
| ENSG000001 | 23.2272895 | -0.843626601 | 0.29907112 | -2.8208227 | 0.00479007 | 0.01982466 | TMEM196 | 256130 |
| ENSG000002 | 3.27713645 | -1.061532591 | 0.37632755 | -2.8207677 | 0.00479089 | 0.01982604 | NA | NA |
| ENSG00000C | 144.099493 | 0.542725992 | 0.19242379 | 2.82047245 | 0.0047953 | 0.01983823 | CACNA2D2 | 9254 |

| | | | | | | | | |
|---|---|---|---|---|---|---|---|---|
| ENSG000001 | 11.7136288 | -0.930440762 | 0.32988699 | -2.8204833 | 0.00479514 | 0.01983823 | NA | NA |
| ENSG000001 | 4.70297563 | -1.067776115 | 0.37858941 | -2.8204067 | 0.00479628 | 0.01983823 | NKX6-2 | 84504 |
| ENSG000001 | 9462.76424 | 0.335509494 | 0.11895622 | 2.82044524 | 0.00479571 | 0.01983823 | ZNF121 | 7675 |
| ENSG000002 | 12.3163179 | -0.944296015 | 0.33480538 | -2.8204326 | 0.00479589 | 0.01983823 | NA | NA |
| ENSG000002 | 7.53638903 | -1.021716035 | 0.36227885 | -2.8202475 | 0.00479866 | 0.01984605 | LOC105373643 | 105373643 |
| ENSG000001 | 15.1705333 | -0.889409683 | 0.31541664 | -2.8197932 | 0.00480546 | 0.01987215 | PGLYRP4 | 57115 |
| ENSG000001 | 197.876271 | 0.418820728 | 0.14853081 | 2.8197565 | 0.00480601 | 0.01987239 | C2orf42 | 54980 |
| ENSG000001 | 543.155482 | 0.332659025 | 0.11798509 | 2.8195005 | 0.00480985 | 0.01988371 | RAB2B | 84932 |
| ENSG000001 | 1066.23368 | 0.348317354 | 0.12353977 | 2.8194756 | 0.00481022 | 0.01988371 | UBR1 | 197131 |
| ENSG000001 | 5962.26689 | 0.319875248 | 0.11345013 | 2.81952307 | 0.00480951 | 0.01988371 | NA | NA |
| ENSG000001 | 388.661108 | 0.424435202 | 0.15054988 | 2.81923302 | 0.00481386 | 0.01989469 | ADAMTS1 | 9510 |
| ENSG000002 | 11.1250359 | -0.941785383 | 0.33405349 | -2.8192652 | 0.00481337 | 0.01989469 | LINC00640 | 283553 |
| ENSG00000C | 304.047232 | 0.377120771 | 0.13377032 | 2.81916618 | 0.00481486 | 0.0198968 | TGDS | 23483 |
| ENSG000001 | 23.5382734 | -0.770562613 | 0.27334273 | -2.8190346 | 0.00481683 | 0.01990293 | NA | NA |
| ENSG000001 | 2414.46399 | 0.34199211 | 0.12132288 | 2.81885925 | 0.00481946 | 0.01991178 | CLINT1 | 9685 |
| ENSG000002 | 8.08748957 | -1.005364576 | 0.35667993 | -2.8186743 | 0.00482224 | 0.01992122 | NA | NA |
| ENSG000002 | 1426.09412 | 0.315461291 | 0.11192572 | 2.81848789 | 0.00482504 | 0.01993077 | SNX2 | 6643 |
| ENSG000002 | 3.95752 | -1.067010842 | 0.37858016 | -2.8184542 | 0.00482555 | 0.01993083 | LINC01372 | 101929736 |
| ENSG000001 | 6.46366211 | -1.040739282 | 0.36928945 | -2.818221 | 0.00482906 | 0.01994328 | DHH | 50846 |
| ENSG000001 | 128.580049 | 0.500819877 | 0.17772887 | 2.81788705 | 0.00483408 | 0.019962 | PABPC1L | 80336 |
| ENSG000002 | 17.8588296 | -0.845840232 | 0.30018123 | -2.8177652 | 0.00483592 | 0.01996754 | NA | NA |
| ENSG000001 | 1875.50973 | 0.311321464 | 0.1104974 | 2.81745523 | 0.00484059 | 0.01998479 | WNK3 | 65267 |
| ENSG000001 | 31.4212741 | -0.679835935 | 0.24130436 | -2.8173379 | 0.00484235 | 0.01999006 | ZAN | 7455 |
| ENSG000001 | 5633.76259 | 0.484506741 | 0.17198504 | 2.81714461 | 0.00484527 | 0.01999906 | PTGES3 | 10728 |
| ENSG000001 | 43.8849012 | -0.640627031 | 0.2274043 | -2.817128 | 0.00484552 | 0.01999906 | MYO1H | 283446 |
| ENSG000001 | 2950.89961 | 0.362529776 | 0.12869108 | 2.81705443 | 0.00484663 | 0.0200016 | ZMYND8 | 23613 |
| ENSG000001 | 139.647525 | 0.424077954 | 0.15054224 | 2.81700316 | 0.0048474 | 0.02000276 | EVC2 | 132884 |
| ENSG000002 | 11.9245367 | -0.923221101 | 0.32777053 | -2.816669 | 0.00485245 | 0.02002154 | LOC101929754 | 101929754 |
| ENSG000001 | 421.745533 | 0.521710215 | 0.18523101 | 2.81653826 | 0.00485443 | 0.02002766 | PPTC7 | 160760 |

| | | | | | | | | |
|---|---|---|---|---|---|---|---|---|
| ENSG000001 | 647.186077 | 0.453589677 | 0.16106429 | 2.81620263 | 0.0048595 | 0.02004655 | NOM1 | 64434 |
| ENSG000001 | 287.662737 | 0.367499775 | 0.13050245 | 2.81603745 | 0.004862 | 0.02005278 | TMEM254 | 80195 |
| ENSG000002 | 4.6843175 | -1.061852819 | 0.37707184 | -2.8160491 | 0.00486182 | 0.02005278 | OR2W3 | 343171 |
| ENSG000002 | 4.37356871 | -1.065556531 | 0.37840664 | -2.8159034 | 0.00486403 | 0.02005911 | NA | NA |
| ENSG000001 | 561.648799 | 0.396542409 | 0.14082593 | 2.81583381 | 0.00486508 | 0.02006142 | MKKS | 8195 |
| ENSG000001 | 3.47729354 | -1.056170781 | 0.37508795 | -2.815795 | 0.00486567 | 0.0200618 | GPHB5 | 122876 |
| ENSG000001 | 119.533936 | 0.461630049 | 0.1639461 | 2.81574277 | 0.00486646 | 0.02006189 | ECE2 | 9718 |
| ENSG000002 | 10.3205258 | -0.961831295 | 0.34159239 | -2.8157281 | 0.00486668 | 0.02006189 | NA | NA |
| ENSG000002 | 178.676561 | 0.430945249 | 0.15307501 | 2.81525537 | 0.00487385 | 0.02008939 | LOC101927027 | 101927027 |
| ENSG000002 | 5.66721298 | -1.04540673 | 0.37135705 | -2.8150987 | 0.00487623 | 0.02009715 | NA | NA |
| ENSG000001 | 255.438973 | 0.371548292 | 0.13198869 | 2.81500104 | 0.00487771 | 0.02010121 | C11orf74 | 119710 |
| ENSG000001 | 7.36254949 | -1.033405365 | 0.36714152 | -2.814733 | 0.00488178 | 0.02011594 | TRIM42 | 287015 |
| ENSG000001 | 27.7345137 | -0.754734078 | 0.26814711 | -2.8146269 | 0.00488339 | 0.02011849 | LPA | 4018 |
| ENSG000002 | 2.81340378 | -1.052038558 | 0.37377502 | -2.8146304 | 0.00488334 | 0.02011849 | NA | NA |
| ENSG000001 | 140.291435 | 0.459984778 | 0.16343511 | 2.81447954 | 0.00488563 | 0.02012567 | LDLRAP1 | 26119 |
| ENSG00000C | 2537.45228 | 0.431546279 | 0.15333518 | 2.81439831 | 0.00488686 | 0.02012872 | BOD1L1 | 259282 |
| ENSG000002 | 6.32726946 | -1.049013134 | 0.37274018 | -2.814328 | 0.00488793 | 0.02013107 | LINC00671 | 388387 |
| ENSG000002 | 14.5422911 | -0.86664889 | 0.30795723 | -2.8141858 | 0.00489009 | 0.02013793 | NA | NA |
| ENSG000001 | 94.8008399 | 0.491980615 | 0.1748264 | 2.81410933 | 0.00489126 | 0.02014068 | SYNGR3 | 9143 |
| ENSG000001 | 1771.64979 | 0.377204146 | 0.13404488 | 2.81401387 | 0.00489271 | 0.02014462 | EIF3J | 8669 |
| ENSG000001 | 138.468426 | 0.43455003 | 0.15443171 | 2.81386534 | 0.00489497 | 0.02015188 | ZNF558 | 148156 |
| ENSG000001 | 19.5281542 | -0.802432691 | 0.28517608 | -2.8138148 | 0.00489574 | 0.020153 | FGF9 | 2254 |
| ENSG000001 | 109.81935 | 0.527060692 | 0.18734761 | 2.8132769 | 0.00490394 | 0.0201847 | BAG3 | 9531 |
| ENSG00000C | 7.72686768 | -1.043724682 | 0.37101699 | -2.8131452 | 0.00490595 | 0.02018993 | DLX6 | 1750 |
| ENSG000001 | 207.067702 | 0.380075895 | 0.13510791 | 2.81312835 | 0.00490621 | 0.02018993 | TRMT2B | 79979 |
| ENSG000001 | 558.549579 | 0.332867822 | 0.11833064 | 2.81303153 | 0.00490768 | 0.02019396 | COX17 | 10063 |
| ENSG000001 | 121.538295 | 0.522821868 | 0.18587337 | 2.81278518 | 0.00491145 | 0.02020534 | HSCB | 150274 |
| ENSG000002 | 8.14070256 | -1.009051315 | 0.35873397 | -2.8128122 | 0.00491103 | 0.02020534 | NA | NA |
| ENSG000001 | 10.641178 | -0.954750644 | 0.33945381 | -2.8126084 | 0.00491415 | 0.02021439 | FLJ34503 | 285759 |

| | | | | | | | | |
|---|---|---|---|---|---|---|---|---|
| ENSG000001 | 32.5875246 | -0.67640636 | 0.24052279 | -2.8122339 | 0.00491987 | 0.02023589 | MSRB3 | 253827 |
| ENSG000001 | 395.070035 | 0.366283146 | 0.13024837 | 2.81218987 | 0.00492054 | 0.02023661 | TMEM65 | 157378 |
| ENSG000001 | 43.795495 | -0.697304849 | 0.24796334 | -2.8121288 | 0.00492148 | 0.02023841 | TRPM1 | 4308 |
| ENSG000001 | 10.3922011 | -0.944956379 | 0.33603368 | -2.8120883 | 0.0049221 | 0.0202389 | PP2D1 | 151649 |
| ENSG000001 | 360.053408 | 0.416371322 | 0.14808885 | 2.81163175 | 0.00492909 | 0.02026559 | EEF1E1 | 9521 |
| ENSG000002 | 3.19101136 | -1.059479343 | 0.37690467 | -2.8110008 | 0.00493877 | 0.02030332 | RNU6-1323P | 106481598 |
| ENSG000001 | 17.4583822 | -0.8316345 | 0.2958536 | -2.8109663 | 0.0049393 | 0.02030344 | RHOJ | 57381 |
| ENSG000002 | 4.77729865 | -1.061554059 | 0.37765336 | -2.8109218 | 0.00493998 | 0.02030418 | NA | NA |
| ENSG000002 | 2.82268652 | -1.049546803 | 0.37339168 | -2.8108468 | 0.00494113 | 0.02030686 | NA | NA |
| ENSG000001 | 166.712742 | 0.527472057 | 0.18766684 | 2.81068328 | 0.00494364 | 0.02031512 | SLC12A9 | 56996 |
| ENSG000002 | 10.5592302 | -0.953876024 | 0.33941221 | -2.8103763 | 0.00494836 | 0.02033245 | NA | NA |
| ENSG000002 | 4.31018736 | -1.062521053 | 0.37811161 | -2.8100726 | 0.00495303 | 0.02034959 | LOC100506530 | 100506530 |
| ENSG000002 | 8.62592146 | -1.007464231 | 0.35852352 | -2.8100367 | 0.00495359 | 0.0203498 | LINC01118 | 388948 |
| ENSG000002 | 23.2141103 | -0.764098821 | 0.27194857 | -2.8097182 | 0.00495849 | 0.02036788 | KCNMA1-AS1 | 101929328 |
| ENSG000001 | 76.2785573 | 0.548669066 | 0.19529786 | 2.80939613 | 0.00496345 | 0.0203862 | LRRC75A | 388341 |
| ENSG000002 | 4.40484622 | -1.062921254 | 0.37836167 | -2.8092731 | 0.00496535 | 0.0203878 | NA | NA |
| ENSG000002 | 3.99346066 | -1.063592618 | 0.37859259 | -2.8093329 | 0.00496443 | 0.0203878 | NA | NA |
| ENSG000002 | 9.40465376 | -0.959558459 | 0.34156473 | -2.8093019 | 0.00496491 | 0.0203878 | CFAP44-AS1 | 100874029 |
| ENSG000002 | 31.0103718 | -0.712645612 | 0.25374478 | -2.8085134 | 0.00497708 | 0.0204339 | ARHGEF26-AS1 | 100507524 |
| ENSG000001 | 160.896684 | 0.491869622 | 0.1751388 | 2.80845607 | 0.00497797 | 0.02043536 | ATG2A | 23130 |
| ENSG000002 | 3.17450511 | -1.058206046 | 0.37679696 | -2.8084251 | 0.00497845 | 0.02043536 | NA | NA |
| ENSG000002 | 15.3124871 | -0.891018791 | 0.31729236 | -2.8081949 | 0.00498201 | 0.02044791 | LINC00837 | 100507605 |
| ENSG000001 | 51.1113028 | -0.615767227 | 0.21928549 | -2.8080619 | 0.00498407 | 0.02045429 | PDE8B | 8622 |
| ENSG00000C | 253.435774 | 0.358361042 | 0.12765611 | 2.80723775 | 0.00499683 | 0.02050254 | CDC14A | 8556 |
| ENSG000001 | 325.137706 | 0.414196213 | 0.14754557 | 2.80724258 | 0.00499676 | 0.02050254 | TRIM41 | 90933 |
| ENSG000002 | 10.6417202 | -0.935774443 | 0.33335068 | -2.8071772 | 0.00499777 | 0.02050432 | NA | NA |
| ENSG000002 | 4.48149669 | -1.06210955 | 0.37838628 | -2.8069452 | 0.00500137 | 0.02051702 | MIR583 | 693168 |
| ENSG000001 | 747.00227 | 0.321094737 | 0.11440695 | 2.80660167 | 0.00500671 | 0.02053683 | PELI2 | 57161 |
| ENSG000001 | 31.0160682 | 0.70822373 | 0.25235362 | 2.80647345 | 0.0050087 | 0.02054085 | NA | NA |

| | | | | | | | | |
|---|---|---|---|---|---|---|---|---|
| ENSG000001 | 9.77543177 | -0.955565122 | 0.34048347 | -2.8064949 | 0.00500837 | 0.02054085 | FAM180B | 399888 |
| ENSG000001 | 247.888804 | 0.528255443 | 0.18823617 | 2.80634398 | 0.00501072 | 0.02054495 | MEN1 | 4221 |
| ENSG000002 | 208.744524 | 0.381665625 | 0.13600043 | 2.80635604 | 0.00501053 | 0.02054495 | NA | NA |
| ENSG000001 | 2599.46488 | 0.747075972 | 0.26621319 | 2.80630715 | 0.00501129 | 0.02054522 | HNRNPA3 | 220988 |
| ENSG000001 | 47.4859234 | 0.640335337 | 0.2281858 | 2.80620147 | 0.00501293 | 0.02054989 | NA | NA |
| ENSG000002 | 10.7027274 | -0.941929678 | 0.33568155 | -2.8060216 | 0.00501573 | 0.02055928 | LOC100996447 | 100996447 |
| ENSG000001 | 224.963307 | 0.4309905 | 0.1536 | 2.80592767 | 0.0050172 | 0.0205632 | RPP40 | 10799 |
| ENSG000002 | 7.70426286 | -1.006844897 | 0.35883711 | -2.805855 | 0.00501833 | 0.02056576 | NA | NA |
| ENSG000001 | 449.891936 | 0.349802435 | 0.12468914 | 2.80539619 | 0.00502548 | 0.02059298 | PDP2 | 57546 |
| ENSG000002 | 4.46866375 | -1.061639139 | 0.37844793 | -2.8052449 | 0.00502784 | 0.0205985 | NA | NA |
| ENSG000002 | 5.82078152 | -1.048308176 | 0.37369581 | -2.8052447 | 0.00502784 | 0.0205985 | NA | NA |
| ENSG000002 | 7.39659486 | -1.015110091 | 0.36186634 | -2.8052073 | 0.00502842 | 0.02059881 | NA | NA |
| ENSG000001 | 15.4794015 | -0.857326988 | 0.30564088 | -2.8050141 | 0.00503144 | 0.02060699 | PIGR | 5284 |
| ENSG000002 | 4.9233478 | -1.060685872 | 0.37813778 | -2.8050248 | 0.00503127 | 0.02060699 | DPRX | 503834 |
| ENSG000002 | 4.83490538 | -1.059536856 | 0.3777876 | -2.8045835 | 0.00503816 | 0.02063246 | LOC101927358 | 101927358 |
| ENSG00000C | 1491.4342 | 0.32663451 | 0.11647179 | 2.80440882 | 0.00504089 | 0.02064155 | DLG3 | 1741 |
| ENSG000002 | 10.9511508 | -0.927582761 | 0.33076941 | -2.8043184 | 0.00504231 | 0.02064526 | NA | NA |
| ENSG000002 | 10.149674 | -0.952061218 | 0.33950753 | -2.8042418 | 0.00504351 | 0.02064808 | LOC440700 | 440700 |
| ENSG000001 | 14.2662764 | -0.874360213 | 0.31184904 | -2.8037932 | 0.00505053 | 0.02067474 | GCNT7 | 140687 |
| ENSG000001 | 90.5761211 | 0.495910287 | 0.17687517 | 2.80373033 | 0.00505151 | 0.02067669 | ARSG | 22901 |
| ENSG000002 | 5.68286993 | -1.043069961 | 0.3720408 | -2.803644 | 0.00505287 | 0.02068013 | HOXB-AS1 | 100874362 |
| ENSG000001 | 71.6125889 | 0.539804586 | 0.19254351 | 2.80354602 | 0.0050544 | 0.02068433 | FAM109A | 144717 |
| ENSG000001 | 911.608004 | 0.391522173 | 0.13967069 | 2.80318059 | 0.00506013 | 0.0207057 | GABPA | 2551 |
| ENSG000002 | 3.92325019 | -1.061109089 | 0.37856489 | -2.8029781 | 0.00506331 | 0.02071661 | NA | NA |
| ENSG000002 | 19.2174432 | -0.829259278 | 0.29585989 | -2.8028783 | 0.00506488 | 0.02072093 | NA | NA |
| ENSG000002 | 6.09010976 | -1.038351603 | 0.37047697 | -2.8027426 | 0.00506701 | 0.02072547 | NA | NA |
| ENSG000002 | 344.190227 | 0.372363177 | 0.13285634 | 2.80275054 | 0.00506688 | 0.02072547 | YAE1D1 | 57002 |
| ENSG000002 | 38.1423165 | 0.649295837 | 0.23167958 | 2.80255957 | 0.00506989 | 0.02073514 | LOC100129215 | 100129215 |
| ENSG000002 | 19.9708975 | -0.826121237 | 0.29478245 | -2.8024777 | 0.00507117 | 0.02073831 | LOC100506457 | 100506457 |

| | | | | | | | | |
|---|---|---|---|---|---|---|---|---|
| ENSG000002 | 12.0150032 | -0.917728634 | 0.32748661 | -2.8023394 | 0.00507335 | 0.02074511 | NA | NA |
| ENSG000001 | 2790.52255 | 0.399961412 | 0.14272931 | 2.80223744 | 0.00507495 | 0.02074958 | MYSM1 | 114803 |
| ENSG000002 | 4.50488927 | -1.06065573 | 0.37851678 | -2.8021366 | 0.00507654 | 0.02075397 | NA | NA |
| ENSG000002 | 37.7115058 | -0.650628094 | 0.23221933 | -2.8017826 | 0.00508211 | 0.02077466 | POT1-AS1 | 401398 |
| ENSG000001 | 212.873074 | 0.397195051 | 0.14177582 | 2.80157116 | 0.00508544 | 0.02078619 | MED7 | 9443 |
| ENSG000001 | 14.8742184 | -0.874826255 | 0.31227337 | -2.8014757 | 0.00508695 | 0.02079024 | COLEC11 | 78989 |
| ENSG00000C | 287.825242 | 0.423728906 | 0.15125456 | 2.80142895 | 0.00508768 | 0.02079116 | DLGAP4 | 22839 |
| ENSG000001 | 475.993956 | 0.37408269 | 0.13353615 | 2.80135901 | 0.00508879 | 0.02079357 | CETN3 | 1070 |
| ENSG000001 | 38.3384147 | 0.641567797 | 0.22902756 | 2.80126893 | 0.00509021 | 0.02079728 | SH2B2 | 10603 |
| ENSG000001 | 1217.16075 | 0.351378663 | 0.12543793 | 2.8012154 | 0.00509105 | 0.02079838 | RWDD1 | 51389 |
| ENSG000001 | 302.387155 | 0.402454256 | 0.14367276 | 2.80118694 | 0.0050915 | 0.02079838 | ZNRF2 | 223082 |
| ENSG000001 | 40.5905498 | 0.626570699 | 0.22370402 | 2.80089159 | 0.00509616 | 0.02081532 | KCND1 | 3750 |
| ENSG000002 | 4.28735985 | -1.058855412 | 0.3780697 | -2.8006883 | 0.00509937 | 0.02082634 | NA | NA |
| ENSG000002 | 10.7083827 | -0.937556769 | 0.33477982 | -2.8005176 | 0.00510207 | 0.02083526 | NA | NA |
| ENSG000001 | 898.514213 | 0.38982832 | 0.1392123 | 2.80024338 | 0.00510641 | 0.02085087 | URB2 | 9816 |
| ENSG000001 | 11.4671322 | -0.936367205 | 0.33439455 | -2.8001868 | 0.0051073 | 0.02085242 | OR56A4 | 120793 |
| ENSG00000C | 853.140453 | 0.363262934 | 0.12973185 | 2.80010602 | 0.00510858 | 0.02085555 | BTBD7 | 55727 |
| ENSG000001 | 4108.92864 | 0.335598046 | 0.11985464 | 2.80004222 | 0.00510959 | 0.02085757 | PTBP3 | 9991 |
| ENSG000001 | 150.246327 | 0.546681607 | 0.19525359 | 2.79985431 | 0.00511257 | 0.02086551 | FBRSL1 | 57666 |
| ENSG000002 | 17.9586276 | -0.880687773 | 0.31454727 | -2.7998583 | 0.0051125 | 0.02086551 | ZCWPW2 | 152098 |
| ENSG000001 | 22.7455484 | -0.774621341 | 0.27670124 | -2.7994864 | 0.0051184 | 0.0208872 | GOLGA6L2 | 283685 |
| ENSG000001 | 9.68389487 | -0.968444921 | 0.34594767 | -2.7993972 | 0.00511981 | 0.02089087 | KIAA1024L | 100127206 |
| ENSG000001 | 4.55489222 | -1.059734698 | 0.37858599 | -2.7991915 | 0.00512307 | 0.02090208 | C1QTNF8 | 390664 |
| ENSG000002 | 1286.54937 | -0.421779309 | 0.15068786 | -2.7990265 | 0.00512569 | 0.02091066 | NA | NA |
| ENSG00000C | 147.306819 | 0.477979336 | 0.17077354 | 2.79890749 | 0.00512758 | 0.02091456 | DTX2 | 113878 |
| ENSG000001 | 21.8801584 | -0.840051193 | 0.30013606 | -2.7989013 | 0.00512768 | 0.02091456 | CBFA2T3 | 863 |
| ENSG00000C | 2231.33909 | 0.373344887 | 0.13339693 | 2.79875165 | 0.00513006 | 0.02092005 | SEMA3A | 10371 |
| ENSG000002 | 6.6583777 | -1.02462354 | 0.36609834 | -2.7987659 | 0.00512983 | 0.02092005 | LOC101929117 | 101929117 |
| ENSG00000C | 656.483953 | 0.315373862 | 0.11269149 | 2.79855979 | 0.00513311 | 0.02093038 | ANAPC4 | 29945 |

| | | | | | | | | | |
|---|---|---|---|---|---|---|---|---|---|
| ENSG000002 | 164.687412 | 0.527268888 | 0.18841666 | 2.7984196 | 0.00513534 | 0.02093736 | RING1 | | 6015 |
| ENSG000002 | 7.29890238 | -1.006924176 | 0.35987434 | -2.7979882 | 0.0051422 | 0.02096324 | LILRA4 | | 23547 |
| ENSG000002 | 3.90471368 | 1.059144966 | 0.37857854 | 2.79768889 | 0.00514697 | 0.02098056 | NA | NA | |
| ENSG000000 | 2396.72888 | 0.434194025 | 0.15520695 | 2.79751661 | 0.00514971 | 0.02098814 | DSG2 | | 1829 |
| ENSG000002 | 50.9699075 | 0.64547528 | 0.23073229 | 2.79750735 | 0.00514986 | 0.02098814 | LINC00545 | | 440132 |
| ENSG000001 | 265.270734 | 0.421363747 | 0.15063009 | 2.79734106 | 0.00515251 | 0.02099683 | FZD4 | | 8322 |
| ENSG000002 | 14.2553662 | -0.887918645 | 0.31745682 | -2.7969746 | 0.00515836 | 0.02101855 | NA | NA | |
| ENSG000000 | 58.1623571 | -0.600573213 | 0.21473954 | -2.7967519 | 0.00516191 | 0.02103093 | SP100 | | 6672 |
| ENSG000002 | 16.4938204 | -0.832996939 | 0.29789874 | -2.7962419 | 0.00517007 | 0.02106203 | LOC100505635 | | 100505635 |
| ENSG000000 | 1916.83034 | 0.379908135 | 0.13586646 | 2.79618779 | 0.00517093 | 0.02106344 | DCUN1D1 | | 54165 |
| ENSG000001 | 8734.25903 | 0.426102383 | 0.15239526 | 2.79603436 | 0.00517339 | 0.02106772 | NASP | | 4678 |
| ENSG000001 | 64.0641857 | 0.587659915 | 0.21017583 | 2.79603942 | 0.00517331 | 0.02106772 | TOR2A | | 27433 |
| ENSG000002 | 13.7213172 | -0.880617759 | 0.31495349 | -2.7960247 | 0.00517354 | 0.02106772 | GRID1-AS1 | | 100507470 |
| ENSG000002 | 5.44785813 | -1.047992713 | 0.37482683 | -2.7959384 | 0.00517493 | 0.02107124 | NA | NA | |
| ENSG000002 | 4.9568712 | -1.057388117 | 0.3782095 | -2.7957736 | 0.00517757 | 0.02107987 | NA | NA | |
| ENSG000000 | 84.864704 | -0.485846513 | 0.17378154 | -2.7957314 | 0.00517824 | 0.0210805 | COL4A4 | | 1286 |
| ENSG000002 | 10.1021965 | -0.946759784 | 0.33868993 | -2.7953585 | 0.00518422 | 0.02110272 | NA | NA | |
| ENSG000002 | 18.6225247 | 0.833981292 | 0.29835836 | 2.79523356 | 0.00518622 | 0.02110876 | SLC35E2 | | 9906 |
| ENSG000001 | 76.6413814 | -0.571332185 | 0.20439996 | -2.7951678 | 0.00518728 | 0.02111093 | EFCAB13 | | 124989 |
| ENSG000002 | 82.0405616 | 0.609843876 | 0.21818517 | 2.79507486 | 0.00518877 | 0.02111488 | GATSL2 | | 729438 |
| ENSG000002 | 11.8198685 | -0.906411115 | 0.3242957 | -2.7950143 | 0.00518974 | 0.02111672 | LOC729867 | | 729867 |
| ENSG000001 | 624.868264 | 0.438882965 | 0.15704102 | 2.79470275 | 0.00519475 | 0.02113284 | SALL1 | | 6299 |
| ENSG000001 | 5.34283909 | -1.044656008 | 0.37379532 | -2.7947273 | 0.00519435 | 0.02113284 | SPATA31A6 | | 389730 |
| ENSG000002 | 7.64700212 | -1.004011227 | 0.35926494 | -2.7946263 | 0.00519597 | 0.02113572 | NA | NA | |
| ENSG000002 | 7.52964747 | -0.996659843 | 0.35663904 | -2.7945899 | 0.00519656 | 0.02113598 | LINC00656 | | 200261 |
| ENSG000001 | 332.810416 | 0.392205074 | 0.14036714 | 2.79413735 | 0.00520384 | 0.02116345 | NA | NA | |
| ENSG000000 | 158.37371 | 0.559925201 | 0.20039849 | 2.79405898 | 0.0052051 | 0.02116646 | PTPN18 | | 26469 |
| ENSG000002 | 307.403028 | 0.371009077 | 0.1327872 | 2.79401238 | 0.00520585 | 0.02116739 | NA | NA | |
| ENSG000001 | 576.580336 | 0.351054212 | 0.12565329 | 2.79383232 | 0.00520875 | 0.02117493 | THUMPD2 | | 80745 |

| | | | | | | | | |
|---|---|---|---|---|---|---|---|---|
| ENSG000001 | 14.4308549 | -0.886451286 | 0.31728755 | -2.793842 | 0.00520859 | 0.02117493 | CIDEC | 63924 |
| ENSG000001 | 128.482974 | 0.511890442 | 0.18323318 | 2.79365583 | 0.00521159 | 0.02118437 | GPX7 | 2882 |
| ENSG000001 | 386.698364 | 0.365230089 | 0.13074082 | 2.79354293 | 0.00521341 | 0.02118964 | PRDM10 | 56980 |
| ENSG000001 | 1012.74715 | 0.294763358 | 0.10552575 | 2.79328371 | 0.00521759 | 0.0212045 | SMURF2 | 64750 |
| ENSG000001 | 77.055252 | 0.564959263 | 0.20226513 | 2.79316196 | 0.00521956 | 0.02121036 | SNAPC2 | 6618 |
| ENSG000001 | 814.285413 | 0.481753783 | 0.17248478 | 2.79302202 | 0.00522181 | 0.0212145 | GALNT2 | 2590 |
| ENSG000002 | 3.55350279 | -1.056378487 | 0.37822226 | -2.7930098 | 0.00522201 | 0.0212145 | SNORA36C | 100124535 |
| ENSG000002 | 12.1347698 | -0.947579836 | 0.33926934 | -2.7930017 | 0.00522214 | 0.0212145 | HP | 3240 |
| ENSG000001 | 1304.04892 | 0.678960897 | 0.24310985 | 2.7928153 | 0.00522515 | 0.02122404 | WDR6 | 11180 |
| ENSG000002 | 2.85102683 | -1.041381445 | 0.37288193 | -2.7927914 | 0.00522554 | 0.02122404 | LOC101928886 | 101928886 |
| ENSG000002 | 12.5730206 | -0.934103769 | 0.33453201 | -2.7922703 | 0.00523396 | 0.02125612 | LOC100507384 | 100507384 |
| ENSG000001 | 54.4600754 | 0.589412137 | 0.21109868 | 2.79211663 | 0.00523645 | 0.02126409 | CBLN1 | 869 |
| ENSG000001 | 55.4821084 | -0.591298031 | 0.21177687 | -2.7920803 | 0.00523704 | 0.02126435 | CCDC144A | 9720 |
| ENSG000001 | 6758.6495 | 0.3588193 | 0.12851538 | 2.79203382 | 0.00523779 | 0.02126527 | NSD1 | 64324 |
| ENSG00000C | 607.274779 | 0.356164718 | 0.1275822 | 2.79164908 | 0.00524402 | 0.02128844 | RBBP9 | 10741 |
| ENSG000002 | 7.51084511 | -1.016063362 | 0.36400976 | -2.7913081 | 0.00524955 | 0.02130875 | NA | NA |
| ENSG000001 | 250.484138 | 0.378592186 | 0.13566366 | 2.79066768 | 0.00525994 | 0.02134882 | ZFP90 | 146198 |
| ENSG00000C | 397.928837 | 0.355821648 | 0.12750975 | 2.7905446 | 0.00526195 | 0.02135421 | DUSP12 | 11266 |
| ENSG000001 | 66.6379956 | 0.544617164 | 0.19516683 | 2.79052115 | 0.00526233 | 0.02135421 | NA | NA |
| ENSG000001 | 547.750683 | 0.381543413 | 0.13673643 | 2.79035676 | 0.005265 | 0.02136078 | GGPS1 | 9453 |
| ENSG000001 | 325.347136 | 0.376079472 | 0.13477709 | 2.79038138 | 0.0052646 | 0.02136078 | ZNF669 | 79862 |
| ENSG000001 | 16.0084907 | -0.860529212 | 0.30840273 | -2.7902775 | 0.00526629 | 0.02136251 | OGN | 4969 |
| ENSG000001 | 282.441402 | 0.426671409 | 0.15291426 | 2.79026571 | 0.00526648 | 0.02136251 | MARVELD1 | 83742 |
| ENSG000001 | 945.381124 | 0.391253718 | 0.14022464 | 2.79019232 | 0.00526767 | 0.02136522 | NBEA | 26960 |
| ENSG000002 | 24.5549113 | 0.769651995 | 0.27585217 | 2.79008864 | 0.00526936 | 0.02136992 | NA | NA |
| ENSG00000C | 985.741799 | 0.321937819 | 0.11539156 | 2.78995983 | 0.00527146 | 0.02137629 | MYO9A | 4649 |
| ENSG000001 | 509.659793 | 0.339320052 | 0.12163594 | 2.78963656 | 0.00527672 | 0.02138941 | RPGRIP1L | 23322 |
| ENSG000002 | 3.96788073 | -1.056135919 | 0.37859331 | -2.7896317 | 0.0052768 | 0.02138941 | NA | NA |
| ENSG000002 | 4.57428244 | -1.050680292 | 0.37663749 | -2.7896328 | 0.00527679 | 0.02138941 | NA | NA |

| | | | | | | | | | |
|---|---|---|---|---|---|---|---|---|---|
| ENSG000002 | 3.18853173 | -1.050698929 | 0.37664246 | -2.7896455 | 0.00527658 | 0.02138941 | NA | NA | |
| ENSG000001 | 941.857265 | 0.311690488 | 0.11174093 | 2.78940301 | 0.00528053 | 0.02140238 | SYF2 | | 25949 |
| ENSG000002 | 11.4916699 | -0.956426216 | 0.34288542 | -2.7893464 | 0.00528145 | 0.02140398 | NA | NA | |
| ENSG000002 | 11.5716641 | -0.939340867 | 0.33677777 | -2.789201 | 0.00528383 | 0.02141146 | NA | NA | |
| ENSG000001 | 538.788411 | 0.476851771 | 0.17098375 | 2.78887189 | 0.0052892 | 0.02143108 | SPTSSA | | 171546 |
| ENSG000001 | 607.887423 | 0.449629149 | 0.16122637 | 2.78880652 | 0.00529027 | 0.02143327 | CIZ1 | | 25792 |
| ENSG000001 | 373.481974 | 0.359805345 | 0.12902239 | 2.78870478 | 0.00529193 | 0.02143786 | TANGO6 | | 79613 |
| ENSG000001 | 268.982379 | 0.406579021 | 0.14579775 | 2.78865085 | 0.00529281 | 0.02143929 | ZNF367 | | 195828 |
| ENSG000001 | 212.990117 | 0.425029862 | 0.1524192 | 2.7885585 | 0.00529432 | 0.02144326 | CREB3L4 | | 148327 |
| ENSG000001 | 4.94172573 | -1.046836076 | 0.37543859 | -2.7883018 | 0.00529852 | 0.021458 | SPINK4 | | 27290 |
| ENSG000002 | 2.80350007 | -1.040681498 | 0.37323539 | -2.7882713 | 0.00529902 | 0.021458 | NA | NA | |
| ENSG000000 | 165.527417 | 0.496713835 | 0.17816727 | 2.78790727 | 0.00530497 | 0.02147997 | KIAA0556 | | 23247 |
| ENSG000001 | 30.5764061 | -0.698680456 | 0.2506222 | -2.7877836 | 0.005307 | 0.02148204 | FSD2 | | 123722 |
| ENSG000002 | 425.578197 | 0.37239516 | 0.1335808 | 2.78778957 | 0.0053069 | 0.02148204 | ZNF468 | | 90333 |
| ENSG000002 | 3.24491254 | -1.049687571 | 0.37653183 | -2.7877791 | 0.00530707 | 0.02148204 | NA | NA | |
| ENSG000001 | 29.0430623 | -0.765854036 | 0.27474605 | -2.7874978 | 0.00531168 | 0.02149855 | SCN4B | | 6330 |
| ENSG000002 | 3.59704433 | -1.054286224 | 0.37822692 | -2.7874437 | 0.00531257 | 0.0215 | NA | NA | |
| ENSG000001 | 423.568863 | 0.384595512 | 0.13799352 | 2.78705484 | 0.00531895 | 0.02152366 | DDX31 | | 64794 |
| ENSG000001 | 543.985006 | 0.315997594 | 0.11338375 | 2.78697436 | 0.00532027 | 0.02152472 | CDK8 | | 1024 |
| ENSG000002 | 2.95936044 | -1.036151882 | 0.37178266 | -2.7869828 | 0.00532013 | 0.02152472 | NA | NA | |
| ENSG000002 | 70.4074943 | 0.577472499 | 0.20721084 | 2.78688358 | 0.00532176 | 0.0215286 | NA | NA | |
| ENSG000001 | 1600.29275 | 0.720289782 | 0.25846077 | 2.78684377 | 0.00532241 | 0.0215291 | IGF2R | | 3482 |
| ENSG000002 | 6.89144761 | -1.008039228 | 0.36174983 | -2.7865645 | 0.005327 | 0.02154551 | NA | NA | |
| ENSG000002 | 8.31597672 | -0.988935925 | 0.35490568 | -2.7864753 | 0.00532847 | 0.02154929 | NA | NA | |
| ENSG000001 | 506.222905 | 0.356929083 | 0.12810388 | 2.78624722 | 0.00533222 | 0.02156231 | Sep-03 | | 55964 |
| ENSG000001 | 81.3946057 | 0.520355129 | 0.1867771 | 2.78596861 | 0.0053368 | 0.02157654 | BCAR3 | | 8412 |
| ENSG000001 | 3634.28604 | 0.303627465 | 0.10898341 | 2.78599709 | 0.00533634 | 0.02157654 | SMC6 | | 79677 |
| ENSG000001 | 331.807308 | 0.384949221 | 0.13817737 | 2.78590643 | 0.00533783 | 0.02157853 | DAB1 | | 1600 |
| ENSG000001 | 153.714082 | 0.507119488 | 0.18203399 | 2.78585052 | 0.00533875 | 0.0215801 | LRP10 | | 26020 |

| | | | | | | | | |
|---|---|---|---|---|---|---|---|---|
| ENSG000002 | 223.405019 | 0.385663159 | 0.1384391 | 2.78579649 | 0.00533964 | 0.02158155 | ZBED9 | 114821 |
| ENSG000001 | 248.472686 | 0.37222092 | 0.13361664 | 2.78573774 | 0.00534061 | 0.02158331 | HERPUD2 | 64224 |
| ENSG000002 | 20.2468049 | -0.821380439 | 0.29485719 | -2.7856891 | 0.00534141 | 0.0215844 | MTND6P5 | 106480427 |
| ENSG000001 | 1760.27681 | 0.359144384 | 0.12892909 | 2.78559628 | 0.00534294 | 0.02158627 | NOTCH2 | 4853 |
| ENSG000002 | 28.2909046 | 0.710310645 | 0.25499334 | 2.78560468 | 0.0053428 | 0.02158627 | GOLGA2P7 | 388152 |
| ENSG000001 | 357.949198 | 0.403956937 | 0.14502202 | 2.78548689 | 0.00534474 | 0.02159141 | RBM4B | 83759 |
| ENSG000001 | 641.506317 | 0.378639291 | 0.13594914 | 2.78515394 | 0.00535023 | 0.02161144 | FAM217B | 63939 |
| ENSG000001 | 8.15324663 | -1.003587777 | 0.36035193 | -2.7850213 | 0.00535242 | 0.02161813 | ODF4 | 146852 |
| ENSG000002 | 6.72193574 | -1.02031593 | 0.36640993 | -2.7846296 | 0.00535889 | 0.0216421 | NA | NA |
| ENSG000001 | 209.482132 | 0.464296445 | 0.16676466 | 2.78414167 | 0.00536696 | 0.02167253 | Mar-05 | 54708 |
| ENSG000001 | 318.468795 | 0.416776623 | 0.14970112 | 2.78405824 | 0.00536834 | 0.02167595 | UBQLN2 | 29978 |
| ENSG00000C | 36.2208155 | -0.675640714 | 0.24268991 | -2.783967 | 0.00536985 | 0.02167988 | FRMPD1 | 22844 |
| ENSG000001 | 147.137006 | 0.468024141 | 0.16811899 | 2.78388621 | 0.00537119 | 0.02168097 | DPH1 | 1801 |
| ENSG000001 | 103.615935 | 0.514981013 | 0.18498572 | 2.78389598 | 0.00537103 | 0.02168097 | DUSP1 | 1843 |
| ENSG000002 | 7.60549334 | -1.016636525 | 0.3652218 | -2.7836141 | 0.0053757 | 0.02169701 | NA | NA |
| ENSG000002 | 18.4095944 | 0.817955349 | 0.29386575 | 2.78343202 | 0.00537871 | 0.02170703 | PCAT6 | 100506696 |
| ENSG000001 | 1239.90583 | 0.366515584 | 0.13168617 | 2.78325032 | 0.00538173 | 0.02171468 | RALGAPB | 57148 |
| ENSG000002 | 17.8111981 | -0.823126668 | 0.29573954 | -2.7832824 | 0.00538119 | 0.02171468 | LOC101928696 | 101928696 |
| ENSG000002 | 6.68917171 | -1.018263627 | 0.365858 | -2.7832209 | 0.00538222 | 0.02171468 | TARM1 | 441864 |
| ENSG000002 | 28.0968729 | -0.69869681 | 0.25106415 | -2.7829414 | 0.00538685 | 0.02173123 | LACTB2-AS1 | 286190 |
| ENSG000002 | 3.91812152 | -1.053539753 | 0.37859165 | -2.7827866 | 0.00538942 | 0.02173944 | SERPINA3 | 12 |
| ENSG000002 | 8.23854188 | -1.002659281 | 0.36032617 | -2.7826435 | 0.0053918 | 0.02174686 | DCAF8L1 | 139425 |
| ENSG000001 | 45.3706978 | -0.628337216 | 0.22581767 | -2.782498 | 0.00539422 | 0.0217522 | CNTNAP5 | 129684 |
| ENSG000001 | 31.7874318 | -0.693563622 | 0.24925776 | -2.7825157 | 0.00539393 | 0.0217522 | SERPINB13 | 5275 |
| ENSG000002 | 22.0045193 | -0.771034396 | 0.2771046 | -2.782467 | 0.00539474 | 0.0217522 | NA | NA |
| ENSG000001 | 9.16980559 | -0.96417609 | 0.34655963 | -2.7821362 | 0.00540024 | 0.02177222 | SULT2A1 | 6822 |
| ENSG000001 | 149.560447 | 0.476503785 | 0.17131207 | 2.78149569 | 0.0054109 | 0.02181306 | NA | NA |
| ENSG000001 | 177.976503 | 0.521453769 | 0.18748976 | 2.7812386 | 0.00541519 | 0.02182601 | ZMIZ2 | 83637 |
| ENSG000001 | 60.0567319 | 0.562645837 | 0.20229981 | 2.78124747 | 0.00541504 | 0.02182601 | DAGLA | 747 |

| | | | | | | | | |
|---|---|---|---|---|---|---|---|---|
| ENSG000002 | 5.94540572 | -1.028023726 | 0.3696562 | -2.7810266 | 0.00541873 | 0.0218381 | NA | NA |
| ENSG000001 | 69.8956535 | -0.588930855 | 0.21177761 | -2.780893 | 0.00542096 | 0.02184057 | FOXP2 | 93986 |
| ENSG000001 | 2964.1112 | 0.356109536 | 0.12805571 | 2.78089537 | 0.00542092 | 0.02184057 | SRRM1 | 10250 |
| ENSG000001 | 29.2894266 | -0.707771749 | 0.25450849 | -2.7809357 | 0.00542025 | 0.02184057 | KCNC2 | 3747 |
| ENSG000001 | 787.871914 | 0.335303757 | 0.12057954 | 2.78076818 | 0.00542304 | 0.02184463 | LATS1 | 9113 |
| ENSG000002 | 23.2890542 | -0.768114816 | 0.2762211 | -2.780797 | 0.00542256 | 0.02184463 | IL21-AS1 | 100996941 |
| ENSG000001 | 77.7241545 | 0.493681984 | 0.17753705 | 2.7807266 | 0.00542374 | 0.02184526 | HVCN1 | 84329 |
| ENSG000002 | 4.09339558 | -1.052096744 | 0.3783722 | -2.7805868 | 0.00542608 | 0.0218525 | SPATS1 | 221409 |
| ENSG000001 | 1104.34163 | 0.357857261 | 0.1287007 | 2.78053867 | 0.00542688 | 0.02185356 | DPY19L3 | 147991 |
| ENSG000001 | 70.3422479 | 0.590904255 | 0.21254315 | 2.78016138 | 0.00543319 | 0.0218768 | TCTA | 6988 |
| ENSG000001 | 466.818505 | 0.387352134 | 0.13933415 | 2.78002288 | 0.00543551 | 0.02188179 | CHMP7 | 91782 |
| ENSG000002 | 18.4702499 | -0.824142774 | 0.2964499 | -2.7800406 | 0.00543521 | 0.02188179 | FLJ31662 | 440594 |
| ENSG000001 | 1781.49707 | 0.342808648 | 0.12331785 | 2.77987859 | 0.00543792 | 0.02188934 | ZNF708 | 7562 |
| ENSG000002 | 11.1731836 | -0.934728901 | 0.33628129 | -2.7796042 | 0.00544252 | 0.02190566 | IZUMO3 | 100129669 |
| ENSG000001 | 51.6802555 | 0.582275626 | 0.20948578 | 2.77954724 | 0.00544347 | 0.02190733 | NR2C2AP | 126382 |
| ENSG000001 | 69.1872199 | -0.559506468 | 0.20132813 | -2.7790775 | 0.00545135 | 0.02193686 | NEGR1 | 257194 |
| ENSG000002 | 16.8285295 | -0.905254018 | 0.32574281 | -2.7790453 | 0.00545189 | 0.02193686 | PSG11 | 5680 |
| ENSG000002 | 9.6738009 | -0.985496091 | 0.35464213 | -2.7788466 | 0.00545523 | 0.0219481 | NA | NA |
| ENSG000001 | 107.888597 | 0.48293704 | 0.17385564 | 2.77780489 | 0.00547275 | 0.02200774 | ZNF174 | 7727 |
| ENSG000001 | 25.8755766 | -0.752236218 | 0.27080517 | -2.7777764 | 0.00547323 | 0.02200774 | OR4D9 | 390199 |
| ENSG000001 | 36.627144 | 0.654370711 | 0.23556899 | 2.77783046 | 0.00547232 | 0.02200774 | TMEM120A | 83862 |
| ENSG000002 | 2.40008368 | -1.018057999 | 0.36648743 | -2.7778797 | 0.00547149 | 0.02200774 | NA | NA |
| ENSG000002 | 14.3091978 | -0.871964091 | 0.31389979 | -2.7778422 | 0.00547212 | 0.02200774 | LOC101930028 | 101930028 |
| ENSG000002 | 15.9921939 | -0.845657975 | 0.30443755 | -2.7777716 | 0.00547331 | 0.02200774 | NA | NA |
| ENSG000001 | 16.3388531 | -0.877495352 | 0.31590901 | -2.7776838 | 0.00547479 | 0.02200932 | FAM69C | 125704 |
| ENSG000002 | 12.4142864 | -0.917392459 | 0.33027133 | -2.7776933 | 0.00547463 | 0.02200932 | OR5K1 | 26339 |
| ENSG000001 | 2152.44261 | 0.34408313 | 0.12388263 | 2.77749296 | 0.005478 | 0.02202007 | NA | NA |
| ENSG000002 | 6.63253773 | -1.015696601 | 0.36573277 | -2.777155 | 0.0054837 | 0.0220408 | LOC101928937 | 101928937 |
| ENSG000001 | 17.3665484 | -0.82064955 | 0.29550865 | -2.7770745 | 0.00548506 | 0.02204367 | RGS13 | 6003 |

| | | | | | | | | |
|---|---|---|---|---|---|---|---|---|
| ENSG000001 | 156.911104 | 0.407577443 | 0.14676762 | 2.77702556 | 0.00548589 | 0.02204367 | NTM | 50863 |
| ENSG000002 | 2.87988941 | -1.035374416 | 0.37283704 | -2.777016 | 0.00548605 | 0.02204367 | NA | NA |
| ENSG000001 | 246.185332 | 0.421753791 | 0.15187831 | 2.77691917 | 0.00548768 | 0.02204805 | NA | NA |
| ENSG000001 | 500.208264 | 0.379757376 | 0.13678137 | 2.77638227 | 0.00549675 | 0.02208142 | HOOK3 | 84376 |
| ENSG000002 | 5.23043779 | -1.048299883 | 0.37758023 | -2.7763633 | 0.00549707 | 0.02208142 | NA | NA |
| ENSG000001 | 662.335345 | 0.324782704 | 0.11698379 | 2.77630523 | 0.00549806 | 0.02208317 | AFTPH | 54812 |
| ENSG000002 | 9.25320242 | -0.963486387 | 0.34708095 | -2.7759702 | 0.00550372 | 0.02210156 | NA | NA |
| ENSG000002 | 7.32520053 | -1.022414882 | 0.36830521 | -2.7759989 | 0.00550324 | 0.02210156 | NA | NA |
| ENSG000001 | 1058.7036 | 0.350377067 | 0.12622341 | 2.7758485 | 0.00550579 | 0.022106 | NVL | 4931 |
| ENSG000002 | 6.39895538 | -1.02615418 | 0.36967333 | -2.7758405 | 0.00550592 | 0.022106 | NA | NA |
| ENSG00000C | 322.271424 | 0.361770792 | 0.13033892 | 2.77561602 | 0.00550972 | 0.02211908 | PIBF1 | 10464 |
| ENSG000002 | 17.1558264 | -0.825587718 | 0.29745541 | -2.7755008 | 0.00551168 | 0.02212473 | NA | NA |
| ENSG000002 | 9.65611414 | -0.988533368 | 0.35618302 | -2.7753523 | 0.00551419 | 0.02213264 | NA | NA |
| ENSG000002 | 2.39160231 | -1.016922975 | 0.36645634 | -2.7750181 | 0.00551986 | 0.02215321 | NA | NA |
| ENSG000001 | 7.98518544 | -1.009145093 | 0.36367094 | -2.7748852 | 0.00552212 | 0.02215641 | IRG1 | 730249 |
| ENSG000002 | 3.30944401 | -1.042550372 | 0.37571082 | -2.7748745 | 0.0055223 | 0.02215641 | NA | NA |
| ENSG000002 | 3.30944401 | -1.042550372 | 0.37571082 | -2.7748745 | 0.0055223 | 0.02215641 | NA | NA |
| ENSG000002 | 12.1581261 | -0.907202486 | 0.32696476 | -2.7746185 | 0.00552665 | 0.02216967 | LINC01033 | 104355136 |
| ENSG000002 | 13.8884841 | -0.886845811 | 0.31962835 | -2.7746156 | 0.0055267 | 0.02216967 | NA | NA |
| ENSG000001 | 87.856303 | 0.574201021 | 0.20695314 | 2.7745461 | 0.00552788 | 0.02217221 | SYNGR1 | 9145 |
| ENSG000001 | 9995.53875 | 0.315153774 | 0.11359335 | 2.77440328 | 0.00553031 | 0.02217976 | AZIN1 | 51582 |
| ENSG000002 | 10.300584 | -0.955559768 | 0.34443494 | -2.7742823 | 0.00553236 | 0.02218581 | NA | NA |
| ENSG000001 | 209.019854 | 0.532890983 | 0.19209111 | 2.77415742 | 0.00553449 | 0.02219214 | RSAD1 | 55316 |
| ENSG000002 | 2.45541188 | -1.014062307 | 0.36557048 | -2.7739173 | 0.00553857 | 0.02220632 | NA | NA |
| ENSG00000C | 53.111811 | -0.598403459 | 0.21573208 | -2.773827 | 0.00554011 | 0.02221029 | IMPG2 | 50939 |
| ENSG000001 | 5174.56761 | 0.349259309 | 0.12591679 | 2.77373111 | 0.00554175 | 0.02221093 | PTPRZ1 | 5803 |
| ENSG000001 | 30.159092 | -0.733063326 | 0.26429182 | -2.773689 | 0.00554246 | 0.02221093 | ADGRG7 | 84873 |
| ENSG000002 | 9.33947077 | -0.970586827 | 0.34991577 | -2.7737727 | 0.00554104 | 0.02221093 | NA | NA |
| ENSG000002 | 7.51576158 | -1.026375442 | 0.37003755 | -2.7737062 | 0.00554217 | 0.02221093 | NA | NA |

| | | | | | | | | | |
|---|---|---|---|---|---|---|---|---|---|
| ENSG000002 | 22.1733759 | -0.799607032 | 0.28832666 | -2.7732678 | 0.00554964 | 0.02223751 | LINC00906 | | 148145 |
| ENSG000002 | 4.3430602 | -1.049434219 | 0.3784352 | -2.7730883 | 0.00555271 | 0.02224758 | NA | NA | |
| ENSG000001 | 219.537101 | 0.443501958 | 0.15993596 | 2.77299715 | 0.00555426 | 0.02225161 | EFHD2 | | 79180 |
| ENSG000001 | 60123.2961 | 0.305280209 | 0.11009941 | 2.7727687 | 0.00555816 | 0.02226504 | HSPA8 | | 3312 |
| ENSG000001 | 616.45624 | 0.318970249 | 0.11504305 | 2.77261647 | 0.00556076 | 0.02227203 | PAM | | 5066 |
| ENSG000001 | 2048.98447 | 0.355185802 | 0.12810558 | 2.77260216 | 0.00556101 | 0.02227203 | AGFG1 | | 3267 |
| ENSG000002 | 23.1872623 | -0.762949869 | 0.27518088 | -2.7725396 | 0.00556208 | 0.02227203 | LOC729950 | | 729950 |
| ENSG000002 | 8.97685716 | -0.97585171 | 0.35197055 | -2.772538 | 0.0055621 | 0.02227203 | NA | NA | |
| ENSG000002 | 3.60900313 | -1.048078912 | 0.37805409 | -2.7722988 | 0.00556619 | 0.0222862 | NA | NA | |
| ENSG000001 | 30.5902217 | -0.680977515 | 0.24564177 | -2.7722383 | 0.00556723 | 0.02228814 | HMGCLL1 | | 54511 |
| ENSG000001 | 716.849709 | 0.327711115 | 0.11822721 | 2.77187559 | 0.00557343 | 0.02231079 | RNF169 | | 254225 |
| ENSG000001 | 469.007789 | 0.362410614 | 0.1307553 | 2.77167065 | 0.00557694 | 0.02231856 | AKAP10 | | 11216 |
| ENSG000001 | 1859.39771 | 0.303056405 | 0.10933965 | 2.77169726 | 0.00557649 | 0.02231856 | SMAD5 | | 4090 |
| ENSG000002 | 4.98273501 | -1.041283819 | 0.37569081 | -2.771651 | 0.00557728 | 0.02231856 | NA | NA | |
| ENSG000002 | 31.4079897 | -0.726021593 | 0.26194716 | -2.7716338 | 0.00557757 | 0.02231856 | LINC00299 | | 339789 |
| ENSG000002 | 25.0970759 | 0.731170105 | 0.26381164 | 2.77156118 | 0.00557882 | 0.02232133 | NA | NA | |
| ENSG00000C | 184.884449 | 0.454042407 | 0.16383182 | 2.77139338 | 0.00558169 | 0.02233063 | WIZ | | 58525 |
| ENSG000001 | 65.0043412 | 0.55110325 | 0.19887023 | 2.77117017 | 0.00558552 | 0.02234374 | UBOX5 | | 22888 |
| ENSG000001 | 83.3187318 | 0.566401821 | 0.20439928 | 2.77105586 | 0.00558748 | 0.02234805 | ZBTB45 | | 84878 |
| ENSG000001 | 67.7670565 | 0.550437054 | 0.19863893 | 2.77104324 | 0.0055877 | 0.02234805 | ABHD6 | | 57406 |
| ENSG000002 | 27.4053139 | -0.723733938 | 0.26119651 | -2.7708408 | 0.00559118 | 0.02235974 | LINC00939 | | 400084 |
| ENSG000001 | 181.838188 | 0.437936962 | 0.1580594 | 2.77071129 | 0.0055934 | 0.02236643 | LACE1 | | 246269 |
| ENSG000001 | 330.704853 | 0.47099915 | 0.16999792 | 2.77061712 | 0.00559502 | 0.02237069 | DENND4B | | 9909 |
| ENSG000002 | 2.41809627 | -1.015180704 | 0.36641856 | -2.7705494 | 0.00559618 | 0.02237314 | RN7SL271P | | 106479321 |
| ENSG00000C | 220.083352 | 0.422523301 | 0.15253754 | 2.76996273 | 0.00560627 | 0.02241127 | MBTPS2 | | 51360 |
| ENSG000001 | 260.76983 | 0.364721279 | 0.13167423 | 2.76987593 | 0.00560777 | 0.02241503 | ZKSCAN7 | | 55888 |
| ENSG000002 | 19.5143183 | -0.80480102 | 0.29056491 | -2.7697805 | 0.00560941 | 0.02241938 | NA | NA | |
| ENSG000001 | 25.3425141 | -0.740468856 | 0.26737571 | -2.7693946 | 0.00561606 | 0.02244154 | LOC344887 | | 344887 |
| ENSG000002 | 2.48131298 | -1.012443105 | 0.36558247 | -2.7693973 | 0.00561601 | 0.02244154 | NA | NA | |

| | | | | | | | | |
|---|---|---|---|---|---|---|---|---|
| ENSG000001 | 12.8906762 | -0.901281938 | 0.3254666 | -2.7691995 | 0.00561942 | 0.02245277 | CFAP126 | 257177 |
| ENSG000001 | 1337.75787 | 0.410848225 | 0.14836554 | 2.76916208 | 0.00562007 | 0.02245314 | RTF1 | 23168 |
| ENSG000001 | 28.2238116 | -0.755006355 | 0.27265757 | -2.7690643 | 0.00562175 | 0.02245766 | DNAJC22 | 79962 |
| ENSG000002 | 3.55876676 | -1.046443971 | 0.37791615 | -2.7689845 | 0.00562313 | 0.02246095 | NA | NA |
| ENSG000002 | 6.25512603 | -1.017364555 | 0.36741877 | -2.768951 | 0.00562371 | 0.02246105 | NA | NA |
| ENSG000001 | 282.244122 | 0.400368736 | 0.14459816 | 2.76883704 | 0.00562568 | 0.02246669 | NADSYN1 | 55191 |
| ENSG000001 | 1646.36172 | 0.31151566 | 0.11251015 | 2.76877836 | 0.00562669 | 0.02246853 | SH3D19 | 152503 |
| ENSG000001 | 3443.56631 | 0.351874317 | 0.12710809 | 2.76830772 | 0.00563482 | 0.0224965 | CAPZA1 | 829 |
| ENSG000002 | 15.415673 | -0.870653684 | 0.31450445 | -2.768335 | 0.00563435 | 0.0224965 | NA | NA |
| ENSG000002 | 9.40278759 | -0.95277113 | 0.34417482 | -2.7682767 | 0.00563536 | 0.0224965 | NA | NA |
| ENSG000001 | 34.6726504 | -0.672969701 | 0.24310967 | -2.7681733 | 0.00563715 | 0.02249921 | KCND3 | 3752 |
| ENSG000001 | 24.9948911 | -0.719960114 | 0.26008433 | -2.7681796 | 0.00563704 | 0.02249921 | NA | NA |
| ENSG000001 | 163.680681 | 0.429730473 | 0.15525227 | 2.76794961 | 0.00564102 | 0.02251244 | CCDC64 | 92558 |
| ENSG000002 | 6.33234398 | -1.018906574 | 0.36811532 | -2.7679005 | 0.00564187 | 0.02251362 | NA | NA |
| ENSG00000C | 204.14892 | 0.474359581 | 0.17138499 | 2.76780117 | 0.00564359 | 0.02251383 | PKD1 | 5310 |
| ENSG000001 | 29.8532427 | -0.694371612 | 0.25087337 | -2.7678172 | 0.00564331 | 0.02251383 | CLCA2 | 9635 |
| ENSG000002 | 5.57876965 | -1.042240719 | 0.37655452 | -2.7678349 | 0.005643 | 0.02251383 | NA | NA |
| ENSG000002 | 12.3733997 | -0.89127693 | 0.32203305 | -2.7676567 | 0.00564609 | 0.0225216 | NA | NA |
| ENSG000001 | 367.410989 | 0.404661643 | 0.14624249 | 2.76705926 | 0.00565645 | 0.02255848 | KDELC1 | 79070 |
| ENSG000001 | 33.5996177 | -0.696042909 | 0.25154414 | -2.7670806 | 0.00565608 | 0.02255848 | GPR78 | 27201 |
| ENSG000001 | 54.7405588 | -0.571639183 | 0.20659816 | -2.7669132 | 0.00565898 | 0.02256637 | CLVS2 | 134829 |
| ENSG000001 | 47.3298744 | -0.5824399 | 0.21051415 | -2.7667493 | 0.00566183 | 0.02257549 | SPATA17 | 128153 |
| ENSG00000C | 60.634348 | -0.534544579 | 0.19321221 | -2.7666191 | 0.00566409 | 0.02258007 | SPTB | 6710 |
| ENSG000001 | 176.176669 | 0.417306547 | 0.15083553 | 2.76663302 | 0.00566385 | 0.02258007 | CAB39L | 81617 |
| ENSG000001 | 3078.83086 | 0.782777913 | 0.28294105 | 2.76657598 | 0.00566484 | 0.02258084 | MYH9 | 4627 |
| ENSG000002 | 14.5502127 | -0.882347926 | 0.31895353 | -2.766384 | 0.00566818 | 0.02259192 | LINC01016 | 100507584 |
| ENSG000002 | 8.26322412 | -0.981757708 | 0.35493283 | -2.7660381 | 0.00567419 | 0.02261367 | NA | NA |
| ENSG000001 | 26.4183099 | -0.759158951 | 0.27447542 | -2.7658541 | 0.00567739 | 0.02262421 | NPSR1-AS1 | 404744 |
| ENSG000001 | 1810.02286 | 0.377514186 | 0.13649787 | 2.76571477 | 0.00567982 | 0.02262721 | ICE2 | 79664 |

| | | | | | | | | |
|---|---|---|---|---|---|---|---|---|
| ENSG000001 | 7.48810207 | -0.989816663 | 0.35788672 | -2.7657261 | 0.00567962 | 0.02262721 | AQP10 | 89872 |
| ENSG000002 | 21.0189364 | -0.806798245 | 0.29171213 | -2.7657343 | 0.00567948 | 0.02262721 | LINC01581 | 101927112 |
| ENSG000001 | 8.5291371 | -0.974423789 | 0.35234639 | -2.7655279 | 0.00568307 | 0.02263795 | YIPF7 | 285525 |
| ENSG00000C | 53.6272312 | -0.589691395 | 0.21324602 | -2.7653102 | 0.00568687 | 0.02264894 | LCP2 | 3937 |
| ENSG000001 | 964.484543 | 0.370553691 | 0.134001 | 2.76530551 | 0.00568695 | 0.02264894 | MAP9 | 79884 |
| ENSG000001 | 694.633008 | 0.36084654 | 0.13050462 | 2.76500962 | 0.00569211 | 0.02266727 | EIF2D | 1939 |
| ENSG000002 | 91.2694023 | 0.547500052 | 0.1980125 | 2.76497726 | 0.00569268 | 0.02266729 | FAM229B | 619208 |
| ENSG000001 | 914.771401 | 0.370805496 | 0.13413162 | 2.76448978 | 0.00570119 | 0.02269451 | FARP1 | 10160 |
| ENSG000001 | 119.80957 | 0.560407899 | 0.20271378 | 2.76452786 | 0.00570052 | 0.02269451 | RPRM | 56475 |
| ENSG000001 | 22.6783757 | -0.753739653 | 0.27264989 | -2.7644965 | 0.00570107 | 0.02269451 | CCDC60 | 160777 |
| ENSG000001 | 14.9283306 | -0.843540738 | 0.30514461 | -2.7643967 | 0.00570282 | 0.02269875 | FCRLB | 127943 |
| ENSG000002 | 5.94545888 | -1.021039463 | 0.36938396 | -2.7641684 | 0.00570681 | 0.02271242 | NA | NA |
| ENSG000001 | 1838.24811 | 0.344272578 | 0.12455152 | 2.76409778 | 0.00570804 | 0.02271347 | PPP1R15B | 84919 |
| ENSG000002 | 46.3059791 | 0.631716979 | 0.22854436 | 2.76408917 | 0.00570819 | 0.02271347 | BORCS8 | 729991 |
| ENSG000002 | 205.87105 | 0.440200404 | 0.15926157 | 2.76400891 | 0.0057096 | 0.02271683 | NA | NA |
| ENSG000002 | 15.2045752 | -0.857163262 | 0.31012935 | -2.7638895 | 0.00571169 | 0.02272291 | NA | NA |
| ENSG000001 | 23.6892943 | -0.750872612 | 0.27167579 | -2.7638554 | 0.00571228 | 0.02272306 | ZP2 | 7783 |
| ENSG000002 | 237.886985 | 0.430942079 | 0.15592278 | 2.76381736 | 0.00571295 | 0.02272348 | H2AFJ | 55766 |
| ENSG000001 | 132.311954 | 0.575186198 | 0.20811929 | 2.76373322 | 0.00571442 | 0.02272711 | ZNF513 | 130557 |
| ENSG000002 | 862995.327 | 0.859858444 | 0.31121527 | 2.76290574 | 0.00572893 | 0.02278257 | NA | NA |
| ENSG000001 | 12.3605589 | -0.910615072 | 0.32959913 | -2.7627957 | 0.00573086 | 0.02278801 | NA | NA |
| ENSG000001 | 15.6216712 | -0.855390596 | 0.30962933 | -2.7626278 | 0.00573381 | 0.02279751 | ZNF534 | 147658 |
| ENSG000002 | 7.74160136 | -0.994951121 | 0.3601567 | -2.7625506 | 0.00573517 | 0.02280066 | NA | NA |
| ENSG00000C | 10424.0992 | 0.328177117 | 0.11880581 | 2.76229862 | 0.0057396 | 0.0228138 | RIF1 | 55183 |
| ENSG000001 | 1052.75793 | 0.352318367 | 0.12754485 | 2.76230957 | 0.0057394 | 0.0228138 | NPEPPS | 9520 |
| ENSG00000C | 66.2210272 | 0.6025989 | 0.21816045 | 2.7621822 | 0.00574164 | 0.02281882 | ZC3H3 | 23144 |
| ENSG000002 | 6.53979712 | -1.007296277 | 0.36467665 | -2.7621628 | 0.00574198 | 0.02281882 | NA | NA |
| ENSG000002 | 11.1760432 | -0.912280002 | 0.33028877 | -2.7620679 | 0.00574365 | 0.02282321 | NA | NA |
| ENSG000001 | 63.2994397 | 0.611480848 | 0.22139382 | 2.76195984 | 0.00574555 | 0.02282853 | RFNG | 5986 |

| | | | | | | | | |
|---|---|---|---|---|---|---|---|---|
| ENSG000002 | 23.0860275 | -0.755150977 | 0.27341771 | -2.7618949 | 0.0057467 | 0.0228286 | NA | NA |
| ENSG000002 | 5.01047184 | -1.039778185 | 0.37646965 | -2.7619177 | 0.0057463 | 0.0228286 | NA | NA |
| ENSG000001 | 393.821027 | 0.320965877 | 0.11622897 | 2.76149638 | 0.00575372 | 0.02285424 | SPICE1 | 152185 |
| ENSG000001 | 461.73712 | 0.35220163 | 0.12754995 | 2.76128387 | 0.00575746 | 0.02286688 | RMDN1 | 51115 |
| ENSG000001 | 332.747716 | 0.369323218 | 0.13375378 | 2.76121696 | 0.00575864 | 0.02286932 | TUBGCP4 | 27229 |
| ENSG000002 | 34.4189151 | 0.648302577 | 0.23479603 | 2.76113088 | 0.00576016 | 0.02287087 | NA | NA |
| ENSG000002 | 102.427328 | 0.559284486 | 0.20255532 | 2.76114443 | 0.00575992 | 0.02287087 | ZNF853 | 54753 |
| ENSG000002 | 3.14709788 | -1.038734008 | 0.37623608 | -2.760857 | 0.00576499 | 0.02288782 | NA | NA |
| ENSG000001 | 36.9917289 | -0.645141395 | 0.23368371 | -2.7607461 | 0.00576695 | 0.02289335 | ZPLD1 | 131368 |
| ENSG000001 | 75.8318417 | 0.579934504 | 0.21008029 | 2.7605374 | 0.00577063 | 0.02290574 | SULT4A1 | 25830 |
| ENSG000001 | 509.248747 | 0.355617362 | 0.12882662 | 2.76043395 | 0.00577246 | 0.02291075 | CAPN7 | 23473 |
| ENSG000001 | 824.224144 | 0.348393394 | 0.12621349 | 2.76034987 | 0.00577395 | 0.02291441 | RBBP5 | 5929 |
| ENSG00000C | 367.665007 | 0.380052466 | 0.13768613 | 2.76028148 | 0.00577516 | 0.02291696 | ZBTB25 | 7597 |
| ENSG000002 | 8.13014497 | -0.97409776 | 0.352903 | -2.7602422 | 0.00577585 | 0.02291747 | NA | NA |
| ENSG000002 | 7.26397279 | -1.017098254 | 0.3685019 | -2.7600896 | 0.00577855 | 0.02292594 | NA | NA |
| ENSG000001 | 21.7090589 | -0.784909273 | 0.28439424 | -2.7599338 | 0.00578131 | 0.02293239 | ANKRD30A | 91074 |
| ENSG000002 | 4.28644658 | -1.044030745 | 0.37828023 | -2.75994 | 0.0057812 | 0.02293239 | NA | NA |
| ENSG000002 | 5.65902391 | -1.029339274 | 0.37298577 | -2.759728 | 0.00578495 | 0.02294459 | NA | NA |
| ENSG000001 | 17.6490765 | -0.810297428 | 0.29364213 | -2.7594727 | 0.00578947 | 0.02295828 | SULT6B1 | 391365 |
| ENSG000002 | 26.1278206 | -0.721668578 | 0.26152442 | -2.7594692 | 0.00578953 | 0.02295828 | NA | NA |
| ENSG00000C | 140.547846 | 0.452626644 | 0.16403722 | 2.75929229 | 0.00579267 | 0.02296847 | PIGV | 55650 |
| ENSG000001 | 217.134306 | 0.478141089 | 0.17328606 | 2.7592589 | 0.00579326 | 0.02296857 | CLASRP | 11129 |
| ENSG000002 | 10.1203613 | -0.945345397 | 0.34261558 | -2.7592015 | 0.00579428 | 0.02297036 | NA | NA |
| ENSG000001 | 1200.81113 | 0.344282051 | 0.12479334 | 2.75881761 | 0.00580109 | 0.02299511 | DSE | 29940 |
| ENSG000001 | 13.2257226 | -0.874030124 | 0.31683095 | -2.758664 | 0.00580382 | 0.02300368 | C10orf53 | 282966 |
| ENSG000001 | 17.597292 | -0.828659897 | 0.30039874 | -2.7585332 | 0.00580614 | 0.02301025 | OIT3 | 170392 |
| ENSG000002 | 7.86074997 | -1.001647459 | 0.3631122 | -2.7585068 | 0.00580661 | 0.02301025 | NA | NA |
| ENSG000001 | 820.5626 | 0.326247719 | 0.11827249 | 2.75844138 | 0.00580777 | 0.02301073 | ZKSCAN8 | 7745 |
| ENSG000002 | 23.2221517 | -0.744048577 | 0.26973648 | -2.7584277 | 0.00580802 | 0.02301073 | OR14J1 | 442191 |

| | | | | | | | | |
|---|---|---|---|---|---|---|---|---|
| ENSG000002 | 3.23649589 | -1.03712039 | 0.37598567 | -2.7584041 | 0.00580844 | 0.02301073 | RN7SKP299 | 106480925 |
| ENSG000000 | 221.895619 | 0.403375747 | 0.14624238 | 2.75826855 | 0.00581084 | 0.02301803 | ASTE1 | 28990 |
| ENSG000001 | 69.3755215 | 0.583810284 | 0.21167761 | 2.75801628 | 0.00581533 | 0.02303355 | REEP4 | 80346 |
| ENSG000001 | 259.776832 | 0.358118803 | 0.12986608 | 2.75760088 | 0.00582272 | 0.02306058 | PDE3A | 5139 |
| ENSG000002 | 8.01890833 | -0.990632008 | 0.35924401 | -2.7575463 | 0.0058237 | 0.02306218 | NA | NA |
| ENSG000001 | 25.4775358 | -0.727962205 | 0.26401811 | -2.7572435 | 0.00582909 | 0.0230813 | NLRP14 | 338323 |
| ENSG000001 | 67.196636 | 0.559303939 | 0.20286294 | 2.75705329 | 0.00583248 | 0.02309022 | CBX4 | 8535 |
| ENSG000001 | 240.85768 | 0.408386368 | 0.14812365 | 2.75706386 | 0.0058323 | 0.02309022 | BTG3 | 10950 |
| ENSG000000 | 19.1742878 | -0.803299025 | 0.29136818 | -2.7569896 | 0.00583362 | 0.02309246 | SLC11A1 | 6556 |
| ENSG000001 | 525.927933 | 0.340177643 | 0.1233915 | 2.75689691 | 0.00583527 | 0.02309675 | AZI2 | 64343 |
| ENSG000001 | 38.3610288 | -0.61562728 | 0.22331313 | -2.7567895 | 0.00583719 | 0.02309758 | NR1I2 | 8856 |
| ENSG000002 | 4.2587256 | -1.0431339 | 0.37837903 | -2.7568491 | 0.00583613 | 0.02309758 | LRCOL1 | 100507055 |
| ENSG000002 | 66.4836319 | 0.557279902 | 0.20214772 | 2.75679538 | 0.00583709 | 0.02309758 | EBLN2 | 55096 |
| ENSG000001 | 578.453928 | 0.325928088 | 0.11823049 | 2.75671775 | 0.00583847 | 0.02310039 | CAPRIN2 | 65981 |
| ENSG000002 | 11.7315528 | -0.948631723 | 0.34415998 | -2.7563685 | 0.00584471 | 0.02312282 | ONECUT3 | 390874 |
| ENSG000002 | 8.44697526 | -0.969541313 | 0.35177235 | -2.7561613 | 0.00584842 | 0.02313522 | MTND6P6 | 106478944 |
| ENSG000001 | 13.225406 | -0.895670485 | 0.32501561 | -2.7557768 | 0.00585529 | 0.02316017 | EPGN | 255324 |
| ENSG000001 | 91.5525381 | 0.522429568 | 0.18959133 | 2.75555623 | 0.00585924 | 0.02317353 | IQCC | 55721 |
| ENSG000001 | 16.0355175 | -0.823668447 | 0.29892725 | -2.7554144 | 0.00586178 | 0.02318131 | C4BPA | 722 |
| ENSG000001 | 95.3067333 | 0.50890514 | 0.18470047 | 2.75529964 | 0.00586384 | 0.02318719 | EPHB6 | 2051 |
| ENSG000001 | 625.778704 | 0.351770576 | 0.12767455 | 2.75521295 | 0.00586539 | 0.02319107 | BLOC1S2 | 282991 |
| ENSG000002 | 15.1740849 | -0.880471428 | 0.3195826 | -2.7550669 | 0.00586801 | 0.02319779 | NA | NA |
| ENSG000002 | 13.7710981 | -0.859036484 | 0.31180382 | -2.7550543 | 0.00586824 | 0.02319779 | NA | NA |
| ENSG000001 | 3.63677508 | -1.041016625 | 0.37791781 | -2.7546112 | 0.00587619 | 0.02322697 | OR8J1 | 219477 |
| ENSG000001 | 1874.97523 | 0.33647028 | 0.12215635 | 2.75442316 | 0.00587957 | 0.02323805 | SELT | 51714 |
| ENSG000002 | 5.29921432 | -1.030724377 | 0.37422994 | -2.7542542 | 0.00588261 | 0.02324779 | NA | NA |
| ENSG000001 | 76.5426185 | 0.491001462 | 0.17827969 | 2.75410761 | 0.00588524 | 0.02325594 | L3HYPDH | 112849 |
| ENSG000002 | 3.18785455 | -1.036175227 | 0.37627374 | -2.7537803 | 0.00589113 | 0.02327467 | NA | NA |
| ENSG000002 | 13.1113943 | -0.912978579 | 0.33153538 | -2.7537893 | 0.00589097 | 0.02327467 | NA | NA |

| | | | | | | | | |
|---|---|---|---|---|---|---|---|---|
| ENSG000001 | 3047.25472 | 0.350662585 | 0.12735651 | 2.75339341 | 0.0058981 | 0.02329993 | SPIN1 | 10927 |
| ENSG00000C | 4.44603624 | -1.042169343 | 0.37858093 | -2.752831 | 0.00590824 | 0.02333748 | KCNK16 | 83795 |
| ENSG000001 | 6.96510537 | -1.003689097 | 0.36460631 | -2.7528024 | 0.00590876 | 0.02333748 | NA | NA |
| ENSG000002 | 37.5354845 | 0.631965087 | 0.22957625 | 2.75274589 | 0.00590977 | 0.02333924 | NA | NA |
| ENSG000001 | 4545.96769 | 0.336934511 | 0.1224056 | 2.752607 | 0.00591228 | 0.02334686 | PHIP | 55023 |
| ENSG000001 | 21.7461703 | -0.820170468 | 0.29797385 | -2.7524914 | 0.00591437 | 0.02335284 | FBXW10 | 10517 |
| ENSG000002 | 203.634544 | 0.539802511 | 0.19612379 | 2.75235615 | 0.00591681 | 0.02336021 | CHCHD10 | 400916 |
| ENSG00000C | 3864.6099 | 0.310848645 | 0.11294387 | 2.75224014 | 0.00591891 | 0.02336621 | WAC | 51322 |
| ENSG000001 | 104.090264 | 0.48809354 | 0.17737337 | 2.75178589 | 0.00592713 | 0.02339637 | C19orf52 | 90580 |
| ENSG000002 | 17.1960487 | -0.808293008 | 0.29376881 | -2.7514596 | 0.00593303 | 0.02341741 | LINC01254 | 101927350 |
| ENSG000001 | 127.516341 | 0.441194971 | 0.16036149 | 2.75125257 | 0.00593678 | 0.02342993 | CYB5R1 | 51706 |
| ENSG000001 | 130.211917 | 0.539984839 | 0.19627403 | 2.75117819 | 0.00593813 | 0.02343297 | CASKIN2 | 57513 |
| ENSG000001 | 1968.03888 | 0.372094487 | 0.13525261 | 2.75110762 | 0.00593941 | 0.02343574 | AFF1 | 4299 |
| ENSG000002 | 4.33506755 | -1.041277089 | 0.37850766 | -2.7510066 | 0.00594124 | 0.02344069 | NA | NA |
| ENSG000001 | 13.8670033 | -0.864148339 | 0.31417446 | -2.7505366 | 0.00594978 | 0.02346987 | NPSR1 | 387129 |
| ENSG000002 | 5.87345382 | -1.031262113 | 0.37493142 | -2.7505354 | 0.0059498 | 0.02346987 | NA | NA |
| ENSG000001 | 16.06534 | -0.866847038 | 0.31516566 | -2.7504489 | 0.00595137 | 0.02347149 | TDRD10 | 126668 |
| ENSG000001 | 423.841094 | 0.353406059 | 0.12848981 | 2.75045971 | 0.00595117 | 0.02347149 | CEP63 | 80254 |
| ENSG000002 | 19.65813 | 0.806724732 | 0.29333382 | 2.75019341 | 0.00595601 | 0.02348752 | NA | NA |
| ENSG000001 | 6459.11254 | 0.412088002 | 0.14984417 | 2.75011043 | 0.00595752 | 0.02349118 | HDAC2 | 3066 |
| ENSG000002 | 6.57233356 | -1.002954077 | 0.36474858 | -2.7497134 | 0.00596474 | 0.02351737 | NA | NA |
| ENSG000002 | 10.6569496 | -0.922106748 | 0.3353548 | -2.7496453 | 0.00596598 | 0.02351998 | CTB-113P19.1 | 101927096 |
| ENSG000002 | 17.5416827 | -0.810182309 | 0.2946653 | -2.7495003 | 0.00596862 | 0.0235281 | TCAM1P | 146771 |
| ENSG000001 | 476.271654 | 0.387271826 | 0.14085975 | 2.74934335 | 0.00597148 | 0.02353708 | NCS1 | 23413 |
| ENSG000002 | 10.5457713 | -0.916750604 | 0.33345401 | -2.7492565 | 0.00597306 | 0.02354103 | NA | NA |
| ENSG000002 | 11.5228751 | -0.909198887 | 0.33071664 | -2.7491779 | 0.00597449 | 0.02354438 | NA | NA |
| ENSG000001 | 5030.63698 | 0.302033925 | 0.10987327 | 2.74892998 | 0.00597902 | 0.02355991 | SOX2 | 6657 |
| ENSG000002 | 106.728306 | 0.538774922 | 0.19604416 | 2.74823248 | 0.00599175 | 0.02360711 | NA | NA |
| ENSG000002 | 124.13924 | -0.417469814 | 0.15190607 | -2.7482102 | 0.00599216 | 0.02360711 | NBPF26 | 101060684 |

| | | | | | | | | | |
|---|---|---|---|---|---|---|---|---|---|
| ENSG000001 | 50.9225402 | -0.628080369 | 0.22859977 | -2.747511 | 0.00600495 | 0.02365521 | SCN7A | | 6332 |
| ENSG000001 | 29.5698304 | -0.716951336 | 0.26095558 | -2.7474076 | 0.00600684 | 0.02366037 | FAM198B | | 51313 |
| ENSG000001 | 256.115923 | 0.464682078 | 0.16915108 | 2.74714232 | 0.0060117 | 0.02367722 | CLP1 | | 10978 |
| ENSG000002 | 16.2223814 | -0.834818679 | 0.30390475 | -2.7469748 | 0.00601478 | 0.02368701 | AKR7L | | 246181 |
| ENSG00000C | 144.762262 | 0.475859148 | 0.17325251 | 2.74662198 | 0.00602125 | 0.0237102 | CHDH | | 55349 |
| ENSG000001 | 3082.40411 | 0.336173606 | 0.12241491 | 2.74618194 | 0.00602933 | 0.02373972 | SETX | | 23064 |
| ENSG000001 | 199.002233 | -0.417241338 | 0.15193969 | -2.7460983 | 0.00603087 | 0.02374347 | MB21D1 | | 115004 |
| ENSG000002 | 6.89424046 | -0.996972268 | 0.36306205 | -2.7460107 | 0.00603248 | 0.02374751 | LINC00880 | | 339894 |
| ENSG000001 | 898.823899 | 0.341217247 | 0.12428238 | 2.74549978 | 0.00604188 | 0.02378221 | MAGOHB | | 55110 |
| ENSG000001 | 564.207723 | 0.337141004 | 0.12280114 | 2.74542237 | 0.00604331 | 0.0237832 | BTRC | | 8945 |
| ENSG000001 | 877.109168 | 0.330808395 | 0.12049402 | 2.74543424 | 0.00604309 | 0.0237832 | CDC40 | | 51362 |
| ENSG000001 | 324.419599 | 0.370647642 | 0.13500827 | 2.74536999 | 0.00604427 | 0.02378469 | DCK | | 1633 |
| ENSG000002 | 21.3067796 | -0.77102978 | 0.28085106 | -2.7453334 | 0.00604495 | 0.02378504 | NA | NA | |
| ENSG000001 | 417.262334 | 0.370949744 | 0.13512322 | 2.74527017 | 0.00604611 | 0.02378731 | UBE2E2 | | 7325 |
| ENSG000002 | 31.3133533 | -0.721010836 | 0.26266995 | -2.7449308 | 0.00605237 | 0.02380961 | LINC01121 | | 400952 |
| ENSG00000C | 1178.72457 | 0.355577343 | 0.1295434 | 2.74485099 | 0.00605384 | 0.02381309 | ABLIM1 | | 3983 |
| ENSG00000C | 192.334721 | 0.42792184 | 0.15591838 | 2.7445246 | 0.00605986 | 0.02383447 | NDE1 | | 54820 |
| ENSG000001 | 28.0882911 | -0.716524178 | 0.26109002 | -2.7443568 | 0.00606296 | 0.02384435 | KCNV1 | | 27012 |
| ENSG000001 | 360.285437 | 0.375682864 | 0.13689726 | 2.74426862 | 0.00606459 | 0.02384844 | RAB28 | | 9364 |
| ENSG000002 | 4.16691199 | -1.037515105 | 0.37807527 | -2.7442025 | 0.00606581 | 0.02385093 | NA | NA | |
| ENSG000002 | 8.93366163 | -0.969189913 | 0.35319675 | -2.7440511 | 0.00606861 | 0.02385963 | NA | NA | |
| ENSG000001 | 193.013174 | 0.419989834 | 0.15307016 | 2.74377341 | 0.00607374 | 0.0238775 | GDPD2 | | 54857 |
| ENSG000001 | 110.111837 | 0.466670413 | 0.17009085 | 2.74365376 | 0.00607596 | 0.02388389 | ARHGAP27 | | 201176 |
| ENSG000001 | 143.110593 | 0.440286407 | 0.1604904 | 2.74338149 | 0.006081 | 0.02390139 | MTFR2 | | 113115 |
| ENSG000001 | 16.1098916 | -0.824171685 | 0.30043144 | -2.7432937 | 0.00608262 | 0.02390546 | MPPED1 | | 758 |
| ENSG000001 | 424.262406 | 0.407872151 | 0.14868333 | 2.74322714 | 0.00608386 | 0.02390799 | TAOK2 | | 9344 |
| ENSG000002 | 6.85078323 | -0.996758376 | 0.36339759 | -2.7428866 | 0.00609017 | 0.02393048 | NA | NA | |
| ENSG000001 | 5360.46461 | 0.469250499 | 0.17108864 | 2.74273321 | 0.00609302 | 0.0239347 | SON | | 6651 |
| ENSG000002 | 334.024059 | 0.335352894 | 0.12226871 | 2.74275315 | 0.00609265 | 0.0239347 | ZNF585B | | 92285 |

| | | | | | | | | |
|---|---|---|---|---|---|---|---|---|
| ENSG000002 | 12.7224203 | -0.916485793 | 0.33414586 | -2.7427717 | 0.0060923 | 0.0239347 | LOC101928597 | 101928597 |
| ENSG000001 | 674.884635 | 0.413532901 | 0.15077791 | 2.74266229 | 0.00609433 | 0.02393755 | CRBN | 51185 |
| ENSG000001 | 132.342904 | 0.505552666 | 0.1843323 | 2.74261577 | 0.0060952 | 0.02393862 | CCDC9 | 26093 |
| ENSG000001 | 435.911502 | 0.330695124 | 0.12058363 | 2.7424545 | 0.00609819 | 0.02394806 | C2orf49 | 79074 |
| ENSG000001 | 256.301016 | 0.455895466 | 0.16623885 | 2.74241237 | 0.00609897 | 0.02394882 | TP53I3 | 9540 |
| ENSG000001 | 3000.97624 | 0.433936348 | 0.15823675 | 2.74232344 | 0.00610062 | 0.02395298 | PPIG | 9360 |
| ENSG000002 | 17.984483 | -0.812007839 | 0.29610839 | -2.7422656 | 0.0061017 | 0.02395488 | NA | NA |
| ENSG000001 | 81.3390204 | 0.515431765 | 0.18799553 | 2.74172345 | 0.00611178 | 0.02399213 | DIABLO | 56616 |
| ENSG000002 | 3.74228752 | -1.033140144 | 0.3768532 | -2.7414923 | 0.00611608 | 0.02400669 | NA | NA |
| ENSG000001 | 88.1998353 | 0.492261379 | 0.17956253 | 2.74144818 | 0.0061169 | 0.02400759 | IFT43 | 112752 |
| ENSG000001 | 668.228991 | 0.452227418 | 0.1649694 | 2.74128058 | 0.00612002 | 0.02401752 | SMNDC1 | 10285 |
| ENSG000001 | 5.57511808 | -1.020219314 | 0.37218432 | -2.7411668 | 0.00612214 | 0.02402351 | ZNF676 | 163223 |
| ENSG000002 | 11.2370976 | -0.912608757 | 0.33295234 | -2.7409591 | 0.00612601 | 0.02403638 | NA | NA |
| ENSG000001 | 2043.52783 | 0.425537347 | 0.15526322 | 2.74074796 | 0.00612995 | 0.0240495 | UBE2I | 7329 |
| ENSG000002 | 12.1871639 | -0.893744305 | 0.32612891 | -2.7404633 | 0.00613526 | 0.02406801 | NA | NA |
| ENSG000001 | 25.5768741 | -0.733911771 | 0.26784167 | -2.7400955 | 0.00614213 | 0.02409263 | PNLIPRP1 | 5407 |
| ENSG000002 | 4.36376791 | -1.037178282 | 0.37853097 | -2.740009 | 0.00614375 | 0.02409664 | NA | NA |
| ENSG000001 | 4.03555637 | -1.036917177 | 0.37846123 | -2.7398241 | 0.00614721 | 0.02410787 | OR2T8 | 343172 |
| ENSG000001 | 19.3674118 | -0.811058027 | 0.29607299 | -2.7393854 | 0.00615542 | 0.02413773 | TGM5 | 9333 |
| ENSG000002 | 8.85460507 | -0.960584315 | 0.35067314 | -2.7392583 | 0.0061578 | 0.02414473 | NA | NA |
| ENSG000002 | 6.6148852 | -1.004586187 | 0.36677557 | -2.738967 | 0.00616326 | 0.0241638 | NA | NA |
| ENSG000002 | 25.8319476 | -0.709028046 | 0.25887794 | -2.7388508 | 0.00616543 | 0.02417001 | LINC01619 | 256021 |
| ENSG000002 | 59.1203249 | 0.557648982 | 0.20361625 | 2.73872531 | 0.00616779 | 0.02417689 | NA | NA |
| ENSG000001 | 329.871721 | 0.337711778 | 0.12331471 | 2.73861704 | 0.00616982 | 0.02418252 | ZNF606 | 80095 |
| ENSG000002 | 58.634584 | 0.553339295 | 0.20206721 | 2.73839236 | 0.00617404 | 0.02419671 | NA | NA |
| ENSG000001 | 103.822904 | 0.472068736 | 0.17239648 | 2.73827371 | 0.00617626 | 0.0242031 | ZFYVE19 | 84936 |
| ENSG000001 | 2475.35275 | 0.33907459 | 0.12383564 | 2.73810173 | 0.0061795 | 0.02421342 | TLK1 | 9874 |
| ENSG000001 | 9.41751739 | -0.968464285 | 0.35371394 | -2.7379873 | 0.00618165 | 0.02421951 | SBSN | 374897 |
| ENSG000001 | 5970.4738 | 0.300541955 | 0.10977289 | 2.73785217 | 0.00618419 | 0.02422664 | TJP1 | 7082 |

| | | | | | | | | |
|---|---|---|---|---|---|---|---|---|
| ENSG000001 | 9.3165029 | -0.958866772 | 0.35022914 | -2.7378269 | 0.00618466 | 0.02422664 | PHKG1 | 5260 |
| ENSG000002 | 3.97871753 | -1.036314213 | 0.37856381 | -2.7374888 | 0.00619102 | 0.02424922 | NA | NA |
| ENSG000001 | 16.0709154 | -0.822868877 | 0.30060613 | -2.7373655 | 0.00619334 | 0.02425596 | SFRP4 | 6424 |
| ENSG000002 | 7.45446295 | -0.998361605 | 0.36472742 | -2.7372814 | 0.00619493 | 0.02425983 | NA | NA |
| ENSG000002 | 138.298165 | 0.515736182 | 0.18844824 | 2.7367525 | 0.0062049 | 0.02429652 | NA | NA |
| ENSG000002 | 23.4529365 | -0.749953117 | 0.27404847 | -2.7365711 | 0.00620832 | 0.02430757 | NA | NA |
| ENSG000002 | 5.6381592 | -1.022596061 | 0.37368239 | -2.736538 | 0.00620894 | 0.02430766 | NA | NA |
| ENSG000002 | 46.9513326 | 0.600218128 | 0.21936168 | 2.73620324 | 0.00621526 | 0.02432771 | NA | NA |
| ENSG000002 | 2.43537144 | -0.99939448 | 0.36524553 | -2.7362265 | 0.00621482 | 0.02432771 | MIR4307 | 100423019 |
| ENSG000001 | 121.489679 | 0.437063988 | 0.15973628 | 2.73615977 | 0.00621608 | 0.0243283 | NLRP12 | 91662 |
| ENSG000001 | 7.3115852 | -0.992574907 | 0.36276997 | -2.7361 | 0.00621721 | 0.0243283 | TPPP3 | 51673 |
| ENSG000002 | 9.38564392 | -0.942678211 | 0.34453219 | -2.7361107 | 0.00621701 | 0.0243283 | NA | NA |
| ENSG000001 | 545.289146 | 0.322856739 | 0.11801248 | 2.73578468 | 0.00622317 | 0.02434928 | PIK3CA | 5290 |
| ENSG000002 | 6.4992296 | -1.013298451 | 0.37044409 | -2.7353614 | 0.00623118 | 0.02437826 | NA | NA |
| ENSG000001 | 320.827281 | 0.407587399 | 0.14903457 | 2.73485136 | 0.00624085 | 0.02441371 | ADAM19 | 8728 |
| ENSG000001 | 26.689702 | -0.736088613 | 0.26915744 | -2.7347883 | 0.00624204 | 0.02441576 | OR8D1 | 283159 |
| ENSG000001 | 19.3296821 | -0.790258707 | 0.28896819 | -2.7347602 | 0.00624257 | 0.02441576 | PRB3 | 5544 |
| ENSG000002 | 10.0191435 | 0.93607612 | 0.34230843 | 2.73459854 | 0.00624564 | 0.0244254 | NA | NA |
| ENSG000001 | 235.098164 | 0.423137007 | 0.1547523 | 2.73428573 | 0.00625158 | 0.02444626 | INTS12 | 57117 |
| ENSG00000C | 2226.24805 | 0.436284105 | 0.15956804 | 2.7341572 | 0.00625402 | 0.02445109 | PUM2 | 23369 |
| ENSG000002 | 5.71385981 | -1.020027871 | 0.37306551 | -2.734179 | 0.0062536 | 0.02445109 | NA | NA |
| ENSG000002 | 4.26932765 | -1.034694576 | 0.37844376 | -2.7340775 | 0.00625553 | 0.02445465 | NA | NA |
| ENSG000001 | 70.9218366 | -0.503620267 | 0.18420589 | -2.7340074 | 0.00625686 | 0.0244575 | PCDHB15 | 56121 |
| ENSG000001 | 12.5935763 | -0.899606695 | 0.32905424 | -2.7339161 | 0.0062586 | 0.02446192 | LOC93463 | 93463 |
| ENSG000001 | 1081.47942 | 0.493746469 | 0.18060675 | 2.73382066 | 0.00626041 | 0.02446666 | CTPS1 | 1503 |
| ENSG000001 | 650.809442 | 0.346671534 | 0.12682137 | 2.73354186 | 0.00626572 | 0.0244839 | SMAP1 | 60682 |
| ENSG000002 | 8.51477965 | -0.965559908 | 0.35322883 | -2.7335252 | 0.00626603 | 0.0244839 | NA | NA |
| ENSG000001 | 44.5472845 | -0.607752764 | 0.22234644 | -2.7333596 | 0.00626918 | 0.02449385 | NBPF20 | 100288142 |
| ENSG000001 | 15.5487398 | -0.849893231 | 0.31096441 | -2.7330884 | 0.00627435 | 0.02451167 | FAM89A | 375061 |

| | | | | | | | | | |
|---|---|---|---|---|---|---|---|---|---|
| ENSG000001 | 41.8404069 | -0.616074314 | 0.22545274 | -2.7326096 | 0.00628348 | 0.02454496 | ARHGEF37 | | 389337 |
| ENSG000001 | 4139.96853 | 0.321713477 | 0.11773304 | 2.73256739 | 0.00628428 | 0.02454574 | SMC3 | | 9126 |
| ENSG000001 | 34.1202555 | -0.667719468 | 0.24438212 | -2.7322762 | 0.00628984 | 0.02456508 | CHIA | | 27159 |
| ENSG000001 | 46.3300043 | 0.581713924 | 0.21291657 | 2.73212142 | 0.00629279 | 0.02457426 | IL23A | | 51561 |
| ENSG000002 | 4.32157813 | -1.034022188 | 0.37847394 | -2.7320829 | 0.00629353 | 0.02457473 | NA | NA | |
| ENSG000002 | 11.7996855 | -0.898561575 | 0.32889627 | -2.7320516 | 0.00629413 | 0.02457473 | NA | NA | |
| ENSG00000C | 133.660367 | 0.415268568 | 0.15200498 | 2.73194057 | 0.00629625 | 0.02458065 | CYBRD1 | | 79901 |
| ENSG00000C | 1842.70329 | 0.345970678 | 0.1266422 | 2.73187525 | 0.0062975 | 0.02458294 | DHX29 | | 54505 |
| ENSG00000C | 23.4110169 | -0.732607953 | 0.26817317 | -2.7318465 | 0.00629805 | 0.02458294 | ANGPT2 | | 285 |
| ENSG000002 | 11.0908711 | -0.920604814 | 0.3370029 | -2.7317415 | 0.00630005 | 0.0245884 | NA | NA | |
| ENSG000001 | 213.868134 | 0.499192885 | 0.18274622 | 2.73161811 | 0.00630241 | 0.02459525 | SETD1B | | 23067 |
| ENSG000001 | 24.4165672 | 0.739731549 | 0.27083624 | 2.73128716 | 0.00630875 | 0.02461522 | LINC01547 | | 84536 |
| ENSG000001 | 14.1485567 | -0.855772189 | 0.31332181 | -2.7312883 | 0.00630873 | 0.02461522 | GPR132 | | 29933 |
| ENSG000001 | 509.043373 | 0.336083361 | 0.12305465 | 2.73117157 | 0.00631096 | 0.02461715 | RTTN | | 25914 |
| ENSG000002 | 89.6281034 | -0.47233375 | 0.17294415 | -2.7311346 | 0.00631167 | 0.02461715 | PLCXD2 | | 257068 |
| ENSG000002 | 3.44907193 | -1.032976828 | 0.37820959 | -2.7312286 | 0.00630987 | 0.02461715 | NA | NA | |
| ENSG000002 | 5.28196562 | -1.02348162 | 0.37474225 | -2.7311615 | 0.00631115 | 0.02461715 | NA | NA | |
| ENSG000002 | 11.9364227 | -0.901311653 | 0.33003628 | -2.7309472 | 0.00631526 | 0.02462878 | HSD3B1 | | 3283 |
| ENSG000001 | 1857.26182 | 0.337301703 | 0.12351585 | 2.73083735 | 0.00631736 | 0.02463462 | CD46 | | 4179 |
| ENSG000002 | 10.4011196 | -0.929740489 | 0.34054487 | -2.7301556 | 0.00633044 | 0.02468325 | NA | NA | |
| ENSG000001 | 493.398481 | 0.37098172 | 0.13588857 | 2.73004366 | 0.00633259 | 0.02468926 | RPRD1B | | 58490 |
| ENSG000001 | 11.0248301 | -0.919327482 | 0.33675122 | -2.7299901 | 0.00633362 | 0.0246909 | CYP8B1 | | 1582 |
| ENSG00000C | 25.9659567 | -0.729054951 | 0.26706264 | -2.7299024 | 0.00633531 | 0.02469272 | GLP2R | | 9340 |
| ENSG000001 | 2088.48996 | 0.330164652 | 0.1209435 | 2.7299083 | 0.00633519 | 0.02469272 | RBFOX2 | | 23543 |
| ENSG000001 | 650.344771 | 0.320022725 | 0.11723638 | 2.72972201 | 0.00633878 | 0.02469911 | SGTB | | 54557 |
| ENSG000002 | 11.9669602 | -0.902132166 | 0.33048038 | -2.7297601 | 0.00633804 | 0.02469911 | NA | NA | |
| ENSG000002 | 4.5460733 | -1.029665045 | 0.37720251 | -2.7297407 | 0.00633842 | 0.02469911 | NA | NA | |
| ENSG000002 | 11.3920846 | -0.895433282 | 0.32805249 | -2.7295427 | 0.00634222 | 0.02471018 | NA | NA | |
| ENSG000002 | 18.8529037 | -0.815838783 | 0.29891146 | -2.729366 | 0.00634562 | 0.02472104 | NA | NA | |

| | | | | | | | | |
|---|---|---|---|---|---|---|---|---|
| ENSG000002 | 12.876726 | -0.909331166 | 0.33317928 | -2.7292548 | 0.00634776 | 0.02472701 | NA | NA |
| ENSG000002 | 13.0171648 | -0.878295461 | 0.32183514 | -2.7290228 | 0.00635223 | 0.02474203 | LOC101928322 | 101928322 |
| ENSG000002 | 10.0540877 | -0.953970478 | 0.34960022 | -2.7287468 | 0.00635755 | 0.02476038 | NA | NA |
| ENSG000001 | 65.0137365 | -0.544676411 | 0.19961267 | -2.7286665 | 0.0063591 | 0.02476402 | KCNA2 | 3737 |
| ENSG000001 | 16.6755579 | -0.833842475 | 0.30561189 | -2.7284359 | 0.00636354 | 0.02477896 | C8A | 731 |
| ENSG00000C | 80.6992282 | 0.552616053 | 0.20254218 | 2.72839985 | 0.00636424 | 0.02477929 | THAP3 | 90326 |
| ENSG000002 | 9.47332673 | -0.943333146 | 0.34575104 | -2.7283596 | 0.00636502 | 0.02477994 | NA | NA |
| ENSG000002 | 214.590729 | 0.45602674 | 0.16714773 | 2.72828559 | 0.00636645 | 0.02478312 | NA | NA |
| ENSG00000C | 168.281735 | 0.425352123 | 0.155908 | 2.72822515 | 0.00636761 | 0.02478528 | ACAP1 | 9744 |
| ENSG000002 | 47.9963874 | -0.589554981 | 0.21609741 | -2.7281909 | 0.00636828 | 0.02478548 | NA | NA |
| ENSG000002 | 22.502041 | -0.775392296 | 0.28422359 | -2.7281068 | 0.0063699 | 0.02478942 | LIPJ | 142910 |
| ENSG000001 | 2862.47577 | 0.408792078 | 0.1498569 | 2.727883 | 0.00637422 | 0.02480386 | HSPH1 | 10808 |
| ENSG000001 | 29.4896823 | -0.709319471 | 0.26002979 | -2.7278393 | 0.00637507 | 0.02480477 | PLA1A | 51365 |
| ENSG000002 | 2.92736065 | -1.011330318 | 0.37076744 | -2.7276676 | 0.00637838 | 0.0248153 | NA | NA |
| ENSG000001 | 13.7913989 | -0.857393258 | 0.31440633 | -2.7270229 | 0.00639086 | 0.02486145 | AMBN | 258 |
| ENSG00000C | 147.099448 | 0.42013902 | 0.15412121 | 2.72602983 | 0.00641012 | 0.02493398 | ENPP4 | 22875 |
| ENSG000002 | 20.8318936 | -0.799030752 | 0.29313471 | -2.7258142 | 0.00641431 | 0.02494789 | SLC26A4-AS1 | 286002 |
| ENSG000002 | 7.14687178 | -0.984141383 | 0.36105751 | -2.7257192 | 0.00641616 | 0.02495268 | NA | NA |
| ENSG000002 | 11.8373391 | -0.915895943 | 0.33602413 | -2.725685 | 0.00641682 | 0.02495286 | NA | NA |
| ENSG000002 | 16.5049395 | -0.829043429 | 0.30417588 | -2.7255397 | 0.00641965 | 0.02496146 | IQCA1L | 392843 |
| ENSG000002 | 3.82485276 | -1.031851283 | 0.37859234 | -2.7254943 | 0.00642053 | 0.02496249 | NA | NA |
| ENSG000001 | 75.2524159 | 0.511347721 | 0.18762392 | 2.72538663 | 0.00642262 | 0.02496794 | PARL | 55486 |
| ENSG000001 | 19.0114294 | -0.804020012 | 0.29501434 | -2.725359 | 0.00642316 | 0.02496794 | FAM216B | 144809 |
| ENSG000001 | 143.719701 | 0.441064876 | 0.16183934 | 2.72532543 | 0.00642381 | 0.02496809 | ZFP69 | 339559 |
| ENSG00000C | 589.600843 | 0.339270694 | 0.12449127 | 2.72525686 | 0.00642515 | 0.02497088 | HEATR5B | 54497 |
| ENSG000001 | 623.223796 | 0.360521941 | 0.13229807 | 2.72507331 | 0.00642872 | 0.02498237 | DEGS1 | 8560 |
| ENSG000001 | 531.526805 | 0.408213576 | 0.14982674 | 2.72457087 | 0.00643851 | 0.02501802 | TARS2 | 80222 |
| ENSG000002 | 30.3085064 | 0.662557513 | 0.24318892 | 2.724456 | 0.00644075 | 0.02502193 | NA | NA |
| ENSG000002 | 5.22819186 | -1.019293257 | 0.37412478 | -2.724474 | 0.0064404 | 0.02502193 | NA | NA |

| | | | | | | | | |
|---|---|---|---|---|---|---|---|---|
| ENSG000001 | 51.3922412 | 0.56092833 | 0.20589065 | 2.72439921 | 0.00644186 | 0.02502384 | ZFP36 | 7538 |
| ENSG000002 | 12.7156543 | -0.877186077 | 0.32198005 | -2.7243491 | 0.00644284 | 0.02502523 | NA | NA |
| ENSG000002 | 5.34796199 | -1.022468451 | 0.37531823 | -2.7242707 | 0.00644437 | 0.02502878 | NA | NA |
| ENSG000001 | 21.8613011 | -0.760548058 | 0.27918809 | -2.7241422 | 0.00644687 | 0.02503612 | NA | NA |
| ENSG000001 | 41.0925711 | -0.635049113 | 0.23312916 | -2.7240227 | 0.00644921 | 0.02504279 | CYP3A5 | 1577 |
| ENSG000001 | 142.322156 | 0.466400539 | 0.17121975 | 2.7239879 | 0.00644989 | 0.02504303 | GCH1 | 2643 |
| ENSG000001 | 459.868255 | 0.357662793 | 0.13130919 | 2.72382145 | 0.00645314 | 0.02505325 | ZNF83 | 55769 |
| ENSG000002 | 73.110119 | 0.552503271 | 0.20284668 | 2.72374819 | 0.00645457 | 0.02505641 | NA | NA |
| ENSG000002 | 7.41542947 | -0.974181732 | 0.35769115 | -2.7235276 | 0.00645888 | 0.02507075 | NA | NA |
| ENSG000001 | 26.5169932 | -0.733783766 | 0.26944187 | -2.7233472 | 0.00646241 | 0.02508204 | PATE1 | 160065 |
| ENSG000002 | 7.60697478 | -0.981584728 | 0.36048588 | -2.7229492 | 0.0064702 | 0.02510988 | NA | NA |
| ENSG000002 | 5.27994301 | -1.021079467 | 0.37499853 | -2.7228893 | 0.00647137 | 0.02511203 | NA | NA |
| ENSG000001 | 4.03869467 | -1.030402467 | 0.3784303 | -2.7228329 | 0.00647248 | 0.02511391 | LALBA | 3906 |
| ENSG000002 | 7.93652844 | -0.976993692 | 0.35882619 | -2.7227491 | 0.00647412 | 0.02511788 | NA | NA |
| ENSG00000C | 289.496577 | 0.360126798 | 0.13227265 | 2.7226097 | 0.00647685 | 0.02511926 | CASP8 | 841 |
| ENSG000001 | 320.543429 | 0.376881071 | 0.13842523 | 2.72263282 | 0.0064764 | 0.02511926 | SETD6 | 79918 |
| ENSG000002 | 14.0402598 | -0.864538202 | 0.31754455 | -2.7225729 | 0.00647757 | 0.02511926 | LOC151475 | 151475 |
| ENSG000002 | 56.3810983 | 0.575308275 | 0.21130873 | 2.7225959 | 0.00647712 | 0.02511926 | SNHG19 | 100507303 |
| ENSG000002 | 113.63059 | 0.448720208 | 0.16480939 | 2.72266171 | 0.00647583 | 0.02511926 | LINC00909 | 400657 |
| ENSG000001 | 2.78488666 | -1.011108756 | 0.37138578 | -2.7225296 | 0.00647842 | 0.02512015 | NA | NA |
| ENSG000001 | 41.4096997 | -0.643741966 | 0.23647167 | -2.7222795 | 0.00648333 | 0.02513677 | WDR63 | 126820 |
| ENSG000001 | 17.2557458 | -0.808795872 | 0.29711573 | -2.7221577 | 0.00648572 | 0.02514312 | LRRC74A | 145497 |
| ENSG000001 | 31.5476589 | -0.71255174 | 0.2617623 | -2.7221328 | 0.00648621 | 0.02514312 | CLVS1 | 157807 |
| ENSG000001 | 244.045544 | 0.379637234 | 0.13946849 | 2.72202877 | 0.00648825 | 0.02514863 | RIC3 | 79608 |
| ENSG000001 | 4.26669788 | -1.030358704 | 0.37853378 | -2.7219729 | 0.00648935 | 0.02515048 | CYP2G1P | 22952 |
| ENSG000002 | 14.4760359 | -0.888180926 | 0.32635434 | -2.7215233 | 0.00649818 | 0.02518231 | LOC100506274 | 100506274 |
| ENSG000002 | 11.7953977 | -0.89342156 | 0.32829618 | -2.7213889 | 0.00650082 | 0.02519015 | C5orf66-AS2 | 340073 |
| ENSG00000C | 318.026354 | 0.351059835 | 0.12900948 | 2.72119402 | 0.00650466 | 0.02520259 | NEBL | 10529 |
| ENSG000002 | 7.93740249 | -0.977432448 | 0.35921404 | -2.7210307 | 0.00650787 | 0.02521264 | NA | NA |

| | | | | | | | | |
|---|---|---|---|---|---|---|---|---|
| ENSG000001 | 896.850619 | 0.340309819 | 0.12507767 | 2.72078793 | 0.00651265 | 0.02522875 | DOCK4 | 9732 |
| ENSG000001 | 12.1872204 | -0.892493649 | 0.32804072 | -2.7206794 | 0.00651479 | 0.02523462 | IZUMO2 | 126123 |
| ENSG000001 | 237.315524 | 0.400761739 | 0.14730813 | 2.72056763 | 0.00651699 | 0.02524075 | BLVRA | 644 |
| ENSG000002 | 7.17279304 | -0.981458452 | 0.36077344 | -2.7204287 | 0.00651973 | 0.02524894 | NA | NA |
| ENSG000001 | 50.149219 | -0.578650944 | 0.21271334 | -2.7203322 | 0.00652164 | 0.0252539 | CCDC108 | 255101 |
| ENSG000001 | 324.756284 | 0.390920048 | 0.1437093 | 2.72021407 | 0.00652397 | 0.02526051 | PSMC3IP | 29893 |
| ENSG000002 | 6.15964249 | -1.015207242 | 0.37323248 | -2.7200399 | 0.0065274 | 0.02527141 | NA | NA |
| ENSG000002 | 97.0467359 | 0.521573308 | 0.19175697 | 2.71997055 | 0.00652877 | 0.0252743 | NA | NA |
| ENSG000001 | 925.924173 | 0.319048212 | 0.11730179 | 2.7198922 | 0.00653032 | 0.02527787 | CWF19L2 | 143884 |
| ENSG000002 | 16.0115087 | -0.842569 | 0.30980911 | -2.7196392 | 0.00653532 | 0.02529481 | FENDRR | 400550 |
| ENSG000001 | 403.329306 | 0.493977515 | 0.1816718 | 2.71906546 | 0.00654667 | 0.02533631 | CYTH2 | 9266 |
| ENSG000002 | 11.5610718 | -0.898769034 | 0.33062043 | -2.7184316 | 0.00655922 | 0.02538248 | NA | NA |
| ENSG000002 | 24.4808646 | 0.730819374 | 0.2688437 | 2.71838011 | 0.00656024 | 0.02538401 | NA | NA |
| ENSG000002 | 3.6757367 | -1.026043757 | 0.37750799 | -2.7179392 | 0.00656899 | 0.02541543 | NA | NA |
| ENSG000001 | 954.627453 | 0.365294761 | 0.13441358 | 2.71769239 | 0.00657389 | 0.02543197 | WDR89 | 112840 |
| ENSG000001 | 982.226295 | 0.332605294 | 0.12239534 | 2.71746706 | 0.00657837 | 0.02544687 | GEN1 | 348654 |
| ENSG000001 | 23.8621317 | -0.758367709 | 0.27907643 | -2.7174194 | 0.00657932 | 0.02544811 | FCRL2 | 79368 |
| ENSG00000C | 2271.9256 | 0.307428553 | 0.11314379 | 2.71714923 | 0.00658469 | 0.02546646 | DCBLD2 | 131566 |
| ENSG00000C | 123.66101 | 0.47242962 | 0.17390637 | 2.71657451 | 0.00659613 | 0.02550626 | OGFR | 11054 |
| ENSG000002 | 695.738285 | 0.321387951 | 0.11830656 | 2.71656913 | 0.00659624 | 0.02550626 | ERVMER34-1 | 100288413 |
| ENSG000001 | 21.2166152 | 0.750578166 | 0.27631994 | 2.71633733 | 0.00660086 | 0.0255217 | FGF21 | 26291 |
| ENSG000001 | 66.9175159 | 0.518253639 | 0.19082991 | 2.71578836 | 0.00661182 | 0.02556105 | CCDC142 | 84865 |
| ENSG000002 | 2.31250398 | -0.994912257 | 0.36634708 | -2.7157641 | 0.0066123 | 0.02556105 | NA | NA |
| ENSG000002 | 31.3318724 | 0.703566887 | 0.25907089 | 2.71573111 | 0.00661296 | 0.02556116 | NA | NA |
| ENSG000002 | 10.6549978 | -0.918603598 | 0.33830104 | -2.7153437 | 0.0066207 | 0.02558865 | NA | NA |
| ENSG00000C | 1888.70974 | 0.328673709 | 0.12105648 | 2.71504437 | 0.00662669 | 0.02560935 | SUCO | 51430 |
| ENSG00000C | 208.570705 | 0.495769281 | 0.18260456 | 2.71498848 | 0.00662781 | 0.02561123 | CROCC | 9696 |
| ENSG000001 | 5.14895815 | -1.015061332 | 0.37388986 | -2.7148672 | 0.00663023 | 0.02561817 | MAGEC3 | 139081 |
| ENSG000002 | 13.4545244 | -0.857847938 | 0.31598675 | -2.7148225 | 0.00663113 | 0.02561918 | LOC101928887 | 101928887 |

| | | | | | | | | |
|---|---|---|---|---|---|---|---|---|
| ENSG000001 | 229.987876 | 0.408800468 | 0.15059278 | 2.71460867 | 0.00663541 | 0.02563329 | RAD54B | 25788 |
| ENSG000002 | 18.3971829 | -0.788136517 | 0.29038916 | -2.7140701 | 0.00664621 | 0.02567256 | KIAA0125 | 9834 |
| ENSG000001 | 52.2416489 | -0.565530598 | 0.20838504 | -2.7138734 | 0.00665016 | 0.02568536 | SYNPO | 11346 |
| ENSG000002 | 5.38463985 | -1.018837089 | 0.37542956 | -2.7137903 | 0.00665183 | 0.02568935 | NA | NA |
| ENSG000001 | 2173.13318 | 0.327354158 | 0.12062812 | 2.71374657 | 0.0066527 | 0.0256903 | ATM | 472 |
| ENSG000001 | 9653.4112 | 0.333135306 | 0.12276087 | 2.71369287 | 0.00665378 | 0.02569201 | HNRNPH1 | 3187 |
| ENSG000001 | 31.1759114 | -0.675931598 | 0.24909949 | -2.7135005 | 0.00665765 | 0.02570204 | AQP7 | 364 |
| ENSG000002 | 10.9546098 | -0.938010188 | 0.34568122 | -2.7135121 | 0.00665741 | 0.02570204 | NA | NA |
| ENSG000002 | 6.93517417 | -1.006605587 | 0.37096733 | -2.7134615 | 0.00665843 | 0.02570262 | NA | NA |
| ENSG000002 | 10.5864699 | -0.917309924 | 0.33809026 | -2.7132102 | 0.00666348 | 0.02571967 | NA | NA |
| ENSG000002 | 6.25932161 | -1.001063052 | 0.36897054 | -2.7131246 | 0.0066652 | 0.02572387 | LOC100506142 | 100506142 |
| ENSG000002 | 4.95965433 | -1.021029579 | 0.37635506 | -2.7129423 | 0.00666887 | 0.02573557 | NA | NA |
| ENSG000002 | 14.1863766 | -0.874632217 | 0.32239805 | -2.7128955 | 0.00666981 | 0.02573676 | NA | NA |
| ENSG000002 | 4.95117264 | -1.021748966 | 0.37666868 | -2.7125933 | 0.0066759 | 0.02575779 | NA | NA |
| ENSG000002 | 7.51630708 | -0.975400916 | 0.35959691 | -2.7124842 | 0.0066781 | 0.02576382 | OR3A2 | 4995 |
| ENSG000001 | 265.814902 | 0.366285806 | 0.13504893 | 2.71224507 | 0.00668292 | 0.02577506 | GPSM2 | 29899 |
| ENSG000001 | 817.573188 | 0.330527833 | 0.12186463 | 2.71225406 | 0.00668274 | 0.02577506 | RIOK1 | 83732 |
| ENSG000002 | 22.6470273 | -0.761081074 | 0.28060637 | -2.712273 | 0.00668235 | 0.02577506 | LINC00550 | 338862 |
| ENSG000001 | 1225.66652 | 0.294030312 | 0.10841648 | 2.71204437 | 0.00668696 | 0.02578822 | CBLL1 | 79872 |
| ENSG000001 | 279.240493 | 0.357524056 | 0.13185254 | 2.71154464 | 0.00669705 | 0.02582466 | MTHFD2L | 441024 |
| ENSG000002 | 2.86064238 | -1.005379891 | 0.37082604 | -2.7111901 | 0.00670422 | 0.02584984 | NA | NA |
| ENSG000002 | 14.157792 | -0.858369357 | 0.31664134 | -2.7108569 | 0.00671096 | 0.02587337 | NA | NA |
| ENSG000002 | 7.0373113 | -0.989930755 | 0.36517741 | -2.7108214 | 0.00671168 | 0.02587368 | NA | NA |
| ENSG000002 | 4.93713392 | -1.020400357 | 0.37642953 | -2.7107341 | 0.00671344 | 0.02587803 | NA | NA |
| ENSG000001 | 14.9589632 | -0.836577766 | 0.30866674 | -2.7102945 | 0.00672235 | 0.0259099 | OR4K13 | 390433 |
| ENSG000001 | 26.8419987 | -0.712449374 | 0.26287148 | -2.7102574 | 0.0067231 | 0.02591033 | CA5A | 763 |
| ENSG000001 | 7.77682005 | -0.965661348 | 0.35631157 | -2.7101599 | 0.00672508 | 0.02591548 | CATSPER1 | 117144 |
| ENSG000002 | 9.65383394 | -0.949034767 | 0.35019469 | -2.7100204 | 0.00672791 | 0.02592392 | NA | NA |
| ENSG000002 | 4.1628417 | -1.024990669 | 0.37823175 | -2.7099541 | 0.00672925 | 0.02592664 | NA | NA |

| | | | | | | | | |
|---|---|---|---|---|---|---|---|---|
| ENSG000001 | 69.3781865 | 0.581861196 | 0.21471835 | 2.70988115 | 0.00673073 | 0.02592988 | MACROD1 | 28992 |
| ENSG000001 | 24.4178879 | -0.749861733 | 0.2767275 | -2.7097478 | 0.00673344 | 0.02593652 | FGD2 | 221472 |
| ENSG000001 | 160.695918 | 0.494020488 | 0.18231333 | 2.70973325 | 0.00673373 | 0.02593652 | CNNM4 | 26504 |
| ENSG000001 | 7.45633462 | -0.971972793 | 0.35874881 | -2.7093408 | 0.0067417 | 0.02596475 | AMTN | 401138 |
| ENSG000002 | 8.66735152 | -0.944500727 | 0.34864905 | -2.7090299 | 0.00674802 | 0.02598662 | NA | NA |
| ENSG000001 | 28.2851246 | -0.747470928 | 0.27594963 | -2.7087224 | 0.00675428 | 0.02600826 | FAR2P1 | 440905 |
| ENSG000001 | 150.464319 | 0.443050789 | 0.16357931 | 2.70847688 | 0.00675928 | 0.02602256 | PNKD | 25953 |
| ENSG000002 | 13.1432031 | -0.861588871 | 0.31810703 | -2.7084874 | 0.00675907 | 0.02602256 | NA | NA |
| ENSG000001 | 662.104367 | 0.448891919 | 0.16574359 | 2.70835165 | 0.00676183 | 0.02602991 | UTP3 | 57050 |
| ENSG000001 | 773.183215 | 0.414826927 | 0.15317112 | 2.70825807 | 0.00676374 | 0.02603394 | ARHGEF2 | 9181 |
| ENSG000001 | 78.7601577 | 0.556281858 | 0.20540366 | 2.70823738 | 0.00676416 | 0.02603394 | SH2D5 | 400745 |
| ENSG000001 | 69.4737406 | -0.527081754 | 0.19464626 | -2.7078956 | 0.00677113 | 0.02605581 | ST8SIA5 | 29906 |
| ENSG000001 | 23.8637709 | -0.72177586 | 0.26654189 | -2.7079266 | 0.0067705 | 0.02605581 | CILP | 8483 |
| ENSG000001 | 27.8336169 | -0.693129465 | 0.25597002 | -2.707854 | 0.00677198 | 0.0260566 | DUSP15 | 128853 |
| ENSG000002 | 25.8132284 | 0.737464091 | 0.27236313 | 2.70765027 | 0.00677614 | 0.02607013 | CEBPD | 1052 |
| ENSG000001 | 5.72609881 | -1.01183412 | 0.37372731 | -2.7074129 | 0.00678099 | 0.02608631 | NA | NA |
| ENSG000001 | 123.16918 | 0.474171459 | 0.17515847 | 2.70709986 | 0.00678738 | 0.02610693 | SMG1P3 | 100271836 |
| ENSG000002 | 22.1064261 | -0.764040697 | 0.28223715 | -2.7070877 | 0.00678763 | 0.02610693 | NA | NA |
| ENSG000002 | 4.64583616 | -1.022730201 | 0.37787046 | -2.706563 | 0.00679837 | 0.02614574 | LOC105378656 | 105378656 |
| ENSG000001 | 1107.86648 | 0.308257084 | 0.11390638 | 2.70623183 | 0.00680515 | 0.02616934 | FNBP4 | 23360 |
| ENSG000002 | 19.2555329 | -0.781070875 | 0.28863605 | -2.7060753 | 0.00680836 | 0.0261792 | PCDHGA9 | 56107 |
| ENSG00000C | 442.055559 | 0.34104343 | 0.12603683 | 2.70590291 | 0.0068119 | 0.02618686 | BCAP29 | 55973 |
| ENSG000001 | 211.276948 | 0.3896913 | 0.14401628 | 2.70588372 | 0.00681229 | 0.02618686 | PDSS1 | 23590 |
| ENSG000002 | 7.20271983 | -1.009605492 | 0.37311087 | -2.7059128 | 0.00681169 | 0.02618686 | CTGLF12P | 414224 |
| ENSG000001 | 207.100422 | 0.374650985 | 0.1384725 | 2.70559856 | 0.00681814 | 0.02620687 | EGLN1 | 54583 |
| ENSG000002 | 4.95939396 | -1.018520297 | 0.37647306 | -2.7054268 | 0.00682167 | 0.02621794 | NA | NA |
| ENSG000002 | 6.55783032 | -0.991831571 | 0.36661649 | -2.7053654 | 0.00682293 | 0.0262203 | NA | NA |
| ENSG000001 | 37.4661137 | -0.686310262 | 0.25369312 | -2.7052774 | 0.00682474 | 0.02622476 | LEKR1 | 389170 |
| ENSG00000C | 22.1999154 | -0.754762991 | 0.27902225 | -2.705028 | 0.00682986 | 0.02623951 | CAMK2B | 816 |

| | | | | | | | | |
|---|---|---|---|---|---|---|---|---|
| ENSG000001 | 1133.49962 | 0.4279309 | 0.15819834 | 2.70502767 | 0.00682987 | 0.02623951 | SUZ12 | 23512 |
| ENSG000001 | 27.9114076 | -0.697273112 | 0.25779497 | -2.7047584 | 0.00683541 | 0.02625333 | CYTIP | 9595 |
| ENSG000001 | 64.66648 | -0.522349053 | 0.19311816 | -2.7048159 | 0.00683423 | 0.02625333 | SV2B | 9899 |
| ENSG000002 | 18.4322152 | -0.793303736 | 0.29329658 | -2.7047834 | 0.00683489 | 0.02625333 | NA | NA |
| ENSG000001 | 83.1200147 | -0.524081643 | 0.19377252 | -2.7046231 | 0.00683819 | 0.02626153 | CACNG8 | 59283 |
| ENSG000001 | 124.260727 | 0.44056501 | 0.16291362 | 2.704286 | 0.00684513 | 0.02628571 | SENP8 | 123228 |
| ENSG000000 | 3753.72015 | 0.37887989 | 0.14011201 | 2.70412152 | 0.00684852 | 0.02629374 | BAZ1B | 9031 |
| ENSG000002 | 8.97645417 | -0.973632914 | 0.36005251 | -2.7041415 | 0.00684811 | 0.02629374 | NA | NA |
| ENSG000002 | 1121.35955 | 0.323865561 | 0.11977137 | 2.70403164 | 0.00685038 | 0.02629609 | RBMXL1 | 494115 |
| ENSG000002 | 9.52593737 | -0.92183776 | 0.34091268 | -2.704029 | 0.00685043 | 0.02629609 | ANKRD34C-AS1 | 729911 |
| ENSG000002 | 3.15452993 | -1.01525195 | 0.37549332 | -2.7037816 | 0.00685553 | 0.02631317 | NA | NA |
| ENSG000000 | 1256.54255 | 0.361299898 | 0.13363668 | 2.70359838 | 0.00685931 | 0.02632344 | RIOK2 | 55781 |
| ENSG000002 | 2.29761595 | -0.988757593 | 0.36572036 | -2.7035892 | 0.0068595 | 0.02632344 | NA | NA |
| ENSG000002 | 3.60925053 | -1.019903937 | 0.3772465 | -2.7035478 | 0.00686036 | 0.02632422 | NA | NA |
| ENSG000001 | 14.6176128 | -0.838230427 | 0.31005874 | -2.7034569 | 0.00686224 | 0.02632893 | CELF3 | 11189 |
| ENSG000001 | 36.252608 | -0.647068355 | 0.23935518 | -2.7033815 | 0.00686379 | 0.02633242 | PAK7 | 57144 |
| ENSG000002 | 5.99466879 | -1.004525599 | 0.3716198 | -2.7031003 | 0.0068696 | 0.02635221 | NA | NA |
| ENSG000002 | 11.582453 | -0.903004953 | 0.3340753 | -2.7029983 | 0.00687171 | 0.02635781 | NA | NA |
| ENSG000001 | 49.3549231 | -0.588952826 | 0.21789155 | -2.7029631 | 0.00687244 | 0.0263581 | MYLK3 | 91807 |
| ENSG000001 | 14.7617066 | -0.842262032 | 0.31163266 | -2.7027399 | 0.00687705 | 0.02637331 | F13B | 2165 |
| ENSG000002 | 10.7985216 | -0.94127823 | 0.34828329 | -2.7026224 | 0.00687949 | 0.02638014 | NA | NA |
| ENSG000002 | 2.32673858 | -0.986494959 | 0.36502572 | -2.7025355 | 0.00688128 | 0.02638434 | LOC101927526 | 101927526 |
| ENSG000002 | 1535.02898 | 0.308111926 | 0.11400969 | 2.70250657 | 0.00688188 | 0.02638434 | LINC00665 | 100506930 |
| ENSG000002 | 25.6329228 | -0.713019324 | 0.26386441 | -2.7022186 | 0.00688785 | 0.02640471 | LOC101927164 | 101927164 |
| ENSG000001 | 443.675068 | 0.311604472 | 0.11533895 | 2.70164141 | 0.00689981 | 0.02644809 | KLF12 | 11278 |
| ENSG000001 | 64.7522975 | 0.506194603 | 0.18737045 | 2.70157114 | 0.00690127 | 0.02644868 | ZDHHC11 | 79844 |
| ENSG000002 | 17.9236517 | -0.802486922 | 0.29704443 | -2.701572 | 0.00690125 | 0.02644868 | NA | NA |
| ENSG000001 | 1021.23973 | 0.342376266 | 0.12673556 | 2.70150126 | 0.00690272 | 0.02645174 | ZNF730 | 100129543 |
| ENSG000001 | 25.5243453 | -0.72488966 | 0.26834247 | -2.7013601 | 0.00690565 | 0.02645546 | NA | NA |

| | | | | | | | | |
|---|---|---|---|---|---|---|---|---|
| ENSG000001 | 54.882635 | -0.549828422 | 0.20353679 | -2.7013712 | 0.00690542 | 0.02645546 | SYCP2 | 10388 |
| ENSG000002 | 4.88999174 | -1.017059356 | 0.37649382 | -2.7013972 | 0.00690488 | 0.02645546 | NA | NA |
| ENSG000002 | 33.928341 | 0.64229116 | 0.23777695 | 2.70123394 | 0.00690827 | 0.02646301 | NA | NA |
| ENSG000002 | 7.31324832 | -0.983395366 | 0.36407154 | -2.7011048 | 0.00691096 | 0.02647079 | NA | NA |
| ENSG000001 | 67.3041844 | -0.545242274 | 0.2018728 | -2.7009199 | 0.0069148 | 0.026483 | SVOPL | 136306 |
| ENSG000001 | 106.482848 | 0.463433715 | 0.17159426 | 2.70075295 | 0.00691827 | 0.02649379 | VKORC1 | 79001 |
| ENSG000001 | 121.590312 | 0.44243913 | 0.1638242 | 2.70069454 | 0.00691949 | 0.02649595 | SLC19A3 | 80704 |
| ENSG000001 | 119.10827 | 0.438859656 | 0.16250522 | 2.70058802 | 0.0069217 | 0.02650193 | NPR1 | 4881 |
| ENSG000002 | 7.34244785 | -0.984532262 | 0.36456956 | -2.7005334 | 0.00692284 | 0.02650378 | OR10G4 | 390264 |
| ENSG000002 | 3337.26358 | 0.298582824 | 0.11056846 | 2.700434 | 0.00692491 | 0.0265092 | RBM8A | 9939 |
| ENSG000001 | 1519.86119 | 0.332086285 | 0.1229805 | 2.7003165 | 0.00692735 | 0.02651356 | SKA2 | 348235 |
| ENSG000002 | 21.0756188 | -0.789641731 | 0.29242225 | -2.7003476 | 0.00692671 | 0.02651356 | OR5H1 | 26341 |
| ENSG00000C | 38.8458167 | -0.645005301 | 0.23889617 | -2.6999399 | 0.0069352 | 0.02654108 | ADGRF5 | 221395 |
| ENSG000001 | 8244.57184 | 0.297522596 | 0.11020471 | 2.69972675 | 0.00693964 | 0.02655558 | PRPF40A | 55660 |
| ENSG000001 | 681.077991 | 0.300990783 | 0.11149283 | 2.69964247 | 0.0069414 | 0.0265573 | SEC24B | 10427 |
| ENSG000002 | 4.66695835 | -1.020437395 | 0.37798818 | -2.6996542 | 0.00694116 | 0.0265573 | NA | NA |
| ENSG000001 | 46.9406465 | 0.590798699 | 0.21885102 | 2.69954743 | 0.00694339 | 0.02656238 | PPP1R13L | 10848 |
| ENSG000001 | 89.8579124 | -0.462455334 | 0.1713364 | -2.6991073 | 0.00695257 | 0.02659255 | CAPS2 | 84698 |
| ENSG000002 | 27.5146487 | 0.725411461 | 0.26875981 | 2.69910687 | 0.00695258 | 0.02659255 | NA | NA |
| ENSG000001 | 427.254607 | 0.353357495 | 0.13092073 | 2.69901874 | 0.00695443 | 0.02659709 | ZNF347 | 84671 |
| ENSG000002 | 26.1962739 | -0.719622353 | 0.26663021 | -2.6989528 | 0.0069558 | 0.02659985 | LINC01524 | 101927700 |
| ENSG000002 | 6.91541041 | -0.983688624 | 0.36455974 | -2.698292 | 0.00696963 | 0.0266502 | NA | NA |
| ENSG000002 | 2.29926664 | -0.986208073 | 0.36549856 | -2.6982543 | 0.00697042 | 0.0266507 | RNA5SP134 | 106480756 |
| ENSG000001 | 3165.49867 | 0.271485519 | 0.10062418 | 2.69801468 | 0.00697544 | 0.02666738 | TCERG1 | 10915 |
| ENSG000002 | 6.71924127 | -0.996383908 | 0.36933855 | -2.6977522 | 0.00698094 | 0.0266859 | NA | NA |
| ENSG000001 | 273.12819 | 0.407187561 | 0.15094161 | 2.69764943 | 0.00698309 | 0.02669162 | TRIM11 | 81559 |
| ENSG000001 | 198.156984 | 0.39486847 | 0.14638446 | 2.69747525 | 0.00698675 | 0.02670307 | HACD1 | 9200 |
| ENSG000001 | 749.786232 | 0.333157848 | 0.12351303 | 2.6973498 | 0.00698938 | 0.0267101 | MSH3 | 4437 |
| ENSG000001 | 152.517239 | 0.454637327 | 0.16855877 | 2.69720358 | 0.00699245 | 0.0267101 | AMN1 | 196394 |

| | | | | | | | | |
|---|---|---|---|---|---|---|---|---|
| ENSG000001 | 117.957311 | 0.453766076 | 0.16823335 | 2.6972421 | 0.00699164 | 0.0267101 | LYSMD1 | 388695 |
| ENSG000001 | 72.2668318 | -0.531163674 | 0.19693155 | -2.6971994 | 0.00699254 | 0.0267101 | MACROD2 | 140733 |
| ENSG000002 | 262.046115 | 0.395279986 | 0.14654713 | 2.69728923 | 0.00699065 | 0.0267101 | METTL6 | 131965 |
| ENSG000002 | 8.90424063 | -0.951275541 | 0.3526788 | -2.6972859 | 0.00699072 | 0.0267101 | NA | NA |
| ENSG000002 | 6.9091459 | -0.987315692 | 0.36606444 | -2.697109 | 0.00699444 | 0.02671483 | NA | NA |
| ENSG000002 | 8.83088599 | -0.968906371 | 0.35927747 | -2.6968192 | 0.00700053 | 0.02673558 | MIR4461 | 100616209 |
| ENSG000002 | 45.7533068 | 0.592033477 | 0.21953737 | 2.69673209 | 0.00700236 | 0.02674005 | LOC440173 | 440173 |
| ENSG000002 | 22.954717 | -0.724921277 | 0.26881981 | -2.696681 | 0.00700343 | 0.02674163 | NA | NA |
| ENSG000002 | 6.22808286 | -0.99258289 | 0.36814112 | -2.6962022 | 0.00701351 | 0.02677759 | NA | NA |
| ENSG000002 | 6.92862216 | -0.98443127 | 0.3651366 | -2.696063 | 0.00701644 | 0.02678626 | NA | NA |
| ENSG000000 | 810.45282 | 0.320088102 | 0.11873196 | 2.69588827 | 0.00702012 | 0.02679779 | ATP11B | 23200 |
| ENSG000001 | 1472.04744 | 0.675874364 | 0.25072978 | 2.69562862 | 0.00702559 | 0.02681616 | CHAF1A | 10036 |
| ENSG000001 | 55.1147253 | 0.557270986 | 0.20674617 | 2.69543559 | 0.00702967 | 0.02682413 | PABPC3 | 5042 |
| ENSG000001 | 6.91926032 | -0.982913292 | 0.36465343 | -2.6954725 | 0.00702889 | 0.02682413 | CT55 | 54967 |
| ENSG000002 | 20.9789653 | -0.778660189 | 0.2888784 | -2.6954601 | 0.00702915 | 0.02682413 | OR2J3 | 442186 |
| ENSG000001 | 3.06535107 | -1.013644295 | 0.37607122 | -2.695352 | 0.00703143 | 0.02682745 | ASIP | 434 |
| ENSG000001 | 75.8829293 | -0.523584378 | 0.19425601 | -2.6953317 | 0.00703186 | 0.02682745 | CAV2 | 858 |
| ENSG000002 | 13.4020916 | -0.897904313 | 0.33315241 | -2.6951758 | 0.00703515 | 0.02683748 | NA | NA |
| ENSG000000 | 2075.18711 | 0.358775929 | 0.13312329 | 2.69506504 | 0.00703749 | 0.02684387 | MTA3 | 57504 |
| ENSG000002 | 4.94393736 | -1.015056302 | 0.37665502 | -2.6949231 | 0.00704049 | 0.02685279 | OR5H15 | 403274 |
| ENSG000002 | 35.0003037 | -0.644559278 | 0.23918612 | -2.6948022 | 0.00704304 | 0.02686001 | LOC102546299 | 102546299 |
| ENSG000001 | 43.6189721 | -0.601106256 | 0.22306581 | -2.6947485 | 0.00704418 | 0.0268602 | CCDC158 | 339965 |
| ENSG000001 | 26.0894051 | -0.720262311 | 0.2672855 | -2.6947302 | 0.00704456 | 0.0268602 | OR5AU1 | 390445 |
| ENSG000002 | 6.26062335 | -0.997745335 | 0.37026133 | -2.6947057 | 0.00704508 | 0.0268602 | TMEM225 | 338661 |
| ENSG000002 | 8.72824901 | -0.948130653 | 0.35188014 | -2.6944705 | 0.00705006 | 0.02687664 | NA | NA |
| ENSG000001 | 20.779732 | -0.789982568 | 0.29319916 | -2.6943548 | 0.0070525 | 0.02688345 | CCDC83 | 220047 |
| ENSG000002 | 28.004868 | -0.682242905 | 0.25328358 | -2.6935931 | 0.00706864 | 0.02694242 | LOC100287290 | 100287290 |
| ENSG000002 | 7.50468797 | -0.969700893 | 0.36002806 | -2.6934036 | 0.00707266 | 0.02695521 | NA | NA |
| ENSG000002 | 3.94366312 | -1.019034571 | 0.37841524 | -2.6929005 | 0.00708334 | 0.02699338 | NA | NA |

| | | | | | | | | |
|---|---|---|---|---|---|---|---|---|
| ENSG000001 | 34.7485686 | -0.649827724 | 0.2413734 | -2.6922093 | 0.00709804 | 0.02704685 | PTPN22 | 26191 |
| ENSG000002 | 5.49644552 | -1.003397128 | 0.37271926 | -2.6920989 | 0.00710039 | 0.02705326 | NA | NA |
| ENSG000001 | 7392.12076 | 0.33959848 | 0.12614796 | 2.69206475 | 0.00710112 | 0.02705349 | USP44 | 84101 |
| ENSG000002 | 912.268625 | 0.709765835 | 0.26365464 | 2.69202861 | 0.00710189 | 0.02705388 | VARS | 7407 |
| ENSG000002 | 3.19094665 | -1.010238719 | 0.3752978 | -2.6918322 | 0.00710607 | 0.02706727 | NA | NA |
| ENSG00000C | 235.859054 | 0.444160022 | 0.1650155 | 2.69162612 | 0.00711046 | 0.02708146 | SYT13 | 57586 |
| ENSG000001 | 934.423095 | 0.415388437 | 0.15433066 | 2.69154831 | 0.00711212 | 0.02708523 | MSL2 | 55167 |
| ENSG000001 | 26.4969946 | -0.733959428 | 0.27270132 | -2.6914407 | 0.00711441 | 0.02708634 | RS1 | 6247 |
| ENSG000001 | 9.75363882 | -0.928145199 | 0.34484685 | -2.6914707 | 0.00711377 | 0.02708634 | C1orf100 | 200159 |
| ENSG000002 | 168.678606 | 0.396498822 | 0.14731531 | 2.69149771 | 0.0071132 | 0.02708634 | NA | NA |
| ENSG000001 | 30.076672 | 0.666594925 | 0.24769678 | 2.69117313 | 0.00712012 | 0.02710553 | SULT1A1 | 6817 |
| ENSG000002 | 5.27603875 | -1.016936107 | 0.37788824 | -2.6911028 | 0.00712162 | 0.0271087 | NA | NA |
| ENSG000002 | 3.52663222 | -1.016239725 | 0.37765275 | -2.6909369 | 0.00712517 | 0.02711964 | NA | NA |
| ENSG000002 | 2.4529865 | -0.976944006 | 0.36308073 | -2.6907074 | 0.00713007 | 0.0271284 | RNU6-933P | 106479979 |
| ENSG000002 | 4.6428523 | -1.017255364 | 0.37805053 | -2.690792 | 0.00712826 | 0.0271284 | NA | NA |
| ENSG000002 | 13.0552446 | -0.871719262 | 0.32397732 | -2.6906799 | 0.00713066 | 0.0271284 | NA | NA |
| ENSG000002 | 9.55092444 | -0.919511719 | 0.3417372 | -2.6906983 | 0.00713026 | 0.0271284 | NA | NA |
| ENSG000002 | 28.5523353 | 0.723577056 | 0.26892052 | 2.69067256 | 0.00713081 | 0.0271284 | NA | NA |
| ENSG000002 | 92.164094 | 0.49352456 | 0.18343176 | 2.69050774 | 0.00713434 | 0.02713672 | NA | NA |
| ENSG000002 | 2.73384844 | -1.001209655 | 0.37212439 | -2.6905241 | 0.00713399 | 0.02713672 | NA | NA |
| ENSG000002 | 2.78908106 | 0.978803053 | 0.36382896 | 2.69028351 | 0.00713913 | 0.02714987 | RNA5S9 | 100169760 |
| ENSG000002 | 3.6356605 | -1.01516366 | 0.37734083 | -2.6903096 | 0.00713858 | 0.02714987 | NA | NA |
| ENSG000001 | 17.2861639 | -0.825908597 | 0.30700066 | -2.6902503 | 0.00713985 | 0.02715002 | CTSE | 1510 |
| ENSG000002 | 15.5325206 | -0.824407425 | 0.30644812 | -2.6902022 | 0.00714087 | 0.02715139 | IGFBP7-AS1 | 255130 |
| ENSG000001 | 88.8341835 | 0.508003208 | 0.18884289 | 2.6900839 | 0.00714341 | 0.02715847 | TMUB1 | 83590 |
| ENSG00000C | 1068.43439 | 0.34132842 | 0.12688971 | 2.68996147 | 0.00714603 | 0.02716589 | MRE11A | 4361 |
| ENSG00000C | 1686.13901 | 0.32977817 | 0.12260134 | 2.68984157 | 0.00714859 | 0.02717056 | ZNF268 | 10795 |
| ENSG000001 | 384.979886 | 0.376581523 | 0.14000009 | 2.68986632 | 0.00714806 | 0.02717056 | LRAT | 9227 |
| ENSG000001 | 27.6221063 | -0.697941959 | 0.25948264 | -2.6897444 | 0.00715068 | 0.02717338 | DAW1 | 164781 |

| | | | | | | | | |
|---|---|---|---|---|---|---|---|---|
| ENSG000002 | 223.028701 | 0.384003756 | 0.14276448 | 2.68977094 | 0.00715011 | 0.02717338 | ZNF155 | 7711 |
| ENSG000001 | 35.8090925 | -0.631389868 | 0.23476592 | -2.6894443 | 0.00715711 | 0.02719527 | NAT2 | 10 |
| ENSG000002 | 14.8755775 | -0.827071449 | 0.30756934 | -2.6890569 | 0.00716542 | 0.0272243 | NA | NA |
| ENSG000001 | 188.759473 | 0.394839126 | 0.14684723 | 2.68877485 | 0.00717148 | 0.02724476 | PEX14 | 5195 |
| ENSG000001 | 1625.81978 | 0.376223447 | 0.13993334 | 2.68859052 | 0.00717544 | 0.02725725 | FKBP3 | 2287 |
| ENSG000001 | 90.7100573 | 0.504906518 | 0.18781303 | 2.68834657 | 0.00718068 | 0.02727462 | PLBD2 | 196463 |
| ENSG000001 | 7.9671012 | -0.967519952 | 0.35990603 | -2.6882571 | 0.00718261 | 0.02727938 | XKR3 | 150165 |
| ENSG000001 | 29.2834748 | -0.694144496 | 0.258229 | -2.6880966 | 0.00718606 | 0.02728993 | AKR1D1 | 6718 |
| ENSG000002 | 5.55923279 | -1.002597056 | 0.37301978 | -2.6877852 | 0.00719276 | 0.02731284 | NA | NA |
| ENSG000001 | 16.2412427 | -0.809480564 | 0.30122883 | -2.6872612 | 0.00720406 | 0.02735316 | FAM217A | 222826 |
| ENSG000001 | 29.5356619 | -0.699777607 | 0.26042034 | -2.6871081 | 0.00720736 | 0.02736315 | EXD1 | 161829 |
| ENSG000001 | 8.44002678 | -0.948560091 | 0.35303214 | -2.6868944 | 0.00721197 | 0.02737342 | ADCYAP1 | 116 |
| ENSG000001 | 37.829668 | -0.624066425 | 0.23226008 | -2.6869293 | 0.00721122 | 0.02737342 | MEI1 | 150365 |
| ENSG000002 | 5.79985511 | -0.993582179 | 0.36978908 | -2.6868889 | 0.00721209 | 0.02737342 | HOXA10 | 3206 |
| ENSG000001 | 44.9484159 | 0.592686918 | 0.22060045 | 2.68669857 | 0.0072162 | 0.02738646 | MOB3A | 126308 |
| ENSG000002 | 4.93847642 | -1.012672558 | 0.37693852 | -2.6865722 | 0.00721893 | 0.02739425 | SPRR2E | 6704 |
| ENSG000001 | 193.994895 | 0.38009433 | 0.14149291 | 2.68631355 | 0.00722453 | 0.02741291 | TRAPPC2 | 6399 |
| ENSG000002 | 10.2208836 | -0.90881286 | 0.33832224 | -2.6862344 | 0.00722624 | 0.02741684 | ITGB2-AS1 | 100505746 |
| ENSG000001 | 525.111129 | 0.354709013 | 0.13205288 | 2.68611332 | 0.00722886 | 0.02742421 | USP31 | 57478 |
| ENSG000001 | 215.421119 | 0.474895991 | 0.17680183 | 2.68603552 | 0.00723054 | 0.02742803 | SLX4 | 84464 |
| ENSG000001 | 22.2036236 | 0.774492579 | 0.2883598 | 2.68585488 | 0.00723445 | 0.0274389 | CD52 | 1043 |
| ENSG000002 | 24.554236 | -0.749196447 | 0.27894301 | -2.6858406 | 0.00723476 | 0.0274389 | OR2F1 | 26211 |
| ENSG000001 | 301.121152 | 0.364557792 | 0.13574423 | 2.68562284 | 0.00723948 | 0.02745422 | ZFAND4 | 93550 |
| ENSG000002 | 2.30750006 | -0.980109777 | 0.36498931 | -2.6853109 | 0.00724624 | 0.02747729 | NA | NA |
| ENSG000001 | 18.3396407 | -0.782324095 | 0.29136908 | -2.6849936 | 0.00725312 | 0.02750082 | ESM1 | 11082 |
| ENSG000001 | 33.0932365 | -0.662157488 | 0.24662126 | -2.6849165 | 0.00725479 | 0.02750459 | DAO | 1610 |
| ENSG000002 | 3.8455216 | -1.016347209 | 0.37855783 | -2.6847872 | 0.0072576 | 0.02751266 | NA | NA |
| ENSG000001 | 121.118231 | 0.435976193 | 0.16239141 | 2.68472452 | 0.00725896 | 0.02751525 | PCGF5 | 84333 |
| ENSG000002 | 513.405735 | -0.314543345 | 0.11716188 | -2.6846901 | 0.00725971 | 0.02751551 | NA | NA |

| | | | | | | | | | |
|---|---|---|---|---|---|---|---|---|---|
| ENSG000002 | 44.9977085 | 0.671218582 | 0.25004988 | 2.68433876 | 0.00726734 | 0.02754187 | RNU5B-1 | | 26832 |
| ENSG000002 | 3.08859571 | -1.009131209 | 0.37594266 | -2.684269 | 0.00726886 | 0.02754505 | NA | NA | |
| ENSG000001 | 368.156638 | 0.356136217 | 0.13268659 | 2.68404076 | 0.00727382 | 0.02756108 | NDUFAF5 | | 79133 |
| ENSG000001 | 3.54246544 | -1.012788291 | 0.37734121 | -2.6840119 | 0.00727445 | 0.02756108 | CCDC42 | | 146849 |
| ENSG000001 | 212.218579 | 0.393942188 | 0.14679082 | 2.68369771 | 0.00728129 | 0.02758442 | TM2D3 | | 80213 |
| ENSG000001 | 953.959118 | 0.367861941 | 0.13707552 | 2.68364429 | 0.00728245 | 0.02758625 | ID1 | | 3397 |
| ENSG000001 | 9.49818656 | -0.938281369 | 0.3496632 | -2.6833861 | 0.00728808 | 0.0276024 | CLEC10A | | 10462 |
| ENSG000001 | 3.42323086 | -1.014130847 | 0.37792729 | -2.683402 | 0.00728773 | 0.0276024 | ZNF733P | | 643955 |
| ENSG000001 | 4076.04006 | 0.313673449 | 0.11689766 | 2.68331678 | 0.00728959 | 0.02760297 | ARHGAP5 | | 394 |
| ENSG000002 | 330.64056 | 0.391990233 | 0.14608259 | 2.68334662 | 0.00728894 | 0.02760297 | PIGW | | 284098 |
| ENSG000002 | 3.49230027 | -1.013774064 | 0.37783945 | -2.6830815 | 0.00729472 | 0.02761981 | NA | NA | |
| ENSG000001 | 285.704999 | 0.371822794 | 0.13858702 | 2.68295532 | 0.00729747 | 0.02762436 | B4GALT3 | | 8703 |
| ENSG000002 | 7.67917283 | -0.973644431 | 0.3628984 | -2.682967 | 0.00729722 | 0.02762436 | NA | NA | |
| ENSG000002 | 7.45840496 | 0.967806559 | 0.36072709 | 2.68293286 | 0.00729796 | 0.02762436 | NA | NA | |
| ENSG000002 | 43.7973515 | 0.595133446 | 0.22182712 | 2.68287051 | 0.00729932 | 0.02762693 | NA | NA | |
| ENSG000002 | 12.4436728 | -0.862120073 | 0.32137871 | -2.6825675 | 0.00730594 | 0.02764939 | NA | NA | |
| ENSG000002 | 24.7089197 | -0.699606226 | 0.26081449 | -2.6823902 | 0.00730982 | 0.02765889 | NA | NA | |
| ENSG000002 | 295.893038 | 0.351567873 | 0.13106372 | 2.68241936 | 0.00730918 | 0.02765889 | DGKK | | 139189 |
| ENSG000002 | 5.25671851 | -1.006447253 | 0.3752168 | -2.6823086 | 0.0073116 | 0.02766305 | RPL31P11 | | 641311 |
| ENSG000001 | 179.977076 | 0.398804908 | 0.14869063 | 2.68211197 | 0.0073159 | 0.02767674 | ACTRT3 | | 84517 |
| ENSG000002 | 8.01969974 | -0.948748749 | 0.35373893 | -2.6820592 | 0.00731705 | 0.02767852 | NA | NA | |
| ENSG000001 | 501.620504 | 0.329520885 | 0.12287203 | 2.68182173 | 0.00732225 | 0.02769559 | PRUNE | | 58497 |
| ENSG000001 | 15.6864022 | -0.871319929 | 0.32490975 | -2.6817291 | 0.00732427 | 0.02769809 | FGF23 | | 8074 |
| ENSG000002 | 166.865103 | 0.418319742 | 0.15598877 | 2.68172994 | 0.00732426 | 0.02769809 | NA | NA | |
| ENSG000001 | 142.231578 | 0.41756127 | 0.15571459 | 2.68158091 | 0.00732752 | 0.02770778 | FKBP9 | | 11328 |
| ENSG000001 | 2.69998248 | -0.997040962 | 0.37185303 | -2.681277 | 0.00733418 | 0.02773037 | NA | NA | |
| ENSG000002 | 55.3709504 | 0.59390206 | 0.22153332 | 2.68087016 | 0.0073431 | 0.02776152 | NA | NA | |
| ENSG000001 | 12.7922436 | -0.883035452 | 0.3294331 | -2.6804697 | 0.00735189 | 0.02779216 | DSCR4 | | 10281 |
| ENSG000001 | 2730.4871 | 0.330986933 | 0.12348786 | 2.6803196 | 0.00735519 | 0.02780204 | Mar-07 | | 64844 |

| | | | | | | | | |
|---|---|---|---|---|---|---|---|---|
| ENSG000002 | 3.80956775 | -1.014606332 | 0.37855148 | -2.6802335 | 0.00735708 | 0.02780661 | NA | NA |
| ENSG000002 | 26.9106138 | -0.693641935 | 0.25880848 | -2.680136 | 0.00735923 | 0.02781211 | FPGT-TNNI3K | 100526835 |
| ENSG000001 | 15.0395604 | -0.819706341 | 0.30584955 | -2.6800966 | 0.00736009 | 0.02781279 | FCMR | 9214 |
| ENSG00000C | 1342.05354 | 0.32482894 | 0.12120618 | 2.67997003 | 0.00736288 | 0.02782072 | PSMD5 | 5711 |
| ENSG00000C | 115.381481 | 0.506018284 | 0.18883264 | 2.67971836 | 0.00736841 | 0.02782349 | PTPN23 | 25930 |
| ENSG00000C | 75.1097777 | -0.511951475 | 0.1910348 | -2.679886 | 0.00736472 | 0.02782349 | RBFOX1 | 54715 |
| ENSG000001 | 72.0790556 | 0.498387339 | 0.18598363 | 2.67973768 | 0.00736799 | 0.02782349 | HRASLS | 57110 |
| ENSG000001 | 17.836767 | -0.807862629 | 0.30147238 | -2.6797235 | 0.0073683 | 0.02782349 | KLF17 | 128209 |
| ENSG000001 | 188.992536 | 0.387677123 | 0.14466754 | 2.67977961 | 0.00736706 | 0.02782349 | ZSCAN26 | 7741 |
| ENSG000002 | 21.5144448 | -0.736182557 | 0.27470881 | -2.6798651 | 0.00736518 | 0.02782349 | NA | NA |
| ENSG000002 | 13.7242603 | -0.864715144 | 0.32267802 | -2.679808 | 0.00736644 | 0.02782349 | NA | NA |
| ENSG000002 | 11.3303246 | -0.903745977 | 0.33725872 | -2.6796816 | 0.00736922 | 0.02782396 | LINC01085 | 152742 |
| ENSG000002 | 4.57729023 | -1.012602083 | 0.37791636 | -2.6794344 | 0.00737467 | 0.02784192 | NA | NA |
| ENSG000002 | 71.5981981 | 0.521947142 | 0.19480643 | 2.67931161 | 0.00737737 | 0.02784954 | NA | NA |
| ENSG000002 | 266.043394 | 0.39978364 | 0.14921667 | 2.67921565 | 0.00737949 | 0.02785234 | C22orf29 | 79680 |
| ENSG000002 | 4.064946 | -1.012545031 | 0.37792342 | -2.6792333 | 0.0073791 | 0.02785234 | LINC01333 | 101929082 |
| ENSG000002 | 20.0668509 | -0.766279177 | 0.28601812 | -2.6791281 | 0.00738141 | 0.02785703 | NA | NA |
| ENSG000002 | 61.4368261 | -0.539499951 | 0.20138681 | -2.6789239 | 0.00738592 | 0.02787142 | ZRANB2-AS2 | 100852410 |
| ENSG000002 | 6.75310114 | -0.989980938 | 0.36955981 | -2.6788111 | 0.00738841 | 0.02787822 | NA | NA |
| ENSG000001 | 28.2777349 | -0.75578036 | 0.28214003 | -2.678742 | 0.00738993 | 0.02788138 | FCRL1 | 115350 |
| ENSG000001 | 157.180484 | 0.450712421 | 0.16828536 | 2.6782628 | 0.00740051 | 0.02791743 | VAMP8 | 8673 |
| ENSG000001 | 139.165171 | 0.435453115 | 0.16258886 | 2.67824695 | 0.00740086 | 0.02791743 | C19orf47 | 126526 |
| ENSG000001 | 75.089525 | -0.5048005 | 0.18848617 | -2.6781832 | 0.00740227 | 0.02792015 | INHBA | 3624 |
| ENSG000001 | 302.57222 | 0.354628667 | 0.13241773 | 2.67810556 | 0.00740399 | 0.02792079 | DCTN6 | 10671 |
| ENSG000001 | 71.8561316 | 0.486441613 | 0.18163805 | 2.67808212 | 0.00740451 | 0.02792079 | USP18 | 11274 |
| ENSG000002 | 25.4563441 | -0.699567214 | 0.26121704 | -2.6781071 | 0.00740395 | 0.02792079 | LOC101929741 | 101929741 |
| ENSG000002 | 7.40640732 | -0.96272128 | 0.35948904 | -2.6780268 | 0.00740573 | 0.0279228 | NA | NA |
| ENSG000001 | 232.389569 | 0.396913267 | 0.14821515 | 2.67795344 | 0.00740735 | 0.02792632 | CALN1 | 83698 |
| ENSG000002 | 5.83605581 | -0.993143757 | 0.37088507 | -2.6777669 | 0.00741148 | 0.02793928 | NA | NA |

| | | | | | | | | |
|---|---|---|---|---|---|---|---|---|
| ENSG00000C | 153.138753 | 0.455085213 | 0.16995892 | 2.67761892 | 0.00741475 | 0.02794756 | SLC46A1 | 113235 |
| ENSG000002 | 5.85005915 | -0.994027558 | 0.37123752 | -2.6776053 | 0.00741505 | 0.02794756 | NA | NA |
| ENSG000001 | 29.1424212 | -0.668721788 | 0.24975682 | -2.6774916 | 0.00741757 | 0.02795445 | PLEKHD1 | 400224 |
| ENSG000001 | 11.0454793 | -0.886466605 | 0.33111547 | -2.6772129 | 0.00742374 | 0.02797118 | CYP4A11 | 1579 |
| ENSG000002 | 8.87896072 | -0.945220838 | 0.353058 | -2.6772395 | 0.00742315 | 0.02797118 | NA | NA |
| ENSG000002 | 5.64888397 | -1.000403689 | 0.37367568 | -2.6771978 | 0.00742408 | 0.02797118 | NA | NA |
| ENSG000002 | 3.08259398 | -1.006187347 | 0.37585131 | -2.6770888 | 0.00742649 | 0.02797768 | NA | NA |
| ENSG000002 | 6.25834343 | -0.990724415 | 0.3701043 | -2.676879 | 0.00743115 | 0.02799261 | LINC00620 | 285375 |
| ENSG000002 | 224.900696 | 0.396259703 | 0.14805595 | 2.67641872 | 0.00744136 | 0.02802848 | NA | NA |
| ENSG000002 | 3.46771814 | -1.011171732 | 0.3778423 | -2.676174 | 0.0074468 | 0.02804636 | RNF223 | 401934 |
| ENSG000002 | 5.21735506 | -1.003735211 | 0.3750712 | -2.6761191 | 0.00744802 | 0.02804834 | NA | NA |
| ENSG000001 | 39.1846158 | -0.631651788 | 0.23603843 | -2.6760549 | 0.00744944 | 0.02805111 | IL18 | 3606 |
| ENSG000002 | 8.89893795 | -0.927549696 | 0.34662991 | -2.6759078 | 0.00745271 | 0.02805821 | NA | NA |
| ENSG000002 | 6.60220026 | -0.984592721 | 0.36794614 | -2.6759153 | 0.00745255 | 0.02805821 | LOC101060498 | 101060498 |
| ENSG000001 | 281.913019 | 0.42795051 | 0.15993819 | 2.67572438 | 0.00745679 | 0.02806836 | TTC28 | 23331 |
| ENSG000001 | 62.4825619 | -0.524260236 | 0.19593175 | -2.6757289 | 0.00745669 | 0.02806836 | DNAH1 | 25981 |
| ENSG000002 | 8.32034098 | -0.940630834 | 0.35157009 | -2.6755144 | 0.00746147 | 0.02808335 | NA | NA |
| ENSG00000C | 4864.52349 | 0.482730699 | 0.18043403 | 2.67538608 | 0.00746432 | 0.02808911 | HSPA5 | 3309 |
| ENSG000002 | 11.5420969 | 0.887040787 | 0.33155651 | 2.6753834 | 0.00746438 | 0.02808911 | NA | NA |
| ENSG000001 | 220.976811 | 0.426555424 | 0.15945416 | 2.67509747 | 0.00747075 | 0.02811047 | CNTROB | 116840 |
| ENSG000001 | 2949.52732 | 0.31385111 | 0.11732893 | 2.67496772 | 0.00747364 | 0.02811874 | SMAD2 | 4087 |
| ENSG00000C | 75.0571947 | -0.514891788 | 0.1924999 | -2.674764 | 0.00747819 | 0.02813322 | KIAA2022 | 340533 |
| ENSG000001 | 140.365113 | 0.418250826 | 0.15638927 | 2.67442156 | 0.00748583 | 0.02815935 | SCLY | 51540 |
| ENSG000001 | 364.312756 | 0.391700745 | 0.14646598 | 2.67434632 | 0.00748751 | 0.02816306 | TRPC4 | 7223 |
| ENSG000002 | 11.1264787 | -0.910936739 | 0.34062459 | -2.6743129 | 0.00748825 | 0.02816325 | NA | NA |
| ENSG000001 | 45.6648874 | -0.697121175 | 0.26069751 | -2.6740615 | 0.00749387 | 0.02818176 | GABRG3 | 2567 |
| ENSG000001 | 2654.38955 | 0.437877686 | 0.1637596 | 2.67390546 | 0.00749736 | 0.02819226 | LUC7L3 | 51747 |
| ENSG000002 | 13.2280318 | -0.862642086 | 0.32263658 | -2.6737268 | 0.00750135 | 0.02820467 | LOC283299 | 283299 |
| ENSG000002 | 189.752172 | 0.375342819 | 0.14039214 | 2.67353165 | 0.00750572 | 0.02821585 | GTF2H5 | 404672 |

| | | | | | | | | |
|---|---|---|---|---|---|---|---|---|
| ENSG000002 | 3.44931987 | -1.009989752 | 0.37777325 | -2.6735343 | 0.00750566 | 0.02821585 | NA | NA |
| ENSG000002 | 8.92758029 | -0.932872482 | 0.34893297 | -2.6735006 | 0.00750642 | 0.02821585 | NA | NA |
| ENSG000001 | 60.3586071 | 0.579158932 | 0.21663398 | 2.67344456 | 0.00750767 | 0.02821795 | NA | NA |
| ENSG000001 | 246.873217 | 0.414256395 | 0.15496095 | 2.67329531 | 0.00751101 | 0.02822789 | ZBTB4 | 57659 |
| ENSG000002 | 6.91815043 | -0.978892216 | 0.36619113 | -2.6731729 | 0.00751375 | 0.02823558 | NA | NA |
| ENSG000002 | 10.1344594 | -0.905066175 | 0.33859201 | -2.6730287 | 0.00751698 | 0.0282451 | NA | NA |
| ENSG000002 | 28.2866003 | -0.667024463 | 0.24959751 | -2.6724003 | 0.00753107 | 0.02829543 | NA | NA |
| ENSG000002 | 10.6111222 | -0.905831157 | 0.33897427 | -2.6722711 | 0.00753397 | 0.02830371 | NA | NA |
| ENSG000001 | 693.849841 | 0.380075021 | 0.14224541 | 2.67196695 | 0.00754081 | 0.02832675 | PCCA | 5095 |
| ENSG000000 | 3031.22852 | 0.334108096 | 0.1250501 | 2.6717939 | 0.0075447 | 0.02833874 | ANLN | 54443 |
| ENSG000002 | 3.85340214 | -1.011197808 | 0.37850065 | -2.671588 | 0.00754933 | 0.02835275 | NA | NA |
| ENSG000002 | 9.8674801 | -0.912018339 | 0.3413797 | -2.6715658 | 0.00754983 | 0.02835275 | NA | NA |
| ENSG000002 | 260.270685 | 0.430816228 | 0.16126193 | 2.67153084 | 0.00755061 | 0.02835308 | NA | NA |
| ENSG000000 | 297.914332 | 0.456399987 | 0.17084205 | 2.6714733 | 0.00755191 | 0.02835338 | DBNDD1 | 79007 |
| ENSG000001 | 14.4963446 | -0.824338657 | 0.30857174 | -2.6714652 | 0.00755209 | 0.02835338 | SAMD7 | 344658 |
| ENSG000002 | 6.89183219 | -0.978515966 | 0.36631171 | -2.6712658 | 0.00755658 | 0.0283676 | LOC101928988 | 101928988 |
| ENSG000001 | 362.776124 | 0.373255058 | 0.13973261 | 2.67120945 | 0.00755785 | 0.02836974 | PPM1E | 22843 |
| ENSG000002 | 4.79614177 | -1.004898689 | 0.37620074 | -2.6711768 | 0.00755858 | 0.02836987 | NA | NA |
| ENSG000002 | 2.65441831 | -0.992492722 | 0.37159462 | -2.6709018 | 0.00756478 | 0.0283905 | NA | NA |
| ENSG000000 | 957.149425 | 0.328973788 | 0.12317322 | 2.67082233 | 0.00756657 | 0.02839459 | HIVEP1 | 3096 |
| ENSG000002 | 5.62250796 | -1.007904735 | 0.37738188 | -2.670782 | 0.00756748 | 0.02839537 | LOC101929353 | 101929353 |
| ENSG000002 | 3.57273717 | -1.007981804 | 0.37741692 | -2.6707382 | 0.00756846 | 0.02839645 | NA | NA |
| ENSG000001 | 380.26942 | 0.4184987 | 0.15670545 | 2.6706071 | 0.00757142 | 0.02840491 | FASTKD3 | 79072 |
| ENSG000001 | 19.173773 | -0.766845332 | 0.28716471 | -2.6704024 | 0.00757604 | 0.02841961 | ADAM5 | 255926 |
| ENSG000001 | 21.5689105 | -0.780330434 | 0.29224234 | -2.6701485 | 0.00758177 | 0.02843848 | GDNF | 2668 |
| ENSG000000 | 318.068839 | 0.393718162 | 0.14746376 | 2.66993172 | 0.00758667 | 0.02845421 | GLI2 | 2736 |
| ENSG000002 | 2.6777877 | -0.992251804 | 0.37165937 | -2.6697882 | 0.00758991 | 0.02846375 | NA | NA |
| ENSG000002 | 11.011105 | -0.883401795 | 0.33092302 | -2.6695084 | 0.00759624 | 0.02848483 | NA | NA |
| ENSG000001 | 1685.13796 | 0.332857947 | 0.12469593 | 2.66935703 | 0.00759966 | 0.02848977 | UBP1 | 7342 |

| | | | | | | | | |
|---|---|---|---|---|---|---|---|---|
| ENSG000001 | 287.51532 | 0.35853321 | 0.13431365 | 2.66937282 | 0.0075993 | 0.02848977 | CSRP1 | 1465 |
| ENSG000002 | 45.1465234 | 0.591489506 | 0.22158227 | 2.66939003 | 0.00759892 | 0.02848977 | NA | NA |
| ENSG000001 | 48.3624165 | 0.581942176 | 0.21802467 | 2.66915744 | 0.00760418 | 0.02849918 | PRADC1 | 84279 |
| ENSG000001 | 1378.39159 | 0.358711654 | 0.13439156 | 2.66915307 | 0.00760428 | 0.02849918 | LUZP1 | 7798 |
| ENSG000001 | 71.6648801 | 0.497692776 | 0.18645846 | 2.66918855 | 0.00760348 | 0.02849918 | EHBP1L1 | 254102 |
| ENSG000000 | 424.246505 | 0.352511235 | 0.13207963 | 2.66892958 | 0.00760934 | 0.02851551 | TIMM21 | 29090 |
| ENSG000002 | 8.53897133 | -0.949319632 | 0.35572161 | -2.6687151 | 0.0076142 | 0.02853109 | NA | NA |
| ENSG000001 | 173.319121 | 0.772658532 | 0.28953062 | 2.66865915 | 0.00761547 | 0.02853321 | PSKH1 | 5681 |
| ENSG000002 | 7.39281102 | -0.958708559 | 0.35926172 | -2.6685519 | 0.0076179 | 0.02853968 | NA | NA |
| ENSG000001 | 612.96004 | 0.324958618 | 0.12177932 | 2.668422 | 0.00762085 | 0.02854808 | VWDE | 221806 |
| ENSG000001 | 497.260943 | 0.343087026 | 0.12857771 | 2.66832438 | 0.00762306 | 0.02855374 | AHI1 | 54806 |
| ENSG000001 | 8.37619637 | -0.962823842 | 0.36085419 | -2.6681797 | 0.00762635 | 0.0285634 | CRP | 1401 |
| ENSG000002 | 11.6943135 | -0.875107477 | 0.32799959 | -2.6680139 | 0.00763011 | 0.02857486 | NA | NA |
| ENSG000002 | 9.17767322 | -0.973942716 | 0.36507642 | -2.6677776 | 0.00763548 | 0.02859232 | LVCAT1 | 100506827 |
| ENSG000001 | 513.667485 | 0.354046686 | 0.13272872 | 2.66744592 | 0.00764302 | 0.02861792 | DCUN1D3 | 123879 |
| ENSG000001 | 662.052282 | 0.33114979 | 0.12414796 | 2.66737994 | 0.00764452 | 0.0286209 | NEDD1 | 121441 |
| ENSG000001 | 19.3252377 | -0.797819949 | 0.29911508 | -2.6672676 | 0.00764708 | 0.02862782 | BPIFB4 | 149954 |
| ENSG000001 | 873.036993 | 0.304983648 | 0.11434847 | 2.66714232 | 0.00764993 | 0.02863585 | C3orf58 | 205428 |
| ENSG000000 | 396.397949 | 0.354536149 | 0.13294782 | 2.66673151 | 0.00765928 | 0.02866823 | ZZEF1 | 23140 |
| ENSG000002 | 6.8272371 | -0.971308162 | 0.36424197 | -2.6666563 | 0.007661 | 0.028672 | NA | NA |
| ENSG000002 | 1527.14335 | 0.303095088 | 0.1136642 | 2.66658364 | 0.00766265 | 0.02867555 | ZNF134 | 7693 |
| ENSG000001 | 78.4438761 | 0.506059437 | 0.18979639 | 2.66632802 | 0.00766848 | 0.02869472 | TCTE3 | 6991 |
| ENSG000002 | 17.2412552 | 0.785222261 | 0.29451096 | 2.66619029 | 0.00767163 | 0.02870383 | NA | NA |
| ENSG000001 | 44.2471377 | 0.606918911 | 0.22766151 | 2.66588278 | 0.00767865 | 0.02872479 | IL34 | 146433 |
| ENSG000001 | 101.986556 | 0.461974752 | 0.17329098 | 2.6658903 | 0.00767847 | 0.02872479 | C12orf76 | 400073 |
| ENSG000002 | 22.4110705 | -0.794258566 | 0.29795034 | -2.6657414 | 0.00768188 | 0.02873422 | NA | NA |
| ENSG000002 | 6.82347893 | -0.972115534 | 0.36468286 | -2.6656464 | 0.00768405 | 0.0287397 | RFPL4B | 442247 |
| ENSG000002 | 13.4323906 | -0.86260217 | 0.32362855 | -2.6654082 | 0.00768949 | 0.02875741 | DAOA-AS1 | 282706 |
| ENSG000001 | 804.212488 | 0.440573336 | 0.16529902 | 2.66531132 | 0.00769171 | 0.02875945 | CIRH1A | 84916 |

| | | | | | | | | |
|---|---|---|---|---|---|---|---|---|
| ENSG000002 | 10.0129881 | -0.937415537 | 0.35171222 | -2.6652914 | 0.00769216 | 0.02875945 | LOC101928226 | 101928226 |
| ENSG000002 | 14.9784819 | -0.83007666 | 0.3114356 | -2.6653236 | 0.00769143 | 0.02875945 | FGF12-AS1 | 100873986 |
| ENSG000002 | 7.35448124 | -0.989508708 | 0.37127015 | -2.6651987 | 0.00769428 | 0.02876472 | NA | NA |
| ENSG000001 | 53.5612581 | -0.603679942 | 0.22652255 | -2.6649883 | 0.0076991 | 0.02878007 | TTC6 | 319089 |
| ENSG000001 | 8.1052401 | -0.949547556 | 0.35633306 | -2.6647754 | 0.00770398 | 0.02879564 | NA | NA |
| ENSG000001 | 22.7210984 | -0.74300624 | 0.27884919 | -2.6645451 | 0.00770925 | 0.02881271 | ICOS | 29851 |
| ENSG000002 | 8.30812956 | -0.935479335 | 0.35117324 | -2.6638685 | 0.00772477 | 0.02886806 | NA | NA |
| ENSG000002 | 3.02985424 | -1.000795712 | 0.37569988 | -2.663817 | 0.00772596 | 0.02886983 | NA | NA |
| ENSG000002 | 36.400894 | -0.627965947 | 0.23574207 | -2.663784 | 0.00772672 | 0.02887 | LINC00511 | 400619 |
| ENSG000002 | 31.73298 | 0.678973189 | 0.25491045 | 2.66357532 | 0.00773151 | 0.02888259 | NA | NA |
| ENSG000002 | 2.65154224 | -0.990831163 | 0.37199241 | -2.663579 | 0.00773143 | 0.02888259 | NA | NA |
| ENSG000001 | 122.562737 | 0.429912869 | 0.16140707 | 2.6635319 | 0.00773251 | 0.02888365 | NA | NA |
| ENSG000001 | 8.56377581 | -0.950681762 | 0.35693769 | -2.663439 | 0.00773464 | 0.02888897 | LGALS13 | 29124 |
| ENSG000002 | 3.40489786 | -1.006143065 | 0.3777835 | -2.6632795 | 0.00773831 | 0.0289 | LINC01402 | 104797536 |
| ENSG000001 | 555.127805 | 0.357992248 | 0.13443804 | 2.66287906 | 0.00774753 | 0.02893175 | EPG5 | 57724 |
| ENSG000002 | 19.7669404 | 0.76971962 | 0.28906001 | 2.66283672 | 0.0077485 | 0.02893273 | NA | NA |
| ENSG00000C | 891.724182 | 0.28811651 | 0.10821051 | 2.66255571 | 0.00775497 | 0.02894624 | PIAS1 | 8554 |
| ENSG000001 | 1617.40961 | 0.305629553 | 0.11478771 | 2.66256346 | 0.00775479 | 0.02894624 | SMAD4 | 4089 |
| ENSG000001 | 180.475197 | 0.378263635 | 0.14206749 | 2.66256292 | 0.00775481 | 0.02894624 | CREBRF | 153222 |
| ENSG000002 | 19.7441732 | -0.788475747 | 0.29612791 | -2.6626188 | 0.00775352 | 0.02894624 | LOC101928651 | 101928651 |
| ENSG000002 | 29.4694347 | -0.679106733 | 0.25506169 | -2.6625196 | 0.00775581 | 0.02894668 | NA | NA |
| ENSG000001 | 356.301106 | 0.389840079 | 0.14644007 | 2.66211353 | 0.00776517 | 0.02897896 | ING1 | 3621 |
| ENSG000001 | 24.6551963 | -0.707362779 | 0.26573163 | -2.6619442 | 0.00776908 | 0.02898877 | EPHX4 | 253152 |
| ENSG000002 | 8.02166288 | -0.945317905 | 0.35512399 | -2.6619376 | 0.00776923 | 0.02898877 | CASC23 | 103581031 |
| ENSG000002 | 2.39859661 | -0.965717946 | 0.36290914 | -2.6610461 | 0.00778983 | 0.02906297 | NA | NA |
| ENSG000002 | 73.2468268 | 0.583772722 | 0.21939044 | 2.66088499 | 0.00779356 | 0.0290742 | NA | NA |
| ENSG000001 | 492.792036 | 0.369522326 | 0.13888058 | 2.66071991 | 0.00779738 | 0.02908578 | AGAP1 | 116987 |
| ENSG000002 | 11.4252154 | -0.889266463 | 0.33423373 | -2.6606126 | 0.00779986 | 0.02909238 | NA | NA |
| ENSG000002 | 23.1229313 | -0.773748565 | 0.29082252 | -2.6605524 | 0.00780126 | 0.0290949 | NA | NA |

| | | | | | | | | |
|---|---|---|---|---|---|---|---|---|
| ENSG000001 | 125.339283 | 0.457540625 | 0.17198823 | 2.66030194 | 0.00780706 | 0.02911387 | ZBED3 | 84327 |
| ENSG000001 | 3.01449013 | -0.999392942 | 0.37569184 | -2.6601402 | 0.00781081 | 0.02912518 | NA | NA |
| ENSG000001 | 307.263735 | 0.36580038 | 0.13753165 | 2.6597541 | 0.00781977 | 0.0291559 | C5orf24 | 134553 |
| ENSG000002 | 915.453526 | 0.334538175 | 0.12579022 | 2.65949264 | 0.00782584 | 0.02917586 | NA | NA |
| ENSG000001 | 704.203265 | 0.35116863 | 0.13206091 | 2.65914134 | 0.00783401 | 0.02920361 | ZNF181 | 339318 |
| ENSG000001 | 702.322219 | 0.294003344 | 0.11056711 | 2.65904876 | 0.00783616 | 0.02920895 | CNTRL | 11064 |
| ENSG00000C | 214.227967 | 0.369698024 | 0.1390377 | 2.65897692 | 0.00783783 | 0.0292125 | CREM | 1390 |
| ENSG000002 | 17.3963224 | -0.803469384 | 0.30220532 | -2.6586871 | 0.00784458 | 0.02923226 | NA | NA |
| ENSG000002 | 8.97801308 | -0.926421839 | 0.34844756 | -2.6587123 | 0.00784399 | 0.02923226 | NA | NA |
| ENSG000001 | 328.168773 | 0.424853025 | 0.15980915 | 2.65850249 | 0.00784888 | 0.0292456 | CHD3 | 1107 |
| ENSG000001 | 180.34682 | 0.453932873 | 0.17076945 | 2.65816213 | 0.00785681 | 0.02927246 | ARL4C | 10123 |
| ENSG000001 | 52.3634756 | -0.660207505 | 0.24839437 | -2.6579004 | 0.00786291 | 0.02929251 | PLCD4 | 84812 |
| ENSG000001 | 19.5007164 | -0.80283871 | 0.30206789 | -2.6578088 | 0.00786505 | 0.02929778 | RP1L1 | 94137 |
| ENSG00000C | 270.474733 | 0.350760695 | 0.1319757 | 2.65776733 | 0.00786602 | 0.02929869 | SLC25A40 | 55972 |
| ENSG000002 | 30.0357374 | -0.712755523 | 0.26819082 | -2.6576433 | 0.00786891 | 0.02930679 | NA | NA |
| ENSG000001 | 285.406486 | 0.3658392 | 0.13766221 | 2.65751365 | 0.00787194 | 0.02931537 | GNE | 10020 |
| ENSG000001 | 16.9930918 | -0.800661504 | 0.30129457 | -2.6574044 | 0.00787449 | 0.02932218 | DNAI1 | 27019 |
| ENSG000001 | 7.37071515 | -0.954197581 | 0.35908124 | -2.6573307 | 0.00787621 | 0.0293259 | TMEM174 | 134288 |
| ENSG00000C | 21.9363395 | -0.749280229 | 0.28197185 | -2.6572874 | 0.00787723 | 0.02932697 | NME8 | 51314 |
| ENSG000002 | 5.47930551 | -0.989479413 | 0.37237665 | -2.6572005 | 0.00787926 | 0.02933184 | NA | NA |
| ENSG000001 | 367.739171 | 0.338492274 | 0.12739107 | 2.65711156 | 0.00788134 | 0.02933689 | COPS7A | 50813 |
| ENSG000002 | 299.561899 | 0.36431506 | 0.13712351 | 2.65683881 | 0.00788771 | 0.02935794 | ZNF550 | 162972 |
| ENSG000001 | 194.393382 | 0.488194464 | 0.18375813 | 2.65672306 | 0.00789042 | 0.02936532 | DOCK6 | 57572 |
| ENSG000001 | 4.15929204 | -1.005395488 | 0.3785112 | -2.6561842 | 0.00790304 | 0.02940959 | ACTRT1 | 139741 |
| ENSG000001 | 6.55327131 | -0.977147198 | 0.36792487 | -2.6558335 | 0.00791127 | 0.02943749 | AGAP5 | 729092 |
| ENSG000001 | 15.0807088 | -0.843857099 | 0.31775854 | -2.6556552 | 0.00791545 | 0.02945036 | SLC36A3 | 285641 |
| ENSG000002 | 6.68811286 | -0.980206853 | 0.36912557 | -2.6554835 | 0.00791948 | 0.02946265 | NA | NA |
| ENSG000001 | 606.546028 | 0.385581375 | 0.14521587 | 2.65522886 | 0.00792546 | 0.02947679 | KIAA1958 | 158405 |
| ENSG000002 | 2.30704359 | -0.967537359 | 0.36438932 | -2.6552297 | 0.00792544 | 0.02947679 | NA | NA |

| | | | | | | | | |
|---|---|---|---|---|---|---|---|---|
| ENSG000002 | 3.03668491 | -0.997130811 | 0.37553447 | -2.6552311 | 0.00792541 | 0.02947679 | LOC101927577 | 101927577 |
| ENSG000001 | 69.089547 | -0.486993535 | 0.18341729 | -2.6551124 | 0.0079282 | 0.02948426 | CASZ1 | 54897 |
| ENSG000001 | 176.014912 | 0.424108969 | 0.1597377 | 2.65503367 | 0.00793005 | 0.02948844 | SLC27A2 | 11001 |
| ENSG000002 | 8.30994937 | -0.955386042 | 0.35985482 | -2.6549208 | 0.0079327 | 0.02949561 | NA | NA |
| ENSG000001 | 1936.84654 | 0.305665806 | 0.11513375 | 2.6548758 | 0.00793376 | 0.02949683 | F11R | 50848 |
| ENSG000002 | 2.30490364 | -0.967284865 | 0.36437792 | -2.6546198 | 0.00793978 | 0.02951652 | NA | NA |
| ENSG000002 | 9.26296327 | -0.92074691 | 0.3468572 | -2.6545417 | 0.00794162 | 0.02952064 | NA | NA |
| ENSG000001 | 293.798841 | 0.382967201 | 0.14427858 | 2.65435929 | 0.00794592 | 0.0295339 | SLC35F1 | 222553 |
| ENSG000001 | 1034.16627 | 0.312346155 | 0.11767949 | 2.65421057 | 0.00794942 | 0.02954422 | LRRC8B | 23507 |
| ENSG000002 | 23.5783645 | -0.72606585 | 0.27360443 | -2.6537065 | 0.0079613 | 0.02958567 | LOC101929634 | 101929634 |
| ENSG000001 | 1555.75252 | 0.328505034 | 0.12380932 | 2.65331425 | 0.00797056 | 0.02961194 | RICTOR | 253260 |
| ENSG000001 | 6.90996751 | -0.970007524 | 0.36557779 | -2.6533546 | 0.00796961 | 0.02961194 | C2CD4A | 145741 |
| ENSG000002 | 7.62207908 | -0.963991265 | 0.36331218 | -2.6533415 | 0.00796992 | 0.02961194 | LOC101929109 | 101929109 |
| ENSG000002 | 16.884805 | -0.815737822 | 0.30744936 | -2.6532428 | 0.00797225 | 0.02961549 | NA | NA |
| ENSG000001 | 10923.0843 | 0.326472167 | 0.12304877 | 2.6531932 | 0.00797342 | 0.02961713 | CSE1L | 1434 |
| ENSG000002 | 78.1943942 | 0.587097259 | 0.22129508 | 2.65300633 | 0.00797784 | 0.02963082 | ZNF316 | 100131017 |
| ENSG000002 | 10.1536534 | -0.898952143 | 0.33885432 | -2.6529162 | 0.00797997 | 0.02963601 | MKNK1-AS1 | 100507423 |
| ENSG00000C | 1795.15567 | 0.29837645 | 0.1124742 | 2.65284343 | 0.00798169 | 0.02963698 | DLG1 | 1739 |
| ENSG000002 | 8.57533701 | -0.92441386 | 0.34846104 | -2.6528471 | 0.0079816 | 0.02963698 | LOC101927690 | 101927690 |
| ENSG000001 | 35.3020723 | -0.616976048 | 0.23259493 | -2.6525774 | 0.00798798 | 0.02965763 | NA | NA |
| ENSG00000C | 1203.99283 | 0.317661391 | 0.11977586 | 2.65213209 | 0.00799852 | 0.02969406 | UBR2 | 23304 |
| ENSG00000C | 445.757952 | 0.356456853 | 0.13442895 | 2.65163765 | 0.00801025 | 0.02973485 | ZNF184 | 7738 |
| ENSG000001 | 80.1884194 | 0.472437587 | 0.17819258 | 2.65127537 | 0.00801884 | 0.02976404 | CCDC126 | 90693 |
| ENSG000002 | 21.391502 | -0.746967859 | 0.28174318 | -2.6512367 | 0.00801976 | 0.02976472 | NA | NA |
| ENSG000001 | 8.16731763 | -0.950916741 | 0.35870214 | -2.6509927 | 0.00802556 | 0.02978351 | CD300LB | 124599 |
| ENSG000001 | 729.140358 | 0.372209752 | 0.14041599 | 2.65076467 | 0.00803098 | 0.02979817 | ZNF827 | 152485 |
| ENSG000002 | 403.019099 | 0.424110732 | 0.15999547 | 2.65076713 | 0.00803092 | 0.02979817 | ARHGAP23 | 57636 |
| ENSG00000C | 2455.26586 | 0.33332531 | 0.1257613 | 2.65046002 | 0.00803822 | 0.02982233 | ROCK1 | 6093 |
| ENSG000001 | 37.618462 | -0.710413544 | 0.26804792 | -2.6503229 | 0.00804149 | 0.0298317 | ZBTB7C | 201501 |

| | | | | | | | | |
|---|---|---|---|---|---|---|---|---|
| ENSG000002 | 3.40170858 | -1.001191936 | 0.37777206 | -2.650254 | 0.00804313 | 0.02983506 | NA | NA |
| ENSG000000 | 20.387752 | -0.81219472 | 0.30651891 | -2.6497377 | 0.00805543 | 0.02987521 | PAX7 | 5081 |
| ENSG000002 | 17.523458 | -0.79249307 | 0.29908707 | -2.6497069 | 0.00805616 | 0.02987521 | NA | NA |
| ENSG000002 | 84.0627227 | 0.475691936 | 0.17952272 | 2.64975902 | 0.00805492 | 0.02987521 | NA | NA |
| ENSG000002 | 4.8947578 | -0.998457303 | 0.37682948 | -2.6496263 | 0.00805809 | 0.02987961 | NA | NA |
| ENSG000002 | 12.672575 | -0.864690608 | 0.32637613 | -2.6493684 | 0.00806424 | 0.02989968 | NA | NA |
| ENSG000001 | 11.530115 | -0.865098047 | 0.32654533 | -2.6492434 | 0.00806722 | 0.02990802 | NA | NA |
| ENSG000001 | 330.022967 | 0.364399201 | 0.13756214 | 2.64897893 | 0.00807354 | 0.02992869 | FBXO9 | 26268 |
| ENSG000001 | 17.5466228 | -0.830388526 | 0.31350158 | -2.6487539 | 0.00807891 | 0.02994342 | HORMAD2 | 150280 |
| ENSG000002 | 10.6281148 | -0.902098519 | 0.34057507 | -2.6487509 | 0.00807899 | 0.02994342 | NA | NA |
| ENSG000001 | 693.039664 | 0.690309923 | 0.26062367 | 2.64868471 | 0.00808057 | 0.02994655 | GNA12 | 2768 |
| ENSG000000 | 728.562259 | 0.316790171 | 0.11960436 | 2.64865077 | 0.00808138 | 0.02994682 | RAD18 | 56852 |
| ENSG000001 | 10.0338901 | -0.893506894 | 0.3373968 | -2.6482376 | 0.00809126 | 0.02997842 | BCL2L10 | 10017 |
| ENSG000001 | 8.1706416 | -0.946646733 | 0.3574636 | -2.6482325 | 0.00809139 | 0.02997842 | HTR3C | 170572 |
| ENSG000002 | 31.7691703 | -0.641690041 | 0.24231615 | -2.6481522 | 0.00809331 | 0.0299828 | NA | NA |
| ENSG000002 | 40.1949546 | -0.596300875 | 0.22519212 | -2.6479651 | 0.00809779 | 0.02999666 | NA | NA |
| ENSG000001 | 26.0170263 | -0.696423345 | 0.26300892 | -2.6479077 | 0.00809916 | 0.02999901 | COL20A1 | 57642 |
| ENSG000001 | 12.2052504 | -0.863263628 | 0.32604037 | -2.64772 | 0.00810366 | 0.03001293 | FAM26D | 221301 |
| ENSG000000 | 3662.70788 | 0.398248692 | 0.15041456 | 2.6476738 | 0.00810477 | 0.03001354 | CECR2 | 27443 |
| ENSG000001 | 15.0468492 | -0.812224971 | 0.30677484 | -2.6476258 | 0.00810592 | 0.03001354 | CLRN1 | 7401 |
| ENSG000002 | 50.7919828 | 0.563178515 | 0.21271119 | 2.64762056 | 0.00810605 | 0.03001354 | NA | NA |
| ENSG000002 | 9.83713053 | -0.901755682 | 0.34060447 | -2.6475157 | 0.00810856 | 0.0300201 | NA | NA |
| ENSG000001 | 144.341352 | 0.405494959 | 0.15316905 | 2.64736878 | 0.00811208 | 0.03002887 | TRAPPC9 | 83696 |
| ENSG000001 | 15.0356108 | -0.84026531 | 0.31739802 | -2.6473553 | 0.00811241 | 0.03002887 | C1orf105 | 92346 |
| ENSG000002 | 9.21809853 | -0.916148192 | 0.34606749 | -2.6473108 | 0.00811348 | 0.03003008 | NA | NA |
| ENSG000001 | 67.7452078 | 0.518787092 | 0.19599246 | 2.64697469 | 0.00812154 | 0.03005369 | BCL6 | 604 |
| ENSG000001 | 114.062179 | 0.449463464 | 0.16980414 | 2.64695238 | 0.00812208 | 0.03005369 | PAQR8 | 85315 |
| ENSG000002 | 12.6986942 | -0.845773629 | 0.31952465 | -2.6469746 | 0.00812155 | 0.03005369 | NA | NA |
| ENSG000001 | 3694.80547 | 0.285551094 | 0.10791535 | 2.64606555 | 0.0081434 | 0.03012985 | UBE3A | 7337 |

| | | | | | | | | |
|---|---|---|---|---|---|---|---|---|
| ENSG000001 | 1499.19527 | 0.328426507 | 0.12412232 | 2.64599066 | 0.00814521 | 0.03013377 | PRELID3B | 51012 |
| ENSG000001 | 266.858679 | 0.413485551 | 0.15627288 | 2.64592006 | 0.00814691 | 0.03013456 | SOWAHC | 65124 |
| ENSG000002 | 5.1516257 | -0.993304525 | 0.37540857 | -2.6459293 | 0.00814669 | 0.03013456 | NA | NA |
| ENSG000002 | 4.4989423 | -0.999080563 | 0.37761038 | -2.6457974 | 0.00814986 | 0.03014274 | NA | NA |
| ENSG00000C | 240.406586 | 0.444079087 | 0.16785744 | 2.64557292 | 0.00815527 | 0.03016 | HDAC4 | 9759 |
| ENSG000001 | 626.620172 | 0.321232099 | 0.12142512 | 2.64551594 | 0.00815664 | 0.03016233 | RNF111 | 54778 |
| ENSG000001 | 10.4419589 | -0.892453231 | 0.33737492 | -2.6452863 | 0.00816218 | 0.03018006 | IL5 | 3567 |
| ENSG000001 | 16.7040476 | -0.787895348 | 0.29787324 | -2.6450692 | 0.00816742 | 0.03019667 | NA | NA |
| ENSG000001 | 8.46660485 | -0.940124049 | 0.35544072 | -2.6449532 | 0.00817022 | 0.03020427 | C4orf26 | 152816 |
| ENSG000001 | 946.149967 | 0.322967941 | 0.12211804 | 2.64471925 | 0.00817587 | 0.03022241 | CUL2 | 8453 |
| ENSG000002 | 8.79873178 | -0.934797522 | 0.35348395 | -2.6445261 | 0.00818054 | 0.03023691 | LINC01391 | 103344930 |
| ENSG000001 | 16.2259297 | -0.81852835 | 0.30959292 | -2.6438859 | 0.00819603 | 0.03028966 | CPN1 | 1369 |
| ENSG000001 | 25.8560131 | -0.719096742 | 0.27198915 | -2.6438435 | 0.00819705 | 0.03028966 | RSAD2 | 91543 |
| ENSG000002 | 11.4080424 | -0.880205389 | 0.33292484 | -2.6438562 | 0.00819675 | 0.03028966 | NA | NA |
| ENSG000001 | 18.5859071 | -0.753057682 | 0.28484809 | -2.6437168 | 0.00820012 | 0.03029548 | LRRC74B | 400891 |
| ENSG000002 | 4.43273686 | -0.997857501 | 0.37744103 | -2.6437441 | 0.00819946 | 0.03029548 | NA | NA |
| ENSG000001 | 487.781737 | 0.329366966 | 0.12458927 | 2.64362217 | 0.00820242 | 0.03030119 | SCN8A | 6334 |
| ENSG000001 | 408.78018 | 0.377410579 | 0.14277674 | 2.64336183 | 0.00820873 | 0.03032174 | C1orf123 | 54987 |
| ENSG000001 | 542.025749 | 0.367320825 | 0.13896843 | 2.64319615 | 0.00821274 | 0.03033382 | NA | NA |
| ENSG000001 | 8.76579634 | -0.932757578 | 0.35290368 | -2.643094 | 0.00821522 | 0.03034021 | AGXT | 189 |
| ENSG00000C | 181.784774 | 0.383814236 | 0.14521842 | 2.64301342 | 0.00821718 | 0.03034467 | ZNF671 | 79891 |
| ENSG000002 | 40.5091905 | -0.605483775 | 0.22911715 | -2.642682 | 0.00822522 | 0.03037161 | VIT | 5212 |
| ENSG000002 | 18.0073756 | -0.777033556 | 0.29403834 | -2.6426266 | 0.00822657 | 0.03037382 | LOC101927683 | 101927683 |
| ENSG000001 | 4492.54396 | 0.297793334 | 0.11269015 | 2.64258536 | 0.00822757 | 0.03037475 | LMNB1 | 4001 |
| ENSG000001 | 20.7056072 | 0.756053987 | 0.28619534 | 2.64174108 | 0.00824811 | 0.0304478 | CH25H | 9023 |
| ENSG000002 | 25.6165384 | -0.737104718 | 0.27905216 | -2.6414586 | 0.00825499 | 0.03047043 | LOC100506725 | 100506725 |
| ENSG000001 | 18482.0983 | 0.353340123 | 0.13377719 | 2.64125846 | 0.00825987 | 0.03048566 | HNRNPU | 3192 |
| ENSG000001 | 45.5436516 | -0.572702168 | 0.21683518 | -2.6411866 | 0.00826162 | 0.03048935 | NA | NA |
| ENSG000001 | 265.191206 | 0.415767238 | 0.15742866 | 2.6409882 | 0.00826646 | 0.03050166 | NAA20 | 51126 |

| | | | | | | | | |
|---|---|---|---|---|---|---|---|---|
| ENSG000002 | 26.0080099 | -0.698954735 | 0.26465446 | -2.6410087 | 0.00826596 | 0.03050166 | NA | NA |
| ENSG000001 | 554.887152 | 0.35302963 | 0.13368368 | 2.64078334 | 0.00827146 | 0.03051733 | PHF12 | 57649 |
| ENSG000001 | 67.3118968 | 0.515097253 | 0.19506174 | 2.64068825 | 0.00827378 | 0.03052312 | PCED1B | 91523 |
| ENSG000001 | 186.542758 | 0.476897537 | 0.18062243 | 2.6403007 | 0.00828325 | 0.03055249 | MAPK8IP3 | 23162 |
| ENSG000002 | 11.1009553 | -0.886391657 | 0.33571328 | -2.6403235 | 0.00828269 | 0.03055249 | NA | NA |
| ENSG000001 | 15.352496 | -0.80973952 | 0.30669884 | -2.640178 | 0.00828625 | 0.03056078 | C2orf71 | 388939 |
| ENSG000002 | 5.08155803 | -0.987992363 | 0.37425637 | -2.6398812 | 0.00829351 | 0.03058477 | NA | NA |
| ENSG000002 | 408.761702 | 0.336941598 | 0.12764036 | 2.63977317 | 0.00829615 | 0.03059174 | FAM200B | 285550 |
| ENSG000002 | 6.27462548 | -0.980346805 | 0.37144183 | -2.639301 | 0.00830772 | 0.03063159 | NA | NA |
| ENSG000001 | 476.840255 | 0.33420639 | 0.1266328 | 2.63917705 | 0.00831076 | 0.03063723 | PDP1 | 54704 |
| ENSG000002 | 5.46715145 | -0.995241132 | 0.37710262 | -2.6391785 | 0.00831072 | 0.03063723 | LINC00398 | 100874158 |
| ENSG000002 | 6.44661155 | -0.985522412 | 0.37344137 | -2.6390285 | 0.0083144 | 0.03064787 | NA | NA |
| ENSG000002 | 16.6601465 | -0.80175848 | 0.30381531 | -2.6389667 | 0.00831592 | 0.03065067 | TMPRSS4-AS1 | 100526771 |
| ENSG000001 | 5.80159012 | -0.978981537 | 0.37098756 | -2.6388527 | 0.00831871 | 0.03065819 | NA | NA |
| ENSG000002 | 4.52800122 | -0.997422971 | 0.37801335 | -2.6385919 | 0.00832511 | 0.03067899 | LINC01364 | 100505768 |
| ENSG000001 | 222.43736 | 0.405370604 | 0.15363964 | 2.63845065 | 0.00832858 | 0.03068899 | GFOD1 | 54438 |
| ENSG000002 | 3.88880201 | -0.998353482 | 0.37840396 | -2.6383273 | 0.00833161 | 0.03069737 | LINC00587 | 414319 |
| ENSG000001 | 2978.5907 | 0.328114982 | 0.12436884 | 2.63824116 | 0.00833373 | 0.03070238 | RPL36AL | 6166 |
| ENSG000001 | 956.199399 | 0.360835735 | 0.136775 | 2.63817019 | 0.00833547 | 0.03070601 | UBTD2 | 92181 |
| ENSG000001 | 33.1981959 | -0.671606603 | 0.25459651 | -2.6379254 | 0.00834149 | 0.03072539 | A1CF | 29974 |
| ENSG000001 | 702.788167 | 0.317071816 | 0.12019923 | 2.63788561 | 0.00834247 | 0.03072621 | TRAK2 | 66008 |
| ENSG000001 | 2081.10955 | 0.323785031 | 0.1227507 | 2.63774497 | 0.00834593 | 0.03073616 | SMC2 | 10592 |
| ENSG000002 | 3.13160856 | -0.989230896 | 0.3750668 | -2.6374792 | 0.00835247 | 0.03075746 | NA | NA |
| ENSG000001 | 267.661204 | 0.333225113 | 0.12635032 | 2.63731117 | 0.00835661 | 0.03076991 | SGK1 | 6446 |
| ENSG000001 | 114.631203 | 0.517922194 | 0.19639343 | 2.63716666 | 0.00836017 | 0.03078023 | PROCA1 | 147011 |
| ENSG000001 | 21.3248564 | -0.737407904 | 0.27962916 | -2.6370923 | 0.00836201 | 0.03078418 | BTG4 | 54766 |
| ENSG000001 | 1583.33549 | 0.317826095 | 0.12053425 | 2.63681147 | 0.00836893 | 0.03080688 | TGS1 | 96764 |
| ENSG000002 | 12.0642421 | -0.874648505 | 0.33171154 | -2.6367744 | 0.00836985 | 0.03080745 | LEMD1-AS1 | 284576 |
| ENSG000002 | 9.12605048 | -0.908621934 | 0.3446013 | -2.6367339 | 0.00837085 | 0.03080833 | NA | NA |

| | | | | | | | | |
|---|---|---|---|---|---|---|---|---|
| ENSG000001 | 423.935321 | 0.339631614 | 0.12881248 | 2.63663597 | 0.00837326 | 0.03081443 | ARMCX3 | 51566 |
| ENSG000001 | 31.8050457 | -0.674100813 | 0.25567144 | -2.6365902 | 0.00837439 | 0.03081579 | TTLL9 | 164395 |
| ENSG000002 | 31.1273545 | -0.654096866 | 0.24810632 | -2.6363571 | 0.00838015 | 0.03083417 | LINC00271 | 100131814 |
| ENSG000001 | 459.794334 | 0.331296549 | 0.12566714 | 2.6363021 | 0.00838151 | 0.03083637 | CASP2 | 835 |
| ENSG000002 | 13.2185334 | -0.848480485 | 0.32185464 | -2.6362226 | 0.00838347 | 0.0308408 | NA | NA |
| ENSG00000C | 6266.56911 | 0.293683369 | 0.11140933 | 2.63607518 | 0.00838711 | 0.0308514 | QSER1 | 79832 |
| ENSG000001 | 11.9494776 | -0.886546012 | 0.33631983 | -2.6360207 | 0.00838846 | 0.03085317 | C5AR1 | 728 |
| ENSG000002 | 63.4792257 | 0.498698514 | 0.18918801 | 2.63599424 | 0.00838912 | 0.03085317 | NA | NA |
| ENSG000001 | 25.0576877 | -0.691578409 | 0.26237063 | -2.6358835 | 0.00839185 | 0.03086044 | TRPM2 | 7226 |
| ENSG000002 | 29.5319374 | -0.706040551 | 0.26787296 | -2.6357291 | 0.00839567 | 0.03087169 | GPR179 | 440435 |
| ENSG000001 | 680.451902 | 0.319027282 | 0.12104188 | 2.63567694 | 0.00839696 | 0.03087363 | C18orf25 | 147339 |
| ENSG000002 | 3.09253071 | -0.989326913 | 0.37541999 | -2.6352537 | 0.00840744 | 0.03090936 | NA | NA |
| ENSG000001 | 11.0871258 | -0.883724289 | 0.33536351 | -2.6351235 | 0.00841067 | 0.03091841 | FASLG | 356 |
| ENSG000002 | 8.78885137 | -0.91004345 | 0.34535607 | -2.6350874 | 0.00841156 | 0.03091889 | NA | NA |
| ENSG000001 | 101.445033 | 0.454391189 | 0.17244265 | 2.6350279 | 0.00841304 | 0.03092151 | CCDC53 | 51019 |
| ENSG000001 | 242.506025 | 0.405530642 | 0.1539032 | 2.6349721 | 0.00841442 | 0.03092379 | DCAF10 | 79269 |
| ENSG000002 | 3.80486439 | -0.997155795 | 0.3784481 | -2.6348548 | 0.00841733 | 0.03093168 | NA | NA |
| ENSG000001 | 1042.17607 | 0.334327395 | 0.12689011 | 2.63477903 | 0.00841921 | 0.03093578 | KLHL12 | 59349 |
| ENSG000001 | 378.576309 | 0.438707923 | 0.1665104 | 2.6347178 | 0.00842073 | 0.03093856 | LRP4 | 4038 |
| ENSG000001 | 32.6456654 | -0.676133369 | 0.25663025 | -2.6346596 | 0.00842217 | 0.03094106 | NPR3 | 4883 |
| ENSG000002 | 5.53105611 | -0.985112954 | 0.37393874 | -2.6344234 | 0.00842803 | 0.03095979 | NA | NA |
| ENSG000001 | 1385.5168 | 0.323096298 | 0.12264976 | 2.63430031 | 0.00843109 | 0.03096821 | CEP192 | 55125 |
| ENSG000002 | 6.53907897 | -0.972701837 | 0.36928367 | -2.6340235 | 0.00843797 | 0.03099067 | NA | NA |
| ENSG000002 | 39.0020194 | 0.598653389 | 0.22728808 | 2.63389701 | 0.00844111 | 0.0309994 | NA | NA |
| ENSG000001 | 199.299936 | 0.361705475 | 0.1373305 | 2.63383209 | 0.00844272 | 0.03100252 | FAXC | 84553 |
| ENSG000002 | 24.3300766 | -0.70487599 | 0.26762695 | -2.6338005 | 0.00844351 | 0.0310026 | GPR1-AS | 101669764 |
| ENSG000001 | 1268.34544 | 0.303938339 | 0.11540419 | 2.63368555 | 0.00844637 | 0.03101029 | MED13L | 23389 |
| ENSG000001 | 349.427063 | 0.366660262 | 0.13925718 | 2.63297198 | 0.00846413 | 0.0310727 | ZDHHC23 | 254887 |
| ENSG000001 | 1071.35307 | 0.32166951 | 0.12217283 | 2.63290551 | 0.00846579 | 0.03107597 | DNAJC21 | 134218 |

| | | | | | | | | |
|---|---|---|---|---|---|---|---|---|
| ENSG000002 | 43.7852547 | 0.564335161 | 0.21436 | 2.63265138 | 0.00847213 | 0.03109641 | NA | NA |
| ENSG000001 | 343.991651 | 0.384121339 | 0.14592047 | 2.63240207 | 0.00847835 | 0.03111643 | ASAP2 | 8853 |
| ENSG000002 | 7.43139541 | -0.955120993 | 0.36285868 | -2.632212 | 0.00848309 | 0.03113102 | LINC01586 | 101929743 |
| ENSG000001 | 19.9571505 | -0.752979246 | 0.28607041 | -2.6321465 | 0.00848473 | 0.0311324 | ITGB7 | 3695 |
| ENSG000001 | 30.1187008 | -0.64683068 | 0.24574369 | -2.6321355 | 0.008485 | 0.0311324 | SLFN11 | 91607 |
| ENSG000001 | 492.166956 | 0.350489006 | 0.13316877 | 2.63191582 | 0.00849049 | 0.03114972 | ZNF140 | 7699 |
| ENSG000002 | 6.98060027 | -0.951904992 | 0.36168322 | -2.6318749 | 0.00849151 | 0.03115065 | NA | NA |
| ENSG000001 | 36.7538385 | 0.658899856 | 0.25035834 | 2.63182707 | 0.00849271 | 0.03115222 | CDA | 978 |
| ENSG00000C | 590.129172 | 0.337431482 | 0.12822415 | 2.6315751 | 0.00849901 | 0.03117251 | TPD52 | 7163 |
| ENSG000002 | 173.031715 | 0.386829886 | 0.1470123 | 2.63127559 | 0.0085065 | 0.03119718 | FAM155A | 728215 |
| ENSG000001 | 398.542561 | 0.329661308 | 0.1252894 | 2.63119864 | 0.00850843 | 0.03120142 | SERPING1 | 710 |
| ENSG000001 | 244.076993 | 0.385135535 | 0.14638849 | 2.63091403 | 0.00851556 | 0.03122474 | LRRC57 | 255252 |
| ENSG000002 | 9.22114886 | -0.910832238 | 0.34622123 | -2.630781 | 0.00851889 | 0.03123414 | NA | NA |
| ENSG00000C | 19.8527439 | -0.79939576 | 0.30388021 | -2.6306279 | 0.00852273 | 0.03123989 | CD6 | 923 |
| ENSG00000C | 61.1842944 | -0.527732291 | 0.20060749 | -2.6306709 | 0.00852165 | 0.03123989 | ABCB1 | 5243 |
| ENSG000001 | 826.126599 | 0.30486248 | 0.1158897 | 2.63062627 | 0.00852277 | 0.03123989 | ABI1 | 10006 |
| ENSG000002 | 39.9517868 | -0.676380913 | 0.25714494 | -2.6303489 | 0.00852973 | 0.03126257 | BAHCC1 | 57597 |
| ENSG000001 | 191.297096 | 0.39712806 | 0.15100468 | 2.62990565 | 0.00854086 | 0.03130052 | KIF24 | 347240 |
| ENSG00000C | 437.616494 | 0.335034501 | 0.12739979 | 2.62978842 | 0.0085438 | 0.03130849 | AP4E1 | 23431 |
| ENSG00000C | 983.885795 | 0.33371749 | 0.12691202 | 2.62951833 | 0.00855059 | 0.03133054 | CRYBG3 | 131544 |
| ENSG000002 | 38.6021835 | -0.613763446 | 0.23342055 | -2.6294319 | 0.00855277 | 0.03133567 | NA | NA |
| ENSG000001 | 111.454849 | 0.455721993 | 0.17333408 | 2.62915404 | 0.00855976 | 0.03135846 | SFR1 | 119392 |
| ENSG000001 | 233.318382 | 0.441422353 | 0.16790269 | 2.62903687 | 0.00856271 | 0.03136543 | ST14 | 6768 |
| ENSG000001 | 294.152583 | 0.361903457 | 0.13765733 | 2.62901698 | 0.00856321 | 0.03136543 | SNHG12 | 85028 |
| ENSG000001 | 12.7307782 | -0.859572309 | 0.32696536 | -2.62894 | 0.00856515 | 0.03136954 | RXRG | 6258 |
| ENSG000001 | 16.9687522 | -0.775654903 | 0.29505099 | -2.6288843 | 0.00856655 | 0.03136954 | IL22RA2 | 116379 |
| ENSG000002 | 6.71931763 | -0.959698464 | 0.36505979 | -2.6288803 | 0.00856665 | 0.03136954 | NA | NA |
| ENSG000001 | 200.78087 | 0.404569627 | 0.15390617 | 2.62867711 | 0.00857177 | 0.03138262 | DUSP22 | 56940 |
| ENSG000001 | 561.086231 | 0.329707125 | 0.1254261 | 2.62869632 | 0.00857129 | 0.03138262 | UAP1 | 6675 |

| | | | | | | | | | |
|---|---|---|---|---|---|---|---|---|---|
| ENSG000001 | 1100.40141 | 0.336892599 | 0.12816904 | 2.62850219 | 0.00857618 | 0.03139112 | PDCD10 | | 11235 |
| ENSG000001 | 279.550565 | 0.337169418 | 0.12827218 | 2.62854672 | 0.00857506 | 0.03139112 | RAB39A | | 54734 |
| ENSG000001 | 748.608013 | 0.353822848 | 0.13461054 | 2.62849288 | 0.00857642 | 0.03139112 | SATB1 | | 6304 |
| ENSG000001 | 205.032335 | 0.398257319 | 0.15152162 | 2.6283861 | 0.00857911 | 0.03139814 | TTC8 | | 123016 |
| ENSG000001 | 10.3346004 | -0.885482877 | 0.33690411 | -2.6282935 | 0.00858144 | 0.03140102 | LILRB2 | | 10288 |
| ENSG000002 | 16.3106973 | -0.796542759 | 0.30306132 | -2.628322 | 0.00858072 | 0.03140102 | NA | NA | |
| ENSG000002 | 7.14290673 | -0.956577062 | 0.36396387 | -2.6282198 | 0.0085833 | 0.03140499 | NA | NA | |
| ENSG000001 | 446.118395 | 0.384308465 | 0.14623832 | 2.62795998 | 0.00858986 | 0.03142235 | ABCC5 | | 10057 |
| ENSG000001 | 54.2831991 | 0.536091834 | 0.20399696 | 2.62794029 | 0.00859036 | 0.03142235 | HPCAL1 | | 3241 |
| ENSG000001 | 48.1769688 | 0.554583343 | 0.2110335 | 2.6279398 | 0.00859037 | 0.03142235 | TSPAN7 | | 7102 |
| ENSG000001 | 611.175208 | 0.365184949 | 0.13896824 | 2.62783028 | 0.00859314 | 0.0314268 | SAP130 | | 79595 |
| ENSG000002 | 3.43627233 | -0.991905561 | 0.37745762 | -2.6278594 | 0.0085924 | 0.0314268 | NA | NA | |
| ENSG000002 | 3.10456063 | -0.985565733 | 0.37505629 | -2.6277808 | 0.00859439 | 0.03142853 | NA | NA | |
| ENSG000001 | 1777.88176 | 0.327981205 | 0.12481838 | 2.62766755 | 0.00859725 | 0.03143616 | MKLN1 | | 4289 |
| ENSG00000C | 46.3489519 | -0.612305412 | 0.23304202 | -2.6274464 | 0.00860284 | 0.03145094 | TP63 | | 8626 |
| ENSG000001 | 11.9744344 | -0.867466974 | 0.33015561 | -2.6274488 | 0.00860278 | 0.03145094 | TREML4 | | 285852 |
| ENSG000002 | 154.259451 | 0.427990708 | 0.16289956 | 2.62732889 | 0.00860581 | 0.03145896 | NA | NA | |
| ENSG000001 | 17.6686921 | -0.775912572 | 0.29533491 | -2.6272294 | 0.00860833 | 0.03146532 | C4orf22 | | 255119 |
| ENSG000001 | 64.0007647 | -0.525945231 | 0.20019362 | -2.6271828 | 0.00860951 | 0.0314668 | ERBB4 | | 2066 |
| ENSG000001 | 100.723154 | 0.437528009 | 0.1665446 | 2.62709218 | 0.0086118 | 0.03147234 | SH3GL2 | | 6456 |
| ENSG000001 | 94.7521261 | 0.501335386 | 0.19083862 | 2.62701226 | 0.00861382 | 0.03147445 | KRBA1 | | 84626 |
| ENSG000001 | 1558.72983 | 0.311229814 | 0.11847311 | 2.6270081 | 0.00861393 | 0.03147445 | NCOA2 | | 10499 |
| ENSG00000C | 357.612901 | 0.379726645 | 0.14455171 | 2.62692596 | 0.00861601 | 0.03147921 | NSF | | 4905 |
| ENSG000002 | 17.4139153 | -0.780857523 | 0.29725638 | -2.6268823 | 0.00861711 | 0.03148041 | GBA3 | | 57733 |
| ENSG000002 | 12.2530912 | -0.916573341 | 0.3489276 | -2.6268296 | 0.00861845 | 0.03148245 | C1orf140 | | 400804 |
| ENSG000002 | 3.76372492 | -0.984374527 | 0.37481954 | -2.6262626 | 0.00863282 | 0.03153211 | NA | NA | |
| ENSG000001 | 394.602753 | 0.349980065 | 0.13326969 | 2.62610397 | 0.00863684 | 0.03154396 | ARL3 | | 403 |
| ENSG000002 | 2.70041225 | -0.973465637 | 0.37069753 | -2.6260376 | 0.00863853 | 0.03154728 | NA | NA | |
| ENSG000001 | 110.77796 | 0.559169044 | 0.21295399 | 2.62577398 | 0.00864522 | 0.03156887 | HELZ2 | | 85441 |

| | | | | | | | | |
|---|---|---|---|---|---|---|---|---|
| ENSG000001 | 30.9523484 | -0.667801823 | 0.25434914 | -2.625532 | 0.00865137 | 0.03158847 | ALS2CR11 | 151254 |
| ENSG000002 | 20.6492524 | -0.754678386 | 0.28744195 | -2.6254984 | 0.00865222 | 0.03158875 | NA | NA |
| ENSG000001 | 11.5404198 | -0.885477284 | 0.33727884 | -2.6253567 | 0.00865582 | 0.03159905 | ATXN3L | 92552 |
| ENSG000002 | 4.08721284 | -0.993434653 | 0.37843502 | -2.6251129 | 0.00866202 | 0.03161884 | NA | NA |
| ENSG000001 | 54.5651632 | 0.536408288 | 0.2043536 | 2.62490254 | 0.00866738 | 0.03163531 | RBM11 | 54033 |
| ENSG000002 | 16.7197317 | -0.820635091 | 0.31263785 | -2.6248744 | 0.00866809 | 0.03163531 | LINC01516 | 101929025 |
| ENSG000002 | 6.83711772 | -0.975888471 | 0.37180117 | -2.624759 | 0.00867103 | 0.03164318 | NA | NA |
| ENSG000002 | 4.84483049 | -0.98893053 | 0.37678198 | -2.6246758 | 0.00867315 | 0.03164522 | NA | NA |
| ENSG000002 | 4.10559737 | -0.993343593 | 0.37845895 | -2.6247063 | 0.00867237 | 0.03164522 | NA | NA |
| ENSG000002 | 3.50787289 | -0.989641626 | 0.3770751 | -2.6245213 | 0.00867709 | 0.03165673 | NA | NA |
| ENSG000002 | 14.6876382 | -0.812603735 | 0.30964227 | -2.6243307 | 0.00868194 | 0.0316716 | NA | NA |
| ENSG000002 | 110.052887 | 0.44623519 | 0.17006534 | 2.62390443 | 0.00869282 | 0.03170841 | NA | NA |
| ENSG000002 | 4.48251981 | -0.991555553 | 0.37793086 | -2.6236427 | 0.0086995 | 0.03172993 | NA | NA |
| ENSG000002 | 79.7320856 | 0.54238236 | 0.20674257 | 2.62346729 | 0.00870398 | 0.03174342 | FKBPL | 63943 |
| ENSG000002 | 15.537534 | -0.797571199 | 0.30403559 | -2.6232823 | 0.00870871 | 0.03175495 | OR2J2 | 26707 |
| ENSG000002 | 10.0177442 | -0.908230473 | 0.34621708 | -2.6232977 | 0.00870831 | 0.03175495 | CCL5 | 6352 |
| ENSG000002 | 5.45014401 | -0.978344086 | 0.3729698 | -2.6231188 | 0.00871289 | 0.03176734 | NA | NA |
| ENSG000002 | 6.32820952 | -0.975082928 | 0.3717313 | -2.6230853 | 0.00871374 | 0.0317676 | NA | NA |
| ENSG000000 | 27.8485138 | -0.671354404 | 0.25595643 | -2.6229246 | 0.00871786 | 0.03177974 | MOV10L1 | 54456 |
| ENSG000002 | 8.34848486 | -0.929778797 | 0.35449042 | -2.6228602 | 0.00871951 | 0.03178289 | NA | NA |
| ENSG000001 | 508.829506 | 0.676450637 | 0.25791845 | 2.62273068 | 0.00872282 | 0.03179211 | PPP6R1 | 22870 |
| ENSG000001 | 3277.85548 | 0.30738717 | 0.11721089 | 2.62251379 | 0.00872837 | 0.0318095 | KIF11 | 3832 |
| ENSG000001 | 213.196047 | 0.36483318 | 0.13912511 | 2.62233877 | 0.00873286 | 0.03182298 | PTPDC1 | 138639 |
| ENSG000002 | 7.36720892 | -0.942843325 | 0.35955076 | -2.6222815 | 0.00873432 | 0.03182547 | NA | NA |
| ENSG000002 | 5.16044839 | -0.984004082 | 0.37532931 | -2.6217086 | 0.00874902 | 0.03187615 | NA | NA |
| ENSG000002 | 8.49939501 | -0.932270046 | 0.35560545 | -2.6216416 | 0.00875074 | 0.03187955 | NA | NA |
| ENSG000001 | 143.093368 | 0.394690149 | 0.15056382 | 2.62141435 | 0.00875658 | 0.03189508 | ACTR5 | 79913 |
| ENSG000002 | 96.9199318 | 0.437580906 | 0.16692365 | 2.62144343 | 0.00875583 | 0.03189508 | SPON1 | 10418 |
| ENSG000002 | 5.24043608 | -0.985958466 | 0.37612919 | -2.6213293 | 0.00875876 | 0.03190017 | LOC100129175 | 100129175 |

| | | | | | | | | |
|---|---|---|---|---|---|---|---|---|
| ENSG000002 | 417.343659 | 0.327623459 | 0.12498874 | 2.62122389 | 0.00876147 | 0.03190717 | NA | NA |
| ENSG000000 | 813.787629 | 0.34647879 | 0.13218813 | 2.62110365 | 0.00876456 | 0.03191556 | SH3GLB1 | 51100 |
| ENSG000002 | 6.50419981 | -0.96625327 | 0.3686573 | -2.6210067 | 0.00876705 | 0.03192177 | HBB | 3043 |
| ENSG000002 | 8.38580974 | -0.927869165 | 0.3540247 | -2.6209165 | 0.00876937 | 0.03192736 | NA | NA |
| ENSG000002 | 9.78165657 | -0.935826598 | 0.35707123 | -2.6208401 | 0.00877134 | 0.03193164 | NA | NA |
| ENSG000002 | 2.27274044 | -0.951253209 | 0.36297101 | -2.6207415 | 0.00877388 | 0.03193801 | NA | NA |
| ENSG000001 | 1722.58601 | 0.326739634 | 0.1246842 | 2.6205375 | 0.00877913 | 0.03195426 | ADIPOR1 | 51094 |
| ENSG000002 | 10.9006215 | -0.88768766 | 0.3387625 | -2.6203835 | 0.00878309 | 0.03196583 | LOC101927450 | 101927450 |
| ENSG000002 | 11.8845976 | 0.871269398 | 0.33250605 | 2.62031141 | 0.00878495 | 0.03196971 | NA | NA |
| ENSG000000 | 729.2833 | 0.343971239 | 0.1312744 | 2.62024609 | 0.00878663 | 0.03197297 | RSBN1 | 54665 |
| ENSG000001 | 511.524318 | 0.344603465 | 0.13152098 | 2.6201407 | 0.00878935 | 0.03197425 | PMM2 | 5373 |
| ENSG000002 | 5.05612088 | -0.990354597 | 0.37797159 | -2.6201826 | 0.00878827 | 0.03197425 | NA | NA |
| ENSG000002 | 8.20145357 | -0.940094814 | 0.35879321 | -2.6201578 | 0.00878891 | 0.03197425 | NA | NA |
| ENSG000002 | 27.0764345 | 0.682566996 | 0.26052423 | 2.61997515 | 0.00879362 | 0.0319869 | NA | NA |
| ENSG000001 | 4.44924258 | -0.98976383 | 0.37779499 | -2.6198437 | 0.00879701 | 0.03199636 | EVI2A | 2123 |
| ENSG000002 | 19.1074825 | -0.834805729 | 0.3186565 | -2.6197668 | 0.00879899 | 0.03200071 | LOC105375875 | 105375875 |
| ENSG000000 | 31.2196479 | -0.626689401 | 0.23921862 | -2.6197351 | 0.00879981 | 0.03200081 | ACTL6B | 51412 |
| ENSG000000 | 213.500514 | 0.381488528 | 0.14563449 | 2.61949294 | 0.00880606 | 0.0320178 | UPRT | 139596 |
| ENSG000001 | 2789.62587 | 0.321837529 | 0.12286173 | 2.61951001 | 0.00880562 | 0.0320178 | MIS18BP1 | 55320 |
| ENSG000001 | 33.3633696 | -0.661068081 | 0.25237181 | -2.6194212 | 0.00880791 | 0.03202166 | DSG1 | 1828 |
| ENSG000002 | 28.4633497 | -0.702074699 | 0.26804716 | -2.6192208 | 0.00881309 | 0.03203761 | NA | NA |
| ENSG000002 | 8.6516218 | -0.925991893 | 0.35356227 | -2.6190348 | 0.0088179 | 0.03205221 | NA | NA |
| ENSG000000 | 176.255356 | -0.374595099 | 0.14305214 | -2.6185913 | 0.00882937 | 0.03209103 | LIMCH1 | 22998 |
| ENSG000001 | 76.3653833 | 0.494090221 | 0.18870123 | 2.61837306 | 0.00883502 | 0.03210869 | ANKRD16 | 54522 |
| ENSG000002 | 6.4034714 | -0.961918346 | 0.36738814 | -2.6182619 | 0.0088379 | 0.03211627 | NA | NA |
| ENSG000001 | 6.7183719 | -0.956201217 | 0.36521346 | -2.6181982 | 0.00883955 | 0.03211939 | ZNF98 | 148198 |
| ENSG000002 | 96.618276 | 0.437592145 | 0.16714948 | 2.61796891 | 0.00884549 | 0.0321381 | MATR3 | 9782 |
| ENSG000002 | 5.18732747 | -0.983262402 | 0.37564631 | -2.6175218 | 0.00885708 | 0.03217735 | NA | NA |
| ENSG000001 | 955.081963 | 0.342336673 | 0.13079216 | 2.61740975 | 0.00885999 | 0.03218503 | JMY | 133746 |

| | | | | | | | | |
|---|---|---|---|---|---|---|---|---|
| ENSG000002 | 2.30256781 | -0.95015633 | 0.36304321 | -2.617199 | 0.00886546 | 0.03220202 | NA | NA |
| ENSG000002 | 4.42248824 | -0.98841619 | 0.37768939 | -2.6170081 | 0.00887042 | 0.03221716 | NA | NA |
| ENSG000002 | 13.901948 | -0.842543414 | 0.32196959 | -2.6168416 | 0.00887475 | 0.03222999 | LINC00879 | 255025 |
| ENSG000002 | 14.1783527 | -0.837709861 | 0.32014169 | -2.6166847 | 0.00887883 | 0.03224192 | NA | NA |
| ENSG000001 | 10.4566241 | -0.889876621 | 0.34009044 | -2.6165882 | 0.00888134 | 0.03224526 | HOXA2 | 3199 |
| ENSG000002 | 180.371004 | -0.365070103 | 0.13952106 | -2.6165951 | 0.00888116 | 0.03224526 | NA | NA |
| ENSG000001 | 704.925416 | 0.346217245 | 0.13238121 | 2.61530498 | 0.00891478 | 0.03236377 | ATP1B1 | 481 |
| ENSG000001 | 1415.06518 | 0.327164096 | 0.12510779 | 2.61505767 | 0.00892124 | 0.03238431 | TMEM132B | 114795 |
| ENSG000002 | 10.5981701 | -0.89623518 | 0.34274888 | -2.6148449 | 0.0089268 | 0.03239965 | NA | NA |
| ENSG000002 | 7.65915598 | -0.939572724 | 0.35932394 | -2.6148348 | 0.00892706 | 0.03239965 | LOC101927318 | 101927318 |
| ENSG000001 | 70.3340698 | 0.484970761 | 0.18548823 | 2.6145635 | 0.00893416 | 0.03242249 | CCDC102B | 79839 |
| ENSG000001 | 202.845529 | 0.364328117 | 0.13935128 | 2.61445835 | 0.00893691 | 0.03242957 | FAIM | 55179 |
| ENSG000001 | 5.20489515 | -0.983440961 | 0.37618052 | -2.6142793 | 0.00894159 | 0.03244367 | GPR31 | 2853 |
| ENSG000002 | 27.5988038 | -0.678863099 | 0.25969863 | -2.6140418 | 0.00894781 | 0.03246332 | CCDC13 | 152206 |
| ENSG000001 | 16.3532113 | -0.783054039 | 0.29957736 | -2.6138626 | 0.0089525 | 0.03247745 | UCP3 | 7352 |
| ENSG000002 | 12.4033924 | -0.851700514 | 0.32585844 | -2.6137132 | 0.00895642 | 0.03248695 | LOC101928371 | 101928371 |
| ENSG000002 | 20.788083 | -0.742026027 | 0.28389853 | -2.6137016 | 0.00895672 | 0.03248695 | MGC32805 | 153163 |
| ENSG000001 | 280.433516 | 0.370790244 | 0.14188772 | 2.61326529 | 0.00896817 | 0.03252554 | THG1L | 54974 |
| ENSG000001 | 387.005546 | 0.326522677 | 0.12495122 | 2.61320115 | 0.00896985 | 0.03252874 | EIF1AD | 84285 |
| ENSG000001 | 16.9683587 | -0.825992367 | 0.31608975 | -2.6131577 | 0.00897099 | 0.03252996 | ZSWIM2 | 151112 |
| ENSG000001 | 17.7840091 | -0.765601794 | 0.29299083 | -2.6130572 | 0.00897363 | 0.03253663 | KRT77 | 374454 |
| ENSG000002 | 2.67960235 | -0.968816389 | 0.37076505 | -2.6130197 | 0.00897461 | 0.03253718 | NA | NA |
| ENSG000002 | 24.7493136 | -0.690530266 | 0.26426822 | -2.6129902 | 0.00897539 | 0.03253718 | LOC285556 | 285556 |
| ENSG000001 | 240.128329 | 0.403433883 | 0.15440748 | 2.61278718 | 0.00898072 | 0.03255047 | ZBTB5 | 9925 |
| ENSG000002 | 3.41895619 | -0.985058821 | 0.37701629 | -2.6127752 | 0.00898104 | 0.03255047 | NA | NA |
| ENSG000002 | 2.75764651 | -0.964662522 | 0.36921221 | -2.612759 | 0.00898146 | 0.03255047 | NA | NA |
| ENSG00000C | 12.2869909 | -0.842638248 | 0.32252534 | -2.6126265 | 0.00898494 | 0.03255876 | ALX4 | 60529 |
| ENSG000002 | 6.24917536 | -0.969717565 | 0.37117521 | -2.6125602 | 0.00898669 | 0.03255876 | NA | NA |
| ENSG000002 | 100.359654 | 0.4424233 | 0.1693454 | 2.61254982 | 0.00898696 | 0.03255876 | IKBKE | 9641 |

| | | | | | | | | |
|---|---|---|---|---|---|---|---|---|
| ENSG000002 | 7.12285884 | -0.965152067 | 0.36942142 | -2.6126045 | 0.00898552 | 0.03255876 | NA | NA |
| ENSG000001 | 224.777205 | 0.382555284 | 0.14645293 | 2.61213821 | 0.00899779 | 0.03259507 | ZNF136 | 7695 |
| ENSG000001 | 1605.06515 | 0.299266611 | 0.11457148 | 2.61205152 | 0.00900007 | 0.03260043 | MTR | 4548 |
| ENSG000001 | 246.662325 | 0.36625121 | 0.14021981 | 2.61197905 | 0.00900198 | 0.03260181 | PIGF | 5281 |
| ENSG000002 | 10.5625301 | -0.89577662 | 0.34294979 | -2.611976 | 0.00900206 | 0.03260181 | NA | NA |
| ENSG000002 | 46.3175128 | -0.589646683 | 0.22575535 | -2.6118836 | 0.00900449 | 0.03260771 | NA | NA |
| ENSG000002 | 17.3268236 | -0.81551174 | 0.31227115 | -2.6115501 | 0.00901328 | 0.03263662 | NA | NA |
| ENSG000001 | 3939.43747 | 0.280178187 | 0.10729357 | 2.61132326 | 0.00901926 | 0.03265536 | NREP | 9315 |
| ENSG000001 | 132.226328 | 0.526917551 | 0.20180979 | 2.61096128 | 0.00902881 | 0.03268702 | CAPN5 | 726 |
| ENSG000001 | 215.235029 | 0.376036906 | 0.14402456 | 2.61092204 | 0.00902985 | 0.03268785 | DMRTA1 | 63951 |
| ENSG000001 | 39.8658711 | -0.614906167 | 0.23552635 | -2.6107744 | 0.00903375 | 0.03269905 | PDE6A | 5145 |
| ENSG000002 | 5.13968696 | -0.981883635 | 0.37609543 | -2.61073 | 0.00903492 | 0.03270037 | NA | NA |
| ENSG000001 | 162.641045 | 0.451145621 | 0.17280733 | 2.61068569 | 0.00903609 | 0.03270169 | ILKAP | 80895 |
| ENSG000001 | 172.498111 | 0.398833843 | 0.15277881 | 2.61053121 | 0.00904017 | 0.03271355 | NEIL2 | 252969 |
| ENSG000002 | 17.546218 | -0.773078383 | 0.29615554 | -2.6103796 | 0.00904418 | 0.03272513 | NA | NA |
| ENSG000002 | 5.51538513 | -0.97739755 | 0.37444867 | -2.610231 | 0.00904811 | 0.03273643 | NA | NA |
| ENSG000001 | 2.66402918 | -0.967059651 | 0.3705002 | -2.6101461 | 0.00905036 | 0.03273871 | CST6 | 1474 |
| ENSG000002 | 12.602483 | -0.833686672 | 0.31939936 | -2.6101702 | 0.00904972 | 0.03273871 | LINC00486 | 285045 |
| ENSG000002 | 8.12253096 | -0.933560803 | 0.35768659 | -2.6099967 | 0.00905431 | 0.0327501 | NA | NA |
| ENSG000002 | 6.60658587 | -0.966441169 | 0.37031544 | -2.6097782 | 0.00906009 | 0.03276809 | NA | NA |
| ENSG000001 | 943.941442 | 0.293384098 | 0.11242015 | 2.60971089 | 0.00906188 | 0.0327687 | DYNLRB1 | 83658 |
| ENSG000001 | 9.52536467 | -0.901272459 | 0.34535077 | -2.6097305 | 0.00906136 | 0.0327687 | NA | NA |
| ENSG000001 | 1431.58515 | 0.335208246 | 0.12844916 | 2.60965697 | 0.00906331 | 0.03277094 | CBL | 867 |
| ENSG000002 | 3.02223428 | -0.977626031 | 0.3746986 | -2.6090997 | 0.00907808 | 0.03282143 | NA | NA |
| ENSG000001 | 303.420553 | 0.363178068 | 0.13919943 | 2.60904845 | 0.00907944 | 0.03282342 | AGO4 | 192670 |
| ENSG000001 | 11.1332717 | -0.88282613 | 0.33840161 | -2.6088118 | 0.00908572 | 0.0328432 | MSX2 | 4488 |
| ENSG000001 | 824.53108 | 0.319152704 | 0.12234889 | 2.60854604 | 0.00909278 | 0.03286578 | ISG20L2 | 81875 |
| ENSG000002 | 69.1847619 | -0.487414644 | 0.18687774 | -2.6082006 | 0.00910196 | 0.03289604 | CR1 | 1378 |
| ENSG00000C | 866.284507 | 0.347142296 | 0.1330997 | 2.60813725 | 0.00910364 | 0.03289696 | KLHL42 | 57542 |

| | | | | | | | | |
|---|---|---|---|---|---|---|---|---|
| ENSG000002 | 4.75356687 | -0.982615715 | 0.37675104 | -2.60813 | 0.00910384 | 0.03289696 | NA | NA |
| ENSG000002 | 23.9477661 | -0.722289767 | 0.276944 | -2.6080715 | 0.00910539 | 0.03289965 | LOC100507195 | 100507195 |
| ENSG000002 | 23.6909345 | -0.700568554 | 0.26862823 | -2.6079484 | 0.00910867 | 0.03290602 | NA | NA |
| ENSG000002 | 11.8950843 | -0.874306899 | 0.33524754 | -2.6079443 | 0.00910878 | 0.03290602 | NA | NA |
| ENSG000002 | 26.1798321 | -0.689795171 | 0.26450946 | -2.607828 | 0.00911187 | 0.03291427 | LOC101929199 | 101929199 |
| ENSG000001 | 45.6605981 | 0.560352705 | 0.21488295 | 2.60771134 | 0.00911498 | 0.03291669 | C14orf28 | 122525 |
| ENSG000002 | 51.1255261 | -0.552183909 | 0.21174841 | -2.6077358 | 0.00911433 | 0.03291669 | CYP1B1-AS1 | 285154 |
| ENSG000002 | 2.7754908 | -0.963877404 | 0.36961873 | -2.6077613 | 0.00911365 | 0.03291669 | NA | NA |
| ENSG000001 | 18.1620886 | -0.765571541 | 0.2936228 | -2.6073301 | 0.00912514 | 0.03294163 | AK8 | 158067 |
| ENSG000002 | 12.0149054 | -0.859890963 | 0.32979468 | -2.6073524 | 0.00912454 | 0.03294163 | NA | NA |
| ENSG000002 | 7.99787369 | -0.927531013 | 0.35572771 | -2.6074185 | 0.00912278 | 0.03294163 | LOC105376100 | 105376100 |
| ENSG000002 | 5.85001717 | -0.97206188 | 0.37281157 | -2.6073812 | 0.00912377 | 0.03294163 | FCGR1C | 100132417 |
| ENSG00000C | 23.1713507 | -0.713396483 | 0.27362441 | -2.6072107 | 0.00912832 | 0.03294726 | PTPRH | 5794 |
| ENSG000002 | 4.47218796 | -0.985114465 | 0.37784214 | -2.6072118 | 0.00912829 | 0.03294726 | NA | NA |
| ENSG000001 | 941.779918 | 0.331543809 | 0.12716797 | 2.60713294 | 0.00913039 | 0.0329518 | MORC3 | 23515 |
| ENSG000002 | 10.9918371 | -0.876882295 | 0.33634607 | -2.6070835 | 0.00913171 | 0.03295363 | NA | NA |
| ENSG000002 | 9.97886491 | -0.901026766 | 0.3456371 | -2.6068578 | 0.00913773 | 0.03297243 | NA | NA |
| ENSG000001 | 622.234378 | 0.353740721 | 0.13570029 | 2.60677935 | 0.00913982 | 0.03297411 | TLN2 | 83660 |
| ENSG000002 | 19.3817244 | -0.77596325 | 0.29767022 | -2.6067883 | 0.00913959 | 0.03297411 | NA | NA |
| ENSG000001 | 377.43019 | 0.389385236 | 0.14938918 | 2.60651567 | 0.00914686 | 0.03299657 | LPAR4 | 2846 |
| ENSG00000C | 752.018945 | 0.335682099 | 0.12878986 | 2.60643272 | 0.00914908 | 0.03300163 | UFL1 | 23376 |
| ENSG000002 | 3.02513106 | -0.976564025 | 0.37470555 | -2.6062171 | 0.00915484 | 0.03301947 | NA | NA |
| ENSG000001 | 18.1862114 | -0.802562319 | 0.30798702 | -2.6058316 | 0.00916515 | 0.03305372 | NA | NA |
| ENSG000002 | 8.23460956 | -0.936920343 | 0.35958207 | -2.6055813 | 0.00917185 | 0.03307494 | LOC105369860 | 105369860 |
| ENSG000002 | 12.0537984 | -0.848722941 | 0.32574597 | -2.6054749 | 0.0091747 | 0.03308228 | LOC105378966 | 105378966 |
| ENSG000002 | 12.0628608 | -0.872612732 | 0.33495158 | -2.6051907 | 0.00918231 | 0.03310678 | LINC01320 | 104355288 |
| ENSG000002 | 286.381618 | 0.441539689 | 0.16951902 | 2.60466158 | 0.0091965 | 0.033155 | CCDC85C | 317762 |
| ENSG00000C | 3638.6033 | 0.297547027 | 0.11425293 | 2.60428365 | 0.00920665 | 0.03318864 | RB1CC1 | 9821 |
| ENSG000001 | 880.387562 | 0.372659623 | 0.14311279 | 2.60395746 | 0.00921542 | 0.03321433 | PAN3 | 255967 |

| | | | | | | | | |
|---|---|---|---|---|---|---|---|---|
| ENSG000002 | 9.05156117 | -0.915037436 | 0.35139931 | -2.6039819 | 0.00921476 | 0.03321433 | NA | NA |
| ENSG000002 | 11.2515734 | -0.868477045 | 0.33353004 | -2.6038946 | 0.00921711 | 0.03321748 | NA | NA |
| ENSG000002 | 12.5555939 | -0.847539349 | 0.32553889 | -2.6034965 | 0.00922782 | 0.03325312 | LINC01426 | 100506385 |
| ENSG000001 | 8.62694284 | -0.955650358 | 0.36707286 | -2.6034351 | 0.00922947 | 0.03325322 | SLC26A1 | 10861 |
| ENSG000002 | 3.70400793 | -0.985426002 | 0.37850999 | -2.6034346 | 0.00922949 | 0.03325322 | NA | NA |
| ENSG000002 | 14.4245947 | -0.825779926 | 0.31720584 | -2.6032936 | 0.00923329 | 0.03326394 | NA | NA |
| ENSG000002 | 5.80837535 | -0.968319545 | 0.37199205 | -2.6030651 | 0.00923944 | 0.03328316 | LOC102467079 | 102467079 |
| ENSG000001 | 10.899243 | -0.86364445 | 0.33178403 | -2.603032 | 0.00924033 | 0.03328342 | NA | NA |
| ENSG000002 | 58.632718 | -0.512002337 | 0.19670503 | -2.6028939 | 0.00924405 | 0.03329387 | NA | NA |
| ENSG000001 | 211.652175 | 0.349310389 | 0.13420372 | 2.60283677 | 0.0092456 | 0.03329551 | ENPP5 | 59084 |
| ENSG000001 | 3546.52925 | 0.430693699 | 0.16547347 | 2.60279609 | 0.00924669 | 0.03329551 | MKI67 | 4288 |
| ENSG000001 | 70.5398262 | 0.549583571 | 0.21115207 | 2.60278564 | 0.00924698 | 0.03329551 | CCDC24 | 149473 |
| ENSG000002 | 12.3578043 | -0.839885218 | 0.3226947 | -2.602724 | 0.00924864 | 0.03329855 | NA | NA |
| ENSG000001 | 5340.29641 | 0.301720984 | 0.11593084 | 2.60259472 | 0.00925213 | 0.03330814 | HELLS | 3070 |
| ENSG000001 | 1489.22145 | 0.410712948 | 0.15781929 | 2.60242557 | 0.00925669 | 0.03332162 | ARF6 | 382 |
| ENSG000001 | 445.778695 | 0.343817594 | 0.1321277 | 2.6021614 | 0.00926382 | 0.03334138 | VPS54 | 51542 |
| ENSG000002 | 20.8938698 | 0.72899236 | 0.28014875 | 2.60216179 | 0.00926381 | 0.03334138 | NA | NA |
| ENSG000001 | 2324.00214 | 0.361907032 | 0.13908417 | 2.60207202 | 0.00926624 | 0.03334711 | INO80D | 54891 |
| ENSG000001 | 133.398353 | -0.401732389 | 0.15441154 | -2.6016993 | 0.00927632 | 0.03337745 | SRGAP3 | 9901 |
| ENSG000002 | 43.0855594 | 0.62599663 | 0.2406079 | 2.60172935 | 0.0092755 | 0.03337745 | NA | NA |
| ENSG00000C | 2.3041711 | -0.940247025 | 0.36141107 | -2.6015999 | 0.009279 | 0.03337823 | NA | NA |
| ENSG000002 | 5.67825246 | -0.964134764 | 0.37058872 | -2.6016301 | 0.00927819 | 0.03337823 | LOC100996351 | 100996351 |
| ENSG000002 | 3.80582221 | -0.983843206 | 0.37815974 | -2.6016604 | 0.00927737 | 0.03337823 | NA | NA |
| ENSG000001 | 176.238672 | 0.401754573 | 0.15443405 | 2.60146372 | 0.00928269 | 0.03338853 | TBC1D10B | 26000 |
| ENSG000001 | 22.5836229 | -0.738478965 | 0.28389529 | -2.6012371 | 0.00928882 | 0.03340763 | NA | NA |
| ENSG000001 | 936.183619 | 0.294700934 | 0.11329953 | 2.60107808 | 0.00929313 | 0.03341762 | DENND4C | 55667 |
| ENSG000002 | 10.9749792 | -0.866977861 | 0.33331538 | -2.6010737 | 0.00929325 | 0.03341762 | LIPN | 643418 |
| ENSG000001 | 13.9274304 | -0.818876249 | 0.31483961 | -2.6009315 | 0.0092971 | 0.03342852 | MAB21L3 | 126868 |
| ENSG000001 | 23.5395747 | -0.740698532 | 0.28479853 | -2.6007807 | 0.00930119 | 0.03344024 | ANKDD1B | 728780 |

| | | | | | | | | |
|---|---|---|---|---|---|---|---|---|
| ENSG000002 | 3.68962953 | -0.984455935 | 0.37855584 | -2.6005567 | 0.00930726 | 0.03345912 | NA | NA |
| ENSG000001 | 719.090633 | 0.439563703 | 0.16905367 | 2.60014297 | 0.00931849 | 0.03349652 | CNOT6 | 57472 |
| ENSG000002 | 19.5263011 | -0.816747767 | 0.31413405 | -2.5999975 | 0.00932244 | 0.03350775 | NA | NA |
| ENSG000001 | 125.24329 | 0.503347979 | 0.19360152 | 2.59991749 | 0.00932462 | 0.03351259 | TMEM201 | 199953 |
| ENSG000002 | 17.9680016 | -0.792844054 | 0.3049586 | -2.5998416 | 0.00932668 | 0.03351704 | NA | NA |
| ENSG000001 | 4.41161898 | -0.982263114 | 0.37782416 | -2.5997891 | 0.00932811 | 0.03351919 | LINC00308 | 54143 |
| ENSG000002 | 10.4047155 | -0.878295247 | 0.33783741 | -2.5997572 | 0.00932897 | 0.03351933 | NA | NA |
| ENSG000002 | 30.2625884 | -0.636076553 | 0.2446842 | -2.5995817 | 0.00933375 | 0.03353351 | NA | NA |
| ENSG000002 | 3.34333584 | -0.981650115 | 0.37766722 | -2.5992463 | 0.00934287 | 0.03356332 | NA | NA |
| ENSG000001 | 11.4207604 | -0.912744943 | 0.35116834 | -2.5991663 | 0.00934505 | 0.03356672 | MMEL1 | 79258 |
| ENSG000001 | 9.42921929 | -0.891042106 | 0.3428245 | -2.5991203 | 0.0093463 | 0.03356672 | GOT1L1 | 137362 |
| ENSG000002 | 18.6367343 | 0.829996743 | 0.31933414 | 2.5991482 | 0.00934554 | 0.03356672 | NA | NA |
| ENSG000001 | 15.7981913 | -0.80718834 | 0.31061913 | -2.5986434 | 0.00935929 | 0.03361041 | HMGCS2 | 3158 |
| ENSG000002 | 815.367411 | -0.365911654 | 0.14081555 | -2.5985174 | 0.00936273 | 0.03361977 | LOC101927721 | 101927721 |
| ENSG000002 | 201.197488 | -0.367269734 | 0.14134666 | -2.5983616 | 0.00936698 | 0.03363205 | NEAT1 | 283131 |
| ENSG000001 | 151.24578 | 0.427124449 | 0.16440917 | 2.5979357 | 0.00937861 | 0.03367081 | AP4B1 | 10717 |
| ENSG000002 | 11.7710651 | -0.847774884 | 0.3263469 | -2.5977721 | 0.00938307 | 0.03368089 | LRRC37A5P | 652972 |
| ENSG000002 | 4.04845506 | -0.98319526 | 0.37847198 | -2.597802 | 0.00938226 | 0.03368089 | NA | NA |
| ENSG000001 | 6.10331622 | -0.960170125 | 0.36965613 | -2.5974684 | 0.00939138 | 0.03370771 | LINC01561 | 404216 |
| ENSG000002 | 6.00816443 | -0.958602436 | 0.36906056 | -2.5974123 | 0.00939291 | 0.03371024 | THEM7P | 106480297 |
| ENSG000001 | 28.1310008 | -0.728360234 | 0.28044167 | -2.5971898 | 0.009399 | 0.0337291 | GSTM5 | 2949 |
| ENSG000001 | 191.073359 | 0.505576841 | 0.19467055 | 2.59708954 | 0.00940174 | 0.03373595 | SNAPC4 | 6621 |
| ENSG000001 | 60.8653494 | -0.49452364 | 0.19041756 | -2.5970486 | 0.00940286 | 0.03373699 | DYNC1I1 | 1780 |
| ENSG000002 | 5.07535192 | -0.975129291 | 0.3755077 | -2.596829 | 0.00940887 | 0.03375557 | NA | NA |
| ENSG000002 | 18.2730826 | -0.789373393 | 0.30400237 | -2.5966028 | 0.00941507 | 0.03377483 | LOC100288181 | 100288181 |
| ENSG000001 | 1361.02419 | 0.307319758 | 0.11835686 | 2.59655224 | 0.00941646 | 0.03377681 | BAZ1A | 11177 |
| ENSG000002 | 21.3414819 | -0.730404673 | 0.2813543 | -2.5960317 | 0.00943074 | 0.03382504 | LOC101927973 | 101927973 |
| ENSG000001 | 177.849871 | 0.392893286 | 0.1513682 | 2.59561316 | 0.00944223 | 0.03386326 | MED19 | 219541 |
| ENSG000001 | 6.69661644 | -0.948105465 | 0.36530642 | -2.5953704 | 0.0094489 | 0.03388175 | CPN2 | 1370 |

| | | | | | | | | |
|---|---|---|---|---|---|---|---|---|
| ENSG000002 | 7.13532903 | -0.951105847 | 0.36646327 | -2.5953648 | 0.00944906 | 0.03388175 | NA | NA |
| ENSG000001 | 8.88845043 | -0.906452596 | 0.34927486 | -2.5952415 | 0.00945245 | 0.03389091 | OPN4 | 94233 |
| ENSG000002 | 23.7782326 | 0.701941258 | 0.27047688 | 2.59519875 | 0.00945362 | 0.03389213 | NA | NA |
| ENSG000001 | 26.6515139 | -0.66313611 | 0.25554347 | -2.5950032 | 0.009459 | 0.03390585 | ASB2 | 51676 |
| ENSG000001 | 5.07757151 | -0.973468999 | 0.37513272 | -2.5949989 | 0.00945912 | 0.03390585 | C15orf53 | 400359 |
| ENSG000002 | 116.987581 | 0.454087057 | 0.1750161 | 2.59454446 | 0.00947164 | 0.0339477 | NA | NA |
| ENSG000001 | 13.7112418 | -0.809621749 | 0.31208622 | -2.5942246 | 0.00948045 | 0.0339763 | MS4A8 | 83661 |
| ENSG000001 | 40.4403927 | 0.613695079 | 0.23664089 | 2.59336021 | 0.00950432 | 0.0340528 | PRR5 | 55615 |
| ENSG000001 | 230.510123 | 0.405829585 | 0.15648519 | 2.59340571 | 0.00950306 | 0.0340528 | CACNA1H | 8912 |
| ENSG000002 | 9.41550478 | -0.892487078 | 0.34414151 | -2.5933723 | 0.00950398 | 0.0340528 | NA | NA |
| ENSG000002 | 16.7189632 | -0.779701216 | 0.30066714 | -2.5932372 | 0.00950772 | 0.03406197 | NA | NA |
| ENSG000001 | 42.062778 | -0.58026319 | 0.22376541 | -2.5931765 | 0.0095094 | 0.03406498 | SLC30A10 | 55532 |
| ENSG000002 | 2.17831025 | -0.943321825 | 0.36382288 | -2.5928051 | 0.00951967 | 0.03409877 | NA | NA |
| ENSG000001 | 5.10648507 | -0.973797007 | 0.37559793 | -2.5926581 | 0.00952374 | 0.03411033 | DIRC1 | 116093 |
| ENSG000002 | 33.3506355 | -0.620104429 | 0.23919943 | -2.592416 | 0.00953044 | 0.03413133 | CPB2-AS1 | 100509894 |
| ENSG000001 | 9.35012164 | -0.888650402 | 0.3428015 | -2.5923177 | 0.00953317 | 0.03413807 | WFDC10B | 280664 |
| ENSG000001 | 25.8192333 | -0.665954771 | 0.25691962 | -2.5920744 | 0.00953991 | 0.03415921 | SLC5A11 | 115584 |
| ENSG000001 | 116.802153 | 0.772106044 | 0.29787963 | 2.59200684 | 0.00954179 | 0.0341629 | TELO2 | 9894 |
| ENSG000001 | 92.5522784 | 0.439844176 | 0.16969818 | 2.59192046 | 0.00954418 | 0.03416846 | STRA6 | 64220 |
| ENSG000001 | 1008.03775 | 0.376327241 | 0.1452045 | 2.59170504 | 0.00955016 | 0.03418404 | SHOC2 | 8036 |
| ENSG000001 | 134.984618 | 0.483071311 | 0.18639366 | 2.59167249 | 0.00955106 | 0.03418404 | PER1 | 5187 |
| ENSG000001 | 215.491048 | 0.344313222 | 0.13285225 | 2.5917003 | 0.00955029 | 0.03418404 | ZNF860 | 344787 |
| ENSG000001 | 474.843923 | 0.403859896 | 0.15584236 | 2.59146418 | 0.00955685 | 0.03420084 | KIF21B | 23046 |
| ENSG000002 | 11.5692756 | -0.857683536 | 0.33096759 | -2.5914427 | 0.00955745 | 0.03420084 | NA | NA |
| ENSG000002 | 25.1013573 | -0.72953979 | 0.28153129 | -2.5913276 | 0.00956064 | 0.03420927 | NA | NA |
| ENSG000002 | 816.533397 | 0.412908087 | 0.15935947 | 2.59104838 | 0.0095684 | 0.03423402 | OTUD7B | 56957 |
| ENSG000001 | 4943.4779 | 0.315719533 | 0.121853 | 2.59098698 | 0.00957011 | 0.0342371 | RBPJ | 3516 |
| ENSG000002 | 4.75914383 | -0.977220449 | 0.37716651 | -2.5909524 | 0.00957107 | 0.03423753 | HSPE1P21 | 106480258 |
| ENSG000002 | 214.696782 | 0.35181258 | 0.13580372 | 2.59059608 | 0.00958099 | 0.03426997 | FMN1 | 342184 |

| | | | | | | | | |
|---|---|---|---|---|---|---|---|---|
| ENSG000001 | 27.9419655 | -0.658176742 | 0.25407048 | -2.5905282 | 0.00958288 | 0.03427371 | PADI1 | 29943 |
| ENSG000001 | 545.818102 | 0.31366287 | 0.12110902 | 2.58992168 | 0.00959978 | 0.03433024 | FBXO3 | 26273 |
| ENSG000002 | 15.0175696 | -0.796930482 | 0.30770703 | -2.5899001 | 0.00960038 | 0.03433024 | NA | NA |
| ENSG000001 | 20.0318217 | -0.771864614 | 0.29808614 | -2.5894012 | 0.0096143 | 0.03437493 | ZNF536 | 9745 |
| ENSG000002 | 11.4920029 | -0.855223422 | 0.33027969 | -2.5893915 | 0.00961457 | 0.03437493 | LOC101928823 | 101928823 |
| ENSG000001 | 78.9221119 | 0.454418719 | 0.17549976 | 2.58928401 | 0.00961757 | 0.03438263 | MSRB2 | 22921 |
| ENSG00000C | 215.577344 | 0.449563564 | 0.17364134 | 2.58903534 | 0.00962452 | 0.03440444 | CELSR1 | 9620 |
| ENSG000001 | 7.53218631 | -0.922900519 | 0.35647234 | -2.5889821 | 0.00962601 | 0.03440673 | TEKT5 | 146279 |
| ENSG000001 | 311.604564 | 0.317488638 | 0.12264909 | 2.58859343 | 0.00963688 | 0.03444255 | DTD2 | 112487 |
| ENSG000001 | 752.466853 | 0.330503287 | 0.12768012 | 2.58852582 | 0.00963877 | 0.03444347 | PLEKHA1 | 59338 |
| ENSG000001 | 231.893173 | 0.392118305 | 0.15148339 | 2.58852346 | 0.00963884 | 0.03444347 | PLEKHA2 | 59339 |
| ENSG000001 | 196.442939 | 0.407573081 | 0.15747064 | 2.588248 | 0.00964655 | 0.03446799 | HAUS8 | 93323 |
| ENSG000001 | 3889.24671 | 0.362101285 | 0.13992624 | 2.58780113 | 0.00965907 | 0.0345097 | SLTM | 79811 |
| ENSG000002 | 15.0389486 | -0.796685056 | 0.30787183 | -2.5877166 | 0.00966144 | 0.03451512 | NA | NA |
| ENSG000002 | 4799.94533 | 0.331293658 | 0.12803271 | 2.58757054 | 0.00966554 | 0.03452672 | PWAR5 | 8123 |
| ENSG000001 | 38.9630837 | -0.581511525 | 0.22473948 | -2.5874916 | 0.00966775 | 0.03453004 | MFAP5 | 8076 |
| ENSG000002 | 7.52988356 | -0.944030781 | 0.36484609 | -2.5874768 | 0.00966817 | 0.03453004 | NA | NA |
| ENSG000001 | 4375.66771 | 0.375345272 | 0.14506609 | 2.58740876 | 0.00967008 | 0.03453077 | ACVR2B | 93 |
| ENSG000002 | 11.8478668 | -0.873758975 | 0.33769259 | -2.587439 | 0.00966923 | 0.03453077 | LOC101929237 | 101929237 |
| ENSG000001 | 18.2546462 | 0.771676837 | 0.29825522 | 2.58730374 | 0.00967303 | 0.03453826 | NA | NA |
| ENSG000001 | 1499.19985 | 0.318866394 | 0.1232494 | 2.58716379 | 0.00967696 | 0.03454925 | INTS8 | 55656 |
| ENSG000001 | 1320.01753 | 0.707407141 | 0.27343299 | 2.58713168 | 0.00967786 | 0.03454942 | UPF3B | 65109 |
| ENSG000001 | 15.1617594 | -0.789798072 | 0.30531253 | -2.5868512 | 0.00968574 | 0.03457452 | ZNF662 | 389114 |
| ENSG000002 | 6.68637991 | -0.947385383 | 0.36623931 | -2.5867933 | 0.00968737 | 0.03457728 | NA | NA |
| ENSG000001 | 331.640512 | 0.34043295 | 0.13160952 | 2.58668934 | 0.00969029 | 0.03458467 | HEATR5A | 25938 |
| ENSG00000C | 18.8882627 | -0.764975161 | 0.29574895 | -2.5865694 | 0.00969367 | 0.03459367 | STMN4 | 81551 |
| ENSG000001 | 294.651239 | 0.417519408 | 0.16142846 | 2.5864052 | 0.00969828 | 0.03460406 | MARVELD3 | 91862 |
| ENSG000002 | 5.09485643 | -0.972170079 | 0.37587628 | -2.5864098 | 0.00969816 | 0.03460406 | NA | NA |
| ENSG000001 | 85.0389747 | -0.471107662 | 0.18215479 | -2.586304 | 0.00970113 | 0.03461117 | COL24A1 | 255631 |

| | | | | | | | | | |
|---|---|---|---|---|---|---|---|---|---|
| ENSG00000C | 2130.52312 | 0.330596642 | 0.12783039 | 2.58621314 | 0.00970369 | 0.03461372 | NUP133 | | 55746 |
| ENSG000001 | 32.3050473 | -0.667345172 | 0.25803717 | -2.5862366 | 0.00970303 | 0.03461372 | GALNT5 | | 11227 |
| ENSG000001 | 169.693718 | 0.381374484 | 0.14746589 | 2.58618771 | 0.00970441 | 0.03461372 | DHRS3 | | 9249 |
| ENSG000002 | 6.01975075 | -0.955731716 | 0.36958644 | -2.5859491 | 0.00971113 | 0.03463465 | LOC102723427 | | 102723427 |
| ENSG000002 | 5.38447883 | -0.967219095 | 0.3740392 | -2.5858763 | 0.00971318 | 0.03463892 | NA | NA | |
| ENSG000001 | 576.322315 | 0.409452501 | 0.15834836 | 2.58577044 | 0.00971616 | 0.03464401 | SASH1 | | 23328 |
| ENSG000001 | 11.2644094 | -0.860940757 | 0.33295399 | -2.5857649 | 0.00971632 | 0.03464401 | GATA6 | | 2627 |
| ENSG000002 | 25.9148056 | -0.691592155 | 0.26746642 | -2.5857158 | 0.0097177 | 0.0346459 | NA | NA | |
| ENSG000001 | 454.155167 | 0.358957751 | 0.13883368 | 2.58552352 | 0.00972312 | 0.03466219 | KPNA5 | | 3841 |
| ENSG00000C | 11.1963935 | -0.8563149 | 0.33125475 | -2.5850645 | 0.00973608 | 0.03470532 | ARX | | 170302 |
| ENSG000001 | 385.890338 | 0.416395921 | 0.16108307 | 2.58497624 | 0.00973857 | 0.03471046 | TLE3 | | 7090 |
| ENSG000002 | 3.2865575 | -0.976059847 | 0.3775929 | -2.5849528 | 0.00973923 | 0.03471046 | NA | NA | |
| ENSG000002 | 24.2429446 | -0.704928167 | 0.27272194 | -2.5847872 | 0.00974391 | 0.03472409 | LOC101927292 | | 101927292 |
| ENSG000002 | 70.9525138 | 0.531409679 | 0.20559624 | 2.5847247 | 0.00974568 | 0.03472733 | NA | NA | |
| ENSG000001 | 1014.13508 | 0.383555265 | 0.14841018 | 2.58442696 | 0.0097541 | 0.03475427 | TAF1 | | 6872 |
| ENSG000001 | 817.721852 | 0.360397543 | 0.13945376 | 2.58435153 | 0.00975623 | 0.03475882 | ZIK1 | | 284307 |
| ENSG000002 | 3.68222506 | -0.978209918 | 0.37852389 | -2.5842752 | 0.00975839 | 0.03476346 | NA | NA | |
| ENSG000002 | 20.1608321 | 0.809022996 | 0.31307915 | 2.58408456 | 0.00976379 | 0.0347772 | NA | NA | |
| ENSG000002 | 12.8628158 | -0.82063885 | 0.31757506 | -2.5840784 | 0.00976396 | 0.0347772 | NA | NA | |
| ENSG000001 | 945.806614 | 0.356161542 | 0.13783166 | 2.58403285 | 0.00976525 | 0.03477873 | ZFP36L1 | | 677 |
| ENSG00000C | 470.030623 | 0.325729552 | 0.12606324 | 2.58385838 | 0.00977019 | 0.03478716 | ING3 | | 54556 |
| ENSG00000C | 13.4571929 | -0.830160446 | 0.32128361 | -2.5838867 | 0.00976939 | 0.03478716 | NOX3 | | 50508 |
| ENSG000001 | 37.7576631 | -0.601766589 | 0.23289016 | -2.5839073 | 0.00976881 | 0.03478716 | SLCO2B1 | | 11309 |
| ENSG000001 | 6562.84399 | 0.300110217 | 0.11615398 | 2.58372728 | 0.0097739 | 0.03479733 | SSB | | 6741 |
| ENSG00000C | 269.17354 | 0.383268548 | 0.14837035 | 2.58318829 | 0.00978919 | 0.03484563 | BID | | 637 |
| ENSG000001 | 4444.01809 | 0.283569077 | 0.10977461 | 2.58319362 | 0.00978904 | 0.03484563 | ACSL3 | | 2181 |
| ENSG00000C | 246.643535 | 0.33507647 | 0.12971745 | 2.58312567 | 0.00979097 | 0.03484728 | PCSK5 | | 5125 |
| ENSG000001 | 9.96987577 | -0.893175713 | 0.34577879 | -2.5830842 | 0.00979214 | 0.03484728 | NA | NA | |
| ENSG000002 | 450.152633 | 0.368444243 | 0.14263751 | 2.58308109 | 0.00979223 | 0.03484728 | LYN | | 4067 |

| | | | | | | | | |
|---|---|---|---|---|---|---|---|---|
| ENSG000002 | 8.41455499 | -0.919235807 | 0.35587685 | -2.5830166 | 0.00979406 | 0.03485074 | NA | NA |
| ENSG000001 | 9.43041352 | -0.887013158 | 0.34346708 | -2.5825275 | 0.00980796 | 0.03489711 | OR6C75 | 390323 |
| ENSG000001 | 39.4021056 | 0.694828025 | 0.26906199 | 2.58240867 | 0.00981133 | 0.03490607 | NPDC1 | 56654 |
| ENSG000002 | 2.18425991 | -0.938330384 | 0.36337126 | -2.5822912 | 0.00981467 | 0.03491489 | NA | NA |
| ENSG000001 | 1496.187 | 0.356620045 | 0.13810648 | 2.58221083 | 0.00981696 | 0.03491928 | PCNX | 22990 |
| ENSG000001 | 562.142139 | 0.429916043 | 0.16649298 | 2.58218722 | 0.00981763 | 0.03491928 | PVRL1 | 5818 |
| ENSG00000C | 6.46420291 | -0.969447754 | 0.37544195 | -2.5821509 | 0.00981867 | 0.0349199 | TACR2 | 6865 |
| ENSG000001 | 2372.11665 | 0.320053325 | 0.12395713 | 2.58196778 | 0.00982388 | 0.0349323 | NANOG | 79923 |
| ENSG000001 | 406.508116 | 0.311575215 | 0.12067285 | 2.58198267 | 0.00982345 | 0.0349323 | NA | NA |
| ENSG000002 | 22.8490525 | -0.69255387 | 0.26826514 | -2.5816022 | 0.00983429 | 0.03496626 | LINC01553 | 283025 |
| ENSG000001 | 94.8077932 | 0.459276234 | 0.17790846 | 2.58153117 | 0.00983631 | 0.03496732 | KIAA1161 | 57462 |
| ENSG000001 | 1137.48912 | 0.338407569 | 0.13108727 | 2.5815442 | 0.00983594 | 0.03496732 | XRCC2 | 7516 |
| ENSG000001 | 295.946937 | 0.41576749 | 0.16105655 | 2.58150002 | 0.0098372 | 0.03496741 | ZNF17 | 7565 |
| ENSG000002 | 96.4020163 | 0.44967571 | 0.17419889 | 2.5813925 | 0.00984026 | 0.03497524 | NA | NA |
| ENSG000001 | 88.5053333 | 0.443385432 | 0.17177215 | 2.58124167 | 0.00984456 | 0.03498746 | NMRK1 | 54981 |
| ENSG000001 | 80.3934003 | -0.455646016 | 0.17655546 | -2.5807529 | 0.00985851 | 0.03503396 | TMEM232 | 642987 |
| ENSG000002 | 6.07044805 | -0.954640745 | 0.36994485 | -2.5804947 | 0.00986589 | 0.03505709 | NA | NA |
| ENSG000001 | 56.7424602 | -0.513661782 | 0.19905822 | -2.58046 | 0.00986688 | 0.03505755 | NCKAP5 | 344148 |
| ENSG000001 | 851.626219 | 0.298020458 | 0.11549582 | 2.58035716 | 0.00986982 | 0.03506492 | LRIF1 | 55791 |
| ENSG000001 | 14.1045324 | -0.862452268 | 0.33425281 | -2.5802394 | 0.00987319 | 0.03506767 | MKRN3 | 7681 |
| ENSG000002 | 17.4203094 | -0.763323029 | 0.29583318 | -2.5802482 | 0.00987293 | 0.03506767 | SAMD5 | 389432 |
| ENSG000002 | 8.76518251 | -0.898287581 | 0.34813755 | -2.5802662 | 0.00987242 | 0.03506767 | NA | NA |
| ENSG000001 | 2505.80147 | 0.295618594 | 0.11457191 | 2.58020138 | 0.00987427 | 0.03506845 | NET1 | 10276 |
| ENSG000001 | 11.2748657 | -0.86500525 | 0.33525241 | -2.5801612 | 0.00987542 | 0.03506947 | CYP17A1 | 1586 |
| ENSG000001 | 190.540217 | 0.427538357 | 0.16570751 | 2.58007829 | 0.00987779 | 0.03507481 | IGLON5 | 402665 |
| ENSG000002 | 13.9603167 | -0.846279973 | 0.32801797 | -2.5799805 | 0.00988059 | 0.03508167 | NA | NA |
| ENSG000002 | 24.7304268 | -0.719572028 | 0.27892794 | -2.5797775 | 0.0098864 | 0.03509923 | NA | NA |
| ENSG000002 | 17.0659119 | -0.776953813 | 0.30118476 | -2.5796584 | 0.00988981 | 0.03510826 | NA | NA |
| ENSG000001 | 41.6233966 | 0.590201974 | 0.22879446 | 2.57961654 | 0.00989101 | 0.03510944 | ASPHD1 | 253982 |

| | | | | | | | | | |
|---|---|---|---|---|---|---|---|---|---|
| ENSG000002 | 4.08421945 | -0.97647527 | 0.37854621 | -2.5795405 | 0.00989319 | 0.0351141 | NA | NA | |
| ENSG000002 | 10.3965307 | -0.897079344 | 0.34777253 | -2.5795003 | 0.00989434 | 0.03511512 | LINC01296 | | 642477 |
| ENSG000001 | 13.7397551 | -0.83591844 | 0.3241024 | -2.57918 | 0.00990352 | 0.03514153 | ADGRE3 | | 84658 |
| ENSG000001 | 1011.92667 | 0.303645727 | 0.11772859 | 2.57920122 | 0.00990291 | 0.03514153 | FLVCR1 | | 28982 |
| ENSG000001 | 10.8931503 | -0.887944197 | 0.34430147 | -2.578973 | 0.00990945 | 0.03515953 | ZNF541 | | 84215 |
| ENSG00000C | 4898.22391 | 0.29253857 | 0.11344448 | 2.57869381 | 0.00991746 | 0.03517954 | ZNF638 | | 27332 |
| ENSG000002 | 2.53792506 | -0.956995128 | 0.37111739 | -2.5786857 | 0.0099177 | 0.03517954 | NA | NA | |
| ENSG000002 | 4.10219261 | -0.976203326 | 0.37856344 | -2.5787047 | 0.00991715 | 0.03517954 | NA | NA | |
| ENSG000001 | 70.1610325 | 0.501447168 | 0.19446533 | 2.57859425 | 0.00992032 | 0.03518076 | SRPX | | 8406 |
| ENSG000001 | 459.311065 | 0.34276057 | 0.13292449 | 2.57861115 | 0.00991984 | 0.03518076 | CRLF3 | | 51379 |
| ENSG000002 | 9.49384684 | -0.895013347 | 0.34709502 | -2.5785831 | 0.00992064 | 0.03518076 | NA | NA | |
| ENSG000002 | 3.36289132 | -0.972428266 | 0.37713095 | -2.5784897 | 0.00992333 | 0.0351872 | NA | NA | |
| ENSG000001 | 379.491534 | 0.340661108 | 0.13211903 | 2.57844082 | 0.00992473 | 0.0351891 | TUB | | 7275 |
| ENSG000001 | 21.9167449 | -0.713868695 | 0.27689389 | -2.5781309 | 0.00993364 | 0.03521759 | NTRK1 | | 4914 |
| ENSG000002 | 11.26561 | -0.857246404 | 0.3325206 | -2.5780249 | 0.00993668 | 0.03522532 | NXPE2 | | 120406 |
| ENSG000002 | 2.7295302 | -0.952323198 | 0.36942547 | -2.5778493 | 0.00994173 | 0.03524015 | NA | NA | |
| ENSG000001 | 334.220853 | 0.366259519 | 0.14210104 | 2.57745844 | 0.00995298 | 0.03527694 | NA | NA | |
| ENSG000001 | 4354.68804 | 0.303781173 | 0.1178647 | 2.57737207 | 0.00995547 | 0.03528268 | CHD1 | | 1105 |
| ENSG000002 | 27.5124686 | -0.667352168 | 0.25893197 | -2.5773263 | 0.00995679 | 0.03528426 | NA | NA | |
| ENSG000001 | 2.94369689 | -0.967332638 | 0.37534921 | -2.5771538 | 0.00996176 | 0.03529867 | FBXL8 | | 55336 |
| ENSG000001 | 1523.74983 | 0.421057817 | 0.16338278 | 2.57712477 | 0.0099626 | 0.03529867 | UBXN7 | | 26043 |
| ENSG000002 | 6.13802619 | -0.956838405 | 0.37132238 | -2.5768401 | 0.00997081 | 0.03532467 | CHEK2P2 | | 646096 |
| ENSG000001 | 274.610493 | 0.377272633 | 0.14641102 | 2.57680488 | 0.00997182 | 0.03532518 | KATNA1 | | 11104 |
| ENSG000002 | 5.57733822 | -0.966298948 | 0.37502405 | -2.576632 | 0.00997681 | 0.03533769 | NA | NA | |
| ENSG000002 | 5.64536503 | -0.956414438 | 0.37118925 | -2.5766222 | 0.00997709 | 0.03533769 | GPIHBP1 | | 338328 |
| ENSG000001 | 422.1167 | 0.308011061 | 0.11954616 | 2.57650309 | 0.00998053 | 0.03534614 | ZNF354A | | 6940 |
| ENSG000002 | 5.4371348 | -0.964215078 | 0.3742375 | -2.5764791 | 0.00998123 | 0.03534614 | NA | NA | |
| ENSG000002 | 8.42700658 | -0.919807678 | 0.35704786 | -2.5761467 | 0.00999082 | 0.03537444 | NA | NA | |
| ENSG000002 | 10.3442438 | -0.898515957 | 0.34878356 | -2.576142 | 0.00999096 | 0.03537444 | NA | NA | |

| | | | | | | | | |
|---|---|---|---|---|---|---|---|---|
| ENSG000002 | 2.55755588 | -0.956187843 | 0.37118408 | -2.5760475 | 0.00999369 | 0.03538102 | NA | NA |
| ENSG000001 | 2273.00341 | 0.395380218 | 0.15349482 | 2.57585374 | 0.00999929 | 0.03539776 | CTCF | 10664 |
| ENSG000001 | 130.603348 | 0.417999072 | 0.16228895 | 2.57564716 | 0.01000527 | 0.03541582 | ZNF71 | 58491 |
| ENSG00000C | 53.5277678 | 0.632688578 | 0.24566824 | 2.57537794 | 0.01001306 | 0.03544031 | OCEL1 | 79629 |
| ENSG000001 | 8.196777 | -0.915319847 | 0.35541671 | -2.5753428 | 0.01001408 | 0.03544082 | PRL | 5617 |
| ENSG000001 | 91.5716626 | 0.481079005 | 0.1868215 | 2.57507301 | 0.01002189 | 0.03545919 | MCEE | 84693 |
| ENSG000001 | 18.7165953 | -0.796752375 | 0.30940723 | -2.575093 | 0.01002132 | 0.03545919 | PADI4 | 23569 |
| ENSG000001 | 534.174838 | 0.29152144 | 0.11320838 | 2.57508705 | 0.01002149 | 0.03545919 | SMAD1 | 4086 |
| ENSG000002 | 10.0398316 | -0.897870703 | 0.34869453 | -2.5749492 | 0.01002548 | 0.03546879 | NA | NA |
| ENSG000001 | 244.182114 | 0.375440618 | 0.1458078 | 2.57490072 | 0.01002689 | 0.03547067 | ATF7IP2 | 80063 |
| ENSG000001 | 60.4463209 | 0.536809365 | 0.20848563 | 2.57480277 | 0.01002973 | 0.03547762 | PKP3 | 11187 |
| ENSG000002 | 14.7355351 | -0.819906303 | 0.31844939 | -2.5746832 | 0.01003319 | 0.03548679 | LOC101927356 | 101927356 |
| ENSG000002 | 90.1904101 | 0.491355729 | 0.19085826 | 2.5744536 | 0.01003986 | 0.03550725 | LOC100287896 | 100287896 |
| ENSG000001 | 1797.20233 | 0.323990385 | 0.12585219 | 2.57437234 | 0.01004221 | 0.03551027 | RBBP6 | 5930 |
| ENSG000001 | 28.7668874 | -0.635554243 | 0.24687817 | -2.5743639 | 0.01004246 | 0.03551027 | ADGRF2 | 222611 |
| ENSG000002 | 12.9986414 | -0.832685288 | 0.32346243 | -2.5742875 | 0.01004468 | 0.03551501 | NA | NA |
| ENSG000002 | 7.76610857 | -0.909694687 | 0.35338349 | -2.5742422 | 0.01004599 | 0.03551657 | NA | NA |
| ENSG000002 | 3.35631204 | -0.971399761 | 0.37736375 | -2.5741735 | 0.01004799 | 0.03552052 | NA | NA |
| ENSG000002 | 11.2984713 | -0.859340317 | 0.33387026 | -2.573875 | 0.01005666 | 0.03554808 | ZBTB20-AS4 | 100874131 |
| ENSG000001 | 102.103018 | 0.438071226 | 0.17021094 | 2.57369606 | 0.01006186 | 0.03556337 | IFNLR1 | 163702 |
| ENSG00000C | 740.411924 | 0.299755432 | 0.11648077 | 2.57343281 | 0.01006952 | 0.03558734 | PRKAR2B | 5577 |
| ENSG000001 | 12.5600538 | -0.848092268 | 0.32961243 | -2.5729984 | 0.01008217 | 0.03562893 | OR2V2 | 285659 |
| ENSG000002 | 14.721804 | -0.797570805 | 0.31000682 | -2.5727524 | 0.01008934 | 0.03565115 | LOC101927229 | 101927229 |
| ENSG000002 | 5.24816345 | -0.968175087 | 0.37632799 | -2.5726896 | 0.01009117 | 0.03565452 | NA | NA |
| ENSG000001 | 6.98735725 | -0.933723037 | 0.36295056 | -2.5725902 | 0.01009407 | 0.03565787 | H2BFM | 286436 |
| ENSG000001 | 603.371967 | 0.319355232 | 0.12413685 | 2.57260619 | 0.0100936 | 0.03565787 | ODF2L | 57489 |
| ENSG000002 | 10.0953672 | -0.879864302 | 0.34201809 | -2.5725666 | 0.01009475 | 0.03565787 | NA | NA |
| ENSG000001 | 760.682834 | 0.30140552 | 0.11717292 | 2.57231372 | 0.01010213 | 0.0356626 | RPS6KB1 | 6198 |
| ENSG000001 | 87.7429872 | 0.46493634 | 0.18073488 | 2.57247703 | 0.01009737 | 0.0356626 | BIVM | 54841 |

| | | | | | | | | |
|---|---|---|---|---|---|---|---|---|
| ENSG000002 | 5.87841522 | -0.961848891 | 0.37392298 | -2.5723182 | 0.010102 | 0.0356626 | FLJ26850 | 400710 |
| ENSG000002 | 6.31316467 | -0.943783283 | 0.36688222 | -2.5724421 | 0.01009838 | 0.0356626 | NA | NA |
| ENSG000002 | 137.434946 | 0.404782563 | 0.15736152 | 2.57230964 | 0.01010225 | 0.0356626 | NA | NA |
| ENSG000002 | 4.44425902 | -0.972504761 | 0.37805723 | -2.5723744 | 0.01010036 | 0.0356626 | NA | NA |
| ENSG000002 | 23.9546722 | -0.693789236 | 0.26970197 | -2.5724292 | 0.01009876 | 0.0356626 | LOC105374989 | 105374989 |
| ENSG000002 | 3.73502952 | -0.973378653 | 0.37842902 | -2.5721565 | 0.01010672 | 0.03567528 | NA | NA |
| ENSG000002 | 35.0652988 | 0.621178431 | 0.24153809 | 2.57176183 | 0.01011825 | 0.03571286 | LOC102724023 | 102724023 |
| ENSG000002 | 5.50236225 | -0.962979569 | 0.37444858 | -2.5717272 | 0.01011926 | 0.03571333 | NA | NA |
| ENSG000001 | 295.175666 | 0.362750466 | 0.14105954 | 2.57161251 | 0.01012261 | 0.03571994 | MFSD8 | 256471 |
| ENSG000002 | 9.147248 | -0.893547774 | 0.34747133 | -2.5715727 | 0.01012378 | 0.03571994 | CCDC26 | 137196 |
| ENSG000002 | 5.70635005 | -0.95570066 | 0.37163679 | -2.5715986 | 0.01012302 | 0.03571994 | NA | NA |
| ENSG000001 | 93.210453 | 0.468976487 | 0.18237588 | 2.57148301 | 0.0101264 | 0.03572608 | SLC43A3 | 29015 |
| ENSG000001 | 2452.50294 | 0.329699695 | 0.12822374 | 2.57128426 | 0.01013221 | 0.03574348 | EPT1 | 85465 |
| ENSG000002 | 49.926604 | -0.538209451 | 0.20932788 | -2.5711312 | 0.01013669 | 0.03575618 | NA | NA |
| ENSG000001 | 6.4489332 | -0.950515617 | 0.36972248 | -2.5708894 | 0.01014377 | 0.03577492 | NA | NA |
| ENSG000001 | 14.5048066 | -0.810151875 | 0.31512384 | -2.5709 | 0.01014346 | 0.03577492 | CLEC17A | 388512 |
| ENSG000001 | 6.53435532 | -0.950174759 | 0.36960596 | -2.5707777 | 0.01014704 | 0.03578336 | MAGEB10 | 139422 |
| ENSG000001 | 6.27547582 | -0.942441181 | 0.36662712 | -2.5705714 | 0.01015309 | 0.03580156 | SFTA2 | 389376 |
| ENSG000001 | 282.785895 | 0.322589512 | 0.12549664 | 2.57050318 | 0.01015509 | 0.0358055 | GLTSCR1L | 23506 |
| ENSG000002 | 5.16239879 | -0.967963905 | 0.37657278 | -2.5704564 | 0.01015646 | 0.03580722 | NA | NA |
| ENSG000001 | 4555.2112 | 0.341006441 | 0.1326665 | 2.57040361 | 0.01015801 | 0.03580956 | SALL2 | 6297 |
| ENSG000001 | 231.105997 | 0.369463232 | 0.14374092 | 2.57034133 | 0.01015984 | 0.03581289 | HDAC8 | 55869 |
| ENSG000002 | 2.74487516 | -0.944648685 | 0.36752498 | -2.5702979 | 0.01016111 | 0.03581427 | NA | NA |
| ENSG000002 | 62.919641 | -0.515799586 | 0.20069601 | -2.570054 | 0.01016827 | 0.03583638 | LOC101927687 | 101927687 |
| ENSG000002 | 7.50752024 | -0.936403754 | 0.36435756 | -2.5700133 | 0.01016946 | 0.03583748 | NA | NA |
| ENSG000001 | 258.518692 | 0.338220859 | 0.13161308 | 2.56981186 | 0.01017538 | 0.0358552 | MTERF3 | 51001 |
| ENSG000001 | 399.115654 | 0.316248103 | 0.12306664 | 2.56973053 | 0.01017776 | 0.0358605 | SLC26A2 | 1836 |
| ENSG000001 | 2365.14649 | 0.300514035 | 0.11694796 | 2.56963891 | 0.01018046 | 0.03586687 | MRPS30 | 10884 |
| ENSG00000C | 103.347925 | 0.461977527 | 0.17978855 | 2.56956039 | 0.01018276 | 0.03586983 | ADCK1 | 57143 |

| | | | | | | | | |
|---|---|---|---|---|---|---|---|---|
| ENSG000001 | 235.249315 | 0.44689183 | 0.17392035 | 2.56952006 | 0.01018395 | 0.03586983 | LENG8 | 114823 |
| ENSG000001 | 21.5094046 | 0.733237033 | 0.28535667 | 2.5695458 | 0.01018319 | 0.03586983 | CRYBA4 | 1413 |
| ENSG000002 | 14.2126643 | -0.794872855 | 0.30935102 | -2.5694852 | 0.01018497 | 0.03587032 | NA | NA |
| ENSG000002 | 19.8143565 | 0.734411469 | 0.28582849 | 2.56941314 | 0.01018709 | 0.03587467 | NA | NA |
| ENSG000001 | 184.519239 | 0.376825187 | 0.14666914 | 2.56921923 | 0.0101928 | 0.03589163 | ZNF597 | 146434 |
| ENSG000001 | 119.001392 | 0.469842903 | 0.18289407 | 2.56893462 | 0.01020117 | 0.035918 | CCDC85B | 11007 |
| ENSG000001 | 844.668248 | 0.302958669 | 0.11793383 | 2.56888688 | 0.01020258 | 0.03591983 | ZNF660 | 285349 |
| ENSG000002 | 141.678498 | 0.397958092 | 0.15493888 | 2.56848441 | 0.01021443 | 0.03595845 | RNF103 | 7844 |
| ENSG000002 | 45.4184246 | 0.65390669 | 0.25466531 | 2.56771012 | 0.01023727 | 0.03603573 | SNORA23 | 677808 |
| ENSG000002 | 10.0069807 | -0.8750349 | 0.34079209 | -2.5676502 | 0.01023904 | 0.03603883 | NA | NA |
| ENSG000002 | 4.52613092 | -0.971450866 | 0.37834743 | -2.5676158 | 0.01024006 | 0.03603928 | NA | NA |
| ENSG000002 | 7.59469297 | -0.926006499 | 0.36068832 | -2.5673315 | 0.01024846 | 0.03606571 | NA | NA |
| ENSG00000C | 1954.00115 | 0.325112975 | 0.12664415 | 2.56713763 | 0.01025419 | 0.03608043 | USP48 | 84196 |
| ENSG000001 | 36.0120998 | -0.589788897 | 0.22974642 | -2.5671299 | 0.01025442 | 0.03608043 | AZIN2 | 113451 |
| ENSG000001 | 8.03471231 | -0.921551015 | 0.358999 | -2.5670016 | 0.01025821 | 0.03609065 | NA | NA |
| ENSG000001 | 72.5701914 | -0.476777162 | 0.18574041 | -2.5669006 | 0.0102612 | 0.03609803 | NRG1 | 3084 |
| ENSG000001 | 41.0757438 | 0.606694112 | 0.23635845 | 2.56683911 | 0.01026302 | 0.0361013 | RTN4RL2 | 349667 |
| ENSG000001 | 13.8894986 | -0.813099044 | 0.31684989 | -2.5661964 | 0.01028206 | 0.03616513 | DEFB128 | 245939 |
| ENSG000002 | 6.55556774 | -0.933216667 | 0.36367654 | -2.5660623 | 0.01028603 | 0.03617597 | LINC00484 | 100129347 |
| ENSG00000C | 18.45844 | -0.736002336 | 0.28682941 | -2.5659932 | 0.01028808 | 0.03617691 | CASS4 | 57091 |
| ENSG000002 | 5.24171391 | -0.967090031 | 0.37688345 | -2.5660188 | 0.01028733 | 0.03617691 | NA | NA |
| ENSG000002 | 24.0331392 | -0.734936244 | 0.28644127 | -2.5657484 | 0.01029535 | 0.03619931 | LOC101928404 | 101928404 |
| ENSG00000C | 1819.9528 | 0.346745524 | 0.13514997 | 2.56563525 | 0.01029871 | 0.03620045 | G2E3 | 55632 |
| ENSG000001 | 49.3572419 | -0.566835197 | 0.22092897 | -2.5656898 | 0.01029709 | 0.03620045 | CTCFL | 140690 |
| ENSG000001 | 14.9424668 | -0.83388732 | 0.32502407 | -2.5656171 | 0.01029924 | 0.03620045 | C9orf153 | 389766 |
| ENSG000002 | 17.2241423 | -0.804437454 | 0.3135443 | -2.5656261 | 0.01029898 | 0.03620045 | LOC254028 | 254028 |
| ENSG000001 | 30.5590468 | -0.644957378 | 0.2514097 | -2.5653639 | 0.01030676 | 0.03622374 | EFCAB12 | 90288 |
| ENSG000002 | 11.5809818 | -0.868594356 | 0.33859635 | -2.5652797 | 0.01030927 | 0.0362294 | OR5V1 | 81696 |
| ENSG000001 | 313.330452 | 0.335568561 | 0.1308232 | 2.56505399 | 0.01031598 | 0.03624355 | NA | NA |

| | | | | | | | | | |
|---|---|---|---|---|---|---|---|---|---|
| ENSG000001 | 142.429095 | 0.552904852 | 0.21555111 | 2.56507536 | 0.01031534 | 0.03624355 | HIRIP3 | | 8479 |
| ENSG000001 | 711.724532 | 0.321979377 | 0.12552435 | 2.56507503 | 0.01031535 | 0.03624355 | BNC2 | | 54796 |
| ENSG000001 | 1313.96955 | 0.332187693 | 0.12951103 | 2.56493756 | 0.01031944 | 0.03624482 | KANSL1 | | 284058 |
| ENSG000001 | 76.9231649 | 0.520172978 | 0.20279695 | 2.56499404 | 0.01031776 | 0.03624482 | NAIF1 | | 203245 |
| ENSG000002 | 6.71776952 | -0.939757139 | 0.36638825 | -2.5649216 | 0.01031991 | 0.03624482 | NA | NA | |
| ENSG000002 | 3.66652127 | -0.970775998 | 0.37847577 | -2.5649621 | 0.01031871 | 0.03624482 | NA | NA | |
| ENSG000001 | 2184.44514 | 0.336936336 | 0.13136886 | 2.56481126 | 0.01032319 | 0.03625321 | PAIP2 | | 51247 |
| ENSG000001 | 35.4380931 | -0.603696345 | 0.23538811 | -2.564685 | 0.01032695 | 0.03626327 | RNF212 | | 285498 |
| ENSG000002 | 3.31205365 | -0.968084454 | 0.37747218 | -2.5646511 | 0.01032796 | 0.03626367 | NA | NA | |
| ENSG000001 | 15.7671104 | -0.786258376 | 0.30658455 | -2.5645727 | 0.01033029 | 0.03626872 | SLC23A1 | | 9963 |
| ENSG000001 | 33.3940907 | -0.649583504 | 0.25329831 | -2.5644999 | 0.01033246 | 0.03627319 | GSG1 | | 83445 |
| ENSG000001 | 7.70042914 | -0.925690824 | 0.36099037 | -2.5643089 | 0.01033815 | 0.03629002 | SIGLEC9 | | 27180 |
| ENSG00000C | 136.782353 | 0.433406479 | 0.16903231 | 2.56404525 | 0.01034601 | 0.03631445 | PARP12 | | 64761 |
| ENSG000001 | 451.011141 | 0.327606319 | 0.12778674 | 2.56369562 | 0.01035643 | 0.0363479 | ZNF143 | | 7702 |
| ENSG000002 | 11.7785189 | -0.86120629 | 0.3359809 | -2.5632596 | 0.01036945 | 0.03639043 | NA | NA | |
| ENSG000002 | 3.84243052 | -0.967449676 | 0.37753201 | -2.5625633 | 0.01039026 | 0.03645718 | NA | NA | |
| ENSG000002 | 3.69386324 | -0.969691262 | 0.37840427 | -2.5625801 | 0.01038976 | 0.03645718 | NA | NA | |
| ENSG000002 | 4.11946488 | -0.970132941 | 0.37859246 | -2.562473 | 0.01039297 | 0.03646351 | NA | NA | |
| ENSG000002 | 9.24855506 | -0.896019202 | 0.34970634 | -2.5622046 | 0.010401 | 0.03648624 | BOD1L2 | | 284257 |
| ENSG000002 | 27.1070063 | -0.663309223 | 0.25888305 | -2.5621964 | 0.01040125 | 0.03648624 | NA | NA | |
| ENSG000001 | 114.975809 | 0.490716184 | 0.19152668 | 2.56212959 | 0.01040325 | 0.03649011 | CNPY4 | | 245812 |
| ENSG000001 | 26.7548467 | -0.715863146 | 0.2794216 | -2.5619463 | 0.01040874 | 0.03650305 | CNGA1 | | 1259 |
| ENSG000002 | 8.02607504 | -0.91803636 | 0.3583348 | -2.5619514 | 0.01040859 | 0.03650305 | NA | NA | |
| ENSG000001 | 48.3158739 | 0.610456079 | 0.23830277 | 2.56168272 | 0.01041664 | 0.03652445 | RHPN1 | | 114822 |
| ENSG000002 | 150.583419 | 0.407616955 | 0.15911895 | 2.56171221 | 0.01041576 | 0.03652445 | NA | NA | |
| ENSG000001 | 134.737801 | 0.387948464 | 0.15145607 | 2.56145867 | 0.01042336 | 0.03654486 | ZNF514 | | 84874 |
| ENSG000001 | 1431.32515 | 0.321828734 | 0.12564672 | 2.56137792 | 0.01042579 | 0.0365502 | EI24 | | 9538 |
| ENSG000002 | 14.2552753 | -0.833632706 | 0.32550522 | -2.5610425 | 0.01043586 | 0.03658234 | NA | NA | |
| ENSG000001 | 2107.33992 | 0.295053865 | 0.11522124 | 2.56075942 | 0.01044437 | 0.03660899 | KMT2A | | 4297 |

| | | | | | | | | |
|---|---|---|---|---|---|---|---|---|
| ENSG000001 | 89.1218137 | 0.46463979 | 0.18150191 | 2.55997191 | 0.01046806 | 0.03668888 | ZNF696 | 79943 |
| ENSG000001 | 13.5831188 | -0.817020603 | 0.31917624 | -2.5597789 | 0.01047388 | 0.03670074 | KRT17 | 3872 |
| ENSG000002 | 5.98650022 | -0.959036458 | 0.37465318 | -2.559798 | 0.0104733 | 0.03670074 | NA | NA |
| ENSG000002 | 10.0408979 | -0.897925836 | 0.35078386 | -2.5597695 | 0.01047416 | 0.03670074 | NA | NA |
| ENSG000001 | 229.345584 | 0.34357976 | 0.13422795 | 2.55967375 | 0.01047705 | 0.03670768 | ZNF573 | 126231 |
| ENSG000001 | 1742.09036 | 0.340424837 | 0.13301915 | 2.55921666 | 0.01049083 | 0.03674964 | TMED7 | 51014 |
| ENSG000001 | 695.822612 | 0.40159706 | 0.15692164 | 2.55922035 | 0.01049072 | 0.03674964 | BRD4 | 23476 |
| ENSG000001 | 15.6218773 | -0.769325547 | 0.30065557 | -2.5588269 | 0.0105026 | 0.03678769 | IP6K3 | 117283 |
| ENSG000001 | 3.6843124 | -0.968065033 | 0.37835365 | -2.5586248 | 0.01050871 | 0.03679664 | FOXF2 | 2295 |
| ENSG000002 | 128.259206 | 0.434060298 | 0.16964563 | 2.5586294 | 0.01050857 | 0.03679664 | DNASE1 | 1773 |
| ENSG000002 | 3.6843124 | -0.968065033 | 0.37835365 | -2.5586248 | 0.01050871 | 0.03679664 | NA | NA |
| ENSG000002 | 7.59582669 | -0.919634839 | 0.3594258 | -2.5586222 | 0.01050879 | 0.03679664 | MAPT-IT1 | 100130148 |
| ENSG000001 | 472.146312 | 0.316877732 | 0.12385255 | 2.55850788 | 0.01051224 | 0.03680218 | CCDC174 | 51244 |
| ENSG000002 | 10.1034095 | -0.915525019 | 0.35783946 | -2.5584798 | 0.01051309 | 0.03680218 | MROH5 | 389690 |
| ENSG000002 | 5.33543361 | -0.9652639 | 0.37727599 | -2.5585087 | 0.01051222 | 0.03680218 | NA | NA |
| ENSG000001 | 499.480873 | 0.328109666 | 0.12825788 | 2.55820274 | 0.01052147 | 0.03682833 | SHQ1 | 55164 |
| ENSG000001 | 169.711182 | 0.419178905 | 0.16386447 | 2.55808299 | 0.0105251 | 0.03683784 | COG7 | 91949 |
| ENSG000002 | 116.595292 | 0.466465485 | 0.18236 | 2.5579375 | 0.0105295 | 0.03685007 | ARPC1A | 10552 |
| ENSG000002 | 2.186928 | -0.92813641 | 0.36285303 | -2.5578852 | 0.01053108 | 0.03685243 | NA | NA |
| ENSG000001 | 28.3268042 | -0.674236123 | 0.26361455 | -2.557659 | 0.01053794 | 0.03687324 | VNN2 | 8875 |
| ENSG000001 | 3168.50797 | 0.313170574 | 0.12245001 | 2.55753822 | 0.0105416 | 0.03688286 | ADGRV1 | 84059 |
| ENSG000001 | 336.161839 | 0.352411633 | 0.13780333 | 2.55735204 | 0.01054724 | 0.03689306 | NCK1 | 4690 |
| ENSG000001 | 410.5822 | 0.395773752 | 0.1547579 | 2.55737342 | 0.01054659 | 0.03689306 | MTCL1 | 23255 |
| ENSG000001 | 523.026231 | 0.354609617 | 0.13866073 | 2.5573903 | 0.01054608 | 0.03689306 | CMTR2 | 55783 |
| ENSG000002 | 32.535382 | -0.660116326 | 0.25813103 | -2.5572916 | 0.01054907 | 0.0368931 | NA | NA |
| ENSG000002 | 34.7396023 | -0.619581415 | 0.24227991 | -2.5572959 | 0.01054894 | 0.0368931 | FCGBP | 8857 |
| ENSG000002 | 13.4438751 | 0.822196674 | 0.32151986 | 2.55721895 | 0.01055128 | 0.03689712 | RPL13P5 | 283345 |
| ENSG000002 | 14.9210347 | -0.815283582 | 0.31881963 | -2.5571938 | 0.01055204 | 0.03689712 | DSG1-AS1 | 101927718 |
| ENSG000002 | 50.3036431 | 0.568251392 | 0.22222908 | 2.5570524 | 0.01055633 | 0.03690893 | NA | NA |

| | | | | | | | | |
|---|---|---|---|---|---|---|---|---|
| ENSG000002 | 26.3985214 | -0.674894038 | 0.26394713 | -2.5569289 | 0.01056008 | 0.03691885 | NA | NA |
| ENSG000002 | 2.63497127 | -0.943947393 | 0.36917909 | -2.5568821 | 0.0105615 | 0.03692064 | NA | NA |
| ENSG000001 | 4863.69838 | 0.293551147 | 0.11482028 | 2.55661406 | 0.01056964 | 0.03694591 | GLUL | 2752 |
| ENSG000001 | 69.655596 | 0.501155852 | 0.19602953 | 2.5565324 | 0.01057212 | 0.0369514 | DMPK | 1760 |
| ENSG000002 | 558.113676 | 0.298728036 | 0.11686771 | 2.55612115 | 0.01058463 | 0.03699191 | DCP1A | 55802 |
| ENSG000001 | 167.312829 | -0.377756632 | 0.14778801 | -2.5560709 | 0.01058616 | 0.03699407 | ABCA5 | 23461 |
| ENSG000001 | 29.7286034 | -0.721918598 | 0.28245624 | -2.55586 | 0.01059257 | 0.03701331 | FPR2 | 2358 |
| ENSG000001 | 29.6880552 | -0.660012942 | 0.25825321 | -2.5556814 | 0.01059801 | 0.03702672 | MYO7B | 4648 |
| ENSG000001 | 40.3417161 | -0.563827529 | 0.22062053 | -2.5556439 | 0.01059915 | 0.03702672 | PIWIL3 | 440822 |
| ENSG000002 | 7.01764163 | -0.92892889 | 0.36347978 | -2.5556549 | 0.01059882 | 0.03702672 | NA | NA |
| ENSG000001 | 9.44261916 | -0.884306044 | 0.34603748 | -2.555521 | 0.0106029 | 0.037034 | CLEC18B | 497190 |
| ENSG000001 | 51.7889178 | 0.545449071 | 0.21343993 | 2.55551555 | 0.01060306 | 0.037034 | MIEF2 | 125170 |
| ENSG000002 | 3.13954804 | -0.95252208 | 0.37275897 | -2.5553297 | 0.01060873 | 0.03704959 | RNU6-889P | 106480402 |
| ENSG000002 | 5.67313175 | -0.950124675 | 0.37182377 | -2.5553092 | 0.01060935 | 0.03704959 | SIGLEC12 | 89858 |
| ENSG000002 | 18.734462 | -0.737118797 | 0.2884948 | -2.5550505 | 0.01061724 | 0.03707394 | NA | NA |
| ENSG000002 | 4.70546976 | -0.962875527 | 0.37686042 | -2.5549925 | 0.01061901 | 0.03707693 | LINC00501 | 100820709 |
| ENSG000002 | 10.6173656 | -0.88311045 | 0.34571562 | -2.5544419 | 0.01063582 | 0.03713242 | NA | NA |
| ENSG000001 | 707.175791 | 0.287649575 | 0.11261289 | 2.55432189 | 0.01063948 | 0.03714202 | FBXW7 | 55294 |
| ENSG000002 | 4.77614677 | -0.963945218 | 0.37747949 | -2.553636 | 0.01066046 | 0.03721205 | NA | NA |
| ENSG00000C | 274.800359 | 0.330337314 | 0.12936209 | 2.55358668 | 0.01066197 | 0.03721412 | PAK3 | 5063 |
| ENSG000001 | 95.2371126 | 0.484282646 | 0.18966589 | 2.55334604 | 0.01066934 | 0.03723664 | WRAP73 | 49856 |
| ENSG000001 | 258.803979 | 0.349376125 | 0.13683611 | 2.55324505 | 0.01067244 | 0.03724424 | ZNF419 | 79744 |
| ENSG000001 | 2266.93007 | 0.369251738 | 0.14462765 | 2.55311987 | 0.01067627 | 0.03725442 | MED1 | 5469 |
| ENSG000001 | 63.5482081 | 0.502518955 | 0.19683122 | 2.55304495 | 0.01067857 | 0.03725923 | CLCN2 | 1181 |
| ENSG00000C | 142.500521 | 0.437089517 | 0.17120875 | 2.55296257 | 0.0106811 | 0.03726163 | SERPINB1 | 1992 |
| ENSG000002 | 37.1267839 | 0.593671589 | 0.23254154 | 2.5529701 | 0.01068087 | 0.03726163 | LOC642361 | 642361 |
| ENSG000001 | 728.622147 | 0.414221607 | 0.16225806 | 2.55285689 | 0.01068434 | 0.03726973 | VKORC1L1 | 154807 |
| ENSG000002 | 5.08331442 | 0.956804553 | 0.37482023 | 2.55270254 | 0.01068907 | 0.03728304 | NA | NA |
| ENSG000001 | 1153.95129 | 0.329627807 | 0.12913496 | 2.55258385 | 0.01069272 | 0.03729254 | GAPVD1 | 26130 |

| | | | | | | | | |
|---|---|---|---|---|---|---|---|---|
| ENSG00000( | 30.4954625 | -0.648631994 | 0.25411581 | -2.5525055 | 0.01069512 | 0.03729452 | ROPN1 | 54763 |
| ENSG000002 | 60.2720402 | 0.562433773 | 0.22034435 | 2.55252185 | 0.01069462 | 0.03729452 | NA | NA |
| ENSG000001 | 13.0552701 | -0.831823692 | 0.3259229 | -2.55221 | 0.0107042 | 0.03732295 | SEC14L4 | 284904 |
| ENSG000002 | 10.1072368 | -0.856193565 | 0.33553124 | -2.5517551 | 0.01071818 | 0.0373685 | LOC101929529 | 101929529 |
| ENSG000001 | 4.30084671 | -0.963718975 | 0.37767806 | -2.5516944 | 0.01072005 | 0.0373718 | NA | NA |
| ENSG000001 | 288.024101 | 0.39530733 | 0.15493704 | 2.5514063 | 0.01072892 | 0.03739949 | NSUN4 | 387338 |
| ENSG000002 | 3.32890427 | -0.961628248 | 0.37692857 | -2.5512214 | 0.01073461 | 0.03741613 | NA | NA |
| ENSG000001 | 501.876236 | 0.30199904 | 0.11838133 | 2.55106988 | 0.01073928 | 0.03742142 | NA | NA |
| ENSG000001 | 240.656136 | 0.361514444 | 0.14171189 | 2.55105226 | 0.01073982 | 0.03742142 | SFT2D3 | 84826 |
| ENSG000002 | 4.76048927 | -0.962986615 | 0.37747625 | -2.5511184 | 0.01073778 | 0.03742142 | NA | NA |
| ENSG000002 | 23.9469148 | -0.686065171 | 0.26893217 | -2.5510714 | 0.01073923 | 0.03742142 | PCDHGB5 | 56101 |
| ENSG000001 | 4.82725564 | -0.962947538 | 0.37750341 | -2.5508314 | 0.01074663 | 0.03744193 | NA | NA |
| ENSG000001 | 985.430424 | 0.388696573 | 0.15238399 | 2.55077034 | 0.01074851 | 0.03744379 | CHTOP | 26097 |
| ENSG000001 | 164.429822 | 0.39600189 | 0.15524894 | 2.55075421 | 0.01074901 | 0.03744379 | TMEM133 | 83935 |
| ENSG000002 | 11.568566 | -0.850829317 | 0.33356633 | -2.550705 | 0.01075053 | 0.03744585 | NA | NA |
| ENSG000001 | 388.054344 | 0.318908564 | 0.12502933 | 2.55066999 | 0.01075161 | 0.0374464 | BBS10 | 79738 |
| ENSG00000( | 47.0721369 | 0.580669385 | 0.22768596 | 2.55030822 | 0.01076277 | 0.03748207 | DFNB31 | 25861 |
| ENSG000001 | 13.5374814 | -0.817540148 | 0.32057781 | -2.5502081 | 0.01076586 | 0.03748961 | KCNN4 | 3783 |
| ENSG000002 | 30.4863582 | -0.630305886 | 0.24716549 | -2.5501371 | 0.01076806 | 0.03749403 | LINC00923 | 91948 |
| ENSG00000( | 20.2154209 | -0.755544983 | 0.29631047 | -2.5498424 | 0.01077716 | 0.03752041 | INSRR | 3645 |
| ENSG000001 | 354.487174 | 0.362750751 | 0.14226457 | 2.54983199 | 0.01077748 | 0.03752041 | SLC38A4 | 55089 |
| ENSG000002 | 9.46661367 | -0.88520565 | 0.34719065 | -2.5496241 | 0.01078391 | 0.03753957 | LINC01104 | 150577 |
| ENSG000001 | 31.2055181 | 0.625640245 | 0.24540154 | 2.5494553 | 0.01078913 | 0.0375513 | U2AF1L4 | 199746 |
| ENSG000001 | 559.735827 | 0.351747782 | 0.13796836 | 2.54948143 | 0.01078833 | 0.0375513 | THBS2 | 7058 |
| ENSG000002 | 8.726957 | -0.892094996 | 0.34992794 | -2.5493677 | 0.01079184 | 0.03755751 | NA | NA |
| ENSG000002 | 6.58892517 | -0.92830947 | 0.36420047 | -2.548897 | 0.01080642 | 0.03760501 | NA | NA |
| ENSG000002 | 4.10449564 | -0.964912454 | 0.37859175 | -2.5486885 | 0.01081288 | 0.03762427 | NA | NA |
| ENSG000001 | 20.6982485 | -0.739405451 | 0.29016665 | -2.5482096 | 0.01082774 | 0.03767273 | BPIFC | 254240 |
| ENSG000001 | 678.841868 | 0.347985256 | 0.13658334 | 2.5477869 | 0.01084087 | 0.03771517 | TIMM17A | 10440 |

| | | | | | | | | |
|---|---|---|---|---|---|---|---|---|
| ENSG000002 | 4.31908743 | -0.962409625 | 0.37777131 | -2.5475985 | 0.01084672 | 0.0377323 | NA | NA |
| ENSG000001 | 13.8703977 | -0.812596266 | 0.31901871 | -2.5471743 | 0.01085992 | 0.03777496 | LSP1 | 4046 |
| ENSG000000 | 28273.4103 | 0.35594957 | 0.13975009 | 2.54704356 | 0.01086399 | 0.0377812 | HSP90AA1 | 3320 |
| ENSG000001 | 20.4021467 | -0.769305108 | 0.30204044 | -2.5470269 | 0.01086451 | 0.0377812 | OR4N2 | 390429 |
| ENSG000002 | 8.33224358 | -0.909270825 | 0.35699133 | -2.5470389 | 0.01086413 | 0.0377812 | MTND6P14 | 106478948 |
| ENSG000001 | 1011.03296 | 0.428484478 | 0.16823309 | 2.54696906 | 0.01086631 | 0.03778422 | VAPB | 9217 |
| ENSG000002 | 60.5790356 | -0.529810532 | 0.20805479 | -2.5464953 | 0.01088107 | 0.03783231 | MALRD1 | 340895 |
| ENSG000002 | 11.1120074 | -0.847041595 | 0.33263921 | -2.5464274 | 0.01088318 | 0.03783354 | NA | NA |
| ENSG000002 | 7.14894118 | -0.915282379 | 0.35943832 | -2.5464241 | 0.01088329 | 0.03783354 | NA | NA |
| ENSG000002 | 8.57747676 | -0.901313591 | 0.35399225 | -2.5461394 | 0.01089217 | 0.03786117 | ZEB2-AS1 | 100303491 |
| ENSG000002 | 8.24908915 | -0.924469444 | 0.36309178 | -2.5461041 | 0.01089327 | 0.03786175 | NA | NA |
| ENSG000002 | 37.3409062 | 0.676278308 | 0.26563432 | 2.54589959 | 0.01089965 | 0.03788069 | NA | NA |
| ENSG000002 | 22.6318564 | -0.68901639 | 0.27065146 | -2.5457701 | 0.0109037 | 0.03789133 | NA | NA |
| ENSG000002 | 2.22000469 | -0.920642315 | 0.36164012 | -2.5457416 | 0.01090459 | 0.03789133 | NA | NA |
| ENSG000001 | 79.0673725 | 0.490202514 | 0.19257128 | 2.54556397 | 0.01091014 | 0.03790737 | TMEM178A | 130733 |
| ENSG000002 | 8.37305849 | -0.893570394 | 0.35104272 | -2.5454748 | 0.01091292 | 0.03791381 | NA | NA |
| ENSG000000 | 743.384872 | 0.290758222 | 0.11423126 | 2.5453472 | 0.01091691 | 0.03792442 | EPB41L4B | 54566 |
| ENSG000002 | 5.0862493 | -0.957105926 | 0.37603851 | -2.5452338 | 0.01092046 | 0.03793336 | NA | NA |
| ENSG000002 | 2.54882657 | -0.942698219 | 0.37038202 | -2.5452051 | 0.01092136 | 0.03793336 | NA | NA |
| ENSG000001 | 8.817633 | -0.894775196 | 0.35156778 | -2.5451001 | 0.01092464 | 0.03793519 | NA | NA |
| ENSG000002 | 10.1448946 | -0.877224129 | 0.34467197 | -2.5450986 | 0.01092469 | 0.03793519 | LINC01375 | 101926924 |
| ENSG000002 | 5.19274187 | -0.959140468 | 0.37685564 | -2.5451138 | 0.01092421 | 0.03793519 | NA | NA |
| ENSG000001 | 51.7930448 | 0.547515583 | 0.2151618 | 2.54466908 | 0.01093813 | 0.03797862 | LRP5L | 91355 |
| ENSG000002 | 3.28066046 | -0.960226972 | 0.37737165 | -2.5445128 | 0.01094303 | 0.03799237 | NA | NA |
| ENSG000001 | 5.80416812 | -0.950886981 | 0.37371014 | -2.5444506 | 0.01094498 | 0.03799588 | UTF1 | 8433 |
| ENSG000001 | 5.07718031 | -0.956157549 | 0.37578618 | -2.5444192 | 0.01094596 | 0.03799605 | PROP1 | 5626 |
| ENSG000001 | 557.880366 | 0.318972126 | 0.12537173 | 2.54421093 | 0.01095249 | 0.03801545 | CHRAC1 | 54108 |
| ENSG000001 | 5.6419997 | -0.945580478 | 0.37167992 | -2.544072 | 0.01095685 | 0.03802732 | DEFB125 | 245938 |
| ENSG000002 | 5.78392384 | -0.950013953 | 0.37343569 | -2.5439828 | 0.01095965 | 0.03803378 | NA | NA |

| | | | | | | | | |
|---|---|---|---|---|---|---|---|---|
| ENSG000001 | 14.7111657 | -0.775615359 | 0.30489251 | -2.5438977 | 0.01096232 | 0.03803666 | CRTAM | 56253 |
| ENSG000002 | 4.32384286 | -0.961316566 | 0.37789137 | -2.5438966 | 0.01096235 | 0.03803666 | NA | NA |
| ENSG000001 | 24.3793051 | -0.695721676 | 0.27350886 | -2.5436897 | 0.01096885 | 0.03805593 | OR6Y1 | 391112 |
| ENSG000002 | 18.3513776 | -0.759590371 | 0.29864391 | -2.5434651 | 0.0109759 | 0.03807389 | NA | NA |
| ENSG000002 | 6.95327525 | -0.926126807 | 0.36411782 | -2.5434811 | 0.0109754 | 0.03807389 | NA | NA |
| ENSG000000 | 4101.94169 | 0.322816237 | 0.12692651 | 2.54333188 | 0.01098009 | 0.03808516 | ATRX | 546 |
| ENSG000002 | 90.5306638 | 0.518176046 | 0.20375865 | 2.54308734 | 0.01098778 | 0.03810714 | FAM127B | 26071 |
| ENSG000002 | 8.97050924 | -0.897330074 | 0.352853 | -2.5430705 | 0.0109883 | 0.03810714 | NA | NA |
| ENSG000001 | 21.8541144 | -0.685955461 | 0.26973965 | -2.5430279 | 0.01098964 | 0.03810852 | TLR4 | 7099 |
| ENSG000002 | 10.8080161 | -0.854405474 | 0.33599887 | -2.542882 | 0.01099423 | 0.03811792 | MYBPHL | 343263 |
| ENSG000002 | 6.30200203 | -0.936295582 | 0.36820225 | -2.5428839 | 0.01099417 | 0.03811792 | NA | NA |
| ENSG000001 | 678.310552 | 0.315529191 | 0.12409872 | 2.54256603 | 0.01100418 | 0.03814914 | CEP55 | 55165 |
| ENSG000002 | 12.6206729 | -0.828446402 | 0.32583692 | -2.5425185 | 0.01100568 | 0.03815107 | NA | NA |
| ENSG000001 | 46.7297258 | 0.547638736 | 0.21541481 | 2.54225198 | 0.01101407 | 0.03817691 | SNTA1 | 6640 |
| ENSG000002 | 1449.28848 | 0.303517273 | 0.11939282 | 2.54217362 | 0.01101654 | 0.03818221 | GTF2I | 2969 |
| ENSG000002 | 36.8282402 | -0.572871157 | 0.22535307 | -2.5421049 | 0.01101871 | 0.03818645 | NA | NA |
| ENSG000002 | 14.40403 | -0.806058046 | 0.3170904 | -2.5420449 | 0.0110206 | 0.03818975 | ANTXRL | 195977 |
| ENSG000002 | 7.74960142 | -0.935562653 | 0.36804887 | -2.5419523 | 0.01102352 | 0.03819533 | LOC101927972 | 101927972 |
| ENSG000002 | 53.2951749 | 0.567184989 | 0.22313389 | 2.54190425 | 0.01102504 | 0.03819533 | NA | NA |
| ENSG000002 | 8.7948275 | -0.891243909 | 0.35061697 | -2.5419303 | 0.01102421 | 0.03819533 | NA | NA |
| ENSG000002 | 19.5393612 | 0.752036807 | 0.2958615 | 2.54185425 | 0.01102661 | 0.03819753 | NA | NA |
| ENSG000002 | 3.6356631 | -0.96168247 | 0.37834443 | -2.5418174 | 0.01102778 | 0.03819829 | NA | NA |
| ENSG000001 | 636.975523 | 0.33654117 | 0.13242101 | 2.5414484 | 0.01103942 | 0.03823537 | ACVR2A | 92 |
| ENSG000002 | 21.4800316 | -0.762836112 | 0.30020266 | -2.5410705 | 0.01105136 | 0.03827345 | NA | NA |
| ENSG000001 | 360.114022 | 0.371675358 | 0.1462817 | 2.54081933 | 0.01105931 | 0.03829768 | GALNT11 | 63917 |
| ENSG000001 | 1236.82222 | 0.294494253 | 0.11592633 | 2.54035682 | 0.01107394 | 0.03833855 | GON4L | 54856 |
| ENSG000001 | 12.7242157 | -0.833683504 | 0.32817197 | -2.5403861 | 0.01107302 | 0.03833855 | ALAS2 | 212 |
| ENSG000002 | 3.73596206 | -0.960660471 | 0.37815366 | -2.5403971 | 0.01107267 | 0.03833855 | NA | NA |
| ENSG000001 | 140.306115 | 0.394440424 | 0.15527517 | 2.54026728 | 0.01107678 | 0.03834509 | PMS2CL | 441194 |

| | | | | | | | | |
|---|---|---|---|---|---|---|---|---|
| ENSG000002 | 6.74766628 | -0.934853976 | 0.36802646 | -2.5401814 | 0.0110795 | 0.03835124 | NA | NA |
| ENSG000001 | 31.4366519 | -0.65251744 | 0.25689491 | -2.540017 | 0.01108471 | 0.03836406 | GCNT3 | 9245 |
| ENSG000002 | 8.46198941 | -0.897846022 | 0.35348202 | -2.5400048 | 0.0110851 | 0.03836406 | NA | NA |
| ENSG000001 | 7.34958753 | -0.922151879 | 0.36306597 | -2.5399017 | 0.01108836 | 0.03837209 | TRIM51 | 84767 |
| ENSG000001 | 7.21483395 | -0.914833187 | 0.36020369 | -2.5397663 | 0.01109266 | 0.03838368 | FAM71B | 153745 |
| ENSG000002 | 28.4778444 | -0.637100271 | 0.25088898 | -2.5393713 | 0.01110519 | 0.03842377 | NA | NA |
| ENSG000002 | 2.92805204 | -0.950797161 | 0.37443255 | -2.5393015 | 0.0111074 | 0.03842487 | NA | NA |
| ENSG000002 | 4.3912272 | -0.959686667 | 0.37793266 | -2.539306 | 0.01110726 | 0.03842487 | NA | NA |
| ENSG000002 | 15.4398903 | -0.775597386 | 0.30545231 | -2.5391767 | 0.01111137 | 0.03843531 | IPO9-AS1 | 100873949 |
| ENSG000001 | 14.5746249 | -0.80669925 | 0.31771503 | -2.5390654 | 0.0111149 | 0.03844269 | B3GALT2 | 8707 |
| ENSG000002 | 10.3186177 | -0.861369743 | 0.33924886 | -2.5390498 | 0.0111154 | 0.03844269 | LOC100129931 | 100129931 |
| ENSG000001 | 539.991333 | 0.314601473 | 0.12390869 | 2.53897829 | 0.01111767 | 0.03844633 | MTERF4 | 130916 |
| ENSG000001 | 12.08121 | -0.83338578 | 0.32824601 | -2.538906 | 0.01111997 | 0.03844633 | BPIFB3 | 359710 |
| ENSG000001 | 20.0876067 | -0.713124639 | 0.28087966 | -2.5388974 | 0.01112024 | 0.03844633 | HRH1 | 3269 |
| ENSG000002 | 18.8347584 | -0.75608273 | 0.29779315 | -2.5389528 | 0.01111848 | 0.03844633 | LINC01225 | 149086 |
| ENSG000002 | 10.9932466 | -0.856860516 | 0.33749781 | -2.5388624 | 0.01112135 | 0.03844689 | KIR3DL2 | 3812 |
| ENSG000001 | 312.746387 | 0.448952925 | 0.1768552 | 2.53853398 | 0.0111318 | 0.03847972 | EPHX1 | 2052 |
| ENSG000001 | 4.28007429 | -0.958886627 | 0.37774845 | -2.5384264 | 0.01113522 | 0.03848828 | NA | NA |
| ENSG000001 | 233.84071 | 0.338340275 | 0.13329373 | 2.53830604 | 0.01113905 | 0.03849823 | DNAJC17 | 55192 |
| ENSG000002 | 5.0259092 | -0.953796909 | 0.37576799 | -2.5382601 | 0.01114051 | 0.0385 | NA | NA |
| ENSG000002 | 12.5587663 | -0.820632122 | 0.32332388 | -2.5381117 | 0.01114524 | 0.03851305 | LOC284865 | 284865 |
| ENSG00000C | 5234.76237 | 0.362909944 | 0.14299094 | 2.53799263 | 0.01114903 | 0.03851866 | PAPOLA | 10914 |
| ENSG000002 | 1597.52184 | 0.422552766 | 0.16649203 | 2.53797596 | 0.01114956 | 0.03851866 | GIGYF2 | 26058 |
| ENSG000002 | 7.94390943 | 0.907061232 | 0.35739617 | 2.53797131 | 0.01114971 | 0.03851866 | NA | NA |
| ENSG000001 | 128.123889 | 0.467240694 | 0.1841067 | 2.53787995 | 0.01115262 | 0.03852543 | NINJ1 | 4814 |
| ENSG000001 | 151.039676 | 0.433693917 | 0.17090419 | 2.53764362 | 0.01116016 | 0.03854817 | B4GALNT4 | 338707 |
| ENSG000002 | 8.38002665 | -0.889735838 | 0.35062027 | -2.5376053 | 0.01116138 | 0.03854911 | LOC100294362 | 100294362 |
| ENSG000002 | 11.0044302 | -0.858901634 | 0.33849784 | -2.5373918 | 0.01116819 | 0.03856934 | ADAMTS7P1 | 390660 |
| ENSG000002 | 15.4305532 | -0.77673105 | 0.30611815 | -2.5373571 | 0.0111693 | 0.03856988 | OOSP1 | 255649 |

| | | | | | | | | |
|---|---|---|---|---|---|---|---|---|
| ENSG000002 | 4.30301892 | -0.958786186 | 0.37787757 | -2.5372932 | 0.01117133 | 0.03857364 | NA | NA |
| ENSG000002 | 18.9222908 | -0.722488258 | 0.28477299 | -2.5370673 | 0.01117854 | 0.03859525 | IGFL4 | 444882 |
| ENSG000002 | 3.26191066 | -0.957065749 | 0.37724841 | -2.5369643 | 0.01118183 | 0.03860003 | NA | NA |
| ENSG000002 | 29.8762695 | -0.631252909 | 0.24881939 | -2.5369924 | 0.01118094 | 0.03860003 | NA | NA |
| ENSG000001 | 319.142447 | 0.337059466 | 0.13286677 | 2.53682282 | 0.01118636 | 0.03861235 | CMC2 | 56942 |
| ENSG000001 | 200.261346 | 0.362726592 | 0.14298821 | 2.53675879 | 0.0111884 | 0.03861613 | DNAJB9 | 4189 |
| ENSG000001 | 117.023857 | 0.400289438 | 0.15780357 | 2.53663101 | 0.01119249 | 0.03862036 | ZDHHC15 | 158866 |
| ENSG000002 | 3.70773539 | -0.948511558 | 0.37391792 | -2.5366839 | 0.0111908 | 0.03862036 | NA | NA |
| ENSG000002 | 39.241232 | 0.607455065 | 0.2394729 | 2.53663381 | 0.0111924 | 0.03862036 | SNORA13 | 654322 |
| ENSG000001 | 88.2089754 | 0.498315479 | 0.1964516 | 2.53658144 | 0.01119407 | 0.03862254 | TCEAL1 | 9338 |
| ENSG000001 | 588.336464 | 0.340078897 | 0.13407777 | 2.53643019 | 0.01119891 | 0.03863594 | ZNF621 | 285268 |
| ENSG000002 | 4.44369841 | -0.959356911 | 0.37824385 | -2.5363451 | 0.01120163 | 0.03864205 | MIR124-2 | 406908 |
| ENSG000001 | 355.102597 | 0.36370511 | 0.14340693 | 2.53617533 | 0.01120706 | 0.03865377 | PGPEP1 | 54858 |
| ENSG000002 | 4.08600598 | -0.960140137 | 0.3785894 | -2.5360988 | 0.01120951 | 0.03865377 | NA | NA |
| ENSG000002 | 7.57547571 | -0.914022549 | 0.36040571 | -2.5360934 | 0.01120968 | 0.03865377 | NA | NA |
| ENSG000002 | 5.69854915 | -0.943776799 | 0.37212652 | -2.5361719 | 0.01120717 | 0.03865377 | NA | NA |
| ENSG000002 | 5.9555211 | -0.9411215 | 0.37109153 | -2.53609 | 0.01120979 | 0.03865377 | NA | NA |
| ENSG000002 | 92.1040625 | 0.447378732 | 0.17640984 | 2.53601919 | 0.01121206 | 0.03865829 | AIF1 | 199 |
| ENSG000001 | 247.126876 | 0.338480658 | 0.1334724 | 2.53595987 | 0.01121396 | 0.03866156 | ZNF135 | 7694 |
| ENSG000001 | 4.70605794 | -0.956190545 | 0.37709687 | -2.5356629 | 0.01122347 | 0.03868905 | NA | NA |
| ENSG000002 | 2.9183123 | -0.949545932 | 0.37447812 | -2.5356513 | 0.01122384 | 0.03868905 | NA | NA |
| ENSG000001 | 1433.65724 | 0.286812666 | 0.11311958 | 2.53548209 | 0.01122927 | 0.03870446 | SERINC3 | 10955 |
| ENSG000001 | 40.8618049 | 0.568316354 | 0.22415986 | 2.5353172 | 0.01123455 | 0.03871611 | SDF2L1 | 23753 |
| ENSG000001 | 1053.33169 | 0.331107375 | 0.13059705 | 2.53533587 | 0.01123396 | 0.03871611 | VPS13C | 54832 |
| ENSG000002 | 12.7475579 | -0.825160597 | 0.32551973 | -2.5349019 | 0.01124788 | 0.03875869 | LOC101930452 | 101930452 |
| ENSG000002 | 216.337191 | 0.515379948 | 0.20331592 | 2.53487259 | 0.01124882 | 0.03875869 | KMT2B | 9757 |
| ENSG000001 | 578.208977 | 0.282502063 | 0.11145415 | 2.53469308 | 0.01125459 | 0.03877526 | C5orf42 | 65250 |
| ENSG000001 | 221.960902 | 0.330376028 | 0.13034689 | 2.53459074 | 0.01125788 | 0.03877671 | SLC38A6 | 145389 |
| ENSG000002 | 2.60534377 | -0.936904699 | 0.36964355 | -2.5346166 | 0.01125704 | 0.03877671 | NA | NA |

| | | | | | | | | |
|---|---|---|---|---|---|---|---|---|
| ENSG000002 | 14.8847115 | -0.80778163 | 0.31869802 | -2.5346302 | 0.01125661 | 0.03877671 | NA | NA |
| ENSG000002 | 4.50357749 | -0.959259823 | 0.37849902 | -2.5343786 | 0.01126469 | 0.03879689 | NA | NA |
| ENSG000001 | 52.005718 | 0.557080188 | 0.21981546 | 2.53430849 | 0.01126695 | 0.03880136 | SRR | 63826 |
| ENSG000001 | 5329.85871 | 0.275492913 | 0.10871797 | 2.53401445 | 0.01127641 | 0.03883028 | KIAA0368 | 23392 |
| ENSG000002 | 10.8682598 | -0.85134842 | 0.33597177 | -2.5339879 | 0.01127726 | 0.03883028 | NA | NA |
| ENSG000002 | 5.33017168 | -0.947460118 | 0.37391677 | -2.5338797 | 0.01128074 | 0.03883898 | NLGN1-AS1 | 100874010 |
| ENSG000001 | 62.5811758 | 0.514313575 | 0.20297949 | 2.53382041 | 0.01128265 | 0.03884225 | ZDHHC24 | 254359 |
| ENSG000002 | 6.24538786 | -0.93140939 | 0.36760139 | -2.5337483 | 0.01128498 | 0.03884695 | LOC339807 | 339807 |
| ENSG000001 | 58.9755643 | 0.54713181 | 0.21597192 | 2.53334701 | 0.0112979 | 0.03888815 | TMEM198 | 130612 |
| ENSG000001 | 228.081548 | 0.376970256 | 0.1488179 | 2.53309749 | 0.01130595 | 0.03891254 | MLYCD | 23417 |
| ENSG000001 | 232.455982 | 0.355530992 | 0.14035786 | 2.53303232 | 0.01130805 | 0.03891386 | IPPK | 64768 |
| ENSG000001 | 367.019034 | 0.334377386 | 0.13200994 | 2.53297123 | 0.01131002 | 0.03891386 | EGLN3 | 112399 |
| ENSG000001 | 6.32269122 | -0.931345091 | 0.36768837 | -2.5329741 | 0.01130993 | 0.03891386 | C16orf78 | 123970 |
| ENSG000002 | 2.20722436 | -0.914933705 | 0.36121033 | -2.5329666 | 0.01131017 | 0.03891386 | NA | NA |
| ENSG000002 | 7.04270762 | -0.926690245 | 0.36586248 | -2.5328923 | 0.01131257 | 0.03891881 | NA | NA |
| ENSG000001 | 2.55781702 | -0.937583504 | 0.37023123 | -2.5324269 | 0.0113276 | 0.03896721 | NA | NA |
| ENSG000002 | 5.35911404 | -0.949101702 | 0.37478465 | -2.5323922 | 0.01132872 | 0.03896776 | NA | NA |
| ENSG000002 | 6.56382545 | -0.925397693 | 0.36552256 | -2.5317116 | 0.01135073 | 0.03904016 | NA | NA |
| ENSG000002 | 7.10816515 | -0.926146599 | 0.36586011 | -2.5314228 | 0.01136008 | 0.03906902 | NA | NA |
| ENSG000001 | 30.8696177 | -0.655001313 | 0.25879296 | -2.5309858 | 0.01137425 | 0.03911196 | ADTRP | 84830 |
| ENSG000002 | 5.89201555 | -0.934953791 | 0.36940417 | -2.5309779 | 0.0113745 | 0.03911196 | DEFB122 | 245935 |
| ENSG000002 | 6.16901499 | -0.943741332 | 0.37290831 | -2.5307597 | 0.01138158 | 0.03913298 | NA | NA |
| ENSG000001 | 369.859545 | 0.338353953 | 0.13370252 | 2.53064762 | 0.01138522 | 0.03913885 | NT5C3A | 51251 |
| ENSG000002 | 14.8534444 | -0.782094188 | 0.30904658 | -2.5306677 | 0.01138457 | 0.03913885 | CLLU1 | 574028 |
| ENSG000002 | 3.6996765 | -0.956943681 | 0.37816138 | -2.5305167 | 0.01138947 | 0.03915014 | MIR517B | 574483 |
| ENSG000001 | 92.8104763 | 0.424313764 | 0.16768379 | 2.53043999 | 0.01139196 | 0.03915539 | SRRM4 | 84530 |
| ENSG000002 | 8.34486254 | -0.887957736 | 0.35095939 | -2.5300868 | 0.01140343 | 0.0391915 | NA | NA |
| ENSG000001 | 7.0470165 | -0.924121181 | 0.36530223 | -2.5297441 | 0.01141457 | 0.03922647 | NRN1L | 123904 |
| ENSG000001 | 1958.42814 | 0.331849932 | 0.1311936 | 2.52946735 | 0.01142358 | 0.03925077 | LBR | 3930 |

| | | | | | | | | |
|---|---|---|---|---|---|---|---|---|
| ENSG000002 | 30.6570625 | 0.635692221 | 0.25131438 | 2.52947017 | 0.01142349 | 0.03925077 | NA | NA |
| ENSG000000 | 61.2490496 | -0.517780132 | 0.20471423 | -2.5292826 | 0.0114296 | 0.03926812 | ABCC2 | 1244 |
| ENSG000002 | 5.78302776 | -0.94325184 | 0.37295811 | -2.5291093 | 0.01143524 | 0.03928418 | NA | NA |
| ENSG000001 | 12.3103683 | -0.82427204 | 0.32593318 | -2.5289602 | 0.0114401 | 0.03929277 | SLC5A4 | 6527 |
| ENSG000001 | 12.4563316 | -0.811585834 | 0.32091894 | -2.5289434 | 0.01144065 | 0.03929277 | PTGIR | 5739 |
| ENSG000002 | 6.53104049 | 0.93074197 | 0.36803075 | 2.52897878 | 0.01143949 | 0.03929277 | NA | NA |
| ENSG000001 | 103.629788 | 0.43511831 | 0.17206311 | 2.52882967 | 0.01144436 | 0.03930217 | BNIP1 | 662 |
| ENSG000001 | 415.72197 | 0.342399646 | 0.13540401 | 2.52872604 | 0.01144773 | 0.03930828 | MTMR3 | 8897 |
| ENSG000001 | 1138.04602 | 0.308168998 | 0.12186779 | 2.52871564 | 0.01144807 | 0.03930828 | RNF219 | 79596 |
| ENSG000001 | 9.51723437 | -0.878276879 | 0.34734421 | -2.5285491 | 0.01145351 | 0.03932028 | CCKAR | 886 |
| ENSG000001 | 171.553156 | 0.456819461 | 0.18066385 | 2.52856041 | 0.01145314 | 0.03932028 | SEMA4B | 10509 |
| ENSG000001 | 58.9007994 | 0.571138162 | 0.22588291 | 2.52847 | 0.01145609 | 0.03932581 | LRFN3 | 79414 |
| ENSG000001 | 22.9210607 | -0.684522238 | 0.27074813 | -2.5282621 | 0.01146287 | 0.03933921 | IRF5 | 3663 |
| ENSG000002 | 18.2133167 | 0.760858229 | 0.30093537 | 2.52831107 | 0.01146128 | 0.03933921 | NA | NA |
| ENSG000002 | 3.89942537 | -0.956951995 | 0.37850202 | -2.5282613 | 0.0114629 | 0.03933921 | NA | NA |
| ENSG000001 | 12.3276124 | -0.848341555 | 0.33554909 | -2.5282189 | 0.01146429 | 0.03934063 | NA | NA |
| ENSG000001 | 3017.72998 | 0.297244607 | 0.11757548 | 2.52811721 | 0.01146761 | 0.03934733 | SCAF11 | 9169 |
| ENSG000001 | 195.187177 | 0.492660783 | 0.19487395 | 2.52809968 | 0.01146818 | 0.03934733 | FBN3 | 84467 |
| ENSG000002 | 11.8572346 | -0.849838752 | 0.33619697 | -2.5278002 | 0.01147797 | 0.03937758 | NA | NA |
| ENSG000000 | 47.995429 | -0.529863292 | 0.20963018 | -2.5276098 | 0.01148419 | 0.03938824 | FER1L4 | 80307 |
| ENSG000001 | 9.51551405 | -0.860488729 | 0.34043891 | -2.5275863 | 0.01148496 | 0.03938824 | COMP | 1311 |
| ENSG000001 | 50.3426484 | 0.56029108 | 0.2216669 | 2.52762623 | 0.01148365 | 0.03938824 | ZNF77 | 58492 |
| ENSG000002 | 6.05450185 | -0.939800977 | 0.37181469 | -2.5276058 | 0.01148432 | 0.03938824 | NA | NA |
| ENSG000002 | 12.8361664 | -0.811137593 | 0.32092484 | -2.5275002 | 0.01148778 | 0.03939457 | LOC102724201 | 102724201 |
| ENSG000000 | 12.9316614 | -0.817312111 | 0.32337161 | -2.5274702 | 0.01148876 | 0.0393946 | PTHLH | 5744 |
| ENSG000002 | 17.6710982 | -0.760487899 | 0.30090138 | -2.5273659 | 0.01149217 | 0.03940297 | NA | NA |
| ENSG000001 | 110.388393 | 0.425161848 | 0.16823136 | 2.52724488 | 0.01149613 | 0.03941154 | C5orf28 | 64417 |
| ENSG000002 | 5.35177849 | -0.946194177 | 0.37439968 | -2.5272302 | 0.01149661 | 0.03941154 | NA | NA |
| ENSG000002 | 36.3741003 | -0.592707549 | 0.23453426 | -2.5271683 | 0.01149864 | 0.03941516 | LRRC37A6P | 387646 |

| | | | | | | | | |
|---|---|---|---|---|---|---|---|---|
| ENSG000002 | 12.3207319 | -0.822058587 | 0.32529902 | -2.5270859 | 0.01150134 | 0.03942107 | NA | NA |
| ENSG000001 | 124.321777 | 0.464406303 | 0.18378153 | 2.52694765 | 0.01150587 | 0.03943326 | C3 | 718 |
| ENSG000001 | 10.3393827 | -0.859087418 | 0.33998288 | -2.526855 | 0.0115089 | 0.03944034 | APOBEC1 | 339 |
| ENSG000001 | 12.2325482 | -0.820149362 | 0.32459954 | -2.5266498 | 0.01151563 | 0.03946005 | AQP9 | 366 |
| ENSG000001 | 21.8777421 | 0.68854839 | 0.27252382 | 2.52656226 | 0.0115185 | 0.03946655 | LAMP5 | 24141 |
| ENSG000001 | 105.235329 | 0.47390937 | 0.18757958 | 2.52644441 | 0.01152236 | 0.03947312 | PC | 5091 |
| ENSG000001 | 30.351367 | 0.622685161 | 0.24646461 | 2.52646886 | 0.01152156 | 0.03947312 | BORCS6 | 54785 |
| ENSG000001 | 99.5490239 | 0.536376056 | 0.21230784 | 2.52640716 | 0.01152358 | 0.03947397 | HES3 | 390992 |
| ENSG000001 | 23.4931558 | -0.681948431 | 0.26998274 | -2.5258964 | 0.01154035 | 0.03952473 | ZIC1 | 7545 |
| ENSG000002 | 8.5227845 | -0.890619043 | 0.35259315 | -2.5259114 | 0.01153986 | 0.03952473 | NA | NA |
| ENSG000001 | 775.348328 | 0.277508134 | 0.10987594 | 2.52564961 | 0.01154846 | 0.03954916 | CCDC112 | 153733 |
| ENSG000001 | 204.629236 | 0.348423826 | 0.13795753 | 2.52558763 | 0.0115505 | 0.0395528 | CLUAP1 | 23059 |
| ENSG000002 | 11.0234468 | -0.835585589 | 0.33085904 | -2.5255033 | 0.01155327 | 0.03955895 | NA | NA |
| ENSG000001 | 9.63266577 | -0.869740733 | 0.34439033 | -2.5254505 | 0.01155501 | 0.03956155 | FCGR3B | 2215 |
| ENSG00000C | 681.55463 | 0.335252506 | 0.13275439 | 2.5253591 | 0.01155801 | 0.03956851 | CDK14 | 5218 |
| ENSG000002 | 6.86224671 | -0.918043207 | 0.3635498 | -2.5252199 | 0.01156259 | 0.03958084 | NA | NA |
| ENSG000002 | 6.32402221 | -0.934556316 | 0.37011219 | -2.5250623 | 0.01156778 | 0.03959526 | LOC105378052 | 105378052 |
| ENSG000001 | 8.98706851 | -0.876720871 | 0.34721493 | -2.5250091 | 0.01156953 | 0.03959772 | KLRB1 | 3820 |
| ENSG000001 | 250.623037 | 0.352044515 | 0.13942841 | 2.5249123 | 0.01157272 | 0.03959772 | PRKCA | 5578 |
| ENSG000001 | 5.43986293 | -0.946898245 | 0.37502162 | -2.5249164 | 0.01157258 | 0.03959772 | CALML3 | 810 |
| ENSG000002 | 8.19288892 | -0.897609558 | 0.3554976 | -2.5249384 | 0.01157186 | 0.03959772 | GACAT3 | 104797537 |
| ENSG000002 | 54.8487121 | -0.508511899 | 0.20139945 | -2.5248922 | 0.01157338 | 0.03959772 | KIAA1456 | 57604 |
| ENSG000001 | 10.7294999 | -0.84213279 | 0.33355479 | -2.524721 | 0.01157902 | 0.03961366 | RGL4 | 266747 |
| ENSG000001 | 1242.59065 | 0.262832992 | 0.10410791 | 2.52462075 | 0.01158232 | 0.03962162 | PAPOLG | 64895 |
| ENSG000001 | 11.3347857 | -0.848665815 | 0.33617011 | -2.524513 | 0.01158587 | 0.03963043 | LRRC66 | 339977 |
| ENSG000002 | 5.47705401 | -0.948242777 | 0.37565203 | -2.5242583 | 0.01159427 | 0.0396558 | NA | NA |
| ENSG00000C | 4825.51389 | 0.284227804 | 0.11260166 | 2.52418832 | 0.01159658 | 0.03965701 | PIK3C2A | 5286 |
| ENSG000002 | 2.92594489 | -0.944991266 | 0.37437124 | -2.524209 | 0.0115959 | 0.03965701 | NA | NA |
| ENSG000001 | 1093.98197 | 0.343084158 | 0.13592179 | 2.52412911 | 0.01159853 | 0.03965774 | SUV420H1 | 51111 |

| | | | | | | | | | |
|---|---|---|---|---|---|---|---|---|---|
| ENSG000001 | 8.45592618 | -0.888695679 | 0.35208243 | -2.5241126 | 0.01159908 | 0.03965774 | NA | NA | |
| ENSG000002 | 510.730394 | -0.409207308 | 0.16212054 | -2.5240929 | 0.01159973 | 0.03965774 | NA | NA | |
| ENSG000002 | 3.40092508 | -0.949553109 | 0.37621533 | -2.5239618 | 0.01160405 | 0.0396692 | NA | NA | |
| ENSG000001 | 29.1859587 | -0.625892812 | 0.24801508 | -2.5236079 | 0.01161574 | 0.03970465 | HIVEP3 | | 59269 |
| ENSG000002 | 12.9885787 | -0.800072901 | 0.31703779 | -2.5235884 | 0.01161638 | 0.03970465 | NA | NA | |
| ENSG000002 | 27.0678125 | -0.644010624 | 0.2552097 | -2.5234567 | 0.01162074 | 0.03971619 | NA | NA | |
| ENSG000002 | 5.27497694 | -0.952368207 | 0.37743193 | -2.5232847 | 0.01162642 | 0.03973227 | NA | NA | |
| ENSG000001 | 278.720088 | 0.333819776 | 0.13231703 | 2.52287839 | 0.01163986 | 0.0397715 | FKBP14 | | 55033 |
| ENSG000002 | 9.80179439 | -0.877530215 | 0.34782756 | -2.5228887 | 0.01163952 | 0.0397715 | NA | NA | |
| ENSG000001 | 624.174015 | 0.333234228 | 0.13209029 | 2.52277609 | 0.01164325 | 0.03977972 | SOCS5 | | 9655 |
| ENSG000001 | 9.52717932 | -0.880149617 | 0.34889621 | -2.5226689 | 0.0116468 | 0.03978849 | GFI1 | | 2672 |
| ENSG000002 | 12.9930988 | -0.818580925 | 0.32449711 | -2.5226139 | 0.01164862 | 0.03979136 | NA | NA | |
| ENSG000001 | 17.6381916 | -0.747872765 | 0.29647182 | -2.5225762 | 0.01164987 | 0.03979227 | KRT2 | | 3849 |
| ENSG000001 | 4098.561 | 0.297536495 | 0.11797261 | 2.52208115 | 0.01166628 | 0.03984497 | SACS | | 26278 |
| ENSG000001 | 40.6997828 | -0.581943468 | 0.23074762 | -2.5219912 | 0.01166926 | 0.0398518 | DISP2 | | 85455 |
| ENSG000001 | 192.487302 | 0.402050119 | 0.15942239 | 2.52191759 | 0.01167171 | 0.03985678 | PSMA2 | | 5683 |
| ENSG000001 | 349.693311 | 0.332855459 | 0.13199407 | 2.5217456 | 0.01167741 | 0.03986383 | LPCAT3 | | 10162 |
| ENSG000001 | 8.77948377 | -0.886823941 | 0.35166015 | -2.521821 | 0.01167491 | 0.03986383 | DKFZP434K028 | | 26070 |
| ENSG000001 | 17.5678574 | 0.76262573 | 0.30241797 | 2.52176065 | 0.01167691 | 0.03986383 | SMPD1 | | 6609 |
| ENSG000001 | 373.307706 | 0.332143934 | 0.13171236 | 2.52173694 | 0.0116777 | 0.03986383 | HIBCH | | 26275 |
| ENSG000001 | 31.2312531 | -0.640411869 | 0.25396014 | -2.5217023 | 0.01167885 | 0.0398644 | MAP3K7CL | | 56911 |
| ENSG000001 | 341.56705 | 0.469275886 | 0.1861022 | 2.52160316 | 0.01168214 | 0.03987228 | ECE1 | | 1889 |
| ENSG000001 | 24.4010561 | -0.694452465 | 0.27540749 | -2.5215454 | 0.01168406 | 0.03987547 | Sep-14 | | 346288 |
| ENSG000002 | 8.691553 | -0.883312736 | 0.3503145 | -2.521485 | 0.01168607 | 0.03987896 | NA | NA | |
| ENSG000001 | 207.309235 | 0.371537822 | 0.14736148 | 2.52126827 | 0.01169327 | 0.03990017 | TTC30A | | 92104 |
| ENSG000002 | 8.93117642 | -0.876010643 | 0.34746369 | -2.5211574 | 0.01169695 | 0.03990939 | SLC9A9-AS1 | | 100885796 |
| ENSG000002 | 3.44372419 | -0.948489661 | 0.37628157 | -2.5206913 | 0.01171246 | 0.03995893 | NA | NA | |
| ENSG000002 | 10.7792718 | -0.839542958 | 0.3330812 | -2.5205354 | 0.01171764 | 0.03997326 | MTND6P16 | | 106478950 |
| ENSG000002 | 11.3413225 | -0.835581065 | 0.33154592 | -2.5202574 | 0.0117269 | 0.04000148 | NA | NA | |

| | | | | | | | | |
|---|---|---|---|---|---|---|---|---|
| ENSG000002 | 4.34721171 | -0.953045055 | 0.37819925 | -2.5199549 | 0.01173699 | 0.04003251 | NA | NA |
| ENSG000001 | 685.347866 | 0.327957956 | 0.13014779 | 2.51988872 | 0.01173919 | 0.04003331 | UGGT2 | 55757 |
| ENSG000002 | 172.530669 | 0.422130819 | 0.1675185 | 2.51990564 | 0.01173863 | 0.04003331 | AFG3L1P | 172 |
| ENSG000001 | 704.515902 | 0.278824949 | 0.11065688 | 2.51972534 | 0.01174464 | 0.04004276 | CCSER2 | 54462 |
| ENSG000001 | 1320.78557 | 0.356766829 | 0.14159005 | 2.51971681 | 0.01174493 | 0.04004276 | DCUN1D4 | 23142 |
| ENSG000002 | 2.65356541 | -0.927919864 | 0.36826065 | -2.5197367 | 0.01174426 | 0.04004276 | NA | NA |
| ENSG000001 | 666.185967 | 0.378156716 | 0.15009776 | 2.51940287 | 0.01175541 | 0.04007512 | FAM160A1 | 729830 |
| ENSG000002 | 4.26845518 | -0.952114695 | 0.37794928 | -2.51916 | 0.01176352 | 0.0400994 | NA | NA |
| ENSG000002 | 29.2095909 | -0.654095756 | 0.25966161 | -2.5190314 | 0.01176782 | 0.04011068 | FLJ46284 | 441369 |
| ENSG000001 | 45.9576301 | -0.527980094 | 0.20961362 | -2.5188253 | 0.01177471 | 0.04013078 | MYRF | 745 |
| ENSG000001 | 13.7821668 | -0.797257527 | 0.31652408 | -2.5187895 | 0.0117759 | 0.0401315 | NR1I3 | 9970 |
| ENSG000001 | 441.570675 | 0.363134779 | 0.14417788 | 2.51865805 | 0.0117803 | 0.04014044 | SVIL | 6840 |
| ENSG000002 | 6.23690595 | -0.928032204 | 0.36846387 | -2.5186518 | 0.01178051 | 0.04014044 | LINC01467 | 101928559 |
| ENSG000002 | 772.823832 | 0.330689508 | 0.13130008 | 2.51857812 | 0.01178297 | 0.04014186 | NA | NA |
| ENSG000002 | 10.7846333 | -0.840859862 | 0.3338613 | -2.5185904 | 0.01178256 | 0.04014186 | NA | NA |
| ENSG000002 | 5.66641765 | -0.93874013 | 0.3727303 | -2.5185506 | 0.01178389 | 0.04014186 | NA | NA |
| ENSG000002 | 2.89992777 | -0.943429452 | 0.37460045 | -2.5184953 | 0.01178575 | 0.0401448 | NA | NA |
| ENSG000001 | 81.5167305 | -0.452138627 | 0.17953355 | -2.5184075 | 0.01178869 | 0.04015143 | WDR66 | 144406 |
| ENSG000001 | 2942.94465 | 0.360248144 | 0.14305168 | 2.51830762 | 0.01179203 | 0.04015945 | YAP1 | 10413 |
| ENSG000002 | 12.0726415 | -0.810642709 | 0.32191519 | -2.5181872 | 0.01179606 | 0.04016981 | NA | NA |
| ENSG000002 | 113.299467 | 0.475131299 | 0.18868478 | 2.51812201 | 0.01179824 | 0.04017387 | NA | NA |
| ENSG000002 | 30.07429 | -0.650034242 | 0.25816743 | -2.5178785 | 0.0118064 | 0.04019827 | LINC00925 | 254559 |
| ENSG000002 | 5.02278464 | -0.946883095 | 0.3760862 | -2.5177289 | 0.01181142 | 0.04021198 | NA | NA |
| ENSG000002 | 4.29037498 | -0.951557899 | 0.37795737 | -2.5176329 | 0.01181464 | 0.04021619 | NA | NA |
| ENSG000002 | 6.18548257 | -0.938840822 | 0.37290227 | -2.5176592 | 0.01181376 | 0.04021619 | NA | NA |
| ENSG000001 | 1609.12203 | 0.635080464 | 0.25225997 | 2.5175634 | 0.01181697 | 0.04022049 | PRRC2B | 84726 |
| ENSG000001 | 119.538838 | 0.43974236 | 0.17467172 | 2.51753607 | 0.01181789 | 0.04022049 | RRNAD1 | 51093 |
| ENSG000001 | 219.759712 | 0.357464439 | 0.14199562 | 2.5174329 | 0.01182135 | 0.04022552 | DLG4 | 1742 |
| ENSG000001 | 184.103094 | 0.396943971 | 0.15767657 | 2.51745689 | 0.01182054 | 0.04022552 | E2F6 | 1876 |

| | | | | | | | | |
|---|---|---|---|---|---|---|---|---|
| ENSG000002 | 15.0208048 | -0.768534293 | 0.30535959 | -2.5168173 | 0.01184202 | 0.04029249 | PCDHA1 | 56147 |
| ENSG000001 | 49.4360829 | -0.522039197 | 0.20742686 | -2.5167386 | 0.01184467 | 0.0402981 | SCUBE3 | 222663 |
| ENSG000001 | 33.5484109 | -0.652803522 | 0.25938788 | -2.5167079 | 0.0118457 | 0.04029824 | SCEL | 8796 |
| ENSG000001 | 270.653764 | 0.354760739 | 0.14096623 | 2.51663627 | 0.01184811 | 0.04029827 | COCH | 1690 |
| ENSG000001 | 1291.30168 | 0.34749629 | 0.13808062 | 2.51661891 | 0.01184869 | 0.04029827 | SLC5A3 | 6526 |
| ENSG000002 | 163.376803 | 0.401949667 | 0.15971594 | 2.51665335 | 0.01184753 | 0.04029827 | LINC00094 | 266655 |
| ENSG000002 | 290.689837 | 0.387792378 | 0.15411979 | 2.51617503 | 0.01186362 | 0.04034568 | PSMD5-AS1 | 253039 |
| ENSG000002 | 16.5800843 | 0.745766588 | 0.29639918 | 2.51608857 | 0.01186654 | 0.04035219 | MRPL38 | 64978 |
| ENSG00000C | 13.9956217 | -0.812422409 | 0.32290192 | -2.5160036 | 0.0118694 | 0.04035853 | AOC1 | 26 |
| ENSG000002 | 3.39380844 | -0.945772456 | 0.37592787 | -2.5158349 | 0.01187508 | 0.04037447 | NA | NA |
| ENSG000002 | 12.5371586 | -0.817890293 | 0.3251384 | -2.5155143 | 0.01188589 | 0.04040783 | LOC102724434 | 102724434 |
| ENSG000002 | 6.85917477 | -0.914073051 | 0.36338632 | -2.5154305 | 0.01188871 | 0.04041404 | NA | NA |
| ENSG000002 | 12905.4054 | 0.356401773 | 0.14170097 | 2.51516817 | 0.01189756 | 0.04044074 | TRIM71 | 131405 |
| ENSG000002 | 6.55773199 | -0.920698 | 0.36608024 | -2.5150169 | 0.01190267 | 0.0404547 | NA | NA |
| ENSG000002 | 8.10460659 | -0.890621868 | 0.35415507 | -2.5147794 | 0.01191069 | 0.04047857 | NA | NA |
| ENSG000001 | 85.8331395 | -0.433675437 | 0.17245934 | -2.5146533 | 0.01191495 | 0.04048966 | GRIN2A | 2903 |
| ENSG000002 | 32.1270905 | -0.614327727 | 0.2443123 | -2.5145182 | 0.01191952 | 0.04050178 | RNF212B | 100507650 |
| ENSG000002 | 3.01887397 | -0.937750711 | 0.37294126 | -2.5144729 | 0.01192105 | 0.04050359 | NA | NA |
| ENSG00000C | 21.273907 | -0.748600023 | 0.29773731 | -2.514297 | 0.01192699 | 0.04051513 | VNN3 | 55350 |
| ENSG000001 | 19.6975041 | -0.766377476 | 0.30480289 | -2.514338 | 0.01192561 | 0.04051513 | LMNTD1 | 160492 |
| ENSG000001 | 31.9291876 | -0.632072011 | 0.25139247 | -2.5142838 | 0.01192744 | 0.04051513 | TOPAZ1 | 375337 |
| ENSG000002 | 10.4328105 | -0.845423403 | 0.33629402 | -2.5139412 | 0.01193903 | 0.04055111 | LILRA2 | 11027 |
| ENSG000001 | 74.947885 | 0.475272926 | 0.18906565 | 2.51379839 | 0.01194387 | 0.04056414 | DCTN3 | 11258 |
| ENSG000001 | 7.22846305 | -0.909160125 | 0.36167482 | -2.5137501 | 0.0119455 | 0.04056629 | ANKRD2 | 26287 |
| ENSG000002 | 4.69758181 | -0.948481161 | 0.37735282 | -2.5135128 | 0.01195354 | 0.04059019 | NA | NA |
| ENSG000002 | 9.87826641 | -0.858421398 | 0.34156588 | -2.5131942 | 0.01196435 | 0.04062348 | LINC00211 | 101929559 |
| ENSG000002 | 5.66459887 | -0.936937977 | 0.37281782 | -2.5131255 | 0.01196668 | 0.04062799 | NA | NA |
| ENSG00000C | 219.8549 | 0.347841914 | 0.13841991 | 2.51294711 | 0.01197273 | 0.04064007 | ARHGAP44 | 9912 |
| ENSG00000C | 323.829734 | 0.334104256 | 0.13295241 | 2.51296125 | 0.01197225 | 0.04064007 | KCNQ2 | 3785 |

| | | | | | | | | |
|---|---|---|---|---|---|---|---|---|
| ENSG000002 | 14.1486394 | -0.781219029 | 0.31087949 | -2.512932 | 0.01197324 | 0.04064007 | NA | NA |
| ENSG000002 | 6.86697513 | -0.916137193 | 0.36457677 | -2.5128787 | 0.01197505 | 0.04064281 | NA | NA |
| ENSG000002 | 951.082358 | 0.340971691 | 0.13569793 | 2.51272589 | 0.01198024 | 0.04065701 | NUDT3 | 11165 |
| ENSG000001 | 11.1547006 | -0.839432158 | 0.33408822 | -2.5126063 | 0.0119843 | 0.04066739 | TCN1 | 6947 |
| ENSG000000 | 141.253431 | 0.465276623 | 0.18517979 | 2.51256697 | 0.01198564 | 0.04066852 | MNT | 4335 |
| ENSG000001 | 20.5202484 | -0.703160668 | 0.27986745 | -2.5124774 | 0.01198868 | 0.04067544 | PLSCR2 | 57047 |
| ENSG000001 | 51.6799538 | -0.5398281 | 0.21488819 | -2.5121348 | 0.01200032 | 0.04070814 | PLK3 | 1263 |
| ENSG000001 | 141.336279 | 0.453297421 | 0.18044264 | 2.51214139 | 0.0120001 | 0.04070814 | TPCN1 | 53373 |
| ENSG000001 | 82.9227717 | 0.542683087 | 0.21602942 | 2.51207953 | 0.0120022 | 0.04071111 | VEGFB | 7423 |
| ENSG000001 | 14.8991448 | 0.826584149 | 0.32908822 | 2.5117403 | 0.01201375 | 0.04074345 | PAOX | 196743 |
| ENSG000002 | 13.8207028 | -0.798786003 | 0.31802008 | -2.511747 | 0.01201352 | 0.04074345 | NA | NA |
| ENSG000001 | 60.1123383 | 0.513276248 | 0.20435335 | 2.51170952 | 0.01201479 | 0.0407436 | NFKBID | 84807 |
| ENSG000002 | 6.80311941 | 0.910494173 | 0.36252372 | 2.51154372 | 0.01202044 | 0.0407587 | NA | NA |
| ENSG000002 | 5.44655844 | -0.943911951 | 0.37583299 | -2.5115197 | 0.01202126 | 0.0407587 | OVAAL | 148756 |
| ENSG000001 | 1279.66517 | 0.333757081 | 0.13289279 | 2.51147626 | 0.01202274 | 0.04076031 | TFG | 10342 |
| ENSG000002 | 8.29237159 | -0.898347139 | 0.35770434 | -2.5114236 | 0.01202453 | 0.04076298 | NA | NA |
| ENSG000001 | 30.7500037 | -0.641969189 | 0.25564188 | -2.5112051 | 0.01203198 | 0.04078481 | DDC | 1644 |
| ENSG000002 | 12.9826391 | -0.81445806 | 0.32435061 | -2.5110422 | 0.01203753 | 0.04080023 | LINC01250 | 101927554 |
| ENSG000001 | 756.914373 | 0.316007863 | 0.12585358 | 2.51091674 | 0.01204181 | 0.04081132 | GTF2A2 | 2958 |
| ENSG000001 | 10.4754098 | -0.850604753 | 0.3387881 | -2.5107279 | 0.01204825 | 0.04082974 | C8orf4 | 56892 |
| ENSG000002 | 7.06444202 | -0.918682085 | 0.36592028 | -2.5106072 | 0.01205237 | 0.04084029 | NA | NA |
| ENSG000001 | 19.8653277 | -0.717104647 | 0.28563492 | -2.5105636 | 0.01205386 | 0.04084192 | DCSTAMP | 81501 |
| ENSG000002 | 28.344518 | -0.64604971 | 0.25733975 | -2.5104933 | 0.01205626 | 0.04084664 | NA | NA |
| ENSG000001 | 16.0846453 | -0.772979097 | 0.30791441 | -2.51037 | 0.01206047 | 0.04085749 | SPDYE1 | 285955 |
| ENSG000001 | 14.7719505 | -0.771696274 | 0.3074304 | -2.5101496 | 0.012068 | 0.04087959 | CLDN16 | 10686 |
| ENSG000002 | 4.58037123 | -0.950297458 | 0.37858853 | -2.5101063 | 0.01206948 | 0.04088119 | AMY2A | 279 |
| ENSG000002 | 2.96980753 | -0.935281302 | 0.3726388 | -2.509887 | 0.01207698 | 0.04090316 | NA | NA |
| ENSG000000 | 79.2041462 | -0.470725156 | 0.187559 | -2.5097445 | 0.01208185 | 0.04091626 | COL17A1 | 1308 |
| ENSG000001 | 887.615143 | 0.309754441 | 0.12342258 | 2.50970644 | 0.01208316 | 0.04091725 | MBD4 | 8930 |

| | | | | | | | | |
|---|---|---|---|---|---|---|---|---|
| ENSG000002 | 4.2668364 | -0.949009277 | 0.37815555 | -2.5095738 | 0.01208769 | 0.0409292 | CRAT8 | 105370070 |
| ENSG000000 | 659.017006 | 0.30540685 | 0.12169947 | 2.50951672 | 0.01208965 | 0.0409324 | PHTF2 | 57157 |
| ENSG000001 | 14.1992073 | -0.789559946 | 0.31463289 | -2.5094642 | 0.01209145 | 0.04093508 | SPDYE7P | 441251 |
| ENSG000002 | 10.1934743 | -0.853513347 | 0.34013126 | -2.5093646 | 0.01209486 | 0.0409432 | NA | NA |
| ENSG000001 | 25.0556944 | -0.650654204 | 0.25931496 | -2.5091272 | 0.01210299 | 0.04096731 | PARVG | 64098 |
| ENSG000001 | 11.7379892 | -0.829191675 | 0.33048452 | -2.5090182 | 0.01210672 | 0.04097311 | UGT2A3 | 79799 |
| ENSG000001 | 43.5887357 | 0.594080613 | 0.23677785 | 2.50902108 | 0.01210663 | 0.04097311 | ZNF517 | 340385 |
| ENSG000002 | 2.95326834 | -0.938645818 | 0.37411958 | -2.508946 | 0.0121092 | 0.04097807 | NA | NA |
| ENSG000001 | 447.388687 | 0.382369076 | 0.15240613 | 2.50888245 | 0.01211138 | 0.04098202 | UNC5B | 219699 |
| ENSG000001 | 5.78341531 | -0.939070562 | 0.3743391 | -2.5086093 | 0.01212075 | 0.04101029 | LCN10 | 414332 |
| ENSG000002 | 3.63041177 | -0.948696189 | 0.37820964 | -2.5083872 | 0.01212837 | 0.04103267 | NA | NA |
| ENSG000001 | 12.8250253 | -0.811405375 | 0.32350261 | -2.508188 | 0.01213521 | 0.04105237 | PDX1 | 3651 |
| ENSG000002 | 5.98115296 | -0.929158599 | 0.37045476 | -2.5081567 | 0.01213628 | 0.04105259 | LOC100996286 | 100996286 |
| ENSG000001 | 12.9373065 | -0.812292452 | 0.32387684 | -2.5080288 | 0.01214068 | 0.0410606 | ZPBP2 | 124626 |
| ENSG000002 | 10.2636357 | -0.871412389 | 0.34744905 | -2.5080292 | 0.01214066 | 0.0410606 | LOC100287944 | 100287944 |
| ENSG000002 | 16.937274 | -0.742825782 | 0.29620043 | -2.5078484 | 0.01214688 | 0.04107405 | GPX5 | 2880 |
| ENSG000002 | 35.0100268 | -0.598232272 | 0.23854414 | -2.5078473 | 0.01214691 | 0.04107405 | NA | NA |
| ENSG000002 | 2.49677196 | -0.926321726 | 0.36937694 | -2.5077952 | 0.0121487 | 0.04107405 | NA | NA |
| ENSG000002 | 6.38249155 | -0.927933816 | 0.37001814 | -2.5078063 | 0.01214832 | 0.04107405 | NA | NA |
| ENSG000001 | 1377.10183 | 0.318400939 | 0.12697266 | 2.50763382 | 0.01215425 | 0.04108939 | GNB4 | 59345 |
| ENSG000001 | 132.658149 | 0.399031649 | 0.15913619 | 2.50748523 | 0.01215937 | 0.04110324 | IRX2 | 153572 |
| ENSG000001 | 429.798924 | 0.32548677 | 0.12981125 | 2.50738497 | 0.01216282 | 0.04111148 | PLCB4 | 5332 |
| ENSG000001 | 17.1866574 | -0.736464814 | 0.29372911 | -2.5072926 | 0.012166 | 0.04111194 | SPDYE5 | 442590 |
| ENSG000002 | 21.9997704 | -0.707494477 | 0.28217241 | -2.5073127 | 0.0121653 | 0.04111194 | LOC401242 | 401242 |
| ENSG000002 | 10.8623231 | -0.845945558 | 0.33738821 | -2.5073359 | 0.01216451 | 0.04111194 | NA | NA |
| ENSG000002 | 8.88482687 | -0.884957064 | 0.35297431 | -2.507143 | 0.01217115 | 0.04112592 | FAM196B | 100131897 |
| ENSG000002 | 27.8185727 | -0.633069363 | 0.25251187 | -2.5070876 | 0.01217305 | 0.04112894 | LINC00426 | 100188949 |
| ENSG000000 | 88.176337 | 0.459561602 | 0.18330729 | 2.50705583 | 0.01217415 | 0.04112921 | AAMDC | 28971 |
| ENSG000002 | 25.7845879 | -0.666093187 | 0.26569073 | -2.5070245 | 0.01217523 | 0.04112942 | NA | NA |

| | | | | | | | | |
|---|---|---|---|---|---|---|---|---|
| ENSG00000( | 311.64215 | 0.324475102 | 0.12942914 | 2.506971 | 0.01217707 | 0.04113223 | EVI5 | 7813 |
| ENSG000001 | 1516.72742 | 0.29344883 | 0.11706236 | 2.50677367 | 0.01218387 | 0.04115177 | STYX | 6815 |
| ENSG000002 | 9.07315032 | -0.876066602 | 0.34949445 | -2.5066682 | 0.01218751 | 0.04116062 | NA | NA |
| ENSG000001 | 26.6772708 | -0.650781948 | 0.25962631 | -2.5066102 | 0.01218951 | 0.04116395 | NWD2 | 57495 |
| ENSG000002 | 2.97970226 | -0.935923801 | 0.3734237 | -2.5063321 | 0.0121991 | 0.04119291 | NA | NA |
| ENSG000001 | 18.2263147 | -0.719784267 | 0.28719913 | -2.5062202 | 0.01220296 | 0.04120251 | TBX18 | 9096 |
| ENSG000001 | 444.800381 | 0.316468836 | 0.12627875 | 2.50611317 | 0.01220665 | 0.04120813 | AGO3 | 192669 |
| ENSG000001 | 852.91158 | 0.356970531 | 0.14243937 | 2.50612268 | 0.01220633 | 0.04120813 | NFXL1 | 152518 |
| ENSG000002 | 5.31253442 | -0.946495214 | 0.37771421 | -2.5058502 | 0.01221574 | 0.04123536 | NA | NA |
| ENSG000001 | 6.16786091 | -0.918520147 | 0.36656462 | -2.5057523 | 0.01221912 | 0.04124334 | LGALS9B | 284194 |
| ENSG000001 | 34.2235874 | -0.613128683 | 0.24469472 | -2.5056882 | 0.01222133 | 0.04124738 | GBP4 | 115361 |
| ENSG000001 | 1447.32642 | 0.270103167 | 0.10779907 | 2.50561694 | 0.0122238 | 0.04124883 | PARD3 | 56288 |
| ENSG000002 | 10.5882764 | -0.851141426 | 0.3396916 | -2.5056299 | 0.01222335 | 0.04124883 | NA | NA |
| ENSG000002 | 11.7149072 | -0.83016785 | 0.33134436 | -2.5054534 | 0.01222945 | 0.04126448 | LOC101929595 | 101929595 |
| ENSG000002 | 22.3981942 | -0.690907671 | 0.27576853 | -2.5053898 | 0.01223165 | 0.04126846 | LOC574538 | 574538 |
| ENSG00000( | 293.717266 | 0.316306195 | 0.12626065 | 2.50518428 | 0.01223876 | 0.04128902 | GALC | 2581 |
| ENSG000001 | 31.387195 | -0.640680259 | 0.25575495 | -2.5050552 | 0.01224323 | 0.04129722 | FILIP1L | 11259 |
| ENSG000002 | 3.30749409 | -0.944088737 | 0.37687037 | -2.5050755 | 0.01224252 | 0.04129722 | NA | NA |
| ENSG00000( | 11.2975539 | -0.850379942 | 0.33952734 | -2.5045993 | 0.01225902 | 0.04134704 | FXYD3 | 5349 |
| ENSG000001 | 264.8822 | 0.431490915 | 0.17229016 | 2.50444324 | 0.01226443 | 0.04136185 | POM121 | 9883 |
| ENSG000002 | 9.2667872 | -0.868009692 | 0.34661356 | -2.5042577 | 0.01227086 | 0.0413801 | NA | NA |
| ENSG000001 | 116.540267 | 0.403204653 | 0.16100999 | 2.50422133 | 0.01227212 | 0.04138092 | SLC15A2 | 6565 |
| ENSG000002 | 13.1108923 | -0.798177101 | 0.31874737 | -2.5041057 | 0.01227614 | 0.041391 | PLCG1-AS1 | 101927117 |
| ENSG000001 | 192.031654 | 0.365436107 | 0.14594313 | 2.50396233 | 0.01228111 | 0.04140433 | ZC3H6 | 376940 |
| ENSG000002 | 4.71870366 | -0.945999479 | 0.37781816 | -2.5038486 | 0.01228506 | 0.0414142 | NA | NA |
| ENSG000002 | 10.5137925 | -0.869993264 | 0.34749309 | -2.5036275 | 0.01229274 | 0.04143664 | NA | NA |
| ENSG000001 | 76.9849476 | -0.474904432 | 0.18969201 | -2.5035553 | 0.01229525 | 0.04144166 | C5 | 727 |
| ENSG000001 | 642.198324 | 0.316259353 | 0.1263317 | 2.50340451 | 0.01230049 | 0.04145587 | TIAM1 | 7074 |
| ENSG000001 | 4.37729656 | -0.947398492 | 0.37846628 | -2.5032573 | 0.0123056 | 0.04146967 | LHX3 | 8022 |

| | | | | | | | | |
|---|---|---|---|---|---|---|---|---|
| ENSG000001 | 26.9321882 | -0.652720325 | 0.26077561 | -2.502996 | 0.01231469 | 0.04149685 | TBR1 | 10716 |
| ENSG000002 | 4.38384273 | -0.947008751 | 0.37836345 | -2.5029076 | 0.01231777 | 0.04150377 | NA | NA |
| ENSG000001 | 216.941677 | 0.347854098 | 0.13898716 | 2.50277861 | 0.01232226 | 0.04151545 | C16orf70 | 80262 |
| ENSG000001 | 23.0257002 | -0.736203518 | 0.29419269 | -2.5024535 | 0.01233358 | 0.04155015 | CYP4B1 | 1580 |
| ENSG000002 | 2.88783258 | -0.936744432 | 0.37433869 | -2.5023981 | 0.01233551 | 0.04155319 | SLC25A18P1 | 106480776 |
| ENSG000001 | 435.684324 | 0.297955723 | 0.11907235 | 2.50230832 | 0.01233864 | 0.04155683 | STAM2 | 10254 |
| ENSG000001 | 25.4899326 | -0.674467478 | 0.26953509 | -2.5023364 | 0.01233766 | 0.04155683 | NA | NA |
| ENSG000002 | 17.1686725 | 0.73641209 | 0.29430207 | 2.50223208 | 0.0123413 | 0.04156233 | NA | NA |
| ENSG000001 | 88.3448251 | 0.509536203 | 0.20364327 | 2.50210188 | 0.01234584 | 0.04157417 | ALDH4A1 | 8659 |
| ENSG000001 | 3336.31241 | 0.292990841 | 0.11710499 | 2.50195004 | 0.01235114 | 0.04158855 | TROVE2 | 6738 |
| ENSG000001 | 16.2380266 | -0.792710177 | 0.31684237 | -2.5019071 | 0.01235263 | 0.04159013 | ZNF208 | 7757 |
| ENSG000001 | 3262.63705 | 0.324767914 | 0.12981488 | 2.50177727 | 0.01235716 | 0.04159506 | TPM1 | 7168 |
| ENSG000001 | 17.4321987 | -0.731327688 | 0.29231755 | -2.5018261 | 0.01235546 | 0.04159506 | CD3G | 917 |
| ENSG000002 | 2.88851772 | -0.936516998 | 0.37434071 | -2.5017771 | 0.01235717 | 0.04159506 | NA | NA |
| ENSG00000C | 529.431007 | 0.332261731 | 0.13281766 | 2.50163818 | 0.01236202 | 0.04160447 | ARNTL2 | 56938 |
| ENSG000002 | 13.0171989 | -0.799623861 | 0.3196373 | -2.50166 | 0.01236126 | 0.04160447 | NA | NA |
| ENSG000001 | 542.805428 | 0.340849971 | 0.13625869 | 2.50149162 | 0.01236714 | 0.04161819 | CCDC82 | 79780 |
| ENSG000002 | 20.6117337 | -0.705827967 | 0.2821661 | -2.5014626 | 0.01236815 | 0.04161819 | LINC01192 | 647107 |
| ENSG000002 | 8.74554701 | -0.878394091 | 0.35118437 | -2.5012335 | 0.01237616 | 0.04164168 | NA | NA |
| ENSG000002 | 6.94376363 | -0.914862657 | 0.36578328 | -2.5011058 | 0.01238062 | 0.04165324 | NA | NA |
| ENSG000002 | 7.92367911 | -0.899605807 | 0.35968947 | -2.5010624 | 0.01238214 | 0.04165489 | NA | NA |
| ENSG000002 | 8.44104844 | -0.885273211 | 0.35400106 | -2.5007643 | 0.01239256 | 0.0416865 | NA | NA |
| ENSG000001 | 27.4756584 | -0.629278383 | 0.25166116 | -2.5004986 | 0.01240186 | 0.04171432 | HCK | 3055 |
| ENSG000002 | 6.84368436 | -0.908049015 | 0.36315211 | -2.5004647 | 0.01240305 | 0.04171486 | NA | NA |
| ENSG000002 | 7.78072228 | -0.893483792 | 0.3573661 | -2.5001918 | 0.01241261 | 0.04174354 | LOC101927650 | 101927650 |
| ENSG000001 | 20.9188075 | -0.685116435 | 0.27403787 | -2.5000794 | 0.01241655 | 0.04175333 | BEGAIN | 57596 |
| ENSG000002 | 20.522907 | -0.719401825 | 0.28778033 | -2.4998298 | 0.0124253 | 0.0417793 | RDH10-AS1 | 101926926 |
| ENSG000002 | 9.68299326 | 0.86699824 | 0.34682746 | 2.49979701 | 0.01242645 | 0.0417797 | SNORA74B | 677841 |
| ENSG000001 | 275.274012 | 0.332573073 | 0.13304365 | 2.49972897 | 0.01242884 | 0.04178426 | CCDC84 | 338657 |

| | | | | | | | | |
|---|---|---|---|---|---|---|---|---|
| ENSG000001 | 434.730533 | 0.441696361 | 0.17670898 | 2.49956939 | 0.01243443 | 0.04179961 | SEC14L1 | 6397 |
| ENSG000001 | 45.6297048 | -0.547206472 | 0.21892633 | -2.4995005 | 0.01243685 | 0.04180428 | MYCBPAP | 84073 |
| ENSG00000C | 566.51972 | 0.308338648 | 0.12336541 | 2.49939298 | 0.01244063 | 0.04181003 | FAM13B | 51306 |
| ENSG000002 | 4.33997719 | -0.945170804 | 0.3781596 | -2.4993966 | 0.0124405 | 0.04181003 | NA | NA |
| ENSG000001 | 155.978871 | 0.385742786 | 0.15434081 | 2.49929215 | 0.01244417 | 0.04181847 | LGI2 | 55203 |
| ENSG000002 | 8.03513515 | -0.884773759 | 0.35401649 | -2.4992445 | 0.01244584 | 0.04182063 | NA | NA |
| ENSG000001 | 42.024902 | -0.577023705 | 0.23088918 | -2.4991371 | 0.01244961 | 0.04182246 | FEZF1 | 389549 |
| ENSG000001 | 4.7073673 | -0.944039878 | 0.3777502 | -2.4991115 | 0.01245051 | 0.04182246 | RNU6-486P | 106481898 |
| ENSG000002 | 10.6594429 | -0.853785286 | 0.34162989 | -2.4991528 | 0.01244906 | 0.04182246 | NA | NA |
| ENSG000002 | 10.883958 | -0.844925172 | 0.33808279 | -2.4991665 | 0.01244858 | 0.04182246 | NA | NA |
| ENSG000002 | 3.35651314 | -0.940385537 | 0.37630626 | -2.4989899 | 0.01245479 | 0.04183335 | NA | NA |
| ENSG000001 | 879.105287 | 0.314158591 | 0.12572426 | 2.49879059 | 0.01246179 | 0.04185342 | ZCRB1 | 85437 |
| ENSG000001 | 7.98930593 | -0.882472886 | 0.35320315 | -2.4984853 | 0.01247253 | 0.04188601 | LINC00615 | 439916 |
| ENSG00000C | 286.97333 | 0.370484161 | 0.14829378 | 2.49831222 | 0.01247862 | 0.04189855 | GPATCH1 | 55094 |
| ENSG000001 | 2728.76842 | 0.368567927 | 0.14752801 | 2.49829118 | 0.01247936 | 0.04189855 | PPP4R3A | 55671 |
| ENSG000001 | 8481.30101 | 0.368396778 | 0.14745701 | 2.49833344 | 0.01247788 | 0.04189855 | PARP1 | 142 |
| ENSG000001 | 11.5172187 | -0.839908108 | 0.33626086 | -2.4977873 | 0.01249711 | 0.04195421 | DGAT2L6 | 347516 |
| ENSG000002 | 5.95921478 | -0.928774617 | 0.37184274 | -2.4977619 | 0.01249801 | 0.04195421 | NA | NA |
| ENSG00000C | 27.2735099 | -0.666942973 | 0.26702625 | -2.4976682 | 0.01250131 | 0.04196182 | PKD2L2 | 27039 |
| ENSG000001 | 18.0671347 | -0.762904879 | 0.30545747 | -2.4975814 | 0.01250438 | 0.04196863 | TBX4 | 9496 |
| ENSG000001 | 137.532603 | 0.414932187 | 0.16614057 | 2.49747657 | 0.01250807 | 0.04197756 | ELL | 8178 |
| ENSG000002 | 3.93719332 | -0.945462522 | 0.37857651 | -2.4974146 | 0.01251026 | 0.04198142 | NA | NA |
| ENSG000002 | 71.8520784 | 0.500234345 | 0.20033642 | 2.4969716 | 0.0125259 | 0.04203043 | SH3D21 | 79729 |
| ENSG000001 | 994.574857 | 0.626010527 | 0.25072197 | 2.49683158 | 0.01253085 | 0.04204355 | TNPO2 | 30000 |
| ENSG000001 | 828.847119 | 0.393041995 | 0.15741834 | 2.49679926 | 0.01253199 | 0.0420439 | ZFC3H1 | 196441 |
| ENSG000002 | 2.67561583 | -0.917044251 | 0.36730456 | -2.4966863 | 0.01253598 | 0.04205382 | NA | NA |
| ENSG000002 | 8.46616316 | -0.88336963 | 0.35387725 | -2.49626 | 0.01255106 | 0.04210091 | NA | NA |
| ENSG00000C | 200.552638 | 0.360585309 | 0.14446627 | 2.49598261 | 0.01256088 | 0.04213036 | ZNF800 | 168850 |
| ENSG000001 | 809.606734 | 0.415827098 | 0.16660656 | 2.49586272 | 0.01256512 | 0.04213763 | SNRPA1 | 6627 |

| | | | | | | | | |
|---|---|---|---|---|---|---|---|---|
| ENSG000001 | 30.59152 | -0.60553486 | 0.24261392 | -2.4958785 | 0.01256456 | 0.04213763 | TTBK1 | 84630 |
| ENSG000002 | 4.85464192 | -0.943817921 | 0.37816028 | -2.4958145 | 0.01256683 | 0.04213987 | LOC101929164 | 101929164 |
| ENSG000001 | 13.3611455 | -0.797776342 | 0.31966854 | -2.495636 | 0.01257315 | 0.04215411 | NA | NA |
| ENSG000002 | 4.26894389 | -0.943949154 | 0.37823884 | -2.4956431 | 0.0125729 | 0.04215411 | LOC105378738 | 105378738 |
| ENSG000001 | 208.993705 | 0.387177127 | 0.15514839 | 2.49552778 | 0.01257699 | 0.04215876 | TMEM218 | 219854 |
| ENSG000001 | 383.47767 | 0.319589715 | 0.12806355 | 2.49555573 | 0.012576 | 0.04215876 | TRIM36 | 55521 |
| ENSG000001 | 182.589521 | 0.399461624 | 0.16007409 | 2.49547955 | 0.0125787 | 0.04215876 | ZFPL1 | 7542 |
| ENSG000002 | 8.09809475 | -0.904980309 | 0.36264432 | -2.4955039 | 0.01257784 | 0.04215876 | NA | NA |
| ENSG000001 | 4.32170291 | -0.944053276 | 0.37833994 | -2.4952514 | 0.01258679 | 0.0421824 | LRRC38 | 126755 |
| ENSG000001 | 2465.89051 | 0.316094409 | 0.12668506 | 2.49511982 | 0.01259146 | 0.04219455 | ZRANB2 | 9406 |
| ENSG000001 | 14.2711426 | -0.788951317 | 0.31622714 | -2.4948881 | 0.01259969 | 0.04221863 | CXCR6 | 10663 |
| ENSG000002 | 2.19233577 | -0.896009824 | 0.35914752 | -2.4948239 | 0.01260197 | 0.04222278 | NA | NA |
| ENSG000001 | 5.30413474 | -0.934525215 | 0.37460345 | -2.4947053 | 0.01260618 | 0.0422334 | AGT | 183 |
| ENSG000002 | 6.27678159 | -0.91999462 | 0.36881068 | -2.4944902 | 0.01261382 | 0.04225552 | ECM1P2 | 106481698 |
| ENSG000002 | 8.97040615 | -0.869860362 | 0.3487329 | -2.4943456 | 0.01261896 | 0.04226925 | NA | NA |
| ENSG000001 | 182.088107 | 0.403228816 | 0.1616724 | 2.49411041 | 0.01262733 | 0.04229377 | MVK | 4598 |
| ENSG000002 | 6.14708739 | -0.91697462 | 0.36767657 | -2.4939708 | 0.01263229 | 0.04230692 | LOC105372918 | 105372918 |
| ENSG000002 | 7.11242178 | -0.916810533 | 0.3676255 | -2.4938709 | 0.01263585 | 0.04231533 | NA | NA |
| ENSG000001 | 8.32709796 | -0.89765165 | 0.36000351 | -2.4934525 | 0.01265075 | 0.04236174 | HOXC11 | 3227 |
| ENSG000002 | 6.94294733 | -0.913377655 | 0.36634343 | -2.4932279 | 0.01265876 | 0.04238504 | NA | NA |
| ENSG00000C | 110.805037 | 0.397404572 | 0.15941854 | 2.49283787 | 0.01267267 | 0.04242812 | NID2 | 22795 |
| ENSG000001 | 169.688237 | 0.405493778 | 0.16267574 | 2.49265055 | 0.01267936 | 0.04244 | RHOC | 389 |
| ENSG000002 | 4992.93949 | 0.279604691 | 0.11216905 | 2.49270807 | 0.0126773 | 0.04244 | RBM12 | 10137 |
| ENSG000002 | 3.95637235 | -0.943691757 | 0.37858827 | -2.4926598 | 0.01267903 | 0.04244 | NA | NA |
| ENSG000002 | 3.29590027 | -0.93798485 | 0.37631098 | -2.492579 | 0.01268191 | 0.04244505 | NA | NA |
| ENSG000002 | 7.45305573 | -0.893888888 | 0.35863751 | -2.4924579 | 0.01268624 | 0.04245602 | NA | NA |
| ENSG000001 | 338.696249 | 0.436981633 | 0.17533821 | 2.49222135 | 0.01269469 | 0.0424808 | ZNF706 | 51123 |
| ENSG00000C | 3385.12952 | 0.43854462 | 0.17598707 | 2.49191394 | 0.01270568 | 0.042511 | MAP4K4 | 9448 |
| ENSG000002 | 11.8112653 | -0.832144257 | 0.33393828 | -2.4919104 | 0.01270581 | 0.042511 | FAM183CP | 286135 |

| | | | | | | | | |
|---|---|---|---|---|---|---|---|---|
| ENSG000001 | 545.443825 | 0.319002242 | 0.12802192 | 2.49177826 | 0.01271054 | 0.0425233 | TYW5 | 129450 |
| ENSG000002 | 6.46224175 | -0.923897441 | 0.37078285 | -2.491748 | 0.01271162 | 0.04252342 | NA | NA |
| ENSG000002 | 5.35039536 | -0.9346263 | 0.37509474 | -2.4917073 | 0.01271308 | 0.04252478 | NA | NA |
| ENSG000001 | 22.3725484 | -0.69437408 | 0.27868942 | -2.4915696 | 0.012718 | 0.04253776 | GRM3 | 2913 |
| ENSG000002 | 21.5006896 | 0.732412369 | 0.29397702 | 2.49139329 | 0.01272432 | 0.04255537 | NA | NA |
| ENSG000001 | 231.644158 | 0.375353467 | 0.15067128 | 2.4912078 | 0.01273096 | 0.04257408 | CDCP1 | 64866 |
| ENSG000002 | 6.93909361 | -0.908436534 | 0.36471027 | -2.4908444 | 0.01274399 | 0.04261414 | LINC01271 | 101927586 |
| ENSG000002 | 5.17303387 | -0.929268443 | 0.3730973 | -2.4906866 | 0.01274965 | 0.04262955 | NA | NA |
| ENSG000002 | 52.0110813 | 0.586441954 | 0.23545714 | 2.49065271 | 0.01275087 | 0.0426301 | RNVU1-19 | 101954277 |
| ENSG000001 | 189.427154 | 0.39526631 | 0.15870935 | 2.49050417 | 0.0127562 | 0.04264441 | PFKFB3 | 5209 |
| ENSG000002 | 21.9889311 | 0.690002445 | 0.27706102 | 2.49043492 | 0.01275869 | 0.0426492 | RNU6-2 | 103625684 |
| ENSG000002 | 4.89243307 | -0.935013105 | 0.37555141 | -2.4897073 | 0.01278483 | 0.04273308 | NA | NA |
| ENSG000002 | 7.81681252 | -0.891891233 | 0.35823987 | -2.4896482 | 0.01278696 | 0.04273667 | NA | NA |
| ENSG000002 | 2.97458678 | -0.928702675 | 0.37304014 | -2.4895516 | 0.01279044 | 0.04274477 | NA | NA |
| ENSG00000C | 20.6049348 | -0.69924151 | 0.28087752 | -2.489489 | 0.01279269 | 0.04274876 | CRTAC1 | 55118 |
| ENSG000002 | 8.47981709 | -0.881473443 | 0.35408701 | -2.4894261 | 0.01279495 | 0.04275281 | NA | NA |
| ENSG000002 | 13.5384692 | -0.78225383 | 0.31423666 | -2.489378 | 0.01279668 | 0.04275507 | LOC100996478 | 100996478 |
| ENSG000001 | 244.824968 | 0.343249769 | 0.13789122 | 2.48927945 | 0.01280023 | 0.04275988 | MRGBP | 55257 |
| ENSG000001 | 25.4328295 | -0.66085089 | 0.26547659 | -2.4893 | 0.01279949 | 0.04275988 | RGS4 | 5999 |
| ENSG000001 | 239.996607 | 0.401687441 | 0.16137397 | 2.48917122 | 0.01280413 | 0.04276702 | TAB1 | 10454 |
| ENSG000001 | 60.505094 | 0.514091868 | 0.20653215 | 2.4891615 | 0.01280448 | 0.04276702 | NOS3 | 4846 |
| ENSG000002 | 13.6754722 | -0.794749705 | 0.31929128 | -2.4891056 | 0.01280649 | 0.04277023 | NA | NA |
| ENSG000002 | 4.3332377 | -0.941580607 | 0.37832212 | -2.4888331 | 0.01281631 | 0.04279949 | NA | NA |
| ENSG000001 | 263.82431 | -0.317788094 | 0.12770556 | -2.4884437 | 0.01283035 | 0.04284286 | ITPR1 | 3708 |
| ENSG00000C | 3552.20893 | 0.317812919 | 0.12772343 | 2.48828981 | 0.01283591 | 0.04285788 | CCAR1 | 55749 |
| ENSG000002 | 6.29035011 | -0.920454097 | 0.36993324 | -2.4881627 | 0.0128405 | 0.04286967 | NA | NA |
| ENSG00000C | 2066.57925 | 0.333151313 | 0.13389807 | 2.48809637 | 0.01284289 | 0.04287414 | MSMO1 | 6307 |
| ENSG000002 | 10.1779703 | 0.841985892 | 0.33841685 | 2.48801411 | 0.01284586 | 0.04288053 | SNORD116-15 | 100033427 |
| ENSG000002 | 56.3922249 | 0.497483534 | 0.19995596 | 2.48796547 | 0.01284762 | 0.04288286 | NA | NA |

| | | | | | | | | |
|---|---|---|---|---|---|---|---|---|
| ENSG000001 | 9.96330074 | -0.854986153 | 0.34368543 | -2.4876997 | 0.01285722 | 0.04291138 | SPZ1 | 84654 |
| ENSG000002 | 16.9388101 | -0.734317466 | 0.29521078 | -2.4874345 | 0.01286682 | 0.04293638 | LOC100506085 | 100506085 |
| ENSG000002 | 293.704133 | 0.334986982 | 0.13467171 | 2.48743402 | 0.01286683 | 0.04293638 | NA | NA |
| ENSG000001 | 31.3990863 | -0.646649077 | 0.25997268 | -2.4873732 | 0.01286903 | 0.04293666 | LIPH | 200879 |
| ENSG000002 | 4.74143078 | -0.939101109 | 0.37754688 | -2.4873761 | 0.01286893 | 0.04293666 | LOC101929019 | 101929019 |
| ENSG000001 | 90.5506913 | 0.439096922 | 0.17654224 | 2.48720602 | 0.01287508 | 0.04295112 | SLC35G2 | 80723 |
| ENSG000002 | 11.6109243 | -0.822833009 | 0.33082773 | -2.4871948 | 0.01287549 | 0.04295112 | LOC101929524 | 101929524 |
| ENSG000002 | 15.576009 | -0.758012989 | 0.30477667 | -2.4871096 | 0.01287857 | 0.04295788 | NA | NA |
| ENSG000001 | 15.296983 | -0.757035517 | 0.30440202 | -2.4869596 | 0.012884 | 0.04297247 | BFSP2 | 8419 |
| ENSG000002 | 3.92446451 | -0.941449301 | 0.37858736 | -2.4867426 | 0.01289186 | 0.04299217 | NA | NA |
| ENSG000002 | 22.7379318 | 0.685334394 | 0.27559574 | 2.48673795 | 0.01289203 | 0.04299217 | NA | NA |
| ENSG000002 | 5.50998664 | 0.9311448 | 0.37446947 | 2.48657065 | 0.0128981 | 0.04300667 | SCARNA3 | 677679 |
| ENSG000002 | 3.36773005 | -0.935475378 | 0.37621275 | -2.4865595 | 0.0128985 | 0.04300667 | NA | NA |
| ENSG000002 | 9.44329949 | -0.85041596 | 0.34208679 | -2.4859655 | 0.01292005 | 0.04307497 | PRSS58 | 136541 |
| ENSG000001 | 18.6881617 | -0.739396929 | 0.29743436 | -2.4859163 | 0.01292184 | 0.04307739 | POF1B | 79983 |
| ENSG000002 | 5.26036617 | -0.930607414 | 0.37436037 | -2.4858599 | 0.01292388 | 0.04308068 | NA | NA |
| ENSG000001 | 296.824735 | 0.352923439 | 0.14197452 | 2.48582229 | 0.01292525 | 0.04308168 | ST5 | 6764 |
| ENSG000001 | 222.343131 | 0.337143734 | 0.13562976 | 2.48576517 | 0.01292732 | 0.04308305 | HELQ | 113510 |
| ENSG000002 | 3.7572243 | -0.93913121 | 0.37780559 | -2.4857525 | 0.01292778 | 0.04308305 | LINC00636 | 285205 |
| ENSG000002 | 11.1461137 | -0.83767019 | 0.33703762 | -2.4853908 | 0.01294093 | 0.04312331 | NA | NA |
| ENSG000002 | 14.6085723 | -0.776293979 | 0.3123641 | -2.4852215 | 0.01294708 | 0.04313674 | LINC00558 | 100861552 |
| ENSG000002 | 19.0533214 | -0.72673097 | 0.29241768 | -2.4852497 | 0.01294606 | 0.04313674 | NA | NA |
| ENSG000001 | 3590.23567 | 0.331803427 | 0.13351258 | 2.48518466 | 0.01294842 | 0.04313766 | ATF7IP | 55729 |
| ENSG000001 | 380.322098 | 0.348498584 | 0.1402333 | 2.48513429 | 0.01295026 | 0.04313976 | SLC7A6 | 9057 |
| ENSG000002 | 4.30921651 | -0.939851794 | 0.37819342 | -2.4851088 | 0.01295118 | 0.04313976 | NA | NA |
| ENSG000002 | 17.2509234 | -0.739221782 | 0.29747766 | -2.4849657 | 0.01295639 | 0.04315356 | PCAT5 | 102578074 |
| ENSG000002 | 22.2298924 | -0.679922233 | 0.27364156 | -2.4847185 | 0.01296539 | 0.04318 | RPL34-AS1 | 285456 |
| ENSG000002 | 13.2663416 | -0.827608584 | 0.33308492 | -2.4846775 | 0.01296689 | 0.04318142 | NA | NA |
| ENSG000002 | 4.62375175 | -0.937751536 | 0.37742841 | -2.4845812 | 0.01297039 | 0.04318955 | NA | NA |

| | | | | | | | | | |
|---|---|---|---|---|---|---|---|---|---|
| ENSG000001 | 111.465432 | -0.399869185 | 0.16096586 | -2.4841863 | 0.01298479 | 0.04323394 | PLAT | | 5327 |
| ENSG000001 | 14.3876289 | -0.790295555 | 0.31813739 | -2.4841329 | 0.01298673 | 0.04323686 | TXNDC2 | | 84203 |
| ENSG000002 | 6.15354838 | -0.927159074 | 0.37324213 | -2.4840686 | 0.01298908 | 0.04324112 | NA | NA | |
| ENSG000002 | 9.45885687 | -0.870164785 | 0.35030856 | -2.4839952 | 0.01299176 | 0.04324648 | NA | NA | |
| ENSG000001 | 13.2251227 | -0.802982011 | 0.32327728 | -2.4838801 | 0.01299596 | 0.04325691 | PODN | | 127435 |
| ENSG000002 | 4.33686161 | -0.939725616 | 0.37843936 | -2.4831604 | 0.01302225 | 0.04334086 | NA | NA | |
| ENSG000001 | 762.758688 | 0.282190802 | 0.11367333 | 2.48247154 | 0.01304745 | 0.04342118 | ZNF626 | | 199777 |
| ENSG000002 | 15.6662584 | -0.759957304 | 0.30615435 | -2.4822685 | 0.01305489 | 0.04344236 | NA | NA | |
| ENSG000001 | 526.326202 | 0.301352398 | 0.12140432 | 2.48222134 | 0.01305661 | 0.04344455 | YY1AP1 | | 55249 |
| ENSG000002 | 9.73229787 | -0.84071199 | 0.33872294 | -2.4820049 | 0.01306455 | 0.04346739 | NA | NA | |
| ENSG000002 | 4.74625523 | -0.938022594 | 0.37793831 | -2.4819463 | 0.0130667 | 0.04347097 | NA | NA | |
| ENSG000001 | 663.863647 | 0.305229964 | 0.12298598 | 2.48182733 | 0.01307106 | 0.0434792 | SGOL1 | | 151648 |
| ENSG000001 | 40.5328262 | -0.564770287 | 0.22756544 | -2.4817929 | 0.01307232 | 0.0434792 | KIAA0895 | | 23366 |
| ENSG000002 | 12.6043396 | -0.814501738 | 0.32819109 | -2.4817911 | 0.01307239 | 0.0434792 | RNF157-AS1 | | 100507218 |
| ENSG000001 | 3842.1833 | 0.306838842 | 0.12364155 | 2.4816807 | 0.01307644 | 0.04348554 | PRDX6 | | 9588 |
| ENSG000002 | 20.3724277 | -0.70343728 | 0.28344889 | -2.4817077 | 0.01307545 | 0.04348554 | NA | NA | |
| ENSG000002 | 9.80713845 | -0.871119754 | 0.35102425 | -2.4816512 | 0.01307752 | 0.04348557 | NA | NA | |
| ENSG000001 | 340.036606 | 0.341005456 | 0.13742629 | 2.48136994 | 0.01308785 | 0.04351138 | MBOAT2 | | 129642 |
| ENSG000001 | 18.7324733 | -0.7739463 | 0.31189814 | -2.4814072 | 0.01308648 | 0.04351138 | BOLL | | 66037 |
| ENSG000002 | 18.3250065 | -0.72299452 | 0.29137119 | -2.4813521 | 0.0130885 | 0.04351138 | NA | NA | |
| ENSG000001 | 3022.87641 | 0.307147311 | 0.12379135 | 2.48116933 | 0.01309521 | 0.04352656 | ZMYM4 | | 9202 |
| ENSG000002 | 89.586456 | -0.432270997 | 0.17422018 | -2.4811764 | 0.01309496 | 0.04352656 | LINC01108 | | 102216342 |
| ENSG00000C | 174.100522 | 0.391964639 | 0.15798548 | 2.48101693 | 0.01310082 | 0.04354161 | AP1M1 | | 8907 |
| ENSG000001 | 140.419583 | 0.386368781 | 0.15573247 | 2.48097771 | 0.01310226 | 0.04354283 | SRSF12 | | 135295 |
| ENSG000002 | 2.98180975 | -0.926013326 | 0.37329487 | -2.4806484 | 0.01311436 | 0.0435795 | NA | NA | |
| ENSG000001 | 54.7480449 | -0.515757282 | 0.20791605 | -2.4806035 | 0.01311602 | 0.04358142 | SLC9C1 | | 285335 |
| ENSG000001 | 1194.10363 | 0.29138563 | 0.11746719 | 2.48057031 | 0.01311724 | 0.04358191 | MAN1A2 | | 10905 |
| ENSG000001 | 554.300126 | 0.322770456 | 0.13012531 | 2.4804587 | 0.01312135 | 0.04359198 | NEK2 | | 4751 |
| ENSG000001 | 1616.05992 | 0.342046953 | 0.13790267 | 2.48035047 | 0.01312533 | 0.04359807 | CASC3 | | 22794 |

| | | | | | | | | |
|---|---|---|---|---|---|---|---|---|
| ENSG000002 | 3.62032281 | -0.938322949 | 0.37829886 | -2.4803748 | 0.01312443 | 0.04359807 | NA | NA |
| ENSG000002 | 10.1969988 | -0.846904891 | 0.34146359 | -2.4802202 | 0.01313013 | 0.04361043 | NA | NA |
| ENSG000002 | 10.0253988 | -0.833081122 | 0.3358988 | -2.4801551 | 0.01313252 | 0.04361483 | NA | NA |
| ENSG000002 | 20.5653852 | -0.705112473 | 0.28432438 | -2.4799579 | 0.01313979 | 0.04363383 | NA | NA |
| ENSG000002 | 43.9406975 | 0.53441818 | 0.2154963 | 2.47994137 | 0.0131404 | 0.04363383 | NA | NA |
| ENSG000002 | 6.34230222 | -0.916433125 | 0.36956893 | -2.4797353 | 0.013148 | 0.04365548 | NA | NA |
| ENSG000001 | 98.4152189 | 0.419695096 | 0.16925864 | 2.47960804 | 0.01315269 | 0.04366748 | MAF | 4094 |
| ENSG000002 | 21.1927591 | -0.694758909 | 0.28019672 | -2.4795398 | 0.0131552 | 0.04367109 | OTX2-AS1 | 100309464 |
| ENSG000002 | 5.70765474 | -0.928840455 | 0.37460492 | -2.4795202 | 0.01315593 | 0.04367109 | NA | NA |
| ENSG000002 | 8.5302833 | -0.861586453 | 0.3474934 | -2.4794325 | 0.01315916 | 0.04367825 | LINC00410 | 144776 |
| ENSG000001 | 144.061814 | 0.385231355 | 0.15538334 | 2.47923197 | 0.01316656 | 0.04369924 | NRARP | 441478 |
| ENSG000002 | 3.64810374 | -0.9370281 | 0.37801113 | -2.4788373 | 0.01318114 | 0.04374404 | LOC105370024 | 105370024 |
| ENSG000001 | 92.087213 | 0.422452581 | 0.17043466 | 2.47867764 | 0.01318704 | 0.04376005 | ETFB | 2109 |
| ENSG000002 | 49.8088771 | 0.533216462 | 0.2151335 | 2.47853762 | 0.01319222 | 0.04377364 | BTBD17 | 388419 |
| ENSG000000 | 481.908744 | 0.291999952 | 0.1178209 | 2.47833745 | 0.01319962 | 0.04379463 | WDR76 | 79968 |
| ENSG000002 | 2.84496774 | -0.928242516 | 0.37460405 | -2.4779297 | 0.01321472 | 0.04384112 | C6orf15 | 29113 |
| ENSG000001 | 15.2397618 | -0.777151748 | 0.31365175 | -2.4777536 | 0.01322124 | 0.04385917 | KL | 9365 |
| ENSG000002 | 6.21279201 | -0.911188754 | 0.36775721 | -2.4776911 | 0.01322356 | 0.04386327 | NA | NA |
| ENSG000001 | 648.351875 | 0.331522432 | 0.13380987 | 2.47756332 | 0.01322829 | 0.0438718 | SH3BP4 | 23677 |
| ENSG000001 | 975.221868 | 0.326071699 | 0.1316087 | 2.47758469 | 0.0132275 | 0.0438718 | FOXN2 | 3344 |
| ENSG000002 | 8.06362791 | -0.885144658 | 0.35728411 | -2.4774252 | 0.01323341 | 0.04388519 | NA | NA |
| ENSG000001 | 8.2688563 | -0.869460256 | 0.3509723 | -2.4772903 | 0.01323842 | 0.0438951 | KIR2DL4 | 3805 |
| ENSG000002 | 3.94729813 | -0.937884053 | 0.37859332 | -2.4772863 | 0.01323856 | 0.0438951 | NA | NA |
| ENSG000001 | 20.46228 | -0.688910382 | 0.27809802 | -2.4772215 | 0.01324097 | 0.04389948 | OR5AN1 | 390195 |
| ENSG000002 | 57.8669485 | 0.519831186 | 0.20984982 | 2.47715815 | 0.01324332 | 0.04390369 | NA | NA |
| ENSG000002 | 16.5964003 | -0.752204384 | 0.30367796 | -2.4769805 | 0.01324991 | 0.04392196 | NA | NA |
| ENSG000001 | 3.56633731 | -0.93716174 | 0.37836391 | -2.4768793 | 0.01325367 | 0.04393082 | GSX1 | 219409 |
| ENSG000001 | 58.3357883 | -0.516567153 | 0.20856183 | -2.4768058 | 0.0132564 | 0.04393443 | SLC8A3 | 6547 |
| ENSG000002 | 10.9488597 | -0.824962496 | 0.33307707 | -2.4767916 | 0.01325693 | 0.04393443 | NA | NA |

| | | | | | | | | |
|---|---|---|---|---|---|---|---|---|
| ENSG000002 | 8.929036 | -0.861518367 | 0.34784586 | -2.4767245 | 0.01325942 | 0.0439391 | NA | NA |
| ENSG000001 | 1080.42806 | 0.269483384 | 0.10881056 | 2.47662899 | 0.01326297 | 0.04394727 | AGGF1 | 55109 |
| ENSG000001 | 3.2832839 | -0.932289434 | 0.37644574 | -2.4765572 | 0.01326564 | 0.04394897 | NA | NA |
| ENSG000002 | 23.0657477 | -0.695988853 | 0.28103084 | -2.4765568 | 0.01326565 | 0.04394897 | LOC100506869 | 100506869 |
| ENSG000001 | 342.368689 | 0.323095556 | 0.13046605 | 2.47647232 | 0.01326879 | 0.04395578 | ENTPD7 | 57089 |
| ENSG000002 | 3.35802277 | -0.931289704 | 0.37609463 | -2.4762111 | 0.0132785 | 0.04398141 | LINC00582 | 100287814 |
| ENSG000002 | 8.15656707 | -0.888186205 | 0.35868834 | -2.4762059 | 0.0132787 | 0.04398141 | NA | NA |
| ENSG000000 | 695.872461 | 0.296366845 | 0.11969222 | 2.47607451 | 0.01328359 | 0.04399401 | CNOT4 | 4850 |
| ENSG000001 | 811.976442 | 0.44119333 | 0.17818512 | 2.4760392 | 0.0132849 | 0.04399477 | ZNF326 | 284695 |
| ENSG000002 | 14.1232093 | 0.796394378 | 0.32164586 | 2.47599762 | 0.01328645 | 0.0439963 | NA | NA |
| ENSG000001 | 11.7786913 | -0.804016428 | 0.32474942 | -2.4758056 | 0.01329359 | 0.04401638 | LAPTM5 | 7805 |
| ENSG000001 | 9.80962504 | -0.866400509 | 0.35001294 | -2.4753385 | 0.01331099 | 0.04407039 | CD37 | 951 |
| ENSG000002 | 110.850995 | -0.407921674 | 0.1647994 | -2.4752619 | 0.01331385 | 0.04407625 | CCDC7 | 79741 |
| ENSG000002 | 327.794425 | 0.380106034 | 0.15358686 | 2.4748604 | 0.01332883 | 0.04412222 | SRSF8 | 10929 |
| ENSG000000 | 400.069604 | 0.327655294 | 0.13239535 | 2.47482477 | 0.01333016 | 0.04412272 | NUCB2 | 4925 |
| ENSG000002 | 27.4082304 | -0.726062709 | 0.29338261 | -2.474798 | 0.01333115 | 0.04412272 | LINC01252 | 338817 |
| ENSG000001 | 150.489076 | 0.454632485 | 0.18371823 | 2.47461829 | 0.01333786 | 0.04414133 | UCP2 | 7351 |
| ENSG000002 | 33.0972144 | 0.646841985 | 0.26140263 | 2.47450454 | 0.01334211 | 0.04415179 | NA | NA |
| ENSG000001 | 139.220181 | 0.426252209 | 0.17226229 | 2.47443721 | 0.01334463 | 0.04415651 | GSTK1 | 373156 |
| ENSG000002 | 18.526048 | -0.728190078 | 0.29429163 | -2.4743826 | 0.01334667 | 0.04415965 | LINC00222 | 387111 |
| ENSG000000 | 62.7734098 | -0.51246066 | 0.20711539 | -2.4742761 | 0.01335065 | 0.04416922 | PTGER3 | 5733 |
| ENSG000002 | 5.57485961 | -0.921147322 | 0.37229454 | -2.4742435 | 0.01335187 | 0.04416965 | NA | NA |
| ENSG000001 | 29.1188688 | -0.628638452 | 0.25409143 | -2.474064 | 0.01335858 | 0.04418824 | OAS2 | 4939 |
| ENSG000002 | 13.2060143 | -0.820284411 | 0.33159807 | -2.4737309 | 0.01337104 | 0.04422585 | NA | NA |
| ENSG000002 | 4.30154395 | -0.936048969 | 0.37842595 | -2.4735327 | 0.01337846 | 0.04424678 | MIR3156-1 | 100422988 |
| ENSG000001 | 112.816305 | -0.407533546 | 0.16476589 | -2.4734097 | 0.01338306 | 0.04425841 | MYOF | 26509 |
| ENSG000002 | 3.14666261 | -0.933159896 | 0.377285 | -2.4733554 | 0.0133851 | 0.04426152 | NA | NA |
| ENSG000002 | 2.43779317 | -0.914572395 | 0.36982253 | -2.4730035 | 0.01339828 | 0.04430152 | NA | NA |
| ENSG000001 | 9.49381605 | -0.84662472 | 0.3423602 | -2.4729064 | 0.01340192 | 0.04430994 | SPATA32 | 124783 |

| | | | | | | | | |
|---|---|---|---|---|---|---|---|---|
| ENSG000001 | 811.663384 | 0.286525191 | 0.11586919 | 2.47283321 | 0.01340467 | 0.04431179 | ELL2 | 22936 |
| ENSG000002 | 18.70748 | -0.719311015 | 0.29088454 | -2.4728403 | 0.0134044 | 0.04431179 | NA | NA |
| ENSG000001 | 44.624987 | 0.571082322 | 0.23095966 | 2.47264963 | 0.01341156 | 0.04433094 | METRN | 79006 |
| ENSG000001 | 182.817691 | 0.356867482 | 0.14432955 | 2.47258785 | 0.01341387 | 0.04433499 | KIAA0226 | 9711 |
| ENSG000001 | 38.1303712 | 0.606598644 | 0.24535462 | 2.47233434 | 0.01342339 | 0.04436283 | MTG1 | 92170 |
| ENSG000001 | 5.2692919 | -0.925600712 | 0.37440193 | -2.4722114 | 0.01342801 | 0.04437448 | OR5J2 | 282775 |
| ENSG000002 | 5.60747241 | -0.920557702 | 0.37237981 | -2.4720935 | 0.01343244 | 0.0443855 | NA | NA |
| ENSG000002 | 10.9559103 | -0.819799858 | 0.33166076 | -2.4718024 | 0.01344338 | 0.04441803 | NA | NA |
| ENSG000002 | 20.7876737 | -0.726065736 | 0.29374444 | -2.4717599 | 0.01344498 | 0.04441969 | TMPRSS11BNL | 401136 |
| ENSG000001 | 10.3519792 | -0.852140206 | 0.34479418 | -2.471446 | 0.01345679 | 0.04445509 | SUN5 | 140732 |
| ENSG000002 | 8.29658937 | -0.869110716 | 0.35167701 | -2.4713322 | 0.01346107 | 0.04446561 | NA | NA |
| ENSG000001 | 349.919437 | 0.31461514 | 0.12731799 | 2.47109738 | 0.01346991 | 0.0444813 | SLC24A2 | 25769 |
| ENSG000001 | 88.9370545 | -0.42790992 | 0.17316518 | -2.4711084 | 0.0134695 | 0.0444813 | SLITRK5 | 26050 |
| ENSG000001 | 1269.43152 | 0.300700884 | 0.12168709 | 2.47109936 | 0.01346984 | 0.0444813 | TEAD1 | 7003 |
| ENSG000002 | 6.56120577 | -0.911238299 | 0.36875971 | -2.4710896 | 0.01347021 | 0.0444813 | NA | NA |
| ENSG000001 | 1534.26464 | 0.294727355 | 0.11927578 | 2.47097393 | 0.01347456 | 0.04449206 | CWC22 | 57703 |
| ENSG000002 | 7.20061843 | -0.897743341 | 0.36335303 | -2.4707193 | 0.01348416 | 0.04452013 | LINC00690 | 100996597 |
| ENSG000002 | 386.468937 | 0.314808767 | 0.12743834 | 2.47028295 | 0.01350062 | 0.04457085 | SPIN3 | 169981 |
| ENSG000002 | 8.57789459 | -0.862040509 | 0.34900991 | -2.4699599 | 0.01351282 | 0.04460748 | NA | NA |
| ENSG000001 | 53.1012807 | 0.540540702 | 0.2188636 | 2.46976061 | 0.01352035 | 0.04462871 | GNB1L | 54584 |
| ENSG000001 | 286.250196 | 0.32616362 | 0.13207083 | 2.46961143 | 0.01352599 | 0.04464369 | PDE9A | 5152 |
| ENSG000001 | 11.3930212 | -0.821268838 | 0.33255474 | -2.4695749 | 0.01352737 | 0.04464414 | IL33 | 90865 |
| ENSG000001 | 761.003822 | 0.29554554 | 0.1196773 | 2.46952044 | 0.01352943 | 0.04464414 | ESCO2 | 157570 |
| ENSG000002 | 8.53483076 | -0.858181476 | 0.34750582 | -2.4695456 | 0.01352848 | 0.04464414 | LOC101928460 | 101928460 |
| ENSG00000( | 4448.86832 | 0.319442838 | 0.12935935 | 2.46942207 | 0.01353315 | 0.04465279 | SYNE2 | 23224 |
| ENSG000001 | 1843.11788 | 0.259841688 | 0.1052263 | 2.46936061 | 0.01353547 | 0.04465682 | MPHOSPH10 | 10199 |
| ENSG000002 | 26.7175632 | -0.622112745 | 0.25200177 | -2.468684 | 0.01356109 | 0.0447377 | LINC01359 | 101927084 |
| ENSG000001 | 6.1306015 | -0.908463144 | 0.368022 | -2.4685023 | 0.01356798 | 0.0447495 | LINC00477 | 144360 |
| ENSG000002 | 20.5989563 | -0.679783239 | 0.27537676 | -2.4685571 | 0.0135659 | 0.0447495 | NA | NA |

| | | | | | | | | |
|---|---|---|---|---|---|---|---|---|
| ENSG000002 | 3.95597592 | -0.934565549 | 0.37859192 | -2.4685301 | 0.01356692 | 0.0447495 | NA | NA |
| ENSG000001 | 97.7880828 | 0.414318678 | 0.16784627 | 2.46844143 | 0.01357029 | 0.04475347 | CRACR2B | 283229 |
| ENSG000000 | 1283.02843 | 0.616777862 | 0.24987253 | 2.46837006 | 0.01357299 | 0.04475875 | CAD | 790 |
| ENSG000001 | 16.4942235 | -0.734074873 | 0.29742233 | -2.4681229 | 0.01358237 | 0.04478603 | CLEC4E | 26253 |
| ENSG000002 | 200.814131 | 0.338791713 | 0.13727209 | 2.46803049 | 0.01358587 | 0.04479295 | ADSL | 158 |
| ENSG000002 | 2.50077754 | -0.908356861 | 0.36805243 | -2.4680094 | 0.01358667 | 0.04479295 | NA | NA |
| ENSG000000 | 582.915038 | 0.661716458 | 0.26812633 | 2.46792796 | 0.01358977 | 0.0447995 | UPF1 | 5976 |
| ENSG000001 | 85.0150787 | 0.442171047 | 0.17917647 | 2.46779636 | 0.01359476 | 0.0448034 | HERC2P3 | 283755 |
| ENSG000002 | 5.86475422 | -0.927981831 | 0.37603907 | -2.4677804 | 0.01359537 | 0.0448034 | NA | NA |
| ENSG000002 | 19.8283424 | -0.693633501 | 0.2810717 | -2.4678169 | 0.01359398 | 0.0448034 | ERVW-1 | 30816 |
| ENSG000002 | 4.17521867 | -0.932676488 | 0.37793054 | -2.4678516 | 0.01359267 | 0.0448034 | NA | NA |
| ENSG000002 | 35.6596268 | -0.585564094 | 0.23728853 | -2.4677303 | 0.01359727 | 0.04480603 | NA | NA |
| ENSG000000 | 37.8663766 | -0.574847642 | 0.23297997 | -2.4673694 | 0.01361099 | 0.04484426 | CYP46A1 | 10858 |
| ENSG000002 | 5.28141934 | -0.924534641 | 0.37470499 | -2.4673667 | 0.01361109 | 0.04484426 | NA | NA |
| ENSG000002 | 8.5923542 | -0.865895699 | 0.35096444 | -2.4671892 | 0.01361784 | 0.04486286 | LOC105371849 | 105371849 |
| ENSG000002 | 10.6953247 | -0.826123554 | 0.33485562 | -2.4671038 | 0.01362109 | 0.04486384 | NA | NA |
| ENSG000002 | 18.2235172 | 0.746785594 | 0.30269846 | 2.46709411 | 0.01362146 | 0.04486384 | NA | NA |
| ENSG000002 | 16.2769313 | -0.739402878 | 0.29970076 | -2.4671372 | 0.01361982 | 0.04486384 | NA | NA |
| ENSG000001 | 31.6363287 | -0.611731807 | 0.24799788 | -2.4666816 | 0.01363716 | 0.04490462 | NA | NA |
| ENSG000002 | 32.2558572 | -0.626256608 | 0.25388156 | -2.4667274 | 0.01363541 | 0.04490462 | NA | NA |
| ENSG000002 | 19.3241102 | -0.699041939 | 0.28339312 | -2.4666864 | 0.01363697 | 0.04490462 | LOC286059 | 286059 |
| ENSG000001 | 414.642863 | 0.318638674 | 0.12918048 | 2.46661629 | 0.01363964 | 0.04490916 | WASF1 | 8936 |
| ENSG000002 | 48.9210081 | 0.580871683 | 0.23549837 | 2.46656353 | 0.01364165 | 0.04491213 | NA | NA |
| ENSG000002 | 3.56172514 | -0.932976349 | 0.37826237 | -2.4664794 | 0.01364486 | 0.04491903 | MIR3912 | 100500831 |
| ENSG000001 | 7.42118571 | -0.886333502 | 0.35939287 | -2.4661966 | 0.01365563 | 0.04494997 | TMEM150B | 284417 |
| ENSG000002 | 10.6831457 | -0.823372984 | 0.33386645 | -2.4661746 | 0.01365647 | 0.04494997 | NA | NA |
| ENSG000000 | 35.7470464 | -0.591881411 | 0.24000365 | -2.466135 | 0.01365798 | 0.0449513 | DLEC1 | 9940 |
| ENSG000002 | 4.69053892 | -0.933346905 | 0.37847691 | -2.4660604 | 0.01366083 | 0.04495702 | NA | NA |
| ENSG000001 | 9.81523861 | -0.847222459 | 0.3435581 | -2.4660238 | 0.01366222 | 0.04495795 | FAM87A | 157693 |

| | | | | | | | | | |
|---|---|---|---|---|---|---|---|---|---|
| ENSG000001 | 39.7612469 | -0.556778719 | 0.22579862 | -2.4658198 | 0.01367001 | 0.04497992 | SIM1 | | 6492 |
| ENSG000001 | 56.3196712 | 0.518139278 | 0.21013656 | 2.46572642 | 0.01367357 | 0.04498453 | C3orf62 | | 375341 |
| ENSG000002 | 305.946277 | 0.319455418 | 0.12955841 | 2.46572499 | 0.01367363 | 0.04498453 | ZNF280B | | 140883 |
| ENSG000001 | 177.001022 | 0.508533005 | 0.20625442 | 2.46556173 | 0.01367986 | 0.04500138 | TRIP4 | | 9325 |
| ENSG000002 | 7.2849416 | -0.899498452 | 0.36486656 | -2.4652806 | 0.0136906 | 0.04503306 | C1orf146 | | 388649 |
| ENSG000001 | 2787.06382 | 0.383259971 | 0.15546501 | 2.46524906 | 0.0136918 | 0.04503337 | ERBB2 | | 2064 |
| ENSG000000 | 103.07868 | 0.433600375 | 0.17590071 | 2.46502915 | 0.01370021 | 0.04505529 | LOC81691 | | 81691 |
| ENSG000002 | 4.76893894 | -0.932242653 | 0.37818921 | -2.4650165 | 0.01370069 | 0.04505529 | NA | NA | |
| ENSG000001 | 328.505522 | 0.404825399 | 0.16423419 | 2.46492773 | 0.01370409 | 0.0450628 | KANK2 | | 25959 |
| ENSG000002 | 13.0063497 | 0.792191164 | 0.32139657 | 2.4648401 | 0.01370744 | 0.04507017 | NA | NA | |
| ENSG000002 | 128.836439 | 0.428908378 | 0.17402857 | 2.46458595 | 0.01371717 | 0.04509849 | NA | NA | |
| ENSG000002 | 9.47842417 | -0.847102076 | 0.34371887 | -2.4645202 | 0.01371968 | 0.04510312 | NA | NA | |
| ENSG000001 | 36.0686051 | 0.618729192 | 0.25107583 | 2.46431206 | 0.01372765 | 0.04511232 | C21orf2 | | 755 |
| ENSG000001 | 12.1668099 | -0.822071269 | 0.33358366 | -2.4643631 | 0.0137257 | 0.04511232 | GABRB1 | | 2560 |
| ENSG000001 | 22.5673565 | -0.666609916 | 0.27050078 | -2.4643548 | 0.01372601 | 0.04511232 | TMEM100 | | 55273 |
| ENSG000002 | 10.2682172 | -0.841855698 | 0.34162037 | -2.4643018 | 0.01372805 | 0.04511232 | NA | NA | |
| ENSG000002 | 5.54876318 | -0.917147123 | 0.37216777 | -2.4643379 | 0.01372666 | 0.04511232 | NA | NA | |
| ENSG000002 | 16.4313438 | -0.724624595 | 0.29406627 | -2.4641541 | 0.01373371 | 0.04512726 | NA | NA | |
| ENSG000002 | 7.81400911 | -0.881811359 | 0.35790937 | -2.463784 | 0.01374789 | 0.04517022 | KLHL41 | | 10324 |
| ENSG000002 | 8.65167797 | -0.863071527 | 0.35031662 | -2.4636899 | 0.0137515 | 0.0451775 | NA | NA | |
| ENSG000002 | 6.57842494 | -0.906942148 | 0.36812675 | -2.4636682 | 0.01375234 | 0.0451775 | NA | NA | |
| ENSG000000 | 182.554991 | 0.493289283 | 0.20023005 | 2.46361259 | 0.01375447 | 0.04518084 | ACAA1 | | 30 |
| ENSG000001 | 3.27852847 | -0.927325239 | 0.37641702 | -2.4635582 | 0.01375655 | 0.04518403 | OR5B3 | | 441608 |
| ENSG000002 | 23.5541581 | -0.663468587 | 0.2693404 | -2.4633088 | 0.01376613 | 0.04521182 | GUCY2GP | | 390003 |
| ENSG000001 | 35.949446 | 0.590343331 | 0.23966782 | 2.46317311 | 0.01377134 | 0.04522527 | SWSAP1 | | 126074 |
| ENSG000002 | 4.53390094 | -0.928554778 | 0.3769924 | -2.4630597 | 0.0137757 | 0.04523592 | NA | NA | |
| ENSG000002 | 14.9663284 | -0.791844581 | 0.32149988 | -2.4629701 | 0.01377914 | 0.04524356 | LOC101927502 | | 101927502 |
| ENSG000002 | 6.27373395 | -0.9114943 | 0.3700862 | -2.4629243 | 0.0137809 | 0.04524567 | NA | NA | |
| ENSG000002 | 10.1589071 | -0.840185906 | 0.34114284 | -2.4628567 | 0.0137835 | 0.04525055 | LINC01095 | | 100505545 |

| | | | | | | | | |
|---|---|---|---|---|---|---|---|---|
| ENSG000002 | 7.47835525 | -0.886698159 | 0.36004774 | -2.4627239 | 0.01378861 | 0.04526364 | LGALS16 | 148003 |
| ENSG000002 | 2.85725881 | -0.921498735 | 0.37418412 | -2.462688 | 0.01378999 | 0.04526451 | NA | NA |
| ENSG000002 | 7.88197067 | -0.867191993 | 0.3521592 | -2.4624999 | 0.01379722 | 0.04528459 | PRAMEF20 | 645425 |
| ENSG000001 | 10.2415357 | -0.894104225 | 0.36309666 | -2.4624413 | 0.01379948 | 0.04528832 | KRT27 | 342574 |
| ENSG000002 | 13.2106747 | -0.781313621 | 0.31730858 | -2.4623149 | 0.01380434 | 0.04530063 | NA | NA |
| ENSG000000 | 1215.67608 | 0.371614211 | 0.15092444 | 2.4622533 | 0.01380671 | 0.04530474 | KDM2B | 84678 |
| ENSG000001 | 3861.86691 | 0.305319105 | 0.12400347 | 2.46218194 | 0.01380946 | 0.04530642 | BRWD1 | 54014 |
| ENSG000002 | 2.81069739 | -0.92209729 | 0.37450171 | -2.4621978 | 0.01380885 | 0.04530642 | KRTAP4-9 | 100132386 |
| ENSG000001 | 15.4943597 | -0.7725154 | 0.31376177 | -2.4621081 | 0.0138123 | 0.04531208 | SLC22A1 | 6580 |
| ENSG000001 | 6.47490359 | -0.899033656 | 0.36515393 | -2.4620676 | 0.01381386 | 0.04531354 | KRT83 | 3889 |
| ENSG000002 | 37.4243565 | 0.567459453 | 0.23048378 | 2.46203642 | 0.01381506 | 0.04531381 | LOC100507437 | 100507437 |
| ENSG000001 | 78.8371542 | -0.482455853 | 0.19597384 | -2.4618381 | 0.01382271 | 0.04533521 | SLC12A5 | 57468 |
| ENSG000001 | 21.2986852 | -0.681562741 | 0.2768581 | -2.4617765 | 0.01382508 | 0.04533933 | GIMAP8 | 155038 |
| ENSG000001 | 844.61637 | 0.432603964 | 0.17573188 | 2.46172732 | 0.01382697 | 0.04534188 | KMT2D | 8085 |
| ENSG000001 | 36.426244 | 0.583960778 | 0.23725107 | 2.46136204 | 0.01384106 | 0.04538073 | TMEM52 | 339456 |
| ENSG000001 | 3.14231157 | -0.928284682 | 0.37714059 | -2.4613757 | 0.01384053 | 0.04538073 | IFNA1 | 3439 |
| ENSG000002 | 22.4994006 | -0.678451516 | 0.27566401 | -2.4611538 | 0.0138491 | 0.04540341 | NA | NA |
| ENSG000001 | 143.053081 | 0.377601255 | 0.15343042 | 2.46105859 | 0.01385277 | 0.04541179 | LPPR3 | 79948 |
| ENSG000002 | 11.3908146 | -0.821662193 | 0.33387976 | -2.4609524 | 0.01385688 | 0.04542156 | NA | NA |
| ENSG000001 | 224.589094 | 0.374418692 | 0.15216478 | 2.46061344 | 0.01386997 | 0.04546082 | CPT1C | 126129 |
| ENSG000000 | 554.993459 | 0.291203843 | 0.11836319 | 2.46025681 | 0.01388376 | 0.04550234 | CTTNBP2 | 83992 |
| ENSG000002 | 3.14780476 | -0.92804112 | 0.37723107 | -2.4601397 | 0.01388829 | 0.04551351 | NA | NA |
| ENSG000002 | 3.55736669 | -0.93033653 | 0.37817846 | -2.4600463 | 0.01389191 | 0.04552168 | NA | NA |
| ENSG000002 | 3.02533114 | -0.916070834 | 0.37238973 | -2.4599788 | 0.01389452 | 0.04552657 | NA | NA |
| ENSG000001 | 668.806791 | 0.398101794 | 0.16183363 | 2.45994472 | 0.01389584 | 0.04552721 | ARHGEF7 | 8874 |
| ENSG000002 | 861.899733 | 0.287291829 | 0.11679962 | 2.45969832 | 0.01390538 | 0.0455548 | CEP95 | 90799 |
| ENSG000001 | 639.425638 | 0.297206551 | 0.12083929 | 2.45951925 | 0.01391232 | 0.04557385 | RAB11FIP2 | 22841 |
| ENSG000001 | 17.9903065 | -0.732985893 | 0.29803196 | -2.4594204 | 0.01391615 | 0.04558271 | NA | NA |
| ENSG000001 | 11.4374251 | -0.80559506 | 0.32759002 | -2.4591563 | 0.0139264 | 0.04561258 | PLN | 5350 |

| | | | | | | | | |
|---|---|---|---|---|---|---|---|---|
| ENSG000001 | 146.691347 | 0.44321433 | 0.18025314 | 2.45884393 | 0.01393852 | 0.0456486 | SLC27A3 | 11000 |
| ENSG000002 | 8.12312649 | -0.874205251 | 0.35557083 | -2.4585966 | 0.01394812 | 0.04567636 | NA | NA |
| ENSG000002 | 4.23058364 | -0.930176839 | 0.37834663 | -2.4585308 | 0.01395068 | 0.04568105 | NA | NA |
| ENSG000001 | 50.1791102 | 0.545911019 | 0.22205645 | 2.45843348 | 0.01395446 | 0.04568974 | C19orf70 | 125988 |
| ENSG000001 | 1260.83005 | 0.30749751 | 0.12508235 | 2.45836058 | 0.0139573 | 0.04569533 | RASSF8 | 11228 |
| ENSG00000C | 1373.2248 | 0.28547131 | 0.11613017 | 2.45820099 | 0.0139635 | 0.04571195 | MLF2 | 8079 |
| ENSG00000C | 240.496793 | 0.717645476 | 0.29194917 | 2.45811788 | 0.01396673 | 0.04571884 | CCDC124 | 115098 |
| ENSG000001 | 2856.27535 | 0.251076385 | 0.10214527 | 2.4580323 | 0.01397006 | 0.04572235 | AKAP9 | 10142 |
| ENSG000002 | 3.29348067 | -0.92399835 | 0.37590862 | -2.4580398 | 0.01396977 | 0.04572235 | NA | NA |
| ENSG000001 | 22.5807168 | -0.708870682 | 0.28840975 | -2.4578597 | 0.01397678 | 0.04574065 | ADGRE1 | 2015 |
| ENSG000001 | 31.2846111 | -0.637405691 | 0.25935542 | -2.4576532 | 0.01398481 | 0.04575956 | SNED1 | 25992 |
| ENSG000001 | 626.691385 | 0.300985924 | 0.1224687 | 2.45765586 | 0.01398471 | 0.04575956 | DDIAS | 220042 |
| ENSG000001 | 74.208629 | 0.467980232 | 0.19044758 | 2.45726535 | 0.01399992 | 0.0458048 | INPP5E | 56623 |
| ENSG000001 | 11.0049488 | -0.819798972 | 0.33362589 | -2.4572403 | 0.0140009 | 0.0458048 | KRT20 | 54474 |
| ENSG000002 | 13.1883776 | -0.797233648 | 0.32445292 | -2.4571628 | 0.01400392 | 0.04581099 | NA | NA |
| ENSG000001 | 8.48162187 | -0.874331967 | 0.35584887 | -2.4570317 | 0.01400903 | 0.04582401 | OR10G3 | 26533 |
| ENSG000001 | 417.123149 | 0.326847149 | 0.13303195 | 2.45690712 | 0.01401389 | 0.04583459 | UBQLN4 | 56893 |
| ENSG000002 | 4.93742813 | -0.924909997 | 0.37645811 | -2.4568736 | 0.0140152 | 0.04583459 | NA | NA |
| ENSG000002 | 156.384736 | 0.384867953 | 0.15665022 | 2.45686186 | 0.01401566 | 0.04583459 | NA | NA |
| ENSG000001 | 4490.49717 | 0.330194667 | 0.13440211 | 2.45676707 | 0.01401936 | 0.04584299 | ZNF462 | 58499 |
| ENSG000002 | 7.78326696 | -0.881597878 | 0.35885188 | -2.456718 | 0.01402127 | 0.04584555 | NA | NA |
| ENSG000002 | 11.177666 | -0.81491591 | 0.33175404 | -2.4563858 | 0.01403424 | 0.04588426 | NA | NA |
| ENSG000002 | 2.81724786 | -0.919932421 | 0.37452193 | -2.4562845 | 0.0140382 | 0.0458935 | NA | NA |
| ENSG000002 | 5.23690348 | -0.919460063 | 0.37436056 | -2.4560816 | 0.01404613 | 0.04591572 | NA | NA |
| ENSG000002 | 4.06691072 | -0.9291542 | 0.37832955 | -2.4559387 | 0.01405171 | 0.04593027 | NA | NA |
| ENSG000001 | 31.5801117 | -0.62918968 | 0.25620198 | -2.4558346 | 0.01405578 | 0.04593988 | CASC1 | 55259 |
| ENSG000002 | 34.9883321 | -0.644051133 | 0.26226174 | -2.4557571 | 0.01405881 | 0.04594608 | SEMA6A-AS1 | 101927233 |
| ENSG000002 | 3.21682154 | -0.925218816 | 0.37676291 | -2.4557057 | 0.01406083 | 0.04594895 | NA | NA |
| ENSG000002 | 25.9696334 | -0.691221087 | 0.28149277 | -2.4555554 | 0.01406671 | 0.04596447 | NA | NA |

| | | | | | | | | | |
|---|---|---|---|---|---|---|---|---|---|
| ENSG000001 | 54.8943085 | 0.54380626 | 0.22146519 | 2.45549317 | 0.01406914 | 0.04596872 | PRX | | 57716 |
| ENSG000001 | 2615.51114 | -0.307849509 | 0.12537598 | -2.4554107 | 0.01407237 | 0.04597557 | MDM2 | | 4193 |
| ENSG000002 | 6.28353351 | -0.908480553 | 0.37001488 | -2.4552541 | 0.0140785 | 0.04599189 | DSCR9 | | 257203 |
| ENSG000001 | 172.894368 | 0.40026003 | 0.1630338 | 2.45507386 | 0.01408556 | 0.04599652 | BCAM | | 4059 |
| ENSG000002 | 9.88808983 | -0.863737381 | 0.35182151 | -2.4550443 | 0.01408672 | 0.04599652 | NA | NA | |
| ENSG000002 | 4.19533875 | -0.92810124 | 0.37801839 | -2.4551749 | 0.01408161 | 0.04599652 | NA | NA | |
| ENSG000002 | 6.42273633 | -0.898148993 | 0.36583796 | -2.4550459 | 0.01408666 | 0.04599652 | NA | NA | |
| ENSG000002 | 9.46547259 | -0.860321425 | 0.35041552 | -2.4551465 | 0.01408272 | 0.04599652 | NA | NA | |
| ENSG000002 | 3.56083148 | -0.928628384 | 0.37824764 | -2.4550804 | 0.01408531 | 0.04599652 | NA | NA | |
| ENSG000002 | 15.4820001 | -0.749467531 | 0.30529145 | -2.4549248 | 0.01409141 | 0.04600811 | NA | NA | |
| ENSG000001 | 16.098139 | -0.747843588 | 0.30463916 | -2.4548504 | 0.01409432 | 0.04601391 | CD300LF | | 146722 |
| ENSG000002 | 8.99399523 | -0.856699403 | 0.34900419 | -2.4546966 | 0.01410035 | 0.0460299 | NA | NA | |
| ENSG000002 | 629.873423 | 0.294924354 | 0.12015734 | 2.45448462 | 0.01410867 | 0.04605334 | NA | NA | |
| ENSG000002 | 22.1481908 | -0.692193679 | 0.28203051 | -2.454322 | 0.01411505 | 0.04607047 | LOC400541 | | 400541 |
| ENSG000002 | 10.0048599 | -0.846093651 | 0.3447611 | -2.4541448 | 0.01412201 | 0.04608947 | NA | NA | |
| ENSG000002 | 9.30817858 | -0.852989402 | 0.34757613 | -2.4541081 | 0.01412345 | 0.04609046 | NA | NA | |
| ENSG000001 | 19.7225024 | -0.68447598 | 0.27895803 | -2.453688 | 0.01413996 | 0.04614062 | PSD2 | | 84249 |
| ENSG000001 | 141.721563 | 0.413117681 | 0.1683686 | 2.45365041 | 0.01414144 | 0.04614173 | LYSMD4 | | 145748 |
| ENSG000002 | 7.75047504 | 0.875217516 | 0.35670947 | 2.45358642 | 0.01414396 | 0.04614623 | NA | NA | |
| ENSG000001 | 21.4006399 | -0.699796895 | 0.28526407 | -2.4531547 | 0.01416094 | 0.04619455 | ITGAD | | 3681 |
| ENSG000002 | 7.83632809 | -0.881065671 | 0.35915655 | -2.4531522 | 0.01416104 | 0.04619455 | NA | NA | |
| ENSG000002 | 5.60183121 | -0.916743143 | 0.37372184 | -2.4530093 | 0.01416667 | 0.04620918 | NA | NA | |
| ENSG000002 | 16.6167679 | -0.735781934 | 0.29998988 | -2.4526892 | 0.01417928 | 0.0462466 | SCGB2B2 | | 284402 |
| ENSG000002 | 6.37862284 | -0.894608248 | 0.36475391 | -2.4526351 | 0.01418141 | 0.04624983 | NA | NA | |
| ENSG000002 | 13.9804061 | -0.786869011 | 0.32087775 | -2.4522393 | 0.01419702 | 0.04629702 | NA | NA | |
| ENSG000002 | 6.11459192 | -0.904506643 | 0.36885986 | -2.4521688 | 0.0141998 | 0.04630236 | NA | NA | |
| ENSG000001 | 24.6655054 | 0.641851312 | 0.26177034 | 2.45196347 | 0.01420791 | 0.04632135 | DUSP26 | | 78986 |
| ENSG000002 | 3.21622073 | -0.923365468 | 0.37658076 | -2.451972 | 0.01420757 | 0.04632135 | NA | NA | |
| ENSG000001 | 48.3595708 | 0.571080334 | 0.23292198 | 2.45180957 | 0.01421399 | 0.04633744 | HDHD3 | | 81932 |

| | | | | | | | | |
|---|---|---|---|---|---|---|---|---|
| ENSG000002 | 3.62932168 | -0.926311191 | 0.37781556 | -2.451755 | 0.01421614 | 0.04634074 | RN7SL344P | 106480503 |
| ENSG000001 | 198.2992 | 0.349058967 | 0.14238302 | 2.45154918 | 0.01422427 | 0.04636352 | TBCCD1 | 55171 |
| ENSG000002 | 9.25890919 | -0.850433691 | 0.34694207 | -2.4512268 | 0.01423702 | 0.04640134 | NA | NA |
| ENSG000001 | 15.8657353 | -0.750153775 | 0.30603682 | -2.451188 | 0.01423856 | 0.04640262 | MGAT4EP | 641515 |
| ENSG000001 | 730.523151 | 0.303946151 | 0.12400173 | 2.45114435 | 0.01424028 | 0.04640451 | XRN1 | 54464 |
| ENSG000001 | 179.059809 | 0.349369067 | 0.14255186 | 2.45082078 | 0.01425309 | 0.04643878 | TGFBR2 | 7048 |
| ENSG000002 | 9.21642466 | -0.867326438 | 0.35388985 | -2.4508372 | 0.01425244 | 0.04643878 | LINC01365 | 101927007 |
| ENSG000002 | 731.322971 | 0.347162124 | 0.14166346 | 2.45061165 | 0.01426137 | 0.04646203 | PHACTR4 | 65979 |
| ENSG000001 | 4314.75312 | 0.286236848 | 0.11680728 | 2.45050521 | 0.01426559 | 0.04647204 | UQCRB | 7381 |
| ENSG00000C | 36.4682366 | -0.579094593 | 0.2363244 | -2.4504224 | 0.01426887 | 0.04647526 | RASGRF1 | 5923 |
| ENSG000001 | 135.523819 | 0.484469268 | 0.19770804 | 2.45042773 | 0.01426866 | 0.04647526 | CPPED1 | 55313 |
| ENSG000001 | 1082.22824 | 0.265693355 | 0.10844101 | 2.45011878 | 0.01428091 | 0.04650864 | SEC24A | 10802 |
| ENSG000002 | 4.17149328 | -0.926163044 | 0.37800936 | -2.4501061 | 0.01428141 | 0.04650864 | NA | NA |
| ENSG000001 | 2.88693174 | -0.913621446 | 0.37291651 | -2.4499356 | 0.01428818 | 0.04652693 | NA | NA |
| ENSG000002 | 34.2057132 | 0.599759772 | 0.24482172 | 2.44978173 | 0.01429428 | 0.04654308 | SNORA65 | 26783 |
| ENSG000002 | 12.9369605 | -0.804545825 | 0.32842214 | -2.4497308 | 0.01429631 | 0.04654593 | EHHADH-AS1 | 339926 |
| ENSG000002 | 4.89527518 | -0.926818425 | 0.37842575 | -2.4491421 | 0.01431969 | 0.04661833 | LOC105378138 | 105378138 |
| ENSG000002 | 4.30569443 | -0.926887059 | 0.3784586 | -2.4491108 | 0.01432094 | 0.04661864 | NA | NA |
| ENSG000001 | 17.7209822 | -0.752689957 | 0.30736914 | -2.4488144 | 0.01433273 | 0.04665328 | HR | 55806 |
| ENSG000002 | 7.86560877 | -0.86415442 | 0.35289673 | -2.4487459 | 0.01433545 | 0.0466584 | NA | NA |
| ENSG000001 | 18.4563465 | -0.704366752 | 0.28766869 | -2.4485347 | 0.01434386 | 0.04668202 | TTC29 | 83894 |
| ENSG000002 | 9.16817104 | -0.84620247 | 0.34560779 | -2.4484474 | 0.01434734 | 0.04668959 | NA | NA |
| ENSG000001 | 18.8542667 | -0.703584473 | 0.28736954 | -2.4483613 | 0.01435077 | 0.04669324 | TCHH | 7062 |
| ENSG000002 | 11.7666776 | -0.801484572 | 0.32735331 | -2.4483778 | 0.01435011 | 0.04669324 | NA | NA |
| ENSG000002 | 12.149617 | -0.816596302 | 0.33354577 | -2.4482286 | 0.01435605 | 0.0467067 | PSG2 | 5670 |
| ENSG000002 | 8.56621846 | -0.875938497 | 0.35779775 | -2.4481387 | 0.01435964 | 0.04671461 | LINC01447 | 101929086 |
| ENSG00000C | 37.4801176 | 0.626413815 | 0.25589056 | 2.44797544 | 0.01436614 | 0.04672079 | MCOLN1 | 57192 |
| ENSG000001 | 289.257556 | 0.309544528 | 0.12644766 | 2.44800514 | 0.01436496 | 0.04672079 | TAOK3 | 51347 |
| ENSG000001 | 43.8146888 | -0.541124412 | 0.22104588 | -2.4480185 | 0.01436443 | 0.04672079 | GLRA3 | 8001 |

| | | | | | | | | |
|---|---|---|---|---|---|---|---|---|
| ENSG000001 | 67.264052 | 0.478409601 | 0.19542972 | 2.44798793 | 0.01436565 | 0.04672079 | POU6F1 | 5463 |
| ENSG000001 | 122.661301 | 0.445991512 | 0.18219324 | 2.44790378 | 0.014369 | 0.04672634 | NEK6 | 10783 |
| ENSG000001 | 868.838434 | 0.31869538 | 0.13020085 | 2.44772125 | 0.01437628 | 0.04674252 | ATP5E | 514 |
| ENSG000002 | 8.31043918 | -0.863529444 | 0.35278533 | -2.4477476 | 0.01437523 | 0.04674252 | NA | NA |
| ENSG000002 | 2.81084119 | -0.915774566 | 0.37417058 | -2.4474788 | 0.01438596 | 0.04677022 | NA | NA |
| ENSG000001 | 9.82008818 | -0.863264464 | 0.35272701 | -2.4474011 | 0.01438906 | 0.04677657 | HSD17B13 | 345275 |
| ENSG000001 | 838.615397 | 0.332767192 | 0.13598676 | 2.4470559 | 0.01440285 | 0.04681696 | UBE2Z | 65264 |
| ENSG000002 | 2.82339285 | -0.91474937 | 0.37381992 | -2.4470322 | 0.0144038 | 0.04681696 | LINC00396 | 104355146 |
| ENSG000001 | 15.0570162 | -0.741736772 | 0.30313104 | -2.4469179 | 0.01440836 | 0.04682683 | CYP4F2 | 8529 |
| ENSG000002 | 73.8655347 | 0.460801265 | 0.18832055 | 2.44689841 | 0.01440914 | 0.04682683 | TMEM185A | 84548 |
| ENSG000001 | 87.6904265 | 0.457069567 | 0.18680529 | 2.44676996 | 0.01441428 | 0.04683977 | FJX1 | 24147 |
| ENSG000002 | 11.6006575 | -0.810343355 | 0.33123056 | -2.4464631 | 0.01442655 | 0.0468759 | NA | NA |
| ENSG000002 | 6.82406126 | -0.891464172 | 0.36440014 | -2.4463881 | 0.01442956 | 0.0468819 | NA | NA |
| ENSG000001 | 10.2438002 | -0.819552606 | 0.33501597 | -2.446309 | 0.01443272 | 0.04688467 | IFI44 | 10561 |
| ENSG000001 | 335.09768 | 0.360064834 | 0.14718621 | 2.44632179 | 0.01443221 | 0.04688467 | PRMT9 | 90826 |
| ENSG000001 | 35.8396675 | -0.606283972 | 0.24784939 | -2.446179 | 0.01443793 | 0.04689407 | CCDC146 | 57639 |
| ENSG000002 | 4.81853608 | -0.925610642 | 0.37838734 | -2.4461987 | 0.01443714 | 0.04689407 | NA | NA |
| ENSG000002 | 40.7153904 | 0.56456385 | 0.23080073 | 2.44610946 | 0.01444072 | 0.04689767 | NA | NA |
| ENSG000002 | 9.28557486 | -0.855202436 | 0.34961967 | -2.4460936 | 0.01444135 | 0.04689767 | NA | NA |
| ENSG000002 | 12.6147906 | -0.822476442 | 0.33624855 | -2.4460371 | 0.01444361 | 0.04690126 | NA | NA |
| ENSG000002 | 5.21852684 | -0.918494486 | 0.37554102 | -2.44579 | 0.01445352 | 0.04692966 | NA | NA |
| ENSG000002 | 10.9980459 | -0.819223815 | 0.33495908 | -2.445743 | 0.0144554 | 0.04693203 | NA | NA |
| ENSG000001 | 636.674135 | 0.320117702 | 0.13089574 | 2.44559304 | 0.01446141 | 0.04694778 | ASAP1 | 50807 |
| ENSG000002 | 5.79487237 | -0.905024093 | 0.37006907 | -2.4455546 | 0.01446296 | 0.04694904 | NA | NA |
| ENSG000002 | 3.90153341 | -0.925680481 | 0.37859121 | -2.445066 | 0.01448256 | 0.04700892 | NA | NA |
| ENSG00000C | 27.6903995 | -0.627693208 | 0.25673103 | -2.4449449 | 0.01448743 | 0.0470184 | SLC6A16 | 28968 |
| ENSG000001 | 18.5039576 | -0.730043004 | 0.29859397 | -2.4449355 | 0.0144878 | 0.0470184 | GRIN2C | 2905 |
| ENSG000002 | 5.80731192 | -0.90617984 | 0.37065183 | -2.4448276 | 0.01449214 | 0.04702871 | NA | NA |
| ENSG000002 | 4.29349377 | -0.925306701 | 0.3784964 | -2.4446909 | 0.01449763 | 0.04704277 | NA | NA |

| | | | | | | | | |
|---|---|---|---|---|---|---|---|---|
| ENSG000001 | 11.87284 | -0.807880426 | 0.3304763 | -2.4445941 | 0.01450152 | 0.04705163 | HLX | 3142 |
| ENSG000002 | 14.48898 | -0.793767613 | 0.324708 | -2.4445583 | 0.01450297 | 0.04705254 | NA | NA |
| ENSG000001 | 82.1859945 | 0.466521911 | 0.19086007 | 2.44431382 | 0.0145128 | 0.04708067 | NA | NA |
| ENSG000002 | 8.34522604 | -0.865175597 | 0.35396996 | -2.4442063 | 0.01451712 | 0.04709095 | NA | NA |
| ENSG000001 | 244.062776 | 0.3325277 | 0.13605334 | 2.44409804 | 0.01452148 | 0.04709754 | ZSCAN30 | 100101467 |
| ENSG000002 | 2.25799676 | -0.870984867 | 0.3563594 | -2.4441193 | 0.01452063 | 0.04709754 | NA | NA |
| ENSG000002 | 18.6204893 | -0.71507097 | 0.29260399 | -2.4438182 | 0.01453275 | 0.0471266 | NA | NA |
| ENSG000002 | 16.0211743 | -0.756436542 | 0.30953066 | -2.4438178 | 0.01453276 | 0.0471266 | NA | NA |
| ENSG000001 | 46.7331234 | -0.550885128 | 0.22544529 | -2.4435424 | 0.01454386 | 0.04715881 | OVCH1 | 341350 |
| ENSG000002 | 4.21168049 | -0.924191409 | 0.37822737 | -2.4434811 | 0.01454633 | 0.04716306 | NA | NA |
| ENSG000001 | 78.3117781 | 0.481772083 | 0.19721066 | 2.44293125 | 0.01456851 | 0.04723119 | HEXDC | 284004 |
| ENSG000002 | 7.76311883 | -0.874776168 | 0.35809525 | -2.4428589 | 0.01457143 | 0.04723689 | LOC101928418 | 101928418 |
| ENSG000001 | 19.5200643 | -0.688989316 | 0.28205089 | -2.4427837 | 0.01457447 | 0.04724029 | TRIM7 | 81786 |
| ENSG000002 | 5.17638656 | -0.914629898 | 0.37442246 | -2.4427752 | 0.01457481 | 0.04724029 | NA | NA |
| ENSG000002 | 25.6355135 | -0.663798437 | 0.27176346 | -2.4425595 | 0.01458352 | 0.04726474 | NA | NA |
| ENSG000001 | 4.5859101 | -0.921671173 | 0.37737318 | -2.4423336 | 0.01459266 | 0.04728029 | NA | NA |
| ENSG000001 | 46.1446805 | 0.549170277 | 0.22484958 | 2.44238958 | 0.01459039 | 0.04728029 | RIPK4 | 54101 |
| ENSG000002 | 42.1774708 | 0.556196509 | 0.22773235 | 2.44232547 | 0.01459298 | 0.04728029 | DNAJC3-AS1 | 100289274 |
| ENSG000002 | 3.52945038 | -0.923590947 | 0.37815809 | -2.4423409 | 0.01459236 | 0.04728029 | NA | NA |
| ENSG000002 | 5.42310234 | -0.907155235 | 0.37143912 | -2.4422717 | 0.01459516 | 0.04728356 | NA | NA |
| ENSG00000C | 416.124978 | 0.361184838 | 0.14790234 | 2.44204945 | 0.01460415 | 0.04730889 | CDIP1 | 29965 |
| ENSG000001 | 240.372396 | 0.34546002 | 0.141466 | 2.44200029 | 0.01460613 | 0.04731156 | PPP4R4 | 57718 |
| ENSG00000C | 9.43430905 | -0.843179537 | 0.34531362 | -2.441779 | 0.01461509 | 0.047333 | OTC | 5009 |
| ENSG000001 | 20.9880372 | -0.696376987 | 0.28518941 | -2.4418052 | 0.01461403 | 0.047333 | GOLGA7B | 401647 |
| ENSG000001 | 513.591025 | 0.327360068 | 0.1340745 | 2.44162812 | 0.0146212 | 0.04734901 | CREB3L2 | 64764 |
| ENSG000001 | 489.439907 | 0.301601989 | 0.12353658 | 2.44139826 | 0.01463051 | 0.04737538 | CEP162 | 22832 |
| ENSG000001 | 13.0319253 | -0.806842016 | 0.33050774 | -2.4412197 | 0.01463775 | 0.04739502 | OR51E1 | 143503 |
| ENSG000002 | 8.74720879 | -0.865536911 | 0.3545615 | -2.4411475 | 0.01464067 | 0.04740072 | HSPE1P24 | 106480259 |
| ENSG000001 | 827.002067 | 0.307631696 | 0.1260409 | 2.44072916 | 0.01465764 | 0.04745086 | PSD3 | 23362 |

| | | | | | | | | |
|---|---|---|---|---|---|---|---|---|
| ENSG000002 | 8.34431214 | -0.871123627 | 0.35691432 | -2.440708 | 0.0146585 | 0.04745086 | DBIL5P2 | 100169989 |
| ENSG000001 | 12.0751631 | -0.802594066 | 0.32885607 | -2.4405633 | 0.01466438 | 0.04746608 | CAPSL | 133690 |
| ENSG000001 | 17.5233414 | -0.728733589 | 0.29862012 | -2.4403365 | 0.01467359 | 0.0474921 | NA | NA |
| ENSG000002 | 14.1623138 | -0.751014306 | 0.30776715 | -2.4402029 | 0.01467901 | 0.04750588 | NA | NA |
| ENSG000001 | 5.48891126 | -0.910736847 | 0.37323545 | -2.4401135 | 0.01468265 | 0.04751385 | MAB21L1 | 4081 |
| ENSG000000 | 4679.14614 | 0.30883636 | 0.12657073 | 2.44002976 | 0.01468605 | 0.04751607 | JADE1 | 79960 |
| ENSG000001 | 32.9980966 | -0.579843762 | 0.2376427 | -2.4399813 | 0.01468802 | 0.04751607 | ALK | 238 |
| ENSG000002 | 11.926152 | -0.787982625 | 0.32294507 | -2.4399896 | 0.01468768 | 0.04751607 | LOC101929406 | 101929406 |
| ENSG000002 | 4.23479773 | -0.923494162 | 0.37847888 | -2.4400151 | 0.01468665 | 0.04751607 | NA | NA |
| ENSG000002 | 14.6053382 | -0.7544367 | 0.30920152 | -2.4399515 | 0.01468924 | 0.04751621 | NA | NA |
| ENSG000002 | 7.99987517 | -0.869164924 | 0.35623999 | -2.4398298 | 0.01469418 | 0.04752842 | NA | NA |
| ENSG000002 | 20.2973975 | -0.71539023 | 0.29323602 | -2.4396397 | 0.01470192 | 0.04754965 | LINC01024 | 100505636 |
| ENSG000002 | 2528.34249 | 0.641082704 | 0.26280118 | 2.43942097 | 0.01471082 | 0.04757464 | PRRC2A | 7916 |
| ENSG000001 | 125.453061 | 0.445568591 | 0.18265702 | 2.43937291 | 0.01471278 | 0.04757718 | ABCC10 | 89845 |
| ENSG000002 | 4.65547079 | -0.922068522 | 0.37800083 | -2.4393294 | 0.01471455 | 0.04757912 | NA | NA |
| ENSG000002 | 4.69373909 | -0.922216239 | 0.3780758 | -2.4392364 | 0.01471834 | 0.04758757 | OSTN-AS1 | 106480738 |
| ENSG000002 | 5.53895121 | -0.910067461 | 0.37311091 | -2.4391339 | 0.01472251 | 0.04759727 | NA | NA |
| ENSG000002 | 7.31410602 | -0.873452049 | 0.35810722 | -2.4390797 | 0.01472472 | 0.04760062 | LOC729652 | 729652 |
| ENSG000001 | 69.2551174 | -0.521940463 | 0.21400787 | -2.4388844 | 0.01473268 | 0.04761875 | ZNF835 | 90485 |
| ENSG000001 | 18.2992762 | -0.735617319 | 0.30161724 | -2.43891 | 0.01473164 | 0.04761875 | HRH4 | 59340 |
| ENSG000002 | 5.45056163 | -0.906622152 | 0.37175138 | -2.4387863 | 0.01473668 | 0.04762789 | NA | NA |
| ENSG000000 | 24.3626315 | -0.6754612 | 0.27697217 | -2.4387331 | 0.01473885 | 0.04763111 | TGFB2 | 7042 |
| ENSG000001 | 3518.80888 | 0.28163955 | 0.11549979 | 2.43844202 | 0.01475072 | 0.04766188 | SNRPD1 | 6632 |
| ENSG000002 | 6.59623103 | -0.900395068 | 0.36924807 | -2.4384557 | 0.01475017 | 0.04766188 | NA | NA |
| ENSG000002 | 3.93203294 | -0.923036084 | 0.37856143 | -2.4382729 | 0.01475763 | 0.04768039 | NA | NA |
| ENSG000001 | 1728.56786 | 0.329378962 | 0.13509087 | 2.43820304 | 0.01476048 | 0.04768581 | ROCK2 | 9475 |
| ENSG000002 | 61.8001699 | 0.482651611 | 0.1979691 | 2.43801491 | 0.01476816 | 0.0477036 | NA | NA |
| ENSG000002 | 33.1368443 | 0.600562281 | 0.24633293 | 2.43801057 | 0.01476834 | 0.0477036 | NA | NA |
| ENSG000002 | 2.93302873 | -0.90753286 | 0.37224793 | -2.4379796 | 0.01476961 | 0.04770389 | NA | NA |

| | | | | | | | | |
|---|---|---|---|---|---|---|---|---|
| ENSG000002 | 3.15850985 | -0.918478148 | 0.37675562 | -2.4378618 | 0.01477442 | 0.04771183 | NA | NA |
| ENSG000002 | 17.070309 | -0.717232332 | 0.2942023 | -2.4378883 | 0.01477334 | 0.04771183 | NA | NA |
| ENSG000002 | 36.6890732 | -0.583594198 | 0.23939307 | -2.4378074 | 0.01477665 | 0.04771522 | NA | NA |
| ENSG000002 | 73.8216497 | 0.51322472 | 0.21053418 | 2.43772641 | 0.01477996 | 0.04772211 | AP5B1 | 91056 |
| ENSG000000 | 2705.39258 | 0.352648411 | 0.14467029 | 2.43760071 | 0.0147851 | 0.0477349 | WNK1 | 65125 |
| ENSG000002 | 19.3260191 | -0.700643736 | 0.28748045 | -2.4371874 | 0.01480201 | 0.04778375 | GS1-600G8.3 | 100093698 |
| ENSG000002 | 7.00819285 | -0.879106477 | 0.36070741 | -2.4371733 | 0.01480258 | 0.04778375 | NA | NA |
| ENSG000002 | 9.45489431 | -0.839066204 | 0.34428665 | -2.4371151 | 0.01480497 | 0.04778764 | NA | NA |
| ENSG000002 | 7.29887831 | -0.873069107 | 0.35825852 | -2.4369807 | 0.01481047 | 0.0478016 | NA | NA |
| ENSG000001 | 19.0040846 | -0.745734508 | 0.30604247 | -2.4367027 | 0.01482186 | 0.04783456 | KLK2 | 3817 |
| ENSG000001 | 29.1801074 | 0.648283707 | 0.26607155 | 2.43650145 | 0.01483011 | 0.04785738 | WASH1 | 100287171 |
| ENSG000001 | 592.564589 | 0.311299791 | 0.12776666 | 2.43647116 | 0.01483135 | 0.04785758 | RTKN2 | 219790 |
| ENSG000001 | 8.59309622 | -0.875102014 | 0.3592023 | -2.4362372 | 0.01484095 | 0.04788473 | NA | NA |
| ENSG000002 | 7.73208514 | -0.871507708 | 0.35773324 | -2.4361944 | 0.01484271 | 0.0478866 | NA | NA |
| ENSG000002 | 8.88763341 | -0.852679043 | 0.35002275 | -2.4360675 | 0.01484791 | 0.04789958 | LOC101927412 | 101927412 |
| ENSG000002 | 22.3924371 | -0.652026161 | 0.26766612 | -2.4359683 | 0.01485199 | 0.04790891 | LOC392232 | 392232 |
| ENSG000002 | 166.8317 | 0.382220136 | 0.15690888 | 2.43593698 | 0.01485327 | 0.04790925 | COX19 | 90639 |
| ENSG000002 | 148.119404 | 0.388888246 | 0.15964838 | 2.43590471 | 0.0148546 | 0.04790971 | RAB30-AS1 | 100506233 |
| ENSG000002 | 10.5941104 | -0.835343542 | 0.34298543 | -2.4355074 | 0.01487092 | 0.04795854 | NA | NA |
| ENSG000001 | 26.6339177 | 0.668911471 | 0.27465866 | 2.43542826 | 0.01487417 | 0.04796522 | SNAI1 | 6615 |
| ENSG000001 | 2.95371216 | -0.90676241 | 0.3723378 | -2.4353219 | 0.01487855 | 0.0479755 | MBD3L1 | 85509 |
| ENSG000001 | 383.472732 | 0.320610315 | 0.13165424 | 2.43524497 | 0.01488171 | 0.04798189 | TAF13 | 6884 |
| ENSG000002 | 15.7433859 | -0.779028133 | 0.31994045 | -2.434916 | 0.01489525 | 0.04802172 | MTND6P18 | 106478951 |
| ENSG000002 | 5.27982018 | -0.914542571 | 0.37562462 | -2.4347248 | 0.01490312 | 0.04804328 | NA | NA |
| ENSG000002 | 8.8247936 | -0.84467873 | 0.34695248 | -2.434566 | 0.01490966 | 0.04806054 | NA | NA |
| ENSG000001 | 1114.56458 | 0.373104083 | 0.15326554 | 2.43436375 | 0.01491799 | 0.04807977 | RAB1A | 5861 |
| ENSG000002 | 3.52529577 | -0.92048605 | 0.37812004 | -2.4343752 | 0.01491752 | 0.04807977 | NA | NA |
| ENSG000002 | 16.2830573 | -0.728061318 | 0.29910649 | -2.4341208 | 0.01492801 | 0.04810823 | NA | NA |
| ENSG000002 | 6.57090195 | -0.899306423 | 0.36947888 | -2.433986 | 0.01493357 | 0.04812232 | NA | NA |

| | | | | | | | | |
|---|---|---|---|---|---|---|---|---|
| ENSG000001 | 9.47273353 | -0.83674714 | 0.343782 | -2.4339469 | 0.01493518 | 0.04812369 | WFDC10A | 140832 |
| ENSG000002 | 53.6874849 | 0.493598244 | 0.20280368 | 2.43387218 | 0.01493827 | 0.04812981 | ZNF324B | 388569 |
| ENSG000001 | 467.570133 | 0.306580582 | 0.12596597 | 2.43383665 | 0.01493974 | 0.04813071 | NTAN1 | 123803 |
| ENSG000001 | 14.1202864 | -0.768309702 | 0.31570074 | -2.4336645 | 0.01494684 | 0.04814977 | CD300LG | 146894 |
| ENSG000001 | 289.560828 | 0.422095835 | 0.17345138 | 2.43351099 | 0.01495318 | 0.04816637 | SOGA1 | 140710 |
| ENSG000001 | 1012.72585 | 0.35959797 | 0.1477868 | 2.43322123 | 0.01496515 | 0.04820111 | COL4A1 | 1282 |
| ENSG000002 | 9.1695271 | -0.863925246 | 0.35506426 | -2.4331518 | 0.01496802 | 0.04820653 | SERPINB10 | 5273 |
| ENSG000001 | 93.4618334 | -0.419135667 | 0.17226816 | -2.4330421 | 0.01497256 | 0.04821348 | INPP4B | 8821 |
| ENSG000002 | 10.8301031 | -0.806099221 | 0.33131004 | -2.4330661 | 0.01497157 | 0.04821348 | LOC105375483 | 105375483 |
| ENSG000001 | 8.68361062 | -0.860758353 | 0.35379723 | -2.4329143 | 0.01497784 | 0.04822666 | CRISP2 | 7180 |
| ENSG000002 | 2.4487462 | -0.895249229 | 0.36802128 | -2.4326018 | 0.01499078 | 0.04826447 | NA | NA |
| ENSG000001 | 533.934052 | 0.321041434 | 0.13197613 | 2.43257199 | 0.01499201 | 0.04826461 | CEP83 | 51134 |
| ENSG00000C | 414.155449 | 0.44234541 | 0.18185602 | 2.43239358 | 0.0149994 | 0.04828456 | HCCS | 3052 |
| ENSG000002 | 9.10905742 | -0.840075952 | 0.34541334 | -2.4320889 | 0.01501202 | 0.04831752 | NA | NA |
| ENSG000002 | 871.181518 | 0.290818484 | 0.11957435 | 2.43211426 | 0.01501097 | 0.04831752 | ATP5J2 | 9551 |
| ENSG000002 | 10.4996624 | -0.808559287 | 0.33247706 | -2.431925 | 0.01501882 | 0.04833362 | NA | NA |
| ENSG000002 | 310.818107 | 0.427118696 | 0.1756309 | 2.43191081 | 0.01501941 | 0.04833362 | NA | NA |
| ENSG000002 | 10.2625511 | -0.815217105 | 0.33526227 | -2.4315802 | 0.01503312 | 0.04837391 | NA | NA |
| ENSG000002 | 19.3859077 | -0.682688882 | 0.28080038 | -2.4312249 | 0.01504787 | 0.04841753 | NA | NA |
| ENSG000001 | 73.783705 | -0.459134498 | 0.18887335 | -2.4309121 | 0.01506087 | 0.04845551 | LRRIQ1 | 84125 |
| ENSG000002 | 8.54960146 | -0.850397828 | 0.34985231 | -2.4307338 | 0.01506828 | 0.04847551 | LINC01203 | 100133123 |
| ENSG000002 | 7.14511276 | -0.88453914 | 0.36391116 | -2.4306458 | 0.01507194 | 0.04848344 | NA | NA |
| ENSG000002 | 12.2084321 | -0.801840789 | 0.32989604 | -2.4305863 | 0.01507442 | 0.04848756 | NA | NA |
| ENSG000002 | 7.80516303 | -0.874853822 | 0.3599754 | -2.4303156 | 0.01508568 | 0.04851994 | NA | NA |
| ENSG000002 | 56.0932597 | -0.511566884 | 0.21050686 | -2.4301673 | 0.01509186 | 0.04853596 | NA | NA |
| ENSG000002 | 2.81203487 | -0.907681538 | 0.37351975 | -2.4300764 | 0.01509564 | 0.04854427 | NA | NA |
| ENSG000002 | 23.0771462 | -0.705633383 | 0.29037996 | -2.4300347 | 0.01509738 | 0.04854456 | NA | NA |
| ENSG000002 | 14.1132579 | -0.806709034 | 0.33197674 | -2.4300168 | 0.01509812 | 0.04854456 | NA | NA |
| ENSG000002 | 22.2749789 | -0.682210921 | 0.28077374 | -2.4297533 | 0.0151091 | 0.04857601 | TSPAN19 | 144448 |

| | | | | | | | | |
|---|---|---|---|---|---|---|---|---|
| ENSG00000C | 6.33934802 | -0.888997448 | 0.36591925 | -2.4294908 | 0.01512005 | 0.04860735 | POU1F1 | 5449 |
| ENSG000001 | 62.8987155 | 0.461157766 | 0.18985271 | 2.4290292 | 0.01513931 | 0.04866542 | SAMD15 | 161394 |
| ENSG000001 | 2367.68661 | 0.323604254 | 0.13323616 | 2.428802 | 0.0151488 | 0.04869207 | CALD1 | 800 |
| ENSG000002 | 9.34126951 | -0.831341784 | 0.3423056 | -2.4286538 | 0.015155 | 0.04870811 | NA | NA |
| ENSG000002 | 6.05950824 | -0.892129567 | 0.36736439 | -2.4284596 | 0.01516311 | 0.04873034 | NA | NA |
| ENSG000002 | 2.41172398 | -0.895859431 | 0.36894046 | -2.4281951 | 0.01517418 | 0.04875816 | NA | NA |
| ENSG000002 | 10.1113585 | -0.827933527 | 0.34096333 | -2.4282187 | 0.01517319 | 0.04875816 | NA | NA |
| ENSG00000C | 908.584182 | 0.426400646 | 0.17562733 | 2.42787187 | 0.01518771 | 0.04879778 | CREBBP | 1387 |
| ENSG000002 | 12.2899411 | -0.780480207 | 0.3214779 | -2.4277881 | 0.01519122 | 0.04880518 | PLSCR5 | 389158 |
| ENSG000002 | 5.9365406 | -0.905164468 | 0.37286125 | -2.4276174 | 0.01519837 | 0.04882429 | NA | NA |
| ENSG000002 | 11.4641147 | -0.823478307 | 0.33921867 | -2.4275737 | 0.0152002 | 0.04882631 | NA | NA |
| ENSG000002 | 310.34013 | 0.414116731 | 0.17060972 | 2.42727516 | 0.01521271 | 0.04886263 | PI4KA | 5297 |
| ENSG000001 | 463.899531 | 0.280817463 | 0.11569847 | 2.42714942 | 0.01521799 | 0.0488757 | PAPD5 | 64282 |
| ENSG000001 | 401.786866 | 0.284044534 | 0.11703484 | 2.4270084 | 0.0152239 | 0.04888881 | HOMER2 | 9455 |
| ENSG000001 | 209.899498 | 0.351387963 | 0.14478316 | 2.42699464 | 0.01522448 | 0.04888881 | CD47 | 961 |
| ENSG000001 | 16.4150636 | -0.735508261 | 0.30306392 | -2.426908 | 0.01522812 | 0.04889662 | FSHR | 2492 |
| ENSG000002 | 4.56698424 | -0.916110598 | 0.37749628 | -2.426807 | 0.01523236 | 0.04890636 | NA | NA |
| ENSG000001 | 26.8487857 | -0.626843154 | 0.25831748 | -2.4266385 | 0.01523943 | 0.04891482 | IDO2 | 169355 |
| ENSG000002 | 3.90436458 | -0.918641654 | 0.37856694 | -2.4266293 | 0.01523982 | 0.04891482 | NA | NA |
| ENSG000002 | 7.8935058 | -0.85853542 | 0.35379039 | -2.4266782 | 0.01523777 | 0.04891482 | NA | NA |
| ENSG000002 | 4.93223495 | -0.914894814 | 0.37701212 | -2.4266987 | 0.0152369 | 0.04891482 | NA | NA |
| ENSG000001 | 78.2585204 | -0.46788095 | 0.19281701 | -2.4265543 | 0.01524297 | 0.04892106 | FDXR | 2232 |
| ENSG00000C | 1288.73192 | 0.248090242 | 0.10224303 | 2.42647579 | 0.01524627 | 0.04892525 | KMT2E | 55904 |
| ENSG000001 | 17.3041594 | -0.712327895 | 0.29356601 | -2.4264658 | 0.01524669 | 0.04892525 | NA | NA |
| ENSG000001 | 739.592044 | 0.42307898 | 0.17437916 | 2.42620148 | 0.0152578 | 0.04895703 | AUTS2 | 26053 |
| ENSG000001 | 126.23899 | 0.415049346 | 0.17107879 | 2.42607126 | 0.01526327 | 0.04896685 | FCHO1 | 23149 |
| ENSG000001 | 2917.48625 | 0.299910478 | 0.12361876 | 2.42609188 | 0.01526241 | 0.04896685 | BRCA2 | 675 |
| ENSG000001 | 1800.42388 | 0.262264515 | 0.10810494 | 2.42601776 | 0.01526552 | 0.04897019 | SEC31A | 22872 |
| ENSG000002 | 17.3604419 | -0.769395309 | 0.31714952 | -2.4259703 | 0.01526752 | 0.04897273 | LOC284798 | 284798 |

| | | | | | | | | |
|---|---|---|---|---|---|---|---|---|
| ENSG00000( | 927.589766 | 0.365464899 | 0.15065469 | 2.42584476 | 0.0152728 | 0.04898579 | FKBP5 | 2289 |
| ENSG000001 | 47.7882422 | 0.517884124 | 0.21350788 | 2.42559722 | 0.01528322 | 0.0490052 | PMM1 | 5372 |
| ENSG000001 | 69.6431583 | 0.463161559 | 0.19094241 | 2.42566097 | 0.01528054 | 0.0490052 | FBXO10 | 26267 |
| ENSG000001 | 22.2459708 | -0.663663427 | 0.2736095 | -2.4255862 | 0.01528369 | 0.0490052 | CYP4F11 | 57834 |
| ENSG000001 | 42.2944378 | -0.553863184 | 0.22833851 | -2.4256232 | 0.01528213 | 0.0490052 | RGS6 | 9628 |
| ENSG000002 | 117.558206 | 0.39391562 | 0.16240767 | 2.42547423 | 0.0152884 | 0.04901644 | NA | NA |
| ENSG000002 | 17.2006278 | -0.726600199 | 0.29959388 | -2.4252838 | 0.01529642 | 0.04903828 | LINC00276 | 100499171 |
| ENSG000001 | 1351.50834 | 0.343998075 | 0.1418537 | 2.42502012 | 0.01530754 | 0.04907004 | TCEAL4 | 79921 |
| ENSG000001 | 44.9865107 | -0.540963904 | 0.22309275 | -2.4248385 | 0.0153152 | 0.04909071 | C11orf65 | 160140 |
| ENSG000001 | 30.1183572 | -0.640387877 | 0.26412371 | -2.4245755 | 0.0153263 | 0.04912241 | TMEM255B | 348013 |
| ENSG000002 | 3.69552767 | -0.914863663 | 0.37734015 | -2.4245065 | 0.01532921 | 0.04912397 | NA | NA |
| ENSG000002 | 6.23104069 | -0.895849874 | 0.36949514 | -2.4245241 | 0.01532847 | 0.04912397 | NA | NA |
| ENSG000001 | 947.370653 | 0.430263059 | 0.17747677 | 2.42433445 | 0.01533647 | 0.04914337 | UPF3A | 65110 |
| ENSG000002 | 11.2023013 | -0.800429489 | 0.33016983 | -2.4242963 | 0.01533809 | 0.04914465 | NA | NA |
| ENSG000002 | 5.20006806 | -0.909194485 | 0.37503933 | -2.4242644 | 0.01533943 | 0.04914509 | NA | NA |
| ENSG000002 | 23.2541813 | -0.6634662 | 0.27369684 | -2.4240915 | 0.01534674 | 0.04916417 | NA | NA |
| ENSG000002 | 4.48035502 | -0.917564209 | 0.37852278 | -2.424066 | 0.01534782 | 0.04916417 | NA | NA |
| ENSG000001 | 716.585452 | 0.271280899 | 0.11191379 | 2.42401675 | 0.0153499 | 0.04916696 | STIM2 | 57620 |
| ENSG000002 | 19.1254087 | -0.695970016 | 0.28713399 | -2.423851 | 0.01535691 | 0.04918552 | MGAT4D | 152586 |
| ENSG000002 | 6.63674208 | -0.897310672 | 0.37021425 | -2.4237605 | 0.01536073 | 0.04919389 | NA | NA |
| ENSG000001 | 390.44297 | 0.326348789 | 0.13464762 | 2.42372485 | 0.01536224 | 0.04919483 | NOCT | 25819 |
| ENSG000001 | 236.796514 | 0.34655419 | 0.14298952 | 2.42363352 | 0.0153661 | 0.04920256 | B3GALNT1 | 8706 |
| ENSG000002 | 9.1560753 | -0.840848418 | 0.34694041 | -2.4236105 | 0.01536708 | 0.04920256 | NA | NA |
| ENSG000002 | 5.24914301 | -0.908427459 | 0.37485083 | -2.4234373 | 0.01537441 | 0.04922214 | NA | NA |
| ENSG000001 | 842.23058 | 0.331972513 | 0.13698596 | 2.42340539 | 0.01537576 | 0.04922257 | NHLRC2 | 374354 |
| ENSG00000( | 339.983122 | 0.334461706 | 0.13801609 | 2.42335302 | 0.01537797 | 0.04922579 | TFPI | 7035 |
| ENSG000001 | 32.3951611 | -0.61688519 | 0.25457941 | -2.4231543 | 0.01538639 | 0.04924884 | CCDC170 | 80129 |
| ENSG000001 | 506.262946 | 0.289810838 | 0.11960708 | 2.42302407 | 0.01539191 | 0.04926261 | CBLB | 868 |
| ENSG000001 | 3.937879 | -0.917246742 | 0.37856119 | -2.4229815 | 0.01539371 | 0.04926449 | SATL1 | 340562 |

| | | | | | | | | |
|---|---|---|---|---|---|---|---|---|
| ENSG000002 | 112.175414 | 0.394978106 | 0.16301613 | 2.42293882 | 0.01539552 | 0.04926639 | ATP6V1E2 | 90423 |
| ENSG000001 | 35.902035 | 0.579737868 | 0.23928209 | 2.42282187 | 0.01540048 | 0.04927836 | EMILIN1 | 11117 |
| ENSG000002 | 4.56845204 | -0.914646874 | 0.37752048 | -2.4227742 | 0.0154025 | 0.04928094 | NA | NA |
| ENSG000002 | 13.1654326 | -0.793702992 | 0.32761447 | -2.4226738 | 0.01540676 | 0.04929068 | NA | NA |
| ENSG000001 | 13.1886511 | -0.774239629 | 0.31959939 | -2.4225316 | 0.01541279 | 0.04930379 | NR2E1 | 7101 |
| ENSG000001 | 35.0722076 | -0.570067836 | 0.2353202 | -2.4225198 | 0.01541329 | 0.04930379 | DRC1 | 92749 |
| ENSG000002 | 9.23573899 | -0.842090824 | 0.34764298 | -2.4222863 | 0.0154232 | 0.04932771 | NA | NA |
| ENSG000002 | 6.4142505 | -0.887425965 | 0.36635798 | -2.4222919 | 0.01542296 | 0.04932771 | LOC101928791 | 101928791 |
| ENSG000001 | 262.603958 | 0.321557469 | 0.13275174 | 2.42224671 | 0.01542487 | 0.04932919 | ZNF548 | 147694 |
| ENSG000002 | 19.3327842 | -0.698608722 | 0.28842322 | -2.4221654 | 0.01542833 | 0.04933634 | NA | NA |
| ENSG000001 | 9.358316 | -0.831436844 | 0.34326996 | -2.4221078 | 0.01543077 | 0.04934027 | RGPD2 | 729857 |
| ENSG000001 | 14.4773288 | -0.745269872 | 0.30770167 | -2.4220534 | 0.01543308 | 0.04934376 | PLIN1 | 5346 |
| ENSG000001 | 45.5776135 | -0.526970742 | 0.21759453 | -2.4218014 | 0.01544379 | 0.0493741 | NPAS2 | 4862 |
| ENSG00000C | 3.21672981 | -0.911140566 | 0.37622884 | -2.4217723 | 0.01544503 | 0.04937416 | MAGEC2 | 51438 |
| ENSG000001 | 60.2180721 | 0.537046653 | 0.22176621 | 2.42167934 | 0.01544898 | 0.0493829 | KHK | 3795 |
| ENSG000002 | 5.42583047 | -0.900047852 | 0.37167028 | -2.4216299 | 0.01545108 | 0.04938572 | NA | NA |
| ENSG000001 | 16.2484435 | -0.753628373 | 0.31122595 | -2.4214831 | 0.01545732 | 0.04939789 | IAPP | 3375 |
| ENSG000002 | 34.2243634 | 0.697047422 | 0.28785668 | 2.4215086 | 0.01545624 | 0.04939789 | NA | NA |
| ENSG000001 | 982.587581 | 0.289916861 | 0.1197308 | 2.42140588 | 0.0154606 | 0.04940449 | YOD1 | 55432 |
| ENSG000001 | 38.1056711 | 0.613045919 | 0.25318928 | 2.42129488 | 0.01546533 | 0.04941568 | NBR2 | 10230 |
| ENSG000001 | 8.79931705 | -0.840868923 | 0.34728488 | -2.4212655 | 0.01546657 | 0.04941578 | TMEM132D | 121256 |
| ENSG000002 | 4.59684386 | -0.91434099 | 0.37763884 | -2.4212049 | 0.01546916 | 0.04942014 | NA | NA |
| ENSG000001 | 248.980811 | 0.339553341 | 0.14024374 | 2.42116574 | 0.01547082 | 0.04942156 | FRMD6 | 122786 |
| ENSG000002 | 15.1017203 | -0.738473321 | 0.3050126 | -2.421124 | 0.0154726 | 0.04942334 | OR2H1 | 26716 |
| ENSG000001 | 13.8256837 | 0.781893801 | 0.32295967 | 2.42102612 | 0.01547677 | 0.04943276 | SPATA25 | 128497 |
| ENSG000002 | 77.9525321 | 0.453400699 | 0.18728597 | 2.42090051 | 0.01548211 | 0.04944595 | NA | NA |
| ENSG000002 | 7.76442652 | -0.870026972 | 0.3594367 | -2.4205291 | 0.01549794 | 0.0494887 | NA | NA |
| ENSG000002 | 5.46303849 | -0.902935664 | 0.37303148 | -2.4205348 | 0.0154977 | 0.0494887 | MIR4289 | 100423015 |
| ENSG000001 | 106125.409 | 0.292545436 | 0.12087003 | 2.42033068 | 0.0155064 | 0.04951181 | COX1 | 4512 |

| | | | | | | | | |
|---|---|---|---|---|---|---|---|---|
| ENSG000001 | 52.3377827 | -0.512458873 | 0.21173804 | -2.4202494 | 0.01550987 | 0.04951558 | IL15 | 3600 |
| ENSG000001 | 1222.53738 | 0.315629358 | 0.13041211 | 2.42024577 | 0.01551002 | 0.04951558 | SNRPE | 6635 |
| ENSG000001 | 65.2659552 | 0.473563412 | 0.19567916 | 2.42010146 | 0.01551618 | 0.04953133 | IL17RC | 84818 |
| ENSG000001 | 1450.00292 | 0.295585105 | 0.12215342 | 2.41978567 | 0.01552966 | 0.04956266 | NKTR | 4820 |
| ENSG000001 | 43.09407 | -0.586380998 | 0.24232279 | -2.4198343 | 0.01552758 | 0.04956266 | Dec-01 | 50514 |
| ENSG000002 | 59.4669697 | -0.470009485 | 0.19423484 | -2.4198001 | 0.01552904 | 0.04956266 | NA | NA |
| ENSG000002 | 7.70727013 | -0.870415253 | 0.35971542 | -2.4197329 | 0.01553191 | 0.04956594 | OSTCP1 | 202459 |
| ENSG000002 | 8.13229504 | -0.848370294 | 0.35061006 | -2.4196975 | 0.01553342 | 0.04956687 | KCNIP2-AS1 | 100289509 |
| ENSG000001 | 31.8660539 | -0.584549199 | 0.24159185 | -2.4195734 | 0.01553873 | 0.04957989 | PCDH12 | 51294 |
| ENSG000002 | 5.44780125 | -0.90122152 | 0.37250619 | -2.4193464 | 0.01554843 | 0.04960694 | NA | NA |
| ENSG000002 | 13.2086814 | -0.772072025 | 0.31914144 | -2.4192159 | 0.01555401 | 0.04962084 | METTL21EP | 121952 |
| ENSG000001 | 1007.83176 | 0.290194433 | 0.11996431 | 2.41900648 | 0.01556296 | 0.04964551 | ERRFI1 | 54206 |
| ENSG000002 | 7.62793273 | -0.888924119 | 0.36748451 | -2.4189431 | 0.01556567 | 0.04965025 | LOC101928514 | 101928514 |
| ENSG000002 | 13.8514471 | -0.766926259 | 0.31705434 | -2.4189111 | 0.01556704 | 0.04965072 | CLRN1-AS1 | 116933 |
| ENSG000002 | 49.4462878 | 0.504526021 | 0.20858595 | 2.41879195 | 0.01557214 | 0.04966308 | NA | NA |
| ENSG000002 | 3.2482679 | -0.910239752 | 0.3763392 | -2.4186685 | 0.01557743 | 0.04967603 | NA | NA |
| ENSG000001 | 86.0406372 | 0.436160401 | 0.18033454 | 2.41861825 | 0.01557958 | 0.04967898 | RPS6KL1 | 83694 |
| ENSG000001 | 4107.9444 | 0.270290255 | 0.11175694 | 2.41855448 | 0.01558231 | 0.04968378 | KDM5B | 10765 |
| ENSG000002 | 67.7109448 | 0.465832971 | 0.19262215 | 2.41837703 | 0.01558991 | 0.04970411 | NA | NA |
| ENSG000001 | 10.0861826 | -0.830784894 | 0.34354193 | -2.4182926 | 0.01559353 | 0.04971174 | MAB21L2 | 10586 |
| ENSG000002 | 4.21607892 | -0.914760543 | 0.37828574 | -2.4181735 | 0.01559864 | 0.04972411 | NA | NA |
| ENSG000002 | 9.07642877 | -0.831612988 | 0.34390698 | -2.4181335 | 0.01560035 | 0.04972567 | NA | NA |
| ENSG000002 | 50.3549116 | 0.537757371 | 0.22238866 | 2.41809708 | 0.01560191 | 0.04972673 | RNU6-8 | 101954278 |
| ENSG000001 | 4.75595558 | -0.908358395 | 0.3756718 | -2.4179574 | 0.0156079 | 0.04974192 | SPATC1 | 375686 |
| ENSG000002 | 27.3465178 | -0.643487068 | 0.26613191 | -2.4179252 | 0.01560928 | 0.0497424 | NA | NA |
| ENSG000001 | 352.20059 | 0.317059114 | 0.13113114 | 2.41787804 | 0.01561131 | 0.04974494 | PM20D2 | 135293 |
| ENSG000001 | 4.03555795 | -0.91441544 | 0.37820423 | -2.4177821 | 0.01561542 | 0.04975415 | NA | NA |
| ENSG000001 | 44.054584 | -0.530730027 | 0.21952214 | -2.4176606 | 0.01562064 | 0.04976686 | KCNH7 | 90134 |
| ENSG000001 | 11.6331299 | -0.811497973 | 0.33566987 | -2.4175479 | 0.01562548 | 0.04977837 | CXCL11 | 6373 |

| | | | | | | | | |
|---|---|---|---|---|---|---|---|---|
| ENSG000001 | 23.5540282 | -0.661971235 | 0.27384124 | -2.4173541 | 0.0156338 | 0.04980096 | PLEKHG7 | 440107 |
| ENSG000002 | 19.1797762 | -0.707333806 | 0.29262399 | -2.4172106 | 0.01563996 | 0.04981668 | NA | NA |
| ENSG000001 | 23.6937811 | -0.638355739 | 0.26409896 | -2.4171081 | 0.01564437 | 0.0498268 | SBSPON | 157869 |
| ENSG000002 | 10.284103 | -0.833075606 | 0.34469426 | -2.4168538 | 0.01565531 | 0.04985772 | NA | NA |
| ENSG000001 | 106.894253 | 0.441383547 | 0.18263846 | 2.41670649 | 0.01566164 | 0.04987397 | CDAN1 | 146059 |
| ENSG000002 | 3.27899017 | -0.906343696 | 0.37505193 | -2.4165819 | 0.015667 | 0.04988713 | NA | NA |
| ENSG000002 | 2.77540962 | -0.903507186 | 0.37388801 | -2.4165182 | 0.01566974 | 0.04989194 | NA | NA |
| ENSG000001 | 12.032826 | -0.785593689 | 0.32511277 | -2.4163729 | 0.015676 | 0.04990793 | ESAM | 90952 |
| ENSG000002 | 9.88747333 | -0.837327924 | 0.34654175 | -2.4162397 | 0.01568174 | 0.04992228 | NA | NA |
| ENSG000002 | 2.77925158 | -0.902826298 | 0.37365438 | -2.4162069 | 0.01568315 | 0.04992286 | IGHV3OR16-6 | 647187 |
| ENSG000001 | 12.2569281 | -0.781922993 | 0.32362061 | -2.4161718 | 0.01568466 | 0.04992375 | CDH22 | 64405 |
| ENSG000001 | 114.666622 | 0.40673134 | 0.16834675 | 2.41603325 | 0.01569063 | 0.04993883 | EPHX3 | 79852 |
| ENSG000002 | 10.7263859 | -0.817802791 | 0.33849741 | -2.4159795 | 0.01569294 | 0.04994229 | LINC00854 | 100874261 |
| ENSG000001 | 53.628336 | -0.509945845 | 0.21108174 | -2.4158691 | 0.0156977 | 0.04995351 | SH3RF2 | 153769 |
| ENSG000001 | 10.7362341 | -0.840162845 | 0.34777965 | -2.4157907 | 0.01570108 | 0.04995857 | KRT39 | 390792 |
| ENSG000002 | 129.416686 | 0.40997872 | 0.16970898 | 2.41577509 | 0.01570176 | 0.04995857 | NA | NA |
| ENSG000001 | 320.602987 | 0.358263199 | 0.14830374 | 2.41573947 | 0.01570329 | 0.04995953 | NAGK | 55577 |
| ENSG000001 | 99.5740001 | -0.400143541 | 0.1656428 | -2.4157014 | 0.01570493 | 0.04996043 | GABRG2 | 2566 |
| ENSG000001 | 6.43427167 | -0.886549681 | 0.36699862 | -2.4156758 | 0.01570604 | 0.04996043 | ODF3 | 113746 |
| ENSG000001 | 35.0569933 | -0.568558693 | 0.23538535 | -2.4154379 | 0.0157163 | 0.04998915 | CDH18 | 1016 |
| ENSG000002 | 17.3119422 | -0.749863095 | 0.31045129 | -2.415397 | 0.01571807 | 0.04999086 | FZD10-AS1 | 440119 |
| ENSG000002 | 4.19841179 | -0.913488432 | 0.37821559 | -2.4152585 | 0.01572405 | 0.05000408 | LOC105374698 | 105374698 |
| ENSG000002 | 38.5719214 | -0.565322887 | 0.23406455 | -2.4152435 | 0.01572469 | 0.05000408 | NA | NA |
| ENSG000001 | 46.4482704 | -0.552271987 | 0.22866661 | -2.4151842 | 0.01572725 | 0.0500083 | TMEFF2 | 23671 |
| ENSG000001 | 6.00215687 | -0.88702877 | 0.36727776 | -2.4151442 | 0.01572898 | 0.05000987 | C14orf177 | 283598 |
| ENSG000001 | 18.2753337 | -0.713115348 | 0.29527186 | -2.4151145 | 0.01573026 | 0.05001003 | UBASH3A | 53347 |
| ENSG000002 | 755.861065 | 0.616828108 | 0.25542595 | 2.41489989 | 0.01573954 | 0.05003558 | TAF15 | 8148 |
| ENSG000001 | 3.51472653 | -0.912710981 | 0.37798507 | -2.4146747 | 0.01574927 | 0.0500626 | NA | NA |
| ENSG00000C | 682.307378 | 0.309173877 | 0.12804705 | 2.4145334 | 0.01575538 | 0.0500781 | NEO1 | 4756 |

| | | | | | | | | |
|---|---|---|---|---|---|---|---|---|
| ENSG000001 | 390.860779 | 0.308795569 | 0.12789775 | 2.41439401 | 0.01576141 | 0.05008941 | GAS2L3 | 283431 |
| ENSG000002 | 11.6783911 | -0.793712442 | 0.32874175 | -2.414395 | 0.01576136 | 0.05008941 | NA | NA |
| ENSG000002 | 36.1203174 | -0.564696413 | 0.23390002 | -2.414264 | 0.01576703 | 0.05010336 | NA | NA |
| ENSG000002 | 2.79920302 | -0.902169159 | 0.37369302 | -2.4141986 | 0.01576987 | 0.05010843 | NA | NA |
| ENSG000001 | 60.7231834 | -0.509418498 | 0.21102483 | -2.4140216 | 0.01577753 | 0.05012884 | PKD1L1 | 168507 |
| ENSG000002 | 35.9694694 | 0.578856372 | 0.23982606 | 2.41365084 | 0.01579359 | 0.05017595 | CAPN10-AS1 | 101752400 |
| ENSG000001 | 418.19853 | 0.307320859 | 0.12733173 | 2.41354493 | 0.01579818 | 0.0501866 | NDRG3 | 57446 |
| ENSG000002 | 2.43753258 | -0.888759836 | 0.36826315 | -2.4133825 | 0.01580522 | 0.05020505 | NA | NA |
| ENSG000002 | 6.721484 | -0.881612341 | 0.36531423 | -2.4132987 | 0.01580886 | 0.05021266 | HERC6 | 55008 |
| ENSG000001 | 5.14201337 | -0.903451552 | 0.37440263 | -2.4130481 | 0.01581973 | 0.05023625 | NA | NA |
| ENSG000002 | 2.39569434 | -0.889499893 | 0.36861949 | -2.4130572 | 0.01581934 | 0.05023625 | NA | NA |
| ENSG000002 | 31.5576575 | 0.595803546 | 0.24690974 | 2.41304188 | 0.01582 | 0.05023625 | NA | NA |
| ENSG000001 | 161.251225 | 0.410518371 | 0.17014584 | 2.41274413 | 0.01583293 | 0.05027328 | OSGEPL1 | 64172 |
| ENSG000001 | 94.6989828 | 0.459302801 | 0.19036752 | 2.41271622 | 0.01583414 | 0.05027328 | MIB2 | 142678 |
| ENSG00000C | 12.081294 | -0.805574814 | 0.33389368 | -2.4126687 | 0.01583621 | 0.0502759 | CRISP3 | 10321 |
| ENSG000002 | 5.21367102 | -0.90428123 | 0.37486169 | -2.4123063 | 0.01585196 | 0.05032196 | NA | NA |
| ENSG000001 | 399.52236 | 0.295616563 | 0.12255633 | 2.41208732 | 0.01586148 | 0.05034826 | PON2 | 5445 |
| ENSG000002 | 49.6105525 | 0.534607661 | 0.22164236 | 2.4120284 | 0.01586405 | 0.05035245 | NA | NA |
| ENSG000002 | 6.98869897 | -0.871935637 | 0.36152132 | -2.4118512 | 0.01587176 | 0.05037298 | LINC01300 | 731779 |
| ENSG000001 | 50.9245079 | 0.56266866 | 0.23329834 | 2.41179886 | 0.01587404 | 0.05037627 | CAMK2N2 | 94032 |
| ENSG000001 | 32.3799732 | 0.572591479 | 0.23744018 | 2.41151885 | 0.01588623 | 0.05041103 | HYAL3 | 8372 |
| ENSG000001 | 6.07119849 | -0.888105363 | 0.36830757 | -2.4113145 | 0.01589514 | 0.05043534 | OR8S1 | 341568 |
| ENSG000001 | 14.3447316 | -0.761425796 | 0.31577727 | -2.4112749 | 0.01589686 | 0.05043687 | ELF5 | 2001 |
| ENSG000002 | 48.8419611 | -0.550181913 | 0.22818291 | -2.4111442 | 0.01590256 | 0.050451 | COL6A4P1 | 344875 |
| ENSG000001 | 5.56312915 | -0.900998122 | 0.37370407 | -2.4109936 | 0.01590913 | 0.05046789 | TUBA8 | 51807 |
| ENSG00000C | 66.7537808 | 0.462600948 | 0.19189099 | 2.41074868 | 0.01591981 | 0.05049389 | NA | NA |
| ENSG000002 | 9.40067183 | -0.827698326 | 0.34333634 | -2.4107507 | 0.01591973 | 0.05049389 | USP17L18 | 100287364 |
| ENSG000001 | 10.7549709 | -0.796678357 | 0.33053464 | -2.4102719 | 0.01594063 | 0.05055597 | NA | NA |
| ENSG000001 | 217.884909 | 0.469090012 | 0.19463414 | 2.41011171 | 0.01594764 | 0.05057423 | DCHS1 | 8642 |

| | | | | | | | | |
|---|---|---|---|---|---|---|---|---|
| ENSG000002 | 6.0562888 | -0.900045886 | 0.37345625 | -2.4100437 | 0.01595061 | 0.05057969 | NA | NA |
| ENSG000002 | 2.79661073 | -0.898520264 | 0.37282875 | -2.410008 | 0.01595217 | 0.0505807 | NA | NA |
| ENSG000001 | 2.39590286 | -0.887066954 | 0.36810402 | -2.4098269 | 0.01596009 | 0.05060185 | MAL | 4118 |
| ENSG000001 | 593.309159 | 0.346673121 | 0.14386256 | 2.40975222 | 0.01596336 | 0.05060429 | BCL9 | 607 |
| ENSG000002 | 24.5938539 | -0.653409849 | 0.27115122 | -2.4097618 | 0.01596294 | 0.05060429 | SMIM2-AS1 | 101929212 |
| ENSG000001 | 620.777955 | 0.314188689 | 0.13038676 | 2.40966712 | 0.01596708 | 0.05061214 | SLC25A44 | 9673 |
| ENSG000001 | 12.9194805 | -0.776305956 | 0.3221751 | -2.4095778 | 0.01597099 | 0.05061662 | ADGRA1 | 84435 |
| ENSG000002 | 9.4478474 | -0.836478079 | 0.34714379 | -2.4096012 | 0.01596997 | 0.05061662 | LOC101929544 | 101929544 |
| ENSG000001 | 1157.39378 | 0.405589769 | 0.1683365 | 2.40939891 | 0.01597882 | 0.05063748 | IGF1R | 3480 |
| ENSG000001 | 150.543074 | 0.752466731 | 0.31235188 | 2.40903537 | 0.01599475 | 0.05068399 | MSRB1 | 51734 |
| ENSG000001 | 13.742741 | -0.761953577 | 0.31630278 | -2.4089373 | 0.01599905 | 0.05069365 | RD3 | 343035 |
| ENSG000002 | 15.5387113 | -0.724138698 | 0.30065258 | -2.4085564 | 0.01601575 | 0.05074261 | NA | NA |
| ENSG000001 | 353.950768 | 0.308624417 | 0.12814253 | 2.40844633 | 0.01602058 | 0.05075395 | SPRYD4 | 283377 |
| ENSG000002 | 12.7683919 | -0.788729276 | 0.32750432 | -2.4083019 | 0.01602692 | 0.05077007 | PCAT29 | 104472713 |
| ENSG00000C | 81.8882364 | 0.434474069 | 0.18042098 | 2.40811274 | 0.01603523 | 0.0507774 | PYGM | 5837 |
| ENSG000001 | 294.531799 | 0.685174034 | 0.28452811 | 2.40810669 | 0.01603549 | 0.0507774 | NA | NA |
| ENSG000001 | 17.7515965 | -0.698791979 | 0.29017875 | -2.4081432 | 0.01603389 | 0.0507774 | NA | NA |
| ENSG000002 | 21.4300434 | -0.674285472 | 0.27999341 | -2.4082191 | 0.01603056 | 0.0507774 | CYP2B7P | 1556 |
| ENSG000002 | 8.61887483 | -0.853205024 | 0.35429699 | -2.4081633 | 0.01603301 | 0.0507774 | LOC100129935 | 100129935 |
| ENSG000001 | 95.7724588 | 0.407814441 | 0.16935982 | 2.40797637 | 0.01604122 | 0.05079156 | MINOS1 | 440574 |
| ENSG000002 | 4.20706942 | -0.911061094 | 0.37837258 | -2.4078412 | 0.01604716 | 0.0508064 | NA | NA |
| ENSG000002 | 10.5399545 | -0.81300828 | 0.33765962 | -2.4077747 | 0.01605008 | 0.05081169 | MGC16275 | 85001 |
| ENSG000001 | 42.1933426 | -0.534548426 | 0.2220278 | -2.4075743 | 0.01605889 | 0.05083562 | OPCML | 4978 |
| ENSG000001 | 2.87476287 | -0.897317492 | 0.37272949 | -2.4074228 | 0.01606556 | 0.0508448 | NA | NA |
| ENSG000002 | 7.78348286 | -0.868413916 | 0.36071532 | -2.4074772 | 0.01606317 | 0.0508448 | TCEAL6 | 158931 |
| ENSG000002 | 6.9888216 | 0.874173155 | 0.36311482 | 2.40742899 | 0.01606529 | 0.0508448 | ADM5 | 199800 |
| ENSG000002 | 7.14240445 | -0.878944204 | 0.36510718 | -2.4073594 | 0.01606835 | 0.05084966 | NA | NA |
| ENSG000002 | 10.8796781 | -0.802695492 | 0.3334401 | -2.4073154 | 0.01607028 | 0.05085182 | NA | NA |
| ENSG00000C | 101.954487 | 0.465554565 | 0.19341008 | 2.40708536 | 0.01608041 | 0.0508799 | H6PD | 9563 |

| | | | | | | | | |
|---|---|---|---|---|---|---|---|---|
| ENSG000002 | 7.38000778 | -0.870304745 | 0.3615791 | -2.4069553 | 0.01608614 | 0.05089404 | NA | NA |
| ENSG000001 | 544.107611 | 0.300358285 | 0.12479067 | 2.40689694 | 0.01608871 | 0.05089776 | DTWD1 | 56986 |
| ENSG000002 | 1633.70485 | 0.322575444 | 0.13402347 | 2.40685778 | 0.01609043 | 0.05089776 | NA | NA |
| ENSG000002 | 17.0756233 | -0.729188335 | 0.30296984 | -2.4068017 | 0.0160929 | 0.05089776 | NA | NA |
| ENSG000002 | 4.81022183 | -0.904940792 | 0.37600176 | -2.4067462 | 0.01609535 | 0.05089776 | NA | NA |
| ENSG000002 | 8.74570199 | -0.853033985 | 0.35443703 | -2.4067293 | 0.0160961 | 0.05089776 | NA | NA |
| ENSG000002 | 3.848702 | 0.910613981 | 0.37835026 | 2.40680151 | 0.01609291 | 0.05089776 | NA | NA |
| ENSG000002 | 7.54013773 | -0.873706143 | 0.36302117 | -2.4067636 | 0.01609458 | 0.05089776 | NA | NA |
| ENSG000002 | 4.17134425 | -0.910481743 | 0.37831194 | -2.4066958 | 0.01609757 | 0.05089846 | NA | NA |
| ENSG000002 | 2.81863796 | -0.899136486 | 0.37361454 | -2.4065886 | 0.0161023 | 0.05090943 | NA | NA |
| ENSG000001 | 15.7345845 | -0.737994624 | 0.30667234 | -2.4064597 | 0.01610798 | 0.0509155 | SERPINI2 | 5276 |
| ENSG000001 | 25.336429 | -0.622083861 | 0.25850084 | -2.4065061 | 0.01610593 | 0.0509155 | C8orf46 | 254778 |
| ENSG000002 | 8.74737642 | -0.855736251 | 0.35559968 | -2.4064596 | 0.01610799 | 0.0509155 | LOC399715 | 399715 |
| ENSG000002 | 17.3399753 | -0.711431731 | 0.29564505 | -2.4063712 | 0.01611188 | 0.05092385 | TMEM72-AS1 | 220980 |
| ENSG000002 | 3396.79801 | 0.289607133 | 0.12035945 | 2.40618524 | 0.01612009 | 0.05094581 | NA | NA |
| ENSG000001 | 82.3289545 | 0.430301547 | 0.17883671 | 2.40611416 | 0.01612323 | 0.05094778 | LOC728673 | 728673 |
| ENSG000002 | 31.655824 | -0.600306695 | 0.24949185 | -2.4061174 | 0.01612308 | 0.05094778 | PCDHGA6 | 56109 |
| ENSG000001 | 1031.20954 | 0.318800065 | 0.13250013 | 2.40603584 | 0.01612668 | 0.05095077 | SP4 | 6671 |
| ENSG000001 | 4.19924095 | -0.910275343 | 0.37832988 | -2.4060361 | 0.01612667 | 0.05095077 | NA | NA |
| ENSG000001 | 68.2914234 | 0.451586035 | 0.18770906 | 2.40577645 | 0.01613814 | 0.05098298 | TAC3 | 6866 |
| ENSG000002 | 3.95390122 | -0.91060937 | 0.37852674 | -2.4056672 | 0.01614296 | 0.05099426 | NA | NA |
| ENSG000002 | 2.84656636 | -0.896911139 | 0.37284932 | -2.4055592 | 0.01614773 | 0.05100535 | NA | NA |
| ENSG000001 | 197.741355 | 0.338530732 | 0.14073136 | 2.40551023 | 0.0161499 | 0.05100822 | MTFR1L | 56181 |
| ENSG000002 | 166.640767 | 0.341813142 | 0.14210022 | 2.40543715 | 0.01615313 | 0.05101445 | NA | NA |
| ENSG000001 | 2729.17082 | 0.5799676 | 0.24112534 | 2.40525359 | 0.01616125 | 0.05103611 | NES | 10763 |
| ENSG000002 | 5.50460839 | -0.898550848 | 0.3735938 | -2.4051546 | 0.01616563 | 0.05104597 | NA | NA |
| ENSG000002 | 2.39951668 | -0.88481743 | 0.36789624 | -2.4050733 | 0.01616922 | 0.05105335 | NA | NA |
| ENSG000002 | 2.49476738 | -0.883538447 | 0.36737259 | -2.40502 | 0.01617158 | 0.05105681 | NA | NA |
| ENSG000002 | 10.6278035 | -0.809919766 | 0.33678426 | -2.4048623 | 0.01617856 | 0.05106693 | LOC100506321 | 100506321 |

| | | | | | | | | |
|---|---|---|---|---|---|---|---|---|
| ENSG000002 | 4.25115222 | -0.910376185 | 0.37855284 | -2.4048854 | 0.01617754 | 0.05106693 | NA | NA |
| ENSG000002 | 11.3402207 | -0.811049764 | 0.3372472 | -2.4049118 | 0.01617637 | 0.05106693 | NA | NA |
| ENSG000001 | 225.378251 | 0.334129078 | 0.13894352 | 2.40478339 | 0.01618205 | 0.05107001 | C9orf156 | 51531 |
| ENSG000002 | 3.1794787 | -0.904578697 | 0.37616253 | -2.4047549 | 0.01618331 | 0.05107001 | NA | NA |
| ENSG000002 | 8.13857752 | -0.844448762 | 0.35115654 | -2.4047645 | 0.01618289 | 0.05107001 | NA | NA |
| ENSG000001 | 741.358221 | 0.300297802 | 0.12488207 | 2.40465114 | 0.01618791 | 0.05108054 | COIL | 8161 |
| ENSG000001 | 12.8205849 | -0.795826123 | 0.33102852 | -2.4041014 | 0.01621228 | 0.05115344 | SIRPB1 | 10326 |
| ENSG00000C | 395.003729 | 0.280623949 | 0.11673364 | 2.40396814 | 0.01621819 | 0.05116413 | ATP6V0A1 | 535 |
| ENSG000001 | 386.901823 | 0.278350297 | 0.11578749 | 2.4039755 | 0.01621786 | 0.05116413 | DUSP11 | 8446 |
| ENSG000002 | 22.4722372 | 0.671006734 | 0.2791347 | 2.40388148 | 0.01622203 | 0.05117228 | NA | NA |
| ENSG000002 | 5.81445167 | -0.891628886 | 0.37091974 | -2.4038324 | 0.01622421 | 0.05117517 | C2-AS1 | 102060414 |
| ENSG00000C | 295.549257 | 0.314508642 | 0.13083964 | 2.4037718 | 0.0162269 | 0.05117967 | HSF2 | 3298 |
| ENSG000001 | 862.867039 | 0.293832598 | 0.12225648 | 2.40341131 | 0.01624291 | 0.05122617 | ZNF85 | 7639 |
| ENSG000002 | 2.49435126 | -0.87790638 | 0.36531048 | -2.4031787 | 0.01625324 | 0.05125478 | OR2AE1 | 81392 |
| ENSG000002 | 5.42061803 | -0.894237218 | 0.37213464 | -2.4029937 | 0.01626147 | 0.05127672 | NA | NA |
| ENSG000001 | 29.048855 | -0.595707349 | 0.24792156 | -2.4028057 | 0.01626983 | 0.05129911 | ST8SIA4 | 7903 |
| ENSG000002 | 10.9584095 | -0.806330241 | 0.33558616 | -2.4027518 | 0.01627223 | 0.05130268 | NA | NA |
| ENSG000002 | 12.2207546 | -0.774998872 | 0.32255776 | -2.4026669 | 0.016276 | 0.0513106 | NA | NA |
| ENSG000001 | 36.4779739 | -0.610394683 | 0.2540562 | -2.4025971 | 0.01627911 | 0.05131142 | SLIT1 | 6585 |
| ENSG000002 | 7.04700216 | -0.873065089 | 0.36338118 | -2.4026151 | 0.01627831 | 0.05131142 | PRAMEF14 | 729528 |
| ENSG000002 | 269.760661 | 0.363342189 | 0.15123027 | 2.4025758 | 0.01628006 | 0.05131142 | PRR3 | 80742 |
| ENSG00000C | 451.856644 | 0.322302365 | 0.13417647 | 2.40207817 | 0.01630222 | 0.05137728 | WDR7 | 23335 |
| ENSG000001 | 148.763477 | 0.392218874 | 0.16329299 | 2.40193335 | 0.01630868 | 0.05138565 | RP2 | 6102 |
| ENSG000001 | 233.426539 | 0.365555821 | 0.15219144 | 2.40194734 | 0.01630806 | 0.05138565 | PIK3R3 | 8503 |
| ENSG000002 | 14.9057067 | -0.765582306 | 0.31873041 | -2.4019745 | 0.01630685 | 0.05138565 | CALR3 | 125972 |
| ENSG000002 | 8.91126428 | -0.843636102 | 0.35124615 | -2.4018373 | 0.01631296 | 0.05139514 | NA | NA |
| ENSG000001 | 21.8786342 | -0.678385006 | 0.28248593 | -2.4014824 | 0.01632879 | 0.05142505 | KCNJ5 | 3762 |
| ENSG000001 | 86.4910384 | 0.439632806 | 0.1830672 | 2.40148319 | 0.01632876 | 0.05142505 | ACAP3 | 116983 |
| ENSG000001 | 240.686317 | 0.349367305 | 0.14547856 | 2.40150366 | 0.01632785 | 0.05142505 | SAP30L | 79685 |

| | | | | | | | | |
|---|---|---|---|---|---|---|---|---|
| ENSG000002 | 2.40525727 | -0.882228246 | 0.36735732 | -2.4015534 | 0.01632562 | 0.05142505 | NA | NA |
| ENSG000002 | 14.1032646 | -0.778630202 | 0.32421824 | -2.401562 | 0.01632524 | 0.05142505 | CSMD2-AS1 | 402779 |
| ENSG00000C | 1708.30297 | 0.801928814 | 0.33395003 | 2.40134374 | 0.01633498 | 0.05143655 | FUS | 2521 |
| ENSG000001 | 15.9887905 | -0.761683932 | 0.317187 | -2.4013718 | 0.01633373 | 0.05143655 | ZNRF3-AS1 | 100874123 |
| ENSG000002 | 2.41869049 | -0.882717296 | 0.36760073 | -2.4012936 | 0.01633722 | 0.0514396 | NA | NA |
| ENSG000002 | 10.8473777 | -0.803770102 | 0.33475427 | -2.401075 | 0.01634699 | 0.05146635 | NA | NA |
| ENSG000001 | 331.244417 | 0.3501147 | 0.14582179 | 2.40097659 | 0.01635138 | 0.0514762 | DENND6A | 201627 |
| ENSG000001 | 979.811845 | 0.295191285 | 0.12295048 | 2.40089576 | 0.01635499 | 0.05148357 | WDFY2 | 115825 |
| ENSG000001 | 53.5802724 | 0.548571123 | 0.22849369 | 2.40081517 | 0.0163586 | 0.05149092 | FRAT1 | 10023 |
| ENSG000001 | 9.19412421 | -0.836480408 | 0.34842193 | -2.4007686 | 0.01636068 | 0.05149239 | KRT79 | 338785 |
| ENSG000002 | 69.0675418 | 0.480968322 | 0.20034104 | 2.40074786 | 0.01636161 | 0.05149239 | NA | NA |
| ENSG000002 | 10.7462937 | -0.794715036 | 0.33103707 | -2.4006829 | 0.01636451 | 0.05149753 | NA | NA |
| ENSG000002 | 17.4922192 | -0.7424629 | 0.30927923 | -2.4006232 | 0.01636718 | 0.05150194 | SPDYE3 | 441272 |
| ENSG000002 | 2.38861572 | -0.883974344 | 0.36823555 | -2.4005676 | 0.01636967 | 0.05150577 | NA | NA |
| ENSG000001 | 1813.02653 | 0.613820175 | 0.25571746 | 2.40038429 | 0.01637787 | 0.05152568 | HK2 | 3099 |
| ENSG000001 | 702.14936 | 0.292391821 | 0.12181118 | 2.40036936 | 0.01637854 | 0.05152568 | NAA16 | 79612 |
| ENSG000002 | 4.62270154 | -0.907733696 | 0.37823106 | -2.3999449 | 0.01639754 | 0.05157036 | NA | NA |
| ENSG000002 | 17.8603364 | -0.702620138 | 0.29275694 | -2.4000119 | 0.01639454 | 0.05157036 | MTND6P8 | 106480726 |
| ENSG000002 | 5.22228556 | -0.902079302 | 0.375876 | -2.3999385 | 0.01639783 | 0.05157036 | DKFZP434H168 | 26077 |
| ENSG000002 | 9.05057295 | -0.851320115 | 0.35472434 | -2.3999484 | 0.01639738 | 0.05157036 | NA | NA |
| ENSG000001 | 26.5936835 | -0.628790842 | 0.26203821 | -2.3996151 | 0.01641232 | 0.05161193 | ASIC1 | 41 |
| ENSG000002 | 28.8098465 | 0.601758375 | 0.25079985 | 2.39935703 | 0.01642389 | 0.05164433 | NA | NA |
| ENSG000001 | 204.752759 | 0.333534191 | 0.13901548 | 2.39925942 | 0.01642827 | 0.05165409 | SRP19 | 6728 |
| ENSG000002 | 8.72057228 | -0.835762505 | 0.34835945 | -2.3991383 | 0.01643371 | 0.05166717 | NA | NA |
| ENSG000001 | 12.9635107 | -0.776168183 | 0.32355934 | -2.3988434 | 0.01644695 | 0.0517048 | FBXO39 | 162517 |
| ENSG000001 | 22.2353529 | -0.65629177 | 0.27368587 | -2.3979746 | 0.01648601 | 0.05182357 | GBP1 | 2633 |
| ENSG000001 | 218.474537 | 0.409159015 | 0.17063618 | 2.39784446 | 0.01649187 | 0.05183797 | FXN | 2395 |
| ENSG000002 | 5.0727061 | -0.896304845 | 0.37383955 | -2.3975656 | 0.01650442 | 0.05187342 | NA | NA |
| ENSG000001 | 1016.35268 | 0.295209674 | 0.12313196 | 2.39750647 | 0.01650709 | 0.05187778 | RFWD2 | 64326 |

| | | | | | | | | |
|---|---|---|---|---|---|---|---|---|
| ENSG000002 | 4.934533 | -0.904056471 | 0.37708901 | -2.3974617 | 0.01650911 | 0.05188009 | NA | NA |
| ENSG000001 | 127.396979 | 0.396717721 | 0.16547622 | 2.39743034 | 0.01651052 | 0.05188052 | GSTM4 | 2948 |
| ENSG000001 | 38.9331106 | -0.546621367 | 0.22800768 | -2.3973814 | 0.01651272 | 0.05188343 | SYTL4 | 94121 |
| ENSG000001 | 285.580741 | 0.337890026 | 0.14094519 | 2.39731512 | 0.01651571 | 0.0518888 | CHST6 | 4166 |
| ENSG000002 | 4.86353748 | 0.90191555 | 0.37623594 | 2.39720731 | 0.01652057 | 0.05190005 | MIR7111 | 102465668 |
| ENSG000002 | 14.8786715 | -0.765798914 | 0.31946555 | -2.3971252 | 0.01652428 | 0.05190766 | NA | NA |
| ENSG000002 | 5.67709261 | -0.885753136 | 0.36952716 | -2.3969906 | 0.01653035 | 0.0519227 | NA | NA |
| ENSG000001 | 35.9041028 | 0.597684309 | 0.24935089 | 2.39696081 | 0.01653169 | 0.05192291 | CD72 | 971 |
| ENSG000002 | 24.4254011 | -0.636959014 | 0.26574256 | -2.3969025 | 0.01653432 | 0.05192714 | SLC5A8 | 160728 |
| ENSG000002 | 11.4662329 | -0.81260341 | 0.33903414 | -2.3968188 | 0.0165381 | 0.05193096 | NA | NA |
| ENSG000002 | 3.63074302 | -0.904976254 | 0.37756981 | -2.3968449 | 0.01653692 | 0.05193096 | SPDYE6 | 729597 |
| ENSG000001 | 310.786587 | 0.401303671 | 0.1674383 | 2.39672569 | 0.0165423 | 0.05194014 | PTPRJ | 5795 |
| ENSG000001 | 1183.72423 | 0.282114851 | 0.11772632 | 2.39636172 | 0.01655874 | 0.05198514 | KIF13A | 63971 |
| ENSG000002 | 852.856409 | 0.295028411 | 0.12311566 | 2.39635155 | 0.0165592 | 0.05198514 | ZNF595 | 152687 |
| ENSG000001 | 24.485837 | -0.654837361 | 0.27326821 | -2.3963173 | 0.01656075 | 0.05198597 | HRG | 3273 |
| ENSG000002 | 12.1453159 | -0.789671416 | 0.32958384 | -2.3959652 | 0.01657666 | 0.05203191 | TAS2R13 | 50838 |
| ENSG000002 | 3.14319259 | -0.901244065 | 0.37616174 | -2.3958951 | 0.01657983 | 0.05203784 | NA | NA |
| ENSG000001 | 620.379538 | 0.368121816 | 0.15365535 | 2.39576302 | 0.01658581 | 0.05205157 | ZNF423 | 23090 |
| ENSG000001 | 103.373167 | 0.425148946 | 0.17746027 | 2.39574162 | 0.01658678 | 0.05205157 | ZNF554 | 115196 |
| ENSG000002 | 10.9647193 | -0.804474082 | 0.33581216 | -2.3956073 | 0.01659286 | 0.05206662 | LINC00645 | 100505967 |
| ENSG000002 | 6.07575357 | -0.882832621 | 0.36852673 | -2.3955729 | 0.01659442 | 0.05206748 | NA | NA |
| ENSG000002 | 4.4128991 | -0.90296705 | 0.37697182 | -2.3953171 | 0.016606 | 0.05209979 | NA | NA |
| ENSG000001 | 9109.66428 | 0.295363726 | 0.12331048 | 2.39528494 | 0.01660745 | 0.05210033 | EPRS | 2058 |
| ENSG000001 | 683.01315 | 0.644787499 | 0.26919781 | 2.39521822 | 0.01661048 | 0.05210578 | INCENP | 3619 |
| ENSG000002 | 24.6808559 | -0.642840221 | 0.26840416 | -2.3950456 | 0.0166183 | 0.05212628 | NA | NA |
| ENSG000001 | 206.514123 | 0.334401461 | 0.1396262 | 2.39497653 | 0.01662143 | 0.05213207 | ZNF391 | 346157 |
| ENSG000001 | 1155.47485 | 0.388169748 | 0.16209026 | 2.39477525 | 0.01663056 | 0.05215666 | PEA15 | 8682 |
| ENSG00000C | 1212.92371 | 0.294079415 | 0.12280521 | 2.39468189 | 0.01663479 | 0.05216591 | FNDC3B | 64778 |
| ENSG00000C | 9.63852843 | -0.843154891 | 0.35210629 | -2.3946033 | 0.01663836 | 0.0521709 | WISP2 | 8839 |

| | | | | | | | | |
|---|---|---|---|---|---|---|---|---|
| ENSG000002 | 16.1710081 | -0.741080362 | 0.30948109 | -2.3945901 | 0.01663895 | 0.0521709 | NA | NA |
| ENSG000000 | 22.7408912 | -0.650144449 | 0.27153623 | -2.3943194 | 0.01665124 | 0.05220539 | TNFRSF9 | 3604 |
| ENSG000002 | 15.2652461 | -0.730486452 | 0.30510071 | -2.394247 | 0.01665453 | 0.05221166 | NA | NA |
| ENSG000002 | 7.90190338 | -0.84934057 | 0.35475902 | -2.3941338 | 0.01665967 | 0.05222374 | NA | NA |
| ENSG000001 | 229.691901 | 0.332358155 | 0.1388262 | 2.39405922 | 0.01666306 | 0.05223032 | PTS | 5805 |
| ENSG000002 | 3.22100394 | -0.899378356 | 0.37568813 | -2.3939494 | 0.01666805 | 0.05224193 | NA | NA |
| ENSG000002 | 10.4333008 | -0.800705123 | 0.33448541 | -2.3938417 | 0.01667294 | 0.05225323 | NA | NA |
| ENSG000001 | 5.43551073 | -0.892192763 | 0.37273124 | -2.3936624 | 0.01668109 | 0.05227252 | CXCR1 | 3577 |
| ENSG000002 | 5.42915614 | -0.892469907 | 0.37284902 | -2.3936496 | 0.01668168 | 0.05227252 | NA | NA |
| ENSG000002 | 2.86881321 | -0.89125844 | 0.37239709 | -2.3933013 | 0.01669752 | 0.05231813 | NA | NA |
| ENSG000002 | 15.099136 | -0.757338229 | 0.31645904 | -2.3931635 | 0.01670379 | 0.05233374 | NA | NA |
| ENSG000002 | 5.17627109 | -0.897164072 | 0.37490046 | -2.3930727 | 0.01670793 | 0.0523386 | NA | NA |
| ENSG000002 | 20.0704249 | -0.677357391 | 0.28304846 | -2.3930792 | 0.01670763 | 0.0523386 | IL12A-AS1 | 101928376 |
| ENSG000002 | 6.9117712 | -0.864323123 | 0.36119323 | -2.392966 | 0.01671279 | 0.05234454 | NA | NA |
| ENSG000002 | 4.59200302 | -0.90469019 | 0.37805922 | -2.3929854 | 0.01671191 | 0.05234454 | NA | NA |
| ENSG000002 | 2.38340273 | -0.87890206 | 0.3672887 | -2.3929461 | 0.0167137 | 0.05234454 | NA | NA |
| ENSG000001 | 87.4425102 | 0.444404423 | 0.18573607 | 2.39266618 | 0.01672645 | 0.05238044 | RAD50 | 10111 |
| ENSG000002 | 7.3459333 | -0.865503312 | 0.36179944 | -2.3922185 | 0.01674687 | 0.05243196 | SMYD3-IT1 | 106478975 |
| ENSG000002 | 18.0998752 | -0.728656307 | 0.30459774 | -2.3921921 | 0.01674807 | 0.05243196 | NA | NA |
| ENSG000002 | 5.37881204 | -0.890562147 | 0.37226895 | -2.3922547 | 0.01674522 | 0.05243196 | NA | NA |
| ENSG000002 | 14.8537814 | -0.766143385 | 0.32026663 | -2.3922049 | 0.01674749 | 0.05243196 | NA | NA |
| ENSG000002 | 5.38293118 | -0.903198297 | 0.37760182 | -2.3919331 | 0.0167599 | 0.05246492 | NA | NA |
| ENSG000002 | 3.32918556 | -0.897018447 | 0.37503089 | -2.3918522 | 0.01676359 | 0.05247244 | NA | NA |
| ENSG000000 | 9229.81052 | 0.281721506 | 0.11779693 | 2.39158612 | 0.01677575 | 0.05249947 | FAT1 | 2195 |
| ENSG000001 | 18.0668275 | -0.692180592 | 0.28942763 | -2.3915498 | 0.01677741 | 0.05249947 | ARHGAP40 | 343578 |
| ENSG000001 | 17.2373922 | -0.71758368 | 0.30004779 | -2.3915646 | 0.01677673 | 0.05249947 | FGL2 | 10875 |
| ENSG000001 | 46.1699433 | 0.528666064 | 0.22105024 | 2.39161047 | 0.01677463 | 0.05249947 | ZNF25 | 219749 |
| ENSG000002 | 6.66136195 | -0.873372563 | 0.36520753 | -2.3914418 | 0.01678234 | 0.05251071 | NA | NA |
| ENSG000002 | 3.83538371 | -0.905352296 | 0.37858876 | -2.3913871 | 0.01678484 | 0.05251071 | LOC102723778 | 102723778 |

| | | | | | | | | | |
|---|---|---|---|---|---|---|---|---|---|
| ENSG000002 | 6.69026396 | -0.872265049 | 0.36475289 | -2.3913863 | 0.01678488 | 0.05251071 | NA | NA | |
| ENSG000001 | 65.5821175 | 0.46075751 | 0.19268243 | 2.39127928 | 0.01678978 | 0.05252196 | GPR37 | | 2861 |
| ENSG000002 | 6.65348965 | -0.887374634 | 0.37110915 | -2.3911419 | 0.01679606 | 0.05253352 | NA | NA | |
| ENSG000002 | 5.22811413 | -0.898659386 | 0.37582441 | -2.3911682 | 0.01679486 | 0.05253352 | LOC340357 | | 340357 |
| ENSG000002 | 6.65039737 | -0.869301467 | 0.36355659 | -2.3911036 | 0.01679781 | 0.05253494 | NA | NA | |
| ENSG000001 | 285.004287 | 0.30652596 | 0.12821342 | 2.39074784 | 0.0168141 | 0.05258182 | C3orf52 | | 79669 |
| ENSG000002 | 2.40773746 | -0.879114001 | 0.3677308 | -2.3906455 | 0.01681878 | 0.05259242 | NA | NA | |
| ENSG000001 | 52.3132924 | -0.487912539 | 0.20409574 | -2.3906062 | 0.01682058 | 0.05259399 | ZBBX | | 79740 |
| ENSG000001 | 12.5259252 | -0.789529949 | 0.33027145 | -2.3905486 | 0.01682322 | 0.05259819 | ZBED2 | | 79413 |
| ENSG000002 | 16.19948 | -0.738494602 | 0.3089392 | -2.3904205 | 0.01682909 | 0.05261248 | LINC00668 | | 400643 |
| ENSG000002 | 4.98754895 | -0.902917201 | 0.3777592 | -2.3901925 | 0.01683954 | 0.0526411 | NA | NA | |
| ENSG000002 | 3.16001205 | -0.899136848 | 0.37629632 | -2.3894383 | 0.01687416 | 0.05274524 | NA | NA | |
| ENSG000002 | 19.4161002 | -0.711692429 | 0.29785622 | -2.3893824 | 0.01687672 | 0.05274919 | NA | NA | |
| ENSG000001 | 91.0777863 | 0.422352115 | 0.17676793 | 2.38930284 | 0.01688038 | 0.05275656 | ANKRD13D | | 338692 |
| ENSG000001 | 68.4366924 | 0.452650088 | 0.18946603 | 2.38908305 | 0.01689048 | 0.05278289 | NUDT7 | | 283927 |
| ENSG000002 | 6.0685183 | -0.880563286 | 0.36858103 | -2.3890629 | 0.01689141 | 0.05278289 | NA | NA | |
| ENSG000002 | 8.68152667 | -0.832040977 | 0.34828025 | -2.3889984 | 0.01689438 | 0.05278809 | NA | NA | |
| ENSG000002 | 10.5148032 | -0.819826175 | 0.34318726 | -2.3888596 | 0.01690076 | 0.05280397 | LOC101927740 | | 101927740 |
| ENSG00000C | 772.763755 | 0.288514797 | 0.12078644 | 2.38863559 | 0.01691107 | 0.05281372 | DIMT1 | | 27292 |
| ENSG000001 | 28.6426774 | -0.616749469 | 0.25819813 | -2.3886674 | 0.0169096 | 0.05281372 | SLC35F4 | | 341880 |
| ENSG000001 | 6.63961951 | -0.870319522 | 0.3643524 | -2.3886751 | 0.01690925 | 0.05281372 | C3AR1 | | 719 |
| ENSG000002 | 4.537385 | -0.902487827 | 0.37781907 | -2.3886773 | 0.01690915 | 0.05281372 | NA | NA | |
| ENSG000002 | 98.0969406 | 0.409923606 | 0.17161101 | 2.388679 | 0.01690907 | 0.05281372 | NA | NA | |
| ENSG000002 | 4.87420298 | -0.899394824 | 0.37653294 | -2.3886219 | 0.0169117 | 0.05281372 | NA | NA | |
| ENSG000001 | 212.953414 | 0.338210894 | 0.14160716 | 2.38837427 | 0.0169231 | 0.05284526 | ZCCHC4 | | 29063 |
| ENSG000001 | 201.733381 | 0.350700417 | 0.1468506 | 2.38814426 | 0.01693369 | 0.05287427 | KCTD6 | | 200845 |
| ENSG000002 | 8.33294487 | -0.869335102 | 0.36404546 | -2.387985 | 0.01694103 | 0.05289312 | LOC101928622 | | 101928622 |
| ENSG000002 | 7.87697471 | -0.849857268 | 0.35589745 | -2.3879274 | 0.01694369 | 0.05289734 | LOC101928769 | | 101928769 |
| ENSG000002 | 8.14572771 | -0.842456187 | 0.35282036 | -2.3877765 | 0.01695065 | 0.05291498 | NA | NA | |

| | | | | | | | | |
|---|---|---|---|---|---|---|---|---|
| ENSG000002 | 5.18353895 | -0.895469839 | 0.37503184 | -2.3877168 | 0.0169534 | 0.05291951 | NA | NA |
| ENSG000002 | 4.77548281 | -0.898136648 | 0.37616404 | -2.3876197 | 0.01695788 | 0.05292942 | NA | NA |
| ENSG000002 | 5.546507 | -0.89265331 | 0.3738719 | -2.3875914 | 0.01695919 | 0.05292942 | NA | NA |
| ENSG000001 | 2522.62297 | 0.277368329 | 0.11619242 | 2.3871465 | 0.01697972 | 0.05298944 | RBM26 | 64062 |
| ENSG000001 | 20.3508121 | -0.723166971 | 0.30295316 | -2.3870587 | 0.01698378 | 0.05299634 | C7orf57 | 136288 |
| ENSG000002 | 4.7734919 | -0.898492248 | 0.37640403 | -2.3870421 | 0.01698455 | 0.05299634 | NA | NA |
| ENSG000002 | 5.75359473 | -0.886383577 | 0.37139289 | -2.3866466 | 0.01700283 | 0.05304929 | NA | NA |
| ENSG000002 | 7.74165604 | -0.860249714 | 0.36047411 | -2.3864397 | 0.0170124 | 0.05307507 | LOC102725079 | 102725079 |
| ENSG000001 | 4.13733449 | -0.902955174 | 0.37840957 | -2.3861848 | 0.01702419 | 0.05310778 | NA | NA |
| ENSG000002 | 9.0432474 | -0.826041122 | 0.34621643 | -2.3859097 | 0.01703694 | 0.05314344 | LOC100507600 | 100507600 |
| ENSG000001 | 19.5344908 | -0.687968487 | 0.28838692 | -2.3855745 | 0.01705247 | 0.05318781 | KLHL10 | 317719 |
| ENSG000001 | 4014.89997 | 0.299412941 | 0.12551852 | 2.38540847 | 0.01706017 | 0.05320772 | TOMM20 | 9804 |
| ENSG000002 | 60.9858762 | 0.491162037 | 0.20592337 | 2.38516903 | 0.01707128 | 0.05323828 | CDPF1 | 150383 |
| ENSG000002 | 21.6523177 | -0.720573368 | 0.30211821 | -2.3850709 | 0.01707583 | 0.05324838 | MIR8078 | 102466878 |
| ENSG000001 | 938.826477 | 0.41425841 | 0.17369517 | 2.38497372 | 0.01708034 | 0.05325836 | GSE1 | 23199 |
| ENSG000001 | 13.0180787 | -0.779550379 | 0.32688934 | -2.3847531 | 0.01709059 | 0.0532862 | C22orf23 | 84645 |
| ENSG000002 | 6.03098699 | -0.87788104 | 0.3681359 | -2.3846656 | 0.01709465 | 0.05329478 | NA | NA |
| ENSG000001 | 264.756077 | 0.321245181 | 0.13472334 | 2.38448049 | 0.01710325 | 0.0533175 | SLC35D1 | 23169 |
| ENSG000002 | 3.83526693 | -0.90268981 | 0.37857387 | -2.3844483 | 0.01710475 | 0.05331807 | NA | NA |
| ENSG000002 | 51.067507 | 0.499645486 | 0.20954766 | 2.38440019 | 0.01710699 | 0.05332094 | NA | NA |
| ENSG000002 | 9.07158847 | -0.824713362 | 0.34589362 | -2.3842977 | 0.01711176 | 0.0533317 | NA | NA |
| ENSG000002 | 25.1803301 | 0.628160861 | 0.26351375 | 2.38378777 | 0.01713548 | 0.05340154 | NA | NA |
| ENSG000001 | 62.3641349 | 0.464863734 | 0.19501411 | 2.38374405 | 0.01713752 | 0.05340378 | MYOZ1 | 58529 |
| ENSG000002 | 154.528988 | 0.37864113 | 0.15887112 | 2.38332253 | 0.01715715 | 0.05345768 | LOC220729 | 220729 |
| ENSG000002 | 9.25293244 | -0.816296737 | 0.34250459 | -2.3833162 | 0.01715745 | 0.05345768 | LOC101929007 | 101929007 |
| ENSG000001 | 16.7087475 | -0.72705628 | 0.30508331 | -2.3831401 | 0.01716566 | 0.05347915 | SERPINA10 | 51156 |
| ENSG000001 | 42.0188445 | 0.569365168 | 0.23891705 | 2.38310818 | 0.01716715 | 0.05347968 | FBXL6 | 26233 |
| ENSG000002 | 6.04744531 | -0.879419848 | 0.36903002 | -2.3830577 | 0.0171695 | 0.0534829 | LOC401286 | 401286 |
| ENSG000001 | 42.6885166 | 0.556382257 | 0.23350254 | 2.3827675 | 0.01718304 | 0.05352098 | CIDECP | 152302 |

| | | | | | | | | |
|---|---|---|---|---|---|---|---|---|
| ENSG000002 | 6.71394727 | -0.874898191 | 0.36718989 | -2.3826859 | 0.01718685 | 0.05352872 | DISC1-IT1 | 104472714 |
| ENSG000002 | 10.4133235 | -0.812677275 | 0.34109519 | -2.3825527 | 0.01719307 | 0.05354398 | NA | NA |
| ENSG000001 | 63.3646997 | 0.551619201 | 0.23152747 | 2.38252163 | 0.01719452 | 0.05354439 | CTXN1 | 404217 |
| ENSG000001 | 481.910781 | 0.385532322 | 0.16182012 | 2.38247461 | 0.01719672 | 0.05354712 | RNF214 | 257160 |
| ENSG000002 | 6.78882993 | -0.87565885 | 0.367553 | -2.3824016 | 0.01720013 | 0.05355362 | LOC100132813 | 100132813 |
| ENSG000001 | 6996.97426 | 0.610174077 | 0.25612833 | 2.38229827 | 0.01720496 | 0.05356454 | ILF3 | 3609 |
| ENSG000001 | 12.9686886 | -0.790854637 | 0.33198221 | -2.3822199 | 0.01720862 | 0.05357183 | OR51G2 | 81282 |
| ENSG00000C | 18.3163805 | -0.68691662 | 0.28838169 | -2.3819703 | 0.01722029 | 0.05360404 | CLDN18 | 51208 |
| ENSG000001 | 488.300688 | 0.389105662 | 0.16336303 | 2.38184652 | 0.01722608 | 0.05361794 | TRERF1 | 55809 |
| ENSG000001 | 13.6285275 | -0.769584451 | 0.32313078 | -2.3816501 | 0.01723527 | 0.05364243 | OR6P1 | 128366 |
| ENSG000001 | 356.476069 | 0.333635416 | 0.14009029 | 2.38157419 | 0.01723882 | 0.05364903 | TFB2M | 64216 |
| ENSG000002 | 2.7536383 | -0.886860386 | 0.37238816 | -2.3815483 | 0.01724003 | 0.05364903 | NA | NA |
| ENSG000001 | 66.5865932 | 0.487227316 | 0.2045936 | 2.38143965 | 0.01724512 | 0.05366074 | PROSER3 | 148137 |
| ENSG000001 | 3.57756587 | -0.89910644 | 0.37758001 | -2.3812342 | 0.01725474 | 0.05368655 | LOC100286922 | 100286922 |
| ENSG000002 | 7.48257249 | -0.849679033 | 0.3568466 | -2.3810764 | 0.01726213 | 0.05370543 | NA | NA |
| ENSG000002 | 14.1946222 | -0.742804947 | 0.31198721 | -2.3808827 | 0.01727121 | 0.05372907 | NA | NA |
| ENSG000002 | 4.42021904 | -0.898017574 | 0.37718236 | -2.3808578 | 0.01727238 | 0.05372907 | NA | NA |
| ENSG000001 | 447.67265 | 0.371120114 | 0.15589152 | 2.38063053 | 0.01728304 | 0.05375465 | PRDM2 | 7799 |
| ENSG000001 | 25.5844215 | -0.635033307 | 0.26675216 | -2.3806116 | 0.01728392 | 0.05375465 | ADCY4 | 196883 |
| ENSG000001 | 17.2439354 | -0.717194403 | 0.3012739 | -2.3805395 | 0.01728731 | 0.05375465 | CD1E | 913 |
| ENSG000001 | 95.6884616 | 0.433320671 | 0.1820281 | 2.38051532 | 0.01728844 | 0.05375465 | SIRT7 | 51547 |
| ENSG000002 | 43.5294852 | -0.524968787 | 0.22052803 | -2.3805082 | 0.01728878 | 0.05375465 | RBM20 | 282996 |
| ENSG000002 | 14.3410155 | -0.730486681 | 0.30686469 | -2.3804847 | 0.01728988 | 0.05375465 | NA | NA |
| ENSG000002 | 6.04330683 | -0.891982397 | 0.3747001 | -2.3805235 | 0.01728806 | 0.05375465 | NA | NA |
| ENSG000001 | 3033.68015 | 0.295934461 | 0.1243216 | 2.38039457 | 0.01729411 | 0.05376368 | ZNF518A | 9849 |
| ENSG000001 | 40.0512177 | 0.57791938 | 0.24281313 | 2.38009937 | 0.01730797 | 0.05380264 | SGSH | 6448 |
| ENSG000001 | 27.4154739 | -0.623384617 | 0.26194862 | -2.3797973 | 0.01732216 | 0.05383438 | TMEM229B | 161145 |
| ENSG000002 | 21.259666 | -0.653411473 | 0.27456567 | -2.3798003 | 0.01732202 | 0.05383438 | NA | NA |
| ENSG000002 | 10.1172883 | -0.816794691 | 0.34321895 | -2.3798065 | 0.01732173 | 0.05383438 | HNRNPKP3 | 399881 |

| | | | | | | | | |
|---|---|---|---|---|---|---|---|---|
| ENSG00000C | 83.7921557 | -0.427288778 | 0.17958009 | -2.3793773 | 0.01734192 | 0.05388751 | NOS1 | 4842 |
| ENSG000001 | 80.2723021 | 0.423326258 | 0.17791391 | 2.37938822 | 0.0173414 | 0.05388751 | CD58 | 965 |
| ENSG000002 | 276.765764 | -0.424447437 | 0.17839233 | -2.379292 | 0.01734593 | 0.05389586 | NA | NA |
| ENSG00000C | 65.3686475 | -0.459585971 | 0.19316537 | -2.3792358 | 0.01734857 | 0.05389994 | PLXNA2 | 5362 |
| ENSG000002 | 15.7587789 | -0.748579468 | 0.31463981 | -2.3791632 | 0.01735199 | 0.05390643 | LINC01307 | 104355286 |
| ENSG000001 | 15.3677993 | -0.777557717 | 0.3268349 | -2.3790535 | 0.01735715 | 0.05391834 | KIR3DX1 | 90011 |
| ENSG000001 | 4.11303201 | -0.900333623 | 0.37844702 | -2.3790216 | 0.01735866 | 0.05391888 | ERAS | 3266 |
| ENSG000001 | 124.792911 | 0.398133046 | 0.16736615 | 2.37881456 | 0.01736841 | 0.05394504 | UBE2F | 140739 |
| ENSG000002 | 12.5090359 | -0.770579874 | 0.32395092 | -2.3786933 | 0.01737413 | 0.05395866 | NA | NA |
| ENSG000002 | 12.69995 | -0.757340746 | 0.31839622 | -2.378611 | 0.017378 | 0.05396657 | MOXD2P | 100289017 |
| ENSG000001 | 22.5452489 | -0.636724445 | 0.2677298 | -2.3782352 | 0.01739572 | 0.05401746 | TMEM71 | 137835 |
| ENSG000001 | 420.597096 | 0.4095891 | 0.1722373 | 2.37805107 | 0.01740442 | 0.05404031 | AKIRIN2 | 55122 |
| ENSG000001 | 21.7536342 | -0.661528285 | 0.27819242 | -2.3779523 | 0.01740908 | 0.05405065 | NA | NA |
| ENSG000002 | 7.55852539 | -0.848859378 | 0.35697571 | -2.3779192 | 0.01741064 | 0.05405137 | MIR548K | 100313770 |
| ENSG000001 | 219.282949 | 0.338411675 | 0.14232138 | 2.37779928 | 0.0174163 | 0.0540648 | SPATA6 | 54558 |
| ENSG000002 | 16.0025343 | -0.711483961 | 0.29924684 | -2.3775822 | 0.01742656 | 0.05409249 | SPACA6P-AS | 102238594 |
| ENSG000002 | 17.7905226 | -0.711619696 | 0.29931126 | -2.3775239 | 0.01742931 | 0.0540969 | CEACAM20 | 125931 |
| ENSG000001 | 185.078011 | 0.343873437 | 0.14465917 | 2.37712849 | 0.01744801 | 0.0541425 | UEVLD | 55293 |
| ENSG000002 | 8.92740437 | -0.840968585 | 0.35376929 | -2.3771668 | 0.0174462 | 0.0541425 | NA | NA |
| ENSG000002 | 12.3359368 | -0.756556178 | 0.3182613 | -2.3771542 | 0.01744679 | 0.0541425 | NA | NA |
| ENSG000002 | 15.2107647 | -0.739700341 | 0.31117766 | -2.3770998 | 0.01744937 | 0.05414257 | NA | NA |
| ENSG000002 | 13.0504428 | -0.753263456 | 0.31693114 | -2.3767417 | 0.01746631 | 0.05419101 | LOC101928995 | 101928995 |
| ENSG000002 | 15.4249248 | -0.734966167 | 0.30923688 | -2.3767093 | 0.01746785 | 0.05419163 | LOC729224 | 729224 |
| ENSG000001 | 2091.35987 | 0.303344967 | 0.12763575 | 2.37664576 | 0.01747086 | 0.05419681 | SPARC | 6678 |
| ENSG000002 | 3.21226254 | -0.893518439 | 0.37596508 | -2.3765996 | 0.01747304 | 0.05419751 | NA | NA |
| ENSG000002 | 2.48028429 | -0.870965309 | 0.36647773 | -2.3765845 | 0.01747376 | 0.05419751 | NA | NA |
| ENSG000001 | 23.2412194 | -0.708375132 | 0.29807485 | -2.3765008 | 0.01747772 | 0.05420567 | SPAM1 | 6677 |
| ENSG000001 | 1241.91406 | 0.277214895 | 0.11666281 | 2.37620632 | 0.01749168 | 0.05424481 | SIPA1L1 | 26037 |
| ENSG000002 | 18.7337079 | -0.708569382 | 0.29821504 | -2.376035 | 0.0174998 | 0.05426585 | C1QTNF9B | 387911 |

| | | | | | | | | |
|---|---|---|---|---|---|---|---|---|
| ENSG000001 | 238.392616 | 0.422092178 | 0.17765514 | 2.37590752 | 0.01750585 | 0.05428045 | TMEM214 | 54867 |
| ENSG000001 | 10305.7864 | 0.290580518 | 0.12230945 | 2.37578136 | 0.01751184 | 0.05429486 | SFPQ | 6421 |
| ENSG000002 | 6.13124971 | -0.879991044 | 0.37043368 | -2.3755697 | 0.01752188 | 0.05432185 | LOC101059948 | 101059948 |
| ENSG000001 | 2336.52871 | 0.258044382 | 0.10862716 | 2.37550525 | 0.01752494 | 0.05432719 | LSM3 | 27258 |
| ENSG000002 | 11.2053759 | -0.774751164 | 0.32615713 | -2.3753924 | 0.0175303 | 0.05433965 | NA | NA |
| ENSG000002 | 97.3133573 | 0.414018069 | 0.1743048 | 2.37525339 | 0.01753691 | 0.05435597 | SETSIP | 646817 |
| ENSG000001 | 188.718528 | 0.335079132 | 0.14108051 | 2.37509163 | 0.01754459 | 0.05437564 | C7orf60 | 154743 |
| ENSG000002 | 3.87030696 | -0.898932336 | 0.37852144 | -2.3748518 | 0.017556 | 0.05440682 | NA | NA |
| ENSG000002 | 6.04839766 | -0.876686325 | 0.36918435 | -2.3746573 | 0.01756525 | 0.05443133 | NA | NA |
| ENSG000001 | 152.282967 | 0.394435558 | 0.16610648 | 2.37459459 | 0.01756823 | 0.05443642 | HS6ST2 | 90161 |
| ENSG000001 | 843.355288 | 0.253298783 | 0.10667366 | 2.37452038 | 0.01757177 | 0.05444321 | FIP1L1 | 81608 |
| ENSG000002 | 6.26052988 | -0.868932708 | 0.36597512 | -2.3742945 | 0.01758252 | 0.05447237 | NA | NA |
| ENSG000001 | 9.11861643 | -0.82753874 | 0.34857228 | -2.3740808 | 0.0175927 | 0.05449558 | ELOVL3 | 83401 |
| ENSG000002 | 17.4254835 | -0.690469821 | 0.29083453 | -2.3740985 | 0.01759186 | 0.05449558 | NA | NA |
| ENSG000002 | 2.385627 | -0.871076375 | 0.36696218 | -2.3737498 | 0.01760848 | 0.05454029 | NA | NA |
| ENSG000002 | 19.1213893 | -0.694314986 | 0.29250797 | -2.3736618 | 0.01761267 | 0.05454624 | NA | NA |
| ENSG000002 | 6.83743124 | -0.876035626 | 0.36906642 | -2.373653 | 0.01761309 | 0.05454624 | NA | NA |
| ENSG000001 | 2817.03565 | 0.296946955 | 0.12510777 | 2.3735293 | 0.01761899 | 0.05455619 | ZNF292 | 23036 |
| ENSG000002 | 24.5086915 | 0.668939339 | 0.28183019 | 2.37355462 | 0.01761779 | 0.05455619 | NA | NA |
| ENSG000001 | 27.1939225 | -0.67745953 | 0.28543308 | -2.3734443 | 0.01762305 | 0.05456064 | ECM2 | 1842 |
| ENSG000001 | 314.015192 | 0.35932681 | 0.15139656 | 2.37341456 | 0.01762447 | 0.05456064 | NT5DC3 | 51559 |
| ENSG000001 | 1354.14749 | 0.282968338 | 0.11922365 | 2.37342446 | 0.017624 | 0.05456064 | ACBD3 | 64746 |
| ENSG000002 | 3.5934506 | -0.895120248 | 0.37715671 | -2.3733377 | 0.01762814 | 0.05456783 | NA | NA |
| ENSG000002 | 454.189306 | 0.28154216 | 0.11863933 | 2.37309302 | 0.01763982 | 0.05459983 | NA | NA |
| ENSG000001 | 16.7636219 | -0.712742503 | 0.30034778 | -2.3730573 | 0.01764152 | 0.05460094 | IL1R2 | 7850 |
| ENSG000002 | 10.3163002 | -0.810055514 | 0.34136789 | -2.3729693 | 0.01764573 | 0.05460979 | FLJ43879 | 401039 |
| ENSG000002 | 9.06848819 | -0.822251798 | 0.34652502 | -2.3728497 | 0.01765144 | 0.0546233 | TH2LCRR | 101927761 |
| ENSG000001 | 13.9926288 | -0.748781901 | 0.31558263 | -2.3726968 | 0.01765875 | 0.05462924 | SERPINB7 | 8710 |
| ENSG000001 | 157.840229 | 0.354171177 | 0.14926508 | 2.37276651 | 0.01765542 | 0.05462924 | ZNF610 | 162963 |

| | | | | | | | | |
|---|---|---|---|---|---|---|---|---|
| ENSG000001 | 447.321021 | 0.302799625 | 0.12761784 | 2.37270611 | 0.01765831 | 0.05462924 | ZNF566 | 84924 |
| ENSG000002 | 46.8373568 | -0.516819736 | 0.217819 | -2.3727027 | 0.01765847 | 0.05462924 | NA | NA |
| ENSG000001 | 476.248521 | 0.304160048 | 0.12819695 | 2.37259981 | 0.01766339 | 0.05463942 | KIF3A | 11127 |
| ENSG000002 | 6.09943432 | -0.878116347 | 0.37022134 | -2.3718686 | 0.01769839 | 0.0547435 | NA | NA |
| ENSG000000 | 873.510387 | 0.655929877 | 0.27655573 | 2.37178187 | 0.01770254 | 0.05474799 | TACC3 | 10460 |
| ENSG000002 | 14.032088 | -0.772377103 | 0.32565129 | -2.3717919 | 0.01770206 | 0.05474799 | NA | NA |
| ENSG000001 | 12.7315456 | -0.765139621 | 0.32263617 | -2.3715246 | 0.01771487 | 0.05477776 | SDR9C7 | 121214 |
| ENSG000002 | 38.9411408 | 0.563929748 | 0.23778994 | 2.37154582 | 0.01771385 | 0.05477776 | NA | NA |
| ENSG000002 | 5.02005013 | -0.887973611 | 0.37445186 | -2.3713959 | 0.01772104 | 0.05479266 | NA | NA |
| ENSG000002 | 8.98374866 | -0.820278941 | 0.3459239 | -2.37127 | 0.01772708 | 0.05480716 | NA | NA |
| ENSG000000 | 181.126355 | 0.337823903 | 0.14247633 | 2.37108795 | 0.01773581 | 0.05482609 | SEC61A2 | 55176 |
| ENSG000001 | 179.115153 | 0.329222116 | 0.13884991 | 2.37106467 | 0.01773693 | 0.05482609 | CPNE2 | 221184 |
| ENSG000001 | 30.0753598 | -0.595303799 | 0.25107191 | -2.3710489 | 0.01773768 | 0.05482609 | CNDP1 | 84735 |
| ENSG000001 | 14.3619997 | -0.753126625 | 0.31764283 | -2.3709857 | 0.01774072 | 0.05482609 | C1orf204 | 284677 |
| ENSG000001 | 61.0884422 | 0.474231446 | 0.20001272 | 2.3710064 | 0.01773972 | 0.05482609 | SMIM10L2B | 644596 |
| ENSG000002 | 12.9293571 | -0.750078279 | 0.3163588 | -2.3709733 | 0.01774131 | 0.05482609 | NA | NA |
| ENSG000001 | 9.25809978 | -0.813158853 | 0.34297102 | -2.3709258 | 0.01774359 | 0.05482895 | DMRT2 | 10655 |
| ENSG000002 | 5.42151169 | -0.884770955 | 0.37319473 | -2.3708024 | 0.01774952 | 0.05484309 | NA | NA |
| ENSG000001 | 19.4721682 | -0.697602658 | 0.29425361 | -2.3707531 | 0.01775189 | 0.05484623 | C9 | 735 |
| ENSG000001 | 453.435064 | 0.321938767 | 0.13580274 | 2.37063536 | 0.01775754 | 0.05485534 | ZNF430 | 80264 |
| ENSG000002 | 35.2691509 | -0.574729524 | 0.24243448 | -2.3706592 | 0.0177564 | 0.05485534 | STARD4-AS1 | 100505678 |
| ENSG000002 | 4.06592027 | -0.896466913 | 0.37817449 | -2.3705113 | 0.0177635 | 0.05486957 | NA | NA |
| ENSG000001 | 13.5979273 | -0.743237866 | 0.31355297 | -2.3703742 | 0.01777009 | 0.05488574 | ARG1 | 383 |
| ENSG000001 | 189.794267 | 0.358778176 | 0.15137089 | 2.37019275 | 0.01777881 | 0.05490851 | R3HDM2 | 22864 |
| ENSG000002 | 9.39189425 | -0.823167624 | 0.3473274 | -2.3700049 | 0.01778785 | 0.05493223 | LOC101929745 | 101929745 |
| ENSG000002 | 7.52192663 | -0.846351247 | 0.35714629 | -2.3697607 | 0.0177996 | 0.05496433 | NA | NA |
| ENSG000002 | 9.64458069 | -0.814913726 | 0.34388739 | -2.369711 | 0.01780199 | 0.05496754 | NA | NA |
| ENSG000001 | 147.394968 | 0.366282212 | 0.15457543 | 2.36960175 | 0.01780725 | 0.05497775 | HILPDA | 29923 |
| ENSG000002 | 4.791356 | -0.891960509 | 0.3764204 | -2.369586 | 0.01780801 | 0.05497775 | NA | NA |

| | | | | | | | | |
|---|---|---|---|---|---|---|---|---|
| ENSG000000 | 109.930899 | 0.454838274 | 0.19196083 | 2.36943276 | 0.01781539 | 0.05499635 | FLYWCH1 | 84256 |
| ENSG000002 | 4.26347085 | -0.897009047 | 0.37859156 | -2.3693319 | 0.01782025 | 0.05500716 | NA | NA |
| ENSG000001 | 3.41081663 | -0.895723217 | 0.37807231 | -2.369185 | 0.01782734 | 0.05501646 | C1QC | 714 |
| ENSG000002 | 53.2532198 | 0.484117276 | 0.20433886 | 2.36918855 | 0.01782716 | 0.05501646 | NA | NA |
| ENSG000002 | 4.49166517 | -0.895701378 | 0.37805821 | -2.3692155 | 0.01782586 | 0.05501646 | NA | NA |
| ENSG000001 | 99.2084352 | 0.442080834 | 0.18660148 | 2.36911751 | 0.01783059 | 0.05502231 | MAP7D2 | 256714 |
| ENSG000002 | 3.48632495 | -0.895291052 | 0.37795175 | -2.3687972 | 0.01784604 | 0.05506579 | NA | NA |
| ENSG000002 | 4.13411709 | -0.89663389 | 0.37853726 | -2.3686807 | 0.01785166 | 0.05507895 | LOC101928992 | 101928992 |
| ENSG000001 | 390.748795 | 0.388162869 | 0.16389226 | 2.36840275 | 0.01786508 | 0.05511196 | WIPF2 | 147179 |
| ENSG000001 | 451.99821 | 0.324802455 | 0.13713885 | 2.3684204 | 0.01786422 | 0.05511196 | ZNF239 | 8187 |
| ENSG000001 | 207.709627 | 0.399499401 | 0.16869028 | 2.36824203 | 0.01787284 | 0.05513171 | AFAP1 | 60312 |
| ENSG000002 | 30.4357911 | 0.579080146 | 0.24452677 | 2.36816668 | 0.01787648 | 0.05513875 | NAT6 | 24142 |
| ENSG000001 | 64.0976203 | 0.466257453 | 0.19689121 | 2.36809683 | 0.01787986 | 0.05514497 | OVOL2 | 58495 |
| ENSG000001 | 3.41027639 | -0.895617861 | 0.37820843 | -2.3680537 | 0.01788194 | 0.0551472 | ODF3B | 440836 |
| ENSG000002 | 3.41229105 | -0.895230803 | 0.37807292 | -2.3678787 | 0.0178904 | 0.0551691 | NA | NA |
| ENSG000002 | 3.64966092 | -0.892571079 | 0.37700687 | -2.3675194 | 0.01790778 | 0.05521849 | NA | NA |
| ENSG000001 | 9.85423416 | -0.802930549 | 0.33916413 | -2.3673805 | 0.0179145 | 0.05523502 | CXCR5 | 643 |
| ENSG000001 | 7.84503963 | -0.844053267 | 0.35655568 | -2.3672411 | 0.01792125 | 0.05525163 | P2RY13 | 53829 |
| ENSG000002 | 12.7437965 | -0.759522383 | 0.3208726 | -2.3670528 | 0.01793037 | 0.05527555 | NA | NA |
| ENSG000002 | 24.2785393 | -0.650125958 | 0.27466599 | -2.3669692 | 0.01793442 | 0.05528383 | NA | NA |
| ENSG000002 | 5.6777783 | -0.875917398 | 0.37007415 | -2.36687 | 0.01793923 | 0.05529446 | LINC01537 | 101928555 |
| ENSG000001 | 741.869837 | 0.324874445 | 0.13726745 | 2.36672596 | 0.01794622 | 0.05530757 | PRCC | 5546 |
| ENSG000002 | 7.69247886 | -0.85860116 | 0.36277715 | -2.3667454 | 0.01794527 | 0.05530757 | NA | NA |
| ENSG000001 | 69.4639563 | 0.448249291 | 0.18940628 | 2.3666021 | 0.01795222 | 0.05532187 | TMEM74 | 157753 |
| ENSG000000 | 47.6974915 | -0.529449199 | 0.2237314 | -2.3664502 | 0.01795959 | 0.05534038 | GABRA1 | 2554 |
| ENSG000001 | 64.0687584 | -0.444887002 | 0.18802464 | -2.3661102 | 0.01797609 | 0.05538702 | POU2F3 | 25833 |
| ENSG000002 | 5.06387928 | -0.886201497 | 0.37454929 | -2.3660477 | 0.01797913 | 0.05538795 | NA | NA |
| ENSG000002 | 3.09711145 | -0.891122705 | 0.37662545 | -2.3660714 | 0.01797798 | 0.05538795 | NA | NA |
| ENSG000001 | 10.3261956 | -0.809153543 | 0.34200708 | -2.3658971 | 0.01798644 | 0.05540628 | SKOR1 | 390598 |

| | | | | | | | | |
|---|---|---|---|---|---|---|---|---|
| ENSG000001 | 463.835764 | 0.421239168 | 0.17805755 | 2.36574726 | 0.01799372 | 0.0554245 | ZNF618 | 114991 |
| ENSG000001 | 172.62976 | 0.365562399 | 0.15452509 | 2.36571542 | 0.01799527 | 0.05542505 | MBD2 | 8932 |
| ENSG000001 | 159.682229 | 0.350723701 | 0.14825934 | 2.3656095 | 0.01800042 | 0.05543364 | ZNF790 | 388536 |
| ENSG000002 | 4.20164736 | -0.895516648 | 0.37855764 | -2.3656018 | 0.01800079 | 0.05543364 | NA | NA |
| ENSG000002 | 4.55469263 | -0.894357038 | 0.37808073 | -2.3655187 | 0.01800483 | 0.05543766 | NA | NA |
| ENSG000002 | 4.33011315 | -0.895437615 | 0.3785361 | -2.3655277 | 0.0180044 | 0.05543766 | LOC102723655 | 102723655 |
| ENSG000001 | 15.5941332 | -0.718909405 | 0.30392168 | -2.365443 | 0.01800851 | 0.05544479 | BTC | 685 |
| ENSG000001 | 207.618027 | 0.381650503 | 0.16135415 | 2.36529709 | 0.01801561 | 0.05546243 | FAM155B | 27112 |
| ENSG000002 | 11.5401054 | -0.794535811 | 0.33592586 | -2.3652118 | 0.01801976 | 0.05547099 | LOC100132078 | 100132078 |
| ENSG000002 | 4.77215332 | -0.891324092 | 0.37690075 | -2.3648775 | 0.01803604 | 0.05551688 | NA | NA |
| ENSG000001 | 166.75991 | 0.364024342 | 0.15395397 | 2.36450122 | 0.01805437 | 0.05556065 | PCBD2 | 84105 |
| ENSG000002 | 11.9540401 | -0.770908226 | 0.32603402 | -2.3645024 | 0.01805431 | 0.05556065 | NA | NA |
| ENSG000002 | 8.45017917 | -0.850993488 | 0.35989905 | -2.3645339 | 0.01805278 | 0.05556065 | CHL1-AS1 | 101927193 |
| ENSG000001 | 149.514356 | 0.385922397 | 0.16322647 | 2.36433709 | 0.01806237 | 0.05558105 | BACH2 | 60468 |
| ENSG000001 | 99.421985 | 0.430926784 | 0.18228081 | 2.36408205 | 0.01807481 | 0.05561219 | SLC1A1 | 6505 |
| ENSG000002 | 3.8623617 | -0.894907863 | 0.37854487 | -2.3640734 | 0.01807523 | 0.05561219 | NA | NA |
| ENSG000002 | 142.832137 | 0.399146139 | 0.16884164 | 2.36402668 | 0.01807751 | 0.05561499 | NA | NA |
| ENSG000002 | 3.05957163 | -0.889939791 | 0.37647044 | -2.3639035 | 0.01808352 | 0.05562926 | NA | NA |
| ENSG000002 | 5.64102776 | -0.875174269 | 0.37023912 | -2.3638082 | 0.01808817 | 0.05563935 | NA | NA |
| ENSG000001 | 1843.481 | 0.291125714 | 0.12316257 | 2.36375155 | 0.01809094 | 0.05564364 | RNF2 | 6045 |
| ENSG00000C | 1257.21925 | 0.326090988 | 0.13796738 | 2.36353689 | 0.01810142 | 0.05567166 | FAM120A | 23196 |
| ENSG000002 | 53.6203371 | 0.515671198 | 0.21819584 | 2.36334112 | 0.01811099 | 0.05569686 | NA | NA |
| ENSG000001 | 87.8290408 | 0.415218558 | 0.17571132 | 2.36307236 | 0.01812413 | 0.05572882 | ZMAT5 | 55954 |
| ENSG000002 | 5.82582925 | -0.879483715 | 0.37217496 | -2.3630921 | 0.01812316 | 0.05572882 | NA | NA |
| ENSG00000C | 186.314011 | 0.372496401 | 0.15763891 | 2.36297248 | 0.01812902 | 0.05573961 | PRDM11 | 56981 |
| ENSG000002 | 3.93483788 | -0.893388577 | 0.37808393 | -2.3629372 | 0.01813074 | 0.05574069 | NA | NA |
| ENSG00000C | 25.454236 | -0.669915135 | 0.2835249 | -2.3628089 | 0.01813702 | 0.05575577 | CLCA1 | 1179 |
| ENSG000001 | 9281.37207 | 0.304949633 | 0.12906769 | 2.3627108 | 0.01814182 | 0.0557663 | DST | 667 |
| ENSG000002 | 9.26055349 | -0.80940451 | 0.34258591 | -2.3626322 | 0.01814567 | 0.05577225 | LOC101929488 | 101929488 |

| | | | | | | | | |
|---|---|---|---|---|---|---|---|---|
| ENSG000002 | 21.2876491 | -0.667865227 | 0.28268051 | -2.362615 | 0.01814651 | 0.05577225 | NA | NA |
| ENSG000001 | 69.1504894 | 0.467001813 | 0.19769063 | 2.36228607 | 0.01816262 | 0.05581754 | CTSK | 1513 |
| ENSG000002 | 6.89745228 | -0.854322133 | 0.36165821 | -2.3622362 | 0.01816506 | 0.05582081 | NA | NA |
| ENSG000002 | 14.0128453 | -0.750094978 | 0.317589 | -2.3618418 | 0.0181844 | 0.05587601 | NA | NA |
| ENSG000001 | 7.63426505 | -0.850762262 | 0.36022381 | -2.3617602 | 0.0181884 | 0.05587983 | ZSCAN4 | 201516 |
| ENSG000002 | 3.69518056 | -0.887331773 | 0.37570731 | -2.3617634 | 0.01818825 | 0.05587983 | NA | NA |
| ENSG000002 | 23.7424928 | -0.619715474 | 0.26243782 | -2.3613802 | 0.01820705 | 0.05593142 | ALOX12-AS1 | 100506713 |
| ENSG000002 | 2.6977331 | -0.882604037 | 0.37376907 | -2.3613619 | 0.01820795 | 0.05593142 | NA | NA |
| ENSG000001 | 353.232841 | 0.623681122 | 0.26419152 | 2.36071586 | 0.0182397 | 0.0560247 | STX16 | 8675 |
| ENSG000001 | 126.091326 | 0.401518139 | 0.17008597 | 2.36067753 | 0.01824158 | 0.05602624 | CEP131 | 22994 |
| ENSG000001 | 71.5375204 | -0.454486223 | 0.19252876 | -2.3606147 | 0.01824468 | 0.05603149 | FMN2 | 56776 |
| ENSG000001 | 14.3451701 | 0.75438564 | 0.31959132 | 2.36046971 | 0.01825181 | 0.05604915 | PXMP2 | 5827 |
| ENSG000002 | 27.1831087 | -0.638384812 | 0.27045399 | -2.3604193 | 0.01825429 | 0.05605253 | NA | NA |
| ENSG000002 | 7.94121267 | 0.859343058 | 0.36406809 | 2.36039105 | 0.01825568 | 0.05605255 | SNORA71D | 677840 |
| ENSG000002 | 3.81762703 | -0.893463445 | 0.37855243 | -2.3602106 | 0.01826456 | 0.05607558 | NA | NA |
| ENSG000002 | 156.792993 | 0.373096384 | 0.15809948 | 2.35988366 | 0.01828067 | 0.05612078 | MRPL46 | 26589 |
| ENSG00000C | 180.029928 | 0.391990426 | 0.16610857 | 2.35984475 | 0.01828258 | 0.05612241 | RGS17 | 26575 |
| ENSG000002 | 79.86345 | 0.427696738 | 0.1812436 | 2.35978947 | 0.01828531 | 0.05612653 | NA | NA |
| ENSG000001 | 14.8815022 | -0.752114461 | 0.31872738 | -2.3597423 | 0.01828764 | 0.05612942 | PAX1 | 5075 |
| ENSG000002 | 67.5462364 | 0.527763036 | 0.22365765 | 2.35969142 | 0.01829014 | 0.05613287 | NA | NA |
| ENSG000001 | 32.1251912 | -0.593573633 | 0.25155395 | -2.3596276 | 0.01829329 | 0.05613828 | STEAP4 | 79689 |
| ENSG000001 | 12.419089 | -0.78021719 | 0.3306723 | -2.3594876 | 0.01830019 | 0.05615521 | RSPO1 | 284654 |
| ENSG000002 | 11.2087291 | -0.772779048 | 0.32754184 | -2.3593292 | 0.018308 | 0.05617493 | NA | NA |
| ENSG000001 | 68.693837 | -0.434982574 | 0.18438102 | -2.3591505 | 0.01831682 | 0.05619774 | ANO4 | 121601 |
| ENSG000001 | 17.8112113 | -0.695346451 | 0.29474963 | -2.3591088 | 0.01831888 | 0.05619915 | TPTEP1 | 387590 |
| ENSG000002 | 19.7673472 | -0.739187912 | 0.31333669 | -2.3590851 | 0.01832006 | 0.05619915 | NA | NA |
| ENSG000001 | 142.484412 | 0.36584152 | 0.15508238 | 2.35901414 | 0.01832356 | 0.05620564 | NME6 | 10201 |
| ENSG000001 | 19.7688705 | -0.660953519 | 0.28018756 | -2.3589681 | 0.01832583 | 0.05620837 | RIN2 | 54453 |
| ENSG000002 | 3.54561228 | -0.890627936 | 0.37762332 | -2.3585088 | 0.01834852 | 0.05627371 | NA | NA |

| | | | | | | | | |
|---|---|---|---|---|---|---|---|---|
| ENSG000001 | 991.596641 | 0.278991423 | 0.11830489 | 2.35824081 | 0.01836178 | 0.0563101 | PARP14 | 54625 |
| ENSG000001 | 3731.45141 | 0.347427174 | 0.14735219 | 2.35780124 | 0.01838353 | 0.05637255 | ATP2A2 | 488 |
| ENSG000002 | 62.6016139 | -0.524590065 | 0.22251668 | -2.3575314 | 0.0183969 | 0.05640928 | LINC01410 | 103352539 |
| ENSG000002 | 7.29624362 | -0.870043063 | 0.36906909 | -2.3573989 | 0.01840347 | 0.05642514 | NA | NA |
| ENSG000002 | 6.68096196 | -0.861415367 | 0.3654254 | -2.3572947 | 0.01840863 | 0.05643671 | NA | NA |
| ENSG000002 | 29.953653 | 0.593549549 | 0.25182526 | 2.3569897 | 0.01842376 | 0.05647882 | NA | NA |
| ENSG000002 | 2.45042914 | -0.861608774 | 0.36556157 | -2.3569457 | 0.01842594 | 0.05648124 | NA | NA |
| ENSG000002 | 107.667778 | 0.423249476 | 0.17958201 | 2.356859 | 0.01843024 | 0.05649016 | NA | NA |
| ENSG000002 | 22.7825502 | -0.665250615 | 0.28228168 | -2.3566907 | 0.0184386 | 0.0565115 | CCDC79 | 283847 |
| ENSG000001 | 18.1030407 | -0.70771565 | 0.30031372 | -2.3565878 | 0.01844371 | 0.05651925 | OR4M1 | 441670 |
| ENSG000002 | 8.34129822 | -0.844545359 | 0.358377 | -2.3565836 | 0.01844392 | 0.05651925 | NA | NA |
| ENSG000001 | 162.183236 | 0.389597908 | 0.16532721 | 2.3565263 | 0.01844676 | 0.05651943 | ZNF185 | 7739 |
| ENSG000001 | 21.9959046 | -0.637472258 | 0.27051032 | -2.3565543 | 0.01844537 | 0.05651943 | TRIM55 | 84675 |
| ENSG000002 | 6.85864031 | -0.871597429 | 0.36988304 | -2.3564136 | 0.01845236 | 0.05653231 | HTR5A-AS1 | 100128264 |
| ENSG000002 | 2.81149354 | -0.875347975 | 0.37149562 | -2.3562807 | 0.01845896 | 0.05654826 | NA | NA |
| ENSG000002 | 8.68798073 | -0.821836582 | 0.34879325 | -2.3562285 | 0.01846156 | 0.05655194 | LOC101927460 | 101927460 |
| ENSG000001 | 27.4788986 | -0.640642692 | 0.27189725 | -2.3561941 | 0.01846327 | 0.05655291 | TMPRSS11B | 132724 |
| ENSG000002 | 6.97728859 | -0.857765534 | 0.3641008 | -2.3558464 | 0.01848056 | 0.0566016 | NA | NA |
| ENSG000001 | 17.4161185 | -0.749844878 | 0.3183088 | -2.3557152 | 0.01848709 | 0.05661731 | HTR2A | 3356 |
| ENSG000001 | 4254.02078 | 0.597692151 | 0.25372646 | 2.35565556 | 0.01849005 | 0.0566205 | PRPF8 | 10594 |
| ENSG000001 | 13.3904152 | 0.748051304 | 0.31755781 | 2.35563816 | 0.01849092 | 0.0566205 | NA | NA |
| ENSG00000C | 1064.64162 | 0.279012064 | 0.11844645 | 2.3555967 | 0.01849298 | 0.05662254 | MLH1 | 4292 |
| ENSG000001 | 14.1518039 | -0.740409352 | 0.31433469 | -2.3554809 | 0.01849875 | 0.05662737 | ATP6V0D2 | 245972 |
| ENSG000001 | 122.919015 | 0.419318048 | 0.1780161 | 2.35550637 | 0.01849748 | 0.05662737 | PNMA3 | 29944 |
| ENSG000002 | 428.536659 | 0.315765705 | 0.13405267 | 2.35553466 | 0.01849607 | 0.05662737 | ZNF37BP | 100129482 |
| ENSG000002 | 24.6497031 | -0.617098816 | 0.26200152 | -2.3553253 | 0.0185065 | 0.05664357 | NA | NA |
| ENSG000002 | 6.69267281 | -0.861740839 | 0.3658702 | -2.3553185 | 0.01850684 | 0.05664357 | NA | NA |
| ENSG000001 | 12.5482229 | -0.764878433 | 0.32474931 | -2.3552889 | 0.01850831 | 0.05664381 | VGLL2 | 245806 |
| ENSG000002 | 18.0312334 | -0.684292285 | 0.2905566 | -2.3551084 | 0.0185173 | 0.05666706 | LOC401134 | 401134 |

| | | | | | | | | |
|---|---|---|---|---|---|---|---|---|
| ENSG000002 | 4.16686847 | -0.891450736 | 0.37853364 | -2.3550106 | 0.01852218 | 0.0566777 | NA | NA |
| ENSG000002 | 7.30287258 | -0.851356704 | 0.36152113 | -2.3549293 | 0.01852623 | 0.05668582 | NA | NA |
| ENSG000001 | 6.03806506 | -0.870480075 | 0.36965316 | -2.3548563 | 0.01852987 | 0.05669268 | SECTM1 | 6398 |
| ENSG000002 | 17.4565878 | -0.692887264 | 0.29425364 | -2.3547279 | 0.01853627 | 0.05670799 | NA | NA |
| ENSG000001 | 4.78500085 | -0.887131803 | 0.37676964 | -2.3545735 | 0.01854398 | 0.05671872 | CDX1 | 1044 |
| ENSG000001 | 137.27646 | 0.384609328 | 0.16334562 | 2.35457386 | 0.01854396 | 0.05671872 | RAB29 | 8934 |
| ENSG000002 | 6.6932017 | -0.859530313 | 0.36503953 | -2.3546226 | 0.01854153 | 0.05671872 | NA | NA |
| ENSG000001 | 46.2990976 | 0.493734537 | 0.20971298 | 2.35433464 | 0.0185559 | 0.05674662 | CEP19 | 84984 |
| ENSG000002 | 9.88691799 | -0.80266479 | 0.34092924 | -2.3543442 | 0.01855542 | 0.05674662 | BHLHE40-AS1 | 100507582 |
| ENSG000001 | 96.168198 | 0.426630935 | 0.18122706 | 2.35412382 | 0.01856643 | 0.05677453 | ZSCAN20 | 7579 |
| ENSG000002 | 8.05523354 | -0.83173581 | 0.35332103 | -2.3540513 | 0.01857005 | 0.05677705 | NA | NA |
| ENSG000002 | 9.43892054 | -0.818692484 | 0.34777957 | -2.3540557 | 0.01856983 | 0.05677705 | NA | NA |
| ENSG000002 | 18.1839249 | -0.755850688 | 0.32108935 | -2.3540198 | 0.01857162 | 0.05677758 | LOC101927040 | 101927040 |
| ENSG000001 | 13.9506907 | -0.748451167 | 0.31798817 | -2.3537076 | 0.01858722 | 0.056821 | NHEJ1 | 79840 |
| ENSG000002 | 7.88484449 | 0.832506098 | 0.35370536 | 2.35367113 | 0.01858905 | 0.05682229 | NA | NA |
| ENSG000002 | 4.76991695 | -0.886861789 | 0.37682184 | -2.3535308 | 0.01859607 | 0.05683947 | NA | NA |
| ENSG000002 | 2.7563064 | -0.875003646 | 0.37184909 | -2.3531149 | 0.01861688 | 0.05689878 | SMARCE1P7 | 106481716 |
| ENSG000002 | 27.6228962 | -0.594603034 | 0.25270807 | -2.3529246 | 0.01862641 | 0.05692362 | LOC101928865 | 101928865 |
| ENSG000002 | 13.6342723 | -0.745418909 | 0.31682056 | -2.352811 | 0.0186321 | 0.05693672 | NA | NA |
| ENSG000001 | 1499.57639 | 0.659981087 | 0.2805128 | 2.35276642 | 0.01863434 | 0.05693776 | PTPRF | 5792 |
| ENSG000002 | 5.65645135 | -0.871806589 | 0.3705482 | -2.3527482 | 0.01863525 | 0.05693776 | NA | NA |
| ENSG000002 | 3.15563268 | -0.882856552 | 0.37525119 | -2.3527082 | 0.01863725 | 0.05693959 | LOC105373278 | 105373278 |
| ENSG00000C | 56.09362 | -0.494455829 | 0.21017366 | -2.352606 | 0.01864237 | 0.05694487 | REEP1 | 65055 |
| ENSG000001 | 410.640413 | 0.302800153 | 0.12870929 | 2.35258973 | 0.01864319 | 0.05694487 | TBCEL | 219899 |
| ENSG000001 | 51.5162675 | -0.48349605 | 0.20551329 | -2.3526266 | 0.01864134 | 0.05694487 | CCDC73 | 493860 |
| ENSG000001 | 8.32277327 | -0.82483689 | 0.35061702 | -2.3525295 | 0.01864621 | 0.05694979 | FAM90A1 | 55138 |
| ENSG000001 | 18.8369231 | -0.69493263 | 0.29542393 | -2.3523234 | 0.01865655 | 0.05697278 | Mar-10 | 162333 |
| ENSG000001 | 304.215759 | 0.42749786 | 0.18173346 | 2.35233431 | 0.018656 | 0.05697278 | SCRIB | 23513 |
| ENSG000001 | 8.69839697 | -0.817432473 | 0.34750583 | -2.3522842 | 0.01865851 | 0.0569745 | SPNS3 | 201305 |

| | | | | | | | | |
|---|---|---|---|---|---|---|---|---|
| ENSG000002 | 51.1846491 | 0.527238721 | 0.22414972 | 2.35217215 | 0.01866413 | 0.05698738 | NA | NA |
| ENSG000002 | 9.2679204 | -0.813600165 | 0.34590289 | -2.3521057 | 0.01866747 | 0.05699327 | LOC102723927 | 102723927 |
| ENSG000001 | 3.07678285 | -0.885366017 | 0.37649362 | -2.3516096 | 0.01869238 | 0.05706503 | OR10H2 | 26538 |
| ENSG000002 | 3.41073286 | -0.888900993 | 0.37800341 | -2.3515687 | 0.01869444 | 0.05706701 | NA | NA |
| ENSG000002 | 46.3586049 | 0.538317165 | 0.22892664 | 2.35148328 | 0.01869873 | 0.05707235 | LOC100506603 | 100506603 |
| ENSG000002 | 46.5007959 | -0.543993696 | 0.2313412 | -2.3514778 | 0.018699 | 0.05707235 | MRC1 | 4360 |
| ENSG000002 | 5.84170658 | -0.878214831 | 0.37352701 | -2.3511414 | 0.01871592 | 0.05711806 | LOC105372257 | 105372257 |
| ENSG000002 | 4.10814018 | -0.889884186 | 0.37849309 | -2.351124 | 0.0187168 | 0.05711806 | NA | NA |
| ENSG000002 | 27.7840011 | -0.591250845 | 0.25148525 | -2.3510359 | 0.01872123 | 0.05712614 | ITGA9-AS1 | 101928153 |
| ENSG000002 | 4.08223065 | -0.88934874 | 0.37828283 | -2.3510154 | 0.01872226 | 0.05712614 | NA | NA |
| ENSG000001 | 39.7797771 | -0.562363703 | 0.23921114 | -2.3509093 | 0.0187276 | 0.05713509 | EPHA5 | 2044 |
| ENSG000002 | 15.5488516 | -0.715720004 | 0.30444497 | -2.3509011 | 0.01872801 | 0.05713509 | LOC101929260 | 101929260 |
| ENSG000001 | 5.11227012 | -0.883016319 | 0.3756166 | -2.3508448 | 0.01873085 | 0.05713944 | COX7A1 | 1346 |
| ENSG000001 | 303.151569 | 0.437257903 | 0.18601253 | 2.35069053 | 0.01873861 | 0.05715453 | PRR12 | 57479 |
| ENSG000002 | 4.89384645 | -0.887100456 | 0.37737701 | -2.3507009 | 0.01873809 | 0.05715453 | LOC101928326 | 101928326 |
| ENSG000001 | 14.5109907 | -0.73896346 | 0.31437342 | -2.3505914 | 0.0187436 | 0.05716545 | ACTA2-AS1 | 100132116 |
| ENSG000001 | 135.879484 | 0.476021771 | 0.20251743 | 2.35052252 | 0.01874707 | 0.05717173 | IGHMBP2 | 3508 |
| ENSG000002 | 3.15784432 | -0.882017396 | 0.37525371 | -2.3504562 | 0.01875042 | 0.05717763 | NA | NA |
| ENSG000002 | 10.0311872 | -0.807923513 | 0.34374869 | -2.3503319 | 0.01875668 | 0.05718382 | NA | NA |
| ENSG000002 | 5.95383714 | -0.881381003 | 0.37499923 | -2.3503541 | 0.01875556 | 0.05718382 | NA | NA |
| ENSG000002 | 4.05257934 | -0.888979574 | 0.37823439 | -2.3503404 | 0.01875625 | 0.05718382 | NA | NA |
| ENSG000002 | 5.16626214 | -0.884310649 | 0.37627456 | -2.3501739 | 0.01876464 | 0.0572038 | NA | NA |
| ENSG000002 | 2.3724497 | -0.860514524 | 0.36622529 | -2.3496862 | 0.01878924 | 0.05727449 | NA | NA |
| ENSG000002 | 4.16235435 | -0.889352167 | 0.37851857 | -2.3495602 | 0.0187956 | 0.05728958 | LINC01048 | 103695431 |
| ENSG000002 | 5.05719765 | -0.881483555 | 0.37518081 | -2.34949 | 0.01879915 | 0.05729608 | NA | NA |
| ENSG00000C | 204.757534 | 0.438994984 | 0.18685759 | 2.34935591 | 0.01880592 | 0.05730531 | PLXND1 | 23129 |
| ENSG00000C | 14.1947353 | -0.74251793 | 0.31605196 | -2.3493539 | 0.01880602 | 0.05730531 | TKTL1 | 8277 |
| ENSG000001 | 326.038885 | 0.320743441 | 0.13652456 | 2.3493461 | 0.01880642 | 0.05730531 | LAD1 | 3898 |
| ENSG000002 | 4.0386574 | -0.888380157 | 0.37816019 | -2.3492165 | 0.01881296 | 0.05732095 | NA | NA |

| | | | | | | | | |
|---|---|---|---|---|---|---|---|---|
| ENSG000001 | 8.70447926 | -0.822810505 | 0.35026218 | -2.3491274 | 0.01881746 | 0.05733035 | NA | NA |
| ENSG000002 | 17.8810461 | 0.716193389 | 0.30490319 | 2.34892058 | 0.01882792 | 0.0573579 | NA | NA |
| ENSG000001 | 26.0944776 | -0.633691031 | 0.26978439 | -2.3488796 | 0.01883 | 0.05735991 | CERKL | 375298 |
| ENSG000001 | 156.767782 | 0.365805618 | 0.15574302 | 2.34877698 | 0.01883518 | 0.05736032 | PEX12 | 5193 |
| ENSG000001 | 22.0315221 | -0.655862792 | 0.27923447 | -2.3487888 | 0.01883459 | 0.05736032 | PIWIL1 | 9271 |
| ENSG000001 | 784.835833 | 0.628516057 | 0.26759426 | 2.34876507 | 0.01883579 | 0.05736032 | UBE2O | 63893 |
| ENSG000002 | 7.95172053 | -0.840729009 | 0.35794491 | -2.3487665 | 0.01883572 | 0.05736032 | NA | NA |
| ENSG000002 | 3.94507854 | -0.88742998 | 0.37784647 | -2.3486523 | 0.01884149 | 0.05737338 | NA | NA |
| ENSG000001 | 497.166953 | 0.340099811 | 0.14480964 | 2.34859923 | 0.01884418 | 0.05737725 | ST6GALNAC3 | 256435 |
| ENSG000002 | 12.8691208 | -0.75976081 | 0.32350028 | -2.3485631 | 0.01884601 | 0.05737851 | LOC101928580 | 101928580 |
| ENSG000001 | 111.0634 | 0.396192709 | 0.16869851 | 2.3485252 | 0.01884792 | 0.05738004 | DPP4 | 1803 |
| ENSG000001 | 18.5545843 | -0.684025711 | 0.29127399 | -2.3483927 | 0.01885463 | 0.05738323 | NEUROD4 | 58158 |
| ENSG000001 | 48.3168684 | 0.522600766 | 0.22252788 | 2.34847324 | 0.01885055 | 0.05738323 | STAR | 6770 |
| ENSG000001 | 117.840523 | 0.384694705 | 0.16381108 | 2.34840468 | 0.01885402 | 0.05738323 | ENDOD1 | 23052 |
| ENSG000002 | 172.282647 | 0.366139708 | 0.15590788 | 2.34843623 | 0.01885243 | 0.05738323 | NA | NA |
| ENSG000002 | 211.690265 | 0.316327853 | 0.13470488 | 2.34830283 | 0.01885918 | 0.05739277 | UBA6-AS1 | 550112 |
| ENSG000001 | 35.2634617 | 0.560434138 | 0.238683 | 2.34802706 | 0.01887315 | 0.05743097 | FBF1 | 85302 |
| ENSG000002 | 9.18724864 | -0.822326286 | 0.35023295 | -2.3479409 | 0.01887751 | 0.05743994 | LINC00221 | 338005 |
| ENSG000001 | 2094.46777 | 0.276838884 | 0.11790899 | 2.34790314 | 0.01887943 | 0.05744146 | ZC3HAV1 | 56829 |
| ENSG000001 | 13.8805743 | -0.742137948 | 0.3160912 | -2.3478602 | 0.01888161 | 0.05744378 | RANBP3L | 202151 |
| ENSG000002 | 8.50007059 | -0.830391561 | 0.35370481 | -2.3476966 | 0.0188899 | 0.05746039 | ZNF503-AS2 | 100131213 |
| ENSG000002 | 28.7676885 | -0.608189195 | 0.25905775 | -2.3476974 | 0.01888986 | 0.05746039 | NA | NA |
| ENSG000001 | 85.5644577 | 0.431966842 | 0.1840083 | 2.34753993 | 0.01889785 | 0.05747829 | TANGO2 | 128989 |
| ENSG000002 | 3.06870643 | -0.884082563 | 0.37660204 | -2.3475246 | 0.01889862 | 0.05747829 | RN7SKP186 | 106480692 |
| ENSG000001 | 244.953896 | 0.330366376 | 0.14073319 | 2.34746597 | 0.0189016 | 0.05748303 | LRRC1 | 55227 |
| ENSG000002 | 14.6589611 | -0.727439845 | 0.3099017 | -2.3473245 | 0.01890878 | 0.05750055 | LINC00243 | 401247 |
| ENSG000001 | 409.677553 | 0.306954237 | 0.13077597 | 2.34717622 | 0.01891631 | 0.05751611 | TMEM263 | 90488 |
| ENSG000002 | 16.8257464 | -0.692889797 | 0.29520249 | -2.3471678 | 0.01891673 | 0.05751611 | NA | NA |
| ENSG000002 | 18.8711713 | -0.739351363 | 0.31500285 | -2.3471259 | 0.01891886 | 0.05751827 | NA | NA |

| | | | | | | | | |
|---|---|---|---|---|---|---|---|---|
| ENSG000001 | 13.7197856 | -0.748343921 | 0.31885353 | -2.3469834 | 0.0189261 | 0.05753597 | INSL4 | 3641 |
| ENSG000001 | 578.08308 | 0.430084659 | 0.18325916 | 2.34686582 | 0.01893207 | 0.05754707 | EP400 | 57634 |
| ENSG000002 | 5.00978278 | -0.878759572 | 0.37444127 | -2.3468556 | 0.01893259 | 0.05754707 | NA | NA |
| ENSG000002 | 34.3501114 | -0.594244166 | 0.25325919 | -2.3463874 | 0.0189564 | 0.05761511 | NA | NA |
| ENSG000002 | 4.46121771 | -0.887160674 | 0.37810636 | -2.3463257 | 0.01895953 | 0.05762032 | NA | NA |
| ENSG000002 | 5.02538575 | -0.887829507 | 0.37841611 | -2.3461726 | 0.01896732 | 0.05763968 | NA | NA |
| ENSG000002 | 44.3730513 | 0.53389915 | 0.22760137 | 2.34576421 | 0.01898812 | 0.05769854 | HIST2H2AA4 | 723790 |
| ENSG000001 | 202.760035 | 0.38155193 | 0.16267251 | 2.34552185 | 0.01900046 | 0.05773174 | RNF24 | 11237 |
| ENSG000001 | 14.7113751 | -0.729824297 | 0.31117004 | -2.3454195 | 0.01900568 | 0.05774327 | CXCL9 | 4283 |
| ENSG000000 | 25.5381709 | -0.610260512 | 0.26021524 | -2.3452143 | 0.01901615 | 0.05777073 | CDKL3 | 51265 |
| ENSG000001 | 25.0689434 | -0.618562883 | 0.26378461 | -2.3449544 | 0.01902941 | 0.05780669 | PART1 | 25859 |
| ENSG000000 | 539.51002 | 0.330059321 | 0.14076345 | 2.34477992 | 0.01903831 | 0.05782538 | PHLDB1 | 23187 |
| ENSG000001 | 11.4974216 | -0.783826743 | 0.33428613 | -2.344778 | 0.01903841 | 0.05782538 | WNT9B | 7484 |
| ENSG000001 | 54.6745266 | 0.490323216 | 0.20912123 | 2.34468409 | 0.01904321 | 0.05783245 | PMS2P3 | 5387 |
| ENSG000002 | 3.12369871 | -0.880096547 | 0.37535947 | -2.3446765 | 0.01904359 | 0.05783245 | LINC00500 | 106480440 |
| ENSG000000 | 19.2183967 | -0.665508613 | 0.28385229 | -2.3445596 | 0.01904957 | 0.05784626 | TBX15 | 6913 |
| ENSG000002 | 6.62017302 | -0.856495453 | 0.36533577 | -2.3444062 | 0.0190574 | 0.05786573 | NA | NA |
| ENSG000002 | 9.0885531 | -0.819117294 | 0.34941389 | -2.3442608 | 0.01906483 | 0.05788395 | NA | NA |
| ENSG000001 | 5.72646703 | -0.869760606 | 0.37102795 | -2.3441916 | 0.01906837 | 0.05789037 | TBC1D21 | 161514 |
| ENSG000002 | 51.7131028 | 0.505293663 | 0.21556159 | 2.34408027 | 0.01907407 | 0.05790331 | LIX1L | 128077 |
| ENSG000002 | 5.62008334 | -0.880440564 | 0.37560923 | -2.3440334 | 0.01907647 | 0.05790626 | NA | NA |
| ENSG000002 | 251.897888 | 0.336911623 | 0.14373923 | 2.34390864 | 0.01908285 | 0.05792129 | ZSWIM7 | 125150 |
| ENSG000001 | 139.608637 | 0.367814485 | 0.15693948 | 2.34367087 | 0.01909501 | 0.05794521 | CAND2 | 23066 |
| ENSG000002 | 2.77817 | -0.872651709 | 0.372343 | -2.343677 | 0.0190947 | 0.05794521 | NA | NA |
| ENSG000002 | 14.1847153 | -0.739275553 | 0.31543081 | -2.3437011 | 0.01909346 | 0.05794521 | LOC105369443 | 105369443 |
| ENSG000001 | 27.3563706 | -0.609779775 | 0.26019109 | -2.3435844 | 0.01909944 | 0.05795431 | SLC30A4 | 7782 |
| ENSG000002 | 3.87141665 | -0.886869528 | 0.37844255 | -2.343472 | 0.01910519 | 0.05796743 | NA | NA |
| ENSG000002 | 2.31455302 | -0.861914429 | 0.36784713 | -2.3431321 | 0.01912261 | 0.05801159 | NA | NA |
| ENSG000002 | 6.36803277 | -0.861919975 | 0.36784544 | -2.3431579 | 0.01912129 | 0.05801159 | LINC01476 | 101927728 |

| | | | | | | | | |
|---|---|---|---|---|---|---|---|---|
| ENSG000001 | 558.266886 | 0.286019935 | 0.12207236 | 2.34303601 | 0.01912754 | 0.0580222 | KIAA2026 | 158358 |
| ENSG000001 | 117.832123 | 0.410011427 | 0.17499638 | 2.34297088 | 0.01913088 | 0.05802799 | INO80C | 125476 |
| ENSG000001 | 25.9410949 | -0.618653888 | 0.26405806 | -2.3428707 | 0.01913601 | 0.0580354 | LHX8 | 431707 |
| ENSG000001 | 16.9233474 | -0.70046391 | 0.29897719 | -2.3428674 | 0.01913618 | 0.0580354 | MFSD4 | 148808 |
| ENSG000002 | 3.14298407 | -0.87927392 | 0.37536703 | -2.3424378 | 0.01915823 | 0.05809791 | NA | NA |
| ENSG000001 | 22.7429893 | -0.642804429 | 0.27443993 | -2.3422409 | 0.01916834 | 0.05812422 | FAM83A | 84985 |
| ENSG000002 | 3.43964437 | -0.873923203 | 0.37312544 | -2.34217 | 0.01917198 | 0.05813093 | NA | NA |
| ENSG000001 | 1314.36477 | 0.265285974 | 0.11327264 | 2.34201283 | 0.01918006 | 0.05815106 | FAM111B | 374393 |
| ENSG000002 | 3.0971109 | -0.880219586 | 0.37586984 | -2.3418202 | 0.01918996 | 0.05817673 | NA | NA |
| ENSG000001 | 5.73638331 | -0.873506556 | 0.37302168 | -2.3417045 | 0.0191959 | 0.05819041 | NA | NA |
| ENSG000001 | 176.220191 | 0.356719169 | 0.15234118 | 2.34158077 | 0.01920227 | 0.05820536 | MED18 | 54797 |
| ENSG000001 | 5.76398751 | -0.872941492 | 0.37280619 | -2.3415424 | 0.01920425 | 0.05820699 | FAM151A | 338094 |
| ENSG000001 | 11.7454719 | -0.794611329 | 0.33935804 | -2.3415132 | 0.01920575 | 0.0582072 | SPATA19 | 219938 |
| ENSG000002 | 4.45411637 | -0.885126455 | 0.37801975 | -2.3414821 | 0.01920735 | 0.0582077 | NA | NA |
| ENSG000001 | 131.869036 | 0.382170202 | 0.16322912 | 2.34131136 | 0.01921613 | 0.05822186 | ATXN7L2 | 127002 |
| ENSG000001 | 28.8375548 | 0.58939542 | 0.25174281 | 2.34126015 | 0.01921877 | 0.05822186 | DLL1 | 28514 |
| ENSG000002 | 4.84531582 | 0.882944282 | 0.37712488 | 2.34125176 | 0.0192192 | 0.05822186 | RNU6-1 | 26827 |
| ENSG000002 | 52.3403735 | 0.483821104 | 0.20664832 | 2.34127774 | 0.01921786 | 0.05822186 | SNORD116-2 | 100033414 |
| ENSG000002 | 2.77274153 | -0.870229073 | 0.37168984 | -2.3412775 | 0.01921788 | 0.05822186 | NA | NA |
| ENSG000002 | 4.43376794 | -0.884681055 | 0.37789905 | -2.3410513 | 0.01922952 | 0.05824529 | NA | NA |
| ENSG000002 | 6.81811597 | -0.865052528 | 0.36951543 | -2.3410458 | 0.01922981 | 0.05824529 | NA | NA |
| ENSG000001 | 19.5018896 | -0.660563625 | 0.28217308 | -2.3409874 | 0.01923281 | 0.05824569 | IL13RA2 | 3598 |
| ENSG000001 | 1084.0728 | 0.287432426 | 0.12278231 | 2.34099218 | 0.01923257 | 0.05824569 | PRKAB2 | 5565 |
| ENSG00000C | 326.707474 | 0.297730083 | 0.12718895 | 2.34084872 | 0.01923996 | 0.05826299 | EML1 | 2009 |
| ENSG000001 | 10.7858941 | -0.786841927 | 0.3361425 | -2.3407987 | 0.01924254 | 0.05826645 | KCNG4 | 93107 |
| ENSG000002 | 5.66787643 | -0.882159779 | 0.37686858 | -2.3407624 | 0.01924441 | 0.05826776 | NA | NA |
| ENSG00000C | 96.4365834 | 0.457655017 | 0.19552454 | 2.34065257 | 0.01925007 | 0.05828002 | PLEKHG6 | 55200 |
| ENSG000002 | 8.96950008 | -0.834387557 | 0.3564802 | -2.3406281 | 0.01925133 | 0.05828002 | NA | NA |
| ENSG000001 | 262.245014 | 0.327908799 | 0.14010408 | 2.34046578 | 0.0192597 | 0.058301 | RAB5C | 5878 |

| | | | | | | | | |
|---|---|---|---|---|---|---|---|---|
| ENSG000001 | 7.85214729 | -0.834495365 | 0.35656445 | -2.3403773 | 0.01926427 | 0.05831046 | GPR4 | 2828 |
| ENSG000002 | 14.8746738 | -0.713874848 | 0.30503912 | -2.3402731 | 0.01926964 | 0.05832238 | LOC101929227 | 101929227 |
| ENSG000002 | 2.75337826 | -0.870828167 | 0.37212849 | -2.3401276 | 0.01927715 | 0.05834075 | NA | NA |
| ENSG000002 | 6.62291792 | -0.856212365 | 0.36590279 | -2.3399996 | 0.01928376 | 0.0583564 | NA | NA |
| ENSG000001 | 33.1069089 | -0.559469515 | 0.23910153 | -2.3398826 | 0.0192898 | 0.05836838 | TGFB3 | 7043 |
| ENSG000002 | 12.7764694 | -0.79369814 | 0.33920649 | -2.3398672 | 0.0192906 | 0.05836838 | LOC101929123 | 101929123 |
| ENSG000002 | 7.84051287 | -0.835745005 | 0.35725795 | -2.3393321 | 0.01931825 | 0.05844769 | NA | NA |
| ENSG000001 | 16.8445798 | -0.715634782 | 0.30592252 | -2.3392681 | 0.01932156 | 0.05844899 | ANGPT4 | 51378 |
| ENSG000002 | 5.64109418 | -0.879276279 | 0.37587477 | -2.3392799 | 0.01932095 | 0.05844899 | NA | NA |
| ENSG000001 | 20.0661441 | -0.653777487 | 0.27949865 | -2.3391078 | 0.01932985 | 0.05846971 | GUCY1A3 | 2982 |
| ENSG000001 | 18.3301404 | -0.682585533 | 0.29182954 | -2.3389871 | 0.0193361 | 0.05848424 | TBX19 | 9095 |
| ENSG000002 | 8.91801172 | -0.826279283 | 0.35327823 | -2.338891 | 0.01934107 | 0.05849492 | LOC101928118 | 101928118 |
| ENSG000001 | 19.0729037 | -0.665872382 | 0.28470126 | -2.338846 | 0.01934341 | 0.05849761 | CASP5 | 838 |
| ENSG000002 | 64.3230361 | 0.458910345 | 0.19621698 | 2.33879021 | 0.01934629 | 0.05850198 | SNORA26 | 677810 |
| ENSG000002 | 2.36803917 | -0.856475002 | 0.36623241 | -2.3386106 | 0.0193556 | 0.05852575 | NA | NA |
| ENSG000001 | 4.75575972 | -0.882066394 | 0.37718499 | -2.3385511 | 0.01935868 | 0.05852633 | GOLGA6L4 | 643707 |
| ENSG000002 | 4.8377826 | -0.882947856 | 0.37755814 | -2.3385746 | 0.01935746 | 0.05852633 | NA | NA |
| ENSG000002 | 6.10472805 | -0.865649269 | 0.37017463 | -2.3384889 | 0.0193619 | 0.05853171 | NA | NA |
| ENSG000002 | 10.7723708 | -0.784646382 | 0.33555911 | -2.3383254 | 0.01937037 | 0.05855297 | LOC101928335 | 101928335 |
| ENSG000001 | 297.360021 | 0.313804542 | 0.13420763 | 2.33820196 | 0.01937677 | 0.05856795 | KLHL8 | 57563 |
| ENSG000001 | 131.119581 | 0.366203454 | 0.15662031 | 2.33816067 | 0.01937892 | 0.05857005 | ANKRA2 | 57763 |
| ENSG000001 | 16.7971537 | -0.689030721 | 0.29469439 | -2.3381196 | 0.01938104 | 0.05857212 | CXorf58 | 254158 |
| ENSG000001 | 180.929835 | 0.371179166 | 0.15878691 | 2.33759299 | 0.01940837 | 0.05865028 | C19orf66 | 55337 |
| ENSG000001 | 3.74513226 | -0.884887779 | 0.378551 | -2.3375655 | 0.0194098 | 0.05865028 | GPR21 | 2844 |
| ENSG000002 | 14.0385326 | -0.749719845 | 0.32074789 | -2.3374117 | 0.01941779 | 0.05867005 | TMEM213 | 155006 |
| ENSG000001 | 2013.81639 | 0.296244391 | 0.12674749 | 2.33728012 | 0.01942462 | 0.05868633 | PVRL3 | 25945 |
| ENSG000001 | 20.9912571 | 0.683487835 | 0.29243859 | 2.33720126 | 0.01942872 | 0.05869434 | TSSK6 | 83983 |
| ENSG000001 | 60.2409013 | 0.472088386 | 0.20200641 | 2.33699704 | 0.01943934 | 0.05872203 | PACSIN1 | 29993 |
| ENSG000001 | 36.5801 | -0.564374725 | 0.24150603 | -2.3368971 | 0.01944454 | 0.05872899 | SULT1B1 | 27284 |

| | | | | | | | | |
|---|---|---|---|---|---|---|---|---|
| ENSG000002 | 3.77513645 | -0.884637385 | 0.37854848 | -2.3369197 | 0.01944336 | 0.05872899 | LACTBL1 | 646262 |
| ENSG000002 | 22.4453279 | -0.676376814 | 0.28948367 | -2.3364939 | 0.01946552 | 0.0587871 | NA | NA |
| ENSG000002 | 68.0950927 | 0.449984385 | 0.19259142 | 2.33647158 | 0.01946668 | 0.0587871 | NA | NA |
| ENSG000002 | 7.28424653 | -0.846419818 | 0.36229373 | -2.3362806 | 0.01947662 | 0.05881276 | NA | NA |
| ENSG000002 | 8.51035998 | -0.828854174 | 0.35479582 | -2.3361441 | 0.01948373 | 0.05882984 | NA | NA |
| ENSG000002 | 14.2542767 | -0.737795569 | 0.3158275 | -2.3360713 | 0.01948752 | 0.05883691 | LINC00282 | 283521 |
| ENSG000002 | 2.66936033 | -0.872204665 | 0.37336924 | -2.3360378 | 0.01948927 | 0.05883781 | RNU6-1090P | 106480045 |
| ENSG000002 | 3.39480019 | -0.882389559 | 0.37774286 | -2.335953 | 0.01949369 | 0.05884677 | NA | NA |
| ENSG000001 | 1852.34095 | 0.265764236 | 0.11377277 | 2.33592123 | 0.01949535 | 0.05884739 | APPL1 | 26060 |
| ENSG000001 | 9.75228816 | -0.7892704 | 0.33789461 | -2.3358479 | 0.01949917 | 0.05885455 | ISX | 91464 |
| ENSG000002 | 3.84819128 | -0.883719636 | 0.37834367 | -2.335759 | 0.01950381 | 0.05886416 | NA | NA |
| ENSG000002 | 3.39089352 | -0.882727368 | 0.37792411 | -2.3357265 | 0.0195055 | 0.05886488 | NA | NA |
| ENSG000002 | 2.90376871 | -0.864997508 | 0.37036275 | -2.3355413 | 0.01951516 | 0.05888966 | NA | NA |
| ENSG000001 | 5.38815536 | -0.872296958 | 0.37350969 | -2.3354065 | 0.0195222 | 0.05890651 | OR3A1 | 4994 |
| ENSG000002 | 11.6908519 | -0.777643211 | 0.33299724 | -2.3352842 | 0.01952858 | 0.05892138 | NA | NA |
| ENSG000002 | 27.2618759 | -0.589176107 | 0.25229658 | -2.3352521 | 0.01953026 | 0.05892206 | NA | NA |
| ENSG000002 | 9.45544552 | -0.816766629 | 0.34976251 | -2.3352035 | 0.01953279 | 0.05892533 | NA | NA |
| ENSG000001 | 777.002322 | 0.292603918 | 0.12530917 | 2.33505596 | 0.0195405 | 0.05894371 | EHBP1 | 23301 |
| ENSG000001 | 21.2404119 | -0.669812695 | 0.2868541 | -2.3350292 | 0.0195419 | 0.05894371 | KIAA1755 | 85449 |
| ENSG000002 | 339.001054 | 0.337772402 | 0.14465607 | 2.33500334 | 0.01954325 | 0.05894371 | LINC01224 | 104472717 |
| ENSG000001 | 93.1421884 | 0.451480737 | 0.19335568 | 2.33497528 | 0.01954471 | 0.05894375 | PTCD1 | 26024 |
| ENSG000001 | 32.9979407 | -0.564665162 | 0.24183533 | -2.3349159 | 0.01954781 | 0.05894872 | SLCO1C1 | 53919 |
| ENSG000002 | 18.6322561 | -0.708346834 | 0.30338113 | -2.3348414 | 0.01955171 | 0.05895607 | NA | NA |
| ENSG000001 | 4.46241029 | -0.883246183 | 0.37836439 | -2.3343798 | 0.01957584 | 0.05902447 | KLKP1 | 606293 |
| ENSG000002 | 3.37848804 | 0.87894479 | 0.37654996 | 2.33420496 | 0.01958499 | 0.05904766 | RNU1-73P | 106481616 |
| ENSG000002 | 7.58321279 | -0.838622751 | 0.35928659 | -2.3341332 | 0.01958875 | 0.05905459 | NA | NA |
| ENSG000001 | 579.107715 | 0.310557301 | 0.13305253 | 2.33409545 | 0.01959072 | 0.05905616 | CTTNBP2NL | 55917 |
| ENSG000001 | 686.807741 | 0.26730987 | 0.1145297 | 2.33397867 | 0.01959684 | 0.0590702 | MCPH1 | 79648 |
| ENSG000002 | 2.4110116 | -0.853288531 | 0.36561026 | -2.3338747 | 0.01960229 | 0.05908223 | NA | NA |

| | | | | | | | | |
|---|---|---|---|---|---|---|---|---|
| ENSG000002 | 51.792608 | 0.499856507 | 0.21418071 | 2.33380727 | 0.01960582 | 0.05908848 | NA | NA |
| ENSG000001 | 8.72427872 | -0.819261876 | 0.35109094 | -2.3334748 | 0.01962324 | 0.05913659 | GZMB | 3002 |
| ENSG000002 | 16.9937936 | -0.700690935 | 0.30028299 | -2.3334353 | 0.01962531 | 0.05913844 | NA | NA |
| ENSG000001 | 13.4393989 | -0.754732979 | 0.32348808 | -2.3331091 | 0.01964242 | 0.0591856 | FTCD | 10841 |
| ENSG000001 | 7.9719428 | -0.836964634 | 0.35874892 | -2.3330095 | 0.01964765 | 0.05919695 | TKTL2 | 84076 |
| ENSG000002 | 16.9728626 | -0.720731975 | 0.30893706 | -2.3329412 | 0.01965123 | 0.05920336 | GGTA1P | 2681 |
| ENSG000001 | 21.0342661 | -0.674277932 | 0.28902928 | -2.3329053 | 0.01965312 | 0.05920463 | ASGR2 | 433 |
| ENSG000001 | 98.924335 | 0.44976082 | 0.19281363 | 2.33261942 | 0.01966813 | 0.05924105 | TSC22D4 | 81628 |
| ENSG000002 | 10.9196962 | -0.792326456 | 0.33966901 | -2.3326428 | 0.0196669 | 0.05924105 | NA | NA |
| ENSG000002 | 3.03810939 | -0.877866017 | 0.37637315 | -2.3324353 | 0.0196778 | 0.05926579 | LOC101929552 | 101929552 |
| ENSG000002 | 6.29559061 | -0.855984962 | 0.36699922 | -2.332389 | 0.01968023 | 0.05926871 | NA | NA |
| ENSG000002 | 2.41337466 | -0.852381222 | 0.36546109 | -2.3323447 | 0.01968257 | 0.05927133 | NA | NA |
| ENSG000002 | 3.25558784 | -0.873796502 | 0.37465965 | -2.3322407 | 0.01968803 | 0.05928338 | NA | NA |
| ENSG000001 | 2725.47081 | 0.281297584 | 0.12062281 | 2.332043 | 0.01969843 | 0.05930265 | SUGT1 | 10910 |
| ENSG000001 | 131.148081 | 0.373880391 | 0.16032362 | 2.33203562 | 0.01969882 | 0.05930265 | UCKL1 | 54963 |
| ENSG000002 | 26.1144398 | -0.65228478 | 0.27970392 | -2.3320544 | 0.01969783 | 0.05930265 | LINC01539 | 100505474 |
| ENSG000002 | 6.5365029 | 0.852151684 | 0.36541714 | 2.33199703 | 0.01970085 | 0.05930436 | NA | NA |
| ENSG000002 | 8.68888557 | -0.8127397 | 0.34853789 | -2.3318547 | 0.01970834 | 0.05932251 | NA | NA |
| ENSG000001 | 6.3436338 | -0.862051211 | 0.36969061 | -2.331818 | 0.01971027 | 0.05932392 | METTL21C | 196541 |
| ENSG000002 | 248.562235 | 0.330073131 | 0.1415545 | 2.33177422 | 0.01971257 | 0.05932644 | NA | NA |
| ENSG000002 | 10.6008504 | -0.79202056 | 0.33969979 | -2.3315309 | 0.01972538 | 0.05936041 | GIMAP1 | 170575 |
| ENSG000002 | 27.2372172 | -0.588344266 | 0.25234535 | -2.3315043 | 0.01972678 | 0.05936041 | LOC101928303 | 101928303 |
| ENSG000001 | 485.424959 | 0.344555924 | 0.14779456 | 2.33131669 | 0.01973667 | 0.05937693 | CLK2 | 1196 |
| ENSG000002 | 513.124704 | 0.312886616 | 0.13420769 | 2.33136134 | 0.01973431 | 0.05937693 | CSNK2A3 | 283106 |
| ENSG000002 | 9.73894097 | -0.808867448 | 0.34695602 | -2.3313256 | 0.0197362 | 0.05937693 | NA | NA |
| ENSG000002 | 13.0277381 | -0.748543341 | 0.3210888 | -2.3312658 | 0.01973935 | 0.05938058 | NA | NA |
| ENSG000002 | 5.00001024 | -0.873853109 | 0.37487186 | -2.3310715 | 0.01974959 | 0.05940699 | NA | NA |
| ENSG000002 | 8.36303359 | -0.818348453 | 0.35106724 | -2.3310305 | 0.01975175 | 0.05940908 | NA | NA |
| ENSG000001 | 13.63278 | -0.760452691 | 0.32623896 | -2.3309683 | 0.01975503 | 0.05941454 | NNMT | 4837 |

| | | | | | | | | |
|---|---|---|---|---|---|---|---|---|
| ENSG00000C | 251.599175 | 0.330020759 | 0.1415873 | 2.33086419 | 0.01976052 | 0.05942406 | PLEKHO1 | 51177 |
| ENSG000002 | 2.40798486 | -0.849850898 | 0.36460943 | -2.3308527 | 0.01976113 | 0.05942406 | NA | NA |
| ENSG000001 | 25.7463561 | 0.624975502 | 0.26814167 | 2.33076604 | 0.0197657 | 0.0594334 | CITED4 | 163732 |
| ENSG000001 | 1069.36908 | 0.263149154 | 0.11291123 | 2.33058435 | 0.01977529 | 0.05945782 | SBF2 | 81846 |
| ENSG000002 | 12.8391065 | -0.792077255 | 0.33988412 | -2.3304332 | 0.01978327 | 0.0594774 | NA | NA |
| ENSG000001 | 222.669994 | 0.423452607 | 0.18171003 | 2.3303755 | 0.01978631 | 0.05948216 | WDR60 | 55112 |
| ENSG000001 | 52.7822914 | 0.500656692 | 0.21485144 | 2.33024588 | 0.01979316 | 0.05949833 | TMEM25 | 84866 |
| ENSG000002 | 6.73413388 | -0.854304121 | 0.36662581 | -2.33018 | 0.01979664 | 0.05950438 | NA | NA |
| ENSG000002 | 7.51674493 | -0.850107422 | 0.36485222 | -2.3300048 | 0.0198059 | 0.05952779 | LOC101928809 | 101928809 |
| ENSG000001 | 1484.08803 | 0.295227894 | 0.12671117 | 2.32992802 | 0.01980996 | 0.05953116 | UBN2 | 254048 |
| ENSG000001 | 51.8052641 | 0.493217692 | 0.21168706 | 2.32993786 | 0.01980944 | 0.05953116 | GLUD2 | 2747 |
| ENSG000002 | 42.2366349 | 0.525689825 | 0.2256399 | 2.32977339 | 0.01981813 | 0.05955132 | NA | NA |
| ENSG000001 | 170.592587 | 0.383522509 | 0.16463171 | 2.32957855 | 0.01982844 | 0.05955846 | DYNLT3 | 6990 |
| ENSG000001 | 9.30194214 | -0.805556356 | 0.34578561 | -2.3296411 | 0.01982513 | 0.05955846 | MS4A14 | 84689 |
| ENSG000001 | 7.70386626 | -0.84447908 | 0.3625016 | -2.3295872 | 0.01982798 | 0.05955846 | C2orf54 | 79919 |
| ENSG000001 | 15.6400834 | -0.700221904 | 0.30058095 | -2.3295619 | 0.01982932 | 0.05955846 | ZNF366 | 167465 |
| ENSG000002 | 3.39050122 | -0.880236105 | 0.37784561 | -2.3296184 | 0.01982633 | 0.05955846 | NA | NA |
| ENSG000002 | 36.7780815 | 0.547233887 | 0.23490245 | 2.32962188 | 0.01982614 | 0.05955846 | LOC102723897 | 102723897 |
| ENSG000001 | 18.9492625 | -0.675968691 | 0.290181 | -2.3294726 | 0.01983404 | 0.05956823 | CWH43 | 80157 |
| ENSG00000C | 191.31798 | 0.389389196 | 0.16718961 | 2.32902742 | 0.01985761 | 0.05963461 | PHPT1 | 29085 |
| ENSG000001 | 6.77447511 | -0.86087692 | 0.36964706 | -2.3289159 | 0.01986352 | 0.05964793 | SOHLH1 | 402381 |
| ENSG000002 | 3.0691825 | -0.876011472 | 0.37615621 | -2.3288502 | 0.019867 | 0.05965194 | NA | NA |
| ENSG000002 | 12.7692477 | -0.790146757 | 0.3392884 | -2.3288352 | 0.0198678 | 0.05965194 | NA | NA |
| ENSG000001 | 2560.27081 | 0.587368367 | 0.2522313 | 2.32868944 | 0.01987552 | 0.05967072 | SLC7A1 | 6541 |
| ENSG000001 | 61.3373675 | -0.453739697 | 0.19488094 | -2.3282918 | 0.01989662 | 0.05972796 | KIAA0226L | 80183 |
| ENSG000002 | 15.8039753 | 0.706875723 | 0.30360499 | 2.3282744 | 0.01989754 | 0.05972796 | NA | NA |
| ENSG000002 | 2.65677587 | -0.868968163 | 0.3732421 | -2.3281622 | 0.01990349 | 0.05974141 | NA | NA |
| ENSG00000C | 7.39544927 | -0.830828049 | 0.35687236 | -2.3280818 | 0.01990776 | 0.05974538 | BPIFB2 | 80341 |
| ENSG000001 | 8104.74693 | 0.313966795 | 0.1348606 | 2.32808395 | 0.01990765 | 0.05974538 | NA | NA |

| | | | | | | | | |
|---|---|---|---|---|---|---|---|---|
| ENSG000002 | 24.4271301 | -0.62018246 | 0.26641561 | -2.3278758 | 0.0199187 | 0.05977378 | PLCE1-AS2 | 101927049 |
| ENSG000001 | 23.5157822 | -0.645616714 | 0.27734599 | -2.3278386 | 0.01992067 | 0.05977529 | ACKR2 | 1238 |
| ENSG000001 | 16.7997695 | -0.688928799 | 0.29596567 | -2.3277321 | 0.01992633 | 0.05978785 | IL7 | 3574 |
| ENSG000001 | 60.1595098 | -0.471710995 | 0.20265242 | -2.327685 | 0.01992883 | 0.05979093 | SPTBN4 | 57731 |
| ENSG000002 | 11.2296116 | -0.782952392 | 0.33636948 | -2.3276559 | 0.01993038 | 0.05979115 | LOC101926955 | 101926955 |
| ENSG000001 | 541.35514 | 0.274822424 | 0.1180727 | 2.32756963 | 0.01993497 | 0.05980048 | PLEKHA3 | 65977 |
| ENSG000001 | 131.788862 | 0.385047878 | 0.16545682 | 2.32718039 | 0.01995567 | 0.05985374 | ZSWIM3 | 140831 |
| ENSG000001 | 10.2463677 | -0.801092184 | 0.34423652 | -2.3271563 | 0.01995695 | 0.05985374 | LPAR5 | 57121 |
| ENSG000001 | 205.173476 | 0.369404203 | 0.15873657 | 2.32715249 | 0.01995715 | 0.05985374 | ITPRIPL1 | 150771 |
| ENSG000002 | 3.07325334 | -0.875580004 | 0.37625165 | -2.3271127 | 0.01995927 | 0.05985566 | NA | NA |
| ENSG000002 | 5.68935644 | -0.865236999 | 0.37183542 | -2.3269354 | 0.01996871 | 0.05987068 | NA | NA |
| ENSG000002 | 6.28042926 | -0.855796735 | 0.3677783 | -2.3269364 | 0.01996865 | 0.05987068 | NA | NA |
| ENSG000002 | 3.46719003 | -0.878348689 | 0.3774701 | -2.3269358 | 0.01996868 | 0.05987068 | SNORA59B | 677882 |
| ENSG000001 | 1206.70119 | 0.27111164 | 0.11651716 | 2.32679583 | 0.01997613 | 0.05988848 | VAT1L | 57687 |
| ENSG000002 | 5.34467056 | -0.870503977 | 0.37412575 | -2.3267684 | 0.01997759 | 0.05988848 | NA | NA |
| ENSG000002 | 9.49051626 | -0.794549782 | 0.34148897 | -2.3267217 | 0.01998008 | 0.05989151 | LOC105370306 | 105370306 |
| ENSG000002 | 8.90183638 | -0.822290042 | 0.35342556 | -2.3266287 | 0.01998504 | 0.05990193 | LOC101928733 | 101928733 |
| ENSG000002 | 4.66012321 | -0.875115014 | 0.37615607 | -2.3264679 | 0.0199936 | 0.05992318 | NA | NA |
| ENSG000001 | 10.6829279 | -0.778460904 | 0.33464508 | -2.3262284 | 0.02000637 | 0.05994372 | CCRL2 | 9034 |
| ENSG000001 | 13.6356545 | -0.741677596 | 0.31882931 | -2.3262528 | 0.02000507 | 0.05994372 | HCAR1 | 27198 |
| ENSG000002 | 3.60428043 | -0.875152813 | 0.37620384 | -2.3262729 | 0.020004 | 0.05994372 | NA | NA |
| ENSG000002 | 13.70846 | -0.749848517 | 0.32234188 | -2.3262523 | 0.0200051 | 0.05994372 | NA | NA |
| ENSG000001 | 335.252419 | -0.347432716 | 0.14937414 | -2.3259227 | 0.02002267 | 0.05997927 | NR5A2 | 2494 |
| ENSG000002 | 866.010487 | 0.346847212 | 0.14912147 | 2.32593753 | 0.02002188 | 0.05997927 | C7orf73 | 647087 |
| ENSG000002 | 4.06119654 | -0.880224531 | 0.37843843 | -2.3259386 | 0.02002183 | 0.05997927 | LOC729506 | 729506 |
| ENSG000002 | 19.7910276 | -0.668001859 | 0.28720858 | -2.3258422 | 0.02002697 | 0.05998328 | LOC101927282 | 101927282 |
| ENSG000002 | 2.46161902 | -0.847004218 | 0.3641681 | -2.3258605 | 0.02002599 | 0.05998328 | NA | NA |
| ENSG000002 | 32.5300629 | 0.583783089 | 0.25100325 | 2.3257989 | 0.02002928 | 0.05998577 | LINC01484 | 101928136 |
| ENSG00000C | 14.3037501 | -0.746146907 | 0.3208354 | -2.3256378 | 0.02003788 | 0.06000186 | CETP | 1071 |

| | | | | | | | | | |
|---|---|---|---|---|---|---|---|---|---|
| ENSG000001 | 988.387429 | 0.294291145 | 0.12654336 | 2.32561516 | 0.02003909 | 0.06000186 | GOLIM4 | | 27333 |
| ENSG000002 | 9.58599764 | -0.822454837 | 0.35364417 | -2.3256565 | 0.02003689 | 0.06000186 | NA | NA | |
| ENSG000001 | 77.8106374 | -0.424179816 | 0.18241285 | -2.3253834 | 0.02005147 | 0.06003449 | CRACR2A | | 84766 |
| ENSG000001 | 76.9274282 | -0.420521675 | 0.18085447 | -2.3251937 | 0.02006161 | 0.06005597 | THSD7B | | 80731 |
| ENSG000002 | 2.37426903 | -0.849384929 | 0.36529367 | -2.3252112 | 0.02006067 | 0.06005597 | NA | NA | |
| ENSG000001 | 10.8509925 | -0.786304944 | 0.33820074 | -2.3249652 | 0.02007382 | 0.06008025 | TTR | | 7276 |
| ENSG000001 | 21.3825817 | -0.641392133 | 0.27587247 | -2.3249588 | 0.02007416 | 0.06008025 | CLEC4D | | 338339 |
| ENSG000002 | 2.78899078 | -0.861025756 | 0.37033305 | -2.3250038 | 0.02007176 | 0.06008025 | NA | NA | |
| ENSG000001 | 25.618532 | -0.657720942 | 0.28292119 | -2.3247496 | 0.02008535 | 0.06010931 | OPN5 | | 221391 |
| ENSG000001 | 159.314597 | 0.373578951 | 0.1607221 | 2.32437825 | 0.02010523 | 0.06016436 | GRAMD3 | | 65983 |
| ENSG00000C | 404.531037 | 0.340385444 | 0.14646033 | 2.32407944 | 0.02012124 | 0.06020205 | PIK3CB | | 5291 |
| ENSG000002 | 4.52801077 | -0.879398123 | 0.37838494 | -2.3240833 | 0.02012103 | 0.06020205 | NA | NA | |
| ENSG000002 | 9.87324571 | -0.815923672 | 0.35107686 | -2.32406 | 0.02012228 | 0.06020205 | NA | NA | |
| ENSG000001 | 6.68702371 | -0.850820847 | 0.36612153 | -2.3238755 | 0.02013217 | 0.0602233 | SLN | | 6588 |
| ENSG000002 | 7.43626295 | -0.851378793 | 0.36636217 | -2.323872 | 0.02013235 | 0.0602233 | NA | NA | |
| ENSG000002 | 59.7071533 | 0.487197028 | 0.20965472 | 2.32380664 | 0.02013586 | 0.0602249 | NA | NA | |
| ENSG000002 | 5.11648889 | -0.87504446 | 0.37655516 | -2.3238148 | 0.02013542 | 0.0602249 | NA | NA | |
| ENSG000002 | 3.84886845 | -0.878871566 | 0.37820873 | -2.3237739 | 0.02013762 | 0.06022571 | NA | NA | |
| ENSG000002 | 9.16241142 | -0.79557934 | 0.34237744 | -2.3236909 | 0.02014207 | 0.06023458 | NA | NA | |
| ENSG00000C | 506.838344 | 0.305970051 | 0.13168719 | 2.32346102 | 0.0201544 | 0.06026702 | VRK2 | | 7444 |
| ENSG000002 | 8.04821294 | -0.818766469 | 0.3524206 | -2.3232651 | 0.02016492 | 0.06028514 | NA | NA | |
| ENSG000002 | 8.65987918 | -0.811334074 | 0.34921865 | -2.3232839 | 0.02016391 | 0.06028514 | NA | NA | |
| ENSG000002 | 3.92256752 | -0.878701329 | 0.37821785 | -2.3232677 | 0.02016477 | 0.06028514 | NA | NA | |
| ENSG000002 | 5.16710382 | -0.874268376 | 0.37632227 | -2.3231906 | 0.02016891 | 0.06029264 | NA | NA | |
| ENSG000001 | 9.71324283 | -0.808644504 | 0.34807986 | -2.323158 | 0.02017067 | 0.06029343 | IL20 | | 50604 |
| ENSG000002 | 3.06326531 | -0.873544805 | 0.37604063 | -2.3230064 | 0.02017881 | 0.06031332 | NA | NA | |
| ENSG000001 | 27.5265714 | -0.599006759 | 0.25786517 | -2.3229456 | 0.02018207 | 0.06031864 | TRPC6 | | 7225 |
| ENSG000001 | 2178.1785 | 0.318117425 | 0.13695026 | 2.3228684 | 0.02018622 | 0.06032659 | SIN3A | | 25942 |
| ENSG000001 | 88.145072 | 0.405023141 | 0.17439004 | 2.32251301 | 0.02020533 | 0.06037479 | RTP1 | | 132112 |

| | | | | | | | | |
|---|---|---|---|---|---|---|---|---|
| ENSG000002 | 7.22216257 | -0.843946976 | 0.36337597 | -2.3225173 | 0.0202051 | 0.06037479 | PQLC1P1 | 106479032 |
| ENSG000002 | 3.06705574 | -0.873718273 | 0.37620372 | -2.3224605 | 0.02020815 | 0.06037878 | NA | NA |
| ENSG000002 | 4.38891725 | -0.877384133 | 0.37786804 | -2.3219327 | 0.02023656 | 0.06045921 | RFPL4A | 342931 |
| ENSG000001 | 220.049581 | 0.359930259 | 0.15506861 | 2.32110323 | 0.02028127 | 0.06058832 | CTC1 | 80169 |
| ENSG000001 | 7.28881326 | -0.841997145 | 0.36279446 | -2.3208655 | 0.0202941 | 0.06061285 | PNOC | 5368 |
| ENSG000002 | 8.9391591 | -0.806668385 | 0.3475746 | -2.3208497 | 0.02029496 | 0.06061285 | NA | NA |
| ENSG000002 | 10.835629 | -0.765110063 | 0.3296627 | -2.3208876 | 0.02029291 | 0.06061285 | NA | NA |
| ENSG000002 | 6.55980413 | -0.846273538 | 0.364641 | -2.3208404 | 0.02029546 | 0.06061285 | LOC101929497 | 101929497 |
| ENSG000002 | 5.25032818 | -0.874995758 | 0.37703254 | -2.3207433 | 0.0203007 | 0.06062403 | NA | NA |
| ENSG000002 | 4.00780445 | -0.87811414 | 0.37838096 | -2.3207144 | 0.02030226 | 0.06062424 | NA | NA |
| ENSG000002 | 201.47964 | 0.34507295 | 0.14870153 | 2.3205743 | 0.02030983 | 0.06064236 | PPAPDC2 | 403313 |
| ENSG000001 | 10.3882308 | -0.799940981 | 0.34474495 | -2.3203849 | 0.02032007 | 0.06066846 | AQP2 | 359 |
| ENSG000002 | 2.37125602 | -0.84817691 | 0.36554279 | -2.3203218 | 0.02032348 | 0.06067417 | NA | NA |
| ENSG000002 | 12.7767649 | -0.749949952 | 0.32321536 | -2.3202794 | 0.02032577 | 0.0606746 | HEPN1 | 641654 |
| ENSG000002 | 15.6400841 | -0.722751766 | 0.31149552 | -2.3202638 | 0.02032661 | 0.0606746 | NA | NA |
| ENSG00000( | 573.624725 | 0.278447594 | 0.12001138 | 2.32017652 | 0.02033133 | 0.06068223 | PPP2R3A | 5523 |
| ENSG000002 | 4.75926994 | -0.875275138 | 0.37724756 | -2.3201612 | 0.02033216 | 0.06068223 | NA | NA |
| ENSG000001 | 29.4359632 | -0.580502122 | 0.25020481 | -2.3201077 | 0.02033505 | 0.06068639 | OPALIN | 93377 |
| ENSG000001 | 8.83379907 | -0.801209978 | 0.34539218 | -2.3197108 | 0.02035653 | 0.06074601 | SLC52A1 | 55065 |
| ENSG000002 | 4.68303558 | -0.872457697 | 0.37617902 | -2.319262 | 0.02038083 | 0.06081407 | NA | NA |
| ENSG000002 | 56.9947654 | -0.469443703 | 0.20245375 | -2.3187701 | 0.02040751 | 0.06088917 | PCDHGA11 | 56105 |
| ENSG000001 | 49.7870149 | 0.508558494 | 0.21933586 | 2.31862906 | 0.02041516 | 0.06090753 | SLC2A4RG | 56731 |
| ENSG000001 | 317.013304 | 0.293092397 | 0.1264149 | 2.3184956 | 0.0204224 | 0.0609213 | AUH | 549 |
| ENSG000002 | 10.632071 | -0.770529206 | 0.33234116 | -2.3184886 | 0.02042278 | 0.0609213 | NA | NA |
| ENSG000001 | 33.5861853 | -0.565216883 | 0.24382224 | -2.3181515 | 0.02044109 | 0.06096961 | LCT | 3938 |
| ENSG000002 | 28.5282482 | 0.594579744 | 0.25649055 | 2.31813507 | 0.02044198 | 0.06096961 | MIR762HG | 101928736 |
| ENSG000002 | 14.608571 | 0.739784819 | 0.31916749 | 2.31785769 | 0.02045706 | 0.06101008 | NA | NA |
| ENSG000002 | 13.4529716 | -0.751130723 | 0.3241117 | -2.3175057 | 0.0204762 | 0.06106268 | LINC01440 | 102723578 |
| ENSG000001 | 90.5321617 | 0.416583371 | 0.17976202 | 2.31741592 | 0.02048109 | 0.06107276 | ARSE | 415 |

| | | | | | | | | |
|---|---|---|---|---|---|---|---|---|
| ENSG000002 | 10.2249082 | 0.78848693 | 0.34025559 | 2.31733717 | 0.02048537 | 0.06108105 | LOC554206 | 554206 |
| ENSG000002 | 3.81687717 | -0.876894622 | 0.3784344 | -2.3171641 | 0.02049479 | 0.06110465 | LOC105375671 | 105375671 |
| ENSG000002 | 4.0616213 | -0.87694418 | 0.37847743 | -2.3170316 | 0.02050201 | 0.06112167 | NA | NA |
| ENSG000001 | 491.107861 | 0.334890009 | 0.14456451 | 2.31654377 | 0.0205286 | 0.06119321 | NEK1 | 4750 |
| ENSG000001 | 1622.27504 | 0.282562591 | 0.12197635 | 2.31653596 | 0.02052902 | 0.06119321 | WDFY3 | 23001 |
| ENSG000002 | 3.65800903 | -0.876975537 | 0.37859301 | -2.3164071 | 0.02053605 | 0.06120516 | NA | NA |
| ENSG000002 | 2.74066264 | -0.859862187 | 0.37120436 | -2.3164119 | 0.02053579 | 0.06120516 | MIR4314 | 100422983 |
| ENSG000001 | 160.032081 | 0.338434743 | 0.14610826 | 2.31632859 | 0.02054033 | 0.06121343 | C3orf14 | 57415 |
| ENSG000001 | 460.49027 | 0.357450547 | 0.15432493 | 2.31622039 | 0.02054624 | 0.06122468 | TNS3 | 64759 |
| ENSG000002 | 18.7223705 | -0.6622833 | 0.2859382 | -2.3161764 | 0.02054864 | 0.06122468 | LOC100652999 | 100652999 |
| ENSG000002 | 14.3191941 | -0.719342582 | 0.31056975 | -2.316203 | 0.02054719 | 0.06122468 | NA | NA |
| ENSG000001 | 204.539146 | 0.355933248 | 0.15367494 | 2.31614378 | 0.02055042 | 0.06122548 | CCNG2 | 901 |
| ENSG000001 | 1280.01304 | 0.262114072 | 0.11317865 | 2.31593218 | 0.02056197 | 0.0612554 | SEC24D | 9871 |
| ENSG000002 | 14.5211876 | -0.752452198 | 0.32491874 | -2.3158166 | 0.02056829 | 0.06126971 | NA | NA |
| ENSG000002 | 9.77486373 | -0.791862408 | 0.34196243 | -2.3156415 | 0.02057785 | 0.0612937 | LOC101927431 | 101927431 |
| ENSG000001 | 82.9858654 | 0.414224478 | 0.17889178 | 2.31550316 | 0.02058541 | 0.06131172 | ZNF767P | 79970 |
| ENSG000002 | 4.40162545 | -0.875105039 | 0.37793935 | -2.3154642 | 0.02058754 | 0.06131355 | NA | NA |
| ENSG000001 | 141.479914 | 0.352333028 | 0.15217382 | 2.3153327 | 0.02059473 | 0.06133047 | ZNF442 | 79973 |
| ENSG000002 | 19.9363101 | -0.67863669 | 0.2931277 | -2.3151572 | 0.02060433 | 0.06135455 | PATE4 | 399968 |
| ENSG000001 | 41.8343102 | -0.520755509 | 0.22494755 | -2.3150086 | 0.02061246 | 0.06137425 | APBA1 | 320 |
| ENSG000001 | 157.978191 | 0.336561989 | 0.14538612 | 2.31495268 | 0.02061552 | 0.06137886 | NA | NA |
| ENSG000001 | 9.2782577 | -0.819672125 | 0.35409323 | -2.3148484 | 0.02062123 | 0.06139135 | OR6N2 | 81442 |
| ENSG000002 | 6.03141099 | -0.858015684 | 0.37066358 | -2.3148098 | 0.02062335 | 0.06139314 | NA | NA |
| ENSG000001 | 464.822358 | 0.326594563 | 0.14109831 | 2.3146596 | 0.02063157 | 0.0614131 | EXOC8 | 149371 |
| ENSG000001 | 44.1743292 | 0.515213574 | 0.22261077 | 2.31441441 | 0.020645 | 0.06144858 | APBB3 | 10307 |
| ENSG000002 | 2.89013906 | -0.854772726 | 0.3693494 | -2.3142659 | 0.02065314 | 0.06146829 | NA | NA |
| ENSG000000 | 210.204836 | 0.463090692 | 0.20010921 | 2.31418984 | 0.02065731 | 0.06147619 | SETD1A | 9739 |
| ENSG000001 | 122.029402 | -0.38463259 | 0.16621423 | -2.3140774 | 0.02066348 | 0.061481 | NA | NA |
| ENSG000002 | 5.58790313 | -0.860321571 | 0.37177714 | -2.3140787 | 0.0206634 | 0.061481 | TCP11X3P | 106481605 |

| | | | | | | | | |
|---|---|---|---|---|---|---|---|---|
| ENSG000002 | 34.0105953 | 0.580926671 | 0.25103564 | 2.31412034 | 0.02066112 | 0.061481 | NA | NA |
| ENSG000001 | 992.016728 | 0.278184694 | 0.12022825 | 2.31380469 | 0.02067844 | 0.06152101 | METTL8 | 79828 |
| ENSG000001 | 385.50667 | 0.337891405 | 0.14603839 | 2.31371626 | 0.02068329 | 0.06153093 | ADAM23 | 8745 |
| ENSG000002 | 27.4242563 | -0.583888852 | 0.2523653 | -2.3136654 | 0.02068608 | 0.06153299 | NA | NA |
| ENSG000002 | 7.5273879 | -0.830633559 | 0.3590146 | -2.3136484 | 0.02068702 | 0.06153299 | NA | NA |
| ENSG000001 | 9.71763955 | -0.784702343 | 0.33917866 | -2.3135369 | 0.02069314 | 0.06154668 | CPLX1 | 10815 |
| ENSG000001 | 135.967655 | 0.361227676 | 0.15614692 | 2.31338331 | 0.02070157 | 0.06155371 | CARF | 79800 |
| ENSG000001 | 14.6434618 | 0.719034638 | 0.31080919 | 2.31342787 | 0.02069913 | 0.06155371 | CSDC2 | 27254 |
| ENSG000001 | 132.938468 | 0.377719553 | 0.1632716 | 2.31344312 | 0.02069829 | 0.06155371 | MORN2 | 729967 |
| ENSG000002 | 4.04022356 | -0.875346664 | 0.37838342 | -2.3133853 | 0.02070146 | 0.06155371 | NA | NA |
| ENSG000001 | 252.520883 | 0.36510468 | 0.15783246 | 2.31324209 | 0.02070933 | 0.06157227 | ACTR3B | 57180 |
| ENSG000001 | 847.640518 | 0.239048715 | 0.1033447 | 2.31312022 | 0.02071603 | 0.06158676 | TTLL5 | 23093 |
| ENSG000002 | 3.36494035 | -0.873631847 | 0.37768906 | -2.3130981 | 0.02071725 | 0.06158676 | RNU6-1261P | 106480116 |
| ENSG000002 | 13.8584107 | -0.737377097 | 0.31880338 | -2.3129526 | 0.02072525 | 0.06160152 | NA | NA |
| ENSG000002 | 27.6913363 | -0.623415891 | 0.26952964 | -2.3129771 | 0.0207239 | 0.06160152 | LOC101928595 | 101928595 |
| ENSG000001 | 234.685457 | 0.34451082 | 0.14895184 | 2.31290075 | 0.0207281 | 0.06160547 | FAM118A | 55007 |
| ENSG000002 | 18.2887114 | -0.681122437 | 0.29449324 | -2.3128627 | 0.02073019 | 0.06160717 | NA | NA |
| ENSG000002 | 3.0591752 | -0.869409711 | 0.37592438 | -2.312725 | 0.02073776 | 0.06162517 | NA | NA |
| ENSG000001 | 1442.66285 | 0.257811551 | 0.11148443 | 2.31253415 | 0.02074827 | 0.06165186 | LARP1B | 55132 |
| ENSG000001 | 176.487188 | 0.333297296 | 0.14413125 | 2.31245688 | 0.02075252 | 0.06165998 | OSTF1 | 26578 |
| ENSG000001 | 135.214202 | 0.418887519 | 0.18118489 | 2.31193409 | 0.02078132 | 0.06173649 | LRSAM1 | 90678 |
| ENSG000001 | 13.1318891 | -0.774809062 | 0.33513148 | -2.3119555 | 0.02078014 | 0.06173649 | NA | NA |
| ENSG000001 | 4.24900151 | -0.875169103 | 0.37855337 | -2.3118777 | 0.02078442 | 0.0617412 | NA | NA |
| ENSG000002 | 35.1266391 | -0.54029189 | 0.23372197 | -2.3116864 | 0.02079497 | 0.06176802 | EFCAB8 | 388795 |
| ENSG000002 | 3.0768151 | -0.868228043 | 0.37562391 | -2.3114291 | 0.02080917 | 0.06180564 | NA | NA |
| ENSG000002 | 2.721467 | -0.861088239 | 0.37254856 | -2.3113449 | 0.02081381 | 0.06181491 | NA | NA |
| ENSG000002 | 4.36603712 | -0.873565261 | 0.37796609 | -2.3112265 | 0.02082035 | 0.06182979 | LOC100505515 | 100505515 |
| ENSG000002 | 11.063042 | -0.77853258 | 0.33687077 | -2.3110719 | 0.02082888 | 0.06185062 | NA | NA |
| ENSG000002 | 2.65851033 | -0.860640553 | 0.37244552 | -2.3107824 | 0.02084488 | 0.06189358 | NA | NA |

| | | | | | | | | |
|---|---|---|---|---|---|---|---|---|
| ENSG000002 | 4.17915414 | -0.874526852 | 0.37848152 | -2.3106197 | 0.02085387 | 0.06191575 | NA | NA |
| ENSG000002 | 2.71742863 | -0.86000621 | 0.37222448 | -2.3104504 | 0.02086323 | 0.061939 | NA | NA |
| ENSG000002 | 29.8100532 | 0.597799309 | 0.25875562 | 2.31028528 | 0.02087237 | 0.06196159 | NA | NA |
| ENSG000001 | 347.516984 | 0.308181446 | 0.13340001 | 2.31020559 | 0.02087677 | 0.06197014 | ZNF449 | 203523 |
| ENSG000002 | 3.68522038 | -0.874531595 | 0.37858951 | -2.3099732 | 0.02088964 | 0.06200379 | NA | NA |
| ENSG00000C | 384.557298 | 0.331162124 | 0.14336415 | 2.30993676 | 0.02089166 | 0.06200523 | BAIAP2L1 | 55971 |
| ENSG00000C | 80.0876223 | 0.454643991 | 0.19684509 | 2.30965375 | 0.02090733 | 0.06204485 | SIRT2 | 22933 |
| ENSG000001 | 83.0729384 | 0.400866029 | 0.17356529 | 2.30959796 | 0.02091042 | 0.06204485 | UNC93B1 | 81622 |
| ENSG000001 | 55.5290847 | -0.491379581 | 0.21275663 | -2.3095853 | 0.02091113 | 0.06204485 | SPTLC3 | 55304 |
| ENSG000001 | 11.1420513 | -0.783055954 | 0.3390452 | -2.3095917 | 0.02091077 | 0.06204485 | TMEM207 | 131920 |
| ENSG000001 | 8.41983862 | -0.816174997 | 0.35340445 | -2.3094644 | 0.02091783 | 0.0620602 | FPR3 | 2359 |
| ENSG000002 | 828.615895 | 0.281406825 | 0.1218509 | 2.3094358 | 0.02091941 | 0.06206036 | SMIM10L1 | 100129361 |
| ENSG000001 | 48.8646195 | -0.51676014 | 0.22376644 | -2.3093729 | 0.0209229 | 0.06206163 | KIAA1324 | 57535 |
| ENSG000002 | 5.92946466 | -0.85263693 | 0.36920445 | -2.3093896 | 0.02092197 | 0.06206163 | NA | NA |
| ENSG000002 | 3.02778146 | -0.868304543 | 0.37603173 | -2.3091257 | 0.02093661 | 0.06209667 | NA | NA |
| ENSG000002 | 43.5301021 | 0.500314245 | 0.21667024 | 2.30910464 | 0.02093778 | 0.06209667 | NA | NA |
| ENSG000002 | 20.0801567 | -0.650623056 | 0.28177154 | -2.3090446 | 0.02094111 | 0.06209746 | MIR646HG | 284757 |
| ENSG000002 | 4.04870492 | -0.873787191 | 0.37841802 | -2.3090528 | 0.02094065 | 0.06209746 | NA | NA |
| ENSG000001 | 15.6578553 | -0.717142282 | 0.31061809 | -2.3087589 | 0.02095696 | 0.06213994 | FAM19A1 | 407738 |
| ENSG000001 | 77.2235582 | 0.466364819 | 0.20201831 | 2.30852743 | 0.02096982 | 0.06217351 | NUDT2 | 318 |
| ENSG000002 | 3.86165729 | -0.873221849 | 0.37826605 | -2.3084859 | 0.02097212 | 0.0621758 | NA | NA |
| ENSG000001 | 100.474029 | 0.455608143 | 0.19736673 | 2.30843434 | 0.02097499 | 0.06217672 | MON1A | 84315 |
| ENSG000001 | 21.1069304 | -0.66668794 | 0.28880639 | -2.3084252 | 0.0209755 | 0.06217672 | C11orf85 | 283129 |
| ENSG00000C | 12.6427581 | -0.750103467 | 0.32498051 | -2.3081491 | 0.02099085 | 0.06221766 | IBSP | 3381 |
| ENSG000002 | 17.872923 | -0.670805276 | 0.29064078 | -2.3080218 | 0.02099792 | 0.06223408 | NA | NA |
| ENSG000001 | 12.0501796 | -0.746115397 | 0.32330845 | -2.307751 | 0.02101299 | 0.06227419 | ZAP70 | 7535 |
| ENSG000002 | 6.94826115 | -0.842381837 | 0.3650465 | -2.3076014 | 0.02102131 | 0.06228976 | NA | NA |
| ENSG000002 | 8.68408904 | -0.831905461 | 0.36050515 | -2.3076105 | 0.02102081 | 0.06228976 | NA | NA |
| ENSG000002 | 12.5137471 | -0.735558642 | 0.31877174 | -2.3074776 | 0.02102821 | 0.06230564 | TNFRSF13B | 23495 |

| | | | | | | | | |
|---|---|---|---|---|---|---|---|---|
| ENSG000001 | 952.715231 | 0.327066896 | 0.14175089 | 2.30733578 | 0.02103611 | 0.06232448 | FGFBP3 | 143282 |
| ENSG000001 | 691.076957 | 0.278301241 | 0.12061901 | 2.30727522 | 0.02103948 | 0.06232606 | CDC37L1 | 55664 |
| ENSG000002 | 3.21283633 | -0.864157184 | 0.37453648 | -2.3072711 | 0.02103971 | 0.06232606 | NA | NA |
| ENSG000001 | 128.018047 | 0.367200651 | 0.15916248 | 2.30708046 | 0.02105033 | 0.06235297 | MAN1C1 | 57134 |
| ENSG000002 | 17.1965792 | 0.697084205 | 0.30215443 | 2.30704613 | 0.02105225 | 0.06235408 | NA | NA |
| ENSG000002 | 16.0384182 | -0.720921735 | 0.3124964 | -2.3069761 | 0.02105615 | 0.06236109 | NA | NA |
| ENSG000002 | 2.41499289 | -0.840906159 | 0.36451594 | -2.3069119 | 0.02105973 | 0.06236714 | LINC01124 | 440925 |
| ENSG000001 | 16.3532827 | -0.731446668 | 0.31708749 | -2.3067661 | 0.02106786 | 0.0623821 | SLC22A24 | 283238 |
| ENSG000002 | 7.78088173 | -0.822164458 | 0.35641034 | -2.3067918 | 0.02106643 | 0.0623821 | NA | NA |
| ENSG000002 | 14.9813271 | -0.697714908 | 0.30246967 | -2.3067268 | 0.02107005 | 0.06238404 | C8orf37-AS1 | 100616530 |
| ENSG000002 | 8.70223391 | -0.807894443 | 0.35027368 | -2.3064663 | 0.02108459 | 0.06242252 | NA | NA |
| ENSG000002 | 4.42829492 | -0.872325224 | 0.37821698 | -2.3064148 | 0.02108747 | 0.06242314 | NA | NA |
| ENSG000002 | 6.50489774 | -0.84030344 | 0.36433435 | -2.3064074 | 0.02108788 | 0.06242314 | NA | NA |
| ENSG000001 | 156.90126 | 0.393813549 | 0.1707548 | 2.30631026 | 0.0210933 | 0.06243463 | CLSTN3 | 9746 |
| ENSG000002 | 203.258294 | 0.352146558 | 0.15269274 | 2.30624297 | 0.02109706 | 0.0624412 | ZNF286B | 729288 |
| ENSG000002 | 5.84194669 | 0.860238954 | 0.37305518 | 2.30592954 | 0.02111457 | 0.06248846 | NA | NA |
| ENSG000001 | 39.0760352 | -0.55642729 | 0.24131146 | -2.3058469 | 0.02111919 | 0.06249757 | FAM78B | 149297 |
| ENSG000001 | 5.30605925 | -0.861758406 | 0.37375089 | -2.3057026 | 0.02112725 | 0.06250988 | KCNE2 | 9992 |
| ENSG000001 | 1202.37078 | 0.361349127 | 0.15672061 | 2.30568985 | 0.02112797 | 0.06250988 | CYP2S1 | 29785 |
| ENSG000002 | 3.75140199 | -0.872337591 | 0.37834287 | -2.30568 | 0.02112852 | 0.06250988 | NA | NA |
| ENSG000002 | 17.7867562 | -0.68600217 | 0.29752935 | -2.3056622 | 0.02112951 | 0.06250988 | NA | NA |
| ENSG000002 | 21.7533114 | 0.635317605 | 0.27556109 | 2.30554177 | 0.02113625 | 0.06252069 | NA | NA |
| ENSG000002 | 7.67263266 | -0.816859537 | 0.35430014 | -2.3055581 | 0.02113534 | 0.06252069 | NA | NA |
| ENSG000002 | 7.85950632 | -0.825339917 | 0.35800566 | -2.3053823 | 0.02114517 | 0.06253795 | NA | NA |
| ENSG000002 | 10.3646536 | -0.777530446 | 0.33726719 | -2.3053842 | 0.02114507 | 0.06253795 | CLSTN2-AS1 | 101927808 |
| ENSG000002 | 2.32775756 | -0.844145074 | 0.36617974 | -2.3052752 | 0.02115117 | 0.06255112 | NA | NA |
| ENSG00000C | 911.975607 | 0.326581304 | 0.14167669 | 2.30511676 | 0.02116003 | 0.06257203 | JADE2 | 23338 |
| ENSG000002 | 2.27405226 | -0.844614426 | 0.36641218 | -2.3050938 | 0.02116132 | 0.06257203 | PRAMEF27 | 101929983 |
| ENSG000002 | 5.10580294 | -0.865818189 | 0.37561825 | -2.3050483 | 0.02116387 | 0.062575 | NA | NA |

| | | | | | | | | |
|---|---|---|---|---|---|---|---|---|
| ENSG000001 | 558.281838 | 0.274086491 | 0.11892297 | 2.30473971 | 0.02118116 | 0.06262154 | STEAP2 | 261729 |
| ENSG000002 | 11.8504557 | -0.761623699 | 0.3304645 | -2.3047066 | 0.02118301 | 0.06262247 | ZNF99 | 7652 |
| ENSG000001 | 85.6055834 | 0.446272496 | 0.1936535 | 2.30448971 | 0.02119517 | 0.06265383 | SPR | 6697 |
| ENSG000001 | 20.380891 | -0.690344196 | 0.29959801 | -2.3042349 | 0.02120946 | 0.06268694 | REG4 | 83998 |
| ENSG000002 | 5.0529268 | -0.865496689 | 0.3756073 | -2.3042595 | 0.02120808 | 0.06268694 | NA | NA |
| ENSG00000C | 1168.52289 | 0.418589583 | 0.18167572 | 2.30404805 | 0.02121995 | 0.06270879 | BAZ2A | 11176 |
| ENSG00000C | 238.557983 | 0.748866163 | 0.32502167 | 2.30404993 | 0.02121984 | 0.06270879 | CHERP | 10523 |
| ENSG000001 | 814.300471 | 0.278578717 | 0.12091372 | 2.30394625 | 0.02122566 | 0.06272111 | NUP214 | 8021 |
| ENSG000002 | 11.5929423 | -0.763219094 | 0.33127381 | -2.303892 | 0.02122871 | 0.06272554 | NA | NA |
| ENSG000001 | 1040.33558 | 0.268949313 | 0.11673966 | 2.30383846 | 0.02123171 | 0.06272985 | BCL2L1 | 598 |
| ENSG000002 | 2.26081547 | -0.845350181 | 0.3669424 | -2.3037681 | 0.02123567 | 0.06273696 | NA | NA |
| ENSG000001 | 9.45830286 | -0.808044831 | 0.35076867 | -2.3036403 | 0.02124284 | 0.06275105 | HOXB13 | 10481 |
| ENSG000002 | 9.26795515 | -0.815243541 | 0.35389547 | -2.303628 | 0.02124353 | 0.06275105 | NA | NA |
| ENSG000002 | 12.3137883 | -0.746177972 | 0.32392312 | -2.303565 | 0.02124707 | 0.06275694 | NA | NA |
| ENSG000002 | 6.28546488 | -0.849335334 | 0.36872184 | -2.3034582 | 0.02125308 | 0.0627701 | NA | NA |
| ENSG000002 | 7.40267486 | -0.840156336 | 0.36477232 | -2.3032349 | 0.02126563 | 0.0628026 | NA | NA |
| ENSG000002 | 5.06538596 | -0.865933495 | 0.37597404 | -2.3031736 | 0.02126908 | 0.06280821 | NA | NA |
| ENSG000001 | 9.03728566 | -0.807333487 | 0.35054103 | -2.303107 | 0.02127283 | 0.0628147 | GUCA1B | 2979 |
| ENSG000002 | 3.26571935 | -0.859884649 | 0.37336942 | -2.3030398 | 0.0212766 | 0.06282128 | NA | NA |
| ENSG000001 | 96.5874899 | 0.430572268 | 0.18699057 | 2.30264167 | 0.02129901 | 0.06288287 | HNRNPA1L2 | 144983 |
| ENSG000002 | 251.730807 | 0.330711114 | 0.14362794 | 2.30255412 | 0.02130394 | 0.06289285 | ZNF234 | 10780 |
| ENSG000001 | 2324.72284 | 0.273201066 | 0.11865512 | 2.30248014 | 0.02130811 | 0.06289599 | UPF2 | 26019 |
| ENSG000002 | 11.5037694 | -0.756008874 | 0.32834294 | -2.3024977 | 0.02130712 | 0.06289599 | NA | NA |
| ENSG000001 | 2677.72763 | 0.320479106 | 0.13920567 | 2.30219869 | 0.02132397 | 0.0629355 | UBAP2L | 9898 |
| ENSG000002 | 11.1435533 | -0.755103582 | 0.32799395 | -2.3021876 | 0.0213246 | 0.0629355 | NA | NA |
| ENSG000001 | 13.9483775 | -0.776672405 | 0.3374598 | -2.3015257 | 0.02136193 | 0.06303651 | ZSCAN5B | 342933 |
| ENSG000002 | 2.66207319 | -0.857981831 | 0.37278606 | -2.3015395 | 0.02136116 | 0.06303651 | NA | NA |
| ENSG000001 | 1491.85669 | 0.284020956 | 0.12344546 | 2.3007808 | 0.02140402 | 0.06315612 | SGOL2 | 151246 |
| ENSG000002 | 10.654627 | -0.788265305 | 0.34261923 | -2.3007036 | 0.02140839 | 0.06316441 | LOC100133077 | 100133077 |

| | | | | | | | | |
|---|---|---|---|---|---|---|---|---|
| ENSG000001 | 11.2374746 | -0.76408276 | 0.3321241 | -2.3005942 | 0.02141458 | 0.06317808 | NA | NA |
| ENSG000001 | 18.7546718 | -0.647077273 | 0.2812691 | -2.300563 | 0.02141635 | 0.06317869 | SCNN1B | 6338 |
| ENSG000001 | 143.848267 | 0.353621449 | 0.15372244 | 2.30038929 | 0.02142618 | 0.06320191 | MED11 | 400569 |
| ENSG000002 | 4.86143357 | 0.867398786 | 0.37706945 | 2.30036878 | 0.02142734 | 0.06320191 | NA | NA |
| ENSG000001 | 13.2146088 | -0.74980852 | 0.32596507 | -2.3002726 | 0.02143278 | 0.06321266 | HEMGN | 55363 |
| ENSG000002 | 104.038927 | 0.401887044 | 0.17471456 | 2.30024935 | 0.0214341 | 0.06321266 | NA | NA |
| ENSG000001 | 113.969808 | 0.40331554 | 0.17534663 | 2.30010437 | 0.02144231 | 0.06323227 | PARD6G | 84552 |
| ENSG000002 | 10.8963292 | -0.762373114 | 0.33146671 | -2.299999 | 0.02144828 | 0.06324528 | NA | NA |
| ENSG000001 | 356.441579 | 0.31076651 | 0.13519167 | 2.29871042 | 0.02152139 | 0.06345625 | CENPL | 91687 |
| ENSG000002 | 18.489698 | 0.673284577 | 0.29290045 | 2.29868058 | 0.02152308 | 0.06345664 | NA | NA |
| ENSG000002 | 7.12321031 | -0.832025971 | 0.36197123 | -2.298597 | 0.02152783 | 0.06346603 | NA | NA |
| ENSG000001 | 13.6088012 | -0.752339041 | 0.32731669 | -2.298505 | 0.02153306 | 0.06347683 | PRAME | 23532 |
| ENSG000001 | 76.1112768 | 0.416319718 | 0.18112987 | 2.29845973 | 0.02153564 | 0.0634798 | TMEM198B | 440104 |
| ENSG000002 | 5.94765386 | -0.84975046 | 0.36971801 | -2.2983746 | 0.02154048 | 0.06348946 | NA | NA |
| ENSG000001 | 23.9024512 | -0.634645629 | 0.27614269 | -2.2982525 | 0.02154742 | 0.06350043 | KNG1 | 3827 |
| ENSG000002 | 2.28634387 | -0.843128502 | 0.36685348 | -2.2982704 | 0.0215464 | 0.06350043 | NA | NA |
| ENSG000002 | 10.9111172 | -0.765092011 | 0.33290539 | -2.2982266 | 0.0215489 | 0.06350043 | NA | NA |
| ENSG000001 | 50.1454398 | 0.480122484 | 0.20892295 | 2.29808401 | 0.02155701 | 0.06351971 | FAM167A | 83648 |
| ENSG000002 | 6.22648004 | -0.858191231 | 0.37344701 | -2.2980268 | 0.02156026 | 0.06352469 | NA | NA |
| ENSG000002 | 11.9806523 | -0.781778922 | 0.34020041 | -2.2979952 | 0.02156206 | 0.06352537 | NA | NA |
| ENSG00000C | 229.324334 | 0.320322192 | 0.13940982 | 2.2977018 | 0.02157877 | 0.06356557 | CCDC109B | 55013 |
| ENSG000002 | 2.73105772 | -0.854801174 | 0.37202463 | -2.2977005 | 0.02157884 | 0.06356557 | NA | NA |
| ENSG000002 | 10.241655 | -0.766876216 | 0.33376605 | -2.297646 | 0.02158195 | 0.0635701 | MILR1 | 284021 |
| ENSG000001 | 17.4231518 | -0.713808415 | 0.31068214 | -2.2975521 | 0.02158729 | 0.06358122 | SLC22A8 | 9376 |
| ENSG000001 | 192.187012 | 0.347452609 | 0.15123594 | 2.29742086 | 0.02159477 | 0.06359864 | BANP | 54971 |
| ENSG000002 | 8.64758756 | -0.804254392 | 0.35007614 | -2.2973699 | 0.02159767 | 0.06360256 | NA | NA |
| ENSG000002 | 3.09186621 | -0.861868943 | 0.37517681 | -2.297234 | 0.02160542 | 0.06362076 | NA | NA |
| ENSG000001 | 11.1464189 | -0.760373319 | 0.33104556 | -2.2968842 | 0.02162538 | 0.06367489 | NEURL1 | 9148 |
| ENSG000001 | 18.8318687 | -0.669901842 | 0.29166821 | -2.2967942 | 0.02163051 | 0.0636854 | WNK4 | 65266 |

| | | | | | | | | |
|---|---|---|---|---|---|---|---|---|
| ENSG000001 | 6.62476537 | -0.842386217 | 0.36678474 | -2.2966774 | 0.02163718 | 0.0637004 | C16orf92 | 146378 |
| ENSG000002 | 3.44291533 | -0.866442089 | 0.3772748 | -2.2965809 | 0.02164269 | 0.06371199 | NA | NA |
| ENSG000002 | 8.73142103 | -0.807407761 | 0.35158042 | -2.2965094 | 0.02164677 | 0.06371938 | NA | NA |
| ENSG000002 | 3.45561691 | -0.866412627 | 0.37727866 | -2.2964793 | 0.02164849 | 0.06371982 | EIF3IP1 | 442720 |
| ENSG000001 | 545.806361 | 0.27888755 | 0.12144736 | 2.29636568 | 0.02165498 | 0.0637343 | MPHOSPH6 | 10200 |
| ENSG000002 | 7.94297421 | -0.810683953 | 0.35306922 | -2.2961049 | 0.02166988 | 0.06377353 | NA | NA |
| ENSG000001 | 137.173591 | 0.367826204 | 0.16020997 | 2.29590078 | 0.02168156 | 0.06380325 | APOL2 | 23780 |
| ENSG000002 | 120.126641 | 0.429013781 | 0.18687747 | 2.29569551 | 0.0216933 | 0.06383317 | VPS16 | 64601 |
| ENSG00000C | 79.9077533 | 0.420855383 | 0.18335677 | 2.29528136 | 0.02171701 | 0.06389829 | FLT4 | 2324 |
| ENSG000002 | 13.9287898 | -0.731106232 | 0.31853052 | -2.295247 | 0.02171897 | 0.06389944 | ARNTL2-AS1 | 101928646 |
| ENSG000002 | 11.6641554 | -0.765165812 | 0.33337781 | -2.2951912 | 0.02172217 | 0.06390421 | NA | NA |
| ENSG000002 | 3.32090166 | -0.867738598 | 0.37807885 | -2.295126 | 0.02172591 | 0.06391057 | RNU6-1047P | 106481511 |
| ENSG000002 | 31.3684859 | 0.590286671 | 0.25719704 | 2.29507565 | 0.02172879 | 0.06391442 | NA | NA |
| ENSG000002 | 8.31433479 | -0.808005426 | 0.35211206 | -2.2947394 | 0.02174806 | 0.06396646 | NA | NA |
| ENSG000002 | 21.1164775 | -0.641441919 | 0.27954653 | -2.2945802 | 0.0217572 | 0.06398869 | NA | NA |
| ENSG000001 | 21.1056525 | -0.653869964 | 0.2849702 | -2.2945205 | 0.02176062 | 0.06399412 | PCK1 | 5105 |
| ENSG000002 | 2.32661595 | -0.840212054 | 0.36620392 | -2.294383 | 0.02176851 | 0.06401267 | RNU6-602P | 106481371 |
| ENSG000002 | 4.00848904 | -0.868500766 | 0.37854789 | -2.2942956 | 0.02177352 | 0.06402278 | NA | NA |
| ENSG000001 | 22.9163222 | -0.645279123 | 0.2812807 | -2.2940754 | 0.02178617 | 0.06404409 | NOXRED1 | 122945 |
| ENSG000002 | 9.39946753 | -0.806128002 | 0.35138587 | -2.2941389 | 0.02178252 | 0.06404409 | NA | NA |
| ENSG000002 | 113.011517 | 0.407251043 | 0.17752419 | 2.29405938 | 0.02178709 | 0.06404409 | EIF5AL1 | 143244 |
| ENSG000002 | 2.31139566 | -0.838555902 | 0.36553168 | -2.2940717 | 0.02178638 | 0.06404409 | NA | NA |
| ENSG000002 | 12.2924288 | -0.741039342 | 0.32303068 | -2.2940215 | 0.02178926 | 0.06404584 | NA | NA |
| ENSG000001 | 40.3741538 | 0.545015018 | 0.23758875 | 2.29394282 | 0.02179378 | 0.06405448 | CXorf40A | 91966 |
| ENSG000001 | 76.5304627 | 0.449084607 | 0.19577928 | 2.29383108 | 0.0218002 | 0.06406787 | GNAZ | 2781 |
| ENSG000002 | 3.67958283 | -0.868213888 | 0.3785032 | -2.2938086 | 0.0218015 | 0.06406787 | NA | NA |
| ENSG000001 | 43.8280299 | 0.544128072 | 0.23722581 | 2.29371363 | 0.02180695 | 0.06407926 | NFATC1 | 4772 |
| ENSG000002 | 4.44979758 | -0.867716443 | 0.37832239 | -2.29359 | 0.02181406 | 0.0640955 | NA | NA |
| ENSG000001 | 9.46742064 | -0.786678038 | 0.34300307 | -2.2935015 | 0.02181915 | 0.06410581 | SFTPA2 | 729238 |

| | | | | | | | | |
|---|---|---|---|---|---|---|---|---|
| ENSG000002 | 11.0930973 | -0.752454741 | 0.32810524 | -2.2933335 | 0.02182881 | 0.06412955 | LOC101929718 | 101929718 |
| ENSG000001 | 2963.19908 | 0.304347566 | 0.13272082 | 2.29314102 | 0.02183989 | 0.06415744 | TAF1D | 79101 |
| ENSG000002 | 29.8609803 | 0.616524579 | 0.26888389 | 2.29290261 | 0.02185361 | 0.06419311 | NA | NA |
| ENSG000002 | 40.4240923 | 0.519780651 | 0.2267055 | 2.2927571 | 0.02186199 | 0.06421307 | NA | NA |
| ENSG000002 | 6.55869945 | -0.838745444 | 0.3658596 | -2.2925337 | 0.02187487 | 0.06424623 | NA | NA |
| ENSG000002 | 3.70670986 | -0.867687199 | 0.37849471 | -2.2924685 | 0.02187862 | 0.06425261 | NA | NA |
| ENSG000002 | 93.0691121 | 0.423196685 | 0.18462946 | 2.29214058 | 0.02189753 | 0.06430349 | METTL12 | 751071 |
| ENSG000001 | 110.717477 | 0.436223085 | 0.19031534 | 2.29210678 | 0.02189948 | 0.06430456 | EEFSEC | 60678 |
| ENSG000002 | 3.70903359 | -0.867687761 | 0.37856627 | -2.2920367 | 0.02190353 | 0.06431178 | NA | NA |
| ENSG000001 | 62.0088871 | 0.451594467 | 0.19704355 | 2.29185108 | 0.02191424 | 0.06433856 | YPEL1 | 29799 |
| ENSG000002 | 6.58060193 | -0.837664804 | 0.36555792 | -2.2914694 | 0.02193628 | 0.06439861 | NA | NA |
| ENSG000002 | 22.2549466 | -0.628844612 | 0.27444092 | -2.2913661 | 0.02194225 | 0.06441148 | WDR11-AS1 | 283089 |
| ENSG000002 | 23.2252043 | 0.625629533 | 0.27304237 | 2.29132762 | 0.02194447 | 0.06441172 | SNORA71C | 677839 |
| ENSG000002 | 16.4026288 | -0.679056282 | 0.29636163 | -2.2913097 | 0.02194551 | 0.06441172 | NA | NA |
| ENSG000001 | 177.877275 | 0.347226806 | 0.15154797 | 2.29120061 | 0.02195181 | 0.06442556 | EVL | 51466 |
| ENSG00000C | 157.012242 | 0.372006364 | 0.16236774 | 2.29113471 | 0.02195562 | 0.06442742 | JMJD6 | 23210 |
| ENSG000002 | 5.51802545 | -0.861244594 | 0.37589906 | -2.2911592 | 0.02195421 | 0.06442742 | ACOT6 | 641372 |
| ENSG000001 | 9.93172604 | -0.809091377 | 0.35316031 | -2.2910032 | 0.02196323 | 0.06444508 | TBX2 | 6909 |
| ENSG000001 | 42.2651004 | -0.495658483 | 0.21636438 | -2.2908507 | 0.02197205 | 0.06446629 | DCLK3 | 85443 |
| ENSG000001 | 27.3267464 | -0.588316895 | 0.25681623 | -2.2908088 | 0.02197447 | 0.06446873 | HIC1 | 3090 |
| ENSG000002 | 15.7326875 | -0.695239282 | 0.30350556 | -2.290697 | 0.02198094 | 0.06448305 | LOC101927277 | 101927277 |
| ENSG000002 | 9.0657193 | -0.836557341 | 0.36520381 | -2.2906588 | 0.02198315 | 0.06448487 | NA | NA |
| ENSG000002 | 11.1262458 | -0.751959066 | 0.32831137 | -2.2903838 | 0.02199908 | 0.06452691 | LINC00323 | 284835 |
| ENSG000002 | 4.42610344 | -0.866199182 | 0.37821653 | -2.2902203 | 0.02200855 | 0.06455003 | NA | NA |
| ENSG000002 | 26.4280957 | -0.600381466 | 0.26216367 | -2.2901017 | 0.02201542 | 0.06456552 | NA | NA |
| ENSG000001 | 32.0533579 | -0.579746948 | 0.25316798 | -2.2899695 | 0.02202309 | 0.06458332 | PRMT8 | 56341 |
| ENSG000001 | 32.0231107 | -0.561839281 | 0.24535556 | -2.2898983 | 0.02202721 | 0.06459076 | MYT1 | 4661 |
| ENSG000001 | 46.5030053 | -0.49499658 | 0.2161833 | -2.2897077 | 0.02203827 | 0.06461099 | FLT3 | 2322 |
| ENSG000001 | 430.333457 | 0.36940337 | 0.16133481 | 2.28966941 | 0.02204049 | 0.06461099 | EIF4ENIF1 | 56478 |

| | | | | | | | | |
|---|---|---|---|---|---|---|---|---|
| ENSG000002 | 8.26657969 | -0.804438836 | 0.35133371 | -2.2896716 | 0.02204036 | 0.06461099 | LINC00606 | 285370 |
| ENSG000002 | 6.82333064 | -0.830120404 | 0.36254435 | -2.2897072 | 0.02203829 | 0.06461099 | LINC01506 | 101927015 |
| ENSG000001 | 25.7791768 | -0.609908968 | 0.26639845 | -2.2894614 | 0.02205256 | 0.06464169 | EVPL | 2125 |
| ENSG000002 | 13.7018428 | -0.735874555 | 0.32144057 | -2.2893021 | 0.02206181 | 0.06466414 | NA | NA |
| ENSG000001 | 843.905615 | 0.313939777 | 0.1371433 | 2.28913676 | 0.02207141 | 0.0646876 | DIAPH3 | 81624 |
| ENSG000002 | 6.8016608 | -0.834147581 | 0.36440581 | -2.2890623 | 0.02207574 | 0.06469561 | NA | NA |
| ENSG000001 | 103.936371 | 0.425061687 | 0.18570574 | 2.28889904 | 0.02208522 | 0.06471873 | PQLC3 | 130814 |
| ENSG000002 | 8.23478863 | -0.802861424 | 0.35078771 | -2.2887388 | 0.02209453 | 0.06474134 | LOC105376609 | 105376609 |
| ENSG000001 | 397.577947 | 0.28594068 | 0.12496198 | 2.2882215 | 0.02212462 | 0.06482483 | AXL | 558 |
| ENSG000002 | 26.0240457 | 0.608978908 | 0.26615138 | 2.28809227 | 0.02213215 | 0.06484219 | NA | NA |
| ENSG000001 | 11.8876128 | -0.75893278 | 0.33171388 | -2.2879138 | 0.02214254 | 0.06486795 | OR4K1 | 79544 |
| ENSG000002 | 4.68457362 | -0.862709935 | 0.37708238 | -2.2878554 | 0.02214595 | 0.06487324 | NA | NA |
| ENSG000002 | 8.61775411 | -0.801582137 | 0.35037332 | -2.2877944 | 0.0221495 | 0.06487895 | ALG1L13P | 106479038 |
| ENSG000001 | 725.573599 | 0.279746626 | 0.12229274 | 2.28751624 | 0.02216571 | 0.06491706 | MED21 | 9412 |
| ENSG000002 | 14.7920598 | -0.73822947 | 0.32271886 | -2.2875312 | 0.02216483 | 0.06491706 | NA | NA |
| ENSG00000C | 340.31192 | 0.350925721 | 0.15341598 | 2.28741312 | 0.02217172 | 0.06492806 | PLEKHH1 | 57475 |
| ENSG000001 | 35.4831496 | -0.551769167 | 0.24122143 | -2.2873969 | 0.02217266 | 0.06492806 | ART3 | 419 |
| ENSG00000C | 446.97638 | 0.296941644 | 0.12982887 | 2.28717735 | 0.02218547 | 0.06495618 | ICA1 | 3382 |
| ENSG000002 | 35.1082897 | 0.534093528 | 0.23351372 | 2.28720411 | 0.02218391 | 0.06495618 | NA | NA |
| ENSG000001 | 62.7793713 | 0.436668697 | 0.19093921 | 2.28695138 | 0.02219866 | 0.0649901 | ZNF490 | 57474 |
| ENSG000001 | 7.24243497 | -0.832597696 | 0.36408289 | -2.2868356 | 0.02220542 | 0.06500521 | SLITRK2 | 84631 |
| ENSG000001 | 29.4243148 | -0.568888129 | 0.24882923 | -2.2862592 | 0.0222391 | 0.06509909 | TNFRSF19 | 55504 |
| ENSG000002 | 7.12946326 | -0.825937603 | 0.36127283 | -2.286188 | 0.02224326 | 0.06510658 | NA | NA |
| ENSG000002 | 4.07168201 | -0.865397652 | 0.37855628 | -2.2860475 | 0.02225148 | 0.06512592 | NA | NA |
| ENSG000002 | 2.31301443 | -0.83695058 | 0.36616301 | -2.2857322 | 0.02226993 | 0.06517523 | NA | NA |
| ENSG000002 | 13.7082571 | -0.7180467 | 0.31416052 | -2.2856045 | 0.02227741 | 0.0651924 | NA | NA |
| ENSG000002 | 2.76045688 | -0.846750613 | 0.37047689 | -2.2855693 | 0.02227947 | 0.06519372 | NA | NA |
| ENSG000001 | 112.057899 | 0.428024124 | 0.18728187 | 2.2854541 | 0.02228621 | 0.06520248 | FCGRT | 2217 |
| ENSG000001 | 1070.37238 | 0.274089646 | 0.119926 | 2.28548974 | 0.02228413 | 0.06520248 | MAP3K2 | 10746 |

| | | | | | | | | |
|---|---|---|---|---|---|---|---|---|
| ENSG000001 | 28.9398687 | -0.602202583 | 0.26350206 | -2.285381 | 0.0222905 | 0.06520248 | ARHGAP30 | 257106 |
| ENSG000002 | 4.52671718 | -0.865040018 | 0.37850957 | -2.2853848 | 0.02229028 | 0.06520248 | NA | NA |
| ENSG000002 | 2.33532599 | -0.837083473 | 0.36627177 | -2.2854163 | 0.02228843 | 0.06520248 | NA | NA |
| ENSG000002 | 125.401642 | 0.363744968 | 0.15917463 | 2.28519436 | 0.02230143 | 0.06522976 | NA | NA |
| ENSG000002 | 2.64783805 | -0.850965218 | 0.37244195 | -2.2848265 | 0.02232301 | 0.06528815 | NA | NA |
| ENSG000002 | 5.54725047 | -0.846803772 | 0.37063898 | -2.2847132 | 0.02232965 | 0.06530288 | GCGR | 2642 |
| ENSG000002 | 13.7164453 | -0.716624274 | 0.31367364 | -2.2846174 | 0.02233527 | 0.0653146 | NA | NA |
| ENSG000002 | 4.02084014 | -0.864838607 | 0.37856011 | -2.2845476 | 0.02233937 | 0.06532187 | LOC101927592 | 101927592 |
| ENSG00000C | 118.393086 | -0.365897137 | 0.16017645 | -2.2843379 | 0.02235168 | 0.06535317 | NAV3 | 89795 |
| ENSG000001 | 1238.99751 | 0.559651002 | 0.24500907 | 2.28420525 | 0.02235947 | 0.06536651 | NUS1 | 116150 |
| ENSG000002 | 63.8029076 | 0.456883396 | 0.20001699 | 2.28422297 | 0.02235843 | 0.06536651 | LOC100499484 | 100499484 |
| ENSG000001 | 92.7412532 | 0.38889077 | 0.17026586 | 2.28402085 | 0.02237031 | 0.06539347 | SPRYD7 | 57213 |
| ENSG000002 | 8.71506894 | -0.820986289 | 0.35946085 | -2.283938 | 0.02237518 | 0.065403 | NA | NA |
| ENSG00000C | 25.2516921 | -0.591900092 | 0.25917699 | -2.2837679 | 0.02238518 | 0.0654228 | NTN4 | 59277 |
| ENSG000002 | 9.14109924 | -0.791783114 | 0.34669781 | -2.2837846 | 0.0223842 | 0.0654228 | NA | NA |
| ENSG000002 | 12.2797665 | -0.759003779 | 0.3323581 | -2.2836927 | 0.0223896 | 0.06543099 | LOC101927914 | 101927914 |
| ENSG000002 | 4.44714168 | -0.864181813 | 0.37843814 | -2.2835484 | 0.02239808 | 0.06545108 | NA | NA |
| ENSG000001 | 304.550674 | 0.290775244 | 0.1273391 | 2.28347182 | 0.02240259 | 0.06545954 | SFXN5 | 94097 |
| ENSG000001 | 531.629341 | -0.325080314 | 0.14237134 | -2.2833269 | 0.02241112 | 0.06547502 | PTCH1 | 5727 |
| ENSG000002 | 19.9562648 | -0.662898189 | 0.29031982 | -2.2833377 | 0.02241048 | 0.06547502 | NA | NA |
| ENSG000001 | 153.042628 | 0.367725668 | 0.16105217 | 2.28327045 | 0.02241444 | 0.06548001 | NKD1 | 85407 |
| ENSG000001 | 19.8625341 | -0.667827566 | 0.29249442 | -2.2832147 | 0.02241772 | 0.06548488 | C5orf64 | 285668 |
| ENSG000001 | 105.188178 | 0.380906199 | 0.16683606 | 2.28311667 | 0.0224235 | 0.06549231 | MPPE1 | 65258 |
| ENSG000002 | 16.9969288 | 0.708200542 | 0.31018813 | 2.28313231 | 0.02242258 | 0.06549231 | NA | NA |
| ENSG000002 | 2.67441577 | -0.848198334 | 0.37154534 | -2.2828932 | 0.02243666 | 0.06552603 | NA | NA |
| ENSG00000C | 92.3664797 | 0.38101897 | 0.16692016 | 2.28264199 | 0.02245147 | 0.06556455 | GLRX2 | 51022 |
| ENSG000001 | 55.9196784 | 0.45219363 | 0.19813158 | 2.28228953 | 0.02247225 | 0.06561581 | TINAGL1 | 64129 |
| ENSG000002 | 2.3043372 | -0.835613975 | 0.36612912 | -2.2822931 | 0.02247204 | 0.06561581 | NA | NA |
| ENSG000001 | 33.2452896 | -0.544831128 | 0.23873181 | -2.2821891 | 0.02247818 | 0.06562838 | ACAN | 176 |

| | | | | | | | | |
|---|---|---|---|---|---|---|---|---|
| ENSG000001 | 47.2363041 | 0.535080657 | 0.23447109 | 2.2820752 | 0.0224849 | 0.06564328 | NPTX1 | 4884 |
| ENSG000000 | 102.704508 | 0.391281522 | 0.17146816 | 2.28194862 | 0.02249237 | 0.06566037 | CCDC28A | 25901 |
| ENSG000001 | 16.1735182 | -0.687041757 | 0.30110714 | -2.2817186 | 0.02250596 | 0.06568585 | SGK2 | 10110 |
| ENSG000002 | 15.1634959 | -0.700511804 | 0.3070046 | -2.2817632 | 0.02250333 | 0.06568585 | DLEU7-AS1 | 100874074 |
| ENSG000002 | 5.21820834 | -0.852292395 | 0.37352816 | -2.2817353 | 0.02250497 | 0.06568585 | NA | NA |
| ENSG000002 | 7.93830029 | -0.8229744 | 0.36069133 | -2.2816584 | 0.02250951 | 0.0656915 | NA | NA |
| ENSG000000 | 82.6388202 | 0.442279452 | 0.19385831 | 2.28145732 | 0.0225214 | 0.06572145 | AIFM2 | 84883 |
| ENSG000002 | 13.4236808 | -0.717404896 | 0.31451015 | -2.281023 | 0.02254708 | 0.06579167 | NA | NA |
| ENSG000002 | 4.90350481 | -0.854814406 | 0.37476236 | -2.2809505 | 0.02255137 | 0.06579945 | NA | NA |
| ENSG000002 | 203.018661 | 0.326654764 | 0.14322153 | 2.28076578 | 0.02256231 | 0.06582662 | NA | NA |
| ENSG000000 | 403.56717 | 0.320260096 | 0.14042771 | 2.28060475 | 0.02257185 | 0.06583035 | PPP2R3C | 55012 |
| ENSG000001 | 32.1132171 | -0.561918384 | 0.2463923 | -2.2805842 | 0.02257306 | 0.06583035 | PTPRE | 5791 |
| ENSG000001 | 190.917463 | 0.313316122 | 0.13738132 | 2.28063111 | 0.02257028 | 0.06583035 | NA | NA |
| ENSG000002 | 17.5392579 | -0.705066248 | 0.30914672 | -2.2806849 | 0.0225671 | 0.06583035 | OR11A1 | 26531 |
| ENSG000002 | 4.52301802 | -0.862906149 | 0.37836817 | -2.2805992 | 0.02257217 | 0.06583035 | NA | NA |
| ENSG000002 | 11.6064925 | -0.757984444 | 0.33236481 | -2.2805797 | 0.02257333 | 0.06583035 | NA | NA |
| ENSG000002 | 5.40497351 | -0.855317666 | 0.37505229 | -2.2805291 | 0.02257633 | 0.06583436 | LOC101928944 | 101928944 |
| ENSG000001 | 472.591349 | 0.298407907 | 0.1308612 | 2.28033909 | 0.02258759 | 0.06585772 | ZDHHC21 | 340481 |
| ENSG000002 | 4.35541636 | -0.862385634 | 0.37818296 | -2.2803397 | 0.02258755 | 0.06585772 | NA | NA |
| ENSG000002 | 2.72633399 | -0.845841612 | 0.37095687 | -2.2801616 | 0.02259811 | 0.06588366 | NA | NA |
| ENSG000002 | 7.79501882 | -0.817584064 | 0.35857011 | -2.2801233 | 0.02260038 | 0.06588554 | LOC105372672 | 105372672 |
| ENSG000002 | 2.62111542 | -0.848088095 | 0.3719566 | -2.2800727 | 0.02260337 | 0.06588954 | IL31 | 386653 |
| ENSG000002 | 44.1272952 | 0.527141692 | 0.23120752 | 2.27995047 | 0.02261063 | 0.06590594 | LINC01481 | 101928062 |
| ENSG000001 | 10.2691513 | -0.784478373 | 0.34408226 | -2.2799152 | 0.02261272 | 0.06590731 | TM4SF4 | 7104 |
| ENSG000001 | 217.369624 | 0.372496577 | 0.16340551 | 2.27958393 | 0.02263238 | 0.06595986 | GPR27 | 2850 |
| ENSG000001 | 141.05901 | 0.349381108 | 0.15327361 | 2.27946023 | 0.02263972 | 0.06597178 | RBP1 | 5947 |
| ENSG000002 | 10.1893613 | -0.783413354 | 0.34368207 | -2.2794711 | 0.02263907 | 0.06597178 | NA | NA |
| ENSG000001 | 136.788316 | 0.394603796 | 0.17312288 | 2.27932777 | 0.02264759 | 0.06598523 | ANKRD13B | 124930 |
| ENSG000002 | 8.06881433 | -0.809308549 | 0.35506458 | -2.2793277 | 0.02264759 | 0.06598523 | NA | NA |

| | | | | | | | | |
|---|---|---|---|---|---|---|---|---|
| ENSG000002 | 10.4129859 | -0.775129233 | 0.34007468 | -2.2792912 | 0.02264976 | 0.06598681 | LOC729987 | 729987 |
| ENSG000002 | 13.2579647 | -0.734403316 | 0.32222545 | -2.2791599 | 0.02265756 | 0.06600055 | FBXO47 | 494188 |
| ENSG000002 | 4.67981929 | -0.858812292 | 0.3768159 | -2.2791297 | 0.02265936 | 0.06600055 | NA | NA |
| ENSG000002 | 15.8544973 | -0.684227909 | 0.30021197 | -2.2791493 | 0.02265819 | 0.06600055 | PRCAT47 | 100996425 |
| ENSG000001 | 454.064298 | 0.307089128 | 0.13474238 | 2.27908349 | 0.0226621 | 0.0660038 | TTBK2 | 146057 |
| ENSG000001 | 1423.01801 | 0.292975643 | 0.1285582 | 2.27893389 | 0.022671 | 0.06602496 | ZNF431 | 170959 |
| ENSG00000C | 156.292074 | 0.41866586 | 0.18371515 | 2.27888584 | 0.02267385 | 0.06602854 | CBFA2T2 | 9139 |
| ENSG000001 | 118.705229 | 0.378190098 | 0.16596584 | 2.27872254 | 0.02268356 | 0.0660517 | NA | NA |
| ENSG000002 | 8.2184845 | -0.816268765 | 0.35821729 | -2.2786973 | 0.02268507 | 0.0660517 | NA | NA |
| ENSG000001 | 34.4043262 | -0.568552148 | 0.24952343 | -2.2785521 | 0.0226937 | 0.06607211 | ABCC12 | 94160 |
| ENSG000002 | 9.79033074 | -0.77915215 | 0.34197202 | -2.2784091 | 0.02270222 | 0.06608741 | NA | NA |
| ENSG000002 | 2.66728563 | -0.847719688 | 0.37206442 | -2.2784218 | 0.02270146 | 0.06608741 | NA | NA |
| ENSG000001 | 290.494629 | 0.353205435 | 0.15503276 | 2.27826317 | 0.0227109 | 0.06610509 | CBS | 875 |
| ENSG000001 | 27.0596659 | -0.587681301 | 0.25795269 | -2.2782523 | 0.02271155 | 0.06610509 | WT1 | 7490 |
| ENSG000001 | 64.7111578 | 0.443932831 | 0.19486173 | 2.27819401 | 0.02271502 | 0.06611044 | ZNF276 | 92822 |
| ENSG000002 | 122.004669 | 0.35935896 | 0.15774522 | 2.27809731 | 0.02272078 | 0.06611772 | SLC25A53 | 401612 |
| ENSG000002 | 21.8988138 | 0.63700096 | 0.27961834 | 2.27810862 | 0.02272011 | 0.06611772 | SNORD91B | 692208 |
| ENSG00000C | 212.9487 | 0.300963042 | 0.1321311 | 2.27776084 | 0.02274083 | 0.06617131 | TRHDE | 29953 |
| ENSG000002 | 334.228208 | 0.32708976 | 0.14361243 | 2.27758677 | 0.02275121 | 0.06619676 | MIR302A | 407028 |
| ENSG00000C | 602.020407 | 0.291673866 | 0.1280657 | 2.27753315 | 0.02275441 | 0.06620132 | NCOA1 | 8648 |
| ENSG000001 | 16.8522265 | -0.701049378 | 0.3078162 | -2.2774935 | 0.02275678 | 0.06620346 | OR4K17 | 390436 |
| ENSG000001 | 487.591767 | 0.284639078 | 0.12498109 | 2.27745709 | 0.02275894 | 0.06620502 | RWDD4 | 201965 |
| ENSG000002 | 9.67507784 | -0.775436163 | 0.34049748 | -2.2773624 | 0.02276459 | 0.0662167 | LOC105377105 | 105377105 |
| ENSG000001 | 8.95376433 | -0.812756695 | 0.35688939 | -2.277335 | 0.02276623 | 0.06621671 | HOXB9 | 3219 |
| ENSG000002 | 8.00891143 | -0.807483386 | 0.35458189 | -2.2772832 | 0.02276932 | 0.06622094 | LINC01068 | 103724388 |
| ENSG000001 | 42.6717383 | -0.494809542 | 0.21729261 | -2.2771577 | 0.02277681 | 0.06623798 | LTBP2 | 4053 |
| ENSG000001 | 18.222947 | -0.672390519 | 0.29529737 | -2.2769946 | 0.02278655 | 0.06626067 | OVCH2 | 341277 |
| ENSG000002 | 2.32576719 | -0.831331118 | 0.36510374 | -2.2769723 | 0.02278788 | 0.06626067 | NA | NA |
| ENSG000002 | 3.78381726 | -0.86056514 | 0.37795623 | -2.276891 | 0.02279274 | 0.06627004 | NA | NA |

| | | | | | | | | |
|---|---|---|---|---|---|---|---|---|
| ENSG000002 | 6.0122282 | -0.843700735 | 0.37062595 | -2.2764211 | 0.02282082 | 0.06634693 | NA | NA |
| ENSG000001 | 10.1870166 | -0.761735215 | 0.33463486 | -2.2763176 | 0.02282701 | 0.06636017 | PLD4 | 122618 |
| ENSG000002 | 3.39604071 | -0.858353717 | 0.37709169 | -2.2762467 | 0.02283125 | 0.06636774 | NA | NA |
| ENSG000001 | 256.823865 | 0.339947129 | 0.14934999 | 2.27617782 | 0.02283537 | 0.06637496 | ELK3 | 2004 |
| ENSG000001 | 50.3855113 | 0.491735151 | 0.21604057 | 2.27612407 | 0.02283859 | 0.06637955 | NA | NA |
| ENSG000001 | 329.572774 | 0.346241121 | 0.15213146 | 2.27593378 | 0.02284997 | 0.06640789 | SMAD7 | 4092 |
| ENSG000002 | 11.3679564 | -0.744860894 | 0.32728135 | -2.2759039 | 0.02285176 | 0.06640833 | LOC101927901 | 101927901 |
| ENSG000001 | 13.9654563 | -0.711443464 | 0.31260644 | -2.2758439 | 0.02285535 | 0.066414 | OXGR1 | 27199 |
| ENSG000002 | 111.627197 | 0.382069569 | 0.16788881 | 2.27572983 | 0.02286218 | 0.0664291 | STX18-AS1 | 100507266 |
| ENSG000002 | 7.0661471 | -0.824274816 | 0.36226494 | -2.275337 | 0.02288572 | 0.06649272 | NA | NA |
| ENSG000001 | 11.4198817 | -0.775689152 | 0.34092823 | -2.2752271 | 0.02289231 | 0.0665071 | LILRB4 | 11006 |
| ENSG000002 | 5.41406037 | -0.854112294 | 0.37541233 | -2.2751312 | 0.02289806 | 0.06651904 | NA | NA |
| ENSG000001 | 70.7361813 | 0.449919569 | 0.1977877 | 2.27476008 | 0.02292033 | 0.06657895 | TESK1 | 7016 |
| ENSG000002 | 26.0685222 | 0.59148688 | 0.26002964 | 2.27469023 | 0.02292452 | 0.06658636 | SNORD3C | 780853 |
| ENSG000002 | 10.1541582 | 0.778561177 | 0.34231009 | 2.27443248 | 0.02294 | 0.06662654 | TMEFF1 | 8577 |
| ENSG000001 | 50.777596 | 0.469067441 | 0.20627403 | 2.27400147 | 0.0229659 | 0.066697 | ST20 | 400410 |
| ENSG000001 | 166.976367 | 0.364728812 | 0.16039498 | 2.27394161 | 0.0229695 | 0.06670267 | GAREML | 150946 |
| ENSG000002 | 76.3741027 | 0.435979686 | 0.19173507 | 2.27386507 | 0.0229741 | 0.06671126 | LOC100287015 | 100287015 |
| ENSG000001 | 9.40722436 | -0.782515473 | 0.34415151 | -2.2737528 | 0.02298085 | 0.06671653 | CCDC36 | 339834 |
| ENSG000002 | 18.1287023 | 0.68344816 | 0.30057967 | 2.27376712 | 0.02297999 | 0.06671653 | NA | NA |
| ENSG000002 | 2.31901561 | -0.831072036 | 0.36550651 | -2.2737544 | 0.02298076 | 0.06671653 | RN7SKP164 | 106480885 |
| ENSG000002 | 3.68404542 | -0.860477126 | 0.37844526 | -2.2737162 | 0.02298306 | 0.06671815 | NA | NA |
| ENSG000002 | 21.603009 | 0.636456414 | 0.27992771 | 2.27364559 | 0.0229873 | 0.06672571 | NA | NA |
| ENSG000001 | 4.89504326 | -0.851960603 | 0.37474461 | -2.2734432 | 0.02299948 | 0.06675629 | NA | NA |
| ENSG000002 | 2.33931251 | -0.831452617 | 0.36572859 | -2.2734143 | 0.02300122 | 0.06675656 | NA | NA |
| ENSG000002 | 56.8463965 | 0.462347306 | 0.20337401 | 2.27338445 | 0.02300302 | 0.066757 | LOC101929494 | 101929494 |
| ENSG000001 | 5.86747333 | -0.842178989 | 0.37049065 | -2.2731451 | 0.02301744 | 0.06679406 | FER1L6-AS1 | 439941 |
| ENSG00000C | 7.96942056 | -0.807536587 | 0.35526954 | -2.2730251 | 0.02302467 | 0.06680145 | MYOC | 4653 |
| ENSG000002 | 4.98910927 | -0.854310231 | 0.37584796 | -2.2730208 | 0.02302493 | 0.06680145 | NA | NA |

| | | | | | | | | |
|---|---|---|---|---|---|---|---|---|
| ENSG000002 | 7.3319857 | -0.813473881 | 0.35788182 | -2.2730238 | 0.02302475 | 0.06680145 | NA | NA |
| ENSG000002 | 8.41522083 | -0.811088576 | 0.35689282 | -2.2726391 | 0.02304794 | 0.06686342 | NA | NA |
| ENSG000001 | 1464.08582 | 0.248193616 | 0.10921482 | 2.27252694 | 0.0230547 | 0.06686935 | EXOSC9 | 5393 |
| ENSG000002 | 14.65755 | 0.714551237 | 0.31443077 | 2.27252322 | 0.02305493 | 0.06686935 | NA | NA |
| ENSG000002 | 3.68626969 | -0.860109272 | 0.37847355 | -2.2725743 | 0.02305184 | 0.06686935 | NA | NA |
| ENSG000002 | 13.6789619 | 0.729708712 | 0.32117781 | 2.27197734 | 0.02308788 | 0.06696013 | FAM27E3 | 100131997 |
| ENSG000000 | 202.189785 | 0.354194914 | 0.15590661 | 2.27184027 | 0.02309616 | 0.06696838 | DGKD | 8527 |
| ENSG000001 | 106.746877 | -0.367559932 | 0.16179089 | -2.271821 | 0.02309733 | 0.06696838 | LAMA4 | 3910 |
| ENSG000002 | 47.5913633 | 0.489867826 | 0.21562506 | 2.27185018 | 0.02309556 | 0.06696838 | SNORD116-3 | 100033415 |
| ENSG000002 | 241.55706 | 0.409854393 | 0.18040378 | 2.27187251 | 0.02309421 | 0.06696838 | POM121C | 100101267 |
| ENSG000001 | 154.95906 | 0.352874957 | 0.1553313 | 2.27175699 | 0.02310119 | 0.06697479 | PIP5K1B | 8395 |
| ENSG000002 | 2.44692687 | -0.821187478 | 0.36150701 | -2.2715672 | 0.02311266 | 0.06700326 | NA | NA |
| ENSG000001 | 109.168097 | 0.379937798 | 0.16727646 | 2.27131665 | 0.02312782 | 0.06704124 | SLC27A6 | 28965 |
| ENSG000002 | 38.8194174 | 0.505826247 | 0.22270381 | 2.27129588 | 0.02312907 | 0.06704124 | DGCR11 | 25786 |
| ENSG000001 | 10.2050384 | -0.780703524 | 0.34375147 | -2.2711278 | 0.02313924 | 0.06706593 | NPHS2 | 7827 |
| ENSG000000 | 348.906936 | 0.379509596 | 0.1671222 | 2.27085092 | 0.023156 | 0.06710083 | SSH1 | 54434 |
| ENSG000001 | 1813.70103 | 0.374733941 | 0.16502145 | 2.27081957 | 0.0231579 | 0.06710083 | YLPM1 | 56252 |
| ENSG000001 | 9.80483094 | -0.780988658 | 0.34392152 | -2.2708339 | 0.02315703 | 0.06710083 | IGFL3 | 388555 |
| ENSG000001 | 369.768539 | 0.297366871 | 0.13094698 | 2.27089524 | 0.02315332 | 0.06710083 | L3MBTL3 | 84456 |
| ENSG000002 | 4.26927828 | -0.857263382 | 0.37757094 | -2.2704697 | 0.0231791 | 0.06715745 | NEURL1-AS1 | 102724341 |
| ENSG000001 | 1063.98554 | 0.361998152 | 0.15943976 | 2.27043844 | 0.02318099 | 0.06715814 | COL4A3BP | 10087 |
| ENSG000002 | 9.08902324 | -0.783590884 | 0.34516871 | -2.2701678 | 0.0231974 | 0.06720088 | NA | NA |
| ENSG000001 | 248.901204 | 0.327229067 | 0.14414547 | 2.2701307 | 0.02319965 | 0.06720095 | B3GNT5 | 84002 |
| ENSG000002 | 5.54681327 | -0.842497789 | 0.37112596 | -2.2701128 | 0.02320074 | 0.06720095 | LOC339622 | 339622 |
| ENSG000001 | 63.4527376 | -0.441902257 | 0.1946637 | -2.2700805 | 0.0232027 | 0.06720182 | CFAP43 | 80217 |
| ENSG000002 | 2.27753025 | -0.829491683 | 0.36542245 | -2.2699527 | 0.02321045 | 0.06721947 | LOC105372894 | 105372894 |
| ENSG000001 | 35.8768733 | 0.53802924 | 0.23707874 | 2.26941158 | 0.02324331 | 0.06730982 | CORO2B | 10391 |
| ENSG000001 | 12.0268254 | -0.738984221 | 0.32564939 | -2.2692633 | 0.02325232 | 0.0673311 | SLC28A1 | 9154 |
| ENSG000002 | 3.70980305 | -0.858727776 | 0.37847894 | -2.2688919 | 0.0232749 | 0.06739169 | RN7SL282P | 106480374 |

| | | | | | | | | | |
|---|---|---|---|---|---|---|---|---|---|
| ENSG000001 | 21.2218606 | 0.630515089 | 0.27790898 | 2.26878271 | 0.02328154 | 0.0674061 | RASD2 | | 23551 |
| ENSG000002 | 5.48824786 | -0.841718435 | 0.37102845 | -2.2686089 | 0.02329212 | 0.06743192 | PTX4 | | 390667 |
| ENSG000002 | 4.18868413 | -0.858685865 | 0.37852937 | -2.2684788 | 0.02330004 | 0.06745002 | NA | NA | |
| ENSG000002 | 5.49905182 | -0.854051887 | 0.3765051 | -2.2683674 | 0.02330682 | 0.06746484 | NA | NA | |
| ENSG000001 | 68.2359496 | -0.434002117 | 0.19135852 | -2.2680052 | 0.02332889 | 0.06751908 | ZFHX3 | | 463 |
| ENSG000002 | 6.45393321 | -0.828557875 | 0.36532289 | -2.2680152 | 0.02332828 | 0.06751908 | NA | NA | |
| ENSG000001 | 89.3858964 | 0.393432402 | 0.17347415 | 2.26795985 | 0.02333165 | 0.06752226 | TMEM108 | | 66000 |
| ENSG000002 | 7.86276598 | -0.798440403 | 0.35205945 | -2.2679136 | 0.02333448 | 0.06752561 | NA | NA | |
| ENSG000002 | 3.7277403 | -0.857969438 | 0.37831557 | -2.2678671 | 0.02333731 | 0.06752898 | NA | NA | |
| ENSG000002 | 27.8101512 | 0.567659494 | 0.25031471 | 2.26778323 | 0.02334242 | 0.06753896 | NA | NA | |
| ENSG000002 | 7.37302725 | -0.813959164 | 0.3589341 | -2.267712 | 0.02334677 | 0.06754671 | NA | NA | |
| ENSG000002 | 3.28216816 | -0.856766538 | 0.37784824 | -2.2674885 | 0.0233604 | 0.06758134 | NA | NA | |
| ENSG000001 | 1162.48698 | 0.284489409 | 0.12546943 | 2.26740011 | 0.02336579 | 0.06759173 | EPC2 | | 26122 |
| ENSG000002 | 9.64457574 | -0.767516046 | 0.33850424 | -2.267375 | 0.02336733 | 0.06759173 | NA | NA | |
| ENSG000002 | 52.0798007 | 0.532705456 | 0.23496371 | 2.26718185 | 0.02337912 | 0.06762102 | LTB4R | | 1241 |
| ENSG000001 | 10.9075185 | -0.7559605 | 0.33346953 | -2.2669553 | 0.02339296 | 0.06765622 | KCNK9 | | 51305 |
| ENSG000001 | 282.515924 | 0.326483852 | 0.14402939 | 2.26678626 | 0.02340329 | 0.06768126 | PLEKHG4B | | 153478 |
| ENSG000001 | 36.8590148 | 0.55884445 | 0.24655071 | 2.26665116 | 0.02341154 | 0.06769792 | ZNF775 | | 285971 |
| ENSG000002 | 3.2884258 | -0.856591647 | 0.37791296 | -2.2666374 | 0.02341238 | 0.06769792 | NA | NA | |
| ENSG000001 | 5.21832044 | -0.847092192 | 0.37373008 | -2.2665882 | 0.02341539 | 0.06769957 | IGFBP1 | | 3484 |
| ENSG000002 | 3.74591997 | -0.857692089 | 0.37840912 | -2.2665735 | 0.02341629 | 0.06769957 | NA | NA | |
| ENSG000001 | 16.1192923 | -0.720497765 | 0.31788718 | -2.2665203 | 0.02341954 | 0.06770414 | OR52K1 | | 390036 |
| ENSG000002 | 287.450104 | 0.341750112 | 0.15078857 | 2.26641922 | 0.02342573 | 0.06771719 | NA | NA | |
| ENSG000001 | 743.648549 | 0.286056123 | 0.12624971 | 2.26579631 | 0.02346386 | 0.06782257 | ANKRD26 | | 22852 |
| ENSG000001 | 1694.78203 | 0.26693419 | 0.11781974 | 2.26561513 | 0.02347496 | 0.06784499 | SHPRH | | 257218 |
| ENSG000002 | 3.09594548 | -0.845522076 | 0.37319406 | -2.2656365 | 0.02347365 | 0.06784499 | NA | NA | |
| ENSG000002 | 16.9126677 | -0.678163171 | 0.29936825 | -2.2653143 | 0.0234934 | 0.06789345 | NA | NA | |
| ENSG000002 | 2.94185191 | -0.852150003 | 0.37619908 | -2.265157 | 0.02350304 | 0.06791123 | NA | NA | |
| ENSG000002 | 15.396574 | 0.696098772 | 0.30730436 | 2.26517701 | 0.02350182 | 0.06791123 | NA | NA | |

| | | | | | | | | |
|---|---|---|---|---|---|---|---|---|
| ENSG000002 | 3.61512093 | -0.857420336 | 0.37853396 | -2.2651081 | 0.02350605 | 0.06791123 | SLIT1-AS1 | 100505540 |
| ENSG000002 | 37.4709894 | 0.557513535 | 0.24613146 | 2.26510477 | 0.02350625 | 0.06791123 | NA | NA |
| ENSG00000C | 1275.26969 | 0.38568031 | 0.1702758 | 2.265033 | 0.02351065 | 0.06791911 | SPEN | 23013 |
| ENSG000001 | 52.3990549 | -0.459178736 | 0.20272988 | -2.264978 | 0.02351403 | 0.06792089 | SAMD3 | 154075 |
| ENSG000002 | 50.5991746 | 0.501446927 | 0.22139246 | 2.26496837 | 0.02351462 | 0.06792089 | LRRC37A15P | 106481796 |
| ENSG000001 | 147.444554 | 0.362686924 | 0.16014816 | 2.26469623 | 0.02353133 | 0.0679643 | ZNF789 | 285989 |
| ENSG000001 | 22.4061733 | -0.618584057 | 0.27314792 | -2.2646486 | 0.02353425 | 0.06796791 | C6orf222 | 389384 |
| ENSG000001 | 11.5788379 | -0.753951333 | 0.33294916 | -2.2644638 | 0.0235456 | 0.06799585 | KRT35 | 3886 |
| ENSG00000C | 209.39706 | 0.309918869 | 0.1368689 | 2.26434843 | 0.02355269 | 0.06801148 | SAMD4A | 23034 |
| ENSG000001 | 28.9180425 | -0.58774271 | 0.25957586 | -2.2642426 | 0.0235592 | 0.06802543 | TMEM163 | 81615 |
| ENSG000002 | 32.0689153 | -0.543566874 | 0.24007013 | -2.2642003 | 0.0235618 | 0.06802646 | NA | NA |
| ENSG000002 | 4.34029176 | -0.855401282 | 0.37779702 | -2.2641822 | 0.02356291 | 0.06802646 | NA | NA |
| ENSG000002 | 4.0777028 | -0.857186314 | 0.37859209 | -2.2641422 | 0.02356537 | 0.06802872 | LINC01180 | 101927017 |
| ENSG000002 | 2.92127536 | -0.851835544 | 0.37629242 | -2.2637595 | 0.02358891 | 0.06809183 | NA | NA |
| ENSG000001 | 27.6617568 | -0.587262354 | 0.25942827 | -2.2636791 | 0.02359386 | 0.06809642 | GRIK1 | 2897 |
| ENSG000001 | 220.890423 | 0.37810986 | 0.16703229 | 2.26369316 | 0.02359299 | 0.06809642 | ZNF335 | 63925 |
| ENSG000002 | 6.85924701 | -0.823097642 | 0.36364086 | -2.2634905 | 0.02360547 | 0.06812507 | LOC100506498 | 100506498 |
| ENSG000002 | 17.6471924 | -0.65243717 | 0.28824888 | -2.2634508 | 0.02360791 | 0.06812727 | WTAPP1 | 100288077 |
| ENSG000002 | 7.70412336 | -0.808106663 | 0.3570683 | -2.2631711 | 0.02362514 | 0.06817214 | NA | NA |
| ENSG000002 | 2.31360674 | -0.824180747 | 0.36421963 | -2.2628675 | 0.02364386 | 0.06821645 | NA | NA |
| ENSG000002 | 4.94513453 | -0.849174981 | 0.37526427 | -2.2628719 | 0.02364359 | 0.06821645 | NA | NA |
| ENSG000002 | 19.6180065 | -0.636884248 | 0.28146611 | -2.2627387 | 0.0236518 | 0.06823451 | NA | NA |
| ENSG000001 | 968.174702 | 0.350278538 | 0.15480677 | 2.26268231 | 0.02365528 | 0.06823968 | MOB1B | 92597 |
| ENSG000002 | 2.65313481 | -0.84049161 | 0.37147239 | -2.2625951 | 0.02366066 | 0.06825035 | NA | NA |
| ENSG000002 | 17.2383637 | -0.667276703 | 0.29492295 | -2.2625459 | 0.0236637 | 0.06825425 | NA | NA |
| ENSG000001 | 575.186378 | 0.296379117 | 0.13099933 | 2.26244762 | 0.02366976 | 0.06826689 | LYSMD3 | 116068 |
| ENSG000002 | 7.88297038 | -0.797677364 | 0.35259925 | -2.2622775 | 0.02368026 | 0.06829232 | NA | NA |
| ENSG00000C | 4839.19032 | 0.24871844 | 0.10994415 | 2.26222523 | 0.02368349 | 0.06829678 | NOP58 | 51602 |
| ENSG000002 | 5.35239364 | -0.847548648 | 0.37465961 | -2.2621831 | 0.02368609 | 0.06829942 | NA | NA |

| | | | | | | | | |
|---|---|---|---|---|---|---|---|---|
| ENSG000001 | 37.7584472 | -0.526845014 | 0.23289772 | -2.2621305 | 0.02368934 | 0.06830394 | HPSE2 | 60495 |
| ENSG000002 | 2.72694004 | -0.838348503 | 0.37062973 | -2.261957 | 0.02370006 | 0.06832998 | NA | NA |
| ENSG000001 | 8.23522528 | -0.799452954 | 0.35346041 | -2.2617892 | 0.02371043 | 0.06835501 | GSTA5 | 221357 |
| ENSG000002 | 17.1135389 | -0.679429044 | 0.30039938 | -2.2617524 | 0.0237127 | 0.06835671 | NA | NA |
| ENSG000002 | 10.4389825 | -0.766878661 | 0.33908972 | -2.2615804 | 0.02372334 | 0.06838252 | NA | NA |
| ENSG000001 | 60.9505894 | 0.471954379 | 0.20869311 | 2.26147558 | 0.02372982 | 0.06839634 | DBP | 1628 |
| ENSG000001 | 7.79745933 | -0.810574893 | 0.35844943 | -2.2613368 | 0.02373841 | 0.06841135 | CLDN22 | 53842 |
| ENSG000001 | 11.9988502 | -0.75953588 | 0.33587835 | -2.2613422 | 0.02373808 | 0.06841135 | TPPP2 | 122664 |
| ENSG00000C | 2114.95585 | 0.272014312 | 0.12030001 | 2.26113297 | 0.02375102 | 0.06844285 | RASAL2 | 9462 |
| ENSG000001 | 6.14955651 | -0.830898414 | 0.3674803 | -2.2610693 | 0.02375497 | 0.06844935 | NA | NA |
| ENSG000001 | 839.271948 | 0.280791627 | 0.12418898 | 2.26100281 | 0.02375908 | 0.06845309 | CACNA2D1 | 781 |
| ENSG000001 | 21.9318951 | -0.652416282 | 0.28855594 | -2.2609699 | 0.02376112 | 0.06845309 | OLR1 | 4973 |
| ENSG000002 | 5.2595584 | -0.844847895 | 0.3736667 | -2.2609665 | 0.02376133 | 0.06845309 | NA | NA |
| ENSG000002 | 3.0331303 | -0.847439465 | 0.37483795 | -2.2608156 | 0.02377068 | 0.06847516 | NA | NA |
| ENSG000002 | 31.5983206 | -0.562007137 | 0.24859189 | -2.2607622 | 0.02377399 | 0.06847982 | ECT2L | 345930 |
| ENSG000002 | 7.42488041 | -0.812905189 | 0.3595921 | -2.2606314 | 0.02378209 | 0.06849831 | ANXA8 | 653145 |
| ENSG000001 | 25.311054 | -0.62141024 | 0.27488887 | -2.2605872 | 0.02378483 | 0.06850134 | BST1 | 683 |
| ENSG000001 | 123.081709 | 0.398452087 | 0.17626817 | 2.26048806 | 0.02379098 | 0.06851416 | WDR13 | 64743 |
| ENSG000002 | 11.7022522 | -0.740952497 | 0.32778903 | -2.2604554 | 0.023793 | 0.06851512 | TSGA13 | 114960 |
| ENSG000002 | 10.4559395 | -0.7685976 | 0.34006424 | -2.2601542 | 0.02381169 | 0.06855955 | NA | NA |
| ENSG000002 | 15.0009613 | -0.688526885 | 0.3046374 | -2.2601522 | 0.02381181 | 0.06855955 | NA | NA |
| ENSG000002 | 8.96070389 | -0.793496792 | 0.35109267 | -2.2600779 | 0.02381642 | 0.06856794 | NA | NA |
| ENSG000001 | 246.338055 | 0.303801152 | 0.1344331 | 2.25986861 | 0.02382941 | 0.06860047 | OGG1 | 4968 |
| ENSG000001 | 176.171421 | 0.317935219 | 0.14069631 | 2.2597267 | 0.02383822 | 0.06862097 | RGS9 | 8787 |
| ENSG000002 | 17.5812851 | -0.66806055 | 0.29565497 | -2.2595952 | 0.02384638 | 0.0686396 | NA | NA |
| ENSG000002 | 2.91088033 | -0.832315217 | 0.36837085 | -2.2594492 | 0.02385546 | 0.06866085 | NA | NA |
| ENSG000002 | 3.09374502 | -0.846128503 | 0.37449607 | -2.2593789 | 0.02385983 | 0.06866855 | NA | NA |
| ENSG00000C | 289.494253 | 0.390607073 | 0.17288653 | 2.25932622 | 0.0238631 | 0.0686731 | MCAM | 4162 |
| ENSG000001 | 4.38399765 | -0.854945133 | 0.37842804 | -2.2592013 | 0.02387087 | 0.06868866 | NCF1 | 653361 |

| | | | | | | | | |
|---|---|---|---|---|---|---|---|---|
| ENSG000002 | 8.10714215 | -0.814356303 | 0.36046468 | -2.2591847 | 0.0238719 | 0.06868866 | NA | NA |
| ENSG000001 | 3.32959154 | -0.852993522 | 0.37758176 | -2.2590962 | 0.0238774 | 0.06869741 | OR11H6 | 122748 |
| ENSG000001 | 7.75039249 | -0.807247252 | 0.3573343 | -2.2590813 | 0.02387833 | 0.06869741 | GP5 | 2814 |
| ENSG000001 | 49.3835037 | -0.498518791 | 0.22067928 | -2.2590195 | 0.02388217 | 0.06870361 | FGF1 | 2246 |
| ENSG000001 | 23.7213735 | -0.607569585 | 0.26896643 | -2.2589049 | 0.0238893 | 0.06871923 | BTLA | 151888 |
| ENSG000001 | 38.8302096 | -0.518387413 | 0.22950778 | -2.2586921 | 0.02390254 | 0.06874928 | MYLK4 | 340156 |
| ENSG000002 | 72.7236869 | 0.423496523 | 0.18749714 | 2.25868257 | 0.02390314 | 0.06874928 | NA | NA |
| ENSG000002 | 2.30800255 | -0.822526741 | 0.36416963 | -2.2586363 | 0.02390602 | 0.06875269 | NA | NA |
| ENSG000001 | 604.619223 | 0.416672575 | 0.18449516 | 2.25844714 | 0.0239178 | 0.06877681 | SAFB2 | 9667 |
| ENSG000002 | 4.91787166 | -0.8466464 | 0.37487757 | -2.2584611 | 0.02391693 | 0.06877681 | NA | NA |
| ENSG000001 | 92.339064 | -0.387285373 | 0.1714923 | -2.2583251 | 0.0239254 | 0.06879379 | MICAL2 | 9645 |
| ENSG000002 | 28.0303373 | 0.582102979 | 0.25776805 | 2.25824329 | 0.0239305 | 0.06880357 | NA | NA |
| ENSG000001 | 19.9647568 | -0.654285843 | 0.28974211 | -2.2581662 | 0.0239353 | 0.06880762 | RSPH10B | 222967 |
| ENSG000001 | 25.1543656 | -0.590309294 | 0.26141025 | -2.258172 | 0.02393494 | 0.06880762 | TCP10L2 | 401285 |
| ENSG000001 | 13.025557 | -0.715298578 | 0.31678292 | -2.2580087 | 0.02394512 | 0.06882652 | SPATA45 | 149643 |
| ENSG000002 | 10.6123002 | 0.755203217 | 0.33445577 | 2.25800628 | 0.02394527 | 0.06882652 | NA | NA |
| ENSG000002 | 5.51778707 | -0.839539144 | 0.3718396 | -2.2577992 | 0.02395818 | 0.06885876 | NA | NA |
| ENSG000002 | 5.89837127 | -0.836124406 | 0.37035332 | -2.2576398 | 0.02396812 | 0.06888245 | NA | NA |
| ENSG000001 | 255.394674 | 0.432237783 | 0.19146019 | 2.2575857 | 0.0239715 | 0.06888727 | NEFH | 4744 |
| ENSG000002 | 6.84646668 | -0.821202726 | 0.36376731 | -2.2574946 | 0.02397719 | 0.06889874 | LOC441178 | 441178 |
| ENSG000002 | 8.53818448 | -0.79090427 | 0.35035394 | -2.2574437 | 0.02398037 | 0.06890299 | NA | NA |
| ENSG000002 | 28.969278 | 0.595169716 | 0.2636754 | 2.25720612 | 0.0239952 | 0.06894072 | NA | NA |
| ENSG000001 | 11.2046462 | -0.753530581 | 0.33384559 | -2.2571231 | 0.02400039 | 0.06894754 | IL22RA1 | 58985 |
| ENSG000002 | 6.37327712 | -0.838409721 | 0.37145215 | -2.2571137 | 0.02400097 | 0.06894754 | NA | NA |
| ENSG000002 | 1128.175 | 0.282222112 | 0.12504997 | 2.25687466 | 0.02401591 | 0.06898555 | ZNF525 | 170958 |
| ENSG000002 | 3.70192196 | -0.853911493 | 0.37838724 | -2.2567133 | 0.024026 | 0.06900965 | NA | NA |
| ENSG000001 | 310.853554 | 0.305226059 | 0.13526066 | 2.25657681 | 0.02403453 | 0.0690285 | ZNF420 | 147923 |
| ENSG000002 | 3.96893511 | -0.85426123 | 0.37856895 | -2.2565539 | 0.02403597 | 0.0690285 | NA | NA |
| ENSG000001 | 63.2378999 | 0.435290871 | 0.19290407 | 2.2565147 | 0.02403842 | 0.06903065 | CRELD1 | 78987 |

| | | | | | | | | |
|---|---|---|---|---|---|---|---|---|
| ENSG000002 | 8.48151879 | -0.787191121 | 0.34888292 | -2.2563189 | 0.02405067 | 0.06906094 | LINC00293 | 497634 |
| ENSG000001 | 16.3261517 | -0.7296579 | 0.32342683 | -2.2560215 | 0.02406928 | 0.0691095 | BHMT2 | 23743 |
| ENSG000001 | 573.117855 | 0.269333322 | 0.11939081 | 2.25589665 | 0.0240771 | 0.06912706 | PFKFB2 | 5208 |
| ENSG000002 | 8.15068623 | -0.8084656 | 0.35840453 | -2.2557349 | 0.02408724 | 0.06915126 | LOC387810 | 387810 |
| ENSG000002 | 2.97857066 | -0.845359797 | 0.37479786 | -2.2555086 | 0.02410142 | 0.06918709 | RNU6-1262P | 106480409 |
| ENSG000002 | 8.80734407 | -0.779939341 | 0.34586676 | -2.2550283 | 0.02413155 | 0.06926867 | NA | NA |
| ENSG000001 | 17.3207485 | -0.68973083 | 0.30587045 | -2.254977 | 0.02413477 | 0.06927302 | ZBP1 | 81030 |
| ENSG000002 | 7.67412088 | -0.823270146 | 0.36510763 | -2.2548697 | 0.02414151 | 0.06928744 | LOC101928203 | 101928203 |
| ENSG000001 | 153.724206 | 0.340626608 | 0.15108529 | 2.2545319 | 0.02416273 | 0.06934186 | HEMK1 | 51409 |
| ENSG000001 | 12.1256986 | -0.763924675 | 0.33884238 | -2.2545134 | 0.02416389 | 0.06934186 | OR9G1 | 390174 |
| ENSG000001 | 18.3149141 | -0.653620965 | 0.28994147 | -2.2543204 | 0.02417602 | 0.06937177 | NA | NA |
| ENSG000002 | 18.9651409 | -0.636120279 | 0.28221094 | -2.2540596 | 0.02419242 | 0.06941391 | NA | NA |
| ENSG000001 | 767.826658 | 0.287471221 | 0.12753906 | 2.25398566 | 0.02419707 | 0.06942234 | CEP97 | 79598 |
| ENSG000001 | 56.9105809 | -0.467860049 | 0.20761551 | -2.2534928 | 0.02422809 | 0.06950459 | C2CD2L | 9854 |
| ENSG000002 | 5.48890074 | -0.836770523 | 0.37132439 | -2.2534758 | 0.02422917 | 0.06950459 | LOC101929541 | 101929541 |
| ENSG000001 | 13.8318289 | -0.725733827 | 0.3220552 | -2.2534454 | 0.02423108 | 0.06950516 | TCF21 | 6943 |
| ENSG000001 | 161.230011 | 0.350418462 | 0.1555114 | 2.25332978 | 0.02423836 | 0.06951735 | AADAT | 51166 |
| ENSG000002 | 7.65456855 | -0.803270631 | 0.3564826 | -2.2533235 | 0.02423876 | 0.06951735 | NA | NA |
| ENSG000001 | 1007.109 | 0.292744363 | 0.12992227 | 2.25322696 | 0.02424484 | 0.06952989 | LEO1 | 123169 |
| ENSG000001 | 3913.01762 | 0.25868487 | 0.11481774 | 2.25300434 | 0.02425888 | 0.06956086 | SFRP1 | 6422 |
| ENSG000001 | 12.8610691 | -0.724403441 | 0.3215285 | -2.2529991 | 0.0242592 | 0.06956086 | LGI3 | 203190 |
| ENSG000002 | 6.75317061 | -0.820972521 | 0.36439503 | -2.252974 | 0.02426079 | 0.06956086 | NA | NA |
| ENSG000001 | 229.016095 | 0.412043035 | 0.18289427 | 2.25290295 | 0.02426527 | 0.06956879 | DOT1L | 84444 |
| ENSG000002 | 2.39670478 | -0.813983018 | 0.36132704 | -2.2527598 | 0.0242743 | 0.06958976 | RN7SL62P | 106479246 |
| ENSG000001 | 15.5310608 | -0.682724566 | 0.30313498 | -2.2522131 | 0.02430881 | 0.06966406 | DNAJC5G | 285126 |
| ENSG000001 | 229.284615 | 0.332068875 | 0.14743765 | 2.25226643 | 0.02430544 | 0.06966406 | LIG4 | 3981 |
| ENSG000002 | 13.6740462 | -0.712610086 | 0.31639491 | -2.2522805 | 0.02430455 | 0.06966406 | NA | NA |
| ENSG000002 | 7.07747371 | -0.816055975 | 0.36233353 | -2.2522232 | 0.02430817 | 0.06966406 | NA | NA |
| ENSG000002 | 5.0267772 | -0.848172313 | 0.37659073 | -2.2522389 | 0.02430718 | 0.06966406 | NA | NA |

| | | | | | | | | |
|---|---|---|---|---|---|---|---|---|
| ENSG000002 | 4.02505534 | -0.852652503 | 0.37859199 | -2.2521673 | 0.0243117 | 0.06966743 | NA | NA |
| ENSG000001 | 189.613163 | 0.316702379 | 0.14062608 | 2.25208856 | 0.02431668 | 0.06967676 | STS | 412 |
| ENSG000002 | 4.98158518 | -0.848441117 | 0.37675621 | -2.2519632 | 0.0243246 | 0.06969422 | NA | NA |
| ENSG000002 | 4.26535648 | -0.851195376 | 0.37798353 | -2.2519378 | 0.02432621 | 0.06969422 | RN7SL517P | 106481064 |
| ENSG000002 | 7.29585925 | -0.804328761 | 0.35717815 | -2.251898 | 0.02432872 | 0.06969649 | NA | NA |
| ENSG000002 | 403.181813 | 0.314246164 | 0.13955539 | 2.25176652 | 0.02433703 | 0.06970845 | ZNF407 | 55628 |
| ENSG000002 | 83.0942179 | -0.391308678 | 0.17377586 | -2.2518011 | 0.02433484 | 0.06970845 | INTS6-AS1 | 100507398 |
| ENSG000002 | 27.763184 | 0.597704459 | 0.26543992 | 2.25175041 | 0.02433805 | 0.06970845 | NA | NA |
| ENSG000002 | 4.86941072 | -0.844782888 | 0.37522509 | -2.251403 | 0.02436003 | 0.06976647 | NA | NA |
| ENSG000002 | 20.3992802 | -0.63968926 | 0.28414244 | -2.2512978 | 0.02436668 | 0.0697806 | NA | NA |
| ENSG000001 | 44.4872476 | -0.503645916 | 0.2237231 | -2.2512021 | 0.02437274 | 0.06979302 | ERAP2 | 64167 |
| ENSG000002 | 15.6175359 | -0.684993375 | 0.30431111 | -2.2509641 | 0.02438781 | 0.06983125 | NA | NA |
| ENSG000001 | 13.1943846 | -0.720005756 | 0.31988044 | -2.250859 | 0.02439447 | 0.06984539 | HEPACAM | 220296 |
| ENSG000002 | 9.45258896 | -0.798094092 | 0.35457946 | -2.2508188 | 0.02439702 | 0.06984775 | CASC17 | 101928165 |
| ENSG000002 | 8.26838215 | -0.795125432 | 0.35327365 | -2.2507352 | 0.02440231 | 0.06985798 | PIRT | 644139 |
| ENSG000002 | 4.64893353 | -0.849074727 | 0.37724997 | -2.2506953 | 0.02440485 | 0.06986029 | NA | NA |
| ENSG000001 | 6.41399706 | -0.820308177 | 0.3645123 | -2.2504266 | 0.02442188 | 0.06990411 | HMHB1 | 57824 |
| ENSG000001 | 8.23578285 | -0.791960061 | 0.35192027 | -2.2503963 | 0.0244238 | 0.06990468 | DDI1 | 414301 |
| ENSG000002 | 26.9745151 | -0.583698492 | 0.25939134 | -2.250262 | 0.02443232 | 0.06992412 | FAM71F2 | 346653 |
| ENSG00000C | 30.6231772 | -0.546863997 | 0.24303595 | -2.2501362 | 0.0244403 | 0.06994203 | MGLL | 11343 |
| ENSG00000C | 625.997733 | 0.304271989 | 0.13522678 | 2.25008674 | 0.02444344 | 0.06994608 | MON2 | 23041 |
| ENSG000001 | 8911.44114 | 0.554862363 | 0.24660486 | 2.25000577 | 0.02444858 | 0.06994928 | EIF3A | 8661 |
| ENSG000001 | 748.908414 | 0.267192195 | 0.11875118 | 2.25001727 | 0.02444785 | 0.06994928 | LPGAT1 | 9926 |
| ENSG000002 | 7.371371 | -0.81033092 | 0.36014906 | -2.2499876 | 0.02444973 | 0.06994928 | NA | NA |
| ENSG000002 | 10.4266603 | -0.78360878 | 0.34827816 | -2.249951 | 0.02445206 | 0.069951 | NA | NA |
| ENSG000001 | 10.4581211 | -0.792671473 | 0.35235303 | -2.2496514 | 0.02447108 | 0.07000049 | CLDN14 | 23562 |
| ENSG000002 | 11.5944531 | -0.750097861 | 0.33344421 | -2.2495453 | 0.02447782 | 0.07001482 | NA | NA |
| ENSG000001 | 26.4673258 | -0.618840736 | 0.27511262 | -2.2494088 | 0.0244865 | 0.07003471 | CCDC38 | 120935 |
| ENSG000002 | 13.6202657 | -0.727663184 | 0.32355392 | -2.2489704 | 0.02451438 | 0.07010951 | MTND6P12 | 106481743 |

| | | | | | | | | |
|---|---|---|---|---|---|---|---|---|
| ENSG00000C | 17.9440336 | -0.666292515 | 0.29627686 | -2.2488848 | 0.02451983 | 0.07011025 | MPO | 4353 |
| ENSG000001 | 149.244456 | 0.383745082 | 0.17063392 | 2.24893782 | 0.02451645 | 0.07011025 | STIM1 | 6786 |
| ENSG000002 | 8.69692515 | -0.776857857 | 0.34543847 | -2.2489037 | 0.02451862 | 0.07011025 | NA | NA |
| ENSG000001 | 20.0614581 | -0.638399305 | 0.28388361 | -2.2488065 | 0.02452481 | 0.07011955 | P2RY14 | 9934 |
| ENSG000002 | 3.0988933 | -0.842087924 | 0.37449052 | -2.2486228 | 0.0245365 | 0.07014804 | NA | NA |
| ENSG000002 | 189.027239 | 0.354644929 | 0.15771924 | 2.24858387 | 0.02453898 | 0.07015018 | NA | NA |
| ENSG00000C | 174.190752 | 0.354287029 | 0.1575681 | 2.24846932 | 0.02454628 | 0.07016609 | NR1H3 | 10062 |
| ENSG000002 | 2.31987702 | -0.817172225 | 0.36346868 | -2.2482604 | 0.02455959 | 0.0701992 | RN7SKP165 | 106480367 |
| ENSG000002 | 5.97949698 | -0.834098041 | 0.37102953 | -2.2480638 | 0.02457212 | 0.07023007 | NA | NA |
| ENSG000001 | 33.5011026 | 0.53994903 | 0.24019244 | 2.24798514 | 0.02457714 | 0.07023945 | CCL2 | 6347 |
| ENSG000001 | 16.8106417 | -0.68119657 | 0.303068 | -2.2476691 | 0.0245973 | 0.07029211 | NR2F2 | 7026 |
| ENSG000002 | 3.27061431 | -0.848775433 | 0.3776817 | -2.24733 | 0.02461894 | 0.07034405 | NA | NA |
| ENSG000002 | 12.2605586 | -0.750738162 | 0.33405439 | -2.2473531 | 0.02461747 | 0.07034405 | NA | NA |
| ENSG000002 | 5.20933054 | -0.840411097 | 0.37396502 | -2.2472987 | 0.02462094 | 0.07034421 | NA | NA |
| ENSG000002 | 3.30903515 | -0.848919881 | 0.37775526 | -2.2472748 | 0.02462247 | 0.07034421 | NA | NA |
| ENSG000001 | 15.2566541 | -0.702529252 | 0.31264443 | -2.2470551 | 0.02463651 | 0.07037441 | ZNF843 | 283933 |
| ENSG000002 | 23.1452964 | -0.640653229 | 0.28510503 | -2.247078 | 0.02463504 | 0.07037441 | KIZ-AS1 | 101929591 |
| ENSG000001 | 2057.60948 | 0.357382158 | 0.15905487 | 2.24691108 | 0.02464571 | 0.07039573 | OTUD4 | 54726 |
| ENSG000002 | 158.200649 | -0.333170098 | 0.14830486 | -2.2465217 | 0.02467061 | 0.07046189 | NA | NA |
| ENSG000002 | 5.60992584 | -0.836977499 | 0.37262342 | -2.2461752 | 0.02469279 | 0.07052026 | NA | NA |
| ENSG000002 | 9.04358234 | -0.770835657 | 0.34318387 | -2.2461302 | 0.02469567 | 0.07052353 | NA | NA |
| ENSG00000C | 112.222284 | 0.422646411 | 0.18818295 | 2.24593366 | 0.02470826 | 0.07053961 | FUZ | 80199 |
| ENSG000001 | 42.19543 | -0.502660937 | 0.22380646 | -2.2459626 | 0.0247064 | 0.07053961 | Mar-01 | 55016 |
| ENSG000001 | 210.023758 | 0.363132008 | 0.16167907 | 2.24600505 | 0.02470368 | 0.07053961 | PRPSAP1 | 5635 |
| ENSG000001 | 31.1121 | 0.553987816 | 0.24665972 | 2.24595983 | 0.02470658 | 0.07053961 | TMEM145 | 284339 |
| ENSG00000C | 185.214541 | 0.31243463 | 0.1391279 | 2.24566475 | 0.02472549 | 0.07058383 | FAM214A | 56204 |
| ENSG000001 | 60.2388261 | -0.482220524 | 0.21474361 | -2.245564 | 0.02473195 | 0.07059729 | CDKL1 | 8814 |
| ENSG000001 | 353.267255 | 0.306427736 | 0.13646934 | 2.24539626 | 0.02474271 | 0.07061806 | ZFP1 | 162239 |
| ENSG000002 | 10.8789577 | -0.778993557 | 0.34692658 | -2.2454133 | 0.02474161 | 0.07061806 | NA | NA |

| | | | | | | | | |
|---|---|---|---|---|---|---|---|---|
| ENSG000002 | 4.71100396 | -0.847592387 | 0.37752432 | -2.2451332 | 0.02475958 | 0.07066126 | NA | NA |
| ENSG000002 | 8.12814443 | -0.802610249 | 0.35751263 | -2.2449843 | 0.02476914 | 0.07068355 | NA | NA |
| ENSG000002 | 3.09298909 | -0.840664764 | 0.37447206 | -2.2449332 | 0.02477242 | 0.07068794 | NA | NA |
| ENSG000001 | 17.5378506 | -0.648031225 | 0.28866844 | -2.244898 | 0.02477468 | 0.07068942 | SLC5A9 | 200010 |
| ENSG000002 | 3.90753671 | -0.849389712 | 0.37838043 | -2.2448035 | 0.02478075 | 0.07070177 | NA | NA |
| ENSG000002 | 7.8051316 | -0.808421767 | 0.36013707 | -2.2447613 | 0.02478346 | 0.07070452 | NA | NA |
| ENSG000001 | 126.817154 | 0.363487452 | 0.16193015 | 2.24471754 | 0.02478627 | 0.07070757 | WWOX | 51741 |
| ENSG000002 | 3.2932661 | -0.84774731 | 0.37768102 | -2.2446119 | 0.02479306 | 0.07072196 | NA | NA |
| ENSG00000C | 200.723252 | 0.316294955 | 0.14092046 | 2.2444927 | 0.02480072 | 0.07073883 | TASP1 | 55617 |
| ENSG000001 | 7.15237963 | -0.832068994 | 0.37075367 | -2.2442636 | 0.02481544 | 0.07077585 | MMP26 | 56547 |
| ENSG000001 | 33.1719693 | 0.552317761 | 0.2461207 | 2.24409307 | 0.02482641 | 0.07080216 | TMEM160 | 54958 |
| ENSG000002 | 3.90425461 | -0.849140426 | 0.37842374 | -2.2438878 | 0.02483962 | 0.07083484 | NA | NA |
| ENSG000001 | 379.318973 | 0.317326682 | 0.14143293 | 2.24365493 | 0.02485461 | 0.07086763 | SCCPDH | 51097 |
| ENSG000002 | 5.29578722 | -0.839830362 | 0.37431163 | -2.2436662 | 0.02485388 | 0.07086763 | NA | NA |
| ENSG000002 | 9.63223315 | -0.761056426 | 0.33921706 | -2.2435677 | 0.02486023 | 0.07087866 | NA | NA |
| ENSG000002 | 7.14919956 | -0.814102741 | 0.36288659 | -2.2434081 | 0.02487051 | 0.07090298 | NA | NA |
| ENSG000002 | 7.68971107 | -0.805019616 | 0.35887825 | -2.2431552 | 0.02488681 | 0.07094446 | NA | NA |
| ENSG000001 | 42.8294513 | -0.511321863 | 0.22797329 | -2.2429025 | 0.0249031 | 0.07098592 | CACNA1I | 8911 |
| ENSG000002 | 4.35842779 | -0.848154915 | 0.37819571 | -2.242635 | 0.02492036 | 0.07103013 | NA | NA |
| ENSG000002 | 11.9270861 | -0.745950842 | 0.33263725 | -2.2425355 | 0.02492678 | 0.07104344 | LINC01087 | 101927994 |
| ENSG000002 | 9.21673825 | -0.777878784 | 0.34693803 | -2.242126 | 0.02495323 | 0.07111381 | NA | NA |
| ENSG000002 | 71.0227255 | -0.408983093 | 0.18241649 | -2.2420291 | 0.02495949 | 0.07112165 | ATP6AP1L | 92270 |
| ENSG000002 | 16.0859678 | -0.693737558 | 0.30942261 | -2.242039 | 0.02495885 | 0.07112165 | NA | NA |
| ENSG000002 | 23.172545 | -0.618902724 | 0.27605063 | -2.2419899 | 0.02496203 | 0.07112388 | NA | NA |
| ENSG000002 | 31.2530545 | -0.558395341 | 0.2491336 | -2.241349 | 0.02500348 | 0.07123699 | NA | NA |
| ENSG000002 | 18.4259037 | -0.655192525 | 0.29238046 | -2.2408903 | 0.02503318 | 0.07131661 | LOC101927787 | 101927787 |
| ENSG000001 | 76.7728423 | -0.415115104 | 0.18527598 | -2.240523 | 0.02505699 | 0.07137942 | IQCH | 64799 |
| ENSG000002 | 58.0510297 | 0.489120879 | 0.21832589 | 2.24032469 | 0.02506985 | 0.07141104 | NA | NA |
| ENSG000002 | 6.5087482 | -0.821085505 | 0.36652617 | -2.2401825 | 0.02507908 | 0.0714323 | NA | NA |

| | | | | | | | | | |
|---|---|---|---|---|---|---|---|---|---|
| ENSG000001 | 14.8737104 | -0.703918223 | 0.31423214 | -2.2401217 | 0.02508302 | 0.07143853 | FCAMR | | 83953 |
| ENSG000001 | 345.220712 | 0.298767045 | 0.13339642 | 2.2396932 | 0.02511085 | 0.07151208 | LRP12 | | 29967 |
| ENSG000002 | 11.00337 | -0.756971496 | 0.33798355 | -2.2396696 | 0.02511238 | 0.07151208 | NA | NA | |
| ENSG000002 | 17.3949164 | -0.677725369 | 0.30261745 | -2.2395449 | 0.02512048 | 0.07152555 | ARSH | | 347527 |
| ENSG000002 | 3.29409525 | -0.845841125 | 0.37768477 | -2.2395426 | 0.02512063 | 0.07152555 | NA | NA | |
| ENSG000002 | 371.017456 | 0.330195276 | 0.14744477 | 2.23945054 | 0.02512662 | 0.07153756 | NA | NA | |
| ENSG000002 | 13.0881468 | -0.75645648 | 0.33782146 | -2.2392197 | 0.02514162 | 0.07157525 | GOLGA6L22 | | 440243 |
| ENSG000002 | 57.5481842 | 0.465122005 | 0.20774515 | 2.23890665 | 0.02516199 | 0.07162821 | UBL7-AS1 | | 440288 |
| ENSG000001 | 14.7084253 | -0.693638643 | 0.30982946 | -2.2387756 | 0.02517052 | 0.07164746 | ASPN | | 54829 |
| ENSG000002 | 4.28214293 | -0.846247148 | 0.37800806 | -2.2387013 | 0.02517535 | 0.07165392 | NA | NA | |
| ENSG000002 | 4.90844402 | -0.840428418 | 0.3754114 | -2.2386865 | 0.02517632 | 0.07165392 | NA | NA | |
| ENSG000001 | 15.1675736 | -0.716070905 | 0.31989219 | -2.2384757 | 0.02519005 | 0.0716779 | NA | NA | |
| ENSG000001 | 18.9078462 | -0.648879033 | 0.28987371 | -2.2384887 | 0.0251892 | 0.0716779 | KLHL1 | | 57626 |
| ENSG000001 | 30.159595 | 0.562191144 | 0.25114716 | 2.2384929 | 0.02518893 | 0.0716779 | LBHD1 | | 79081 |
| ENSG000002 | 2.34426327 | -0.811786524 | 0.36267076 | -2.2383567 | 0.0251978 | 0.07169493 | NA | NA | |
| ENSG000002 | 128.124111 | 0.403451609 | 0.18024691 | 2.23832748 | 0.02519971 | 0.07169532 | ZNF783 | | 100289678 |
| ENSG000002 | 3.66330112 | -0.847012853 | 0.37844298 | -2.2381519 | 0.02521115 | 0.07172285 | NA | NA | |
| ENSG000001 | 13.0787093 | -0.714518507 | 0.31925456 | -2.238084 | 0.02521558 | 0.07172537 | FOXG1 | | 2290 |
| ENSG000001 | 29.6964093 | -0.555501449 | 0.24820218 | -2.2381006 | 0.0252145 | 0.07172537 | BCO2 | | 83875 |
| ENSG000001 | 89.1325066 | 0.390817705 | 0.17463143 | 2.23795738 | 0.02522383 | 0.07174383 | CSRNP1 | | 64651 |
| ENSG000001 | 1194.06608 | 0.246626093 | 0.11020442 | 2.23789666 | 0.02522779 | 0.07175006 | CLOCK | | 9575 |
| ENSG000001 | 122.748965 | 0.479122044 | 0.21409911 | 2.23785159 | 0.02523074 | 0.07175339 | UBALD1 | | 124402 |
| ENSG000002 | 7.02744883 | 0.806393581 | 0.36035084 | 2.23780132 | 0.02523401 | 0.07175768 | NA | NA | |
| ENSG000001 | 37.9158285 | -0.523018386 | 0.23372317 | -2.2377687 | 0.02523614 | 0.07175871 | CCDC33 | | 80125 |
| ENSG000001 | 25.4822861 | -0.600839164 | 0.26850578 | -2.2377141 | 0.02523971 | 0.0717638 | FGF7 | | 2252 |
| ENSG000002 | 4.32020283 | -0.84644486 | 0.37826786 | -2.2376864 | 0.02524151 | 0.0717639 | NA | NA | |
| ENSG000002 | 5.50237089 | -0.833109214 | 0.37234372 | -2.2374735 | 0.02525541 | 0.07179838 | NA | NA | |
| ENSG000002 | 18.3422181 | -0.685510636 | 0.30640554 | -2.2372658 | 0.02526897 | 0.07183191 | NA | NA | |
| ENSG000002 | 6.02685978 | -0.8332549 | 0.37245652 | -2.237187 | 0.02527412 | 0.07183646 | NA | NA | |

| | | | | | | | | |
|---|---|---|---|---|---|---|---|---|
| ENSG000002 | 3.6297525 | -0.846318698 | 0.37829418 | -2.2371973 | 0.02527345 | 0.07183646 | NA | NA |
| ENSG000001 | 319.732168 | 0.331888747 | 0.14835339 | 2.2371498 | 0.02527655 | 0.07183834 | TRIM6 | 117854 |
| ENSG000001 | 9.09278197 | -0.775218629 | 0.34658279 | -2.2367488 | 0.02530276 | 0.07190566 | P2RY8 | 286530 |
| ENSG000001 | 5.20433725 | -0.834981033 | 0.37330834 | -2.2367061 | 0.02530556 | 0.07190566 | NA | NA |
| ENSG000002 | 7.85926301 | -0.809136135 | 0.3617532 | -2.2367076 | 0.02530546 | 0.07190566 | DLX2-AS1 | 104326193 |
| ENSG000001 | 102.204396 | 0.365573352 | 0.16344864 | 2.23662521 | 0.02531085 | 0.07191565 | SLC2A12 | 154091 |
| ENSG000002 | 2.31182076 | -0.815538129 | 0.36464124 | -2.2365493 | 0.02531581 | 0.07192471 | NA | NA |
| ENSG000002 | 2.27089545 | -0.81419942 | 0.3640586 | -2.2364516 | 0.02532221 | 0.07193785 | NA | NA |
| ENSG000002 | 3.27013301 | -0.844568516 | 0.3776536 | -2.2363576 | 0.02532835 | 0.07195027 | NA | NA |
| ENSG000001 | 218.804663 | 0.321255789 | 0.14365393 | 2.23631738 | 0.02533099 | 0.07195271 | CNTNAP3 | 79937 |
| ENSG000002 | 2.27563824 | -0.813964634 | 0.36401537 | -2.2360722 | 0.02534704 | 0.07199327 | LARP7P3 | 106481700 |
| ENSG000002 | 14.3029889 | 0.709480589 | 0.31730101 | 2.23598591 | 0.02535269 | 0.07200428 | NA | NA |
| ENSG000002 | 3.35278466 | -0.843690675 | 0.37732969 | -2.235951 | 0.02535498 | 0.07200574 | NA | NA |
| ENSG000002 | 6.21515028 | 0.841950939 | 0.3765637 | 2.23587917 | 0.02535969 | 0.07201405 | NA | NA |
| ENSG00000C | 16.1417844 | -0.682524539 | 0.30527121 | -2.2357972 | 0.02536506 | 0.07202425 | CDH10 | 1008 |
| ENSG000002 | 22.6034416 | -0.605123098 | 0.27066217 | -2.2357136 | 0.02537054 | 0.07202973 | NA | NA |
| ENSG000002 | 10.5575196 | -0.749521541 | 0.33524863 | -2.2357185 | 0.02537022 | 0.07202973 | NA | NA |
| ENSG000002 | 7.04365084 | -0.808979645 | 0.36185151 | -2.2356675 | 0.02537356 | 0.07203327 | NA | NA |
| ENSG000001 | 24.5828612 | -0.60348695 | 0.26994558 | -2.2355875 | 0.02537881 | 0.07204312 | KANK4 | 163782 |
| ENSG000001 | 18.6947972 | -0.636633002 | 0.28477687 | -2.2355503 | 0.02538125 | 0.072045 | HSF5 | 124535 |
| ENSG000001 | 126.520631 | 0.374264457 | 0.1674415 | 2.23519534 | 0.02540453 | 0.07210198 | ZNF821 | 55565 |
| ENSG000001 | 8.14087987 | -0.800488213 | 0.35813547 | -2.2351548 | 0.02540719 | 0.07210198 | SERPINB12 | 89777 |
| ENSG000002 | 41.9514504 | -0.497565981 | 0.22261107 | -2.2351358 | 0.02540843 | 0.07210198 | NA | NA |
| ENSG000002 | 7.47837698 | -0.807225704 | 0.36114897 | -2.2351599 | 0.02540685 | 0.07210198 | NA | NA |
| ENSG000001 | 3.33575723 | -0.843635791 | 0.37750902 | -2.2347434 | 0.0254342 | 0.07217005 | FXYD4 | 53828 |
| ENSG000002 | 8.19986125 | -0.78730482 | 0.3523096 | -2.2346959 | 0.02543732 | 0.07217386 | NA | NA |
| ENSG000001 | 6.00115941 | -0.832662239 | 0.37262363 | -2.2345933 | 0.02544406 | 0.07218145 | EVX1 | 2128 |
| ENSG000002 | 8.5014871 | -0.783136558 | 0.35046773 | -2.2345468 | 0.02544712 | 0.07218145 | LOC101928233 | 101928233 |
| ENSG000002 | 6.70935793 | -0.813589371 | 0.36409393 | -2.234559 | 0.02544632 | 0.07218145 | SIRPG-AS1 | 101929010 |

| | | | | | | | | |
|---|---|---|---|---|---|---|---|---|
| ENSG000002 | 9.53895041 | -0.776793999 | 0.34762314 | -2.2345866 | 0.0254445 | 0.07218145 | C2orf61 | 285051 |
| ENSG000002 | 4.74635121 | -0.844156859 | 0.37778039 | -2.2345174 | 0.02544905 | 0.07218188 | LOC102724913 | 102724913 |
| ENSG000001 | 945.060971 | -0.269757225 | 0.12073854 | -2.2342263 | 0.02546819 | 0.07223112 | HMCN1 | 83872 |
| ENSG000002 | 18.0746587 | -0.668623225 | 0.29929671 | -2.2339812 | 0.02548431 | 0.07227179 | NA | NA |
| ENSG000002 | 4.54647432 | -0.840629874 | 0.37631849 | -2.2338256 | 0.02549455 | 0.07229577 | NA | NA |
| ENSG000001 | 1188.08267 | 0.282384411 | 0.12642156 | 2.23367293 | 0.0255046 | 0.07231922 | PRKX | 5613 |
| ENSG000001 | 214.01545 | 0.329500758 | 0.14752011 | 2.23359898 | 0.02550947 | 0.07232796 | MATN2 | 4147 |
| ENSG000001 | 306.488079 | 0.326142107 | 0.14602021 | 2.23354093 | 0.02551329 | 0.07233375 | PIGM | 93183 |
| ENSG000002 | 1658.81972 | 0.268035831 | 0.12000928 | 2.23345917 | 0.02551868 | 0.0723397 | ZNF814 | 730051 |
| ENSG000002 | 11.5034885 | -0.744252058 | 0.33322905 | -2.2334549 | 0.02551896 | 0.0723397 | LOC101927876 | 101927876 |
| ENSG00000C | 552.544561 | 0.406199568 | 0.18187468 | 2.2334036 | 0.02552234 | 0.07234063 | FBLN1 | 2192 |
| ENSG000002 | 7.88060285 | -0.790307212 | 0.35385901 | -2.2333958 | 0.02552286 | 0.07234063 | NA | NA |
| ENSG000001 | 134.376059 | 0.374885873 | 0.16786397 | 2.23327178 | 0.02553103 | 0.07234862 | PPFIA3 | 8541 |
| ENSG000002 | 6.40443357 | -0.814413954 | 0.36467268 | -2.2332738 | 0.02553089 | 0.07234862 | LOC400800 | 400800 |
| ENSG000002 | 12.1257868 | -0.720885504 | 0.32278841 | -2.2333067 | 0.02552873 | 0.07234862 | NA | NA |
| ENSG00000C | 336.327429 | 0.322174277 | 0.14426947 | 2.2331425 | 0.02553955 | 0.07236771 | FARP2 | 9855 |
| ENSG000002 | 5.55140095 | -0.830321639 | 0.37184725 | -2.2329643 | 0.0255513 | 0.07239594 | NA | NA |
| ENSG000001 | 9.98814961 | -0.759717913 | 0.34023998 | -2.2328885 | 0.0255563 | 0.07240506 | PRKAG3 | 53632 |
| ENSG000002 | 2.2001131 | 0.789998486 | 0.35381241 | 2.23281737 | 0.02556099 | 0.07241328 | NA | NA |
| ENSG000002 | 5.34999043 | -0.838916865 | 0.37575283 | -2.2326295 | 0.02557338 | 0.07244333 | NA | NA |
| ENSG000001 | 417.693312 | 0.283464951 | 0.1269679 | 2.23257179 | 0.0255772 | 0.07244401 | EBAG9 | 9166 |
| ENSG000002 | 11.8512493 | -0.724037326 | 0.32430437 | -2.2325858 | 0.02557627 | 0.07244401 | RAB6C-AS1 | 100131320 |
| ENSG000002 | 2.92107948 | -0.838699178 | 0.37569372 | -2.2324014 | 0.02558845 | 0.07247081 | LOC101928225 | 101928225 |
| ENSG000001 | 1439.69467 | 0.304849941 | 0.13655863 | 2.23237402 | 0.02559025 | 0.07247087 | AKAP13 | 11214 |
| ENSG000001 | 150.985148 | 0.357377855 | 0.16009378 | 2.23230312 | 0.02559494 | 0.07247907 | SIMC1 | 375484 |
| ENSG000001 | 308.930398 | 0.326932418 | 0.14645982 | 2.2322328 | 0.02559958 | 0.07248716 | TCP11L1 | 55346 |
| ENSG000001 | 7.92602266 | -0.789770277 | 0.35380911 | -2.2321932 | 0.0256022 | 0.07248951 | LOC541472 | 541472 |
| ENSG000001 | 13.3307734 | -0.70962601 | 0.31793174 | -2.2320074 | 0.02561447 | 0.0725192 | ANP32D | 23519 |
| ENSG000002 | 4.55495514 | -0.840677122 | 0.3766532 | -2.2319659 | 0.02561721 | 0.0725219 | NA | NA |

| | | | | | | | | |
|---|---|---|---|---|---|---|---|---|
| ENSG000001 | 247.850902 | 0.340133803 | 0.15241102 | 2.23168776 | 0.02563561 | 0.0725689 | COA6 | 388753 |
| ENSG000001 | 98.6576407 | 0.38378358 | 0.17197534 | 2.2316198 | 0.0256401 | 0.07257323 | FZD8 | 8325 |
| ENSG000002 | 4.90817592 | -0.838848193 | 0.37589363 | -2.2316105 | 0.02564071 | 0.07257323 | NA | NA |
| ENSG000001 | 5.76871226 | -0.824154917 | 0.36931901 | -2.2315529 | 0.02564453 | 0.07257533 | REG3G | 130120 |
| ENSG000002 | 3.34965159 | -0.841579598 | 0.37712864 | -2.2315452 | 0.02564504 | 0.07257533 | NA | NA |
| ENSG000001 | 270.978506 | 0.312617344 | 0.14010604 | 2.23129105 | 0.02566186 | 0.07261786 | LYPLAL1 | 127018 |
| ENSG000001 | 4.2649267 | -0.84362824 | 0.3781208 | -2.2311077 | 0.025674 | 0.07264714 | NA | NA |
| ENSG000001 | 1674.47337 | 0.291638663 | 0.13073671 | 2.23073283 | 0.02569883 | 0.07271234 | DNAJC2 | 27000 |
| ENSG000001 | 6.37739829 | -0.81453819 | 0.36515076 | -2.2306901 | 0.02570166 | 0.07271528 | HOXA7 | 3204 |
| ENSG000001 | 11.1484803 | -0.742550085 | 0.33290123 | -2.2305417 | 0.0257115 | 0.07273631 | DCAF8L2 | 347442 |
| ENSG000002 | 14.6588834 | -0.750202552 | 0.33633468 | -2.2305239 | 0.02571268 | 0.07273631 | NA | NA |
| ENSG000002 | 4.28640195 | -0.842762935 | 0.37783978 | -2.230477 | 0.02571579 | 0.07274004 | NA | NA |
| ENSG000002 | 4.93711937 | -0.839484101 | 0.37638322 | -2.2303972 | 0.02572108 | 0.07274992 | NA | NA |
| ENSG000001 | 149.342313 | 0.339604063 | 0.15227189 | 2.23024788 | 0.02573099 | 0.07276975 | ELOVL7 | 79993 |
| ENSG000002 | 5.29991427 | -0.837376868 | 0.37546534 | -2.2302375 | 0.02573168 | 0.07276975 | NA | NA |
| ENSG000001 | 3384.68498 | 0.284711412 | 0.12766457 | 2.23015211 | 0.02573735 | 0.07278069 | FGF2 | 2247 |
| ENSG000001 | 110.919819 | 0.3611359 | 0.16194197 | 2.23003276 | 0.02574527 | 0.07279802 | SLC41A2 | 84102 |
| ENSG000002 | 5.5547861 | -0.830791437 | 0.37256536 | -2.2299213 | 0.02575267 | 0.07281386 | NA | NA |
| ENSG000001 | 4700.58591 | 0.267398705 | 0.1199186 | 2.22983516 | 0.02575839 | 0.07282496 | TAOK1 | 57551 |
| ENSG000001 | 1328.71297 | 0.250808238 | 0.11248005 | 2.22980206 | 0.02576059 | 0.0728261 | MMS22L | 253714 |
| ENSG000002 | 7.29795431 | -0.797777961 | 0.35779386 | -2.2297139 | 0.02576644 | 0.07283757 | CCL16 | 6360 |
| ENSG000002 | 9.59668425 | -0.757155242 | 0.33959411 | -2.2295889 | 0.02577475 | 0.07285598 | NA | NA |
| ENSG000002 | 17.064826 | 0.674291462 | 0.30246299 | 2.22933542 | 0.0257916 | 0.07289492 | ZNF321P | 399669 |
| ENSG000002 | 7.67186408 | -0.7961142 | 0.35710957 | -2.2293275 | 0.02579212 | 0.07289492 | LOC105370622 | 105370622 |
| ENSG000001 | 6.1809836 | -0.820875667 | 0.36825939 | -2.2290692 | 0.0258093 | 0.07293838 | CNTD2 | 79935 |
| ENSG000001 | 30.3554778 | -0.572492275 | 0.25683712 | -2.2290091 | 0.0258133 | 0.07294131 | SLC16A4 | 9122 |
| ENSG000002 | 10.1533873 | -0.76667075 | 0.34395285 | -2.2289996 | 0.02581393 | 0.07294131 | NA | NA |
| ENSG000002 | 3.94133741 | -0.843659843 | 0.37850061 | -2.2289524 | 0.02581707 | 0.0729451 | NA | NA |
| ENSG000001 | 158.882066 | 0.368475335 | 0.16533275 | 2.22868932 | 0.02583458 | 0.07298949 | HTR7 | 3363 |

| | | | | | | | | |
|---|---|---|---|---|---|---|---|---|
| ENSG000002 | 30.5612456 | -0.558221641 | 0.25049022 | -2.2285167 | 0.02584608 | 0.07301689 | DOK6 | 220164 |
| ENSG000002 | 3.9414859 | -0.843537779 | 0.37855596 | -2.228304 | 0.02586025 | 0.07305182 | NA | NA |
| ENSG000002 | 10.0966254 | -0.768479891 | 0.34488183 | -2.2282412 | 0.02586444 | 0.07305856 | NA | NA |
| ENSG000001 | 168.434483 | 0.334875842 | 0.15029375 | 2.22814215 | 0.02587104 | 0.07306343 | MAP3K12 | 7786 |
| ENSG000002 | 10.5877529 | -0.752980678 | 0.33793656 | -2.2281717 | 0.02586907 | 0.07306343 | LOC101928241 | 101928241 |
| ENSG000002 | 3.93564835 | -0.843409209 | 0.37852711 | -2.2281342 | 0.02587156 | 0.07306343 | NA | NA |
| ENSG000002 | 9.19142129 | -0.795748466 | 0.35715176 | -2.2280401 | 0.02587784 | 0.07307606 | NA | NA |
| ENSG000001 | 29.3490758 | -0.561172103 | 0.25187608 | -2.227969 | 0.02588258 | 0.07308437 | MLIP | 90523 |
| ENSG000001 | 14.0408263 | -0.686845454 | 0.30835791 | -2.2274293 | 0.02591859 | 0.07317674 | OR5AS1 | 219447 |
| ENSG000002 | 19.6536385 | -0.636340166 | 0.28568426 | -2.2274247 | 0.02591891 | 0.07317674 | NA | NA |
| ENSG000002 | 2.71530132 | -0.822138702 | 0.36915583 | -2.2270777 | 0.02594208 | 0.07323708 | NA | NA |
| ENSG000002 | 6.50022652 | -0.819700789 | 0.36806604 | -2.2270481 | 0.02594406 | 0.07323755 | LOC100128554 | 100128554 |
| ENSG000002 | 5.86861637 | -0.826226665 | 0.37105707 | -2.2266835 | 0.02596843 | 0.07329616 | NA | NA |
| ENSG000002 | 3.64371386 | -0.842128798 | 0.37819788 | -2.2266883 | 0.02596811 | 0.07329616 | NA | NA |
| ENSG000002 | 200.36399 | 0.306068638 | 0.13746058 | 2.22659208 | 0.02597455 | 0.07330832 | LINC00599 | 157627 |
| ENSG00000C | 587.997523 | 0.251187955 | 0.11282281 | 2.22639339 | 0.02598784 | 0.07334074 | NECAP1 | 25977 |
| ENSG000001 | 20.9026381 | -0.621364342 | 0.279101 | -2.2263064 | 0.02599367 | 0.07334696 | BMP3 | 651 |
| ENSG000002 | 14.7742914 | 0.694940237 | 0.31214616 | 2.22632957 | 0.02599211 | 0.07334696 | SCARNA1 | 677774 |
| ENSG000002 | 32.5899375 | 0.545243003 | 0.24493592 | 2.22606386 | 0.0260099 | 0.07338767 | NA | NA |
| ENSG000002 | 17.4197403 | -0.681438119 | 0.30613748 | -2.2259219 | 0.02601941 | 0.0734094 | LOC101928002 | 101928002 |
| ENSG00000C | 95.5215497 | 0.383940408 | 0.17249478 | 2.22580888 | 0.02602699 | 0.0734233 | SPA17 | 53340 |
| ENSG000001 | 235.295271 | 0.382358124 | 0.17178502 | 2.22579431 | 0.02602796 | 0.0734233 | C10orf76 | 79591 |
| ENSG000001 | 65.8968709 | -0.417270556 | 0.18748018 | -2.2256782 | 0.02603574 | 0.07344014 | ADAMTSL1 | 92949 |
| ENSG000001 | 536.558849 | 0.573697977 | 0.25778886 | 2.22545684 | 0.02605059 | 0.07347219 | COPS6 | 10980 |
| ENSG000002 | 10.6231604 | -0.753863669 | 0.33874591 | -2.2254547 | 0.02605073 | 0.07347219 | NA | NA |
| ENSG000001 | 5.27547181 | -0.833288846 | 0.37444718 | -2.2253842 | 0.02605546 | 0.07348041 | LINC01565 | 23434 |
| ENSG000001 | 231.63358 | 0.297621352 | 0.13375937 | 2.22505056 | 0.02607784 | 0.07353843 | ADAMTS20 | 80070 |
| ENSG000002 | 12.5068368 | -0.736861913 | 0.33117409 | -2.2249987 | 0.02608133 | 0.07354314 | NA | NA |
| ENSG000001 | 17.8446581 | -0.666399627 | 0.29952513 | -2.2248538 | 0.02609105 | 0.07356545 | ADH4 | 127 |

| | | | | | | | | |
|---|---|---|---|---|---|---|---|---|
| ENSG000001 | 2011.49207 | 0.662887403 | 0.2979675 | 2.22469703 | 0.02610158 | 0.07359003 | FASN | 2194 |
| ENSG000000 | 425.234926 | 0.276287184 | 0.12419291 | 2.22466154 | 0.02610397 | 0.07359163 | RHOBTB1 | 9886 |
| ENSG000001 | 5.10893547 | -0.838982286 | 0.37713668 | -2.2246107 | 0.02610739 | 0.07359615 | SLC16A8 | 23539 |
| ENSG000001 | 34.0846567 | -0.549609303 | 0.24707267 | -2.2244844 | 0.02611587 | 0.07361193 | PKNOX2 | 63876 |
| ENSG000002 | 6.40980944 | -0.813118744 | 0.36553315 | -2.2244733 | 0.02611662 | 0.07361193 | NA | NA |
| ENSG000002 | 10.1734011 | -0.773301934 | 0.3476534 | -2.2243474 | 0.02612508 | 0.07363067 | FAM95B1 | 100133036 |
| ENSG000001 | 860.675748 | 0.299794562 | 0.13480061 | 2.22398522 | 0.02614944 | 0.0736942 | PLS1 | 5357 |
| ENSG000001 | 2.93544524 | -0.834426658 | 0.375223 | -2.2238153 | 0.02616087 | 0.07371742 | S100A8 | 6279 |
| ENSG000002 | 9.04112283 | -0.785650073 | 0.35329562 | -2.2237753 | 0.02616357 | 0.07371742 | NA | NA |
| ENSG000002 | 4.66047608 | -0.839709496 | 0.37760886 | -2.2237547 | 0.02616495 | 0.07371742 | NA | NA |
| ENSG000002 | 4.71871586 | -0.840018097 | 0.37773966 | -2.2238017 | 0.02616179 | 0.07371742 | NA | NA |
| ENSG000002 | 5.68252311 | -0.834312223 | 0.37519255 | -2.2236908 | 0.02616926 | 0.07372443 | NA | NA |
| ENSG000001 | 46.1525011 | -0.509328025 | 0.22905029 | -2.2236515 | 0.0261719 | 0.07372675 | LRP2BP | 55805 |
| ENSG000001 | 71.0873757 | 0.420533655 | 0.18913812 | 2.22342092 | 0.02618743 | 0.07376538 | SDHAF2 | 54949 |
| ENSG000001 | 462.749092 | 0.267302005 | 0.12022576 | 2.22333383 | 0.0261933 | 0.07377679 | SOS2 | 6655 |
| ENSG000001 | 513.032001 | 0.257487895 | 0.11581469 | 2.22327482 | 0.02619727 | 0.07378286 | GAB1 | 2549 |
| ENSG000001 | 5.8463479 | -0.8224001 | 0.36991514 | -2.2232129 | 0.02620145 | 0.0737895 | SIRPD | 128646 |
| ENSG000001 | 693.328729 | 0.359766704 | 0.16182697 | 2.22315667 | 0.02620524 | 0.07379504 | ZC3H7A | 29066 |
| ENSG000002 | 4.71142851 | -0.839252186 | 0.37752322 | -2.2230479 | 0.02621257 | 0.07381057 | NA | NA |
| ENSG000001 | 2.2707707 | -0.808040891 | 0.36349481 | -2.2229778 | 0.0262173 | 0.07381875 | NA | NA |
| ENSG000002 | 6.54982962 | -0.818265728 | 0.36816154 | -2.2225725 | 0.02624464 | 0.07388197 | C11orf39 | 101929637 |
| ENSG000002 | 5.59897226 | -0.829911673 | 0.37340233 | -2.2225669 | 0.02624502 | 0.07388197 | EDRF1-AS1 | 101927983 |
| ENSG000002 | 8.44006913 | -0.813650625 | 0.36608648 | -2.222564 | 0.02624522 | 0.07388197 | NA | NA |
| ENSG000002 | 9.98136827 | -0.775241694 | 0.34881281 | -2.2225149 | 0.02624853 | 0.07388315 | NA | NA |
| ENSG000002 | 7.06377481 | 0.801159257 | 0.360476 | 2.22250375 | 0.02624928 | 0.07388315 | NA | NA |
| ENSG000001 | 578.180345 | 0.328684799 | 0.14789219 | 2.22246212 | 0.02625209 | 0.07388594 | SLAIN2 | 57606 |
| ENSG000002 | 22.8662654 | 0.599166722 | 0.26960148 | 2.22241629 | 0.02625519 | 0.07388952 | NA | NA |
| ENSG000002 | 3.34185933 | -0.836713153 | 0.37649992 | -2.2223462 | 0.02625992 | 0.07389771 | NA | NA |
| ENSG000001 | 12.2039612 | -0.726232196 | 0.3267905 | -2.2223174 | 0.02626187 | 0.07389806 | ALDH3A1 | 218 |

| | | | | | | | | |
|---|---|---|---|---|---|---|---|---|
| ENSG000002 | 12.0255804 | -0.727376729 | 0.32732599 | -2.2221783 | 0.02627126 | 0.07391935 | NA | NA |
| ENSG000002 | 2.28509017 | -0.806312037 | 0.36285335 | -2.222143 | 0.02627364 | 0.07392094 | RNU6-231P | 106481251 |
| ENSG000001 | 13.0052844 | -0.707474094 | 0.3184321 | -2.2217424 | 0.02630072 | 0.07399199 | SLC36A2 | 153201 |
| ENSG000001 | 901.119663 | 0.252405321 | 0.11360982 | 2.22168579 | 0.02630455 | 0.07399762 | PDCD5 | 9141 |
| ENSG000002 | 15.1677211 | -0.674596016 | 0.30367111 | -2.2214692 | 0.0263192 | 0.0740337 | LOC728084 | 728084 |
| ENSG000002 | 2.40427321 | -0.801850409 | 0.36098057 | -2.2213119 | 0.02632985 | 0.07405564 | NA | NA |
| ENSG000002 | 5.70883923 | -0.83229982 | 0.37469042 | -2.2213 | 0.02633065 | 0.07405564 | NA | NA |
| ENSG000001 | 15.8473498 | -0.716174736 | 0.32241966 | -2.2212502 | 0.02633402 | 0.07405998 | SLC22A7 | 10864 |
| ENSG000002 | 236.07818 | 0.334508783 | 0.15061772 | 2.22091254 | 0.02635689 | 0.07411915 | NA | NA |
| ENSG000001 | 245.222465 | 0.366633131 | 0.16509511 | 2.22073895 | 0.02636865 | 0.07414708 | ACVR1B | 91 |
| ENSG000002 | 18.6536079 | -0.630315603 | 0.28385349 | -2.2205667 | 0.02638033 | 0.07417477 | LOC101929457 | 101929457 |
| ENSG000001 | 25.9426708 | -0.57540492 | 0.25913364 | -2.2204949 | 0.02638519 | 0.07418332 | NA | NA |
| ENSG000002 | 5.3223644 | -0.834148865 | 0.375674 | -2.2204061 | 0.02639121 | 0.07419509 | PGM5-AS1 | 572558 |
| ENSG000002 | 3.60219398 | -0.832988686 | 0.37515809 | -2.2203671 | 0.02639386 | 0.07419739 | NA | NA |
| ENSG000002 | 8.4677448 | -0.782391083 | 0.35237603 | -2.22033 | 0.02639637 | 0.07419932 | GOLGA8IP | 283796 |
| ENSG000002 | 58.9579169 | 0.518463643 | 0.23352981 | 2.22011763 | 0.02641078 | 0.07422889 | NPTXR | 23467 |
| ENSG000002 | 2.97119056 | -0.833280764 | 0.3753358 | -2.220094 | 0.02641238 | 0.07422889 | NA | NA |
| ENSG000002 | 7.38526177 | -0.800740017 | 0.36067076 | -2.2201412 | 0.02640918 | 0.07422889 | NA | NA |
| ENSG000001 | 30.2369085 | -0.560664031 | 0.25255805 | -2.2199413 | 0.02642276 | 0.07424784 | CPNE5 | 57699 |
| ENSG000001 | 11.0411009 | -0.758822572 | 0.3418211 | -2.2199407 | 0.02642279 | 0.07424784 | OR2M4 | 26245 |
| ENSG000002 | 9.36965683 | -0.765122255 | 0.34468086 | -2.2197991 | 0.02643241 | 0.07426972 | LINC01205 | 401082 |
| ENSG000002 | 6.94285543 | -0.802029744 | 0.36134151 | -2.2195893 | 0.02644666 | 0.07430461 | NA | NA |
| ENSG000001 | 9.61133348 | -0.801262612 | 0.36101637 | -2.2194634 | 0.02645521 | 0.0743235 | SGCA | 6442 |
| ENSG000001 | 52.7832972 | 0.471063178 | 0.21226628 | 2.21920871 | 0.02647253 | 0.07436699 | GNG11 | 2791 |
| ENSG000002 | 3.33060895 | -0.836355458 | 0.37692264 | -2.2189048 | 0.0264932 | 0.07441991 | NA | NA |
| ENSG000002 | 9.82350907 | -0.771943624 | 0.34792078 | -2.2187339 | 0.02650483 | 0.07444743 | LOC101927342 | 101927342 |
| ENSG000001 | 9.65100147 | -0.759002083 | 0.3421439 | -2.2183709 | 0.02652955 | 0.07451135 | CRYBA2 | 1412 |
| ENSG000002 | 5.78547966 | -0.819663373 | 0.36949306 | -2.2183458 | 0.02653127 | 0.07451135 | NA | NA |
| ENSG000002 | 190.356804 | 0.341258426 | 0.15384683 | 2.21817009 | 0.02654324 | 0.07453981 | POLG2 | 11232 |

| | | | | | | | | |
|---|---|---|---|---|---|---|---|---|
| ENSG000002 | 6.27686069 | -0.83384099 | 0.37592775 | -2.2180884 | 0.02654881 | 0.07455028 | CBWD6 | 644019 |
| ENSG000001 | 641.51543 | 0.268176065 | 0.12091146 | 2.21795412 | 0.02655796 | 0.07457083 | MED12L | 116931 |
| ENSG000001 | 187.266067 | 0.371939494 | 0.1677077 | 2.21778422 | 0.02656955 | 0.0745982 | COG1 | 9382 |
| ENSG000002 | 9.11173949 | -0.77019078 | 0.34728615 | -2.2177411 | 0.02657249 | 0.07460128 | NA | NA |
| ENSG000001 | 29.4152499 | -0.59445321 | 0.2680557 | -2.2176481 | 0.02657884 | 0.07461394 | DMGDH | 29958 |
| ENSG000002 | 12.9757175 | -0.714259538 | 0.32211026 | -2.217438 | 0.02659317 | 0.07464902 | HCG25 | 414765 |
| ENSG000002 | 4.34487835 | -0.838857956 | 0.37832542 | -2.2172921 | 0.02660314 | 0.07467182 | NA | NA |
| ENSG000002 | 12.5441934 | -0.718801992 | 0.32419359 | -2.2171999 | 0.02660943 | 0.07467476 | MRLN | 100507027 |
| ENSG000002 | 11.7081896 | -0.732246536 | 0.33025691 | -2.2172028 | 0.02660924 | 0.07467476 | NA | NA |
| ENSG000002 | 10.6845994 | -0.756461316 | 0.34117929 | -2.2171959 | 0.02660971 | 0.07467476 | WFDC21P | 645638 |
| ENSG000002 | 17.4424012 | -0.67362165 | 0.30382325 | -2.2171498 | 0.02661286 | 0.07467843 | NA | NA |
| ENSG000001 | 19.088696 | -0.649896172 | 0.29313529 | -2.2170519 | 0.02661954 | 0.07468089 | KAZALD1 | 81621 |
| ENSG000002 | 3.96926581 | -0.839341352 | 0.37858833 | -2.2170291 | 0.0266211 | 0.07468089 | LINGO4 | 339398 |
| ENSG000002 | 4.53961119 | -0.834851254 | 0.37655476 | -2.2170779 | 0.02661777 | 0.07468089 | NA | NA |
| ENSG000002 | 3.96459525 | -0.839195699 | 0.37851478 | -2.2170751 | 0.02661796 | 0.07468089 | NA | NA |
| ENSG000001 | 8.7629821 | -0.770878542 | 0.34777634 | -2.2165928 | 0.02665093 | 0.07474388 | LYL1 | 4066 |
| ENSG000002 | 22.6456244 | -0.620086624 | 0.27974648 | -2.2166021 | 0.0266503 | 0.07474388 | OR6B1 | 135946 |
| ENSG000002 | 4.87369024 | -0.839060729 | 0.37853116 | -2.2166226 | 0.02664889 | 0.07474388 | NA | NA |
| ENSG000002 | 34.6540161 | -0.512454609 | 0.23118963 | -2.2165986 | 0.02665054 | 0.07474388 | LOC101927056 | 101927056 |
| ENSG000001 | 13.5489875 | -0.696960897 | 0.31443893 | -2.2165223 | 0.02665575 | 0.07474945 | NXNL2 | 158046 |
| ENSG000001 | 7.08800639 | -0.804454938 | 0.36293767 | -2.2165099 | 0.0266566 | 0.07474945 | NA | NA |
| ENSG000002 | 8.93471053 | 0.765024657 | 0.34516358 | 2.21641188 | 0.02666331 | 0.07476309 | NA | NA |
| ENSG000002 | 2.27068693 | -0.806115169 | 0.36373251 | -2.2162308 | 0.0266757 | 0.07479267 | NA | NA |
| ENSG000002 | 66.0401765 | 0.432473555 | 0.19514189 | 2.21620046 | 0.02667778 | 0.07479332 | NA | NA |
| ENSG000001 | 131.242016 | 0.354180913 | 0.15982812 | 2.21601121 | 0.02669073 | 0.07482447 | ANKRD42 | 338699 |
| ENSG000001 | 821.080685 | 0.279186765 | 0.12598903 | 2.2159609 | 0.02669418 | 0.07482896 | ZNF148 | 7707 |
| ENSG000002 | 8.08471017 | -0.776779948 | 0.35059433 | -2.215609 | 0.02671829 | 0.07489137 | NA | NA |
| ENSG000002 | 11.3214069 | -0.7485136 | 0.33788074 | -2.2153189 | 0.02673818 | 0.07492641 | NA | NA |
| ENSG000002 | 2.99083402 | -0.83092355 | 0.37506975 | -2.2153841 | 0.02673371 | 0.07492641 | NA | NA |

| | | | | | | | | |
|---|---|---|---|---|---|---|---|---|
| ENSG000002 | 13.0152412 | -0.703769956 | 0.31768294 | -2.2153219 | 0.02673798 | 0.07492641 | NA | NA |
| ENSG000002 | 3.37023767 | -0.833928665 | 0.37642859 | -2.2153702 | 0.02673467 | 0.07492641 | NA | NA |
| ENSG000002 | 2.34506574 | -0.802137588 | 0.36213907 | -2.2149987 | 0.02676015 | 0.07498279 | LOC101928087 | 101928087 |
| ENSG000001 | 7.73795686 | -0.796686627 | 0.35969127 | -2.2149179 | 0.0267657 | 0.07499315 | LGALS2 | 3957 |
| ENSG000002 | 4.01485823 | -0.838447338 | 0.37855928 | -2.2148376 | 0.02677121 | 0.07500341 | LOC440040 | 440040 |
| ENSG000002 | 24.6169752 | -0.594917649 | 0.26861053 | -2.2147964 | 0.02677404 | 0.07500615 | NA | NA |
| ENSG00000C | 1399.04091 | 0.619197486 | 0.27959126 | 2.21465249 | 0.02678393 | 0.07502867 | CAPZB | 832 |
| ENSG000001 | 18.5022823 | -0.675138183 | 0.30486071 | -2.2145792 | 0.02678896 | 0.07503237 | CPA6 | 57094 |
| ENSG000001 | 7.40949697 | -0.807400524 | 0.36458857 | -2.2145525 | 0.0267908 | 0.07503237 | ZAR1 | 326340 |
| ENSG000002 | 2.35294106 | -0.802856569 | 0.36253631 | -2.2145549 | 0.02679063 | 0.07503237 | NA | NA |
| ENSG000002 | 4.30210647 | -0.83766281 | 0.37827905 | -2.2144044 | 0.02680097 | 0.07505249 | NA | NA |
| ENSG000002 | 2.89246539 | -0.831520258 | 0.37550691 | -2.2143941 | 0.02680168 | 0.07505249 | NA | NA |
| ENSG000002 | 3.53724592 | -0.831668925 | 0.37558491 | -2.21433 | 0.02680609 | 0.07505446 | NA | NA |
| ENSG000002 | 20.5711792 | -0.630255473 | 0.28462503 | -2.2143361 | 0.02680567 | 0.07505446 | LINC01146 | 283587 |
| ENSG000001 | 2527.97756 | 0.244545699 | 0.11044911 | 2.21410303 | 0.02682169 | 0.07509296 | THOC2 | 57187 |
| ENSG000001 | 11.7976023 | -0.74193288 | 0.33510782 | -2.2140124 | 0.02682792 | 0.07510522 | CLEC3A | 10143 |
| ENSG000002 | 11.5655992 | -0.723955649 | 0.32700004 | -2.2139314 | 0.0268335 | 0.07511564 | NA | NA |
| ENSG000002 | 221.905812 | 0.350654611 | 0.15839222 | 2.21383741 | 0.02683997 | 0.07512856 | NA | NA |
| ENSG000002 | 34.9356146 | -0.523423755 | 0.23645374 | -2.2136413 | 0.02685347 | 0.07516116 | BAALC-AS1 | 100499183 |
| ENSG000001 | 55.6452038 | 0.463148679 | 0.20925126 | 2.21336153 | 0.02687273 | 0.07520989 | CELF5 | 60680 |
| ENSG000002 | 2.92764956 | -0.830991871 | 0.37547968 | -2.2131474 | 0.02688749 | 0.075246 | NA | NA |
| ENSG000001 | 5.54085176 | -0.824860507 | 0.37277369 | -2.2127648 | 0.02691386 | 0.07531342 | PRSS33 | 260429 |
| ENSG000002 | 145.998183 | 0.345033117 | 0.15592997 | 2.21274407 | 0.0269153 | 0.07531342 | NA | NA |
| ENSG000001 | 5246.75901 | 0.252869985 | 0.11429423 | 2.21244757 | 0.02693575 | 0.07536547 | CCND2 | 894 |
| ENSG000001 | 188.670094 | 0.31846949 | 0.14394908 | 2.21237605 | 0.02694069 | 0.07537409 | SCFD2 | 152579 |
| ENSG000001 | 142.031583 | 0.651023771 | 0.29427628 | 2.2122876 | 0.0269468 | 0.07538597 | WDR4 | 10785 |
| ENSG000001 | 2.55599109 | -0.822076805 | 0.37163152 | -2.212075 | 0.02696148 | 0.07541664 | TAAR1 | 134864 |
| ENSG000002 | 158.920059 | 0.319529525 | 0.14444771 | 2.21207751 | 0.02696131 | 0.07541664 | LOC645513 | 645513 |
| ENSG000002 | 305.792476 | 0.333120099 | 0.15061127 | 2.21178729 | 0.02698137 | 0.07546706 | ALG11 | 440138 |

| | | | | | | | | |
|---|---|---|---|---|---|---|---|---|
| ENSG000001 | 178.437569 | 0.30794717 | 0.13923353 | 2.21173146 | 0.02698523 | 0.07547265 | ADAL | 161823 |
| ENSG000002 | 2.5321977 | -0.822310744 | 0.37182128 | -2.2115752 | 0.02699603 | 0.07549765 | PRKX-AS1 | 100873944 |
| ENSG000002 | 8.14222654 | -0.798049113 | 0.36090941 | -2.2112173 | 0.0270208 | 0.07556171 | NA | NA |
| ENSG000002 | 2.9271381 | -0.809150498 | 0.36593697 | -2.2111745 | 0.02702376 | 0.07556477 | NA | NA |
| ENSG000002 | 9.83949729 | -0.747822858 | 0.33820959 | -2.2111225 | 0.02702735 | 0.07556962 | LINC00367 | 101930748 |
| ENSG000001 | 1417.15176 | 0.270143623 | 0.12218484 | 2.21094221 | 0.02703984 | 0.07559238 | SPATS2L | 26010 |
| ENSG000002 | 12.4396303 | -0.733126363 | 0.33159271 | -2.2109242 | 0.02704108 | 0.07559238 | LOC101927701 | 101927701 |
| ENSG000002 | 30.2979849 | -0.55031176 | 0.24890098 | -2.2109666 | 0.02703815 | 0.07559238 | LOC105369748 | 105369748 |
| ENSG000001 | 27.4297432 | -0.609554808 | 0.27572354 | -2.2107463 | 0.02705341 | 0.07562162 | SPATA16 | 83893 |
| ENSG000002 | 4.2199967 | -0.835847363 | 0.37809313 | -2.2106918 | 0.02705719 | 0.07562697 | NA | NA |
| ENSG000002 | 2.55713324 | -0.822042908 | 0.37189006 | -2.210446 | 0.02707422 | 0.07566936 | NA | NA |
| ENSG00000C | 21.1992516 | -0.62012525 | 0.28056933 | -2.2102389 | 0.02708858 | 0.07570429 | CDH19 | 28513 |
| ENSG000001 | 530.060345 | 0.384069499 | 0.1737839 | 2.2100407 | 0.02710234 | 0.0757346 | CTGF | 1490 |
| ENSG000002 | 3.94654931 | -0.83669108 | 0.37858833 | -2.2100287 | 0.02710317 | 0.0757346 | NA | NA |
| ENSG000001 | 22.4938545 | -0.606865439 | 0.27460292 | -2.2099745 | 0.02710693 | 0.07573991 | EBF3 | 253738 |
| ENSG000002 | 4.51784042 | -0.832447717 | 0.3766905 | -2.2098983 | 0.02711222 | 0.07574945 | NA | NA |
| ENSG000002 | 33.4364309 | -0.542409317 | 0.24546454 | -2.2097258 | 0.0271242 | 0.07577771 | LOC100507557 | 100507557 |
| ENSG000002 | 18.3946928 | -0.632275919 | 0.2861528 | -2.2095745 | 0.02713471 | 0.07580185 | LOC649133 | 649133 |
| ENSG000001 | 40.5128603 | -0.508450167 | 0.23012575 | -2.2094449 | 0.02714371 | 0.07581132 | VWA5B1 | 127731 |
| ENSG000002 | 3.92438788 | -0.836340687 | 0.37852417 | -2.2094777 | 0.02714143 | 0.07581132 | NA | NA |
| ENSG000002 | 7.14077446 | -0.82259899 | 0.37230701 | -2.2094642 | 0.02714237 | 0.07581132 | NA | NA |
| ENSG000001 | 1015.54233 | 0.369226975 | 0.16713206 | 2.20919293 | 0.02716122 | 0.07584979 | TRRAP | 8295 |
| ENSG000002 | 9.21027045 | -0.7749789 | 0.35079541 | -2.2092048 | 0.0271604 | 0.07584979 | NA | NA |
| ENSG000001 | 90.4997642 | 0.386991919 | 0.17519808 | 2.20888216 | 0.02718284 | 0.07590492 | NA | NA |
| ENSG00000C | 281.81068 | 0.348892232 | 0.15795584 | 2.20879598 | 0.02718883 | 0.07591644 | EPN2 | 22905 |
| ENSG000001 | 96.0382571 | -0.364168601 | 0.16488788 | -2.2085832 | 0.02720364 | 0.07595256 | LRRK1 | 79705 |
| ENSG000002 | 3.90212785 | -0.836063153 | 0.37856848 | -2.2084859 | 0.02721042 | 0.07596625 | NA | NA |
| ENSG000001 | 12.072418 | -0.710766738 | 0.32184283 | -2.2084281 | 0.02721445 | 0.07597226 | ANKRD55 | 79722 |
| ENSG000002 | 3.15261069 | -0.822283729 | 0.37235615 | -2.2083259 | 0.02722156 | 0.07598685 | NA | NA |

| | | | | | | | | |
|---|---|---|---|---|---|---|---|---|
| ENSG000002 | 6.59097646 | -0.827136359 | 0.3745581 | -2.2082992 | 0.02722342 | 0.07598685 | NA | NA |
| ENSG000002 | 3.08173361 | -0.822450277 | 0.37246759 | -2.2081123 | 0.02723645 | 0.07601274 | NA | NA |
| ENSG000002 | 5.64315497 | -0.827845779 | 0.37490795 | -2.2081307 | 0.02723516 | 0.07601274 | NA | NA |
| ENSG000001 | 234.81739 | 0.29962663 | 0.13570593 | 2.20791108 | 0.02725047 | 0.07603609 | TEP1 | 7011 |
| ENSG000001 | 559.744277 | 0.282773155 | 0.12807099 | 2.20794079 | 0.0272484 | 0.07603609 | OXR1 | 55074 |
| ENSG000002 | 5.47023581 | -0.822412283 | 0.37248324 | -2.2079175 | 0.02725002 | 0.07603609 | NA | NA |
| ENSG000002 | 16.6504966 | 0.670822021 | 0.30383019 | 2.20788471 | 0.02725231 | 0.07603609 | NA | NA |
| ENSG00000C | 108.404848 | -0.354931618 | 0.16075957 | -2.2078412 | 0.02725534 | 0.07603931 | TNS1 | 7145 |
| ENSG000002 | 3.25222455 | -0.833372865 | 0.37748173 | -2.2077171 | 0.027264 | 0.07605824 | NA | NA |
| ENSG000002 | 3.30557035 | -0.832885834 | 0.37726892 | -2.2076715 | 0.02726718 | 0.07606188 | NA | NA |
| ENSG000002 | 3.70095663 | -0.834342449 | 0.37794315 | -2.2075871 | 0.02727307 | 0.07607306 | NA | NA |
| ENSG000002 | 21.1271556 | -0.620829952 | 0.28124034 | -2.2074712 | 0.02728116 | 0.07609039 | NA | NA |
| ENSG000001 | 72.1863913 | 0.42034074 | 0.19042497 | 2.20738248 | 0.02728735 | 0.07610243 | NMNAT1 | 64802 |
| ENSG000002 | 3.91951512 | -0.835523366 | 0.37852428 | -2.2073178 | 0.02729187 | 0.0761098 | NA | NA |
| ENSG000001 | 489.870844 | 0.389334974 | 0.17639091 | 2.2072281 | 0.02729813 | 0.07612094 | RAB14 | 51552 |
| ENSG000001 | 220.840779 | 0.297047217 | 0.13458061 | 2.20720673 | 0.02729962 | 0.07612094 | NDRG2 | 57447 |
| ENSG000001 | 172.534171 | 0.356684342 | 0.16160334 | 2.20715951 | 0.02730292 | 0.0761249 | C1orf216 | 127703 |
| ENSG000001 | 15.0339109 | -0.665942534 | 0.30174954 | -2.206938 | 0.02731839 | 0.07615234 | HTN1 | 3346 |
| ENSG000001 | 737.082472 | 0.389769555 | 0.17660801 | 2.20697551 | 0.02731577 | 0.07615234 | NSRP1 | 84081 |
| ENSG000002 | 6.38506225 | -0.810240844 | 0.36713326 | -2.2069394 | 0.0273183 | 0.07615234 | LOC101928251 | 101928251 |
| ENSG000002 | 3.26318968 | -0.831719371 | 0.37688122 | -2.2068475 | 0.02732472 | 0.07616187 | NA | NA |
| ENSG000002 | 7.13050073 | -0.806774484 | 0.36557983 | -2.2068353 | 0.02732557 | 0.07616187 | LOC644090 | 644090 |
| ENSG000002 | 2.70524946 | -0.81600341 | 0.36982117 | -2.2064811 | 0.02735033 | 0.07622566 | NA | NA |
| ENSG000001 | 8.44312232 | -0.771106411 | 0.34948294 | -2.2064208 | 0.02735455 | 0.07622694 | NA | NA |
| ENSG000002 | 14.6615197 | -0.683783451 | 0.30990422 | -2.2064348 | 0.02735357 | 0.07622694 | NA | NA |
| ENSG000002 | 2.9522647 | -0.827693308 | 0.37515181 | -2.206289 | 0.02736377 | 0.07624739 | NA | NA |
| ENSG000001 | 26.4757843 | 0.578479693 | 0.26221353 | 2.20613975 | 0.02737422 | 0.07626601 | EID2B | 126272 |
| ENSG000002 | 3.06700422 | -0.824541744 | 0.37374436 | -2.206165 | 0.02737245 | 0.07626601 | NA | NA |
| ENSG000001 | 91.9929552 | -0.401592044 | 0.18205427 | -2.2058919 | 0.02739157 | 0.07629863 | AK9 | 221264 |

| | | | | | | | | |
|---|---|---|---|---|---|---|---|---|
| ENSG000002 | 16.2648911 | -0.674898394 | 0.30594992 | -2.2059113 | 0.02739021 | 0.07629863 | LINC01204 | 101927528 |
| ENSG000002 | 4.01745554 | -0.835108017 | 0.37857864 | -2.2059037 | 0.02739074 | 0.07629863 | NA | NA |
| ENSG000002 | 11.0341716 | -0.728270295 | 0.33017116 | -2.2057356 | 0.02740252 | 0.07631864 | NA | NA |
| ENSG000002 | 10.8603815 | -0.744026869 | 0.33731254 | -2.2057492 | 0.02740157 | 0.07631864 | NA | NA |
| ENSG00000C | 263.252889 | 0.432017041 | 0.19587669 | 2.20555615 | 0.02741509 | 0.07634842 | IKZF5 | 64376 |
| ENSG000002 | 2.20232594 | -0.805505045 | 0.36523754 | -2.2054278 | 0.02742409 | 0.07636299 | NA | NA |
| ENSG000002 | 2.20232594 | -0.805505045 | 0.36523754 | -2.2054278 | 0.02742409 | 0.07636299 | RN7SL397P | 106479369 |
| ENSG000001 | 2.63995751 | -0.81546168 | 0.36976254 | -2.2053659 | 0.02742843 | 0.07636984 | OR14C36 | 127066 |
| ENSG000001 | 11.6029404 | -0.728718321 | 0.33044087 | -2.2052911 | 0.02743367 | 0.07637918 | C10orf107 | 219621 |
| ENSG000001 | 99.4966566 | 0.379900061 | 0.17227772 | 2.20516069 | 0.02744282 | 0.07639941 | ZSCAN23 | 222696 |
| ENSG000002 | 2.6964747 | -0.815284204 | 0.36980884 | -2.2046099 | 0.02748148 | 0.0765018 | NPS | 594857 |
| ENSG00000C | 211.756528 | 0.304267534 | 0.13802375 | 2.20445783 | 0.02749216 | 0.07651757 | RAD52 | 5893 |
| ENSG000002 | 11.6168524 | -0.722742327 | 0.32785627 | -2.2044487 | 0.02749281 | 0.07651757 | LINC00658 | 100507629 |
| ENSG000002 | 3.29919092 | -0.831316565 | 0.37711322 | -2.2044217 | 0.0274947 | 0.07651757 | NA | NA |
| ENSG000002 | 7.29948767 | -0.792318967 | 0.35942163 | -2.2044276 | 0.02749429 | 0.07651757 | NA | NA |
| ENSG000002 | 11.5488271 | -0.721682547 | 0.32738509 | -2.2043843 | 0.02749733 | 0.07651964 | NA | NA |
| ENSG000001 | 13.7435637 | -0.702699739 | 0.31877953 | -2.204344 | 0.02750016 | 0.07652227 | AGTR1 | 185 |
| ENSG000001 | 84.7175283 | 0.436392332 | 0.19799761 | 2.20402829 | 0.02752235 | 0.07657352 | CCDC130 | 81576 |
| ENSG000001 | 14.0335641 | -0.715224918 | 0.32450471 | -2.204051 | 0.02752076 | 0.07657352 | CDH5 | 1003 |
| ENSG000002 | 12.9710384 | -0.723776617 | 0.32841672 | -2.2038361 | 0.02753587 | 0.07660588 | NA | NA |
| ENSG000001 | 161.399735 | 0.370686275 | 0.16820938 | 2.20371944 | 0.02754408 | 0.07661448 | LSM11 | 134353 |
| ENSG000002 | 40.7360016 | 0.498081405 | 0.22601933 | 2.20371156 | 0.02754464 | 0.07661448 | NA | NA |
| ENSG000002 | 7.17706325 | -0.805915438 | 0.36570401 | -2.2037369 | 0.02754285 | 0.07661448 | NA | NA |
| ENSG000001 | 323.776757 | 0.308419188 | 0.13996522 | 2.20354158 | 0.0275566 | 0.07663906 | RGS2 | 5997 |
| ENSG000001 | 198.618486 | 0.326266113 | 0.14806649 | 2.2035108 | 0.02755877 | 0.07663906 | NDUFAF1 | 51103 |
| ENSG000001 | 16.0740224 | -0.668231201 | 0.30325824 | -2.2035055 | 0.02755914 | 0.07663906 | SLC10A6 | 345274 |
| ENSG000001 | 207.731917 | 0.612964571 | 0.27819996 | 2.20332372 | 0.02757194 | 0.07665888 | PAK4 | 10298 |
| ENSG000001 | 4.6601324 | -0.833259529 | 0.37817906 | -2.2033465 | 0.02757033 | 0.07665888 | MYADML2 | 255275 |
| ENSG000002 | 9.32501333 | -0.762204879 | 0.34593089 | -2.2033443 | 0.02757049 | 0.07665888 | NA | NA |

| | | | | | | | | |
|---|---|---|---|---|---|---|---|---|
| ENSG000001 | 222.289813 | 0.306908192 | 0.13930173 | 2.20319008 | 0.02758135 | 0.07667722 | UBE3D | 90025 |
| ENSG000002 | 19.5384929 | -0.642334679 | 0.29155292 | -2.2031496 | 0.02758421 | 0.07667722 | NA | NA |
| ENSG000002 | 5.56159771 | -0.821864282 | 0.37303737 | -2.2031688 | 0.02758285 | 0.07667722 | NA | NA |
| ENSG000002 | 6.00577002 | -0.821677314 | 0.37298731 | -2.2029632 | 0.02759734 | 0.07670847 | NA | NA |
| ENSG000001 | 282.303783 | 0.327882624 | 0.14884182 | 2.20289321 | 0.02760228 | 0.07671166 | CEP164 | 22897 |
| ENSG000002 | 3.30838226 | -0.830676883 | 0.37708215 | -2.2029069 | 0.02760131 | 0.07671166 | NA | NA |
| ENSG000001 | 143.86792 | 0.3685021 | 0.16729376 | 2.20272466 | 0.02761416 | 0.07673418 | SLC25A19 | 60386 |
| ENSG000001 | 69.1377252 | -0.420110783 | 0.19072266 | -2.2027314 | 0.02761369 | 0.07673418 | GPM6A | 2823 |
| ENSG000001 | 361.482467 | 0.301424678 | 0.13684515 | 2.20266974 | 0.02761803 | 0.07673968 | SHCBP1 | 79801 |
| ENSG000001 | 10.2247856 | -0.746949064 | 0.33911551 | -2.2026391 | 0.0276202 | 0.07674043 | CDH16 | 1014 |
| ENSG000002 | 19.3134685 | -0.657637942 | 0.29861699 | -2.2022791 | 0.0276456 | 0.07680575 | NA | NA |
| ENSG000002 | 3.90935082 | -0.83344885 | 0.37855121 | -2.2016806 | 0.02768788 | 0.07691794 | NA | NA |
| ENSG000001 | 6.21965541 | -0.816046897 | 0.37067583 | -2.2015109 | 0.02769987 | 0.07694599 | LRTM1 | 57408 |
| ENSG000002 | 9.38196249 | -0.761123922 | 0.3457563 | -2.2013306 | 0.02771263 | 0.07697615 | NA | NA |
| ENSG000002 | 10.1696939 | -0.761893089 | 0.34611187 | -2.2012914 | 0.0277154 | 0.07697857 | LOC100506895 | 100506895 |
| ENSG000002 | 2.58132362 | -0.817433647 | 0.37134907 | -2.201254 | 0.02771804 | 0.07698065 | RNU6-354P | 106481290 |
| ENSG000002 | 2.91580232 | -0.826063449 | 0.3753055 | -2.2010428 | 0.027733 | 0.0770169 | NA | NA |
| ENSG000001 | 4.49154255 | -0.829179898 | 0.37673552 | -2.2009602 | 0.02773884 | 0.07702525 | NA | NA |
| ENSG000001 | 29.7648474 | -0.551066261 | 0.25037903 | -2.2009281 | 0.02774111 | 0.07702525 | IGSF22 | 283284 |
| ENSG000002 | 6.46204032 | -0.808629714 | 0.36740535 | -2.2009198 | 0.0277417 | 0.07702525 | NA | NA |
| ENSG000002 | 8.20442191 | -0.778813534 | 0.35386967 | -2.2008485 | 0.02774675 | 0.07703399 | NA | NA |
| ENSG000002 | 165.483349 | 0.363595466 | 0.16521137 | 2.20078956 | 0.02775093 | 0.07704031 | PABPC4L | 132430 |
| ENSG000001 | 1282.98414 | 0.287680086 | 0.13072062 | 2.20072455 | 0.02775553 | 0.07704781 | KIAA1429 | 25962 |
| ENSG000002 | 13.4868083 | -0.707186043 | 0.32138715 | -2.2004179 | 0.02777726 | 0.07710286 | NA | NA |
| ENSG000002 | 8.39108759 | -0.789159722 | 0.3586564 | -2.2003225 | 0.02778402 | 0.07711636 | LOC101929584 | 101929584 |
| ENSG000002 | 2.27666829 | -0.79728661 | 0.36237929 | -2.200144 | 0.02779668 | 0.07714621 | NA | NA |
| ENSG000002 | 8.00160001 | -0.788093653 | 0.35820713 | -2.200106 | 0.02779938 | 0.07714841 | LOC151121 | 151121 |
| ENSG000001 | 56.4317745 | 0.455331178 | 0.20698669 | 2.19980902 | 0.02782045 | 0.07719451 | MAP4K1 | 11184 |
| ENSG000002 | 2.25746278 | -0.798846264 | 0.36314637 | -2.1997914 | 0.0278217 | 0.07719451 | RNU4-78P | 106479594 |

| | | | | | | | | |
|---|---|---|---|---|---|---|---|---|
| ENSG000002 | 5.81291117 | -0.815756181 | 0.37083148 | -2.199803 | 0.02782088 | 0.07719451 | NA | NA |
| ENSG000002 | 82.4240892 | 0.383996685 | 0.17458316 | 2.19950582 | 0.02784198 | 0.07724548 | NA | NA |
| ENSG000001 | 31.617768 | -0.531217958 | 0.24152231 | -2.1994571 | 0.02784543 | 0.07724816 | C1orf87 | 127795 |
| ENSG000002 | 7.47929361 | -0.781295725 | 0.35522507 | -2.1994386 | 0.02784675 | 0.07724816 | NA | NA |
| ENSG000002 | 2.63756164 | -0.811979919 | 0.36918075 | -2.1994102 | 0.02784877 | 0.07724847 | NA | NA |
| ENSG000002 | 6.19232893 | -0.812092862 | 0.36925059 | -2.1993001 | 0.02785659 | 0.07726488 | NA | NA |
| ENSG000001 | 12.7251604 | -0.708667026 | 0.32224506 | -2.1991556 | 0.02786686 | 0.07728808 | ZNF683 | 257101 |
| ENSG00000C | 21.6671551 | -0.627279237 | 0.285249 | -2.1990585 | 0.02787377 | 0.07729137 | SLC26A3 | 1811 |
| ENSG000001 | 3.28631864 | -0.829593462 | 0.37724518 | -2.199083 | 0.02787202 | 0.07729137 | S100A7L2 | 645922 |
| ENSG000002 | 6.32262754 | -0.817933696 | 0.37193921 | -2.1991058 | 0.0278704 | 0.07729137 | NA | NA |
| ENSG000002 | 2.56988132 | -0.817343406 | 0.37169222 | -2.198979 | 0.02787941 | 0.07730175 | NA | NA |
| ENSG000002 | 7.23357564 | -0.789922404 | 0.3592798 | -2.1986274 | 0.02790443 | 0.07736582 | NA | NA |
| ENSG000002 | 41.0857189 | 0.534142598 | 0.24295064 | 2.19856426 | 0.02790892 | 0.07737298 | SNORD116-24 | 100033435 |
| ENSG000001 | 511.825022 | 0.295139151 | 0.13425355 | 2.19837134 | 0.02792265 | 0.07740577 | GMEB1 | 10691 |
| ENSG000002 | 3.95377806 | -0.832242839 | 0.37858382 | -2.1983053 | 0.02792735 | 0.0774135 | NA | NA |
| ENSG000002 | 15.2858863 | -0.67633199 | 0.307689 | -2.1981026 | 0.02794179 | 0.07744824 | NA | NA |
| ENSG000002 | 7.90058449 | -0.796149066 | 0.36222218 | -2.1979578 | 0.02795212 | 0.07747155 | MSMB | 4477 |
| ENSG000001 | 21.680712 | -0.625382029 | 0.28453529 | -2.1979067 | 0.02795575 | 0.07747634 | ADAD1 | 132612 |
| ENSG000001 | 13.1077681 | -0.706987788 | 0.32168886 | -2.1977379 | 0.02796779 | 0.07749145 | TMEM236 | 653567 |
| ENSG000001 | 8.18153191 | -0.776273074 | 0.35322698 | -2.1976608 | 0.02797329 | 0.07749145 | SOX17 | 64321 |
| ENSG000001 | 207.444788 | 0.297885319 | 0.13554616 | 2.19766703 | 0.02797284 | 0.07749145 | ZSWIM1 | 90204 |
| ENSG000002 | 3.36272029 | -0.827733881 | 0.376646 | -2.1976442 | 0.02797447 | 0.07749145 | NA | NA |
| ENSG000002 | 13.8400191 | -0.708487208 | 0.32238306 | -2.1976564 | 0.0279736 | 0.07749145 | LINC01354 | 100506795 |
| ENSG000002 | 4.25334093 | -0.831217751 | 0.37820732 | -2.1977833 | 0.02796455 | 0.07749145 | LINC01287 | 103724390 |
| ENSG000002 | 4.60674916 | -0.828970652 | 0.37720904 | -2.1976426 | 0.02797459 | 0.07749145 | NA | NA |
| ENSG000002 | 6.02850784 | -0.808287104 | 0.36781422 | -2.1975418 | 0.02798178 | 0.07750589 | LOC101929771 | 101929771 |
| ENSG000002 | 3.93533623 | -0.831855218 | 0.37854342 | -2.197516 | 0.02798362 | 0.07750589 | NA | NA |
| ENSG000001 | 49.6758239 | -0.496809266 | 0.22613025 | -2.1970049 | 0.0280201 | 0.07759633 | STAT4 | 6775 |
| ENSG000001 | 90.6482458 | 0.370887156 | 0.16881384 | 2.19701865 | 0.02801911 | 0.07759633 | AFAP1L1 | 134265 |

| | | | | | | | | | |
|---|---|---|---|---|---|---|---|---|---|
| ENSG000001 | 13.6712427 | 0.688567084 | 0.3134494 | 2.19674078 | 0.02803896 | 0.07764327 | NA | NA | |
| ENSG000002 | 3.31370681 | -0.828329821 | 0.37709353 | -2.1966163 | 0.02804786 | 0.07766259 | NA | NA | |
| ENSG000001 | 62.0768007 | -0.469342074 | 0.21367852 | -2.1964869 | 0.02805711 | 0.07768291 | ADGRD1 | | 283383 |
| ENSG000002 | 655.599512 | 0.341449357 | 0.15545989 | 2.19638228 | 0.02806459 | 0.07769832 | NA | NA | |
| ENSG000002 | 5.48833239 | -0.819791117 | 0.37325667 | -2.1963201 | 0.02806904 | 0.07770377 | NA | NA | |
| ENSG000002 | 31.2397318 | -0.549763934 | 0.25031354 | -2.1963012 | 0.02807039 | 0.07770377 | NA | NA | |
| ENSG000001 | 27.1531758 | -0.576877241 | 0.2626854 | -2.1960765 | 0.02808647 | 0.07774295 | ADGRF3 | | 165082 |
| ENSG000001 | 173.222962 | 0.305122964 | 0.13894694 | 2.19596743 | 0.02809427 | 0.07775926 | ABAT | | 18 |
| ENSG000001 | 3239.42352 | 0.258121781 | 0.11755764 | 2.19570396 | 0.02811314 | 0.07777962 | TET1 | | 80312 |
| ENSG000001 | 74.2844768 | 0.43345369 | 0.1974088 | 2.1957161 | 0.02811227 | 0.07777962 | JUNB | | 3726 |
| ENSG000001 | 2.17745042 | -0.801702605 | 0.36512116 | -2.1957166 | 0.02811223 | 0.07777962 | KRTAP1-1 | | 81851 |
| ENSG000002 | 4.52137047 | -0.827371236 | 0.37681352 | -2.1957047 | 0.02811308 | 0.07777962 | NA | NA | |
| ENSG000002 | 8.18803729 | -0.775729543 | 0.35328285 | -2.1957747 | 0.02810807 | 0.07777962 | LOC100506272 | | 100506272 |
| ENSG000002 | 5.45434355 | -0.828665973 | 0.37738134 | -2.1958319 | 0.02810398 | 0.07777962 | NA | NA | |
| ENSG000001 | 564.125064 | 0.315972695 | 0.14391704 | 2.19551969 | 0.02812634 | 0.07781084 | CEPT1 | | 10390 |
| ENSG000002 | 11.8051259 | -0.714252074 | 0.32533662 | -2.1954248 | 0.02813314 | 0.07782434 | NA | NA | |
| ENSG00000C | 17.6466902 | -0.646711699 | 0.2946141 | -2.1951146 | 0.02815538 | 0.07785136 | ALDH3B1 | | 221 |
| ENSG000001 | 4.29581245 | -0.830373607 | 0.37828013 | -2.1951288 | 0.02815436 | 0.07785136 | LBX1 | | 10660 |
| ENSG000001 | 161.264738 | 0.38742967 | 0.17649741 | 2.19510111 | 0.02815635 | 0.07785136 | C2CD2 | | 25966 |
| ENSG000002 | 9.53747309 | -0.769341963 | 0.35046193 | -2.1952227 | 0.02814763 | 0.07785136 | NA | NA | |
| ENSG000002 | 7.76624791 | -0.774824143 | 0.35297765 | -2.1951082 | 0.02815584 | 0.07785136 | NA | NA | |
| ENSG000002 | 12.8443757 | -0.692268033 | 0.31536749 | -2.1951154 | 0.02815532 | 0.07785136 | ASIC5 | | 51802 |
| ENSG000002 | 177.311909 | 0.320286052 | 0.1458994 | 2.19525266 | 0.02814548 | 0.07785136 | NA | NA | |
| ENSG000002 | 314.659987 | 0.30618367 | 0.13949415 | 2.19495708 | 0.02816668 | 0.07787016 | NA | NA | |
| ENSG000002 | 14.8800781 | -0.677125399 | 0.30849201 | -2.1949528 | 0.02816699 | 0.07787016 | NA | NA | |
| ENSG000002 | 2.40037864 | -0.788137555 | 0.35908863 | -2.1948274 | 0.02817598 | 0.07788971 | NA | NA | |
| ENSG000002 | 5.13481138 | -0.820535996 | 0.37386039 | -2.1947658 | 0.0281804 | 0.07789663 | NA | NA | |
| ENSG000002 | 18.7222444 | -0.632492198 | 0.28821238 | -2.1945352 | 0.02819696 | 0.07793708 | ANKRD34A | | 284615 |
| ENSG000001 | 185.460533 | 0.327940136 | 0.14945278 | 2.19427264 | 0.02821581 | 0.07798388 | DIP2C | | 22982 |

| | | | | | | | | |
|---|---|---|---|---|---|---|---|---|
| ENSG000001 | 38.1160069 | -0.497286788 | 0.22664216 | -2.1941495 | 0.02822466 | 0.07799842 | KIAA1257 | 57501 |
| ENSG000002 | 4.08251936 | -0.830537561 | 0.37852431 | -2.1941459 | 0.02822492 | 0.07799842 | NA | NA |
| ENSG000001 | 276.871843 | 0.307983076 | 0.14037209 | 2.19404778 | 0.02823197 | 0.07801259 | LPIN1 | 23175 |
| ENSG000002 | 4.55736264 | -0.827220746 | 0.37703442 | -2.1940192 | 0.02823403 | 0.07801296 | LOC105372292 | 105372292 |
| ENSG000001 | 24.447514 | -0.573900112 | 0.2615826 | -2.1939537 | 0.02823874 | 0.07802066 | GPR85 | 54329 |
| ENSG000002 | 22.1623844 | -0.618376882 | 0.28187124 | -2.1938275 | 0.02824781 | 0.07804041 | FLJ37035 | 399821 |
| ENSG000002 | 15.5179669 | -0.672757824 | 0.30666772 | -2.193768 | 0.02825209 | 0.07804691 | LOC256880 | 256880 |
| ENSG000001 | 47.3334204 | 0.458218496 | 0.20887794 | 2.19371412 | 0.02825597 | 0.07804698 | CCDC107 | 203260 |
| ENSG000002 | 2.52312348 | -0.814725444 | 0.37138924 | -2.1937239 | 0.02825527 | 0.07804698 | NA | NA |
| ENSG000001 | 911.467357 | 0.259811204 | 0.11844834 | 2.19345587 | 0.02827455 | 0.07809299 | SASS6 | 163786 |
| ENSG000002 | 2.91521415 | -0.821863447 | 0.37469713 | -2.1934074 | 0.02827804 | 0.07809432 | NA | NA |
| ENSG000002 | 2.91483644 | -0.822216688 | 0.37486017 | -2.1933957 | 0.02827888 | 0.07809432 | NA | NA |
| ENSG000001 | 366.665555 | 0.311895771 | 0.14219989 | 2.19336161 | 0.02828134 | 0.07809578 | AMOTL2 | 51421 |
| ENSG000002 | 14.2755776 | -0.6952736 | 0.3170011 | -2.1932845 | 0.02828689 | 0.07810579 | NA | NA |
| ENSG000002 | 21.7036912 | -0.633671215 | 0.28892595 | -2.1931959 | 0.02829327 | 0.07811809 | NA | NA |
| ENSG00000C | 14.0608949 | -0.681968496 | 0.31101842 | -2.1926949 | 0.02832937 | 0.07820222 | KCNH4 | 23415 |
| ENSG000001 | 11.0768735 | -0.753225067 | 0.34350852 | -2.1927406 | 0.02832608 | 0.07820222 | ARL5C | 390790 |
| ENSG000002 | 4.03724007 | -0.829935935 | 0.3785008 | -2.1926927 | 0.02832953 | 0.07820222 | NA | NA |
| ENSG000002 | 11.2094164 | -0.717870717 | 0.32742572 | -2.192469 | 0.02834566 | 0.07823611 | NA | NA |
| ENSG000002 | 6.74405228 | -0.802698201 | 0.36611575 | -2.1924711 | 0.02834551 | 0.07823611 | NA | NA |
| ENSG00000C | 389.997745 | 0.299446228 | 0.13658557 | 2.19237096 | 0.02835273 | 0.0782503 | C4orf27 | 54969 |
| ENSG000001 | 25.2347873 | -0.569374404 | 0.25973561 | -2.1921307 | 0.02837007 | 0.07829039 | NPHS1 | 4868 |
| ENSG000002 | 10.0214227 | -0.752060454 | 0.34307509 | -2.1921162 | 0.02837112 | 0.07829039 | LOC101927311 | 101927311 |
| ENSG000002 | 14.0902203 | -0.703108502 | 0.32076957 | -2.1919427 | 0.02838365 | 0.07831963 | NA | NA |
| ENSG000001 | 22.5085935 | 0.595944444 | 0.27189623 | 2.19180839 | 0.02839335 | 0.07834106 | ZNF497 | 162968 |
| ENSG000002 | 3.30803735 | -0.826653063 | 0.377173 | -2.1917079 | 0.0284006 | 0.07835357 | NA | NA |
| ENSG000002 | 3.66526392 | -0.828217058 | 0.37788933 | -2.1916921 | 0.02840175 | 0.07835357 | NA | NA |
| ENSG000001 | 16.0204239 | -0.68420145 | 0.31218505 | -2.1916535 | 0.02840454 | 0.07835473 | STARD6 | 147323 |
| ENSG000002 | 4.74206013 | -0.829363074 | 0.37842245 | -2.1916329 | 0.02840603 | 0.07835473 | NA | NA |

| | | | | | | | | |
|---|---|---|---|---|---|---|---|---|
| ENSG000002 | 3.56408738 | -0.829390964 | 0.37844899 | -2.1915528 | 0.02841181 | 0.07836102 | NA | NA |
| ENSG000002 | 14.0537815 | -0.705278799 | 0.32181767 | -2.1915478 | 0.02841218 | 0.07836102 | THRA1/BTR | 105371807 |
| ENSG000002 | 3.88313727 | -0.829576355 | 0.37854174 | -2.1915056 | 0.02841522 | 0.0783641 | NA | NA |
| ENSG000001 | 23.6414969 | -0.629667906 | 0.28736367 | -2.1911883 | 0.02843817 | 0.07842055 | LYVE1 | 10894 |
| ENSG000002 | 2.56447245 | -0.812626873 | 0.37086452 | -2.1911691 | 0.02843956 | 0.07842055 | RN7SKP21 | 106480839 |
| ENSG000001 | 31.5019843 | -0.553569981 | 0.25266067 | -2.1909622 | 0.02845453 | 0.07845649 | DSC1 | 1823 |
| ENSG000001 | 89.4706686 | 0.416265248 | 0.19003057 | 2.19051726 | 0.02848674 | 0.07852949 | AGFG2 | 3268 |
| ENSG000002 | 5.38623714 | -0.813261777 | 0.37126354 | -2.1905242 | 0.02848624 | 0.07852949 | NA | NA |
| ENSG000002 | 3.86563321 | -0.829257526 | 0.37856715 | -2.1905163 | 0.02848681 | 0.07852949 | NA | NA |
| ENSG000001 | 2901.65764 | 0.535008787 | 0.2442448 | 2.19046135 | 0.02849079 | 0.07852979 | ACTR2 | 10097 |
| ENSG000002 | 189.090312 | 0.625143287 | 0.28539016 | 2.19048647 | 0.02848898 | 0.07852979 | ZBTB12 | 221527 |
| ENSG000001 | 5.83124383 | -0.813088303 | 0.37122229 | -2.1903004 | 0.02850246 | 0.0785566 | CRH | 1392 |
| ENSG000001 | 71.5937405 | 0.464551357 | 0.2121202 | 2.1900383 | 0.02852146 | 0.07860362 | MXRA5 | 25878 |
| ENSG000001 | 97.0686438 | 0.412162451 | 0.18821308 | 2.18987141 | 0.02853356 | 0.07863164 | USP43 | 124739 |
| ENSG000002 | 2.94606655 | -0.819737409 | 0.37434005 | -2.1898202 | 0.02853728 | 0.07863653 | NA | NA |
| ENSG000002 | 3.54330231 | -0.828838454 | 0.37850774 | -2.189753 | 0.02854216 | 0.07864463 | SEC62-AS1 | 106478967 |
| ENSG000001 | 21.8645731 | -0.614647885 | 0.28070087 | -2.18969 | 0.02854673 | 0.07865188 | AGXT2 | 64902 |
| ENSG000002 | 52.0749821 | -0.499785978 | 0.2282498 | -2.1896447 | 0.02855001 | 0.07865558 | NA | NA |
| ENSG000002 | 15.0923012 | -0.691871453 | 0.31599567 | -2.1894966 | 0.02856076 | 0.07867986 | NA | NA |
| ENSG000002 | 4.77644232 | -0.820992688 | 0.37497378 | -2.1894669 | 0.02856292 | 0.07868046 | NA | NA |
| ENSG000002 | 23.9686896 | -0.592458678 | 0.27061467 | -2.1893073 | 0.02857451 | 0.07870703 | NA | NA |
| ENSG000001 | 26.6299356 | -0.560910046 | 0.25621052 | -2.1892545 | 0.02857835 | 0.07871225 | DUSP27 | 92235 |
| ENSG000002 | 2.95437164 | -0.820208333 | 0.37473711 | -2.1887567 | 0.02861453 | 0.07880657 | NA | NA |
| ENSG000002 | 26.575702 | -0.578084472 | 0.26413002 | -2.188636 | 0.0286233 | 0.07882537 | LOC101929567 | 101929567 |
| ENSG000002 | 8.1426077 | -0.789563857 | 0.36088506 | -2.1878541 | 0.02868023 | 0.07897678 | NA | NA |
| ENSG000001 | 20.973145 | -0.623384781 | 0.28493726 | -2.1877966 | 0.02868442 | 0.07898295 | OR1A1 | 8383 |
| ENSG000002 | 2.58117981 | -0.811459718 | 0.37091474 | -2.1877257 | 0.02868959 | 0.07898645 | NA | NA |
| ENSG000002 | 6.13382136 | -0.808279474 | 0.36945722 | -2.1877485 | 0.02868793 | 0.07898645 | NA | NA |
| ENSG000002 | 2.52690093 | -0.812464842 | 0.37140935 | -2.1875185 | 0.02870469 | 0.07902267 | NA | NA |

| | | | | | | | | |
|---|---|---|---|---|---|---|---|---|
| ENSG000002 | 15.7325514 | -0.660992727 | 0.30217645 | -2.1874396 | 0.02871045 | 0.07903316 | NA | NA |
| ENSG000001 | 1248.06612 | 0.290091795 | 0.13265253 | 2.18685457 | 0.02875314 | 0.07913662 | ARL5A | 26225 |
| ENSG000001 | 91.4182659 | 0.375574736 | 0.17174528 | 2.18681256 | 0.02875621 | 0.07913662 | MROH1 | 727957 |
| ENSG000002 | 36.5289947 | 0.528358634 | 0.2416137 | 2.18679088 | 0.02875779 | 0.07913662 | NA | NA |
| ENSG000002 | 4.94456204 | -0.823014314 | 0.37634945 | -2.1868355 | 0.02875454 | 0.07913662 | NA | NA |
| ENSG000002 | 8.17730533 | -0.779063367 | 0.35624491 | -2.1868758 | 0.02875159 | 0.07913662 | NA | NA |
| ENSG000000 | 168.75605 | -0.304816172 | 0.1393932 | -2.1867363 | 0.02876178 | 0.07913843 | DGKA | 1606 |
| ENSG000002 | 7.22846812 | -0.786165565 | 0.35951678 | -2.1867284 | 0.02876235 | 0.07913843 | TAT-AS1 | 100132529 |
| ENSG000002 | 889.406168 | 0.257517456 | 0.11777868 | 2.18645226 | 0.02878253 | 0.07918858 | NA | NA |
| ENSG000002 | 4.21056961 | -0.826905759 | 0.378203 | -2.1864072 | 0.02878583 | 0.07919227 | NA | NA |
| ENSG000001 | 14.057188 | -0.682984463 | 0.31238653 | -2.1863441 | 0.02879044 | 0.07919958 | KRT76 | 51350 |
| ENSG000002 | 19.9360686 | -0.676575624 | 0.30947629 | -2.1861954 | 0.02880132 | 0.07922413 | LCMT1-AS2 | 100506655 |
| ENSG000001 | 961.871218 | 0.272367864 | 0.12459253 | 2.18606904 | 0.02881055 | 0.07924417 | SPPL2A | 84888 |
| ENSG000001 | 82.2750087 | 0.411818044 | 0.18839278 | 2.18595451 | 0.02881893 | 0.07926184 | BVES | 11149 |
| ENSG000002 | 8.91582705 | -0.770577959 | 0.35251985 | -2.1859137 | 0.02882192 | 0.07926469 | C3orf36 | 80111 |
| ENSG000002 | 14.6693713 | -0.680305408 | 0.31123368 | -2.1858348 | 0.02882769 | 0.07927275 | LINC01554 | 202299 |
| ENSG000002 | 8.17964923 | -0.776441245 | 0.35521735 | -2.1858201 | 0.02882877 | 0.07927275 | NA | NA |
| ENSG000001 | 157.024859 | 0.336603221 | 0.15400949 | 2.18560053 | 0.02884484 | 0.07931158 | C12orf66 | 144577 |
| ENSG000001 | 19.5894814 | -0.645541064 | 0.29537292 | -2.185512 | 0.02885133 | 0.07932404 | SLC45A2 | 51151 |
| ENSG000002 | 2.57202073 | -0.809694128 | 0.37054637 | -2.1851358 | 0.02887889 | 0.07939444 | NA | NA |
| ENSG000001 | 132.987719 | 0.433731549 | 0.19850058 | 2.18503923 | 0.02888597 | 0.07940851 | MED25 | 81857 |
| ENSG000002 | 14.5985576 | 0.696353824 | 0.31869776 | 2.18499753 | 0.02888903 | 0.07941153 | HIST2H3A | 333932 |
| ENSG000002 | 53.9585443 | 0.444872039 | 0.20362204 | 2.18479317 | 0.02890402 | 0.07944175 | NA | NA |
| ENSG000002 | 4.28429085 | -0.826624453 | 0.37835808 | -2.1847675 | 0.0289059 | 0.07944175 | NA | NA |
| ENSG000002 | 4.08046373 | -0.826989257 | 0.37852122 | -2.1847897 | 0.02890427 | 0.07944175 | NA | NA |
| ENSG000001 | 19.1364258 | -0.675922189 | 0.30938949 | -2.1846967 | 0.0289111 | 0.07945065 | SCRG1 | 11341 |
| ENSG000002 | 6.02834476 | -0.803970748 | 0.36802536 | -2.1845526 | 0.02892167 | 0.07947432 | UGT1A7 | 54577 |
| ENSG000002 | 9.33404069 | -0.757571099 | 0.34679747 | -2.184477 | 0.02892722 | 0.07948418 | NA | NA |
| ENSG000002 | 13.7307428 | -0.685226777 | 0.31368911 | -2.1844137 | 0.02893187 | 0.07949157 | LINC00538 | 100861504 |

| | | | | | | | | |
|---|---|---|---|---|---|---|---|---|
| ENSG000002 | 4.08364777 | -0.826731697 | 0.37848191 | -2.1843361 | 0.02893756 | 0.07950184 | LOC102724452 | 102724452 |
| ENSG000002 | 8.4917282 | -0.773128874 | 0.35396796 | -2.1841775 | 0.02894921 | 0.07952844 | NA | NA |
| ENSG000002 | 2.54770615 | -0.81129469 | 0.37147152 | -2.1840024 | 0.02896207 | 0.07955839 | NA | NA |
| ENSG000001 | 179.110774 | 0.339842625 | 0.15561042 | 2.18393225 | 0.02896723 | 0.07956716 | SLC36A1 | 206358 |
| ENSG000001 | 3234.83196 | 0.235865439 | 0.10800866 | 2.18376406 | 0.02897959 | 0.07959573 | CACYBP | 27101 |
| ENSG000001 | 10.700954 | -0.73079411 | 0.33465521 | -2.1837225 | 0.02898265 | 0.07959874 | NA | NA |
| ENSG000002 | 4.27811197 | -0.8262688 | 0.37841897 | -2.1834762 | 0.02900076 | 0.07964309 | LOC101929297 | 101929297 |
| ENSG000002 | 4.63319152 | -0.825787701 | 0.37821775 | -2.1833658 | 0.02900888 | 0.07965999 | MED15P9 | 285103 |
| ENSG000002 | 2.56393634 | -0.808967284 | 0.37052416 | -2.1833051 | 0.02901335 | 0.07966687 | NA | NA |
| ENSG000001 | 24.7966098 | -0.581722553 | 0.26645153 | -2.1832209 | 0.02901955 | 0.07967849 | CFAP52 | 146845 |
| ENSG000001 | 186.08342 | 0.364063063 | 0.16676724 | 2.18306107 | 0.02903131 | 0.0797054 | IL17RA | 23765 |
| ENSG000001 | 36.7921936 | -0.514211252 | 0.23555397 | -2.182987 | 0.02903677 | 0.07971498 | CDK15 | 65061 |
| ENSG000001 | 6.06804359 | -0.803385211 | 0.36803378 | -2.1829116 | 0.02904232 | 0.07971943 | OCM | 654231 |
| ENSG000002 | 7.06271151 | -0.810976295 | 0.3715109 | -2.1829139 | 0.02904215 | 0.07971943 | NA | NA |
| ENSG000001 | 9.28320665 | -0.749731519 | 0.34346987 | -2.182816 | 0.02904937 | 0.07973337 | LINC00839 | 84856 |
| ENSG000001 | 53.3011382 | -0.451259556 | 0.20674829 | -2.1826519 | 0.02906145 | 0.07975575 | CPNE4 | 131034 |
| ENSG000002 | 5.01189333 | 0.820999946 | 0.37614529 | 2.1826671 | 0.02906033 | 0.07975575 | HIST2H4A | 8370 |
| ENSG000001 | 141.27861 | 0.363919617 | 0.16675012 | 2.18242489 | 0.02907819 | 0.07979628 | FIZ1 | 84922 |
| ENSG000001 | 3.32099339 | -0.821962767 | 0.37663974 | -2.182358 | 0.02908312 | 0.07980441 | SPINT3 | 10816 |
| ENSG000001 | 243.368672 | -0.304826575 | 0.13968094 | -2.1823061 | 0.02908695 | 0.07980952 | KALRN | 8997 |
| ENSG000001 | 109.73595 | 0.349703865 | 0.16025452 | 2.18217791 | 0.02909641 | 0.07982209 | CIART | 148523 |
| ENSG000002 | 13.6793961 | -0.68095535 | 0.31205509 | -2.1821639 | 0.02909744 | 0.07982209 | NA | NA |
| ENSG000002 | 13.7582646 | -0.702465085 | 0.32190718 | -2.1821976 | 0.02909495 | 0.07982209 | LOC101929657 | 101929657 |
| ENSG000002 | 4.89362921 | -0.821108169 | 0.37629176 | -2.1821051 | 0.02910178 | 0.0798286 | OR6X1 | 390260 |
| ENSG000002 | 4.61248021 | -0.824041471 | 0.37765298 | -2.1820071 | 0.02910901 | 0.07984304 | NA | NA |
| ENSG000001 | 17.9106758 | -0.659838878 | 0.30240525 | -2.181969 | 0.02911182 | 0.07984534 | NIM1K | 167359 |
| ENSG000001 | 11.263275 | -0.721909761 | 0.33085941 | -2.181923 | 0.02911521 | 0.07984925 | ZG16 | 653808 |
| ENSG000002 | 5.93159973 | -0.813782586 | 0.37298113 | -2.1818331 | 0.02912185 | 0.07986205 | NA | NA |
| ENSG000002 | 11.3497268 | -0.722180201 | 0.33101579 | -2.1817092 | 0.029131 | 0.07988174 | NA | NA |

| | | | | | | | | |
|---|---|---|---|---|---|---|---|---|
| ENSG000002 | 5.76578412 | -0.808352067 | 0.37056051 | -2.1814307 | 0.02915158 | 0.07993276 | NA | NA |
| ENSG000002 | 2.8590887 | -0.81760434 | 0.37481551 | -2.1813514 | 0.02915743 | 0.07994341 | NA | NA |
| ENSG000001 | 7.05320213 | -0.79588178 | 0.36486293 | -2.1813172 | 0.02915997 | 0.07994495 | NA | NA |
| ENSG000001 | 259.48502 | 0.297505953 | 0.13639617 | 2.18118997 | 0.02916937 | 0.07995456 | MSANTD2 | 79684 |
| ENSG000001 | 313.013707 | 0.3479871 | 0.15954115 | 2.18117464 | 0.0291705 | 0.07995456 | IFNAR1 | 3454 |
| ENSG000002 | 2.53632857 | -0.810208864 | 0.37144861 | -2.1812139 | 0.0291676 | 0.07995456 | NA | NA |
| ENSG000002 | 6.96095179 | -0.792787872 | 0.36347025 | -2.181163 | 0.02917136 | 0.07995456 | LOC101929470 | 101929470 |
| ENSG000001 | 693.653544 | 0.284662566 | 0.13052549 | 2.18089641 | 0.02919108 | 0.08000321 | CNST | 163882 |
| ENSG000002 | 3.56912356 | -0.825204348 | 0.3783925 | -2.1808158 | 0.02919704 | 0.08001414 | LOC389273 | 389273 |
| ENSG000002 | 5.00226574 | -0.821914013 | 0.37689708 | -2.1807386 | 0.02920275 | 0.08002439 | NA | NA |
| ENSG000001 | 229.526429 | 0.310158106 | 0.14223554 | 2.180595 | 0.02921339 | 0.0800427 | DUSP5 | 1847 |
| ENSG000001 | 774.866421 | 0.277833135 | 0.12741045 | 2.18061497 | 0.02921191 | 0.0800427 | UBR3 | 130507 |
| ENSG000002 | 14.1576554 | 0.695527789 | 0.31897139 | 2.18053348 | 0.02921794 | 0.08004977 | NA | NA |
| ENSG000002 | 76.5138028 | 0.436650043 | 0.20025422 | 2.18047865 | 0.029222 | 0.08005549 | ZNF674-AS1 | 401588 |
| ENSG000001 | 73.7971435 | 0.427026381 | 0.1958623 | 2.18023779 | 0.02923984 | 0.08009354 | CD40 | 958 |
| ENSG000002 | 2.97532191 | -0.813260303 | 0.37301159 | -2.1802548 | 0.02923858 | 0.08009354 | NA | NA |
| ENSG000001 | 55.4389552 | -0.480392874 | 0.2203573 | -2.1800633 | 0.02925277 | 0.08012354 | FAM26E | 254228 |
| ENSG000001 | 16.97311 | -0.659800091 | 0.30265552 | -2.1800365 | 0.02925475 | 0.08012357 | UNC93A | 54346 |
| ENSG000002 | 9.07195545 | -0.757394112 | 0.34744596 | -2.1798904 | 0.02926559 | 0.08014782 | NA | NA |
| ENSG000001 | 637.482525 | 0.275684319 | 0.12647203 | 2.17980467 | 0.02927194 | 0.08015982 | RIC1 | 57589 |
| ENSG000002 | 2.1758518 | -0.794426301 | 0.36448106 | -2.1796093 | 0.02928644 | 0.08019409 | NA | NA |
| ENSG000001 | 217.359981 | 0.294715689 | 0.13522075 | 2.17951534 | 0.02929341 | 0.08020776 | TRIM21 | 6737 |
| ENSG000001 | 1203.56003 | 0.279743769 | 0.1283557 | 2.17944177 | 0.02929887 | 0.08021188 | DSC2 | 1824 |
| ENSG000002 | 3.33408693 | -0.820323758 | 0.37639091 | -2.1794462 | 0.02929853 | 0.08021188 | TCEB1P32 | 106481964 |
| ENSG000002 | 5.12574847 | -0.816313189 | 0.37455639 | -2.1794133 | 0.02930098 | 0.08021224 | NA | NA |
| ENSG000002 | 18.1284068 | -0.631658771 | 0.28983405 | -2.1793809 | 0.02930339 | 0.08021342 | NA | NA |
| ENSG000002 | 3.30784113 | -0.821104542 | 0.37677959 | -2.1792702 | 0.0293116 | 0.08023049 | NA | NA |
| ENSG000002 | 6.17881531 | -0.808356723 | 0.37094207 | -2.1791994 | 0.02931686 | 0.08023947 | NA | NA |
| ENSG000001 | 2.53086763 | -0.807996812 | 0.37079443 | -2.1790964 | 0.02932451 | 0.08025498 | NA | NA |

| | | | | | | | | |
|---|---|---|---|---|---|---|---|---|
| ENSG000002 | 7.49519905 | -0.775676502 | 0.35599112 | -2.1789209 | 0.02933754 | 0.0802744 | PPP1R3G | 648791 |
| ENSG000002 | 2.25928953 | -0.788094444 | 0.36168249 | -2.1789677 | 0.02933407 | 0.0802744 | NA | NA |
| ENSG000002 | 43.7842773 | 0.546579355 | 0.25084698 | 2.17893537 | 0.02933647 | 0.0802744 | NA | NA |
| ENSG000002 | 7.49517868 | -0.777524305 | 0.35686963 | -2.1787349 | 0.02935137 | 0.08029738 | LOC102724849 | 102724849 |
| ENSG000002 | 53.6126893 | 0.464808009 | 0.21333916 | 2.17872801 | 0.02935188 | 0.08029738 | NA | NA |
| ENSG000002 | 8.32674165 | -0.760957866 | 0.34926337 | -2.1787509 | 0.02935018 | 0.08029738 | NA | NA |
| ENSG000001 | 12.1327701 | -0.724607461 | 0.332592 | -2.1786677 | 0.02935636 | 0.08030422 | CHRNA6 | 8973 |
| ENSG000002 | 51.8023071 | 0.446528023 | 0.20496086 | 2.17860143 | 0.02936129 | 0.08031229 | NA | NA |
| ENSG000002 | 5.20733495 | -0.818036556 | 0.37551394 | -2.1784452 | 0.0293729 | 0.08033864 | SCEL-AS1 | 104355296 |
| ENSG000002 | 4.79721474 | -0.817711476 | 0.37537401 | -2.1783913 | 0.02937692 | 0.08034065 | CSHL1 | 1444 |
| ENSG000002 | 3.59473127 | -0.82423042 | 0.37837236 | -2.1783579 | 0.0293794 | 0.08034065 | NA | NA |
| ENSG000002 | 11.4604483 | -0.726205081 | 0.3333786 | -2.1783194 | 0.02938226 | 0.08034065 | NA | NA |
| ENSG000002 | 3.16686178 | -0.8086698 | 0.37123857 | -2.1783022 | 0.02938355 | 0.08034065 | UGT1A10 | 54575 |
| ENSG000002 | 2.57341083 | -0.807408866 | 0.37065889 | -2.178307 | 0.02938319 | 0.08034065 | NA | NA |
| ENSG000001 | 68.7428306 | 0.412549571 | 0.18939618 | 2.17823591 | 0.02938848 | 0.08034872 | RASL12 | 51285 |
| ENSG000001 | 253.103258 | 0.313079564 | 0.14374443 | 2.17802922 | 0.02940386 | 0.08038536 | ANXA3 | 306 |
| ENSG000002 | 7.65456964 | -0.782319265 | 0.35919485 | -2.1779802 | 0.02940751 | 0.08038992 | NA | NA |
| ENSG000002 | 5.30221261 | -0.818671132 | 0.37590391 | -2.1778734 | 0.02941546 | 0.08040623 | NA | NA |
| ENSG000002 | 8.19951095 | -0.774441528 | 0.35560989 | -2.177784 | 0.02942212 | 0.08041901 | NA | NA |
| ENSG00000C | 366.628877 | 0.605754916 | 0.27816007 | 2.17772061 | 0.02942684 | 0.08042447 | CA11 | 770 |
| ENSG000002 | 3.17887945 | -0.822622803 | 0.37774777 | -2.1777039 | 0.02942809 | 0.08042447 | NA | NA |
| ENSG000001 | 8.18599108 | -0.776231238 | 0.35647412 | -2.1775248 | 0.02944143 | 0.08045553 | HMGB4 | 127540 |
| ENSG000002 | 7.47850891 | -0.791150872 | 0.36333666 | -2.1774595 | 0.0294463 | 0.08045901 | NA | NA |
| ENSG000002 | 6.17145704 | -0.806319488 | 0.37030372 | -2.1774545 | 0.02944667 | 0.08045901 | NA | NA |
| ENSG000002 | 6.4755584 | -0.801143441 | 0.36795439 | -2.1772901 | 0.02945893 | 0.08048707 | NA | NA |
| ENSG000002 | 6.89269081 | -0.785577417 | 0.36086199 | -2.176947 | 0.02948452 | 0.08055156 | NA | NA |
| ENSG000002 | 3.11729157 | -0.811346483 | 0.37275505 | -2.176621 | 0.02950885 | 0.08061261 | NA | NA |
| ENSG000002 | 12.4787857 | -0.707483429 | 0.32507559 | -2.1763659 | 0.02952791 | 0.08065924 | NA | NA |
| ENSG000001 | 7.60938349 | -0.783147442 | 0.3598551 | -2.1762855 | 0.02953392 | 0.0806702 | SAMD11 | 148398 |

| | | | | | | | | |
|---|---|---|---|---|---|---|---|---|
| ENSG000002 | 11.878766 | -0.732548443 | 0.33662113 | -2.1761808 | 0.02954174 | 0.08068614 | NA | NA |
| ENSG000001 | 231.521336 | 0.29780952 | 0.13686007 | 2.17601474 | 0.02955416 | 0.08071462 | NIT1 | 4817 |
| ENSG000002 | 4.77254829 | -0.816249223 | 0.37511932 | -2.1759722 | 0.02955734 | 0.08071787 | NA | NA |
| ENSG000001 | 24.2433836 | -0.592943253 | 0.27250864 | -2.1758695 | 0.02956502 | 0.0807334 | CHODL | 140578 |
| ENSG000002 | 48.0179742 | 0.506753156 | 0.23290939 | 2.17575239 | 0.02957378 | 0.08075189 | RNU1-85P | 106480177 |
| ENSG000002 | 3.5206781 | -0.823364608 | 0.37848733 | -2.1754086 | 0.02959951 | 0.0808167 | NA | NA |
| ENSG000002 | 36.8531986 | 0.522682334 | 0.2402766 | 2.175336 | 0.02960495 | 0.08082611 | NA | NA |
| ENSG000002 | 6.99789821 | -0.793214431 | 0.36465112 | -2.1752694 | 0.02960993 | 0.08082883 | LOC100289650 | 100289650 |
| ENSG000002 | 7.23204228 | -0.783411412 | 0.36014144 | -2.1752882 | 0.02960853 | 0.08082883 | NA | NA |
| ENSG000002 | 7.54783377 | -0.778010899 | 0.35773754 | -2.1748092 | 0.02964442 | 0.08091753 | FLJ41941 | 100192420 |
| ENSG000002 | 9.2133546 | -0.746434681 | 0.34323029 | -2.1747343 | 0.02965003 | 0.08092741 | NA | NA |
| ENSG000001 | 6.84598985 | -0.798698185 | 0.36728772 | -2.1745845 | 0.02966126 | 0.08095261 | RTN4RL1 | 146760 |
| ENSG00000C | 403.225436 | 0.297992685 | 0.13704379 | 2.17443406 | 0.02967255 | 0.08096923 | SLC23A2 | 9962 |
| ENSG000002 | 3.1843721 | -0.821045595 | 0.37758949 | -2.1744398 | 0.02967212 | 0.08096923 | NA | NA |
| ENSG000002 | 21.4465081 | -0.605861878 | 0.27863103 | -2.1744235 | 0.02967334 | 0.08096923 | NA | NA |
| ENSG000002 | 14.5646308 | -0.673576849 | 0.30978398 | -2.1743437 | 0.02967933 | 0.08098013 | HSD3B2 | 3284 |
| ENSG000002 | 9.8490087 | -0.742452329 | 0.34147071 | -2.1742782 | 0.02968425 | 0.08098808 | LOC101929662 | 101929662 |
| ENSG000001 | 68.6355214 | 0.456974506 | 0.21017765 | 2.17422978 | 0.02968788 | 0.08098944 | NA | NA |
| ENSG000001 | 3.18809803 | -0.821168514 | 0.37768447 | -2.1742184 | 0.02968874 | 0.08098944 | ACER1 | 125981 |
| ENSG000002 | 283.760575 | 0.285569331 | 0.13135585 | 2.17401302 | 0.02970416 | 0.08102605 | NA | NA |
| ENSG00000C | 21.9014595 | -0.628558734 | 0.28914327 | -2.173866 | 0.0297152 | 0.08104526 | OAS1 | 4938 |
| ENSG000002 | 5.81262629 | -0.80660714 | 0.37104419 | -2.1738843 | 0.02971382 | 0.08104526 | NA | NA |
| ENSG000002 | 3.35121795 | -0.81790437 | 0.37626103 | -2.1737685 | 0.02972253 | 0.0810598 | NA | NA |
| ENSG000002 | 3.30107028 | -0.818917785 | 0.37678185 | -2.1734534 | 0.02974621 | 0.08111894 | NA | NA |
| ENSG000001 | 12.8722341 | -0.713335565 | 0.32822595 | -2.1733064 | 0.02975726 | 0.08114266 | ALOX15B | 247 |
| ENSG000002 | 7.04495053 | -0.790388801 | 0.36368401 | -2.1732845 | 0.02975891 | 0.08114266 | NA | NA |
| ENSG000002 | 4.49137838 | -0.820304797 | 0.37746479 | -2.1731955 | 0.02976561 | 0.08115545 | LOC101927334 | 101927334 |
| ENSG000002 | 14.6752761 | -0.678594143 | 0.31226284 | -2.1731505 | 0.029769 | 0.08115924 | LOC100505817 | 100505817 |
| ENSG00000C | 512.36304 | 0.296511642 | 0.13645319 | 2.17299172 | 0.02978094 | 0.08118089 | CENPQ | 55166 |

| | | | | | | | | |
|---|---|---|---|---|---|---|---|---|
| ENSG000002 | 1957.28361 | 0.244965639 | 0.1127313 | 2.17300472 | 0.02977996 | 0.08118089 | NA | NA |
| ENSG000002 | 5.94488824 | -0.811554609 | 0.37348333 | -2.172934 | 0.02978529 | 0.08118728 | NA | NA |
| ENSG000001 | 8.80198112 | -0.768147651 | 0.35352287 | -2.1728372 | 0.02979257 | 0.08120168 | RPL13AP3 | 645683 |
| ENSG000001 | 4391.02846 | 0.368324216 | 0.16952051 | 2.17274135 | 0.02979979 | 0.08120573 | ANP32E | 81611 |
| ENSG000002 | 15.3117596 | -0.657144313 | 0.30244829 | -2.1727493 | 0.02979919 | 0.08120573 | NA | NA |
| ENSG000002 | 2.25125199 | -0.785443496 | 0.36149946 | -2.1727377 | 0.02980007 | 0.08120573 | NA | NA |
| ENSG000002 | 11.6985865 | -0.704835974 | 0.32442689 | -2.1725572 | 0.02981366 | 0.08123731 | GRXCR2 | 643226 |
| ENSG00000C | 187.838188 | 0.334611951 | 0.15403028 | 2.17237779 | 0.02982718 | 0.08126323 | ZNF416 | 55659 |
| ENSG000002 | 4.36001955 | -0.822313052 | 0.37853115 | -2.1723788 | 0.0298271 | 0.08126323 | NA | NA |
| ENSG000001 | 26.6173633 | 0.556528977 | 0.25621778 | 2.17209351 | 0.02984861 | 0.08131616 | S1PR5 | 53637 |
| ENSG000002 | 3.49207665 | -0.822063543 | 0.37847295 | -2.1720536 | 0.02985162 | 0.08131889 | NA | NA |
| ENSG000001 | 7.38325855 | -0.787894605 | 0.36275912 | -2.1719498 | 0.02985945 | 0.08133046 | XIRP1 | 165904 |
| ENSG000002 | 5.82668373 | -0.808759846 | 0.37236678 | -2.1719441 | 0.02985988 | 0.08133046 | NA | NA |
| ENSG000002 | 5.54685871 | -0.812192021 | 0.37398087 | -2.1717475 | 0.02987471 | 0.08136541 | NA | NA |
| ENSG000001 | 19.4079471 | -0.638625786 | 0.2940718 | -2.1716662 | 0.02988085 | 0.08137285 | SIM2 | 6493 |
| ENSG000001 | 184.645935 | 0.31792467 | 0.1463972 | 2.17165811 | 0.02988146 | 0.08137285 | SLC9A8 | 23315 |
| ENSG000002 | 3.18263764 | -0.820166283 | 0.37768495 | -2.171562 | 0.02988872 | 0.08138715 | NA | NA |
| ENSG000001 | 40.7735506 | -0.512175824 | 0.23587802 | -2.1713588 | 0.02990406 | 0.08142345 | ABLIM3 | 22885 |
| ENSG000002 | 4.61038372 | -0.821378962 | 0.37828714 | -2.1713108 | 0.02990768 | 0.08142785 | NA | NA |
| ENSG000001 | 29.6885993 | -0.567666747 | 0.26147171 | -2.1710446 | 0.0299278 | 0.08147169 | ADH6 | 130 |
| ENSG000002 | 7.84541892 | -0.776105258 | 0.35747816 | -2.1710564 | 0.02992691 | 0.08147169 | NA | NA |
| ENSG000001 | 4908.23456 | 0.261202322 | 0.1203465 | 2.17041887 | 0.02997513 | 0.08157531 | HEATR1 | 55127 |
| ENSG000002 | 12.5979652 | -0.6952585 | 0.32033159 | -2.1704337 | 0.029974 | 0.08157531 | TXNDC8 | 255220 |
| ENSG000002 | 23.7941523 | 0.587285385 | 0.27057707 | 2.17049211 | 0.02996959 | 0.08157531 | NA | NA |
| ENSG000002 | 31.2028756 | -0.541033598 | 0.24926886 | -2.1704821 | 0.02997034 | 0.08157531 | NA | NA |
| ENSG000002 | 94.5819314 | 0.459332344 | 0.21163407 | 2.17040834 | 0.02997593 | 0.08157531 | NA | NA |
| ENSG000002 | 4.56121944 | 0.819225227 | 0.37745771 | 2.17037619 | 0.02997836 | 0.08157646 | NA | NA |
| ENSG000001 | 13.6924522 | -0.712304245 | 0.32821212 | -2.1702558 | 0.02998748 | 0.08158606 | CBLN2 | 147381 |
| ENSG000002 | 6.03619305 | -0.800151212 | 0.36869082 | -2.1702499 | 0.02998792 | 0.08158606 | CYP4F62P | 646802 |

| | | | | | | | | |
|---|---|---|---|---|---|---|---|---|
| ENSG000002 | 4.2651044 | -0.821137691 | 0.37836019 | -2.1702539 | 0.02998761 | 0.08158606 | NA | NA |
| ENSG000001 | 8.83750119 | -0.773377964 | 0.35636625 | -2.1701773 | 0.02999341 | 0.08159553 | OR4F6 | 390648 |
| ENSG000002 | 5.4427075 | -0.808204245 | 0.37246602 | -2.1698737 | 0.03001641 | 0.08165262 | NA | NA |
| ENSG000002 | 3.31632751 | -0.816842459 | 0.37646321 | -2.1697803 | 0.03002349 | 0.0816664 | NA | NA |
| ENSG000001 | 13.4106911 | -0.702433585 | 0.32375626 | -2.1696371 | 0.03003435 | 0.08169045 | C14orf105 | 55195 |
| ENSG000001 | 103.027318 | 0.386914902 | 0.17834334 | 2.16949451 | 0.03004516 | 0.08169794 | NEK8 | 284086 |
| ENSG000001 | 11.632744 | -0.721443922 | 0.33253622 | -2.1695198 | 0.03004324 | 0.08169794 | S100A7A | 338324 |
| ENSG000002 | 3.51524962 | -0.820903833 | 0.37837752 | -2.1695365 | 0.03004198 | 0.08169794 | NA | NA |
| ENSG000002 | 3.30114664 | -0.817258394 | 0.37670172 | -2.1695107 | 0.03004394 | 0.08169794 | NA | NA |
| ENSG000002 | 6.00840641 | -0.809693297 | 0.37322512 | -2.1694502 | 0.03004852 | 0.0817016 | NA | NA |
| ENSG000001 | 10.0624128 | -0.736088945 | 0.3393233 | -2.169285 | 0.03006106 | 0.08172471 | NXF5 | 55998 |
| ENSG000002 | 20.9132143 | 0.612976831 | 0.28256826 | 2.16930532 | 0.03005951 | 0.08172471 | NA | NA |
| ENSG000001 | 144.961655 | 0.327527109 | 0.15098783 | 2.1692285 | 0.03006534 | 0.08172541 | ERMARD | 55780 |
| ENSG000002 | 6.03360873 | -0.800889569 | 0.36920473 | -2.1692289 | 0.03006531 | 0.08172541 | NA | NA |
| ENSG000002 | 3.90088265 | -0.821201212 | 0.37858871 | -2.1691117 | 0.0300742 | 0.08174402 | NA | NA |
| ENSG000002 | 10.440437 | -0.731354402 | 0.33717711 | -2.1690512 | 0.0300788 | 0.08175103 | NA | NA |
| ENSG000002 | 6.10572322 | -0.802330125 | 0.36990833 | -2.1689972 | 0.0300829 | 0.08175669 | NA | NA |
| ENSG000001 | 1042.83152 | 0.261658072 | 0.12064566 | 2.16881467 | 0.03009676 | 0.0817832 | EPC1 | 80314 |
| ENSG000001 | 14.0457938 | -0.703136543 | 0.32420835 | -2.1687799 | 0.0300994 | 0.0817832 | DUOXA1 | 90527 |
| ENSG000001 | 7.27167003 | -0.798971042 | 0.36840402 | -2.168736 | 0.03010274 | 0.0817832 | ADAMTSL5 | 339366 |
| ENSG000002 | 10.7012585 | -0.730113447 | 0.33664684 | -2.1687815 | 0.03009927 | 0.0817832 | LINC01208 | 100505547 |
| ENSG000002 | 5.4214932 | -0.80649667 | 0.37187113 | -2.1687531 | 0.03010143 | 0.0817832 | NA | NA |
| ENSG000002 | 2.91668857 | -0.812003403 | 0.37453281 | -2.1680435 | 0.03015538 | 0.08191908 | NA | NA |
| ENSG000002 | 10.6668806 | -0.724923341 | 0.33437039 | -2.1680249 | 0.03015679 | 0.08191908 | OR4F13P | 390651 |
| ENSG000002 | 8.78996814 | -0.765615129 | 0.35315973 | -2.1679004 | 0.03016626 | 0.08193933 | NA | NA |
| ENSG000001 | 194.809068 | 0.326882981 | 0.15078711 | 2.16784435 | 0.03017053 | 0.08193994 | PPP1R18 | 170954 |
| ENSG000002 | 5.42286349 | -0.806166187 | 0.37187382 | -2.1678487 | 0.0301702 | 0.08193994 | NA | NA |
| ENSG000002 | 11.2571954 | -0.720239209 | 0.3322549 | -2.1677309 | 0.03017917 | 0.0819579 | NA | NA |
| ENSG000002 | 2.58297376 | -0.800604832 | 0.36939388 | -2.1673473 | 0.03020838 | 0.08203175 | NA | NA |

| | | | | | | | | |
|---|---|---|---|---|---|---|---|---|
| ENSG000002 | 59.6846872 | 0.425930823 | 0.19654492 | 2.16709149 | 0.03022788 | 0.0820792 | NA | NA |
| ENSG000001 | 16.9608146 | -0.640256205 | 0.29548318 | -2.166811 | 0.03024927 | 0.08213178 | KRT72 | 140807 |
| ENSG000002 | 19.8123779 | 0.622380978 | 0.28723857 | 2.16677368 | 0.03025211 | 0.08213401 | NA | NA |
| ENSG000001 | 8.14611653 | -0.772215438 | 0.35641126 | -2.1666415 | 0.0302622 | 0.08214679 | IL17F | 112744 |
| ENSG000002 | 9.22205518 | 0.74949903 | 0.34592811 | 2.16663232 | 0.0302629 | 0.08214679 | NA | NA |
| ENSG000002 | 3.56838635 | -0.819656891 | 0.37830078 | -2.1666804 | 0.03025923 | 0.08214679 | NA | NA |
| ENSG000001 | 50.3274285 | -0.450723237 | 0.20803323 | -2.1665925 | 0.03026594 | 0.08214954 | LRRC4C | 57689 |
| ENSG000002 | 3.21388702 | -0.81776951 | 0.3774686 | -2.166457 | 0.03027628 | 0.08217211 | NA | NA |
| ENSG000001 | 14.4351985 | 0.670227236 | 0.30937272 | 2.16640704 | 0.0302801 | 0.08217696 | IPO4 | 79711 |
| ENSG000002 | 10.0571045 | -0.730887984 | 0.33738582 | -2.1663269 | 0.03028622 | 0.08218807 | MIR3663HG | 101927704 |
| ENSG000002 | 5.79993083 | -0.803490268 | 0.37092591 | -2.1661746 | 0.03029785 | 0.08221261 | NA | NA |
| ENSG000002 | 6.31586452 | -0.795205742 | 0.36710466 | -2.1661554 | 0.03029931 | 0.08221261 | NA | NA |
| ENSG000002 | 9.3355213 | -0.749797486 | 0.34620595 | -2.1657556 | 0.03032986 | 0.08229 | CDRT1 | 374286 |
| ENSG000002 | 7.68921856 | -0.780961793 | 0.3606027 | -2.1657125 | 0.03033316 | 0.08229344 | NA | NA |
| ENSG000001 | 76.4214505 | 0.425512093 | 0.19649767 | 2.16548161 | 0.03035082 | 0.08233129 | TUT1 | 64852 |
| ENSG000001 | 157.551381 | 0.360512269 | 0.16648169 | 2.16547699 | 0.03035118 | 0.08233129 | RASGEF1A | 221002 |
| ENSG000002 | 2.94956882 | -0.809678567 | 0.37391399 | -2.1654139 | 0.030356 | 0.08233887 | NA | NA |
| ENSG00000C | 5.5213065 | -0.812321749 | 0.37514616 | -2.1653473 | 0.0303611 | 0.08234719 | IZUMO4 | 113177 |
| ENSG000001 | 10.8541341 | -0.734927926 | 0.33941362 | -2.1652871 | 0.03036571 | 0.08234868 | BEAN1 | 146227 |
| ENSG000002 | 12.3400418 | -0.700900695 | 0.32369794 | -2.1652925 | 0.0303653 | 0.08234868 | NA | NA |
| ENSG000001 | 2356.9003 | 0.53241932 | 0.24589844 | 2.16520007 | 0.03037237 | 0.08236123 | HNRNPD | 3184 |
| ENSG000001 | 10.8529886 | -0.730594332 | 0.33746218 | -2.1649666 | 0.03039024 | 0.0824042 | CCER1 | 196477 |
| ENSG000002 | 5.43101671 | -0.806927329 | 0.37273497 | -2.1648823 | 0.0303967 | 0.08241069 | DIAPH3-AS1 | 100874195 |
| ENSG000002 | 5.57938884 | -0.810892789 | 0.37456433 | -2.1648959 | 0.03039566 | 0.08241069 | NA | NA |
| ENSG000001 | 373.109498 | 0.282564067 | 0.13053282 | 2.16469751 | 0.03041086 | 0.0824235 | ZNF549 | 256051 |
| ENSG000002 | 6.56845673 | -0.788725225 | 0.36435978 | -2.164688 | 0.03041159 | 0.0824235 | NA | NA |
| ENSG000002 | 7.77729432 | -0.766067952 | 0.35388119 | -2.1647603 | 0.03040605 | 0.0824235 | HCG22 | 285834 |
| ENSG000002 | 12.2713869 | -0.713555998 | 0.3296277 | -2.1647331 | 0.03040813 | 0.0824235 | LINC01487 | 101928190 |
| ENSG000002 | 11.6312633 | -0.74012715 | 0.34189491 | -2.1647796 | 0.03040456 | 0.0824235 | TBX5-AS1 | 255480 |

| | | | | | | | | |
|---|---|---|---|---|---|---|---|---|
| ENSG000002 | 25.2237226 | 0.559665029 | 0.25860635 | 2.16415809 | 0.03045222 | 0.0825281 | NA | NA |
| ENSG000001 | 85.623469 | 0.382994858 | 0.17698645 | 2.1639784 | 0.03046601 | 0.08255995 | RABGGTA | 5875 |
| ENSG000002 | 6.86962188 | -0.782522017 | 0.36162647 | -2.1638958 | 0.03047235 | 0.08257161 | NA | NA |
| ENSG000000 | 2142.62313 | 0.328816454 | 0.15196455 | 2.16377084 | 0.03048194 | 0.08259209 | EIF2AK2 | 5610 |
| ENSG000001 | 20.3735075 | -0.612608708 | 0.28312457 | -2.1637426 | 0.03048411 | 0.08259244 | ASZ1 | 136991 |
| ENSG000001 | 175.852843 | 0.384257786 | 0.17759527 | 2.16367125 | 0.03048959 | 0.08259625 | RANBP10 | 57610 |
| ENSG000002 | 10.2738282 | -0.725035145 | 0.33509197 | -2.16369 | 0.03048815 | 0.08259625 | NA | NA |
| ENSG000002 | 2.56079859 | -0.799554251 | 0.36955582 | -2.1635547 | 0.03049854 | 0.08261467 | NA | NA |
| ENSG000002 | 10.5630269 | -0.719916422 | 0.33275088 | -2.1635297 | 0.03050046 | 0.08261467 | NA | NA |
| ENSG000000 | 157.412204 | 0.37984955 | 0.1755745 | 2.16346645 | 0.03050532 | 0.08262231 | TTC7A | 57217 |
| ENSG000002 | 6.99446871 | -0.785855208 | 0.36325354 | -2.1633793 | 0.03051202 | 0.08263492 | NA | NA |
| ENSG000002 | 4.65237437 | -0.817679221 | 0.37797776 | -2.1632998 | 0.03051813 | 0.08264596 | NA | NA |
| ENSG000002 | 7.92905717 | 0.765041624 | 0.35365083 | 2.16326829 | 0.03052055 | 0.082647 | NA | NA |
| ENSG000001 | 28.3036354 | 0.560422361 | 0.25907718 | 2.16314834 | 0.03052977 | 0.08266645 | FIBCD1 | 84929 |
| ENSG000002 | 17.3061122 | -0.64488095 | 0.29812845 | -2.1630977 | 0.03053367 | 0.08267148 | STARD13-AS | 100874241 |
| ENSG000002 | 6.02079648 | -0.796665152 | 0.36833998 | -2.1628528 | 0.03055251 | 0.08271362 | NA | NA |
| ENSG000002 | 2.18268248 | -0.786546576 | 0.36367076 | -2.1627985 | 0.03055668 | 0.08271362 | RN7SL806P | 106479515 |
| ENSG000002 | 11.5227189 | -0.735549098 | 0.34008933 | -2.1628114 | 0.03055569 | 0.08271362 | NA | NA |
| ENSG000002 | 9.79559354 | -0.768641724 | 0.35539372 | -2.1627893 | 0.03055739 | 0.08271362 | NA | NA |
| ENSG000002 | 2.97867412 | -0.809181672 | 0.37414932 | -2.1627239 | 0.03056242 | 0.08272172 | NA | NA |
| ENSG000001 | 3.15284252 | -0.816792355 | 0.37768177 | -2.162647 | 0.03056834 | 0.08273221 | DEFA5 | 1670 |
| ENSG000001 | 12.5683037 | -0.689629161 | 0.31891393 | -2.1624304 | 0.03058502 | 0.08276078 | UMOD | 7369 |
| ENSG000002 | 19.703647 | -0.631278005 | 0.29192894 | -2.1624372 | 0.03058449 | 0.08276078 | PIK3R6 | 146850 |
| ENSG000002 | 13.9690772 | -0.690594775 | 0.31935666 | -2.1624561 | 0.03058304 | 0.08276078 | NA | NA |
| ENSG000001 | 54.8230524 | -0.434820699 | 0.20108237 | -2.1624009 | 0.03058729 | 0.08276141 | COL13A1 | 1305 |
| ENSG000001 | 103.632313 | 0.395868701 | 0.18310374 | 2.16199129 | 0.03061885 | 0.08283232 | SSBP4 | 170463 |
| ENSG000001 | 128.871941 | 0.345696529 | 0.15989803 | 2.16198118 | 0.03061963 | 0.08283232 | KCNK1 | 3775 |
| ENSG000001 | 134.637503 | -0.336379725 | 0.15558577 | -2.1620211 | 0.03061655 | 0.08283232 | ANKS1B | 56899 |
| ENSG000001 | 19.9002193 | -0.628081807 | 0.29052203 | -2.1619077 | 0.03062529 | 0.08284129 | PNLDC1 | 154197 |

| | | | | | | | | |
|---|---|---|---|---|---|---|---|---|
| ENSG000001 | 24.7357107 | 0.560997962 | 0.2594948 | 2.16188516 | 0.03062703 | 0.08284129 | CES3 | 23491 |
| ENSG000002 | 2.18015077 | -0.786164004 | 0.36365708 | -2.1618278 | 0.03063145 | 0.08284772 | NA | NA |
| ENSG000001 | 1094.86177 | 0.265306273 | 0.12272937 | 2.16171786 | 0.03063993 | 0.08286513 | TMCO1 | 54499 |
| ENSG000001 | 9.88605184 | -0.739249959 | 0.34198171 | -2.1616652 | 0.03064399 | 0.08287059 | APOL5 | 80831 |
| ENSG000002 | 4.59921298 | -0.816817282 | 0.37792527 | -2.1613196 | 0.03067066 | 0.08293718 | RN7SKP256 | 106480911 |
| ENSG000001 | 464.069585 | 0.247694841 | 0.11460871 | 2.16122175 | 0.03067821 | 0.08295207 | PITPNC1 | 26207 |
| ENSG000002 | 9.49486833 | -0.739612412 | 0.34222558 | -2.1611839 | 0.03068114 | 0.08295444 | NA | NA |
| ENSG000002 | 13.5171168 | -0.70970959 | 0.32841536 | -2.1610121 | 0.0306944 | 0.08298478 | CASC16 | 643714 |
| ENSG000001 | 8.72712409 | -0.779415233 | 0.36071779 | -2.1607341 | 0.03071588 | 0.08303732 | PROZ | 8858 |
| ENSG000002 | 7.36954513 | -0.783613934 | 0.36266917 | -2.1606853 | 0.03071966 | 0.08304199 | KRT25 | 147183 |
| ENSG000001 | 166.052487 | 0.330536327 | 0.15298301 | 2.16060812 | 0.03072562 | 0.08305258 | DMC1 | 11144 |
| ENSG000001 | 4.99304351 | -0.815550388 | 0.377537 | -2.1601866 | 0.03075822 | 0.08313516 | NA | NA |
| ENSG000002 | 6.2877439 | -0.788859086 | 0.36518644 | -2.1601544 | 0.03076072 | 0.08313636 | NA | NA |
| ENSG00000C | 60.205638 | -0.455734382 | 0.21098268 | -2.1600559 | 0.03076834 | 0.08315142 | SLC7A14 | 57709 |
| ENSG000001 | 7.86203119 | -0.789637326 | 0.36562437 | -2.1596955 | 0.03079625 | 0.08322128 | OOSP2 | 219990 |
| ENSG000002 | 2.9716991 | -0.806757929 | 0.3735767 | -2.159551 | 0.03080745 | 0.08324433 | NA | NA |
| ENSG000002 | 3.88744943 | -0.817584259 | 0.37859318 | -2.1595325 | 0.03080888 | 0.08324433 | HNRNPA1P65 | 106481664 |
| ENSG000001 | 28.9710029 | -0.563057074 | 0.26074844 | -2.1593881 | 0.03082007 | 0.08326902 | SYNPO2L | 79933 |
| ENSG000001 | 2.1880908 | -0.784602964 | 0.36336947 | -2.1592429 | 0.03083132 | 0.08329387 | RNF186 | 54546 |
| ENSG000001 | 25.7117609 | 0.575348656 | 0.26651907 | 2.15875232 | 0.03086939 | 0.08338559 | TCF15 | 6939 |
| ENSG000001 | 20.6524498 | -0.622819883 | 0.28850909 | -2.1587531 | 0.03086933 | 0.08338559 | SLC25A45 | 283130 |
| ENSG000001 | 54.288804 | -0.429415617 | 0.19892279 | -2.158705 | 0.03087306 | 0.08338996 | ZNF385B | 151126 |
| ENSG000002 | 8.57102835 | -0.768295709 | 0.35594279 | -2.1584809 | 0.03089046 | 0.08343141 | NA | NA |
| ENSG000001 | 78.4303871 | -0.398965833 | 0.18486326 | -2.1581672 | 0.03091483 | 0.08349083 | PDLIM2 | 64236 |
| ENSG000001 | 16.7163925 | -0.667980653 | 0.30952003 | -2.1581177 | 0.03091869 | 0.08349083 | BTBD16 | 118663 |
| ENSG000001 | 1471.32624 | 0.246080811 | 0.11402505 | 2.15812935 | 0.03091778 | 0.08349083 | EXO1 | 9156 |
| ENSG000002 | 47.1623118 | 0.462104119 | 0.21412626 | 2.15809173 | 0.0309207 | 0.08349083 | NA | NA |
| ENSG000001 | 1080.16448 | 0.273660274 | 0.12681334 | 2.15797698 | 0.03092962 | 0.08350936 | UBAP2 | 55833 |
| ENSG000002 | 5.37940702 | -0.802394867 | 0.37183301 | -2.1579441 | 0.03093217 | 0.08351069 | LOC105371115 | 105371115 |

| | | | | | | | | |
|---|---|---|---|---|---|---|---|---|
| ENSG000001 | 32.9159085 | -0.516602838 | 0.23940252 | -2.1578839 | 0.03093686 | 0.08351223 | FAM151B | 167555 |
| ENSG000002 | 3.52053429 | -0.81655951 | 0.37840685 | -2.1578878 | 0.03093656 | 0.08351223 | NA | NA |
| ENSG000001 | 357.33373 | 0.315177136 | 0.14606424 | 2.157798 | 0.03094354 | 0.0835247 | MRPS14 | 63931 |
| ENSG000001 | 11.7013926 | -0.701196813 | 0.32496659 | -2.1577505 | 0.03094724 | 0.08352912 | DRD1 | 1812 |
| ENSG000002 | 6.88575566 | -0.781440888 | 0.36216914 | -2.1576684 | 0.03095363 | 0.0835408 | NA | NA |
| ENSG000002 | 5.76326318 | -0.80318095 | 0.37225041 | -2.1576362 | 0.03095613 | 0.08354199 | LOC102723927 | 102723927 |
| ENSG000001 | 14.4143294 | -0.661101834 | 0.3064155 | -2.1575339 | 0.03096409 | 0.08355791 | ABO | 28 |
| ENSG000001 | 278.662999 | 0.280876031 | 0.13020994 | 2.1571013 | 0.03099777 | 0.08363768 | ZNF501 | 115560 |
| ENSG000002 | 4.17427506 | -0.815389558 | 0.37799961 | -2.1571175 | 0.03099651 | 0.08363768 | NA | NA |
| ENSG000002 | 5.78132497 | -0.805095436 | 0.37323906 | -2.1570503 | 0.03100174 | 0.08364283 | NA | NA |
| ENSG000002 | 4.8537972 | -0.811083847 | 0.37604338 | -2.156889 | 0.03101431 | 0.08367117 | NA | NA |
| ENSG000001 | 20.3463695 | 0.636453034 | 0.29509521 | 2.15677185 | 0.03102345 | 0.08369024 | PRPH2 | 5961 |
| ENSG000002 | 7.13680962 | -0.773047716 | 0.35844541 | -2.1566679 | 0.03103155 | 0.08370653 | NA | NA |
| ENSG000002 | 8.29931406 | -0.757201928 | 0.35111284 | -2.1565772 | 0.03103863 | 0.08372006 | NA | NA |
| ENSG000002 | 4.82084527 | -0.810214864 | 0.37571564 | -2.1564577 | 0.03104795 | 0.08372849 | NA | NA |
| ENSG000002 | 2.17619671 | -0.783572343 | 0.3633597 | -2.1564646 | 0.0310474 | 0.08372849 | NA | NA |
| ENSG000002 | 7.35194668 | -0.783002129 | 0.36309575 | -2.1564618 | 0.03104762 | 0.08372849 | NA | NA |
| ENSG000002 | 14.721171 | -0.656303884 | 0.30435154 | -2.1564007 | 0.03105239 | 0.0837349 | LOC101927244 | 101927244 |
| ENSG000002 | 89.6622818 | 0.389500694 | 0.18062767 | 2.15637334 | 0.03105452 | 0.08373509 | NA | NA |
| ENSG000002 | 7.66422507 | -0.777947504 | 0.3607782 | -2.1563041 | 0.03105993 | 0.0837441 | NA | NA |
| ENSG000002 | 4.11826619 | -0.814991768 | 0.37796722 | -2.1562499 | 0.03106416 | 0.08374993 | RNASE13 | 440163 |
| ENSG000001 | 145.802832 | 0.343511644 | 0.15933379 | 2.15592462 | 0.03108955 | 0.08380355 | STAMBPL1 | 57559 |
| ENSG000002 | 6.30606894 | -0.788702369 | 0.36583173 | -2.1559157 | 0.03109025 | 0.08380355 | MIR4325 | 100422883 |
| ENSG000002 | 16.0948653 | -0.666496213 | 0.30914585 | -2.155928 | 0.03108928 | 0.08380355 | NA | NA |
| ENSG000001 | 12.6154873 | -0.69440828 | 0.32211069 | -2.1558064 | 0.03109879 | 0.083821 | NA | NA |
| ENSG000001 | 2.17960943 | -0.783229368 | 0.36332561 | -2.155723 | 0.0311053 | 0.08382772 | OR2T3 | 343173 |
| ENSG000002 | 41.6831255 | 0.498194927 | 0.23110356 | 2.15572152 | 0.03110542 | 0.08382772 | NA | NA |
| ENSG000002 | 4.16642622 | -0.815124253 | 0.37812743 | -2.1556867 | 0.03110814 | 0.08382949 | NA | NA |
| ENSG000001 | 16.1435104 | -0.651209463 | 0.30212542 | -2.1554276 | 0.03112839 | 0.08387741 | TPO | 7173 |

| | | | | | | | | |
|---|---|---|---|---|---|---|---|---|
| ENSG000002 | 7.09111886 | -0.786621457 | 0.36495276 | -2.1554062 | 0.03113006 | 0.08387741 | NA | NA |
| ENSG000001 | 7.32043029 | -0.780527826 | 0.36214454 | -2.1552937 | 0.03113886 | 0.08389555 | PI16 | 221476 |
| ENSG000002 | 12.8517016 | -0.706580646 | 0.32787149 | -2.1550537 | 0.03115764 | 0.08394056 | NA | NA |
| ENSG000001 | 219.847116 | 0.359072066 | 0.16663125 | 2.15489028 | 0.03117042 | 0.08396385 | PIAS3 | 10401 |
| ENSG000002 | 3.15531419 | -0.813383069 | 0.37745783 | -2.1548979 | 0.03116983 | 0.08396385 | LOC100506393 | 100506393 |
| ENSG000001 | 35.527274 | -0.514090491 | 0.23858047 | -2.1547887 | 0.03117838 | 0.08397524 | PTH2R | 5746 |
| ENSG000001 | 9.59381505 | -0.742784817 | 0.34471439 | -2.1547833 | 0.03117879 | 0.08397524 | NAP1L5 | 266812 |
| ENSG000002 | 5.23057262 | -0.810557159 | 0.37619416 | -2.1546245 | 0.03119123 | 0.08400316 | NA | NA |
| ENSG000002 | 3.15423262 | -0.813758888 | 0.3776869 | -2.154586 | 0.03119425 | 0.08400571 | NA | NA |
| ENSG000000 | 17.2428865 | -0.636767126 | 0.29554917 | -2.1545218 | 0.03119927 | 0.08400685 | TMEM40 | 55287 |
| ENSG000001 | 318.680458 | 0.274734441 | 0.12751547 | 2.15451847 | 0.03119954 | 0.08400685 | MPV17L | 255027 |
| ENSG000002 | 5.45053184 | -0.802200851 | 0.37233715 | -2.1545012 | 0.03120089 | 0.08400685 | NA | NA |
| ENSG000002 | 3.85841079 | -0.815659771 | 0.37859093 | -2.154462 | 0.03120396 | 0.08400955 | NA | NA |
| ENSG000002 | 10.5928509 | -0.715006083 | 0.33187668 | -2.154433 | 0.03120623 | 0.08401007 | NA | NA |
| ENSG000002 | 6.89002325 | -0.778004975 | 0.36113575 | -2.1543283 | 0.03121443 | 0.08402373 | NA | NA |
| ENSG000002 | 12.7113137 | -0.704906315 | 0.32720665 | -2.1543154 | 0.03121545 | 0.08402373 | NA | NA |
| ENSG000002 | 3.93578419 | -0.815402966 | 0.37852516 | -2.1541579 | 0.03122779 | 0.08405138 | NA | NA |
| ENSG000001 | 20.0112229 | -0.610824686 | 0.28366424 | -2.1533369 | 0.03129221 | 0.08421918 | KCNIP4 | 80333 |
| ENSG000002 | 3.85422317 | -0.81521948 | 0.37859179 | -2.1532941 | 0.03129557 | 0.08422183 | NA | NA |
| ENSG000002 | 17.7621356 | 0.738884169 | 0.34314492 | 2.15327148 | 0.03129735 | 0.08422183 | RNA5-8S5 | 100008587 |
| ENSG000002 | 2.52119259 | -0.797423791 | 0.37036169 | -2.1530947 | 0.03131124 | 0.08425361 | NA | NA |
| ENSG000001 | 8.57985831 | -0.747226629 | 0.34709303 | -2.1528137 | 0.03133332 | 0.08430602 | RFPL1 | 5988 |
| ENSG000002 | 2.92682804 | -0.804702891 | 0.37379465 | -2.152794 | 0.03133487 | 0.08430602 | NA | NA |
| ENSG000001 | 4.82909775 | -0.809567595 | 0.37610857 | -2.1524838 | 0.03135927 | 0.08436607 | GK2 | 2712 |
| ENSG000001 | 12.0826082 | -0.705219776 | 0.32766492 | -2.152259 | 0.03137696 | 0.08438646 | C8orf86 | 389649 |
| ENSG000001 | 3.05476662 | 0.810723316 | 0.37668548 | 2.15225526 | 0.03137726 | 0.08438646 | C9orf173 | 441476 |
| ENSG000002 | 17.4751797 | -0.643099977 | 0.29880226 | -2.1522594 | 0.03137693 | 0.08438646 | NA | NA |
| ENSG000002 | 7.50985614 | -0.784529474 | 0.36451122 | -2.152278 | 0.03137547 | 0.08438646 | NA | NA |
| ENSG000002 | 12.9540989 | -0.694340997 | 0.32260691 | -2.1522818 | 0.03137516 | 0.08438646 | NA | NA |

| | | | | | | | | |
|---|---|---|---|---|---|---|---|---|
| ENSG000001 | 87.9871933 | -0.38786223 | 0.18021658 | -2.1522006 | 0.03138156 | 0.08439244 | TRIM66 | 9866 |
| ENSG000001 | 15.5275308 | -0.66406711 | 0.30858292 | -2.1519892 | 0.03139821 | 0.08443161 | FAM83C | 128876 |
| ENSG000001 | 2.56880443 | -0.796405811 | 0.37011803 | -2.1517617 | 0.03141613 | 0.0844686 | CCDC182 | 101927581 |
| ENSG000002 | 25.688686 | -0.562690551 | 0.26150187 | -2.1517649 | 0.03141588 | 0.0844686 | LOC101927533 | 101927533 |
| ENSG000002 | 960.776173 | 0.267662983 | 0.12441207 | 2.15142295 | 0.03144283 | 0.08453021 | NA | NA |
| ENSG000002 | 710.301427 | 0.284375162 | 0.13218033 | 2.15141814 | 0.03144321 | 0.08453021 | RNF115 | 27246 |
| ENSG000002 | 6.42051582 | -0.791123162 | 0.36774288 | -2.1512943 | 0.03145298 | 0.08454525 | NA | NA |
| ENSG000002 | 7.73364785 | -0.763514744 | 0.35490713 | -2.1513086 | 0.03145185 | 0.08454525 | NA | NA |
| ENSG000002 | 9.14087839 | -0.756193438 | 0.35152466 | -2.1511817 | 0.03146186 | 0.08456352 | NA | NA |
| ENSG000001 | 292.957455 | 0.38896516 | 0.18082149 | 2.15110031 | 0.03146828 | 0.08456958 | ECH1 | 1891 |
| ENSG000002 | 2.99429882 | -0.803303938 | 0.37343856 | -2.1511007 | 0.03146825 | 0.08456958 | RNU2-56P | 106480221 |
| ENSG000002 | 2.57249735 | -0.79542424 | 0.36978184 | -2.1510636 | 0.03147118 | 0.08457176 | NA | NA |
| ENSG000002 | 46.1917082 | 0.472023051 | 0.21944741 | 2.15096207 | 0.03147919 | 0.08458208 | NA | NA |
| ENSG000002 | 32.193851 | -0.518726371 | 0.2411578 | -2.1509832 | 0.03147753 | 0.08458208 | NA | NA |
| ENSG000002 | 4.50056579 | -0.811477283 | 0.37729877 | -2.1507552 | 0.03149553 | 0.08462036 | RN7SL288P | 106481000 |
| ENSG000001 | 9.47795767 | -0.735644524 | 0.34207358 | -2.1505447 | 0.03151216 | 0.08465943 | CEACAM5 | 1048 |
| ENSG000002 | 2.17848047 | -0.780637073 | 0.36304119 | -2.1502713 | 0.03153376 | 0.08471185 | NA | NA |
| ENSG000002 | 9.0272188 | -0.75417917 | 0.35075161 | -2.1501802 | 0.03154096 | 0.08472559 | NA | NA |
| ENSG000001 | 801.820534 | 0.255009795 | 0.1186042 | 2.15009068 | 0.03154804 | 0.084739 | COG2 | 22796 |
| ENSG000002 | 5.21540238 | -0.806807234 | 0.37527329 | -2.1499192 | 0.03156161 | 0.08476983 | NA | NA |
| ENSG000002 | 4.61753932 | -0.812199052 | 0.37779752 | -2.1498263 | 0.03156896 | 0.08478394 | NA | NA |
| ENSG000001 | 176.899978 | 0.328019275 | 0.15259839 | 2.1495592 | 0.0315901 | 0.0848351 | ORAOV1 | 220064 |
| ENSG000002 | 3.59615417 | -0.812989836 | 0.37830482 | -2.1490338 | 0.03163172 | 0.08494126 | NA | NA |
| ENSG000002 | 2.19871266 | -0.781802901 | 0.3638095 | -2.1489348 | 0.03163957 | 0.0849567 | NA | NA |
| ENSG000001 | 16.7570765 | -0.638132208 | 0.29696995 | -2.1488107 | 0.03164941 | 0.08495967 | BMP5 | 653 |
| ENSG000001 | 164.061347 | 0.355076758 | 0.16524711 | 2.14876232 | 0.03165325 | 0.08495967 | GRB7 | 2886 |
| ENSG000001 | 4.16898222 | -0.812911236 | 0.37830008 | -2.1488529 | 0.03164606 | 0.08495967 | IFNA5 | 3442 |
| ENSG000002 | 6.97186649 | -0.784380098 | 0.36503544 | -2.1487779 | 0.03165201 | 0.08495967 | NA | NA |
| ENSG000002 | 2.52510643 | -0.795420922 | 0.37016723 | -2.1488151 | 0.03164906 | 0.08495967 | NA | NA |

| | | | | | | | | |
|---|---|---|---|---|---|---|---|---|
| ENSG000002 | 6.71596796 | -0.800304644 | 0.37244733 | -2.1487726 | 0.03165243 | 0.08495967 | NA | NA |
| ENSG000002 | 4.34664059 | -0.81344507 | 0.37857841 | -2.1486832 | 0.03165952 | 0.08497089 | NA | NA |
| ENSG000001 | 6.08898166 | -0.79700301 | 0.37094074 | -2.1485993 | 0.03166618 | 0.08498314 | PSG9 | 5678 |
| ENSG000001 | 106.405376 | 0.358239048 | 0.16675308 | 2.14832042 | 0.03168831 | 0.08503128 | TJP3 | 27134 |
| ENSG000002 | 48.7809309 | 0.493381371 | 0.22965906 | 2.14832094 | 0.03168827 | 0.08503128 | NA | NA |
| ENSG000002 | 30.9202586 | -0.523676781 | 0.24377685 | -2.1481809 | 0.03169939 | 0.08504975 | NA | NA |
| ENSG000002 | 2.1904787 | -0.779436874 | 0.36283464 | -2.1481876 | 0.03169886 | 0.08504975 | NA | NA |
| ENSG000002 | 3.36005916 | -0.807222534 | 0.37578792 | -2.1480801 | 0.0317074 | 0.0850656 | NA | NA |
| ENSG000002 | 17.6175467 | -0.635382356 | 0.29583246 | -2.1477777 | 0.03173142 | 0.08512443 | MTND6P3 | 106478942 |
| ENSG000001 | 1180.20944 | 0.267475736 | 0.12454422 | 2.14763673 | 0.03174263 | 0.08514885 | LPP | 4026 |
| ENSG000001 | 5.3555291 | -0.799460173 | 0.37226941 | -2.1475312 | 0.03175102 | 0.08516009 | XCL1 | 6375 |
| ENSG000002 | 7.70471359 | -0.763240673 | 0.35540235 | -2.1475398 | 0.03175033 | 0.08516009 | NA | NA |
| ENSG000002 | 3.82980207 | -0.808871136 | 0.37668637 | -2.1473332 | 0.03176677 | 0.0851967 | NA | NA |
| ENSG000002 | 3.86111114 | -0.812922637 | 0.37858485 | -2.1472667 | 0.03177206 | 0.08520526 | NA | NA |
| ENSG000002 | 9.26032674 | -0.743077772 | 0.34610986 | -2.1469419 | 0.03179791 | 0.08526894 | LOC102724917 | 102724917 |
| ENSG000001 | 15.6813519 | -0.653716075 | 0.30450153 | -2.14684 | 0.03180602 | 0.08528506 | C4orf17 | 84103 |
| ENSG000002 | 7.75805471 | -0.763320292 | 0.35556777 | -2.1467646 | 0.03181203 | 0.08529553 | NA | NA |
| ENSG000001 | 7.70618802 | -0.762927133 | 0.35541373 | -2.1465888 | 0.03182603 | 0.08532501 | TBATA | 219793 |
| ENSG000002 | 37.8209503 | -0.49015969 | 0.22834514 | -2.1465737 | 0.03182723 | 0.08532501 | NA | NA |
| ENSG000002 | 11.5078274 | -0.727500271 | 0.33892423 | -2.1464983 | 0.03183325 | 0.08533549 | NA | NA |
| ENSG000002 | 18.7787923 | -0.643620806 | 0.29985554 | -2.1464363 | 0.03183819 | 0.08534309 | PAQR9-AS1 | 101927832 |
| ENSG000001 | 7.71545053 | -0.776693023 | 0.36188402 | -2.1462485 | 0.03185316 | 0.08536066 | EVX2 | 344191 |
| ENSG000002 | 25.7529835 | -0.588683042 | 0.27428131 | -2.1462747 | 0.03185107 | 0.08536066 | NA | NA |
| ENSG000002 | 3.58468409 | -0.811850465 | 0.37825462 | -2.1463068 | 0.03184851 | 0.08536066 | NA | NA |
| ENSG000002 | 22.6131333 | -0.603437205 | 0.28115707 | -2.1462637 | 0.03185195 | 0.08536066 | NA | NA |
| ENSG000002 | 4.24363731 | -0.811942645 | 0.37832555 | -2.146148 | 0.03186117 | 0.08537233 | NA | NA |
| ENSG000002 | 6.98389587 | -0.785726188 | 0.36611566 | -2.1461147 | 0.03186383 | 0.08537233 | NA | NA |
| ENSG000002 | 21.1056374 | -0.604226631 | 0.28154355 | -2.1461214 | 0.0318633 | 0.08537233 | NA | NA |
| ENSG000002 | 98.804297 | 0.382636866 | 0.1783036 | 2.1459851 | 0.03187417 | 0.08538874 | ACTN3 | 89 |

| | | | | | | | | |
|---|---|---|---|---|---|---|---|---|
| ENSG000002 | 9.75707288 | -0.73667314 | 0.34327921 | -2.1459882 | 0.03187392 | 0.08538874 | NA | NA |
| ENSG000002 | 6.94544983 | -0.779289628 | 0.36316342 | -2.1458374 | 0.03188596 | 0.08541468 | NA | NA |
| ENSG000001 | 12.3972203 | -0.705088825 | 0.32861202 | -2.1456575 | 0.03190032 | 0.08544186 | GIMAP6 | 474344 |
| ENSG000002 | 9.28552984 | -0.741491215 | 0.34557401 | -2.1456799 | 0.03189852 | 0.08544186 | NA | NA |
| ENSG000001 | 249.823765 | 0.283048917 | 0.13192221 | 2.14557434 | 0.03190695 | 0.08545001 | RFK | 55312 |
| ENSG000002 | 2.99671897 | -0.801437383 | 0.37353182 | -2.1455666 | 0.03190757 | 0.08545001 | NA | NA |
| ENSG000002 | 6.89592029 | -0.777149185 | 0.36224313 | -2.14538 | 0.03192247 | 0.08548427 | NA | NA |
| ENSG000001 | 210.502419 | -0.302907713 | 0.14122691 | -2.1448301 | 0.03196644 | 0.08559634 | TANC2 | 26115 |
| ENSG000001 | 21.7760009 | 0.589250567 | 0.27474094 | 2.14474974 | 0.03197286 | 0.08560789 | PSPN | 5623 |
| ENSG000001 | 7.75115191 | -0.76126414 | 0.35496998 | -2.1445874 | 0.03198585 | 0.08563701 | C3orf30 | 152405 |
| ENSG000001 | 62.8546099 | 0.459203042 | 0.21414709 | 2.14433473 | 0.03200608 | 0.08568551 | CORO1A | 11151 |
| ENSG00000C | 79.5602119 | 0.41864777 | 0.19527573 | 2.14388021 | 0.03204249 | 0.08577166 | STXBP2 | 6813 |
| ENSG000002 | 2.53242637 | -0.793198098 | 0.36997842 | -2.1439037 | 0.03204061 | 0.08577166 | NA | NA |
| ENSG000002 | 7.07214555 | -0.797040488 | 0.37180654 | -2.1436968 | 0.03205719 | 0.08580535 | NA | NA |
| ENSG000002 | 20.4769675 | -0.605028603 | 0.28224704 | -2.1436136 | 0.03206386 | 0.08581754 | NA | NA |
| ENSG000001 | 1170.73977 | 0.286174636 | 0.13350996 | 2.14347027 | 0.03207536 | 0.08584264 | RALGPS2 | 55103 |
| ENSG000001 | 240.566623 | 0.337492767 | 0.15745523 | 2.14342049 | 0.03207935 | 0.08584767 | B3GAT1 | 27087 |
| ENSG000001 | 408.611555 | 0.338234251 | 0.15782417 | 2.14310809 | 0.03210442 | 0.08590342 | C1orf106 | 55765 |
| ENSG000002 | 2.61177163 | -0.791613981 | 0.36937383 | -2.1431241 | 0.03210313 | 0.08590342 | NA | NA |
| ENSG000002 | 6.75863795 | -0.787011298 | 0.36724313 | -2.1430252 | 0.03211107 | 0.08591555 | NA | NA |
| ENSG00000C | 8.05608375 | -0.777007769 | 0.36259516 | -2.1429072 | 0.03212055 | 0.08592734 | VSIG2 | 23584 |
| ENSG000001 | 83.3869878 | 0.475046842 | 0.22168047 | 2.14293505 | 0.03211831 | 0.08592734 | TRAPPC1 | 58485 |
| ENSG000002 | 8.71033729 | -0.748862862 | 0.34946378 | -2.1428912 | 0.03212184 | 0.08592734 | NA | NA |
| ENSG000002 | 3.21001261 | -0.808433938 | 0.37729576 | -2.1427061 | 0.0321367 | 0.08596145 | NA | NA |
| ENSG000002 | 6.21946323 | -0.783909499 | 0.3658725 | -2.1425756 | 0.03214719 | 0.08598382 | NA | NA |
| ENSG000002 | 7.14378376 | -0.770639389 | 0.35974317 | -2.1421932 | 0.03217794 | 0.08606038 | SMIM9 | 100132963 |
| ENSG000001 | 267.873178 | 0.319589891 | 0.14919199 | 2.14213835 | 0.03218235 | 0.08606549 | RARG | 5916 |
| ENSG000002 | 6.23657031 | -0.783087239 | 0.36556703 | -2.1421167 | 0.03218409 | 0.08606549 | NA | NA |
| ENSG000001 | 242.672722 | 0.317867871 | 0.14839229 | 2.14207814 | 0.03218719 | 0.08606811 | MIR302C | 442895 |

| | | | | | | | | | |
|---|---|---|---|---|---|---|---|---|---|
| ENSG000001 | 8.89309839 | -0.740018865 | 0.34549376 | -2.1419168 | 0.03220018 | 0.08609716 | VSX2 | | 338917 |
| ENSG000002 | 6.75386013 | -0.786452532 | 0.36718622 | -2.1418357 | 0.03220671 | 0.08610894 | NA | NA | |
| ENSG000002 | 12.867376 | -0.689481244 | 0.32193406 | -2.1416846 | 0.03221887 | 0.08613576 | NA | NA | |
| ENSG00000C | 6.51981905 | -0.777457821 | 0.36302851 | -2.1415889 | 0.03222658 | 0.08615071 | TMEM176A | | 55365 |
| ENSG00000C | 637.289158 | 0.277232279 | 0.12946488 | 2.1413706 | 0.03224416 | 0.08618019 | KIZ | | 55857 |
| ENSG000001 | 562.254577 | 0.290830644 | 0.13581204 | 2.14142016 | 0.03224017 | 0.08618019 | SERPINE2 | | 5270 |
| ENSG000001 | 12.3369499 | -0.708959603 | 0.33107543 | -2.1413839 | 0.03224309 | 0.08618019 | C9orf131 | | 138724 |
| ENSG000002 | 17.395339 | -0.621542586 | 0.29025784 | -2.1413464 | 0.03224611 | 0.08618019 | NA | NA | |
| ENSG000001 | 98.9339301 | 0.369290579 | 0.17246293 | 2.14127511 | 0.03225186 | 0.08618987 | RPS10 | | 6204 |
| ENSG000002 | 2.94526463 | -0.799262484 | 0.37329395 | -2.1411075 | 0.03226537 | 0.08622029 | NA | NA | |
| ENSG000002 | 3.59615307 | -0.809148701 | 0.37793611 | -2.1409669 | 0.03227671 | 0.08623922 | NA | NA | |
| ENSG000002 | 3.20555002 | -0.807528246 | 0.37717622 | -2.140984 | 0.03227533 | 0.08623922 | NA | NA | |
| ENSG000001 | 834.309749 | 0.264666992 | 0.12362406 | 2.14090195 | 0.03228195 | 0.08624754 | BRAF | | 673 |
| ENSG000002 | 9.07366774 | -0.750309984 | 0.35049902 | -2.1406907 | 0.03229899 | 0.08628739 | NA | NA | |
| ENSG000002 | 5.17821653 | -0.802797828 | 0.37503661 | -2.1405852 | 0.0323075 | 0.08630444 | NA | NA | |
| ENSG000001 | 6.63153237 | -0.782739497 | 0.36568309 | -2.1404859 | 0.03231552 | 0.08632017 | GPER1 | | 2852 |
| ENSG000001 | 98.6182071 | 0.419956643 | 0.19624035 | 2.14001174 | 0.03235382 | 0.08641679 | OLFML2A | | 169611 |
| ENSG000001 | 19.9053927 | -0.610834538 | 0.28543943 | -2.1399795 | 0.03235642 | 0.08641805 | SLC6A3 | | 6531 |
| ENSG000001 | 17.2701639 | -0.637432403 | 0.29788302 | -2.1398749 | 0.03236488 | 0.08643494 | CDKN2A | | 1029 |
| ENSG000002 | 9.49173689 | -0.731711131 | 0.34199432 | -2.1395418 | 0.03239182 | 0.08650119 | NA | NA | |
| ENSG000002 | 6.32172032 | -0.785422903 | 0.36710455 | -2.1395074 | 0.0323946 | 0.08650292 | CAPNS2 | | 84290 |
| ENSG000001 | 16.2902178 | -0.635323376 | 0.29697885 | -2.1392883 | 0.03241233 | 0.08654456 | NT5DC4 | | 284958 |
| ENSG000001 | 169.423777 | 0.336385774 | 0.15724818 | 2.13920298 | 0.03241923 | 0.0865573 | ACSL1 | | 2180 |
| ENSG000002 | 32.644248 | 0.529041133 | 0.24732438 | 2.13905774 | 0.03243099 | 0.086583 | NA | NA | |
| ENSG000002 | 18.5913932 | 0.630054512 | 0.29456079 | 2.13896262 | 0.0324387 | 0.08659786 | NA | NA | |
| ENSG000002 | 4.43993576 | -0.806070125 | 0.37686352 | -2.1388914 | 0.03244447 | 0.08660757 | NA | NA | |
| ENSG000001 | 26.517439 | -0.575188452 | 0.26893273 | -2.1387819 | 0.03245334 | 0.0866171 | TNNI1 | | 7135 |
| ENSG000002 | 3.15027801 | -0.806834125 | 0.3772339 | -2.1388166 | 0.03245053 | 0.0866171 | NA | NA | |
| ENSG000002 | 6.63816771 | -0.781235949 | 0.36527378 | -2.1387682 | 0.03245445 | 0.0866171 | NA | NA | |

| | | | | | | | | |
|---|---|---|---|---|---|---|---|---|
| ENSG000002 | 3.18374699 | -0.806767482 | 0.37722188 | -2.1387081 | 0.03245932 | 0.08662441 | NA | NA |
| ENSG000001 | 1528.51384 | 0.674521171 | 0.31539987 | 2.13862221 | 0.03246628 | 0.08663203 | LRP1 | 4035 |
| ENSG000002 | 13.5051831 | 0.672547988 | 0.31447754 | 2.13862009 | 0.03246645 | 0.08663203 | NA | NA |
| ENSG000001 | 14.8978588 | -0.667533304 | 0.31214544 | -2.1385329 | 0.03247352 | 0.08664152 | NA | NA |
| ENSG000002 | 8.93136049 | -0.740906252 | 0.34645691 | -2.1385235 | 0.03247428 | 0.08664152 | USP17L2 | 377630 |
| ENSG000001 | 76.500847 | 0.429275421 | 0.20075536 | 2.13830121 | 0.0324923 | 0.08666873 | MAT1A | 4143 |
| ENSG000001 | 326.398443 | 0.308219936 | 0.14414481 | 2.13826597 | 0.03249516 | 0.08666873 | ATP9B | 374868 |
| ENSG000002 | 11.2041529 | -0.714750869 | 0.33426538 | -2.1382737 | 0.03249453 | 0.08666873 | LIPK | 643414 |
| ENSG000002 | 2.27439304 | -0.770205122 | 0.36018581 | -2.138355 | 0.03248794 | 0.08666873 | NA | NA |
| ENSG000002 | 3.57902727 | -0.808438533 | 0.37806782 | -2.1383426 | 0.03248895 | 0.08666873 | NA | NA |
| ENSG000001 | 628.036773 | 0.270637593 | 0.12657217 | 2.13820772 | 0.03249989 | 0.08667563 | ERI1 | 90459 |
| ENSG000001 | 515.742005 | 0.301381233 | 0.14096402 | 2.13800105 | 0.03251666 | 0.08671465 | SIK3 | 23387 |
| ENSG000002 | 20.4529926 | -0.604534282 | 0.28276879 | -2.1379102 | 0.03252403 | 0.08672862 | DIAPH2-AS1 | 10824 |
| ENSG000001 | 16.2275362 | -0.64419578 | 0.30133369 | -2.1378153 | 0.03253174 | 0.08674345 | SPATA3 | 130560 |
| ENSG000001 | 14.4364674 | -0.67713548 | 0.31675436 | -2.1377306 | 0.03253862 | 0.08675609 | SLC1A7 | 6512 |
| ENSG000002 | 22.5075023 | -0.583719646 | 0.27306245 | -2.1376782 | 0.03254287 | 0.08676174 | RAPGEF4-AS1 | 91149 |
| ENSG000001 | 25.6093436 | -0.56622203 | 0.2648871 | -2.1375976 | 0.03254942 | 0.08677349 | MAK | 4117 |
| ENSG000001 | 122.382574 | 0.374694003 | 0.17529591 | 2.13749425 | 0.03255781 | 0.08677875 | POLE4 | 56655 |
| ENSG000002 | 6.88564847 | -0.789111076 | 0.36917348 | -2.1375075 | 0.03255674 | 0.08677875 | NA | NA |
| ENSG000002 | 9.738822 | -0.742349298 | 0.34729554 | -2.1375146 | 0.03255616 | 0.08677875 | NA | NA |
| ENSG000002 | 6.44661149 | -0.788812919 | 0.36904978 | -2.1374161 | 0.03256417 | 0.08678998 | PRSS56 | 646960 |
| ENSG000001 | 17.3700561 | -0.643845305 | 0.30124221 | -2.1373011 | 0.03257351 | 0.08680918 | CNGA3 | 1261 |
| ENSG000000 | 213.889016 | 0.352771539 | 0.16506224 | 2.13720309 | 0.03258148 | 0.0868247 | MATR3 | 9782 |
| ENSG000002 | 6.32729729 | -0.785881581 | 0.36773555 | -2.1370835 | 0.0325912 | 0.0868449 | GAS2L2 | 246176 |
| ENSG000001 | 18.1461952 | -0.650203227 | 0.3042771 | -2.1368786 | 0.03260787 | 0.08688361 | FA2H | 79152 |
| ENSG000002 | 12.014293 | -0.693024283 | 0.32432208 | -2.1368397 | 0.03261103 | 0.08688634 | NA | NA |
| ENSG000001 | 53.1853467 | 0.446114939 | 0.20878269 | 2.13674297 | 0.0326189 | 0.08690159 | FAM189A1 | 23359 |
| ENSG000001 | 546.904547 | -0.274176939 | 0.12831838 | -2.1366927 | 0.03262299 | 0.08690678 | CCDC152 | 100129792 |
| ENSG000002 | 3.3332533 | -0.803898907 | 0.376263 | -2.1365345 | 0.03263587 | 0.08693537 | NA | NA |

| | | | | | | | | |
|---|---|---|---|---|---|---|---|---|
| ENSG000002 | 110.698065 | -0.372314358 | 0.17426828 | -2.1364436 | 0.03264327 | 0.08694366 | NA | NA |
| ENSG000002 | 62.850716 | 0.415690592 | 0.19457013 | 2.13645631 | 0.03264224 | 0.08694366 | B3GNT9 | 84752 |
| ENSG000002 | 4.9497565 | -0.808460197 | 0.37842291 | -2.1363934 | 0.03264736 | 0.08694885 | NA | NA |
| ENSG000002 | 25.4171092 | -0.567196977 | 0.26550429 | -2.1363006 | 0.03265492 | 0.08696223 | SLCO6A1 | 133482 |
| ENSG000002 | 16.5557006 | 0.656046885 | 0.30709793 | 2.13627907 | 0.03265667 | 0.08696223 | NA | NA |
| ENSG000002 | 10.7707818 | -0.720498963 | 0.33733186 | -2.1358758 | 0.03268954 | 0.08704402 | NA | NA |
| ENSG000002 | 4.99480179 | -0.806406537 | 0.37759659 | -2.1356298 | 0.0327096 | 0.087086 | NA | NA |
| ENSG000002 | 10.6509662 | -0.71412372 | 0.33438358 | -2.1356423 | 0.03270858 | 0.087086 | NA | NA |
| ENSG000002 | 5.10717603 | -0.80142556 | 0.37527709 | -2.1355568 | 0.03271555 | 0.08709614 | NA | NA |
| ENSG000001 | 16.7973456 | -0.629724303 | 0.29488065 | -2.1355226 | 0.03271835 | 0.08709785 | ANKRD30BL | 554226 |
| ENSG000002 | 4.61973346 | -0.808139143 | 0.37845968 | -2.1353375 | 0.03273345 | 0.08712217 | KRT16P3 | 644945 |
| ENSG000002 | 9.92537929 | -0.738376533 | 0.34578405 | -2.135369 | 0.03273088 | 0.08712217 | NA | NA |
| ENSG000002 | 3.80645854 | -0.808357054 | 0.37856547 | -2.1353164 | 0.03273517 | 0.08712217 | NA | NA |
| ENSG000002 | 24.9121622 | -0.579182778 | 0.27124119 | -2.1353054 | 0.03273607 | 0.08712217 | LOC101927666 | 101927666 |
| ENSG000001 | 36.6650165 | -0.511321495 | 0.23946942 | -2.1352266 | 0.0327425 | 0.08713356 | OR56A1 | 120796 |
| ENSG000002 | 11.059734 | -0.701154783 | 0.32838819 | -2.13514 | 0.03274957 | 0.08714094 | NA | NA |
| ENSG000002 | 3.58196392 | -0.807310201 | 0.37810342 | -2.1351571 | 0.03274818 | 0.08714094 | NA | NA |
| ENSG000002 | 12.9067788 | -0.693629018 | 0.32488301 | -2.1350117 | 0.03276006 | 0.08716312 | NA | NA |
| ENSG000002 | 2.87575858 | -0.798413229 | 0.37399305 | -2.1348344 | 0.03277454 | 0.08719593 | NA | NA |
| ENSG00000C | 23.2253569 | -0.581401637 | 0.27234954 | -2.1347627 | 0.0327804 | 0.08720008 | HES2 | 54626 |
| ENSG000002 | 8.14393459 | -0.762194469 | 0.35703565 | -2.1347853 | 0.03277855 | 0.08720008 | NA | NA |
| ENSG00000C | 446.45019 | 0.268749129 | 0.12589589 | 2.13469337 | 0.03278606 | 0.08720943 | HACE1 | 57531 |
| ENSG000002 | 3.49555464 | -0.807538195 | 0.37831175 | -2.1345839 | 0.03279501 | 0.0872275 | NA | NA |
| ENSG00000C | 971.297666 | 0.271061008 | 0.12700121 | 2.13431826 | 0.03281674 | 0.08727957 | R3HDM1 | 23518 |
| ENSG000001 | 13.2189503 | -0.687373383 | 0.32209332 | -2.1340815 | 0.03283611 | 0.08731964 | MLC1 | 23209 |
| ENSG000001 | 4.86628561 | -0.805310497 | 0.37735249 | -2.1341067 | 0.03283404 | 0.08731964 | SLC51B | 123264 |
| ENSG000002 | 9.81196375 | -0.728271493 | 0.34128018 | -2.1339402 | 0.03284768 | 0.08734466 | NA | NA |
| ENSG000001 | 30.2993706 | -0.5235928 | 0.24538377 | -2.1337711 | 0.03286152 | 0.08737576 | TAL1 | 6886 |
| ENSG00000C | 14.5862131 | -0.670212607 | 0.31412804 | -2.133565 | 0.0328784 | 0.08741491 | AQP6 | 363 |

| | | | | | | | | |
|---|---|---|---|---|---|---|---|---|
| ENSG000002 | 18.2876376 | -0.627856442 | 0.2943265 | -2.1331971 | 0.03290856 | 0.08748935 | LINC01114 | 284998 |
| ENSG000002 | 7.54799719 | -0.768807249 | 0.36041325 | -2.133127 | 0.03291431 | 0.08749315 | LOC101927257 | 101927257 |
| ENSG000002 | 4.17453599 | -0.806725343 | 0.3781861 | -2.1331438 | 0.03291293 | 0.08749315 | SPANXA2-OT1 | 619455 |
| ENSG000001 | 156.094263 | 0.351750933 | 0.16490509 | 2.13305081 | 0.03292056 | 0.08749826 | PLAU | 5328 |
| ENSG000001 | 10.8016412 | -0.70587243 | 0.33091853 | -2.1330701 | 0.03291897 | 0.08749826 | LOC100130264 | 100130264 |
| ENSG000002 | 4.2011732 | -0.806956559 | 0.37832028 | -2.1329984 | 0.03292486 | 0.08749826 | NA | NA |
| ENSG000002 | 13.3092796 | -0.688336297 | 0.32270817 | -2.1329993 | 0.03292478 | 0.08749826 | NA | NA |
| ENSG000001 | 92.249589 | 0.377533289 | 0.17700414 | 2.13290656 | 0.03293239 | 0.08751254 | ZADH2 | 284273 |
| ENSG000001 | 32.270407 | -0.528298958 | 0.24769806 | -2.1328344 | 0.03293831 | 0.08752254 | TMPRSS7 | 344805 |
| ENSG000002 | 8.04223818 | -0.750783985 | 0.35203727 | -2.1326832 | 0.03295072 | 0.08754977 | NA | NA |
| ENSG000001 | 56.0548888 | 0.437397843 | 0.20510044 | 2.13260312 | 0.0329573 | 0.08755796 | BCL6B | 255877 |
| ENSG000001 | 30.8044939 | -0.534875464 | 0.2508099 | -2.1325931 | 0.03295812 | 0.08755796 | GDPD4 | 220032 |
| ENSG000001 | 248.469927 | 0.290564379 | 0.13625598 | 2.13248903 | 0.03296667 | 0.08757493 | TSPAN12 | 23554 |
| ENSG000002 | 4.21707864 | -0.806771732 | 0.37834639 | -2.1323627 | 0.03297704 | 0.08759147 | NA | NA |
| ENSG000002 | 15.636439 | -0.655847264 | 0.30756864 | -2.1323606 | 0.03297721 | 0.08759147 | LOC100506551 | 100506551 |
| ENSG000002 | 13.7899939 | -0.658758636 | 0.30899986 | -2.131906 | 0.03301458 | 0.08768497 | NA | NA |
| ENSG000002 | 4.42035701 | -0.803606556 | 0.37700012 | -2.1315817 | 0.03304125 | 0.08775006 | NA | NA |
| ENSG000002 | 5.38798012 | -0.795212239 | 0.37307879 | -2.1314861 | 0.03304912 | 0.08776522 | NA | NA |
| ENSG000002 | 8.3075376 | -0.780102884 | 0.36602115 | -2.1313055 | 0.03306398 | 0.08779894 | NA | NA |
| ENSG000001 | 19.613424 | -0.611208293 | 0.28679648 | -2.1311569 | 0.03307622 | 0.08782567 | GYPA | 2993 |
| ENSG000001 | 157.317106 | 0.360825231 | 0.16931583 | 2.13107799 | 0.03308272 | 0.08783718 | ITGA7 | 3679 |
| ENSG000002 | 5.6979237 | -0.790971056 | 0.3711779 | -2.1309756 | 0.03309115 | 0.08785383 | NA | NA |
| ENSG000001 | 23.7421317 | -0.580789067 | 0.27255974 | -2.1308689 | 0.03309995 | 0.08787142 | GBX2 | 2637 |
| ENSG000002 | 412.425068 | 0.35704411 | 0.16756429 | 2.13078876 | 0.03310655 | 0.08788319 | MRPS6 | 64968 |
| ENSG00000C | 322.66038 | 0.382621106 | 0.179581 | 2.13063242 | 0.03311944 | 0.08791165 | HDAC7 | 51564 |
| ENSG000001 | 241.572447 | 0.294894939 | 0.1384115 | 2.13056672 | 0.03312485 | 0.08792027 | SLC25A12 | 8604 |
| ENSG000001 | 662.15587 | 0.318948764 | 0.14970447 | 2.13052263 | 0.03312849 | 0.08792417 | KIT | 3815 |
| ENSG000001 | 3.22264858 | -0.802625108 | 0.37675763 | -2.1303487 | 0.03314284 | 0.08795649 | OR4K15 | 81127 |
| ENSG00000C | 179.933291 | 0.305122719 | 0.14323545 | 2.13021789 | 0.03315363 | 0.08797937 | CECR1 | 51816 |

| | | | | | | | | |
|---|---|---|---|---|---|---|---|---|
| ENSG000002 | 13.9571197 | -0.671148116 | 0.31506813 | -2.1301682 | 0.03315773 | 0.0879845 | STMND1 | 401236 |
| ENSG000001 | 9.54596207 | -0.73612749 | 0.34559512 | -2.1300286 | 0.03316925 | 0.08800931 | ANGPTL7 | 10218 |
| ENSG000002 | 4.21247115 | -0.806127561 | 0.37848543 | -2.1298774 | 0.03318174 | 0.08803093 | NA | NA |
| ENSG000002 | 5.48566429 | -0.798985016 | 0.3751305 | -2.1298855 | 0.03318107 | 0.08803093 | NA | NA |
| ENSG000001 | 17.2617649 | -0.629245261 | 0.2955228 | -2.1292613 | 0.03323265 | 0.08815445 | GPR55 | 9290 |
| ENSG000002 | 12.0897037 | -0.70368311 | 0.33048009 | -2.1292754 | 0.03323148 | 0.08815445 | NA | NA |
| ENSG000002 | 9.3277613 | -0.744523684 | 0.34970582 | -2.129 | 0.03325426 | 0.08820602 | NA | NA |
| ENSG000002 | 2.80625598 | -0.799040893 | 0.37534083 | -2.1288409 | 0.03326743 | 0.08823517 | NA | NA |
| ENSG000002 | 5.33168308 | -0.794153916 | 0.37305256 | -2.128799 | 0.03327089 | 0.08823859 | NA | NA |
| ENSG000002 | 14.9465714 | -0.651011208 | 0.30581972 | -2.1287418 | 0.03327563 | 0.08824034 | NA | NA |
| ENSG000002 | 14.13454 | 0.664910692 | 0.31234964 | 2.12873845 | 0.03327591 | 0.08824034 | LOC100129534 | 100129534 |
| ENSG000001 | 10.4003772 | -0.724641508 | 0.3404165 | -2.1286909 | 0.03327984 | 0.08824195 | MUC15 | 143662 |
| ENSG000002 | 17.0565675 | -0.637194869 | 0.29933823 | -2.1286785 | 0.03328087 | 0.08824195 | LOC101928539 | 101928539 |
| ENSG000001 | 15.3305496 | -0.65167187 | 0.3061486 | -2.1286129 | 0.0332863 | 0.08825058 | PTK6 | 5753 |
| ENSG000002 | 8.06663342 | -0.755975996 | 0.3551606 | -2.1285469 | 0.03329176 | 0.08825931 | NA | NA |
| ENSG000001 | 40.1545459 | -0.498563418 | 0.23424339 | -2.1283991 | 0.03330401 | 0.08828599 | PEX11A | 8800 |
| ENSG000001 | 13.4258254 | -0.677721745 | 0.31844296 | -2.1282359 | 0.03331753 | 0.08829875 | GABRG1 | 2565 |
| ENSG000002 | 210.181358 | 0.346630793 | 0.16286755 | 2.12829875 | 0.03331232 | 0.08829875 | NA | NA |
| ENSG000002 | 4.77898721 | -0.800575328 | 0.37616002 | -2.1282839 | 0.03331355 | 0.08829875 | NA | NA |
| ENSG000002 | 2.25119992 | -0.764706352 | 0.35931067 | -2.1282595 | 0.03331557 | 0.08829875 | NA | NA |
| ENSG000002 | 13.6017402 | 0.725285858 | 0.34079849 | 2.12819565 | 0.03332087 | 0.08830182 | NA | NA |
| ENSG00000C | 96.0018808 | 0.371557273 | 0.17459863 | 2.12806517 | 0.03333168 | 0.08832471 | GMIP | 51291 |
| ENSG000002 | 40.9084079 | 0.479640096 | 0.22540774 | 2.12787762 | 0.03334723 | 0.08836014 | SLC2A1-AS1 | 440584 |
| ENSG000001 | 15.3393145 | 0.65965435 | 0.3100207 | 2.12777515 | 0.03335573 | 0.08837689 | PLAC9 | 219348 |
| ENSG000002 | 5.47430611 | -0.798081974 | 0.37518787 | -2.1271529 | 0.03340738 | 0.08850794 | NA | NA |
| ENSG000002 | 22.5065299 | -0.608641777 | 0.28614601 | -2.1270322 | 0.03341741 | 0.08852873 | LOC100129029 | 100129029 |
| ENSG000001 | 343.481975 | 0.625815203 | 0.29423351 | 2.1269338 | 0.03342558 | 0.08853882 | PLEKHM2 | 23207 |
| ENSG000002 | 3.53089447 | -0.804220634 | 0.37811192 | -2.1269381 | 0.03342523 | 0.08853882 | NA | NA |
| ENSG000001 | 233.056248 | 0.321537576 | 0.15117834 | 2.12687601 | 0.03343039 | 0.08854576 | AMOT | 154796 |

| | | | | | | | | |
|---|---|---|---|---|---|---|---|---|
| ENSG000002 | 5.76557451 | -0.793126308 | 0.37295073 | -2.1266249 | 0.03345126 | 0.08859527 | NA | NA |
| ENSG000002 | 11.5454876 | -0.706849802 | 0.33239502 | -2.1265355 | 0.0334587 | 0.08860917 | HSD52 | 729467 |
| ENSG000002 | 6.59781599 | -0.777516448 | 0.3656379 | -2.1264657 | 0.0334645 | 0.08861876 | NA | NA |
| ENSG000001 | 260.988947 | 0.296670243 | 0.13952178 | 2.12633639 | 0.03347526 | 0.08862988 | LINS1 | 55180 |
| ENSG000001 | 2536.13529 | 0.354895716 | 0.16690294 | 2.12635988 | 0.03347331 | 0.08862988 | ZNF217 | 7764 |
| ENSG000002 | 4.16447528 | -0.804356134 | 0.3782794 | -2.1263546 | 0.03347374 | 0.08862988 | NA | NA |
| ENSG000001 | 862.972377 | 0.255339192 | 0.12010522 | 2.12596241 | 0.03350639 | 0.08870072 | HELB | 92797 |
| ENSG000001 | 1809.06276 | 0.265302873 | 0.12479039 | 2.12598801 | 0.03350426 | 0.08870072 | SREK1 | 140890 |
| ENSG000002 | 9.79272158 | -0.730023603 | 0.34339139 | -2.1259229 | 0.03350968 | 0.08870364 | NA | NA |
| ENSG000002 | 19.7962029 | -0.61908606 | 0.29121216 | -2.1258936 | 0.03351212 | 0.08870432 | NA | NA |
| ENSG000002 | 16.6454589 | -0.630952559 | 0.29680226 | -2.1258347 | 0.03351702 | 0.0887115 | GOLGA6L7P | 728310 |
| ENSG00000C | 137.171246 | 0.338758968 | 0.15937068 | 2.12560415 | 0.03353623 | 0.08875655 | RHOBTB2 | 23221 |
| ENSG000002 | 2.50835223 | -0.784525958 | 0.36909231 | -2.1255549 | 0.03354034 | 0.08876162 | NA | NA |
| ENSG000002 | 11.5921797 | -0.710463725 | 0.33429322 | -2.1252711 | 0.03356399 | 0.08881263 | MTND6P11 | 106480727 |
| ENSG000002 | 4.23714631 | -0.804290125 | 0.37844049 | -2.125275 | 0.03356367 | 0.08881263 | NA | NA |
| ENSG000002 | 16.1124739 | -0.655665285 | 0.30851383 | -2.1252379 | 0.03356677 | 0.08881418 | LOC100505625 | 100505625 |
| ENSG000002 | 8.599428 | -0.744894951 | 0.35053809 | -2.1250043 | 0.03358625 | 0.08885993 | NA | NA |
| ENSG000001 | 11.0908675 | -0.698790549 | 0.32884925 | -2.1249571 | 0.03359019 | 0.08886456 | FCGR1B | 2210 |
| ENSG000002 | 68.0997585 | 0.427527628 | 0.20119619 | 2.12492904 | 0.03359253 | 0.08886495 | NA | NA |
| ENSG000002 | 6.63111338 | -0.778369435 | 0.36631501 | -2.1248636 | 0.033598 | 0.0888736 | KCNAB1-AS1 | 100874084 |
| ENSG000001 | 185.592922 | 0.298540475 | 0.14050108 | 2.12482696 | 0.03360105 | 0.0888759 | TMEM5 | 10329 |
| ENSG000002 | 4.17488035 | -0.803886577 | 0.37836729 | -2.1246196 | 0.03361836 | 0.08891588 | NA | NA |
| ENSG000002 | 6.01253878 | -0.784461939 | 0.36925574 | -2.1244407 | 0.03363331 | 0.0889496 | NA | NA |
| ENSG000001 | 82.3181194 | -0.37482512 | 0.17644215 | -2.1243514 | 0.03364077 | 0.08895971 | NRXN2 | 9379 |
| ENSG000002 | 4.17043235 | -0.803758891 | 0.37835655 | -2.1243425 | 0.03364152 | 0.08895971 | NA | NA |
| ENSG00000C | 96.8182744 | 0.362687165 | 0.17073332 | 2.12429044 | 0.03364586 | 0.08896541 | PDK2 | 5164 |
| ENSG000002 | 74.5651389 | -0.715211021 | 0.33671064 | -2.1241117 | 0.0336608 | 0.0889933 | NA | NA |
| ENSG000002 | 20.0419711 | -0.636454407 | 0.29963074 | -2.1241292 | 0.03365934 | 0.0889933 | NA | NA |
| ENSG000001 | 17.5296002 | -0.647338077 | 0.30477951 | -2.1239554 | 0.03367388 | 0.08902206 | MMRN1 | 22915 |

| | | | | | | | | |
|---|---|---|---|---|---|---|---|---|
| ENSG000002 | 4.11545428 | -0.803224989 | 0.37819181 | -2.1238561 | 0.03368218 | 0.0890382 | NA | NA |
| ENSG000002 | 9.91569919 | -0.737827805 | 0.34745256 | -2.1235354 | 0.03370901 | 0.08910333 | NA | NA |
| ENSG000002 | 3.25103034 | -0.797627281 | 0.37562137 | -2.1234875 | 0.03371302 | 0.08910812 | NA | NA |
| ENSG000001 | 19.9632731 | -0.586742969 | 0.27632155 | -2.1234065 | 0.0337198 | 0.08911442 | CD28 | 940 |
| ENSG000002 | 8.50686038 | -0.755096002 | 0.35560478 | -2.1234135 | 0.03371921 | 0.08911442 | NA | NA |
| ENSG00000C | 81.7339442 | 0.421244842 | 0.19840225 | 2.12318583 | 0.03373828 | 0.08915745 | ABCA7 | 10347 |
| ENSG000001 | 15.2658196 | -0.678147391 | 0.31940778 | -2.1231399 | 0.03374213 | 0.08916182 | C3orf49 | 132200 |
| ENSG000002 | 16.6199101 | -0.633479487 | 0.29838788 | -2.1230068 | 0.03375328 | 0.08917966 | LGALS17A | 400696 |
| ENSG000002 | 10.829002 | -0.728457946 | 0.34312341 | -2.1230202 | 0.03375216 | 0.08917966 | NA | NA |
| ENSG000001 | 71.3790305 | -0.384880702 | 0.1812952 | -2.1229503 | 0.03375801 | 0.08918053 | C5orf63 | 401207 |
| ENSG000002 | 25.9146756 | -0.551814324 | 0.25992563 | -2.12297 | 0.03375637 | 0.08918053 | TMEM253 | 643382 |
| ENSG000002 | 16.853948 | -0.64039452 | 0.30167245 | -2.1228141 | 0.03376944 | 0.0892049 | RNF217-AS1 | 7955 |
| ENSG000002 | 4.0394253 | -0.80292486 | 0.37827041 | -2.1226214 | 0.03378559 | 0.08924176 | NA | NA |
| ENSG000002 | 7.23813472 | -0.765921076 | 0.3608673 | -2.1224452 | 0.03380037 | 0.08927497 | NA | NA |
| ENSG000001 | 94.2328215 | 0.39445886 | 0.18586657 | 2.12226904 | 0.03381515 | 0.08930821 | CDH24 | 64403 |
| ENSG000002 | 2.21396466 | -0.767880555 | 0.36182671 | -2.1222329 | 0.03381818 | 0.08931039 | NA | NA |
| ENSG000001 | 705.156011 | 0.267442637 | 0.12602528 | 2.12213477 | 0.03382642 | 0.08932312 | GPR180 | 160897 |
| ENSG000002 | 23.707052 | 0.561800013 | 0.26473489 | 2.12212305 | 0.03382741 | 0.08932312 | LOC105377371 | 105377371 |
| ENSG000001 | 81.9237481 | 0.400972665 | 0.18895515 | 2.12205202 | 0.03383337 | 0.08932965 | C1QTNF6 | 114904 |
| ENSG000001 | 1626.46521 | 0.29053613 | 0.13691353 | 2.12204112 | 0.03383429 | 0.08932965 | LCOR | 84458 |
| ENSG000001 | 31.2822003 | -0.586795541 | 0.27654485 | -2.121882 | 0.03384765 | 0.0893533 | EHF | 26298 |
| ENSG000002 | 7.78213093 | -0.774101253 | 0.36481711 | -2.1218886 | 0.0338471 | 0.0893533 | NA | NA |
| ENSG000001 | 382.274667 | 0.554975983 | 0.26157338 | 2.12168369 | 0.03386431 | 0.08939146 | SLC6A8 | 6535 |
| ENSG000001 | 265.481202 | 0.400236529 | 0.18864397 | 2.12165024 | 0.03386712 | 0.08939306 | ZRSR2 | 8233 |
| ENSG000001 | 81.4808158 | 0.378508587 | 0.17842049 | 2.1214412 | 0.03388469 | 0.08941618 | GRIK2 | 2898 |
| ENSG000001 | 8.60720606 | -0.743202342 | 0.350329 | -2.1214411 | 0.0338847 | 0.08941618 | CXCL8 | 3576 |
| ENSG000002 | 5.45056266 | -0.794536504 | 0.37452378 | -2.1214581 | 0.03388327 | 0.08941618 | NA | NA |
| ENSG000002 | 7.69564576 | -0.774015687 | 0.36484352 | -2.1215004 | 0.03387971 | 0.08941618 | NA | NA |
| ENSG00000C | 124.529997 | 0.324317826 | 0.15288412 | 2.12133095 | 0.03389396 | 0.0894348 | LXN | 56925 |

| | | | | | | | | |
|---|---|---|---|---|---|---|---|---|
| ENSG000001 | 3999.85989 | 0.237198391 | 0.11182543 | 2.12114893 | 0.03390927 | 0.08946938 | KIF20B | 9585 |
| ENSG000001 | 37.8488302 | 0.532826901 | 0.2512054 | 2.12108056 | 0.03391502 | 0.08947873 | PPP1R14A | 94274 |
| ENSG00000C | 12.4079723 | -0.692894418 | 0.32671915 | -2.1207646 | 0.03394161 | 0.08952261 | ETV7 | 51513 |
| ENSG000001 | 17.2770048 | -0.629754865 | 0.29694174 | -2.1208027 | 0.03393841 | 0.08952261 | CSF2RB | 1439 |
| ENSG000001 | 99.9992241 | -0.382354383 | 0.18029222 | -2.1207481 | 0.03394301 | 0.08952261 | KCNC4 | 3749 |
| ENSG000002 | 7.37156137 | -0.755345587 | 0.3561621 | -2.1207916 | 0.03393935 | 0.08952261 | HBE1 | 3046 |
| ENSG000002 | 12.300818 | -0.687920947 | 0.32436706 | -2.1208101 | 0.03393778 | 0.08952261 | NA | NA |
| ENSG000002 | 12.9901406 | 0.674299989 | 0.3179572 | 2.12072563 | 0.0339449 | 0.08952261 | NA | NA |
| ENSG000002 | 11.1158353 | 0.710408276 | 0.33499611 | 2.12064634 | 0.03395158 | 0.08953439 | NA | NA |
| ENSG000002 | 3.23867822 | -0.798323042 | 0.37647074 | -2.1205447 | 0.03396013 | 0.08955071 | NA | NA |
| ENSG000002 | 5.96586581 | -0.782577354 | 0.36904966 | -2.1205205 | 0.03396218 | 0.08955071 | NA | NA |
| ENSG000001 | 5.40807297 | -0.79164175 | 0.3733348 | -2.1204606 | 0.03396722 | 0.08955234 | DEFB118 | 117285 |
| ENSG000001 | 10.6474078 | -0.716280707 | 0.33779215 | -2.1204776 | 0.03396579 | 0.08955234 | WNT10B | 7480 |
| ENSG000002 | 3.49802685 | -0.802067102 | 0.37826649 | -2.1203758 | 0.03397437 | 0.08956538 | NA | NA |
| ENSG000002 | 3.69301192 | -0.800062758 | 0.37735961 | -2.12016 | 0.03399255 | 0.08960749 | NA | NA |
| ENSG000001 | 16.5443207 | -0.626281321 | 0.29540487 | -2.1200778 | 0.03399948 | 0.08961411 | AKNAD1 | 254268 |
| ENSG000001 | 6.84963764 | -0.771933177 | 0.36410156 | -2.120104 | 0.03399727 | 0.08961411 | VAMP5 | 10791 |
| ENSG000002 | 4.39235639 | -0.799399411 | 0.37706716 | -2.1200452 | 0.03400224 | 0.08961554 | NA | NA |
| ENSG000001 | 12.530855 | -0.684707972 | 0.32298048 | -2.119967 | 0.03400883 | 0.08961669 | TSPAN1 | 10103 |
| ENSG000002 | 3.61449131 | -0.800565325 | 0.37763203 | -2.1199614 | 0.0340093 | 0.08961669 | CDSN | 1041 |
| ENSG000002 | 11.5681568 | -0.710192486 | 0.33499718 | -2.1199954 | 0.03400643 | 0.08961669 | LINC01492 | 101928496 |
| ENSG000002 | 2.23158027 | -0.767297072 | 0.36196234 | -2.1198257 | 0.03402075 | 0.08964103 | NA | NA |
| ENSG000001 | 24.690685 | -0.560157041 | 0.26425753 | -2.1197392 | 0.03402804 | 0.08965442 | C5orf47 | 133491 |
| ENSG000001 | 93.2694853 | 0.389433357 | 0.18372977 | 2.11959856 | 0.03403991 | 0.08967987 | CNN1 | 1264 |
| ENSG000002 | 3.25930263 | -0.796210364 | 0.37565177 | -2.1195438 | 0.03404454 | 0.08968623 | NA | NA |
| ENSG000002 | 3.3625718 | -0.795892149 | 0.3755161 | -2.1194621 | 0.03405143 | 0.08969856 | NA | NA |
| ENSG000001 | 149.302013 | 0.348874552 | 0.16461012 | 2.11939924 | 0.03405674 | 0.08970672 | CYTH1 | 9267 |
| ENSG00000C | 975.797263 | 0.294910368 | 0.13915493 | 2.11929517 | 0.03406553 | 0.08972379 | LAMB1 | 3912 |
| ENSG000001 | 272.558961 | 0.399592082 | 0.18855175 | 2.11927011 | 0.03406765 | 0.08972379 | MYD88 | 4615 |

| | | | | | | | | |
|---|---|---|---|---|---|---|---|---|
| ENSG000001 | 37.8978831 | -0.52839797 | 0.24937194 | -2.1189151 | 0.03409764 | 0.08979695 | CCDC81 | 60494 |
| ENSG000001 | 10.0438801 | -0.721790421 | 0.34066256 | -2.1187841 | 0.03410871 | 0.08982028 | LINC00523 | 283601 |
| ENSG000002 | 6.22697905 | -0.777423437 | 0.36694902 | -2.1186143 | 0.03412308 | 0.08985226 | NA | NA |
| ENSG000001 | 675.826958 | 0.250694149 | 0.11833322 | 2.11854417 | 0.03412901 | 0.08986205 | ZNF35 | 7584 |
| ENSG000002 | 7.89663786 | -0.776534074 | 0.36656643 | -2.1183993 | 0.03414126 | 0.08988848 | LOC105374516 | 105374516 |
| ENSG000000 | 54.7554714 | -0.42842061 | 0.2022537 | -2.1182337 | 0.03415528 | 0.08991954 | YAF2 | 10138 |
| ENSG000002 | 12.0156874 | -0.692293388 | 0.32683261 | -2.1181895 | 0.03415903 | 0.08992357 | LOC101928174 | 101928174 |
| ENSG000002 | 8.41036661 | -0.750029669 | 0.35411331 | -2.1180499 | 0.03417084 | 0.08994883 | NA | NA |
| ENSG000002 | 6.56077151 | -0.776521854 | 0.36662632 | -2.1180199 | 0.03417338 | 0.08994968 | LOC100996291 | 100996291 |
| ENSG000000 | 12.3799614 | -0.693524266 | 0.32744932 | -2.1179591 | 0.03417853 | 0.0899574 | SLC13A2 | 9058 |
| ENSG000002 | 4.76269699 | -0.796450325 | 0.37610681 | -2.1176174 | 0.03420748 | 0.09002775 | NA | NA |
| ENSG000001 | 191.996 | 0.313309865 | 0.14796891 | 2.11740328 | 0.03422564 | 0.09006384 | TRAF3 | 7187 |
| ENSG000002 | 28.9388195 | -0.522313343 | 0.24667511 | -2.117414 | 0.03422473 | 0.09006384 | NA | NA |
| ENSG000002 | 5.94229897 | -0.778845999 | 0.36787922 | -2.1171242 | 0.03424931 | 0.09010929 | NA | NA |
| ENSG000002 | 12.9388184 | -0.681547179 | 0.32192168 | -2.1171211 | 0.03424957 | 0.09010929 | NA | NA |
| ENSG000002 | 13.8685196 | -0.691042189 | 0.32640586 | -2.1171255 | 0.0342492 | 0.09010929 | LOC101928894 | 101928894 |
| ENSG000002 | 9.73862295 | -0.722035928 | 0.34106236 | -2.1170203 | 0.03425813 | 0.0901201 | NA | NA |
| ENSG000002 | 4.47613762 | 0.798649979 | 0.3772481 | 2.11704178 | 0.0342563 | 0.0901201 | NA | NA |
| ENSG000002 | 7.7415501 | -0.753602154 | 0.35598922 | -2.1169241 | 0.03426629 | 0.09013573 | NA | NA |
| ENSG000002 | 4.251349 | -0.801191604 | 0.37849059 | -2.1168072 | 0.03427622 | 0.09015598 | NA | NA |
| ENSG000000 | 22.0170905 | -0.580579022 | 0.27428321 | -2.1167137 | 0.03428416 | 0.09017068 | C8B | 732 |
| ENSG000002 | 4.20174334 | -0.801260408 | 0.37854423 | -2.116689 | 0.03428625 | 0.09017068 | LOC105376365 | 105376365 |
| ENSG000002 | 13.5311566 | 0.671935813 | 0.31748636 | 2.11642419 | 0.03430875 | 0.09022399 | NA | NA |
| ENSG000002 | 2.60989281 | -0.78043391 | 0.36875708 | -2.1163903 | 0.03431163 | 0.09022571 | NA | NA |
| ENSG000002 | 3.48325506 | -0.799838416 | 0.37797303 | -2.1161256 | 0.03433412 | 0.09027421 | NA | NA |
| ENSG000002 | 18.31969 | 0.648673462 | 0.30653893 | 2.11612095 | 0.03433452 | 0.09027421 | NA | NA |
| ENSG000002 | 3.47369213 | -0.800219132 | 0.37816417 | -2.1160628 | 0.03433947 | 0.09027552 | NA | NA |
| ENSG000002 | 6.03288644 | -0.783375136 | 0.37020447 | -2.1160607 | 0.03433964 | 0.09027552 | NA | NA |
| ENSG000002 | 10.0413672 | -0.715344178 | 0.33805851 | -2.1160366 | 0.0343417 | 0.09027552 | LOC730668 | 730668 |

| | | | | | | | | |
|---|---|---|---|---|---|---|---|---|
| ENSG000002 | 14.536642 | -0.678013691 | 0.32042961 | -2.1159521 | 0.03434888 | 0.09028856 | NA | NA |
| ENSG000001 | 7.64866458 | -0.748682046 | 0.35384137 | -2.1158692 | 0.03435594 | 0.09030124 | KCNJ13 | 3769 |
| ENSG000002 | 18.8724809 | -0.629063833 | 0.29732537 | -2.1157422 | 0.03436674 | 0.09031793 | NA | NA |
| ENSG000002 | 5.96848128 | -0.781798954 | 0.36951348 | -2.1157522 | 0.03436589 | 0.09031793 | NA | NA |
| ENSG00000C | 11.5655139 | -0.706332867 | 0.33385877 | -2.1156637 | 0.03437342 | 0.09032964 | SPP2 | 6694 |
| ENSG000001 | 13.6684754 | -0.689014302 | 0.32569308 | -2.1155325 | 0.03438459 | 0.09035312 | OR10A2 | 341276 |
| ENSG000001 | 4.25472297 | -0.800876768 | 0.37857926 | -2.1154798 | 0.03438908 | 0.09035906 | OR2T1 | 26696 |
| ENSG000001 | 11.286173 | -0.68947921 | 0.32593353 | -2.1153982 | 0.03439602 | 0.0903656 | RHCG | 51458 |
| ENSG000002 | 8.94314732 | -0.738413387 | 0.34906523 | -2.1154023 | 0.03439568 | 0.0903656 | NA | NA |
| ENSG000002 | 12.3774609 | -0.709665971 | 0.33556383 | -2.1148464 | 0.03444304 | 0.09048326 | LINC00924 | 145820 |
| ENSG00000C | 74.7629338 | 0.415427405 | 0.19645028 | 2.11466944 | 0.03445813 | 0.09051705 | TMEM14A | 28978 |
| ENSG000001 | 664.241395 | 0.247875509 | 0.11722226 | 2.1145771 | 0.03446601 | 0.09052108 | FER | 2241 |
| ENSG000001 | 24.4145433 | -0.601427709 | 0.28442042 | -2.1145729 | 0.03446636 | 0.09052108 | ASB5 | 140458 |
| ENSG000002 | 2.23045076 | -0.762290155 | 0.36049278 | -2.1145782 | 0.03446592 | 0.09052108 | NA | NA |
| ENSG00000C | 483.295474 | 0.372842506 | 0.17632834 | 2.11447861 | 0.03447441 | 0.09052336 | NOTCH3 | 4854 |
| ENSG000002 | 10.5098781 | -0.707571581 | 0.33462288 | -2.1145344 | 0.03446965 | 0.09052336 | NA | NA |
| ENSG000002 | 79.5133981 | 0.381429802 | 0.18038934 | 2.11448088 | 0.03447422 | 0.09052336 | NA | NA |
| ENSG000002 | 3.15551061 | -0.797441928 | 0.37713773 | -2.1144581 | 0.03447616 | 0.09052336 | NA | NA |
| ENSG000002 | 2.84840688 | -0.791621014 | 0.37441676 | -2.1142777 | 0.03449155 | 0.09055791 | NA | NA |
| ENSG000002 | 14.3737918 | -0.67100252 | 0.31737681 | -2.1142141 | 0.03449699 | 0.09056632 | NA | NA |
| ENSG000001 | 3.5618642 | -0.799381634 | 0.37811155 | -2.1141424 | 0.03450311 | 0.09057653 | PSAPL1 | 768239 |
| ENSG000001 | 16.8569093 | -0.641711477 | 0.30353991 | -2.1140926 | 0.03450736 | 0.09058181 | ACVRL1 | 94 |
| ENSG000002 | 35.1245337 | 0.519196185 | 0.24559592 | 2.11402607 | 0.03451304 | 0.090585 | SPTY2D1-AS1 | 100506540 |
| ENSG000002 | 3.51256833 | -0.798949655 | 0.3779235 | -2.1140512 | 0.03451089 | 0.090585 | NA | NA |
| ENSG000002 | 8.40570147 | -0.749602863 | 0.35461627 | -2.1138423 | 0.03452874 | 0.09061448 | NA | NA |
| ENSG000002 | 3.81918735 | -0.800265545 | 0.37858105 | -2.1138553 | 0.03452763 | 0.09061448 | NA | NA |
| ENSG000001 | 3.7986108 | -0.800177322 | 0.37857008 | -2.1136835 | 0.03454231 | 0.09064422 | IFNW1 | 3467 |
| ENSG000002 | 12.7071655 | -0.694629654 | 0.32864444 | -2.1136206 | 0.03454769 | 0.09065247 | LOC100505658 | 100505658 |
| ENSG000002 | 7.62241649 | -0.779168636 | 0.36869872 | -2.1132936 | 0.03457565 | 0.09071997 | NA | NA |

| | | | | | | | | |
|---|---|---|---|---|---|---|---|---|
| ENSG000002 | 3.54145466 | -0.797949052 | 0.37760978 | -2.1131578 | 0.03458727 | 0.09074458 | NA | NA |
| ENSG000002 | 7.51104366 | -0.778365597 | 0.36836788 | -2.1130116 | 0.03459978 | 0.09076566 | NA | NA |
| ENSG000002 | 14.6543865 | -0.647289386 | 0.30633372 | -2.1130203 | 0.03459903 | 0.09076566 | NA | NA |
| ENSG000002 | 68.3861502 | 0.397808019 | 0.18829734 | 2.11265879 | 0.03462998 | 0.09083902 | STON1 | 11037 |
| ENSG000002 | 7.11773418 | -0.761629954 | 0.3605614 | -2.1123447 | 0.0346569 | 0.09090374 | NA | NA |
| ENSG000002 | 41.2527987 | 0.460542515 | 0.21803794 | 2.1122127 | 0.03466821 | 0.09092753 | FAM218A | 152756 |
| ENSG000001 | 422.349345 | 0.26564298 | 0.12577396 | 2.11206668 | 0.03468073 | 0.09095449 | ZFP14 | 57677 |
| ENSG000002 | 4.80729581 | -0.794459315 | 0.37623033 | -2.1116302 | 0.03471818 | 0.09104682 | NA | NA |
| ENSG000002 | 35.5040559 | 0.499338559 | 0.23651401 | 2.11124308 | 0.03475142 | 0.0911281 | NA | NA |
| ENSG000001 | 6.53934518 | -0.770653782 | 0.36504129 | -2.1111414 | 0.03476016 | 0.09114512 | RNASE7 | 84659 |
| ENSG000002 | 5.10795953 | -0.792835671 | 0.37555777 | -2.1110884 | 0.03476471 | 0.09115116 | NA | NA |
| ENSG000001 | 232.332679 | 0.302924435 | 0.14350264 | 2.11093287 | 0.03477808 | 0.09118032 | SH3BGRL3 | 83442 |
| ENSG000001 | 5.02611435 | -0.789641513 | 0.37408218 | -2.1108771 | 0.03478288 | 0.091187 | OR51Q1 | 390061 |
| ENSG000001 | 10.0085225 | -0.717245757 | 0.33981183 | -2.1107145 | 0.03479686 | 0.09121775 | APOD | 347 |
| ENSG000001 | 1280.11391 | 0.255770191 | 0.12118266 | 2.11061708 | 0.03480524 | 0.09122798 | FAM20B | 9917 |
| ENSG000002 | 2.54157326 | -0.77790947 | 0.36856973 | -2.1106168 | 0.03480526 | 0.09122798 | LOC645188 | 645188 |
| ENSG000002 | 7.51338121 | -0.766681314 | 0.36326489 | -2.1105296 | 0.03481276 | 0.09124174 | LINC01352 | 101929730 |
| ENSG000002 | 251.531781 | 0.328148969 | 0.15550694 | 2.11018852 | 0.03484212 | 0.09131279 | NA | NA |
| ENSG000000 | 282.554917 | 0.312683775 | 0.14818003 | 2.11016136 | 0.03484446 | 0.09131302 | IL4R | 3566 |
| ENSG000002 | 19.5050197 | -0.601632759 | 0.28513338 | -2.1100046 | 0.03485796 | 0.09134249 | NA | NA |
| ENSG000001 | 83.4931244 | 0.370484456 | 0.17559656 | 2.10986169 | 0.03487027 | 0.09136885 | BFSP1 | 631 |
| ENSG000001 | 45.2895316 | -0.474524654 | 0.22491728 | -2.1097741 | 0.03487781 | 0.09137091 | FGF14 | 2259 |
| ENSG000001 | 7.16387214 | -0.761771621 | 0.3610638 | -2.1097978 | 0.03487577 | 0.09137091 | KRT6B | 3854 |
| ENSG000002 | 4.90054798 | 0.79693687 | 0.37773454 | 2.10978025 | 0.03487729 | 0.09137091 | NA | NA |
| ENSG000001 | 3.18966941 | -0.794789378 | 0.37674425 | -2.1096258 | 0.0348906 | 0.09139259 | HAND1 | 9421 |
| ENSG000002 | 3.49908359 | -0.79746703 | 0.3780133 | -2.1096269 | 0.03489051 | 0.09139259 | NA | NA |
| ENSG000001 | 21.9517033 | -0.593739789 | 0.28144831 | -2.1095873 | 0.03489392 | 0.09139538 | PPP1R42 | 286187 |
| ENSG000002 | 5.69928944 | -0.787118047 | 0.37314731 | -2.109403 | 0.03490981 | 0.0914311 | AATK-AS1 | 388428 |
| ENSG000001 | 185.605106 | 0.294950594 | 0.1398441 | 2.10913867 | 0.03493261 | 0.09148491 | RNF150 | 57484 |

| | | | | | | | | |
|---|---|---|---|---|---|---|---|---|
| ENSG000002 | 7.12619681 | -0.758924539 | 0.35990498 | -2.1086803 | 0.03497219 | 0.09158264 | DENND5B-AS1 | 100874249 |
| ENSG000002 | 4.87567303 | -0.796275928 | 0.37764933 | -2.1085061 | 0.03498723 | 0.09161613 | NA | NA |
| ENSG000002 | 2.7953024 | -0.79006323 | 0.37471938 | -2.108413 | 0.03499528 | 0.09163127 | NA | NA |
| ENSG000002 | 2.50166591 | -0.777094521 | 0.36858195 | -2.1083358 | 0.03500196 | 0.09164284 | FAM225A | 286333 |
| ENSG000000 | 1481.21635 | 0.301554123 | 0.14303442 | 2.10826262 | 0.03500828 | 0.09164756 | TAB2 | 23118 |
| ENSG000002 | 10.5386891 | -0.704583658 | 0.33419748 | -2.1082854 | 0.03500631 | 0.09164756 | NA | NA |
| ENSG000002 | 5.94779162 | -0.776958706 | 0.3685492 | -2.1081546 | 0.03501762 | 0.09166609 | NA | NA |
| ENSG000002 | 9.16780136 | -0.746575556 | 0.3541712 | -2.107951 | 0.03503522 | 0.09170626 | NA | NA |
| ENSG000002 | 5.035853 | -0.790442446 | 0.37501993 | -2.1077345 | 0.03505396 | 0.09174938 | NA | NA |
| ENSG000001 | 8.13334215 | -0.751456255 | 0.35655147 | -2.1075674 | 0.03506843 | 0.09177552 | SSX5 | 6758 |
| ENSG000002 | 2.6723681 | -0.771295052 | 0.36596469 | -2.1075669 | 0.03506847 | 0.09177552 | NA | NA |
| ENSG000001 | 4.37157132 | -0.794606701 | 0.37703344 | -2.1075232 | 0.03507226 | 0.09177949 | CLC | 1178 |
| ENSG000002 | 20.1669229 | -0.581795064 | 0.27611057 | -2.107109 | 0.03510814 | 0.09186746 | NA | NA |
| ENSG000001 | 7.51270888 | -0.761925505 | 0.36161619 | -2.1070005 | 0.03511753 | 0.09188612 | OTP | 23440 |
| ENSG000002 | 26.0624175 | -0.551260134 | 0.26164343 | -2.1069138 | 0.03512506 | 0.09189987 | NA | NA |
| ENSG000001 | 28.3715528 | 0.534049219 | 0.25348842 | 2.10679925 | 0.03513498 | 0.09191399 | SHC2 | 25759 |
| ENSG000001 | 5970.10763 | 0.263575345 | 0.1251066 | 2.10680605 | 0.0351344 | 0.09191399 | BPTF | 2186 |
| ENSG000001 | 95.9503802 | 0.350343688 | 0.16630551 | 2.10662707 | 0.03514992 | 0.09194713 | TMEM50B | 757 |
| ENSG000002 | 4.7266635 | -0.793478985 | 0.37666388 | -2.106597 | 0.03515253 | 0.09194803 | NA | NA |
| ENSG000002 | 4.97551346 | -0.788664831 | 0.37439217 | -2.1065206 | 0.03515915 | 0.09195349 | NA | NA |
| ENSG000002 | 12.2547815 | -0.688446215 | 0.32681668 | -2.106521 | 0.03515912 | 0.09195349 | NA | NA |
| ENSG000002 | 2.9125138 | -0.784093169 | 0.37225601 | -2.1063278 | 0.03517589 | 0.09199133 | NA | NA |
| ENSG000002 | 5.92826439 | -0.776602925 | 0.36870974 | -2.1062718 | 0.03518075 | 0.0919981 | NA | NA |
| ENSG000001 | 1516.66768 | 0.510703652 | 0.24247779 | 2.10618731 | 0.03518809 | 0.09201135 | CORO1C | 23603 |
| ENSG000002 | 19.7043227 | -0.618026777 | 0.29348862 | -2.1057947 | 0.03522219 | 0.0920946 | NA | NA |
| ENSG000002 | 3.19237738 | -0.793002129 | 0.37661455 | -2.1056067 | 0.03523853 | 0.09213137 | LOC102723604 | 102723604 |
| ENSG000001 | 2165.63806 | 0.316989543 | 0.15055148 | 2.10552257 | 0.03524584 | 0.09214456 | PRR14L | 253143 |
| ENSG000002 | 2.81527868 | -0.789024446 | 0.37474712 | -2.105485 | 0.03524911 | 0.09214716 | PCDH9-AS4 | 100874087 |
| ENSG000001 | 852.196103 | 0.322459054 | 0.15316335 | 2.10532779 | 0.03526279 | 0.09217696 | RBM33 | 155435 |

| | | | | | | | | | |
|---|---|---|---|---|---|---|---|---|---|
| ENSG000001 | 10.538143 | -0.725876979 | 0.34479186 | -2.1052613 | 0.03526856 | 0.09218613 | PRPH | | 5630 |
| ENSG000002 | 6.04832846 | -0.781365925 | 0.37116718 | -2.105159 | 0.03527747 | 0.09220346 | NA | NA | |
| ENSG000002 | 4.17817705 | -0.796548345 | 0.37839505 | -2.1050707 | 0.03528515 | 0.0922176 | NA | NA | |
| ENSG000001 | 23.4849535 | -0.56283601 | 0.2674019 | -2.1048317 | 0.03530596 | 0.09226602 | C3orf20 | | 84077 |
| ENSG000002 | 5.73125318 | -0.783870457 | 0.37246005 | -2.1045759 | 0.03532824 | 0.0923183 | NA | NA | |
| ENSG000001 | 209.098982 | 0.396879366 | 0.18859657 | 2.10438275 | 0.03534507 | 0.09235634 | HSPG2 | | 3339 |
| ENSG000002 | 5.51117264 | -0.789952162 | 0.37540621 | -2.1042597 | 0.0353558 | 0.09236773 | NA | NA | |
| ENSG000002 | 11.6257432 | 0.691351288 | 0.32854755 | 2.10426557 | 0.03535529 | 0.09236773 | LOC105374366 | | 105374366 |
| ENSG000002 | 6.01489422 | -0.778479697 | 0.36995513 | -2.1042543 | 0.03535627 | 0.09236773 | NA | NA | |
| ENSG000002 | 4.23238378 | 0.796527382 | 0.37854913 | 2.10415851 | 0.03536462 | 0.09238361 | RN7SL449P | | 106481045 |
| ENSG000002 | 3.15665277 | -0.792963171 | 0.37686205 | -2.1041205 | 0.03536794 | 0.09238632 | NA | NA | |
| ENSG00000C | 17.4466066 | -0.636914056 | 0.30270958 | -2.1040433 | 0.03537467 | 0.09239795 | SPI1 | | 6688 |
| ENSG000001 | 2077.20541 | 0.255486597 | 0.1214393 | 2.10382137 | 0.03539403 | 0.09244257 | KCTD3 | | 51133 |
| ENSG000002 | 4.84006397 | -0.793197518 | 0.37706726 | -2.1035969 | 0.03541363 | 0.09248779 | NA | NA | |
| ENSG000001 | 55.5751343 | -0.42806485 | 0.20349591 | -2.103555 | 0.03541728 | 0.09249138 | RHOU | | 58480 |
| ENSG000002 | 3.13172463 | -0.792828909 | 0.37693347 | -2.1033656 | 0.03543382 | 0.09252862 | NA | NA | |
| ENSG000002 | 3.16541432 | -0.79281343 | 0.37694171 | -2.1032786 | 0.03544142 | 0.09254251 | NA | NA | |
| ENSG000002 | 2.98909956 | -0.783212245 | 0.37238388 | -2.1032388 | 0.0354449 | 0.09254562 | NA | NA | |
| ENSG000001 | 166.155732 | -0.311644193 | 0.14818103 | -2.1031315 | 0.03545427 | 0.0925595 | ARHGAP29 | | 9411 |
| ENSG000002 | 11.2246378 | -0.710495916 | 0.33782855 | -2.1031257 | 0.03545478 | 0.0925595 | NA | NA | |
| ENSG000002 | 25.1854887 | 0.54829462 | 0.26071591 | 2.10303476 | 0.03546273 | 0.0925743 | NME2 | | 4831 |
| ENSG000002 | 3.83760434 | -0.796143543 | 0.37858184 | -2.1029628 | 0.03546902 | 0.09258475 | NA | NA | |
| ENSG000002 | 500.138786 | 0.281366688 | 0.13381861 | 2.10259758 | 0.03550096 | 0.09266217 | GABBR1 | | 2550 |
| ENSG000001 | 100.228163 | 0.362492391 | 0.17240971 | 2.10250566 | 0.035509 | 0.09266898 | HGSNAT | | 138050 |
| ENSG000002 | 4.78267924 | -0.792583778 | 0.37697394 | -2.1024896 | 0.03551041 | 0.09266898 | NA | NA | |
| ENSG000002 | 2.54976069 | -0.772053814 | 0.36720937 | -2.1024894 | 0.03551042 | 0.09266898 | NA | NA | |
| ENSG000001 | 83.5857136 | -0.438103738 | 0.2083848 | -2.1023785 | 0.03552013 | 0.09268834 | NA | NA | |
| ENSG000002 | 5.94329779 | -0.778690094 | 0.37041023 | -2.102237 | 0.03553252 | 0.09270277 | NA | NA | |
| ENSG000002 | 2.79630075 | -0.787939905 | 0.37480468 | -2.1022681 | 0.0355298 | 0.09270277 | NA | NA | |

| | | | | | | | | |
|---|---|---|---|---|---|---|---|---|
| ENSG000002 | 206.841907 | 0.301151693 | 0.14325122 | 2.10226267 | 0.03553027 | 0.09270277 | LINC00294 | 283267 |
| ENSG000000 | 85.6523314 | 0.449067168 | 0.21362451 | 2.10213315 | 0.03554161 | 0.09272053 | STX1B | 112755 |
| ENSG000001 | 9.36745723 | -0.759333643 | 0.36122758 | -2.1020921 | 0.03554521 | 0.09272395 | APOBEC4 | 403314 |
| ENSG000001 | 1600.99322 | 0.265574895 | 0.12634335 | 2.10200923 | 0.03555247 | 0.09273095 | GPD2 | 2820 |
| ENSG000001 | 376.596485 | 0.396568438 | 0.18865963 | 2.10203122 | 0.03555054 | 0.09273095 | BCL9L | 283149 |
| ENSG000002 | 9.76280722 | -0.717242918 | 0.34122229 | -2.1019814 | 0.0355549 | 0.09273134 | NA | NA |
| ENSG000001 | 62.2540202 | -0.399670007 | 0.19014837 | -2.101885 | 0.03556335 | 0.09274675 | NFIA | 4774 |
| ENSG000001 | 14.7672522 | -0.648989766 | 0.308769 | -2.1018618 | 0.03556539 | 0.09274675 | RNF222 | 643904 |
| ENSG000002 | 8.7261065 | -0.747650084 | 0.35572663 | -2.1017546 | 0.03557478 | 0.09276528 | LOC105369364 | 105369364 |
| ENSG000001 | 30.0750259 | -0.518012812 | 0.24647714 | -2.1016668 | 0.03558247 | 0.09277938 | SCN10A | 6336 |
| ENSG000002 | 26.6843762 | -0.547323458 | 0.26046178 | -2.1013581 | 0.03560955 | 0.092844 | NA | NA |
| ENSG000001 | 1514.07045 | 0.303546925 | 0.14446426 | 2.10119053 | 0.03562424 | 0.09287635 | TAF7 | 6879 |
| ENSG000001 | 49.2287449 | 0.469251137 | 0.22333052 | 2.10115092 | 0.03562772 | 0.09287944 | GPR3 | 2827 |
| ENSG000002 | 6.35024246 | -0.774678348 | 0.36872395 | -2.100971 | 0.03564351 | 0.09291463 | NA | NA |
| ENSG000000 | 32.1482971 | -0.533333347 | 0.25385539 | -2.1009337 | 0.03564678 | 0.09291719 | NOS2 | 4843 |
| ENSG000001 | 66.7847861 | 0.396496384 | 0.18873079 | 2.10085689 | 0.03565353 | 0.09292283 | TMEM51 | 55092 |
| ENSG000002 | 39.9429693 | 0.469671849 | 0.22355997 | 2.10087632 | 0.03565182 | 0.09292283 | NA | NA |
| ENSG000001 | 160.914608 | 0.370539359 | 0.17639984 | 2.10056516 | 0.03567915 | 0.09298364 | PECR | 55825 |
| ENSG000002 | 4.85443082 | -0.791423576 | 0.37681497 | -2.1002976 | 0.03570267 | 0.09303895 | LINC00351 | 100874137 |
| ENSG000001 | 3.1728177 | -0.790006091 | 0.37615056 | -2.1002391 | 0.03570782 | 0.09304638 | OR4S1 | 256148 |
| ENSG000001 | 357.778328 | 0.276337292 | 0.13159111 | 2.09996926 | 0.03573155 | 0.09309495 | BCL10 | 8915 |
| ENSG000001 | 10.196445 | -0.706932676 | 0.33664282 | -2.0999488 | 0.03573335 | 0.09309495 | B3GNT6 | 192134 |
| ENSG000001 | 2599.94476 | 0.258254672 | 0.12298132 | 2.09995033 | 0.03573321 | 0.09309495 | IPO9 | 55705 |
| ENSG000001 | 28.0176982 | -0.53040873 | 0.25260682 | -2.0997404 | 0.03575169 | 0.09312591 | SAMD9L | 219285 |
| ENSG000002 | 3.4757153 | -0.793919068 | 0.37810096 | -2.0997542 | 0.03575047 | 0.09312591 | LINC00160 | 54064 |
| ENSG000002 | 6.78264994 | -0.776283425 | 0.36970535 | -2.0997355 | 0.03575212 | 0.09312591 | OR8K3 | 219473 |
| ENSG000001 | 565.177291 | 0.271520445 | 0.12931465 | 2.09968819 | 0.03575628 | 0.09313076 | SCAI | 286205 |
| ENSG000000 | 19.4969131 | -0.614917785 | 0.29290408 | -2.0993828 | 0.03578317 | 0.09319135 | SLC6A13 | 6540 |
| ENSG000002 | 12.0618178 | -0.6693022 | 0.3188107 | -2.0993718 | 0.03578414 | 0.09319135 | NA | NA |

| | | | | | | | | |
|---|---|---|---|---|---|---|---|---|
| ENSG000002 | 9.16430992 | -0.741164713 | 0.35306386 | -2.099237 | 0.03579602 | 0.09321307 | MTND6P15 | 106478949 |
| ENSG000002 | 9.16063811 | -0.72480222 | 0.34527134 | -2.099225 | 0.03579707 | 0.09321307 | NA | NA |
| ENSG000002 | 3.15665222 | -0.790218151 | 0.37644382 | -2.0991662 | 0.03580225 | 0.09322057 | NA | NA |
| ENSG000002 | 8.45482849 | 0.737154641 | 0.35117716 | 2.09909622 | 0.03580842 | 0.09323064 | RNU1-28P | 26866 |
| ENSG000001 | 76.0938329 | -0.38891713 | 0.18528871 | -2.0989791 | 0.03581874 | 0.09325153 | LOXHD1 | 125336 |
| ENSG000001 | 22.3895412 | -0.576018637 | 0.27443171 | -2.0989507 | 0.03582124 | 0.09325206 | NOV | 4856 |
| ENSG000001 | 810.941628 | 0.257334018 | 0.12261764 | 2.09867047 | 0.03584596 | 0.09330442 | BMP2K | 55589 |
| ENSG000002 | 3.27868605 | -0.788433464 | 0.3756809 | -2.0986786 | 0.03584524 | 0.09330442 | NA | NA |
| ENSG000002 | 2.96772109 | -0.782288048 | 0.37275888 | -2.0986436 | 0.03584833 | 0.0933046 | NA | NA |
| ENSG000002 | 8.59760314 | -0.736171846 | 0.35078964 | -2.0986134 | 0.03585099 | 0.09330554 | LOC105374060 | 105374060 |
| ENSG000002 | 4.45417035 | -0.792313482 | 0.37756643 | -2.0984744 | 0.03586326 | 0.09333148 | NA | NA |
| ENSG000001 | 881.224547 | 0.279411989 | 0.13315616 | 2.0983782 | 0.03587175 | 0.09334758 | DARS2 | 55157 |
| ENSG000002 | 5.63850505 | -0.779922323 | 0.37170673 | -2.0982196 | 0.03588576 | 0.09337803 | NA | NA |
| ENSG000001 | 101.030964 | 0.361851865 | 0.17247801 | 2.09795935 | 0.03590874 | 0.09341385 | FUCA1 | 2517 |
| ENSG000002 | 27.8351104 | -0.524763678 | 0.25012442 | -2.0980106 | 0.03590421 | 0.09341385 | LINC00856 | 100132987 |
| ENSG000002 | 11.2201777 | -0.708431945 | 0.33767545 | -2.097967 | 0.03590806 | 0.09341385 | NA | NA |
| ENSG000002 | 3.40023024 | -0.794003077 | 0.37845418 | -2.0980164 | 0.0359037 | 0.09341385 | LOC101927539 | 101927539 |
| ENSG000002 | 31.2669211 | 0.528440288 | 0.25188629 | 2.0979319 | 0.03591116 | 0.09341417 | NA | NA |
| ENSG000002 | 10.4252593 | -0.697152701 | 0.33233223 | -2.0977583 | 0.0359265 | 0.09344808 | C17orf78 | 284099 |
| ENSG000002 | 3.41397612 | -0.793807772 | 0.3784261 | -2.097656 | 0.03593555 | 0.09346562 | RN7SKP135 | 106480875 |
| ENSG000002 | 8.2327627 | -0.736705803 | 0.35120905 | -2.0976276 | 0.03593805 | 0.09346613 | NA | NA |
| ENSG00000C | 175.455316 | 0.301890844 | 0.14392796 | 2.09751356 | 0.03594814 | 0.09348637 | MFSD11 | 79157 |
| ENSG000002 | 5.4694577 | -0.784650298 | 0.3741429 | -2.0971941 | 0.03597639 | 0.09354185 | NA | NA |
| ENSG000002 | 17.2229188 | -0.622830421 | 0.29697809 | -2.0972268 | 0.0359735 | 0.09354185 | LOC102724612 | 102724612 |
| ENSG000002 | 4.774115 | -0.789453448 | 0.37643316 | -2.0971942 | 0.03597639 | 0.09354185 | NA | NA |
| ENSG000001 | 11.5180858 | -0.694987419 | 0.33139593 | -2.0971513 | 0.03598018 | 0.0935457 | SPATA4 | 132851 |
| ENSG000002 | 10.2190577 | -0.704888272 | 0.33616465 | -2.0968542 | 0.03600648 | 0.09360808 | NA | NA |
| ENSG000001 | 31.9737754 | -0.527588389 | 0.25161575 | -2.0968019 | 0.03601111 | 0.09361412 | GSDMC | 56169 |
| ENSG000001 | 7.1150383 | -0.759404742 | 0.3621827 | -2.0967449 | 0.03601616 | 0.09361524 | ZNF80 | 7634 |

| | | | | | | | | |
|---|---|---|---|---|---|---|---|---|
| ENSG000002 | 6.516463 | -0.778858628 | 0.3714594 | -2.096753 | 0.03601544 | 0.09361524 | NA | NA |
| ENSG000001 | 1405.82279 | 0.316664129 | 0.15105333 | 2.096373 | 0.03604912 | 0.09369489 | C2CD5 | 9847 |
| ENSG000001 | 200.486174 | 0.319565638 | 0.1524443 | 2.09627808 | 0.03605753 | 0.09371075 | ZNF443 | 10224 |
| ENSG000001 | 21.420257 | -0.580316731 | 0.27684292 | -2.096195 | 0.0360649 | 0.0937239 | ANKS4B | 257629 |
| ENSG000001 | 386.696745 | 0.331540099 | 0.15816914 | 2.09611115 | 0.03607233 | 0.09373721 | B3GALT5 | 10317 |
| ENSG000002 | 10.4302099 | 0.734961481 | 0.35067202 | 2.09586575 | 0.0360941 | 0.09378777 | HIST2H4B | 554313 |
| ENSG000002 | 3.20862196 | -0.787861274 | 0.37597632 | -2.0955077 | 0.03612589 | 0.09386434 | NA | NA |
| ENSG000002 | 5.88616556 | -0.772850906 | 0.36883014 | -2.0954115 | 0.03613443 | 0.09388053 | NA | NA |
| ENSG000002 | 10.0103895 | -0.712997463 | 0.34027163 | -2.0953773 | 0.03613746 | 0.09388239 | NA | NA |
| ENSG000002 | 8.18173814 | -0.735124763 | 0.3508371 | -2.095345 | 0.03614034 | 0.09388384 | NA | NA |
| ENSG00000C | 228.128823 | 0.281708127 | 0.13445283 | 2.09521897 | 0.03615153 | 0.09390691 | IFT80 | 57560 |
| ENSG000002 | 14.4185476 | -0.649802427 | 0.31014477 | -2.0951584 | 0.03615691 | 0.09391487 | LOC101927924 | 101927924 |
| ENSG00000C | 27.4854838 | -0.542369941 | 0.25887615 | -2.0950943 | 0.03616261 | 0.09391827 | TNIP3 | 79931 |
| ENSG000001 | 6.83437754 | -0.759655683 | 0.36258829 | -2.0950916 | 0.03616285 | 0.09391827 | IL37 | 27178 |
| ENSG000002 | 6.77334735 | -0.772729696 | 0.36885571 | -2.0949376 | 0.03617654 | 0.0939478 | LOC105375614 | 105375614 |
| ENSG000001 | 47.11128 | -0.468688716 | 0.22380066 | -2.094224 | 0.03624003 | 0.09410665 | IQUB | 154865 |
| ENSG000002 | 3.15451227 | -0.788095938 | 0.37635332 | -2.0940321 | 0.03625712 | 0.094145 | NA | NA |
| ENSG000002 | 152.496292 | 0.312464001 | 0.14922009 | 2.09398079 | 0.03626169 | 0.09415084 | C12orf75 | 387882 |
| ENSG000002 | 13.5117345 | -0.657906481 | 0.31419487 | -2.0939441 | 0.03626496 | 0.0941533 | NA | NA |
| ENSG00000C | 1896.67341 | 0.287674878 | 0.13738894 | 2.09387218 | 0.03627137 | 0.09416391 | PDS5B | 23047 |
| ENSG000002 | 10.680508 | -0.690075752 | 0.32957524 | -2.0938337 | 0.03627479 | 0.09416678 | LOC100506675 | 100506675 |
| ENSG000001 | 656.535088 | 0.243662331 | 0.11637854 | 2.09370506 | 0.03628626 | 0.09418492 | DAAM1 | 23002 |
| ENSG000001 | 53.7324739 | 0.443018028 | 0.21159651 | 2.09369256 | 0.03628738 | 0.09418492 | GCSH | 2653 |
| ENSG000002 | 2.57865607 | -0.770260378 | 0.36789836 | -2.0936771 | 0.03628875 | 0.09418492 | NA | NA |
| ENSG000002 | 3.49976077 | -0.790387191 | 0.37755133 | -2.0934562 | 0.03630845 | 0.09423001 | NA | NA |
| ENSG000001 | 837.648294 | 0.562902589 | 0.26890485 | 2.09331509 | 0.03632104 | 0.09425665 | BANF1 | 8815 |
| ENSG000002 | 9.39773889 | -0.715693136 | 0.34192432 | -2.0931332 | 0.03633727 | 0.09429274 | NA | NA |
| ENSG000002 | 115.489932 | 0.36428535 | 0.17404702 | 2.09302835 | 0.03634662 | 0.09430758 | RNU5D-1 | 26830 |
| ENSG000002 | 3.75831115 | -0.792396815 | 0.37859073 | -2.0930169 | 0.03634764 | 0.09430758 | NA | NA |

| | | | | | | | | |
|---|---|---|---|---|---|---|---|---|
| ENSG00000C | 745.830213 | 0.365237214 | 0.17452388 | 2.09276353 | 0.03637027 | 0.09435691 | NFYA | 4800 |
| ENSG000002 | 6.83692171 | -0.775985228 | 0.37079657 | -2.0927519 | 0.03637131 | 0.09435691 | NA | NA |
| ENSG000001 | 162.975929 | 0.310928784 | 0.14859611 | 2.09244235 | 0.03639896 | 0.09442262 | RGMA | 56963 |
| ENSG000001 | 508.908619 | 0.275619322 | 0.13172483 | 2.09238703 | 0.03640391 | 0.0944294 | ZNF732 | 654254 |
| ENSG000001 | 11.4343769 | -0.692154258 | 0.33081003 | -2.0923013 | 0.03641157 | 0.09444324 | TTLL10 | 254173 |
| ENSG000001 | 7784.57376 | 0.232291029 | 0.11102515 | 2.092238 | 0.03641723 | 0.09445188 | USO1 | 8615 |
| ENSG000002 | 5.66917487 | -0.776277637 | 0.37106315 | -2.0920365 | 0.03643525 | 0.09449258 | LOC105372941 | 105372941 |
| ENSG000001 | 169.162684 | 0.326438515 | 0.15605436 | 2.09182564 | 0.03645412 | 0.09453545 | PSEN2 | 5664 |
| ENSG000001 | 23.9121839 | -0.585399626 | 0.2798988 | -2.0914689 | 0.03648605 | 0.09460191 | IL18R1 | 8809 |
| ENSG000001 | 3.50362996 | -0.789727747 | 0.37759645 | -2.0914597 | 0.03648688 | 0.09460191 | OR5P3 | 120066 |
| ENSG000002 | 15.8257231 | -0.636470607 | 0.30432242 | -2.0914351 | 0.03648907 | 0.09460191 | CDKL4 | 344387 |
| ENSG000002 | 5.6672297 | -0.77798269 | 0.37197714 | -2.0914798 | 0.03648507 | 0.09460191 | NA | NA |
| ENSG000001 | 10.66406 | -0.704765395 | 0.33698194 | -2.091404 | 0.03649186 | 0.09460309 | FPR1 | 2357 |
| ENSG000001 | 665.509284 | 0.312629115 | 0.14948965 | 2.09130948 | 0.03650033 | 0.09461652 | LSM12 | 124801 |
| ENSG000002 | 10.2754378 | -0.706048735 | 0.33761332 | -2.0912941 | 0.03650171 | 0.09461652 | NA | NA |
| ENSG000001 | 25.381149 | -0.547879618 | 0.26199307 | -2.0911989 | 0.03651024 | 0.09462529 | PITX2 | 5308 |
| ENSG000002 | 6.00602852 | -0.774891209 | 0.37055245 | -2.0911782 | 0.03651209 | 0.09462529 | NA | NA |
| ENSG000002 | 15.0251425 | -0.652028482 | 0.31179456 | -2.0912118 | 0.03650908 | 0.09462529 | NA | NA |
| ENSG000001 | 68.6725399 | -0.401950443 | 0.19221938 | -2.0911026 | 0.03651886 | 0.09463679 | IFI16 | 3428 |
| ENSG000001 | 71.2131469 | 0.409424023 | 0.19581638 | 2.09085685 | 0.0365409 | 0.09468784 | MMP17 | 4326 |
| ENSG000002 | 2.8455717 | -0.779927891 | 0.37302679 | -2.0908093 | 0.03654516 | 0.09469283 | NA | NA |
| ENSG000001 | 9.43129164 | -0.7163911 | 0.34270193 | -2.0904204 | 0.03658006 | 0.09477719 | MUC7 | 4589 |
| ENSG000001 | 398.472597 | 0.314517589 | 0.15047258 | 2.09019867 | 0.03659996 | 0.0948227 | ELP5 | 23587 |
| ENSG000002 | 4.46930688 | -0.789207784 | 0.37762076 | -2.0899481 | 0.03662246 | 0.09487494 | TRIM34 | 53840 |
| ENSG000002 | 30.0442884 | -0.52660774 | 0.25201263 | -2.0896085 | 0.03665298 | 0.09494794 | NA | NA |
| ENSG00000C | 200.355537 | 0.301931907 | 0.14450143 | 2.08947352 | 0.03666512 | 0.09497332 | CTNNA2 | 1496 |
| ENSG000001 | 548.888224 | 0.269652716 | 0.1290704 | 2.08919095 | 0.03669054 | 0.09502702 | B3GALNT2 | 148789 |
| ENSG000001 | 8.71788557 | -0.73859046 | 0.35352878 | -2.0891947 | 0.0366902 | 0.09502702 | OSR2 | 116039 |
| ENSG000001 | 55.3978686 | 0.456861255 | 0.21868719 | 2.08910849 | 0.03669796 | 0.0950341 | ESPN | 83715 |

| | | | | | | | | |
|---|---|---|---|---|---|---|---|---|
| ENSG000002 | 7.17690096 | -0.754513946 | 0.36116403 | -2.0891171 | 0.03669718 | 0.0950341 | FLJ46066 | 401103 |
| ENSG000001 | 133.366464 | 0.377743079 | 0.1808213 | 2.08904087 | 0.03670404 | 0.09504229 | TMEM39B | 55116 |
| ENSG000001 | 82.4041237 | 0.382106311 | 0.1829162 | 2.08896918 | 0.0367105 | 0.09504229 | SH2D3A | 10045 |
| ENSG000002 | 5.27950271 | -0.778245269 | 0.37254652 | -2.0889882 | 0.03670879 | 0.09504229 | PP14571 | 100130449 |
| ENSG000002 | 5.26199975 | -0.77691923 | 0.37191029 | -2.0889963 | 0.03670805 | 0.09504229 | NA | NA |
| ENSG000002 | 9.19594282 | -0.7279444 | 0.34848384 | -2.08889 | 0.03671763 | 0.09504861 | NA | NA |
| ENSG000002 | 173.856736 | 0.348347731 | 0.16676155 | 2.08889714 | 0.03671698 | 0.09504861 | NA | NA |
| ENSG000002 | 17.079218 | -0.634514145 | 0.30377482 | -2.0887648 | 0.0367289 | 0.09507172 | NA | NA |
| ENSG000002 | 16.5880497 | -0.641543957 | 0.30717702 | -2.0885155 | 0.03675136 | 0.0951238 | LOC101928940 | 101928940 |
| ENSG000001 | 20.51055 | 0.632177711 | 0.30269698 | 2.08848374 | 0.03675422 | 0.09512512 | SOWAHD | 347454 |
| ENSG000002 | 4.50264103 | -0.789308874 | 0.37795333 | -2.0883765 | 0.03676388 | 0.09514405 | NA | NA |
| ENSG000000 | 14.4138391 | 0.652055023 | 0.31224223 | 2.08829862 | 0.03677091 | 0.09515616 | DDTL | 100037417 |
| ENSG000002 | 5.71939503 | -0.78701528 | 0.37691509 | -2.088044 | 0.03679387 | 0.0952095 | NA | NA |
| ENSG000002 | 2.40921346 | -0.774104926 | 0.37074646 | -2.0879631 | 0.03680116 | 0.09521623 | NA | NA |
| ENSG000002 | 5.65623778 | -0.775776729 | 0.3715471 | -2.0879633 | 0.03680114 | 0.09521623 | NA | NA |
| ENSG000002 | 3.21733254 | -0.785830125 | 0.37637012 | -2.0879184 | 0.03680519 | 0.09522058 | NA | NA |
| ENSG000001 | 9.63171246 | -0.708298993 | 0.33925689 | -2.0877955 | 0.03681629 | 0.09523374 | DAOA | 267012 |
| ENSG000001 | 12.7911331 | -0.689223324 | 0.33011402 | -2.0878342 | 0.0368128 | 0.09523374 | NA | NA |
| ENSG000002 | 6.48168638 | -0.763246897 | 0.36557752 | -2.087784 | 0.03681733 | 0.09523374 | NA | NA |
| ENSG000002 | 2.47701797 | -0.767427457 | 0.36758685 | -2.0877446 | 0.03682088 | 0.09523687 | NA | NA |
| ENSG000002 | 7.1100751 | -0.753987113 | 0.36115433 | -2.0877145 | 0.0368236 | 0.09523782 | NA | NA |
| ENSG000002 | 15.9358092 | -0.637802248 | 0.30552274 | -2.087577 | 0.03683601 | 0.09526384 | NA | NA |
| ENSG000002 | 5.88199113 | -0.769885776 | 0.36881863 | -2.0874373 | 0.03684862 | 0.0952843 | LOC101928516 | 101928516 |
| ENSG000002 | 19.7503892 | -0.600557466 | 0.2876981 | -2.0874572 | 0.03684683 | 0.0952843 | TMEM229A | 730130 |
| ENSG000001 | 4.37339861 | -0.787390713 | 0.37722591 | -2.0873187 | 0.03685933 | 0.09529377 | XAGE5 | 170627 |
| ENSG000002 | 8.57628737 | -0.729019641 | 0.34925951 | -2.0873294 | 0.03685837 | 0.09529377 | LOC101929128 | 101929128 |
| ENSG000002 | 3.23161976 | -0.78541562 | 0.37627247 | -2.0873587 | 0.03685572 | 0.09529377 | NA | NA |
| ENSG000001 | 16.1143085 | -0.637843561 | 0.30564106 | -2.0869041 | 0.03689681 | 0.09538458 | LINC01558 | 26238 |
| ENSG000002 | 17.5189154 | -0.62017272 | 0.29720248 | -2.086701 | 0.03691517 | 0.09542596 | NA | NA |

| | | | | | | | | |
|---|---|---|---|---|---|---|---|---|
| ENSG000001 | 106.533412 | 0.352470216 | 0.16891529 | 2.08666847 | 0.03691811 | 0.09542749 | PPOX | 5498 |
| ENSG000002 | 8.50811162 | -0.727815212 | 0.34880483 | -2.0865973 | 0.03692455 | 0.09543804 | NA | NA |
| ENSG000002 | 13.3489445 | -0.702392848 | 0.33663034 | -2.0865405 | 0.03692969 | 0.09544524 | NA | NA |
| ENSG000002 | 9.4015179 | -0.717699786 | 0.34401265 | -2.0862599 | 0.03695509 | 0.0955048 | CARD16 | 114769 |
| ENSG000002 | 4.80260379 | -0.786556414 | 0.37703374 | -2.0861698 | 0.03696324 | 0.09551978 | NA | NA |
| ENSG00000C | 72.6439091 | -0.406479654 | 0.19484825 | -2.0861345 | 0.03696645 | 0.09552197 | ABCB4 | 5244 |
| ENSG000001 | 601.550011 | 0.240420448 | 0.11525887 | 2.08591708 | 0.03698613 | 0.09556676 | TAF10 | 6881 |
| ENSG000002 | 6.75066806 | -0.755952988 | 0.36242182 | -2.0858374 | 0.03699336 | 0.09557933 | NA | NA |
| ENSG000001 | 13.5426773 | -0.660202013 | 0.31653646 | -2.085706 | 0.03700526 | 0.0955927 | TRIM63 | 84676 |
| ENSG000002 | 2.42225437 | -0.772009267 | 0.37013822 | -2.0857324 | 0.03700287 | 0.0955927 | NA | NA |
| ENSG000002 | 4.85836051 | -0.787744026 | 0.3777061 | -2.0856005 | 0.03701483 | 0.0955927 | NA | NA |
| ENSG000002 | 5.94323151 | -0.77093847 | 0.36963711 | -2.085663 | 0.03700916 | 0.0955927 | NA | NA |
| ENSG000002 | 3.74962868 | -0.789626081 | 0.37859307 | -2.0856855 | 0.03700712 | 0.0955927 | NA | NA |
| ENSG000002 | 3.16309114 | -0.785165589 | 0.37647021 | -2.0855982 | 0.03701503 | 0.0955927 | NA | NA |
| ENSG000002 | 2.2160071 | -0.750614292 | 0.3599028 | -2.0856028 | 0.03701462 | 0.0955927 | NA | NA |
| ENSG000002 | 60.9519107 | 0.399690816 | 0.19164682 | 2.08555938 | 0.03701855 | 0.09559571 | LOC148709 | 148709 |
| ENSG000001 | 2.40306792 | -0.773118014 | 0.37073154 | -2.085385 | 0.03703436 | 0.09562626 | NKX2-1 | 7080 |
| ENSG000002 | 24.6384925 | 0.590831323 | 0.28332396 | 2.085356 | 0.037037 | 0.09562626 | SNORD14A | 26822 |
| ENSG000002 | 13.3077596 | -0.659837241 | 0.31641545 | -2.0853509 | 0.03703746 | 0.09562626 | NA | NA |
| ENSG000002 | 4.7784525 | -0.786086197 | 0.37696199 | -2.0853195 | 0.03704031 | 0.09562752 | NA | NA |
| ENSG000001 | 29.4446895 | -0.549088338 | 0.26331723 | -2.0852731 | 0.03704452 | 0.0956323 | LMOD3 | 56203 |
| ENSG000002 | 4.67056929 | -0.783842989 | 0.37591552 | -2.0851573 | 0.03705502 | 0.09565334 | NA | NA |
| ENSG000001 | 79.3528731 | 0.378380404 | 0.18146726 | 2.08511666 | 0.03705871 | 0.09565677 | THNSL2 | 55258 |
| ENSG000002 | 14.6218033 | -0.659056101 | 0.31612576 | -2.0847909 | 0.03708829 | 0.09572701 | LINC00968 | 100507632 |
| ENSG000002 | 4.10857044 | -0.789178832 | 0.37855895 | -2.0846921 | 0.03709726 | 0.09574409 | NA | NA |
| ENSG000002 | 4.09310993 | -0.789008001 | 0.37848694 | -2.0846373 | 0.03710224 | 0.09575083 | RNU6-456P | 106481323 |
| ENSG000002 | 2.769774 | -0.779764678 | 0.37407266 | -2.0845273 | 0.03711223 | 0.09577052 | NA | NA |
| ENSG000001 | 254.241008 | 0.267195766 | 0.12819137 | 2.08435067 | 0.03712828 | 0.09580585 | KLHL2 | 11275 |
| ENSG000002 | 6.57476989 | -0.764114336 | 0.36663035 | -2.0841546 | 0.0371461 | 0.09584574 | NA | NA |

| | | | | | | | | |
|---|---|---|---|---|---|---|---|---|
| ENSG000001 | 543.614316 | 0.287528378 | 0.1379644 | 2.0840766 | 0.0371532 | 0.09585795 | TRIAP1 | 51499 |
| ENSG000002 | 4.36070151 | -0.78682998 | 0.37755123 | -2.084035 | 0.03715698 | 0.09586161 | NA | NA |
| ENSG000002 | 2.7667015 | -0.780571591 | 0.37456394 | -2.0839475 | 0.03716494 | 0.09587604 | NA | NA |
| ENSG000001 | 15.4424061 | -0.631031153 | 0.30281761 | -2.0838654 | 0.03717241 | 0.0958891 | SLC6A1 | 6529 |
| ENSG000002 | 9.57792581 | -0.723813201 | 0.34734588 | -2.0838399 | 0.03717473 | 0.0958891 | NA | NA |
| ENSG000001 | 109.747952 | -0.33761937 | 0.16202739 | -2.0837179 | 0.03718584 | 0.09590554 | HHIP | 64399 |
| ENSG000002 | 2.39820039 | -0.772475245 | 0.37071685 | -2.0837339 | 0.03718438 | 0.09590554 | NA | NA |
| ENSG000002 | 2.42745418 | -0.771725051 | 0.37036896 | -2.0836656 | 0.0371906 | 0.09591172 | NA | NA |
| ENSG00000C | 2034.75594 | 0.29046834 | 0.1394429 | 2.08306293 | 0.03724549 | 0.09603496 | FOXN3 | 1112 |
| ENSG000001 | 87.5135197 | 0.3548075 | 0.17032929 | 2.0830681 | 0.03724502 | 0.09603496 | ST3GAL4 | 6484 |
| ENSG000001 | 7.68527076 | -0.741249407 | 0.35583877 | -2.0831047 | 0.03724168 | 0.09603496 | OPN1SW | 611 |
| ENSG000001 | 10.5011993 | -0.697608283 | 0.33491633 | -2.082933 | 0.03725733 | 0.09605058 | GJB2 | 2706 |
| ENSG000002 | 3.09979011 | -0.784818 | 0.37678766 | -2.0829185 | 0.03725865 | 0.09605058 | NA | NA |
| ENSG000002 | 17.7788418 | -0.617356694 | 0.29638495 | -2.0829556 | 0.03725527 | 0.09605058 | AKR1C8P | 340811 |
| ENSG000002 | 3.39240177 | -0.788190088 | 0.37842852 | -2.0827978 | 0.03726966 | 0.09607284 | NA | NA |
| ENSG000002 | 3.15318688 | -0.783146001 | 0.37606673 | -2.0824655 | 0.03729997 | 0.09614487 | NA | NA |
| ENSG000002 | 339.334359 | 0.284322654 | 0.1365383 | 2.08236554 | 0.03730909 | 0.09615166 | NA | NA |
| ENSG000002 | 3.14168096 | -0.784779021 | 0.37687024 | -2.0823587 | 0.03730972 | 0.09615166 | NA | NA |
| ENSG000002 | 5.6750074 | -0.7736804 | 0.37153786 | -2.082373 | 0.03730841 | 0.09615166 | NA | NA |
| ENSG000002 | 78.4698653 | 0.385631151 | 0.18519945 | 2.08224787 | 0.03731984 | 0.09617162 | NA | NA |
| ENSG000001 | 357.146161 | 0.300528316 | 0.14433618 | 2.08214132 | 0.03732956 | 0.09618824 | FAM122A | 116224 |
| ENSG000002 | 11.8730246 | -0.679583311 | 0.32638925 | -2.0821253 | 0.03733103 | 0.09618824 | NA | NA |
| ENSG000001 | 5.12266467 | -0.782922088 | 0.37606694 | -2.081869 | 0.03735444 | 0.09623375 | NA | NA |
| ENSG000002 | 19.5397634 | -0.593291955 | 0.28497895 | -2.0818799 | 0.03735344 | 0.09623375 | ADIPOQ-AS1 | 100874095 |
| ENSG000002 | 2.18726981 | -0.749776969 | 0.36014868 | -2.081854 | 0.03735581 | 0.09623375 | NA | NA |
| ENSG000002 | 8.51010667 | -0.728468689 | 0.34992271 | -2.0817989 | 0.03736085 | 0.09624061 | ZNF728 | 388523 |
| ENSG000002 | 19.3651919 | -0.619505197 | 0.29759578 | -2.0817002 | 0.03736986 | 0.09625771 | NA | NA |
| ENSG000001 | 347.585968 | 0.308051876 | 0.14798488 | 2.08164425 | 0.03737498 | 0.09626478 | ASXL1 | 171023 |
| ENSG000001 | 14.7933185 | -0.68717792 | 0.33012636 | -2.0815603 | 0.03738265 | 0.09627843 | IL1RL2 | 8808 |

| | | | | | | | | |
|---|---|---|---|---|---|---|---|---|
| ENSG000001 | 7.45136777 | -0.748776097 | 0.35976798 | -2.081275 | 0.03740875 | 0.09633952 | GPR88 | 54112 |
| ENSG000001 | 2702.06544 | 0.261433315 | 0.12561787 | 2.08117929 | 0.0374175 | 0.09635594 | RAB3GAP2 | 25782 |
| ENSG000001 | 61.8593597 | -0.426667355 | 0.20502285 | -2.0810722 | 0.03742729 | 0.09637504 | PCDHB3 | 56132 |
| ENSG000001 | 2.78420556 | -0.779047311 | 0.37437188 | -2.0809451 | 0.03743894 | 0.09639278 | BECN2 | 441925 |
| ENSG000002 | 4.45024442 | -0.785852636 | 0.37764165 | -2.080948 | 0.03743866 | 0.09639278 | NA | NA |
| ENSG000001 | 11.6202865 | -0.678278088 | 0.32597704 | -2.0807542 | 0.03745641 | 0.09640716 | MEOX2 | 4223 |
| ENSG000001 | 18.9933118 | -0.603171223 | 0.28987408 | -2.0808043 | 0.03745182 | 0.09640716 | NA | NA |
| ENSG000002 | 7.97712173 | -0.738815581 | 0.35506535 | -2.0807876 | 0.03745335 | 0.09640716 | NA | NA |
| ENSG000002 | 22.7888829 | 0.590308338 | 0.28369598 | 2.08077794 | 0.03745424 | 0.09640716 | LOC101928707 | 101928707 |
| ENSG000002 | 5.86799492 | -0.779029885 | 0.37439676 | -2.0807602 | 0.03745586 | 0.09640716 | NA | NA |
| ENSG000002 | 2.9609268 | -0.772250336 | 0.37117513 | -2.0805552 | 0.03747464 | 0.09644797 | NA | NA |
| ENSG000002 | 31.3454742 | 0.547396072 | 0.26311742 | 2.08042504 | 0.03748656 | 0.09647253 | NA | NA |
| ENSG000001 | 12.1089402 | -0.668193883 | 0.32118639 | -2.0803929 | 0.03748951 | 0.096474 | GPRC6A | 222545 |
| ENSG000002 | 19.2984831 | -0.598274223 | 0.28758276 | -2.080355 | 0.03749298 | 0.09647477 | PCAT4 | 118425 |
| ENSG000002 | 3.15201357 | -0.783075702 | 0.37641759 | -2.0803377 | 0.03749457 | 0.09647477 | LOC101929504 | 101929504 |
| ENSG000001 | 290.983338 | 0.31210718 | 0.15002947 | 2.08030584 | 0.03749749 | 0.09647616 | SNAPC5 | 10302 |
| ENSG000001 | 49.1025719 | -0.431719618 | 0.2075454 | -2.0801214 | 0.0375144 | 0.09651354 | CASC2 | 255082 |
| ENSG000001 | 123.184887 | 0.337592699 | 0.16230341 | 2.08000989 | 0.03752463 | 0.09653022 | CDYL2 | 124359 |
| ENSG000002 | 71.4319706 | 0.466413609 | 0.22423744 | 2.07999877 | 0.03752565 | 0.09653022 | LOC652276 | 652276 |
| ENSG000001 | 223.033625 | 0.296017972 | 0.14233059 | 2.0797917 | 0.03754464 | 0.09657297 | ZNF14 | 7561 |
| ENSG000001 | 20.1936709 | -0.61030683 | 0.29345446 | -2.0797327 | 0.03755006 | 0.09657928 | DRD2 | 1813 |
| ENSG000001 | 163.377899 | 0.339524517 | 0.1632556 | 2.07971125 | 0.03755203 | 0.09657928 | DACT1 | 51339 |
| ENSG000002 | 23.7963211 | -0.587142812 | 0.2823277 | -2.0796501 | 0.03755764 | 0.09657928 | SAMD13 | 148418 |
| ENSG000002 | 2.54003468 | -0.765005078 | 0.36785112 | -2.0796595 | 0.03755677 | 0.09657928 | NA | NA |
| ENSG000002 | 11.795702 | -0.689921403 | 0.33175116 | -2.0796352 | 0.037559 | 0.09657928 | LOC100505851 | 100505851 |
| ENSG000002 | 5.92626988 | -0.766682785 | 0.36867015 | -2.0795901 | 0.03756315 | 0.0965838 | NA | NA |
| ENSG000001 | 475.731642 | 0.611509738 | 0.29408367 | 2.07937335 | 0.03758305 | 0.09662885 | RMI1 | 80010 |
| ENSG000002 | 5.71449918 | -0.7759235 | 0.37317791 | -2.0792322 | 0.03759602 | 0.09665607 | NA | NA |
| ENSG000002 | 7.68356689 | -0.763674201 | 0.36731386 | -2.0790781 | 0.03761017 | 0.09668633 | NA | NA |

| | | | | | | | | |
|---|---|---|---|---|---|---|---|---|
| ENSG000001 | 149.489152 | 0.36163157 | 0.17394959 | 2.07894459 | 0.03762244 | 0.09670561 | NFIL3 | 4783 |
| ENSG000002 | 8.28412366 | -0.735905462 | 0.35397845 | -2.0789555 | 0.03762144 | 0.09670561 | NA | NA |
| ENSG000002 | 10.6317084 | -0.702673981 | 0.33800309 | -2.0788981 | 0.03762671 | 0.09671046 | LINC00534 | 100874052 |
| ENSG000001 | 326.615749 | 0.365704876 | 0.1759166 | 2.07885367 | 0.0376308 | 0.09671483 | TRIM56 | 81844 |
| ENSG000002 | 6.32356238 | -0.770192316 | 0.37052008 | -2.078679 | 0.03764686 | 0.09674998 | NA | NA |
| ENSG000002 | 25.3728241 | -0.547370405 | 0.26333399 | -2.0786166 | 0.0376526 | 0.0967586 | KLF3-AS1 | 79667 |
| ENSG000002 | 23.8074001 | -0.560496335 | 0.26965361 | -2.078579 | 0.03765606 | 0.09676136 | NA | NA |
| ENSG000002 | 3.8209022 | -0.786738502 | 0.37850698 | -2.078531 | 0.03766048 | 0.0967661 | NA | NA |
| ENSG000002 | 5.13765575 | -0.783339498 | 0.37687604 | -2.078507 | 0.03766268 | 0.0967661 | NA | NA |
| ENSG000002 | 2.56017349 | -0.761885037 | 0.36656777 | -2.0784289 | 0.03766987 | 0.09677845 | NA | NA |
| ENSG000002 | 3.91680213 | -0.786381073 | 0.37835899 | -2.0783994 | 0.03767259 | 0.0967793 | NA | NA |
| ENSG00000C | 570.188658 | 0.254736492 | 0.12258362 | 2.07806304 | 0.03770355 | 0.0968527 | CYLD | 1540 |
| ENSG000001 | 7.72629621 | -0.744046484 | 0.35806648 | -2.0779563 | 0.03771339 | 0.09687183 | FCAR | 2204 |
| ENSG000002 | 2.82509752 | -0.776262529 | 0.3736534 | -2.0774935 | 0.03775603 | 0.09697522 | LINC00317 | 378828 |
| ENSG000001 | 590.00556 | 0.284383419 | 0.13689447 | 2.07739161 | 0.03776543 | 0.09698976 | ZNFX1 | 57169 |
| ENSG000001 | 2028.45828 | 0.288538575 | 0.1388954 | 2.07738028 | 0.03776647 | 0.09698976 | DAG1 | 1605 |
| ENSG000001 | 5.04693775 | -0.779994182 | 0.37548434 | -2.0773015 | 0.03777374 | 0.09700228 | GZMA | 3001 |
| ENSG000002 | 8.76171732 | -0.717187954 | 0.34525573 | -2.0772659 | 0.03777702 | 0.09700424 | GRXCR1 | 389207 |
| ENSG000002 | 17.4506597 | -0.619812535 | 0.29838253 | -2.0772414 | 0.03777929 | 0.09700424 | NA | NA |
| ENSG000002 | 4.56619845 | -0.786237553 | 0.37851374 | -2.0771704 | 0.03778583 | 0.0970149 | NA | NA |
| ENSG000001 | 113.101363 | 0.339901737 | 0.16365644 | 2.07692246 | 0.03780872 | 0.09706751 | F3 | 2152 |
| ENSG000001 | 38.2177219 | -0.500449881 | 0.24096263 | -2.0768776 | 0.03781286 | 0.09707164 | NA | NA |
| ENSG000001 | 288.674064 | 0.275288043 | 0.13255056 | 2.07685317 | 0.03781511 | 0.09707164 | PIGC | 5279 |
| ENSG000001 | 144.296964 | 0.346856898 | 0.1670172 | 2.0767735 | 0.03782247 | 0.09708054 | ZSCAN9 | 7746 |
| ENSG000002 | 50.5028453 | 0.441246675 | 0.2124684 | 2.07676374 | 0.03782337 | 0.09708054 | NA | NA |
| ENSG000002 | 11.8859792 | -0.674960592 | 0.32503268 | -2.076593 | 0.03783914 | 0.09711204 | IGSF11-AS1 | 100506765 |
| ENSG000002 | 12.7072226 | -0.687173329 | 0.33091605 | -2.076579 | 0.03784043 | 0.09711204 | NA | NA |
| ENSG000001 | 118.207702 | 0.391148677 | 0.18836992 | 2.07649227 | 0.03784845 | 0.09712474 | MAPK13 | 5603 |
| ENSG000002 | 3.55508375 | -0.783127156 | 0.37714284 | -2.0764736 | 0.03785017 | 0.09712474 | NA | NA |

| | | | | | | | | |
|---|---|---|---|---|---|---|---|---|
| ENSG000002 | 5.19380843 | -0.782802184 | 0.3769935 | -2.0764341 | 0.03785382 | 0.09712787 | NA | NA |
| ENSG000002 | 17.295454 | -0.60844653 | 0.29302833 | -2.0764086 | 0.03785618 | 0.09712787 | ERC2-IT1 | 711 |
| ENSG000001 | 13.9312501 | -0.658333884 | 0.317068 | -2.0763176 | 0.03786458 | 0.09713647 | NA | NA |
| ENSG000002 | 2.42057143 | -0.768539279 | 0.37014945 | -2.0762945 | 0.03786672 | 0.09713647 | NA | NA |
| ENSG000002 | 3.53117934 | -0.783625158 | 0.37741122 | -2.0763165 | 0.03786469 | 0.09713647 | NA | NA |
| ENSG000002 | 4.03016706 | -0.784472074 | 0.37785858 | -2.0760997 | 0.03788473 | 0.09717654 | NA | NA |
| ENSG000002 | 3.63584238 | -0.782406529 | 0.37686961 | -2.0760669 | 0.03788776 | 0.09717816 | NA | NA |
| ENSG000001 | 8.07607113 | -0.739424186 | 0.35617292 | -2.0760258 | 0.03789156 | 0.09718176 | NA | NA |
| ENSG000001 | 7792.04278 | -0.296924819 | 0.14303159 | -2.0759387 | 0.03789962 | 0.09719628 | ND6 | 4541 |
| ENSG000001 | 22.1875847 | -0.584156721 | 0.28141998 | -2.0757472 | 0.03791734 | 0.09723339 | HNF4A | 3172 |
| ENSG000002 | 356.282393 | 0.280801954 | 0.13527862 | 2.07573044 | 0.03791889 | 0.09723339 | THUMPD3-AS1 | 440944 |
| ENSG000002 | 2.81157998 | -0.775648164 | 0.3736811 | -2.0756955 | 0.03792213 | 0.09723555 | NA | NA |
| ENSG000001 | 230.277885 | 0.324156099 | 0.15617934 | 2.07553758 | 0.03793674 | 0.09724899 | CCDC136 | 64753 |
| ENSG000001 | 24.6449496 | -0.570648903 | 0.27493979 | -2.0755414 | 0.03793639 | 0.09724899 | TCL6 | 27004 |
| ENSG000002 | 5.38268892 | -0.776597322 | 0.37416962 | -2.0755221 | 0.03793817 | 0.09724899 | STEAP3-AS1 | 100874111 |
| ENSG000002 | 15.6498757 | 0.643746627 | 0.31015867 | 2.07553965 | 0.03793655 | 0.09724899 | RNF148 | 378925 |
| ENSG000002 | 4.19427781 | -0.785674824 | 0.37854556 | -2.0755093 | 0.03793936 | 0.09724899 | NA | NA |
| ENSG00000C | 18.5216521 | -0.607307023 | 0.29261575 | -2.075442 | 0.03794558 | 0.09725406 | LAMP3 | 27074 |
| ENSG000002 | 4.35862627 | -0.784013238 | 0.37775831 | -2.0754361 | 0.03794613 | 0.09725406 | NA | NA |
| ENSG000002 | 6.75273373 | -0.767455737 | 0.3699076 | -2.0747228 | 0.03801223 | 0.0974173 | NA | NA |
| ENSG000001 | 129.957265 | 0.329983533 | 0.15906211 | 2.07455769 | 0.03802755 | 0.09743808 | PLCB2 | 5330 |
| ENSG000002 | 4.44915488 | -0.783412184 | 0.37761992 | -2.074605 | 0.03802316 | 0.09743808 | NA | NA |
| ENSG000002 | 5.80488269 | -0.775741926 | 0.37392971 | -2.0745662 | 0.03802676 | 0.09743808 | NA | NA |
| ENSG000001 | 776.585037 | 0.247892039 | 0.11949551 | 2.07448828 | 0.03803399 | 0.09744842 | FAM175A | 84142 |
| ENSG000001 | 13.4180801 | -0.672196794 | 0.32404548 | -2.0743903 | 0.03804308 | 0.09746555 | OR2T12 | 127064 |
| ENSG000002 | 7.13527158 | -0.749312656 | 0.36123913 | -2.0742843 | 0.03805292 | 0.0974823 | NA | NA |
| ENSG000002 | 19.9831068 | 0.579793334 | 0.27951707 | 2.07426807 | 0.03805442 | 0.0974823 | NA | NA |
| ENSG000002 | 4.64819824 | -0.779873464 | 0.37598457 | -2.0742167 | 0.03805919 | 0.09748837 | LOC102723544 | 102723544 |
| ENSG000002 | 57.1850397 | -0.457113819 | 0.22038543 | -2.0741562 | 0.0380648 | 0.09749658 | LINC00907 | 284260 |

| | | | | | | | | |
|---|---|---|---|---|---|---|---|---|
| ENSG000001 | 12.2065807 | -0.692185334 | 0.33372856 | -2.0740968 | 0.03807032 | 0.09750455 | LINC00670 | 284034 |
| ENSG000000 | 55.3139658 | 0.447794395 | 0.21593143 | 2.07378051 | 0.0380997 | 0.09757363 | RASSF1 | 11186 |
| ENSG000002 | 11.0166447 | -0.707223204 | 0.34105443 | -2.0736373 | 0.03811301 | 0.09760155 | NA | NA |
| ENSG000002 | 5.019368 | -0.77718639 | 0.37483926 | -2.0733858 | 0.03813639 | 0.09765205 | NA | NA |
| ENSG000002 | 3.80947951 | -0.784924192 | 0.37857349 | -2.0733734 | 0.03813754 | 0.09765205 | NA | NA |
| ENSG000002 | 2.42250287 | -0.767990135 | 0.3704356 | -2.0732083 | 0.0381529 | 0.09768521 | NA | NA |
| ENSG000001 | 297.952635 | 0.279928652 | 0.13503436 | 2.07301803 | 0.0381706 | 0.09771202 | ZC3H8 | 84524 |
| ENSG000001 | 920.688891 | 0.3158787 | 0.15237283 | 2.0730645 | 0.03816628 | 0.09771202 | PTMS | 5763 |
| ENSG000002 | 4.00984621 | -0.783801449 | 0.37809541 | -2.0730255 | 0.0381699 | 0.09771202 | NA | NA |
| ENSG000002 | 63.690802 | 0.452262229 | 0.21817788 | 2.07290593 | 0.03818104 | 0.09773256 | NA | NA |
| ENSG000002 | 3.83592187 | -0.783809497 | 0.37812977 | -2.0728585 | 0.03818545 | 0.0977377 | SENCR | 100507392 |
| ENSG000001 | 10.4607509 | -0.714738669 | 0.34481677 | -2.0728072 | 0.03819023 | 0.09774375 | FABP2 | 2169 |
| ENSG000002 | 12.0980519 | -0.665323475 | 0.32100328 | -2.0726376 | 0.03820602 | 0.09777799 | NA | NA |
| ENSG000002 | 3.38465762 | -0.784181248 | 0.37838957 | -2.0724177 | 0.0382265 | 0.09782424 | OR1B1 | 347169 |
| ENSG000001 | 4919.24363 | 0.281783974 | 0.13597574 | 2.07231058 | 0.03823649 | 0.09784361 | SMC4 | 10051 |
| ENSG000002 | 10.1838204 | -0.73210138 | 0.3532914 | -2.072231 | 0.0382439 | 0.09785642 | NA | NA |
| ENSG000001 | 307.252356 | 0.302002504 | 0.14575133 | 2.0720395 | 0.03826176 | 0.09788855 | POP1 | 10940 |
| ENSG000001 | 7.16234544 | -0.752546145 | 0.36319468 | -2.0720186 | 0.0382637 | 0.09788855 | DEFB119 | 245932 |
| ENSG000002 | 5.66554122 | -0.770551815 | 0.37187562 | -2.0720686 | 0.03825904 | 0.09788855 | NA | NA |
| ENSG000001 | 9.06230921 | -0.734337018 | 0.35441393 | -2.0719756 | 0.03826772 | 0.09789265 | NA | NA |
| ENSG000002 | 14.2841371 | -0.657708125 | 0.31744209 | -2.0718996 | 0.03827481 | 0.09790461 | NA | NA |
| ENSG000001 | 6.77946559 | -0.7521356 | 0.36305054 | -2.0717104 | 0.03829245 | 0.09794357 | PAGE5 | 90737 |
| ENSG000002 | 5.04304482 | -0.776903444 | 0.3750677 | -2.0713685 | 0.03832437 | 0.09801901 | NA | NA |
| ENSG000001 | 73.4258082 | 0.387516534 | 0.18709241 | 2.07125736 | 0.03833475 | 0.09803939 | NA | NA |
| ENSG000002 | 10.4587134 | -0.718631268 | 0.34697179 | -2.0711518 | 0.03834461 | 0.09805841 | NA | NA |
| ENSG000001 | 3.79058645 | -0.783999802 | 0.37859018 | -2.0708403 | 0.03837372 | 0.09812049 | BMP15 | 9210 |
| ENSG000002 | 5.83915509 | -0.763620996 | 0.36874666 | -2.0708553 | 0.03837232 | 0.09812049 | NA | NA |
| ENSG000001 | 90.646209 | 0.42881235 | 0.20709379 | 2.07061911 | 0.0383944 | 0.09816099 | C10orf82 | 143379 |
| ENSG000001 | 8.45479679 | -0.73810013 | 0.35646032 | -2.0706376 | 0.03839268 | 0.09816099 | NA | NA |

| | | | | | | | | |
|---|---|---|---|---|---|---|---|---|
| ENSG000001 | 7.59642205 | -0.735195519 | 0.35507802 | -2.0705183 | 0.03840384 | 0.09817888 | RAX | 30062 |
| ENSG000002 | 7.93561998 | -0.731959508 | 0.35351951 | -2.0704925 | 0.03840624 | 0.09817888 | LOC102467225 | 102467225 |
| ENSG000002 | 23.4465169 | -0.605454181 | 0.2924337 | -2.0703981 | 0.03841508 | 0.09819528 | NA | NA |
| ENSG000001 | 6.59733992 | -0.76236068 | 0.3682377 | -2.070295 | 0.03842473 | 0.09820716 | IFNG | 3458 |
| ENSG000002 | 9.8635673 | -0.716771159 | 0.34621643 | -2.070298 | 0.03842445 | 0.09820716 | NA | NA |
| ENSG000002 | 5.0695836 | -0.780360269 | 0.37693632 | -2.0702708 | 0.03842699 | 0.09820716 | NA | NA |
| ENSG000002 | 12.003166 | -0.679786756 | 0.32837112 | -2.0701782 | 0.03843566 | 0.09822271 | CHIAP2 | 149620 |
| ENSG000002 | 6.98380219 | 0.75591089 | 0.36514717 | 2.07015405 | 0.03843792 | 0.09822271 | NA | NA |
| ENSG000001 | 46.3363056 | -0.463706169 | 0.22400792 | -2.0700437 | 0.03844826 | 0.09824293 | CEP170P1 | 645455 |
| ENSG000001 | 8.41094019 | -0.738125951 | 0.35659873 | -2.0699063 | 0.03846112 | 0.0982696 | PRELP | 5549 |
| ENSG000002 | 51.7182032 | 0.447668665 | 0.2162862 | 2.06979763 | 0.0384713 | 0.09828692 | SLC25A5-AS1 | 100303728 |
| ENSG000002 | 7.15915849 | -0.75225297 | 0.36344547 | -2.0697822 | 0.03847275 | 0.09828692 | LOC101929607 | 101929607 |
| ENSG000001 | 696.710378 | 0.257375728 | 0.12435411 | 2.06970025 | 0.03848042 | 0.09829415 | HBP1 | 26959 |
| ENSG000002 | 5.62802864 | -0.770045441 | 0.37205631 | -2.0697013 | 0.03848032 | 0.09829415 | NA | NA |
| ENSG000001 | 5838.81125 | 0.502084749 | 0.24262542 | 2.06938233 | 0.03851022 | 0.09836408 | FTH1 | 2495 |
| ENSG000002 | 4.37472758 | -0.781938526 | 0.37787154 | -2.0693237 | 0.03851572 | 0.09837192 | NA | NA |
| ENSG000001 | 25.7947953 | -0.582603271 | 0.2815613 | -2.069188 | 0.03852845 | 0.09839823 | RFX6 | 222546 |
| ENSG00000C | 566.446119 | 0.235516722 | 0.113834 | 2.06894885 | 0.03855089 | 0.09844934 | LRRFIP2 | 9209 |
| ENSG000001 | 210.241329 | 0.299966474 | 0.14500034 | 2.06872945 | 0.03857148 | 0.09849573 | SLC16A10 | 117247 |
| ENSG000002 | 3.85018424 | -0.782265403 | 0.37817379 | -2.0685341 | 0.03858982 | 0.09853636 | NA | NA |
| ENSG000001 | 9.74738987 | -0.719246912 | 0.34773473 | -2.0683782 | 0.03860448 | 0.0985627 | LILRB1 | 10859 |
| ENSG000001 | 6.49515184 | -0.757242611 | 0.3661066 | -2.0683665 | 0.03860558 | 0.0985627 | FFAR2 | 2867 |
| ENSG000001 | 18.7516322 | -0.596266036 | 0.28828147 | -2.0683467 | 0.03860743 | 0.0985627 | CD200R1 | 131450 |
| ENSG000002 | 4.03490114 | -0.782188014 | 0.3781772 | -2.0683109 | 0.0386108 | 0.09856395 | NA | NA |
| ENSG000002 | 2.45585974 | -0.76502048 | 0.3698807 | -2.0682898 | 0.03861278 | 0.09856395 | NA | NA |
| ENSG000001 | 20.8344619 | 0.563997012 | 0.27269587 | 2.06822717 | 0.03861867 | 0.09857277 | RARRES3 | 5920 |
| ENSG000002 | 13.14058 | -0.672979739 | 0.32542169 | -2.0680236 | 0.0386378 | 0.0986154 | LINC00330 | 144817 |
| ENSG000001 | 12.1711882 | -0.690660995 | 0.33398974 | -2.0679108 | 0.03864841 | 0.09863627 | ADAMDEC1 | 27299 |
| ENSG000002 | 16.2233794 | 0.662770425 | 0.32052165 | 2.06778676 | 0.03866008 | 0.09865984 | SNORD116-9 | 100033421 |

| | | | | | | | | | |
|---|---|---|---|---|---|---|---|---|---|
| ENSG000002 | 26.5577461 | 0.534610252 | 0.25855496 | 2.06768515 | 0.03866964 | 0.09867803 | NA | NA | |
| ENSG000001 | 8.5744379 | -0.726238283 | 0.35124627 | -2.0676043 | 0.03867725 | 0.09869124 | CRYBB3 | | 1417 |
| ENSG000001 | 7.10414584 | -0.747944156 | 0.36175101 | -2.0675662 | 0.03868084 | 0.09869418 | NA | NA | |
| ENSG000001 | 1327.69622 | 0.241698904 | 0.11691264 | 2.06734621 | 0.03870154 | 0.0987408 | CCDC186 | | 55088 |
| ENSG000001 | 10.0353332 | -0.714181752 | 0.34548821 | -2.0671668 | 0.03871844 | 0.09877769 | LOC388882 | | 388882 |
| ENSG000002 | 3.43244497 | -0.781576519 | 0.37810577 | -2.0670843 | 0.03872621 | 0.0987913 | NA | NA | |
| ENSG000001 | 15.1032232 | -0.628135186 | 0.30391234 | -2.0668302 | 0.03875016 | 0.09884618 | CNBD1 | | 168975 |
| ENSG000002 | 2.84006932 | -0.771210095 | 0.3731467 | -2.0667745 | 0.0387554 | 0.09885334 | FOXN3-AS2 | | 29018 |
| ENSG000001 | 4.0858136 | -0.781878217 | 0.37832107 | -2.0667055 | 0.03876192 | 0.09886373 | NCF4 | | 4689 |
| ENSG00000C | 13.4429675 | -0.696536274 | 0.33706331 | -2.066485 | 0.03878271 | 0.0988981 | LTK | | 4058 |
| ENSG000001 | 33.2016405 | -0.514013517 | 0.24873642 | -2.0664988 | 0.0387814 | 0.0988981 | PI15 | | 51050 |
| ENSG000001 | 1965.09739 | 0.613821767 | 0.29703204 | 2.06651701 | 0.03877969 | 0.0988981 | ANKRD11 | | 29123 |
| ENSG000002 | 4.5817706 | -0.78178405 | 0.37833842 | -2.0663618 | 0.03879433 | 0.09892152 | NA | NA | |
| ENSG000001 | 192.040202 | 0.321912117 | 0.15579238 | 2.0662892 | 0.03880118 | 0.09893276 | STXBP1 | | 6812 |
| ENSG000002 | 6.53824853 | -0.770590115 | 0.37296121 | -2.0661401 | 0.03881525 | 0.09896242 | NA | NA | |
| ENSG000001 | 43.7647588 | 0.464556373 | 0.22487098 | 2.06587961 | 0.03883985 | 0.09900826 | CRYBB1 | | 1414 |
| ENSG000001 | 8.25432245 | -0.730502567 | 0.35360493 | -2.0658721 | 0.03884056 | 0.09900826 | RARRES1 | | 5918 |
| ENSG000001 | 10.520334 | -0.692445068 | 0.33518215 | -2.0658769 | 0.0388401 | 0.09900826 | RAG2 | | 5897 |
| ENSG000002 | 19.7360622 | -0.608857071 | 0.29476504 | -2.0655675 | 0.03886934 | 0.09907007 | DBNDD2 | | 55861 |
| ENSG000002 | 6.77688215 | -0.749594389 | 0.36290063 | -2.0655637 | 0.03886969 | 0.09907007 | NA | NA | |
| ENSG000002 | 11.8656077 | 0.676536254 | 0.32755729 | 2.06539826 | 0.03888533 | 0.09910371 | NA | NA | |
| ENSG000001 | 23.1740466 | -0.555817378 | 0.26912891 | -2.0652459 | 0.03889974 | 0.09913419 | SLC17A7 | | 57030 |
| ENSG000001 | 6.85641321 | -0.753574624 | 0.36491247 | -2.0650832 | 0.03891513 | 0.09916718 | OTOP2 | | 92736 |
| ENSG000002 | 3.36453754 | -0.781237737 | 0.37832697 | -2.0649803 | 0.03892486 | 0.09918576 | NA | NA | |
| ENSG000001 | 8.43506103 | -0.735009797 | 0.35597952 | -2.064753 | 0.03894637 | 0.09923433 | REC114 | | 283677 |
| ENSG000001 | 17.9401283 | -0.593375692 | 0.28740364 | -2.0646074 | 0.03896016 | 0.09925698 | CCDC114 | | 93233 |
| ENSG000002 | 4.02849208 | -0.780274832 | 0.37792446 | -2.0646317 | 0.03895786 | 0.09925698 | NA | NA | |
| ENSG000001 | 415.210738 | 0.330853455 | 0.16025334 | 2.06456517 | 0.03896416 | 0.09926094 | PARP2 | | 10038 |
| ENSG000002 | 2.67831027 | 0.766132063 | 0.37110561 | 2.06445833 | 0.03897428 | 0.09928049 | NA | NA | |

| | | | | | | | | |
|---|---|---|---|---|---|---|---|---|
| ENSG000001 | 776.346559 | 0.292838015 | 0.14185133 | 2.06440084 | 0.03897973 | 0.09928303 | FAM129A | 116496 |
| ENSG000002 | 13.032894 | -0.66157465 | 0.32046886 | -2.0643961 | 0.03898018 | 0.09928303 | NA | NA |
| ENSG000001 | 29.9438726 | -0.522112682 | 0.25292621 | -2.0642886 | 0.03899036 | 0.09929027 | GSG1L | 146395 |
| ENSG000001 | 7.7352723 | -0.742822086 | 0.35984079 | -2.0643076 | 0.03898856 | 0.09929027 | TEX26 | 122046 |
| ENSG000002 | 2.77062799 | -0.771471891 | 0.37371772 | -2.0643171 | 0.03898766 | 0.09929027 | NA | NA |
| ENSG000002 | 13.1203052 | -0.666395922 | 0.32283767 | -2.0641827 | 0.0390004 | 0.09930335 | UGT2B11 | 10720 |
| ENSG000002 | 3.45632344 | -0.780582024 | 0.37815431 | -2.0641892 | 0.03899978 | 0.09930335 | NA | NA |
| ENSG000001 | 106.340642 | 0.391246959 | 0.18956337 | 2.06393757 | 0.03902364 | 0.09933246 | RAPGEFL1 | 51195 |
| ENSG000001 | 13.9022604 | -0.670134861 | 0.32469236 | -2.0639071 | 0.03902653 | 0.09933246 | GATA4 | 2626 |
| ENSG000002 | 11.6728665 | -0.681673342 | 0.33027066 | -2.063984 | 0.03901924 | 0.09933246 | CD200R1L | 344807 |
| ENSG000002 | 5.07048843 | -0.776484588 | 0.37621691 | -2.0639279 | 0.03902455 | 0.09933246 | NA | NA |
| ENSG000002 | 3.89378185 | -0.781081353 | 0.37844542 | -2.0639207 | 0.03902523 | 0.09933246 | NA | NA |
| ENSG000002 | 11.0241447 | -0.679860754 | 0.32939114 | -2.0639923 | 0.03901845 | 0.09933246 | NA | NA |
| ENSG00000C | 1282.24294 | 0.236537553 | 0.11461615 | 2.06373659 | 0.0390427 | 0.09935491 | FRYL | 285527 |
| ENSG000001 | 15.2478496 | -0.634815505 | 0.30760302 | -2.0637493 | 0.03904149 | 0.09935491 | WNT7A | 7476 |
| ENSG000001 | 10.696548 | -0.703069121 | 0.34067736 | -2.0637389 | 0.03904248 | 0.09935491 | WFDC11 | 259239 |
| ENSG000001 | 26.7426759 | -0.539433075 | 0.26139442 | -2.0636748 | 0.03904856 | 0.09935735 | SLC14A1 | 6563 |
| ENSG000001 | 24.5003232 | -0.590150427 | 0.28597031 | -2.0636772 | 0.03904833 | 0.09935735 | DPCR1 | 135656 |
| ENSG000002 | 4.35780528 | -0.779648637 | 0.37780646 | -2.0636191 | 0.03905384 | 0.09936457 | OR6C65 | 403282 |
| ENSG000001 | 15.3689641 | -0.640874684 | 0.31057515 | -2.0635092 | 0.03906428 | 0.09937641 | VNN1 | 8876 |
| ENSG000001 | 50.5154916 | 0.444196186 | 0.21526245 | 2.06350982 | 0.03906422 | 0.09937641 | GAS1 | 2619 |
| ENSG000001 | 484.815351 | 0.324504154 | 0.15725967 | 2.06349257 | 0.03906585 | 0.09937641 | HIST3H2BB | 128312 |
| ENSG000001 | 7.52928128 | -0.74780715 | 0.36240535 | -2.0634551 | 0.03906941 | 0.09937923 | OR4N5 | 390437 |
| ENSG000001 | 28.7806623 | -0.521627101 | 0.25280688 | -2.0633422 | 0.03908013 | 0.09940026 | RSPH1 | 89765 |
| ENSG000001 | 509.239463 | 0.248486206 | 0.12043341 | 2.06326631 | 0.03908733 | 0.09941235 | ZNF772 | 400720 |
| ENSG000002 | 19.9842886 | -0.597593522 | 0.28964617 | -2.0631846 | 0.03909509 | 0.09942584 | LINC01043 | 101928752 |
| ENSG000002 | 4.85415979 | 0.778174308 | 0.37718164 | 2.06312882 | 0.03910039 | 0.09942685 | NA | NA |
| ENSG000002 | 24.5161198 | -0.534147972 | 0.25890133 | -2.0631333 | 0.03909996 | 0.09942685 | GRM5-AS1 | 100873989 |
| ENSG000002 | 6.89855201 | -0.752950302 | 0.36497144 | -2.063039 | 0.03910893 | 0.09944232 | NA | NA |

| | | | | | | | | |
|---|---|---|---|---|---|---|---|---|
| ENSG000001 | 589.84686 | 0.277349944 | 0.13444227 | 2.06296685 | 0.03911578 | 0.0994535 | PPFIBP1 | 8496 |
| ENSG000001 | 166.107125 | 0.326851563 | 0.15844318 | 2.06289451 | 0.03912265 | 0.09946474 | ACAD10 | 80724 |
| ENSG000002 | 4.66472377 | -0.776199362 | 0.37634277 | -2.0624798 | 0.03916208 | 0.09955874 | NA | NA |
| ENSG000001 | 8.80953713 | -0.717473441 | 0.34787979 | -2.0624177 | 0.03916799 | 0.09956751 | GUCA1C | 9626 |
| ENSG000001 | 6.40408845 | -0.765165168 | 0.371031 | -2.0622675 | 0.03918228 | 0.0995976 | KRT6C | 286887 |
| ENSG000002 | 39.0614888 | 0.493996346 | 0.23955621 | 2.06213121 | 0.03919525 | 0.09962432 | NA | NA |
| ENSG000001 | 4.05607109 | -0.77979263 | 0.37816625 | -2.0620365 | 0.03920426 | 0.09964098 | RAET1L | 154064 |
| ENSG000002 | 4.66100635 | -0.775368767 | 0.37603027 | -2.061985 | 0.03920917 | 0.09964721 | NA | NA |
| ENSG000002 | 3.71485242 | -0.775502314 | 0.37614218 | -2.0617265 | 0.03923378 | 0.09970351 | NA | NA |
| ENSG000002 | 7.15082613 | 0.746861692 | 0.36226093 | 2.06166779 | 0.03923938 | 0.09971149 | NA | NA |
| ENSG000001 | 3.79478845 | -0.780436898 | 0.37858032 | -2.061483 | 0.03925698 | 0.09974998 | GAL3ST2 | 64090 |
| ENSG000001 | 671.323033 | 0.278635299 | 0.13517196 | 2.06133952 | 0.03927066 | 0.09977848 | FEM1C | 56929 |
| ENSG000001 | 618.049063 | 0.304319129 | 0.14764291 | 2.06118347 | 0.03928554 | 0.09979133 | C18orf54 | 162681 |
| ENSG000002 | 4.03370746 | -0.779482579 | 0.37816929 | -2.0612001 | 0.03928396 | 0.09979133 | BRWD1P1 | 106481724 |
| ENSG000002 | 2.44471029 | -0.76132884 | 0.36936495 | -2.0611832 | 0.03928556 | 0.09979133 | NA | NA |
| ENSG000002 | 33.1580268 | 0.501934144 | 0.24351265 | 2.0612241 | 0.03928167 | 0.09979133 | NA | NA |
| ENSG000001 | 16.5815055 | -0.620072899 | 0.30084302 | -2.0611178 | 0.03929181 | 0.09980094 | PDZK1 | 5174 |
| ENSG000001 | 79.5923939 | -0.375918633 | 0.18239796 | -2.0609804 | 0.03930491 | 0.09982179 | PSTPIP2 | 9050 |
| ENSG000001 | 58.1381975 | 0.479366689 | 0.23259452 | 2.06095433 | 0.0393074 | 0.09982179 | FES | 2242 |
| ENSG000002 | 9.48021803 | -0.714899585 | 0.34687506 | -2.0609714 | 0.03930577 | 0.09982179 | NA | NA |
| ENSG000002 | 8.2778497 | -0.730162867 | 0.35438034 | -2.0603933 | 0.03936096 | 0.09994529 | NA | NA |
| ENSG000002 | 4.1210783 | -0.779996835 | 0.37856665 | -2.060395 | 0.03936079 | 0.09994529 | NA | NA |
| ENSG000002 | 6.60891447 | -0.757598003 | 0.36771169 | -2.0603044 | 0.03936945 | 0.0999606 | LINC01519 | 101929624 |
| ENSG000001 | 464.393145 | 0.29320693 | 0.14231481 | 2.06027 | 0.03937274 | 0.09996108 | ETV6 | 2120 |
| ENSG000002 | 10.6188998 | -0.706155955 | 0.34275243 | -2.0602508 | 0.03937458 | 0.09996108 | NA | NA |
| ENSG000001 | 2.77858817 | -0.770626748 | 0.37410912 | -2.0598983 | 0.03940826 | 0.10003872 | NA | NA |
| ENSG000002 | 11.0610108 | -0.683465319 | 0.33179874 | -2.0598792 | 0.03941009 | 0.10003872 | NA | NA |
| ENSG000001 | 9.89439674 | -0.700439664 | 0.34005363 | -2.0597918 | 0.03941845 | 0.10004742 | CASP14 | 23581 |
| ENSG000001 | 158.171405 | 0.29490732 | 0.14317312 | 2.05979527 | 0.03941812 | 0.10004742 | TNFRSF11A | 8792 |

| | | | | | | | | |
|---|---|---|---|---|---|---|---|---|
| ENSG000002 | 5.90910551 | -0.761508246 | 0.3697236 | -2.059669 | 0.03943019 | 0.10007096 | NA | NA |
| ENSG000001 | 16.5717189 | 0.612621949 | 0.29745229 | 2.05956373 | 0.03944027 | 0.100084 | PROKR2 | 128674 |
| ENSG000001 | 65.7063449 | 0.40120893 | 0.1948022 | 2.05957084 | 0.03943959 | 0.100084 | DGAT1 | 8694 |
| ENSG000002 | 3.72671685 | -0.779694128 | 0.37858544 | -2.0594932 | 0.03944702 | 0.10009487 | NA | NA |
| ENSG000001 | 651.901437 | 0.275959378 | 0.13400135 | 2.05937758 | 0.03945808 | 0.10011667 | THBS1 | 7057 |
| ENSG000001 | 21.4124509 | -0.590137174 | 0.28659391 | -2.0591407 | 0.03948076 | 0.10016169 | SLC34A1 | 6569 |
| ENSG000002 | 10.8646709 | -0.691234638 | 0.3356908 | -2.0591408 | 0.03948075 | 0.10016169 | NA | NA |
| ENSG000002 | 7.8727435 | -0.726401493 | 0.35277739 | -2.0590931 | 0.03948532 | 0.10016698 | NA | NA |
| ENSG000001 | 631.220115 | 0.281803765 | 0.13686387 | 2.05900772 | 0.0394935 | 0.1001752 | DNAJB5 | 25822 |
| ENSG000002 | 3.37708919 | -0.778675522 | 0.37817829 | -2.059017 | 0.03949261 | 0.1001752 | NA | NA |
| ENSG000001 | 18.9205606 | -0.592714485 | 0.28787472 | -2.058932 | 0.03950075 | 0.10018732 | LMOD1 | 25802 |
| ENSG000001 | 17.2660866 | -0.641634443 | 0.31167116 | -2.0586905 | 0.0395239 | 0.10023977 | NA | NA |
| ENSG000002 | 16.2622001 | -0.639508674 | 0.31065807 | -2.0585613 | 0.03953628 | 0.10026491 | LINC01241 | 101929582 |
| ENSG000002 | 8.99036217 | -0.725396853 | 0.35240832 | -2.0583988 | 0.03955187 | 0.10029816 | LOC101927851 | 101927851 |
| ENSG000002 | 51.970498 | -0.437179518 | 0.21240965 | -2.0581905 | 0.03957185 | 0.10034255 | PPP3CB-AS1 | 101929145 |
| ENSG000001 | 20.212964 | -0.595382784 | 0.28931423 | -2.0579105 | 0.03959873 | 0.10040443 | PAX3 | 5077 |
| ENSG000002 | 11.5547778 | -0.674419529 | 0.32776028 | -2.057661 | 0.03962269 | 0.1004589 | NA | NA |
| ENSG000002 | 2.88442178 | -0.764164519 | 0.3713946 | -2.0575542 | 0.03963295 | 0.10047863 | NA | NA |
| ENSG000001 | 5.62802919 | -0.765386899 | 0.37201733 | -2.0573958 | 0.03964817 | 0.10051093 | HOXB2 | 3212 |
| ENSG000002 | 13.8230555 | -0.66732649 | 0.32436426 | -2.0573367 | 0.03965385 | 0.10051906 | NA | NA |
| ENSG000002 | 2.40092797 | -0.760584642 | 0.36972139 | -2.0571833 | 0.0396686 | 0.10055016 | LOC284933 | 284933 |
| ENSG000002 | 4.06543347 | -0.778041466 | 0.37822601 | -2.0570808 | 0.03967845 | 0.10056885 | NA | NA |
| ENSG000002 | 5.0439815 | -0.774159574 | 0.37636294 | -2.0569495 | 0.03969108 | 0.10059457 | NA | NA |
| ENSG000001 | 71.6254487 | 0.396585443 | 0.19281472 | 2.05682142 | 0.03970341 | 0.10061911 | TERT | 7015 |
| ENSG000001 | 3.49013678 | -0.777214798 | 0.37787622 | -2.0567973 | 0.03970573 | 0.10061911 | C15orf48 | 84419 |
| ENSG000002 | 4.21529678 | -0.77867241 | 0.37859086 | -2.056765 | 0.03970884 | 0.1006207 | NA | NA |
| ENSG000002 | 13.4912478 | -0.65542014 | 0.31869265 | -2.0565901 | 0.03972567 | 0.10065707 | NA | NA |
| ENSG000002 | 5.93760825 | -0.761351415 | 0.3702177 | -2.0564965 | 0.03973468 | 0.1006736 | NA | NA |
| ENSG000002 | 16.6091754 | -0.622134497 | 0.30252732 | -2.0564572 | 0.03973847 | 0.1006769 | STRC | 161497 |

| | | | | | | | | |
|---|---|---|---|---|---|---|---|---|
| ENSG000002 | 8.80676653 | -0.73706818 | 0.35842539 | -2.0564062 | 0.03974338 | 0.10068307 | TUB-AS1 | 101927917 |
| ENSG000002 | 11.0010235 | -0.680520977 | 0.33093357 | -2.0563673 | 0.03974712 | 0.10068625 | NA | NA |
| ENSG000001 | 198.462215 | 0.364880906 | 0.1774447 | 2.05630768 | 0.03975287 | 0.10069452 | ABHD17C | 58489 |
| ENSG000001 | 363.020278 | 0.256572178 | 0.12478459 | 2.0561207 | 0.03977088 | 0.10073386 | RAB22A | 57403 |
| ENSG000001 | 72.4618314 | 0.396792606 | 0.19298512 | 2.05607875 | 0.03977493 | 0.10073781 | PCSK1 | 5122 |
| ENSG000002 | 9.16483479 | -0.713711064 | 0.34712874 | -2.0560414 | 0.03977852 | 0.10074063 | NA | NA |
| ENSG000002 | 5.68055932 | -0.766790676 | 0.37297984 | -2.0558502 | 0.03979696 | 0.10078102 | NA | NA |
| ENSG000002 | 5.53637069 | -0.760956106 | 0.37014876 | -2.0558116 | 0.03980068 | 0.10078415 | NA | NA |
| ENSG000002 | 3.08021136 | -0.773414795 | 0.37624283 | -2.0556267 | 0.03981851 | 0.10082301 | NA | NA |
| ENSG000002 | 3.52533656 | -0.775991728 | 0.3775138 | -2.0555321 | 0.03982764 | 0.10083983 | NA | NA |
| ENSG000002 | 3.78930736 | -0.77787322 | 0.37845414 | -2.0553963 | 0.03984075 | 0.10086671 | NA | NA |
| ENSG000002 | 14.1458019 | -0.647460079 | 0.31502296 | -2.055279 | 0.03985206 | 0.10088906 | SERPINB5 | 5268 |
| ENSG000001 | 68.761132 | -0.39044775 | 0.18998532 | -2.055147 | 0.03986481 | 0.10090243 | EFCAB6 | 64800 |
| ENSG000002 | 2.83224085 | -0.766879902 | 0.37314737 | -2.0551663 | 0.03986294 | 0.10090243 | LOC101929153 | 101929153 |
| ENSG000002 | 8.03702621 | -0.739206791 | 0.35967676 | -2.0551975 | 0.03985993 | 0.10090243 | NA | NA |
| ENSG000001 | 47.1731099 | 0.491615103 | 0.23923592 | 2.05493852 | 0.03988494 | 0.10094079 | KLHL26 | 55295 |
| ENSG000002 | 33.0125858 | 0.765517985 | 0.37252217 | 2.05495954 | 0.03988291 | 0.10094079 | RNA5-8S5 | 100008587 |
| ENSG000001 | 8.1511374 | -0.742952534 | 0.36156234 | -2.0548394 | 0.03989452 | 0.10095873 | TAAR6 | 319100 |
| ENSG000002 | 17.9565999 | -0.624937611 | 0.30415681 | -2.054656 | 0.03991224 | 0.10099728 | LINC01235 | 401492 |
| ENSG000002 | 5.99926919 | -0.764326538 | 0.3720208 | -2.0545264 | 0.03992477 | 0.10102268 | NA | NA |
| ENSG000001 | 198.878248 | 0.284557591 | 0.13850812 | 2.05444695 | 0.03993245 | 0.10103581 | ZNF641 | 121274 |
| ENSG000002 | 7.07330428 | -0.7403893 | 0.36039158 | -2.0544023 | 0.03993677 | 0.10104043 | NA | NA |
| ENSG000001 | 330.733623 | 0.342141757 | 0.16656995 | 2.05404251 | 0.03997158 | 0.10112218 | HIST3H2A | 92815 |
| ENSG000002 | 22.7961816 | 0.59483877 | 0.28961185 | 2.05391725 | 0.0399837 | 0.10114023 | NA | NA |
| ENSG000002 | 6.97628611 | -0.738351959 | 0.35948141 | -2.0539364 | 0.03998185 | 0.10114023 | NA | NA |
| ENSG000001 | 120.957385 | 0.340718921 | 0.16589422 | 2.05383233 | 0.03999192 | 0.10114363 | SLC39A11 | 201266 |
| ENSG000001 | 11.6999661 | -0.70407185 | 0.34280987 | -2.0538261 | 0.03999253 | 0.10114363 | AGR3 | 155465 |
| ENSG000002 | 14.1396036 | -0.66766134 | 0.3250751 | -2.053868 | 0.03998847 | 0.10114363 | NA | NA |
| ENSG000001 | 129.186331 | 0.318876855 | 0.15527101 | 2.05367924 | 0.04000675 | 0.10116581 | HSPBAP1 | 79663 |

| | | | | | | | | |
|---|---|---|---|---|---|---|---|---|
| ENSG000001 | 17.6231372 | -0.611383126 | 0.29770201 | -2.0536748 | 0.04000717 | 0.10116581 | SLC47A2 | 146802 |
| ENSG000002 | 3.43358767 | -0.776719816 | 0.37821279 | -2.0536582 | 0.04000878 | 0.10116581 | NA | NA |
| ENSG000001 | 85.2649541 | 0.465158691 | 0.226512 | 2.05357197 | 0.04001714 | 0.10117896 | PLEKHA4 | 57664 |
| ENSG000002 | 3.3292841 | -0.767247507 | 0.37361953 | -2.053553 | 0.04001897 | 0.10117896 | NA | NA |
| ENSG000001 | 24.7216115 | 0.53575346 | 0.26089794 | 2.05349825 | 0.04002428 | 0.10118521 | RAB33A | 9363 |
| ENSG000002 | 3.87241291 | -0.776936461 | 0.37835186 | -2.053476 | 0.04002643 | 0.10118521 | NA | NA |
| ENSG000001 | 5.98928651 | -0.763164812 | 0.37165721 | -2.0534105 | 0.04003278 | 0.10119495 | NA | NA |
| ENSG000001 | 26.4527386 | -0.524523882 | 0.25545865 | -2.0532633 | 0.04004705 | 0.10122469 | NOD2 | 64127 |
| ENSG000002 | 3.78389849 | -0.77696326 | 0.37844472 | -2.0530429 | 0.04006842 | 0.10127239 | NA | NA |
| ENSG000002 | 7.35824473 | -0.739008329 | 0.35997877 | -2.052922 | 0.04008014 | 0.10129573 | NA | NA |
| ENSG000002 | 5.26917533 | -0.766682082 | 0.37349106 | -2.0527454 | 0.04009727 | 0.1013327 | NA | NA |
| ENSG000002 | 3.51554942 | -0.774458424 | 0.3772872 | -2.0527026 | 0.04010143 | 0.10133688 | RNU6-1183P | 106481941 |
| ENSG000002 | 7.65096794 | -0.732568772 | 0.35697369 | -2.0521646 | 0.04015367 | 0.10146257 | LOC102724710 | 102724710 |
| ENSG000001 | 6.2104364 | -0.756166676 | 0.36848079 | -2.0521197 | 0.04015803 | 0.10146728 | RRAD | 6236 |
| ENSG000002 | 3.40321896 | -0.776098456 | 0.37820173 | -2.0520754 | 0.04016234 | 0.10147184 | MTRNR2L3 | 100462983 |
| ENSG000002 | 5.7399229 | -0.768806527 | 0.37467662 | -2.0519202 | 0.04017742 | 0.10150361 | LOC284632 | 284632 |
| ENSG000001 | 70.7817209 | 0.407198925 | 0.19848536 | 2.05153132 | 0.04021523 | 0.10158342 | SNHG11 | 128439 |
| ENSG000002 | 4.00956861 | -0.775901945 | 0.37819955 | -2.0515676 | 0.0402117 | 0.10158342 | NA | NA |
| ENSG000002 | 5.93173174 | -0.762953228 | 0.37189691 | -2.0515181 | 0.04021652 | 0.10158342 | NA | NA |
| ENSG000002 | 167.435469 | 0.303721872 | 0.14808526 | 2.05099333 | 0.0402676 | 0.10170238 | NA | NA |
| ENSG000002 | 7.72400507 | -0.74001684 | 0.36081087 | -2.0509827 | 0.04026863 | 0.10170238 | NA | NA |
| ENSG000001 | 6.65791163 | -0.769591137 | 0.37523961 | -2.0509326 | 0.04027352 | 0.10170837 | NA | NA |
| ENSG000002 | 6.50948665 | -0.751075851 | 0.36634303 | -2.0501983 | 0.04034509 | 0.10188278 | NA | NA |
| ENSG000001 | 15.3610288 | -0.644171918 | 0.31420691 | -2.050152 | 0.0403496 | 0.10188783 | LDHC | 3948 |
| ENSG000002 | 7.17984895 | -0.747284971 | 0.36457512 | -2.0497421 | 0.0403896 | 0.10198249 | FAM41C | 284593 |
| ENSG000002 | 4.06079427 | -0.775384133 | 0.37828863 | -2.0497156 | 0.04039219 | 0.10198267 | NA | NA |
| ENSG000002 | 11.1234034 | -0.683718958 | 0.33359295 | -2.0495606 | 0.04040733 | 0.1020086 | LOC101927189 | 101927189 |
| ENSG000002 | 9.28980281 | -0.699549686 | 0.34131718 | -2.049559 | 0.04040749 | 0.1020086 | NA | NA |
| ENSG000002 | 5.44948142 | -0.769278715 | 0.37539116 | -2.0492723 | 0.04043549 | 0.10207294 | NA | NA |

| | | | | | | | | |
|---|---|---|---|---|---|---|---|---|
| ENSG000002 | 26.3913558 | -0.550552656 | 0.26867821 | -2.0491154 | 0.04045083 | 0.1021053 | NA | NA |
| ENSG000001 | 84.7434339 | 0.369305522 | 0.18023298 | 2.04904525 | 0.04045769 | 0.10211626 | SMCO4 | 56935 |
| ENSG000001 | 47.5880988 | -0.44716544 | 0.21823873 | -2.0489738 | 0.04046468 | 0.10212753 | DPYS | 1807 |
| ENSG00000C | 15.5445345 | -0.635920319 | 0.31043374 | -2.0484897 | 0.04051204 | 0.10223434 | NFIX | 4784 |
| ENSG000002 | 10.7234635 | -0.68324144 | 0.33353012 | -2.048515 | 0.04050957 | 0.10223434 | NA | NA |
| ENSG000002 | 7.88117074 | -0.739295866 | 0.36091998 | -2.048365 | 0.04052425 | 0.10225879 | NA | NA |
| ENSG000001 | 134.193444 | 0.330535867 | 0.16137069 | 2.04830178 | 0.04053044 | 0.10226805 | PEX7 | 5191 |
| ENSG000001 | 450.353599 | 0.312451743 | 0.15254993 | 2.04819325 | 0.04054107 | 0.10228343 | PPARGC1B | 133522 |
| ENSG000002 | 6.53792426 | -0.751425087 | 0.36687771 | -2.0481623 | 0.0405441 | 0.10228343 | LOC100506271 | 100506271 |
| ENSG000002 | 2.4032638 | -0.756508359 | 0.36935799 | -2.0481711 | 0.04054324 | 0.10228343 | LINC01444 | 101927642 |
| ENSG000001 | 265.737326 | 0.278549264 | 0.13600617 | 2.04806349 | 0.04055378 | 0.10230148 | AHSA2 | 130872 |
| ENSG000001 | 2.38778814 | -0.756107812 | 0.36918769 | -2.0480309 | 0.04055697 | 0.10230257 | RNU6-545P | 106479808 |
| ENSG000002 | 2.52138902 | -0.750362657 | 0.36638666 | -2.0480076 | 0.04055926 | 0.10230257 | NA | NA |
| ENSG000002 | 2.78702214 | -0.764052713 | 0.37308975 | -2.047906 | 0.04056921 | 0.10232132 | FAM222A-AS1 | 84983 |
| ENSG000001 | 8.44871045 | -0.737519279 | 0.36014041 | -2.0478659 | 0.04057314 | 0.10232486 | SLC34A3 | 142680 |
| ENSG000001 | 183.176187 | 0.581226608 | 0.28384136 | 2.04771638 | 0.0405878 | 0.10235546 | MANBAL | 63905 |
| ENSG000001 | 18.1520384 | -0.597705534 | 0.29190927 | -2.047573 | 0.04060186 | 0.10238456 | SCARA5 | 286133 |
| ENSG000002 | 8.94949444 | -0.704369538 | 0.34406178 | -2.0472182 | 0.04063667 | 0.10246595 | NA | NA |
| ENSG000002 | 20.5234815 | -0.576822635 | 0.28176448 | -2.0471801 | 0.04064041 | 0.10246902 | ZMIZ1-AS1 | 283050 |
| ENSG000002 | 2.41931773 | -0.753744691 | 0.36826684 | -2.0467352 | 0.04068409 | 0.10257279 | NA | NA |
| ENSG000001 | 25.4163352 | -0.554273216 | 0.27082666 | -2.0465977 | 0.0406976 | 0.10260046 | NA | NA |
| ENSG000001 | 14.2853319 | -0.632993768 | 0.30930452 | -2.0465067 | 0.04070655 | 0.10261663 | F2 | 2147 |
| ENSG000002 | 9.02580158 | -0.709249567 | 0.34657189 | -2.0464717 | 0.04070998 | 0.10261891 | NA | NA |
| ENSG000001 | 113.928079 | -0.335444118 | 0.16391738 | -2.046422 | 0.04071487 | 0.10262393 | MEIS3 | 56917 |
| ENSG000002 | 8.91723027 | -0.720052552 | 0.35186306 | -2.0464 | 0.04071703 | 0.10262393 | NA | NA |
| ENSG000001 | 204.333979 | 0.290155968 | 0.14179453 | 2.04631284 | 0.0407256 | 0.10263915 | BTN2A2 | 10385 |
| ENSG000002 | 8.98533359 | -0.724186066 | 0.35392135 | -2.0461779 | 0.04073887 | 0.10265982 | NA | NA |
| ENSG000002 | 5.95120434 | -0.759025697 | 0.37094409 | -2.0461998 | 0.04073672 | 0.10265982 | PCAT18 | 728606 |
| ENSG000001 | 315.537951 | 0.29504539 | 0.14419888 | 2.04610046 | 0.04074649 | 0.10267264 | GSK3A | 2931 |

| | | | | | | | | |
|---|---|---|---|---|---|---|---|---|
| ENSG000002 | 3.39829881 | -0.7731769 | 0.37799032 | -2.0454939 | 0.04080619 | 0.10281669 | NA | NA |
| ENSG000001 | 24.4914911 | -0.541016194 | 0.26449815 | -2.0454441 | 0.04081109 | 0.10282265 | NA | NA |
| ENSG000002 | 5.87506872 | -0.757552243 | 0.37036909 | -2.0453981 | 0.04081562 | 0.1028243 | NA | NA |
| ENSG000002 | 9.30029562 | -0.704879028 | 0.34461907 | -2.045386 | 0.04081682 | 0.1028243 | NA | NA |
| ENSG000002 | 3.55417899 | -0.770913644 | 0.37693884 | -2.0451956 | 0.04083558 | 0.10286518 | NA | NA |
| ENSG00000C | 4.62875066 | -0.769826791 | 0.37644009 | -2.0450181 | 0.04085308 | 0.10290286 | TNFRSF17 | 608 |
| ENSG000002 | 3.42572639 | -0.773293411 | 0.37814043 | -2.04499 | 0.04085584 | 0.10290343 | NA | NA |
| ENSG000001 | 27.9629415 | -0.515375769 | 0.25202663 | -2.0449259 | 0.04086217 | 0.10291009 | GLIPR1L2 | 144321 |
| ENSG000002 | 3.1838908 | -0.76775019 | 0.37544417 | -2.0449117 | 0.04086356 | 0.10291009 | NA | NA |
| ENSG00000C | 10.3737837 | -0.700888632 | 0.34276048 | -2.044835 | 0.04087113 | 0.10291636 | TFAP2D | 83741 |
| ENSG000002 | 23.0011297 | -0.594111807 | 0.29054196 | -2.0448399 | 0.04087064 | 0.10291636 | UG0898H09 | 643763 |
| ENSG000002 | 4.18267135 | -0.774143904 | 0.37858979 | -2.0448092 | 0.04087367 | 0.10291637 | NA | NA |
| ENSG000002 | 4.90824972 | -0.765015177 | 0.37413098 | -2.044779 | 0.04087665 | 0.10291749 | NA | NA |
| ENSG000001 | 168.58001 | 0.315864563 | 0.15447571 | 2.0447523 | 0.04087928 | 0.10291773 | ZMYM6 | 9204 |
| ENSG000001 | 115.745418 | 0.328956487 | 0.16088268 | 2.04469789 | 0.04088465 | 0.10292485 | C15orf57 | 90416 |
| ENSG000001 | 224.408241 | 0.288525421 | 0.14111152 | 2.04466243 | 0.04088815 | 0.10292727 | DTX3L | 151636 |
| ENSG000001 | 1539.56939 | 0.516127593 | 0.25245609 | 2.04442518 | 0.04091156 | 0.10297981 | HCFC1 | 3054 |
| ENSG000002 | 3.78969225 | -0.773663044 | 0.37844975 | -2.0442953 | 0.04092439 | 0.1030057 | NA | NA |
| ENSG000002 | 16.2168845 | -0.65303722 | 0.31946201 | -2.044178 | 0.04093597 | 0.10302845 | NA | NA |
| ENSG000002 | 4.70910691 | -0.770242161 | 0.37681582 | -2.0440813 | 0.04094552 | 0.1030461 | NA | NA |
| ENSG000001 | 1367.8525 | 0.30171108 | 0.14761131 | 2.04395642 | 0.04095785 | 0.1030704 | LRRFIP1 | 9208 |
| ENSG000002 | 3.40375975 | -0.772545295 | 0.37797014 | -2.0439321 | 0.04096026 | 0.1030704 | NA | NA |
| ENSG000001 | 5.82542849 | -0.756038721 | 0.36990662 | -2.0438637 | 0.04096701 | 0.10308099 | NA | NA |
| ENSG00000C | 4.09094227 | -0.773415306 | 0.37842613 | -2.0437682 | 0.04097645 | 0.10308905 | HOXA11 | 3207 |
| ENSG000001 | 72.3036877 | 0.390135873 | 0.1908938 | 2.0437325 | 0.04097998 | 0.10308905 | IGSF11 | 152404 |
| ENSG000001 | 248.403295 | 0.290887177 | 0.14233453 | 2.04368664 | 0.04098451 | 0.10308905 | ZNF182 | 7569 |
| ENSG000001 | 26.1092009 | -0.550581806 | 0.26940745 | -2.043677 | 0.04098547 | 0.10308905 | ABCD2 | 225 |
| ENSG000002 | 15.3064706 | -0.622513342 | 0.30459707 | -2.0437273 | 0.0409805 | 0.10308905 | LOC101926892 | 101926892 |
| ENSG000002 | 4.28214451 | -0.77067339 | 0.37709953 | -2.043687 | 0.04098448 | 0.10308905 | NA | NA |

| | | | | | | | | |
|---|---|---|---|---|---|---|---|---|
| ENSG000001 | 2.7393786 | -0.76325223 | 0.37348577 | -2.0435912 | 0.04099395 | 0.1030976 | CCL8 | 6355 |
| ENSG000001 | 898.020261 | 0.270330742 | 0.13228062 | 2.04361557 | 0.04099154 | 0.1030976 | TAF9B | 51616 |
| ENSG000002 | 2.82606362 | -0.762592847 | 0.37318635 | -2.0434639 | 0.04100653 | 0.10312285 | NA | NA |
| ENSG000002 | 19.6382033 | -0.581509284 | 0.28458232 | -2.0433781 | 0.04101502 | 0.1031378 | NA | NA |
| ENSG000002 | 8.78000956 | -0.712264486 | 0.34858846 | -2.0432819 | 0.04102454 | 0.10315535 | NA | NA |
| ENSG000001 | 3.85393398 | -0.773071907 | 0.37839459 | -2.0430311 | 0.04104936 | 0.10321135 | NA | NA |
| ENSG000002 | 10.2610401 | -0.694054717 | 0.33975057 | -2.0428361 | 0.04106867 | 0.1032535 | NA | NA |
| ENSG000001 | 8.22353781 | -0.721854103 | 0.35339684 | -2.0426162 | 0.04109045 | 0.10329544 | FOLR2 | 2350 |
| ENSG000002 | 15.7660676 | -0.630068509 | 0.30846068 | -2.0426218 | 0.04108989 | 0.10329544 | NA | NA |
| ENSG000002 | 3.36769435 | -0.772238169 | 0.37807052 | -2.0425771 | 0.04109432 | 0.10329878 | NA | NA |
| ENSG000002 | 4.42463169 | -0.77203543 | 0.378002 | -2.042411 | 0.04111078 | 0.10333375 | NA | NA |
| ENSG000002 | 22.5107794 | -0.564789916 | 0.27654914 | -2.0422769 | 0.04112407 | 0.10336075 | ANKRD18DP | 348840 |
| ENSG000002 | 7.72337043 | -0.733848543 | 0.35934653 | -2.0421751 | 0.04113416 | 0.1033797 | NA | NA |
| ENSG000001 | 22.4405946 | -0.592655301 | 0.29021981 | -2.0420912 | 0.04114249 | 0.10339422 | MYLK2 | 85366 |
| ENSG000002 | 10.1334676 | -0.687405612 | 0.33663407 | -2.0419965 | 0.04115187 | 0.10341139 | NA | NA |
| ENSG000001 | 835.454028 | 0.297169704 | 0.14554422 | 2.04178295 | 0.04117307 | 0.10345823 | ARHGEF11 | 9826 |
| ENSG000001 | 247.25027 | 0.293181461 | 0.14361173 | 2.04148687 | 0.04120246 | 0.10352567 | FGF19 | 9965 |
| ENSG000001 | 835.444618 | 0.254359774 | 0.12460875 | 2.04126735 | 0.04122426 | 0.10357404 | CMSS1 | 84319 |
| ENSG000001 | 16.3151487 | -0.622600231 | 0.30501071 | -2.0412405 | 0.04122692 | 0.10357431 | NA | NA |
| ENSG000002 | 13.9875092 | 0.632172209 | 0.30972452 | 2.04107897 | 0.04124298 | 0.10360181 | NA | NA |
| ENSG000002 | 2.36598512 | -0.753408481 | 0.36911993 | -2.041094 | 0.04124148 | 0.10360181 | NA | NA |
| ENSG000002 | 8.86919529 | -0.718629505 | 0.35209731 | -2.0409969 | 0.04125114 | 0.10361589 | NA | NA |
| ENSG000002 | 14.3441066 | -0.64135733 | 0.31424268 | -2.0409619 | 0.04125462 | 0.10361821 | LOC101929473 | 101929473 |
| ENSG000001 | 29.4641203 | -0.535161191 | 0.26226094 | -2.0405676 | 0.04129382 | 0.10371025 | KLB | 152831 |
| ENSG000002 | 67.1317524 | 0.401002946 | 0.19651938 | 2.04052623 | 0.04129794 | 0.10371418 | NA | NA |
| ENSG000002 | 8.84773915 | -0.71403761 | 0.34993773 | -2.0404705 | 0.04130349 | 0.10372169 | NA | NA |
| ENSG000001 | 16.304886 | -0.622601996 | 0.30514295 | -2.0403617 | 0.04131431 | 0.10373602 | TCF23 | 150921 |
| ENSG000002 | 3.0255836 | -0.769266521 | 0.3770208 | -2.0403822 | 0.04131228 | 0.10373602 | NA | NA |
| ENSG000002 | 6.87140462 | -0.747125246 | 0.36620053 | -2.040208 | 0.04132961 | 0.10376636 | NA | NA |

| | | | | | | | | |
|---|---|---|---|---|---|---|---|---|
| ENSG000002 | 8.4618304 | -0.715385661 | 0.35064678 | -2.0401889 | 0.04133152 | 0.10376636 | LOC158434 | 158434 |
| ENSG000002 | 4.91727187 | -0.764345973 | 0.37469497 | -2.0399152 | 0.04135877 | 0.10382836 | NA | NA |
| ENSG000002 | 3.37831621 | -0.771359096 | 0.37817327 | -2.0396975 | 0.04138046 | 0.10387638 | NA | NA |
| ENSG000001 | 21.7184547 | 0.553646121 | 0.27144773 | 2.03960488 | 0.0413897 | 0.10389313 | FBP1 | 2203 |
| ENSG000002 | 11.6620079 | -0.700319693 | 0.34338307 | -2.0394706 | 0.04140308 | 0.1039203 | NA | NA |
| ENSG000002 | 8.65110679 | -0.726395334 | 0.35618483 | -2.0393775 | 0.04141236 | 0.10393716 | GSN-AS1 | 57000 |
| ENSG000001 | 22.0186769 | -0.564587233 | 0.27685209 | -2.03931 | 0.0414191 | 0.10394763 | EPS8L3 | 79574 |
| ENSG000002 | 10.9466904 | -0.677565508 | 0.33227034 | -2.0391995 | 0.04143012 | 0.10396887 | MIR4458HG | 100505738 |
| ENSG000002 | 4.64236069 | -0.767716374 | 0.37649758 | -2.0391004 | 0.04144001 | 0.10398724 | NA | NA |
| ENSG000002 | 10.6862888 | -0.68259482 | 0.33479807 | -2.0388254 | 0.04146745 | 0.10404824 | NA | NA |
| ENSG000002 | 11.0026106 | -0.675122174 | 0.33113614 | -2.0388054 | 0.04146945 | 0.10404824 | NA | NA |
| ENSG000002 | 2.93724083 | -0.755015506 | 0.37043696 | -2.0381754 | 0.04153239 | 0.10419972 | NA | NA |
| ENSG000002 | 222.112126 | 0.272720732 | 0.13380991 | 2.03812062 | 0.04153787 | 0.10420702 | ZNF225 | 7768 |
| ENSG000002 | 8.26237671 | -0.72650937 | 0.3564927 | -2.0379361 | 0.04155632 | 0.10424685 | NA | NA |
| ENSG000001 | 6.01139642 | -0.759314623 | 0.37260662 | -2.0378452 | 0.04156541 | 0.10426322 | CTRC | 11330 |
| ENSG000001 | 24.0803492 | -0.557017597 | 0.27334503 | -2.0377821 | 0.04157173 | 0.10427261 | KLHDC1 | 122773 |
| ENSG000002 | 52.5478758 | 0.43079305 | 0.21142247 | 2.03759351 | 0.0415906 | 0.10431349 | NA | NA |
| ENSG000001 | 191.495682 | 0.284245972 | 0.13950918 | 2.03747142 | 0.04160282 | 0.10433489 | SLC35G1 | 159371 |
| ENSG000001 | 9.69519427 | -0.703351154 | 0.34521033 | -2.0374569 | 0.04160428 | 0.10433489 | NTF3 | 4908 |
| ENSG000002 | 20.0512627 | -0.568293589 | 0.27892782 | -2.0374217 | 0.0416078 | 0.10433726 | LOC553103 | 553103 |
| ENSG000002 | 12.4152264 | -0.657042429 | 0.32258366 | -2.0368125 | 0.04166883 | 0.10448385 | LINC00487 | 400941 |
| ENSG000002 | 6.22208042 | -0.765036627 | 0.3756142 | -2.0367617 | 0.04167392 | 0.10449016 | NA | NA |
| ENSG000001 | 805.057067 | 0.504112491 | 0.24754405 | 2.03645566 | 0.04170462 | 0.10454773 | PITPNB | 23760 |
| ENSG000002 | 11.6565008 | -0.671578718 | 0.32977377 | -2.0364831 | 0.04170186 | 0.10454773 | NA | NA |
| ENSG000002 | 8.71386963 | -0.709818234 | 0.34855182 | -2.0364784 | 0.04170234 | 0.10454773 | NA | NA |
| ENSG000001 | 5.06449387 | -0.766093949 | 0.37619967 | -2.0364025 | 0.04170995 | 0.10455373 | SPDYE2B | 100310812 |
| ENSG000002 | 5.58226804 | -0.759512096 | 0.37297162 | -2.0363804 | 0.04171217 | 0.10455373 | NA | NA |
| ENSG000001 | 31.0837586 | -0.502257083 | 0.24664937 | -2.0363201 | 0.04171822 | 0.10456243 | SLC25A48 | 153328 |
| ENSG000001 | 331.010778 | 0.256520011 | 0.12598438 | 2.03612549 | 0.04173776 | 0.10459846 | MPP7 | 143098 |

| | | | | | | | | |
|---|---|---|---|---|---|---|---|---|
| ENSG000002 | 4.42988356 | -0.769672315 | 0.37800529 | -2.0361417 | 0.04173613 | 0.10459846 | NA | NA |
| ENSG000002 | 8.123385 | -0.715325126 | 0.35132434 | -2.0360819 | 0.04174213 | 0.10460297 | NA | NA |
| ENSG000001 | 39.0603813 | 0.484709377 | 0.23807424 | 2.03595892 | 0.04175448 | 0.10460805 | ELMO3 | 79767 |
| ENSG000001 | 8.11630885 | -0.733197382 | 0.36011342 | -2.0360179 | 0.04174856 | 0.10460805 | SPRR3 | 6707 |
| ENSG000002 | 5.55279621 | -0.757485555 | 0.37204576 | -2.036001 | 0.04175025 | 0.10460805 | LOC101928298 | 101928298 |
| ENSG000002 | 23.1070698 | -0.549390499 | 0.26984102 | -2.0359784 | 0.04175252 | 0.10460805 | NA | NA |
| ENSG000002 | 3.77649566 | -0.770576963 | 0.37853518 | -2.0356813 | 0.04178237 | 0.10467146 | NA | NA |
| ENSG000002 | 6.82540692 | -0.744778515 | 0.36593849 | -2.035256 | 0.04182512 | 0.10476401 | KRT81 | 3887 |
| ENSG000002 | 33.5599197 | -0.507420423 | 0.24931765 | -2.0352367 | 0.04182707 | 0.10476401 | PATL2 | 197135 |
| ENSG000002 | 10.6901596 | -0.677109168 | 0.33268734 | -2.0352718 | 0.04182353 | 0.10476401 | NA | NA |
| ENSG00000C | 526.257368 | 0.278950598 | 0.13706417 | 2.03518252 | 0.04183251 | 0.10477118 | WWC3 | 55841 |
| ENSG000002 | 12.1632481 | -0.667925479 | 0.32819585 | -2.035143 | 0.04183649 | 0.10477468 | LINC01255 | 101927433 |
| ENSG000002 | 2.75493803 | -0.759133188 | 0.37302657 | -2.0350647 | 0.04184437 | 0.10478793 | MAS1L | 116511 |
| ENSG000002 | 4.04903877 | -0.770144982 | 0.37844283 | -2.0350365 | 0.0418472 | 0.10478854 | NA | NA |
| ENSG000002 | 4.87720281 | -0.769525048 | 0.37815442 | -2.0349493 | 0.04185598 | 0.10480406 | NA | NA |
| ENSG000002 | 2.91131215 | -0.75377761 | 0.37044303 | -2.0348003 | 0.04187097 | 0.10483512 | NA | NA |
| ENSG000001 | 397.4684 | 0.324693131 | 0.15959653 | 2.03446233 | 0.041905 | 0.10490424 | WDR59 | 79726 |
| ENSG000001 | 82.5866438 | 0.373261175 | 0.18347039 | 2.03444918 | 0.04190633 | 0.10490424 | LPAR6 | 10161 |
| ENSG000002 | 3.07428497 | -0.766186416 | 0.37660633 | -2.0344491 | 0.04190634 | 0.10490424 | NA | NA |
| ENSG00000C | 40.6073763 | -0.456120651 | 0.22421367 | -2.0343124 | 0.04192011 | 0.10492577 | PTGS1 | 5742 |
| ENSG000001 | 244.14223 | 0.300340782 | 0.14763718 | 2.03431667 | 0.04191968 | 0.10492577 | ANKRD52 | 283373 |
| ENSG000002 | 3.07075546 | -0.766472726 | 0.37678674 | -2.0342349 | 0.04192792 | 0.10493884 | NA | NA |
| ENSG00000C | 125.511146 | 0.328065207 | 0.16128049 | 2.03412835 | 0.04193866 | 0.10494516 | SNCAIP | 9627 |
| ENSG000001 | 187.066378 | 0.624818742 | 0.30716832 | 2.03412495 | 0.041939 | 0.10494516 | NEFM | 4741 |
| ENSG000001 | 30.191484 | -0.532227416 | 0.26164508 | -2.034158 | 0.04193567 | 0.10494516 | ZNF540 | 163255 |
| ENSG000002 | 3.07095134 | -0.766413945 | 0.37678151 | -2.0341071 | 0.0419408 | 0.10494516 | NA | NA |
| ENSG000002 | 4.38586594 | -0.76881287 | 0.37797777 | -2.0340161 | 0.04194997 | 0.10496164 | CBY3 | 646019 |
| ENSG000002 | 8.98761502 | -0.700770734 | 0.34453207 | -2.0339782 | 0.04195379 | 0.10496472 | NA | NA |
| ENSG000001 | 29.9175208 | -0.52198996 | 0.25664319 | -2.0339131 | 0.04196036 | 0.10497467 | CCDC178 | 374864 |

| | | | | | | | | |
|---|---|---|---|---|---|---|---|---|
| ENSG000002 | 5.86937388 | -0.754580766 | 0.37103465 | -2.0337205 | 0.04197979 | 0.1050168 | NA | NA |
| ENSG000001 | 61.0498962 | -0.410809912 | 0.20201 | -2.0336117 | 0.04199076 | 0.10503265 | NA | NA |
| ENSG000002 | 6.50158475 | -0.747199175 | 0.36742567 | -2.0336063 | 0.0419913 | 0.10503265 | NA | NA |
| ENSG000002 | 3.45942764 | -0.7676288 | 0.37748736 | -2.0335219 | 0.04199983 | 0.10504101 | NA | NA |
| ENSG000002 | 17.2950581 | -0.634995493 | 0.31226173 | -2.0335361 | 0.04199839 | 0.10504101 | NA | NA |
| ENSG000002 | 7.06347303 | -0.735313376 | 0.36160447 | -2.0334743 | 0.04200463 | 0.10504655 | GPR52 | 9293 |
| ENSG000002 | 49.4980931 | -0.434925925 | 0.21391474 | -2.0331741 | 0.04203493 | 0.10511585 | FER1L5 | 90342 |
| ENSG000002 | 23.6077313 | -0.572758162 | 0.28172168 | -2.0330639 | 0.04204607 | 0.10513508 | NA | NA |
| ENSG000002 | 3.37171312 | -0.768245135 | 0.37787875 | -2.0330467 | 0.04204781 | 0.10513508 | RN7SL291P | 106479328 |
| ENSG000002 | 4.04913518 | -0.769433077 | 0.37852855 | -2.032695 | 0.04208335 | 0.10521745 | NA | NA |
| ENSG00000C | 6.47408767 | -0.74445598 | 0.36626466 | -2.032563 | 0.0420967 | 0.10523785 | CPA1 | 1357 |
| ENSG000002 | 16.9740735 | 0.644135292 | 0.31690435 | 2.03258581 | 0.04209439 | 0.10523785 | SNORA47 | 677828 |
| ENSG000001 | 87.0665225 | 0.352111167 | 0.17323749 | 2.03253447 | 0.04209958 | 0.10523856 | KHDRBS2 | 202559 |
| ENSG000002 | 3.56408793 | -0.765519956 | 0.37666898 | -2.0323414 | 0.04211911 | 0.10528089 | NA | NA |
| ENSG000002 | 13.4391051 | -0.645527214 | 0.31764989 | -2.0321972 | 0.04213369 | 0.10531086 | NA | NA |
| ENSG000001 | 165.234188 | 0.320818854 | 0.15787351 | 2.03212592 | 0.04214091 | 0.10532241 | EPHX2 | 2053 |
| ENSG000001 | 292.204486 | 0.272726731 | 0.13422591 | 2.03184865 | 0.04216898 | 0.10538607 | OGFRL1 | 79627 |
| ENSG000002 | 5.01732054 | -0.763266983 | 0.37566909 | -2.0317535 | 0.04217862 | 0.10540366 | NA | NA |
| ENSG000001 | 12.5588974 | -0.646334997 | 0.31813251 | -2.0316534 | 0.04218875 | 0.10542249 | DLX4 | 1748 |
| ENSG000002 | 2.7466531 | -0.757947611 | 0.3730802 | -2.0315943 | 0.04219475 | 0.10543097 | NA | NA |
| ENSG000002 | 3.86137016 | -0.768076854 | 0.37808957 | -2.031468 | 0.04220754 | 0.10545645 | NA | NA |
| ENSG000001 | 112.254815 | 0.347448772 | 0.17104283 | 2.03135537 | 0.04221896 | 0.10547846 | TMCC2 | 9911 |
| ENSG000001 | 161.863073 | 0.312314928 | 0.15375233 | 2.03128585 | 0.04222601 | 0.10548958 | ZNF44 | 51710 |
| ENSG000002 | 10.0100663 | -0.697393518 | 0.34334775 | -2.031158 | 0.04223897 | 0.10551546 | LOC388942 | 388942 |
| ENSG000002 | 3.43631525 | -0.767743938 | 0.37799996 | -2.0310688 | 0.04224801 | 0.10553154 | NA | NA |
| ENSG000001 | 91.8000954 | 0.393736878 | 0.19388561 | 2.03076897 | 0.04227844 | 0.10560104 | CTIF | 9811 |
| ENSG000002 | 3.42507405 | -0.767800101 | 0.37809989 | -2.0306805 | 0.04228741 | 0.10561695 | NA | NA |
| ENSG000001 | 8.59999552 | 0.704169889 | 0.34678967 | 2.03053881 | 0.0423018 | 0.10563742 | BGN | 633 |
| ENSG000002 | 162.779096 | 0.323222743 | 0.15918404 | 2.03049717 | 0.04230603 | 0.10563742 | NA | NA |

| | | | | | | | | |
|---|---|---|---|---|---|---|---|---|
| ENSG000002 | 14.9058159 | -0.62861976 | 0.30958742 | -2.0305081 | 0.04230492 | 0.10563742 | NA | NA |
| ENSG000002 | 5.59329229 | -0.755208394 | 0.37192085 | -2.0305621 | 0.04229943 | 0.10563742 | NA | NA |
| ENSG000002 | 4.08007246 | -0.768604611 | 0.37853999 | -2.030445 | 0.04231133 | 0.10564416 | NA | NA |
| ENSG000002 | 6.83284418 | -0.741980101 | 0.36543589 | -2.0303975 | 0.04231615 | 0.10564678 | LOC102724919 | 102724919 |
| ENSG000002 | 6.42308238 | -0.743741006 | 0.36630571 | -2.0303833 | 0.04231759 | 0.10564678 | NA | NA |
| ENSG000002 | 6.16521558 | -0.747923357 | 0.36837617 | -2.030325 | 0.04232351 | 0.10565506 | NA | NA |
| ENSG000001 | 50.6619069 | 0.435865996 | 0.21468772 | 2.03023252 | 0.04233291 | 0.10567201 | GAMT | 2593 |
| ENSG000002 | 3.01106826 | -0.765473465 | 0.37705093 | -2.0301593 | 0.04234035 | 0.10568407 | NA | NA |
| ENSG000001 | 9.38658577 | -0.706510744 | 0.34802191 | -2.0300755 | 0.04234887 | 0.10569233 | UGT2B7 | 7364 |
| ENSG000002 | 3.4157387 | -0.767496818 | 0.37805865 | -2.0300999 | 0.04234639 | 0.10569233 | NA | NA |
| ENSG000002 | 2.72533934 | -0.757606901 | 0.37321633 | -2.0299404 | 0.0423626 | 0.10572009 | NA | NA |
| ENSG000001 | 15.7709097 | -0.605393174 | 0.29825069 | -2.0298132 | 0.04237553 | 0.10573915 | DKK2 | 27123 |
| ENSG000001 | 7.63247982 | -0.72521817 | 0.35729987 | -2.0297185 | 0.04238516 | 0.10573915 | PGLYRP3 | 114771 |
| ENSG000001 | 14.2561236 | -0.651287545 | 0.32086516 | -2.0297858 | 0.04237832 | 0.10573915 | TMEM31 | 203562 |
| ENSG000002 | 5.50389483 | -0.752505681 | 0.37074457 | -2.0297146 | 0.04238556 | 0.10573915 | NA | NA |
| ENSG000002 | 10.1026873 | -0.685424977 | 0.33768012 | -2.0298055 | 0.04237631 | 0.10573915 | NA | NA |
| ENSG000002 | 13.4962894 | -0.66721399 | 0.32872358 | -2.0297114 | 0.04238588 | 0.10573915 | OR8G5 | 219865 |
| ENSG000002 | 6.56667871 | -0.748820532 | 0.36894622 | -2.0296197 | 0.04239521 | 0.10575006 | NA | NA |
| ENSG000002 | 3.01482644 | -0.765274198 | 0.37705401 | -2.0296143 | 0.04239576 | 0.10575006 | NA | NA |
| ENSG000002 | 2.40916139 | -0.747987667 | 0.36854099 | -2.0295915 | 0.04239808 | 0.10575006 | NA | NA |
| ENSG000002 | 7.83091531 | -0.736047373 | 0.36267351 | -2.0295041 | 0.04240697 | 0.10576573 | NA | NA |
| ENSG000001 | 97.6082995 | -0.380598085 | 0.18754887 | -2.0293275 | 0.04242495 | 0.10580297 | EPS8L2 | 64787 |
| ENSG000002 | 15.7258349 | -0.62653095 | 0.30874146 | -2.0293062 | 0.04242712 | 0.10580297 | LOC101929570 | 101929570 |
| ENSG000001 | 12.7514421 | -0.660710994 | 0.32561803 | -2.0290983 | 0.04244828 | 0.10584923 | PCDHB18P | 54660 |
| ENSG000001 | 9.67012322 | -0.699693832 | 0.34486425 | -2.0288964 | 0.04246885 | 0.1058906 | ZIM3 | 114026 |
| ENSG000002 | 9.92044465 | -0.695703143 | 0.3428994 | -2.0288841 | 0.04247009 | 0.1058906 | OR13A1 | 79290 |
| ENSG000002 | 3.01061125 | -0.764542412 | 0.37688709 | -2.0285715 | 0.04250195 | 0.10596353 | NA | NA |
| ENSG000000 | 236.76008 | 0.271099165 | 0.13364658 | 2.02847807 | 0.04251148 | 0.10598076 | FRY | 10129 |
| ENSG000001 | 584.225952 | 0.249543982 | 0.12302872 | 2.02833922 | 0.04252564 | 0.10600954 | CEP152 | 22995 |

| | | | | | | | | |
|---|---|---|---|---|---|---|---|---|
| ENSG000002 | 21.8739365 | -0.598688044 | 0.29517405 | -2.0282543 | 0.0425343 | 0.10602461 | ZNF469 | 84627 |
| ENSG000002 | 3.41332323 | -0.766448123 | 0.37792599 | -2.0280376 | 0.04255641 | 0.10606977 | NA | NA |
| ENSG000002 | 3.5260217 | -0.76481621 | 0.37712357 | -2.0280255 | 0.04255765 | 0.10606977 | NA | NA |
| ENSG000002 | 5.54941051 | -0.754519083 | 0.37207563 | -2.0278648 | 0.04257404 | 0.10610412 | NA | NA |
| ENSG000002 | 3.22230422 | -0.761288905 | 0.37543012 | -2.0277779 | 0.04258292 | 0.10611971 | MIR1273G | 100616145 |
| ENSG000001 | 260.949784 | 0.319629532 | 0.15764269 | 2.02755691 | 0.04260549 | 0.10616944 | SMAD3 | 4088 |
| ENSG000001 | 6.64089359 | -0.733651943 | 0.3618558 | -2.0274704 | 0.04261433 | 0.10618493 | MS4A10 | 341116 |
| ENSG000002 | 3.09269884 | -0.762133366 | 0.37595821 | -2.0271758 | 0.04264444 | 0.10625344 | NA | NA |
| ENSG000002 | 6.36500162 | -0.751838297 | 0.37088723 | -2.0271345 | 0.04264866 | 0.10625742 | NA | NA |
| ENSG000002 | 6.83096013 | -0.739399229 | 0.36477158 | -2.0270198 | 0.04266038 | 0.1062801 | NA | NA |
| ENSG000001 | 12.9017511 | -0.663404001 | 0.3273061 | -2.0268611 | 0.04267662 | 0.10630749 | GPR87 | 53836 |
| ENSG000002 | 3.86130124 | -0.765813775 | 0.37782959 | -2.0268761 | 0.04267509 | 0.10630749 | NA | NA |
| ENSG000002 | 11.5948932 | -0.670507242 | 0.33082593 | -2.0267675 | 0.04268619 | 0.10632481 | NA | NA |
| ENSG000002 | 16.0942643 | -0.606435205 | 0.29921962 | -2.0267227 | 0.04269078 | 0.10632969 | NA | NA |
| ENSG000002 | 2.89303796 | -0.753115673 | 0.37160254 | -2.0266699 | 0.04269618 | 0.1063301 | NA | NA |
| ENSG000002 | 5.47976807 | -0.752656313 | 0.37137216 | -2.0266902 | 0.04269411 | 0.1063301 | NA | NA |
| ENSG000001 | 196.225459 | 0.315698896 | 0.15579286 | 2.02640159 | 0.04272365 | 0.10637889 | ZC3H10 | 84872 |
| ENSG000002 | 5.67620883 | -0.756723166 | 0.37343044 | -2.0264099 | 0.04272279 | 0.10637889 | NA | NA |
| ENSG000002 | 6.10917445 | -0.745890679 | 0.36808086 | -2.0264316 | 0.04272058 | 0.10637889 | NA | NA |
| ENSG000002 | 43.4977285 | 0.518199063 | 0.2557441 | 2.02624053 | 0.04274014 | 0.10641343 | NA | NA |
| ENSG000002 | 7.11176655 | -0.735697383 | 0.36309128 | -2.026205 | 0.04274378 | 0.10641594 | NA | NA |
| ENSG000001 | 30.8629317 | -0.495964258 | 0.24480706 | -2.0259394 | 0.042771 | 0.10647137 | CCNA1 | 8900 |
| ENSG000002 | 17.8222385 | 0.609403975 | 0.3008037 | 2.02591913 | 0.04277307 | 0.10647137 | NA | NA |
| ENSG000002 | 37.1362345 | 0.487172108 | 0.24047199 | 2.02589957 | 0.04277508 | 0.10647137 | DPP10-AS1 | 389023 |
| ENSG000002 | 12.9378655 | -0.667733247 | 0.32960072 | -2.0258853 | 0.04277654 | 0.10647137 | ZFHX4-AS1 | 100192378 |
| ENSG000001 | 6.76633263 | -0.740889832 | 0.36571752 | -2.0258527 | 0.04277988 | 0.10647314 | GIMAP7 | 168537 |
| ENSG000002 | 2.46614028 | -0.743540913 | 0.36705127 | -2.0257141 | 0.04279409 | 0.10650199 | NA | NA |
| ENSG000002 | 2.21169749 | 0.73914674 | 0.36489063 | 2.02566657 | 0.04279897 | 0.10650757 | NA | NA |
| ENSG000002 | 17.1493011 | -0.594379569 | 0.29344318 | -2.0255354 | 0.04281241 | 0.1065345 | LINC00052 | 145978 |

| | | | | | | | | |
|---|---|---|---|---|---|---|---|---|
| ENSG000002 | 6.98605309 | -0.731760301 | 0.36129284 | -2.0253939 | 0.04282693 | 0.10656409 | NA | NA |
| ENSG000001 | 223.004648 | 0.322800084 | 0.15938384 | 2.02529996 | 0.04283657 | 0.10658154 | ARHGAP18 | 93663 |
| ENSG000001 | 17.848294 | -0.608977593 | 0.30069011 | -2.0252664 | 0.04284001 | 0.10658356 | SLC39A4 | 55630 |
| ENSG000001 | 41.0361315 | -0.457839635 | 0.22609455 | -2.024992 | 0.04286819 | 0.10664712 | SYBU | 55638 |
| ENSG000002 | 5.25232417 | -0.757999669 | 0.37432791 | -2.0249617 | 0.0428713 | 0.10664831 | NA | NA |
| ENSG000001 | 46.397752 | 0.448803056 | 0.22164707 | 2.02485448 | 0.04288231 | 0.10666915 | PRRT2 | 112476 |
| ENSG000002 | 16.3586569 | -0.607527233 | 0.30007609 | -2.0245773 | 0.04291079 | 0.10673345 | NA | NA |
| ENSG000001 | 78.3700196 | 0.370796699 | 0.18316442 | 2.02439266 | 0.04292977 | 0.1067741 | NA | NA |
| ENSG000002 | 3.38960947 | -0.765355823 | 0.37807622 | -2.0243427 | 0.04293491 | 0.10678034 | NA | NA |
| ENSG000002 | 3.44345913 | -0.765044916 | 0.37799435 | -2.0239586 | 0.04297441 | 0.10687204 | NA | NA |
| ENSG000002 | 55.3998236 | 0.401744714 | 0.19850598 | 2.02384188 | 0.04298642 | 0.10689535 | NA | NA |
| ENSG000002 | 4.08240883 | -0.765977899 | 0.37850845 | -2.0236745 | 0.04300366 | 0.10693164 | NA | NA |
| ENSG000002 | 13.1856561 | -0.664602577 | 0.32842735 | -2.0235909 | 0.04301227 | 0.10694649 | MTND6P9 | 106478946 |
| ENSG000001 | 28.3495379 | -0.542329271 | 0.26802307 | -2.0234425 | 0.04302754 | 0.10697793 | CLCA3P | 9629 |
| ENSG000002 | 8.52883164 | -0.71571999 | 0.35374086 | -2.023289 | 0.04304336 | 0.10701069 | LINC00692 | 285326 |
| ENSG00000C | 25.049975 | -0.524532111 | 0.25925404 | -2.023236 | 0.04304882 | 0.1070177 | CYP24A1 | 1591 |
| ENSG000001 | 7.28829329 | -0.725431617 | 0.35861926 | -2.0228462 | 0.043089 | 0.10711102 | TEX36 | 387718 |
| ENSG000002 | 83.5282468 | -0.382613607 | 0.18916729 | -2.0226203 | 0.04311231 | 0.10714437 | LAYN | 143903 |
| ENSG000002 | 5.53372186 | -0.751257736 | 0.37141793 | -2.0226749 | 0.04310667 | 0.10714437 | NA | NA |
| ENSG000002 | 11.426129 | -0.661642382 | 0.3271163 | -2.0226518 | 0.04310906 | 0.10714437 | LINC01119 | 100134259 |
| ENSG000002 | 11.5951135 | -0.672713675 | 0.33259622 | -2.0226137 | 0.04311299 | 0.10714437 | NA | NA |
| ENSG000002 | 5.46308555 | -0.751408106 | 0.37154168 | -2.0224059 | 0.04313443 | 0.10718817 | POU6F2-AS1 | 100861520 |
| ENSG000002 | 8.13562905 | -0.733880251 | 0.3628774 | -2.0223917 | 0.0431359 | 0.10718817 | LINC00457 | 100874179 |
| ENSG000001 | 7.86510427 | -0.717012219 | 0.35457569 | -2.0221697 | 0.04315882 | 0.10723857 | ADAM30 | 11085 |
| ENSG000001 | 18.3370015 | -0.587175927 | 0.29038831 | -2.0220371 | 0.04317252 | 0.10725289 | ABCG5 | 64240 |
| ENSG000002 | 8.29827035 | -0.721014271 | 0.35657011 | -2.0220827 | 0.04316781 | 0.10725289 | SPATA31B1P | 404770 |
| ENSG000002 | 2.4249825 | -0.745041443 | 0.36845998 | -2.0220417 | 0.04317205 | 0.10725289 | NA | NA |
| ENSG000002 | 3.7897002 | -0.765234449 | 0.37845645 | -2.0219881 | 0.04317758 | 0.10725888 | NA | NA |
| ENSG000001 | 745.990045 | 0.305327332 | 0.15100786 | 2.02193004 | 0.04318358 | 0.10726722 | PEAK1 | 79834 |

| | | | | | | | | |
|---|---|---|---|---|---|---|---|---|
| ENSG000002 | 3.08136359 | -0.760825949 | 0.37632274 | -2.0217379 | 0.04320343 | 0.10730996 | NA | NA |
| ENSG000001 | 26.1600908 | -0.525098342 | 0.25972986 | -2.0217096 | 0.04320636 | 0.10731066 | AMZ1 | 155185 |
| ENSG000002 | 554.876642 | -0.336027063 | 0.16621284 | -2.0216673 | 0.04321073 | 0.10731495 | NA | NA |
| ENSG000001 | 109.536059 | -0.341531168 | 0.16893763 | -2.0216406 | 0.04321349 | 0.10731523 | ADCY8 | 114 |
| ENSG000002 | 5.76051322 | -0.744754635 | 0.3684076 | -2.0215507 | 0.04322279 | 0.10732964 | NA | NA |
| ENSG000002 | 8.78322876 | -0.710379151 | 0.35140611 | -2.0215333 | 0.04322459 | 0.10732964 | NA | NA |
| ENSG000002 | 6.79222774 | -0.737099532 | 0.36463932 | -2.0214483 | 0.04323338 | 0.10734489 | NA | NA |
| ENSG000002 | 2.87507399 | -0.750300571 | 0.37118252 | -2.0213791 | 0.04324054 | 0.1073561 | NA | NA |
| ENSG000002 | 3.11716942 | -0.758970241 | 0.37547795 | -2.0213444 | 0.04324413 | 0.10735843 | NA | NA |
| ENSG000001 | 55.9445419 | 0.407697878 | 0.20172549 | 2.02105289 | 0.04327429 | 0.10742674 | HARBI1 | 283254 |
| ENSG000002 | 8.12537503 | -0.713090214 | 0.35287429 | -2.0208053 | 0.04329993 | 0.1074838 | LOC105374524 | 105374524 |
| ENSG000002 | 4.50412551 | 0.763952532 | 0.37808011 | 2.02061022 | 0.04332013 | 0.10752737 | NA | NA |
| ENSG000001 | 412.219565 | 0.26336985 | 0.13035047 | 2.0204748 | 0.04333416 | 0.10754245 | KIAA1715 | 80856 |
| ENSG000001 | 395.599145 | 0.260829615 | 0.12909094 | 2.02051068 | 0.04333044 | 0.10754245 | CCDC150 | 284992 |
| ENSG000002 | 5.0165878 | -0.760364551 | 0.37632959 | -2.0204751 | 0.04333413 | 0.10754245 | NA | NA |
| ENSG000001 | 107.095206 | 0.356640188 | 0.17652216 | 2.02037067 | 0.04334495 | 0.10756235 | KIF17 | 57576 |
| ENSG000002 | 4.65157817 | -0.762009104 | 0.37716758 | -2.0203462 | 0.04334748 | 0.10756235 | NA | NA |
| ENSG000001 | 20.4152762 | -0.62429241 | 0.30900962 | -2.0203009 | 0.04335219 | 0.10756743 | FGF5 | 2250 |
| ENSG000002 | 2.40895287 | -0.744740001 | 0.36863727 | -2.0202515 | 0.04335731 | 0.10757355 | NA | NA |
| ENSG000001 | 405.007182 | 0.286481942 | 0.14182238 | 2.02000516 | 0.04338285 | 0.10761168 | PDCD4 | 27250 |
| ENSG000001 | 102.809402 | 0.34375637 | 0.17017628 | 2.02000161 | 0.04338322 | 0.10761168 | TMEM144 | 55314 |
| ENSG000001 | 10.5793686 | -0.706139369 | 0.34957378 | -2.020001 | 0.04338329 | 0.10761168 | MRGPRX3 | 117195 |
| ENSG000002 | 9.7561541 | -0.718200323 | 0.35553478 | -2.0200565 | 0.04337753 | 0.10761168 | NA | NA |
| ENSG000002 | 5.56472419 | -0.750398296 | 0.3714973 | -2.0199293 | 0.04339072 | 0.10762353 | NA | NA |
| ENSG000002 | 164.007101 | 0.294301106 | 0.14571169 | 2.01974951 | 0.04340938 | 0.10766322 | ZNF658 | 26149 |
| ENSG000002 | 6.5524039 | -0.745601516 | 0.36916467 | -2.019699 | 0.04341462 | 0.10766964 | NA | NA |
| ENSG000001 | 98.0248511 | 0.346424282 | 0.17152951 | 2.01961916 | 0.04342291 | 0.1076836 | ADAMTS10 | 81794 |
| ENSG000001 | 13.183318 | -0.657198121 | 0.32543615 | -2.019438 | 0.04344172 | 0.10771049 | IL2RG | 3561 |
| ENSG000002 | 2.38437019 | -0.743333138 | 0.36808822 | -2.0194429 | 0.04344121 | 0.10771049 | RNU6ATAC31P | 106481169 |

| | | | | | | | | |
|---|---|---|---|---|---|---|---|---|
| ENSG000002 | 7.67739544 | -0.726365817 | 0.35968018 | -2.0194769 | 0.04343767 | 0.10771049 | NA | NA |
| ENSG000002 | 2.4168455 | -0.742121912 | 0.36749614 | -2.0194006 | 0.0434456 | 0.10771354 | NA | NA |
| ENSG000001 | 11.8143535 | -0.686554677 | 0.33999333 | -2.0193181 | 0.04345417 | 0.10772161 | CES1 | 1066 |
| ENSG000002 | 5.93280717 | -0.749230771 | 0.37102999 | -2.0193267 | 0.04345327 | 0.10772161 | NA | NA |
| ENSG000001 | 40.7806453 | -0.477579399 | 0.2365483 | -2.0189509 | 0.04349232 | 0.10780538 | CLCN1 | 1180 |
| ENSG000002 | 15.3036208 | -0.612544745 | 0.30339893 | -2.0189417 | 0.04349328 | 0.10780538 | NA | NA |
| ENSG000001 | 340.835308 | 0.253986594 | 0.12581324 | 2.01875884 | 0.04351229 | 0.10784584 | TEX9 | 374618 |
| ENSG000002 | 16.6653777 | -0.619403459 | 0.30682774 | -2.0187335 | 0.04351492 | 0.10784584 | NA | NA |
| ENSG00000C | 2927.4714 | 0.275725764 | 0.13658734 | 2.01867729 | 0.04352077 | 0.10785375 | RSF1 | 51773 |
| ENSG000001 | 21.5956501 | -0.547793921 | 0.27137007 | -2.0186232 | 0.04352639 | 0.10786109 | CCDC63 | 160762 |
| ENSG000001 | 9.49064066 | -0.687771396 | 0.34072594 | -2.0185472 | 0.0435343 | 0.10787409 | TREML3P | 340206 |
| ENSG000002 | 345.860579 | 0.287038864 | 0.14220338 | 2.01850944 | 0.04353823 | 0.10787724 | ZNF10 | 7556 |
| ENSG000002 | 6.87715894 | -0.740121595 | 0.36668039 | -2.0184379 | 0.04354568 | 0.10788909 | CFHR1 | 3078 |
| ENSG000001 | 171.877891 | 0.314936747 | 0.15603371 | 2.01838918 | 0.04355075 | 0.10789506 | CYB561A3 | 220002 |
| ENSG000002 | 2.24328371 | -0.719085859 | 0.35639791 | -2.0176489 | 0.04362784 | 0.10807945 | NA | NA |
| ENSG000002 | 10.1714533 | -0.689502683 | 0.34177041 | -2.0174441 | 0.04364919 | 0.10812574 | NA | NA |
| ENSG000002 | 13.053098 | -0.653567244 | 0.32397106 | -2.0173631 | 0.04365764 | 0.10814006 | NA | NA |
| ENSG000001 | 8.81817684 | -0.709948386 | 0.35194635 | -2.0172063 | 0.04367399 | 0.10817396 | SIGLEC17P | 284367 |
| ENSG000002 | 3.77206389 | -0.762781618 | 0.37815329 | -2.0171228 | 0.0436827 | 0.10818893 | LOC101927418 | 101927418 |
| ENSG000002 | 5.26443254 | -0.754625754 | 0.37414431 | -2.0169377 | 0.04370202 | 0.10821693 | NA | NA |
| ENSG000002 | 3.85097753 | -0.762251584 | 0.37791786 | -2.0169769 | 0.04369793 | 0.10821693 | NA | NA |
| ENSG000002 | 6.41598935 | -0.736772004 | 0.36529222 | -2.0169387 | 0.04370192 | 0.10821693 | NA | NA |
| ENSG000002 | 3.98714106 | 0.763198771 | 0.37843948 | 2.01669968 | 0.04372686 | 0.10827185 | NA | NA |
| ENSG000002 | 10.1581348 | -0.690478356 | 0.34245384 | -2.016267 | 0.04377207 | 0.10837716 | LOC101928837 | 101928837 |
| ENSG000001 | 144.809608 | 0.304183444 | 0.15087958 | 2.01606769 | 0.0437929 | 0.10842211 | PDZRN3 | 23024 |
| ENSG000001 | 37.9284145 | -0.453827176 | 0.22514521 | -2.0157087 | 0.04383044 | 0.10850844 | CPM | 1368 |
| ENSG000001 | 7.02137833 | -0.730321328 | 0.36235508 | -2.0154853 | 0.04385383 | 0.10855308 | GCG | 2641 |
| ENSG000002 | 4.32671556 | -0.761302803 | 0.37772547 | -2.0154923 | 0.04385309 | 0.10855308 | NA | NA |
| ENSG000001 | 212.479001 | 0.272196796 | 0.13506375 | 2.01532089 | 0.04387104 | 0.10858905 | ZFP37 | 7539 |

| | | | | | | | | |
|---|---|---|---|---|---|---|---|---|
| ENSG000001 | 15.0762879 | -0.651613896 | 0.32338998 | -2.0149477 | 0.04391013 | 0.10867918 | GRM6 | 2916 |
| ENSG000002 | 9.88665204 | -0.691727871 | 0.34331156 | -2.0148691 | 0.04391836 | 0.10868629 | LINC01486 | 101928138 |
| ENSG000002 | 7.22690845 | -0.735618774 | 0.3650914 | -2.0148893 | 0.04391625 | 0.10868629 | NA | NA |
| ENSG000002 | 15.7894565 | -0.625836037 | 0.31061391 | -2.0148358 | 0.04392185 | 0.1086883 | NA | NA |
| ENSG000002 | 2.67985847 | -0.754244837 | 0.37436536 | -2.0147292 | 0.04393303 | 0.10870931 | NA | NA |
| ENSG000002 | 3.42139965 | -0.760361251 | 0.37742034 | -2.0146271 | 0.04394373 | 0.10872916 | NA | NA |
| ENSG000001 | 38.2476381 | -0.46361413 | 0.23012777 | -2.0145944 | 0.04394716 | 0.10873101 | BAALC | 79870 |
| ENSG000002 | 13.5020788 | -0.63932408 | 0.31735282 | -2.014553 | 0.04395151 | 0.10873513 | MIR4500HG | 642345 |
| ENSG000002 | 34.2097141 | 0.477240449 | 0.23690498 | 2.01448045 | 0.04395912 | 0.10874732 | SPRY4-IT1 | 100642175 |
| ENSG000002 | 3.0964475 | -0.757723226 | 0.37623179 | -2.0139798 | 0.04401165 | 0.10887064 | LOC100509620 | 100509620 |
| ENSG000002 | 4.10445925 | -0.762398115 | 0.37856389 | -2.0139219 | 0.04401773 | 0.10887905 | NA | NA |
| ENSG000002 | 4.41994739 | -0.761575739 | 0.37818021 | -2.0137906 | 0.04403153 | 0.10889988 | NA | NA |
| ENSG000002 | 3.00541145 | -0.758165207 | 0.3764837 | -2.0138062 | 0.04402989 | 0.10889988 | NA | NA |
| ENSG00000C | 12.5474472 | -0.643288922 | 0.31947915 | -2.0135553 | 0.04405625 | 0.10894739 | PCDHGA2 | 56113 |
| ENSG000002 | 7.1811631 | -0.721021437 | 0.35808468 | -2.0135501 | 0.04405679 | 0.10894739 | NA | NA |
| ENSG000002 | 2.72509139 | -0.75047867 | 0.37271771 | -2.013531 | 0.04405879 | 0.10894739 | NA | NA |
| ENSG000001 | 6.30644959 | -0.74664993 | 0.37084432 | -2.0133784 | 0.04407484 | 0.10898042 | ADRB3 | 155 |
| ENSG000002 | 7.29344157 | -0.722413046 | 0.35884837 | -2.0131429 | 0.0440996 | 0.10903499 | NA | NA |
| ENSG000002 | 7.33113596 | -0.725073038 | 0.36019084 | -2.0130247 | 0.04411203 | 0.10905244 | NA | NA |
| ENSG000002 | 10.4595572 | -0.6809444 | 0.33826924 | -2.0130249 | 0.04411201 | 0.10905244 | NA | NA |
| ENSG000002 | 13.5496252 | -0.640146508 | 0.31804551 | -2.0127513 | 0.0441408 | 0.10911423 | NA | NA |
| ENSG000002 | 8.13281285 | -0.709336382 | 0.35242396 | -2.012736 | 0.04414241 | 0.10911423 | NA | NA |
| ENSG000001 | 8.11154627 | -0.70754845 | 0.35155446 | -2.0126283 | 0.04415375 | 0.1091356 | C10orf71 | 118461 |
| ENSG000002 | 6.33344596 | -0.735746232 | 0.36557551 | -2.0125698 | 0.04415991 | 0.10914417 | NA | NA |
| ENSG000002 | 9.91147355 | -0.685706801 | 0.34073246 | -2.0124493 | 0.0441726 | 0.10916888 | LINC00581 | 100874531 |
| ENSG000002 | 6.92946376 | -0.726592592 | 0.36106536 | -2.0123575 | 0.04418227 | 0.10918612 | NA | NA |
| ENSG000001 | 29.3771185 | -0.510303464 | 0.25359551 | -2.0122733 | 0.04419114 | 0.10919092 | FRMD7 | 90167 |
| ENSG000001 | 16.0075259 | 0.599371751 | 0.29785323 | 2.01230568 | 0.04418772 | 0.10919092 | NA | NA |
| ENSG000002 | 3.47494132 | -0.759343204 | 0.37735795 | -2.0122624 | 0.04419229 | 0.10919092 | NA | NA |

| | | | | | | | | |
|---|---|---|---|---|---|---|---|---|
| ENSG000002 | 3.07787498 | -0.754205434 | 0.37483964 | -2.0120749 | 0.04421204 | 0.10923307 | NA | NA |
| ENSG000002 | 2.39005926 | -0.740774879 | 0.36817915 | -2.0119957 | 0.04422039 | 0.10924704 | NA | NA |
| ENSG000001 | 37.8555881 | -0.538264875 | 0.26755836 | -2.0117662 | 0.04424459 | 0.10930017 | GUCY2F | 2986 |
| ENSG000001 | 3.7418134 | -0.76119039 | 0.37839666 | -2.0116203 | 0.04425998 | 0.10931822 | NA | NA |
| ENSG000001 | 128.084982 | 0.335778698 | 0.16691709 | 2.01164963 | 0.04425688 | 0.10931822 | IMMP2L | 83943 |
| ENSG000002 | 6.38302945 | -0.735546206 | 0.36564518 | -2.0116393 | 0.04425797 | 0.10931822 | NA | NA |
| ENSG000001 | 1657.98341 | 0.23187147 | 0.11527427 | 2.0114763 | 0.04427517 | 0.10934908 | GCNT2 | 2651 |
| ENSG000001 | 21.7536 | -0.58729133 | 0.29199061 | -2.0113364 | 0.04428994 | 0.10937889 | MMP20 | 9313 |
| ENSG000001 | 93.6267451 | 0.339783873 | 0.16894502 | 2.01120972 | 0.04430331 | 0.10939858 | ALDH5A1 | 7915 |
| ENSG000001 | 202.632196 | 0.314220193 | 0.1562334 | 2.01122289 | 0.04430192 | 0.10939858 | ZNF165 | 7718 |
| ENSG000002 | 8.41777148 | -0.712353183 | 0.35421666 | -2.0110663 | 0.04431846 | 0.10942932 | NA | NA |
| ENSG000002 | 122.439994 | 0.337031369 | 0.16759648 | 2.01096926 | 0.0443287 | 0.10944129 | DHRS4-AS1 | 55449 |
| ENSG000002 | 4.92405102 | -0.75436281 | 0.37512061 | -2.0109874 | 0.04432679 | 0.10944129 | NA | NA |
| ENSG000001 | 717.994345 | 0.334436904 | 0.16631977 | 2.01080666 | 0.04434588 | 0.10947704 | CHAMP1 | 283489 |
| ENSG000002 | 3.04243476 | -0.757130756 | 0.37654656 | -2.0107228 | 0.04435474 | 0.10949225 | RN7SL240P | 106479310 |
| ENSG000001 | 28.3593916 | 0.521210623 | 0.25922021 | 2.01068667 | 0.04435856 | 0.10949501 | CCDC167 | 154467 |
| ENSG000001 | 35.9917042 | 0.469343187 | 0.23343191 | 2.01062136 | 0.04436547 | 0.10950539 | EFEMP2 | 30008 |
| ENSG000002 | 6.46716294 | -0.735960538 | 0.36604934 | -2.0105501 | 0.044373 | 0.10951731 | NA | NA |
| ENSG000002 | 6.35069425 | -0.74844267 | 0.37226275 | -2.0105226 | 0.0443759 | 0.10951782 | NA | NA |
| ENSG000001 | 9.26995825 | -0.690943831 | 0.34368852 | -2.010378 | 0.0443912 | 0.10954485 | CATSPER3 | 347732 |
| ENSG000001 | 5.29007967 | -0.752228826 | 0.3741747 | -2.010368 | 0.04439226 | 0.10954485 | FCN2 | 2220 |
| ENSG000002 | 4.03316558 | -0.761010247 | 0.37855337 | -2.0103116 | 0.04439822 | 0.10955288 | OR1F2P | 26184 |
| ENSG000002 | 106.65962 | 0.335152188 | 0.16673086 | 2.01013893 | 0.04441649 | 0.1095913 | NA | NA |
| ENSG000002 | 5.09229145 | -0.757977454 | 0.3770906 | -2.0100672 | 0.04442408 | 0.10960335 | NA | NA |
| ENSG000001 | 2161.30966 | 0.218062113 | 0.10849343 | 2.00991068 | 0.04444064 | 0.10963756 | RPL26 | 6154 |
| ENSG000002 | 3.03355643 | -0.756937258 | 0.37660904 | -2.0098754 | 0.04444437 | 0.10964009 | NA | NA |
| ENSG000001 | 28.1044385 | -0.50654324 | 0.25203191 | -2.0098377 | 0.04444837 | 0.10964327 | VIPR2 | 7434 |
| ENSG000002 | 281.70227 | 0.258895396 | 0.12882169 | 2.00971893 | 0.04446095 | 0.10966763 | NEFL | 4747 |
| ENSG000001 | 351.991126 | 0.258586435 | 0.12867434 | 2.00961923 | 0.0444715 | 0.109687 | KLHL3 | 26249 |

| | | | | | | | | |
|---|---|---|---|---|---|---|---|---|
| ENSG000002 | 4.2722736 | -0.758498356 | 0.37746291 | -2.0094646 | 0.04448788 | 0.10971405 | NA | NA |
| ENSG000002 | 93.0327832 | -0.340474399 | 0.16943426 | -2.0094779 | 0.04448647 | 0.10971405 | NA | NA |
| ENSG000001 | 701.898797 | 0.240114721 | 0.11950233 | 2.00928898 | 0.0445065 | 0.10973626 | KIAA0922 | 23240 |
| ENSG000001 | 2.38080843 | -0.740419661 | 0.36850045 | -2.0092775 | 0.04450771 | 0.10973626 | KRTAP19-5 | 337972 |
| ENSG000002 | 4.97266066 | -0.754592953 | 0.37554896 | -2.0093065 | 0.04450464 | 0.10973626 | LOC105377671 | 105377671 |
| ENSG000002 | 4.04102652 | -0.760688537 | 0.378581 | -2.0093151 | 0.04450372 | 0.10973626 | NA | NA |
| ENSG000001 | 15.5556563 | -0.610741541 | 0.30397141 | -2.0092072 | 0.04451516 | 0.10974795 | C1orf64 | 149563 |
| ENSG000002 | 2.24580008 | 0.73009706 | 0.36339368 | 2.00910777 | 0.04452571 | 0.10976728 | LOC105370400 | 105370400 |
| ENSG000001 | 12.0612322 | -0.689628303 | 0.34326588 | -2.0090208 | 0.04453493 | 0.10978333 | NA | NA |
| ENSG000001 | 14.6177081 | -0.652709624 | 0.3248946 | -2.0089888 | 0.04453832 | 0.10978502 | ANXA13 | 312 |
| ENSG000002 | 2.39581304 | -0.739183323 | 0.36796991 | -2.0088146 | 0.0445568 | 0.10981723 | NA | NA |
| ENSG000002 | 6.38218068 | -0.734280976 | 0.36552653 | -2.0088309 | 0.04455507 | 0.10981723 | HNRNPA1P62 | 106480235 |
| ENSG000002 | 10.937107 | -0.683977491 | 0.34050446 | -2.0087182 | 0.04456703 | 0.10983576 | NA | NA |
| ENSG000002 | 5.45585844 | -0.746129735 | 0.37146497 | -2.008614 | 0.04457809 | 0.10985634 | NA | NA |
| ENSG000002 | 9.03968808 | -0.717392378 | 0.35717946 | -2.0084928 | 0.04459095 | 0.10988135 | NA | NA |
| ENSG000001 | 27.4886208 | -0.516831134 | 0.25732951 | -2.008441 | 0.04459645 | 0.10988169 | ZNF781 | 163115 |
| ENSG000002 | 2.4032966 | -0.739681634 | 0.36828656 | -2.0084405 | 0.0445965 | 0.10988169 | DEFA10P | 449493 |
| ENSG000002 | 3.66105119 | -0.760327844 | 0.37859206 | -2.0083037 | 0.04461103 | 0.1099108 | LOC105371508 | 105371508 |
| ENSG000002 | 9.5157563 | -0.707014829 | 0.35205622 | -2.0082441 | 0.04461736 | 0.10991971 | LYPD8 | 646627 |
| ENSG000002 | 240.139207 | 0.26358353 | 0.13126181 | 2.00807481 | 0.04463534 | 0.10995734 | NA | NA |
| ENSG000001 | 436.88071 | 0.258553868 | 0.1287681 | 2.00790312 | 0.04465359 | 0.10998892 | FAM126B | 285172 |
| ENSG000001 | 245.561272 | 0.356859733 | 0.17772606 | 2.00792007 | 0.04465179 | 0.10998892 | MYOZ3 | 91977 |
| ENSG000002 | 25.6718443 | -0.560084508 | 0.27894665 | -2.0078553 | 0.04465867 | 0.10999476 | NA | NA |
| ENSG000002 | 15.9264481 | -0.613517711 | 0.30557894 | -2.0077225 | 0.04467279 | 0.11002285 | PCAT19 | 100505495 |
| ENSG000001 | 95.6423256 | 0.346909911 | 0.17279685 | 2.00761712 | 0.04468399 | 0.11004377 | RCAN1 | 1827 |
| ENSG000002 | 58.9063418 | -0.411249618 | 0.2048484 | -2.0075803 | 0.04468791 | 0.11004672 | ANKRD36BP2 | 645784 |
| ENSG000002 | 3.97444949 | -0.759702619 | 0.37845631 | -2.0073721 | 0.04471006 | 0.1100879 | NA | NA |
| ENSG000002 | 111.926206 | 0.392845901 | 0.19570148 | 2.00737321 | 0.04470994 | 0.1100879 | NA | NA |
| ENSG000001 | 11.5243421 | -0.65754554 | 0.32758974 | -2.0072226 | 0.04472596 | 0.11011702 | NA | NA |

| | | | | | | | | |
|---|---|---|---|---|---|---|---|---|
| ENSG000002 | 5.82832947 | -0.741699781 | 0.36952249 | -2.0071844 | 0.04473003 | 0.11011702 | NA | NA |
| ENSG000002 | 4.88844318 | -0.752163914 | 0.37473173 | -2.0072064 | 0.04472769 | 0.11011702 | NA | NA |
| ENSG000002 | 9.13674228 | -0.721019208 | 0.35924327 | -2.00705 | 0.04474434 | 0.11013218 | PSG3 | 5671 |
| ENSG000002 | 3.04556729 | -0.755901027 | 0.3766158 | -2.0070879 | 0.0447403 | 0.11013218 | RN7SL317P | 106479338 |
| ENSG000002 | 5.28951907 | -0.751422483 | 0.37438763 | -2.0070708 | 0.04474212 | 0.11013218 | FLJ42969 | 441374 |
| ENSG00000C | 43.0093548 | -0.442179914 | 0.22032233 | -2.0069682 | 0.04475305 | 0.11014693 | F11 | 2160 |
| ENSG000002 | 57.5318232 | -0.413438439 | 0.2060102 | -2.0068833 | 0.04476209 | 0.11016249 | NA | NA |
| ENSG000001 | 187.269726 | 0.314568358 | 0.15675406 | 2.00676365 | 0.04477484 | 0.11018718 | INSIG2 | 51141 |
| ENSG000002 | 3.07012981 | -0.75464464 | 0.37605573 | -2.0067362 | 0.04477776 | 0.1101877 | LOC102724357 | 102724357 |
| ENSG000001 | 167.8042 | 0.300764922 | 0.1498839 | 2.00665265 | 0.04478666 | 0.11020291 | C18orf8 | 29919 |
| ENSG000002 | 3.38183253 | -0.758129148 | 0.3778157 | -2.006611 | 0.04479109 | 0.11020713 | NA | NA |
| ENSG000002 | 7.77435925 | -0.707037254 | 0.35248334 | -2.0058742 | 0.04486967 | 0.11038706 | LOC101927062 | 101927062 |
| ENSG000002 | 4.92171519 | -0.752674362 | 0.37523477 | -2.0058759 | 0.04486949 | 0.11038706 | NA | NA |
| ENSG000001 | 48.7602756 | -0.425851908 | 0.21233061 | -2.0056077 | 0.04489812 | 0.11045036 | RGS7 | 6000 |
| ENSG000002 | 7.06689098 | -0.728256013 | 0.36311769 | -2.0055647 | 0.04490271 | 0.11045496 | NA | NA |
| ENSG000001 | 4.08907188 | -0.759248446 | 0.37858806 | -2.0054738 | 0.04491241 | 0.11046819 | MAGEE2 | 139599 |
| ENSG000002 | 16.7598643 | -0.604701579 | 0.30152712 | -2.0054633 | 0.04491354 | 0.11046819 | NA | NA |
| ENSG000001 | 22.2779233 | -0.534321928 | 0.26643858 | -2.0054225 | 0.04491789 | 0.11046871 | PENK | 5179 |
| ENSG000002 | 24.7872562 | 0.521394535 | 0.25999394 | 2.00541031 | 0.0449192 | 0.11046871 | WBP1 | 23559 |
| ENSG000001 | 21.5833007 | -0.562331178 | 0.2804151 | -2.0053527 | 0.04492536 | 0.11047715 | DACT2 | 168002 |
| ENSG000001 | 142.047318 | 0.30160399 | 0.1504019 | 2.00532031 | 0.04492881 | 0.11047895 | C12orf65 | 91574 |
| ENSG00000C | 351.081044 | 0.276798567 | 0.13804225 | 2.00517275 | 0.04494458 | 0.11051102 | SPRTN | 83932 |
| ENSG000001 | 4.25695634 | -0.756823245 | 0.3774517 | -2.0050864 | 0.04495382 | 0.11051637 | NA | NA |
| ENSG000002 | 6.73959955 | -0.731377038 | 0.36476277 | -2.0050759 | 0.04495493 | 0.11051637 | NA | NA |
| ENSG000002 | 2.67464603 | -0.750264318 | 0.37417608 | -2.0051103 | 0.04495126 | 0.11051637 | NA | NA |
| ENSG000001 | 9.41751385 | -0.696752111 | 0.34752614 | -2.0048912 | 0.04497468 | 0.11055821 | OR7D2 | 162998 |
| ENSG000002 | 44.3715648 | 0.448790523 | 0.2238555 | 2.00482238 | 0.04498204 | 0.1105629 | NA | NA |
| ENSG000002 | 20.2648052 | -0.56100625 | 0.27982646 | -2.0048364 | 0.04498054 | 0.1105629 | MIR4477B | 100616194 |
| ENSG000002 | 8.14338669 | -0.740584837 | 0.36941493 | -2.0047507 | 0.0449897 | 0.11057504 | NA | NA |

| | | | | | | | | |
|---|---|---|---|---|---|---|---|---|
| ENSG000002 | 4.91488506 | -0.751473688 | 0.37486792 | -2.0046359 | 0.04500199 | 0.11059852 | NA | NA |
| ENSG000002 | 10.3002382 | -0.686922352 | 0.34267532 | -2.0045866 | 0.04500726 | 0.11060478 | LOC101927438 | 101927438 |
| ENSG000002 | 4.67150237 | -0.756022482 | 0.37718767 | -2.0043669 | 0.04503077 | 0.11065585 | NA | NA |
| ENSG000002 | 3.39754951 | -0.756895638 | 0.37763963 | -2.0042802 | 0.04504005 | 0.11067194 | NA | NA |
| ENSG000002 | 12.2935807 | -0.648496474 | 0.3235623 | -2.0042399 | 0.04504437 | 0.11067585 | TBC1D3P5 | 440419 |
| ENSG00000C | 17.8549565 | -0.57704645 | 0.28792463 | -2.004158 | 0.04505314 | 0.11067728 | EDN1 | 1906 |
| ENSG000001 | 3.69083175 | -0.751850901 | 0.37514501 | -2.0041607 | 0.04505285 | 0.11067728 | C19orf35 | 374872 |
| ENSG000002 | 6.15222618 | -0.739758707 | 0.3691033 | -2.0042051 | 0.0450481 | 0.11067728 | KIR2DS4 | 3809 |
| ENSG000002 | 3.69233337 | -0.758734898 | 0.37858681 | -2.004124 | 0.04505678 | 0.11067953 | NA | NA |
| ENSG000001 | 20.1340451 | -0.605422853 | 0.30210832 | -2.0039927 | 0.04507085 | 0.11070737 | PTPN7 | 5778 |
| ENSG00000C | 252.629303 | 0.309347818 | 0.15437511 | 2.00387107 | 0.04508387 | 0.11072865 | GNB5 | 10681 |
| ENSG000002 | 7.03929506 | -0.725848348 | 0.36222493 | -2.0038608 | 0.04508497 | 0.11072865 | NA | NA |
| ENSG000002 | 3.12158462 | -0.749879256 | 0.37425815 | -2.0036417 | 0.04510845 | 0.1107796 | NA | NA |
| ENSG000002 | 3.65621557 | -0.758473168 | 0.37855835 | -2.0035832 | 0.04511472 | 0.11078829 | NA | NA |
| ENSG000002 | 11.9054007 | -0.684805893 | 0.34179527 | -2.0035558 | 0.04511766 | 0.1107888 | IQCJ | 654502 |
| ENSG000002 | 5.63819652 | -0.750968657 | 0.37484161 | -2.0034293 | 0.04513123 | 0.11081542 | NA | NA |
| ENSG000001 | 20.1475019 | -0.552608024 | 0.27585559 | -2.0032511 | 0.04515034 | 0.11085053 | SP6 | 80320 |
| ENSG000002 | 12.1304901 | -0.656754896 | 0.32784552 | -2.003245 | 0.045151 | 0.11085053 | LINC00570 | 100874055 |
| ENSG000002 | 3.62946157 | -0.758300578 | 0.37859338 | -2.002942 | 0.04518351 | 0.11092364 | NA | NA |
| ENSG000001 | 4.85568658 | -0.750595308 | 0.37479678 | -2.0026728 | 0.04521242 | 0.11098789 | NA | NA |
| ENSG000002 | 2.46730204 | -0.734052438 | 0.36661201 | -2.0022596 | 0.04525682 | 0.11109014 | CNTN4-AS2 | 100873976 |
| ENSG000002 | 3.11262305 | -0.751784101 | 0.37549568 | -2.0021112 | 0.04527278 | 0.11112259 | NA | NA |
| ENSG000002 | 7.03797687 | -0.726507051 | 0.36287617 | -2.0020798 | 0.04527615 | 0.11112414 | NA | NA |
| ENSG000002 | 52.6824331 | 0.413799494 | 0.20671575 | 2.00178018 | 0.04530838 | 0.11119652 | NA | NA |
| ENSG000001 | 105.892995 | 0.322553024 | 0.16113725 | 2.00172844 | 0.04531395 | 0.11120345 | NRG3 | 10718 |
| ENSG00000C | 185.706514 | 0.325098699 | 0.16241455 | 2.00165995 | 0.04532132 | 0.11121481 | RNF31 | 55072 |
| ENSG000002 | 13.4149016 | -0.657343633 | 0.32841251 | -2.0015792 | 0.04533001 | 0.11122942 | NA | NA |
| ENSG000001 | 695.467453 | -0.269257959 | 0.13452546 | -2.001539 | 0.04533434 | 0.11123329 | TTC39C | 125488 |
| ENSG000002 | 10.4060788 | 0.683541606 | 0.34152306 | 2.0014508 | 0.04534383 | 0.11124986 | NA | NA |

| | | | | | | | | |
|---|---|---|---|---|---|---|---|---|
| ENSG000002 | 21.2087632 | 0.572249374 | 0.28592894 | 2.00136917 | 0.04535262 | 0.11126469 | NA | NA |
| ENSG000001 | 12.0258692 | -0.669874149 | 0.33475886 | -2.0010647 | 0.04538541 | 0.11133841 | LGALS9 | 3965 |
| ENSG000001 | 291.229009 | 0.275534883 | 0.13770408 | 2.0009203 | 0.04540098 | 0.11136986 | DIRC2 | 84925 |
| ENSG000002 | 3.43232894 | -0.755276348 | 0.37753063 | -2.0005697 | 0.04543878 | 0.11145584 | NA | NA |
| ENSG000002 | 36.3840774 | -0.472312988 | 0.23610391 | -2.0004455 | 0.04545218 | 0.11146175 | NA | NA |
| ENSG000002 | 6.0805852 | -0.737728682 | 0.36877811 | -2.0004677 | 0.04544979 | 0.11146175 | NA | NA |
| ENSG000002 | 9.91122451 | -0.68504639 | 0.34244303 | -2.0004682 | 0.04544973 | 0.11146175 | NA | NA |
| ENSG000002 | 2.55559466 | -0.728594166 | 0.36421332 | -2.0004599 | 0.04545062 | 0.11146175 | NA | NA |
| ENSG000001 | 327.708429 | 0.259191669 | 0.12957905 | 2.00025912 | 0.04547229 | 0.11150432 | ARSK | 153642 |
| ENSG000002 | 2.39037192 | -0.734443459 | 0.36718567 | -2.0001964 | 0.04547906 | 0.11151417 | NA | NA |
| ENSG000002 | 7.34986641 | -0.7277594 | 0.36386286 | -2.0000925 | 0.04549027 | 0.11153492 | NA | NA |
| ENSG000002 | 5.5516312 | -0.743634459 | 0.37181972 | -1.9999866 | 0.04550171 | 0.11155622 | NA | NA |
| ENSG000001 | 100.530805 | 0.340571415 | 0.17030846 | 1.99973282 | 0.04552912 | 0.11161668 | NBPF3 | 84224 |
| ENSG000002 | 773.56754 | 0.318257329 | 0.15915656 | 1.99964951 | 0.04553812 | 0.11163199 | SYNRG | 11276 |
| ENSG000001 | 5.51861924 | -0.743709798 | 0.37193192 | -1.9995858 | 0.045545 | 0.11164211 | RCVRN | 5957 |
| ENSG000002 | 5.94362491 | -0.742549593 | 0.37136733 | -1.9995016 | 0.0455541 | 0.11165767 | NA | NA |
| ENSG000002 | 5.57177203 | -0.747661371 | 0.37392899 | -1.9994742 | 0.04555707 | 0.11165819 | NA | NA |
| ENSG000002 | 7.29223884 | -0.721592647 | 0.36094594 | -1.999171 | 0.04558986 | 0.1117318 | NA | NA |
| ENSG000001 | 31.8588674 | -0.489157414 | 0.24469299 | -1.9990659 | 0.04560122 | 0.11175289 | SLC6A4 | 6532 |
| ENSG000002 | 5.98260837 | -0.745547382 | 0.37295423 | -1.9990318 | 0.04560491 | 0.11175518 | NA | NA |
| ENSG000001 | 12.3910453 | -0.675063042 | 0.33770766 | -1.9989569 | 0.04561302 | 0.1117683 | ANKRD20A4 | 728747 |
| ENSG000000 | 18.3356343 | -0.621553347 | 0.31095646 | -1.9988437 | 0.04562527 | 0.11179156 | CHAT | 1103 |
| ENSG000001 | 9.3321175 | -0.699876971 | 0.35016498 | -1.9987063 | 0.04564014 | 0.11182123 | TNFSF8 | 944 |
| ENSG000001 | 140.158868 | 0.3458561 | 0.17304643 | 1.99863177 | 0.04564821 | 0.11183425 | CTSZ | 1522 |
| ENSG000001 | 13.0469701 | -0.655918714 | 0.32822786 | -1.998364 | 0.04567722 | 0.11189856 | OLFML1 | 283298 |
| ENSG000001 | 407.068809 | 0.309820252 | 0.15506036 | 1.99806224 | 0.04570991 | 0.11196513 | DSTYK | 25778 |
| ENSG000002 | 5.6342913 | -0.747702824 | 0.37421257 | -1.9980698 | 0.04570909 | 0.11196513 | NA | NA |
| ENSG000002 | 89.0968415 | 0.348314493 | 0.17434091 | 1.9978931 | 0.04572825 | 0.11200328 | LOXL1-AS1 | 100287616 |
| ENSG000001 | 29.8948854 | -0.501003701 | 0.25077539 | -1.9978184 | 0.04573635 | 0.11201635 | SLC9B1 | 150159 |

| | | | | | | | | |
|---|---|---|---|---|---|---|---|---|
| ENSG000001 | 199.773542 | 0.273664999 | 0.1369894 | 1.99770934 | 0.04574818 | 0.11203206 | ZNF700 | 90592 |
| ENSG000002 | 4.58571874 | -0.753001643 | 0.37693272 | -1.9977084 | 0.04574829 | 0.11203206 | NA | NA |
| ENSG000002 | 8.7868933 | -0.702753812 | 0.35181447 | -1.9975125 | 0.04576953 | 0.11207732 | LOC647323 | 647323 |
| ENSG000002 | 3.00441364 | -0.751691085 | 0.37634244 | -1.9973593 | 0.04578616 | 0.11211126 | NA | NA |
| ENSG000002 | 9.12377986 | -0.701294207 | 0.35112402 | -1.9972835 | 0.0457944 | 0.11212466 | NA | NA |
| ENSG000002 | 11.1401919 | -0.66015584 | 0.3305571 | -1.9971008 | 0.04581424 | 0.11216647 | NA | NA |
| ENSG000002 | 5.78071784 | -0.73778956 | 0.36943815 | -1.9970584 | 0.04581884 | 0.11217095 | NA | NA |
| ENSG000002 | 4.08216697 | -0.75598444 | 0.37859329 | -1.9968247 | 0.04584423 | 0.11222634 | NA | NA |
| ENSG000001 | 8.15956428 | -0.708676322 | 0.35494224 | -1.9965962 | 0.04586907 | 0.11228036 | TFDP3 | 51270 |
| ENSG000002 | 2.43894854 | -0.731242602 | 0.36624964 | -1.9965688 | 0.04587205 | 0.11228088 | NA | NA |
| ENSG000002 | 101.612574 | 0.378526388 | 0.18959282 | 1.99652282 | 0.04587704 | 0.11228634 | RNVU1-15 | 101954267 |
| ENSG000002 | 21.0764642 | -0.565470039 | 0.28324152 | -1.9964236 | 0.04588784 | 0.11230598 | OR7E122P | 81428 |
| ENSG000002 | 13.5164083 | -0.670115297 | 0.33569642 | -1.9961943 | 0.04591277 | 0.11236022 | NA | NA |
| ENSG000002 | 16.7029371 | -0.583585236 | 0.29236759 | -1.9960668 | 0.04592665 | 0.1123874 | NA | NA |
| ENSG000002 | 239.439512 | 0.273095977 | 0.13682108 | 1.99600803 | 0.04593305 | 0.11239628 | NA | NA |
| ENSG000001 | 151.73666 | 0.294480164 | 0.14753902 | 1.99594772 | 0.04593961 | 0.11239877 | GRIA4 | 2893 |
| ENSG000001 | 146.279002 | 0.302956433 | 0.1517855 | 1.9959511 | 0.04593925 | 0.11239877 | PIGP | 51227 |
| ENSG000001 | 239.187208 | 0.273162394 | 0.13686146 | 1.99590438 | 0.04594433 | 0.11240354 | NSUN3 | 63899 |
| ENSG000002 | 13.0480861 | -0.657032869 | 0.32920149 | -1.9958381 | 0.04595155 | 0.11240435 | LINC01020 | 340094 |
| ENSG000002 | 7.34630455 | -0.727031435 | 0.364267 | -1.9958751 | 0.04594752 | 0.11240435 | LOC146880 | 146880 |
| ENSG000002 | 3.00987459 | -0.751038388 | 0.37630474 | -1.9958249 | 0.04595298 | 0.11240435 | NA | NA |
| ENSG000001 | 11.395284 | -0.655381486 | 0.3283953 | -1.9957091 | 0.0459656 | 0.11240808 | NA | NA |
| ENSG000002 | 2.65199424 | -0.74604529 | 0.37382382 | -1.9957136 | 0.0459651 | 0.11240808 | NA | NA |
| ENSG000002 | 2.65199424 | -0.74604529 | 0.37382382 | -1.9957136 | 0.0459651 | 0.11240808 | NA | NA |
| ENSG000002 | 5.06122028 | -0.752234077 | 0.37691554 | -1.9957629 | 0.04595973 | 0.11240808 | NA | NA |
| ENSG000002 | 7.65358975 | -0.71923667 | 0.36041547 | -1.9955766 | 0.04598003 | 0.11243661 | LUCAT1 | 100505994 |
| ENSG000002 | 14.6364776 | -0.612475246 | 0.30694833 | -1.9953692 | 0.04600263 | 0.11248349 | TDRD15 | 100129278 |
| ENSG000002 | 10.4953994 | -0.69869051 | 0.35015943 | -1.9953497 | 0.04600475 | 0.11248349 | NA | NA |
| ENSG000001 | 17.3514453 | -0.606572818 | 0.30399944 | -1.995309 | 0.04600919 | 0.11248757 | RGPD4 | 285190 |

| | | | | | | | | |
|---|---|---|---|---|---|---|---|---|
| ENSG000002 | 23.0763875 | 0.588291887 | 0.2948558 | 1.99518509 | 0.0460227 | 0.1125138 | CNTNAP3 | 79937 |
| ENSG000001 | 233.438763 | 0.295848761 | 0.1482892 | 1.99507966 | 0.04603419 | 0.11253512 | GALNT12 | 79695 |
| ENSG000001 | 15.9958228 | -0.594909746 | 0.29820372 | -1.9949776 | 0.04604532 | 0.1125489 | DQX1 | 165545 |
| ENSG000002 | 2.65521163 | -0.746819374 | 0.37434986 | -1.9949771 | 0.04604538 | 0.1125489 | NA | NA |
| ENSG000002 | 17.5310858 | 0.587818135 | 0.29466792 | 1.99484945 | 0.0460593 | 0.11257166 | NA | NA |
| ENSG000002 | 5.49385975 | -0.742637562 | 0.37227911 | -1.9948408 | 0.04606024 | 0.11257166 | NA | NA |
| ENSG000001 | 2.43067102 | -0.731582025 | 0.3667881 | -1.9945631 | 0.04609055 | 0.11263893 | ELANE | 1991 |
| ENSG000001 | 5.46891952 | -0.74156069 | 0.37180135 | -1.9945078 | 0.04609659 | 0.1126469 | LINC00301 | 283197 |
| ENSG000001 | 62.9718262 | 0.39925418 | 0.20021467 | 1.99413048 | 0.0461378 | 0.11274081 | TTC32 | 130502 |
| ENSG000002 | 4.06697947 | -0.754936763 | 0.37858889 | -1.9940806 | 0.04614325 | 0.11274734 | NA | NA |
| ENSG000000 | 230.451085 | 0.279909033 | 0.14037472 | 1.99401305 | 0.04615063 | 0.11275787 | CLEC16A | 23274 |
| ENSG000002 | 7.00647353 | -0.722739198 | 0.36245874 | -1.9939903 | 0.04615312 | 0.11275787 | HOXB7 | 3217 |
| ENSG000002 | 79.3962688 | 0.358732252 | 0.1799256 | 1.99378107 | 0.04617598 | 0.11280694 | NA | NA |
| ENSG000001 | 76.1151414 | 0.373985175 | 0.18757908 | 1.99374672 | 0.04617974 | 0.11280932 | TRIM16L | 147166 |
| ENSG000002 | 4.6965875 | -0.752281297 | 0.37733399 | -1.9936749 | 0.0461876 | 0.11282172 | NA | NA |
| ENSG000001 | 766.996065 | 0.233655617 | 0.11723295 | 1.9930882 | 0.04625179 | 0.1129492 | PLCH1 | 23007 |
| ENSG000001 | 14.7619256 | -0.616945146 | 0.30953742 | -1.9931198 | 0.04624834 | 0.1129492 | GJB6 | 10804 |
| ENSG000001 | 16.4148551 | -0.619995924 | 0.31108254 | -1.9930271 | 0.04625848 | 0.1129492 | NA | NA |
| ENSG000002 | 13969.7556 | 0.208547536 | 0.10463398 | 1.9931149 | 0.04624887 | 0.1129492 | TDGF1 | 6997 |
| ENSG000002 | 4.7385661 | -0.753340579 | 0.37798952 | -1.9930197 | 0.04625928 | 0.1129492 | NA | NA |
| ENSG000002 | 12.5292474 | -0.662510824 | 0.33239073 | -1.9931688 | 0.04624297 | 0.1129492 | LOC101929468 | 101929468 |
| ENSG000002 | 148.081224 | 0.318287171 | 0.15969961 | 1.99303663 | 0.04625744 | 0.1129492 | NA | NA |
| ENSG000002 | 2.38556442 | -0.732249719 | 0.36741758 | -1.9929632 | 0.04626548 | 0.11295752 | NA | NA |
| ENSG000001 | 134.30202 | 0.310224234 | 0.15566703 | 1.99287057 | 0.04627562 | 0.11297548 | STEAP1 | 26872 |
| ENSG000001 | 7.59717887 | -0.715149467 | 0.35886479 | -1.9928103 | 0.04628222 | 0.11297798 | MTHFS | 10588 |
| ENSG000002 | 3.4945801 | -0.751765186 | 0.37723659 | -1.9928215 | 0.046281 | 0.11297798 | NA | NA |
| ENSG000002 | 14.8369329 | -0.60271673 | 0.30245185 | -1.9927692 | 0.04628673 | 0.11298219 | C6orf165 | 154313 |
| ENSG000002 | 10.6564194 | -0.691018337 | 0.34678675 | -1.9926319 | 0.04630177 | 0.11301211 | NA | NA |
| ENSG000001 | 10.3295834 | -0.669684324 | 0.33608992 | -1.9925748 | 0.04630802 | 0.11302056 | ICAM2 | 3384 |

| | | | | | | | | |
|---|---|---|---|---|---|---|---|---|
| ENSG000002 | 268.062721 | 0.276770123 | 0.13890411 | 1.99252654 | 0.04631332 | 0.11302668 | NA | NA |
| ENSG000002 | 4.35421107 | -0.753957803 | 0.37840492 | -1.992463 | 0.04632028 | 0.11303234 | NA | NA |
| ENSG000002 | 105.21168 | 0.331875233 | 0.16656603 | 1.99245453 | 0.04632121 | 0.11303234 | NA | NA |
| ENSG000002 | 3.00466214 | -0.75022289 | 0.37659754 | -1.9921078 | 0.04635924 | 0.11311832 | NA | NA |
| ENSG000002 | 4.10382674 | -0.754174833 | 0.37859222 | -1.9920505 | 0.04636552 | 0.11312684 | NA | NA |
| ENSG000001 | 7.03621016 | -0.722607544 | 0.36276566 | -1.9919403 | 0.04637761 | 0.11314574 | C2orf70 | 339778 |
| ENSG000002 | 4.94889428 | -0.748216481 | 0.37562406 | -1.991929 | 0.04637884 | 0.11314574 | HSPE1P18 | 106480257 |
| ENSG000002 | 7.47502621 | -0.72378569 | 0.36338969 | -1.9917618 | 0.0463972 | 0.11318372 | NA | NA |
| ENSG000002 | 5.80096191 | -0.736615679 | 0.36983772 | -1.9917268 | 0.04640105 | 0.11318629 | NA | NA |
| ENSG000002 | 5.76689211 | -0.736821615 | 0.36995415 | -1.9916566 | 0.04640875 | 0.11319827 | NA | NA |
| ENSG000001 | 14.7527121 | -0.618201751 | 0.31040621 | -1.9915895 | 0.04641612 | 0.11320943 | TMPRSS11F | 389208 |
| ENSG000001 | 8.61477551 | -0.689860041 | 0.34644494 | -1.9912545 | 0.04645291 | 0.11329236 | KRT80 | 144501 |
| ENSG000001 | 214.26257 | 0.297330849 | 0.14932834 | 1.99112136 | 0.04646755 | 0.11332123 | PLEKHA7 | 144100 |
| ENSG000001 | 2072.5566 | 0.615997427 | 0.30938115 | 1.99106324 | 0.04647394 | 0.11333 | HNRNPA0 | 10949 |
| ENSG000001 | 43.1433698 | 0.439832021 | 0.22093281 | 1.99079544 | 0.04650338 | 0.11338807 | FAAP24 | 91442 |
| ENSG000002 | 4.23023426 | -0.751578667 | 0.3775291 | -1.9907834 | 0.04650471 | 0.11338807 | NA | NA |
| ENSG000002 | 12.3599896 | 0.641611811 | 0.32229323 | 1.99077038 | 0.04650614 | 0.11338807 | LINC00622 | 644242 |
| ENSG000001 | 13.9443435 | -0.631548895 | 0.3172494 | -1.9907016 | 0.0465137 | 0.11339969 | SLC5A10 | 125206 |
| ENSG000002 | 2.76590013 | -0.740971636 | 0.37223426 | -1.9906057 | 0.04652425 | 0.1134186 | NA | NA |
| ENSG000001 | 10.0253761 | -0.669543506 | 0.33636887 | -1.9905038 | 0.04653547 | 0.11343534 | SDPR | 8436 |
| ENSG000002 | 8.1022158 | -0.70266814 | 0.35301221 | -1.9904925 | 0.04653671 | 0.11343534 | NA | NA |
| ENSG000002 | 3.68043406 | -0.753525046 | 0.37857197 | -1.9904407 | 0.04654241 | 0.11344241 | NA | NA |
| ENSG000002 | 20.8477748 | 0.560668423 | 0.28170951 | 1.99023608 | 0.04656494 | 0.11349048 | NA | NA |
| ENSG000002 | 3.36334387 | -0.751268751 | 0.37748754 | -1.9901816 | 0.04657094 | 0.11349829 | NA | NA |
| ENSG000001 | 17.9266285 | -0.59693167 | 0.29994563 | -1.9901329 | 0.04657629 | 0.11350453 | KIF19 | 124602 |
| ENSG000002 | 24.3590857 | -0.53936577 | 0.27102972 | -1.9900613 | 0.04658418 | 0.11351692 | NA | NA |
| ENSG000002 | 3.17009066 | -0.744090055 | 0.3739325 | -1.9899048 | 0.04660143 | 0.11355213 | NA | NA |
| ENSG000001 | 9.69048954 | -0.671373647 | 0.33739818 | -1.9898556 | 0.04660685 | 0.11355378 | CA6 | 765 |
| ENSG000002 | 3.38724085 | -0.750857562 | 0.37734422 | -1.9898478 | 0.04660771 | 0.11355378 | NA | NA |

| | | | | | | | | |
|---|---|---|---|---|---|---|---|---|
| ENSG000001 | 320.584106 | 0.246319923 | 0.12379176 | 1.9897926 | 0.04661379 | 0.11356177 | DTNA | 1837 |
| ENSG000002 | 16.0788103 | -0.601232551 | 0.30218368 | -1.9896262 | 0.04663213 | 0.11359963 | LINC01473 | 101927217 |
| ENSG000002 | 14.4778087 | -0.640429723 | 0.32190634 | -1.9894908 | 0.04664705 | 0.11362916 | NA | NA |
| ENSG000001 | 47.3393917 | -0.419698617 | 0.21096093 | -1.9894614 | 0.0466503 | 0.11363024 | SHE | 126669 |
| ENSG000001 | 3.5761512 | -0.748883074 | 0.37647743 | -1.9891845 | 0.04668084 | 0.1136978 | RNU6-552P | 106481356 |
| ENSG000001 | 156.907358 | 0.310433384 | 0.15606875 | 1.98908105 | 0.04669226 | 0.11371552 | SLC25A30 | 253512 |
| ENSG000002 | 9.70694319 | -0.673798808 | 0.33875106 | -1.9890678 | 0.04669372 | 0.11371552 | NA | NA |
| ENSG000001 | 586.382274 | 0.25772135 | 0.12959311 | 1.98869638 | 0.04673473 | 0.11380854 | ZNF318 | 24149 |
| ENSG000001 | 9.21612123 | -0.684923801 | 0.34441646 | -1.98865 | 0.04673985 | 0.11381418 | LOC728175 | 728175 |
| ENSG000002 | 3.58082232 | -0.747126734 | 0.37570625 | -1.9885928 | 0.04674617 | 0.11382275 | NA | NA |
| ENSG000002 | 52.7069068 | 0.426932822 | 0.21470918 | 1.98842373 | 0.04676485 | 0.11386138 | NA | NA |
| ENSG000001 | 8.89511719 | -0.690297312 | 0.34716258 | -1.9883978 | 0.04676771 | 0.11386151 | PSG8 | 440533 |
| ENSG000001 | 3.01471434 | -0.74776858 | 0.37609447 | -1.9882467 | 0.04678442 | 0.11389535 | NA | NA |
| ENSG00000C | 204.822126 | 0.31761578 | 0.15975477 | 1.98814581 | 0.04679557 | 0.11390882 | SNAP29 | 9342 |
| ENSG000001 | 8.1034487 | -0.70250876 | 0.35334458 | -1.9881691 | 0.04679299 | 0.11390882 | GGT6 | 124975 |
| ENSG000002 | 11.5863595 | -0.657748317 | 0.33084998 | -1.9880561 | 0.04680549 | 0.11392614 | NA | NA |
| ENSG000002 | 4.99219687 | -0.748451752 | 0.37650525 | -1.9878919 | 0.04682364 | 0.11396348 | NA | NA |
| ENSG000002 | 2.83053068 | -0.73681285 | 0.3706552 | -1.9878659 | 0.04682652 | 0.11396364 | NA | NA |
| ENSG000001 | 14.2183403 | -0.630823094 | 0.3173436 | -1.9878236 | 0.0468312 | 0.1139682 | DIRAS2 | 54769 |
| ENSG000002 | 7.4562388 | -0.708210633 | 0.35630588 | -1.9876479 | 0.04685065 | 0.11400868 | NA | NA |
| ENSG000002 | 6.99124493 | -0.723276695 | 0.36389045 | -1.9876221 | 0.0468535 | 0.11400879 | NA | NA |
| ENSG000002 | 7.29078211 | -0.719983372 | 0.36224168 | -1.9875774 | 0.04685845 | 0.11401397 | LOC100507091 | 100507091 |
| ENSG000002 | 128.216566 | 0.354823447 | 0.17853163 | 1.98745421 | 0.04687208 | 0.11404031 | HERC2P9 | 440248 |
| ENSG000002 | 62.4752507 | 0.385265468 | 0.19386317 | 1.98730612 | 0.04688848 | 0.1140704 | KCTD11 | 147040 |
| ENSG000002 | 5.59065644 | -0.741918109 | 0.37333125 | -1.9872917 | 0.04689008 | 0.1140704 | NA | NA |
| ENSG000002 | 11.1488618 | -0.659540872 | 0.3319127 | -1.9870914 | 0.04691227 | 0.11411754 | NA | NA |
| ENSG000002 | 3.68835605 | -0.751711285 | 0.37830272 | -1.9870629 | 0.04691543 | 0.11411839 | NA | NA |
| ENSG000001 | 6.38460028 | -0.728978211 | 0.36687771 | -1.9869788 | 0.04692475 | 0.11413421 | PRNT | 149830 |
| ENSG000001 | 13.8927874 | -0.617798748 | 0.31095681 | -1.986767 | 0.04694822 | 0.11417937 | NLRP9 | 338321 |

| | | | | | | | | |
|---|---|---|---|---|---|---|---|---|
| ENSG000002 | 5.61677459 | -0.743436109 | 0.37419514 | -1.9867605 | 0.04694895 | 0.11417937 | NA | NA |
| ENSG000002 | 6.00434162 | -0.741554326 | 0.37325664 | -1.9867144 | 0.04695406 | 0.11418495 | NA | NA |
| ENSG000001 | 5.55331414 | -0.739999251 | 0.372483 | -1.9866658 | 0.04695944 | 0.11419119 | NA | NA |
| ENSG000002 | 3.02917314 | -0.747527458 | 0.37631583 | -1.9864364 | 0.04698489 | 0.11424622 | AA06 | 100506677 |
| ENSG000001 | 1248.42962 | 0.357077927 | 0.17976664 | 1.98634141 | 0.04699543 | 0.11425994 | ZNF507 | 22847 |
| ENSG000002 | 6.20372129 | -0.734486323 | 0.3697782 | -1.9862889 | 0.04700126 | 0.11425994 | NA | NA |
| ENSG000002 | 11.9609694 | -0.640532412 | 0.32246986 | -1.9863326 | 0.0469964 | 0.11425994 | LOC102723536 | 102723536 |
| ENSG000002 | 4.68115725 | -0.75048808 | 0.37783524 | -1.986284 | 0.0470018 | 0.11425994 | NA | NA |
| ENSG000002 | 8.48645674 | -0.707351778 | 0.35613447 | -1.986193 | 0.0470119 | 0.11427764 | NA | NA |
| ENSG000002 | 4.45477227 | -0.751671336 | 0.37847977 | -1.9860278 | 0.04703024 | 0.11431537 | NA | NA |
| ENSG000002 | 11.1478802 | -0.653908086 | 0.32926672 | -1.9859526 | 0.04703859 | 0.11432881 | NA | NA |
| ENSG000002 | 25.0154289 | -0.548984011 | 0.27643948 | -1.9859103 | 0.04704328 | 0.11433336 | NA | NA |
| ENSG000002 | 2.61976373 | -0.719510464 | 0.36232279 | -1.9858272 | 0.04705251 | 0.11434894 | NA | NA |
| ENSG000002 | 4.70335422 | -0.749121391 | 0.37726047 | -1.9856875 | 0.04706804 | 0.11437982 | NA | NA |
| ENSG000002 | 11.3707114 | -0.678808976 | 0.34189024 | -1.9854588 | 0.04709345 | 0.11443472 | NA | NA |
| ENSG000002 | 4.88627098 | -0.744619165 | 0.3751412 | -1.9849037 | 0.04715518 | 0.11457786 | LINC00866 | 100505561 |
| ENSG000001 | 57.4711632 | 0.396835125 | 0.19993666 | 1.98480421 | 0.04716626 | 0.11459583 | CA13 | 377677 |
| ENSG000001 | 6.31471481 | -0.73754073 | 0.37159984 | -1.9847714 | 0.04716991 | 0.11459583 | LINC00207 | 388910 |
| ENSG000002 | 3.6386796 | -0.751289311 | 0.37852884 | -1.9847611 | 0.04717106 | 0.11459583 | NA | NA |
| ENSG00000C | 10.1633439 | -0.682103752 | 0.34367829 | -1.9847159 | 0.04717609 | 0.1146012 | SIRPG | 55423 |
| ENSG000002 | 7.22615347 | -0.714496614 | 0.36003411 | -1.9845248 | 0.04719737 | 0.11464601 | NA | NA |
| ENSG000001 | 9.99432313 | -0.665563132 | 0.33538682 | -1.9844642 | 0.04720412 | 0.11465554 | HAS1 | 3036 |
| ENSG000002 | 8.54231634 | 0.696281443 | 0.35087277 | 1.9844271 | 0.04720825 | 0.1146587 | SNHG9 | 735301 |
| ENSG00000C | 14.3270742 | -0.624784663 | 0.31486096 | -1.9843192 | 0.04722026 | 0.11468102 | RDH8 | 50700 |
| ENSG00000C | 6.39456403 | -0.727768061 | 0.36677462 | -1.9842378 | 0.04722934 | 0.11468931 | TREM2 | 54209 |
| ENSG000002 | 7.78640237 | -0.702605294 | 0.35409107 | -1.9842503 | 0.04722795 | 0.11468931 | RGPD4-AS1 | 729121 |
| ENSG000002 | 9.7884808 | -0.672987414 | 0.33917991 | -1.9841606 | 0.04723794 | 0.11470333 | RNU6-531P | 106479802 |
| ENSG000002 | 17.0649451 | 0.582456591 | 0.29358558 | 1.98394144 | 0.04726237 | 0.11475578 | NA | NA |
| ENSG000002 | 4.13764007 | -0.751032697 | 0.37858977 | -1.9837638 | 0.04728217 | 0.11479012 | NA | NA |

| | | | | | | | | |
|---|---|---|---|---|---|---|---|---|
| ENSG000002 | 7.94651736 | -0.709800261 | 0.357802 | -1.9837795 | 0.04728043 | 0.11479012 | NA | NA |
| ENSG000002 | 2.86295 | -0.734835921 | 0.3704328 | -1.9837226 | 0.04728677 | 0.11479441 | NA | NA |
| ENSG000001 | 643.756348 | 0.259184262 | 0.13066718 | 1.98354523 | 0.04730656 | 0.11483558 | PIPOX | 51268 |
| ENSG000002 | 10.0114997 | -0.691369854 | 0.34855933 | -1.9835069 | 0.04731083 | 0.11483908 | NA | NA |
| ENSG000002 | 2.66684981 | -0.741121628 | 0.37364861 | -1.9834722 | 0.04731471 | 0.11484162 | NA | NA |
| ENSG000001 | 3.66013792 | -0.750828865 | 0.37856716 | -1.9833439 | 0.04732903 | 0.11486949 | NA | NA |
| ENSG000002 | 2.46821607 | -0.727120727 | 0.36664436 | -1.9831772 | 0.04734764 | 0.11490091 | DIAPH3-AS2 | 100874196 |
| ENSG000002 | 7.88138653 | -0.708667497 | 0.35733597 | -1.9831966 | 0.04734547 | 0.11490091 | NA | NA |
| ENSG000001 | 441.068633 | 0.252799555 | 0.12747792 | 1.98308503 | 0.04735793 | 0.11491901 | CREG1 | 8804 |
| ENSG000000 | 102.400801 | -0.32219823 | 0.16248395 | -1.9829541 | 0.04737255 | 0.1149476 | PLD1 | 5337 |
| ENSG000001 | 17.5984057 | -0.595051613 | 0.30009579 | -1.9828722 | 0.0473817 | 0.11496161 | CTSS | 1520 |
| ENSG000002 | 2.7328229 | -0.737553097 | 0.37196583 | -1.9828517 | 0.04738399 | 0.11496161 | NA | NA |
| ENSG000002 | 6.18171208 | -0.745632283 | 0.37607885 | -1.9826488 | 0.04740667 | 0.11500975 | NA | NA |
| ENSG000002 | 7.25945589 | -0.729869169 | 0.36813815 | -1.9825958 | 0.04741259 | 0.11501723 | NA | NA |
| ENSG000002 | 52.0581515 | 0.424271293 | 0.21400399 | 1.98253916 | 0.04741893 | 0.11502572 | NUDT19 | 390916 |
| ENSG000000 | 54.3591021 | -0.401800257 | 0.2026794 | -1.9824425 | 0.04742974 | 0.11504505 | PITPNM2 | 57605 |
| ENSG000002 | 5.18584219 | -0.741126977 | 0.37387869 | -1.9822659 | 0.04744949 | 0.11508609 | NA | NA |
| ENSG000002 | 11.4534982 | 0.664175544 | 0.3350634 | 1.98223843 | 0.04745256 | 0.11508665 | FLJ20021 | 90024 |
| ENSG000001 | 53.7363583 | -0.40014923 | 0.20188647 | -1.9820508 | 0.04747356 | 0.11513069 | FAXDC2 | 10826 |
| ENSG000002 | 3.682117 | -0.750186679 | 0.37852632 | -1.9818613 | 0.04749476 | 0.11517523 | NA | NA |
| ENSG000002 | 3.61414954 | -0.750221622 | 0.37857976 | -1.9816739 | 0.04751575 | 0.11521923 | NA | NA |
| ENSG000001 | 53.2418185 | -0.403140549 | 0.20346014 | -1.9814228 | 0.04754388 | 0.11528056 | CHD5 | 26038 |
| ENSG000002 | 36.0729858 | 0.488578589 | 0.24659275 | 1.98131777 | 0.04755565 | 0.1152953 | LOC101927752 | 101927752 |
| ENSG000002 | 12.4272332 | 0.64010971 | 0.32306966 | 1.98133652 | 0.04755355 | 0.1152953 | NA | NA |
| ENSG000002 | 7.03241497 | -0.72420349 | 0.3655563 | -1.9810997 | 0.04758009 | 0.11534765 | HOXB-AS3 | 404266 |
| ENSG000000 | 161.000914 | 0.344659499 | 0.17398295 | 1.98099579 | 0.04759175 | 0.11536902 | SNX11 | 29916 |
| ENSG000002 | 13.2677778 | -0.640069229 | 0.3231207 | -1.9808983 | 0.04760268 | 0.11538863 | LOC101927378 | 101927378 |
| ENSG000001 | 9.83873682 | -0.680196507 | 0.34338381 | -1.9808636 | 0.04760657 | 0.11539115 | MNDA | 4332 |
| ENSG000001 | 104.730391 | -0.367518573 | 0.18554756 | -1.9807244 | 0.04762219 | 0.11542213 | DGKI | 9162 |

| | | | | | | | | |
|---|---|---|---|---|---|---|---|---|
| ENSG000002 | 4.89211704 | -0.743203116 | 0.37524853 | -1.9805624 | 0.04764037 | 0.11545929 | NA | NA |
| ENSG000002 | 3.65126426 | -0.749607004 | 0.37851728 | -1.9803773 | 0.04766115 | 0.11550274 | NA | NA |
| ENSG000002 | 7.4851546 | -0.723267492 | 0.36523835 | -1.9802617 | 0.04767413 | 0.1155273 | NA | NA |
| ENSG000002 | 3.68604294 | -0.749246904 | 0.37837894 | -1.9801496 | 0.04768672 | 0.1155509 | NA | NA |
| ENSG000001 | 21.9227832 | 0.576930406 | 0.29137943 | 1.97999702 | 0.04770386 | 0.11558553 | MYPOP | 339344 |
| ENSG000001 | 5.72289018 | -0.731297866 | 0.36936991 | -1.9798523 | 0.04772013 | 0.11561141 | ATP6V1G3 | 127124 |
| ENSG000002 | 12.8039285 | -0.646498445 | 0.32654727 | -1.9798005 | 0.04772595 | 0.11561141 | NA | NA |
| ENSG000002 | 11.2840154 | -0.643522314 | 0.32503966 | -1.9798271 | 0.04772297 | 0.11561141 | NA | NA |
| ENSG000002 | 4.32659953 | -0.748316806 | 0.37797435 | -1.9798084 | 0.04772506 | 0.11561141 | NA | NA |
| ENSG000002 | 3.61499776 | -0.749440219 | 0.37855565 | -1.9797359 | 0.04773321 | 0.11562209 | NA | NA |
| ENSG000000 | 518.800946 | 0.252236283 | 0.12742381 | 1.97950663 | 0.04775899 | 0.11567763 | TDRD3 | 81550 |
| ENSG000002 | 22.101296 | -0.552241042 | 0.27898317 | -1.979478 | 0.04776221 | 0.11567853 | GSTM2 | 2946 |
| ENSG000002 | 4.31424376 | -0.747904997 | 0.37783696 | -1.9794384 | 0.04776667 | 0.1156824 | NA | NA |
| ENSG000002 | 5.52954057 | -0.736915294 | 0.37232437 | -1.9792293 | 0.0477902 | 0.11573248 | NA | NA |
| ENSG000002 | 7.23438002 | -0.711503197 | 0.35950545 | -1.9791166 | 0.04780288 | 0.11575628 | NA | NA |
| ENSG000001 | 315.943803 | -0.26981729 | 0.13634054 | -1.9789953 | 0.04781654 | 0.11578217 | ENC1 | 8507 |
| ENSG000002 | 3.02290286 | -0.744515067 | 0.37621325 | -1.9789709 | 0.04781928 | 0.11578217 | NA | NA |
| ENSG000002 | 4.50936063 | -0.744319213 | 0.37612624 | -1.9789079 | 0.04782637 | 0.11579242 | NA | NA |
| ENSG000002 | 6.56696899 | -0.732873972 | 0.37035215 | -1.9788571 | 0.0478321 | 0.11579937 | ZNF385D-AS2 | 100874221 |
| ENSG000002 | 5.05539483 | 0.745190553 | 0.3765846 | 1.97881314 | 0.04783705 | 0.11580444 | NA | NA |
| ENSG000001 | 57.4883226 | 0.394633063 | 0.19945068 | 1.9785997 | 0.04786109 | 0.11585573 | ZC3H12A | 80149 |
| ENSG000002 | 4.00453188 | -0.748979188 | 0.37859245 | -1.9783257 | 0.04789197 | 0.11592356 | NA | NA |
| ENSG000001 | 29.2131916 | -0.539730081 | 0.27283458 | -1.9782319 | 0.04790255 | 0.11594224 | MROH2A | 339766 |
| ENSG000001 | 135.431589 | 0.294547303 | 0.14889952 | 1.9781615 | 0.04791049 | 0.11595454 | KIAA0391 | 9692 |
| ENSG000001 | 16.7125065 | -0.598256713 | 0.3024495 | -1.9780384 | 0.04792438 | 0.11598123 | LINC00994 | 100287879 |
| ENSG000001 | 31.641493 | -0.486887276 | 0.24615546 | -1.9779666 | 0.04793248 | 0.11599391 | WBP2NL | 164684 |
| ENSG000001 | 1823.46011 | 0.269267929 | 0.1361443 | 1.97781269 | 0.04794984 | 0.116029 | IL17RD | 54756 |
| ENSG000002 | 2.75815543 | -0.734192881 | 0.37122162 | -1.9777751 | 0.04795408 | 0.11603233 | NA | NA |
| ENSG000000 | 65.655148 | 0.398709948 | 0.20162234 | 1.9775088 | 0.04798415 | 0.11609816 | STYK1 | 55359 |

| | | | | | | | | |
|---|---|---|---|---|---|---|---|---|
| ENSG000000 | 1191.03748 | 0.262026759 | 0.13251512 | 1.97733474 | 0.0480038 | 0.11613563 | ARID1B | 57492 |
| ENSG000002 | 4.24074292 | -0.747456057 | 0.37801798 | -1.9773029 | 0.0480074 | 0.11613563 | NA | NA |
| ENSG000002 | 5.15854686 | -0.738153826 | 0.37331485 | -1.9772956 | 0.04800822 | 0.11613563 | NA | NA |
| ENSG000002 | 4.600523 | -0.745626832 | 0.37710335 | -1.977248 | 0.04801361 | 0.11614172 | NA | NA |
| ENSG000001 | 23.7793484 | -0.532779408 | 0.26947995 | -1.9770651 | 0.04803427 | 0.11618476 | LRRC31 | 79782 |
| ENSG000002 | 2.99965767 | -0.742842139 | 0.37574105 | -1.9770055 | 0.048041 | 0.11619413 | LMF1-AS1 | 101929387 |
| ENSG000002 | 2.41624001 | -0.72385986 | 0.36620185 | -1.9766691 | 0.04807905 | 0.11627921 | NA | NA |
| ENSG000002 | 7.57591056 | -0.711534838 | 0.35997763 | -1.9766085 | 0.0480859 | 0.11628885 | NA | NA |
| ENSG000001 | 16.8423745 | -0.586842794 | 0.29689877 | -1.9765754 | 0.04808965 | 0.11629097 | LBP | 3929 |
| ENSG000002 | 9.94207155 | -0.665743396 | 0.33684368 | -1.9764164 | 0.04810763 | 0.11632752 | NA | NA |
| ENSG000001 | 27.9331582 | -0.512162839 | 0.25916038 | -1.9762389 | 0.04812772 | 0.11636916 | ENKUR | 219670 |
| ENSG000002 | 19.0168343 | -0.568084312 | 0.28746271 | -1.9762018 | 0.04813193 | 0.1163724 | ZNF812 | 729648 |
| ENSG000002 | 8.70456582 | -0.689027875 | 0.34872738 | -1.9758353 | 0.04817343 | 0.11646579 | NA | NA |
| ENSG000002 | 15.6395132 | -0.605274013 | 0.30634383 | -1.9757996 | 0.04817748 | 0.11646865 | NR2E3 | 10002 |
| ENSG000002 | 15.1410448 | -0.611668609 | 0.3095916 | -1.9757274 | 0.04818566 | 0.1164785 | NA | NA |
| ENSG000002 | 9.25970299 | -0.68225456 | 0.3453207 | -1.9757129 | 0.0481873 | 0.1164785 | NA | NA |
| ENSG000001 | 6.79719798 | -0.725828086 | 0.36738346 | -1.9756689 | 0.04819229 | 0.1164836 | CLEC14A | 161198 |
| ENSG000002 | 19.6772322 | -0.557247699 | 0.28206327 | -1.9756124 | 0.04819869 | 0.11649215 | NA | NA |
| ENSG000001 | 965.588029 | 0.245270997 | 0.12415195 | 1.97557106 | 0.04820338 | 0.11649653 | DDAH1 | 23576 |
| ENSG000002 | 5.21916838 | -0.74093389 | 0.37507854 | -1.97541 | 0.04822164 | 0.11653371 | NA | NA |
| ENSG000002 | 2.63582876 | -0.738227515 | 0.37371349 | -1.9753836 | 0.04822463 | 0.11653399 | NA | NA |
| ENSG000000 | 87.1090519 | 0.341430047 | 0.17284624 | 1.97533971 | 0.04822961 | 0.11653908 | FSTL4 | 23105 |
| ENSG000002 | 2.36740802 | -0.725651095 | 0.36737228 | -1.9752473 | 0.04824009 | 0.11655747 | WBP1LP1 | 106479029 |
| ENSG000002 | 3.07724178 | -0.742631807 | 0.37598447 | -1.9751662 | 0.04824929 | 0.11657275 | NA | NA |
| ENSG000002 | 9.65756593 | -0.686892808 | 0.34778418 | -1.9750548 | 0.04826193 | 0.11659634 | LINC00840 | 100506835 |
| ENSG000002 | 5.7711536 | -0.731510872 | 0.37040802 | -1.9748786 | 0.04828193 | 0.1166377 | NA | NA |
| ENSG000002 | 4.58557337 | -0.743936201 | 0.37670959 | -1.9748268 | 0.04828781 | 0.11664495 | NA | NA |
| ENSG000002 | 2.74467069 | -0.733844395 | 0.37163554 | -1.9746346 | 0.04830963 | 0.11669073 | NA | NA |
| ENSG000002 | 3.37085967 | -0.745356355 | 0.37747303 | -1.974595 | 0.04831413 | 0.11669463 | NA | NA |

| | | | | | | | | |
|---|---|---|---|---|---|---|---|---|
| ENSG000002 | 4.39043192 | -0.747350323 | 0.37851191 | -1.9744434 | 0.04833135 | 0.11672928 | NA | NA |
| ENSG000002 | 38.0923436 | 0.465825688 | 0.23593425 | 1.97438773 | 0.04833767 | 0.1167376 | NA | NA |
| ENSG000002 | 2.63875744 | -0.737559546 | 0.37361165 | -1.9741342 | 0.04836648 | 0.11680021 | RN7SL819P | 106481145 |
| ENSG000001 | 69.1039377 | -0.381700087 | 0.19337448 | -1.9738907 | 0.04839417 | 0.11686011 | PDE1B | 5153 |
| ENSG000001 | 57.1925833 | -0.389130609 | 0.19714279 | -1.9738516 | 0.04839862 | 0.11686391 | ASXL3 | 80816 |
| ENSG000002 | 4.51675246 | -0.742361746 | 0.37613166 | -1.9736753 | 0.04841868 | 0.11690538 | NA | NA |
| ENSG000000 | 40.2772221 | -0.438335536 | 0.2220978 | -1.973615 | 0.04842554 | 0.11691007 | HFE | 3077 |
| ENSG000002 | 96.1501623 | 0.345283273 | 0.17495032 | 1.97360752 | 0.04842639 | 0.11691007 | NA | NA |
| ENSG000001 | 4.70138944 | -0.745681129 | 0.37787173 | -1.973371 | 0.0484533 | 0.11696809 | MAGEB18 | 286514 |
| ENSG000002 | 9.04047476 | -0.693977646 | 0.35168128 | -1.9733142 | 0.04845978 | 0.11697676 | LOC101928139 | 101928139 |
| ENSG000002 | 13.2945033 | -0.62237857 | 0.3154056 | -1.9732642 | 0.04846547 | 0.11698353 | NA | NA |
| ENSG000002 | 8.97137766 | -0.68722192 | 0.34833477 | -1.9728777 | 0.04850949 | 0.11708282 | NA | NA |
| ENSG000002 | 171.860315 | 0.315425548 | 0.15988304 | 1.97285189 | 0.04851244 | 0.11708296 | SLC16A1-AS1 | 100506392 |
| ENSG000002 | 3.68134754 | -0.746699616 | 0.37851561 | -1.972705 | 0.04852919 | 0.11711641 | NA | NA |
| ENSG000001 | 19.3962527 | -0.558697058 | 0.28322227 | -1.9726452 | 0.048536 | 0.11712587 | LRRC36 | 55282 |
| ENSG000002 | 16.0294519 | -0.596104423 | 0.30219688 | -1.9725697 | 0.0485446 | 0.11713968 | NA | NA |
| ENSG000002 | 8.3830312 | -0.695692533 | 0.35269178 | -1.9725227 | 0.04854997 | 0.11714566 | NA | NA |
| ENSG000002 | 3.26989829 | -0.745793403 | 0.37810663 | -1.972442 | 0.04855917 | 0.11716088 | NCOR1P1 | 149934 |
| ENSG000002 | 6.9439439 | 0.709355847 | 0.35964722 | 1.97236572 | 0.04856787 | 0.11716096 | NA | NA |
| ENSG000002 | 10.057176 | -0.673045307 | 0.34123539 | -1.9723784 | 0.04856643 | 0.11716096 | PRAMEF11 | 440560 |
| ENSG000002 | 5.73893943 | -0.729402526 | 0.3698099 | -1.9723716 | 0.04856721 | 0.11716096 | NA | NA |
| ENSG000002 | 3.62871835 | 0.746195246 | 0.37833519 | 1.97231254 | 0.04857394 | 0.11716863 | NA | NA |
| ENSG000002 | 6.06069332 | -0.724241484 | 0.36722711 | -1.9721896 | 0.04858797 | 0.11719329 | LINC00272 | 388719 |
| ENSG000002 | 6.72963402 | -0.720519288 | 0.36534297 | -1.9721723 | 0.04858994 | 0.11719329 | LINC00200 | 399706 |
| ENSG000001 | 12.486502 | -0.633996834 | 0.32153278 | -1.9717953 | 0.04863297 | 0.1172901 | LMOD2 | 442721 |
| ENSG000002 | 14.2000211 | -0.617767727 | 0.31331638 | -1.9717058 | 0.0486432 | 0.1173008 | NA | NA |
| ENSG000002 | 3.65024664 | -0.746274173 | 0.37848787 | -1.9717255 | 0.04864095 | 0.1173008 | NA | NA |
| ENSG000001 | 269.760798 | 0.265542979 | 0.13468825 | 1.97153785 | 0.04866239 | 0.11733734 | MTUS1 | 57509 |
| ENSG000002 | 4.25720951 | -0.745031531 | 0.37789653 | -1.9715225 | 0.04866414 | 0.11733734 | NA | NA |

| | | | | | | | | |
|---|---|---|---|---|---|---|---|---|
| ENSG000002 | 2.69027127 | -0.735974847 | 0.37331921 | -1.9714358 | 0.04867405 | 0.11735427 | NA | NA |
| ENSG000001 | 27.4980615 | 0.520655989 | 0.26411138 | 1.97135008 | 0.04868385 | 0.11737091 | PXDC1 | 221749 |
| ENSG000002 | 10.9357678 | -0.657811586 | 0.33369225 | -1.9713121 | 0.04868818 | 0.11737438 | LST1 | 7940 |
| ENSG000001 | 4452.4821 | 0.229163036 | 0.1162522 | 1.97125771 | 0.04869441 | 0.11738241 | IQGAP1 | 8826 |
| ENSG000001 | 4.60050721 | -0.746234635 | 0.3785885 | -1.971097 | 0.04871279 | 0.11741974 | IL17A | 3605 |
| ENSG000002 | 2.39133726 | -0.722039066 | 0.36636542 | -1.9708167 | 0.04874485 | 0.11748323 | NA | NA |
| ENSG000002 | 2.36131401 | -0.721943362 | 0.36631697 | -1.970816 | 0.04874492 | 0.11748323 | NA | NA |
| ENSG00000C | 41.9401004 | 0.446088413 | 0.22635367 | 1.97075844 | 0.04875151 | 0.11748813 | FSTL3 | 10272 |
| ENSG000002 | 13.6473276 | -0.624202072 | 0.31673364 | -1.9707476 | 0.04875275 | 0.11748813 | PSG4 | 5672 |
| ENSG000001 | 24.1503022 | 0.532725487 | 0.27033299 | 1.97062697 | 0.04876656 | 0.11751442 | TNFAIP6 | 7130 |
| ENSG000001 | 3.62698935 | -0.745892659 | 0.37851976 | -1.9705514 | 0.04877521 | 0.11752828 | R3HDML | 140902 |
| ENSG000002 | 16.6536255 | -0.595656695 | 0.30229001 | -1.9704809 | 0.04878328 | 0.11754076 | TMEM211 | 255349 |
| ENSG00000C | 17.4606151 | -0.597282133 | 0.30312926 | -1.9703876 | 0.04879397 | 0.11755253 | PON1 | 5444 |
| ENSG000002 | 4.26950909 | -0.74449451 | 0.37784093 | -1.9703914 | 0.04879353 | 0.11755253 | LINC00706 | 100652997 |
| ENSG000001 | 11.7666606 | -0.647338737 | 0.3285678 | -1.9701831 | 0.04881739 | 0.11760196 | SERPINC1 | 462 |
| ENSG00000C | 58.3331665 | 0.395588131 | 0.2007976 | 1.97008393 | 0.04882875 | 0.11762236 | C3orf18 | 51161 |
| ENSG000002 | 2.7942718 | -0.729503078 | 0.37030684 | -1.9699962 | 0.0488388 | 0.11763957 | LOC105374731 | 105374731 |
| ENSG000002 | 2.44009866 | -0.721036405 | 0.36603636 | -1.9698492 | 0.04885566 | 0.1176732 | LOC100506384 | 100506384 |
| ENSG000002 | 43.6809238 | -0.452544497 | 0.22974813 | -1.9697418 | 0.04886797 | 0.11769584 | DARS-AS1 | 101928243 |
| ENSG000001 | 2.37065678 | -0.720950839 | 0.36602773 | -1.9696618 | 0.04887714 | 0.11771095 | OR13F1 | 138805 |
| ENSG000001 | 712.829058 | 0.330112583 | 0.16762125 | 1.96939582 | 0.04890766 | 0.11777095 | HMGXB4 | 10042 |
| ENSG000002 | 4.00376689 | -0.745559246 | 0.37857292 | -1.969394 | 0.04890786 | 0.11777095 | TSSK2 | 23617 |
| ENSG000001 | 89.2282336 | 0.332844324 | 0.16901441 | 1.96932507 | 0.04891577 | 0.117783 | FAM69A | 388650 |
| ENSG000002 | 3.00859659 | -0.740625446 | 0.37610697 | -1.9691883 | 0.04893147 | 0.1178138 | NA | NA |
| ENSG000002 | 3.03110347 | -0.739798932 | 0.37569605 | -1.9691422 | 0.04893677 | 0.11781956 | NA | NA |
| ENSG000002 | 2.44083587 | -0.720744969 | 0.36603994 | -1.9690337 | 0.04894922 | 0.11784255 | NA | NA |
| ENSG000002 | 34.0309162 | 0.465988276 | 0.23666249 | 1.96899931 | 0.04895317 | 0.11784506 | SNHG18 | 100505806 |
| ENSG000001 | 47.4616064 | -0.417081739 | 0.21183973 | -1.9688552 | 0.04896973 | 0.11787792 | SPTBN5 | 51332 |
| ENSG000001 | 28.5191023 | 0.490425809 | 0.24910677 | 1.96873737 | 0.04898326 | 0.11789649 | FGF11 | 2256 |

| | | | | | | | | |
|---|---|---|---|---|---|---|---|---|
| ENSG000002 | 6.10058044 | -0.730600526 | 0.37109766 | -1.9687554 | 0.04898118 | 0.11789649 | TM4SF1-AS1 | 100874091 |
| ENSG000002 | 8.37497528 | -0.692532307 | 0.35177584 | -1.968675 | 0.04899042 | 0.11790674 | NA | NA |
| ENSG000002 | 11.1024037 | -0.648224247 | 0.32930828 | -1.968442 | 0.04901721 | 0.1179642 | NA | NA |
| ENSG000002 | 2.65686232 | -0.735262581 | 0.37355182 | -1.9683014 | 0.04903336 | 0.11799608 | NA | NA |
| ENSG000001 | 252.21195 | 0.286919981 | 0.1457828 | 1.9681333 | 0.0490527 | 0.11803561 | GK | 2710 |
| ENSG000001 | 40.005205 | -0.456175469 | 0.23178957 | -1.9680586 | 0.04906129 | 0.11804528 | DACH2 | 117154 |
| ENSG000002 | 3.26255908 | -0.744254196 | 0.37816877 | -1.9680478 | 0.04906254 | 0.11804528 | NA | NA |
| ENSG000001 | 72.4192159 | -0.36456342 | 0.18526972 | -1.9677443 | 0.04909747 | 0.11810129 | TRIM3 | 10612 |
| ENSG000002 | 5.80865375 | -0.729259192 | 0.37059861 | -1.9677872 | 0.04909253 | 0.11810129 | NA | NA |
| ENSG000002 | 4.67754431 | -0.743263846 | 0.37771901 | -1.9677692 | 0.0490946 | 0.11810129 | NA | NA |
| ENSG000002 | 26.7673504 | 0.498121494 | 0.25313884 | 1.96777983 | 0.04909338 | 0.11810129 | DSTNP2 | 171220 |
| ENSG000001 | 793.022176 | 0.213541958 | 0.10852838 | 1.96761407 | 0.04911246 | 0.11812333 | LRRCC1 | 85444 |
| ENSG000002 | 3.27556774 | -0.743973956 | 0.37810893 | -1.967618 | 0.049112 | 0.11812333 | NA | NA |
| ENSG000002 | 3.30148845 | -0.743983587 | 0.3781241 | -1.9675646 | 0.04911816 | 0.11813003 | NA | NA |
| ENSG000001 | 19.2192473 | -0.576992993 | 0.2932574 | -1.9675309 | 0.04912204 | 0.11813236 | GLRB | 2743 |
| ENSG000002 | 6.1772698 | -0.731316089 | 0.37173257 | -1.9673178 | 0.04914659 | 0.11818439 | NA | NA |
| ENSG000001 | 794.838454 | 0.342113101 | 0.17390916 | 1.96719424 | 0.04916082 | 0.1182116 | RAB13 | 5872 |
| ENSG000002 | 2.4311008 | -0.720048563 | 0.36606748 | -1.9669831 | 0.04918516 | 0.11825609 | NA | NA |
| ENSG000002 | 4.92356286 | -0.739187183 | 0.37579489 | -1.9669963 | 0.04918363 | 0.11825609 | NA | NA |
| ENSG000002 | 3.97035959 | -0.744611457 | 0.3785695 | -1.9669082 | 0.04919379 | 0.11826985 | NA | NA |
| ENSG000001 | 42.6409177 | -0.470922493 | 0.23944398 | -1.9667335 | 0.04921394 | 0.11831127 | SLC3A1 | 6519 |
| ENSG000002 | 10.8569864 | -0.651685034 | 0.33136536 | -1.9666661 | 0.04922171 | 0.11832294 | MTHFD2P1 | 100287639 |
| ENSG000002 | 3.75095773 | -0.743853964 | 0.37830025 | -1.9663058 | 0.0492633 | 0.11841589 | LOC101928386 | 101928386 |
| ENSG000002 | 95.3808395 | 0.355101953 | 0.18059665 | 1.96627099 | 0.04926732 | 0.11841852 | NA | NA |
| ENSG000002 | 2.69456969 | -0.732369251 | 0.37247409 | -1.9662287 | 0.0492722 | 0.11842324 | NA | NA |
| ENSG000002 | 4.23686851 | -0.743263911 | 0.37803192 | -1.9661406 | 0.04928237 | 0.11843365 | MANSC4 | 100287284 |
| ENSG000002 | 6.73847536 | -0.723498346 | 0.36797471 | -1.9661632 | 0.04927976 | 0.11843365 | NA | NA |
| ENSG000002 | 3.66298208 | -0.74406604 | 0.37844899 | -1.9660933 | 0.04928783 | 0.11843975 | NA | NA |
| ENSG000002 | 8.41710188 | -0.691585695 | 0.35178354 | -1.965941 | 0.04930542 | 0.11847099 | NA | NA |

| | | | | | | | | |
|---|---|---|---|---|---|---|---|---|
| ENSG000002 | 18.1605229 | 0.60972497 | 0.31014578 | 1.96593022 | 0.04930667 | 0.11847099 | NA | NA |
| ENSG000002 | 19.0575956 | 0.5755557 | 0.29278998 | 1.965763 | 0.04932599 | 0.11851039 | NA | NA |
| ENSG000002 | 4.91947288 | -0.741308942 | 0.37714 | -1.9656068 | 0.04934405 | 0.11854674 | NA | NA |
| ENSG000002 | 6.84046789 | -0.724837333 | 0.3687653 | -1.965579 | 0.04934726 | 0.11854743 | NA | NA |
| ENSG000001 | 142.80154 | 0.400683256 | 0.20385666 | 1.96551464 | 0.0493547 | 0.11855829 | SLCO3A1 | 28232 |
| ENSG000002 | 57.2149719 | 0.412255561 | 0.20975598 | 1.9654055 | 0.04936732 | 0.11858159 | NA | NA |
| ENSG000001 | 53.4109238 | 0.441608147 | 0.22470215 | 1.96530453 | 0.049379 | 0.11860261 | MAGEE1 | 57692 |
| ENSG000001 | 97.5370445 | -0.323380942 | 0.1645581 | -1.9651475 | 0.04939716 | 0.11863921 | ITGA4 | 3676 |
| ENSG000002 | 3.05944901 | -0.737659091 | 0.37537899 | -1.9651049 | 0.0494021 | 0.11864326 | LOC101927553 | 101927553 |
| ENSG000002 | 7.87498042 | -0.705622371 | 0.3590803 | -1.9650824 | 0.0494047 | 0.11864326 | NA | NA |
| ENSG000001 | 4.50617604 | -0.739872985 | 0.3765318 | -1.9649681 | 0.04941792 | 0.11866799 | NA | NA |
| ENSG00000C | 531.00119 | 0.226503016 | 0.11527855 | 1.96483222 | 0.04943366 | 0.11869171 | PTCD2 | 79810 |
| ENSG000002 | 2.6321549 | -0.733126919 | 0.37312194 | -1.9648454 | 0.04943214 | 0.11869171 | NA | NA |
| ENSG000002 | 17.4670581 | -0.600554315 | 0.30566734 | -1.9647317 | 0.0494453 | 0.11871262 | LINC01299 | 286186 |
| ENSG000001 | 24.9107658 | 0.540921712 | 0.27533208 | 1.96461566 | 0.04945874 | 0.11873786 | EMID1 | 129080 |
| ENSG000002 | 5.99424999 | -0.734097954 | 0.37366876 | -1.9645687 | 0.04946418 | 0.1187439 | NA | NA |
| ENSG000001 | 82.7778217 | -0.346265864 | 0.176273 | -1.9643727 | 0.04948688 | 0.11879137 | GRHL1 | 29841 |
| ENSG000002 | 7.70511895 | -0.71103543 | 0.36198019 | -1.9642938 | 0.04949604 | 0.11880383 | NA | NA |
| ENSG000002 | 4.37387889 | -0.742875126 | 0.37819258 | -1.9642774 | 0.04949793 | 0.11880383 | NA | NA |
| ENSG000001 | 26.7469004 | -0.510760653 | 0.26003267 | -1.9642173 | 0.0495049 | 0.11881352 | GAL3ST4 | 79690 |
| ENSG000002 | 5.33175319 | -0.738783702 | 0.37618093 | -1.9639053 | 0.04954108 | 0.11889332 | NA | NA |
| ENSG000002 | 2.35724317 | -0.719730719 | 0.3664886 | -1.9638557 | 0.04954683 | 0.11890008 | NA | NA |
| ENSG000002 | 11.1086975 | -0.683535666 | 0.34808127 | -1.9637243 | 0.04956207 | 0.11892961 | NA | NA |
| ENSG000001 | 6.15482451 | -0.730521491 | 0.37202176 | -1.9636526 | 0.0495704 | 0.11894255 | Sep-12 | 124404 |
| ENSG000002 | 3.66958517 | -0.743283453 | 0.37853381 | -1.9635854 | 0.0495782 | 0.11895423 | RPS23P5 | 100271093 |
| ENSG000001 | 357.749874 | 0.276642464 | 0.14089529 | 1.96346142 | 0.04959258 | 0.11897467 | AR | 367 |
| ENSG000001 | 26.0181308 | 0.554242427 | 0.28227761 | 1.96346578 | 0.04959208 | 0.11897467 | NA | NA |
| ENSG000002 | 5.17157514 | -0.734011674 | 0.37384841 | -1.9633939 | 0.04960043 | 0.11898645 | NA | NA |
| ENSG000002 | 20.6726342 | -0.549608933 | 0.27994082 | -1.963304 | 0.04961086 | 0.11900444 | LOC102723766 | 102723766 |

| | | | | | | | | |
|---|---|---|---|---|---|---|---|---|
| ENSG000001 | 56.7748829 | -0.39445073 | 0.20092617 | -1.9631625 | 0.04962729 | 0.11903682 | AGBL2 | 79841 |
| ENSG000001 | 1955.16667 | 0.250909786 | 0.12781093 | 1.96313251 | 0.04963078 | 0.11903813 | GCC2 | 9648 |
| ENSG000002 | 3.70593435 | -0.742411553 | 0.37818523 | -1.9630897 | 0.04963575 | 0.11904301 | NA | NA |
| ENSG000000 | 9.0742875 | -0.672113885 | 0.34238323 | -1.9630456 | 0.04964088 | 0.11904536 | DLX3 | 1747 |
| ENSG000002 | 2.36945046 | -0.719982811 | 0.36677214 | -1.9630248 | 0.04964329 | 0.11904536 | NA | NA |
| ENSG000002 | 41.4970235 | 0.443553611 | 0.22595929 | 1.96298022 | 0.04964847 | 0.11904536 | NA | NA |
| ENSG000002 | 14.7462419 | -0.613741246 | 0.31265413 | -1.9630038 | 0.04964573 | 0.11904536 | NA | NA |
| ENSG000002 | 8.57487558 | -0.683793741 | 0.34835119 | -1.9629436 | 0.04965273 | 0.11904853 | LOC101927416 | 101927416 |
| ENSG000000 | 22.9435644 | -0.524339311 | 0.26712889 | -1.9628701 | 0.04966127 | 0.11906196 | SEMA3B | 7869 |
| ENSG000001 | 168.82862 | 0.346234109 | 0.17640582 | 1.96271366 | 0.04967946 | 0.11909853 | RGS12 | 6002 |
| ENSG000001 | 58.1822532 | 0.460672943 | 0.23472606 | 1.96259817 | 0.04969288 | 0.11912368 | KRT10 | 3858 |
| ENSG000002 | 11.3024074 | -0.676225761 | 0.34457276 | -1.962505 | 0.04970372 | 0.11914261 | NA | NA |
| ENSG000001 | 5.37396776 | 0.733115636 | 0.37358606 | 1.96237419 | 0.04971894 | 0.11917204 | NA | NA |
| ENSG000001 | 32.364333 | -0.478147162 | 0.24366929 | -1.9622791 | 0.04973 | 0.11919151 | C2orf80 | 389073 |
| ENSG000002 | 17.6700459 | -0.563175302 | 0.28702863 | -1.9620875 | 0.0497523 | 0.1192379 | NA | NA |
| ENSG000002 | 3.38722103 | -0.739713693 | 0.37702809 | -1.9619591 | 0.04976724 | 0.11926668 | MIR8066 | 102465868 |
| ENSG000002 | 3.33327917 | -0.74143313 | 0.37794242 | -1.9617621 | 0.04979019 | 0.11931461 | HCAR3 | 8843 |
| ENSG000002 | 3.71250553 | -0.742390208 | 0.3784675 | -1.9615693 | 0.04981265 | 0.11935435 | NA | NA |
| ENSG000002 | 163.440744 | 0.368848145 | 0.18803579 | 1.96158479 | 0.04981084 | 0.11935435 | NA | NA |
| ENSG000002 | 24.9039802 | -0.550927106 | 0.28088066 | -1.9614277 | 0.04982915 | 0.11938682 | LOC101927359 | 101927359 |
| ENSG000002 | 6.83677066 | -0.722655104 | 0.36844402 | -1.9613702 | 0.04983586 | 0.11939583 | NA | NA |
| ENSG000001 | 27.1494958 | 0.502142185 | 0.25605571 | 1.96106618 | 0.0498713 | 0.11947369 | RIN1 | 9610 |
| ENSG000002 | 65.8915624 | -0.374635488 | 0.19104542 | -1.960976 | 0.04988182 | 0.11949183 | LINC00882 | 100302640 |
| ENSG000001 | 1000.1276 | 0.315796051 | 0.16107676 | 1.96053144 | 0.04993371 | 0.11960906 | PDLIM1 | 9124 |
| ENSG000002 | 6.92911223 | -0.712933728 | 0.36365725 | -1.9604552 | 0.04994261 | 0.11962333 | NA | NA |
| ENSG000002 | 4.64277693 | -0.740464734 | 0.37771568 | -1.9603759 | 0.04995187 | 0.11963842 | FOXD2-AS1 | 84793 |
| ENSG000002 | 6.62291913 | -0.726063527 | 0.37040087 | -1.9602101 | 0.04997124 | 0.11967775 | TUBB4BP4 | 106480434 |
| ENSG000002 | 200.082964 | 0.291366141 | 0.14866257 | 1.95991598 | 0.05000561 | 0.119753 | RDH14 | 57665 |
| ENSG000001 | 20.9203256 | -0.541842538 | 0.27647656 | -1.9598136 | 0.05001758 | 0.11976791 | GPR83 | 10888 |

| | | | | | | | | |
|---|---|---|---|---|---|---|---|---|
| ENSG000001 | 187.693488 | 0.299803402 | 0.15297558 | 1.95981219 | 0.05001775 | 0.11976791 | TRMT10A | 93587 |
| ENSG000001 | 9.52223568 | -0.677197653 | 0.34556077 | -1.9597064 | 0.05003012 | 0.11978339 | HTR3D | 200909 |
| ENSG000002 | 3.32732951 | -0.740660035 | 0.37794277 | -1.9597148 | 0.05002913 | 0.11978339 | NA | NA |
| ENSG000002 | 37.0917442 | 0.456654095 | 0.23303246 | 1.95961581 | 0.05004071 | 0.11979685 | LOC101929448 | 101929448 |
| ENSG000002 | 3.68664485 | -0.741687166 | 0.37849243 | -1.9595826 | 0.0500446 | 0.11979685 | NA | NA |
| ENSG000002 | 12.5445094 | -0.639804714 | 0.32650005 | -1.9595853 | 0.05004428 | 0.11979685 | LINC01227 | 101928276 |
| ENSG000002 | 2.41325183 | -0.717212498 | 0.3660105 | -1.9595408 | 0.05004949 | 0.11980147 | NA | NA |
| ENSG000002 | 7.89872118 | -0.704004023 | 0.35928206 | -1.9594745 | 0.05005724 | 0.11981297 | SMIM5 | 643008 |
| ENSG000002 | 6.95780407 | -0.712501815 | 0.36363898 | -1.9593659 | 0.05006995 | 0.11983631 | NA | NA |
| ENSG000001 | 73.4258165 | 0.364824292 | 0.1862182 | 1.95912262 | 0.05009843 | 0.11987434 | ITLN2 | 142683 |
| ENSG000002 | 14.6416995 | -0.604638064 | 0.30862302 | -1.9591477 | 0.05009549 | 0.11987434 | NA | NA |
| ENSG000002 | 4.22334355 | -0.740535893 | 0.37799551 | -1.9591129 | 0.05009956 | 0.11987434 | NA | NA |
| ENSG000002 | 5.29012112 | -0.737220142 | 0.37629769 | -1.9591408 | 0.05009631 | 0.11987434 | KIAA0196-AS1 | 106479020 |
| ENSG000002 | 7.27086832 | -0.709492011 | 0.3621513 | -1.9591039 | 0.05010062 | 0.11987434 | LINC01169 | 102723165 |
| ENSG000001 | 1259.19087 | 0.316625188 | 0.16162133 | 1.95905571 | 0.05010626 | 0.11988076 | RYBP | 23429 |
| ENSG000002 | 2.82864684 | -0.725239084 | 0.370204 | -1.9590255 | 0.05010979 | 0.11988214 | NA | NA |
| ENSG000002 | 3.07885967 | -0.735247436 | 0.37532171 | -1.9589792 | 0.05011522 | 0.1198862 | NA | NA |
| ENSG000002 | 6.29590622 | -0.714709309 | 0.36484109 | -1.9589606 | 0.0501174 | 0.1198862 | NA | NA |
| ENSG000001 | 6.98474765 | -0.714611793 | 0.36479954 | -1.9589164 | 0.05012258 | 0.11989152 | FAM150A | 389658 |
| ENSG000002 | 3.69375593 | -0.741291665 | 0.37847051 | -1.9586511 | 0.05015367 | 0.11995879 | LOC101927926 | 101927926 |
| ENSG000002 | 2.65998688 | -0.730876268 | 0.37317964 | -1.9585106 | 0.05017013 | 0.1199911 | NA | NA |
| ENSG000002 | 2.67886067 | -0.730473774 | 0.37298565 | -1.9584501 | 0.05017722 | 0.12000098 | NA | NA |
| ENSG000002 | 7.31005829 | -0.710515185 | 0.36281563 | -1.9583368 | 0.05019051 | 0.12002567 | HP09025 | 100652929 |
| ENSG000002 | 13.5245969 | -0.621537652 | 0.31739183 | -1.9582661 | 0.0501988 | 0.12003842 | NA | NA |
| ENSG000002 | 14.0882547 | 0.606902911 | 0.3099363 | 1.9581537 | 0.05021198 | 0.12006286 | LOC100506405 | 100506405 |
| ENSG000002 | 4.66492816 | -0.739534111 | 0.37769267 | -1.9580314 | 0.05022633 | 0.12009008 | NA | NA |
| ENSG000001 | 464.194418 | -0.265132612 | 0.13541728 | -1.9578936 | 0.0502425 | 0.12011459 | PDE4B | 5142 |
| ENSG000002 | 3.06489068 | -0.735127456 | 0.37546778 | -1.9578976 | 0.05024203 | 0.12011459 | NA | NA |
| ENSG000002 | 3.74470009 | -0.740401073 | 0.37817042 | -1.9578503 | 0.05024758 | 0.12011965 | NA | NA |

| | | | | | | | | | |
|---|---|---|---|---|---|---|---|---|---|
| ENSG000002 | 6.62501572 | -0.714313725 | 0.36486028 | -1.9577734 | 0.05025661 | 0.12013416 | NA | NA | |
| ENSG000002 | 14.5747341 | -0.61983003 | 0.31664651 | -1.9574826 | 0.05029076 | 0.1202087 | NA | NA | |
| ENSG000002 | 11.2623559 | 0.677490519 | 0.34610883 | 1.95744941 | 0.05029465 | 0.12021092 | SNORD94 | | 692225 |
| ENSG000002 | 3.93294195 | -0.740860451 | 0.378488 | -1.9574212 | 0.05029797 | 0.12021175 | NA | NA | |
| ENSG000002 | 12.6536744 | -0.619035602 | 0.31632355 | -1.9569697 | 0.05035103 | 0.12033149 | NA | NA | |
| ENSG000001 | 132.659576 | 0.320284592 | 0.16367043 | 1.95688738 | 0.05036071 | 0.12034065 | PARVB | | 29780 |
| ENSG000002 | 4.89276993 | -0.73512473 | 0.37566036 | -1.9568866 | 0.0503608 | 0.12034065 | NA | NA | |
| ENSG000001 | 112.5728 | 0.355578309 | 0.18171759 | 1.95676329 | 0.05037531 | 0.12036822 | CHST3 | | 9469 |
| ENSG000002 | 4.58079812 | -0.737651769 | 0.37698977 | -1.9566891 | 0.05038403 | 0.12038198 | NA | NA | |
| ENSG000002 | 7.75443614 | -0.693124585 | 0.35424559 | -1.9566216 | 0.05039197 | 0.12039385 | NA | NA | |
| ENSG000001 | 49.5194313 | 0.413307346 | 0.21124051 | 1.95657235 | 0.05039777 | 0.12040061 | CD1D | | 912 |
| ENSG000001 | 21.2742575 | 0.542059099 | 0.27705236 | 1.95652221 | 0.05040367 | 0.12040761 | HRH3 | | 11255 |
| ENSG000001 | 61.150703 | 0.393052796 | 0.20090233 | 1.95643726 | 0.05041367 | 0.1204244 | SNPH | | 9751 |
| ENSG000002 | 3.93053748 | 0.740244613 | 0.37837012 | 1.95640349 | 0.05041764 | 0.1204268 | NA | NA | |
| ENSG000002 | 4.21199354 | -0.739244654 | 0.3778867 | -1.95626 | 0.05043453 | 0.12045295 | NA | NA | |
| ENSG000002 | 20.610591 | -0.540830006 | 0.27645838 | -1.95628 | 0.05043218 | 0.12045295 | NA | NA | |
| ENSG000002 | 6.06579421 | -0.720345504 | 0.36826007 | -1.9560782 | 0.05045594 | 0.12049698 | NA | NA | |
| ENSG000001 | 14.140549 | -0.615398164 | 0.31461572 | -1.9560312 | 0.05046148 | 0.12050214 | CAT | | 847 |
| ENSG000002 | 3.02720183 | -0.734094751 | 0.37530225 | -1.9560094 | 0.05046404 | 0.12050214 | NA | NA | |
| ENSG000001 | 20.7331415 | -0.556999127 | 0.28478713 | -1.9558437 | 0.05048357 | 0.12054167 | ITGB2 | | 3689 |
| ENSG000002 | 11.7168696 | -0.644072407 | 0.32931152 | -1.955815 | 0.05048695 | 0.12054265 | NA | NA | |
| ENSG000002 | 3.44783036 | -0.73409553 | 0.37537908 | -1.9556112 | 0.05051098 | 0.12058511 | NA | NA | |
| ENSG000002 | 6.48568468 | -0.73410386 | 0.37537836 | -1.9556371 | 0.05050792 | 0.12058511 | NA | NA | |
| ENSG000002 | 4.92424745 | -0.735985323 | 0.3763498 | -1.9555884 | 0.05051366 | 0.12058511 | SALRNA3 | | 104548972 |
| ENSG000002 | 13.4896194 | -0.627792886 | 0.32105082 | -1.9554315 | 0.05053216 | 0.12062219 | NA | NA | |
| ENSG000002 | 5.37202958 | -0.737088593 | 0.37695116 | -1.9553955 | 0.05053641 | 0.12062522 | NA | NA | |
| ENSG00000C | 51.0910499 | -0.416816283 | 0.21317462 | -1.955281 | 0.05054991 | 0.12063615 | RFX2 | | 5990 |
| ENSG000002 | 4.96093741 | -0.736676099 | 0.37676137 | -1.9552856 | 0.05054937 | 0.12063615 | GS1-594A7.3 | | 104798195 |
| ENSG000002 | 5.08500018 | -0.73815357 | 0.37751002 | -1.9553218 | 0.0505451 | 0.12063615 | NA | NA | |

| | | | | | | | | |
|---|---|---|---|---|---|---|---|---|
| ENSG000002 | 13.6186067 | -0.624190754 | 0.31923793 | -1.9552525 | 0.05055328 | 0.12063708 | NA | NA |
| ENSG000001 | 21.4502423 | -0.551077079 | 0.28185052 | -1.9552105 | 0.05055823 | 0.12064181 | SMYD1 | 150572 |
| ENSG000002 | 20.7821265 | -0.552766645 | 0.28271986 | -1.9551744 | 0.05056248 | 0.12064485 | NA | NA |
| ENSG000002 | 38.1304257 | -0.459141126 | 0.23486394 | -1.9549239 | 0.05059205 | 0.12070502 | EBF2 | 64641 |
| ENSG000002 | 15.5303406 | -0.614339316 | 0.31425447 | -1.9549104 | 0.05059365 | 0.12070502 | NA | NA |
| ENSG000001 | 126.545061 | 0.369502522 | 0.18902681 | 1.95476254 | 0.0506111 | 0.12073955 | C19orf54 | 284325 |
| ENSG000002 | 157.262707 | 0.279354375 | 0.14291503 | 1.95468859 | 0.05061984 | 0.12075328 | NA | NA |
| ENSG000002 | 3.27197353 | -0.738881001 | 0.37801594 | -1.9546292 | 0.05062685 | 0.1207558 | NA | NA |
| ENSG000002 | 8.33266097 | -0.688579561 | 0.35228005 | -1.9546368 | 0.05062595 | 0.1207558 | NA | NA |
| ENSG00000C | 15.9965585 | -0.602911029 | 0.30847447 | -1.9544925 | 0.050643 | 0.12078012 | COPZ2 | 51226 |
| ENSG000002 | 7.69602584 | -0.694622203 | 0.35539678 | -1.9544978 | 0.05064238 | 0.12078012 | NA | NA |
| ENSG000001 | 1057.01512 | 0.232056693 | 0.11873783 | 1.95436193 | 0.05065843 | 0.1208027 | RAP2B | 5912 |
| ENSG000001 | 4.54633133 | -0.736509884 | 0.37684996 | -1.9543849 | 0.05065571 | 0.1208027 | NA | NA |
| ENSG000002 | 59.9379189 | 0.406027672 | 0.20775814 | 1.95432859 | 0.05066237 | 0.12080499 | NA | NA |
| ENSG000001 | 23.7268665 | -0.527857331 | 0.27013075 | -1.9540809 | 0.05069165 | 0.12086061 | LAIR1 | 3903 |
| ENSG000002 | 4.91302586 | -0.73501372 | 0.37614272 | -1.954082 | 0.05069152 | 0.12086061 | NA | NA |
| ENSG000002 | 12.8351059 | -0.635212077 | 0.32508303 | -1.9539995 | 0.05070128 | 0.12087645 | NA | NA |
| ENSG000002 | 27.5695883 | -0.498152493 | 0.2549557 | -1.9538786 | 0.05071557 | 0.12090343 | NA | NA |
| ENSG000001 | 76.144132 | -0.38391598 | 0.19649975 | -1.9537734 | 0.05072802 | 0.12092598 | CDH13 | 1012 |
| ENSG000001 | 797.531974 | 0.235483835 | 0.12053734 | 1.95361738 | 0.05074648 | 0.12096289 | RIMS2 | 9699 |
| ENSG000001 | 11.0688998 | -0.664509474 | 0.34016286 | -1.9535039 | 0.05075991 | 0.12098068 | A4GNT | 51146 |
| ENSG000002 | 15.9505246 | -0.604734931 | 0.30956276 | -1.9535132 | 0.05075882 | 0.12098068 | BTBD18 | 643376 |
| ENSG000002 | 3.89461394 | -0.739091377 | 0.37835323 | -1.9534427 | 0.05076716 | 0.12099083 | NA | NA |
| ENSG000001 | 234.275914 | 0.267145589 | 0.13676305 | 1.95334624 | 0.05077858 | 0.12101095 | RNPC3 | 55599 |
| ENSG000001 | 9.96802505 | -0.659429367 | 0.33761248 | -1.9532138 | 0.05079426 | 0.12104121 | LILRA1 | 11024 |
| ENSG000002 | 5.33745179 | -0.736105971 | 0.37688827 | -1.9531146 | 0.05080602 | 0.12106211 | NA | NA |
| ENSG000002 | 17.4923823 | 0.58181657 | 0.29789601 | 1.95308616 | 0.05080939 | 0.12106302 | NA | NA |
| ENSG000001 | 118.018197 | 0.318179695 | 0.16292196 | 1.95295771 | 0.05082461 | 0.12109216 | EID2 | 163126 |
| ENSG000002 | 9.01004326 | -0.686439632 | 0.35150106 | -1.9528807 | 0.05083374 | 0.1211068 | ELMO1-AS1 | 100861514 |

| | | | | | | | | |
|---|---|---|---|---|---|---|---|---|
| ENSG000001 | 15.0115433 | -0.587627857 | 0.30093307 | -1.9526862 | 0.05085679 | 0.12115461 | LDHAL6A | 160287 |
| ENSG000001 | 20.1691547 | -0.562254581 | 0.28796373 | -1.9525188 | 0.05087665 | 0.12119479 | RFX8 | 731220 |
| ENSG000001 | 6.14207431 | -0.724737416 | 0.3712062 | -1.952385 | 0.05089251 | 0.12121123 | ZNF729 | 100287226 |
| ENSG000002 | 2.614515 | -0.728654793 | 0.37320983 | -1.9523998 | 0.05089076 | 0.12121123 | NA | NA |
| ENSG000002 | 9.77045578 | -0.665599823 | 0.34091405 | -1.9523977 | 0.050891 | 0.12121123 | NA | NA |
| ENSG000001 | 6.60631854 | -0.711659376 | 0.36451669 | -1.9523369 | 0.05089822 | 0.1212177 | MYL1 | 4632 |
| ENSG000001 | 905.026019 | 0.292512753 | 0.14983317 | 1.95225629 | 0.05090778 | 0.12123335 | POLR2A | 5430 |
| ENSG000002 | 4.49705693 | -0.734850465 | 0.37643384 | -1.9521371 | 0.05092193 | 0.12125993 | LOC105375844 | 105375844 |
| ENSG000002 | 4.25880345 | -0.737774651 | 0.37794352 | -1.9520765 | 0.05092912 | 0.12126993 | NA | NA |
| ENSG000001 | 21.8299952 | -0.549835471 | 0.28168669 | -1.9519398 | 0.05094535 | 0.12130145 | ART4 | 420 |
| ENSG000002 | 4.84621155 | -0.73471808 | 0.37643841 | -1.9517617 | 0.0509665 | 0.12134468 | NA | NA |
| ENSG000001 | 15.6013352 | -0.588730628 | 0.30165301 | -1.9516816 | 0.05097602 | 0.12135309 | KIAA1875 | 340390 |
| ENSG000001 | 8.26625755 | -0.681936274 | 0.349406 | -1.9517017 | 0.05097363 | 0.12135309 | CRYGB | 1419 |
| ENSG000001 | 173.326039 | 0.577296403 | 0.29580581 | 1.95160601 | 0.050985 | 0.12136734 | RAB11B | 9230 |
| ENSG000001 | 9.6822017 | -0.682738597 | 0.34990579 | -1.9512069 | 0.05103244 | 0.12147313 | C11orf21 | 29125 |
| ENSG000001 | 672.22519 | 0.250423939 | 0.12834709 | 1.95114625 | 0.05103965 | 0.12148318 | SLF1 | 84250 |
| ENSG000002 | 3.67454967 | -0.738444122 | 0.37847324 | -1.9511132 | 0.05104358 | 0.12148541 | NA | NA |
| ENSG000002 | 3.8962837 | -0.738393731 | 0.37845644 | -1.9510666 | 0.05104912 | 0.12149146 | NA | NA |
| ENSG000002 | 8.03244896 | -0.691405718 | 0.35438497 | -1.9510018 | 0.05105683 | 0.12150267 | CYS1 | 192668 |
| ENSG000002 | 2.68693764 | -0.727711491 | 0.37300825 | -1.950926 | 0.05106585 | 0.121517 | NA | NA |
| ENSG000002 | 8.87067795 | -0.674399764 | 0.34570041 | -1.9508214 | 0.05107829 | 0.12153948 | LOC100507144 | 100507144 |
| ENSG000002 | 2.63697091 | -0.72812067 | 0.37325518 | -1.9507316 | 0.05108898 | 0.12155778 | NA | NA |
| ENSG000001 | 50.5029189 | -0.402484509 | 0.20633634 | -1.9506235 | 0.05110185 | 0.12156701 | GALR1 | 2587 |
| ENSG000001 | 285.607984 | 0.279162955 | 0.1431143 | 1.95062933 | 0.05110116 | 0.12156701 | PLEKHF2 | 79666 |
| ENSG000002 | 13.6478684 | -0.622630885 | 0.31919238 | -1.9506446 | 0.05109934 | 0.12156701 | NA | NA |
| ENSG000001 | 102.392679 | 0.324948148 | 0.16659972 | 1.9504724 | 0.05111984 | 0.12160266 | C8orf37 | 157657 |
| ENSG000001 | 200.698523 | 0.290280851 | 0.14883155 | 1.95039866 | 0.05112862 | 0.12160384 | ATG7 | 10533 |
| ENSG000002 | 3.33072419 | -0.735898072 | 0.3773002 | -1.9504312 | 0.05112475 | 0.12160384 | NA | NA |
| ENSG000002 | 31.4505619 | -0.48653209 | 0.2494534 | -1.9503927 | 0.05112933 | 0.12160384 | NA | NA |

| | | | | | | | | |
|---|---|---|---|---|---|---|---|---|
| ENSG00000( | 405.154349 | 0.257214968 | 0.13189691 | 1.95012123 | 0.05116167 | 0.12166649 | CDON | 50937 |
| ENSG000002 | 14.685357 | -0.603854 | 0.30964699 | -1.9501368 | 0.05115981 | 0.12166649 | NA | NA |
| ENSG000002 | 7.41108394 | -0.711911819 | 0.36506863 | -1.9500767 | 0.05116698 | 0.12167198 | NA | NA |
| ENSG000001 | 716.110615 | 0.333484826 | 0.17101381 | 1.95004615 | 0.05117062 | 0.12167349 | SF3B4 | 10262 |
| ENSG000002 | 14.2930911 | -0.62063155 | 0.31827438 | -1.9499891 | 0.05117742 | 0.12168254 | LOC101060019 | 101060019 |
| ENSG000001 | 3.93968885 | -0.73774759 | 0.37842149 | -1.9495393 | 0.05123105 | 0.12176589 | IL9R | 3581 |
| ENSG000001 | 6.88112589 | -0.704393813 | 0.3613199 | -1.9495019 | 0.05123552 | 0.12176589 | SPINK7 | 84651 |
| ENSG000001 | 6.81860696 | -0.703597631 | 0.36091601 | -1.9494775 | 0.05123843 | 0.12176589 | ZNF503 | 84858 |
| ENSG000002 | 35.0321279 | 0.475220988 | 0.24376802 | 1.94948045 | 0.05123808 | 0.12176589 | RN7SL767P | 106481131 |
| ENSG000002 | 2.4730508 | -0.710352151 | 0.36438247 | -1.9494685 | 0.0512395 | 0.12176589 | RN7SL482P | 106480513 |
| ENSG000002 | 128.324976 | -0.302197531 | 0.1550083 | -1.9495571 | 0.05122893 | 0.12176589 | NA | NA |
| ENSG000002 | 6.63266758 | -0.71250983 | 0.36548149 | -1.9495101 | 0.05123454 | 0.12176589 | NA | NA |
| ENSG000002 | 3.30371272 | -0.736943178 | 0.37802012 | -1.9494814 | 0.05123796 | 0.12176589 | NA | NA |
| ENSG000002 | 6.03870809 | -0.72238139 | 0.37052882 | -1.9495957 | 0.05122433 | 0.12176589 | NA | NA |
| ENSG000001 | 446.228118 | 0.251012304 | 0.12876424 | 1.94939459 | 0.05124832 | 0.12176668 | DDHD1 | 80821 |
| ENSG000001 | 338.813452 | 0.266206013 | 0.13655861 | 1.94939019 | 0.05124885 | 0.12176668 | TAF1B | 9014 |
| ENSG000002 | 10.2077845 | -0.655945644 | 0.33648112 | -1.9494278 | 0.05124436 | 0.12176668 | LOC645949 | 645949 |
| ENSG000002 | 13.8976483 | 0.624888838 | 0.32066947 | 1.94870074 | 0.05133117 | 0.12195515 | NA | NA |
| ENSG000002 | 8.64950212 | -0.683328631 | 0.3506647 | -1.9486667 | 0.05133524 | 0.12195766 | NA | NA |
| ENSG000001 | 7.53661411 | -0.701967006 | 0.36024628 | -1.9485753 | 0.05134616 | 0.12197647 | P2RY12 | 64805 |
| ENSG000002 | 5.30864081 | -0.732606516 | 0.37597894 | -1.9485307 | 0.05135149 | 0.12198198 | NA | NA |
| ENSG000001 | 15.3083397 | -0.641071273 | 0.32902742 | -1.9483825 | 0.05136921 | 0.12201693 | GLRA1 | 2741 |
| ENSG000002 | 3.31507015 | -0.735953898 | 0.377743 | -1.9482926 | 0.05137996 | 0.1220353 | NA | NA |
| ENSG000002 | 3.72812014 | -0.736534151 | 0.37808182 | -1.9480814 | 0.05140523 | 0.12208817 | TXNDC12-AS1 | 104355143 |
| ENSG000001 | 3.96249811 | -0.737412137 | 0.37859336 | -1.9477683 | 0.0514427 | 0.12216285 | TICAM1 | 148022 |
| ENSG000002 | 6.00306642 | -0.716623654 | 0.36791707 | -1.9477858 | 0.05144059 | 0.12216285 | NA | NA |
| ENSG000002 | 5.80066797 | -0.724572021 | 0.37202773 | -1.9476291 | 0.05145936 | 0.12218335 | NA | NA |
| ENSG000002 | 4.58188024 | -0.734567062 | 0.37716019 | -1.9476262 | 0.05145971 | 0.12218335 | NA | NA |
| ENSG000002 | 2.65506783 | -0.726380395 | 0.37295784 | -1.9476207 | 0.05146037 | 0.12218335 | NA | NA |

| | | | | | | | | |
|---|---|---|---|---|---|---|---|---|
| ENSG000001 | 7.13388251 | -0.698411627 | 0.35862734 | -1.9474579 | 0.05147987 | 0.12221533 | HSF4 | 3299 |
| ENSG000001 | 733.578209 | 0.248337842 | 0.12751769 | 1.94747762 | 0.0514775 | 0.12221533 | VPS13B | 157680 |
| ENSG000002 | 144.668653 | 0.376942022 | 0.1935834 | 1.94718153 | 0.05151297 | 0.12228677 | NA | NA |
| ENSG000001 | 274.209057 | 0.305840756 | 0.15707329 | 1.94712128 | 0.0515202 | 0.12229675 | CABLES1 | 91768 |
| ENSG000002 | 6.59528871 | 0.706160979 | 0.36267662 | 1.94708159 | 0.05152495 | 0.12230089 | NA | NA |
| ENSG000002 | 6.2531545 | -0.710224108 | 0.36478249 | -1.9469797 | 0.05153716 | 0.12232271 | NA | NA |
| ENSG000002 | 2.62981907 | -0.725335705 | 0.37255579 | -1.9469183 | 0.05154453 | 0.12232587 | RN7SKP107 | 106480464 |
| ENSG000002 | 6.25440767 | -0.711370613 | 0.3653821 | -1.9469224 | 0.05154403 | 0.12232587 | NA | NA |
| ENSG000002 | 6.4092202 | 0.727082763 | 0.37347486 | 1.9468051 | 0.05155811 | 0.12235093 | NA | NA |
| ENSG000001 | 8.8780409 | -0.676096806 | 0.34731197 | -1.9466557 | 0.05157603 | 0.12237379 | GPR171 | 29909 |
| ENSG000001 | 617.116312 | 0.332201633 | 0.17065254 | 1.9466551 | 0.0515761 | 0.12237379 | MYO6 | 4646 |
| ENSG000002 | 9.30426249 | -0.675446792 | 0.34699263 | -1.9465739 | 0.05158584 | 0.12237379 | NA | NA |
| ENSG000002 | 2.98662939 | -0.730144072 | 0.37509106 | -1.9465782 | 0.05158533 | 0.12237379 | NA | NA |
| ENSG000002 | 7.07869483 | -0.698947197 | 0.35905209 | -1.9466457 | 0.05157723 | 0.12237379 | NA | NA |
| ENSG000002 | 6.92596603 | -0.703808896 | 0.36155521 | -1.9466153 | 0.05158087 | 0.12237379 | NA | NA |
| ENSG000001 | 19.2676608 | -0.569465957 | 0.29257254 | -1.9464094 | 0.05160558 | 0.12241344 | OR2AG2 | 338755 |
| ENSG000002 | 56.7623722 | 0.434979654 | 0.22350736 | 1.94615361 | 0.05163629 | 0.12247913 | ARHGEF35 | 445328 |
| ENSG000001 | 42.8021245 | -0.453041397 | 0.23282427 | -1.9458512 | 0.05167261 | 0.12255812 | KIAA1614 | 57710 |
| ENSG000001 | 19.6419639 | -0.551633833 | 0.28351402 | -1.9457021 | 0.05169053 | 0.12259344 | ACSM1 | 116285 |
| ENSG000002 | 2.63076535 | -0.725400649 | 0.3728369 | -1.9456246 | 0.05169985 | 0.12260837 | LOC105371506 | 105371506 |
| ENSG000002 | 5.85627748 | -0.724921821 | 0.37261287 | -1.9455094 | 0.0517137 | 0.12263406 | NA | NA |
| ENSG000001 | 20.6530008 | -0.557037072 | 0.2863439 | -1.9453429 | 0.05173372 | 0.12267435 | CTLA4 | 1493 |
| ENSG000001 | 34.5496941 | -0.462331899 | 0.23767318 | -1.9452422 | 0.05174584 | 0.12269591 | NRG4 | 145957 |
| ENSG000002 | 8.28154256 | -0.681753495 | 0.35050105 | -1.9450826 | 0.05176503 | 0.12273425 | NA | NA |
| ENSG000002 | 6.37929169 | -0.728441078 | 0.37455491 | -1.9448179 | 0.0517969 | 0.12280263 | NA | NA |
| ENSG000002 | 3.01219722 | -0.729890699 | 0.375316 | -1.9447364 | 0.05180671 | 0.1228187 | NA | NA |
| ENSG000000 | 51.5017332 | 0.456210126 | 0.23459677 | 1.94465647 | 0.05181634 | 0.12283435 | LIPE | 3991 |
| ENSG000002 | 4.26257373 | -0.735482783 | 0.37824308 | -1.9444712 | 0.05183865 | 0.12287288 | NA | NA |
| ENSG000002 | 8.04622607 | -0.686040774 | 0.35281363 | -1.9444849 | 0.05183701 | 0.12287288 | NA | NA |

| | | | | | | | | |
|---|---|---|---|---|---|---|---|---|
| ENSG000002 | 8.13465343 | -0.694180724 | 0.35700798 | -1.9444404 | 0.05184237 | 0.12287451 | NA | NA |
| ENSG000001 | 23.9603197 | -0.511106126 | 0.26286454 | -1.9443708 | 0.05185075 | 0.1228872 | ELFN2 | 114794 |
| ENSG000002 | 6.62498457 | -0.708821868 | 0.36459605 | -1.9441293 | 0.05187986 | 0.1229418 | NA | NA |
| ENSG000002 | 4.24710524 | -0.734883316 | 0.37799734 | -1.9441494 | 0.05187743 | 0.1229418 | NA | NA |
| ENSG000001 | 8.89767162 | -0.678706515 | 0.34911196 | -1.9440941 | 0.0518841 | 0.12294467 | MSTN | 2660 |
| ENSG000002 | 45.6792339 | 0.416713497 | 0.21435165 | 1.94406478 | 0.05188764 | 0.12294587 | NA | NA |
| ENSG000002 | 3.86153672 | -0.73574934 | 0.37847649 | -1.9439764 | 0.0518983 | 0.1229618 | NA | NA |
| ENSG000002 | 4.42347081 | -0.735869929 | 0.37854195 | -1.9439587 | 0.05190043 | 0.1229618 | NA | NA |
| ENSG000002 | 3.08847522 | -0.728672177 | 0.37484601 | -1.943924 | 0.05190461 | 0.12296453 | NA | NA |
| ENSG000002 | 2.62435868 | -0.724155611 | 0.3725497 | -1.9437826 | 0.05192167 | 0.12299777 | LINC01082 | 100506542 |
| ENSG000001 | 18.916794 | -0.576509299 | 0.29661494 | -1.9436287 | 0.05194025 | 0.12303386 | LIPM | 340654 |
| ENSG000002 | 10.9529759 | -0.655341648 | 0.33717823 | -1.943606 | 0.05194298 | 0.12303386 | NA | NA |
| ENSG000002 | 3.3064403 | -0.73453505 | 0.37794616 | -1.9434913 | 0.05195683 | 0.12305948 | PFN1P2 | 767846 |
| ENSG000001 | 7.97299375 | -0.684293044 | 0.3521063 | -1.9434274 | 0.05196454 | 0.12307055 | TRAT1 | 50852 |
| ENSG00000C | 48.9965782 | 0.423213207 | 0.2177809 | 1.94329806 | 0.05198016 | 0.12310035 | RTN4R | 65078 |
| ENSG000001 | 11.6362575 | -0.634255757 | 0.32640913 | -1.9431312 | 0.05200031 | 0.1231409 | LGALS4 | 3960 |
| ENSG00000C | 1366.67898 | 0.245519511 | 0.12636349 | 1.94296234 | 0.05202071 | 0.12317746 | KMT2C | 58508 |
| ENSG000001 | 8.8027413 | -0.673675064 | 0.3467274 | -1.9429531 | 0.05202183 | 0.12317746 | DOK2 | 9046 |
| ENSG000001 | 3.27406141 | -0.734160307 | 0.37787937 | -1.9428431 | 0.05203512 | 0.12319455 | SPRR4 | 163778 |
| ENSG000002 | 11.9559199 | -0.644178099 | 0.33156241 | -1.9428563 | 0.05203353 | 0.12319455 | NA | NA |
| ENSG000002 | 12.1058698 | -0.621521714 | 0.31991533 | -1.9427694 | 0.05204403 | 0.12320844 | LINC00499 | 100874047 |
| ENSG00000C | 64.6311857 | 0.442560869 | 0.22781692 | 1.94261632 | 0.05206253 | 0.12324506 | MPND | 84954 |
| ENSG000002 | 47.2408417 | 0.437944245 | 0.22549797 | 1.94212055 | 0.05212251 | 0.12337983 | PGAM4 | 441531 |
| ENSG000001 | 80.8554041 | 0.369840906 | 0.19043799 | 1.9420542 | 0.05213054 | 0.12338743 | RAB11FIP5 | 26056 |
| ENSG000001 | 144.398213 | 0.303324687 | 0.15618839 | 1.94204376 | 0.0521318 | 0.12338743 | NAPEPLD | 222236 |
| ENSG000001 | 17.073826 | -0.575621902 | 0.29641301 | -1.9419589 | 0.05214207 | 0.12339732 | INSC | 387755 |
| ENSG000002 | 2.29559628 | -0.714211992 | 0.36777545 | -1.9419784 | 0.05213971 | 0.12339732 | NA | NA |
| ENSG000001 | 17.5388795 | -0.585626791 | 0.30158969 | -1.9417998 | 0.05216135 | 0.12342853 | TLR6 | 10333 |
| ENSG000002 | 138.810401 | 0.346198992 | 0.17828608 | 1.9418173 | 0.05215922 | 0.12342853 | NA | NA |

| | | | | | | | | |
|---|---|---|---|---|---|---|---|---|
| ENSG000001 | 23.2525096 | -0.542559907 | 0.27941971 | -1.9417381 | 0.05216881 | 0.12343899 | TRPC3 | 7222 |
| ENSG000001 | 17.497987 | -0.571793151 | 0.29448898 | -1.9416453 | 0.05218005 | 0.12345838 | CATSPER4 | 378807 |
| ENSG000002 | 16.4794992 | -0.593059177 | 0.30545057 | -1.9415881 | 0.05218699 | 0.1234676 | CRAT37 | 101926928 |
| ENSG000001 | 3.64315593 | -0.734584431 | 0.37839635 | -1.9413095 | 0.05222075 | 0.12354025 | RTP2 | 344892 |
| ENSG000002 | 7.98848597 | -0.687753779 | 0.35429746 | -1.9411761 | 0.05223692 | 0.1235713 | NA | NA |
| ENSG000001 | 3.01778793 | -0.728802546 | 0.37545916 | -1.9410967 | 0.05224655 | 0.12358687 | FEV | 54738 |
| ENSG000002 | 3.6154355 | -0.73440032 | 0.37839826 | -1.9408132 | 0.05228094 | 0.12364659 | NA | NA |
| ENSG000002 | 12.3073741 | -0.638123186 | 0.32878808 | -1.9408343 | 0.05227838 | 0.12364659 | LOC439933 | 439933 |
| ENSG000002 | 3.9527955 | -0.734726391 | 0.37857116 | -1.940788 | 0.05228399 | 0.12364659 | NA | NA |
| ENSG000002 | 35.3559676 | 0.503725918 | 0.25953865 | 1.94085127 | 0.05227632 | 0.12364659 | NA | NA |
| ENSG000001 | 258.736375 | 0.27754151 | 0.14300695 | 1.94075543 | 0.05228795 | 0.12364874 | CD109 | 135228 |
| ENSG000002 | 3.03867191 | -0.72775045 | 0.37503594 | -1.9404819 | 0.05232116 | 0.12372005 | NA | NA |
| ENSG000002 | 22.3816956 | 0.563734485 | 0.29053884 | 1.94030681 | 0.05234241 | 0.12376311 | NA | NA |
| ENSG000001 | 143.857854 | 0.364793777 | 0.18801749 | 1.9402119 | 0.05235394 | 0.12378314 | TRABD | 80305 |
| ENSG000002 | 28.6302966 | -0.479791802 | 0.24729539 | -1.9401567 | 0.05236065 | 0.12379179 | NA | NA |
| ENSG000001 | 12.5924354 | -0.645085985 | 0.33250032 | -1.9401064 | 0.05236677 | 0.12379181 | HES7 | 84667 |
| ENSG000002 | 3.96974456 | -0.734421089 | 0.37854358 | -1.940123 | 0.05236475 | 0.12379181 | NA | NA |
| ENSG000001 | 38.67198 | -0.481497787 | 0.24818869 | -1.9400472 | 0.05237395 | 0.12380157 | HLF | 3131 |
| ENSG000001 | 89.1967329 | 0.414773626 | 0.21381508 | 1.93987074 | 0.0523954 | 0.12384506 | FAM20C | 56975 |
| ENSG000002 | 17.1430665 | 0.58631535 | 0.30225317 | 1.93981538 | 0.05240213 | 0.12385375 | NA | NA |
| ENSG000002 | 42.0870794 | 0.474702083 | 0.24472423 | 1.93974291 | 0.05241094 | 0.12386736 | NA | NA |
| ENSG000002 | 2.98903745 | -0.728245865 | 0.37544535 | -1.9396854 | 0.05241793 | 0.12387666 | NA | NA |
| ENSG000001 | 698.453478 | 0.219683776 | 0.11326551 | 1.93954703 | 0.05243476 | 0.12390798 | C1orf27 | 54953 |
| ENSG000002 | 4.66944087 | -0.733199937 | 0.37803044 | -1.9395262 | 0.0524373 | 0.12390798 | NA | NA |
| ENSG000002 | 6.83295106 | -0.716026731 | 0.36918226 | -1.9394939 | 0.05244123 | 0.12391005 | NA | NA |
| ENSG000002 | 16.919406 | -0.585993231 | 0.30215798 | -1.9393604 | 0.05245746 | 0.12393397 | NA | NA |
| ENSG000002 | 2.45535121 | -0.707704663 | 0.36491422 | -1.9393727 | 0.05245598 | 0.12393397 | NA | NA |
| ENSG000002 | 83.8743913 | 0.389341958 | 0.20076775 | 1.93926547 | 0.05246902 | 0.12395405 | RPL32P3 | 132241 |
| ENSG000001 | 7.72603036 | -0.689203765 | 0.35542845 | -1.9390788 | 0.05249175 | 0.12400051 | POU3F4 | 5456 |

| | | | | | | | | |
|---|---|---|---|---|---|---|---|---|
| ENSG000002 | 3.88046313 | -0.733830015 | 0.37845074 | -1.9390371 | 0.05249683 | 0.12400529 | NA | NA |
| ENSG000000 | 72.8606199 | 0.412572315 | 0.21278244 | 1.93893967 | 0.05250868 | 0.12402608 | USE1 | 55850 |
| ENSG000001 | 20.9729434 | -0.550370377 | 0.28385843 | -1.9388904 | 0.05251469 | 0.12403304 | USP26 | 83844 |
| ENSG000001 | 212.890134 | 0.319212823 | 0.16464689 | 1.93877229 | 0.05252907 | 0.12404658 | MICALL1 | 85377 |
| ENSG000001 | 11.9821668 | -0.643904376 | 0.33212039 | -1.938768 | 0.0525296 | 0.12404658 | ESRRB | 2103 |
| ENSG000002 | 2.73481849 | -0.717924601 | 0.37029453 | -1.9387934 | 0.0525265 | 0.12404658 | NA | NA |
| ENSG000002 | 6.68869068 | -0.71501772 | 0.36881125 | -1.9387091 | 0.05253677 | 0.12405628 | GLYCAM1 | 644076 |
| ENSG000002 | 103.167228 | 0.343494453 | 0.1771802 | 1.93867286 | 0.05254119 | 0.12405949 | NA | NA |
| ENSG000002 | 9.67686424 | 0.667299886 | 0.34421363 | 1.93862131 | 0.05254747 | 0.12406709 | SNORD91A | 692207 |
| ENSG000001 | 10.3577541 | -0.669460871 | 0.34536132 | -1.9384362 | 0.05257003 | 0.12411314 | IL24 | 11009 |
| ENSG000002 | 7.71018112 | -0.691413022 | 0.3567069 | -1.9383225 | 0.05258389 | 0.12413862 | NA | NA |
| ENSG000002 | 7.75849434 | -0.688822941 | 0.35537627 | -1.9382919 | 0.05258762 | 0.12414022 | IQCF5-AS1 | 101928999 |
| ENSG000002 | 4.31978792 | -0.733284624 | 0.37833535 | -1.9381869 | 0.05260042 | 0.12416321 | NA | NA |
| ENSG000002 | 17.3147021 | 0.582338759 | 0.3004685 | 1.93810255 | 0.05261071 | 0.12418027 | YY2 | 404281 |
| ENSG000001 | 32.7506263 | -0.51797243 | 0.26728014 | -1.9379383 | 0.05263075 | 0.12422033 | SLC26A10 | 65012 |
| ENSG000001 | 31.1038933 | -0.517348159 | 0.26696272 | -1.9379041 | 0.05263492 | 0.12422295 | UNC45B | 146862 |
| ENSG000002 | 2.71332745 | -0.720757589 | 0.37193795 | -1.9378436 | 0.0526423 | 0.12423314 | NA | NA |
| ENSG000001 | 6.66847851 | -0.708726854 | 0.36573767 | -1.9378011 | 0.05264749 | 0.12423816 | STX19 | 415117 |
| ENSG000002 | 2.64331232 | -0.722307195 | 0.372765 | -1.9377012 | 0.05265968 | 0.12425969 | NA | NA |
| ENSG000002 | 26.1624219 | -0.502730711 | 0.25945217 | -1.9376624 | 0.05266443 | 0.12426365 | LOC283194 | 283194 |
| ENSG000001 | 13.357303 | -0.618783503 | 0.31936214 | -1.9375606 | 0.05267685 | 0.12428574 | SUCNR1 | 56670 |
| ENSG000001 | 32.4099552 | -0.468829424 | 0.24199778 | -1.9373295 | 0.05270508 | 0.1243403 | ZC3H12B | 340554 |
| ENSG000001 | 8.26739228 | -0.677488659 | 0.34970386 | -1.9373211 | 0.05270611 | 0.1243403 | DRGX | 644168 |
| ENSG000002 | 4.55726531 | -0.730984159 | 0.37733769 | -1.9372148 | 0.05271909 | 0.12436369 | NA | NA |
| ENSG000002 | 2.67083687 | -0.722057516 | 0.37276299 | -1.9370418 | 0.05274023 | 0.12440632 | NA | NA |
| ENSG000001 | 337.77557 | 0.24628856 | 0.12715786 | 1.93687247 | 0.05276093 | 0.12444792 | ZC3H12C | 85463 |
| ENSG000002 | 7.9990941 | -0.687720928 | 0.35510735 | -1.9366564 | 0.05278736 | 0.12449576 | NA | NA |
| ENSG000002 | 9.31789722 | -0.661493152 | 0.34156133 | -1.9366746 | 0.05278513 | 0.12449576 | NA | NA |
| ENSG000002 | 9.78675444 | -0.669574096 | 0.34575821 | -1.9365386 | 0.05280178 | 0.12452252 | NA | NA |

| | | | | | | | | |
|---|---|---|---|---|---|---|---|---|
| ENSG000002 | 11.6102779 | -0.656242104 | 0.33888118 | -1.9364962 | 0.05280696 | 0.12452751 | LINC00592 | 283404 |
| ENSG000002 | 5.24813012 | -0.728541357 | 0.37632425 | -1.9359405 | 0.05287499 | 0.12468069 | NA | NA |
| ENSG000001 | 36.97754 | -0.459312147 | 0.23726726 | -1.9358429 | 0.05288695 | 0.12469437 | SHC4 | 399694 |
| ENSG000002 | 8.93149302 | -0.675089819 | 0.34872942 | -1.9358557 | 0.05288538 | 0.12469437 | NA | NA |
| ENSG000002 | 3.40771436 | -0.729108336 | 0.37664775 | -1.9357831 | 0.05289427 | 0.1247044 | NA | NA |
| ENSG000002 | 36.5341199 | -0.456006194 | 0.23557104 | -1.9357481 | 0.05289856 | 0.12470725 | SRCIN1 | 80725 |
| ENSG000002 | 4.27552933 | -0.732342684 | 0.37833823 | -1.9356825 | 0.0529066 | 0.12471895 | ENPP7P5 | 106480674 |
| ENSG000002 | 2.32872502 | -0.711151667 | 0.36739641 | -1.9356522 | 0.05291031 | 0.12471932 | NA | NA |
| ENSG000002 | 3.62140443 | -0.732204168 | 0.37827672 | -1.935631 | 0.0529129 | 0.12471932 | NA | NA |
| ENSG000001 | 25.0157065 | 0.52249479 | 0.26998073 | 1.93530404 | 0.052953 | 0.12480657 | NR1D1 | 9572 |
| ENSG000002 | 3.05231419 | -0.726164894 | 0.3752418 | -1.9351919 | 0.05296675 | 0.12483172 | NA | NA |
| ENSG000002 | 11.983322 | -0.631768037 | 0.32648083 | -1.9350846 | 0.05297991 | 0.12485549 | PAPPA-AS1 | 493913 |
| ENSG00000C | 150.214442 | 0.314879278 | 0.16272396 | 1.93505173 | 0.05298395 | 0.12485775 | ULK2 | 9706 |
| ENSG000002 | 2.65967422 | -0.721299562 | 0.37277089 | -1.9349675 | 0.05299428 | 0.12487484 | RN7SKP14 | 106479101 |
| ENSG000002 | 3.25868466 | -0.731248693 | 0.37794508 | -1.9348015 | 0.05301466 | 0.1249156 | GRM7-AS1 | 100873937 |
| ENSG000002 | 10.2733878 | -0.676066318 | 0.34946325 | -1.9345849 | 0.05304126 | 0.12497101 | NA | NA |
| ENSG000002 | 3.89956526 | -0.732304989 | 0.3785572 | -1.9344633 | 0.0530562 | 0.12499894 | NA | NA |
| ENSG000001 | 104.290108 | 0.318964022 | 0.16490463 | 1.93423323 | 0.05308446 | 0.1250302 | CLDN1 | 9076 |
| ENSG000001 | 8.37851329 | -0.682255435 | 0.35272719 | -1.9342298 | 0.05308488 | 0.1250302 | CCR3 | 1232 |
| ENSG000002 | 5.97576204 | -0.714544953 | 0.36941582 | -1.9342565 | 0.0530816 | 0.1250302 | NA | NA |
| ENSG000002 | 3.01741021 | -0.726226097 | 0.37545595 | -1.9342511 | 0.05308226 | 0.1250302 | NA | NA |
| ENSG000002 | 6.41947681 | 0.715494086 | 0.3699025 | 1.9342775 | 0.05307902 | 0.1250302 | NA | NA |
| ENSG000002 | 5.53321312 | -0.724350966 | 0.37449856 | -1.9341889 | 0.0530899 | 0.12503477 | NA | NA |
| ENSG00000C | 351.564576 | 0.274035299 | 0.14169587 | 1.93396816 | 0.05311704 | 0.1250822 | SYDE2 | 84144 |
| ENSG000002 | 3.24402497 | -0.73079914 | 0.37787474 | -1.9339719 | 0.05311658 | 0.1250822 | NA | NA |
| ENSG000002 | 3.69703384 | -0.730942642 | 0.37795668 | -1.9339323 | 0.05312145 | 0.1250822 | NA | NA |
| ENSG000002 | 3.61918517 | -0.731555926 | 0.37827528 | -1.9339247 | 0.05312238 | 0.1250822 | NA | NA |
| ENSG000002 | 2.6505915 | -0.719079114 | 0.37184349 | -1.933822 | 0.05313502 | 0.12510469 | NA | NA |
| ENSG000001 | 20.5659519 | -0.533541966 | 0.27593153 | -1.9336028 | 0.05316199 | 0.12516092 | SLC14A2 | 8170 |

| | | | | | | | | |
|---|---|---|---|---|---|---|---|---|
| ENSG000001 | 18.638715 | -0.592231521 | 0.30629777 | -1.9335156 | 0.05317271 | 0.12517891 | GRK7 | 131890 |
| ENSG000002 | 34.0260529 | 0.461330053 | 0.23864425 | 1.93312875 | 0.05322034 | 0.12528375 | NA | NA |
| ENSG000002 | 3.03162554 | -0.725050521 | 0.37511132 | -1.9328943 | 0.05324923 | 0.12534448 | NA | NA |
| ENSG000001 | 460.30925 | 0.225053541 | 0.11643722 | 1.93283168 | 0.05325694 | 0.12535535 | MPV17 | 4358 |
| ENSG000002 | 17.6360705 | 0.560173985 | 0.28983653 | 1.93272387 | 0.05327022 | 0.12537935 | NA | NA |
| ENSG000002 | 8.35564424 | -0.702002512 | 0.36323378 | -1.9326466 | 0.05327975 | 0.1253945 | NA | NA |
| ENSG000002 | 7.94293804 | -0.681156238 | 0.352593 | -1.9318485 | 0.05337821 | 0.12561894 | OR4D2 | 124538 |
| ENSG000001 | 17.0967601 | -0.576190496 | 0.29827179 | -1.9317633 | 0.05338873 | 0.12563639 | NRG2 | 9542 |
| ENSG000002 | 8.04339503 | -0.68992363 | 0.35716629 | -1.9316594 | 0.05340156 | 0.12565929 | NA | NA |
| ENSG000002 | 4.54729689 | -0.729137785 | 0.37747642 | -1.9316115 | 0.05340748 | 0.12566269 | NA | NA |
| ENSG000002 | 20.8386415 | -0.562791516 | 0.29136065 | -1.9315975 | 0.0534092 | 0.12566269 | NA | NA |
| ENSG000002 | 9.39545705 | -0.659012701 | 0.34118316 | -1.9315511 | 0.05341494 | 0.1256689 | LINC01207 | 100505989 |
| ENSG000002 | 8.27496813 | -0.675620922 | 0.34979529 | -1.9314752 | 0.05342432 | 0.12568368 | NA | NA |
| ENSG000002 | 5.98034972 | -0.712638349 | 0.36896872 | -1.9314329 | 0.05342953 | 0.12568866 | NA | NA |
| ENSG000001 | 61.5644302 | -0.372007894 | 0.19261886 | -1.931316 | 0.05344398 | 0.12571535 | KIAA0319 | 9856 |
| ENSG000002 | 6.93732814 | -0.702783045 | 0.36392185 | -1.9311372 | 0.05346608 | 0.12575275 | LOC101928909 | 101928909 |
| ENSG000002 | 4.01091873 | -0.73093801 | 0.37849845 | -1.9311519 | 0.05346427 | 0.12575275 | NA | NA |
| ENSG000002 | 12.3590509 | -0.623803835 | 0.32303807 | -1.9310536 | 0.05347642 | 0.12576976 | NA | NA |
| ENSG000002 | 3.29247043 | -0.728844014 | 0.37747376 | -1.9308468 | 0.053502 | 0.12582263 | NA | NA |
| ENSG000001 | 218.45955 | 0.521756178 | 0.27031945 | 1.93014665 | 0.05358867 | 0.12601915 | FAM102A | 399665 |
| ENSG000002 | 2.61356927 | -0.719566187 | 0.37282154 | -1.9300553 | 0.05359999 | 0.12603845 | NA | NA |
| ENSG000002 | 6.9971678 | -0.709129028 | 0.36743579 | -1.92994 | 0.05361427 | 0.12606473 | NA | NA |
| ENSG000002 | 4.54997349 | -0.727628764 | 0.37702779 | -1.9299075 | 0.0536183 | 0.12606689 | NA | NA |
| ENSG000002 | 3.04156171 | -0.723517911 | 0.37492913 | -1.9297458 | 0.05363834 | 0.12610671 | LOC105378420 | 105378420 |
| ENSG000001 | 25.2529347 | -0.528086937 | 0.27371871 | -1.9293052 | 0.05369299 | 0.12622785 | VDR | 7421 |
| ENSG000001 | 17.231995 | -0.602304937 | 0.3121986 | -1.9292365 | 0.0537015 | 0.12624056 | TRPC2 | 7221 |
| ENSG000001 | 644.562343 | 0.260920199 | 0.13526692 | 1.92892833 | 0.05373976 | 0.12632317 | MYO5C | 55930 |
| ENSG000001 | 11.9013771 | -0.627850574 | 0.3255209 | -1.9287565 | 0.05376109 | 0.12635134 | GKN2 | 200504 |
| ENSG000002 | 107.896461 | 0.343301214 | 0.17798724 | 1.92879672 | 0.0537561 | 0.12635134 | AP4M1 | 9179 |

| | | | | | | | | |
|---|---|---|---|---|---|---|---|---|
| ENSG000002 | 58.5740074 | 0.378453738 | 0.19621399 | 1.92878057 | 0.05375811 | 0.12635134 | HYPK | 25764 |
| ENSG000002 | 42.7118676 | 0.438283136 | 0.22725071 | 1.92863266 | 0.05377648 | 0.12637013 | ZNF579 | 163033 |
| ENSG000002 | 2.69209856 | -0.717944639 | 0.37226365 | -1.9285918 | 0.05378155 | 0.12637013 | NA | NA |
| ENSG000002 | 3.28037057 | -0.728567556 | 0.37776401 | -1.9286315 | 0.05377663 | 0.12637013 | NA | NA |
| ENSG000002 | 20.2337226 | 0.566229311 | 0.29359373 | 1.92861516 | 0.05377866 | 0.12637013 | LENG9 | 94059 |
| ENSG000002 | 2.27317594 | 0.692718798 | 0.35920165 | 1.9284956 | 0.05379351 | 0.12639091 | PLCH1-AS2 | 100874036 |
| ENSG000002 | 14.0181719 | -0.621864776 | 0.32247178 | -1.9284316 | 0.05380146 | 0.12640226 | NA | NA |
| ENSG000002 | 6.04459578 | 0.717563236 | 0.37211723 | 1.92832575 | 0.05381462 | 0.12642585 | INAFM1 | 255783 |
| ENSG000002 | 3.29273139 | 0.726684635 | 0.37686616 | 1.92823002 | 0.05382652 | 0.12644649 | NA | NA |
| ENSG000001 | 377.261764 | 0.34281252 | 0.17778989 | 1.92818904 | 0.05383162 | 0.12645113 | TSC22D2 | 9819 |
| ENSG000001 | 57.0962209 | 0.391110131 | 0.20284329 | 1.92813932 | 0.0538378 | 0.12645116 | GPR75 | 10936 |
| ENSG000001 | 13.2943819 | -0.614385982 | 0.31864199 | -1.9281388 | 0.05383786 | 0.12645116 | GTF2IRD2 | 84163 |
| ENSG000002 | 6.28705668 | -0.708880682 | 0.3676662 | -1.9280551 | 0.05384827 | 0.12646828 | NA | NA |
| ENSG000002 | 4.10123773 | -0.729646489 | 0.37850313 | -1.9277159 | 0.05389048 | 0.12656007 | NA | NA |
| ENSG000002 | 2.30720274 | -0.709071823 | 0.36785066 | -1.9276078 | 0.05390392 | 0.12658432 | RNU6ATAC22P | 106479553 |
| ENSG000002 | 2.66418226 | -0.717414762 | 0.37221342 | -1.9274285 | 0.05392625 | 0.12662942 | NA | NA |
| ENSG000002 | 15.5554435 | -0.626343501 | 0.32496792 | -1.927401 | 0.05392967 | 0.12663012 | LINC00689 | 154822 |
| ENSG000002 | 2.7013485 | -0.715448833 | 0.37122059 | -1.9272876 | 0.0539438 | 0.12665595 | NA | NA |
| ENSG000002 | 11.2734875 | -0.631408606 | 0.32765135 | -1.9270746 | 0.05397033 | 0.12671091 | NA | NA |
| ENSG000002 | 2.98160541 | -0.722407937 | 0.3748919 | -1.9269766 | 0.05398254 | 0.12673224 | NA | NA |
| ENSG000002 | 6.67621757 | -0.711267192 | 0.36911619 | -1.9269466 | 0.05398628 | 0.12673369 | NA | NA |
| ENSG000002 | 2.26064078 | -0.708369831 | 0.36764897 | -1.9267559 | 0.05401005 | 0.12678215 | NA | NA |
| ENSG000002 | 60.2946984 | 0.381925929 | 0.19823454 | 1.92663667 | 0.05402493 | 0.12680972 | NA | NA |
| ENSG000001 | 12.1716029 | -0.63141781 | 0.32776121 | -1.9264568 | 0.05404736 | 0.12685123 | ANXA10 | 11199 |
| ENSG000001 | 7.94810049 | 0.695905545 | 0.36123826 | 1.92644475 | 0.05404886 | 0.12685123 | HCST | 10870 |
| ENSG000002 | 18.926725 | 0.558094754 | 0.28974431 | 1.92616293 | 0.05408403 | 0.12692642 | NA | NA |
| ENSG000001 | 59.2498911 | 0.401750033 | 0.20858406 | 1.92608218 | 0.05409411 | 0.12694273 | AP5S1 | 55317 |
| ENSG000001 | 4.32901969 | -0.728898812 | 0.37846325 | -1.9259434 | 0.05411144 | 0.12697604 | PRH2 | 5555 |
| ENSG000002 | 5.66066208 | -0.711209618 | 0.36928896 | -1.9258892 | 0.05411821 | 0.12698458 | NA | NA |

| | | | | | | | | |
|---|---|---|---|---|---|---|---|---|
| ENSG000002 | 27.9664863 | 0.480090698 | 0.24929058 | 1.92582771 | 0.05412589 | 0.1269881 | LOC101929726 | 101929726 |
| ENSG000002 | 7.15044139 | -0.692703974 | 0.35969168 | -1.9258271 | 0.05412597 | 0.1269881 | NA | NA |
| ENSG000001 | 12.0998998 | -0.620845702 | 0.32240693 | -1.9256587 | 0.05414701 | 0.12700086 | KIR2DL1 | 3802 |
| ENSG000002 | 3.88706568 | -0.728963979 | 0.37854404 | -1.9257045 | 0.05414128 | 0.12700086 | HIGD1C | 613227 |
| ENSG000002 | 10.4919561 | -0.643688792 | 0.33426778 | -1.9256681 | 0.05414583 | 0.12700086 | LINC00682 | 101927074 |
| ENSG000002 | 7.31572066 | -0.700219711 | 0.36361218 | -1.9257323 | 0.05413781 | 0.12700086 | NA | NA |
| ENSG000002 | 4.22740861 | -0.728604506 | 0.37836648 | -1.9256582 | 0.05414707 | 0.12700086 | NA | NA |
| ENSG000002 | 7.7760863 | -0.689808902 | 0.35822475 | -1.9256316 | 0.05415039 | 0.12700129 | NA | NA |
| ENSG000001 | 1006.11401 | 0.225693489 | 0.11720938 | 1.92555824 | 0.05415956 | 0.12701044 | SLF2 | 55719 |
| ENSG000001 | 8.7368579 | -0.681413061 | 0.35388315 | -1.9255313 | 0.05416293 | 0.12701044 | PDCL2 | 132954 |
| ENSG000002 | 12.7134851 | -0.618858941 | 0.32139747 | -1.9255252 | 0.05416368 | 0.12701044 | NA | NA |
| ENSG000002 | 4.54561504 | -0.725803643 | 0.37696675 | -1.9253784 | 0.05418203 | 0.12704612 | NA | NA |
| ENSG000001 | 129.861982 | 0.297161438 | 0.15434398 | 1.92531923 | 0.05418943 | 0.12704878 | EIF4A1 | 1973 |
| ENSG000002 | 3.25752235 | -0.726704496 | 0.37744577 | -1.9253216 | 0.05418914 | 0.12704878 | NA | NA |
| ENSG000001 | 26.0110167 | -0.514271459 | 0.26713068 | -1.9251681 | 0.05420834 | 0.12705689 | SLC28A3 | 64078 |
| ENSG000002 | 2.27729552 | -0.706171475 | 0.36680076 | -1.9252181 | 0.05420208 | 0.12705689 | NA | NA |
| ENSG000002 | 3.94488271 | -0.728767105 | 0.37853518 | -1.9252295 | 0.05420065 | 0.12705689 | NA | NA |
| ENSG000002 | 13.7765128 | -0.627114643 | 0.32574009 | -1.9251995 | 0.05420441 | 0.12705689 | LOC101927476 | 101927476 |
| ENSG000002 | 2.65663398 | -0.716653636 | 0.37225545 | -1.9251663 | 0.05420856 | 0.12705689 | NA | NA |
| ENSG000001 | 3.24332001 | -0.727046332 | 0.37766367 | -1.9251159 | 0.05421486 | 0.12706432 | NA | NA |
| ENSG000002 | 7.07883863 | -0.693635928 | 0.36032754 | -1.925015 | 0.05422748 | 0.1270792 | SCRT2 | 85508 |
| ENSG000002 | 3.88453397 | -0.728705743 | 0.3785439 | -1.9250231 | 0.05422647 | 0.1270792 | NA | NA |
| ENSG000001 | 25.4166851 | -0.493577297 | 0.25643534 | -1.9247632 | 0.054259 | 0.12714261 | ARHGDIB | 397 |
| ENSG000001 | 77.4030681 | 0.355693962 | 0.18480021 | 1.92474869 | 0.05426081 | 0.12714261 | CXXC4 | 80319 |
| ENSG000001 | 326.569716 | 0.233350709 | 0.12124113 | 1.92468277 | 0.05426906 | 0.12714674 | STK3 | 6788 |
| ENSG000001 | 42.5667458 | -0.414311968 | 0.21526508 | -1.9246594 | 0.05427198 | 0.12714674 | ZNF438 | 220929 |
| ENSG000002 | 13.026682 | -0.618468178 | 0.3213388 | -1.9246608 | 0.05427181 | 0.12714674 | LOC100507389 | 100507389 |
| ENSG000002 | 59.6636909 | 0.394785798 | 0.20512894 | 1.92457391 | 0.05428269 | 0.12716448 | NA | NA |
| ENSG000001 | 61.172915 | 0.419729027 | 0.21809315 | 1.92454015 | 0.05428691 | 0.12716704 | ARID5A | 10865 |

| | | | | | | | | |
|---|---|---|---|---|---|---|---|---|
| ENSG000002 | 9.09196523 | -0.6830383 | 0.35492249 | -1.9244717 | 0.05429548 | 0.12717976 | NA | NA |
| ENSG000002 | 3.93047027 | -0.728429286 | 0.37852887 | -1.9243691 | 0.05430833 | 0.12720251 | NA | NA |
| ENSG000002 | 16.4007342 | -0.57784515 | 0.30029183 | -1.9242786 | 0.05431967 | 0.12722097 | NA | NA |
| ENSG000002 | 3.87932097 | -0.728417756 | 0.37854511 | -1.9242562 | 0.05432248 | 0.12722097 | NA | NA |
| ENSG000001 | 68.8746371 | -0.366878863 | 0.19066682 | -1.9241883 | 0.05433099 | 0.12723354 | NALCN | 259232 |
| ENSG000001 | 8.13372144 | -0.688935535 | 0.3580702 | -1.9240237 | 0.05435162 | 0.12727449 | SFTPB | 6439 |
| ENSG000002 | 21.1647671 | -0.567769848 | 0.29511927 | -1.9238657 | 0.05437141 | 0.1273135 | NA | NA |
| ENSG000001 | 210.491988 | -0.331342067 | 0.17223505 | -1.9237784 | 0.05438237 | 0.12732896 | TRANK1 | 9881 |
| ENSG000002 | 13.5902532 | 0.614118515 | 0.31922774 | 1.92376299 | 0.0543843 | 0.12732896 | NA | NA |
| ENSG000002 | 2.91762323 | -0.724291934 | 0.3765059 | -1.92372 | 0.05438969 | 0.12733423 | NA | NA |
| ENSG000001 | 114.734326 | 0.385562706 | 0.20042829 | 1.92369405 | 0.05439294 | 0.12733451 | P3H3 | 10536 |
| ENSG000002 | 16.0866888 | -0.595701237 | 0.30967598 | -1.9236275 | 0.05440129 | 0.12734669 | NA | NA |
| ENSG000001 | 58.415562 | 0.393998059 | 0.20483889 | 1.92345339 | 0.05442314 | 0.12739047 | C1R | 715 |
| ENSG000002 | 2.95216316 | -0.720887801 | 0.37482271 | -1.9232767 | 0.0544453 | 0.12743501 | NA | NA |
| ENSG000002 | 5.19694068 | -0.721161653 | 0.37501544 | -1.9230185 | 0.05447772 | 0.12750352 | NA | NA |
| ENSG000002 | 13.4963052 | -0.627137975 | 0.32614068 | -1.9229063 | 0.05449182 | 0.1275218 | NA | NA |
| ENSG000002 | 13.6032371 | -0.611786273 | 0.31815346 | -1.9229283 | 0.05448906 | 0.1275218 | NA | NA |
| ENSG000001 | 1960.35791 | 0.223902109 | 0.11644383 | 1.92283367 | 0.05450094 | 0.12753579 | DTL | 51514 |
| ENSG000002 | 4.56032516 | -0.726119823 | 0.37766209 | -1.9226707 | 0.05452142 | 0.12757635 | OR7E91P | 79315 |
| ENSG000001 | 20.3347463 | 0.549347566 | 0.285758 | 1.92242236 | 0.05455263 | 0.12764202 | FAM83E | 54854 |
| ENSG000002 | 15.3675553 | -0.592357818 | 0.30814402 | -1.9223408 | 0.05456289 | 0.12765865 | NA | NA |
| ENSG000001 | 13.6579942 | -0.604340695 | 0.31440374 | -1.9221804 | 0.05458306 | 0.12769848 | EGR3 | 1960 |
| ENSG000001 | 8.65223025 | -0.676167143 | 0.35178245 | -1.9221173 | 0.05459099 | 0.12770967 | AADAC | 13 |
| ENSG000002 | 24.6519803 | -0.527726514 | 0.27456733 | -1.9220295 | 0.05460204 | 0.12772814 | NA | NA |
| ENSG000002 | 3.37864325 | -0.724046801 | 0.37671708 | -1.9219909 | 0.0546069 | 0.12773215 | LINC00557 | 100861544 |
| ENSG000001 | 24.6742102 | -0.507449227 | 0.26404185 | -1.9218515 | 0.05462445 | 0.12776582 | FAM129C | 199786 |
| ENSG000002 | 5.53072184 | -0.720046032 | 0.37467023 | -1.9218128 | 0.05462932 | 0.12776985 | NA | NA |
| ENSG000002 | 18.7996369 | -0.552021371 | 0.28724823 | -1.9217572 | 0.05463631 | 0.12777882 | ROR1-AS1 | 101927034 |
| ENSG00000C | 101.668359 | 0.326588492 | 0.16995423 | 1.92162609 | 0.05465282 | 0.12781007 | SARM1 | 23098 |

| | | | | | | | | |
|---|---|---|---|---|---|---|---|---|
| ENSG000001 | 412.983438 | 0.303394231 | 0.15788897 | 1.92156701 | 0.05466026 | 0.12781273 | SDC1 | 6382 |
| ENSG000002 | 2.36809316 | -0.701352506 | 0.3649862 | -1.9215864 | 0.05465782 | 0.12781273 | NA | NA |
| ENSG000001 | 144.921115 | -0.319128432 | 0.16608435 | -1.9214841 | 0.0546707 | 0.12782331 | ID2 | 3398 |
| ENSG000002 | 10.6570567 | -0.634182467 | 0.33004878 | -1.921481 | 0.05467109 | 0.12782331 | LOC101927571 | 101927571 |
| ENSG000001 | 365.682394 | 0.239928145 | 0.12488246 | 1.92123168 | 0.05470251 | 0.12788938 | FIGN | 55137 |
| ENSG00000C | 812.652511 | 0.60181154 | 0.31326639 | 1.92108557 | 0.05472092 | 0.12792506 | PSMC5 | 5705 |
| ENSG000002 | 97.3311312 | 0.32713633 | 0.17029103 | 1.92104259 | 0.05472634 | 0.12793035 | NA | NA |
| ENSG000002 | 2.67587195 | -0.713480277 | 0.37140938 | -1.9210077 | 0.05473073 | 0.12793325 | NA | NA |
| ENSG00000C | 35.3656561 | -0.459697025 | 0.23930566 | -1.9209617 | 0.05473653 | 0.12793942 | KCNK2 | 3776 |
| ENSG000002 | 9.73225088 | -0.660986995 | 0.34410729 | -1.9208747 | 0.0547475 | 0.12795769 | NA | NA |
| ENSG000002 | 3.29722228 | -0.725502399 | 0.37772775 | -1.9207019 | 0.05476931 | 0.12800127 | NA | NA |
| ENSG000002 | 55.5167551 | 0.396251761 | 0.20631721 | 1.92059482 | 0.05478281 | 0.12802353 | NA | NA |
| ENSG000002 | 2.59828949 | -0.715198343 | 0.37238736 | -1.9205763 | 0.05478514 | 0.12802353 | NA | NA |
| ENSG000002 | 16.783125 | 0.568856925 | 0.29619512 | 1.92054795 | 0.05478872 | 0.12802452 | NA | NA |
| ENSG000001 | 14.1171863 | -0.602110348 | 0.31352022 | -1.9204833 | 0.05479688 | 0.1280362 | NA | NA |
| ENSG000002 | 7.18992052 | -0.694651031 | 0.36175954 | -1.920201 | 0.05483251 | 0.12811208 | NA | NA |
| ENSG000002 | 3.90594022 | -0.726790633 | 0.37851143 | -1.9201286 | 0.05484166 | 0.12812606 | NA | NA |
| ENSG000002 | 7.87728624 | -0.705791209 | 0.36758985 | -1.9200509 | 0.05485147 | 0.12812863 | NA | NA |
| ENSG000002 | 12.0842156 | -0.619211542 | 0.32249728 | -1.920052 | 0.05485134 | 0.12812863 | LOC102724555 | 102724555 |
| ENSG000002 | 3.05111997 | -0.719251206 | 0.37460125 | -1.9200448 | 0.05485223 | 0.12812863 | NA | NA |
| ENSG000001 | 315.037234 | 0.288833905 | 0.15043477 | 1.9199943 | 0.05485862 | 0.1281349 | XYLT1 | 64131 |
| ENSG000001 | 120.16378 | 0.322632891 | 0.16804028 | 1.91997357 | 0.05486124 | 0.1281349 | DNAJC27 | 51277 |
| ENSG000001 | 4625.37558 | 0.212659556 | 0.11076891 | 1.91984885 | 0.05487699 | 0.12815717 | GOLGA4 | 2803 |
| ENSG000002 | 3.86307117 | -0.726624805 | 0.37848037 | -1.9198481 | 0.05487709 | 0.12815717 | NA | NA |
| ENSG000001 | 26.5498189 | 0.517421062 | 0.26953382 | 1.91968879 | 0.05489722 | 0.1281894 | C1QL1 | 10882 |
| ENSG000002 | 4.73318309 | -0.720466203 | 0.37530266 | -1.9196938 | 0.05489658 | 0.1281894 | NA | NA |
| ENSG000002 | 4.92361327 | -0.723933382 | 0.37711825 | -1.9196456 | 0.05490268 | 0.12819478 | NA | NA |
| ENSG000002 | 7.73624673 | -0.684346047 | 0.35650772 | -1.9195827 | 0.05491063 | 0.12820595 | NA | NA |
| ENSG000002 | 5.3508394 | -0.714595067 | 0.37227529 | -1.919534 | 0.05491679 | 0.12821295 | NA | NA |

| | | | | | | | | |
|---|---|---|---|---|---|---|---|---|
| ENSG000001 | 6.20551096 | -0.701345846 | 0.36538212 | -1.919486 | 0.05492286 | 0.12821975 | NA | NA |
| ENSG000002 | 4.03226396 | -0.72626234 | 0.37840478 | -1.9192737 | 0.05494971 | 0.12827503 | NA | NA |
| ENSG000002 | 9.71084584 | -0.663713981 | 0.34582136 | -1.9192394 | 0.05495404 | 0.12827777 | NA | NA |
| ENSG000001 | 222.87205 | 0.271368029 | 0.14140291 | 1.91911205 | 0.05497016 | 0.12830799 | MLLT3 | 4300 |
| ENSG000001 | 6.0480568 | -0.709759856 | 0.36985319 | -1.9190313 | 0.05498037 | 0.12832445 | NA | NA |
| ENSG000000 | 51.9233347 | -0.397168624 | 0.20696978 | -1.9189691 | 0.05498824 | 0.12832805 | SNAP91 | 9892 |
| ENSG000002 | 9.63119573 | 0.663395847 | 0.34570243 | 1.91897944 | 0.05498694 | 0.12832805 | NA | NA |
| ENSG000002 | 9.28991403 | 0.676202309 | 0.35238933 | 1.91890688 | 0.05499612 | 0.12833905 | PRR22 | 163154 |
| ENSG000002 | 13.2764647 | -0.613573019 | 0.31975734 | -1.9188708 | 0.05500069 | 0.12834233 | NA | NA |
| ENSG000001 | 118.908443 | 0.328520412 | 0.17121362 | 1.91877496 | 0.05501282 | 0.12835781 | BACE1 | 23621 |
| ENSG000002 | 12.1310082 | -0.636781092 | 0.33186971 | -1.9187684 | 0.05501366 | 0.12835781 | NA | NA |
| ENSG000002 | 12.3491518 | 0.630547727 | 0.32863899 | 1.91866377 | 0.0550269 | 0.12838133 | NA | NA |
| ENSG000002 | 7.83686266 | -0.690339288 | 0.35982605 | -1.9185362 | 0.05504306 | 0.12841164 | NA | NA |
| ENSG000002 | 12.9624712 | -0.611086832 | 0.31853359 | -1.9184377 | 0.05505554 | 0.12843337 | NA | NA |
| ENSG000002 | 7.10718472 | -0.692606779 | 0.36108982 | -1.9181011 | 0.05509819 | 0.12852547 | NA | NA |
| ENSG000002 | 9.38346599 | -0.654636335 | 0.34130459 | -1.9180414 | 0.05510576 | 0.12853573 | NA | NA |
| ENSG000001 | 166.402803 | 0.277356889 | 0.14462928 | 1.91770913 | 0.0551479 | 0.12862663 | MTRF1 | 9617 |
| ENSG000002 | 5.63021407 | -0.708596079 | 0.36951873 | -1.9176188 | 0.05515936 | 0.12864595 | NA | NA |
| ENSG000001 | 15.5607613 | -0.576874243 | 0.30083502 | -1.9175768 | 0.0551647 | 0.12864781 | TMEM139 | 135932 |
| ENSG000002 | 9.3056541 | -0.654236556 | 0.34118133 | -1.9175626 | 0.0551665 | 0.12864781 | NA | NA |
| ENSG000002 | 6.26527671 | -0.702971068 | 0.36660607 | -1.9175107 | 0.05517308 | 0.12865575 | NA | NA |
| ENSG000001 | 143.432017 | 0.298988628 | 0.15593902 | 1.91734331 | 0.05519433 | 0.12869791 | GALNT14 | 79623 |
| ENSG000002 | 3.21591279 | -0.724029348 | 0.37767464 | -1.9170717 | 0.05522883 | 0.12877094 | NA | NA |
| ENSG000002 | 12.0055425 | -0.616440297 | 0.32156317 | -1.9170115 | 0.05523647 | 0.12878135 | LINC00838 | 100505583 |
| ENSG000002 | 15.7601652 | -0.569781521 | 0.29723231 | -1.9169569 | 0.05524341 | 0.12879014 | ADPGK-AS1 | 100287559 |
| ENSG000002 | 6.03585053 | -0.712449156 | 0.37169321 | -1.9167667 | 0.05526758 | 0.12883907 | NA | NA |
| ENSG000002 | 43.6430252 | -0.414842793 | 0.21643407 | -1.9167167 | 0.05527393 | 0.12884647 | NA | NA |
| ENSG000002 | 2.68916137 | -0.710606271 | 0.37077331 | -1.9165519 | 0.05529489 | 0.12888792 | NA | NA |
| ENSG000002 | 4.79715478 | -0.725252396 | 0.37842383 | -1.9165083 | 0.05530043 | 0.12889343 | LINC01395 | 101929557 |

| | | | | | | | | |
|---|---|---|---|---|---|---|---|---|
| ENSG00000C | 280.450736 | 0.256453074 | 0.13382748 | 1.91629602 | 0.05532743 | 0.12894893 | RNF19A | 25897 |
| ENSG000001 | 5.75920277 | -0.71348858 | 0.37236468 | -1.9161017 | 0.05535216 | 0.12899915 | BDKRB1 | 623 |
| ENSG000002 | 4.15369192 | -0.723784031 | 0.37775864 | -1.915996 | 0.05536561 | 0.12902309 | NA | NA |
| ENSG000002 | 3.64514685 | -0.724852973 | 0.37833191 | -1.9159182 | 0.05537551 | 0.12903876 | NA | NA |
| ENSG000002 | 64.1670635 | 0.383753261 | 0.20030773 | 1.91581853 | 0.0553882 | 0.1290609 | ARHGEF25 | 115557 |
| ENSG000001 | 15.1708606 | -0.602693874 | 0.31460752 | -1.9157008 | 0.0554032 | 0.12908843 | TGM7 | 116179 |
| ENSG000002 | 43.4194384 | 0.421690252 | 0.22013874 | 1.91556587 | 0.05542038 | 0.12912104 | NA | NA |
| ENSG000001 | 124.61305 | 0.31510159 | 0.1645104 | 1.91539014 | 0.05544277 | 0.12916579 | TRIM45 | 80263 |
| ENSG000002 | 15.556769 | -0.597918533 | 0.31219189 | -1.9152276 | 0.05546348 | 0.12919284 | OR51A4 | 401666 |
| ENSG000002 | 437.412512 | 0.236800695 | 0.12364125 | 1.91522406 | 0.05546394 | 0.12919284 | HAUS3 | 79441 |
| ENSG000002 | 7.39985328 | -0.702252654 | 0.36666069 | -1.9152657 | 0.05545862 | 0.12919284 | NA | NA |
| ENSG000001 | 124.184274 | 0.306369134 | 0.15998201 | 1.91502237 | 0.05548965 | 0.12924531 | ZMAT4 | 79698 |
| ENSG00000C | 40.7855842 | -0.449047896 | 0.23449554 | -1.9149528 | 0.05549852 | 0.12924928 | EPHA3 | 2042 |
| ENSG000001 | 230.472886 | 0.265979437 | 0.13889925 | 1.91490903 | 0.05550411 | 0.12924928 | GFOD2 | 81577 |
| ENSG000002 | 10.4784683 | -0.657891698 | 0.34355147 | -1.9149727 | 0.05549598 | 0.12924928 | NA | NA |
| ENSG000002 | 10.3701905 | -0.656340313 | 0.34274986 | -1.9149251 | 0.05550206 | 0.12924928 | NA | NA |
| ENSG000002 | 6.87681865 | -0.697867922 | 0.36445755 | -1.9148126 | 0.0555164 | 0.1292705 | NA | NA |
| ENSG000002 | 4.48360444 | -0.721847944 | 0.37702824 | -1.9145726 | 0.05554703 | 0.12933438 | NA | NA |
| ENSG000001 | 17.1731278 | -0.579197366 | 0.30253737 | -1.9144655 | 0.0555607 | 0.12935879 | SRL | 6345 |
| ENSG000001 | 1324.69522 | 0.273093535 | 0.14266297 | 1.91425665 | 0.05558737 | 0.12941345 | VASH2 | 79805 |
| ENSG000002 | 4.1935155 | -0.72310212 | 0.37775185 | -1.9142252 | 0.05559139 | 0.12941537 | SFTA3 | 253970 |
| ENSG000001 | 6.00457997 | -0.708840486 | 0.37033332 | -1.9140608 | 0.05561239 | 0.12945683 | NA | NA |
| ENSG000001 | 114.462579 | 0.3891545 | 0.20332839 | 1.91392113 | 0.05563023 | 0.12947805 | NUMBL | 9253 |
| ENSG000001 | 13.3036262 | -0.60926683 | 0.31833152 | -1.9139381 | 0.05562806 | 0.12947805 | SIGLEC6 | 946 |
| ENSG000001 | 36.2786005 | 0.449735719 | 0.23498214 | 1.91391447 | 0.05563109 | 0.12947805 | DNAJC12 | 56521 |
| ENSG000002 | 5.11815876 | -0.71656828 | 0.37440551 | -1.9138829 | 0.05563512 | 0.12948002 | NA | NA |
| ENSG000002 | 4.51913263 | -0.722517368 | 0.37752099 | -1.9138469 | 0.05563972 | 0.1294833 | NA | NA |
| ENSG000002 | 2.92229489 | -0.720064988 | 0.37627254 | -1.9136794 | 0.05566114 | 0.1295257 | NA | NA |
| ENSG000002 | 2.3134326 | -0.70158376 | 0.36670028 | -1.9132348 | 0.055718 | 0.12965057 | NA | NA |

| | | | | | | | | |
|---|---|---|---|---|---|---|---|---|
| ENSG000002 | 289.118137 | 0.240476095 | 0.12569378 | 1.91319009 | 0.05572372 | 0.12965509 | VGLL3 | 389136 |
| ENSG000002 | 2.92015494 | -0.719862579 | 0.3762699 | -1.9131548 | 0.05572824 | 0.12965509 | SPATA17-AS1 | 103752555 |
| ENSG000002 | 2.35428223 | -0.69763488 | 0.36465348 | -1.9131447 | 0.05572953 | 0.12965509 | NA | NA |
| ENSG000002 | 8.21660348 | -0.673142317 | 0.3518581 | -1.9131074 | 0.05573431 | 0.12965877 | NA | NA |
| ENSG000001 | 198.41706 | 0.278382882 | 0.14552976 | 1.9128931 | 0.05576174 | 0.12968865 | MYBL1 | 4603 |
| ENSG000002 | 13.9187366 | -0.613714408 | 0.320832 | -1.912884 | 0.0557629 | 0.12968865 | NA | NA |
| ENSG000002 | 6.03722034 | -0.707155289 | 0.36966264 | -1.9129747 | 0.05575129 | 0.12968865 | NA | NA |
| ENSG000002 | 4.14990179 | -0.723128016 | 0.37801949 | -1.9129384 | 0.05575593 | 0.12968865 | NA | NA |
| ENSG000002 | 12.8159532 | 0.66730843 | 0.34884974 | 1.91288213 | 0.05576314 | 0.12968865 | NA | NA |
| ENSG000002 | 7.16870926 | -0.695747526 | 0.36375077 | -1.9127039 | 0.05578596 | 0.12972684 | NA | NA |
| ENSG000002 | 13.1653525 | 0.613829256 | 0.32091956 | 1.91272 | 0.05578391 | 0.12972684 | STX17-AS1 | 441461 |
| ENSG000002 | 7.97675248 | -0.675890545 | 0.35339712 | -1.9125525 | 0.05580537 | 0.12976451 | NA | NA |
| ENSG000002 | 2.30797221 | -0.701469206 | 0.36680342 | -1.9123846 | 0.05582688 | 0.12980711 | NA | NA |
| ENSG000001 | 131.422197 | 0.310402912 | 0.16231909 | 1.91230076 | 0.05583763 | 0.12982465 | IGFBPL1 | 347252 |
| ENSG000002 | 4.20016957 | -0.722658197 | 0.37790825 | -1.9122583 | 0.05584307 | 0.12982984 | DEFB116 | 245930 |
| ENSG000001 | 38.5876648 | 0.431858788 | 0.22584136 | 1.91222185 | 0.05584774 | 0.12983041 | PPP1R3C | 5507 |
| ENSG000002 | 33.5725328 | 0.464925413 | 0.24313557 | 1.91220652 | 0.05584971 | 0.12983041 | MTRNR2L8 | 100463486 |
| ENSG000001 | 15.5097401 | -0.589722658 | 0.3084228 | -1.9120592 | 0.0558686 | 0.12986687 | TTC16 | 158248 |
| ENSG000002 | 15.3388329 | -0.584265176 | 0.30557946 | -1.911991 | 0.05587735 | 0.12987977 | LOC100507373 | 100507373 |
| ENSG000001 | 59.4016415 | -0.376014906 | 0.19666583 | -1.9119483 | 0.05588282 | 0.12988504 | KLHL14 | 57565 |
| ENSG000002 | 3.85931354 | -0.723671193 | 0.37850762 | -1.9119065 | 0.05588819 | 0.12989006 | NA | NA |
| ENSG000001 | 19.7825159 | -0.572583493 | 0.29948803 | -1.9118744 | 0.05589231 | 0.12989221 | MMP10 | 4319 |
| ENSG000002 | 2.70929064 | -0.705980578 | 0.36928355 | -1.9117575 | 0.05590731 | 0.12991873 | NA | NA |
| ENSG000002 | 3.28398384 | -0.721168137 | 0.37723218 | -1.9117355 | 0.05591013 | 0.12991873 | NA | NA |
| ENSG000002 | 5.16260397 | -0.718025802 | 0.37559414 | -1.9117066 | 0.05591384 | 0.1299199 | NA | NA |
| ENSG000001 | 10.4597786 | -0.641803221 | 0.33577099 | -1.9114314 | 0.05594916 | 0.12999452 | EMX2 | 2018 |
| ENSG000002 | 74.4122382 | 0.35573226 | 0.18611586 | 1.9113484 | 0.05595982 | 0.13001185 | NA | NA |
| ENSG000002 | 5.61413202 | -0.707071156 | 0.36994492 | -1.9112877 | 0.05596762 | 0.13002252 | NA | NA |
| ENSG000002 | 5.62017396 | -0.707080129 | 0.36995697 | -1.9112496 | 0.05597251 | 0.13002642 | NA | NA |

| | | | | | | | | |
|---|---|---|---|---|---|---|---|---|
| ENSG000002 | 5.7873606 | -0.711810794 | 0.37244851 | -1.9111656 | 0.0559833 | 0.13004404 | NA | NA |
| ENSG000002 | 2.91788382 | -0.718622714 | 0.37601928 | -1.9111326 | 0.05598754 | 0.13004644 | NA | NA |
| ENSG000002 | 8.27161434 | -0.671926883 | 0.3516288 | -1.9108983 | 0.05601765 | 0.13010893 | NA | NA |
| ENSG000002 | 5.16401679 | -0.718671251 | 0.37610498 | -1.9108262 | 0.05602692 | 0.130123 | NA | NA |
| ENSG000001 | 43.2084722 | -0.430109585 | 0.2250971 | -1.9107735 | 0.05603369 | 0.13013128 | CCDC3 | 83643 |
| ENSG000001 | 150.09353 | 0.292553972 | 0.15312181 | 1.91059641 | 0.05605647 | 0.13017671 | TXNDC11 | 51061 |
| ENSG000002 | 2.38214592 | -0.6944191 | 0.36347422 | -1.9105044 | 0.0560683 | 0.13019673 | NA | NA |
| ENSG000002 | 3.24662139 | -0.721024515 | 0.37741785 | -1.9104145 | 0.05607987 | 0.13021615 | NA | NA |
| ENSG000002 | 10.1707817 | -0.644840081 | 0.33755017 | -1.9103533 | 0.05608774 | 0.13022696 | NA | NA |
| ENSG000002 | 4.68666052 | -0.722954226 | 0.37846093 | -1.910248 | 0.05610129 | 0.13025097 | NA | NA |
| ENSG000002 | 7.23391778 | -0.694269185 | 0.36348048 | -1.9100591 | 0.05612561 | 0.13029997 | FLJ22763 | 401081 |
| ENSG000001 | 72.821388 | 0.378779864 | 0.19833713 | 1.9097779 | 0.05616182 | 0.13037656 | NA | NA |
| ENSG000002 | 6.92252929 | -0.699418947 | 0.36624524 | -1.9097011 | 0.05617171 | 0.13039206 | LOC102724651 | 102724651 |
| ENSG000001 | 79.9270849 | 0.365464709 | 0.19137794 | 1.90964909 | 0.05617841 | 0.13039369 | SYDE1 | 85360 |
| ENSG000001 | 8.23522573 | -0.688739702 | 0.3606636 | -1.9096457 | 0.05617884 | 0.13039369 | REG1A | 5967 |
| ENSG000001 | 134.195645 | 0.29536551 | 0.154679 | 1.90953848 | 0.05619266 | 0.13041831 | ADARB1 | 104 |
| ENSG000002 | 6.49033123 | -0.70736037 | 0.37044402 | -1.9094933 | 0.05619849 | 0.13042436 | NA | NA |
| ENSG000001 | 11.3907996 | -0.637640007 | 0.33394622 | -1.9094093 | 0.05620932 | 0.13043456 | GNAT2 | 2780 |
| ENSG000002 | 2.59348144 | -0.70974439 | 0.37170853 | -1.9094111 | 0.05620908 | 0.13043456 | NA | NA |
| ENSG000001 | 62.5550925 | 0.383610696 | 0.20091602 | 1.90930867 | 0.05622228 | 0.13044973 | SDHAP1 | 255812 |
| ENSG000001 | 10.5759852 | -0.641194675 | 0.33582332 | -1.9093215 | 0.05622063 | 0.13044973 | ZNF257 | 113835 |
| ENSG000001 | 60.642754 | 0.40739775 | 0.21338153 | 1.90924564 | 0.05623041 | 0.13046112 | LPPR2 | 64748 |
| ENSG000002 | 2.26980783 | -0.70205149 | 0.36772351 | -1.909183 | 0.05623848 | 0.13047238 | NA | NA |
| ENSG000002 | 2.31645357 | -0.699659997 | 0.36650791 | -1.9089902 | 0.05626336 | 0.13051803 | NA | NA |
| ENSG000002 | 5.3857498 | -0.711500386 | 0.37271222 | -1.9089806 | 0.0562646 | 0.13051803 | NA | NA |
| ENSG000002 | 27.5518757 | -0.489114983 | 0.25623354 | -1.908864 | 0.05627964 | 0.13054545 | NA | NA |
| ENSG000001 | 10.2314471 | -0.64649626 | 0.33869524 | -1.9087846 | 0.05628988 | 0.13055236 | CEACAM3 | 1084 |
| ENSG000002 | 8.0836643 | -0.699491098 | 0.36646245 | -1.9087661 | 0.05629227 | 0.13055236 | NA | NA |
| ENSG000002 | 5.8501723 | -0.716793705 | 0.3755215 | -1.9087954 | 0.0562885 | 0.13055236 | NA | NA |

| | | | | | | | | |
|---|---|---|---|---|---|---|---|---|
| ENSG000002 | 86.2369601 | 0.394429025 | 0.20664402 | 1.90873671 | 0.05629607 | 0.13055369 | NA | NA |
| ENSG000002 | 3.54748663 | -0.722528501 | 0.37857997 | -1.9085228 | 0.05632368 | 0.13061026 | NA | NA |
| ENSG000001 | 162.417698 | 0.338553444 | 0.17739395 | 1.90848358 | 0.05632874 | 0.13061304 | FBLN2 | 2199 |
| ENSG000002 | 12.7114054 | -0.614992406 | 0.32224477 | -1.9084636 | 0.05633132 | 0.13061304 | NA | NA |
| ENSG00000C | 62.7057256 | 0.413887005 | 0.21688787 | 1.90829944 | 0.05635253 | 0.13064324 | KCNH2 | 3757 |
| ENSG000001 | 353.944799 | 0.230988849 | 0.12104522 | 1.90828559 | 0.05635431 | 0.13064324 | DIXDC1 | 85458 |
| ENSG000002 | 3.62841225 | -0.721824276 | 0.37826247 | -1.908263 | 0.05635723 | 0.13064324 | NA | NA |
| ENSG000002 | 8.69956449 | -0.657334872 | 0.34445498 | -1.9083332 | 0.05634817 | 0.13064324 | NA | NA |
| ENSG000001 | 10.6715225 | -0.650972394 | 0.34114412 | -1.9082035 | 0.05636492 | 0.1306536 | WFDC8 | 90199 |
| ENSG000001 | 11.4650673 | -0.656153438 | 0.34387158 | -1.9081351 | 0.05637376 | 0.1306666 | REN | 5972 |
| ENSG000002 | 5.69131861 | -0.706807053 | 0.37042842 | -1.90808 | 0.05638088 | 0.13067564 | NA | NA |
| ENSG00000C | 8.78224384 | -0.664775686 | 0.34844101 | -1.9078572 | 0.05640968 | 0.13073492 | LYZ | 4069 |
| ENSG000002 | 4.82136361 | -0.719152993 | 0.37695325 | -1.9078042 | 0.05641653 | 0.13074333 | NA | NA |
| ENSG000001 | 4.15094005 | -0.720495741 | 0.37766843 | -1.9077468 | 0.05642395 | 0.13075304 | IFNA10 | 3446 |
| ENSG000001 | 7.45104695 | -0.687272421 | 0.36030059 | -1.9074974 | 0.05645621 | 0.13082034 | NA | NA |
| ENSG000001 | 13.4708376 | -0.598856258 | 0.31395681 | -1.9074479 | 0.05646261 | 0.13082768 | HTR2B | 3357 |
| ENSG000002 | 2.2914665 | -0.700535813 | 0.36728296 | -1.9073464 | 0.05647575 | 0.13085065 | NA | NA |
| ENSG000001 | 14.8378639 | -0.608410556 | 0.3190084 | -1.9071929 | 0.05649561 | 0.1308892 | FFAR4 | 338557 |
| ENSG000001 | 26.6435876 | -0.503553464 | 0.26406074 | -1.9069607 | 0.05652568 | 0.13094676 | CXCL16 | 58191 |
| ENSG000002 | 4.18984109 | -0.72101019 | 0.37809579 | -1.9069511 | 0.05652692 | 0.13094676 | NA | NA |
| ENSG000002 | 3.00192933 | -0.714533504 | 0.37470927 | -1.9069011 | 0.0565334 | 0.13094854 | NA | NA |
| ENSG000002 | 5.81224721 | -0.711440397 | 0.37308833 | -1.9068953 | 0.05653414 | 0.13094854 | NA | NA |
| ENSG00000C | 659.428032 | 0.218700453 | 0.11469256 | 1.9068408 | 0.05654121 | 0.13094994 | SORBS1 | 10580 |
| ENSG000002 | 10.4596641 | -0.63678734 | 0.33394468 | -1.9068648 | 0.0565381 | 0.13094994 | NA | NA |
| ENSG000001 | 24.5538374 | -0.502327537 | 0.26344815 | -1.9067416 | 0.05655406 | 0.13095727 | CACNA1F | 778 |
| ENSG000001 | 148.634638 | 0.315447445 | 0.16543588 | 1.90676557 | 0.05655095 | 0.13095727 | ADCY9 | 115 |
| ENSG000001 | 1298.09529 | 0.590958526 | 0.30992517 | 1.90677807 | 0.05654933 | 0.13095727 | PCBP1 | 5093 |
| ENSG000001 | 12.6942794 | -0.621905823 | 0.32618407 | -1.90661 | 0.0565711 | 0.13098926 | IFI27 | 3429 |
| ENSG000001 | 5.37120723 | -0.710792953 | 0.37282483 | -1.9065065 | 0.05658453 | 0.13101285 | VIP | 7432 |

| | | | | | | | | |
|---|---|---|---|---|---|---|---|---|
| ENSG000002 | 4.8187593 | -0.716014877 | 0.37559634 | -1.9063414 | 0.05660592 | 0.13104742 | NA | NA |
| ENSG000002 | 6.94385514 | -0.698798731 | 0.36656298 | -1.9063538 | 0.05660432 | 0.13104742 | NA | NA |
| ENSG000001 | 4.50910473 | -0.72009396 | 0.37775801 | -1.9062308 | 0.05662026 | 0.13105293 | CST9L | 128821 |
| ENSG000001 | 2.69113213 | -0.705646176 | 0.37018109 | -1.9062188 | 0.05662182 | 0.13105293 | SMR3A | 26952 |
| ENSG000002 | 15.5460542 | -0.576135386 | 0.30224296 | -1.9061995 | 0.05662432 | 0.13105293 | INHBA-AS1 | 285954 |
| ENSG000002 | 3.35488156 | -0.716782938 | 0.37600868 | -1.9062936 | 0.05661212 | 0.13105293 | SLC6A1-AS1 | 100874090 |
| ENSG000002 | 6.90831796 | -0.693196874 | 0.36365409 | -1.9061985 | 0.05662446 | 0.13105293 | NA | NA |
| ENSG000001 | 25.2316973 | 0.508184069 | 0.26661262 | 1.9060766 | 0.05664027 | 0.13107456 | KREMEN2 | 79412 |
| ENSG000002 | 3.02379762 | -0.714354409 | 0.37477619 | -1.9060827 | 0.05663948 | 0.13107456 | NA | NA |
| ENSG000000 | 30.0703178 | -0.496685391 | 0.26059584 | -1.9059605 | 0.05665533 | 0.13109752 | ABCC8 | 6833 |
| ENSG000002 | 2.65893756 | -0.708738195 | 0.37185555 | -1.9059503 | 0.05665665 | 0.13109752 | NA | NA |
| ENSG000002 | 5.03288657 | -0.711528416 | 0.37332909 | -1.9059013 | 0.05666301 | 0.13110474 | NMS | 129521 |
| ENSG000002 | 18.6612606 | -0.541400023 | 0.28410127 | -1.9056586 | 0.05669452 | 0.13115939 | NA | NA |
| ENSG000002 | 10.7335052 | -0.634125897 | 0.33275508 | -1.9056836 | 0.05669127 | 0.13115939 | NA | NA |
| ENSG000002 | 8.85878051 | -0.68202252 | 0.35789597 | -1.9056446 | 0.05669633 | 0.13115939 | NA | NA |
| ENSG000002 | 3.97539054 | -0.721297167 | 0.37852522 | -1.9055459 | 0.05670915 | 0.13118017 | AGBL4-IT1 | 100874313 |
| ENSG000002 | 4.28700904 | 0.720578974 | 0.37815235 | 1.90552559 | 0.05671178 | 0.13118017 | NA | NA |
| ENSG000002 | 6.30616226 | -0.700882581 | 0.36782917 | -1.9054568 | 0.05672072 | 0.13118063 | LOC101929452 | 101929452 |
| ENSG000002 | 4.45491812 | -0.718943709 | 0.37730912 | -1.9054501 | 0.05672159 | 0.13118063 | NA | NA |
| ENSG000002 | 4.76027842 | -0.715545208 | 0.37552571 | -1.9054493 | 0.05672169 | 0.13118063 | NA | NA |
| ENSG000002 | 3.90713369 | -0.721355376 | 0.37858513 | -1.9053981 | 0.05672835 | 0.13118854 | NA | NA |
| ENSG000000 | 335.502617 | 0.527688926 | 0.27695338 | 1.90533485 | 0.05673656 | 0.13120005 | KLF6 | 1316 |
| ENSG000001 | 113.751583 | 0.339390203 | 0.17813511 | 1.90524041 | 0.05674883 | 0.13120731 | ATP7B | 540 |
| ENSG000001 | 1679.71348 | 0.316927763 | 0.16634896 | 1.90519839 | 0.05675429 | 0.13120731 | KRAS | 3845 |
| ENSG000001 | 40.996813 | -0.428611517 | 0.22496894 | -1.9052031 | 0.05675368 | 0.13120731 | FAM161B | 145483 |
| ENSG000002 | 3.88470969 | -0.721314324 | 0.37859052 | -1.9052625 | 0.05674595 | 0.13120731 | NA | NA |
| ENSG000002 | 9.78916389 | -0.67186703 | 0.35265164 | -1.9051862 | 0.05675587 | 0.13120731 | NA | NA |
| ENSG000001 | 3.6079393 | -0.720361892 | 0.37814661 | -1.9049804 | 0.05678261 | 0.1312436 | NPBWR1 | 2831 |
| ENSG000001 | 9.03000461 | -0.65733474 | 0.3450638 | -1.9049658 | 0.05678452 | 0.1312436 | SULT1C3 | 442038 |

| | | | | | | | | |
|---|---|---|---|---|---|---|---|---|
| ENSG000002 | 2.70479402 | -0.706419422 | 0.37082765 | -1.9049804 | 0.05678261 | 0.1312436 | MIR320C1 | 100302135 |
| ENSG000002 | 2.34622908 | -0.696698554 | 0.36571858 | -1.9050128 | 0.05677841 | 0.1312436 | NA | NA |
| ENSG000002 | 15.9976488 | -0.57616413 | 0.3024704 | -1.9048612 | 0.05679812 | 0.13126755 | NA | NA |
| ENSG000001 | 17.0635003 | -0.556277582 | 0.29205763 | -1.9046843 | 0.05682112 | 0.13131324 | OR8B8 | 26493 |
| ENSG000002 | 8.91392392 | -0.66931458 | 0.35144336 | -1.9044735 | 0.05684854 | 0.13136911 | NA | NA |
| ENSG000002 | 2.88565331 | -0.699131401 | 0.36715758 | -1.9041726 | 0.05688771 | 0.13145213 | LOC728093 | 728093 |
| ENSG000001 | 13.0318135 | -0.635468421 | 0.3337436 | -1.9040617 | 0.05690214 | 0.13147092 | NLRP13 | 126204 |
| ENSG000001 | 4.79257801 | -0.715473297 | 0.37577069 | -1.9040157 | 0.05690814 | 0.13147092 | NA | NA |
| ENSG000002 | 6.90102974 | -0.695234326 | 0.36513934 | -1.9040247 | 0.05690697 | 0.13147092 | NA | NA |
| ENSG000002 | 2.65145346 | -0.705907635 | 0.37074775 | -1.9040106 | 0.05690881 | 0.13147092 | NA | NA |
| ENSG000002 | 5.30618233 | -0.717027695 | 0.37659812 | -1.9039598 | 0.05691542 | 0.1314787 | NA | NA |
| ENSG000002 | 8.47806764 | -0.680359819 | 0.3573658 | -1.9038191 | 0.05693375 | 0.13151356 | NA | NA |
| ENSG000002 | 12.1660506 | -0.619743688 | 0.32553097 | -1.9037933 | 0.05693711 | 0.13151382 | NA | NA |
| ENSG000002 | 9.6145758 | -0.646429772 | 0.33955628 | -1.9037485 | 0.05694294 | 0.13151981 | NA | NA |
| ENSG000001 | 5.24952719 | -0.716507276 | 0.37638839 | -1.9036381 | 0.05695734 | 0.13154556 | NA | NA |
| ENSG000002 | 5.66220067 | -0.706169926 | 0.37097945 | -1.9035284 | 0.05697163 | 0.13157107 | NA | NA |
| ENSG000001 | 440.914491 | 0.250816264 | 0.13176853 | 1.903461 | 0.05698042 | 0.13157639 | CSTF2 | 1478 |
| ENSG000002 | 6.182512 | -0.707869717 | 0.3718819 | -1.9034799 | 0.05697796 | 0.13157639 | NA | NA |
| ENSG000002 | 8.0464542 | -0.677186535 | 0.35579089 | -1.9033274 | 0.05699784 | 0.13160912 | NA | NA |
| ENSG000001 | 47.1227181 | 0.423406695 | 0.22247831 | 1.903137 | 0.05702267 | 0.13164397 | GLT1D1 | 144423 |
| ENSG000002 | 8.22900494 | -0.684268218 | 0.3595384 | -1.9031854 | 0.05701636 | 0.13164397 | NA | NA |
| ENSG000002 | 2.32812475 | -0.697423567 | 0.36645772 | -1.9031488 | 0.05702114 | 0.13164397 | NA | NA |
| ENSG000002 | 4.28540601 | -0.720301869 | 0.37850239 | -1.9030312 | 0.05703647 | 0.13166833 | LOC102724551 | 102724551 |
| ENSG000002 | 9.0324168 | -0.658322563 | 0.34594776 | -1.9029537 | 0.05704659 | 0.1316842 | NA | NA |
| ENSG000002 | 3.08637603 | -0.712288965 | 0.37432281 | -1.9028735 | 0.05705705 | 0.13170085 | NA | NA |
| ENSG000002 | 10.9523611 | -0.645019981 | 0.33900297 | -1.9026971 | 0.05708008 | 0.1317465 | NA | NA |
| ENSG000002 | 9.06247937 | -0.660973636 | 0.34739409 | -1.9026623 | 0.05708463 | 0.1317495 | NA | NA |
| ENSG000002 | 7.76613381 | -0.683076753 | 0.35901763 | -1.9026273 | 0.05708919 | 0.13175254 | NA | NA |
| ENSG000002 | 30.7047883 | 0.479418797 | 0.25198799 | 1.90254623 | 0.05709978 | 0.13176947 | NA | NA |

| | | | | | | | | |
|---|---|---|---|---|---|---|---|---|
| ENSG000001 | 9.55583357 | -0.646577685 | 0.33986268 | -1.9024675 | 0.05711007 | 0.13178055 | CLECL1 | 160365 |
| ENSG000002 | 5.20572926 | -0.714513314 | 0.37557342 | -1.9024597 | 0.05711108 | 0.13178055 | NA | NA |
| ENSG000002 | 18.1009594 | -0.564881988 | 0.29693802 | -1.9023566 | 0.05712456 | 0.13180415 | LINC00589 | 619351 |
| ENSG000002 | 2.30267544 | -0.697913748 | 0.36687949 | -1.902297 | 0.05713234 | 0.1318146 | NA | NA |
| ENSG000001 | 225.921986 | 0.266012917 | 0.13984538 | 1.90219314 | 0.05714591 | 0.13183841 | MYCL | 4610 |
| ENSG00000C | 31.3126621 | 0.466180442 | 0.24508261 | 1.90213597 | 0.05715338 | 0.13184261 | CCL26 | 10344 |
| ENSG000002 | 3.25425398 | -0.717798413 | 0.3773657 | -1.9021295 | 0.05715423 | 0.13184261 | NA | NA |
| ENSG000002 | 2.93493086 | -0.714952889 | 0.37587518 | -1.9021019 | 0.05715784 | 0.13184342 | NA | NA |
| ENSG000001 | 20.9295746 | -0.538573398 | 0.28316325 | -1.9019891 | 0.05717258 | 0.13186994 | UMODL1 | 89766 |
| ENSG000001 | 10.2516861 | -0.645325635 | 0.33930252 | -1.9019182 | 0.05718185 | 0.13188382 | NIPSNAP3B | 55335 |
| ENSG000001 | 3110.54274 | 0.510194882 | 0.26827087 | 1.90179008 | 0.0571986 | 0.13191495 | PNN | 5411 |
| ENSG000002 | 6.94037723 | -0.693023325 | 0.36441518 | -1.9017411 | 0.05720501 | 0.13192222 | NA | NA |
| ENSG000002 | 3.52388911 | -0.719946124 | 0.37858043 | -1.9016993 | 0.05721048 | 0.13192734 | NA | NA |
| ENSG000001 | 2.66923716 | -0.703581932 | 0.37000195 | -1.9015628 | 0.05722834 | 0.13196101 | OR51I1 | 390063 |
| ENSG000002 | 3.56419344 | -0.719726824 | 0.3785215 | -1.901416 | 0.05724755 | 0.1319903 | NA | NA |
| ENSG000002 | 14.8836699 | -0.58681164 | 0.3086151 | -1.9014353 | 0.05724502 | 0.1319903 | NA | NA |
| ENSG000001 | 7.23328276 | -0.693195753 | 0.36457425 | -1.9013843 | 0.0572517 | 0.13199236 | MYEOV | 26579 |
| ENSG000001 | 119.428079 | 0.354580092 | 0.18651161 | 1.90111538 | 0.0572869 | 0.13204349 | CARD10 | 29775 |
| ENSG000002 | 3.57193704 | -0.719437073 | 0.37841856 | -1.9011675 | 0.05728008 | 0.13204349 | USP32P2 | 220594 |
| ENSG000002 | 4.74519506 | -0.713532492 | 0.37531932 | -1.9011345 | 0.0572844 | 0.13204349 | NA | NA |
| ENSG000002 | 3.60060009 | -0.719036469 | 0.37821246 | -1.9011443 | 0.05728311 | 0.13204349 | NA | NA |
| ENSG000002 | 14.7438969 | 0.594257896 | 0.31258971 | 1.90107953 | 0.0572916 | 0.13204681 | NA | NA |
| ENSG000002 | 13.0479414 | -0.63340853 | 0.33320042 | -1.9009836 | 0.05730416 | 0.13206827 | LOC102724933 | 102724933 |
| ENSG000002 | 4.38657435 | -0.71966505 | 0.37858565 | -1.9009306 | 0.0573111 | 0.13207675 | NA | NA |
| ENSG000002 | 8.49914258 | -0.664478592 | 0.3496026 | -1.9006683 | 0.05734547 | 0.13214844 | NA | NA |
| ENSG000002 | 2.29585741 | -0.696208225 | 0.36631877 | -1.900553 | 0.05736059 | 0.13217578 | RNU6ATAC8P | 106481161 |
| ENSG000002 | 5.61214907 | -0.704291877 | 0.37058294 | -1.9004973 | 0.05736789 | 0.13217757 | NA | NA |
| ENSG000002 | 2.29217091 | -0.696366066 | 0.36641142 | -1.9005032 | 0.05736712 | 0.13217757 | NA | NA |
| ENSG000002 | 6.48865611 | -0.690635898 | 0.36344049 | -1.9002723 | 0.05739739 | 0.13223052 | NA | NA |

| | | | | | | | | |
|---|---|---|---|---|---|---|---|---|
| ENSG000002 | 32.4786314 | 0.502908237 | 0.26464873 | 1.90028582 | 0.05739562 | 0.13223052 | NA | NA |
| ENSG000002 | 9.59874652 | -0.648357519 | 0.34121849 | -1.9001242 | 0.05741682 | 0.13226777 | LLPHP1 | 106480685 |
| ENSG000002 | 3.85316746 | -0.719338997 | 0.378581 | -1.9000927 | 0.05742095 | 0.13226978 | NA | NA |
| ENSG000001 | 385.512673 | 0.243603885 | 0.12821145 | 1.90001656 | 0.05743095 | 0.13228528 | TRMT13 | 54482 |
| ENSG000002 | 2.64259994 | -0.705701274 | 0.37148261 | -1.8996886 | 0.057474 | 0.13237693 | NA | NA |
| ENSG000002 | 4.50331814 | -0.7175655 | 0.37776819 | -1.8994863 | 0.05750057 | 0.1324306 | NA | NA |
| ENSG000002 | 5.45805337 | -0.710821637 | 0.37422481 | -1.8994508 | 0.05750522 | 0.1324338 | LOC101929261 | 101929261 |
| ENSG000002 | 3.3309513 | -0.716460937 | 0.37721086 | -1.8993646 | 0.05751656 | 0.13245238 | NA | NA |
| ENSG000001 | 3.55577157 | -0.718881122 | 0.37849085 | -1.8993355 | 0.05752038 | 0.13245365 | NA | NA |
| ENSG000001 | 353.579448 | 0.533720092 | 0.2810167 | 1.89924688 | 0.05753202 | 0.13246719 | ZFAND3 | 60685 |
| ENSG000001 | 90.7510019 | 0.345628452 | 0.18198479 | 1.8992162 | 0.05753606 | 0.13246719 | CHRNB1 | 1140 |
| ENSG000002 | 2.30743087 | -0.69536637 | 0.36613131 | -1.8992267 | 0.05753467 | 0.13246719 | CDRT15L2 | 256223 |
| ENSG000001 | 249.598824 | 0.240202731 | 0.12648789 | 1.89901759 | 0.05756216 | 0.13251977 | SCAPER | 49855 |
| ENSG000002 | 3.94468105 | -0.71858331 | 0.37840447 | -1.8989821 | 0.05756683 | 0.13252299 | NA | NA |
| ENSG000002 | 2.91330499 | -0.714387167 | 0.37623086 | -1.8988 | 0.05759077 | 0.13257059 | NA | NA |
| ENSG000002 | 83.7700492 | 0.344133195 | 0.18123972 | 1.89877361 | 0.05759425 | 0.13257106 | LINC01278 | 92249 |
| ENSG000002 | 2.33706258 | -0.694012328 | 0.36551465 | -1.8987264 | 0.05760045 | 0.13257781 | NA | NA |
| ENSG000002 | 3.64224245 | -0.718045657 | 0.37820326 | -1.8985708 | 0.05762093 | 0.13261743 | NA | NA |
| ENSG000001 | 33.9645967 | -0.464923949 | 0.24489662 | -1.8984498 | 0.05763685 | 0.13264653 | ARHGAP20 | 57569 |
| ENSG000002 | 6.5238262 | -0.692997237 | 0.36511838 | -1.898007 | 0.05769516 | 0.1327732 | NA | NA |
| ENSG000002 | 5.42052127 | -0.70887362 | 0.37348842 | -1.8979802 | 0.05769869 | 0.13277379 | NA | NA |
| ENSG000002 | 5.05568339 | -0.711533457 | 0.3749018 | -1.8979196 | 0.05770668 | 0.13278463 | NA | NA |
| ENSG000002 | 2.65543997 | -0.704336063 | 0.37112452 | -1.897843 | 0.05771677 | 0.13279576 | NA | NA |
| ENSG000002 | 7.31576754 | -0.699297405 | 0.36847149 | -1.8978332 | 0.05771806 | 0.13279576 | PRR23C | 389152 |
| ENSG000001 | 276.999422 | 0.314931574 | 0.16595165 | 1.89773094 | 0.05773153 | 0.13280595 | FAM161A | 84140 |
| ENSG000002 | 3.63555579 | -0.717558883 | 0.37811532 | -1.897725 | 0.05773232 | 0.13280595 | NA | NA |
| ENSG000002 | 3.6447223 | -0.717525281 | 0.37809332 | -1.8977465 | 0.05772948 | 0.13280595 | NA | NA |
| ENSG000002 | 3.0362523 | -0.710871966 | 0.37463257 | -1.8975178 | 0.05775963 | 0.13285446 | NA | NA |
| ENSG000002 | 5.61522266 | -0.702420812 | 0.37017926 | -1.8975153 | 0.05775996 | 0.13285446 | MGC39584 | 441058 |

| | | | | | | | | |
|---|---|---|---|---|---|---|---|---|
| ENSG000001 | 128.726971 | 0.303071798 | 0.15973128 | 1.89738536 | 0.0577771 | 0.13288634 | CDC42EP4 | 23580 |
| ENSG000002 | 6.09052124 | -0.70282488 | 0.37044021 | -1.8972694 | 0.05779239 | 0.13291397 | NA | NA |
| ENSG000002 | 2.27709497 | -0.695431341 | 0.36656644 | -1.8971495 | 0.05780821 | 0.13294282 | NA | NA |
| ENSG000001 | 2.26920646 | -0.695440401 | 0.366612 | -1.8969385 | 0.05783606 | 0.13299178 | SPRR2G | 6706 |
| ENSG000002 | 7.80811585 | -0.681883243 | 0.35946426 | -1.8969431 | 0.05783545 | 0.13299178 | FAM170B-AS1 | 100506733 |
| ENSG000001 | 19.3594917 | -0.544968726 | 0.2873056 | -1.896826 | 0.05785091 | 0.13301648 | SPATA21 | 374955 |
| ENSG000001 | 88.1347332 | 0.341020573 | 0.17978661 | 1.8968074 | 0.05785336 | 0.13301648 | NA | NA |
| ENSG000001 | 371.506119 | 0.305295139 | 0.16095897 | 1.89672652 | 0.05786404 | 0.13303349 | NID1 | 4811 |
| ENSG000002 | 4.60902654 | -0.717254534 | 0.37817243 | -1.8966336 | 0.05787631 | 0.13305416 | CYP4F44P | 106480733 |
| ENSG000002 | 7.93572227 | -0.674075484 | 0.35542294 | -1.8965447 | 0.05788806 | 0.13307362 | LOC101927987 | 101927987 |
| ENSG000002 | 3.8996779 | -0.717852345 | 0.37855554 | -1.8962933 | 0.05792127 | 0.13313901 | NA | NA |
| ENSG000002 | 17.6789094 | 0.568866506 | 0.29999082 | 1.89627969 | 0.05792307 | 0.13313901 | NA | NA |
| ENSG000002 | 18.9500235 | -0.539193919 | 0.28435004 | -1.896233 | 0.05792924 | 0.13314111 | NA | NA |
| ENSG000002 | 5.88232497 | -0.696826447 | 0.36748126 | -1.8962231 | 0.05793055 | 0.13314111 | NA | NA |
| ENSG000002 | 3.91466953 | -0.717826345 | 0.37858397 | -1.8960823 | 0.05794917 | 0.13317635 | NA | NA |
| ENSG000002 | 3.2418982 | -0.715715022 | 0.37748649 | -1.8960017 | 0.05795982 | 0.13318813 | NA | NA |
| ENSG000002 | 6.02597915 | -0.702385525 | 0.37045771 | -1.8959938 | 0.05796086 | 0.13318813 | LINC00642 | 400238 |
| ENSG000002 | 6.51158496 | -0.705099931 | 0.37190284 | -1.8959251 | 0.05796995 | 0.13320147 | NA | NA |
| ENSG000001 | 26.3387456 | -0.505425947 | 0.26659034 | -1.8958899 | 0.0579746 | 0.1332046 | B4GALNT1 | 2583 |
| ENSG000002 | 16.3894838 | -0.590686741 | 0.31156887 | -1.8958464 | 0.05798035 | 0.13321027 | NA | NA |
| ENSG000002 | 18.1361988 | 0.583413858 | 0.30777995 | 1.8955551 | 0.0580189 | 0.13329128 | NA | NA |
| ENSG000002 | 58.3726044 | 0.382646632 | 0.20187833 | 1.8954319 | 0.05803521 | 0.13331364 | NA | NA |
| ENSG000002 | 4.54177288 | -0.715350021 | 0.3774039 | -1.8954495 | 0.05803288 | 0.13331364 | NA | NA |
| ENSG000002 | 6.7457799 | -0.701289922 | 0.37003544 | -1.8951966 | 0.05806636 | 0.13337764 | LINC01475 | 101927324 |
| ENSG000002 | 4.77707751 | -0.71207598 | 0.37574493 | -1.8951047 | 0.05807853 | 0.13339117 | NA | NA |
| ENSG000002 | 6.35641028 | -0.69798449 | 0.36830963 | -1.8951025 | 0.05807882 | 0.13339117 | NA | NA |
| ENSG000001 | 153.357439 | 0.301460677 | 0.15908194 | 1.89500253 | 0.05809207 | 0.13340647 | EGF | 1950 |
| ENSG000002 | 7.62120775 | -0.671494633 | 0.35434676 | -1.8950212 | 0.05808959 | 0.13340647 | NA | NA |
| ENSG000002 | 7.17583012 | -0.687709014 | 0.36292039 | -1.8949308 | 0.05810157 | 0.13342074 | NA | NA |

| | | | | | | | | |
|---|---|---|---|---|---|---|---|---|
| ENSG000002 | 3.17575545 | -0.703188761 | 0.37110194 | -1.8948669 | 0.05811004 | 0.13342508 | LOC283352 | 283352 |
| ENSG000002 | 4.35839004 | -0.71733392 | 0.3785631 | -1.894886 | 0.05810751 | 0.13342508 | NA | NA |
| ENSG000001 | 10.2234509 | -0.641704923 | 0.33866746 | -1.8947936 | 0.05811976 | 0.13343983 | SRGN | 5552 |
| ENSG000002 | 14.9231554 | -0.592966963 | 0.31300825 | -1.8944132 | 0.05817018 | 0.13354804 | NA | NA |
| ENSG000001 | 72.0204586 | 0.388634678 | 0.2051743 | 1.89416846 | 0.05820265 | 0.1336117 | PLCD1 | 5333 |
| ENSG000002 | 12.7026119 | -0.611445253 | 0.32280642 | -1.8941546 | 0.0582045 | 0.1336117 | NA | NA |
| ENSG000002 | 4.77247002 | -0.712612811 | 0.3762569 | -1.8939528 | 0.05823127 | 0.13366559 | NA | NA |
| ENSG000002 | 9.08532172 | -0.657068438 | 0.34693599 | -1.8939183 | 0.05823585 | 0.13366854 | LOC101928405 | 101928405 |
| ENSG000001 | 142.959021 | 0.367553641 | 0.19407412 | 1.89388279 | 0.05824057 | 0.1336718 | HSD11B2 | 3291 |
| ENSG000001 | 27.9947998 | -0.483250557 | 0.25520724 | -1.8935613 | 0.05828326 | 0.13376221 | CLEC7A | 64581 |
| ENSG000002 | 3.32258672 | -0.713847166 | 0.37699252 | -1.8935314 | 0.05828724 | 0.13376376 | NA | NA |
| ENSG000002 | 4.4584997 | -0.71462271 | 0.3774235 | -1.893424 | 0.05830151 | 0.13378895 | NA | NA |
| ENSG000002 | 2.92132934 | -0.71164169 | 0.3758567 | -1.8933857 | 0.0583066 | 0.13379306 | NA | NA |
| ENSG000002 | 34.0798209 | 0.470133342 | 0.24830993 | 1.89333282 | 0.05831362 | 0.13379403 | RNU6ATAC | 100151684 |
| ENSG000002 | 3.24258335 | -0.714189135 | 0.37720983 | -1.8933471 | 0.05831173 | 0.13379403 | NA | NA |
| ENSG000001 | 4.78802373 | -0.716775575 | 0.3785872 | -1.8932906 | 0.05831924 | 0.13379934 | LINC00982 | 440556 |
| ENSG000002 | 2.2886535 | -0.690815352 | 0.36494631 | -1.8929232 | 0.05836808 | 0.13389718 | NA | NA |
| ENSG000002 | 2.25577324 | -0.693073096 | 0.36613963 | -1.8929202 | 0.05836848 | 0.13389718 | NA | NA |
| ENSG000002 | 2.71239382 | -0.699057239 | 0.3693349 | -1.8927463 | 0.05839162 | 0.13394268 | LOC101927869 | 101927869 |
| ENSG000001 | 285.807598 | 0.296920632 | 0.15687989 | 1.89266215 | 0.05840281 | 0.13396077 | GLI3 | 2737 |
| ENSG000001 | 32.5360117 | -0.448013841 | 0.23671729 | -1.8926114 | 0.05840957 | 0.13396548 | CD101 | 9398 |
| ENSG000002 | 369.634634 | 0.272229812 | 0.14383929 | 1.89259708 | 0.05841147 | 0.13396548 | NA | NA |
| ENSG000001 | 32.9400557 | -0.447695355 | 0.23656171 | -1.8925098 | 0.05842308 | 0.13398453 | CCDC62 | 84660 |
| ENSG000002 | 3.55909255 | -0.716313193 | 0.37852244 | -1.8923929 | 0.05843865 | 0.13401265 | RNU6-252P | 106480564 |
| ENSG000002 | 22.0715201 | -0.530096392 | 0.28013635 | -1.8922799 | 0.05845369 | 0.13403957 | NA | NA |
| ENSG000002 | 6.42666478 | -0.688435641 | 0.36382442 | -1.8922194 | 0.05846175 | 0.13405046 | NA | NA |
| ENSG000001 | 2.66987132 | -0.702479617 | 0.3712738 | -1.8920797 | 0.05848036 | 0.13405761 | CCL13 | 6357 |
| ENSG000002 | 6.22187287 | -0.694745036 | 0.36717406 | -1.8921409 | 0.05847222 | 0.13405761 | SPDYC | 387778 |
| ENSG000002 | 2.93427797 | -0.710315248 | 0.37542445 | -1.8920325 | 0.05848666 | 0.13405761 | NA | NA |

| | | | | | | | | |
|---|---|---|---|---|---|---|---|---|
| ENSG000002 | 9.63472041 | -0.647659862 | 0.34230628 | -1.8920479 | 0.0584846 | 0.13405761 | NA | NA |
| ENSG000002 | 5.40439567 | -0.706730773 | 0.37352833 | -1.8920406 | 0.05848557 | 0.13405761 | NA | NA |
| ENSG000002 | 24.2204894 | 0.496699134 | 0.26252287 | 1.89202232 | 0.05848801 | 0.13405761 | NA | NA |
| ENSG000002 | 7.31133963 | -0.67652692 | 0.35754853 | -1.8921262 | 0.05847417 | 0.13405761 | NA | NA |
| ENSG000002 | 8.43347793 | -0.661558824 | 0.34968705 | -1.8918596 | 0.05850968 | 0.13409086 | NA | NA |
| ENSG000002 | 5.05599537 | -0.706751962 | 0.37357433 | -1.8918644 | 0.05850906 | 0.13409086 | NA | NA |
| ENSG000002 | 11.2242585 | 0.631158572 | 0.33362172 | 1.891839 | 0.05851244 | 0.13409086 | NA | NA |
| ENSG000002 | 2.90755122 | -0.711498616 | 0.3761185 | -1.8916874 | 0.05853265 | 0.1341296 | NA | NA |
| ENSG000002 | 6.8855179 | -0.688716632 | 0.36409473 | -1.8915864 | 0.05854611 | 0.13413772 | DSCAM-AS1 | 100506492 |
| ENSG000002 | 22.6170107 | -0.554575748 | 0.29317815 | -1.8915998 | 0.05854432 | 0.13413772 | NA | NA |
| ENSG000002 | 11.1678149 | -0.633825164 | 0.33507467 | -1.8915938 | 0.05854513 | 0.13413772 | NA | NA |
| ENSG000001 | 5.50646245 | -0.708355491 | 0.37449567 | -1.8914918 | 0.05855872 | 0.13414568 | THEG | 51298 |
| ENSG000001 | 294.360993 | 0.248148638 | 0.13119032 | 1.89151644 | 0.05855544 | 0.13414568 | DUS4L | 11062 |
| ENSG000001 | 95.8758416 | 0.32111174 | 0.16976692 | 1.89148588 | 0.05855951 | 0.13414568 | COQ6 | 51004 |
| ENSG000001 | 3.68091068 | -0.71498888 | 0.37801384 | -1.8914357 | 0.0585662 | 0.13415342 | OR10T2 | 128360 |
| ENSG000001 | 8.86848592 | -0.660534988 | 0.34924356 | -1.8913305 | 0.05858024 | 0.134178 | CSNK1A1L | 122011 |
| ENSG000001 | 18.5729838 | -0.550633298 | 0.29114504 | -1.891268 | 0.05858858 | 0.13418952 | CFAP74 | 85452 |
| ENSG000001 | 7.48515187 | -0.679879177 | 0.35950959 | -1.8911295 | 0.05860706 | 0.13422427 | PGK2 | 5232 |
| ENSG000001 | 571.987831 | 0.279471198 | 0.14779136 | 1.89098468 | 0.05862639 | 0.13426095 | CCP110 | 9738 |
| ENSG000001 | 657.026525 | 0.230472082 | 0.12188533 | 1.89089278 | 0.05863866 | 0.13428147 | CSPP1 | 79848 |
| ENSG000002 | 14.4451547 | 0.595480795 | 0.31493307 | 1.89081695 | 0.05864878 | 0.13429707 | SNORD116-16 | 100033428 |
| ENSG000002 | 83.551015 | 0.348778663 | 0.18447442 | 1.89066137 | 0.05866956 | 0.13433706 | FBXO48 | 554251 |
| ENSG000002 | 6.60798993 | -0.692482536 | 0.36627818 | -1.8905918 | 0.05867885 | 0.13435075 | ITPK1-AS1 | 319085 |
| ENSG000002 | 10.3596048 | -0.632936772 | 0.33483011 | -1.8903221 | 0.05871489 | 0.13442567 | NA | NA |
| ENSG000002 | 36.7747705 | 0.453661617 | 0.24000333 | 1.89023051 | 0.05872714 | 0.13444612 | AARD | 441376 |
| ENSG000002 | 3.53787575 | -0.71534373 | 0.37847125 | -1.8900873 | 0.05874628 | 0.13448235 | NA | NA |
| ENSG000002 | 11.5586941 | -0.623673661 | 0.32998727 | -1.8899931 | 0.05875888 | 0.13450361 | LOC101927207 | 101927207 |
| ENSG000002 | 3.36966655 | -0.710975051 | 0.37618493 | -1.8899616 | 0.0587631 | 0.13450567 | NA | NA |
| ENSG000001 | 6.5613722 | -0.704717434 | 0.37289094 | -1.8898754 | 0.05877462 | 0.13452445 | OR2L13 | 284521 |

| | | | | | | | | |
|---|---|---|---|---|---|---|---|---|
| ENSG000002 | 23.8188674 | -0.496943933 | 0.26296588 | -1.8897658 | 0.05878929 | 0.13455042 | NA | NA |
| ENSG000002 | 2.77134125 | -0.696996086 | 0.36885482 | -1.8896217 | 0.05880857 | 0.13458696 | NA | NA |
| ENSG000002 | 10.0479801 | -0.652909368 | 0.34553838 | -1.8895423 | 0.0588192 | 0.1345961 | NA | NA |
| ENSG000002 | 8.48374656 | -0.661214548 | 0.34993018 | -1.8895614 | 0.05881665 | 0.1345961 | NA | NA |
| ENSG000002 | 15.161688 | 0.587710914 | 0.3110669 | 1.88933929 | 0.05884638 | 0.13465069 | PNMA6A | 84968 |
| ENSG000002 | 4.83028934 | -0.712315937 | 0.37707254 | -1.8890687 | 0.05888262 | 0.13472324 | SCRT1 | 83482 |
| ENSG000002 | 4.87044789 | 0.709119727 | 0.37538372 | 1.88905298 | 0.05888473 | 0.13472324 | NA | NA |
| ENSG000002 | 6.24621446 | -0.694061681 | 0.36746743 | -1.8887706 | 0.05892257 | 0.13480222 | LOC101928751 | 101928751 |
| ENSG000002 | 2.9513528 | -0.70907519 | 0.37544409 | -1.8886306 | 0.05894135 | 0.13482379 | NA | NA |
| ENSG000002 | 2.32733492 | -0.688578113 | 0.36459165 | -1.8886283 | 0.05894165 | 0.13482379 | NA | NA |
| ENSG000002 | 2.24923073 | -0.691461569 | 0.36611887 | -1.8886259 | 0.05894198 | 0.13482379 | NA | NA |
| ENSG000001 | 6.92251528 | -0.689188533 | 0.36493033 | -1.8885483 | 0.05895239 | 0.13483999 | HOXD13 | 3239 |
| ENSG000002 | 21.858434 | 0.51662808 | 0.27356635 | 1.88849282 | 0.05895982 | 0.13484939 | NA | NA |
| ENSG000002 | 4.48020261 | -0.714688095 | 0.37845601 | -1.8884311 | 0.0589681 | 0.13486072 | NA | NA |
| ENSG000002 | 2.94548692 | -0.708581937 | 0.37522862 | -1.8884006 | 0.0589722 | 0.13486248 | NA | NA |
| ENSG000002 | 4.85780491 | -0.710689262 | 0.37635422 | -1.8883521 | 0.0589787 | 0.13486975 | NA | NA |
| ENSG000001 | 645.258332 | 0.272478307 | 0.14429842 | 1.88829726 | 0.05898606 | 0.13487136 | TBCE | 6905 |
| ENSG000001 | 682.675637 | 0.534386569 | 0.28299902 | 1.88829831 | 0.05898591 | 0.13487136 | ZC3H18 | 124245 |
| ENSG000002 | 8.80467363 | -0.660063264 | 0.34957704 | -1.8881768 | 0.05900221 | 0.1349007 | NA | NA |
| ENSG000001 | 15.6969929 | -0.567315802 | 0.30047361 | -1.888072 | 0.05901628 | 0.13492526 | CD1B | 910 |
| ENSG000002 | 2.91088485 | -0.709484947 | 0.37580433 | -1.8879105 | 0.05903796 | 0.13496722 | NA | NA |
| ENSG000002 | 3.48322473 | -0.714627621 | 0.37854317 | -1.8878365 | 0.0590479 | 0.13498232 | NA | NA |
| ENSG000001 | 10.0845449 | -0.634014874 | 0.33584756 | -1.8878055 | 0.05905207 | 0.13498424 | NAT16 | 375607 |
| ENSG000001 | 4.85172098 | -0.711324279 | 0.37681588 | -1.8877237 | 0.05906305 | 0.13499425 | SLC35G3 | 146861 |
| ENSG000001 | 27.0704519 | -0.493843066 | 0.26160776 | -1.8877233 | 0.0590631 | 0.13499425 | GLDN | 342035 |
| ENSG000001 | 16.4748538 | -0.573599073 | 0.30387471 | -1.887617 | 0.05907739 | 0.13501928 | OR4D1 | 26689 |
| ENSG000002 | 5.40817291 | -0.705210545 | 0.37367649 | -1.8872222 | 0.05913045 | 0.13513294 | NA | NA |
| ENSG000002 | 5.6998985 | -0.703737902 | 0.3729259 | -1.8870717 | 0.05915069 | 0.13517019 | NA | NA |
| ENSG000002 | 4.11640856 | -0.713056665 | 0.37786818 | -1.8870514 | 0.05915341 | 0.13517019 | TEX36-AS1 | 100169752 |

| | | | | | | | | |
|---|---|---|---|---|---|---|---|---|
| ENSG000001 | 4.55787364 | -0.712550537 | 0.37761438 | -1.8869794 | 0.0591631 | 0.1351847 | PDHA2 | 5161 |
| ENSG000001 | 70.1972917 | 0.356497866 | 0.18893111 | 1.88691991 | 0.0591711 | 0.13519046 | RILPL2 | 196383 |
| ENSG000002 | 23.7647725 | 0.511206498 | 0.27092597 | 1.88688629 | 0.05917562 | 0.13519046 | NA | NA |
| ENSG000002 | 2.61283151 | -0.699136207 | 0.37051932 | -1.8869089 | 0.05917258 | 0.13519046 | LOC105370605 | 105370605 |
| ENSG000002 | 3.2991011 | -0.711490381 | 0.377083 | -1.886827 | 0.0591836 | 0.13520107 | NA | NA |
| ENSG000002 | 2.69309582 | -0.697118112 | 0.36947525 | -1.8867789 | 0.05919007 | 0.13520823 | NA | NA |
| ENSG000002 | 11.200102 | -0.652782149 | 0.34598237 | -1.8867497 | 0.05919399 | 0.13520958 | NBAT1 | 729177 |
| ENSG000002 | 6.45734092 | -0.687740607 | 0.36452929 | -1.8866539 | 0.05920689 | 0.13523142 | NA | NA |
| ENSG000002 | 2.91116504 | -0.709294991 | 0.3759694 | -1.8865764 | 0.05921733 | 0.13524764 | NA | NA |
| ENSG000002 | 4.19232182 | -0.713087575 | 0.37799101 | -1.8865199 | 0.05922493 | 0.13525738 | NA | NA |
| ENSG000002 | 18.9988015 | -0.552507495 | 0.292906 | -1.8862963 | 0.05925504 | 0.13531853 | NA | NA |
| ENSG000002 | 3.06585046 | -0.703179787 | 0.37280299 | -1.8861968 | 0.05926844 | 0.13534152 | NA | NA |
| ENSG000001 | 4.15241392 | -0.71286695 | 0.37795225 | -1.8861296 | 0.05927749 | 0.1353466 | MUC20 | 200958 |
| ENSG000002 | 6.54272483 | -0.690241942 | 0.36596204 | -1.8861026 | 0.05928113 | 0.1353466 | NA | NA |
| ENSG000002 | 8.27628084 | -0.682655655 | 0.36193565 | -1.8861244 | 0.05927819 | 0.1353466 | NA | NA |
| ENSG000002 | 14.0012067 | -0.58907144 | 0.31232903 | -1.8860605 | 0.0592868 | 0.1353466 | NA | NA |
| ENSG000002 | 6.71128783 | -0.696765442 | 0.3694298 | -1.8860564 | 0.05928736 | 0.1353466 | NA | NA |
| ENSG000002 | 770.896996 | 0.244254987 | 0.12951471 | 1.88592472 | 0.0593051 | 0.13537948 | ATXN2 | 6311 |
| ENSG000002 | 6.1867532 | -0.692597212 | 0.36729806 | -1.8856544 | 0.05934154 | 0.13545504 | NA | NA |
| ENSG000002 | 2.86932625 | -0.708604441 | 0.37583551 | -1.8854111 | 0.05937436 | 0.13552232 | NA | NA |
| ENSG000002 | 5.89997339 | -0.69302431 | 0.36762569 | -1.8851357 | 0.05941152 | 0.13559951 | NA | NA |
| ENSG000002 | 1076.34199 | 0.25055058 | 0.13291658 | 1.88502124 | 0.05942697 | 0.13562714 | MEX3A | 92312 |
| ENSG000002 | 12.6207032 | -0.605404851 | 0.32117543 | -1.8849663 | 0.05943439 | 0.13563645 | HOXA-AS3 | 100133311 |
| ENSG000001 | 7.05213747 | -0.680687197 | 0.36112906 | -1.8848862 | 0.0594452 | 0.13563993 | TMIE | 259236 |
| ENSG000002 | 10.3290805 | -0.679034459 | 0.36025329 | -1.8848807 | 0.05944595 | 0.13563993 | NA | NA |
| ENSG000002 | 6.45782814 | -0.700093984 | 0.37142344 | -1.8848945 | 0.05944409 | 0.13563993 | NA | NA |
| ENSG000002 | 2.31264387 | -0.688745707 | 0.36541882 | -1.8848118 | 0.05945525 | 0.13565351 | NA | NA |
| ENSG000001 | 92.6534187 | -0.31929909 | 0.16940956 | -1.8847761 | 0.05946007 | 0.13565688 | ONECUT2 | 9480 |
| ENSG000002 | 2.89431388 | -0.708391505 | 0.3758576 | -1.8847338 | 0.05946579 | 0.1356623 | UGT1A2P | 54580 |

| | | | | | | | | |
|---|---|---|---|---|---|---|---|---|
| ENSG000001 | 72.724919 | 0.352388524 | 0.18697649 | 1.88466757 | 0.05947473 | 0.13567506 | PDE6B | 5158 |
| ENSG000002 | 5.62965367 | -0.698937624 | 0.37087155 | -1.8845814 | 0.05948638 | 0.13569399 | LOC105373496 | 105373496 |
| ENSG000002 | 8.32008814 | -0.66701306 | 0.3539451 | -1.88451 | 0.05949603 | 0.13570837 | NA | NA |
| ENSG000002 | 2.70744251 | -0.696339907 | 0.36951855 | -1.8844518 | 0.05950389 | 0.13571103 | RNU6-35P | 100873750 |
| ENSG000002 | 3.8862315 | -0.71344461 | 0.37859226 | -1.884467 | 0.05950184 | 0.13571103 | NA | NA |
| ENSG000002 | 10.7645844 | -0.630397113 | 0.3345396 | -1.8843722 | 0.05951465 | 0.13572794 | LOC338963 | 338963 |
| ENSG000001 | 270.403167 | 0.285111052 | 0.15131601 | 1.8842094 | 0.05953666 | 0.13576939 | IFT122 | 55764 |
| ENSG000002 | 2.86665871 | -0.708554151 | 0.37605274 | -1.8841882 | 0.05953952 | 0.13576939 | NA | NA |
| ENSG000001 | 209.342623 | 0.276889794 | 0.14696461 | 1.88405758 | 0.05955719 | 0.13580204 | PRPF40B | 25766 |
| ENSG000001 | 16.685547 | -0.568462005 | 0.30172882 | -1.8840163 | 0.05956277 | 0.13580321 | BLK | 640 |
| ENSG000002 | 13.0602288 | -0.593560837 | 0.31505281 | -1.8840043 | 0.0595644 | 0.13580321 | NA | NA |
| ENSG000002 | 2.30362172 | -0.690510489 | 0.36652506 | -1.883938 | 0.05957336 | 0.13581602 | RN7SL211P | 106479299 |
| ENSG000002 | 3.28187745 | -0.710330838 | 0.37707456 | -1.8837941 | 0.05959283 | 0.13585276 | NA | NA |
| ENSG000002 | 32.7563435 | 0.468161363 | 0.24853481 | 1.88368527 | 0.05960756 | 0.13587106 | FAHD2CP | 729234 |
| ENSG000002 | 2.26560528 | -0.6884459 | 0.36547605 | -1.8836963 | 0.05960606 | 0.13587106 | NA | NA |
| ENSG000002 | 8.52205495 | -0.66163167 | 0.35124968 | -1.8836506 | 0.05961225 | 0.13587412 | NA | NA |
| ENSG000001 | 117.627151 | 0.301454219 | 0.16006486 | 1.88332547 | 0.05965627 | 0.13596681 | GLT8D2 | 83468 |
| ENSG000002 | 11.8178776 | -0.615362414 | 0.32676353 | -1.8832041 | 0.05967272 | 0.13598117 | NA | NA |
| ENSG000002 | 3.89582778 | -0.712950667 | 0.37858871 | -1.8831799 | 0.05967599 | 0.13598117 | NA | NA |
| ENSG000002 | 3.52235465 | -0.712639499 | 0.37841515 | -1.8832214 | 0.05967037 | 0.13598117 | NA | NA |
| ENSG000002 | 66.7757313 | 0.369463123 | 0.19618782 | 1.8832113 | 0.05967174 | 0.13598117 | MRPS17 | 51373 |
| ENSG000001 | 35.0649326 | -0.493317178 | 0.26197641 | -1.8830596 | 0.0596923 | 0.1360107 | CLNK | 116449 |
| ENSG000002 | 3.4834206 | -0.71271774 | 0.3785254 | -1.8828796 | 0.05971669 | 0.13605863 | ZNF735 | 730291 |
| ENSG000002 | 3.98340591 | -0.71270801 | 0.37854714 | -1.8827457 | 0.05973483 | 0.13609232 | NA | NA |
| ENSG000002 | 4.14100387 | -0.71152952 | 0.37793849 | -1.8826596 | 0.05974652 | 0.13610365 | NA | NA |
| ENSG000002 | 4.80251308 | -0.708686772 | 0.37642639 | -1.8826703 | 0.05974507 | 0.13610365 | NA | NA |
| ENSG000002 | 16.7792489 | -0.551361763 | 0.29287791 | -1.8825652 | 0.05975931 | 0.13612515 | NA | NA |
| ENSG000002 | 3.28368493 | -0.709864373 | 0.37708428 | -1.8825085 | 0.059767 | 0.13612886 | FAM66E | 100132103 |
| ENSG000002 | 2.86506009 | -0.707903391 | 0.37604356 | -1.8825037 | 0.05976766 | 0.13612886 | NA | NA |

| | | | | | | | | |
|---|---|---|---|---|---|---|---|---|
| ENSG000001 | 8.1970851 | -0.66332773 | 0.35237794 | -1.8824326 | 0.0597773 | 0.13614319 | EFCC1 | 79825 |
| ENSG000001 | 9.58139444 | -0.64442175 | 0.34235449 | -1.8823231 | 0.05979216 | 0.13616939 | SPERT | 220082 |
| ENSG000002 | 5.21487615 | -0.708002579 | 0.3761474 | -1.8822477 | 0.05980239 | 0.13618503 | B4GALT4-AS1 | 100874201 |
| ENSG000002 | 4.23316762 | -0.712489599 | 0.37857247 | -1.8820428 | 0.0598302 | 0.1362407 | NA | NA |
| ENSG000001 | 102.898467 | 0.309019869 | 0.16419621 | 1.88201589 | 0.05983386 | 0.13624138 | GRIA3 | 2892 |
| ENSG000002 | 37.9426093 | 0.451288511 | 0.23980343 | 1.88191014 | 0.05984822 | 0.13626643 | NA | NA |
| ENSG000002 | 2.66701398 | -0.698537715 | 0.37119595 | -1.881857 | 0.05985543 | 0.1362752 | NA | NA |
| ENSG000002 | 4.99859606 | -0.704190613 | 0.37421011 | -1.8818054 | 0.05986244 | 0.13628351 | NA | NA |
| ENSG000002 | 2.9034856 | -0.706969339 | 0.37570621 | -1.8817079 | 0.05987569 | 0.13630602 | NA | NA |
| ENSG000000 | 26.520942 | -0.483380918 | 0.25691348 | -1.881493 | 0.05990488 | 0.13636482 | CAMK1G | 57172 |
| ENSG000001 | 9.40488631 | -0.671736911 | 0.35704651 | -1.8813709 | 0.05992148 | 0.13639495 | HAPLN2 | 60484 |
| ENSG000001 | 582.061277 | 0.226177619 | 0.12022667 | 1.88126001 | 0.05993656 | 0.13642161 | NA | NA |
| ENSG000001 | 108.371123 | 0.356263046 | 0.18938184 | 1.88118902 | 0.05994621 | 0.13643592 | SLC18A2 | 6571 |
| ENSG000002 | 16.8317556 | -0.560338459 | 0.29789245 | -1.8810093 | 0.05997066 | 0.13648391 | LOC101927087 | 101927087 |
| ENSG000001 | 47.2262057 | -0.401014341 | 0.21320069 | -1.8809243 | 0.05998222 | 0.13649695 | KCNJ2 | 3759 |
| ENSG000001 | 22.934065 | -0.511088377 | 0.27172289 | -1.8809176 | 0.05998312 | 0.13649695 | LHCGR | 3973 |
| ENSG000001 | 8.64558141 | -0.665927548 | 0.35406206 | -1.8808215 | 0.0599962 | 0.13651905 | APOA5 | 116519 |
| ENSG000002 | 2.63779244 | -0.698079259 | 0.37122038 | -1.8804982 | 0.06004021 | 0.13661152 | NA | NA |
| ENSG000002 | 5.89474078 | -0.692734644 | 0.36838901 | -1.8804433 | 0.06004769 | 0.13662088 | NA | NA |
| ENSG000002 | 9.38681041 | -0.65225458 | 0.34687818 | -1.8803563 | 0.06005953 | 0.13664015 | NA | NA |
| ENSG000001 | 261.535297 | 0.26108717 | 0.13885905 | 1.88023159 | 0.06007652 | 0.13667114 | IRF6 | 3664 |
| ENSG000002 | 2.29345632 | -0.685930444 | 0.36483063 | -1.8801339 | 0.06008983 | 0.13669374 | NA | NA |
| ENSG000002 | 7.63144394 | -0.667210709 | 0.354901 | -1.8799911 | 0.0601093 | 0.13673036 | NA | NA |
| ENSG000002 | 3.53456194 | -0.711540527 | 0.37849884 | -1.8799015 | 0.0601215 | 0.13675046 | NA | NA |
| ENSG000001 | 78.8526375 | 0.358244726 | 0.19058041 | 1.8797563 | 0.0601413 | 0.13678781 | TMEM143 | 55260 |
| ENSG000001 | 411.721016 | 0.255378438 | 0.13587012 | 1.87957765 | 0.06016566 | 0.13683555 | DPH6 | 89978 |
| ENSG000001 | 9.82038457 | -0.637477252 | 0.33923922 | -1.8791379 | 0.06022567 | 0.13696435 | KLK3 | 354 |
| ENSG000002 | 8.56747777 | -0.662502988 | 0.35256376 | -1.8791012 | 0.06023067 | 0.13696803 | NA | NA |
| ENSG000002 | 10.6562548 | -0.626059596 | 0.33318884 | -1.8789933 | 0.06024541 | 0.13699387 | NA | NA |

| | | | | | | | | |
|---|---|---|---|---|---|---|---|---|
| ENSG000001 | 32.2372924 | -0.458839599 | 0.24420327 | -1.8789249 | 0.06025475 | 0.13699974 | SYNPR | 132204 |
| ENSG000002 | 8.7959301 | -0.652051267 | 0.34703156 | -1.8789394 | 0.06025277 | 0.13699974 | NA | NA |
| ENSG000002 | 3.92770968 | -0.711268438 | 0.37856519 | -1.8788533 | 0.06026453 | 0.1370143 | NA | NA |
| ENSG000002 | 4.10110862 | -0.710493242 | 0.37816667 | -1.8787834 | 0.06027408 | 0.13702833 | NA | NA |
| ENSG000001 | 10.0540796 | -0.629539849 | 0.33509718 | -1.8786784 | 0.06028841 | 0.13705322 | CLEC4C | 170482 |
| ENSG000002 | 2.29084139 | -0.688393407 | 0.36644654 | -1.8785644 | 0.060304 | 0.13708098 | NA | NA |
| ENSG000001 | 804.972779 | 0.561978406 | 0.29916131 | 1.87851299 | 0.06031102 | 0.13708157 | PELP1 | 27043 |
| ENSG000002 | 3.42598787 | -0.706109108 | 0.37588337 | -1.8785325 | 0.06030836 | 0.13708157 | LINC01047 | 105616982 |
| ENSG000002 | 2.32644181 | -0.68709314 | 0.36577064 | -1.8784809 | 0.06031541 | 0.13708386 | RN7SL654P | 106479461 |
| ENSG000001 | 81.1596723 | -0.342824325 | 0.18251221 | -1.8783638 | 0.06033142 | 0.13711255 | NPL | 80896 |
| ENSG000001 | 32.8835756 | -0.460669048 | 0.24525408 | -1.8783339 | 0.0603355 | 0.13711415 | ENOX1 | 55068 |
| ENSG000001 | 8.26944825 | -0.662887168 | 0.35292475 | -1.8782677 | 0.06034455 | 0.13712702 | NA | NA |
| ENSG000002 | 76.7385045 | 0.343599993 | 0.18294992 | 1.87810953 | 0.06036618 | 0.1371685 | NA | NA |
| ENSG000002 | 7.38948842 | -0.690045561 | 0.36742023 | -1.8780826 | 0.06036986 | 0.13716917 | GABARAPL3 | 23766 |
| ENSG000002 | 3.2361965 | -0.708448085 | 0.37723806 | -1.8779868 | 0.06038298 | 0.13719129 | NA | NA |
| ENSG000002 | 6.62180605 | -0.692121113 | 0.36857097 | -1.8778503 | 0.06040165 | 0.13722601 | NA | NA |
| ENSG000002 | 37.8266644 | 0.469868458 | 0.25022395 | 1.87779167 | 0.06040967 | 0.13723656 | NA | NA |
| ENSG000001 | 452.338482 | 0.239250797 | 0.1274206 | 1.87764607 | 0.0604296 | 0.13727414 | CIPC | 85457 |
| ENSG000001 | 47.0989698 | -0.401589053 | 0.21389415 | -1.8775131 | 0.06044781 | 0.13730781 | RFTN2 | 130132 |
| ENSG000002 | 166.031197 | 0.341519422 | 0.18191161 | 1.8773921 | 0.06046437 | 0.13733775 | NA | NA |
| ENSG000002 | 7.6060211 | 0.677398593 | 0.36084427 | 1.87726024 | 0.06048243 | 0.137348 | LOC101928402 | 101928402 |
| ENSG000002 | 6.5606297 | -0.685322091 | 0.36506447 | -1.8772632 | 0.06048203 | 0.137348 | LINC00911 | 100996280 |
| ENSG000002 | 3.56580059 | 0.70950742 | 0.37794613 | 1.87727129 | 0.06048092 | 0.137348 | NA | NA |
| ENSG000002 | 9.16614084 | -0.654722603 | 0.34875351 | -1.8773219 | 0.06047399 | 0.137348 | NA | NA |
| ENSG000001 | 62.0328262 | 0.413552276 | 0.22030394 | 1.87718967 | 0.0604921 | 0.13736226 | CXCR2 | 3579 |
| ENSG000002 | 5.87315848 | -0.692205812 | 0.36875594 | -1.8771381 | 0.06049917 | 0.13737062 | NA | NA |
| ENSG000001 | 316.238419 | 0.273493315 | 0.14570935 | 1.87697851 | 0.06052104 | 0.13741229 | ZNF256 | 10172 |
| ENSG000001 | 16.9741918 | -0.566122268 | 0.30161743 | -1.8769547 | 0.0605243 | 0.13741229 | CLEC1B | 51266 |
| ENSG000002 | 2.62621789 | -0.693555473 | 0.36953397 | -1.8768382 | 0.06054027 | 0.13742546 | NA | NA |

| | | | | | | | | | |
|---|---|---|---|---|---|---|---|---|---|
| ENSG000002 | 87.5291513 | 0.3457958 | 0.18424214 | 1.87685513 | 0.06053795 | 0.13742546 | C8orf58 | | 541565 |
| ENSG000002 | 72.8117896 | 0.373448771 | 0.19897533 | 1.8768597 | 0.06053733 | 0.13742546 | NA | NA | |
| ENSG000001 | 6.5798761 | -0.690762436 | 0.36805522 | -1.8767902 | 0.06054685 | 0.1374327 | WNT7B | | 7477 |
| ENSG000002 | 7.2186215 | -0.684355106 | 0.36464827 | -1.876754 | 0.06055181 | 0.13743628 | NA | NA | |
| ENSG000001 | 7.36773398 | -0.673614527 | 0.35893454 | -1.8767058 | 0.06055843 | 0.13744361 | NA | NA | |
| ENSG000002 | 5.41446912 | -0.700318725 | 0.37316917 | -1.8766789 | 0.06056211 | 0.13744426 | NA | NA | |
| ENSG000001 | 69.4207413 | 0.354964818 | 0.18915033 | 1.87662805 | 0.06056909 | 0.13745241 | IFI27L1 | | 122509 |
| ENSG000001 | 73.1846558 | 0.353692245 | 0.1884796 | 1.8765545 | 0.06057918 | 0.13745992 | KLHDC8B | | 200942 |
| ENSG000002 | 6.46668423 | -0.684225058 | 0.36461423 | -1.8765726 | 0.0605767 | 0.13745992 | NA | NA | |
| ENSG00000C | 80.725463 | 0.338103667 | 0.18018036 | 1.87647352 | 0.06059029 | 0.13746889 | CAMKK1 | | 84254 |
| ENSG000001 | 11.7490235 | -0.613974582 | 0.32719189 | -1.8764969 | 0.06058708 | 0.13746889 | LECT2 | | 3950 |
| ENSG000002 | 5.25306098 | 0.701307418 | 0.37374575 | 1.87642915 | 0.06059637 | 0.13746889 | NA | NA | |
| ENSG000002 | 3.23025948 | -0.707568103 | 0.3770827 | -1.8764269 | 0.06059669 | 0.13746889 | NA | NA | |
| ENSG000002 | 328.065831 | 0.235289934 | 0.12539771 | 1.87634955 | 0.0606073 | 0.13748526 | NA | NA | |
| ENSG000002 | 11.8044793 | -0.620944741 | 0.33094313 | -1.8762884 | 0.06061569 | 0.13749661 | NA | NA | |
| ENSG000001 | 102.457911 | 0.321408573 | 0.17131097 | 1.87617044 | 0.06063188 | 0.13752564 | TOMM5 | | 401505 |
| ENSG000002 | 2.89243451 | -0.704213998 | 0.37540189 | -1.8758936 | 0.06066989 | 0.13760416 | NA | NA | |
| ENSG000001 | 335.655453 | 0.236880006 | 0.12628322 | 1.87578363 | 0.060685 | 0.13762303 | HMBOX1 | | 79618 |
| ENSG000001 | 10.8064812 | -0.651643849 | 0.34739497 | -1.8758011 | 0.0606826 | 0.13762303 | SPDYE4 | | 388333 |
| ENSG000001 | 15.8883004 | 0.58319527 | 0.3109325 | 1.87563302 | 0.06070569 | 0.13765554 | TRIM61 | | 391712 |
| ENSG000002 | 3.56855189 | -0.709590127 | 0.37832098 | -1.8756299 | 0.06070612 | 0.13765554 | NA | NA | |
| ENSG000001 | 532.594211 | 0.586019337 | 0.3124494 | 1.87556559 | 0.06071495 | 0.13766787 | CSNK1D | | 1453 |
| ENSG000002 | 11.1812276 | -0.617718742 | 0.3293667 | -1.8754742 | 0.06072752 | 0.13768866 | NA | NA | |
| ENSG000001 | 186.456469 | 0.276114244 | 0.14723001 | 1.87539372 | 0.06073858 | 0.13769438 | IRF2 | | 3660 |
| ENSG000001 | 13.5986432 | -0.60592927 | 0.32309394 | -1.8753966 | 0.06073819 | 0.13769438 | GABRD | | 2563 |
| ENSG000002 | 4.82559018 | -0.705625115 | 0.3762568 | -1.8753817 | 0.06074023 | 0.13769438 | NA | NA | |
| ENSG000002 | 4.38250666 | -0.706184328 | 0.37658252 | -1.8752446 | 0.06075908 | 0.1377273 | NA | NA | |
| ENSG000002 | 31.2617662 | 0.493674745 | 0.26326137 | 1.8752267 | 0.06076154 | 0.1377273 | NA | NA | |
| ENSG000002 | 3.57628808 | -0.709212158 | 0.37821178 | -1.875172 | 0.06076906 | 0.13773664 | NA | NA | |

| | | | | | | | | |
|---|---|---|---|---|---|---|---|---|
| ENSG000002 | 3.60102485 | -0.708924629 | 0.37810866 | -1.874923 | 0.06080331 | 0.13779961 | NA | NA |
| ENSG000002 | 2.89258354 | -0.703726513 | 0.37534163 | -1.874896 | 0.06080704 | 0.13779961 | LCA10 | 100996465 |
| ENSG000002 | 19.4826351 | -0.533580415 | 0.28459042 | -1.8749065 | 0.06080559 | 0.13779961 | NA | NA |
| ENSG000001 | 9.63454921 | -0.645017662 | 0.34404235 | -1.8748205 | 0.06081742 | 0.13781544 | TMCO2 | 127391 |
| ENSG000002 | 2.22538527 | -0.685636418 | 0.36578306 | -1.8744346 | 0.06087055 | 0.13792812 | RNU6-59P | 100873767 |
| ENSG000001 | 18.0427917 | -0.551396422 | 0.29418743 | -1.8743031 | 0.06088865 | 0.13795386 | TNFAIP8L3 | 388121 |
| ENSG000002 | 70.3656539 | -0.349008712 | 0.18620723 | -1.8743027 | 0.06088871 | 0.13795386 | NA | NA |
| ENSG000002 | 3.56871606 | -0.709319968 | 0.37846441 | -1.8742052 | 0.06090215 | 0.13796889 | NA | NA |
| ENSG000002 | 2.98992212 | -0.702201988 | 0.37466524 | -1.8742117 | 0.06090125 | 0.13796889 | NA | NA |
| ENSG000000 | 8.17768053 | -0.664168054 | 0.35441198 | -1.874 | 0.06093042 | 0.13800981 | LHX5 | 64211 |
| ENSG000001 | 22.4181978 | 0.52463926 | 0.27995102 | 1.8740395 | 0.06092498 | 0.13800981 | CEND1 | 51286 |
| ENSG000002 | 2.88558457 | -0.704020206 | 0.3756734 | -1.874022 | 0.0609274 | 0.13800981 | NA | NA |
| ENSG000002 | 5.90295622 | -0.7023226 | 0.37478939 | -1.8739127 | 0.06094246 | 0.13802935 | NA | NA |
| ENSG000002 | 4.7909868 | -0.705138985 | 0.37630899 | -1.8738298 | 0.06095389 | 0.13804754 | NA | NA |
| ENSG000002 | 29.9637522 | -0.473917352 | 0.25293007 | -1.873709 | 0.06097054 | 0.13807753 | NA | NA |
| ENSG000001 | 123.622439 | 0.319899035 | 0.17074211 | 1.87358024 | 0.0609883 | 0.13811005 | ADGRB2 | 576 |
| ENSG000001 | 520.854097 | 0.594281972 | 0.31719885 | 1.87353132 | 0.06099505 | 0.13811346 | SEPHS2 | 22928 |
| ENSG000002 | 3.88134436 | -0.709248417 | 0.37856963 | -1.8734953 | 0.06100002 | 0.13811346 | NA | NA |
| ENSG000002 | 8.06439411 | -0.671336789 | 0.35833098 | -1.8735103 | 0.06099795 | 0.13811346 | NA | NA |
| ENSG000002 | 8.79121902 | -0.652780093 | 0.34844789 | -1.8733937 | 0.06101403 | 0.13813746 | C12orf74 | 338809 |
| ENSG000001 | 10.4409148 | -0.629128379 | 0.33584408 | -1.8732752 | 0.0610304 | 0.13814366 | NA | NA |
| ENSG000002 | 4.17609631 | -0.708226372 | 0.37806841 | -1.8732757 | 0.06103032 | 0.13814366 | NA | NA |
| ENSG000002 | 11.2312005 | -0.65441251 | 0.34933166 | -1.8733273 | 0.06102321 | 0.13814366 | NA | NA |
| ENSG000002 | 34.6414214 | -0.443869588 | 0.23694469 | -1.8733047 | 0.06102633 | 0.13814366 | RASSF5 | 83593 |
| ENSG000002 | 7.5936515 | -0.665415973 | 0.35522974 | -1.873199 | 0.06104092 | 0.13815975 | NA | NA |
| ENSG000001 | 27.3596367 | -0.487233295 | 0.26015112 | -1.8728857 | 0.06108418 | 0.13822982 | CERS3 | 204219 |
| ENSG000002 | 2.5397677 | -0.699059779 | 0.37324864 | -1.8729064 | 0.0610813 | 0.13822982 | SNORD115-17 | 100033455 |
| ENSG000002 | 16.6497193 | -0.56483835 | 0.30157967 | -1.8729324 | 0.06107771 | 0.13822982 | NA | NA |
| ENSG000002 | 11.3821697 | -0.629275563 | 0.33599425 | -1.872876 | 0.06108551 | 0.13822982 | NA | NA |

| | | | | | | | | |
|---|---|---|---|---|---|---|---|---|
| ENSG00000( | 257.715515 | 0.262127716 | 0.13996626 | 1.87279218 | 0.06109709 | 0.13824821 | KIF16B | 55614 |
| ENSG000002 | 3.25973344 | -0.705062994 | 0.37648179 | -1.8727678 | 0.06110045 | 0.13824821 | NA | NA |
| ENSG000002 | 4.40615625 | -0.70557116 | 0.37677184 | -1.8726749 | 0.06111329 | 0.13826955 | NA | NA |
| ENSG000002 | 8.60425205 | -0.663543243 | 0.35438741 | -1.8723669 | 0.06115587 | 0.13835815 | CH17-351M24.1 | 644767 |
| ENSG000001 | 9.13786049 | -0.658198884 | 0.35153962 | -1.872332 | 0.06116068 | 0.13835971 | CD3D | 915 |
| ENSG000002 | 7.77647492 | -0.674233241 | 0.36010721 | -1.8723125 | 0.06116338 | 0.13835971 | LINC01335 | 102503429 |
| ENSG000002 | 6.22515497 | -0.687685088 | 0.36734378 | -1.8720477 | 0.0612 | 0.13843483 | NA | NA |
| ENSG000001 | 2.26766788 | -0.685177005 | 0.36603379 | -1.8718955 | 0.06122107 | 0.13847476 | NA | NA |
| ENSG000001 | 115.533718 | 0.356728559 | 0.19057358 | 1.87186785 | 0.06122489 | 0.13847568 | SIDT2 | 51092 |
| ENSG000002 | 4.49094255 | -0.707608195 | 0.37802788 | -1.8718413 | 0.06122856 | 0.13847627 | LOC105372069 | 105372069 |
| ENSG000002 | 4.70908763 | -0.708544527 | 0.37855724 | -1.8716972 | 0.06124851 | 0.13851365 | NA | NA |
| ENSG000002 | 226.861627 | 0.251899722 | 0.13460679 | 1.8713746 | 0.06129318 | 0.13860694 | CSNK2B | 1460 |
| ENSG000001 | 17.9695871 | -0.549004941 | 0.29338025 | -1.8713085 | 0.06130234 | 0.13861992 | WISP3 | 8838 |
| ENSG000002 | 134.57044 | 0.286346087 | 0.15304726 | 1.87096519 | 0.06134991 | 0.13871653 | FAM185A | 222234 |
| ENSG000002 | 18.1666561 | 0.571829753 | 0.30563591 | 1.8709508 | 0.0613519 | 0.13871653 | NA | NA |
| ENSG000001 | 12.5550867 | -0.620174496 | 0.33155187 | -1.8705203 | 0.0614116 | 0.13884377 | CABP5 | 56344 |
| ENSG000002 | 7.87966061 | -0.658868175 | 0.35228357 | -1.8702779 | 0.06144524 | 0.13891207 | AGBL1-AS1 | 727915 |
| ENSG000002 | 6.35334988 | -0.691435186 | 0.36971048 | -1.8702072 | 0.06145506 | 0.13891878 | NA | NA |
| ENSG000002 | 11.4443804 | -0.648201811 | 0.34658982 | -1.8702275 | 0.06145223 | 0.13891878 | NA | NA |
| ENSG000001 | 46.1423279 | 0.42320959 | 0.22629802 | 1.8701427 | 0.061464 | 0.13893126 | GAL3ST1 | 9514 |
| ENSG000001 | 8.46095848 | -0.654533491 | 0.35002688 | -1.8699521 | 0.06149047 | 0.13897472 | SLC22A12 | 116085 |
| ENSG000002 | 5.00137571 | -0.700453394 | 0.37458799 | -1.8699302 | 0.06149351 | 0.13897472 | NA | NA |
| ENSG000002 | 3.30020194 | -0.704008947 | 0.37648883 | -1.8699332 | 0.0614931 | 0.13897472 | NA | NA |
| ENSG000001 | 51.077249 | 0.402405601 | 0.21521935 | 1.86974639 | 0.06151904 | 0.13900943 | SPIN2B | 474343 |
| ENSG000002 | 10.9552084 | -0.620467313 | 0.33184208 | -1.8697668 | 0.0615162 | 0.13900943 | NA | NA |
| ENSG000002 | 3.58491434 | -0.707529856 | 0.37840969 | -1.8697456 | 0.06151915 | 0.13900943 | LOC101929422 | 101929422 |
| ENSG000002 | 5.79457485 | -0.699658872 | 0.37421355 | -1.8696781 | 0.06152854 | 0.13902289 | NA | NA |
| ENSG000002 | 5.19630293 | -0.702880899 | 0.37597685 | -1.8694792 | 0.06155618 | 0.13907759 | NA | NA |
| ENSG000002 | 10.9214498 | -0.634066251 | 0.33918609 | -1.8693757 | 0.06157056 | 0.13910233 | LOC101927272 | 101927272 |

| | | | | | | | | |
|---|---|---|---|---|---|---|---|---|
| ENSG000002 | 3.58790197 | -0.706925268 | 0.37819225 | -1.869222 | 0.06159194 | 0.13914288 | NA | NA |
| ENSG000001 | 1000.23732 | 0.222907957 | 0.11925434 | 1.8691811 | 0.06159762 | 0.13914797 | ATAD2B | 54454 |
| ENSG000002 | 8.32193159 | -0.665245286 | 0.35595107 | -1.8689234 | 0.06163347 | 0.1392212 | NA | NA |
| ENSG000002 | 55.0389485 | 0.408876058 | 0.21878147 | 1.8688788 | 0.06163968 | 0.13922746 | NA | NA |
| ENSG000001 | 107.544561 | 0.325491427 | 0.17417377 | 1.86877412 | 0.06165425 | 0.13925261 | EPHB1 | 2047 |
| ENSG000000 | 126.769526 | 0.323590591 | 0.17316509 | 1.86868265 | 0.06166698 | 0.13927361 | PHF21B | 112885 |
| ENSG000002 | 7.86993317 | -0.659645997 | 0.35302516 | -1.8685524 | 0.06168512 | 0.13930681 | NA | NA |
| ENSG000002 | 6.20369163 | -0.687181536 | 0.36777398 | -1.8684887 | 0.06169398 | 0.13931907 | GS1-279B7.1 | 100288079 |
| ENSG000002 | 9.90617141 | -0.638856417 | 0.34193014 | -1.8683829 | 0.06170872 | 0.1393446 | LOC285766 | 285766 |
| ENSG000002 | 8.32044127 | -0.668792626 | 0.3579665 | -1.8683107 | 0.06171878 | 0.13935956 | NA | NA |
| ENSG000002 | 11.2531567 | -0.618090719 | 0.33083859 | -1.8682546 | 0.06172659 | 0.13936944 | BHMG1 | 388553 |
| ENSG000002 | 6.44019625 | -0.695924289 | 0.37250851 | -1.8682104 | 0.06173275 | 0.13937162 | NA | NA |
| ENSG000002 | 3.85478569 | -0.707287393 | 0.37859332 | -1.8681983 | 0.06173444 | 0.13937162 | NA | NA |
| ENSG000002 | 2.88162996 | -0.700585379 | 0.37501863 | -1.8681349 | 0.06174328 | 0.13938382 | NA | NA |
| ENSG000001 | 5.30269796 | -0.694347834 | 0.37168508 | -1.8681079 | 0.06174704 | 0.13938455 | ASB16 | 92591 |
| ENSG000002 | 3.36060262 | -0.703191366 | 0.37644262 | -1.8679908 | 0.06176335 | 0.13941362 | NA | NA |
| ENSG000002 | 2.52331092 | -0.67192632 | 0.35972221 | -1.8679033 | 0.06177555 | 0.13943338 | NA | NA |
| ENSG000001 | 19.5816962 | -0.535340434 | 0.2866054 | -1.8678658 | 0.06178077 | 0.13943742 | TGM4 | 7047 |
| ENSG000002 | 3.9450779 | -0.707059398 | 0.37854548 | -1.8678321 | 0.06178548 | 0.13943989 | NA | NA |
| ENSG000002 | 24.3019356 | 0.50937093 | 0.27271404 | 1.86778401 | 0.06179218 | 0.13943989 | NA | NA |
| ENSG000002 | 2.86457082 | -0.701697229 | 0.37568357 | -1.8677879 | 0.06179164 | 0.13943989 | NA | NA |
| ENSG000001 | 8.70662295 | -0.649572786 | 0.34778422 | -1.8677466 | 0.0617974 | 0.13944391 | SP7 | 121340 |
| ENSG000002 | 69.06425 | 0.366955775 | 0.19647904 | 1.86765861 | 0.06180967 | 0.13945607 | NA | NA |
| ENSG000002 | 5.61883538 | -0.69474529 | 0.37198275 | -1.8676815 | 0.06180649 | 0.13945607 | ANKRD20A3 | 441425 |
| ENSG000001 | 279.78638 | 0.24827357 | 0.13294255 | 1.86752522 | 0.06182828 | 0.13947632 | FAM149B1 | 317662 |
| ENSG000001 | 58.9591292 | -0.36130973 | 0.19347031 | -1.8675203 | 0.06182897 | 0.13947632 | HEPHL1 | 341208 |
| ENSG000002 | 5.03627227 | -0.699843293 | 0.37473788 | -1.8675542 | 0.06182424 | 0.13947632 | NA | NA |
| ENSG000001 | 11.5969409 | -0.621811332 | 0.33299709 | -1.8673176 | 0.06185724 | 0.13953235 | TMEM140 | 55281 |
| ENSG000002 | 14.0206688 | -0.586627606 | 0.3142203 | -1.866931 | 0.06191123 | 0.13964636 | ANHX | 647589 |

| | | | | | | | | |
|---|---|---|---|---|---|---|---|---|
| ENSG000001 | 2.22062984 | -0.683303913 | 0.36603426 | -1.8667758 | 0.0619329 | 0.13968747 | MC3R | 4159 |
| ENSG000002 | 22.0283574 | -0.534371711 | 0.28628317 | -1.8665844 | 0.06195965 | 0.13974002 | LINC00922 | 283867 |
| ENSG000001 | 6.8034303 | -0.677707769 | 0.3631214 | -1.8663394 | 0.0619939 | 0.1398095 | C1QA | 712 |
| ENSG000002 | 2.54064872 | -0.696205629 | 0.37304513 | -1.8662772 | 0.06200259 | 0.13982132 | NA | NA |
| ENSG000001 | 2814.48995 | 0.455049954 | 0.2438361 | 1.86621237 | 0.06201166 | 0.13982623 | KHDRBS1 | 10657 |
| ENSG000001 | 1130.15943 | 0.209349438 | 0.11218175 | 1.86616303 | 0.06201856 | 0.13982623 | VEZF1 | 7716 |
| ENSG000002 | 7.16038889 | -0.679195699 | 0.3639518 | -1.8661693 | 0.06201768 | 0.13982623 | NA | NA |
| ENSG000002 | 8.93907168 | -0.641912226 | 0.34396956 | -1.866189 | 0.06201492 | 0.13982623 | NA | NA |
| ENSG000001 | 9.16910438 | -0.636887988 | 0.3412958 | -1.866088 | 0.06202906 | 0.13984213 | SPACA3 | 124912 |
| ENSG000002 | 8.25155986 | -0.661382616 | 0.35443006 | -1.8660455 | 0.062035 | 0.13984774 | NA | NA |
| ENSG000002 | 6.28384349 | -0.688753468 | 0.36910663 | -1.8660014 | 0.06204118 | 0.13985389 | NA | NA |
| ENSG000001 | 5.39841355 | -0.698199431 | 0.37423614 | -1.8656655 | 0.06208819 | 0.13995209 | CIB4 | 130106 |
| ENSG000001 | 64.899852 | 0.38664908 | 0.2072523 | 1.86559611 | 0.06209789 | 0.13996619 | KPTN | 11133 |
| ENSG000002 | 7.30869224 | -0.667810044 | 0.35797007 | -1.8655471 | 0.06210475 | 0.13997387 | RPSAP52 | 204010 |
| ENSG000001 | 4.49208635 | -0.70442078 | 0.37760392 | -1.8655017 | 0.06211111 | 0.13998041 | LACRT | 90070 |
| ENSG000002 | 5.11420918 | -0.702145518 | 0.37640445 | -1.8654017 | 0.06212511 | 0.1400042 | NA | NA |
| ENSG000001 | 100.790092 | 0.319802592 | 0.1714463 | 1.86532222 | 0.06213625 | 0.14000677 | ZNF221 | 7638 |
| ENSG000002 | 2.53495965 | -0.695254368 | 0.37272402 | -1.8653329 | 0.06213476 | 0.14000677 | NA | NA |
| ENSG000002 | 3.2559769 | -0.703370115 | 0.37707753 | -1.8653196 | 0.06213662 | 0.14000677 | NA | NA |
| ENSG000002 | 7.15926243 | -0.676502765 | 0.36269684 | -1.8652017 | 0.06215314 | 0.14003621 | NA | NA |
| ENSG000001 | 8.15103221 | -0.657268345 | 0.35240241 | -1.8651074 | 0.06216635 | 0.14005821 | OR10G7 | 390265 |
| ENSG000002 | 6.85120812 | -0.681189084 | 0.3652367 | -1.865062 | 0.06217272 | 0.14006477 | LOC101060542 | 101060542 |
| ENSG000002 | 7.57297494 | -0.665561911 | 0.35687245 | -1.8649854 | 0.06218345 | 0.14008116 | LINC01514 | 101927396 |
| ENSG000002 | 6.92541598 | -0.684324285 | 0.36694003 | -1.8649486 | 0.06218861 | 0.14008502 | NA | NA |
| ENSG000002 | 11.2836236 | -0.65231359 | 0.34979383 | -1.8648516 | 0.0622022 | 0.14010785 | NA | NA |
| ENSG000002 | 4.81862236 | -0.702472722 | 0.37671733 | -1.864721 | 0.06222052 | 0.14014131 | NA | NA |
| ENSG000002 | 2.90194702 | -0.700280927 | 0.3755619 | -1.8646219 | 0.06223443 | 0.14016485 | NA | NA |
| ENSG000001 | 501.261319 | 0.216362354 | 0.11605245 | 1.8643497 | 0.06227261 | 0.14024307 | CRIM1 | 51232 |
| ENSG000002 | 4.38754284 | -0.702489705 | 0.37681069 | -1.8643041 | 0.06227902 | 0.1402497 | NA | NA |

| | | | | | | | | |
|---|---|---|---|---|---|---|---|---|
| ENSG000001 | 184.518713 | 0.351433654 | 0.18851804 | 1.86419108 | 0.06229488 | 0.14027763 | ZSWIM4 | 65249 |
| ENSG000002 | 6.47597937 | -0.681794089 | 0.36575732 | -1.8640613 | 0.06231309 | 0.14031085 | GS1-120K12.4 | 101929069 |
| ENSG000002 | 2.51258861 | -0.695255089 | 0.37298548 | -1.8640272 | 0.06231788 | 0.14031384 | NA | NA |
| ENSG000002 | 9.31618314 | -0.645249635 | 0.3461783 | -1.8639228 | 0.06233254 | 0.14032348 | NA | NA |
| ENSG000002 | 7.88723317 | -0.659048824 | 0.35358096 | -1.8639262 | 0.06233206 | 0.14032348 | NA | NA |
| ENSG000002 | 3.3184319 | -0.702142446 | 0.3766953 | -1.8639533 | 0.06232827 | 0.14032348 | NA | NA |
| ENSG000002 | 24.9262835 | -0.505972929 | 0.27146029 | -1.8638929 | 0.06233674 | 0.14032514 | NA | NA |
| ENSG000001 | 250.2063 | 0.304610279 | 0.16343824 | 1.86376378 | 0.06235488 | 0.14034387 | HSPA2 | 3306 |
| ENSG000002 | 5.23836529 | -0.702617509 | 0.37699051 | -1.8637538 | 0.06235628 | 0.14034387 | NA | NA |
| ENSG000002 | 2.23625453 | -0.68044209 | 0.36508636 | -1.8637839 | 0.06235205 | 0.14034387 | NA | NA |
| ENSG000002 | 3.34466472 | -0.701658337 | 0.37647964 | -1.8637352 | 0.0623589 | 0.14034387 | NA | NA |
| ENSG000002 | 7.57405259 | -0.663492436 | 0.35602953 | -1.8635882 | 0.06237956 | 0.14038256 | NA | NA |
| ENSG000002 | 2.96057137 | -0.699101671 | 0.37516061 | -1.863473 | 0.06239575 | 0.1404112 | NA | NA |
| ENSG000002 | 11.1978388 | -0.614857666 | 0.32996412 | -1.8634077 | 0.06240493 | 0.14042408 | NA | NA |
| ENSG000002 | 4.11060848 | -0.704533597 | 0.37809749 | -1.8633649 | 0.06241094 | 0.14042981 | NA | NA |
| ENSG000001 | 23.2256359 | 0.523037565 | 0.28069901 | 1.86333955 | 0.06241451 | 0.14043005 | NRROS | 375387 |
| ENSG000002 | 6.43489669 | -0.691477972 | 0.37111671 | -1.8632359 | 0.06242908 | 0.14045505 | NA | NA |
| ENSG000002 | 12.5144866 | 0.607382344 | 0.32599232 | 1.86317994 | 0.06243695 | 0.14046496 | NA | NA |
| ENSG000001 | 4.56021767 | 0.701353975 | 0.37643358 | 1.86315466 | 0.06244051 | 0.14046517 | NRGN | 4900 |
| ENSG000001 | 363.931213 | 0.236271585 | 0.12682124 | 1.86302852 | 0.06245825 | 0.1404895 | LUC7L2 | 51631 |
| ENSG000002 | 3.24229574 | -0.702299538 | 0.37696412 | -1.8630408 | 0.06245653 | 0.1404895 | NA | NA |
| ENSG000002 | 7.8223734 | -0.657322016 | 0.35283629 | -1.862966 | 0.06246705 | 0.14050149 | NA | NA |
| ENSG000001 | 9.02989258 | -0.642394658 | 0.3448326 | -1.8629174 | 0.06247389 | 0.14050907 | NA | NA |
| ENSG000002 | 4.9947418 | -0.69676736 | 0.37404338 | -1.8627983 | 0.06249066 | 0.14053899 | NA | NA |
| ENSG000002 | 12.9827402 | -0.608552939 | 0.32671414 | -1.8626465 | 0.06251202 | 0.14057924 | PCDHB17P | 54661 |
| ENSG000002 | 6.21246357 | -0.696192156 | 0.37377889 | -1.8625775 | 0.06252173 | 0.14059328 | NA | NA |
| ENSG000002 | 6.521348 | -0.680174866 | 0.36519432 | -1.8625012 | 0.06253247 | 0.14060964 | NA | NA |
| ENSG000002 | 5.15844327 | -0.700856 | 0.37632622 | -1.8623629 | 0.06255195 | 0.14064564 | NA | NA |
| ENSG000001 | 237.616351 | 0.265741285 | 0.14269832 | 1.86225935 | 0.06256654 | 0.14067065 | STXBP5L | 9515 |

| | | | | | | | | |
|---|---|---|---|---|---|---|---|---|
| ENSG000001 | 15.4109117 | -0.556572382 | 0.29888853 | -1.8621403 | 0.06258331 | 0.14068496 | TMEM74B | 55321 |
| ENSG000001 | 51.3521035 | 0.384070424 | 0.20625208 | 1.86214088 | 0.06258324 | 0.14068496 | VWA2 | 340706 |
| ENSG000002 | 7.41108463 | -0.672921491 | 0.36136553 | -1.8621629 | 0.06258013 | 0.14068496 | NA | NA |
| ENSG000002 | 3.54703016 | -0.704681451 | 0.37844799 | -1.8620298 | 0.06259888 | 0.14071216 | NA | NA |
| ENSG000002 | 4.49208491 | -0.703849208 | 0.3780064 | -1.8620034 | 0.06260261 | 0.14071273 | NA | NA |
| ENSG000001 | 2.90727047 | -0.698449972 | 0.37512371 | -1.861919 | 0.0626145 | 0.14073167 | MYOG | 4656 |
| ENSG000002 | 8.70893951 | -0.647542304 | 0.34779919 | -1.8618281 | 0.06262732 | 0.14075267 | NA | NA |
| ENSG000002 | 5.89260606 | -0.688324182 | 0.36973002 | -1.861694 | 0.06264623 | 0.14078737 | NA | NA |
| ENSG000002 | 5.02355176 | -0.696902258 | 0.37435458 | -1.8616101 | 0.06265807 | 0.14080617 | LINC00921 | 283876 |
| ENSG000002 | 4.09649754 | -0.703833136 | 0.37809168 | -1.8615409 | 0.06266782 | 0.14082029 | NA | NA |
| ENSG000002 | 18.492561 | -0.576722416 | 0.30985766 | -1.8612495 | 0.06270895 | 0.1409049 | NA | NA |
| ENSG000001 | 5403.26596 | 0.508361706 | 0.27314301 | 1.86115585 | 0.06272217 | 0.14092679 | LARP1 | 23367 |
| ENSG000002 | 2.23479972 | -0.67926864 | 0.36500528 | -1.860983 | 0.06274657 | 0.14097381 | NA | NA |
| ENSG000001 | 21.9519992 | -0.500446778 | 0.26892603 | -1.8609087 | 0.06275707 | 0.14098863 | XPNPEP2 | 7512 |
| ENSG000001 | 159.482981 | 0.279589603 | 0.15024533 | 1.86088711 | 0.06276012 | 0.14098863 | ZNF337 | 26152 |
| ENSG000001 | 157.837458 | 0.355264028 | 0.19092738 | 1.86072854 | 0.06278252 | 0.14102464 | GRIK5 | 2901 |
| ENSG000002 | 7.58376685 | -0.663452095 | 0.35655581 | -1.8607244 | 0.06278311 | 0.14102464 | LINC01422 | 101929539 |
| ENSG000001 | 56.4680972 | 0.397076706 | 0.21342292 | 1.86051577 | 0.06281259 | 0.14107112 | CCDC120 | 90060 |
| ENSG000002 | 2.53792059 | -0.692446103 | 0.37218198 | -1.8605041 | 0.06281424 | 0.14107112 | RN7SL645P | 106479457 |
| ENSG000002 | 26.0933223 | -0.521255071 | 0.28016448 | -1.8605323 | 0.06281025 | 0.14107112 | NA | NA |
| ENSG000001 | 96.9817551 | 0.396815322 | 0.21328918 | 1.86045686 | 0.06282092 | 0.14107831 | REC8 | 9985 |
| ENSG000002 | 11.9240326 | -0.596378052 | 0.3205841 | -1.8602858 | 0.0628451 | 0.14112481 | LOC100128770 | 100128770 |
| ENSG000001 | 11.4754539 | 0.612618406 | 0.32932216 | 1.86024045 | 0.06285151 | 0.14113139 | NA | NA |
| ENSG000002 | 6.00710528 | -0.694901263 | 0.37357038 | -1.8601616 | 0.06286267 | 0.14114862 | NA | NA |
| ENSG000002 | 4.72395188 | -0.700200556 | 0.37646807 | -1.8599202 | 0.06289682 | 0.14121748 | LINC01541 | 100505776 |
| ENSG000002 | 8.67611188 | -0.644901724 | 0.34675542 | -1.8598173 | 0.06291138 | 0.14124235 | NA | NA |
| ENSG000001 | 14.803523 | -0.579514782 | 0.31164216 | -1.8595519 | 0.06294894 | 0.14131104 | SLC6A19 | 340024 |
| ENSG000002 | 6.80178108 | -0.680250788 | 0.36581332 | -1.8595572 | 0.0629482 | 0.14131104 | NA | NA |
| ENSG000002 | 2.52052864 | -0.693019821 | 0.3726978 | -1.8594685 | 0.06296076 | 0.14131409 | NA | NA |

| | | | | | | | | |
|---|---|---|---|---|---|---|---|---|
| ENSG000002 | 69.8394985 | 0.352432525 | 0.18953297 | 1.85947872 | 0.06295931 | 0.14131409 | NA | NA |
| ENSG000002 | 2.27654621 | -0.679105075 | 0.36521172 | -1.8594833 | 0.06295867 | 0.14131409 | RN7SL303P | 106481005 |
| ENSG000002 | 2.89582021 | -0.697452962 | 0.37513175 | -1.8592213 | 0.06299577 | 0.14138485 | NA | NA |
| ENSG000002 | 2.90556084 | -0.697908961 | 0.3753868 | -1.8591729 | 0.06300264 | 0.14139244 | RNU6ATAC25P | 106480384 |
| ENSG000002 | 15.1624188 | -0.564048991 | 0.30341562 | -1.8589979 | 0.06302744 | 0.14142742 | LINC00473 | 90632 |
| ENSG000002 | 18.059664 | -0.550704774 | 0.29623193 | -1.8590325 | 0.06302253 | 0.14142742 | MGC27382 | 149047 |
| ENSG000002 | 25.3920781 | -0.479280019 | 0.25781755 | -1.8589891 | 0.06302869 | 0.14142742 | NA | NA |
| ENSG000002 | 5.02334738 | -0.697077888 | 0.37505216 | -1.8586158 | 0.06308161 | 0.14153834 | NA | NA |
| ENSG000002 | 3.20762786 | -0.700206906 | 0.37680276 | -1.858285 | 0.06312856 | 0.14163584 | NA | NA |
| ENSG000002 | 21.7410885 | 0.525942514 | 0.28303173 | 1.85824574 | 0.06313413 | 0.14164049 | NA | NA |
| ENSG000002 | 3.20729559 | -0.700374477 | 0.37693753 | -1.8580651 | 0.06315977 | 0.14169018 | NA | NA |
| ENSG000001 | 81.4754621 | -0.343286008 | 0.18476419 | -1.8579683 | 0.06317352 | 0.14171167 | PRKD1 | 5587 |
| ENSG000002 | 7.01260605 | -0.6684455 | 0.35977613 | -1.8579484 | 0.06317634 | 0.14171167 | NA | NA |
| ENSG000002 | 13.1482571 | -0.591420021 | 0.31833543 | -1.8578517 | 0.06319008 | 0.14173466 | LOC101928295 | 101928295 |
| ENSG000002 | 28.1939385 | -0.488093013 | 0.26273331 | -1.8577508 | 0.06320442 | 0.14175898 | NA | NA |
| ENSG000002 | 5.98861146 | -0.689259134 | 0.37103843 | -1.8576489 | 0.06321889 | 0.1417836 | NA | NA |
| ENSG000001 | 3.28684229 | -0.699913549 | 0.37678497 | -1.8575941 | 0.06322667 | 0.14179321 | SCGB1A1 | 7356 |
| ENSG000002 | 4.77428467 | -0.700206066 | 0.37695084 | -1.8575527 | 0.06323256 | 0.14179856 | HNRNPA1P49 | 106481663 |
| ENSG000002 | 5.085542 | -0.697004812 | 0.37526423 | -1.8573707 | 0.06325843 | 0.14184873 | NA | NA |
| ENSG000002 | 2.54292452 | -0.691577439 | 0.37235491 | -1.8573072 | 0.06326747 | 0.14186116 | NA | NA |
| ENSG000002 | 48.985851 | -0.384991236 | 0.20730523 | -1.8571227 | 0.06329371 | 0.14191215 | NOTCH4 | 4855 |
| ENSG000002 | 8.85325353 | -0.651603971 | 0.35087304 | -1.8570933 | 0.06329788 | 0.14191366 | NA | NA |
| ENSG000002 | 12.0042521 | -0.60568718 | 0.32615646 | -1.8570449 | 0.06330477 | 0.14192126 | LINC01215 | 101929623 |
| ENSG000001 | 12.4573311 | -0.610084653 | 0.32855219 | -1.8568881 | 0.06332707 | 0.14193987 | NR2F1 | 7025 |
| ENSG000002 | 3.30126011 | -0.69914242 | 0.3765018 | -1.8569431 | 0.06331925 | 0.14193987 | NA | NA |
| ENSG000002 | 3.31557455 | -0.699505831 | 0.37670142 | -1.8569238 | 0.06332199 | 0.14193987 | NA | NA |
| ENSG000002 | 14.5634052 | -0.581574629 | 0.31319563 | -1.8569053 | 0.06332463 | 0.14193987 | NA | NA |
| ENSG000001 | 4.96942191 | -0.695157486 | 0.37440629 | -1.8566928 | 0.06335487 | 0.14199432 | CLDN5 | 7122 |
| ENSG000002 | 7.78069653 | -0.670135609 | 0.36093506 | -1.8566653 | 0.06335879 | 0.14199526 | CTAGE11P | 647288 |

| | | | | | | | | |
|---|---|---|---|---|---|---|---|---|
| ENSG000002 | 40.1775909 | -0.413995771 | 0.2229828 | -1.8566264 | 0.06336432 | 0.14199981 | LOC349160 | 349160 |
| ENSG000001 | 1047.57228 | 0.207926525 | 0.11200111 | 1.85646846 | 0.06338681 | 0.14204237 | ZNF605 | 100289635 |
| ENSG000002 | 66.7347641 | 0.388899476 | 0.20951067 | 1.8562275 | 0.06342114 | 0.14211143 | SNORD14E | 85391 |
| ENSG000001 | 28.7281407 | -0.484418152 | 0.26100686 | -1.8559594 | 0.06345934 | 0.14216561 | CRYAB | 1410 |
| ENSG000002 | 24.8332413 | -0.51211061 | 0.27592137 | -1.856002 | 0.06345327 | 0.14216561 | NA | NA |
| ENSG000002 | 8.61573296 | -0.655131067 | 0.35297626 | -1.8560202 | 0.06345068 | 0.14216561 | NA | NA |
| ENSG000002 | 2.66745118 | -0.685764144 | 0.36949092 | -1.8559702 | 0.06345781 | 0.14216561 | RNU1-68P | 106481615 |
| ENSG000001 | 40.2255747 | -0.424832995 | 0.22892425 | -1.8557798 | 0.06348496 | 0.14219167 | KCP | 375616 |
| ENSG000002 | 9.07513158 | -0.647898197 | 0.34912109 | -1.855798 | 0.06348236 | 0.14219167 | NA | NA |
| ENSG000002 | 8.22134715 | -0.65677243 | 0.35390346 | -1.8557954 | 0.06348272 | 0.14219167 | NA | NA |
| ENSG000002 | 4.21261021 | -0.702139797 | 0.37835304 | -1.8557795 | 0.063485 | 0.14219167 | NA | NA |
| ENSG000001 | 3.5193618 | -0.702230255 | 0.37841822 | -1.8556989 | 0.06349649 | 0.14220956 | OR6C6 | 283365 |
| ENSG000002 | 5.28206934 | -0.690139461 | 0.37190862 | -1.8556694 | 0.06350069 | 0.14221063 | NA | NA |
| ENSG000002 | 6.12565505 | -0.679551959 | 0.36620769 | -1.8556463 | 0.06350398 | 0.14221063 | NA | NA |
| ENSG000001 | 64.5312496 | 0.405009623 | 0.21829372 | 1.85534252 | 0.06354733 | 0.14229984 | IGSF21 | 84966 |
| ENSG000002 | 4.45203253 | -0.700170017 | 0.37739283 | -1.8552817 | 0.06355601 | 0.14231141 | NA | NA |
| ENSG000002 | 2.64342965 | -0.686647592 | 0.37012271 | -1.855189 | 0.06356924 | 0.14233318 | PRAMEF5 | 343068 |
| ENSG000002 | 3.24343266 | -0.699379918 | 0.37699802 | -1.8551289 | 0.06357782 | 0.14234454 | NA | NA |
| ENSG000002 | 44.3609675 | -0.405402495 | 0.21853563 | -1.8550865 | 0.06358388 | 0.14235023 | NA | NA |
| ENSG000002 | 8.83594105 | -0.649226271 | 0.34997629 | -1.8550579 | 0.06358796 | 0.14235152 | NA | NA |
| ENSG000001 | 11.2424841 | -0.613508712 | 0.33075849 | -1.854854 | 0.06361708 | 0.14238724 | NMUR2 | 56923 |
| ENSG000002 | 5.6607585 | -0.691925657 | 0.37303635 | -1.8548478 | 0.06361796 | 0.14238724 | NA | NA |
| ENSG000002 | 16.3046724 | -0.55637645 | 0.29994702 | -1.8549158 | 0.06360826 | 0.14238724 | LOC440910 | 440910 |
| ENSG000002 | 2.87854538 | -0.696501358 | 0.37549801 | -1.8548736 | 0.06361427 | 0.14238724 | NA | NA |
| ENSG000002 | 4.36534751 | -0.699531899 | 0.37714997 | -1.8547844 | 0.06362702 | 0.14239964 | NA | NA |
| ENSG000002 | 6.7256237 | -0.690842526 | 0.37247347 | -1.8547429 | 0.06363295 | 0.14240506 | NA | NA |
| ENSG000001 | 7.04474683 | -0.687977404 | 0.37094302 | -1.8546714 | 0.06364317 | 0.14242007 | NA | NA |
| ENSG000001 | 11.7215555 | -0.628851998 | 0.33907074 | -1.8546337 | 0.06364856 | 0.14242427 | LRTM2 | 654429 |
| ENSG000002 | 2.71483412 | -0.68447736 | 0.36911292 | -1.8543847 | 0.06368414 | 0.14248031 | LCN8 | 138307 |

| | | | | | | | | |
|---|---|---|---|---|---|---|---|---|
| ENSG000002 | 2.9693971 | -0.692964441 | 0.37368626 | -1.8544017 | 0.06368171 | 0.14248031 | NA | NA |
| ENSG000002 | 4.04688064 | -0.70150772 | 0.37828679 | -1.8544335 | 0.06367716 | 0.14248031 | NA | NA |
| ENSG000002 | 7.69946292 | -0.663464164 | 0.35780095 | -1.8542829 | 0.0636987 | 0.14250502 | NA | NA |
| ENSG000002 | 8.19269921 | -0.671085419 | 0.36194555 | -1.854106 | 0.063724 | 0.14255375 | LOC101928797 | 101928797 |
| ENSG000001 | 1549.25814 | 0.232724094 | 0.12553807 | 1.85381285 | 0.06376594 | 0.14263971 | MPDZ | 8777 |
| ENSG000002 | 4.74261018 | -0.698318768 | 0.37675442 | -1.8535118 | 0.06380903 | 0.14272823 | NA | NA |
| ENSG000000 | 370.730112 | 0.241314637 | 0.13020209 | 1.85338521 | 0.06382717 | 0.14275342 | CELF2 | 10659 |
| ENSG000002 | 3.93236925 | -0.701622158 | 0.37856275 | -1.853384 | 0.06382733 | 0.14275342 | NA | NA |
| ENSG000002 | 5.05889126 | -0.695661853 | 0.37535728 | -1.8533325 | 0.06383471 | 0.14276205 | KRT128P | 106480238 |
| ENSG000002 | 3.92524666 | -0.700796257 | 0.37816669 | -1.8531412 | 0.06386213 | 0.14281548 | NA | NA |
| ENSG000001 | 4.52155975 | -0.700806662 | 0.37821107 | -1.8529512 | 0.06388935 | 0.14286061 | FAM181A | 90050 |
| ENSG000001 | 31.0337999 | -0.447871454 | 0.2417057 | -1.8529619 | 0.06388782 | 0.14286061 | TNFSF15 | 9966 |
| ENSG000001 | 221.587479 | 0.273242847 | 0.14746975 | 1.85287384 | 0.06390044 | 0.14287208 | TRIM32 | 22954 |
| ENSG000001 | 6.74374151 | -0.686900046 | 0.37072295 | -1.8528663 | 0.06390153 | 0.14287208 | SHH | 6469 |
| ENSG000001 | 28.5713584 | -0.479759659 | 0.25894982 | -1.8527128 | 0.06392353 | 0.14291339 | OTOA | 146183 |
| ENSG000002 | 160.019281 | 0.336340505 | 0.18154766 | 1.85262923 | 0.06393552 | 0.14293232 | ARPIN | 348110 |
| ENSG000002 | 17.8401651 | 0.54830884 | 0.29598084 | 1.85251467 | 0.06395195 | 0.14296117 | NA | NA |
| ENSG000002 | 3.15124547 | -0.699828846 | 0.37779248 | -1.852416 | 0.06396611 | 0.14298493 | NA | NA |
| ENSG000002 | 2.86084489 | -0.694783824 | 0.37507574 | -1.8523827 | 0.06397088 | 0.14298773 | NA | NA |
| ENSG000002 | 16.616708 | -0.571740878 | 0.30865976 | -1.8523337 | 0.06397792 | 0.14299556 | NA | NA |
| ENSG000001 | 97.6347211 | 0.360566178 | 0.1946604 | 1.85228311 | 0.06398518 | 0.14299603 | ENO3 | 2027 |
| ENSG000002 | 6.27894459 | -0.683389943 | 0.36894376 | -1.8522876 | 0.06398454 | 0.14299603 | NA | NA |
| ENSG000001 | 1033.56103 | 0.24606663 | 0.13285239 | 1.85218068 | 0.06399988 | 0.14300896 | UCHL5 | 51377 |
| ENSG000001 | 40.9883187 | 0.415905109 | 0.22454499 | 1.85221281 | 0.06399527 | 0.14300896 | ACBD4 | 79777 |
| ENSG000002 | 13.3374334 | -0.589451054 | 0.31824905 | -1.8521691 | 0.06400154 | 0.14300896 | NA | NA |
| ENSG000001 | 17.9454807 | -0.528736269 | 0.28549185 | -1.8520188 | 0.06402313 | 0.14304931 | THEMIS | 387357 |
| ENSG000002 | 16.477373 | 0.546227014 | 0.2949416 | 1.85198365 | 0.06402817 | 0.14305269 | LOC102724094 | 102724094 |
| ENSG000002 | 58.8576156 | 0.372740316 | 0.20127188 | 1.85192443 | 0.06403668 | 0.14306381 | NA | NA |
| ENSG000001 | 275.921153 | 0.241803412 | 0.13057119 | 1.85188951 | 0.06404169 | 0.1430641 | PXYLP1 | 92370 |

| | | | | | | | | |
|---|---|---|---|---|---|---|---|---|
| ENSG000002 | 48.1984334 | 0.401880882 | 0.21701303 | 1.8518744 | 0.06404386 | 0.1430641 | PTENP1 | 11191 |
| ENSG000002 | 2.54278868 | -0.689842363 | 0.37253418 | -1.8517559 | 0.06406089 | 0.14309425 | NA | NA |
| ENSG000002 | 5.69123428 | -0.689535432 | 0.37238116 | -1.8516926 | 0.06406998 | 0.14310669 | NA | NA |
| ENSG000002 | 8.22811574 | -0.671182396 | 0.36248497 | -1.8516144 | 0.06408121 | 0.14312389 | NA | NA |
| ENSG000002 | 5.88206907 | -0.685575821 | 0.37026455 | -1.8515837 | 0.06408562 | 0.14312584 | NA | NA |
| ENSG000002 | 8.64257763 | -0.643397684 | 0.34749813 | -1.8515141 | 0.06409563 | 0.14314032 | NA | NA |
| ENSG000002 | 23.7720689 | -0.493300842 | 0.26646818 | -1.8512561 | 0.06413271 | 0.14321525 | NA | NA |
| ENSG000001 | 3.79100462 | -0.700729168 | 0.37852419 | -1.8512137 | 0.06413882 | 0.14322099 | NA | NA |
| ENSG000001 | 6.48356024 | -0.678073932 | 0.36636694 | -1.8508055 | 0.06419754 | 0.14334423 | TEX33 | 339669 |
| ENSG000002 | 4.16663642 | -0.700247615 | 0.37836936 | -1.8506985 | 0.06421294 | 0.14337071 | NA | NA |
| ENSG00000C | 15.4211376 | -0.55960371 | 0.30240009 | -1.8505408 | 0.06423564 | 0.14341351 | SLC6A7 | 6534 |
| ENSG000001 | 9.4426391 | -0.644521617 | 0.34829776 | -1.8504903 | 0.06424292 | 0.14342186 | KCTD4 | 386618 |
| ENSG000002 | 5.06751752 | -0.693402191 | 0.37473482 | -1.850381 | 0.06425865 | 0.14344909 | NA | NA |
| ENSG000001 | 42.6701875 | 0.404133076 | 0.21843187 | 1.85015622 | 0.06429104 | 0.14350409 | CLDN15 | 24146 |
| ENSG000001 | 14.9600879 | -0.575973122 | 0.31131389 | -1.8501363 | 0.06429391 | 0.14350409 | NA | NA |
| ENSG000001 | 12.2441994 | -0.596792899 | 0.3225598 | -1.8501776 | 0.06428796 | 0.14350409 | CYP51A1-AS1 | 613126 |
| ENSG000002 | 45.8700457 | 0.409804678 | 0.22151105 | 1.85004171 | 0.06430754 | 0.14352661 | RBM26-AS1 | 100505538 |
| ENSG000002 | 6.19537735 | -0.678843003 | 0.36696777 | -1.8498709 | 0.06433216 | 0.14357365 | NA | NA |
| ENSG000001 | 15.6719622 | -0.558978138 | 0.30218445 | -1.8497912 | 0.06434365 | 0.14359139 | USP32P1 | 162632 |
| ENSG000002 | 7.3457415 | -0.666770522 | 0.3604636 | -1.8497583 | 0.0643484 | 0.1435941 | CACNA2D3-AS1 | 100874237 |
| ENSG000001 | 7.66118681 | -0.669473973 | 0.36198411 | -1.8494568 | 0.06439188 | 0.14368322 | MTNR1B | 4544 |
| ENSG000002 | 2.60824417 | -0.681750581 | 0.36863585 | -1.8493877 | 0.06440186 | 0.14369757 | UBE2Q2L | 100505679 |
| ENSG000002 | 7.25220674 | -0.681066686 | 0.36834719 | -1.8489803 | 0.06446066 | 0.14382087 | NA | NA |
| ENSG000001 | 4.68960985 | -0.696557226 | 0.3767469 | -1.8488732 | 0.06447613 | 0.14384746 | NA | NA |
| ENSG000001 | 16.8830028 | -0.555625494 | 0.30055695 | -1.848653 | 0.06450794 | 0.14391051 | EDAR | 10913 |
| ENSG000001 | 373.809424 | 0.243271096 | 0.13160676 | 1.84846957 | 0.06453444 | 0.14396172 | PCDHB2 | 56133 |
| ENSG000001 | 135.415091 | 0.286025919 | 0.15476736 | 1.84810228 | 0.06458755 | 0.14407226 | C11orf54 | 28970 |
| ENSG000001 | 56.566888 | -0.376645321 | 0.20380931 | -1.8480281 | 0.06459828 | 0.14408036 | SPATA6L | 55064 |
| ENSG000001 | 131.452921 | 0.281216483 | 0.15217006 | 1.84804077 | 0.06459644 | 0.14408036 | C1S | 716 |

| | | | | | | | | |
|---|---|---|---|---|---|---|---|---|
| ENSG000002 | 10.8939717 | -0.610004678 | 0.33009434 | -1.8479707 | 0.06460658 | 0.14409094 | NA | NA |
| ENSG000002 | 6.0300453 | -0.685891916 | 0.37116907 | -1.8479231 | 0.06461346 | 0.14409836 | DNAH10OS | 642797 |
| ENSG000002 | 9.37727151 | 0.63813574 | 0.34534168 | 1.84783873 | 0.06462568 | 0.14411768 | NA | NA |
| ENSG00000( | 27.9843415 | 0.490102479 | 0.26527664 | 1.84751463 | 0.06467259 | 0.14421436 | HSD3B7 | 80270 |
| ENSG000002 | 145.399555 | 0.293575168 | 0.15890602 | 1.84747666 | 0.06467809 | 0.14421869 | NA | NA |
| ENSG00000( | 15.1160071 | -0.594252776 | 0.32166998 | -1.8473989 | 0.06468934 | 0.14422 | DHRS9 | 10170 |
| ENSG000001 | 12.3199843 | -0.603282591 | 0.32654974 | -1.8474447 | 0.06468271 | 0.14422 | SERHL2 | 253190 |
| ENSG000002 | 6.17668016 | -0.67700638 | 0.36646017 | -1.8474215 | 0.06468608 | 0.14422 | NA | NA |
| ENSG000002 | 3.90926099 | -0.699332781 | 0.37856711 | -1.8473152 | 0.06470147 | 0.14423883 | DNM3-IT1 | 100874284 |
| ENSG000002 | 4.85045954 | -0.698069055 | 0.3778929 | -1.8472669 | 0.06470846 | 0.14423883 | LINC01511 | 100506791 |
| ENSG000002 | 2.83738095 | 0.688645726 | 0.37279115 | 1.84726953 | 0.06470809 | 0.14423883 | NA | NA |
| ENSG000002 | 7.90159894 | -0.657527003 | 0.35597732 | -1.8471036 | 0.06473212 | 0.14428364 | ADORA3 | 140 |
| ENSG000002 | 5.55797466 | -0.686506295 | 0.37167716 | -1.84705 | 0.0647399 | 0.14429304 | NA | NA |
| ENSG00000( | 54.231085 | -0.384669022 | 0.20827504 | -1.8469281 | 0.06475756 | 0.14430862 | CROCCP3 | 114819 |
| ENSG000002 | 2.86927418 | -0.692314913 | 0.37484358 | -1.8469435 | 0.06475533 | 0.14430862 | NA | NA |
| ENSG000002 | 12.4085316 | -0.585944797 | 0.31725072 | -1.8469455 | 0.06475504 | 0.14430862 | NA | NA |
| ENSG000001 | 12.7247295 | -0.607637915 | 0.32901431 | -1.8468434 | 0.06476984 | 0.14431598 | OTOL1 | 131149 |
| ENSG000002 | 2.50524886 | -0.687241454 | 0.3721229 | -1.8468131 | 0.06477423 | 0.14431598 | NA | NA |
| ENSG000002 | 2.50524886 | -0.687241454 | 0.3721229 | -1.8468131 | 0.06477423 | 0.14431598 | NA | NA |
| ENSG000002 | 6.7775456 | -0.675329677 | 0.36567418 | -1.8468071 | 0.0647751 | 0.14431598 | PACRG-AS1 | 285796 |
| ENSG000002 | 5.82768686 | -0.692872658 | 0.37518903 | -1.8467295 | 0.06478635 | 0.14433311 | NA | NA |
| ENSG000002 | 2.29881367 | -0.672621521 | 0.36423482 | -1.84667 | 0.06479498 | 0.14434441 | NA | NA |
| ENSG000002 | 8.30929315 | -0.640715209 | 0.34696646 | -1.84662 | 0.06480223 | 0.14435264 | RN7SL138P | 106480489 |
| ENSG000001 | 9.73722323 | -0.642489289 | 0.34794457 | -1.8465277 | 0.06481562 | 0.14437308 | NLRP6 | 171389 |
| ENSG000002 | 7.02874724 | -0.66674836 | 0.36108617 | -1.8465076 | 0.06481853 | 0.14437308 | NA | NA |
| ENSG000002 | 2.74569121 | -0.677935144 | 0.36717228 | -1.846368 | 0.0648388 | 0.14441028 | NA | NA |
| ENSG000001 | 406.762382 | 0.236787962 | 0.1282534 | 1.84625098 | 0.06485577 | 0.14442125 | STK38 | 11329 |
| ENSG000001 | 43.6378287 | -0.431150064 | 0.23352115 | -1.8462998 | 0.06484868 | 0.14442125 | MCMDC2 | 157777 |
| ENSG000002 | 240.628046 | 0.271647503 | 0.14713651 | 1.84622773 | 0.06485915 | 0.14442125 | HSPA1A | 3303 |

| | | | | | | | | |
|---|---|---|---|---|---|---|---|---|
| ENSG000002 | 14.3917078 | -0.585291266 | 0.3170229 | -1.8462113 | 0.06486153 | 0.14442125 | NA | NA |
| ENSG000002 | 248.112263 | 0.629295624 | 0.3408521 | 1.84624246 | 0.06485701 | 0.14442125 | HMBS | 3145 |
| ENSG000002 | 4.19158461 | -0.698645754 | 0.37845037 | -1.8460697 | 0.06488209 | 0.1444591 | NA | NA |
| ENSG000002 | 3.54313504 | -0.698003267 | 0.37812265 | -1.8459706 | 0.06489648 | 0.14448321 | NA | NA |
| ENSG000002 | 3.60491731 | -0.697831738 | 0.37807333 | -1.8457576 | 0.0649274 | 0.14454412 | NA | NA |
| ENSG000002 | 3.22592806 | -0.695027985 | 0.37656194 | -1.8457202 | 0.06493284 | 0.14454828 | NA | NA |
| ENSG000002 | 4.09059941 | -0.69831234 | 0.37834852 | -1.8456854 | 0.0649379 | 0.14455161 | NA | NA |
| ENSG000002 | 10.0028343 | -0.62227052 | 0.33719279 | -1.8454443 | 0.06497293 | 0.14462166 | NA | NA |
| ENSG000001 | 510.377137 | 0.264821478 | 0.14350915 | 1.84532816 | 0.06498982 | 0.1446513 | GNG4 | 2786 |
| ENSG000001 | 2.60287682 | -0.682380201 | 0.36983614 | -1.8450879 | 0.06502475 | 0.14472111 | CRNN | 49860 |
| ENSG000001 | 416.725117 | 0.262430878 | 0.14224221 | 1.84495781 | 0.06504368 | 0.14475529 | DDX58 | 23586 |
| ENSG000002 | 5.42079017 | -0.692085619 | 0.37514313 | -1.8448575 | 0.06505827 | 0.14477983 | NA | NA |
| ENSG000002 | 3.7910641 | -0.698263711 | 0.37852542 | -1.8446944 | 0.06508201 | 0.14482471 | NA | NA |
| ENSG000002 | 8.43091542 | -0.649136463 | 0.3519043 | -1.8446392 | 0.06509004 | 0.14483463 | NA | NA |
| ENSG000002 | 4.20157285 | -0.697983552 | 0.37839829 | -1.8445738 | 0.06509957 | 0.14484712 | MIR4682 | 100616322 |
| ENSG000002 | 6.95778323 | -0.678044253 | 0.36759301 | -1.8445515 | 0.0651028 | 0.14484712 | LOC63930 | 63930 |
| ENSG000001 | 95.4005053 | 0.331500913 | 0.17973299 | 1.84440771 | 0.06512374 | 0.14488577 | RABIF | 5877 |
| ENSG000002 | 3.8911949 | -0.697762327 | 0.37831809 | -1.84438 | 0.06512777 | 0.14488679 | NA | NA |
| ENSG000002 | 5.5520382 | -0.685081566 | 0.37148047 | -1.8441927 | 0.06515506 | 0.14493954 | NA | NA |
| ENSG000001 | 16.7793846 | -0.550858142 | 0.29872912 | -1.8440055 | 0.06518234 | 0.14499227 | CA7 | 766 |
| ENSG000002 | 4.5099865 | -0.697388454 | 0.3782276 | -1.8438328 | 0.06520751 | 0.14503236 | NA | NA |
| ENSG000002 | 3.76823605 | -0.697941132 | 0.37852387 | -1.8438497 | 0.06520504 | 0.14503236 | NA | NA |
| ENSG000001 | 14.5401519 | -0.600354296 | 0.32566591 | -1.8434668 | 0.06526088 | 0.14513514 | TLR1 | 7096 |
| ENSG000002 | 7.699547 | -0.677811071 | 0.36768067 | -1.8434776 | 0.06525931 | 0.14513514 | LAMP5-AS1 | 101929329 |
| ENSG000002 | 9.80209398 | -0.625281391 | 0.33923902 | -1.8431883 | 0.06530152 | 0.14521755 | NA | NA |
| ENSG000001 | 7.05333046 | -0.664153999 | 0.36034951 | -1.8430828 | 0.06531692 | 0.14523586 | CIB3 | 117286 |
| ENSG000001 | 5.81152111 | -0.679845864 | 0.36886323 | -1.8430839 | 0.06531677 | 0.14523586 | ZXDA | 7789 |
| ENSG000002 | 3.72457946 | -0.692757957 | 0.3758848 | -1.8430061 | 0.06532813 | 0.14524485 | NA | NA |
| ENSG000002 | 54.372438 | 0.390223363 | 0.21172972 | 1.84302592 | 0.06532523 | 0.14524485 | NA | NA |

| | | | | | | | | |
|---|---|---|---|---|---|---|---|---|
| ENSG000002 | 7.55241127 | -0.671505653 | 0.36436235 | -1.8429611 | 0.06533469 | 0.14524968 | NA | NA |
| ENSG000002 | 4.11921404 | -0.697218287 | 0.3783231 | -1.8429176 | 0.06534104 | 0.14524968 | OR2F2 | 135948 |
| ENSG000002 | 2.95350174 | -0.687986931 | 0.37331233 | -1.8429258 | 0.06533984 | 0.14524968 | NA | NA |
| ENSG000001 | 153.706645 | 0.293940309 | 0.15953222 | 1.8425137 | 0.06540004 | 0.14537287 | FAM173B | 134145 |
| ENSG00000C | 7.06112668 | -0.66429271 | 0.36054955 | -1.842445 | 0.06541009 | 0.14537926 | MEOX1 | 4222 |
| ENSG000002 | 3.78565523 | -0.697501416 | 0.37857229 | -1.8424524 | 0.06540901 | 0.14537926 | RN7SL177P | 106480969 |
| ENSG000002 | 5.42372702 | -0.690780764 | 0.37494832 | -1.8423359 | 0.06542603 | 0.14539296 | NA | NA |
| ENSG000002 | 12.5982387 | -0.595560464 | 0.32326495 | -1.8423292 | 0.06542701 | 0.14539296 | NA | NA |
| ENSG000002 | 7.3269307 | -0.675652678 | 0.36673544 | -1.8423436 | 0.06542491 | 0.14539296 | NA | NA |
| ENSG000001 | 10.1396642 | -0.629044108 | 0.3414638 | -1.8421985 | 0.06544611 | 0.14541947 | NA | NA |
| ENSG000001 | 16.3283846 | -0.558850503 | 0.3033593 | -1.8422066 | 0.06544494 | 0.14541947 | RNASE9 | 390443 |
| ENSG000002 | 8.47076983 | 0.646622424 | 0.35106035 | 1.84191242 | 0.06548796 | 0.14550448 | NA | NA |
| ENSG000002 | 4.7358269 | -0.693728499 | 0.37663993 | -1.8418878 | 0.06549156 | 0.14550451 | NA | NA |
| ENSG000002 | 6.52354867 | -0.67830563 | 0.36830787 | -1.841681 | 0.06552182 | 0.14556376 | NA | NA |
| ENSG000002 | 4.42052255 | -0.694814359 | 0.37728266 | -1.8416281 | 0.06552956 | 0.14557299 | PGA5 | 5222 |
| ENSG000001 | 74.1549878 | 0.389260731 | 0.21137794 | 1.84153902 | 0.06554261 | 0.14559136 | ABHD11 | 83451 |
| ENSG000001 | 116.116509 | 0.290837171 | 0.15793299 | 1.84152259 | 0.06554501 | 0.14559136 | EFNB3 | 1949 |
| ENSG000002 | 5.09584269 | -0.691954303 | 0.37575669 | -1.8414956 | 0.06554896 | 0.14559215 | LOC100506585 | 100506585 |
| ENSG000002 | 8.33143327 | -0.658307903 | 0.35749244 | -1.8414596 | 0.06555423 | 0.14559588 | NA | NA |
| ENSG000002 | 5.50546725 | -0.680897992 | 0.36981873 | -1.8411669 | 0.06559711 | 0.14567516 | MIR670HG | 100507261 |
| ENSG000002 | 5.02466166 | -0.689688287 | 0.374589 | -1.8411867 | 0.0655942 | 0.14567516 | SMIM22 | 440335 |
| ENSG000002 | 3.2732925 | -0.692853212 | 0.37633866 | -1.8410365 | 0.06561621 | 0.14570959 | NA | NA |
| ENSG000001 | 23.7173975 | -0.513788898 | 0.2791136 | -1.8407878 | 0.06565266 | 0.14577969 | CD300E | 342510 |
| ENSG000002 | 15.4157073 | -0.561878683 | 0.30524076 | -1.8407721 | 0.06565496 | 0.14577969 | NA | NA |
| ENSG000001 | 49.5687971 | -0.396825155 | 0.2156061 | -1.8405099 | 0.06569342 | 0.14584909 | PHEX | 5251 |
| ENSG000002 | 2.8567746 | -0.690342013 | 0.37508079 | -1.8405155 | 0.06569259 | 0.14584909 | RN7SKP35 | 106479111 |
| ENSG000002 | 7.63302883 | -0.660994072 | 0.35914211 | -1.8404806 | 0.06569771 | 0.14585065 | NA | NA |
| ENSG000002 | 8.71312508 | -0.658755304 | 0.35794201 | -1.8403968 | 0.06571001 | 0.14586996 | NA | NA |
| ENSG000002 | 12.2817956 | -0.597227028 | 0.32452358 | -1.8403193 | 0.06572137 | 0.1458792 | NA | NA |

| | | | | | | | | |
|---|---|---|---|---|---|---|---|---|
| ENSG000002 | 26.312353 | 0.491731172 | 0.26719546 | 1.84034251 | 0.06571797 | 0.1458792 | TXNDC5 | 81567 |
| ENSG000002 | 7.84416433 | -0.650650523 | 0.35357815 | -1.8401887 | 0.06574053 | 0.14591376 | NA | NA |
| ENSG000000 | 58.4179277 | 0.392263315 | 0.21319866 | 1.83989575 | 0.06578354 | 0.14599995 | CHRNA3 | 1136 |
| ENSG000002 | 45.0683926 | -0.409157462 | 0.22238327 | -1.8398752 | 0.06578657 | 0.14599995 | NA | NA |
| ENSG000002 | 15.9471903 | 0.550717453 | 0.29933311 | 1.83981468 | 0.06579545 | 0.14601167 | RPL23P8 | 222901 |
| ENSG000002 | 3.5480395 | -0.695247984 | 0.37791796 | -1.8396796 | 0.06581528 | 0.1460477 | NA | NA |
| ENSG000002 | 13.9829115 | -0.582137966 | 0.31644564 | -1.8396144 | 0.06582486 | 0.14606096 | CASC8 | 727677 |
| ENSG000002 | 2.52349012 | -0.684927252 | 0.37233204 | -1.8395603 | 0.06583281 | 0.1460706 | NA | NA |
| ENSG000001 | 16.3498313 | -0.55604938 | 0.30228902 | -1.8394627 | 0.06584716 | 0.14609444 | DMRTC2 | 63946 |
| ENSG000002 | 15.2141676 | -0.564515134 | 0.30689827 | -1.8394211 | 0.06585328 | 0.14610003 | NA | NA |
| ENSG000002 | 275.939089 | 0.261505925 | 0.14218671 | 1.83917272 | 0.06588979 | 0.14616503 | NA | NA |
| ENSG000002 | 8.02189339 | -0.645699369 | 0.351079 | -1.8391854 | 0.06588793 | 0.14616503 | NA | NA |
| ENSG000002 | 2.51139493 | -0.685009942 | 0.37247507 | -1.839076 | 0.06590402 | 0.1461886 | NA | NA |
| ENSG000002 | 9.68880852 | -0.618401216 | 0.33630359 | -1.8388183 | 0.06594191 | 0.14626046 | NA | NA |
| ENSG000002 | 2.89167463 | 0.686043809 | 0.37309186 | 1.83880667 | 0.06594363 | 0.14626046 | NA | NA |
| ENSG000002 | 2.28592179 | -0.668461596 | 0.36354503 | -1.8387312 | 0.06595473 | 0.14627709 | NA | NA |
| ENSG000002 | 4.14416068 | -0.695743458 | 0.37840971 | -1.8385983 | 0.06597429 | 0.14631246 | LOC100129027 | 100129027 |
| ENSG000002 | 6.81081184 | -0.671652018 | 0.36532416 | -1.8385097 | 0.06598733 | 0.14632581 | NA | NA |
| ENSG000002 | 5.26706975 | -0.68535503 | 0.37277775 | -1.8385084 | 0.06598752 | 0.14632581 | NA | NA |
| ENSG000001 | 3.1690818 | -0.694256392 | 0.3776408 | -1.8384041 | 0.06600288 | 0.14635187 | ELL3 | 80237 |
| ENSG000001 | 16263.7235 | 0.463372516 | 0.25209048 | 1.83811989 | 0.06604474 | 0.14643667 | HNRNPA1 | 3178 |
| ENSG000001 | 9.33334408 | -0.643023645 | 0.34985055 | -1.8379953 | 0.0660631 | 0.14646137 | FHL5 | 9457 |
| ENSG000002 | 4.42320384 | -0.693146773 | 0.37711843 | -1.8380082 | 0.0660612 | 0.14646137 | NA | NA |
| ENSG000001 | 65.0549633 | 0.405257273 | 0.22049618 | 1.83793333 | 0.06607223 | 0.14647359 | UBXN11 | 91544 |
| ENSG000001 | 79.3056888 | -0.329163795 | 0.17909761 | -1.8379017 | 0.0660769 | 0.14647593 | XRRA1 | 143570 |
| ENSG000001 | 441.010412 | 0.254721009 | 0.13859863 | 1.83783211 | 0.06608715 | 0.14649065 | ZNF567 | 163081 |
| ENSG000001 | 49.1626042 | -0.389935783 | 0.2121908 | -1.8376659 | 0.06611166 | 0.14653268 | TMEM255A | 55026 |
| ENSG000002 | 9.72609875 | -0.62136784 | 0.33813094 | -1.8376545 | 0.06611334 | 0.14653268 | NA | NA |
| ENSG000002 | 7.25074241 | -0.659167959 | 0.35871284 | -1.8375923 | 0.0661225 | 0.14654212 | LOC727993 | 727993 |

| | | | | | | | | |
|---|---|---|---|---|---|---|---|---|
| ENSG000002 | 5.02594379 | -0.690615309 | 0.3758294 | -1.8375766 | 0.06612482 | 0.14654212 | NA | NA |
| ENSG000001 | 14.2662422 | -0.577684867 | 0.3143878 | -1.8374913 | 0.06613739 | 0.14656197 | PKD2L1 | 9033 |
| ENSG000002 | 3.52580072 | -0.694995504 | 0.37827759 | -1.8372632 | 0.06617105 | 0.14662853 | NA | NA |
| ENSG000002 | 8.12352406 | -0.646011099 | 0.35162808 | -1.8371999 | 0.06618038 | 0.14664121 | NA | NA |
| ENSG000001 | 5.0532548 | -0.689281263 | 0.37523971 | -1.8369092 | 0.0662233 | 0.14672829 | OR6S1 | 341799 |
| ENSG000002 | 11.0453002 | 0.602960686 | 0.32826456 | 1.83681323 | 0.06623747 | 0.14675167 | NA | NA |
| ENSG000002 | 20.3967396 | -0.512066108 | 0.27881183 | -1.8366011 | 0.06626881 | 0.14681307 | NA | NA |
| ENSG000002 | 66.5530625 | 0.35432762 | 0.19292942 | 1.83656602 | 0.06627399 | 0.14681652 | NA | NA |
| ENSG000002 | 4.1156719 | -0.694823625 | 0.37834435 | -1.8364848 | 0.066286 | 0.1468351 | NA | NA |
| ENSG000000 | 195.11091 | 0.277583626 | 0.15116308 | 1.8363189 | 0.06631051 | 0.14687335 | IDS | 3423 |
| ENSG000001 | 8.41408752 | -0.642271704 | 0.34975643 | -1.8363399 | 0.0663074 | 0.14687335 | OR4K5 | 79317 |
| ENSG000002 | 6.19779749 | -0.674609868 | 0.36738018 | -1.8362718 | 0.06631747 | 0.14688075 | LINC01567 | 400511 |
| ENSG000002 | 15.9543087 | -0.550779669 | 0.29997428 | -1.8360896 | 0.0663444 | 0.14693238 | NA | NA |
| ENSG000001 | 10.5836066 | -0.615406703 | 0.33519952 | -1.8359415 | 0.06636631 | 0.14697288 | ACKR4 | 51554 |
| ENSG000002 | 14.4569526 | -0.589435191 | 0.32110444 | -1.8356495 | 0.06640952 | 0.14705249 | TMPRSS11GP | 644759 |
| ENSG000002 | 3.81693796 | -0.694905907 | 0.37855632 | -1.8356738 | 0.06640592 | 0.14705249 | NA | NA |
| ENSG000001 | 25.8794666 | -0.507880194 | 0.27669588 | -1.8355177 | 0.06642902 | 0.14708765 | CLEC12A | 160364 |
| ENSG000002 | 4.58520809 | -0.694669732 | 0.37850163 | -1.835315 | 0.06645903 | 0.14714605 | NA | NA |
| ENSG000001 | 517.823409 | 0.253434145 | 0.13809368 | 1.83523348 | 0.0664711 | 0.14716476 | C2CD3 | 26005 |
| ENSG000001 | 202.869065 | 0.345618615 | 0.18833621 | 1.83511508 | 0.06648864 | 0.14719555 | AEBP1 | 165 |
| ENSG000001 | 33.703638 | 0.439979378 | 0.239763 | 1.83505952 | 0.06649687 | 0.14720573 | ZBTB47 | 92999 |
| ENSG000002 | 22.8814515 | 0.503100415 | 0.27416947 | 1.83499798 | 0.06650599 | 0.14721788 | NA | NA |
| ENSG000002 | 5.36805041 | -0.687090268 | 0.37446802 | -1.8348436 | 0.06652886 | 0.14726047 | LINC00564 | 100861546 |
| ENSG000000 | 6.44892014 | -0.671678385 | 0.36607888 | -1.8347914 | 0.0665366 | 0.14726957 | PITX1 | 5307 |
| ENSG000002 | 8.1926726 | -0.656592516 | 0.35787946 | -1.834675 | 0.06655385 | 0.14729971 | NA | NA |
| ENSG000001 | 6.15691397 | -0.676823327 | 0.36892771 | -1.834569 | 0.06656958 | 0.14732648 | KERA | 11081 |
| ENSG000002 | 6.73937542 | -0.679837944 | 0.37057926 | -1.8345278 | 0.06657569 | 0.14733196 | NA | NA |
| ENSG000002 | 3.23523095 | -0.690559038 | 0.37645799 | -1.8343588 | 0.06660075 | 0.1473714 | NA | NA |
| ENSG000002 | 7.64910424 | -0.657919766 | 0.35866475 | -1.8343586 | 0.06660078 | 0.1473714 | WBP11P1 | 441818 |

| | | | | | | | | |
|---|---|---|---|---|---|---|---|---|
| ENSG000002 | 19.9451837 | -0.528117637 | 0.28791544 | -1.8342803 | 0.06661239 | 0.14738101 | NA | NA |
| ENSG000002 | 2.844119 | 0.684359197 | 0.37309194 | 1.83429102 | 0.06661081 | 0.14738101 | NA | NA |
| ENSG000002 | 6.65427788 | -0.667238703 | 0.36377393 | -1.8342126 | 0.06662245 | 0.14739521 | NA | NA |
| ENSG000002 | 5.16438692 | -0.690999563 | 0.37681018 | -1.8338134 | 0.0666817 | 0.14750215 | NA | NA |
| ENSG000002 | 4.64674555 | -0.688464178 | 0.37542186 | -1.8338415 | 0.06667753 | 0.14750215 | NA | NA |
| ENSG000002 | 4.84677257 | -0.691690293 | 0.37718448 | -1.8338249 | 0.06667999 | 0.14750215 | NA | NA |
| ENSG000002 | 12.5640556 | -0.614383051 | 0.33504054 | -1.8337573 | 0.06669002 | 0.14750448 | NA | NA |
| ENSG000002 | 4.3962716 | -0.692502321 | 0.37763875 | -1.8337692 | 0.06668826 | 0.14750448 | CCAT2 | 101805488 |
| ENSG000001 | 112.268602 | 0.335506339 | 0.18296406 | 1.8337281 | 0.06669437 | 0.14750604 | UCHL3 | 7347 |
| ENSG000002 | 8.57544333 | -0.632935492 | 0.34517138 | -1.8336847 | 0.0667008 | 0.14751223 | NA | NA |
| ENSG000001 | 165.710156 | 0.512323718 | 0.27940842 | 1.83360156 | 0.06671316 | 0.14752347 | SLC12A7 | 10723 |
| ENSG000002 | 8.8418071 | -0.652076032 | 0.35562407 | -1.8336105 | 0.06671183 | 0.14752347 | NA | NA |
| ENSG000002 | 4.18446055 | -0.69400411 | 0.37850845 | -1.8335234 | 0.06672477 | 0.14754109 | NA | NA |
| ENSG000002 | 4.49184814 | -0.692666429 | 0.37780922 | -1.8333762 | 0.06674665 | 0.14756638 | NA | NA |
| ENSG000002 | 6.06027728 | -0.682666951 | 0.37236159 | -1.8333442 | 0.0667514 | 0.14756638 | NA | NA |
| ENSG000002 | 2.91323973 | -0.686263316 | 0.3743235 | -1.8333429 | 0.0667516 | 0.14756638 | NA | NA |
| ENSG000002 | 2.25437846 | -0.666910119 | 0.36377101 | -1.833324 | 0.0667544 | 0.14756638 | RN7SKP43 | 106481806 |
| ENSG000002 | 3.57961757 | -0.693404004 | 0.37820612 | -1.8334024 | 0.06674275 | 0.14756638 | NA | NA |
| ENSG000001 | 769.339182 | 0.218804645 | 0.11935187 | 1.83327376 | 0.06676187 | 0.14757485 | TRMT1L | 81627 |
| ENSG000002 | 2.4901327 | -0.681115263 | 0.37174577 | -1.8322071 | 0.06692056 | 0.14791757 | NA | NA |
| ENSG000001 | 158.294931 | 0.311794761 | 0.17020734 | 1.83185261 | 0.06697338 | 0.14801206 | PNPLA3 | 80339 |
| ENSG000001 | 33.7055503 | 0.46852861 | 0.25577463 | 1.83180248 | 0.06698085 | 0.14801206 | SFRP5 | 6425 |
| ENSG000002 | 7.90875133 | -0.651407115 | 0.35561432 | -1.8317798 | 0.06698424 | 0.14801206 | NA | NA |
| ENSG000002 | 9.94801124 | -0.611525009 | 0.33384091 | -1.8317857 | 0.06698335 | 0.14801206 | LOC100507140 | 100507140 |
| ENSG000002 | 6.66248177 | -0.680655828 | 0.37158302 | -1.8317732 | 0.06698521 | 0.14801206 | LOC101927606 | 101927606 |
| ENSG000002 | 2.51506825 | -0.68109265 | 0.3718091 | -1.8318343 | 0.06697611 | 0.14801206 | NA | NA |
| ENSG000002 | 7.7868816 | -0.650631737 | 0.35520511 | -1.8317071 | 0.06699507 | 0.14801974 | NA | NA |
| ENSG000002 | 18.5999308 | -0.541853456 | 0.29581983 | -1.8317009 | 0.06699599 | 0.14801974 | LOC100506606 | 100506606 |
| ENSG000002 | 14.6116424 | -0.566597015 | 0.30936159 | -1.8315041 | 0.06702534 | 0.14806846 | MLK7-AS1 | 339751 |

| | | | | | | | | |
|---|---|---|---|---|---|---|---|---|
| ENSG000002 | 6.08924529 | -0.670754174 | 0.36622721 | -1.8315247 | 0.06702227 | 0.14806846 | NA | NA |
| ENSG000001 | 42.7852389 | -0.392683853 | 0.21442105 | -1.831368 | 0.06704563 | 0.14808927 | CCDC176 | 80127 |
| ENSG000002 | 11.4670284 | -0.607024124 | 0.33145768 | -1.8313775 | 0.06704422 | 0.14808927 | NA | NA |
| ENSG000002 | 2.58095305 | -0.679206826 | 0.37087414 | -1.8313674 | 0.06704572 | 0.14808927 | NA | NA |
| ENSG000002 | 7.28271028 | -0.656681007 | 0.35859025 | -1.8312852 | 0.06705799 | 0.14810831 | LOC101929144 | 101929144 |
| ENSG000002 | 4.62483097 | -0.68760658 | 0.37548571 | -1.8312457 | 0.06706388 | 0.14811326 | NA | NA |
| ENSG000002 | 2.88574819 | -0.686204442 | 0.37475017 | -1.8310984 | 0.06708585 | 0.14815371 | NA | NA |
| ENSG000002 | 3.13462298 | -0.69139719 | 0.37760321 | -1.8310151 | 0.06709829 | 0.1481731 | NA | NA |
| ENSG000002 | 3.5879445 | -0.691412045 | 0.37763444 | -1.8309031 | 0.06711501 | 0.14820197 | NA | NA |
| ENSG000002 | 7.25196372 | -0.671938096 | 0.36703102 | -1.8307392 | 0.06713947 | 0.14824719 | NA | NA |
| ENSG000002 | 4.99607983 | -0.686136072 | 0.37479092 | -1.8307169 | 0.06714281 | 0.14824719 | NA | NA |
| ENSG000002 | 9.15441223 | -0.630041191 | 0.3441636 | -1.8306445 | 0.06715363 | 0.14826302 | NA | NA |
| ENSG000002 | 14.4491737 | -0.587791726 | 0.32110298 | -1.8305396 | 0.06716929 | 0.14828952 | NA | NA |
| ENSG000001 | 21.5952174 | -0.531552801 | 0.29039677 | -1.8304363 | 0.06718472 | 0.14831551 | C1QTNF1 | 114897 |
| ENSG000001 | 24.4631382 | -0.510411252 | 0.27886939 | -1.8302878 | 0.06720691 | 0.14835643 | CYP3A4 | 1576 |
| ENSG000001 | 83.3966298 | 0.323122493 | 0.17657223 | 1.82997342 | 0.06725391 | 0.14844719 | RNLS | 55328 |
| ENSG000002 | 6.7792293 | -0.669796185 | 0.36601608 | -1.8299638 | 0.06725535 | 0.14844719 | NA | NA |
| ENSG000002 | 7.34376985 | -0.663596126 | 0.36264098 | -1.8298984 | 0.06726514 | 0.14846071 | NA | NA |
| ENSG000002 | 16.2502758 | -0.56429734 | 0.30839816 | -1.8297688 | 0.06728451 | 0.14849539 | LOC101929505 | 101929505 |
| ENSG000002 | 3.29820026 | -0.688048236 | 0.37608714 | -1.8294915 | 0.06732601 | 0.14856272 | NA | NA |
| ENSG000002 | 2.53167809 | -0.680245919 | 0.37182017 | -1.8295025 | 0.06732437 | 0.14856272 | NA | NA |
| ENSG000002 | 3.74376547 | -0.692508558 | 0.37852037 | -1.8295146 | 0.06732255 | 0.14856272 | IMMP1LP3 | 106481715 |
| ENSG000001 | 695.839074 | 0.574871812 | 0.31422914 | 1.82946691 | 0.06732969 | 0.14856276 | BRD3 | 8019 |
| ENSG000001 | 85.3078382 | 0.335436408 | 0.18336881 | 1.82929917 | 0.0673548 | 0.14861008 | MED26 | 9441 |
| ENSG000002 | 6.98679222 | -0.660425081 | 0.36104496 | -1.8292045 | 0.06736898 | 0.14863326 | NA | NA |
| ENSG000002 | 6.10305478 | -0.672376113 | 0.36760013 | -1.8290965 | 0.06738515 | 0.14866086 | NA | NA |
| ENSG000002 | 29.9922553 | -0.450774886 | 0.24645656 | -1.8290237 | 0.06739606 | 0.14867685 | NA | NA |
| ENSG000002 | 3.07529736 | -0.678833894 | 0.37116977 | -1.8289041 | 0.06741398 | 0.14870828 | NA | NA |
| ENSG000002 | 30.3528999 | 0.458068524 | 0.25046686 | 1.82885884 | 0.06742076 | 0.14871514 | SNORD101 | 594837 |

| | | | | | | | | |
|---|---|---|---|---|---|---|---|---|
| ENSG000002 | 6.70559005 | -0.666749871 | 0.36457957 | -1.8288185 | 0.0674268 | 0.14872037 | NA | NA |
| ENSG000001 | 11.2518698 | -0.617145583 | 0.33750478 | -1.8285536 | 0.06746651 | 0.14879987 | DLK1 | 8788 |
| ENSG000002 | 4.14734528 | -0.69196035 | 0.37843335 | -1.8284867 | 0.06747654 | 0.1488058 | NA | NA |
| ENSG000002 | 33.4364417 | 0.437282672 | 0.23914862 | 1.82849758 | 0.06747491 | 0.1488058 | NA | NA |
| ENSG000001 | 15.9603989 | -0.569011072 | 0.31121645 | -1.8283451 | 0.06749778 | 0.14884455 | XCR1 | 2829 |
| ENSG000002 | 4.08226708 | -0.691837488 | 0.37845138 | -1.828075 | 0.0675383 | 0.14892582 | PACERR | 103752588 |
| ENSG000002 | 3.55078724 | -0.691004902 | 0.37800363 | -1.8280377 | 0.06754389 | 0.14893003 | NA | NA |
| ENSG000001 | 240.904395 | 0.250940971 | 0.13728063 | 1.82794154 | 0.06755833 | 0.14895165 | RAB3C | 115827 |
| ENSG000002 | 9.50370434 | -0.633245784 | 0.34642905 | -1.8279235 | 0.06756104 | 0.14895165 | NA | NA |
| ENSG000002 | 2.2839909 | -0.6645028 | 0.36353459 | -1.8278943 | 0.06756542 | 0.1489532 | NA | NA |
| ENSG000002 | 3.75958604 | -0.691968393 | 0.3785679 | -1.8278581 | 0.06757085 | 0.14895709 | NA | NA |
| ENSG000001 | 352.352875 | 0.254255037 | 0.13911035 | 1.82772197 | 0.06759129 | 0.14895823 | PCSK7 | 9159 |
| ENSG000002 | 61.5479819 | 0.356529939 | 0.19506944 | 1.82770784 | 0.06759341 | 0.14895823 | NA | NA |
| ENSG000002 | 2.52667472 | -0.680273995 | 0.3721961 | -1.82773 | 0.06759008 | 0.14895823 | NA | NA |
| ENSG000002 | 3.13937842 | -0.68977245 | 0.37738541 | -1.8277666 | 0.06758459 | 0.14895823 | NA | NA |
| ENSG000002 | 9.74184076 | -0.618897216 | 0.33860236 | -1.8277995 | 0.06757966 | 0.14895823 | NA | NA |
| ENSG000002 | 2.55146591 | -0.679080062 | 0.37153807 | -1.8277536 | 0.06758653 | 0.14895823 | MIR7975 | 102466872 |
| ENSG000002 | 3.18112492 | -0.689188284 | 0.37713695 | -1.8274218 | 0.06763637 | 0.14904029 | NA | NA |
| ENSG000002 | 4.44892812 | -0.68990587 | 0.37753186 | -1.827411 | 0.067638 | 0.14904029 | NA | NA |
| ENSG000002 | 7.601707 | -0.657286993 | 0.35972757 | -1.8271799 | 0.06767272 | 0.1491087 | NA | NA |
| ENSG000001 | 269.570716 | 0.294256019 | 0.16104805 | 1.82713183 | 0.06767995 | 0.14911653 | SYNGAP1 | 8831 |
| ENSG000001 | 13.2567625 | -0.576149052 | 0.3153435 | -1.8270523 | 0.06769191 | 0.14913478 | MOGAT1 | 116255 |
| ENSG000002 | 7.69141345 | 0.650035757 | 0.35578915 | 1.82702525 | 0.06769597 | 0.14913562 | NA | NA |
| ENSG000002 | 4.6595774 | -0.687016396 | 0.37604921 | -1.8269321 | 0.06770997 | 0.14915836 | NA | NA |
| ENSG000001 | 40.3622165 | 0.420145315 | 0.23000796 | 1.82665555 | 0.06775158 | 0.14923851 | CXCL6 | 6372 |
| ENSG000002 | 2.4926644 | -0.678603073 | 0.37150319 | -1.8266413 | 0.06775372 | 0.14923851 | NA | NA |
| ENSG000002 | 12.9323266 | -0.581821351 | 0.31854343 | -1.8265056 | 0.06777414 | 0.1492754 | NA | NA |
| ENSG000002 | 2.90877769 | -0.683978911 | 0.37449082 | -1.8264237 | 0.06778647 | 0.14929442 | NA | NA |
| ENSG000002 | 6.49344629 | -0.669501826 | 0.36659182 | -1.8262869 | 0.06780706 | 0.14932914 | LOC102723968 | 102723968 |

| | | | | | | | | |
|---|---|---|---|---|---|---|---|---|
| ENSG000002 | 8.68587124 | -0.639317945 | 0.35006758 | -1.8262701 | 0.06780959 | 0.14932914 | NA | NA |
| ENSG000000 | 90.4505054 | 0.313765217 | 0.17181533 | 1.82617705 | 0.0678236 | 0.14935187 | HSD17B6 | 8630 |
| ENSG000001 | 14.7952324 | -0.557691828 | 0.30540486 | -1.8260738 | 0.06783914 | 0.14936987 | ADAM21 | 8747 |
| ENSG000002 | 135.442699 | 0.275376361 | 0.15080109 | 1.82609001 | 0.0678367 | 0.14936987 | EMBP1 | 647121 |
| ENSG000001 | 198.85494 | 0.295571514 | 0.16188599 | 1.82580051 | 0.06788032 | 0.14945242 | FOXRED2 | 80020 |
| ENSG000002 | 5.29132016 | -0.681461177 | 0.37325045 | -1.8257478 | 0.06788826 | 0.14945369 | FABP12P1 | 106479057 |
| ENSG000002 | 26.4387554 | -0.474192951 | 0.25972187 | -1.8257721 | 0.06788459 | 0.14945369 | NA | NA |
| ENSG000002 | 6.40180043 | -0.664780025 | 0.36414909 | -1.825571 | 0.06791491 | 0.14950423 | NA | NA |
| ENSG000001 | 73.2332096 | -0.344630898 | 0.1887851 | -1.8255196 | 0.06792265 | 0.14951316 | NA | NA |
| ENSG000001 | 22.7769456 | -0.498170234 | 0.27290795 | -1.8254149 | 0.06793844 | 0.1495398 | SPHKAP | 80309 |
| ENSG000001 | 160.675027 | 0.292312606 | 0.16013776 | 1.82538216 | 0.06794338 | 0.14954255 | SNRNP35 | 11066 |
| ENSG000002 | 7.47330723 | -0.64690863 | 0.35443171 | -1.8251996 | 0.06797091 | 0.14959503 | LOC101927815 | 101927815 |
| ENSG000002 | 3.82305125 | -0.69096959 | 0.37859164 | -1.8251053 | 0.06798514 | 0.14961823 | NA | NA |
| ENSG000002 | 2.8914499 | -0.682573838 | 0.37400983 | -1.8250158 | 0.06799865 | 0.14963628 | NA | NA |
| ENSG000002 | 21.2860916 | -0.492495144 | 0.26986006 | -1.825002 | 0.06800073 | 0.14963628 | NA | NA |
| ENSG000002 | 13.03309 | -0.584433588 | 0.32030824 | -1.8245974 | 0.0680618 | 0.14976255 | NA | NA |
| ENSG000002 | 12.2502334 | -0.593783835 | 0.32545741 | -1.8244594 | 0.06808264 | 0.14980029 | DLGAP1-AS5 | 284215 |
| ENSG000002 | 6.39061873 | -0.664214493 | 0.36407651 | -1.8243816 | 0.0680944 | 0.14981802 | NA | NA |
| ENSG000002 | 12.9407029 | -0.595166439 | 0.32623817 | -1.8243311 | 0.06810203 | 0.14982669 | NA | NA |
| ENSG000002 | 110.642792 | 0.301183834 | 0.16509761 | 1.82427732 | 0.06811016 | 0.14983644 | RXRB | 6257 |
| ENSG000002 | 6.92261485 | -0.658513997 | 0.36100482 | -1.8241141 | 0.06813482 | 0.14988255 | NA | NA |
| ENSG000002 | 8.60542231 | 0.636122975 | 0.34874132 | 1.82405391 | 0.06814392 | 0.14989055 | NA | NA |
| ENSG000002 | 3.2365095 | -0.686332087 | 0.37627006 | -1.8240412 | 0.06814585 | 0.14989055 | NA | NA |
| ENSG000002 | 3.500299 | -0.689654705 | 0.37812992 | -1.8238565 | 0.06817377 | 0.14994384 | NA | NA |
| ENSG000002 | 4.63370463 | -0.684632844 | 0.37538808 | -1.8238002 | 0.06818229 | 0.14995444 | LOC105377247 | 105377247 |
| ENSG000002 | 7.9118328 | -0.653531723 | 0.35834563 | -1.8237469 | 0.06819035 | 0.14996403 | NA | NA |
| ENSG000002 | 11.6244655 | 0.594125683 | 0.3257829 | 1.82368593 | 0.06819957 | 0.14997617 | SNORD116-11 | 100033423 |
| ENSG000002 | 4.1546277 | 0.689997326 | 0.37838467 | 1.82353402 | 0.06822255 | 0.14997783 | NA | NA |
| ENSG000002 | 3.49180968 | -0.689428973 | 0.37806321 | -1.8235812 | 0.06821541 | 0.14997783 | NA | NA |

| | | | | | | | | | |
|---|---|---|---|---|---|---|---|---|---|
| ENSG000002 | 7.38523693 | -0.661035779 | 0.36250599 | -1.8235168 | 0.06822515 | 0.14997783 | LINC01153 | | 101927889 |
| ENSG000002 | 3.14879809 | -0.687921093 | 0.37725445 | -1.8234937 | 0.06822865 | 0.14997783 | NA | NA | |
| ENSG000002 | 8.37073464 | -0.639511104 | 0.3506947 | -1.8235551 | 0.06821936 | 0.14997783 | NA | NA | |
| ENSG000002 | 6.74835506 | -0.664549915 | 0.36444353 | -1.8234647 | 0.06823304 | 0.14997783 | NA | NA | |
| ENSG000002 | 4.97755836 | -0.68388983 | 0.37505044 | -1.8234609 | 0.06823361 | 0.14997783 | NA | NA | |
| ENSG000002 | 4.11283381 | 0.689975792 | 0.37837471 | 1.82352514 | 0.06822389 | 0.14997783 | NA | NA | |
| ENSG000002 | 8.40845843 | 0.644084873 | 0.35320917 | 1.82352253 | 0.06822429 | 0.14997783 | NA | NA | |
| ENSG000001 | 7.80834733 | -0.647090437 | 0.3549088 | -1.8232584 | 0.06826427 | 0.1500371 | SLCO1B3 | | 28234 |
| ENSG000001 | 280.3273 | 0.231851047 | 0.1271681 | 1.82318552 | 0.0682753 | 0.15004507 | SLC35E3 | | 55508 |
| ENSG000002 | 5.61601118 | -0.680350081 | 0.37316298 | -1.8231982 | 0.06827338 | 0.15004507 | NA | NA | |
| ENSG000000 | 61.2468541 | 0.384104207 | 0.21068458 | 1.82312447 | 0.06828454 | 0.15005725 | TSPAN15 | | 23555 |
| ENSG000002 | 18.6236535 | -0.529961736 | 0.29070676 | -1.8230114 | 0.06830166 | 0.15008674 | LINC00309 | | 150992 |
| ENSG000002 | 5.22491906 | -0.679012434 | 0.37248215 | -1.8229395 | 0.06831255 | 0.15010254 | NA | NA | |
| ENSG000002 | 4.43433314 | -0.688261841 | 0.37758725 | -1.8227889 | 0.06833537 | 0.1501364 | NA | NA | |
| ENSG000002 | 4.09936151 | -0.689688829 | 0.37836838 | -1.8227972 | 0.06833411 | 0.1501364 | NA | NA | |
| ENSG000002 | 52.0110477 | 0.382060823 | 0.20960544 | 1.82276194 | 0.06833945 | 0.15013724 | ASB16-AS1 | | 339201 |
| ENSG000001 | 50.7094438 | -0.38729101 | 0.21250088 | -1.8225384 | 0.06837333 | 0.15020353 | CCDC144B | | 284047 |
| ENSG000002 | 2.89797922 | -0.680563009 | 0.37350124 | -1.8221171 | 0.06843721 | 0.15033573 | NA | NA | |
| ENSG000001 | 640.594126 | 0.234079507 | 0.12847354 | 1.82200563 | 0.06845413 | 0.15036474 | GXYLT1 | | 283464 |
| ENSG000002 | 8.53118931 | -0.645691929 | 0.35440203 | -1.8219194 | 0.06846722 | 0.15038535 | NA | NA | |
| ENSG000001 | 72.1912649 | 0.351207372 | 0.19277264 | 1.82187352 | 0.06847418 | 0.15039248 | ZNF771 | | 51333 |
| ENSG000002 | 8.04438155 | -0.642435618 | 0.35263375 | -1.8218211 | 0.06848213 | 0.1504018 | NA | NA | |
| ENSG000001 | 922.350075 | 0.245395112 | 0.13471038 | 1.82164966 | 0.06850816 | 0.15045082 | NDUFS2 | | 4720 |
| ENSG000001 | 12.1488115 | -0.588136912 | 0.32288294 | -1.8215174 | 0.06852824 | 0.15047861 | ZAR1L | | 646799 |
| ENSG000002 | 3.78172133 | -0.689593737 | 0.3785783 | -1.8215353 | 0.06852552 | 0.15047861 | NA | NA | |
| ENSG000001 | 105.418269 | 0.320034562 | 0.17570227 | 1.82145943 | 0.06853705 | 0.15048981 | IQCE | | 23288 |
| ENSG000001 | 505.532332 | 0.228293133 | 0.12535163 | 1.82122184 | 0.06857314 | 0.15054601 | VIPAS39 | | 63894 |
| ENSG000002 | 7.46443637 | -0.66217673 | 0.36358867 | -1.8212249 | 0.06857268 | 0.15054601 | CLLU1OS | | 574016 |
| ENSG000002 | 7.66268671 | 0.647002185 | 0.35525803 | 1.82121763 | 0.06857378 | 0.15054601 | NA | NA | |

| | | | | | | | | |
|---|---|---|---|---|---|---|---|---|
| ENSG000002 | 2.51979143 | -0.67608505 | 0.37124976 | -1.8211057 | 0.06859079 | 0.1505752 | NA | NA |
| ENSG000001 | 864.284376 | 0.232674356 | 0.12776717 | 1.82108099 | 0.06859455 | 0.1505753 | PANK1 | 53354 |
| ENSG000002 | 11.0878162 | 0.598712761 | 0.32877335 | 1.82105015 | 0.06859923 | 0.15057744 | NA | NA |
| ENSG00000C | 321.454387 | 0.233237269 | 0.12808573 | 1.8209465 | 0.06861499 | 0.15060387 | OSBPL3 | 26031 |
| ENSG000002 | 2.5142393 | -0.676209772 | 0.37137389 | -1.8208328 | 0.06863227 | 0.15063365 | NA | NA |
| ENSG000001 | 8.1819224 | -0.650192909 | 0.35710045 | -1.8207564 | 0.0686439 | 0.15065102 | LY6K | 54742 |
| ENSG000001 | 10.5331775 | -0.629213816 | 0.34559976 | -1.8206431 | 0.06866113 | 0.15067251 | SLC10A1 | 6554 |
| ENSG000001 | 57.628441 | -0.366361367 | 0.2012254 | -1.8206517 | 0.06865981 | 0.15067251 | IKZF3 | 22806 |
| ENSG000001 | 15.7783638 | -0.554229006 | 0.30442801 | -1.8205585 | 0.06867399 | 0.15068545 | SLC10A2 | 6555 |
| ENSG000001 | 69.5419989 | -0.357290288 | 0.19625359 | -1.8205542 | 0.06867465 | 0.15068545 | SLC4A10 | 57282 |
| ENSG000002 | 3.11857374 | -0.687261327 | 0.37750596 | -1.8205311 | 0.06867817 | 0.15068545 | RN7SKP234 | 106480368 |
| ENSG000002 | 6.17425946 | -0.670127278 | 0.36811161 | -1.8204459 | 0.06869113 | 0.15070573 | NA | NA |
| ENSG000001 | 12.2653982 | -0.587309308 | 0.32263375 | -1.8203591 | 0.06870433 | 0.15072653 | LHFPL5 | 222662 |
| ENSG000002 | 2.50479239 | -0.676281439 | 0.3715401 | -1.8202112 | 0.06872685 | 0.15076779 | NA | NA |
| ENSG000001 | 252.94936 | 0.244027375 | 0.13408045 | 1.82000718 | 0.06875791 | 0.15079447 | ATP5S | 27109 |
| ENSG000001 | 9.20551243 | -0.643264702 | 0.3534245 | -1.8200909 | 0.06874517 | 0.15079447 | KRT84 | 3890 |
| ENSG000001 | 4.65524673 | -0.683767753 | 0.37569605 | -1.8200025 | 0.06875863 | 0.15079447 | OR2T29 | 343563 |
| ENSG000002 | 3.51509977 | -0.687687504 | 0.37785344 | -1.8199848 | 0.06876133 | 0.15079447 | NA | NA |
| ENSG000002 | 16.5702392 | -0.564397717 | 0.31009544 | -1.8200774 | 0.06874721 | 0.15079447 | NA | NA |
| ENSG000002 | 8.35573834 | -0.635924985 | 0.34940489 | -1.8200231 | 0.06875548 | 0.15079447 | NA | NA |
| ENSG000002 | 3.12445868 | -0.687218605 | 0.37760542 | -1.8199384 | 0.06876838 | 0.15080179 | NA | NA |
| ENSG000002 | 3.15716267 | -0.686641131 | 0.37730362 | -1.8198636 | 0.06877978 | 0.15081862 | NA | NA |
| ENSG000001 | 7.31010972 | 0.648188 | 0.35621251 | 1.81966661 | 0.06880979 | 0.15087628 | TMEM88 | 92162 |
| ENSG000002 | 4.47770997 | -0.687994976 | 0.37809643 | -1.8196283 | 0.06881563 | 0.15088091 | NA | NA |
| ENSG000001 | 226.634998 | 0.335746526 | 0.18451848 | 1.81958216 | 0.06882266 | 0.15088818 | NBEAL2 | 23218 |
| ENSG000001 | 321.443664 | 0.278014017 | 0.15281866 | 1.81924132 | 0.06887462 | 0.15099393 | ZCCHC14 | 23174 |
| ENSG000002 | 2.56162225 | -0.674874315 | 0.37098545 | -1.8191396 | 0.06889014 | 0.15101978 | NA | NA |
| ENSG000002 | 10.667825 | -0.61405347 | 0.33762516 | -1.8187432 | 0.06895061 | 0.15114419 | LOC100507388 | 100507388 |
| ENSG000001 | 174.364605 | 0.255262494 | 0.14035406 | 1.81870405 | 0.06895659 | 0.15114912 | RP9 | 6100 |

| | | | | | | | | |
|---|---|---|---|---|---|---|---|---|
| ENSG000002 | 5.1125969 | -0.68418807 | 0.37625662 | -1.8184081 | 0.06900179 | 0.15124001 | NA | NA |
| ENSG000001 | 48.1897524 | 0.390514906 | 0.21477392 | 1.81826034 | 0.06902435 | 0.15127351 | NA | NA |
| ENSG000002 | 3.78303976 | -0.688378649 | 0.37859215 | -1.8182592 | 0.06902453 | 0.15127351 | NA | NA |
| ENSG000002 | 22.1561267 | 0.49270837 | 0.27101166 | 1.81803382 | 0.06905896 | 0.15134079 | NA | NA |
| ENSG000001 | 43.4173241 | 0.435338769 | 0.23946647 | 1.81795296 | 0.06907132 | 0.1513597 | ZNF319 | 57567 |
| ENSG000001 | 11.4209698 | 0.595193094 | 0.32740466 | 1.8179127 | 0.06907748 | 0.151365 | NDUFA4L2 | 56901 |
| ENSG000002 | 2.5201237 | -0.673998119 | 0.37078342 | -1.8177677 | 0.06909965 | 0.15140541 | LOC105372897 | 105372897 |
| ENSG000001 | 580.741232 | 0.223959729 | 0.12321012 | 1.81770558 | 0.06910914 | 0.15141803 | POLE2 | 5427 |
| ENSG000001 | 561.054539 | 0.238555181 | 0.1312459 | 1.81762009 | 0.06912222 | 0.15143849 | MCC | 4163 |
| ENSG000002 | 3.92314958 | -0.687134751 | 0.37813643 | -1.817161 | 0.06919247 | 0.15158209 | NA | NA |
| ENSG000002 | 10.2035378 | -0.620802034 | 0.34163633 | -1.8171429 | 0.06919524 | 0.15158209 | NA | NA |
| ENSG000001 | 20.0207166 | -0.529169856 | 0.29124118 | -1.8169472 | 0.0692252 | 0.15163245 | LOXL4 | 84171 |
| ENSG000002 | 10.8530341 | -0.602896063 | 0.33181876 | -1.8169439 | 0.06922571 | 0.15163245 | NA | NA |
| ENSG000001 | 4.9605356 | -0.681822704 | 0.37528922 | -1.8167927 | 0.06924886 | 0.15167437 | KLHL38 | 340359 |
| ENSG000001 | 49.7687787 | 0.395129725 | 0.21749314 | 1.8167457 | 0.06925606 | 0.15167437 | TMEM80 | 283232 |
| ENSG000002 | 6.93011857 | -0.654857422 | 0.36045479 | -1.8167533 | 0.0692549 | 0.15167437 | NA | NA |
| ENSG000001 | 10.8659798 | -0.607542643 | 0.33443466 | -1.8166258 | 0.06927443 | 0.151697 | AANAT | 15 |
| ENSG000002 | 5.47982206 | -0.673039275 | 0.37049879 | -1.8165762 | 0.06928203 | 0.151697 | NA | NA |
| ENSG000002 | 8.56910082 | -0.63083358 | 0.34725454 | -1.8166316 | 0.06927354 | 0.151697 | NA | NA |
| ENSG000002 | 7.90189287 | -0.6477419 | 0.35658165 | -1.8165318 | 0.06928884 | 0.151697 | LINC01370 | 100505663 |
| ENSG000002 | 2.5827153 | -0.673116351 | 0.37054465 | -1.8165594 | 0.06928461 | 0.151697 | NA | NA |
| ENSG000002 | 2.54393028 | -0.673577748 | 0.37080303 | -1.8165379 | 0.0692879 | 0.151697 | NA | NA |
| ENSG000002 | 8.64072209 | -0.632383443 | 0.3481648 | -1.8163336 | 0.06931921 | 0.15175529 | LOC101929210 | 101929210 |
| ENSG000001 | 187.190497 | 0.286387748 | 0.15768227 | 1.816233 | 0.06933464 | 0.15177154 | CPT1A | 1374 |
| ENSG000001 | 36.4064235 | -0.430947778 | 0.23727835 | -1.816212 | 0.06933786 | 0.15177154 | RERG | 85004 |
| ENSG000002 | 10.3898805 | -0.614620001 | 0.33839935 | -1.8162565 | 0.06933104 | 0.15177154 | ID2-AS1 | 100506299 |
| ENSG000001 | 257.632854 | 0.293796137 | 0.16176718 | 1.81616656 | 0.06934483 | 0.1517786 | ZSCAN18 | 65982 |
| ENSG000002 | 21.6645925 | -0.528600273 | 0.29105751 | -1.8161369 | 0.06934938 | 0.15178037 | LOC730100 | 730100 |
| ENSG000002 | 2.59837279 | -0.672602894 | 0.37037166 | -1.8160215 | 0.06936708 | 0.15181091 | NA | NA |

| | | | | | | | | |
|---|---|---|---|---|---|---|---|---|
| ENSG00000( | 139.436162 | -0.281325213 | 0.15492473 | -1.8158832 | 0.06938829 | 0.15184095 | HECW1 | 23072 |
| ENSG000002 | 4.99081497 | -0.681697292 | 0.37540429 | -1.8159017 | 0.06938545 | 0.15184095 | NA | NA |
| ENSG000002 | 4.61435402 | -0.682268404 | 0.37575796 | -1.8157125 | 0.06941449 | 0.15189009 | NA | NA |
| ENSG000001 | 9.20263752 | -0.627022316 | 0.3453847 | -1.8154316 | 0.0694576 | 0.15197621 | TMEM262 | 100130348 |
| ENSG000001 | 162.015985 | 0.258641548 | 0.14247743 | 1.81531589 | 0.06947538 | 0.15199871 | CHST11 | 50515 |
| ENSG000002 | 18.0454025 | -0.527241295 | 0.29043762 | -1.8153341 | 0.06947259 | 0.15199871 | NA | NA |
| ENSG000001 | 177.996311 | 0.280886719 | 0.15474008 | 1.81521633 | 0.06949067 | 0.15202397 | RBM4 | 5936 |
| ENSG000002 | 4.05154482 | -0.686207687 | 0.37806781 | -1.8150386 | 0.06951797 | 0.15207549 | NA | NA |
| ENSG000001 | 666.934825 | 0.485954449 | 0.26774736 | 1.81497379 | 0.06952794 | 0.15208908 | SCAF4 | 57466 |
| ENSG000002 | 2.57405767 | -0.672827429 | 0.37072045 | -1.8149186 | 0.06953642 | 0.15209944 | NA | NA |
| ENSG000001 | 4.77290971 | -0.684604559 | 0.37722212 | -1.8148579 | 0.06954575 | 0.15211163 | FOXR2 | 139628 |
| ENSG000001 | 9.52392396 | -0.619604853 | 0.34142765 | -1.8147471 | 0.06956278 | 0.15214067 | NXPH1 | 30010 |
| ENSG000002 | 6.82327432 | -0.66578829 | 0.36692978 | -1.8144842 | 0.06960321 | 0.15222089 | NA | NA |
| ENSG000002 | 8.77575327 | -0.636717201 | 0.35091602 | -1.8144432 | 0.06960951 | 0.15222647 | NA | NA |
| ENSG000002 | 6.9863156 | -0.657369937 | 0.36230659 | -1.8144024 | 0.0696158 | 0.15223201 | NA | NA |
| ENSG00000( | 4.67108028 | -0.682326505 | 0.37607286 | -1.8143466 | 0.06962438 | 0.15224257 | CDHR5 | 53841 |
| ENSG000002 | 4.380335 | -0.684941776 | 0.37752269 | -1.8143063 | 0.06963059 | 0.15224792 | NA | NA |
| ENSG000002 | 2.49534514 | -0.672936754 | 0.37092451 | -1.8142149 | 0.06964465 | 0.15227046 | NA | NA |
| ENSG000002 | 17.8072305 | -0.531810985 | 0.29314206 | -1.8141749 | 0.0696508 | 0.15227569 | NA | NA |
| ENSG000002 | 2.63099402 | -0.667992396 | 0.3682155 | -1.8141344 | 0.06965704 | 0.15228113 | RN7SKP158 | 106480883 |
| ENSG000002 | 6.38266004 | -0.6765219 | 0.3729548 | -1.8139514 | 0.0696852 | 0.15233449 | NA | NA |
| ENSG000002 | 2.30965514 | -0.656951594 | 0.36220485 | -1.813757 | 0.06971514 | 0.1523835 | NA | NA |
| ENSG000002 | 2.4983014 | -0.673521481 | 0.37133956 | -1.8137617 | 0.06971442 | 0.1523835 | NA | NA |
| ENSG000002 | 2.50791695 | -0.673196073 | 0.37118904 | -1.8136206 | 0.06973616 | 0.15242123 | NA | NA |
| ENSG000001 | 8.18418853 | -0.644901264 | 0.35562548 | -1.8134282 | 0.06976581 | 0.15247781 | FANCD2OS | 115795 |
| ENSG000002 | 2.51706384 | -0.672737222 | 0.37098166 | -1.8133975 | 0.06977053 | 0.15247992 | NA | NA |
| ENSG000002 | 6.19241046 | -0.672137088 | 0.37065726 | -1.8133655 | 0.06977546 | 0.15248248 | LINC01158 | 100506421 |
| ENSG000002 | 8.74333451 | -0.6369509 | 0.35127726 | -1.8132426 | 0.06979441 | 0.15250745 | NA | NA |
| ENSG000002 | 5.60689282 | -0.675108444 | 0.37231941 | -1.8132507 | 0.06979316 | 0.15250745 | PPP4R1-AS1 | 101927323 |

| | | | | | | | | |
|---|---|---|---|---|---|---|---|---|
| ENSG000002 | 4.70839986 | -0.683803069 | 0.37713878 | -1.8131338 | 0.06981119 | 0.15253589 | NA | NA |
| ENSG000002 | 9.47011288 | -0.622266905 | 0.34321635 | -1.8130456 | 0.06982478 | 0.15255737 | LINC00114 | 400866 |
| ENSG000002 | 3.5083567 | -0.68557109 | 0.37818629 | -1.8127867 | 0.06986472 | 0.1526364 | NA | NA |
| ENSG000002 | 7.59176004 | -0.649499807 | 0.35833141 | -1.8125673 | 0.06989858 | 0.15270086 | PGM5-AS1 | 572558 |
| ENSG000002 | 4.64525849 | -0.681691818 | 0.37609613 | -1.8125468 | 0.06990176 | 0.15270086 | MCM8-AS1 | 101929225 |
| ENSG000002 | 5.25752664 | -0.675869664 | 0.37290328 | -1.812453 | 0.06991623 | 0.15272426 | NA | NA |
| ENSG000001 | 3.80597174 | -0.686157164 | 0.37859049 | -1.8123994 | 0.06992451 | 0.15273412 | NA | NA |
| ENSG000001 | 66.4475183 | 0.375544788 | 0.20721335 | 1.81235804 | 0.06993089 | 0.15273672 | ABCA2 | 20 |
| ENSG000001 | 1837.37569 | 0.228237002 | 0.12593478 | 1.81234284 | 0.06993324 | 0.15273672 | PRKD3 | 23683 |
| ENSG000002 | 3.23208623 | -0.680496065 | 0.3754997 | -1.8122413 | 0.06994892 | 0.15276275 | NA | NA |
| ENSG000001 | 1676.20573 | 0.221050308 | 0.12201219 | 1.81170675 | 0.07003152 | 0.1529349 | DCLK1 | 9201 |
| ENSG000001 | 361.466431 | 0.223491698 | 0.12337148 | 1.8115346 | 0.07005814 | 0.1529745 | ARHGAP26 | 23092 |
| ENSG000001 | 8.61301335 | -0.644129742 | 0.35557491 | -1.8115163 | 0.07006097 | 0.1529745 | TRHR | 7201 |
| ENSG000002 | 11.5034098 | -0.618241711 | 0.34127892 | -1.8115438 | 0.07005672 | 0.1529745 | NA | NA |
| ENSG000002 | 3.53132115 | -0.684861158 | 0.37809573 | -1.8113433 | 0.07008773 | 0.15301679 | NA | NA |
| ENSG000002 | 3.13045986 | -0.683131954 | 0.37714128 | -1.8113423 | 0.07008788 | 0.15301679 | OR5K2 | 402135 |
| ENSG000002 | 7.7497103 | -0.656977778 | 0.3627209 | -1.8112488 | 0.07010235 | 0.1530319 | LOC100287792 | 100287792 |
| ENSG000002 | 5.26999376 | -0.676117639 | 0.37328433 | -1.8112671 | 0.07009951 | 0.1530319 | LOC101929420 | 101929420 |
| ENSG000002 | 3.52944233 | -0.684957535 | 0.37817687 | -1.8112095 | 0.07010843 | 0.15303692 | NA | NA |
| ENSG000001 | 3.41027192 | -0.685539783 | 0.37854425 | -1.8109898 | 0.07014243 | 0.1531029 | CELA2A | 63036 |
| ENSG000001 | 2.51984404 | -0.673000714 | 0.37164034 | -1.8108925 | 0.07015749 | 0.15312753 | NA | NA |
| ENSG000002 | 20.3900105 | -0.522186776 | 0.28836935 | -1.8108262 | 0.07016776 | 0.1531417 | LOC105372038 | 105372038 |
| ENSG000002 | 167.788156 | 0.264260213 | 0.14593708 | 1.81078185 | 0.07017463 | 0.15314845 | LINC00630 | 100287765 |
| ENSG000001 | 12.1999702 | -0.582963453 | 0.32194837 | -1.8107358 | 0.07018175 | 0.15315576 | C2orf82 | 389084 |
| ENSG000001 | 56.5149495 | -0.397837559 | 0.21971413 | -1.8107054 | 0.07018646 | 0.15315779 | MUC4 | 4585 |
| ENSG000002 | 6.73594666 | -0.660289909 | 0.36467122 | -1.8106444 | 0.07019591 | 0.15317017 | NA | NA |
| ENSG000001 | 666.748899 | 0.284050108 | 0.15688868 | 1.81052007 | 0.07021517 | 0.15320396 | KIRREL | 55243 |
| ENSG000001 | 73.719629 | 0.339753559 | 0.18768053 | 1.81027601 | 0.070253 | 0.15326992 | MTMR8 | 55613 |
| ENSG000002 | 9.565607 | -0.625460836 | 0.34551108 | -1.8102483 | 0.0702573 | 0.15326992 | NA | NA |

| | | | | | | | | |
|---|---|---|---|---|---|---|---|---|
| ENSG000002 | 2.54478372 | -0.669800336 | 0.36999766 | -1.8102826 | 0.07025197 | 0.15326992 | NA | NA |
| ENSG000002 | 2.85441916 | -0.677316284 | 0.37416088 | -1.8102274 | 0.07026053 | 0.15326992 | NA | NA |
| ENSG000002 | 7.28510159 | -0.655044743 | 0.3618842 | -1.8100949 | 0.07028108 | 0.1533065 | LRP4-AS1 | 100507401 |
| ENSG000001 | 18.3673647 | -0.53329498 | 0.29463359 | -1.8100277 | 0.07029148 | 0.15332096 | PAX9 | 5083 |
| ENSG000002 | 4.38540364 | -0.683418656 | 0.37762564 | -1.8097782 | 0.07033019 | 0.15339713 | MYADML | 151325 |
| ENSG000002 | 7.53893942 | -0.644097237 | 0.35592171 | -1.8096599 | 0.07034855 | 0.15342892 | LOC286114 | 286114 |
| ENSG000002 | 4.6355512 | -0.680009688 | 0.37577895 | -1.8096003 | 0.0703578 | 0.15344084 | NA | NA |
| ENSG000002 | 5.21680909 | -0.67590037 | 0.37353028 | -1.8094929 | 0.07037447 | 0.15346895 | CST13P | 164380 |
| ENSG000002 | 5.77048962 | -0.679847239 | 0.3757255 | -1.8094253 | 0.07038495 | 0.15348355 | NA | NA |
| ENSG000002 | 9.54892436 | -0.628323274 | 0.34725644 | -1.8093927 | 0.07039001 | 0.15348633 | PBOV1 | 59351 |
| ENSG000000 | 113.165424 | -0.312519894 | 0.17273366 | -1.8092588 | 0.0704108 | 0.15350691 | SLC12A1 | 6557 |
| ENSG000001 | 6.92725872 | -0.654946004 | 0.36198782 | -1.809304 | 0.07040379 | 0.15350691 | NAPSB | 256236 |
| ENSG000002 | 4.7646261 | 0.68006919 | 0.37587805 | 1.80928147 | 0.07040729 | 0.15350691 | NA | NA |
| ENSG000002 | 2.52198345 | -0.670737021 | 0.37075564 | -1.8091081 | 0.07043422 | 0.1535497 | NA | NA |
| ENSG000002 | 6.44985688 | -0.661721362 | 0.36579586 | -1.8089908 | 0.07045243 | 0.15358114 | NA | NA |
| ENSG000002 | 3.16871707 | -0.682081629 | 0.37705859 | -1.808954 | 0.07045816 | 0.15358537 | NA | NA |
| ENSG000002 | 3.94726678 | -0.675461075 | 0.37340783 | -1.8089098 | 0.07046502 | 0.15359208 | NA | NA |
| ENSG000002 | 10.1266984 | -0.619914059 | 0.34271786 | -1.8088175 | 0.07047936 | 0.15361508 | NA | NA |
| ENSG000002 | 3.1915139 | -0.680168173 | 0.37604736 | -1.8087301 | 0.07049295 | 0.15363644 | IFNA4 | 3441 |
| ENSG000002 | 3.01618951 | -0.6721641 | 0.37163172 | -1.8086833 | 0.07050021 | 0.15364401 | NA | NA |
| ENSG000002 | 7.36665489 | -0.659159139 | 0.36445258 | -1.808628 | 0.07050881 | 0.1536545 | NA | NA |
| ENSG000001 | 915.811676 | 0.259695554 | 0.14359644 | 1.80850971 | 0.07052721 | 0.15368632 | RBPMS | 11030 |
| ENSG000002 | 205.779059 | -0.260797969 | 0.14421714 | -1.8083701 | 0.07054891 | 0.15370911 | ZNF880 | 400713 |
| ENSG000002 | 3.41622158 | -0.684534142 | 0.37852726 | -1.8084144 | 0.07054203 | 0.15370911 | ZNF295-AS1 | 150142 |
| ENSG000002 | 4.76253026 | -0.682175241 | 0.37723225 | -1.8083693 | 0.07054904 | 0.15370911 | NA | NA |
| ENSG000002 | 8.4419188 | 0.635933558 | 0.35166822 | 1.80833386 | 0.07055455 | 0.15371287 | NA | NA |
| ENSG000002 | 49.9157243 | 0.393322605 | 0.21751504 | 1.80825477 | 0.07056686 | 0.15373142 | DANT2 | 642776 |
| ENSG000002 | 4.61239033 | -0.679840388 | 0.37598231 | -1.8081712 | 0.07057986 | 0.15375148 | RN7SKP195 | 106481816 |
| ENSG000001 | 472.853622 | 0.619117229 | 0.34241744 | 1.80807738 | 0.07059446 | 0.15377502 | ANP32B | 10541 |

| | | | | | | | | |
|---|---|---|---|---|---|---|---|---|
| ENSG000002 | 17.1162197 | -0.568890036 | 0.31464742 | -1.8080238 | 0.07060279 | 0.15378491 | LOC646652 | 646652 |
| ENSG000002 | 12.8907862 | -0.584748289 | 0.32344352 | -1.8078837 | 0.0706246 | 0.15382415 | LOC101928012 | 101928012 |
| ENSG000001 | 9.45396722 | -0.619156564 | 0.34248978 | -1.8078104 | 0.07063602 | 0.15384077 | DRICH1 | 51233 |
| ENSG000002 | 4.00258548 | -0.683498824 | 0.37810705 | -1.807686 | 0.07065538 | 0.15386641 | NA | NA |
| ENSG000002 | 9.2022618 | -0.622043662 | 0.34410851 | -1.8076963 | 0.07065379 | 0.15386641 | NA | NA |
| ENSG000002 | 9.50479768 | -0.623658803 | 0.34502292 | -1.8075866 | 0.07067087 | 0.15389186 | NA | NA |
| ENSG000001 | 36.6076939 | 0.489273532 | 0.27068992 | 1.80750557 | 0.07068349 | 0.15391108 | ZNF414 | 84330 |
| ENSG000002 | 3.57919281 | -0.682736863 | 0.37773814 | -1.8074343 | 0.0706946 | 0.153927 | NA | NA |
| ENSG000001 | 6.75509079 | -0.66316485 | 0.36691996 | -1.8073829 | 0.0707026 | 0.15393617 | RTP4 | 64108 |
| ENSG000002 | 5.88263405 | -0.668675861 | 0.36998882 | -1.8072867 | 0.07071759 | 0.15396055 | NA | NA |
| ENSG000002 | 3.50459055 | -0.683109287 | 0.37803022 | -1.807023 | 0.0707587 | 0.15404176 | NA | NA |
| ENSG000002 | 4.74087627 | -0.681548781 | 0.37717574 | -1.8069794 | 0.0707655 | 0.1540483 | PHGR1 | 644844 |
| ENSG000002 | 9.65197759 | -0.630668649 | 0.34908359 | -1.8066408 | 0.0708183 | 0.15415088 | NA | NA |
| ENSG000002 | 20.0714923 | -0.503992044 | 0.27896828 | -1.8066285 | 0.07082022 | 0.15415088 | NA | NA |
| ENSG000002 | 8.42776868 | -0.635612125 | 0.35183981 | -1.8065384 | 0.07083428 | 0.15417321 | NA | NA |
| ENSG000001 | 98.3406332 | 0.309040957 | 0.17109045 | 1.8063016 | 0.07087124 | 0.15422882 | ZNF132 | 7691 |
| ENSG000001 | 10.4651099 | 0.606875883 | 0.33597471 | 1.80631418 | 0.07086928 | 0.15422882 | CDC42EP5 | 148170 |
| ENSG000002 | 2.53043235 | -0.669478881 | 0.37062531 | -1.8063496 | 0.07086374 | 0.15422882 | LOC101929523 | 101929523 |
| ENSG000002 | 5.21175822 | -0.681461347 | 0.37736991 | -1.8058179 | 0.07094678 | 0.15438492 | NA | NA |
| ENSG000002 | 3.11911507 | -0.681288697 | 0.37728052 | -1.8057881 | 0.07095144 | 0.15438677 | NA | NA |
| ENSG000001 | 8.35894688 | -0.63458593 | 0.35145693 | -1.8055866 | 0.07098294 | 0.15444702 | NEUROD6 | 63974 |
| ENSG000001 | 1471.77862 | 0.182917467 | 0.10131058 | 1.80551194 | 0.07099461 | 0.15446413 | ZCCHC11 | 23318 |
| ENSG000002 | 13.3787537 | -0.616251443 | 0.34133114 | -1.8054357 | 0.07100652 | 0.15448176 | NA | NA |
| ENSG000002 | 2.8684532 | -0.674924223 | 0.37388803 | -1.8051506 | 0.07105111 | 0.15457048 | LOC102724502 | 102724502 |
| ENSG000001 | 18.0653812 | -0.551339413 | 0.30543123 | -1.805118 | 0.07105622 | 0.15457329 | ZC3H12D | 340152 |
| ENSG000002 | 3.73610564 | -0.683299231 | 0.37858912 | -1.804857 | 0.07109706 | 0.15465384 | OR4A47 | 403253 |
| ENSG000001 | 178.399197 | 0.270217126 | 0.14971908 | 1.80482759 | 0.07110166 | 0.15465556 | KIF27 | 55582 |
| ENSG000001 | 99.2286148 | 0.295543415 | 0.16376383 | 1.80469285 | 0.07112276 | 0.15468485 | C14orf93 | 60686 |
| ENSG000001 | 561.43584 | 0.265107831 | 0.1468991 | 1.80469336 | 0.07112268 | 0.15468485 | FAM222B | 55731 |

| | | | | | | | | |
|---|---|---|---|---|---|---|---|---|
| ENSG000002 | 6.15001014 | -0.663403593 | 0.36761241 | -1.8046278 | 0.07113294 | 0.15469869 | NA | NA |
| ENSG000002 | 64.8732017 | 0.353113769 | 0.1956781 | 1.80456456 | 0.07114285 | 0.15471194 | NA | NA |
| ENSG000002 | 3.75566369 | -0.683034539 | 0.37855471 | -1.8043219 | 0.07118087 | 0.15478632 | NA | NA |
| ENSG000002 | 53.3859416 | 0.377314543 | 0.20915055 | 1.80403322 | 0.0712261 | 0.15487638 | NA | NA |
| ENSG000001 | 25.9302726 | -0.461122927 | 0.25561348 | -1.8039852 | 0.07123363 | 0.15488446 | PAX5 | 5079 |
| ENSG000001 | 19.7366234 | -0.509447704 | 0.28240688 | -1.8039493 | 0.07123925 | 0.15488836 | CASQ2 | 845 |
| ENSG000002 | 5.78345318 | -0.677811586 | 0.37576894 | -1.8037989 | 0.07126284 | 0.15491718 | NA | NA |
| ENSG000002 | 10.2225242 | -0.625472553 | 0.34675431 | -1.8037917 | 0.07126397 | 0.15491718 | LOC643542 | 643542 |
| ENSG000002 | 9.58734809 | -0.623045266 | 0.34540414 | -1.8038153 | 0.07126027 | 0.15491718 | NA | NA |
| ENSG000002 | 6.69692054 | -0.657455205 | 0.36452335 | -1.8036024 | 0.07129366 | 0.15495439 | NA | NA |
| ENSG000002 | 2.88890501 | -0.672801153 | 0.37302605 | -1.8036305 | 0.07128926 | 0.15495439 | NA | NA |
| ENSG000002 | 3.12326446 | -0.680344704 | 0.37720579 | -1.8036433 | 0.07128724 | 0.15495439 | NA | NA |
| ENSG000002 | 16.6295918 | 0.539236231 | 0.29898019 | 1.80358513 | 0.07129637 | 0.15495439 | NA | NA |
| ENSG000002 | 4.62535912 | -0.678658404 | 0.37631571 | -1.8034283 | 0.07132098 | 0.15499957 | NA | NA |
| ENSG000002 | 169.662005 | 0.266101364 | 0.14756438 | 1.80328994 | 0.07134269 | 0.15503106 | NA | NA |
| ENSG000002 | 20.5686474 | -0.50120308 | 0.27793857 | -1.8032873 | 0.07134311 | 0.15503106 | LOC102546294 | 102546294 |
| ENSG000002 | 5.9383868 | -0.668598106 | 0.37086074 | -1.8028279 | 0.07141524 | 0.15517948 | NA | NA |
| ENSG000002 | 6.64248148 | -0.653365936 | 0.36243034 | -1.8027352 | 0.07142981 | 0.15520281 | LOC101927596 | 101927596 |
| ENSG000002 | 2.17423549 | -0.659635367 | 0.365919 | -1.8026814 | 0.07143827 | 0.15521288 | NA | NA |
| ENSG000001 | 14.1916814 | -0.573065665 | 0.31791952 | -1.8025495 | 0.07145899 | 0.15524843 | DEFB132 | 400830 |
| ENSG000002 | 175.813688 | 0.279001842 | 0.15478491 | 1.80251314 | 0.07146471 | 0.15524843 | NA | NA |
| ENSG000002 | 9.15628316 | -0.617277146 | 0.34245532 | -1.8025042 | 0.07146612 | 0.15524843 | LOC102723526 | 102723526 |
| ENSG000002 | 12.6768313 | -0.577991455 | 0.32066727 | -1.8024648 | 0.07147232 | 0.15525358 | NA | NA |
| ENSG000002 | 4.80062904 | -0.681083478 | 0.37789108 | -1.8023275 | 0.0714939 | 0.15529214 | NA | NA |
| ENSG000002 | 15.4638093 | 0.553791171 | 0.30731518 | 1.80203002 | 0.07154068 | 0.15538544 | NA | NA |
| ENSG000001 | 214.886412 | 0.25405262 | 0.14101383 | 1.80161491 | 0.07160601 | 0.15551901 | C6orf203 | 51250 |
| ENSG000002 | 5.13994935 | -0.678990515 | 0.37691129 | -1.8014597 | 0.07163045 | 0.15556375 | LINC00264 | 645528 |
| ENSG000002 | 5.30571783 | -0.674662898 | 0.37452297 | -1.8013926 | 0.07164103 | 0.15557839 | LINC01451 | 401561 |
| ENSG00000C | 41.8792802 | -0.411581874 | 0.22848594 | -1.8013444 | 0.07164862 | 0.15558654 | BTK | 695 |

| | | | | | | | | |
|---|---|---|---|---|---|---|---|---|
| ENSG000002 | 3.21390821 | -0.678215327 | 0.37651119 | -1.8013152 | 0.07165322 | 0.1555882 | NA | NA |
| ENSG000001 | 4.00185829 | -0.680917492 | 0.37803318 | -1.8012109 | 0.07166964 | 0.15561296 | CLDN8 | 9073 |
| ENSG000001 | 25.2949902 | 0.476802249 | 0.26471453 | 1.80119406 | 0.0716723 | 0.15561296 | BRICD5 | 283870 |
| ENSG000002 | 17.2725212 | -0.51856749 | 0.28790812 | -1.8011562 | 0.07167826 | 0.15561758 | MIR497HG | 100506755 |
| ENSG000002 | 6.25296693 | -0.667641469 | 0.37068224 | -1.8011154 | 0.0716847 | 0.15562323 | LOC400661 | 400661 |
| ENSG000001 | 16.6844304 | -0.530515635 | 0.29455711 | -1.8010621 | 0.0716931 | 0.15563313 | BPIFB1 | 92747 |
| ENSG000002 | 6.87407483 | -0.662489738 | 0.36784277 | -1.8010133 | 0.07170078 | 0.15564147 | NA | NA |
| ENSG000002 | 8.37685642 | -0.630865235 | 0.35029015 | -1.8009791 | 0.07170618 | 0.15564485 | NA | NA |
| ENSG000002 | 4.81317226 | 0.681825388 | 0.37859246 | 1.80094816 | 0.07171105 | 0.1556471 | PANO1 | 101927423 |
| ENSG000002 | 7.50097538 | -0.641883258 | 0.35644527 | -1.8007905 | 0.07173591 | 0.15569272 | NA | NA |
| ENSG000002 | 4.45196836 | -0.68039892 | 0.37785223 | -1.8007011 | 0.07175001 | 0.15571499 | NA | NA |
| ENSG000002 | 27.0893989 | 0.48424747 | 0.26892562 | 1.80067434 | 0.07175422 | 0.15571581 | SNORD3B-2 | 780852 |
| ENSG000002 | 7.47615139 | -0.641209015 | 0.35610396 | -1.8006231 | 0.0717623 | 0.15572501 | NA | NA |
| ENSG000001 | 12.9457326 | -0.577682446 | 0.32083213 | -1.8005754 | 0.07176983 | 0.155733 | PAX4 | 5078 |
| ENSG000002 | 2.71420847 | -0.659095303 | 0.36607248 | -1.8004503 | 0.07178956 | 0.15576749 | NA | NA |
| ENSG000002 | 36.2515878 | 0.457963764 | 0.25437782 | 1.80032897 | 0.07180871 | 0.15580069 | TRG-AS1 | 100506776 |
| ENSG000002 | 2.18276892 | -0.658250755 | 0.36564316 | -1.8002545 | 0.07182046 | 0.15581785 | NA | NA |
| ENSG000002 | 4.14943426 | -0.68131739 | 0.37849015 | -1.8000928 | 0.07184599 | 0.1558649 | NA | NA |
| ENSG000001 | 41.4139923 | 0.424436943 | 0.23580646 | 1.79993773 | 0.07187047 | 0.15590967 | NMB | 4828 |
| ENSG000002 | 9.41205043 | -0.615317882 | 0.34187343 | -1.7998412 | 0.07188572 | 0.15593441 | LOC101929631 | 101929631 |
| ENSG000002 | 9.28404244 | -0.628429796 | 0.34916653 | -1.7997996 | 0.07189228 | 0.1559403 | LINC01347 | 731275 |
| ENSG000001 | 24.8378752 | -0.522434826 | 0.29030235 | -1.7996231 | 0.07192017 | 0.15599244 | CSF1R | 1436 |
| ENSG000002 | 17.4055706 | 0.526524275 | 0.29260894 | 1.7994128 | 0.07195341 | 0.15605619 | NA | NA |
| ENSG000001 | 7.49614718 | -0.643546755 | 0.35766945 | -1.799278 | 0.07197472 | 0.15609407 | OR9G4 | 283189 |
| ENSG000001 | 9.08951733 | -0.614103377 | 0.34131284 | -1.799239 | 0.07198088 | 0.15609907 | KRT73 | 319101 |
| ENSG000002 | 8.59226999 | -0.624808773 | 0.34727347 | -1.7991837 | 0.07198962 | 0.15610969 | NA | NA |
| ENSG000001 | 49.604224 | 0.380170177 | 0.21130721 | 1.79913494 | 0.07199734 | 0.15611807 | CERCAM | 51148 |
| ENSG000000 | 11548.973 | 0.459138748 | 0.25520567 | 1.79909305 | 0.07200396 | 0.15612409 | PABPC1 | 26986 |
| ENSG000000 | 16.293422 | -0.531657073 | 0.29552653 | -1.7990164 | 0.07201609 | 0.15614203 | CPB2 | 1361 |

| | | | | | | | | |
|---|---|---|---|---|---|---|---|---|
| ENSG000002 | 6.51817138 | -0.663282424 | 0.3687023 | -1.7989647 | 0.07202426 | 0.15615141 | NA | NA |
| ENSG000002 | 43.5859191 | 0.39779146 | 0.22113715 | 1.79884505 | 0.0720432 | 0.1561841 | ZNF582-AS1 | 386758 |
| ENSG000001 | 330.971099 | 0.263621299 | 0.14655755 | 1.79875616 | 0.07205726 | 0.15618463 | NLGN1 | 22871 |
| ENSG000002 | 7.48136329 | -0.643531091 | 0.35776082 | -1.7987746 | 0.07205434 | 0.15618463 | NA | NA |
| ENSG000002 | 8.63831546 | -0.626324543 | 0.34819914 | -1.7987538 | 0.07205763 | 0.15618463 | LOC101927058 | 101927058 |
| ENSG000002 | 7.22741289 | -0.649118683 | 0.36087287 | -1.7987462 | 0.07205884 | 0.15618463 | NA | NA |
| ENSG000001 | 8.913378 | -0.631203508 | 0.35093597 | -1.7986287 | 0.07207743 | 0.15621657 | TP53TG5 | 27296 |
| ENSG000002 | 2.80400605 | -0.675116738 | 0.37536666 | -1.7985527 | 0.07208946 | 0.15623429 | RNU6-321P | 106479709 |
| ENSG000002 | 5.20652562 | -0.679229413 | 0.37770655 | -1.7982993 | 0.07212959 | 0.15631291 | LOC283585 | 283585 |
| ENSG000002 | 3.10437085 | -0.678103844 | 0.3770904 | -1.7982528 | 0.07213696 | 0.15632053 | NA | NA |
| ENSG000002 | 2.50917121 | -0.667821228 | 0.37138683 | -1.7981823 | 0.07214812 | 0.15633634 | NA | NA |
| ENSG000002 | 5.5307477 | -0.668250244 | 0.37168151 | -1.7979109 | 0.07219113 | 0.15642119 | NA | NA |
| ENSG000001 | 19.038913 | -0.512775193 | 0.28522747 | -1.7977763 | 0.07221246 | 0.15645905 | KLHL35 | 283212 |
| ENSG000002 | 6.18730093 | -0.664343722 | 0.3695705 | -1.7976103 | 0.07223879 | 0.15650773 | NA | NA |
| ENSG000001 | 2.51654212 | -0.665717119 | 0.37039415 | -1.7973208 | 0.0722847 | 0.15659883 | PALM2-AKAP2 | 445815 |
| ENSG000002 | 9.11507901 | -0.621488132 | 0.34581072 | -1.7971916 | 0.0723052 | 0.15663488 | NA | NA |
| ENSG000002 | 14.4801656 | -0.59032952 | 0.32850559 | -1.7970151 | 0.07233321 | 0.15668719 | NA | NA |
| ENSG000002 | 8.33459323 | -0.631300098 | 0.35133442 | -1.7968638 | 0.07235725 | 0.15673088 | PATE3 | 100169851 |
| ENSG000001 | 2821.09321 | 0.251315851 | 0.13987726 | 1.79668836 | 0.07238511 | 0.15677463 | POU2F1 | 5451 |
| ENSG000002 | 3.38949895 | -0.679926223 | 0.37843312 | -1.7966879 | 0.07238518 | 0.15677463 | NA | NA |
| ENSG000001 | 14.9322799 | -0.578447471 | 0.3219893 | -1.7964804 | 0.07241814 | 0.1568361 | MLANA | 2315 |
| ENSG000002 | 7.53683137 | -0.64244195 | 0.35762023 | -1.7964363 | 0.07242516 | 0.1568361 | NA | NA |
| ENSG000002 | 2.88455452 | -0.670943675 | 0.37348225 | -1.796454 | 0.07242235 | 0.1568361 | NA | NA |
| ENSG000002 | 11.0715367 | -0.616332805 | 0.34310653 | -1.7963307 | 0.07244194 | 0.15685567 | LINC01490 | 101928420 |
| ENSG000002 | 14.2558634 | -0.560361916 | 0.31194537 | -1.7963463 | 0.07243945 | 0.15685567 | NA | NA |
| ENSG000002 | 2.27449855 | -0.650287693 | 0.36202104 | -1.7962704 | 0.07245152 | 0.15686804 | NA | NA |
| ENSG000002 | 3.19442174 | -0.676716184 | 0.37674919 | -1.7961981 | 0.07246302 | 0.15688456 | NA | NA |
| ENSG000002 | 2.78457166 | -0.674894792 | 0.37574057 | -1.7961723 | 0.07246713 | 0.15688508 | NA | NA |
| ENSG000001 | 46.6078022 | 0.417093365 | 0.2322219 | 1.79609832 | 0.07247888 | 0.15689378 | GLIS2 | 84662 |

| | | | | | | | | |
|---|---|---|---|---|---|---|---|---|
| ENSG000002 | 6.4898804 | -0.666091784 | 0.37085021 | -1.7961208 | 0.0724753 | 0.15689378 | NA | NA |
| ENSG000001 | 5.07095398 | -0.676234417 | 0.37651054 | -1.7960571 | 0.07248544 | 0.1568996 | FAM3D | 131177 |
| ENSG000002 | 23.4389539 | -0.488802829 | 0.27217646 | -1.7959041 | 0.07250977 | 0.15694388 | LOC101927032 | 101927032 |
| ENSG000001 | 7.28803062 | -0.651647317 | 0.3628933 | -1.7956995 | 0.07254232 | 0.15700596 | TSPYL6 | 388951 |
| ENSG000002 | 3.17716618 | -0.676762757 | 0.376889 | -1.7956554 | 0.07254934 | 0.15701277 | NA | NA |
| ENSG000002 | 3.73206726 | -0.679805387 | 0.37859156 | -1.7956169 | 0.07255547 | 0.15701766 | NA | NA |
| ENSG000002 | 30.5826229 | 0.460259406 | 0.25639405 | 1.79512513 | 0.07263377 | 0.15717539 | NA | NA |
| ENSG000002 | 43.3201952 | 0.43796705 | 0.24397778 | 1.79511042 | 0.07263611 | 0.15717539 | LOC100129917 | 100129917 |
| ENSG000002 | 26.8865869 | 0.473657251 | 0.26386495 | 1.79507454 | 0.07264183 | 0.15717937 | SNORA49 | 677829 |
| ENSG000002 | 3.50258689 | -0.677061464 | 0.37720521 | -1.794942 | 0.07266295 | 0.15721669 | NA | NA |
| ENSG000002 | 4.41290314 | -0.678003444 | 0.37774952 | -1.7948493 | 0.07267772 | 0.15724026 | NA | NA |
| ENSG000001 | 5.22463756 | -0.671766345 | 0.37429014 | -1.7947743 | 0.07268966 | 0.15724932 | PANX3 | 116337 |
| ENSG000002 | 5.48593534 | -0.666330303 | 0.37125652 | -1.7947976 | 0.07268596 | 0.15724932 | NA | NA |
| ENSG000002 | 3.14616942 | -0.676710497 | 0.3770555 | -1.7947238 | 0.07269771 | 0.15725834 | NA | NA |
| ENSG000001 | 32.5390984 | 0.446696512 | 0.2489023 | 1.79466607 | 0.07270692 | 0.15726987 | NA | NA |
| ENSG000001 | 39.5732368 | -0.419378482 | 0.23370317 | -1.7944921 | 0.07273466 | 0.15732149 | FBXO32 | 114907 |
| ENSG000002 | 6.74681647 | -0.655020748 | 0.3650365 | -1.7943979 | 0.07274967 | 0.15734557 | NA | NA |
| ENSG000002 | 3.76453735 | -0.679273891 | 0.37856435 | -1.7943419 | 0.07275861 | 0.1573565 | NA | NA |
| ENSG000001 | 10.828296 | -0.597482523 | 0.33300698 | -1.7942042 | 0.07278058 | 0.15739563 | CD14 | 929 |
| ENSG000001 | 6.13582668 | 0.660568059 | 0.36817398 | 1.79417368 | 0.07278545 | 0.15739776 | NA | NA |
| ENSG000002 | 3.40646222 | -0.679007044 | 0.37847989 | -1.7940373 | 0.07280721 | 0.15743643 | NA | NA |
| ENSG000001 | 5.78386771 | -0.678562257 | 0.37827257 | -1.7938447 | 0.07283795 | 0.1574861 | HSPB7 | 27129 |
| ENSG000002 | 4.03239117 | -0.678150248 | 0.37804202 | -1.7938488 | 0.0728373 | 0.1574861 | NA | NA |
| ENSG000001 | 9.2026188 | -0.621504586 | 0.34647909 | -1.7937723 | 0.07284952 | 0.15750272 | NA | NA |
| ENSG000001 | 100.632796 | -0.302802869 | 0.16881657 | -1.7936798 | 0.07286429 | 0.15752625 | CDH6 | 1004 |
| ENSG000002 | 5.2164047 | -0.671145258 | 0.37418318 | -1.7936276 | 0.07287263 | 0.15753589 | NA | NA |
| ENSG000001 | 30.8181403 | 0.476875224 | 0.26589375 | 1.7934804 | 0.07289614 | 0.1575624 | ZNF296 | 162979 |
| ENSG000002 | 10.0217251 | -0.606650743 | 0.33825842 | -1.7934535 | 0.07290044 | 0.1575624 | NA | NA |
| ENSG000002 | 2.54912954 | -0.662264909 | 0.36926543 | -1.7934658 | 0.07289848 | 0.1575624 | NA | NA |

| | | | | | | | | |
|---|---|---|---|---|---|---|---|---|
| ENSG000002 | 7.73484612 | -0.632688424 | 0.35277444 | -1.7934645 | 0.07289869 | 0.1575624 | NA | NA |
| ENSG000001 | 2.98613203 | -0.66732123 | 0.37210568 | -1.7933648 | 0.07291461 | 0.1575803 | DHRS7C | 201140 |
| ENSG000002 | 3.41829682 | -0.678772611 | 0.37849872 | -1.7933287 | 0.07292038 | 0.1575803 | NA | NA |
| ENSG000002 | 2.18837312 | -0.654740003 | 0.36509567 | -1.7933382 | 0.07291887 | 0.1575803 | NA | NA |
| ENSG000002 | 7.50674843 | -0.639932571 | 0.35685117 | -1.7932758 | 0.07292883 | 0.15759017 | NA | NA |
| ENSG000001 | 9.90708571 | -0.603743811 | 0.33668421 | -1.793205 | 0.07294015 | 0.15759782 | KRT74 | 121391 |
| ENSG000002 | 3.11430703 | -0.675896446 | 0.37692006 | -1.793209 | 0.07293951 | 0.15759782 | NA | NA |
| ENSG000002 | 15.0949364 | -0.549039849 | 0.30619391 | -1.7931116 | 0.07295508 | 0.15760283 | NA | NA |
| ENSG000002 | 3.93867123 | -0.677985412 | 0.37809903 | -1.7931424 | 0.07295015 | 0.15760283 | NA | NA |
| ENSG000002 | 3.13746735 | -0.67610194 | 0.37705899 | -1.7930933 | 0.07295801 | 0.15760283 | NA | NA |
| ENSG000002 | 12.2235134 | -0.58865487 | 0.32828811 | -1.7931044 | 0.07295623 | 0.15760283 | NA | NA |
| ENSG000002 | 10.6942416 | -0.609474879 | 0.3399397 | -1.7928911 | 0.07299033 | 0.15766425 | NA | NA |
| ENSG000001 | 108.032338 | -0.293822753 | 0.16388704 | -1.792837 | 0.07299899 | 0.15766615 | RMDN2 | 151393 |
| ENSG000001 | 8.93630265 | -0.621868506 | 0.34686151 | -1.7928438 | 0.0729979 | 0.15766615 | SLFNL1 | 200172 |
| ENSG000002 | 4.11444541 | -0.678612832 | 0.37852036 | -1.792804 | 0.07300426 | 0.15766915 | PAGE3 | 139793 |
| ENSG000002 | 7.29958145 | 0.644163819 | 0.35931126 | 1.7927738 | 0.0730091 | 0.1576712 | APOC1P1 | 342 |
| ENSG000001 | 10.3213152 | -0.605801045 | 0.33791906 | -1.7927401 | 0.07301449 | 0.15767445 | S100Z | 170591 |
| ENSG000002 | 5.57863506 | -0.666696207 | 0.37190827 | -1.7926362 | 0.07303112 | 0.15770195 | NA | NA |
| ENSG000001 | 184.33414 | 0.3052207 | 0.17026622 | 1.79260862 | 0.07303553 | 0.15770307 | JRKL | 8690 |
| ENSG000002 | 6.72662471 | -0.655710356 | 0.36579246 | -1.7925748 | 0.07304094 | 0.15770636 | NA | NA |
| ENSG00000C | 14.5086292 | -0.550285755 | 0.30699211 | -1.7925078 | 0.07305166 | 0.15771117 | WNT16 | 51384 |
| ENSG000002 | 13.6663997 | -0.590395154 | 0.32937636 | -1.7924637 | 0.07305872 | 0.15771117 | NA | NA |
| ENSG000002 | 426.431424 | 0.240933366 | 0.13441441 | 1.79246675 | 0.07305823 | 0.15771117 | NA | NA |
| ENSG000002 | 4.12680098 | -0.678598931 | 0.37857465 | -1.7925102 | 0.07305127 | 0.15771117 | NA | NA |
| ENSG000001 | 14.2822464 | -0.597105466 | 0.33312764 | -1.7924225 | 0.07306531 | 0.157717 | ABHD1 | 84696 |
| ENSG000002 | 6.99189177 | -0.652326915 | 0.36396944 | -1.7922574 | 0.07309175 | 0.15775727 | NA | NA |
| ENSG000002 | 5.26340181 | -0.677085126 | 0.37778252 | -1.7922617 | 0.07309106 | 0.15775727 | NA | NA |
| ENSG000002 | 17.5159701 | -0.531160966 | 0.29637062 | -1.7922187 | 0.07309794 | 0.15776222 | NA | NA |
| ENSG000002 | 3.16034726 | -0.675482391 | 0.37692941 | -1.792066 | 0.07312239 | 0.1577982 | NA | NA |

| | | | | | | | | |
|---|---|---|---|---|---|---|---|---|
| ENSG000002 | 3.4104678 | -0.678231832 | 0.37845924 | -1.7920869 | 0.07311905 | 0.1577982 | NA | NA |
| ENSG000002 | 36.0764412 | -0.424110428 | 0.23671641 | -1.7916393 | 0.07319076 | 0.15793735 | NA | NA |
| ENSG000002 | 8.69644082 | -0.629810525 | 0.35153719 | -1.79159 | 0.07319866 | 0.15794599 | NA | NA |
| ENSG000002 | 16.3862546 | -0.554026406 | 0.309243 | -1.7915568 | 0.07320398 | 0.15794906 | LINC01468 | 101928687 |
| ENSG000001 | 36.6452004 | -0.419916286 | 0.23439038 | -1.7915252 | 0.07320905 | 0.1579514 | CNBD2 | 140894 |
| ENSG000002 | 9.74990533 | -0.612826548 | 0.34207426 | -1.7915015 | 0.07321285 | 0.1579514 | TMEM233 | 387890 |
| ENSG000002 | 65.668612 | 0.355677646 | 0.1985411 | 1.79145599 | 0.07322015 | 0.15795873 | SNHG20 | 654434 |
| ENSG000002 | 4.86587002 | -0.676894117 | 0.37790371 | -1.7911814 | 0.07326418 | 0.15804421 | HPYR1 | 93668 |
| ENSG000002 | 6.09902502 | -0.659147364 | 0.3680002 | -1.7911604 | 0.07326757 | 0.15804421 | NA | NA |
| ENSG000002 | 3.82502703 | -0.677962474 | 0.37851023 | -1.7911338 | 0.07327182 | 0.15804498 | NA | NA |
| ENSG000000 | 33.0695293 | -0.433920727 | 0.24229473 | -1.7908798 | 0.07331259 | 0.1581245 | DCT | 1638 |
| ENSG000002 | 3.13838062 | -0.674896164 | 0.37688113 | -1.7907401 | 0.07333501 | 0.15816445 | NA | NA |
| ENSG000001 | 48.8022005 | -0.393342538 | 0.2197023 | -1.7903433 | 0.07339874 | 0.15826835 | MTUS2 | 23281 |
| ENSG000001 | 53.8547808 | 0.394694278 | 0.22045736 | 1.79034293 | 0.0733988 | 0.15826835 | S100A14 | 57402 |
| ENSG000002 | 12.0449426 | -0.574523271 | 0.32089508 | -1.7903773 | 0.07339328 | 0.15826835 | LINC01531 | 100128682 |
| ENSG000002 | 2.20334547 | -0.652972756 | 0.36470759 | -1.7904008 | 0.07338951 | 0.15826835 | NA | NA |
| ENSG000001 | 5.05847555 | -0.673218388 | 0.37603377 | -1.7903137 | 0.0734035 | 0.15827008 | NA | NA |
| ENSG000001 | 147.255176 | 0.267089477 | 0.14919043 | 1.79025881 | 0.07341232 | 0.15828067 | LRRC28 | 123355 |
| ENSG000002 | 5.24970209 | -0.670249924 | 0.37441346 | -1.7901331 | 0.07343252 | 0.15830034 | NA | NA |
| ENSG000002 | 7.21198875 | -0.647205945 | 0.36154148 | -1.7901291 | 0.07343315 | 0.15830034 | LOC343052 | 343052 |
| ENSG000002 | 2.79210729 | -0.672662663 | 0.37575238 | -1.7901754 | 0.07342572 | 0.15830034 | NA | NA |
| ENSG000001 | 16.2497418 | -0.552168503 | 0.30846552 | -1.7900494 | 0.07344598 | 0.15831957 | UBA7 | 7318 |
| ENSG000002 | 4.55559675 | -0.677631045 | 0.37858166 | -1.7899204 | 0.07346671 | 0.1583545 | NA | NA |
| ENSG000002 | 3.19208591 | -0.674130865 | 0.37663046 | -1.7899 | 0.07346999 | 0.1583545 | NA | NA |
| ENSG000001 | 5490.56198 | 0.229945463 | 0.1284755 | 1.78980016 | 0.07348604 | 0.15838068 | IVNS1ABP | 10625 |
| ENSG000002 | 9.24758844 | -0.62330792 | 0.34826279 | -1.7897632 | 0.07349199 | 0.15838508 | LOC286177 | 286177 |
| ENSG000001 | 39.0155052 | 0.404056767 | 0.2257776 | 1.78962293 | 0.07351455 | 0.15842528 | RADIL | 55698 |
| ENSG000001 | 83.6584196 | 0.326241801 | 0.18230076 | 1.78958003 | 0.07352145 | 0.15843173 | Mar-03 | 115123 |
| ENSG000001 | 352.659019 | 0.25147924 | 0.14052842 | 1.7895258 | 0.07353018 | 0.15844212 | NUDT10 | 170685 |

| | | | | | | | | |
|---|---|---|---|---|---|---|---|---|
| ENSG000002 | 14.707556 | -0.546498611 | 0.3053998 | -1.7894531 | 0.07354188 | 0.15845892 | NA | NA |
| ENSG000002 | 23.3633331 | -0.493288378 | 0.27567359 | -1.7893929 | 0.07355156 | 0.15846802 | NA | NA |
| ENSG000002 | 3.98272391 | -0.676391914 | 0.37800388 | -1.7893782 | 0.07355392 | 0.15846802 | NA | NA |
| ENSG000001 | 48.9135189 | -0.388614026 | 0.21719675 | -1.7892258 | 0.07357846 | 0.15851203 | ACYP2 | 98 |
| ENSG000002 | 2.19592139 | -0.652459229 | 0.36466973 | -1.7891785 | 0.07358607 | 0.15851203 | NA | NA |
| ENSG000002 | 9.60097828 | -0.618223568 | 0.34553082 | -1.7891995 | 0.07358269 | 0.15851203 | SHANK2-AS1 | 100874198 |
| ENSG000001 | 698.006257 | 0.237872128 | 0.13295963 | 1.78905524 | 0.07360592 | 0.15853793 | ETV3 | 2117 |
| ENSG000001 | 57.2458553 | 0.350752262 | 0.19605389 | 1.78906041 | 0.07360509 | 0.15853793 | LYPD1 | 116372 |
| ENSG000001 | 31.8667543 | -0.428875164 | 0.23973554 | -1.7889511 | 0.07362269 | 0.15856563 | C15orf52 | 388115 |
| ENSG000001 | 1513.61602 | 0.233225537 | 0.13037424 | 1.78889276 | 0.07363209 | 0.15857745 | ANKRD12 | 23253 |
| ENSG000001 | 657.6996 | 0.227918298 | 0.12742798 | 1.78860477 | 0.07367849 | 0.15866896 | EPM2AIP1 | 9852 |
| ENSG000002 | 6.00830679 | -0.668668862 | 0.37387139 | -1.7884997 | 0.07369543 | 0.158697 | NA | NA |
| ENSG000002 | 4.92953783 | -0.670254302 | 0.37477115 | -1.7884363 | 0.07370565 | 0.15871059 | NA | NA |
| ENSG000002 | 2.81501967 | -0.671526689 | 0.37549555 | -1.7883746 | 0.0737156 | 0.15872358 | NA | NA |
| ENSG000001 | 20.063799 | -0.518320642 | 0.28984067 | -1.7882951 | 0.07372841 | 0.15874274 | PRSS42 | 339906 |
| ENSG000002 | 3.7770901 | -0.676959256 | 0.37856229 | -1.7882374 | 0.07373771 | 0.15875434 | NA | NA |
| ENSG000002 | 5.62748071 | -0.669259908 | 0.37428388 | -1.7881078 | 0.07375862 | 0.15879093 | PP2672 | 100130249 |
| ENSG000001 | 11.2558712 | -0.620957775 | 0.34731739 | -1.7878684 | 0.07379725 | 0.15885431 | OR1D2 | 4991 |
| ENSG000002 | 4.4023984 | -0.675733826 | 0.37795839 | -1.7878524 | 0.07379982 | 0.15885431 | NA | NA |
| ENSG000002 | 2.50302458 | -0.66174344 | 0.37012868 | -1.787874 | 0.07379634 | 0.15885431 | NA | NA |
| ENSG000002 | 7.87013014 | -0.639211652 | 0.35755589 | -1.7877251 | 0.07382036 | 0.1588901 | LOC105377494 | 105377494 |
| ENSG000002 | 8.06923332 | -0.632838199 | 0.35400068 | -1.7876751 | 0.07382844 | 0.15889906 | NA | NA |
| ENSG000001 | 2119.06887 | 0.204602172 | 0.1145025 | 1.78687946 | 0.07395697 | 0.15916724 | TOR1AIP2 | 163590 |
| ENSG000002 | 4.00217149 | -0.675550427 | 0.37811422 | -1.7866306 | 0.07399722 | 0.1592454 | NA | NA |
| ENSG000002 | 4.4738678 | -0.675978236 | 0.37836783 | -1.7865637 | 0.07400803 | 0.15926022 | NA | NA |
| ENSG000002 | 3.79636142 | -0.676262397 | 0.37854049 | -1.7864995 | 0.07401841 | 0.15927412 | NA | NA |
| ENSG000001 | 368.684287 | 0.234938996 | 0.13151754 | 1.78637006 | 0.07403936 | 0.15930227 | ZNF529 | 57711 |
| ENSG000002 | 26.845868 | -0.458034362 | 0.25640469 | -1.7863728 | 0.07403892 | 0.15930227 | NA | NA |
| ENSG000002 | 8.66535817 | -0.624706335 | 0.34971662 | -1.7863215 | 0.07404721 | 0.15930227 | NA | NA |

| | | | | | | | | |
|---|---|---|---|---|---|---|---|---|
| ENSG000002 | 7.81258268 | -0.636236248 | 0.35616804 | -1.7863373 | 0.07404466 | 0.15930227 | NA | NA |
| ENSG000002 | 32.5049691 | -0.478889849 | 0.26812075 | -1.7860977 | 0.07408344 | 0.15937176 | CCDC144NL | 339184 |
| ENSG000001 | 84.2250004 | 0.32815732 | 0.18374171 | 1.78597071 | 0.074104 | 0.15938653 | RINL | 126432 |
| ENSG000002 | 56.4053112 | -0.353536643 | 0.19794828 | -1.7860052 | 0.07409842 | 0.15938653 | ISPD | 729920 |
| ENSG000002 | 21.3239972 | -0.509242832 | 0.28513704 | -1.7859582 | 0.07410603 | 0.15938653 | NA | NA |
| ENSG000002 | 9.23284511 | -0.621414129 | 0.34793574 | -1.7860026 | 0.07409884 | 0.15938653 | LOC400620 | 400620 |
| ENSG000002 | 5.51786986 | -0.664426395 | 0.37204354 | -1.7858834 | 0.07411814 | 0.15940411 | NA | NA |
| ENSG000002 | 4.61853641 | -0.671858543 | 0.37622003 | -1.7858128 | 0.07412957 | 0.15942025 | MIR4518 | 100616405 |
| ENSG000002 | 9.72455071 | -0.627273108 | 0.35129713 | -1.7855913 | 0.07416545 | 0.15948896 | NA | NA |
| ENSG000002 | 3.73937401 | -0.675975886 | 0.37858382 | -1.7855382 | 0.07417405 | 0.159499 | NA | NA |
| ENSG000001 | 22.1522263 | 0.482547534 | 0.27026055 | 1.78549008 | 0.07418186 | 0.15950732 | ACOT4 | 122970 |
| ENSG000002 | 6.22961571 | -0.664442522 | 0.37215481 | -1.7853928 | 0.07419762 | 0.15953276 | NA | NA |
| ENSG000002 | 2.29943932 | -0.645467141 | 0.36153466 | -1.7853534 | 0.07420401 | 0.15953802 | NA | NA |
| ENSG000002 | 4.88824834 | -0.668841301 | 0.37464878 | -1.7852489 | 0.07422096 | 0.15956602 | NA | NA |
| ENSG000001 | 149.573981 | 0.304450138 | 0.17053973 | 1.78521531 | 0.0742264 | 0.15956722 | TPP1 | 1200 |
| ENSG000001 | 236.066789 | 0.264675001 | 0.14826096 | 1.78519685 | 0.07422939 | 0.15956722 | NANP | 140838 |
| ENSG000002 | 14.0548188 | -0.565987589 | 0.3170876 | -1.7849565 | 0.07426837 | 0.15964254 | NA | NA |
| ENSG000002 | 224.040834 | 0.23813961 | 0.13341863 | 1.78490519 | 0.0742767 | 0.15965199 | TATDN3 | 128387 |
| ENSG000002 | 4.09090397 | -0.675253483 | 0.37833111 | -1.7848215 | 0.07429028 | 0.15967272 | NA | NA |
| ENSG00000C | 15.7015413 | -0.544085884 | 0.30489104 | -1.7845257 | 0.07433829 | 0.15976743 | APBB1IP | 54518 |
| ENSG000002 | 4.08174488 | -0.675132532 | 0.37837422 | -1.7842984 | 0.07437518 | 0.15983825 | NA | NA |
| ENSG00000C | 3477.03713 | 0.210029411 | 0.11771405 | 1.78423406 | 0.07438564 | 0.15984459 | N4BP2 | 55728 |
| ENSG000001 | 226.871651 | 0.249269805 | 0.13970725 | 1.78422953 | 0.07438638 | 0.15984459 | HSD17B7 | 51478 |
| ENSG000001 | 23.0133059 | -0.486231335 | 0.2725195 | -1.7842075 | 0.07438995 | 0.15984459 | NA | NA |
| ENSG000001 | 45.8899191 | -0.373086565 | 0.20911063 | -1.7841588 | 0.07439786 | 0.15984464 | TRPV3 | 162514 |
| ENSG000002 | 3.83743917 | -0.675014336 | 0.37833353 | -1.7841779 | 0.07439476 | 0.15984464 | NA | NA |
| ENSG000002 | 3.17871741 | -0.671126873 | 0.37618331 | -1.7840421 | 0.07441682 | 0.1598769 | NA | NA |
| ENSG000002 | 3.80585964 | -0.675200193 | 0.37847892 | -1.7839836 | 0.07442633 | 0.15988886 | NA | NA |
| ENSG000002 | 4.02795994 | -0.674784559 | 0.37837058 | -1.7833959 | 0.07452187 | 0.16008563 | LOC101927637 | 101927637 |

| | | | | | | | | |
|---|---|---|---|---|---|---|---|---|
| ENSG000002 | 13.4755745 | -0.569205068 | 0.31922412 | -1.7830891 | 0.07457179 | 0.16018438 | NA | NA |
| ENSG000000 | 598.358187 | 0.451003831 | 0.25294567 | 1.78300671 | 0.07458521 | 0.16020472 | SNX1 | 6642 |
| ENSG000002 | 15.0938007 | -0.54363757 | 0.30490535 | -1.7829716 | 0.07459093 | 0.16020853 | NA | NA |
| ENSG000001 | 29.2116407 | -0.45475261 | 0.255059 | -1.7829311 | 0.07459752 | 0.16021419 | RHBDL2 | 54933 |
| ENSG000002 | 6.90275678 | -0.662084194 | 0.37136246 | -1.7828517 | 0.07461045 | 0.16023347 | NA | NA |
| ENSG000002 | 9.74034011 | -0.623319808 | 0.34964415 | -1.7827263 | 0.07463088 | 0.16026885 | LINC00871 | 100506412 |
| ENSG000002 | 6.4130595 | -0.654647177 | 0.36723017 | -1.7826617 | 0.07464138 | 0.16027444 | NA | NA |
| ENSG000002 | 47.8126675 | 0.381278372 | 0.21388047 | 1.78267033 | 0.07463998 | 0.16027444 | TRIM52-AS1 | 100507602 |
| ENSG000001 | 22.1035321 | -0.486315647 | 0.27280924 | -1.7826216 | 0.07464792 | 0.16027999 | RXFP2 | 122042 |
| ENSG000001 | 65.5980572 | 0.35101767 | 0.19691843 | 1.7825537 | 0.07465898 | 0.16028676 | MYO1F | 4542 |
| ENSG000001 | 13.9775661 | -0.569757287 | 0.31962734 | -1.7825674 | 0.07465675 | 0.16028676 | NA | NA |
| ENSG000001 | 5.51604346 | -0.663070169 | 0.37199459 | -1.7824726 | 0.0746722 | 0.16030664 | OR5B2 | 390190 |
| ENSG000002 | 3.76476657 | -0.67478682 | 0.37858943 | -1.782371 | 0.07468875 | 0.1603337 | LOC102724163 | 102724163 |
| ENSG000001 | 26.8733735 | -0.473677967 | 0.26578898 | -1.7821581 | 0.07472346 | 0.16039621 | DCLK2 | 166614 |
| ENSG000002 | 5.18324369 | -0.666236451 | 0.37384498 | -1.7821195 | 0.07472974 | 0.16039621 | NA | NA |
| ENSG000002 | 12.7008772 | -0.591250789 | 0.33176651 | -1.7821292 | 0.07472817 | 0.16039621 | LINC00908 | 284276 |
| ENSG000002 | 5.51884894 | -0.662228227 | 0.37163549 | -1.7819295 | 0.07476073 | 0.16045425 | NA | NA |
| ENSG000001 | 9.94606716 | -0.603645751 | 0.33879601 | -1.7817381 | 0.07479195 | 0.16051276 | RASL10A | 10633 |
| ENSG000002 | 13.8444815 | -0.560569196 | 0.31463157 | -1.7816686 | 0.07480329 | 0.16052859 | LMLN-AS1 | 100873947 |
| ENSG000002 | 5.00370412 | -0.670593774 | 0.37639547 | -1.7816202 | 0.07481119 | 0.16052931 | NA | NA |
| ENSG000002 | 6.4075346 | -0.65321805 | 0.36664316 | -1.781618 | 0.07481155 | 0.16052931 | NA | NA |
| ENSG000001 | 2.5439829 | -0.659398725 | 0.3701981 | -1.781205 | 0.07487896 | 0.16066547 | OR10H3 | 26532 |
| ENSG000002 | 5.20095572 | -0.666708477 | 0.37432841 | -1.781079 | 0.07489954 | 0.16070113 | NA | NA |
| ENSG000002 | 3.72338425 | -0.674147862 | 0.37856701 | -1.7807887 | 0.07494697 | 0.16078587 | NA | NA |
| ENSG000002 | 5.69607718 | -0.656087312 | 0.36842192 | -1.7808042 | 0.07494444 | 0.16078587 | NA | NA |
| ENSG000001 | 57.2875498 | -0.375289542 | 0.21074853 | -1.7807457 | 0.07495401 | 0.16079246 | NEK11 | 79858 |
| ENSG000001 | 7.67067466 | -0.64284694 | 0.36101081 | -1.7806861 | 0.07496374 | 0.16080484 | RHO | 6010 |
| ENSG000002 | 2.51446742 | -0.659044943 | 0.37011631 | -1.7806428 | 0.07497082 | 0.16081152 | NA | NA |
| ENSG000001 | 8.81788716 | -0.614915276 | 0.34535462 | -1.7805329 | 0.07498879 | 0.16083456 | OR9K2 | 441639 |

| | | | | | | | | |
|---|---|---|---|---|---|---|---|---|
| ENSG000002 | 2.68072685 | -0.651056596 | 0.36565355 | -1.7805286 | 0.07498949 | 0.16083456 | NA | NA |
| ENSG000001 | 3.18098798 | -0.669227039 | 0.3758639 | -1.7805036 | 0.07499358 | 0.16083481 | NA | NA |
| ENSG000002 | 15.444032 | -0.551986874 | 0.3100325 | -1.7804162 | 0.07500788 | 0.16085697 | NA | NA |
| ENSG000002 | 7.96196886 | -0.625556568 | 0.35140771 | -1.7801447 | 0.07505229 | 0.1609388 | LOC644669 | 644669 |
| ENSG000002 | 15.1191077 | -0.564570642 | 0.31715058 | -1.7801344 | 0.07505397 | 0.1609388 | NA | NA |
| ENSG000002 | 6.77241646 | -0.653519157 | 0.3671361 | -1.780046 | 0.07506843 | 0.16096129 | SRGAP2-AS1 | 100873165 |
| ENSG000002 | 3.42712254 | -0.673304199 | 0.37826552 | -1.7799777 | 0.07507961 | 0.16097674 | RN7SL266P | 106480994 |
| ENSG000002 | 5.44820074 | -0.660891103 | 0.37129897 | -1.7799433 | 0.07508524 | 0.16098032 | NA | NA |
| ENSG000002 | 3.7005892 | -0.673824927 | 0.37859074 | -1.7798241 | 0.07510475 | 0.16101363 | NA | NA |
| ENSG000001 | 38.8382617 | 0.417593504 | 0.23463418 | 1.77976412 | 0.07511457 | 0.16102617 | CYP4F22 | 126410 |
| ENSG000002 | 15.8421376 | -0.538458801 | 0.30255326 | -1.7797157 | 0.07512249 | 0.16103464 | CTB-12O2.1 | 101927115 |
| ENSG000002 | 3.4160257 | -0.673311253 | 0.37836667 | -1.7795205 | 0.07515446 | 0.16109465 | NA | NA |
| ENSG000002 | 28.0250041 | 0.48283144 | 0.27133176 | 1.77948737 | 0.07515989 | 0.16109778 | NA | NA |
| ENSG000002 | 3.38676669 | -0.673418721 | 0.37844183 | -1.7794511 | 0.07516584 | 0.16110201 | NA | NA |
| ENSG000001 | 10.6204182 | -0.599960054 | 0.33716682 | -1.779416 | 0.07517158 | 0.16110579 | HNF1A | 6927 |
| ENSG000001 | 25.6833502 | 0.468825367 | 0.26348261 | 1.77934081 | 0.0751839 | 0.16110666 | CLEC11A | 6320 |
| ENSG000002 | 2.17827353 | -0.648454871 | 0.36442746 | -1.7793798 | 0.07517751 | 0.16110666 | NA | NA |
| ENSG000002 | 17.5417557 | -0.530644028 | 0.29822289 | -1.7793538 | 0.07518177 | 0.16110666 | LOC100131347 | 100131347 |
| ENSG000002 | 3.77313638 | -0.673562002 | 0.37856874 | -1.779233 | 0.07520156 | 0.16113599 | LINC00261 | 140828 |
| ENSG000002 | 4.64205483 | -0.670192899 | 0.3766865 | -1.7791795 | 0.07521034 | 0.16114628 | LOC105374132 | 105374132 |
| ENSG000001 | 3.76606439 | -0.673489484 | 0.3785706 | -1.7790327 | 0.0752344 | 0.16118931 | NA | NA |
| ENSG000002 | 89.7291692 | 0.30400988 | 0.17088879 | 1.77899251 | 0.07524099 | 0.16119492 | NA | NA |
| ENSG000001 | 275.913187 | -0.342744332 | 0.19267777 | -1.7788473 | 0.0752648 | 0.16123136 | COL1A1 | 1277 |
| ENSG000002 | 6.60745485 | -0.64501971 | 0.36260688 | -1.7788403 | 0.07526595 | 0.16123136 | NA | NA |
| ENSG000002 | 4.29566418 | -0.670484986 | 0.37694586 | -1.7787302 | 0.075284 | 0.1612615 | NA | NA |
| ENSG00000C | 164.153942 | 0.354376433 | 0.19925931 | 1.77846861 | 0.07532692 | 0.16132788 | GPC1 | 2817 |
| ENSG000002 | 45.8810863 | 0.379761496 | 0.2135298 | 1.77849416 | 0.07532273 | 0.16132788 | NA | NA |
| ENSG000002 | 2.60590788 | -0.654589473 | 0.36806266 | -1.7784729 | 0.07532622 | 0.16132788 | NA | NA |
| ENSG000002 | 2.54828186 | -0.658048514 | 0.37003783 | -1.7783277 | 0.07535005 | 0.16135378 | NA | NA |

| | | | | | | | | |
|---|---|---|---|---|---|---|---|---|
| ENSG000002 | 10.1451433 | -0.596213687 | 0.33526753 | -1.7783222 | 0.07535096 | 0.16135378 | NA | NA |
| ENSG000002 | 47.4207165 | 0.382011729 | 0.21481326 | 1.77834334 | 0.07534748 | 0.16135378 | TMEM220-AS1 | 101101775 |
| ENSG000001 | 430.929173 | 0.274589384 | 0.15441472 | 1.77825917 | 0.0753613 | 0.16136524 | TAGLN2 | 8407 |
| ENSG000001 | 291.488339 | 0.334718906 | 0.18823033 | 1.77824112 | 0.07536426 | 0.16136524 | KIAA1522 | 57648 |
| ENSG000001 | 11.2462824 | -0.583778568 | 0.32829816 | -1.7781963 | 0.07537163 | 0.16137248 | ANGPTL6 | 83854 |
| ENSG000001 | 36.8887572 | -0.419290021 | 0.23580904 | -1.7780914 | 0.07538885 | 0.16140083 | AMER2 | 219287 |
| ENSG000001 | 59.2907605 | 0.36265759 | 0.20397642 | 1.77793877 | 0.07541391 | 0.16143745 | TRMT10B | 158234 |
| ENSG000002 | 3.80338742 | -0.67293226 | 0.37848553 | -1.7779604 | 0.07541037 | 0.16143745 | NA | NA |
| ENSG000001 | 20.3209174 | -0.538363401 | 0.30281566 | -1.7778585 | 0.0754271 | 0.16145715 | TPD52L3 | 89882 |
| ENSG000001 | 12.5793067 | -0.585645977 | 0.3294445 | -1.7776772 | 0.07545689 | 0.16151159 | PCDHB6 | 56130 |
| ENSG000001 | 230.896302 | -0.24278874 | 0.13657808 | -1.7776552 | 0.0754605 | 0.16151159 | HSPA4L | 22824 |
| ENSG000002 | 67.4761075 | 0.341215208 | 0.19197258 | 1.77741641 | 0.07549975 | 0.16158708 | ZNF2 | 7549 |
| ENSG000002 | 6.48014338 | -0.655152617 | 0.36861157 | -1.7773523 | 0.07551029 | 0.16160111 | NA | NA |
| ENSG000001 | 326.016356 | 0.234846499 | 0.13213497 | 1.77732286 | 0.07551513 | 0.16160295 | ADAT2 | 134637 |
| ENSG000002 | 47.7177022 | 0.3953478 | 0.22244944 | 1.77724792 | 0.07552746 | 0.16162079 | NA | NA |
| ENSG000002 | 6.02682941 | -0.656015054 | 0.369127 | -1.7772069 | 0.07553421 | 0.1616267 | NA | NA |
| ENSG000001 | 19.0499137 | -0.5263328 | 0.29617501 | -1.7771006 | 0.07555168 | 0.16165557 | NA | NA |
| ENSG000002 | 4.45352171 | -0.672484811 | 0.37848557 | -1.7767779 | 0.07560479 | 0.16176066 | LINC01284 | 101926971 |
| ENSG00000C | 31.5056041 | -0.442975671 | 0.24932937 | -1.7766686 | 0.07562278 | 0.16178208 | ATP12A | 479 |
| ENSG000002 | 21.6012289 | 0.490222574 | 0.27592089 | 1.77667801 | 0.07562123 | 0.16178208 | LOC101928069 | 101928069 |
| ENSG000002 | 9.51240868 | -0.615641985 | 0.34652814 | -1.7766003 | 0.07563403 | 0.16179762 | NA | NA |
| ENSG000001 | 3.50388057 | -0.670857045 | 0.37761308 | -1.7765726 | 0.07563859 | 0.16179883 | HNRNPCL1 | 343069 |
| ENSG000001 | 5.73870525 | -0.654808581 | 0.36858991 | -1.7765234 | 0.07564669 | 0.1617991 | VTN | 7448 |
| ENSG000001 | 77.3177344 | -0.338488554 | 0.19053191 | -1.7765452 | 0.07564309 | 0.1617991 | PROX1 | 5629 |
| ENSG000002 | 3.13382519 | -0.667880432 | 0.37595842 | -1.7764742 | 0.07565479 | 0.16180788 | NA | NA |
| ENSG000001 | 11.8167285 | -0.579426862 | 0.3261887 | -1.7763548 | 0.07567445 | 0.1618414 | HAVCR1 | 26762 |
| ENSG000001 | 5.95738096 | -0.66410665 | 0.37387267 | -1.776291 | 0.07568497 | 0.16185536 | SH2D7 | 646892 |
| ENSG000001 | 120.996806 | 0.290662844 | 0.16363812 | 1.77625387 | 0.07569108 | 0.16185989 | ALKBH2 | 121642 |
| ENSG000001 | 4.61868076 | -0.668554291 | 0.37639568 | -1.7762007 | 0.07569984 | 0.16187007 | OR52N4 | 390072 |

| | | | | | | | | |
|---|---|---|---|---|---|---|---|---|
| ENSG000002 | 3.19621781 | -0.666034449 | 0.37498633 | -1.7761566 | 0.07570711 | 0.16187708 | NA | NA |
| ENSG000001 | 3.24694703 | -0.665376758 | 0.37462724 | -1.7761035 | 0.07571586 | 0.16187872 | C11orf52 | 91894 |
| ENSG000002 | 4.91291588 | -0.665171546 | 0.37450894 | -1.7761166 | 0.0757137 | 0.16187872 | NA | NA |
| ENSG000002 | 9.97236558 | -0.601950354 | 0.33893216 | -1.7760202 | 0.0757296 | 0.1618974 | C5orf49 | 134121 |
| ENSG000002 | 21.3685389 | -0.494513166 | 0.27844178 | -1.7760021 | 0.07573258 | 0.1618974 | NA | NA |
| ENSG000002 | 88.2580012 | 0.305307641 | 0.17191078 | 1.77596562 | 0.07573858 | 0.1619017 | NA | NA |
| ENSG000001 | 3.7613216 | -0.672312593 | 0.37858996 | -1.7758332 | 0.07576042 | 0.16193629 | NA | NA |
| ENSG000002 | 15.2506376 | -0.541834387 | 0.30511802 | -1.775819 | 0.07576275 | 0.16193629 | NA | NA |
| ENSG000001 | 90.9988873 | 0.348486224 | 0.19626151 | 1.77562187 | 0.07579526 | 0.16199724 | WFDC2 | 10406 |
| ENSG000002 | 9.62123852 | -0.619862377 | 0.34910716 | -1.7755648 | 0.07580468 | 0.16200883 | NA | NA |
| ENSG000002 | 6.65061052 | -0.645849066 | 0.36375301 | -1.7755154 | 0.07581283 | 0.16201771 | NA | NA |
| ENSG000001 | 252.327022 | 0.234973157 | 0.13234737 | 1.77542746 | 0.07582733 | 0.16204017 | RBM45 | 129831 |
| ENSG000001 | 4.03745925 | -0.671553189 | 0.37827158 | -1.7753202 | 0.07584503 | 0.16206943 | NA | NA |
| ENSG000002 | 5.25824403 | -0.663815081 | 0.37392129 | -1.7752803 | 0.07585162 | 0.16207499 | HOXA10-AS | 100874323 |
| ENSG000002 | 2.1737983 | -0.646847178 | 0.36440961 | -1.7750552 | 0.07588877 | 0.16214582 | NA | NA |
| ENSG00000C | 211.74026 | 0.248508354 | 0.14001524 | 1.77486642 | 0.07591995 | 0.16219956 | MAGI3 | 260425 |
| ENSG000002 | 4.6719285 | -0.66930071 | 0.37710174 | -1.7748545 | 0.07592192 | 0.16219956 | NA | NA |
| ENSG000002 | 2.56364542 | -0.655896465 | 0.3695648 | -1.7747807 | 0.07593411 | 0.16221704 | NA | NA |
| ENSG000002 | 15.255632 | -0.539137964 | 0.30380251 | -1.774633 | 0.07595851 | 0.16226062 | NA | NA |
| ENSG00000C | 24.2562037 | -0.484500152 | 0.27302972 | -1.7745326 | 0.07597509 | 0.16227895 | TXK | 7294 |
| ENSG000002 | 8.98843307 | 0.639290319 | 0.36025644 | 1.77454239 | 0.07597348 | 0.16227895 | NA | NA |
| ENSG000002 | 22.6109824 | -0.488128347 | 0.2750879 | -1.774445 | 0.07598957 | 0.16230133 | DPPA5 | 340168 |
| ENSG000001 | 9.26536118 | -0.618720971 | 0.34875016 | -1.7741095 | 0.07604504 | 0.16241125 | ANKRD22 | 118932 |
| ENSG000002 | 56.0917786 | 0.377943763 | 0.21304002 | 1.77405057 | 0.07605479 | 0.16242351 | NA | NA |
| ENSG000002 | 9.48924902 | -0.617549132 | 0.34818135 | -1.7736422 | 0.07612236 | 0.16255925 | NA | NA |
| ENSG000002 | 2.80727552 | -0.665392844 | 0.37519159 | -1.7734748 | 0.07615006 | 0.16260985 | NA | NA |
| ENSG000001 | 130.102887 | 0.364586808 | 0.20560099 | 1.77327364 | 0.07618337 | 0.16263426 | ADAMTS7 | 11173 |
| ENSG000001 | 253.323207 | 0.270082572 | 0.15230846 | 1.77326047 | 0.07618556 | 0.16263426 | SMARCAL1 | 50485 |
| ENSG000002 | 7.41036409 | -0.632430251 | 0.35663115 | -1.7733455 | 0.07617147 | 0.16263426 | NA | NA |

| | | | | | | | | |
|---|---|---|---|---|---|---|---|---|
| ENSG000002 | 4.04881778 | -0.670885228 | 0.3783106 | -1.7733715 | 0.07616717 | 0.16263426 | LOC105371620 | 105371620 |
| ENSG000002 | 9.84535299 | 0.611882505 | 0.34504686 | 1.77333164 | 0.07617377 | 0.16263426 | NA | NA |
| ENSG000002 | 3.49074109 | -0.669773317 | 0.37770573 | -1.7732676 | 0.07618438 | 0.16263426 | NA | NA |
| ENSG000002 | 8.09007743 | -0.630018572 | 0.35530525 | -1.7731755 | 0.07619963 | 0.16265574 | NA | NA |
| ENSG000001 | 5.26102368 | -0.663170743 | 0.37401511 | -1.7731121 | 0.07621013 | 0.16265804 | GALP | 85569 |
| ENSG000002 | 2.21940336 | -0.64002259 | 0.36095727 | -1.7731257 | 0.07620789 | 0.16265804 | NA | NA |
| ENSG000002 | 11.4151969 | -0.591820066 | 0.33377772 | -1.7730964 | 0.07621274 | 0.16265804 | NA | NA |
| ENSG000002 | 167.54754 | 0.324495068 | 0.18301507 | 1.77305108 | 0.07622024 | 0.16266198 | RNY1 | 6084 |
| ENSG000002 | 3.52636929 | -0.669343641 | 0.37751255 | -1.7730368 | 0.07622261 | 0.16266198 | NA | NA |
| ENSG000001 | 116.76899 | -0.296038221 | 0.1669721 | -1.7729802 | 0.07623199 | 0.16267345 | ASAH2 | 56624 |
| ENSG000002 | 14.7140259 | -0.579381393 | 0.3268081 | -1.7728489 | 0.07625374 | 0.16270273 | NA | NA |
| ENSG000002 | 2.19526905 | -0.646124831 | 0.36445436 | -1.7728553 | 0.07625269 | 0.16270273 | NA | NA |
| ENSG000001 | 1437.43097 | 0.221747372 | 0.12509035 | 1.7726977 | 0.07627881 | 0.16274766 | BRWD3 | 254065 |
| ENSG000002 | 3.37219187 | -0.670543637 | 0.37842829 | -1.7719173 | 0.07640829 | 0.16300244 | NA | NA |
| ENSG000002 | 12.6473004 | -0.583121858 | 0.32909313 | -1.7719053 | 0.07641028 | 0.16300244 | NA | NA |
| ENSG000002 | 17.1384458 | -0.525295297 | 0.29644986 | -1.7719533 | 0.07640231 | 0.16300244 | SSTR3 | 6753 |
| ENSG000001 | 46.1157895 | 0.403010094 | 0.22746946 | 1.77171082 | 0.07644257 | 0.16306275 | RUNDC3A | 10900 |
| ENSG000002 | 11.0635946 | -0.589850977 | 0.33294305 | -1.7716273 | 0.07645645 | 0.16308377 | NA | NA |
| ENSG000002 | 2.92429083 | -0.65983366 | 0.37249275 | -1.7714 | 0.07649421 | 0.16315573 | NA | NA |
| ENSG000002 | 2.49923449 | -0.65465313 | 0.36960454 | -1.7712259 | 0.07652314 | 0.16320886 | NA | NA |
| ENSG000002 | 11.108058 | -0.594443809 | 0.33563024 | -1.7711271 | 0.07653957 | 0.1632283 | NA | NA |
| ENSG000002 | 21.7492114 | -0.512762275 | 0.28951257 | -1.7711227 | 0.07654031 | 0.1632283 | NA | NA |
| ENSG000001 | 9.41567147 | -0.610813323 | 0.34487905 | -1.7710943 | 0.07654502 | 0.16322976 | CELF2-AS1 | 414196 |
| ENSG000001 | 15.791912 | -0.568771146 | 0.32118032 | -1.770878 | 0.076581 | 0.1632979 | TRIM73 | 375593 |
| ENSG000002 | 9.63131671 | -0.606929214 | 0.3427442 | -1.7707936 | 0.07659504 | 0.16331924 | NA | NA |
| ENSG000001 | 17.4079793 | 0.514971856 | 0.29082147 | 1.77074907 | 0.07660244 | 0.16332643 | ISL2 | 64843 |
| ENSG00000C | 12.2669992 | -0.562795806 | 0.31783557 | -1.7707137 | 0.07660831 | 0.16332822 | MARCO | 8685 |
| ENSG000001 | 197.446841 | 0.249813815 | 0.1410823 | 1.77069563 | 0.07661133 | 0.16332822 | NKIRAS1 | 28512 |
| ENSG000002 | 4.86211443 | -0.662937605 | 0.37441249 | -1.7706076 | 0.07662597 | 0.16335085 | NA | NA |

| | | | | | | | | |
|---|---|---|---|---|---|---|---|---|
| ENSG000001 | 11.0692327 | -0.586152833 | 0.33105488 | -1.7705609 | 0.07663375 | 0.16335187 | GDF5 | 8200 |
| ENSG000002 | 7.00517729 | -0.644582529 | 0.36405649 | -1.7705563 | 0.07663451 | 0.16335187 | NA | NA |
| ENSG000002 | 3.40956697 | -0.669694347 | 0.37824692 | -1.7705216 | 0.07664029 | 0.1633556 | LOC101927845 | 101927845 |
| ENSG000002 | 8.80444632 | -0.614002411 | 0.34679887 | -1.7704856 | 0.07664628 | 0.16335978 | NA | NA |
| ENSG000002 | 8.63670465 | -0.620992075 | 0.35075493 | -1.7704443 | 0.07665315 | 0.16336584 | NA | NA |
| ENSG000001 | 23.0254911 | -0.469099197 | 0.26499472 | -1.7702209 | 0.07669035 | 0.16343653 | TCP11 | 6954 |
| ENSG000001 | 115.387279 | -0.286951214 | 0.16210603 | -1.7701452 | 0.07670295 | 0.1634548 | ST3GAL5 | 8869 |
| ENSG000001 | 110.472912 | 0.348228388 | 0.19674451 | 1.76995228 | 0.07673509 | 0.1635147 | NA | NA |
| ENSG000002 | 6.58508381 | -0.642989687 | 0.363289 | -1.7699123 | 0.07674175 | 0.16352028 | NA | NA |
| ENSG000001 | 153.893772 | 0.30600065 | 0.17290656 | 1.76974571 | 0.07676951 | 0.16357085 | IBA57 | 200205 |
| ENSG000002 | 3.11507648 | -0.666520033 | 0.37662491 | -1.7697184 | 0.07677407 | 0.16357196 | NA | NA |
| ENSG000001 | 459.182906 | 0.293632662 | 0.16593386 | 1.76957653 | 0.07679771 | 0.16360515 | SGCB | 6443 |
| ENSG000002 | 4.0423464 | -0.669460074 | 0.37831395 | -1.7695886 | 0.07679569 | 0.16360515 | NA | NA |
| ENSG000002 | 4.37843527 | -0.669541295 | 0.37837667 | -1.76951 | 0.07680881 | 0.16362019 | NA | NA |
| ENSG000002 | 10.268079 | -0.602710974 | 0.34065175 | -1.7692878 | 0.07684586 | 0.16369052 | NA | NA |
| ENSG000001 | 6.14295169 | -0.652865214 | 0.36901993 | -1.7691869 | 0.07686269 | 0.16371778 | NA | NA |
| ENSG000002 | 6.9190767 | -0.639922969 | 0.36175291 | -1.7689504 | 0.07690215 | 0.16379322 | NA | NA |
| ENSG000002 | 5.11488011 | -0.66781542 | 0.37754322 | -1.768845 | 0.07691975 | 0.1638221 | NA | NA |
| ENSG000002 | 3.07031751 | 0.665491917 | 0.37623892 | 1.76880136 | 0.07692703 | 0.163829 | NA | NA |
| ENSG000002 | 3.13055269 | -0.666270874 | 0.37670781 | -1.7686675 | 0.07694938 | 0.16386798 | NA | NA |
| ENSG000000 | 2.54016 | -0.654556136 | 0.37009743 | -1.7686049 | 0.07695983 | 0.16388164 | CEBPE | 1053 |
| ENSG000001 | 4.68763861 | -0.6662923 | 0.37675383 | -1.7685084 | 0.07697596 | 0.16390737 | ACTL7B | 10880 |
| ENSG000002 | 10.0792475 | 0.600394423 | 0.33949751 | 1.76847961 | 0.07698076 | 0.16390899 | NA | NA |
| ENSG000002 | 2.65447967 | 0.633364659 | 0.35815466 | 1.76841106 | 0.07699221 | 0.16392477 | NA | NA |
| ENSG000001 | 17.2238457 | -0.543057007 | 0.30714633 | -1.7680726 | 0.07704877 | 0.16403657 | EREG | 2069 |
| ENSG000002 | 6.57963246 | -0.654838275 | 0.37043737 | -1.7677436 | 0.07710378 | 0.16413646 | NA | NA |
| ENSG000002 | 5.30227264 | -0.66368746 | 0.37544238 | -1.7677478 | 0.07710307 | 0.16413646 | NA | NA |
| ENSG000002 | 4.86635392 | -0.662305045 | 0.37469961 | -1.7675627 | 0.07713404 | 0.16419226 | NA | NA |
| ENSG000002 | 2.49415146 | -0.652884451 | 0.36938745 | -1.7674787 | 0.07714809 | 0.16421354 | NA | NA |

| | | | | | | | | |
|---|---|---|---|---|---|---|---|---|
| ENSG000001 | 218.881822 | 0.285295517 | 0.16142937 | 1.7673086 | 0.07717656 | 0.16426552 | THNSL1 | 79896 |
| ENSG000001 | 55.9503409 | -0.345447007 | 0.19546949 | -1.7672681 | 0.07718333 | 0.16427132 | KLRG1 | 10219 |
| ENSG000002 | 5.47064291 | -0.657558649 | 0.37210865 | -1.7671147 | 0.07720903 | 0.16431738 | NA | NA |
| ENSG000002 | 9.94525806 | -0.6015357 | 0.34042223 | -1.7670282 | 0.0772235 | 0.16433955 | NA | NA |
| ENSG000001 | 6.21990898 | -0.659196044 | 0.37306945 | -1.7669526 | 0.07723616 | 0.16435788 | NA | NA |
| ENSG000002 | 9.73558508 | -0.606260695 | 0.34311941 | -1.7669088 | 0.07724351 | 0.16436487 | NA | NA |
| ENSG000001 | 39.4580574 | -0.42176939 | 0.23871363 | -1.7668425 | 0.07725461 | 0.16437987 | PTPRR | 5801 |
| ENSG000002 | 2.79552524 | -0.662717294 | 0.37509149 | -1.7668151 | 0.0772592 | 0.16438102 | NA | NA |
| ENSG000001 | 13.3897172 | -0.568169112 | 0.32159837 | -1.766704 | 0.07727782 | 0.164412 | CABP4 | 57010 |
| ENSG000002 | 13.1867258 | -0.561494595 | 0.3178565 | -1.7665034 | 0.07731143 | 0.16447488 | CASP12 | 100506742 |
| ENSG000002 | 4.85977806 | -0.662306128 | 0.37494117 | -1.7664268 | 0.07732428 | 0.16449359 | NA | NA |
| ENSG000002 | 2.18391108 | -0.643662764 | 0.36440321 | -1.7663477 | 0.07733754 | 0.16451317 | RNU4-75P | 106479593 |
| ENSG000002 | 2.76607035 | -0.663015692 | 0.37540325 | -1.7661427 | 0.07737192 | 0.16457766 | NA | NA |
| ENSG00000C | 128.799037 | -0.27301027 | 0.15458253 | -1.7661133 | 0.07737683 | 0.16457949 | TNC | 3371 |
| ENSG000001 | 2.64182526 | -0.646544197 | 0.36609455 | -1.766058 | 0.07738613 | 0.16458199 | TRADD | 8717 |
| ENSG000002 | 3.50375638 | -0.666675263 | 0.37749171 | -1.766066 | 0.07738478 | 0.16458199 | NA | NA |
| ENSG000002 | 6.17497795 | -0.652816634 | 0.36965792 | -1.766002 | 0.07739551 | 0.16458563 | NA | NA |
| ENSG000002 | 2.24223643 | -0.636012005 | 0.36014282 | -1.7659994 | 0.07739595 | 0.16458563 | NA | NA |
| ENSG000002 | 4.05652893 | -0.668337386 | 0.37847837 | -1.7658536 | 0.07742042 | 0.16462901 | NA | NA |
| ENSG000002 | 12.2721662 | -0.562447097 | 0.31852786 | -1.7657705 | 0.07743436 | 0.16465002 | NA | NA |
| ENSG000002 | 4.00252044 | -0.666130727 | 0.37725362 | -1.7657371 | 0.07743996 | 0.16465331 | NA | NA |
| ENSG000002 | 12.5454424 | -0.578634131 | 0.32771511 | -1.765662 | 0.07745257 | 0.16467148 | NA | NA |
| ENSG000002 | 5.48624909 | -0.656136217 | 0.37164841 | -1.7654757 | 0.07748386 | 0.16472509 | TAAR9 | 134860 |
| ENSG000002 | 11.9589352 | -0.584500475 | 0.33107637 | -1.7654551 | 0.07748731 | 0.16472509 | NA | NA |
| ENSG000002 | 4.03666943 | -0.66826856 | 0.37852821 | -1.7654393 | 0.07748997 | 0.16472509 | NA | NA |
| ENSG000001 | 318.418558 | 0.257493819 | 0.14585748 | 1.76537957 | 0.07749999 | 0.16473776 | TMEM63B | 55362 |
| ENSG000002 | 3.77199477 | -0.668278556 | 0.37855473 | -1.765342 | 0.07750631 | 0.16474254 | NA | NA |
| ENSG000002 | 3.73154609 | -0.668240448 | 0.37856854 | -1.7651769 | 0.07753404 | 0.16479284 | NA | NA |
| ENSG000001 | 52.0894272 | 0.365901562 | 0.20729632 | 1.76511368 | 0.07754466 | 0.16480679 | AMIGO1 | 57463 |

| | | | | | | | | |
|---|---|---|---|---|---|---|---|---|
| ENSG000002 | 18.4784026 | 0.509321914 | 0.28855769 | 1.76506098 | 0.07755352 | 0.16481697 | NA | NA |
| ENSG000002 | 6.45324258 | -0.650191457 | 0.36838074 | -1.7649985 | 0.07756401 | 0.16483064 | NA | NA |
| ENSG000001 | 707.299679 | 0.303922744 | 0.1722181 | 1.76475495 | 0.07760496 | 0.16490901 | AIDA | 64853 |
| ENSG000002 | 4.31766308 | -0.665755044 | 0.3773265 | -1.7644005 | 0.07766458 | 0.16502706 | HCCAT5 | 283902 |
| ENSG000001 | 42.9515158 | 0.438289477 | 0.2484223 | 1.76429203 | 0.07768283 | 0.16505718 | ARL4D | 379 |
| ENSG000001 | 183.712183 | 0.28175278 | 0.15970654 | 1.7641906 | 0.0776999 | 0.1650848 | MAOA | 4128 |
| ENSG000002 | 4.86071582 | -0.662304339 | 0.37542497 | -1.7641457 | 0.07770745 | 0.1650922 | NA | NA |
| ENSG000001 | 5.58267364 | -0.657315361 | 0.37260814 | -1.7640929 | 0.07771634 | 0.16510244 | SEMG2 | 6407 |
| ENSG000001 | 31.8601587 | 0.431170306 | 0.24446814 | 1.76370754 | 0.07778123 | 0.16522298 | NA | NA |
| ENSG000001 | 20.7121312 | -0.499845546 | 0.2834043 | -1.763719 | 0.0777793 | 0.16522298 | TMC3 | 342125 |
| ENSG000002 | 4.07414508 | -0.667329914 | 0.37838644 | -1.7636201 | 0.07779597 | 0.16523442 | KIRREL3-AS3 | 283165 |
| ENSG000002 | 7.91118042 | -0.649191135 | 0.36811007 | -1.7635788 | 0.07780291 | 0.16523442 | LINC01268 | 285758 |
| ENSG000002 | 9.92096535 | 0.604676195 | 0.34285893 | 1.76362972 | 0.07779434 | 0.16523442 | LOC100505771 | 100505771 |
| ENSG000002 | 6.70754219 | -0.645989457 | 0.36629081 | -1.7635972 | 0.07779981 | 0.16523442 | LOC101927286 | 101927286 |
| ENSG000002 | 7.6978228 | -0.625887078 | 0.35491837 | -1.7634677 | 0.07782164 | 0.16526554 | NA | NA |
| ENSG000002 | 6.91731732 | -0.642535394 | 0.36437359 | -1.7633972 | 0.07783351 | 0.16528209 | LINC01185 | 400957 |
| ENSG000002 | 4.26160234 | -0.665059936 | 0.37715793 | -1.763346 | 0.07784215 | 0.16528958 | LINC01494 | 101928537 |
| ENSG000002 | 15.0910169 | 0.58117417 | 0.32958938 | 1.76332796 | 0.07784519 | 0.16528958 | NA | NA |
| ENSG000002 | 197.068787 | 0.575252357 | 0.32626155 | 1.76316321 | 0.07787297 | 0.1653399 | HNRNPUL2 | 221092 |
| ENSG000002 | 8.83051408 | -0.616815685 | 0.34986236 | -1.7630239 | 0.07789647 | 0.16538113 | NA | NA |
| ENSG000001 | 2.56444212 | -0.649790772 | 0.3686036 | -1.7628443 | 0.07792675 | 0.16542809 | UNCX | 340260 |
| ENSG000002 | 26.8050118 | -0.46710113 | 0.26496867 | -1.7628542 | 0.07792509 | 0.16542809 | PSORS1C1 | 170679 |
| ENSG000002 | 717.819947 | 0.196807411 | 0.11165019 | 1.76271451 | 0.07794865 | 0.16546594 | FTX | 100302692 |
| ENSG000002 | 2.49784493 | -0.651638977 | 0.36970193 | -1.7626064 | 0.0779669 | 0.16549602 | NA | NA |
| ENSG000002 | 2.99621201 | -0.653344407 | 0.3706816 | -1.7625488 | 0.07797662 | 0.16550798 | NA | NA |
| ENSG000001 | 55.9499072 | 0.401705184 | 0.22793037 | 1.76240309 | 0.07800122 | 0.16555152 | RUSC2 | 9853 |
| ENSG000001 | 45.6177371 | -0.393273396 | 0.22319093 | -1.7620492 | 0.07806099 | 0.16566971 | KIFC3 | 3801 |
| ENSG000002 | 7.78670452 | -0.628740302 | 0.3568445 | -1.7619448 | 0.07807863 | 0.16569848 | NA | NA |
| ENSG000002 | 7.46991491 | -0.632206877 | 0.35882179 | -1.7618966 | 0.07808678 | 0.16570709 | LOC102724890 | 102724890 |

| | | | | | | | | |
|---|---|---|---|---|---|---|---|---|
| ENSG000001 | 22.9396282 | 0.485925352 | 0.27580676 | 1.76183265 | 0.07809758 | 0.16572134 | SUSD3 | 203328 |
| ENSG000001 | 22.099467 | -0.483734208 | 0.27457159 | -1.7617781 | 0.0781068 | 0.16573058 | SYT5 | 6861 |
| ENSG000002 | 5.76262284 | -0.651244774 | 0.36965609 | -1.7617585 | 0.0781101 | 0.16573058 | NA | NA |
| ENSG000001 | 72.2820908 | 0.333204472 | 0.18914658 | 1.76162042 | 0.07813345 | 0.16576277 | SMPDL3A | 10924 |
| ENSG000001 | 4.15561812 | -0.666940256 | 0.37859045 | -1.7616405 | 0.07813006 | 0.16576277 | UMODL1-AS1 | 150147 |
| ENSG000001 | 103.063871 | 0.335243917 | 0.19032021 | 1.76147308 | 0.07815837 | 0.16580695 | ATPAF2 | 91647 |
| ENSG000002 | 4.55088786 | -0.666786498 | 0.37854704 | -1.7614363 | 0.07816459 | 0.16581147 | LOC101927798 | 101927798 |
| ENSG000002 | 7.38859128 | -0.644714054 | 0.36602996 | -1.7613696 | 0.07817587 | 0.16582673 | NA | NA |
| ENSG000001 | 2.54068152 | -0.650724085 | 0.36945567 | -1.7613049 | 0.07818681 | 0.16582705 | NA | NA |
| ENSG000002 | 3.90267817 | -0.664810019 | 0.37745498 | -1.7612962 | 0.07818829 | 0.16582705 | NA | NA |
| ENSG000002 | 4.89102799 | -0.660536403 | 0.37502641 | -1.7613064 | 0.07818656 | 0.16582705 | NA | NA |
| ENSG000001 | 102.671537 | 0.293054277 | 0.16640637 | 1.76107612 | 0.07822552 | 0.16589637 | C21orf59 | 56683 |
| ENSG000002 | 46.6375152 | 0.38024491 | 0.21591886 | 1.76105465 | 0.07822915 | 0.16589637 | NA | NA |
| ENSG000002 | 3.17648159 | -0.663076105 | 0.37654002 | -1.7609711 | 0.07824329 | 0.16591767 | NA | NA |
| ENSG000001 | 12.8924036 | -0.566353918 | 0.32163442 | -1.7608623 | 0.07826172 | 0.16594807 | DLX1 | 1745 |
| ENSG000002 | 6.88961384 | -0.636396632 | 0.36142741 | -1.7607869 | 0.07827448 | 0.16595946 | NA | NA |
| ENSG000002 | 4.23512821 | -0.663875531 | 0.37703444 | -1.7607822 | 0.07827527 | 0.16595946 | NA | NA |
| ENSG000002 | 4.06808388 | -0.666289102 | 0.37843326 | -1.7606515 | 0.0782974 | 0.16599769 | NA | NA |
| ENSG000001 | 13.742236 | -0.555106688 | 0.31531992 | -1.7604555 | 0.07833061 | 0.16605601 | NCR1 | 9437 |
| ENSG000002 | 5.83335192 | -0.654871593 | 0.37199296 | -1.7604408 | 0.0783331 | 0.16605601 | NA | NA |
| ENSG000002 | 2.17554072 | -0.639889927 | 0.36349053 | -1.7604033 | 0.07833945 | 0.16606081 | MIR944 | 100126340 |
| ENSG000002 | 13.4903511 | -0.556750668 | 0.31627342 | -1.760346 | 0.07834915 | 0.16607268 | LINC00691 | 152024 |
| ENSG000002 | 4.95882042 | -0.663447409 | 0.37689827 | -1.7602824 | 0.07835993 | 0.16608684 | NA | NA |
| ENSG000001 | 18.8644352 | -0.513772681 | 0.29188253 | -1.7602036 | 0.07837329 | 0.16610649 | PLA2G4E | 123745 |
| ENSG000002 | 3.19952263 | -0.661224112 | 0.3756623 | -1.7601556 | 0.07838142 | 0.16610789 | NA | NA |
| ENSG000002 | 2.83628136 | -0.659145036 | 0.37448202 | -1.7601513 | 0.07838215 | 0.16610789 | NA | NA |
| ENSG000002 | 6.08524026 | -0.649374785 | 0.36894233 | -1.7600983 | 0.07839114 | 0.16611826 | NA | NA |
| ENSG000002 | 2.50325381 | -0.650415976 | 0.36956647 | -1.7599432 | 0.07841745 | 0.16616533 | NA | NA |
| ENSG000002 | 8.39777245 | -0.624016131 | 0.35457295 | -1.759909 | 0.07842323 | 0.1661689 | NA | NA |

| | | | | | | | | |
|---|---|---|---|---|---|---|---|---|
| ENSG000001 | 7.9486483 | -0.619020599 | 0.35181153 | -1.7595234 | 0.07848866 | 0.16629015 | IL4I1 | 259307 |
| ENSG000002 | 6.65115981 | -0.639912494 | 0.36368217 | -1.7595377 | 0.07848622 | 0.16629015 | CADM2-AS1 | 100874038 |
| ENSG000001 | 12.6555637 | -0.584432069 | 0.33220049 | -1.7592752 | 0.07853078 | 0.16637071 | C9orf24 | 84688 |
| ENSG000001 | 55.5273677 | 0.376564804 | 0.21405129 | 1.759227 | 0.07853896 | 0.16637893 | TMEM220 | 388335 |
| ENSG000002 | 4.28009155 | -0.663363596 | 0.37708168 | -1.759204 | 0.07854287 | 0.16637893 | NA | NA |
| ENSG000002 | 3.99741606 | -0.665464914 | 0.37828343 | -1.7591701 | 0.07854861 | 0.16638241 | IQCF6 | 440956 |
| ENSG000001 | 39.8608046 | -0.43383971 | 0.24664425 | -1.7589695 | 0.07858269 | 0.16642916 | MORC1 | 27136 |
| ENSG000001 | 47.7692007 | 0.391859031 | 0.22277784 | 1.75896769 | 0.078583 | 0.16642916 | PHOSPHO2 | 493911 |
| ENSG000002 | 6.62199899 | -0.643696193 | 0.36594464 | -1.7589988 | 0.07857771 | 0.16642916 | CST2P1 | 101410546 |
| ENSG000000 | 33.7696067 | -0.413285838 | 0.23500375 | -1.7586351 | 0.07863951 | 0.16645321 | PPEF1 | 5475 |
| ENSG000001 | 295.316881 | 0.246712134 | 0.14026972 | 1.75884104 | 0.07860451 | 0.16645321 | SPATA13 | 221178 |
| ENSG000001 | 44.2912329 | -0.372842069 | 0.21200266 | -1.7586669 | 0.0786341 | 0.16645321 | C2orf88 | 84281 |
| ENSG000002 | 7.20894548 | -0.635501135 | 0.36134272 | -1.7587213 | 0.07862486 | 0.16645321 | NA | NA |
| ENSG000002 | 2.78601383 | -0.65944104 | 0.37497118 | -1.7586446 | 0.07863789 | 0.16645321 | NA | NA |
| ENSG000002 | 5.50680418 | -0.657460816 | 0.37384426 | -1.7586489 | 0.07863716 | 0.16645321 | NA | NA |
| ENSG000002 | 14.2041538 | 0.562715352 | 0.3199555 | 1.75873007 | 0.07862337 | 0.16645321 | NA | NA |
| ENSG000002 | 4.06404549 | -0.665513868 | 0.3784064 | -1.7587278 | 0.07862375 | 0.16645321 | NA | NA |
| ENSG000002 | 2.48811804 | -0.649982528 | 0.36959018 | -1.7586575 | 0.07863571 | 0.16645321 | NA | NA |
| ENSG000002 | 9.11118843 | -0.60817049 | 0.3458107 | -1.7586804 | 0.07863181 | 0.16645321 | NA | NA |
| ENSG000002 | 7.11521064 | -0.631745633 | 0.35918399 | -1.7588357 | 0.07860542 | 0.16645321 | NA | NA |
| ENSG000002 | 11.4507912 | -0.604034194 | 0.34348069 | -1.7585681 | 0.07865089 | 0.16646862 | NA | NA |
| ENSG000002 | 13.4202332 | -0.589048201 | 0.33498169 | -1.758449 | 0.07867115 | 0.16650281 | NA | NA |
| ENSG000001 | 23.6195245 | -0.515313302 | 0.29307257 | -1.758313 | 0.07869428 | 0.16653668 | GP2 | 2813 |
| ENSG000002 | 3.99313615 | -0.665139004 | 0.37828387 | -1.7583065 | 0.07869537 | 0.16653668 | NA | NA |
| ENSG000002 | 7.8686236 | -0.631009895 | 0.35887886 | -1.758281 | 0.07869971 | 0.16653717 | NA | NA |
| ENSG000002 | 12.0134897 | -0.567894991 | 0.32300645 | -1.7581537 | 0.07872136 | 0.16657429 | MTND6P10 | 106478947 |
| ENSG000001 | 59.5323497 | 0.345098596 | 0.19631698 | 1.75786425 | 0.07877061 | 0.16665242 | C3orf33 | 285315 |
| ENSG000001 | 3.96304102 | -0.665039729 | 0.37832017 | -1.7578754 | 0.07876871 | 0.16665242 | NA | NA |
| ENSG000002 | 7.97296754 | -0.621238698 | 0.35340441 | -1.7578691 | 0.07876978 | 0.16665242 | TCONS_000291 | 101928399 |

| | | | | | | | | |
|---|---|---|---|---|---|---|---|---|
| ENSG000001 | 7.9616245 | -0.619057329 | 0.35218616 | -1.757756 | 0.07878904 | 0.16668271 | TMEM202 | 338949 |
| ENSG000002 | 89.0059337 | 0.304881434 | 0.17345421 | 1.75770561 | 0.07879761 | 0.16669216 | NA | NA |
| ENSG000002 | 11.5354363 | -0.590391772 | 0.33589825 | -1.7576506 | 0.07880698 | 0.16670327 | NA | NA |
| ENSG000002 | 5.1888849 | -0.655964791 | 0.37324142 | -1.7574812 | 0.07883582 | 0.16674372 | NA | NA |
| ENSG000002 | 4.97763425 | -0.659895285 | 0.37548114 | -1.7574659 | 0.07883843 | 0.16674372 | LOC105369911 | 105369911 |
| ENSG000002 | 2.78738411 | -0.659017832 | 0.3749736 | -1.7575046 | 0.07883184 | 0.16674372 | NA | NA |
| ENSG000001 | 3.38502372 | -0.664540725 | 0.37813251 | -1.7574282 | 0.07884485 | 0.1667486 | GPR174 | 84636 |
| ENSG00000C | 348.294308 | 0.227184389 | 0.12927658 | 1.75735142 | 0.07885793 | 0.16676755 | EED | 8726 |
| ENSG000002 | 5.45189366 | -0.653666036 | 0.37197163 | -1.7573008 | 0.07886656 | 0.1667771 | NA | NA |
| ENSG000002 | 6.31858002 | 0.644315301 | 0.36666219 | 1.75724502 | 0.07887605 | 0.16678849 | NA | NA |
| ENSG000001 | 12.3482142 | -0.567934816 | 0.32321103 | -1.7571641 | 0.07888985 | 0.16680896 | SERPINA1 | 5265 |
| ENSG000002 | 6.76490119 | -0.644953922 | 0.36706305 | -1.7570658 | 0.0789066 | 0.16682392 | NA | NA |
| ENSG000002 | 2.528045 | -0.649097827 | 0.3694293 | -1.7570285 | 0.07891294 | 0.16682392 | NA | NA |
| ENSG000002 | 35.9368438 | 0.410446225 | 0.23360282 | 1.757026 | 0.07891338 | 0.16682392 | NA | NA |
| ENSG000002 | 4.04101634 | -0.664886347 | 0.37840043 | -1.7570972 | 0.07890124 | 0.16682392 | NA | NA |
| ENSG000001 | 6.61948423 | -0.641367263 | 0.36506881 | -1.7568394 | 0.07894519 | 0.16687806 | ZNF703 | 80139 |
| ENSG000001 | 15.2573407 | -0.533883634 | 0.30389076 | -1.7568275 | 0.07894722 | 0.16687806 | SLC38A3 | 10991 |
| ENSG00000C | 7.15581086 | -0.630477205 | 0.3588876 | -1.7567539 | 0.07895976 | 0.16689587 | TSPAN32 | 10077 |
| ENSG000002 | 6.20094944 | 0.653942577 | 0.37225741 | 1.7566946 | 0.07896988 | 0.16690856 | NA | NA |
| ENSG000001 | 165.782311 | 0.285444014 | 0.16249649 | 1.75661651 | 0.0789832 | 0.16692801 | CDK19 | 23097 |
| ENSG000002 | 10.9505187 | -0.577673787 | 0.3288754 | -1.7565126 | 0.07900092 | 0.16695676 | LOC101927657 | 101927657 |
| ENSG000002 | 3.726835 | -0.66478048 | 0.37851255 | -1.756297 | 0.0790377 | 0.16702579 | NA | NA |
| ENSG000002 | 3.14054093 | -0.661581117 | 0.37669719 | -1.7562677 | 0.0790427 | 0.16702685 | NA | NA |
| ENSG000002 | 4.19456516 | -0.664797423 | 0.37853324 | -1.7562458 | 0.07904644 | 0.16702685 | NA | NA |
| ENSG000002 | 9.38650917 | -0.604630999 | 0.34431783 | -1.7560258 | 0.079084 | 0.16709751 | NA | NA |
| ENSG000002 | 13.6137033 | -0.565533086 | 0.32208871 | -1.7558302 | 0.07911741 | 0.16715938 | ANKRD34C | 390616 |
| ENSG000002 | 17.686764 | -0.516645667 | 0.29427734 | -1.755642 | 0.07914955 | 0.16721858 | F11-AS1 | 285441 |
| ENSG000002 | 6.69256499 | -0.643531584 | 0.36657319 | -1.7555337 | 0.07916806 | 0.16724898 | NA | NA |
| ENSG000002 | 25.3105004 | -0.457820674 | 0.26081886 | -1.7553204 | 0.07920452 | 0.16731727 | WEE2-AS1 | 285962 |

| | | | | | | | | |
|---|---|---|---|---|---|---|---|---|
| ENSG000002 | 17.7363685 | -0.516335039 | 0.29418574 | -1.7551328 | 0.07923659 | 0.16737631 | NA | NA |
| ENSG000002 | 28.4441023 | -0.44632751 | 0.25431898 | -1.7549909 | 0.07926086 | 0.16741885 | FAM66C | 440078 |
| ENSG000002 | 29.4702285 | -0.452009186 | 0.25756372 | -1.7549412 | 0.07926936 | 0.16742809 | TCEA3 | 6920 |
| ENSG000002 | 4.04568058 | -0.663986279 | 0.3783716 | -1.7548523 | 0.07928457 | 0.16745149 | NA | NA |
| ENSG000001 | 10.3913969 | -0.586955471 | 0.33449717 | -1.7547397 | 0.07930384 | 0.16748346 | CHI3L1 | 1116 |
| ENSG000002 | 12.2461766 | -0.581224365 | 0.33131539 | -1.7542933 | 0.07938027 | 0.16762741 | NA | NA |
| ENSG000002 | 6.16425171 | -0.649607202 | 0.37029502 | -1.7542964 | 0.07937974 | 0.16762741 | NA | NA |
| ENSG000001 | 436.661414 | 0.245177221 | 0.13976049 | 1.75426709 | 0.07938475 | 0.16762815 | FAM117B | 150864 |
| ENSG00000C | 759.185942 | 0.211907498 | 0.1208041 | 1.7541416 | 0.07940625 | 0.16766481 | OFD1 | 8481 |
| ENSG000002 | 96.1573711 | 0.315579693 | 0.17990804 | 1.7541167 | 0.07941051 | 0.16766509 | LOC101929378 | 101929378 |
| ENSG000002 | 2.77161506 | -0.657359974 | 0.37476454 | -1.7540613 | 0.07942001 | 0.1676764 | NA | NA |
| ENSG000001 | 23.4762944 | -0.476559135 | 0.2717147 | -1.7538953 | 0.07944845 | 0.16772771 | C11orf87 | 399947 |
| ENSG000001 | 4.43460866 | -0.663426683 | 0.37828382 | -1.7537802 | 0.07946817 | 0.16776055 | SMR3B | 10879 |
| ENSG000002 | 32.5555452 | -0.420807789 | 0.23994656 | -1.7537563 | 0.07947227 | 0.16776055 | NA | NA |
| ENSG000002 | 3.71316101 | -0.663847883 | 0.37854953 | -1.7536619 | 0.07948846 | 0.16777725 | RNU6-919P | 106479972 |
| ENSG000002 | 4.70431717 | 0.661194088 | 0.37703597 | 1.75366316 | 0.07948824 | 0.16777725 | RHPN1-AS1 | 78998 |
| ENSG000001 | 732.694252 | 0.48461348 | 0.27635052 | 1.75361887 | 0.07949583 | 0.16778408 | SSBP3 | 23648 |
| ENSG000002 | 8.32820692 | -0.618825829 | 0.35291298 | -1.7534799 | 0.07951966 | 0.16782564 | NA | NA |
| ENSG000001 | 1187.76619 | 0.198187551 | 0.11302872 | 1.75342655 | 0.07952882 | 0.16783623 | CERS6 | 253782 |
| ENSG000002 | 4.16249424 | -0.663785219 | 0.37857814 | -1.7533638 | 0.07953958 | 0.16785019 | NA | NA |
| ENSG000002 | 4.07076407 | -0.663725544 | 0.37855263 | -1.7533243 | 0.07954635 | 0.16785576 | NA | NA |
| ENSG000002 | 3.44169792 | -0.662627786 | 0.37794365 | -1.7532449 | 0.07955997 | 0.16787577 | NA | NA |
| ENSG000001 | 68.6163488 | 0.352923425 | 0.20132331 | 1.75301815 | 0.07959889 | 0.16794915 | APOBEC3F | 200316 |
| ENSG000002 | 189.028458 | 0.247381746 | 0.14112447 | 1.75293307 | 0.0796135 | 0.16797122 | RPL36A | 6173 |
| ENSG000002 | 2.81997097 | -0.655714269 | 0.3740974 | -1.7527902 | 0.07963802 | 0.16801422 | NA | NA |
| ENSG000002 | 15.5397162 | -0.54564809 | 0.31130776 | -1.752761 | 0.07964304 | 0.16801607 | NA | NA |
| ENSG000001 | 41.2111607 | 0.388612184 | 0.22173753 | 1.75257743 | 0.07967457 | 0.16807384 | KDM4D | 55693 |
| ENSG000002 | 2.93963064 | -0.652408907 | 0.37230155 | -1.7523669 | 0.07971074 | 0.1681414 | NA | NA |
| ENSG000001 | 25.8065862 | -0.451993589 | 0.25795604 | -1.7522117 | 0.07973741 | 0.16818017 | CYP2C8 | 1558 |

| | | | | | | | | |
|---|---|---|---|---|---|---|---|---|
| ENSG000002 | 4.31935521 | -0.662594637 | 0.37814528 | -1.7522224 | 0.07973557 | 0.16818017 | NA | NA |
| ENSG000001 | 16.8081734 | -0.542553953 | 0.30968636 | -1.7519466 | 0.07978299 | 0.16826755 | GPX6 | 257202 |
| ENSG000002 | 15.7373798 | 0.534041851 | 0.30483268 | 1.75191799 | 0.07978791 | 0.16826917 | NA | NA |
| ENSG000002 | 5.14860967 | -0.661758937 | 0.37777414 | -1.7517317 | 0.07981995 | 0.16832799 | NA | NA |
| ENSG000002 | 10.8659737 | -0.574813281 | 0.32816217 | -1.7516135 | 0.07984029 | 0.16835361 | CTXN3 | 613212 |
| ENSG000002 | 4.81802469 | -0.656403048 | 0.37474209 | -1.7516128 | 0.0798404 | 0.16835361 | NA | NA |
| ENSG000001 | 12.351609 | -0.561517362 | 0.32059128 | -1.7515054 | 0.07985889 | 0.16838383 | CD180 | 4064 |
| ENSG000001 | 19.5722281 | 0.507013705 | 0.28949581 | 1.751368 | 0.07988254 | 0.16840743 | RSG1 | 79363 |
| ENSG000002 | 3.80691768 | -0.662722743 | 0.37840104 | -1.7513766 | 0.07988105 | 0.16840743 | NA | NA |
| ENSG000002 | 10.0928696 | -0.608895867 | 0.34766093 | -1.7514072 | 0.07987579 | 0.16840743 | NA | NA |
| ENSG000002 | 11.8769406 | -0.571706857 | 0.32644943 | -1.7512877 | 0.07989637 | 0.16842761 | NA | NA |
| ENSG000002 | 3.42811567 | -0.662103174 | 0.37807156 | -1.7512642 | 0.07990042 | 0.16842761 | NA | NA |
| ENSG000001 | 14.8246612 | -0.546875022 | 0.31228122 | -1.7512261 | 0.07990697 | 0.16843266 | NA | NA |
| ENSG000002 | 5.05129125 | -0.660779551 | 0.37733487 | -1.7511754 | 0.0799157 | 0.1684423 | NA | NA |
| ENSG000001 | 199.341875 | 0.252598056 | 0.14427551 | 1.75080346 | 0.07997977 | 0.16856859 | PARVA | 55742 |
| ENSG000002 | 5.51406105 | -0.652060391 | 0.37244424 | -1.7507598 | 0.0799873 | 0.1685757 | NA | NA |
| ENSG000001 | 176.570487 | 0.251323112 | 0.14355864 | 1.75066518 | 0.0800036 | 0.16860129 | TIPARP | 25976 |
| ENSG000002 | 2.18245626 | -0.63649235 | 0.36359549 | -1.7505507 | 0.08002333 | 0.16862534 | NA | NA |
| ENSG000002 | 9.62053687 | -0.611890097 | 0.34953949 | -1.7505607 | 0.0800216 | 0.16862534 | LOC102724804 | 102724804 |
| ENSG000002 | 5.25299824 | -0.657477787 | 0.37559739 | -1.7504855 | 0.08003458 | 0.16864028 | NA | NA |
| ENSG000001 | 14.5985901 | -0.531750236 | 0.30379021 | -1.7503863 | 0.08005167 | 0.16865798 | UPK1B | 7348 |
| ENSG000001 | 11.0165231 | -0.59778104 | 0.34151805 | -1.7503644 | 0.08005546 | 0.16865798 | NYX | 60506 |
| ENSG000002 | 5.21460573 | -0.655584943 | 0.37453544 | -1.750395 | 0.08005018 | 0.16865798 | LOC105373100 | 105373100 |
| ENSG00000C | 6.01341533 | -0.643425493 | 0.36761928 | -1.7502496 | 0.08007526 | 0.16869094 | CBLN4 | 140689 |
| ENSG000002 | 2.60049955 | -0.643536335 | 0.36773635 | -1.7499938 | 0.08011938 | 0.16877512 | NA | NA |
| ENSG000002 | 7.17229696 | -0.630067503 | 0.36010129 | -1.7496952 | 0.08017092 | 0.16887491 | NA | NA |
| ENSG000002 | 16.2367237 | -0.523843157 | 0.29941187 | -1.7495738 | 0.08019189 | 0.16891031 | NA | NA |
| ENSG000001 | 14.6649367 | -0.552084329 | 0.315572 | -1.7494719 | 0.08020949 | 0.1689309 | TRAPPC3L | 100128327 |
| ENSG000001 | 9.39474515 | -0.601442307 | 0.34379513 | -1.7494207 | 0.08021833 | 0.1689309 | OR52N1 | 79473 |

| | | | | | | | | |
|---|---|---|---|---|---|---|---|---|
| ENSG000002 | 3.71962607 | -0.662039874 | 0.37842769 | -1.7494488 | 0.08021347 | 0.1689309 | NA | NA |
| ENSG000002 | 17.7832628 | 0.502100462 | 0.28700898 | 1.74942423 | 0.08021772 | 0.1689309 | NA | NA |
| ENSG000002 | 25.2698264 | 0.449596954 | 0.25702808 | 1.74921334 | 0.08025415 | 0.16899756 | LOC731157 | 731157 |
| ENSG000002 | 12.8224632 | -0.571628818 | 0.32681237 | -1.749104 | 0.08027304 | 0.16902857 | LOC101928283 | 101928283 |
| ENSG000002 | 27.9913759 | 0.438902954 | 0.25094885 | 1.74897379 | 0.08029555 | 0.16906718 | TMEM9B-AS1 | 493900 |
| ENSG000001 | 289.332659 | 0.27297878 | 0.15608493 | 1.7489118 | 0.08030627 | 0.16908097 | S100A10 | 6281 |
| ENSG00000C | 394.078002 | 0.230715768 | 0.13192477 | 1.7488434 | 0.08031809 | 0.16909709 | FAM168A | 23201 |
| ENSG000001 | 11.9849957 | -0.563700667 | 0.32234887 | -1.7487285 | 0.08033796 | 0.16912563 | FLJ42102 | 399923 |
| ENSG000002 | 9.75246419 | -0.610828979 | 0.34930608 | -1.7486926 | 0.08034416 | 0.16912563 | FLJ31104 | 441072 |
| ENSG000002 | 16.9590797 | 0.523489269 | 0.2993588 | 1.74870179 | 0.08034258 | 0.16912563 | NA | NA |
| ENSG00000C | 7.8208357 | -0.627111328 | 0.35865003 | -1.7485327 | 0.08037182 | 0.16917507 | MISP | 126353 |
| ENSG000002 | 6.91636371 | -0.634013439 | 0.36260881 | -1.7484778 | 0.08038133 | 0.16917753 | NA | NA |
| ENSG000002 | 16.4158935 | 0.525410593 | 0.30049197 | 1.74850128 | 0.08037726 | 0.16917753 | NA | NA |
| ENSG000002 | 4.8540623 | -0.655997915 | 0.3751902 | -1.7484409 | 0.0803877 | 0.16918215 | NA | NA |
| ENSG000001 | 1349.50471 | 0.214858002 | 0.12288776 | 1.7484085 | 0.08039332 | 0.16918519 | TEX15 | 56154 |
| ENSG000001 | 895.929224 | 0.444151036 | 0.25406344 | 1.74818946 | 0.08043123 | 0.16925618 | DLG5 | 9231 |
| ENSG000002 | 6.88834381 | -0.632428739 | 0.36176809 | -1.7481606 | 0.08043621 | 0.1692579 | LOC643201 | 643201 |
| ENSG000002 | 45.5106772 | -0.386221219 | 0.22095243 | -1.7479836 | 0.08046686 | 0.16931361 | SAMD9 | 54809 |
| ENSG000002 | 6.63379683 | -0.63910744 | 0.36564913 | -1.7478708 | 0.0804864 | 0.16934592 | NA | NA |
| ENSG000002 | 25.8815874 | 0.449293007 | 0.25706089 | 1.7478077 | 0.08049733 | 0.16936014 | SNAI3-AS1 | 197187 |
| ENSG000001 | 204.247679 | 0.298766285 | 0.17094161 | 1.74776801 | 0.08050421 | 0.16936582 | KCTD12 | 115207 |
| ENSG000002 | 4.08786769 | -0.661554269 | 0.3785931 | -1.7474018 | 0.08056766 | 0.1694879 | LINC00092 | 100188953 |
| ENSG000002 | 12.2185644 | -0.572664145 | 0.32772639 | -1.7473849 | 0.0805706 | 0.1694879 | NA | NA |
| ENSG000002 | 12.1453226 | -0.576720386 | 0.33005387 | -1.7473523 | 0.08057625 | 0.16949099 | NA | NA |
| ENSG000001 | 298.060364 | 0.233245065 | 0.13348991 | 1.74728605 | 0.08058773 | 0.16950636 | CAMKMT | 79823 |
| ENSG000002 | 7.68162954 | -0.622017201 | 0.35600406 | -1.7472194 | 0.08059929 | 0.16952187 | NA | NA |
| ENSG000002 | 2.68468291 | 0.650336451 | 0.37222833 | 1.7471439 | 0.08061238 | 0.16954062 | NA | NA |
| ENSG000002 | 4.39297131 | -0.660866033 | 0.37827028 | -1.7470736 | 0.08062457 | 0.16955746 | NA | NA |
| ENSG000002 | 5.31540661 | -0.65699917 | 0.37608883 | -1.7469255 | 0.08065026 | 0.1696027 | NA | NA |

| | | | | | | | | |
|---|---|---|---|---|---|---|---|---|
| ENSG000002 | 13.6388863 | -0.547433794 | 0.31340072 | -1.7467535 | 0.08068011 | 0.16965666 | NA | NA |
| ENSG000002 | 11.1745839 | -0.607003897 | 0.34751389 | -1.746704 | 0.0806887 | 0.16966593 | NA | NA |
| ENSG000001 | 9.58430577 | -0.593108366 | 0.33956985 | -1.7466461 | 0.08069874 | 0.16967825 | NA | NA |
| ENSG000001 | 33.6947611 | -0.425431211 | 0.24357888 | -1.746585 | 0.08070935 | 0.16968296 | NA | NA |
| ENSG000002 | 3.39231195 | -0.660653825 | 0.37825194 | -1.7465973 | 0.08070721 | 0.16968296 | NA | NA |
| ENSG000002 | 5.35267842 | -0.656108038 | 0.37567112 | -1.7464958 | 0.08072484 | 0.16970672 | NA | NA |
| ENSG000002 | 2.73206004 | -0.653795674 | 0.37437447 | -1.7463682 | 0.08074699 | 0.1697445 | NA | NA |
| ENSG000002 | 5.64364315 | -0.654171685 | 0.37460556 | -1.7462946 | 0.08075977 | 0.16976255 | SPANXN5 | 494197 |
| ENSG000002 | 13.7607459 | -0.575395731 | 0.3295232 | -1.7461463 | 0.08078553 | 0.16980789 | NA | NA |
| ENSG000001 | 32.2428312 | -0.446230499 | 0.25561008 | -1.7457469 | 0.08085495 | 0.16993407 | LGR5 | 8549 |
| ENSG000002 | 6.59582832 | -0.635867831 | 0.36423631 | -1.7457563 | 0.08085331 | 0.16993407 | NA | NA |
| ENSG000002 | 44.4646339 | -0.383994653 | 0.21996649 | -1.7456962 | 0.08086376 | 0.16993407 | DUBR | 344595 |
| ENSG000002 | 5.76848975 | -0.649407403 | 0.37199859 | -1.7457254 | 0.08085867 | 0.16993407 | NA | NA |
| ENSG000002 | 5.75659402 | -0.651057351 | 0.37295335 | -1.7456804 | 0.08086651 | 0.16993407 | NA | NA |
| ENSG000002 | 2.50462375 | -0.642884951 | 0.36828349 | -1.7456252 | 0.08087609 | 0.16994541 | NA | NA |
| ENSG000001 | 42.7695929 | 0.424313931 | 0.2430981 | 1.74544321 | 0.08090775 | 0.16999431 | C9orf116 | 138162 |
| ENSG000002 | 2.49098201 | -0.644772115 | 0.36939905 | -1.7454623 | 0.08090443 | 0.16999431 | NA | NA |
| ENSG00000C | 49.1803809 | 0.372261897 | 0.2132936 | 1.74530274 | 0.08093219 | 0.17003684 | TBC1D25 | 4943 |
| ENSG000002 | 16.3417277 | -0.525204052 | 0.3009442 | -1.7451875 | 0.08095224 | 0.17007017 | LINC01550 | 388011 |
| ENSG000001 | 64.3896298 | 0.327854505 | 0.18787111 | 1.74510336 | 0.08096688 | 0.17008329 | ZNF584 | 201514 |
| ENSG000002 | 14.5666089 | -0.542270679 | 0.31073593 | -1.7451174 | 0.08096444 | 0.17008329 | NA | NA |
| ENSG000002 | 5.15863222 | -0.652735542 | 0.37405864 | -1.7450086 | 0.08098337 | 0.17010913 | NA | NA |
| ENSG000001 | 15.3903504 | -0.527354967 | 0.30222457 | -1.744911 | 0.08100036 | 0.17013601 | TRPV5 | 56302 |
| ENSG000001 | 199.276356 | 0.246321605 | 0.14116768 | 1.74488666 | 0.0810046 | 0.17013609 | BCAS3 | 54828 |
| ENSG000002 | 3.12656584 | -0.655867961 | 0.3759006 | -1.744791 | 0.08102126 | 0.17016228 | RFPL4AL1 | 729974 |
| ENSG000002 | 9.90115147 | 0.609975872 | 0.34962292 | 1.74466786 | 0.0810427 | 0.17019848 | NA | NA |
| ENSG000002 | 17.6850944 | -0.515704666 | 0.29559741 | -1.7446184 | 0.08105132 | 0.17020778 | FAM21EP | 100421577 |
| ENSG000002 | 4.60727594 | -0.657650868 | 0.37697583 | -1.7445439 | 0.0810643 | 0.17022621 | NA | NA |
| ENSG000001 | 1124.95582 | 0.4817529 | 0.27618666 | 1.74430187 | 0.08110647 | 0.1702883 | HIF1AN | 55662 |

| | | | | | | | | |
|---|---|---|---|---|---|---|---|---|
| ENSG000002 | 7.37147258 | -0.620252121 | 0.35558639 | -1.7443078 | 0.08110543 | 0.1702883 | LINC01548 | 728409 |
| ENSG000002 | 5.99492667 | -0.641242141 | 0.36761365 | -1.7443372 | 0.08110032 | 0.1702883 | NA | NA |
| ENSG000002 | 4.04645657 | -0.660261149 | 0.37856284 | -1.7441256 | 0.08113718 | 0.17034398 | NA | NA |
| ENSG000002 | 6.54416365 | -0.647493396 | 0.37127223 | -1.7439855 | 0.08116162 | 0.17038645 | NA | NA |
| ENSG000002 | 6.01148444 | -0.641113113 | 0.36763156 | -1.7439012 | 0.08117632 | 0.17040516 | LINC01166 | 101927590 |
| ENSG000002 | 8.72623931 | 0.618684131 | 0.35477322 | 1.74388621 | 0.08117893 | 0.17040516 | NA | NA |
| ENSG000002 | 2.7671797 | -0.653450747 | 0.37471632 | -1.7438545 | 0.08118447 | 0.17040795 | NA | NA |
| ENSG000002 | 2.77102166 | -0.652879858 | 0.37441956 | -1.7437119 | 0.08120934 | 0.17045133 | NA | NA |
| ENSG000002 | 5.98536731 | -0.642060153 | 0.36825068 | -1.743541 | 0.08123916 | 0.17050511 | NA | NA |
| ENSG000001 | 87.3074833 | 0.338759765 | 0.19431233 | 1.74337757 | 0.08126768 | 0.17054732 | HIATL2 | 84278 |
| ENSG000002 | 3.97512824 | -0.659869642 | 0.3784956 | -1.7434011 | 0.08126358 | 0.17054732 | NA | NA |
| ENSG000001 | 12.1238375 | -0.566599047 | 0.32501443 | -1.7433043 | 0.08128048 | 0.17056534 | NA | NA |
| ENSG000001 | 27.6045271 | -0.43909531 | 0.25191892 | -1.7430025 | 0.08133317 | 0.17066708 | AIM2 | 9447 |
| ENSG000001 | 1377.36587 | 0.195689532 | 0.11227756 | 1.74290873 | 0.08134956 | 0.17069196 | ASH1L | 55870 |
| ENSG000002 | 2.40576985 | -0.647876229 | 0.371726 | -1.7428865 | 0.08135344 | 0.17069196 | RNF126P1 | 376412 |
| ENSG000001 | 16.1988093 | -0.528356765 | 0.30321001 | -1.7425439 | 0.08141331 | 0.17080874 | TMPRSS12 | 283471 |
| ENSG000002 | 9.91990421 | -0.610110786 | 0.3501347 | -1.7425031 | 0.08142046 | 0.17081489 | POLR2J4 | 84820 |
| ENSG000001 | 9.03570844 | -0.599911985 | 0.34431422 | -1.7423387 | 0.0814492 | 0.17086634 | HAO1 | 54363 |
| ENSG000002 | 4.70440546 | -0.657828092 | 0.37756708 | -1.7422814 | 0.08145922 | 0.17087854 | NA | NA |
| ENSG000001 | 29.0416765 | 0.463941118 | 0.26631099 | 1.74210279 | 0.08149046 | 0.17093522 | GDPGP1 | 390637 |
| ENSG000002 | 41.9368379 | 0.414068694 | 0.23768988 | 1.74205435 | 0.08149893 | 0.17094415 | NA | NA |
| ENSG000002 | 3.81773411 | -0.658895067 | 0.3782397 | -1.742004 | 0.08150775 | 0.17095379 | NA | NA |
| ENSG000002 | 8.20741177 | -0.624215864 | 0.35839404 | -1.7417027 | 0.08156048 | 0.17105555 | NA | NA |
| ENSG000001 | 161.238527 | 0.26233464 | 0.15063114 | 1.74156978 | 0.08158375 | 0.17109551 | SH2D4A | 63898 |
| ENSG000001 | 9.679923 | -0.599360807 | 0.34415666 | -1.7415348 | 0.08158987 | 0.17109605 | HCAR2 | 338442 |
| ENSG000002 | 24.3938719 | -0.47792783 | 0.27443141 | -1.7415201 | 0.08159245 | 0.17109605 | NA | NA |
| ENSG000002 | 3.11927904 | -0.655526911 | 0.37642004 | -1.7414772 | 0.08159997 | 0.17110298 | NA | NA |
| ENSG000002 | 9.71049388 | -0.60330237 | 0.34644482 | -1.7414097 | 0.08161179 | 0.1711189 | NA | NA |
| ENSG000002 | 6.1817354 | -0.647450817 | 0.37180711 | -1.7413621 | 0.08162013 | 0.17112386 | NA | NA |

| | | | | | | | | |
|---|---|---|---|---|---|---|---|---|
| ENSG000002 | 4.56751131 | -0.657291598 | 0.37746136 | -1.741348 | 0.08162259 | 0.17112386 | NA | NA |
| ENSG000002 | 10.4650547 | -0.583147359 | 0.3348889 | -1.7413159 | 0.08162823 | 0.17112683 | LOC105372751 | 105372751 |
| ENSG000001 | 1602.1298 | 0.507462561 | 0.29145284 | 1.74114813 | 0.08165762 | 0.17115968 | TOP1 | 7150 |
| ENSG000002 | 2.54663118 | -0.641488537 | 0.36843584 | -1.7411133 | 0.08166373 | 0.17115968 | NA | NA |
| ENSG000002 | 2.19734429 | -0.632586961 | 0.36332878 | -1.7410869 | 0.08166835 | 0.17115968 | NA | NA |
| ENSG000002 | 2.55488441 | -0.64195373 | 0.36870186 | -1.7411188 | 0.08166276 | 0.17115968 | NA | NA |
| ENSG000002 | 7.14429688 | -0.625813924 | 0.35943968 | -1.7410819 | 0.08166922 | 0.17115968 | LOC101927430 | 101927430 |
| ENSG000002 | 2.92136235 | -0.647100912 | 0.37164604 | -1.7411753 | 0.08165285 | 0.17115968 | MLECP1 | 106480317 |
| ENSG000002 | 7.20292116 | -0.63422637 | 0.36429074 | -1.7409895 | 0.08168541 | 0.17117591 | NA | NA |
| ENSG000002 | 2.83513975 | -0.651476523 | 0.37419413 | -1.7410121 | 0.08168145 | 0.17117591 | NA | NA |
| ENSG000002 | 2.78149807 | -0.650219692 | 0.3734869 | -1.7409438 | 0.08169343 | 0.17118388 | NA | NA |
| ENSG000002 | 2.81129319 | -0.649505845 | 0.37311171 | -1.7407812 | 0.08172194 | 0.17123477 | WWC2-AS1 | 101928734 |
| ENSG000002 | 9.73606693 | -0.600255511 | 0.34483596 | -1.7406987 | 0.0817364 | 0.17125622 | SPATA31D5P | 347127 |
| ENSG000001 | 44.9485752 | 0.370604737 | 0.21291567 | 1.74061746 | 0.08175065 | 0.17126986 | H3F3C | 440093 |
| ENSG000002 | 20.1973004 | -0.492459355 | 0.28292287 | -1.7406134 | 0.08175136 | 0.17126986 | LOC442028 | 442028 |
| ENSG000002 | 24.9945367 | 0.462362633 | 0.265647 | 1.74051519 | 0.08176859 | 0.17128024 | NA | NA |
| ENSG000002 | 4.27124546 | -0.656956023 | 0.37744966 | -1.740513 | 0.08176899 | 0.17128024 | NA | NA |
| ENSG000002 | 3.08224775 | -0.654524563 | 0.3760494 | -1.7405281 | 0.08176632 | 0.17128024 | NA | NA |
| ENSG000002 | 5.29045373 | -0.653152716 | 0.37533419 | -1.7401898 | 0.0818257 | 0.17138757 | NA | NA |
| ENSG000002 | 5.72385709 | -0.644102917 | 0.37013733 | -1.7401728 | 0.08182868 | 0.17138757 | NA | NA |
| ENSG000001 | 85.8919745 | 0.306838979 | 0.17633865 | 1.74005513 | 0.08184934 | 0.17141314 | THAP10 | 56906 |
| ENSG000002 | 32.2295161 | 0.49061665 | 0.28195099 | 1.74007776 | 0.08184536 | 0.17141314 | UGDH-AS1 | 100885776 |
| ENSG000001 | 945.502607 | 0.217022413 | 0.12473195 | 1.73991044 | 0.08187475 | 0.17145749 | SNX27 | 81609 |
| ENSG000002 | 6.26758289 | -0.639248995 | 0.36743752 | -1.7397488 | 0.08190313 | 0.17149554 | NA | NA |
| ENSG000002 | 2.80881355 | -0.649122073 | 0.3731081 | -1.7397695 | 0.08189951 | 0.17149554 | NA | NA |
| ENSG000002 | 2.49485642 | -0.641176977 | 0.3685487 | -1.7397348 | 0.0819056 | 0.17149554 | NA | NA |
| ENSG000002 | 6.8828553 | -0.631006365 | 0.36271227 | -1.7396885 | 0.08191373 | 0.17150369 | NA | NA |
| ENSG000002 | 4.26629361 | -0.65680825 | 0.37755396 | -1.7396407 | 0.08192212 | 0.17151241 | NA | NA |
| ENSG000002 | 23.8622828 | -0.495608911 | 0.28491518 | -1.7394964 | 0.08194749 | 0.17155668 | LOC101927598 | 101927598 |

| | | | | | | | | |
|---|---|---|---|---|---|---|---|---|
| ENSG000002 | 4.08762982 | -0.658486 | 0.37856587 | -1.7394225 | 0.08196047 | 0.17157499 | LOC101929445 | 101929445 |
| ENSG000002 | 28.1142193 | -0.451807772 | 0.25975601 | -1.7393544 | 0.08197244 | 0.17159118 | NA | NA |
| ENSG000002 | 11.1669641 | -0.591203605 | 0.3399139 | -1.7392746 | 0.08198648 | 0.17161171 | NA | NA |
| ENSG000001 | 68.5898273 | -0.321716677 | 0.18497752 | -1.7392204 | 0.08199601 | 0.1716228 | CDH23 | 64072 |
| ENSG000002 | 27.3141405 | -0.445522642 | 0.25618031 | -1.7390979 | 0.08201754 | 0.17165858 | XKR9 | 389668 |
| ENSG000002 | 2.83818842 | -0.646924643 | 0.37199352 | -1.739075 | 0.08202157 | 0.17165858 | NA | NA |
| ENSG000001 | 591.515791 | 0.49778744 | 0.28625928 | 1.73893905 | 0.08204548 | 0.17169977 | CGN | 57530 |
| ENSG000002 | 3.74373301 | -0.658044612 | 0.37843821 | -1.7388429 | 0.08206239 | 0.17172629 | NA | NA |
| ENSG000001 | 25.1388386 | -0.461256046 | 0.2652784 | -1.7387622 | 0.08207661 | 0.17174717 | EOMES | 8320 |
| ENSG000002 | 3.08211136 | -0.653970545 | 0.37617656 | -1.738467 | 0.08212856 | 0.17184701 | NA | NA |
| ENSG00000C | 17.5459047 | -0.556938739 | 0.32042094 | -1.7381471 | 0.08218489 | 0.17195174 | DAPP1 | 27071 |
| ENSG000002 | 3.4489856 | -0.656706545 | 0.37782259 | -1.7381347 | 0.08218709 | 0.17195174 | NA | NA |
| ENSG000002 | 3.68444645 | -0.657955441 | 0.37858852 | -1.737917 | 0.08222543 | 0.17201421 | NA | NA |
| ENSG000002 | 3.9691871 | -0.657581211 | 0.37836958 | -1.7379336 | 0.08222251 | 0.17201421 | NA | NA |
| ENSG000002 | 4.37385754 | -0.65727706 | 0.37822487 | -1.7377944 | 0.08224705 | 0.17205056 | NA | NA |
| ENSG000002 | 8.04110006 | -0.614639535 | 0.3537104 | -1.7376914 | 0.0822652 | 0.17207965 | NA | NA |
| ENSG000002 | 28.7103908 | -0.454959996 | 0.26184078 | -1.7375444 | 0.08229112 | 0.17211611 | LOC100128164 | 100128164 |
| ENSG000002 | 10.0596009 | -0.598088808 | 0.34421315 | -1.7375536 | 0.0822895 | 0.17211611 | NA | NA |
| ENSG000001 | 39.3246357 | -0.40132734 | 0.23098897 | -1.7374308 | 0.08231116 | 0.17214916 | FBLN5 | 10516 |
| ENSG000001 | 6.88113935 | -0.631950364 | 0.36374971 | -1.737322 | 0.08233035 | 0.17217153 | NA | NA |
| ENSG000002 | 8.70516114 | -0.612733729 | 0.35268472 | -1.7373413 | 0.08232694 | 0.17217153 | KLRF2 | 100431172 |
| ENSG000002 | 6.9027027 | -0.628800056 | 0.36197609 | -1.7371315 | 0.08236396 | 0.17223294 | NA | NA |
| ENSG000002 | 3.23343599 | -0.652234783 | 0.3754924 | -1.7370119 | 0.08238506 | 0.17226817 | NA | NA |
| ENSG000002 | 5.25085165 | -0.652402685 | 0.37560015 | -1.7369607 | 0.08239411 | 0.17227821 | FAM24A | 118670 |
| ENSG000002 | 2.23144757 | -0.625406634 | 0.36007849 | -1.7368619 | 0.08241154 | 0.17230577 | NA | NA |
| ENSG000001 | 382.39536 | 0.220920722 | 0.12720292 | 1.73675829 | 0.08242984 | 0.17233423 | HECTD2 | 143279 |
| ENSG000001 | 4.58926943 | -0.655050272 | 0.37717304 | -1.7367367 | 0.08243365 | 0.17233423 | OR4A16 | 81327 |
| ENSG000002 | 4.30676156 | -0.655789526 | 0.3776292 | -1.7365965 | 0.08245843 | 0.17237713 | NA | NA |
| ENSG000001 | 14.9791022 | -0.536756047 | 0.30910025 | -1.7365112 | 0.08247349 | 0.17239973 | RNF183 | 138065 |

| | | | | | | | | |
|---|---|---|---|---|---|---|---|---|
| ENSG000001 | 22.0803948 | 0.477859612 | 0.27522673 | 1.73623982 | 0.08252144 | 0.17248492 | FAM50B | 26240 |
| ENSG000001 | 318.345432 | 0.248617755 | 0.14319779 | 1.73618428 | 0.08253126 | 0.17248492 | C1D | 10438 |
| ENSG000002 | 14.108318 | -0.536475164 | 0.30898833 | -1.7362312 | 0.08252297 | 0.17248492 | HAGLR | 401022 |
| ENSG000002 | 3.41087329 | -0.656413206 | 0.37807609 | -1.7361934 | 0.08252965 | 0.17248492 | NA | NA |
| ENSG000001 | 41.3487458 | -0.389153023 | 0.22415739 | -1.7360705 | 0.08255138 | 0.17251808 | EPSTI1 | 94240 |
| ENSG000001 | 3.3952486 | -0.656580563 | 0.37821744 | -1.735987 | 0.08256613 | 0.17254002 | OR2T27 | 403239 |
| ENSG000001 | 97.9049471 | 0.306281607 | 0.17643872 | 1.73590926 | 0.08257988 | 0.17255986 | ALPK2 | 115701 |
| ENSG000002 | 3.82173282 | -0.656602884 | 0.37826268 | -1.7358384 | 0.08259241 | 0.17257715 | NA | NA |
| ENSG000002 | 15.6869993 | -0.523855142 | 0.3018084 | -1.7357209 | 0.0826132 | 0.17261169 | LOC100506071 | 100506071 |
| ENSG000002 | 3.83230208 | -0.656214937 | 0.37807394 | -1.7356788 | 0.08262064 | 0.17261447 | NA | NA |
| ENSG000002 | 3.21833247 | -0.651400782 | 0.37530323 | -1.7356653 | 0.08262304 | 0.17261447 | NA | NA |
| ENSG000001 | 89.1442163 | 0.304170489 | 0.17525929 | 1.73554556 | 0.08264422 | 0.17263655 | BBS12 | 166379 |
| ENSG000002 | 4.38937068 | -0.656555816 | 0.37829387 | -1.7355709 | 0.08263974 | 0.17263655 | NA | NA |
| ENSG000002 | 7.70207024 | -0.61808996 | 0.35613834 | -1.7355333 | 0.08264638 | 0.17263655 | NA | NA |
| ENSG000001 | 10.1890761 | -0.588762857 | 0.33925922 | -1.7354366 | 0.08266351 | 0.17266343 | NA | NA |
| ENSG000002 | 3.95201476 | -0.656791722 | 0.37849151 | -1.7352879 | 0.08268983 | 0.17269173 | NA | NA |
| ENSG000002 | 4.60183537 | -0.654191281 | 0.37698506 | -1.7353242 | 0.08268341 | 0.17269173 | NA | NA |
| ENSG000002 | 12.7534404 | -0.553202768 | 0.31879536 | -1.7352912 | 0.08268925 | 0.17269173 | NA | NA |
| ENSG000002 | 9.12181189 | 0.5994063 | 0.34543285 | 1.73523247 | 0.08269964 | 0.17270332 | NA | NA |
| ENSG000001 | 9.33517115 | -0.593820478 | 0.34226436 | -1.7349761 | 0.08274504 | 0.17278923 | INHA | 3623 |
| ENSG000002 | 8.61557522 | -0.608911109 | 0.35098639 | -1.7348568 | 0.08276618 | 0.17282448 | NA | NA |
| ENSG000002 | 4.24859313 | -0.655353728 | 0.37778031 | -1.7347482 | 0.08278541 | 0.17285573 | NA | NA |
| ENSG000002 | 4.31624127 | -0.654968658 | 0.37758759 | -1.7346138 | 0.08280923 | 0.17289656 | NA | NA |
| ENSG000002 | 2.93749069 | -0.64461441 | 0.37164172 | -1.7345049 | 0.08282853 | 0.17291905 | NA | NA |
| ENSG000002 | 3.02512871 | -0.654634168 | 0.37741713 | -1.734511 | 0.08282745 | 0.17291905 | NA | NA |
| ENSG000001 | 2.75824078 | -0.647946363 | 0.3736053 | -1.7343072 | 0.08286359 | 0.17298334 | SPATA31D3 | 389762 |
| ENSG000002 | 4.30579568 | -0.65546674 | 0.37797974 | -1.7341319 | 0.08289467 | 0.17303211 | NA | NA |
| ENSG000002 | 2.76152289 | -0.648396358 | 0.37390354 | -1.7341274 | 0.08289549 | 0.17303211 | NA | NA |
| ENSG000001 | 9.82090876 | -0.584215055 | 0.33691368 | -1.73402 | 0.08291453 | 0.17304891 | MPL | 4352 |

| | | | | | | | | |
|---|---|---|---|---|---|---|---|---|
| ENSG000001 | 27.733368 | -0.43260704 | 0.24948308 | -1.7340136 | 0.08291568 | 0.17304891 | ERN2 | 10595 |
| ENSG000002 | 17.2442418 | -0.549762203 | 0.31704676 | -1.7340098 | 0.08291634 | 0.17304891 | NA | NA |
| ENSG000002 | 3.69492029 | -0.656296881 | 0.37849868 | -1.7339476 | 0.08292738 | 0.17306306 | NA | NA |
| ENSG000002 | 6.31416868 | -0.634638022 | 0.36604074 | -1.7337907 | 0.08295523 | 0.17311226 | RAI1-AS1 | 100861516 |
| ENSG000002 | 3.70338184 | -0.656089284 | 0.37844514 | -1.7336444 | 0.0829812 | 0.17315755 | NA | NA |
| ENSG000001 | 21.3067316 | -0.489280001 | 0.28225043 | -1.7334961 | 0.08300754 | 0.1732036 | SULT1C4 | 27233 |
| ENSG000002 | 3.70022448 | -0.655853553 | 0.37839279 | -1.7332612 | 0.08304925 | 0.17328172 | NA | NA |
| ENSG000001 | 114.968333 | -0.285169478 | 0.1645321 | -1.7332148 | 0.08305749 | 0.17328999 | SLC8A1 | 6546 |
| ENSG000001 | 2.55001165 | -0.638540151 | 0.3684495 | -1.7330466 | 0.08308739 | 0.17334345 | APOA4 | 337 |
| ENSG000001 | 10.6078705 | -0.594686107 | 0.34316447 | -1.7329478 | 0.08310495 | 0.17337117 | CST8 | 10047 |
| ENSG000002 | 2.75229635 | -0.647413072 | 0.37359735 | -1.7329166 | 0.08311049 | 0.1733738 | NA | NA |
| ENSG000001 | 342.035667 | 0.232479478 | 0.13416049 | 1.73284601 | 0.08312304 | 0.17339108 | DNAL1 | 83544 |
| ENSG000002 | 3.03062136 | -0.653603308 | 0.37720337 | -1.7327611 | 0.08313815 | 0.17341366 | NA | NA |
| ENSG000001 | 4.42319065 | -0.655842435 | 0.37852655 | -1.7326194 | 0.08316335 | 0.17345618 | NA | NA |
| ENSG000002 | 7.58708971 | -0.628357159 | 0.36266753 | -1.7325983 | 0.08316708 | 0.17345618 | NA | NA |
| ENSG000001 | 8.57920848 | -0.610310651 | 0.35228575 | -1.7324307 | 0.08319691 | 0.17350945 | ZMYND15 | 84225 |
| ENSG000002 | 3.4596458 | -0.654473146 | 0.3777929 | -1.7323596 | 0.08320956 | 0.17352692 | NA | NA |
| ENSG000002 | 4.43908444 | -0.655581948 | 0.37844725 | -1.7322941 | 0.08322121 | 0.17354228 | NA | NA |
| ENSG000001 | 701.423221 | 0.211783296 | 0.12226308 | 1.73219332 | 0.08323915 | 0.17357077 | MLH3 | 27030 |
| ENSG000002 | 7.13908498 | -0.622069363 | 0.35912917 | -1.7321605 | 0.08324499 | 0.17357402 | NA | NA |
| ENSG00000( | 438.198358 | 0.269926824 | 0.15583671 | 1.73211318 | 0.08325341 | 0.17358266 | BRPF3 | 27154 |
| ENSG000002 | 3.35428287 | -0.654773115 | 0.37805702 | -1.7319428 | 0.08328375 | 0.17363699 | PRAMEF9 | 343070 |
| ENSG000002 | 2.41191538 | -0.642825924 | 0.37117187 | -1.7318821 | 0.08329455 | 0.17364164 | NA | NA |
| ENSG000002 | 5.20773597 | -0.649388868 | 0.37495737 | -1.7319005 | 0.08329128 | 0.17364164 | NA | NA |
| ENSG000002 | 6.08019434 | -0.63942294 | 0.36921872 | -1.731827 | 0.08330438 | 0.17364428 | NA | NA |
| ENSG000002 | 9.68736276 | -0.594000059 | 0.34298733 | -1.7318426 | 0.08330159 | 0.17364428 | NA | NA |
| ENSG000002 | 8.37080298 | -0.616306685 | 0.35589712 | -1.731699 | 0.08332717 | 0.17368286 | NA | NA |
| ENSG000002 | 2.43463188 | -0.642116788 | 0.3708077 | -1.7316706 | 0.08333223 | 0.17368448 | NA | NA |
| ENSG000001 | 50.8692816 | -0.362545764 | 0.20937383 | -1.7315715 | 0.08334988 | 0.17371234 | SORCS3 | 22986 |

| | | | | | | | | |
|---|---|---|---|---|---|---|---|---|
| ENSG000002 | 5.67242759 | -0.641161256 | 0.37036523 | -1.7311594 | 0.08342334 | 0.17384853 | NA | NA |
| ENSG000002 | 5.15139302 | -0.64612326 | 0.37323209 | -1.7311568 | 0.0834238 | 0.17384853 | MEG9 | 100507257 |
| ENSG000001 | 38.8310556 | 0.396928417 | 0.22929287 | 1.73109793 | 0.0834343 | 0.17386147 | NLRP7 | 199713 |
| ENSG000001 | 60.8277097 | 0.366937979 | 0.21197412 | 1.73105086 | 0.08344269 | 0.17387003 | ADRA1A | 148 |
| ENSG000002 | 6.02776203 | -0.637921909 | 0.3685442 | -1.7309238 | 0.08346536 | 0.17390833 | LOC101928443 | 101928443 |
| ENSG000002 | 11.7652621 | 0.56340616 | 0.32550953 | 1.73084381 | 0.08347963 | 0.17392911 | NA | NA |
| ENSG000002 | 21.4546613 | -0.483574052 | 0.2793962 | -1.7307825 | 0.08349056 | 0.17394296 | NA | NA |
| ENSG000002 | 3.63515167 | -0.655250911 | 0.37859325 | -1.7307517 | 0.08349606 | 0.17394547 | NA | NA |
| ENSG000001 | 193.055934 | 0.502511193 | 0.29035945 | 1.7306521 | 0.08351383 | 0.17396463 | C22orf46 | 79640 |
| ENSG000002 | 8.24429131 | -0.623578498 | 0.36030962 | -1.730674 | 0.08350992 | 0.17396463 | NA | NA |
| ENSG000001 | 21.4576445 | -0.49067616 | 0.28359278 | -1.7302139 | 0.08359207 | 0.17411561 | SLC9A4 | 389015 |
| ENSG000002 | 3.40054537 | -0.654113159 | 0.37805681 | -1.730198 | 0.0835949 | 0.17411561 | NA | NA |
| ENSG000001 | 30.6711846 | 0.450367021 | 0.26035328 | 1.7298304 | 0.08366058 | 0.17424346 | ARTN | 9048 |
| ENSG000001 | 91.0516961 | 0.303550575 | 0.17548694 | 1.72976164 | 0.08367287 | 0.17424919 | EML3 | 256364 |
| ENSG000001 | 3.0502001 | -0.652395482 | 0.37717076 | -1.7297085 | 0.08368236 | 0.17424919 | FOXR1 | 283150 |
| ENSG000002 | 3.67800807 | -0.654850927 | 0.37859334 | -1.7296948 | 0.08368482 | 0.17424919 | NA | NA |
| ENSG000002 | 4.01435897 | -0.654767256 | 0.37853827 | -1.7297254 | 0.08367935 | 0.17424919 | NA | NA |
| ENSG000002 | 4.8446947 | -0.650285872 | 0.3759343 | -1.7297859 | 0.08366853 | 0.17424919 | NA | NA |
| ENSG000001 | 132.265988 | 0.315580332 | 0.18245775 | 1.7296077 | 0.08370039 | 0.17426391 | ALAD | 210 |
| ENSG000002 | 11.8613853 | -0.579780599 | 0.3352094 | -1.7296072 | 0.08370048 | 0.17426391 | NA | NA |
| ENSG000001 | 25.9134317 | -0.474902185 | 0.27458548 | -1.729524 | 0.08371536 | 0.17428593 | SLC17A8 | 246213 |
| ENSG000002 | 11.4089851 | -0.57666301 | 0.33346843 | -1.7292882 | 0.08375753 | 0.17436478 | NA | NA |
| ENSG000001 | 6.01954071 | -0.638857446 | 0.36945417 | -1.7291927 | 0.08377462 | 0.17439004 | NA | NA |
| ENSG000002 | 3.40867385 | -0.653789182 | 0.37809372 | -1.7291723 | 0.08377826 | 0.17439004 | NA | NA |
| ENSG000002 | 20.4157513 | 0.535034274 | 0.3094715 | 1.72886442 | 0.08383336 | 0.17449577 | NA | NA |
| ENSG000001 | 37.8313573 | -0.396557977 | 0.22938964 | -1.7287528 | 0.08385334 | 0.17452841 | ZFYVE28 | 57732 |
| ENSG000002 | 3.42919834 | -0.653040262 | 0.37780412 | -1.7285155 | 0.08389585 | 0.17460791 | NA | NA |
| ENSG000001 | 3.39317227 | -0.652603497 | 0.37756375 | -1.7284591 | 0.08390595 | 0.17461101 | OR4D11 | 219986 |
| ENSG000002 | 2.2022956 | -0.625357929 | 0.36180075 | -1.7284594 | 0.08390589 | 0.17461101 | NA | NA |

| | | | | | | | | |
|---|---|---|---|---|---|---|---|---|
| ENSG000002 | 10.9134035 | -0.572080775 | 0.33098373 | -1.7284257 | 0.08391194 | 0.17461451 | NA | NA |
| ENSG000002 | 12.7745655 | 0.569480001 | 0.32950142 | 1.72830818 | 0.08393299 | 0.17464935 | NA | NA |
| ENSG000002 | 5.98162874 | -0.63852676 | 0.36946057 | -1.7282677 | 0.08394024 | 0.17465549 | NA | NA |
| ENSG000002 | 9.94287126 | -0.592248401 | 0.34270197 | -1.7281733 | 0.08395715 | 0.17467275 | NA | NA |
| ENSG000002 | 9.64607052 | 0.604566367 | 0.34982829 | 1.72818031 | 0.0839559 | 0.17467275 | NA | NA |
| ENSG000001 | 42.416074 | -0.371431333 | 0.21495455 | -1.7279529 | 0.08399667 | 0.17473762 | SCD5 | 79966 |
| ENSG000002 | 7.39175014 | -0.623052151 | 0.36057275 | -1.7279513 | 0.08399695 | 0.17473762 | SEPT4-AS1 | 101927688 |
| ENSG000002 | 2.18431547 | -0.624563359 | 0.36145346 | -1.7279219 | 0.08400222 | 0.17473962 | NA | NA |
| ENSG000002 | 4.00179935 | -0.65403268 | 0.37852676 | -1.7278374 | 0.08401738 | 0.17476219 | NA | NA |
| ENSG000002 | 2.39619155 | 0.635050384 | 0.36755048 | 1.72779093 | 0.08402571 | 0.17477055 | NA | NA |
| ENSG000002 | 6.25071666 | -0.631201789 | 0.36536497 | -1.7275925 | 0.08406131 | 0.17483562 | NA | NA |
| ENSG000002 | 2.41359887 | -0.641619244 | 0.37142815 | -1.7274384 | 0.08408896 | 0.17488417 | NA | NA |
| ENSG000002 | 6.8832793 | -0.628312477 | 0.36375502 | -1.7272957 | 0.08411457 | 0.17492846 | NA | NA |
| ENSG000001 | 18.8691689 | -0.510507294 | 0.29558646 | -1.7270997 | 0.08414975 | 0.17499265 | PASD1 | 139135 |
| ENSG000002 | 7.24683717 | -0.626526429 | 0.36277 | -1.7270624 | 0.08415645 | 0.17499761 | LINC01493 | 101928536 |
| ENSG000001 | 8.816858 | -0.601142145 | 0.34808087 | -1.7270186 | 0.08416432 | 0.17500463 | CPA5 | 93979 |
| ENSG000002 | 6.27263069 | -0.632267881 | 0.36611301 | -1.7269747 | 0.08417222 | 0.17500463 | NA | NA |
| ENSG000002 | 4.24818928 | -0.651997348 | 0.37754183 | -1.726954 | 0.08417593 | 0.17500463 | NA | NA |
| ENSG000002 | 10.5469367 | -0.583603129 | 0.33793913 | -1.7269475 | 0.08417709 | 0.17500463 | LINC01478 | 101927921 |
| ENSG000002 | 3.98508403 | -0.653730935 | 0.3785649 | -1.7268662 | 0.08419169 | 0.17502504 | NA | NA |
| ENSG000002 | 4.6437114 | -0.652651816 | 0.37794469 | -1.7268448 | 0.08419554 | 0.17502504 | NA | NA |
| ENSG000002 | 7.65011813 | -0.617669101 | 0.35770873 | -1.7267376 | 0.08421481 | 0.17505611 | NA | NA |
| ENSG000000 | 8.29593534 | -0.630793091 | 0.36533211 | -1.7266292 | 0.08423428 | 0.17508761 | HSD17B7P2 | 158160 |
| ENSG000001 | 2.79532827 | -0.643013032 | 0.37242287 | -1.726567 | 0.08424546 | 0.17510188 | ACY3 | 91703 |
| ENSG000002 | 4.80895047 | -0.648015548 | 0.37534698 | -1.726444 | 0.08426756 | 0.17513884 | NA | NA |
| ENSG000002 | 16.638785 | -0.518084166 | 0.30009643 | -1.7263923 | 0.08427686 | 0.1751492 | LOC102723661 | 102723661 |
| ENSG000002 | 16.3287531 | 0.525643932 | 0.30448236 | 1.72635267 | 0.08428399 | 0.17515502 | NA | NA |
| ENSG000002 | 2.535612 | -0.631983478 | 0.36609241 | -1.726295 | 0.08429436 | 0.17516183 | NA | NA |
| ENSG000002 | 3.09962806 | -0.648186456 | 0.37548025 | -1.7262864 | 0.0842959 | 0.17516183 | NA | NA |

| | | | | | | | | |
|---|---|---|---|---|---|---|---|---|
| ENSG000002 | 4.86075713 | -0.648437539 | 0.37563591 | -1.7262395 | 0.08430434 | 0.17517038 | NA | NA |
| ENSG000002 | 8.66500734 | -0.595373021 | 0.34493588 | -1.7260397 | 0.08434028 | 0.17523608 | NA | NA |
| ENSG000001 | 23.4948464 | -0.537462672 | 0.31139312 | -1.725994 | 0.08434848 | 0.17524415 | MUC5B | 727897 |
| ENSG000001 | 265.181742 | 0.22485034 | 0.13028526 | 1.72583104 | 0.08437781 | 0.17529611 | ERC2 | 26059 |
| ENSG00000C | 127.229992 | 0.300400808 | 0.17407928 | 1.7256552 | 0.08440946 | 0.17535288 | RASGRP2 | 10235 |
| ENSG000002 | 9.0806914 | -0.596572837 | 0.34574634 | -1.7254639 | 0.0844439 | 0.17541543 | PCDHGA3 | 56112 |
| ENSG000002 | 5.1047195 | -0.643761814 | 0.37312814 | -1.7253103 | 0.08447157 | 0.17546392 | NA | NA |
| ENSG000002 | 5.14761447 | -0.645436693 | 0.37411682 | -1.7252277 | 0.08448645 | 0.17548583 | NA | NA |
| ENSG000002 | 6.42903881 | -0.638035963 | 0.36983587 | -1.7251868 | 0.08449382 | 0.17549215 | NA | NA |
| ENSG000001 | 23.1917061 | -0.456005458 | 0.26433208 | -1.7251234 | 0.08450524 | 0.17550688 | PDGFB | 5155 |
| ENSG000002 | 5.24993674 | -0.650013321 | 0.37680298 | -1.7250748 | 0.08451399 | 0.17551608 | NA | NA |
| ENSG000002 | 4.03700966 | -0.653085148 | 0.37858996 | -1.7250461 | 0.08451916 | 0.17551782 | NA | NA |
| ENSG000002 | 6.62627657 | -0.632183838 | 0.36648254 | -1.725004 | 0.08452676 | 0.1755246 | NA | NA |
| ENSG000001 | 9.5505566 | -0.589118366 | 0.34154023 | -1.7248872 | 0.0845478 | 0.17555932 | FNDC9 | 408263 |
| ENSG000002 | 6.24629349 | -0.62919168 | 0.36479243 | -1.7247937 | 0.08456466 | 0.17558534 | NA | NA |
| ENSG000001 | 99.3432592 | 0.306085818 | 0.17747871 | 1.72463396 | 0.08459346 | 0.17563613 | THAP2 | 83591 |
| ENSG000002 | 5.80210198 | -0.64320204 | 0.37296882 | -1.7245464 | 0.08460925 | 0.17565992 | NA | NA |
| ENSG000001 | 47.4380117 | 0.384286186 | 0.22284747 | 1.72443594 | 0.08462917 | 0.1756923 | GNG3 | 2785 |
| ENSG000002 | 15.3182471 | -0.538691949 | 0.3123999 | -1.7243666 | 0.08464168 | 0.17570927 | GRM7-AS3 | 101927347 |
| ENSG000002 | 4.50850135 | -0.647972397 | 0.37582022 | -1.7241552 | 0.08467982 | 0.17577944 | NA | NA |
| ENSG000002 | 4.26813221 | -0.651487004 | 0.37791724 | -1.723888 | 0.08472806 | 0.17587058 | NA | NA |
| ENSG000002 | 12.7737345 | -0.554238242 | 0.32153331 | -1.723735 | 0.08475569 | 0.17591891 | LINC01099 | 101928656 |
| ENSG000001 | 104.395727 | 0.282464736 | 0.1638748 | 1.72366185 | 0.08476891 | 0.17592981 | MPZL3 | 196264 |
| ENSG000002 | 56.9116748 | 0.351559618 | 0.20396136 | 1.72365793 | 0.08476961 | 0.17592981 | MKLN1-AS | 100506881 |
| ENSG000002 | 13.7857088 | -0.569397461 | 0.33036325 | -1.7235497 | 0.08478917 | 0.1759614 | NA | NA |
| ENSG000002 | 2.80081732 | -0.644679885 | 0.37406874 | -1.7234262 | 0.08481147 | 0.17599867 | NA | NA |
| ENSG000002 | 3.38818904 | -0.651303568 | 0.37793328 | -1.7233295 | 0.08482896 | 0.17602595 | NA | NA |
| ENSG000001 | 78.6994523 | 0.313114382 | 0.18170408 | 1.72321053 | 0.08485046 | 0.17604524 | PEX11B | 8799 |
| ENSG000002 | 3.68643736 | -0.65239451 | 0.37858873 | -1.7232275 | 0.08484739 | 0.17604524 | NA | NA |

| | | | | | | | | |
|---|---|---|---|---|---|---|---|---|
| ENSG000002 | 9.22790794 | -0.591838706 | 0.34345209 | -1.723206 | 0.08485128 | 0.17604524 | NA | NA |
| ENSG000002 | 14.4008802 | 0.548077717 | 0.31806308 | 1.72317302 | 0.08485724 | 0.1760486 | MRPL53 | 116540 |
| ENSG000001 | 80.8193005 | 0.339657949 | 0.19712066 | 1.72309665 | 0.08487105 | 0.17606824 | KIAA0895L | 653319 |
| ENSG000001 | 11.5563205 | -0.569330681 | 0.33042541 | -1.7230233 | 0.08488432 | 0.17608675 | CYP4Z2P | 163720 |
| ENSG000001 | 833.581009 | 0.238650829 | 0.13852082 | 1.72285171 | 0.08491534 | 0.17613858 | PBDC1 | 51260 |
| ENSG000001 | 6.10356062 | -0.638256482 | 0.37046827 | -1.7228371 | 0.08491799 | 0.17613858 | NA | NA |
| ENSG000001 | 408.852198 | 0.305624323 | 0.17740781 | 1.72272192 | 0.08493883 | 0.17617279 | PPL | 5493 |
| ENSG000001 | 29.3788271 | -0.440571806 | 0.25575208 | -1.7226519 | 0.08495149 | 0.17619005 | VCAM1 | 7412 |
| ENSG00000C | 154.982661 | 0.325086409 | 0.1887287 | 1.72250644 | 0.08497782 | 0.1762086 | HEBP1 | 50865 |
| ENSG00000C | 48.1824431 | 0.380011832 | 0.22061459 | 1.72251448 | 0.08497636 | 0.1762086 | MMP11 | 4320 |
| ENSG000002 | 15.4332012 | 0.535519676 | 0.31089207 | 1.722526 | 0.08497428 | 0.1762086 | SERF1B | 728492 |
| ENSG000002 | 6.72641476 | -0.637358832 | 0.37001207 | -1.7225353 | 0.0849726 | 0.1762086 | NA | NA |
| ENSG000001 | 30.0443318 | -0.443496872 | 0.2575056 | -1.7222805 | 0.08501872 | 0.17628173 | SEZ6L | 23544 |
| ENSG000002 | 161.425588 | 0.267833942 | 0.15551275 | 1.7222636 | 0.08502178 | 0.17628173 | NA | NA |
| ENSG000002 | 4.26654156 | -0.650675572 | 0.37781648 | -1.7222001 | 0.08503328 | 0.17629658 | RN7SL258P | 106480373 |
| ENSG000002 | 10.2533247 | -0.587122918 | 0.34092736 | -1.7221349 | 0.08504508 | 0.17631201 | NA | NA |
| ENSG00000C | 440.295738 | 0.232632872 | 0.13510771 | 1.7218327 | 0.08509983 | 0.1764165 | VPS50 | 55610 |
| ENSG000002 | 8.47167379 | -0.605226455 | 0.35157324 | -1.7214805 | 0.08516368 | 0.17653983 | LINC00106 | 751580 |
| ENSG00000C | 104.833974 | 0.319971745 | 0.18588504 | 1.72134209 | 0.08518877 | 0.17657379 | KCNK6 | 9424 |
| ENSG000001 | 6.06453795 | -0.634919011 | 0.36884771 | -1.7213582 | 0.08518585 | 0.17657379 | C9orf152 | 401546 |
| ENSG000002 | 22.6375051 | 0.480216063 | 0.27899033 | 1.72126417 | 0.0852029 | 0.17658503 | CEBPA-AS1 | 80054 |
| ENSG000002 | 6.8915855 | -0.625420856 | 0.36334935 | -1.7212659 | 0.08520258 | 0.17658503 | NA | NA |
| ENSG000002 | 8.89403552 | -0.604616938 | 0.35127829 | -1.7211907 | 0.08521622 | 0.17660362 | NA | NA |
| ENSG000001 | 181.695913 | 0.327235623 | 0.19012651 | 1.72114673 | 0.0852242 | 0.17660966 | JUN | 3725 |
| ENSG000002 | 9.84961489 | -0.58067362 | 0.33737995 | -1.7211266 | 0.08522785 | 0.17660966 | C16orf95 | 100506581 |
| ENSG000002 | 3.01203572 | -0.64956346 | 0.37741784 | -1.7210725 | 0.08523767 | 0.176621 | NA | NA |
| ENSG000002 | 13.9209448 | -0.539960444 | 0.31374736 | -1.721004 | 0.08525011 | 0.17663773 | NA | NA |
| ENSG000001 | 18.9958971 | -0.487285007 | 0.28315191 | -1.7209314 | 0.08526328 | 0.17664697 | CD86 | 942 |
| ENSG000001 | 76.7090036 | 0.339771512 | 0.19743364 | 1.72094032 | 0.08526165 | 0.17664697 | IFNAR2 | 3455 |

| | | | | | | | | |
|---|---|---|---|---|---|---|---|---|
| ENSG000002 | 13.2865659 | -0.542097876 | 0.31501406 | -1.7208688 | 0.08527463 | 0.17666146 | NA | NA |
| ENSG000002 | 5.11174625 | -0.642244036 | 0.37321482 | -1.7208428 | 0.08527935 | 0.17666222 | NA | NA |
| ENSG000002 | 2.45994114 | -0.633928353 | 0.3683878 | -1.720818 | 0.08528386 | 0.17666253 | RN7SL335P | 106480375 |
| ENSG000001 | 16.7371689 | -0.528552931 | 0.30715926 | -1.7207781 | 0.0852911 | 0.17666494 | LRRC43 | 254050 |
| ENSG000002 | 19.4307245 | 0.525335463 | 0.30529206 | 1.72076358 | 0.08529373 | 0.17666494 | NA | NA |
| ENSG000001 | 714.922307 | 0.227268388 | 0.13208127 | 1.72067083 | 0.08531057 | 0.17669079 | RERE | 473 |
| ENSG000001 | 18.8827512 | -0.502399325 | 0.29199373 | -1.7205826 | 0.0853266 | 0.17671496 | NA | NA |
| ENSG000001 | 16.4686944 | -0.510813651 | 0.29689678 | -1.7205092 | 0.08533992 | 0.17673353 | HABP2 | 3026 |
| ENSG000001 | 112.310574 | 0.298011004 | 0.17321626 | 1.72045628 | 0.08534954 | 0.17673538 | CRELD2 | 79174 |
| ENSG000002 | 3.39945528 | -0.650373071 | 0.37802189 | -1.7204641 | 0.08534812 | 0.17673538 | RN7SL246P | 106480989 |
| ENSG000002 | 4.09928873 | -0.650670987 | 0.37826281 | -1.7201559 | 0.08540411 | 0.17683936 | NA | NA |
| ENSG000002 | 13.0078504 | -0.56923141 | 0.33097255 | -1.719875 | 0.08545517 | 0.17693606 | NA | NA |
| ENSG000002 | 5.79259634 | -0.640699362 | 0.37256435 | -1.7197012 | 0.08548677 | 0.17699245 | NA | NA |
| ENSG000001 | 14.7370498 | -0.532321164 | 0.30958452 | -1.7194696 | 0.0855289 | 0.17707064 | PCDHB7 | 56129 |
| ENSG000002 | 5.02429007 | -0.648335968 | 0.37710063 | -1.7192652 | 0.0855661 | 0.1771386 | NA | NA |
| ENSG000001 | 74.9397565 | -0.327661616 | 0.19059151 | -1.7191827 | 0.08558111 | 0.17716064 | ANKRD31 | 256006 |
| ENSG000001 | 4.54606491 | -0.647390853 | 0.37657781 | -1.7191423 | 0.08558846 | 0.1771668 | ADIG | 149685 |
| ENSG000001 | 213.674005 | 0.236886623 | 0.13780276 | 1.71902667 | 0.08560952 | 0.1771923 | LACTB | 114294 |
| ENSG000002 | 4.59252376 | -0.64944103 | 0.37779191 | -1.7190443 | 0.08560631 | 0.1771923 | NA | NA |
| ENSG000002 | 3.6758483 | -0.650773443 | 0.37857675 | -1.719 | 0.08561437 | 0.1771933 | NA | NA |
| ENSG000001 | 74.8021752 | -0.330166752 | 0.19208912 | -1.7188207 | 0.08564702 | 0.17724278 | DCC | 1630 |
| ENSG000002 | 6.23948985 | -0.627831783 | 0.36526509 | -1.7188387 | 0.08564374 | 0.17724278 | NA | NA |
| ENSG000002 | 3.04658682 | -0.648534417 | 0.37732548 | -1.7187666 | 0.08565689 | 0.17725415 | NA | NA |
| ENSG000001 | 55.5797562 | 0.360687295 | 0.20987887 | 1.71854979 | 0.08569638 | 0.1773215 | PPM1M | 132160 |
| ENSG000002 | 12.650473 | -0.558519787 | 0.32499669 | -1.7185399 | 0.08569818 | 0.1773215 | NA | NA |
| ENSG000001 | 6.89401327 | -0.624312074 | 0.36337899 | -1.7180742 | 0.08578308 | 0.17748811 | TMEM119 | 338773 |
| ENSG000002 | 6.0604916 | -0.634033605 | 0.36910013 | -1.7177821 | 0.08583636 | 0.17758931 | LOC102723809 | 102723809 |
| ENSG000001 | 11.1001433 | -0.576782686 | 0.33578748 | -1.7177016 | 0.08585106 | 0.1775952 | GPR84 | 53831 |
| ENSG000001 | 42.644973 | -0.37813621 | 0.22014171 | -1.7176945 | 0.08585235 | 0.1775952 | SCN3B | 55800 |

| | | | | | | | | |
|---|---|---|---|---|---|---|---|---|
| ENSG000002 | 3.64847388 | -0.650319525 | 0.37859325 | -1.7177262 | 0.08584657 | 0.1775952 | NA | NA |
| ENSG000001 | 145.451498 | 0.295286081 | 0.17191276 | 1.7176507 | 0.08586035 | 0.17760268 | HENMT1 | 113802 |
| ENSG000002 | 2.75636306 | -0.64255942 | 0.37409839 | -1.7176214 | 0.08586569 | 0.17760466 | NA | NA |
| ENSG000002 | 3.05025217 | -0.647391581 | 0.37693158 | -1.7175308 | 0.08588224 | 0.17762007 | NA | NA |
| ENSG000002 | 3.35870548 | -0.648476822 | 0.37756847 | -1.7175079 | 0.08588642 | 0.17762007 | NA | NA |
| ENSG000002 | 10.0664499 | -0.576150397 | 0.33546341 | -1.7174761 | 0.08589221 | 0.17762007 | NA | NA |
| ENSG000002 | 2.70980054 | -0.642303979 | 0.37398468 | -1.7174607 | 0.08589503 | 0.17762007 | NA | NA |
| ENSG000002 | 3.68088469 | -0.650222432 | 0.37858192 | -1.7175211 | 0.08588401 | 0.17762007 | NA | NA |
| ENSG000001 | 25.5656118 | -0.450388239 | 0.26224476 | -1.7174347 | 0.08589978 | 0.17762083 | SYTL3 | 94120 |
| ENSG000002 | 4.73710944 | -0.649120283 | 0.37798521 | -1.7173166 | 0.08592133 | 0.17765633 | NA | NA |
| ENSG000002 | 2.37859076 | -0.636583243 | 0.37070495 | -1.7172235 | 0.08593835 | 0.17768246 | NA | NA |
| ENSG000001 | 97.175713 | 0.293846784 | 0.17113255 | 1.71707126 | 0.08596615 | 0.17773088 | CLHC1 | 130162 |
| ENSG000001 | 2.74967619 | -0.642385333 | 0.3741282 | -1.7170193 | 0.08597565 | 0.17773907 | RNU1-94P | 106481956 |
| ENSG000002 | 3.03836605 | -0.647607059 | 0.37717324 | -1.7170016 | 0.08597888 | 0.17773907 | NA | NA |
| ENSG000001 | 4.06466593 | -0.649994048 | 0.37858886 | -1.7168864 | 0.08599994 | 0.17777354 | OR4D6 | 219983 |
| ENSG00000C | 4.87885295 | -0.646210036 | 0.37644175 | -1.7166269 | 0.08604736 | 0.17786042 | NCR2 | 9436 |
| ENSG000002 | 28.634151 | 0.435205844 | 0.25352657 | 1.71660843 | 0.08605074 | 0.17786042 | NA | NA |
| ENSG000002 | 6.31947332 | -0.634056396 | 0.36937992 | -1.7165427 | 0.08606276 | 0.1778762 | NA | NA |
| ENSG000002 | 6.22537204 | -0.626221387 | 0.36484686 | -1.7163952 | 0.08608974 | 0.17792289 | NA | NA |
| ENSG000002 | 4.76012972 | -0.649297117 | 0.37830168 | -1.7163474 | 0.08609848 | 0.17793188 | LINC01282 | 101927449 |
| ENSG000002 | 8.21745525 | -0.602676345 | 0.35116904 | -1.7162001 | 0.08612542 | 0.17797849 | NA | NA |
| ENSG000002 | 5.40623619 | -0.638218535 | 0.37189351 | -1.7161325 | 0.0861378 | 0.177995 | LOC102724265 | 102724265 |
| ENSG000002 | 142.502236 | 0.300221685 | 0.1749546 | 1.71599762 | 0.08616248 | 0.17803692 | LYRM5 | 144363 |
| ENSG000002 | 4.08333663 | -0.649596214 | 0.37858161 | -1.7158684 | 0.08618614 | 0.17807673 | NA | NA |
| ENSG000002 | 13.2832198 | -0.539322435 | 0.31436364 | -1.7156005 | 0.08623518 | 0.17816899 | NA | NA |
| ENSG000001 | 73.0597465 | 0.351816568 | 0.20509495 | 1.7153839 | 0.08627487 | 0.1782419 | ADA | 100 |
| ENSG000001 | 1314.12591 | 0.190234142 | 0.11090377 | 1.71530818 | 0.08628874 | 0.17826144 | UHMK1 | 127933 |
| ENSG000002 | 3.35494895 | -0.648454986 | 0.37804519 | -1.7152843 | 0.08629311 | 0.17826144 | NA | NA |
| ENSG000001 | 8.35570974 | -0.608427452 | 0.35473791 | -1.7151464 | 0.08631839 | 0.17829548 | CYP2C18 | 1562 |

| | | | | | | | | |
|---|---|---|---|---|---|---|---|---|
| ENSG000001 | 1077.25075 | 0.208371639 | 0.1214885 | 1.71515524 | 0.08631677 | 0.17829548 | NSL1 | 25936 |
| ENSG000002 | 4.05068422 | 0.649081815 | 0.37845223 | 1.71509576 | 0.08632767 | 0.17830558 | SNORA14A | 677801 |
| ENSG000002 | 6.06729744 | -0.633275501 | 0.36937887 | -1.7144335 | 0.08644914 | 0.17854738 | NA | NA |
| ENSG000002 | 4.53152975 | -0.645601352 | 0.37657884 | -1.7143856 | 0.08645793 | 0.17855643 | NA | NA |
| ENSG000002 | 4.54333265 | -0.645098831 | 0.37631051 | -1.7142727 | 0.08647866 | 0.17859014 | NA | NA |
| ENSG000001 | 3.59911146 | -0.643965022 | 0.37566098 | -1.7142186 | 0.08648859 | 0.17860157 | APCS | 325 |
| ENSG000001 | 17.2057309 | -0.505788883 | 0.29507722 | -1.7140899 | 0.08651222 | 0.17864125 | ALOXE3 | 59344 |
| ENSG000001 | 11.8746784 | -0.562939481 | 0.32845788 | -1.7138864 | 0.0865496 | 0.17870934 | SLC7A13 | 157724 |
| ENSG000002 | 4.02512276 | -0.648828444 | 0.37859335 | -1.7137872 | 0.08656781 | 0.17873785 | NA | NA |
| ENSG000002 | 6.01213733 | -0.632087885 | 0.36884041 | -1.7137165 | 0.08658081 | 0.17875558 | NA | NA |
| ENSG000002 | 4.58519217 | -0.647032846 | 0.37756995 | -1.7136768 | 0.08658811 | 0.17876155 | LOC105379374 | 105379374 |
| ENSG000002 | 11.3639967 | -0.575255091 | 0.33574 | -1.7133946 | 0.08663998 | 0.17885953 | NA | NA |
| ENSG000001 | 5.35015563 | -0.637006409 | 0.37180898 | -1.7132626 | 0.08666424 | 0.17890051 | MYL3 | 4634 |
| ENSG000002 | 4.18919781 | -0.64834295 | 0.3784509 | -1.7131495 | 0.08668505 | 0.17893435 | NA | NA |
| ENSG000001 | 256.02045 | 0.240417537 | 0.14034943 | 1.71299263 | 0.0867139 | 0.17898479 | CEP41 | 95681 |
| ENSG00000C | 104.928161 | -0.299190358 | 0.1746944 | -1.71265 | 0.08677695 | 0.17910583 | RIMS1 | 22999 |
| ENSG000002 | 4.8665255 | -0.643646823 | 0.37586842 | -1.7124259 | 0.0868182 | 0.17918185 | NA | NA |
| ENSG000001 | 6.79236308 | -0.620831961 | 0.36257108 | -1.7123041 | 0.08684063 | 0.17920975 | FAM182B | 728882 |
| ENSG000001 | 22.5313493 | -0.470591846 | 0.27483003 | -1.7123014 | 0.08684113 | 0.17920975 | TNP2 | 7142 |
| ENSG000002 | 6.08384548 | -0.634742378 | 0.37069999 | -1.7122806 | 0.08684498 | 0.17920975 | NA | NA |
| ENSG000002 | 5.99844352 | -0.631245027 | 0.36867916 | -1.7121799 | 0.08686353 | 0.1792389 | SMCR5 | 140771 |
| ENSG000002 | 2.75343382 | -0.640161212 | 0.37390583 | -1.7120921 | 0.0868797 | 0.17924592 | NA | NA |
| ENSG000002 | 22.4222377 | -0.466337419 | 0.27237913 | -1.7120894 | 0.08688019 | 0.17924592 | NA | NA |
| ENSG000002 | 4.93630509 | -0.643991063 | 0.37613993 | -1.712105 | 0.08687731 | 0.17924592 | NA | NA |
| ENSG000002 | 85.1512177 | 0.301944484 | 0.17636705 | 1.71202319 | 0.0868924 | 0.17926198 | NA | NA |
| ENSG000001 | 18.3653002 | -0.509678015 | 0.29773062 | -1.7118764 | 0.08691945 | 0.17930867 | GPR65 | 8477 |
| ENSG000002 | 4.87989815 | -0.644747667 | 0.37665436 | -1.7117754 | 0.08693806 | 0.17933795 | NA | NA |
| ENSG000002 | 5.09749185 | -0.639292528 | 0.37351042 | -1.7115788 | 0.08697431 | 0.17940359 | EFCAB9 | 285588 |
| ENSG000002 | 6.51560735 | -0.623445076 | 0.36425917 | -1.7115426 | 0.086981 | 0.17940827 | CACNA1C-IT2 | 100874369 |

| | | | | | | | | |
|---|---|---|---|---|---|---|---|---|
| ENSG000002 | 7.75398159 | -0.612086677 | 0.35764606 | -1.7114313 | 0.08700152 | 0.17944146 | LOC105378622 | 105378622 |
| ENSG000001 | 517.468149 | 0.257862518 | 0.15068575 | 1.71126013 | 0.0870331 | 0.17948835 | KDM3A | 55818 |
| ENSG000002 | 3.94908662 | -0.647751036 | 0.37851927 | -1.7112762 | 0.08703013 | 0.17948835 | NA | NA |
| ENSG000001 | 441.279716 | 0.240636687 | 0.14062676 | 1.71117279 | 0.08704922 | 0.17951246 | FAM193A | 8603 |
| ENSG000002 | 2.40625911 | -0.633740484 | 0.37040974 | -1.7109174 | 0.08709636 | 0.17960053 | NA | NA |
| ENSG000002 | 2.77179187 | -0.639697391 | 0.37392866 | -1.7107472 | 0.0871278 | 0.17965622 | NA | NA |
| ENSG000001 | 11.3484464 | -0.588711838 | 0.34415799 | -1.710586 | 0.08715756 | 0.17969932 | OMD | 4958 |
| ENSG000001 | 275.625928 | 0.235505375 | 0.1376738 | 1.71060416 | 0.08715421 | 0.17969932 | NA | NA |
| ENSG000001 | 37.848214 | 0.403957862 | 0.23617181 | 1.71044064 | 0.08718442 | 0.17974416 | PWWP2B | 170394 |
| ENSG000002 | 6.25096351 | -0.627850631 | 0.36707388 | -1.7104203 | 0.08718818 | 0.17974416 | NA | NA |
| ENSG000002 | 5.0848589 | -0.640400449 | 0.37445791 | -1.7102067 | 0.08722765 | 0.1798164 | NA | NA |
| ENSG000002 | 15.3193616 | 0.528513095 | 0.30904745 | 1.71013577 | 0.08724077 | 0.17983431 | RPS2P32 | 256355 |
| ENSG000002 | 95.7743737 | 0.306932975 | 0.1794822 | 1.71010256 | 0.08724691 | 0.17983782 | NA | NA |
| ENSG000001 | 2.38661565 | -0.633944614 | 0.37073856 | -1.7099506 | 0.08727501 | 0.17987746 | NA | NA |
| ENSG000002 | 5.16100598 | 0.646333471 | 0.37797848 | 1.70997427 | 0.08727063 | 0.17987746 | NA | NA |
| ENSG000001 | 18.4492344 | -0.499714738 | 0.29224639 | -1.709909 | 0.0872827 | 0.17988415 | KCNA4 | 3739 |
| ENSG000002 | 14.3269285 | -0.528535836 | 0.30913762 | -1.7097105 | 0.08731942 | 0.1799507 | LOC153910 | 153910 |
| ENSG000002 | 2.72289408 | -0.639214107 | 0.37388841 | -1.7096387 | 0.0873327 | 0.17996891 | NA | NA |
| ENSG000002 | 5.14442988 | -0.640753229 | 0.37479732 | -1.7095993 | 0.08734 | 0.17997482 | HAO2-IT1 | 100874270 |
| ENSG000002 | 5.13125312 | -0.63964702 | 0.3741993 | -1.7093752 | 0.08738147 | 0.18005111 | NA | NA |
| ENSG000001 | 104.650814 | 0.27959795 | 0.16357476 | 1.70929765 | 0.08739583 | 0.18006552 | TMEM56 | 148534 |
| ENSG000002 | 13.5710253 | -0.55501667 | 0.32470607 | -1.7092895 | 0.08739734 | 0.18006552 | NA | NA |
| ENSG000001 | 9.91470519 | -0.580963818 | 0.33991029 | -1.7091681 | 0.08741982 | 0.18007842 | TGM6 | 343641 |
| ENSG000001 | 55.6205939 | 0.371055908 | 0.21709844 | 1.70915973 | 0.08742137 | 0.18007842 | ENTHD2 | 146705 |
| ENSG000002 | 4.58066487 | -0.645714574 | 0.37778717 | -1.709202 | 0.08741354 | 0.18007842 | NA | NA |
| ENSG000002 | 3.3697202 | -0.645956079 | 0.37793388 | -1.7091775 | 0.08741808 | 0.18007842 | NA | NA |
| ENSG000002 | 3.47955031 | -0.642463462 | 0.37590173 | -1.7091261 | 0.08742759 | 0.1800821 | NA | NA |
| ENSG000002 | 5.38693819 | -0.635548666 | 0.37186403 | -1.7090889 | 0.08743449 | 0.18008716 | NA | NA |
| ENSG000002 | 9.67677371 | -0.590753561 | 0.34566334 | -1.7090431 | 0.08744296 | 0.18009545 | NA | NA |

| | | | | | | | | |
|---|---|---|---|---|---|---|---|---|
| ENSG000002 | 5.74000385 | -0.635032369 | 0.37158625 | -1.7089771 | 0.0874552 | 0.18011152 | LINC00930 | 100144604 |
| ENSG000001 | 15.3264408 | -0.55067218 | 0.32224321 | -1.7088713 | 0.08747479 | 0.18013356 | GBP6 | 163351 |
| ENSG000002 | 6.56947817 | -0.625598494 | 0.36608376 | -1.7088944 | 0.08747051 | 0.18013356 | NA | NA |
| ENSG000002 | 3.67704307 | -0.646939363 | 0.37859312 | -1.7087985 | 0.08748828 | 0.18015219 | NA | NA |
| ENSG000002 | 7.33484788 | -0.609122071 | 0.35651015 | -1.7085687 | 0.08753088 | 0.18023077 | NA | NA |
| ENSG000002 | 10.9113329 | -0.573540698 | 0.33570946 | -1.7084437 | 0.08755405 | 0.18026933 | NA | NA |
| ENSG000002 | 3.71316102 | -0.645951822 | 0.37814154 | -1.7082276 | 0.08759412 | 0.18034267 | NA | NA |
| ENSG000002 | 3.68547182 | -0.646646954 | 0.37856236 | -1.7081649 | 0.08760574 | 0.18035743 | NA | NA |
| ENSG000001 | 2241.37412 | 0.1977467 | 0.11577937 | 1.70796138 | 0.08764351 | 0.18042603 | ZNF480 | 147657 |
| ENSG000001 | 183.501158 | 0.256332422 | 0.15008703 | 1.70789191 | 0.0876564 | 0.18044341 | HECW2 | 57520 |
| ENSG000001 | 9.29612465 | -0.587900077 | 0.34423076 | -1.7078662 | 0.08766118 | 0.18044408 | KRT23 | 25984 |
| ENSG000002 | 20.07114 | 0.487939782 | 0.28571722 | 1.70777171 | 0.08767871 | 0.18047101 | NA | NA |
| ENSG000002 | 2.4157898 | -0.631002301 | 0.36951505 | -1.7076498 | 0.08770134 | 0.1804901 | RNU6ATAC35P | 106481171 |
| ENSG000002 | 4.0819811 | -0.645869108 | 0.37821201 | -1.7076906 | 0.08769376 | 0.1804901 | LINC00210 | 100885798 |
| ENSG000002 | 9.36033136 | -0.600847004 | 0.35185197 | -1.7076699 | 0.08769762 | 0.1804901 | RN7SL582P | 106479435 |
| ENSG000002 | 9.85566097 | -0.581235551 | 0.34037974 | -1.7076091 | 0.0877089 | 0.18049649 | NA | NA |
| ENSG000002 | 3.64585786 | -0.646416686 | 0.37857835 | -1.7074846 | 0.08773202 | 0.18053493 | NA | NA |
| ENSG000002 | 2.40943575 | -0.632008965 | 0.37014559 | -1.7074605 | 0.0877365 | 0.18053497 | NA | NA |
| ENSG000001 | 7.81662622 | -0.599629465 | 0.35122826 | -1.7072358 | 0.08777823 | 0.18061168 | NA | NA |
| ENSG000002 | 4.9051786 | -0.642670752 | 0.37651577 | -1.7068893 | 0.08784262 | 0.180735 | NA | NA |
| ENSG000001 | 264.327211 | 0.253081038 | 0.14827541 | 1.70683084 | 0.0878535 | 0.18074821 | WDR72 | 256764 |
| ENSG000001 | 2.3861911 | -0.631628846 | 0.37007723 | -1.7067487 | 0.08786877 | 0.18077045 | AQP12A | 375318 |
| ENSG000002 | 2.42048107 | -0.63063555 | 0.36953183 | -1.7065798 | 0.08790019 | 0.18082591 | NA | NA |
| ENSG000001 | 3.5324754 | -0.641461034 | 0.37589016 | -1.7065119 | 0.08791281 | 0.18084271 | DUPD1 | 338599 |
| ENSG000002 | 2.19760488 | -0.615139108 | 0.3605106 | -1.7062996 | 0.08795231 | 0.18091478 | NA | NA |
| ENSG000001 | 23.8385273 | -0.446311282 | 0.26158867 | -1.7061568 | 0.0879789 | 0.1809603 | NR4A3 | 8013 |
| ENSG000001 | 18.4098555 | 0.5083589 | 0.29796393 | 1.70610884 | 0.08798782 | 0.18096946 | GRASP | 160622 |
| ENSG000002 | 14.2943202 | -0.528309813 | 0.30966474 | -1.7060703 | 0.08799499 | 0.18097504 | LINC00559 | 100874187 |
| ENSG000002 | 2.47643944 | -0.626768682 | 0.36740907 | -1.7059151 | 0.08802389 | 0.1810253 | NA | NA |

| | | | | | | | | |
|---|---|---|---|---|---|---|---|---|
| ENSG000002 | 2.86166541 | -0.634245124 | 0.37181127 | -1.7058255 | 0.08804058 | 0.18105044 | NA | NA |
| ENSG000000 | 122.971921 | 0.278926136 | 0.16352112 | 1.70574992 | 0.08805465 | 0.18107019 | OPHN1 | 4983 |
| ENSG000002 | 3.73403781 | -0.644870433 | 0.3780879 | -1.7056098 | 0.08808075 | 0.18111468 | NA | NA |
| ENSG000001 | 30.2146923 | 0.42704388 | 0.25038821 | 1.70552711 | 0.08809616 | 0.18113718 | LRFN1 | 57622 |
| ENSG000001 | 6.86755929 | -0.618824434 | 0.36285183 | -1.7054466 | 0.08811116 | 0.18115883 | CYP2A7 | 1549 |
| ENSG000002 | 2.41200025 | -0.631905385 | 0.37055178 | -1.7053093 | 0.08813676 | 0.18120229 | NA | NA |
| ENSG000002 | 5.98220056 | 0.628562063 | 0.36860153 | 1.70526168 | 0.08814563 | 0.18121133 | NA | NA |
| ENSG000002 | 4.36590798 | -0.644946191 | 0.37825708 | -1.7050473 | 0.08818561 | 0.18128433 | NA | NA |
| ENSG000002 | 2.4242717 | -0.631148334 | 0.37019561 | -1.704905 | 0.08821215 | 0.18132969 | NA | NA |
| ENSG000001 | 13.9779571 | -0.559377703 | 0.32810852 | -1.7048558 | 0.08822133 | 0.18133018 | HPGDS | 27306 |
| ENSG000002 | 2.19332497 | -0.614567706 | 0.36048014 | -1.7048587 | 0.08822078 | 0.18133018 | NA | NA |
| ENSG000002 | 169.001987 | -0.247942374 | 0.14545289 | -1.7046232 | 0.08826472 | 0.18139273 | NA | NA |
| ENSG000002 | 3.36542013 | -0.643666089 | 0.37760076 | -1.7046208 | 0.08826518 | 0.18139273 | NA | NA |
| ENSG000002 | 8.42871721 | -0.624081187 | 0.36610431 | -1.704654 | 0.08825897 | 0.18139273 | NA | NA |
| ENSG000002 | 3.42533656 | -0.643826607 | 0.37770218 | -1.704588 | 0.08827129 | 0.1813961 | NA | NA |
| ENSG000002 | 5.95498761 | -0.627923219 | 0.36838088 | -1.7045489 | 0.08827859 | 0.1814019 | NA | NA |
| ENSG000001 | 86.4001324 | 0.295530628 | 0.17338378 | 1.7044883 | 0.0882899 | 0.18141595 | LHFP | 10186 |
| ENSG000001 | 30.9100987 | -0.420961994 | 0.24699703 | -1.7043201 | 0.08832131 | 0.18147129 | TMEM117 | 84216 |
| ENSG000001 | 3.63168687 | -0.645131684 | 0.37855404 | -1.7041997 | 0.08834379 | 0.18150828 | NA | NA |
| ENSG000001 | 3375.58565 | 0.215463702 | 0.12643837 | 1.70410066 | 0.08836228 | 0.18152958 | ZNF281 | 23528 |
| ENSG000001 | 16.6877519 | -0.518999813 | 0.30456322 | -1.7040791 | 0.08836631 | 0.18152958 | LIPC | 3990 |
| ENSG000001 | 66.2155999 | 0.39421558 | 0.23133736 | 1.70407227 | 0.08836759 | 0.18152958 | LYNX1 | 66004 |
| ENSG000001 | 142.256457 | 0.250057619 | 0.14675135 | 1.7039545 | 0.08838959 | 0.18154826 | AP1S3 | 130340 |
| ENSG000002 | 2.17423604 | -0.616268419 | 0.36167012 | -1.7039517 | 0.08839011 | 0.18154826 | NA | NA |
| ENSG000002 | 9.92677015 | -0.585298645 | 0.34348682 | -1.7039916 | 0.08838266 | 0.18154826 | NA | NA |
| ENSG000002 | 5.11847074 | -0.643795257 | 0.37787928 | -1.7037062 | 0.08843599 | 0.1816333 | NA | NA |
| ENSG000002 | 18.7024619 | -0.512618981 | 0.30089828 | -1.7036288 | 0.08845046 | 0.18164461 | SNAP25-AS1 | 100131208 |
| ENSG000002 | 3.0397884 | -0.642159227 | 0.3769352 | -1.7036329 | 0.08844968 | 0.18164461 | NA | NA |
| ENSG000002 | 11.2198378 | -0.569616937 | 0.33436527 | -1.7035769 | 0.08846017 | 0.18165535 | NA | NA |

| | | | | | | | | |
|---|---|---|---|---|---|---|---|---|
| ENSG000002 | 7.32715361 | -0.611744591 | 0.35910289 | -1.7035357 | 0.08846786 | 0.18166196 | LOC101927660 | 101927660 |
| ENSG000002 | 2.17815675 | -0.612037423 | 0.35936019 | -1.7031308 | 0.08854359 | 0.18180824 | NA | NA |
| ENSG000002 | 5.1180356 | -0.63746118 | 0.37429999 | -1.7030756 | 0.08855392 | 0.18182026 | NA | NA |
| ENSG000001 | 2.79105797 | -0.636101787 | 0.37352238 | -1.7029817 | 0.08857149 | 0.18184701 | TM4SF5 | 9032 |
| ENSG000002 | 3.67650174 | -0.644663884 | 0.37856053 | -1.7029348 | 0.08858028 | 0.18184701 | OR2L1P | 26247 |
| ENSG000002 | 17.1486396 | 0.507563542 | 0.29805237 | 1.70293409 | 0.0885804 | 0.18184701 | SLC7A5P1 | 81893 |
| ENSG000001 | 17.3645747 | -0.50273548 | 0.29524009 | -1.7028022 | 0.08860509 | 0.18188305 | SLC22A9 | 114571 |
| ENSG000002 | 6.94531906 | -0.626284313 | 0.3677984 | -1.7027924 | 0.08860693 | 0.18188305 | NA | NA |
| ENSG000002 | 6.74719674 | -0.628748179 | 0.36925406 | -1.7027523 | 0.08861444 | 0.18188926 | RNU7-94P | 106481799 |
| ENSG000002 | 7.83498372 | -0.600077477 | 0.35242559 | -1.7027069 | 0.08862293 | 0.18189748 | NA | NA |
| ENSG000002 | 4.65272333 | -0.643054971 | 0.37768143 | -1.7026386 | 0.08863573 | 0.18191454 | NA | NA |
| ENSG000002 | 11.9424637 | -0.548738636 | 0.32230089 | -1.7025663 | 0.08864926 | 0.1819331 | NA | NA |
| ENSG000002 | 2.41658704 | -0.629570091 | 0.36981366 | -1.7023982 | 0.08868075 | 0.18198853 | NA | NA |
| ENSG000001 | 8.03065536 | -0.606457996 | 0.35624904 | -1.7023428 | 0.08869113 | 0.1820006 | TSSK4 | 283629 |
| ENSG000001 | 25.449969 | -0.44983919 | 0.26427511 | -1.7021625 | 0.08872491 | 0.18205149 | NA | NA |
| ENSG000001 | 4.31317777 | -0.644103746 | 0.37840133 | -1.7021709 | 0.08872333 | 0.18205149 | SLPI | 6590 |
| ENSG000002 | 4.53412617 | -0.641741681 | 0.37702944 | -1.7020997 | 0.08873669 | 0.18206645 | NA | NA |
| ENSG000001 | 6.0437221 | -0.632258069 | 0.37147052 | -1.702041 | 0.08874767 | 0.18207978 | NA | NA |
| ENSG000002 | 7.2889498 | -0.610139852 | 0.35848993 | -1.7019721 | 0.08876061 | 0.1820971 | NA | NA |
| ENSG000001 | 101.988732 | 0.279867415 | 0.16443985 | 1.70194396 | 0.08876587 | 0.1820987 | KDELR3 | 11015 |
| ENSG000001 | 28.6335728 | -0.458937837 | 0.26967492 | -1.7018188 | 0.08878935 | 0.18212947 | BMX | 660 |
| ENSG000002 | 14.1004996 | -0.527233741 | 0.30980654 | -1.701816 | 0.08878986 | 0.18212947 | NA | NA |
| ENSG000000 | 6.55457905 | -0.623110975 | 0.36617318 | -1.7016838 | 0.08881465 | 0.18216246 | METTL24 | 728464 |
| ENSG000002 | 2.17634319 | -0.61374344 | 0.36066862 | -1.7016824 | 0.08881493 | 0.18216246 | NA | NA |
| ENSG000002 | 45.0313718 | 0.39549408 | 0.23242036 | 1.70163268 | 0.08882425 | 0.18217237 | ZBTB11-AS1 | 100009676 |
| ENSG000002 | 8.96737813 | -0.607793198 | 0.35718901 | -1.7016011 | 0.08883018 | 0.18217531 | NA | NA |
| ENSG000001 | 7.82035215 | -0.597783566 | 0.35131394 | -1.7015652 | 0.08883691 | 0.18217991 | SNX20 | 124460 |
| ENSG000001 | 21.3170375 | -0.468803891 | 0.27553577 | -1.7014266 | 0.08886291 | 0.18221292 | MAP6 | 4135 |
| ENSG000001 | 4.2338172 | -0.643090378 | 0.3779684 | -1.7014396 | 0.08886048 | 0.18221292 | PRAMEF10 | 343071 |

| | | | | | | | | |
|---|---|---|---|---|---|---|---|---|
| ENSG000002 | 4.84350048 | -0.6410465 | 0.37677422 | -1.7014075 | 0.08886648 | 0.18221292 | NA | NA |
| ENSG000002 | 23.7852228 | 0.449922563 | 0.26445741 | 1.70130441 | 0.08888584 | 0.18224339 | NA | NA |
| ENSG000002 | 3.3924684 | -0.642292506 | 0.37756716 | -1.7011345 | 0.08891774 | 0.18229625 | NA | NA |
| ENSG000002 | 2.38047012 | -0.630200073 | 0.37046204 | -1.7011192 | 0.08892061 | 0.18229625 | LLPHP2 | 106480300 |
| ENSG00000C | 246.952086 | 0.236301602 | 0.13891867 | 1.70100682 | 0.08894171 | 0.18233027 | DNAJC25 | 548645 |
| ENSG000002 | 2.18484361 | -0.613067223 | 0.36043034 | -1.7009312 | 0.0889559 | 0.18234918 | NA | NA |
| ENSG000002 | 6.58369206 | -0.626267159 | 0.36819539 | -1.7009098 | 0.08895992 | 0.18234918 | NA | NA |
| ENSG000002 | 6.01683438 | -0.628223198 | 0.36937897 | -1.7007552 | 0.08898896 | 0.1823981 | NA | NA |
| ENSG000002 | 10.5796622 | -0.564083915 | 0.33167069 | -1.7007349 | 0.08899278 | 0.1823981 | NA | NA |
| ENSG000002 | 3.71626529 | -0.64374124 | 0.37853081 | -1.700631 | 0.08901229 | 0.18242886 | NA | NA |
| ENSG000001 | 5.11239839 | -0.636531255 | 0.3743002 | -1.7005902 | 0.08901996 | 0.18243536 | PRR35 | 146325 |
| ENSG000002 | 3.66850965 | -0.643720325 | 0.37856372 | -1.7004279 | 0.08905046 | 0.18248864 | NA | NA |
| ENSG000002 | 15.0534454 | -0.530223736 | 0.31183311 | -1.7003446 | 0.08906613 | 0.18250232 | LINC00547 | 400121 |
| ENSG000002 | 51.1110834 | 0.368924346 | 0.21696902 | 1.70035492 | 0.08906419 | 0.18250232 | NA | NA |
| ENSG000002 | 3.7013571 | -0.643404737 | 0.37841098 | -1.7002803 | 0.08907821 | 0.18251525 | NA | NA |
| ENSG000002 | 6.30233386 | -0.625032708 | 0.3676094 | -1.7002631 | 0.08908145 | 0.18251525 | NA | NA |
| ENSG000002 | 8.96545654 | -0.583329615 | 0.34311246 | -1.700112 | 0.08910986 | 0.18256423 | NA | NA |
| ENSG000001 | 12.5596702 | -0.550264666 | 0.32367151 | -1.7000714 | 0.0891175 | 0.18257066 | GPR37L1 | 9283 |
| ENSG000002 | 32.1673265 | 0.414206345 | 0.24366904 | 1.69987266 | 0.08915488 | 0.18263802 | NA | NA |
| ENSG000001 | 63.9066816 | -0.349240058 | 0.20545658 | -1.6998242 | 0.089164 | 0.18264747 | PRKCE | 5581 |
| ENSG00000C | 83.9319003 | 0.360842047 | 0.21229263 | 1.69973889 | 0.08918005 | 0.18265267 | CAMSAP3 | 57662 |
| ENSG000001 | 3.08077443 | -0.63827497 | 0.37551279 | -1.6997423 | 0.08917941 | 0.18265267 | OR52W1 | 120787 |
| ENSG000002 | 10.6220619 | -0.573794135 | 0.33757112 | -1.6997726 | 0.08917371 | 0.18265267 | NA | NA |
| ENSG000002 | 8.40622809 | -0.606576938 | 0.35689929 | -1.6995745 | 0.08921099 | 0.1827068 | NA | NA |
| ENSG00000C | 1958.96605 | 0.20982629 | 0.12346354 | 1.69949997 | 0.08922502 | 0.18272632 | PLEKHA5 | 54477 |
| ENSG000002 | 4.80345728 | -0.639325912 | 0.37620023 | -1.6994299 | 0.08923821 | 0.18274411 | NA | NA |
| ENSG000002 | 2.82606499 | -0.632323736 | 0.37214855 | -1.6991165 | 0.08929724 | 0.18284511 | NA | NA |
| ENSG000002 | 3.65894561 | -0.643220823 | 0.37856402 | -1.6991071 | 0.089299 | 0.18284511 | NA | NA |
| ENSG000002 | 36.0907773 | 0.425338462 | 0.25033219 | 1.69909618 | 0.08930106 | 0.18284511 | A1BG-AS1 | 503538 |

| | | | | | | | | |
|---|---|---|---|---|---|---|---|---|
| ENSG000001 | 31.4288093 | -0.437560156 | 0.25754206 | -1.6989852 | 0.08932197 | 0.18287261 | CACNA2D4 | 93589 |
| ENSG000002 | 29.6847557 | 0.425449398 | 0.25041504 | 1.69897701 | 0.08932352 | 0.18287261 | NA | NA |
| ENSG000002 | 73.7095499 | 0.324051047 | 0.19073985 | 1.69891631 | 0.08933495 | 0.1828868 | NA | NA |
| ENSG000001 | 17.3000905 | -0.504214258 | 0.29683697 | -1.6986235 | 0.08939014 | 0.18299054 | CD80 | 941 |
| ENSG000001 | 44.2803695 | 0.377687923 | 0.22237936 | 1.6983947 | 0.08943329 | 0.18306963 | PI4KAP2 | 375133 |
| ENSG000002 | 7.88404094 | -0.604142941 | 0.35572726 | -1.6983319 | 0.08944515 | 0.18308465 | NA | NA |
| ENSG000001 | 11.1209203 | -0.556270764 | 0.32755736 | -1.6982393 | 0.08946261 | 0.18311116 | SIGLEC15 | 284266 |
| ENSG000001 | 7.15095973 | -0.619674939 | 0.36490421 | -1.6981852 | 0.08947281 | 0.18312279 | ANKRD33 | 341405 |
| ENSG000001 | 633.577484 | 0.219277449 | 0.12912924 | 1.69812392 | 0.08948438 | 0.18313524 | ZNF569 | 148266 |
| ENSG000002 | 2.76896678 | -0.63423741 | 0.37349714 | -1.6981051 | 0.08948792 | 0.18313524 | NA | NA |
| ENSG000002 | 2.41693195 | -0.627458401 | 0.36953021 | -1.6979895 | 0.08950975 | 0.18317067 | NA | NA |
| ENSG000001 | 6.00657348 | -0.631222278 | 0.37178616 | -1.6978101 | 0.08954362 | 0.18323072 | MAGEA8 | 4107 |
| ENSG00000C | 20.8354602 | -0.476447505 | 0.28065638 | -1.6976186 | 0.08957976 | 0.18323311 | CX3CL1 | 6376 |
| ENSG00000C | 3436.23465 | 0.515196772 | 0.30346279 | 1.69772635 | 0.08955942 | 0.18323311 | HNRNPM | 4670 |
| ENSG000001 | 95.1194871 | 0.282431098 | 0.16636587 | 1.69765044 | 0.08957376 | 0.18323311 | FKBP1B | 2281 |
| ENSG000001 | 855.241561 | 0.198354219 | 0.11684114 | 1.69764017 | 0.0895757 | 0.18323311 | NA | NA |
| ENSG000001 | 242.291886 | 0.242346436 | 0.14274729 | 1.69773055 | 0.08955863 | 0.18323311 | NA | NA |
| ENSG000002 | 6.26925773 | -0.62276605 | 0.36683051 | -1.6976943 | 0.08956548 | 0.18323311 | OR7E24 | 26648 |
| ENSG000002 | 6.77056351 | -0.61340639 | 0.36133476 | -1.6976124 | 0.08958093 | 0.18323311 | NA | NA |
| ENSG000002 | 16.0512658 | -0.515058199 | 0.30340094 | -1.6976157 | 0.08958032 | 0.18323311 | NA | NA |
| ENSG000002 | 12.9213157 | -0.551087487 | 0.3246484 | -1.6974902 | 0.08960402 | 0.18327109 | LOC101929099 | 101929099 |
| ENSG00000C | 622.031053 | 0.222399476 | 0.1310365 | 1.69723297 | 0.08965262 | 0.18336125 | C1orf112 | 55732 |
| ENSG000002 | 5.78920146 | -0.631925495 | 0.37233428 | -1.6971993 | 0.08965898 | 0.18336501 | NA | NA |
| ENSG000002 | 41.3901228 | 0.40985453 | 0.24149278 | 1.69717097 | 0.08966434 | 0.18336672 | NA | NA |
| ENSG000001 | 117.001885 | 0.275828011 | 0.16252659 | 1.6971254 | 0.08967295 | 0.18337509 | TGFBI | 7045 |
| ENSG000002 | 2.46070162 | -0.626695203 | 0.36927654 | -1.6970891 | 0.08967981 | 0.18337987 | RNU2-12P | 106480202 |
| ENSG000002 | 8.78050699 | -0.592713315 | 0.34927834 | -1.6969656 | 0.08970317 | 0.18341837 | NA | NA |
| ENSG000002 | 6.2625518 | -0.621427144 | 0.36621513 | -1.696891 | 0.08971728 | 0.18343797 | NA | NA |
| ENSG000001 | 47.8276513 | 0.3733292 | 0.22001999 | 1.69679675 | 0.08973509 | 0.1834559 | NA | NA |

| | | | | | | | | |
|---|---|---|---|---|---|---|---|---|
| ENSG000002 | 22.7377592 | 0.468354849 | 0.27602261 | 1.69679884 | 0.0897347 | 0.1834559 | SNORD6 | 692075 |
| ENSG000002 | 2.39756869 | -0.626842024 | 0.36946848 | -1.6966049 | 0.08977139 | 0.18352085 | NA | NA |
| ENSG000001 | 121.517291 | -0.274336324 | 0.16171297 | -1.6964399 | 0.08980261 | 0.18354767 | GADD45A | 1647 |
| ENSG000002 | 129.700941 | 0.26623615 | 0.15693175 | 1.69650918 | 0.08978949 | 0.18354767 | TAP2 | 6891 |
| ENSG000002 | 9.87229915 | -0.576277627 | 0.33969368 | -1.6964626 | 0.08979831 | 0.18354767 | NA | NA |
| ENSG000002 | 4.76206686 | -0.635486517 | 0.37459234 | -1.6964749 | 0.08979597 | 0.18354767 | NA | NA |
| ENSG000002 | 2.7559055 | -0.631604634 | 0.3723183 | -1.6964104 | 0.08980819 | 0.18354981 | NA | NA |
| ENSG000002 | 18.6683605 | -0.506067548 | 0.29834491 | -1.69625 | 0.08983855 | 0.18360261 | GUSBP2 | 387036 |
| ENSG000001 | 38.544772 | 0.386344061 | 0.22778367 | 1.69610081 | 0.08986679 | 0.18362163 | PKLR | 5313 |
| ENSG000001 | 3.88669456 | -0.641666139 | 0.37832128 | -1.6960879 | 0.08986924 | 0.18362163 | ZNF474 | 133923 |
| ENSG000002 | 2.41331812 | -0.626762325 | 0.36952275 | -1.69614 | 0.08985936 | 0.18362163 | NA | NA |
| ENSG000002 | 35.9863762 | -0.395188526 | 0.23299289 | -1.6961398 | 0.0898594 | 0.18362163 | LINC00461 | 645323 |
| ENSG000002 | 6.57946064 | -0.622431875 | 0.36698235 | -1.6960812 | 0.0898705 | 0.18362163 | NA | NA |
| ENSG000002 | 5.40283143 | -0.631591566 | 0.37240212 | -1.6959935 | 0.08988711 | 0.18364633 | NA | NA |
| ENSG000002 | 12.9305267 | -0.552422321 | 0.32575501 | -1.6958214 | 0.08991971 | 0.18370366 | LOC642929 | 642929 |
| ENSG000002 | 15.7932447 | -0.521505032 | 0.30754583 | -1.6956986 | 0.08994298 | 0.18374195 | NA | NA |
| ENSG000002 | 11.3662262 | 0.5563682 | 0.3281293 | 1.69557608 | 0.08996619 | 0.18378011 | NA | NA |
| ENSG000001 | 9.73891 | -0.574418303 | 0.33878884 | -1.6955054 | 0.08997959 | 0.18379823 | IL2RA | 3559 |
| ENSG000002 | 3.37627012 | -0.640023605 | 0.37755906 | -1.6951616 | 0.09004478 | 0.18392212 | LOC729083 | 729083 |
| ENSG000002 | 11.8153964 | -0.582315239 | 0.34352592 | -1.695113 | 0.09005399 | 0.18393168 | NA | NA |
| ENSG000002 | 4.10613222 | -0.641743372 | 0.3785908 | -1.6950844 | 0.0900594 | 0.18393346 | NA | NA |
| ENSG000002 | 2.38334674 | -0.627285186 | 0.37007974 | -1.6950001 | 0.09007541 | 0.18395688 | NA | NA |
| ENSG000002 | 5.48371609 | -0.63644899 | 0.37550283 | -1.6949246 | 0.09008973 | 0.18397686 | NA | NA |
| ENSG000002 | 2.3724973 | -0.62673502 | 0.36981155 | -1.6947416 | 0.09012445 | 0.18403052 | NA | NA |
| ENSG000002 | 3.71865209 | -0.641259361 | 0.37838253 | -1.6947383 | 0.09012508 | 0.18403052 | NA | NA |
| ENSG000002 | 2.40596627 | -0.627964384 | 0.37054662 | -1.6946974 | 0.09013285 | 0.18403711 | LOC105369509 | 105369509 |
| ENSG000001 | 88.6173486 | 0.333307084 | 0.19668443 | 1.69462874 | 0.09014587 | 0.18405444 | SP2 | 6668 |
| ENSG000002 | 4.75056453 | -0.634693134 | 0.3745649 | -1.6944811 | 0.09017391 | 0.18410241 | RN7SL364P | 106479356 |
| ENSG000001 | 50.2893226 | 0.351069524 | 0.20718897 | 1.69444119 | 0.09018148 | 0.1841086 | KIFC2 | 90990 |

| | | | | | | | | |
|---|---|---|---|---|---|---|---|---|
| ENSG000002 | 5.39176629 | -0.630873213 | 0.37233562 | -1.6943671 | 0.09019555 | 0.18412806 | NA | NA |
| ENSG000001 | 29.0613191 | -0.425698916 | 0.25125528 | -1.6942885 | 0.09021048 | 0.18414 | AVIL | 10677 |
| ENSG000002 | 5.34186328 | -0.630523882 | 0.3721446 | -1.6942981 | 0.09020866 | 0.18414 | NA | NA |
| ENSG000002 | 9.01097277 | -0.586064508 | 0.34598535 | -1.6938998 | 0.09028433 | 0.18428147 | NA | NA |
| ENSG000002 | 2.99601818 | -0.637622889 | 0.37643843 | -1.6938305 | 0.0902975 | 0.18429908 | BRWD1P3 | 106480688 |
| ENSG000002 | 3.08110513 | -0.637771501 | 0.37654244 | -1.6937573 | 0.09031141 | 0.18431819 | NA | NA |
| ENSG000002 | 4.24610936 | -0.640048915 | 0.37790152 | -1.6936924 | 0.09032375 | 0.1843341 | NA | NA |
| ENSG000001 | 3.64029199 | -0.641046438 | 0.37850943 | -1.6936076 | 0.09033988 | 0.18435772 | OR1L8 | 138881 |
| ENSG000002 | 4.78745091 | -0.635376869 | 0.37517013 | -1.69357 | 0.09034703 | 0.18436304 | MIRLET7DHG | 158257 |
| ENSG000001 | 183.73114 | 0.255884245 | 0.15112029 | 1.69324873 | 0.09040813 | 0.18445589 | NA | NA |
| ENSG000002 | 2.45355774 | -0.624288405 | 0.3687034 | -1.6931995 | 0.0904175 | 0.18445589 | NA | NA |
| ENSG000002 | 10.3584031 | -0.569879266 | 0.3365719 | -1.6931873 | 0.09041982 | 0.18445589 | LOC105375166 | 105375166 |
| ENSG000002 | 2.43488038 | -0.626523385 | 0.37000805 | -1.6932696 | 0.09040416 | 0.18445589 | NA | NA |
| ENSG000002 | 7.44547504 | -0.61408021 | 0.36267626 | -1.6931911 | 0.0904191 | 0.18445589 | NA | NA |
| ENSG000002 | 11.836891 | -0.548489587 | 0.32392978 | -1.6932361 | 0.09041054 | 0.18445589 | NA | NA |
| ENSG000002 | 2.41691931 | -0.626147445 | 0.36982065 | -1.6931111 | 0.09043432 | 0.18447619 | NA | NA |
| ENSG000001 | 12.079145 | -0.55424733 | 0.32739946 | -1.692878 | 0.0904787 | 0.18455742 | GFRAL | 389400 |
| ENSG000001 | 6.50822482 | -0.620543041 | 0.36658119 | -1.6927847 | 0.09049645 | 0.18457932 | SSMEM1 | 136263 |
| ENSG000001 | 2.75180241 | -0.631183381 | 0.3728693 | -1.6927738 | 0.09049853 | 0.18457932 | PRSS57 | 400668 |
| ENSG000002 | 4.30715332 | -0.64039841 | 0.37832075 | -1.6927393 | 0.0905051 | 0.18458343 | NA | NA |
| ENSG000002 | 3.0168953 | -0.637548446 | 0.37665236 | -1.6926708 | 0.09051815 | 0.18459148 | LOC105378349 | 105378349 |
| ENSG000002 | 20.1677499 | -0.475485937 | 0.28090505 | -1.6926928 | 0.09051396 | 0.18459148 | NA | NA |
| ENSG000002 | 4.24647408 | -0.63955899 | 0.37784932 | -1.6926297 | 0.09052597 | 0.18459814 | NA | NA |
| ENSG000002 | 2.8497991 | -0.629924101 | 0.37217042 | -1.692569 | 0.09053755 | 0.18461246 | NA | NA |
| ENSG000001 | 81.8564729 | -0.296958578 | 0.1754556 | -1.6924999 | 0.0905507 | 0.18463 | CRTC3 | 64784 |
| ENSG000002 | 107.272081 | 0.294408582 | 0.17395906 | 1.6924015 | 0.09056945 | 0.18465894 | XKR7 | 343702 |
| ENSG000002 | 12.6799963 | -0.539878619 | 0.31901111 | -1.6923505 | 0.09057916 | 0.18466017 | NA | NA |
| ENSG000002 | 3.88373362 | -0.640185113 | 0.37828046 | -1.6923558 | 0.09057816 | 0.18466017 | NA | NA |
| ENSG000001 | 1416.30376 | 0.241392715 | 0.14264388 | 1.69227532 | 0.09059349 | 0.18468011 | ANXA2 | 302 |

| | | | | | | | | |
|---|---|---|---|---|---|---|---|---|
| ENSG000002 | 9.08209524 | -0.588030027 | 0.3474948 | -1.692198 | 0.09060822 | 0.18470084 | NA | NA |
| ENSG000002 | 4.4924519 | -0.637263067 | 0.37659682 | -1.6921626 | 0.09061497 | 0.18470532 | NA | NA |
| ENSG000001 | 229.628334 | 0.227863037 | 0.13466318 | 1.69209608 | 0.09062765 | 0.18471478 | DNALI1 | 7802 |
| ENSG000002 | 5.36594429 | -0.63035348 | 0.37252941 | -1.6920905 | 0.09062872 | 0.18471478 | NA | NA |
| ENSG000000 | 35.0712437 | -0.424178588 | 0.25071109 | -1.691902 | 0.09066467 | 0.18477876 | ARHGAP15 | 55843 |
| ENSG000000 | 105.896903 | 0.314642147 | 0.18597828 | 1.69182197 | 0.09067992 | 0.18479668 | CRAMP1 | 57585 |
| ENSG000002 | 20.2309127 | -0.484711374 | 0.28650494 | -1.6918081 | 0.09068258 | 0.18479668 | APCDD1L-AS1 | 149773 |
| ENSG000001 | 22.2662024 | -0.481474347 | 0.28459883 | -1.691765 | 0.09069078 | 0.18480412 | MYO1G | 64005 |
| ENSG000002 | 9.49129394 | -0.578265494 | 0.34182399 | -1.6917054 | 0.09070216 | 0.18481754 | INMT | 11185 |
| ENSG000002 | 3.44514255 | -0.637413929 | 0.37679284 | -1.6916827 | 0.09070649 | 0.18481754 | NA | NA |
| ENSG000002 | 13.81533 | 0.61838981 | 0.36556685 | 1.6915916 | 0.09072387 | 0.18484367 | RNA5-8S5 | 100008587 |
| ENSG000002 | 2.44504413 | -0.624514741 | 0.36919539 | -1.6915562 | 0.09073063 | 0.1848456 | NA | NA |
| ENSG000002 | 6.09029882 | -0.628577312 | 0.37160087 | -1.6915389 | 0.09073393 | 0.1848456 | NA | NA |
| ENSG000002 | 2.36823058 | -0.626088934 | 0.37013676 | -1.691507 | 0.09074 | 0.18484868 | NA | NA |
| ENSG000002 | 2.7378733 | -0.630274918 | 0.37264834 | -1.6913397 | 0.09077195 | 0.18490447 | EFCAB14-AS1 | 100130197 |
| ENSG000002 | 9.73092149 | -0.577701623 | 0.34159962 | -1.6911659 | 0.09080512 | 0.18496276 | NA | NA |
| ENSG000002 | 10.4917697 | -0.574389848 | 0.33965251 | -1.6911103 | 0.09081575 | 0.18496581 | LOC101927196 | 101927196 |
| ENSG000002 | 4.36153422 | -0.640062307 | 0.37848262 | -1.6911273 | 0.09081248 | 0.18496581 | HOXC13-AS | 100874366 |
| ENSG000001 | 17.0926131 | -0.524518519 | 0.31018441 | -1.6909893 | 0.09083885 | 0.18500014 | LINC00483 | 55018 |
| ENSG000002 | 10.8421235 | -0.561963185 | 0.33233102 | -1.6909742 | 0.09084173 | 0.18500014 | NA | NA |
| ENSG000001 | 477.257705 | 0.22291034 | 0.13183659 | 1.69080782 | 0.09087351 | 0.18505558 | TSNAX | 7257 |
| ENSG000002 | 4.7752714 | -0.633910269 | 0.3749584 | -1.6906149 | 0.09091037 | 0.18511753 | NA | NA |
| ENSG000002 | 8.13262743 | -0.590305432 | 0.34916901 | -1.6906009 | 0.09091306 | 0.18511753 | NA | NA |
| ENSG000001 | 585.216768 | 0.47568544 | 0.28138142 | 1.6905361 | 0.09092544 | 0.18513343 | SRP68 | 6730 |
| ENSG000002 | 6.02869511 | -0.626172814 | 0.3704387 | -1.6903548 | 0.0909601 | 0.18519472 | NA | NA |
| ENSG000002 | 2.51102987 | -0.616353047 | 0.36464096 | -1.6903012 | 0.09097035 | 0.18519863 | DSCR10 | 259234 |
| ENSG000002 | 4.2156608 | -0.639094737 | 0.37809613 | -1.6902969 | 0.09097116 | 0.18519863 | NA | NA |
| ENSG000001 | 53.8334935 | 0.341649823 | 0.20213274 | 1.69022504 | 0.09098491 | 0.18521732 | SNTG1 | 54212 |
| ENSG000002 | 2.42543292 | -0.623895225 | 0.36913467 | -1.6901561 | 0.09099809 | 0.18523485 | NA | NA |

| | | | | | | | | |
|---|---|---|---|---|---|---|---|---|
| ENSG000002 | 24.0505643 | -0.462605518 | 0.27371932 | -1.6900726 | 0.09101407 | 0.18525809 | RGAG1 | 57529 |
| ENSG000001 | 7.05964614 | -0.612529041 | 0.36243998 | -1.6900151 | 0.09102507 | 0.18527117 | LUM | 4060 |
| ENSG000001 | 70.9689672 | 0.341344976 | 0.20200551 | 1.68978051 | 0.09106995 | 0.18534393 | TMEM53 | 79639 |
| ENSG000001 | 70.5285173 | -0.324336805 | 0.19193815 | -1.6897985 | 0.0910665 | 0.18534393 | GRIK3 | 2899 |
| ENSG000001 | 12.0415345 | -0.558887401 | 0.33079183 | -1.6895441 | 0.0911152 | 0.18541548 | AHSG | 197 |
| ENSG000001 | 52.7727206 | 0.369541045 | 0.21872657 | 1.68951148 | 0.09112145 | 0.18541548 | IL1RAPL2 | 26280 |
| ENSG000002 | 2.76324526 | -0.629052213 | 0.37233012 | -1.6895013 | 0.0911234 | 0.18541548 | NA | NA |
| ENSG000002 | 7.64892179 | -0.602430469 | 0.35657158 | -1.6895078 | 0.09112216 | 0.18541548 | C2orf74 | 339804 |
| ENSG000002 | 5.70037147 | -0.628219174 | 0.37184428 | -1.6894684 | 0.0911297 | 0.18541898 | NA | NA |
| ENSG000002 | 6.60806069 | -0.621681571 | 0.36798668 | -1.6894133 | 0.09114026 | 0.18543118 | NA | NA |
| ENSG000002 | 5.58726805 | -0.623888912 | 0.36929914 | -1.6893863 | 0.09114542 | 0.18543237 | LOC101927210 | 101927210 |
| ENSG000002 | 4.52751208 | -0.636197638 | 0.37659825 | -1.6893271 | 0.09115676 | 0.18544613 | NA | NA |
| ENSG000002 | 3.60362209 | -0.639434737 | 0.37856695 | -1.6890929 | 0.09120163 | 0.18552811 | DEFB115 | 245929 |
| ENSG000002 | 2.44379839 | -0.622730132 | 0.36870356 | -1.6889724 | 0.09122472 | 0.18556577 | NA | NA |
| ENSG000002 | 15.4090265 | -0.516325306 | 0.30573806 | -1.6887832 | 0.09126098 | 0.18562239 | DIO1 | 1733 |
| ENSG000002 | 5.66996655 | -0.625686046 | 0.37049601 | -1.6887794 | 0.09126171 | 0.18562239 | NA | NA |
| ENSG000001 | 8.5716248 | -0.605131731 | 0.35834191 | -1.6886993 | 0.09127707 | 0.18564201 | CD27 | 939 |
| ENSG000001 | 74.0373164 | -0.314792279 | 0.18641307 | -1.6886814 | 0.09128051 | 0.18564201 | KCNH5 | 27133 |
| ENSG000001 | 5.15118863 | -0.633080149 | 0.37494514 | -1.6884607 | 0.09132282 | 0.18571874 | MYL2 | 4633 |
| ENSG000002 | 5.25604012 | -0.634835586 | 0.37599808 | -1.6884011 | 0.09133426 | 0.18572337 | NA | NA |
| ENSG000002 | 4.85707859 | -0.635443963 | 0.37635457 | -1.6884183 | 0.09133095 | 0.18572337 | NA | NA |
| ENSG000002 | 2.41333828 | -0.625054283 | 0.37022873 | -1.6882922 | 0.09135515 | 0.18575654 | NA | NA |
| ENSG000002 | 3.88561299 | -0.638528718 | 0.37824566 | -1.6881323 | 0.09138584 | 0.18580962 | NA | NA |
| ENSG000002 | 6.02495046 | -0.625211595 | 0.3703626 | -1.6881067 | 0.09139074 | 0.18581028 | NA | NA |
| ENSG000002 | 6.30718897 | -0.620803312 | 0.3677727 | -1.6880081 | 0.09140967 | 0.18583944 | NA | NA |
| ENSG000001 | 180.196644 | 0.573531297 | 0.33977429 | 1.68797732 | 0.09141558 | 0.18584215 | QDPR | 5860 |
| ENSG000002 | 8.21461463 | 0.600901307 | 0.35599438 | 1.68795169 | 0.0914205 | 0.18584283 | NA | NA |
| ENSG000002 | 6.75696321 | -0.613494204 | 0.3634748 | -1.687859 | 0.0914383 | 0.1858697 | NA | NA |
| ENSG000002 | 3.07806434 | -0.634993633 | 0.37622041 | -1.6878235 | 0.09144512 | 0.18587425 | NA | NA |

| | | | | | | | | |
|---|---|---|---|---|---|---|---|---|
| ENSG000001 | 8.33704005 | -0.604458744 | 0.35820308 | -1.687475 | 0.09151205 | 0.18599728 | HOXC4 | 3221 |
| ENSG000002 | 11.4831148 | -0.556333483 | 0.3296868 | -1.6874606 | 0.09151482 | 0.18599728 | LOC730338 | 730338 |
| ENSG000002 | 2.38052219 | -0.62404057 | 0.36984598 | -1.6872985 | 0.09154597 | 0.18605124 | TUBB4BP5 | 106480747 |
| ENSG000002 | 11.1426874 | -0.560160539 | 0.33199668 | -1.6872474 | 0.09155579 | 0.18606189 | NA | NA |
| ENSG000001 | 57.9841641 | 0.341668767 | 0.20251253 | 1.68714879 | 0.09157475 | 0.18608176 | NA | NA |
| ENSG000002 | 7.63220975 | -0.600512547 | 0.35593041 | -1.6871628 | 0.09157205 | 0.18608176 | NA | NA |
| ENSG000001 | 7129.16093 | 0.236632169 | 0.14026104 | 1.68708412 | 0.09158718 | 0.18609769 | ND3 | 4537 |
| ENSG000001 | 193.588896 | 0.269437291 | 0.15970927 | 1.68704857 | 0.09159401 | 0.18610226 | ZBTB7A | 51341 |
| ENSG000002 | 8.43917056 | -0.590043884 | 0.34975424 | -1.6870242 | 0.09159869 | 0.18610244 | NA | NA |
| ENSG000002 | 7.07187458 | -0.610243665 | 0.36173593 | -1.6869866 | 0.09160593 | 0.18610782 | LOC101927835 | 101927835 |
| ENSG000002 | 4.49236758 | -0.635733565 | 0.37685596 | -1.6869405 | 0.0916148 | 0.18611652 | NA | NA |
| ENSG000001 | 28.0005743 | -0.424002818 | 0.25141942 | -1.6864362 | 0.09171181 | 0.1862856 | ACTR3C | 653857 |
| ENSG000002 | 6.70214549 | -0.621193231 | 0.36834612 | -1.6864389 | 0.09171129 | 0.1862856 | NA | NA |
| ENSG000002 | 6.56350368 | -0.619280268 | 0.3672084 | -1.6864545 | 0.09170829 | 0.1862856 | CCL4 | 6351 |
| ENSG000001 | 13.5078373 | -0.54555413 | 0.3235074 | -1.686373 | 0.09172398 | 0.18629613 | SERPINA9 | 327657 |
| ENSG000001 | 60.1005279 | 0.325632393 | 0.19309762 | 1.68636155 | 0.09172618 | 0.18629613 | B3GNTL1 | 146712 |
| ENSG000002 | 3.86745604 | -0.637873208 | 0.37827944 | -1.6862487 | 0.09174791 | 0.18633093 | NA | NA |
| ENSG000001 | 79.2674399 | -0.298234342 | 0.17686911 | -1.6861867 | 0.09175985 | 0.1863365 | RNF152 | 220441 |
| ENSG000001 | 180.731589 | 0.265412276 | 0.15740168 | 1.68620989 | 0.09175538 | 0.1863365 | ZNF552 | 79818 |
| ENSG000002 | 36.7480179 | -0.39922923 | 0.23677777 | -1.6860925 | 0.09177798 | 0.18636399 | NA | NA |
| ENSG000002 | 5.3685418 | -0.627211149 | 0.37201776 | -1.685971 | 0.09180139 | 0.18640219 | NA | NA |
| ENSG000002 | 2.41855072 | -0.623577426 | 0.36996581 | -1.6855001 | 0.09189212 | 0.18657707 | NA | NA |
| ENSG000002 | 57.3200388 | 0.413720002 | 0.24546712 | 1.68543956 | 0.0919038 | 0.18659144 | NA | NA |
| ENSG000001 | 83.3950285 | 0.298440151 | 0.17707887 | 1.68535159 | 0.09192076 | 0.18659785 | CPXM2 | 119587 |
| ENSG000002 | 36.8657838 | 0.388555468 | 0.23054528 | 1.68537594 | 0.09191606 | 0.18659785 | NA | NA |
| ENSG000002 | 6.71938195 | -0.609854363 | 0.36185563 | -1.6853527 | 0.09192054 | 0.18659785 | NA | NA |
| ENSG000002 | 3.63159101 | -0.638017423 | 0.37858753 | -1.6852574 | 0.09193893 | 0.18662539 | NA | NA |
| ENSG000002 | 3.02948771 | -0.634667298 | 0.37660973 | -1.6852122 | 0.09194764 | 0.18663374 | NA | NA |
| ENSG000001 | 95.2512745 | 0.305835051 | 0.18148777 | 1.68515518 | 0.09195864 | 0.18663954 | STOX1 | 219736 |

| | | | | | | | | |
|---|---|---|---|---|---|---|---|---|
| ENSG000002 | 2.48409982 | -0.618015627 | 0.36674229 | -1.6851496 | 0.09195971 | 0.18663954 | NA | NA |
| ENSG000002 | 3.41889819 | -0.636006729 | 0.37745389 | -1.6849919 | 0.09199014 | 0.18669197 | NA | NA |
| ENSG000002 | 5.35737916 | -0.627495206 | 0.37241717 | -1.6849256 | 0.09200293 | 0.18670859 | LINC01602 | 100505477 |
| ENSG000002 | 3.55983827 | -0.633519555 | 0.37601141 | -1.6848413 | 0.09201919 | 0.18673223 | NA | NA |
| ENSG000002 | 5.33520399 | -0.62688912 | 0.37212254 | -1.6846309 | 0.09205981 | 0.18680532 | NA | NA |
| ENSG000001 | 5.45132462 | -0.628025728 | 0.37283397 | -1.6844649 | 0.09209186 | 0.186861 | FOXD4L4 | 349334 |
| ENSG000002 | 36.5732443 | -0.386130912 | 0.22927074 | -1.6841701 | 0.09214881 | 0.18696719 | NA | NA |
| ENSG000002 | 7.93588631 | -0.614063556 | 0.36463352 | -1.6840568 | 0.0921707 | 0.18700225 | RBP3 | 5949 |
| ENSG000001 | 517.835268 | 0.506212829 | 0.30059971 | 1.68400968 | 0.0921798 | 0.18701136 | NCOR2 | 9612 |
| ENSG000001 | 2.40845059 | -0.621382957 | 0.36900767 | -1.6839296 | 0.09219527 | 0.18703338 | NA | NA |
| ENSG000002 | 2.78171486 | -0.627222873 | 0.37249766 | -1.6838304 | 0.09221445 | 0.18706295 | NA | NA |
| ENSG000002 | 6.59980299 | -0.619781253 | 0.36810508 | -1.6837074 | 0.09223823 | 0.18710182 | NA | NA |
| ENSG000001 | 257.470045 | 0.257230717 | 0.15282358 | 1.68318734 | 0.09233883 | 0.18729652 | PTRF | 284119 |
| ENSG000002 | 5.80565475 | -0.628204439 | 0.37322986 | -1.683157 | 0.09234471 | 0.18729906 | NA | NA |
| ENSG000002 | 15.084457 | -0.562429952 | 0.33416773 | -1.6830768 | 0.09236023 | 0.18732117 | NA | NA |
| ENSG000002 | 3.88441931 | -0.636528076 | 0.37821852 | -1.6829638 | 0.0923821 | 0.18735616 | NA | NA |
| ENSG000002 | 6.4207495 | -0.612304514 | 0.36383105 | -1.6829364 | 0.09238741 | 0.18735756 | NA | NA |
| ENSG000002 | 3.88026828 | -0.63689992 | 0.37846706 | -1.6828411 | 0.09240586 | 0.18738561 | NA | NA |
| ENSG000002 | 5.66556467 | -0.626114322 | 0.37208237 | -1.6827304 | 0.09242729 | 0.18741969 | NA | NA |
| ENSG000001 | 29.113599 | -0.423879381 | 0.251939 | -1.6824683 | 0.09247807 | 0.18751329 | HGD | 3081 |
| ENSG000002 | 24.1187788 | 0.454849261 | 0.27038635 | 1.68221975 | 0.09252624 | 0.18760157 | MAFB | 9935 |
| ENSG000002 | 3.00515298 | -0.633671002 | 0.37669986 | -1.6821642 | 0.09253701 | 0.18761404 | NA | NA |
| ENSG00000C | 4.47819714 | -0.633723088 | 0.37675717 | -1.6820465 | 0.09255982 | 0.18764409 | KRT33A | 3883 |
| ENSG000001 | 374.026089 | 0.221689194 | 0.13181073 | 1.6818752 | 0.09259304 | 0.18764409 | KANSL1L | 151050 |
| ENSG000001 | 11.0478828 | 0.574762557 | 0.34172325 | 1.68195333 | 0.09257789 | 0.18764409 | ATOH1 | 474 |
| ENSG000002 | 13.43982 | 0.528334852 | 0.31411585 | 1.68197453 | 0.09257378 | 0.18764409 | NA | NA |
| ENSG000002 | 2.75770054 | -0.626507161 | 0.37247245 | -1.6820228 | 0.09256441 | 0.18764409 | LINC00442 | 348021 |
| ENSG000002 | 3.6776439 | -0.636607952 | 0.37851132 | -1.681873 | 0.09259348 | 0.18764409 | NA | NA |
| ENSG000002 | 3.01615863 | -0.633730356 | 0.37679732 | -1.6818866 | 0.09259083 | 0.18764409 | NA | NA |

| | | | | | | | | |
|---|---|---|---|---|---|---|---|---|
| ENSG000002 | 15349.2721 | 0.561915781 | 0.33410092 | 1.68187441 | 0.0925932 | 0.18764409 | MALAT1 | 378938 |
| ENSG000002 | 3.77645682 | -0.634912366 | 0.37748283 | -1.6819636 | 0.09257591 | 0.18764409 | NA | NA |
| ENSG000002 | 11.1074677 | -0.561694092 | 0.33398084 | -1.6818153 | 0.09260466 | 0.18765736 | NA | NA |
| ENSG000002 | 6.30071096 | -0.618710181 | 0.36791361 | -1.6816725 | 0.09263237 | 0.18770415 | NA | NA |
| ENSG000002 | 3.6665465 | -0.636578784 | 0.37855139 | -1.6816179 | 0.09264296 | 0.18771623 | NFIA-AS1 | 645030 |
| ENSG000001 | 20.4633143 | 0.48689184 | 0.28964629 | 1.68098767 | 0.09276531 | 0.18795473 | CCDC78 | 124093 |
| ENSG000002 | 6.22077305 | -0.61640955 | 0.36670954 | -1.6809204 | 0.09277838 | 0.18796243 | FLJ16171 | 441116 |
| ENSG000002 | 3.06611474 | -0.630034765 | 0.37481298 | -1.680931 | 0.09277631 | 0.18796243 | NA | NA |
| ENSG000002 | 8.33726498 | -0.602011259 | 0.35820241 | -1.6806455 | 0.09283179 | 0.18806125 | SPINK9 | 643394 |
| ENSG000001 | 348.891889 | 0.200266886 | 0.1191666 | 1.68056217 | 0.09284799 | 0.18808467 | ZNF26 | 7574 |
| ENSG000001 | 195.008689 | 0.25707655 | 0.15298261 | 1.68042992 | 0.0928737 | 0.18809916 | LOC105372481 | 105372481 |
| ENSG000001 | 12.9196129 | -0.548356542 | 0.32631148 | -1.6804697 | 0.09286597 | 0.18809916 | HRNR | 388697 |
| ENSG000002 | 10.488027 | -0.57700329 | 0.34336427 | -1.6804407 | 0.0928716 | 0.18809916 | NA | NA |
| ENSG000002 | 5.94816937 | -0.620810399 | 0.3694213 | -1.6804943 | 0.09286117 | 0.18809916 | NA | NA |
| ENSG000002 | 4.89792371 | -0.633511757 | 0.37700498 | -1.6803803 | 0.09288334 | 0.18810929 | NA | NA |
| ENSG000002 | 4.8046227 | -0.630993109 | 0.37553638 | -1.6802449 | 0.09290967 | 0.18815321 | LOC105370027 | 105370027 |
| ENSG000002 | 9.46598593 | 0.59155668 | 0.35208418 | 1.68015695 | 0.09292678 | 0.18816908 | NA | NA |
| ENSG000002 | 33.1226094 | 0.410579329 | 0.24436703 | 1.68017482 | 0.09292331 | 0.18816908 | FGF14-AS2 | 283481 |
| ENSG000002 | 4.11273478 | -0.635939182 | 0.37853313 | -1.6800093 | 0.0929555 | 0.18821783 | NA | NA |
| ENSG000002 | 11.3972776 | 0.578889222 | 0.34462433 | 1.67976885 | 0.0930023 | 0.18830319 | RN7SL648P | 106479459 |
| ENSG000001 | 158.113613 | 0.264299966 | 0.15735611 | 1.67962952 | 0.09302942 | 0.18833611 | C16orf45 | 89927 |
| ENSG000001 | 383.128149 | 0.216867636 | 0.12912123 | 1.67956604 | 0.09304178 | 0.18833611 | ZNF81 | 347344 |
| ENSG000002 | 4.57042742 | -0.633222933 | 0.3770064 | -1.6796079 | 0.09303363 | 0.18833611 | NA | NA |
| ENSG000002 | 3.01020898 | -0.632860589 | 0.37679608 | -1.6795838 | 0.09303832 | 0.18833611 | NA | NA |
| ENSG000002 | 10.8180281 | -0.580136723 | 0.34539182 | -1.6796481 | 0.0930258 | 0.18833611 | LOC101928100 | 101928100 |
| ENSG000002 | 2.98599878 | -0.632893899 | 0.37683383 | -1.6795039 | 0.09305387 | 0.18835119 | LINC01262 | 101928971 |
| ENSG000002 | 3.10432595 | -0.629088351 | 0.37459716 | -1.679373 | 0.09307937 | 0.18839339 | NA | NA |
| ENSG000002 | 3.63618282 | -0.635659642 | 0.37852761 | -1.6792953 | 0.0930945 | 0.18841461 | NA | NA |
| ENSG000001 | 1068.41548 | 0.514536288 | 0.30644901 | 1.6790274 | 0.0931467 | 0.18851085 | RPA1 | 6117 |

| | | | | | | | | |
|---|---|---|---|---|---|---|---|---|
| ENSG000001 | 66.4771121 | -0.313956579 | 0.1869982 | -1.6789283 | 0.09316601 | 0.18853111 | MCF2 | 4168 |
| ENSG000002 | 5.2138444 | -0.631926982 | 0.37638698 | -1.6789289 | 0.0931659 | 0.18853111 | NA | NA |
| ENSG000002 | 12.4684592 | -0.541108539 | 0.32236033 | -1.6785829 | 0.09323336 | 0.18865798 | NA | NA |
| ENSG000002 | 11.9847557 | -0.550278101 | 0.32784544 | -1.6784681 | 0.09325576 | 0.18869389 | NA | NA |
| ENSG000002 | 2.76554828 | -0.624475453 | 0.37207789 | -1.678346 | 0.09327956 | 0.18872259 | NA | NA |
| ENSG000002 | 10.4236662 | 0.557337854 | 0.33208005 | 1.67832381 | 0.0932839 | 0.18872259 | NA | NA |
| ENSG000002 | 6.42720824 | -0.610737028 | 0.3638875 | -1.6783677 | 0.09327534 | 0.18872259 | NA | NA |
| ENSG000002 | 19.6081694 | 0.481384031 | 0.28682874 | 1.67829778 | 0.09328898 | 0.18872345 | NA | NA |
| ENSG000002 | 8.60247556 | -0.614941199 | 0.36642185 | -1.6782329 | 0.09330164 | 0.18873964 | ANKRD65 | 441869 |
| ENSG000002 | 65.612705 | -0.311181502 | 0.1854257 | -1.6782005 | 0.09330796 | 0.18874301 | MAGI2-AS3 | 100505881 |
| ENSG000001 | 28.0607844 | -0.44057247 | 0.26253776 | -1.6781299 | 0.09332174 | 0.18875205 | AKR1C2 | 1646 |
| ENSG000002 | 3.89417867 | -0.63483557 | 0.3782988 | -1.6781327 | 0.0933212 | 0.18875205 | NA | NA |
| ENSG000002 | 2.75585453 | -0.626126107 | 0.37311694 | -1.6780962 | 0.09332832 | 0.18875596 | NA | NA |
| ENSG000001 | 12.1885664 | -0.540431489 | 0.32206306 | -1.67803 | 0.09334124 | 0.18876324 | NA | NA |
| ENSG000001 | 11.7924087 | -0.559213676 | 0.33325573 | -1.6780317 | 0.09334091 | 0.18876324 | MCEMP1 | 199675 |
| ENSG00000C | 11.7779551 | -0.561348605 | 0.33457355 | -1.6778033 | 0.09338551 | 0.18882452 | FCN1 | 2219 |
| ENSG000001 | 11.5270619 | -0.569295663 | 0.33930802 | -1.6778138 | 0.09338346 | 0.18882452 | SHISA7 | 729956 |
| ENSG000002 | 165.740854 | 0.254937024 | 0.15194563 | 1.67781737 | 0.09338276 | 0.18882452 | MICB | 4277 |
| ENSG000002 | 4.76434321 | -0.629122681 | 0.37499233 | -1.6776948 | 0.0934067 | 0.18885795 | LINC01474 | 101927258 |
| ENSG000002 | 11.901911 | -0.542315882 | 0.32325528 | -1.6776706 | 0.09341142 | 0.18885808 | NA | NA |
| ENSG000002 | 17.2392649 | -0.501776057 | 0.29912166 | -1.6774982 | 0.0934451 | 0.18891675 | NA | NA |
| ENSG000002 | 6.69935122 | -0.621758617 | 0.37070849 | -1.6772171 | 0.09350004 | 0.18900898 | NA | NA |
| ENSG000002 | 12.8251592 | -0.547562614 | 0.3264665 | -1.6772398 | 0.0934956 | 0.18900898 | NA | NA |
| ENSG000002 | 3.33698733 | -0.632009488 | 0.37683308 | -1.6771603 | 0.09351114 | 0.18902199 | KRTAP3-3 | 85293 |
| ENSG000001 | 249.889178 | 0.221548128 | 0.13210546 | 1.67705505 | 0.09353172 | 0.18903532 | AP4S1 | 11154 |
| ENSG000001 | 13.5318841 | -0.530275909 | 0.31618815 | -1.6770898 | 0.09352493 | 0.18903532 | UROC1 | 131669 |
| ENSG000001 | 42.1184327 | 0.389949206 | 0.2325171 | 1.67707751 | 0.09352733 | 0.18903532 | SLC35A1 | 10559 |
| ENSG000002 | 232.799768 | 0.305120242 | 0.18194318 | 1.67700842 | 0.09354084 | 0.18904433 | NA | NA |
| ENSG000002 | 6.06209872 | -0.633595411 | 0.37783912 | -1.676892 | 0.0935636 | 0.18908091 | NA | NA |

| | | | | | | | | |
|---|---|---|---|---|---|---|---|---|
| ENSG000002 | 2.2501173 | 0.614071825 | 0.36620551 | 1.67685034 | 0.09357175 | 0.18908796 | NA | NA |
| ENSG000002 | 3.69441019 | -0.634531137 | 0.37843818 | -1.6767101 | 0.09359918 | 0.18913228 | NA | NA |
| ENSG000002 | 4.49345024 | -0.631739612 | 0.3767777 | -1.6766905 | 0.09360301 | 0.18913228 | NA | NA |
| ENSG000002 | 4.54048849 | -0.631835976 | 0.37687115 | -1.6765305 | 0.09363433 | 0.18915786 | NA | NA |
| ENSG000002 | 24.1025881 | 0.443968491 | 0.26481275 | 1.67653745 | 0.09363297 | 0.18915786 | NA | NA |
| ENSG000002 | 23.3807338 | -0.467545428 | 0.27887119 | -1.6765641 | 0.09362775 | 0.18915786 | NA | NA |
| ENSG000002 | 6.00958581 | -0.62244809 | 0.37126918 | -1.6765412 | 0.09363222 | 0.18915786 | NA | NA |
| ENSG000001 | 10.1047982 | -0.592428201 | 0.35341902 | -1.6762771 | 0.09368393 | 0.18924864 | CEACAM8 | 1088 |
| ENSG000002 | 2.41376249 | -0.61941298 | 0.36954423 | -1.6761538 | 0.09370807 | 0.18928796 | NA | NA |
| ENSG000001 | 258.170023 | 0.492086806 | 0.29360134 | 1.67603734 | 0.09373088 | 0.18931876 | FYCO1 | 79443 |
| ENSG000002 | 73.095009 | 0.33951758 | 0.2025727 | 1.6760283 | 0.09373265 | 0.18931876 | TECPR1 | 25851 |
| ENSG000002 | 7.58164525 | -0.601624643 | 0.35897629 | -1.6759453 | 0.0937489 | 0.18934215 | HLA-DOB | 3112 |
| ENSG000001 | 47.268112 | 0.37326263 | 0.22273514 | 1.67581386 | 0.09377466 | 0.18938474 | SARS2 | 54938 |
| ENSG000001 | 65.6318242 | -0.332447712 | 0.1983982 | -1.6756589 | 0.09380503 | 0.18940353 | RNASEL | 6041 |
| ENSG000001 | 29.6503918 | -0.410428113 | 0.24494055 | -1.6756234 | 0.09381199 | 0.18940353 | SLA | 6503 |
| ENSG000001 | 212.94812 | 0.450902576 | 0.26908495 | 1.67568856 | 0.09379921 | 0.18940353 | WNK2 | 65268 |
| ENSG000002 | 3.65799988 | -0.634262461 | 0.37850864 | -1.6756882 | 0.09379928 | 0.18940353 | NA | NA |
| ENSG000002 | 5.33474718 | -0.624289374 | 0.3725611 | -1.6756698 | 0.09380289 | 0.18940353 | NA | NA |
| ENSG000002 | 2.19731095 | -0.597889746 | 0.35681283 | -1.6756397 | 0.09380878 | 0.18940353 | NA | NA |
| ENSG000002 | 2.39623917 | -0.619057892 | 0.36949127 | -1.6754331 | 0.09384929 | 0.18946942 | NA | NA |
| ENSG000002 | 3.95149779 | -0.634124885 | 0.37850502 | -1.6753407 | 0.0938674 | 0.18949653 | NA | NA |
| ENSG000001 | 17.2300326 | -0.502066882 | 0.29970258 | -1.6752171 | 0.09389165 | 0.18951718 | DMTN | 2039 |
| ENSG000001 | 38.0168383 | 0.379634215 | 0.22661746 | 1.67522052 | 0.09389097 | 0.18951718 | ECEL1 | 9427 |
| ENSG000002 | 6.2780071 | -0.616877656 | 0.36823576 | -1.6752247 | 0.09389015 | 0.18951718 | NA | NA |
| ENSG000002 | 11.6936206 | -0.551229471 | 0.32907115 | -1.6751073 | 0.09391319 | 0.18955123 | NA | NA |
| ENSG000002 | 11.7687032 | -0.554816779 | 0.33125711 | -1.6748826 | 0.09395726 | 0.18963023 | NA | NA |
| ENSG000002 | 4.67189713 | -0.633299259 | 0.37812069 | -1.6748601 | 0.09396168 | 0.18963023 | NA | NA |
| ENSG000001 | 29.0067974 | -0.418961046 | 0.2501747 | -1.6746739 | 0.09399823 | 0.18969098 | RSPO2 | 340419 |
| ENSG000002 | 5.85965088 | -0.626813066 | 0.37429293 | -1.6746591 | 0.09400114 | 0.18969098 | NA | NA |

| | | | | | | | | |
|---|---|---|---|---|---|---|---|---|
| ENSG000001 | 92.0478863 | 0.341193693 | 0.20374927 | 1.67457629 | 0.09401739 | 0.18971434 | NOMO3 | 408050 |
| ENSG000001 | 398.139046 | 0.227143018 | 0.13564522 | 1.67453754 | 0.094025 | 0.18972025 | TERF2IP | 54386 |
| ENSG000002 | 3.06018835 | -0.62926605 | 0.37580199 | -1.6744617 | 0.09403989 | 0.18973373 | NA | NA |
| ENSG000002 | 6.88665703 | -0.614359246 | 0.36690083 | -1.6744559 | 0.09404104 | 0.18973373 | NA | NA |
| ENSG000001 | 19.168172 | -0.47372036 | 0.28294901 | -1.6742252 | 0.09408636 | 0.18973459 | RNF207 | 388591 |
| ENSG000001 | 9.46762327 | -0.568079118 | 0.33930188 | -1.6742587 | 0.09407977 | 0.18973459 | ATP4B | 496 |
| ENSG000002 | 53.0227946 | 0.568417489 | 0.33949802 | 1.67428808 | 0.094074 | 0.18973459 | SMG1P2 | 440354 |
| ENSG000002 | 3.2784661 | -0.633201805 | 0.37820807 | -1.6742155 | 0.09408825 | 0.18973459 | NA | NA |
| ENSG000002 | 4.76155799 | -0.628425276 | 0.37534557 | -1.6742579 | 0.09407993 | 0.18973459 | NA | NA |
| ENSG000002 | 4.80854552 | -0.63038529 | 0.37650879 | -1.674291 | 0.09407342 | 0.18973459 | NA | NA |
| ENSG000002 | 6.27247458 | -0.616274112 | 0.36806222 | -1.6743748 | 0.09405696 | 0.18973459 | SPESP1 | 246777 |
| ENSG000002 | 3.87840962 | -0.633500949 | 0.37834928 | -1.6743813 | 0.09405569 | 0.18973459 | NA | NA |
| ENSG000002 | 24.9407728 | 0.454966451 | 0.27174022 | 1.67426986 | 0.09407758 | 0.18973459 | NA | NA |
| ENSG000002 | 5.69671797 | -0.622692981 | 0.37192804 | -1.6742297 | 0.09408546 | 0.18973459 | LOC286083 | 286083 |
| ENSG000001 | 12.7370985 | -0.535424537 | 0.31982748 | -1.6741042 | 0.09411012 | 0.18976107 | C1orf162 | 128346 |
| ENSG000002 | 3.87832585 | -0.633277932 | 0.3782794 | -1.674101 | 0.09411075 | 0.18976107 | NA | NA |
| ENSG000002 | 22.445103 | 0.456584442 | 0.27274028 | 1.6740631 | 0.0941182 | 0.18976406 | SNORA14B | 677802 |
| ENSG000002 | 8.73888032 | -0.583985603 | 0.34884786 | -1.6740409 | 0.09412255 | 0.18976406 | NA | NA |
| ENSG000002 | 5.57470912 | -0.628090125 | 0.37519824 | -1.674022 | 0.09412627 | 0.18976406 | NA | NA |
| ENSG000001 | 84.5670351 | 0.309397141 | 0.18482968 | 1.67395813 | 0.09413883 | 0.18977687 | EMILIN2 | 84034 |
| ENSG000002 | 6.77725991 | -0.610066336 | 0.36444889 | -1.6739421 | 0.09414198 | 0.18977687 | NA | NA |
| ENSG000001 | 60.1400364 | 0.331648339 | 0.19812823 | 1.67390758 | 0.09414876 | 0.18978111 | GPR153 | 387509 |
| ENSG000001 | 6.8990051 | -0.610304404 | 0.36461102 | -1.6738507 | 0.09415995 | 0.18979399 | OR11G2 | 390439 |
| ENSG000002 | 4.74203165 | -0.627781989 | 0.37506245 | -1.6738066 | 0.09416862 | 0.18979399 | NA | NA |
| ENSG000002 | 10.0376598 | -0.56885745 | 0.33985913 | -1.6738037 | 0.09416919 | 0.18979399 | NA | NA |
| ENSG000002 | 6.90754987 | -0.612751995 | 0.36609732 | -1.6737407 | 0.09418157 | 0.18980949 | LOC101928708 | 101928708 |
| ENSG000002 | 5.11807939 | -0.629391306 | 0.37604513 | -1.6737122 | 0.09418718 | 0.18981137 | NA | NA |
| ENSG000002 | 3.44850898 | -0.629893887 | 0.37639269 | -1.6735019 | 0.09422853 | 0.18988527 | NA | NA |
| ENSG000002 | 3.30265647 | -0.632670064 | 0.3780957 | -1.6733067 | 0.09426693 | 0.18995321 | NA | NA |

| | | | | | | | | |
|---|---|---|---|---|---|---|---|---|
| ENSG000002 | 25.2949553 | -0.432446549 | 0.25844419 | -1.6732686 | 0.09427443 | 0.18995887 | NA | NA |
| ENSG000002 | 3.8929048 | -0.633341901 | 0.37853453 | -1.6731417 | 0.09429941 | 0.18999976 | NA | NA |
| ENSG000002 | 3.68731893 | -0.633233005 | 0.37848141 | -1.6730888 | 0.09430982 | 0.1900113 | NA | NA |
| ENSG000001 | 53.2354034 | 0.335691865 | 0.20065267 | 1.67299974 | 0.09432735 | 0.19002773 | KIRREL2 | 84063 |
| ENSG000002 | 3.14512806 | -0.62669366 | 0.37458884 | -1.6730174 | 0.09432387 | 0.19002773 | LOC105375734 | 105375734 |
| ENSG000002 | 32.9444625 | -0.421491807 | 0.25195083 | -1.672913 | 0.09434443 | 0.19004327 | NA | NA |
| ENSG000002 | 3.25435949 | -0.632692183 | 0.3781948 | -1.6729267 | 0.09434172 | 0.19004327 | NA | NA |
| ENSG000002 | 9.04270048 | -0.598454276 | 0.35774426 | -1.672855 | 0.09435584 | 0.19005681 | SMIM17 | 147670 |
| ENSG000002 | 3.62091653 | -0.63301876 | 0.37842022 | -1.6727932 | 0.09436802 | 0.19007191 | NUTF2P3 | 106480298 |
| ENSG000000 | 54.3034378 | 0.35296805 | 0.21102369 | 1.67264656 | 0.09439689 | 0.19010173 | MAN2B2 | 23324 |
| ENSG000002 | 7.58519132 | -0.59484332 | 0.35562102 | -1.6726889 | 0.09438856 | 0.19010173 | TRIM77 | 390231 |
| ENSG000002 | 13.6701678 | 0.545309673 | 0.3260146 | 1.67265414 | 0.0943954 | 0.19010173 | NA | NA |
| ENSG000002 | 3.40563327 | -0.630942562 | 0.37723155 | -1.6725604 | 0.09441387 | 0.19012647 | NA | NA |
| ENSG000002 | 36.5917339 | 0.446551728 | 0.26701096 | 1.67240971 | 0.09444356 | 0.19017682 | NA | NA |
| ENSG000002 | 3.89584145 | -0.632981458 | 0.37851023 | -1.6722968 | 0.09446581 | 0.19021217 | NA | NA |
| ENSG000002 | 24.2045296 | -0.431863131 | 0.25827145 | -1.6721288 | 0.09449892 | 0.19024975 | ERICH2 | 285141 |
| ENSG000002 | 8.1917169 | -0.594186644 | 0.35533964 | -1.6721654 | 0.09449171 | 0.19024975 | STX18-IT1 | 104472519 |
| ENSG000002 | 8.47339243 | -0.592325142 | 0.35423879 | -1.6721069 | 0.09450324 | 0.19024975 | NA | NA |
| ENSG000002 | 6.54368816 | -0.610677491 | 0.36520829 | -1.6721348 | 0.09449774 | 0.19024975 | NA | NA |
| ENSG000001 | 2793.41357 | 0.434723118 | 0.25999996 | 1.67201223 | 0.09452191 | 0.1902779 | SRRM2 | 23524 |
| ENSG000002 | 5.41090022 | -0.624506986 | 0.37355891 | -1.6717764 | 0.09456842 | 0.19035758 | NA | NA |
| ENSG000002 | 8.24297068 | -0.593066349 | 0.35475484 | -1.6717639 | 0.09457088 | 0.19035758 | NA | NA |
| ENSG000001 | 55.7706107 | 0.333120978 | 0.19927458 | 1.67166819 | 0.09458977 | 0.19038615 | EGFL6 | 25975 |
| ENSG000002 | 16.5171528 | 0.510805111 | 0.30558466 | 1.67156662 | 0.09460981 | 0.19041704 | SNORA54 | 677833 |
| ENSG000001 | 403.585241 | -0.246677287 | 0.1475763 | -1.6715237 | 0.09461829 | 0.19042464 | PDE12 | 201626 |
| ENSG000001 | 7.58793732 | -0.595194249 | 0.35612898 | -1.6712884 | 0.09466472 | 0.19050863 | FSHB | 2488 |
| ENSG000002 | 5.77287113 | -0.625423253 | 0.37422782 | -1.6712366 | 0.09467494 | 0.19051976 | ALDH1L1-AS2 | 100862662 |
| ENSG000002 | 5.7332589 | -0.62296415 | 0.37276626 | -1.6711924 | 0.09468368 | 0.19052788 | MIR320B2 | 100313769 |
| ENSG000001 | 1130.47789 | 0.190500878 | 0.11399966 | 1.67106529 | 0.09470878 | 0.19056894 | ZNF614 | 80110 |

| | | | | | | | | |
|---|---|---|---|---|---|---|---|---|
| ENSG000001 | 347.987622 | 0.207929181 | 0.12444019 | 1.67091656 | 0.09473816 | 0.19060914 | PATL1 | 219988 |
| ENSG000002 | 2.97116283 | -0.629253521 | 0.37658842 | -1.6709317 | 0.09473517 | 0.19060914 | NA | NA |
| ENSG000002 | 3.01746833 | -0.628000299 | 0.37588805 | -1.670711 | 0.09477878 | 0.19067449 | RNU6-21P | 100302738 |
| ENSG000002 | 11.6744275 | -0.547595499 | 0.32776322 | -1.6707046 | 0.09478005 | 0.19067449 | NA | NA |
| ENSG000001 | 961.574173 | 0.218598075 | 0.13085608 | 1.67052294 | 0.09481595 | 0.19070997 | RPRD2 | 23248 |
| ENSG000002 | 8.07483113 | -0.585430868 | 0.35044272 | -1.6705465 | 0.09481128 | 0.19070997 | NA | NA |
| ENSG000002 | 4.46584605 | -0.630307567 | 0.37731216 | -1.6705202 | 0.09481649 | 0.19070997 | LOC100133920 | 100133920 |
| ENSG000002 | 5.88509161 | -0.615585179 | 0.36849766 | -1.6705267 | 0.0948152 | 0.19070997 | PAUPAR | 103157000 |
| ENSG000002 | 6.56192889 | -0.611779087 | 0.36623644 | -1.6704484 | 0.09483067 | 0.19072903 | NA | NA |
| ENSG000002 | 2.46496779 | -0.613420177 | 0.36722897 | -1.6704025 | 0.09483976 | 0.19073784 | NA | NA |
| ENSG000001 | 3.56961232 | -0.632259556 | 0.3785157 | -1.6703655 | 0.09484707 | 0.19074309 | SMCP | 4184 |
| ENSG000002 | 2.78236003 | -0.619090474 | 0.37065506 | -1.6702604 | 0.09486785 | 0.19077541 | NA | NA |
| ENSG000001 | 288.09072 | 0.222006173 | 0.13291912 | 1.67023505 | 0.09487287 | 0.19077604 | CLK3 | 1198 |
| ENSG000001 | 62.667249 | 0.332247612 | 0.19895788 | 1.66993948 | 0.09493134 | 0.19086385 | GALNT18 | 374378 |
| ENSG000001 | 20.0339739 | -0.482033824 | 0.28865006 | -1.6699592 | 0.09492744 | 0.19086385 | KCTD19 | 146212 |
| ENSG000002 | 3.93453507 | -0.632069044 | 0.37850278 | -1.6699191 | 0.09493536 | 0.19086385 | CT47B1 | 643311 |
| ENSG000002 | 13.8996008 | -0.533910083 | 0.31971351 | -1.6699641 | 0.09492647 | 0.19086385 | NA | NA |
| ENSG000002 | 5.59190746 | -0.617857712 | 0.37002247 | -1.6697843 | 0.09496205 | 0.19090804 | NA | NA |
| ENSG000002 | 3.61790875 | -0.631895765 | 0.37845125 | -1.6696887 | 0.09498098 | 0.19093662 | NA | NA |
| ENSG000001 | 12.5928844 | -0.532768516 | 0.31910214 | -1.6695861 | 0.09500128 | 0.19096796 | C11orf45 | 219833 |
| ENSG000002 | 4.49742357 | -0.629143876 | 0.3768415 | -1.6695185 | 0.09501466 | 0.19098466 | NA | NA |
| ENSG000002 | 4.59821181 | -0.630595322 | 0.37771585 | -1.6694966 | 0.095019 | 0.19098466 | NA | NA |
| ENSG000002 | 11.5533778 | -0.561573183 | 0.33638508 | -1.6694355 | 0.09503111 | 0.19099953 | NA | NA |
| ENSG000001 | 3.65521172 | -0.631264496 | 0.3781473 | -1.6693614 | 0.09504579 | 0.19101956 | CHAD | 1101 |
| ENSG000002 | 2.34591902 | -0.615954888 | 0.36907333 | -1.6689228 | 0.09513269 | 0.19117526 | NA | NA |
| ENSG000002 | 6.78162178 | -0.604574306 | 0.36225117 | -1.6689368 | 0.0951299 | 0.19117526 | DUX4 | 100288687 |
| ENSG000001 | 77.2618327 | -0.328082933 | 0.19660571 | -1.6687355 | 0.0951698 | 0.19124037 | GFRA1 | 2674 |
| ENSG000002 | 3.60242045 | -0.631546871 | 0.37848587 | -1.6686141 | 0.09519387 | 0.19127405 | NA | NA |
| ENSG000002 | 3.69560441 | -0.631463812 | 0.37843852 | -1.6686034 | 0.095196 | 0.19127405 | NA | NA |

| | | | | | | | | | |
|---|---|---|---|---|---|---|---|---|---|
| ENSG000001 | 17.380692 | -0.493606745 | 0.29583324 | -1.6685304 | 0.09521049 | 0.19129369 | RHAG | | 6005 |
| ENSG000002 | 16.7215623 | 0.511905006 | 0.30681613 | 1.66844228 | 0.09522796 | 0.19131931 | SNORA72 | | 26775 |
| ENSG000002 | 19.5741591 | -0.483358698 | 0.28971253 | -1.668408 | 0.09523476 | 0.1913235 | NA | NA | |
| ENSG000002 | 460.742193 | 0.287697246 | 0.17246128 | 1.66818456 | 0.0952791 | 0.19139895 | APOBEC3C | | 27350 |
| ENSG000002 | 6.47777612 | -0.608628409 | 0.36484771 | -1.6681711 | 0.09528176 | 0.19139895 | LOC388406 | | 388406 |
| ENSG000002 | 3.07781294 | -0.625245977 | 0.37482475 | -1.6681022 | 0.09529544 | 0.19141478 | NA | NA | |
| ENSG000002 | 7.85814383 | -0.592297682 | 0.35507668 | -1.6680839 | 0.09529908 | 0.19141478 | NA | NA | |
| ENSG000002 | 56.3111332 | 0.349079753 | 0.20928802 | 1.66793946 | 0.09532775 | 0.19146287 | NA | NA | |
| ENSG000002 | 7.0284161 | -0.599169832 | 0.35925496 | -1.6678123 | 0.09535299 | 0.1915041 | NA | NA | |
| ENSG000002 | 2.67308179 | -0.624898488 | 0.3746884 | -1.6677818 | 0.09535905 | 0.19150678 | NA | NA | |
| ENSG000002 | 8.12246019 | -0.601056559 | 0.36041418 | -1.6676829 | 0.0953787 | 0.19153675 | NA | NA | |
| ENSG000001 | 2.74073727 | -0.62042125 | 0.37204521 | -1.6675964 | 0.09539587 | 0.19155227 | NA | NA | |
| ENSG000001 | 29.9919706 | -0.410506454 | 0.24616439 | -1.667611 | 0.09539297 | 0.19155227 | CD247 | | 919 |
| ENSG000002 | 5.44494566 | -0.625633963 | 0.37519052 | -1.66751 | 0.09541303 | 0.19157723 | NA | NA | |
| ENSG000001 | 269.37963 | 0.253832494 | 0.15222783 | 1.6674513 | 0.0954247 | 0.19159118 | BRMS1L | | 84312 |
| ENSG000002 | 4.52848505 | -0.628984276 | 0.37722735 | -1.6673878 | 0.09543732 | 0.19160703 | NA | NA | |
| ENSG000001 | 2.37216503 | -0.615076252 | 0.36890761 | -1.6672908 | 0.0954566 | 0.19162624 | FFAR3 | | 2865 |
| ENSG000002 | 8.64393456 | -0.610908926 | 0.36641309 | -1.6672683 | 0.09546107 | 0.19162624 | NA | NA | |
| ENSG000002 | 23.0722507 | -0.460112382 | 0.27596171 | -1.6673052 | 0.09545374 | 0.19162624 | NA | NA | |
| ENSG000002 | 2.9930774 | -0.627652142 | 0.37651789 | -1.6669915 | 0.09551609 | 0.1917272 | NA | NA | |
| ENSG000001 | 162.42098 | 0.297265626 | 0.17833475 | 1.66689682 | 0.09553492 | 0.19173653 | IQSEC1 | | 9922 |
| ENSG000001 | 34.4995368 | 0.410259882 | 0.24612189 | 1.6668972 | 0.09553485 | 0.19173653 | HAAO | | 23498 |
| ENSG000002 | 5.97391814 | -0.61762358 | 0.37051719 | -1.6669229 | 0.09552974 | 0.19173653 | NA | NA | |
| ENSG000002 | 8.97798783 | -0.579865484 | 0.34789449 | -1.6667855 | 0.09555706 | 0.19177148 | NA | NA | |
| ENSG000002 | 9.32332789 | 0.577535734 | 0.34651275 | 1.66670846 | 0.09557239 | 0.19179275 | NA | NA | |
| ENSG000001 | 22.2937024 | -0.448863223 | 0.26934527 | -1.6664975 | 0.09561436 | 0.19184849 | HSD17B1 | | 3292 |
| ENSG000001 | 3.88937062 | -0.630661653 | 0.3784336 | -1.6665055 | 0.09561278 | 0.19184849 | NKX3-2 | | 579 |
| ENSG000002 | 3.19177347 | -0.621583741 | 0.37297933 | -1.6665367 | 0.09560656 | 0.19184849 | NA | NA | |
| ENSG000001 | 18.4683691 | -0.485039011 | 0.29105966 | -1.6664591 | 0.09562202 | 0.19185436 | FAM20A | | 54757 |

| | | | | | | | | |
|---|---|---|---|---|---|---|---|---|
| ENSG000001 | 8.61588095 | -0.574633851 | 0.34483036 | -1.6664248 | 0.09562884 | 0.19185856 | MCHR2 | 84539 |
| ENSG000000 | 481.624008 | 0.216558929 | 0.12996309 | 1.66631103 | 0.09565148 | 0.19188499 | ATP2B4 | 493 |
| ENSG000001 | 202.719629 | 0.244092764 | 0.14648559 | 1.66632608 | 0.09564848 | 0.19188499 | TTC26 | 79989 |
| ENSG000002 | 5.44423302 | -0.622381666 | 0.37354112 | -1.6661664 | 0.09568027 | 0.19193325 | NA | NA |
| ENSG000002 | 3.59161177 | -0.630693243 | 0.37854008 | -1.6661201 | 0.09568949 | 0.19194226 | NA | NA |
| ENSG000001 | 3.06995512 | -0.625042966 | 0.37517475 | -1.6660049 | 0.09571244 | 0.19197879 | OR56B4 | 196335 |
| ENSG000000 | 10.0459244 | 0.566320097 | 0.33993881 | 1.66594717 | 0.09572394 | 0.19198489 | CDK11A | 728642 |
| ENSG000002 | 6.77383352 | -0.602851802 | 0.36186841 | -1.6659421 | 0.09572495 | 0.19198489 | NA | NA |
| ENSG000001 | 7.95146276 | -0.600717797 | 0.36061022 | -1.6658369 | 0.09574591 | 0.19201743 | ANKRD53 | 79998 |
| ENSG000001 | 92.0512262 | 0.290186728 | 0.17420938 | 1.66573534 | 0.09576614 | 0.1920485 | B3GALT5-AS1 | 114041 |
| ENSG000001 | 22.3187534 | -0.478942282 | 0.28753058 | -1.665709 | 0.09577138 | 0.19204951 | PRLHR | 2834 |
| ENSG000001 | 230.877054 | 0.236271121 | 0.14185443 | 1.66558855 | 0.09579539 | 0.19208817 | WIPF3 | 644150 |
| ENSG000002 | 2.63004932 | -0.624246496 | 0.3748084 | -1.6655083 | 0.09581139 | 0.19211075 | NA | NA |
| ENSG000001 | 62.2506745 | 0.354481369 | 0.21286772 | 1.66526597 | 0.0958597 | 0.19219811 | MIR22HG | 84981 |
| ENSG000002 | 5.29949167 | -0.62033341 | 0.37252069 | -1.6652321 | 0.09586646 | 0.19220215 | NA | NA |
| ENSG000002 | 11.6240175 | -0.546552751 | 0.32822133 | -1.6651957 | 0.09587372 | 0.1922072 | NA | NA |
| ENSG000002 | 2.41143876 | -0.611725441 | 0.36740289 | -1.664999 | 0.09591296 | 0.19227636 | NA | NA |
| ENSG000002 | 6.45268403 | -0.609521237 | 0.36608582 | -1.6649682 | 0.09591911 | 0.19227919 | NA | NA |
| ENSG000001 | 2.72915116 | -0.620112362 | 0.37247785 | -1.6648302 | 0.09594664 | 0.19231233 | NA | NA |
| ENSG000002 | 3.87892612 | -0.630002616 | 0.37842222 | -1.664814 | 0.09594987 | 0.19231233 | NA | NA |
| ENSG000002 | 11.8811165 | -0.555280281 | 0.33353051 | -1.664856 | 0.09594148 | 0.19231233 | NA | NA |
| ENSG000001 | 2.83312868 | -0.615061853 | 0.36945457 | -1.6647835 | 0.09595597 | 0.19231505 | NA | NA |
| ENSG000001 | 11.6068203 | -0.542828006 | 0.32607789 | -1.6647188 | 0.09596887 | 0.1923314 | APOBR | 55911 |
| ENSG000001 | 13.2489206 | -0.564352359 | 0.33903843 | -1.6645675 | 0.09599907 | 0.1923729 | HSD17B3 | 3293 |
| ENSG000002 | 5.60063711 | -0.615833096 | 0.36996254 | -1.6645823 | 0.09599612 | 0.1923729 | NA | NA |
| ENSG000002 | 4.56500341 | -0.628969583 | 0.37786766 | -1.6645235 | 0.09600787 | 0.19238103 | NA | NA |
| ENSG000002 | 7.17092794 | -0.613260539 | 0.36846324 | -1.6643737 | 0.09603777 | 0.19243144 | NA | NA |
| ENSG000002 | 16.745346 | -0.506030049 | 0.30404808 | -1.6643093 | 0.09605063 | 0.19244771 | LOC101928858 | 101928858 |
| ENSG000002 | 68.1464796 | 0.309294797 | 0.18584726 | 1.66424193 | 0.09606409 | 0.19246517 | PINLYP | 390940 |

| | | | | | | | | |
|---|---|---|---|---|---|---|---|---|
| ENSG000002 | 2.83888731 | -0.614703243 | 0.36937606 | -1.6641665 | 0.09607917 | 0.19247636 | NA | NA |
| ENSG000002 | 2.38158002 | -0.615119498 | 0.36962212 | -1.6641848 | 0.09607551 | 0.19247636 | RN7SL237P | 106479308 |
| ENSG000001 | 27.18185 | 0.448330413 | 0.26942964 | 1.66399809 | 0.09611281 | 0.19252473 | C16orf74 | 404550 |
| ENSG000001 | 48.1776824 | -0.344756112 | 0.20718429 | -1.664007 | 0.09611103 | 0.19252473 | OTUD7A | 161725 |
| ENSG000001 | 118.505762 | 0.273812449 | 0.16455965 | 1.66391002 | 0.09613041 | 0.19253531 | CPNE8 | 144402 |
| ENSG000002 | 3.03619384 | -0.625937597 | 0.37618667 | -1.6639016 | 0.0961321 | 0.19253531 | NA | NA |
| ENSG000002 | 2.48173621 | -0.606551551 | 0.36454118 | -1.6638766 | 0.09613709 | 0.19253531 | NA | NA |
| ENSG000002 | 2.71919593 | -0.619975316 | 0.37260538 | -1.6638926 | 0.0961339 | 0.19253531 | NA | NA |
| ENSG000002 | 5.72855991 | -0.622980114 | 0.37442543 | -1.6638296 | 0.09614648 | 0.19254462 | NA | NA |
| ENSG000001 | 3.85843389 | -0.629613411 | 0.37842222 | -1.6637855 | 0.0961553 | 0.19255277 | NA | NA |
| ENSG000002 | 2.67100655 | -0.622634933 | 0.37423396 | -1.6637585 | 0.0961607 | 0.19255407 | NA | NA |
| ENSG000002 | 12.1159232 | 0.556918878 | 0.33474506 | 1.66371054 | 0.09617029 | 0.19256377 | MBLAC1 | 255374 |
| ENSG000001 | 270.289959 | 0.271600824 | 0.16325619 | 1.66364796 | 0.0961828 | 0.1925698 | JPH1 | 56704 |
| ENSG000001 | 221.328698 | 0.228845279 | 0.13755554 | 1.66365731 | 0.09618093 | 0.1925698 | ZNF672 | 79894 |
| ENSG000002 | 4.87118899 | -0.627574743 | 0.37724705 | -1.6635643 | 0.09619952 | 0.19259377 | LOC101928201 | 101928201 |
| ENSG000001 | 42.856351 | 0.371417343 | 0.22329621 | 1.66333918 | 0.09624456 | 0.19265891 | FAM78A | 286336 |
| ENSG000001 | 184.123165 | 0.247963166 | 0.14907465 | 1.66334899 | 0.0962426 | 0.19265891 | NEMP2 | 100131211 |
| ENSG000002 | 3.31361747 | -0.628602338 | 0.3779179 | -1.6633304 | 0.09624631 | 0.19265891 | LOC101928417 | 101928417 |
| ENSG000002 | 3.87010452 | -0.629436132 | 0.37842512 | -1.6633043 | 0.09625154 | 0.19265986 | NA | NA |
| ENSG000002 | 3.87508864 | -0.629482177 | 0.37850836 | -1.6630602 | 0.09630039 | 0.19273862 | NA | NA |
| ENSG000002 | 3.01955154 | -0.625616562 | 0.37618229 | -1.6630675 | 0.09629892 | 0.19273862 | NA | NA |
| ENSG00000C | 99.5457311 | -0.279530666 | 0.16809306 | -1.6629518 | 0.09632208 | 0.19277251 | ANK1 | 286 |
| ENSG000002 | 17.6089688 | -0.483117064 | 0.29053992 | -1.6628251 | 0.09634746 | 0.19281379 | NA | NA |
| ENSG000002 | 17.978381 | -0.489028928 | 0.29410965 | -1.6627436 | 0.09636378 | 0.19282121 | NA | NA |
| ENSG000002 | 13.6926933 | -0.521225532 | 0.31347475 | -1.6627353 | 0.09636543 | 0.19282121 | LINC01411 | 101928176 |
| ENSG000002 | 5.61571991 | -0.61634469 | 0.37067251 | -1.6627742 | 0.09635764 | 0.19282121 | NA | NA |
| ENSG000001 | 580.619896 | 0.211074448 | 0.12695165 | 1.66263652 | 0.09638522 | 0.19285128 | DCAF8 | 50717 |
| ENSG000002 | 17.8922302 | -0.482969524 | 0.29050251 | -1.6625313 | 0.09640628 | 0.19288392 | NA | NA |
| ENSG000002 | 3.61991983 | -0.629227468 | 0.37851622 | -1.6623527 | 0.09644207 | 0.19293647 | NA | NA |

| | | | | | | | | |
|---|---|---|---|---|---|---|---|---|
| ENSG000002 | 28.8642744 | -0.446438155 | 0.26855717 | -1.662358 | 0.09644101 | 0.19293647 | NA | NA |
| ENSG000002 | 6.23911661 | -0.614105301 | 0.36943876 | -1.6622655 | 0.09645956 | 0.19296194 | CRLF2 | 64109 |
| ENSG000002 | 2.38914791 | -0.614147185 | 0.36948199 | -1.6621844 | 0.09647581 | 0.19298494 | NA | NA |
| ENSG000000 | 47.0929221 | 0.378713405 | 0.22785295 | 1.66209566 | 0.0964936 | 0.19301099 | EPHA8 | 2046 |
| ENSG000001 | 12.0660498 | -0.530256086 | 0.31907336 | -1.6618626 | 0.09654033 | 0.19309494 | ASMT | 438 |
| ENSG000002 | 3.71517466 | -0.628152259 | 0.37798786 | -1.6618319 | 0.09654649 | 0.19309775 | LOC105375929 | 105375929 |
| ENSG000001 | 50.6401373 | -0.338268545 | 0.20355686 | -1.6617889 | 0.0965551 | 0.19310544 | WDR31 | 114987 |
| ENSG000001 | 202.505298 | -0.264658128 | 0.15926337 | -1.6617639 | 0.09656012 | 0.19310596 | OPTN | 10133 |
| ENSG000001 | 770.265845 | 0.203558947 | 0.12250109 | 1.66169095 | 0.09657476 | 0.1931257 | VWA8 | 23078 |
| ENSG000001 | 5.99132393 | -0.617639262 | 0.37172578 | -1.6615454 | 0.09660396 | 0.19317457 | SLC6A18 | 348932 |
| ENSG000001 | 984.570523 | 0.214174331 | 0.12890788 | 1.66145258 | 0.09662258 | 0.19320229 | AGL | 178 |
| ENSG000001 | 490.484187 | 0.189885781 | 0.11429635 | 1.66134601 | 0.09664397 | 0.19322745 | CCDC34 | 91057 |
| ENSG000002 | 10.624186 | -0.560578831 | 0.33742522 | -1.6613424 | 0.0966447 | 0.19322745 | NA | NA |
| ENSG000002 | 17.4094314 | 0.491060331 | 0.29558564 | 1.66131323 | 0.09665055 | 0.19322963 | NA | NA |
| ENSG000001 | 3.71814512 | -0.627503275 | 0.3777305 | -1.661246 | 0.09666405 | 0.19324709 | NA | NA |
| ENSG000001 | 5.88862112 | -0.613529791 | 0.36934308 | -1.6611379 | 0.09668575 | 0.19328094 | BSPH1 | 100131137 |
| ENSG000001 | 281.694213 | 0.237567444 | 0.14302262 | 1.66104799 | 0.09670381 | 0.19329798 | TMEM168 | 64418 |
| ENSG000001 | 4.77407827 | -0.623241245 | 0.3752064 | -1.6610624 | 0.09670091 | 0.19329798 | GPR15 | 2838 |
| ENSG000001 | 119.439227 | 0.272456942 | 0.16405058 | 1.6608106 | 0.09675149 | 0.19338376 | FBXO8 | 26269 |
| ENSG000002 | 3.97533209 | -0.628488347 | 0.37843466 | -1.6607579 | 0.09676208 | 0.1933954 | NA | NA |
| ENSG000002 | 11.0972509 | -0.551171221 | 0.33190738 | -1.6606176 | 0.09679028 | 0.19344221 | LOC101927228 | 101927228 |
| ENSG000002 | 2.82367763 | -0.616756888 | 0.37141319 | -1.6605681 | 0.09680022 | 0.19345256 | NA | NA |
| ENSG000001 | 9.80740421 | -0.570709973 | 0.34370366 | -1.660471 | 0.09681973 | 0.193482 | TMPRSS6 | 164656 |
| ENSG000002 | 23.2120705 | -0.446363668 | 0.26884913 | -1.6602756 | 0.09685901 | 0.19355097 | NA | NA |
| ENSG000001 | 13.9296359 | -0.519008701 | 0.31262744 | -1.6601508 | 0.09688411 | 0.19358256 | NAALADL1 | 10004 |
| ENSG000002 | 3.97535692 | -0.628255522 | 0.3784385 | -1.6601258 | 0.09688914 | 0.19358256 | NA | NA |
| ENSG000002 | 2.68449183 | -0.621319456 | 0.37425582 | -1.6601464 | 0.096885 | 0.19358256 | RPL34P6 | 100270998 |
| ENSG000001 | 3.01200794 | -0.623601715 | 0.37567685 | -1.6599418 | 0.09692615 | 0.19364697 | EDDM3B | 64184 |
| ENSG000002 | 12.3590235 | -0.531442752 | 0.32021033 | -1.6596677 | 0.09698132 | 0.19374763 | NA | NA |

| | | | | | | | | |
|---|---|---|---|---|---|---|---|---|
| ENSG000002 | 3.26631884 | -0.627494521 | 0.37809813 | -1.6596076 | 0.09699342 | 0.19375272 | RNU6-73P | 106478989 |
| ENSG000002 | 4.23818861 | -0.627881368 | 0.37832665 | -1.6596277 | 0.09698938 | 0.19375272 | LEUTX | 342900 |
| ENSG000002 | 57.0090988 | 0.351356387 | 0.21171841 | 1.65954576 | 0.09700587 | 0.19376804 | NA | NA |
| ENSG000002 | 55.9738316 | -0.34113569 | 0.20558571 | -1.6593356 | 0.09704818 | 0.19384301 | NA | NA |
| ENSG000001 | 34.5908559 | -0.412535323 | 0.24863362 | -1.6592098 | 0.09707353 | 0.1938841 | SYTL5 | 94122 |
| ENSG000002 | 3.73891573 | 0.627483347 | 0.37821565 | 1.65906237 | 0.09710322 | 0.19393385 | NA | NA |
| ENSG000002 | 6.59526195 | -0.613097747 | 0.36957519 | -1.6589256 | 0.09713078 | 0.19397933 | LOC101928009 | 101928009 |
| ENSG000002 | 15.2420298 | 0.514103069 | 0.30995483 | 1.6586387 | 0.09718862 | 0.19408528 | NA | NA |
| ENSG000002 | 7.96047125 | -0.594502983 | 0.35848628 | -1.6583703 | 0.09724276 | 0.19418383 | NA | NA |
| ENSG00000C | 52.8389175 | -0.34409006 | 0.20749184 | -1.6583306 | 0.09725076 | 0.19419024 | ELN | 2006 |
| ENSG000001 | 9.8675104 | -0.572675129 | 0.34534229 | -1.6582827 | 0.09726042 | 0.19419042 | SERPINA7 | 6906 |
| ENSG000001 | 56.5364604 | 0.328236809 | 0.19793769 | 1.6582835 | 0.09726026 | 0.19419042 | FAM133A | 286499 |
| ENSG000002 | 11.8433095 | -0.558434134 | 0.33677502 | -1.6581816 | 0.09728082 | 0.19421202 | NEU4 | 129807 |
| ENSG000002 | 5.28261699 | -0.626495161 | 0.37781971 | -1.6581855 | 0.09728003 | 0.19421202 | RN7SL220P | 106479303 |
| ENSG000002 | 13.1896944 | -0.52574169 | 0.3170804 | -1.6580706 | 0.09730321 | 0.19424716 | NA | NA |
| ENSG000002 | 6.81192567 | -0.603945278 | 0.36428395 | -1.657897 | 0.09733825 | 0.19430754 | NA | NA |
| ENSG000002 | 2.66739272 | -0.62018707 | 0.3740883 | -1.6578628 | 0.09734517 | 0.19431179 | NA | NA |
| ENSG000001 | 7.26305167 | -0.611138185 | 0.36866212 | -1.657719 | 0.0973742 | 0.19436018 | REG3A | 5068 |
| ENSG000001 | 338.083972 | 0.203812399 | 0.12295399 | 1.65763139 | 0.09739189 | 0.19438459 | SCLT1 | 132320 |
| ENSG000001 | 1515.66896 | 0.486643382 | 0.29358118 | 1.65761094 | 0.09739602 | 0.19438459 | ALDH7A1 | 501 |
| ENSG000002 | 8.44493127 | -0.599839131 | 0.36189856 | -1.6574786 | 0.09742275 | 0.19442836 | NA | NA |
| ENSG000001 | 22.9926656 | -0.440006691 | 0.26549635 | -1.6572985 | 0.09745915 | 0.19449143 | IGIP | 492311 |
| ENSG000002 | 11.8931207 | 0.533934787 | 0.32226392 | 1.65682461 | 0.09755494 | 0.19467302 | LINC01162 | 104355138 |
| ENSG000002 | 3.0670785 | -0.622011596 | 0.37545269 | -1.6566977 | 0.09758061 | 0.19470509 | NA | NA |
| ENSG000002 | 3.30453583 | -0.62623818 | 0.3780009 | -1.6567108 | 0.09757796 | 0.19470509 | NA | NA |
| ENSG000002 | 11.8678199 | -0.536555694 | 0.32388397 | -1.6566294 | 0.09759443 | 0.19472307 | NA | NA |
| ENSG000002 | 2.79265659 | -0.615779917 | 0.37172068 | -1.6565662 | 0.09760721 | 0.19473899 | NA | NA |
| ENSG000001 | 38.3724908 | 0.398841336 | 0.24077696 | 1.65647636 | 0.09762539 | 0.19475609 | NA | NA |
| ENSG000002 | 13.3670263 | -0.53553867 | 0.32329678 | -1.6564924 | 0.09762214 | 0.19475609 | OR2A42 | 402317 |

| | | | | | | | | |
|---|---|---|---|---|---|---|---|---|
| ENSG000002 | 8.8977758 | -0.574439205 | 0.34680778 | -1.6563619 | 0.09764855 | 0.19479271 | NA | NA |
| ENSG000002 | 5.58037322 | -0.613426044 | 0.37035431 | -1.6563221 | 0.0976566 | 0.1947992 | NA | NA |
| ENSG000002 | 5.99021066 | -0.613690061 | 0.3705592 | -1.6561188 | 0.09769776 | 0.19487171 | NA | NA |
| ENSG000002 | 5.83050117 | -0.60824047 | 0.36732367 | -1.6558706 | 0.09774802 | 0.19496237 | NA | NA |
| ENSG000002 | 11.0084586 | -0.560983579 | 0.33882668 | -1.6556653 | 0.09778961 | 0.19503574 | NA | NA |
| ENSG000002 | 9.02718619 | -0.577814312 | 0.34900723 | -1.6555941 | 0.09780405 | 0.19505493 | LINC00347 | 338864 |
| ENSG000001 | 15.3213588 | -0.507157971 | 0.306335 | -1.6555665 | 0.09780963 | 0.19505647 | RIIAD1 | 284485 |
| ENSG000002 | 6.03564291 | -0.615724581 | 0.37192211 | -1.6555203 | 0.09781901 | 0.19506559 | NA | NA |
| ENSG000002 | 2.42104963 | -0.608971683 | 0.36787011 | -1.6553987 | 0.09784366 | 0.19510514 | RN7SKP272 | 106479217 |
| ENSG000002 | 3.03513656 | -0.622418489 | 0.3760229 | -1.6552675 | 0.09787024 | 0.19514856 | NA | NA |
| ENSG000002 | 42.7961217 | -0.364268534 | 0.2200861 | -1.6551183 | 0.0979005 | 0.19516565 | MUC12 | 10071 |
| ENSG000002 | 4.9931574 | -0.618056457 | 0.37341787 | -1.6551336 | 0.0978974 | 0.19516565 | RN7SKP137 | 106479156 |
| ENSG000002 | 3.87693465 | -0.626575183 | 0.37855472 | -1.6551773 | 0.09788854 | 0.19516565 | NA | NA |
| ENSG000002 | 9.49983855 | -0.566162712 | 0.34206858 | -1.6551146 | 0.09790125 | 0.19516565 | NA | NA |
| ENSG000002 | 12.4050046 | -0.531945507 | 0.32139653 | -1.6551066 | 0.09790288 | 0.19516565 | NA | NA |
| ENSG00000C | 853.187476 | 0.252823405 | 0.15275967 | 1.65504031 | 0.09791632 | 0.19518285 | ERC1 | 23085 |
| ENSG000002 | 15.0299175 | -0.540862439 | 0.32680225 | -1.6550144 | 0.09792158 | 0.19518373 | LINC01182 | 101929071 |
| ENSG000002 | 2.72363925 | -0.616158943 | 0.3723159 | -1.6549359 | 0.0979375 | 0.19520589 | NA | NA |
| ENSG000001 | 215.703518 | 0.243883512 | 0.14737892 | 1.65480593 | 0.09796387 | 0.19521851 | ZC2HC1A | 51101 |
| ENSG000001 | 602.715536 | 0.202692437 | 0.12248411 | 1.65484685 | 0.09795557 | 0.19521851 | NUF2 | 83540 |
| ENSG000001 | 107.578497 | 0.291041121 | 0.17587481 | 1.65481984 | 0.09796105 | 0.19521851 | LPL | 4023 |
| ENSG000002 | 3.07505235 | -0.618717773 | 0.37389594 | -1.654786 | 0.09796791 | 0.19521851 | NA | NA |
| ENSG000002 | 15.3679694 | -0.511990468 | 0.30938744 | -1.6548521 | 0.09795449 | 0.19521851 | NA | NA |
| ENSG000002 | 4.76350903 | -0.621582964 | 0.37564514 | -1.6547078 | 0.09798377 | 0.19524053 | LOC729307 | 729307 |
| ENSG00000C | 17.6388247 | -0.505019957 | 0.3052182 | -1.6546194 | 0.09800173 | 0.1952667 | SIRT4 | 23409 |
| ENSG000002 | 3.89115769 | -0.626175458 | 0.3784693 | -1.6544947 | 0.09802703 | 0.19530753 | NA | NA |
| ENSG000002 | 4.74832153 | -0.620897352 | 0.37529122 | -1.6544415 | 0.09803785 | 0.19530988 | NA | NA |
| ENSG000002 | 3.03169934 | -0.622587923 | 0.37630857 | -1.6544612 | 0.09803384 | 0.19530988 | NA | NA |
| ENSG000002 | 3.94316078 | -0.626178578 | 0.37851641 | -1.654297 | 0.09806718 | 0.19535126 | NA | NA |

| | | | | | | | | |
|---|---|---|---|---|---|---|---|---|
| ENSG000002 | 2.3922204 | -0.61040875 | 0.36898494 | -1.6542917 | 0.09806825 | 0.19535126 | NA | NA |
| ENSG000002 | 4.72464383 | -0.622134907 | 0.37609426 | -1.6541994 | 0.098087 | 0.19537901 | NA | NA |
| ENSG000001 | 2.99247659 | -0.621957352 | 0.37600955 | -1.6540999 | 0.09810722 | 0.19540969 | DMRTA2 | 63950 |
| ENSG000002 | 5.18715099 | -0.621603738 | 0.37580289 | -1.6540686 | 0.09811358 | 0.19541277 | NA | NA |
| ENSG000001 | 8.60058827 | -0.590341367 | 0.356941 | -1.6538906 | 0.09814974 | 0.19547519 | ZNF716 | 441234 |
| ENSG000001 | 8.2834457 | -0.575942673 | 0.34825796 | -1.6537818 | 0.09817186 | 0.19550963 | RSPH6A | 81492 |
| ENSG000001 | 2.40506544 | -0.60888363 | 0.36819892 | -1.6536812 | 0.0981923 | 0.19553114 | NA | NA |
| ENSG000002 | 2.96928401 | -0.622439199 | 0.37639449 | -1.6536884 | 0.09819084 | 0.19553114 | NA | NA |
| ENSG000002 | 4.76198309 | -0.620616129 | 0.37529954 | -1.6536554 | 0.09819754 | 0.19553197 | LOC101930100 | 101930100 |
| ENSG000002 | 3.88590528 | -0.625986202 | 0.37856652 | -1.6535699 | 0.09821493 | 0.19555699 | NA | NA |
| ENSG000002 | 8.01872998 | -0.582180368 | 0.35209813 | -1.6534605 | 0.09823717 | 0.19559167 | NA | NA |
| ENSG000001 | 16.8861668 | -0.508311682 | 0.30742851 | -1.6534305 | 0.09824328 | 0.19559422 | HAVCR2 | 84868 |
| ENSG000001 | 4.72874279 | -0.620559306 | 0.37532688 | -1.6533836 | 0.09825282 | 0.19559697 | MIOX | 55586 |
| ENSG000002 | 5.32312089 | -0.61754516 | 0.37351079 | -1.6533529 | 0.09825906 | 0.19559697 | RN7SKP121 | 106481811 |
| ENSG000002 | 6.4728515 | -0.604113428 | 0.36538693 | -1.6533526 | 0.09825913 | 0.19559697 | NA | NA |
| ENSG00000C | 513.494425 | 0.447470867 | 0.27065454 | 1.65329153 | 0.09827155 | 0.19560249 | PAF1 | 54623 |
| ENSG000001 | 5.84893776 | -0.608050812 | 0.36777957 | -1.6533023 | 0.09826936 | 0.19560249 | NA | NA |
| ENSG000001 | 11.77184 | -0.530222017 | 0.32071498 | -1.6532499 | 0.09828001 | 0.19560973 | GIMAP2 | 26157 |
| ENSG000001 | 79.4175114 | 0.293123181 | 0.17730622 | 1.65320305 | 0.09828955 | 0.19561911 | USP30 | 84749 |
| ENSG000002 | 4.06912797 | -0.62539795 | 0.37830213 | -1.6531706 | 0.09829615 | 0.19562266 | NA | NA |
| ENSG000002 | 17.2583408 | -0.478769375 | 0.28961523 | -1.6531222 | 0.09830599 | 0.19563264 | C4orf51 | 646603 |
| ENSG000001 | 108.688749 | 0.281411237 | 0.17024421 | 1.65298564 | 0.09833379 | 0.19565914 | DET1 | 55070 |
| ENSG000002 | 193.081563 | 0.226797862 | 0.1372047 | 1.65298903 | 0.0983331 | 0.19565914 | TCTN1 | 79600 |
| ENSG000002 | 41.3167412 | 0.361760249 | 0.2188526 | 1.65298587 | 0.09833374 | 0.19565914 | NA | NA |
| ENSG000002 | 6.2796457 | -0.609886766 | 0.36896803 | -1.6529529 | 0.09834045 | 0.19566281 | NA | NA |
| ENSG000002 | 113.487867 | 0.270536916 | 0.16369133 | 1.65272596 | 0.09838665 | 0.19573856 | NA | NA |
| ENSG000002 | 3.16405392 | -0.617638062 | 0.37371039 | -1.6527184 | 0.09838818 | 0.19573856 | NA | NA |
| ENSG000002 | 4.44041656 | -0.623553357 | 0.37735968 | -1.6524112 | 0.09845076 | 0.19585345 | RNU6-940P | 106479982 |
| ENSG000001 | 2.40689936 | -0.609045066 | 0.36861572 | -1.6522493 | 0.09848374 | 0.19590945 | TSPO2 | 222642 |

| | | | | | | | | |
|---|---|---|---|---|---|---|---|---|
| ENSG000002 | 4.55836503 | -0.623329375 | 0.37728357 | -1.6521509 | 0.0985038 | 0.19593974 | NA | NA |
| ENSG000002 | 2.79385026 | -0.613073243 | 0.37109378 | -1.6520709 | 0.09852009 | 0.19596254 | NA | NA |
| ENSG000002 | 2.65538185 | -0.618359747 | 0.37434711 | -1.6518352 | 0.09856814 | 0.1960485 | NA | NA |
| ENSG000001 | 593.616801 | 0.203722335 | 0.12333649 | 1.65176048 | 0.09858339 | 0.1960692 | PNRC1 | 10957 |
| ENSG000002 | 3.87121443 | -0.625102034 | 0.37849231 | -1.6515581 | 0.09862467 | 0.19614169 | NA | NA |
| ENSG000002 | 7.06437001 | 0.591601359 | 0.35823977 | 1.65141174 | 0.09865453 | 0.19619145 | NPIPB11 | 728888 |
| ENSG000000 | 155.421964 | 0.249211186 | 0.15093607 | 1.65110425 | 0.09871729 | 0.19629366 | CLTCL1 | 8218 |
| ENSG000001 | 3.57168756 | -0.624862604 | 0.37845929 | -1.6510695 | 0.09872438 | 0.19629366 | IL17C | 27189 |
| ENSG000001 | 40.0203808 | 0.363033742 | 0.21987822 | 1.65106729 | 0.09872484 | 0.19629366 | OXSM | 54995 |
| ENSG000001 | 10.7256462 | -0.563120345 | 0.3410649 | -1.6510651 | 0.09872529 | 0.19629366 | OR52E4 | 390081 |
| ENSG000001 | 11.730118 | -0.561668963 | 0.34019206 | -1.6510349 | 0.09873145 | 0.19629628 | RLBP1 | 6017 |
| ENSG000002 | 13.2625746 | -0.525393309 | 0.31825664 | -1.6508479 | 0.09876964 | 0.19636258 | NA | NA |
| ENSG000002 | 4.43387872 | -0.622983664 | 0.37738025 | -1.6508115 | 0.09877706 | 0.19636772 | NA | NA |
| ENSG000002 | 7.53687921 | -0.591602177 | 0.35844111 | -1.6504864 | 0.09884349 | 0.19649014 | NA | NA |
| ENSG000002 | 3.97447919 | -0.624815507 | 0.37857187 | -1.6504541 | 0.09885009 | 0.19649362 | NA | NA |
| ENSG000000 | 9.44540922 | -0.561871463 | 0.34045631 | -1.6503482 | 0.09887174 | 0.19652702 | UTS2 | 10911 |
| ENSG000002 | 3.89612274 | -0.624612202 | 0.37849436 | -1.650255 | 0.0988908 | 0.19655528 | LOC101927038 | 101927038 |
| ENSG000002 | 2.98132712 | -0.620807089 | 0.3761955 | -1.6502246 | 0.098897 | 0.19655797 | NA | NA |
| ENSG000001 | 22.9908849 | -0.464349038 | 0.28139769 | -1.6501523 | 0.0989118 | 0.19657775 | IRF8 | 3394 |
| ENSG000000 | 41.3834469 | 0.367771236 | 0.22287485 | 1.65012447 | 0.09891748 | 0.19657941 | P2RX5 | 5026 |
| ENSG000001 | 12.3148917 | -0.524859417 | 0.3180843 | -1.6500639 | 0.09892987 | 0.19659438 | TUBB1 | 81027 |
| ENSG000002 | 8.11466582 | -0.584115492 | 0.35400357 | -1.6500271 | 0.09893739 | 0.19659971 | NA | NA |
| ENSG000002 | 20.4046558 | 0.507352136 | 0.30748818 | 1.64998904 | 0.09894518 | 0.19660554 | SNORD83A | 116937 |
| ENSG000002 | 63.2964053 | 0.338970726 | 0.20544382 | 1.64994364 | 0.09895446 | 0.19661436 | NA | NA |
| ENSG000002 | 2.36440769 | -0.606722883 | 0.36774 | -1.6498691 | 0.0989697 | 0.19663501 | NA | NA |
| ENSG000002 | 4.1432818 | -0.623069273 | 0.37766606 | -1.6497889 | 0.09898612 | 0.19665798 | NA | NA |
| ENSG000001 | 17.915522 | -0.509271318 | 0.30872784 | -1.6495801 | 0.09902884 | 0.19672358 | OLFM4 | 10562 |
| ENSG000002 | 2.73592977 | -0.612104313 | 0.3710644 | -1.6495905 | 0.09902672 | 0.19672358 | NA | NA |
| ENSG000002 | 2.39470471 | -0.606332386 | 0.36758141 | -1.6495187 | 0.09904143 | 0.19673895 | NA | NA |

| | | | | | | | | | |
|---|---|---|---|---|---|---|---|---|---|
| ENSG000002 | 3.59865814 | -0.624297221 | 0.37848781 | -1.6494513 | 0.09905521 | 0.19675668 | NA | NA | |
| ENSG000001 | 10.5372446 | -0.551501187 | 0.33439202 | -1.6492654 | 0.09909328 | 0.19682267 | MMP7 | | 4316 |
| ENSG000001 | 9.7857525 | -0.565355906 | 0.34284182 | -1.6490284 | 0.09914181 | 0.19690942 | PRDM13 | | 59336 |
| ENSG000001 | 353.355591 | 0.244441914 | 0.1482428 | 1.64892947 | 0.09916208 | 0.19693866 | AP1S1 | | 1174 |
| ENSG000002 | 3.67076005 | -0.623456746 | 0.37810254 | -1.6489092 | 0.09916625 | 0.19693866 | NA | NA | |
| ENSG000002 | 9.09006453 | 0.574568203 | 0.34847769 | 1.64879478 | 0.09918968 | 0.19697556 | NA | NA | |
| ENSG000002 | 3.1805921 | -0.614845984 | 0.37293433 | -1.6486709 | 0.09921507 | 0.19701632 | NA | NA | |
| ENSG00000C | 354.572848 | 0.200651788 | 0.12172597 | 1.64838925 | 0.09927282 | 0.19712135 | CC2D2A | | 57545 |
| ENSG000001 | 6.76925772 | -0.600705399 | 0.36443806 | -1.6483059 | 0.09928992 | 0.19713599 | CYP2F1 | | 1572 |
| ENSG000002 | 4.13998042 | -0.622498529 | 0.37765689 | -1.6483177 | 0.0992875 | 0.19713599 | C6orf165 | | 154313 |
| ENSG000002 | 2.6556298 | -0.616791138 | 0.37422104 | -1.6482001 | 0.09931162 | 0.19716943 | MTHFD2P2 | | 106480237 |
| ENSG000002 | 4.7842624 | -0.61916216 | 0.37568217 | -1.648101 | 0.09933194 | 0.19720012 | NA | NA | |
| ENSG000002 | 3.87974766 | -0.623857209 | 0.37854145 | -1.6480552 | 0.09934134 | 0.19720913 | NA | NA | |
| ENSG000001 | 140.31358 | 0.251041782 | 0.15234949 | 1.64780192 | 0.09939332 | 0.19730266 | PXK | | 54899 |
| ENSG000002 | 5.08257029 | 0.620147474 | 0.37638822 | 1.64762721 | 0.09942919 | 0.1973642 | NA | NA | |
| ENSG000002 | 4.48197569 | -0.621164764 | 0.37703433 | -1.6475019 | 0.09945494 | 0.19740299 | NA | NA | |
| ENSG000002 | 8.28735979 | -0.576788426 | 0.3501075 | -1.6474609 | 0.09946334 | 0.19740299 | NA | NA | |
| ENSG000002 | 10.521689 | -0.568325129 | 0.34496693 | -1.6474771 | 0.09946002 | 0.19740299 | NA | NA | |
| ENSG00000C | 3.59365421 | -0.623402911 | 0.37843881 | -1.6473017 | 0.09949604 | 0.19745823 | IGFALS | | 3483 |
| ENSG000001 | 60.2727627 | 0.354377396 | 0.2151291 | 1.64727781 | 0.09950096 | 0.19745832 | MAST1 | | 22983 |
| ENSG000002 | 47.6007428 | -0.342116884 | 0.20769767 | -1.6471869 | 0.09951964 | 0.19748444 | NA | NA | |
| ENSG000002 | 23.5338351 | -0.441556924 | 0.26807063 | -1.6471664 | 0.09952386 | 0.19748444 | NA | NA | |
| ENSG000002 | 7.66449008 | -0.591071556 | 0.35884898 | -1.6471318 | 0.09953096 | 0.19748887 | NA | NA | |
| ENSG000001 | 13.2416459 | -0.523975661 | 0.31812629 | -1.6470681 | 0.09954405 | 0.19750215 | C2 | | 717 |
| ENSG000002 | 48.7333889 | 0.355687296 | 0.21595392 | 1.64705183 | 0.09954739 | 0.19750215 | C2orf15 | | 150590 |
| ENSG000001 | 3.63810847 | -0.623171942 | 0.37838368 | -1.6469313 | 0.09957216 | 0.19754162 | CSAG1 | | 158511 |
| ENSG00000C | 351.190362 | 0.209382894 | 0.12714004 | 1.64686821 | 0.09958514 | 0.1975577 | FAM234B | | 57613 |
| ENSG000001 | 11.9376641 | -0.538095037 | 0.32679615 | -1.646577 | 0.09964502 | 0.19766681 | ALMS1P | | 200420 |
| ENSG000001 | 59.30661 | 0.339408511 | 0.20615202 | 1.64639917 | 0.0996816 | 0.19772972 | PRKCDBP | | 112464 |

| | | | | | | | | |
|---|---|---|---|---|---|---|---|---|
| ENSG000001 | 34.8745975 | -0.401298195 | 0.2437729 | -1.6461969 | 0.09972322 | 0.1978026 | FGD3 | 89846 |
| ENSG000002 | 6.08773257 | -0.601240568 | 0.36526257 | -1.6460503 | 0.0997534 | 0.19785279 | NA | NA |
| ENSG000002 | 7.65834454 | -0.590550537 | 0.35880164 | -1.645897 | 0.09978497 | 0.19789604 | NA | NA |
| ENSG000002 | 3.65808366 | -0.622249555 | 0.37805926 | -1.6459048 | 0.09978336 | 0.19789604 | NA | NA |
| ENSG000002 | 5.40874807 | -0.616028948 | 0.37434199 | -1.6456314 | 0.09983966 | 0.19796578 | NA | NA |
| ENSG000002 | 4.79419836 | -0.618701038 | 0.37595818 | -1.6456645 | 0.09983285 | 0.19796578 | NA | NA |
| ENSG000002 | 8.1997654 | -0.584217529 | 0.35500755 | -1.6456482 | 0.09983621 | 0.19796578 | DCTN1-AS1 | 100189589 |
| ENSG000002 | 3.86418657 | -0.62301277 | 0.3785843 | -1.6456381 | 0.09983828 | 0.19796578 | NA | NA |
| ENSG000001 | 113.217484 | 0.275020889 | 0.16713312 | 1.64551998 | 0.09986262 | 0.19800163 | C1RL | 51279 |
| ENSG000002 | 3.91777564 | -0.622933736 | 0.37858694 | -1.6454179 | 0.09988366 | 0.19803292 | NA | NA |
| ENSG000002 | 3.88085756 | -0.622765178 | 0.37848954 | -1.645396 | 0.09988817 | 0.19803292 | NA | NA |
| ENSG000001 | 2.68202016 | -0.615529243 | 0.37411974 | -1.6452734 | 0.09991345 | 0.19807335 | ODF1 | 4956 |
| ENSG000002 | 9.97675806 | -0.553996763 | 0.33672754 | -1.6452375 | 0.09992085 | 0.19807833 | NA | NA |
| ENSG000002 | 18.066235 | 0.49245216 | 0.29932927 | 1.64518544 | 0.09993157 | 0.19808991 | DPY19L2P4 | 442523 |
| ENSG000002 | 8.13938167 | -0.586819714 | 0.35672482 | -1.6450207 | 0.09996554 | 0.19814756 | FAM95B1 | 100133036 |
| ENSG000002 | 4.35113261 | -0.622628706 | 0.3785211 | -1.6448983 | 0.09999079 | 0.19818791 | NA | NA |
| ENSG00000C | 604.459319 | 0.208537901 | 0.12678631 | 1.64479823 | 0.10001143 | 0.19820944 | ATXN3 | 4287 |
| ENSG000002 | 32.0575631 | 0.39057998 | 0.23746109 | 1.64481676 | 0.1000076 | 0.19820944 | LOC150051 | 150051 |
| ENSG000001 | 3.90064482 | -0.622552844 | 0.37853083 | -1.6446556 | 0.10004085 | 0.19825807 | GPR17 | 2840 |
| ENSG00000C | 186.47356 | 0.261997906 | 0.15930739 | 1.64460607 | 0.10005107 | 0.19826101 | DKK3 | 27122 |
| ENSG000002 | 3.88883397 | -0.622630583 | 0.37859105 | -1.6445993 | 0.10005247 | 0.19826101 | NA | NA |
| ENSG000002 | 4.72344602 | -0.617282574 | 0.37534421 | -1.6445773 | 0.100057 | 0.19826101 | CTD-3080P12.3 | 101928857 |
| ENSG000001 | 4.98774908 | -0.61457298 | 0.37371176 | -1.6445107 | 0.10007076 | 0.19827858 | MZB1 | 51237 |
| ENSG000002 | 14.8237292 | -0.511664712 | 0.31114367 | -1.6444645 | 0.10008029 | 0.19828778 | MS4A4E | 643680 |
| ENSG000002 | 16.6399574 | 0.486545717 | 0.29588466 | 1.64437628 | 0.1000985 | 0.19831416 | NA | NA |
| ENSG000001 | 11.0532145 | -0.558594988 | 0.33971309 | -1.644314 | 0.10011136 | 0.19832994 | C8orf74 | 203076 |
| ENSG000002 | 4.13087395 | -0.620771094 | 0.3775541 | -1.6441911 | 0.10013673 | 0.19837052 | NA | NA |
| ENSG000001 | 80.809787 | 0.299451041 | 0.18213294 | 1.64413443 | 0.10014844 | 0.19838402 | FICD | 11153 |
| ENSG000002 | 4.67650877 | -0.62226907 | 0.3785311 | -1.6439047 | 0.10019588 | 0.19846332 | NA | NA |

| | | | | | | | | |
|---|---|---|---|---|---|---|---|---|
| ENSG000002 | 17.4130393 | -0.510164112 | 0.31033896 | -1.6438932 | 0.10019826 | 0.19846332 | NA | NA |
| ENSG000002 | 3.23796589 | -0.621558027 | 0.37812608 | -1.6437851 | 0.1002206 | 0.1984953 | NA | NA |
| ENSG000002 | 3.03708806 | -0.617164079 | 0.37545699 | -1.6437677 | 0.10022419 | 0.1984953 | NA | NA |
| ENSG000002 | 85.9363564 | 0.300423359 | 0.18277511 | 1.64367765 | 0.10024281 | 0.19852247 | LINC01234 | 100506465 |
| ENSG000001 | 927.01949 | 0.414820271 | 0.25238025 | 1.64363207 | 0.10025222 | 0.19853142 | MORC2 | 22880 |
| ENSG000002 | 7.23033195 | -0.587091359 | 0.35720094 | -1.6435885 | 0.10026123 | 0.19853956 | NA | NA |
| ENSG000002 | 20.387488 | 0.496378707 | 0.30201968 | 1.643531 | 0.10027312 | 0.1985534 | NA | NA |
| ENSG000002 | 2.61721692 | -0.615027644 | 0.37422084 | -1.6434885 | 0.10028191 | 0.19856111 | MIR1255A | 100302193 |
| ENSG000002 | 3.28928328 | -0.621184469 | 0.37797982 | -1.6434329 | 0.1002934 | 0.19857418 | NA | NA |
| ENSG000001 | 305.190335 | 0.199732177 | 0.12154724 | 1.64324728 | 0.10033178 | 0.19864047 | LMO7 | 4008 |
| ENSG000001 | 3.90149881 | -0.621956529 | 0.37851152 | -1.643164 | 0.100349 | 0.19866485 | TMEM252 | 169693 |
| ENSG000002 | 3.85592201 | -0.621927524 | 0.37850067 | -1.6431345 | 0.1003551 | 0.19866724 | NA | NA |
| ENSG000001 | 5.87550541 | -0.609042367 | 0.37070197 | -1.6429434 | 0.10039464 | 0.19873581 | TAS2R60 | 338398 |
| ENSG000001 | 10.2361617 | -0.549445541 | 0.33446275 | -1.6427705 | 0.10043042 | 0.19878723 | SLC25A41 | 284427 |
| ENSG000002 | 2.19938673 | -0.586522436 | 0.35702792 | -1.6427915 | 0.10042608 | 0.19878723 | NA | NA |
| ENSG000002 | 3.23938824 | -0.621138622 | 0.37812638 | -1.6426747 | 0.10045026 | 0.1988168 | NA | NA |
| ENSG000001 | 44.5123373 | 0.389886502 | 0.23739637 | 1.64234399 | 0.10051874 | 0.19893291 | APC2 | 10297 |
| ENSG000001 | 868.005183 | 0.211166022 | 0.12857447 | 1.6423636 | 0.10051467 | 0.19893291 | PTPRD | 5789 |
| ENSG000001 | 15.6337649 | -0.510889678 | 0.31112061 | -1.6420953 | 0.10057026 | 0.1989887 | KCNMB2 | 10242 |
| ENSG000002 | 22.7474331 | -0.445700965 | 0.27141392 | -1.6421448 | 0.10055999 | 0.1989887 | C1orf95 | 375057 |
| ENSG000002 | 40.8162721 | 0.390203353 | 0.23762121 | 1.64212343 | 0.10056443 | 0.1989887 | LOC729603 | 729603 |
| ENSG000002 | 5.57559762 | -0.608876703 | 0.37077922 | -1.6421543 | 0.10055802 | 0.1989887 | LGALS8-AS1 | 100287902 |
| ENSG000002 | 3.04962761 | -0.617083032 | 0.37579136 | -1.6420895 | 0.10057147 | 0.1989887 | NA | NA |
| ENSG000002 | 3.31282077 | -0.620222689 | 0.37771544 | -1.6420369 | 0.10058236 | 0.19900054 | NA | NA |
| ENSG000001 | 11.0521848 | -0.540983354 | 0.32947732 | -1.6419441 | 0.10060158 | 0.19902888 | MBL2 | 4153 |
| ENSG000001 | 8.79713366 | -0.578008148 | 0.35207125 | -1.6417363 | 0.10064466 | 0.19910439 | NA | NA |
| ENSG000002 | 6.78838056 | -0.598106309 | 0.36434957 | -1.6415727 | 0.10067859 | 0.19916178 | NA | NA |
| ENSG000001 | 60.6642974 | -0.328622733 | 0.20019715 | -1.6414956 | 0.10069459 | 0.19918372 | STK32A | 202374 |
| ENSG000002 | 5.61793568 | -0.611783381 | 0.37270534 | -1.6414666 | 0.10070059 | 0.19918587 | NA | NA |

| | | | | | | | | |
|---|---|---|---|---|---|---|---|---|
| ENSG000001 | 359.243842 | 0.201039622 | 0.12249254 | 1.64123973 | 0.10074766 | 0.19926926 | BDNF | 627 |
| ENSG000001 | 68.9506421 | 0.335130083 | 0.20422 | 1.64102477 | 0.10079227 | 0.19934778 | BEST1 | 7439 |
| ENSG000002 | 3.65206232 | -0.62023087 | 0.37799439 | -1.6408468 | 0.10082923 | 0.19940142 | NA | NA |
| ENSG000002 | 2.34542976 | -0.603209127 | 0.36761652 | -1.6408651 | 0.10082542 | 0.19940142 | NA | NA |
| ENSG000002 | 2.40306188 | -0.602364008 | 0.36711222 | -1.6408171 | 0.10083539 | 0.19940387 | NA | NA |
| ENSG000001 | 11.5650127 | -0.533507252 | 0.32515852 | -1.6407605 | 0.10084714 | 0.19941739 | TRIML1 | 339976 |
| ENSG000002 | 6.82284836 | -0.602663474 | 0.36731479 | -1.6407275 | 0.100854 | 0.19942122 | CTAGE7P | 119437 |
| ENSG000002 | 11.5027862 | -0.551194338 | 0.33595268 | -1.6406904 | 0.1008617 | 0.19942673 | NA | NA |
| ENSG000002 | 2.42176757 | -0.603426099 | 0.36779455 | -1.6406608 | 0.10086785 | 0.19942916 | RN7SL606P | 106479443 |
| ENSG000002 | 5.90589738 | -0.605920816 | 0.36932874 | -1.6406002 | 0.10088043 | 0.19943459 | NA | NA |
| ENSG000002 | 4.24418923 | -0.621060942 | 0.37855519 | -1.6406087 | 0.10087867 | 0.19943459 | LOC101928845 | 101928845 |
| ENSG000001 | 71.291278 | 0.330605629 | 0.20152852 | 1.64049055 | 0.10090321 | 0.1994699 | TMEM186 | 25880 |
| ENSG000002 | 10.5914734 | -0.550466056 | 0.33555613 | -1.640459 | 0.10090978 | 0.19947316 | ALG1L9P | 285407 |
| ENSG000001 | 388.452484 | 0.300354254 | 0.18309609 | 1.64041873 | 0.10091813 | 0.19947995 | MGRN1 | 23295 |
| ENSG000002 | 19.6472076 | -0.485281445 | 0.2958514 | -1.6402878 | 0.10094534 | 0.19951678 | NA | NA |
| ENSG000002 | 109.170355 | 0.279869045 | 0.17062255 | 1.64028171 | 0.10094661 | 0.19951678 | NA | NA |
| ENSG000002 | 2.65745709 | -0.613918672 | 0.37428327 | -1.6402514 | 0.1009529 | 0.1995195 | NA | NA |
| ENSG000002 | 14.3210036 | -0.499919172 | 0.30478933 | -1.6402122 | 0.10096105 | 0.19952588 | NA | NA |
| ENSG000002 | 2.62295806 | -0.613939355 | 0.37437187 | -1.6399185 | 0.10102212 | 0.19963684 | NA | NA |
| ENSG000002 | 4.717352 | -0.615709581 | 0.37547403 | -1.6398193 | 0.10104273 | 0.19966785 | NA | NA |
| ENSG000002 | 2.37973346 | -0.604139996 | 0.3685158 | -1.639387 | 0.10113269 | 0.19983588 | NA | NA |
| ENSG000002 | 3.26730344 | -0.619356574 | 0.37780424 | -1.6393585 | 0.10113863 | 0.19983786 | NA | NA |
| ENSG000002 | 7.68338368 | -0.58921218 | 0.35943206 | -1.6392867 | 0.10115357 | 0.19985765 | LOC283575 | 283575 |
| ENSG000002 | 2.35366372 | -0.603973992 | 0.36844959 | -1.639231 | 0.10116516 | 0.19987081 | NA | NA |
| ENSG000002 | 4.28861726 | -0.620561881 | 0.37858163 | -1.6391759 | 0.10117663 | 0.19988374 | NA | NA |
| ENSG000002 | 10.2233287 | -0.549845326 | 0.33544749 | -1.6391398 | 0.10118415 | 0.19988885 | NA | NA |
| ENSG000001 | 5.62850024 | -0.609486165 | 0.37187484 | -1.6389551 | 0.10122261 | 0.19993562 | CXCL17 | 284340 |
| ENSG000002 | 9.71131138 | -0.565445003 | 0.34500305 | -1.6389565 | 0.10122231 | 0.19993562 | NA | NA |
| ENSG000002 | 16.9485401 | -0.477908215 | 0.29159039 | -1.6389711 | 0.10121928 | 0.19993562 | NA | NA |

| | | | | | | | | |
|---|---|---|---|---|---|---|---|---|
| ENSG000001 | 25.1750539 | 0.432959674 | 0.26418887 | 1.63882635 | 0.10124943 | 0.19997884 | NA | NA |
| ENSG000001 | 7.59212384 | -0.589059139 | 0.35947196 | -1.6386789 | 0.10128014 | 0.20002002 | CCR1 | 1230 |
| ENSG000002 | 4.49991475 | -0.619475406 | 0.37803022 | -1.6386928 | 0.10127726 | 0.20002002 | NA | NA |
| ENSG000002 | 9.61495432 | -0.570933585 | 0.3484488 | -1.6385007 | 0.1013173 | 0.20008365 | TM4SF19-AS1 | 100874214 |
| ENSG000002 | 16.1845531 | 0.494858702 | 0.3020554 | 1.63830444 | 0.1013582 | 0.20015468 | RNU6-798P | 106479919 |
| ENSG00000C | 89.1675598 | -0.292104017 | 0.17830033 | -1.6382696 | 0.10136546 | 0.20015927 | NLRP1 | 22861 |
| ENSG000001 | 991.343745 | 0.196015698 | 0.11965563 | 1.63816533 | 0.10138721 | 0.20019247 | ERCC6L2 | 375748 |
| ENSG000002 | 5.88530786 | -0.605737895 | 0.36978204 | -1.6380944 | 0.101402 | 0.20021193 | NA | NA |
| ENSG000002 | 3.92364152 | -0.620069067 | 0.37858819 | -1.6378458 | 0.10145387 | 0.20030458 | NA | NA |
| ENSG000002 | 2.65150744 | -0.612964647 | 0.37427645 | -1.6377323 | 0.10147755 | 0.20032207 | FRG2IP | 106480351 |
| ENSG000002 | 82.0331596 | 0.323646463 | 0.19761335 | 1.6377763 | 0.10146837 | 0.20032207 | NA | NA |
| ENSG000002 | 3.56961287 | -0.619764895 | 0.37842808 | -1.6377349 | 0.101477 | 0.20032207 | NA | NA |
| ENSG000002 | 2.38155224 | -0.601565579 | 0.36734212 | -1.6376167 | 0.10150168 | 0.20035996 | NA | NA |
| ENSG000002 | 3.07634969 | -0.614702481 | 0.37537452 | -1.6375711 | 0.10151119 | 0.20036897 | NA | NA |
| ENSG000002 | 3.36960341 | -0.617114201 | 0.37689517 | -1.637363 | 0.10155465 | 0.200445 | NA | NA |
| ENSG000002 | 4.235004 | -0.619495608 | 0.37843884 | -1.6369768 | 0.10163531 | 0.20059444 | NA | NA |
| ENSG000002 | 2.66685193 | -0.61218072 | 0.37399386 | -1.6368737 | 0.10165686 | 0.20062721 | NA | NA |
| ENSG000002 | 4.52907824 | -0.618049315 | 0.37760267 | -1.6367716 | 0.10167821 | 0.20065958 | NA | NA |
| ENSG000002 | 5.37844518 | -0.612591458 | 0.37427944 | -1.6367222 | 0.10168854 | 0.20067019 | NA | NA |
| ENSG000002 | 3.01038559 | -0.614770211 | 0.3756192 | -1.6366847 | 0.10169636 | 0.20067587 | NA | NA |
| ENSG000002 | 11.300891 | -0.535418357 | 0.32715016 | -1.6366135 | 0.10171126 | 0.20068982 | NA | NA |
| ENSG000002 | 4.40957061 | -0.619592091 | 0.3785841 | -1.6366036 | 0.10171333 | 0.20068982 | NA | NA |
| ENSG000002 | 7.05099621 | -0.591549965 | 0.3614869 | -1.6364354 | 0.1017485 | 0.20072991 | LINC00376 | 104355293 |
| ENSG000002 | 4.74681486 | -0.615294941 | 0.37599652 | -1.6364378 | 0.10174798 | 0.20072991 | LOC613038 | 613038 |
| ENSG000002 | 3.91622985 | -0.619495102 | 0.37855772 | -1.6364614 | 0.10174305 | 0.20072991 | NA | NA |
| ENSG000001 | 18.9232635 | -0.473379638 | 0.28928277 | -1.6363907 | 0.10175784 | 0.20073857 | NXF3 | 56000 |
| ENSG000001 | 408.049012 | 0.250093607 | 0.1528447 | 1.63625964 | 0.10178526 | 0.20077928 | FNBP1 | 23048 |
| ENSG000002 | 7.68300449 | -0.594567958 | 0.36337349 | -1.6362447 | 0.10178837 | 0.20077928 | NA | NA |
| ENSG000002 | 52.0061524 | 0.340781363 | 0.20829542 | 1.63604828 | 0.10182948 | 0.20084083 | NAP1L6 | 645996 |

| | | | | | | | | |
|---|---|---|---|---|---|---|---|---|
| ENSG000002 | 3.61460847 | -0.61902857 | 0.37836724 | -1.6360522 | 0.10182866 | 0.20084083 | MIR4263 | 100422965 |
| ENSG000000 | 4.18505532 | -0.618298008 | 0.37796176 | -1.6358745 | 0.10186585 | 0.20090279 | REM1 | 28954 |
| ENSG000002 | 16.1234062 | -0.495088093 | 0.30264881 | -1.6358501 | 0.10187096 | 0.20090308 | NA | NA |
| ENSG000002 | 4.39328555 | -0.615267717 | 0.37612995 | -1.635785 | 0.1018846 | 0.20092021 | NA | NA |
| ENSG000001 | 130.977726 | 0.272048483 | 0.16632592 | 1.63563489 | 0.10191602 | 0.20097241 | TMEM154 | 201799 |
| ENSG000002 | 2.99123084 | -0.614613316 | 0.37577046 | -1.6356084 | 0.10192157 | 0.20097359 | NA | NA |
| ENSG000002 | 4.47421172 | -0.617113137 | 0.37731493 | -1.6355386 | 0.10193619 | 0.20099263 | NA | NA |
| ENSG000002 | 2.40439991 | -0.600831845 | 0.3673676 | -1.6355058 | 0.10194306 | 0.2009964 | NA | NA |
| ENSG000002 | 246.951112 | 0.313188538 | 0.19149754 | 1.63547033 | 0.10195049 | 0.20100128 | TRAF3IP1 | 26146 |
| ENSG000001 | 58.8567847 | 0.324148384 | 0.19821615 | 1.6353278 | 0.10198035 | 0.2010406 | PAMR1 | 25891 |
| ENSG000002 | 3.67952271 | -0.618245481 | 0.37805298 | -1.6353408 | 0.10197762 | 0.2010406 | NA | NA |
| ENSG000002 | 4.72312506 | -0.614668585 | 0.37588277 | -1.6352667 | 0.10199315 | 0.20105606 | NA | NA |
| ENSG000002 | 3.54449948 | -0.619082808 | 0.3785896 | -1.6352346 | 0.10199988 | 0.20105956 | NA | NA |
| ENSG000002 | 5.44925836 | -0.61406374 | 0.37555872 | -1.6350672 | 0.10203496 | 0.20111893 | NA | NA |
| ENSG000002 | 11.2848481 | -0.536412951 | 0.32807638 | -1.6350246 | 0.10204389 | 0.20112675 | CSTF3-AS1 | 338739 |
| ENSG000002 | 21.5504065 | -0.454403763 | 0.27793185 | -1.6349467 | 0.10206022 | 0.20114917 | SMC2-AS1 | 101928550 |
| ENSG000001 | 36.1827375 | 0.382322217 | 0.23385451 | 1.63487213 | 0.10207586 | 0.20116515 | TNNC1 | 7134 |
| ENSG000001 | 2.43809668 | -0.598246464 | 0.36593116 | -1.6348607 | 0.10207826 | 0.20116515 | IVL | 3713 |
| ENSG000001 | 12.1197233 | -0.525727433 | 0.32159833 | -1.6347331 | 0.10210501 | 0.20120339 | OR7A17 | 26333 |
| ENSG000002 | 5.88031636 | -0.605094445 | 0.37015155 | -1.6347208 | 0.10210758 | 0.20120339 | LOC102546229 | 102546229 |
| ENSG000001 | 36.0943229 | -0.373482385 | 0.22847864 | -1.634649 | 0.10212265 | 0.20121079 | RSPH4A | 345895 |
| ENSG000001 | 12.2305882 | -0.533424687 | 0.32632709 | -1.634632 | 0.10212622 | 0.20121079 | CXXC1P1 | 392459 |
| ENSG000002 | 2.35974867 | -0.598872299 | 0.366362 | -1.6346463 | 0.10212321 | 0.20121079 | LINC01425 | 101927821 |
| ENSG000002 | 10.9165099 | -0.555342432 | 0.33975096 | -1.6345574 | 0.10214187 | 0.20123185 | SALRNA2 | 104548973 |
| ENSG000002 | 3.02685904 | -0.614153439 | 0.37580167 | -1.6342488 | 0.10220661 | 0.20133004 | NA | NA |
| ENSG000002 | 8.53529165 | -0.584431771 | 0.35760732 | -1.6342836 | 0.10219932 | 0.20133004 | NA | NA |
| ENSG000002 | 4.50739996 | -0.617002395 | 0.37754444 | -1.6342511 | 0.10220614 | 0.20133004 | NA | NA |
| ENSG000002 | 9.21323263 | -0.569276242 | 0.3483575 | -1.6341725 | 0.10222263 | 0.20135183 | LINC00365 | 100874146 |
| ENSG000002 | 2.63901995 | -0.61048464 | 0.37365015 | -1.6338402 | 0.1022924 | 0.20147947 | B4GALNT2P1 | 106481717 |

| | | | | | | | | |
|---|---|---|---|---|---|---|---|---|
| ENSG000002 | 2.66497257 | -0.610232873 | 0.37352047 | -1.6337334 | 0.10231484 | 0.20151388 | NA | NA |
| ENSG000001 | 353.686866 | 0.240571649 | 0.14725973 | 1.63365535 | 0.10233124 | 0.20152659 | DTWD2 | 285605 |
| ENSG000002 | 7.55634302 | -0.583702935 | 0.35729651 | -1.6336654 | 0.10232913 | 0.20152659 | NA | NA |
| ENSG000002 | 3.25422365 | -0.617118544 | 0.37781402 | -1.6333924 | 0.1023865 | 0.20161584 | RNU6-481P | 106479779 |
| ENSG000002 | 2.42910733 | -0.600804881 | 0.3678226 | -1.6334094 | 0.10238293 | 0.20161584 | NA | NA |
| ENSG000002 | 2.36857136 | -0.601827269 | 0.36848359 | -1.6332539 | 0.10241561 | 0.20165357 | LOC105374580 | 105374580 |
| ENSG000002 | 3.91866188 | -0.618342917 | 0.37859261 | -1.6332672 | 0.1024128 | 0.20165357 | NA | NA |
| ENSG000001 | 4.50503635 | -0.616945752 | 0.37777757 | -1.6330926 | 0.10244952 | 0.2016636 | MYL10 | 93408 |
| ENSG000002 | 12.8323613 | -0.517856947 | 0.31708541 | -1.6331781 | 0.10243154 | 0.2016636 | IGFL2 | 147920 |
| ENSG000002 | 13.0033404 | -0.526601549 | 0.32245759 | -1.6330878 | 0.10245054 | 0.2016636 | OR2A12 | 346525 |
| ENSG000002 | 3.03545698 | -0.612838451 | 0.37525283 | -1.6331348 | 0.10244065 | 0.2016636 | NA | NA |
| ENSG000002 | 5.29379396 | -0.607310671 | 0.37185745 | -1.6331814 | 0.10243085 | 0.2016636 | NA | NA |
| ENSG000002 | 9.91881642 | 0.554923618 | 0.33979545 | 1.63311078 | 0.1024457 | 0.2016636 | CCND2-AS1 | 103752584 |
| ENSG000001 | 15.8794756 | -0.516838466 | 0.3165185 | -1.6328855 | 0.10249309 | 0.20173756 | CDR1 | 1038 |
| ENSG000001 | 5.11866866 | -0.613914453 | 0.37600449 | -1.6327317 | 0.10252543 | 0.20179143 | ZG16B | 124220 |
| ENSG000002 | 22.3182955 | 0.456693575 | 0.27974706 | 1.63252322 | 0.10256931 | 0.20185894 | RNU1-4 | 6060 |
| ENSG000002 | 7.73511968 | -0.592776401 | 0.36310482 | -1.6325214 | 0.10256969 | 0.20185894 | NA | NA |
| ENSG000002 | 7.51774381 | -0.580828947 | 0.35580373 | -1.632442 | 0.10258641 | 0.20188206 | NA | NA |
| ENSG000001 | 5.91186218 | -0.603061497 | 0.36943954 | -1.6323686 | 0.10260187 | 0.20190267 | NA | NA |
| ENSG000002 | 3.82556573 | -0.617852372 | 0.37853798 | -1.6322071 | 0.10263587 | 0.20195609 | LINC00395 | 100874157 |
| ENSG000002 | 4.83229051 | -0.615582154 | 0.3771505 | -1.6321923 | 0.10263897 | 0.20195609 | NA | NA |
| ENSG000001 | 51.5593251 | 0.333397385 | 0.20427285 | 1.63211796 | 0.10265464 | 0.20197711 | HSPA12B | 116835 |
| ENSG000002 | 7.06647744 | -0.590631002 | 0.36189667 | -1.6320432 | 0.10267038 | 0.20199829 | NA | NA |
| ENSG000002 | 2.60873556 | -0.610260243 | 0.37393157 | -1.6320105 | 0.10267728 | 0.20200205 | NA | NA |
| ENSG000002 | 2.70828523 | -0.608575452 | 0.37290845 | -1.6319701 | 0.10268578 | 0.20200898 | MIR1299 | 100302167 |
| ENSG000002 | 10.4134103 | -0.539578262 | 0.33063508 | -1.631945 | 0.10269108 | 0.2020096 | ETV3L | 440695 |
| ENSG000001 | 30.315121 | -0.422473193 | 0.25890407 | -1.631775 | 0.1027269 | 0.20207026 | CYBB | 1536 |
| ENSG000001 | 4.21577813 | -0.617477334 | 0.37843271 | -1.6316701 | 0.102749 | 0.20209413 | HOXC13 | 3229 |
| ENSG000002 | 2.63967229 | -0.608878082 | 0.37315768 | -1.6316911 | 0.10274457 | 0.20209413 | NA | NA |

| | | | | | | | | |
|---|---|---|---|---|---|---|---|---|
| ENSG000001 | 12.4192873 | -0.525624334 | 0.32215637 | -1.6315814 | 0.10276771 | 0.20211132 | LRRC63 | 220416 |
| ENSG000002 | 7.40410565 | -0.593622679 | 0.36383215 | -1.6315839 | 0.10276718 | 0.20211132 | SLC5A4P1 | 100422550 |
| ENSG000002 | 4.78213433 | -0.614703643 | 0.37676096 | -1.6315481 | 0.10277471 | 0.20211529 | NA | NA |
| ENSG000001 | 91.2982719 | 0.280796504 | 0.17212793 | 1.63132443 | 0.10282188 | 0.20219825 | NUDT6 | 11162 |
| ENSG000002 | 194.937722 | 0.238857585 | 0.14642225 | 1.6312929 | 0.10282853 | 0.20220152 | SLC12A8 | 84561 |
| ENSG000002 | 6.32661933 | -0.592061874 | 0.36294673 | -1.6312639 | 0.10283466 | 0.20220376 | CLPSL1 | 340204 |
| ENSG000001 | 7.23227602 | -0.58289486 | 0.35744742 | -1.630715 | 0.10295047 | 0.20242168 | HBG2 | 3048 |
| ENSG000002 | 5.04366601 | -0.61013213 | 0.37415814 | -1.6306798 | 0.1029579 | 0.20242646 | NA | NA |
| ENSG000002 | 2.36796999 | -0.59805934 | 0.36676557 | -1.6306311 | 0.10296819 | 0.20243687 | NA | NA |
| ENSG000002 | 3.09195512 | -0.611497152 | 0.3750402 | -1.6304843 | 0.10299919 | 0.202488 | NA | NA |
| ENSG000001 | 16.4725935 | -0.482084257 | 0.29569227 | -1.630358 | 0.10302585 | 0.2025219 | TMC8 | 147138 |
| ENSG000002 | 13.0149327 | 0.539606882 | 0.33097502 | 1.63035533 | 0.10302642 | 0.2025219 | NA | NA |
| ENSG000002 | 5.68334715 | -0.608851494 | 0.37345382 | -1.6303261 | 0.1030326 | 0.20252424 | NA | NA |
| ENSG00000C | 21.3571531 | 0.470671818 | 0.28872298 | 1.63018481 | 0.10306244 | 0.20257308 | SNCB | 6620 |
| ENSG000002 | 6.2286635 | 0.601851108 | 0.3692323 | 1.63000668 | 0.10310009 | 0.20262742 | NA | NA |
| ENSG000002 | 5.69534162 | -0.608855664 | 0.37352693 | -1.6300181 | 0.10309767 | 0.20262742 | LINC01213 | 101927992 |
| ENSG000002 | 10.2887101 | -0.55137269 | 0.3382818 | -1.6299212 | 0.10311815 | 0.2026531 | LOC283332 | 283332 |
| ENSG000002 | 28.0196603 | -0.430908407 | 0.2644023 | -1.6297453 | 0.10315534 | 0.20271636 | NA | NA |
| ENSG000001 | 7.82904899 | -0.580312652 | 0.35608337 | -1.6297101 | 0.10316279 | 0.20272117 | HSPA6 | 3310 |
| ENSG000002 | 3.3029967 | -0.61513485 | 0.37747695 | -1.6295958 | 0.10318696 | 0.20275884 | NA | NA |
| ENSG000001 | 193.439784 | 0.225561459 | 0.13843278 | 1.62939338 | 0.10322977 | 0.20283314 | CCDC15 | 80071 |
| ENSG000001 | 2.42397832 | -0.596863259 | 0.36633623 | -1.6292772 | 0.10325435 | 0.20287162 | NA | NA |
| ENSG000002 | 7.84392962 | -0.58173277 | 0.35707436 | -1.6291642 | 0.10327826 | 0.20290875 | ACCSL | 390110 |
| ENSG000002 | 6.25816851 | -0.601743018 | 0.36936642 | -1.6291222 | 0.10328716 | 0.20291641 | OR7E47P | 26628 |
| ENSG000001 | 136.312044 | 0.252819964 | 0.15519405 | 1.6290571 | 0.10330093 | 0.20293363 | ARMC4 | 55130 |
| ENSG000002 | 7.78316425 | -0.59210603 | 0.3634925 | -1.628936 | 0.10332656 | 0.20297416 | NA | NA |
| ENSG000002 | 10.3333909 | -0.574441301 | 0.35265858 | -1.6288879 | 0.10333675 | 0.2029755 | LINC01289 | 286184 |
| ENSG000002 | 11.2711344 | -0.537271214 | 0.32983977 | -1.6288855 | 0.10333726 | 0.2029755 | NA | NA |
| ENSG000002 | 7.93217572 | -0.58518686 | 0.35929461 | -1.6287104 | 0.10337435 | 0.20303851 | NA | NA |

| | | | | | | | | | |
|---|---|---|---|---|---|---|---|---|---|
| ENSG000002 | 5.33314029 | -0.611063044 | 0.3751952 | -1.6286537 | 0.10338635 | 0.20305225 | C18orf42 | | 642597 |
| ENSG000002 | 7.14281162 | -0.599808278 | 0.36832677 | -1.6284678 | 0.10342573 | 0.20311976 | TRIM49C | | 642612 |
| ENSG000001 | 18.9488801 | -0.513284892 | 0.3152295 | -1.6282895 | 0.10346351 | 0.20314477 | TM4SF20 | | 79853 |
| ENSG000001 | 202.384178 | 0.22996055 | 0.14122633 | 1.62831216 | 0.10345871 | 0.20314477 | ZFP3 | | 124961 |
| ENSG000002 | 9.74146586 | -0.554815648 | 0.34072627 | -1.6283324 | 0.10345441 | 0.20314477 | NA | NA | |
| ENSG000002 | 5.03225596 | -0.609119125 | 0.37408027 | -1.6283113 | 0.1034589 | 0.20314477 | NA | NA | |
| ENSG000002 | 3.27294567 | -0.614643063 | 0.3774749 | -1.6283018 | 0.1034609 | 0.20314477 | NA | NA | |
| ENSG000001 | 231.539972 | 0.260706397 | 0.16011981 | 1.62819579 | 0.10348338 | 0.2031641 | ARNT2 | | 9915 |
| ENSG000002 | 3.39823732 | -0.613228686 | 0.37663075 | -1.628196 | 0.10348333 | 0.2031641 | NA | NA | |
| ENSG000002 | 4.23421528 | -0.616313127 | 0.37854454 | -1.6281126 | 0.10350101 | 0.20318888 | NA | NA | |
| ENSG000001 | 7.18266084 | -0.582432823 | 0.35775634 | -1.6280154 | 0.10352162 | 0.2032195 | CCDC96 | | 257236 |
| ENSG000002 | 5.62639329 | -0.605314904 | 0.37182998 | -1.6279347 | 0.10353874 | 0.2032236 | CCDC148-AS1 | | 554201 |
| ENSG000002 | 2.35896049 | -0.597896959 | 0.36727214 | -1.6279398 | 0.10353764 | 0.2032236 | NA | NA | |
| ENSG000002 | 8.20473045 | -0.579017072 | 0.3556687 | -1.6279675 | 0.10353178 | 0.2032236 | NA | NA | |
| ENSG000001 | 195.245146 | 0.245480223 | 0.15079939 | 1.62785954 | 0.10355467 | 0.20324503 | CXorf23 | | 256643 |
| ENSG000001 | 20.5449879 | -0.481209675 | 0.29561784 | -1.62781 | 0.10356518 | 0.20325581 | AMPD1 | | 270 |
| ENSG000001 | 68.5128479 | 0.365335124 | 0.22444809 | 1.62770429 | 0.1035876 | 0.20328926 | ZNF784 | | 147808 |
| ENSG000002 | 3.38270904 | -0.612202832 | 0.37611935 | -1.6276824 | 0.10359225 | 0.20328926 | NA | NA | |
| ENSG000002 | 10.6941494 | -0.556915793 | 0.34220757 | -1.627421 | 0.10364771 | 0.20338826 | NA | NA | |
| ENSG000001 | 116.496669 | 0.278881012 | 0.17137698 | 1.6272956 | 0.10367433 | 0.20343065 | FOS | | 2353 |
| ENSG000002 | 226.32127 | 0.240347116 | 0.14770147 | 1.62724928 | 0.10368416 | 0.2034401 | NA | NA | |
| ENSG000002 | 3.96063259 | -0.616040997 | 0.37858921 | -1.6272017 | 0.10369427 | 0.20345008 | NA | NA | |
| ENSG000002 | 4.22558379 | -0.616000147 | 0.37858439 | -1.6271145 | 0.10371278 | 0.20347656 | NA | NA | |
| ENSG000002 | 17.7591116 | -0.481825958 | 0.29616811 | -1.6268664 | 0.10376547 | 0.20357008 | NA | NA | |
| ENSG000000 | 283.096426 | 0.220109492 | 0.13529957 | 1.62683065 | 0.10377307 | 0.20357514 | NA | NA | |
| ENSG000002 | 11.3510089 | -0.535866194 | 0.32941404 | -1.6267254 | 0.10379542 | 0.2035993 | LOC400958 | | 400958 |
| ENSG000002 | 77.9196064 | 0.333332369 | 0.20490903 | 1.62673339 | 0.10379373 | 0.2035993 | NA | NA | |
| ENSG000002 | 9.68062595 | -0.572114325 | 0.35174954 | -1.6264821 | 0.10384714 | 0.20369088 | NA | NA | |
| ENSG000000 | 6.96547296 | -0.58770397 | 0.36135992 | -1.6263674 | 0.10387153 | 0.20372887 | FNDC8 | | 54752 |

| | | | | | | | | |
|---|---|---|---|---|---|---|---|---|
| ENSG000001 | 53.7505465 | -0.345356498 | 0.21238674 | -1.6260737 | 0.10393398 | 0.20383163 | COLEC12 | 81035 |
| ENSG000002 | 2.64367842 | -0.607502342 | 0.37360057 | -1.6260744 | 0.10393382 | 0.20383163 | NA | NA |
| ENSG000002 | 4.69988075 | -0.611082585 | 0.37582545 | -1.6259745 | 0.10395509 | 0.20386318 | NA | NA |
| ENSG000001 | 198.236193 | 0.223816334 | 0.1376596 | 1.62586799 | 0.10397774 | 0.20389774 | MFSD1 | 64747 |
| ENSG000001 | 7.78308409 | -0.575127146 | 0.35374681 | -1.6258158 | 0.10398885 | 0.20390966 | C7orf33 | 202865 |
| ENSG000002 | 4.45829255 | -0.614180421 | 0.37777804 | -1.6257706 | 0.10399847 | 0.20391866 | NA | NA |
| ENSG000001 | 86.4857121 | 0.281730179 | 0.17330375 | 1.62564387 | 0.10402544 | 0.20394875 | GKAP1 | 80318 |
| ENSG000002 | 71.1741766 | 0.326375373 | 0.20076796 | 1.62563474 | 0.10402738 | 0.20394875 | ZNF580 | 51157 |
| ENSG000002 | 2.62524127 | -0.60658299 | 0.37313773 | -1.6256276 | 0.1040289 | 0.20394875 | NA | NA |
| ENSG000002 | 5.82201981 | -0.600292078 | 0.36929456 | -1.6255102 | 0.10405389 | 0.20398787 | NA | NA |
| ENSG000001 | 91.1315509 | 0.309423781 | 0.19036117 | 1.62545635 | 0.10406536 | 0.2040005 | MEX3B | 84206 |
| ENSG000002 | 6.43678976 | -0.593375838 | 0.36508195 | -1.6253223 | 0.1040939 | 0.20404409 | NA | NA |
| ENSG000002 | 20.5339894 | -0.464167778 | 0.28558817 | -1.6253047 | 0.10409766 | 0.20404409 | LOC400655 | 400655 |
| ENSG000001 | 7.81998763 | -0.579127138 | 0.35636757 | -1.6250837 | 0.10414472 | 0.20412647 | CNGA2 | 1260 |
| ENSG000002 | 6.7080655 | -0.590788369 | 0.36357035 | -1.624963 | 0.10417044 | 0.20416702 | NA | NA |
| ENSG000001 | 348.981511 | 0.475591428 | 0.29269002 | 1.62489798 | 0.1041843 | 0.20417444 | HEXIM1 | 10614 |
| ENSG000002 | 3.54355375 | -0.615168863 | 0.37858718 | -1.6249067 | 0.10418243 | 0.20417444 | NA | NA |
| ENSG000002 | 6.51235446 | -0.597183501 | 0.36752637 | -1.6248725 | 0.10418973 | 0.20417521 | LINC00102 | 100359394 |
| ENSG000002 | 8.35243854 | -0.570678912 | 0.35122584 | -1.6248204 | 0.10420083 | 0.2041871 | NA | NA |
| ENSG000002 | 4.3709386 | -0.615062187 | 0.37855968 | -1.624743 | 0.10421734 | 0.20420958 | NA | NA |
| ENSG000002 | 11.4638263 | 0.574668697 | 0.35371455 | 1.62466796 | 0.10423333 | 0.20423104 | NA | NA |
| ENSG000002 | 8.44999859 | -0.571556304 | 0.35181487 | -1.6245939 | 0.10424911 | 0.20425211 | NA | NA |
| ENSG000001 | 81.5226914 | 0.32102872 | 0.19760853 | 1.62456911 | 0.1042544 | 0.2042526 | VAMP1 | 6843 |
| ENSG000002 | 3.06812871 | -0.608490194 | 0.37457796 | -1.6244687 | 0.10427581 | 0.20427479 | LOC105375553 | 105375553 |
| ENSG000002 | 4.43701235 | -0.613147243 | 0.37744271 | -1.6244776 | 0.10427391 | 0.20427479 | NA | NA |
| ENSG000002 | 6.11366592 | -0.595009189 | 0.36631482 | -1.624311 | 0.10430945 | 0.20433084 | NA | NA |
| ENSG00000C | 31.7703825 | 0.403007241 | 0.24811716 | 1.6242619 | 0.10431993 | 0.20434148 | ZNF446 | 55663 |
| ENSG000001 | 230.058198 | -0.232041998 | 0.14286812 | -1.6241691 | 0.10433971 | 0.20437037 | ADAMTS4 | 9507 |
| ENSG000002 | 37.5412041 | 0.383340241 | 0.2360451 | 1.62401268 | 0.1043731 | 0.20441602 | NA | NA |

| | | | | | | | | | |
|---|---|---|---|---|---|---|---|---|---|
| ENSG000002 | 2.44265733 | -0.596034763 | 0.36701313 | -1.6240148 | 0.10437265 | 0.20441602 | NA | NA | |
| ENSG000002 | 2.64206074 | -0.60668761 | 0.37358038 | -1.6239814 | 0.10437977 | 0.2044192 | NA | NA | |
| ENSG000001 | 28.8657088 | -0.448622443 | 0.27625968 | -1.6239158 | 0.10439379 | 0.20443679 | BEST3 | | 144453 |
| ENSG000002 | 41.2505537 | -0.371729499 | 0.228922 | -1.623826 | 0.10441294 | 0.20446443 | NA | NA | |
| ENSG000002 | 4.16626585 | -0.613678545 | 0.37794615 | -1.6237195 | 0.10443568 | 0.20449908 | NA | NA | |
| ENSG000001 | 13.0653105 | -0.513414143 | 0.31621414 | -1.623628 | 0.10445522 | 0.2045077 | NA | NA | |
| ENSG000002 | 8.56772723 | -0.56612989 | 0.34867883 | -1.6236428 | 0.10445206 | 0.2045077 | KRT6A | | 3853 |
| ENSG000002 | 9.71604846 | -0.554489007 | 0.34150796 | -1.6236488 | 0.10445078 | 0.2045077 | NA | NA | |
| ENSG000002 | 9.26052659 | -0.568553214 | 0.35023359 | -1.6233543 | 0.10451369 | 0.2046123 | NA | NA | |
| ENSG000001 | 74.2915677 | 0.303264603 | 0.18682102 | 1.62328956 | 0.10452752 | 0.20462884 | GPRASP1 | | 9737 |
| ENSG000002 | 55.3751133 | 0.335028947 | 0.20639243 | 1.62326181 | 0.10453345 | 0.20462884 | NA | NA | |
| ENSG000002 | 3.06024817 | -0.608566 | 0.37490731 | -1.6232439 | 0.10453728 | 0.20462884 | NA | NA | |
| ENSG000001 | 150.837216 | 0.253745319 | 0.15633401 | 1.6230974 | 0.10456859 | 0.20468025 | TEX30 | | 93081 |
| ENSG000002 | 22.7383775 | -0.438138176 | 0.2699569 | -1.6229931 | 0.10459089 | 0.20471403 | SIAH3 | | 283514 |
| ENSG000002 | 5.95581204 | -0.602450937 | 0.37120303 | -1.6229688 | 0.10459608 | 0.2047143 | NA | NA | |
| ENSG000001 | 24.2089457 | -0.45312791 | 0.27923677 | -1.6227373 | 0.10464558 | 0.20477378 | TNFSF4 | | 7292 |
| ENSG000001 | 50.6304051 | -0.352526996 | 0.21724537 | -1.6227135 | 0.10465066 | 0.20477378 | N4BP2L1 | | 90634 |
| ENSG000001 | 8472.41903 | 0.186138895 | 0.1147092 | 1.62270235 | 0.10465305 | 0.20477378 | ENAH | | 55740 |
| ENSG000002 | 11.3028127 | 0.545885028 | 0.33639282 | 1.62276064 | 0.10464058 | 0.20477378 | NA | NA | |
| ENSG000002 | 2.39717226 | -0.595297984 | 0.36686361 | -1.6226684 | 0.10466031 | 0.20477378 | NA | NA | |
| ENSG000002 | 2.43553327 | -0.595517235 | 0.3669853 | -1.6227278 | 0.10464761 | 0.20477378 | NA | NA | |
| ENSG000002 | 2.66966384 | -0.605491774 | 0.37314734 | -1.6226614 | 0.10466182 | 0.20477378 | NA | NA | |
| ENSG000002 | 93.1991866 | 0.308309637 | 0.19001116 | 1.62258703 | 0.10467772 | 0.204795 | LOC283922 | | 283922 |
| ENSG00000C | 165.530976 | 0.243286928 | 0.14994393 | 1.62251936 | 0.10469219 | 0.20480356 | SNX29 | | 92017 |
| ENSG000002 | 8.06358526 | -0.573452492 | 0.35343204 | -1.6225255 | 0.10469088 | 0.20480356 | NA | NA | |
| ENSG000002 | 4.72358187 | -0.610783056 | 0.37646916 | -1.6223986 | 0.10471802 | 0.20484421 | NA | NA | |
| ENSG000002 | 2.44524001 | -0.592967935 | 0.36551235 | -1.6222925 | 0.10474074 | 0.20487877 | NA | NA | |
| ENSG000002 | 2.68702409 | -0.605932322 | 0.37356091 | -1.6220442 | 0.10479388 | 0.20497282 | OR6Q1 | | 219952 |
| ENSG000002 | 12.3450311 | 0.525685962 | 0.3240954 | 1.62200994 | 0.10480122 | 0.20497729 | NA | NA | |

| | | | | | | | | | |
|---|---|---|---|---|---|---|---|---|---|
| ENSG000001 | 118.65036 | 0.280717601 | 0.17308142 | 1.6218818 | 0.10482866 | 0.20502108 | HIAT1 | | 64645 |
| ENSG000002 | 4.11175112 | -0.613090936 | 0.37804385 | -1.6217455 | 0.10485784 | 0.20506826 | NA | NA | |
| ENSG000001 | 5.82036967 | -0.598079935 | 0.36880543 | -1.6216679 | 0.10487447 | 0.20508101 | PLA2G2D | | 26279 |
| ENSG000002 | 6.18957966 | -0.600974539 | 0.37058622 | -1.6216862 | 0.10487056 | 0.20508101 | NA | NA | |
| ENSG000002 | 12.0256588 | -0.536933831 | 0.33112039 | -1.6215668 | 0.10489614 | 0.20511348 | NA | NA | |
| ENSG000002 | 68.5251674 | 0.31234732 | 0.19264162 | 1.62139064 | 0.10493389 | 0.2051774 | NA | NA | |
| ENSG000002 | 3.25480386 | -0.612448088 | 0.37774929 | -1.6213084 | 0.10495152 | 0.20520199 | NA | NA | |
| ENSG000002 | 6.51044339 | -0.595163243 | 0.36712113 | -1.6211631 | 0.10498266 | 0.20525296 | NA | NA | |
| ENSG000002 | 5.85388907 | -0.599512687 | 0.36981255 | -1.6211259 | 0.10499065 | 0.20525869 | NA | NA | |
| ENSG000002 | 3.5016957 | -0.613712724 | 0.37859143 | -1.6210423 | 0.10500857 | 0.20528383 | NA | NA | |
| ENSG000001 | 2.41655513 | -0.594778948 | 0.3669262 | -1.6209771 | 0.10502256 | 0.20529376 | KCNG2 | | 26251 |
| ENSG000002 | 16.2246844 | -0.495405369 | 0.30562252 | -1.6209714 | 0.10502377 | 0.20529376 | NA | NA | |
| ENSG000001 | 46.4825445 | 0.369333517 | 0.22785418 | 1.62092049 | 0.10503469 | 0.2053052 | ZNF764 | | 92595 |
| ENSG000001 | 173.268047 | 0.266416803 | 0.16437162 | 1.62081992 | 0.10505627 | 0.20533748 | PPP2R3B | | 28227 |
| ENSG000001 | 5.14439974 | -0.611800515 | 0.37749495 | -1.6206853 | 0.10508514 | 0.2053657 | PLGLB1 | | 5343 |
| ENSG000001 | 2.61249375 | -0.605395443 | 0.37354738 | -1.6206657 | 0.10508935 | 0.2053657 | NA | NA | |
| ENSG000002 | 3.54923165 | -0.612932419 | 0.37820519 | -1.6206346 | 0.10509603 | 0.2053657 | NA | NA | |
| ENSG000002 | 6.14843272 | -0.594905175 | 0.3670609 | -1.6207261 | 0.10507639 | 0.2053657 | NA | NA | |
| ENSG000002 | 2.37238106 | -0.596006196 | 0.36775766 | -1.6206493 | 0.10509287 | 0.2053657 | NA | NA | |
| ENSG000001 | 127.218288 | 0.268486596 | 0.16567049 | 1.62060605 | 0.10510215 | 0.20536777 | CDK5R1 | | 8851 |
| ENSG000001 | 1053.30416 | 0.193591384 | 0.11946184 | 1.62052908 | 0.10511867 | 0.20539016 | BROX | | 148362 |
| ENSG000001 | 67.0066901 | 0.324020904 | 0.19996248 | 1.62040854 | 0.10514455 | 0.20542091 | DYRK1B | | 9149 |
| ENSG000002 | 3.08503957 | -0.606695595 | 0.37440707 | -1.620417 | 0.10514273 | 0.20542091 | NA | NA | |
| ENSG000002 | 3.02612128 | -0.607326663 | 0.37482103 | -1.6203111 | 0.10516547 | 0.20545189 | NA | NA | |
| ENSG000001 | 113.438309 | 0.274537637 | 0.16944746 | 1.6201933 | 0.10519076 | 0.2054914 | TRIM38 | | 10475 |
| ENSG000002 | 11.9001991 | -0.531059996 | 0.32780883 | -1.6200296 | 0.10522593 | 0.2055502 | NA | NA | |
| ENSG000001 | 3.66648179 | -0.612674987 | 0.37819776 | -1.6199858 | 0.10523533 | 0.20555867 | CYP11B2 | | 1585 |
| ENSG000001 | 51.8123919 | 0.334315904 | 0.20638002 | 1.61990439 | 0.10525282 | 0.20557122 | CRYM | | 1428 |
| ENSG000001 | 4.14638599 | -0.612857548 | 0.37833948 | -1.6198615 | 0.10526204 | 0.20557122 | NT5C1A | | 84618 |

| | | | | | | | | | |
|---|---|---|---|---|---|---|---|---|---|
| ENSG000002 | 58.7808055 | 0.337253297 | 0.20819815 | 1.61986691 | 0.10526087 | 0.20557122 | PCDHGC3 | | 5098 |
| ENSG000002 | 6.15173355 | -0.594989837 | 0.36730632 | -1.6198737 | 0.10525942 | 0.20557122 | NA | NA | |
| ENSG000002 | 4.48184322 | -0.612246217 | 0.37797968 | -1.6197861 | 0.10527824 | 0.20559297 | NA | NA | |
| ENSG000001 | 71.9579941 | -0.316833071 | 0.19561785 | -1.6196532 | 0.10530679 | 0.2056362 | DRAM1 | | 55332 |
| ENSG000002 | 4.44238766 | -0.61186484 | 0.37777926 | -1.6196359 | 0.10531052 | 0.2056362 | NA | NA | |
| ENSG000001 | 150.931025 | 0.254272847 | 0.15700649 | 1.61950535 | 0.10533858 | 0.20568108 | DCLRE1C | | 64421 |
| ENSG000002 | 3.53689914 | -0.612997025 | 0.37856467 | -1.6192663 | 0.10538997 | 0.20577152 | NA | NA | |
| ENSG000001 | 1152.65971 | 0.293493668 | 0.18127793 | 1.61902596 | 0.10544168 | 0.20586257 | KAT6B | | 23522 |
| ENSG000002 | 88.6255701 | 0.280474181 | 0.17324552 | 1.61894042 | 0.10546008 | 0.20588859 | C15orf59 | | 388135 |
| ENSG000002 | 4.45314537 | -0.611705459 | 0.37792823 | -1.6185757 | 0.10553858 | 0.20603192 | NA | NA | |
| ENSG000002 | 6.41768309 | -0.594805859 | 0.36749783 | -1.6185289 | 0.10554866 | 0.20604168 | NA | NA | |
| ENSG000002 | 4.10798084 | -0.611707003 | 0.37795745 | -1.6184547 | 0.10556465 | 0.2060588 | NA | NA | |
| ENSG000002 | 3.27572587 | -0.611114036 | 0.37759427 | -1.618441 | 0.1055676 | 0.2060588 | NA | NA | |
| ENSG000002 | 10.7363173 | -0.539330699 | 0.33324676 | -1.6184124 | 0.10557374 | 0.20606088 | NA | NA | |
| ENSG000002 | 2.39287274 | -0.593172892 | 0.36652685 | -1.6183614 | 0.10558474 | 0.20607243 | NA | NA | |
| ENSG000002 | 4.26140069 | -0.612644172 | 0.37859313 | -1.6182126 | 0.10561679 | 0.20612506 | SP5 | | 389058 |
| ENSG000002 | 4.94855812 | 0.606695729 | 0.37494335 | 1.61809971 | 0.10564111 | 0.2061626 | PDXP | | 57026 |
| ENSG000001 | 67.7623529 | -0.313658283 | 0.19385822 | -1.6179777 | 0.1056674 | 0.20620398 | LTBP3 | | 4054 |
| ENSG000002 | 4.87232916 | -0.611973575 | 0.37824491 | -1.6179294 | 0.10567781 | 0.20621437 | NA | NA | |
| ENSG000002 | 4.70199945 | -0.60969093 | 0.37685047 | -1.617859 | 0.10569299 | 0.20623407 | NA | NA | |
| ENSG000002 | 3.2743958 | -0.61126287 | 0.37783318 | -1.6178115 | 0.10570322 | 0.20624411 | NA | NA | |
| ENSG000001 | 93.7264682 | 0.270069578 | 0.16695226 | 1.61764556 | 0.105739 | 0.20629495 | ENTPD1 | | 953 |
| ENSG000001 | 12.7548934 | -0.51640769 | 0.31923683 | -1.6176319 | 0.10574194 | 0.20629495 | CARNS1 | | 57571 |
| ENSG000002 | 4.99191883 | -0.605070787 | 0.37405004 | -1.6176199 | 0.10574454 | 0.20629495 | NA | NA | |
| ENSG000002 | 4.74472753 | -0.608371316 | 0.37610108 | -1.617574 | 0.10575445 | 0.20630435 | NA | NA | |
| ENSG000002 | 3.84684779 | -0.612347873 | 0.37857656 | -1.6175008 | 0.10577023 | 0.20632522 | NA | NA | |
| ENSG000002 | 3.93612901 | -0.612226958 | 0.37859335 | -1.6171097 | 0.10585461 | 0.20647989 | NA | NA | |
| ENSG000001 | 3.88482425 | -0.612132671 | 0.37857552 | -1.6169367 | 0.10589194 | 0.20653282 | KLK8 | | 11202 |
| ENSG000002 | 6.07362218 | -0.592887582 | 0.36666997 | -1.6169516 | 0.10588873 | 0.20653282 | NA | NA | |

| | | | | | | | | |
|---|---|---|---|---|---|---|---|---|
| ENSG000001 | 18.5600378 | 0.461968807 | 0.2857122 | 1.61690261 | 0.1058993 | 0.20653726 | NDUFA13 | 51079 |
| ENSG00000C | 1460.73623 | 0.422834711 | 0.26152926 | 1.61677786 | 0.10592624 | 0.20657986 | RAB7A | 7879 |
| ENSG000002 | 3.24270868 | -0.610693265 | 0.37775017 | -1.6166591 | 0.10595188 | 0.20660999 | NA | NA |
| ENSG000002 | 3.51405147 | -0.612001611 | 0.3785552 | -1.6166773 | 0.10594795 | 0.20660999 | NRG1-IT3 | 100874286 |
| ENSG000002 | 3.02643483 | -0.606840498 | 0.37538538 | -1.61658 | 0.10596897 | 0.20663337 | MIR544A | 664613 |
| ENSG000001 | 2.71993204 | -0.597457452 | 0.36961682 | -1.6164239 | 0.1060027 | 0.20668921 | CCNO | 10309 |
| ENSG000002 | 2.63947696 | -0.603430007 | 0.37332435 | -1.6163693 | 0.1060145 | 0.20670228 | NA | NA |
| ENSG000001 | 8.57174599 | -0.564086662 | 0.34901262 | -1.6162357 | 0.10604335 | 0.2067486 | DEGS2 | 123099 |
| ENSG000002 | 23.0792535 | -0.426173309 | 0.26370527 | -1.6160971 | 0.10607333 | 0.20679709 | NA | NA |
| ENSG000001 | 479.54188 | 0.249440607 | 0.15435809 | 1.6159866 | 0.10609721 | 0.20682098 | VPS53 | 55275 |
| ENSG000001 | 228.11583 | 0.272701598 | 0.16875909 | 1.61592244 | 0.10611108 | 0.20682098 | NA | NA |
| ENSG000002 | 15.9658008 | -0.497785576 | 0.30804477 | -1.615952 | 0.10610468 | 0.20682098 | LINC00933 | 100506874 |
| ENSG000002 | 6.19483106 | -0.594282921 | 0.36776566 | -1.6159283 | 0.10610982 | 0.20682098 | NA | NA |
| ENSG000002 | 9.26548706 | -0.562456771 | 0.34805951 | -1.6159788 | 0.10609891 | 0.20682098 | NA | NA |
| ENSG000001 | 41.8951785 | 0.362676852 | 0.22445576 | 1.61580554 | 0.10613636 | 0.20685037 | C6orf1 | 221491 |
| ENSG000002 | 2.3789629 | -0.590363984 | 0.36536619 | -1.6158145 | 0.10613443 | 0.20685037 | NA | NA |
| ENSG000001 | 115.301766 | -0.270758724 | 0.16757515 | -1.6157451 | 0.10614944 | 0.20685598 | LEFTY2 | 7044 |
| ENSG000002 | 12.5068176 | -0.528642709 | 0.32717881 | -1.6157608 | 0.10614603 | 0.20685598 | NA | NA |
| ENSG000002 | 4.44902096 | -0.611308527 | 0.37835266 | -1.6157109 | 0.10615682 | 0.20686042 | NA | NA |
| ENSG000002 | 3.82863932 | -0.611559186 | 0.37854066 | -1.6155707 | 0.10618717 | 0.20690961 | NA | NA |
| ENSG000002 | 3.26224799 | -0.609972115 | 0.37759588 | -1.6154099 | 0.10622196 | 0.20696747 | NA | NA |
| ENSG000001 | 288.49818 | 0.224093038 | 0.13873638 | 1.61524351 | 0.10625797 | 0.20702767 | CSNK1G1 | 53944 |
| ENSG000002 | 15.2437534 | -0.516694338 | 0.31991032 | -1.6151225 | 0.10628418 | 0.20706878 | NA | NA |
| ENSG000001 | 311.803657 | 0.22772976 | 0.14101709 | 1.61490898 | 0.10633041 | 0.20713876 | ZNF710 | 374655 |
| ENSG000002 | 15.1230748 | -0.505247258 | 0.31286275 | -1.6149166 | 0.10632875 | 0.20713876 | NA | NA |
| ENSG000002 | 4.82848744 | -0.609176914 | 0.37722599 | -1.6148858 | 0.10633542 | 0.20713876 | NA | NA |
| ENSG000002 | 5.62678615 | -0.600329892 | 0.37175356 | -1.6148598 | 0.10634105 | 0.20713978 | MTRNR2L4 | 100463285 |
| ENSG000002 | 3.32584905 | -0.608450377 | 0.37680134 | -1.6147776 | 0.10635886 | 0.20716451 | NA | NA |
| ENSG000002 | 18.7810519 | -0.471703977 | 0.29215856 | -1.6145478 | 0.10640865 | 0.20725155 | NA | NA |

| | | | | | | | | |
|---|---|---|---|---|---|---|---|---|
| ENSG000002 | 6.25832452 | -0.598548996 | 0.37075072 | -1.6144244 | 0.10643541 | 0.20729371 | NA | NA |
| ENSG000002 | 5.02124942 | -0.60605728 | 0.37542838 | -1.6143087 | 0.10646049 | 0.2073326 | NA | NA |
| ENSG000001 | 13.9497283 | -0.50498427 | 0.31283287 | -1.6142302 | 0.1064775 | 0.20733656 | TMEM78 | 677790 |
| ENSG000002 | 2.40173236 | -0.592233661 | 0.3668834 | -1.6142285 | 0.10647786 | 0.20733656 | NA | NA |
| ENSG000002 | 18.5446708 | -0.464594617 | 0.28780611 | -1.6142625 | 0.10647049 | 0.20733656 | DYTN | 391475 |
| ENSG000001 | 45.8892271 | 0.377346834 | 0.23378444 | 1.61408019 | 0.10651003 | 0.20737341 | CCDC157 | 550631 |
| ENSG000001 | 91.4947388 | -0.272362624 | 0.16874518 | -1.6140468 | 0.10651728 | 0.20737341 | SEMA4A | 64218 |
| ENSG000002 | 6.85830542 | -0.596373207 | 0.36948452 | -1.6140682 | 0.10651264 | 0.20737341 | NA | NA |
| ENSG000002 | 4.41827317 | -0.607972757 | 0.37668151 | -1.6140234 | 0.10652236 | 0.20737341 | NA | NA |
| ENSG000002 | 26.9825691 | -0.418534582 | 0.25930719 | -1.6140493 | 0.10651674 | 0.20737341 | NA | NA |
| ENSG000001 | 55.2659676 | 0.338071701 | 0.20946767 | 1.61395645 | 0.10653687 | 0.20739171 | CUEDC1 | 404093 |
| ENSG000002 | 8.07349344 | -0.57234596 | 0.35464437 | -1.6138588 | 0.10655805 | 0.20740306 | NA | NA |
| ENSG000002 | 3.91964257 | -0.611007954 | 0.37859338 | -1.6138897 | 0.10655136 | 0.20740306 | NA | NA |
| ENSG000002 | 2.66918722 | -0.60089824 | 0.37233303 | -1.613873 | 0.10655497 | 0.20740306 | NA | NA |
| ENSG000002 | 4.99393404 | -0.604510849 | 0.37459998 | -1.6137504 | 0.10658158 | 0.20743891 | NA | NA |
| ENSG000002 | 6.17044392 | -0.593872981 | 0.36803525 | -1.6136307 | 0.10660755 | 0.2074795 | NA | NA |
| ENSG000001 | 10.6792355 | -0.540605941 | 0.33503447 | -1.613583 | 0.1066179 | 0.20748968 | PCDHB8 | 56128 |
| ENSG000002 | 4.52489151 | -0.61063353 | 0.37844525 | -1.613532 | 0.10662897 | 0.20750127 | NA | NA |
| ENSG000001 | 5.14242218 | -0.607767122 | 0.3767531 | -1.6131709 | 0.10670739 | 0.20763431 | DIRAS3 | 9077 |
| ENSG000001 | 14.9906247 | -0.507690088 | 0.3147158 | -1.61317 | 0.10670758 | 0.20763431 | TMEM217 | 221468 |
| ENSG000001 | 286.390993 | 0.207592063 | 0.12869091 | 1.61310594 | 0.10672149 | 0.20765142 | GAREM | 64762 |
| ENSG000001 | 4.91113619 | -0.609595262 | 0.37790723 | -1.6130818 | 0.10672674 | 0.20765167 | NA | NA |
| ENSG000001 | 23.5486048 | -0.423048071 | 0.26227518 | -1.6129931 | 0.106746 | 0.20767917 | PCSK2 | 5126 |
| ENSG000000 | 6.18463444 | -0.598778046 | 0.37123461 | -1.612937 | 0.10675819 | 0.20769293 | PRSS3 | 5646 |
| ENSG000001 | 13.0268993 | -0.525679213 | 0.32592854 | -1.6128664 | 0.10677353 | 0.2077128 | TSPEAR | 54084 |
| ENSG000002 | 7.28515216 | -0.580158759 | 0.35972212 | -1.612797 | 0.10678862 | 0.20773219 | NA | NA |
| ENSG000001 | 70.3812819 | 0.295947426 | 0.18352736 | 1.61255207 | 0.10684186 | 0.20781582 | DUSP10 | 11221 |
| ENSG000002 | 3.0597393 | -0.602890888 | 0.37387078 | -1.6125649 | 0.10683908 | 0.20781582 | NA | NA |
| ENSG000002 | 3.55192356 | -0.610349422 | 0.37852266 | -1.6124515 | 0.10686373 | 0.20783842 | NA | NA |

| | | | | | | | | |
|---|---|---|---|---|---|---|---|---|
| ENSG000002 | 5.02216324 | -0.605026457 | 0.37521853 | -1.6124642 | 0.10686096 | 0.20783842 | KCNAB1-AS2 | 100874083 |
| ENSG000001 | 533.802723 | 0.226512046 | 0.14048195 | 1.61239258 | 0.10687654 | 0.20785211 | UXT | 8409 |
| ENSG000002 | 5.91811311 | -0.600245983 | 0.37227512 | -1.612372 | 0.10688102 | 0.20785211 | NA | NA |
| ENSG000002 | 2.66158742 | -0.601720783 | 0.37320535 | -1.6123048 | 0.10689563 | 0.20787055 | NA | NA |
| ENSG000002 | 86.3171507 | 0.33871706 | 0.21009042 | 1.61224422 | 0.10690881 | 0.20788621 | NA | NA |
| ENSG000001 | 189.867436 | 0.226341304 | 0.14040137 | 1.61210182 | 0.10693978 | 0.20791654 | MMAA | 166785 |
| ENSG000002 | 5.85528027 | -0.595252403 | 0.36923891 | -1.6121064 | 0.10693879 | 0.20791654 | LOC101928855 | 101928855 |
| ENSG000002 | 2.6853734 | -0.601700605 | 0.37323845 | -1.6121078 | 0.10693849 | 0.20791654 | NA | NA |
| ENSG000002 | 4.08388688 | -0.609567994 | 0.37813251 | -1.6120486 | 0.10695136 | 0.20792908 | NA | NA |
| ENSG000002 | 6.38535613 | -0.586914847 | 0.3640986 | -1.6119668 | 0.10696917 | 0.20795373 | LTA | 4049 |
| ENSG000002 | 10.5562876 | -0.547769918 | 0.33982183 | -1.6119327 | 0.10697659 | 0.20795818 | NA | NA |
| ENSG000001 | 13.9552289 | -0.503522213 | 0.31240293 | -1.6117718 | 0.10701162 | 0.20800633 | RBM24 | 221662 |
| ENSG000002 | 5.01653898 | -0.60393016 | 0.37469542 | -1.6117895 | 0.10700775 | 0.20800633 | NA | NA |
| ENSG000002 | 7.08747976 | -0.58803582 | 0.36484394 | -1.6117462 | 0.10701719 | 0.20800719 | NA | NA |
| ENSG000002 | 3.52065402 | -0.610118575 | 0.3785552 | -1.611703 | 0.10702657 | 0.20801546 | NA | NA |
| ENSG000001 | 13.9909536 | -0.505310429 | 0.31354959 | -1.6115806 | 0.10705324 | 0.20804735 | CPA3 | 1359 |
| ENSG000001 | 25.1641299 | -0.437478045 | 0.27145573 | -1.6116 | 0.10704901 | 0.20804735 | RBM44 | 375316 |
| ENSG000002 | 2.67780442 | -0.598840424 | 0.37162119 | -1.611427 | 0.10708669 | 0.20810238 | NA | NA |
| ENSG000002 | 2.63867504 | -0.601832992 | 0.37349533 | -1.6113535 | 0.10710271 | 0.20811356 | NA | NA |
| ENSG000002 | 6.37555655 | -0.586564931 | 0.36401917 | -1.6113573 | 0.10710188 | 0.20811356 | NA | NA |
| ENSG000002 | 12.2693689 | -0.514944659 | 0.3195902 | -1.6112655 | 0.10712187 | 0.20814083 | NA | NA |
| ENSG00000C | 95.5704648 | 0.274989011 | 0.17067109 | 1.61122199 | 0.10713135 | 0.20814679 | CTNS | 1497 |
| ENSG000001 | 2522.61151 | 0.186150117 | 0.11553646 | 1.6111807 | 0.10714034 | 0.20814679 | SPG20 | 23111 |
| ENSG000001 | 3.61993301 | -0.609354964 | 0.37820057 | -1.6111952 | 0.10713718 | 0.20814679 | RNASE3 | 6037 |
| ENSG00000C | 134.992067 | 0.255816949 | 0.15878383 | 1.61110197 | 0.1071575 | 0.20815031 | MGAT4A | 11320 |
| ENSG000002 | 5.61572156 | -0.598848533 | 0.37169047 | -1.6111485 | 0.10714737 | 0.20815031 | NA | NA |
| ENSG000002 | 9.24989448 | -0.561329635 | 0.34841353 | -1.6111017 | 0.10715755 | 0.20815031 | NA | NA |
| ENSG000001 | 621.158066 | 0.420851976 | 0.26123514 | 1.61100831 | 0.10717791 | 0.20817989 | PLEC | 5339 |
| ENSG000002 | 8.6460303 | -0.567540608 | 0.35233607 | -1.6107934 | 0.10722476 | 0.20826091 | NA | NA |

| | | | | | | | | |
|---|---|---|---|---|---|---|---|---|
| ENSG000002 | 2.38475665 | -0.590870141 | 0.36683328 | -1.6107321 | 0.10723813 | 0.2082769 | NA | NA |
| ENSG000002 | 7.26535161 | -0.582252657 | 0.36149729 | -1.6106695 | 0.10725177 | 0.20829343 | NA | NA |
| ENSG000002 | 6.01033111 | -0.603936287 | 0.37497545 | -1.6106022 | 0.10726645 | 0.20831195 | NA | NA |
| ENSG000002 | 3.25256032 | -0.608148858 | 0.37762073 | -1.6104753 | 0.10729413 | 0.20835573 | NA | NA |
| ENSG000002 | 5.59070093 | -0.599366957 | 0.37218035 | -1.6104207 | 0.10730604 | 0.20836887 | NA | NA |
| ENSG000002 | 24.5283428 | 0.422401368 | 0.26229873 | 1.61038281 | 0.10731431 | 0.20837496 | NA | NA |
| ENSG000002 | 12.6161825 | -0.524450181 | 0.3257472 | -1.6099914 | 0.10739974 | 0.20853085 | LOC101927239 | 101927239 |
| ENSG000002 | 2.38909584 | -0.590526681 | 0.3668517 | -1.609715 | 0.10746009 | 0.20863804 | NA | NA |
| ENSG000002 | 6.39740392 | -0.587963697 | 0.36526491 | -1.6096911 | 0.1074653 | 0.20863817 | NA | NA |
| ENSG000002 | 5.27415104 | -0.598773514 | 0.37199231 | -1.6096395 | 0.10747658 | 0.20865008 | NA | NA |
| ENSG000002 | 31.4729836 | -0.389631813 | 0.24207039 | -1.6095807 | 0.10748943 | 0.20866504 | NA | NA |
| ENSG000001 | 6.63870718 | -0.586829091 | 0.36459671 | -1.6095293 | 0.10750066 | 0.20867683 | SELPLG | 6404 |
| ENSG000002 | 6.42198681 | -0.588188603 | 0.36544846 | -1.6094981 | 0.10750747 | 0.20868007 | RPL23AP87 | 388574 |
| ENSG000002 | 3.52392239 | -0.609106036 | 0.37846356 | -1.6094179 | 0.10752499 | 0.2087041 | NA | NA |
| ENSG000002 | 5.59442632 | -0.59955009 | 0.37253552 | -1.609377 | 0.10753394 | 0.20871146 | NA | NA |
| ENSG000002 | 4.98925019 | -0.6034711 | 0.37498046 | -1.6093401 | 0.10754199 | 0.2087171 | NA | NA |
| ENSG000002 | 3.25839808 | -0.607476159 | 0.37748665 | -1.6092653 | 0.10755835 | 0.20873886 | NA | NA |
| ENSG000001 | 4.00425415 | -0.608272405 | 0.37807277 | -1.6088765 | 0.10764334 | 0.20888206 | COL18A1-AS1 | 378832 |
| ENSG000002 | 3.64497161 | -0.60771518 | 0.37773098 | -1.6088571 | 0.10764758 | 0.20888206 | NA | NA |
| ENSG000002 | 4.28324389 | -0.609087451 | 0.37857611 | -1.6088904 | 0.10764032 | 0.20888206 | MIR4312 | 100422971 |
| ENSG00000C | 85.6346413 | 0.292386719 | 0.18173996 | 1.60881909 | 0.10765591 | 0.20888821 | MRVI1 | 10335 |
| ENSG000002 | 4.38952792 | -0.606088024 | 0.37675255 | -1.6087165 | 0.10767835 | 0.20892177 | NA | NA |
| ENSG000002 | 21.8490542 | -0.438645767 | 0.27268674 | -1.6086069 | 0.10770233 | 0.20895828 | NA | NA |
| ENSG000002 | 2.75572923 | -0.594108286 | 0.36933968 | -1.6085688 | 0.10771066 | 0.20896445 | RNU6-246P | 106481884 |
| ENSG000002 | 4.10488853 | -0.607958432 | 0.3779699 | -1.6084837 | 0.10772928 | 0.20899057 | NA | NA |
| ENSG000002 | 5.62725559 | -0.598705151 | 0.37222553 | -1.6084473 | 0.10773725 | 0.20899603 | NA | NA |
| ENSG000001 | 376.730712 | 0.188947754 | 0.11748697 | 1.60824436 | 0.10778167 | 0.20907221 | MOCS2 | 4338 |
| ENSG000001 | 17.5863316 | -0.483441143 | 0.30061907 | -1.6081519 | 0.10780191 | 0.20909228 | TLR5 | 7100 |
| ENSG000001 | 9.55945639 | -0.565656683 | 0.35174373 | -1.60815 | 0.10780233 | 0.20909228 | LRRIQ4 | 344657 |

| | | | | | | | | |
|---|---|---|---|---|---|---|---|---|
| ENSG000001 | 73.8270229 | 0.317988584 | 0.19774134 | 1.60810368 | 0.10781247 | 0.20910195 | FAM174A | 345757 |
| ENSG000002 | 12.4034972 | -0.525940709 | 0.32706701 | -1.6080518 | 0.10782383 | 0.20911397 | PPAPDC1A | 196051 |
| ENSG000002 | 3.68320125 | -0.606953115 | 0.37747386 | -1.6079342 | 0.10784959 | 0.20915394 | RNU6-183P | 106481234 |
| ENSG000001 | 12.585457 | 0.518115185 | 0.32225136 | 1.6077983 | 0.10787936 | 0.20919166 | NRTN | 4902 |
| ENSG000002 | 5.00580055 | -0.603237535 | 0.375192 | -1.6078102 | 0.10787675 | 0.20919166 | SPATA13 | 221178 |
| ENSG000001 | 8.32106985 | -0.56765062 | 0.35307572 | -1.6077306 | 0.1078942 | 0.20921043 | EPX | 8288 |
| ENSG000002 | 7.88385647 | -0.577652105 | 0.35932051 | -1.6076235 | 0.10791766 | 0.20924591 | LCNL1 | 401562 |
| ENSG000001 | 279.562926 | 0.223595043 | 0.13910252 | 1.60741188 | 0.10796405 | 0.20932584 | RNF217 | 154214 |
| ENSG000002 | 7.86506426 | -0.578260898 | 0.35976386 | -1.6073346 | 0.10798099 | 0.20934868 | NA | NA |
| ENSG000001 | 18.166763 | -0.45636647 | 0.28397698 | -1.6070544 | 0.10804243 | 0.20944777 | WT1-AS | 51352 |
| ENSG000002 | 29.0062919 | -0.418986336 | 0.26071591 | -1.6070608 | 0.10804102 | 0.20944777 | PCDHGA1 | 56114 |
| ENSG000001 | 172.930104 | 0.256759485 | 0.15977324 | 1.60702436 | 0.10804902 | 0.20945054 | THEM4 | 117145 |
| ENSG000001 | 43.5268321 | 0.344740825 | 0.21453643 | 1.60691044 | 0.10807402 | 0.20948898 | TNFAIP3 | 7128 |
| ENSG000002 | 3.33465048 | -0.604824455 | 0.37645626 | -1.6066261 | 0.10813641 | 0.2095999 | NA | NA |
| ENSG000001 | 9.44091936 | -0.556144747 | 0.34619108 | -1.6064676 | 0.10817121 | 0.20965734 | ADRA1B | 147 |
| ENSG000001 | 6.75302753 | -0.587652594 | 0.36584141 | -1.6063042 | 0.10820709 | 0.20968031 | ACTBL2 | 345651 |
| ENSG000001 | 2.74589875 | -0.593406865 | 0.36942563 | -1.6062959 | 0.10820892 | 0.20968031 | OR5AK2 | 390181 |
| ENSG000002 | 6.63314083 | -0.580570615 | 0.36142852 | -1.6063221 | 0.10820316 | 0.20968031 | NA | NA |
| ENSG000002 | 4.7004212 | -0.605442051 | 0.37690162 | -1.6063663 | 0.10819346 | 0.20968031 | NA | NA |
| ENSG000002 | 70.8613165 | 0.303089938 | 0.18868801 | 1.60630205 | 0.10820756 | 0.20968031 | LINC00869 | 57234 |
| ENSG000002 | 2.6802846 | -0.597880796 | 0.37222454 | -1.6062369 | 0.10822186 | 0.20969537 | NA | NA |
| ENSG000001 | 132.195574 | 0.300221771 | 0.18691827 | 1.60616604 | 0.10823743 | 0.20969548 | STK11IP | 114790 |
| ENSG000002 | 6.69222401 | -0.587529975 | 0.36579009 | -1.6061943 | 0.10823122 | 0.20969548 | LOC151484 | 151484 |
| ENSG000002 | 3.28604607 | -0.605340815 | 0.3768842 | -1.6061719 | 0.10823614 | 0.20969548 | NA | NA |
| ENSG000001 | 21.5413639 | -0.432848407 | 0.26950694 | -1.6060752 | 0.1082574 | 0.20972414 | BRDT | 676 |
| ENSG000001 | 228.427118 | 0.227373416 | 0.14159079 | 1.60584894 | 0.1083071 | 0.20981041 | DIP2A | 23181 |
| ENSG000002 | 2.70254464 | -0.597816081 | 0.37230001 | -1.6057375 | 0.10833161 | 0.20984785 | NA | NA |
| ENSG000001 | 217.023051 | 0.232473518 | 0.14478595 | 1.60563594 | 0.10835392 | 0.20988106 | TMEM200A | 114801 |
| ENSG000002 | 11.9411356 | -0.516550722 | 0.32171914 | -1.6055952 | 0.10836287 | 0.20988837 | NALCN-AS1 | 100885778 |

| | | | | | | | | |
|---|---|---|---|---|---|---|---|---|
| ENSG000002 | 3.30320577 | -0.605529052 | 0.37716912 | -1.6054577 | 0.10839312 | 0.2099269 | NA | NA |
| ENSG000002 | 2.64532965 | -0.599655185 | 0.37350959 | -1.6054613 | 0.10839233 | 0.2099269 | NA | NA |
| ENSG000002 | 2.67466776 | -0.598450284 | 0.37277008 | -1.6054139 | 0.10840274 | 0.2099355 | NA | NA |
| ENSG000002 | 2.77284916 | -0.591915475 | 0.36871301 | -1.6053555 | 0.10841559 | 0.20995035 | NA | NA |
| ENSG000001 | 209.91624 | 0.272702282 | 0.16987937 | 1.60527015 | 0.10843437 | 0.20997669 | CELSR2 | 1952 |
| ENSG000001 | 48.5692861 | 0.364567711 | 0.22711817 | 1.60518954 | 0.1084521 | 0.20999962 | VGF | 7425 |
| ENSG000001 | 201.94343 | 0.281349495 | 0.17527715 | 1.60516926 | 0.10845656 | 0.20999962 | PGBD2 | 267002 |
| ENSG000002 | 2.35849083 | -0.587605257 | 0.36612118 | -1.6049475 | 0.10850537 | 0.21007406 | NA | NA |
| ENSG000002 | 3.26400337 | -0.606188832 | 0.37769492 | -1.6049695 | 0.10850051 | 0.21007406 | NA | NA |
| ENSG000002 | 3.83671519 | -0.607595194 | 0.3785918 | -1.6048821 | 0.10851976 | 0.21009189 | NA | NA |
| ENSG000001 | 80.0681317 | -0.286176641 | 0.17833825 | -1.6046846 | 0.10856323 | 0.21015632 | FAM122C | 159091 |
| ENSG000002 | 10.4663061 | -0.569108929 | 0.35465487 | -1.6046838 | 0.1085634 | 0.21015632 | NA | NA |
| ENSG000002 | 7.27882838 | -0.581401426 | 0.3623339 | -1.6046012 | 0.10858159 | 0.21018149 | NA | NA |
| ENSG000001 | 748.075611 | 0.232832254 | 0.14513599 | 1.60423512 | 0.10866224 | 0.21032756 | GOLPH3L | 55204 |
| ENSG000001 | 42.9178813 | -0.344620171 | 0.21482608 | -1.6041822 | 0.10867391 | 0.21034011 | CPQ | 10404 |
| ENSG000002 | 5.20969809 | -0.60519969 | 0.3773073 | -1.6039967 | 0.10871477 | 0.21040915 | NA | NA |
| ENSG000002 | 2.62736913 | -0.598955001 | 0.37343103 | -1.603924 | 0.1087308 | 0.21043013 | NA | NA |
| ENSG000002 | 4.41482765 | -0.604797897 | 0.37709059 | -1.6038531 | 0.10874645 | 0.21044606 | NA | NA |
| ENSG000002 | 3.85892315 | -0.607200662 | 0.37859188 | -1.6038396 | 0.10874941 | 0.21044606 | NA | NA |
| ENSG000001 | 36.7621783 | -0.366265262 | 0.22837828 | -1.6037658 | 0.10876569 | 0.21045557 | DCDC2 | 51473 |
| ENSG000001 | 9.57359375 | -0.543381207 | 0.33881984 | -1.6037467 | 0.10876989 | 0.21045557 | C1orf111 | 284680 |
| ENSG000002 | 2.6474366 | -0.597853492 | 0.37277507 | -1.6037915 | 0.10876001 | 0.21045557 | NA | NA |
| ENSG000002 | 5.84629003 | -0.592756325 | 0.36962609 | -1.6036647 | 0.10878797 | 0.21047093 | NA | NA |
| ENSG000002 | 9.83597374 | -0.557737573 | 0.34778962 | -1.6036637 | 0.10878821 | 0.21047093 | AATBC | 284837 |
| ENSG000001 | 219.72911 | 0.430138795 | 0.26825798 | 1.60345203 | 0.10883489 | 0.2105359 | SCAF1 | 58506 |
| ENSG000001 | 5.61872486 | -0.600929431 | 0.37477533 | -1.6034391 | 0.10883774 | 0.2105359 | ANXA2R | 389289 |
| ENSG000002 | 2.91671133 | -0.603760259 | 0.37655146 | -1.6033937 | 0.10884775 | 0.2105359 | NA | NA |
| ENSG000002 | 2.69613797 | -0.596363596 | 0.37192311 | -1.6034594 | 0.10883327 | 0.2105359 | NA | NA |
| ENSG000002 | 3.02069369 | -0.600858705 | 0.37473915 | -1.6034052 | 0.10884523 | 0.2105359 | NA | NA |

| | | | | | | | | |
|---|---|---|---|---|---|---|---|---|
| ENSG000001 | 4.16545642 | -0.606812838 | 0.37846434 | -1.6033554 | 0.10885622 | 0.21054224 | OR6K2 | 81448 |
| ENSG000001 | 16.4702393 | -0.478174144 | 0.2982436 | -1.6033006 | 0.1088683 | 0.21055557 | LEP | 3952 |
| ENSG000002 | 687.795723 | 0.214999667 | 0.13410168 | 1.60325851 | 0.10887759 | 0.21056349 | NA | NA |
| ENSG000001 | 107.535816 | 0.259742433 | 0.16201833 | 1.60316691 | 0.10889781 | 0.2105825 | FAM159B | 100132916 |
| ENSG000001 | 57.6225602 | 0.342199786 | 0.21345187 | 1.60317071 | 0.10889697 | 0.2105825 | DPP7 | 29952 |
| ENSG000002 | 3.22513349 | -0.605381819 | 0.37762276 | -1.6031391 | 0.10890394 | 0.21058431 | NA | NA |
| ENSG000002 | 5.25562958 | -0.596866147 | 0.37233086 | -1.6030531 | 0.10892293 | 0.21061099 | NA | NA |
| ENSG000002 | 3.22481286 | -0.605294794 | 0.37762278 | -1.6029086 | 0.10895483 | 0.21066072 | NA | NA |
| ENSG000002 | 40.5945486 | 0.361854263 | 0.22575121 | 1.60288956 | 0.10895904 | 0.21066072 | ZNF488 | 118738 |
| ENSG000002 | 6.42070915 | -0.585449435 | 0.36527966 | -1.6027431 | 0.10899139 | 0.21070318 | NA | NA |
| ENSG000002 | 2.81103728 | -0.586623095 | 0.36600785 | -1.6027609 | 0.10898744 | 0.21070318 | NA | NA |
| ENSG000001 | 4.16143853 | -0.606078257 | 0.37816518 | -1.6026813 | 0.10900504 | 0.21071952 | F2RL3 | 9002 |
| ENSG000002 | 7.06354977 | -0.585779208 | 0.36550737 | -1.6026468 | 0.10901266 | 0.2107242 | NA | NA |
| ENSG000001 | 141.011389 | 0.258108571 | 0.16107547 | 1.60240768 | 0.10906549 | 0.21079618 | KLHDC7A | 127707 |
| ENSG000001 | 75.0322998 | 0.311857176 | 0.19461574 | 1.60242527 | 0.1090616 | 0.21079618 | LRRC16B | 90668 |
| ENSG000002 | 3.52431991 | -0.606528631 | 0.37851022 | -1.6024102 | 0.10906494 | 0.21079618 | NA | NA |
| ENSG000001 | 66.4806871 | 0.303615844 | 0.18950365 | 1.60216355 | 0.10911945 | 0.21089042 | KTN1-AS1 | 100129075 |
| ENSG000002 | 5.31178253 | -0.600848153 | 0.37503578 | -1.6021089 | 0.10913154 | 0.21090373 | NA | NA |
| ENSG000002 | 2.87531115 | 0.601489315 | 0.37546214 | 1.60199724 | 0.10915622 | 0.21093133 | NA | NA |
| ENSG000002 | 2.660302 | -0.597882166 | 0.37320973 | -1.6020005 | 0.1091555 | 0.21093133 | NA | NA |
| ENSG000002 | 6.08130739 | -0.587373484 | 0.36665732 | -1.6019685 | 0.10916257 | 0.21093355 | NA | NA |
| ENSG000002 | 16.0640217 | 0.485032174 | 0.30278433 | 1.60190646 | 0.10917629 | 0.21095001 | NA | NA |
| ENSG000002 | 6.10489695 | -0.588499481 | 0.36739547 | -1.6018148 | 0.10919658 | 0.21097915 | NA | NA |
| ENSG000002 | 6.33580544 | -0.584334751 | 0.36482253 | -1.6016959 | 0.10922287 | 0.2110199 | OR4G2P | 26680 |
| ENSG000002 | 7.52033913 | -0.575881448 | 0.359555 | -1.6016505 | 0.10923291 | 0.21102924 | NA | NA |
| ENSG000001 | 91.3198717 | 0.287424621 | 0.17948672 | 1.60136981 | 0.10929504 | 0.21113878 | MAN1A1 | 4121 |
| ENSG000002 | 2.68335819 | -0.597039725 | 0.37283588 | -1.6013473 | 0.10930002 | 0.21113878 | NA | NA |
| ENSG000002 | 13.5221107 | -0.513544151 | 0.32070551 | -1.6012951 | 0.10931158 | 0.21115105 | FAM24B | 196792 |
| ENSG000002 | 13.446428 | -0.507351968 | 0.31686536 | -1.6011595 | 0.1093416 | 0.21119898 | LINC01102 | 150568 |

| | | | | | | | | |
|---|---|---|---|---|---|---|---|---|
| ENSG000002 | 4.10369431 | -0.605619888 | 0.37829015 | -1.6009401 | 0.10939018 | 0.21127804 | NA | NA |
| ENSG000002 | 6.13788408 | -0.589108238 | 0.36797931 | -1.6009276 | 0.10939296 | 0.21127804 | NA | NA |
| ENSG000002 | 58.1732164 | 0.315596635 | 0.19713875 | 1.60088588 | 0.1094022 | 0.21128583 | NA | NA |
| ENSG000002 | 4.1510778 | -0.605501656 | 0.37823787 | -1.6008489 | 0.1094104 | 0.21129161 | NA | NA |
| ENSG000001 | 55.9494376 | 0.322315862 | 0.20135109 | 1.60076545 | 0.10942888 | 0.21131723 | C2orf76 | 130355 |
| ENSG000002 | 34.3162896 | -0.411186919 | 0.25688772 | -1.6006484 | 0.10945482 | 0.21135725 | ARHGEF38 | 54848 |
| ENSG000002 | 3.49056606 | -0.605853815 | 0.3785562 | -1.6004329 | 0.10950257 | 0.2114394 | NA | NA |
| ENSG000002 | 11.5541063 | -0.532974931 | 0.33308093 | -1.6001364 | 0.10956832 | 0.21155628 | LINC01247 | 101929390 |
| ENSG000002 | 12.3553928 | -0.518583114 | 0.32413387 | -1.5999041 | 0.10961985 | 0.2116457 | NA | NA |
| ENSG000001 | 22.6035369 | -0.435850676 | 0.27244758 | -1.5997598 | 0.10965189 | 0.21169747 | TSPAN8 | 7103 |
| ENSG000002 | 3.255646 | -0.603496358 | 0.37725547 | -1.5997021 | 0.10966469 | 0.21171211 | RNU1-106P | 106481624 |
| ENSG000002 | 8.88029225 | -0.561503974 | 0.35101797 | -1.5996445 | 0.10967746 | 0.21172669 | NA | NA |
| ENSG000002 | 6.39056959 | -0.587957248 | 0.36756047 | -1.5996204 | 0.10968281 | 0.21172693 | NA | NA |
| ENSG000002 | 5.78521775 | -0.591765573 | 0.36996019 | -1.5995385 | 0.109701 | 0.21175197 | NA | NA |
| ENSG000001 | 79.7059015 | 0.303565305 | 0.18980346 | 1.59936653 | 0.10973918 | 0.21177875 | SEC22A | 26984 |
| ENSG000001 | 10.0293633 | -0.550034863 | 0.34390859 | -1.5993635 | 0.10973985 | 0.21177875 | SGPP2 | 130367 |
| ENSG000001 | 14.5269425 | -0.520424669 | 0.32539933 | -1.5993415 | 0.10974474 | 0.21177875 | DNASE1L3 | 1776 |
| ENSG000002 | 1107.56441 | 0.185160297 | 0.11576792 | 1.5994094 | 0.10972967 | 0.21177875 | ZNF90 | 7643 |
| ENSG000002 | 2.68720014 | -0.595514784 | 0.37235153 | -1.5993349 | 0.10974621 | 0.21177875 | NA | NA |
| ENSG000002 | 11.5063364 | -0.546123216 | 0.34146549 | -1.5993511 | 0.1097426 | 0.21177875 | NA | NA |
| ENSG000002 | 6.10449853 | -0.589663718 | 0.36874186 | -1.5991234 | 0.1097932 | 0.21185933 | NA | NA |
| ENSG000001 | 23.4738609 | -0.452226484 | 0.28284904 | -1.5988263 | 0.1098592 | 0.21197662 | CDC20B | 166979 |
| ENSG000002 | 20.6217875 | -0.445882835 | 0.27888952 | -1.5987794 | 0.10986962 | 0.21198663 | LOC104968399 | 104968399 |
| ENSG000002 | 40.531611 | 0.364490735 | 0.22799023 | 1.59871206 | 0.1098846 | 0.21200544 | NA | NA |
| ENSG000002 | 6.77834101 | -0.586690919 | 0.36698835 | -1.5986636 | 0.10989537 | 0.21201613 | NA | NA |
| ENSG000002 | 2.38947211 | -0.580867134 | 0.36336553 | -1.5985752 | 0.10991502 | 0.21204396 | NA | NA |
| ENSG000001 | 262.179565 | 0.253876899 | 0.1588178 | 1.59854188 | 0.10992243 | 0.21204584 | PRDM15 | 63977 |
| ENSG000002 | 5.21045716 | -0.595000101 | 0.37221848 | -1.5985238 | 0.10992645 | 0.21204584 | NA | NA |
| ENSG000002 | 4.39391862 | -0.602649337 | 0.37701132 | -1.5984914 | 0.10993366 | 0.21204966 | LOC101929439 | 101929439 |

| | | | | | | | | |
|---|---|---|---|---|---|---|---|---|
| ENSG000001 | 3.60508387 | -0.60409058 | 0.37792618 | -1.5984354 | 0.10994611 | 0.21206075 | OR51F2 | 119694 |
| ENSG000002 | 13.8828716 | -0.499482557 | 0.31248547 | -1.5984185 | 0.10994987 | 0.21206075 | RPLP0P2 | 113157 |
| ENSG000002 | 15.1897111 | -0.521173987 | 0.32611173 | -1.5981455 | 0.11001061 | 0.2121678 | NANOGNB | 360030 |
| ENSG000002 | 5.51166731 | -0.591153566 | 0.36993458 | -1.5979949 | 0.11004412 | 0.21222006 | NA | NA |
| ENSG000002 | 22.3012663 | -0.43439445 | 0.2718403 | -1.5979767 | 0.11004817 | 0.21222006 | NA | NA |
| ENSG000001 | 1285.06584 | 0.188097503 | 0.11772007 | 1.59783715 | 0.11007923 | 0.21226985 | TTC3 | 7267 |
| ENSG000001 | 305.348244 | 0.461594035 | 0.28891754 | 1.59766704 | 0.1101171 | 0.21233278 | SPCS2 | 9789 |
| ENSG000002 | 21.901313 | -0.444679065 | 0.27835571 | -1.5975209 | 0.11014965 | 0.21238545 | NA | NA |
| ENSG000002 | 4.8805444 | -0.604125499 | 0.37821592 | -1.5973032 | 0.11019813 | 0.21246883 | NA | NA |
| ENSG000002 | 8.86551277 | -0.555606544 | 0.34787178 | -1.597159 | 0.11023027 | 0.21252001 | CASC20 | 101929244 |
| ENSG000002 | 2.62731652 | -0.594763018 | 0.37239322 | -1.5971371 | 0.11023516 | 0.21252001 | MIR3686 | 100500839 |
| ENSG000002 | 3.51270877 | -0.604507982 | 0.37851495 | -1.5970518 | 0.11025416 | 0.21254654 | NA | NA |
| ENSG000002 | 3.22748838 | -0.602881071 | 0.37750509 | -1.5970144 | 0.11026249 | 0.2125525 | NA | NA |
| ENSG000002 | 4.25954271 | -0.604139084 | 0.37830822 | -1.5969494 | 0.11027699 | 0.21257034 | NA | NA |
| ENSG000002 | 7.91782745 | -0.573273045 | 0.35899716 | -1.5968735 | 0.11029391 | 0.21259285 | TNF | 7124 |
| ENSG000002 | 10.9605411 | -0.525906801 | 0.32934332 | -1.5968346 | 0.11030258 | 0.21259946 | VIM-AS1 | 100507347 |
| ENSG000002 | 6.35715908 | -0.583288006 | 0.36529355 | -1.5967651 | 0.11031807 | 0.21261921 | NA | NA |
| ENSG000002 | 24.3397949 | -0.423968294 | 0.26552719 | -1.5967039 | 0.11033173 | 0.21263542 | NA | NA |
| ENSG000002 | 9.8025975 | -0.533968159 | 0.33443041 | -1.5966495 | 0.11034386 | 0.21264869 | NA | NA |
| ENSG000001 | 8.78216008 | -0.556349386 | 0.34850301 | -1.5963977 | 0.11040004 | 0.21274685 | DNASE2B | 58511 |
| ENSG000001 | 2.26448486 | -0.590592444 | 0.36996918 | -1.5963288 | 0.11041541 | 0.21276636 | PI3 | 5266 |
| ENSG000002 | 4.1238791 | -0.603794846 | 0.37827593 | -1.5961757 | 0.11044958 | 0.21282208 | NA | NA |
| ENSG000001 | 80.1355738 | 0.284087996 | 0.17799801 | 1.59601779 | 0.11048482 | 0.21287988 | CDK5 | 1020 |
| ENSG000002 | 6.91799827 | -0.577829555 | 0.36207263 | -1.5958941 | 0.11051245 | 0.212923 | NA | NA |
| ENSG000002 | 8.65051224 | -0.558019578 | 0.34967635 | -1.5958173 | 0.11052958 | 0.21293588 | OR2A1 | 346528 |
| ENSG000002 | 2.40573794 | -0.583661221 | 0.36574983 | -1.5957936 | 0.11053489 | 0.21293588 | TRIM49D1 | 399939 |
| ENSG000002 | 3.94233816 | -0.603843753 | 0.37839426 | -1.5958058 | 0.11053215 | 0.21293588 | NA | NA |
| ENSG000002 | 3.50120698 | -0.604057724 | 0.37857734 | -1.5955993 | 0.11057829 | 0.21300938 | NA | NA |
| ENSG000001 | 67.3737237 | -0.330950135 | 0.20746059 | -1.5952434 | 0.11065782 | 0.21315245 | OTOGL | 283310 |

| | | | | | | | | |
|---|---|---|---|---|---|---|---|---|
| ENSG000002 | 4.39929524 | -0.601339419 | 0.37696685 | -1.595205 | 0.1106664 | 0.21315884 | NA | NA |
| ENSG000001 | 45.1786946 | -0.342945574 | 0.21499158 | -1.5951581 | 0.11067688 | 0.2131689 | TTC21A | 199223 |
| ENSG000002 | 5.56443607 | -0.593453398 | 0.3720405 | -1.5951312 | 0.1106829 | 0.21317038 | LOC102724596 | 102724596 |
| ENSG000001 | 14.1908577 | 0.499825539 | 0.31336303 | 1.59503673 | 0.11070402 | 0.21320093 | PEX11G | 92960 |
| ENSG000002 | 6.38604127 | -0.58231815 | 0.36511611 | -1.5948848 | 0.11073799 | 0.21325622 | NA | NA |
| ENSG000002 | 13.0195768 | -0.50050843 | 0.31384261 | -1.5947752 | 0.1107625 | 0.2132933 | NA | NA |
| ENSG000001 | 10.4910573 | -0.538812915 | 0.33789365 | -1.5946228 | 0.11079662 | 0.21334886 | OR8B12 | 219858 |
| ENSG000002 | 3.46887961 | -0.603498082 | 0.37850112 | -1.594442 | 0.11083708 | 0.21341663 | NA | NA |
| ENSG000000 | 567.216629 | 0.195923879 | 0.12288256 | 1.59439933 | 0.11084662 | 0.21342488 | FBXL3 | 26224 |
| ENSG000001 | 2094.21994 | 0.44853678 | 0.28132588 | 1.59436731 | 0.11085379 | 0.21342855 | DNMT3A | 1788 |
| ENSG000002 | 3.53305719 | -0.603372051 | 0.37849632 | -1.5941292 | 0.11090709 | 0.21352104 | NA | NA |
| ENSG000002 | 4.15896597 | -0.603177342 | 0.37841022 | -1.5939774 | 0.1109411 | 0.21357636 | NA | NA |
| ENSG000002 | 6.22967951 | -0.589998394 | 0.37018197 | -1.5938064 | 0.11097941 | 0.21363997 | NA | NA |
| ENSG000002 | 11.9549727 | -0.516807827 | 0.32428333 | -1.5936922 | 0.11100499 | 0.21367907 | BPESC1 | 60467 |
| ENSG000002 | 2.9805252 | -0.59654547 | 0.37433228 | -1.5936255 | 0.11101994 | 0.2136977 | LOC101927950 | 101927950 |
| ENSG000002 | 6.10618899 | -0.586825986 | 0.36824284 | -1.5935842 | 0.11102919 | 0.21370537 | TMEM92-AS1 | 103752589 |
| ENSG000001 | 8.15153403 | -0.569355884 | 0.35732722 | -1.5933739 | 0.11107633 | 0.21376566 | DUOXA2 | 405753 |
| ENSG000002 | 4.42743171 | -0.601255754 | 0.3773422 | -1.5933965 | 0.11107127 | 0.21376566 | NA | NA |
| ENSG000002 | 2.63117829 | -0.593007931 | 0.37216103 | -1.5934176 | 0.11106654 | 0.21376566 | NA | NA |
| ENSG000002 | 2.69380269 | -0.593291556 | 0.37236465 | -1.5933079 | 0.11109113 | 0.21378399 | RN7SKP219 | 106479192 |
| ENSG000002 | 75.324681 | -0.294446863 | 0.184805 | -1.5932841 | 0.11109648 | 0.21378415 | NA | NA |
| ENSG000002 | 7.0762359 | -0.581846987 | 0.36523124 | -1.5930921 | 0.11113954 | 0.21385686 | NA | NA |
| ENSG000002 | 5.19395146 | -0.593287026 | 0.37245071 | -1.5929276 | 0.11117643 | 0.21391769 | NA | NA |
| ENSG000001 | 2.89567832 | -0.600064259 | 0.37672576 | -1.5928411 | 0.11119585 | 0.21394451 | CCDC105 | 126402 |
| ENSG000001 | 12.9743575 | -0.499313122 | 0.31347773 | -1.5928185 | 0.11120092 | 0.21394451 | CDX2 | 1045 |
| ENSG000002 | 3.26278988 | -0.601316876 | 0.3775249 | -1.5927873 | 0.11120791 | 0.21394781 | NA | NA |
| ENSG000001 | 11.938957 | -0.514618291 | 0.32310811 | -1.5927124 | 0.11122473 | 0.21395986 | VSX1 | 30813 |
| ENSG000001 | 15.928629 | -0.494393005 | 0.31040758 | -1.5927221 | 0.11122255 | 0.21395986 | CD48 | 962 |
| ENSG000002 | 6.15849343 | -0.58670172 | 0.36837278 | -1.5926848 | 0.11123092 | 0.21396163 | NA | NA |

| | | | | | | | | |
|---|---|---|---|---|---|---|---|---|
| ENSG000001 | 12.1506736 | -0.519379774 | 0.32612584 | -1.5925747 | 0.11125562 | 0.21399006 | PDCD1LG2 | 80380 |
| ENSG000002 | 5.03331379 | -0.597603512 | 0.37524428 | -1.5925719 | 0.11125626 | 0.21399006 | NA | NA |
| ENSG000002 | 3.5284693 | -0.602497984 | 0.37832357 | -1.5925468 | 0.1112619 | 0.21399077 | NA | NA |
| ENSG000001 | 6.60750774 | -0.578049241 | 0.36299013 | -1.5924655 | 0.11128016 | 0.21401573 | C5orf46 | 389336 |
| ENSG000002 | 7.84929327 | -0.571414141 | 0.35888841 | -1.5921778 | 0.11134477 | 0.2141278 | PRSS1 | 5644 |
| ENSG000002 | 301.956558 | 0.226761261 | 0.14242376 | 1.59215896 | 0.11134899 | 0.2141278 | NA | NA |
| ENSG000002 | 4.7725386 | -0.600832272 | 0.3773869 | -1.5920857 | 0.11136546 | 0.21414931 | NA | NA |
| ENSG00000C | 8.79819689 | -0.553482943 | 0.34767283 | -1.5919649 | 0.1113926 | 0.21415596 | ZMYND10 | 51364 |
| ENSG000002 | 3.87488863 | -0.602728294 | 0.37859293 | -1.5920221 | 0.11137975 | 0.21415596 | NA | NA |
| ENSG000002 | 2.27845941 | -0.589077573 | 0.37002676 | -1.5919864 | 0.11138776 | 0.21415596 | NA | NA |
| ENSG000002 | 6.96190215 | -0.581692962 | 0.36539587 | -1.5919527 | 0.11139532 | 0.21415596 | NA | NA |
| ENSG000002 | 3.48798228 | -0.602628726 | 0.37852669 | -1.5920376 | 0.11137626 | 0.21415596 | NA | NA |
| ENSG000002 | 23.9637162 | -0.424761117 | 0.2668222 | -1.5919257 | 0.1114014 | 0.21415749 | NA | NA |
| ENSG000002 | 2.66069376 | -0.593449015 | 0.37280371 | -1.5918538 | 0.11141755 | 0.21417838 | NA | NA |
| ENSG000001 | 1181.69299 | 0.215159699 | 0.13517478 | 1.59171475 | 0.11144881 | 0.21422833 | FZD5 | 7855 |
| ENSG000002 | 12.7073178 | -0.505317001 | 0.31747653 | -1.5916673 | 0.11145949 | 0.2142387 | LOC101927406 | 101927406 |
| ENSG000002 | 3.77606664 | -0.602139922 | 0.37835729 | -1.5914585 | 0.11150643 | 0.21431876 | NA | NA |
| ENSG000001 | 94.8079246 | -0.281240555 | 0.17672539 | -1.5913987 | 0.11151987 | 0.21433443 | DNHD1 | 144132 |
| ENSG000001 | 3.31449958 | -0.600421278 | 0.37732088 | -1.591275 | 0.11154771 | 0.21436841 | NA | NA |
| ENSG000002 | 9.4071529 | -0.548257984 | 0.34454046 | -1.5912731 | 0.11154812 | 0.21436841 | MYCBP2-AS1 | 100874212 |
| ENSG000001 | 15.5911602 | -0.499065376 | 0.31363115 | -1.5912494 | 0.11155346 | 0.21436851 | VAX1 | 11023 |
| ENSG000002 | 10.9202889 | -0.52386422 | 0.3292484 | -1.5910912 | 0.11158906 | 0.21442677 | FLJ45079 | 400624 |
| ENSG000001 | 21.7372577 | -0.444533443 | 0.27946818 | -1.5906407 | 0.11169047 | 0.21457079 | IMPG1 | 3617 |
| ENSG000001 | 7.11600987 | -0.582754403 | 0.3663625 | -1.5906497 | 0.11168843 | 0.21457079 | NINJ2 | 4815 |
| ENSG000002 | 4.955501 | -0.596944414 | 0.37527852 | -1.5906703 | 0.11168379 | 0.21457079 | NA | NA |
| ENSG000002 | 2.63633921 | -0.591762609 | 0.37202228 | -1.5906644 | 0.11168512 | 0.21457079 | NA | NA |
| ENSG000002 | 2.74346595 | -0.58798764 | 0.36963973 | -1.5907046 | 0.11167606 | 0.21457079 | NA | NA |
| ENSG000001 | 274.140938 | 0.27995167 | 0.17601496 | 1.59049931 | 0.1117223 | 0.21460144 | TAF3 | 83860 |
| ENSG000002 | 5.2045181 | -0.600493769 | 0.3775428 | -1.5905316 | 0.11171502 | 0.21460144 | NA | NA |

| | | | | | | | | |
|---|---|---|---|---|---|---|---|---|
| ENSG000002 | 10.0315596 | -0.56340437 | 0.35422655 | -1.5905199 | 0.11171767 | 0.21460144 | NA | NA |
| ENSG000001 | 7.80341809 | -0.567829186 | 0.35704165 | -1.5903724 | 0.11175089 | 0.21464285 | PLD6 | 201164 |
| ENSG000002 | 8.56049786 | -0.57123675 | 0.35918784 | -1.5903566 | 0.11175444 | 0.21464285 | NA | NA |
| ENSG000002 | 8.19514404 | -0.571800066 | 0.35956064 | -1.5902743 | 0.11177298 | 0.21466828 | NA | NA |
| ENSG000002 | 147.442478 | 0.319406688 | 0.20088115 | 1.59002813 | 0.11182846 | 0.21476467 | NA | NA |
| ENSG000002 | 4.93904661 | -0.596993287 | 0.37549051 | -1.5899024 | 0.1118568 | 0.21480891 | NA | NA |
| ENSG000002 | 6.17638128 | -0.586874174 | 0.36913401 | -1.5898675 | 0.11186467 | 0.21481386 | LOC105376382 | 105376382 |
| ENSG000002 | 11.0837782 | -0.537206777 | 0.33791976 | -1.5897466 | 0.11189194 | 0.21485605 | NA | NA |
| ENSG000002 | 3.27661485 | -0.599692793 | 0.37724639 | -1.5896581 | 0.11191189 | 0.21488418 | LOC105372267 | 105372267 |
| ENSG000001 | 388.643719 | 0.201167461 | 0.12656234 | 1.58947331 | 0.11195358 | 0.21495405 | VANGL1 | 81839 |
| ENSG000002 | 4.78007979 | -0.599442384 | 0.37715578 | -1.5893761 | 0.11197551 | 0.21498597 | NA | NA |
| ENSG000001 | 5.60445477 | -0.592702915 | 0.37295266 | -1.5892175 | 0.11201129 | 0.21504449 | FAM183BP | 340286 |
| ENSG000002 | 6.38046953 | -0.580586515 | 0.36540928 | -1.5888664 | 0.11209057 | 0.21518651 | NA | NA |
| ENSG000001 | 5.6927481 | -0.594694595 | 0.3743092 | -1.588779 | 0.1121103 | 0.2152142 | OR10W1 | 81341 |
| ENSG000001 | 15.3646651 | -0.47977404 | 0.30199227 | -1.5886964 | 0.11212895 | 0.21523804 | MICALCL | 84953 |
| ENSG000002 | 18.3194242 | -0.477943038 | 0.30084343 | -1.588677 | 0.11213333 | 0.21523804 | NA | NA |
| ENSG000002 | 6.77387677 | -0.585638794 | 0.36865302 | -1.5885908 | 0.11215281 | 0.21526523 | LOC101929080 | 101929080 |
| ENSG000002 | 16.8750746 | -0.470615208 | 0.29627964 | -1.5884156 | 0.11219239 | 0.21532082 | NA | NA |
| ENSG000002 | 7.75359804 | -0.562098824 | 0.35387276 | -1.5884207 | 0.11219123 | 0.21532082 | NA | NA |
| ENSG00000C | 79.9108997 | -0.283405943 | 0.17842936 | -1.5883369 | 0.11221017 | 0.21532877 | TDRD1 | 56165 |
| ENSG000001 | 190.084525 | 0.268506422 | 0.16904986 | 1.58832679 | 0.11221246 | 0.21532877 | MKL1 | 57591 |
| ENSG000002 | 10.9534923 | 0.540475645 | 0.34027795 | 1.58833577 | 0.11221043 | 0.21532877 | LOC102724428 | 102724428 |
| ENSG000001 | 560.67625 | 0.199881834 | 0.12586201 | 1.588103 | 0.11226305 | 0.21541565 | CDC42EP3 | 10602 |
| ENSG000002 | 2.92081442 | -0.597160264 | 0.37612029 | -1.5876842 | 0.11235776 | 0.21558718 | NA | NA |
| ENSG00000C | 14.0473587 | -0.524361876 | 0.33030956 | -1.5874862 | 0.11240256 | 0.21566295 | TYR | 7299 |
| ENSG000002 | 2.28460494 | -0.586841063 | 0.36967784 | -1.5874391 | 0.11241322 | 0.2156728 | NA | NA |
| ENSG000002 | 6.09339735 | -0.583575106 | 0.36762569 | -1.5874165 | 0.11241834 | 0.2156728 | NA | NA |
| ENSG000002 | 9.10154972 | -0.547142666 | 0.34469445 | -1.5873266 | 0.1124387 | 0.21570166 | NA | NA |
| ENSG000001 | 9.14195102 | -0.550764066 | 0.34698451 | -1.5872872 | 0.11244762 | 0.21570305 | IL36B | 27177 |

| | | | | | | | | |
|---|---|---|---|---|---|---|---|---|
| ENSG000002 | 6.12123436 | -0.584610332 | 0.36831036 | -1.5872764 | 0.11245006 | 0.21570305 | NA | NA |
| ENSG000002 | 25.7803345 | 0.423177927 | 0.26664228 | 1.58706235 | 0.11249853 | 0.2157858 | NA | NA |
| ENSG000002 | 7.34819692 | 0.570553681 | 0.35953875 | 1.58690456 | 0.11253426 | 0.21584414 | ANG | 283 |
| ENSG000002 | 9.33610919 | -0.546428812 | 0.34435322 | -1.5868265 | 0.11255195 | 0.21585764 | LOC101927310 | 101927310 |
| ENSG000002 | 5.5909882 | -0.592699882 | 0.37351116 | -1.5868331 | 0.11255045 | 0.21585764 | NA | NA |
| ENSG00000C | 14.516783 | -0.515662686 | 0.32498787 | -1.5867137 | 0.11257751 | 0.21589454 | PIH1D3 | 139212 |
| ENSG000002 | 28.4197399 | -0.401371573 | 0.25296369 | -1.5866766 | 0.11258591 | 0.21589454 | MOG | 4340 |
| ENSG000002 | 7.93222793 | -0.557343335 | 0.35126583 | -1.5866711 | 0.11258716 | 0.21589454 | NA | NA |
| ENSG000002 | 2.86867549 | -0.597471352 | 0.37657569 | -1.5865904 | 0.11260545 | 0.21591941 | NA | NA |
| ENSG000002 | 3.70796053 | -0.598672305 | 0.37740802 | -1.5862734 | 0.1126773 | 0.21602653 | NA | NA |
| ENSG000002 | 5.0510975 | -0.595583796 | 0.3754562 | -1.5862937 | 0.1126727 | 0.21602653 | NA | NA |
| ENSG000002 | 5.50901904 | -0.587711257 | 0.37049194 | -1.5862997 | 0.11267134 | 0.21602653 | NA | NA |
| ENSG000002 | 2.28372338 | -0.584500572 | 0.36851867 | -1.5860813 | 0.11272086 | 0.21608815 | NA | NA |
| ENSG000002 | 2.68121824 | -0.590403609 | 0.37223643 | -1.5860984 | 0.11271699 | 0.21608815 | NA | NA |
| ENSG000002 | 2.86936063 | -0.597273252 | 0.37657642 | -1.5860612 | 0.11272542 | 0.21608815 | NA | NA |
| ENSG00000C | 82.3613797 | 0.29764431 | 0.18768402 | 1.58587989 | 0.11276656 | 0.21613635 | ANO8 | 57719 |
| ENSG000002 | 2.59028578 | -0.590082478 | 0.37207983 | -1.5859029 | 0.11276134 | 0.21613635 | NA | NA |
| ENSG000002 | 4.71565 | -0.597366208 | 0.37667533 | -1.5858915 | 0.11276392 | 0.21613635 | KRTAP7-1 | 337878 |
| ENSG000001 | 237.81647 | 0.243456311 | 0.15353041 | 1.58572048 | 0.11280273 | 0.21619546 | ING2 | 3622 |
| ENSG000001 | 463.085627 | 0.201207388 | 0.12691877 | 1.58532411 | 0.11289271 | 0.21635769 | ARHGAP32 | 9743 |
| ENSG000002 | 4.63993661 | -0.594552669 | 0.37505202 | -1.5852539 | 0.11290867 | 0.21637804 | NA | NA |
| ENSG000002 | 5.26308537 | -0.592592665 | 0.37383929 | -1.5851535 | 0.11293146 | 0.21641149 | NA | NA |
| ENSG000002 | 2.2905546 | -0.585553058 | 0.36943584 | -1.5849926 | 0.11296802 | 0.21647133 | FGF14-AS1 | 100874081 |
| ENSG000001 | 13.8292131 | -0.494744464 | 0.31215083 | -1.5849532 | 0.11297696 | 0.21647823 | NA | NA |
| ENSG000001 | 33.204228 | -0.382313077 | 0.24122697 | -1.5848687 | 0.11299616 | 0.21650479 | SPACA6P | 147650 |
| ENSG000002 | 2.62464045 | -0.589229236 | 0.37179969 | -1.5848029 | 0.11301112 | 0.21652322 | NA | NA |
| ENSG000002 | 3.53578979 | -0.59964101 | 0.37838991 | -1.5847172 | 0.1130306 | 0.2165503 | NA | NA |
| ENSG000001 | 68.7299884 | 0.30514162 | 0.19256205 | 1.5846405 | 0.11304804 | 0.21657348 | ZNF34 | 80778 |
| ENSG000002 | 9.06191594 | -0.546231794 | 0.34471648 | -1.5845828 | 0.11306117 | 0.2165884 | NA | NA |

| | | | | | | | | |
|---|---|---|---|---|---|---|---|---|
| ENSG000002 | 5.01893266 | -0.596732944 | 0.37659349 | -1.5845546 | 0.11306756 | 0.21659043 | NA | NA |
| ENSG000002 | 3.8068352 | -0.599766276 | 0.37853082 | -1.5844582 | 0.11308948 | 0.21662218 | NA | NA |
| ENSG000002 | 9.01854988 | -0.546800812 | 0.34511099 | -1.5844202 | 0.11309814 | 0.21662853 | NA | NA |
| ENSG000002 | 6.15527182 | -0.587760593 | 0.37097629 | -1.5843616 | 0.11311146 | 0.21664381 | NA | NA |
| ENSG000002 | 4.33676835 | -0.597134394 | 0.37691323 | -1.5842755 | 0.11313104 | 0.21665062 | NA | NA |
| ENSG000002 | 314.512646 | 0.229725907 | 0.14500172 | 1.58429776 | 0.11312597 | 0.21665062 | GABRQ | 55879 |
| ENSG000002 | 2.34932378 | -0.576289071 | 0.36374719 | -1.5843121 | 0.11312271 | 0.21665062 | NA | NA |
| ENSG000001 | 3.10017534 | -0.590826132 | 0.37297693 | -1.5840822 | 0.11317501 | 0.21671828 | FZD10 | 11211 |
| ENSG000002 | 3.74706026 | -0.599493821 | 0.37845083 | -1.5840732 | 0.11317706 | 0.21671828 | LOC101927346 | 101927346 |
| ENSG000002 | 4.39590157 | -0.597436534 | 0.37719999 | -1.5838721 | 0.11322283 | 0.21679569 | NA | NA |
| ENSG000002 | 3.22919826 | 0.597464665 | 0.3772284 | 1.58382736 | 0.11323301 | 0.21680494 | RNA5SP448 | 106479015 |
| ENSG000002 | 12.0114395 | -0.513067654 | 0.32396601 | -1.5837083 | 0.11326012 | 0.21683636 | NA | NA |
| ENSG000002 | 50.4009584 | 0.33567002 | 0.211949 | 1.58373016 | 0.11325514 | 0.21683636 | NA | NA |
| ENSG000002 | 50.0181268 | 0.335597125 | 0.21191299 | 1.58365529 | 0.11327218 | 0.21684923 | TNFRSF25 | 8718 |
| ENSG000002 | 9.88137474 | -0.533969717 | 0.33718922 | -1.5835907 | 0.11328689 | 0.21686654 | LOC100505716 | 100505716 |
| ENSG000002 | 5.2779285 | -0.591236873 | 0.37335728 | -1.5835686 | 0.11329192 | 0.21686654 | NA | NA |
| ENSG000002 | 3.4867239 | -0.599269259 | 0.37849947 | -1.5832763 | 0.11335849 | 0.21698373 | NA | NA |
| ENSG000002 | 6.5414191 | -0.574217057 | 0.36269016 | -1.5832165 | 0.11337212 | 0.21699957 | LINC00358 | 100874143 |
| ENSG000001 | 4.64681936 | -0.59394801 | 0.37517496 | -1.5831227 | 0.11339349 | 0.21703023 | TRIM49B | 283116 |
| ENSG000001 | 77.1696857 | -0.289663296 | 0.18297373 | -1.5830868 | 0.11340167 | 0.21703565 | PDGFC | 56034 |
| ENSG000002 | 2.95901611 | -0.593400238 | 0.37485199 | -1.5830255 | 0.11341566 | 0.21705217 | NA | NA |
| ENSG000002 | 6.30413151 | -0.57883875 | 0.36570427 | -1.5828056 | 0.11346579 | 0.21713786 | NA | NA |
| ENSG000002 | 3.31188355 | -0.596620345 | 0.37699618 | -1.5825634 | 0.11352101 | 0.21723329 | NA | NA |
| ENSG000001 | 3.25530109 | -0.5969962 | 0.37724797 | -1.5825034 | 0.11353469 | 0.21724921 | IQCF1 | 132141 |
| ENSG000002 | 3.53075306 | -0.598481149 | 0.37821422 | -1.5823867 | 0.11356132 | 0.21728991 | NA | NA |
| ENSG000001 | 2019.342 | 0.178143543 | 0.11258776 | 1.58226384 | 0.11358935 | 0.21732304 | CDC42BPA | 8476 |
| ENSG000002 | 3.23724883 | -0.597351784 | 0.3775268 | -1.5822765 | 0.11358647 | 0.21732304 | NA | NA |
| ENSG000002 | 24.6301897 | -0.415379328 | 0.26254017 | -1.5821553 | 0.11361411 | 0.21736015 | PCDHB16 | 57717 |
| ENSG000002 | 6.32811357 | -0.57840146 | 0.36561354 | -1.5820023 | 0.11364904 | 0.21741672 | NA | NA |

| | | | | | | | | |
|---|---|---|---|---|---|---|---|---|
| ENSG000001 | 367.543789 | 0.189498415 | 0.11978928 | 1.58193138 | 0.11366524 | 0.21743745 | C15orf41 | 84529 |
| ENSG000002 | 5.54445053 | -0.591700176 | 0.3740684 | -1.5817967 | 0.11369598 | 0.217486 | NA | NA |
| ENSG00000C | 59.41286 | 0.319053906 | 0.20173826 | 1.58152405 | 0.11375826 | 0.21759488 | IFFO1 | 25900 |
| ENSG000002 | 5.14769521 | -0.596222368 | 0.37701619 | -1.5814238 | 0.11378117 | 0.21762535 | NA | NA |
| ENSG000002 | 3.75369505 | -0.598528697 | 0.37847851 | -1.5814073 | 0.11378493 | 0.21762535 | NA | NA |
| ENSG000001 | 5.940724 | -0.588108101 | 0.37190229 | -1.5813511 | 0.11379778 | 0.21763966 | KCNV2 | 169522 |
| ENSG000001 | 1345.69942 | -0.168849268 | 0.10678924 | -1.5811449 | 0.1138449 | 0.21771317 | TRDN | 10345 |
| ENSG000002 | 6.03179433 | -0.592087636 | 0.37447534 | -1.5811125 | 0.11385232 | 0.21771317 | NA | NA |
| ENSG000002 | 7.1452872 | 0.571809754 | 0.36164719 | 1.58112596 | 0.11384924 | 0.21771317 | NA | NA |
| ENSG000001 | 14.331385 | -0.505617387 | 0.3198494 | -1.5807983 | 0.11392417 | 0.21784028 | C11orf16 | 56673 |
| ENSG000002 | 2.60708487 | -0.587939302 | 0.37197191 | -1.5806013 | 0.11396922 | 0.21791615 | NA | NA |
| ENSG000002 | 4.40815376 | -0.596571896 | 0.37746853 | -1.5804546 | 0.11400279 | 0.21797006 | NA | NA |
| ENSG000002 | 5.24520469 | -0.589550109 | 0.37305115 | -1.5803466 | 0.11402752 | 0.21800706 | NA | NA |
| ENSG000001 | 222.286427 | 0.214628433 | 0.13583075 | 1.58011668 | 0.11408015 | 0.21809741 | NA | NA |
| ENSG000001 | 82.391284 | 0.271744341 | 0.17198029 | 1.58009003 | 0.11408625 | 0.21809879 | SLC25A51 | 92014 |
| ENSG000002 | 5.2138432 | -0.589115888 | 0.37288479 | -1.5798871 | 0.11413272 | 0.21817734 | ASIC3 | 9311 |
| ENSG000001 | 60.864029 | -0.315787563 | 0.19988895 | -1.579815 | 0.11414924 | 0.21819863 | IKZF1 | 10320 |
| ENSG000002 | 4.96617363 | -0.593716425 | 0.37588201 | -1.5795287 | 0.11421484 | 0.21831373 | NA | NA |
| ENSG000002 | 13.3303769 | -0.507421932 | 0.32129405 | -1.579307 | 0.11426566 | 0.21839093 | NA | NA |
| ENSG000002 | 14.6317628 | -0.505039293 | 0.31978568 | -1.5793055 | 0.11426599 | 0.21839093 | NA | NA |
| ENSG000001 | 51.7385198 | 0.35262566 | 0.2232937 | 1.57920113 | 0.11428993 | 0.21842149 | D2HGDH | 728294 |
| ENSG000002 | 5.88446541 | -0.590017115 | 0.37362037 | -1.5791888 | 0.11429276 | 0.21842149 | HOXC-AS3 | 100874365 |
| ENSG000002 | 7.44349497 | -0.56495463 | 0.357764 | -1.5791266 | 0.11430703 | 0.21843848 | NA | NA |
| ENSG000002 | 3.27757244 | -0.594932172 | 0.37678193 | -1.5789827 | 0.11434001 | 0.21849121 | NA | NA |
| ENSG000002 | 5.96975919 | -0.589027242 | 0.37309231 | -1.5787708 | 0.11438864 | 0.21857383 | NA | NA |
| ENSG000001 | 18.428935 | -0.451352265 | 0.28592133 | -1.578589 | 0.11443036 | 0.21864324 | ADAM20 | 8748 |
| ENSG000001 | 21.2845229 | -0.473125842 | 0.29971887 | -1.5785654 | 0.11443577 | 0.21864328 | SLC24A5 | 283652 |
| ENSG000002 | 20.1070013 | -0.47071591 | 0.29819688 | -1.5785407 | 0.11444145 | 0.21864382 | FLJ27354 | 400761 |
| ENSG000002 | 2.95244493 | -0.589824642 | 0.37366336 | -1.578492 | 0.11445261 | 0.21865486 | NA | NA |

| | | | | | | | | |
|---|---|---|---|---|---|---|---|---|
| ENSG000001 | 1589.0436 | 0.188342897 | 0.11932421 | 1.57841306 | 0.11447075 | 0.21867282 | SUMO3 | 6612 |
| ENSG000002 | 10.4976225 | -0.53301554 | 0.33769692 | -1.5783844 | 0.11447733 | 0.21867282 | NA | NA |
| ENSG000002 | 33.3358831 | 0.376194348 | 0.23834197 | 1.57838062 | 0.11447819 | 0.21867282 | NA | NA |
| ENSG000002 | 7.17190292 | -0.569661888 | 0.36093402 | -1.5782993 | 0.11449688 | 0.2186982 | NA | NA |
| ENSG000002 | 4.46792925 | -0.596948014 | 0.37823056 | -1.5782649 | 0.11450477 | 0.21870298 | NA | NA |
| ENSG000002 | 3.57398195 | -0.596985129 | 0.37827382 | -1.5781825 | 0.11452369 | 0.21872881 | NA | NA |
| ENSG000001 | 1006.36017 | 0.197336973 | 0.12505091 | 1.57805304 | 0.11455342 | 0.21872914 | FRS2 | 10818 |
| ENSG000001 | 420.511622 | 0.211616946 | 0.13409488 | 1.57811358 | 0.11453952 | 0.21872914 | KCNG3 | 170850 |
| ENSG000001 | 3.55357425 | -0.596912865 | 0.37826199 | -1.5780408 | 0.11455623 | 0.21872914 | IFNB1 | 3456 |
| ENSG000002 | 2.28079469 | -0.581098314 | 0.3682193 | -1.578131 | 0.11453551 | 0.21872914 | NA | NA |
| ENSG000002 | 2.62331039 | -0.586322141 | 0.37154445 | -1.5780673 | 0.11455015 | 0.21872914 | NA | NA |
| ENSG000002 | 12.8606054 | 0.504624847 | 0.31976666 | 1.57810339 | 0.11454186 | 0.21872914 | GS1-124K5.4 | 100289098 |
| ENSG000002 | 6.23841992 | -0.586693951 | 0.37184176 | -1.5778054 | 0.11461031 | 0.21882211 | NA | NA |
| ENSG000002 | 14.7919595 | -0.495891849 | 0.31430025 | -1.5777647 | 0.11461967 | 0.21882967 | SCART1 | 619207 |
| ENSG000001 | 86.4897304 | -0.286142589 | 0.18136757 | -1.5776944 | 0.11463584 | 0.21885024 | KCNAB1 | 7881 |
| ENSG000002 | 2.30916801 | -0.581547369 | 0.36864398 | -1.5775311 | 0.11467336 | 0.21890126 | NA | NA |
| ENSG000002 | 2.60973391 | -0.587172181 | 0.37220492 | -1.5775508 | 0.11466884 | 0.21890126 | NA | NA |
| ENSG000001 | 9.78052616 | -0.531861082 | 0.3371617 | -1.5774659 | 0.11468836 | 0.21891959 | RGR | 5995 |
| ENSG00000C | 8.41240875 | -0.545750899 | 0.3459825 | -1.5773945 | 0.11470478 | 0.21892781 | ZBTB32 | 27033 |
| ENSG000002 | 10.0031455 | -0.543930291 | 0.34483007 | -1.5773865 | 0.11470662 | 0.21892781 | NA | NA |
| ENSG000002 | 2.66246102 | -0.587111661 | 0.37220763 | -1.5773767 | 0.11470886 | 0.21892781 | NA | NA |
| ENSG000002 | 2.84997776 | -0.593554534 | 0.3763231 | -1.5772471 | 0.11473867 | 0.21897439 | NA | NA |
| ENSG000002 | 14.5696501 | -0.486739638 | 0.30864504 | -1.5770208 | 0.11479075 | 0.21906347 | NA | NA |
| ENSG000002 | 3.47513724 | -0.59691442 | 0.37851691 | -1.5769822 | 0.11479962 | 0.21907009 | NA | NA |
| ENSG000002 | 5.53969931 | -0.587491503 | 0.37257697 | -1.5768326 | 0.11483405 | 0.21912549 | NA | NA |
| ENSG000002 | 5.73115801 | -0.580849888 | 0.36839376 | -1.5767093 | 0.11486242 | 0.2191693 | NA | NA |
| ENSG000002 | 4.39223568 | -0.594834858 | 0.37731705 | -1.5764855 | 0.11491396 | 0.21924735 | LOC101927884 | 101927884 |
| ENSG000002 | 7.55956244 | -0.568961508 | 0.36091056 | -1.5764612 | 0.11491954 | 0.21924735 | NA | NA |
| ENSG000002 | 4.39117751 | -0.595503788 | 0.37774549 | -1.5764683 | 0.11491792 | 0.21924735 | NA | NA |

| | | | | | | | | |
|---|---|---|---|---|---|---|---|---|
| ENSG000002 | 5.76551268 | -0.58124334 | 0.36873929 | -1.5762989 | 0.11495693 | 0.21930013 | NA | NA |
| ENSG000002 | 4.97866167 | -0.59266969 | 0.37598927 | -1.5762942 | 0.11495802 | 0.21930013 | NA | NA |
| ENSG000001 | 12.9789026 | -0.513996447 | 0.32609355 | -1.5762239 | 0.11497422 | 0.2193207 | TBC1D10C | 374403 |
| ENSG000002 | 4.75064383 | -0.594715209 | 0.3773249 | -1.5761356 | 0.11499456 | 0.21934918 | NA | NA |
| ENSG000002 | 7.43988756 | -0.563378413 | 0.35747944 | -1.5759743 | 0.11503172 | 0.21940975 | NA | NA |
| ENSG000002 | 7.71070034 | -0.563005114 | 0.35726469 | -1.5758767 | 0.11505422 | 0.21944234 | NA | NA |
| ENSG000002 | 5.71052939 | -0.580012473 | 0.36808719 | -1.5757475 | 0.115084 | 0.21947449 | RN7SKP200 | 106480895 |
| ENSG000002 | 9.08973675 | -0.550241667 | 0.34919724 | -1.5757332 | 0.11508731 | 0.21947449 | LOC101929282 | 101929282 |
| ENSG000002 | 4.87091245 | -0.595767286 | 0.37807944 | -1.5757728 | 0.11507818 | 0.21947449 | NA | NA |
| ENSG000002 | 4.15968337 | -0.596478672 | 0.37854702 | -1.5757056 | 0.11509366 | 0.21947628 | NA | NA |
| ENSG000001 | 22.4518199 | -0.43331829 | 0.27500901 | -1.5756512 | 0.1151062 | 0.21948457 | Mar-04 | 57574 |
| ENSG000002 | 2.90075437 | -0.592429497 | 0.37599857 | -1.5756163 | 0.11511425 | 0.21948457 | NA | NA |
| ENSG000002 | 7.93778116 | -0.553607155 | 0.35135481 | -1.5756356 | 0.1151098 | 0.21948457 | NA | NA |
| ENSG000002 | 5.31968654 | -0.590144923 | 0.37456422 | -1.5755507 | 0.11512939 | 0.21950311 | NA | NA |
| ENSG000002 | 7.6579215 | -0.56148669 | 0.3563827 | -1.5755161 | 0.11513736 | 0.21950799 | DKFZP434A062 | 26102 |
| ENSG000002 | 7.81171854 | -0.562579164 | 0.35710346 | -1.5753954 | 0.11516519 | 0.21955074 | NA | NA |
| ENSG000002 | 3.52087005 | -0.596195507 | 0.378472 | -1.5752698 | 0.11519419 | 0.21958656 | LPP-AS1 | 100873917 |
| ENSG000002 | 4.38269649 | -0.594713367 | 0.37753177 | -1.5752671 | 0.11519481 | 0.21958656 | NA | NA |
| ENSG000002 | 6.86917527 | -0.568706398 | 0.36102904 | -1.5752373 | 0.11520168 | 0.21958933 | KCNQ5-AS1 | 100873997 |
| ENSG000002 | 2.29035817 | -0.579338239 | 0.36781459 | -1.5750823 | 0.11523746 | 0.2196472 | RNU6-700P | 106480398 |
| ENSG000002 | 8.4927451 | -0.555498399 | 0.35270742 | -1.5749552 | 0.11526678 | 0.21969276 | NA | NA |
| ENSG000001 | 23.0471345 | 0.441181965 | 0.28013342 | 1.57489946 | 0.11527965 | 0.21970697 | RAB3IL1 | 5866 |
| ENSG000002 | 2.96823469 | -0.589764499 | 0.37450781 | -1.5747722 | 0.11530903 | 0.21975262 | NA | NA |
| ENSG000002 | 6.84131954 | -0.566289856 | 0.35962641 | -1.5746615 | 0.11533461 | 0.21979104 | NA | NA |
| ENSG000002 | 12.1169599 | -0.520469438 | 0.33055098 | -1.5745512 | 0.11536007 | 0.21982924 | LOC339298 | 339298 |
| ENSG000002 | 2.8977662 | -0.592408013 | 0.37626152 | -1.5744581 | 0.11538158 | 0.21985989 | NA | NA |
| ENSG000002 | 3.79234423 | -0.595907115 | 0.37851469 | -1.5743302 | 0.11541112 | 0.21990586 | NA | NA |
| ENSG000001 | 5.5895199 | -0.588826965 | 0.37404427 | -1.5742173 | 0.11543722 | 0.21993492 | SMCO3 | 440087 |
| ENSG000002 | 2.26587496 | -0.581452982 | 0.36935584 | -1.5742353 | 0.11543307 | 0.21993492 | NA | NA |

| | | | | | | | | |
|---|---|---|---|---|---|---|---|---|
| ENSG000002 | 3.50784178 | -0.595649465 | 0.37839556 | -1.574145 | 0.11545394 | 0.21994611 | NA | NA |
| ENSG000002 | 4.39691202 | -0.594509648 | 0.37766947 | -1.5741533 | 0.11545201 | 0.21994611 | NA | NA |
| ENSG000001 | 116.736763 | 0.245778555 | 0.15613812 | 1.57410984 | 0.11546206 | 0.21994611 | SLC7A7 | 9056 |
| ENSG000002 | 4.37387022 | -0.594185529 | 0.37747683 | -1.574098 | 0.11546479 | 0.21994611 | NA | NA |
| ENSG000001 | 19.1124355 | -0.458732743 | 0.29145723 | -1.5739282 | 0.11550405 | 0.22001056 | GCSAM | 257144 |
| ENSG000001 | 91.6402518 | 0.275759469 | 0.17520899 | 1.5738888 | 0.11551316 | 0.22001757 | LIPT1 | 51601 |
| ENSG000001 | 8.60854895 | -0.555035965 | 0.35269145 | -1.5737154 | 0.11555326 | 0.22008362 | NA | NA |
| ENSG000001 | 2793.91319 | 0.211180968 | 0.13419517 | 1.57368529 | 0.11556022 | 0.22008654 | SNRPF | 6636 |
| ENSG000001 | 24.0808695 | 0.445426365 | 0.28306315 | 1.57359361 | 0.11558143 | 0.22011089 | NACAD | 23148 |
| ENSG000001 | 186.801101 | 0.235334199 | 0.14955452 | 1.57356798 | 0.11558736 | 0.22011089 | ZNF521 | 25925 |
| ENSG000002 | 3.56407879 | -0.59523221 | 0.37827115 | -1.5735596 | 0.11558929 | 0.22011089 | NA | NA |
| ENSG000002 | 2.26590721 | -0.580458935 | 0.36889049 | -1.5735264 | 0.11559697 | 0.22011517 | NA | NA |
| ENSG000001 | 3.27483245 | -0.592678345 | 0.37666464 | -1.5734908 | 0.11560522 | 0.22012054 | S1PR4 | 8698 |
| ENSG00000C | 7.7407779 | -0.575949732 | 0.36607575 | -1.5733075 | 0.11564763 | 0.22019096 | SMPX | 23676 |
| ENSG000002 | 4.09625814 | -0.59518691 | 0.37831492 | -1.5732578 | 0.11565913 | 0.22020252 | NA | NA |
| ENSG000002 | 3.23366616 | -0.592830471 | 0.37683283 | -1.5731922 | 0.11567432 | 0.22022109 | NA | NA |
| ENSG000002 | 7.08739367 | -0.575802341 | 0.36601505 | -1.5731657 | 0.11568045 | 0.22022242 | NA | NA |
| ENSG000001 | 45.7915157 | 0.349663668 | 0.2222759 | 1.57310655 | 0.11569415 | 0.22023817 | LINGO2 | 158038 |
| ENSG000002 | 3.55562967 | -0.595015489 | 0.3783607 | -1.5726144 | 0.11580813 | 0.2204448 | NA | NA |
| ENSG000002 | 3.222485 | -0.59326839 | 0.37725761 | -1.5725816 | 0.11581572 | 0.2204489 | GOLGA8O | 728047 |
| ENSG000002 | 3.89826215 | -0.594843164 | 0.37835465 | -1.5721841 | 0.11590786 | 0.22061392 | NA | NA |
| ENSG000001 | 13.866417 | -0.510753176 | 0.32488859 | -1.5720871 | 0.11593034 | 0.22064636 | SLC22A25 | 387601 |
| ENSG000002 | 9.23432893 | -0.53736274 | 0.34182573 | -1.5720372 | 0.11594194 | 0.22064771 | ISPD-AS1 | 100506025 |
| ENSG000002 | 2.88825479 | -0.591568106 | 0.37630574 | -1.5720411 | 0.11594101 | 0.22064771 | HAVCR1P1 | 100101266 |
| ENSG000002 | 6.7267544 | -0.579791249 | 0.36882693 | -1.5719873 | 0.1159535 | 0.22065936 | NA | NA |
| ENSG000002 | 2.47318911 | -0.570992765 | 0.36323851 | -1.57195 | 0.11596215 | 0.22066548 | NA | NA |
| ENSG000002 | 2.90963217 | -0.589734393 | 0.37517958 | -1.5718723 | 0.11598017 | 0.2206894 | NA | NA |
| ENSG000001 | 2.87764612 | -0.591227041 | 0.37613823 | -1.5718345 | 0.11598894 | 0.22069574 | GJA8 | 2703 |
| ENSG000002 | 2.91358732 | -0.59139795 | 0.37627289 | -1.5717262 | 0.11601407 | 0.22072386 | NA | NA |

| | | | | | | | | |
|---|---|---|---|---|---|---|---|---|
| ENSG000002 | 2.2583914 | -0.579751808 | 0.36886365 | -1.5717239 | 0.11601461 | 0.22072386 | NA | NA |
| ENSG000002 | 21.942062 | 0.463596502 | 0.29497274 | 1.57165882 | 0.1160297 | 0.22074221 | NA | NA |
| ENSG000001 | 30.1757088 | -0.387743234 | 0.24672372 | -1.5715685 | 0.11605065 | 0.22077172 | RALYL | 138046 |
| ENSG000002 | 3.85650356 | -0.594858226 | 0.37851944 | -1.5715394 | 0.1160574 | 0.2207742 | LOC101927143 | 101927143 |
| ENSG000001 | 13.0549472 | -0.503099704 | 0.32015404 | -1.5714301 | 0.11608278 | 0.2207914 | C11orf88 | 399949 |
| ENSG000002 | 35.4572707 | 0.372032003 | 0.23674422 | 1.57145123 | 0.11607787 | 0.2207914 | NA | NA |
| ENSG000002 | 2.86354737 | -0.590992917 | 0.37607528 | -1.571475 | 0.11607235 | 0.2207914 | NA | NA |
| ENSG000002 | 6.14455113 | -0.581172947 | 0.36987923 | -1.5712506 | 0.11612446 | 0.22086032 | NA | NA |
| ENSG000001 | 22.9406167 | -0.416416033 | 0.26504134 | -1.5711361 | 0.11615103 | 0.22090049 | PLAC8 | 51316 |
| ENSG000002 | 2.23776278 | -0.580111485 | 0.36925022 | -1.5710525 | 0.11617045 | 0.22090644 | NA | NA |
| ENSG000002 | 2.65740502 | -0.583804924 | 0.37159884 | -1.5710623 | 0.11616818 | 0.22090644 | NA | NA |
| ENSG000002 | 24.0981206 | -0.437039804 | 0.27818285 | -1.5710523 | 0.1161705 | 0.22090644 | NA | NA |
| ENSG000000 | 2.2777222 | -0.579605336 | 0.36893943 | -1.571004 | 0.11618171 | 0.2209174 | MLN | 4295 |
| ENSG000001 | 267.836659 | 0.231912693 | 0.14762898 | 1.5709158 | 0.11620221 | 0.2209253 | CNNM2 | 54805 |
| ENSG000001 | 67.3112862 | 0.291283535 | 0.18541725 | 1.5709624 | 0.11619138 | 0.2209253 | PLA2G6 | 8398 |
| ENSG000002 | 3.57757562 | -0.593692282 | 0.37792663 | -1.5709194 | 0.11620137 | 0.2209253 | NA | NA |
| ENSG000002 | 11.2496635 | -0.527522806 | 0.33581395 | -1.5708782 | 0.11621095 | 0.22093156 | NA | NA |
| ENSG000001 | 162.105705 | 0.231881649 | 0.1476208 | 1.57079256 | 0.11623084 | 0.22095902 | LRRC2 | 79442 |
| ENSG000002 | 142.224252 | 0.246802408 | 0.15712447 | 1.57074456 | 0.116242 | 0.22096986 | TMPO-AS1 | 100128191 |
| ENSG000001 | 31.26865 | -0.419334096 | 0.2669772 | -1.5706738 | 0.11625843 | 0.22099074 | TEKT3 | 64518 |
| ENSG000002 | 5.27685051 | -0.588723975 | 0.37484548 | -1.5705778 | 0.11628076 | 0.22101661 | NA | NA |
| ENSG000002 | 6.80260532 | -0.565395662 | 0.3599943 | -1.5705684 | 0.11628294 | 0.22101661 | LOC101928052 | 101928052 |
| ENSG000002 | 46.2180311 | 0.333666846 | 0.21245365 | 1.5705395 | 0.11628965 | 0.22101902 | NA | NA |
| ENSG000002 | 2.34690418 | -0.571097274 | 0.36365653 | -1.5704304 | 0.11631502 | 0.22105686 | NA | NA |
| ENSG000002 | 4.43461525 | -0.593042549 | 0.37767576 | -1.5702425 | 0.11635869 | 0.22112714 | NA | NA |
| ENSG000002 | 8.20563443 | -0.547595533 | 0.34873711 | -1.5702244 | 0.1163629 | 0.22112714 | NA | NA |
| ENSG000001 | 7.67011669 | -0.556725152 | 0.35457897 | -1.570102 | 0.11639138 | 0.22117089 | POM121L2 | 94026 |
| ENSG000001 | 62.7849164 | 0.315803816 | 0.20114116 | 1.57006063 | 0.11640101 | 0.22117882 | CABLES2 | 81928 |
| ENSG000001 | 2.92314971 | -0.58874579 | 0.37499384 | -1.5700146 | 0.11641172 | 0.22118882 | OR6C70 | 390327 |

| | | | | | | | | |
|---|---|---|---|---|---|---|---|---|
| ENSG00000C | 78.2036058 | -0.284189418 | 0.18101835 | -1.5699481 | 0.11642718 | 0.22119747 | P4HA2 | 8974 |
| ENSG000002 | 4.61960747 | -0.589893731 | 0.37573577 | -1.5699696 | 0.11642219 | 0.22119747 | NA | NA |
| ENSG000001 | 13.2630982 | -0.497880977 | 0.31714173 | -1.5699006 | 0.11643825 | 0.22120812 | C12orf54 | 121273 |
| ENSG000002 | 2.90299792 | -0.589868532 | 0.37575137 | -1.5698373 | 0.11645296 | 0.22122071 | NA | NA |
| ENSG000002 | 12.4914737 | -0.52768912 | 0.33614515 | -1.5698252 | 0.11645579 | 0.22122071 | NA | NA |
| ENSG00000C | 76.7084959 | 0.277584886 | 0.17683115 | 1.5697737 | 0.11646777 | 0.22122275 | ACOX3 | 8310 |
| ENSG000001 | 19.2697715 | 0.444539674 | 0.28318632 | 1.56977805 | 0.11646676 | 0.22122275 | CFHR3 | 10878 |
| ENSG000002 | 27.0349428 | -0.404278984 | 0.25755871 | -1.5696576 | 0.1164948 | 0.22125336 | COLQ | 8292 |
| ENSG000002 | 5.521622 | -0.583987892 | 0.37204593 | -1.5696661 | 0.11649281 | 0.22125336 | RN7SL541P | 106481069 |
| ENSG000002 | 11.0876421 | 0.526796103 | 0.33564955 | 1.5694825 | 0.11653555 | 0.2213204 | TEX22 | 647310 |
| ENSG000001 | 28.5398856 | -0.38708716 | 0.24664001 | -1.5694419 | 0.11654501 | 0.221328 | FAM186B | 84070 |
| ENSG000002 | 4.60139955 | -0.588972519 | 0.3753079 | -1.5693049 | 0.11657691 | 0.22137821 | NA | NA |
| ENSG000002 | 2.2741927 | -0.578275083 | 0.36850249 | -1.5692569 | 0.11658809 | 0.22138906 | NA | NA |
| ENSG000002 | 3.55559742 | -0.593751957 | 0.3783883 | -1.5691604 | 0.11661056 | 0.22142137 | NA | NA |
| ENSG000001 | 14.5461953 | -0.489010518 | 0.31164542 | -1.5691247 | 0.1166189 | 0.22142683 | KRT78 | 196374 |
| ENSG000002 | 11.2047205 | 0.518438028 | 0.33046707 | 1.56880391 | 0.11669364 | 0.22155837 | RNASEK | 440400 |
| ENSG000001 | 62.3254711 | -0.310491416 | 0.19792365 | -1.5687434 | 0.11670774 | 0.22157477 | LOC400499 | 400499 |
| ENSG000002 | 157.101726 | -0.268390386 | 0.17110019 | -1.5686153 | 0.1167376 | 0.22162108 | NA | NA |
| ENSG000002 | 3.82809854 | -0.593793081 | 0.37857189 | -1.5685081 | 0.1167626 | 0.22165816 | NA | NA |
| ENSG000002 | 7.20922039 | -0.566475539 | 0.36117237 | -1.5684354 | 0.11677955 | 0.22167997 | NA | NA |
| ENSG000001 | 1852.53753 | 0.1901865 | 0.12126709 | 1.56832739 | 0.11680475 | 0.22169978 | RC3H1 | 149041 |
| ENSG000002 | 9.56600816 | -0.539678813 | 0.34411261 | -1.5683203 | 0.11680639 | 0.22169978 | NA | NA |
| ENSG000002 | 13.8459772 | 0.492841747 | 0.31424228 | 1.56834956 | 0.11679957 | 0.22169978 | NA | NA |
| ENSG000002 | 5.45246086 | -0.582379877 | 0.37135933 | -1.5682382 | 0.11682556 | 0.22172579 | NA | NA |
| ENSG000002 | 3.46740573 | -0.593453374 | 0.37843619 | -1.5681729 | 0.11684079 | 0.22174432 | NA | NA |
| ENSG000002 | 6.65454688 | -0.57113112 | 0.36421954 | -1.5680958 | 0.11685878 | 0.22176808 | NA | NA |
| ENSG000002 | 3.44642424 | -0.593399463 | 0.37843724 | -1.5680261 | 0.11687505 | 0.22177819 | NA | NA |
| ENSG000002 | 3.50022183 | -0.593278003 | 0.37835646 | -1.5680398 | 0.11687184 | 0.22177819 | NA | NA |
| ENSG000002 | 2.34111207 | -0.573121176 | 0.36554142 | -1.5678693 | 0.11691164 | 0.22183724 | LOC101929207 | 101929207 |

| | | | | | | | | |
|---|---|---|---|---|---|---|---|---|
| ENSG000001 | 43.3988099 | -0.340172835 | 0.21697388 | -1.5678055 | 0.11692654 | 0.22185514 | CCDC67 | 159989 |
| ENSG000001 | 18.3914189 | -0.451660985 | 0.28810106 | -1.5677172 | 0.11694715 | 0.22188387 | RGN | 9104 |
| ENSG000002 | 4.59402754 | -0.588340084 | 0.37532554 | -1.5675461 | 0.1169871 | 0.22194193 | NA | NA |
| ENSG000002 | 3.17692083 | -0.590948846 | 0.37699142 | -1.5675392 | 0.1169887 | 0.22194193 | NA | NA |
| ENSG000002 | 7.41664292 | -0.561102278 | 0.35798361 | -1.5673965 | 0.11702203 | 0.22199478 | NA | NA |
| ENSG000001 | 4.09812148 | -0.593316161 | 0.37858224 | -1.5672055 | 0.11706667 | 0.22206907 | COX4I2 | 84701 |
| ENSG000001 | 49.9364525 | 0.330220986 | 0.21074737 | 1.56690442 | 0.11713703 | 0.22218397 | AQP3 | 360 |
| ENSG000001 | 38.1959993 | -0.375175622 | 0.23944181 | -1.566876 | 0.11714368 | 0.22218397 | LAMB3 | 3914 |
| ENSG000002 | 2.86791878 | 0.587613882 | 0.37501713 | 1.56689877 | 0.11713835 | 0.22218397 | NA | NA |
| ENSG000001 | 103.069849 | 0.286925231 | 0.18313952 | 1.56670302 | 0.11718413 | 0.22224679 | NOD1 | 10392 |
| ENSG000002 | 2.41894248 | -0.567665733 | 0.36233502 | -1.5666875 | 0.11718776 | 0.22224679 | LOC105373051 | 105373051 |
| ENSG000002 | 3.72093899 | -0.592762872 | 0.37845808 | -1.5662577 | 0.11728831 | 0.22242707 | RNASE4 | 6038 |
| ENSG000002 | 18.338373 | -0.46288392 | 0.29554268 | -1.5662169 | 0.11729786 | 0.22243478 | KRT42P | 284116 |
| ENSG000002 | 3.22288088 | -0.589270861 | 0.37624934 | -1.5661711 | 0.11730856 | 0.22244467 | NA | NA |
| ENSG000002 | 4.61807411 | -0.58827113 | 0.37562088 | -1.56613 | 0.11731818 | 0.22245252 | NA | NA |
| ENSG000002 | 13.9807434 | 0.492523946 | 0.3145024 | 1.56604192 | 0.1173388 | 0.2224645 | NA | NA |
| ENSG000002 | 3.44735733 | -0.592627946 | 0.37842628 | -1.5660327 | 0.11734096 | 0.2224645 | NA | NA |
| ENSG000002 | 7.37137104 | -0.562475517 | 0.35916569 | -1.5660614 | 0.11733424 | 0.2224645 | KCNJ2-AS1 | 400617 |
| ENSG000002 | 7.89033742 | -0.552808768 | 0.35301627 | -1.5659583 | 0.11735838 | 0.22247672 | NA | NA |
| ENSG000002 | 5.20730503 | -0.585324936 | 0.37377661 | -1.5659753 | 0.1173544 | 0.22247672 | NA | NA |
| ENSG000002 | 2.29496512 | -0.575962225 | 0.36784421 | -1.5657776 | 0.11740069 | 0.22253613 | NA | NA |
| ENSG000002 | 7.20092967 | -0.564425442 | 0.36047146 | -1.5657979 | 0.11739595 | 0.22253613 | NA | NA |
| ENSG000002 | 4.07528187 | -0.592452228 | 0.37838149 | -1.5657537 | 0.11740629 | 0.22253633 | NA | NA |
| ENSG000002 | 6.561827 | -0.570329943 | 0.36427758 | -1.5656466 | 0.11743138 | 0.22257349 | NA | NA |
| ENSG000002 | 2.58621549 | -0.581923274 | 0.37174213 | -1.565395 | 0.11749032 | 0.22267479 | NA | NA |
| ENSG000002 | 6.69027205 | -0.571584523 | 0.3651563 | -1.5653147 | 0.11750913 | 0.22268964 | NA | NA |
| ENSG000002 | 2.85538608 | -0.588429093 | 0.37591447 | -1.5653271 | 0.11750624 | 0.22268964 | NA | NA |
| ENSG000002 | 3.1328811 | -0.580678086 | 0.37097514 | -1.5652749 | 0.11751846 | 0.2226969 | NA | NA |
| ENSG000001 | 5.72518908 | -0.576785322 | 0.36851017 | -1.5651815 | 0.11754035 | 0.22272343 | HOXD8 | 3234 |

| | | | | | | | | |
|---|---|---|---|---|---|---|---|---|
| ENSG000002 | 43.0458682 | -0.335231322 | 0.21418228 | -1.5651684 | 0.11754344 | 0.22272343 | NA | NA |
| ENSG000002 | 2.64912009 | -0.581588231 | 0.37159211 | -1.5651254 | 0.11755352 | 0.22273211 | NA | NA |
| ENSG000001 | 38.3036441 | 0.352385787 | 0.22516672 | 1.56499944 | 0.11758304 | 0.22277764 | CAPN12 | 147968 |
| ENSG000002 | 5.98943067 | 0.57360852 | 0.36654249 | 1.56491683 | 0.11760241 | 0.22280393 | SNORA5A | 654319 |
| ENSG000001 | 14.2850502 | -0.479745937 | 0.30656956 | -1.5648845 | 0.11761 | 0.22280791 | SPG20-AS1 | 100507135 |
| ENSG000001 | 7.76102212 | -0.556710414 | 0.3557808 | -1.5647568 | 0.11763995 | 0.22285423 | FABP1 | 2168 |
| ENSG000000 | 50.8251035 | -0.332293788 | 0.21237911 | -1.5646256 | 0.11767072 | 0.22290211 | MSH4 | 4438 |
| ENSG000002 | 7.99088598 | -0.553261928 | 0.35363655 | -1.564493 | 0.11770183 | 0.22295061 | LINC01594 | 102724774 |
| ENSG000002 | 11.7740081 | -0.512377433 | 0.32751142 | -1.5644567 | 0.11771035 | 0.22295635 | NA | NA |
| ENSG000002 | 4.64619893 | -0.588168019 | 0.37596841 | -1.5644081 | 0.11772176 | 0.22296755 | NA | NA |
| ENSG000002 | 3.87559359 | -0.592178951 | 0.37855636 | -1.5643085 | 0.11774513 | 0.22300092 | NA | NA |
| ENSG000002 | 5.85751141 | -0.582054154 | 0.37208931 | -1.5642862 | 0.11775038 | 0.22300092 | NA | NA |
| ENSG000001 | 455.646241 | 0.435049426 | 0.27812964 | 1.56419656 | 0.11777142 | 0.22303034 | SKI | 6497 |
| ENSG000002 | 2.98860162 | -0.584134641 | 0.37344958 | -1.5641593 | 0.11778015 | 0.22303647 | LOC101927694 | 101927694 |
| ENSG000001 | 24.9822325 | -0.480566239 | 0.30725761 | -1.5640499 | 0.11780585 | 0.22307471 | AATK | 9625 |
| ENSG000002 | 10.3295567 | -0.529019185 | 0.33824485 | -1.5640126 | 0.11781462 | 0.22308091 | LOC101929084 | 101929084 |
| ENSG000001 | 12.8085067 | -0.502112264 | 0.32107228 | -1.5638605 | 0.11785033 | 0.22313809 | LILRB5 | 10990 |
| ENSG000002 | 4.09601019 | -0.591726007 | 0.37839278 | -1.5637878 | 0.11786741 | 0.22316002 | NA | NA |
| ENSG000002 | 6.86513276 | -0.563740943 | 0.36052742 | -1.5636562 | 0.11789832 | 0.22320813 | NA | NA |
| ENSG000000 | 4.6384939 | -0.58830167 | 0.37631132 | -1.5633377 | 0.11797319 | 0.22333388 | PLA2G10 | 8399 |
| ENSG000001 | 40.6876532 | 0.3638428 | 0.23273625 | 1.56332674 | 0.11797576 | 0.22333388 | ZNF284 | 342909 |
| ENSG000002 | 2.292694 | -0.573848023 | 0.36709609 | -1.5632093 | 0.11800337 | 0.22334177 | NA | NA |
| ENSG000002 | 6.62874637 | -0.57360565 | 0.36693533 | -1.5632337 | 0.11799765 | 0.22334177 | NA | NA |
| ENSG000002 | 7.658288 | -0.567868205 | 0.36327479 | -1.5631919 | 0.11800747 | 0.22334177 | MTND6P17 | 106480728 |
| ENSG000002 | 2.28104319 | -0.576118172 | 0.36853542 | -1.563264 | 0.1179905 | 0.22334177 | RNU6-572P | 106481362 |
| ENSG000002 | 7.07437504 | 0.573879803 | 0.36710913 | 1.56324034 | 0.11799608 | 0.22334177 | NA | NA |
| ENSG000002 | 11.9792119 | -0.507859747 | 0.32490106 | -1.5631212 | 0.11802408 | 0.2233628 | NA | NA |
| ENSG000000 | 10.7442877 | -0.522802891 | 0.33449812 | -1.5629472 | 0.11806502 | 0.22342983 | C1QTNF3 | 114899 |
| ENSG000001 | 41.9202082 | 0.359186283 | 0.22982678 | 1.56285653 | 0.11808634 | 0.22345977 | RLN2 | 6019 |

| | | | | | | | | |
|---|---|---|---|---|---|---|---|---|
| ENSG000002 | 16.9839308 | 0.464739984 | 0.29737879 | 1.56278793 | 0.11810248 | 0.22347988 | NA | NA |
| ENSG000002 | 8.02692756 | -0.557854791 | 0.35701821 | -1.5625387 | 0.11816113 | 0.22358041 | NA | NA |
| ENSG000002 | 2.68383427 | -0.579358086 | 0.37079166 | -1.5624895 | 0.11817271 | 0.22359191 | NA | NA |
| ENSG000001 | 2.66841013 | -0.577426185 | 0.36963018 | -1.5621727 | 0.1182473 | 0.22372259 | RNY1P16 | 106480312 |
| ENSG000002 | 3.30155563 | -0.588351646 | 0.37663552 | -1.5621247 | 0.11825862 | 0.22372475 | NA | NA |
| ENSG000002 | 5.7307338 | -0.5765432 | 0.36907716 | -1.562121 | 0.11825948 | 0.22372475 | NA | NA |
| ENSG000002 | 19.0875066 | -0.457664903 | 0.29302505 | -1.5618627 | 0.11832033 | 0.22381898 | NDUFA6-AS1 | 100132273 |
| ENSG000002 | 5.48675844 | -0.580571112 | 0.37171561 | -1.5618691 | 0.11831884 | 0.22381898 | NA | NA |
| ENSG000002 | 10.3711255 | -0.532793951 | 0.34114124 | -1.5617987 | 0.11833541 | 0.22383708 | LOC101929710 | 101929710 |
| ENSG000001 | 2.90294585 | -0.586424832 | 0.37550437 | -1.5616991 | 0.1183589 | 0.22387106 | OR4F21 | 441308 |
| ENSG000002 | 3.09341892 | -0.578948538 | 0.37077107 | -1.5614717 | 0.11841249 | 0.22396198 | NA | NA |
| ENSG000001 | 85.492641 | 0.277683654 | 0.17784095 | 1.56141572 | 0.11842569 | 0.22397651 | PAPLN | 89932 |
| ENSG000002 | 14.2528667 | -0.500109409 | 0.32031683 | -1.5612961 | 0.11845391 | 0.22400898 | NA | NA |
| ENSG000002 | 4.61535284 | -0.587644588 | 0.37638009 | -1.5613063 | 0.11845151 | 0.22400898 | OR52B2 | 255725 |
| ENSG000002 | 2.62125585 | -0.579960345 | 0.37147162 | -1.5612508 | 0.1184646 | 0.22400962 | NA | NA |
| ENSG000002 | 4.73230455 | -0.589765227 | 0.37775248 | -1.5612478 | 0.1184653 | 0.22400962 | NA | NA |
| ENSG000001 | 310.264773 | 0.248185439 | 0.15899042 | 1.56100881 | 0.11852167 | 0.22410039 | FAM131B | 9715 |
| ENSG000002 | 2.88756965 | -0.586928865 | 0.37599604 | -1.5609975 | 0.11852435 | 0.22410039 | NA | NA |
| ENSG000002 | 7.19462409 | -0.56613142 | 0.36271757 | -1.5608051 | 0.11856976 | 0.22417578 | COX6CP10 | 106481681 |
| ENSG000002 | 2.26595928 | -0.574708465 | 0.36821854 | -1.5607809 | 0.11857545 | 0.2241761 | LOC105373525 | 105373525 |
| ENSG000002 | 3.5916829 | -0.58853831 | 0.37709157 | -1.5607305 | 0.11858736 | 0.2241777 | NA | NA |
| ENSG000002 | 2.88235665 | -0.586836553 | 0.37600048 | -1.5607335 | 0.11858664 | 0.2241777 | NA | NA |
| ENSG000002 | 2.92895142 | -0.586559292 | 0.37583505 | -1.5606828 | 0.11859862 | 0.22418069 | NA | NA |
| ENSG000002 | 5.39088094 | -0.587349002 | 0.37634246 | -1.560677 | 0.11859999 | 0.22418069 | NA | NA |
| ENSG000001 | 6.87560896 | -0.563991157 | 0.36140057 | -1.5605707 | 0.11862507 | 0.22421098 | FOXA2 | 3170 |
| ENSG000001 | 126.515346 | 0.250430763 | 0.1604747 | 1.56056223 | 0.11862708 | 0.22421098 | FRRS1 | 391059 |
| ENSG000001 | 6.99339534 | -0.570870118 | 0.36582937 | -1.5604819 | 0.11864604 | 0.22423638 | NTN5 | 126147 |
| ENSG000002 | 5.62119383 | -0.583958816 | 0.37426837 | -1.5602676 | 0.11869666 | 0.22432158 | SPINK8 | 646424 |
| ENSG000002 | 2.90000974 | -0.586283207 | 0.37578989 | -1.5601356 | 0.11872783 | 0.22435958 | RNU1-36P | 106481783 |

| | | | | | | | | |
|---|---|---|---|---|---|---|---|---|
| ENSG000002 | 6.89157443 | -0.563593881 | 0.36124605 | -1.5601385 | 0.11872715 | 0.22435958 | NA | NA |
| ENSG000002 | 9.74337865 | -0.528438174 | 0.3387333 | -1.560042 | 0.11874995 | 0.22438047 | NA | NA |
| ENSG000002 | 5.25866906 | -0.584278085 | 0.37452234 | -1.560062 | 0.11874522 | 0.22438047 | NA | NA |
| ENSG000001 | 10.0541599 | -0.524892454 | 0.3364761 | -1.5599695 | 0.11876709 | 0.22439195 | FCER2 | 2208 |
| ENSG000002 | 47.3387591 | 0.352233124 | 0.22579221 | 1.55998795 | 0.11876273 | 0.22439195 | NA | NA |
| ENSG000002 | 5.71622313 | -0.576200013 | 0.36939607 | -1.5598434 | 0.1187969 | 0.22443781 | CCL14 | 6358 |
| ENSG000001 | 216.185719 | 0.224046524 | 0.14364447 | 1.5597295 | 0.11882382 | 0.2244782 | ZNF701 | 55762 |
| ENSG000002 | 8.23877746 | 0.555927198 | 0.35645113 | 1.55961688 | 0.11885045 | 0.22451805 | NA | NA |
| ENSG000001 | 164.717768 | -0.22751682 | 0.14589094 | -1.5594993 | 0.11887825 | 0.22455849 | UNC79 | 57578 |
| ENSG000002 | 2.30161974 | -0.573696824 | 0.36787711 | -1.5594795 | 0.11888293 | 0.22455849 | NA | NA |
| ENSG000002 | 7.75005333 | -0.564607603 | 0.36208473 | -1.5593245 | 0.11891959 | 0.22461728 | NA | NA |
| ENSG000002 | 23.2852215 | 0.4166008 | 0.26717282 | 1.55929335 | 0.11892696 | 0.22462074 | C2orf72 | 257407 |
| ENSG000002 | 2.93019662 | -0.585380086 | 0.375432 | -1.5592174 | 0.11894494 | 0.22464424 | RN7SL269P | 106479320 |
| ENSG000001 | 2781.7302 | 0.214806097 | 0.13776826 | 1.55918417 | 0.1189528 | 0.22464861 | ZNF483 | 158399 |
| ENSG000002 | 8.16944418 | -0.546880046 | 0.3507914 | -1.5589893 | 0.1189989 | 0.22472521 | NA | NA |
| ENSG000002 | 5.73070209 | -0.574121984 | 0.36830183 | -1.5588355 | 0.11903531 | 0.2247835 | NA | NA |
| ENSG000002 | 2.69086549 | -0.575826062 | 0.36942653 | -1.5587025 | 0.1190668 | 0.2248325 | NA | NA |
| ENSG000001 | 20.2567532 | -0.442237401 | 0.28372911 | -1.5586607 | 0.11907672 | 0.22484058 | DSG4 | 147409 |
| ENSG000002 | 2.67670358 | -0.577777842 | 0.37069971 | -1.5586142 | 0.11908772 | 0.22484058 | NA | NA |
| ENSG000002 | 25.7900685 | -0.399454525 | 0.25628783 | -1.5586168 | 0.1190871 | 0.22484058 | NA | NA |
| ENSG000002 | 8.28899791 | -0.548363449 | 0.35185192 | -1.5585063 | 0.11911326 | 0.22487834 | NA | NA |
| ENSG000002 | 6.55032576 | -0.565754504 | 0.36303072 | -1.5584205 | 0.11913361 | 0.22490628 | NA | NA |
| ENSG000001 | 4.67483153 | -0.587155061 | 0.37677454 | -1.5583725 | 0.11914498 | 0.22490681 | WBSCR28 | 135886 |
| ENSG000002 | 7.93654381 | -0.568178947 | 0.36459241 | -1.5583949 | 0.11913966 | 0.22490681 | NA | NA |
| ENSG000002 | 8.14365519 | -0.546883924 | 0.3509381 | -1.5583487 | 0.11915061 | 0.22490697 | GSTA2 | 2939 |
| ENSG000002 | 18.0316643 | -0.448656371 | 0.28792301 | -1.5582512 | 0.11917371 | 0.2249401 | NA | NA |
| ENSG000001 | 8.3970014 | 0.555013075 | 0.35619366 | 1.55817788 | 0.11919109 | 0.22496243 | NA | NA |
| ENSG000001 | 242.808068 | 0.24675643 | 0.15836678 | 1.55813247 | 0.11920185 | 0.22497227 | ARL8A | 127829 |
| ENSG000002 | 15.2518996 | -0.490345057 | 0.31473304 | -1.5579714 | 0.11924004 | 0.22503387 | LINC01450 | 101928744 |

| | | | | | | | | |
|---|---|---|---|---|---|---|---|---|
| ENSG000002 | 7.53399023 | -0.560313233 | 0.35965024 | -1.5579393 | 0.11924764 | 0.22503775 | NA | NA |
| ENSG000001 | 12.3854648 | -0.507457475 | 0.32575428 | -1.5577922 | 0.11928251 | 0.22506166 | FMO6P | 388714 |
| ENSG000002 | 3.65255986 | -0.588129137 | 0.3775255 | -1.5578527 | 0.11926816 | 0.22506166 | MIR1253 | 100302208 |
| ENSG000002 | 3.23492565 | -0.586531002 | 0.37650565 | -1.5578279 | 0.11927405 | 0.22506166 | NA | NA |
| ENSG000002 | 5.51945034 | -0.579858618 | 0.37222656 | -1.557811 | 0.11927806 | 0.22506166 | NA | NA |
| ENSG000002 | 3.38733599 | 0.584860717 | 0.37546464 | 1.55769851 | 0.11930473 | 0.22507389 | LOC101060389 | 101060389 |
| ENSG000002 | 3.77253714 | -0.589637629 | 0.3785211 | -1.5577404 | 0.11929479 | 0.22507389 | NA | NA |
| ENSG000002 | 2.30276189 | -0.573742964 | 0.36832826 | -1.5576947 | 0.11930565 | 0.22507389 | NA | NA |
| ENSG000002 | 2.94036991 | -0.581390583 | 0.37325022 | -1.557643 | 0.11931791 | 0.22508655 | NA | NA |
| ENSG000002 | 4.38770042 | -0.588653047 | 0.377939 | -1.5575345 | 0.11934363 | 0.22509318 | NA | NA |
| ENSG000002 | 9.05461418 | -0.544237936 | 0.34941997 | -1.5575467 | 0.11934073 | 0.22509318 | NA | NA |
| ENSG000002 | 2.29171602 | -0.573001996 | 0.36787537 | -1.5575982 | 0.11932854 | 0.22509318 | NA | NA |
| ENSG000002 | 7.74921557 | -0.560306977 | 0.35973857 | -1.5575393 | 0.11934249 | 0.22509318 | NA | NA |
| ENSG000002 | 2.90129571 | -0.584777343 | 0.37548918 | -1.5573747 | 0.11938155 | 0.22515424 | NA | NA |
| ENSG000002 | 4.69536575 | -0.587723764 | 0.37745262 | -1.5570796 | 0.1194516 | 0.22527587 | LOC101927635 | 101927635 |
| ENSG000002 | 2.62017318 | -0.577973736 | 0.37119848 | -1.5570477 | 0.11945916 | 0.22527965 | NA | NA |
| ENSG000002 | 3.76428356 | -0.589384896 | 0.37857695 | -1.5568431 | 0.11950775 | 0.22535295 | NA | NA |
| ENSG000002 | 3.57801999 | -0.588435768 | 0.37796872 | -1.5568372 | 0.11950914 | 0.22535295 | NA | NA |
| ENSG000001 | 4.39493658 | -0.587758152 | 0.37764337 | -1.5563841 | 0.11961677 | 0.22549299 | HOXD12 | 3238 |
| ENSG000002 | 5.26746364 | -0.583555711 | 0.37493327 | -1.5564255 | 0.11960694 | 0.22549299 | NA | NA |
| ENSG000002 | 4.03279816 | -0.589016482 | 0.37844798 | -1.5564001 | 0.11961298 | 0.22549299 | NA | NA |
| ENSG000002 | 3.2995 | -0.585722043 | 0.3763229 | -1.5564348 | 0.11960474 | 0.22549299 | NA | NA |
| ENSG000002 | 3.50852658 | -0.588748928 | 0.37826532 | -1.5564444 | 0.11960247 | 0.22549299 | NA | NA |
| ENSG000002 | 3.77708405 | -0.589229327 | 0.37857754 | -1.5564297 | 0.11960595 | 0.22549299 | NA | NA |
| ENSG000001 | 11.8802254 | -0.503824005 | 0.3237252 | -1.5563324 | 0.11962908 | 0.2255057 | TTLL2 | 83887 |
| ENSG000002 | 14.7806513 | 0.494165111 | 0.31752867 | 1.556285 | 0.11964034 | 0.22551645 | NA | NA |
| ENSG000002 | 10.1970631 | -0.531614156 | 0.34160295 | -1.5562341 | 0.11965243 | 0.22552876 | NA | NA |
| ENSG000002 | 35.4974328 | 0.361961663 | 0.23260047 | 1.55615192 | 0.11967197 | 0.22554575 | NA | NA |
| ENSG000002 | 4.63609445 | -0.584847384 | 0.37582984 | -1.5561494 | 0.11967257 | 0.22554575 | NA | NA |

| | | | | | | | | |
|---|---|---|---|---|---|---|---|---|
| ENSG000002 | 50.7981747 | 0.335738068 | 0.21577924 | 1.55593315 | 0.11972399 | 0.22563217 | NA | NA |
| ENSG000002 | 5.43354483 | -0.586610332 | 0.37702441 | -1.5558948 | 0.1197331 | 0.22563701 | NA | NA |
| ENSG000002 | 7.19144166 | -0.563741365 | 0.36233063 | -1.5558756 | 0.11973769 | 0.22563701 | NA | NA |
| ENSG000002 | 2.61578193 | -0.577315158 | 0.3710814 | -1.5557642 | 0.11976417 | 0.22566596 | NA | NA |
| ENSG000002 | 3.48345553 | -0.588764298 | 0.37843595 | -1.5557832 | 0.11975965 | 0.22566596 | NA | NA |
| ENSG000001 | 74.0506537 | 0.28611646 | 0.18392706 | 1.55559738 | 0.11980386 | 0.22571499 | TRIM68 | 55128 |
| ENSG000002 | 8.91885994 | 0.555695072 | 0.35721657 | 1.55562514 | 0.11979726 | 0.22571499 | NA | NA |
| ENSG000002 | 3.74372662 | -0.588870168 | 0.37855232 | -1.5555846 | 0.11980689 | 0.22571499 | UGT1A5 | 54579 |
| ENSG000001 | 18.8443184 | -0.463078185 | 0.29770341 | -1.5555018 | 0.11982661 | 0.22574165 | GALNTL5 | 168391 |
| ENSG000002 | 2.63408791 | -0.577272776 | 0.37114785 | -1.5553715 | 0.11985762 | 0.22578958 | EFCAB14P1 | 106479047 |
| ENSG000002 | 4.32578197 | -0.587450039 | 0.37770439 | -1.555317 | 0.1198706 | 0.22579336 | NA | NA |
| ENSG000002 | 7.35625598 | -0.556821391 | 0.35801168 | -1.5553163 | 0.11987077 | 0.22579336 | NA | NA |
| ENSG000001 | 216.348055 | 0.447415238 | 0.28767787 | 1.55526472 | 0.11988303 | 0.22580598 | EPS15L1 | 58513 |
| ENSG000001 | 1624.82783 | 0.183335712 | 0.11788551 | 1.55520142 | 0.11989811 | 0.22582388 | HERC1 | 8925 |
| ENSG000002 | 6.33349466 | -0.571154116 | 0.3672762 | -1.5551079 | 0.11992037 | 0.22585533 | NA | NA |
| ENSG000002 | 6.87693994 | -0.569480245 | 0.36624363 | -1.5549219 | 0.11996467 | 0.22592826 | LINC01497 | 102723487 |
| ENSG000001 | 9.54080687 | -0.527596533 | 0.33933549 | -1.5547933 | 0.11999532 | 0.2259545 | PRSS37 | 136242 |
| ENSG000002 | 2.38154722 | -0.564264485 | 0.36291563 | -1.554809 | 0.11999157 | 0.2259545 | RNU6-660P | 106480599 |
| ENSG000002 | 2.61561886 | -0.576507689 | 0.37078948 | -1.5548113 | 0.11999101 | 0.2259545 | NA | NA |
| ENSG000002 | 6.32748108 | -0.570734711 | 0.36711414 | -1.5546519 | 0.12002899 | 0.22600741 | LOC101929239 | 101929239 |
| ENSG000001 | 18.914283 | -0.442093743 | 0.28438398 | -1.5545663 | 0.1200494 | 0.22603535 | RHBG | 57127 |
| ENSG000001 | 199.232009 | 0.249388937 | 0.16043607 | 1.55444434 | 0.12007846 | 0.22606318 | CSGALNACT2 | 55454 |
| ENSG000002 | 99.5650327 | 0.279804289 | 0.18000396 | 1.55443409 | 0.12008091 | 0.22606318 | ERV3-1 | 2086 |
| ENSG000002 | 24.5068726 | -0.408341893 | 0.26268711 | -1.5544802 | 0.12006993 | 0.22606318 | NA | NA |
| ENSG000002 | 7.75971112 | -0.555313373 | 0.35726526 | -1.5543447 | 0.12010221 | 0.22609279 | NA | NA |
| ENSG000002 | 2.88652391 | -0.582675035 | 0.37489198 | -1.5542478 | 0.12012532 | 0.22612068 | NA | NA |
| ENSG000002 | 5.98570225 | -0.569229759 | 0.36624414 | -1.5542358 | 0.12012818 | 0.22612068 | NA | NA |
| ENSG000002 | 20.5673273 | -0.439604355 | 0.28284849 | -1.5542043 | 0.12013568 | 0.22612431 | NA | NA |
| ENSG000001 | 4.21963413 | -0.587984363 | 0.37832955 | -1.5541592 | 0.12014644 | 0.22613407 | NA | NA |

| | | | | | | | | |
|---|---|---|---|---|---|---|---|---|
| ENSG000001 | 4.11208338 | -0.588226061 | 0.37854606 | -1.5539088 | 0.12020616 | 0.22622208 | PF4V1 | 5197 |
| ENSG000001 | 368.460737 | 0.232838107 | 0.14984179 | 1.55389301 | 0.12020994 | 0.22622208 | KIF5A | 3798 |
| ENSG000002 | 3.77589038 | -0.588250574 | 0.37856209 | -1.5539078 | 0.12020641 | 0.22622208 | MIR548B | 693128 |
| ENSG000002 | 15.3609667 | -0.493922101 | 0.31786816 | -1.5538584 | 0.1202182 | 0.22622713 | NA | NA |
| ENSG000002 | 5.32495991 | -0.582255038 | 0.37473862 | -1.5537631 | 0.12024093 | 0.2262489 | NA | NA |
| ENSG000002 | 4.02157555 | -0.586813454 | 0.37767026 | -1.5537719 | 0.12023883 | 0.2262489 | NA | NA |
| ENSG000001 | 22.1956969 | -0.44155661 | 0.28421224 | -1.5536158 | 0.1202761 | 0.22630457 | FREM3 | 166752 |
| ENSG000002 | 5.91930686 | -0.578834441 | 0.37258258 | -1.5535735 | 0.12028617 | 0.22631303 | NA | NA |
| ENSG000002 | 2.29101161 | -0.572138603 | 0.36828539 | -1.5535197 | 0.12029903 | 0.22632671 | NA | NA |
| ENSG000001 | 4.98378979 | -0.584368571 | 0.37619084 | -1.5533833 | 0.12033159 | 0.22636697 | FABP12 | 646486 |
| ENSG000001 | 2.30817021 | -0.571504716 | 0.36790966 | -1.5533833 | 0.12033159 | 0.22636697 | RNU6-1120P | 106480059 |
| ENSG000002 | 2.40412439 | -0.561594812 | 0.36156506 | -1.5532331 | 0.12036746 | 0.22642394 | NA | NA |
| ENSG000002 | 3.01976778 | -0.580302041 | 0.3736183 | -1.5531949 | 0.12037657 | 0.22643058 | NA | NA |
| ENSG000001 | 7.15480005 | -0.559332508 | 0.36012841 | -1.5531474 | 0.12038792 | 0.22643092 | TAS2R9 | 50835 |
| ENSG000002 | 2.31059036 | -0.57060811 | 0.36738626 | -1.5531558 | 0.12038591 | 0.22643092 | NA | NA |
| ENSG000001 | 9.52031714 | -0.534437103 | 0.34413456 | -1.5529888 | 0.12042582 | 0.22649169 | PLA2G12B | 84647 |
| ENSG000002 | 20.8461672 | -0.45130711 | 0.29063523 | -1.55283 | 0.12046374 | 0.22653871 | NA | NA |
| ENSG000002 | 2.62666472 | -0.57737851 | 0.37183157 | -1.5527958 | 0.12047192 | 0.22653871 | NA | NA |
| ENSG000002 | 35.8419509 | 0.366364952 | 0.2359397 | 1.55279062 | 0.12047317 | 0.22653871 | NA | NA |
| ENSG000002 | 25.5394888 | -0.402326886 | 0.25908858 | -1.5528546 | 0.12045788 | 0.22653871 | NA | NA |
| ENSG000002 | 3.14987711 | -0.586952882 | 0.37800552 | -1.5527627 | 0.12047983 | 0.22654074 | KRT18P32 | 391179 |
| ENSG000002 | 6.3332817 | -0.570268733 | 0.36728135 | -1.5526754 | 0.12050069 | 0.22656946 | NA | NA |
| ENSG00000C | 11.3781289 | -0.512246617 | 0.32992808 | -1.5526008 | 0.12051852 | 0.22658197 | CEACAM7 | 1087 |
| ENSG000002 | 3.73599512 | -0.58778533 | 0.37857583 | -1.5526224 | 0.12051336 | 0.22658197 | NA | NA |
| ENSG00000C | 53.6539007 | -0.308783329 | 0.19891434 | -1.5523432 | 0.12058012 | 0.22661373 | SLC15A1 | 6564 |
| ENSG000001 | 104.30138 | 0.262062755 | 0.16880251 | 1.5524814 | 0.12054708 | 0.22661373 | METAP1D | 254042 |
| ENSG000001 | 6.91548461 | -0.562991901 | 0.36266092 | -1.552392 | 0.12056847 | 0.22661373 | CTAGE10P | 220429 |
| ENSG000001 | 18.0111811 | -0.458044624 | 0.29506545 | -1.5523492 | 0.12057869 | 0.22661373 | QRFPR | 84109 |
| ENSG000002 | 25.5050714 | -0.422883428 | 0.27239775 | -1.5524483 | 0.12055499 | 0.22661373 | NA | NA |

| | | | | | | | | |
|---|---|---|---|---|---|---|---|---|
| ENSG000002 | 187.404156 | 0.250084477 | 0.16108942 | 1.55245749 | 0.1205528 | 0.22661373 | NA | NA |
| ENSG000002 | 5.7303172 | -0.57289041 | 0.36904389 | -1.5523639 | 0.12057518 | 0.22661373 | NA | NA |
| ENSG000002 | 3.13851913 | -0.586929156 | 0.37807162 | -1.5524285 | 0.12055973 | 0.22661373 | NA | NA |
| ENSG000001 | 304.185279 | 0.205659891 | 0.13249197 | 1.55224416 | 0.12060381 | 0.22664775 | ADCK3 | 56997 |
| ENSG000002 | 3.44830361 | -0.587312105 | 0.3783842 | -1.5521581 | 0.1206244 | 0.22667594 | NA | NA |
| ENSG000001 | 3.76946355 | -0.587573071 | 0.3785911 | -1.5519992 | 0.12066242 | 0.22672638 | CSN3 | 1448 |
| ENSG000002 | 3.99331174 | -0.586694839 | 0.37802176 | -1.5520134 | 0.12065902 | 0.22672638 | TMEM257 | 9142 |
| ENSG000002 | 7.13041325 | -0.55947333 | 0.36049377 | -1.5519639 | 0.12067086 | 0.22673172 | NA | NA |
| ENSG000002 | 2.85244943 | -0.582421762 | 0.37528841 | -1.5519311 | 0.12067871 | 0.22673596 | NA | NA |
| ENSG000002 | 6.37967208 | -0.571325746 | 0.36815309 | -1.55187 | 0.12069333 | 0.22675293 | LOC101929608 | 101929608 |
| ENSG000002 | 3.75647471 | -0.587469241 | 0.37858884 | -1.5517342 | 0.12072584 | 0.22680351 | NA | NA |
| ENSG000002 | 2.29223754 | -0.569627563 | 0.36710481 | -1.5516756 | 0.12073987 | 0.22681935 | NA | NA |
| ENSG000001 | 5.99099379 | -0.569074343 | 0.36679709 | -1.5514691 | 0.12078931 | 0.22688808 | WNT3A | 89780 |
| ENSG000001 | 6.56957204 | -0.564814242 | 0.36405509 | -1.5514527 | 0.12079324 | 0.22688808 | LILRP2 | 79166 |
| ENSG000002 | 2.58693309 | -0.575642994 | 0.37103192 | -1.5514649 | 0.12079031 | 0.22688808 | RNU6-168P | 106479640 |
| ENSG000002 | 4.203193 | -0.587305967 | 0.37856721 | -1.5513915 | 0.12080788 | 0.22690507 | NA | NA |
| ENSG000001 | 60.2308489 | 0.297905457 | 0.19203526 | 1.55130604 | 0.12082836 | 0.22692251 | PELI3 | 246330 |
| ENSG000002 | 2.25266987 | -0.569965643 | 0.3674089 | -1.5513115 | 0.12082706 | 0.22692251 | RN7SL635P | 106479454 |
| ENSG000002 | 3.72420791 | -0.587205525 | 0.37853816 | -1.5512453 | 0.12084292 | 0.22693934 | NA | NA |
| ENSG000002 | 10.2746542 | -0.533406555 | 0.34388584 | -1.5511152 | 0.12087408 | 0.22698735 | NA | NA |
| ENSG000001 | 7.0669354 | -0.557077215 | 0.35916266 | -1.5510443 | 0.12089106 | 0.22700872 | FGF17 | 8822 |
| ENSG000002 | 2.25526628 | -0.570355277 | 0.36773411 | -1.5509991 | 0.1209019 | 0.22701857 | NA | NA |
| ENSG000002 | 6.59761677 | -0.566036301 | 0.36496287 | -1.5509421 | 0.12091555 | 0.22702688 | PGA3 | 643834 |
| ENSG000002 | 9.21620546 | -0.544668688 | 0.35118755 | -1.5509339 | 0.12091752 | 0.22702688 | NA | NA |
| ENSG000002 | 2.25144394 | -0.571415025 | 0.3684439 | -1.5508875 | 0.12092866 | 0.22703727 | NA | NA |
| ENSG000001 | 86.7572488 | -0.266147376 | 0.17161354 | -1.550853 | 0.12093692 | 0.22704227 | PKD1L2 | 114780 |
| ENSG000002 | 7.31790525 | -0.551661896 | 0.35573187 | -1.5507801 | 0.1209544 | 0.22706457 | NA | NA |
| ENSG000001 | 120.719018 | 0.254080249 | 0.16386497 | 1.55054649 | 0.1210104 | 0.2270939 | SLC15A4 | 121260 |
| ENSG000001 | 15.5191264 | -0.466936245 | 0.3011427 | -1.5505481 | 0.12101001 | 0.2270939 | TEKT1 | 83659 |

| | | | | | | | | |
|---|---|---|---|---|---|---|---|---|
| ENSG000001 | 16.2777734 | -0.4720835 | 0.30444955 | -1.5506132 | 0.1209944 | 0.2270939 | P2RY6 | 5031 |
| ENSG000002 | 3.83023629 | -0.586444009 | 0.37819467 | -1.5506406 | 0.12098783 | 0.2270939 | NA | NA |
| ENSG000002 | 2.25119599 | -0.57126898 | 0.36843081 | -1.5505461 | 0.12101049 | 0.2270939 | NA | NA |
| ENSG000002 | 109.381303 | 0.256572517 | 0.16547428 | 1.55052805 | 0.12101483 | 0.2270939 | TTC28-AS1 | 284900 |
| ENSG000002 | 78.9574847 | -0.284576227 | 0.18353243 | -1.5505501 | 0.12100954 | 0.2270939 | LINC00669 | 647946 |
| ENSG000002 | 8.91857553 | -0.529283109 | 0.34134398 | -1.5505857 | 0.12100099 | 0.2270939 | NA | NA |
| ENSG000001 | 2.85324723 | -0.583128193 | 0.37609417 | -1.5504845 | 0.12102527 | 0.22710299 | IL9 | 3578 |
| ENSG000002 | 6.47919151 | -0.575827495 | 0.3713965 | -1.5504387 | 0.12103626 | 0.22711311 | NA | NA |
| ENSG000001 | 25.3677936 | 0.413583981 | 0.26676591 | 1.55036295 | 0.12105443 | 0.22713668 | BNC1 | 646 |
| ENSG000001 | 4.17059707 | -0.586797426 | 0.37858208 | -1.5499873 | 0.12114456 | 0.22729528 | GOLGA8R | 101059918 |
| ENSG000002 | 3.23022228 | -0.584231691 | 0.37694286 | -1.5499211 | 0.12116046 | 0.2273146 | NA | NA |
| ENSG000002 | 13.4329306 | -0.498835538 | 0.32186018 | -1.5498517 | 0.12117712 | 0.22733534 | OR2L2 | 26246 |
| ENSG000002 | 15.8367364 | -0.471567467 | 0.30429612 | -1.5496993 | 0.12121371 | 0.22739347 | NA | NA |
| ENSG000002 | 2.24202894 | -0.569222023 | 0.36736499 | -1.5494727 | 0.12126813 | 0.22748503 | RN7SKP108 | 106480866 |
| ENSG000002 | 7.3572765 | -0.553464116 | 0.35720502 | -1.5494298 | 0.12127842 | 0.22749382 | NA | NA |
| ENSG000001 | 44.4732894 | -0.346558666 | 0.22368955 | -1.5492841 | 0.12131344 | 0.22753456 | FBXO40 | 51725 |
| ENSG000002 | 3.12343523 | -0.58561911 | 0.37800265 | -1.549246 | 0.12132259 | 0.22753456 | NA | NA |
| ENSG000002 | 3.74939608 | -0.586489161 | 0.37855891 | -1.5492679 | 0.12131734 | 0.22753456 | NA | NA |
| ENSG000002 | 13.935972 | -0.480025887 | 0.30983891 | -1.5492757 | 0.12131546 | 0.22753456 | NA | NA |
| ENSG000002 | 4.61381426 | -0.583178727 | 0.37647117 | -1.5490661 | 0.12136583 | 0.22760514 | FTH1P9 | 106478936 |
| ENSG000001 | 23.6648719 | -0.415464625 | 0.2682167 | -1.5489887 | 0.12138445 | 0.22762952 | GLRA4 | 441509 |
| ENSG000001 | 12.2418885 | -0.498186539 | 0.32163034 | -1.5489414 | 0.12139581 | 0.22764031 | COL10A1 | 1300 |
| ENSG000002 | 2.33683582 | -0.566332699 | 0.36571075 | -1.5485809 | 0.12148249 | 0.22779231 | NA | NA |
| ENSG000002 | 21.6477485 | -0.422629464 | 0.27292869 | -1.5484977 | 0.12150251 | 0.2278193 | LINC00346 | 283487 |
| ENSG000002 | 42.8117304 | -0.354893448 | 0.22921063 | -1.5483289 | 0.12154312 | 0.22788493 | NA | NA |
| ENSG000001 | 729.484315 | 0.236564888 | 0.15279426 | 1.54825773 | 0.12156026 | 0.22790651 | VANGL2 | 57216 |
| ENSG000002 | 2.90524976 | -0.581435762 | 0.37568938 | -1.5476502 | 0.12170653 | 0.22817021 | NA | NA |
| ENSG000002 | 11.4668735 | -0.522147881 | 0.33739659 | -1.547579 | 0.12172368 | 0.22819181 | LINC01343 | 339442 |
| ENSG000001 | 337.797801 | 0.221979337 | 0.14344318 | 1.54750706 | 0.12174102 | 0.22821377 | SLC36A4 | 120103 |

| | | | | | | | | |
|---|---|---|---|---|---|---|---|---|
| ENSG000002 | 19.1550193 | 0.445269315 | 0.28773893 | 1.5474768 | 0.12174832 | 0.22821689 | NA | NA |
| ENSG000002 | 4.00763325 | -0.584672958 | 0.37792395 | -1.5470651 | 0.12184755 | 0.22839234 | NA | NA |
| ENSG000002 | 6.39820889 | -0.567877123 | 0.36713154 | -1.5467947 | 0.12191276 | 0.22850261 | NA | NA |
| ENSG000002 | 10.2785705 | -0.516677666 | 0.33403556 | -1.5467744 | 0.12191764 | 0.22850261 | NA | NA |
| ENSG000002 | 7.34601081 | -0.550988993 | 0.35622893 | -1.5467272 | 0.12192903 | 0.22850283 | NA | NA |
| ENSG000002 | 2.29741773 | -0.567858254 | 0.3671309 | -1.546746 | 0.12192451 | 0.22850283 | NA | NA |
| ENSG000001 | 14.9063138 | -0.500230929 | 0.3234317 | -1.5466354 | 0.12195118 | 0.22853377 | AKR1B10 | 57016 |
| ENSG000001 | 4.61556997 | -0.58135391 | 0.37589619 | -1.546581 | 0.12196431 | 0.22854781 | NA | NA |
| ENSG000002 | 3.15841054 | -0.584385583 | 0.37787263 | -1.5465147 | 0.12198031 | 0.22856724 | NA | NA |
| ENSG000002 | 3.21460968 | -0.582343747 | 0.37656546 | -1.5464609 | 0.1219933 | 0.22858102 | NA | NA |
| ENSG000002 | 6.15701791 | -0.574409285 | 0.37147327 | -1.5463005 | 0.12203202 | 0.228643 | NA | NA |
| ENSG000001 | 30.3518468 | -0.396033496 | 0.25619304 | -1.5458402 | 0.12214318 | 0.22883331 | PPP1R36 | 145376 |
| ENSG000002 | 4.38046579 | -0.584345068 | 0.37801709 | -1.5458165 | 0.12214888 | 0.22883331 | NA | NA |
| ENSG000002 | 3.51276083 | -0.584683231 | 0.37823751 | -1.5458097 | 0.12215052 | 0.22883331 | NA | NA |
| ENSG000001 | 102.388381 | 0.274341577 | 0.17747987 | 1.54576162 | 0.12216215 | 0.22883852 | ELAC1 | 55520 |
| ENSG000002 | 4.34814559 | -0.584106119 | 0.37787841 | -1.5457515 | 0.12216459 | 0.22883852 | NA | NA |
| ENSG000002 | 5.72049836 | -0.571075925 | 0.36946595 | -1.5456795 | 0.122182 | 0.22884958 | NA | NA |
| ENSG000002 | 9.04003881 | -0.541334671 | 0.35023532 | -1.5456313 | 0.12219364 | 0.22884958 | NA | NA |
| ENSG000002 | 3.21428439 | -0.582134944 | 0.37663711 | -1.5456123 | 0.12219823 | 0.22884958 | NA | NA |
| ENSG000002 | 6.3317152 | -0.568177933 | 0.36761242 | -1.5455896 | 0.12220371 | 0.22884958 | LOC340107 | 340107 |
| ENSG000002 | 4.87719229 | -0.578000295 | 0.37396815 | -1.545587 | 0.12220435 | 0.22884958 | NA | NA |
| ENSG000002 | 2.86409612 | -0.580477799 | 0.37556019 | -1.5456319 | 0.1221935 | 0.22884958 | NA | NA |
| ENSG000002 | 3.75326637 | -0.585023769 | 0.37854009 | -1.5454737 | 0.12223172 | 0.22887968 | NA | NA |
| ENSG000002 | 2.96028458 | -0.577925295 | 0.37394694 | -1.5454741 | 0.12223164 | 0.22887968 | NA | NA |
| ENSG000002 | 14.3670659 | -0.484020009 | 0.31320211 | -1.5453919 | 0.12225149 | 0.22890614 | TCP10 | 6953 |
| ENSG000002 | 6.4170549 | -0.569896624 | 0.3687844 | -1.5453382 | 0.12226448 | 0.22891988 | NA | NA |
| ENSG000002 | 11.4765783 | -0.511598623 | 0.33108513 | -1.5452178 | 0.1222936 | 0.22896384 | NA | NA |
| ENSG000002 | 12.3174471 | -0.508505397 | 0.3291215 | -1.5450385 | 0.12233695 | 0.22903443 | POTEJ | 653781 |
| ENSG000002 | 92.3327639 | 0.281423508 | 0.18215354 | 1.54497966 | 0.12235118 | 0.2290505 | NA | NA |

| | | | | | | | | |
|---|---|---|---|---|---|---|---|---|
| ENSG000002 | 3.98477295 | -0.58372821 | 0.37783367 | -1.5449343 | 0.12236215 | 0.22906045 | MTRNR2L6 | 100463482 |
| ENSG000001 | 44.0052088 | 0.338370632 | 0.21903759 | 1.54480619 | 0.12239315 | 0.2290746 | EFHD1 | 80303 |
| ENSG000002 | 10.3301172 | -0.516743396 | 0.334508 | -1.5447864 | 0.12239795 | 0.2290746 | NA | NA |
| ENSG000002 | 2.65441652 | -0.570614211 | 0.369366 | -1.5448477 | 0.12238311 | 0.2290746 | NA | NA |
| ENSG000002 | 10.107661 | -0.526067837 | 0.34053457 | -1.5448294 | 0.12238752 | 0.2290746 | LINC00929 | 503519 |
| ENSG000002 | 5.75975945 | -0.571011143 | 0.36962546 | -1.5448372 | 0.12238566 | 0.2290746 | NA | NA |
| ENSG000001 | 390.673681 | 0.192931653 | 0.12489995 | 1.54468958 | 0.12242137 | 0.22910784 | DMD | 1756 |
| ENSG000002 | 5.76990305 | -0.57372804 | 0.37146648 | -1.5444948 | 0.12246851 | 0.2291855 | LINC00375 | 104355140 |
| ENSG000002 | 8.47116389 | -0.546288599 | 0.35372158 | -1.5444028 | 0.12249078 | 0.2292166 | NA | NA |
| ENSG000001 | 209.025827 | 0.211505665 | 0.1369597 | 1.54429118 | 0.12251781 | 0.22923544 | A2ML1 | 144568 |
| ENSG000002 | 7.1399398 | -0.557126178 | 0.36076295 | -1.5442999 | 0.1225157 | 0.22923544 | NA | NA |
| ENSG000002 | 9.62308961 | 0.520903558 | 0.33730615 | 1.54430494 | 0.12251448 | 0.22923544 | NA | NA |
| ENSG000001 | 425.640768 | 0.247416878 | 0.16022739 | 1.54416097 | 0.12254934 | 0.22926565 | MED12 | 9968 |
| ENSG000002 | 6.62745919 | -0.565282383 | 0.36607891 | -1.5441545 | 0.12255092 | 0.22926565 | NA | NA |
| ENSG000002 | 4.92955956 | -0.578592325 | 0.37469472 | -1.5441699 | 0.12254717 | 0.22926565 | NA | NA |
| ENSG000002 | 8.69235775 | -0.533273649 | 0.34535767 | -1.5441199 | 0.12255929 | 0.22927073 | NA | NA |
| ENSG000002 | 6.37465159 | -0.566394425 | 0.36681876 | -1.5440716 | 0.12257099 | 0.22928204 | NA | NA |
| ENSG000001 | 1134.34663 | 0.186180566 | 0.12058579 | 1.54396772 | 0.12259615 | 0.22931036 | C12orf45 | 121053 |
| ENSG000002 | 6.78551348 | -0.572262194 | 0.37064516 | -1.5439624 | 0.12259743 | 0.22931036 | TLR8-AS1 | 349408 |
| ENSG000001 | 394.568559 | -0.206776782 | 0.13393083 | -1.5439073 | 0.1226108 | 0.22932478 | HERC5 | 51191 |
| ENSG000001 | 3.47332293 | -0.58391165 | 0.3782232 | -1.5438282 | 0.12262995 | 0.22935002 | NA | NA |
| ENSG000002 | 3.145206 | -0.583449683 | 0.37793201 | -1.5437953 | 0.12263792 | 0.22935435 | NA | NA |
| ENSG000002 | 3.79548245 | -0.58392178 | 0.37827145 | -1.5436581 | 0.12267118 | 0.22940598 | NA | NA |
| ENSG000002 | 5.97292172 | -0.566290355 | 0.36687109 | -1.5435677 | 0.1226931 | 0.22943638 | KDM4E | 390245 |
| ENSG000001 | 12.1829171 | -0.499970925 | 0.32394668 | -1.5433741 | 0.12274003 | 0.22951355 | C4orf45 | 152940 |
| ENSG000002 | 4.13812066 | -0.584255702 | 0.37859329 | -1.5432279 | 0.1227755 | 0.2295693 | RNU6-534P | 106481978 |
| ENSG000002 | 2.89165955 | -0.57897085 | 0.37517652 | -1.5431959 | 0.12278326 | 0.22957322 | NA | NA |
| ENSG000002 | 9.33881969 | -0.531062767 | 0.34414531 | -1.5431353 | 0.12279796 | 0.22958478 | SMIM21 | 284274 |
| ENSG000002 | 4.79727031 | -0.583864717 | 0.37836547 | -1.5431237 | 0.12280077 | 0.22958478 | NA | NA |

| | | | | | | | | |
|---|---|---|---|---|---|---|---|---|
| ENSG000002 | 12.0149371 | -0.506839868 | 0.3284653 | -1.5430545 | 0.12281755 | 0.22960557 | OR7E47P | 26628 |
| ENSG000002 | 3.16458887 | -0.583158931 | 0.37793433 | -1.5430166 | 0.12282676 | 0.2296122 | LOC339666 | 339666 |
| ENSG000002 | 3.11298868 | -0.573618742 | 0.37178097 | -1.5428943 | 0.12285642 | 0.22965623 | NA | NA |
| ENSG000002 | 3.78830619 | -0.584119416 | 0.37859207 | -1.5428728 | 0.12286164 | 0.22965623 | NA | NA |
| ENSG000002 | 2.85472055 | -0.57900182 | 0.37529249 | -1.5428015 | 0.12287895 | 0.22966768 | NA | NA |
| ENSG000002 | 2.92422715 | -0.578110799 | 0.3747151 | -1.5428009 | 0.12287909 | 0.22966768 | NA | NA |
| ENSG000001 | 128.150247 | 0.234605135 | 0.15206705 | 1.54277432 | 0.12288554 | 0.22966915 | TSGA10 | 80705 |
| ENSG000002 | 10.0329063 | -0.517017151 | 0.33513979 | -1.542691 | 0.12290576 | 0.22967519 | IDI2-AS1 | 55853 |
| ENSG000002 | 9.37655484 | -0.532430721 | 0.345129 | -1.5427006 | 0.12290343 | 0.22967519 | NA | NA |
| ENSG000002 | 6.00095013 | -0.565699519 | 0.36668849 | -1.5427251 | 0.1228975 | 0.22967519 | NA | NA |
| ENSG000002 | 3.1256267 | -0.582920876 | 0.37786776 | -1.5426584 | 0.12291368 | 0.2296794 | LOC100508046 | 100508046 |
| ENSG000002 | 5.43653113 | -0.573488986 | 0.37179095 | -1.5425039 | 0.12295119 | 0.22973891 | NA | NA |
| ENSG000002 | 2.92565025 | -0.579186516 | 0.3755241 | -1.5423418 | 0.12299056 | 0.22980188 | LOC103171574 | 103171574 |
| ENSG000002 | 2.90143951 | -0.578291778 | 0.37499229 | -1.5421431 | 0.12303882 | 0.22988147 | NA | NA |
| ENSG000002 | 7.38670744 | -0.55059772 | 0.35709556 | -1.5418778 | 0.12310329 | 0.22998258 | LOC400002 | 400002 |
| ENSG000002 | 3.38886099 | -0.576104342 | 0.37363912 | -1.5418737 | 0.12310429 | 0.22998258 | NA | NA |
| ENSG000002 | 5.74999213 | -0.571176534 | 0.37051691 | -1.5415667 | 0.12317892 | 0.23010081 | NA | NA |
| ENSG000002 | 44.3573615 | 0.349435085 | 0.22667477 | 1.54157024 | 0.12317806 | 0.23010081 | NA | NA |
| ENSG000002 | 2.3122733 | -0.566029015 | 0.36719526 | -1.5414932 | 0.12319679 | 0.23012359 | NA | NA |
| ENSG000002 | 6.84873839 | -0.556871216 | 0.36127647 | -1.5413991 | 0.12321969 | 0.23015576 | NA | NA |
| ENSG000001 | 5.07890987 | -0.580935656 | 0.37693532 | -1.5412078 | 0.12326622 | 0.2302271 | GJB4 | 127534 |
| ENSG000002 | 2.90768918 | -0.578569947 | 0.37540337 | -1.5411954 | 0.12326924 | 0.2302271 | NA | NA |
| ENSG000001 | 30.6175695 | -0.378414816 | 0.24553851 | -1.5411628 | 0.12327718 | 0.23023133 | PIH1D2 | 120379 |
| ENSG000001 | 8.73721139 | -0.533986502 | 0.34649073 | -1.5411278 | 0.12328569 | 0.23023661 | GPAT2 | 150763 |
| ENSG000002 | 3.53242412 | -0.582512424 | 0.37798844 | -1.5410853 | 0.12329603 | 0.23024533 | RN7SKP279 | 106480479 |
| ENSG000002 | 17.5262266 | -0.491048766 | 0.31865078 | -1.5410249 | 0.12331074 | 0.23026219 | LOC284825 | 284825 |
| ENSG000002 | 9.53833715 | -0.532883879 | 0.34583004 | -1.5408837 | 0.12334508 | 0.23031572 | NA | NA |
| ENSG000000 | 101.993077 | -0.260204058 | 0.16887753 | -1.5407855 | 0.12336899 | 0.23033942 | ADAMTS6 | 11174 |
| ENSG000002 | 6.72392733 | 0.562123439 | 0.36483119 | 1.54077681 | 0.12337112 | 0.23033942 | NA | NA |

| | | | | | | | | |
|---|---|---|---|---|---|---|---|---|
| ENSG000002 | 2.32861084 | -0.557809557 | 0.36203495 | -1.5407616 | 0.12337482 | 0.23033942 | NA | NA |
| ENSG000002 | 6.39568351 | -0.568749596 | 0.36914897 | -1.5407048 | 0.12338864 | 0.23035462 | MYT1L-AS1 | 730811 |
| ENSG000002 | 3.12081865 | -0.581954723 | 0.37772981 | -1.540664 | 0.12339858 | 0.23036216 | RNU1-44P | 106480160 |
| ENSG000002 | 3.90413335 | -0.582299043 | 0.3779588 | -1.5406416 | 0.12340404 | 0.23036216 | NA | NA |
| ENSG000001 | 80.7116032 | -0.268130473 | 0.17404479 | -1.5405831 | 0.12341828 | 0.23037814 | TMEM35 | 59353 |
| ENSG000002 | 6.35604143 | -0.569765973 | 0.36989079 | -1.5403627 | 0.12347196 | 0.23045713 | NA | NA |
| ENSG000002 | 3.83551308 | -0.583171191 | 0.37859305 | -1.5403642 | 0.12347159 | 0.23045713 | NA | NA |
| ENSG000001 | 19.2267661 | 0.443795625 | 0.28812716 | 1.54027695 | 0.12349286 | 0.23048552 | PDYN | 5173 |
| ENSG000002 | 4.97520296 | -0.580585214 | 0.37696568 | -1.5401541 | 0.12352281 | 0.23053081 | FDPSP2 | 619190 |
| ENSG000002 | 8.97874698 | -0.52899966 | 0.34348563 | -1.5400925 | 0.1235378 | 0.23054817 | SCHLAP1 | 101669767 |
| ENSG000002 | 2.87069866 | -0.578394026 | 0.37556651 | -1.5400575 | 0.12354634 | 0.2305535 | NA | NA |
| ENSG000002 | 3.9921127 | -0.58181397 | 0.37783588 | -1.539859 | 0.12359474 | 0.23062329 | NA | NA |
| ENSG000002 | 2.63707643 | -0.569236306 | 0.36966819 | -1.5398574 | 0.12359511 | 0.23062329 | NA | NA |
| ENSG000001 | 17.0899821 | -0.462942124 | 0.30067353 | -1.5396837 | 0.12363748 | 0.23069173 | ANPEP | 290 |
| ENSG00000C | 59.2244813 | 0.331741037 | 0.21547015 | 1.53961483 | 0.12365427 | 0.23071244 | MAP3K14 | 9020 |
| ENSG000002 | 8.91259027 | -0.544573934 | 0.35373959 | -1.5394769 | 0.12368791 | 0.23075449 | LINC00619 | 414260 |
| ENSG000002 | 9.69037102 | -0.535744815 | 0.34800471 | -1.5394758 | 0.12368819 | 0.23075449 | NA | NA |
| ENSG000002 | 5.20715578 | -0.57639216 | 0.37441763 | -1.5394365 | 0.12369778 | 0.23076177 | NA | NA |
| ENSG000002 | 4.31228458 | -0.581289615 | 0.37763017 | -1.5393092 | 0.12372882 | 0.23080906 | NA | NA |
| ENSG000002 | 10.6076243 | -0.523190109 | 0.33991246 | -1.5391908 | 0.12375773 | 0.23085238 | NA | NA |
| ENSG000002 | 4.88101464 | -0.577083591 | 0.37494558 | -1.539113 | 0.12377672 | 0.23087718 | NA | NA |
| ENSG000002 | 3.10831907 | -0.581614437 | 0.37792711 | -1.5389593 | 0.12381423 | 0.23093652 | NA | NA |
| ENSG000001 | 306.617584 | 0.225993285 | 0.14685114 | 1.53892767 | 0.12382195 | 0.2309403 | DEDD | 9191 |
| ENSG000002 | 2.9232946 | -0.577377046 | 0.37519707 | -1.5388634 | 0.12383764 | 0.23095894 | NA | NA |
| ENSG000002 | 2.61049019 | -0.568770169 | 0.36961686 | -1.5388101 | 0.12385065 | 0.23097258 | NA | NA |
| ENSG000002 | 2.27271031 | -0.563249911 | 0.36603738 | -1.538777 | 0.12385874 | 0.23097704 | NA | NA |
| ENSG000001 | 68.9580034 | 0.288669116 | 0.18760315 | 1.53872208 | 0.12387216 | 0.23099144 | BMI1 | 648 |
| ENSG000002 | 6.29545283 | -0.568333838 | 0.36936493 | -1.5386784 | 0.12388283 | 0.23100071 | NA | NA |
| ENSG000001 | 6.26590608 | -0.561221795 | 0.36474886 | -1.5386526 | 0.12388912 | 0.23100183 | MNX1 | 3110 |

| | | | | | | | | |
|---|---|---|---|---|---|---|---|---|
| ENSG000002 | 6.77736989 | -0.569278094 | 0.37002705 | -1.538477 | 0.12393203 | 0.23107122 | LINC01304 | 100505964 |
| ENSG000002 | 2.93197184 | -0.576999618 | 0.37507078 | -1.5383753 | 0.12395688 | 0.23110692 | NA | NA |
| ENSG000002 | 2.63181157 | -0.569013909 | 0.36989813 | -1.5382989 | 0.12397554 | 0.23113108 | NA | NA |
| ENSG000001 | 570.554658 | 0.177032262 | 0.11508523 | 1.53827092 | 0.12398239 | 0.23113322 | MAGI2 | 9863 |
| ENSG000001 | 1729.77955 | 0.171563736 | 0.11153411 | 1.53821768 | 0.1239954 | 0.23114685 | POGK | 57645 |
| ENSG000002 | 4.10122263 | -0.582286031 | 0.378552 | -1.538193 | 0.12400144 | 0.23114748 | NA | NA |
| ENSG000002 | 7.66617439 | -0.552014758 | 0.35894396 | -1.5378856 | 0.12407658 | 0.23126628 | NA | NA |
| ENSG000002 | 19.3114402 | -0.46219205 | 0.30053697 | -1.5378875 | 0.12407612 | 0.23126628 | NA | NA |
| ENSG000002 | 17.8073756 | 0.448927049 | 0.29193008 | 1.53778965 | 0.12410006 | 0.23129941 | NA | NA |
| ENSG000002 | 2.88820217 | -0.576145253 | 0.37469826 | -1.5376246 | 0.12414044 | 0.23135341 | NA | NA |
| ENSG000002 | 4.02664365 | -0.581472892 | 0.37816012 | -1.5376368 | 0.12413746 | 0.23135341 | NA | NA |
| ENSG000002 | 2.93857439 | -0.576105327 | 0.37469199 | -1.5375437 | 0.12416021 | 0.23137964 | LINC00864 | 728218 |
| ENSG000002 | 3.76544587 | -0.582007747 | 0.37855969 | -1.5374266 | 0.12418887 | 0.23142239 | NA | NA |
| ENSG000001 | 8.69691621 | -0.534458369 | 0.34765971 | -1.5373031 | 0.12421909 | 0.23145923 | PPP3R2 | 5535 |
| ENSG000002 | 3.52568497 | -0.580915787 | 0.37788648 | -1.5372759 | 0.12422576 | 0.23145923 | NA | NA |
| ENSG000002 | 19.1716871 | -0.436425484 | 0.28389391 | -1.5372837 | 0.12422385 | 0.23145923 | NA | NA |
| ENSG000001 | 29.7888864 | 0.395263474 | 0.25712736 | 1.53722835 | 0.1242374 | 0.23147029 | PRRG2 | 5639 |
| ENSG000002 | 23.4102597 | 0.42692529 | 0.27778783 | 1.53687544 | 0.12432382 | 0.23162065 | NA | NA |
| ENSG000002 | 66.3652486 | 0.297534102 | 0.19360382 | 1.53681937 | 0.12433755 | 0.23162495 | C6orf136 | 221545 |
| ENSG000002 | 7.6912946 | -0.547675343 | 0.35636455 | -1.5368401 | 0.12433247 | 0.23162495 | NA | NA |
| ENSG000001 | 8.02585951 | -0.564614277 | 0.36739885 | -1.5367883 | 0.12434515 | 0.23162847 | FOXN1 | 8456 |
| ENSG000002 | 4.5991805 | -0.578459999 | 0.37642676 | -1.5367133 | 0.12436354 | 0.23165208 | HNF4A-AS1 | 101927219 |
| ENSG000002 | 104.723772 | -0.309091713 | 0.20114628 | -1.5366514 | 0.12437869 | 0.23166966 | NA | NA |
| ENSG000001 | 17.0793254 | -0.48072853 | 0.31292422 | -1.5362459 | 0.12447809 | 0.23182287 | LRRC39 | 127495 |
| ENSG000001 | 9.44632475 | -0.524052732 | 0.34112199 | -1.536262 | 0.12447414 | 0.23182287 | HAND2 | 9464 |
| ENSG000001 | 20.0394789 | -0.437810135 | 0.28498495 | -1.536257 | 0.12447535 | 0.23182287 | PCDHB11 | 56125 |
| ENSG000002 | 3.13613123 | -0.580451642 | 0.37785261 | -1.5361853 | 0.12449293 | 0.23183986 | NA | NA |
| ENSG000001 | 19.8727881 | -0.437318478 | 0.28470052 | -1.5360649 | 0.12452246 | 0.2318629 | AZGP1 | 563 |
| ENSG000002 | 5.16623048 | -0.575408357 | 0.37459179 | -1.5360944 | 0.12451523 | 0.2318629 | NA | NA |

| | | | | | | | | |
|---|---|---|---|---|---|---|---|---|
| ENSG000002 | 3.45900402 | -0.581083004 | 0.37828984 | -1.5360788 | 0.12451906 | 0.2318629 | NA | NA |
| ENSG000002 | 197.917881 | 0.225323325 | 0.14669824 | 1.53596474 | 0.12454703 | 0.231898 | NA | NA |
| ENSG000001 | 38.6250581 | 0.359832398 | 0.2342865 | 1.53586487 | 0.12457153 | 0.23193296 | HS3ST4 | 9951 |
| ENSG000002 | 44.9166526 | 0.327745344 | 0.21340264 | 1.53580735 | 0.12458564 | 0.23194859 | FDXACB1 | 91893 |
| ENSG000002 | 2.53225326 | -0.573436013 | 0.37341189 | -1.5356662 | 0.12462028 | 0.23200243 | NA | NA |
| ENSG000001 | 917.502663 | 0.237257897 | 0.15451932 | 1.53545782 | 0.12467141 | 0.23208697 | TBC1D15 | 64786 |
| ENSG000000 | 20.7205421 | -0.432555391 | 0.28173095 | -1.5353492 | 0.12469807 | 0.23211529 | LIMS2 | 55679 |
| ENSG000002 | 5.56809223 | -0.574328981 | 0.37406911 | -1.5353553 | 0.12469658 | 0.23211529 | NA | NA |
| ENSG000001 | 11.8332589 | -0.507897867 | 0.33081167 | -1.5353082 | 0.12470814 | 0.23212338 | CCR9 | 10803 |
| ENSG000002 | 39.8179012 | 0.36769081 | 0.23949745 | 1.53525979 | 0.12472003 | 0.23213484 | NA | NA |
| ENSG000002 | 4.47225812 | -0.580961527 | 0.37844559 | -1.5351257 | 0.12475297 | 0.23218549 | LOC101926933 | 101926933 |
| ENSG000001 | 21.121975 | -0.430582112 | 0.28051138 | -1.5349898 | 0.12478633 | 0.23223694 | MSANTD1 | 345222 |
| ENSG000001 | 20.0194454 | -0.444763492 | 0.28976607 | -1.5349054 | 0.12480707 | 0.23225422 | LOC101929106 | 101929106 |
| ENSG000002 | 11.881367 | -0.501282839 | 0.32658485 | -1.5349237 | 0.12480257 | 0.23225422 | NA | NA |
| ENSG000001 | 176.977364 | 0.21925781 | 0.14285148 | 1.53486548 | 0.12481687 | 0.2322618 | COL27A1 | 85301 |
| ENSG000002 | 7.08416036 | -0.551551026 | 0.3593623 | -1.5348049 | 0.12483176 | 0.23227883 | NA | NA |
| ENSG000002 | 59.3737972 | 0.322246079 | 0.20996444 | 1.534765 | 0.12484156 | 0.23228642 | NA | NA |
| ENSG000002 | 2320.7641 | -0.210488491 | 0.13715164 | -1.5347137 | 0.12485418 | 0.23229924 | NA | NA |
| ENSG000002 | 7.05159682 | -0.565155148 | 0.36825957 | -1.5346652 | 0.12486609 | 0.23231073 | C8orf89 | 100130301 |
| ENSG000002 | 11.6160316 | -0.512094395 | 0.33372617 | -1.5344748 | 0.12491288 | 0.23238714 | NA | NA |
| ENSG000001 | 11.4478288 | -0.512022271 | 0.33372389 | -1.5342692 | 0.12496345 | 0.23247054 | PNPLA1 | 285848 |
| ENSG000002 | 5.00943589 | 0.573106239 | 0.37355396 | 1.53419934 | 0.12498062 | 0.23249182 | PDF | 64146 |
| ENSG000002 | 4.43840489 | 0.57836516 | 0.37699523 | 1.53414451 | 0.1249941 | 0.23250624 | NA | NA |
| ENSG000001 | 83.7230805 | 0.268563883 | 0.17507742 | 1.53397215 | 0.1250365 | 0.23257444 | CYP2B6 | 1555 |
| ENSG000002 | 3.74846245 | -0.580668372 | 0.37858335 | -1.5337927 | 0.12508066 | 0.2326459 | NA | NA |
| ENSG000002 | 6.55553025 | -0.560995018 | 0.36577774 | -1.5337046 | 0.12510233 | 0.23267554 | NA | NA |
| ENSG000001 | 224.796462 | -0.21771254 | 0.14195603 | -1.5336618 | 0.12511287 | 0.23268446 | ZDHHC4 | 55146 |
| ENSG000001 | 32.7630614 | -0.379530351 | 0.24748581 | -1.5335439 | 0.12514191 | 0.23271712 | SMOC1 | 64093 |
| ENSG000002 | 3.5147292 | -0.579494587 | 0.37787612 | -1.533557 | 0.12513866 | 0.23271712 | NA | NA |

| | | | | | | | | |
|---|---|---|---|---|---|---|---|---|
| ENSG000001 | 19.9330907 | -0.435058701 | 0.28370797 | -1.5334737 | 0.12515919 | 0.2327279 | SLC5A7 | 60482 |
| ENSG000002 | 2.94411855 | -0.573601006 | 0.37405163 | -1.5334809 | 0.12515742 | 0.2327279 | MIR3692 | 100500899 |
| ENSG000002 | 5.15091035 | -0.570878714 | 0.37231975 | -1.5333023 | 0.12520139 | 0.23279571 | NA | NA |
| ENSG000002 | 2.96985603 | -0.572948823 | 0.37369394 | -1.5332034 | 0.12522574 | 0.23283031 | NA | NA |
| ENSG000001 | 13.457078 | -0.482226963 | 0.31453703 | -1.5331326 | 0.1252432 | 0.23285209 | TMEM173 | 340061 |
| ENSG000002 | 3.9974695 | -0.579412355 | 0.37794781 | -1.5330486 | 0.12526388 | 0.23287914 | NA | NA |
| ENSG000002 | 3.19276522 | -0.578321354 | 0.3772415 | -1.5330269 | 0.12526923 | 0.23287914 | RN7SL700P | 106479478 |
| ENSG000001 | 4.60991282 | -0.5776501 | 0.37682175 | -1.5329532 | 0.1252874 | 0.23289155 | NA | NA |
| ENSG000002 | 9.73511799 | -0.522911356 | 0.34110964 | -1.5329715 | 0.12528288 | 0.23289155 | LINC00348 | 100885781 |
| ENSG000002 | 3.70243769 | -0.580256125 | 0.37854826 | -1.5328458 | 0.12531384 | 0.2329198 | NA | NA |
| ENSG000002 | 69.6242664 | 0.296085508 | 0.19316078 | 1.53284487 | 0.12531408 | 0.2329198 | LRRC37A2 | 474170 |
| ENSG000002 | 432.900165 | 0.207304877 | 0.13524612 | 1.53279726 | 0.12532582 | 0.23293094 | LOC644554 | 644554 |
| ENSG000002 | 4.66273344 | -0.578980366 | 0.37775189 | -1.5327001 | 0.12534976 | 0.23295805 | NA | NA |
| ENSG000002 | 2.24672076 | -0.563543214 | 0.3676821 | -1.5326915 | 0.12535189 | 0.23295805 | MIR4282 | 100423005 |
| ENSG000001 | 189.522262 | 0.224784851 | 0.14666572 | 1.53263385 | 0.1253661 | 0.23297377 | ZNF571 | 51276 |
| ENSG000001 | 5.78207843 | -0.5688192 | 0.37120114 | -1.5323746 | 0.12543002 | 0.23306053 | PRDM12 | 59335 |
| ENSG000002 | 5.71013763 | -0.566653423 | 0.36978477 | -1.5323871 | 0.12542693 | 0.23306053 | NA | NA |
| ENSG000002 | 5.44955995 | -0.56977433 | 0.37182386 | -1.532377 | 0.12542943 | 0.23306053 | NA | NA |
| ENSG000002 | 12.7274967 | -0.490403878 | 0.32003695 | -1.5323352 | 0.12543974 | 0.23306791 | NA | NA |
| ENSG000002 | 3.7031749 | -0.580044092 | 0.3785483 | -1.5322856 | 0.12545198 | 0.23307997 | NA | NA |
| ENSG000001 | 29.5795814 | -0.37788363 | 0.24663301 | -1.5321697 | 0.12548055 | 0.23311169 | HTRA4 | 203100 |
| ENSG000002 | 2.86539448 | -0.575226041 | 0.37543088 | -1.5321756 | 0.12547911 | 0.23311169 | LOC338694 | 338694 |
| ENSG000002 | 100.285741 | 0.274560121 | 0.17921285 | 1.5320337 | 0.12551412 | 0.23316336 | FPGT | 8790 |
| ENSG000002 | 5.77818997 | -0.567553719 | 0.37046487 | -1.5320042 | 0.12552141 | 0.23316622 | NA | NA |
| ENSG000001 | 59.1104217 | 0.306684618 | 0.20020473 | 1.53185498 | 0.12555822 | 0.23318567 | PTPN5 | 84867 |
| ENSG000001 | 756.730239 | 0.18728359 | 0.1222568 | 1.53188686 | 0.12555036 | 0.23318567 | NCSTN | 23385 |
| ENSG000001 | 18.9831305 | 0.447611636 | 0.2922008 | 1.53186313 | 0.12555621 | 0.23318567 | NA | NA |
| ENSG000002 | 5.70389568 | -0.565239274 | 0.36899241 | -1.5318452 | 0.12556063 | 0.23318567 | NA | NA |
| ENSG000002 | 3.7220685 | -0.579907773 | 0.37854976 | -1.5319196 | 0.12554229 | 0.23318567 | NA | NA |

| | | | | | | | | |
|---|---|---|---|---|---|---|---|---|
| ENSG000002 | 2.23841567 | -0.563129727 | 0.36764916 | -1.5317041 | 0.12559547 | 0.23323969 | LINC00307 | 266919 |
| ENSG000002 | 11.0343156 | -0.515687293 | 0.33670006 | -1.5315925 | 0.12562302 | 0.23328018 | ERICH4 | 100170765 |
| ENSG000002 | 2.51167671 | -0.572160974 | 0.37358029 | -1.5315609 | 0.12563082 | 0.23328397 | NA | NA |
| ENSG000001 | 6.80570593 | -0.553629954 | 0.361493 | -1.5315095 | 0.12564353 | 0.233295 | SERPINA4 | 5267 |
| ENSG000002 | 4.01010493 | -0.579268255 | 0.37823828 | -1.5314903 | 0.12564827 | 0.233295 | NA | NA |
| ENSG000002 | 63.7105583 | 0.30338608 | 0.19811284 | 1.53138021 | 0.12567545 | 0.23332411 | SNORD116-1 | 100033413 |
| ENSG000002 | 6.92673811 | -0.558841578 | 0.36492247 | -1.5313981 | 0.12567103 | 0.23332411 | NA | NA |
| ENSG000002 | 8.69629961 | -0.530262661 | 0.3462769 | -1.5313255 | 0.12568896 | 0.2333385 | NA | NA |
| ENSG000002 | 3.28567447 | -0.574892865 | 0.37543312 | -1.531279 | 0.12570044 | 0.23334226 | NA | NA |
| ENSG000002 | 54.1340638 | -0.315742412 | 0.20619633 | -1.5312707 | 0.12570249 | 0.23334226 | NA | NA |
| ENSG000002 | 5.00215426 | -0.577025756 | 0.37686079 | -1.5311377 | 0.12573537 | 0.23338193 | NA | NA |
| ENSG000002 | 3.24306045 | -0.574156963 | 0.3749844 | -1.5311489 | 0.12573259 | 0.23338193 | NA | NA |
| ENSG000002 | 13.7486311 | -0.481924572 | 0.3147895 | -1.5309423 | 0.12578365 | 0.23346086 | NA | NA |
| ENSG000002 | 442.319923 | 0.457530563 | 0.29886941 | 1.53087117 | 0.12580124 | 0.23348281 | HNRNPA1P10 | 664709 |
| ENSG000001 | 63.7104574 | 0.308635921 | 0.2016129 | 1.53083418 | 0.12581038 | 0.23348909 | RPS6KA4 | 8986 |
| ENSG000002 | 2.8383592 | -0.574744242 | 0.37545157 | -1.5308079 | 0.12581687 | 0.23349046 | NA | NA |
| ENSG000002 | 3.98189633 | -0.578631865 | 0.37801018 | -1.5307309 | 0.12583592 | 0.23350991 | NA | NA |
| ENSG000002 | 3.77030545 | -0.579517844 | 0.37859193 | -1.5307189 | 0.12583887 | 0.23350991 | NA | NA |
| ENSG000002 | 9.34384343 | -0.527424487 | 0.34457364 | -1.5306582 | 0.12585388 | 0.23352707 | NA | NA |
| ENSG000001 | 4.63302142 | -0.576765065 | 0.37682125 | -1.5306065 | 0.12586667 | 0.23353285 | OR2Z1 | 284383 |
| ENSG000002 | 5.26936206 | -0.574810379 | 0.37554601 | -1.5305991 | 0.12586851 | 0.23353285 | NA | NA |
| ENSG000002 | 4.93329704 | -0.575610985 | 0.37610089 | -1.5304696 | 0.12590053 | 0.23358156 | NA | NA |
| ENSG000001 | 322.345166 | 0.202479022 | 0.13230078 | 1.53044464 | 0.12590671 | 0.23358234 | ZXDB | 158586 |
| ENSG000002 | 7.40953238 | -0.5502841 | 0.35960083 | -1.5302637 | 0.12595146 | 0.23365468 | NA | NA |
| ENSG000000 | 117.032574 | 0.259055621 | 0.16929477 | 1.53020451 | 0.12596612 | 0.23366049 | RETSAT | 54884 |
| ENSG000001 | 5.16813366 | -0.570780764 | 0.37300704 | -1.5302145 | 0.12596365 | 0.23366049 | FOXA1 | 3169 |
| ENSG000002 | 3.12977664 | -0.577849142 | 0.37766591 | -1.5300538 | 0.12600342 | 0.23370961 | NA | NA |
| ENSG000002 | 3.7594447 | -0.579228947 | 0.37856841 | -1.5300509 | 0.12600412 | 0.23370961 | NA | NA |
| ENSG000002 | 4.08895118 | -0.57916555 | 0.37855712 | -1.5299291 | 0.12603428 | 0.23375485 | NA | NA |

| | | | | | | | | |
|---|---|---|---|---|---|---|---|---|
| ENSG000001 | 40.9581677 | 0.381644795 | 0.24947415 | 1.52979694 | 0.126067 | 0.23380484 | Mar-09 | 92979 |
| ENSG000002 | 2.28468927 | -0.559943201 | 0.36603962 | -1.5297339 | 0.12608262 | 0.23382311 | NA | NA |
| ENSG000001 | 38.8469201 | 0.347074297 | 0.2268985 | 1.52964565 | 0.12610446 | 0.23385294 | RWDD3 | 25950 |
| ENSG000002 | 6.20542217 | -0.55647324 | 0.36381342 | -1.5295567 | 0.1261265 | 0.23388311 | NA | NA |
| ENSG000002 | 4.76029939 | -0.578369558 | 0.37814725 | -1.5294824 | 0.1261449 | 0.23390253 | NA | NA |
| ENSG000002 | 2.26234546 | -0.560621932 | 0.36654706 | -1.5294678 | 0.12614851 | 0.23390253 | NA | NA |
| ENSG000001 | 18.9702325 | -0.44056539 | 0.28808252 | -1.5293028 | 0.12618941 | 0.23395453 | PIANP | 196500 |
| ENSG000001 | 208.990711 | 0.239827942 | 0.15682947 | 1.52922753 | 0.12620805 | 0.23395453 | MRPL24 | 79590 |
| ENSG000002 | 3.11019789 | -0.577488767 | 0.37763762 | -1.5292141 | 0.12621138 | 0.23395453 | NA | NA |
| ENSG000002 | 5.40285364 | -0.569463041 | 0.37240049 | -1.5291684 | 0.12622271 | 0.23395453 | NA | NA |
| ENSG000002 | 12.2027368 | -0.497614313 | 0.32541454 | -1.5291705 | 0.12622218 | 0.23395453 | LOC285740 | 285740 |
| ENSG000002 | 5.7182939 | -0.565433066 | 0.36975328 | -1.5292172 | 0.12621062 | 0.23395453 | RN7SL754P | 106479497 |
| ENSG000002 | 4.02749241 | -0.578354063 | 0.37818119 | -1.5293042 | 0.12618906 | 0.23395453 | LOC101928738 | 101928738 |
| ENSG000002 | 13.2694338 | 0.488609988 | 0.31951508 | 1.52922356 | 0.12620903 | 0.23395453 | NA | NA |
| ENSG000002 | 4.19515546 | -0.578919998 | 0.37859329 | -1.5291344 | 0.12623114 | 0.23395946 | NA | NA |
| ENSG000001 | 694.883127 | 0.177690665 | 0.11620572 | 1.52910425 | 0.1262386 | 0.23396259 | FOXP1 | 27086 |
| ENSG000001 | 91.1966307 | 0.294464298 | 0.19258298 | 1.52902555 | 0.12625811 | 0.23396667 | PPP1R9B | 84687 |
| ENSG000001 | 58.9305525 | 0.333654263 | 0.21821131 | 1.52904203 | 0.12625402 | 0.23396667 | NA | NA |
| ENSG000001 | 60.6519256 | -0.297226568 | 0.19438414 | -1.529068 | 0.12624759 | 0.23396667 | PCBP3 | 54039 |
| ENSG000002 | 2.35506591 | -0.551125164 | 0.36046807 | -1.5289153 | 0.12628543 | 0.2340066 | NA | NA |
| ENSG000001 | 15.2024647 | -0.460738745 | 0.30137416 | -1.5287931 | 0.12631574 | 0.23405207 | KCNK13 | 56659 |
| ENSG000002 | 3.21027716 | -0.574207126 | 0.37560535 | -1.5287512 | 0.12632613 | 0.23406064 | NA | NA |
| ENSG000002 | 3.75448378 | -0.578673889 | 0.37857129 | -1.5285731 | 0.12637031 | 0.23413179 | LOC100506679 | 100506679 |
| ENSG000002 | 4.01964412 | -0.577960582 | 0.37815226 | -1.5283806 | 0.12641807 | 0.23420957 | RN7SL143P | 106479273 |
| ENSG000001 | 18.8068785 | -0.436583935 | 0.28566265 | -1.52832 | 0.12643311 | 0.23422674 | CCDC155 | 147872 |
| ENSG000002 | 11.2183623 | -0.525366393 | 0.34379076 | -1.5281574 | 0.12647347 | 0.2342908 | NA | NA |
| ENSG000001 | 52.929584 | 0.305072899 | 0.19964052 | 1.52811109 | 0.12648496 | 0.23430138 | DLEU7 | 220107 |
| ENSG000001 | 5.96569461 | -0.560608908 | 0.36689255 | -1.5279921 | 0.12651451 | 0.23434542 | SPACA7 | 122258 |
| ENSG000002 | 11.0578842 | -0.518996361 | 0.33968013 | -1.5278973 | 0.12653804 | 0.23436759 | NA | NA |

| | | | | | | | | |
|---|---|---|---|---|---|---|---|---|
| ENSG000002 | 2.2765795 | -0.55702364 | 0.36456773 | -1.5279017 | 0.12653696 | 0.23436759 | NA | NA |
| ENSG000000 | 121.304593 | 0.237477488 | 0.15543388 | 1.5278361 | 0.12655324 | 0.23438504 | AMPH | 273 |
| ENSG000002 | 3.5705135 | -0.575743191 | 0.37690637 | -1.5275496 | 0.1266244 | 0.23450611 | NA | NA |
| ENSG000000 | 7.23085285 | 0.554402392 | 0.3629618 | 1.52744005 | 0.12665162 | 0.23453594 | RTEL1-TNFRSF6 | 100533107 |
| ENSG000002 | 3.76343067 | -0.578262652 | 0.37858332 | -1.5274383 | 0.12665207 | 0.23453594 | NA | NA |
| ENSG000002 | 17.6087699 | 0.454860924 | 0.2978054 | 1.52737636 | 0.12666745 | 0.23455372 | NA | NA |
| ENSG000002 | 7.30413861 | -0.557808926 | 0.3652254 | -1.5273005 | 0.12668632 | 0.23457794 | NA | NA |
| ENSG000002 | 3.45124026 | -0.577622872 | 0.37821622 | -1.5272292 | 0.12670402 | 0.23460001 | NA | NA |
| ENSG000002 | 2.21672922 | -0.56017974 | 0.36680597 | -1.5271827 | 0.12671558 | 0.2346107 | NA | NA |
| ENSG000002 | 7.35879113 | -0.546762287 | 0.3580257 | -1.5271593 | 0.1267214 | 0.23461076 | NA | NA |
| ENSG000002 | 5.97259663 | -0.561093321 | 0.36747626 | -1.5268832 | 0.12679006 | 0.23472716 | NA | NA |
| ENSG000002 | 2.91901525 | -0.572424327 | 0.37491893 | -1.5267949 | 0.12681201 | 0.23475708 | NA | NA |
| ENSG000002 | 7.91803748 | -0.544777713 | 0.35686024 | -1.5265856 | 0.12686409 | 0.23483205 | LINC00332 | 100874127 |
| ENSG000002 | 13.7857479 | -0.496854859 | 0.32546649 | -1.526593 | 0.12686225 | 0.23483205 | VWA8-AS1 | 100507240 |
| ENSG000002 | 2.49984051 | 0.563339184 | 0.36906265 | 1.52640529 | 0.12690896 | 0.23490439 | TBC1D3L | 101060376 |
| ENSG000002 | 6.22870648 | -0.556832552 | 0.3648105 | -1.5263611 | 0.12691996 | 0.23491402 | NA | NA |
| ENSG000002 | 3.73530475 | -0.57784011 | 0.37858357 | -1.5263211 | 0.12692991 | 0.23492171 | LOC105377480 | 105377480 |
| ENSG000001 | 28.84339 | 0.394033464 | 0.25819031 | 1.52613575 | 0.12697606 | 0.23498568 | SOX21 | 11166 |
| ENSG000002 | 2.3012795 | -0.558259483 | 0.36579654 | -1.5261475 | 0.12697313 | 0.23498568 | MIR4428 | 100616141 |
| ENSG000002 | 12.7336949 | -0.488194806 | 0.31998219 | -1.5256937 | 0.12708617 | 0.23517871 | NA | NA |
| ENSG000002 | 7.65535706 | -0.542509019 | 0.35561557 | -1.5255491 | 0.12712219 | 0.2352308 | NA | NA |
| ENSG000002 | 5.50438649 | -0.570097061 | 0.37370324 | -1.5255342 | 0.12712591 | 0.2352308 | NA | NA |
| ENSG000002 | 2.69104177 | -0.558820724 | 0.36632433 | -1.5254808 | 0.12713923 | 0.2352447 | NA | NA |
| ENSG000001 | 27.8652699 | 0.396214356 | 0.25974379 | 1.52540452 | 0.12715823 | 0.23526913 | SYNDIG1 | 79953 |
| ENSG000002 | 2.83816277 | -0.571496759 | 0.37466482 | -1.5253547 | 0.12717064 | 0.23528136 | NA | NA |
| ENSG000002 | 3.16043371 | -0.576060513 | 0.37767054 | -1.525299 | 0.12718453 | 0.2352936 | NA | NA |
| ENSG000002 | 10.5356442 | -0.524668413 | 0.34398134 | -1.5252816 | 0.12718887 | 0.2352936 | NA | NA |
| ENSG000002 | 2.61468577 | -0.563251832 | 0.36928876 | -1.5252342 | 0.1272007 | 0.23529402 | NA | NA |
| ENSG000002 | 11.4071358 | -0.517190465 | 0.33908826 | -1.5252385 | 0.12719962 | 0.23529402 | NA | NA |

| | | | | | | | | |
|---|---|---|---|---|---|---|---|---|
| ENSG000002 | 3.16473268 | -0.575909663 | 0.37764535 | -1.5250013 | 0.12725876 | 0.23537995 | NA | NA |
| ENSG000002 | 10.7046741 | -0.502729533 | 0.32965469 | -1.5250186 | 0.12725446 | 0.23537995 | LOC101927237 | 101927237 |
| ENSG000002 | 2.87872356 | -0.572291243 | 0.37531715 | -1.5248204 | 0.1273039 | 0.2354527 | NA | NA |
| ENSG000002 | 46.3759831 | 0.321772078 | 0.21103175 | 1.52475671 | 0.12731979 | 0.23547135 | LOC100288748 | 100288748 |
| ENSG000001 | 12.7828829 | -0.50960732 | 0.33429674 | -1.5244161 | 0.12740481 | 0.23559664 | FAM9A | 171482 |
| ENSG000002 | 4.64103746 | -0.57515647 | 0.37729641 | -1.5244155 | 0.12740496 | 0.23559664 | NA | NA |
| ENSG000002 | 2.62590681 | -0.5613887 | 0.36825622 | -1.5244514 | 0.127396 | 0.23559664 | NA | NA |
| ENSG000001 | 17.7544416 | -0.43975308 | 0.28849558 | -1.5242975 | 0.12743441 | 0.23561378 | EXOC3L1 | 283849 |
| ENSG000002 | 13.1300223 | -0.479829261 | 0.3147872 | -1.5242973 | 0.12743447 | 0.23561378 | FAM13A-AS1 | 285512 |
| ENSG000002 | 3.74104643 | -0.577073067 | 0.37858601 | -1.5242852 | 0.12743747 | 0.23561378 | NA | NA |
| ENSG000002 | 8.75193559 | -0.531386955 | 0.34860689 | -1.5243157 | 0.12742987 | 0.23561378 | NA | NA |
| ENSG000001 | 18.6516365 | -0.438620502 | 0.28776352 | -1.5242394 | 0.12744892 | 0.23562419 | NA | NA |
| ENSG000002 | 2.24644056 | -0.560434389 | 0.36769106 | -1.5241991 | 0.12745898 | 0.23563206 | NA | NA |
| ENSG000001 | 2.54320684 | -0.568716459 | 0.37314156 | -1.5241306 | 0.1274761 | 0.23565296 | NA | NA |
| ENSG000002 | 2.26809814 | -0.555522654 | 0.36453133 | -1.5239367 | 0.12752453 | 0.23573175 | NA | NA |
| ENSG000002 | 14.4397387 | -0.48709455 | 0.31966302 | -1.5237751 | 0.1275649 | 0.23579562 | NA | NA |
| ENSG00000C | 10.5188796 | -0.509665598 | 0.3345038 | -1.5236467 | 0.127597 | 0.23582749 | SCGN | 10590 |
| ENSG000001 | 2587.34376 | 0.173430319 | 0.1138244 | 1.52366563 | 0.12759226 | 0.23582749 | PHACTR2 | 9749 |
| ENSG000001 | 42.0440814 | -0.352069839 | 0.2310721 | -1.5236363 | 0.12759959 | 0.23582749 | GLIPR1 | 11010 |
| ENSG000002 | 5.9957892 | -0.560323135 | 0.36776219 | -1.5236018 | 0.12760822 | 0.23583269 | NA | NA |
| ENSG000001 | 1672.77725 | -0.219913407 | 0.14434526 | -1.5235235 | 0.12762778 | 0.23585809 | PTPN14 | 5784 |
| ENSG00000C | 273.262757 | 0.21013952 | 0.13797017 | 1.52307933 | 0.12773886 | 0.23605261 | CFLAR | 8837 |
| ENSG000001 | 6.11949935 | -0.566179977 | 0.37174048 | -1.5230517 | 0.12774577 | 0.23605463 | KLK12 | 43849 |
| ENSG000002 | 57.8377265 | 0.306965328 | 0.2015615 | 1.52293633 | 0.12777463 | 0.2360972 | NA | NA |
| ENSG000002 | 5.15454511 | 0.573307523 | 0.37646382 | 1.5228755 | 0.12778986 | 0.23611457 | NA | NA |
| ENSG000001 | 73.6556252 | 0.284541511 | 0.1868615 | 1.52274019 | 0.12782372 | 0.23616095 | PRRG3 | 79057 |
| ENSG000002 | 38.7732071 | 0.362707836 | 0.23819597 | 1.52272866 | 0.12782661 | 0.23616095 | NA | NA |
| ENSG000002 | 9.6663657 | -0.528024725 | 0.34676834 | -1.5227017 | 0.12783336 | 0.23616268 | NA | NA |
| ENSG000001 | 70.2450912 | -0.318540816 | 0.20921807 | -1.5225302 | 0.12787629 | 0.23623122 | CCSER1 | 401145 |

| | | | | | | | | |
|---|---|---|---|---|---|---|---|---|
| ENSG000002 | 3.98363079 | -0.575835243 | 0.37822924 | -1.5224504 | 0.12789627 | 0.23625737 | NA | NA |
| ENSG000002 | 16.794275 | -0.451519749 | 0.29666043 | -1.5220087 | 0.12800691 | 0.23645097 | VWA7 | 80737 |
| ENSG000002 | 5.47619422 | -0.569022881 | 0.37394839 | -1.5216615 | 0.12809391 | 0.23659012 | NA | NA |
| ENSG000002 | 4.40111293 | -0.576037199 | 0.37855227 | -1.5216847 | 0.1280881 | 0.23659012 | NA | NA |
| ENSG000002 | 3.12384017 | -0.57471718 | 0.37769996 | -1.5216236 | 0.12810342 | 0.23659691 | NA | NA |
| ENSG00000C | 19.7855186 | -0.422961276 | 0.27798397 | -1.5215312 | 0.12812659 | 0.23662893 | SEZ6 | 124925 |
| ENSG000002 | 4.34578527 | -0.575216084 | 0.37807 | -1.5214539 | 0.12814597 | 0.23665393 | NA | NA |
| ENSG000002 | 11.0558808 | -0.505638981 | 0.33238331 | -1.5212526 | 0.12819646 | 0.2367364 | LOC101927377 | 101927377 |
| ENSG000002 | 3.13757286 | -0.574113516 | 0.37743386 | -1.521097 | 0.12823549 | 0.2367977 | NA | NA |
| ENSG000002 | 2.8523332 | -0.569907847 | 0.37467561 | -1.52107 | 0.12824226 | 0.23679942 | NA | NA |
| ENSG000001 | 69.512798 | 0.31058595 | 0.20421768 | 1.52085728 | 0.12829566 | 0.23688722 | RNF128 | 79589 |
| ENSG000001 | 102.468742 | -0.252984137 | 0.16634836 | -1.5208093 | 0.12830769 | 0.23688787 | ERN1 | 2081 |
| ENSG000002 | 4.52868181 | -0.575735196 | 0.37856832 | -1.5208224 | 0.12830441 | 0.23688787 | NA | NA |
| ENSG000002 | 2.87120665 | -0.568530369 | 0.37389204 | -1.5205736 | 0.12836688 | 0.23698635 | NA | NA |
| ENSG000002 | 4.04306558 | -0.575147138 | 0.37825451 | -1.5205295 | 0.12837795 | 0.23699601 | NA | NA |
| ENSG000001 | 20.0168375 | -0.427910793 | 0.28143858 | -1.5204411 | 0.12840014 | 0.23702618 | TRPV4 | 59341 |
| ENSG000002 | 333.148769 | 0.212220011 | 0.13958195 | 1.52039728 | 0.12841116 | 0.23703573 | MICAL3 | 57553 |
| ENSG000001 | 12.6629361 | -0.483983593 | 0.31834367 | -1.5203179 | 0.12843108 | 0.23703614 | CPA2 | 1358 |
| ENSG000001 | 27.648475 | 0.402322129 | 0.26463281 | 1.52030331 | 0.12843476 | 0.23703614 | FLJ37453 | 729614 |
| ENSG000002 | 2.94441249 | -0.569078042 | 0.37431118 | -1.5203341 | 0.12842704 | 0.23703614 | RNU6-392P | 106480577 |
| ENSG000002 | 2.26661162 | -0.55514557 | 0.36514024 | -1.5203626 | 0.12841987 | 0.23703614 | NA | NA |
| ENSG000001 | 3.769139 | -0.57553863 | 0.37859328 | -1.520203 | 0.12845998 | 0.23704835 | RIPK3 | 11035 |
| ENSG000001 | 237.278439 | 0.282129098 | 0.18558563 | 1.52020985 | 0.12845824 | 0.23704835 | TAF4 | 6874 |
| ENSG000001 | 44.107505 | 0.345560254 | 0.22731477 | 1.52018391 | 0.12846476 | 0.23704835 | NA | NA |
| ENSG000002 | 2.59067808 | -0.562433995 | 0.36996147 | -1.5202502 | 0.12844811 | 0.23704835 | NA | NA |
| ENSG000002 | 11.2731687 | 0.503153594 | 0.33099537 | 1.52012276 | 0.12848013 | 0.23706591 | NA | NA |
| ENSG000002 | 2.24456065 | -0.555643724 | 0.36555523 | -1.5199994 | 0.12851113 | 0.23710154 | NA | NA |
| ENSG000002 | 15.0430224 | 0.4695843 | 0.30893568 | 1.52000669 | 0.12850929 | 0.23710154 | NA | NA |
| ENSG000002 | 39.5726913 | -0.335373088 | 0.22064778 | -1.5199477 | 0.12852412 | 0.23711471 | NA | NA |

| | | | | | | | | |
|---|---|---|---|---|---|---|---|---|
| ENSG000001 | 43.636512 | -0.37549317 | 0.24705057 | -1.5199041 | 0.12853508 | 0.23712415 | LYPD6B | 130576 |
| ENSG000001 | 116.990618 | 0.256191457 | 0.16856642 | 1.519825 | 0.12855497 | 0.23714654 | WDR5B | 54554 |
| ENSG000002 | 43.3023851 | 0.334091034 | 0.21982431 | 1.5198093 | 0.12855891 | 0.23714654 | PPP1R26-AS1 | 100506599 |
| ENSG000002 | 4.75994652 | -0.574946718 | 0.37832162 | -1.51973 | 0.12857885 | 0.23717253 | NA | NA |
| ENSG000002 | 3.49434275 | -0.57366868 | 0.37749273 | -1.5196814 | 0.12859107 | 0.23718429 | RN7SL162P | 106481828 |
| ENSG000002 | 6.87191047 | -0.553624337 | 0.36431474 | -1.519632 | 0.12860349 | 0.2371964 | NA | NA |
| ENSG000002 | 4.89559001 | -0.570275054 | 0.37530409 | -1.5195013 | 0.12863636 | 0.23724624 | NA | NA |
| ENSG000002 | 141.21841 | -0.23883375 | 0.15719986 | -1.5193 | 0.128687 | 0.23732884 | SOX2-OT | 347689 |
| ENSG000002 | 23.9196577 | -0.413490957 | 0.27217175 | -1.5192281 | 0.1287051 | 0.23735143 | NA | NA |
| ENSG000001 | 8.95856273 | -0.522561597 | 0.3439866 | -1.5191336 | 0.12872888 | 0.23738448 | TNFRSF14 | 8764 |
| ENSG000002 | 7.24864511 | -0.55528911 | 0.36554813 | -1.5190588 | 0.1287477 | 0.23740839 | NA | NA |
| ENSG000002 | 2.5343285 | -0.566790506 | 0.37312952 | -1.5190181 | 0.12875793 | 0.23741647 | NA | NA |
| ENSG000001 | 27.9252984 | -0.392075535 | 0.25813847 | -1.5188575 | 0.12879838 | 0.23748025 | XAF1 | 54739 |
| ENSG000002 | 18.517059 | 0.437629528 | 0.28813979 | 1.51880974 | 0.12881039 | 0.2374916 | FOXN3-AS1 | 400236 |
| ENSG000002 | 2.86896833 | -0.567832109 | 0.37389112 | -1.5187098 | 0.12883556 | 0.23752721 | NA | NA |
| ENSG000001 | 14.5219784 | -0.492278439 | 0.32416034 | -1.5186264 | 0.12885657 | 0.23755514 | ZNF491 | 126069 |
| ENSG000002 | 4.87751292 | -0.569368582 | 0.37493059 | -1.5185973 | 0.12886389 | 0.23755784 | NA | NA |
| ENSG000002 | 10.9145198 | -0.513485264 | 0.33814705 | -1.5185265 | 0.12888171 | 0.23757989 | NA | NA |
| ENSG000002 | 20.4731127 | 0.420196599 | 0.27673683 | 1.5183978 | 0.12891415 | 0.23762887 | ATP6V0E2-AS1 | 401431 |
| ENSG000002 | 2.509145 | -0.566467272 | 0.37307767 | -1.5183629 | 0.12892295 | 0.2376343 | NA | NA |
| ENSG000002 | 2.28321539 | -0.556510297 | 0.36653855 | -1.5182859 | 0.12894235 | 0.23765925 | NA | NA |
| ENSG000001 | 15.7496006 | -0.477889195 | 0.31477523 | -1.5181918 | 0.12896604 | 0.23769212 | CFHR4 | 10877 |
| ENSG000002 | 12.4008636 | -0.487807075 | 0.32132228 | -1.518124 | 0.12898313 | 0.23771281 | NA | NA |
| ENSG000001 | 3.09529079 | -0.573247602 | 0.3776641 | -1.5178769 | 0.12904544 | 0.23781684 | NA | NA |
| ENSG000001 | 183.199598 | 0.252875427 | 0.16660155 | 1.51784559 | 0.12905333 | 0.23782057 | ARMCX4 | 100131755 |
| ENSG000002 | 5.97365947 | -0.557964758 | 0.36761677 | -1.5177892 | 0.12906755 | 0.23783596 | NA | NA |
| ENSG000002 | 5.7309897 | -0.562232231 | 0.37044028 | -1.5177405 | 0.12907982 | 0.23784776 | NA | NA |
| ENSG000002 | 9.73181636 | -0.513185549 | 0.33814653 | -1.5176425 | 0.12910454 | 0.23788251 | NA | NA |
| ENSG000001 | 774.114939 | 0.446354515 | 0.29411526 | 1.51761769 | 0.1291108 | 0.23788324 | MIDN | 90007 |

| | | | | | | | | |
|---|---|---|---|---|---|---|---|---|
| ENSG000002 | 10.5407742 | -0.508544986 | 0.33510637 | -1.5175629 | 0.12912463 | 0.23789767 | DIP2A-IT1 | 100862692 |
| ENSG000002 | 2.27481333 | -0.556347895 | 0.36661165 | -1.5175401 | 0.12913037 | 0.23789767 | NA | NA |
| ENSG000002 | 6.08328878 | -0.56103326 | 0.36971101 | -1.5174913 | 0.12914268 | 0.23790828 | LOC105375448 | 105375448 |
| ENSG000002 | 3.12248895 | -0.572568497 | 0.37731764 | -1.5174708 | 0.12914786 | 0.23790828 | NA | NA |
| ENSG000002 | 81.5057476 | 0.271425331 | 0.17891128 | 1.51709457 | 0.12924281 | 0.23807237 | NA | NA |
| ENSG000002 | 5.94497371 | -0.557094948 | 0.36723995 | -1.5169781 | 0.12927222 | 0.23810491 | NA | NA |
| ENSG000002 | 7.38637384 | -0.545988531 | 0.35991845 | -1.5169785 | 0.12927212 | 0.23810491 | NA | NA |
| ENSG000002 | 11.8138639 | -0.504809149 | 0.3328032 | -1.5168398 | 0.12930713 | 0.23815839 | LOC101928270 | 101928270 |
| ENSG000002 | 22.7551703 | 0.419483372 | 0.27655785 | 1.51680154 | 0.1293168 | 0.23816539 | NA | NA |
| ENSG000001 | 12.1970046 | -0.500783197 | 0.33016475 | -1.5167676 | 0.12932537 | 0.23817035 | OR4F17 | 81099 |
| ENSG00000C | 276.330926 | 0.214105907 | 0.14116775 | 1.51667724 | 0.12934819 | 0.23820157 | FAM114A2 | 10827 |
| ENSG000002 | 3.14716914 | -0.572491043 | 0.37747069 | -1.5166503 | 0.12935499 | 0.23820327 | NA | NA |
| ENSG000002 | 2.89508601 | -0.569013253 | 0.37519003 | -1.5166001 | 0.12936767 | 0.23821581 | RN7SL756P | 106481128 |
| ENSG000001 | 10.2767865 | 0.518373095 | 0.34181584 | 1.5165274 | 0.12938605 | 0.23823601 | CD19 | 930 |
| ENSG000002 | 10.7801295 | -0.506252705 | 0.33382743 | -1.5165102 | 0.1293904 | 0.23823601 | NA | NA |
| ENSG000001 | 9.5024093 | -0.515615489 | 0.34001214 | -1.516462 | 0.12940257 | 0.23824762 | SPACA1 | 81833 |
| ENSG000002 | 2.51141612 | -0.566163562 | 0.37338446 | -1.5163019 | 0.12944304 | 0.23831129 | NA | NA |
| ENSG000002 | 6.48165212 | -0.54968416 | 0.36257179 | -1.5160699 | 0.12950169 | 0.23840845 | NA | NA |
| ENSG000002 | 5.9350689 | -0.559730991 | 0.36920922 | -1.5160266 | 0.12951263 | 0.23841297 | NA | NA |
| ENSG000002 | 2.46823054 | 0.559908384 | 0.36932938 | 1.51601365 | 0.1295159 | 0.23841297 | MIR3176 | 100423037 |
| ENSG000001 | 2002.75228 | 0.446176693 | 0.29433475 | 1.51588182 | 0.12954924 | 0.23846351 | ATIC | 471 |
| ENSG000002 | 7.33505777 | -0.542558602 | 0.35792253 | -1.5158549 | 0.12955606 | 0.23846524 | NA | NA |
| ENSG000002 | 6.03348547 | -0.558658226 | 0.36855349 | -1.5158132 | 0.1295666 | 0.23847382 | NA | NA |
| ENSG000001 | 18.2241157 | -0.43777094 | 0.28883613 | -1.5156377 | 0.12961098 | 0.23854141 | PLAC8L1 | 153770 |
| ENSG000002 | 2.28548596 | -0.554321131 | 0.3657385 | -1.5156215 | 0.12961509 | 0.23854141 | EHMT2-AS1 | 106478954 |
| ENSG000002 | 31.2612685 | 0.375557458 | 0.24779795 | 1.51557933 | 0.12962576 | 0.23855022 | NA | NA |
| ENSG000002 | 6.85934457 | -0.552199846 | 0.36438214 | -1.5154416 | 0.12966062 | 0.23860354 | NA | NA |
| ENSG000002 | 4.9226253 | -0.569061774 | 0.37553036 | -1.5153549 | 0.12968256 | 0.23863309 | NA | NA |
| ENSG000002 | 4.90250101 | -0.571066108 | 0.37687216 | -1.515278 | 0.12970201 | 0.23864723 | NA | NA |

| | | | | | | | | |
|---|---|---|---|---|---|---|---|---|
| ENSG000002 | 3.16971645 | -0.571568659 | 0.37720097 | -1.5152895 | 0.12969911 | 0.23864723 | NA | NA |
| ENSG000001 | 207.154897 | 0.205263249 | 0.13546612 | 1.51523683 | 0.12971244 | 0.23865532 | PBXIP1 | 57326 |
| ENSG000002 | 4.4311151 | -0.5735381 | 0.3785195 | -1.5152141 | 0.12971818 | 0.23865532 | NA | NA |
| ENSG000002 | 6.00668375 | -0.559301034 | 0.36916889 | -1.5150275 | 0.12976543 | 0.23873142 | LOC102477328 | 102477328 |
| ENSG000002 | 6.19123192 | -0.553246516 | 0.36520106 | -1.5149094 | 0.12979535 | 0.23877564 | NA | NA |
| ENSG000002 | 2.26904496 | -0.554415082 | 0.36599059 | -1.5148343 | 0.12981438 | 0.2387998 | NA | NA |
| ENSG000002 | 16.9510668 | -0.440577037 | 0.29085906 | -1.5147441 | 0.12983723 | 0.238831 | NA | NA |
| ENSG000001 | 4.34227613 | -0.572428446 | 0.37793335 | -1.514628 | 0.12986663 | 0.23887425 | H1FOO | 132243 |
| ENSG000002 | 59.3374433 | 0.30850682 | 0.20368801 | 1.51460475 | 0.12987253 | 0.23887427 | NA | NA |
| ENSG000002 | 2.30661249 | -0.5466101 | 0.36090841 | -1.5145396 | 0.12988903 | 0.23889377 | NA | NA |
| ENSG000001 | 19.2893692 | -0.429242442 | 0.28345365 | -1.5143303 | 0.12994209 | 0.23898053 | PSTPIP1 | 9051 |
| ENSG000001 | 16.615592 | -0.464188564 | 0.30654762 | -1.5142462 | 0.12996341 | 0.23899806 | OSTN | 344901 |
| ENSG000002 | 8.69629733 | -0.539899861 | 0.35654633 | -1.5142488 | 0.12996274 | 0.23899806 | NA | NA |
| ENSG000001 | 4.91395469 | -0.568573702 | 0.3754965 | -1.5141917 | 0.12997722 | 0.23900892 | RNU6-212P | 106481243 |
| ENSG000002 | 4.91125413 | -0.568760111 | 0.37562341 | -1.5141764 | 0.1299811 | 0.23900892 | NA | NA |
| ENSG000001 | 3.16162739 | -0.571271844 | 0.37728915 | -1.5141486 | 0.12998815 | 0.23901105 | NA | NA |
| ENSG000001 | 137.838369 | 0.251050318 | 0.16581415 | 1.51404643 | 0.13001406 | 0.23904785 | LMBR1L | 55716 |
| ENSG000001 | 51.5994343 | -0.317738989 | 0.20987211 | -1.5139648 | 0.13003477 | 0.23907507 | LETM2 | 137994 |
| ENSG000002 | 3.78642628 | -0.57281676 | 0.37838414 | -1.5138498 | 0.13006394 | 0.23911786 | NA | NA |
| ENSG000001 | 88.7585761 | 0.255332491 | 0.16868002 | 1.51370917 | 0.13009962 | 0.23916177 | CA8 | 767 |
| ENSG000001 | 61.1675512 | -0.307392171 | 0.20306982 | -1.5137265 | 0.13009522 | 0.23916177 | AHNAK2 | 113146 |
| ENSG000002 | 2.32516519 | -0.551890198 | 0.36461478 | -1.5136254 | 0.13012087 | 0.23919 | NA | NA |
| ENSG000002 | 4.24104698 | -0.570419295 | 0.3768873 | -1.513501 | 0.13015246 | 0.23923722 | NA | NA |
| ENSG000002 | 2.56131172 | -0.564235122 | 0.3728082 | -1.5134729 | 0.13015957 | 0.23923945 | LOC102724323 | 102724323 |
| ENSG000002 | 2.51921234 | -0.564601026 | 0.3731026 | -1.5132594 | 0.13021378 | 0.23932823 | NA | NA |
| ENSG000002 | 4.05125912 | -0.572353911 | 0.3782661 | -1.5130986 | 0.13025461 | 0.23939244 | NA | NA |
| ENSG000002 | 7.62363803 | -0.538190296 | 0.35569828 | -1.5130529 | 0.13026623 | 0.23940295 | NA | NA |
| ENSG000002 | 3.43762967 | -0.571529639 | 0.37774825 | -1.5129908 | 0.13028199 | 0.23942106 | NA | NA |
| ENSG000002 | 6.04940301 | -0.559747633 | 0.36998799 | -1.5128806 | 0.13031 | 0.23946168 | NA | NA |

| | | | | | | | | |
|---|---|---|---|---|---|---|---|---|
| ENSG000002 | 4.84480076 | -0.567178855 | 0.37493405 | -1.512743 | 0.13034495 | 0.23951506 | NA | NA |
| ENSG000001 | 61.1079637 | 0.312356196 | 0.20650047 | 1.51261739 | 0.13037687 | 0.23956285 | SNX22 | 79856 |
| ENSG000001 | 106.474901 | 0.252641698 | 0.16702745 | 1.51257588 | 0.13038743 | 0.23957139 | RHBDL3 | 162494 |
| ENSG000001 | 410.936166 | 0.200740742 | 0.13271727 | 1.51254424 | 0.13039547 | 0.2395753 | NAF1 | 92345 |
| ENSG000002 | 4.86424725 | -0.568760257 | 0.37607367 | -1.5123639 | 0.1304413 | 0.23964568 | LINC01257 | 116437 |
| ENSG000002 | 11.2760359 | -0.492439317 | 0.32561263 | -1.5123471 | 0.13044559 | 0.23964568 | ST8SIA6-AS1 | 100128098 |
| ENSG000002 | 3.76704504 | -0.572549047 | 0.37859266 | -1.5123089 | 0.1304553 | 0.23965267 | NA | NA |
| ENSG000002 | 4.85329421 | -0.568214928 | 0.37573948 | -1.5122577 | 0.13046832 | 0.23966572 | NA | NA |
| ENSG000001 | 333.210337 | 0.278359421 | 0.18411693 | 1.51186215 | 0.13056893 | 0.23983968 | AGRN | 375790 |
| ENSG000002 | 8.45212873 | -0.530448644 | 0.35087567 | -1.5117852 | 0.13058852 | 0.23986479 | NA | NA |
| ENSG000001 | 55.2860757 | 0.322398363 | 0.21327576 | 1.51165031 | 0.13062285 | 0.23991698 | POPDC3 | 64208 |
| ENSG000002 | 3.76936659 | -0.572151072 | 0.37851694 | -1.51156 | 0.13064583 | 0.23994832 | NA | NA |
| ENSG000002 | 3.75329917 | -0.572152424 | 0.37859317 | -1.5112592 | 0.13072242 | 0.24007811 | NA | NA |
| ENSG000002 | 2.70876206 | -0.556070505 | 0.36796009 | -1.511225 | 0.13073113 | 0.24008323 | RN7SL760P | 106481129 |
| ENSG000002 | 3.74336245 | -0.572101066 | 0.37858436 | -1.5111587 | 0.13074801 | 0.24010337 | NA | NA |
| ENSG000002 | 3.10277381 | -0.570122309 | 0.3773122 | -1.5110095 | 0.13078604 | 0.24014058 | NA | NA |
| ENSG000002 | 6.47041037 | -0.548772317 | 0.36317184 | -1.5110542 | 0.13077465 | 0.24014058 | NA | NA |
| ENSG000002 | 3.983247 | -0.571561302 | 0.37826072 | -1.5110247 | 0.13078215 | 0.24014058 | NA | NA |
| ENSG000002 | 3.13518496 | -0.569916137 | 0.37723562 | -1.5107697 | 0.13084714 | 0.2402383 | NA | NA |
| ENSG000002 | 12.3422051 | -0.485213426 | 0.321173 | -1.5107541 | 0.13085111 | 0.2402383 | NA | NA |
| ENSG000001 | 3.48051365 | -0.57087537 | 0.37789378 | -1.5106768 | 0.13087082 | 0.2402636 | FAM47B | 170062 |
| ENSG000002 | 5.48491222 | -0.564763026 | 0.37390488 | -1.5104457 | 0.13092973 | 0.24036088 | NA | NA |
| ENSG000002 | 4.02573813 | -0.571454704 | 0.37834503 | -1.5104063 | 0.13093979 | 0.24036847 | NA | NA |
| ENSG000001 | 17.9481665 | -0.464119535 | 0.30730464 | -1.5102913 | 0.13096911 | 0.2404114 | LINC01559 | 283422 |
| ENSG000002 | 2.2268092 | -0.551918515 | 0.36548509 | -1.5100986 | 0.13101828 | 0.24049077 | NA | NA |
| ENSG000002 | 5.81846375 | -0.570073723 | 0.37751368 | -1.5100743 | 0.13102446 | 0.24049123 | NA | NA |
| ENSG000001 | 97.906875 | 0.293240313 | 0.19420763 | 1.50993201 | 0.13106077 | 0.240547 | PLCD3 | 113026 |
| ENSG000002 | 6.503798 | 0.559040426 | 0.37028276 | 1.50976627 | 0.13110308 | 0.24061375 | NA | NA |
| ENSG000001 | 22.0188405 | 0.420548021 | 0.27857446 | 1.50964311 | 0.13113452 | 0.24066056 | CYB561D2 | 11068 |

| | | | | | | | | |
|---|---|---|---|---|---|---|---|---|
| ENSG000002 | 3.73743261 | -0.571486793 | 0.37859249 | -1.5095038 | 0.13117009 | 0.24071495 | NA | NA |
| ENSG000001 | 10.5236071 | 0.500769037 | 0.33175481 | 1.50945523 | 0.13118249 | 0.24072681 | DLL4 | 54567 |
| ENSG000002 | 3.19076962 | -0.569173002 | 0.37711635 | -1.5092769 | 0.13122803 | 0.2407876 | NA | NA |
| ENSG000002 | 3.73980069 | -0.571380124 | 0.37858402 | -1.5092558 | 0.13123343 | 0.2407876 | NA | NA |
| ENSG000002 | 4.26692029 | -0.569507237 | 0.37733704 | -1.50928 | 0.13122725 | 0.2407876 | NA | NA |
| ENSG000002 | 12.4039568 | -0.484811328 | 0.32124167 | -1.5091795 | 0.13125293 | 0.24081248 | PSG7 | 5676 |
| ENSG000002 | 2.24438493 | -0.552916975 | 0.36637868 | -1.5091407 | 0.13126283 | 0.24081975 | NA | NA |
| ENSG000002 | 22.287857 | 0.417231379 | 0.27648476 | 1.5090574 | 0.13128412 | 0.24084791 | NA | NA |
| ENSG000002 | 13.8945707 | -0.48705517 | 0.3227869 | -1.5089063 | 0.13132274 | 0.24090787 | NA | NA |
| ENSG000002 | 3.09551892 | -0.569377018 | 0.37736014 | -1.5088425 | 0.13133903 | 0.24091998 | NA | NA |
| ENSG000002 | 4.04772962 | -0.570871075 | 0.37835249 | -1.5088339 | 0.13134122 | 0.24091998 | NA | NA |
| ENSG000002 | 13.6457772 | -0.48639511 | 0.32237537 | -1.5087849 | 0.13135375 | 0.24093206 | NA | NA |
| ENSG000001 | 35.2218692 | -0.376066513 | 0.24927168 | -1.5086612 | 0.13138538 | 0.24097164 | NFATC2 | 4773 |
| ENSG000002 | 3.11702856 | -0.569060123 | 0.37719723 | -1.508654 | 0.13138722 | 0.24097164 | NA | NA |
| ENSG000001 | 3.79317474 | -0.570519234 | 0.37820177 | -1.5085049 | 0.13142534 | 0.24103067 | ANKRD60 | 140731 |
| ENSG000002 | 5.07272597 | -0.562867502 | 0.3731429 | -1.5084503 | 0.13143933 | 0.24104541 | CCER2 | 643669 |
| ENSG000001 | 125.385331 | 0.249389791 | 0.16533214 | 1.50841691 | 0.13144786 | 0.24105016 | ZNF684 | 127396 |
| ENSG000002 | 5.39265394 | -0.570048818 | 0.37794363 | -1.5082906 | 0.13148017 | 0.24109851 | LINC01216 | 100874275 |
| ENSG000001 | 172.125292 | 0.210913439 | 0.13985211 | 1.50811766 | 0.13152442 | 0.24115902 | FGGY | 55277 |
| ENSG000002 | 4.85092723 | -0.565774599 | 0.37515345 | -1.5081151 | 0.13152506 | 0.24115902 | NA | NA |
| ENSG000002 | 2.82467803 | -0.564744973 | 0.37456935 | -1.507718 | 0.13162671 | 0.24133447 | MIR454 | 768216 |
| ENSG000001 | 16273.1756 | 0.524416301 | 0.34785665 | 1.50756441 | 0.13166605 | 0.24139569 | NCL | 4691 |
| ENSG000002 | 25.099982 | 0.390588848 | 0.25909809 | 1.50749414 | 0.13168405 | 0.24141777 | NA | NA |
| ENSG000002 | 4.31274139 | -0.569919412 | 0.37806702 | -1.507456 | 0.1316938 | 0.24142475 | NA | NA |
| ENSG000001 | 25.1233369 | -0.401484451 | 0.26637024 | -1.5072422 | 0.1317486 | 0.24150833 | PDZRN4 | 29951 |
| ENSG000002 | 11.3862594 | -0.502513766 | 0.33341725 | -1.5071619 | 0.13176917 | 0.24150833 | NA | NA |
| ENSG000002 | 11.9295295 | -0.492573149 | 0.32681519 | -1.5071917 | 0.13176152 | 0.24150833 | NA | NA |
| ENSG000002 | 2.86970499 | -0.562719583 | 0.37335128 | -1.5072121 | 0.13175629 | 0.24150833 | NA | NA |
| ENSG000002 | 48.3854825 | 0.353660092 | 0.23465279 | 1.50716338 | 0.13176878 | 0.24150833 | NA | NA |

| | | | | | | | | |
|---|---|---|---|---|---|---|---|---|
| ENSG000001 | 13.9435344 | -0.485859164 | 0.32238044 | -1.5070988 | 0.13178534 | 0.24152704 | NA | NA |
| ENSG000002 | 2.27786656 | -0.550741043 | 0.36545396 | -1.5070052 | 0.13180931 | 0.24154914 | NA | NA |
| ENSG000002 | 3.99622376 | -0.569926861 | 0.37818251 | -1.5070154 | 0.13180671 | 0.24154914 | NA | NA |
| ENSG000002 | 11.2476435 | -0.490487085 | 0.32550949 | -1.5068289 | 0.13185453 | 0.24162109 | NA | NA |
| ENSG000002 | 3.52449184 | -0.569261614 | 0.37782113 | -1.5066961 | 0.13188858 | 0.24167256 | LOC400553 | 400553 |
| ENSG000001 | 231.487089 | 0.209160832 | 0.13883286 | 1.50656572 | 0.13192201 | 0.24172078 | CEP57L1 | 285753 |
| ENSG000002 | 6.53213746 | -0.551392384 | 0.36599747 | -1.506547 | 0.13192682 | 0.24172078 | NA | NA |
| ENSG000001 | 358.486797 | 0.209150144 | 0.13883424 | 1.50647384 | 0.13194558 | 0.24173331 | SNTB2 | 6645 |
| ENSG000002 | 2.4826188 | -0.561963074 | 0.37303125 | -1.5064772 | 0.13194472 | 0.24173331 | NA | NA |
| ENSG000002 | 13.4345041 | -0.488868357 | 0.32454361 | -1.5063256 | 0.1319836 | 0.24179204 | PAPOLB | 56903 |
| ENSG000002 | 6.73019976 | -0.554375901 | 0.36806128 | -1.5062054 | 0.13201445 | 0.24183764 | NA | NA |
| ENSG000001 | 7.78036653 | -0.531784159 | 0.35308475 | -1.5061091 | 0.13203917 | 0.24185014 | KRT32 | 3882 |
| ENSG000002 | 5.40892783 | -0.561974921 | 0.3731284 | -1.5061167 | 0.13203722 | 0.24185014 | NA | NA |
| ENSG000002 | 5.09946069 | -0.563154624 | 0.37391224 | -1.5061144 | 0.1320378 | 0.24185014 | NA | NA |
| ENSG000001 | 17.6848976 | -0.445994083 | 0.29615551 | -1.5059456 | 0.13208114 | 0.2418724 | PFKFB1 | 5207 |
| ENSG000001 | 3.38739338 | -0.569973709 | 0.37848073 | -1.5059517 | 0.13207957 | 0.2418724 | ODF3L2 | 284451 |
| ENSG000002 | 6.63728875 | 0.549166431 | 0.36466394 | 1.50595211 | 0.13207947 | 0.2418724 | NA | NA |
| ENSG000002 | 5.07613924 | -0.561290978 | 0.37269882 | -1.5060176 | 0.13206266 | 0.2418724 | NA | NA |
| ENSG000002 | 5.92743631 | -0.556175916 | 0.36931839 | -1.5059524 | 0.1320794 | 0.2418724 | NA | NA |
| ENSG000001 | 38.5761433 | -0.335953597 | 0.22309104 | -1.5059036 | 0.13209193 | 0.24188123 | STPG2 | 285555 |
| ENSG000002 | 6.20511365 | -0.552809241 | 0.36714889 | -1.5056814 | 0.13214899 | 0.24197479 | NA | NA |
| ENSG000001 | 23.3747896 | 0.409992188 | 0.27231298 | 1.50559177 | 0.132172 | 0.242006 | TM9SF1 | 10548 |
| ENSG000001 | 119.17841 | 0.251893643 | 0.16733281 | 1.50534519 | 0.13223535 | 0.24211106 | ZNF233 | 353355 |
| ENSG000002 | 7.58368411 | 0.549559099 | 0.36509537 | 1.50524805 | 0.13226032 | 0.24213648 | IDH1-AS1 | 100507475 |
| ENSG000002 | 10.0902331 | -0.51258517 | 0.34053279 | -1.5052447 | 0.13226118 | 0.24213648 | NA | NA |
| ENSG000002 | 7.20867742 | -0.55413064 | 0.36816361 | -1.5051206 | 0.13229307 | 0.24218393 | NA | NA |
| ENSG000001 | 24.359889 | -0.393650223 | 0.26156124 | -1.5050021 | 0.13232353 | 0.24222876 | NA | NA |
| ENSG000001 | 15.1146248 | -0.469322569 | 0.31192314 | -1.5046097 | 0.13242446 | 0.24239707 | NA | NA |
| ENSG000002 | 7.85481179 | -0.533499771 | 0.35457958 | -1.5045981 | 0.13242743 | 0.24239707 | PLA2G4E-AS1 | 101928388 |

| | | | | | | | | |
|---|---|---|---|---|---|---|---|---|
| ENSG000002 | 4.63491287 | -0.567861815 | 0.37743988 | -1.5045093 | 0.13245028 | 0.24242795 | NA | NA |
| ENSG000001 | 9.10276814 | -0.534248726 | 0.35514058 | -1.5043303 | 0.13249635 | 0.24250132 | DYNLRB2 | 83657 |
| ENSG000002 | 3.16337449 | -0.566382659 | 0.37656757 | -1.5040665 | 0.13256425 | 0.24261466 | NA | NA |
| ENSG000002 | 5.32666417 | -0.557287711 | 0.37056349 | -1.5038926 | 0.13260901 | 0.24267467 | NA | NA |
| ENSG000002 | 240.67886 | 0.225479698 | 0.14992849 | 1.50391499 | 0.13260326 | 0.24267467 | NA | NA |
| ENSG000002 | 2.2617526 | -0.550761145 | 0.36623229 | -1.5038574 | 0.13261809 | 0.242679 | NA | NA |
| ENSG000002 | 5.37723226 | -0.557513477 | 0.37072734 | -1.503837 | 0.13262335 | 0.242679 | NA | NA |
| ENSG000002 | 6.48917217 | -0.558948362 | 0.37168962 | -1.5038041 | 0.13263183 | 0.24268357 | NA | NA |
| ENSG00000C | 284.100649 | 0.26626143 | 0.17706864 | 1.50371871 | 0.13265381 | 0.24271284 | THOC3 | 84321 |
| ENSG000002 | 9.37935386 | -0.510412375 | 0.33944509 | -1.503667 | 0.13266714 | 0.24272627 | NA | NA |
| ENSG000002 | 8.37696532 | -0.527873094 | 0.35108062 | -1.5035666 | 0.132693 | 0.24276263 | NA | NA |
| ENSG00000C | 1988.42512 | 0.486675802 | 0.3237043 | 1.50345795 | 0.13272099 | 0.24279194 | HNRNPH3 | 3189 |
| ENSG000002 | 2.89571112 | -0.562945875 | 0.37443378 | -1.5034591 | 0.1327207 | 0.24279194 | NA | NA |
| ENSG000002 | 2.58047435 | -0.553597619 | 0.36823147 | -1.5033957 | 0.13273703 | 0.24281032 | NA | NA |
| ENSG000002 | 10.0851649 | -0.511067415 | 0.34000884 | -1.5031004 | 0.13281316 | 0.24293863 | ASAP1-IT2 | 100507117 |
| ENSG000002 | 4.35830556 | -0.568464445 | 0.37821275 | -1.5030282 | 0.13283176 | 0.2429617 | LINC00935 | 255411 |
| ENSG000002 | 2.53567121 | -0.558955313 | 0.37191207 | -1.5029233 | 0.13285882 | 0.24298927 | NA | NA |
| ENSG000002 | 4.17759933 | -0.568947203 | 0.37855464 | -1.502946 | 0.13285296 | 0.24298927 | HLX-AS1 | 100873924 |
| ENSG000001 | 727.251888 | -0.173919822 | 0.11572467 | -1.502876 | 0.13287103 | 0.24300064 | NTPCR | 84284 |
| ENSG000002 | 6.45534161 | -0.544905154 | 0.36258439 | -1.5028368 | 0.13288114 | 0.24300817 | NA | NA |
| ENSG000002 | 20.2274637 | -0.413464303 | 0.27512776 | -1.5028084 | 0.13288846 | 0.24301059 | NA | NA |
| ENSG000002 | 10.736505 | -0.498854358 | 0.33199397 | -1.5026007 | 0.13294205 | 0.24309763 | NA | NA |
| ENSG000002 | 4.84276594 | -0.563884281 | 0.37531681 | -1.5024221 | 0.13298812 | 0.24317091 | NA | NA |
| ENSG00000C | 51.0313001 | -0.308838342 | 0.20558593 | -1.5022348 | 0.13303648 | 0.24324702 | TLL1 | 7092 |
| ENSG000002 | 10.1721058 | -0.516253295 | 0.34366152 | -1.5022144 | 0.13304174 | 0.24324702 | NA | NA |
| ENSG000002 | 11.7934282 | -0.505270412 | 0.3363696 | -1.5021286 | 0.13306389 | 0.24327655 | NA | NA |
| ENSG000001 | 15.6331024 | -0.449919494 | 0.29954126 | -1.5020284 | 0.13308976 | 0.24331288 | Sep-01 | 1731 |
| ENSG000002 | 5.09504549 | -0.561245705 | 0.37366701 | -1.5019943 | 0.13309859 | 0.24331804 | NA | NA |
| ENSG000002 | 2.92907507 | -0.560141946 | 0.37296666 | -1.5018553 | 0.13313449 | 0.2433727 | NA | NA |

| | | | | | | | | |
|---|---|---|---|---|---|---|---|---|
| ENSG000002 | 2.2988599 | -0.548242167 | 0.36505185 | -1.50182 | 0.1331436 | 0.24337839 | NA | NA |
| ENSG000002 | 3.83088163 | -0.568388194 | 0.37847885 | -1.50177 | 0.13315651 | 0.24338032 | NA | NA |
| ENSG000002 | 5.7895148 | -0.560016622 | 0.37290452 | -1.5017695 | 0.13315666 | 0.24338032 | LOC100505853 | 100505853 |
| ENSG000002 | 2.51255827 | -0.560206535 | 0.37315183 | -1.5012831 | 0.13328235 | 0.24359908 | RNU6-5P | 106478913 |
| ENSG000002 | 36.7090637 | 0.339710203 | 0.22629286 | 1.50119716 | 0.13330458 | 0.24362871 | NA | NA |
| ENSG000001 | 36.1672081 | 0.344363311 | 0.22940442 | 1.5011189 | 0.13332481 | 0.24363432 | EFNA3 | 1944 |
| ENSG000001 | 3.47457664 | -0.566861118 | 0.37762656 | -1.5011156 | 0.13332567 | 0.24363432 | CRYGC | 1420 |
| ENSG000001 | 4.92859456 | -0.564385149 | 0.37597531 | -1.5011229 | 0.13332378 | 0.24363432 | EDDM3A | 10876 |
| ENSG000002 | 5.18959256 | 0.564836128 | 0.37630448 | 1.50100825 | 0.13335343 | 0.24367406 | NA | NA |
| ENSG000002 | 2.59583955 | -0.555295274 | 0.36998647 | -1.5008529 | 0.13339361 | 0.2437365 | NA | NA |
| ENSG000002 | 9.95352664 | -0.504277237 | 0.3360221 | -1.5007264 | 0.13342634 | 0.24378532 | NA | NA |
| ENSG000002 | 5.11778215 | -0.559635809 | 0.37292525 | -1.5006648 | 0.13344228 | 0.24379247 | NA | NA |
| ENSG000002 | 4.95562876 | -0.565536469 | 0.37685604 | -1.5006698 | 0.133441 | 0.24379247 | NA | NA |
| ENSG000001 | 170.774761 | 0.22657953 | 0.15099024 | 1.50062371 | 0.13345292 | 0.24380093 | C9orf85 | 138241 |
| ENSG000002 | 7.81708627 | -0.545051824 | 0.36326382 | -1.5004297 | 0.13350314 | 0.24387766 | NA | NA |
| ENSG000002 | 6.50484414 | -0.547026265 | 0.36458332 | -1.500415 | 0.13350695 | 0.24387766 | NA | NA |
| ENSG000001 | 592.989887 | 0.187759324 | 0.12514801 | 1.50029817 | 0.13353718 | 0.24389994 | RB1 | 5925 |
| ENSG000001 | 67.9092669 | 0.279573446 | 0.18634371 | 1.50031058 | 0.13353397 | 0.24389994 | GPRC5C | 55890 |
| ENSG000002 | 2.48990648 | -0.559693072 | 0.37305173 | -1.5003096 | 0.13353423 | 0.24389994 | NA | NA |
| ENSG000002 | 2.98493197 | -0.557977789 | 0.37193534 | -1.5002011 | 0.13356233 | 0.24393487 | NA | NA |
| ENSG000002 | 5.53074111 | -0.563196132 | 0.37542333 | -1.5001628 | 0.13357223 | 0.24394197 | NA | NA |
| ENSG000002 | 289.938538 | 0.513672784 | 0.34243927 | 1.50004051 | 0.13360391 | 0.24398884 | CDK11B | 984 |
| ENSG000002 | 2.54008173 | -0.556834717 | 0.37125752 | -1.4998611 | 0.13365039 | 0.24406272 | RN7SL589P | 106479438 |
| ENSG000001 | 149.150622 | 0.261308487 | 0.1742381 | 1.49972075 | 0.13368675 | 0.24411814 | LEPR | 3953 |
| ENSG000001 | 3.42905556 | -0.567409719 | 0.3783703 | -1.4996148 | 0.1337142 | 0.24414627 | NA | NA |
| ENSG000002 | 6.21883612 | -0.548580814 | 0.36580904 | -1.4996371 | 0.13370843 | 0.24414627 | NA | NA |
| ENSG000002 | 5.07708552 | -0.558857031 | 0.37269768 | -1.4994916 | 0.13374616 | 0.24419362 | NA | NA |
| ENSG000002 | 9.74797459 | -0.511021095 | 0.34081497 | -1.4994092 | 0.13376751 | 0.24421125 | NA | NA |
| ENSG000002 | 6.80232023 | -0.545599719 | 0.36387679 | -1.4994079 | 0.13376786 | 0.24421125 | NA | NA |

| | | | | | | | | |
|---|---|---|---|---|---|---|---|---|
| ENSG000001 | 140.165686 | 0.225662313 | 0.15051282 | 1.49928966 | 0.1337985 | 0.24424657 | ZNF773 | 374928 |
| ENSG000001 | 3.11313508 | -0.566038999 | 0.37753884 | -1.4992868 | 0.13379925 | 0.24424657 | THEM5 | 284486 |
| ENSG000002 | 2.27529463 | -0.546984124 | 0.36486528 | -1.49914 | 0.13383732 | 0.2442988 | RPPH1-2P | 106481202 |
| ENSG000002 | 93.2314751 | -0.564408718 | 0.37649084 | -1.49913 | 0.13383991 | 0.2442988 | U2AF1L5 | 102724594 |
| ENSG000002 | 4.22428013 | -0.565281018 | 0.3770896 | -1.4990629 | 0.13385732 | 0.24431958 | NA | NA |
| ENSG000002 | 2.83286655 | -0.561794813 | 0.37477611 | -1.4990145 | 0.13386988 | 0.24433151 | NA | NA |
| ENSG000002 | 9.0027724 | -0.52306618 | 0.34895264 | -1.4989604 | 0.13388392 | 0.24434613 | NA | NA |
| ENSG000002 | 4.02740898 | -0.567044311 | 0.37829782 | -1.4989362 | 0.13389018 | 0.24434656 | LOC101929125 | 101929125 |
| ENSG000002 | 2.26297056 | -0.545514249 | 0.36395661 | -1.4988442 | 0.13391406 | 0.24436815 | RNU6-362P | 106481292 |
| ENSG000002 | 2.87932851 | -0.559454317 | 0.37325289 | -1.4988613 | 0.13390961 | 0.24436815 | NA | NA |
| ENSG000001 | 26.9144536 | -0.378905262 | 0.25281439 | -1.4987488 | 0.13393882 | 0.24439134 | EGR2 | 1959 |
| ENSG000002 | 2.30225335 | -0.547983843 | 0.36562653 | -1.498753 | 0.13393773 | 0.24439134 | NA | NA |
| ENSG000001 | 20.4278903 | -0.426075053 | 0.28430649 | -1.4986469 | 0.13396525 | 0.24442856 | ANO7 | 50636 |
| ENSG000002 | 3.12554238 | -0.564726169 | 0.37684215 | -1.4985748 | 0.13398396 | 0.24445171 | NA | NA |
| ENSG000002 | 5.88722949 | -0.552412251 | 0.36864789 | -1.4984821 | 0.13400805 | 0.24448465 | NA | NA |
| ENSG000002 | 4.0456169 | -0.567013588 | 0.37844198 | -1.4982841 | 0.13405946 | 0.24456745 | DSCAS | 101927698 |
| ENSG000002 | 5.33924189 | -0.563580233 | 0.3762115 | -1.4980409 | 0.13412261 | 0.24467165 | EGOT | 100126791 |
| ENSG00000C | 58.8979825 | -0.319834051 | 0.21353513 | -1.4978053 | 0.13418384 | 0.24477233 | POU2F2 | 5452 |
| ENSG000001 | 6.8011075 | -0.54449413 | 0.36356868 | -1.4976376 | 0.13422743 | 0.24484083 | HOXC10 | 3226 |
| ENSG000002 | 26.9645416 | 0.389288035 | 0.2599581 | 1.49750299 | 0.13426243 | 0.24489365 | NA | NA |
| ENSG000002 | 6.74095842 | -0.540416853 | 0.3609051 | -1.4973932 | 0.13429097 | 0.24493469 | NA | NA |
| ENSG000001 | 12.5296898 | -0.48896306 | 0.3265638 | -1.4972972 | 0.13431594 | 0.24496923 | ELSPBP1 | 64100 |
| ENSG000002 | 44.6426053 | 0.366965334 | 0.24510659 | 1.49716633 | 0.13434998 | 0.24502029 | VAMP2 | 6844 |
| ENSG00000C | 255.956164 | 0.2057006 | 0.13740157 | 1.49707602 | 0.13437348 | 0.24505212 | SLC24A1 | 9187 |
| ENSG000002 | 2.22834778 | -0.544628118 | 0.36380412 | -1.4970367 | 0.13438371 | 0.24505976 | NA | NA |
| ENSG000001 | 11.2753956 | -0.49139371 | 0.32825772 | -1.4969753 | 0.13439967 | 0.24507785 | RSPH14 | 27156 |
| ENSG000001 | 60.0880288 | -0.290386485 | 0.19399135 | -1.4969043 | 0.13441816 | 0.24510054 | RHOH | 399 |
| ENSG000002 | 3.39071491 | -0.566583783 | 0.37852078 | -1.4968367 | 0.13443577 | 0.24512162 | NA | NA |
| ENSG00000C | 139.417935 | -0.223801646 | 0.1495231 | -1.4967697 | 0.13445319 | 0.24514236 | NRXN3 | 9369 |

| | | | | | | | | |
|---|---|---|---|---|---|---|---|---|
| ENSG000001 | 10.783939 | -0.50343662 | 0.33638975 | -1.4965873 | 0.13450068 | 0.24521792 | SLAMF8 | 56833 |
| ENSG000002 | 4.34726546 | -0.56596918 | 0.37818339 | -1.4965469 | 0.13451119 | 0.24522607 | RN7SL153P | 106480490 |
| ENSG000001 | 22.4852183 | -0.42033439 | 0.28091315 | -1.4963144 | 0.13457175 | 0.24529235 | PTGFR | 5737 |
| ENSG000001 | 15.64794 | -0.465715443 | 0.31123253 | -1.4963585 | 0.13456026 | 0.24529235 | ALLC | 55821 |
| ENSG000002 | 2.3411162 | -0.535234679 | 0.35770026 | -1.4963218 | 0.13456983 | 0.24529235 | NA | NA |
| ENSG000002 | 7.02873966 | -0.539926676 | 0.36082207 | -1.4963793 | 0.13455484 | 0.24529235 | NA | NA |
| ENSG000002 | 3.69233809 | -0.566464261 | 0.37858536 | -1.4962656 | 0.13458444 | 0.24530447 | NA | NA |
| ENSG000002 | 5.38945165 | -0.555668625 | 0.37140485 | -1.4961265 | 0.1346207 | 0.24535952 | NA | NA |
| ENSG000002 | 5.34137593 | -0.554767939 | 0.37081034 | -1.4960962 | 0.13462858 | 0.24536286 | NA | NA |
| ENSG000002 | 16.2750794 | -0.447654207 | 0.29922502 | -1.4960454 | 0.13464184 | 0.24537599 | CLEC6A | 93978 |
| ENSG000001 | 144.951255 | 0.236336214 | 0.15797975 | 1.49599054 | 0.13465612 | 0.24539099 | FRMD5 | 84978 |
| ENSG000001 | 140.818966 | 0.248447409 | 0.16609358 | 1.49582787 | 0.13469852 | 0.24545722 | GALNT6 | 11226 |
| ENSG000002 | 2.25895975 | -0.546251188 | 0.36519583 | -1.4957761 | 0.13471202 | 0.24547079 | NA | NA |
| ENSG000001 | 68.2253269 | 0.289910444 | 0.19382728 | 1.4957154 | 0.13472784 | 0.24548859 | PABPC5 | 140886 |
| ENSG000002 | 2.91190438 | -0.558635593 | 0.37352941 | -1.4955599 | 0.13476839 | 0.2455462 | NA | NA |
| ENSG000002 | 3.14907574 | -0.564131227 | 0.37720712 | -1.4955477 | 0.13477157 | 0.2455462 | NA | NA |
| ENSG000002 | 4.95403864 | -0.562281536 | 0.37603961 | -1.4952721 | 0.13484345 | 0.2456596 | NA | NA |
| ENSG000002 | 13.5675742 | -0.480402001 | 0.3212827 | -1.4952626 | 0.13484593 | 0.2456596 | C15orf65 | 145788 |
| ENSG000002 | 3.93738945 | -0.565635308 | 0.37834652 | -1.4950192 | 0.13490943 | 0.24576426 | NA | NA |
| ENSG000002 | 3.09551947 | -0.564250612 | 0.37743293 | -1.4949692 | 0.13492247 | 0.24577697 | NA | NA |
| ENSG000002 | 3.38713279 | -0.565776518 | 0.37846731 | -1.4949152 | 0.13493658 | 0.24579163 | NA | NA |
| ENSG000002 | 2.29942771 | -0.54410091 | 0.36398469 | -1.4948456 | 0.13495475 | 0.24581368 | NA | NA |
| ENSG000000 | 928.521921 | 0.187883766 | 0.12570157 | 1.49468119 | 0.13499767 | 0.24588081 | NT5C2 | 22978 |
| ENSG000001 | 120.214389 | 0.270977928 | 0.1813066 | 1.49458389 | 0.13502307 | 0.24591604 | WDR91 | 29062 |
| ENSG000002 | 4.94232028 | -0.563054553 | 0.3767561 | -1.4944803 | 0.13505013 | 0.24594324 | NA | NA |
| ENSG000002 | 3.82379037 | -0.565665347 | 0.37850033 | -1.494491 | 0.13504733 | 0.24594324 | NA | NA |
| ENSG000001 | 187.185069 | 0.214398032 | 0.14346507 | 1.49442671 | 0.13506413 | 0.24595731 | LRRC37B | 114659 |
| ENSG000002 | 4.01613477 | -0.565438602 | 0.37837058 | -1.4944042 | 0.13506999 | 0.24595731 | NA | NA |
| ENSG000002 | 7.82454776 | -0.528216303 | 0.35347602 | -1.4943483 | 0.1350846 | 0.2459682 | NA | NA |

| | | | | | | | | |
|---|---|---|---|---|---|---|---|---|
| ENSG000002 | 2.26369613 | -0.544345342 | 0.36427266 | -1.4943349 | 0.13508811 | 0.2459682 | OR7E2P | 8587 |
| ENSG000002 | 2.60682428 | -0.550318058 | 0.36828427 | -1.4942752 | 0.13510369 | 0.24598554 | NA | NA |
| ENSG000001 | 245.066636 | 0.210589287 | 0.14093569 | 1.49422253 | 0.13511747 | 0.24599957 | MESDC1 | 59274 |
| ENSG000002 | 7.70419482 | -0.536277812 | 0.35896043 | -1.4939747 | 0.13518223 | 0.24610643 | NA | NA |
| ENSG000000 | 11.0067175 | -0.496391451 | 0.3322785 | -1.4939018 | 0.13520129 | 0.24611903 | CAPN3 | 825 |
| ENSG000002 | 3.97040251 | -0.564910634 | 0.37814255 | -1.4939092 | 0.13519936 | 0.24611903 | NA | NA |
| ENSG000001 | 237.90354 | 0.196831241 | 0.13176095 | 1.49385113 | 0.13521454 | 0.24613209 | NDUFA9 | 4704 |
| ENSG000002 | 13.338803 | -0.475284642 | 0.31818628 | -1.4937308 | 0.135246 | 0.24617413 | NA | NA |
| ENSG000002 | 3.95471724 | -0.565200643 | 0.37838552 | -1.4937164 | 0.13524977 | 0.24617413 | NA | NA |
| ENSG000002 | 4.95470072 | -0.562917349 | 0.37686692 | -1.4936767 | 0.13526013 | 0.24618194 | NA | NA |
| ENSG000002 | 5.10266175 | -0.565174016 | 0.37838599 | -1.4936441 | 0.13526866 | 0.2461864 | NA | NA |
| ENSG000002 | 28.8624444 | 0.383954841 | 0.25709223 | 1.49345177 | 0.13531897 | 0.24626692 | SLC25A21-AS1 | 100129794 |
| ENSG000002 | 3.75254822 | -0.565169506 | 0.37845829 | -1.4933469 | 0.13534641 | 0.2463058 | NA | NA |
| ENSG000002 | 2.5239157 | -0.555036867 | 0.37167888 | -1.4933237 | 0.13535249 | 0.24630581 | NA | NA |
| ENSG000002 | 3.75417964 | -0.56519837 | 0.37850926 | -1.493222 | 0.13537908 | 0.24634315 | NA | NA |
| ENSG000001 | 10.2082168 | -0.517725669 | 0.34674604 | -1.4930976 | 0.13541165 | 0.24639135 | UGT2B15 | 7366 |
| ENSG000001 | 146.484003 | 0.238549442 | 0.15977956 | 1.49299094 | 0.13543956 | 0.24643108 | FUT4 | 2526 |
| ENSG000000 | 40.1925048 | -0.340365999 | 0.22799055 | -1.4928952 | 0.13546461 | 0.24645929 | TBC1D10A | 83874 |
| ENSG000002 | 6.75839777 | -0.539108622 | 0.36111859 | -1.4928853 | 0.13546722 | 0.24645929 | NA | NA |
| ENSG000001 | 13.1865385 | -0.491674104 | 0.32936631 | -1.4927881 | 0.13549266 | 0.24647589 | FXYD2 | 486 |
| ENSG000001 | 1911.84692 | 0.160767624 | 0.1076957 | 1.49279518 | 0.13549081 | 0.24647589 | HELZ | 9931 |
| ENSG000002 | 2.27780707 | -0.545181005 | 0.3652117 | -1.4927808 | 0.13549458 | 0.24647589 | NA | NA |
| ENSG000001 | 84.2437861 | 0.275395837 | 0.18449103 | 1.49273294 | 0.13550711 | 0.24648763 | DPYSL5 | 56896 |
| ENSG000002 | 2.85956085 | -0.557482076 | 0.37347691 | -1.4926815 | 0.13552058 | 0.24650107 | NA | NA |
| ENSG000002 | 3.1931049 | -0.561527624 | 0.37621523 | -1.4925701 | 0.13554974 | 0.24654306 | NA | NA |
| ENSG000002 | 9.2616957 | -0.517589881 | 0.34679701 | -1.4924866 | 0.13557163 | 0.24657181 | NA | NA |
| ENSG000002 | 4.87634407 | -0.560376559 | 0.37552075 | -1.4922652 | 0.13562962 | 0.24666623 | SAMSN1-AS1 | 100874190 |
| ENSG000001 | 7.63116703 | -0.532805852 | 0.35706285 | -1.4921907 | 0.13564917 | 0.24669071 | CCDC140 | 151278 |
| ENSG000001 | 19.8939996 | -0.420470036 | 0.28183428 | -1.4919052 | 0.13572399 | 0.24681571 | LOC494141 | 494141 |

| | | | | | | | | |
|---|---|---|---|---|---|---|---|---|
| ENSG000002 | 95.2330998 | 0.272361579 | 0.18257702 | 1.49176262 | 0.13576139 | 0.24687265 | LINC-PINT | 378805 |
| ENSG000002 | 9.9764604 | -0.50607318 | 0.33926856 | -1.4916595 | 0.13578843 | 0.24691075 | NA | NA |
| ENSG000002 | 3.37628335 | -0.564445035 | 0.37841703 | -1.4915952 | 0.1358053 | 0.24693034 | NA | NA |
| ENSG000002 | 2.57266969 | -0.554180129 | 0.3715492 | -1.491539 | 0.13582004 | 0.24694608 | NA | NA |
| ENSG000001 | 96.7795743 | 0.260310844 | 0.17453812 | 1.4914269 | 0.13584945 | 0.24697619 | RNU1-120P | 106481627 |
| ENSG000002 | 7.81940038 | -0.52820334 | 0.35416463 | -1.4914062 | 0.13585487 | 0.24697619 | LINC01449 | 101928773 |
| ENSG000002 | 19.7737813 | -0.420820161 | 0.28216079 | -1.4914197 | 0.13585134 | 0.24697619 | NA | NA |
| ENSG00000C | 183.705065 | 0.236713856 | 0.1587295 | 1.49130346 | 0.13588184 | 0.24700902 | CLCN6 | 1185 |
| ENSG00000C | 413.655981 | 0.213495359 | 0.14316144 | 1.49129099 | 0.13588511 | 0.24700902 | AACS | 65985 |
| ENSG000002 | 2.8869964 | -0.558329069 | 0.37443276 | -1.4911331 | 0.13592656 | 0.24707329 | NA | NA |
| ENSG000001 | 151.496594 | 0.222647672 | 0.14931974 | 1.49107995 | 0.1359405 | 0.24708028 | USB1 | 79650 |
| ENSG000002 | 3.69341499 | -0.564482991 | 0.37857528 | -1.491072 | 0.1359426 | 0.24708028 | SPATA31A3 | 727830 |
| ENSG000002 | 4.02090251 | -0.564440367 | 0.37856597 | -1.490996 | 0.13596254 | 0.24710545 | NA | NA |
| ENSG000002 | 3.1432104 | -0.562543544 | 0.37732559 | -1.4908704 | 0.13599553 | 0.24715434 | NA | NA |
| ENSG000002 | 100.743198 | 0.260746524 | 0.17490334 | 1.49080354 | 0.13601308 | 0.24717516 | NA | NA |
| ENSG000002 | 69.0211571 | 0.308020354 | 0.20663348 | 1.49066042 | 0.13605067 | 0.24722716 | FSBP | 100861412 |
| ENSG000002 | 5.45185526 | -0.558816876 | 0.37488695 | -1.4906277 | 0.13605926 | 0.24722716 | NA | NA |
| ENSG000002 | 3.36959703 | -0.564238786 | 0.37852498 | -1.490625 | 0.13605999 | 0.24722716 | NA | NA |
| ENSG000001 | 56.9687194 | 0.348011395 | 0.2334718 | 1.49059287 | 0.13606842 | 0.24723008 | RFXAP | 5994 |
| ENSG000002 | 4.0455048 | -0.564302946 | 0.37858137 | -1.4905724 | 0.13607379 | 0.24723008 | NA | NA |
| ENSG000001 | 16.988443 | -0.445737629 | 0.29905369 | -1.4904937 | 0.13609448 | 0.24725331 | ADRB2 | 154 |
| ENSG000002 | 111.765319 | 0.248448543 | 0.16669059 | 1.49047734 | 0.13609877 | 0.24725331 | EPHA1-AS1 | 285965 |
| ENSG000002 | 8.30166988 | -0.536144382 | 0.35973236 | -1.490398 | 0.13611962 | 0.24728011 | EXOC3L4 | 91828 |
| ENSG000001 | 27.212466 | -0.393217955 | 0.26384262 | -1.4903504 | 0.13613213 | 0.24729176 | ZNF648 | 127665 |
| ENSG000001 | 13.1603167 | -0.501418194 | 0.33644971 | -1.4903214 | 0.13613976 | 0.24729454 | DRC7 | 84229 |
| ENSG00000C | 8.96186301 | -0.518481658 | 0.34791007 | -1.4902749 | 0.13615196 | 0.24730563 | HOXC8 | 3224 |
| ENSG000002 | 127.235063 | -0.260755504 | 0.17498916 | -1.4901237 | 0.1361917 | 0.24736166 | NA | NA |
| ENSG000002 | 5.39890357 | -0.55447432 | 0.37210845 | -1.490088 | 0.13620111 | 0.24736166 | NA | NA |
| ENSG000002 | 56.7998754 | -0.315938147 | 0.2120254 | -1.4900957 | 0.13619907 | 0.24736166 | NA | NA |

| | | | | | | | | |
|---|---|---|---|---|---|---|---|---|
| ENSG000002 | 4.88143198 | -0.56008756 | 0.3759183 | -1.4899183 | 0.13624572 | 0.2474316 | NA | NA |
| ENSG000002 | 3.15417663 | -0.561402061 | 0.37684887 | -1.4897273 | 0.13629595 | 0.24751173 | NA | NA |
| ENSG000002 | 4.00255721 | -0.563868289 | 0.37852573 | -1.4896432 | 0.13631807 | 0.24754082 | NA | NA |
| ENSG00000C | 376.578024 | -0.183936191 | 0.12349024 | -1.4894796 | 0.13636111 | 0.24757463 | ZNF506 | 440515 |
| ENSG000001 | 28.8654466 | 0.376793219 | 0.25296915 | 1.4894829 | 0.13636025 | 0.24757463 | ABI3 | 51225 |
| ENSG000001 | 3.42407124 | -0.563359882 | 0.37822506 | -1.4894832 | 0.13636017 | 0.24757463 | RNU6-874P | 106479952 |
| ENSG000002 | 29.9579197 | 0.374338725 | 0.25132 | 1.48949041 | 0.13635828 | 0.24757463 | NA | NA |
| ENSG000002 | 2.34798582 | -0.541646492 | 0.36368198 | -1.489341 | 0.13639759 | 0.24760761 | NA | NA |
| ENSG000002 | 4.23751692 | -0.562291315 | 0.37754362 | -1.4893413 | 0.13639751 | 0.24760761 | ADAM6 | 8755 |
| ENSG000002 | 2.53387204 | -0.554520696 | 0.37231856 | -1.4893716 | 0.13638954 | 0.24760761 | NA | NA |
| ENSG000002 | 4.63500327 | -0.562730825 | 0.37786931 | -1.4892208 | 0.13642924 | 0.24765398 | NA | NA |
| ENSG000002 | 4.32104559 | -0.563510221 | 0.37840242 | -1.4891824 | 0.13643935 | 0.24766124 | NA | NA |
| ENSG000002 | 3.3732431 | -0.563484932 | 0.37846803 | -1.4888574 | 0.13652492 | 0.24780547 | NA | NA |
| ENSG00000C | 1548.29035 | 0.174900253 | 0.11747467 | 1.48883379 | 0.13653114 | 0.24780567 | RRP15 | 51018 |
| ENSG000002 | 2.84994605 | -0.55734273 | 0.37440968 | -1.4885906 | 0.13659521 | 0.24791085 | NA | NA |
| ENSG000001 | 277.127995 | 0.193775021 | 0.13018846 | 1.48841934 | 0.13664034 | 0.24798167 | ZNF570 | 148268 |
| ENSG000002 | 5.79690676 | -0.555223372 | 0.37305053 | -1.4883329 | 0.13666313 | 0.24801193 | NA | NA |
| ENSG000001 | 23.3466976 | -0.398932012 | 0.26804782 | -1.4882867 | 0.13667529 | 0.24801475 | GRID1 | 2894 |
| ENSG000002 | 3.1928575 | -0.56048456 | 0.37659872 | -1.4882806 | 0.13667691 | 0.24801475 | NA | NA |
| ENSG000002 | 4.26469656 | -0.561532436 | 0.3773352 | -1.4881528 | 0.1367106 | 0.24806477 | NA | NA |
| ENSG000002 | 8.56020647 | -0.512561795 | 0.34443871 | -1.4881074 | 0.13672257 | 0.24807538 | ZNF385D-AS1 | 100874216 |
| ENSG000001 | 30.3430976 | 0.410138718 | 0.27562198 | 1.48804792 | 0.13673826 | 0.24808165 | SLC16A13 | 201232 |
| ENSG000002 | 4.2682776 | -0.5621544 | 0.37777408 | -1.4880703 | 0.13673235 | 0.24808165 | NA | NA |
| ENSG000001 | 18.4716878 | -0.452391296 | 0.30409135 | -1.4876822 | 0.13683471 | 0.24822934 | SLITRK1 | 114798 |
| ENSG000002 | 4.56984926 | -0.561776246 | 0.3776267 | -1.4876497 | 0.1368433 | 0.24822934 | MIR609 | 693194 |
| ENSG000002 | 2.52635622 | -0.553858129 | 0.37230493 | -1.4876465 | 0.13684415 | 0.24822934 | RN7SL683P | 106481109 |
| ENSG000002 | 6.77998985 | 0.538132283 | 0.361718 | 1.48771222 | 0.1368268 | 0.24822934 | NA | NA |
| ENSG000002 | 9.71732914 | -0.503918294 | 0.33878572 | -1.4874248 | 0.13690264 | 0.24832434 | NA | NA |
| ENSG000002 | 5.99861429 | -0.549099791 | 0.3691691 | -1.4873937 | 0.13691087 | 0.24832815 | NA | NA |

| | | | | | | | | |
|---|---|---|---|---|---|---|---|---|
| ENSG000001 | 13.8794425 | -0.461311479 | 0.31016085 | -1.4873298 | 0.13692773 | 0.24834763 | IL1RN | 3557 |
| ENSG000002 | 11.0799462 | -0.515505815 | 0.34660446 | -1.4873029 | 0.13693483 | 0.2483494 | NIFKP7 | 106479049 |
| ENSG000001 | 11.3236399 | -0.49975905 | 0.33603768 | -1.4872113 | 0.136959 | 0.24838213 | DNAI2 | 64446 |
| ENSG000002 | 2.63468852 | -0.546991048 | 0.36782603 | -1.4870917 | 0.13699059 | 0.24842831 | NA | NA |
| ENSG000001 | 72.9579342 | -0.283272833 | 0.19050628 | -1.4869475 | 0.13702868 | 0.24848197 | GRIK4 | 2900 |
| ENSG000001 | 177.35431 | 0.259372689 | 0.17443956 | 1.4868914 | 0.13704349 | 0.24848197 | ZNF852 | 285346 |
| ENSG000002 | 8.93473992 | -0.521972154 | 0.35104576 | -1.4869063 | 0.13703956 | 0.24848197 | LDLRAD1 | 388633 |
| ENSG000002 | 2.34087075 | -0.535798799 | 0.3603494 | -1.4868869 | 0.13704469 | 0.24848197 | NA | NA |
| ENSG000002 | 18.2327026 | -0.431463806 | 0.29018452 | -1.4868602 | 0.13705175 | 0.24848366 | SLFN14 | 342618 |
| ENSG000002 | 7.18516353 | -0.543599169 | 0.36562308 | -1.4867748 | 0.1370743 | 0.24851344 | NA | NA |
| ENSG000001 | 118.23723 | 0.242942421 | 0.16340746 | 1.48672788 | 0.1370867 | 0.2485248 | ZNF19 | 7567 |
| ENSG000001 | 668.614412 | 0.190384371 | 0.12807705 | 1.48648308 | 0.13715139 | 0.24862069 | NR3C1 | 2908 |
| ENSG000002 | 4.51060246 | -0.558595242 | 0.37578356 | -1.4864813 | 0.13715186 | 0.24862069 | NA | NA |
| ENSG000002 | 2.52347906 | -0.553307903 | 0.37223964 | -1.4864293 | 0.13716561 | 0.24863451 | NA | NA |
| ENSG000001 | 86.0163476 | 0.286363656 | 0.192664 | 1.48633713 | 0.13718997 | 0.24864051 | VSIG10L | 147645 |
| ENSG000002 | 5.08220458 | -0.556610339 | 0.3744879 | -1.486324 | 0.13719345 | 0.24864051 | NA | NA |
| ENSG000002 | 6.31543294 | -0.550457215 | 0.37034306 | -1.4863441 | 0.13718814 | 0.24864051 | LOC100420587 | 100420587 |
| ENSG000002 | 2.73194846 | 0.557267512 | 0.37491276 | 1.48639248 | 0.13717534 | 0.24864051 | NA | NA |
| ENSG000002 | 2.72516363 | -0.54315491 | 0.36547689 | -1.4861539 | 0.13723841 | 0.24871088 | NA | NA |
| ENSG000002 | 12.2978155 | -0.472526967 | 0.31798101 | -1.4860226 | 0.13727314 | 0.24876271 | NA | NA |
| ENSG000002 | 2.28810274 | -0.542418558 | 0.36502747 | -1.4859664 | 0.137288 | 0.24877853 | NA | NA |
| ENSG000001 | 535.719023 | 0.251340923 | 0.16915205 | 1.4858875 | 0.13730888 | 0.24880524 | SHANK2 | 22941 |
| ENSG000002 | 2.29770589 | -0.54001803 | 0.36346829 | -1.4857363 | 0.13734889 | 0.24886661 | LRP1-AS | 105751187 |
| ENSG000001 | 48.4801264 | 0.313356403 | 0.21092125 | 1.48565594 | 0.13737015 | 0.24889402 | SLC9A5 | 6553 |
| ENSG000002 | 11.0451012 | -0.494500319 | 0.33287249 | -1.4855548 | 0.13739692 | 0.24893139 | NA | NA |
| ENSG000002 | 4.61067815 | -0.560049186 | 0.37705718 | -1.4853163 | 0.13746005 | 0.2490124 | NA | NA |
| ENSG000002 | 3.10001397 | -0.559866913 | 0.37692929 | -1.4853367 | 0.13745465 | 0.2490124 | NA | NA |
| ENSG000002 | 9.10283053 | -0.510517358 | 0.34370013 | -1.4853569 | 0.13744931 | 0.2490124 | LINC01258 | 101928776 |
| ENSG000002 | 50.0715942 | 0.3241607 | 0.21824903 | 1.48527898 | 0.13746994 | 0.24901918 | NA | NA |

| | | | | | | | | |
|---|---|---|---|---|---|---|---|---|
| ENSG000002 | 4.09895903 | -0.562284709 | 0.37858421 | -1.4852302 | 0.13748285 | 0.24903144 | NA | NA |
| ENSG000001 | 5.1363754 | -0.556043579 | 0.37441999 | -1.4850798 | 0.13752268 | 0.24909246 | SMAD5-AS1 | 9597 |
| ENSG000001 | 116.641237 | 0.233221841 | 0.15704607 | 1.48505366 | 0.13752961 | 0.24909389 | KCTD1 | 284252 |
| ENSG000002 | 48.341372 | 0.317416535 | 0.21374574 | 1.48501921 | 0.13753874 | 0.2490993 | NA | NA |
| ENSG000002 | 4.54997801 | -0.560081668 | 0.37720073 | -1.4848372 | 0.13758697 | 0.24917552 | NA | NA |
| ENSG000002 | 2.5499532 | -0.550515564 | 0.37080134 | -1.4846645 | 0.13763273 | 0.24924727 | NA | NA |
| ENSG000002 | 6.4761074 | -0.54069874 | 0.3642334 | -1.4844842 | 0.13768051 | 0.24932266 | NA | NA |
| ENSG000001 | 7.99820763 | -0.520111583 | 0.35037611 | -1.4844379 | 0.13769279 | 0.24933376 | KLRF1 | 51348 |
| ENSG000001 | 10.7610866 | -0.499495855 | 0.3365204 | -1.4842959 | 0.13773043 | 0.24939079 | ZNF804A | 91752 |
| ENSG00000C | 18.930786 | -0.430866361 | 0.29030234 | -1.4841987 | 0.13775621 | 0.24940861 | SPAG6 | 9576 |
| ENSG000002 | 3.47811724 | -0.559359194 | 0.37687862 | -1.4841892 | 0.13775872 | 0.24940861 | LOC105373635 | 105373635 |
| ENSG000002 | 3.98310319 | -0.56159818 | 0.37838519 | -1.484197 | 0.13775665 | 0.24940861 | NA | NA |
| ENSG000002 | 7.99898327 | 0.524598989 | 0.35358666 | 1.4836504 | 0.13790169 | 0.2496563 | PET117 | 100303755 |
| ENSG000001 | 15.1008443 | -0.466223178 | 0.31429559 | -1.4833908 | 0.13797062 | 0.24976016 | HEPACAM2 | 253012 |
| ENSG000002 | 318.811395 | 0.216642138 | 0.14604551 | 1.4833879 | 0.13797138 | 0.24976016 | NA | NA |
| ENSG000002 | 3.74731726 | -0.561476977 | 0.37856947 | -1.4831544 | 0.13803339 | 0.24983896 | NA | NA |
| ENSG000002 | 5.10644344 | -0.554368554 | 0.37377063 | -1.4831785 | 0.138027 | 0.24983896 | NA | NA |
| ENSG000002 | 6.80078944 | -0.539879189 | 0.36400182 | -1.4831772 | 0.13802734 | 0.24983896 | NA | NA |
| ENSG000002 | 4.4939279 | -0.557151101 | 0.37565901 | -1.4831299 | 0.13803991 | 0.24983961 | C10orf142 | 100130539 |
| ENSG000002 | 5.31556733 | -0.550330395 | 0.37107169 | -1.4830837 | 0.13805216 | 0.24985063 | NA | NA |
| ENSG000001 | 63.2864248 | 0.286490232 | 0.1931875 | 1.48296466 | 0.1380838 | 0.24989674 | CABP1 | 9478 |
| ENSG000001 | 5.50470004 | -0.555580214 | 0.37470643 | -1.482708 | 0.13815201 | 0.25000717 | NA | NA |
| ENSG000002 | 21.0311509 | -0.408783709 | 0.27570866 | -1.4826654 | 0.13816332 | 0.25000717 | CLEC5A | 23601 |
| ENSG000002 | 2.4941732 | -0.551313041 | 0.37183648 | -1.482676 | 0.1381605 | 0.25000717 | NA | NA |
| ENSG000002 | 3.45125088 | -0.560885981 | 0.37830183 | -1.4826415 | 0.1381697 | 0.25000756 | NA | NA |
| ENSG000002 | 3.20680878 | -0.556613556 | 0.37544249 | -1.4825535 | 0.13819309 | 0.25003873 | NA | NA |
| ENSG000001 | 7.15455696 | -0.542985642 | 0.36626568 | -1.4824912 | 0.13820965 | 0.25005753 | PRG3 | 10394 |
| ENSG000002 | 38.9064818 | 0.33835028 | 0.22827345 | 1.48221477 | 0.13828316 | 0.25017937 | NA | NA |
| ENSG000002 | 6.79858788 | -0.541076248 | 0.36505155 | -1.4821913 | 0.13828939 | 0.25017948 | NA | NA |

| | | | | | | | | |
|---|---|---|---|---|---|---|---|---|
| ENSG000002 | 2.51929667 | -0.550362583 | 0.37134126 | -1.4820938 | 0.13831533 | 0.25021525 | NA | NA |
| ENSG000002 | 3.18761226 | -0.557562616 | 0.37621647 | -1.4820261 | 0.13833336 | 0.2502367 | NA | NA |
| ENSG000001 | 4.91136623 | -0.55721106 | 0.3759972 | -1.4819553 | 0.13835218 | 0.25025958 | POTEH | 23784 |
| ENSG000000 | 413.244971 | 0.395902044 | 0.26716872 | 1.48184282 | 0.13838212 | 0.25030258 | CNOT3 | 4849 |
| ENSG000001 | 5.06750016 | -0.55980205 | 0.37779596 | -1.4817576 | 0.1384048 | 0.25032126 | OR2T33 | 391195 |
| ENSG000002 | 5.27717148 | -0.558785847 | 0.37710853 | -1.481764 | 0.1384031 | 0.25032126 | CST12P | 106478911 |
| ENSG000001 | 31.6527934 | 0.351523315 | 0.23724193 | 1.48170818 | 0.13841796 | 0.25032282 | LRRC37A3 | 374819 |
| ENSG000002 | 9.89955044 | -0.517669587 | 0.34937355 | -1.481708 | 0.138418 | 0.25032282 | LINGO1-AS1 | 253044 |
| ENSG000002 | 4.20659079 | -0.560253951 | 0.37817008 | -1.4814867 | 0.13847692 | 0.25041819 | NA | NA |
| ENSG000001 | 15.0478176 | -0.444750331 | 0.30021523 | -1.4814383 | 0.13848982 | 0.25043036 | LURAP1L | 286343 |
| ENSG000002 | 3.97863538 | -0.560612186 | 0.37843895 | -1.4813808 | 0.13850514 | 0.25044689 | NA | NA |
| ENSG000002 | 38.0494665 | 0.337856361 | 0.22807398 | 1.48134548 | 0.13851454 | 0.25045271 | NA | NA |
| ENSG000000 | 15.8092218 | -0.454974673 | 0.30716374 | -1.4812122 | 0.13855005 | 0.25049944 | ALOX5 | 240 |
| ENSG000001 | 19.1304469 | -0.442673722 | 0.29886113 | -1.4812021 | 0.13855273 | 0.25049944 | GIMAP4 | 55303 |
| ENSG000002 | 3.46734157 | -0.559495484 | 0.37773977 | -1.4811665 | 0.13856222 | 0.25050542 | NA | NA |
| ENSG000001 | 18.6915962 | -0.425542642 | 0.28731568 | -1.4810979 | 0.13858049 | 0.25051612 | NA | NA |
| ENSG000002 | 10.2303379 | -0.495015673 | 0.33421844 | -1.4811142 | 0.13857614 | 0.25051612 | NA | NA |
| ENSG000002 | 4.3027251 | -0.55935898 | 0.37771606 | -1.4808981 | 0.13863374 | 0.25060121 | NA | NA |
| ENSG000000 | 264.95214 | 0.217314082 | 0.14675087 | 1.48083671 | 0.13865009 | 0.25061959 | BAMBI | 25805 |
| ENSG000001 | 220.619971 | 0.223770654 | 0.15111432 | 1.48080378 | 0.13865887 | 0.25062428 | ZSCAN16 | 80345 |
| ENSG000002 | 2.84685265 | -0.552391939 | 0.37304746 | -1.4807551 | 0.13867183 | 0.25063655 | NA | NA |
| ENSG000002 | 19.9687046 | -0.413845476 | 0.27949943 | -1.4806666 | 0.13869543 | 0.25066802 | NA | NA |
| ENSG000002 | 2.2881339 | -0.536617776 | 0.36242318 | -1.4806387 | 0.13870288 | 0.25067032 | VTA1P1 | 106481708 |
| ENSG000002 | 3.34702956 | -0.560275251 | 0.37841869 | -1.4805697 | 0.13872127 | 0.25069238 | NA | NA |
| ENSG000002 | 10.9800715 | -0.488699542 | 0.33008923 | -1.4805074 | 0.13873789 | 0.25071124 | NA | NA |
| ENSG000001 | 70.862881 | 0.296433776 | 0.20023759 | 1.48041023 | 0.1387638 | 0.25074689 | IL18BP | 10068 |
| ENSG000002 | 3.98725204 | -0.56041968 | 0.37857722 | -1.4803312 | 0.13878488 | 0.2507738 | NA | NA |
| ENSG000001 | 7.64120334 | -0.532290816 | 0.35958312 | -1.4802998 | 0.13879327 | 0.25077778 | MAG | 4099 |
| ENSG000001 | 10.3376806 | -0.503024227 | 0.33985747 | -1.4801035 | 0.13884563 | 0.2508501 | NA | NA |

| | | | | | | | | |
|---|---|---|---|---|---|---|---|---|
| ENSG000002 | 13.3344709 | -0.465130751 | 0.31425559 | -1.4801034 | 0.13884566 | 0.2508501 | NA | NA |
| ENSG000002 | 4.00026235 | -0.560119776 | 0.37844728 | -1.480047 | 0.1388607 | 0.25086608 | NA | NA |
| ENSG000001 | 12.9301582 | -0.481484342 | 0.32533455 | -1.4799668 | 0.1388821 | 0.25089358 | G6PC | 2538 |
| ENSG000002 | 24.8808739 | -0.386872152 | 0.26141274 | -1.4799285 | 0.13889233 | 0.25090089 | NA | NA |
| ENSG000001 | 6.45894029 | -0.540760244 | 0.36541501 | -1.4798523 | 0.13891267 | 0.25092644 | CACNG2 | 10369 |
| ENSG000002 | 3.43190026 | -0.559630276 | 0.37818576 | -1.4797762 | 0.13893299 | 0.25095197 | NA | NA |
| ENSG000002 | 4.52568636 | -0.556304075 | 0.37595373 | -1.4797142 | 0.13894953 | 0.25097067 | NA | NA |
| ENSG000002 | 2.51275415 | -0.550772111 | 0.37222229 | -1.479686 | 0.13895707 | 0.2509731 | NA | NA |
| ENSG000002 | 3.76952881 | -0.559778978 | 0.37833267 | -1.4795946 | 0.13898146 | 0.25100598 | NA | NA |
| ENSG000001 | 32.5848613 | -0.351927578 | 0.23787011 | -1.4794948 | 0.13900813 | 0.25104296 | NT5E | 4907 |
| ENSG000001 | 65.2918866 | -0.283970236 | 0.19194431 | -1.4794408 | 0.13902254 | 0.25104663 | KAT2B | 8850 |
| ENSG000001 | 7.78738119 | -0.523450551 | 0.3538118 | -1.4794604 | 0.1390173 | 0.25104663 | GBX1 | 2636 |
| ENSG000002 | 58.4496787 | 0.295411711 | 0.19969848 | 1.47928873 | 0.13906316 | 0.2511088 | NA | NA |
| ENSG000002 | 2.24503782 | -0.540132589 | 0.36514432 | -1.4792304 | 0.13907874 | 0.25112574 | NA | NA |
| ENSG000002 | 61.3175484 | 0.287312341 | 0.19423461 | 1.47920261 | 0.13908617 | 0.25112798 | NA | NA |
| ENSG000002 | 12.5467099 | -0.468710319 | 0.316931 | -1.4789033 | 0.13916615 | 0.2512612 | NA | NA |
| ENSG000002 | 6.04882289 | -0.548093863 | 0.37062094 | -1.4788529 | 0.13917963 | 0.25127435 | NA | NA |
| ENSG000001 | 582.049175 | 0.197742577 | 0.13371774 | 1.47880589 | 0.1391922 | 0.25128586 | EOGT | 285203 |
| ENSG000002 | 44.2460639 | 0.335278875 | 0.22672657 | 1.47878073 | 0.13919893 | 0.25128682 | NA | NA |
| ENSG000002 | 3.42974124 | -0.559349832 | 0.37833508 | -1.4784509 | 0.13928714 | 0.25143486 | NA | NA |
| ENSG000002 | 8.08706287 | 0.517768598 | 0.35023321 | 1.47835381 | 0.1393131 | 0.25147054 | NA | NA |
| ENSG000001 | 22.9843315 | -0.412139688 | 0.27880844 | -1.4782181 | 0.13934941 | 0.25152488 | KCNJ3 | 3760 |
| ENSG000002 | 6.63768884 | -0.543869421 | 0.36795075 | -1.4781038 | 0.13937999 | 0.25156887 | NA | NA |
| ENSG000001 | 2.53967734 | -0.54789563 | 0.37071417 | -1.4779463 | 0.13942216 | 0.25161551 | UBQLN3 | 50613 |
| ENSG000002 | 3.17617552 | -0.55684039 | 0.3767685 | -1.4779377 | 0.13942444 | 0.25161551 | LOC105376935 | 105376935 |
| ENSG000002 | 2.49471344 | -0.547611095 | 0.3705207 | -1.4779501 | 0.13942114 | 0.25161551 | RN7SL284P | 106480999 |
| ENSG000001 | 634.041983 | 0.187930599 | 0.12716213 | 1.47788187 | 0.1394394 | 0.25163131 | SLX4IP | 128710 |
| ENSG000001 | 8.94407525 | -0.542223074 | 0.36692864 | -1.4777344 | 0.13947888 | 0.25169136 | ATP4A | 495 |
| ENSG000002 | 2.4944534 | -0.549724459 | 0.37201444 | -1.4776966 | 0.13948901 | 0.25169843 | PPIAL4C | 653598 |

| | | | | | | | | |
|---|---|---|---|---|---|---|---|---|
| ENSG000002 | 11.0805008 | -0.494200332 | 0.3344555 | -1.4776266 | 0.13950776 | 0.25172029 | NA | NA |
| ENSG000002 | 3.36418871 | -0.559228516 | 0.37846956 | -1.477605 | 0.13951354 | 0.25172029 | NA | NA |
| ENSG000002 | 4.29013323 | -0.557894799 | 0.37757707 | -1.4775654 | 0.13952415 | 0.25172823 | NA | NA |
| ENSG000002 | 31.2917409 | 0.37527968 | 0.25399921 | 1.47748362 | 0.13954605 | 0.25174534 | LOC101927043 | 101927043 |
| ENSG000002 | 3.3826702 | -0.558894893 | 0.37827143 | -1.477497 | 0.13954247 | 0.25174534 | NA | NA |
| ENSG000002 | 4.84688919 | -0.554979263 | 0.37563142 | -1.477457 | 0.13955318 | 0.25174702 | NA | NA |
| ENSG000002 | 9.45964204 | -0.524388852 | 0.35493821 | -1.4774088 | 0.13956608 | 0.25175908 | NA | NA |
| ENSG000001 | 7.49973326 | -0.528149742 | 0.35748975 | -1.4773843 | 0.13957266 | 0.25175975 | ABRA | 137735 |
| ENSG000002 | 3.25928651 | -0.55144934 | 0.37337869 | -1.476917 | 0.13969788 | 0.25195605 | NA | NA |
| ENSG000002 | 3.4514594 | -0.558476288 | 0.37813038 | -1.4769411 | 0.13969144 | 0.25195605 | NA | NA |
| ENSG000002 | 12.2809061 | -0.478461408 | 0.32396141 | -1.4769086 | 0.13970013 | 0.25195605 | LOC101929162 | 101929162 |
| ENSG000002 | 2.21911766 | -0.539088236 | 0.36502799 | -1.4768408 | 0.13971831 | 0.25197764 | NA | NA |
| ENSG000002 | 7.25590695 | -0.533171924 | 0.36108012 | -1.4766028 | 0.13978213 | 0.25208153 | NA | NA |
| ENSG000002 | 2.52645264 | -0.547419017 | 0.37074819 | -1.4765251 | 0.13980297 | 0.2521079 | NA | NA |
| ENSG000001 | 106.115329 | 0.238182547 | 0.16132172 | 1.4764444 | 0.13982463 | 0.25213574 | WHAMM | 123720 |
| ENSG000001 | 32.5828478 | 0.363098781 | 0.24596639 | 1.47621298 | 0.13988673 | 0.2522365 | CCDC61 | 729440 |
| ENSG000002 | 9.18117833 | -0.509366499 | 0.34505501 | -1.4761893 | 0.13989309 | 0.25223677 | NA | NA |
| ENSG000001 | 12.9792832 | -0.461935502 | 0.31293615 | -1.4761334 | 0.13990808 | 0.25225095 | DPEP2 | 64174 |
| ENSG000002 | 8.07216417 | -0.520186857 | 0.35240853 | -1.4760904 | 0.13991962 | 0.25225095 | NA | NA |
| ENSG000002 | 2.57053029 | -0.547567998 | 0.37095313 | -1.476111 | 0.13991409 | 0.25225095 | NA | NA |
| ENSG000002 | 3.41450831 | -0.558385889 | 0.3783054 | -1.4760188 | 0.13993884 | 0.25227438 | NA | NA |
| ENSG000002 | 5.1421039 | -0.552434475 | 0.37428019 | -1.4759918 | 0.13994611 | 0.25227626 | NA | NA |
| ENSG000002 | 3.99236861 | -0.558541106 | 0.37845903 | -1.4758298 | 0.13998959 | 0.25234344 | MIR633 | 693218 |
| ENSG000001 | 93.6692043 | 0.270992881 | 0.18362741 | 1.47577573 | 0.14000411 | 0.25234717 | SUV420H2 | 84787 |
| ENSG000001 | 11.7176244 | -0.494990028 | 0.335405 | -1.475798 | 0.13999813 | 0.25234717 | WNT4 | 54361 |
| ENSG000002 | 159.031173 | 0.259257757 | 0.17567846 | 1.47575154 | 0.1400106 | 0.25234766 | NA | NA |
| ENSG000001 | 16.3040821 | -0.448813877 | 0.30413488 | -1.4757067 | 0.14002265 | 0.25235815 | GATA2 | 2624 |
| ENSG000001 | 185.307881 | 0.2371144 | 0.16068305 | 1.47566525 | 0.14003378 | 0.252367 | HUNK | 30811 |
| ENSG000001 | 16.6974138 | -0.460885715 | 0.31235074 | -1.475539 | 0.14006768 | 0.25241688 | ALDOB | 229 |

| | | | | | | | | |
|---|---|---|---|---|---|---|---|---|
| ENSG000002 | 6.48952743 | -0.538700502 | 0.36509552 | -1.4755056 | 0.14007665 | 0.25242182 | NA | NA |
| ENSG000001 | 9.56646003 | -0.513038869 | 0.34771101 | -1.4754749 | 0.14008492 | 0.2524255 | SLC25A31 | 83447 |
| ENSG000002 | 2.30244923 | -0.537210655 | 0.36414058 | -1.4752837 | 0.14013629 | 0.25250685 | NA | NA |
| ENSG000001 | 91.9783452 | 0.269616282 | 0.18276597 | 1.47519959 | 0.14015889 | 0.25253636 | PORCN | 64840 |
| ENSG000002 | 2.52995023 | -0.547572893 | 0.37124737 | -1.4749543 | 0.14022483 | 0.25264394 | NA | NA |
| ENSG000002 | 3.37507006 | -0.55795429 | 0.37832791 | -1.4747902 | 0.14026895 | 0.25271221 | NA | NA |
| ENSG00000C | 31.1967771 | -0.365421388 | 0.24779084 | -1.4747171 | 0.14028862 | 0.25271988 | STAB1 | 23166 |
| ENSG000002 | 11.289507 | 0.487091833 | 0.33030156 | 1.47468826 | 0.14029638 | 0.25271988 | MYLK-AS1 | 100506826 |
| ENSG000002 | 5.41449188 | -0.548347967 | 0.37184158 | -1.4746817 | 0.14029814 | 0.25271988 | LINC01532 | 100505835 |
| ENSG000002 | 5.74538901 | -0.548346653 | 0.37183066 | -1.4747215 | 0.14028744 | 0.25271988 | NA | NA |
| ENSG000002 | 3.36371208 | -0.557958582 | 0.37837552 | -1.474616 | 0.14031582 | 0.2527405 | NA | NA |
| ENSG000002 | 3.39120418 | -0.557902004 | 0.37837456 | -1.4744702 | 0.14035504 | 0.25279991 | COX6CP13 | 106480267 |
| ENSG000001 | 11.3226701 | -0.509794348 | 0.34575683 | -1.4744303 | 0.14036577 | 0.252808 | HYAL4 | 23553 |
| ENSG000002 | 3.41348239 | -0.557391622 | 0.37805215 | -1.4743776 | 0.14037995 | 0.25282233 | NA | NA |
| ENSG000002 | 7.39170723 | -0.530949509 | 0.3601377 | -1.4742958 | 0.14040196 | 0.25284204 | NA | NA |
| ENSG000002 | 7.37375935 | -0.531510458 | 0.36051947 | -1.4742906 | 0.14040337 | 0.25284204 | NA | NA |
| ENSG000002 | 18.0797157 | -0.43068574 | 0.29215104 | -1.4741886 | 0.14043081 | 0.25287961 | TRIM10 | 10107 |
| ENSG000002 | 17.076503 | 0.434695807 | 0.2948756 | 1.47416673 | 0.1404367 | 0.25287961 | NA | NA |
| ENSG000002 | 5.60471646 | -0.554945579 | 0.3764731 | -1.4740644 | 0.14046426 | 0.25291801 | NA | NA |
| ENSG000002 | 8.59766589 | -0.512588134 | 0.34774472 | -1.4740357 | 0.14047197 | 0.25292066 | NA | NA |
| ENSG000002 | 45.7081675 | -0.320878855 | 0.21771247 | -1.4738653 | 0.14051786 | 0.25299203 | GVINP1 | 387751 |
| ENSG000001 | 4.25450001 | -0.556433713 | 0.37755027 | -1.4738003 | 0.14053536 | 0.25301231 | EDN2 | 1907 |
| ENSG000001 | 72.2258924 | 0.271659804 | 0.18434129 | 1.47367855 | 0.14056816 | 0.25306013 | SYT9 | 143425 |
| ENSG000001 | 12.013588 | -0.493317475 | 0.33477301 | -1.4735879 | 0.14059257 | 0.25309284 | GAPT | 202309 |
| ENSG000002 | 6.99658011 | -0.528991097 | 0.35898937 | -1.4735564 | 0.14060106 | 0.25309688 | NA | NA |
| ENSG000002 | 4.43593057 | -0.557810341 | 0.37858208 | -1.4734198 | 0.14063787 | 0.25315157 | NA | NA |
| ENSG000002 | 5.80932202 | -0.551634023 | 0.37439596 | -1.4733974 | 0.14064392 | 0.25315157 | NA | NA |
| ENSG000001 | 10.9853994 | -0.488115393 | 0.33132241 | -1.4732339 | 0.14068799 | 0.25321964 | NA | NA |
| ENSG000002 | 3.75104766 | -0.557475709 | 0.37844877 | -1.4730546 | 0.14073631 | 0.25329537 | PCDH9-AS3 | 100874086 |

| | | | | | | | | |
|---|---|---|---|---|---|---|---|---|
| ENSG000002 | 4.27507656 | -0.556353447 | 0.3777012 | -1.4729989 | 0.14075134 | 0.25329673 | RNU6ATAC27P | 106481167 |
| ENSG000002 | 9.69506964 | -0.499631254 | 0.33920238 | -1.4729592 | 0.14076205 | 0.25329673 | NA | NA |
| ENSG000002 | 5.07384383 | -0.550599486 | 0.37380314 | -1.4729664 | 0.14076009 | 0.25329673 | NA | NA |
| ENSG000002 | 4.06549802 | -0.557588607 | 0.37854227 | -1.472989 | 0.140754 | 0.25329673 | NA | NA |
| ENSG000002 | 13.3921492 | -0.468835943 | 0.31832683 | -1.4728132 | 0.14080142 | 0.25335633 | NA | NA |
| ENSG000002 | 6.45834125 | -0.547299698 | 0.3716105 | -1.4727778 | 0.14081095 | 0.25336225 | LINC00507 | 100862680 |
| ENSG000002 | 7.03001656 | -0.536315132 | 0.36415885 | -1.4727505 | 0.14081833 | 0.25336427 | NA | NA |
| ENSG00000C | 271.910395 | 0.19704135 | 0.13379736 | 1.47268485 | 0.14083603 | 0.25338489 | USP40 | 55230 |
| ENSG000002 | 4.77824345 | -0.553347543 | 0.3757525 | -1.4726383 | 0.14084858 | 0.25339622 | NA | NA |
| ENSG000002 | 2.49784651 | -0.547240735 | 0.37163615 | -1.4725175 | 0.14088119 | 0.25344364 | NA | NA |
| ENSG000001 | 68.8926393 | 0.280225625 | 0.19032663 | 1.47234061 | 0.14092892 | 0.25350001 | HRH2 | 3274 |
| ENSG000001 | 4.47930046 | -0.55397596 | 0.37625582 | -1.4723386 | 0.14092947 | 0.25350001 | PMCHL1 | 5369 |
| ENSG000002 | 5.12026199 | -0.550567593 | 0.37394259 | -1.4723319 | 0.14093128 | 0.25350001 | NA | NA |
| ENSG000002 | 8.85218218 | -0.510038605 | 0.34645172 | -1.472178 | 0.14097282 | 0.25356348 | NA | NA |
| ENSG00000C | 3059.40276 | 0.176868271 | 0.12015303 | 1.472025 | 0.14101413 | 0.25361959 | LRP6 | 4040 |
| ENSG000001 | 35.7999452 | -0.359033198 | 0.24390575 | -1.4720161 | 0.14101652 | 0.25361959 | FBXO15 | 201456 |
| ENSG000001 | 209.915555 | 0.213972874 | 0.14536279 | 1.47199203 | 0.14102303 | 0.25362005 | C3orf67 | 200844 |
| ENSG000002 | 13.0002198 | -0.460209348 | 0.31265376 | -1.4719457 | 0.14103555 | 0.25362527 | TAS2R46 | 259292 |
| ENSG000002 | 3.66068322 | -0.557027946 | 0.37843244 | -1.471935 | 0.14103844 | 0.25362527 | NA | NA |
| ENSG000002 | 4.52913189 | -0.553294405 | 0.37595663 | -1.4716974 | 0.1411026 | 0.2537294 | NA | NA |
| ENSG000002 | 8.52689341 | -0.529392211 | 0.35972757 | -1.4716476 | 0.14111606 | 0.25374235 | NA | NA |
| ENSG000001 | 11.564545 | -0.480648068 | 0.32662754 | -1.4715479 | 0.141143 | 0.25377954 | TAC4 | 255061 |
| ENSG000002 | 2.30319908 | -0.534360733 | 0.36315387 | -1.4714444 | 0.14117096 | 0.25381856 | NA | NA |
| ENSG000002 | 3.43018507 | -0.556574309 | 0.37826113 | -1.4714023 | 0.14118234 | 0.25382776 | NA | NA |
| ENSG000001 | 118.535007 | 0.250793554 | 0.17045033 | 1.47135857 | 0.14119417 | 0.25383777 | MYRFL | 196446 |
| ENSG000002 | 7.67975011 | -0.537686034 | 0.36546225 | -1.4712492 | 0.14122374 | 0.25386554 | LOC101928778 | 101928778 |
| ENSG000002 | 3.34410088 | -0.556660853 | 0.37835933 | -1.4712492 | 0.14122374 | 0.25386554 | CDRT7 | 94150 |
| ENSG000002 | 38.9559335 | 0.34768426 | 0.23632185 | 1.47123197 | 0.14122839 | 0.25386554 | LOC100506083 | 100506083 |
| ENSG000002 | 6.53134599 | -0.544734809 | 0.3702793 | -1.4711457 | 0.14125172 | 0.2538962 | NA | NA |

| | | | | | | | | | |
|---|---|---|---|---|---|---|---|---|---|
| ENSG000001 | 31.2480121 | -0.362030084 | 0.24610504 | -1.4710389 | 0.1412806 | 0.25392705 | EPHA10 | | 284656 |
| ENSG000002 | 6.41208494 | -0.538113055 | 0.36580552 | -1.4710359 | 0.1412814 | 0.25392705 | OR2A25 | | 392138 |
| ENSG000002 | 4.3011791 | -0.555432793 | 0.37760428 | -1.4709388 | 0.14130767 | 0.253963 | NA | NA | |
| ENSG000001 | 558.357285 | 0.224274821 | 0.15247838 | 1.47086306 | 0.14132816 | 0.25398857 | UFC1 | | 51506 |
| ENSG000002 | 2.52764218 | -0.547072293 | 0.37195998 | -1.4707827 | 0.1413499 | 0.2540164 | NA | NA | |
| ENSG000002 | 10.7364857 | -0.505003867 | 0.3433832 | -1.4706714 | 0.14137999 | 0.25405921 | LINC00927 | | 283688 |
| ENSG000001 | 8.07265453 | -0.516115617 | 0.35096337 | -1.4705684 | 0.14140789 | 0.25408682 | LINC00905 | | 148231 |
| ENSG000002 | 2.86406442 | -0.548913746 | 0.37326229 | -1.4705845 | 0.14140351 | 0.25408682 | NA | NA | |
| ENSG000001 | 11.7121717 | -0.478259656 | 0.32524402 | -1.4704641 | 0.1414361 | 0.25412035 | RSPH10B2 | | 728194 |
| ENSG000001 | 12.8670899 | -0.473235473 | 0.32183984 | -1.4704068 | 0.14145162 | 0.25412035 | C2orf73 | | 129852 |
| ENSG000002 | 3.4486763 | -0.555094376 | 0.37750672 | -1.4704225 | 0.14144736 | 0.25412035 | NA | NA | |
| ENSG000002 | 23.6155076 | -0.413926097 | 0.28149966 | -1.470432 | 0.1414448 | 0.25412035 | SERPINE3 | | 647174 |
| ENSG000001 | 295.193173 | 0.203795563 | 0.13860817 | 1.4702998 | 0.14148057 | 0.25416111 | RIT1 | | 6016 |
| ENSG000002 | 2.91050632 | -0.546171053 | 0.37153133 | -1.4700538 | 0.14154718 | 0.25425904 | MIR601 | | 693186 |
| ENSG000002 | 8.174178 | -0.528569847 | 0.35955857 | -1.4700522 | 0.14154763 | 0.25425904 | NA | NA | |
| ENSG000002 | 7003705.45 | 0.170944891 | 0.11629174 | 1.46996592 | 0.14157098 | 0.25428973 | NA | NA | |
| ENSG000002 | 2.29078327 | -0.53421113 | 0.36343334 | -1.4699013 | 0.14158849 | 0.25430992 | NA | NA | |
| ENSG000002 | 4.05032136 | -0.556425544 | 0.37855283 | -1.4698756 | 0.14159546 | 0.25431117 | NA | NA | |
| ENSG000002 | 48.141237 | 0.327879764 | 0.22307226 | 1.46983654 | 0.14160603 | 0.25431888 | NA | NA | |
| ENSG000001 | 79.3184286 | 0.269535593 | 0.18341306 | 1.46955509 | 0.14168229 | 0.25443331 | AFAP1L2 | | 84632 |
| ENSG000002 | 12.9189545 | -0.486291862 | 0.33090997 | -1.4695594 | 0.14168112 | 0.25443331 | NA | NA | |
| ENSG000002 | 3.4836842 | -0.554694354 | 0.37747307 | -1.4694938 | 0.14169889 | 0.25445186 | NA | NA | |
| ENSG00000C | 23.8228763 | -0.417049973 | 0.28383042 | -1.4693632 | 0.14173431 | 0.25450418 | PALMD | | 54873 |
| ENSG000002 | 4.47147253 | -0.551866477 | 0.37565623 | -1.4690731 | 0.14181295 | 0.25462285 | NA | NA | |
| ENSG000002 | 10.4599826 | 0.494335686 | 0.33649218 | 1.46908523 | 0.14180967 | 0.25462285 | NA | NA | |
| ENSG000001 | 8.620346 | -0.508802471 | 0.34637428 | -1.4689384 | 0.1418495 | 0.25465465 | PRR27 | | 401137 |
| ENSG000002 | 3.44143945 | -0.55555029 | 0.37819754 | -1.4689421 | 0.1418485 | 0.25465465 | NA | NA | |
| ENSG000002 | 7.26855052 | -0.524183391 | 0.35683625 | -1.4689746 | 0.14183968 | 0.25465465 | NA | NA | |
| ENSG000002 | 3.15471273 | -0.553353923 | 0.37675977 | -1.4687182 | 0.14190924 | 0.25475062 | NA | NA | |

| | | | | | | | | | |
|---|---|---|---|---|---|---|---|---|---|
| ENSG000001 | 5.1221089 | -0.548890712 | 0.37375226 | -1.468595 | 0.14194267 | 0.25477678 | FEZF2 | | 55079 |
| ENSG000002 | 2.52493387 | -0.545391645 | 0.371362 | -1.4686253 | 0.14193444 | 0.25477678 | NA | NA | |
| ENSG000002 | 206.934206 | 0.21683443 | 0.1476471 | 1.46859927 | 0.14194151 | 0.25477678 | NA | NA | |
| ENSG000002 | 2.76922874 | -0.551951694 | 0.37585002 | -1.4685424 | 0.14195695 | 0.25479114 | NA | NA | |
| ENSG000001 | 17.4368997 | -0.430283173 | 0.29302014 | -1.4684423 | 0.14198412 | 0.25482863 | PATE2 | | 399967 |
| ENSG000002 | 3.08973344 | -0.553701715 | 0.37707963 | -1.4683947 | 0.14199705 | 0.25484056 | NA | NA | |
| ENSG000001 | 249.105845 | -0.195303883 | 0.13300869 | -1.4683543 | 0.142008 | 0.25484893 | MAPK10 | | 5602 |
| ENSG000002 | 4.0760781 | -0.555864619 | 0.37857111 | -1.4683229 | 0.14201655 | 0.25485299 | NA | NA | |
| ENSG000002 | 3.98506111 | -0.555832412 | 0.3785577 | -1.4682898 | 0.14202552 | 0.2548561 | NA | NA | |
| ENSG000002 | 9.643887 | -0.501004589 | 0.34122098 | -1.4682702 | 0.14203085 | 0.2548561 | NA | NA | |
| ENSG000001 | 15.5691573 | -0.485048547 | 0.33037501 | -1.4681757 | 0.14205651 | 0.25489087 | LIPF | | 8513 |
| ENSG000001 | 7.59029489 | -0.524408546 | 0.35721021 | -1.468067 | 0.14208601 | 0.25493252 | MC2R | | 4158 |
| ENSG000001 | 227.802472 | 0.24761507 | 0.16867899 | 1.46796625 | 0.14211339 | 0.25497036 | PER2 | | 8864 |
| ENSG000002 | 2.51162519 | -0.545581777 | 0.37167016 | -1.4679192 | 0.14212616 | 0.25498199 | NA | NA | |
| ENSG000001 | 181.374787 | 0.206030989 | 0.14038558 | 1.46760796 | 0.14221074 | 0.25507644 | MBP | | 4155 |
| ENSG000002 | 5.94131195 | -0.545644606 | 0.3717931 | -1.4676028 | 0.14221214 | 0.25507644 | NA | NA | |
| ENSG000002 | 8.17596149 | -0.519575246 | 0.35400678 | -1.4676986 | 0.14218612 | 0.25507644 | NA | NA | |
| ENSG000002 | 3.63890401 | -0.55557725 | 0.37856097 | -1.4676031 | 0.14221206 | 0.25507644 | NA | NA | |
| ENSG000002 | 3.38400823 | -0.555114301 | 0.37824977 | -1.4675866 | 0.14221655 | 0.25507644 | NA | NA | |
| ENSG000002 | 3.36300713 | -0.55509018 | 0.37823181 | -1.4675925 | 0.14221494 | 0.25507644 | NA | NA | |
| ENSG000002 | 4.20785874 | -0.554115036 | 0.37759861 | -1.4674711 | 0.14224795 | 0.25512148 | NA | NA | |
| ENSG000002 | 3.26742709 | -0.545686244 | 0.37186322 | -1.4674381 | 0.14225693 | 0.2551263 | NA | NA | |
| ENSG000002 | 6.71972486 | -0.531131821 | 0.3619614 | -1.4673714 | 0.14227504 | 0.2551475 | NA | NA | |
| ENSG000001 | 24.7190546 | -0.388233487 | 0.26459277 | -1.4672869 | 0.14229804 | 0.25517745 | NA | NA | |
| ENSG000002 | 16.1842572 | -0.450438555 | 0.30699522 | -1.4672494 | 0.14230823 | 0.25518444 | NA | NA | |
| ENSG000002 | 5.88055379 | -0.53847173 | 0.36700428 | -1.4672083 | 0.14231939 | 0.25519318 | NA | NA | |
| ENSG000001 | 6.70986494 | -0.533470736 | 0.36360833 | -1.4671576 | 0.14233318 | 0.25520662 | FAM182A | | 284800 |
| ENSG000001 | 12.4089278 | -0.47891799 | 0.32646973 | -1.4669599 | 0.14238697 | 0.25529177 | CLEC4F | | 165530 |
| ENSG000002 | 3.95606736 | -0.555276059 | 0.37856239 | -1.466802 | 0.14242994 | 0.25534623 | NA | NA | |

| | | | | | | | | |
|---|---|---|---|---|---|---|---|---|
| ENSG000002 | 77.0167645 | 0.283933154 | 0.1935707 | 1.46681889 | 0.14242533 | 0.25534623 | NA | NA |
| ENSG000001 | 10.0284623 | -0.497294605 | 0.33904919 | -1.4667329 | 0.14244873 | 0.25536864 | CD69 | 969 |
| ENSG000002 | 4.31244479 | -0.554415954 | 0.37800118 | -1.4667043 | 0.14245651 | 0.2553713 | LOC105370473 | 105370473 |
| ENSG000001 | 29.4535757 | -0.373839781 | 0.25488956 | -1.4666736 | 0.14246487 | 0.25537499 | VSTM2A | 222008 |
| ENSG000002 | 6.21882652 | -0.543137204 | 0.3703481 | -1.4665586 | 0.14249616 | 0.25541979 | NA | NA |
| ENSG000002 | 2.5007752 | -0.543119549 | 0.37035085 | -1.4665001 | 0.1425121 | 0.25543707 | RNU6-1270P | 106481581 |
| ENSG00000C | 712.451121 | 0.246641922 | 0.16818707 | 1.46647371 | 0.14251928 | 0.25543864 | ZIC2 | 7546 |
| ENSG000002 | 2.76239807 | -0.55111333 | 0.37584543 | -1.4663297 | 0.14255848 | 0.25546374 | NA | NA |
| ENSG000002 | 11.2294137 | -0.499794613 | 0.34084619 | -1.4663347 | 0.14255712 | 0.25546374 | TARP | 445347 |
| ENSG000002 | 4.01810608 | -0.555001984 | 0.37848575 | -1.4663748 | 0.1425462 | 0.25546374 | HBG1 | 3047 |
| ENSG000002 | 3.70051337 | 0.55514853 | 0.37858672 | 1.46637084 | 0.14254729 | 0.25546374 | NA | NA |
| ENSG000002 | 3.94601516 | -0.554945125 | 0.37847262 | -1.4662755 | 0.14257325 | 0.25547892 | NA | NA |
| ENSG00000C | 31.7780188 | -0.363190964 | 0.24773508 | -1.4660457 | 0.14263583 | 0.25557374 | PDE4A | 5141 |
| ENSG000001 | 29.3170624 | 0.394479166 | 0.26907897 | 1.46603491 | 0.14263878 | 0.25557374 | GOLT1A | 127845 |
| ENSG000001 | 7.77850319 | -0.518929843 | 0.35402136 | -1.4658151 | 0.14269867 | 0.25562459 | RETNLB | 84666 |
| ENSG000002 | 4.46270988 | -0.551641073 | 0.37633678 | -1.4658176 | 0.14269799 | 0.25562459 | NA | NA |
| ENSG000002 | 27.0548735 | 0.382153601 | 0.26070736 | 1.46583356 | 0.14269363 | 0.25562459 | NA | NA |
| ENSG000002 | 5.36434835 | -0.545701262 | 0.37228181 | -1.4658284 | 0.14269504 | 0.25562459 | NA | NA |
| ENSG000002 | 5.79837008 | -0.547013325 | 0.3731706 | -1.4658532 | 0.14268828 | 0.25562459 | NA | NA |
| ENSG00000C | 462.75014 | 0.178422362 | 0.12172655 | 1.46576376 | 0.14271266 | 0.25563835 | GCFC2 | 6936 |
| ENSG000002 | 4.5128336 | -0.552278834 | 0.3768031 | -1.4656961 | 0.1427311 | 0.25566009 | NA | NA |
| ENSG00000C | 507.371226 | 0.193138584 | 0.13177559 | 1.4656628 | 0.14274017 | 0.25566505 | RPS6KA6 | 27330 |
| ENSG000002 | 12.5049145 | 0.488871953 | 0.33355836 | 1.46562646 | 0.14275008 | 0.2556715 | NA | NA |
| ENSG000002 | 2.47640856 | -0.544880831 | 0.37180353 | -1.4655074 | 0.14278253 | 0.25571833 | NA | NA |
| ENSG000002 | 23.9641326 | -0.38898118 | 0.26545389 | -1.4653437 | 0.14282717 | 0.25575309 | PPP5D1 | 100506012 |
| ENSG000002 | 3.35690102 | -0.55440452 | 0.3783293 | -1.465402 | 0.14281126 | 0.25575309 | NA | NA |
| ENSG000002 | 12.0195492 | -0.474873538 | 0.32406572 | -1.4653618 | 0.14282223 | 0.25575309 | LINC01229 | 101928248 |
| ENSG000002 | 2.78980529 | -0.550216011 | 0.37548061 | -1.4653647 | 0.14282145 | 0.25575309 | NA | NA |
| ENSG000001 | 32.7872389 | 0.356654762 | 0.24340488 | 1.46527366 | 0.14284627 | 0.255776 | HEYL | 26508 |

| | | | | | | | | |
|---|---|---|---|---|---|---|---|---|
| ENSG000001 | 3.50523685 | -0.551090869 | 0.37611667 | -1.4652126 | 0.14286293 | 0.25579454 | CCL1 | 6346 |
| ENSG000002 | 3.38762205 | -0.554033573 | 0.3782003 | -1.464921 | 0.14294247 | 0.25592565 | NA | NA |
| ENSG000001 | 62.688578 | 0.314892092 | 0.21498193 | 1.46473748 | 0.14299256 | 0.25600402 | C20orf96 | 140680 |
| ENSG000002 | 3.85436671 | -0.553126809 | 0.37767197 | -1.4645694 | 0.14303844 | 0.25607486 | WSPAR | 105664404 |
| ENSG000001 | 417.643578 | 0.181415992 | 0.12389346 | 1.46429034 | 0.14311464 | 0.25618574 | GTDC1 | 79712 |
| ENSG000001 | 19.0978405 | -0.439141583 | 0.29989855 | -1.4643004 | 0.14311188 | 0.25618574 | NA | NA |
| ENSG000002 | 15.762456 | -0.445303685 | 0.30411244 | -1.4642732 | 0.14311933 | 0.25618574 | LINC01482 | 101928104 |
| ENSG000002 | 11.7189555 | -0.493172389 | 0.33685661 | -1.4640425 | 0.14318234 | 0.25628723 | LOC105378311 | 105378311 |
| ENSG000001 | 46.761071 | 0.337317346 | 0.23043788 | 1.46381034 | 0.14324578 | 0.25638945 | PP7080 | 25845 |
| ENSG000002 | 2.23529056 | -0.530140784 | 0.3621708 | -1.4637867 | 0.14325224 | 0.25638971 | NA | NA |
| ENSG000002 | 7.66410232 | -0.529652145 | 0.36187607 | -1.4636286 | 0.14329546 | 0.25645575 | LOC101929268 | 101929268 |
| ENSG000001 | 5.31873156 | -0.545352673 | 0.37263625 | -1.4634987 | 0.14333096 | 0.25650796 | C1orf189 | 388701 |
| ENSG00000C | 116.591549 | -0.245938539 | 0.16806835 | -1.4633245 | 0.14337861 | 0.2565819 | SERTAD4 | 56256 |
| ENSG000002 | 8.67778512 | -0.515956933 | 0.35259931 | -1.4632953 | 0.14338658 | 0.25658485 | NA | NA |
| ENSG000001 | 139.398354 | 0.231962498 | 0.15852697 | 1.4632368 | 0.14340259 | 0.25660218 | PCOLCE2 | 26577 |
| ENSG000002 | 7.65234385 | -0.527243157 | 0.36033776 | -1.4631915 | 0.14341498 | 0.25661302 | SPATA8-AS1 | 100652749 |
| ENSG000002 | 2.77303899 | -0.549666597 | 0.37569847 | -1.4630525 | 0.14345301 | 0.25666974 | NA | NA |
| ENSG000001 | 13.8287699 | -0.457038365 | 0.31240492 | -1.4629679 | 0.14347617 | 0.25668923 | LYG2 | 254773 |
| ENSG000002 | 11.6349381 | -0.500032826 | 0.34179378 | -1.4629664 | 0.14347656 | 0.25668923 | NA | NA |
| ENSG000002 | 14.1266019 | -0.45373119 | 0.31016034 | -1.4628923 | 0.14349684 | 0.25670286 | C21orf91-OT1 | 246312 |
| ENSG000002 | 2.51249824 | -0.542651964 | 0.37094412 | -1.4628941 | 0.14349636 | 0.25670286 | NA | NA |
| ENSG000002 | 4.51134763 | -0.55039643 | 0.37624762 | -1.4628569 | 0.14350654 | 0.25670889 | NA | NA |
| ENSG000002 | 2.89952103 | -0.544524999 | 0.37225206 | -1.4627857 | 0.14352602 | 0.25673241 | NA | NA |
| ENSG000002 | 2.27455274 | -0.532333696 | 0.3639507 | -1.4626533 | 0.14356227 | 0.25678592 | NA | NA |
| ENSG00000C | 35.5075429 | 0.342073169 | 0.23389544 | 1.46250468 | 0.14360296 | 0.25684738 | SH2D2A | 9047 |
| ENSG000001 | 19.3989588 | -0.42708593 | 0.29203044 | -1.4624706 | 0.1436123 | 0.25685274 | MST1 | 4485 |
| ENSG000002 | 7.20523428 | 0.532695724 | 0.36428702 | 1.46229676 | 0.1436599 | 0.25692656 | LOC101928659 | 101928659 |
| ENSG000002 | 9.12871214 | -0.504471895 | 0.34500352 | -1.4622225 | 0.14368026 | 0.25695163 | NA | NA |
| ENSG000001 | 11.041697 | -0.486792258 | 0.33294588 | -1.4620762 | 0.14372032 | 0.25701194 | IL36RN | 26525 |

| | | | | | | | | |
|---|---|---|---|---|---|---|---|---|
| ENSG000002 | 4.25959981 | -0.553062305 | 0.37830397 | -1.4619522 | 0.14375432 | 0.25705006 | NA | NA |
| ENSG000002 | 4.91004119 | -0.550658738 | 0.37665812 | -1.461959 | 0.14375244 | 0.25705006 | NA | NA |
| ENSG000002 | 6.74980017 | -0.530075828 | 0.36260444 | -1.461857 | 0.14378041 | 0.25708539 | NA | NA |
| ENSG000001 | 6.06844096 | -0.543728231 | 0.37195423 | -1.4618149 | 0.14379194 | 0.25709467 | HOXB5 | 3215 |
| ENSG000001 | 239.95449 | -0.2016802 | 0.13796934 | -1.4617755 | 0.14380274 | 0.25710263 | DCP1B | 196513 |
| ENSG000002 | 3.10352366 | -0.54931709 | 0.37580177 | -1.4617203 | 0.14381786 | 0.25711834 | USP17L15 | 100288520 |
| ENSG000001 | 35.0616337 | 0.346625973 | 0.23715095 | 1.46162594 | 0.14384374 | 0.25715327 | NMNAT3 | 349565 |
| ENSG000002 | 5.29780269 | -0.543784171 | 0.37206332 | -1.4615366 | 0.14386823 | 0.25717565 | NA | NA |
| ENSG000002 | 3.16193984 | -0.549499676 | 0.37597461 | -1.461534 | 0.14386895 | 0.25717565 | NA | NA |
| ENSG000002 | 7.59900327 | -0.524016506 | 0.35855249 | -1.4614778 | 0.14388437 | 0.25717769 | NA | NA |
| ENSG000002 | 5.15051337 | -0.547605061 | 0.37469714 | -1.4614605 | 0.14388912 | 0.25717769 | NA | NA |
| ENSG000002 | 3.0866921 | -0.549922211 | 0.37627402 | -1.461494 | 0.14387993 | 0.25717769 | NA | NA |
| ENSG000002 | 5.63538979 | -0.541484291 | 0.37054829 | -1.4613056 | 0.14393159 | 0.25724228 | NA | NA |
| ENSG000002 | 6.98627125 | -0.527800878 | 0.36120812 | -1.4612099 | 0.14395785 | 0.25727485 | RN7SL792P | 106479509 |
| ENSG000002 | 5.05986111 | -0.547615092 | 0.37477262 | -1.4611929 | 0.14396251 | 0.25727485 | NA | NA |
| ENSG000001 | 100.46016 | 0.257200617 | 0.17603191 | 1.46110219 | 0.1439874 | 0.25728532 | AAED1 | 195827 |
| ENSG000002 | 22.0067385 | -0.395663308 | 0.2707925 | -1.461131 | 0.1439795 | 0.25728532 | NA | NA |
| ENSG000002 | 8.77894013 | -0.504011894 | 0.34494907 | -1.4611198 | 0.14398258 | 0.25728532 | NA | NA |
| ENSG000002 | 3.39857563 | -0.552499997 | 0.37819947 | -1.4608693 | 0.14405131 | 0.25737684 | NA | NA |
| ENSG000002 | 2.46571611 | -0.543112988 | 0.37177174 | -1.4608775 | 0.14404905 | 0.25737684 | OR52R1 | 119695 |
| ENSG000001 | 5.12184776 | -0.546774987 | 0.37429179 | -1.4608255 | 0.14406334 | 0.257387 | NA | NA |
| ENSG000002 | 9.50012766 | -0.51298781 | 0.35117998 | -1.4607547 | 0.14408277 | 0.25740347 | NA | NA |
| ENSG000002 | 18.9699463 | -0.423737249 | 0.29008284 | -1.4607456 | 0.14408526 | 0.25740347 | NA | NA |
| ENSG000002 | 53.1087871 | 0.5159238 | 0.35321663 | 1.46064414 | 0.14411313 | 0.25744191 | NA | NA |
| ENSG000002 | 2.57912788 | -0.539254765 | 0.36921589 | -1.4605405 | 0.14414159 | 0.25748062 | NA | NA |
| ENSG000002 | 2.33956395 | -0.527488426 | 0.36116506 | -1.460519 | 0.14414749 | 0.25748062 | NA | NA |
| ENSG000002 | 5.04025657 | -0.552175164 | 0.37807737 | -1.4604819 | 0.14415767 | 0.25748674 | NA | NA |
| ENSG000002 | 7.18482142 | -0.52058684 | 0.35645396 | -1.4604603 | 0.14416362 | 0.25748674 | NA | NA |
| ENSG000002 | 4.47352761 | -0.54935758 | 0.37618328 | -1.4603455 | 0.14419513 | 0.25753168 | NA | NA |

| | | | | | | | | |
|---|---|---|---|---|---|---|---|---|
| ENSG000002 | 2.49767025 | -0.542045404 | 0.37118313 | -1.460318 | 0.1442027 | 0.25753385 | NA | NA |
| ENSG000001 | 165.424737 | 0.234167027 | 0.16037212 | 1.46014793 | 0.14424942 | 0.25760595 | FAM120C | 54954 |
| ENSG000002 | 2.27678115 | -0.527493495 | 0.36129076 | -1.4600249 | 0.14428324 | 0.25765499 | NA | NA |
| ENSG000001 | 12.5553077 | -0.46687705 | 0.3197797 | -1.4599959 | 0.1442912 | 0.25765786 | GDF9 | 2661 |
| ENSG000002 | 3.38234456 | -0.552045117 | 0.37813654 | -1.4599095 | 0.14431496 | 0.25768894 | NA | NA |
| ENSG000002 | 46.0207867 | -0.320824181 | 0.21976367 | -1.45986 | 0.14432857 | 0.25770189 | LINC00240 | 100133205 |
| ENSG000002 | 8.44891775 | -0.519237804 | 0.35570002 | -1.4597632 | 0.14435517 | 0.25773804 | OR4D10 | 390197 |
| ENSG000002 | 6.27925531 | -0.53805672 | 0.3686127 | -1.4596804 | 0.14437794 | 0.25776736 | LOC101928162 | 101928162 |
| ENSG000002 | 3.47351037 | -0.551380354 | 0.37776141 | -1.4595995 | 0.14440019 | 0.25779572 | NA | NA |
| ENSG000001 | 105.299483 | 0.239127171 | 0.16383877 | 1.45952739 | 0.14442001 | 0.25781975 | ARHGAP22 | 58504 |
| ENSG000001 | 156.346698 | 0.212653391 | 0.14571808 | 1.45934799 | 0.14446935 | 0.25789649 | SHROOM4 | 57477 |
| ENSG000002 | 2.48495408 | -0.539570663 | 0.36975892 | -1.4592499 | 0.14449634 | 0.25792631 | LOC100128531 | 100128531 |
| ENSG000002 | 2.77295467 | -0.548397853 | 0.37581033 | -1.459241 | 0.14449878 | 0.25792631 | CACNA1C-AS3 | 106478964 |
| ENSG000001 | 56.869473 | 0.282671087 | 0.19373184 | 1.45908427 | 0.14454192 | 0.25799092 | DDIT4L | 115265 |
| ENSG000002 | 7.25369611 | -0.537808445 | 0.36860417 | -1.4590406 | 0.14455394 | 0.25799092 | NA | NA |
| ENSG000002 | 8.73140644 | -0.514844252 | 0.35286504 | -1.4590401 | 0.14455406 | 0.25799092 | LOC440390 | 440390 |
| ENSG000002 | 4.35355013 | -0.552226999 | 0.37849817 | -1.4589952 | 0.14456644 | 0.25800166 | LOC101927616 | 101927616 |
| ENSG000001 | 2497.86429 | 0.20176025 | 0.13829251 | 1.45893837 | 0.14458207 | 0.2580182 | CCND1 | 595 |
| ENSG000002 | 2.90430349 | -0.54140211 | 0.37115057 | -1.4587129 | 0.14464415 | 0.25810627 | NA | NA |
| ENSG000002 | 2.48737478 | -0.541508083 | 0.37122219 | -1.4587169 | 0.14464304 | 0.25810627 | NA | NA |
| ENSG000001 | 240.307898 | 0.250621968 | 0.17181918 | 1.45863794 | 0.14466479 | 0.25813173 | NA | NA |
| ENSG000002 | 3.68306746 | -0.551768232 | 0.37830333 | -1.4585339 | 0.14469344 | 0.25817149 | FABP5P3 | 220832 |
| ENSG000002 | 9.24776952 | -0.526290115 | 0.36086024 | -1.458432 | 0.14472151 | 0.25821022 | NA | NA |
| ENSG000002 | 15.4990306 | 0.459066419 | 0.3147752 | 1.45839452 | 0.14473183 | 0.25821728 | NA | NA |
| ENSG000002 | 2.48614885 | -0.541419259 | 0.37126221 | -1.4583204 | 0.14475225 | 0.25824234 | NA | NA |
| ENSG000001 | 405.465754 | 0.177151017 | 0.12148452 | 1.45821887 | 0.14478022 | 0.25828089 | ZBTB18 | 10472 |
| ENSG000001 | 14.0877445 | -0.461349976 | 0.3164076 | -1.4580875 | 0.14481642 | 0.25831136 | PROC | 5624 |
| ENSG000001 | 552.40901 | 0.184893256 | 0.1268021 | 1.45812451 | 0.14480623 | 0.25831136 | IL6ST | 3572 |
| ENSG000002 | 3.15943536 | -0.548043764 | 0.37585954 | -1.4581079 | 0.1448108 | 0.25831136 | NA | NA |

| | | | | | | | | |
|---|---|---|---|---|---|---|---|---|
| ENSG000002 | 2.49511893 | -0.540900328 | 0.37098607 | -1.4580071 | 0.14483859 | 0.25833955 | NA | NA |
| ENSG000001 | 5.86873935 | -0.53489414 | 0.36687747 | -1.457964 | 0.14485047 | 0.25834937 | C17orf99 | 100141515 |
| ENSG000002 | 4.50909766 | -0.551484903 | 0.3782834 | -1.4578618 | 0.14487865 | 0.25837691 | NA | NA |
| ENSG000002 | 52.5650109 | 0.31231401 | 0.21422479 | 1.45787991 | 0.14487365 | 0.25837691 | NA | NA |
| ENSG000002 | 3.13616403 | -0.548763871 | 0.37643108 | -1.457807 | 0.14489374 | 0.25839246 | NA | NA |
| ENSG000002 | 8.50739387 | -0.517246227 | 0.35485667 | -1.4576201 | 0.14494528 | 0.25847299 | NA | NA |
| ENSG000002 | 7.58450013 | -0.522365999 | 0.35839998 | -1.4574945 | 0.14497993 | 0.25852342 | NA | NA |
| ENSG000001 | 140.742581 | -0.220387159 | 0.15122108 | -1.4573838 | 0.14501046 | 0.2585665 | CDH8 | 1006 |
| ENSG000002 | 5.82208274 | -0.546774621 | 0.3752017 | -1.4572818 | 0.1450386 | 0.25860529 | NA | NA |
| ENSG000001 | 3.14431921 | -0.547911099 | 0.37598932 | -1.4572518 | 0.1450469 | 0.25860872 | FSCN2 | 25794 |
| ENSG000002 | 311.690211 | 0.189006884 | 0.12971155 | 1.45713222 | 0.14507989 | 0.25864826 | FGFR1OP | 11116 |
| ENSG000002 | 26.6503427 | 0.375337442 | 0.25758765 | 1.45712517 | 0.14508183 | 0.25864826 | TAF1A-AS1 | 100506161 |
| ENSG000002 | 3.42037309 | -0.550565566 | 0.37789624 | -1.4569226 | 0.14513775 | 0.25871933 | NA | NA |
| ENSG000002 | 10.1851578 | -0.493860264 | 0.3389763 | -1.4569168 | 0.14513935 | 0.25871933 | NA | NA |
| ENSG000002 | 16.3400637 | 0.444265416 | 0.30493647 | 1.4569114 | 0.14514084 | 0.25871933 | NA | NA |
| ENSG000002 | 3.13142101 | 0.54787392 | 0.37605948 | 1.45688103 | 0.14514922 | 0.25872291 | SNORA11 | 677799 |
| ENSG000001 | 205.369928 | 0.24480701 | 0.16803838 | 1.45685172 | 0.14515732 | 0.25872596 | MOV10 | 4343 |
| ENSG000001 | 6.97759009 | -0.523283 | 0.35928616 | -1.4564519 | 0.14526774 | 0.2589114 | HOXA3 | 3200 |
| ENSG000001 | 9.44419433 | -0.505976792 | 0.34743157 | -1.4563351 | 0.1453 | 0.25895752 | MAGEA11 | 4110 |
| ENSG000001 | 9.44070018 | -0.507979138 | 0.34886484 | -1.4560915 | 0.14536731 | 0.25904094 | CNGA4 | 1262 |
| ENSG000001 | 239.568851 | 0.189976827 | 0.13047202 | 1.45607328 | 0.14537236 | 0.25904094 | NA | NA |
| ENSG000002 | 13.0438942 | 0.469687813 | 0.32256457 | 1.45610478 | 0.14536366 | 0.25904094 | USP27X-AS1 | 158572 |
| ENSG000002 | 2.53760951 | -0.537893804 | 0.36940569 | -1.4561059 | 0.14536335 | 0.25904094 | NA | NA |
| ENSG000001 | 4.49184001 | -0.548117242 | 0.37644306 | -1.4560429 | 0.14538076 | 0.25904452 | IRX1 | 79192 |
| ENSG000001 | 433.715314 | 0.205198223 | 0.1409351 | 1.45597672 | 0.14539906 | 0.25905671 | IQGAP3 | 128239 |
| ENSG000002 | 3.75564631 | -0.550588955 | 0.37815905 | -1.4559719 | 0.14540038 | 0.25905671 | NA | NA |
| ENSG00000C | 383.418231 | 0.195856249 | 0.13452627 | 1.45589596 | 0.14542138 | 0.25905999 | DOCK9 | 23348 |
| ENSG000001 | 5.61713643 | -0.538882446 | 0.37013269 | -1.4559169 | 0.14541559 | 0.25905999 | OR8H2 | 390151 |
| ENSG000002 | 5.5574098 | -0.538763314 | 0.37005168 | -1.4559137 | 0.14541647 | 0.25905999 | NA | NA |

| | | | | | | | | | |
|---|---|---|---|---|---|---|---|---|---|
| ENSG000001 | 68.6224665 | 0.301448801 | 0.20707847 | 1.45572253 | 0.14546934 | 0.25912101 | TMEM92 | | 162461 |
| ENSG000002 | 2.56173572 | -0.537961623 | 0.36956045 | -1.4556796 | 0.1454812 | 0.25912101 | NA | NA | |
| ENSG000002 | 9.13039454 | -0.506433644 | 0.34789179 | -1.4557218 | 0.14546955 | 0.25912101 | NA | NA | |
| ENSG000002 | 4.27169141 | -0.549986034 | 0.37781886 | -1.4556871 | 0.14547914 | 0.25912101 | NA | NA | |
| ENSG000001 | 7.80750957 | -0.51621935 | 0.35468232 | -1.4554415 | 0.14554708 | 0.25922697 | AKR1C4 | | 1109 |
| ENSG000002 | 6.1665402 | -0.535340561 | 0.36786232 | -1.4552742 | 0.14559336 | 0.259298 | NA | NA | |
| ENSG000002 | 7.59966194 | -0.521810458 | 0.35857861 | -1.4552191 | 0.14560861 | 0.25931377 | NA | NA | |
| ENSG000002 | 2.22440865 | -0.527754841 | 0.36268711 | -1.4551243 | 0.14563485 | 0.25934911 | NA | NA | |
| ENSG000002 | 5.63367494 | -0.538909733 | 0.37036756 | -1.4550673 | 0.14565063 | 0.25936582 | NA | NA | |
| ENSG000002 | 5.02694163 | -0.544784437 | 0.37443099 | -1.4549662 | 0.14567862 | 0.25940427 | NA | NA | |
| ENSG000002 | 3.18162529 | -0.543930828 | 0.37389695 | -1.4547613 | 0.14573535 | 0.2594939 | NA | NA | |
| ENSG000002 | 4.270987 | -0.54940967 | 0.37769454 | -1.4546402 | 0.14576889 | 0.25954223 | NA | NA | |
| ENSG000002 | 3.65133998 | -0.550647573 | 0.37856096 | -1.454581 | 0.1457853 | 0.25956005 | NA | NA | |
| ENSG000002 | 2.47214184 | -0.539465709 | 0.37093414 | -1.4543437 | 0.14585105 | 0.25966571 | NA | NA | |
| ENSG000001 | 22.2231861 | -0.397554243 | 0.27337676 | -1.4542357 | 0.14588098 | 0.25969914 | ZNF876P | | 642280 |
| ENSG000002 | 5.14519405 | -0.546151615 | 0.37556076 | -1.4542297 | 0.14588263 | 0.25969914 | NA | NA | |
| ENSG000002 | 4.80330633 | -0.54830442 | 0.37707058 | -1.4541162 | 0.1459141 | 0.25974376 | NA | NA | |
| ENSG000002 | 7.08147282 | -0.543372957 | 0.3737152 | -1.4539761 | 0.14595294 | 0.25979836 | NA | NA | |
| ENSG000002 | 3.20423475 | -0.545279473 | 0.37503077 | -1.4539593 | 0.14595759 | 0.25979836 | NA | NA | |
| ENSG000001 | 198.358898 | 0.205147383 | 0.14110303 | 1.45388362 | 0.14597858 | 0.25981709 | PLCL2 | | 23228 |
| ENSG000002 | 17.1904894 | -0.447286211 | 0.30765105 | -1.4538751 | 0.14598093 | 0.25981709 | NA | NA | |
| ENSG000002 | 4.25737697 | -0.549482223 | 0.37795919 | -1.4538136 | 0.14599799 | 0.25983605 | NA | NA | |
| ENSG000002 | 5.14675628 | -0.544241664 | 0.37441563 | -1.4535763 | 0.14606382 | 0.25994181 | NA | NA | |
| ENSG000002 | 4.22879534 | -0.549193264 | 0.37783193 | -1.4535385 | 0.14607429 | 0.25994904 | NA | NA | |
| ENSG000002 | 4.57979295 | -0.549197547 | 0.37784801 | -1.453488 | 0.14608831 | 0.25996258 | NA | NA | |
| ENSG000002 | 188.774004 | -0.221421617 | 0.15234226 | -1.4534484 | 0.14609929 | 0.25996414 | NHSL2 | | 340527 |
| ENSG000002 | 2.773496 | -0.54599183 | 0.37565524 | -1.4534386 | 0.14610201 | 0.25996414 | NA | NA | |
| ENSG000001 | 2.54468868 | -0.538087596 | 0.3702449 | -1.4533288 | 0.14613247 | 0.26000342 | SPPL2C | | 162540 |
| ENSG000002 | 3.64445724 | -0.550164277 | 0.37855874 | -1.4533128 | 0.14613691 | 0.26000342 | NA | NA | |

| | | | | | | | | |
|---|---|---|---|---|---|---|---|---|
| ENSG000001 | 344.979568 | 0.205789757 | 0.14160634 | 1.45325239 | 0.14615369 | 0.26002186 | RCAN3 | 11123 |
| ENSG000001 | 14.3868973 | -0.464408373 | 0.31959643 | -1.4531088 | 0.14619355 | 0.26008137 | NRSN1 | 140767 |
| ENSG000002 | 4.25757773 | -0.549402468 | 0.37813848 | -1.4529134 | 0.1462478 | 0.26015994 | NA | NA |
| ENSG000002 | 2.68494417 | -0.53149307 | 0.36581443 | -1.4529035 | 0.14625055 | 0.26015994 | NA | NA |
| ENSG000002 | 8.66841431 | -0.507463949 | 0.34929807 | -1.4528106 | 0.14627634 | 0.26017895 | NA | NA |
| ENSG000002 | 31.0937823 | 0.369212822 | 0.25413953 | 1.45279572 | 0.14628048 | 0.26017895 | ZNF625 | 90589 |
| ENSG000002 | 2.49579143 | -0.539409996 | 0.37128729 | -1.4528103 | 0.14627644 | 0.26017895 | NA | NA |
| ENSG000001 | 8.32080413 | -0.517446595 | 0.35618572 | -1.4527438 | 0.1462949 | 0.26018177 | STRA8 | 346673 |
| ENSG000002 | 8.62573451 | -0.506324823 | 0.34852878 | -1.4527489 | 0.14629347 | 0.26018177 | NA | NA |
| ENSG000002 | 8.21878201 | -0.52020459 | 0.35809912 | -1.4526832 | 0.14631172 | 0.26020028 | NA | NA |
| ENSG000002 | 5.85787009 | -0.534081665 | 0.36765903 | -1.4526548 | 0.14631962 | 0.26020291 | NA | NA |
| ENSG000002 | 46.6996602 | 0.309870033 | 0.21333691 | 1.45249142 | 0.14636501 | 0.26026408 | HSPA1L | 3305 |
| ENSG000002 | 9.44644837 | -0.500310228 | 0.34445127 | -1.4524848 | 0.14636686 | 0.26026408 | FLJ31356 | 403150 |
| ENSG000001 | 252.285359 | 0.196797151 | 0.13549853 | 1.45239325 | 0.14639229 | 0.26028648 | RIMKLA | 284716 |
| ENSG000002 | 3.96261097 | -0.549824125 | 0.37856217 | -1.4524011 | 0.1463901 | 0.26028648 | NA | NA |
| ENSG000002 | 7.58908857 | -0.519204828 | 0.3575209 | -1.4522363 | 0.14643591 | 0.26035262 | NA | NA |
| ENSG00000C | 796.966659 | 0.213394423 | 0.14694761 | 1.45218026 | 0.14645149 | 0.26036011 | NCBP3 | 55421 |
| ENSG000002 | 3.64846281 | -0.549699386 | 0.37856931 | -1.4520442 | 0.1464893 | 0.26036011 | RNU1-129P | 106481630 |
| ENSG000002 | 13.2504976 | -0.454182657 | 0.31278677 | -1.452052 | 0.14648715 | 0.26036011 | LINC01249 | 727982 |
| ENSG000002 | 191.051867 | 0.204350213 | 0.14072201 | 1.45215533 | 0.14645842 | 0.26036011 | NA | NA |
| ENSG000002 | 2.76145233 | -0.545523004 | 0.37569101 | -1.4520523 | 0.14648706 | 0.26036011 | NA | NA |
| ENSG000002 | 4.25263418 | -0.548822696 | 0.37796414 | -1.4520497 | 0.1464878 | 0.26036011 | NA | NA |
| ENSG000002 | 13.5615041 | -0.456391566 | 0.31430969 | -1.4520442 | 0.14648931 | 0.26036011 | NA | NA |
| ENSG000002 | 28.1395764 | -0.36073399 | 0.24843316 | -1.4520364 | 0.14649149 | 0.26036011 | NA | NA |
| ENSG000002 | 6.46958973 | -0.531496384 | 0.36612258 | -1.4516897 | 0.1465879 | 0.26052004 | NA | NA |
| ENSG000002 | 9.9261203 | -0.490389168 | 0.3378144 | -1.4516527 | 0.14659821 | 0.26052695 | LOC100505920 | 100505920 |
| ENSG000001 | 7.7932474 | -0.514356098 | 0.354355 | -1.4515277 | 0.14663298 | 0.26057733 | LINC01164 | 399827 |
| ENSG000002 | 3.68214188 | -0.549340786 | 0.37846861 | -1.4514831 | 0.14664539 | 0.26058796 | NA | NA |
| ENSG000001 | 34.2803746 | 0.365890226 | 0.25209886 | 1.45137598 | 0.14667519 | 0.26061808 | KCNC3 | 3748 |

| | | | | | | | | |
|---|---|---|---|---|---|---|---|---|
| ENSG000002 | 23.8485403 | -0.396460476 | 0.27316007 | -1.4513852 | 0.14667264 | 0.26061808 | HEATR9 | 256957 |
| ENSG000002 | 6.58273623 | -0.533019524 | 0.36729185 | -1.4512152 | 0.14671994 | 0.26068617 | NA | NA |
| ENSG000001 | 75.3778664 | -0.265492399 | 0.18297251 | -1.4509961 | 0.14678094 | 0.26078312 | ADGRB3 | 577 |
| ENSG000001 | 149.723375 | 0.213619209 | 0.14722712 | 1.45095011 | 0.14679375 | 0.26079446 | CLMN | 79789 |
| ENSG000002 | 6.9636795 | -0.524445278 | 0.36146721 | -1.4508793 | 0.14681349 | 0.26080666 | NA | NA |
| ENSG000002 | 11.6942697 | -0.475447438 | 0.32769147 | -1.4508996 | 0.14680781 | 0.26080666 | NA | NA |
| ENSG000002 | 7.42521431 | -0.514486756 | 0.35461088 | -1.4508488 | 0.14682198 | 0.26081032 | NA | NA |
| ENSG000002 | 2.83647928 | -0.538308346 | 0.37103649 | -1.4508232 | 0.1468291 | 0.26081155 | NA | NA |
| ENSG000001 | 2.54477355 | -0.537855163 | 0.37073501 | -1.4507806 | 0.14684096 | 0.26082119 | NA | NA |
| ENSG000002 | 6.12140369 | -0.530624292 | 0.36577507 | -1.4506847 | 0.14686768 | 0.26085723 | NA | NA |
| ENSG000002 | 4.22235034 | -0.54766397 | 0.37758471 | -1.45044 | 0.14693586 | 0.26096689 | NA | NA |
| ENSG000002 | 2.54275834 | -0.537716263 | 0.37073532 | -1.4504047 | 0.14694569 | 0.26097292 | NA | NA |
| ENSG000002 | 4.17083542 | -0.548121348 | 0.37791909 | -1.4503669 | 0.14695622 | 0.26098019 | LOC101928495 | 101928495 |
| ENSG000002 | 6.95849223 | -0.535044145 | 0.36891273 | -1.4503271 | 0.14696731 | 0.26098846 | NA | NA |
| ENSG000001 | 2.52531845 | -0.535700268 | 0.36938335 | -1.4502556 | 0.14698726 | 0.26101246 | NA | NA |
| ENSG000001 | 15.1837739 | 0.457148813 | 0.31530537 | 1.44986055 | 0.14709741 | 0.26119661 | SAMD10 | 140700 |
| ENSG000002 | 3.4069121 | -0.547123737 | 0.37739772 | -1.4497272 | 0.14713461 | 0.26125123 | NA | NA |
| ENSG000002 | 9.41008337 | -0.497568248 | 0.34323657 | -1.4496364 | 0.14715993 | 0.26128474 | NA | NA |
| ENSG000002 | 6.97958438 | -0.523609408 | 0.36122362 | -1.4495436 | 0.14718582 | 0.26131484 | NA | NA |
| ENSG000002 | 10.4463609 | -0.490731363 | 0.33854527 | -1.4495295 | 0.14718976 | 0.26131484 | NA | NA |
| ENSG000002 | 2.51047039 | -0.537479466 | 0.37082844 | -1.4494019 | 0.14722537 | 0.2613666 | NA | NA |
| ENSG000002 | 39.7128711 | 0.332145593 | 0.2291672 | 1.44935922 | 0.14723729 | 0.26137633 | ZNF688 | 146542 |
| ENSG000001 | 3.70131535 | -0.548280616 | 0.37829802 | -1.4493352 | 0.14724401 | 0.26137681 | EGR4 | 1961 |
| ENSG000001 | 5.87692069 | -0.532711228 | 0.36756701 | -1.4492901 | 0.14725659 | 0.26138196 | NKX6-3 | 157848 |
| ENSG000002 | 3.40005713 | -0.547460977 | 0.37774724 | -1.4492786 | 0.1472598 | 0.26138196 | NA | NA |
| ENSG000002 | 3.44394338 | -0.547264347 | 0.37763607 | -1.4491845 | 0.14728606 | 0.26139425 | NA | NA |
| ENSG000002 | 4.5087316 | -0.54554159 | 0.37644561 | -1.449191 | 0.14728425 | 0.26139425 | NA | NA |
| ENSG000002 | 2.5371208 | -0.534945502 | 0.36912657 | -1.4492197 | 0.14727624 | 0.26139425 | NA | NA |
| ENSG000002 | 3.82225124 | -0.547699639 | 0.37797539 | -1.4490352 | 0.14732776 | 0.2614568 | NA | NA |

| | | | | | | | | |
|---|---|---|---|---|---|---|---|---|
| ENSG000001 | 417.585231 | 0.175164985 | 0.12089324 | 1.4489229 | 0.14735912 | 0.26145751 | SNRNP27 | 11017 |
| ENSG000002 | 4.48843939 | -0.546497814 | 0.37717641 | -1.4489183 | 0.14736039 | 0.26145751 | POLHP1 | 106480782 |
| ENSG000002 | 11.6127524 | -0.48560871 | 0.3351471 | -1.448942 | 0.14735377 | 0.26145751 | HNF1A-AS1 | 283460 |
| ENSG000002 | 2.19308018 | -0.525322593 | 0.36253976 | -1.4490068 | 0.14733568 | 0.26145751 | NA | NA |
| ENSG000002 | 17.7896394 | -0.44245896 | 0.30536039 | -1.448973 | 0.14734512 | 0.26145751 | NA | NA |
| ENSG000001 | 55.4209014 | 0.299514123 | 0.20673162 | 1.44880655 | 0.14739161 | 0.26150146 | TIGD4 | 201798 |
| ENSG000002 | 12.3068515 | -0.487245961 | 0.3363322 | -1.4487045 | 0.14742013 | 0.26152917 | NA | NA |
| ENSG000002 | 5.10360649 | -0.543265291 | 0.37499499 | -1.4487268 | 0.14741389 | 0.26152917 | NA | NA |
| ENSG000001 | 4.09943877 | -0.548214286 | 0.37848315 | -1.448451 | 0.14749096 | 0.26162049 | ZNF479 | 90827 |
| ENSG000002 | 5.38748906 | -0.540227526 | 0.37296863 | -1.448453 | 0.1474904 | 0.26162049 | NA | NA |
| ENSG000002 | 6.13703118 | -0.541214521 | 0.37364089 | -1.4484885 | 0.14748049 | 0.26162049 | NA | NA |
| ENSG000002 | 4.54598217 | -0.546322332 | 0.37721015 | -1.4483235 | 0.14752659 | 0.26167225 | NA | NA |
| ENSG000002 | 8.34825085 | -0.507713276 | 0.35059828 | -1.4481339 | 0.14757959 | 0.26174918 | LOC101929528 | 101929528 |
| ENSG000002 | 9.29762856 | -0.508707221 | 0.35128749 | -1.4481222 | 0.14758287 | 0.26174918 | NA | NA |
| ENSG000002 | 2.75077895 | -0.543187221 | 0.37514897 | -1.4479241 | 0.14763829 | 0.261836 | NA | NA |
| ENSG000001 | 74.8393461 | -0.267301133 | 0.18461695 | -1.4478689 | 0.14765372 | 0.26185193 | ADCY10P1 | 221442 |
| ENSG000002 | 8.90539392 | -0.504423018 | 0.34840169 | -1.4478202 | 0.14766734 | 0.26186461 | NA | NA |
| ENSG000002 | 4.89622377 | -0.547733736 | 0.37833286 | -1.4477562 | 0.14768526 | 0.26187349 | NA | NA |
| ENSG000002 | 3.38405974 | -0.547198017 | 0.37796209 | -1.447759 | 0.14768446 | 0.26187349 | NA | NA |
| ENSG000001 | 125.725761 | 0.236401747 | 0.16330593 | 1.44760052 | 0.1477288 | 0.26193925 | JAM2 | 58494 |
| ENSG000002 | 117.562167 | 0.271046891 | 0.18724766 | 1.44753154 | 0.14774811 | 0.26196203 | NA | NA |
| ENSG000001 | 23.9708949 | 0.381924112 | 0.26386253 | 1.44743598 | 0.14777485 | 0.261998 | RIPPLY3 | 53820 |
| ENSG000002 | 2.52826729 | -0.53465031 | 0.36938998 | -1.4473871 | 0.14778853 | 0.26199932 | NA | NA |
| ENSG000002 | 4.75452071 | -0.541952938 | 0.37443389 | -1.4473928 | 0.14778693 | 0.26199932 | LOC101929064 | 101929064 |
| ENSG000002 | 4.04548262 | -0.547769423 | 0.37849024 | -1.4472485 | 0.14782734 | 0.26205668 | NA | NA |
| ENSG000001 | 7.49446031 | -0.514764956 | 0.35569707 | -1.4472004 | 0.14784079 | 0.26206907 | OCM2 | 4951 |
| ENSG000001 | 199.106952 | 0.20958675 | 0.14484497 | 1.44697296 | 0.1479045 | 0.26215907 | EPHA7 | 2045 |
| ENSG000002 | 4.22092799 | -0.546470468 | 0.37766357 | -1.4469769 | 0.14790339 | 0.26215907 | NA | NA |
| ENSG000002 | 25.606713 | 0.389305908 | 0.26909363 | 1.4467303 | 0.14797248 | 0.2622681 | NA | NA |

| | | | | | | | | |
|---|---|---|---|---|---|---|---|---|
| ENSG000001 | 94.2632759 | 0.246846852 | 0.17063706 | 1.44661926 | 0.14800359 | 0.26231178 | ASB8 | 140461 |
| ENSG000001 | 22.6246579 | -0.398431956 | 0.27543459 | -1.4465574 | 0.14802092 | 0.26233102 | POU3F2 | 5454 |
| ENSG000001 | 17.8081107 | 0.428443328 | 0.29618681 | 1.44653073 | 0.1480284 | 0.26233282 | NPY | 4852 |
| ENSG000001 | 6.68579996 | -0.523462529 | 0.36188755 | -1.4464784 | 0.14804306 | 0.26234734 | NA | NA |
| ENSG000001 | 34.331345 | 0.342841614 | 0.23703365 | 1.4463837 | 0.14806961 | 0.26237613 | ZNF628 | 89887 |
| ENSG000002 | 16.3724778 | -0.42853795 | 0.29628427 | -1.4463743 | 0.14807225 | 0.26237613 | NA | NA |
| ENSG000002 | 2.50686976 | -0.536849369 | 0.37121612 | -1.4461909 | 0.14812366 | 0.26245577 | NA | NA |
| ENSG000001 | 17.6463108 | 0.418601938 | 0.28947176 | 1.44608906 | 0.14815222 | 0.26249276 | C8orf48 | 157773 |
| ENSG000002 | 2.7971445 | -0.541301314 | 0.37432573 | -1.4460703 | 0.14815749 | 0.26249276 | CYP17A1-AS1 | 102724307 |
| ENSG000002 | 2.74989848 | -0.543214542 | 0.37568423 | -1.4459339 | 0.14819575 | 0.26254909 | NA | NA |
| ENSG000001 | 4.54193203 | -0.544546818 | 0.37662948 | -1.4458423 | 0.14822145 | 0.26258314 | ASCL4 | 121549 |
| ENSG000001 | 81.5406105 | 0.267690458 | 0.18515477 | 1.44576595 | 0.14824286 | 0.26260959 | TRIM46 | 80128 |
| ENSG000002 | 4.46864743 | -0.544066793 | 0.37634515 | -1.4456591 | 0.14827284 | 0.26265122 | NA | NA |
| ENSG000002 | 3.34248209 | -0.546483172 | 0.37802468 | -1.4456283 | 0.14828148 | 0.26265506 | LOC100505609 | 100505609 |
| ENSG000002 | 7.06548989 | 0.517488548 | 0.35798107 | 1.44557518 | 0.14829639 | 0.26265853 | RNU1-2 | 26870 |
| ENSG000002 | 6.21324487 | -0.534063657 | 0.36944345 | -1.4455897 | 0.14829231 | 0.26265853 | NA | NA |
| ENSG000001 | 8.07066458 | 0.515037079 | 0.35632149 | 1.44542804 | 0.14833769 | 0.2627202 | NKX2-4 | 644524 |
| ENSG000002 | 2.76928136 | -0.543024466 | 0.3757018 | -1.4453603 | 0.14835671 | 0.26274241 | NA | NA |
| ENSG000002 | 3.45461676 | -0.545920529 | 0.37776762 | -1.4451226 | 0.14842345 | 0.26284913 | NA | NA |
| ENSG000002 | 9.53181541 | -0.501700643 | 0.34719217 | -1.4450229 | 0.14845144 | 0.26288722 | NA | NA |
| ENSG000001 | 11.1253701 | -0.502505592 | 0.34776564 | -1.4449547 | 0.14847062 | 0.26289821 | UGT2A1 | 10941 |
| ENSG000001 | 31.304268 | -0.363906679 | 0.25184584 | -1.4449581 | 0.14846966 | 0.26289821 | GPC5 | 2262 |
| ENSG000002 | 4.7935465 | -0.542290429 | 0.37530757 | -1.4449227 | 0.14847959 | 0.26290263 | NA | NA |
| ENSG000001 | 110.908422 | -0.231000489 | 0.15988779 | -1.4447663 | 0.14852355 | 0.26295694 | HMCN2 | 256158 |
| ENSG000002 | 18.5857194 | -0.4118294 | 0.28504925 | -1.4447658 | 0.14852369 | 0.26295694 | NA | NA |
| ENSG000002 | 3.34943042 | 0.546595401 | 0.37833366 | 1.4447443 | 0.14852972 | 0.26295694 | NA | NA |
| ENSG000002 | 7.64288994 | -0.521958947 | 0.36130266 | -1.4446584 | 0.14855385 | 0.26298817 | NA | NA |
| ENSG000002 | 46.5901057 | 0.308875626 | 0.21381345 | 1.44460335 | 0.14856933 | 0.2630041 | LOC257396 | 257396 |
| ENSG000002 | 2.54289419 | -0.532781142 | 0.36886698 | -1.444372 | 0.14863436 | 0.26310465 | NA | NA |

| | | | | | | | | |
|---|---|---|---|---|---|---|---|---|
| ENSG000002 | 2.45677774 | -0.535423584 | 0.37070079 | -1.4443551 | 0.14863911 | 0.26310465 | NA | NA |
| ENSG000002 | 16.0651293 | -0.427769912 | 0.29620241 | -1.4441811 | 0.14868805 | 0.26317979 | NA | NA |
| ENSG000002 | 6.77037727 | -0.524899468 | 0.36346954 | -1.4441361 | 0.14870071 | 0.26319071 | GPR158-AS1 | 100128811 |
| ENSG000002 | 13.340785 | -0.474791311 | 0.32878614 | -1.4440734 | 0.14871833 | 0.26321042 | NA | NA |
| ENSG000002 | 2.57824666 | -0.531188951 | 0.36786372 | -1.443983 | 0.14874376 | 0.26324393 | NA | NA |
| ENSG000001 | 11.841506 | -0.481173474 | 0.33323313 | -1.4439545 | 0.14875178 | 0.26324663 | GATA3 | 2625 |
| ENSG000002 | 8.31341214 | -0.506173499 | 0.35056778 | -1.4438677 | 0.14877619 | 0.26327834 | NA | NA |
| ENSG000002 | 32.2194353 | -0.368305789 | 0.25509142 | -1.4438188 | 0.14878995 | 0.2632912 | LOC285389 | 285389 |
| ENSG000002 | 4.01157667 | 0.546432401 | 0.37852497 | 1.44358349 | 0.14885617 | 0.2633969 | NA | NA |
| ENSG000002 | 177.047405 | 0.225908142 | 0.15650389 | 1.44346667 | 0.14888905 | 0.26344358 | TIGD1 | 200765 |
| ENSG000002 | 4.06334963 | -0.546465804 | 0.37859154 | -1.4434179 | 0.14890278 | 0.26345638 | NA | NA |
| ENSG000002 | 2.50894913 | -0.533238863 | 0.36944033 | -1.4433694 | 0.14891644 | 0.26346906 | CSTP1 | 1480 |
| ENSG000001 | 12.9301652 | -0.486918978 | 0.33736539 | -1.4432986 | 0.14893639 | 0.26348135 | MEPE | 56955 |
| ENSG000002 | 2.54935183 | -0.530858909 | 0.36780403 | -1.44332 | 0.14893035 | 0.26348135 | NA | NA |
| ENSG000001 | 8.18432842 | -0.516241075 | 0.35770539 | -1.4432018 | 0.14896362 | 0.26351803 | CDKN2C | 1031 |
| ENSG000002 | 4.50883554 | -0.543658181 | 0.37671286 | -1.4431633 | 0.14897447 | 0.26352572 | NA | NA |
| ENSG000001 | 53.7206509 | 0.29384594 | 0.20361675 | 1.44313246 | 0.14898316 | 0.26352934 | RDH12 | 145226 |
| ENSG000002 | 5.32103514 | -0.538739882 | 0.37331868 | -1.4431099 | 0.14898951 | 0.26352934 | NA | NA |
| ENSG000002 | 5.38707383 | -0.540718919 | 0.37470853 | -1.4430387 | 0.14900957 | 0.26354183 | NA | NA |
| ENSG000002 | 5.41455694 | -0.538483749 | 0.37315592 | -1.443053 | 0.14900555 | 0.26354183 | NA | NA |
| ENSG00000C | 3.32224578 | -0.545544011 | 0.37812191 | -1.4427728 | 0.14908448 | 0.26364133 | TYROBP | 7305 |
| ENSG000001 | 4.22210185 | -0.545309196 | 0.37795995 | -1.4427698 | 0.14908533 | 0.26364133 | IL2 | 3558 |
| ENSG000002 | 2.57141131 | -0.532686275 | 0.36920514 | -1.4427921 | 0.14907903 | 0.26364133 | NA | NA |
| ENSG000002 | 4.71559134 | -0.540101935 | 0.37436987 | -1.4426961 | 0.14910611 | 0.26365759 | NA | NA |
| ENSG000002 | 2.75764209 | -0.540724795 | 0.37480291 | -1.442691 | 0.14910753 | 0.26365759 | NA | NA |
| ENSG000002 | 3.40437571 | -0.544799634 | 0.37764447 | -1.4426258 | 0.14912593 | 0.26367862 | NA | NA |
| ENSG000001 | 8.58017089 | -0.502941629 | 0.34867391 | -1.4424412 | 0.14917794 | 0.26372458 | CATIP | 375307 |
| ENSG000001 | 14.6984939 | -0.451157398 | 0.3127614 | -1.4424971 | 0.1491622 | 0.26372458 | SHOX2 | 6474 |
| ENSG000002 | 3.99248574 | -0.546095483 | 0.37858597 | -1.442461 | 0.14917237 | 0.26372458 | NA | NA |

| | | | | | | | | |
|---|---|---|---|---|---|---|---|---|
| ENSG000002 | 4.50630383 | -0.543394718 | 0.37671096 | -1.4424712 | 0.14916948 | 0.26372458 | NA | NA |
| ENSG000002 | 2.52175724 | -0.533925139 | 0.37016867 | -1.4423834 | 0.14919425 | 0.26373826 | NA | NA |
| ENSG000002 | 13.0770599 | 0.46310122 | 0.32107016 | 1.44236767 | 0.14919868 | 0.26373826 | NA | NA |
| ENSG000002 | 3.43008093 | -0.545008856 | 0.37786945 | -1.4423205 | 0.14921198 | 0.26375027 | NA | NA |
| ENSG000002 | 4.80116768 | -0.543508838 | 0.37684615 | -1.4422566 | 0.14923 | 0.26377063 | LOC105377245 | 105377245 |
| ENSG000002 | 16.1104054 | -0.431165295 | 0.29896443 | -1.442196 | 0.1492471 | 0.26378935 | LOC101927543 | 101927543 |
| ENSG000002 | 7.41266211 | -0.512618643 | 0.35547104 | -1.4420827 | 0.14927904 | 0.2638343 | NA | NA |
| ENSG000002 | 13.2458527 | -0.455490234 | 0.31587035 | -1.4420164 | 0.14929774 | 0.26385586 | LOC101929538 | 101929538 |
| ENSG000002 | 4.23913625 | -0.544818722 | 0.37783383 | -1.4419533 | 0.14931556 | 0.26387585 | NA | NA |
| ENSG000002 | 5.22038104 | -0.543929131 | 0.37722351 | -1.441928 | 0.1493227 | 0.26387697 | NA | NA |
| ENSG000001 | 86.7732817 | 0.243856912 | 0.16917563 | 1.44144229 | 0.14945977 | 0.26410768 | SOBP | 55084 |
| ENSG000001 | 2.473284 | -0.534201378 | 0.37074418 | -1.4408894 | 0.14961592 | 0.26434906 | CACNG1 | 786 |
| ENSG000002 | 5.75293708 | -0.538979461 | 0.37405874 | -1.4408952 | 0.14961428 | 0.26434906 | NA | NA |
| ENSG000002 | 3.96326386 | -0.545528559 | 0.37859335 | -1.4409354 | 0.14960294 | 0.26434906 | NA | NA |
| ENSG000002 | 3.43884303 | -0.544552538 | 0.37795324 | -1.4407934 | 0.14964306 | 0.26437397 | NA | NA |
| ENSG000002 | 4.7333428 | -0.540500252 | 0.3751402 | -1.4407953 | 0.14964251 | 0.26437397 | NA | NA |
| ENSG000002 | 158.278456 | 0.208934922 | 0.14501789 | 1.44075276 | 0.14965454 | 0.26438274 | TMEM161B-AS1 | 100505894 |
| ENSG000002 | 4.51858276 | -0.545223784 | 0.37843985 | -1.4407145 | 0.14966535 | 0.26439031 | NA | NA |
| ENSG000001 | 41.3211251 | 0.318191044 | 0.22086143 | 1.44068183 | 0.14967459 | 0.26439512 | BEND5 | 79656 |
| ENSG000002 | 5.14865416 | -0.541001714 | 0.37553173 | -1.4406285 | 0.14968966 | 0.26441022 | NA | NA |
| ENSG000002 | 3.44569306 | 0.54390033 | 0.37755642 | 1.44058025 | 0.1497033 | 0.2644228 | NA | NA |
| ENSG000002 | 7.50411274 | 0.514867121 | 0.35743198 | 1.44046184 | 0.14973678 | 0.26447041 | NA | NA |
| ENSG000002 | 2.82555061 | -0.539814097 | 0.37478212 | -1.4403411 | 0.14977091 | 0.26451918 | NA | NA |
| ENSG000002 | 26.1273924 | -0.388182475 | 0.26959924 | -1.43985 | 0.14990983 | 0.264753 | NA | NA |
| ENSG000001 | 10.4117707 | -0.485615943 | 0.33728119 | -1.4397955 | 0.14992527 | 0.26475899 | TLDC2 | 140711 |
| ENSG000002 | 12.9089741 | -0.455480678 | 0.31635174 | -1.4397919 | 0.14992628 | 0.26475899 | NA | NA |
| ENSG000002 | 3.64133158 | -0.545065536 | 0.37858417 | -1.4397473 | 0.14993891 | 0.26476976 | NA | NA |
| ENSG000002 | 4.25658823 | -0.544122162 | 0.37795201 | -1.4396594 | 0.14996379 | 0.26480216 | NA | NA |
| ENSG000001 | 126.931147 | 0.222714856 | 0.15470338 | 1.43962505 | 0.14997351 | 0.26480779 | PTGR2 | 145482 |

| | | | | | | | | |
|---|---|---|---|---|---|---|---|---|
| ENSG000001 | 8.03433339 | -0.508444678 | 0.35319151 | -1.4395722 | 0.14998847 | 0.26482268 | FOLR3 | 2352 |
| ENSG000001 | 10.5247932 | -0.488490086 | 0.33934056 | -1.4395276 | 0.15000111 | 0.26483346 | GNLY | 10578 |
| ENSG000002 | 4.59106584 | -0.544308468 | 0.37812432 | -1.4394961 | 0.15001002 | 0.26483767 | RTL1 | 388015 |
| ENSG000001 | 7.27557683 | -0.532912794 | 0.37023627 | -1.4393857 | 0.15004127 | 0.26484671 | MGC45922 | 284365 |
| ENSG000002 | 15.3689109 | 0.436191539 | 0.30303943 | 1.43938873 | 0.15004042 | 0.26484671 | NA | NA |
| ENSG000002 | 17.2548901 | -0.417920551 | 0.29033738 | -1.4394307 | 0.15002852 | 0.26484671 | NA | NA |
| ENSG000002 | 7.41576795 | -0.511203624 | 0.35515155 | -1.4393957 | 0.15003843 | 0.26484671 | NA | NA |
| ENSG000002 | 7.31504955 | -0.519913396 | 0.36122329 | -1.4393131 | 0.15006185 | 0.2648715 | NA | NA |
| ENSG000002 | 4.49733266 | -0.542706244 | 0.37709465 | -1.4391778 | 0.15010015 | 0.26492759 | LINC01546 | 100129464 |
| ENSG000002 | 6.14057859 | -0.525678847 | 0.36532374 | -1.4389397 | 0.1501676 | 0.26502653 | NA | NA |
| ENSG000002 | 9.0761923 | -0.493352748 | 0.34285994 | -1.4389338 | 0.15016928 | 0.26502653 | NA | NA |
| ENSG000002 | 7.48810681 | -0.51047753 | 0.35477873 | -1.4388617 | 0.15018971 | 0.26505104 | NA | NA |
| ENSG000001 | 2.72451389 | -0.540223172 | 0.37549157 | -1.4387092 | 0.15023294 | 0.26511579 | OR13C9 | 286362 |
| ENSG000001 | 9.57825991 | -0.487039312 | 0.3385429 | -1.438634 | 0.15025426 | 0.26514188 | GSDMA | 284110 |
| ENSG000001 | 12.0607331 | -0.470444386 | 0.32703561 | -1.4385112 | 0.15028907 | 0.26519177 | NUTM2G | 441457 |
| ENSG000002 | 6.21461679 | -0.529727506 | 0.36827667 | -1.4383955 | 0.15032187 | 0.26523811 | NA | NA |
| ENSG000002 | 7.84438136 | -0.514664429 | 0.35782178 | -1.4383262 | 0.15034154 | 0.26524972 | NA | NA |
| ENSG000002 | 3.21656835 | -0.536933226 | 0.37330099 | -1.4383386 | 0.150338 | 0.26524972 | SUMO2P15 | 106480417 |
| ENSG000002 | 37.2726307 | 0.340471237 | 0.23673018 | 1.43822491 | 0.15037025 | 0.26527104 | GPSM3 | 63940 |
| ENSG000002 | 3.64311811 | -0.54443611 | 0.37855004 | -1.4382144 | 0.15037324 | 0.26527104 | RNU6-370P | 106481295 |
| ENSG000002 | 2.2205995 | -0.522333355 | 0.3631789 | -1.438226 | 0.15036993 | 0.26527104 | NA | NA |
| ENSG000002 | 3.66160787 | -0.54444408 | 0.37858342 | -1.4381086 | 0.15040324 | 0.26528933 | HNRNPA1P33 | 728643 |
| ENSG000002 | 11.4040711 | -0.488273056 | 0.33951471 | -1.4381499 | 0.15039152 | 0.26528933 | NA | NA |
| ENSG000002 | 4.38140888 | 0.543791546 | 0.37812884 | 1.43811178 | 0.15040234 | 0.26528933 | NA | NA |
| ENSG000002 | 4.55983252 | -0.542613106 | 0.37732587 | -1.438049 | 0.15042014 | 0.26530761 | NA | NA |
| ENSG000002 | 2.79341966 | -0.538529393 | 0.37452674 | -1.437893 | 0.15046441 | 0.26537414 | NA | NA |
| ENSG000002 | 13.7998597 | -0.470689078 | 0.32736117 | -1.4378281 | 0.15048284 | 0.26539511 | LOC101928307 | 101928307 |
| ENSG000001 | 8.39890273 | -0.509023693 | 0.3540419 | -1.4377499 | 0.15050504 | 0.26542271 | PRND | 23627 |
| ENSG000002 | 27.5260013 | 0.370685494 | 0.25785381 | 1.43758004 | 0.15055325 | 0.2654862 | NA | NA |

| | | | | | | | | |
|---|---|---|---|---|---|---|---|---|
| ENSG000002 | 10.9163941 | -0.474899793 | 0.3303474 | -1.4375769 | 0.15055413 | 0.2654862 | LINC01361 | 101927498 |
| ENSG000002 | 3.61533407 | -0.544237284 | 0.37858855 | -1.4375429 | 0.15056379 | 0.26549167 | NA | NA |
| ENSG000002 | 12.6189714 | 0.46881445 | 0.32613796 | 1.43747282 | 0.15058369 | 0.26551028 | NA | NA |
| ENSG000002 | 4.42028517 | -0.543658214 | 0.37820764 | -1.4374596 | 0.15058743 | 0.26551028 | NA | NA |
| ENSG000002 | 20.3619057 | -0.42616095 | 0.29647555 | -1.4374236 | 0.15059767 | 0.26551678 | FLJ21408 | 400512 |
| ENSG000002 | 230.936099 | 0.192998457 | 0.13427616 | 1.43732477 | 0.15062573 | 0.26554317 | ZNF611 | 81856 |
| ENSG000002 | 6.10451435 | -0.538723506 | 0.37480559 | -1.4373412 | 0.15062107 | 0.26554317 | NA | NA |
| ENSG000002 | 2.7853306 | -0.53936022 | 0.37528426 | -1.4372045 | 0.1506599 | 0.26559186 | NA | NA |
| ENSG000002 | 6.17121164 | -0.52963598 | 0.36856567 | -1.4370193 | 0.15071251 | 0.26567305 | NA | NA |
| ENSG000001 | 291.021738 | 0.210269987 | 0.14632763 | 1.43698078 | 0.15072346 | 0.2656808 | NSMCE2 | 286053 |
| ENSG000002 | 3.4563602 | -0.540736451 | 0.37631496 | -1.4369252 | 0.15073924 | 0.26569027 | NA | NA |
| ENSG000002 | 3.8748817 | -0.543325787 | 0.37811944 | -1.4369158 | 0.15074193 | 0.26569027 | NA | NA |
| ENSG000002 | 3.62296721 | -0.54394646 | 0.37857898 | -1.436811 | 0.1507717 | 0.26573118 | NA | NA |
| ENSG000002 | 3.69659562 | -0.543802782 | 0.37849651 | -1.4367445 | 0.15079061 | 0.26575296 | NA | NA |
| ENSG000002 | 5.02470872 | -0.536049304 | 0.37311765 | -1.4366763 | 0.15080998 | 0.26577555 | NA | NA |
| ENSG000002 | 5.04532882 | 0.539505736 | 0.37553204 | 1.43664368 | 0.15081927 | 0.26578037 | MIR1270 | 100302179 |
| ENSG000002 | 5.23454398 | 0.537212999 | 0.37395856 | 1.43655756 | 0.15084375 | 0.26581196 | NA | NA |
| ENSG000002 | 7.70399482 | -0.520419784 | 0.36228569 | -1.43649 | 0.15086295 | 0.26583425 | NA | NA |
| ENSG000001 | 160.618965 | 0.264733745 | 0.18429856 | 1.43643955 | 0.15087731 | 0.26583867 | ATN1 | 1822 |
| ENSG000001 | 49.6706485 | 0.306338107 | 0.21326276 | 1.43643509 | 0.15087858 | 0.26583867 | RABL2A | 11159 |
| ENSG000002 | 5.42044101 | 0.539506569 | 0.37559799 | 1.43639365 | 0.15089036 | 0.26584789 | NA | NA |
| ENSG000001 | 4.80330798 | -0.54110979 | 0.3767485 | -1.4362626 | 0.15092763 | 0.265902 | FUT6 | 2528 |
| ENSG000002 | 20.912804 | -0.398559038 | 0.27751077 | -1.436193 | 0.15094742 | 0.26591476 | LOC102723493 | 102723493 |
| ENSG000002 | 5.62117481 | -0.533433883 | 0.37142265 | -1.436191 | 0.15094799 | 0.26591476 | NA | NA |
| ENSG000001 | 404.013955 | 0.192375913 | 0.13396801 | 1.43598391 | 0.15100692 | 0.26600702 | TEX2 | 55852 |
| ENSG000002 | 4.20525013 | -0.542925462 | 0.378102 | -1.4359233 | 0.15102417 | 0.2660143 | NA | NA |
| ENSG000002 | 6.14440093 | -0.525632679 | 0.36605795 | -1.4359275 | 0.15102297 | 0.2660143 | LOC105370616 | 105370616 |
| ENSG000002 | 4.74291424 | -0.538666112 | 0.37518719 | -1.4357263 | 0.15108024 | 0.2661015 | NA | NA |
| ENSG000001 | 6.41232267 | -0.524937846 | 0.36563486 | -1.4356887 | 0.15109094 | 0.26610536 | OR4C11 | 219429 |

| | | | | | | | | |
|---|---|---|---|---|---|---|---|---|
| ENSG000001 | 85.9475629 | 0.291353248 | 0.20293895 | 1.43566942 | 0.15109643 | 0.26610536 | NA | NA |
| ENSG000002 | 9.63614114 | -0.507908973 | 0.35378342 | -1.4356494 | 0.15110212 | 0.26610536 | C9orf172 | 389813 |
| ENSG000002 | 2.75955371 | -0.53776032 | 0.37460963 | -1.4355219 | 0.15113844 | 0.26615777 | NA | NA |
| ENSG000001 | 80.8280009 | -0.254168557 | 0.17707586 | -1.4353654 | 0.15118301 | 0.26622469 | ASTN1 | 460 |
| ENSG000001 | 3.63878723 | -0.543381039 | 0.37859109 | -1.4352716 | 0.15120972 | 0.26626016 | ALX3 | 257 |
| ENSG000001 | 160.959391 | 0.214009508 | 0.14911811 | 1.43516776 | 0.1512393 | 0.26630069 | TMCO6 | 55374 |
| ENSG000002 | 2.45128509 | -0.53216764 | 0.37081572 | -1.435127 | 0.15125092 | 0.26630959 | NA | NA |
| ENSG000002 | 9.08396904 | -0.492513852 | 0.34320979 | -1.4350227 | 0.15128062 | 0.26634402 | NA | NA |
| ENSG000002 | 21.023972 | 0.408927197 | 0.28496426 | 1.43501222 | 0.15128362 | 0.26634402 | SNORD93 | 692210 |
| ENSG000002 | 4.51113856 | -0.540739725 | 0.37683957 | -1.4349335 | 0.15130605 | 0.26637196 | NA | NA |
| ENSG000002 | 4.22058198 | -0.542525556 | 0.37809792 | -1.4348811 | 0.15132099 | 0.26638435 | NA | NA |
| ENSG000002 | 7.21669124 | -0.514975041 | 0.35890196 | -1.4348627 | 0.15132623 | 0.26638435 | NA | NA |
| ENSG000001 | 201.68832 | 0.204854671 | 0.1427788 | 1.43476957 | 0.15135277 | 0.26641639 | TMOD2 | 29767 |
| ENSG000002 | 9.57452951 | -0.484915532 | 0.33798252 | -1.4347355 | 0.15136247 | 0.26641639 | NA | NA |
| ENSG000002 | 2.49883222 | -0.532006658 | 0.37080619 | -1.4347297 | 0.15136414 | 0.26641639 | NA | NA |
| ENSG000002 | 92.3908499 | 0.259862056 | 0.18112561 | 1.43470632 | 0.1513708 | 0.26641655 | NA | NA |
| ENSG000002 | 6.10219797 | -0.524425694 | 0.36553849 | -1.4346661 | 0.15138226 | 0.26642515 | NA | NA |
| ENSG000002 | 6.68446139 | -0.52312093 | 0.36463901 | -1.4346269 | 0.15139344 | 0.26643328 | NA | NA |
| ENSG000002 | 3.3596735 | -0.541695251 | 0.3776556 | -1.4343631 | 0.15146868 | 0.26655411 | NA | NA |
| ENSG000001 | 91.0062548 | 0.28648796 | 0.19973945 | 1.43430834 | 0.15148429 | 0.26657002 | TBKBP1 | 9755 |
| ENSG000001 | 2.20790295 | -0.517767988 | 0.36101931 | -1.4341836 | 0.15151987 | 0.26662106 | CCL17 | 6361 |
| ENSG000002 | 7.46105137 | -0.511681532 | 0.35680381 | -1.4340697 | 0.15155237 | 0.26666667 | USP17L7 | 392197 |
| ENSG000002 | 2.36121432 | 0.53030935 | 0.36979957 | 1.43404535 | 0.15155932 | 0.26666733 | NA | NA |
| ENSG000002 | 8.78869948 | 0.517937499 | 0.36120123 | 1.4339306 | 0.15159207 | 0.2667029 | NA | NA |
| ENSG000002 | 2.51366708 | -0.528738656 | 0.36873434 | -1.4339284 | 0.15159269 | 0.2667029 | NA | NA |
| ENSG000002 | 3.45007883 | -0.539902399 | 0.37655591 | -1.4337908 | 0.15163197 | 0.26674544 | NA | NA |
| ENSG000002 | 4.74484567 | -0.537049543 | 0.37457042 | -1.4337746 | 0.1516366 | 0.26674544 | NA | NA |
| ENSG000002 | 5.15104729 | -0.53881157 | 0.37579601 | -1.4337874 | 0.15163296 | 0.26674544 | PRSS46 | 100287362 |
| ENSG000001 | 192.573905 | 0.216716894 | 0.15115612 | 1.43372891 | 0.15164964 | 0.26674785 | GLCCI1 | 113263 |

| | | | | | | | | |
|---|---|---|---|---|---|---|---|---|
| ENSG000002 | 23.7063438 | -0.394494177 | 0.27515356 | -1.4337237 | 0.15165113 | 0.26674785 | NA | NA |
| ENSG00000C | 91.1137052 | 0.257964143 | 0.17993072 | 1.43368597 | 0.1516619 | 0.26675522 | SLC8B1 | 80024 |
| ENSG000002 | 4.23061413 | -0.542297446 | 0.37830643 | -1.4334873 | 0.15171864 | 0.26684345 | NA | NA |
| ENSG000002 | 6.43378184 | 0.524446933 | 0.36587677 | 1.43339774 | 0.1517442 | 0.26687684 | NA | NA |
| ENSG000002 | 4.50958539 | -0.540541713 | 0.37713333 | -1.4332908 | 0.15177476 | 0.26688428 | NA | NA |
| ENSG000002 | 2.81815479 | -0.533077553 | 0.37191318 | -1.4333387 | 0.15176107 | 0.26688428 | NA | NA |
| ENSG000002 | 2.54401563 | -0.526304359 | 0.36719855 | -1.4332964 | 0.15177316 | 0.26688428 | NA | NA |
| ENSG000002 | 3.88950914 | -0.54183503 | 0.37803466 | -1.4332946 | 0.15177367 | 0.26688428 | NA | NA |
| ENSG000002 | 2.48677341 | -0.527775239 | 0.36824708 | -1.4332096 | 0.15179796 | 0.26691349 | NA | NA |
| ENSG000002 | 2.53674866 | -0.527926257 | 0.36836863 | -1.4331466 | 0.15181594 | 0.26693354 | NA | NA |
| ENSG000002 | 5.4049612 | -0.534811679 | 0.37319607 | -1.4330582 | 0.15184121 | 0.2669664 | NA | NA |
| ENSG000002 | 8.8756864 | -0.498477607 | 0.34785791 | -1.432992 | 0.15186012 | 0.26698807 | NA | NA |
| ENSG000001 | 7.51682504 | -0.51010224 | 0.35600191 | -1.4328638 | 0.15189677 | 0.2670348 | WFDC13 | 164237 |
| ENSG000002 | 5.84756939 | -0.527329449 | 0.36802762 | -1.4328529 | 0.15189987 | 0.2670348 | NA | NA |
| ENSG000001 | 2.19353664 | -0.516484187 | 0.36047096 | -1.4328039 | 0.15191389 | 0.26704787 | PHOX2A | 401 |
| ENSG000002 | 5.53830767 | -0.531343324 | 0.37086415 | -1.4327169 | 0.15193876 | 0.26708001 | NA | NA |
| ENSG000001 | 65.2662824 | 0.29387228 | 0.20513648 | 1.43256958 | 0.15198088 | 0.26714246 | NA | NA |
| ENSG000002 | 9.09034345 | -0.491012637 | 0.34276294 | -1.4325138 | 0.15199683 | 0.26715069 | NA | NA |
| ENSG000002 | 24.7851828 | -0.376659361 | 0.26293716 | -1.4325071 | 0.15199874 | 0.26715069 | NA | NA |
| ENSG000002 | 4.77957195 | -0.538251205 | 0.37582984 | -1.4321673 | 0.15209594 | 0.26730996 | LINC00637 | 145216 |
| ENSG000002 | 2.75485034 | -0.537707425 | 0.37546207 | -1.4321218 | 0.15210895 | 0.26732124 | NA | NA |
| ENSG000001 | 174.28783 | 0.209762274 | 0.14647627 | 1.43205633 | 0.1521277 | 0.2673426 | CHRM3 | 1131 |
| ENSG000002 | 9.80194598 | -0.481101183 | 0.33596968 | -1.431978 | 0.15215012 | 0.26736774 | NA | NA |
| ENSG000002 | 2.48176536 | -0.529984059 | 0.37011087 | -1.4319603 | 0.1521552 | 0.26736774 | NA | NA |
| ENSG000002 | 3.66691206 | -0.541962409 | 0.37848472 | -1.4319268 | 0.15216477 | 0.26737298 | NA | NA |
| ENSG000001 | 401.685494 | 0.174999275 | 0.12222023 | 1.43183556 | 0.15219089 | 0.26740662 | MTBP | 27085 |
| ENSG000002 | 12.8647952 | -0.46444017 | 0.3243719 | -1.4318138 | 0.15219711 | 0.26740662 | NA | NA |
| ENSG000002 | 2.81038238 | -0.536421036 | 0.37465544 | -1.4317717 | 0.15220918 | 0.26741625 | DEFA8P | 449491 |
| ENSG000002 | 8.87490987 | -0.497193513 | 0.34726583 | -1.4317375 | 0.15221896 | 0.26742184 | NA | NA |

| | | | | | | | | |
|---|---|---|---|---|---|---|---|---|
| ENSG000002 | 3.12560743 | -0.537567733 | 0.37547256 | -1.4317098 | 0.1522269 | 0.26742421 | NA | NA |
| ENSG000000 | 10.1563685 | -0.484801747 | 0.33864097 | -1.4316099 | 0.1522555 | 0.26746287 | BCL3 | 602 |
| ENSG000001 | 68.0944731 | 0.269668222 | 0.18838268 | 1.43149158 | 0.15228938 | 0.2674992 | PPP1R3D | 5509 |
| ENSG000002 | 4.97598773 | -0.533440144 | 0.37264217 | -1.4315077 | 0.15228475 | 0.2674992 | NA | NA |
| ENSG000001 | 7.1469695 | -0.512716671 | 0.35819742 | -1.4313801 | 0.1523213 | 0.26754369 | MMD2 | 221938 |
| ENSG000001 | 9.08105853 | -0.494087927 | 0.34522764 | -1.4311946 | 0.15237445 | 0.26761683 | NA | NA |
| ENSG000002 | 70.8105302 | 0.295114725 | 0.20620253 | 1.43118869 | 0.15237614 | 0.26761683 | HLA-DMA | 3108 |
| ENSG000002 | 9.47149329 | 0.48927131 | 0.34188749 | 1.43108867 | 0.1524048 | 0.26765557 | NA | NA |
| ENSG000001 | 3.13513453 | -0.538366618 | 0.37622088 | -1.4309855 | 0.15243438 | 0.26768646 | NA | NA |
| ENSG000002 | 4.71599574 | -0.5367371 | 0.37508326 | -1.4309812 | 0.1524356 | 0.26768646 | LOC441025 | 441025 |
| ENSG000002 | 5.83987732 | -0.526497009 | 0.36796084 | -1.4308506 | 0.15247303 | 0.2677406 | NA | NA |
| ENSG000001 | 13.6238507 | -0.470023885 | 0.32849812 | -1.4308267 | 0.15247989 | 0.26774106 | ANGPTL4 | 51129 |
| ENSG000002 | 5.64278329 | 0.534855267 | 0.37384361 | 1.43069256 | 0.15251835 | 0.26776551 | RNU2-64P | 106481654 |
| ENSG000002 | 4.81531673 | -0.538324369 | 0.37625214 | -1.4307543 | 0.15250064 | 0.26776551 | NA | NA |
| ENSG000002 | 2.57135127 | -0.526304172 | 0.36785934 | -1.4307212 | 0.15251013 | 0.26776551 | NA | NA |
| ENSG000002 | 11.3940774 | -0.474989892 | 0.3320015 | -1.430686 | 0.15252023 | 0.26776551 | NA | NA |
| ENSG000001 | 66.5452306 | 0.27619979 | 0.19307232 | 1.43055091 | 0.15255896 | 0.26782192 | TRIM72 | 493829 |
| ENSG000000 | 75.0597509 | -0.270144772 | 0.18884634 | -1.4305005 | 0.15257344 | 0.26783573 | CROT | 54677 |
| ENSG000001 | 5.28206725 | -0.540866567 | 0.37810856 | -1.4304531 | 0.15258701 | 0.26784797 | LINC00304 | 283860 |
| ENSG000001 | 2.7343055 | -0.536049945 | 0.37478235 | -1.4302967 | 0.15263189 | 0.26791515 | NA | NA |
| ENSG000002 | 23.2177821 | -0.414188623 | 0.28958854 | -1.4302659 | 0.15264071 | 0.26791904 | NA | NA |
| ENSG000001 | 3.77567076 | -0.540880576 | 0.37817462 | -1.4302403 | 0.15264807 | 0.26792035 | LRRC25 | 126364 |
| ENSG000002 | 4.48098394 | -0.538971141 | 0.37685072 | -1.430198 | 0.15266021 | 0.26793006 | NA | NA |
| ENSG000002 | 36.6232855 | -0.341028921 | 0.23846279 | -1.4301138 | 0.15268437 | 0.26796088 | NA | NA |
| ENSG000002 | 7.57800142 | -0.517824896 | 0.36209377 | -1.4300851 | 0.15269261 | 0.26796373 | NA | NA |
| ENSG000002 | 12.3758841 | 0.464766915 | 0.32502775 | 1.42992997 | 0.15273712 | 0.26803025 | NA | NA |
| ENSG000002 | 6.91678526 | -0.517137162 | 0.36166091 | -1.4298951 | 0.15274713 | 0.26803622 | NA | NA |
| ENSG000002 | 11.2078391 | -0.471192063 | 0.32953935 | -1.4298507 | 0.15275988 | 0.26804698 | NA | NA |
| ENSG000002 | 2.86774241 | -0.528633238 | 0.36974613 | -1.4297195 | 0.15279755 | 0.26810148 | NA | NA |

| | | | | | | | | |
|---|---|---|---|---|---|---|---|---|
| ENSG000002 | 8.22844526 | -0.501111096 | 0.35051285 | -1.4296511 | 0.15281717 | 0.26812432 | NA | NA |
| ENSG000002 | 6.73325558 | -0.519952373 | 0.36375173 | -1.4294155 | 0.15288484 | 0.26823143 | B4GALT1-AS1 | 101929639 |
| ENSG000002 | 2.72332021 | -0.536691145 | 0.37549406 | -1.4292933 | 0.15291995 | 0.26827251 | LOC344967 | 344967 |
| ENSG000002 | 18.931911 | -0.421302613 | 0.29476398 | -1.429288 | 0.15292148 | 0.26827251 | NA | NA |
| ENSG000002 | 23.2698603 | 0.377657229 | 0.26427187 | 1.42904817 | 0.15299039 | 0.26838178 | NA | NA |
| ENSG000002 | 13.2101826 | -0.465951625 | 0.32606325 | -1.4290222 | 0.15299786 | 0.26838327 | LOC100996251 | 100996251 |
| ENSG000002 | 7.93515909 | -0.498685117 | 0.34898498 | -1.4289587 | 0.1530161 | 0.26840337 | NA | NA |
| ENSG000002 | 11.4785735 | -0.469716616 | 0.32871768 | -1.4289363 | 0.15302255 | 0.26840337 | NA | NA |
| ENSG000002 | 2.7483916 | -0.536045664 | 0.3751534 | -1.4288706 | 0.15304143 | 0.2684186 | NA | NA |
| ENSG000002 | 28.2290462 | -0.355908309 | 0.2490855 | -1.42886 | 0.15304447 | 0.2684186 | NA | NA |
| ENSG000002 | 24.8557917 | 0.399631264 | 0.2796982 | 1.42879453 | 0.1530633 | 0.2684284 | NA | NA |
| ENSG000002 | 6.12191311 | -0.523344808 | 0.36627933 | -1.4288134 | 0.15305788 | 0.2684284 | NA | NA |
| ENSG000002 | 2.80204483 | -0.53438046 | 0.37401737 | -1.4287584 | 0.15307368 | 0.268435 | NA | NA |
| ENSG000002 | 12.3341438 | -0.470879662 | 0.32962716 | -1.428522 | 0.15314166 | 0.26854259 | PCDHGC5 | 56097 |
| ENSG000002 | 29.3552265 | 0.355475367 | 0.24885359 | 1.42845183 | 0.15316185 | 0.26856639 | NA | NA |
| ENSG000001 | 27.809971 | -0.377011192 | 0.26395874 | -1.4282959 | 0.15320671 | 0.26863342 | PHYHIP | 9796 |
| ENSG000002 | 4.7051791 | -0.535565514 | 0.37498575 | -1.428229 | 0.15322597 | 0.26865557 | NA | NA |
| ENSG000002 | 4.05565037 | -0.54064238 | 0.37854898 | -1.4281966 | 0.15323527 | 0.26866027 | NA | NA |
| ENSG000001 | 67.5065461 | 0.293951287 | 0.20583074 | 1.42812142 | 0.15325691 | 0.2686866 | GAS6 | 2621 |
| ENSG000001 | 65.345503 | 0.291576658 | 0.20420321 | 1.42787498 | 0.15332785 | 0.26879934 | ARHGEF10 | 9639 |
| ENSG000002 | 2.74045103 | -0.534842384 | 0.37458942 | -1.4278097 | 0.15334665 | 0.26882069 | NA | NA |
| ENSG000001 | 52.3949911 | 0.302145051 | 0.2116195 | 1.4277751 | 0.1533566 | 0.2688265 | FAM160B2 | 64760 |
| ENSG000002 | 9.4019272 | -0.496671966 | 0.3479022 | -1.4276195 | 0.1534014 | 0.26889341 | NA | NA |
| ENSG000001 | 52.1955049 | -0.313367717 | 0.21952768 | -1.4274634 | 0.15344637 | 0.2689386 | KCNQ5 | 56479 |
| ENSG000002 | 9.57385735 | -0.483339728 | 0.33860102 | -1.4274609 | 0.15344708 | 0.2689386 | NA | NA |
| ENSG000002 | 3.65087234 | -0.53989001 | 0.37821092 | -1.4274839 | 0.15344045 | 0.2689386 | NA | NA |
| ENSG000002 | 2.53498194 | -0.528394346 | 0.37019094 | -1.4273562 | 0.15347724 | 0.26897984 | RNU6-603P | 106480594 |
| ENSG000001 | 143.040002 | 0.225529482 | 0.15801159 | 1.42729712 | 0.15349427 | 0.26899806 | PRELID2 | 153768 |
| ENSG00000C | 11.9138652 | -0.459593124 | 0.32203247 | -1.4271639 | 0.15353265 | 0.2690537 | SULT2B1 | 6820 |

| | | | | | | | | |
|---|---|---|---|---|---|---|---|---|
| ENSG000002 | 5.8914881 | -0.526290888 | 0.36883264 | -1.4269097 | 0.15360591 | 0.26915882 | NA | NA |
| ENSG000002 | 6.12886002 | -0.523758251 | 0.36705621 | -1.4269157 | 0.1536042 | 0.26915882 | LOC100652911 | 100652911 |
| ENSG000002 | 12.4975117 | -0.462624746 | 0.32423619 | -1.426814 | 0.15363351 | 0.26919554 | NA | NA |
| ENSG000002 | 3.4315876 | -0.538557629 | 0.37747838 | -1.4267244 | 0.15365934 | 0.26922917 | NA | NA |
| ENSG000002 | 4.49124661 | -0.538133218 | 0.37719735 | -1.4266622 | 0.15367728 | 0.26924896 | NA | NA |
| ENSG000002 | 4.54143614 | -0.53794589 | 0.37710137 | -1.4265286 | 0.15371581 | 0.26930484 | NA | NA |
| ENSG000002 | 11.3770421 | -0.464780489 | 0.32582907 | -1.426455 | 0.15373706 | 0.26933043 | NA | NA |
| ENSG000002 | 2.74551499 | -0.535122666 | 0.37515302 | -1.4264117 | 0.15374953 | 0.26934065 | NA | NA |
| ENSG000002 | 2.46525965 | -0.52859755 | 0.37063088 | -1.4262102 | 0.15380767 | 0.26943085 | RN7SL790P | 106481136 |
| ENSG000002 | 5.31013662 | 0.531570071 | 0.37276033 | 1.42603714 | 0.15385762 | 0.26950671 | NA | NA |
| ENSG000001 | 4.76959666 | -0.535349591 | 0.37543047 | -1.425962 | 0.15387931 | 0.26953305 | NA | NA |
| ENSG000002 | 2.75695729 | -0.534232815 | 0.37465708 | -1.4259248 | 0.15389006 | 0.26954025 | NA | NA |
| ENSG000001 | 7.49550551 | -0.508997308 | 0.35698848 | -1.4258088 | 0.15392354 | 0.26958076 | NEUROG2 | 63973 |
| ENSG000002 | 3.33017784 | -0.538454781 | 0.37765136 | -1.4257986 | 0.15392649 | 0.26958076 | LOC101928942 | 101928942 |
| ENSG000002 | 3.01413683 | -0.538226995 | 0.37749846 | -1.4257727 | 0.15393397 | 0.26958223 | NA | NA |
| ENSG000001 | 4.47087858 | -0.537810993 | 0.37723121 | -1.42568 | 0.15396073 | 0.26961745 | CAV3 | 859 |
| ENSG000000 | 257.429166 | 0.213373915 | 0.14967003 | 1.42562886 | 0.1539755 | 0.26963167 | EPB41L1 | 2036 |
| ENSG000001 | 5.60081529 | -0.528743425 | 0.37096861 | -1.425305 | 0.15406905 | 0.26977028 | NA | NA |
| ENSG000002 | 2.78470963 | -0.532620162 | 0.37369963 | -1.4252627 | 0.15408127 | 0.26977028 | NA | NA |
| ENSG000002 | 3.37330369 | -0.538899581 | 0.37810168 | -1.4252769 | 0.15407718 | 0.26977028 | NA | NA |
| ENSG000002 | 5.11898433 | -0.534853095 | 0.37525335 | -1.425312 | 0.15406703 | 0.26977028 | NA | NA |
| ENSG000002 | 3.10998185 | -0.533936523 | 0.37465204 | -1.4251531 | 0.15411294 | 0.26981409 | NA | NA |
| ENSG000001 | 15.3678201 | -0.434337999 | 0.30477709 | -1.4251005 | 0.15412814 | 0.26982905 | KIF25 | 3834 |
| ENSG000002 | 8.2610541 | 0.50032451 | 0.35109344 | 1.4250466 | 0.15414373 | 0.26984469 | NEXN-AS1 | 374987 |
| ENSG000002 | 37.7223358 | 0.333386662 | 0.23395248 | 1.42501874 | 0.15415178 | 0.26984714 | DECR2 | 26063 |
| ENSG000001 | 17.2624597 | -0.426521046 | 0.29932036 | -1.424965 | 0.15416731 | 0.26986267 | LRRC15 | 131578 |
| ENSG000000 | 3.7004751 | -0.53893546 | 0.37823488 | -1.4248698 | 0.15419484 | 0.26988756 | MAGEB2 | 4113 |
| ENSG000002 | 3.45709215 | -0.444923046 | 0.31225076 | -1.4248902 | 0.15418894 | 0.26988756 | NA | NA |
| ENSG000001 | 34.3243876 | 0.342638038 | 0.24047674 | 1.42482822 | 0.15420686 | 0.26988833 | KHDC1 | 80759 |

| | | | | | | | | |
|---|---|---|---|---|---|---|---|---|
| ENSG000002 | 3.62718076 | -0.53933634 | 0.37852886 | -1.4248222 | 0.15420859 | 0.26988833 | NA | NA |
| ENSG000002 | 8.11110762 | -0.505056131 | 0.35449663 | -1.4247135 | 0.15424004 | 0.26993172 | NA | NA |
| ENSG000002 | 5.55729356 | -0.528014502 | 0.37066771 | -1.4244955 | 0.15430308 | 0.27003038 | NA | NA |
| ENSG000002 | 2.77955721 | -0.533888698 | 0.37483051 | -1.4243469 | 0.15434606 | 0.27009396 | NA | NA |
| ENSG000002 | 25.5337052 | 0.371273838 | 0.26067203 | 1.42429486 | 0.15436113 | 0.27010867 | NA | NA |
| ENSG000001 | 2543.03945 | 0.177583781 | 0.12469829 | 1.42410756 | 0.15441534 | 0.27018021 | ZNF678 | 339500 |
| ENSG000002 | 3.3093539 | -0.538050643 | 0.37781135 | -1.4241251 | 0.15441025 | 0.27018021 | NA | NA |
| ENSG000002 | 19.6556795 | -0.401395394 | 0.28187181 | -1.4240353 | 0.15443627 | 0.27020516 | NA | NA |
| ENSG000002 | 6.30541012 | -0.526914717 | 0.37007338 | -1.4238115 | 0.15450105 | 0.27029738 | RAET1G | 353091 |
| ENSG000002 | 3.5978432 | -0.539030862 | 0.37858873 | -1.4237901 | 0.15450726 | 0.27029738 | NA | NA |
| ENSG000002 | 3.61106735 | -0.539017202 | 0.37858071 | -1.4237841 | 0.15450897 | 0.27029738 | NA | NA |
| ENSG000002 | 6.42549476 | -0.52289691 | 0.36729948 | -1.4236255 | 0.15455491 | 0.27036608 | NA | NA |
| ENSG000001 | 15.3154976 | -0.451529891 | 0.31718284 | -1.4235634 | 0.15457289 | 0.27038588 | OTOR | 56914 |
| ENSG000002 | 4.71579931 | -0.534974932 | 0.37580919 | -1.4235281 | 0.15458314 | 0.27039213 | NA | NA |
| ENSG000001 | 548.259151 | 0.204019555 | 0.14332295 | 1.42349535 | 0.15459261 | 0.27039704 | STOM | 2040 |
| ENSG000002 | 6.13589474 | -0.521983345 | 0.36676397 | -1.4232133 | 0.15467435 | 0.27052833 | NA | NA |
| ENSG000001 | 10.4776686 | -0.503300664 | 0.35366024 | -1.4231192 | 0.15470161 | 0.27055269 | AMN | 81693 |
| ENSG000002 | 3.58681694 | -0.538767265 | 0.37858138 | -1.4231214 | 0.15470096 | 0.27055269 | LOC102467212 | 102467212 |
| ENSG000002 | 6.84389843 | -0.523982232 | 0.36823034 | -1.4229741 | 0.15474367 | 0.27061456 | NA | NA |
| ENSG000002 | 11.2320224 | 0.468928204 | 0.32954633 | 1.42295079 | 0.15475042 | 0.2706147 | NA | NA |
| ENSG000001 | 84.222957 | 0.267555112 | 0.18803561 | 1.42289594 | 0.15476632 | 0.27063084 | IFIT1 | 3434 |
| ENSG000002 | 9.12988997 | -0.491357606 | 0.34539068 | -1.422614 | 0.1548481 | 0.27076216 | NA | NA |
| ENSG000002 | 7.1262504 | 0.51272736 | 0.36042653 | 1.42255721 | 0.15486456 | 0.27077926 | TM6SF2 | 53345 |
| ENSG000002 | 11.2952357 | -0.478831657 | 0.33660879 | -1.4225168 | 0.15487627 | 0.27078807 | NA | NA |
| ENSG000001 | 4.20737002 | -0.537803775 | 0.37808224 | -1.4224518 | 0.15489513 | 0.27080936 | IL3RA | 3563 |
| ENSG000002 | 4.72683254 | -0.534020115 | 0.37544142 | -1.4223793 | 0.15491616 | 0.27083445 | NA | NA |
| ENSG000001 | 103.271146 | 0.248013265 | 0.1743923 | 1.42215717 | 0.15498063 | 0.27093548 | MED31 | 51003 |
| ENSG000002 | 8.80061254 | -0.494770676 | 0.34793482 | -1.4220212 | 0.1550201 | 0.27099281 | NA | NA |
| ENSG000002 | 5.93816163 | -0.526514524 | 0.37027076 | -1.4219717 | 0.15503447 | 0.27100625 | NA | NA |

| | | | | | | | | |
|---|---|---|---|---|---|---|---|---|
| ENSG000002 | 3.35625609 | -0.537470837 | 0.37800303 | -1.4218691 | 0.15506425 | 0.27104661 | NA | NA |
| ENSG000002 | 61.1276541 | 0.298349997 | 0.20988308 | 1.42150573 | 0.15516979 | 0.2712194 | NA | NA |
| ENSG000002 | 3.60588736 | -0.538142483 | 0.37859336 | -1.4214261 | 0.15519292 | 0.27124815 | NA | NA |
| ENSG000002 | 3.62979734 | -0.538055297 | 0.37855322 | -1.4213465 | 0.15521605 | 0.27127688 | NA | NA |
| ENSG000001 | 2474.52646 | 0.373980757 | 0.26313322 | 1.42126011 | 0.15524115 | 0.27129736 | NIPBL | 25836 |
| ENSG000002 | 4.27948046 | -0.537875843 | 0.37844741 | -1.4212697 | 0.15523837 | 0.27129736 | GABRG3-AS1 | 101928869 |
| ENSG000002 | 2.26308157 | -0.507483157 | 0.35709887 | -1.4211279 | 0.15527958 | 0.27135283 | NA | NA |
| ENSG000002 | 2.77253046 | -0.532258244 | 0.37458966 | -1.4209102 | 0.15534288 | 0.27145174 | NA | NA |
| ENSG000002 | 5.00493354 | -0.531215594 | 0.37387493 | -1.4208377 | 0.15536395 | 0.27146517 | NA | NA |
| ENSG000002 | 3.86071127 | -0.537564973 | 0.37834102 | -1.4208477 | 0.15536104 | 0.27146517 | NA | NA |
| ENSG000002 | 12.057441 | 0.458326588 | 0.32258855 | 1.42077761 | 0.15538142 | 0.271484 | NA | NA |
| ENSG000002 | 3.64189307 | -0.537856397 | 0.37858419 | -1.4207049 | 0.15540258 | 0.27150926 | NA | NA |
| ENSG000002 | 3.36794689 | -0.536748726 | 0.37782818 | -1.4206159 | 0.15542845 | 0.27153107 | NA | NA |
| ENSG000002 | 5.26428163 | -0.530892668 | 0.37370083 | -1.4206355 | 0.15542275 | 0.27153107 | LOC101929181 | 101929181 |
| ENSG000001 | 243.31061 | 0.18452162 | 0.12990813 | 1.42040091 | 0.155491 | 0.27161626 | POLK | 51426 |
| ENSG000001 | 498.679279 | 0.202203584 | 0.14235459 | 1.42042196 | 0.15548487 | 0.27161626 | HECTD4 | 283450 |
| ENSG000002 | 4.70727361 | -0.533075741 | 0.37530523 | -1.4203792 | 0.15549731 | 0.27161626 | NA | NA |
| ENSG000001 | 12.0275614 | -0.466309311 | 0.32841599 | -1.419874 | 0.15564438 | 0.27183949 | EQTN | 54586 |
| ENSG000001 | 77.646541 | -0.268464336 | 0.18907656 | -1.4198711 | 0.15564522 | 0.27183949 | GDPD5 | 81544 |
| ENSG000002 | 3.0335197 | -0.535856996 | 0.37739142 | -1.4198972 | 0.15563763 | 0.27183949 | NA | NA |
| ENSG000001 | 17.7678864 | -0.421486755 | 0.29687478 | -1.4197459 | 0.15568167 | 0.27189145 | ARPC1B | 10095 |
| ENSG000001 | 5.54150322 | 0.526458836 | 0.37084433 | 1.41962219 | 0.1557177 | 0.27194266 | BHLHA15 | 168620 |
| ENSG000002 | 5.28310411 | -0.530577832 | 0.37376137 | -1.4195631 | 0.15573492 | 0.27196102 | LOC100506388 | 100506388 |
| ENSG000002 | 10.4929465 | -0.479293501 | 0.33764413 | -1.4195227 | 0.15574669 | 0.27196986 | NA | NA |
| ENSG000002 | 9.69536228 | -0.490854921 | 0.34579496 | -1.4194971 | 0.15575415 | 0.27197118 | NA | NA |
| ENSG000002 | 7.2630344 | -0.515065778 | 0.36286471 | -1.419443 | 0.1557699 | 0.27198697 | NA | NA |
| ENSG000002 | 8.51149324 | -0.496053231 | 0.34947789 | -1.4194123 | 0.15577884 | 0.27199086 | FAM92A1P2 | 403315 |
| ENSG000002 | 5.04386346 | -0.5302829 | 0.37363966 | -1.4192361 | 0.15583019 | 0.27206175 | LOC101929340 | 101929340 |
| ENSG000002 | 3.68621061 | -0.537100748 | 0.37844599 | -1.419227 | 0.15583286 | 0.27206175 | NA | NA |

| | | | | | | | | | |
|---|---|---|---|---|---|---|---|---|---|
| ENSG000002 | 2.79335495 | -0.531041446 | 0.37418782 | -1.4191842 | 0.15584532 | 0.27207179 | LINC00316 | | 388830 |
| ENSG000000 | 48.4016279 | -0.29606877 | 0.20862249 | -1.4191604 | 0.15585226 | 0.2720722 | PAG1 | | 55824 |
| ENSG000001 | 19.2640838 | -0.417426712 | 0.29415528 | -1.4190693 | 0.15587882 | 0.27210328 | GJA10 | | 84694 |
| ENSG000002 | 2.73724214 | -0.53163752 | 0.3746424 | -1.4190533 | 0.15588349 | 0.27210328 | NA | NA | |
| ENSG000002 | 6.70232159 | -0.529960463 | 0.37350553 | -1.4188825 | 0.15593328 | 0.27217848 | NA | NA | |
| ENSG000001 | 151.289445 | 0.214441203 | 0.15114835 | 1.41874652 | 0.15597293 | 0.27223597 | GLS2 | | 27165 |
| ENSG000002 | 3.0454785 | -0.534930671 | 0.37706325 | -1.4186762 | 0.15599343 | 0.27224832 | NA | NA | |
| ENSG000002 | 2.7350231 | -0.530974031 | 0.37427226 | -1.4186839 | 0.1559912 | 0.27224832 | NA | NA | |
| ENSG000001 | 336.035851 | 0.198768717 | 0.14011397 | 1.41862164 | 0.15600935 | 0.27226438 | KLF7 | | 8609 |
| ENSG000002 | 3.18042926 | -0.531265207 | 0.37450537 | -1.4185783 | 0.15602198 | 0.27227471 | NA | NA | |
| ENSG000001 | 775.688578 | 0.215052952 | 0.15160672 | 1.41849217 | 0.15604712 | 0.27230686 | UXS1 | | 80146 |
| ENSG000002 | 3.34217053 | -0.536210156 | 0.37804511 | -1.4183761 | 0.15608098 | 0.27235423 | NA | NA | |
| ENSG000002 | 7.47720551 | -0.505500837 | 0.35642597 | -1.4182492 | 0.15611803 | 0.27240715 | NA | NA | |
| ENSG000002 | 74.367986 | 0.300281294 | 0.21177402 | 1.41793266 | 0.15621043 | 0.27255664 | NA | NA | |
| ENSG000002 | 2.73976643 | -0.531532205 | 0.37489228 | -1.4178265 | 0.15624143 | 0.27259901 | NA | NA | |
| ENSG000001 | 43.3709266 | -0.308756941 | 0.21783249 | -1.4174054 | 0.15636445 | 0.27277982 | LANCL3 | | 347404 |
| ENSG000001 | 176.651359 | 0.221822538 | 0.1565044 | 1.4173566 | 0.1563787 | 0.27277982 | EFNA5 | | 1946 |
| ENSG000001 | 435.595821 | 0.170559633 | 0.12033166 | 1.41741277 | 0.15636228 | 0.27277982 | ZNF615 | | 284370 |
| ENSG000002 | 4.57290122 | -0.535356713 | 0.37770615 | -1.4173894 | 0.1563691 | 0.27277982 | NA | NA | |
| ENSG000002 | 24.272207 | -0.36880144 | 0.26020339 | -1.4173583 | 0.1563782 | 0.27277982 | LOC389641 | | 389641 |
| ENSG000002 | 6.37624774 | -0.517154669 | 0.3648863 | -1.4173036 | 0.15639418 | 0.2727951 | NA | NA | |
| ENSG000002 | 4.72133989 | -0.531936659 | 0.37532865 | -1.4172557 | 0.15640819 | 0.27279616 | NA | NA | |
| ENSG000002 | 16.165563 | 0.4236724 | 0.29893862 | 1.4172555 | 0.15640824 | 0.27279616 | NA | NA | |
| ENSG000002 | 4.58623443 | -0.535921268 | 0.37816871 | -1.4171486 | 0.15643948 | 0.2728389 | NA | NA | |
| ENSG000000 | 74.2213198 | 0.266512747 | 0.18807868 | 1.41702797 | 0.15647475 | 0.27288138 | PHF7 | | 51533 |
| ENSG000001 | 89.4906358 | -0.262219652 | 0.18505017 | -1.4170193 | 0.15647729 | 0.27288138 | IER5 | | 51278 |
| ENSG000002 | 13.2862638 | -0.458015942 | 0.32326272 | -1.4168536 | 0.15652574 | 0.27295412 | NA | NA | |
| ENSG000002 | 2.73973363 | -0.530519578 | 0.3744414 | -1.4168294 | 0.15653282 | 0.27295473 | NA | NA | |
| ENSG000002 | 12.0717267 | -0.469620986 | 0.33148768 | -1.4167072 | 0.15656855 | 0.27299357 | NA | NA | |

| | | | | | | | | |
|---|---|---|---|---|---|---|---|---|
| ENSG000002 | 4.7274254 | -0.532779588 | 0.37606421 | -1.4167251 | 0.15656333 | 0.27299357 | NA | NA |
| ENSG000002 | 6.22407579 | -0.524399424 | 0.37017979 | -1.4166074 | 0.15659775 | 0.273021 | NA | NA |
| ENSG000002 | 2.53352658 | -0.522070342 | 0.3685342 | -1.416613 | 0.15659611 | 0.273021 | NA | NA |
| ENSG000002 | 4.52744895 | -0.534737178 | 0.37748829 | -1.4165663 | 0.15660978 | 0.27303023 | NA | NA |
| ENSG000002 | 4.32820984 | -0.536274505 | 0.37858271 | -1.416532 | 0.15661981 | 0.27303599 | NA | NA |
| ENSG000002 | 9.58594044 | -0.479535723 | 0.33854664 | -1.416454 | 0.15664264 | 0.27306404 | NA | NA |
| ENSG000001 | 3.60807918 | -0.536210464 | 0.37858526 | -1.4163532 | 0.15667211 | 0.27309674 | CEACAM4 | 1089 |
| ENSG000001 | 294.864454 | 0.216155168 | 0.1526149 | 1.41634382 | 0.15667487 | 0.27309674 | TP53I11 | 9537 |
| ENSG000002 | 3.60016584 | -0.536132502 | 0.37858581 | -1.4161453 | 0.15673298 | 0.27318629 | NA | NA |
| ENSG000001 | 7.67055546 | -0.499337166 | 0.35262023 | -1.4160763 | 0.15675316 | 0.27320972 | HOXA1 | 3198 |
| ENSG000001 | 15.3519935 | -0.437831743 | 0.30921919 | -1.4159268 | 0.15679693 | 0.27327426 | FAM178B | 51252 |
| ENSG000001 | 4.99547475 | -0.528578958 | 0.37334801 | -1.4157808 | 0.15683968 | 0.27332528 | GJB1 | 2705 |
| ENSG000002 | 11.3066668 | -0.475462799 | 0.33582977 | -1.4157851 | 0.15683842 | 0.27332528 | LOC100505685 | 100505685 |
| ENSG000002 | 5.39361532 | -0.530269682 | 0.37455982 | -1.4157143 | 0.15685918 | 0.27334751 | NA | NA |
| ENSG000001 | 25.7201873 | -0.370765638 | 0.26193727 | -1.4154749 | 0.15692929 | 0.27341094 | KCNIP1 | 30820 |
| ENSG000002 | 27.8869842 | 0.361278758 | 0.25523126 | 1.41549574 | 0.1569232 | 0.27341094 | NA | NA |
| ENSG000002 | 2.53107919 | -0.520953946 | 0.36802891 | -1.4155245 | 0.15691477 | 0.27341094 | NA | NA |
| ENSG000002 | 9.40481653 | -0.488151229 | 0.34486495 | -1.4154852 | 0.15692629 | 0.27341094 | NA | NA |
| ENSG000002 | 5.34200956 | -0.529862911 | 0.37431222 | -1.415564 | 0.15690319 | 0.27341094 | NA | NA |
| ENSG000002 | 5.43378138 | -0.531372966 | 0.37548807 | -1.4151527 | 0.15702372 | 0.2735637 | NA | NA |
| ENSG000002 | 2.99970581 | -0.534062266 | 0.37739478 | -1.4151289 | 0.1570307 | 0.27356411 | NA | NA |
| ENSG000002 | 3.42709242 | -0.533412965 | 0.37695722 | -1.4150491 | 0.15705411 | 0.27359314 | LOC101929337 | 101929337 |
| ENSG000001 | 17.5759637 | -0.408832549 | 0.28892501 | -1.4150127 | 0.15706478 | 0.27359997 | SLC22A18 | 5002 |
| ENSG000002 | 8.40732016 | -0.508847369 | 0.35961487 | -1.4149787 | 0.15707475 | 0.2736056 | NA | NA |
| ENSG000002 | 2.81106698 | -0.528039977 | 0.37319541 | -1.4149155 | 0.15709327 | 0.27361435 | ACTP1 | 106480335 |
| ENSG000002 | 3.33870465 | -0.533979688 | 0.37739011 | -1.4149276 | 0.15708972 | 0.27361435 | NA | NA |
| ENSG000002 | 4.470964 | -0.53372729 | 0.3772286 | -1.4148643 | 0.15710828 | 0.27362188 | NA | NA |
| ENSG000002 | 3.89637227 | -0.535252559 | 0.3783092 | -1.4148547 | 0.15711109 | 0.27362188 | NA | NA |
| ENSG000001 | 33.6974816 | 0.343532841 | 0.24282331 | 1.41474406 | 0.15714355 | 0.27366666 | CTU1 | 90353 |

| | | | | | | | | |
|---|---|---|---|---|---|---|---|---|
| ENSG000001 | 100.615637 | 0.270190191 | 0.19099064 | 1.41467768 | 0.15716302 | 0.27368881 | POLM | 27434 |
| ENSG000002 | 3.59936969 | -0.535574805 | 0.37859323 | -1.4146444 | 0.15717277 | 0.27369404 | NA | NA |
| ENSG000002 | 5.1475849 | -0.532464578 | 0.37641383 | -1.4145723 | 0.15719394 | 0.27371914 | NA | NA |
| ENSG000001 | 3060.75012 | 0.484894466 | 0.3428299 | 1.41438792 | 0.15724803 | 0.27380158 | RPL28 | 6158 |
| ENSG000002 | 3.7406817 | -0.534473195 | 0.37790584 | -1.4143025 | 0.15727309 | 0.27383346 | NA | NA |
| ENSG000002 | 2.75297098 | -0.528687602 | 0.37383762 | -1.4142172 | 0.15729813 | 0.27386528 | NA | NA |
| ENSG000001 | 14001.2008 | 0.159407139 | 0.11272773 | 1.41408982 | 0.15733553 | 0.27390689 | CENPF | 1063 |
| ENSG000001 | 3.46247301 | -0.533485719 | 0.37726164 | -1.4141001 | 0.15733252 | 0.27390689 | GJD4 | 219770 |
| ENSG000002 | 2.78696147 | -0.528964199 | 0.37409196 | -1.4139951 | 0.15736334 | 0.27394287 | NA | NA |
| ENSG000002 | 4.50796145 | -0.534800595 | 0.37822905 | -1.4139596 | 0.15737377 | 0.27394287 | NA | NA |
| ENSG000002 | 7.68440493 | -0.499614756 | 0.35334674 | -1.4139504 | 0.15737647 | 0.27394287 | LOC101929511 | 101929511 |
| ENSG000001 | 63.9592349 | 0.277953433 | 0.19659061 | 1.41386935 | 0.15740027 | 0.27397254 | NA | NA |
| ENSG000002 | 10.9883519 | -0.474704835 | 0.33576821 | -1.4137873 | 0.15742436 | 0.27399303 | NA | NA |
| ENSG000002 | 4.83685971 | -0.531990861 | 0.37628884 | -1.4137832 | 0.15742555 | 0.27399303 | NA | NA |
| ENSG000002 | 17.4647151 | -0.421885862 | 0.29842613 | -1.4137028 | 0.15744917 | 0.27401061 | NA | NA |
| ENSG000002 | 9.96219059 | -0.485519522 | 0.34343595 | -1.413712 | 0.15744648 | 0.27401061 | NA | NA |
| ENSG000002 | 3.63150064 | -0.535163252 | 0.37857712 | -1.4136175 | 0.15747423 | 0.27404247 | NA | NA |
| ENSG000002 | 3.02811138 | -0.533358891 | 0.37732354 | -1.413532 | 0.15749937 | 0.27407444 | NA | NA |
| ENSG000002 | 3.71813772 | -0.534505411 | 0.37817961 | -1.4133639 | 0.15754876 | 0.27414863 | NA | NA |
| ENSG000001 | 135.931557 | 0.230269471 | 0.16292816 | 1.41331907 | 0.15756193 | 0.27415979 | HECA | 51696 |
| ENSG000002 | 4.36396658 | -0.534973587 | 0.37853947 | -1.4132571 | 0.15758015 | 0.27416185 | NA | NA |
| ENSG000002 | 6.89779992 | -0.512742633 | 0.36280974 | -1.4132549 | 0.15758079 | 0.27416185 | NA | NA |
| ENSG000002 | 3.67789789 | -0.53461516 | 0.37829497 | -1.413223 | 0.15759016 | 0.27416185 | NA | NA |
| ENSG000002 | 11.9023744 | -0.460965501 | 0.326175 | -1.413246 | 0.15758342 | 0.27416185 | NA | NA |
| ENSG000001 | 222.222864 | 0.185526845 | 0.13129702 | 1.41303165 | 0.15764642 | 0.27422819 | SERAC1 | 84947 |
| ENSG000001 | 5.60400292 | -0.527565517 | 0.3733586 | -1.4130263 | 0.15764799 | 0.27422819 | DNM1P46 | 196968 |
| ENSG000001 | 5.48878721 | 0.524157974 | 0.37095364 | 1.41300129 | 0.15765534 | 0.27422819 | OXCT2 | 64064 |
| ENSG000002 | 2.81331053 | -0.525864991 | 0.37215703 | -1.4130191 | 0.1576501 | 0.27422819 | NA | NA |
| ENSG000001 | 13.2737785 | -0.479564182 | 0.33946757 | -1.4126951 | 0.15774539 | 0.27437305 | NA | NA |

| | | | | | | | | |
|---|---|---|---|---|---|---|---|---|
| ENSG000001 | 15.6334188 | -0.433219504 | 0.30666847 | -1.412664 | 0.15775455 | 0.27437721 | TMPRSS5 | 80975 |
| ENSG000001 | 69.8238753 | 0.273825275 | 0.19385875 | 1.41249891 | 0.15780311 | 0.27444991 | C19orf44 | 84167 |
| ENSG000002 | 2.87585959 | -0.523009447 | 0.37029003 | -1.4124319 | 0.15782283 | 0.27447243 | NA | NA |
| ENSG000001 | 166.383264 | 0.216161641 | 0.15305269 | 1.41233483 | 0.15785139 | 0.27451033 | MCF2L | 23263 |
| ENSG000001 | 64.9569266 | -0.278648 | 0.19731867 | -1.4121725 | 0.15789918 | 0.27456988 | SYT12 | 91683 |
| ENSG000002 | 2.49981075 | -0.522126608 | 0.36973141 | -1.4121781 | 0.15789753 | 0.27456988 | NA | NA |
| ENSG000002 | 10.8965478 | -0.472757714 | 0.33478082 | -1.412141 | 0.15790845 | 0.27457423 | THEGL | 100506564 |
| ENSG000001 | 803.921032 | -0.184030593 | 0.1303397 | -1.4119304 | 0.15797045 | 0.27467025 | CCDC141 | 285025 |
| ENSG000001 | 5.42575525 | -0.530774529 | 0.37592847 | -1.411903 | 0.15797853 | 0.27467252 | KRT75 | 9119 |
| ENSG000001 | 37.5176993 | -0.340003966 | 0.24081813 | -1.4118703 | 0.15798815 | 0.27467747 | TEX11 | 56159 |
| ENSG000001 | 3.50156703 | -0.530550819 | 0.37582025 | -1.4117143 | 0.1580341 | 0.27474558 | UTS2R | 2837 |
| ENSG000002 | 4.5596689 | -0.533503879 | 0.37794039 | -1.4116085 | 0.15806526 | 0.27478798 | NA | NA |
| ENSG000002 | 35.0897085 | 0.343641998 | 0.24344902 | 1.4115563 | 0.15808065 | 0.27480295 | NA | NA |
| ENSG000002 | 5.6092644 | -0.524403992 | 0.37153166 | -1.4114651 | 0.15810751 | 0.27483785 | NA | NA |
| ENSG000001 | 108.344653 | 0.240005586 | 0.17004654 | 1.41141115 | 0.15812342 | 0.27484195 | FAM149A | 25854 |
| ENSG000001 | 8.61164044 | -0.500089322 | 0.35431855 | -1.4114116 | 0.15812327 | 0.27484195 | NA | NA |
| ENSG000001 | 182.738531 | 0.247881705 | 0.17563105 | 1.41137744 | 0.15813335 | 0.27484743 | TXNL4B | 54957 |
| ENSG000002 | 6.15933635 | -0.517585581 | 0.36674388 | -1.4112999 | 0.15815622 | 0.2748754 | NA | NA |
| ENSG000002 | 2.7563837 | -0.528312165 | 0.37435807 | -1.4112482 | 0.15817144 | 0.27489007 | NA | NA |
| ENSG000002 | 7.51556101 | -0.511600706 | 0.36253689 | -1.4111687 | 0.15819488 | 0.27491902 | EPN2-AS1 | 100874018 |
| ENSG000002 | 6.35210999 | -0.513476703 | 0.36390406 | -1.4110222 | 0.15823808 | 0.27498231 | NA | NA |
| ENSG000002 | 3.46611543 | -0.530623178 | 0.37606549 | -1.4109861 | 0.15824871 | 0.27498901 | NA | NA |
| ENSG000001 | 211.100905 | 0.237306033 | 0.16819774 | 1.41087531 | 0.15828138 | 0.27499779 | FANCC | 2176 |
| ENSG000002 | 7.69444449 | -0.501049358 | 0.35514485 | -1.410831 | 0.15829446 | 0.27499779 | NA | NA |
| ENSG000002 | 3.96846998 | -0.533807247 | 0.37836102 | -1.410841 | 0.15829149 | 0.27499779 | NA | NA |
| ENSG000002 | 2.99236606 | -0.532165651 | 0.37717025 | -1.4109428 | 0.15826148 | 0.27499779 | RN7SL293P | 106481835 |
| ENSG000002 | 6.0614559 | -0.517383154 | 0.36671167 | -1.4108718 | 0.15828242 | 0.27499779 | NA | NA |
| ENSG000002 | 57.1849997 | 0.304394227 | 0.2157526 | 1.41084845 | 0.15828931 | 0.27499779 | NA | NA |
| ENSG000002 | 3.88684049 | -0.533790841 | 0.37839399 | -1.4106747 | 0.15834055 | 0.27506608 | NA | NA |

| | | | | | | | | |
|---|---|---|---|---|---|---|---|---|
| ENSG000001 | 45.2123882 | -0.299661186 | 0.21242863 | -1.4106441 | 0.15834958 | 0.27507 | SLC9A9 | 285195 |
| ENSG000002 | 6.59827715 | -0.520970803 | 0.36933837 | -1.4105515 | 0.15837689 | 0.27510564 | NA | NA |
| ENSG000002 | 6.30657305 | -0.512615036 | 0.3634359 | -1.4104689 | 0.15840127 | 0.2751362 | NA | NA |
| ENSG000002 | 55.3051865 | 0.288802608 | 0.20476371 | 1.41041893 | 0.15841602 | 0.27515004 | NA | NA |
| ENSG000002 | 3.60230634 | -0.533963972 | 0.37859334 | -1.4103892 | 0.15842478 | 0.27515347 | NA | NA |
| ENSG000001 | 62.6245562 | 0.278609419 | 0.1975543 | 1.41029289 | 0.15845322 | 0.27519108 | S100A13 | 6284 |
| ENSG000001 | 710.76728 | 0.217245082 | 0.15404791 | 1.41024358 | 0.15846777 | 0.27520457 | TCF7L2 | 6934 |
| ENSG000002 | 12.7252756 | -0.452208069 | 0.32071928 | -1.4099809 | 0.15854531 | 0.27531355 | NA | NA |
| ENSG000002 | 7.96751707 | -0.497928243 | 0.35314417 | -1.4099857 | 0.1585439 | 0.27531355 | HNCAT21 | 100505811 |
| ENSG000002 | 2.45253084 | -0.520164282 | 0.36892077 | -1.409962 | 0.15855089 | 0.27531355 | SNRPGP20 | 106480271 |
| ENSG000002 | 31.8147698 | 0.341866401 | 0.24250481 | 1.40973038 | 0.15861931 | 0.27542056 | NA | NA |
| ENSG000001 | 7.73824686 | -0.502534678 | 0.35648603 | -1.4096897 | 0.15863133 | 0.27542964 | SERINC4 | 619189 |
| ENSG000001 | 3.0402666 | -0.531850019 | 0.37732952 | -1.4095108 | 0.15868419 | 0.27550962 | RN7SKP149 | 106479161 |
| ENSG000002 | 10.7627935 | 0.477148532 | 0.33856653 | 1.40931985 | 0.15874061 | 0.27559261 | NA | NA |
| ENSG000002 | 3.84904063 | -0.533169421 | 0.37832135 | -1.409303 | 0.15874559 | 0.27559261 | NA | NA |
| ENSG000000 | 874.484345 | 0.148871548 | 0.10564085 | 1.40922324 | 0.15876917 | 0.27562175 | CNTLN | 54875 |
| ENSG000001 | 7.26242459 | -0.508074246 | 0.36059284 | -1.408997 | 0.15883604 | 0.27572604 | AIRE | 326 |
| ENSG000001 | 36.0524021 | -0.324075233 | 0.23002291 | -1.4088824 | 0.15886993 | 0.27576416 | LPO | 4025 |
| ENSG000002 | 3.60426893 | -0.533253918 | 0.37850196 | -1.4088538 | 0.1588784 | 0.27576416 | NA | NA |
| ENSG000002 | 3.86293086 | -0.532863936 | 0.37822365 | -1.4088594 | 0.15887676 | 0.27576416 | NA | NA |
| ENSG000002 | 6.26408762 | -0.521901396 | 0.37047781 | -1.4087251 | 0.15891647 | 0.27581843 | NA | NA |
| ENSG000002 | 3.69099087 | -0.532821985 | 0.37825445 | -1.4086338 | 0.15894349 | 0.27585352 | NA | NA |
| ENSG000002 | 2.75220131 | -0.526332486 | 0.37366893 | -1.408553 | 0.15896739 | 0.27588319 | NA | NA |
| ENSG000001 | 2.79563871 | -0.526340152 | 0.37371307 | -1.4084071 | 0.15901055 | 0.27594629 | C7orf34 | 135927 |
| ENSG000001 | 19.5839455 | -0.393686445 | 0.27954888 | -1.408292 | 0.15904463 | 0.27599361 | RIN3 | 79890 |
| ENSG000001 | 8.34535572 | -0.497153396 | 0.35303772 | -1.4082161 | 0.15906709 | 0.27601312 | AGTR2 | 186 |
| ENSG000002 | 6.10446929 | -0.516309766 | 0.3666491 | -1.408185 | 0.15907629 | 0.27601312 | ASPRV1 | 151516 |
| ENSG000002 | 10.6287366 | 0.477290398 | 0.33893799 | 1.40819388 | 0.15907367 | 0.27601312 | NA | NA |
| ENSG000002 | 7.35690484 | 0.503996245 | 0.35794348 | 1.40803305 | 0.15912128 | 0.27607938 | NA | NA |

| | | | | | | | | |
|---|---|---|---|---|---|---|---|---|
| ENSG000002 | 7.19120827 | -0.506814663 | 0.36003228 | -1.4076923 | 0.15922221 | 0.27624266 | CARD18 | 59082 |
| ENSG000002 | 9.33690224 | -0.482269477 | 0.34261849 | -1.4075991 | 0.15924981 | 0.27627874 | NA | NA |
| ENSG000002 | 15.3392526 | -0.450684052 | 0.32020644 | -1.4074796 | 0.15928521 | 0.27632832 | NA | NA |
| ENSG000002 | 3.45943223 | -0.530498803 | 0.37693357 | -1.4074066 | 0.15930685 | 0.27634221 | UGT1A11P | 54574 |
| ENSG000002 | 3.88413252 | -0.532560096 | 0.37839695 | -1.4074112 | 0.1593055 | 0.27634221 | NA | NA |
| ENSG000002 | 11.1597256 | -0.482176858 | 0.34261901 | -1.4073266 | 0.15933055 | 0.27636636 | LOC101928476 | 101928476 |
| ENSG000002 | 2.44986219 | -0.518018444 | 0.36809025 | -1.4073137 | 0.15933439 | 0.27636636 | NA | NA |
| ENSG000002 | 3.18019374 | -0.526400475 | 0.37409172 | -1.4071428 | 0.15938505 | 0.27644239 | NA | NA |
| ENSG000001 | 13.9949617 | -0.454084265 | 0.32271932 | -1.4070563 | 0.15941069 | 0.27647504 | PSMA8 | 143471 |
| ENSG000002 | 6.52919511 | -0.518060065 | 0.36819609 | -1.4070222 | 0.15942082 | 0.27648078 | NA | NA |
| ENSG000002 | 12.2430011 | -0.457504904 | 0.32516714 | -1.4069838 | 0.15943218 | 0.27648866 | AFAP1-AS1 | 84740 |
| ENSG000002 | 2.27518527 | -0.504141627 | 0.35833279 | -1.406909 | 0.15945438 | 0.27651533 | NA | NA |
| ENSG000002 | 3.62885161 | -0.532521829 | 0.37852595 | -1.4068305 | 0.15947767 | 0.27653207 | NA | NA |
| ENSG000002 | 4.26752112 | -0.532618838 | 0.37858873 | -1.4068534 | 0.15947087 | 0.27653207 | NA | NA |
| ENSG000002 | 6.40558661 | -0.514227038 | 0.36553108 | -1.4067943 | 0.15948839 | 0.27653571 | NA | NA |
| ENSG000002 | 11.072328 | -0.465725181 | 0.33105819 | -1.4067774 | 0.15949341 | 0.27653571 | NA | NA |
| ENSG000002 | 3.40604676 | -0.530890511 | 0.37739758 | -1.4067141 | 0.15951218 | 0.27655643 | NA | NA |
| ENSG000002 | 3.93096427 | -0.532415476 | 0.37854284 | -1.4064867 | 0.15957965 | 0.27664975 | NA | NA |
| ENSG000002 | 7.56554978 | -0.509204751 | 0.36203446 | -1.4065091 | 0.15957301 | 0.27664975 | NA | NA |
| ENSG000002 | 94.5281276 | 0.254827125 | 0.18119293 | 1.40638561 | 0.15960965 | 0.27667811 | TRIL | 9865 |
| ENSG000002 | 4.45386434 | -0.53087274 | 0.37747288 | -1.4063864 | 0.15960941 | 0.27667811 | NA | NA |
| ENSG000002 | 5.26817464 | -0.523623613 | 0.37232938 | -1.4063451 | 0.15962166 | 0.2766871 | MIR4755 | 100616434 |
| ENSG000002 | 17.4132131 | -0.418660946 | 0.2976995 | -1.4063206 | 0.15962893 | 0.27668787 | NA | NA |
| ENSG000002 | 5.82693391 | -0.518495709 | 0.36873997 | -1.4061283 | 0.15968603 | 0.27677502 | NA | NA |
| ENSG000001 | 49.7384806 | -0.301607401 | 0.21452673 | -1.4059199 | 0.15974791 | 0.27685834 | FAM134B | 54463 |
| ENSG000001 | 14.396074 | -0.44161619 | 0.31411694 | -1.4058974 | 0.15975459 | 0.27685834 | PLK5 | 126520 |
| ENSG000002 | 4.29143792 | -0.532244925 | 0.37856834 | -1.4059415 | 0.1597415 | 0.27685834 | NA | NA |
| ENSG00000C | 6.78152741 | -0.519577319 | 0.36958305 | -1.4058473 | 0.15976949 | 0.27687233 | PRSS21 | 10942 |
| ENSG000002 | 3.63492491 | -0.531909696 | 0.3783642 | -1.405814 | 0.15977935 | 0.27687759 | NA | NA |

| | | | | | | | | |
|---|---|---|---|---|---|---|---|---|
| ENSG000001 | 105.011262 | 0.229076827 | 0.16296613 | 1.40567143 | 0.15982171 | 0.27693916 | AK5 | 26289 |
| ENSG000002 | 3.62221681 | -0.532044534 | 0.37852878 | -1.405559 | 0.15985512 | 0.2769767 | NA | NA |
| ENSG000002 | 131.982006 | 0.22661301 | 0.16122699 | 1.40555254 | 0.15985704 | 0.2769767 | NA | NA |
| ENSG000002 | 4.70117242 | -0.528746559 | 0.3762119 | -1.4054488 | 0.15988787 | 0.27701828 | NA | NA |
| ENSG000002 | 3.05500973 | -0.529542091 | 0.37680062 | -1.4053642 | 0.15991301 | 0.27705001 | CEACAM16 | 388551 |
| ENSG000002 | 2.53754535 | -0.515997676 | 0.36720061 | -1.4052201 | 0.15995585 | 0.27710054 | LOC101927839 | 101927839 |
| ENSG000002 | 12.1547228 | -0.472482811 | 0.33623396 | -1.4052204 | 0.15995576 | 0.27710054 | NA | NA |
| ENSG000002 | 11.1922744 | -0.463607216 | 0.32992493 | -1.40519 | 0.15996479 | 0.27710419 | NA | NA |
| ENSG000001 | 4.50476469 | -0.530326301 | 0.37742175 | -1.4051291 | 0.15998288 | 0.2771237 | C10orf120 | 399814 |
| ENSG000002 | 9.26933373 | -0.480316581 | 0.34186262 | -1.4049988 | 0.16002162 | 0.27717897 | NA | NA |
| ENSG000002 | 22.4657792 | 0.38225457 | 0.27207907 | 1.40493925 | 0.16003934 | 0.27719782 | NA | NA |
| ENSG000002 | 3.35183293 | -0.530736296 | 0.3777954 | -1.4048247 | 0.16007343 | 0.27724501 | NA | NA |
| ENSG000002 | 3.81852586 | -0.529588504 | 0.37701099 | -1.4047031 | 0.16010958 | 0.27729579 | NA | NA |
| ENSG000002 | 3.03599934 | -0.529568104 | 0.37701915 | -1.4046186 | 0.16013473 | 0.27730382 | NA | NA |
| ENSG000002 | 15.1145756 | 0.438191965 | 0.31196148 | 1.40463485 | 0.16012989 | 0.27730382 | LOC374443 | 374443 |
| ENSG000002 | 4.73710153 | -0.527969969 | 0.37587186 | -1.4046542 | 0.16012415 | 0.27730382 | NA | NA |
| ENSG000002 | 2.79358274 | -0.523269002 | 0.37257823 | -1.404454 | 0.16018369 | 0.27736786 | NA | NA |
| ENSG000002 | 3.03610348 | -0.52950368 | 0.37701899 | -1.4044483 | 0.1601854 | 0.27736786 | NA | NA |
| ENSG000002 | 3.66619555 | -0.531582022 | 0.37852521 | -1.4043504 | 0.16021454 | 0.27740234 | NA | NA |
| ENSG000002 | 48.572311 | -0.307271507 | 0.21880207 | -1.4043354 | 0.16021899 | 0.27740234 | MYO15B | 80022 |
| ENSG000001 | 13.6982032 | -0.439062077 | 0.3126755 | -1.4042101 | 0.16025631 | 0.27745511 | SCIMP | 388325 |
| ENSG000002 | 2.51432553 | -0.517161986 | 0.3683214 | -1.4041052 | 0.16028753 | 0.27749732 | NA | NA |
| ENSG000001 | 90.8245672 | 0.269901612 | 0.19223992 | 1.4039832 | 0.16032385 | 0.27754835 | RAB6B | 51560 |
| ENSG000002 | 3.61926906 | -0.53148752 | 0.37858074 | -1.4038948 | 0.16035019 | 0.27757782 | NA | NA |
| ENSG000002 | 3.01254618 | -0.529619759 | 0.37725428 | -1.4038801 | 0.16035457 | 0.27757782 | NA | NA |
| ENSG000002 | 5.31080612 | -0.525397524 | 0.37425346 | -1.4038548 | 0.16036208 | 0.27757899 | TMEM235 | 283999 |
| ENSG000002 | 2.19865212 | -0.504738277 | 0.35960539 | -1.4035893 | 0.16044118 | 0.27770404 | NA | NA |
| ENSG000001 | 434.730834 | 0.199650225 | 0.14228101 | 1.40321065 | 0.16055403 | 0.27788752 | NEK7 | 140609 |
| ENSG000001 | 10.0823984 | -0.488548677 | 0.34820946 | -1.403031 | 0.16060758 | 0.27796833 | CXorf21 | 80231 |

| | | | | | | | | |
|---|---|---|---|---|---|---|---|---|
| ENSG000002 | 23.3006797 | -0.371037998 | 0.26446268 | -1.4029881 | 0.16062038 | 0.27797862 | NA | NA |
| ENSG000002 | 23.2310324 | 0.386892865 | 0.27578735 | 1.40286659 | 0.16065662 | 0.27801762 | NA | NA |
| ENSG000002 | 5.63458951 | -0.521525923 | 0.37175385 | -1.4028797 | 0.16065272 | 0.27801762 | RN7SL822P | 106479521 |
| ENSG000001 | 1170.511 | 0.174882861 | 0.12466871 | 1.40278067 | 0.16068225 | 0.27803847 | TP53BP2 | 7159 |
| ENSG000001 | 31.5840138 | -0.342091358 | 0.24387024 | -1.4027598 | 0.16068848 | 0.27803847 | ENTPD3 | 956 |
| ENSG000002 | 5.04657279 | -0.527253925 | 0.37586969 | -1.4027572 | 0.16068925 | 0.27803847 | NA | NA |
| ENSG000001 | 12.884567 | -0.443583817 | 0.31622995 | -1.4027255 | 0.16069871 | 0.27804299 | NA | NA |
| ENSG000002 | 8.87920115 | 0.490446586 | 0.34965648 | 1.40265264 | 0.16072045 | 0.27806873 | NA | NA |
| ENSG000002 | 15.788375 | -0.420427148 | 0.29974787 | -1.4026026 | 0.16073537 | 0.27808268 | NA | NA |
| ENSG000002 | 34.5167352 | 0.329879365 | 0.23520697 | 1.40250679 | 0.16076397 | 0.27810597 | NA | NA |
| ENSG000002 | 46.0718234 | 0.310672492 | 0.22151517 | 1.40248857 | 0.1607694 | 0.27810597 | NA | NA |
| ENSG000002 | 91.7260608 | -0.24121611 | 0.17198922 | -1.4025072 | 0.16076385 | 0.27810597 | NA | NA |
| ENSG000001 | 64.2010286 | 0.273318212 | 0.19489778 | 1.40236697 | 0.16080569 | 0.27813315 | CDKN1C | 1028 |
| ENSG000002 | 9.30711377 | -0.481432967 | 0.34329522 | -1.4023876 | 0.16079952 | 0.27813315 | NA | NA |
| ENSG000002 | 3.49813727 | 0.525953019 | 0.37504039 | 1.40239033 | 0.16079872 | 0.27813315 | NA | NA |
| ENSG000002 | 9.5165903 | -0.476834401 | 0.3400396 | -1.4022908 | 0.16082842 | 0.2781606 | NA | NA |
| ENSG000002 | 3.33613601 | -0.52951263 | 0.37765893 | -1.4020922 | 0.16088772 | 0.27822402 | UNC5B-AS1 | 728978 |
| ENSG000002 | 4.70457305 | -0.526692814 | 0.37564832 | -1.4020901 | 0.16088833 | 0.27822402 | NA | NA |
| ENSG000002 | 4.51916868 | -0.529245579 | 0.37748518 | -1.4020301 | 0.16090626 | 0.27822402 | NA | NA |
| ENSG000002 | 4.9854548 | -0.523608278 | 0.37346425 | -1.4020305 | 0.16090613 | 0.27822402 | LOC101928710 | 101928710 |
| ENSG000002 | 3.42313402 | -0.529121858 | 0.3773757 | -1.402109 | 0.1608827 | 0.27822402 | NA | NA |
| ENSG000002 | 33.7494021 | -0.339644478 | 0.24224907 | -1.4020466 | 0.16090133 | 0.27822402 | NA | NA |
| ENSG000001 | 5816.13629 | 0.147866906 | 0.10547797 | 1.40187476 | 0.16095265 | 0.27829237 | TRIM24 | 8805 |
| ENSG000002 | 9.81125683 | -0.474391343 | 0.3384746 | -1.4015567 | 0.16104767 | 0.27844479 | LINC01491 | 101928442 |
| ENSG000002 | 24.1142602 | -0.367685023 | 0.26237247 | -1.4013857 | 0.16109877 | 0.27852126 | PCDHA13 | 56136 |
| ENSG000002 | 20.0107006 | -0.388808472 | 0.27745925 | -1.4013174 | 0.16111919 | 0.27854469 | NA | NA |
| ENSG000002 | 6.69007801 | -0.514240166 | 0.36700928 | -1.4011639 | 0.16116506 | 0.27861212 | NA | NA |
| ENSG000002 | 3.60688571 | -0.530415572 | 0.37858105 | -1.4010621 | 0.16119549 | 0.27865284 | NA | NA |
| ENSG000002 | 2.18112036 | -0.511291711 | 0.36496058 | -1.4009505 | 0.16122887 | 0.27869867 | NA | NA |

| | | | | | | | | | |
|---|---|---|---|---|---|---|---|---|---|
| ENSG000002 | 17.9919147 | -0.41990095 | 0.29974278 | -1.400871 | 0.16125266 | 0.27872791 | NA | NA | |
| ENSG000001 | 209.459562 | 0.215238704 | 0.15370851 | 1.40030439 | 0.16142219 | 0.27896148 | CMTM7 | | 112616 |
| ENSG000001 | 6.16793429 | -0.514718608 | 0.36757602 | -1.4003052 | 0.16142195 | 0.27896148 | TRIM74 | | 378108 |
| ENSG000002 | 3.16110621 | -0.524504571 | 0.37455227 | -1.4003508 | 0.1614083 | 0.27896148 | NA | NA | |
| ENSG000002 | 2.7358394 | -0.523654735 | 0.37395452 | -1.4003166 | 0.16141853 | 0.27896148 | NDFIP2-AS1 | | 100874208 |
| ENSG000002 | 3.61830819 | -0.530033373 | 0.37850488 | -1.4003343 | 0.16141322 | 0.27896148 | NA | NA | |
| ENSG000001 | 178.960305 | 0.215730396 | 0.15407902 | 1.40012831 | 0.1614749 | 0.27904068 | ZNF444 | | 55311 |
| ENSG000002 | 4.45497424 | -0.528254918 | 0.37733252 | -1.3999719 | 0.16152174 | 0.27910974 | NA | NA | |
| ENSG000001 | 16.0849423 | -0.424550446 | 0.30326975 | -1.3999103 | 0.16154018 | 0.27912971 | DMP1 | | 1758 |
| ENSG000002 | 3.8146046 | -0.52793236 | 0.37713383 | -1.3998542 | 0.16155699 | 0.27913741 | NA | NA | |
| ENSG000002 | 2.45179418 | -0.517166185 | 0.36945595 | -1.3998047 | 0.1615718 | 0.27913741 | MIR5585 | | 100847018 |
| ENSG000002 | 4.27115585 | -0.52962649 | 0.37835775 | -1.3998035 | 0.16157218 | 0.27913741 | NA | NA | |
| ENSG000002 | 11.6445584 | -0.456302299 | 0.32597279 | -1.3998171 | 0.16156808 | 0.27913741 | NA | NA | |
| ENSG000002 | 3.63337892 | -0.52954255 | 0.3783398 | -1.399648 | 0.16161875 | 0.27920598 | NA | NA | |
| ENSG000002 | 20.0199528 | -0.386500655 | 0.27614773 | -1.3996155 | 0.16162848 | 0.2792109 | NA | NA | |
| ENSG000002 | 5.02248554 | -0.523147259 | 0.37379832 | -1.3995442 | 0.16164986 | 0.27923593 | NA | NA | |
| ENSG000002 | 3.64794632 | -0.529504458 | 0.37835907 | -1.3994761 | 0.16167027 | 0.2792474 | NA | NA | |
| ENSG000002 | 2.74950095 | -0.523363967 | 0.3739703 | -1.39948 | 0.16166908 | 0.2792474 | NA | NA | |
| ENSG000001 | 381.815087 | 0.195636899 | 0.13981661 | 1.39923938 | 0.16174121 | 0.27935803 | PPM1L | | 151742 |
| ENSG000002 | 6.61928553 | -0.505779006 | 0.36148131 | -1.3991844 | 0.1617577 | 0.27937461 | NA | NA | |
| ENSG000001 | 46.8094577 | 0.316665283 | 0.22636562 | 1.39891065 | 0.16183978 | 0.27948825 | AQP4 | | 361 |
| ENSG000001 | 10.2260021 | -0.463743269 | 0.33149749 | -1.3989345 | 0.16183263 | 0.27948825 | MROH7 | | 374977 |
| ENSG000002 | 3.59394066 | -0.529458065 | 0.37848279 | -1.398896 | 0.16184417 | 0.27948825 | NA | NA | |
| ENSG000002 | 2.17448556 | -0.510376432 | 0.36492893 | -1.3985639 | 0.16194379 | 0.27964837 | NA | NA | |
| ENSG000001 | 6.58955949 | -0.517629382 | 0.37015487 | -1.398413 | 0.16198908 | 0.27971466 | PTCHD3 | | 374308 |
| ENSG000002 | 3.00418105 | -0.527039703 | 0.37691618 | -1.3982942 | 0.16202474 | 0.27976433 | NA | NA | |
| ENSG000002 | 4.23197386 | -0.528883853 | 0.37827869 | -1.3981328 | 0.16207318 | 0.27983605 | NA | NA | |
| ENSG000002 | 7.44303212 | -0.500574899 | 0.35808669 | -1.3979154 | 0.16213848 | 0.27993687 | NA | NA | |
| ENSG000002 | 7.46736664 | -0.501640393 | 0.35887803 | -1.3978019 | 0.16217256 | 0.27997188 | NA | NA | |

| | | | | | | | | |
|---|---|---|---|---|---|---|---|---|
| ENSG000002 | 5.56508297 | -0.528197394 | 0.37787134 | -1.3978234 | 0.16216612 | 0.27997188 | NA | NA |
| ENSG000002 | 4.776656 | -0.525747812 | 0.37613248 | -1.397773 | 0.16218126 | 0.27997498 | NA | NA |
| ENSG000002 | 14.3005209 | -0.433910303 | 0.31044988 | -1.3976823 | 0.16220849 | 0.28001006 | NA | NA |
| ENSG000002 | 102.923434 | 0.242297328 | 0.17336069 | 1.39764858 | 0.16221862 | 0.28001564 | ZYG11A | 440590 |
| ENSG000002 | 9.02613342 | -0.483177871 | 0.34577201 | -1.3973886 | 0.16229674 | 0.28013014 | NA | NA |
| ENSG000002 | 5.32222915 | -0.524063974 | 0.37503276 | -1.3973819 | 0.16229877 | 0.28013014 | NA | NA |
| ENSG000002 | 3.89882467 | -0.528841353 | 0.37846211 | -1.3973429 | 0.16231048 | 0.28013842 | NA | NA |
| ENSG000002 | 11.8844661 | -0.457840679 | 0.32766144 | -1.397298 | 0.16232397 | 0.28014979 | NA | NA |
| ENSG000002 | 9.16705177 | -0.489128715 | 0.35007545 | -1.3972094 | 0.16235061 | 0.28018383 | NA | NA |
| ENSG000002 | 3.03164034 | -0.526149057 | 0.37660352 | -1.3970901 | 0.16238647 | 0.28023379 | NA | NA |
| ENSG000002 | 3.04322754 | -0.526392158 | 0.37679907 | -1.3970102 | 0.16241049 | 0.28026332 | NA | NA |
| ENSG000001 | 8.18955843 | -0.489629605 | 0.35050102 | -1.396942 | 0.162431 | 0.28028679 | OR2D2 | 120776 |
| ENSG000001 | 4.19644925 | -0.528564567 | 0.37843348 | -1.3967173 | 0.16249861 | 0.2803823 | OR1N2 | 138882 |
| ENSG000002 | 18.7724938 | -0.413724653 | 0.29621328 | -1.396712 | 0.16250018 | 0.2803823 | CHMP4A | 29082 |
| ENSG000002 | 3.70067442 | -0.528325089 | 0.37827472 | -1.3966703 | 0.16251273 | 0.28039202 | RNU2-58P | 106481652 |
| ENSG000001 | 5.22932566 | -0.51841172 | 0.37119611 | -1.3965979 | 0.16253451 | 0.28041768 | P2RX2 | 22953 |
| ENSG000002 | 40.5152179 | 0.338345705 | 0.24236803 | 1.39599975 | 0.16271457 | 0.28070444 | NA | NA |
| ENSG000002 | 5.87114023 | -0.517055239 | 0.37038094 | -1.3960093 | 0.16271169 | 0.28070444 | FER1L6-AS2 | 157376 |
| ENSG000002 | 3.62467684 | -0.528300369 | 0.3784516 | -1.3959523 | 0.16272887 | 0.28071716 | NA | NA |
| ENSG000002 | 15.6233473 | 0.42729769 | 0.30610457 | 1.39592067 | 0.16273839 | 0.28072164 | NA | NA |
| ENSG000002 | 2.90230789 | -0.514833845 | 0.36883664 | -1.3958316 | 0.1627652 | 0.28075595 | NA | NA |
| ENSG000001 | 157.977951 | 0.211896674 | 0.15181509 | 1.39575504 | 0.16278827 | 0.2807838 | MIR302D | 442896 |
| ENSG000002 | 13.2461945 | -0.44178036 | 0.31654502 | -1.395632 | 0.16282533 | 0.28083578 | LINC01305 | 285084 |
| ENSG000002 | 3.04197725 | -0.522948148 | 0.37472559 | -1.3955496 | 0.16285017 | 0.28086668 | NA | NA |
| ENSG000002 | 6.82128481 | -0.501298119 | 0.35922179 | -1.3955114 | 0.16286167 | 0.28087455 | NA | NA |
| ENSG000002 | 7.79679707 | -0.501151229 | 0.35916077 | -1.3953396 | 0.16291347 | 0.28095195 | NA | NA |
| ENSG000001 | 54.2422687 | 0.277683236 | 0.19901304 | 1.3953017 | 0.16292488 | 0.28095954 | CCKBR | 887 |
| ENSG000001 | 24.7525276 | 0.368740226 | 0.26427706 | 1.39527898 | 0.16293173 | 0.28095954 | S1PR1 | 1901 |
| ENSG000002 | 5.84352395 | -0.516612756 | 0.37026734 | -1.3952426 | 0.1629427 | 0.28096651 | NA | NA |

| | | | | | | | | |
|---|---|---|---|---|---|---|---|---|
| ENSG000002 | 2.75214944 | -0.520844995 | 0.37332088 | -1.3951671 | 0.16296547 | 0.28099383 | NA | NA |
| ENSG000002 | 4.02401029 | -0.527997414 | 0.37845677 | -1.3951327 | 0.16297583 | 0.28099974 | NA | NA |
| ENSG000001 | 424.273275 | 0.228008709 | 0.16343635 | 1.39509176 | 0.16298817 | 0.28100907 | ANKRD33B | 651746 |
| ENSG000002 | 3.89395191 | -0.527979225 | 0.37846136 | -1.3950677 | 0.16299542 | 0.28100963 | RN7SL299P | 106479331 |
| ENSG000002 | 66.7597356 | -0.282246155 | 0.20233903 | -1.394917 | 0.16304087 | 0.28107603 | UBE2Q2P2 | 100134869 |
| ENSG000002 | 6.15315836 | -0.512689698 | 0.36755435 | -1.3948677 | 0.16305574 | 0.28108971 | LOC100506489 | 100506489 |
| ENSG000002 | 4.42926957 | -0.526291824 | 0.37735493 | -1.3946865 | 0.1631104 | 0.28117198 | NA | NA |
| ENSG000002 | 4.99261002 | -0.522559707 | 0.37468732 | -1.3946554 | 0.16311977 | 0.28117619 | NA | NA |
| ENSG000002 | 2.7462522 | -0.52156988 | 0.37398852 | -1.3946147 | 0.16313206 | 0.28118543 | MIR7849 | 102465837 |
| ENSG000001 | 262.553204 | 0.244504952 | 0.17532694 | 1.39456578 | 0.16314682 | 0.28119891 | MAP1A | 4130 |
| ENSG000002 | 2.55748883 | -0.51021312 | 0.3658867 | -1.3944566 | 0.16317977 | 0.28124375 | NA | NA |
| ENSG00000C | 407.175754 | 0.224397592 | 0.16092386 | 1.3944333 | 0.1631868 | 0.28124391 | CCNK | 8812 |
| ENSG000002 | 7.55345028 | -0.502230967 | 0.36018258 | -1.3943789 | 0.16320323 | 0.28125292 | NA | NA |
| ENSG000002 | 6.38180062 | -0.50975195 | 0.36557868 | -1.39437 | 0.1632059 | 0.28125292 | NA | NA |
| ENSG000002 | 48.1902404 | -0.301423909 | 0.21619258 | -1.394238 | 0.16324575 | 0.28130964 | NA | NA |
| ENSG000002 | 8.3282457 | -0.500545962 | 0.35902274 | -1.3941901 | 0.16326019 | 0.28132257 | RGS21 | 431704 |
| ENSG000001 | 80.2094458 | 0.256770109 | 0.18420048 | 1.39397089 | 0.1633264 | 0.28142469 | IQCK | 124152 |
| ENSG000002 | 12.1450869 | -0.450085706 | 0.3229209 | -1.3937955 | 0.16337936 | 0.28150399 | DLG1-AS1 | 100507086 |
| ENSG000001 | 14.9079957 | -0.421897392 | 0.30270437 | -1.3937605 | 0.16338994 | 0.28151026 | DBX2 | 440097 |
| ENSG000002 | 5.28866741 | -0.518967631 | 0.3723869 | -1.3936248 | 0.16343093 | 0.28156891 | NA | NA |
| ENSG000002 | 12.8092194 | -0.473590009 | 0.33983542 | -1.3935864 | 0.16344254 | 0.28157696 | PROSER2-AS1 | 219731 |
| ENSG000002 | 5.69400027 | -0.521727719 | 0.37441312 | -1.3934547 | 0.16348233 | 0.28162158 | NA | NA |
| ENSG000002 | 3.44400396 | -0.525358417 | 0.37701279 | -1.3934764 | 0.16347578 | 0.28162158 | MIR4713 | 100616369 |
| ENSG000001 | 223.248613 | 0.221495088 | 0.15895953 | 1.39340555 | 0.16349719 | 0.28163521 | TMEM126A | 84233 |
| ENSG000002 | 17.0766277 | -0.419474349 | 0.30105236 | -1.3933601 | 0.16351092 | 0.2816469 | FAM166B | 730112 |
| ENSG000002 | 6.9190495 | -0.505299937 | 0.36265604 | -1.3933311 | 0.16351969 | 0.28165004 | NA | NA |
| ENSG000002 | 2.99082802 | -0.525772828 | 0.37739855 | -1.3931501 | 0.1635744 | 0.28173231 | NA | NA |
| ENSG000002 | 2.74107112 | -0.519603898 | 0.37300982 | -1.3930033 | 0.1636188 | 0.28179682 | NA | NA |
| ENSG000001 | 46.7611037 | 0.311521238 | 0.2236448 | 1.39292857 | 0.16364139 | 0.28182375 | GOLGA6L9 | 440295 |

| | | | | | | | | |
|---|---|---|---|---|---|---|---|---|
| ENSG000001 | 5.5731673 | -0.517442451 | 0.37153337 | -1.3927213 | 0.16370408 | 0.28191974 | ASCL1 | 429 |
| ENSG000002 | 4.12073551 | -0.525821576 | 0.37758391 | -1.3925953 | 0.16374221 | 0.28197342 | NA | NA |
| ENSG000002 | 3.59565172 | -0.5270887 | 0.37854984 | -1.3923892 | 0.16380458 | 0.28206886 | NA | NA |
| ENSG000002 | 4.96268568 | -0.520915456 | 0.37412343 | -1.3923626 | 0.16381263 | 0.28207075 | DEFB133 | 403339 |
| ENSG000002 | 2.81314222 | -0.51864692 | 0.37258784 | -1.3920125 | 0.16391863 | 0.28224127 | NA | NA |
| ENSG000002 | 12.6546702 | -0.445041489 | 0.31972869 | -1.3919348 | 0.16394215 | 0.28226979 | NA | NA |
| ENSG000002 | 3.92758865 | -0.526921107 | 0.37859264 | -1.3917891 | 0.16398627 | 0.28233377 | NA | NA |
| ENSG000002 | 47.1238025 | -0.289254375 | 0.20783766 | -1.3917322 | 0.1640035 | 0.282348 | NA | NA |
| ENSG000002 | 3.01493352 | -0.524520008 | 0.37688728 | -1.3917159 | 0.16400846 | 0.282348 | NA | NA |
| ENSG000002 | 4.09682534 | -0.525366939 | 0.37754298 | -1.3915421 | 0.16406111 | 0.28242664 | LINC01455 | 105616916 |
| ENSG000002 | 9.16055426 | -0.485823894 | 0.349137 | -1.3914993 | 0.16407407 | 0.28242897 | NA | NA |
| ENSG000002 | 6.01927049 | -0.520670938 | 0.37418186 | -1.3914917 | 0.16407639 | 0.28242897 | LINC01237 | 101927289 |
| ENSG000002 | 3.64018255 | -0.526556088 | 0.37842855 | -1.391428 | 0.16409567 | 0.28245016 | NA | NA |
| ENSG000002 | 4.2263361 | -0.526566836 | 0.37845211 | -1.3913698 | 0.16411331 | 0.28246855 | NA | NA |
| ENSG000001 | 7.37682764 | -0.512900743 | 0.36864553 | -1.3913114 | 0.164131 | 0.28247502 | SST | 6750 |
| ENSG000002 | 5.29147986 | -0.517786538 | 0.37215472 | -1.3913206 | 0.16412822 | 0.28247502 | NA | NA |
| ENSG000001 | 2.19556036 | -0.500700048 | 0.35989594 | -1.3912356 | 0.16415399 | 0.28250259 | NA | NA |
| ENSG000002 | 10.5634682 | -0.467998165 | 0.33644581 | -1.3910061 | 0.16422358 | 0.28261036 | SPATA31D1 | 389763 |
| ENSG000002 | 12.6495669 | -0.446410265 | 0.32093795 | -1.3909551 | 0.16423905 | 0.28262498 | NA | NA |
| ENSG000002 | 2.57016556 | -0.510044495 | 0.3667173 | -1.3908384 | 0.16427444 | 0.28266436 | NA | NA |
| ENSG000002 | 3.94731752 | -0.526509115 | 0.3785565 | -1.3908337 | 0.16427588 | 0.28266436 | NA | NA |
| ENSG00000C | 52.4597751 | -0.304812619 | 0.21916312 | -1.3908025 | 0.16428532 | 0.28266861 | EPHA6 | 285220 |
| ENSG000001 | 7.40870501 | -0.496196195 | 0.3567849 | -1.3907432 | 0.1643033 | 0.28267557 | OR12D3 | 81797 |
| ENSG000002 | 3.56764368 | -0.526475634 | 0.3785547 | -1.3907518 | 0.1643007 | 0.28267557 | NA | NA |
| ENSG000002 | 5.23262574 | -0.51808951 | 0.37253568 | -1.3907111 | 0.16431305 | 0.28268035 | NA | NA |
| ENSG000002 | 3.86227852 | -0.526034393 | 0.37832859 | -1.3904167 | 0.16440238 | 0.28282203 | NA | NA |
| ENSG000002 | 2.69594469 | -0.519184825 | 0.37341352 | -1.390375 | 0.16441504 | 0.2828318 | NA | NA |
| ENSG000002 | 8.44091982 | -0.482897011 | 0.34733501 | -1.3902918 | 0.16444029 | 0.28286325 | NA | NA |
| ENSG000002 | 8.75010191 | -0.48233989 | 0.34694921 | -1.390232 | 0.16445844 | 0.28288247 | NA | NA |

| | | | | | | | | |
|---|---|---|---|---|---|---|---|---|
| ENSG000001 | 8.60799313 | 0.484996724 | 0.34887979 | 1.39015426 | 0.16448204 | 0.28289906 | CLPSL2 | 389383 |
| ENSG000002 | 7.95097247 | -0.491570142 | 0.35360581 | -1.3901642 | 0.16447901 | 0.28289906 | NA | NA |
| ENSG000002 | 2.81010164 | -0.516438138 | 0.37151692 | -1.3900797 | 0.16450466 | 0.28291398 | RNU6-1048P | 106480027 |
| ENSG000002 | 2.78174869 | -0.517906468 | 0.37257054 | -1.3900897 | 0.16450164 | 0.28291398 | NA | NA |
| ENSG000002 | 11.0169832 | -0.454068049 | 0.32666464 | -1.3900128 | 0.16452498 | 0.28293691 | LOC101929723 | 101929723 |
| ENSG000002 | 2.71518376 | -0.519083739 | 0.37346256 | -1.3899218 | 0.16455264 | 0.28296048 | GPR142 | 350383 |
| ENSG000002 | 6.09760595 | -0.511247918 | 0.36782188 | -1.3899334 | 0.1645491 | 0.28296048 | NA | NA |
| ENSG000002 | 6.15008533 | -0.512190762 | 0.36851366 | -1.3898827 | 0.1645645 | 0.28296887 | NA | NA |
| ENSG000002 | 44.67326 | 0.321906792 | 0.23163006 | 1.38974529 | 0.16460624 | 0.28302169 | NA | NA |
| ENSG000002 | 2.7096385 | -0.518496415 | 0.37308996 | -1.3897356 | 0.16460917 | 0.28302169 | NA | NA |
| ENSG000001 | 279.517115 | 0.271751147 | 0.19556159 | 1.38959365 | 0.16465231 | 0.28305985 | ZNHIT6 | 54680 |
| ENSG000001 | 15.7541279 | -0.421431725 | 0.30327618 | -1.3895972 | 0.16465123 | 0.28305985 | PODNL1 | 79883 |
| ENSG000001 | 703.649268 | 0.212539272 | 0.15295004 | 1.3895993 | 0.16465059 | 0.28305985 | SOX13 | 9580 |
| ENSG000002 | 5.24136878 | -0.516602451 | 0.3717741 | -1.3895601 | 0.16466251 | 0.28306538 | NA | NA |
| ENSG000002 | 3.01115662 | -0.524021819 | 0.37713747 | -1.3894717 | 0.16468937 | 0.28309957 | NA | NA |
| ENSG000002 | 3.71553864 | -0.524199088 | 0.37729473 | -1.3893623 | 0.16472259 | 0.28314467 | NA | NA |
| ENSG000002 | 7.00369263 | -0.505535249 | 0.36387058 | -1.3893271 | 0.16473331 | 0.28315109 | TLX1NB | 100038246 |
| ENSG000002 | 5.79728225 | -0.513622467 | 0.36969884 | -1.3892997 | 0.16474165 | 0.28315341 | NA | NA |
| ENSG00000C | 35.3944509 | -0.327540852 | 0.23579033 | -1.3891191 | 0.16479654 | 0.28323575 | HS3ST1 | 9957 |
| ENSG000002 | 14.1856306 | -0.435164586 | 0.31336568 | -1.3886798 | 0.16493014 | 0.28343772 | TRIM31 | 11074 |
| ENSG000002 | 744.63552 | 0.228511359 | 0.16454951 | 1.38870888 | 0.16492129 | 0.28343772 | SNORD116-6 | 100033418 |
| ENSG000002 | 130.469607 | 0.212116222 | 0.15274844 | 1.38866376 | 0.16493502 | 0.28343772 | NA | NA |
| ENSG000002 | 11.0268745 | -0.452861416 | 0.3261418 | -1.3885415 | 0.16497223 | 0.28348965 | LINC00967 | 100505659 |
| ENSG000001 | 69.2218064 | 0.462862971 | 0.33335085 | 1.38851595 | 0.16497999 | 0.28349098 | PRKCD | 5580 |
| ENSG000002 | 5.29577718 | 0.519929898 | 0.37445736 | 1.38848892 | 0.16498822 | 0.2834931 | NA | NA |
| ENSG000002 | 3.64229658 | -0.524794129 | 0.37797107 | -1.3884505 | 0.1649999 | 0.28350117 | NA | NA |
| ENSG000002 | 15.1285084 | -0.421864921 | 0.30385001 | -1.3883986 | 0.1650157 | 0.28351599 | NA | NA |
| ENSG000002 | 3.35353514 | -0.524005147 | 0.37742302 | -1.3883762 | 0.16502251 | 0.28351599 | NA | NA |
| ENSG00000C | 17.1264225 | -0.423457102 | 0.30501392 | -1.3883206 | 0.16503945 | 0.28353308 | GGT5 | 2687 |

| | | | | | | | | |
|---|---|---|---|---|---|---|---|---|
| ENSG000002 | 36.1347693 | -0.331396793 | 0.2387586 | -1.3879994 | 0.16513724 | 0.28368905 | LOC101928673 | 101928673 |
| ENSG000000 | 232.446623 | 0.232778578 | 0.16772184 | 1.38788474 | 0.16517214 | 0.28372247 | MED15 | 51586 |
| ENSG000002 | 2.76236581 | -0.517924172 | 0.37318009 | -1.3878666 | 0.16517768 | 0.28372247 | NA | NA |
| ENSG000002 | 5.89595921 | -0.513949408 | 0.37031421 | -1.3878739 | 0.16517546 | 0.28372247 | NA | NA |
| ENSG000000 | 57.1122716 | 0.302340058 | 0.21787638 | 1.38766792 | 0.16523819 | 0.28376455 | VASH1 | 22846 |
| ENSG000002 | 5.26173682 | -0.516812554 | 0.37241811 | -1.3877213 | 0.16522192 | 0.28376455 | NA | NA |
| ENSG000002 | 16.7520376 | 0.420855118 | 0.30328314 | 1.38766408 | 0.16523936 | 0.28376455 | SNORD116-8 | 100033420 |
| ENSG000002 | 3.6963992 | -0.524050069 | 0.37763754 | -1.3877065 | 0.16522643 | 0.28376455 | NA | NA |
| ENSG000002 | 16.2097639 | -0.416710374 | 0.30030467 | -1.3876253 | 0.16525116 | 0.28376455 | NA | NA |
| ENSG000002 | 7.88688674 | -0.502094217 | 0.3618343 | -1.3876358 | 0.16524799 | 0.28376455 | MTOR-AS1 | 100873935 |
| ENSG000002 | 9.33418076 | -0.479695263 | 0.34568236 | -1.3876764 | 0.16523559 | 0.28376455 | NA | NA |
| ENSG000002 | 8.4378787 | -0.48389879 | 0.34877645 | -1.3874182 | 0.16531428 | 0.28386092 | NA | NA |
| ENSG000001 | 8.8309627 | 0.479944577 | 0.345939 | 1.38736765 | 0.16532968 | 0.28387534 | KIAA0408 | 9729 |
| ENSG000002 | 3.03628008 | -0.522790139 | 0.376845 | -1.3872816 | 0.16535591 | 0.28390835 | NA | NA |
| ENSG000002 | 8.67403354 | -0.478358036 | 0.34485839 | -1.3871144 | 0.16540688 | 0.28398385 | NA | NA |
| ENSG000002 | 2.96276736 | -0.522691742 | 0.37690695 | -1.3867925 | 0.16550503 | 0.28414033 | LINC00162 | 378825 |
| ENSG000001 | 11.3844861 | -0.469781818 | 0.33879977 | -1.3866061 | 0.16556192 | 0.28421393 | ADAM3A | 1587 |
| ENSG000002 | 12.8264708 | -0.439924427 | 0.31726184 | -1.3866289 | 0.16555496 | 0.28421393 | LINC00284 | 121838 |
| ENSG000002 | 3.01428119 | -0.522134474 | 0.37660428 | -1.3864273 | 0.16561645 | 0.28428348 | NA | NA |
| ENSG000002 | 51.0818347 | 0.298552595 | 0.21533766 | 1.3864393 | 0.1656128 | 0.28428348 | NA | NA |
| ENSG000002 | 3.01496632 | -0.522683319 | 0.37701668 | -1.3863666 | 0.165635 | 0.28430328 | NA | NA |
| ENSG000002 | 14.2730338 | -0.43265203 | 0.31209464 | -1.3862847 | 0.16565998 | 0.28433412 | NA | NA |
| ENSG000002 | 6.84154576 | -0.513087751 | 0.37016272 | -1.386114 | 0.16571208 | 0.28441151 | NA | NA |
| ENSG000002 | 5.4777429 | -0.512609801 | 0.36983695 | -1.3860427 | 0.16573387 | 0.28443685 | NA | NA |
| ENSG000002 | 3.35462522 | -0.523264312 | 0.37754064 | -1.3859814 | 0.16575258 | 0.28445694 | NA | NA |
| ENSG000002 | 4.09580793 | -0.523253308 | 0.37754121 | -1.3859502 | 0.16576212 | 0.28446118 | NA | NA |
| ENSG000002 | 3.0525056 | -0.52195431 | 0.3766136 | -1.3859147 | 0.16577297 | 0.28446118 | NA | NA |
| ENSG000002 | 117.680147 | 0.223748923 | 0.16144614 | 1.3859044 | 0.1657761 | 0.28446118 | NA | NA |
| ENSG000002 | 8.77032436 | -0.483328565 | 0.3487533 | -1.3858753 | 0.165785 | 0.28446441 | NA | NA |

| | | | | | | | | |
|---|---|---|---|---|---|---|---|---|
| ENSG000002 | 4.22720468 | -0.524352261 | 0.3783938 | -1.3857316 | 0.16582887 | 0.28450784 | NA | NA |
| ENSG000002 | 10.5029568 | -0.458544833 | 0.33090645 | -1.3857235 | 0.16583135 | 0.28450784 | LINC01504 | 100507540 |
| ENSG000002 | 3.15806618 | -0.518254252 | 0.37398631 | -1.3857573 | 0.16582103 | 0.28450784 | NA | NA |
| ENSG000002 | 2.41956971 | -0.514958856 | 0.37167589 | -1.3855052 | 0.16589806 | 0.28461024 | MIR3922 | 100500843 |
| ENSG000002 | 5.79082765 | -0.514773948 | 0.37154863 | -1.385482 | 0.16590512 | 0.28461032 | NA | NA |
| ENSG000001 | 5.23817896 | -0.515604107 | 0.37219508 | -1.3853061 | 0.1659589 | 0.28464988 | SSTR5 | 6755 |
| ENSG000002 | 10.1353669 | -0.467620823 | 0.33754992 | -1.3853383 | 0.16594906 | 0.28464988 | LSAMP-AS1 | 101926903 |
| ENSG000002 | 5.13216906 | -0.5219908 | 0.3768018 | -1.3853193 | 0.16595486 | 0.28464988 | NA | NA |
| ENSG000002 | 5.21389685 | -0.514997683 | 0.37175445 | -1.3853168 | 0.16595562 | 0.28464988 | NA | NA |
| ENSG000002 | 2.97384514 | -0.522131844 | 0.37691111 | -1.3852917 | 0.16596328 | 0.28464988 | NA | NA |
| ENSG000002 | 3.31555891 | -0.522383762 | 0.37711229 | -1.3852207 | 0.16598498 | 0.28467505 | NA | NA |
| ENSG000002 | 2.80011394 | -0.515963118 | 0.37248861 | -1.3851783 | 0.16599797 | 0.28468529 | NA | NA |
| ENSG000002 | 3.01069961 | -0.52195801 | 0.37683405 | -1.3851137 | 0.1660177 | 0.28470708 | NA | NA |
| ENSG000002 | 8.96115105 | -0.487937325 | 0.35227849 | -1.3850898 | 0.16602502 | 0.28470761 | NA | NA |
| ENSG000002 | 158.110963 | 0.214636478 | 0.1549655 | 1.38505976 | 0.1660342 | 0.28471129 | NA | NA |
| ENSG000002 | 7.1064931 | -0.493817092 | 0.35655797 | -1.384956 | 0.16606593 | 0.28475366 | NA | NA |
| ENSG000002 | 6.38735365 | -0.507414473 | 0.36640691 | -1.3848387 | 0.16610179 | 0.28480312 | NA | NA |
| ENSG000001 | 6.85947697 | -0.499398259 | 0.36062885 | -1.3847984 | 0.16611412 | 0.28481222 | PIGZ | 80235 |
| ENSG000001 | 11.8520937 | -0.455059661 | 0.32864609 | -1.3846495 | 0.16615965 | 0.28485415 | NPAS4 | 266743 |
| ENSG000002 | 13.9794744 | -0.446760573 | 0.32264948 | -1.3846623 | 0.16615575 | 0.28485415 | NA | NA |
| ENSG000002 | 7.19568406 | -0.502136592 | 0.36264046 | -1.3846679 | 0.16615404 | 0.28485415 | NA | NA |
| ENSG000002 | 9.46516607 | 0.473794047 | 0.34219964 | 1.38455451 | 0.16618873 | 0.28489196 | NA | NA |
| ENSG000002 | 5.29697251 | -0.516249599 | 0.37287352 | -1.3845166 | 0.16620031 | 0.28489977 | NA | NA |
| ENSG000001 | 515.782512 | 0.182007609 | 0.13146916 | 1.38441295 | 0.16623205 | 0.28494212 | ZNF354C | 30832 |
| ENSG000001 | 28.8296601 | 0.364875763 | 0.26357071 | 1.38435626 | 0.1662494 | 0.28494777 | KLHDC9 | 126823 |
| ENSG000002 | 9.59086397 | -0.476952349 | 0.34452899 | -1.3843606 | 0.16624807 | 0.28494777 | NA | NA |
| ENSG000002 | 6.08271704 | -0.511535699 | 0.3695361 | -1.3842645 | 0.16627749 | 0.28498388 | TPM3P3 | 106478937 |
| ENSG000002 | 2.38482274 | -0.514650291 | 0.37182146 | -1.3841328 | 0.16631779 | 0.2850409 | NA | NA |
| ENSG000002 | 4.93724215 | -0.517687117 | 0.37403084 | -1.384076 | 0.16633518 | 0.28504661 | NA | NA |

| | | | | | | | | |
|---|---|---|---|---|---|---|---|---|
| ENSG000002 | 4.94914887 | -0.51763444 | 0.37398799 | -1.3840937 | 0.16632975 | 0.28504661 | NA | NA |
| ENSG000002 | 2.45168208 | -0.50763785 | 0.36680248 | -1.3839542 | 0.16637247 | 0.28508641 | NA | NA |
| ENSG000002 | 71.1552847 | 0.268937305 | 0.19432344 | 1.3839674 | 0.16636844 | 0.28508641 | LINC01232 | 102725509 |
| ENSG000002 | 3.02423697 | -0.521567449 | 0.3768901 | -1.3838714 | 0.16639782 | 0.28511781 | NA | NA |
| ENSG000002 | 3.96333452 | -0.523880107 | 0.37857358 | -1.3838264 | 0.16641162 | 0.28512941 | NA | NA |
| ENSG000001 | 8.04762227 | -0.497406247 | 0.35949423 | -1.3836279 | 0.16647242 | 0.28519743 | PLET1 | 349633 |
| ENSG000002 | 5.30478137 | -0.51664973 | 0.37339866 | -1.3836411 | 0.16646838 | 0.28519743 | NA | NA |
| ENSG000002 | 4.511368 | -0.522520201 | 0.37763403 | -1.3836682 | 0.16646007 | 0.28519743 | NA | NA |
| ENSG000001 | 26.5316001 | -0.364999508 | 0.26380662 | -1.3835874 | 0.16648484 | 0.28520665 | KRT15 | 3866 |
| ENSG000002 | 4.25437661 | -0.523769731 | 0.37857892 | -1.3835153 | 0.16650692 | 0.28523243 | NA | NA |
| ENSG000001 | 63.5121654 | -0.275080077 | 0.19886118 | -1.3832769 | 0.16657998 | 0.28534553 | CEMIP | 57214 |
| ENSG000002 | 88.3352919 | 0.241269097 | 0.17442301 | 1.38324119 | 0.16659092 | 0.28535222 | WDR92 | 116143 |
| ENSG000001 | 9.71336449 | -0.46791111 | 0.33828509 | -1.3831857 | 0.16660793 | 0.28535724 | FLJ33534 | 285150 |
| ENSG000001 | 384.057379 | 0.216233927 | 0.15632998 | 1.3831891 | 0.16660689 | 0.28535724 | SPRY1 | 10252 |
| ENSG000001 | 3.8903579 | -0.523504813 | 0.3784869 | -1.3831517 | 0.16661834 | 0.28535948 | PDIA2 | 64714 |
| ENSG000002 | 4.10402815 | -0.522520957 | 0.37778513 | -1.3831168 | 0.16662906 | 0.28535948 | NA | NA |
| ENSG000002 | 6.11924556 | -0.50993239 | 0.36868466 | -1.3831126 | 0.16663035 | 0.28535948 | NA | NA |
| ENSG000002 | 180.458542 | 0.278583514 | 0.20143944 | 1.38296409 | 0.16667588 | 0.2854254 | NA | NA |
| ENSG000001 | 39.3060264 | 0.326519146 | 0.23611317 | 1.38289258 | 0.16669781 | 0.28544658 | ARRDC2 | 27106 |
| ENSG000002 | 4.74695207 | -0.521375091 | 0.37702434 | -1.3828685 | 0.16670518 | 0.28544658 | NA | NA |
| ENSG000002 | 6.3181267 | -0.506025486 | 0.36592812 | -1.3828549 | 0.16670936 | 0.28544658 | NA | NA |
| ENSG000001 | 26.1720162 | -0.369702584 | 0.2674152 | -1.382504 | 0.16681701 | 0.28559858 | ACADL | 33 |
| ENSG000002 | 3.01965726 | -0.520639479 | 0.37660617 | -1.3824507 | 0.16683335 | 0.28559858 | NA | NA |
| ENSG000002 | 2.41342418 | -0.513812304 | 0.37166265 | -1.3824696 | 0.16682756 | 0.28559858 | NA | NA |
| ENSG000002 | 3.668784 | -0.523038176 | 0.37832957 | -1.3824935 | 0.16682022 | 0.28559858 | NA | NA |
| ENSG000002 | 7.49966808 | -0.498071001 | 0.36027527 | -1.3824735 | 0.16682638 | 0.28559858 | NA | NA |
| ENSG000002 | 4.53617771 | -0.522761258 | 0.37815296 | -1.3824069 | 0.1668468 | 0.28560954 | NA | NA |
| ENSG000002 | 184.193635 | 0.197985602 | 0.14322058 | 1.38238234 | 0.16685434 | 0.28561039 | NA | NA |
| ENSG000002 | 101.505907 | 0.253332203 | 0.18326679 | 1.38231371 | 0.1668754 | 0.28563439 | KCTD7 | 154881 |

| | | | | | | | | |
|---|---|---|---|---|---|---|---|---|
| ENSG000001 | 46.4951169 | 0.297821311 | 0.21545825 | 1.38226922 | 0.16688905 | 0.2856457 | SLC27A1 | 376497 |
| ENSG000002 | 4.86281169 | 0.519972108 | 0.37618375 | 1.38222906 | 0.16690138 | 0.28565474 | NA | NA |
| ENSG000002 | 21.6580637 | 0.38903981 | 0.28146645 | 1.38218895 | 0.16691369 | 0.28566376 | NA | NA |
| ENSG000001 | 158.211335 | 0.234050883 | 0.16934769 | 1.38207305 | 0.16694927 | 0.28571259 | IRS2 | 8660 |
| ENSG000002 | 2.73317185 | -0.516388495 | 0.37365802 | -1.3819816 | 0.16697736 | 0.2857486 | NA | NA |
| ENSG000001 | 1791.95752 | 0.160937098 | 0.11646016 | 1.38190693 | 0.16700028 | 0.28576137 | STRN | 6801 |
| ENSG000001 | 821.091635 | 0.169059602 | 0.12233955 | 1.38188842 | 0.16700596 | 0.28576137 | EDEM3 | 80267 |
| ENSG000002 | 2.75206546 | -0.516384699 | 0.37367614 | -1.3819044 | 0.16700107 | 0.28576137 | NA | NA |
| ENSG000002 | 2.96657707 | -0.520557952 | 0.37672432 | -1.3818008 | 0.16703289 | 0.28578333 | MRGPRG-AS1 | 283303 |
| ENSG000002 | 3.86893203 | -0.523021406 | 0.37850196 | -1.3818195 | 0.16702712 | 0.28578333 | NA | NA |
| ENSG000002 | 2.17613604 | -0.501792485 | 0.36318857 | -1.3816307 | 0.16708511 | 0.28586062 | NA | NA |
| ENSG000000 | 239.913915 | 0.186567496 | 0.13503723 | 1.3816004 | 0.16709443 | 0.2858645 | ARHGAP10 | 79658 |
| ENSG000002 | 3.63765881 | -0.522806295 | 0.37842605 | -1.3815283 | 0.16711659 | 0.28589035 | NA | NA |
| ENSG000002 | 3.93678849 | -0.522940751 | 0.37857 | -1.3813582 | 0.16716887 | 0.28596771 | NA | NA |
| ENSG000001 | 124.365383 | 0.225850753 | 0.16350311 | 1.38132393 | 0.16717938 | 0.28597364 | ANGPTL2 | 23452 |
| ENSG000002 | 3.89916436 | -0.52290807 | 0.37858971 | -1.3811999 | 0.1672175 | 0.28602679 | NA | NA |
| ENSG000002 | 2.75379992 | -0.51536386 | 0.37313632 | -1.3811678 | 0.16722738 | 0.28603161 | NA | NA |
| ENSG000002 | 4.40680467 | -0.51992929 | 0.37645445 | -1.3811214 | 0.16724164 | 0.28604394 | NA | NA |
| ENSG000002 | 6.06581991 | -0.508851755 | 0.36851636 | -1.3808118 | 0.16733682 | 0.28619466 | PPP2R2B-IT1 | 100874361 |
| ENSG000001 | 84.9685416 | 0.240950174 | 0.17452093 | 1.38063767 | 0.16739039 | 0.28625805 | EAF2 | 55840 |
| ENSG000002 | 10.3690798 | 0.476676703 | 0.34527112 | 1.38058666 | 0.16740609 | 0.28625805 | NA | NA |
| ENSG000002 | 3.61396324 | -0.522108449 | 0.37816513 | -1.3806361 | 0.16739087 | 0.28625805 | LOC102723883 | 102723883 |
| ENSG000002 | 3.8396777 | -0.522594585 | 0.37853357 | -1.3805766 | 0.16740918 | 0.28625805 | NA | NA |
| ENSG000002 | 6.59868772 | -0.503100364 | 0.36440796 | -1.3805965 | 0.16740306 | 0.28625805 | NA | NA |
| ENSG000002 | 4.08418882 | -0.52134078 | 0.37768479 | -1.3803595 | 0.16747599 | 0.28636022 | NA | NA |
| ENSG000001 | 3.03680072 | -0.518112243 | 0.37544213 | -1.3800056 | 0.16758492 | 0.28653439 | HOXD9 | 3235 |
| ENSG000002 | 10.8326978 | 0.454767898 | 0.32954852 | 1.37997252 | 0.16759511 | 0.28653972 | NA | NA |
| ENSG000000 | 173.893314 | 0.190885538 | 0.13833067 | 1.37992199 | 0.16761067 | 0.28655424 | WDR47 | 22911 |
| ENSG000002 | 2.82174091 | -0.513555695 | 0.37226559 | -1.3795411 | 0.16772798 | 0.28674271 | HNRNPCL2 | 440563 |

| | | | | | | | | |
|---|---|---|---|---|---|---|---|---|
| ENSG000001 | 14.9427069 | -0.442739929 | 0.32097061 | -1.3793784 | 0.16777811 | 0.28676225 | F2RL2 | 2151 |
| ENSG000002 | 2.39785101 | -0.512208512 | 0.37132326 | -1.379414 | 0.16776714 | 0.28676225 | NA | NA |
| ENSG000002 | 88.3650899 | 0.264735675 | 0.1919141 | 1.37944877 | 0.16775643 | 0.28676225 | NA | NA |
| ENSG000002 | 3.62872099 | -0.521941103 | 0.37839194 | -1.3793663 | 0.16778183 | 0.28676225 | NA | NA |
| ENSG000002 | 3.83384015 | -0.522100861 | 0.37847807 | -1.3794745 | 0.16774849 | 0.28676225 | LINC01151 | 104266958 |
| ENSG000002 | 7.69887671 | -0.491738377 | 0.35648476 | -1.3794092 | 0.16776861 | 0.28676225 | NA | NA |
| ENSG000002 | 5.22033523 | -0.514162625 | 0.37277392 | -1.3792881 | 0.16780595 | 0.2867793 | NA | NA |
| ENSG000002 | 5.23188853 | -0.51461599 | 0.37310033 | -1.3792965 | 0.16780334 | 0.2867793 | NA | NA |
| ENSG000001 | 17.127327 | -0.399747439 | 0.28984589 | -1.3791724 | 0.16784162 | 0.28681607 | OLAH | 55301 |
| ENSG000001 | 4.1750937 | -0.521528836 | 0.3781452 | -1.3791762 | 0.16784045 | 0.28681607 | NPPA | 4878 |
| ENSG000002 | 2.777698 | -0.515061258 | 0.37350249 | -1.3790035 | 0.16789366 | 0.28689292 | NA | NA |
| ENSG000002 | 5.87674519 | -0.511778493 | 0.37114182 | -1.3789297 | 0.16791642 | 0.28691971 | NA | NA |
| ENSG000002 | 3.43541049 | -0.519058658 | 0.37646005 | -1.3787881 | 0.16796008 | 0.28698223 | NA | NA |
| ENSG000002 | 7.42438157 | -0.494779704 | 0.35888226 | -1.3786686 | 0.16799695 | 0.28703313 | LINC00974 | 147093 |
| ENSG000002 | 3.60432896 | -0.521735696 | 0.37845419 | -1.3785967 | 0.16801914 | 0.28705894 | NA | NA |
| ENSG000002 | 14.6855869 | -0.440170332 | 0.31933866 | -1.378381 | 0.16808569 | 0.28716055 | SKOR2 | 652991 |
| ENSG000001 | 987.278974 | 0.161014624 | 0.11682339 | 1.37827384 | 0.16811876 | 0.28719284 | USP15 | 9958 |
| ENSG000002 | 3.32407307 | -0.519777124 | 0.37711841 | -1.3782863 | 0.16811493 | 0.28719284 | LOC221122 | 221122 |
| ENSG000002 | 3.23475041 | -0.52144692 | 0.37834762 | -1.3782217 | 0.16813484 | 0.28720821 | NA | NA |
| ENSG000002 | 3.60195237 | -0.521329592 | 0.37828682 | -1.3781331 | 0.16816219 | 0.28724284 | NA | NA |
| ENSG000001 | 4.7641487 | -0.519677406 | 0.37710189 | -1.3780822 | 0.16817791 | 0.28725759 | CD300C | 10871 |
| ENSG000001 | 3.03406846 | -0.518250886 | 0.37607741 | -1.3780431 | 0.16818997 | 0.28726609 | NA | NA |
| ENSG000002 | 11.7693971 | 0.462914899 | 0.33597025 | 1.37784493 | 0.16825117 | 0.28735852 | SNORA45B | 677826 |
| ENSG000002 | 15.9188738 | -0.409247002 | 0.29703926 | -1.3777539 | 0.16827927 | 0.28739441 | NA | NA |
| ENSG000002 | 10.0626481 | -0.480852495 | 0.34902631 | -1.377697 | 0.16829686 | 0.28740023 | LOC101927285 | 101927285 |
| ENSG000002 | 4.76801898 | -0.518811523 | 0.37657435 | -1.3777134 | 0.16829179 | 0.28740023 | NA | NA |
| ENSG000001 | 3.58853266 | 0.521537808 | 0.37859309 | 1.37756822 | 0.16833664 | 0.28745606 | NA | NA |
| ENSG000002 | 7.05180723 | -0.504989974 | 0.36659707 | -1.3775069 | 0.1683556 | 0.28747633 | NA | NA |
| ENSG000002 | 2.1798661 | -0.497926323 | 0.36157591 | -1.3771004 | 0.16848122 | 0.28767872 | NA | NA |

| | | | | | | | | |
|---|---|---|---|---|---|---|---|---|
| ENSG00000C | 36.9180866 | 0.311277525 | 0.22606044 | 1.37696592 | 0.16852278 | 0.28773758 | DLL3 | 10683 |
| ENSG000002 | 7.2319908 | -0.503818767 | 0.36594829 | -1.3767485 | 0.16859001 | 0.28783677 | NA | NA |
| ENSG000002 | 8.45756292 | -0.480868686 | 0.34928267 | -1.3767321 | 0.16859508 | 0.28783677 | NA | NA |
| ENSG000002 | 57.3993241 | 0.28548755 | 0.20737146 | 1.37669644 | 0.16860612 | 0.28784351 | NA | NA |
| ENSG000002 | 3.08703343 | -0.515852224 | 0.37478791 | -1.3763844 | 0.16870266 | 0.2879962 | NA | NA |
| ENSG000001 | 115.625658 | 0.219513359 | 0.15951188 | 1.37615676 | 0.16877311 | 0.28810432 | XRCC6BP1 | 91419 |
| ENSG000002 | 9.74006559 | -0.481092906 | 0.34964083 | -1.3759632 | 0.16883303 | 0.28819449 | ARHGEF38-IT1 | 100874374 |
| ENSG000002 | 22.6888322 | -0.368859983 | 0.26808244 | -1.37592 | 0.16884641 | 0.28820519 | PCDHA5 | 56143 |
| ENSG000002 | 4.14002142 | -0.519871486 | 0.37787084 | -1.3757915 | 0.16888619 | 0.28826096 | EGFLAM-AS2 | 100506475 |
| ENSG000002 | 3.86350411 | -0.520790699 | 0.37857992 | -1.3756427 | 0.16893228 | 0.2883275 | LOC105377390 | 105377390 |
| ENSG000002 | 5.22018619 | -0.512204829 | 0.37235566 | -1.3755796 | 0.16895184 | 0.28834874 | NA | NA |
| ENSG000001 | 51.6927077 | 0.279568086 | 0.20324701 | 1.37550899 | 0.1689737 | 0.28835178 | ZNF547 | 284306 |
| ENSG000001 | 15.1085544 | -0.426503243 | 0.31007031 | -1.375505 | 0.16897495 | 0.28835178 | TLR10 | 81793 |
| ENSG000002 | 18.4927903 | -0.407194084 | 0.29602613 | -1.3755343 | 0.16896587 | 0.28835178 | NA | NA |
| ENSG000002 | 7.95496705 | -0.48474973 | 0.35243991 | -1.375411 | 0.16900406 | 0.28838932 | NA | NA |
| ENSG000002 | 2.21707 | -0.492394096 | 0.35801875 | -1.3753305 | 0.16902901 | 0.28841125 | NA | NA |
| ENSG000002 | 2.96785425 | 0.517409318 | 0.37620914 | 1.37532361 | 0.16903114 | 0.28841125 | NA | NA |
| ENSG000002 | 2.38621284 | -0.511292406 | 0.37183227 | -1.375062 | 0.16911224 | 0.28853748 | NA | NA |
| ENSG000002 | 45.636872 | 0.29121908 | 0.21179348 | 1.3750144 | 0.16912698 | 0.2885505 | NA | NA |
| ENSG000001 | 95.7704797 | -0.238759434 | 0.17364544 | -1.3749825 | 0.16913687 | 0.28855524 | HHIPL1 | 84439 |
| ENSG000002 | 12.3629855 | -0.443766848 | 0.32276169 | -1.3749056 | 0.16916072 | 0.28858378 | NA | NA |
| ENSG000002 | 5.62426124 | -0.513258254 | 0.37331208 | -1.3748772 | 0.16916953 | 0.28858667 | NA | NA |
| ENSG000002 | 11.8946376 | -0.453884005 | 0.3301376 | -1.3748328 | 0.16918329 | 0.28859799 | NA | NA |
| ENSG000002 | 2.49641152 | -0.504169882 | 0.36675072 | -1.3746936 | 0.16922645 | 0.28865221 | NA | NA |
| ENSG000002 | 2.38294392 | -0.51113916 | 0.37182291 | -1.3746844 | 0.16922931 | 0.28865221 | NA | NA |
| ENSG000002 | 4.10334301 | -0.519529568 | 0.37793849 | -1.3746405 | 0.16924291 | 0.28866328 | NA | NA |
| ENSG000002 | 162.57399 | 0.228426309 | 0.16618017 | 1.37457024 | 0.16926472 | 0.28866405 | CRYZL1 | 9946 |
| ENSG000002 | 10.2497757 | -0.460981499 | 0.33536225 | -1.3745778 | 0.16926238 | 0.28866405 | NA | NA |
| ENSG000002 | 3.02688436 | -0.516409908 | 0.37567955 | -1.3746021 | 0.16925483 | 0.28866405 | NA | NA |

| | | | | | | | | |
|---|---|---|---|---|---|---|---|---|
| ENSG000001 | 3.82223423 | -0.520183842 | 0.37847582 | -1.3744176 | 0.16931206 | 0.28873264 | NA | NA |
| ENSG000002 | 3.06638787 | -0.516553481 | 0.37584365 | -1.374384 | 0.16932251 | 0.28873831 | NA | NA |
| ENSG000001 | 3.59786281 | -0.51991989 | 0.37831446 | -1.3743062 | 0.16934665 | 0.28876037 | OR4X1 | 390113 |
| ENSG000001 | 4.13047113 | -0.519075126 | 0.37770246 | -1.3742964 | 0.16934969 | 0.28876037 | SRPK3 | 26576 |
| ENSG000002 | 5.2330516 | -0.511247045 | 0.37204338 | -1.3741598 | 0.16939208 | 0.28882052 | NA | NA |
| ENSG000002 | 5.15853936 | -0.517715784 | 0.3767593 | -1.3741288 | 0.16940171 | 0.28882479 | NA | NA |
| ENSG000001 | 145.97319 | 0.220096469 | 0.16017455 | 1.37410384 | 0.16940945 | 0.28882584 | SH3GL3 | 6457 |
| ENSG000002 | 5.25981948 | -0.519177623 | 0.37800391 | -1.3734715 | 0.1696058 | 0.28914844 | NA | NA |
| ENSG000001 | 6.83274225 | -0.494922876 | 0.36036718 | -1.373385 | 0.16963269 | 0.28915803 | MMP21 | 118856 |
| ENSG000002 | 22.2502434 | 0.395559794 | 0.28801812 | 1.37338511 | 0.16963266 | 0.28915803 | C19orf81 | 342918 |
| ENSG000002 | 19.704891 | 0.382512177 | 0.27852111 | 1.37336869 | 0.16963776 | 0.28915803 | NA | NA |
| ENSG000002 | 36.0181841 | 0.340833903 | 0.24817491 | 1.37336164 | 0.16963995 | 0.28915803 | NA | NA |
| ENSG000001 | 185.785381 | 0.227217888 | 0.16545183 | 1.37331749 | 0.16965367 | 0.28916926 | CCBE1 | 147372 |
| ENSG000002 | 2.74491362 | -0.511580287 | 0.37253101 | -1.3732556 | 0.16967291 | 0.28918991 | NA | NA |
| ENSG000001 | 5.28982829 | -0.513563739 | 0.37403579 | -1.3730337 | 0.16974187 | 0.28929528 | PGLYRP2 | 114770 |
| ENSG000001 | 6.22051492 | -0.509650585 | 0.37122895 | -1.372874 | 0.16979152 | 0.28936773 | HOXC6 | 3223 |
| ENSG000002 | 5.24865647 | -0.511980276 | 0.37293865 | -1.3728271 | 0.16980611 | 0.28938044 | NA | NA |
| ENSG000001 | 29.3340139 | -0.356763798 | 0.25993831 | -1.3724941 | 0.16990968 | 0.28953865 | RORC | 6097 |
| ENSG000002 | 12.6072685 | -0.439680955 | 0.32035548 | -1.3724783 | 0.16991458 | 0.28953865 | NA | NA |
| ENSG000002 | 24.6781668 | -0.356844399 | 0.26000355 | -1.3724597 | 0.16992037 | 0.28953865 | NA | NA |
| ENSG000002 | 11.435244 | 0.458118873 | 0.33380013 | 1.37243468 | 0.16992816 | 0.28953976 | PAM16 | 51025 |
| ENSG000001 | 10.7207765 | -0.454112082 | 0.3309014 | -1.3723486 | 0.16995493 | 0.28956861 | CCDC27 | 148870 |
| ENSG000002 | 3.94486491 | -0.519545587 | 0.37859161 | -1.3723114 | 0.16996651 | 0.28956861 | GNA14-AS1 | 101927422 |
| ENSG000002 | 3.64362808 | -0.519205621 | 0.37834239 | -1.3723168 | 0.16996484 | 0.28956861 | NA | NA |
| ENSG000002 | 2.79133919 | -0.510438099 | 0.37197369 | -1.3722425 | 0.16998796 | 0.28959298 | NA | NA |
| ENSG000002 | 3.58494609 | -0.519221419 | 0.37839903 | -1.3721531 | 0.17001577 | 0.28962819 | SNORD59A | 26789 |
| ENSG000001 | 268.231134 | -0.185998096 | 0.1355583 | -1.3720893 | 0.17003563 | 0.28964309 | FAM65B | 9750 |
| ENSG000001 | 8.45658494 | -0.478558707 | 0.3487836 | -1.3720791 | 0.1700388 | 0.28964309 | DMBX1 | 127343 |
| ENSG000001 | 2.44060327 | -0.508981424 | 0.37096605 | -1.3720431 | 0.17005001 | 0.28965002 | NLGN4Y | 22829 |

| | | | | | | | | |
|---|---|---|---|---|---|---|---|---|
| ENSG000001 | 243.509118 | 0.218482624 | 0.15925061 | 1.37194216 | 0.17008145 | 0.28967923 | DIS3L2 | 129563 |
| ENSG000002 | 12.9442501 | -0.438665831 | 0.31973667 | -1.3719597 | 0.17007598 | 0.28967923 | NA | NA |
| ENSG000001 | 181.040928 | 0.186182063 | 0.13571125 | 1.37189852 | 0.17009503 | 0.2896902 | TIGD7 | 91151 |
| ENSG000002 | 7.36435724 | -0.488546126 | 0.35614772 | -1.3717514 | 0.17014085 | 0.28974389 | NA | NA |
| ENSG000002 | 21.1420091 | -0.382382985 | 0.27875177 | -1.3717688 | 0.17013541 | 0.28974389 | NA | NA |
| ENSG000002 | 5.21930462 | -0.511159886 | 0.37264682 | -1.3717006 | 0.17015665 | 0.28975863 | NA | NA |
| ENSG000002 | 2.7518101 | -0.511414466 | 0.3728993 | -1.3714546 | 0.17023329 | 0.28987696 | NA | NA |
| ENSG000001 | 225.49944 | -0.191563362 | 0.13968173 | -1.3714275 | 0.17024173 | 0.28987917 | DOCK2 | 1794 |
| ENSG000001 | 9.41581092 | 0.470863469 | 0.34334717 | 1.37139173 | 0.17025288 | 0.28988597 | NA | NA |
| ENSG000001 | 25.4991333 | -0.357735113 | 0.26089679 | -1.3711748 | 0.17032047 | 0.28998889 | SPOCD1 | 90853 |
| ENSG000001 | 87.9256744 | 0.242384937 | 0.17677792 | 1.37112678 | 0.17033544 | 0.2900022 | FBRS | 64319 |
| ENSG000002 | 6.08047762 | -0.50875698 | 0.37105712 | -1.3711015 | 0.17034331 | 0.29000343 | NA | NA |
| ENSG000002 | 15.4684649 | -0.427881329 | 0.31211573 | -1.370906 | 0.17040426 | 0.29009502 | CPEB1-AS1 | 283692 |
| ENSG000002 | 3.27837573 | -0.518397994 | 0.3781513 | -1.3708745 | 0.17041407 | 0.29009953 | NA | NA |
| ENSG000002 | 9.81537533 | -0.481581934 | 0.35130889 | -1.370822 | 0.17043046 | 0.29011526 | CES1P1 | 51716 |
| ENSG000002 | 3.00986957 | -0.515108098 | 0.37577836 | -1.3707764 | 0.17044467 | 0.29012727 | NA | NA |
| ENSG000001 | 1789.54662 | 0.160682664 | 0.11722718 | 1.37069465 | 0.17047016 | 0.29015848 | RFX7 | 64864 |
| ENSG000002 | 9.30562671 | -0.475522958 | 0.34693804 | -1.3706279 | 0.17049098 | 0.29016956 | NA | NA |
| ENSG000002 | 2.39776669 | -0.508889584 | 0.37128046 | -1.3706339 | 0.17048911 | 0.29016956 | NA | NA |
| ENSG000001 | 32.0496172 | -0.33147056 | 0.24184946 | -1.3705657 | 0.17051039 | 0.29019041 | CARD6 | 84674 |
| ENSG000002 | 8.68485073 | -0.475371713 | 0.34687776 | -1.3704301 | 0.17055269 | 0.29025021 | NA | NA |
| ENSG000002 | 10.5856036 | -0.456886154 | 0.33341094 | -1.3703394 | 0.17058097 | 0.29028616 | NA | NA |
| ENSG000002 | 6.541762 | -0.496477605 | 0.36231296 | -1.3703004 | 0.17059314 | 0.29029469 | NA | NA |
| ENSG000002 | 9.5823861 | -0.471038465 | 0.34378033 | -1.3701728 | 0.17063296 | 0.29035026 | KRT73-AS1 | 100127967 |
| ENSG000001 | 1281.43145 | 0.181346708 | 0.13235787 | 1.37012413 | 0.17064816 | 0.29036394 | WDR26 | 80232 |
| ENSG000002 | 2.39845183 | -0.508682383 | 0.37128285 | -1.370067 | 0.17066599 | 0.2903821 | NA | NA |
| ENSG000001 | 67.4616966 | 0.302045582 | 0.2204657 | 1.37003435 | 0.17067618 | 0.29038725 | LMTK3 | 114783 |
| ENSG000002 | 6.6016312 | 0.508006619 | 0.37082382 | 1.36994063 | 0.17070544 | 0.29042484 | MIR25 | 407014 |
| ENSG000002 | 10.9497344 | -0.460061818 | 0.33584993 | -1.3698434 | 0.17073579 | 0.29046429 | NA | NA |

| | | | | | | | | |
|---|---|---|---|---|---|---|---|---|
| ENSG000001 | 802.321274 | 0.181158111 | 0.13225429 | 1.36977112 | 0.17075836 | 0.29046873 | ZNF124 | 7678 |
| ENSG000002 | 16.554709 | 0.40354474 | 0.29460847 | 1.36976624 | 0.17075988 | 0.29046873 | MTFP1 | 51537 |
| ENSG000002 | 5.0778492 | -0.516737179 | 0.37723541 | -1.3698003 | 0.17074924 | 0.29046873 | NA | NA |
| ENSG000001 | 7.53365273 | -0.498031369 | 0.36360355 | -1.3697099 | 0.17077749 | 0.29048649 | GPR18 | 2841 |
| ENSG000001 | 34.0910599 | -0.334572595 | 0.24427496 | -1.3696557 | 0.1707944 | 0.29050085 | SPIN2A | 54466 |
| ENSG000002 | 3.87056236 | -0.518530994 | 0.3785901 | -1.369637 | 0.17080026 | 0.29050085 | NA | NA |
| ENSG000002 | 3.91325101 | -0.51844065 | 0.37855337 | -1.3695312 | 0.17083329 | 0.29054484 | NA | NA |
| ENSG000002 | 10.303032 | 0.459056707 | 0.33520904 | 1.369464 | 0.17085428 | 0.29056836 | NA | NA |
| ENSG000002 | 7.72876879 | -0.489332107 | 0.35732273 | -1.3694402 | 0.17086172 | 0.29056882 | NA | NA |
| ENSG000002 | 2.73917303 | -0.510621322 | 0.37288129 | -1.3693938 | 0.17087621 | 0.29058128 | NA | NA |
| ENSG000002 | 2.95850065 | -0.515793807 | 0.37669028 | -1.3692782 | 0.17091232 | 0.2906305 | NA | NA |
| ENSG000002 | 4.18981288 | -0.517767034 | 0.37815205 | -1.3692033 | 0.17093573 | 0.29065812 | LOC100421746 | 100421746 |
| ENSG000002 | 4.08747038 | -0.517524938 | 0.37802205 | -1.3690337 | 0.17098873 | 0.29072789 | LINC01549 | 100505929 |
| ENSG000002 | 4.76268996 | -0.515320693 | 0.37641518 | -1.3690221 | 0.17099237 | 0.29072789 | NA | NA |
| ENSG000002 | 6.89216346 | -0.499287382 | 0.3647087 | -1.3690032 | 0.17099827 | 0.29072789 | NA | NA |
| ENSG000002 | 7.36638236 | 0.490646437 | 0.35843622 | 1.36885283 | 0.17104528 | 0.29079563 | INSL3 | 3640 |
| ENSG000001 | 24.3584917 | -0.364168939 | 0.26604771 | -1.3688107 | 0.17105846 | 0.29080584 | GRAMD2 | 196996 |
| ENSG000002 | 42.7457067 | 0.30587592 | 0.2234766 | 1.36871566 | 0.17108817 | 0.29083197 | NA | NA |
| ENSG000002 | 3.22999498 | -0.517870212 | 0.37835662 | -1.3687357 | 0.17108191 | 0.29083197 | NA | NA |
| ENSG00000C | 1323.55623 | 0.198220356 | 0.14483063 | 1.36863561 | 0.17111321 | 0.29086233 | CLSPN | 63967 |
| ENSG000002 | 2.21521445 | -0.49494859 | 0.36164374 | -1.3686082 | 0.17112178 | 0.29086471 | NA | NA |
| ENSG000002 | 3.16439824 | -0.373889376 | 0.27320516 | -1.3685297 | 0.17114633 | 0.29089425 | RN7SL413P | 106479374 |
| ENSG000002 | 3.24957373 | -0.51777965 | 0.37835677 | -1.3684958 | 0.17115694 | 0.29090008 | NA | NA |
| ENSG000002 | 8.8867062 | -0.470599394 | 0.3438925 | -1.3684491 | 0.17117153 | 0.29091269 | NA | NA |
| ENSG000002 | 7.44505583 | -0.491417687 | 0.35911197 | -1.3684247 | 0.17117917 | 0.29091349 | NA | NA |
| ENSG000002 | 2.79509792 | -0.510069611 | 0.37275823 | -1.3683658 | 0.1711976 | 0.29092042 | NA | NA |
| ENSG000002 | 2.37178182 | -0.50875947 | 0.37179747 | -1.368378 | 0.17119379 | 0.29092042 | NA | NA |
| ENSG000001 | 11.1184841 | -0.451291665 | 0.32981056 | -1.368336 | 0.17120693 | 0.29092408 | MRVI1-AS1 | 100129827 |
| ENSG000002 | 14.3405981 | -0.420049719 | 0.30705188 | -1.3680089 | 0.17130928 | 0.2910858 | NA | NA |

| | | | | | | | | |
|---|---|---|---|---|---|---|---|---|
| ENSG000002 | 5.22213027 | -0.509865664 | 0.37273453 | -1.3679057 | 0.17134161 | 0.29112854 | OR7E162P | 442101 |
| ENSG000001 | 15.0722644 | 0.418809562 | 0.30617475 | 1.36787755 | 0.17135041 | 0.29113129 | UGT2B4 | 7363 |
| ENSG000002 | 5.45027096 | -0.506062052 | 0.36999486 | -1.3677543 | 0.171389 | 0.2911742 | NA | NA |
| ENSG000002 | 14.2713146 | 0.420414113 | 0.3073762 | 1.36775102 | 0.17139002 | 0.2911742 | NA | NA |
| ENSG000002 | 13.9235491 | -0.43460244 | 0.31781296 | -1.3674786 | 0.17147533 | 0.29130691 | NA | NA |
| ENSG000001 | 80.6336675 | -0.238708782 | 0.17456419 | -1.3674556 | 0.17148253 | 0.29130695 | AGBL3 | 340351 |
| ENSG000002 | 3.01045829 | -0.514553945 | 0.37631449 | -1.3673509 | 0.17151535 | 0.29134738 | NA | NA |
| ENSG000002 | 5.2190168 | -0.510034413 | 0.37301383 | -1.3673338 | 0.1715207 | 0.29134738 | NA | NA |
| ENSG000001 | 40.0023428 | -0.318972303 | 0.23329762 | -1.3672334 | 0.17155215 | 0.29138859 | SFXN3 | 81855 |
| ENSG000002 | 16.8310896 | -0.417995252 | 0.30575353 | -1.3670987 | 0.17159437 | 0.2914481 | NA | NA |
| ENSG000001 | 29.9395467 | -0.365114695 | 0.26708971 | -1.3670115 | 0.17162171 | 0.29148232 | KIAA1024 | 23251 |
| ENSG000002 | 3.03935258 | -0.513538491 | 0.37568444 | -1.3669411 | 0.17164378 | 0.2915076 | NA | NA |
| ENSG000002 | 4.07332011 | -0.516432024 | 0.37782571 | -1.3668525 | 0.17167154 | 0.29154253 | LOC100506937 | 100506937 |
| ENSG000000 | 38.1087854 | -0.308820232 | 0.22594852 | -1.3667726 | 0.17169661 | 0.29157289 | EYA2 | 2139 |
| ENSG000002 | 6.38834198 | 0.50731941 | 0.37120235 | 1.36669236 | 0.17172175 | 0.29159118 | NA | NA |
| ENSG000002 | 4.14772615 | -0.516869239 | 0.37818564 | -1.3667077 | 0.17171693 | 0.29159118 | NA | NA |
| ENSG000002 | 8.91962643 | -0.471370955 | 0.34496276 | -1.3664401 | 0.17180086 | 0.29171329 | NA | NA |
| ENSG000002 | 2.55798673 | -0.493306948 | 0.36104351 | -1.3663366 | 0.17183335 | 0.29175623 | NA | NA |
| ENSG000002 | 2.43761456 | -0.505721345 | 0.37018115 | -1.3661456 | 0.17189328 | 0.29184577 | NA | NA |
| ENSG000001 | 14.2179968 | -0.431294217 | 0.31570761 | -1.3661191 | 0.17190157 | 0.29184763 | EPHB3 | 2049 |
| ENSG000002 | 3.01087588 | -0.513919192 | 0.37622402 | -1.3659925 | 0.17194132 | 0.29190289 | LINC01115 | 339822 |
| ENSG000002 | 26.1489679 | 0.35325991 | 0.2586775 | 1.36563834 | 0.17205251 | 0.29207942 | RNVU1-3 | 101954272 |
| ENSG000002 | 9.22714628 | -0.467033532 | 0.34201599 | -1.3655313 | 0.17208613 | 0.29208916 | NA | NA |
| ENSG000002 | 7.06642954 | -0.488386562 | 0.35765314 | -1.3655313 | 0.17208612 | 0.29208916 | NA | NA |
| ENSG000002 | 7.17550732 | -0.493320591 | 0.3612672 | -1.3655283 | 0.17208705 | 0.29208916 | NA | NA |
| ENSG000002 | 4.40108369 | -0.514273047 | 0.37659986 | -1.3655689 | 0.17207432 | 0.29208916 | NA | NA |
| ENSG000002 | 10.8238443 | 0.452923395 | 0.33170999 | 1.3654198 | 0.17212114 | 0.29213479 | NA | NA |
| ENSG000002 | 3.0127426 | -0.514401496 | 0.37674404 | -1.3653872 | 0.17213138 | 0.29213994 | NA | NA |
| ENSG000001 | 63.0816581 | 0.262169119 | 0.19203211 | 1.36523586 | 0.17217893 | 0.29220679 | SARDH | 1757 |

| | | | | | | | | |
|---|---|---|---|---|---|---|---|---|
| ENSG000002 | 3.02643586 | -0.513354196 | 0.37602416 | -1.365216 | 0.17218518 | 0.29220679 | NA | NA |
| ENSG000002 | 31.6688744 | -0.329216632 | 0.24116602 | -1.3651037 | 0.17222046 | 0.29225443 | LOC101926941 | 101926941 |
| ENSG000002 | 11.7220902 | -0.451154807 | 0.33049991 | -1.3650679 | 0.17223172 | 0.2922613 | NA | NA |
| ENSG000002 | 3.01215332 | -0.512638452 | 0.37555144 | -1.3650286 | 0.17224407 | 0.29226991 | NA | NA |
| ENSG000002 | 3.28424106 | -0.515844008 | 0.37790607 | -1.3650059 | 0.17225121 | 0.29226991 | NA | NA |
| ENSG000001 | 14.1834175 | 0.435206599 | 0.31887735 | 1.36480877 | 0.17231316 | 0.29233888 | BARX1 | 56033 |
| ENSG000002 | 42.295657 | -0.312252495 | 0.22878863 | -1.3648078 | 0.17231348 | 0.29233888 | NA | NA |
| ENSG000002 | 3.28410467 | -0.515918079 | 0.37801469 | -1.3648096 | 0.1723129 | 0.29233888 | NA | NA |
| ENSG000002 | 12.9731509 | -0.454498026 | 0.33302321 | -1.3647638 | 0.1723273 | 0.29235009 | NA | NA |
| ENSG000002 | 5.08519371 | -0.51296609 | 0.37589792 | -1.364642 | 0.17236559 | 0.29240283 | NA | NA |
| ENSG000002 | 14.346067 | -0.41768269 | 0.306104 | -1.3645124 | 0.17240636 | 0.29245155 | NA | NA |
| ENSG000002 | 6.53820438 | -0.497291958 | 0.36444867 | -1.3645048 | 0.17240874 | 0.29245155 | NA | NA |
| ENSG000001 | 16.9567017 | -0.417014397 | 0.30563076 | -1.3644386 | 0.17242958 | 0.29247467 | RAET1E | 135250 |
| ENSG000001 | 63.0191378 | 0.275000882 | 0.20157526 | 1.36425907 | 0.17248604 | 0.2925582 | CDH4 | 1002 |
| ENSG000002 | 11.7924938 | -0.44451055 | 0.32584176 | -1.3641915 | 0.17250731 | 0.29257718 | LOC101928820 | 101928820 |
| ENSG000002 | 10.3230793 | -0.460628892 | 0.33766049 | -1.3641776 | 0.17251166 | 0.29257718 | LINC01584 | 101929701 |
| ENSG000002 | 4.10226646 | -0.515489405 | 0.37793839 | -1.3639509 | 0.17258302 | 0.29268597 | NA | NA |
| ENSG000002 | 4.8264038 | -0.515674544 | 0.37808673 | -1.3639054 | 0.17259733 | 0.292698 | NA | NA |
| ENSG000002 | 2.46659767 | -0.501540061 | 0.36773157 | -1.3638755 | 0.17260677 | 0.29270175 | NA | NA |
| ENSG000001 | 118.731229 | -0.217053854 | 0.15915455 | -1.3637929 | 0.17263274 | 0.29273356 | GARNL3 | 84253 |
| ENSG000001 | 99.1856597 | 0.226348823 | 0.16597827 | 1.36372564 | 0.17265393 | 0.29275725 | SERGEF | 26297 |
| ENSG000001 | 13.6757126 | -0.427642398 | 0.31368972 | -1.3632656 | 0.17279883 | 0.29299068 | RRH | 10692 |
| ENSG000001 | 759.607393 | 0.158898671 | 0.11656869 | 1.36313341 | 0.17284047 | 0.29304123 | RPAP2 | 79871 |
| ENSG000002 | 2.69682626 | -0.508552441 | 0.37307834 | -1.3631251 | 0.17284309 | 0.29304123 | NA | NA |
| ENSG000001 | 16.5310801 | -0.407837002 | 0.29920249 | -1.3630802 | 0.17285722 | 0.29304344 | MATN4 | 8785 |
| ENSG000002 | 6.82645236 | -0.49227534 | 0.36115056 | -1.3630751 | 0.17285884 | 0.29304344 | NA | NA |
| ENSG000002 | 5.20577946 | -0.508819095 | 0.37330906 | -1.3629969 | 0.17288348 | 0.29307295 | NA | NA |
| ENSG000002 | 6.42047706 | 0.503004928 | 0.36905491 | 1.36295415 | 0.17289696 | 0.29308355 | NA | NA |
| ENSG000002 | 8.14299681 | -0.480630695 | 0.35267949 | -1.3627974 | 0.17294637 | 0.29314653 | LINC01529 | 644050 |

| | | | | | | | | |
|---|---|---|---|---|---|---|---|---|
| ENSG000002 | 6.25514355 | -0.499325118 | 0.36639906 | -1.3627904 | 0.17294858 | 0.29314653 | NA | NA |
| ENSG000002 | 3.9645553 | 0.515523507 | 0.37831539 | 1.36268181 | 0.17298281 | 0.29319231 | NPIPA5 | 100288332 |
| ENSG000002 | 51.5362927 | 0.282514783 | 0.20735216 | 1.3624878 | 0.17304399 | 0.29327225 | NA | NA |
| ENSG000002 | 4.50385884 | -0.515244526 | 0.3781649 | -1.3624864 | 0.17304444 | 0.29327225 | NA | NA |
| ENSG000001 | 707.667978 | 0.157010726 | 0.11524777 | 1.36237543 | 0.17307943 | 0.29330704 | TAF5L | 27097 |
| ENSG000002 | 2.17494237 | -0.494007718 | 0.36260326 | -1.3623918 | 0.17307426 | 0.29330704 | NA | NA |
| ENSG000002 | 6.30952214 | -0.500099507 | 0.36711709 | -1.3622343 | 0.17312394 | 0.2933592 | NA | NA |
| ENSG000002 | 19.5272301 | -0.396756569 | 0.29125967 | -1.3622091 | 0.17313192 | 0.2933592 | NA | NA |
| ENSG000002 | 2.3790695 | -0.50650657 | 0.37182222 | -1.3622278 | 0.17312601 | 0.2933592 | NA | NA |
| ENSG000001 | 705.170505 | 0.168037538 | 0.12336359 | 1.36213241 | 0.1731561 | 0.29336612 | NFAT5 | 10725 |
| ENSG000002 | 4.08852079 | -0.514610081 | 0.37779881 | -1.3621273 | 0.1731577 | 0.29336612 | NA | NA |
| ENSG000002 | 5.69974657 | -0.500715583 | 0.36759744 | -1.3621302 | 0.17315679 | 0.29336612 | NA | NA |
| ENSG000002 | 3.31850775 | -0.513485139 | 0.37698206 | -1.3620944 | 0.17316811 | 0.29337148 | NA | NA |
| ENSG000002 | 2.76837584 | -0.508017561 | 0.37298804 | -1.3620211 | 0.17319124 | 0.29339842 | NA | NA |
| ENSG000001 | 2.80283357 | -0.503487318 | 0.36968569 | -1.3619335 | 0.17321888 | 0.29342109 | TAS2R8 | 50836 |
| ENSG000002 | 5.27386371 | -0.510235271 | 0.37464056 | -1.3619328 | 0.17321909 | 0.29342109 | NA | NA |
| ENSG000002 | 10.029613 | 0.471666704 | 0.34632774 | 1.36190853 | 0.17322675 | 0.29342181 | NA | NA |
| ENSG000001 | 1180.55261 | 0.402340748 | 0.2954361 | 1.3618537 | 0.17324406 | 0.29342661 | PRPF38B | 55119 |
| ENSG000002 | 9.05464302 | -0.476543909 | 0.34991775 | -1.3618741 | 0.17323763 | 0.29342661 | LINC00575 | 439934 |
| ENSG000001 | 18.9027241 | -0.384217857 | 0.28214572 | -1.361771 | 0.17327017 | 0.29344632 | SLC15A5 | 729025 |
| ENSG000002 | 4.0958928 | -0.51451812 | 0.37782774 | -1.3617796 | 0.17326744 | 0.29344632 | NA | NA |
| ENSG000002 | 2.48282787 | -0.49794275 | 0.36568189 | -1.3616828 | 0.173298 | 0.2934812 | NA | NA |
| ENSG00000C | 190.873701 | 0.183378914 | 0.13467914 | 1.36159846 | 0.17332464 | 0.29350179 | IFT88 | 8100 |
| ENSG000002 | 5.54000975 | -0.504973267 | 0.37086353 | -1.3616148 | 0.17331949 | 0.29350179 | NA | NA |
| ENSG000002 | 14.4521686 | 0.423694665 | 0.3111818 | 1.36156636 | 0.17333478 | 0.29350671 | NA | NA |
| ENSG000002 | 3.08475333 | -0.511461395 | 0.37567236 | -1.361456 | 0.17336962 | 0.29355345 | NA | NA |
| ENSG000002 | 2.36310403 | -0.504480709 | 0.37056757 | -1.3613731 | 0.17339582 | 0.29358555 | NA | NA |
| ENSG000001 | 99.1937618 | 0.238964053 | 0.17554296 | 1.3612853 | 0.17342355 | 0.29361372 | L3MBTL1 | 26013 |
| ENSG000002 | 2.37702652 | -0.504799941 | 0.3708289 | -1.3612746 | 0.17342694 | 0.29361372 | NA | NA |

| | | | | | | | | |
|---|---|---|---|---|---|---|---|---|
| ENSG000002 | 2.98934509 | -0.511464601 | 0.37574689 | -1.3611945 | 0.17345223 | 0.29364427 | NA | NA |
| ENSG000002 | 2.40010252 | -0.504589364 | 0.37072652 | -1.3610825 | 0.17348763 | 0.29369195 | NA | NA |
| ENSG000002 | 3.05638056 | -0.511793903 | 0.37606632 | -1.3609139 | 0.17354089 | 0.29374531 | NA | NA |
| ENSG000002 | 43.6080752 | 0.295024734 | 0.2167822 | 1.36092694 | 0.17353678 | 0.29374531 | NA | NA |
| ENSG000002 | 3.38870004 | -0.512457309 | 0.376553 | -1.3609168 | 0.17353997 | 0.29374531 | NA | NA |
| ENSG000002 | 2.3857043 | -0.505765246 | 0.37165519 | -1.3608454 | 0.17356256 | 0.29376975 | NA | NA |
| ENSG000000 | 24.4047095 | 0.369321834 | 0.27139823 | 1.36081151 | 0.17357326 | 0.29377559 | IFI35 | 3430 |
| ENSG000002 | 2.79475659 | -0.503709798 | 0.37018617 | -1.3606932 | 0.17361067 | 0.29382663 | NA | NA |
| ENSG000002 | 7.29934604 | -0.482855147 | 0.35488869 | -1.360582 | 0.17364584 | 0.29387389 | NA | NA |
| ENSG000002 | 4.39567537 | -0.512459521 | 0.37666153 | -1.3605305 | 0.17366209 | 0.29388914 | NA | NA |
| ENSG000002 | 5.22796314 | -0.507252573 | 0.37285352 | -1.3604607 | 0.17368417 | 0.29391424 | NA | NA |
| ENSG000001 | 34.8304423 | 0.318696595 | 0.23427005 | 1.36038131 | 0.17370929 | 0.29394447 | BSCL2 | 26580 |
| ENSG000002 | 9.45621181 | 0.481245773 | 0.35379403 | 1.36024278 | 0.17375311 | 0.29400636 | NA | NA |
| ENSG000001 | 281.470718 | 0.183010734 | 0.13455634 | 1.36010489 | 0.17379673 | 0.2940311 | MDM1 | 56890 |
| ENSG000001 | 4.946553 | -0.509421921 | 0.37454381 | -1.3601131 | 0.17379415 | 0.2940311 | NA | NA |
| ENSG000002 | 3.4337281 | -0.511721079 | 0.37623425 | -1.360113 | 0.17379418 | 0.2940311 | NA | NA |
| ENSG000002 | 3.76085326 | 0.514501761 | 0.37827389 | 1.36013025 | 0.17378871 | 0.2940311 | NA | NA |
| ENSG000002 | 10.4856533 | -0.456327208 | 0.33553981 | -1.3599793 | 0.17383647 | 0.29408605 | NA | NA |
| ENSG000001 | 14.847322 | 0.416319706 | 0.30614089 | 1.3598958 | 0.1738629 | 0.294113 | UCP1 | 7350 |
| ENSG000002 | 3.89590316 | -0.514807251 | 0.37856727 | -1.3598831 | 0.17386691 | 0.294113 | NA | NA |
| ENSG000002 | 9.5026169 | -0.485239851 | 0.35685453 | -1.3597694 | 0.17390292 | 0.29416166 | NA | NA |
| ENSG000002 | 4.07537574 | -0.513675235 | 0.37785615 | -1.3594466 | 0.17400513 | 0.29432226 | NA | NA |
| ENSG000002 | 27.8205251 | 0.359689411 | 0.2645978 | 1.35938171 | 0.17402567 | 0.29434472 | NA | NA |
| ENSG000001 | 1230.64158 | 0.170026155 | 0.12508956 | 1.35923542 | 0.174072 | 0.29438626 | EEA1 | 8411 |
| ENSG000002 | 8.44275508 | -0.47483768 | 0.34934162 | -1.3592359 | 0.17407185 | 0.29438626 | NA | NA |
| ENSG000002 | 4.0922269 | -0.513591812 | 0.37784399 | -1.3592695 | 0.1740612 | 0.29438626 | NA | NA |
| ENSG000001 | 12.5035596 | -0.442957213 | 0.32592365 | -1.3590828 | 0.17412036 | 0.29445577 | IL12B | 3593 |
| ENSG000001 | 63.1585036 | 0.262118906 | 0.19287503 | 1.35900904 | 0.17414373 | 0.29448299 | OTUD1 | 220213 |
| ENSG000001 | 23.1021066 | 0.370838591 | 0.27290116 | 1.35887509 | 0.17418617 | 0.29453021 | MAGEA4 | 4103 |

| | | | | | | | | |
|---|---|---|---|---|---|---|---|---|
| ENSG000002 | 8.42297957 | -0.475865773 | 0.35019083 | -1.3588756 | 0.17418601 | 0.29453021 | NA | NA |
| ENSG000002 | 32.7539809 | -0.329337478 | 0.24237355 | -1.3588012 | 0.17420958 | 0.29454522 | NA | NA |
| ENSG000002 | 6.34231708 | -0.501050151 | 0.36874241 | -1.3588081 | 0.17420742 | 0.29454522 | LOC105370424 | 105370424 |
| ENSG000002 | 6.10156468 | -0.502497431 | 0.36984775 | -1.3586603 | 0.17425427 | 0.29460851 | NA | NA |
| ENSG000002 | 5.31932394 | -0.508703921 | 0.37448695 | -1.3584023 | 0.17433607 | 0.29473451 | NA | NA |
| ENSG000002 | 11.3018294 | -0.460532834 | 0.33903834 | -1.3583503 | 0.17435256 | 0.2947501 | NA | NA |
| ENSG000001 | 18.128933 | -0.399493038 | 0.29411562 | -1.3582857 | 0.17437304 | 0.29477244 | GOLGA6A | 342096 |
| ENSG000002 | 2.9872142 | -0.511517125 | 0.37660069 | -1.358248 | 0.17438501 | 0.29478038 | H3F3AP1 | 654505 |
| ENSG000001 | 6.86506245 | -0.492001386 | 0.36227408 | -1.3580916 | 0.17443462 | 0.29483581 | SLC46A2 | 57864 |
| ENSG000002 | 3.01456159 | -0.510822749 | 0.37615616 | -1.3580071 | 0.17446143 | 0.29483581 | NA | NA |
| ENSG000002 | 23.70594 | -0.37014686 | 0.27255415 | -1.3580672 | 0.17444235 | 0.29483581 | NA | NA |
| ENSG000002 | 28.0136791 | -0.350158832 | 0.25784247 | -1.358034 | 0.1744529 | 0.29483581 | RAD21L1 | 642636 |
| ENSG000002 | 2.39395699 | -0.503432303 | 0.37071203 | -1.3580145 | 0.1744591 | 0.29483581 | RNU6-1053P | 106481513 |
| ENSG000002 | 4.77432704 | -0.512905935 | 0.37766042 | -1.3581141 | 0.17442749 | 0.29483581 | NA | NA |
| ENSG000000 | 54.9146747 | 0.271163654 | 0.19968279 | 1.35797211 | 0.17447254 | 0.2948423 | OSBPL5 | 114879 |
| ENSG000001 | 3.80669386 | -0.513960605 | 0.37853059 | -1.3577783 | 0.17453405 | 0.29493397 | NA | NA |
| ENSG000002 | 43.2532002 | -0.291549011 | 0.21472921 | -1.357752 | 0.17454239 | 0.29493576 | UBAP1L | 390595 |
| ENSG000002 | 5.46743011 | -0.502547307 | 0.37015825 | -1.3576553 | 0.17457309 | 0.29497535 | NA | NA |
| ENSG000002 | 7.64993849 | -0.485434443 | 0.35756919 | -1.3575958 | 0.17459196 | 0.29499495 | NA | NA |
| ENSG000002 | 2.95807609 | -0.510472933 | 0.37604061 | -1.3574942 | 0.17462424 | 0.2950372 | GTF2H4 | 2968 |
| ENSG000002 | 4.72837963 | -0.511705388 | 0.37697327 | -1.357405 | 0.17465257 | 0.29507276 | NA | NA |
| ENSG000002 | 5.87308875 | -0.508145139 | 0.37436198 | -1.3573631 | 0.17466587 | 0.29508293 | NA | NA |
| ENSG000002 | 6.94724044 | -0.496183567 | 0.36556218 | -1.3573165 | 0.17468068 | 0.29509566 | NA | NA |
| ENSG000001 | 98.8170076 | -0.231534776 | 0.17059439 | -1.3572238 | 0.1747101 | 0.2951306 | CEP112 | 201134 |
| ENSG000002 | 6.57132824 | -0.494955502 | 0.36468722 | -1.3572055 | 0.17471592 | 0.2951306 | NA | NA |
| ENSG000001 | 991.509888 | 0.196690528 | 0.14493435 | 1.35710083 | 0.17474917 | 0.29517448 | TNIK | 23043 |
| ENSG000001 | 56.1183329 | -0.268700311 | 0.19801131 | -1.3569948 | 0.17478288 | 0.29521911 | BNIPL | 149428 |
| ENSG000002 | 4.10896095 | -0.513079859 | 0.37811345 | -1.3569468 | 0.1747981 | 0.29523253 | NA | NA |
| ENSG000002 | 6.30357303 | -0.497303529 | 0.36653131 | -1.3567832 | 0.1748501 | 0.29530805 | NA | NA |

| | | | | | | | | |
|---|---|---|---|---|---|---|---|---|
| ENSG000002 | 3.24214224 | -0.513055279 | 0.37816364 | -1.3567018 | 0.17487599 | 0.29533947 | NA | NA |
| ENSG000002 | 6.87506589 | 0.505116641 | 0.37237603 | 1.35646928 | 0.1749499 | 0.29545201 | NA | NA |
| ENSG000002 | 7.58008805 | -0.497536344 | 0.36680763 | -1.3563958 | 0.17497326 | 0.29546684 | ANKRD36BP1 | 84832 |
| ENSG000002 | 3.57815952 | -0.512929342 | 0.37815587 | -1.3563966 | 0.17497301 | 0.29546684 | NA | NA |
| ENSG000002 | 8.43175408 | -0.474769075 | 0.35003809 | -1.3563355 | 0.17499246 | 0.29548696 | NA | NA |
| ENSG000002 | 3.21275151 | -0.512886798 | 0.37817719 | -1.3562076 | 0.17503311 | 0.29553099 | NA | NA |
| ENSG000002 | 5.76972891 | -0.50166354 | 0.36989665 | -1.3562262 | 0.1750272 | 0.29553099 | NA | NA |
| ENSG000000 | 189.168025 | 0.234530418 | 0.17295647 | 1.35600837 | 0.1750965 | 0.29557649 | TEAD3 | 7005 |
| ENSG000002 | 19.4402045 | -0.383315419 | 0.28267857 | -1.3560116 | 0.17509547 | 0.29557649 | NA | NA |
| ENSG000002 | 4.36647311 | -0.511074253 | 0.37688548 | -1.3560465 | 0.17508436 | 0.29557649 | NA | NA |
| ENSG000002 | 8.93613473 | -0.484470711 | 0.35726114 | -1.3560689 | 0.17507725 | 0.29557649 | NA | NA |
| ENSG000002 | 4.72922119 | -0.511073567 | 0.37688655 | -1.3560409 | 0.17508616 | 0.29557649 | NA | NA |
| ENSG000002 | 3.85002579 | -0.51332813 | 0.37859337 | -1.3558825 | 0.17513655 | 0.29563179 | MIR4803 | 100616377 |
| ENSG000002 | 8.8615758 | -0.48025726 | 0.35424466 | -1.3557219 | 0.17518765 | 0.29570573 | CARD17 | 440068 |
| ENSG000002 | 1854.24592 | 0.152987655 | 0.11285261 | 1.35564123 | 0.17521335 | 0.29573679 | ZNF260 | 339324 |
| ENSG000001 | 3.00315134 | -0.509836875 | 0.37609723 | -1.3555986 | 0.17522692 | 0.2957474 | NA | NA |
| ENSG000002 | 3.07496151 | -0.509250955 | 0.37569901 | -1.3554759 | 0.17526598 | 0.29580101 | RNU4-64P | 106481872 |
| ENSG000002 | 11.9483931 | -0.444518485 | 0.32794984 | -1.3554466 | 0.17527532 | 0.29580447 | NA | NA |
| ENSG000002 | 22.3416384 | -0.37468901 | 0.27645064 | -1.3553559 | 0.17530418 | 0.29584086 | NA | NA |
| ENSG000002 | 9.14655005 | -0.461581865 | 0.34058136 | -1.3552764 | 0.17532952 | 0.29587131 | DIO2-AS1 | 100628307 |
| ENSG000002 | 2.38026318 | -0.502700149 | 0.37093154 | -1.355237 | 0.17534206 | 0.29588014 | NA | NA |
| ENSG000002 | 2.37033965 | -0.502201119 | 0.37059299 | -1.3551285 | 0.17537663 | 0.29592616 | NA | NA |
| ENSG000002 | 3.8135766 | -0.512973188 | 0.37855126 | -1.3550957 | 0.17538707 | 0.29593147 | NA | NA |
| ENSG000001 | 25.6453886 | -0.357153659 | 0.26358007 | -1.3550101 | 0.17541433 | 0.29596516 | C10orf90 | 118611 |
| ENSG000002 | 4.13266294 | -0.512027805 | 0.37789956 | -1.3549309 | 0.17543956 | 0.29599541 | LINC00940 | 100271702 |
| ENSG000002 | 4.42286345 | -0.511818902 | 0.37791142 | -1.3543356 | 0.17562932 | 0.29630323 | NA | NA |
| ENSG000001 | 15.5894953 | 0.417080967 | 0.30799067 | 1.35419999 | 0.17567258 | 0.29636389 | TMEM187 | 8269 |
| ENSG000001 | 108.040699 | -0.215167794 | 0.15890065 | -1.3541027 | 0.17570362 | 0.29636808 | CDHR3 | 222256 |
| ENSG000001 | 2.82925327 | -0.500004114 | 0.36925183 | -1.3541006 | 0.1757043 | 0.29636808 | NA | NA |

| | | | | | | | | |
|---|---|---|---|---|---|---|---|---|
| ENSG000002 | 2.75215033 | -0.505047633 | 0.37297136 | -1.3541191 | 0.17569839 | 0.29636808 | NA | NA |
| ENSG000002 | 3.27189738 | -0.511523079 | 0.37775813 | -1.3541021 | 0.17570381 | 0.29636808 | NA | NA |
| ENSG000001 | 94.4498517 | 0.262615275 | 0.1939529 | 1.3540157 | 0.17573137 | 0.29640141 | ERMAP | 114625 |
| ENSG000001 | 2.41846901 | -0.495030017 | 0.36568124 | -1.35372 | 0.17582571 | 0.2965482 | KISS1 | 3814 |
| ENSG000001 | 2.38640871 | -0.501783708 | 0.37069177 | -1.3536413 | 0.17585083 | 0.29657823 | WFDC12 | 128488 |
| ENSG000002 | 8.25076223 | -0.480572579 | 0.35505215 | -1.3535267 | 0.17588742 | 0.2966276 | NA | NA |
| ENSG000002 | 3.32540369 | -0.510121033 | 0.37692184 | -1.3533868 | 0.17593211 | 0.29669063 | NA | NA |
| ENSG000002 | 15.6066506 | -0.406796282 | 0.30059928 | -1.3532843 | 0.17596482 | 0.29673346 | LOC100506797 | 100506797 |
| ENSG000000 | 120.655043 | -0.209714921 | 0.15497501 | -1.3532177 | 0.1759861 | 0.29675699 | IRAK3 | 11213 |
| ENSG000001 | 91.7203115 | 0.26701618 | 0.1973369 | 1.35309809 | 0.17602429 | 0.29676862 | PPARD | 5467 |
| ENSG000002 | 2.36305251 | -0.502275949 | 0.3711964 | -1.3531272 | 0.176015 | 0.29676862 | KPRP | 448834 |
| ENSG000002 | 16.8975781 | -0.427984668 | 0.31630368 | -1.3530815 | 0.17602958 | 0.29676862 | RPA4 | 29935 |
| ENSG000002 | 11.2758612 | -0.441758096 | 0.32648254 | -1.3530834 | 0.176029 | 0.29676862 | LOC101928205 | 101928205 |
| ENSG000002 | 9.17681482 | -0.461191823 | 0.3408242 | -1.3531663 | 0.1760025 | 0.29676862 | NA | NA |
| ENSG000002 | 4.62913802 | -0.512233777 | 0.37858232 | -1.3530314 | 0.17604559 | 0.29678326 | NA | NA |
| ENSG000002 | 24.9186678 | -0.386500604 | 0.28569041 | -1.3528652 | 0.17609871 | 0.29686047 | FANK1 | 92565 |
| ENSG000002 | 3.62588271 | -0.511264524 | 0.37796814 | -1.3526657 | 0.17616246 | 0.29695003 | NA | NA |
| ENSG000002 | 2.3867021 | -0.501421433 | 0.37069477 | -1.3526531 | 0.17616648 | 0.29695003 | CYP4F35P | 284233 |
| ENSG000002 | 186.321808 | 0.207340257 | 0.15328763 | 1.35262225 | 0.17617634 | 0.29695431 | FAM200A | 221786 |
| ENSG000002 | 26.3663988 | -0.370030306 | 0.27357797 | -1.3525588 | 0.17619661 | 0.29697613 | NA | NA |
| ENSG000002 | 3.47498766 | -0.504751833 | 0.37319432 | -1.3525175 | 0.17620983 | 0.29698607 | LOC105369785 | 105369785 |
| ENSG000000 | 20.4136929 | -0.383315077 | 0.28342969 | -1.3524168 | 0.17624202 | 0.29702152 | AHRR | 57491 |
| ENSG000001 | 8.5724172 | -0.478530165 | 0.35383621 | -1.3524059 | 0.17624551 | 0.29702152 | TRIM60 | 166655 |
| ENSG000002 | 3.58527836 | -0.511328403 | 0.37809863 | -1.3523678 | 0.1762577 | 0.29702971 | NA | NA |
| ENSG000002 | 5.05340245 | -0.508954295 | 0.3763574 | -1.3523164 | 0.17627411 | 0.29704502 | NA | NA |
| ENSG000001 | 39.6499878 | -0.313500627 | 0.2318748 | -1.3520254 | 0.17636719 | 0.29718952 | CSF2RA | 1438 |
| ENSG000002 | 6.09479763 | -0.502012 | 0.37133203 | -1.3519222 | 0.17640021 | 0.29722305 | NA | NA |
| ENSG000002 | 2.37599592 | -0.500486834 | 0.37020519 | -1.3519174 | 0.17640174 | 0.29722305 | NA | NA |
| ENSG000001 | 128.23688 | 0.245314295 | 0.18146135 | 1.3518818 | 0.17641313 | 0.29722988 | NA | NA |

| | | | | | | | | |
|---|---|---|---|---|---|---|---|---|
| ENSG000002 | 2.40531496 | -0.500201674 | 0.37001451 | -1.3518434 | 0.17642542 | 0.29723824 | NA | NA |
| ENSG000000 | 4.8197402 | -0.510579065 | 0.37772497 | -1.3517218 | 0.17646434 | 0.2972791 | CEACAM21 | 90273 |
| ENSG000002 | 3.21536808 | -0.51127121 | 0.37823583 | -1.351726 | 0.17646298 | 0.2972791 | NA | NA |
| ENSG000002 | 2.98958173 | -0.508346588 | 0.3760952 | -1.3516434 | 0.17648943 | 0.29730902 | NA | NA |
| ENSG000000 | 22.1697492 | -0.386737853 | 0.28613905 | -1.3515732 | 0.1765119 | 0.29733453 | OVGP1 | 5016 |
| ENSG000001 | 3.93458336 | 0.511486469 | 0.37845291 | 1.35151946 | 0.17652909 | 0.29735113 | ADRA1D | 146 |
| ENSG000002 | 4.96776283 | -0.506744062 | 0.37498541 | -1.3513701 | 0.17657692 | 0.29741934 | CCDC179 | 100500938 |
| ENSG000002 | 3.27675227 | -0.510549685 | 0.37784585 | -1.3512116 | 0.17662767 | 0.29749247 | NA | NA |
| ENSG000002 | 5.24788735 | -0.504339328 | 0.37326265 | -1.3511647 | 0.17664267 | 0.29750538 | NA | NA |
| ENSG000002 | 2.79619986 | -0.502993901 | 0.37228125 | -1.3511126 | 0.17665936 | 0.29752112 | NA | NA |
| ENSG000002 | 4.92046833 | -0.507205205 | 0.37542545 | -1.3510144 | 0.1766908 | 0.29756173 | NA | NA |
| ENSG000002 | 3.20688618 | -0.510721902 | 0.3780416 | -1.3509675 | 0.17670585 | 0.29757471 | NA | NA |
| ENSG000001 | 4.09509143 | -0.510431617 | 0.3778491 | -1.3508875 | 0.17673148 | 0.29760551 | KNCN | 148930 |
| ENSG000002 | 4.10172513 | -0.511030269 | 0.37830643 | -1.3508369 | 0.17674769 | 0.29762045 | NA | NA |
| ENSG000001 | 12.8488823 | -0.433833415 | 0.32119581 | -1.3506821 | 0.1767973 | 0.29762199 | FIGF | 2277 |
| ENSG000002 | 2.21404004 | -0.48635114 | 0.36007755 | -1.3506844 | 0.17679654 | 0.29762199 | NA | NA |
| ENSG000002 | 3.232611 | -0.510917872 | 0.37823573 | -1.3507922 | 0.17676201 | 0.29762199 | NA | NA |
| ENSG000002 | 6.3113894 | -0.494668578 | 0.36621308 | -1.3507671 | 0.17677005 | 0.29762199 | NA | NA |
| ENSG000002 | 7.1601525 | -0.501284667 | 0.37113677 | -1.3506737 | 0.17679998 | 0.29762199 | TMEM30C | 644444 |
| ENSG000002 | 5.52606799 | -0.501333944 | 0.37116416 | -1.3507068 | 0.17678938 | 0.29762199 | NA | NA |
| ENSG000002 | 136.981194 | 0.23403117 | 0.17326013 | 1.35075029 | 0.17677544 | 0.29762199 | NA | NA |
| ENSG000001 | 164.690358 | 0.210211392 | 0.1556391 | 1.35063352 | 0.17681286 | 0.29763131 | GPHN | 10243 |
| ENSG000002 | 10.3583558 | -0.457259874 | 0.33858156 | -1.3505162 | 0.17685047 | 0.29768226 | NA | NA |
| ENSG000001 | 11.7864613 | -0.439836854 | 0.32572466 | -1.3503333 | 0.17690909 | 0.29774095 | NOTUM | 147111 |
| ENSG000002 | 14.1412078 | -0.418866936 | 0.3102045 | -1.3502929 | 0.17692205 | 0.29774095 | NA | NA |
| ENSG000002 | 43.1119282 | 0.307976551 | 0.22807668 | 1.35032021 | 0.17691329 | 0.29774095 | NA | NA |
| ENSG000002 | 6.62026297 | -0.495222396 | 0.36674651 | -1.3503125 | 0.17691577 | 0.29774095 | NA | NA |
| ENSG000002 | 5.89175501 | -0.505953746 | 0.37468693 | -1.3503373 | 0.17690781 | 0.29774095 | NA | NA |
| ENSG000001 | 40.2081604 | 0.312421857 | 0.23138548 | 1.35022237 | 0.17694466 | 0.2977619 | CYB5D1 | 124637 |

| | | | | | | | | |
|---|---|---|---|---|---|---|---|---|
| ENSG000002 | 7.01029558 | -0.484749223 | 0.35901811 | -1.3502083 | 0.17694918 | 0.2977619 | NA | NA |
| ENSG000001 | 108.149575 | 0.228224482 | 0.16903965 | 1.35012398 | 0.17697622 | 0.29777933 | CST3 | 1471 |
| ENSG000001 | 49.1983418 | -0.282714834 | 0.20940403 | -1.3500926 | 0.17698628 | 0.29777933 | PSD4 | 23550 |
| ENSG000001 | 15.4699003 | -0.428514592 | 0.31739837 | -1.3500844 | 0.17698891 | 0.29777933 | NA | NA |
| ENSG000002 | 6.27569636 | -0.494279113 | 0.36610805 | -1.3500908 | 0.17698686 | 0.29777933 | NA | NA |
| ENSG000002 | 2.43261063 | -0.497782004 | 0.36872832 | -1.3499967 | 0.17701705 | 0.29781431 | NA | NA |
| ENSG000002 | 4.35285142 | -0.508870267 | 0.37695139 | -1.3499626 | 0.17702799 | 0.29782037 | NA | NA |
| ENSG000002 | 3.0756389 | -0.506702217 | 0.3753671 | -1.3498845 | 0.17705304 | 0.29785015 | NA | NA |
| ENSG000002 | 4.78161582 | -0.508982434 | 0.37709206 | -1.3497564 | 0.17709413 | 0.29790691 | NA | NA |
| ENSG000002 | 4.72344793 | -0.508692144 | 0.37688399 | -1.3497314 | 0.17710216 | 0.29790807 | NA | NA |
| ENSG000002 | 2.41911325 | -0.499821346 | 0.37033328 | -1.3496528 | 0.17712738 | 0.29793813 | NA | NA |
| ENSG000002 | 7.30723454 | -0.480419919 | 0.35601326 | -1.3494439 | 0.17719443 | 0.29803854 | NA | NA |
| ENSG000001 | 22.0563299 | -0.383125851 | 0.28395101 | -1.3492674 | 0.17725108 | 0.29811155 | BAAT | 570 |
| ENSG000002 | 2.38273485 | -0.499156426 | 0.36994749 | -1.3492629 | 0.17725253 | 0.29811155 | NA | NA |
| ENSG000002 | 6.95095021 | -0.492078403 | 0.36473122 | -1.3491535 | 0.17728767 | 0.29812118 | NA | NA |
| ENSG000002 | 2.74121657 | -0.503113243 | 0.37289189 | -1.3492201 | 0.17726627 | 0.29812118 | NA | NA |
| ENSG000002 | 23.6244851 | -0.363111251 | 0.26913787 | -1.3491645 | 0.17728415 | 0.29812118 | NA | NA |
| ENSG000002 | 6.17376292 | -0.509653298 | 0.37774631 | -1.3491947 | 0.17727445 | 0.29812118 | NA | NA |
| ENSG000002 | 3.59828593 | -0.509400219 | 0.37758973 | -1.3490839 | 0.17731002 | 0.2981464 | NA | NA |
| ENSG000001 | 55.6059433 | 0.276367846 | 0.20487648 | 1.34894861 | 0.17735347 | 0.29818795 | RAB4B | 53916 |
| ENSG000002 | 2.38755087 | -0.500397673 | 0.37095668 | -1.3489383 | 0.17735679 | 0.29818795 | RNU6-640P | 106479850 |
| ENSG000002 | 3.25293906 | -0.509690146 | 0.37783498 | -1.3489755 | 0.17734483 | 0.29818795 | NA | NA |
| ENSG000001 | 4.35375281 | -0.507693623 | 0.37641314 | -1.348767 | 0.17741182 | 0.29826389 | S100A3 | 6274 |
| ENSG000002 | 7.17201715 | -0.490161928 | 0.36341889 | -1.3487519 | 0.17741667 | 0.29826389 | NA | NA |
| ENSG000002 | 188.655046 | 0.213777885 | 0.15851042 | 1.34866775 | 0.17744371 | 0.29829698 | NA | NA |
| ENSG000001 | 20.1726754 | 0.379694152 | 0.28154337 | 1.34861692 | 0.17746004 | 0.29829971 | CEACAM19 | 56971 |
| ENSG000002 | 4.60341578 | -0.506026109 | 0.3752177 | -1.34862 | 0.17745904 | 0.29829971 | NA | NA |
| ENSG000002 | 11.480567 | -0.450277132 | 0.33388677 | -1.3485923 | 0.17746795 | 0.29830064 | NA | NA |
| ENSG000001 | 266.796579 | 0.181333103 | 0.13446698 | 1.34853258 | 0.17748715 | 0.29832054 | PRTG | 283659 |

| | | | | | | | | |
|---|---|---|---|---|---|---|---|---|
| ENSG000002 | 5.35110257 | -0.505652687 | 0.37499749 | -1.3484162 | 0.17752456 | 0.29835869 | NA | NA |
| ENSG000002 | 4.11334369 | -0.510323732 | 0.37845832 | -1.3484278 | 0.17752082 | 0.29835869 | NA | NA |
| ENSG000001 | 10.8088258 | -0.448669127 | 0.33277226 | -1.3482768 | 0.17756936 | 0.29840127 | WNT2 | 7472 |
| ENSG000002 | 2.80925287 | -0.498401106 | 0.36964825 | -1.3483118 | 0.17755811 | 0.29840127 | NA | NA |
| ENSG000002 | 46.4460306 | 0.297939461 | 0.22097929 | 1.34826871 | 0.17757197 | 0.29840127 | NA | NA |
| ENSG000002 | 5.19988984 | -0.503617039 | 0.37355785 | -1.3481635 | 0.17760581 | 0.29844577 | NA | NA |
| ENSG000002 | 4.10807973 | -0.509965891 | 0.37828593 | -1.3480964 | 0.17762738 | 0.29846965 | NA | NA |
| ENSG000001 | 39.9896859 | 0.341360903 | 0.25322429 | 1.34805749 | 0.17763989 | 0.2984783 | NA | NA |
| ENSG000001 | 6.38197358 | -0.500317071 | 0.37116231 | -1.3479738 | 0.17766681 | 0.29851116 | FCN3 | 8547 |
| ENSG000001 | 81.4855592 | -0.255472621 | 0.18954185 | -1.3478428 | 0.17770895 | 0.2985696 | SEMA4F | 10505 |
| ENSG000002 | 5.25546388 | -0.506103229 | 0.37550231 | -1.3478032 | 0.17772167 | 0.2985786 | NA | NA |
| ENSG000001 | 144.063159 | 0.209423858 | 0.155397 | 1.34766993 | 0.17776457 | 0.29863829 | C11orf49 | 79096 |
| ENSG000001 | 17.8432891 | -0.390339806 | 0.28965289 | -1.3476123 | 0.17778311 | 0.29864616 | CD93 | 22918 |
| ENSG000002 | 8.01841853 | -0.481811653 | 0.35753282 | -1.3476012 | 0.17778669 | 0.29864616 | NA | NA |
| ENSG000002 | 21.1532252 | -0.369074199 | 0.27387789 | -1.3475867 | 0.17779135 | 0.29864616 | NA | NA |
| ENSG000002 | 2.99806309 | -0.506487848 | 0.37585577 | -1.3475591 | 0.17780025 | 0.29864873 | NA | NA |
| ENSG000002 | 2.3511532 | -0.50011733 | 0.37116648 | -1.3474205 | 0.17784483 | 0.29871125 | NA | NA |
| ENSG000001 | 6.765128 | -0.485769156 | 0.36059152 | -1.3471453 | 0.17793345 | 0.29884772 | OR5T2 | 219464 |
| ENSG00000C | 8.25351519 | -0.477715221 | 0.35464739 | -1.3470146 | 0.17797552 | 0.29886886 | CD209 | 30835 |
| ENSG000002 | 2.43738643 | -0.497801473 | 0.36955541 | -1.347028 | 0.17797122 | 0.29886886 | NA | NA |
| ENSG000002 | 32.4260302 | 0.321489403 | 0.23866529 | 1.34703041 | 0.17797044 | 0.29886886 | NA | NA |
| ENSG000002 | 11.5053205 | -0.4375265 | 0.32479948 | -1.3470665 | 0.17795881 | 0.29886886 | NA | NA |
| ENSG000001 | 2.37480279 | -0.499256405 | 0.3706646 | -1.3469223 | 0.17800527 | 0.29890645 | NA | NA |
| ENSG000001 | 28.5434434 | 0.336319906 | 0.24971699 | 1.34680427 | 0.17804328 | 0.29895789 | FAM162B | 221303 |
| ENSG000001 | 499.749559 | 0.166697986 | 0.12378062 | 1.34672117 | 0.17807005 | 0.29899046 | TAB3 | 257397 |
| ENSG000002 | 2.74589236 | -0.499137461 | 0.37064487 | -1.3466731 | 0.17808555 | 0.29900411 | CYMP | 643160 |
| ENSG000001 | 9.09556918 | -0.474568051 | 0.35241035 | -1.3466348 | 0.17809787 | 0.29901242 | NA | NA |
| ENSG000001 | 65.3203886 | -0.257243571 | 0.19103327 | -1.3465905 | 0.17811217 | 0.29901347 | TEX14 | 56155 |
| ENSG000002 | 5.97320349 | -0.493882929 | 0.36677265 | -1.3465642 | 0.17812062 | 0.29901347 | LY6G6C | 80740 |

| | | | | | | | | |
|---|---|---|---|---|---|---|---|---|
| ENSG000002 | 7.7566483 | -0.489531603 | 0.36354006 | -1.3465685 | 0.17811924 | 0.29901347 | NA | NA |
| ENSG000001 | 9.38873742 | -0.45704359 | 0.33943047 | -1.3465014 | 0.17814088 | 0.29903243 | MUCL1 | 118430 |
| ENSG000002 | 66.8266293 | 0.258673592 | 0.19211049 | 1.34648341 | 0.17814667 | 0.29903243 | BMS1P5 | 399761 |
| ENSG000002 | 33.6057773 | -0.318740642 | 0.23674188 | -1.3463636 | 0.17818529 | 0.29908488 | LOC284577 | 284577 |
| ENSG000002 | 14.8644331 | -0.409452605 | 0.30412499 | -1.34633 | 0.17819611 | 0.29909066 | NA | NA |
| ENSG000002 | 9.16242268 | -0.459953957 | 0.34164394 | -1.3462963 | 0.17820699 | 0.29909655 | NA | NA |
| ENSG000002 | 3.37088765 | 0.507683751 | 0.37712041 | 1.34621127 | 0.17823439 | 0.29913015 | NA | NA |
| ENSG000001 | 162.900242 | 0.202456673 | 0.15040112 | 1.34611151 | 0.17826656 | 0.29917175 | LOC100128398 | 100128398 |
| ENSG000002 | 4.14444513 | -0.508961836 | 0.37814527 | -1.3459426 | 0.17832102 | 0.29925078 | NA | NA |
| ENSG000001 | 131.361856 | 0.212428694 | 0.15784867 | 1.34577432 | 0.17837531 | 0.29932949 | GPLD1 | 2822 |
| ENSG000002 | 7.55465397 | -0.476435146 | 0.35404046 | -1.3457082 | 0.17839665 | 0.29935291 | NA | NA |
| ENSG000002 | 5.22896093 | -0.502351406 | 0.37332023 | -1.3456314 | 0.17842143 | 0.29938211 | NA | NA |
| ENSG00000C | 17.3772398 | -0.394241499 | 0.29304404 | -1.3453319 | 0.17851807 | 0.29953187 | ARSF | 416 |
| ENSG000002 | 9.743003 | -0.473201126 | 0.35179962 | -1.3450871 | 0.17859711 | 0.29965209 | TRIM40 | 135644 |
| ENSG000001 | 121.240998 | -0.209917613 | 0.15607197 | -1.3450052 | 0.17862355 | 0.29967539 | SEMA6D | 80031 |
| ENSG000002 | 6.97856394 | -0.492726869 | 0.36634161 | -1.3449929 | 0.17862752 | 0.29967539 | NA | NA |
| ENSG000002 | 14.2507512 | -0.412290622 | 0.30654138 | -1.3449754 | 0.17863317 | 0.29967539 | NA | NA |
| ENSG000002 | 14.8905939 | -0.411709495 | 0.30612108 | -1.3449237 | 0.17864988 | 0.29969103 | NA | NA |
| ENSG000001 | 37.2385918 | -0.306864855 | 0.22818023 | -1.3448354 | 0.17867839 | 0.29972645 | CPLX2 | 10814 |
| ENSG00000C | 1106.04277 | 0.169634729 | 0.12614606 | 1.34474858 | 0.17870644 | 0.29974514 | TBC1D23 | 55773 |
| ENSG000002 | 21.5792907 | 0.367467234 | 0.27326423 | 1.34473229 | 0.1787117 | 0.29974514 | NA | NA |
| ENSG000002 | 5.43626471 | -0.498902436 | 0.37100044 | -1.3447489 | 0.17870633 | 0.29974514 | NA | NA |
| ENSG000002 | 2.3323842 | 0.489158015 | 0.36379715 | 1.34459002 | 0.17875767 | 0.29980873 | MIR6835 | 102465502 |
| ENSG000002 | 2.70387262 | -0.500834113 | 0.37248668 | -1.3445692 | 0.1787644 | 0.29980873 | NA | NA |
| ENSG000001 | 2854.12367 | 0.150315562 | 0.11182209 | 1.34423852 | 0.17887127 | 0.29997166 | SKA3 | 221150 |
| ENSG000002 | 4.65757809 | -0.504930711 | 0.37563022 | -1.3442228 | 0.17887635 | 0.29997166 | NA | NA |
| ENSG000001 | 28.3058325 | -0.354511597 | 0.2637451 | -1.3441448 | 0.17890157 | 0.29999716 | SLC38A11 | 151258 |
| ENSG000001 | 8.88167091 | -0.464362567 | 0.34547444 | -1.34413 | 0.17890635 | 0.29999716 | RESP18 | 389075 |
| ENSG000002 | 4.08034741 | -0.507916628 | 0.37792032 | -1.3439781 | 0.17895547 | 0.30006713 | NA | NA |

| | | | | | | | | | |
|---|---|---|---|---|---|---|---|---|---|
| ENSG000002 | 3.84866849 | -0.508785197 | 0.37858228 | -1.3439224 | 0.17897349 | 0.30008493 | NA | NA | |
| ENSG000002 | 4.07516612 | -0.508247969 | 0.37822268 | -1.3437797 | 0.17901963 | 0.30014988 | NA | NA | |
| ENSG000002 | 3.51640904 | -0.508673354 | 0.37859338 | -1.3435876 | 0.17908179 | 0.30024169 | NA | NA | |
| ENSG000002 | 30.6472583 | 0.3354918 | 0.24974134 | 1.34335711 | 0.17915637 | 0.3003543 | NA | NA | |
| ENSG000001 | 77.0412378 | -0.246645047 | 0.18361558 | -1.3432687 | 0.179185 | 0.30038988 | ACVR1C | | 130399 |
| ENSG000001 | 7.75555753 | -0.48324707 | 0.35977948 | -1.3431758 | 0.17921507 | 0.30042787 | PAPL | | 390928 |
| ENSG000001 | 5.73691336 | -0.496063003 | 0.36933534 | -1.3431236 | 0.17923196 | 0.30044377 | C9orf117 | | 286207 |
| ENSG000002 | 15.1148689 | 0.406643415 | 0.30277473 | 1.34305599 | 0.17925385 | 0.30046803 | NA | NA | |
| ENSG000000 | 13.8065813 | -0.433278075 | 0.32262309 | -1.3429853 | 0.17927672 | 0.30049396 | ITIH1 | | 3697 |
| ENSG000002 | 4.4029429 | -0.506841019 | 0.37740973 | -1.3429463 | 0.17928935 | 0.30050271 | NA | NA | |
| ENSG000002 | 13.0136039 | -0.426674009 | 0.31772302 | -1.3429118 | 0.17930052 | 0.30050901 | NA | NA | |
| ENSG000002 | 16.6567509 | 0.395626784 | 0.29463804 | 1.34275527 | 0.17935123 | 0.30058157 | ABHD14A | | 25864 |
| ENSG000002 | 2.71164957 | -0.500431107 | 0.37273904 | -1.3425777 | 0.17940877 | 0.30066558 | NA | NA | |
| ENSG000002 | 4.11453323 | -0.507641458 | 0.3781194 | -1.3425427 | 0.17942008 | 0.30067211 | NA | NA | |
| ENSG000001 | 9.42603032 | -0.46212193 | 0.34425955 | -1.3423649 | 0.17947772 | 0.30070962 | SERPINB2 | | 5055 |
| ENSG000002 | 12.508196 | -0.444366623 | 0.33102406 | -1.3423998 | 0.17946641 | 0.30070962 | NA | NA | |
| ENSG000002 | 2.40371689 | -0.497610428 | 0.37069406 | -1.342375 | 0.17947445 | 0.30070962 | NA | NA | |
| ENSG000002 | 3.68101832 | -0.507043276 | 0.37772547 | -1.3423592 | 0.17947954 | 0.30070962 | NA | NA | |
| ENSG000002 | 5.21301563 | -0.502003594 | 0.37395094 | -1.3424317 | 0.17945606 | 0.30070962 | NA | NA | |
| ENSG000002 | 6.15788547 | -0.499938308 | 0.37244411 | -1.3423177 | 0.17949301 | 0.30071977 | NA | NA | |
| ENSG000000 | 112.76881 | 0.240291309 | 0.17903387 | 1.34215562 | 0.17954554 | 0.30079535 | TNK2 | | 10188 |
| ENSG000002 | 13.3643208 | -0.422907186 | 0.31511326 | -1.34208 | 0.17957007 | 0.30082401 | LOC101929653 | | 101929653 |
| ENSG000001 | 45.1874327 | 0.323387131 | 0.24099028 | 1.34190943 | 0.17962536 | 0.30090302 | METRNL | | 284207 |
| ENSG000002 | 6.61623692 | -0.501568165 | 0.37377776 | -1.3418887 | 0.17963208 | 0.30090302 | NA | NA | |
| ENSG000002 | 13.3296744 | 0.423622746 | 0.3157044 | 1.34183354 | 0.17964997 | 0.30092057 | NA | NA | |
| ENSG000002 | 24.9926746 | -0.363306111 | 0.27079161 | -1.3416446 | 0.17971124 | 0.30101077 | LOC644838 | | 644838 |
| ENSG000002 | 26.1318498 | -0.347273234 | 0.25884788 | -1.3416113 | 0.17972207 | 0.30101647 | NA | NA | |
| ENSG000002 | 14.3667521 | -0.417616681 | 0.31134791 | -1.3413184 | 0.1798171 | 0.30116003 | SNCA-AS1 | | 644248 |
| ENSG000002 | 3.04549811 | -0.502469651 | 0.37461355 | -1.3413013 | 0.17982264 | 0.30116003 | OR2S2 | | 56656 |

| | | | | | | | | |
|---|---|---|---|---|---|---|---|---|
| ENSG000002 | 3.30035034 | -0.506728145 | 0.37782215 | -1.3411817 | 0.17986147 | 0.30121263 | NA | NA |
| ENSG000002 | 3.22971478 | -0.507100545 | 0.37811463 | -1.3411291 | 0.17987853 | 0.30121631 | NA | NA |
| ENSG000002 | 5.5346933 | -0.503534266 | 0.37544915 | -1.3411517 | 0.1798712 | 0.30121631 | NA | NA |
| ENSG000002 | 12.7735206 | -0.435521734 | 0.32477723 | -1.3409861 | 0.17992498 | 0.30128165 | NA | NA |
| ENSG000002 | 2.4179711 | -0.495062512 | 0.36923905 | -1.3407642 | 0.17999701 | 0.30138982 | NA | NA |
| ENSG000002 | 4.69842861 | -0.505307312 | 0.37688691 | -1.3407399 | 0.18000492 | 0.30139062 | NA | NA |
| ENSG000002 | 5.98472565 | -0.492033525 | 0.36703441 | -1.3405651 | 0.1800617 | 0.30147324 | NA | NA |
| ENSG000001 | 62.3129269 | 0.431678899 | 0.32204278 | 1.34043961 | 0.18010247 | 0.30152905 | GNG7 | 2788 |
| ENSG000002 | 4.32423788 | -0.50468408 | 0.37654074 | -1.3403173 | 0.1801422 | 0.30157067 | NA | NA |
| ENSG000002 | 2.48190195 | -0.489970598 | 0.36556058 | -1.3403267 | 0.18013916 | 0.30157067 | NA | NA |
| ENSG000002 | 4.35780328 | -0.505353547 | 0.37706858 | -1.3402165 | 0.18017496 | 0.30161306 | NA | NA |
| ENSG000002 | 29.4123873 | 0.329465427 | 0.24583626 | 1.34018239 | 0.18018605 | 0.30161918 | LOC100652758 | 100652758 |
| ENSG000002 | 9.38397458 | -0.454203347 | 0.33892292 | -1.3401376 | 0.18020062 | 0.30163111 | NA | NA |
| ENSG000002 | 17.4017018 | 0.393154088 | 0.2933825 | 1.34007341 | 0.18022148 | 0.30165358 | NA | NA |
| ENSG000002 | 9.16417502 | -0.461755731 | 0.34458956 | -1.3400166 | 0.18023995 | 0.30167204 | NA | NA |
| ENSG00000C | 96.4122217 | 0.234521644 | 0.17503442 | 1.33986016 | 0.18029081 | 0.3017367 | PIGB | 9488 |
| ENSG000001 | 9.16011269 | -0.458699193 | 0.34235649 | -1.3398291 | 0.1803009 | 0.3017367 | HAMP | 57817 |
| ENSG000002 | 3.83957466 | -0.507254127 | 0.37859301 | -1.3398402 | 0.18029731 | 0.3017367 | NA | NA |
| ENSG000002 | 15.3385937 | -0.429668532 | 0.32074727 | -1.3395859 | 0.18038 | 0.30185661 | LOC284395 | 284395 |
| ENSG000002 | 3.41025057 | -0.500846013 | 0.37394129 | -1.3393707 | 0.18045002 | 0.30196132 | TCEB1P22 | 100287579 |
| ENSG000001 | 6.87691631 | -0.486579381 | 0.36329684 | -1.3393438 | 0.18045876 | 0.30196349 | NA | NA |
| ENSG000002 | 6.33122819 | -0.492243442 | 0.36754931 | -1.3392582 | 0.18048662 | 0.30199764 | NA | NA |
| ENSG000002 | 7.03981621 | -0.4939019 | 0.36880506 | -1.339195 | 0.18050719 | 0.3020196 | NA | NA |
| ENSG000002 | 3.50202955 | -0.506946017 | 0.37857012 | -1.3391073 | 0.18053574 | 0.30204846 | NA | NA |
| ENSG000002 | 11.0501969 | -0.446801277 | 0.33365882 | -1.3390963 | 0.18053933 | 0.30204846 | NA | NA |
| ENSG000002 | 3.22170895 | -0.505708022 | 0.37772841 | -1.3388138 | 0.18063129 | 0.30218983 | LINC00565 | 100861555 |
| ENSG000002 | 4.63623187 | -0.502673423 | 0.37552011 | -1.3386059 | 0.18069901 | 0.30229066 | NA | NA |
| ENSG000002 | 7.35346871 | -0.479545539 | 0.35826613 | -1.3385177 | 0.18072775 | 0.30232626 | LOC101928517 | 101928517 |
| ENSG000001 | 29.6689409 | -0.331012076 | 0.24730216 | -1.3384924 | 0.18073596 | 0.30232753 | ST8SIA2 | 8128 |

| | | | | | | | | |
|---|---|---|---|---|---|---|---|---|
| ENSG000002 | 5.97540193 | 0.491236517 | 0.36702302 | 1.33843517 | 0.18075462 | 0.30234627 | LOC100131655 | 100131655 |
| ENSG000001 | 108.492063 | -0.222612406 | 0.16634256 | -1.3382769 | 0.1808062 | 0.30242006 | LRRC8C | 84230 |
| ENSG000002 | 11.6535242 | -0.45966946 | 0.34350074 | -1.3381906 | 0.18083431 | 0.30245461 | LINC01416 | 101927273 |
| ENSG000002 | 3.54800413 | 0.506421445 | 0.37851074 | 1.33793151 | 0.18091877 | 0.30257091 | NA | NA |
| ENSG000002 | 13.6291796 | -0.420450656 | 0.3142506 | -1.337947 | 0.1809137 | 0.30257091 | NA | NA |
| ENSG000002 | 2.39204571 | -0.495323254 | 0.37026152 | -1.3377659 | 0.18097275 | 0.30264871 | RNU6-610P | 106479835 |
| ENSG000001 | 3.55161248 | -0.506335908 | 0.37855167 | -1.3375609 | 0.18103962 | 0.30274806 | TMEM95 | 339168 |
| ENSG000001 | 3.90420108 | -0.506299474 | 0.37854204 | -1.3374987 | 0.18105992 | 0.30276952 | FCER1A | 2205 |
| ENSG000002 | 2.96331611 | -0.502438448 | 0.37566836 | -1.3374521 | 0.18107509 | 0.30278075 | NA | NA |
| ENSG000002 | 3.53515884 | -0.506311809 | 0.37858949 | -1.3373636 | 0.18110397 | 0.30278075 | NA | NA |
| ENSG000002 | 3.06952094 | -0.50212487 | 0.37544819 | -1.3374012 | 0.18109171 | 0.30278075 | NA | NA |
| ENSG000002 | 2.71358046 | -0.497585177 | 0.37204825 | -1.3374211 | 0.18108523 | 0.30278075 | FBXO3-AS1 | 101928440 |
| ENSG000002 | 10.0145487 | -0.458781934 | 0.34304874 | -1.3373666 | 0.181103 | 0.30278075 | NA | NA |
| ENSG000002 | 5.70056756 | -0.495082134 | 0.37021202 | -1.3372935 | 0.18112685 | 0.30280651 | NA | NA |
| ENSG000001 | 5.42964234 | -0.496451675 | 0.37129994 | -1.3370637 | 0.18120185 | 0.30291941 | LY96 | 23643 |
| ENSG000001 | 30.9762772 | -0.336823087 | 0.25192347 | -1.3370056 | 0.18122081 | 0.30293862 | TCTEX1D1 | 200132 |
| ENSG000001 | 126.104335 | -0.227167954 | 0.16993233 | -1.3368142 | 0.18128328 | 0.3029997 | SCN1A | 6323 |
| ENSG000001 | 36.816366 | -0.322764684 | 0.24143575 | -1.3368554 | 0.18126984 | 0.3029997 | PPIL6 | 285755 |
| ENSG000002 | 7.79570408 | 0.47586399 | 0.3559719 | 1.33680212 | 0.18128724 | 0.3029997 | NA | NA |
| ENSG000002 | 5.26395552 | -0.50041924 | 0.37433997 | -1.3368042 | 0.18128656 | 0.3029997 | LOC284661 | 284661 |
| ENSG000001 | 115.797864 | 0.209584699 | 0.15679923 | 1.33664364 | 0.18133899 | 0.30304237 | C1orf54 | 79630 |
| ENSG000001 | 53.7890398 | 0.28246522 | 0.21132603 | 1.33663239 | 0.18134266 | 0.30304237 | RHEBL1 | 121268 |
| ENSG000002 | 3.28108424 | -0.505048061 | 0.3778345 | -1.3366912 | 0.18132345 | 0.30304237 | NA | NA |
| ENSG000002 | 5.43954661 | -0.496336359 | 0.37132457 | -1.3366645 | 0.18133219 | 0.30304237 | LOC146513 | 146513 |
| ENSG000002 | 9.7893874 | 0.464535625 | 0.34764495 | 1.3362358 | 0.18147221 | 0.30324637 | MIR3153 | 100422936 |
| ENSG000002 | 5.3172694 | -0.500375577 | 0.37447759 | -1.3361963 | 0.18148511 | 0.30325543 | SIX3-AS1 | 100506108 |
| ENSG000002 | 8.02992076 | -0.480986814 | 0.36002524 | -1.3359808 | 0.18155554 | 0.30336062 | RAB11AP2 | 106480447 |
| ENSG000002 | 22.2820712 | -0.353669416 | 0.26475601 | -1.3358315 | 0.18160436 | 0.30341718 | LINC00858 | 170425 |
| ENSG000002 | 4.32825556 | -0.503995489 | 0.37728794 | -1.3358378 | 0.18160229 | 0.30341718 | UGT1A1 | 54658 |

| | | | | | | | | |
|---|---|---|---|---|---|---|---|---|
| ENSG000002 | 7.87352007 | -0.476825495 | 0.35697164 | -1.3357518 | 0.18163043 | 0.30344823 | NA | NA |
| ENSG000001 | 28.6479337 | 0.327949186 | 0.24555478 | 1.33554388 | 0.1816984 | 0.30354811 | CYP27B1 | 1594 |
| ENSG000002 | 4.47218435 | -0.50539692 | 0.37842618 | -1.3355232 | 0.18170518 | 0.30354811 | NA | NA |
| ENSG000002 | 4.1556217 | 0.504282903 | 0.37764488 | 1.33533627 | 0.18176631 | 0.30363772 | C11orf94 | 143678 |
| ENSG000001 | 13.3491239 | -0.444358671 | 0.33281698 | -1.3351442 | 0.18182914 | 0.30373017 | CPNE6 | 9362 |
| ENSG000002 | 4.674521 | -0.502436711 | 0.37634055 | -1.3350587 | 0.18185715 | 0.30373331 | NA | NA |
| ENSG000002 | 6.49952996 | -0.496294136 | 0.37172725 | -1.3351029 | 0.18184266 | 0.30373331 | DLGAP1-AS4 | 101410534 |
| ENSG000002 | 4.32281553 | -0.502927773 | 0.37671168 | -1.3350469 | 0.18186098 | 0.30373331 | NA | NA |
| ENSG000002 | 4.64636447 | -0.501024294 | 0.37528045 | -1.3350663 | 0.18185463 | 0.30373331 | NA | NA |
| ENSG000002 | 2.40767095 | -0.494371276 | 0.3703117 | -1.3350139 | 0.1818718 | 0.30373886 | LOC105377924 | 105377924 |
| ENSG000001 | 5.42016264 | -0.496855401 | 0.37218875 | -1.3349554 | 0.18189093 | 0.3037583 | KRT16 | 3868 |
| ENSG000002 | 3.86362123 | -0.505310248 | 0.37857737 | -1.3347608 | 0.18195464 | 0.30384718 | NA | NA |
| ENSG000002 | 9.44292044 | -0.453854154 | 0.34003008 | -1.3347471 | 0.18195914 | 0.30384718 | LOC728040 | 728040 |
| ENSG000001 | 13.065942 | -0.422972671 | 0.31692885 | -1.3345982 | 0.18200789 | 0.30390356 | SSTR1 | 6751 |
| ENSG000002 | 4.11045388 | -0.50514554 | 0.37849773 | -1.3346065 | 0.18200517 | 0.30390356 | NA | NA |
| ENSG000002 | 4.38573831 | -0.504148175 | 0.37778586 | -1.3344813 | 0.18204616 | 0.30395495 | NA | NA |
| ENSG000002 | 2.40672935 | -0.491567785 | 0.36837296 | -1.3344296 | 0.1820631 | 0.30396724 | LOC149950 | 149950 |
| ENSG000002 | 2.79301004 | -0.495201528 | 0.37110062 | -1.3344131 | 0.18206852 | 0.30396724 | NA | NA |
| ENSG000002 | 2.36697432 | -0.494587616 | 0.37064953 | -1.3343808 | 0.18207909 | 0.30397238 | NA | NA |
| ENSG000002 | 2.62987464 | -0.500077774 | 0.37479238 | -1.3342795 | 0.18211229 | 0.30401529 | FAM66B | 100128890 |
| ENSG000001 | 5.95494216 | 0.490407792 | 0.36756647 | 1.33420165 | 0.18213778 | 0.30403281 | PPBP | 5473 |
| ENSG000002 | 7.14951292 | -0.481565309 | 0.3609382 | -1.3342043 | 0.1821369 | 0.30403281 | NA | NA |
| ENSG000002 | 3.53112026 | -0.505035051 | 0.37854355 | -1.3341531 | 0.18215369 | 0.30404685 | MIR644A | 693229 |
| ENSG000001 | 690.32382 | 0.152152546 | 0.11405704 | 1.33400402 | 0.18220254 | 0.30406581 | C11orf30 | 56946 |
| ENSG000001 | 8.1274988 | -0.470331291 | 0.35256703 | -1.3340195 | 0.18219747 | 0.30406581 | NA | NA |
| ENSG000002 | 4.34334001 | -0.503266757 | 0.3772518 | -1.3340341 | 0.18219269 | 0.30406581 | NA | NA |
| ENSG000002 | 4.39200206 | -0.503131792 | 0.37713555 | -1.3340874 | 0.18217521 | 0.30406581 | NA | NA |
| ENSG000002 | 5.32270943 | -0.50083401 | 0.37543475 | -1.3340108 | 0.18220031 | 0.30406581 | NA | NA |
| ENSG000002 | 6.6289526 | -0.488995417 | 0.36660311 | -1.3338551 | 0.18225136 | 0.30413476 | NA | NA |

| | | | | | | | | |
|---|---|---|---|---|---|---|---|---|
| ENSG000002 | 7.36859946 | -0.478247103 | 0.35857166 | -1.333756 | 0.18228383 | 0.30417644 | NA | NA |
| ENSG000002 | 22.8440293 | 0.376418311 | 0.28223431 | 1.33370856 | 0.18229939 | 0.30418988 | NA | NA |
| ENSG000002 | 2.39732286 | -0.492327953 | 0.36918574 | -1.3335508 | 0.18235111 | 0.30426366 | NA | NA |
| ENSG000001 | 20.3507554 | -0.371620966 | 0.27872294 | -1.3332988 | 0.18243377 | 0.30433983 | WFDC3 | 140686 |
| ENSG000001 | 9.99343858 | -0.452953862 | 0.33973739 | -1.333247 | 0.18245076 | 0.30433983 | NA | NA |
| ENSG000002 | 4.93862507 | -0.500928358 | 0.37573066 | -1.3332113 | 0.18246248 | 0.30433983 | PCDH9-AS2 | 100874064 |
| ENSG000002 | 15.783967 | -0.407202881 | 0.3054322 | -1.3332022 | 0.18246546 | 0.30433983 | NA | NA |
| ENSG000002 | 2.6495181 | -0.499135464 | 0.37436667 | -1.3332797 | 0.18244004 | 0.30433983 | NA | NA |
| ENSG000002 | 3.00782712 | -0.499234803 | 0.37441712 | -1.3333653 | 0.18241195 | 0.30433983 | NA | NA |
| ENSG000002 | 2.76231463 | -0.49508415 | 0.37133825 | -1.3332431 | 0.18245205 | 0.30433983 | NA | NA |
| ENSG000002 | 3.08057992 | -0.496898378 | 0.37270084 | -1.3332366 | 0.18245419 | 0.30433983 | NA | NA |
| ENSG000002 | 3.89132277 | 0.504727662 | 0.37858847 | 1.33318285 | 0.18247181 | 0.30433983 | NA | NA |
| ENSG000002 | 10.1631475 | 0.443272861 | 0.33248431 | 1.33321437 | 0.18246147 | 0.30433983 | NA | NA |
| ENSG000002 | 6.3002396 | -0.489775542 | 0.36739945 | -1.3330873 | 0.18250315 | 0.30437959 | DEFB121 | 245934 |
| ENSG000002 | 60.070939 | 0.286215588 | 0.21474081 | 1.33284206 | 0.18258364 | 0.3045013 | NA | NA |
| ENSG000002 | 23.4158779 | -0.36017985 | 0.27024427 | -1.3327937 | 0.18259952 | 0.30451526 | NA | NA |
| ENSG000002 | 12.4914675 | -0.429448074 | 0.32225375 | -1.3326395 | 0.18265014 | 0.30458715 | NA | NA |
| ENSG000002 | 7.51802872 | -0.470814796 | 0.35331043 | -1.3325811 | 0.18266932 | 0.3046066 | LINC00242 | 401288 |
| ENSG000002 | 86.108421 | 0.246382844 | 0.18491218 | 1.33243169 | 0.18271838 | 0.30467589 | NA | NA |
| ENSG000002 | 2.39874522 | -0.491859021 | 0.36919059 | -1.3322631 | 0.18277373 | 0.30475566 | RN7SL104P | 106480949 |
| ENSG000002 | 2.54660084 | -0.485150673 | 0.36419532 | -1.3321167 | 0.18282184 | 0.30481081 | NA | NA |
| ENSG000002 | 2.97491517 | -0.500444325 | 0.37567063 | -1.3321359 | 0.18281554 | 0.30481081 | RN7SL283P | 106479325 |
| ENSG00000C | 211.832541 | 0.212643438 | 0.15963162 | 1.33208848 | 0.18283112 | 0.30481118 | TIMP2 | 7077 |
| ENSG000001 | 330.830635 | 0.171500309 | 0.12874938 | 1.33204764 | 0.18284453 | 0.30481118 | CADPS | 8618 |
| ENSG000002 | 2.21065435 | -0.476672999 | 0.35784988 | -1.3320474 | 0.18284461 | 0.30481118 | NA | NA |
| ENSG000002 | 9.055453 | -0.467010731 | 0.35060701 | -1.3320063 | 0.18285813 | 0.30482118 | NA | NA |
| ENSG000002 | 6.2359283 | -0.486108432 | 0.36504419 | -1.3316427 | 0.18297762 | 0.30500783 | NA | NA |
| ENSG000002 | 3.06280574 | -0.497780454 | 0.37382724 | -1.3315789 | 0.18299861 | 0.30503027 | NA | NA |
| ENSG000002 | 3.32191672 | -0.502008306 | 0.37701871 | -1.3315209 | 0.18301766 | 0.30504949 | NA | NA |

| | | | | | | | | |
|---|---|---|---|---|---|---|---|---|
| ENSG000002 | 3.28438596 | -0.503181717 | 0.37791504 | -1.3314678 | 0.18303513 | 0.30505353 | RNU6-922P | 106479974 |
| ENSG000002 | 7.14287116 | -0.481534465 | 0.36165493 | -1.3314749 | 0.18303279 | 0.30505353 | NA | NA |
| ENSG000002 | 7.59983394 | -0.480775528 | 0.36109897 | -1.3314231 | 0.18304982 | 0.30506548 | NA | NA |
| ENSG000002 | 6.60992263 | 0.481207943 | 0.36144521 | 1.33134409 | 0.18307582 | 0.30509627 | NA | NA |
| ENSG000002 | 6.79438123 | -0.48283376 | 0.36268185 | -1.3312873 | 0.18309448 | 0.30510621 | NA | NA |
| ENSG000002 | 3.00657691 | -0.49933304 | 0.37507734 | -1.3312802 | 0.18309683 | 0.30510621 | NA | NA |
| ENSG000002 | 5.22440138 | -0.497320887 | 0.37358596 | -1.3312087 | 0.18312034 | 0.30513285 | NA | NA |
| ENSG000002 | 3.89815467 | -0.503960387 | 0.37858906 | -1.3311541 | 0.1831383 | 0.30515024 | NA | NA |
| ENSG000002 | 15.3551309 | -0.4028658 | 0.30267688 | -1.3310095 | 0.18318588 | 0.30521698 | NA | NA |
| ENSG000002 | 8.79084156 | -0.483044912 | 0.36294411 | -1.3309071 | 0.18321957 | 0.30526056 | NA | NA |
| ENSG000002 | 3.02740195 | -0.499626075 | 0.37542156 | -1.3308401 | 0.18324162 | 0.30527222 | NA | NA |
| ENSG000002 | 2.98528331 | -0.500470041 | 0.37605355 | -1.3308478 | 0.1832391 | 0.30527222 | NA | NA |
| ENSG000002 | 2.38291146 | -0.491707859 | 0.36953783 | -1.3306022 | 0.18331993 | 0.30536503 | UCA1 | 652995 |
| ENSG000002 | 5.58984872 | -0.498873618 | 0.37492142 | -1.3306085 | 0.18331788 | 0.30536503 | NA | NA |
| ENSG000002 | 120.216559 | 0.226055805 | 0.16988479 | 1.33064185 | 0.18330689 | 0.30536503 | NA | NA |
| ENSG000002 | 11.4510201 | -0.446704772 | 0.33574392 | -1.3304925 | 0.18335605 | 0.30541266 | NA | NA |
| ENSG000002 | 5.96369318 | -0.486873067 | 0.36594926 | -1.3304387 | 0.18337376 | 0.30542961 | TRIM31-AS1 | 104533120 |
| ENSG000001 | 3.31614367 | -0.501972804 | 0.37730907 | -1.3304022 | 0.1833858 | 0.30543712 | OR1L1 | 26737 |
| ENSG000002 | 30.2918064 | 0.334133026 | 0.25115756 | 1.33037217 | 0.18339568 | 0.30544103 | NA | NA |
| ENSG000002 | 2.4027191 | -0.49166711 | 0.36960138 | -1.3302632 | 0.18343157 | 0.30548826 | NA | NA |
| ENSG000001 | 8.93381811 | -0.459279957 | 0.34526589 | -1.3302211 | 0.18344545 | 0.30549882 | FOLR1 | 2348 |
| ENSG000002 | 9.24366497 | -0.46109426 | 0.34670081 | -1.3299486 | 0.1835352 | 0.30562866 | KCCAT211 | 102724550 |
| ENSG000002 | 3.24550343 | -0.502738274 | 0.37801614 | -1.3299386 | 0.18353849 | 0.30562866 | NA | NA |
| ENSG000002 | 4.9115768 | -0.499782098 | 0.37582294 | -1.3298339 | 0.183573 | 0.30567358 | LOC101928327 | 101928327 |
| ENSG000002 | 3.286348 | -0.501506128 | 0.37713748 | -1.3297701 | 0.18359404 | 0.30569607 | NA | NA |
| ENSG000001 | 88.603034 | 0.232549645 | 0.17490077 | 1.32960905 | 0.18364712 | 0.30577188 | TMEM182 | 130827 |
| ENSG000002 | 9.67979489 | 0.461612558 | 0.34723354 | 1.32940082 | 0.18371577 | 0.30587363 | NA | NA |
| ENSG000002 | 2.17440103 | -0.478440444 | 0.35992889 | -1.3292638 | 0.18376095 | 0.30592374 | RN7SL392P | 106480506 |
| ENSG000002 | 12.5350081 | -0.431746139 | 0.32479889 | -1.3292722 | 0.18375818 | 0.30592374 | NA | NA |

| | | | | | | | | |
|---|---|---|---|---|---|---|---|---|
| ENSG00000C | 5.61251295 | -0.498965307 | 0.37540689 | -1.329132 | 0.18380443 | 0.30596637 | CYP2W1 | 54905 |
| ENSG000002 | 2.61590063 | -0.498621748 | 0.37514745 | -1.3291354 | 0.18380331 | 0.30596637 | NA | NA |
| ENSG000002 | 10.9597415 | -0.453211502 | 0.34098678 | -1.3291175 | 0.1838092 | 0.30596637 | NA | NA |
| ENSG000002 | 8.12122671 | -0.483023929 | 0.36344739 | -1.3290065 | 0.18384584 | 0.3060148 | NA | NA |
| ENSG000002 | 9.45569083 | 0.459216067 | 0.34554261 | 1.32897086 | 0.18385758 | 0.30602179 | NA | NA |
| ENSG000002 | 6.20849658 | -0.483750597 | 0.36402113 | -1.328908 | 0.18387832 | 0.30604373 | NA | NA |
| ENSG00000C | 49.6050366 | 0.287624557 | 0.21644473 | 1.32885915 | 0.18389444 | 0.30604545 | FAM65A | 79567 |
| ENSG000002 | 7.43920855 | -0.478189705 | 0.35984969 | -1.3288596 | 0.1838943 | 0.30604545 | NA | NA |
| ENSG000001 | 19.5165501 | 0.389049772 | 0.29278897 | 1.32877196 | 0.18392321 | 0.30605641 | ZNF837 | 116412 |
| ENSG000002 | 422.551011 | 0.161810673 | 0.12177607 | 1.32875594 | 0.1839285 | 0.30605641 | RANBP17 | 64901 |
| ENSG000002 | 11.8984714 | -0.426404876 | 0.32092942 | -1.3286562 | 0.18396141 | 0.30605641 | NA | NA |
| ENSG000002 | 5.28808566 | -0.499442514 | 0.37588617 | -1.3287068 | 0.18394471 | 0.30605641 | NA | NA |
| ENSG000002 | 3.47530747 | -0.503012869 | 0.37857116 | -1.3287142 | 0.18394229 | 0.30605641 | NA | NA |
| ENSG000002 | 8.92289432 | 0.475160416 | 0.35760727 | 1.32872134 | 0.18393992 | 0.30605641 | NA | NA |
| ENSG000002 | 5.18751444 | 0.496183107 | 0.37344494 | 1.32866469 | 0.18395862 | 0.30605641 | NA | NA |
| ENSG000002 | 3.60060358 | -0.502244861 | 0.37800417 | -1.3286755 | 0.18395506 | 0.30605641 | NA | NA |
| ENSG000002 | 69.1856089 | 0.262368485 | 0.19749138 | 1.32850602 | 0.18401099 | 0.30612518 | CKMT2-AS1 | 100131067 |
| ENSG000002 | 2.63881301 | -0.497859267 | 0.37475708 | -1.3284853 | 0.18401784 | 0.30612518 | NA | NA |
| ENSG000002 | 2.37053607 | -0.491787543 | 0.37020878 | -1.328406 | 0.18404403 | 0.30614898 | NA | NA |
| ENSG000002 | 3.29250383 | -0.500004627 | 0.3763972 | -1.3283962 | 0.18404724 | 0.30614898 | NA | NA |
| ENSG000002 | 3.01630436 | -0.498091638 | 0.37497805 | -1.3283221 | 0.18407172 | 0.30617713 | NA | NA |
| ENSG000001 | 7.29130983 | -0.474151699 | 0.35699791 | -1.3281638 | 0.184124 | 0.30625154 | LILRA5 | 353514 |
| ENSG00000C | 7.02176676 | -0.47755904 | 0.35961596 | -1.3279695 | 0.18418817 | 0.30632094 | HOXA9 | 3205 |
| ENSG000001 | 44.1357343 | -0.32870503 | 0.24752465 | -1.3279689 | 0.18418839 | 0.30632094 | EFHB | 151651 |
| ENSG000002 | 130.457416 | 0.218651197 | 0.16464948 | 1.3279799 | 0.18418475 | 0.30632094 | ITPRIPL2 | 162073 |
| ENSG000002 | 4.61166902 | -0.49901748 | 0.37586554 | -1.3276489 | 0.18429411 | 0.30648419 | NA | NA |
| ENSG000002 | 3.49414955 | -0.502620198 | 0.37859339 | -1.327599 | 0.18431062 | 0.30649908 | NA | NA |
| ENSG000001 | 35.4411759 | 0.3459943 | 0.26067223 | 1.32731552 | 0.18440433 | 0.30664234 | BLOC1S3 | 388552 |
| ENSG000001 | 104.555758 | 0.240705131 | 0.18138886 | 1.32701163 | 0.18450483 | 0.30678399 | SRGAP2B | 647135 |

| | | | | | | | | |
|---|---|---|---|---|---|---|---|---|
| ENSG000002 | 4.10466178 | -0.502229032 | 0.37847008 | -1.326998 | 0.18450936 | 0.30678399 | NA | NA |
| ENSG000002 | 2.97323964 | -0.498274431 | 0.37549242 | -1.3269893 | 0.18451221 | 0.30678399 | NA | NA |
| ENSG000001 | 175.474126 | -0.196385466 | 0.14801273 | -1.3268147 | 0.18456999 | 0.30684231 | NABP1 | 64859 |
| ENSG000001 | 2.65779094 | -0.496736349 | 0.3743797 | -1.326825 | 0.18456657 | 0.30684231 | OR10S1 | 219873 |
| ENSG000002 | 5.46433691 | -0.494336639 | 0.37256686 | -1.3268401 | 0.18456159 | 0.30684231 | NA | NA |
| ENSG000001 | 129.380452 | 0.208489211 | 0.15713867 | 1.32678491 | 0.18457984 | 0.30684611 | PRSS12 | 8492 |
| ENSG000002 | 5.97835232 | -0.486743259 | 0.3668778 | -1.3267177 | 0.18460209 | 0.30685794 | NA | NA |
| ENSG000002 | 3.46489521 | -0.502285452 | 0.37859028 | -1.3267257 | 0.18459944 | 0.30685794 | NA | NA |
| ENSG000002 | 9.71396726 | -0.467689131 | 0.35256895 | -1.3265182 | 0.18466811 | 0.30694252 | NA | NA |
| ENSG000002 | 4.77285481 | -0.500595243 | 0.37736986 | -1.3265374 | 0.18466175 | 0.30694252 | NA | NA |
| ENSG000002 | 10.3386011 | 0.449621975 | 0.33897492 | 1.32641663 | 0.18470173 | 0.30697324 | NA | NA |
| ENSG000002 | 3.51239027 | -0.502170717 | 0.37858571 | -1.3264386 | 0.18469445 | 0.30697324 | NA | NA |
| ENSG000002 | 8.16734558 | -0.473716683 | 0.35716956 | -1.3263075 | 0.18473785 | 0.30702068 | NA | NA |
| ENSG000001 | 9.1439592 | -0.469169913 | 0.35375008 | -1.3262751 | 0.1847486 | 0.30702596 | LRRC19 | 64922 |
| ENSG000002 | 3.01105911 | -0.497613121 | 0.37520688 | -1.3262367 | 0.18476132 | 0.30703452 | LINC01040 | 105370308 |
| ENSG000001 | 160.250163 | 0.199103067 | 0.15014725 | 1.32605207 | 0.18482245 | 0.30712352 | GABPB2 | 126626 |
| ENSG000002 | 4.96539181 | -0.499132652 | 0.37644351 | -1.3259165 | 0.18486736 | 0.30717297 | NA | NA |
| ENSG000002 | 5.71505881 | -0.490450038 | 0.36989453 | -1.3259186 | 0.18486665 | 0.30717297 | NA | NA |
| ENSG000002 | 2.38877624 | -0.488347875 | 0.36832217 | -1.3258715 | 0.18488226 | 0.30718513 | NA | NA |
| ENSG000002 | 4.84460646 | -0.494844781 | 0.37323821 | -1.3258149 | 0.18490101 | 0.30720371 | NA | NA |
| ENSG000002 | 3.62676472 | -0.500330839 | 0.37741512 | -1.3256778 | 0.18494645 | 0.30726661 | NA | NA |
| ENSG000001 | 13.2905441 | -0.433375987 | 0.32695953 | -1.3254729 | 0.18501437 | 0.30732908 | OR9Q2 | 219957 |
| ENSG000002 | 6.24512415 | -0.486951943 | 0.36736176 | -1.3255379 | 0.1849928 | 0.30732908 | LINC01353 | 100506775 |
| ENSG000002 | 3.21428541 | -0.500805228 | 0.37782421 | -1.325498 | 0.18500604 | 0.30732908 | NA | NA |
| ENSG000002 | 57.3263645 | 0.291296047 | 0.21976553 | 1.32548563 | 0.18501013 | 0.30732908 | NA | NA |
| ENSG000002 | 26.2751552 | 0.344187569 | 0.25968533 | 1.32540242 | 0.18503771 | 0.30735527 | TCEB3-AS1 | 100506963 |
| ENSG000002 | 2.62794375 | -0.496738033 | 0.37479332 | -1.3253652 | 0.18505004 | 0.30736316 | NA | NA |
| ENSG000002 | 2.65039966 | -0.496749743 | 0.37482348 | -1.3252898 | 0.18507505 | 0.30739211 | NA | NA |
| ENSG000001 | 9.90532654 | 0.447835262 | 0.33793682 | 1.32520412 | 0.18510346 | 0.3074178 | BOLA2B | 654483 |

| | | | | | | | | |
|---|---|---|---|---|---|---|---|---|
| ENSG000002 | 4.43166003 | -0.501229959 | 0.37823047 | -1.3251972 | 0.18510576 | 0.3074178 | NA | NA |
| ENSG000002 | 8.30242713 | -0.459572658 | 0.3468016 | -1.3251746 | 0.18511326 | 0.3074178 | NA | NA |
| ENSG000001 | 56.7199401 | 0.277100254 | 0.2091248 | 1.32504729 | 0.18515547 | 0.30745012 | SUN2 | 25777 |
| ENSG000002 | 3.26258294 | -0.500777478 | 0.37792562 | -1.3250689 | 0.18514831 | 0.30745012 | NA | NA |
| ENSG000002 | 5.67822485 | -0.488906209 | 0.36896786 | -1.3250645 | 0.18514976 | 0.30745012 | NA | NA |
| ENSG000002 | 5.6849712 | -0.488615513 | 0.36877332 | -1.3249752 | 0.18517936 | 0.30747721 | NA | NA |
| ENSG000002 | 4.90514243 | -0.495394516 | 0.37397249 | -1.3246817 | 0.18527674 | 0.30761371 | MIR155HG | 114614 |
| ENSG000002 | 3.19618109 | -0.500503537 | 0.37782478 | -1.3246975 | 0.1852715 | 0.30761371 | NA | NA |
| ENSG000002 | 2.36230266 | -0.490308077 | 0.37018081 | -1.3245097 | 0.18533382 | 0.30769587 | RNU6-857P | 106480615 |
| ENSG000002 | 10.3996403 | -0.43798254 | 0.33070305 | -1.3243983 | 0.18537081 | 0.30774469 | NA | NA |
| ENSG000002 | 2.7828782 | -0.49022592 | 0.37016918 | -1.3243294 | 0.18539368 | 0.30777005 | NA | NA |
| ENSG000001 | 86.8929743 | 0.273531898 | 0.20655106 | 1.3242822 | 0.18540934 | 0.30778345 | CMBL | 134147 |
| ENSG000002 | 4.32676904 | -0.499685895 | 0.37734854 | -1.3242025 | 0.18543579 | 0.30781475 | NA | NA |
| ENSG000001 | 204.045817 | -0.201314838 | 0.1520338 | -1.3241453 | 0.1854548 | 0.30783371 | NSMCE1 | 197370 |
| ENSG000001 | 3.00931006 | -0.496784809 | 0.37521994 | -1.323983 | 0.1855087 | 0.30791057 | NA | NA |
| ENSG000001 | 70.4765471 | -0.261874988 | 0.19780832 | -1.3238826 | 0.18554205 | 0.30793154 | DGKB | 1607 |
| ENSG000001 | 2.4201105 | -0.487123491 | 0.36795241 | -1.3238764 | 0.18554411 | 0.30793154 | OR1Q1 | 158131 |
| ENSG000002 | 2.80738899 | -0.490072926 | 0.37018001 | -1.3238773 | 0.18554379 | 0.30793154 | NA | NA |
| ENSG000002 | 2.97398208 | -0.496925526 | 0.37536497 | -1.3238463 | 0.18555409 | 0.30793551 | NA | NA |
| ENSG000001 | 19.1459329 | 0.381243603 | 0.28806485 | 1.32346452 | 0.18568095 | 0.30799379 | LINC00324 | 284029 |
| ENSG000001 | 54.2904183 | -0.263763009 | 0.19930515 | -1.3234129 | 0.18569809 | 0.30799379 | C14orf39 | 317761 |
| ENSG000001 | 7.08017576 | -0.477666766 | 0.3608757 | -1.3236324 | 0.18562516 | 0.30799379 | BPIFA1 | 51297 |
| ENSG000002 | 10.3631197 | -0.449596645 | 0.3397169 | -1.323445 | 0.18568743 | 0.30799379 | NA | NA |
| ENSG000002 | 202.233706 | 0.194993843 | 0.14731859 | 1.32362003 | 0.18562927 | 0.30799379 | NA | NA |
| ENSG000002 | 4.33806753 | -0.499389705 | 0.3773243 | -1.3235026 | 0.18566828 | 0.30799379 | NA | NA |
| ENSG000002 | 4.03188626 | -0.500696155 | 0.37833881 | -1.3234068 | 0.18570013 | 0.30799379 | NA | NA |
| ENSG000002 | 2.74238191 | -0.489602105 | 0.36994237 | -1.3234551 | 0.18568409 | 0.30799379 | NA | NA |
| ENSG000002 | 4.34069152 | -0.499385654 | 0.37729779 | -1.3235849 | 0.18564094 | 0.30799379 | NA | NA |
| ENSG000002 | 5.71468722 | -0.48855864 | 0.36916711 | -1.3234078 | 0.18569979 | 0.30799379 | LINC01177 | 104355218 |

| | | | | | | | | |
|---|---|---|---|---|---|---|---|---|
| ENSG000002 | 3.55146813 | -0.500671734 | 0.37831227 | -1.3234351 | 0.18569072 | 0.30799379 | NA | NA |
| ENSG000002 | 26.0024246 | -0.350194984 | 0.26458699 | -1.3235533 | 0.18565144 | 0.30799379 | NA | NA |
| ENSG000002 | 2.3876671 | -0.488032179 | 0.36870159 | -1.3236509 | 0.18561903 | 0.30799379 | NA | NA |
| ENSG000002 | 4.31268294 | -0.499659721 | 0.37755824 | -1.3233977 | 0.18570314 | 0.30799379 | NA | NA |
| ENSG000002 | 11.7800248 | -0.438837675 | 0.33152206 | -1.3237058 | 0.18560077 | 0.30799379 | NA | NA |
| ENSG000001 | 93.0061895 | -0.234090235 | 0.17689598 | -1.3233214 | 0.18572853 | 0.30799782 | HCFC2 | 29915 |
| ENSG000002 | 2.3779268 | -0.486752631 | 0.36782373 | -1.3233312 | 0.18572525 | 0.30799782 | PRR20D | 729246 |
| ENSG000002 | 15.9204662 | -0.402492406 | 0.30415818 | -1.3232996 | 0.18573575 | 0.30799782 | NA | NA |
| ENSG000002 | 8.38020413 | 0.464884371 | 0.35132159 | 1.32324454 | 0.18575407 | 0.30799782 | NA | NA |
| ENSG000002 | 9.58745679 | -0.445870061 | 0.33695247 | -1.3232432 | 0.18575451 | 0.30799782 | NA | NA |
| ENSG000002 | 9.84756426 | -0.445276848 | 0.3365074 | -1.3232305 | 0.18575874 | 0.30799782 | GFY | 100507003 |
| ENSG000002 | 2.62848508 | -0.495890061 | 0.37474408 | -1.3232766 | 0.18574342 | 0.30799782 | NA | NA |
| ENSG000002 | 4.64079197 | -0.497354391 | 0.37587335 | -1.3231967 | 0.18576996 | 0.30800383 | NA | NA |
| ENSG000002 | 2.38561943 | -0.488010112 | 0.36885097 | -1.323055 | 0.1858171 | 0.30806938 | NA | NA |
| ENSG000002 | 2.66066755 | -0.494940746 | 0.37410357 | -1.3230046 | 0.18583385 | 0.30808456 | NA | NA |
| ENSG000001 | 335.529204 | 0.1867769 | 0.14117961 | 1.32297366 | 0.18584414 | 0.30808902 | GNG2 | 54331 |
| ENSG000002 | 6.28289466 | -0.484789515 | 0.36648336 | -1.3228145 | 0.18589706 | 0.30816416 | NA | NA |
| ENSG000002 | 7.33560655 | -0.47569137 | 0.35961689 | -1.3227726 | 0.18591101 | 0.30817469 | NA | NA |
| ENSG000001 | 12.1765428 | -0.442314852 | 0.33439576 | -1.3227286 | 0.18592564 | 0.3081859 | TMEM132C | 92293 |
| ENSG000002 | 6.50520016 | -0.484829077 | 0.36654318 | -1.3227066 | 0.18593297 | 0.3081859 | NA | NA |
| ENSG00000C | 82.4846792 | 0.237216692 | 0.17935293 | 1.32262516 | 0.18596006 | 0.3082182 | PLEKHB1 | 58473 |
| ENSG000002 | 2.64445 | -0.495727974 | 0.37481711 | -1.3225863 | 0.18597299 | 0.30822702 | OR10G9 | 219870 |
| ENSG000001 | 29.3600936 | 0.332491324 | 0.25141029 | 1.32250481 | 0.18600011 | 0.30824678 | BMP6 | 654 |
| ENSG000002 | 2.99095167 | -0.495433904 | 0.37461175 | -1.3225263 | 0.18599294 | 0.30824678 | NA | NA |
| ENSG000001 | 144.028901 | 0.253890669 | 0.19199536 | 1.32237919 | 0.18604192 | 0.30830346 | NEXN | 91624 |
| ENSG000002 | 3.24365687 | -0.499710519 | 0.37791645 | -1.3222778 | 0.18607567 | 0.30834681 | NA | NA |
| ENSG000002 | 7.55413747 | -0.470992268 | 0.35622146 | -1.3221895 | 0.18610507 | 0.30838291 | NA | NA |
| ENSG000002 | 2.34802068 | -0.488258391 | 0.36929892 | -1.3221224 | 0.18612738 | 0.30840729 | NA | NA |
| ENSG000002 | 2.41951819 | -0.488182244 | 0.36925242 | -1.3220827 | 0.18614061 | 0.30841659 | NA | NA |

| | | | | | | | | |
|---|---|---|---|---|---|---|---|---|
| ENSG000002 | 2.64683791 | -0.495449933 | 0.37477066 | -1.3220083 | 0.18616538 | 0.30844504 | NA | NA |
| ENSG000001 | 295.361848 | 0.173395502 | 0.13116901 | 1.32192432 | 0.18619335 | 0.30847878 | SSH2 | 85464 |
| ENSG000001 | 14.5623868 | -0.406237192 | 0.30732003 | -1.3218702 | 0.18621138 | 0.30849604 | IL6 | 3569 |
| ENSG000001 | 7.52950328 | -0.469937104 | 0.35552688 | -1.3218047 | 0.18623319 | 0.30851958 | GHRL | 51738 |
| ENSG000001 | 8.64569097 | -0.477762876 | 0.36151657 | -1.3215518 | 0.18631745 | 0.30864655 | SDS | 10993 |
| ENSG000002 | 8.12382798 | -0.473396317 | 0.35823208 | -1.3214794 | 0.18634156 | 0.30867388 | LINC01538 | 400654 |
| ENSG000002 | 14.5136985 | 0.40867603 | 0.30929006 | 1.32133579 | 0.18638943 | 0.30872795 | NA | NA |
| ENSG000002 | 3.20850495 | -0.49926976 | 0.37785227 | -1.3213359 | 0.1863894 | 0.30872795 | RN7SL408P | 106481841 |
| ENSG000002 | 2.37995052 | -0.488226034 | 0.36960623 | -1.3209356 | 0.18652283 | 0.30889844 | NA | NA |
| ENSG000002 | 7.50328973 | -0.469467847 | 0.35539962 | -1.3209577 | 0.18651548 | 0.30889844 | NA | NA |
| ENSG000002 | 2.42522653 | -0.487397341 | 0.36896699 | -1.3209782 | 0.18650864 | 0.30889844 | RN7SL91P | 106480946 |
| ENSG000002 | 2.40280341 | -0.488231166 | 0.36960137 | -1.3209669 | 0.18651241 | 0.30889844 | NA | NA |
| ENSG000002 | 9.18930252 | -0.454701852 | 0.3442438 | -1.3208716 | 0.18654419 | 0.3089212 | NA | NA |
| ENSG000001 | 16.5905571 | -0.393288526 | 0.29777584 | -1.3207536 | 0.18658353 | 0.30897372 | BAIAP2L2 | 80115 |
| ENSG000002 | 6.30577003 | -0.488523162 | 0.36989832 | -1.3206958 | 0.18660281 | 0.30899305 | NA | NA |
| ENSG000002 | 6.90292315 | -0.486404724 | 0.36833318 | -1.3205564 | 0.18664933 | 0.30905745 | NA | NA |
| ENSG000001 | 4.89429299 | -0.494184711 | 0.37427674 | -1.3203725 | 0.18671069 | 0.30914643 | GPR101 | 83550 |
| ENSG000002 | 53.9607596 | 0.270045423 | 0.20452688 | 1.32034196 | 0.18672087 | 0.30915067 | PGAM1P5 | 100132594 |
| ENSG000002 | 2.98408854 | -0.494643614 | 0.3746953 | -1.3201223 | 0.1867942 | 0.30924682 | NA | NA |
| ENSG000002 | 3.20299215 | -0.498559538 | 0.37765823 | -1.3201342 | 0.18679022 | 0.30924682 | NA | NA |
| ENSG000002 | 2.42551992 | -0.487031096 | 0.36897022 | -1.319974 | 0.1868437 | 0.30931614 | RNU4ATAC17P | 106480696 |
| ENSG00000C | 12.8978679 | -0.432211767 | 0.32745892 | -1.3198961 | 0.18686969 | 0.30934655 | SERPINB3 | 6317 |
| ENSG000001 | 12.182626 | -0.424603579 | 0.32171143 | -1.3198275 | 0.18689263 | 0.30936841 | MAPK11 | 5600 |
| ENSG000002 | 2.96856853 | -0.496135295 | 0.37591392 | -1.3198109 | 0.18689816 | 0.30936841 | NA | NA |
| ENSG000002 | 11.2768403 | -0.435869272 | 0.33026637 | -1.3197507 | 0.18691826 | 0.30938906 | NA | NA |
| ENSG000002 | 10.1569752 | -0.442224938 | 0.33512608 | -1.3195778 | 0.18697602 | 0.30947202 | NA | NA |
| ENSG000002 | 2.6380758 | -0.493967238 | 0.37435801 | -1.3195049 | 0.18700036 | 0.30948717 | NA | NA |
| ENSG000002 | 16.8180206 | 0.388813087 | 0.29466593 | 1.31950472 | 0.18700043 | 0.30948717 | NA | NA |
| ENSG000002 | 4.13676102 | -0.499181778 | 0.37832168 | -1.3194638 | 0.18701409 | 0.30949714 | NA | NA |

| | | | | | | | | |
|---|---|---|---|---|---|---|---|---|
| ENSG000002 | 4.33453857 | -0.4976467 | 0.3771759 | -1.3194022 | 0.1870347 | 0.30949972 | NA | NA |
| ENSG000002 | 10.3219914 | -0.448287302 | 0.33976157 | -1.3194173 | 0.18702963 | 0.30949972 | NA | NA |
| ENSG000002 | 2.36484646 | -0.485388189 | 0.36788815 | -1.3193906 | 0.18703855 | 0.30949972 | RN7SL96P | 106480486 |
| ENSG000002 | 6.36385309 | -0.489228727 | 0.37081193 | -1.3193446 | 0.18705392 | 0.30951253 | NA | NA |
| ENSG000001 | 566.268711 | 0.168856357 | 0.12798911 | 1.31930257 | 0.18706798 | 0.30952316 | KIAA1804 | 84451 |
| ENSG000001 | 5.73370427 | -0.489674449 | 0.37118067 | -1.3192348 | 0.18709064 | 0.30954802 | SLAMF9 | 89886 |
| ENSG000002 | 11.7066808 | -0.42983949 | 0.32583625 | -1.3191887 | 0.18710604 | 0.30956088 | NA | NA |
| ENSG000002 | 2.45384061 | -0.48547106 | 0.36806047 | -1.3189981 | 0.18716975 | 0.30965364 | RNU7-97P | 106479063 |
| ENSG000002 | 4.15736241 | -0.499154064 | 0.37849946 | -1.3187709 | 0.18724573 | 0.30976265 | NA | NA |
| ENSG000002 | 14.1468468 | -0.413870754 | 0.31383437 | -1.3187554 | 0.18725091 | 0.30976265 | SULT1C2P1 | 151234 |
| ENSG000001 | 13.7274413 | -0.428841931 | 0.32519637 | -1.3187169 | 0.18726379 | 0.30977131 | RSPH9 | 221421 |
| ENSG000001 | 263.021915 | 0.362683174 | 0.27507537 | 1.31848654 | 0.18734083 | 0.30988611 | PFDN2 | 5202 |
| ENSG000002 | 4.85624518 | -0.492175122 | 0.37346893 | -1.3178476 | 0.18755468 | 0.31022719 | NA | NA |
| ENSG000001 | 64.5493228 | -0.24988643 | 0.18962729 | -1.3177767 | 0.18757841 | 0.31025378 | AFF3 | 3899 |
| ENSG000002 | 8.54888232 | -0.452936361 | 0.34373283 | -1.3176989 | 0.18760447 | 0.31028424 | NA | NA |
| ENSG000001 | 4.15942593 | -0.49860607 | 0.37842586 | -1.3175793 | 0.18764453 | 0.31033784 | OR9A2 | 135924 |
| ENSG000002 | 4.66420944 | -0.497228706 | 0.3774188 | -1.3174455 | 0.18768933 | 0.31039927 | SPRN | 503542 |
| ENSG00000C | 65.2182111 | -0.271506017 | 0.20616266 | -1.3169505 | 0.18785522 | 0.31063561 | GUCY2C | 2984 |
| ENSG000001 | 18.7301239 | -0.379818099 | 0.28839849 | -1.3169906 | 0.18784177 | 0.31063561 | IL23R | 149233 |
| ENSG000002 | 9.95613506 | -0.44913506 | 0.34103885 | -1.3169616 | 0.1878515 | 0.31063561 | NA | NA |
| ENSG000002 | 2.78491401 | -0.486792732 | 0.36968594 | -1.3167737 | 0.18791449 | 0.31072094 | NA | NA |
| ENSG000002 | 8.06508747 | -0.461520554 | 0.35051496 | -1.3166929 | 0.18794158 | 0.31075307 | NA | NA |
| ENSG000002 | 6.19215863 | -0.481409316 | 0.36563704 | -1.3166317 | 0.18796211 | 0.31077435 | NA | NA |
| ENSG000002 | 5.16122181 | -0.492662691 | 0.37419688 | -1.3165868 | 0.18797715 | 0.31078654 | NA | NA |
| ENSG000001 | 49.0447721 | 0.312927102 | 0.23769327 | 1.31651644 | 0.18800076 | 0.31080012 | NA | NA |
| ENSG000002 | 7.13964997 | -0.475467123 | 0.36116169 | -1.3164938 | 0.18800836 | 0.31080012 | LINC00996 | 285972 |
| ENSG000002 | 4.38436892 | -0.497508176 | 0.37789501 | -1.3165249 | 0.18799793 | 0.31080012 | NA | NA |
| ENSG000001 | 70.0346847 | 0.266062998 | 0.20211137 | 1.31641775 | 0.18803387 | 0.31082962 | PIGH | 5283 |
| ENSG000002 | 9.38879869 | -0.455504925 | 0.34607131 | -1.3162169 | 0.18810124 | 0.31092831 | NA | NA |

| | | | | | | | | |
|---|---|---|---|---|---|---|---|---|
| ENSG000002 | 5.73297478 | -0.486889553 | 0.36993114 | -1.3161627 | 0.18811945 | 0.31094574 | NA | NA |
| ENSG000002 | 6.91238068 | -0.481340694 | 0.36572234 | -1.316137 | 0.18812808 | 0.31094732 | LOC101929586 | 101929586 |
| ENSG000001 | 7.06562762 | -0.476554211 | 0.36211372 | -1.3160347 | 0.18816241 | 0.31097069 | OR7G2 | 390882 |
| ENSG000002 | 3.94968038 | -0.49769978 | 0.37817938 | -1.3160415 | 0.1881601 | 0.31097069 | NA | NA |
| ENSG000002 | 2.6314738 | -0.493184759 | 0.37475297 | -1.3160263 | 0.18816522 | 0.31097069 | NA | NA |
| ENSG000001 | 11.5412 | -0.437758823 | 0.33264652 | -1.315988 | 0.18817807 | 0.31097925 | POLR2F | 5435 |
| ENSG000002 | 4.75319165 | 0.494164633 | 0.3755288 | 1.31591675 | 0.18820199 | 0.3110061 | NA | NA |
| ENSG000001 | 561.625934 | 0.422004105 | 0.32076699 | 1.3156095 | 0.18830514 | 0.31112904 | HAS3 | 3038 |
| ENSG000002 | 23.5311037 | 0.361750522 | 0.27496497 | 1.31562404 | 0.18830026 | 0.31112904 | C4orf47 | 441054 |
| ENSG000002 | 3.50137665 | -0.497937106 | 0.37848562 | -1.3156038 | 0.18830707 | 0.31112904 | RN7SL228P | 106480983 |
| ENSG000002 | 3.46842472 | -0.497990777 | 0.37852314 | -1.3156151 | 0.18830325 | 0.31112904 | NA | NA |
| ENSG000002 | 2.32875 | 0.476505757 | 0.36220212 | 1.31557969 | 0.18831516 | 0.31112972 | NA | NA |
| ENSG000002 | 2.98056013 | -0.494195538 | 0.37568134 | -1.3154647 | 0.18835378 | 0.31118085 | NA | NA |
| ENSG000002 | 2.37485486 | -0.485127414 | 0.36882304 | -1.3153392 | 0.18839591 | 0.31123779 | RN7SL79P | 106480943 |
| ENSG000002 | 2.36270705 | -0.48529452 | 0.36897358 | -1.3152555 | 0.18842405 | 0.31127159 | NA | NA |
| ENSG000002 | 9.88713139 | -0.447517829 | 0.34029024 | -1.3151063 | 0.18847419 | 0.31134174 | NA | NA |
| ENSG000002 | 6.26130767 | -0.481813513 | 0.36638359 | -1.3150521 | 0.18849239 | 0.31135912 | NA | NA |
| ENSG000001 | 9.90935394 | -0.444258489 | 0.33793848 | -1.3146135 | 0.18863984 | 0.31158998 | ARL13A | 392509 |
| ENSG000002 | 3.79935431 | -0.497637184 | 0.37855883 | -1.3145571 | 0.18865881 | 0.31160863 | NA | NA |
| ENSG000002 | 2.40822383 | -0.482273079 | 0.36691317 | -1.3144066 | 0.18870942 | 0.31167952 | LOC105373580 | 105373580 |
| ENSG000002 | 4.04017229 | -0.497200983 | 0.37830263 | -1.3142943 | 0.18874719 | 0.31172921 | NA | NA |
| ENSG000002 | 8.12012072 | -0.465345829 | 0.35407897 | -1.3142431 | 0.18876442 | 0.31174497 | NA | NA |
| ENSG000001 | 3.52046723 | -0.497499845 | 0.37856965 | -1.3141567 | 0.18879348 | 0.31176757 | NA | NA |
| ENSG000002 | 3.46608944 | -0.497517672 | 0.37858241 | -1.3141595 | 0.18879253 | 0.31176757 | NA | NA |
| ENSG000002 | 3.47664604 | -0.497441365 | 0.37859009 | -1.3139313 | 0.18886933 | 0.31186742 | NA | NA |
| ENSG000002 | 2.6698346 | -0.491937568 | 0.37440023 | -1.313935 | 0.18886807 | 0.31186742 | NA | NA |
| ENSG000002 | 3.21981086 | -0.496124618 | 0.37760342 | -1.3138774 | 0.18888745 | 0.31188465 | NA | NA |
| ENSG000002 | 2.64564423 | -0.49242004 | 0.3748233 | -1.3137392 | 0.188934 | 0.31194881 | NA | NA |
| ENSG000001 | 40.0878866 | 0.297421536 | 0.22640668 | 1.31366062 | 0.18896044 | 0.31197741 | GJB7 | 375519 |

| | | | | | | | | |
|---|---|---|---|---|---|---|---|---|
| ENSG000002 | 8.09837188 | -0.461730272 | 0.35149353 | -1.3136238 | 0.18897283 | 0.31197741 | LOC100506175 | 100506175 |
| ENSG000002 | 7.87650483 | -0.468346155 | 0.35653116 | -1.3136191 | 0.18897441 | 0.31197741 | NA | NA |
| ENSG000002 | 4.65823132 | -0.494043015 | 0.37612215 | -1.3135175 | 0.18900864 | 0.31202123 | NA | NA |
| ENSG000002 | 4.11033069 | -0.497240555 | 0.37858608 | -1.3134148 | 0.18904322 | 0.3120656 | NA | NA |
| ENSG000001 | 57.5795674 | 0.261904913 | 0.19941311 | 1.3133786 | 0.1890554 | 0.31207302 | FAM124B | 79843 |
| ENSG000001 | 4.63490181 | -0.494265263 | 0.37634197 | -1.3133408 | 0.18906814 | 0.31208133 | DLX5 | 1749 |
| ENSG000002 | 3.5041891 | -0.497085476 | 0.37852514 | -1.3132166 | 0.18910999 | 0.3121377 | NA | NA |
| ENSG000002 | 8.01785246 | -0.46124598 | 0.35124154 | -1.3131874 | 0.18911979 | 0.31214118 | NA | NA |
| ENSG000002 | 7.38989863 | -0.474482817 | 0.36134799 | -1.3130911 | 0.18915226 | 0.31218206 | NA | NA |
| ENSG000002 | 4.62687746 | -0.493768012 | 0.37604882 | -1.3130423 | 0.18916869 | 0.31219647 | NA | NA |
| ENSG000002 | 5.97780246 | 0.484398086 | 0.36893722 | 1.31295533 | 0.189198 | 0.31223214 | RNU6-4P | 106478914 |
| ENSG000002 | 4.04010703 | -0.496789684 | 0.37839014 | -1.3129034 | 0.18921551 | 0.31224833 | NA | NA |
| ENSG000002 | 4.37976993 | -0.496170762 | 0.37793754 | -1.312838 | 0.18923754 | 0.31227197 | NA | NA |
| ENSG000002 | 10.5048031 | 0.442675133 | 0.3372059 | 1.31277397 | 0.18925912 | 0.31229488 | NA | NA |
| ENSG000002 | 7.13729378 | -0.476155068 | 0.36276109 | -1.3125858 | 0.18932255 | 0.31238682 | NA | NA |
| ENSG000001 | 156.103145 | -0.201396463 | 0.15344303 | -1.3125162 | 0.18934603 | 0.31241287 | FSIP2 | 401024 |
| ENSG000002 | 4.60068407 | -0.49272043 | 0.37545664 | -1.3123231 | 0.18941113 | 0.31250756 | NA | NA |
| ENSG000001 | 87.4530522 | -0.221743794 | 0.16897443 | -1.312292 | 0.18942164 | 0.31250936 | SEMA3E | 9723 |
| ENSG000002 | 14.3216737 | -0.408106898 | 0.31099209 | -1.3122742 | 0.18942764 | 0.31250936 | NA | NA |
| ENSG000002 | 5.7525714 | -0.486345933 | 0.37063242 | -1.3122056 | 0.18945078 | 0.31253483 | NA | NA |
| ENSG000002 | 10.667044 | -0.432848714 | 0.32987593 | -1.3121561 | 0.18946747 | 0.31254965 | NA | NA |
| ENSG000002 | 2.62729086 | -0.490500217 | 0.37387064 | -1.3119517 | 0.18953643 | 0.31264973 | NA | NA |
| ENSG000002 | 6.41733275 | -0.484773888 | 0.36951185 | -1.3119306 | 0.18954356 | 0.31264973 | EPHA5-AS1 | 100144602 |
| ENSG000002 | 10.6516157 | -0.431191617 | 0.32871291 | -1.3117575 | 0.18960198 | 0.31273336 | NA | NA |
| ENSG000001 | 18.9769147 | -0.389266697 | 0.29680194 | -1.3115369 | 0.18967644 | 0.3128261 | ADAMTSL2 | 9719 |
| ENSG000002 | 58.8826505 | 0.267207354 | 0.20374517 | 1.31147821 | 0.18969625 | 0.3128261 | NA | NA |
| ENSG000002 | 12.8262467 | 0.419741946 | 0.32005293 | 1.31147667 | 0.18969677 | 0.3128261 | NA | NA |
| ENSG000002 | 11.2691354 | -0.433895251 | 0.33084211 | -1.3114874 | 0.18969316 | 0.3128261 | NA | NA |
| ENSG000002 | 4.63672169 | -0.493591354 | 0.37634328 | -1.3115456 | 0.18967351 | 0.3128261 | OR10G2 | 26534 |

| | | | | | | | | |
|---|---|---|---|---|---|---|---|---|
| ENSG000002 | 16.3547671 | -0.391644778 | 0.29863805 | -1.3114363 | 0.18971041 | 0.31283586 | LINC01393 | 102724386 |
| ENSG000002 | 5.72637279 | -0.484772284 | 0.36966679 | -1.3113763 | 0.18973065 | 0.31285651 | OR1S1 | 219959 |
| ENSG000001 | 529.796753 | 0.164091302 | 0.12514275 | 1.31123296 | 0.18977907 | 0.31291215 | MBNL1 | 4154 |
| ENSG000002 | 15.9820832 | 0.402770681 | 0.30716995 | 1.31123074 | 0.18977982 | 0.31291215 | NA | NA |
| ENSG00000C | 1153.59296 | -0.192009776 | 0.14644086 | -1.3111763 | 0.18979821 | 0.31292698 | RRM2B | 50484 |
| ENSG000001 | 13.2493729 | -0.418668449 | 0.31931312 | -1.3111533 | 0.18980599 | 0.31292698 | KCNS2 | 3788 |
| ENSG000002 | 6.48852434 | -0.474689521 | 0.36204457 | -1.3111356 | 0.18981196 | 0.31292698 | NA | NA |
| ENSG000002 | 5.68995476 | -0.485632385 | 0.37040544 | -1.3110833 | 0.18982963 | 0.31294338 | NA | NA |
| ENSG000002 | 4.41005486 | -0.49508393 | 0.37763146 | -1.3110241 | 0.18984963 | 0.31296363 | NA | NA |
| ENSG000001 | 36.1884641 | 0.305956401 | 0.23340329 | 1.31084873 | 0.18990888 | 0.31303648 | SLC2A4 | 6517 |
| ENSG000002 | 12.8158697 | -0.417827134 | 0.31874578 | -1.3108476 | 0.18990926 | 0.31303648 | LINC00887 | 100131551 |
| ENSG000002 | 4.34376457 | -0.494912775 | 0.37758361 | -1.3107369 | 0.18994666 | 0.3130854 | NA | NA |
| ENSG000002 | 4.10083092 | -0.496167554 | 0.37859245 | -1.3105585 | 0.19000696 | 0.31317207 | NA | NA |
| ENSG000002 | 2.65090874 | -0.490878071 | 0.37456991 | -1.3105112 | 0.19002294 | 0.31317295 | NA | NA |
| ENSG000002 | 4.19781347 | 0.495536069 | 0.37811859 | 1.31053084 | 0.19001632 | 0.31317295 | NA | NA |
| ENSG000001 | 13.8380135 | -0.408293745 | 0.31156593 | -1.310457 | 0.19004129 | 0.31319046 | KLF1 | 10661 |
| ENSG000001 | 7.78951328 | -0.477681084 | 0.36452843 | -1.3104083 | 0.19005775 | 0.31319212 | LINC00305 | 221241 |
| ENSG000002 | 2.9722741 | -0.491339813 | 0.3749499 | -1.3104146 | 0.19005563 | 0.31319212 | NA | NA |
| ENSG000002 | 4.90284859 | -0.492449151 | 0.37583407 | -1.3102834 | 0.19009997 | 0.31324898 | NA | NA |
| ENSG000001 | 119.522589 | 0.268388951 | 0.20485461 | 1.3101436 | 0.19014726 | 0.31330143 | C1orf35 | 79169 |
| ENSG000002 | 7.02023264 | -0.472926921 | 0.36096964 | -1.310157 | 0.19014271 | 0.31330143 | NA | NA |
| ENSG000002 | 9.97273571 | -0.443386605 | 0.33843324 | -1.3101154 | 0.19015679 | 0.31330441 | NA | NA |
| ENSG000002 | 3.06234982 | -0.49019372 | 0.37417176 | -1.3100768 | 0.19016987 | 0.31331322 | NA | NA |
| ENSG000002 | 13.4426455 | -0.421978472 | 0.3221684 | -1.3098072 | 0.19026108 | 0.31345077 | NA | NA |
| ENSG000001 | 60.6357902 | 0.252805315 | 0.19306925 | 1.30940225 | 0.19039813 | 0.31361851 | DHRS12 | 79758 |
| ENSG000001 | 70.2200097 | -0.247386673 | 0.18892576 | -1.3094385 | 0.19038585 | 0.31361851 | TNXB | 7148 |
| ENSG000002 | 6.01092746 | -0.487632788 | 0.37241013 | -1.3093972 | 0.19039983 | 0.31361851 | NA | NA |
| ENSG000002 | 5.19323632 | -0.490000924 | 0.37422017 | -1.3093921 | 0.19040157 | 0.31361851 | NA | NA |
| ENSG000002 | 6.72416124 | -0.471406946 | 0.36000867 | -1.3094322 | 0.19038799 | 0.31361851 | NA | NA |

| | | | | | | | | |
|---|---|---|---|---|---|---|---|---|
| ENSG000001 | 145.314512 | 0.212124994 | 0.16201136 | 1.30932172 | 0.1904254 | 0.31364501 | SHC3 | 53358 |
| ENSG000001 | 23.621016 | -0.351160903 | 0.26823281 | -1.3091646 | 0.19047859 | 0.31371989 | XKRX | 402415 |
| ENSG000001 | 15.815292 | -0.390829081 | 0.29855315 | -1.3090771 | 0.19050824 | 0.31375598 | AMHR2 | 269 |
| ENSG000002 | 30.3411029 | 0.319822259 | 0.24432721 | 1.30899156 | 0.19053721 | 0.3137782 | NA | NA |
| ENSG000002 | 2.37144159 | -0.482068974 | 0.36827377 | -1.3089962 | 0.19053563 | 0.3137782 | LOC100507071 | 100507071 |
| ENSG000002 | 2.63850035 | -0.489996697 | 0.37436175 | -1.3088856 | 0.19057311 | 0.31379961 | NA | NA |
| ENSG000002 | 5.4761407 | -0.48741379 | 0.3723822 | -1.3089073 | 0.19056574 | 0.31379961 | NA | NA |
| ENSG000002 | 20.2626829 | 0.376633138 | 0.28775623 | 1.30886182 | 0.19058116 | 0.31379961 | NA | NA |
| ENSG000002 | 3.83162145 | -0.495228518 | 0.37836506 | -1.3088643 | 0.19058033 | 0.31379961 | NA | NA |
| ENSG000002 | 3.76123281 | -0.495176644 | 0.37837154 | -1.3087048 | 0.19063437 | 0.31387448 | NA | NA |
| ENSG00000C | 252.2831 | 0.246702035 | 0.18851372 | 1.30866884 | 0.19064655 | 0.31388178 | CIC | 23152 |
| ENSG000002 | 3.85299578 | -0.495408777 | 0.37856897 | -1.3086355 | 0.19065786 | 0.31388766 | NA | NA |
| ENSG000002 | 6.5383446 | -0.478674788 | 0.36580117 | -1.3085655 | 0.19068158 | 0.31391396 | NA | NA |
| ENSG00000C | 11.4110262 | -0.424387968 | 0.32433954 | -1.3084682 | 0.19071456 | 0.31393897 | FGF20 | 26281 |
| ENSG000001 | 95.8818102 | 0.225646464 | 0.17245297 | 1.30845214 | 0.19072 | 0.31393897 | LPAR1 | 1902 |
| ENSG000002 | 7.52526489 | -0.464631944 | 0.35509538 | -1.3084708 | 0.19071367 | 0.31393897 | NA | NA |
| ENSG000002 | 3.49915238 | -0.494988705 | 0.37836157 | -1.3082425 | 0.19079106 | 0.3140432 | NA | NA |
| ENSG000001 | 5.18474198 | -0.489580761 | 0.37425504 | -1.3081474 | 0.19082332 | 0.31408355 | MAGEA1 | 4100 |
| ENSG000001 | 229.527624 | 0.221171299 | 0.16907518 | 1.30812398 | 0.19083126 | 0.31408388 | ZMIZ1 | 57178 |
| ENSG000002 | 3.23024237 | -0.493521467 | 0.37732009 | -1.307965 | 0.19088518 | 0.31413536 | NA | NA |
| ENSG000002 | 2.77946526 | -0.483851953 | 0.36992779 | -1.3079632 | 0.19088578 | 0.31413536 | LOC100507464 | 100507464 |
| ENSG000002 | 8.6204279 | -0.457102193 | 0.34946451 | -1.3080075 | 0.19087078 | 0.31413536 | NA | NA |
| ENSG000002 | 2.64396074 | -0.489353095 | 0.37417361 | -1.3078236 | 0.19093313 | 0.31419456 | RN7SKP178 | 106479175 |
| ENSG000002 | 2.41590436 | -0.481918137 | 0.36849205 | -1.3078115 | 0.19093725 | 0.31419456 | NA | NA |
| ENSG000002 | 3.50451027 | -0.495048132 | 0.37857063 | -1.3076771 | 0.19098286 | 0.31425685 | NA | NA |
| ENSG000001 | 8.30134501 | -0.455798378 | 0.34860551 | -1.3074905 | 0.19104619 | 0.31433555 | TBC1D28 | 254272 |
| ENSG000002 | 2.40097612 | -0.48133297 | 0.36812858 | -1.3075132 | 0.19103848 | 0.31433555 | MIR5739 | 102466081 |
| ENSG000002 | 87.5835429 | 0.253281046 | 0.19373319 | 1.30737042 | 0.19108694 | 0.31437708 | NA | NA |
| ENSG000002 | 9.15801196 | -0.452974004 | 0.34647209 | -1.3073896 | 0.19108043 | 0.31437708 | NA | NA |

| | | | | | | | | |
|---|---|---|---|---|---|---|---|---|
| ENSG000002 | 8.5998985 | 0.468016445 | 0.35800353 | 1.30729561 | 0.19111233 | 0.31440611 | SNORD116-12 | 100033424 |
| ENSG000002 | 5.24882098 | -0.489539206 | 0.37455702 | -1.3069818 | 0.1912189 | 0.31456866 | NA | NA |
| ENSG000000 | 31.8851026 | -0.327751419 | 0.25079322 | -1.3068591 | 0.19126055 | 0.31461166 | PARP3 | 10039 |
| ENSG000002 | 2.65139691 | -0.487023261 | 0.37266639 | -1.3068612 | 0.19125984 | 0.31461166 | NA | NA |
| ENSG000002 | 14.0130036 | -0.412106447 | 0.31537004 | -1.3067394 | 0.19130124 | 0.31466582 | FLJ22447 | 400221 |
| ENSG000002 | 10.5901659 | -0.429370179 | 0.32860208 | -1.3066569 | 0.19132926 | 0.31469915 | NA | NA |
| ENSG000002 | 2.36225004 | -0.480534885 | 0.36782245 | -1.3064316 | 0.19140581 | 0.31481229 | NA | NA |
| ENSG000001 | 4.65979769 | -0.491444069 | 0.37620014 | -1.3063368 | 0.19143806 | 0.31483979 | CPXCR1 | 53336 |
| ENSG000002 | 4.3121099 | -0.493130422 | 0.37749027 | -1.3063394 | 0.19143715 | 0.31483979 | NA | NA |
| ENSG000001 | 8.75730683 | -0.465271274 | 0.35623503 | -1.3060795 | 0.19152552 | 0.31495701 | SIGLEC11 | 114132 |
| ENSG000002 | 3.21517006 | 0.492850662 | 0.37736384 | 1.30603575 | 0.1915404 | 0.31495701 | NA | NA |
| ENSG000002 | 3.19539235 | -0.493409999 | 0.37778703 | -1.3060533 | 0.19153443 | 0.31495701 | NA | NA |
| ENSG000002 | 10.3839702 | -0.434811421 | 0.33291261 | -1.3060828 | 0.19152441 | 0.31495701 | NA | NA |
| ENSG000002 | 19.7892343 | 0.364022943 | 0.27875341 | 1.30589593 | 0.19158795 | 0.31502242 | NA | NA |
| ENSG000002 | 2.94896996 | -0.489855155 | 0.37514264 | -1.3057837 | 0.19162611 | 0.31505962 | MIR708 | 100126333 |
| ENSG000002 | 3.24738225 | -0.492813048 | 0.37740563 | -1.3057915 | 0.19162348 | 0.31505962 | NA | NA |
| ENSG000002 | 3.68816165 | -0.492210848 | 0.37698153 | -1.305663 | 0.19166717 | 0.31511436 | NA | NA |
| ENSG000002 | 2.9753992 | -0.490050103 | 0.37536074 | -1.3055444 | 0.19170754 | 0.31516794 | NA | NA |
| ENSG000002 | 4.61035856 | -0.491003651 | 0.37611158 | -1.3054734 | 0.19173171 | 0.31519489 | NA | NA |
| ENSG000002 | 5.15853188 | -0.486485898 | 0.37265859 | -1.3054466 | 0.19174081 | 0.31519708 | LINC01159 | 102682016 |
| ENSG000002 | 3.23772649 | -0.492715469 | 0.37744363 | -1.3054015 | 0.19175618 | 0.31520957 | NA | NA |
| ENSG000001 | 57.9013405 | -0.257554307 | 0.19730913 | -1.3053339 | 0.19177916 | 0.31523457 | LHX4 | 89884 |
| ENSG000002 | 6.24299162 | -0.481616402 | 0.36897595 | -1.3052786 | 0.19179801 | 0.31525277 | NA | NA |
| ENSG000002 | 22.7059115 | 0.354090955 | 0.27131792 | 1.30507767 | 0.1918664 | 0.3153524 | C19orf84 | 147646 |
| ENSG000001 | 28.6988604 | -0.331797767 | 0.25426544 | -1.3049267 | 0.19191781 | 0.31542411 | C11orf63 | 79864 |
| ENSG000002 | 3.48382053 | -0.493643877 | 0.37830872 | -1.3048705 | 0.19193696 | 0.31544281 | PRAMEF4 | 400735 |
| ENSG000002 | 4.83421348 | -0.488131625 | 0.37412662 | -1.304723 | 0.19198717 | 0.31551254 | NA | NA |
| ENSG000002 | 4.62127216 | -0.491500494 | 0.37672652 | -1.3046613 | 0.19200822 | 0.31552351 | LINC00161 | 118421 |
| ENSG000002 | 2.41365286 | -0.479852788 | 0.36779974 | -1.3046578 | 0.19200941 | 0.31552351 | NA | NA |

| | | | | | | | | |
|---|---|---|---|---|---|---|---|---|
| ENSG000002 | 8.7285126 | -0.46399448 | 0.35571138 | -1.3044128 | 0.19209287 | 0.31564786 | GSTTP1 | 25774 |
| ENSG000002 | 2.36352913 | -0.481407208 | 0.36909467 | -1.304292 | 0.19213405 | 0.31568995 | NA | NA |
| ENSG000002 | 17.0059899 | -0.383126711 | 0.29374014 | -1.3043049 | 0.19212963 | 0.31568995 | NA | NA |
| ENSG000002 | 6.47251599 | -0.477795086 | 0.36639185 | -1.3040549 | 0.19221486 | 0.31580679 | TMED11P | 100379220 |
| ENSG000002 | 3.49078835 | -0.493484072 | 0.37842778 | -1.3040377 | 0.19222074 | 0.31580679 | NA | NA |
| ENSG000002 | 4.29583177 | -0.492526543 | 0.37771076 | -1.3039781 | 0.19224106 | 0.31582738 | NA | NA |
| ENSG000001 | 50.1399587 | -0.277950557 | 0.21316075 | -1.3039481 | 0.19225128 | 0.31583137 | MYH3 | 4621 |
| ENSG000002 | 3.5107911 | -0.493343123 | 0.3783597 | -1.3038998 | 0.19226776 | 0.31584566 | NA | NA |
| ENSG000002 | 10.6492063 | -0.43364819 | 0.33259264 | -1.3038418 | 0.19228752 | 0.31585253 | MACC1-AS1 | 100874041 |
| ENSG000002 | 4.16851983 | -0.493531474 | 0.37852003 | -1.3038451 | 0.19228641 | 0.31585253 | NA | NA |
| ENSG000002 | 4.65741392 | -0.491693503 | 0.37712937 | -1.3037794 | 0.19230881 | 0.3158747 | NA | NA |
| ENSG000002 | 12.4296581 | -0.416204866 | 0.31927936 | -1.3035759 | 0.19237824 | 0.31597593 | NA | NA |
| ENSG000001 | 23.9483009 | -0.361137083 | 0.27707404 | -1.3033956 | 0.19243974 | 0.31606415 | MIP | 4284 |
| ENSG00000C | 17.0645313 | -0.383587214 | 0.29433361 | -1.3032396 | 0.19249298 | 0.31613879 | TRPC7 | 57113 |
| ENSG000002 | 7.59384895 | -0.467286488 | 0.35857511 | -1.303176 | 0.19251467 | 0.31616161 | NA | NA |
| ENSG000002 | 2.27784322 | 0.476992787 | 0.36604731 | 1.30309053 | 0.19254386 | 0.31619673 | NA | NA |
| ENSG000001 | 1159.25044 | 0.145342785 | 0.1115394 | 1.30306224 | 0.19255351 | 0.31619979 | TOR1AIP1 | 26092 |
| ENSG000002 | 4.35634902 | -0.49223422 | 0.37781862 | -1.3028321 | 0.1926321 | 0.31631602 | NA | NA |
| ENSG000002 | 5.56488581 | -0.48738204 | 0.37410799 | -1.3027844 | 0.19264839 | 0.31632997 | NA | NA |
| ENSG000002 | 2.38874509 | -0.480549955 | 0.36887167 | -1.3027565 | 0.19265792 | 0.3163328 | NA | NA |
| ENSG000002 | 6.23829399 | -0.48796664 | 0.37461251 | -1.3025903 | 0.19271465 | 0.31641315 | NA | NA |
| ENSG000001 | 108.086956 | 0.216642958 | 0.1663241 | 1.30253495 | 0.19273358 | 0.31643141 | RGS7BP | 401190 |
| ENSG000001 | 174.92415 | 0.20480586 | 0.15724108 | 1.30249588 | 0.19274693 | 0.31644051 | GPATCH11 | 253635 |
| ENSG000002 | 8.09860001 | -0.45984142 | 0.35308695 | -1.3023461 | 0.19279809 | 0.3165117 | LOC102724601 | 102724601 |
| ENSG000002 | 14.0751607 | 0.405769858 | 0.31159072 | 1.30225269 | 0.19283002 | 0.3165513 | NA | NA |
| ENSG000002 | 5.92424229 | -0.478073498 | 0.36712075 | -1.3022241 | 0.19283978 | 0.3165545 | NA | NA |
| ENSG000002 | 9.17576864 | -0.452772975 | 0.34773325 | -1.3020698 | 0.19289251 | 0.31662825 | NA | NA |
| ENSG000002 | 5.40963491 | -0.484420782 | 0.37207988 | -1.3019268 | 0.19294139 | 0.31669567 | NA | NA |
| ENSG000002 | 5.52391506 | -0.486485523 | 0.37370262 | -1.3017985 | 0.19298526 | 0.31675485 | NA | NA |

| | | | | | | | | |
|---|---|---|---|---|---|---|---|---|
| ENSG000002 | 3.49747054 | -0.492723462 | 0.37852187 | -1.3017041 | 0.19301756 | 0.31679504 | NA | NA |
| ENSG000001 | 4.40852041 | -0.492236337 | 0.37817033 | -1.301626 | 0.19304426 | 0.31682605 | HOXD3 | 3232 |
| ENSG000002 | 3.83874603 | 0.492663518 | 0.37853792 | 1.30149052 | 0.19309061 | 0.31688928 | NA | NA |
| ENSG000002 | 2.60201095 | -0.487143104 | 0.37431341 | -1.3014311 | 0.19311094 | 0.31690983 | NA | NA |
| ENSG000001 | 17.6671977 | -0.386745607 | 0.29719008 | -1.3013409 | 0.1931418 | 0.31692458 | YPEL4 | 219539 |
| ENSG000001 | 114.528437 | 0.203755407 | 0.15657058 | 1.30136458 | 0.19313369 | 0.31692458 | ZNF461 | 92283 |
| ENSG000002 | 98.1241847 | 0.258045019 | 0.19829234 | 1.30133627 | 0.19314338 | 0.31692458 | PPM1N | 147699 |
| ENSG000002 | 9.33675471 | -0.446171073 | 0.34290614 | -1.3011464 | 0.19320836 | 0.31701838 | NA | NA |
| ENSG000002 | 3.06484105 | 0.487026547 | 0.37434444 | 1.30101184 | 0.1932544 | 0.31708109 | MIR8082 | 102465878 |
| ENSG000001 | 267.844253 | -0.173881253 | 0.13369424 | -1.300589 | 0.19339919 | 0.31730582 | USP3 | 9960 |
| ENSG000002 | 6.49656188 | -0.472188209 | 0.36308261 | -1.300498 | 0.19343034 | 0.31732501 | NA | NA |
| ENSG000002 | 134.424071 | 0.200071387 | 0.15384352 | 1.30048625 | 0.19343436 | 0.31732501 | ZNF345 | 25850 |
| ENSG000002 | 5.64481535 | -0.481057295 | 0.36990354 | -1.3004939 | 0.19343175 | 0.31732501 | NA | NA |
| ENSG000002 | 7.82920098 | -0.463366583 | 0.35632552 | -1.3004024 | 0.19346307 | 0.31735926 | HRAT17 | 101928036 |
| ENSG000002 | 4.36609485 | -0.49179352 | 0.37820916 | -1.3003216 | 0.19349078 | 0.31739188 | NA | NA |
| ENSG000001 | 121.835774 | -0.208196269 | 0.16012911 | -1.3001775 | 0.19354014 | 0.31746 | ACER2 | 340485 |
| ENSG000002 | 5.99043508 | -0.478762416 | 0.36823954 | -1.3001385 | 0.1935535 | 0.31746907 | NA | NA |
| ENSG00000C | 53.3687838 | -0.272011435 | 0.20923149 | -1.3000502 | 0.19358378 | 0.31749285 | MTMR11 | 10903 |
| ENSG000002 | 4.36938108 | -0.49186899 | 0.37835271 | -1.3000277 | 0.19359148 | 0.31749285 | NA | NA |
| ENSG000002 | 9.34613061 | -0.444578921 | 0.34196703 | -1.3000637 | 0.19357914 | 0.31749285 | ADH1C | 126 |
| ENSG000001 | 4.56519988 | -0.488436843 | 0.37573005 | -1.2999675 | 0.19361212 | 0.31750139 | NA | NA |
| ENSG000002 | 3.79606411 | -0.492071225 | 0.37852753 | -1.2999615 | 0.19361415 | 0.31750139 | NA | NA |
| ENSG000002 | 4.63868483 | -0.48969827 | 0.37671217 | -1.2999269 | 0.19362604 | 0.31750139 | NA | NA |
| ENSG000002 | 2.72469104 | -0.482049085 | 0.37082949 | -1.2999211 | 0.19362801 | 0.31750139 | NA | NA |
| ENSG000001 | 181.522456 | 0.190616285 | 0.14664209 | 1.2998743 | 0.19364405 | 0.31751486 | EXOC6 | 54536 |
| ENSG000002 | 4.19216056 | -0.492102992 | 0.37858548 | -1.2998465 | 0.1936536 | 0.31751767 | NA | NA |
| ENSG000002 | 3.4903721 | -0.491939188 | 0.37848119 | -1.2997718 | 0.19367918 | 0.31752543 | NA | NA |
| ENSG000002 | 7.92495624 | -0.466318877 | 0.35876677 | -1.2997828 | 0.19367542 | 0.31752543 | NA | NA |
| ENSG000002 | 2.61297195 | -0.486176653 | 0.3740499 | -1.2997641 | 0.19368182 | 0.31752543 | NA | NA |

| | | | | | | | | |
|---|---|---|---|---|---|---|---|---|
| ENSG000002 | 3.19475928 | -0.490822742 | 0.37765856 | -1.2996468 | 0.19372205 | 0.31757855 | NA | NA |
| ENSG000002 | 3.75844051 | -0.491723058 | 0.37837144 | -1.2995776 | 0.19374577 | 0.3176046 | NA | NA |
| ENSG000002 | 6.74836136 | -0.473476108 | 0.3643679 | -1.2994452 | 0.1937912 | 0.31766622 | LOC101927188 | 101927188 |
| ENSG000002 | 9.13621594 | -0.460327554 | 0.35431 | -1.2992226 | 0.19386755 | 0.31777852 | HPR | 3250 |
| ENSG000002 | 4.613417 | -0.489155981 | 0.37652715 | -1.2991254 | 0.1939009 | 0.31782034 | UBD | 10537 |
| ENSG000001 | 330.976025 | 0.203975759 | 0.15701602 | 1.29907606 | 0.19391783 | 0.31783525 | KIAA0430 | 9665 |
| ENSG000002 | 5.74562209 | 0.483508821 | 0.37220989 | 1.29902196 | 0.19393639 | 0.31785283 | SNORD8 | 319103 |
| ENSG000001 | 8.32412677 | -0.452931308 | 0.34873613 | -1.2987794 | 0.19401964 | 0.31796637 | GLIS1 | 148979 |
| ENSG000002 | 4.63270312 | -0.488831634 | 0.37637917 | -1.2987744 | 0.19402136 | 0.31796637 | NA | NA |
| ENSG000001 | 9.96333039 | -0.453833803 | 0.34944026 | -1.298745 | 0.19403145 | 0.31797006 | CHRFAM7A | 89832 |
| ENSG000002 | 2.63487562 | 0.481512585 | 0.37080202 | 1.29857055 | 0.19409135 | 0.31802967 | NA | NA |
| ENSG000002 | 88.0959038 | -0.219215659 | 0.1688119 | -1.2985794 | 0.1940883 | 0.31802967 | LOC730102 | 730102 |
| ENSG000002 | 18.2215237 | -0.372927618 | 0.28717485 | -1.2986082 | 0.19407842 | 0.31802967 | NA | NA |
| ENSG000002 | 54.0383179 | 0.269145047 | 0.20727968 | 1.29846326 | 0.19412819 | 0.31807341 | NA | NA |
| ENSG000002 | 7.51389317 | -0.462583883 | 0.35625931 | -1.2984471 | 0.19413373 | 0.31807341 | NA | NA |
| ENSG000001 | 126.171328 | 0.230503402 | 0.17753275 | 1.29837118 | 0.19415982 | 0.3181033 | CHST2 | 9435 |
| ENSG000002 | 9.52109156 | -0.454914966 | 0.35039647 | -1.2982864 | 0.19418895 | 0.31813817 | NA | NA |
| ENSG000002 | 8.02022597 | -0.482698589 | 0.3718046 | -1.2982588 | 0.19419842 | 0.31814084 | NA | NA |
| ENSG000001 | 2.65515509 | -0.484293058 | 0.37309019 | -1.298059 | 0.19426708 | 0.31823102 | LCE1E | 353135 |
| ENSG000002 | 3.29950212 | -0.489506297 | 0.37710813 | -1.2980529 | 0.19426917 | 0.31823102 | NA | NA |
| ENSG000002 | 3.72249931 | -0.49118207 | 0.37840868 | -1.2980201 | 0.19428043 | 0.31823661 | NA | NA |
| ENSG00000C | 6.24403284 | -0.475854287 | 0.36661848 | -1.297955 | 0.1943028 | 0.31824869 | CACNG3 | 10368 |
| ENSG000002 | 7.76003929 | -0.462196958 | 0.35610189 | -1.2979346 | 0.19430982 | 0.31824869 | ZNF32-AS3 | 414201 |
| ENSG000002 | 13.3907884 | -0.416997548 | 0.32127889 | -1.2979301 | 0.19431135 | 0.31824869 | NA | NA |
| ENSG000002 | 2.39273085 | -0.477458081 | 0.36787794 | -1.2978709 | 0.1943317 | 0.31825632 | NA | NA |
| ENSG000002 | 2.34688703 | -0.479080279 | 0.36912199 | -1.2978915 | 0.19432464 | 0.31825632 | RNU2-20P | 106481640 |
| ENSG000002 | 3.75144567 | -0.491279072 | 0.37853402 | -1.2978465 | 0.19434007 | 0.31825718 | NA | NA |
| ENSG000002 | 4.32804704 | -0.49069071 | 0.37811332 | -1.2977345 | 0.19437858 | 0.31830738 | NA | NA |
| ENSG000002 | 4.68236963 | -0.489485181 | 0.37721493 | -1.2976294 | 0.19441472 | 0.3183537 | NA | NA |

| | | | | | | | | |
|---|---|---|---|---|---|---|---|---|
| ENSG000002 | 3.75055943 | -0.491046573 | 0.37853358 | -1.2972338 | 0.19455074 | 0.31856358 | NA | NA |
| ENSG000001 | 6.98861819 | -0.468642286 | 0.36129027 | -1.2971351 | 0.19458471 | 0.31860633 | ASB17 | 127247 |
| ENSG000002 | 11.592646 | 0.42360077 | 0.32658411 | 1.29706485 | 0.19460887 | 0.31863304 | NA | NA |
| ENSG000002 | 6.42828121 | -0.469742616 | 0.36216806 | -1.2970294 | 0.19462108 | 0.31864015 | OR5AM1P | 81226 |
| ENSG000002 | 2.61231906 | -0.484472257 | 0.37354512 | -1.2969578 | 0.1946457 | 0.3186676 | NA | NA |
| ENSG000001 | 191.688401 | 0.206539922 | 0.1592657 | 1.29682613 | 0.19469102 | 0.31872892 | MIR367 | 442912 |
| ENSG000001 | 24.5394398 | -0.343525096 | 0.26491581 | -1.2967331 | 0.19472303 | 0.31876847 | CHRNA1 | 1134 |
| ENSG000002 | 23.1505687 | -0.350839687 | 0.27056285 | -1.2967031 | 0.19473336 | 0.3187725 | NA | NA |
| ENSG000002 | 24.5382265 | -0.337982483 | 0.26068048 | -1.2965393 | 0.19478976 | 0.31885196 | NA | NA |
| ENSG000002 | 3.21462977 | -0.488349257 | 0.37669783 | -1.2963952 | 0.19483937 | 0.31892029 | NA | NA |
| ENSG000002 | 9.89894515 | 0.440529894 | 0.33987843 | 1.2961396 | 0.1949274 | 0.3190515 | NA | NA |
| ENSG000002 | 18.2281321 | 0.386086266 | 0.29788165 | 1.29610624 | 0.19493888 | 0.31905742 | LINC01126 | 100129726 |
| ENSG000001 | 90.3846812 | 0.244851745 | 0.18892441 | 1.29603026 | 0.19496506 | 0.31906526 | NA | NA |
| ENSG000001 | 6.39799683 | -0.46965532 | 0.36237361 | -1.2960528 | 0.1949573 | 0.31906526 | GPBAR1 | 151306 |
| ENSG000002 | 14.4827162 | -0.402862057 | 0.31084464 | -1.2960238 | 0.19496728 | 0.31906526 | NA | NA |
| ENSG000002 | 11.2187751 | -0.434567951 | 0.33533771 | -1.2959114 | 0.19500602 | 0.31911579 | NA | NA |
| ENSG000002 | 2.64342464 | -0.48295485 | 0.37272917 | -1.2957259 | 0.19506993 | 0.31920748 | NA | NA |
| ENSG000001 | 128.46046 | 0.234232085 | 0.18077978 | 1.29567636 | 0.19508701 | 0.31922255 | NA | NA |
| ENSG000001 | 17.6160086 | -0.397352485 | 0.30668626 | -1.2956318 | 0.19510235 | 0.31923478 | ACOT12 | 134526 |
| ENSG000001 | 139.933339 | 0.205623831 | 0.15871845 | 1.29552571 | 0.19513894 | 0.31927729 | PARP11 | 57097 |
| ENSG000002 | 2.97878958 | -0.484192292 | 0.37374624 | -1.2955108 | 0.19514408 | 0.31927729 | NA | NA |
| ENSG000002 | 8.8287223 | -0.451818036 | 0.34878243 | -1.2954151 | 0.19517706 | 0.31931836 | NA | NA |
| ENSG000001 | 4.30574812 | -0.489697883 | 0.37804336 | -1.2953485 | 0.19520005 | 0.31933021 | OR51B4 | 79339 |
| ENSG000002 | 4.37117233 | -0.489363996 | 0.37778423 | -1.2953532 | 0.19519843 | 0.31933021 | NA | NA |
| ENSG000002 | 4.06583186 | -0.490087802 | 0.37843179 | -1.2950493 | 0.19530324 | 0.31948613 | NA | NA |
| ENSG000001 | 194.589494 | 0.179409807 | 0.13854044 | 1.29499953 | 0.19532039 | 0.3195013 | DENND2C | 163259 |
| ENSG000002 | 7.02412512 | 0.469590297 | 0.36264116 | 1.29491725 | 0.19534878 | 0.31953484 | NA | NA |
| ENSG000001 | 30.179432 | -0.333642147 | 0.25768466 | -1.2947692 | 0.19539987 | 0.31956712 | CD274 | 29126 |
| ENSG000001 | 1518.07994 | 0.153425563 | 0.11849775 | 1.29475506 | 0.19540474 | 0.31956712 | MIER1 | 57708 |

| | | | | | | | | |
|---|---|---|---|---|---|---|---|---|
| ENSG000002 | 3.87704977 | -0.489921118 | 0.378369 | -1.2948236 | 0.19538108 | 0.31956712 | NA | NA |
| ENSG000002 | 6.91093962 | -0.475692393 | 0.36740213 | -1.2947459 | 0.19540791 | 0.31956712 | NA | NA |
| ENSG000002 | 30.1174642 | -0.322856477 | 0.24935879 | -1.2947467 | 0.19540761 | 0.31956712 | LOC100129550 | 100129550 |
| ENSG000002 | 3.64861217 | 0.489911247 | 0.3783946 | 1.29470994 | 0.19542031 | 0.3195745 | NA | NA |
| ENSG000001 | 17.640519 | -0.379898501 | 0.29343292 | -1.294669 | 0.19543444 | 0.3195821 | TOX2 | 84969 |
| ENSG000002 | 11.7691157 | -0.426756908 | 0.3296309 | -1.2946508 | 0.19544071 | 0.3195821 | NA | NA |
| ENSG000002 | 3.23831272 | -0.487752553 | 0.37677137 | -1.2945584 | 0.1954726 | 0.31962135 | NA | NA |
| ENSG000002 | 3.22959745 | -0.4880356 | 0.37702363 | -1.294443 | 0.19551244 | 0.31967361 | RN7SL782P | 106481133 |
| ENSG000001 | 62.1559829 | 0.246261845 | 0.19025921 | 1.29434917 | 0.19554484 | 0.31969158 | ACYP1 | 97 |
| ENSG000001 | 288.11821 | 0.157690578 | 0.12183051 | 1.29434393 | 0.19554665 | 0.31969158 | TIMP4 | 7079 |
| ENSG000002 | 11.1840402 | -0.445923497 | 0.34451735 | -1.2943427 | 0.19554709 | 0.31969158 | LOC101928583 | 101928583 |
| ENSG000002 | 15.8610662 | -0.404978662 | 0.31290155 | -1.2942686 | 0.19557265 | 0.31970759 | HLA-G | 3135 |
| ENSG000002 | 7.71063941 | -0.469716406 | 0.36291402 | -1.2942912 | 0.19556487 | 0.31970759 | NA | NA |
| ENSG000001 | 84.8085636 | 0.223816943 | 0.17294388 | 1.29415939 | 0.19561037 | 0.31975636 | PIK3R5 | 23533 |
| ENSG000001 | 28.2613861 | -0.329607247 | 0.25470151 | -1.2940922 | 0.19563356 | 0.3197685 | LRRC4B | 94030 |
| ENSG000002 | 8.76840636 | -0.442808676 | 0.34217468 | -1.2941012 | 0.19563046 | 0.3197685 | NA | NA |
| ENSG000000 | 11.6791155 | -0.417847289 | 0.32293093 | -1.2939216 | 0.19569252 | 0.3198385 | WAS | 7454 |
| ENSG000001 | 25.9486688 | -0.345905024 | 0.26733526 | -1.2938997 | 0.19570005 | 0.3198385 | KCNMB1 | 3779 |
| ENSG000002 | 2.2374915 | 0.471987913 | 0.36477374 | 1.29391966 | 0.19569317 | 0.3198385 | ST20-MTHFS | 100528021 |
| ENSG000002 | 4.13170659 | -0.489735622 | 0.37850518 | -1.2938677 | 0.19571113 | 0.31984371 | NA | NA |
| ENSG000002 | 56.8514217 | -0.254474518 | 0.19671096 | -1.2936469 | 0.19578742 | 0.31995549 | NA | NA |
| ENSG000002 | 2.97876563 | -0.485016962 | 0.3749525 | -1.2935424 | 0.19582353 | 0.32000161 | NA | NA |
| ENSG000002 | 7.84137711 | -0.458203747 | 0.35425518 | -1.2934285 | 0.1958629 | 0.32005305 | NA | NA |
| ENSG000002 | 2.60403357 | -0.483124475 | 0.37353256 | -1.2933932 | 0.1958751 | 0.32006008 | RNU1-33P | 106480156 |
| ENSG000002 | 9.97158058 | -0.441146087 | 0.34116135 | -1.2930717 | 0.19598626 | 0.32022882 | SCX | 642658 |
| ENSG000002 | 2.88981456 | -0.487267331 | 0.37683781 | -1.2930426 | 0.19599634 | 0.32023239 | NA | NA |
| ENSG000002 | 18.6379925 | -0.37557588 | 0.29047169 | -1.2929862 | 0.19601585 | 0.32025135 | RPL13AP17 | 399670 |
| ENSG000002 | 2.65656501 | -0.483110857 | 0.3736547 | -1.292934 | 0.19603389 | 0.32026793 | RN7SL679P | 106481855 |
| ENSG000002 | 2.60107731 | -0.482204315 | 0.37301535 | -1.2927197 | 0.196108 | 0.32037611 | NA | NA |

| | | | | | | | | |
|---|---|---|---|---|---|---|---|---|
| ENSG000002 | 6.80085295 | -0.472016607 | 0.365149 | -1.2926685 | 0.19612574 | 0.32039218 | NA | NA |
| ENSG000001 | 3.51279466 | -0.488997436 | 0.37830404 | -1.2926043 | 0.19614795 | 0.32041554 | ANKRD20A2 | 441430 |
| ENSG000001 | 103.707619 | 0.218005117 | 0.16867326 | 1.29246992 | 0.19619446 | 0.3204786 | ZNF70 | 7621 |
| ENSG000001 | 3.0759223 | -0.478290499 | 0.37010081 | -1.2923249 | 0.19624464 | 0.32053476 | KLHDC7B | 113730 |
| ENSG000002 | 3.51204536 | -0.489031335 | 0.37841099 | -1.2923286 | 0.19624338 | 0.32053476 | NA | NA |
| ENSG000001 | 137.530996 | -0.214452067 | 0.16595268 | -1.2922483 | 0.19627119 | 0.32054954 | MAD2L1BP | 9587 |
| ENSG000002 | 2.59760784 | -0.483248613 | 0.37396478 | -1.2922303 | 0.19627741 | 0.32054954 | NA | NA |
| ENSG000002 | 3.74691837 | -0.489183715 | 0.37854521 | -1.2922729 | 0.19626266 | 0.32054954 | NA | NA |
| ENSG000001 | 58.3013935 | -0.250857549 | 0.19413309 | -1.2921937 | 0.19629008 | 0.32055733 | SLC13A5 | 284111 |
| ENSG000002 | 7.25475626 | -0.462756198 | 0.35813905 | -1.2921132 | 0.19631794 | 0.32058991 | NA | NA |
| ENSG000002 | 3.24232879 | 0.488142603 | 0.37781524 | 1.29201406 | 0.19635228 | 0.32063307 | MIR98 | 407054 |
| ENSG000001 | 6.25196535 | -0.476260232 | 0.36871182 | -1.2916869 | 0.19646559 | 0.32078049 | IL21 | 59067 |
| ENSG000002 | 9.58450795 | -0.441227871 | 0.34159873 | -1.2916555 | 0.19647648 | 0.32078049 | NA | NA |
| ENSG000002 | 3.77062799 | -0.488840653 | 0.37846531 | -1.2916393 | 0.1964821 | 0.32078049 | KRT8P22 | 342419 |
| ENSG000002 | 18.8952824 | 0.383541563 | 0.29693828 | 1.29165415 | 0.19647694 | 0.32078049 | NA | NA |
| ENSG000002 | 8.23023317 | -0.450248486 | 0.34857003 | -1.2917017 | 0.19646048 | 0.32078049 | NA | NA |
| ENSG000002 | 3.473997 | -0.488812525 | 0.37846464 | -1.2915672 | 0.19650706 | 0.32080831 | NA | NA |
| ENSG000001 | 12.3112308 | -0.420867638 | 0.32591075 | -1.2913586 | 0.19657937 | 0.32091345 | BHLHE22 | 27319 |
| ENSG000002 | 4.5807519 | -0.486051035 | 0.37642104 | -1.291243 | 0.19661943 | 0.32096592 | NA | NA |
| ENSG000002 | 3.00682574 | -0.483890607 | 0.37476063 | -1.2911992 | 0.19663463 | 0.32097781 | LOC105376606 | 105376606 |
| ENSG000002 | 3.49002664 | -0.488528806 | 0.37840274 | -1.2910287 | 0.19669372 | 0.32106134 | NA | NA |
| ENSG000002 | 3.78038734 | -0.488637658 | 0.37849965 | -1.2909858 | 0.19670861 | 0.32107273 | NA | NA |
| ENSG000002 | 3.23115619 | -0.486366191 | 0.37676467 | -1.2909018 | 0.19673773 | 0.32110734 | NA | NA |
| ENSG000002 | 5.98702902 | -0.477872389 | 0.37019902 | -1.2908527 | 0.19675477 | 0.32112222 | NA | NA |
| ENSG000002 | 37.2096986 | -0.302363926 | 0.23424578 | -1.2907977 | 0.19677383 | 0.3211404 | FAM228B | 375190 |
| ENSG000002 | 4.36616512 | -0.488275709 | 0.37830572 | -1.2906908 | 0.19681092 | 0.32118801 | NA | NA |
| ENSG000002 | 14.3785632 | -0.408456245 | 0.31648197 | -1.2906146 | 0.19683736 | 0.32121823 | NA | NA |
| ENSG000002 | 4.01919026 | -0.488543781 | 0.37857322 | -1.2904869 | 0.19688166 | 0.3212776 | NA | NA |
| ENSG000001 | 2.63944608 | -0.481411426 | 0.3730737 | -1.2903923 | 0.19691448 | 0.32131822 | CCL21 | 6366 |

| | | | | | | | | |
|---|---|---|---|---|---|---|---|---|
| ENSG000002 | 3.75545921 | -0.48841759 | 0.37854081 | -1.290264 | 0.196959 | 0.32137793 | NA | NA |
| ENSG000002 | 7.12955485 | -0.4700291 | 0.36434457 | -1.2900675 | 0.19702721 | 0.32146845 | NA | NA |
| ENSG000002 | 4.39608086 | -0.487388912 | 0.37780371 | -1.2900586 | 0.19703033 | 0.32146845 | NA | NA |
| ENSG000002 | 4.61132803 | -0.487022343 | 0.37753062 | -1.2900208 | 0.19704344 | 0.32147691 | NA | NA |
| ENSG000002 | 6.67045649 | -0.463901702 | 0.35970281 | -1.2896805 | 0.19716161 | 0.32165676 | NA | NA |
| ENSG000002 | 73.289679 | 0.242503698 | 0.18804332 | 1.28961613 | 0.19718397 | 0.3216803 | NA | NA |
| ENSG000002 | 354.926722 | 0.180519472 | 0.14000401 | 1.28938787 | 0.19726328 | 0.32179673 | BMPR2 | 659 |
| ENSG000002 | 5.4228158 | -0.483190139 | 0.37481872 | -1.2891302 | 0.19735282 | 0.32192325 | FOXO6 | 100132074 |
| ENSG000002 | 2.35203422 | -0.474076027 | 0.36775194 | -1.289119 | 0.19735671 | 0.32192325 | NA | NA |
| ENSG000001 | 57.4916124 | -0.257816395 | 0.2000079 | -1.2890311 | 0.19738729 | 0.32194631 | SYCP1 | 6847 |
| ENSG000002 | 2.38015904 | -0.47287904 | 0.36684726 | -1.2890352 | 0.19738584 | 0.32194631 | MIR138-1 | 406929 |
| ENSG000002 | 12.9752033 | -0.40821833 | 0.3166914 | -1.2890098 | 0.19739467 | 0.32194631 | NA | NA |
| ENSG000002 | 15.8327493 | -0.398512943 | 0.30920238 | -1.2888417 | 0.19745311 | 0.32202868 | NA | NA |
| ENSG000002 | 14.2181637 | -0.395633325 | 0.30701107 | -1.2886614 | 0.19751582 | 0.32210504 | NA | NA |
| ENSG000002 | 19.1435943 | -0.368221643 | 0.28573661 | -1.288675 | 0.19751108 | 0.32210504 | NA | NA |
| ENSG000002 | 178.179338 | 0.198452994 | 0.15400481 | 1.28861555 | 0.19753178 | 0.32211811 | CASC10 | 399726 |
| ENSG000002 | 4.7983306 | -0.487633589 | 0.37842817 | -1.2885763 | 0.19754541 | 0.3221274 | NA | NA |
| ENSG000002 | 3.6859313 | -0.484934076 | 0.37636573 | -1.288465 | 0.19758414 | 0.32217759 | NA | NA |
| ENSG000002 | 6.23721013 | -0.471502238 | 0.36596056 | -1.2883963 | 0.19760804 | 0.3222036 | LINC01499 | 102723644 |
| ENSG000002 | 2.99479527 | -0.482437396 | 0.37445878 | -1.2883591 | 0.19762098 | 0.32221174 | NA | NA |
| ENSG000001 | 12.5259145 | -0.414538911 | 0.32177592 | -1.2882844 | 0.19764697 | 0.32224117 | KRT86 | 3892 |
| ENSG000002 | 9151.22902 | 0.465392849 | 0.36130108 | 1.28810257 | 0.19771026 | 0.32233139 | NA | NA |
| ENSG000002 | 6.01324222 | -0.477900729 | 0.37105808 | -1.2879405 | 0.19776668 | 0.32241042 | NA | NA |
| ENSG000002 | 5.72954887 | -0.478892637 | 0.37183633 | -1.2879125 | 0.19777644 | 0.32241336 | NA | NA |
| ENSG00000C | 18.7421382 | 0.37795831 | 0.29348211 | 1.28784105 | 0.1978013 | 0.32244093 | BEST2 | 54831 |
| ENSG000001 | 2.66616816 | -0.479470252 | 0.3723306 | -1.2877541 | 0.19783158 | 0.32246437 | KRTAP21-1 | 337977 |
| ENSG000002 | 3.55064748 | -0.487168674 | 0.37830381 | -1.287771 | 0.19782569 | 0.32246437 | NA | NA |
| ENSG000001 | 131.895435 | -0.201655578 | 0.15660485 | -1.2876713 | 0.1978604 | 0.32249837 | TRPM6 | 140803 |
| ENSG000001 | 7.06012039 | -0.468965664 | 0.36423424 | -1.2875387 | 0.19790661 | 0.32252221 | HTR1E | 3354 |

| | | | | | | | | |
|---|---|---|---|---|---|---|---|---|
| ENSG000002 | 6.84933969 | -0.473341195 | 0.3676328 | -1.287538 | 0.19790684 | 0.32252221 | NA | NA |
| ENSG000002 | 2.98928888 | -0.482067847 | 0.37440998 | -1.2875401 | 0.19790611 | 0.32252221 | NA | NA |
| ENSG000002 | 2.71443391 | -0.47302774 | 0.36737597 | -1.2875849 | 0.19789051 | 0.32252221 | NA | NA |
| ENSG000002 | 6.31490883 | -0.475271389 | 0.36916851 | -1.2874104 | 0.19795128 | 0.32258167 | NA | NA |
| ENSG000002 | 3.46864021 | -0.486897818 | 0.37829369 | -1.2870895 | 0.19806311 | 0.32275094 | NA | NA |
| ENSG000001 | 4.59163414 | -0.484671539 | 0.37657595 | -1.2870486 | 0.19807736 | 0.32276119 | NA | NA |
| ENSG000001 | 60.3757074 | 0.272927258 | 0.21206442 | 1.28700163 | 0.19809372 | 0.32276193 | ANKLE1 | 126549 |
| ENSG000002 | 7.35343288 | -0.463483854 | 0.36012687 | -1.2870016 | 0.19809373 | 0.32276193 | SMAD1-AS2 | 101927659 |
| ENSG000002 | 3.77141672 | -0.487178502 | 0.37857239 | -1.2868833 | 0.19813496 | 0.32281613 | NA | NA |
| ENSG000002 | 8.5946233 | -0.446392398 | 0.34694932 | -1.2866214 | 0.1982263 | 0.32295197 | NA | NA |
| ENSG000002 | 9.4008348 | -0.441260286 | 0.34297497 | -1.286567 | 0.19824524 | 0.32296986 | AKAP2 | 11217 |
| ENSG000002 | 2.62427044 | -0.479603335 | 0.37278488 | -1.2865418 | 0.19825404 | 0.32297122 | NA | NA |
| ENSG000002 | 3.72779553 | -0.487063968 | 0.37859204 | -1.286514 | 0.19826373 | 0.32297403 | NA | NA |
| ENSG00000C | 117.764814 | 0.21550654 | 0.167524 | 1.28642189 | 0.19829586 | 0.3229814 | GEMIN8 | 54960 |
| ENSG000001 | 45.0670194 | -0.296213 | 0.23026334 | -1.2864097 | 0.19830011 | 0.3229814 | SLC51A | 200931 |
| ENSG000002 | 3.84794444 | 0.486989176 | 0.37856406 | 1.28641155 | 0.19829947 | 0.3229814 | NA | NA |
| ENSG000002 | 17.3032881 | -0.382149613 | 0.29705182 | -1.2864746 | 0.19827749 | 0.3229814 | LOC283140 | 283140 |
| ENSG000001 | 3.73285895 | -0.486993276 | 0.37859313 | -1.2863236 | 0.19833016 | 0.32301736 | RHOXF2 | 84528 |
| ENSG000002 | 9.48755207 | -0.443843144 | 0.34508146 | -1.2861982 | 0.19837391 | 0.32306266 | NA | NA |
| ENSG000002 | 5.53603641 | -0.481304855 | 0.3742025 | -1.286215 | 0.19836805 | 0.32306266 | NA | NA |
| ENSG000002 | 14.5344113 | -0.405658875 | 0.31540696 | -1.2861443 | 0.19839269 | 0.32308028 | NA | NA |
| ENSG000002 | 2.58863721 | -0.480762914 | 0.37381221 | -1.2861081 | 0.19840533 | 0.32308789 | NA | NA |
| ENSG000002 | 3.76592353 | -0.486843704 | 0.37854783 | -1.2860824 | 0.19841431 | 0.32308953 | NA | NA |
| ENSG000002 | 2.32915375 | -0.471480721 | 0.36662078 | -1.2860174 | 0.19843697 | 0.32311346 | NA | NA |
| ENSG000002 | 8.55491216 | -0.444348454 | 0.34553042 | -1.2859894 | 0.19844677 | 0.32311643 | NA | NA |
| ENSG000002 | 18.2520719 | 0.376801267 | 0.29302757 | 1.28589016 | 0.1984814 | 0.32315985 | NA | NA |
| ENSG000001 | 24.4868746 | -0.344718041 | 0.26808853 | -1.2858366 | 0.1985001 | 0.32316814 | HTR3B | 9177 |
| ENSG000002 | 85.336754 | 0.262046834 | 0.20379588 | 1.28582991 | 0.19850243 | 0.32316814 | MICA | 100507436 |
| ENSG000002 | 9.40723907 | -0.445593589 | 0.34655365 | -1.2857853 | 0.19851801 | 0.32318053 | MAFTRR | 102467146 |

| | | | | | | | | |
|---|---|---|---|---|---|---|---|---|
| ENSG000001 | 40.2744979 | 0.309389616 | 0.24064197 | 1.28568436 | 0.19855324 | 0.32322491 | ULBP3 | 79465 |
| ENSG000002 | 2.66522243 | -0.479161221 | 0.37273231 | -1.2855371 | 0.19860465 | 0.32328265 | NA | NA |
| ENSG000002 | 6.22901419 | -0.471007653 | 0.36638937 | -1.2855385 | 0.19860416 | 0.32328265 | NA | NA |
| ENSG000001 | 905.65365 | 0.143118456 | 0.11133788 | 1.28544266 | 0.19863764 | 0.32330547 | PHC3 | 80012 |
| ENSG000002 | 9.59865856 | -0.440401402 | 0.34260503 | -1.2854493 | 0.19863532 | 0.32330547 | NA | NA |
| ENSG000002 | 2.8959601 | -0.484000597 | 0.37652861 | -1.2854285 | 0.19864259 | 0.32330547 | IL21R-AS1 | 283888 |
| ENSG000002 | 4.85533115 | -0.482475301 | 0.37536678 | -1.2853436 | 0.19867222 | 0.32334072 | NA | NA |
| ENSG000002 | 50.8790115 | 0.301993333 | 0.23498263 | 1.28517299 | 0.19873184 | 0.32342476 | PCDHGB6 | 56100 |
| ENSG000002 | 9.29583373 | -0.436238418 | 0.33951617 | -1.2848826 | 0.19883332 | 0.32355415 | FLJ33581 | 400839 |
| ENSG000002 | 6.8897498 | -0.47002745 | 0.36581514 | -1.284877 | 0.19883528 | 0.32355415 | NA | NA |
| ENSG000002 | 20.83817 | -0.359310283 | 0.27964542 | -1.2848781 | 0.19883488 | 0.32355415 | NA | NA |
| ENSG000002 | 10.144012 | -0.436400936 | 0.3396552 | -1.2848351 | 0.1988499 | 0.32356262 | NA | NA |
| ENSG000002 | 13.2578134 | -0.409859748 | 0.31900258 | -1.2848164 | 0.19885644 | 0.32356262 | NA | NA |
| ENSG000002 | 6.29188179 | -0.476554977 | 0.37094083 | -1.2847197 | 0.19889027 | 0.32360467 | NA | NA |
| ENSG000002 | 4.10984902 | 0.486207303 | 0.37851944 | 1.28449757 | 0.19896791 | 0.32371802 | RN7SL316P | 106479337 |
| ENSG000002 | 3.7403436 | -0.486051112 | 0.37844427 | -1.28434 | 0.19902302 | 0.32379468 | NA | NA |
| ENSG000002 | 2.3817376 | -0.472417863 | 0.36784678 | -1.2842789 | 0.19904439 | 0.32381645 | RN7SL502P | 106479407 |
| ENSG000001 | 244.788943 | 0.20132757 | 0.15679255 | 1.28403787 | 0.19912871 | 0.32394063 | PKDCC | 91461 |
| ENSG000002 | 2.86616497 | -0.483830023 | 0.37682688 | -1.2839584 | 0.19915653 | 0.32397289 | NA | NA |
| ENSG000002 | 12.2624767 | -0.45844493 | 0.35706643 | -1.2839205 | 0.19916976 | 0.32398142 | NA | NA |
| ENSG000001 | 12.0091537 | 0.412403666 | 0.32124072 | 1.28378389 | 0.19921758 | 0.32404622 | CSTA | 1475 |
| ENSG000001 | 7.99012048 | -0.450840015 | 0.35120769 | -1.2836849 | 0.19925222 | 0.32408295 | NPVF | 64111 |
| ENSG000002 | 9.43112496 | -0.445442851 | 0.34700629 | -1.2836737 | 0.19925615 | 0.32408295 | NA | NA |
| ENSG000001 | 1248.05466 | 0.147201615 | 0.11467806 | 1.28360742 | 0.19927935 | 0.32410769 | SESN3 | 143686 |
| ENSG000001 | 8.54275694 | -0.444097237 | 0.34600192 | -1.2835109 | 0.19931314 | 0.32414964 | KCNH3 | 23416 |
| ENSG000002 | 5.36121739 | -0.47516978 | 0.37023565 | -1.2834252 | 0.19934314 | 0.32418542 | NA | NA |
| ENSG000002 | 3.48419431 | -0.48567079 | 0.37842655 | -1.2833951 | 0.19935368 | 0.32418956 | NA | NA |
| ENSG000001 | 107.742243 | -0.21294137 | 0.16592497 | -1.2833594 | 0.1993662 | 0.32419693 | ATXN1 | 6310 |
| ENSG000001 | 8.50070103 | -0.448319971 | 0.3493631 | -1.2832494 | 0.19940473 | 0.32421583 | NA | NA |

| | | | | | | | | |
|---|---|---|---|---|---|---|---|---|
| ENSG000002 | 5.37492018 | -0.478667834 | 0.37302318 | -1.283212 | 0.1994178 | 0.32421583 | MIR548U | 100422884 |
| ENSG000002 | 3.48772327 | -0.485471472 | 0.37830928 | -1.2832661 | 0.19939886 | 0.32421583 | NA | NA |
| ENSG000002 | 2.86665424 | -0.483567981 | 0.37682783 | -1.2832597 | 0.1994011 | 0.32421583 | NA | NA |
| ENSG000002 | 5.99100703 | -0.477234971 | 0.37190511 | -1.283217 | 0.19941605 | 0.32421583 | NA | NA |
| ENSG000002 | 5.99629931 | -0.475595016 | 0.37070646 | -1.2829424 | 0.19951227 | 0.32435641 | NA | NA |
| ENSG000002 | 4.85565123 | -0.481766468 | 0.37554242 | -1.282855 | 0.19954288 | 0.32439318 | NA | NA |
| ENSG000001 | 34.3015893 | -0.308985582 | 0.24087422 | -1.2827673 | 0.1995736 | 0.32443011 | ADAMTS17 | 170691 |
| ENSG000002 | 3.24470673 | -0.483882712 | 0.37724612 | -1.2826711 | 0.19960732 | 0.32447192 | NA | NA |
| ENSG000002 | 8.96251891 | -0.449752485 | 0.3506459 | -1.2826401 | 0.1996182 | 0.32447659 | NA | NA |
| ENSG000002 | 28.4671161 | 0.317912835 | 0.24788631 | 1.28249454 | 0.19966922 | 0.32454652 | LINC00847 | 729678 |
| ENSG000002 | 3.47810031 | -0.485108454 | 0.37827249 | -1.2824312 | 0.19969142 | 0.32456959 | NA | NA |
| ENSG000002 | 5.16102071 | -0.481518285 | 0.37549529 | -1.2823551 | 0.19971812 | 0.32459998 | NA | NA |
| ENSG000001 | 26.9593159 | 0.330019559 | 0.2574155 | 1.28205007 | 0.19982508 | 0.32474779 | C8orf76 | 84933 |
| ENSG000002 | 3.77768802 | -0.485179784 | 0.37843812 | -1.2820584 | 0.19982215 | 0.32474779 | NA | NA |
| ENSG000001 | 11.937353 | -0.435790288 | 0.33992365 | -1.282024 | 0.19983422 | 0.32474963 | OAF | 220323 |
| ENSG000001 | 257.647213 | 0.168346306 | 0.13133328 | 1.28182519 | 0.19990398 | 0.32484997 | DEPDC5 | 9681 |
| ENSG000001 | 910.864349 | 0.178519385 | 0.13928026 | 1.28172781 | 0.19993814 | 0.32488273 | EZH2 | 2146 |
| ENSG000002 | 4.90860722 | -0.481595101 | 0.37574066 | -1.2817221 | 0.19994016 | 0.32488273 | NA | NA |
| ENSG000002 | 2.91862453 | -0.482036444 | 0.37609701 | -1.2816811 | 0.19995453 | 0.32489306 | NA | NA |
| ENSG000001 | 61.2597898 | -0.244721129 | 0.1909494 | -1.281602 | 0.1999823 | 0.32492516 | ANKAR | 150709 |
| ENSG000001 | 2.61040744 | -0.477581741 | 0.37268364 | -1.2814669 | 0.20002973 | 0.3249892 | OTUD6A | 139562 |
| ENSG000002 | 6.25956448 | 0.467540564 | 0.3648672 | 1.28139926 | 0.20005346 | 0.32501475 | NA | NA |
| ENSG000002 | 38.636162 | 0.313727577 | 0.24483775 | 1.28136928 | 0.20006399 | 0.32501882 | NA | NA |
| ENSG000001 | 124.894807 | -0.193568252 | 0.1510724 | -1.2812946 | 0.20009022 | 0.32504841 | THSD4 | 79875 |
| ENSG00000C | 21.8332978 | -0.36085951 | 0.28165639 | -1.2812048 | 0.20012175 | 0.32508661 | ASIC4 | 55515 |
| ENSG000002 | 7.58119258 | 0.470077084 | 0.36691083 | 1.28117526 | 0.20013211 | 0.32509042 | NA | NA |
| ENSG000002 | 2.60805234 | -0.478195429 | 0.3732641 | -1.2811182 | 0.20015215 | 0.32510995 | NA | NA |
| ENSG000002 | 4.69767445 | -0.483525448 | 0.3774702 | -1.2809632 | 0.20020659 | 0.32518534 | NA | NA |
| ENSG000001 | 118.076916 | -0.202072823 | 0.1577742 | -1.2807723 | 0.20027366 | 0.32528126 | ZHX3 | 23051 |

| | | | | | | | | |
|---|---|---|---|---|---|---|---|---|
| ENSG000002 | 3.54897318 | -0.483173651 | 0.37727692 | -1.280687 | 0.20030363 | 0.3253169 | NA | NA |
| ENSG000001 | 100.525465 | 0.214844554 | 0.16781766 | 1.28022612 | 0.20046562 | 0.32556696 | WIPF1 | 7456 |
| ENSG000002 | 8.38144101 | -0.450677386 | 0.35206391 | -1.2801011 | 0.2005096 | 0.32562534 | NA | NA |
| ENSG000002 | 4.80982702 | -0.479936517 | 0.37494535 | -1.2800173 | 0.20053907 | 0.32566016 | NA | NA |
| ENSG000002 | 4.04105386 | -0.484413355 | 0.37847023 | -1.2799246 | 0.20057165 | 0.32570003 | NA | NA |
| ENSG000002 | 2.62341699 | -0.478435316 | 0.37382602 | -1.2798342 | 0.20060346 | 0.32573864 | NA | NA |
| ENSG000002 | 4.96205449 | -0.482643518 | 0.37712529 | -1.2797962 | 0.20061682 | 0.32574728 | NA | NA |
| ENSG000001 | 49.1572866 | 0.281435744 | 0.21992443 | 1.27969293 | 0.20065315 | 0.32579323 | FBXL16 | 146330 |
| ENSG000001 | 360.533247 | -0.170473935 | 0.13321876 | -1.2796541 | 0.20066681 | 0.32580237 | TSPYL5 | 85453 |
| ENSG000002 | 4.34686985 | -0.483705215 | 0.37803295 | -1.2795319 | 0.20070982 | 0.32585915 | NA | NA |
| ENSG000002 | 3.86137768 | -0.483740864 | 0.37814202 | -1.2792571 | 0.20080654 | 0.32600312 | NA | NA |
| ENSG000001 | 22.9266577 | -0.348124769 | 0.27213937 | -1.279215 | 0.20082135 | 0.32601411 | GC | 2638 |
| ENSG000002 | 2.67919698 | -0.47657349 | 0.37262725 | -1.278955 | 0.2009129 | 0.32614967 | NA | NA |
| ENSG000002 | 3.87893271 | -0.483396423 | 0.37797782 | -1.2789015 | 0.20093173 | 0.32616719 | NA | NA |
| ENSG000002 | 3.4376318 | -0.483874153 | 0.3783914 | -1.2787663 | 0.20097938 | 0.32623147 | NA | NA |
| ENSG000002 | 3.44124508 | -0.483566848 | 0.3781748 | -1.2786861 | 0.20100763 | 0.32626428 | NA | NA |
| ENSG000002 | 6.3332743 | -0.473277102 | 0.37015651 | -1.2785865 | 0.20104272 | 0.32630009 | NA | NA |
| ENSG000002 | 10.9567045 | -0.427718017 | 0.3345264 | -1.2785778 | 0.20104579 | 0.32630009 | NA | NA |
| ENSG000002 | 7.20588119 | -0.455995634 | 0.35665841 | -1.278522 | 0.20106542 | 0.3263189 | NA | NA |
| ENSG000002 | 14.9680585 | 0.394853029 | 0.30885127 | 1.27845687 | 0.20108838 | 0.3263431 | MIR1282 | 100302254 |
| ENSG000002 | 4.00224667 | -0.482973295 | 0.37780487 | -1.278367 | 0.20112004 | 0.32637723 | NA | NA |
| ENSG000002 | 10.943086 | -0.422648338 | 0.33061981 | -1.2783515 | 0.20112551 | 0.32637723 | NA | NA |
| ENSG000002 | 2.94514707 | -0.477740062 | 0.37373682 | -1.2782794 | 0.20115094 | 0.32638373 | NA | NA |
| ENSG000002 | 2.90043533 | -0.480849315 | 0.37617146 | -1.2782716 | 0.20115366 | 0.32638373 | NA | NA |
| ENSG000002 | 13.158956 | -0.404098695 | 0.31612275 | -1.2782968 | 0.20114481 | 0.32638373 | NA | NA |
| ENSG000001 | 28.1772887 | -0.326179661 | 0.25518974 | -1.2781849 | 0.20118425 | 0.32642029 | FAIM2 | 23017 |
| ENSG000001 | 7.57191173 | -0.457030301 | 0.35759466 | -1.278068 | 0.20122544 | 0.32647407 | CNR2 | 1269 |
| ENSG00000C | 19.2004553 | -0.358240068 | 0.28035681 | -1.2778005 | 0.20131978 | 0.32659542 | LAT2 | 7462 |
| ENSG000001 | 2.38788258 | -0.467349606 | 0.36575565 | -1.2777646 | 0.20133245 | 0.32659542 | NA | NA |

| | | | | | | | | |
|---|---|---|---|---|---|---|---|---|
| ENSG000002 | 3.55130084 | -0.481503966 | 0.37682994 | -1.2777752 | 0.20132868 | 0.32659542 | NA | NA |
| ENSG000002 | 2.99897602 | -0.475218055 | 0.37190127 | -1.277807 | 0.20131747 | 0.32659542 | NA | NA |
| ENSG000002 | 3.8634066 | -0.482644487 | 0.37775386 | -1.2776692 | 0.20136607 | 0.32663689 | NA | NA |
| ENSG000002 | 2.36527624 | -0.467232196 | 0.36572144 | -1.2775631 | 0.20140353 | 0.32668459 | NA | NA |
| ENSG000002 | 8.5372315 | -0.444014367 | 0.34755927 | -1.2775213 | 0.20141826 | 0.32669541 | NA | NA |
| ENSG000002 | 14.5626673 | 0.410092458 | 0.32104867 | 1.27735292 | 0.20147768 | 0.32677193 | NA | NA |
| ENSG000002 | 4.3856028 | -0.483285885 | 0.37835279 | -1.2773419 | 0.20148156 | 0.32677193 | NA | NA |
| ENSG000002 | 5.37481871 | -0.474996165 | 0.3718741 | -1.2773037 | 0.20149505 | 0.32678074 | NA | NA |
| ENSG000001 | 11.7829045 | -0.428841499 | 0.33574775 | -1.2772729 | 0.20150592 | 0.3267853 | MS4A7 | 58475 |
| ENSG000002 | 3.27197811 | -0.481614595 | 0.3770854 | -1.277203 | 0.20153059 | 0.32679952 | OR2B3 | 442184 |
| ENSG000002 | 3.77034669 | -0.483534025 | 0.37858841 | -1.2772024 | 0.2015308 | 0.32679952 | NA | NA |
| ENSG000001 | 11.7996747 | -0.426490639 | 0.33394975 | -1.2771102 | 0.20156334 | 0.32681642 | ANGPTL3 | 27329 |
| ENSG000002 | 3.45872827 | 0.483493264 | 0.37858533 | 1.27710511 | 0.20156514 | 0.32681642 | NA | NA |
| ENSG000002 | 2.62108061 | -0.476134379 | 0.37282339 | -1.2771044 | 0.2015654 | 0.32681642 | NA | NA |
| ENSG000001 | 6.93657366 | -0.459003192 | 0.3594495 | -1.2769615 | 0.20161583 | 0.32687562 | OR7A5 | 26659 |
| ENSG000002 | 2.64093977 | -0.475960243 | 0.37273055 | -1.2769553 | 0.20161804 | 0.32687562 | NA | NA |
| ENSG000002 | 10.0280398 | 0.434606523 | 0.3403521 | 1.27693211 | 0.20162622 | 0.32687581 | NA | NA |
| ENSG000001 | 94.2796764 | 0.244163596 | 0.19121565 | 1.27690171 | 0.20163695 | 0.32688015 | SLC43A2 | 124935 |
| ENSG000002 | 12.655018 | -0.411876051 | 0.32257821 | -1.2768254 | 0.2016639 | 0.32691076 | CTAGE1 | 64693 |
| ENSG000002 | 5.14468153 | -0.477852447 | 0.37426608 | -1.276772 | 0.20168276 | 0.32692827 | NA | NA |
| ENSG000002 | 4.03894636 | -0.483208247 | 0.37850169 | -1.2766343 | 0.20173138 | 0.32699014 | NA | NA |
| ENSG000002 | 2.6174987 | -0.474551368 | 0.371732 | -1.2765954 | 0.20174512 | 0.32699014 | NA | NA |
| ENSG000002 | 11.6015653 | -0.412345826 | 0.32300163 | -1.276606 | 0.20174137 | 0.32699014 | NA | NA |
| ENSG000000 | 246.130598 | 0.16241603 | 0.12723883 | 1.27646596 | 0.20179085 | 0.3270456 | MAP3K13 | 9175 |
| ENSG000002 | 9.3568384 | -0.447762938 | 0.35078689 | -1.2764529 | 0.20179547 | 0.3270456 | NA | NA |
| ENSG000002 | 3.46597266 | -0.482832449 | 0.37828473 | -1.2763731 | 0.20182366 | 0.32707822 | TMEM5-AS1 | 104169670 |
| ENSG000002 | 8.77803606 | -0.459153137 | 0.35974985 | -1.2763122 | 0.20184517 | 0.32708918 | LOC284898 | 284898 |
| ENSG000002 | 21.8471894 | -0.350173442 | 0.27436431 | -1.2763083 | 0.20184656 | 0.32708918 | NA | NA |
| ENSG000001 | 34.1322481 | -0.308488723 | 0.24170968 | -1.2762779 | 0.2018573 | 0.32709352 | PCDHB13 | 56123 |

| | | | | | | | | | |
|---|---|---|---|---|---|---|---|---|---|
| ENSG000002 | 16.4228704 | 0.388582438 | 0.30447616 | 1.27623272 | 0.20187326 | 0.32710632 | NA | NA | |
| ENSG000001 | 36.0459498 | -0.318333369 | 0.24945005 | -1.2761408 | 0.20190577 | 0.32714591 | KCNA1 | | 3736 |
| ENSG000002 | 8.49346808 | -0.454891067 | 0.35647234 | -1.2760908 | 0.20192343 | 0.32716146 | NA | NA | |
| ENSG000002 | 5.42249058 | -0.474056602 | 0.3715142 | -1.2760121 | 0.20195126 | 0.32719091 | NA | NA | |
| ENSG000002 | 19.3418994 | -0.367934504 | 0.28835135 | -1.2759937 | 0.20195775 | 0.32719091 | NA | NA | |
| ENSG000002 | 9.34966316 | -0.446278349 | 0.34976729 | -1.2759294 | 0.20198049 | 0.32721469 | NA | NA | |
| ENSG000001 | 14.7823542 | -0.391323868 | 0.30680888 | -1.2754646 | 0.20214485 | 0.32746787 | FOXL1 | | 2300 |
| ENSG000002 | 3.52454549 | -0.482429175 | 0.37825633 | -1.2754028 | 0.20216672 | 0.32749021 | NA | NA | |
| ENSG000002 | 6.69818901 | -0.461458493 | 0.36182166 | -1.2753755 | 0.20217635 | 0.32749274 | NA | NA | |
| ENSG000002 | 2.90438939 | -0.479613544 | 0.3760779 | -1.2753037 | 0.20220176 | 0.32752081 | NA | NA | |
| ENSG000002 | 2.23614083 | 0.463112772 | 0.36315614 | 1.27524423 | 0.20222281 | 0.32754183 | NA | NA | |
| ENSG000002 | 2.65531871 | -0.474773989 | 0.37231442 | -1.2751964 | 0.20223975 | 0.32755618 | NA | NA | |
| ENSG000001 | 81.4045455 | -0.228126587 | 0.17891148 | -1.2750808 | 0.20228065 | 0.32759143 | POU6F2 | | 11281 |
| ENSG000002 | 5.60883325 | -0.469763523 | 0.36841941 | -1.2750781 | 0.20228159 | 0.32759143 | NA | NA | |
| ENSG000002 | 55.7209706 | 0.266796487 | 0.20924125 | 1.27506638 | 0.20228575 | 0.32759143 | NA | NA | |
| ENSG000002 | 5.4915027 | 0.481054843 | 0.37729488 | 1.27501028 | 0.20230561 | 0.32760304 | NA | NA | |
| ENSG000002 | 13.5088775 | -0.429891177 | 0.33716942 | -1.2750005 | 0.20230907 | 0.32760304 | NA | NA | |
| ENSG000001 | 111.464377 | 0.22746747 | 0.17842125 | 1.27489002 | 0.20234817 | 0.3276271 | LFNG | | 3955 |
| ENSG000001 | 16.0855627 | 0.387074589 | 0.30360592 | 1.2749244 | 0.202336 | 0.3276271 | GJA4 | | 2701 |
| ENSG000002 | 2.63303996 | -0.475494565 | 0.37296324 | -1.27491 | 0.2023411 | 0.3276271 | NA | NA | |
| ENSG000002 | 2.3617098 | -0.468156277 | 0.36723312 | -1.2748204 | 0.20237282 | 0.32765392 | NA | NA | |
| ENSG000001 | 194.516028 | 0.184316318 | 0.14459937 | 1.27466891 | 0.20242646 | 0.32772768 | ZNF415 | | 55786 |
| ENSG000002 | 109.695696 | -0.202809728 | 0.15911458 | -1.2746144 | 0.20244577 | 0.32774586 | PCDHA4 | | 56144 |
| ENSG000002 | 12.2339029 | -0.413647246 | 0.32456948 | -1.274449 | 0.20250433 | 0.32782759 | NA | NA | |
| ENSG000002 | 2.63477497 | -0.474682048 | 0.37247461 | -1.2744011 | 0.20252131 | 0.32784198 | NA | NA | |
| ENSG000001 | 3.46062795 | -0.481859847 | 0.37811756 | -1.2743652 | 0.20253404 | 0.32784234 | CARTPT | | 9607 |
| ENSG000002 | 8.34397543 | -0.453032374 | 0.35549941 | -1.2743548 | 0.2025377 | 0.32784234 | NA | NA | |
| ENSG000002 | 2.91721116 | -0.475990525 | 0.37354843 | -1.2742405 | 0.20257821 | 0.32788173 | NA | NA | |
| ENSG000002 | 4.69680111 | -0.482294889 | 0.3784902 | -1.2742599 | 0.20257132 | 0.32788173 | NA | NA | |

| | | | | | | | | |
|---|---|---|---|---|---|---|---|---|
| ENSG000002 | 5.29118757 | -0.479013798 | 0.37593006 | -1.2742099 | 0.20258904 | 0.32788618 | NA | NA |
| ENSG000002 | 9.26939205 | -0.43198661 | 0.33906258 | -1.2740616 | 0.20264159 | 0.32795813 | NA | NA |
| ENSG000002 | 31.5857725 | -0.330118214 | 0.25912169 | -1.2739891 | 0.20266727 | 0.3279866 | NA | NA |
| ENSG000002 | 5.91251959 | -0.46998044 | 0.36891275 | -1.2739609 | 0.20267726 | 0.32798968 | NA | NA |
| ENSG000002 | 3.23666223 | -0.479379044 | 0.37633542 | -1.2738079 | 0.20273149 | 0.32806435 | NA | NA |
| ENSG000001 | 7.33826301 | -0.463335021 | 0.363773 | -1.2736927 | 0.20277234 | 0.32811737 | RHD | 6007 |
| ENSG000002 | 3.81977496 | -0.481922335 | 0.37841108 | -1.2735418 | 0.20282584 | 0.32819084 | NA | NA |
| ENSG000002 | 2.58208674 | -0.474908894 | 0.37294333 | -1.2734077 | 0.20287341 | 0.32825472 | NA | NA |
| ENSG000002 | 6.64311344 | -0.460779907 | 0.36187553 | -1.2733105 | 0.20290789 | 0.32828277 | NA | NA |
| ENSG000002 | 5.0555122 | -0.474755116 | 0.37285251 | -1.2733054 | 0.20290969 | 0.32828277 | NA | NA |
| ENSG000002 | 4.95647654 | -0.480267419 | 0.3771861 | -1.2732903 | 0.20291504 | 0.32828277 | NA | NA |
| ENSG000001 | 9.37011247 | -0.44051521 | 0.346025 | -1.2730734 | 0.20299201 | 0.3283811 | AKR7A3 | 22977 |
| ENSG000002 | 6.24140964 | -0.469444352 | 0.36874747 | -1.2730782 | 0.20299031 | 0.3283811 | NA | NA |
| ENSG000001 | 7.93972513 | -0.456233387 | 0.35843146 | -1.2728609 | 0.20306741 | 0.32848997 | PRAMEF12 | 390999 |
| ENSG000002 | 6.86885178 | -0.469728627 | 0.36905657 | -1.2727822 | 0.20309535 | 0.32852206 | NA | NA |
| ENSG000001 | 641.521268 | 0.145254182 | 0.11412809 | 1.27272943 | 0.20311407 | 0.32852614 | ITSN2 | 50618 |
| ENSG000002 | 27.5269831 | 0.339706796 | 0.26691066 | 1.27273596 | 0.20311175 | 0.32852614 | NA | NA |
| ENSG000002 | 6.00894936 | -0.470853203 | 0.36997201 | -1.2726725 | 0.20313428 | 0.32854572 | NA | NA |
| ENSG000002 | 2.66292804 | -0.469469993 | 0.36893605 | -1.2724969 | 0.20319661 | 0.32863343 | NA | NA |
| ENSG000001 | 20.2166357 | -0.356387062 | 0.28010828 | -1.2723189 | 0.20325983 | 0.32872256 | LRIT3 | 345193 |
| ENSG000002 | 4.77297255 | -0.477231857 | 0.37510039 | -1.2722777 | 0.20327448 | 0.32873313 | LOC105370791 | 105370791 |
| ENSG000002 | 11.3334313 | -0.415253805 | 0.32642686 | -1.272119 | 0.20333085 | 0.32881119 | NA | NA |
| ENSG000001 | 67.8984903 | 0.262918121 | 0.20672935 | 1.27179872 | 0.20344464 | 0.3288954 | TM4SF18 | 116441 |
| ENSG000001 | 7.24971056 | -0.454320817 | 0.3572263 | -1.2718011 | 0.20344378 | 0.3288954 | OR9I1 | 219954 |
| ENSG000002 | 2.60656583 | -0.474856239 | 0.37337353 | -1.2717994 | 0.2034444 | 0.3288954 | RNU6-379P | 106479734 |
| ENSG000002 | 3.7042782 | -0.481448768 | 0.37852958 | -1.271892 | 0.20341148 | 0.3288954 | RNU6-987P | 106480002 |
| ENSG000002 | 2.65935764 | -0.474066249 | 0.37275519 | -1.2717898 | 0.20344781 | 0.3288954 | NA | NA |
| ENSG000002 | 6.99812834 | -0.456817311 | 0.35917311 | -1.2718583 | 0.20342346 | 0.3288954 | TMEM114 | 283953 |
| ENSG000002 | 2.34306827 | -0.464414232 | 0.36516188 | -1.2718037 | 0.20344287 | 0.3288954 | NA | NA |

| | | | | | | | | |
|---|---|---|---|---|---|---|---|---|
| ENSG000002 | 5.43804495 | -0.476236533 | 0.37444638 | -1.2718417 | 0.20342935 | 0.3288954 | NA | NA |
| ENSG000002 | 3.44103766 | -0.481196087 | 0.37841444 | -1.2716113 | 0.20351126 | 0.32897174 | NA | NA |
| ENSG000002 | 5.35543583 | -0.47245522 | 0.37153896 | -1.2716169 | 0.20350928 | 0.32897174 | NA | NA |
| ENSG000002 | 2.91287129 | -0.478653948 | 0.37643712 | -1.2715376 | 0.20353747 | 0.32900099 | NA | NA |
| ENSG000002 | 4.28763115 | -0.480664457 | 0.37807123 | -1.2713595 | 0.20360077 | 0.3290902 | NA | NA |
| ENSG000002 | 5.10036365 | -0.475230846 | 0.37381194 | -1.27131 | 0.2036184 | 0.32910558 | NA | NA |
| ENSG000002 | 2.59609355 | -0.473737869 | 0.37266295 | -1.2712234 | 0.20364917 | 0.32914219 | NA | NA |
| ENSG000002 | 7.16321524 | -0.46381306 | 0.36486831 | -1.2711794 | 0.20366485 | 0.32915441 | NA | NA |
| ENSG000002 | 2.4581594 | -0.464252249 | 0.36524055 | -1.2710863 | 0.20369796 | 0.3291948 | NA | NA |
| ENSG000002 | 4.01152995 | -0.480297064 | 0.37789976 | -1.2709642 | 0.20374139 | 0.32925187 | LOC101929592 | 101929592 |
| ENSG000002 | 5.34528005 | -0.470911421 | 0.37053743 | -1.2708876 | 0.20376864 | 0.32926966 | NA | NA |
| ENSG000002 | 113.336272 | 0.224003457 | 0.17625722 | 1.27088953 | 0.20376795 | 0.32926966 | NA | NA |
| ENSG000002 | 3.4668578 | -0.48025309 | 0.37790228 | -1.2708394 | 0.2037858 | 0.32928427 | NA | NA |
| ENSG000001 | 28.1182069 | 0.312503571 | 0.24595091 | 1.27059325 | 0.20387339 | 0.32941267 | CCNB3 | 85417 |
| ENSG000002 | 8.61574456 | -0.448600182 | 0.35307121 | -1.2705657 | 0.20388318 | 0.32941537 | LINC00704 | 100216001 |
| ENSG000001 | 314.053922 | 0.194550209 | 0.15314844 | 1.2703375 | 0.20396443 | 0.329529 | KIAA1549 | 57670 |
| ENSG000002 | 12.2969138 | -0.422582014 | 0.33265726 | -1.2703225 | 0.20396976 | 0.329529 | NA | NA |
| ENSG000001 | 6.62095114 | -0.468498652 | 0.36886136 | -1.2701212 | 0.20404145 | 0.32961511 | DAPL1 | 92196 |
| ENSG000001 | 8.18629424 | -0.455582382 | 0.35869625 | -1.2701063 | 0.20404676 | 0.32961511 | OR51E2 | 81285 |
| ENSG000002 | 5.67223412 | -0.474748679 | 0.37378714 | -1.2701044 | 0.20404745 | 0.32961511 | NA | NA |
| ENSG000001 | 8.64062816 | -0.453448905 | 0.35708759 | -1.2698535 | 0.20413683 | 0.32974637 | BMP10 | 27302 |
| ENSG000002 | 3.72594483 | -0.480666671 | 0.37854329 | -1.2697799 | 0.20416304 | 0.32977555 | NA | NA |
| ENSG000002 | 2.48643756 | -0.45781277 | 0.36058549 | -1.2696373 | 0.20421387 | 0.32983159 | NA | NA |
| ENSG000002 | 5.41203415 | -0.475874204 | 0.37481126 | -1.2696369 | 0.204214 | 0.32983159 | NA | NA |
| ENSG00000C | 69.6917992 | -0.2403744 | 0.18933138 | -1.2695962 | 0.2042285 | 0.32984188 | STAG3 | 10734 |
| ENSG000002 | 11.6984746 | -0.435942173 | 0.34338831 | -1.2695312 | 0.20425167 | 0.32986617 | TRIM15 | 89870 |
| ENSG000002 | 24.782179 | -0.339431222 | 0.26737598 | -1.2694903 | 0.20426623 | 0.32987654 | LINC00937 | 389634 |
| ENSG000002 | 6.22676508 | -0.467033899 | 0.36792055 | -1.2693879 | 0.20430274 | 0.32992236 | NA | NA |
| ENSG000002 | 5.06854102 | -0.472940179 | 0.37259059 | -1.2693294 | 0.20432362 | 0.32994294 | LOC105373373 | 105373373 |

| | | | | | | | | |
|---|---|---|---|---|---|---|---|---|
| ENSG000001 | 3.50073227 | -0.480039616 | 0.37820086 | -1.2692716 | 0.2043442 | 0.3299499 | IL17B | 27190 |
| ENSG000002 | 20.3240341 | 0.352163467 | 0.27745301 | 1.26927245 | 0.20434391 | 0.3299499 | NA | NA |
| ENSG000002 | 7.45745132 | -0.450420693 | 0.35488399 | -1.2692054 | 0.2043678 | 0.32997152 | LOC101929159 | 101929159 |
| ENSG000002 | 10.8594314 | -0.425453308 | 0.33521682 | -1.2691884 | 0.20437387 | 0.32997152 | NA | NA |
| ENSG000002 | 2.86325785 | 0.477145197 | 0.37597007 | 1.26910422 | 0.2044039 | 0.33000686 | NA | NA |
| ENSG000002 | 5.05499625 | -0.473282794 | 0.37295737 | -1.2689997 | 0.20444117 | 0.3300539 | NA | NA |
| ENSG000002 | 2.36674598 | -0.465409562 | 0.36679327 | -1.2688607 | 0.20449076 | 0.33012082 | NA | NA |
| ENSG000001 | 8.75434673 | -0.439217146 | 0.34616378 | -1.2688131 | 0.20450773 | 0.33012193 | ROPN1L | 83853 |
| ENSG000002 | 2.60068088 | -0.473100144 | 0.37286327 | -1.2688301 | 0.20450166 | 0.33012193 | NA | NA |
| ENSG000002 | 2.46807466 | -0.457187747 | 0.36033756 | -1.2687763 | 0.20452087 | 0.33012999 | NA | NA |
| ENSG000001 | 31.9833791 | 0.327227288 | 0.25791317 | 1.26874983 | 0.2045303 | 0.33013209 | NA | NA |
| ENSG000002 | 32.7430999 | -0.317873191 | 0.25055668 | -1.2686678 | 0.20455958 | 0.33016619 | ENTPD3-AS1 | 285266 |
| ENSG000001 | 68.4887238 | -0.234561915 | 0.18491175 | -1.2685074 | 0.20461681 | 0.33024543 | MEGF9 | 1955 |
| ENSG000002 | 8.12023603 | -0.448780343 | 0.35380085 | -1.2684547 | 0.20463562 | 0.33026264 | NA | NA |
| ENSG000001 | 5.46636152 | -0.474173682 | 0.373836 | -1.2684003 | 0.20465505 | 0.33026771 | TAS2R16 | 50833 |
| ENSG000002 | 2.30504825 | -0.465558611 | 0.36703876 | -1.2684181 | 0.20464867 | 0.33026771 | NA | NA |
| ENSG000002 | 5.08678587 | -0.475392553 | 0.37482109 | -1.2683186 | 0.2046842 | 0.33030161 | RNY1P12 | 106480309 |
| ENSG000001 | 22.656207 | 0.345535113 | 0.27244255 | 1.26828616 | 0.20469578 | 0.33030714 | ACKR3 | 57007 |
| ENSG000002 | 4.13437469 | -0.479783547 | 0.37835288 | -1.2680848 | 0.20476766 | 0.33040999 | NA | NA |
| ENSG000001 | 6.21686082 | -0.467517484 | 0.36872875 | -1.2679171 | 0.20482757 | 0.3304935 | RBMXL3 | 139804 |
| ENSG000002 | 3.81698236 | -0.479161034 | 0.37794561 | -1.2678042 | 0.20486788 | 0.33054539 | NA | NA |
| ENSG000002 | 4.70860555 | -0.479557391 | 0.37832963 | -1.267565 | 0.20495334 | 0.33067013 | NA | NA |
| ENSG000001 | 15.8651886 | -0.375030702 | 0.2958864 | -1.267482 | 0.20498298 | 0.33070479 | RDH16 | 8608 |
| ENSG000002 | 6.91050501 | -0.453782596 | 0.35809923 | -1.2671979 | 0.20508454 | 0.33085547 | NA | NA |
| ENSG000001 | 7.06434938 | -0.460962823 | 0.36378986 | -1.2671129 | 0.20511493 | 0.33086501 | VSIG4 | 11326 |
| ENSG000002 | 2.33122899 | -0.463820107 | 0.36603603 | -1.2671433 | 0.20510405 | 0.33086501 | NA | NA |
| ENSG000002 | 3.28258624 | -0.477032281 | 0.37646548 | -1.2671342 | 0.20510731 | 0.33086501 | TBX2-AS1 | 103689912 |
| ENSG000000 | 54.186937 | 0.255560572 | 0.20169122 | 1.26708825 | 0.20512374 | 0.33086606 | PITPNM3 | 83394 |
| ENSG000001 | 71.7088548 | 0.263938291 | 0.20835631 | 1.26676409 | 0.20523966 | 0.33101354 | CNTNAP1 | 8506 |

| | | | | | | | | |
|---|---|---|---|---|---|---|---|---|
| ENSG000002 | 4.87047509 | -0.475985816 | 0.37574255 | -1.2667871 | 0.20523143 | 0.33101354 | NA | NA |
| ENSG000002 | 4.03486633 | -0.47909926 | 0.37820683 | -1.2667652 | 0.20523925 | 0.33101354 | NA | NA |
| ENSG000001 | 41.4626028 | 0.27987366 | 0.22094257 | 1.26672582 | 0.20525334 | 0.33102245 | NA | NA |
| ENSG000002 | 2.4433314 | -0.462959745 | 0.36550206 | -1.2666406 | 0.20528383 | 0.33105723 | NA | NA |
| ENSG000002 | 7.29720195 | -0.457234892 | 0.36098825 | -1.2666199 | 0.20529124 | 0.33105723 | NA | NA |
| ENSG000002 | 5.15551942 | -0.473132255 | 0.37355596 | -1.2665633 | 0.2053115 | 0.33106357 | LOC101927494 | 101927494 |
| ENSG000002 | 4260.00987 | 0.477360444 | 0.37688892 | 1.26658126 | 0.20530506 | 0.33106357 | MTRNR2L2 | 100462981 |
| ENSG000002 | 5.65584802 | -0.470986119 | 0.37191334 | -1.2663867 | 0.20537465 | 0.33115224 | NA | NA |
| ENSG000002 | 19.2490998 | 0.358905461 | 0.28341829 | 1.26634544 | 0.20538944 | 0.3311629 | ABCA11P | 79963 |
| ENSG000002 | 13.9132847 | -0.394632115 | 0.31164399 | -1.2662915 | 0.20540876 | 0.33118089 | LOC101927410 | 101927410 |
| ENSG000002 | 4.56590463 | -0.477014377 | 0.37671869 | -1.266235 | 0.20542898 | 0.33120032 | NA | NA |
| ENSG000001 | 20.7838288 | -0.35179697 | 0.27784446 | -1.2661651 | 0.20545396 | 0.33122743 | SLC6A17 | 388662 |
| ENSG000001 | 11.6315765 | -0.410218701 | 0.32405618 | -1.2658876 | 0.20555333 | 0.33136128 | OR10A3 | 26496 |
| ENSG000002 | 5.54559613 | -0.446531877 | 0.35273668 | -1.2659071 | 0.20554633 | 0.33136128 | SBF1P1 | 100133234 |
| ENSG000002 | 5.34701505 | -0.469059151 | 0.3705497 | -1.2658468 | 0.20556793 | 0.33137165 | NA | NA |
| ENSG000001 | 78.9385177 | 0.259190551 | 0.20480405 | 1.26555386 | 0.20567285 | 0.3315276 | ADPRM | 56985 |
| ENSG000002 | 6.25289987 | -0.465918868 | 0.3681642 | -1.2655192 | 0.20568527 | 0.33153443 | NA | NA |
| ENSG000001 | 6.48799599 | -0.462388045 | 0.36550422 | -1.2650689 | 0.20584661 | 0.33178131 | SPINK6 | 404203 |
| ENSG000001 | 11.1127336 | -0.433924937 | 0.34301742 | -1.265023 | 0.20586307 | 0.33178146 | LEXM | 163747 |
| ENSG000002 | 13.5387428 | 0.394043058 | 0.31149048 | 1.26502439 | 0.20586259 | 0.33178146 | NA | NA |
| ENSG000001 | 4.3692409 | -0.478606407 | 0.3783468 | -1.2649939 | 0.20587351 | 0.3317851 | FOXQ1 | 94234 |
| ENSG000002 | 12.1845092 | -0.412638816 | 0.32621762 | -1.2649188 | 0.20590042 | 0.33181529 | NA | NA |
| ENSG000001 | 70.6342312 | 0.24306353 | 0.1921664 | 1.2648597 | 0.20592163 | 0.33183627 | SGK494 | 124923 |
| ENSG000002 | 3.83999833 | -0.477285262 | 0.37735012 | -1.264834 | 0.20593084 | 0.33183794 | NA | NA |
| ENSG000002 | 3.48967322 | -0.477583519 | 0.37762266 | -1.264711 | 0.20597496 | 0.33189583 | NA | NA |
| ENSG000001 | 2.66141327 | -0.471411756 | 0.37276298 | -1.2646421 | 0.20599965 | 0.33190925 | SPDEF | 25803 |
| ENSG000002 | 22.2675482 | 0.340265566 | 0.26905827 | 1.26465381 | 0.20599546 | 0.33190925 | RNVU1-1 | 101954273 |
| ENSG000001 | 176.825884 | 0.178146636 | 0.14087409 | 1.26458057 | 0.20602172 | 0.33191843 | C1orf115 | 79762 |
| ENSG000001 | 6.40124592 | -0.460873077 | 0.36444621 | -1.2645846 | 0.20602026 | 0.33191843 | SHOX | 6473 |

| | | | | | | | | |
|---|---|---|---|---|---|---|---|---|
| ENSG000002 | 2.95027465 | -0.470838694 | 0.37235084 | -1.2645029 | 0.2060496 | 0.33191901 | NA | NA |
| ENSG000002 | 4.38621657 | -0.478065645 | 0.37805985 | -1.2645237 | 0.20604212 | 0.33191901 | LOC101927755 | 101927755 |
| ENSG000002 | 7.90963191 | -0.462452838 | 0.36572331 | -1.2644883 | 0.20605483 | 0.33191901 | NA | NA |
| ENSG000002 | 5.04449913 | -0.471811063 | 0.37310776 | -1.2645437 | 0.20603495 | 0.33191901 | LINC01028 | 101928141 |
| ENSG000001 | 34.7547935 | 0.300998528 | 0.23805024 | 1.2644328 | 0.20607473 | 0.33193671 | GLB1L | 79411 |
| ENSG000001 | 5.0991056 | -0.472330353 | 0.37355731 | -1.264412 | 0.20608218 | 0.33193671 | NA | NA |
| ENSG000002 | 3.97856483 | -0.477686041 | 0.37780335 | -1.2643775 | 0.20609458 | 0.33194349 | NA | NA |
| ENSG000002 | 2.34319998 | -0.462308217 | 0.36564784 | -1.2643537 | 0.20610309 | 0.33194401 | RNU7-175P | 106479091 |
| ENSG000002 | 3.79669739 | -0.478638126 | 0.3785875 | -1.2642735 | 0.20613189 | 0.33197722 | NA | NA |
| ENSG000001 | 32.9076709 | -0.299853579 | 0.23717913 | -1.2642494 | 0.20614052 | 0.33197793 | CCDC68 | 80323 |
| ENSG000002 | 5.11152141 | -0.471842399 | 0.37322979 | -1.2642142 | 0.20615316 | 0.33197953 | NA | NA |
| ENSG000002 | 5.61378511 | -0.466310832 | 0.36885814 | -1.264201 | 0.20615789 | 0.33197953 | NA | NA |
| ENSG000001 | 145.3449 | 0.200937329 | 0.15896433 | 1.26404032 | 0.20621555 | 0.3320592 | TCTN2 | 79867 |
| ENSG00000C | 63.9699075 | 0.252200886 | 0.1995305 | 1.26397162 | 0.20624021 | 0.33208573 | CRAT | 1384 |
| ENSG000001 | 927.301888 | 0.159553196 | 0.12623681 | 1.26391972 | 0.20625884 | 0.33210253 | GNPAT | 8443 |
| ENSG000002 | 14.5018799 | 0.402912575 | 0.31880792 | 1.26380982 | 0.2062983 | 0.33215287 | LOC101929243 | 101929243 |
| ENSG000001 | 9.04277785 | -0.432427471 | 0.34219542 | -1.2636857 | 0.20634285 | 0.33218504 | HCRTR2 | 3062 |
| ENSG000001 | 47.143756 | -0.265779351 | 0.21031511 | -1.2637197 | 0.20633064 | 0.33218504 | CASQ1 | 844 |
| ENSG000002 | 2.93666937 | -0.474012459 | 0.37510101 | -1.2636928 | 0.20634029 | 0.33218504 | NA | NA |
| ENSG000001 | 75.4558617 | 0.231896968 | 0.18351605 | 1.2636332 | 0.20636171 | 0.33220221 | TTLL6 | 284076 |
| ENSG000002 | 21.0825565 | 0.374769751 | 0.29662684 | 1.26343843 | 0.20643166 | 0.33230163 | NA | NA |
| ENSG000002 | 85.9057197 | 0.231743364 | 0.18343573 | 1.26334912 | 0.20646374 | 0.33234008 | NA | NA |
| ENSG000001 | 372.439854 | 0.409547887 | 0.32420702 | 1.26322955 | 0.2065067 | 0.33239602 | RBPMS2 | 348093 |
| ENSG000002 | 3.44876199 | -0.477727724 | 0.3782031 | -1.2631513 | 0.20653482 | 0.3324281 | LINC00572 | 100861573 |
| ENSG000002 | 3.54797197 | -0.47740611 | 0.37796931 | -1.2630817 | 0.20655983 | 0.33245515 | RNU6-23P | 100873755 |
| ENSG000001 | 8.34548826 | -0.443426465 | 0.3510876 | -1.263008 | 0.20658629 | 0.33247763 | CD40LG | 959 |
| ENSG000001 | 4.0443455 | -0.477898786 | 0.37838468 | -1.2629972 | 0.20659019 | 0.33247763 | LMO7DN | 729420 |
| ENSG000001 | 5.25671417 | -0.475666537 | 0.37668311 | -1.2627764 | 0.20666956 | 0.33258482 | TPRX1 | 284355 |
| ENSG000002 | 382.801357 | -0.174601878 | 0.13826936 | -1.2627662 | 0.2066732 | 0.33258482 | NA | NA |

| | | | | | | | | |
|---|---|---|---|---|---|---|---|---|
| ENSG000002 | 12.0126612 | 0.414962568 | 0.32862583 | 1.26272049 | 0.20668965 | 0.33259809 | NA | NA |
| ENSG000002 | 16.3913316 | -0.36923789 | 0.29242404 | -1.2626797 | 0.20670432 | 0.3326085 | NA | NA |
| ENSG00000C | 14.2289023 | -0.390701078 | 0.30942885 | -1.2626524 | 0.20671412 | 0.33261108 | WNT8A | 7478 |
| ENSG000002 | 7.37682133 | -0.458497858 | 0.3631388 | -1.2625967 | 0.20673415 | 0.33261691 | NA | NA |
| ENSG000002 | 2.8419746 | -0.475378589 | 0.37650529 | -1.262608 | 0.20673009 | 0.33261691 | NA | NA |
| ENSG000002 | 6.92414675 | -0.453167739 | 0.35894917 | -1.2624844 | 0.20677453 | 0.33266868 | NA | NA |
| ENSG000002 | 3.24740654 | -0.474145368 | 0.37562213 | -1.2622935 | 0.2068432 | 0.33276596 | NA | NA |
| ENSG000002 | 2.36424509 | -0.457766704 | 0.36268317 | -1.2621669 | 0.20688872 | 0.33280189 | NA | NA |
| ENSG000002 | 2.90256319 | -0.475237552 | 0.3765263 | -1.262163 | 0.20689016 | 0.33280189 | NA | NA |
| ENSG000002 | 7.11898834 | -0.461566117 | 0.36568357 | -1.2622009 | 0.20687652 | 0.33280189 | NA | NA |
| ENSG000002 | 4.53124219 | -0.47542507 | 0.37674549 | -1.2619264 | 0.20697528 | 0.3329256 | NA | NA |
| ENSG000002 | 9.22579281 | -0.441786646 | 0.35013932 | -1.2617453 | 0.20704045 | 0.3330083 | LINC01209 | 101928684 |
| ENSG000002 | 3.98090011 | -0.477083059 | 0.37811582 | -1.2617379 | 0.20704311 | 0.3330083 | NA | NA |
| ENSG000002 | 24.4382088 | -0.33808701 | 0.26798776 | -1.2615763 | 0.20710129 | 0.3330836 | NA | NA |
| ENSG000002 | 24.0580296 | -0.347808543 | 0.27570468 | -1.2615257 | 0.2071195 | 0.3330836 | PAXBP1-AS1 | 100506215 |
| ENSG000002 | 4.96444451 | -0.475697091 | 0.3770835 | -1.2615166 | 0.20712278 | 0.3330836 | NA | NA |
| ENSG000002 | 3.47712266 | -0.477297695 | 0.37834462 | -1.2615422 | 0.20711358 | 0.3330836 | LOC105372200 | 105372200 |
| ENSG000002 | 3.51795661 | -0.4748926 | 0.37645788 | -1.2614761 | 0.20713738 | 0.33309386 | CORT | 1325 |
| ENSG000002 | 6.65736296 | -0.456917691 | 0.36221846 | -1.2614423 | 0.20714954 | 0.3331002 | LOC101928401 | 101928401 |
| ENSG000002 | 14.9236563 | 0.396914559 | 0.31469454 | 1.26126927 | 0.20721185 | 0.33318719 | NA | NA |
| ENSG000001 | 27.8784512 | 0.315333828 | 0.25011622 | 1.26074921 | 0.20739922 | 0.33340076 | ADAM33 | 80332 |
| ENSG000002 | 4.62673724 | -0.476943241 | 0.37830852 | -1.2607256 | 0.20740773 | 0.33340076 | NA | NA |
| ENSG000002 | 2.86477542 | -0.474549326 | 0.37641194 | -1.2607181 | 0.20741045 | 0.33340076 | NA | NA |
| ENSG000002 | 3.05961702 | -0.468851817 | 0.37188289 | -1.2607512 | 0.20739849 | 0.33340076 | SPANXN2 | 494119 |
| ENSG000002 | 2.90672522 | -0.473464083 | 0.3755187 | -1.2608269 | 0.20737122 | 0.33340076 | NA | NA |
| ENSG000002 | 4.30155363 | -0.476812471 | 0.37818059 | -1.2608063 | 0.20737865 | 0.33340076 | NA | NA |
| ENSG000002 | 11.0587681 | -0.414049404 | 0.32840313 | -1.2607962 | 0.20738229 | 0.33340076 | NA | NA |
| ENSG000002 | 5.69250756 | 0.466954321 | 0.37036317 | 1.26080118 | 0.20738049 | 0.33340076 | NA | NA |
| ENSG000001 | 4.62946157 | -0.475596714 | 0.37728117 | -1.2605896 | 0.20745675 | 0.33346198 | MAGEB4 | 4115 |

| | | | | | | | | |
|---|---|---|---|---|---|---|---|---|
| ENSG000001 | 40.1898436 | 0.280660875 | 0.222661 | 1.26048513 | 0.20749441 | 0.33350928 | C7orf31 | 136895 |
| ENSG000001 | 6.73541562 | -0.458175873 | 0.36351227 | -1.2604138 | 0.20752013 | 0.33353741 | NR0B1 | 190 |
| ENSG000001 | 23.6167689 | -0.330292282 | 0.26206265 | -1.2603562 | 0.2075409 | 0.33355757 | BTN1A1 | 696 |
| ENSG000002 | 6.97938718 | -0.457767401 | 0.3632723 | -1.260122 | 0.20762537 | 0.3336801 | LINC00603 | 102467077 |
| ENSG000002 | 9.84272199 | -0.440106063 | 0.34932034 | -1.2598925 | 0.20770816 | 0.33379993 | LINC00919 | 100505619 |
| ENSG000002 | 2.71639657 | -0.466971742 | 0.37067463 | -1.2597888 | 0.20774556 | 0.33384681 | NA | NA |
| ENSG000001 | 5.13914955 | -0.47071829 | 0.37366874 | -1.2597208 | 0.2077701 | 0.33386254 | TUSC5 | 286753 |
| ENSG000001 | 6.41652516 | -0.460666823 | 0.36569101 | -1.259716 | 0.20777182 | 0.33386254 | NA | NA |
| ENSG000002 | 8.51734064 | -0.437954196 | 0.34768565 | -1.2596269 | 0.20780399 | 0.3339003 | NA | NA |
| ENSG000002 | 2.86619777 | -0.474131779 | 0.37641298 | -1.2596053 | 0.20781179 | 0.3339003 | NA | NA |
| ENSG000002 | 18.0642944 | 0.375608073 | 0.29822424 | 1.25948204 | 0.20785627 | 0.33395854 | NA | NA |
| ENSG000002 | 2.62697786 | -0.467159854 | 0.37093102 | -1.2594251 | 0.20787681 | 0.33397832 | NA | NA |
| ENSG000002 | 8.99994051 | -0.43249081 | 0.34341255 | -1.2593914 | 0.207889 | 0.33398466 | NA | NA |
| ENSG000002 | 4.6739166 | -0.475226423 | 0.37738041 | -1.2592769 | 0.20793035 | 0.33403785 | NA | NA |
| ENSG000001 | 5.25000147 | -0.474367658 | 0.37670764 | -1.2592462 | 0.20794143 | 0.33404242 | GPRC5D | 55507 |
| ENSG000001 | 11.0691102 | 0.416446021 | 0.33072083 | 1.2592071 | 0.20795554 | 0.33405186 | IL17D | 53342 |
| ENSG000002 | 2.84658622 | -0.473483729 | 0.37603214 | -1.2591576 | 0.20797341 | 0.33405779 | NA | NA |
| ENSG000002 | 8.56764873 | -0.44552896 | 0.35383276 | -1.2591512 | 0.20797571 | 0.33405779 | NA | NA |
| ENSG000002 | 7.6645575 | -0.457127321 | 0.36305293 | -1.2591203 | 0.20798688 | 0.3340625 | NA | NA |
| ENSG000002 | 8.18763172 | -0.453471703 | 0.36018969 | -1.2589802 | 0.20803747 | 0.33413053 | NA | NA |
| ENSG000001 | 150.333272 | 0.180250596 | 0.14318646 | 1.2588522 | 0.20808372 | 0.33418342 | SFI1 | 9814 |
| ENSG000002 | 8.85936754 | 0.438193673 | 0.34809227 | 1.25884344 | 0.20808689 | 0.33418342 | NA | NA |
| ENSG000002 | 4.00010672 | -0.475902824 | 0.37813353 | -1.2585576 | 0.20819018 | 0.33433606 | NA | NA |
| ENSG000001 | 3.52784721 | -0.475822076 | 0.37815845 | -1.2582611 | 0.20829734 | 0.3344949 | MC4R | 4160 |
| ENSG000002 | 9.01546161 | -0.432146511 | 0.3434719 | -1.2581714 | 0.20832979 | 0.33453377 | NA | NA |
| ENSG000002 | 6.82098818 | -0.461813811 | 0.36706821 | -1.2581144 | 0.20835038 | 0.33455359 | NA | NA |
| ENSG000002 | 2.84032391 | -0.473507812 | 0.37639879 | -1.257995 | 0.20839355 | 0.33460966 | NA | NA |
| ENSG000002 | 2.44080868 | -0.454383281 | 0.36120805 | -1.2579545 | 0.20840822 | 0.33461996 | NA | NA |
| ENSG000002 | 4.46343452 | -0.476198957 | 0.3785593 | -1.2579243 | 0.20841913 | 0.33462422 | NA | NA |

| | | | | | | | | |
|---|---|---|---|---|---|---|---|---|
| ENSG000002 | 7.01066359 | -0.458649788 | 0.36462627 | -1.2578627 | 0.20844141 | 0.33464675 | NA | NA |
| ENSG000001 | 87.3417702 | 0.21723227 | 0.17271393 | 1.25775763 | 0.20847942 | 0.33469453 | ARL10 | 285598 |
| ENSG000002 | 2.42345209 | -0.45651008 | 0.36301416 | -1.2575545 | 0.20855292 | 0.33479927 | NA | NA |
| ENSG000001 | 28.907739 | 0.314260084 | 0.24992363 | 1.25742445 | 0.20859998 | 0.3348246 | AKR1E2 | 83592 |
| ENSG000002 | 3.8123734 | -0.475664307 | 0.37828998 | -1.2574066 | 0.20860645 | 0.3348246 | NA | NA |
| ENSG000002 | 4.12778267 | -0.475804214 | 0.37840418 | -1.2573968 | 0.20860998 | 0.3348246 | NA | NA |
| ENSG000002 | 3.47300498 | -0.475577328 | 0.37821688 | -1.2574196 | 0.20860173 | 0.3348246 | NA | NA |
| ENSG000002 | 2.86827247 | -0.472835005 | 0.37604253 | -1.2573977 | 0.20860967 | 0.3348246 | FAM231B | 100133301 |
| ENSG000002 | 2.64752359 | -0.46862689 | 0.37274507 | -1.2572316 | 0.20866977 | 0.3349073 | MIA | 8190 |
| ENSG000002 | 2.68908054 | -0.465081866 | 0.36995366 | -1.2571354 | 0.20870459 | 0.33494993 | NA | NA |
| ENSG000002 | 3.54565596 | -0.475149506 | 0.37799697 | -1.2570194 | 0.20874662 | 0.33499087 | NA | NA |
| ENSG000002 | 4.92068335 | -0.473540859 | 0.37671278 | -1.2570342 | 0.20874124 | 0.33499087 | NA | NA |
| ENSG000002 | 3.17903045 | -0.473364084 | 0.3766082 | -1.2569139 | 0.20878481 | 0.33502665 | NA | NA |
| ENSG000002 | 2.86240734 | -0.472651342 | 0.37604166 | -1.2569122 | 0.20878544 | 0.33502665 | NA | NA |
| ENSG000002 | 7.70250376 | -0.44263008 | 0.35218539 | -1.2568099 | 0.20882249 | 0.33507284 | NA | NA |
| ENSG000002 | 10.1876851 | -0.428330192 | 0.34081383 | -1.2567864 | 0.20883098 | 0.33507321 | NA | NA |
| ENSG000002 | 7.01652075 | -0.458986939 | 0.36522951 | -1.2567082 | 0.20885931 | 0.33510539 | NA | NA |
| ENSG000002 | 37.0515755 | 0.308380298 | 0.2454052 | 1.25661681 | 0.20889243 | 0.33514527 | NA | NA |
| ENSG000002 | 5.47199055 | -0.47109504 | 0.37492496 | -1.2565049 | 0.20893298 | 0.33519707 | NA | NA |
| ENSG000001 | 2.85194247 | -0.472187445 | 0.37581112 | -1.2564489 | 0.20895328 | 0.33520312 | PRSS30P | 124221 |
| ENSG000002 | 5.07042038 | -0.468529119 | 0.37289333 | -1.2564696 | 0.20894577 | 0.33520312 | NA | NA |
| ENSG000001 | 153.053583 | 0.192072266 | 0.1528743 | 1.25640655 | 0.20896861 | 0.33521445 | FHOD3 | 80206 |
| ENSG000002 | 2.60255118 | -0.466041213 | 0.37094781 | -1.2563525 | 0.20898819 | 0.3352326 | NA | NA |
| ENSG000002 | 5.24085612 | -0.472716328 | 0.37629243 | -1.2562472 | 0.20902634 | 0.33528054 | NA | NA |
| ENSG000002 | 4.5135417 | -0.473714932 | 0.37714468 | -1.2560562 | 0.20909558 | 0.33533854 | NA | NA |
| ENSG000002 | 6.96077743 | -0.453724169 | 0.36122162 | -1.2560826 | 0.20908604 | 0.33533854 | NA | NA |
| ENSG000002 | 2.85148601 | -0.472464679 | 0.37614877 | -1.256058 | 0.20909494 | 0.33533854 | NA | NA |
| ENSG000002 | 8.67971912 | 0.442489223 | 0.35227593 | 1.25608703 | 0.20908442 | 0.33533854 | NA | NA |
| ENSG000002 | 9.34866652 | -0.430072885 | 0.34241656 | -1.2559933 | 0.20911842 | 0.3353619 | NA | NA |

| | | | | | | | | |
|---|---|---|---|---|---|---|---|---|
| ENSG000002 | 2.8593671 | -0.472687882 | 0.3763665 | -1.2559244 | 0.20914337 | 0.33538864 | NA | NA |
| ENSG000002 | 2.61441356 | -0.467497261 | 0.37224364 | -1.2558905 | 0.20915566 | 0.33539509 | NA | NA |
| ENSG000002 | 45.7639718 | 0.279392023 | 0.22250545 | 1.25566372 | 0.20923792 | 0.33551373 | HLA-DRA | 3122 |
| ENSG000002 | 5.06914651 | -0.469194624 | 0.37367133 | -1.2556345 | 0.2092485 | 0.33551744 | NA | NA |
| ENSG000002 | 23.0191986 | -0.335360837 | 0.26710565 | -1.2555363 | 0.20928414 | 0.33556132 | NA | NA |
| ENSG000002 | 3.75043523 | -0.475243212 | 0.37859196 | -1.2552913 | 0.20937302 | 0.33569055 | NA | NA |
| ENSG000002 | 5.5893666 | -0.462587023 | 0.36853951 | -1.2551898 | 0.20940988 | 0.33573638 | NA | NA |
| ENSG000001 | 209.004609 | 0.198728862 | 0.15832875 | 1.25516602 | 0.2094185 | 0.33573691 | FUT9 | 10690 |
| ENSG000002 | 5.55856572 | -0.462681373 | 0.36865952 | -1.2550371 | 0.2094653 | 0.33579867 | NA | NA |
| ENSG000000 | 178.062381 | 0.182758151 | 0.14563063 | 1.25494304 | 0.20949944 | 0.33582805 | SPATA7 | 55812 |
| ENSG000002 | 3.45116038 | 0.474928999 | 0.37845139 | 1.25492736 | 0.20950513 | 0.33582805 | RNU6-759P | 106479902 |
| ENSG000002 | 5.19351963 | -0.471558847 | 0.37576861 | -1.2549182 | 0.20950847 | 0.33582805 | LOC105369473 | 105369473 |
| ENSG000001 | 41.6646081 | -0.292674362 | 0.23323195 | -1.2548639 | 0.20952816 | 0.33584634 | LDB3 | 11155 |
| ENSG000002 | 3.27946113 | -0.471901064 | 0.37607097 | -1.2548192 | 0.20954441 | 0.3358591 | NA | NA |
| ENSG000001 | 8.48152308 | -0.437366572 | 0.34856474 | -1.2547642 | 0.20956436 | 0.3358778 | BPIFA3 | 128861 |
| ENSG000001 | 28.6441443 | -0.311992205 | 0.24865316 | -1.2547285 | 0.20957733 | 0.33588532 | LYPD5 | 284348 |
| ENSG000002 | 28.5145591 | 0.319204948 | 0.25440886 | 1.25469276 | 0.20959032 | 0.33589286 | SPATA3-AS1 | 348761 |
| ENSG000001 | 2.49125344 | 0.467373603 | 0.37253135 | 1.25458864 | 0.20962813 | 0.33594018 | NA | NA |
| ENSG000002 | 16.4176875 | -0.373663257 | 0.2978573 | -1.2545043 | 0.20965877 | 0.33597601 | DISC1FP1 | 101929222 |
| ENSG000001 | 6.76913255 | -0.456994925 | 0.3643339 | -1.25433 | 0.2097221 | 0.33606421 | NA | NA |
| ENSG000001 | 14.2072421 | -0.39108525 | 0.31181026 | -1.2542411 | 0.20975438 | 0.33610266 | SPAG8 | 26206 |
| ENSG000002 | 5.76014506 | -0.470680752 | 0.37531603 | -1.2540918 | 0.20980865 | 0.33617633 | NA | NA |
| ENSG000002 | 5.56150182 | -0.463031609 | 0.3692346 | -1.2540309 | 0.20983076 | 0.33619848 | NA | NA |
| ENSG000002 | 16.2052144 | 0.379887811 | 0.30294193 | 1.25399546 | 0.20984365 | 0.33620585 | NA | NA |
| ENSG000002 | 41.1633835 | 0.279555986 | 0.22295346 | 1.25387598 | 0.20988709 | 0.33626215 | NA | NA |
| ENSG000002 | 7.63713148 | -0.459072912 | 0.36615938 | -1.2537516 | 0.20993231 | 0.33632131 | NA | NA |
| ENSG000001 | 212.295285 | 0.175395704 | 0.13990541 | 1.25367354 | 0.20996069 | 0.33635349 | TCEANC2 | 127428 |
| ENSG000001 | 5.13087726 | -0.46893359 | 0.37406641 | -1.2536105 | 0.2099836 | 0.33637691 | MBOAT4 | 619373 |
| ENSG000002 | 12.1830784 | 0.410238766 | 0.32725713 | 1.25356709 | 0.2099994 | 0.33638893 | BRWD1-AS1 | 100874093 |

| | | | | | | | | |
|---|---|---|---|---|---|---|---|---|
| ENSG000002 | 3.52755053 | -0.47316035 | 0.37756095 | -1.2532026 | 0.210132 | 0.33657474 | NA | NA |
| ENSG000002 | 2.65714656 | -0.465960274 | 0.37181544 | -1.2532031 | 0.2101318 | 0.33657474 | NA | NA |
| ENSG000002 | 7.74174982 | -0.469340888 | 0.37452267 | -1.2531708 | 0.21014354 | 0.33657994 | NA | NA |
| ENSG000002 | 3.16487065 | -0.473472453 | 0.37783915 | -1.2531059 | 0.21016719 | 0.33659122 | ARRDC5 | 645432 |
| ENSG000002 | 5.11565221 | -0.471602401 | 0.37634117 | -1.2531247 | 0.21016034 | 0.33659122 | NA | NA |
| ENSG00000C | 7054.14144 | 0.131519348 | 0.10495702 | 1.25307814 | 0.21017727 | 0.33659407 | TPR | 7175 |
| ENSG000001 | 22.1301431 | -0.36258804 | 0.28939604 | -1.2529129 | 0.21023739 | 0.33667639 | HLA-DQA1 | 3117 |
| ENSG000002 | 5.3071077 | -0.464652063 | 0.37086384 | -1.2528913 | 0.21024528 | 0.33667639 | NA | NA |
| ENSG000001 | 20.0018096 | -0.352004287 | 0.28101516 | -1.2526167 | 0.21034524 | 0.33682316 | TNS4 | 84951 |
| ENSG000002 | 12.7016281 | 0.409065624 | 0.32658467 | 1.25255611 | 0.21036729 | 0.33684518 | NA | NA |
| ENSG000002 | 8.57572784 | -0.442406219 | 0.35322742 | -1.2524685 | 0.2103992 | 0.33688297 | NA | NA |
| ENSG000001 | 55.1568762 | 0.269729071 | 0.21538777 | 1.25229521 | 0.21046231 | 0.33697071 | TMEM54 | 113452 |
| ENSG000002 | 25.6681653 | -0.323601952 | 0.25842216 | -1.2522221 | 0.21048893 | 0.33698672 | NA | NA |
| ENSG000002 | 2.59006011 | -0.467323594 | 0.37318878 | -1.2522445 | 0.21048078 | 0.33698672 | NA | NA |
| ENSG000002 | 61.7751828 | 0.253924307 | 0.20278435 | 1.25218884 | 0.21050106 | 0.33699283 | NA | NA |
| ENSG000002 | 4.07821302 | -0.474009974 | 0.3785707 | -1.2521042 | 0.21053189 | 0.33702888 | NA | NA |
| ENSG000002 | 5.1283596 | -0.468767159 | 0.37443121 | -1.2519447 | 0.21059003 | 0.33710864 | NA | NA |
| ENSG000002 | 45.2127872 | 0.277272389 | 0.22149944 | 1.25179726 | 0.21064375 | 0.33718133 | NA | NA |
| ENSG000002 | 5.07468656 | -0.468261722 | 0.37410087 | -1.2516991 | 0.21067953 | 0.33722529 | SLC35G6 | 643664 |
| ENSG000002 | 34.4860227 | 0.307490647 | 0.24566719 | 1.25165535 | 0.21069548 | 0.33723751 | NA | NA |
| ENSG000002 | 4.27650551 | -0.472269654 | 0.3773712 | -1.2514724 | 0.21076216 | 0.33733093 | NA | NA |
| ENSG000002 | 2.88298444 | -0.47071239 | 0.37616891 | -1.2513325 | 0.21081318 | 0.3373862 | NA | NA |
| ENSG000002 | 8.1298103 | -0.449119537 | 0.35891314 | -1.2513321 | 0.21081333 | 0.3373862 | NA | NA |
| ENSG000001 | 99.7223283 | 0.237046405 | 0.1894398 | 1.25130204 | 0.2108243 | 0.33739044 | MAN2A2 | 4122 |
| ENSG000002 | 8.63210974 | -0.442122799 | 0.35333746 | -1.2512763 | 0.21083368 | 0.33739214 | NA | NA |
| ENSG000002 | 2.5913976 | -0.466170936 | 0.37262824 | -1.2510349 | 0.21092174 | 0.33751973 | NA | NA |
| ENSG000002 | 2.92630919 | -0.469235772 | 0.37509505 | -1.2509783 | 0.21094238 | 0.33753945 | NA | NA |
| ENSG000002 | 2.87222673 | -0.470369242 | 0.37600859 | -1.2509535 | 0.21095146 | 0.33754065 | NA | NA |
| ENSG000002 | 20.5807721 | 0.346583706 | 0.27706296 | 1.25092038 | 0.21096353 | 0.33754664 | F8A1 | 8263 |

| | | | | | | | | |
|---|---|---|---|---|---|---|---|---|
| ENSG000002 | 2.37483559 | -0.456892641 | 0.36527651 | -1.2508131 | 0.21100268 | 0.33759597 | NA | NA |
| ENSG000001 | 6.68889419 | -0.453206667 | 0.3623565 | -1.2507204 | 0.21103651 | 0.33760935 | FABP4 | 2167 |
| ENSG000002 | 4.92746195 | -0.471474918 | 0.37695956 | -1.2507308 | 0.21103272 | 0.33760935 | NA | NA |
| ENSG000002 | 25.1900348 | -0.322918018 | 0.25819005 | -1.2506989 | 0.21104435 | 0.33760935 | LOC105370404 | 105370404 |
| ENSG000002 | 2.87276697 | -0.469313679 | 0.37524087 | -1.2506998 | 0.21104403 | 0.33760935 | NA | NA |
| ENSG000002 | 5.38069201 | -0.466147954 | 0.37278295 | -1.2504541 | 0.21113373 | 0.33773901 | NA | NA |
| ENSG000002 | 7.68278917 | -0.439153593 | 0.35120893 | -1.2504055 | 0.21115144 | 0.33775402 | NA | NA |
| ENSG000002 | 3.14978674 | -0.472258478 | 0.37774808 | -1.2501943 | 0.2112286 | 0.33786411 | RN7SKP44 | 106479114 |
| ENSG000002 | 5.62610918 | -0.461706941 | 0.36934073 | -1.250084 | 0.21126886 | 0.33791519 | NA | NA |
| ENSG000002 | 7.20765799 | -0.463193284 | 0.37055089 | -1.2500126 | 0.21129494 | 0.33794357 | NA | NA |
| ENSG000002 | 2.32864522 | -0.456006335 | 0.36482886 | -1.2499185 | 0.21132933 | 0.33798524 | FLJ26245 | 400533 |
| ENSG000001 | 85.525038 | -0.227900309 | 0.18235134 | -1.2497869 | 0.21137741 | 0.3380488 | SYT10 | 341359 |
| ENSG000002 | 6.2288891 | -0.461673749 | 0.36942179 | -1.2497199 | 0.2114019 | 0.33807463 | LOC101928111 | 101928111 |
| ENSG000002 | 15.69005 | -0.380167897 | 0.30421588 | -1.2496649 | 0.211422 | 0.33809344 | HCG27 | 253018 |
| ENSG000001 | 27.8359188 | 0.311601733 | 0.24937039 | 1.24955388 | 0.21146256 | 0.33814497 | TMEM130 | 222865 |
| ENSG000002 | 2.60300352 | -0.465102031 | 0.37222712 | -1.2495114 | 0.21147809 | 0.33815647 | RNA5SP57 | 106480754 |
| ENSG000001 | 192.292403 | 0.184767775 | 0.14787586 | 1.24947897 | 0.21148994 | 0.33816208 | MFAP3L | 9848 |
| ENSG000002 | 3.50596488 | -0.472125631 | 0.37791449 | -1.2492922 | 0.21155821 | 0.33825791 | NA | NA |
| ENSG000001 | 4.40193246 | -0.472928958 | 0.37858174 | -1.2492123 | 0.21158744 | 0.33826263 | GPR6 | 2830 |
| ENSG000002 | 2.84465534 | -0.46969887 | 0.37599081 | -1.2492296 | 0.21158109 | 0.33826263 | DBH-AS1 | 138948 |
| ENSG000002 | 2.19210097 | -0.439732508 | 0.35199876 | -1.2492445 | 0.21157566 | 0.33826263 | NA | NA |
| ENSG000002 | 22.907466 | -0.34387002 | 0.27527376 | -1.2491929 | 0.21159453 | 0.33826263 | NA | NA |
| ENSG000002 | 11.3327728 | -0.422896737 | 0.33854275 | -1.2491679 | 0.21160367 | 0.3382639 | NA | NA |
| ENSG000001 | 80.8541884 | 0.226256251 | 0.18115722 | 1.24894966 | 0.21168349 | 0.33835148 | ARSJ | 79642 |
| ENSG000002 | 2.88356489 | -0.468899539 | 0.37543493 | -1.2489502 | 0.21168328 | 0.33835148 | NA | NA |
| ENSG000002 | 3.68475894 | -0.472847681 | 0.37858772 | -1.2489779 | 0.21167317 | 0.33835148 | NA | NA |
| ENSG000002 | 6.99626958 | -0.45168851 | 0.36171012 | -1.2487583 | 0.21175349 | 0.33843716 | NA | NA |
| ENSG000002 | 4.26557229 | -0.470750247 | 0.37697491 | -1.2487575 | 0.21175378 | 0.33843716 | NA | NA |
| ENSG000002 | 2.90566848 | -0.469277311 | 0.37582932 | -1.2486448 | 0.21179503 | 0.33847997 | LOC105375972 | 105375972 |

| | | | | | | | | |
|---|---|---|---|---|---|---|---|---|
| ENSG000002 | 3.54861112 | -0.471097743 | 0.37728909 | -1.2486387 | 0.21179726 | 0.33847997 | NA | NA |
| ENSG000001 | 32.2125348 | 0.311811951 | 0.24973883 | 1.24855212 | 0.21182893 | 0.33851639 | A4GALT | 53947 |
| ENSG000002 | 6.42886197 | -0.456181247 | 0.36537445 | -1.2485308 | 0.21183675 | 0.33851639 | ERVFRD-1 | 405754 |
| ENSG000000 | 213.394792 | 0.184844423 | 0.14805639 | 1.24847315 | 0.21185783 | 0.33852341 | MED17 | 9440 |
| ENSG000002 | 5.58108112 | -0.460456212 | 0.36881077 | -1.2484891 | 0.21185201 | 0.33852341 | NA | NA |
| ENSG000002 | 3.71039148 | -0.472610106 | 0.37859146 | -1.248338 | 0.21190731 | 0.33858912 | LINC01495 | 102723378 |
| ENSG000001 | 77.5106761 | -0.225692179 | 0.18080606 | -1.2482556 | 0.21193747 | 0.33862398 | CEP126 | 57562 |
| ENSG000002 | 2.60284044 | -0.465288593 | 0.37276009 | -1.2482254 | 0.21194854 | 0.33862832 | NA | NA |
| ENSG000001 | 145.478496 | 0.201853606 | 0.1617212 | 1.24815796 | 0.21197322 | 0.3386544 | GDF3 | 9573 |
| ENSG000002 | 5.08186269 | -0.467182 | 0.37431612 | -1.2480948 | 0.21199635 | 0.33867801 | NA | NA |
| ENSG000002 | 27.0056476 | -0.317515628 | 0.25440601 | -1.2480665 | 0.21200669 | 0.3386812 | NA | NA |
| ENSG000002 | 2.5087892 | 0.46600288 | 0.37339244 | 1.24802441 | 0.21202212 | 0.33869249 | NA | NA |
| ENSG000002 | 7.86128204 | -0.446627775 | 0.35789602 | -1.2479261 | 0.21205813 | 0.33873668 | NA | NA |
| ENSG000001 | 5.12510021 | -0.468847559 | 0.37572861 | -1.2478357 | 0.21209125 | 0.33874862 | C1orf194 | 127003 |
| ENSG000002 | 3.13442264 | -0.471466528 | 0.37781367 | -1.2478811 | 0.21207461 | 0.33874862 | NA | NA |
| ENSG000002 | 5.3544143 | -0.463814931 | 0.37169997 | -1.2478207 | 0.21209671 | 0.33874862 | NA | NA |
| ENSG000002 | 2.61999904 | -0.464647918 | 0.3723694 | -1.2478144 | 0.21209902 | 0.33874862 | NA | NA |
| ENSG000002 | 44.5067917 | 0.2844881 | 0.22802227 | 1.24763297 | 0.2121655 | 0.33884145 | PROB1 | 389333 |
| ENSG000002 | 3.00790786 | -0.465071168 | 0.37278893 | -1.2475455 | 0.21219753 | 0.33887925 | ACKR1 | 2532 |
| ENSG000002 | 4.21454664 | -0.470883631 | 0.37746205 | -1.2474993 | 0.21221449 | 0.338893 | NA | NA |
| ENSG000002 | 3.95189427 | -0.471730953 | 0.37816208 | -1.2474306 | 0.21223965 | 0.33891983 | NA | NA |
| ENSG000002 | 7.0809281 | -0.454848001 | 0.36468274 | -1.247243 | 0.2123084 | 0.33901626 | NA | NA |
| ENSG000002 | 3.84894114 | 0.47216364 | 0.37859251 | 1.24715526 | 0.21234057 | 0.33905428 | NA | NA |
| ENSG000002 | 5.30290047 | -0.462789347 | 0.37115817 | -1.2468791 | 0.21244185 | 0.33920264 | NA | NA |
| ENSG000001 | 5.51542741 | -0.468602509 | 0.3758877 | -1.2466556 | 0.21252381 | 0.33932014 | TAAR8 | 83551 |
| ENSG000001 | 70.0993248 | -0.244626282 | 0.1962387 | -1.2465751 | 0.21255332 | 0.3393539 | LONRF2 | 164832 |
| ENSG000001 | 505.557774 | 0.171003807 | 0.13721187 | 1.2462756 | 0.21266323 | 0.33950317 | TRDMT1 | 1787 |
| ENSG000002 | 4.52125394 | -0.469927623 | 0.37706584 | -1.2462747 | 0.21266356 | 0.33950317 | LINC00867 | 100506126 |
| ENSG000002 | 65.9678785 | -0.263700728 | 0.21159745 | -1.2462378 | 0.21267712 | 0.33951145 | LINC01057 | 101928079 |

| | | | | | | | | |
|---|---|---|---|---|---|---|---|---|
| ENSG000002 | 2.92953523 | -0.464488571 | 0.37273328 | -1.2461688 | 0.21270244 | 0.33953851 | LOC101927138 | 101927138 |
| ENSG000002 | 2.35823127 | -0.452016242 | 0.36276887 | -1.2460172 | 0.21275809 | 0.33961398 | NA | NA |
| ENSG000001 | 432.246387 | -0.165032539 | 0.1324539 | -1.2459621 | 0.21277831 | 0.33961544 | ZNF613 | 79898 |
| ENSG000002 | 5.82445476 | -0.457897056 | 0.36750948 | -1.2459462 | 0.21278414 | 0.33961544 | NA | NA |
| ENSG000002 | 6.01330836 | -0.462517725 | 0.37120502 | -1.24599 | 0.21276808 | 0.33961544 | NA | NA |
| ENSG000002 | 12.0787981 | 0.407065953 | 0.32672545 | 1.24589607 | 0.21280255 | 0.33963147 | NA | NA |
| ENSG000002 | 6.28449868 | -0.463451683 | 0.37202276 | -1.2457616 | 0.21285192 | 0.33969689 | NA | NA |
| ENSG00000C | 87.8407291 | 0.217727035 | 0.17479303 | 1.2456277 | 0.21290111 | 0.33975306 | UBE2D4 | 51619 |
| ENSG000001 | 17.6416093 | -0.36153933 | 0.29024845 | -1.2456202 | 0.21290387 | 0.33975306 | NYAP2 | 57624 |
| ENSG000002 | 2.88074089 | -0.468414142 | 0.37607279 | -1.2455412 | 0.21293289 | 0.33978599 | RN7SL218P | 106480981 |
| ENSG000002 | 8.24078078 | -0.434578356 | 0.34897932 | -1.245284 | 0.21302739 | 0.33992341 | NA | NA |
| ENSG000002 | 18.8263143 | 0.35560798 | 0.28557895 | 1.24521777 | 0.21305171 | 0.33994885 | NA | NA |
| ENSG000002 | 6.72562457 | -0.45314965 | 0.36391959 | -1.2451917 | 0.21306129 | 0.33995076 | NA | NA |
| ENSG000002 | 4.39210085 | 0.470194495 | 0.37762406 | 1.24513915 | 0.21308061 | 0.3399682 | NA | NA |
| ENSG000002 | 11.0233087 | -0.41188639 | 0.33080826 | -1.245091 | 0.21309829 | 0.33998304 | NA | NA |
| ENSG000002 | 26.5364912 | 0.322193577 | 0.25878026 | 1.24504698 | 0.21311448 | 0.33999549 | SNURF | 8926 |
| ENSG000002 | 27.2648933 | 0.315211254 | 0.25318307 | 1.24499341 | 0.21313417 | 0.34001353 | ATP6V1G2 | 534 |
| ENSG000002 | 6.29482045 | -0.464176487 | 0.37285742 | -1.2449169 | 0.21316231 | 0.34004504 | NA | NA |
| ENSG000001 | 52.2554748 | -0.254403654 | 0.20438119 | -1.2447508 | 0.21322336 | 0.34012904 | RNF43 | 54894 |
| ENSG000002 | 5.80821168 | -0.467294179 | 0.37548345 | -1.2445134 | 0.21331068 | 0.34025494 | NA | NA |
| ENSG000001 | 2.56920227 | -0.462531731 | 0.37166829 | -1.2444746 | 0.21332496 | 0.34026434 | DEFA4 | 1669 |
| ENSG000001 | 6.94403129 | -0.448524741 | 0.36054992 | -1.2440018 | 0.21349891 | 0.34052841 | INSL6 | 11172 |
| ENSG00000C | 17.199941 | -0.366002682 | 0.29423331 | -1.2439199 | 0.21352903 | 0.34055243 | TFAP2B | 7021 |
| ENSG000001 | 44.7043793 | 0.277376228 | 0.22298644 | 1.24391522 | 0.21353077 | 0.34055243 | C15orf61 | 145853 |
| ENSG000002 | 7.47862868 | -0.443578219 | 0.35662243 | -1.2438315 | 0.21356157 | 0.34057478 | LOC101929694 | 101929694 |
| ENSG000002 | 2.42996269 | -0.452136897 | 0.36350082 | -1.2438401 | 0.21355841 | 0.34057478 | NA | NA |
| ENSG000001 | 8.29764829 | -0.437367005 | 0.35163639 | -1.2438047 | 0.21357143 | 0.34057711 | CLRN3 | 119467 |
| ENSG000002 | 5.30734793 | -0.463649416 | 0.37279684 | -1.2437053 | 0.21360804 | 0.34062208 | NA | NA |
| ENSG000002 | 2.87749104 | -0.46690422 | 0.37543061 | -1.2436498 | 0.21362846 | 0.34064126 | NA | NA |

| | | | | | | | | |
|---|---|---|---|---|---|---|---|---|
| ENSG000002 | 2.89016743 | -0.466892506 | 0.37543876 | -1.2435916 | 0.2136499 | 0.34066204 | LINC01337 | 103689917 |
| ENSG000002 | 7.45174353 | -0.44280521 | 0.35609667 | -1.2434972 | 0.21368468 | 0.3407041 | NA | NA |
| ENSG000002 | 17.1312561 | -0.402621952 | 0.32379943 | -1.2434301 | 0.2137094 | 0.34073012 | CARD8-AS1 | 100505812 |
| ENSG000002 | 7.45485565 | -0.44189082 | 0.355393 | -1.2433864 | 0.21372549 | 0.34074237 | LOC101927051 | 101927051 |
| ENSG000002 | 3.17765098 | -0.469393248 | 0.37752607 | -1.24334 | 0.21374258 | 0.34075623 | NA | NA |
| ENSG000002 | 15.3747353 | -0.372525133 | 0.29965358 | -1.243186 | 0.2137993 | 0.34083325 | NA | NA |
| ENSG000002 | 4.25609204 | -0.469377532 | 0.37759641 | -1.2430667 | 0.21384325 | 0.34087651 | NA | NA |
| ENSG000002 | 3.70917799 | -0.470597006 | 0.37857276 | -1.2430821 | 0.2138376 | 0.34087651 | FTLP10 | 100130017 |
| ENSG000002 | 16.7225249 | -0.37109543 | 0.29858815 | -1.2428337 | 0.2139291 | 0.34099996 | NA | NA |
| ENSG000001 | 33.846278 | 0.296648032 | 0.23872084 | 1.2426566 | 0.2139944 | 0.34109064 | PDLIM4 | 8572 |
| ENSG000002 | 22.9427655 | -0.348494768 | 0.28046913 | -1.2425423 | 0.21403654 | 0.34114439 | NA | NA |
| ENSG000002 | 3.50640836 | -0.469078537 | 0.37752268 | -1.2425175 | 0.21404569 | 0.34114558 | NA | NA |
| ENSG000002 | 4.33454489 | -0.470358452 | 0.37858092 | -1.2424251 | 0.21407975 | 0.34118644 | NA | NA |
| ENSG000002 | 2.34471483 | -0.45480707 | 0.36608866 | -1.2423413 | 0.21411065 | 0.34122229 | NA | NA |
| ENSG000002 | 25.1500205 | -0.332901065 | 0.26797903 | -1.2422653 | 0.21413868 | 0.34125354 | NA | NA |
| ENSG000002 | 12.8400759 | -0.393127328 | 0.31647572 | -1.2422037 | 0.2141614 | 0.34127633 | NA | NA |
| ENSG000001 | 99.8902857 | 0.22196206 | 0.17870922 | 1.24202917 | 0.2142258 | 0.34136555 | MTHFSD | 64779 |
| ENSG000002 | 7.53666595 | -0.444628433 | 0.35802499 | -1.2418922 | 0.21427635 | 0.34140585 | NA | NA |
| ENSG000002 | 3.17055972 | -0.46913768 | 0.37774998 | -1.2419264 | 0.21426371 | 0.34140585 | NA | NA |
| ENSG000002 | 5.07702293 | -0.463746974 | 0.37341544 | -1.2419063 | 0.21427115 | 0.34140585 | NA | NA |
| ENSG000002 | 3.56179639 | -0.468405822 | 0.37719199 | -1.2418234 | 0.21430174 | 0.34143289 | SLCO4A1-AS1 | 100127888 |
| ENSG000001 | 4.28222181 | -0.468748474 | 0.37748953 | -1.2417523 | 0.21432798 | 0.34144718 | TCL1A | 8115 |
| ENSG000002 | 44.6753639 | 0.280799029 | 0.22612921 | 1.24176361 | 0.21432379 | 0.34144718 | CPTP | 80772 |
| ENSG000002 | 2.82148237 | -0.466715163 | 0.37585863 | -1.2417306 | 0.21433597 | 0.34144718 | NA | NA |
| ENSG000001 | 18.4731291 | 0.363900727 | 0.29310636 | 1.24153133 | 0.21440953 | 0.34155095 | NA | NA |
| ENSG000002 | 4.02122405 | -0.469957347 | 0.37853983 | -1.2415004 | 0.21442095 | 0.34155572 | NA | NA |
| ENSG000002 | 11.4895635 | 0.402111133 | 0.32393588 | 1.24132941 | 0.21448409 | 0.34164287 | NA | NA |
| ENSG000001 | 7.76516143 | -0.445282808 | 0.35873475 | -1.2412592 | 0.21451001 | 0.34167074 | PRR15L | 79170 |
| ENSG000001 | 3.13453419 | -0.468943827 | 0.37781411 | -1.2412025 | 0.21453094 | 0.34169065 | OR2B2 | 81697 |

| | | | | | | | | |
|---|---|---|---|---|---|---|---|---|
| ENSG000002 | 5.55117444 | -0.457976909 | 0.36900082 | -1.2411271 | 0.21455881 | 0.34172162 | NA | NA |
| ENSG000002 | 3.15396913 | -0.468837581 | 0.37781523 | -1.2409176 | 0.21463618 | 0.34183142 | NA | NA |
| ENSG000002 | 3.17783677 | -0.467028039 | 0.3764005 | -1.2407742 | 0.21468917 | 0.34190239 | NA | NA |
| ENSG000002 | 10.0055025 | -0.414585053 | 0.33417093 | -1.2406377 | 0.21473962 | 0.34196931 | NA | NA |
| ENSG000002 | 2.58767167 | -0.460929906 | 0.37161042 | -1.2403579 | 0.21484306 | 0.34212059 | NA | NA |
| ENSG000002 | 5.0885173 | -0.46448722 | 0.37448846 | -1.2403245 | 0.21485539 | 0.34212679 | NA | NA |
| ENSG000001 | 10.7687818 | -0.420014088 | 0.33865792 | -1.2402311 | 0.21488993 | 0.34216836 | PSKH2 | 85481 |
| ENSG000001 | 14.3162372 | 0.401280956 | 0.3235691 | 1.24017083 | 0.21491221 | 0.3421904 | DUSP9 | 1852 |
| ENSG000001 | 43.3291241 | -0.26774036 | 0.21590126 | -1.2401056 | 0.21493634 | 0.34221538 | FMO5 | 2330 |
| ENSG000002 | 5.90090413 | -0.456344644 | 0.3680269 | -1.2399763 | 0.21498415 | 0.34227806 | NA | NA |
| ENSG000001 | 76.3328146 | -0.229795127 | 0.18534077 | -1.239852 | 0.21503013 | 0.34233783 | NDRG1 | 10397 |
| ENSG000002 | 18.0482519 | -0.354829733 | 0.28620279 | -1.2397843 | 0.21505518 | 0.34236426 | NA | NA |
| ENSG000002 | 21.176171 | -0.349700662 | 0.28210262 | -1.2396222 | 0.21511518 | 0.34243289 | NA | NA |
| ENSG000002 | 5.96537143 | -0.459410284 | 0.37059836 | -1.2396447 | 0.21510685 | 0.34243289 | NA | NA |
| ENSG000002 | 2.87761523 | -0.465036152 | 0.37516256 | -1.2395591 | 0.21513853 | 0.34245662 | RN7SKP198 | 106479183 |
| ENSG000002 | 20.4087471 | -0.345024003 | 0.27835397 | -1.2395153 | 0.21515473 | 0.34246898 | NA | NA |
| ENSG000001 | 22.8030739 | -0.342914094 | 0.27669482 | -1.2393224 | 0.21522613 | 0.34254228 | SVOP | 55530 |
| ENSG000001 | 41.942706 | 0.280468794 | 0.22630159 | 1.23935846 | 0.21521278 | 0.34254228 | NA | NA |
| ENSG000002 | 7.34142203 | 0.442708318 | 0.35721529 | 1.23933192 | 0.2152226 | 0.34254228 | C19orf73 | 55150 |
| ENSG000002 | 2.32665484 | -0.451221719 | 0.36414814 | -1.2391158 | 0.21530261 | 0.34264446 | NA | NA |
| ENSG000002 | 4.28650117 | -0.467911515 | 0.37762106 | -1.2391033 | 0.21530723 | 0.34264446 | LOC101928435 | 101928435 |
| ENSG000002 | 2.27828472 | -0.458695099 | 0.37021378 | -1.2390006 | 0.21534527 | 0.34269155 | NA | NA |
| ENSG000002 | 5.40834718 | -0.463104334 | 0.37386419 | -1.2386967 | 0.21545785 | 0.34285725 | LOC100130417 | 100130417 |
| ENSG000001 | 22.2770707 | -0.34255309 | 0.27659034 | -1.2384854 | 0.21553614 | 0.34286302 | SLC46A3 | 283537 |
| ENSG000001 | 15.1767812 | 0.382044414 | 0.30847517 | 1.23849324 | 0.21553323 | 0.34286302 | KRBA2 | 124751 |
| ENSG000001 | 7.16614542 | -0.443225246 | 0.3578818 | -1.2384682 | 0.21554249 | 0.34286302 | TMEM239 | 100288797 |
| ENSG000002 | 10.2560981 | -0.413867618 | 0.33416471 | -1.2385138 | 0.2155256 | 0.34286302 | NA | NA |
| ENSG000002 | 3.94638126 | -0.468681734 | 0.37841648 | -1.2385341 | 0.21551807 | 0.34286302 | NA | NA |
| ENSG000002 | 17.7341027 | 0.378454833 | 0.30558535 | 1.23845869 | 0.21554603 | 0.34286302 | FBXW4P1 | 26226 |

| | | | | | | | | |
|---|---|---|---|---|---|---|---|---|
| ENSG000002 | 2.85159811 | -0.465548126 | 0.37587946 | -1.238557 | 0.21550961 | 0.34286302 | NA | NA |
| ENSG000002 | 3.75172174 | -0.468897225 | 0.37858062 | -1.2385664 | 0.21550613 | 0.34286302 | NA | NA |
| ENSG000002 | 3.77343193 | -0.468746534 | 0.37843882 | -1.2386323 | 0.21548171 | 0.34286302 | NA | NA |
| ENSG000002 | 108.253468 | 0.247514782 | 0.1998522 | 1.23848913 | 0.21553475 | 0.34286302 | NA | NA |
| ENSG000002 | 4.00005431 | -0.468724631 | 0.37848332 | -1.2384288 | 0.21555712 | 0.34286722 | NA | NA |
| ENSG000002 | 2.8228204 | -0.465674503 | 0.37603769 | -1.2383719 | 0.2155782 | 0.34287385 | NA | NA |
| ENSG000002 | 5.0680972 | -0.463867926 | 0.37457758 | -1.2383761 | 0.21557663 | 0.34287385 | NA | NA |
| ENSG000002 | 7.84574055 | -0.445056141 | 0.35943735 | -1.2382023 | 0.21564107 | 0.34296038 | NA | NA |
| ENSG000002 | 5.56760555 | 0.463151868 | 0.37407172 | 1.23813655 | 0.21566543 | 0.34298569 | NA | NA |
| ENSG000002 | 6.42774022 | -0.454149677 | 0.36682745 | -1.2380471 | 0.21569858 | 0.3430115 | NA | NA |
| ENSG000002 | 7.15977211 | -0.442528503 | 0.35744055 | -1.2380479 | 0.21569832 | 0.3430115 | NA | NA |
| ENSG000002 | 15.1265354 | -0.378779256 | 0.30596113 | -1.237998 | 0.21571681 | 0.34302703 | NA | NA |
| ENSG000002 | 2.86402612 | -0.46552251 | 0.37604521 | -1.2379429 | 0.21573722 | 0.34303259 | NA | NA |
| ENSG000002 | 4.87332001 | -0.466115104 | 0.37652071 | -1.2379534 | 0.21573333 | 0.34303259 | NA | NA |
| ENSG000002 | 28.0319339 | 0.313964586 | 0.25363252 | 1.23787197 | 0.21576354 | 0.34306098 | NA | NA |
| ENSG000002 | 273.673948 | 0.16250444 | 0.13129124 | 1.23774014 | 0.21581243 | 0.34312527 | LINC00641 | 283624 |
| ENSG000000 | 5.55420179 | -0.466813339 | 0.37715885 | -1.2377102 | 0.21582355 | 0.34312949 | CMA1 | 1215 |
| ENSG000002 | 4.02399728 | -0.468386491 | 0.37844161 | -1.2376718 | 0.2158378 | 0.34313869 | LOC105377652 | 105377652 |
| ENSG000002 | 4.30055584 | -0.468401911 | 0.3784802 | -1.2375863 | 0.21586949 | 0.34317562 | NA | NA |
| ENSG000002 | 5.72038423 | 0.458671658 | 0.37068535 | 1.23736118 | 0.21595302 | 0.34329496 | NA | NA |
| ENSG000002 | 2.29083637 | -0.456536128 | 0.3689908 | -1.2372561 | 0.21599201 | 0.34333002 | NA | NA |
| ENSG000002 | 14.978497 | -0.382007723 | 0.30875091 | -1.2372683 | 0.21598748 | 0.34333002 | NA | NA |
| ENSG000002 | 7.6795076 | -0.435442593 | 0.35195602 | -1.2372074 | 0.2160101 | 0.34334005 | NA | NA |
| ENSG000002 | 119.805473 | -0.204725525 | 0.16547575 | -1.2371935 | 0.21601526 | 0.34334005 | NA | NA |
| ENSG000001 | 8.73915538 | -0.444660073 | 0.35943782 | -1.2370987 | 0.21605043 | 0.34337563 | IL26 | 55801 |
| ENSG000002 | 2.26431017 | -0.457446687 | 0.36979077 | -1.2370419 | 0.21607151 | 0.34337563 | NA | NA |
| ENSG000002 | 4.9810403 | 0.464527384 | 0.3755013 | 1.23708597 | 0.21605517 | 0.34337563 | MFI2-AS1 | 100507057 |
| ENSG000002 | 5.5973111 | -0.458959562 | 0.37101372 | -1.237042 | 0.21607149 | 0.34337563 | NA | NA |
| ENSG000002 | 18.215005 | -0.359433889 | 0.29059858 | -1.2368742 | 0.21613378 | 0.34346113 | NA | NA |

| | | | | | | | | |
|---|---|---|---|---|---|---|---|---|
| ENSG000002 | 5.99714896 | -0.459771925 | 0.37177211 | -1.2367036 | 0.21619713 | 0.34354833 | NA | NA |
| ENSG000002 | 6.73068127 | -0.456071085 | 0.36879657 | -1.2366468 | 0.21621824 | 0.34356842 | NA | NA |
| ENSG000001 | 19.9614012 | -0.355406921 | 0.28741953 | -1.2365441 | 0.21625638 | 0.34356893 | ANGPTL1 | 9068 |
| ENSG000001 | 196.123483 | 0.202582162 | 0.16383093 | 1.23653187 | 0.21626093 | 0.34356893 | ELMSAN1 | 91748 |
| ENSG000002 | 3.7200478 | -0.468159348 | 0.37858894 | -1.2365901 | 0.21623929 | 0.34356893 | RNU6-921P | 106479973 |
| ENSG000002 | 5.6426468 | -0.459735078 | 0.37178388 | -1.2365654 | 0.21624848 | 0.34356893 | NA | NA |
| ENSG000002 | 2.87479069 | -0.464664473 | 0.37575663 | -1.2366102 | 0.21623184 | 0.34356893 | NA | NA |
| ENSG000002 | 2.88316104 | -0.464889627 | 0.37601573 | -1.2363568 | 0.21632595 | 0.34365876 | NA | NA |
| ENSG000002 | 6.64401406 | -0.449866387 | 0.36396216 | -1.2360252 | 0.21644921 | 0.3438411 | NA | NA |
| ENSG000001 | 322.26075 | 0.379829742 | 0.30731539 | 1.23596067 | 0.21647318 | 0.34386572 | PLXNB2 | 23654 |
| ENSG000002 | 3.99834523 | -0.467483133 | 0.37824168 | -1.2359377 | 0.21648174 | 0.34386583 | NA | NA |
| ENSG000000 | 137.296961 | 0.250390446 | 0.20259617 | 1.23590906 | 0.21649237 | 0.34386926 | ATP2A3 | 489 |
| ENSG000002 | 30.9504633 | 0.303142108 | 0.24528756 | 1.2358642 | 0.21650905 | 0.34387965 | NA | NA |
| ENSG000002 | 6.71060318 | -0.454938273 | 0.36811895 | -1.2358458 | 0.21651588 | 0.34387965 | NA | NA |
| ENSG000000 | 3.12224313 | -0.466901767 | 0.37781434 | -1.2357968 | 0.21653409 | 0.34388877 | CST7 | 8530 |
| ENSG000002 | 4.27795564 | -0.466375988 | 0.37739257 | -1.2357848 | 0.21653858 | 0.34388877 | LOC285889 | 285889 |
| ENSG000002 | 6.69845787 | -0.449248964 | 0.36356151 | -1.235689 | 0.21657418 | 0.34393185 | SLED1 | 643036 |
| ENSG000001 | 44.7120249 | -0.270252703 | 0.21873594 | -1.2355204 | 0.21663691 | 0.34396707 | PTAFR | 5724 |
| ENSG000001 | 5.96641013 | -0.461914057 | 0.37385894 | -1.2355303 | 0.2166332 | 0.34396707 | NA | NA |
| ENSG000001 | 5.30557215 | -0.461669146 | 0.3736641 | -1.2355191 | 0.21663736 | 0.34396707 | LINC00336 | 401253 |
| ENSG000002 | 2.66041493 | -0.457913009 | 0.37062504 | -1.2355156 | 0.21663869 | 0.34396707 | NA | NA |
| ENSG000002 | 3.72628886 | -0.467482907 | 0.37837079 | -1.2355153 | 0.21663878 | 0.34396707 | RN7SL713P | 106480381 |
| ENSG000002 | 3.94600967 | -0.467246241 | 0.37825674 | -1.2352622 | 0.21673295 | 0.34409157 | NA | NA |
| ENSG000002 | 8.1789292 | -0.445879253 | 0.36096016 | -1.2352589 | 0.21673416 | 0.34409157 | NA | NA |
| ENSG000002 | 2.90040397 | -0.463749497 | 0.37544425 | -1.235202 | 0.21675533 | 0.3441117 | NA | NA |
| ENSG000002 | 2.84408209 | -0.463944011 | 0.37562529 | -1.2351245 | 0.21678416 | 0.344144 | NA | NA |
| ENSG000002 | 2.90479433 | -0.462938782 | 0.374829 | -1.2350666 | 0.21680572 | 0.34416475 | NA | NA |
| ENSG000001 | 99.6700741 | 0.222650907 | 0.18028506 | 1.23499365 | 0.21683287 | 0.34419438 | AIM1L | 55057 |
| ENSG000001 | 6.2107666 | -0.458178061 | 0.37103875 | -1.2348523 | 0.21688547 | 0.34426439 | ENTPD8 | 377841 |

| | | | | | | | | |
|---|---|---|---|---|---|---|---|---|
| ENSG000002 | 4.94061855 | -0.466916195 | 0.37814202 | -1.2347641 | 0.21691831 | 0.34430305 | NA | NA |
| ENSG000002 | 10.6253874 | 0.414227122 | 0.33550833 | 1.23462546 | 0.21696993 | 0.3443715 | NA | NA |
| ENSG000001 | 27.0784675 | -0.311055844 | 0.25195415 | -1.2345732 | 0.21698938 | 0.3443889 | AVPR1A | 552 |
| ENSG000002 | 7.50058129 | -0.441916016 | 0.35801273 | -1.2343584 | 0.21706939 | 0.3445024 | NA | NA |
| ENSG000002 | 4.42890236 | 0.465708272 | 0.37731213 | 1.23427857 | 0.21709912 | 0.3445361 | NA | NA |
| ENSG000002 | 10.5216847 | -0.414251525 | 0.33563373 | -1.2342369 | 0.21711465 | 0.34454726 | NA | NA |
| ENSG000002 | 3.95509185 | -0.466795299 | 0.37822602 | -1.2341702 | 0.21713948 | 0.34457318 | LINC00113 | 54088 |
| ENSG000002 | 8.47707112 | -0.427937035 | 0.34675458 | -1.2341208 | 0.2171579 | 0.34458893 | NA | NA |
| ENSG00000C | 632.625882 | 0.208230875 | 0.16873126 | 1.23409776 | 0.21716648 | 0.34458906 | PXN | 5829 |
| ENSG000002 | 2.61473975 | -0.45874007 | 0.37174675 | -1.2340123 | 0.21719832 | 0.34462609 | KRTAP4-11 | 653240 |
| ENSG000002 | 2.28602887 | -0.456060007 | 0.36960502 | -1.2339118 | 0.21723576 | 0.34467201 | NA | NA |
| ENSG000002 | 4.29145303 | -0.466257817 | 0.37788673 | -1.233856 | 0.21725656 | 0.34468995 | NA | NA |
| ENSG000002 | 42.3797835 | 0.28945651 | 0.23459887 | 1.23383589 | 0.21726406 | 0.34468995 | NA | NA |
| ENSG000002 | 3.9987298 | -0.466879945 | 0.37843 | -1.2337287 | 0.21730402 | 0.34471287 | NA | NA |
| ENSG000002 | 6.77742251 | -0.453263254 | 0.36738027 | -1.2337713 | 0.21728812 | 0.34471287 | NA | NA |
| ENSG000002 | 6.62635923 | -0.450527876 | 0.36517235 | -1.2337404 | 0.21729965 | 0.34471287 | NA | NA |
| ENSG000001 | 266.671938 | -0.180154626 | 0.14602829 | -1.2336967 | 0.21731593 | 0.34471828 | LMNA | 4000 |
| ENSG000001 | 3.13489926 | -0.465970871 | 0.37772345 | -1.2336297 | 0.21734092 | 0.34473619 | NA | NA |
| ENSG000001 | 3.95203862 | -0.466487821 | 0.37814522 | -1.2336208 | 0.21734422 | 0.34473619 | OR51T1 | 401665 |
| ENSG000002 | 7.18192025 | -0.441800948 | 0.35817713 | -1.2334706 | 0.21740025 | 0.34480878 | NA | NA |
| ENSG000002 | 3.00751088 | -0.459021976 | 0.37214402 | -1.2334525 | 0.21740699 | 0.34480878 | NA | NA |
| ENSG000002 | 3.43900208 | -0.465865957 | 0.37773986 | -1.2332984 | 0.21746446 | 0.34488644 | NA | NA |
| ENSG000002 | 4.22392938 | -0.4653787 | 0.37737262 | -1.2332074 | 0.2174984 | 0.3448998 | NA | NA |
| ENSG000002 | 3.19634871 | -0.465265146 | 0.37726874 | -1.2332459 | 0.21748402 | 0.3448998 | NA | NA |
| ENSG000002 | 4.29511947 | -0.465694116 | 0.37762202 | -1.2332282 | 0.21749065 | 0.3448998 | NA | NA |
| ENSG000002 | 2.64183343 | -0.454338111 | 0.36842786 | -1.2331806 | 0.21750838 | 0.34490213 | NA | NA |
| ENSG000002 | 2.26243135 | -0.456461627 | 0.37016222 | -1.2331394 | 0.21752375 | 0.34491303 | NA | NA |
| ENSG000002 | 2.63477552 | -0.457829097 | 0.37129108 | -1.2330732 | 0.21754844 | 0.3449252 | NA | NA |
| ENSG000002 | 12.1792685 | -0.401414486 | 0.32553637 | -1.2330864 | 0.21754351 | 0.3449252 | NA | NA |

| | | | | | | | | |
|---|---|---|---|---|---|---|---|---|
| ENSG000001 | 126.238856 | -0.187788066 | 0.15230088 | -1.2330071 | 0.21757311 | 0.34495082 | ZNF385D | 79750 |
| ENSG000002 | 2.61106067 | -0.457669509 | 0.37126553 | -1.2327282 | 0.21767717 | 0.34510231 | NA | NA |
| ENSG000001 | 11.0906215 | -0.405098858 | 0.32864951 | -1.2326166 | 0.21771882 | 0.34515485 | IL36G | 56300 |
| ENSG000002 | 4.24487841 | -0.465135456 | 0.37737756 | -1.2325467 | 0.21774495 | 0.34518278 | NA | NA |
| ENSG000002 | 58.7890752 | -0.254051539 | 0.20612965 | -1.2324842 | 0.21776826 | 0.34520624 | PSMD6-AS2 | 100507062 |
| ENSG000001 | 35.0571723 | -0.327230554 | 0.26552983 | -1.2323683 | 0.21781152 | 0.34524782 | SLC6A12 | 6539 |
| ENSG000001 | 6.2130763 | -0.457158181 | 0.37095363 | -1.2323863 | 0.21780481 | 0.34524782 | NA | NA |
| ENSG000002 | 3.56552287 | -0.465207952 | 0.37755772 | -1.2321506 | 0.21789285 | 0.34536323 | NA | NA |
| ENSG000002 | 6.33790868 | -0.446700234 | 0.36255003 | -1.2321065 | 0.21790932 | 0.34537584 | PRO1804 | 100133319 |
| ENSG000001 | 9.60429665 | -0.427310923 | 0.34685926 | -1.2319433 | 0.21797028 | 0.34544545 | C22orf31 | 25770 |
| ENSG000002 | 8.55797057 | 0.426550293 | 0.3462374 | 1.23195903 | 0.21796439 | 0.34544545 | NA | NA |
| ENSG000002 | 3.16725854 | -0.46527082 | 0.37769313 | -1.2318753 | 0.21799568 | 0.34547221 | NA | NA |
| ENSG000001 | 31.127232 | -0.30957262 | 0.25131186 | -1.2318265 | 0.21801389 | 0.34548756 | AKAP3 | 10566 |
| ENSG000002 | 2.26365783 | -0.455938524 | 0.3701682 | -1.2317064 | 0.21805879 | 0.34554522 | NA | NA |
| ENSG000002 | 3.48873136 | 0.466191635 | 0.37852478 | 1.23160137 | 0.21809803 | 0.34559389 | MIR3671 | 100500854 |
| ENSG000001 | 129.511185 | 0.196322109 | 0.15941754 | 1.23149628 | 0.21813731 | 0.34563119 | BCL2 | 596 |
| ENSG000002 | 2.29743892 | -0.454443098 | 0.36901808 | -1.2314928 | 0.21813862 | 0.34563119 | NA | NA |
| ENSG000001 | 26.5107308 | -0.315554647 | 0.25624577 | -1.2314531 | 0.21815345 | 0.34564119 | C9orf47 | 286223 |
| ENSG000001 | 244.232227 | 0.174644486 | 0.14184469 | 1.23123743 | 0.21823408 | 0.3457479 | ARHGEF26 | 26084 |
| ENSG000002 | 25.5230816 | 0.315430251 | 0.25619172 | 1.23122733 | 0.21823785 | 0.3457479 | NA | NA |
| ENSG000002 | 4.57562456 | -0.465179583 | 0.37783764 | -1.2311626 | 0.21826205 | 0.34577272 | NA | NA |
| ENSG000002 | 5.39490655 | -0.461119131 | 0.37456154 | -1.2310904 | 0.21828904 | 0.34580197 | NA | NA |
| ENSG000001 | 3.45512266 | -0.464957667 | 0.37768998 | -1.2310564 | 0.21830177 | 0.34580863 | TMEM8C | 389827 |
| ENSG000002 | 5.87694188 | -0.453503763 | 0.36843859 | -1.2308802 | 0.21836767 | 0.34589951 | NA | NA |
| ENSG000001 | 41.503448 | 0.271576872 | 0.22065015 | 1.23080303 | 0.21839654 | 0.34593173 | MOSPD3 | 64598 |
| ENSG000002 | 2.50207062 | 0.456221896 | 0.37067753 | 1.23077841 | 0.21840575 | 0.34593281 | NA | NA |
| ENSG000002 | 5.2917436 | -0.457633423 | 0.37185071 | -1.2306913 | 0.21843836 | 0.34596597 | NA | NA |
| ENSG000002 | 5.47628449 | 0.460495661 | 0.37418082 | 1.23067683 | 0.21844376 | 0.34596597 | NA | NA |
| ENSG000002 | 30.638607 | 0.355117201 | 0.28856701 | 1.23062299 | 0.2184639 | 0.34597085 | NA | NA |

| | | | | | | | | |
|---|---|---|---|---|---|---|---|---|
| ENSG000002 | 10.2927089 | -0.417065986 | 0.33890477 | -1.2306289 | 0.21846171 | 0.34597085 | NA | NA |
| ENSG000002 | 3.75196162 | -0.465247969 | 0.37810686 | -1.2304669 | 0.21852232 | 0.34602887 | NA | NA |
| ENSG000002 | 12.7357375 | -0.389422182 | 0.31648215 | -1.2304712 | 0.2185207 | 0.34602887 | NA | NA |
| ENSG000002 | 17.2770189 | -0.368777447 | 0.29970778 | -1.2304567 | 0.21852614 | 0.34602887 | LOC101928766 | 101928766 |
| ENSG000002 | 2.34682177 | -0.447510882 | 0.36375255 | -1.2302618 | 0.21859907 | 0.34613084 | NA | NA |
| ENSG000002 | 3.74906573 | -0.465518491 | 0.37843263 | -1.2301225 | 0.21865123 | 0.34619992 | NA | NA |
| ENSG000002 | 14.1102554 | -0.377957653 | 0.3072638 | -1.2300755 | 0.21866885 | 0.3462143 | MAGIX | 79917 |
| ENSG000001 | 14.4310323 | -0.402499391 | 0.32725544 | -1.2299242 | 0.21872549 | 0.34629045 | OR10H5 | 284433 |
| ENSG000002 | 5.6295547 | -0.455264994 | 0.37017786 | -1.2298547 | 0.21875151 | 0.34631812 | NA | NA |
| ENSG000001 | 57.8194568 | 0.240787017 | 0.19579177 | 1.22981171 | 0.21876762 | 0.34633012 | NA | NA |
| ENSG000002 | 3.12320902 | -0.464706856 | 0.37789802 | -1.229715 | 0.21880385 | 0.34637394 | NA | NA |
| ENSG000002 | 3.14130593 | -0.464499072 | 0.37774885 | -1.2296505 | 0.218828 | 0.34639865 | NA | NA |
| ENSG000001 | 8.20528597 | -0.435682713 | 0.35432506 | -1.229613 | 0.21884205 | 0.34640736 | GPR50 | 9248 |
| ENSG000001 | 7.71730781 | -0.434857647 | 0.35370925 | -1.2294212 | 0.21891393 | 0.34649469 | CHRM1 | 1128 |
| ENSG000002 | 41.1209855 | -0.27958587 | 0.227417 | -1.2293974 | 0.21892285 | 0.34649469 | NA | NA |
| ENSG000002 | 4.22225942 | -0.463766993 | 0.37722539 | -1.2294162 | 0.21891582 | 0.34649469 | NA | NA |
| ENSG000002 | 3.47952106 | 0.464771695 | 0.3780722 | 1.22931995 | 0.21895187 | 0.3465271 | NA | NA |
| ENSG000001 | 79.8979012 | 0.231129957 | 0.18802871 | 1.22922697 | 0.21898672 | 0.34655319 | SLC15A3 | 51296 |
| ENSG000001 | 10.2583104 | -0.410170634 | 0.33367952 | -1.2292353 | 0.21898361 | 0.34655319 | SLC22A11 | 55867 |
| ENSG000002 | 4.46213134 | -0.461518257 | 0.37546482 | -1.2291917 | 0.21899992 | 0.34655319 | NA | NA |
| ENSG000002 | 2.6127119 | -0.457350364 | 0.37207617 | -1.2291848 | 0.21900254 | 0.34655319 | NA | NA |
| ENSG000002 | 2.26431072 | -0.455388528 | 0.37053778 | -1.2289935 | 0.21907424 | 0.34665312 | NA | NA |
| ENSG000002 | 3.41989962 | -0.464015044 | 0.37759216 | -1.2288789 | 0.2191172 | 0.34670757 | NA | NA |
| ENSG000002 | 2.87956649 | -0.461299189 | 0.37541571 | -1.228769 | 0.21915842 | 0.34675927 | NA | NA |
| ENSG000002 | 2.63758742 | -0.456462681 | 0.37149002 | -1.2287347 | 0.21917129 | 0.3467661 | NA | NA |
| ENSG000002 | 10.8960407 | 0.408416952 | 0.33247383 | 1.22841833 | 0.21928997 | 0.34694033 | NA | NA |
| ENSG000002 | 2.28488672 | -0.452733014 | 0.36856231 | -1.2283758 | 0.21930594 | 0.34695205 | NA | NA |
| ENSG000002 | 17.1623521 | -0.375538614 | 0.30574757 | -1.2282636 | 0.21934803 | 0.34699954 | HIPK1-AS1 | 101928846 |
| ENSG000002 | 3.76704586 | -0.464802989 | 0.37842697 | -1.2282502 | 0.21935307 | 0.34699954 | NA | NA |

| | | | | | | | | |
|---|---|---|---|---|---|---|---|---|
| ENSG000002 | 7.49391112 | -0.441977911 | 0.35985635 | -1.2282065 | 0.21936946 | 0.34701193 | LOC100129138 | 100129138 |
| ENSG000001 | 6.09741218 | 0.454832034 | 0.37033835 | 1.22815268 | 0.21938966 | 0.34703035 | HIST3H3 | 8290 |
| ENSG000002 | 4.83739896 | -0.4622349 | 0.3763776 | -1.2281148 | 0.21940388 | 0.34703931 | NA | NA |
| ENSG000002 | 3.16183041 | -0.463591378 | 0.3775214 | -1.227987 | 0.21945184 | 0.34710164 | NA | NA |
| ENSG000002 | 10.0870842 | -0.416652959 | 0.33931665 | -1.2279178 | 0.21947782 | 0.34712918 | NA | NA |
| ENSG000001 | 63.1954918 | 0.290703241 | 0.23675356 | 1.22787273 | 0.21949475 | 0.34714242 | ZFP41 | 286128 |
| ENSG000001 | 18.6777832 | -0.348640915 | 0.28394557 | -1.2278442 | 0.21950547 | 0.34714584 | SLC7A10 | 56301 |
| ENSG000002 | 30.1658354 | -0.303481533 | 0.24720167 | -1.2276678 | 0.21957169 | 0.34723702 | NA | NA |
| ENSG000002 | 5.01530483 | -0.45983989 | 0.3746269 | -1.227461 | 0.21964939 | 0.34734635 | NA | NA |
| ENSG000001 | 13.4495864 | -0.390523896 | 0.3181775 | -1.2273775 | 0.21968075 | 0.34738239 | ERP27 | 121506 |
| ENSG000001 | 33.1084128 | -0.290611947 | 0.23680227 | -1.2272346 | 0.21973442 | 0.34744781 | FABP6 | 2172 |
| ENSG000002 | 8.50101785 | -0.431055301 | 0.35124483 | -1.2272218 | 0.21973925 | 0.34744781 | NA | NA |
| ENSG000002 | 2.90197033 | -0.460634588 | 0.37537094 | -1.227145 | 0.21976809 | 0.34747985 | NA | NA |
| ENSG000002 | 37.3242911 | 0.290436434 | 0.23669379 | 1.22705555 | 0.21980171 | 0.34751947 | ZNF529-AS1 | 101927599 |
| ENSG000002 | 6.9082022 | -0.443600819 | 0.3615342 | -1.2269955 | 0.2198243 | 0.34754163 | NA | NA |
| ENSG000001 | 14.3405283 | -0.377809109 | 0.3079524 | -1.2268426 | 0.21988177 | 0.34761893 | AHSP | 51327 |
| ENSG000002 | 10.3697396 | -0.412985003 | 0.33666181 | -1.2267058 | 0.21993318 | 0.34764645 | NA | NA |
| ENSG000002 | 7.99839377 | -0.438680203 | 0.35759531 | -1.2267504 | 0.21991641 | 0.34764645 | GOLGA6B | 55889 |
| ENSG000002 | 4.48826057 | -0.461095561 | 0.37588136 | -1.2267051 | 0.21993346 | 0.34764645 | NA | NA |
| ENSG000002 | 56.8068794 | 0.258215078 | 0.21048629 | 1.22675487 | 0.21991474 | 0.34764645 | STAG3L5P | 101735302 |
| ENSG000001 | 39.9572318 | 0.280526206 | 0.22873137 | 1.22644393 | 0.22003166 | 0.34776271 | LINC00471 | 151477 |
| ENSG000002 | 5.10339572 | -0.461689701 | 0.37644671 | -1.226441 | 0.22003275 | 0.34776271 | NA | NA |
| ENSG000002 | 2.90649743 | -0.459894224 | 0.37497685 | -1.2264603 | 0.22002551 | 0.34776271 | NA | NA |
| ENSG000002 | 53.0079818 | 0.25534758 | 0.20821175 | 1.2263841 | 0.22005417 | 0.34776946 | NA | NA |
| ENSG000002 | 5.53674343 | -0.453318565 | 0.36963549 | -1.2263935 | 0.22005063 | 0.34776946 | NA | NA |
| ENSG000002 | 5.03168579 | -0.460106675 | 0.37518338 | -1.2263514 | 0.22006645 | 0.34777532 | NA | NA |
| ENSG000001 | 295.666334 | 0.163418621 | 0.13327086 | 1.2262142 | 0.22011808 | 0.34784335 | TDRKH | 11022 |
| ENSG000001 | 15.5487854 | -0.386394903 | 0.31514152 | -1.2260996 | 0.22016119 | 0.34789792 | TLE6 | 79816 |
| ENSG000002 | 8.1943111 | -0.428526856 | 0.34953191 | -1.2260021 | 0.22019789 | 0.34794235 | TUBA4B | 80086 |

| | | | | | | | | |
|---|---|---|---|---|---|---|---|---|
| ENSG000002 | 5.81462059 | 0.453772031 | 0.37014455 | 1.22593195 | 0.22022429 | 0.3479705 | NA | NA |
| ENSG000002 | 7.07451227 | -0.453206655 | 0.3697159 | -1.2258241 | 0.22026489 | 0.34802109 | VSTM2A-OT1 | 285878 |
| ENSG000002 | 2.25175852 | -0.453674435 | 0.37013085 | -1.2257136 | 0.22030645 | 0.34805965 | NA | NA |
| ENSG000002 | 23.0127608 | -0.330364454 | 0.26952628 | -1.2257226 | 0.22030308 | 0.34805965 | NA | NA |
| ENSG000000 | 261.61221 | 0.1601241 | 0.13064817 | 1.22561305 | 0.22034432 | 0.34807915 | ADCYAP1R1 | 117 |
| ENSG000001 | 3.1530683 | -0.462426161 | 0.37729643 | -1.2256309 | 0.22033761 | 0.34807915 | FAM47C | 442444 |
| ENSG000002 | 4.356109 | -0.463786505 | 0.37841203 | -1.2256125 | 0.22034455 | 0.34807915 | NA | NA |
| ENSG000002 | 4.6784504 | 0.460615833 | 0.37584828 | 1.22553663 | 0.2203731 | 0.34809713 | NA | NA |
| ENSG000002 | 2.95711772 | -0.454706878 | 0.37102572 | -1.22554 | 0.22037181 | 0.34809713 | NA | NA |
| ENSG000002 | 9.98940844 | 0.413235584 | 0.33720284 | 1.22548074 | 0.22039414 | 0.34811681 | NA | NA |
| ENSG000002 | 2.85401827 | -0.459562395 | 0.37505913 | -1.2253065 | 0.22045976 | 0.34820689 | PTGES3P5 | 106480359 |
| ENSG000002 | 2.2484896 | -0.45349882 | 0.37011871 | -1.2252794 | 0.22046998 | 0.34820947 | NA | NA |
| ENSG000002 | 29.971249 | -0.324067075 | 0.26449177 | -1.2252444 | 0.22048314 | 0.34821669 | NA | NA |
| ENSG000002 | 3.0996115 | -0.463097093 | 0.37797243 | -1.2252139 | 0.22049462 | 0.34822127 | NA | NA |
| ENSG000001 | 169.174096 | -0.175953003 | 0.14361567 | -1.2251657 | 0.22051278 | 0.34823638 | DDX60L | 91351 |
| ENSG000001 | 48.5836555 | -0.258223605 | 0.21078116 | -1.2250792 | 0.2205454 | 0.34826303 | FAM81A | 145773 |
| ENSG000002 | 7.7818919 | -0.438727273 | 0.3581293 | -1.2250527 | 0.22055536 | 0.34826303 | NA | NA |
| ENSG000002 | 2.63799181 | -0.454055419 | 0.37064158 | -1.2250526 | 0.22055542 | 0.34826303 | NA | NA |
| ENSG000001 | 7.11842259 | -0.439138658 | 0.35847443 | -1.2250209 | 0.22056736 | 0.34826832 | NA | NA |
| ENSG000001 | 50.8748134 | -0.258296635 | 0.21085913 | -1.2249725 | 0.22058558 | 0.34828353 | CDKL2 | 8999 |
| ENSG000002 | 20.4320588 | 0.346999087 | 0.28327709 | 1.22494582 | 0.22059564 | 0.34828584 | NA | NA |
| ENSG000002 | 4.28415325 | -0.46263523 | 0.37769655 | -1.2248861 | 0.22061816 | 0.34830784 | NA | NA |
| ENSG000002 | 7.99216747 | -0.432284704 | 0.35297304 | -1.2246961 | 0.22068974 | 0.34840729 | NA | NA |
| ENSG000002 | 11.5688156 | 0.285880544 | 0.2334448 | 1.22461733 | 0.22071944 | 0.34844061 | NA | NA |
| ENSG000002 | 3.10073032 | 0.462183817 | 0.37745698 | 1.22446755 | 0.2207759 | 0.34851618 | NA | NA |
| ENSG000002 | 16.2989999 | 0.37539962 | 0.30658985 | 1.22443589 | 0.22078784 | 0.34852145 | NA | NA |
| ENSG000002 | 15.603075 | -0.381663221 | 0.31176468 | -1.2242029 | 0.2208757 | 0.34864658 | LOC101927604 | 101927604 |
| ENSG000001 | 2.25036842 | -0.452223661 | 0.36941833 | -1.2241506 | 0.22089543 | 0.34866415 | NA | NA |
| ENSG000002 | 11.7680039 | -0.398445141 | 0.32555429 | -1.2238977 | 0.22099082 | 0.34880114 | NA | NA |

| | | | | | | | | |
|---|---|---|---|---|---|---|---|---|
| ENSG000002 | 6.64790864 | -0.447045394 | 0.36528346 | -1.2238315 | 0.22101581 | 0.348827 | NACAP1 | 83955 |
| ENSG000002 | 7.95106644 | -0.430448329 | 0.35173047 | -1.2238016 | 0.22102706 | 0.34883118 | DYX1C1 | 161582 |
| ENSG000002 | 16.1371743 | -0.370901805 | 0.30309354 | -1.2237206 | 0.22105765 | 0.34885993 | CFLAR-AS1 | 65072 |
| ENSG000002 | 4.53852068 | -0.460589399 | 0.37638839 | -1.2237078 | 0.22106248 | 0.34885993 | NA | NA |
| ENSG000002 | 7.40248838 | -0.435163323 | 0.35571186 | -1.223359 | 0.22119412 | 0.34905408 | NA | NA |
| ENSG000002 | 4.28336342 | -0.46241505 | 0.37803052 | -1.2232215 | 0.22124605 | 0.34912245 | RN7SL728P | 106481857 |
| ENSG000002 | 3.12887792 | -0.461609541 | 0.37748966 | -1.2228402 | 0.22139005 | 0.34933607 | LOC101929240 | 101929240 |
| ENSG000002 | 16.1875971 | 0.386767159 | 0.3162988 | 1.22279047 | 0.22140884 | 0.34935214 | NA | NA |
| ENSG000002 | 6.64865925 | -0.44582959 | 0.36463358 | -1.2226783 | 0.22145121 | 0.34939179 | NA | NA |
| ENSG000002 | 10.6663773 | -0.421433076 | 0.3446782 | -1.2226856 | 0.22144847 | 0.34939179 | LOC101929058 | 101929058 |
| ENSG000001 | 35.8112283 | 0.288028655 | 0.23559375 | 1.22256491 | 0.22149407 | 0.34943069 | DHX58 | 79132 |
| ENSG000001 | 130.026165 | 0.191586487 | 0.1567103 | 1.222552 | 0.22149895 | 0.34943069 | CACNA1B | 774 |
| ENSG000002 | 3.98551207 | -0.462756674 | 0.37851923 | -1.2225447 | 0.22150171 | 0.34943069 | NA | NA |
| ENSG000001 | 47.7750449 | -0.258284203 | 0.21129103 | -1.2224097 | 0.22155274 | 0.34949759 | KCNMB3 | 27094 |
| ENSG000002 | 16.5808509 | 0.373832672 | 0.30583479 | 1.22233534 | 0.22158084 | 0.34952833 | NA | NA |
| ENSG000002 | 2.85924977 | -0.458092084 | 0.37478167 | -1.2222905 | 0.22159778 | 0.34953222 | RNU6-883P | 106481921 |
| ENSG000002 | 6.0944433 | -0.445872088 | 0.36478624 | -1.2222832 | 0.22160055 | 0.34953222 | NA | NA |
| ENSG000002 | 6.3482366 | -0.444519459 | 0.3637138 | -1.2221682 | 0.22164401 | 0.34957358 | RNU6-522P | 106479797 |
| ENSG000002 | 4.25874951 | -0.462604466 | 0.37850767 | -1.2221799 | 0.2216396 | 0.34957358 | NA | NA |
| ENSG000002 | 2.84106057 | -0.457414451 | 0.37430669 | -1.2220312 | 0.22169584 | 0.34964172 | NA | NA |
| ENSG000002 | 3.38795645 | -0.46258151 | 0.37856125 | -1.2219463 | 0.22172794 | 0.34967875 | NA | NA |
| ENSG000002 | 7.26942679 | -0.440151241 | 0.3602142 | -1.2219153 | 0.22173967 | 0.34968364 | NA | NA |
| ENSG000002 | 4.47485905 | -0.459860482 | 0.37640379 | -1.2217212 | 0.22181308 | 0.34978581 | NA | NA |
| ENSG000002 | 4.37181582 | -0.462457449 | 0.37854003 | -1.221687 | 0.221826 | 0.34979258 | SPDYE2 | 441273 |
| ENSG000002 | 4.51168147 | -0.45955337 | 0.37620733 | -1.2215428 | 0.22188055 | 0.34985675 | NA | NA |
| ENSG000002 | 9.45694262 | 0.415792105 | 0.34038524 | 1.22153388 | 0.22188394 | 0.34985675 | MIR4482 | 100616323 |
| ENSG000002 | 2.5455918 | 0.455139191 | 0.37262196 | 1.22145026 | 0.22191559 | 0.34989304 | NA | NA |
| ENSG000001 | 3.72499966 | -0.462376824 | 0.37856047 | -1.2214081 | 0.22193154 | 0.34990459 | PRG2 | 5553 |
| ENSG000002 | 5.80660299 | -0.459798395 | 0.37646281 | -1.2213647 | 0.22194796 | 0.34991686 | LINC00297 | 100874059 |

| | | | | | | | | |
|---|---|---|---|---|---|---|---|---|
| ENSG000002 | 4.05305044 | -0.461803118 | 0.37812947 | -1.2212831 | 0.22197886 | 0.34995198 | NA | NA |
| ENSG000000 | 38.7539282 | -0.273578096 | 0.22402728 | -1.2211821 | 0.2220171 | 0.34997144 | STOML1 | 9399 |
| ENSG000002 | 3.00964308 | -0.453289776 | 0.37118656 | -1.2211912 | 0.22201363 | 0.34997144 | NA | NA |
| ENSG000002 | 5.69247748 | -0.455650661 | 0.37311763 | -1.2211984 | 0.2220109 | 0.34997144 | LINC01477 | 101927900 |
| ENSG000002 | 7.25788015 | -0.440399834 | 0.36067431 | -1.2210458 | 0.2220687 | 0.35002557 | NA | NA |
| ENSG000002 | 6.91089731 | -0.440984121 | 0.3611504 | -1.2210539 | 0.2220656 | 0.35002557 | NA | NA |
| ENSG000002 | 3.82681751 | 0.462162415 | 0.37858169 | 1.22077328 | 0.22217187 | 0.35017458 | RNU6-998P | 106481495 |
| ENSG000002 | 2.25275632 | -0.45060254 | 0.36914294 | -1.2206722 | 0.22221014 | 0.35017475 | NA | NA |
| ENSG000002 | 24.1238275 | -0.350733479 | 0.28731762 | -1.2207169 | 0.22219323 | 0.35017475 | NA | NA |
| ENSG000002 | 9.92682848 | -0.419934379 | 0.34400947 | -1.2207059 | 0.2221974 | 0.35017475 | NA | NA |
| ENSG000002 | 3.45703538 | -0.461286111 | 0.37789924 | -1.220659 | 0.22221516 | 0.35017475 | NA | NA |
| ENSG000002 | 19.4482458 | 0.374405121 | 0.30672043 | 1.22067226 | 0.22221013 | 0.35017475 | NA | NA |
| ENSG000002 | 4.79441084 | -0.45996831 | 0.37684066 | -1.220591 | 0.22224093 | 0.35018814 | NA | NA |
| ENSG000002 | 8.74969544 | -0.429236769 | 0.35165922 | -1.2206043 | 0.22223586 | 0.35018814 | NA | NA |
| ENSG000002 | 14.2201036 | 0.377175149 | 0.30901882 | 1.22055722 | 0.22225371 | 0.35019468 | NA | NA |
| ENSG000002 | 6.07927507 | -0.444982937 | 0.3645897 | -1.2205033 | 0.22227414 | 0.35021325 | NA | NA |
| ENSG000002 | 33.9174073 | 0.291392601 | 0.2387562 | 1.2204609 | 0.2222902 | 0.35022496 | ANKRD39 | 51239 |
| ENSG000001 | 1182.0475 | 0.414807054 | 0.33992026 | 1.22030695 | 0.22234853 | 0.35030325 | TARDBP | 23435 |
| ENSG000002 | 4.6907474 | -0.461677432 | 0.37839495 | -1.2200941 | 0.22242922 | 0.35040749 | NA | NA |
| ENSG000002 | 5.3162414 | -0.454630369 | 0.37262134 | -1.2200868 | 0.22243198 | 0.35040749 | NA | NA |
| ENSG000001 | 194.949374 | 0.167571099 | 0.13735699 | 1.21996777 | 0.22247709 | 0.35045924 | NA | NA |
| ENSG000002 | 5.33061926 | -0.457199355 | 0.37476754 | -1.2199545 | 0.22248212 | 0.35045924 | NA | NA |
| ENSG000001 | 8.33141032 | -0.433194977 | 0.35513247 | -1.2198124 | 0.22253601 | 0.35051691 | DMRT1 | 1761 |
| ENSG000002 | 3.1320551 | -0.460784525 | 0.37774858 | -1.219818 | 0.22253387 | 0.35051691 | NA | NA |
| ENSG000002 | 5.8888853 | -0.451227135 | 0.36993159 | -1.2197583 | 0.22255651 | 0.35053558 | LINC01398 | 101929651 |
| ENSG000001 | 4.42166875 | -0.461767394 | 0.37859237 | -1.2196955 | 0.22258034 | 0.35055949 | OR7G3 | 390883 |
| ENSG000002 | 10.6546524 | -0.411177887 | 0.33715152 | -1.2195641 | 0.22263017 | 0.35062436 | NA | NA |
| ENSG000002 | 4.06870059 | -0.461575744 | 0.37851627 | -1.2194344 | 0.22267937 | 0.35068823 | NA | NA |
| ENSG000002 | 13.0993684 | 0.39470183 | 0.32370541 | 1.21932417 | 0.22272118 | 0.35074045 | NA | NA |

| | | | | | | | | |
|---|---|---|---|---|---|---|---|---|
| ENSG000001 | 75.9874889 | 0.386135134 | 0.31670533 | 1.21922526 | 0.22275871 | 0.35075868 | PMVK | 10654 |
| ENSG000002 | 22.2792597 | -0.326979296 | 0.26817748 | -1.2192646 | 0.22274379 | 0.35075868 | TIAF1 | 9220 |
| ENSG000002 | 5.10326873 | -0.457537605 | 0.37526267 | -1.2192463 | 0.22275074 | 0.35075868 | NA | NA |
| ENSG000001 | 149.018743 | 0.180503584 | 0.14805938 | 1.21912966 | 0.22279498 | 0.35078856 | IQSEC2 | 23096 |
| ENSG000001 | 261.634317 | 0.165907032 | 0.13608406 | 1.21915113 | 0.22278684 | 0.35078856 | BPNT1 | 10380 |
| ENSG000002 | 2.83749937 | -0.457142751 | 0.37506876 | -1.2188238 | 0.22291106 | 0.35094111 | NA | NA |
| ENSG000002 | 7.48363763 | -0.438715787 | 0.35994912 | -1.2188272 | 0.22290978 | 0.35094111 | NA | NA |
| ENSG000002 | 3.17608116 | -0.457550087 | 0.37540846 | -1.218806 | 0.22291784 | 0.35094111 | NA | NA |
| ENSG000001 | 83.7443122 | 0.230443169 | 0.18909184 | 1.21868383 | 0.22296422 | 0.3510005 | HPSE | 10855 |
| ENSG000002 | 86.2355243 | 0.214460373 | 0.17598497 | 1.2186289 | 0.22298508 | 0.35101971 | NA | NA |
| ENSG000002 | 8.24585149 | 0.432533696 | 0.35501571 | 1.21835087 | 0.22309067 | 0.3511723 | NA | NA |
| ENSG000001 | 132.477718 | 0.192876559 | 0.15832402 | 1.21823939 | 0.22313301 | 0.35121136 | ADAMTS3 | 9508 |
| ENSG000002 | 3.15539203 | -0.460157438 | 0.37772628 | -1.2182299 | 0.22313663 | 0.35121136 | NA | NA |
| ENSG000002 | 3.07977161 | -0.460104279 | 0.37768659 | -1.2182171 | 0.22314146 | 0.35121136 | NA | NA |
| ENSG000002 | 4.51413333 | -0.458988837 | 0.37679327 | -1.218145 | 0.22316888 | 0.35124088 | NA | NA |
| ENSG000002 | 2.25924676 | -0.449043797 | 0.36864163 | -1.2181039 | 0.2231845 | 0.35125183 | NA | NA |
| ENSG000002 | 32.9775374 | 0.29313485 | 0.24066334 | 1.21802869 | 0.22321307 | 0.35128316 | NA | NA |
| ENSG000002 | 2.63491157 | -0.450767069 | 0.3700913 | -1.2179888 | 0.22322822 | 0.35129337 | NA | NA |
| ENSG000002 | 2.28469139 | -0.449742866 | 0.36925985 | -1.2179577 | 0.22324006 | 0.35129836 | NA | NA |
| ENSG000001 | 2.97531543 | -0.452968901 | 0.37192705 | -1.2178972 | 0.22326306 | 0.35132092 | C1QTNF4 | 114900 |
| ENSG000001 | 601.682026 | 0.155904777 | 0.12803554 | 1.21766796 | 0.22335018 | 0.35140347 | ARNT | 405 |
| ENSG000001 | 48.225125 | 0.267146626 | 0.2193841 | 1.21771189 | 0.22333348 | 0.35140347 | GOLGA8A | 23015 |
| ENSG000002 | 24.0467439 | -0.334969475 | 0.27508729 | -1.2176843 | 0.22334396 | 0.35140347 | USP3-AS1 | 100130855 |
| ENSG000002 | 3.11642986 | -0.459867587 | 0.37765399 | -1.2176956 | 0.22333969 | 0.35140347 | RN7SL118P | 106480952 |
| ENSG000001 | 243.961372 | 0.197830356 | 0.16248276 | 1.21754672 | 0.22339627 | 0.35146235 | ZSWIM6 | 57688 |
| ENSG000002 | 3.44943305 | -0.458662667 | 0.37671837 | -1.2175214 | 0.22340591 | 0.35146389 | NA | NA |
| ENSG000002 | 10.3755436 | 0.412977226 | 0.33920603 | 1.21748198 | 0.22342089 | 0.35147381 | NA | NA |
| ENSG000002 | 7.67803843 | 0.438039752 | 0.35983875 | 1.21732236 | 0.22348159 | 0.35155567 | NA | NA |
| ENSG000002 | 2.24914194 | -0.448994465 | 0.36884953 | -1.2172835 | 0.22349635 | 0.35156525 | NA | NA |

| | | | | | | | | |
|---|---|---|---|---|---|---|---|---|
| ENSG000001 | 6.42435995 | -0.447465971 | 0.36763027 | -1.217163 | 0.2235422 | 0.3516221 | GAPDHS | 26330 |
| ENSG000002 | 5.29694977 | 0.453236081 | 0.37238401 | 1.21712015 | 0.22355851 | 0.3516221 | NA | NA |
| ENSG000002 | 19.9535258 | 0.354275684 | 0.29107346 | 1.21713495 | 0.22355287 | 0.3516221 | NA | NA |
| ENSG000002 | 3.1763135 | -0.458941675 | 0.37708871 | -1.2170655 | 0.22357929 | 0.35164114 | NA | NA |
| ENSG000001 | 25.6174265 | -0.324003289 | 0.26623514 | -1.2169817 | 0.22361118 | 0.35167767 | NLRC3 | 197358 |
| ENSG000002 | 14.6095796 | 0.391253602 | 0.3215199 | 1.21688768 | 0.22364695 | 0.35172029 | NA | NA |
| ENSG000002 | 3.12025221 | -0.458626676 | 0.37695762 | -1.2166531 | 0.22373621 | 0.35184702 | NA | NA |
| ENSG000001 | 2.59764727 | -0.450857479 | 0.37058107 | -1.2166231 | 0.22374766 | 0.35185138 | C11orf86 | 254439 |
| ENSG000001 | 15.3684221 | -0.366801649 | 0.30150251 | -1.2165791 | 0.2237644 | 0.35186405 | SUN3 | 256979 |
| ENSG000001 | 52.7541356 | 0.244365285 | 0.20088021 | 1.21647267 | 0.22380492 | 0.35191412 | CPNE7 | 27132 |
| ENSG000002 | 7.88050392 | -0.441831261 | 0.36322305 | -1.2164186 | 0.22382551 | 0.35193285 | NA | NA |
| ENSG000002 | 7.52116524 | 0.440055007 | 0.36177527 | 1.21637671 | 0.22384145 | 0.35194427 | NA | NA |
| ENSG000001 | 2.8966656 | -0.456067408 | 0.37497216 | -1.21627 | 0.22388207 | 0.35199449 | PRSS38 | 339501 |
| ENSG000002 | 35.1499389 | 0.292886917 | 0.24084139 | 1.21609875 | 0.22394731 | 0.35208341 | NA | NA |
| ENSG000002 | 6.09386199 | -0.445978815 | 0.36675849 | -1.2160013 | 0.22398442 | 0.35212756 | NA | NA |
| ENSG000002 | 15.3848237 | -0.375083001 | 0.30846163 | -1.2159794 | 0.22399276 | 0.35212756 | NA | NA |
| ENSG000002 | 3.07354176 | -0.459353371 | 0.37783082 | -1.2157647 | 0.22407457 | 0.35224252 | NA | NA |
| ENSG000002 | 7.19941832 | -0.43837702 | 0.36060268 | -1.2156788 | 0.22410731 | 0.35228033 | NA | NA |
| ENSG000002 | 4.53214702 | -0.457493826 | 0.37637931 | -1.2155127 | 0.22417061 | 0.35236618 | NA | NA |
| ENSG000002 | 3.21430578 | -0.456382777 | 0.37549865 | -1.2154046 | 0.22421182 | 0.35241728 | NA | NA |
| ENSG000002 | 3.45435932 | -0.458756835 | 0.37746393 | -1.215366 | 0.22422652 | 0.35242673 | NA | NA |
| ENSG000002 | 2.89413389 | -0.455697401 | 0.37496994 | -1.2152905 | 0.22425532 | 0.35245833 | NA | NA |
| ENSG000002 | 2.6164888 | -0.450258251 | 0.37051066 | -1.215237 | 0.22427573 | 0.35246308 | NA | NA |
| ENSG000002 | 3.13080936 | -0.458782462 | 0.37752167 | -1.215248 | 0.22427151 | 0.35246308 | NA | NA |
| ENSG000001 | 7.9118635 | -0.427242761 | 0.35158202 | -1.2152008 | 0.22428951 | 0.35247108 | NA | NA |
| ENSG000002 | 3.3611495 | -0.459975463 | 0.3785351 | -1.2151461 | 0.22431037 | 0.3524902 | NA | NA |
| ENSG000002 | 6.33264471 | -0.440834417 | 0.36280484 | -1.2150731 | 0.2243382 | 0.35252028 | RNU6-769P | 106479906 |
| ENSG000002 | 9.17759099 | -0.424973732 | 0.34976275 | -1.2150343 | 0.22435302 | 0.3525299 | NA | NA |
| ENSG000001 | 5.16488703 | -0.457227525 | 0.37632186 | -1.2149906 | 0.22436968 | 0.35254242 | PRF1 | 5551 |

| | | | | | | | | |
|---|---|---|---|---|---|---|---|---|
| ENSG000002 | 3.94985237 | -0.459640263 | 0.37835831 | -1.214828 | 0.22443171 | 0.35261256 | NA | NA |
| ENSG000002 | 3.09346596 | -0.459006362 | 0.37783338 | -1.214838 | 0.22442787 | 0.35261256 | NA | NA |
| ENSG000001 | 74.7386814 | 0.239129563 | 0.19685152 | 1.2147712 | 0.22445337 | 0.35263293 | MVB12B | 89853 |
| ENSG000002 | 7.65722885 | -0.429182347 | 0.35338912 | -1.2144753 | 0.22456629 | 0.35278301 | NA | NA |
| ENSG000002 | 85.0543225 | 0.225307995 | 0.18551759 | 1.21448319 | 0.22456326 | 0.35278301 | GLIDR | 389741 |
| ENSG000001 | 5.33078377 | -0.45377944 | 0.3736783 | -1.2143586 | 0.22461083 | 0.35280586 | OR10V1 | 390201 |
| ENSG000002 | 18.9585616 | -0.363254926 | 0.29912909 | -1.2143751 | 0.2246045 | 0.35280586 | LOC400997 | 400997 |
| ENSG000002 | 5.8245344 | -0.44738244 | 0.36840877 | -1.2143643 | 0.22460865 | 0.35280586 | NA | NA |
| ENSG000002 | 7.43662269 | -0.43397072 | 0.35736992 | -1.214346 | 0.22461564 | 0.35280586 | NA | NA |
| ENSG000002 | 3.4882768 | -0.458081493 | 0.37724956 | -1.2142665 | 0.22464599 | 0.35283596 | NA | NA |
| ENSG000002 | 2.48023928 | 0.451594396 | 0.37191216 | 1.21425017 | 0.22465221 | 0.35283596 | NA | NA |
| ENSG000002 | 79.3503153 | 0.22960418 | 0.18909527 | 1.21422489 | 0.22466186 | 0.35283745 | SPIB | 6689 |
| ENSG000002 | 2.25674229 | -0.447241314 | 0.3683454 | -1.21419 | 0.22467517 | 0.35284469 | KRT19P2 | 160313 |
| ENSG000002 | 7.54901697 | -0.442277907 | 0.3643067 | -1.2140263 | 0.2247377 | 0.35292922 | NA | NA |
| ENSG000001 | 23.8251828 | -0.323140803 | 0.26619844 | -1.2139095 | 0.2247823 | 0.35297193 | C1orf50 | 79078 |
| ENSG000002 | 3.11760372 | -0.457851345 | 0.37716801 | -1.2139188 | 0.22477873 | 0.35297193 | RNU6-844P | 106479939 |
| ENSG00000C | 44.5923681 | 0.271749423 | 0.22390531 | 1.21368013 | 0.22486989 | 0.3530583 | MAPK8IP2 | 23542 |
| ENSG000001 | 63.6841856 | -0.23931664 | 0.19721109 | -1.213505 | 0.22493681 | 0.3530583 | COL2A1 | 1280 |
| ENSG000001 | 273.244686 | 0.16454693 | 0.13559785 | 1.21349214 | 0.22494172 | 0.3530583 | TNFAIP8 | 25816 |
| ENSG000002 | 4.47923101 | 0.458885117 | 0.37812232 | 1.21358907 | 0.22490468 | 0.3530583 | NA | NA |
| ENSG000002 | 3.089167 | -0.45854009 | 0.37783504 | -1.2135986 | 0.22490103 | 0.3530583 | NA | NA |
| ENSG000002 | 3.69783265 | -0.459283921 | 0.37847122 | -1.213524 | 0.22492953 | 0.3530583 | NA | NA |
| ENSG000002 | 15.5159611 | 0.368650931 | 0.30377969 | 1.21354702 | 0.22492075 | 0.3530583 | PYCARD-AS1 | 100652740 |
| ENSG000002 | 10.5756295 | -0.405547424 | 0.33418442 | -1.2135438 | 0.22492199 | 0.3530583 | NA | NA |
| ENSG000002 | 70.3680455 | 0.245627539 | 0.20240386 | 1.21355167 | 0.22491897 | 0.3530583 | NA | NA |
| ENSG000002 | 7.99290468 | -0.429723531 | 0.35412145 | -1.213492 | 0.22494179 | 0.3530583 | NA | NA |
| ENSG000002 | 3.37076505 | -0.459335522 | 0.37849899 | -1.2135713 | 0.22491146 | 0.3530583 | NA | NA |
| ENSG000002 | 8.69649047 | -0.420152874 | 0.34622139 | -1.2135382 | 0.22492413 | 0.3530583 | RN7SKP23 | 106479105 |
| ENSG000001 | 236.086903 | 0.171797699 | 0.14158189 | 1.21341579 | 0.22497089 | 0.35307758 | PACRGL | 133015 |

| | | | | | | | | |
|---|---|---|---|---|---|---|---|---|
| ENSG000001 | 4.24258748 | -0.458733625 | 0.37805289 | -1.2134112 | 0.22497264 | 0.35307758 | CCL11 | 6356 |
| ENSG000002 | 2.35218325 | -0.439199278 | 0.36196008 | -1.2133915 | 0.22498019 | 0.35307758 | RNU6-1304P | 106480135 |
| ENSG000001 | 24.4133349 | 0.313169489 | 0.25810313 | 1.21335021 | 0.22499595 | 0.35308866 | ILDR1 | 286676 |
| ENSG000002 | 3.13265537 | -0.45786814 | 0.37738023 | -1.2132807 | 0.22502253 | 0.3531167 | NA | NA |
| ENSG000002 | 5.55757589 | -0.449470738 | 0.37047816 | -1.2132179 | 0.22504652 | 0.35314068 | NA | NA |
| ENSG000002 | 5.81392518 | -0.449246126 | 0.37030899 | -1.2131656 | 0.22506651 | 0.35315839 | ERVV-1 | 147664 |
| ENSG000002 | 2.87433422 | -0.45516139 | 0.37524478 | -1.2129719 | 0.22514055 | 0.35321988 | NA | NA |
| ENSG000002 | 4.21449566 | -0.458007675 | 0.37757619 | -1.2130206 | 0.22512196 | 0.35321988 | NA | NA |
| ENSG000002 | 6.414027 | -0.44560397 | 0.36735272 | -1.2130139 | 0.2251245 | 0.35321988 | NA | NA |
| ENSG000002 | 12.2540946 | -0.392709112 | 0.32375711 | -1.2129745 | 0.22513957 | 0.35321988 | NA | NA |
| ENSG000002 | 3.14039211 | -0.457387029 | 0.37708718 | -1.2129477 | 0.22514981 | 0.35322075 | NA | NA |
| ENSG000002 | 3.94543662 | -0.458805392 | 0.3782747 | -1.2128894 | 0.22517209 | 0.35324204 | DOCK9-AS1 | 100874096 |
| ENSG000002 | 3.08091376 | -0.458129565 | 0.37774449 | -1.2128028 | 0.22520523 | 0.35326791 | NA | NA |
| ENSG000002 | 2.87443836 | -0.455096814 | 0.3752445 | -1.2128008 | 0.22520601 | 0.35326791 | NA | NA |
| ENSG000002 | 2.25147777 | -0.446889772 | 0.36848465 | -1.2127772 | 0.22521503 | 0.35326839 | SLC6A10P | 386757 |
| ENSG000001 | 48.0527908 | -0.260653756 | 0.21494645 | -1.2126451 | 0.22526555 | 0.35329298 | CMKLR1 | 1240 |
| ENSG000002 | 3.42976243 | -0.458921656 | 0.37843577 | -1.2126804 | 0.22525202 | 0.35329298 | NA | NA |
| ENSG000002 | 4.76099394 | -0.457523591 | 0.37728224 | -1.2126826 | 0.22525121 | 0.35329298 | NA | NA |
| ENSG000002 | 6.09979049 | -0.446367521 | 0.36809328 | -1.2126478 | 0.22526449 | 0.35329298 | NA | NA |
| ENSG000001 | 65.7403726 | 0.23877063 | 0.1969093 | 1.21259194 | 0.22528587 | 0.35330646 | RGS14 | 10636 |
| ENSG000002 | 4.46153849 | -0.455963178 | 0.37602821 | -1.212577 | 0.22529157 | 0.35330646 | NA | NA |
| ENSG000002 | 35.4552643 | 0.291885472 | 0.24072077 | 1.21254794 | 0.22530271 | 0.35331026 | NA | NA |
| ENSG000002 | 4.23054402 | -0.458509856 | 0.37816308 | -1.212466 | 0.22533407 | 0.35333211 | LINC00634 | 339674 |
| ENSG000002 | 4.20213935 | -0.458186652 | 0.37789464 | -1.212472 | 0.22533177 | 0.35333211 | NA | NA |
| ENSG000002 | 9.05190228 | -0.425737444 | 0.35115822 | -1.2123807 | 0.22536669 | 0.35336961 | NA | NA |
| ENSG000002 | 5.4289612 | -0.455409672 | 0.37566784 | -1.2122669 | 0.22541024 | 0.35342422 | NA | NA |
| ENSG000002 | 3.12103626 | -0.457304267 | 0.37729457 | -1.2120616 | 0.2254888 | 0.35353372 | NA | NA |
| ENSG000002 | 9.79470073 | -0.412671149 | 0.34048961 | -1.2119934 | 0.22551492 | 0.353561 | NA | NA |
| ENSG000002 | 89.2414942 | 0.206342975 | 0.17026578 | 1.21188752 | 0.22555544 | 0.35360789 | LOC283335 | 283335 |

| | | | | | | | | |
|---|---|---|---|---|---|---|---|---|
| ENSG000002 | 5.67481035 | -0.452650346 | 0.37351404 | -1.2118697 | 0.22556226 | 0.35360789 | NA | NA |
| ENSG000002 | 10.5603762 | -0.406002326 | 0.33503847 | -1.2118081 | 0.22558586 | 0.35363121 | NA | NA |
| ENSG000002 | 2.26653444 | -0.446738178 | 0.36867305 | -1.2117462 | 0.22560954 | 0.35364099 | NA | NA |
| ENSG000002 | 3.1025476 | -0.457426637 | 0.37748804 | -1.2117646 | 0.2256025 | 0.35364099 | NA | NA |
| ENSG000002 | 31.8197939 | -0.298393728 | 0.24626563 | -1.2116742 | 0.2256371 | 0.35367052 | LINC00504 | 201853 |
| ENSG000002 | 3.36590493 | -0.458496879 | 0.37843182 | -1.2115706 | 0.22567679 | 0.35371905 | RN7SKP265 | 106480914 |
| ENSG000002 | 9.51085104 | -0.414274488 | 0.34195254 | -1.211497 | 0.22570497 | 0.35374956 | OXCT1-AS1 | 100874002 |
| ENSG000002 | 4.00455024 | -0.458418607 | 0.37846289 | -1.2112643 | 0.22579411 | 0.35386191 | NA | NA |
| ENSG000002 | 6.99833273 | -0.439535967 | 0.36286828 | -1.2112824 | 0.22578718 | 0.35386191 | NA | NA |
| ENSG000001 | 25.0204605 | -0.314784299 | 0.25991036 | -1.2111264 | 0.22584696 | 0.35393106 | TMOD1 | 7111 |
| ENSG000001 | 5.34806472 | -0.452986654 | 0.37404533 | -1.2110475 | 0.22587721 | 0.35396479 | SSBP3-AS1 | 619518 |
| ENSG000002 | 4.24460345 | -0.457745847 | 0.37799042 | -1.2109985 | 0.22589596 | 0.35398049 | NA | NA |
| ENSG000001 | 167.397543 | 0.183441512 | 0.1514883 | 1.21092861 | 0.22592277 | 0.35400882 | INVS | 27130 |
| ENSG000001 | 20.1720708 | 0.352089896 | 0.29076881 | 1.21089292 | 0.22593645 | 0.35401657 | RSPO4 | 343637 |
| ENSG000001 | 49.0366838 | -0.263131091 | 0.21734642 | -1.210653 | 0.22602843 | 0.35414702 | ATP6V0A4 | 50617 |
| ENSG000001 | 2.90672631 | -0.453899035 | 0.37498142 | -1.2104574 | 0.22610343 | 0.35423715 | NA | NA |
| ENSG000002 | 3.70833972 | 0.457987475 | 0.37835547 | 1.21046876 | 0.22609907 | 0.35423715 | RNU6-125P | 106479621 |
| ENSG000002 | 6.78158116 | -0.44477783 | 0.36747247 | -1.2103705 | 0.22613676 | 0.35427568 | NA | NA |
| ENSG000002 | 2.29124132 | -0.445039986 | 0.36771627 | -1.2102809 | 0.22617112 | 0.35430214 | NA | NA |
| ENSG000002 | 5.06121446 | -0.454645571 | 0.37564657 | -1.2103014 | 0.22616326 | 0.35430214 | NA | NA |
| ENSG000002 | 2.89648216 | -0.45264964 | 0.37402603 | -1.2102089 | 0.22619873 | 0.3543317 | NA | NA |
| ENSG000002 | 10.0257096 | 0.410105059 | 0.33890021 | 1.21010565 | 0.22623835 | 0.35438008 | NA | NA |
| ENSG000002 | 9.99624795 | 0.414444232 | 0.34252885 | 1.20995422 | 0.22629646 | 0.3544574 | NA | NA |
| ENSG000002 | 6.0515869 | -0.442301653 | 0.36556124 | -1.2099249 | 0.2263077 | 0.35446132 | NA | NA |
| ENSG000001 | 4.50370899 | -0.455386414 | 0.37642464 | -1.2097678 | 0.226368 | 0.35452793 | TAS2R7 | 50837 |
| ENSG000002 | 61.2909024 | 0.242056976 | 0.2000886 | 1.20974894 | 0.22637525 | 0.35452793 | LOC644656 | 644656 |
| ENSG000002 | 5.43838126 | -0.45434494 | 0.37557059 | -1.2097458 | 0.22637646 | 0.35452793 | NA | NA |
| ENSG000002 | 2.88968285 | -0.451055982 | 0.37288303 | -1.2096447 | 0.22641526 | 0.354575 | NA | NA |
| ENSG000001 | 77.7896605 | 0.261181577 | 0.21592522 | 1.20959273 | 0.22643521 | 0.35459256 | RHBDF2 | 79651 |

| | | | | | | | | |
|---|---|---|---|---|---|---|---|---|
| ENSG000002 | 6.74839184 | -0.446607236 | 0.36925528 | -1.209481 | 0.22647812 | 0.35463256 | LINC00382 | 101927195 |
| ENSG000002 | 2.23118197 | -0.447132976 | 0.36969007 | -1.2094806 | 0.22647825 | 0.35463256 | NA | NA |
| ENSG000002 | 2.28230294 | -0.444153508 | 0.3672394 | -1.2094386 | 0.22649438 | 0.35464412 | NA | NA |
| ENSG000002 | 2.27352929 | -0.44636856 | 0.36909802 | -1.2093496 | 0.22652855 | 0.35468393 | NA | NA |
| ENSG000000 | 376.222409 | 0.156440845 | 0.12936723 | 1.20927719 | 0.22655637 | 0.3547138 | MYLK | 4638 |
| ENSG000002 | 9.23800684 | -0.418119971 | 0.34579684 | -1.2091492 | 0.22660554 | 0.35477708 | LOC101928834 | 101928834 |
| ENSG000002 | 10.5527294 | -0.405464054 | 0.33533717 | -1.2091235 | 0.2266154 | 0.35477883 | SAMD12-AS1 | 552860 |
| ENSG000001 | 146.703513 | 0.194147068 | 0.16057148 | 1.20910055 | 0.22662422 | 0.35477893 | MAP2K5 | 5607 |
| ENSG000001 | 48.4600864 | 0.252545298 | 0.20889346 | 1.20896698 | 0.22667553 | 0.35479078 | MATN3 | 4148 |
| ENSG000001 | 20.1050598 | 0.340287341 | 0.28145649 | 1.20902289 | 0.22665405 | 0.35479078 | FAM86JP | 100125556 |
| ENSG000001 | 6.55248031 | -0.436391911 | 0.36095913 | -1.2089787 | 0.22667102 | 0.35479078 | HIGD2B | 123346 |
| ENSG000002 | 2.29266367 | -0.444577904 | 0.36772218 | -1.2090049 | 0.22666098 | 0.35479078 | NA | NA |
| ENSG000002 | 2.24919401 | -0.445270412 | 0.36828051 | -1.2090523 | 0.22664274 | 0.35479078 | NA | NA |
| ENSG000001 | 318.604734 | 0.153038884 | 0.12658933 | 1.20893979 | 0.22668598 | 0.35479343 | MCM9 | 254394 |
| ENSG000001 | 55.6805649 | 0.243776765 | 0.20165096 | 1.20890457 | 0.22669951 | 0.35479955 | HTR2C | 3358 |
| ENSG000002 | 5.2558815 | -0.451280359 | 0.37330325 | -1.2088841 | 0.22670738 | 0.35479955 | NA | NA |
| ENSG000001 | 38.9535819 | -0.275353353 | 0.22779686 | -1.2087671 | 0.22675234 | 0.35481606 | PLLP | 51090 |
| ENSG000002 | 6.35911884 | -0.440079702 | 0.36407366 | -1.2087655 | 0.22675293 | 0.35481606 | NA | NA |
| ENSG000002 | 3.3789332 | -0.457342553 | 0.3783523 | -1.2087743 | 0.22674956 | 0.35481606 | EVX1-AS | 101410536 |
| ENSG000002 | 3.10570441 | -0.455926422 | 0.37716798 | -1.2088153 | 0.22673382 | 0.35481606 | NA | NA |
| ENSG000002 | 5.81005186 | -0.445221608 | 0.36833936 | -1.2087267 | 0.22676788 | 0.35482575 | NA | NA |
| ENSG000001 | 57.9620215 | 0.241713796 | 0.19998171 | 1.20867952 | 0.22678599 | 0.35482744 | GDPD1 | 284161 |
| ENSG000001 | 1365.00206 | 0.362819464 | 0.30017868 | 1.20867831 | 0.22678646 | 0.35482744 | SAFB | 6294 |
| ENSG000001 | 34.1173838 | -0.292620879 | 0.24211148 | -1.2086204 | 0.22680872 | 0.35483489 | OAZ3 | 51686 |
| ENSG000002 | 10.8462828 | -0.401289512 | 0.33201748 | -1.2086397 | 0.22680129 | 0.35483489 | NA | NA |
| ENSG000001 | 2189.82691 | 0.401804235 | 0.33248997 | 1.20847024 | 0.22686644 | 0.35491149 | EIF5B | 9669 |
| ENSG000002 | 2.26570027 | -0.445292827 | 0.36850296 | -1.2083833 | 0.22689986 | 0.35494201 | MIR4280 | 100422887 |
| ENSG000002 | 2.83170671 | -0.453247644 | 0.37508888 | -1.208374 | 0.22690345 | 0.35494201 | NA | NA |
| ENSG000002 | 3.93235573 | -0.457274156 | 0.37843673 | -1.2083239 | 0.22692268 | 0.35495841 | NA | NA |

| | | | | | | | | |
|---|---|---|---|---|---|---|---|---|
| ENSG000001 | 1016.99379 | 0.153645791 | 0.12718013 | 1.20809586 | 0.22701039 | 0.3550819 | FBXO22 | 26263 |
| ENSG000002 | 10.0551283 | -0.412117337 | 0.34114614 | -1.2080375 | 0.22703283 | 0.3551033 | SPATA31A1 | 647060 |
| ENSG000002 | 4.00856174 | 0.457262579 | 0.37853932 | 1.20796585 | 0.2270604 | 0.35513272 | MIR3657 | 100500889 |
| ENSG000001 | 2.57415358 | -0.44524072 | 0.36860587 | -1.2079046 | 0.22708396 | 0.35515589 | TGIF2LX | 90316 |
| ENSG000002 | 2.29385789 | -0.444907843 | 0.36835476 | -1.2078243 | 0.22711484 | 0.35517679 | NA | NA |
| ENSG000002 | 2.61741526 | -0.447082463 | 0.37015338 | -1.2078303 | 0.22711255 | 0.35517679 | NA | NA |
| ENSG000001 | 60.1361957 | 0.2547619 | 0.21096291 | 1.20761462 | 0.22719553 | 0.35528927 | MAFF | 23764 |
| ENSG000002 | 3.15605702 | -0.455201828 | 0.37698904 | -1.207467 | 0.22725234 | 0.3553644 | NA | NA |
| ENSG000000 | 66.1899398 | -0.22963887 | 0.1902233 | -1.2072068 | 0.2273525 | 0.35550732 | DAPK2 | 23604 |
| ENSG000002 | 2.84322089 | -0.452445796 | 0.37479719 | -1.207175 | 0.22736475 | 0.35551277 | NA | NA |
| ENSG000002 | 2.28323658 | -0.444958039 | 0.36860355 | -1.2071453 | 0.2273762 | 0.35551695 | NA | NA |
| ENSG000001 | 133.078623 | 0.187230271 | 0.1551091 | 1.20708757 | 0.22739842 | 0.35553799 | HS3ST3B1 | 9953 |
| ENSG000002 | 22.7104037 | -0.347287367 | 0.28774224 | -1.2069391 | 0.22745559 | 0.35559995 | C18orf63 | 644041 |
| ENSG000002 | 2.3047266 | -0.44387991 | 0.36776658 | -1.206961 | 0.22744718 | 0.35559995 | NA | NA |
| ENSG000002 | 1306.4206 | 0.130685709 | 0.10828208 | 1.20690064 | 0.22747041 | 0.35560942 | ZNF616 | 90317 |
| ENSG000002 | 13.8665408 | -0.388034646 | 0.32155891 | -1.2067296 | 0.2275363 | 0.35569871 | NA | NA |
| ENSG000001 | 48.443793 | -0.265865402 | 0.22034087 | -1.2066096 | 0.22758254 | 0.35575728 | CFAP46 | 54777 |
| ENSG000002 | 15.7157033 | 0.366059178 | 0.30342193 | 1.20643612 | 0.22764938 | 0.35582061 | TMEM110 | 375346 |
| ENSG000002 | 7.40022447 | -0.431603965 | 0.35774234 | -1.206466 | 0.22763787 | 0.35582061 | NA | NA |
| ENSG000002 | 6.85601576 | -0.446332987 | 0.36995923 | -1.2064383 | 0.22764853 | 0.35582061 | NA | NA |
| ENSG000002 | 3.08588489 | -0.455414015 | 0.37751931 | -1.206333 | 0.22768911 | 0.35586899 | NA | NA |
| ENSG000002 | 45.6272204 | 0.27098683 | 0.22465443 | 1.20623853 | 0.22772553 | 0.3559122 | FAM72D | 728833 |
| ENSG000002 | 9.04145413 | -0.428325056 | 0.35510089 | -1.2062066 | 0.22773782 | 0.35591769 | LOC101927120 | 101927120 |
| ENSG000002 | 8.30185539 | -0.429279894 | 0.35591842 | -1.2061188 | 0.22777169 | 0.35595691 | PCDHA8 | 56140 |
| ENSG000002 | 5.38615707 | -0.452445597 | 0.37518337 | -1.2059319 | 0.22784375 | 0.3560558 | NA | NA |
| ENSG000002 | 4.74726741 | -0.451849285 | 0.37470086 | -1.2058934 | 0.2278586 | 0.35606529 | NA | NA |
| ENSG000002 | 6.30415427 | -0.438869673 | 0.36394507 | -1.2058679 | 0.22786844 | 0.35606694 | NA | NA |
| ENSG000002 | 40.9220843 | -0.274237887 | 0.22742674 | -1.2058295 | 0.22788322 | 0.35607631 | NA | NA |
| ENSG000001 | 68.2689628 | 0.222824345 | 0.18480778 | 1.20570866 | 0.22792984 | 0.35611978 | ENDOV | 284131 |

| | | | | | | | | |
|---|---|---|---|---|---|---|---|---|
| ENSG000001 | 3.34699922 | -0.456419502 | 0.37856203 | -1.2056663 | 0.22794617 | 0.35611978 | NA | NA |
| ENSG000002 | 6.98294716 | -0.440624856 | 0.36545176 | -1.2056991 | 0.22793353 | 0.35611978 | NA | NA |
| ENSG000002 | 8.71906262 | -0.419677516 | 0.3480834 | -1.2056809 | 0.22794054 | 0.35611978 | LOC101928445 | 101928445 |
| ENSG000002 | 4.53815616 | -0.454476032 | 0.37697509 | -1.2055864 | 0.22797701 | 0.35613833 | NA | NA |
| ENSG000002 | 7.9503132 | -0.429624582 | 0.35636031 | -1.2055904 | 0.22797545 | 0.35613833 | NA | NA |
| ENSG000002 | 5.31134216 | -0.449579524 | 0.37291948 | -1.2055673 | 0.22798439 | 0.35613833 | NA | NA |
| ENSG000002 | 3.12294788 | -0.454676833 | 0.37716924 | -1.2054982 | 0.22801104 | 0.35616625 | NA | NA |
| ENSG000002 | 7.53000126 | -0.43857481 | 0.3638349 | -1.2054226 | 0.2280402 | 0.35619807 | NA | NA |
| ENSG000001 | 262.588518 | 0.162998807 | 0.13523623 | 1.20528948 | 0.22809157 | 0.356252 | FBXL20 | 84961 |
| ENSG000001 | 47.3299023 | 0.260876812 | 0.21644362 | 1.20528759 | 0.2280923 | 0.356252 | CSPG4 | 1464 |
| ENSG000001 | 35.9232635 | 0.282443278 | 0.23435341 | 1.20520236 | 0.22812519 | 0.35628966 | DOK3 | 79930 |
| ENSG000002 | 2.23098609 | -0.445210611 | 0.36942471 | -1.2051457 | 0.22814705 | 0.35629634 | RN7SKP210 | 106480898 |
| ENSG000002 | 7.66023574 | -0.428810724 | 0.35581134 | -1.2051632 | 0.22814032 | 0.35629634 | NA | NA |
| ENSG000002 | 4.25577992 | -0.455409611 | 0.37790628 | -1.2050861 | 0.22817006 | 0.35631856 | NA | NA |
| ENSG000002 | 4.05470513 | -0.456034018 | 0.37845485 | -1.2049892 | 0.22820746 | 0.35636325 | NA | NA |
| ENSG000002 | 8.39339871 | -0.435655965 | 0.36156816 | -1.2049069 | 0.22823926 | 0.35639917 | NA | NA |
| ENSG000002 | 2.23892612 | -0.444712053 | 0.36910123 | -1.2048512 | 0.22826076 | 0.35641902 | LOC105374582 | 105374582 |
| ENSG000002 | 10.0247942 | -0.406616927 | 0.33749181 | -1.2048201 | 0.22827275 | 0.35642355 | NA | NA |
| ENSG000002 | 7.18528015 | -0.436302491 | 0.36213742 | -1.2047981 | 0.22828124 | 0.35642355 | NA | NA |
| ENSG000002 | 9.7983624 | -0.411484105 | 0.34159901 | -1.2045823 | 0.22836461 | 0.35653999 | NA | NA |
| ENSG000002 | 2.82282095 | -0.452250999 | 0.37547443 | -1.2044788 | 0.22840457 | 0.35658865 | NA | NA |
| ENSG000001 | 4.49261215 | -0.453089155 | 0.37621227 | -1.2043444 | 0.22845648 | 0.35665596 | OR5H2 | 79310 |
| ENSG000002 | 10.6922111 | -0.413894904 | 0.343719 | -1.2041665 | 0.22852523 | 0.35673582 | PCDHA2 | 56146 |
| ENSG000002 | 2.23966333 | -0.444470769 | 0.36910432 | -1.2041874 | 0.22851716 | 0.35673582 | NA | NA |
| ENSG000002 | 3.35747618 | -0.455760956 | 0.37852509 | -1.2040442 | 0.22857248 | 0.35679585 | NA | NA |
| ENSG000002 | 7.49436527 | -0.434797524 | 0.36112215 | -1.2040179 | 0.22858266 | 0.35679801 | LOC101929315 | 101929315 |
| ENSG000002 | 2.92972137 | -0.447490778 | 0.37168063 | -1.2039658 | 0.22860279 | 0.35681569 | NA | NA |
| ENSG000001 | 31.763082 | 0.30232635 | 0.25112263 | 1.20389925 | 0.22862852 | 0.35683669 | CAPN11 | 11131 |
| ENSG000002 | 13.0143785 | -0.387001642 | 0.32146051 | -1.2038855 | 0.22863384 | 0.35683669 | PCAT1 | 100750225 |

| | | | | | | | | |
|---|---|---|---|---|---|---|---|---|
| ENSG000001 | 27.7073886 | 0.30538439 | 0.2536842 | 1.20379742 | 0.22866789 | 0.35687609 | ARSI | 340075 |
| ENSG000001 | 9.36289958 | -0.423884972 | 0.35215494 | -1.2036889 | 0.22870986 | 0.35690039 | PHOX2B | 8929 |
| ENSG000002 | 5.75083096 | -0.452759154 | 0.37613559 | -1.2037126 | 0.22870067 | 0.35690039 | NA | NA |
| ENSG000002 | 11.007892 | -0.415081933 | 0.34483867 | -1.2036989 | 0.22870597 | 0.35690039 | LOC101929413 | 101929413 |
| ENSG000002 | 9.79521341 | -0.416104987 | 0.34570277 | -1.2036495 | 0.22872506 | 0.35691037 | NA | NA |
| ENSG000001 | 4.73257525 | -0.451132321 | 0.37486156 | -1.2034638 | 0.22879689 | 0.35700872 | NKX2-3 | 159296 |
| ENSG000002 | 4.74660188 | -0.450963267 | 0.37475651 | -1.2033501 | 0.22884088 | 0.35706363 | SLC16A12-AS1 | 101926906 |
| ENSG000001 | 38.289717 | -0.305677743 | 0.2540603 | -1.2031701 | 0.22891052 | 0.35714113 | SCN1B | 6324 |
| ENSG000001 | 17.8345555 | -0.362423098 | 0.30121954 | -1.2031859 | 0.2289044 | 0.35714113 | TAGAP | 117289 |
| ENSG000002 | 5.87462744 | -0.448309942 | 0.37261247 | -1.2031534 | 0.22891698 | 0.35714113 | NA | NA |
| ENSG000001 | 1879.89894 | 0.132965848 | 0.11053046 | 1.20297918 | 0.22898438 | 0.35723255 | XPR1 | 9213 |
| ENSG000002 | 14.8292982 | 0.370057249 | 0.3076333 | 1.20291674 | 0.22900854 | 0.3572565 | NA | NA |
| ENSG000002 | 2.31027132 | -0.440846582 | 0.36653483 | -1.2027413 | 0.22907644 | 0.35733492 | NA | NA |
| ENSG000002 | 7.10320174 | -0.434368914 | 0.3611424 | -1.2027635 | 0.22906783 | 0.35733492 | LOC102725121 | 102725121 |
| ENSG000001 | 17.483685 | -0.360896692 | 0.3000733 | -1.2026951 | 0.22909433 | 0.35734908 | NXPE4 | 54827 |
| ENSG000002 | 16.2692015 | -0.360804098 | 0.30004692 | -1.2024923 | 0.22917286 | 0.35745783 | PTENP1-AS | 101243555 |
| ENSG000002 | 3.73903314 | -0.454908104 | 0.37837712 | -1.2022611 | 0.22926237 | 0.35758369 | NA | NA |
| ENSG000002 | 3.16360163 | -0.45113579 | 0.37525588 | -1.2022085 | 0.22928277 | 0.35760176 | NA | NA |
| ENSG000002 | 2.2589024 | -0.443585443 | 0.36904941 | -1.2019676 | 0.22937608 | 0.35773352 | PAGE2B | 389860 |
| ENSG000002 | 3.12245861 | -0.453181547 | 0.3770502 | -1.201913 | 0.22939723 | 0.35774712 | NA | NA |
| ENSG000002 | 7.49069397 | -0.436940516 | 0.36354162 | -1.2018996 | 0.22940244 | 0.35774712 | NA | NA |
| ENSG000002 | 4.33348769 | -0.454748087 | 0.37838087 | -1.2018263 | 0.22943082 | 0.35777762 | CYP4F29P | 54055 |
| ENSG000002 | 25.4065345 | 0.308680917 | 0.25685763 | 1.20175882 | 0.22945698 | 0.35780466 | NA | NA |
| ENSG000002 | 10.583615 | -0.40250057 | 0.33494317 | -1.2016981 | 0.22948052 | 0.35782759 | LOC101927274 | 101927274 |
| ENSG000002 | 5.84736245 | -0.4439301 | 0.36944239 | -1.201622 | 0.22951002 | 0.35784893 | LINC01527 | 101927988 |
| ENSG000002 | 6.19181304 | -0.44419497 | 0.36967055 | -1.2015969 | 0.22951976 | 0.35784893 | LOC105375380 | 105375380 |
| ENSG000002 | 5.01644644 | -0.451730198 | 0.37594863 | -1.2015743 | 0.22952851 | 0.35784893 | NA | NA |
| ENSG000002 | 18.8461717 | -0.345390713 | 0.2874491 | -1.2015717 | 0.2295295 | 0.35784893 | NA | NA |
| ENSG000002 | 2.29954641 | -0.440910521 | 0.36698447 | -1.2014419 | 0.22957983 | 0.35791363 | NA | NA |

| | | | | | | | | |
|---|---|---|---|---|---|---|---|---|
| ENSG000001 | 68.8970462 | 0.230226277 | 0.19168146 | 1.20108785 | 0.22971713 | 0.35811391 | PARP16 | 54956 |
| ENSG000001 | 30.5717295 | 0.311090129 | 0.25904632 | 1.2009054 | 0.2297879 | 0.35816917 | COL8A2 | 1296 |
| ENSG000002 | 55.3686261 | 0.262111795 | 0.21825082 | 1.20096593 | 0.22976442 | 0.35816917 | NA | NA |
| ENSG000002 | 2.26476664 | -0.439965508 | 0.3663603 | -1.2009093 | 0.22978637 | 0.35816917 | RN7SL339P | 106479348 |
| ENSG000002 | 2.88981456 | -0.449062442 | 0.37392284 | -1.2009495 | 0.2297708 | 0.35816917 | NA | NA |
| ENSG000002 | 9.69948038 | -0.404168083 | 0.33660171 | -1.2007309 | 0.2298556 | 0.35826092 | NA | NA |
| ENSG000002 | 5.49883854 | -0.446612464 | 0.37200752 | -1.2005469 | 0.22992702 | 0.35834487 | NA | NA |
| ENSG000002 | 11.6852516 | -0.385818942 | 0.3213694 | -1.2005466 | 0.22992713 | 0.35834487 | NA | NA |
| ENSG00000C | 31.4169316 | -0.290479599 | 0.24196995 | -1.200478 | 0.22995376 | 0.35834505 | RAB27B | 5874 |
| ENSG000002 | 2.29148981 | -0.44236444 | 0.36848016 | -1.2005109 | 0.22994099 | 0.35834505 | NA | NA |
| ENSG000002 | 4.75225151 | -0.450156697 | 0.37497544 | -1.2004965 | 0.22994658 | 0.35834505 | NA | NA |
| ENSG000002 | 2.28295583 | -0.440846765 | 0.36724606 | -1.2004125 | 0.22997917 | 0.35836194 | KRTAP4-12 | 83755 |
| ENSG000002 | 14.0563596 | -0.372050447 | 0.30993756 | -1.2004045 | 0.22998227 | 0.35836194 | NA | NA |
| ENSG000002 | 4.69683515 | -0.450261603 | 0.37509878 | -1.2003814 | 0.22999125 | 0.35836216 | NA | NA |
| ENSG000001 | 13.785388 | -0.374722506 | 0.31218068 | -1.2003386 | 0.23000788 | 0.35837123 | PRSS48 | 345062 |
| ENSG000002 | 2.89487056 | -0.448243397 | 0.37343631 | -1.2003209 | 0.23001475 | 0.35837123 | NA | NA |
| ENSG000001 | 2.28305997 | -0.440776766 | 0.36724601 | -1.2002221 | 0.2300531 | 0.35841722 | CCT8L2 | 150160 |
| ENSG000002 | 3.33402247 | -0.454175651 | 0.37845623 | -1.2000744 | 0.23011043 | 0.35848251 | NA | NA |
| ENSG000002 | 3.50899247 | -0.452581831 | 0.37712996 | -1.2000686 | 0.23011269 | 0.35848251 | NA | NA |
| ENSG000001 | 24.7579889 | -0.312854726 | 0.26071792 | -1.199974 | 0.23014944 | 0.35852599 | CMPK2 | 129607 |
| ENSG000002 | 4.73999933 | -0.450056977 | 0.37507314 | -1.1999179 | 0.23017122 | 0.35854614 | FOXD1 | 2297 |
| ENSG000001 | 418.813413 | 0.15651761 | 0.13044677 | 1.19985806 | 0.23019447 | 0.35855481 | BLZF1 | 8548 |
| ENSG000002 | 3.63392182 | -0.454143098 | 0.37849199 | -1.199875 | 0.23018787 | 0.35855481 | LOC102724593 | 102724593 |
| ENSG000002 | 3.99425512 | -0.45417384 | 0.3785364 | -1.1998155 | 0.23021101 | 0.3585668 | NA | NA |
| ENSG000001 | 9.95334423 | 0.411700751 | 0.34315177 | 1.19976286 | 0.23023145 | 0.35858487 | NA | NA |
| ENSG000001 | 218.306707 | 0.172247366 | 0.14358092 | 1.19965361 | 0.2302739 | 0.35862342 | RASSF2 | 9770 |
| ENSG000002 | 9.4079933 | 0.415343237 | 0.34621509 | 1.1996682 | 0.23026823 | 0.35862342 | RN7SKP230 | 106479197 |
| ENSG000002 | 8.42794629 | -0.416637274 | 0.34732538 | -1.199559 | 0.23031067 | 0.35866692 | NA | NA |
| ENSG000002 | 2.24278789 | -0.442111485 | 0.36859088 | -1.199464 | 0.23034758 | 0.35871062 | LOC389033 | 389033 |

| | | | | | | | | |
|---|---|---|---|---|---|---|---|---|
| ENSG000001 | 67.0278814 | 0.261581943 | 0.21808718 | 1.19943751 | 0.23035787 | 0.35871287 | RGPD8 | 727851 |
| ENSG000002 | 5.85308398 | -0.44300691 | 0.36937574 | -1.1993395 | 0.23039597 | 0.35875842 | DNAJB8-AS1 | 285224 |
| ENSG000001 | 1501.21156 | 0.174282879 | 0.14532133 | 1.19929318 | 0.23041396 | 0.35877267 | CDC73 | 79577 |
| ENSG000002 | 3.17408977 | -0.451901596 | 0.37686284 | -1.1991142 | 0.23048353 | 0.35886721 | RN7SKP287 | 106480921 |
| ENSG000001 | 114.523931 | 0.193290089 | 0.1612032 | 1.19904623 | 0.23050997 | 0.35888082 | NA | NA |
| ENSG000002 | 2.30188224 | -0.439741972 | 0.36674155 | -1.1990514 | 0.23050795 | 0.35888082 | NA | NA |
| ENSG000001 | 289.67875 | 0.162032755 | 0.13514708 | 1.19893639 | 0.23055268 | 0.35893353 | SLC12A6 | 9990 |
| ENSG000001 | 202.268305 | 0.16202148 | 0.13516741 | 1.19867264 | 0.23065526 | 0.35907945 | ZBTB8A | 653121 |
| ENSG00000C | 78.9496564 | 0.232064977 | 0.19362599 | 1.19852182 | 0.23071393 | 0.35912943 | SLC9A1 | 6548 |
| ENSG000002 | 8.60490568 | -0.415373962 | 0.34657078 | -1.1985256 | 0.23071246 | 0.35912943 | NA | NA |
| ENSG000002 | 6.64455437 | -0.435080702 | 0.36300797 | -1.1985431 | 0.23070565 | 0.35912943 | NA | NA |
| ENSG000001 | 104.3493 | -0.199536775 | 0.16649553 | -1.1984512 | 0.23074139 | 0.3591471 | CFAP69 | 79846 |
| ENSG000001 | 46.0698669 | 0.283812054 | 0.2368165 | 1.19844712 | 0.230743 | 0.3591471 | C1orf159 | 54991 |
| ENSG000001 | 64.1965637 | 0.247727555 | 0.20673586 | 1.19828051 | 0.23080783 | 0.35923422 | BSPRY | 54836 |
| ENSG000002 | 8.96080215 | -0.412158149 | 0.34399138 | -1.1981642 | 0.23085311 | 0.35927954 | NA | NA |
| ENSG000002 | 4.2393458 | -0.453250777 | 0.37828897 | -1.1981602 | 0.23085467 | 0.35927954 | NA | NA |
| ENSG000002 | 4.04552707 | -0.453591304 | 0.3785903 | -1.198106 | 0.23087576 | 0.35929858 | NA | NA |
| ENSG000002 | 21.6571087 | -0.327482685 | 0.27336581 | -1.197965 | 0.23093063 | 0.35934259 | SEPT14P12 | 106480343 |
| ENSG000002 | 2.24794882 | -0.441389449 | 0.36844448 | -1.1979809 | 0.23092446 | 0.35934259 | NA | NA |
| ENSG000002 | 7.16702699 | -0.431863876 | 0.36049442 | -1.1979766 | 0.23092613 | 0.35934259 | LOC105370681 | 105370681 |
| ENSG000002 | 3.52614356 | -0.450288067 | 0.37590096 | -1.1978902 | 0.23095978 | 0.35936799 | NA | NA |
| ENSG000002 | 3.35761999 | -0.453365784 | 0.37848141 | -1.1978548 | 0.23097354 | 0.35936799 | NA | NA |
| ENSG000002 | 7.64084679 | -0.426733279 | 0.3562477 | -1.1978555 | 0.23097326 | 0.35936799 | NA | NA |
| ENSG000002 | 12.7137571 | -0.377347416 | 0.31503705 | -1.1977874 | 0.23099979 | 0.35939504 | NA | NA |
| ENSG000002 | 3.15733501 | -0.45114109 | 0.37665748 | -1.1977489 | 0.23101478 | 0.35940458 | UBE2NL | 389898 |
| ENSG000001 | 206.930267 | -0.177081819 | 0.14785655 | -1.1976596 | 0.23104955 | 0.35944489 | BCL11B | 64919 |
| ENSG000002 | 9.8590061 | -0.417596191 | 0.34874225 | -1.197435 | 0.23113704 | 0.35956719 | SMCO1 | 255798 |
| ENSG000002 | 3.95525547 | -0.453202941 | 0.37852095 | -1.1972995 | 0.23118984 | 0.35958983 | NA | NA |
| ENSG000002 | 5.67289077 | -0.449235671 | 0.37519783 | -1.1973301 | 0.2311779 | 0.35958983 | NA | NA |

| | | | | | | | | |
|---|---|---|---|---|---|---|---|---|
| ENSG000002 | 7.47406947 | -0.4316471 | 0.36051314 | -1.1973131 | 0.23118454 | 0.35958983 | NA | NA |
| ENSG000002 | 23.5481843 | 0.32148167 | 0.26850914 | 1.19728388 | 0.23119593 | 0.35958983 | NA | NA |
| ENSG000002 | 28.5741789 | -0.301402715 | 0.25173052 | -1.1973229 | 0.23118073 | 0.35958983 | NA | NA |
| ENSG000001 | 2.25130151 | -0.440064492 | 0.367568 | -1.1972329 | 0.2312158 | 0.35960694 | OR51V1 | 283111 |
| ENSG000001 | 6.04550608 | -0.439187936 | 0.36685262 | -1.1971782 | 0.23123712 | 0.35962632 | C8orf22 | 492307 |
| ENSG000002 | 8.87092697 | -0.409986083 | 0.34247759 | -1.197118 | 0.23126057 | 0.35964899 | NA | NA |
| ENSG00000C | 56.0639348 | 0.257789858 | 0.21537847 | 1.19691566 | 0.23133943 | 0.35971964 | ITGA2B | 3674 |
| ENSG000001 | 18.0638064 | -0.358669049 | 0.2996499 | -1.1969603 | 0.23132201 | 0.35971964 | ADAP1 | 11033 |
| ENSG000001 | 286.684377 | 0.163873085 | 0.13691337 | 1.19691079 | 0.23134133 | 0.35971964 | BTBD11 | 121551 |
| ENSG000001 | 278.456947 | 0.150170348 | 0.12546499 | 1.19691039 | 0.23134148 | 0.35971964 | ZNF585A | 199704 |
| ENSG00000C | 2.26186279 | -0.43838388 | 0.36634717 | -1.1966351 | 0.23144882 | 0.35987274 | PGLYRP1 | 8993 |
| ENSG000001 | 12.1927555 | -0.380542017 | 0.31805534 | -1.1964648 | 0.23151523 | 0.35994624 | RERGL | 79785 |
| ENSG000001 | 11.0381122 | -0.404169195 | 0.33781468 | -1.1964228 | 0.23153159 | 0.35994624 | CGA | 1081 |
| ENSG000002 | 4.1784892 | -0.452343434 | 0.378074 | -1.1964415 | 0.2315243 | 0.35994624 | NA | NA |
| ENSG000002 | 3.34039668 | -0.452837637 | 0.3784822 | -1.1964569 | 0.23151831 | 0.35994624 | NA | NA |
| ENSG000002 | 4.69048907 | -0.448044357 | 0.37452044 | -1.1963148 | 0.23157372 | 0.35995725 | NA | NA |
| ENSG000002 | 3.77566471 | -0.452601959 | 0.37833757 | -1.1962913 | 0.23158289 | 0.35995725 | NA | NA |
| ENSG000002 | 7.3691313 | -0.427867406 | 0.35764475 | -1.1963475 | 0.23156097 | 0.35995725 | NA | NA |
| ENSG000002 | 4.54982658 | -0.451438624 | 0.37736526 | -1.1962909 | 0.23158306 | 0.35995725 | RPL34P1 | 26514 |
| ENSG000002 | 3.97847288 | -0.452906114 | 0.37858397 | -1.1963162 | 0.23157321 | 0.35995725 | NA | NA |
| ENSG000001 | 12.0209546 | -0.386060732 | 0.32272437 | -1.1962553 | 0.23159695 | 0.35996505 | SPNS2 | 124976 |
| ENSG000002 | 2.24020411 | -0.440614359 | 0.36841046 | -1.1959877 | 0.23170138 | 0.36011355 | MIR7851 | 102467003 |
| ENSG000001 | 191.57867 | 0.17152879 | 0.14343011 | 1.195905 | 0.23173363 | 0.36012227 | RALGPS1 | 9649 |
| ENSG000002 | 3.35946655 | -0.452597922 | 0.37845501 | -1.1959094 | 0.2317319 | 0.36012227 | NA | NA |
| ENSG000002 | 2.22270061 | -0.441342177 | 0.36904356 | -1.1959081 | 0.23173243 | 0.36012227 | NA | NA |
| ENSG000002 | 4.48737158 | -0.450994702 | 0.37713692 | -1.1958381 | 0.23175974 | 0.36014904 | NA | NA |
| ENSG000001 | 12.4393867 | -0.377339702 | 0.31555775 | -1.1957865 | 0.23177988 | 0.36015274 | COLEC10 | 10584 |
| ENSG000002 | 2.58451431 | -0.440327253 | 0.36823094 | -1.195791 | 0.23177811 | 0.36015274 | NA | NA |
| ENSG00000C | 16.4861802 | -0.351191715 | 0.29371595 | -1.1956849 | 0.23181955 | 0.36015922 | FGF10 | 2255 |

| | | | | | | | | |
|---|---|---|---|---|---|---|---|---|
| ENSG000001 | 15.2721058 | -0.362678115 | 0.30332767 | -1.1956645 | 0.23182752 | 0.36015922 | KCNJ14 | 3770 |
| ENSG000002 | 3.16803597 | -0.450563246 | 0.3768316 | -1.1956621 | 0.23182846 | 0.36015922 | NA | NA |
| ENSG000002 | 5.01403004 | -0.446500115 | 0.37341563 | -1.1957189 | 0.23180628 | 0.36015922 | NA | NA |
| ENSG000002 | 3.08542788 | -0.45082615 | 0.37704553 | -1.1956809 | 0.2318211 | 0.36015922 | NA | NA |
| ENSG000002 | 3.99475842 | -0.45256836 | 0.37852773 | -1.1956016 | 0.23185207 | 0.36017457 | NA | NA |
| ENSG000002 | 4.46922151 | -0.451589751 | 0.37771391 | -1.1955868 | 0.23185785 | 0.36017457 | NA | NA |
| ENSG000002 | 2.60522779 | -0.441861285 | 0.36958257 | -1.1955685 | 0.23186499 | 0.36017457 | NA | NA |
| ENSG000001 | 11.2882699 | -0.39625411 | 0.33145455 | -1.1955006 | 0.23189151 | 0.36020196 | PLXNB3 | 5365 |
| ENSG000002 | 6.06487631 | -0.439114296 | 0.36731462 | -1.1954719 | 0.23190272 | 0.36020558 | NA | NA |
| ENSG000002 | 2.27814092 | -0.438996619 | 0.36723131 | -1.1954226 | 0.23192194 | 0.36022165 | NA | NA |
| ENSG000002 | 5.35890417 | 0.447497972 | 0.37436599 | 1.19534891 | 0.23195074 | 0.36025257 | NA | NA |
| ENSG000002 | 3.38308891 | -0.452273896 | 0.37837546 | -1.1953045 | 0.23196808 | 0.3602657 | NA | NA |
| ENSG000002 | 16.3454582 | -0.366605251 | 0.30671157 | -1.1952769 | 0.23197886 | 0.36026866 | NA | NA |
| ENSG000002 | 2.26050102 | -0.440436283 | 0.36849378 | -1.195234 | 0.23199563 | 0.36028089 | NA | NA |
| ENSG000001 | 2.60892092 | -0.441393027 | 0.36934553 | -1.195068 | 0.23206046 | 0.36034018 | NA | NA |
| ENSG000002 | 10.7707745 | -0.39651814 | 0.33179114 | -1.1950836 | 0.23205438 | 0.36034018 | NA | NA |
| ENSG000002 | 4.79553192 | -0.448815682 | 0.3755491 | -1.1950919 | 0.23205113 | 0.36034018 | LOC101928008 | 101928008 |
| ENSG000002 | 2.64735942 | -0.440169968 | 0.36840621 | -1.1947952 | 0.23216706 | 0.3604919 | NA | NA |
| ENSG000001 | 4.2950018 | -0.452187875 | 0.37848487 | -1.1947317 | 0.23219187 | 0.36050282 | OR11H4 | 390442 |
| ENSG000001 | 3.87167424 | -0.450483946 | 0.3770545 | -1.1947449 | 0.23218671 | 0.36050282 | PRR30 | 339779 |
| ENSG000001 | 8.16536474 | -0.419964481 | 0.35159337 | -1.1944607 | 0.23229779 | 0.36065346 | AWAT2 | 158835 |
| ENSG000002 | 6.85401108 | -0.429460788 | 0.35955356 | -1.1944279 | 0.23231065 | 0.36065961 | NA | NA |
| ENSG000001 | 10.4496148 | 0.401255265 | 0.33600055 | 1.19421015 | 0.23239577 | 0.36077796 | RDH5 | 5959 |
| ENSG000001 | 50.0219302 | 0.260219426 | 0.21794341 | 1.19397702 | 0.23248696 | 0.36087807 | SIRPA | 140885 |
| ENSG000002 | 2.29701471 | -0.437590304 | 0.36648733 | -1.1940121 | 0.23247324 | 0.36087807 | RNA5SP140 | 106478996 |
| ENSG000002 | 25.4146604 | -0.314187766 | 0.26313916 | -1.1939985 | 0.23247855 | 0.36087807 | NA | NA |
| ENSG000002 | 4.739491 | -0.447470551 | 0.37483657 | -1.1937751 | 0.23256597 | 0.3609869 | NA | NA |
| ENSG000002 | 36.8064482 | -0.269976853 | 0.22620301 | -1.1935157 | 0.23266745 | 0.36108913 | LOC100130691 | 100130691 |
| ENSG000002 | 12.5873431 | -0.392535009 | 0.32887994 | -1.1935511 | 0.23265361 | 0.36108913 | NA | NA |

| | | | | | | | | |
|---|---|---|---|---|---|---|---|---|
| ENSG000002 | 9.22881449 | -0.413294267 | 0.3462771 | -1.1935362 | 0.23265943 | 0.36108913 | NA | NA |
| ENSG000002 | 11.5579975 | -0.390889565 | 0.32750853 | -1.1935249 | 0.23266389 | 0.36108913 | NA | NA |
| ENSG000002 | 10.0425429 | 0.411373673 | 0.344681 | 1.19349102 | 0.23267713 | 0.36109033 | NA | NA |
| ENSG000002 | 2.55753631 | -0.439410424 | 0.36818798 | -1.1934404 | 0.23269692 | 0.36110723 | NA | NA |
| ENSG000002 | 14.974237 | -0.362077384 | 0.30340773 | -1.193369 | 0.23272488 | 0.36113679 | GNRHR2 | 114814 |
| ENSG000002 | 3.37070033 | -0.451483267 | 0.37839767 | -1.1931449 | 0.23281261 | 0.36125911 | NA | NA |
| ENSG000002 | 8.35827552 | -0.415194003 | 0.34802783 | -1.1929908 | 0.23287297 | 0.36133893 | NA | NA |
| ENSG000002 | 2.25830899 | -0.439426042 | 0.3683625 | -1.1929174 | 0.23290172 | 0.36136972 | RN7SL300P | 106481004 |
| ENSG000002 | 2.27333341 | -0.438918372 | 0.36797045 | -1.1928088 | 0.23294428 | 0.36142192 | SMIM2-IT1 | 100874377 |
| ENSG000001 | 4.44784413 | -0.449604506 | 0.37695259 | -1.1927349 | 0.23297322 | 0.361453 | OR4C3 | 256144 |
| ENSG000002 | 3.0973357 | -0.450257379 | 0.37751954 | -1.1926731 | 0.23299744 | 0.36146348 | NA | NA |
| ENSG000002 | 3.2163324 | -0.448049834 | 0.37567503 | -1.1926527 | 0.23300542 | 0.36146348 | NA | NA |
| ENSG000002 | 6.83382628 | -0.428308758 | 0.35912378 | -1.1926494 | 0.23300672 | 0.36146348 | NA | NA |
| ENSG000001 | 126.772022 | 0.196821213 | 0.16504894 | 1.19250217 | 0.2330644 | 0.36149765 | CRISPLD2 | 83716 |
| ENSG000001 | 22.2253714 | -0.333389301 | 0.27956779 | -1.1925168 | 0.23305865 | 0.36149765 | KLHL34 | 257240 |
| ENSG000001 | 11.7119368 | -0.407415327 | 0.34164491 | -1.1925111 | 0.23306092 | 0.36149765 | INCA1 | 388324 |
| ENSG000002 | 2.30814401 | -0.435077536 | 0.3648234 | -1.1925703 | 0.23303772 | 0.36149765 | NA | NA |
| ENSG000001 | 11.2474947 | -0.395290974 | 0.33151986 | -1.1923599 | 0.23312017 | 0.36156185 | LMX1B | 4010 |
| ENSG000002 | 6.58677879 | -0.431351924 | 0.36176588 | -1.192351 | 0.23312363 | 0.36156185 | NA | NA |
| ENSG000001 | 26.8594872 | -0.312955349 | 0.26247546 | -1.1923223 | 0.23313488 | 0.36156548 | FKBP9P1 | 360132 |
| ENSG000002 | 12.0035087 | -0.385497681 | 0.32333786 | -1.1922442 | 0.23316552 | 0.36159917 | NA | NA |
| ENSG000002 | 20.1892003 | -0.33995548 | 0.28515047 | -1.1921968 | 0.23318408 | 0.36161412 | NA | NA |
| ENSG000002 | 8.38919243 | -0.41625075 | 0.34915736 | -1.192158 | 0.23319928 | 0.36162386 | HLA-DQB2 | 3120 |
| ENSG000002 | 3.70151369 | -0.451138372 | 0.37847838 | -1.1919792 | 0.23326939 | 0.36171875 | NA | NA |
| ENSG000002 | 2.9334803 | -0.443945437 | 0.37245786 | -1.1919347 | 0.23328686 | 0.361732 | NA | NA |
| ENSG000002 | 22.3957213 | -0.327601505 | 0.27487079 | -1.1918382 | 0.2333247 | 0.36177684 | SMG7-AS1 | 284649 |
| ENSG000001 | 2.50842898 | -0.444860451 | 0.37327906 | -1.1917637 | 0.2333539 | 0.36180828 | NA | NA |
| ENSG000002 | 4.20174291 | -0.450151186 | 0.37773038 | -1.1917262 | 0.23336863 | 0.36181729 | NA | NA |
| ENSG000001 | 5.54726778 | -0.443113552 | 0.37186046 | -1.1916124 | 0.23341325 | 0.36187263 | NA | NA |

| | | | | | | | | |
|---|---|---|---|---|---|---|---|---|
| ENSG000002 | 10.4737047 | -0.399604031 | 0.33537305 | -1.191521 | 0.23344911 | 0.3619144 | NA | NA |
| ENSG000002 | 2.52900608 | -0.445308163 | 0.3737665 | -1.1914074 | 0.2334937 | 0.36196969 | NA | NA |
| ENSG000001 | 20.129561 | 0.337598592 | 0.28339167 | 1.19127918 | 0.233544 | 0.36203163 | SDCBP2 | 27111 |
| ENSG000001 | 1121.74856 | 0.15557681 | 0.13060102 | 1.19123729 | 0.23356044 | 0.36203163 | PPP1R3B | 79660 |
| ENSG000002 | 2.51810457 | -0.444988411 | 0.37354528 | -1.1912569 | 0.23355273 | 0.36203163 | NA | NA |
| ENSG000002 | 4.73494388 | -0.446778814 | 0.37506276 | -1.1912108 | 0.23357084 | 0.36203391 | NA | NA |
| ENSG000001 | 10.6290275 | 0.41356197 | 0.34719309 | 1.19115842 | 0.2335914 | 0.36205194 | JMJD7-PLA2G4E | 8681 |
| ENSG000001 | 12.2687082 | -0.380770797 | 0.31968956 | -1.1910642 | 0.23362837 | 0.3620954 | AK1 | 203 |
| ENSG000002 | 6.54694769 | -0.43296275 | 0.36352511 | -1.1910119 | 0.23364889 | 0.36211337 | NA | NA |
| ENSG000002 | 2.83011277 | -0.444764267 | 0.37344425 | -1.190979 | 0.23366182 | 0.36211956 | NA | NA |
| ENSG000002 | 3.53047588 | -0.447880732 | 0.37607419 | -1.1909372 | 0.23367825 | 0.36213119 | C1DP5 | 106481729 |
| ENSG000001 | 7.12255346 | -0.429453909 | 0.36066537 | -1.1907268 | 0.23376087 | 0.36224538 | MOGAT3 | 346606 |
| ENSG000002 | 48.7401967 | 0.284824758 | 0.23922233 | 1.19062782 | 0.23379973 | 0.36229174 | RNVU1-6 | 101954276 |
| ENSG000002 | 2.23786884 | -0.438451423 | 0.36828553 | -1.1905204 | 0.23384193 | 0.3623433 | NA | NA |
| ENSG000002 | 7.56756701 | -0.436213199 | 0.36641704 | -1.1904828 | 0.23385668 | 0.3623523 | NA | NA |
| ENSG000002 | 8.87895207 | -0.407777187 | 0.34254726 | -1.1904261 | 0.23387897 | 0.36237299 | NA | NA |
| ENSG000002 | 5.55981092 | -0.444812424 | 0.37367375 | -1.1903764 | 0.23389848 | 0.36237553 | NA | NA |
| ENSG000002 | 4.21957923 | -0.449651479 | 0.37773682 | -1.190383 | 0.2338959 | 0.36237553 | LOC105379391 | 105379391 |
| ENSG000001 | 66.7909681 | 0.235691495 | 0.1980239 | 1.19021743 | 0.23396094 | 0.36244462 | SLC16A14 | 151473 |
| ENSG000002 | 7.79833545 | -0.428753093 | 0.36023042 | -1.190219 | 0.23396032 | 0.36244462 | NA | NA |
| ENSG000002 | 4.26092766 | -0.450202587 | 0.37826892 | -1.1901654 | 0.23398139 | 0.36246244 | NA | NA |
| ENSG000001 | 2.33097418 | -0.429566334 | 0.36096355 | -1.1900546 | 0.23402493 | 0.36251604 | GSC | 145258 |
| ENSG000002 | 50.8392031 | 0.246955385 | 0.20753534 | 1.18994379 | 0.23406849 | 0.36256966 | NA | NA |
| ENSG000002 | 211.838859 | 0.161495725 | 0.13572491 | 1.18987535 | 0.23409539 | 0.36259748 | ENTPD1-AS1 | 728558 |
| ENSG000002 | 6.2724531 | -0.434043012 | 0.36481786 | -1.1897526 | 0.23414363 | 0.3626445 | NA | NA |
| ENSG000002 | 2.92288704 | -0.438614182 | 0.36865991 | -1.1897529 | 0.23414354 | 0.3626445 | NA | NA |
| ENSG000002 | 2.49587733 | -0.444893776 | 0.37398659 | -1.1895982 | 0.23420435 | 0.36271415 | NA | NA |
| ENSG000002 | 5.1870338 | 0.378499442 | 0.31817844 | 1.1895823 | 0.23421061 | 0.36271415 | NA | NA |
| ENSG000002 | 10.7047826 | -0.395359537 | 0.33235499 | -1.18957 | 0.23421543 | 0.36271415 | NA | NA |

| | | | | | | | | |
|---|---|---|---|---|---|---|---|---|
| ENSG000002 | 4.71268384 | -0.446179764 | 0.3751239 | -1.1894197 | 0.23427454 | 0.36279184 | LINC00489 | 100861522 |
| ENSG000001 | 8.94008811 | -0.411837953 | 0.34633993 | -1.1891148 | 0.23439448 | 0.36293284 | FRG2C | 100288801 |
| ENSG000002 | 112.59424 | 0.196198334 | 0.16499772 | 1.18909726 | 0.2344014 | 0.36293284 | NA | NA |
| ENSG000002 | 3.37734723 | -0.449393681 | 0.37792638 | -1.1891037 | 0.23439884 | 0.36293284 | NA | NA |
| ENSG000002 | 6.48686703 | -0.441690066 | 0.37143732 | -1.1891376 | 0.23438554 | 0.36293284 | NA | NA |
| ENSG000002 | 15.2023465 | -0.371389799 | 0.31234327 | -1.1890437 | 0.23442246 | 0.3629516 | LOC101927575 | 101927575 |
| ENSG000001 | 239.495767 | 0.198527862 | 0.16697979 | 1.18893345 | 0.23446586 | 0.36300493 | MTF1 | 4520 |
| ENSG000001 | 140.429129 | 0.181532581 | 0.15268887 | 1.18890513 | 0.234477 | 0.36300832 | FAM19A4 | 151647 |
| ENSG000001 | 6.25933995 | -0.434085969 | 0.36516457 | -1.1887406 | 0.23454174 | 0.36304179 | MATN1-AS1 | 100129196 |
| ENSG000002 | 4.80738815 | -0.448621521 | 0.37736762 | -1.1888183 | 0.23451119 | 0.36304179 | NA | NA |
| ENSG000002 | 3.38319992 | -0.449222328 | 0.37789601 | -1.1887459 | 0.23453968 | 0.36304179 | NA | NA |
| ENSG000002 | 2.52166632 | -0.443766305 | 0.37330497 | -1.18875 | 0.23453804 | 0.36304179 | UGT1A9 | 54600 |
| ENSG000002 | 2.8086908 | 0.440850844 | 0.37085666 | 1.18873647 | 0.23454338 | 0.36304179 | NA | NA |
| ENSG000002 | 9.31677013 | -0.417330722 | 0.3511024 | -1.1886296 | 0.23458544 | 0.36309038 | NA | NA |
| ENSG000002 | 19.0361985 | 0.34311293 | 0.28866708 | 1.18861123 | 0.23459269 | 0.36309038 | NA | NA |
| ENSG000002 | 2.23049628 | -0.435833478 | 0.36671702 | -1.1884735 | 0.2346469 | 0.36314806 | NA | NA |
| ENSG000002 | 5.79312139 | -0.439345493 | 0.36967285 | -1.1884711 | 0.23464786 | 0.36314806 | LOC101927692 | 101927692 |
| ENSG000002 | 3.73650507 | 0.449914774 | 0.37859336 | 1.18838526 | 0.23468166 | 0.3631865 | NA | NA |
| ENSG000002 | 2.87531938 | -0.44470536 | 0.37421711 | -1.1883619 | 0.23469086 | 0.36318687 | RN7SL357P | 106481021 |
| ENSG000001 | 6.07712651 | -0.440013156 | 0.37029011 | -1.188293 | 0.23471798 | 0.36320113 | LOC613037 | 613037 |
| ENSG000002 | 12.3046625 | -0.38976857 | 0.32800165 | -1.1883128 | 0.23471018 | 0.36320113 | NA | NA |
| ENSG00000C | 27.5935461 | 0.316528433 | 0.26640166 | 1.18816239 | 0.23476944 | 0.36326689 | PANX2 | 56666 |
| ENSG000001 | 18.4869319 | 0.349471427 | 0.29414127 | 1.18810741 | 0.23479109 | 0.36328654 | OR7E14P | 10819 |
| ENSG000001 | 2349.17703 | 0.141790531 | 0.11935251 | 1.18799786 | 0.23483425 | 0.36331507 | CEP170 | 9859 |
| ENSG000001 | 4.21710701 | -0.448957295 | 0.37790293 | -1.1880228 | 0.23482443 | 0.36331507 | MRGPRX4 | 117196 |
| ENSG000001 | 4.25146025 | -0.449682273 | 0.37852286 | -1.1879924 | 0.23483641 | 0.36331507 | OR56B1 | 387748 |
| ENSG000002 | 11.9908928 | -0.382459722 | 0.3219448 | -1.1879668 | 0.2348465 | 0.36331682 | LOC401177 | 401177 |
| ENSG000002 | 3.72786274 | -0.448984216 | 0.3779553 | -1.1879294 | 0.23486122 | 0.36332574 | NA | NA |
| ENSG000001 | 13.1811502 | -0.369205902 | 0.31082957 | -1.1878082 | 0.23490899 | 0.36336071 | KANK3 | 256949 |

| | | | | | | | | |
|---|---|---|---|---|---|---|---|---|
| ENSG000002 | 6.68695314 | 0.431325202 | 0.36311997 | 1.18783111 | 0.23489996 | 0.36336071 | NA | NA |
| ENSG000002 | 11.7264984 | -0.391672536 | 0.32974514 | -1.1878038 | 0.23491071 | 0.36336071 | NA | NA |
| ENSG000002 | 5.62262597 | -0.443343827 | 0.37330321 | -1.1876239 | 0.2349816 | 0.3634565 | NA | NA |
| ENSG000002 | 4.47174166 | -0.44767788 | 0.37697881 | -1.1875412 | 0.2350142 | 0.36349305 | NA | NA |
| ENSG000002 | 2.87501991 | -0.444463746 | 0.37429868 | -1.1874574 | 0.23504725 | 0.36352068 | PIK3IP1-AS1 | 101929760 |
| ENSG000002 | 3.11209844 | -0.447088261 | 0.37651109 | -1.1874505 | 0.23504999 | 0.36352068 | NA | NA |
| ENSG000002 | 11.8407705 | -0.388283049 | 0.32705748 | -1.1872012 | 0.23514825 | 0.36365877 | NA | NA |
| ENSG000002 | 6.16303642 | -0.437566494 | 0.36857717 | -1.1871774 | 0.23515765 | 0.36365944 | NA | NA |
| ENSG000002 | 16.8580778 | 0.374254547 | 0.31526091 | 1.1871264 | 0.23517777 | 0.36367668 | RNVU1-18 | 26863 |
| ENSG000001 | 4.68513227 | -0.445611387 | 0.37539276 | -1.1870538 | 0.23520641 | 0.36367979 | LINC00208 | 83655 |
| ENSG000002 | 3.32875796 | -0.449130409 | 0.3783568 | -1.1870552 | 0.23520585 | 0.36367979 | NA | NA |
| ENSG000002 | 23.5107846 | 0.317808627 | 0.26772908 | 1.18705308 | 0.23520668 | 0.36367979 | NA | NA |
| ENSG00000C | 62.2099333 | 0.234036972 | 0.19717159 | 1.18697105 | 0.23523904 | 0.36371595 | FOSL2 | 2355 |
| ENSG000002 | 5.23211137 | -0.441234514 | 0.37175568 | -1.1868938 | 0.23526951 | 0.3637492 | NA | NA |
| ENSG000002 | 6.85673921 | -0.426370811 | 0.35927446 | -1.1867552 | 0.23532421 | 0.3638199 | LOC101927907 | 101927907 |
| ENSG000002 | 2.85370615 | -0.444582359 | 0.37463512 | -1.1867076 | 0.23534297 | 0.36382115 | NA | NA |
| ENSG000002 | 6.97797469 | 0.426112477 | 0.359069 | 1.18671475 | 0.23534015 | 0.36382115 | NA | NA |
| ENSG000002 | 364.419684 | 0.145673398 | 0.12279123 | 1.18635019 | 0.23548403 | 0.36399849 | LOC101927768 | 101927768 |
| ENSG000002 | 3.73947287 | 0.449027387 | 0.37848124 | 1.18639273 | 0.23546724 | 0.36399849 | NA | NA |
| ENSG000002 | 2.21820576 | -0.437177243 | 0.36850652 | -1.1863487 | 0.23548461 | 0.36399849 | NA | NA |
| ENSG000001 | 5.53945195 | -0.441358699 | 0.37207596 | -1.1862059 | 0.235541 | 0.36404403 | C20orf144 | 128864 |
| ENSG000002 | 2.2949995 | -0.434044645 | 0.36590993 | -1.1862063 | 0.23554086 | 0.36404403 | NA | NA |
| ENSG000002 | 5.08590924 | -0.446125896 | 0.37608568 | -1.1862347 | 0.23552961 | 0.36404403 | NA | NA |
| ENSG000001 | 9.56316925 | -0.417017056 | 0.35158134 | -1.1861183 | 0.23557559 | 0.36408361 | TMEM45B | 120224 |
| ENSG000002 | 3.65966773 | -0.448640068 | 0.37825699 | -1.1860721 | 0.23559383 | 0.36409792 | NA | NA |
| ENSG000002 | 3.60560112 | -0.448920188 | 0.37850552 | -1.1860334 | 0.23560911 | 0.36410766 | NA | NA |
| ENSG000001 | 151.158848 | -0.186906323 | 0.15760262 | -1.1859341 | 0.23564831 | 0.36415436 | FSD1L | 83856 |
| ENSG000002 | 13.7258009 | -0.367159692 | 0.3096186 | -1.1858451 | 0.23568349 | 0.36419201 | LOC653712 | 653712 |
| ENSG000002 | 40.0471323 | -0.267773882 | 0.22581626 | -1.1858042 | 0.23569962 | 0.36419201 | NA | NA |

| | | | | | | | | |
|---|---|---|---|---|---|---|---|---|
| ENSG000002 | 22.9834737 | -0.32623252 | 0.2751146 | -1.1858059 | 0.23569895 | 0.36419201 | NA | NA |
| ENSG000002 | 3.75568844 | -0.448146498 | 0.37795536 | -1.1857128 | 0.23573574 | 0.36423395 | NA | NA |
| ENSG000002 | 9.10474396 | 0.414250441 | 0.34942794 | 1.18551036 | 0.23581572 | 0.36434365 | NA | NA |
| ENSG000002 | 15.0513837 | -0.361816096 | 0.30522721 | -1.1853992 | 0.23585964 | 0.36439762 | NA | NA |
| ENSG000001 | 6.34839251 | -0.436428423 | 0.36821089 | -1.1852675 | 0.23591168 | 0.36446413 | FAM170A | 340069 |
| ENSG000001 | 29.5224376 | 0.323592757 | 0.27305528 | 1.18508149 | 0.23598523 | 0.36456387 | METTL7B | 196410 |
| ENSG000002 | 3.13616561 | -0.446263869 | 0.37658659 | -1.1850232 | 0.23600825 | 0.36457166 | NA | NA |
| ENSG000002 | 6.28747284 | -0.431939782 | 0.36449531 | -1.1850352 | 0.23600352 | 0.36457166 | NA | NA |
| ENSG000002 | 4.77259518 | -0.447550751 | 0.37769585 | -1.1849501 | 0.23603717 | 0.36460244 | NA | NA |
| ENSG000002 | 4.20538123 | -0.447977093 | 0.3780677 | -1.1849124 | 0.2360521 | 0.36461161 | NA | NA |
| ENSG000002 | 6.0376446 | -0.435993503 | 0.36796528 | -1.1848767 | 0.23606619 | 0.36461949 | NA | NA |
| ENSG000001 | 356.908213 | 0.175267789 | 0.14793334 | 1.18477542 | 0.23610625 | 0.36465359 | USP49 | 25862 |
| ENSG000002 | 4.26821555 | -0.448103311 | 0.37821774 | -1.184776 | 0.23610601 | 0.36465359 | NA | NA |
| ENSG000002 | 2.22062482 | -0.43347969 | 0.36592768 | -1.1846048 | 0.23617374 | 0.36474392 | NA | NA |
| ENSG000001 | 4.56918729 | -0.447830011 | 0.37806132 | -1.1845433 | 0.23619806 | 0.36476759 | CTSW | 1521 |
| ENSG000002 | 2.90711806 | -0.442737764 | 0.37379688 | -1.1844341 | 0.23624127 | 0.36479265 | NA | NA |
| ENSG000002 | 11.4783561 | -0.390171485 | 0.3294128 | -1.1844454 | 0.23623678 | 0.36479265 | NA | NA |
| ENSG000002 | 4.78491081 | -0.444973136 | 0.37567237 | -1.1844713 | 0.23622655 | 0.36479265 | NA | NA |
| ENSG000002 | 16.602021 | -0.349950313 | 0.29548171 | -1.1843383 | 0.23627916 | 0.36483726 | NA | NA |
| ENSG000001 | 14.2070922 | -0.361484987 | 0.30524332 | -1.1842519 | 0.23631335 | 0.36486227 | FAM196A | 642938 |
| ENSG000002 | 35.6021757 | -0.281955261 | 0.23808362 | -1.1842699 | 0.23630625 | 0.36486227 | LOC101929295 | 101929295 |
| ENSG000002 | 5.22815263 | -0.439393322 | 0.37106599 | -1.1841379 | 0.23635846 | 0.36491802 | LOC101928557 | 101928557 |
| ENSG000001 | 41.1803794 | -0.267754917 | 0.22613666 | -1.1840403 | 0.23639711 | 0.3649638 | MROH8 | 140699 |
| ENSG000002 | 13.9327814 | 0.365698171 | 0.30890879 | 1.18383866 | 0.23647694 | 0.3650672 | RNU1-122P | 106480190 |
| ENSG000002 | 3.08249974 | -0.446612888 | 0.37726239 | -1.1838256 | 0.23648209 | 0.3650672 | NA | NA |
| ENSG000002 | 5.06970279 | -0.445526409 | 0.37637534 | -1.183729 | 0.23652035 | 0.36510107 | NA | NA |
| ENSG000002 | 2.89431792 | -0.442264143 | 0.37362076 | -1.1837248 | 0.23652204 | 0.36510107 | NA | NA |
| ENSG000002 | 3.89882204 | -0.44809899 | 0.37858116 | -1.1836273 | 0.23656063 | 0.36514675 | LINC00239 | 145200 |
| ENSG000001 | 861.39866 | 0.155527711 | 0.1314037 | 1.183587 | 0.23657659 | 0.36515747 | THADA | 63892 |

| | | | | | | | | |
|---|---|---|---|---|---|---|---|---|
| ENSG000002 | 2.2306927 | -0.435402351 | 0.36787721 | -1.1835535 | 0.23658986 | 0.36516406 | NA | NA |
| ENSG000001 | 5.67778553 | -0.443658798 | 0.37486694 | -1.1835101 | 0.23660706 | 0.3651767 | NA | NA |
| ENSG000001 | 86.751708 | 0.211439807 | 0.17868542 | 1.18330758 | 0.23668727 | 0.3652866 | IPP | 3652 |
| ENSG000000 | 24.4471633 | -0.32416384 | 0.27397503 | -1.1831875 | 0.23673483 | 0.36532264 | PDK4 | 5166 |
| ENSG000002 | 2.50608919 | 0.436101319 | 0.36859246 | 1.18315312 | 0.23674847 | 0.36532264 | NA | NA |
| ENSG000002 | 6.98014408 | -0.431449978 | 0.36465363 | -1.1831775 | 0.2367388 | 0.36532264 | FRMD6-AS2 | 100874185 |
| ENSG000002 | 7.67884228 | -0.422327893 | 0.35695625 | -1.1831363 | 0.23675514 | 0.36532264 | NA | NA |
| ENSG000002 | 2.2276323 | -0.434253629 | 0.36703643 | -1.183135 | 0.23675567 | 0.36532264 | NA | NA |
| ENSG000002 | 3.36478293 | -0.447477194 | 0.37822309 | -1.1831038 | 0.236768 | 0.36532777 | NA | NA |
| ENSG000002 | 22.653187 | 0.333958773 | 0.28229413 | 1.18301707 | 0.23680238 | 0.36536692 | TAS2R19 | 259294 |
| ENSG000001 | 293.966037 | 0.157914211 | 0.13349841 | 1.1828921 | 0.23685191 | 0.3653934 | GALNT13 | 114805 |
| ENSG000002 | 2.25414642 | -0.434289449 | 0.36714494 | -1.1828828 | 0.23685559 | 0.3653934 | NA | NA |
| ENSG000002 | 4.52369289 | -0.446293945 | 0.37729144 | -1.1828891 | 0.23685309 | 0.3653934 | NA | NA |
| ENSG000002 | 2.23593795 | -0.435835444 | 0.36844066 | -1.182919 | 0.23684127 | 0.3653934 | NA | NA |
| ENSG000002 | 7.76311275 | -0.426010473 | 0.36015497 | -1.1828532 | 0.23686732 | 0.36539759 | RPS4XP20 | 100271386 |
| ENSG000002 | 2.5180525 | -0.44125559 | 0.37308637 | -1.182717 | 0.23692133 | 0.36546701 | LOC105376276 | 105376276 |
| ENSG000002 | 8.63822671 | -0.408436108 | 0.34537201 | -1.1825976 | 0.23696867 | 0.36552613 | NA | NA |
| ENSG000002 | 9.97286227 | -0.402342323 | 0.34024385 | -1.1825117 | 0.23700274 | 0.36556479 | NA | NA |
| ENSG000001 | 9.70851892 | -0.407168267 | 0.34436808 | -1.1823636 | 0.23706148 | 0.36559975 | OR51A7 | 119687 |
| ENSG000002 | 6.99007077 | 0.425555518 | 0.35991809 | 1.18236769 | 0.23705984 | 0.36559975 | NA | NA |
| ENSG000002 | 6.85836899 | -0.428487311 | 0.36238303 | -1.1824155 | 0.23704087 | 0.36559975 | NA | NA |
| ENSG000002 | 3.12995482 | -0.44494298 | 0.37630723 | -1.1823929 | 0.23704985 | 0.36559975 | NA | NA |
| ENSG000001 | 1004.32549 | 0.145924747 | 0.12343047 | 1.18224254 | 0.23710948 | 0.36565987 | STX6 | 10228 |
| ENSG000002 | 3.1159734 | -0.44569011 | 0.37701043 | -1.1821692 | 0.23713858 | 0.36569085 | NA | NA |
| ENSG000002 | 2.53817258 | -0.441083973 | 0.37312311 | -1.1821406 | 0.23714992 | 0.36569443 | NA | NA |
| ENSG000002 | 4.22449829 | -0.446778345 | 0.37796368 | -1.1820669 | 0.23717916 | 0.36572562 | NA | NA |
| ENSG000002 | 8.62230067 | -0.409034334 | 0.34604795 | -1.1820163 | 0.23719921 | 0.36574263 | NA | NA |
| ENSG000001 | 79.564189 | 0.235572394 | 0.19930965 | 1.18194173 | 0.23722882 | 0.36577085 | IFFO2 | 126917 |
| ENSG000002 | 3.94772452 | -0.447351935 | 0.37849443 | -1.1819248 | 0.23723556 | 0.36577085 | LOC105755953 | 105755953 |

| | | | | | | | | |
|---|---|---|---|---|---|---|---|---|
| ENSG000001 | 22.7040904 | 0.324779807 | 0.27481264 | 1.18182268 | 0.23727607 | 0.36581016 | TRPV1 | 7442 |
| ENSG000002 | 5.00485216 | -0.445122688 | 0.37664327 | -1.1818151 | 0.23727909 | 0.36581016 | KHDC3L | 154288 |
| ENSG000002 | 3.14466713 | -0.44552397 | 0.37699057 | -1.1817907 | 0.23728874 | 0.36581112 | NA | NA |
| ENSG000002 | 3.93888505 | -0.447408373 | 0.37859261 | -1.1817673 | 0.23729804 | 0.36581155 | NA | NA |
| ENSG000002 | 8.17762465 | -0.415445558 | 0.35160909 | -1.1815552 | 0.23738222 | 0.36591697 | LOC101928682 | 101928682 |
| ENSG000002 | 2.85375713 | -0.440117492 | 0.37249894 | -1.1815268 | 0.2373935 | 0.36591697 | NA | NA |
| ENSG000002 | 2.26641787 | -0.432885583 | 0.36637619 | -1.1815331 | 0.23739103 | 0.36591697 | NA | NA |
| ENSG000002 | 14.6582255 | 0.375748766 | 0.31802935 | 1.18149083 | 0.23740779 | 0.3659251 | NA | NA |
| ENSG000001 | 1706.01771 | 0.143402052 | 0.12137696 | 1.18146024 | 0.23741994 | 0.36592991 | KIF14 | 9928 |
| ENSG000001 | 47.0862611 | 0.249693874 | 0.21134764 | 1.1814368 | 0.23742925 | 0.36593035 | WDFY3-AS2 | 404201 |
| ENSG000001 | 7.5206311 | -0.429975498 | 0.36395806 | -1.1813875 | 0.23744883 | 0.36594662 | POU4F2 | 5458 |
| ENSG000001 | 7.46753272 | -0.425053683 | 0.35981199 | -1.1813216 | 0.23747498 | 0.36594822 | RFPL3 | 10738 |
| ENSG000001 | 16.8609115 | -0.352962379 | 0.29879301 | -1.181294 | 0.23748596 | 0.36594822 | IL12A | 3592 |
| ENSG000002 | 6.53178753 | -0.427803706 | 0.36213197 | -1.1813475 | 0.2374647 | 0.36594822 | NA | NA |
| ENSG000002 | 3.97599369 | -0.447221106 | 0.37858039 | -1.1813108 | 0.23747927 | 0.36594822 | NA | NA |
| ENSG000001 | 7.42403823 | -0.424347771 | 0.3592532 | -1.1811941 | 0.23752562 | 0.36599541 | FAM183A | 440585 |
| ENSG000002 | 4.03505501 | -0.446976833 | 0.37844961 | -1.1810736 | 0.2375735 | 0.36605529 | NA | NA |
| ENSG000001 | 93.157236 | 0.247990752 | 0.20998213 | 1.18100882 | 0.23759922 | 0.36608101 | HTR1D | 3352 |
| ENSG000002 | 4.99013141 | -0.442925082 | 0.37511015 | -1.1807867 | 0.23768746 | 0.36620305 | NA | NA |
| ENSG000001 | 4.80279945 | -0.444571094 | 0.37652399 | -1.1807245 | 0.23771219 | 0.36622724 | HOXA4 | 3201 |
| ENSG000002 | 4.7692663 | -0.443403234 | 0.3755545 | -1.1806628 | 0.2377367 | 0.36625108 | NA | NA |
| ENSG000002 | 4.53580769 | -0.44558054 | 0.37742729 | -1.1805732 | 0.23777232 | 0.36629204 | NA | NA |
| ENSG000002 | 4.16676726 | -0.446225374 | 0.37800736 | -1.1804674 | 0.23781436 | 0.36634289 | NA | NA |
| ENSG000002 | 3.11082099 | -0.445366526 | 0.37729649 | -1.1804152 | 0.2378351 | 0.366347 | NA | NA |
| ENSG000002 | 3.61234693 | -0.44688919 | 0.3785843 | -1.1804219 | 0.23783247 | 0.366347 | NA | NA |
| ENSG000002 | 3.44638492 | -0.445946087 | 0.37781574 | -1.1803269 | 0.23787022 | 0.36638717 | NA | NA |
| ENSG000001 | 8.65079743 | -0.410467758 | 0.34778955 | -1.1802188 | 0.23791322 | 0.36643949 | OR6N1 | 128372 |
| ENSG000001 | 6.83875908 | -0.424715241 | 0.35991634 | -1.1800388 | 0.23798479 | 0.36650082 | NA | NA |
| ENSG000001 | 20.8637009 | -0.345111142 | 0.29246934 | -1.1799908 | 0.23800386 | 0.36650082 | FAM43A | 131583 |

| | | | | | | | | |
|---|---|---|---|---|---|---|---|---|
| ENSG000002 | 2.94985153 | -0.436450576 | 0.3698763 | -1.1799906 | 0.23800393 | 0.36650082 | RNU6-1327P | 106481599 |
| ENSG000002 | 4.7028771 | -0.443607365 | 0.37594409 | -1.1799823 | 0.23800726 | 0.36650082 | ZNF727 | 442319 |
| ENSG000002 | 2.20986821 | -0.43460365 | 0.36830346 | -1.1800151 | 0.2379942 | 0.36650082 | NA | NA |
| ENSG000002 | 2.20986821 | -0.43460365 | 0.36830346 | -1.1800151 | 0.2379942 | 0.36650082 | NA | NA |
| ENSG000002 | 2.51511639 | -0.440723956 | 0.37354287 | -1.1798484 | 0.23806052 | 0.36651324 | UGT1A12P | 54573 |
| ENSG000002 | 3.37301689 | -0.446423743 | 0.37835332 | -1.1799123 | 0.23803509 | 0.36651324 | NA | NA |
| ENSG000002 | 7.30353762 | -0.433127611 | 0.36709401 | -1.179882 | 0.23804716 | 0.36651324 | LINC00700 | 282980 |
| ENSG000002 | 3.37293257 | -0.446291411 | 0.37826036 | -1.1798524 | 0.23805891 | 0.36651324 | NA | NA |
| ENSG000002 | 22.5290872 | 0.327917746 | 0.27792308 | 1.17988669 | 0.23804528 | 0.36651324 | NA | NA |
| ENSG000002 | 4.14140695 | -0.446205999 | 0.37820414 | -1.179802 | 0.23807898 | 0.36652774 | NA | NA |
| ENSG000002 | 3.59364974 | -0.446517428 | 0.37851655 | -1.179651 | 0.23813905 | 0.3666063 | BRD9P1 | 106481721 |
| ENSG000002 | 77.6145223 | 0.238027505 | 0.20179402 | 1.1795568 | 0.23817653 | 0.36665009 | LEFTY1 | 10637 |
| ENSG000002 | 6.06252142 | -0.433644394 | 0.36764415 | -1.1795221 | 0.23819034 | 0.36665743 | NA | NA |
| ENSG000002 | 3.3982651 | -0.446061301 | 0.37822055 | -1.1793682 | 0.23825158 | 0.36672652 | NA | NA |
| ENSG000002 | 3.04869849 | -0.444672118 | 0.37704404 | -1.1793639 | 0.23825332 | 0.36672652 | NA | NA |
| ENSG000001 | 16.2440892 | -0.350619301 | 0.29730447 | -1.1793274 | 0.23826784 | 0.36673099 | NA | NA |
| ENSG000002 | 6.32137126 | -0.430056338 | 0.36466741 | -1.1793111 | 0.23827431 | 0.36673099 | NA | NA |
| ENSG000001 | 16.8228915 | -0.370240884 | 0.31397312 | -1.1792121 | 0.23831375 | 0.36677777 | GRIN1 | 2902 |
| ENSG000002 | 2.52354569 | -0.440187779 | 0.37331314 | -1.1791382 | 0.23834315 | 0.3668091 | NA | NA |
| ENSG000002 | 3.7315694 | -0.446006442 | 0.37829523 | -1.1789904 | 0.238402 | 0.36688575 | NA | NA |
| ENSG000002 | 3.0596 | -0.444557681 | 0.37713231 | -1.1787844 | 0.23848404 | 0.36699808 | NA | NA |
| ENSG000001 | 8.64409095 | -0.410126831 | 0.34793819 | -1.1787347 | 0.23850382 | 0.36701459 | FOXD2 | 2306 |
| ENSG000002 | 2.2382132 | -0.430945547 | 0.3656716 | -1.1785043 | 0.23859563 | 0.36711947 | NA | NA |
| ENSG000002 | 3.33564125 | -0.445971901 | 0.3784177 | -1.1785175 | 0.23859034 | 0.36711947 | NA | NA |
| ENSG000002 | 4.46501482 | -0.445167643 | 0.37774235 | -1.1784955 | 0.23859914 | 0.36711947 | ZIM2-AS1 | 101929059 |
| ENSG000002 | 37.2847796 | 0.272280287 | 0.23105947 | 1.17839919 | 0.2386375 | 0.36716456 | NA | NA |
| ENSG000002 | 3.91578531 | -0.446092683 | 0.37856894 | -1.1783658 | 0.23865082 | 0.36717112 | NA | NA |
| ENSG000002 | 4.71281969 | -0.442843553 | 0.37589028 | -1.1781192 | 0.23874909 | 0.36730838 | NA | NA |
| ENSG000001 | 3.12208723 | -0.444195524 | 0.37708734 | -1.1779646 | 0.23881073 | 0.36732079 | GZMH | 2999 |

| | | | | | | | | |
|---|---|---|---|---|---|---|---|---|
| ENSG000001 | 166.542616 | 0.176273632 | 0.14963748 | 1.17800451 | 0.23879481 | 0.36732079 | GPD1L | 23171 |
| ENSG000002 | 2.30725831 | -0.430491334 | 0.36544856 | -1.1779806 | 0.23880432 | 0.36732079 | SEPT14P24 | 106480345 |
| ENSG000002 | 4.17384588 | -0.445408688 | 0.37808409 | -1.1780678 | 0.23876958 | 0.36732079 | NA | NA |
| ENSG000002 | 3.0833934 | -0.443941344 | 0.37685045 | -1.1780306 | 0.23878443 | 0.36732079 | NA | NA |
| ENSG000002 | 2.30973053 | -0.429571878 | 0.3646736 | -1.1779626 | 0.23881151 | 0.36732079 | NA | NA |
| ENSG000002 | 6.76979272 | -0.423811179 | 0.35980941 | -1.1778769 | 0.23884569 | 0.36735944 | NA | NA |
| ENSG000002 | 7.31662278 | -0.418689225 | 0.35549861 | -1.1777521 | 0.23889547 | 0.36742206 | FLJ36000 | 284124 |
| ENSG000002 | 7.39608205 | -0.423448842 | 0.35957097 | -1.1776502 | 0.23893607 | 0.36747058 | NA | NA |
| ENSG000001 | 2.50985188 | -0.439883719 | 0.37358636 | -1.177462 | 0.23901118 | 0.3675582 | LRRC18 | 474354 |
| ENSG000002 | 3.445583 | -0.444625055 | 0.37761112 | -1.1774681 | 0.23900871 | 0.3675582 | NA | NA |
| ENSG000002 | 7.86432433 | -0.416273719 | 0.35357914 | -1.1773141 | 0.23907015 | 0.36763495 | NA | NA |
| ENSG000001 | 19.0483973 | -0.33307936 | 0.28294797 | -1.1771753 | 0.23912554 | 0.36770619 | NCMAP | 400746 |
| ENSG000002 | 3.16271197 | -0.443269621 | 0.37660711 | -1.1770081 | 0.23919227 | 0.36778119 | OR6K3 | 391114 |
| ENSG000002 | 3.33177097 | -0.445014708 | 0.37808991 | -1.1770076 | 0.23919246 | 0.36778119 | NA | NA |
| ENSG000001 | 79.1254463 | 0.2126538 | 0.18069609 | 1.17685886 | 0.23925184 | 0.36785855 | AK7 | 122481 |
| ENSG000002 | 51.7998995 | 0.253546453 | 0.21545684 | 1.17678536 | 0.23928119 | 0.36788972 | NA | NA |
| ENSG000002 | 3.38166188 | -0.445193735 | 0.37835302 | -1.1766623 | 0.23933032 | 0.36795132 | NA | NA |
| ENSG000002 | 2.91241538 | -0.439828039 | 0.37380619 | -1.1766205 | 0.239347 | 0.36796301 | NA | NA |
| ENSG000001 | 3.46642069 | -0.444208142 | 0.37756682 | -1.1765021 | 0.23939431 | 0.36802179 | OR7A10 | 390892 |
| ENSG000002 | 5.33042284 | -0.443482437 | 0.37696411 | -1.176458 | 0.23941192 | 0.3680349 | NA | NA |
| ENSG000001 | 79.9382792 | -0.209580245 | 0.17816059 | -1.1763558 | 0.23945272 | 0.368056 | RNF165 | 494470 |
| ENSG000001 | 83.2716494 | -0.222456781 | 0.18910856 | -1.1763444 | 0.2394573 | 0.368056 | SLC24A3 | 57419 |
| ENSG000002 | 2.23687049 | -0.431122897 | 0.36648731 | -1.1763651 | 0.23944899 | 0.368056 | NA | NA |
| ENSG000002 | 14.0897116 | -0.367761465 | 0.31263388 | -1.1763327 | 0.23946194 | 0.368056 | IFNG-AS1 | 100885789 |
| ENSG000002 | 23.055006 | -0.327807978 | 0.27872069 | -1.1761164 | 0.23954836 | 0.36817486 | NA | NA |
| ENSG000002 | 2.21755288 | -0.431700549 | 0.36711281 | -1.1759343 | 0.23962113 | 0.36827275 | NA | NA |
| ENSG000002 | 5.08541976 | -0.442354934 | 0.37618514 | -1.1758969 | 0.23963609 | 0.36828178 | NA | NA |
| ENSG000002 | 5.00743601 | -0.439247653 | 0.37356725 | -1.1758195 | 0.23966702 | 0.36831535 | NA | NA |
| ENSG000002 | 7.68039979 | 0.418354345 | 0.3558122 | 1.1757729 | 0.23968564 | 0.36833001 | NA | NA |

| | | | | | | | | |
|---|---|---|---|---|---|---|---|---|
| ENSG000002 | 3.36800053 | -0.444886437 | 0.37839019 | -1.1757346 | 0.23970095 | 0.36833957 | NA | NA |
| ENSG000002 | 2.24601793 | -0.432121369 | 0.36756122 | -1.1756446 | 0.23973692 | 0.36838089 | NA | NA |
| ENSG000002 | 24.8945849 | -0.316391334 | 0.26914429 | -1.1755454 | 0.23977658 | 0.36842788 | NA | NA |
| ENSG000002 | 3.76847505 | -0.443957127 | 0.37768679 | -1.1754638 | 0.23980923 | 0.36846408 | NA | NA |
| ENSG000002 | 2.22544083 | -0.431010907 | 0.36670416 | -1.1753641 | 0.23984908 | 0.36848343 | COL18A1-AS2 | 100874236 |
| ENSG000002 | 3.53026353 | -0.43935313 | 0.37380071 | -1.1753673 | 0.2398478 | 0.36848343 | NA | NA |
| ENSG000002 | 5.89021865 | -0.436647368 | 0.37149965 | -1.1753641 | 0.23984906 | 0.36848343 | NA | NA |
| ENSG000002 | 4.5098944 | -0.443911671 | 0.37769631 | -1.1753138 | 0.23986921 | 0.3685004 | NA | NA |
| ENSG000002 | 2.51343291 | -0.438091671 | 0.37275931 | -1.1752669 | 0.23988796 | 0.36851524 | NA | NA |
| ENSG000002 | 17.3036988 | 0.342918018 | 0.29181852 | 1.1751071 | 0.23995187 | 0.36859945 | NA | NA |
| ENSG000002 | 2.85904648 | -0.43713783 | 0.37201512 | -1.1750539 | 0.23997314 | 0.36861816 | NA | NA |
| ENSG000002 | 3.67130645 | -0.444324898 | 0.37815973 | -1.1749662 | 0.24000824 | 0.36865811 | MTND6P13 | 106480428 |
| ENSG000001 | 18.8905415 | -0.332329845 | 0.28285573 | -1.1749094 | 0.24003098 | 0.36866588 | TTLL11 | 158135 |
| ENSG000002 | 4.03255883 | -0.444808349 | 0.3785899 | -1.1749081 | 0.24003148 | 0.36866588 | NA | NA |
| ENSG000001 | 341.616577 | 0.157317485 | 0.13390102 | 1.17487894 | 0.24004315 | 0.36866984 | LRRC49 | 54839 |
| ENSG000002 | 2.47461564 | -0.438955218 | 0.3736734 | -1.1747029 | 0.24011361 | 0.36876408 | NA | NA |
| ENSG000002 | 5.54167681 | 0.436086924 | 0.37126876 | 1.17458555 | 0.24016057 | 0.36882224 | NA | NA |
| ENSG000002 | 11.6988294 | -0.380950145 | 0.32435699 | -1.174478 | 0.24020362 | 0.36887439 | LINC00961 | 158376 |
| ENSG000002 | 6.02860174 | -0.433984997 | 0.36955489 | -1.1743452 | 0.2402568 | 0.36894208 | NA | NA |
| ENSG000001 | 8.36915309 | -0.409815742 | 0.34899227 | -1.1742831 | 0.24028163 | 0.36896624 | PTCRA | 171558 |
| ENSG000002 | 3.65279561 | -0.444535976 | 0.37858805 | -1.1741944 | 0.24031717 | 0.36900683 | NA | NA |
| ENSG000002 | 4.67406768 | -0.440477891 | 0.37520366 | -1.1739701 | 0.24040699 | 0.36913079 | NA | NA |
| ENSG000002 | 2.51190696 | -0.437588976 | 0.37276143 | -1.1739116 | 0.24043042 | 0.3691388 | NA | NA |
| ENSG000002 | 2.51190696 | -0.437588976 | 0.37276143 | -1.1739116 | 0.24043042 | 0.3691388 | NA | NA |
| ENSG00000C | 4.47326894 | -0.442719249 | 0.37715612 | -1.1738355 | 0.24046092 | 0.36916526 | CREB3L3 | 84699 |
| ENSG000002 | 5.39749178 | -0.443073844 | 0.37746217 | -1.1738232 | 0.24046586 | 0.36916526 | NA | NA |
| ENSG000002 | 3.17215092 | -0.441317014 | 0.37598169 | -1.1737726 | 0.24048611 | 0.3691684 | NA | NA |
| ENSG000002 | 2.87843752 | -0.437657351 | 0.37286183 | -1.1737789 | 0.2404836 | 0.3691684 | NA | NA |
| ENSG000002 | 3.05817765 | -0.442379541 | 0.37695581 | -1.1735581 | 0.24057208 | 0.36927241 | NA | NA |

| | | | | | | | | |
|---|---|---|---|---|---|---|---|---|
| ENSG000002 | 10.4754675 | -0.393186903 | 0.33503582 | -1.1735668 | 0.2405686 | 0.36927241 | NA | NA |
| ENSG000001 | 91.9556625 | -0.20255237 | 0.17260542 | -1.1734995 | 0.24059556 | 0.36929447 | LSAMP | 4045 |
| ENSG000002 | 3.92183138 | -0.444229897 | 0.3785785 | -1.1734155 | 0.24062921 | 0.36933214 | NA | NA |
| ENSG000001 | 434.230513 | 0.143169689 | 0.1220178 | 1.17335085 | 0.24065514 | 0.36935796 | RAB4A | 5867 |
| ENSG000002 | 4.73587697 | -0.440890894 | 0.37578229 | -1.1732615 | 0.24069095 | 0.36939894 | LOC105372179 | 105372179 |
| ENSG000002 | 2.25355301 | -0.42977513 | 0.36632537 | -1.173206 | 0.2407132 | 0.36941911 | NA | NA |
| ENSG000002 | 3.33388608 | -0.443688022 | 0.37822659 | -1.1730746 | 0.24076587 | 0.36948596 | NA | NA |
| ENSG000002 | 4.23649457 | -0.443964479 | 0.37849136 | -1.1729844 | 0.24080205 | 0.36952748 | PPBPP2 | 10895 |
| ENSG000002 | 5.06022438 | -0.44072496 | 0.37575985 | -1.17289 | 0.24083994 | 0.36957165 | NA | NA |
| ENSG000002 | 6.6271682 | -0.427108362 | 0.36416922 | -1.1728294 | 0.24086423 | 0.36959494 | NA | NA |
| ENSG000002 | 2.84558952 | -0.438257025 | 0.37369705 | -1.1727602 | 0.24089199 | 0.36962354 | NA | NA |
| ENSG000002 | 3.34170246 | -0.443763192 | 0.37840034 | -1.1727347 | 0.24090223 | 0.36962527 | NA | NA |
| ENSG000001 | 2187.28194 | 0.35603227 | 0.30361034 | 1.17266186 | 0.24093143 | 0.36965609 | ZC3H13 | 23091 |
| ENSG000002 | 7.07443596 | -0.418674693 | 0.35706079 | -1.1725586 | 0.24097287 | 0.36970567 | NA | NA |
| ENSG000001 | 7.18651728 | -0.431552754 | 0.36809224 | -1.1724038 | 0.24103496 | 0.36973886 | TNNT3 | 7140 |
| ENSG000001 | 10.2629898 | -0.397001775 | 0.33863885 | -1.1723456 | 0.24105832 | 0.36973886 | CD3E | 916 |
| ENSG000002 | 34.5906989 | 0.294341759 | 0.25106903 | 1.1723539 | 0.241055 | 0.36973886 | MXD3 | 83463 |
| ENSG000002 | 3.33581697 | -0.44326976 | 0.37809077 | -1.1723898 | 0.24104059 | 0.36973886 | NA | NA |
| ENSG000002 | 12.939772 | -0.369067867 | 0.31479694 | -1.1723998 | 0.24103658 | 0.36973886 | ALKBH3-AS1 | 100507300 |
| ENSG000002 | 5.23900262 | -0.435569822 | 0.3715321 | -1.1723612 | 0.24105207 | 0.36973886 | NA | NA |
| ENSG000002 | 3.33544537 | -0.443578843 | 0.37835668 | -1.1723828 | 0.24104342 | 0.36973886 | NA | NA |
| ENSG000002 | 4.71173811 | -0.440202766 | 0.37553915 | -1.1721887 | 0.24112129 | 0.36982145 | NA | NA |
| ENSG000002 | 3.37193477 | -0.443332582 | 0.37826861 | -1.1720047 | 0.24119517 | 0.36992078 | NA | NA |
| ENSG000000 | 13.9432894 | -0.363469164 | 0.31014591 | -1.1719296 | 0.24122533 | 0.36995304 | FOXP3 | 50943 |
| ENSG000002 | 3.97795628 | -0.443331007 | 0.37841343 | -1.171552 | 0.24137697 | 0.37017161 | NA | NA |
| ENSG000001 | 26.1700672 | -0.303854124 | 0.25936571 | -1.1715277 | 0.24138671 | 0.37017254 | FOSB | 2354 |
| ENSG000002 | 19.9104608 | 0.328894353 | 0.28075361 | 1.1714697 | 0.24141003 | 0.3701943 | NA | NA |
| ENSG000002 | 165.741079 | -0.176434369 | 0.15061392 | -1.1714347 | 0.24142411 | 0.37019508 | DENND1B | 163486 |
| ENSG000002 | 4.14595314 | -0.388957355 | 0.33203835 | -1.171423 | 0.24142879 | 0.37019508 | NA | NA |

| | | | | | | | | |
|---|---|---|---|---|---|---|---|---|
| ENSG000002 | 8.15141578 | -0.413138656 | 0.35269543 | -1.1713751 | 0.24144803 | 0.37019657 | NA | NA |
| ENSG000002 | 3.48226343 | -0.440067842 | 0.37568438 | -1.1713765 | 0.24144748 | 0.37019657 | NA | NA |
| ENSG000002 | 10.7422614 | -0.405949634 | 0.34669732 | -1.170905 | 0.24163695 | 0.37047223 | NA | NA |
| ENSG000002 | 7.32464713 | -0.417998797 | 0.35702707 | -1.1707762 | 0.24168874 | 0.37050961 | NA | NA |
| ENSG000002 | 3.08972684 | -0.440458015 | 0.37620703 | -1.1707862 | 0.24168471 | 0.37050961 | NA | NA |
| ENSG000002 | 3.63269535 | -0.443257394 | 0.37859184 | -1.1708055 | 0.24167699 | 0.37050961 | NA | NA |
| ENSG000001 | 309.718626 | 0.186984808 | 0.15971995 | 1.17070414 | 0.24171772 | 0.37054002 | USP42 | 84132 |
| ENSG000002 | 3.59954733 | -0.443140452 | 0.37857298 | -1.1705549 | 0.24177774 | 0.37061801 | LOC105373876 | 105373876 |
| ENSG000002 | 5.24378576 | -0.438889013 | 0.3749824 | -1.1704256 | 0.24182973 | 0.37066969 | NA | NA |
| ENSG000002 | 3.09503212 | -0.440969647 | 0.37675521 | -1.1704407 | 0.24182365 | 0.37066969 | NA | NA |
| ENSG000002 | 4.06842018 | -0.442968192 | 0.37852664 | -1.1702431 | 0.24190315 | 0.37076822 | NA | NA |
| ENSG000001 | 4.00028875 | -0.442994173 | 0.37856422 | -1.1701956 | 0.24192228 | 0.37078352 | PDZK1IP1 | 10158 |
| ENSG000002 | 2.6947624 | -0.427539122 | 0.36538141 | -1.1701173 | 0.24195375 | 0.37081774 | NA | NA |
| ENSG000001 | 4.7390987 | -0.439800245 | 0.3758772 | -1.1700636 | 0.24197536 | 0.37083684 | COX6B2 | 125965 |
| ENSG000001 | 3.72603986 | 0.442936505 | 0.37858906 | 1.16996647 | 0.24201446 | 0.37088275 | MAGED4B | 81557 |
| ENSG000002 | 4.76544396 | -0.439367089 | 0.37562176 | -1.1697062 | 0.24211923 | 0.37102929 | LOC102725019 | 102725019 |
| ENSG000002 | 3.34900224 | -0.442192049 | 0.37808924 | -1.1695441 | 0.24218447 | 0.37111524 | LINC00618 | 145249 |
| ENSG000002 | 5.02861907 | -0.440513328 | 0.37668163 | -1.169458 | 0.24221917 | 0.37115111 | NA | NA |
| ENSG000002 | 4.55964597 | -0.441926065 | 0.37789529 | -1.1694405 | 0.24222619 | 0.37115111 | LINC01387 | 100130480 |
| ENSG000002 | 4.81679361 | -0.440529598 | 0.37676178 | -1.1692523 | 0.24230198 | 0.37121535 | NA | NA |
| ENSG000002 | 6.75997483 | -0.435573192 | 0.37252076 | -1.1692589 | 0.24229936 | 0.37121535 | LINC01135 | 100131060 |
| ENSG000002 | 3.16936658 | -0.440253315 | 0.37652769 | -1.1692455 | 0.24230473 | 0.37121535 | NA | NA |
| ENSG000002 | 2.82917486 | 0.43515524 | 0.37215922 | 1.16927168 | 0.24229419 | 0.37121535 | MIR5192 | 100847087 |
| ENSG000002 | 27.5771308 | -0.311824414 | 0.26669739 | -1.1692068 | 0.24232031 | 0.3712252 | NA | NA |
| ENSG000002 | 2.54252362 | -0.435160038 | 0.37229984 | -1.1688429 | 0.24246691 | 0.37143575 | NA | NA |
| ENSG000002 | 2.48056475 | -0.43496788 | 0.37214867 | -1.1688014 | 0.24248366 | 0.37144738 | NA | NA |
| ENSG000002 | 2.54784762 | -0.434235398 | 0.37153535 | -1.1687593 | 0.24250062 | 0.37145932 | NA | NA |
| ENSG000002 | 6.57019343 | -0.424914574 | 0.36358628 | -1.168676 | 0.24253418 | 0.37148968 | NA | NA |
| ENSG000002 | 23.1568695 | 0.319092799 | 0.27304051 | 1.16866468 | 0.24253876 | 0.37148968 | NA | NA |

| | | | | | | | | |
|---|---|---|---|---|---|---|---|---|
| ENSG000002 | 4.97233616 | -0.436997956 | 0.3739672 | -1.1685462 | 0.24258651 | 0.37154878 | NA | NA |
| ENSG000002 | 4.73298074 | -0.43871138 | 0.37548986 | -1.1683708 | 0.24265724 | 0.37162904 | NA | NA |
| ENSG000002 | 3.0557575 | -0.439985472 | 0.37657397 | -1.1683906 | 0.24264926 | 0.37162904 | NA | NA |
| ENSG000002 | 7.07113424 | -0.419474081 | 0.35907446 | -1.1682092 | 0.2427224 | 0.37169208 | TAS2R50 | 259296 |
| ENSG000002 | 7.33570619 | -0.416333132 | 0.35639386 | -1.1681827 | 0.24273309 | 0.37169208 | NA | NA |
| ENSG000002 | 4.7067672 | -0.438533236 | 0.37540267 | -1.1681676 | 0.24273917 | 0.37169208 | NA | NA |
| ENSG000002 | 9.12743023 | -0.399975349 | 0.34240247 | -1.1681439 | 0.24274873 | 0.37169208 | NA | NA |
| ENSG000002 | 108.616133 | 0.194373324 | 0.16638775 | 1.16819493 | 0.24272814 | 0.37169208 | LINC01572 | 101927957 |
| ENSG000002 | 31.7004039 | 0.291239792 | 0.24932089 | 1.16813231 | 0.2427534 | 0.37169208 | NA | NA |
| ENSG000002 | 2.2125484 | -0.427125047 | 0.36565917 | -1.1680961 | 0.24276801 | 0.37170041 | NA | NA |
| ENSG000002 | 10.5521529 | 0.391494029 | 0.33520663 | 1.16791853 | 0.24283963 | 0.37179604 | NA | NA |
| ENSG000002 | 12.810652 | -0.37326032 | 0.31964558 | -1.1677319 | 0.24291494 | 0.3718973 | NA | NA |
| ENSG000002 | 10.1505799 | -0.406661633 | 0.34826861 | -1.1676666 | 0.24294126 | 0.37192355 | LINC01500 | 102723742 |
| ENSG000002 | 10.7591617 | -0.388926867 | 0.33309817 | -1.1676043 | 0.2429664 | 0.371948 | NA | NA |
| ENSG000002 | 6.62472769 | -0.427857084 | 0.36644768 | -1.1675803 | 0.24297611 | 0.37194844 | NA | NA |
| ENSG000002 | 9.9479929 | 0.393858453 | 0.33733519 | 1.16755817 | 0.24298503 | 0.37194844 | NA | NA |
| ENSG000001 | 2.51993186 | -0.435146783 | 0.37277933 | -1.167304 | 0.24308764 | 0.37207741 | NCF1B | 654816 |
| ENSG000002 | 2.23425501 | -0.428108264 | 0.36674341 | -1.1673237 | 0.24307968 | 0.37207741 | PIGPP2 | 106479056 |
| ENSG000002 | 68.0448844 | 0.222310803 | 0.1904523 | 1.16727813 | 0.24309807 | 0.37207933 | EFNA4 | 1945 |
| ENSG000002 | 6.6868586 | -0.430023853 | 0.36840971 | -1.1672435 | 0.24311204 | 0.37208666 | NA | NA |
| ENSG000002 | 4.69191197 | -0.437817515 | 0.37515388 | -1.1670345 | 0.24319644 | 0.3722018 | PP12613 | 100192379 |
| ENSG000002 | 9.38585531 | -0.397530101 | 0.34065363 | -1.1669628 | 0.2432254 | 0.37223208 | NA | NA |
| ENSG000002 | 3.34936731 | -0.441189119 | 0.37809567 | -1.1668717 | 0.24326221 | 0.37227435 | NA | NA |
| ENSG000002 | 4.77783253 | -0.438589513 | 0.37588367 | -1.1668225 | 0.24328207 | 0.37227665 | LOC101928475 | 101928475 |
| ENSG000002 | 9.01121816 | -0.409361681 | 0.35083203 | -1.166831 | 0.24327864 | 0.37227665 | LOC101928132 | 101928132 |
| ENSG000001 | 2.22338575 | -0.428181935 | 0.36702512 | -1.1666284 | 0.24336046 | 0.37237461 | NA | NA |
| ENSG000002 | 2.49331337 | -0.435451676 | 0.37325969 | -1.1666185 | 0.24336445 | 0.37237461 | NA | NA |
| ENSG000002 | 6.78925469 | -0.421466972 | 0.36129343 | -1.1665503 | 0.24339201 | 0.37240272 | NA | NA |
| ENSG000001 | 3.63900952 | -0.441531637 | 0.37851321 | -1.1664894 | 0.24341664 | 0.37242636 | NA | NA |

| | | | | | | | | |
|---|---|---|---|---|---|---|---|---|
| ENSG000001 | 175.481749 | 0.167035414 | 0.14320453 | 1.16641151 | 0.24344811 | 0.37246045 | RNF19B | 127544 |
| ENSG000002 | 4.98395452 | -0.437179614 | 0.37486366 | -1.1662363 | 0.24351891 | 0.37255471 | NA | NA |
| ENSG000002 | 3.37598491 | -0.440277085 | 0.37759597 | -1.1660005 | 0.24361424 | 0.37267245 | NA | NA |
| ENSG000002 | 2.48953537 | -0.43352162 | 0.37180162 | -1.1660025 | 0.24361343 | 0.37267245 | NA | NA |
| ENSG000002 | 2.23514071 | -0.425424076 | 0.36488107 | -1.1659253 | 0.24364464 | 0.37269082 | NA | NA |
| ENSG000002 | 4.90085881 | 0.437784348 | 0.37548113 | 1.16592901 | 0.24364315 | 0.37269082 | NA | NA |
| ENSG000002 | 12.296505 | -0.381359465 | 0.32713376 | -1.16576 | 0.24371148 | 0.372779 | NA | NA |
| ENSG000002 | 2.30418582 | -0.425010384 | 0.36466134 | -1.1654934 | 0.24381934 | 0.37292633 | RN7SKP188 | 106479179 |
| ENSG000002 | 5.26005992 | -0.435325988 | 0.37351762 | -1.1654764 | 0.24382619 | 0.37292633 | LINC00892 | 100128420 |
| ENSG000002 | 3.57534899 | -0.441220012 | 0.37858282 | -1.1654518 | 0.24383618 | 0.37292754 | NA | NA |
| ENSG000002 | 2.53042788 | -0.432624768 | 0.37123272 | -1.1653735 | 0.24386784 | 0.37295664 | NA | NA |
| ENSG000002 | 3.5929974 | -0.441143832 | 0.3785475 | -1.1653593 | 0.2438736 | 0.37295664 | NA | NA |
| ENSG000002 | 9.26234405 | 0.40119557 | 0.34428716 | 1.16529343 | 0.24390024 | 0.37298332 | NA | NA |
| ENSG000002 | 3.09674697 | -0.438290643 | 0.37613872 | -1.1652367 | 0.24392319 | 0.37300435 | NA | NA |
| ENSG000002 | 3.23241099 | -0.435793558 | 0.37402504 | -1.1651454 | 0.24396014 | 0.37304678 | NA | NA |
| ENSG000002 | 4.89024184 | 0.436543171 | 0.37468684 | 1.16508809 | 0.24398334 | 0.37306819 | NA | NA |
| ENSG000001 | 6.8479441 | -0.423540834 | 0.36354179 | -1.1650403 | 0.24400268 | 0.37308369 | BARHL2 | 343472 |
| ENSG000001 | 47.9775205 | -0.241028834 | 0.20691417 | -1.1648735 | 0.2440702 | 0.37314045 | GPR158 | 57512 |
| ENSG000002 | 2.24487632 | -0.428156545 | 0.36756066 | -1.1648596 | 0.24407584 | 0.37314045 | NA | NA |
| ENSG000002 | 6.90401823 | -0.422698127 | 0.36287534 | -1.1648577 | 0.24407661 | 0.37314045 | NA | NA |
| ENSG000002 | 6.58153409 | -0.423481063 | 0.36353909 | -1.1648845 | 0.24406574 | 0.37314045 | NA | NA |
| ENSG000002 | 2.51295628 | -0.432688202 | 0.37146658 | -1.1648106 | 0.24409567 | 0.37315553 | NA | NA |
| ENSG000001 | 76.2596639 | 0.2207004 | 0.18948974 | 1.16470896 | 0.24413682 | 0.3731781 | GPR137C | 283554 |
| ENSG000001 | 2.50820085 | -0.433501654 | 0.37220564 | -1.1646832 | 0.24414725 | 0.3731781 | NA | NA |
| ENSG000002 | 2.24227881 | -0.425425448 | 0.36525668 | -1.1647301 | 0.24412827 | 0.3731781 | NA | NA |
| ENSG000002 | 5.0627909 | -0.437950744 | 0.37602014 | -1.1647002 | 0.24414036 | 0.3731781 | NA | NA |
| ENSG000001 | 82.5533595 | -0.213484143 | 0.18330995 | -1.1646075 | 0.24417792 | 0.37318311 | MLPH | 79083 |
| ENSG000001 | 27.7624411 | 0.302480406 | 0.25972522 | 1.16461701 | 0.24417406 | 0.37318311 | CMTM8 | 152189 |
| ENSG000002 | 3.34889786 | -0.440406742 | 0.37815912 | -1.1646069 | 0.24417813 | 0.37318311 | NA | NA |

| | | | | | | | | |
|---|---|---|---|---|---|---|---|---|
| ENSG000002 | 16.6765554 | -0.351660818 | 0.30204288 | -1.1642745 | 0.24431279 | 0.37337483 | LOC100996345 | 100996345 |
| ENSG000001 | 14.0298708 | -0.36213078 | 0.31104483 | -1.1642398 | 0.24432684 | 0.37338224 | FAM132B | 151176 |
| ENSG000002 | 3.87511195 | -0.440296247 | 0.37820418 | -1.164176 | 0.24435268 | 0.37339358 | NKX2-1-AS1 | 100506237 |
| ENSG000002 | 11.5004109 | -0.382494393 | 0.3285504 | -1.1641879 | 0.24434789 | 0.37339358 | NA | NA |
| ENSG000002 | 11.1948631 | -0.411178733 | 0.3532104 | -1.1641184 | 0.24437603 | 0.37341519 | ZBTB20-AS1 | 100131117 |
| ENSG000002 | 5.09463027 | -0.439649407 | 0.37769811 | -1.1640233 | 0.24441456 | 0.37342964 | NA | NA |
| ENSG000002 | 8.07764968 | -0.410993131 | 0.35308562 | -1.1640042 | 0.24442232 | 0.37342964 | NA | NA |
| ENSG000002 | 4.90342812 | -0.4399352 | 0.3779372 | -1.1640431 | 0.24440654 | 0.37342964 | NA | NA |
| ENSG000002 | 34.9820627 | 0.270025346 | 0.23196761 | 1.16406485 | 0.24439773 | 0.37342964 | NA | NA |
| ENSG000001 | 153.520345 | 0.176996257 | 0.15206726 | 1.16393403 | 0.24445075 | 0.37344241 | SLCO5A1 | 81796 |
| ENSG000002 | 7.59371625 | -0.416955497 | 0.35822183 | -1.1639589 | 0.24444065 | 0.37344241 | NA | NA |
| ENSG000002 | 4.34574032 | 0.439531845 | 0.37763213 | 1.16391538 | 0.24445831 | 0.37344241 | NA | NA |
| ENSG000002 | 3.40415004 | -0.440030465 | 0.37807981 | -1.163856 | 0.24448236 | 0.37346509 | NA | NA |
| ENSG000001 | 2939.43253 | 0.130471828 | 0.11211036 | 1.16378034 | 0.24451304 | 0.37348381 | CEP350 | 9857 |
| ENSG000002 | 2.52800828 | -0.432768098 | 0.3718569 | -1.1638028 | 0.24450394 | 0.37348381 | NA | NA |
| ENSG000001 | 153.427699 | 0.171499013 | 0.1473761 | 1.16368266 | 0.24455264 | 0.37351616 | NFKB1 | 4790 |
| ENSG000002 | 7.90258763 | -0.414999896 | 0.35662157 | -1.1636983 | 0.24454632 | 0.37351616 | NA | NA |
| ENSG000002 | 4.81099299 | -0.438574146 | 0.37690376 | -1.1636237 | 0.24457655 | 0.37353861 | NA | NA |
| ENSG000002 | 3.8789222 | -0.439816746 | 0.37799458 | -1.1635531 | 0.24460518 | 0.37356827 | NA | NA |
| ENSG000001 | 4.97774448 | -0.434904869 | 0.37378086 | -1.163529 | 0.24461495 | 0.37356911 | LY86 | 9450 |
| ENSG000001 | 147.068011 | 0.177647046 | 0.15268876 | 1.16345858 | 0.24464349 | 0.37358457 | EFCAB11 | 90141 |
| ENSG000001 | 9.32456224 | -0.416244059 | 0.35776303 | -1.163463 | 0.24464169 | 0.37358457 | SERPINA12 | 145264 |
| ENSG000002 | 6.84905509 | -0.41952174 | 0.36060538 | -1.1633818 | 0.24467463 | 0.37361805 | NA | NA |
| ENSG00000C | 126.766397 | 0.216114967 | 0.18578382 | 1.16326042 | 0.24472386 | 0.37367915 | EVC | 2121 |
| ENSG000002 | 7.3127372 | -0.420343126 | 0.36135779 | -1.1632325 | 0.24473519 | 0.37368238 | NA | NA |
| ENSG000002 | 2.61245022 | -0.425494243 | 0.36580623 | -1.1631684 | 0.24476119 | 0.37370801 | RPL23AP89 | 106480336 |
| ENSG000002 | 8.83311574 | -0.412319214 | 0.3544928 | -1.1631244 | 0.24477904 | 0.3737212 | NA | NA |
| ENSG000001 | 2.64700132 | -0.425008357 | 0.36543772 | -1.1630117 | 0.24482474 | 0.37377689 | DUSP21 | 63904 |
| ENSG000002 | 11.7274038 | -0.380215824 | 0.32697083 | -1.1628432 | 0.24489311 | 0.37386719 | NA | NA |

| | | | | | | | | |
|---|---|---|---|---|---|---|---|---|
| ENSG000002 | 5.47232406 | -0.429126347 | 0.36906059 | -1.1627531 | 0.24492969 | 0.37390897 | NA | NA |
| ENSG000002 | 5.08666441 | -0.438704283 | 0.37731124 | -1.1627119 | 0.24494639 | 0.37392039 | DLGAP1-AS3 | 201477 |
| ENSG000002 | 4.9807332 | -0.434996222 | 0.37413133 | -1.1626832 | 0.24495805 | 0.37392411 | LINC00977 | 728724 |
| ENSG000001 | 10.9195231 | -0.38643867 | 0.33238944 | -1.1626081 | 0.24498853 | 0.37393486 | C19orf18 | 147685 |
| ENSG000002 | 12.4178115 | -0.373318231 | 0.32110697 | -1.1625977 | 0.24499276 | 0.37393486 | NA | NA |
| ENSG000002 | 3.72830082 | -0.439507418 | 0.3780281 | -1.1626316 | 0.244979 | 0.37393486 | NA | NA |
| ENSG000002 | 7.18307193 | -0.42151625 | 0.36257238 | -1.1625713 | 0.24500347 | 0.37393714 | NA | NA |
| ENSG000002 | 2.49721536 | -0.433684399 | 0.37305286 | -1.162528 | 0.24502106 | 0.3739499 | NA | NA |
| ENSG000002 | 5.50654179 | -0.431051093 | 0.37083076 | -1.162393 | 0.24507586 | 0.37401946 | NA | NA |
| ENSG000002 | 72.0894055 | 0.425463851 | 0.36604298 | 1.16233303 | 0.24510021 | 0.37402848 | MRPL23 | 6150 |
| ENSG000002 | 3.89474277 | -0.439640327 | 0.37823701 | -1.1623408 | 0.24509704 | 0.37402848 | NA | NA |
| ENSG000002 | 17.673534 | -0.339708394 | 0.29227526 | -1.1622893 | 0.24511799 | 0.37404153 | NA | NA |
| ENSG000002 | 2.46866598 | -0.433666832 | 0.37316284 | -1.1621383 | 0.24517928 | 0.37412098 | NA | NA |
| ENSG000002 | 2.30344119 | -0.424317621 | 0.36515591 | -1.1620177 | 0.24522829 | 0.37416761 | RNU6-433P | 106481315 |
| ENSG000002 | 4.78678997 | -0.437376272 | 0.376391 | -1.1620264 | 0.24522475 | 0.37416761 | LOC105379030 | 105379030 |
| ENSG000002 | 16.002658 | -0.352438377 | 0.30333108 | -1.1618934 | 0.24527878 | 0.37421965 | NA | NA |
| ENSG000002 | 6.62556435 | -0.423034147 | 0.36409194 | -1.1618882 | 0.24528086 | 0.37421965 | LINC01111 | 101926978 |
| ENSG000001 | 70.6096149 | 0.220691851 | 0.18998594 | 1.16162199 | 0.24538904 | 0.37434246 | EEPD1 | 80820 |
| ENSG000001 | 218.707533 | 0.161808953 | 0.13929334 | 1.16164167 | 0.24538105 | 0.37434246 | PARP6 | 56965 |
| ENSG000001 | 401.231618 | 0.166857207 | 0.14363634 | 1.16166431 | 0.24537185 | 0.37434246 | MKL2 | 57496 |
| ENSG000002 | 2.51415051 | -0.431911605 | 0.37185076 | -1.1615187 | 0.24543103 | 0.37439241 | NA | NA |
| ENSG000002 | 9.96198502 | -0.392858776 | 0.33839622 | -1.1609432 | 0.24566501 | 0.37473525 | LOC101929089 | 101929089 |
| ENSG000002 | 3.12254434 | 0.438060774 | 0.37734175 | 1.16091255 | 0.24567746 | 0.37474015 | NA | NA |
| ENSG000001 | 3.34746833 | -0.438543153 | 0.37777558 | -1.1608563 | 0.24570036 | 0.37474688 | PRKACG | 5568 |
| ENSG000002 | 3.6331199 | -0.439491239 | 0.37859097 | -1.1608603 | 0.2456987 | 0.37474688 | NA | NA |
| ENSG000002 | 5.53120934 | -0.430548959 | 0.37090473 | -1.1608074 | 0.24572022 | 0.37476308 | NA | NA |
| ENSG000002 | 2.27321663 | -0.425288755 | 0.36640417 | -1.1607094 | 0.24576011 | 0.37478162 | NA | NA |
| ENSG000002 | 3.09580124 | -0.436782056 | 0.3763039 | -1.1607163 | 0.24575731 | 0.37478162 | NA | NA |
| ENSG000002 | 3.61710345 | -0.439431193 | 0.37857937 | -1.1607373 | 0.24574875 | 0.37478162 | NA | NA |

| | | | | | | | | |
|---|---|---|---|---|---|---|---|---|
| ENSG000001 | 49.665218 | 0.252255344 | 0.21735654 | 1.16056016 | 0.24582081 | 0.3748601 | RAB20 | 55647 |
| ENSG000002 | 2.50846178 | -0.43190135 | 0.3721588 | -1.1605297 | 0.2458332 | 0.37486489 | C3orf84 | 646498 |
| ENSG000002 | 5.18771659 | -0.432258315 | 0.3724762 | -1.1604992 | 0.24584563 | 0.37486975 | NA | NA |
| ENSG000002 | 3.38662583 | -0.438310072 | 0.37776082 | -1.1602846 | 0.24593294 | 0.37497871 | LOC340017 | 340017 |
| ENSG000002 | 6.42050434 | -0.427233835 | 0.36821674 | -1.1602781 | 0.24593558 | 0.37497871 | NA | NA |
| ENSG000002 | 38.5656523 | 0.271434825 | 0.23397618 | 1.160096 | 0.24600972 | 0.37507765 | NA | NA |
| ENSG000001 | 3.63614087 | -0.43917121 | 0.37858958 | -1.1600193 | 0.24604096 | 0.37511117 | TAS2R1 | 50834 |
| ENSG000001 | 27.5653386 | 0.297896636 | 0.25681287 | 1.15997548 | 0.24605879 | 0.37512425 | IL11RA | 3590 |
| ENSG000002 | 30.5448995 | 0.290564036 | 0.25050751 | 1.15990151 | 0.24608891 | 0.37515607 | LOC100996579 | 100996579 |
| ENSG000002 | 6.95721113 | -0.426212501 | 0.36748693 | -1.1598032 | 0.24612896 | 0.37520301 | LOC100996583 | 100996583 |
| ENSG000002 | 2.52565284 | -0.430799741 | 0.3714928 | -1.1596449 | 0.2461934 | 0.37528714 | OPA1-AS1 | 100873941 |
| ENSG000002 | 15.7681787 | 0.349562914 | 0.30144601 | 1.15962032 | 0.24620343 | 0.37528832 | NA | NA |
| ENSG000002 | 12.4726973 | 0.37133152 | 0.32023254 | 1.15956837 | 0.24622459 | 0.37530647 | EXOC3-AS1 | 116349 |
| ENSG000002 | 5.02195026 | -0.436050573 | 0.37607355 | -1.1594821 | 0.24625971 | 0.37534589 | NA | NA |
| ENSG000001 | 140.078753 | -0.199759117 | 0.17229851 | -1.1593781 | 0.24630209 | 0.37539639 | NFASC | 23114 |
| ENSG00000C | 34.4790776 | -0.273001621 | 0.23549806 | -1.1592521 | 0.24635344 | 0.37543509 | CD82 | 3732 |
| ENSG000002 | 11.2547765 | -0.397164633 | 0.34260512 | -1.159249 | 0.24635471 | 0.37543509 | NA | NA |
| ENSG000002 | 8.6420887 | -0.404450758 | 0.34889757 | -1.1592249 | 0.24636452 | 0.37543509 | NA | NA |
| ENSG000002 | 3.13454683 | -0.434498495 | 0.37481766 | -1.1592263 | 0.24636394 | 0.37543509 | NA | NA |
| ENSG000002 | 4.95767646 | -0.43350679 | 0.37399653 | -1.1591198 | 0.24640735 | 0.37548624 | MED4-AS1 | 100873965 |
| ENSG000002 | 2.25030838 | -0.421345833 | 0.36351475 | -1.1590887 | 0.24642003 | 0.37549146 | FAM138B | 654412 |
| ENSG000001 | 90.484226 | 0.219649094 | 0.18953007 | 1.15891423 | 0.24649115 | 0.37558548 | DLGAP3 | 58512 |
| ENSG000002 | 4.80512318 | -0.436361971 | 0.37654117 | -1.1588692 | 0.2465095 | 0.37558548 | LOC101928994 | 101928994 |
| ENSG000002 | 3.91507137 | -0.438499173 | 0.3783854 | -1.1588692 | 0.24650952 | 0.37558548 | NA | NA |
| ENSG000002 | 6.67712199 | 0.425941877 | 0.36757562 | 1.15878708 | 0.24654299 | 0.3755965 | NA | NA |
| ENSG000002 | 262.024742 | 0.178527623 | 0.15406001 | 1.15881873 | 0.24653008 | 0.3755965 | NBPF12 | 149013 |
| ENSG000002 | 5.78148924 | 0.426723353 | 0.36825122 | 1.15878327 | 0.24654454 | 0.3755965 | NA | NA |
| ENSG000002 | 3.68971422 | 0.438436568 | 0.37844544 | 1.1585199 | 0.24665194 | 0.375746 | RN7SL370P | 106481024 |
| ENSG000002 | 12.6958693 | -0.3702024 | 0.31959551 | -1.1583467 | 0.24672259 | 0.3758395 | NA | NA |

| | | | | | | | | |
|---|---|---|---|---|---|---|---|---|
| ENSG000001 | 12.4225941 | -0.383350773 | 0.33095893 | -1.1583031 | 0.24674035 | 0.37585244 | WFDC9 | 259240 |
| ENSG000001 | 39.5129123 | 0.269906704 | 0.23303406 | 1.15822858 | 0.24677077 | 0.37587054 | TCIRG1 | 10312 |
| ENSG000001 | 10.395564 | -0.385700362 | 0.33300647 | -1.1582369 | 0.24676739 | 0.37587054 | PDZD9 | 255762 |
| ENSG000001 | 462.066808 | 0.16457935 | 0.14211514 | 1.15807051 | 0.24683526 | 0.37593793 | KCNT2 | 343450 |
| ENSG000002 | 14.226235 | 0.356554696 | 0.30788204 | 1.15808865 | 0.24682786 | 0.37593793 | URB1-AS1 | 84996 |
| ENSG000002 | 2.27928307 | -0.422787776 | 0.36508532 | -1.158052 | 0.24684282 | 0.37593793 | RN7SL498P | 106479405 |
| ENSG000002 | 4.78177808 | -0.436490432 | 0.37694215 | -1.1579772 | 0.24687333 | 0.37597026 | NA | NA |
| ENSG000002 | 4.40496866 | -0.437001793 | 0.37749515 | -1.1576355 | 0.2470128 | 0.37616855 | NA | NA |
| ENSG000002 | 3.14270399 | -0.435510284 | 0.37624137 | -1.157529 | 0.24705632 | 0.37619829 | NA | NA |
| ENSG000002 | 2.30757152 | -0.42197835 | 0.36456588 | -1.1574817 | 0.24707559 | 0.37619829 | NA | NA |
| ENSG000002 | 4.73852546 | -0.435971515 | 0.3766464 | -1.1575088 | 0.24706456 | 0.37619829 | LOC101927523 | 101927523 |
| ENSG000002 | 7.41012443 | -0.416580546 | 0.35988389 | -1.1575415 | 0.24705119 | 0.37619829 | NA | NA |
| ENSG000002 | 5.21623818 | -0.432355487 | 0.37353362 | -1.1574741 | 0.24707872 | 0.37619829 | GOLGA8K | 653125 |
| ENSG000001 | 4.96028064 | -0.433190411 | 0.37431095 | -1.1573009 | 0.24714943 | 0.37626357 | PROL1 | 58503 |
| ENSG000001 | 9.36363624 | -0.393499348 | 0.34001425 | -1.1573025 | 0.24714878 | 0.37626357 | FLJ40194 | 124871 |
| ENSG000002 | 3.58910283 | -0.438160701 | 0.37859221 | -1.1573421 | 0.24713262 | 0.37626357 | MIR181A2HG | 100379345 |
| ENSG000001 | 7.33840152 | -0.41329267 | 0.35714793 | -1.157203 | 0.24718941 | 0.37631032 | C17orf64 | 124773 |
| ENSG000002 | 2.78286098 | -0.431355316 | 0.37282958 | -1.1569772 | 0.24728168 | 0.37643665 | NA | NA |
| ENSG000002 | 8.14472458 | -0.408401932 | 0.35301972 | -1.1568813 | 0.24732084 | 0.37648213 | NA | NA |
| ENSG000001 | 3.84963506 | -0.437636914 | 0.3782989 | -1.1568548 | 0.24733167 | 0.37648448 | TBX10 | 347853 |
| ENSG000001 | 10.2400182 | -0.395615345 | 0.34200996 | -1.1567363 | 0.24738011 | 0.37653417 | TECTB | 6975 |
| ENSG000001 | 173.589548 | 0.16993722 | 0.1469118 | 1.15672953 | 0.24738288 | 0.37653417 | ENTPD5 | 957 |
| ENSG000002 | 3.87391031 | -0.437567583 | 0.37831241 | -1.1566303 | 0.24742345 | 0.37658177 | NA | NA |
| ENSG000002 | 3.65250332 | -0.437861235 | 0.37858805 | -1.1565638 | 0.24745062 | 0.37659052 | NA | NA |
| ENSG000002 | 7.18707628 | -0.420229891 | 0.3633484 | -1.1565481 | 0.24745705 | 0.37659052 | NA | NA |
| ENSG000002 | 23.6607926 | 0.327385055 | 0.28306261 | 1.15658179 | 0.24744327 | 0.37659052 | LOC102724200 | 102724200 |
| ENSG000001 | 216.496325 | 0.161529029 | 0.13968824 | 1.15635385 | 0.24753645 | 0.37669721 | GALNT10 | 55568 |
| ENSG000002 | 6.51047432 | -0.419412931 | 0.36273988 | -1.1562361 | 0.24758461 | 0.37675636 | NA | NA |
| ENSG000001 | 60.7786573 | -0.234902082 | 0.20316558 | -1.15621 | 0.24759525 | 0.37675842 | SGMS2 | 166929 |

| | | | | | | | | |
|---|---|---|---|---|---|---|---|---|
| ENSG000002 | 3.17822907 | -0.432137929 | 0.37378666 | -1.1561085 | 0.24763678 | 0.37680747 | NA | NA |
| ENSG000001 | 14.1892641 | -0.353695736 | 0.3059714 | -1.1559765 | 0.24769079 | 0.37686137 | USH1G | 124590 |
| ENSG000002 | 2.75516437 | -0.434830148 | 0.37615668 | -1.1559815 | 0.24768874 | 0.37686137 | NA | NA |
| ENSG000001 | 4619.82151 | -0.171248013 | 0.14817185 | -1.1557392 | 0.24778785 | 0.37699491 | HIST1H1D | 3007 |
| ENSG000002 | 6.60091358 | -0.425370496 | 0.36806047 | -1.1557082 | 0.24780054 | 0.37700007 | NA | NA |
| ENSG000001 | 237.590966 | 0.344256524 | 0.29788851 | 1.15565561 | 0.24782205 | 0.37701865 | RAB3D | 9545 |
| ENSG000002 | 3.17039609 | -0.433701732 | 0.37529902 | -1.1556165 | 0.24783806 | 0.37702886 | NA | NA |
| ENSG000002 | 6.31154701 | -0.429519409 | 0.37176115 | -1.1553639 | 0.24794145 | 0.37717199 | NA | NA |
| ENSG000002 | 3.96733392 | 0.436353739 | 0.37776613 | 1.15508964 | 0.24805372 | 0.37732862 | RPL13AP6 | 644511 |
| ENSG000001 | 206.895197 | 0.171491812 | 0.14847347 | 1.15503338 | 0.24807676 | 0.37734951 | RNF157 | 114804 |
| ENSG000001 | 1900.69028 | 0.129526874 | 0.112149 | 1.15495344 | 0.24810949 | 0.37738515 | MIA3 | 375056 |
| ENSG000002 | 3.69249466 | -0.437022567 | 0.37841749 | -1.1548688 | 0.24814414 | 0.3774237 | NA | NA |
| ENSG000002 | 2.46820897 | -0.429908483 | 0.37227547 | -1.1548128 | 0.24816709 | 0.37744444 | NA | NA |
| ENSG000002 | 2.50939487 | -0.4293969 | 0.37184121 | -1.1547857 | 0.24817821 | 0.3774472 | NA | NA |
| ENSG000002 | 19.0542459 | -0.345303908 | 0.29907045 | -1.1545905 | 0.24825815 | 0.37755462 | NA | NA |
| ENSG000002 | 2.47410656 | -0.429576401 | 0.3720909 | -1.1544932 | 0.24829803 | 0.37760111 | NA | NA |
| ENSG000002 | 4.44758354 | -0.435728579 | 0.37743682 | -1.1544411 | 0.24831937 | 0.37761939 | NA | NA |
| ENSG000002 | 2.49345773 | -0.430265247 | 0.37273235 | -1.1543545 | 0.24835488 | 0.37765923 | PDCL2P2 | 106480295 |
| ENSG000001 | 4.66894663 | -0.434321526 | 0.3762578 | -1.154319 | 0.24836941 | 0.37766716 | NA | NA |
| ENSG000002 | 2.52280847 | -0.429245397 | 0.37186982 | -1.1542894 | 0.24838153 | 0.37767143 | NA | NA |
| ENSG000002 | 4.67229191 | -0.434811675 | 0.37673142 | -1.1541689 | 0.24843094 | 0.3777324 | NA | NA |
| ENSG000002 | 5.31393681 | -0.43178788 | 0.37418727 | -1.1539353 | 0.24852671 | 0.37786385 | NA | NA |
| ENSG000002 | 4.73538825 | -0.433840707 | 0.37600429 | -1.1538185 | 0.24857459 | 0.37792247 | NA | NA |
| ENSG000002 | 3.10939754 | -0.432775976 | 0.37510413 | -1.1537489 | 0.24860313 | 0.37794566 | NA | NA |
| ENSG000002 | 8.323286 | -0.400445275 | 0.34708575 | -1.1537359 | 0.24860848 | 0.37794566 | NA | NA |
| ENSG000002 | 2.49403076 | -0.429303761 | 0.37213132 | -1.1536351 | 0.2486498 | 0.3779943 | NA | NA |
| ENSG000000 | 67.791424 | -0.2325149 | 0.20156094 | -1.1535712 | 0.24867601 | 0.37800581 | CACNA1G | 8913 |
| ENSG000002 | 24.9876241 | -0.302950841 | 0.2626151 | -1.1535926 | 0.24866723 | 0.37800581 | LOC100128239 | 100128239 |
| ENSG000001 | 12.8663213 | 0.371678682 | 0.32220833 | 1.1535353 | 0.24869074 | 0.37801002 | ALX1 | 8092 |

| | | | | | | | | |
|---|---|---|---|---|---|---|---|---|
| ENSG000001 | 23.7541241 | 0.307088336 | 0.26621871 | 1.15351899 | 0.24869743 | 0.37801002 | SERPINA5 | 5104 |
| ENSG000001 | 206.007174 | 0.161870919 | 0.14033488 | 1.15346176 | 0.24872091 | 0.37803154 | ZNF704 | 619279 |
| ENSG000002 | 5.34567736 | -0.433546605 | 0.3759031 | -1.1533467 | 0.24876811 | 0.37808911 | NA | NA |
| ENSG000002 | 7.99068891 | -0.412936822 | 0.35804664 | -1.1533046 | 0.2487854 | 0.37810122 | NA | NA |
| ENSG000002 | 3.3642416 | -0.435519479 | 0.37765199 | -1.1532297 | 0.24881613 | 0.37813376 | NA | NA |
| ENSG000002 | 8.21630227 | 0.408263077 | 0.35407415 | 1.15304402 | 0.24889232 | 0.37823536 | MIR135A2 | 406926 |
| ENSG000000 | 3.91278816 | -0.436122909 | 0.37825699 | -1.1529804 | 0.24891842 | 0.37824668 | NA | NA |
| ENSG000001 | 17.2025006 | -0.434595674 | 0.37692756 | -1.1529952 | 0.24891234 | 0.37824668 | SLC2A2 | 6514 |
| ENSG000001 | 19.5190307 | -0.333825717 | 0.28955048 | -1.1529103 | 0.24894722 | 0.37827626 | SH3TC1 | 54436 |
| ENSG000002 | 15.1978538 | 0.350652925 | 0.30419599 | 1.1527204 | 0.24902517 | 0.37838053 | NA | NA |
| ENSG000002 | 2.52333717 | -0.42929778 | 0.37243527 | -1.1526776 | 0.24904275 | 0.37839307 | FAM99A | 387742 |
| ENSG000000 | 39.8953018 | -0.26052787 | 0.22606886 | -1.1524271 | 0.24914562 | 0.37853519 | CDHR2 | 54825 |
| ENSG000002 | 3.58747196 | -0.43622287 | 0.37858925 | -1.1522326 | 0.24922551 | 0.37862819 | NA | NA |
| ENSG000002 | 146.39661 | 0.176843408 | 0.15347606 | 1.15225405 | 0.24921669 | 0.37862819 | NA | NA |
| ENSG000001 | 6.09495286 | -0.429779064 | 0.37302789 | -1.1521365 | 0.24926497 | 0.37866052 | RGS9BP | 388531 |
| ENSG000002 | 2.46474418 | -0.428765756 | 0.37214878 | -1.1521353 | 0.24926547 | 0.37866052 | NA | NA |
| ENSG000002 | 4.47256419 | -0.435155321 | 0.37774692 | -1.1519758 | 0.249331 | 0.3787317 | NA | NA |
| ENSG000002 | 3.88819229 | -0.435931114 | 0.37841776 | -1.1519837 | 0.24932775 | 0.3787317 | NA | NA |
| ENSG000002 | 6.34761238 | -0.42298042 | 0.36719315 | -1.1519289 | 0.24935028 | 0.3787326 | NA | NA |
| ENSG000002 | 6.92234381 | -0.41925794 | 0.36395731 | -1.1519426 | 0.24934465 | 0.3787326 | NA | NA |
| ENSG000002 | 2.26566084 | -0.418212739 | 0.3630718 | -1.1518734 | 0.24937311 | 0.3787531 | NA | NA |
| ENSG000002 | 3.58274878 | -0.436051964 | 0.37858319 | -1.1517996 | 0.24940344 | 0.37878497 | NA | NA |
| ENSG000001 | 157.562717 | 0.17182678 | 0.14919874 | 1.15166377 | 0.24945927 | 0.37885558 | ACTC1 | 70 |
| ENSG000001 | 28.88946 | -0.285870024 | 0.24823947 | -1.1515897 | 0.24948971 | 0.37885924 | HEATR4 | 399671 |
| ENSG000002 | 2.48625436 | -0.429228777 | 0.37271436 | -1.1516293 | 0.24947344 | 0.37885924 | NA | NA |
| ENSG000002 | 3.66594863 | -0.435796805 | 0.37842812 | -1.1515973 | 0.24948659 | 0.37885924 | LOC105375401 | 105375401 |
| ENSG000002 | 2.92363642 | -0.426534171 | 0.37040742 | -1.1515271 | 0.24951547 | 0.37888417 | LOC101927139 | 101927139 |
| ENSG000002 | 120.992748 | 0.196680959 | 0.17080671 | 1.1514826 | 0.24953375 | 0.37889774 | CD99P1 | 401577 |
| ENSG000001 | 15.5518227 | -0.369767182 | 0.32115495 | -1.1513669 | 0.24958132 | 0.37895578 | KY | 339855 |

| | | | | | | | | |
|---|---|---|---|---|---|---|---|---|
| ENSG000001 | 679.512215 | 0.138140295 | 0.1199903 | 1.15126215 | 0.2496244 | 0.37897863 | C1orf198 | 84886 |
| ENSG000002 | 5.22645097 | -0.428384477 | 0.37209508 | -1.1512769 | 0.24961833 | 0.37897863 | MRGPRF-AS1 | 101928200 |
| ENSG000002 | 2.58029123 | -0.425857165 | 0.36989104 | -1.1513044 | 0.24960705 | 0.37897863 | NA | NA |
| ENSG000002 | 5.47705967 | -0.426845512 | 0.37077266 | -1.1512324 | 0.24963663 | 0.378983 | NA | NA |
| ENSG000002 | 3.42721091 | -0.434126553 | 0.37714797 | -1.1510775 | 0.24970035 | 0.37906554 | NA | NA |
| ENSG000001 | 90.6563395 | 0.201098207 | 0.17473595 | 1.15086909 | 0.2497861 | 0.37918139 | TRIM9 | 114088 |
| ENSG000002 | 3.33645154 | 0.435529481 | 0.3784427 | 1.15084658 | 0.24979536 | 0.37918139 | RNU6-82P | 100873781 |
| ENSG000002 | 4.9926323 | -0.430953303 | 0.37455274 | -1.1505811 | 0.24990461 | 0.37933303 | NA | NA |
| ENSG000002 | 5.80781409 | -0.427075236 | 0.37119145 | -1.1505524 | 0.24991641 | 0.37933673 | NA | NA |
| ENSG000002 | 3.30852577 | -0.434330284 | 0.37752392 | -1.1504709 | 0.24994998 | 0.37936013 | LINC01419 | 103352670 |
| ENSG000002 | 4.66425953 | -0.435254309 | 0.37832754 | -1.1504695 | 0.24995053 | 0.37936013 | NA | NA |
| ENSG000002 | 2.74678175 | 0.432049813 | 0.37556061 | 1.15041301 | 0.2499738 | 0.37938125 | NA | NA |
| ENSG000001 | 91.296772 | 0.193973875 | 0.16861814 | 1.15037372 | 0.24998998 | 0.3793916 | C1orf116 | 79098 |
| ENSG000002 | 5.27290029 | -0.43039773 | 0.37416428 | -1.1502908 | 0.25002411 | 0.37941501 | NA | NA |
| ENSG000002 | 2.55479507 | -0.426308749 | 0.37060332 | -1.1503101 | 0.25001617 | 0.37941501 | NA | NA |
| ENSG000002 | 3.59191886 | -0.435296601 | 0.37844982 | -1.1502096 | 0.25005757 | 0.37945157 | NA | NA |
| ENSG000002 | 5.21068103 | -0.42886234 | 0.37287862 | -1.1501393 | 0.25008652 | 0.37948131 | NA | NA |
| ENSG000002 | 3.12316412 | -0.432690332 | 0.37622292 | -1.1500903 | 0.25010668 | 0.37949318 | NA | NA |
| ENSG000002 | 8.21788388 | -0.409455498 | 0.3560251 | -1.1500748 | 0.25011306 | 0.37949318 | NA | NA |
| ENSG000001 | 631.464472 | 0.15197785 | 0.13215519 | 1.14999531 | 0.2501458 | 0.37952866 | PRPS2 | 5634 |
| ENSG000002 | 5.22677517 | -0.429882094 | 0.37382895 | -1.1499433 | 0.25016722 | 0.37953964 | NA | NA |
| ENSG000002 | 7.78177231 | -0.404844168 | 0.35205913 | -1.1499323 | 0.25017176 | 0.37953964 | NA | NA |
| ENSG00000C | 26.4312951 | -0.317488101 | 0.276123 | -1.1498068 | 0.25022345 | 0.37960192 | MYH7B | 57644 |
| ENSG000002 | 2.3205279 | -0.416943885 | 0.36262701 | -1.1497872 | 0.25023154 | 0.37960192 | NA | NA |
| ENSG000002 | 2.51927897 | -0.42679563 | 0.3712169 | -1.1497204 | 0.25025907 | 0.37962949 | NA | NA |
| ENSG000001 | 396.32883 | 0.150831991 | 0.13120581 | 1.14958321 | 0.25031558 | 0.37963056 | LNPEP | 4012 |
| ENSG000001 | 5.57939851 | -0.429581794 | 0.37365819 | -1.1496651 | 0.25028182 | 0.37963056 | CST11 | 140880 |
| ENSG000001 | 5.44783312 | -0.425644159 | 0.3702357 | -1.1496573 | 0.25028507 | 0.37963056 | EN2 | 2020 |
| ENSG000002 | 40.8501956 | 0.269344143 | 0.2342974 | 1.14958231 | 0.25031595 | 0.37963056 | TTC25 | 83538 |

| | | | | | | | | |
|---|---|---|---|---|---|---|---|---|
| ENSG000002 | 10.9529234 | 0.379562699 | 0.3301729 | 1.14958768 | 0.25031373 | 0.37963056 | NA | NA |
| ENSG000002 | 3.33804179 | -0.434695216 | 0.3781113 | -1.1496488 | 0.25028853 | 0.37963056 | NA | NA |
| ENSG000002 | 3.58953206 | -0.435179351 | 0.37857019 | -1.1495341 | 0.25033581 | 0.37963229 | NA | NA |
| ENSG000002 | 4.4893708 | -0.433972656 | 0.3775203 | -1.1495346 | 0.2503356 | 0.37963229 | NA | NA |
| ENSG000002 | 3.11046149 | -0.432327044 | 0.37614678 | -1.1493573 | 0.25040868 | 0.3797286 | NA | NA |
| ENSG000002 | 4.10439535 | -0.433307278 | 0.37702041 | -1.149294 | 0.25043479 | 0.37975399 | NA | NA |
| ENSG000002 | 6.83645082 | -0.416028414 | 0.36201225 | -1.1492109 | 0.25046904 | 0.37979173 | LOC102724153 | 102724153 |
| ENSG000002 | 8.38384307 | 0.403423087 | 0.35108123 | 1.14908759 | 0.25051986 | 0.37985458 | NA | NA |
| ENSG000001 | 4.21995138 | -0.434926738 | 0.37851449 | -1.1490359 | 0.25054118 | 0.3798585 | HOXA5 | 3202 |
| ENSG000001 | 18.119043 | -0.339944821 | 0.29584646 | -1.1490583 | 0.25053195 | 0.3798585 | IDUA | 3425 |
| ENSG000001 | 192.290661 | 0.343786715 | 0.29920617 | 1.14899608 | 0.2505576 | 0.37986918 | NAA60 | 79903 |
| ENSG000002 | 4.14786101 | -0.434688407 | 0.37833476 | -1.1489518 | 0.25057585 | 0.37988265 | NA | NA |
| ENSG000002 | 3.07106954 | -0.431565291 | 0.37565625 | -1.1488303 | 0.25062596 | 0.37994441 | RN7SL715P | 106479483 |
| ENSG000001 | 65.7063592 | 0.233011096 | 0.20286549 | 1.14859901 | 0.25072137 | 0.38007484 | KRT7 | 3855 |
| ENSG000002 | 7.36042205 | -0.416706198 | 0.36282018 | -1.1485199 | 0.250754 | 0.38011011 | NA | NA |
| ENSG000002 | 2.86347299 | 0.425598387 | 0.3705999 | 1.14840393 | 0.25080185 | 0.3801573 | NA | NA |
| ENSG000002 | 4.87260606 | -0.434018859 | 0.37793385 | -1.148399 | 0.25080388 | 0.3801573 | NA | NA |
| ENSG000002 | 4.25478322 | -0.434504299 | 0.37837112 | -1.1483548 | 0.25082212 | 0.3801658 | NA | NA |
| ENSG000002 | 7.20851424 | -0.419749639 | 0.36552732 | -1.14834 | 0.25082824 | 0.3801658 | NA | NA |
| ENSG000002 | 2.47709527 | -0.42734446 | 0.3721784 | -1.1482248 | 0.25087579 | 0.38022265 | NA | NA |
| ENSG000002 | 2.52314129 | -0.427350651 | 0.37219064 | -1.1482037 | 0.2508845 | 0.38022265 | NA | NA |
| ENSG000002 | 3.34557742 | -0.433921748 | 0.377923 | -1.148175 | 0.25089631 | 0.38022617 | NA | NA |
| ENSG000002 | 4.9608931 | -0.430243705 | 0.37472694 | -1.1481526 | 0.25090557 | 0.38022617 | NA | NA |
| ENSG000002 | 4.22152297 | -0.434413967 | 0.37849403 | -1.1477432 | 0.25107458 | 0.38046805 | DDX26B-AS1 | 100874118 |
| ENSG000002 | 23.5759193 | -0.308554488 | 0.2688733 | -1.1475832 | 0.25114065 | 0.38055396 | NA | NA |
| ENSG000002 | 3.63036061 | -0.434433778 | 0.3785855 | -1.1475183 | 0.25116749 | 0.3805804 | NA | NA |
| ENSG000001 | 12.100973 | -0.365440108 | 0.31848322 | -1.1474391 | 0.25120018 | 0.3805881 | OR10H1 | 26539 |
| ENSG000002 | 30.0365542 | 0.280718875 | 0.24464844 | 1.14743781 | 0.25120072 | 0.3805881 | PNMAL2 | 57469 |
| ENSG000002 | 4.39469281 | -0.433316758 | 0.37763053 | -1.1474622 | 0.25119065 | 0.3805881 | NA | NA |

| | | | | | | | | |
|---|---|---|---|---|---|---|---|---|
| ENSG000002 | 2.4859221 | -0.426803875 | 0.37197662 | -1.1473943 | 0.25121868 | 0.38060108 | MIR4514 | 100616181 |
| ENSG000002 | 2.48566096 | -0.426118578 | 0.37141726 | -1.1472773 | 0.25126705 | 0.38066015 | NA | NA |
| ENSG000001 | 11.576814 | -0.369291143 | 0.32190129 | -1.1472186 | 0.2512913 | 0.38068266 | LIPI | 149998 |
| ENSG000001 | 242.670682 | 0.151229033 | 0.13183789 | 1.14708322 | 0.25134723 | 0.38075316 | GRIP1 | 23426 |
| ENSG000002 | 3.39577173 | -0.432362566 | 0.37693579 | -1.1470457 | 0.25136275 | 0.38076245 | NA | NA |
| ENSG000002 | 2.86952727 | -0.427607735 | 0.37284718 | -1.1468713 | 0.25143482 | 0.38085739 | NA | NA |
| ENSG000001 | 53.1994287 | 0.232780805 | 0.20297819 | 1.1468267 | 0.25145325 | 0.38087109 | CECR6 | 27439 |
| ENSG000002 | 7.11425878 | -0.414081595 | 0.36112832 | -1.1466328 | 0.2515334 | 0.38097826 | NA | NA |
| ENSG000002 | 21.6536016 | -0.321215208 | 0.28016857 | -1.1465069 | 0.25158547 | 0.38102484 | NA | NA |
| ENSG000002 | 3.39496229 | -0.431298127 | 0.37618995 | -1.1464903 | 0.25159234 | 0.38102484 | NA | NA |
| ENSG000002 | 7.31889391 | -0.411796436 | 0.35917397 | -1.1465097 | 0.25158432 | 0.38102484 | LINC01050 | 103689845 |
| ENSG000002 | 4.62075675 | -0.43401996 | 0.3785903 | -1.1464107 | 0.25162527 | 0.38106047 | NA | NA |
| ENSG000002 | 2.26396526 | -0.417836499 | 0.36448906 | -1.1463622 | 0.2516453 | 0.38107658 | NA | NA |
| ENSG000002 | 2.79489511 | -0.429769153 | 0.37491566 | -1.1463089 | 0.25166736 | 0.38109575 | NA | NA |
| ENSG000002 | 39.3803686 | -0.255275114 | 0.2227201 | -1.1461701 | 0.25172479 | 0.38116849 | NA | NA |
| ENSG000001 | 10.6600466 | -0.379989919 | 0.33154032 | -1.1461349 | 0.25173936 | 0.38117631 | TP53AIP1 | 63970 |
| ENSG000001 | 4.66443948 | -0.430746444 | 0.37586579 | -1.1460113 | 0.25179047 | 0.38123947 | GATA5 | 140628 |
| ENSG000002 | 2.54084068 | -0.424427925 | 0.37037021 | -1.1459559 | 0.2518134 | 0.38125995 | NA | NA |
| ENSG000002 | 3.31441127 | -0.432720768 | 0.37763124 | -1.1458818 | 0.25184405 | 0.38127789 | NA | NA |
| ENSG000002 | 2.74138624 | -0.431202788 | 0.37629919 | -1.1459041 | 0.25183483 | 0.38127789 | NA | NA |
| ENSG000002 | 7.5690673 | -0.408272931 | 0.35637278 | -1.1456344 | 0.25194644 | 0.38141866 | NA | NA |
| ENSG000002 | 2.25193424 | -0.414708495 | 0.36200045 | -1.1456021 | 0.25195981 | 0.38142466 | NA | NA |
| ENSG000001 | 17.61394 | -0.355604072 | 0.31042048 | -1.1455561 | 0.25197888 | 0.38142505 | PACRG | 135138 |
| ENSG000002 | 11.1024706 | -0.376485331 | 0.32864299 | -1.1455754 | 0.25197087 | 0.38142505 | NA | NA |
| ENSG000002 | 2.50532424 | -0.425205846 | 0.37118928 | -1.145523 | 0.25199259 | 0.38143156 | NA | NA |
| ENSG000002 | 2.21078169 | -0.418746799 | 0.36556983 | -1.1454632 | 0.25201732 | 0.38145476 | NA | NA |
| ENSG000002 | 5.70417087 | -0.421729512 | 0.36820139 | -1.1453773 | 0.25205289 | 0.38149436 | NA | NA |
| ENSG000002 | 13.0089725 | 0.367728997 | 0.32106653 | 1.14533583 | 0.25207007 | 0.38150612 | NA | NA |
| ENSG000002 | 2.22538134 | -0.418163506 | 0.36511076 | -1.1453059 | 0.25208246 | 0.38151063 | RNU6-494P | 106479785 |

| | | | | | | | | |
|---|---|---|---|---|---|---|---|---|
| ENSG000002 | 4.98427568 | -0.429060746 | 0.37465755 | -1.1452078 | 0.2521231 | 0.38155789 | LOC100128108 | 100128108 |
| ENSG000002 | 3.81699558 | -0.433274183 | 0.37835851 | -1.1451419 | 0.25215038 | 0.38158495 | NA | NA |
| ENSG000002 | 2.46058902 | -0.425717521 | 0.37179067 | -1.1450463 | 0.25218999 | 0.38163064 | NA | NA |
| ENSG000002 | 2.76703143 | -0.430247054 | 0.37580464 | -1.1448689 | 0.25226346 | 0.38172757 | NA | NA |
| ENSG000002 | 2.26909895 | -0.417162003 | 0.36441084 | -1.1447574 | 0.25230967 | 0.38176902 | NA | NA |
| ENSG000002 | 2.62474183 | -0.418429639 | 0.36551746 | -1.1447597 | 0.25230873 | 0.38176902 | NA | NA |
| ENSG000002 | 2.49695945 | -0.425167667 | 0.37142968 | -1.1446788 | 0.25234225 | 0.38180406 | NA | NA |
| ENSG000002 | 2.28872977 | -0.417156028 | 0.36448658 | -1.1445031 | 0.25241504 | 0.3818857 | NA | NA |
| ENSG000002 | 12.6609988 | 0.370205625 | 0.32346059 | 1.14451538 | 0.25240996 | 0.3818857 | NA | NA |
| ENSG000001 | 40.6271342 | 0.259675037 | 0.22692363 | 1.14432786 | 0.25248769 | 0.38196711 | FAM110A | 83541 |
| ENSG000002 | 2.75997187 | -0.429359067 | 0.37520373 | -1.1443358 | 0.25248439 | 0.38196711 | NA | NA |
| ENSG000002 | 2.56452273 | -0.42394171 | 0.37057729 | -1.1440035 | 0.2526222 | 0.38215633 | NA | NA |
| ENSG000002 | 5.82727551 | -0.425547682 | 0.37198907 | -1.143979 | 0.25263234 | 0.38215743 | NA | NA |
| ENSG000002 | 23.7426067 | 0.320737621 | 0.28038191 | 1.14393122 | 0.25265216 | 0.38217315 | NA | NA |
| ENSG000001 | 5.46614607 | -0.42332201 | 0.37010926 | -1.1437758 | 0.25271663 | 0.38218994 | GATA1 | 2623 |
| ENSG000001 | 249.256007 | 0.167577927 | 0.14651327 | 1.14377307 | 0.25271776 | 0.38218994 | TANK | 10010 |
| ENSG000002 | 4.11763269 | -0.431361563 | 0.37711925 | -1.1438333 | 0.25269276 | 0.38218994 | NA | NA |
| ENSG000002 | 22.5046975 | 0.306010117 | 0.26753181 | 1.14382703 | 0.25269538 | 0.38218994 | NA | NA |
| ENSG000002 | 3.81884269 | -0.432559937 | 0.37818848 | -1.1437681 | 0.25271981 | 0.38218994 | NA | NA |
| ENSG000002 | 3.61431007 | -0.432709093 | 0.37831421 | -1.1437823 | 0.25271394 | 0.38218994 | NA | NA |
| ENSG000002 | 3.83423905 | -0.432710898 | 0.37834602 | -1.1436909 | 0.25275186 | 0.38220989 | NA | NA |
| ENSG000002 | 7.686286 | -0.413575606 | 0.36161024 | -1.1437055 | 0.25274581 | 0.38220989 | NA | NA |
| ENSG000002 | 3.62367354 | -0.432910922 | 0.37856507 | -1.1435575 | 0.2528072 | 0.38227933 | NA | NA |
| ENSG000001 | 2.23539716 | -0.418152499 | 0.3656779 | -1.1434995 | 0.25283125 | 0.38230144 | RNASE2 | 6036 |
| ENSG000001 | 621.975885 | -0.128684021 | 0.11254508 | -1.1433998 | 0.25287265 | 0.382347 | PPM1D | 8493 |
| ENSG000001 | 2.21299332 | -0.417988795 | 0.36557973 | -1.1433588 | 0.25288966 | 0.382347 | OR5W2 | 390148 |
| ENSG000002 | 3.15408646 | -0.42871911 | 0.37496391 | -1.143361 | 0.25288874 | 0.382347 | RN7SL342P | 106479349 |
| ENSG000002 | 2.53470724 | -0.422371782 | 0.36943399 | -1.1432943 | 0.25291642 | 0.38237319 | NA | NA |
| ENSG000001 | 10.4722693 | -0.382668725 | 0.33472449 | -1.1432349 | 0.25294107 | 0.3823962 | LOC101928372 | 101928372 |

| | | | | | | | | |
|---|---|---|---|---|---|---|---|---|
| ENSG000002 | 3.71398502 | -0.432450104 | 0.37829483 | -1.1431563 | 0.25297372 | 0.3824313 | NA | NA |
| ENSG000002 | 5.68524501 | -0.420531899 | 0.36790759 | -1.1430368 | 0.25302333 | 0.38249205 | NA | NA |
| ENSG000002 | 2.21179964 | -0.418795837 | 0.36639809 | -1.1430077 | 0.2530354 | 0.38249603 | NA | NA |
| ENSG000001 | 127.292418 | -0.186416792 | 0.1631457 | -1.14264 | 0.25318811 | 0.3827126 | ST8SIA3 | 51046 |
| ENSG000002 | 2.86315306 | -0.425191848 | 0.37214821 | -1.1425336 | 0.25323228 | 0.3827651 | NA | NA |
| ENSG000002 | 4.99892597 | -0.429665508 | 0.37607555 | -1.1424978 | 0.25324715 | 0.3827733 | NA | NA |
| ENSG000001 | 76.5470474 | 0.219396444 | 0.19204933 | 1.14239627 | 0.25328934 | 0.38278693 | SPRED3 | 399473 |
| ENSG000002 | 4.14620133 | -0.431225214 | 0.37748077 | -1.1423766 | 0.2532975 | 0.38278693 | NA | NA |
| ENSG000002 | 6.42787267 | 0.417000746 | 0.36501395 | 1.14242415 | 0.25327776 | 0.38278693 | NA | NA |
| ENSG000002 | 107.361546 | 0.182321976 | 0.15961032 | 1.14229439 | 0.25333167 | 0.38278693 | HLA-DPA1 | 3113 |
| ENSG000002 | 2.76127765 | -0.429284203 | 0.37580001 | -1.1423209 | 0.25332066 | 0.38278693 | RNU7-148P | 106480823 |
| ENSG000002 | 5.34555556 | -0.429352285 | 0.37586146 | -1.1423153 | 0.253323 | 0.38278693 | NA | NA |
| ENSG000002 | 5.53363206 | 0.42709126 | 0.37387839 | 1.14232669 | 0.25331825 | 0.38278693 | YTHDF3-AS1 | 101410533 |
| ENSG000002 | 3.32822981 | -0.431547394 | 0.37773865 | -1.1424497 | 0.25326716 | 0.38278693 | NA | NA |
| ENSG000002 | 32.0631107 | -0.284357621 | 0.24896325 | -1.1421671 | 0.25338458 | 0.3828526 | MEI4 | 101928601 |
| ENSG000001 | 2.76498376 | -0.429202158 | 0.37580465 | -1.1420885 | 0.25341724 | 0.38288769 | NA | NA |
| ENSG000002 | 2.51130614 | -0.424070397 | 0.37134979 | -1.1419702 | 0.2534664 | 0.38294769 | MIR4524B | 100847008 |
| ENSG000002 | 4.68146747 | -0.429284054 | 0.37600225 | -1.1417061 | 0.25357622 | 0.38309933 | NA | NA |
| ENSG000001 | 192.582392 | 0.182999975 | 0.16029236 | 1.14166372 | 0.25359383 | 0.38310426 | KIAA1841 | 84542 |
| ENSG000002 | 2.46585353 | -0.423980446 | 0.37137425 | -1.1416528 | 0.25359837 | 0.38310426 | LOC101927464 | 101927464 |
| ENSG00000C | 1681.40445 | 0.162846033 | 0.14265257 | 1.14155694 | 0.25363823 | 0.38312166 | ARID4B | 51742 |
| ENSG000002 | 2.48267279 | -0.424432562 | 0.37179192 | -1.1415863 | 0.25362602 | 0.38312166 | NA | NA |
| ENSG000002 | 5.19784897 | -0.425344586 | 0.37259821 | -1.1415637 | 0.25363543 | 0.38312166 | NA | NA |
| ENSG000002 | 3.86310109 | -0.432005036 | 0.37848392 | -1.1414092 | 0.25369967 | 0.38320018 | NA | NA |
| ENSG000002 | 2.2528158 | -0.414979542 | 0.36361738 | -1.1412533 | 0.25376452 | 0.38328386 | NA | NA |
| ENSG000002 | 7.05852976 | -0.412172266 | 0.36118752 | -1.1411587 | 0.2538039 | 0.38332907 | NA | NA |
| ENSG000002 | 7.55945653 | -0.406540472 | 0.35626939 | -1.1411041 | 0.2538266 | 0.38334906 | NA | NA |
| ENSG000002 | 2.22003306 | -0.417554137 | 0.3659601 | -1.1409827 | 0.25387713 | 0.38341111 | MIR1913 | 100302141 |
| ENSG000001 | 1182.93411 | 0.158613918 | 0.13901924 | 1.14094938 | 0.25389099 | 0.38341775 | POGZ | 23126 |

| | | | | | | | | | |
|---|---|---|---|---|---|---|---|---|---|
| ENSG000001 | 59.5637487 | -0.227839513 | 0.19970255 | -1.1408943 | 0.2539139 | 0.38343508 | TBC1D9 | | 23158 |
| ENSG000002 | 2.77188614 | 0.428155592 | 0.37528658 | 1.14087636 | 0.25392138 | 0.38343508 | NA | NA | |
| ENSG000002 | 2.50955849 | -0.423762593 | 0.37145903 | -1.1408058 | 0.25395077 | 0.38346519 | NA | NA | |
| ENSG000002 | 2.5100604 | -0.423036764 | 0.37083474 | -1.140769 | 0.25396608 | 0.38347403 | NA | NA | |
| ENSG000002 | 2.48397856 | -0.425153883 | 0.3727159 | -1.1406916 | 0.2539983 | 0.38350839 | NA | NA | |
| ENSG000002 | 5.43843773 | -0.423643017 | 0.37140585 | -1.1406471 | 0.25401679 | 0.38352204 | NA | NA | |
| ENSG000001 | 2.47877822 | -0.423292795 | 0.37113487 | -1.1405363 | 0.25406294 | 0.38357743 | C9orf50 | | 375759 |
| ENSG000001 | 46.3707046 | 0.252057495 | 0.22101653 | 1.14044635 | 0.25410039 | 0.38360783 | KCNN1 | | 3780 |
| ENSG000002 | 16.1915547 | 0.341169528 | 0.29916135 | 1.14041979 | 0.25411145 | 0.38360783 | NA | NA | |
| ENSG000002 | 10.8628038 | -0.375237137 | 0.32902897 | -1.140438 | 0.25410388 | 0.38360783 | NA | NA | |
| ENSG000002 | 9.43118433 | 0.396823669 | 0.34803829 | 1.14017245 | 0.25421446 | 0.38374905 | NA | NA | |
| ENSG000002 | 5.64101105 | -0.428411603 | 0.3757604 | -1.1401191 | 0.2542367 | 0.38376833 | NA | NA | |
| ENSG000002 | 3.8827319 | -0.431290747 | 0.37837366 | -1.139854 | 0.25434712 | 0.38392071 | NA | NA | |
| ENSG000002 | 5.70439975 | -0.419158422 | 0.36781791 | -1.1395813 | 0.25446077 | 0.38407075 | NA | NA | |
| ENSG000002 | 5.50078856 | -0.423174321 | 0.37134559 | -1.1395701 | 0.25446546 | 0.38407075 | NA | NA | |
| ENSG000002 | 5.49701166 | -0.422456268 | 0.37073895 | -1.1394979 | 0.25449553 | 0.38410184 | MIR542 | | 664617 |
| ENSG000002 | 3.63316534 | -0.43127491 | 0.37850674 | -1.1394114 | 0.25453159 | 0.38414197 | NA | NA | |
| ENSG000001 | 310.412438 | 0.146870999 | 0.12890767 | 1.13935034 | 0.25455706 | 0.38416381 | STK33 | | 65975 |
| ENSG000002 | 4.14822918 | -0.430386446 | 0.3777611 | -1.1393085 | 0.25457449 | 0.38416381 | NA | NA | |
| ENSG000002 | 5.43615507 | -0.422087954 | 0.37047721 | -1.1393088 | 0.25457437 | 0.38416381 | NA | NA | |
| ENSG000002 | 2.55166996 | -0.419838499 | 0.3685443 | -1.1391806 | 0.25462785 | 0.38423004 | GPR33 | | 2856 |
| ENSG00000C | 181.257009 | -0.162454008 | 0.14261785 | -1.1390861 | 0.25466724 | 0.38423392 | DPF1 | | 8193 |
| ENSG000001 | 3.56669953 | -0.431263696 | 0.37859326 | -1.1391214 | 0.25465251 | 0.38423392 | NA | NA | |
| ENSG000002 | 5.67102306 | -0.418514188 | 0.36740424 | -1.1391109 | 0.25465689 | 0.38423392 | NA | NA | |
| ENSG000002 | 4.98944323 | -0.426547506 | 0.37446552 | -1.1390835 | 0.25466832 | 0.38423392 | NA | NA | |
| ENSG000002 | 2.2677559 | -0.41425147 | 0.36368611 | -1.1390357 | 0.25468825 | 0.3842497 | NA | NA | |
| ENSG000002 | 9.79210455 | -0.398868352 | 0.3501904 | -1.1390042 | 0.25470139 | 0.38425521 | NA | NA | |
| ENSG000002 | 5.71840656 | -0.41918883 | 0.36811323 | -1.1387497 | 0.25480757 | 0.3844011 | NA | NA | |
| ENSG000002 | 4.99254056 | -0.427102681 | 0.37507907 | -1.1387004 | 0.25482815 | 0.38440355 | NA | NA | |

| | | | | | | | | | |
|---|---|---|---|---|---|---|---|---|---|
| ENSG000002 | 2.47435505 | -0.423932132 | 0.37229293 | -1.1387058 | 0.25482586 | 0.38440355 | NA | NA | |
| ENSG000002 | 6.16981204 | -0.399863869 | 0.35125156 | -1.1383974 | 0.25495457 | 0.38457995 | NA | NA | |
| ENSG000002 | 5.76670985 | -0.42164413 | 0.37040934 | -1.1383194 | 0.25498714 | 0.38461477 | GOLGA2P9 | | 440518 |
| ENSG000001 | 169.945171 | 0.172241914 | 0.1513233 | 1.13823791 | 0.25502115 | 0.38465177 | ZNF527 | | 84503 |
| ENSG000002 | 7.62620389 | 0.415897862 | 0.3654111 | 1.13816429 | 0.25505189 | 0.38468382 | MIR3140 | | 100422896 |
| ENSG000002 | 18.2434525 | -0.328049058 | 0.28823549 | -1.1381286 | 0.25506678 | 0.38469197 | NA | NA | |
| ENSG000002 | 2.5098717 | -0.42318867 | 0.37184731 | -1.1380711 | 0.2550908 | 0.38469958 | NA | NA | |
| ENSG000002 | 5.50523189 | -0.422689129 | 0.37140725 | -1.1380745 | 0.25508936 | 0.38469958 | NA | NA | |
| ENSG000002 | 3.95121688 | -0.430667153 | 0.37844523 | -1.1379907 | 0.25512437 | 0.38473589 | NA | NA | |
| ENSG000002 | 2.22767173 | -0.414589123 | 0.36433103 | -1.1379462 | 0.25514293 | 0.38474959 | NA | NA | |
| ENSG000001 | 7.32867599 | -0.41327 | 0.36319146 | -1.1378847 | 0.25516864 | 0.38477403 | WDR88 | | 126248 |
| ENSG000001 | 55.5327346 | 0.244186876 | 0.21462389 | 1.13774321 | 0.25522773 | 0.38484883 | PLXNA3 | | 55558 |
| ENSG00000C | 6009.56208 | 0.136015658 | 0.11956801 | 1.13755889 | 0.25530472 | 0.38492359 | ASPM | | 259266 |
| ENSG000002 | 3.86488761 | -0.430557897 | 0.37848531 | -1.1375815 | 0.25529526 | 0.38492359 | NA | NA | |
| ENSG000002 | 4.13229993 | -0.430608856 | 0.37853848 | -1.1375564 | 0.25530578 | 0.38492359 | NA | NA | |
| ENSG000002 | 2.72180749 | -0.427676975 | 0.37598562 | -1.1374823 | 0.25533673 | 0.38495594 | NA | NA | |
| ENSG00000C | 137.391744 | 0.17797849 | 0.15649248 | 1.13729742 | 0.25541397 | 0.38501568 | MKNK1 | | 8569 |
| ENSG000001 | 14.3377675 | -0.366103374 | 0.32190522 | -1.1373017 | 0.25541216 | 0.38501568 | SCNN1D | | 6339 |
| ENSG000002 | 7.83961902 | 0.402702606 | 0.3540876 | 1.13729654 | 0.25541434 | 0.38501568 | NA | NA | |
| ENSG000002 | 2.73504483 | -0.427758712 | 0.37609866 | -1.1373577 | 0.25538878 | 0.38501568 | NA | NA | |
| ENSG000002 | 34.594673 | 0.265068924 | 0.23309682 | 1.13716235 | 0.25547042 | 0.38508591 | NA | NA | |
| ENSG000001 | 20.1050604 | -0.316662848 | 0.27848254 | -1.1371013 | 0.25549596 | 0.38511009 | CFAP53 | | 220136 |
| ENSG000002 | 3.85783656 | -0.430233826 | 0.37836892 | -1.137075 | 0.25550695 | 0.38511234 | NA | NA | |
| ENSG000002 | 2.7497238 | -0.42732512 | 0.37584042 | -1.1369855 | 0.25554433 | 0.38515436 | NA | NA | |
| ENSG000002 | 13.9015675 | 0.358721079 | 0.31552231 | 1.13691194 | 0.2555751 | 0.38517211 | NA | NA | |
| ENSG000002 | 2.30227454 | -0.413075058 | 0.36332522 | -1.1369292 | 0.25556788 | 0.38517211 | NA | NA | |
| ENSG000002 | 3.40133814 | -0.428967361 | 0.37734948 | -1.1367906 | 0.25562582 | 0.38520867 | NA | NA | |
| ENSG000002 | 10.3467457 | -0.38214811 | 0.33615436 | -1.1368233 | 0.25561216 | 0.38520867 | NA | NA | |
| ENSG000002 | 6.86258052 | -0.418656851 | 0.36828123 | -1.1367858 | 0.25562786 | 0.38520867 | NA | NA | |

| | | | | | | | | |
|---|---|---|---|---|---|---|---|---|
| ENSG000002 | 7.86249629 | -0.409907845 | 0.3605932 | -1.1367598 | 0.25563872 | 0.38521072 | NA | NA |
| ENSG000000 | 115.768973 | 0.190664364 | 0.16774349 | 1.13664244 | 0.25568779 | 0.38524172 | SEMA5B | 54437 |
| ENSG000001 | 43.6038989 | -0.255161938 | 0.2244817 | -1.1366715 | 0.25567565 | 0.38524172 | DPY19L2P2 | 349152 |
| ENSG000002 | 2.50743173 | -0.422600988 | 0.37179617 | -1.136647 | 0.2556859 | 0.38524172 | NA | NA |
| ENSG000001 | 2.49394645 | -0.421069336 | 0.37048384 | -1.136539 | 0.25573105 | 0.38529258 | NAT8 | 9027 |
| ENSG000002 | 5.44000465 | -0.421135136 | 0.37054917 | -1.1365162 | 0.25574059 | 0.38529264 | NA | NA |
| ENSG000002 | 4.14316054 | -0.429327849 | 0.37776634 | -1.1364905 | 0.25575134 | 0.38529452 | C10orf113 | 387638 |
| ENSG000002 | 4.93432006 | -0.426596605 | 0.37537391 | -1.1364578 | 0.25576501 | 0.38530081 | NA | NA |
| ENSG000002 | 6.59341303 | -0.416815513 | 0.36679143 | -1.1363829 | 0.25579635 | 0.38533369 | NA | NA |
| ENSG000001 | 269.076028 | 0.350693679 | 0.30862241 | 1.13631956 | 0.25582285 | 0.3853593 | TCF7L1 | 83439 |
| ENSG000002 | 2.78675453 | -0.426626607 | 0.37554076 | -1.1360328 | 0.25594285 | 0.38551143 | OR10G8 | 219869 |
| ENSG000002 | 2.50604163 | -0.42168691 | 0.37118719 | -1.1360492 | 0.25593598 | 0.38551143 | NA | NA |
| ENSG000002 | 8.02871175 | -0.396881862 | 0.34939954 | -1.1358969 | 0.25599971 | 0.38558275 | NA | NA |
| ENSG000002 | 2.77688252 | -0.425287747 | 0.37443494 | -1.1358121 | 0.2560352 | 0.38560755 | NA | NA |
| ENSG000002 | 7.2611524 | 0.407530472 | 0.3588008 | 1.13581259 | 0.25603501 | 0.38560755 | NA | NA |
| ENSG000002 | 3.34059778 | -0.428706367 | 0.37747856 | -1.1357105 | 0.25607774 | 0.38564298 | NA | NA |
| ENSG000002 | 3.85054943 | -0.42958592 | 0.37825225 | -1.1357128 | 0.25607678 | 0.38564298 | NA | NA |
| ENSG000002 | 5.23566954 | -0.423729855 | 0.37312751 | -1.1356168 | 0.25611699 | 0.38568776 | OR6C1 | 390321 |
| ENSG000001 | 33.2091431 | 0.278808165 | 0.2455469 | 1.13545789 | 0.25618352 | 0.38577362 | KCNS1 | 3787 |
| ENSG000001 | 3.67542779 | -0.429416246 | 0.37825718 | -1.1352494 | 0.25627083 | 0.38580534 | OR1M1 | 125963 |
| ENSG000001 | 7.01032528 | -0.409124524 | 0.36034826 | -1.1353587 | 0.25622507 | 0.38580534 | TBX1 | 6899 |
| ENSG000002 | 3.5897483 | -0.429830464 | 0.37859054 | -1.1353439 | 0.25623125 | 0.38580534 | NA | NA |
| ENSG000002 | 3.30653127 | -0.429124167 | 0.3779886 | -1.1352834 | 0.25625661 | 0.38580534 | NA | NA |
| ENSG000002 | 11.6453041 | -0.367846173 | 0.3240219 | -1.135251 | 0.25627018 | 0.38580534 | NA | NA |
| ENSG000002 | 16.5162449 | -0.340241868 | 0.29969846 | -1.1352807 | 0.25625774 | 0.38580534 | NA | NA |
| ENSG000002 | 8.10249649 | 0.397510143 | 0.35015251 | 1.13524858 | 0.25627118 | 0.38580534 | NA | NA |
| ENSG000002 | 7.47198268 | -0.4131439 | 0.36393731 | -1.1352062 | 0.25628892 | 0.38581491 | NA | NA |
| ENSG000002 | 9.56393819 | 0.387773939 | 0.34159447 | 1.13518798 | 0.25629656 | 0.38581491 | NA | NA |
| ENSG000002 | 2.82173828 | -0.422411564 | 0.37211971 | -1.1351497 | 0.25631262 | 0.38582475 | NA | NA |

| | | | | | | | | |
|---|---|---|---|---|---|---|---|---|
| ENSG00000( | 59.4379619 | 0.223539143 | 0.19692979 | 1.13512099 | 0.25632463 | 0.38582851 | IPCEF1 | 26034 |
| ENSG000002 | 52.5520484 | 0.248477321 | 0.21892294 | 1.134999 | 0.25637574 | 0.38589111 | NA | NA |
| ENSG000002 | 6.0211146 | -0.419742146 | 0.36984027 | -1.1349282 | 0.25640541 | 0.38592145 | NA | NA |
| ENSG000001 | 132.331377 | 0.205066821 | 0.18069914 | 1.13485225 | 0.25643723 | 0.38595502 | UBALD2 | 283991 |
| ENSG000002 | 13.7106027 | 0.354315594 | 0.31226488 | 1.13466361 | 0.25651629 | 0.38605968 | NA | NA |
| ENSG000002 | 4.9085109 | -0.425626797 | 0.37517617 | -1.1344718 | 0.25659668 | 0.38616634 | NREP-AS1 | 100873948 |
| ENSG000002 | 5.2441509 | -0.422824816 | 0.37272614 | -1.1344115 | 0.25662199 | 0.38619009 | NA | NA |
| ENSG00000( | 49.0067331 | -0.253673896 | 0.2236268 | -1.1343627 | 0.25664246 | 0.38620656 | TRPC5 | 7224 |
| ENSG000001 | 4.21375433 | -0.429256512 | 0.37849275 | -1.1341208 | 0.25674386 | 0.38634483 | NDP | 4693 |
| ENSG000002 | 9.16198922 | -0.39947262 | 0.35226321 | -1.1340174 | 0.25678724 | 0.38639576 | NA | NA |
| ENSG000002 | 3.83232743 | -0.429157438 | 0.37846397 | -1.1339453 | 0.25681749 | 0.38642694 | NA | NA |
| ENSG000001 | 11.2775418 | 0.367664018 | 0.32426968 | 1.13382177 | 0.25686932 | 0.38649058 | PYY | 5697 |
| ENSG000002 | 5.46635514 | -0.420729607 | 0.37110601 | -1.1337181 | 0.2569128 | 0.38652733 | LOC105369736 | 105369736 |
| ENSG000002 | 3.64521618 | -0.429173678 | 0.37854847 | -1.1337351 | 0.25690569 | 0.38652733 | NA | NA |
| ENSG000002 | 5.4245486 | -0.420205349 | 0.37065166 | -1.1336934 | 0.25692317 | 0.38652859 | NA | NA |
| ENSG000001 | 18.4136196 | -0.32625894 | 0.28781985 | -1.1335526 | 0.25698226 | 0.38660314 | CPLX4 | 339302 |
| ENSG000002 | 10.2151786 | -0.385505362 | 0.34009989 | -1.1335063 | 0.2570017 | 0.38661805 | NA | NA |
| ENSG000001 | 64.4740598 | 0.225103145 | 0.19861994 | 1.13333609 | 0.25707314 | 0.38670382 | NA | NA |
| ENSG000002 | 6.76042601 | -0.411038877 | 0.36268403 | -1.133325 | 0.25707779 | 0.38670382 | LOC102546226 | 102546226 |
| ENSG000001 | 25.5425722 | -0.306266697 | 0.27026058 | -1.1332274 | 0.25711877 | 0.38675111 | ENO4 | 387712 |
| ENSG000002 | 66.5610847 | 0.230922667 | 0.20377928 | 1.13319992 | 0.25713031 | 0.38675412 | NA | NA |
| ENSG000002 | 7.05688779 | -0.40882598 | 0.36080207 | -1.1331032 | 0.25717092 | 0.38680085 | NA | NA |
| ENSG000002 | 3.09043977 | -0.425357842 | 0.37542516 | -1.133003 | 0.25721299 | 0.38683544 | NA | NA |
| ENSG000002 | 3.1782397 | -0.424859988 | 0.37498267 | -1.1330123 | 0.25720907 | 0.38683544 | LOC101928567 | 101928567 |
| ENSG000002 | 2.46588633 | -0.420947324 | 0.37155444 | -1.1329358 | 0.25724121 | 0.38686354 | RNU6-98P | 106479609 |
| ENSG00000( | 9.23874301 | -0.394230404 | 0.34802108 | -1.1327774 | 0.25730776 | 0.38689187 | SPO11 | 23626 |
| ENSG000001 | 7.01517689 | -0.411007963 | 0.36280796 | -1.1328527 | 0.25727613 | 0.38689187 | KLF2 | 10365 |
| ENSG000001 | 56.8815159 | -0.22997008 | 0.20301402 | -1.1327793 | 0.25730695 | 0.38689187 | FRMPD4 | 9758 |
| ENSG000002 | 2.74798878 | -0.425936322 | 0.37600749 | -1.1327868 | 0.2573038 | 0.38689187 | NA | NA |

| | | | | | | | | |
|---|---|---|---|---|---|---|---|---|
| ENSG000002 | 2.78015199 | -0.424984716 | 0.37516629 | -1.1327903 | 0.25730234 | 0.38689187 | NA | NA |
| ENSG000002 | 5.16705605 | -0.422267132 | 0.37279614 | -1.1327025 | 0.2573392 | 0.3869248 | NA | NA |
| ENSG000002 | 64.7657028 | 0.213543647 | 0.18853027 | 1.13267566 | 0.25735048 | 0.38692742 | NA | NA |
| ENSG000002 | 4.81317978 | -0.427485341 | 0.37742318 | -1.132642 | 0.25736463 | 0.38693434 | NA | NA |
| ENSG000002 | 4.93884323 | -0.424053843 | 0.37440515 | -1.1326069 | 0.25737939 | 0.38694219 | NA | NA |
| ENSG000002 | 5.35935233 | 0.425835758 | 0.37599322 | 1.13256233 | 0.2573981 | 0.38695596 | NA | NA |
| ENSG000002 | 5.22530881 | -0.422301414 | 0.37288161 | -1.1325348 | 0.25740965 | 0.38695898 | NA | NA |
| ENSG000002 | 4.70235668 | -0.42673496 | 0.37684539 | -1.1323874 | 0.25747161 | 0.38703779 | NA | NA |
| ENSG000002 | 3.82428381 | -0.428308043 | 0.37826734 | -1.1322893 | 0.25751285 | 0.38708542 | NA | NA |
| ENSG000002 | 2.29041271 | -0.410383894 | 0.3625012 | -1.1320898 | 0.2575967 | 0.38719711 | NA | NA |
| ENSG000002 | 5.01275259 | -0.424756091 | 0.37520473 | -1.1320649 | 0.25760717 | 0.3871985 | HMBOX1-IT1 | 106481795 |
| ENSG000002 | 73.5200384 | 0.208648006 | 0.18431308 | 1.13203039 | 0.25762166 | 0.38720593 | NA | NA |
| ENSG000002 | 4.23624857 | -0.428528637 | 0.37856117 | -1.1319931 | 0.25763733 | 0.38721512 | LINC00563 | 100861554 |
| ENSG000001 | 171.316803 | 0.164306334 | 0.1451691 | 1.13182717 | 0.2577071 | 0.38729128 | ZNF530 | 348327 |
| ENSG000002 | 5.53074833 | -0.424749174 | 0.37527685 | -1.1318289 | 0.25770639 | 0.38729128 | NA | NA |
| ENSG000002 | 5.22332586 | -0.422342167 | 0.37319826 | -1.1316831 | 0.25776768 | 0.38736757 | NA | NA |
| ENSG000002 | 42.3898975 | -0.256579235 | 0.22672799 | -1.131661 | 0.25777697 | 0.38736757 | RRS1-AS1 | 100505676 |
| ENSG000002 | 6.02127426 | -0.416037779 | 0.36764279 | -1.1316359 | 0.25778755 | 0.38736911 | NA | NA |
| ENSG000002 | 2.49033207 | -0.418752515 | 0.37008212 | -1.1315124 | 0.25783947 | 0.38743278 | NA | NA |
| ENSG000002 | 7.08694798 | -0.411699302 | 0.36387367 | -1.1314347 | 0.25787217 | 0.38746218 | NA | NA |
| ENSG000002 | 7.28685524 | -0.405756713 | 0.35862592 | -1.1314205 | 0.25787815 | 0.38746218 | NA | NA |
| ENSG000002 | 15.2186154 | -0.348713399 | 0.30822457 | -1.1313615 | 0.25790298 | 0.38747093 | NA | NA |
| ENSG000002 | 2.49860546 | -0.420644144 | 0.37180357 | -1.1313612 | 0.25790308 | 0.38747093 | RN7SL461P | 106481844 |
| ENSG000001 | 13.0706919 | -0.369594525 | 0.32670957 | -1.1312633 | 0.25794431 | 0.38750407 | TMPRSS9 | 360200 |
| ENSG000002 | 41.2820579 | 0.251526188 | 0.22234542 | 1.13124068 | 0.25795381 | 0.38750407 | PAXIP1-AS1 | 202781 |
| ENSG000002 | 14.9739568 | -0.35326591 | 0.31227695 | -1.1312584 | 0.25794636 | 0.38750407 | NA | NA |
| ENSG000002 | 2.50230361 | -0.419859451 | 0.37118023 | -1.1311471 | 0.25799317 | 0.38754885 | RNU6-463P | 106481325 |
| ENSG000001 | 7.09283291 | -0.410103614 | 0.36259143 | -1.1310351 | 0.25804034 | 0.38759098 | CYP11B1 | 1584 |
| ENSG000002 | 2.76171429 | -0.423565592 | 0.37448669 | -1.1310565 | 0.25803133 | 0.38759098 | NA | NA |

| | | | | | | | | |
|---|---|---|---|---|---|---|---|---|
| ENSG000002 | 5.70808605 | -0.417144912 | 0.36884056 | -1.1309627 | 0.25807079 | 0.38762237 | NA | NA |
| ENSG000002 | 2.23160618 | -0.411104987 | 0.36352092 | -1.1308978 | 0.25809813 | 0.38764907 | NA | NA |
| ENSG000002 | 56.12827 | -0.223040457 | 0.19723552 | -1.1308331 | 0.25812535 | 0.38766935 | BDNF-AS | 497258 |
| ENSG000002 | 3.86647359 | -0.428089217 | 0.37856522 | -1.1308203 | 0.25813075 | 0.38766935 | NA | NA |
| ENSG000002 | 4.66753265 | -0.427743586 | 0.37831405 | -1.1306574 | 0.25819931 | 0.38775795 | NA | NA |
| ENSG000001 | 6.99944916 | -0.407856857 | 0.36076774 | -1.1305248 | 0.25825517 | 0.38782748 | BCL2A1 | 597 |
| ENSG000001 | 45.7417412 | 0.24795492 | 0.21934308 | 1.13044332 | 0.25828947 | 0.38785026 | RAB42 | 115273 |
| ENSG000002 | 3.58959209 | -0.42789589 | 0.37851658 | -1.1304548 | 0.25828463 | 0.38785026 | NA | NA |
| ENSG000002 | 5.15681277 | 0.423178061 | 0.37438821 | 1.13031887 | 0.25834189 | 0.38791461 | RN7SL378P | 106481026 |
| ENSG000002 | 2.73330927 | -0.424342336 | 0.37543042 | -1.1302822 | 0.25835732 | 0.38792342 | NA | NA |
| ENSG000001 | 34.264281 | -0.263413304 | 0.23308744 | -1.1301051 | 0.25843195 | 0.38800674 | PRORSD1P | 344405 |
| ENSG000002 | 2.48502844 | -0.419881895 | 0.37153894 | -1.1301154 | 0.25842758 | 0.38800674 | NA | NA |
| ENSG000002 | 3.5608659 | -0.42772433 | 0.37853101 | -1.1299585 | 0.25849371 | 0.3880851 | NA | NA |
| ENSG000001 | 44.6130824 | -0.246345062 | 0.21805208 | -1.1297533 | 0.25858019 | 0.38814307 | MTTP | 4547 |
| ENSG000002 | 10.1393255 | -0.381616986 | 0.33778245 | -1.1297715 | 0.25857251 | 0.38814307 | NA | NA |
| ENSG000002 | 7.54329921 | -0.40027406 | 0.35428811 | -1.1297982 | 0.25856128 | 0.38814307 | NA | NA |
| ENSG000002 | 2.46548084 | -0.417358255 | 0.36941165 | -1.1297918 | 0.25856396 | 0.38814307 | NA | NA |
| ENSG000002 | 16.1237714 | -0.348545851 | 0.30849898 | -1.129812 | 0.25855545 | 0.38814307 | NA | NA |
| ENSG000001 | 3.29423966 | -0.42659663 | 0.37763494 | -1.1296535 | 0.25862226 | 0.38814876 | SSX7 | 280658 |
| ENSG000002 | 3.02985641 | -0.423872269 | 0.3752087 | -1.1296973 | 0.25860378 | 0.38814876 | PRB4 | 5545 |
| ENSG000002 | 5.50445914 | -0.422014478 | 0.37357216 | -1.1296733 | 0.25861392 | 0.38814876 | LOC105369486 | 105369486 |
| ENSG000002 | 3.39395872 | -0.426351459 | 0.37741702 | -1.1296562 | 0.25862114 | 0.38814876 | NA | NA |
| ENSG000001 | 3.16122436 | -0.422797264 | 0.37429177 | -1.1295927 | 0.25864787 | 0.38817283 | TPSG1 | 25823 |
| ENSG000002 | 5.29543956 | -0.427064377 | 0.3781221 | -1.1294351 | 0.25871432 | 0.38825819 | NA | NA |
| ENSG000002 | 3.96076363 | 0.426995563 | 0.37809344 | 1.12933873 | 0.25875497 | 0.38830482 | NA | NA |
| ENSG000002 | 2.49761289 | -0.419133112 | 0.37117505 | -1.129206 | 0.25881092 | 0.38837441 | LOC101929395 | 101929395 |
| ENSG000001 | 8.82304611 | -0.388409307 | 0.34400658 | -1.1290752 | 0.2588661 | 0.38842846 | FMOD | 2331 |
| ENSG000002 | 2.76029194 | -0.422641697 | 0.37432532 | -1.1290759 | 0.25886582 | 0.38842846 | NA | NA |
| ENSG000002 | 14.6400228 | -0.357261423 | 0.3164403 | -1.129001 | 0.2588974 | 0.38846106 | GMNC | 647309 |

| | | | | | | | | |
|---|---|---|---|---|---|---|---|---|
| ENSG000002 | 11.2165016 | -0.378522784 | 0.33533751 | -1.1287815 | 0.25899002 | 0.38858564 | NA | NA |
| ENSG000002 | 10.3402942 | -0.379700583 | 0.33639535 | -1.1287331 | 0.25901043 | 0.38860189 | LOC101929002 | 101929002 |
| ENSG000002 | 2.75962696 | -0.423549735 | 0.37527573 | -1.1286361 | 0.25905138 | 0.38864895 | NA | NA |
| ENSG000002 | 2.21474042 | -0.408737572 | 0.3621777 | -1.1285553 | 0.25908547 | 0.38868571 | NA | NA |
| ENSG000001 | 4.09176755 | -0.371540987 | 0.32923609 | -1.1284941 | 0.2591113 | 0.38871009 | C9orf171 | 389799 |
| ENSG000002 | 93.1677781 | 0.19457148 | 0.17242065 | 1.1284697 | 0.25912161 | 0.38871117 | NA | NA |
| ENSG000001 | 82.3165242 | 0.193188305 | 0.17121907 | 1.12831067 | 0.25918874 | 0.38877568 | P2RX4 | 5025 |
| ENSG000002 | 5.25678688 | -0.422642221 | 0.37457812 | -1.1283153 | 0.25918678 | 0.38877568 | NA | NA |
| ENSG000002 | 4.57523952 | -0.426918692 | 0.3783811 | -1.128277 | 0.25920296 | 0.38877568 | NA | NA |
| ENSG000002 | 4.56523604 | 0.423731805 | 0.37555201 | 1.1282906 | 0.25919721 | 0.38877568 | NA | NA |
| ENSG000002 | 5.76874167 | -0.421659554 | 0.37374962 | -1.1281873 | 0.25924084 | 0.38881812 | BNIP3P16 | 106480418 |
| ENSG000002 | 2.7979687 | -0.422928376 | 0.37492232 | -1.1280427 | 0.2593019 | 0.3888953 | NA | NA |
| ENSG000002 | 23.3223198 | -0.300312002 | 0.26623136 | -1.1280114 | 0.25931512 | 0.38890076 | NA | NA |
| ENSG000002 | 9.05868011 | -0.389306705 | 0.34517028 | -1.1278686 | 0.25937544 | 0.38897283 | NA | NA |
| ENSG000002 | 5.46341849 | -0.419335421 | 0.37179999 | -1.1278522 | 0.25938236 | 0.38897283 | NA | NA |
| ENSG000001 | 121.281112 | 0.185452615 | 0.1644607 | 1.12764095 | 0.25947158 | 0.38909224 | PACS1 | 55690 |
| ENSG000002 | 2.50230306 | -0.41672867 | 0.36958995 | -1.1275433 | 0.25951285 | 0.38913972 | NA | NA |
| ENSG000001 | 539.456909 | 0.146835143 | 0.13024637 | 1.12736456 | 0.25958837 | 0.38923684 | SESTD1 | 91404 |
| ENSG000002 | 10.9407909 | -0.376923098 | 0.33434595 | -1.1273446 | 0.25959681 | 0.38923684 | KRT40 | 125115 |
| ENSG000002 | 2.52163462 | -0.417352429 | 0.37022108 | -1.127306 | 0.25961314 | 0.38924693 | NA | NA |
| ENSG000002 | 7.64922254 | -0.403421132 | 0.35787624 | -1.1272644 | 0.25963073 | 0.38925295 | NA | NA |
| ENSG000002 | 34.4452041 | -0.271532664 | 0.24088038 | -1.1272511 | 0.25963635 | 0.38925295 | NA | NA |
| ENSG000002 | 2.18675941 | -0.409542422 | 0.36332656 | -1.127202 | 0.25965708 | 0.38926964 | NA | NA |
| ENSG000002 | 5.95423988 | -0.412776421 | 0.36621526 | -1.1271415 | 0.25968266 | 0.38929358 | NA | NA |
| ENSG000001 | 5.47192763 | -0.417624299 | 0.37053929 | -1.1270716 | 0.25971222 | 0.38932351 | NA | NA |
| ENSG000002 | 13.6123283 | -0.349459597 | 0.31009257 | -1.1269525 | 0.25976257 | 0.3893702 | LOC100506688 | 100506688 |
| ENSG000002 | 7.78579959 | -0.397142747 | 0.35240024 | -1.126965 | 0.25975726 | 0.3893702 | NA | NA |
| ENSG000002 | 2.47570572 | -0.418676336 | 0.3715196 | -1.1269293 | 0.25977236 | 0.38937048 | NA | NA |
| ENSG000002 | 2.50079693 | -0.418447509 | 0.37133177 | -1.1268831 | 0.25979189 | 0.38938536 | NA | NA |

| | | | | | | | | |
|---|---|---|---|---|---|---|---|---|
| ENSG000002 | 8.02797487 | -0.393955764 | 0.34963402 | -1.1267661 | 0.25984138 | 0.38944515 | NA | NA |
| ENSG000002 | 2.75155109 | -0.423418301 | 0.37579746 | -1.1267194 | 0.25986112 | 0.38946033 | NA | NA |
| ENSG000002 | 4.13858138 | -0.425584374 | 0.37782278 | -1.1264127 | 0.25999088 | 0.38961562 | LOC153684 | 153684 |
| ENSG000002 | 2.21835012 | -0.411837618 | 0.36560735 | -1.1264479 | 0.25997599 | 0.38961562 | NA | NA |
| ENSG000002 | 64.378722 | 0.214835214 | 0.19072913 | 1.12638911 | 0.26000085 | 0.38961562 | NA | NA |
| ENSG000002 | 5.94433562 | -0.413509512 | 0.36711249 | -1.1263837 | 0.26000316 | 0.38961562 | NA | NA |
| ENSG000002 | 4.10052561 | -0.425474381 | 0.37778204 | -1.126243 | 0.26006268 | 0.38968596 | NA | NA |
| ENSG000002 | 5.79302864 | -0.417144613 | 0.37039114 | -1.1262273 | 0.26006932 | 0.38968596 | NA | NA |
| ENSG000002 | 2.8113493 | -0.42120404 | 0.37403754 | -1.126101 | 0.26012278 | 0.38975165 | RNU6-1277P | 106480123 |
| ENSG000002 | 8.59717978 | -0.394706348 | 0.35054199 | -1.1259888 | 0.26017027 | 0.38980841 | NA | NA |
| ENSG000002 | 7.45765883 | 0.398227431 | 0.35370717 | 1.12586757 | 0.26022158 | 0.38987088 | NA | NA |
| ENSG000002 | 3.88510053 | -0.426029653 | 0.37843189 | -1.1257763 | 0.26026021 | 0.38991436 | NA | NA |
| ENSG000002 | 3.30174304 | -0.425261759 | 0.37781449 | -1.1255835 | 0.26034186 | 0.39002226 | NA | NA |
| ENSG000002 | 2.5062179 | -0.41534311 | 0.36907639 | -1.1253581 | 0.26043732 | 0.39013645 | NA | NA |
| ENSG000002 | 8.73910669 | -0.401005187 | 0.35633061 | -1.125374 | 0.2604306 | 0.39013645 | LOC105373662 | 105373662 |
| ENSG000002 | 16.9221386 | -0.33136624 | 0.29449709 | -1.1251936 | 0.26050701 | 0.39022643 | TNRC6C-AS1 | 100131096 |
| ENSG000002 | 2.4610981 | -0.41749298 | 0.37109719 | -1.1250233 | 0.26057916 | 0.39030568 | RNU4-36P | 106479573 |
| ENSG000002 | 4.48695554 | -0.425388732 | 0.37811297 | -1.1250308 | 0.26057599 | 0.39030568 | NA | NA |
| ENSG000001 | 11.9235969 | -0.362728946 | 0.32246846 | -1.1248509 | 0.26065221 | 0.39037184 | CRHBP | 1393 |
| ENSG000001 | 7.33431141 | -0.405094329 | 0.36012581 | -1.1248689 | 0.26064458 | 0.39037184 | KRT14 | 3861 |
| ENSG000002 | 6.59366152 | -0.412541688 | 0.36674085 | -1.1248861 | 0.2606373 | 0.39037184 | NA | NA |
| ENSG000001 | 20.4636049 | 0.31956288 | 0.28410606 | 1.12480137 | 0.26067321 | 0.39037446 | MRGPRF | 116535 |
| ENSG000001 | 31.1424589 | -0.274519581 | 0.24406008 | -1.1248033 | 0.2606724 | 0.39037446 | FAM109B | 150368 |
| ENSG000002 | 20.3735514 | -0.318120998 | 0.28283393 | -1.1247625 | 0.26068969 | 0.39038472 | NA | NA |
| ENSG000002 | 4.20717029 | -0.425799102 | 0.37859221 | -1.1246906 | 0.26072016 | 0.39041592 | FAM230C | 26080 |
| ENSG000002 | 2.71966754 | -0.422245027 | 0.37548238 | -1.1245402 | 0.26078393 | 0.390497 | NA | NA |
| ENSG00000C | 159.301704 | 0.165373857 | 0.1470656 | 1.1244904 | 0.26080504 | 0.39051419 | CALCOCO1 | 57658 |
| ENSG000002 | 4.10676343 | -0.424840687 | 0.37783641 | -1.1244038 | 0.26084177 | 0.39055477 | NA | NA |
| ENSG000001 | 8.08881985 | -0.399828046 | 0.35561089 | -1.1243414 | 0.26086822 | 0.39056621 | CMTM2 | 146225 |

| | | | | | | | | |
|---|---|---|---|---|---|---|---|---|
| ENSG000002 | 3.45540294 | -0.423560141 | 0.3767188 | -1.1243403 | 0.26086868 | 0.39056621 | NA | NA |
| ENSG000002 | 2.48731165 | -0.415717902 | 0.36978163 | -1.1242254 | 0.26091742 | 0.39062476 | NA | NA |
| ENSG000002 | 5.69222472 | -0.416266975 | 0.37029755 | -1.1241419 | 0.26095285 | 0.39066338 | NA | NA |
| ENSG000001 | 6.94983089 | -0.41156577 | 0.3661238 | -1.1241164 | 0.26096365 | 0.39066513 | ZC2HC1B | 153918 |
| ENSG000002 | 10.9262354 | -0.374661609 | 0.33332686 | -1.1240066 | 0.26101021 | 0.3907204 | MIR133A1HG | 102723167 |
| ENSG000002 | 2.70973119 | 0.420188054 | 0.3738468 | 1.12395788 | 0.26103089 | 0.39073694 | NA | NA |
| ENSG000001 | 30.2372059 | -0.28524581 | 0.25382997 | -1.1237673 | 0.26111177 | 0.39084357 | NA | NA |
| ENSG000001 | 382.4342 | 0.143444154 | 0.12766272 | 1.1236182 | 0.26117503 | 0.39092146 | ANKRD36 | 375248 |
| ENSG000002 | 4.34416312 | -0.422689659 | 0.37619255 | -1.1235992 | 0.26118308 | 0.39092146 | SLIT2-IT1 | 100505893 |
| ENSG000002 | 2.20565411 | -0.41021778 | 0.36511182 | -1.1235401 | 0.26120817 | 0.39094459 | NA | NA |
| ENSG000001 | 10.6028942 | -0.376505873 | 0.33512301 | -1.1234856 | 0.26123132 | 0.39096481 | AQP11 | 282679 |
| ENSG000002 | 116.588296 | 0.183278543 | 0.16315237 | 1.12335816 | 0.26128541 | 0.39103133 | NA | NA |
| ENSG000002 | 12.2955169 | -0.358304142 | 0.31898501 | -1.1232633 | 0.2613257 | 0.39106276 | NA | NA |
| ENSG000002 | 4.09447181 | -0.424373044 | 0.37779818 | -1.1232797 | 0.2613187 | 0.39106276 | NA | NA |
| ENSG000002 | 3.58708 | -0.425121705 | 0.37848508 | -1.1232192 | 0.26134441 | 0.39107632 | NA | NA |
| ENSG000002 | 3.34564855 | -0.423828839 | 0.37734557 | -1.1231849 | 0.26135898 | 0.39108369 | NA | NA |
| ENSG000002 | 3.57137009 | -0.425018693 | 0.37842037 | -1.123139 | 0.26137845 | 0.39109839 | NA | NA |
| ENSG000002 | 2.45820167 | -0.416214465 | 0.37061288 | -1.1230437 | 0.26141893 | 0.39114453 | NA | NA |
| ENSG000002 | 85.1622349 | -0.194791512 | 0.17345492 | -1.1230094 | 0.26143349 | 0.39115189 | LOC728730 | 728730 |
| ENSG000002 | 2.51884177 | -0.416089279 | 0.37053033 | -1.1229561 | 0.26145615 | 0.39117135 | NA | NA |
| ENSG000002 | 8.14890301 | -0.396546497 | 0.35316411 | -1.1228392 | 0.2615058 | 0.3912312 | NA | NA |
| ENSG000001 | 2.73051753 | -0.421686045 | 0.3756664 | -1.1225014 | 0.26164933 | 0.39142395 | COX8C | 341947 |
| ENSG000002 | 14.3649909 | -0.347976078 | 0.31000358 | -1.1224905 | 0.26165394 | 0.39142395 | GBP7 | 388646 |
| ENSG000002 | 2.74092977 | -0.421779945 | 0.37578671 | -1.1223919 | 0.26169587 | 0.39147223 | NA | NA |
| ENSG000002 | 14.3756673 | -0.356787076 | 0.31789326 | -1.1223487 | 0.26171423 | 0.39148525 | NA | NA |
| ENSG000002 | 46.4337079 | -0.235756257 | 0.21006112 | -1.1223222 | 0.26172548 | 0.39148764 | NA | NA |
| ENSG000002 | 2.19865872 | -0.406681805 | 0.36236489 | -1.1222991 | 0.26173527 | 0.39148785 | NA | NA |
| ENSG000002 | 6.47939167 | -0.406063241 | 0.36184067 | -1.1222156 | 0.26177079 | 0.39152653 | MS4A13 | 503497 |
| ENSG000002 | 25.0228012 | 0.296461435 | 0.26418715 | 1.12216447 | 0.26179251 | 0.39154458 | NA | NA |

| | | | | | | | | |
|---|---|---|---|---|---|---|---|---|
| ENSG000001 | 3.56441577 | -0.424788323 | 0.37856143 | -1.122112 | 0.26181483 | 0.39154907 | NUTM2B | 729262 |
| ENSG000002 | 51.8533967 | -0.240640302 | 0.21445012 | -1.1221271 | 0.26180839 | 0.39154907 | NA | NA |
| ENSG000002 | 9.57384942 | -0.38199892 | 0.34046112 | -1.1220045 | 0.26186051 | 0.39158851 | NA | NA |
| ENSG000002 | 10.7640219 | -0.38224402 | 0.34067345 | -1.1220247 | 0.26185194 | 0.39158851 | NA | NA |
| ENSG000002 | 3.02239198 | -0.423740603 | 0.37769788 | -1.1219036 | 0.26190343 | 0.39163825 | NA | NA |
| ENSG000002 | 4.69193123 | -0.422585969 | 0.37668205 | -1.1218638 | 0.26192034 | 0.39163944 | LINC01331 | 104310351 |
| ENSG000002 | 5.2231498 | -0.419601735 | 0.37402449 | -1.1218563 | 0.26192354 | 0.39163944 | NA | NA |
| ENSG000002 | 67.9561856 | 0.231143664 | 0.20605413 | 1.12176184 | 0.26196372 | 0.39167784 | NA | NA |
| ENSG000002 | 3.84901552 | -0.424562504 | 0.37848212 | -1.1217505 | 0.26196855 | 0.39167784 | NA | NA |
| ENSG000002 | 42.7885612 | 0.249805347 | 0.22272877 | 1.12156751 | 0.26204637 | 0.39177976 | ACVR2B-AS1 | 100128640 |
| ENSG000001 | 88.2059558 | 0.194126074 | 0.17309034 | 1.12153038 | 0.26206217 | 0.39178893 | ZDHHC14 | 79683 |
| ENSG000002 | 2.76643857 | -0.420338887 | 0.37485272 | -1.1213441 | 0.26214141 | 0.39187851 | NA | NA |
| ENSG000002 | 2.49726797 | -0.416946177 | 0.37182673 | -1.1213454 | 0.26214089 | 0.39187851 | NA | NA |
| ENSG000002 | 11.4894051 | -0.377087605 | 0.33630061 | -1.1212814 | 0.26216812 | 0.39190399 | NA | NA |
| ENSG000002 | 2.70899398 | 0.419020396 | 0.37372071 | 1.12121269 | 0.26219734 | 0.39193322 | NA | NA |
| ENSG000002 | 2.46448359 | -0.414535416 | 0.36973 | -1.1211842 | 0.26220948 | 0.39193692 | LOC392364 | 392364 |
| ENSG000001 | 170.280261 | 0.173870272 | 0.15508724 | 1.12111268 | 0.2622399 | 0.39196794 | TMEM151B | 441151 |
| ENSG000002 | 54.8951182 | 0.228603406 | 0.20391399 | 1.12107759 | 0.26225484 | 0.39197582 | MINCR | 100507316 |
| ENSG000002 | 5.23424774 | -0.418403759 | 0.37323483 | -1.1210201 | 0.26227929 | 0.39199791 | LOC340074 | 340074 |
| ENSG000002 | 13.0768424 | -0.353591041 | 0.31543048 | -1.1209793 | 0.26229667 | 0.39200944 | NA | NA |
| ENSG000002 | 2.75434283 | -0.420289007 | 0.37496702 | -1.1208693 | 0.26234352 | 0.39206502 | NA | NA |
| ENSG000001 | 13.5884415 | 0.351272078 | 0.3134229 | 1.12076073 | 0.26238973 | 0.39206549 | ABCB6 | 10058 |
| ENSG000001 | 34.0389162 | -0.260362101 | 0.23230039 | -1.1207993 | 0.26237332 | 0.39206549 | TGM2 | 7052 |
| ENSG000002 | 2.70544504 | -0.420476437 | 0.37515915 | -1.1207948 | 0.26237521 | 0.39206549 | NA | NA |
| ENSG000002 | 22.6744506 | -0.30787688 | 0.27469866 | -1.1207804 | 0.26238134 | 0.39206549 | NA | NA |
| ENSG000002 | 5.53012777 | -0.417542896 | 0.37255503 | -1.120755 | 0.26239218 | 0.39206549 | NA | NA |
| ENSG000002 | 3.42793569 | -0.421499114 | 0.37610174 | -1.120705 | 0.26241343 | 0.3920828 | NA | NA |
| ENSG000002 | 4.97948779 | -0.420499433 | 0.37528891 | -1.1204686 | 0.26251413 | 0.39221881 | NA | NA |
| ENSG000001 | 30.7191732 | 0.274333595 | 0.24484435 | 1.12044076 | 0.26252598 | 0.39222206 | SLC30A2 | 7780 |

| | | | | | | | | |
|---|---|---|---|---|---|---|---|---|
| ENSG000002 | 14.6703967 | 0.351270057 | 0.31356971 | 1.12022954 | 0.26261596 | 0.39234085 | SNORA66 | 26782 |
| ENSG000002 | 4.14911075 | -0.423540919 | 0.37809875 | -1.120186 | 0.26263452 | 0.39234085 | RN7SL51P | 106480936 |
| ENSG000002 | 5.96396644 | -0.412803417 | 0.36850845 | -1.1202007 | 0.26262825 | 0.39234085 | NA | NA |
| ENSG000002 | 2.50341351 | -0.416205735 | 0.37157583 | -1.1201098 | 0.26266697 | 0.39237487 | NA | NA |
| ENSG000002 | 6.81141831 | -0.407127209 | 0.36348818 | -1.1200563 | 0.26268979 | 0.3923945 | AADACP1 | 201651 |
| ENSG000002 | 2.53608524 | -0.415040681 | 0.37060132 | -1.1199115 | 0.26275148 | 0.39245774 | NA | NA |
| ENSG000002 | 5.76536027 | -0.414186473 | 0.36983507 | -1.1199221 | 0.26274695 | 0.39245774 | NA | NA |
| ENSG000002 | 29.0926213 | 0.27647288 | 0.24687912 | 1.11987145 | 0.26276855 | 0.39246878 | NA | NA |
| ENSG000001 | 244.073675 | 0.157745508 | 0.14087236 | 1.11977616 | 0.26280916 | 0.39250053 | ADAMTS16 | 170690 |
| ENSG000001 | 5.66580952 | -0.41487543 | 0.37049134 | -1.1197979 | 0.26279988 | 0.39250053 | NA | NA |
| ENSG000001 | 5.5164136 | -0.417344024 | 0.37273153 | -1.1196907 | 0.2628456 | 0.39254049 | S100A5 | 6276 |
| ENSG000002 | 4.6625086 | -0.42157246 | 0.37655426 | -1.119553 | 0.26290429 | 0.39259921 | NA | NA |
| ENSG000002 | 3.59126597 | -0.423044773 | 0.37786238 | -1.1195737 | 0.26289547 | 0.39259921 | NA | NA |
| ENSG000001 | 2.7083222 | -0.419916577 | 0.37510436 | -1.119466 | 0.26294139 | 0.39264017 | CAPZA3 | 93661 |
| ENSG000001 | 3.90518747 | -0.423731576 | 0.37858785 | -1.1192424 | 0.26303674 | 0.39276809 | TMEM82 | 388595 |
| ENSG000002 | 5.4054858 | -0.415980029 | 0.3716722 | -1.1192121 | 0.26304966 | 0.39277292 | NA | NA |
| ENSG000001 | 21.7570258 | 0.314593269 | 0.28112962 | 1.1190328 | 0.26312615 | 0.39283628 | CITED1 | 4435 |
| ENSG000001 | 4.38759256 | -0.421567275 | 0.37671053 | -1.1190748 | 0.26310822 | 0.39283628 | KRTAP13-4 | 284827 |
| ENSG000002 | 3.85924648 | -0.423414576 | 0.3784176 | -1.1189082 | 0.26317929 | 0.39283628 | RN7SKP245 | 106480909 |
| ENSG000002 | 2.55283227 | -0.413435894 | 0.3694648 | -1.1190129 | 0.26313462 | 0.39283628 | NA | NA |
| ENSG000002 | 7.01084601 | -0.404911326 | 0.36184737 | -1.1190114 | 0.26313528 | 0.39283628 | NA | NA |
| ENSG000002 | 2.77198329 | -0.418700575 | 0.37418667 | -1.1189618 | 0.26315645 | 0.39283628 | NA | NA |
| ENSG000002 | 2.74793617 | -0.419161969 | 0.37460667 | -1.1189389 | 0.26316621 | 0.39283628 | NA | NA |
| ENSG000002 | 3.33957624 | -0.422674139 | 0.37772638 | -1.1189955 | 0.26314204 | 0.39283628 | RN7SL362P | 106481022 |
| ENSG000002 | 6.32913358 | -0.415109946 | 0.37099243 | -1.1189176 | 0.26317531 | 0.39283628 | NA | NA |
| ENSG000002 | 5.47148629 | -0.418813832 | 0.3743493 | -1.1187782 | 0.26323478 | 0.39289019 | NA | NA |
| ENSG000002 | 16.5239546 | -0.332281124 | 0.29700259 | -1.1187819 | 0.26323319 | 0.39289019 | NA | NA |
| ENSG000002 | 2.20179124 | -0.405869491 | 0.36280079 | -1.1187117 | 0.26326315 | 0.39291807 | NA | NA |
| ENSG000002 | 2.46355051 | -0.413315303 | 0.36950773 | -1.1185566 | 0.26332935 | 0.39300242 | NA | NA |

| | | | | | | | | |
|---|---|---|---|---|---|---|---|---|
| ENSG000001 | 20.8795154 | 0.320103015 | 0.28618886 | 1.11850271 | 0.26335235 | 0.39302227 | WDR86 | 349136 |
| ENSG000002 | 7.36107395 | -0.409015648 | 0.36569143 | -1.1184721 | 0.26336543 | 0.39302733 | NA | NA |
| ENSG000002 | 4.10638572 | -0.422599914 | 0.37785217 | -1.1184266 | 0.26338484 | 0.39302737 | NA | NA |
| ENSG000002 | 47.4753563 | 0.252615172 | 0.22586343 | 1.11844209 | 0.26337822 | 0.39302737 | SNORD11 | 692058 |
| ENSG000001 | 32.6129843 | -0.273477955 | 0.24454487 | -1.118314 | 0.26343291 | 0.39307934 | ADORA1 | 134 |
| ENSG000002 | 2.48680366 | -0.415359759 | 0.37142082 | -1.1182996 | 0.26343906 | 0.39307934 | NA | NA |
| ENSG000002 | 76.2233617 | 0.200262886 | 0.17908409 | 1.11826173 | 0.26345522 | 0.393089 | LINC00862 | 554279 |
| ENSG000001 | 69.5806534 | 0.21519624 | 0.19247583 | 1.11804291 | 0.26354867 | 0.39320456 | EFHC2 | 80258 |
| ENSG000002 | 2.50097354 | -0.414120748 | 0.37040054 | -1.1180349 | 0.26355207 | 0.39320456 | NA | NA |
| ENSG000002 | 4.608517 | 0.420204982 | 0.37587561 | 1.11793627 | 0.26359421 | 0.3932385 | NA | NA |
| ENSG000002 | 2.5090626 | -0.413611407 | 0.369973 | -1.1179502 | 0.26358824 | 0.3932385 | NA | NA |
| ENSG000002 | 9.67466633 | -0.382682198 | 0.34232788 | -1.1178821 | 0.26361737 | 0.39325858 | NA | NA |
| ENSG000002 | 5.20231267 | -0.417438947 | 0.37346475 | -1.1177466 | 0.26367525 | 0.39333045 | NA | NA |
| ENSG000001 | 5.73835003 | -0.413697671 | 0.3702064 | -1.1174784 | 0.26378982 | 0.39348688 | NA | NA |
| ENSG000001 | 15.5143764 | -0.348543078 | 0.31195088 | -1.1173011 | 0.26386559 | 0.39355544 | HPX | 3263 |
| ENSG000002 | 18.3145686 | -0.319399753 | 0.28586456 | -1.1173115 | 0.26386117 | 0.39355544 | LOC100505824 | 100505824 |
| ENSG000002 | 4.07920717 | -0.421697104 | 0.37743833 | -1.1172609 | 0.26388277 | 0.39355544 | NA | NA |
| ENSG000002 | 5.23869051 | -0.417309708 | 0.37351261 | -1.1172574 | 0.2638843 | 0.39355544 | NA | NA |
| ENSG000002 | 9.93805591 | -0.382207949 | 0.34207515 | -1.1173216 | 0.26385685 | 0.39355544 | NA | NA |
| ENSG000001 | 62.9143353 | 0.212582273 | 0.19027787 | 1.11722018 | 0.26390019 | 0.39356466 | ZNF385C | 201181 |
| ENSG000002 | 3.60561377 | -0.42243973 | 0.37818944 | -1.1170056 | 0.26399193 | 0.39368699 | NA | NA |
| ENSG000002 | 5.10791539 | 0.419986789 | 0.37600765 | 1.1169634 | 0.26400997 | 0.39369942 | MIR590 | 693175 |
| ENSG000001 | 7.72730551 | -0.394642956 | 0.35336103 | -1.1168265 | 0.26406853 | 0.39377226 | SERPINA3 | 12 |
| ENSG000002 | 13.8436409 | -0.355764727 | 0.31858994 | -1.1166854 | 0.26412887 | 0.39384776 | NA | NA |
| ENSG000002 | 12.5819608 | -0.358218244 | 0.32081216 | -1.1165981 | 0.2641662 | 0.39388895 | PRC1-AS1 | 100507118 |
| ENSG000002 | 3.62262388 | -0.422578106 | 0.37847981 | -1.1165143 | 0.26420207 | 0.39392794 | NA | NA |
| ENSG000001 | 6.55851388 | -0.419970094 | 0.37616934 | -1.1164389 | 0.2642343 | 0.39394855 | TNFRSF4 | 7293 |
| ENSG000002 | 35.3412221 | 0.274286664 | 0.24568048 | 1.11643653 | 0.26423532 | 0.39394855 | NA | NA |
| ENSG00000C | 13.2483466 | -0.362138452 | 0.32444422 | -1.1161809 | 0.2643447 | 0.39409712 | IL12RB1 | 3594 |

| | | | | | | | | |
|---|---|---|---|---|---|---|---|---|
| ENSG000002 | 15.6772891 | -0.337489957 | 0.30239253 | -1.1160658 | 0.26439398 | 0.39415611 | LOC101928438 | 101928438 |
| ENSG000002 | 4.98044448 | -0.419963143 | 0.37629856 | -1.1160371 | 0.26440626 | 0.39415993 | NA | NA |
| ENSG000001 | 19.9800234 | 0.313481148 | 0.28095816 | 1.11575741 | 0.26452599 | 0.39428043 | TMEM175 | 84286 |
| ENSG000002 | 9.78025059 | -0.374260279 | 0.33542528 | -1.1157784 | 0.26451701 | 0.39428043 | NA | NA |
| ENSG000002 | 3.32369041 | -0.421563748 | 0.37780654 | -1.1158191 | 0.26449958 | 0.39428043 | NA | NA |
| ENSG000002 | 3.38013594 | -0.420852601 | 0.37718528 | -1.1157716 | 0.26451993 | 0.39428043 | NA | NA |
| ENSG000002 | 2.72949161 | -0.417706952 | 0.37439241 | -1.1156929 | 0.26455361 | 0.3943071 | NA | NA |
| ENSG000002 | 2.76529697 | -0.418397199 | 0.37503517 | -1.1156212 | 0.2645843 | 0.39432385 | MIR603 | 693188 |
| ENSG000002 | 2.2030566 | -0.403577757 | 0.361749 | -1.1156292 | 0.26458088 | 0.39432385 | NA | NA |
| ENSG000001 | 4.44247602 | -0.421966597 | 0.37825641 | -1.115557 | 0.2646118 | 0.39435035 | IRX5 | 10265 |
| ENSG000002 | 5.48830196 | -0.414873449 | 0.37198343 | -1.1153009 | 0.26472151 | 0.39449935 | NA | NA |
| ENSG000002 | 2.50709946 | -0.412432015 | 0.36982631 | -1.1152046 | 0.26476274 | 0.39453835 | NA | NA |
| ENSG000002 | 5.92236485 | -0.408942595 | 0.36670073 | -1.1151944 | 0.26476713 | 0.39453835 | NA | NA |
| ENSG000002 | 31.9726127 | 0.311265047 | 0.27912438 | 1.11514817 | 0.26478693 | 0.39455335 | NA | NA |
| ENSG000002 | 52.2699556 | 0.23428994 | 0.2101201 | 1.11502867 | 0.26483814 | 0.39461515 | ZNF878 | 729747 |
| ENSG000002 | 17.527105 | -0.330990692 | 0.29687519 | -1.1149153 | 0.26488671 | 0.39467303 | NA | NA |
| ENSG000002 | 5.4592435 | -0.414929753 | 0.3721858 | -1.1148457 | 0.26491653 | 0.39470295 | NA | NA |
| ENSG000002 | 2.53276481 | -0.413003962 | 0.37046618 | -1.1148223 | 0.26492658 | 0.39470342 | RN7SKP172 | 106480469 |
| ENSG000002 | 3.37273581 | -0.419416171 | 0.37623779 | -1.1147635 | 0.26495176 | 0.39472644 | LOC102723885 | 102723885 |
| ENSG000001 | 16.9034044 | -0.338881699 | 0.30402078 | -1.1146663 | 0.26499347 | 0.39474063 | OPTC | 26254 |
| ENSG000002 | 13.1681619 | -0.359682633 | 0.32267464 | -1.1146913 | 0.26498271 | 0.39474063 | NPY6R | 4888 |
| ENSG000002 | 2.4775719 | -0.413315773 | 0.37080302 | -1.1146505 | 0.26500022 | 0.39474063 | NA | NA |
| ENSG000002 | 58.7415143 | 0.22214535 | 0.19928493 | 1.11471223 | 0.26497376 | 0.39474063 | TSC22D1-AS1 | 641467 |
| ENSG000002 | 8.49487687 | -0.400056205 | 0.35893412 | -1.1145672 | 0.26503592 | 0.3947793 | NA | NA |
| ENSG000001 | 9.27050278 | -0.379864316 | 0.34083831 | -1.1145 | 0.26506472 | 0.39480164 | LOC401554 | 401554 |
| ENSG000002 | 2.47119769 | -0.412610051 | 0.37022425 | -1.1144868 | 0.26507039 | 0.39480164 | NA | NA |
| ENSG000002 | 6.16839493 | -0.405534244 | 0.36388652 | -1.1144525 | 0.26508511 | 0.39480906 | NA | NA |
| ENSG000002 | 8.64308216 | -0.391657445 | 0.35144372 | -1.1144244 | 0.26509717 | 0.39481253 | NA | NA |
| ENSG000001 | 37.5387009 | 0.268366215 | 0.24082794 | 1.11434836 | 0.26512977 | 0.39484657 | NA | NA |

| | | | | | | | | |
|---|---|---|---|---|---|---|---|---|
| ENSG00000( | 60.4488746 | -0.220990871 | 0.19834213 | -1.1141903 | 0.26519756 | 0.39488705 | IL12RB2 | 3595 |
| ENSG000001 | 11.8732963 | -0.361514613 | 0.32447079 | -1.1141669 | 0.2652076 | 0.39488705 | BMP8B | 656 |
| ENSG000001 | 5.43289302 | -0.415090322 | 0.3725531 | -1.1141776 | 0.265203 | 0.39488705 | PEBP4 | 157310 |
| ENSG000001 | 23.9101232 | 0.305372398 | 0.27406509 | 1.11423312 | 0.26517919 | 0.39488705 | SYNDIG1L | 646658 |
| ENSG000002 | 2.98207538 | 0.420155944 | 0.37710938 | 1.11414874 | 0.26521538 | 0.39488705 | NA | NA |
| ENSG000002 | 3.61353373 | -0.421530833 | 0.37832785 | -1.1141946 | 0.26519572 | 0.39488705 | NA | NA |
| ENSG000002 | 18.4904673 | 0.327925069 | 0.29434417 | 1.11408721 | 0.26524178 | 0.39491111 | RNVU1-7 | 26864 |
| ENSG000002 | 30.6106883 | -0.266462993 | 0.23918069 | -1.1140657 | 0.26525101 | 0.39491111 | LINC00648 | 100506433 |
| ENSG00000( | 305.319252 | 0.150197994 | 0.13486222 | 1.11371436 | 0.26540175 | 0.39509915 | KLHL20 | 27252 |
| ENSG000002 | 5.98238431 | -0.409832337 | 0.36798786 | -1.1137116 | 0.26540294 | 0.39509915 | NA | NA |
| ENSG000002 | 4.10752787 | -0.420757518 | 0.37780041 | -1.1137032 | 0.26540655 | 0.39509915 | NA | NA |
| ENSG000002 | 19.1770895 | -0.318773815 | 0.28623605 | -1.1136746 | 0.26541882 | 0.39510291 | NA | NA |
| ENSG000002 | 4.09556908 | -0.420743904 | 0.37782571 | -1.1135926 | 0.26545401 | 0.39512629 | NA | NA |
| ENSG000002 | 13.9628656 | -0.349587368 | 0.31392478 | -1.1136023 | 0.26544982 | 0.39512629 | NA | NA |
| ENSG000001 | 5.02172157 | -0.419922632 | 0.3771194 | -1.1135005 | 0.26549354 | 0.39517062 | LINC00525 | 84847 |
| ENSG000002 | 6.85931113 | -0.407292002 | 0.36578747 | -1.1134662 | 0.26550824 | 0.395178 | KRBOX1-AS1 | 100506275 |
| ENSG000002 | 5.43357792 | -0.414344535 | 0.3721301 | -1.11344 | 0.26551951 | 0.39518027 | NA | NA |
| ENSG000002 | 3.87547722 | -0.421445737 | 0.3785277 | -1.1133815 | 0.26554461 | 0.39520313 | NA | NA |
| ENSG000001 | 474.647528 | 0.181556479 | 0.16308044 | 1.11329403 | 0.26558217 | 0.39524252 | ARRB1 | 408 |
| ENSG000002 | 10.6809139 | -0.379364775 | 0.34076483 | -1.1132744 | 0.26559058 | 0.39524252 | NA | NA |
| ENSG000002 | 2.50409865 | -0.413213051 | 0.37118869 | -1.1132156 | 0.26561584 | 0.39526561 | NA | NA |
| ENSG000002 | 17.5301368 | -0.322944283 | 0.29024176 | -1.1126734 | 0.26584873 | 0.39559767 | NA | NA |
| ENSG000002 | 6.72236887 | -0.399921742 | 0.35947315 | -1.1125219 | 0.26591384 | 0.39568002 | NA | NA |
| ENSG000002 | 14.0330952 | 0.350829469 | 0.31537844 | 1.11240789 | 0.26596281 | 0.39573838 | NA | NA |
| ENSG000002 | 4.78171826 | -0.420611311 | 0.37813187 | -1.1123403 | 0.26599187 | 0.3957671 | NA | NA |
| ENSG000002 | 12.4608731 | -0.369971705 | 0.33261983 | -1.112296 | 0.2660109 | 0.39578089 | NA | NA |
| ENSG000002 | 2.52943008 | -0.409445569 | 0.36815335 | -1.1121604 | 0.26606921 | 0.39585311 | NA | NA |
| ENSG000002 | 7.58481307 | -0.403471167 | 0.3627987 | -1.1121075 | 0.26609191 | 0.39587237 | NA | NA |
| ENSG000001 | 196.648143 | 0.151923429 | 0.13662046 | 1.1120108 | 0.26613351 | 0.39589174 | CA5B | 11238 |

| | | | | | | | | |
|---|---|---|---|---|---|---|---|---|
| ENSG000002 | 4.31456415 | -0.420531444 | 0.37816775 | -1.1120235 | 0.26612803 | 0.39589174 | PTTG3P | 26255 |
| ENSG000002 | 2.79509099 | -0.414849719 | 0.37306323 | -1.1120091 | 0.26613422 | 0.39589174 | CHCHD4P5 | 106479039 |
| ENSG000002 | 11.6519268 | -0.362121565 | 0.3256564 | -1.1119744 | 0.26614918 | 0.39589946 | NA | NA |
| ENSG000001 | 86.3378697 | 0.220444971 | 0.19826703 | 1.11185893 | 0.26619881 | 0.39594424 | OVOL1 | 5017 |
| ENSG000002 | 5.6026949 | -0.41890196 | 0.37675393 | -1.1118715 | 0.2661934 | 0.39594424 | NA | NA |
| ENSG000001 | 4.10880532 | -0.420392542 | 0.37815129 | -1.1117046 | 0.26626517 | 0.39602842 | CCDC54 | 84692 |
| ENSG000002 | 3.02532012 | -0.418878032 | 0.3768328 | -1.1115753 | 0.2663208 | 0.39609664 | NA | NA |
| ENSG000001 | 4.36335012 | -0.418617916 | 0.37664426 | -1.1114411 | 0.26637852 | 0.39616795 | OR1L3 | 26735 |
| ENSG000001 | 63.2303374 | -0.217199333 | 0.19543813 | -1.1113458 | 0.26641955 | 0.39621444 | NA | NA |
| ENSG000001 | 11.2678341 | -0.375676991 | 0.33811381 | -1.1110962 | 0.26652692 | 0.39635959 | POTEC | 388468 |
| ENSG000002 | 5.59027089 | -0.416403487 | 0.37478341 | -1.111051 | 0.2665464 | 0.39636813 | RN7SKP240 | 106479202 |
| ENSG000002 | 3.16006417 | -0.409221231 | 0.36832352 | -1.1110375 | 0.26655222 | 0.39636813 | NA | NA |
| ENSG000002 | 12.1900496 | 0.362957709 | 0.32669912 | 1.11098464 | 0.26657496 | 0.39638741 | LOC105375666 | 105375666 |
| ENSG000002 | 2.48172706 | -0.411598093 | 0.37048848 | -1.1109606 | 0.26658532 | 0.39638829 | NA | NA |
| ENSG000002 | 5.2309141 | -0.41478693 | 0.37337925 | -1.1108998 | 0.26661148 | 0.39641265 | NA | NA |
| ENSG000002 | 4.63416665 | -0.419604825 | 0.37775215 | -1.110794 | 0.26665702 | 0.39645129 | NA | NA |
| ENSG000002 | 3.82059676 | -0.420516654 | 0.37857276 | -1.1107948 | 0.26665668 | 0.39645129 | NA | NA |
| ENSG000001 | 58.1999116 | 0.226678635 | 0.20409193 | 1.11066928 | 0.26671073 | 0.3965166 | CCDC110 | 256309 |
| ENSG000002 | 6.2932245 | -0.408481842 | 0.36780303 | -1.1105994 | 0.2667408 | 0.39654677 | NA | NA |
| ENSG000002 | 10.7637541 | -0.383126829 | 0.34500148 | -1.1105078 | 0.26678028 | 0.39659092 | LOC101927641 | 101927641 |
| ENSG000002 | 5.92304943 | -0.40774594 | 0.36721144 | -1.1103846 | 0.26683331 | 0.39665522 | NA | NA |
| ENSG000002 | 9.08934118 | -0.380046506 | 0.34228591 | -1.1103189 | 0.26686164 | 0.39668278 | NA | NA |
| ENSG000002 | 11.1472312 | -0.372275546 | 0.33529425 | -1.110295 | 0.26687191 | 0.39668351 | NA | NA |
| ENSG000002 | 3.02378153 | -0.418682548 | 0.37713377 | -1.1101699 | 0.26692583 | 0.39672004 | NA | NA |
| ENSG000002 | 12.08071 | -0.357264836 | 0.32180921 | -1.1101759 | 0.26692322 | 0.39672004 | NA | NA |
| ENSG000002 | 2.53262045 | -0.409372168 | 0.36873564 | -1.110205 | 0.26691068 | 0.39672004 | RN7SL509P | 106479409 |
| ENSG000001 | 48.4170293 | 0.236276066 | 0.21286406 | 1.1099857 | 0.26700519 | 0.39682343 | PRSS35 | 167681 |
| ENSG000002 | 2.99686358 | -0.418959287 | 0.37746937 | -1.109916 | 0.26703523 | 0.39685354 | NA | NA |
| ENSG000002 | 7.74066019 | -0.39224394 | 0.35341147 | -1.1098789 | 0.26705121 | 0.39686274 | NA | NA |

| | | | | | | | | |
|---|---|---|---|---|---|---|---|---|
| ENSG000002 | 95.4587421 | 0.19469139 | 0.17542577 | 1.10982208 | 0.2670757 | 0.39688459 | LOC101929147 | 101929147 |
| ENSG000002 | 3.81086987 | -0.420106407 | 0.37856726 | -1.1097272 | 0.26711658 | 0.39693079 | NA | NA |
| ENSG000001 | 260.902358 | 0.138318481 | 0.12465154 | 1.10964121 | 0.26715367 | 0.39697136 | ZNF782 | 158431 |
| ENSG000000 | 21341.4307 | 0.122805981 | 0.11067617 | 1.1095973 | 0.26717259 | 0.39698494 | NUCKS1 | 64710 |
| ENSG000001 | 13.7276356 | 0.347554822 | 0.31324186 | 1.10954143 | 0.26719668 | 0.39699164 | OGFOD2 | 79676 |
| ENSG000002 | 19.583449 | 0.311708449 | 0.28093031 | 1.1095579 | 0.26718958 | 0.39699164 | THAP7-AS1 | 439931 |
| ENSG000002 | 47.1526919 | 0.247382077 | 0.22296379 | 1.10951681 | 0.2672073 | 0.39699286 | MZF1-AS1 | 100131691 |
| ENSG000002 | 2.70920377 | -0.416566234 | 0.37547807 | -1.1094289 | 0.26724519 | 0.39703461 | NA | NA |
| ENSG000002 | 2.98165062 | 0.418572234 | 0.37738886 | 1.10912717 | 0.26737532 | 0.39721339 | NA | NA |
| ENSG000000 | 13.4700647 | -0.367589031 | 0.33143005 | -1.1090999 | 0.2673871 | 0.39721633 | FGR | 2268 |
| ENSG000002 | 5.71616137 | -0.41242675 | 0.37201617 | -1.1086259 | 0.26759162 | 0.39750559 | RNU6-644P | 106481385 |
| ENSG000002 | 2.72263644 | -0.415375422 | 0.37474504 | -1.1084214 | 0.26767987 | 0.39762212 | RN7SKP266 | 106479214 |
| ENSG000002 | 5.24160601 | -0.415572359 | 0.37495752 | -1.1083185 | 0.26772429 | 0.39767354 | NA | NA |
| ENSG000002 | 4.91516085 | -0.416652786 | 0.37598058 | -1.1081763 | 0.26778567 | 0.39775014 | NA | NA |
| ENSG000001 | 24.5001456 | 0.330821141 | 0.29863405 | 1.10778105 | 0.26795638 | 0.39798913 | TMEM59L | 25789 |
| ENSG000002 | 2.73772034 | -0.415130132 | 0.3747907 | -1.1076319 | 0.26802082 | 0.39807026 | NA | NA |
| ENSG000002 | 9.08871553 | -0.380203559 | 0.3433076 | -1.107472 | 0.2680899 | 0.39815828 | HNRNPA1P76 | 106480675 |
| ENSG000001 | 2.74654661 | -0.414735789 | 0.3745225 | -1.1073722 | 0.26813306 | 0.39820779 | GHSR | 2693 |
| ENSG000002 | 5.84463631 | -0.413661661 | 0.37356615 | -1.1073318 | 0.26815052 | 0.39821913 | NA | NA |
| ENSG000002 | 8.86760551 | -0.385309174 | 0.34802525 | -1.10713 | 0.26823775 | 0.39833409 | NA | NA |
| ENSG000001 | 15.3541605 | 0.335485275 | 0.30306696 | 1.1069675 | 0.26830798 | 0.39842379 | SHD | 56961 |
| ENSG000001 | 475.812473 | 0.128870119 | 0.11644674 | 1.10668725 | 0.26842917 | 0.39855997 | ELP4 | 26610 |
| ENSG000002 | 15.6777957 | -0.349403448 | 0.31570935 | -1.1067251 | 0.26841282 | 0.39855997 | LINC01583 | 101929690 |
| ENSG000002 | 30.4518771 | -0.268005168 | 0.24216844 | -1.1066891 | 0.26842837 | 0.39855997 | LOC102724532 | 102724532 |
| ENSG000002 | 5.65388352 | -0.409625199 | 0.37017367 | -1.1065757 | 0.2684774 | 0.39860239 | NA | NA |
| ENSG000002 | 2.47657877 | -0.408890982 | 0.36950308 | -1.1065969 | 0.26846823 | 0.39860239 | NA | NA |
| ENSG000001 | 79.0651189 | 0.20502412 | 0.18528315 | 1.1065449 | 0.26849074 | 0.3986076 | QRICH2 | 84074 |
| ENSG000002 | 4.17434577 | -0.418662557 | 0.37836107 | -1.1065159 | 0.26850327 | 0.3986116 | NA | NA |
| ENSG000002 | 2.82148135 | -0.412289209 | 0.37260954 | -1.1064913 | 0.26851393 | 0.39861284 | NA | NA |

| | | | | | | | | |
|---|---|---|---|---|---|---|---|---|
| ENSG000002 | 5.22298562 | -0.41443992 | 0.37458441 | -1.1063993 | 0.26855374 | 0.39865734 | NA | NA |
| ENSG000002 | 12.9059417 | -0.353975512 | 0.31994498 | -1.1063637 | 0.26856912 | 0.39866559 | LOC101929151 | 101929151 |
| ENSG000001 | 1882.51862 | 0.190783959 | 0.17244761 | 1.10632997 | 0.26858372 | 0.39867267 | PTP4A1 | 7803 |
| ENSG000002 | 3.83582557 | -0.418726944 | 0.3784918 | -1.1063039 | 0.26859502 | 0.39867484 | NA | NA |
| ENSG000002 | 6.81087001 | -0.405177024 | 0.36625253 | -1.1062777 | 0.26860632 | 0.39867703 | NA | NA |
| ENSG000002 | 5.17017375 | -0.413549688 | 0.37389387 | -1.1060617 | 0.26869981 | 0.39875741 | NA | NA |
| ENSG000002 | 4.12941521 | -0.418060406 | 0.37796459 | -1.1060835 | 0.26869036 | 0.39875741 | NA | NA |
| ENSG000002 | 10.1905657 | -0.376710543 | 0.34058543 | -1.1060677 | 0.26869722 | 0.39875741 | NA | NA |
| ENSG000002 | 13.3896516 | -0.349605635 | 0.3160716 | -1.1060963 | 0.26868484 | 0.39875741 | NA | NA |
| ENSG000002 | 39.0309726 | 0.249267098 | 0.22539253 | 1.1059244 | 0.26875924 | 0.398831 | NA | NA |
| ENSG000002 | 5.43121562 | 0.410347 | 0.37105532 | 1.1058917 | 0.2687734 | 0.39883741 | NA | NA |
| ENSG000002 | 3.80341387 | -0.418525261 | 0.37847484 | -1.1058206 | 0.2688042 | 0.39886852 | NA | NA |
| ENSG000002 | 5.74322478 | -0.410248896 | 0.37102421 | -1.10572 | 0.26884773 | 0.39891852 | ARHGAP26-AS1 | 100874239 |
| ENSG000001 | 2.73800109 | -0.414834341 | 0.37519245 | -1.1056575 | 0.2688748 | 0.39894409 | KRTAP26-1 | 388818 |
| ENSG000002 | 11.0948678 | -0.371886425 | 0.3363656 | -1.1056018 | 0.2688989 | 0.39895065 | NA | NA |
| ENSG000002 | 5.17195233 | 0.412913788 | 0.37347028 | 1.10561351 | 0.26889384 | 0.39895065 | NA | NA |
| ENSG000001 | 1254.64562 | 0.140658397 | 0.12722682 | 1.10557193 | 0.26891185 | 0.39895527 | NF1 | 4763 |
| ENSG000002 | 5.95026132 | -0.406043998 | 0.36732309 | -1.1054138 | 0.26898035 | 0.39897331 | NA | NA |
| ENSG000002 | 4.1451118 | -0.417977981 | 0.37810869 | -1.105444 | 0.26896725 | 0.39897331 | NA | NA |
| ENSG000002 | 2.71649091 | -0.414243703 | 0.37474077 | -1.105414 | 0.26898023 | 0.39897331 | NA | NA |
| ENSG000002 | 4.80167073 | 0.413519443 | 0.37406834 | 1.10546496 | 0.26895817 | 0.39897331 | LOC102724301 | 102724301 |
| ENSG000002 | 5.76500874 | -0.409785216 | 0.37070963 | -1.1054075 | 0.26898304 | 0.39897331 | NA | NA |
| ENSG000002 | 2.71649091 | -0.414243703 | 0.37474077 | -1.105414 | 0.26898023 | 0.39897331 | NA | NA |
| ENSG000002 | 9.21864872 | -0.376919075 | 0.34098748 | -1.1053751 | 0.26899709 | 0.39897705 | LOC101929415 | 101929415 |
| ENSG000002 | 2.22546099 | -0.401137931 | 0.36290374 | -1.1053563 | 0.26900524 | 0.39897705 | NA | NA |
| ENSG000002 | 2.44769768 | -0.409339578 | 0.37034554 | -1.105291 | 0.26903353 | 0.39900441 | RNU6-593P | 106479828 |
| ENSG000002 | 13.7199157 | -0.370654408 | 0.33536197 | -1.1052369 | 0.26905697 | 0.39902458 | NA | NA |
| ENSG000002 | 4.54085034 | 0.417005886 | 0.37733295 | 1.1051404 | 0.26909876 | 0.39907196 | NA | NA |
| ENSG000002 | 9.06453626 | -0.38075417 | 0.34454525 | -1.1050919 | 0.26911976 | 0.39908851 | NA | NA |

| | | | | | | | | |
|---|---|---|---|---|---|---|---|---|
| ENSG000002 | 5.82890406 | -0.416072945 | 0.37651333 | -1.1050683 | 0.26913 | 0.3990891 | NA | NA |
| ENSG000001 | 4.13545682 | -0.417908763 | 0.3781835 | -1.1050423 | 0.26914126 | 0.39909121 | SCGB2A2 | 4250 |
| ENSG000002 | 6.61669911 | 0.401316908 | 0.36320751 | 1.10492457 | 0.26919227 | 0.39915226 | NA | NA |
| ENSG000002 | 378.995896 | -0.145301986 | 0.13150783 | -1.1048923 | 0.26920627 | 0.39915841 | NA | NA |
| ENSG000002 | 2.51434694 | -0.40667259 | 0.36808159 | -1.1048436 | 0.26922736 | 0.39917509 | NA | NA |
| ENSG000002 | 40.5122692 | 0.240733526 | 0.21789828 | 1.10479775 | 0.26924723 | 0.39918996 | SOX21-AS1 | 100507533 |
| ENSG000001 | 3809.30245 | 0.298883242 | 0.27059904 | 1.10452441 | 0.26936572 | 0.39932183 | RBM25 | 58517 |
| ENSG000002 | 5.87993733 | -0.407045402 | 0.36852512 | -1.1045255 | 0.26936524 | 0.39932183 | NA | NA |
| ENSG000002 | 7.984779 | -0.388252565 | 0.35149906 | -1.1045622 | 0.26934934 | 0.39932183 | LOC101928851 | 101928851 |
| ENSG000002 | 8.27701384 | -0.385037915 | 0.34862619 | -1.1044434 | 0.26940082 | 0.39933073 | LOC101928627 | 101928627 |
| ENSG000002 | 2.77514806 | -0.413343583 | 0.37424195 | -1.1044822 | 0.26938401 | 0.39933073 | NA | NA |
| ENSG000002 | 5.89752158 | -0.404743797 | 0.36647486 | -1.1044244 | 0.26940906 | 0.39933073 | NA | NA |
| ENSG000002 | 10.4741841 | 0.373533694 | 0.33821715 | 1.10441972 | 0.26941111 | 0.39933073 | NA | NA |
| ENSG000001 | 55.8581019 | 0.224241893 | 0.2030467 | 1.1043858 | 0.26942582 | 0.39933793 | ATG16L2 | 89849 |
| ENSG000002 | 2.37464562 | 0.409813281 | 0.3711021 | 1.1043141 | 0.26945691 | 0.39936941 | NA | NA |
| ENSG000001 | 6.03611975 | -0.410339563 | 0.37160527 | -1.104235 | 0.26949119 | 0.39938348 | IL4 | 3565 |
| ENSG000001 | 263.018175 | 0.16790465 | 0.15205668 | 1.10422408 | 0.26949595 | 0.39938348 | BCL2L11 | 10018 |
| ENSG000002 | 4.64434322 | -0.414193729 | 0.37509808 | -1.1042278 | 0.26949434 | 0.39938348 | NA | NA |
| ENSG000001 | 12.2500199 | -0.362042793 | 0.32788181 | -1.1041869 | 0.26951208 | 0.39939279 | CIDEB | 27141 |
| ENSG000002 | 7.33735275 | -0.398133501 | 0.36059716 | -1.104095 | 0.26955195 | 0.39943728 | NA | NA |
| ENSG000001 | 14.6322143 | -0.348521302 | 0.31570503 | -1.103946 | 0.26961656 | 0.39951763 | WFIKKN2 | 124857 |
| ENSG000002 | 2.74099339 | -0.411099344 | 0.37240572 | -1.1039018 | 0.26963573 | 0.39951763 | NA | NA |
| ENSG000002 | 3.84344966 | -0.417913298 | 0.37857517 | -1.103911 | 0.26963174 | 0.39951763 | NA | NA |
| ENSG000002 | 2.73620605 | -0.413158523 | 0.37430205 | -1.1038105 | 0.26967537 | 0.39956176 | NA | NA |
| ENSG000002 | 2.48065703 | -0.407416616 | 0.36911291 | -1.1037723 | 0.26969191 | 0.39957167 | NA | NA |
| ENSG000002 | 20.4008974 | -0.310920781 | 0.28169968 | -1.1037314 | 0.26970967 | 0.39958339 | NA | NA |
| ENSG000002 | 6.07601999 | -0.411951967 | 0.37326665 | -1.1036399 | 0.26974937 | 0.39962761 | MEIS1-AS2 | 100873998 |
| ENSG000001 | 17.7981668 | -0.33193929 | 0.30078048 | -1.1035932 | 0.26976964 | 0.39962843 | ANKRD37 | 353322 |
| ENSG000002 | 2.74801563 | 0.411524633 | 0.37289063 | 1.10360679 | 0.26976374 | 0.39962843 | NA | NA |

| | | | | | | | | |
|---|---|---|---|---|---|---|---|---|
| ENSG000002 | 3.61427945 | -0.417598986 | 0.37848039 | -1.103357 | 0.26987217 | 0.39976571 | NA | NA |
| ENSG000002 | 2.49746385 | -0.408993878 | 0.37069769 | -1.1033084 | 0.26989326 | 0.39977126 | CCDC166 | 100130274 |
| ENSG000002 | 4.06791336 | -0.417172791 | 0.37811582 | -1.1032937 | 0.26989965 | 0.39977126 | NA | NA |
| ENSG000002 | 4.32514477 | -0.415805726 | 0.37688134 | -1.1032802 | 0.26990549 | 0.39977126 | NA | NA |
| ENSG000002 | 4.38124538 | -0.416481664 | 0.37753443 | -1.1031621 | 0.26995678 | 0.39983263 | NA | NA |
| ENSG000002 | 3.8263268 | -0.417423518 | 0.37844168 | -1.1030062 | 0.27002448 | 0.39987447 | NA | NA |
| ENSG000002 | 4.62877005 | -0.413869551 | 0.3752143 | -1.1030218 | 0.2700177 | 0.39987447 | NA | NA |
| ENSG000002 | 2.49223627 | -0.406885539 | 0.36887181 | -1.103054 | 0.27000371 | 0.39987447 | NA | NA |
| ENSG000002 | 12.3936439 | -0.360037732 | 0.32641364 | -1.1030107 | 0.27002253 | 0.39987447 | NA | NA |
| ENSG000002 | 4.65213944 | -0.413953926 | 0.3753147 | -1.1029515 | 0.27004821 | 0.39989501 | NA | NA |
| ENSG000002 | 2.47023214 | -0.407567651 | 0.36955304 | -1.1028665 | 0.27008516 | 0.39990591 | NA | NA |
| ENSG000002 | 10.0673509 | 0.366596593 | 0.33240104 | 1.10287439 | 0.27008172 | 0.39990591 | SMARCA5-AS1 | 100128055 |
| ENSG000002 | 2.44957595 | -0.405827721 | 0.36797468 | -1.1028686 | 0.27008423 | 0.39990591 | NA | NA |
| ENSG000002 | 6.16603928 | -0.402828506 | 0.36530845 | -1.1027079 | 0.27015404 | 0.39997869 | FOCAD-AS1 | 101929548 |
| ENSG000002 | 3.02739591 | -0.416132154 | 0.3773678 | -1.102723 | 0.27014747 | 0.39997869 | NA | NA |
| ENSG000002 | 2.45358262 | -0.408702214 | 0.3706769 | -1.1025835 | 0.2702081 | 0.40002952 | NA | NA |
| ENSG000002 | 4.06696818 | -0.416415375 | 0.37767104 | -1.1025875 | 0.27020634 | 0.40002952 | NA | NA |
| ENSG000002 | 6.74104679 | -0.398063355 | 0.36109794 | -1.1023695 | 0.27030106 | 0.40014376 | NA | NA |
| ENSG000002 | 2.52177897 | -0.407708785 | 0.36985832 | -1.1023377 | 0.27031487 | 0.40014376 | NA | NA |
| ENSG000002 | 6.71070182 | -0.400182135 | 0.36302707 | -1.1023479 | 0.27031044 | 0.40014376 | NA | NA |
| ENSG000002 | 2.72858664 | -0.413550659 | 0.37518448 | -1.1022595 | 0.27034887 | 0.40017949 | ALG1L11P | 106480789 |
| ENSG000001 | 9.11063038 | -0.38676542 | 0.35091582 | -1.1021601 | 0.27039206 | 0.40022881 | CCR4 | 1233 |
| ENSG000002 | 2.79518382 | -0.411872581 | 0.37371716 | -1.102097 | 0.27041948 | 0.40025479 | NA | NA |
| ENSG000002 | 7.01847807 | -0.403410245 | 0.36605894 | -1.1020363 | 0.27044589 | 0.40027926 | LINC01589 | 100506737 |
| ENSG000002 | 12.0081507 | 0.36973889 | 0.33552176 | 1.10198185 | 0.27046956 | 0.40029969 | RN7SL16P | 106479231 |
| ENSG000001 | 3.66449491 | -0.416732554 | 0.37823505 | -1.101782 | 0.27055647 | 0.40039755 | NA | NA |
| ENSG000002 | 2.97460409 | -0.416117352 | 0.37768883 | -1.1017465 | 0.27057189 | 0.40039755 | NA | NA |
| ENSG000002 | 2.974095 | -0.415908168 | 0.37749907 | -1.1017462 | 0.27057203 | 0.40039755 | NA | NA |
| ENSG000002 | 6.85749658 | -0.402867875 | 0.36566546 | -1.1017389 | 0.27057518 | 0.40039755 | NA | NA |

| | | | | | | | | |
|---|---|---|---|---|---|---|---|---|
| ENSG000002 | 4.08746433 | -0.4168394 | 0.37836135 | -1.1016966 | 0.27059361 | 0.40041021 | NA | NA |
| ENSG000002 | 4.11022767 | -0.416751453 | 0.37829157 | -1.1016673 | 0.27060634 | 0.40041444 | NA | NA |
| ENSG000002 | 99.6315411 | 0.201138851 | 0.18258147 | 1.10163889 | 0.27061869 | 0.40041811 | NA | NA |
| ENSG000002 | 71.4768716 | 0.207089446 | 0.18798842 | 1.10160747 | 0.27063236 | 0.40042372 | NA | NA |
| ENSG000002 | 8.35625598 | -0.393104601 | 0.35692653 | -1.10136 | 0.27074001 | 0.40056838 | PRICKLE2-AS3 | 100874243 |
| ENSG000001 | 10.0486526 | -0.368576662 | 0.33474927 | -1.1010529 | 0.27087363 | 0.40075145 | LOC100129345 | 100129345 |
| ENSG000001 | 13.1800097 | -0.358084942 | 0.32522972 | -1.1010216 | 0.27088727 | 0.40075701 | LYG1 | 129530 |
| ENSG000002 | 2.69289339 | -0.412786368 | 0.37496788 | -1.1008579 | 0.2709585 | 0.40084776 | NA | NA |
| ENSG000002 | 14.3665805 | -0.346307397 | 0.31460585 | -1.1007659 | 0.27099855 | 0.40089239 | NA | NA |
| ENSG000002 | 3.82015349 | -0.416727929 | 0.37859073 | -1.1007346 | 0.27101216 | 0.4008979 | NA | NA |
| ENSG000002 | 4.08471963 | -0.415941834 | 0.37788759 | -1.1007025 | 0.27102614 | 0.40090395 | NA | NA |
| ENSG000002 | 6.12572274 | -0.413889527 | 0.3760586 | -1.1005985 | 0.27107145 | 0.40095635 | RNU6-549P | 106479810 |
| ENSG000002 | 6.68644783 | -0.395574485 | 0.35942648 | -1.1005713 | 0.27108327 | 0.40095921 | SLC22A18AS | 5003 |
| ENSG000002 | 2.7415953 | -0.412417332 | 0.37476313 | -1.1004747 | 0.27112536 | 0.40100684 | NA | NA |
| ENSG000002 | 2.73587378 | -0.412120186 | 0.37451507 | -1.1004102 | 0.27115345 | 0.40103376 | NA | NA |
| ENSG000002 | 2.73578149 | -0.411962957 | 0.37439648 | -1.1003388 | 0.27118455 | 0.40106513 | NA | NA |
| ENSG000002 | 2.73244841 | -0.412613159 | 0.3750036 | -1.1002912 | 0.27120527 | 0.40108115 | NA | NA |
| ENSG000002 | 5.46855457 | -0.410261676 | 0.3728742 | -1.1002683 | 0.27121522 | 0.40108123 | CXorf66 | 347487 |
| ENSG000002 | 2.48019315 | -0.406020815 | 0.36908943 | -1.1000608 | 0.27130563 | 0.40118849 | NA | NA |
| ENSG000002 | 4.64772782 | -0.41396149 | 0.37630933 | -1.1000564 | 0.27130754 | 0.40118849 | CCL23 | 6368 |
| ENSG000001 | 14.82368 | 0.335735626 | 0.30522562 | 1.09995885 | 0.27135005 | 0.40123672 | LOC645166 | 645166 |
| ENSG000002 | 6.90267532 | -0.405675362 | 0.36892801 | -1.0996058 | 0.27150393 | 0.40144963 | NA | NA |
| ENSG000002 | 6.46070603 | -0.400948509 | 0.36465617 | -1.0995248 | 0.27153922 | 0.40148716 | HCG17 | 414778 |
| ENSG000002 | 6.09618662 | -0.40967008 | 0.3726102 | -1.0994602 | 0.27156739 | 0.40151417 | NA | NA |
| ENSG000002 | 2.80132935 | -0.410509586 | 0.37340026 | -1.0993822 | 0.27160141 | 0.40154983 | NA | NA |
| ENSG000002 | 3.82706869 | -0.416112208 | 0.37857107 | -1.0991654 | 0.27169594 | 0.40166031 | NA | NA |
| ENSG000002 | 5.92609077 | -0.40310755 | 0.3667354 | -1.0991782 | 0.27169036 | 0.40166031 | NA | NA |
| ENSG000001 | 8.13638757 | -0.393040826 | 0.35762767 | -1.0990224 | 0.27175828 | 0.40173782 | CHADL | 150356 |
| ENSG000002 | 6.25635022 | -0.405310711 | 0.36882571 | -1.0989221 | 0.27180207 | 0.4017879 | NA | NA |

| | | | | | | | | |
|---|---|---|---|---|---|---|---|---|
| ENSG000002 | 2.7233742 | -0.41227499 | 0.37518035 | -1.0988715 | 0.27182413 | 0.40180586 | NA | NA |
| ENSG000002 | 4.44719069 | -0.415962536 | 0.37854938 | -1.098833 | 0.27184093 | 0.40181605 | NA | NA |
| ENSG000002 | 4.3779123 | -0.414246855 | 0.3769979 | -1.0988041 | 0.2718535 | 0.40181999 | NA | NA |
| ENSG000001 | 42.2843295 | 0.259126037 | 0.23585097 | 1.09868547 | 0.27190528 | 0.40185258 | NA | NA |
| ENSG000002 | 7.79197181 | -0.392249434 | 0.35700311 | -1.0987283 | 0.27188658 | 0.40185258 | NA | NA |
| ENSG000002 | 2.4932285 | -0.404392213 | 0.36806711 | -1.0986915 | 0.27190264 | 0.40185258 | NA | NA |
| ENSG000002 | 2.95764082 | -0.414571821 | 0.3773513 | -1.0986363 | 0.27192675 | 0.40186966 | NA | NA |
| ENSG000002 | 3.57275999 | -0.415415845 | 0.37813115 | -1.0986026 | 0.27194146 | 0.40187676 | LOC103312105 | 103312105 |
| ENSG000000 | 62.7270164 | 0.228185702 | 0.20772785 | 1.09848393 | 0.27199323 | 0.4019009 | RAP1GAP | 5909 |
| ENSG000000 | 7.757591 | -0.388734241 | 0.35388111 | -1.0984883 | 0.27199134 | 0.4019009 | GP6 | 51206 |
| ENSG000002 | 30.9218044 | -0.290063724 | 0.26406055 | -1.0984743 | 0.27199745 | 0.4019009 | CATSPER2P1 | 440278 |
| ENSG000002 | 2.4695798 | -0.406496316 | 0.37003604 | -1.0985317 | 0.27197239 | 0.4019009 | NA | NA |
| ENSG000002 | 22.8920866 | 0.320342412 | 0.29168607 | 1.09824379 | 0.27209805 | 0.40203491 | NA | NA |
| ENSG000002 | 38.7231114 | -0.245075547 | 0.22315805 | -1.0982151 | 0.27211057 | 0.40203875 | NA | NA |
| ENSG000002 | 4.59282939 | -0.411647403 | 0.37484489 | -1.0981807 | 0.27212561 | 0.40204633 | NA | NA |
| ENSG000001 | 61.397554 | 0.227910387 | 0.20758804 | 1.09789751 | 0.27224925 | 0.4021759 | TNS2 | 23371 |
| ENSG000002 | 2.74349373 | -0.410968914 | 0.37430886 | -1.0979406 | 0.27223041 | 0.4021759 | NA | NA |
| ENSG000002 | 6.72438408 | -0.39635851 | 0.36101876 | -1.097889 | 0.27225298 | 0.4021759 | NA | NA |
| ENSG000002 | 27.1858565 | -0.278099128 | 0.25329794 | -1.0979131 | 0.27224243 | 0.4021759 | SLC6A14 | 11254 |
| ENSG000002 | 2.4772917 | -0.406468921 | 0.37023886 | -1.0978559 | 0.27226742 | 0.40218257 | NA | NA |
| ENSG000002 | 2.84442287 | -0.406826961 | 0.37057811 | -1.097817 | 0.2722844 | 0.402193 | MIMT1 | 100073347 |
| ENSG000001 | 2.72673919 | -0.409913294 | 0.37342285 | -1.0977188 | 0.27232729 | 0.40221241 | GOLGA8J | 653073 |
| ENSG000002 | 2.26063218 | -0.396814896 | 0.36147637 | -1.0977617 | 0.27230858 | 0.40221241 | NA | NA |
| ENSG000002 | 6.01388264 | -0.406445195 | 0.37025712 | -1.0977377 | 0.27231904 | 0.40221241 | NA | NA |
| ENSG000000 | 90.9554617 | -0.195838755 | 0.17842785 | -1.0975795 | 0.27238813 | 0.40228762 | LNX1 | 84708 |
| ENSG000002 | 2.99472417 | -0.414410756 | 0.37760186 | -1.0974807 | 0.27243131 | 0.40233674 | NA | NA |
| ENSG000002 | 6.49549298 | -0.399070741 | 0.36363679 | -1.0974433 | 0.27244767 | 0.40234625 | LOC286178 | 286178 |
| ENSG000001 | 154.481942 | 0.194946265 | 0.1776453 | 1.09739052 | 0.27247072 | 0.40236562 | ZFHX2 | 85446 |
| ENSG000001 | 38.3716605 | 0.249945232 | 0.22777305 | 1.09734329 | 0.27249135 | 0.40238145 | LPIN3 | 64900 |

| | | | | | | | | |
|---|---|---|---|---|---|---|---|---|
| ENSG000002 | 4.11583297 | -0.41493467 | 0.3781398 | -1.0973049 | 0.27250811 | 0.40239154 | NA | NA |
| ENSG000002 | 7.09124168 | -0.399687756 | 0.36426112 | -1.0972561 | 0.27252944 | 0.40240838 | NA | NA |
| ENSG000002 | 9.61322898 | -0.378705912 | 0.34515025 | -1.0972204 | 0.27254504 | 0.40241676 | NA | NA |
| ENSG000001 | 12.2950276 | -0.353192553 | 0.3219568 | -1.0970185 | 0.27263333 | 0.40249502 | LINC01600 | 154386 |
| ENSG000001 | 7.30465426 | -0.401425045 | 0.36592706 | -1.0970083 | 0.27263774 | 0.40249502 | DLK2 | 65989 |
| ENSG000002 | 12.1269516 | -0.355443382 | 0.32400866 | -1.0970182 | 0.27263345 | 0.40249502 | NA | NA |
| ENSG000002 | 4.58935141 | -0.412158289 | 0.37570665 | -1.0970215 | 0.27263197 | 0.40249502 | NA | NA |
| ENSG000001 | 424.630333 | 0.16678909 | 0.15205292 | 1.09691474 | 0.27267866 | 0.40254077 | COBL | 23242 |
| ENSG000002 | 3.59509365 | 0.415242704 | 0.37857841 | 1.09684729 | 0.27270815 | 0.40256965 | NA | NA |
| ENSG000002 | 3.0587891 | -0.412506563 | 0.37610022 | -1.0967996 | 0.27272901 | 0.40257156 | ZNF705B | 100132396 |
| ENSG000002 | 3.81838677 | -0.415088037 | 0.37846193 | -1.0967762 | 0.27273923 | 0.40257156 | NA | NA |
| ENSG000002 | 2.72602214 | -0.409783972 | 0.37362142 | -1.0967893 | 0.27273351 | 0.40257156 | NA | NA |
| ENSG000002 | 8.50711015 | -0.382594741 | 0.34885047 | -1.0967299 | 0.27275948 | 0.4025868 | NA | NA |
| ENSG000002 | 2.47876612 | -0.405982708 | 0.37024205 | -1.0965332 | 0.2728455 | 0.40266979 | NA | NA |
| ENSG000002 | 27.2148436 | -0.275352094 | 0.25110523 | -1.0965606 | 0.27283352 | 0.40266979 | PCP4L1 | 654790 |
| ENSG000002 | 2.44684314 | -0.404636654 | 0.36901158 | -1.0965419 | 0.27284171 | 0.40266979 | NA | NA |
| ENSG000002 | 3.82920397 | -0.415016213 | 0.37849568 | -1.0964886 | 0.27286499 | 0.4026839 | LOC400748 | 400748 |
| ENSG000002 | 19.9071531 | -0.31079958 | 0.2834619 | -1.0964422 | 0.27288532 | 0.40269925 | NA | NA |
| ENSG000001 | 2.71016932 | -0.410943955 | 0.37482194 | -1.0963711 | 0.27291639 | 0.40273044 | PRM1 | 5619 |
| ENSG000002 | 11.5167354 | -0.358838705 | 0.32735455 | -1.0961775 | 0.27300109 | 0.40284077 | NA | NA |
| ENSG000001 | 4.1896976 | -0.414385931 | 0.37805119 | -1.0961106 | 0.27303036 | 0.40285465 | OR52E2 | 119678 |
| ENSG000002 | 3.01263262 | -0.413386009 | 0.37713389 | -1.0961253 | 0.27302393 | 0.40285465 | NA | NA |
| ENSG000002 | 4.15514025 | -0.41440194 | 0.37808527 | -1.0960542 | 0.27305507 | 0.40287644 | NA | NA |
| ENSG000002 | 5.65522951 | -0.405544684 | 0.37002433 | -1.0959946 | 0.27308112 | 0.40290022 | TTC39C-AS1 | 102724246 |
| ENSG000001 | 83.6375613 | -0.190133366 | 0.17349279 | -1.0959151 | 0.27311593 | 0.40291138 | C8orf34 | 116328 |
| ENSG000001 | 10.0313642 | -0.366405087 | 0.33436293 | -1.0958305 | 0.27315297 | 0.40291138 | OR52A5 | 390054 |
| ENSG000002 | 19.1175287 | 0.316009383 | 0.2883775 | 1.09581845 | 0.27315823 | 0.40291138 | NA | NA |
| ENSG000002 | 3.01740787 | -0.413139703 | 0.37700564 | -1.0958449 | 0.27314667 | 0.40291138 | NA | NA |
| ENSG000002 | 3.01055684 | -0.412264769 | 0.37618696 | -1.0959039 | 0.27312081 | 0.40291138 | NA | NA |

| | | | | | | | | |
|---|---|---|---|---|---|---|---|---|
| ENSG000002 | 3.17511405 | -0.406497648 | 0.37092925 | -1.09589 | 0.27312693 | 0.40291138 | LOC105374244 | 105374244 |
| ENSG000002 | 10.5890736 | 0.368119023 | 0.33592638 | 1.09583245 | 0.2731521 | 0.40291138 | NA | NA |
| ENSG000002 | 3.26056114 | -0.414655796 | 0.37843458 | -1.0957133 | 0.27320427 | 0.40296463 | NA | NA |
| ENSG000002 | 6.20094933 | -0.402095747 | 0.3669917 | -1.0956535 | 0.27323043 | 0.40298855 | NA | NA |
| ENSG000001 | 3.05367362 | -0.412081162 | 0.37612613 | -1.095593 | 0.27325692 | 0.40301296 | KRTAP8-1 | 337879 |
| ENSG000002 | 4.84670756 | -0.409357726 | 0.37369417 | -1.0954351 | 0.27332604 | 0.40310025 | FOXP1-AS1 | 104502416 |
| ENSG000002 | 3.83229497 | -0.41470381 | 0.37858429 | -1.0954068 | 0.27333845 | 0.40310389 | NA | NA |
| ENSG000002 | 4.11161026 | -0.414582025 | 0.37854147 | -1.095209 | 0.27342508 | 0.40321698 | NA | NA |
| ENSG000002 | 3.02014047 | -0.412491596 | 0.37673153 | -1.0949219 | 0.27355087 | 0.4033878 | NA | NA |
| ENSG000002 | 4.63327753 | -0.411597792 | 0.37596481 | -1.0947774 | 0.27361415 | 0.40346645 | LOC105378668 | 105378668 |
| ENSG000002 | 3.57688399 | -0.414241436 | 0.37841083 | -1.0946871 | 0.27365376 | 0.40348084 | NA | NA |
| ENSG000002 | 6.45625563 | -0.399324554 | 0.36478117 | -1.0946962 | 0.27364976 | 0.40348084 | NA | NA |
| ENSG000002 | 2.71520439 | -0.408981368 | 0.37360253 | -1.0946965 | 0.27364964 | 0.40348084 | NA | NA |
| ENSG000002 | 4.61246022 | -0.411117441 | 0.37557508 | -1.0946345 | 0.2736768 | 0.40350014 | NA | NA |
| ENSG000001 | 2.80540918 | -0.405289216 | 0.37026634 | -1.0945883 | 0.27369703 | 0.4035014 | RNF113B | 140432 |
| ENSG000002 | 2.31229449 | -0.3889783 | 0.35536531 | -1.0945871 | 0.27369756 | 0.4035014 | NA | NA |
| ENSG000001 | 7.50230202 | -0.39232231 | 0.35843106 | -1.0945544 | 0.27371188 | 0.40350785 | SIGLEC10 | 89790 |
| ENSG000002 | 3.57453692 | 0.4142988 | 0.37856802 | 1.09438405 | 0.27378658 | 0.4036033 | NA | NA |
| ENSG000001 | 161.508841 | 0.223844682 | 0.20458509 | 1.09413977 | 0.27389368 | 0.40373088 | GCNT4 | 51301 |
| ENSG000002 | 10.8268226 | -0.368202752 | 0.33652912 | -1.0941185 | 0.27390299 | 0.40373088 | NA | NA |
| ENSG000002 | 7.78728088 | -0.39025709 | 0.35667649 | -1.0941486 | 0.2738898 | 0.40373088 | NA | NA |
| ENSG000002 | 34.8550757 | -0.268319157 | 0.24527704 | -1.0939432 | 0.27397988 | 0.40382954 | NA | NA |
| ENSG000002 | 3.85536435 | -0.4141062 | 0.37857391 | -1.0938583 | 0.27401714 | 0.40386289 | NA | NA |
| ENSG000002 | 6.99597858 | -0.397385099 | 0.3632991 | -1.0938235 | 0.27403239 | 0.40386289 | NA | NA |
| ENSG000002 | 2.38995875 | 0.402924254 | 0.36835963 | 1.09383392 | 0.27402783 | 0.40386289 | NA | NA |
| ENSG000001 | 7.07498559 | -0.397181444 | 0.36316053 | -1.0936801 | 0.27409531 | 0.40390396 | CRYBA1 | 1411 |
| ENSG000002 | 2.54163737 | -0.401478798 | 0.36707184 | -1.0937336 | 0.27407185 | 0.40390396 | NA | NA |
| ENSG000002 | 3.64214369 | -0.413794618 | 0.37835144 | -1.0936779 | 0.27409626 | 0.40390396 | RN7SL30P | 106479235 |
| ENSG000002 | 4.4809217 | -0.413929728 | 0.37847801 | -1.0936692 | 0.2741001 | 0.40390396 | NA | NA |

| | | | | | | | | |
|---|---|---|---|---|---|---|---|---|
| ENSG000002 | 3.00026458 | 0.411439985 | 0.37624067 | 1.09355532 | 0.27415005 | 0.4039629 | NA | NA |
| ENSG000001 | 139.069987 | 0.169175556 | 0.15471705 | 1.09345125 | 0.27419572 | 0.40401309 | BTN3A2 | 11118 |
| ENSG000002 | 4.59983634 | -0.410879893 | 0.37577077 | -1.0934323 | 0.27420404 | 0.40401309 | NA | NA |
| ENSG000002 | 2.980273 | -0.412331316 | 0.37712723 | -1.093348 | 0.27424102 | 0.4040382 | NA | NA |
| ENSG000002 | 6.46213611 | -0.397931526 | 0.36395487 | -1.093354 | 0.27423839 | 0.4040382 | NA | NA |
| ENSG000002 | 3.05338078 | -0.411179711 | 0.37609919 | -1.0932747 | 0.27427323 | 0.40407098 | ZNF630-AS1 | 100874120 |
| ENSG000002 | 25.2182757 | 0.287844897 | 0.26329383 | 1.09324589 | 0.27428585 | 0.4040749 | NA | NA |
| ENSG000002 | 2.54019575 | -0.402175834 | 0.36789316 | -1.0931865 | 0.27431193 | 0.40409863 | NA | NA |
| ENSG000002 | 7.56668801 | -0.391862394 | 0.35856687 | -1.0928572 | 0.27445649 | 0.4042969 | NA | NA |
| ENSG000002 | 2.27217199 | -0.392858905 | 0.35957927 | -1.0925516 | 0.27459072 | 0.40447994 | NA | NA |
| ENSG000002 | 6.21229577 | -0.399748206 | 0.36591511 | -1.0924616 | 0.27463026 | 0.4045235 | NA | NA |
| ENSG000001 | 16.6509766 | -0.339212772 | 0.31054015 | -1.0923314 | 0.27468744 | 0.40457833 | CXCR4 | 7852 |
| ENSG000002 | 2.51266454 | -0.40349927 | 0.36938749 | -1.0923469 | 0.27468067 | 0.40457833 | NA | NA |
| ENSG000002 | 4.36491627 | -0.412057668 | 0.37723786 | -1.092302 | 0.27470038 | 0.40458269 | NA | NA |
| ENSG000002 | 3.02221086 | 0.412012998 | 0.37726039 | 1.09211835 | 0.27478108 | 0.40468685 | NA | NA |
| ENSG000002 | 2.79538492 | -0.407364769 | 0.37306388 | -1.0919437 | 0.27485784 | 0.40478519 | NA | NA |
| ENSG000002 | 7.24014103 | -0.389401143 | 0.35664677 | -1.0918398 | 0.27490351 | 0.40483775 | NA | NA |
| ENSG000001 | 46576.2552 | 0.13998398 | 0.12821539 | 1.09178766 | 0.27492644 | 0.40484212 | COX2 | 4513 |
| ENSG000002 | 54.6299693 | -0.228515408 | 0.20930037 | -1.091806 | 0.27491837 | 0.40484212 | LINC01125 | 728537 |
| ENSG000002 | 3.04470354 | -0.410701531 | 0.37619338 | -1.0917298 | 0.27495187 | 0.40486486 | NA | NA |
| ENSG000002 | 2.79509208 | -0.408286593 | 0.37403787 | -1.0915649 | 0.27502441 | 0.40494226 | NA | NA |
| ENSG000002 | 2.99617788 | -0.411337871 | 0.37682674 | -1.0915835 | 0.27501619 | 0.40494226 | NA | NA |
| ENSG000002 | 2.78252007 | -0.407648755 | 0.37346504 | -1.0915312 | 0.27503919 | 0.40494933 | MIR5191 | 100847050 |
| ENSG000002 | 37.4772019 | 0.260730079 | 0.23889965 | 1.09137906 | 0.27510612 | 0.40503316 | NA | NA |
| ENSG000002 | 5.00821898 | 0.412147929 | 0.377748 | 1.09106582 | 0.27524392 | 0.40522133 | NA | NA |
| ENSG000002 | 4.57672808 | -0.409675495 | 0.37550159 | -1.0910087 | 0.27526906 | 0.40524362 | NA | NA |
| ENSG000002 | 2.35182607 | 0.400144983 | 0.36688065 | 1.09066799 | 0.275419 | 0.40544964 | RNU6-122P | 106481876 |
| ENSG000002 | 2.50826123 | -0.402378673 | 0.36893767 | -1.0906413 | 0.27543073 | 0.4054522 | NA | NA |
| ENSG000001 | 6.17112306 | -0.398056721 | 0.36500973 | -1.0905373 | 0.27547654 | 0.4055049 | CABP2 | 51475 |

| | | | | | | | | |
|---|---|---|---|---|---|---|---|---|
| ENSG000002 | 5.14259713 | -0.408344408 | 0.37453 | -1.0902849 | 0.27558766 | 0.40565375 | CRYBB2 | 1415 |
| ENSG000001 | 53.7822639 | -0.219799229 | 0.20160496 | -1.0902471 | 0.27560431 | 0.4056615 | TMEM63C | 57156 |
| ENSG000002 | 5.41412426 | -0.405825164 | 0.37223897 | -1.0902275 | 0.27561293 | 0.4056615 | NA | NA |
| ENSG000002 | 59.4535333 | 0.23793912 | 0.21826259 | 1.09015074 | 0.27564675 | 0.40569655 | BAIAP2-AS1 | 440465 |
| ENSG000002 | 2.53617753 | -0.401864509 | 0.36867429 | -1.0900259 | 0.27570175 | 0.40574805 | NA | NA |
| ENSG000002 | 2.47688621 | -0.400597701 | 0.36750568 | -1.0900449 | 0.27569336 | 0.40574805 | NA | NA |
| ENSG000002 | 4.61951923 | -0.409773316 | 0.375941 | -1.0899937 | 0.27571593 | 0.4057523 | NA | NA |
| ENSG000002 | 5.9415709 | -0.402802969 | 0.36955286 | -1.0899739 | 0.27572465 | 0.4057523 | NA | NA |
| ENSG00000C | 116.962131 | 0.178721906 | 0.16399004 | 1.08983388 | 0.27578632 | 0.40580368 | STPG1 | 90529 |
| ENSG000001 | 932.475607 | 0.307628161 | 0.28229093 | 1.08975576 | 0.27582075 | 0.40580368 | SDHC | 6391 |
| ENSG000001 | 182.970789 | -0.155323497 | 0.14253096 | -1.0897527 | 0.27582209 | 0.40580368 | PRICKLE2 | 166336 |
| ENSG000002 | 2.73065337 | -0.407145146 | 0.37361828 | -1.0897356 | 0.27582962 | 0.40580368 | NA | NA |
| ENSG000002 | 5.88979007 | -0.400920098 | 0.36790007 | -1.0897527 | 0.27582209 | 0.40580368 | NA | NA |
| ENSG000002 | 3.0026168 | -0.410500822 | 0.37669702 | -1.0897374 | 0.27582884 | 0.40580368 | NA | NA |
| ENSG000002 | 4.5643723 | -0.408662519 | 0.37500743 | -1.0897451 | 0.27582543 | 0.40580368 | NA | NA |
| ENSG000002 | 2.74524801 | -0.407122646 | 0.37366455 | -1.0895405 | 0.27591562 | 0.40591548 | NA | NA |
| ENSG000002 | 2.96933957 | -0.411235256 | 0.37746355 | -1.0894701 | 0.27594663 | 0.40594638 | LOC105378137 | 105378137 |
| ENSG000002 | 3.30497307 | -0.410826348 | 0.37711066 | -1.0894053 | 0.2759752 | 0.40597368 | NA | NA |
| ENSG000002 | 2.7075729 | -0.408440044 | 0.3749838 | -1.0892205 | 0.27605666 | 0.40607878 | NA | NA |
| ENSG000002 | 5.21135387 | -0.408520834 | 0.37511639 | -1.0890509 | 0.27613146 | 0.40615933 | NA | NA |
| ENSG000002 | 2.21288973 | -0.395107153 | 0.36279879 | -1.0890531 | 0.27613046 | 0.40615933 | MIR4259 | 100422852 |
| ENSG000001 | 17.1768366 | -0.324218827 | 0.29773408 | -1.0889544 | 0.27617401 | 0.4062072 | OSR1 | 130497 |
| ENSG000001 | 20.9458724 | 0.299812722 | 0.27540962 | 1.08860656 | 0.27632742 | 0.40633486 | SLC25A35 | 399512 |
| ENSG000001 | 17.6023922 | -0.319469033 | 0.29345742 | -1.0886385 | 0.27631334 | 0.40633486 | NA | NA |
| ENSG000002 | 16.7567026 | -0.316793728 | 0.29099725 | -1.0886485 | 0.27630891 | 0.40633486 | NA | NA |
| ENSG000002 | 19.6939567 | -0.313213252 | 0.28772754 | -1.0885758 | 0.27634098 | 0.40633486 | NA | NA |
| ENSG000002 | 2.80334477 | -0.406145763 | 0.37306348 | -1.0886774 | 0.27629619 | 0.40633486 | NA | NA |
| ENSG000002 | 4.36209228 | -0.410224586 | 0.37683248 | -1.0886126 | 0.27632476 | 0.40633486 | NA | NA |
| ENSG000002 | 2.95818215 | -0.41048626 | 0.377081 | -1.0885891 | 0.27633513 | 0.40633486 | NA | NA |

| | | | | | | | | |
|---|---|---|---|---|---|---|---|---|
| ENSG000002 | 2.68848287 | -0.407260209 | 0.37408802 | -1.0886748 | 0.2762973 | 0.40633486 | NA | NA |
| ENSG000002 | 2.80836058 | -0.404085242 | 0.37125006 | -1.0884449 | 0.27639874 | 0.40640506 | NA | NA |
| ENSG000001 | 3.94347748 | -0.411914564 | 0.37852408 | -1.0882123 | 0.27650139 | 0.40654126 | OR13H1 | 347468 |
| ENSG00000C | 204.304492 | -0.15347929 | 0.1410559 | -1.0880742 | 0.27656234 | 0.40661613 | MCF2L2 | 23101 |
| ENSG000002 | 4.35024972 | -0.410351121 | 0.37718826 | -1.0879212 | 0.27662988 | 0.40670068 | RFPL4AP3 | 106480810 |
| ENSG000002 | 2.70894263 | -0.406950491 | 0.37407187 | -1.0878939 | 0.27664196 | 0.4067037 | NA | NA |
| ENSG000002 | 3.03058518 | -0.409588597 | 0.37650789 | -1.0878619 | 0.27665606 | 0.40670969 | NA | NA |
| ENSG000002 | 4.57546969 | -0.407896772 | 0.37499295 | -1.0877452 | 0.27670762 | 0.40675598 | NA | NA |
| ENSG000002 | 6.67199721 | -0.393087222 | 0.36137583 | -1.0877518 | 0.27670467 | 0.40675598 | NA | NA |
| ENSG000002 | 3.25207923 | -0.411272729 | 0.37813976 | -1.087621 | 0.27676247 | 0.40682187 | NA | NA |
| ENSG00000C | 35.6087901 | 0.254263602 | 0.23379489 | 1.08754986 | 0.27679388 | 0.40685329 | PPP2R5B | 5526 |
| ENSG000002 | 3.58201047 | -0.410403226 | 0.37739093 | -1.0874751 | 0.27682691 | 0.40687235 | NA | NA |
| ENSG000002 | 4.68104291 | -0.410578856 | 0.37754483 | -1.087497 | 0.27681724 | 0.40687235 | LINC01580 | 101927129 |
| ENSG000002 | 23.0113013 | -0.291704947 | 0.2682625 | -1.0873862 | 0.27686615 | 0.40691527 | NA | NA |
| ENSG000002 | 19.3190667 | -0.318390522 | 0.29281364 | -1.0873487 | 0.27688275 | 0.40692492 | NA | NA |
| ENSG000002 | 4.10584995 | -0.411430884 | 0.37841565 | -1.087246 | 0.27692812 | 0.4069621 | NA | NA |
| ENSG000002 | 14.1008042 | -0.342849023 | 0.31533252 | -1.0872619 | 0.27692109 | 0.4069621 | NA | NA |
| ENSG000001 | 28.1542731 | -0.284675806 | 0.2619628 | -1.0867032 | 0.27716801 | 0.40729987 | IGFBP5 | 3488 |
| ENSG000001 | 85.4888719 | -0.222138191 | 0.20443864 | -1.0865763 | 0.27722408 | 0.40735879 | TNRC18 | 84629 |
| ENSG000002 | 6.17357546 | -0.396988455 | 0.36536029 | -1.0865671 | 0.27722819 | 0.40735879 | NA | NA |
| ENSG000002 | 2.77276071 | -0.405767932 | 0.37346086 | -1.0865072 | 0.27725464 | 0.40738289 | LOC105370829 | 105370829 |
| ENSG000002 | 10.2887454 | -0.362967217 | 0.33412085 | -1.0863351 | 0.27733075 | 0.40746898 | NA | NA |
| ENSG000002 | 2.98268051 | -0.409498125 | 0.3769558 | -1.0863293 | 0.27733333 | 0.40746898 | NA | NA |
| ENSG000002 | 2.47384617 | -0.400372642 | 0.36860591 | -1.0861807 | 0.27739904 | 0.40755076 | NA | NA |
| ENSG000002 | 4.86168733 | -0.406357802 | 0.37418445 | -1.0859826 | 0.27748669 | 0.40766476 | NA | NA |
| ENSG00000C | 12.6337312 | 0.35537041 | 0.32726962 | 1.08586436 | 0.27753901 | 0.40772317 | LAG3 | 3902 |
| ENSG000002 | 3.01143895 | -0.409463146 | 0.37709679 | -1.0858304 | 0.27755405 | 0.40772317 | NA | NA |
| ENSG000002 | 2.23448313 | -0.39148353 | 0.36054031 | -1.0858246 | 0.27755661 | 0.40772317 | NA | NA |
| ENSG000001 | 152.306359 | 0.202803574 | 0.18679635 | 1.08569345 | 0.27761464 | 0.40777887 | USP21 | 27005 |

| | | | | | | | | |
|---|---|---|---|---|---|---|---|---|
| ENSG000002 | 17.3985967 | -0.32095266 | 0.29561968 | -1.0856945 | 0.27761417 | 0.40777887 | LOC100507194 | 100507194 |
| ENSG000002 | 2.74300969 | -0.405540776 | 0.37355679 | -1.0856201 | 0.27764709 | 0.40781176 | NA | NA |
| ENSG000002 | 44.7723803 | -0.241314927 | 0.22232053 | -1.085437 | 0.27772817 | 0.40791607 | NA | NA |
| ENSG000002 | 2.30601311 | -0.388708176 | 0.35812612 | -1.0853947 | 0.2777469 | 0.40792881 | NA | NA |
| ENSG000002 | 85.0463272 | 0.210686022 | 0.19411835 | 1.0853483 | 0.27776742 | 0.40794418 | NA | NA |
| ENSG000002 | 6.68576215 | -0.394095219 | 0.36313448 | -1.0852597 | 0.27780665 | 0.40798701 | NA | NA |
| ENSG000002 | 4.08678825 | -0.410184551 | 0.37802953 | -1.0850596 | 0.27789528 | 0.4081024 | NA | NA |
| ENSG000002 | 2.45890607 | -0.399643398 | 0.36832635 | -1.0850253 | 0.27791046 | 0.40810991 | NA | NA |
| ENSG000001 | 2.70416759 | -0.406181055 | 0.37439668 | -1.0848949 | 0.2779682 | 0.40816514 | OR1E2 | 8388 |
| ENSG000002 | 3.03257555 | -0.408282662 | 0.37633003 | -1.0849059 | 0.27796334 | 0.40816514 | NA | NA |
| ENSG000002 | 6.95785784 | -0.388514308 | 0.35812163 | -1.0848669 | 0.2779806 | 0.40816856 | NA | NA |
| ENSG000001 | 22.1715031 | -0.297825465 | 0.27453519 | -1.0848353 | 0.27799462 | 0.40817437 | SORCS2 | 57537 |
| ENSG000001 | 58.8940888 | -0.227035625 | 0.20930287 | -1.084723 | 0.27804438 | 0.40823265 | MGAT3 | 4248 |
| ENSG000002 | 5.2050837 | -0.408624649 | 0.37673244 | -1.0846548 | 0.27807457 | 0.4082622 | NA | NA |
| ENSG000002 | 32.5578387 | 0.25502022 | 0.23512283 | 1.08462551 | 0.27808755 | 0.40826647 | NA | NA |
| ENSG000002 | 4.30437234 | -0.409096983 | 0.37721029 | -1.0845329 | 0.27812857 | 0.40831191 | RNU6-990P | 106480627 |
| ENSG000001 | 4.07386247 | -0.410517311 | 0.37854475 | -1.0844618 | 0.27816011 | 0.40834343 | SLC25A2 | 83884 |
| ENSG000002 | 2.38228618 | 0.399845903 | 0.36878347 | 1.08422945 | 0.27826308 | 0.40846502 | NA | NA |
| ENSG000002 | 4.09179217 | -0.409784192 | 0.37794432 | -1.0842449 | 0.27825624 | 0.40846502 | NA | NA |
| ENSG000002 | 3.24359787 | -0.409951381 | 0.37813969 | -1.0841268 | 0.27830858 | 0.40851702 | NA | NA |
| ENSG000001 | 852.168331 | 0.124903396 | 0.11521412 | 1.084098 | 0.27832135 | 0.40852098 | Mar-01 | 64757 |
| ENSG000002 | 3.05460725 | -0.408036209 | 0.3763943 | -1.0840659 | 0.2783356 | 0.4085271 | RN7SKP99 | 106479138 |
| ENSG000001 | 24.4031554 | 0.284387581 | 0.26235789 | 1.08396808 | 0.27837895 | 0.40857595 | PTPRN2 | 5799 |
| ENSG000002 | 5.3084128 | -0.407237385 | 0.3757048 | -1.0839292 | 0.27839621 | 0.40858649 | NA | NA |
| ENSG000002 | 4.36910335 | -0.409602867 | 0.3779064 | -1.0838739 | 0.27842073 | 0.40860769 | RN7SL110P | 106480371 |
| ENSG00000C | 5.73289685 | -0.403395387 | 0.37218774 | -1.0838492 | 0.27843166 | 0.40860894 | SLC9A3 | 6550 |
| ENSG000002 | 15.2413948 | -0.337544439 | 0.31144465 | -1.0838023 | 0.27845245 | 0.40862466 | NA | NA |
| ENSG000001 | 22.630828 | -0.291648145 | 0.26910503 | -1.0837707 | 0.27846648 | 0.40862701 | ACRBP | 84519 |
| ENSG000001 | 15.4289001 | -0.330703435 | 0.30515282 | -1.0837306 | 0.27848428 | 0.40862701 | TNFSF10 | 8743 |

| | | | | | | | | | |
|---|---|---|---|---|---|---|---|---|---|
| ENSG000001 | 90.7687539 | 0.18222531 | 0.16814499 | 1.08373916 | 0.27848047 | 0.40862701 | PPARA | | 5465 |
| ENSG000002 | 2.95195175 | -0.408196098 | 0.37666736 | -1.0837045 | 0.27849585 | 0.4086292 | NA | NA | |
| ENSG000002 | 2.76921767 | -0.403749157 | 0.37259271 | -1.0836207 | 0.27853303 | 0.40866897 | NA | NA | |
| ENSG000002 | 5.37253502 | -0.408598854 | 0.37715626 | -1.0833676 | 0.27864531 | 0.40881892 | NA | NA | |
| ENSG000001 | 66.4267252 | 0.215106999 | 0.19857902 | 1.08323125 | 0.27870579 | 0.40887807 | CYB561D1 | | 284613 |
| ENSG000002 | 3.86341408 | -0.410065715 | 0.37855694 | -1.0832339 | 0.27870461 | 0.40887807 | NA | NA | |
| ENSG000002 | 6.23053041 | -0.397549411 | 0.36702564 | -1.0831652 | 0.27873509 | 0.40890625 | NA | NA | |
| ENSG000002 | 5.9389953 | -0.399927088 | 0.36923644 | -1.0831192 | 0.27875551 | 0.40892141 | NA | NA | |
| ENSG000002 | 7.42603988 | -0.38341973 | 0.35402616 | -1.0830266 | 0.27879663 | 0.40896695 | NA | NA | |
| ENSG000001 | 61.6426216 | -0.22368513 | 0.20657635 | -1.0828206 | 0.27888806 | 0.40908627 | GRM8 | | 2918 |
| ENSG000002 | 2.1795323 | -0.391557965 | 0.36163462 | -1.0827447 | 0.27892176 | 0.40912089 | NA | NA | |
| ENSG000002 | 2.97254846 | -0.408168628 | 0.37699223 | -1.0826977 | 0.27894262 | 0.40913669 | NA | NA | |
| ENSG000001 | 13.7692198 | -0.34185625 | 0.31581101 | -1.082471 | 0.27904331 | 0.40926958 | ALOX12B | | 242 |
| ENSG000002 | 2.76562923 | -0.404602597 | 0.3738031 | -1.082395 | 0.27907706 | 0.40930427 | NA | NA | |
| ENSG000002 | 3.24727173 | -0.409342548 | 0.37822678 | -1.0822675 | 0.27913368 | 0.40937251 | NA | NA | |
| ENSG000002 | 15.9403865 | -0.332907375 | 0.30763065 | -1.0821658 | 0.27917886 | 0.40942396 | NA | NA | |
| ENSG000001 | 6.16919108 | -0.397439049 | 0.36727793 | -1.0821207 | 0.2791989 | 0.40943854 | GKN1 | | 56287 |
| ENSG000001 | 5.22069422 | 0.405757315 | 0.37503089 | 1.0819304 | 0.27928346 | 0.40954773 | GJB3 | | 2707 |
| ENSG000002 | 2.56287945 | -0.394887377 | 0.3650083 | -1.0818586 | 0.27931536 | 0.4095797 | NA | NA | |
| ENSG000002 | 7.46496925 | -0.383966187 | 0.35494341 | -1.0817673 | 0.27935593 | 0.40962438 | NA | NA | |
| ENSG000001 | 5.02287921 | -0.408127026 | 0.3772911 | -1.0817298 | 0.2793726 | 0.40963402 | OR8J3 | | 81168 |
| ENSG00000C | 9.2292329 | -0.370978232 | 0.34297144 | -1.0816593 | 0.27940396 | 0.40966518 | LGALS14 | | 56891 |
| ENSG000001 | 4.13233147 | -0.40926381 | 0.37838725 | -1.0816004 | 0.27943012 | 0.409666 | NA | NA | |
| ENSG000002 | 5.12924721 | -0.402669814 | 0.37229141 | -1.0815984 | 0.27943099 | 0.409666 | NA | NA | |
| ENSG000002 | 4.37457922 | -0.407987901 | 0.3772113 | -1.0815898 | 0.27943483 | 0.409666 | NA | NA | |
| ENSG000002 | 2.79051271 | -0.404142195 | 0.37370562 | -1.0814453 | 0.27949907 | 0.40974471 | NA | NA | |
| ENSG000002 | 2.72081024 | -0.40529208 | 0.37477643 | -1.0814236 | 0.27950872 | 0.40974471 | NA | NA | |
| ENSG000001 | 2.97836227 | -0.40785757 | 0.37716407 | -1.0813797 | 0.27952825 | 0.40975852 | PLA2G2E | | 30814 |
| ENSG000002 | 7.57473218 | -0.389280411 | 0.36006261 | -1.0811465 | 0.27963197 | 0.40989575 | LOC400043 | | 400043 |

| | | | | | | | | | |
|---|---|---|---|---|---|---|---|---|---|
| ENSG000001 | 9.58942083 | -0.371013003 | 0.34318593 | -1.0810846 | 0.2796595 | 0.40991836 | TEX37 | | 200523 |
| ENSG000002 | 21.6402359 | -0.307113357 | 0.28408373 | -1.0810663 | 0.27966761 | 0.40991836 | NA | NA | |
| ENSG000002 | 2.47287321 | -0.397538608 | 0.36775766 | -1.0809798 | 0.27970609 | 0.40995994 | NA | NA | |
| ENSG000000 | 50.4982663 | -0.228628315 | 0.21151546 | -1.0809059 | 0.27973896 | 0.40997847 | FAM184B | | 27146 |
| ENSG000002 | 9.66060615 | -0.377850374 | 0.34956551 | -1.0809143 | 0.27973522 | 0.40997847 | NA | NA | |
| ENSG000001 | 4.6355492 | -0.40655219 | 0.37617121 | -1.0807637 | 0.27980223 | 0.41005639 | OR51S1 | | 119692 |
| ENSG000002 | 11.5464504 | -0.349343513 | 0.32325805 | -1.0806955 | 0.2798326 | 0.41008607 | NA | NA | |
| ENSG000002 | 2.70219628 | -0.403506127 | 0.37339647 | -1.0806372 | 0.27985852 | 0.41009442 | NA | NA | |
| ENSG000002 | 2.70231251 | -0.402703483 | 0.37264685 | -1.0806571 | 0.27984965 | 0.41009442 | NA | NA | |
| ENSG000002 | 6.87208003 | 0.390211136 | 0.36112252 | 1.08055055 | 0.27989709 | 0.41013612 | NA | NA | |
| ENSG000001 | 32.2419814 | 0.266274946 | 0.24644836 | 1.08044924 | 0.27994218 | 0.41016175 | ZNF433 | | 163059 |
| ENSG000002 | 4.038856 | -0.408462491 | 0.37805092 | -1.0804431 | 0.27994493 | 0.41016175 | NA | NA | |
| ENSG000002 | 8.27034679 | -0.378991537 | 0.35077063 | -1.080454 | 0.27994005 | 0.41016175 | NA | NA | |
| ENSG000002 | 11.4785221 | -0.349236018 | 0.32328076 | -1.080287 | 0.28001438 | 0.41024868 | NA | NA | |
| ENSG000002 | 16.5552641 | -0.318244309 | 0.29459938 | -1.0802613 | 0.28002584 | 0.41025065 | NA | NA | |
| ENSG000002 | 2.71545344 | -0.403925862 | 0.37397126 | -1.0800987 | 0.28009824 | 0.41034189 | NA | NA | |
| ENSG000002 | 2.99446303 | -0.40705074 | 0.37687945 | -1.0800556 | 0.28011744 | 0.41035521 | NA | NA | |
| ENSG000002 | 2.71093878 | -0.403512141 | 0.37363628 | -1.0799597 | 0.28016011 | 0.41040288 | NA | NA | |
| ENSG000002 | 6.16627458 | -0.395990469 | 0.36671595 | -1.0798289 | 0.28021839 | 0.41046541 | NA | NA | |
| ENSG000002 | 10.0584815 | -0.368120815 | 0.34090992 | -1.0798184 | 0.28022304 | 0.41046541 | NA | NA | |
| ENSG000001 | 10.9760499 | -0.351308172 | 0.32535934 | -1.0797544 | 0.28025156 | 0.41048035 | FCER1G | | 2207 |
| ENSG000002 | 7.68203122 | -0.392021141 | 0.36306655 | -1.0797501 | 0.28025349 | 0.41048035 | OR13C3 | | 138803 |
| ENSG000002 | 7.70418773 | -0.381404755 | 0.35326077 | -1.0796691 | 0.28028954 | 0.41051834 | NA | NA | |
| ENSG000002 | 4.83561017 | -0.403843099 | 0.37405616 | -1.0796323 | 0.28030597 | 0.41052757 | NA | NA | |
| ENSG000002 | 5.79243524 | -0.403802215 | 0.37409054 | -1.0794238 | 0.28039887 | 0.41064879 | NA | NA | |
| ENSG000002 | 2.44456048 | -0.398829682 | 0.36949511 | -1.079391 | 0.28041348 | 0.41065537 | NA | NA | |
| ENSG000001 | 3.52082216 | -0.407841082 | 0.37786226 | -1.079338 | 0.2804371 | 0.41067084 | DGCR6 | | 8214 |
| ENSG000002 | 2.43756508 | -0.396441991 | 0.36730658 | -1.0793218 | 0.2804443 | 0.41067084 | NA | NA | |
| ENSG000001 | 89.4403201 | 0.184694524 | 0.17114046 | 1.07919851 | 0.28049924 | 0.41072505 | CADM3 | | 57863 |

| | | | | | | | | |
|---|---|---|---|---|---|---|---|---|
| ENSG000002 | 3.27334146 | -0.408085173 | 0.3781391 | -1.0791933 | 0.28050158 | 0.41072505 | RN7SL688P | 106479474 |
| ENSG000002 | 17.7780042 | -0.312206486 | 0.28930309 | -1.0791675 | 0.28051307 | 0.41072704 | NA | NA |
| ENSG000002 | 4.61293886 | -0.407588355 | 0.37772074 | -1.0790733 | 0.28055507 | 0.4107737 | NA | NA |
| ENSG000002 | 4.37002489 | -0.406841746 | 0.37704033 | -1.0790404 | 0.28056973 | 0.41078034 | NA | NA |
| ENSG000001 | 155.154735 | 0.160274265 | 0.14855417 | 1.0788944 | 0.28063481 | 0.41084595 | ZNF383 | 163087 |
| ENSG000001 | 2.77684285 | -0.401481929 | 0.37211719 | -1.0789126 | 0.28062668 | 0.41084595 | RNU6-1266P | 106480118 |
| ENSG000002 | 42.2552772 | 0.245031379 | 0.22716972 | 1.07862692 | 0.28075408 | 0.41100573 | NA | NA |
| ENSG000002 | 4.63748029 | -0.405414059 | 0.37588542 | -1.0785576 | 0.28078501 | 0.41103617 | NA | NA |
| ENSG000002 | 3.02147796 | -0.405180987 | 0.37570011 | -1.0784692 | 0.28082444 | 0.41107904 | NA | NA |
| ENSG000002 | 3.87519159 | -0.408237219 | 0.37857427 | -1.0783544 | 0.28087564 | 0.41113916 | NA | NA |
| ENSG000002 | 13.5612399 | -0.340666808 | 0.31592764 | -1.0783064 | 0.28089703 | 0.41114429 | PTCHD1-AS | 100873065 |
| ENSG000002 | 5.24601875 | -0.407056541 | 0.37749804 | -1.0783011 | 0.28089942 | 0.41114429 | RN7SL650P | 106480523 |
| ENSG000002 | 3.24567311 | -0.407719105 | 0.37817943 | -1.0781102 | 0.28098457 | 0.41125407 | NA | NA |
| ENSG000001 | 14.2346788 | -0.330787976 | 0.30684752 | -1.0780207 | 0.28102452 | 0.4112977 | HFE2 | 148738 |
| ENSG000001 | 14.5679024 | -0.33191919 | 0.30799531 | -1.0776761 | 0.28117833 | 0.41150795 | NAPSA | 9476 |
| ENSG000001 | 9.25510677 | -0.371707523 | 0.34494977 | -1.07757 | 0.2812257 | 0.41156243 | NA | NA |
| ENSG000002 | 3.24052592 | -0.407662864 | 0.37833368 | -1.077522 | 0.28124713 | 0.41157894 | NA | NA |
| ENSG000002 | 5.4796525 | -0.402193523 | 0.3732852 | -1.077443 | 0.28128241 | 0.41161571 | NA | NA |
| ENSG000002 | 4.35213649 | -0.40626708 | 0.37711738 | -1.0772961 | 0.281348 | 0.41169684 | NA | NA |
| ENSG000002 | 4.57818179 | -0.405842542 | 0.37674239 | -1.0772415 | 0.28137238 | 0.41170592 | NA | NA |
| ENSG000002 | 2.74551711 | -0.402887725 | 0.3740011 | -1.0772367 | 0.28137451 | 0.41170592 | NA | NA |
| ENSG000002 | 3.2061707 | -0.40760256 | 0.37839494 | -1.0771882 | 0.2813962 | 0.41172279 | NA | NA |
| ENSG000002 | 9.14853643 | -0.378496761 | 0.35141558 | -1.0770631 | 0.28145207 | 0.41177906 | NA | NA |
| ENSG000002 | 4.55961577 | -0.404843255 | 0.37587926 | -1.0770566 | 0.28145496 | 0.41177906 | NA | NA |
| ENSG000002 | 11.4694415 | 0.351726675 | 0.32657805 | 1.07700649 | 0.28147737 | 0.41179698 | NA | NA |
| ENSG000002 | 2.68802586 | -0.401669191 | 0.37296458 | -1.0769634 | 0.28149664 | 0.41181031 | NA | NA |
| ENSG000002 | 7.42556699 | 0.386775981 | 0.35917578 | 1.07684317 | 0.28155034 | 0.41187401 | NA | NA |
| ENSG000002 | 4.10772429 | -0.40719142 | 0.37821136 | -1.076624 | 0.28164829 | 0.41200245 | NA | NA |
| ENSG000002 | 2.27837756 | -0.386285447 | 0.35881348 | -1.0765634 | 0.28167538 | 0.41202721 | NA | NA |

| | | | | | | | | |
|---|---|---|---|---|---|---|---|---|
| ENSG000001 | 6.84865049 | -0.395946341 | 0.36781442 | -1.076484 | 0.28171085 | 0.41206327 | STATH | 6779 |
| ENSG000002 | 2.2366099 | -0.385924098 | 0.35851134 | -1.0764627 | 0.28172035 | 0.41206327 | RN7SKP79 | 106480857 |
| ENSG000002 | 3.02495649 | -0.40543312 | 0.37664478 | -1.0764337 | 0.28173335 | 0.41206742 | NA | NA |
| ENSG000002 | 2.47074177 | -0.397987119 | 0.36974122 | -1.0763937 | 0.28175122 | 0.41207868 | NA | NA |
| ENSG000002 | 16.4744032 | -0.318175096 | 0.29560043 | -1.0763688 | 0.28176233 | 0.41208008 | NA | NA |
| ENSG000002 | 3.85095437 | -0.407473646 | 0.37859097 | -1.0762899 | 0.28179761 | 0.41211682 | NA | NA |
| ENSG000002 | 19.7090367 | -0.299465251 | 0.27827442 | -1.0761508 | 0.28185981 | 0.4121929 | LOC101928053 | 101928053 |
| ENSG000001 | 7.46250171 | -0.383458218 | 0.35644274 | -1.0757919 | 0.28202032 | 0.41241277 | TOMM20L | 387990 |
| ENSG000002 | 9.73974555 | -0.378776628 | 0.35217818 | -1.0755255 | 0.2821395 | 0.41256612 | LOC284294 | 284294 |
| ENSG000002 | 4.01052802 | -0.40589409 | 0.37739613 | -1.0755121 | 0.28214554 | 0.41256612 | NA | NA |
| ENSG000002 | 35.636457 | -0.251381944 | 0.2337858 | -1.0752661 | 0.2822556 | 0.41271218 | PWRN1 | 791114 |
| ENSG000002 | 3.03775663 | -0.405023997 | 0.37668335 | -1.0752373 | 0.28226849 | 0.41271615 | NA | NA |
| ENSG000002 | 6.38242503 | -0.392758536 | 0.36529616 | -1.0751784 | 0.28229487 | 0.41273984 | NA | NA |
| ENSG000001 | 17.4408748 | 0.319223183 | 0.29691147 | 1.07514602 | 0.28230936 | 0.41274614 | C3orf80 | 401097 |
| ENSG000002 | 6.25386699 | -0.395118736 | 0.36762626 | -1.0747838 | 0.28247155 | 0.41296838 | NA | NA |
| ENSG000001 | 82.0213979 | -0.207521266 | 0.19310271 | -1.0746678 | 0.28252348 | 0.41301452 | TNFAIP2 | 7127 |
| ENSG000002 | 5.63014241 | 0.396318672 | 0.36877562 | 1.07468782 | 0.28251452 | 0.41301452 | NA | NA |
| ENSG000002 | 12.0165729 | -0.350310158 | 0.32600694 | -1.0745481 | 0.28257708 | 0.41307798 | NA | NA |
| ENSG000002 | 3.03767122 | -0.403461073 | 0.37549444 | -1.0744795 | 0.28260783 | 0.41310804 | NA | NA |
| ENSG000002 | 16.7656125 | -0.318308107 | 0.29627145 | -1.07438 | 0.28265242 | 0.41315832 | NA | NA |
| ENSG000002 | 8.75588887 | -0.373448003 | 0.34760763 | -1.0743378 | 0.28267132 | 0.41317106 | CLEC2L | 154790 |
| ENSG000001 | 11.7359182 | -0.345095865 | 0.32123123 | -1.0742911 | 0.28269222 | 0.41317182 | PNLIP | 5406 |
| ENSG000002 | 3.21778926 | -0.406235715 | 0.37814155 | -1.0742954 | 0.28269032 | 0.41317182 | PPIAL4D | 645142 |
| ENSG000000 | 213.327073 | -0.152638866 | 0.14209057 | -1.0742364 | 0.28271674 | 0.41317788 | COBLL1 | 22837 |
| ENSG000002 | 5.13155079 | -0.400108277 | 0.372455 | -1.0742459 | 0.28271246 | 0.41317788 | NA | NA |
| ENSG000002 | 5.25971143 | -0.403168167 | 0.37532028 | -1.0741977 | 0.2827341 | 0.41318835 | NA | NA |
| ENSG000002 | 62.3278903 | 0.219510479 | 0.2044042 | 1.07390397 | 0.28286572 | 0.41336582 | HERC2P2 | 400322 |
| ENSG000001 | 4.58977597 | -0.404106832 | 0.37633023 | -1.0738091 | 0.28290824 | 0.41341305 | DBX1 | 120237 |
| ENSG000002 | 6.63411302 | -0.387075243 | 0.3604856 | -1.0737606 | 0.28292998 | 0.41341502 | NA | NA |

| | | | | | | | | |
|---|---|---|---|---|---|---|---|---|
| ENSG000002 | 17.6443735 | -0.320078323 | 0.29808687 | -1.0737753 | 0.2829234 | 0.41341502 | NOTCH2NL | 388677 |
| ENSG000002 | 36.8022292 | 0.251498367 | 0.23423511 | 1.07370054 | 0.28295692 | 0.41343948 | LOC103344931 | 103344931 |
| ENSG000001 | 9.84932555 | -0.367879375 | 0.34265172 | -1.0736248 | 0.28299089 | 0.41345932 | TLX2 | 3196 |
| ENSG000002 | 2.7143187 | -0.401036373 | 0.37352856 | -1.0736431 | 0.28298265 | 0.41345932 | NA | NA |
| ENSG000002 | 8.07651549 | -0.388857815 | 0.36222639 | -1.0735215 | 0.2830372 | 0.41349718 | CD300LD | 100131439 |
| ENSG000002 | 3.34915878 | -0.403930958 | 0.37625994 | -1.0735423 | 0.28302788 | 0.41349718 | NA | NA |
| ENSG000002 | 3.59632525 | -0.405293233 | 0.377552 | -1.0734766 | 0.28305734 | 0.41351172 | NA | NA |
| ENSG000002 | 2.44555773 | -0.393583622 | 0.36666362 | -1.0734188 | 0.28308325 | 0.41351859 | NA | NA |
| ENSG000002 | 2.24783258 | -0.387388405 | 0.36089475 | -1.073411 | 0.28308674 | 0.41351859 | NA | NA |
| ENSG000002 | 2.78984684 | -0.399900791 | 0.37256297 | -1.0733777 | 0.28310167 | 0.41351859 | ART2BP | 106481723 |
| ENSG000002 | 29.156889 | -0.266499437 | 0.24828699 | -1.0733524 | 0.28311303 | 0.41351859 | MRPS31P5 | 100887750 |
| ENSG000002 | 10.7251645 | -0.354474962 | 0.33024535 | -1.0733685 | 0.2831058 | 0.41351859 | NA | NA |
| ENSG000002 | 4.16897025 | -0.40627968 | 0.37854841 | -1.0732569 | 0.28315588 | 0.41356627 | NA | NA |
| ENSG000002 | 2.99153958 | -0.404268895 | 0.37669471 | -1.0732003 | 0.28318124 | 0.41358842 | NA | NA |
| ENSG000002 | 2.99088167 | -0.404179566 | 0.37663806 | -1.0731246 | 0.28321522 | 0.41362314 | NA | NA |
| ENSG000001 | 79.4040084 | -0.197422109 | 0.18401541 | -1.0728564 | 0.28333555 | 0.41378399 | TCP11L2 | 255394 |
| ENSG000002 | 3.58468021 | -0.405791488 | 0.37827267 | -1.0727486 | 0.28338392 | 0.41383972 | CTF2P | 100289419 |
| ENSG000001 | 6.42003065 | -0.390396603 | 0.3639644 | -1.0726231 | 0.28344028 | 0.41390006 | CCIN | 881 |
| ENSG000001 | 3.26748555 | -0.334212779 | 0.31160125 | -1.0725656 | 0.28346607 | 0.41390006 | SLC22A6 | 9356 |
| ENSG000002 | 14.1159105 | -0.340294155 | 0.31725799 | -1.0726102 | 0.28344605 | 0.41390006 | NA | NA |
| ENSG000002 | 11.8910499 | -0.352871652 | 0.32899389 | -1.0725781 | 0.28346044 | 0.41390006 | NA | NA |
| ENSG000002 | 14.221701 | -0.354766705 | 0.33079372 | -1.0724711 | 0.28350849 | 0.4139471 | MIR137HG | 400765 |
| ENSG000002 | 10.3293781 | -0.361299852 | 0.33697089 | -1.072199 | 0.28363066 | 0.41410981 | NA | NA |
| ENSG000002 | 4.42626189 | -0.404910865 | 0.37765286 | -1.0721774 | 0.28364036 | 0.41410981 | NA | NA |
| ENSG000002 | 12.8474217 | -0.347168477 | 0.32386945 | -1.0719396 | 0.28374717 | 0.41425085 | NA | NA |
| ENSG000002 | 3.56654254 | -0.404959146 | 0.37780529 | -1.0718726 | 0.28377725 | 0.41426493 | NA | NA |
| ENSG000002 | 3.24567365 | -0.405525878 | 0.37833312 | -1.0718752 | 0.28377611 | 0.41426493 | NA | NA |
| ENSG000002 | 3.29035681 | -0.405278087 | 0.37817905 | -1.0716566 | 0.28387429 | 0.41439168 | NA | NA |
| ENSG000002 | 3.52832954 | -0.405681485 | 0.37858993 | -1.0715591 | 0.28391812 | 0.41444073 | NA | NA |

| | | | | | | | | |
|---|---|---|---|---|---|---|---|---|
| ENSG000002 | 2.51133371 | -0.393488263 | 0.36723183 | -1.0714982 | 0.28394546 | 0.41446573 | RNU6-712P | 106481404 |
| ENSG000000 | 11.8535689 | 0.353701702 | 0.33015515 | 1.07131967 | 0.28402572 | 0.41456795 | CACNG5 | 27091 |
| ENSG000000 | 70.2382658 | -0.197138878 | 0.18404737 | -1.0711312 | 0.28411044 | 0.4146324 | ZCWPW1 | 55063 |
| ENSG000002 | 2.38706088 | 0.395978712 | 0.36971243 | 1.07104516 | 0.28414913 | 0.4146324 | NA | NA |
| ENSG000002 | 2.73298975 | -0.400126152 | 0.37358881 | -1.0710336 | 0.28415434 | 0.4146324 | NA | NA |
| ENSG000002 | 4.15227608 | -0.405466467 | 0.37852126 | -1.0711855 | 0.28408605 | 0.4146324 | NA | NA |
| ENSG000002 | 4.22789174 | -0.405324393 | 0.37843103 | -1.0710654 | 0.28414001 | 0.4146324 | NA | NA |
| ENSG000002 | 6.87538747 | -0.383917519 | 0.35844377 | -1.0710676 | 0.28413902 | 0.4146324 | NA | NA |
| ENSG000002 | 4.70778495 | -0.405019978 | 0.37811152 | -1.0711654 | 0.28409507 | 0.4146324 | NA | NA |
| ENSG000002 | 16.1338754 | 0.325169539 | 0.30360827 | 1.07101676 | 0.2841619 | 0.4146324 | LOC100289361 | 100289361 |
| ENSG000002 | 5.38016546 | -0.400441037 | 0.37385736 | -1.0711065 | 0.28412155 | 0.4146324 | NA | NA |
| ENSG000001 | 33.9308393 | 0.253961343 | 0.237143 | 1.07092066 | 0.2842051 | 0.41468053 | NYAP1 | 222950 |
| ENSG000002 | 7.13295877 | -0.381708078 | 0.35644134 | -1.0708861 | 0.28422064 | 0.41468827 | NA | NA |
| ENSG000001 | 151.854923 | 0.163105302 | 0.15232635 | 1.07076226 | 0.28427634 | 0.4147397 | TMEM19 | 55266 |
| ENSG000002 | 3.23531348 | -0.405060225 | 0.37828424 | -1.0707827 | 0.28426715 | 0.4147397 | C5orf67 | 101928448 |
| ENSG000002 | 4.4379402 | -0.405329173 | 0.37859003 | -1.0706282 | 0.28433663 | 0.41481273 | NA | NA |
| ENSG000002 | 9.72028126 | -0.375776862 | 0.35102853 | -1.0705023 | 0.28439325 | 0.41488041 | NA | NA |
| ENSG000002 | 3.22193974 | -0.404985279 | 0.37833399 | -1.0704438 | 0.28441959 | 0.4149039 | NA | NA |
| ENSG000001 | 6.93679595 | -0.388351983 | 0.36282583 | -1.0703537 | 0.28446012 | 0.41493318 | RNASE10 | 338879 |
| ENSG000001 | 6.2169732 | -0.392713824 | 0.36689722 | -1.0703647 | 0.28445519 | 0.41493318 | CALHM3 | 119395 |
| ENSG000002 | 3.88861042 | -0.404977637 | 0.37841196 | -1.0702031 | 0.28452792 | 0.41501714 | NA | NA |
| ENSG000002 | 2.19261265 | -0.38516491 | 0.35991199 | -1.0701642 | 0.28454543 | 0.41502775 | NA | NA |
| ENSG000002 | 3.92938615 | -0.40495531 | 0.37841335 | -1.0701401 | 0.28455625 | 0.41502861 | EVPLL | 645027 |
| ENSG000002 | 2.21951154 | -0.3857298 | 0.36047263 | -1.0700668 | 0.28458923 | 0.41506179 | NA | NA |
| ENSG000002 | 2.98238104 | -0.403402643 | 0.37699819 | -1.0700387 | 0.2846019 | 0.41506533 | NA | NA |
| ENSG000002 | 7.08541911 | 0.384305413 | 0.35918244 | 1.06994489 | 0.28464412 | 0.41510592 | FAM209B | 388799 |
| ENSG000002 | 4.88159835 | -0.40157441 | 0.37532726 | -1.0699314 | 0.2846502 | 0.41510592 | NA | NA |
| ENSG000002 | 3.60136252 | -0.404562351 | 0.37816042 | -1.0698168 | 0.2847018 | 0.41515131 | RNU6-1181P | 106480085 |
| ENSG000002 | 6.17183935 | -0.394016248 | 0.36829731 | -1.069832 | 0.28469493 | 0.41515131 | NA | NA |

| | | | | | | | | |
|---|---|---|---|---|---|---|---|---|
| ENSG000001 | 8.87768272 | -0.378865652 | 0.35415762 | -1.0697656 | 0.28472481 | 0.41516993 | CALY | 50632 |
| ENSG000002 | 5.94247717 | -0.396246948 | 0.37042064 | -1.0697216 | 0.28474465 | 0.41518393 | MTND6P7 | 106478945 |
| ENSG000002 | 6.6584155 | -0.386196481 | 0.36108777 | -1.0695363 | 0.28482808 | 0.41529065 | NA | NA |
| ENSG000002 | 6.75407903 | -0.39322767 | 0.36767206 | -1.0695065 | 0.28484149 | 0.41529527 | NA | NA |
| ENSG000001 | 708.016851 | 0.121860146 | 0.11394758 | 1.0694404 | 0.28487127 | 0.41532376 | SNRNP48 | 154007 |
| ENSG000002 | 4.31715267 | -0.403952281 | 0.37773359 | -1.0694105 | 0.28488472 | 0.41532844 | NA | NA |
| ENSG000002 | 6.46134771 | -0.389580464 | 0.36432072 | -1.0693338 | 0.28491928 | 0.41534895 | CTAGE9 | 643854 |
| ENSG000002 | 6.28591271 | 0.390161031 | 0.36486078 | 1.06934221 | 0.2849155 | 0.41534895 | MIR4258 | 100423020 |
| ENSG000002 | 2.94876248 | -0.402618633 | 0.37657674 | -1.0691543 | 0.28500015 | 0.41545191 | NA | NA |
| ENSG000002 | 4.11683599 | -0.404596264 | 0.37845367 | -1.0690774 | 0.2850348 | 0.41548747 | NA | NA |
| ENSG000002 | 3.23907111 | -0.40434218 | 0.37822653 | -1.0690476 | 0.2850482 | 0.41549208 | NA | NA |
| ENSG000002 | 9.43461932 | -0.36355388 | 0.34008244 | -1.0690169 | 0.28506205 | 0.41549732 | LINC00320 | 387486 |
| ENSG000002 | 5.76201206 | -0.400173326 | 0.374387 | -1.0688761 | 0.2851255 | 0.41557487 | NA | NA |
| ENSG000002 | 12.5849856 | -0.345848521 | 0.32361607 | -1.0687001 | 0.28520483 | 0.41567556 | NA | NA |
| ENSG000002 | 6.98753368 | -0.387393668 | 0.36254224 | -1.0685477 | 0.28527354 | 0.41576076 | NA | NA |
| ENSG000002 | 3.002697 | -0.402749594 | 0.37696099 | -1.0684119 | 0.28533477 | 0.41583505 | NA | NA |
| ENSG000001 | 5.64079795 | -0.397819731 | 0.37237006 | -1.0683451 | 0.28536487 | 0.41585269 | ART1 | 417 |
| ENSG000002 | 3.24346202 | -0.403849889 | 0.37801642 | -1.0683395 | 0.28536738 | 0.41585269 | HOXA11-AS | 221883 |
| ENSG000002 | 7.21546641 | -0.38337438 | 0.35886538 | -1.0682958 | 0.2853871 | 0.41586647 | NAIP | 4671 |
| ENSG000002 | 11.0404382 | -0.362943404 | 0.33978362 | -1.0681604 | 0.28544816 | 0.41594051 | NA | NA |
| ENSG000001 | 96.0170058 | -0.175135524 | 0.16398063 | -1.0680257 | 0.28550894 | 0.41601412 | CHRNB2 | 1141 |
| ENSG000001 | 5.34402737 | -0.395856049 | 0.37069367 | -1.0678792 | 0.28557502 | 0.41609545 | PPP1R14D | 54866 |
| ENSG000002 | 7.17377994 | -0.381034586 | 0.35685987 | -1.0677429 | 0.28563651 | 0.41615515 | NA | NA |
| ENSG000002 | 4.36060521 | -0.402826281 | 0.37726223 | -1.067762 | 0.28562788 | 0.41615515 | NA | NA |
| ENSG000002 | 5.41063875 | -0.399753407 | 0.37440513 | -1.0677028 | 0.28565459 | 0.41616654 | KCNQ5-IT1 | 100507381 |
| ENSG000002 | 4.41397083 | -0.403143437 | 0.37764163 | -1.0675291 | 0.28573297 | 0.41626576 | NA | NA |
| ENSG000001 | 3.24825743 | -0.403792434 | 0.37826158 | -1.0674952 | 0.28574827 | 0.4162731 | APOBEC2 | 10930 |
| ENSG000001 | 16.6082593 | -0.319034245 | 0.29888594 | -1.0674114 | 0.28578612 | 0.41629833 | CBLN3 | 643866 |
| ENSG000002 | 3.45324894 | -0.40006048 | 0.37479015 | -1.0674253 | 0.28577984 | 0.41629833 | NA | NA |

| | | | | | | | | |
|---|---|---|---|---|---|---|---|---|
| ENSG000002 | 25.816147 | 0.274305343 | 0.25699512 | 1.06735622 | 0.28581101 | 0.41631964 | PI4KAP1 | 728233 |
| ENSG000002 | 5.41310682 | 0.400486342 | 0.37529128 | 1.06713468 | 0.28591102 | 0.41645036 | NA | NA |
| ENSG000002 | 3.24698356 | -0.403262433 | 0.37790614 | -1.0670968 | 0.28592813 | 0.41646031 | NA | NA |
| ENSG000002 | 3.53194336 | -0.403949199 | 0.37857257 | -1.0670324 | 0.2859572 | 0.4164877 | RN7SL277P | 106479323 |
| ENSG000002 | 3.2706337 | -0.40336632 | 0.3780625 | -1.0669303 | 0.28600333 | 0.41652496 | NA | NA |
| ENSG000002 | 2.98524447 | -0.401094266 | 0.37593294 | -1.0669304 | 0.28600328 | 0.41652496 | RNU6-561P | 106479815 |
| ENSG000002 | 3.56063309 | -0.403312443 | 0.37807926 | -1.0667404 | 0.28608906 | 0.41663485 | LINC01520 | 101929684 |
| ENSG000002 | 3.2696676 | -0.40289551 | 0.37772164 | -1.0666466 | 0.28613144 | 0.41668161 | RN7SL18P | 106480927 |
| ENSG000002 | 4.84910233 | -0.400653833 | 0.37563563 | -1.0666023 | 0.28615146 | 0.41669579 | SYCE3 | 644186 |
| ENSG000002 | 14.2814936 | 0.332283146 | 0.31155709 | 1.06652411 | 0.28618679 | 0.41673228 | NA | NA |
| ENSG000000 | 18.6783416 | -0.310871997 | 0.29148942 | -1.066495 | 0.28619995 | 0.41673648 | NR1H4 | 9971 |
| ENSG000002 | 16.1194353 | -0.325428658 | 0.3051606 | -1.0664177 | 0.28623487 | 0.41677236 | LINC00485 | 283432 |
| ENSG000002 | 5.07299005 | -0.39800651 | 0.3732771 | -1.0662495 | 0.28631088 | 0.41686807 | NA | NA |
| ENSG000002 | 4.5713994 | -0.40037139 | 0.3755428 | -1.0661139 | 0.28637217 | 0.41694233 | NA | NA |
| ENSG000002 | 9.16420314 | -0.375852505 | 0.35262416 | -1.0658728 | 0.28648114 | 0.41708601 | NA | NA |
| ENSG000002 | 3.63449965 | 0.403443276 | 0.37856125 | 1.06572787 | 0.28654668 | 0.41716645 | NA | NA |
| ENSG000002 | 6.89946106 | -0.382806968 | 0.35924279 | -1.065594 | 0.2866072 | 0.41723958 | NTRK3-AS1 | 283738 |
| ENSG000002 | 3.27120206 | -0.402708483 | 0.37793238 | -1.065557 | 0.28662396 | 0.41724501 | NA | NA |
| ENSG000002 | 27.464065 | -0.264900608 | 0.24860685 | -1.0655403 | 0.28663151 | 0.41724501 | IL10RB | 3588 |
| ENSG000002 | 10.2084101 | -0.35539346 | 0.3335726 | -1.0654156 | 0.28668789 | 0.4173121 | NA | NA |
| ENSG000001 | 7.53314361 | 0.384770133 | 0.36116916 | 1.06534604 | 0.28671937 | 0.41731875 | GPR68 | 8111 |
| ENSG000002 | 2.69849868 | -0.398543841 | 0.37410109 | -1.0653373 | 0.28672333 | 0.41731875 | NA | NA |
| ENSG000002 | 3.22748501 | -0.403077727 | 0.37835324 | -1.0653476 | 0.28671864 | 0.41731875 | NA | NA |
| ENSG000002 | 7.98622892 | -0.378244308 | 0.35506656 | -1.0652772 | 0.28675051 | 0.41734333 | UBE2E1-AS1 | 100874092 |
| ENSG000001 | 125.966959 | -0.163975457 | 0.15394321 | -1.0651685 | 0.28679969 | 0.41738838 | RECK | 8434 |
| ENSG000002 | 4.34472406 | -0.402185919 | 0.37758147 | -1.0651633 | 0.28680205 | 0.41738838 | NA | NA |
| ENSG000002 | 2.46253952 | -0.389031752 | 0.36524106 | -1.065137 | 0.28681395 | 0.41739072 | NA | NA |
| ENSG000002 | 5.37954401 | -0.395542853 | 0.3713756 | -1.065075 | 0.28684201 | 0.41741658 | LINC01384 | 101929685 |
| ENSG000001 | 196.446662 | -0.144230669 | 0.13542365 | -1.0650331 | 0.28686095 | 0.41742916 | UBASH3B | 84959 |

| | | | | | | | | |
|---|---|---|---|---|---|---|---|---|
| ENSG000002 | 5.65963783 | 0.393974092 | 0.36999572 | 1.06480715 | 0.28696321 | 0.41754958 | NA | NA |
| ENSG000002 | 4.65162294 | -0.401256392 | 0.37683565 | -1.0648047 | 0.2869643 | 0.41754958 | NA | NA |
| ENSG000002 | 28.831456 | -0.266122112 | 0.25000085 | -1.0644848 | 0.28710912 | 0.41774323 | LINC00877 | 285286 |
| ENSG000002 | 4.55999294 | -0.401369524 | 0.37707018 | -1.0644425 | 0.28712829 | 0.41774323 | NA | NA |
| ENSG000002 | 4.80270817 | 0.400420672 | 0.37617408 | 1.06445578 | 0.28712228 | 0.41774323 | SCGB1B2P | 643719 |
| ENSG000002 | 2.96393015 | -0.400228932 | 0.37607097 | -1.0642378 | 0.287221 | 0.41786312 | RNU6-123P | 106479620 |
| ENSG000001 | 4.56266159 | -0.400996721 | 0.37681366 | -1.0641778 | 0.28724818 | 0.41787267 | SOX14 | 8403 |
| ENSG000002 | 4.32822522 | -0.402064127 | 0.37781553 | -1.0641811 | 0.28724669 | 0.41787267 | EIF1AXP2 | 106479036 |
| ENSG000002 | 4.89925144 | -0.400011116 | 0.37589849 | -1.0641466 | 0.28726228 | 0.4178782 | NA | NA |
| ENSG000002 | 3.01719163 | -0.399366519 | 0.3753694 | -1.0639293 | 0.28736071 | 0.41800638 | NA | NA |
| ENSG000001 | 18.4205061 | 0.309125332 | 0.29057069 | 1.06385586 | 0.287394 | 0.41800983 | TEX19 | 400629 |
| ENSG000002 | 7.74633585 | -0.381753302 | 0.3588323 | -1.0638767 | 0.28738458 | 0.41800983 | NA | NA |
| ENSG000002 | 3.21322971 | -0.40229863 | 0.37814189 | -1.0638828 | 0.28738182 | 0.41800983 | RN7SL664P | 106479464 |
| ENSG000001 | 4.80427647 | -0.397773103 | 0.37391023 | -1.0638198 | 0.28741035 | 0.41801862 | CYP2C19 | 1557 |
| ENSG000002 | 8.24568282 | -0.378151947 | 0.355481 | -1.0637754 | 0.28743045 | 0.41803286 | NA | NA |
| ENSG000002 | 3.65876597 | -0.401609508 | 0.377541 | -1.0637507 | 0.28744165 | 0.41803415 | NA | NA |
| ENSG000002 | 4.34026944 | -0.401532718 | 0.37753803 | -1.0635557 | 0.28753002 | 0.41814768 | NA | NA |
| ENSG000002 | 4.54222905 | -0.40029565 | 0.37640734 | -1.063464 | 0.28757158 | 0.41819312 | NA | NA |
| ENSG000002 | 7.41183081 | -0.388605822 | 0.36542694 | -1.0634296 | 0.28758717 | 0.4182008 | C9orf57 | 138240 |
| ENSG000002 | 11.9745246 | -0.339914909 | 0.31966321 | -1.0633532 | 0.2876218 | 0.41823615 | NA | NA |
| ENSG000001 | 95.6630334 | 0.178664528 | 0.16802955 | 1.06329231 | 0.28764941 | 0.41824067 | PSTK | 118672 |
| ENSG000002 | 30.7872733 | -0.265121249 | 0.2493473 | -1.0632609 | 0.28766363 | 0.41824067 | XACT | 105463123 |
| ENSG000002 | 5.69285441 | -0.394964192 | 0.37145987 | -1.0632755 | 0.28765701 | 0.41824067 | NA | NA |
| ENSG000002 | 4.66659006 | -0.40086275 | 0.37702262 | -1.0632326 | 0.28767647 | 0.41824067 | LINC01195 | 104266959 |
| ENSG000002 | 3.51677513 | -0.402518907 | 0.37857842 | -1.0632379 | 0.2876741 | 0.41824067 | NA | NA |
| ENSG000002 | 4.53044631 | -0.399952728 | 0.37622429 | -1.0630699 | 0.28775024 | 0.41833291 | NA | NA |
| ENSG000001 | 175.576184 | -0.155504028 | 0.14629908 | -1.0629187 | 0.28781882 | 0.41841762 | EML5 | 161436 |
| ENSG000002 | 283.801205 | 0.146967152 | 0.13827399 | 1.06286908 | 0.28784132 | 0.41843533 | ZNRD1-AS1 | 80862 |
| ENSG000002 | 2.77648664 | -0.395960403 | 0.37255591 | -1.0628214 | 0.28786294 | 0.41844119 | NA | NA |

| | | | | | | | | |
|---|---|---|---|---|---|---|---|---|
| ENSG000002 | 7.21391975 | -0.383602533 | 0.36093077 | -1.0628147 | 0.28786599 | 0.41844119 | NA | NA |
| ENSG000001 | 17.4842153 | -0.314101016 | 0.29555901 | -1.0627354 | 0.28790197 | 0.41847848 | KCNJ12 | 3768 |
| ENSG000002 | 8.08338959 | -0.381896806 | 0.35940383 | -1.0625841 | 0.28797059 | 0.41856323 | NA | NA |
| ENSG000002 | 11.0837552 | -0.360576128 | 0.33934976 | -1.0625501 | 0.28798602 | 0.41857064 | NA | NA |
| ENSG000002 | 9.23154259 | -0.364246979 | 0.34286865 | -1.0623514 | 0.2880762 | 0.41868671 | NA | NA |
| ENSG000001 | 7.56472322 | -0.385646822 | 0.36303532 | -1.0622846 | 0.28810652 | 0.41871577 | CXCL10 | 3627 |
| ENSG000001 | 40.1391942 | 0.237259367 | 0.22342025 | 1.06194209 | 0.28826198 | 0.41892669 | NRBP2 | 340371 |
| ENSG000002 | 3.27385953 | -0.399519714 | 0.37639761 | -1.06143 | 0.28849454 | 0.41924964 | NA | NA |
| ENSG000001 | 10.8528914 | -0.356238911 | 0.33563513 | -1.0613874 | 0.28851387 | 0.41926271 | SGCZ | 137868 |
| ENSG000002 | 2.69782508 | -0.394820436 | 0.37199731 | -1.0613529 | 0.28852953 | 0.41927045 | NA | NA |
| ENSG000001 | 91.0584938 | 0.181972072 | 0.17146078 | 1.06130433 | 0.28855162 | 0.41928752 | FBXL17 | 64839 |
| ENSG000002 | 3.20423926 | -0.401158825 | 0.37801608 | -1.0612216 | 0.28858921 | 0.41932711 | NA | NA |
| ENSG000002 | 5.33244071 | -0.39403925 | 0.3713724 | -1.0610354 | 0.28867383 | 0.41943504 | LOC400548 | 400548 |
| ENSG000002 | 3.10383919 | 0.395279475 | 0.37257026 | 1.06095284 | 0.28871134 | 0.4194745 | RN7SKP292 | 106479225 |
| ENSG000001 | 366.519324 | 0.14261235 | 0.1344291 | 1.06087407 | 0.28874713 | 0.4195097 | LGALS8 | 3964 |
| ENSG000002 | 5.11083044 | -0.395494056 | 0.37280724 | -1.060854 | 0.28875625 | 0.4195097 | LOC105376594 | 105376594 |
| ENSG000002 | 2.72566981 | -0.395771366 | 0.37310465 | -1.0607516 | 0.28880278 | 0.41954724 | NA | NA |
| ENSG000002 | 3.00365403 | -0.398380057 | 0.37555695 | -1.0607714 | 0.28879382 | 0.41954724 | NA | NA |
| ENSG000002 | 2.96486902 | -0.398739591 | 0.37599555 | -1.0604902 | 0.28892166 | 0.41968986 | NA | NA |
| ENSG000002 | 2.78951491 | -0.394591905 | 0.37207828 | -1.0605078 | 0.28891367 | 0.41968986 | NA | NA |
| ENSG000002 | 5.0750532 | -0.394419639 | 0.37197116 | -1.0603501 | 0.28898538 | 0.41976738 | NA | NA |
| ENSG000002 | 2.94335416 | -0.39897857 | 0.37637025 | -1.0600694 | 0.28911304 | 0.41993778 | NA | NA |
| ENSG000002 | 6.00404627 | -0.393290126 | 0.37105687 | -1.0599187 | 0.28918158 | 0.42002228 | NA | NA |
| ENSG000002 | 7.17463359 | -0.3807102 | 0.35920539 | -1.0598677 | 0.28920478 | 0.42004094 | NA | NA |
| ENSG000002 | 8.16142501 | -0.367394732 | 0.34668546 | -1.059735 | 0.28926515 | 0.42011358 | NA | NA |
| ENSG000001 | 43.6006631 | -0.246861704 | 0.23298688 | -1.059552 | 0.28934848 | 0.42021955 | RIT2 | 6014 |
| ENSG000002 | 5.86860748 | -0.391946679 | 0.36998413 | -1.0593608 | 0.2894355 | 0.42033087 | NA | NA |
| ENSG000002 | 3.26246032 | -0.400409676 | 0.37814052 | -1.0588912 | 0.28964932 | 0.42062632 | NA | NA |
| ENSG000002 | 3.24453095 | -0.399800865 | 0.37758547 | -1.0588354 | 0.28967475 | 0.42064819 | NA | NA |

| | | | | | | | | |
|---|---|---|---|---|---|---|---|---|
| ENSG000002 | 6.54975517 | 0.382673768 | 0.36145164 | 1.0587136 | 0.28973023 | 0.4207137 | NA | NA |
| ENSG000002 | 6.70001026 | -0.38626123 | 0.36497414 | -1.0583249 | 0.28990733 | 0.42095579 | NA | NA |
| ENSG000001 | 26.4099466 | 0.302297219 | 0.28567773 | 1.05817566 | 0.28997536 | 0.42103949 | GNGT1 | 2792 |
| ENSG000002 | 31.2008793 | 0.269524038 | 0.25473587 | 1.05805294 | 0.29003131 | 0.42110564 | NA | NA |
| ENSG000002 | 10.0819852 | -0.365933752 | 0.34588014 | -1.0579785 | 0.29006525 | 0.42113984 | NA | NA |
| ENSG000001 | 12.1512809 | -0.350290998 | 0.33110281 | -1.0579523 | 0.29007717 | 0.42114207 | TLR8 | 51311 |
| ENSG000001 | 10.3925915 | -0.361260655 | 0.34150953 | -1.0578348 | 0.29013077 | 0.42120482 | PCP4 | 5121 |
| ENSG00000C | 134.418933 | 0.158921001 | 0.15023781 | 1.05779631 | 0.29014832 | 0.42121122 | TTC39A | 22996 |
| ENSG000002 | 3.46565361 | -0.40046834 | 0.37859338 | -1.0577796 | 0.29015595 | 0.42121122 | NA | NA |
| ENSG000002 | 67.8746627 | -0.205699378 | 0.19447014 | -1.0577427 | 0.29017275 | 0.42122052 | NA | NA |
| ENSG000002 | 4.35115899 | -0.400415457 | 0.37857868 | -1.0576809 | 0.29020093 | 0.42124634 | NA | NA |
| ENSG000002 | 4.26447296 | -0.399221934 | 0.37750788 | -1.0575195 | 0.29027458 | 0.42132308 | SUMO2P14 | 106480264 |
| ENSG000002 | 3.5008576 | -0.400372549 | 0.37859328 | -1.0575268 | 0.29027122 | 0.42132308 | NA | NA |
| ENSG000002 | 8.46576713 | -0.366599591 | 0.34672816 | -1.0573113 | 0.29036956 | 0.42144586 | NA | NA |
| ENSG000002 | 2.67396698 | -0.39326637 | 0.37196868 | -1.0572567 | 0.29039447 | 0.42146693 | NA | NA |
| ENSG000002 | 2.98545353 | -0.397903071 | 0.37646409 | -1.0569483 | 0.2905352 | 0.42162925 | NA | NA |
| ENSG000002 | 4.07531816 | -0.400017561 | 0.37845774 | -1.0569676 | 0.29052638 | 0.42162925 | NA | NA |
| ENSG000002 | 3.2323046 | -0.39893042 | 0.37743789 | -1.0569432 | 0.2905375 | 0.42162925 | NA | NA |
| ENSG000002 | 3.23816994 | -0.399018997 | 0.37755666 | -1.0568453 | 0.29058218 | 0.42166707 | NA | NA |
| ENSG000002 | 2.20748141 | -0.380631366 | 0.36016745 | -1.0568178 | 0.29059476 | 0.42166707 | NA | NA |
| ENSG000002 | 2.9678974 | -0.397491016 | 0.37611787 | -1.0568257 | 0.29059116 | 0.42166707 | NA | NA |
| ENSG000002 | 5.45611949 | -0.395812258 | 0.37455167 | -1.0567628 | 0.29061987 | 0.42168842 | NA | NA |
| ENSG000002 | 8.42444952 | -0.366533413 | 0.34689842 | -1.0566016 | 0.29069347 | 0.42178012 | NA | NA |
| ENSG000002 | 5.96373964 | -0.392434242 | 0.37142558 | -1.0565622 | 0.29071142 | 0.42179108 | LDLRAD4-AS1 | 100288122 |
| ENSG000002 | 3.02868186 | -0.397300416 | 0.37608239 | -1.0564186 | 0.29077703 | 0.42185608 | LINC00583 | 100113404 |
| ENSG000002 | 5.57937252 | -0.388536821 | 0.36777919 | -1.0564405 | 0.29076702 | 0.42185608 | NA | NA |
| ENSG000002 | 13.3688767 | 0.339941055 | 0.32180138 | 1.05636916 | 0.29079959 | 0.42187372 | SCARF2 | 91179 |
| ENSG000001 | 101.690297 | 0.187220233 | 0.17723795 | 1.05632136 | 0.29082142 | 0.4218903 | WTIP | 126374 |
| ENSG000002 | 5.33193272 | -0.391107881 | 0.37027778 | -1.0562553 | 0.29085158 | 0.42191895 | NA | NA |

| | | | | | | | | |
|---|---|---|---|---|---|---|---|---|
| ENSG000001 | 31.4234971 | 0.257439238 | 0.24374247 | 1.0561936 | 0.29087977 | 0.42194476 | IRF7 | 3665 |
| ENSG000002 | 4.57278895 | -0.397367917 | 0.37625655 | -1.056109 | 0.29091844 | 0.42198575 | NA | NA |
| ENSG000002 | 3.24701114 | -0.399080576 | 0.37790683 | -1.056029 | 0.29095496 | 0.42202363 | NA | NA |
| ENSG000001 | 16.2610664 | 0.318918442 | 0.30200581 | 1.05600103 | 0.29096775 | 0.42202708 | TCAP | 8557 |
| ENSG000002 | 3.53060533 | -0.399617925 | 0.37843525 | -1.0559744 | 0.29097993 | 0.42202966 | NA | NA |
| ENSG000002 | 2.45467271 | -0.387115597 | 0.36663771 | -1.0558532 | 0.2910353 | 0.42209486 | NA | NA |
| ENSG000002 | 3.25000508 | -0.399071406 | 0.37801627 | -1.055699 | 0.29110575 | 0.42218194 | LINC00397 | 101930749 |
| ENSG000002 | 27.1083453 | 0.276575144 | 0.26201247 | 1.05558007 | 0.29116011 | 0.42224568 | NA | NA |
| ENSG000002 | 22.886417 | 0.294930222 | 0.27950287 | 1.05519568 | 0.29133584 | 0.42247031 | NA | NA |
| ENSG000002 | 30.6406346 | 0.267032515 | 0.25306123 | 1.0552091 | 0.29132971 | 0.42247031 | NA | NA |
| ENSG000001 | 5.4202444 | -0.395053166 | 0.37446891 | -1.0549692 | 0.29143941 | 0.42260538 | OR4S2 | 219431 |
| ENSG000001 | 14.2610106 | 0.33540209 | 0.31798969 | 1.05475776 | 0.29153613 | 0.4226905 | POLD4 | 57804 |
| ENSG000002 | 7.37656109 | -0.376305415 | 0.35677003 | -1.0547562 | 0.29153683 | 0.4226905 | LOC100506790 | 100506790 |
| ENSG000002 | 3.49881516 | -0.399321234 | 0.37859335 | -1.0547497 | 0.2915398 | 0.4226905 | NA | NA |
| ENSG000002 | 3.82563076 | 0.399023995 | 0.37829959 | 1.05478305 | 0.29152456 | 0.4226905 | NA | NA |
| ENSG000002 | 4.10599506 | -0.3990939 | 0.3783879 | -1.0547216 | 0.29155265 | 0.42269402 | NA | NA |
| ENSG000002 | 6.46088209 | -0.389184472 | 0.36900999 | -1.0546719 | 0.2915754 | 0.42269677 | NA | NA |
| ENSG000002 | 4.89617393 | -0.396555757 | 0.37599634 | -1.0546798 | 0.29157178 | 0.42269677 | NA | NA |
| ENSG000002 | 4.83305418 | -0.395692502 | 0.37522911 | -1.0545357 | 0.29163771 | 0.42277199 | NA | NA |
| ENSG000002 | 7.92287097 | -0.371193763 | 0.35202336 | -1.0544577 | 0.2916734 | 0.4228086 | NA | NA |
| ENSG000001 | 37.2632617 | -0.243630525 | 0.23109323 | -1.0542521 | 0.29176751 | 0.4229148 | FLRT3 | 23767 |
| ENSG000001 | 8.24286693 | -0.374261136 | 0.35500129 | -1.0542529 | 0.29176715 | 0.4229148 | KLHL40 | 131377 |
| ENSG000001 | 84.8795851 | 0.188232892 | 0.17856645 | 1.0541336 | 0.29182175 | 0.4229783 | KCNH8 | 131096 |
| ENSG000002 | 4.35454421 | -0.398035132 | 0.37760482 | -1.054105 | 0.29183484 | 0.42298215 | LOC101929698 | 101929698 |
| ENSG000002 | 33.3625135 | 0.250993913 | 0.2381332 | 1.05400639 | 0.29187999 | 0.42301735 | NA | NA |
| ENSG000002 | 11.5887184 | 0.350891501 | 0.33290774 | 1.05402024 | 0.29187365 | 0.42301735 | ABALON | 103021294 |
| ENSG000002 | 18.008693 | -0.313895962 | 0.29783158 | -1.0539378 | 0.29191139 | 0.42303171 | NA | NA |
| ENSG000002 | 7.01616181 | -0.385239897 | 0.36553103 | -1.0539184 | 0.29192026 | 0.42303171 | NA | NA |
| ENSG000002 | 5.08013233 | 0.39306581 | 0.37295729 | 1.0539164 | 0.29192119 | 0.42303171 | NA | NA |

| | | | | | | | | |
|---|---|---|---|---|---|---|---|---|
| ENSG000002 | 2.75257591 | -0.392075886 | 0.37204633 | -1.0538362 | 0.29195792 | 0.42306982 | NA | NA |
| ENSG000002 | 45.726394 | 0.22948642 | 0.21777124 | 1.05379579 | 0.29197642 | 0.4230815 | NA | NA |
| ENSG000001 | 7.55241329 | 0.377404669 | 0.35816743 | 1.0537102 | 0.29201562 | 0.42310806 | ETV2 | 2116 |
| ENSG000001 | 4.28940109 | -0.39774652 | 0.37746493 | -1.0537311 | 0.29200607 | 0.42310806 | MAGEA10 | 4109 |
| ENSG000002 | 13.0103672 | -0.343445775 | 0.32595158 | -1.0536711 | 0.29203351 | 0.42311887 | NA | NA |
| ENSG000002 | 3.22024454 | 0.398415788 | 0.37816499 | 1.05355016 | 0.29208892 | 0.42318402 | NA | NA |
| ENSG000001 | 45.2577051 | 0.221309153 | 0.2100729 | 1.05348738 | 0.29211767 | 0.42319545 | USP35 | 57558 |
| ENSG000002 | 4.64887031 | -0.397521383 | 0.37733046 | -1.0535099 | 0.29210735 | 0.42319545 | NA | NA |
| ENSG000001 | 6.90194091 | -0.379452641 | 0.36023958 | -1.0533341 | 0.29218788 | 0.42328204 | DKK4 | 27121 |
| ENSG000002 | 3.50319343 | -0.398760444 | 0.37859047 | -1.0532765 | 0.29221429 | 0.42330517 | FRG2LP | 106480352 |
| ENSG000002 | 4.28815391 | -0.398306197 | 0.37819048 | -1.0531894 | 0.29225421 | 0.42333986 | NA | NA |
| ENSG000002 | 3.87504833 | -0.39853765 | 0.37841409 | -1.0531787 | 0.29225911 | 0.42333986 | NA | NA |
| ENSG000002 | 5.4566718 | 0.392188738 | 0.37241799 | 1.05308753 | 0.29230088 | 0.42333986 | LINC01545 | 724087 |
| ENSG000002 | 3.27826053 | -0.396995844 | 0.37697218 | -1.0531171 | 0.29228733 | 0.42333986 | NA | NA |
| ENSG000002 | 25.9301716 | 0.283811506 | 0.2695026 | 1.05309375 | 0.29229803 | 0.42333986 | NA | NA |
| ENSG000002 | 3.25600103 | -0.397559629 | 0.37751654 | -1.053092 | 0.29229885 | 0.42333986 | NA | NA |
| ENSG000002 | 5.6695933 | 0.389536858 | 0.36996373 | 1.05290554 | 0.29238429 | 0.42344554 | LOC105379695 | 105379695 |
| ENSG000001 | 184.132682 | 0.157832379 | 0.14991274 | 1.05282835 | 0.29241967 | 0.42348166 | HERC6 | 55008 |
| ENSG000001 | 7.46672856 | -0.379080921 | 0.36012457 | -1.0526383 | 0.2925068 | 0.42359271 | KRT12 | 3859 |
| ENSG000002 | 10.5631088 | -0.357221555 | 0.33936817 | -1.0526077 | 0.29252082 | 0.42359657 | NA | NA |
| ENSG000002 | 3.5310618 | -0.398158767 | 0.37826687 | -1.0525869 | 0.29253035 | 0.42359657 | RN7SL274P | 106479322 |
| ENSG000001 | 5.35036136 | -0.391843175 | 0.37230414 | -1.0524814 | 0.29257875 | 0.42359838 | CHRNG | 1146 |
| ENSG000002 | 7.48356752 | -0.378711796 | 0.35981068 | -1.0525307 | 0.29255612 | 0.42359838 | NA | NA |
| ENSG000002 | 4.14144624 | -0.398436951 | 0.37855843 | -1.0525111 | 0.29256512 | 0.42359838 | NA | NA |
| ENSG000002 | 23.33098 | 0.286022766 | 0.2717483 | 1.05252825 | 0.29255726 | 0.42359838 | NA | NA |
| ENSG000002 | 3.04683085 | -0.395275658 | 0.37556942 | -1.0524703 | 0.29258383 | 0.42359838 | NA | NA |
| ENSG000001 | 106.762795 | 0.175836191 | 0.16709617 | 1.05230533 | 0.29265948 | 0.42369278 | ADCY6 | 112 |
| ENSG000001 | 48.3298116 | 0.225404074 | 0.21423166 | 1.0521511 | 0.29273023 | 0.42378007 | DHRS1 | 115817 |
| ENSG000002 | 12.2453681 | 0.335482625 | 0.31888432 | 1.05205118 | 0.29277607 | 0.4238313 | NA | NA |

| | | | | | | | | |
|---|---|---|---|---|---|---|---|---|
| ENSG000002 | 4.11521288 | -0.398224492 | 0.3785771 | -1.051898 | 0.29284635 | 0.42391791 | NA | NA |
| ENSG000002 | 3.48703208 | -0.398170604 | 0.37857887 | -1.0517507 | 0.29291393 | 0.4240006 | NA | NA |
| ENSG000002 | 7.64975044 | -0.373688618 | 0.35532649 | -1.0516768 | 0.29294787 | 0.42403459 | NA | NA |
| ENSG000002 | 3.21298121 | -0.396942504 | 0.37747858 | -1.051563 | 0.29300009 | 0.42409504 | NA | NA |
| ENSG000001 | 9.47433594 | 0.360797366 | 0.34312579 | 1.05150173 | 0.29302821 | 0.42411223 | CHRM4 | 1132 |
| ENSG000002 | 2.75385446 | -0.391147698 | 0.37199319 | -1.0514916 | 0.29303288 | 0.42411223 | NA | NA |
| ENSG000002 | 3.53086179 | -0.398061013 | 0.37857798 | -1.0514637 | 0.29304566 | 0.42411558 | NA | NA |
| ENSG000001 | 8.55280638 | -0.371643853 | 0.3535455 | -1.051191 | 0.29317087 | 0.42428165 | REG1B | 5968 |
| ENSG000002 | 13.8696776 | -0.348311908 | 0.3314563 | -1.0508532 | 0.29332603 | 0.42449105 | LOC100133286 | 100133286 |
| ENSG000002 | 5.54019456 | -0.395150128 | 0.37606602 | -1.0507467 | 0.29337494 | 0.42454669 | NA | NA |
| ENSG000001 | 116.602803 | -0.171081265 | 0.16284069 | -1.0506051 | 0.29343999 | 0.42462567 | LYPD6 | 130574 |
| ENSG00000C | 17.0409493 | -0.332187467 | 0.31628643 | -1.0502742 | 0.29359208 | 0.42481543 | CYP3A43 | 64816 |
| ENSG000002 | 3.46633876 | -0.397622559 | 0.37858431 | -1.050288 | 0.29358572 | 0.42481543 | NA | NA |
| ENSG000002 | 17.9055642 | 0.315529357 | 0.30047692 | 1.05009517 | 0.29367436 | 0.42491932 | ALKBH6 | 84964 |
| ENSG000001 | 150.535798 | 0.168944951 | 0.16089435 | 1.05003658 | 0.2937013 | 0.42492797 | C17orf104 | 284071 |
| ENSG000002 | 47.7622744 | -0.22043877 | 0.20993289 | -1.050044 | 0.29369788 | 0.42492797 | NA | NA |
| ENSG00000C | 137.73293 | 0.164731104 | 0.15688718 | 1.04999722 | 0.29371939 | 0.42493899 | GAS7 | 8522 |
| ENSG000002 | 5.05537284 | 0.391120931 | 0.37251259 | 1.0499536 | 0.29373945 | 0.42495284 | NA | NA |
| ENSG000002 | 3.05253997 | -0.393596376 | 0.37491098 | -1.0498396 | 0.29379189 | 0.42501235 | LOC102724231 | 102724231 |
| ENSG000002 | 8.46922239 | -0.36769763 | 0.35024874 | -1.0498186 | 0.29380154 | 0.42501235 | NA | NA |
| ENSG000001 | 8.10010522 | -0.376049029 | 0.35826214 | -1.0496477 | 0.29388012 | 0.42509448 | C8G | 733 |
| ENSG000001 | 6.69425704 | -0.38358882 | 0.36543685 | -1.049672 | 0.29386896 | 0.42509448 | NA | NA |
| ENSG000002 | 6.59469439 | -0.381206874 | 0.36318328 | -1.0496267 | 0.29388976 | 0.42509448 | NA | NA |
| ENSG000001 | 6.26801727 | -0.389711314 | 0.37132401 | -1.0495182 | 0.29393967 | 0.42515151 | CCDC13-AS1 | 100874114 |
| ENSG000002 | 52.6978064 | -0.216032184 | 0.20588051 | -1.0493086 | 0.29403613 | 0.42527585 | RGS5 | 8490 |
| ENSG000002 | 2.73672254 | -0.390322309 | 0.37208039 | -1.0490268 | 0.29416577 | 0.42544819 | NA | NA |
| ENSG000001 | 91.0067625 | 0.185694909 | 0.17702676 | 1.04896517 | 0.29419415 | 0.42547405 | ADPRH | 141 |
| ENSG000002 | 7.1065486 | -0.377753043 | 0.36014579 | -1.0488892 | 0.2942291 | 0.42549186 | NA | NA |
| ENSG000002 | 7.3769936 | -0.373446343 | 0.35604635 | -1.04887 | 0.29423794 | 0.42549186 | KLF14 | 136259 |

| | | | | | | | | |
|---|---|---|---|---|---|---|---|---|
| ENSG000002 | 8.14384811 | -0.37693244 | 0.35936013 | -1.0488989 | 0.29422466 | 0.42549186 | NA | NA |
| ENSG000001 | 51.6092844 | -0.220501302 | 0.21026342 | -1.0486908 | 0.29432047 | 0.42559602 | ACSS1 | 84532 |
| ENSG000002 | 3.64980331 | -0.395311874 | 0.37701697 | -1.0485254 | 0.2943966 | 0.42569094 | NA | NA |
| ENSG000002 | 10.0187796 | -0.357651055 | 0.34113129 | -1.0484264 | 0.29444219 | 0.42573497 | NA | NA |
| ENSG000002 | 5.04844735 | -0.390701944 | 0.3726601 | -1.0484137 | 0.29444805 | 0.42573497 | NA | NA |
| ENSG00000C | 125.201228 | 0.168708573 | 0.16094442 | 1.04824117 | 0.2945275 | 0.4257827 | NAALAD2 | 10003 |
| ENSG000001 | 6.65290249 | -0.380610679 | 0.36307866 | -1.0482871 | 0.29450634 | 0.4257827 | CCDC70 | 83446 |
| ENSG000002 | 5.564236 | -0.387354845 | 0.3695081 | -1.0482986 | 0.29450103 | 0.4257827 | NA | NA |
| ENSG000002 | 3.53755764 | -0.396039943 | 0.37781851 | -1.048228 | 0.29453356 | 0.4257827 | NA | NA |
| ENSG000002 | 8.19219811 | -0.366234384 | 0.34937565 | -1.0482539 | 0.29452165 | 0.4257827 | NA | NA |
| ENSG000002 | 4.31435646 | -0.395660023 | 0.37749127 | -1.0481302 | 0.2945786 | 0.42583263 | NA | NA |
| ENSG000002 | 3.29462243 | -0.394903762 | 0.37679718 | -1.0480539 | 0.29461377 | 0.42586828 | NA | NA |
| ENSG000001 | 5.33079112 | -0.388167828 | 0.37038969 | -1.0479985 | 0.29463932 | 0.42588807 | VPREB3 | 29802 |
| ENSG000002 | 6.492753 | -0.387418111 | 0.36968131 | -1.0479786 | 0.29464846 | 0.42588807 | NA | NA |
| ENSG000002 | 4.8188502 | -0.394225934 | 0.37627257 | -1.0477137 | 0.29477052 | 0.4260282 | NA | NA |
| ENSG000002 | 4.03784501 | -0.396536143 | 0.37847945 | -1.0477085 | 0.29477292 | 0.4260282 | NA | NA |
| ENSG000002 | 9.19814028 | -0.361214974 | 0.34476953 | -1.0476998 | 0.29477693 | 0.4260282 | NA | NA |
| ENSG000001 | 3.83873499 | -0.396255377 | 0.37834879 | -1.0473283 | 0.29494821 | 0.42626055 | FFAR1 | 2864 |
| ENSG000002 | 6.92936774 | -0.378680701 | 0.36158633 | -1.047276 | 0.29497229 | 0.42626497 | CTD-2350J17.1 | 101929472 |
| ENSG000002 | 3.22277337 | -0.39536259 | 0.37751142 | -1.0472865 | 0.29496748 | 0.42626497 | LINC01571 | 101927364 |
| ENSG000001 | 90.5489196 | 0.174655586 | 0.16678413 | 1.04719548 | 0.29500943 | 0.42628826 | DHFRL1 | 200895 |
| ENSG000002 | 2.53076866 | -0.380750788 | 0.36358376 | -1.0472162 | 0.2949999 | 0.42628826 | NA | NA |
| ENSG000002 | 19.6485193 | -0.297700266 | 0.28429527 | -1.0471517 | 0.29502964 | 0.42630227 | NA | NA |
| ENSG000002 | 7.09825116 | 0.378259611 | 0.36126282 | 1.04704828 | 0.29507731 | 0.42635596 | NA | NA |
| ENSG000002 | 4.90995185 | -0.39415999 | 0.37646249 | -1.04701 | 0.29509496 | 0.42636627 | NA | NA |
| ENSG000001 | 3.31858386 | 0.391993007 | 0.37440999 | 1.04696192 | 0.29511714 | 0.42638312 | NA | NA |
| ENSG000001 | 261.544386 | 0.142167717 | 0.13583029 | 1.04665696 | 0.29525782 | 0.42655598 | HDX | 139324 |
| ENSG000002 | 3.19214408 | -0.395456078 | 0.37782097 | -1.0466758 | 0.29524911 | 0.42655598 | NA | NA |
| ENSG000001 | 15549.5164 | 0.119679036 | 0.11434873 | 1.04661444 | 0.29527744 | 0.42656913 | CYTB | 4519 |

| | | | | | | | | | |
|---|---|---|---|---|---|---|---|---|---|
| ENSG000001 | 7.73595973 | -0.374182899 | 0.35756294 | -1.0464812 | 0.29533892 | 0.42663869 | MEIG1 | | 644890 |
| ENSG000002 | 20.1500509 | -0.291517563 | 0.27857378 | -1.0464645 | 0.29534663 | 0.42663869 | NA | NA | |
| ENSG000002 | 3.48320974 | -0.396141552 | 0.37859058 | -1.0463587 | 0.29539545 | 0.426694 | NA | NA | |
| ENSG000002 | 31.9446964 | 0.253834019 | 0.24259822 | 1.04631443 | 0.29541589 | 0.42670833 | NA | NA | |
| ENSG000002 | 3.51534069 | -0.395979916 | 0.37854579 | -1.0460555 | 0.2955354 | 0.42685055 | NA | NA | |
| ENSG000002 | 4.06144198 | -0.395998617 | 0.37856166 | -1.0460611 | 0.29553284 | 0.42685055 | NA | NA | |
| ENSG000002 | 12.4239519 | -0.332188549 | 0.3175992 | -1.0459364 | 0.29559041 | 0.4269148 | NA | NA | |
| ENSG000002 | 17.1931547 | -0.306827017 | 0.29337246 | -1.0458617 | 0.2956249 | 0.4269494 | NA | NA | |
| ENSG000002 | 7.46084689 | -0.373516465 | 0.35714663 | -1.0458351 | 0.2956372 | 0.42695195 | LOC101928989 | | 101928989 |
| ENSG000002 | 5.82844264 | -0.383630219 | 0.36683873 | -1.0457735 | 0.29566562 | 0.4269778 | RNU6-1283P | | 106481782 |
| ENSG000002 | 2.18003219 | -0.380983617 | 0.36431635 | -1.0457494 | 0.29567674 | 0.42697865 | RNU2-29P | | 106480209 |
| ENSG000002 | 4.64352203 | -0.39444544 | 0.37724384 | -1.0455981 | 0.29574663 | 0.42706436 | NA | NA | |
| ENSG000001 | 250.555482 | 0.168204074 | 0.16088618 | 1.04548492 | 0.29579891 | 0.42712464 | LATS2 | | 26524 |
| ENSG000002 | 8.14148153 | -0.36351181 | 0.34771015 | -1.0454449 | 0.29581738 | 0.42713611 | NA | NA | |
| ENSG000002 | 3.19203252 | -0.394748348 | 0.37762462 | -1.0453459 | 0.29586313 | 0.42718696 | ROBO2P1 | | 106480790 |
| ENSG000002 | 9.43836506 | -0.356504941 | 0.34105972 | -1.045286 | 0.29589082 | 0.42721057 | NA | NA | |
| ENSG000002 | 3.28697111 | -0.394562499 | 0.37747608 | -1.0452649 | 0.29590055 | 0.42721057 | NA | NA | |
| ENSG000002 | 12.2221544 | 0.333553642 | 0.31911762 | 1.04523731 | 0.2959133 | 0.42721377 | NA | NA | |
| ENSG000002 | 72.0423137 | 0.202303933 | 0.19355524 | 1.04519996 | 0.29593056 | 0.42722348 | NA | NA | |
| ENSG000001 | 6.2956686 | -0.390562637 | 0.37372168 | -1.0450628 | 0.29599393 | 0.42729975 | TTC9B | | 148014 |
| ENSG000002 | 5.07114278 | 0.39087643 | 0.37403931 | 1.04501432 | 0.29601635 | 0.4273169 | NA | NA | |
| ENSG000002 | 7.86793069 | -0.369783565 | 0.35389823 | -1.0448867 | 0.29607532 | 0.42735638 | NA | NA | |
| ENSG000002 | 4.59493599 | -0.39429142 | 0.37734547 | -1.0449083 | 0.29606535 | 0.42735638 | NA | NA | |
| ENSG000002 | 26.3263945 | -0.270900384 | 0.25925173 | -1.0449318 | 0.29605448 | 0.42735638 | NA | NA | |
| ENSG000002 | 2.69402235 | -0.388564253 | 0.371896 | -1.0448197 | 0.29610633 | 0.42738593 | NA | NA | |
| ENSG000002 | 3.19584222 | -0.394463934 | 0.37755353 | -1.0447894 | 0.2961203 | 0.42739089 | NA | NA | |
| ENSG000002 | 54.8947317 | -0.242104917 | 0.23175339 | -1.0446661 | 0.29617731 | 0.42745795 | S1PR3 | | 1903 |
| ENSG000002 | 4.84805356 | -0.392329291 | 0.37558557 | -1.0445803 | 0.29621698 | 0.42749999 | NA | NA | |
| ENSG000002 | 7.11612004 | -0.376329406 | 0.36033479 | -1.0443882 | 0.29630581 | 0.42756733 | RPTN | | 126638 |

| | | | | | | | | |
|---|---|---|---|---|---|---|---|---|
| ENSG000002 | 10.1425328 | -0.360652548 | 0.34530975 | -1.044432 | 0.29628557 | 0.42756733 | NA | NA |
| ENSG000002 | 59.206314 | 0.200275871 | 0.19175391 | 1.04444218 | 0.29628085 | 0.42756733 | NA | NA |
| ENSG000002 | 6.69546475 | -0.381412339 | 0.36519605 | -1.0444043 | 0.29629836 | 0.42756733 | NA | NA |
| ENSG000002 | 2.6962987 | -0.387760041 | 0.3713087 | -1.0443064 | 0.29634367 | 0.42760673 | NA | NA |
| ENSG000001 | 181.424404 | 0.153981368 | 0.14745456 | 1.04426321 | 0.29636363 | 0.42762033 | C15orf40 | 123207 |
| ENSG000002 | 7.14387929 | 0.373856798 | 0.35803481 | 1.04419119 | 0.29639694 | 0.42762274 | LOH12CR2 | 503693 |
| ENSG000002 | 3.26226334 | -0.394025873 | 0.37734343 | -1.0442102 | 0.29638813 | 0.42762274 | NA | NA |
| ENSG000002 | 9.45955507 | -0.35726023 | 0.34213329 | -1.0442136 | 0.29638659 | 0.42762274 | NA | NA |
| ENSG000002 | 6.39308686 | -0.381724917 | 0.36561978 | -1.0440489 | 0.29646279 | 0.42770252 | NA | NA |
| ENSG000002 | 5.08591473 | -0.390487309 | 0.37405715 | -1.0439242 | 0.29652046 | 0.42773019 | NA | NA |
| ENSG000002 | 2.37289906 | 0.378062171 | 0.36215828 | 1.04391421 | 0.29652508 | 0.42773019 | NA | NA |
| ENSG000002 | 2.9830849 | -0.392033665 | 0.37553184 | -1.0439426 | 0.29651197 | 0.42773019 | NA | NA |
| ENSG000002 | 8.44291096 | -0.36592785 | 0.350527 | -1.0439363 | 0.29651487 | 0.42773019 | NA | NA |
| ENSG000002 | 4.31753691 | -0.395145813 | 0.3785308 | -1.0438934 | 0.29653471 | 0.42773019 | NA | NA |
| ENSG000001 | 124.649904 | -0.163827139 | 0.15696576 | -1.0437126 | 0.29661839 | 0.42782997 | ROBO2 | 6092 |
| ENSG000002 | 73.3379238 | 0.21857734 | 0.20942578 | 1.04369834 | 0.29662498 | 0.42782997 | NA | NA |
| ENSG00000C | 17.7325968 | -0.306026966 | 0.29323273 | -1.0436317 | 0.29665584 | 0.42785122 | WNT8B | 7479 |
| ENSG000002 | 3.25996828 | -0.393947191 | 0.37748112 | -1.0436209 | 0.29666081 | 0.42785122 | NA | NA |
| ENSG000002 | 25.3830405 | 0.281354791 | 0.26965937 | 1.0433711 | 0.29677645 | 0.42800277 | NA | NA |
| ENSG000001 | 13.4288084 | -0.324051172 | 0.31061202 | -1.0432667 | 0.29682479 | 0.42805726 | KCNA5 | 3741 |
| ENSG000001 | 79.1929711 | -0.188107133 | 0.18032786 | -1.0431396 | 0.29688363 | 0.42812689 | FCRL5 | 83416 |
| ENSG00000C | 98.9536324 | 0.178777239 | 0.17141573 | 1.04294535 | 0.2969736 | 0.42824141 | COL9A3 | 1299 |
| ENSG000002 | 2.69713994 | -0.386586312 | 0.3706938 | -1.0428724 | 0.29700741 | 0.42827493 | NA | NA |
| ENSG000002 | 2.73120322 | -0.388563177 | 0.37263797 | -1.0427364 | 0.29707039 | 0.42835051 | NA | NA |
| ENSG00000C | 62.4318601 | 0.208386607 | 0.19985644 | 1.04268145 | 0.29709585 | 0.42837199 | TUBG2 | 27175 |
| ENSG000001 | 9.86519428 | -0.360926 | 0.3462307 | -1.0424437 | 0.29720603 | 0.42850038 | NA | NA |
| ENSG000002 | 7.30115162 | 0.371672045 | 0.3565374 | 1.04244896 | 0.29720358 | 0.42850038 | LOC105369635 | 105369635 |
| ENSG000002 | 47.2420702 | 0.228494688 | 0.21919691 | 1.04241748 | 0.29721817 | 0.42850264 | NA | NA |
| ENSG000002 | 5.26185659 | 0.393192698 | 0.37720915 | 1.04237318 | 0.2972387 | 0.42851701 | NA | NA |

| | | | | | | | | |
|---|---|---|---|---|---|---|---|---|
| ENSG000002 | 8.18098394 | -0.364431614 | 0.34963297 | -1.0423262 | 0.29726046 | 0.42853315 | NA | NA |
| ENSG000001 | 150.050393 | 0.164641894 | 0.15796639 | 1.04225901 | 0.29729161 | 0.42856016 | GAS8 | 2622 |
| ENSG000002 | 3.76054958 | -0.39429818 | 0.37831796 | -1.0422402 | 0.29730033 | 0.42856016 | NA | NA |
| ENSG00000C | 15.2489015 | -0.314406954 | 0.30172075 | -1.0420462 | 0.29739027 | 0.42867456 | PILRA | 29992 |
| ENSG00000C | 74.8961533 | -0.189305844 | 0.18167585 | -1.0419979 | 0.29741267 | 0.42869161 | ANO2 | 57101 |
| ENSG000002 | 13.7227377 | -0.323834974 | 0.31080609 | -1.0419196 | 0.29744894 | 0.42871359 | NA | NA |
| ENSG000002 | 6.0860721 | 0.38644708 | 0.37089921 | 1.04191938 | 0.29744906 | 0.42871359 | NA | NA |
| ENSG000002 | 6.66204177 | -0.38102715 | 0.36574071 | -1.0417958 | 0.29750634 | 0.42878091 | NA | NA |
| ENSG000002 | 2.9360144 | -0.391029381 | 0.37535539 | -1.0417577 | 0.29752402 | 0.42879114 | NA | NA |
| ENSG000002 | 7.62858856 | -0.368159901 | 0.35347719 | -1.0415379 | 0.29762596 | 0.42892282 | NA | NA |
| ENSG000002 | 2.72615798 | -0.385829609 | 0.37046462 | -1.0414749 | 0.29765518 | 0.42893445 | NA | NA |
| ENSG000002 | 3.53686352 | -0.394210025 | 0.37850944 | -1.04148 | 0.29765281 | 0.42893445 | NA | NA |
| ENSG000002 | 3.49049742 | -0.39426196 | 0.3785905 | -1.0413942 | 0.29769263 | 0.42897316 | NA | NA |
| ENSG000002 | 33.7333824 | -0.290618577 | 0.27909599 | -1.0412854 | 0.29774312 | 0.42902483 | NA | NA |
| ENSG000002 | 16.9472435 | -0.304097834 | 0.29204476 | -1.0412713 | 0.29774964 | 0.42902483 | NA | NA |
| ENSG000001 | 89.6522675 | 0.178868565 | 0.17180404 | 1.04111965 | 0.29782002 | 0.42911099 | ZSWIM5 | 57643 |
| ENSG000002 | 2.96120367 | -0.39143097 | 0.37601181 | -1.0410071 | 0.29787224 | 0.42917098 | NA | NA |
| ENSG000002 | 4.83509696 | -0.390819836 | 0.37546936 | -1.0408834 | 0.29792965 | 0.42922365 | NA | NA |
| ENSG000002 | 4.12753641 | -0.393981649 | 0.37850723 | -1.0408828 | 0.29792997 | 0.42922365 | NA | NA |
| ENSG000001 | 9.17639782 | -0.365598808 | 0.3512837 | -1.0407508 | 0.2979912 | 0.42928558 | RAMP3 | 10268 |
| ENSG000002 | 3.20264174 | -0.393421147 | 0.37801894 | -1.0407445 | 0.29799413 | 0.42928558 | NA | NA |
| ENSG000002 | 5.83244932 | -0.381083701 | 0.36620079 | -1.0406414 | 0.29804202 | 0.42933932 | NA | NA |
| ENSG000001 | 73.8926043 | -0.192600552 | 0.18508968 | -1.0405796 | 0.29807069 | 0.42936537 | TPH2 | 121278 |
| ENSG000002 | 14.1090727 | -0.331242345 | 0.31835938 | -1.0404667 | 0.29812311 | 0.42942563 | NA | NA |
| ENSG000002 | 4.03996569 | -0.393770212 | 0.37846773 | -1.0404327 | 0.2981389 | 0.42943311 | NA | NA |
| ENSG000002 | 4.13644754 | -0.393715003 | 0.37858965 | -1.0399518 | 0.29836228 | 0.42973961 | NA | NA |
| ENSG000002 | 62.7070387 | 0.207868803 | 0.199903 | 1.03984832 | 0.29841037 | 0.42978235 | NA | NA |
| ENSG000002 | 5.78869058 | -0.39028037 | 0.37532648 | -1.0398423 | 0.29841315 | 0.42978235 | MIR3165 | 100422953 |
| ENSG000002 | 4.0663555 | -0.393547135 | 0.37851039 | -1.0397261 | 0.29846717 | 0.42984488 | TMEM244 | 253582 |

| | | | | | | | | |
|---|---|---|---|---|---|---|---|---|
| ENSG000001 | 7.1486166 | -0.373638888 | 0.35938418 | -1.0396643 | 0.29849591 | 0.429871 | PRAMENP | 649179 |
| ENSG000001 | 124.436104 | 0.176346961 | 0.16966915 | 1.03935781 | 0.29863836 | 0.43006087 | THRA | 7067 |
| ENSG000001 | 28.5807127 | -0.260753937 | 0.25089958 | -1.0392761 | 0.29867634 | 0.43006975 | TSLP | 85480 |
| ENSG000002 | 5.95592556 | -0.386459543 | 0.37184912 | -1.0392913 | 0.29866929 | 0.43006975 | NA | NA |
| ENSG000002 | 16.2601292 | 0.32506934 | 0.31278222 | 1.03928331 | 0.29867299 | 0.43006975 | AMACR | 23600 |
| ENSG000002 | 3.20605859 | -0.392273881 | 0.37748103 | -1.0391883 | 0.29871716 | 0.43011325 | NA | NA |
| ENSG000002 | 23.2661407 | -0.288880841 | 0.27803561 | -1.0390066 | 0.29880165 | 0.43019567 | NA | NA |
| ENSG000002 | 13.4511509 | -0.326599591 | 0.31434129 | -1.0389968 | 0.29880622 | 0.43019567 | LOC101929486 | 101929486 |
| ENSG000002 | 3.01756432 | -0.388671186 | 0.3740734 | -1.0390238 | 0.29879365 | 0.43019567 | NA | NA |
| ENSG000002 | 38.3186222 | 0.250358875 | 0.24098208 | 1.03891074 | 0.29884625 | 0.43023802 | NA | NA |
| ENSG00000C | 18.954632 | -0.315284031 | 0.30349932 | -1.0388294 | 0.29888407 | 0.43027719 | PGC | 5225 |
| ENSG000002 | 2.95445209 | -0.390960018 | 0.37637149 | -1.038761 | 0.29891592 | 0.43030776 | NA | NA |
| ENSG000001 | 11.6677494 | -0.337093963 | 0.32455949 | -1.0386199 | 0.29898152 | 0.43038693 | ZCCHC16 | 340595 |
| ENSG000002 | 12.9612966 | -0.327273441 | 0.31519333 | -1.038326 | 0.29911829 | 0.43056852 | NA | NA |
| ENSG000002 | 5.85088052 | -0.385694472 | 0.37154202 | -1.0380911 | 0.29922763 | 0.43071062 | NA | NA |
| ENSG000002 | 6.67287856 | -0.378233992 | 0.3643768 | -1.0380298 | 0.29925616 | 0.43072283 | NA | NA |
| ENSG000002 | 7.89731065 | -0.364459174 | 0.35110751 | -1.0380273 | 0.29925736 | 0.43072283 | LINC00463 | 101928922 |
| ENSG000002 | 12.4772445 | -0.335057273 | 0.32279179 | -1.0379981 | 0.29927092 | 0.43072365 | NA | NA |
| ENSG000002 | 4.07994405 | -0.392969912 | 0.37859087 | -1.0379804 | 0.29927917 | 0.43072365 | NA | NA |
| ENSG000001 | 13.3643035 | -0.333735397 | 0.32157075 | -1.0378288 | 0.29934975 | 0.43077937 | C2orf50 | 130813 |
| ENSG000002 | 2.73706855 | -0.386744663 | 0.37264241 | -1.0378439 | 0.29934272 | 0.43077937 | NA | NA |
| ENSG000002 | 8.38671024 | -0.359333114 | 0.34622786 | -1.0378515 | 0.29933919 | 0.43077937 | CTBP1-AS | 285463 |
| ENSG000001 | 25.5772892 | 0.270632623 | 0.26078348 | 1.03776752 | 0.2993783 | 0.43080517 | PTRH1 | 138428 |
| ENSG000002 | 5.83968907 | -0.382266702 | 0.36839112 | -1.0376653 | 0.29942588 | 0.43085834 | NA | NA |
| ENSG000001 | 524.576777 | 0.131799829 | 0.12703236 | 1.03752959 | 0.29948911 | 0.43091875 | TADA1 | 117143 |
| ENSG000001 | 4.31641023 | -0.392527979 | 0.37832564 | -1.03754 | 0.29948427 | 0.43091875 | KLK1 | 3816 |
| ENSG000002 | 5.31077497 | -0.385286086 | 0.37137516 | -1.0374579 | 0.29952252 | 0.43095152 | NA | NA |
| ENSG000002 | 7.35211256 | 0.371974654 | 0.35858082 | 1.03735235 | 0.29957168 | 0.43100696 | NA | NA |
| ENSG000002 | 2.73152274 | -0.384644941 | 0.37082528 | -1.0372673 | 0.2996113 | 0.43104706 | NA | NA |

| | | | | | | | | |
|---|---|---|---|---|---|---|---|---|
| ENSG000002 | 2.72024964 | -0.385892781 | 0.37203561 | -1.0372469 | 0.29962081 | 0.43104706 | RNU6-352P | 106480573 |
| ENSG000002 | 4.34077797 | -0.392058431 | 0.37803222 | -1.0371032 | 0.29968775 | 0.43111277 | NA | NA |
| ENSG000002 | 7.14864906 | -0.372085176 | 0.35876706 | -1.0371219 | 0.29967905 | 0.43111277 | NA | NA |
| ENSG000002 | 4.34685137 | -0.392056396 | 0.37804054 | -1.037075 | 0.2997009 | 0.43111639 | NA | NA |
| ENSG000002 | 14.94157 | -0.329594093 | 0.31787626 | -1.0368629 | 0.29979977 | 0.43124332 | NA | NA |
| ENSG000002 | 2.92665257 | -0.389834808 | 0.37606127 | -1.0366258 | 0.2999103 | 0.43135058 | NA | NA |
| ENSG000002 | 6.88638127 | -0.374872857 | 0.36163288 | -1.0366116 | 0.29991689 | 0.43135058 | NA | NA |
| ENSG000002 | 3.4791717 | -0.392445832 | 0.37857756 | -1.0366326 | 0.29990713 | 0.43135058 | NA | NA |
| ENSG000002 | 3.21793361 | -0.391533278 | 0.37769892 | -1.036628 | 0.29990926 | 0.43135058 | NA | NA |
| ENSG000001 | 7.2579983 | -0.374927192 | 0.36176914 | -1.0363714 | 0.30002891 | 0.4313889 | IL22 | 50616 |
| ENSG000001 | 56.9057797 | -0.205363556 | 0.1981454 | -1.0364286 | 0.30000225 | 0.4313889 | M1AP | 130951 |
| ENSG000001 | 8.90259562 | -0.354593932 | 0.34215296 | -1.0363609 | 0.30003382 | 0.4313889 | LOC151174 | 151174 |
| ENSG000001 | 13.5191341 | -0.342382505 | 0.33034284 | -1.036446 | 0.29999414 | 0.4313889 | MMP1 | 4312 |
| ENSG000001 | 2.7271839 | -0.383980976 | 0.3704985 | -1.0363901 | 0.30002019 | 0.4313889 | NA | NA |
| ENSG000002 | 3.34274102 | -0.388514878 | 0.37485489 | -1.0364407 | 0.29999657 | 0.4313889 | NA | NA |
| ENSG000002 | 6.49026798 | 0.377063783 | 0.36380647 | 1.03644056 | 0.29999666 | 0.4313889 | AGAP1-IT1 | 100506749 |
| ENSG000002 | 3.21412337 | -0.391056318 | 0.37727789 | -1.0365206 | 0.29995932 | 0.4313889 | NA | NA |
| ENSG000002 | 2.97708427 | -0.389859925 | 0.3761873 | -1.0363453 | 0.3000411 | 0.4313889 | NA | NA |
| ENSG000002 | 5.919734 | 0.379731439 | 0.36642071 | 1.03632636 | 0.30004991 | 0.4313889 | NA | NA |
| ENSG000002 | 42.5969031 | 0.236838711 | 0.22855306 | 1.03625262 | 0.3000843 | 0.43142305 | NA | NA |
| ENSG000002 | 3.51219542 | -0.391894378 | 0.37822954 | -1.0361284 | 0.30014223 | 0.43149103 | RN7SL285P | 106479326 |
| ENSG000001 | 8.52576246 | -0.372512084 | 0.35954609 | -1.0360621 | 0.30017316 | 0.4315202 | TRIM64B | 642446 |
| ENSG000002 | 5.59585897 | -0.383065242 | 0.36974625 | -1.036022 | 0.30019188 | 0.43153182 | NA | NA |
| ENSG000002 | 11.7612661 | -0.343764214 | 0.33182426 | -1.0359827 | 0.30021019 | 0.43154284 | TMEM108-AS1 | 101927455 |
| ENSG000002 | 11.6657267 | -0.337768778 | 0.32605996 | -1.03591 | 0.30024412 | 0.43157632 | NA | NA |
| ENSG000002 | 3.77927902 | -0.392160173 | 0.37857708 | -1.0358793 | 0.30025844 | 0.43158159 | NA | NA |
| ENSG000002 | 90.3391705 | 0.184863051 | 0.17847834 | 1.03577301 | 0.30030805 | 0.43163761 | ZNF285 | 26974 |
| ENSG000001 | 6.95306885 | -0.377606508 | 0.36458594 | -1.0357133 | 0.30033591 | 0.43166235 | RGCC | 28984 |
| ENSG000002 | 4.0639604 | -0.392041221 | 0.37854251 | -1.0356597 | 0.30036093 | 0.431683 | SMAD1-AS1 | 104326058 |

| | | | | | | | | |
|---|---|---|---|---|---|---|---|---|
| ENSG000002 | 2.67854669 | -0.384394212 | 0.37118271 | -1.035593 | 0.30039207 | 0.43171246 | NA | NA |
| ENSG000002 | 37.2868673 | 0.255352816 | 0.24658942 | 1.03553842 | 0.30041753 | 0.43173375 | NA | NA |
| ENSG000002 | 3.4529697 | -0.391808652 | 0.37837078 | -1.0355151 | 0.30042842 | 0.4317341 | NA | NA |
| ENSG000002 | 5.61064206 | -0.382429779 | 0.369345 | -1.035427 | 0.30046955 | 0.43177791 | NA | NA |
| ENSG000002 | 2.3767675 | -0.384919026 | 0.37182656 | -1.0352112 | 0.30057028 | 0.43187674 | DUXAP10 | 503639 |
| ENSG000002 | 5.83174381 | -0.381208262 | 0.36824058 | -1.0352152 | 0.3005684 | 0.43187674 | NA | NA |
| ENSG000002 | 2.39025279 | -0.385160596 | 0.37204423 | -1.0352548 | 0.30054991 | 0.43187674 | NA | NA |
| ENSG000002 | 34.1882711 | 0.253142058 | 0.24456239 | 1.03508173 | 0.30063074 | 0.43194831 | NA | NA |
| ENSG000002 | 6.12914877 | -0.381075881 | 0.36818588 | -1.0350095 | 0.30066448 | 0.43198148 | NA | NA |
| ENSG000002 | 32.2119481 | 0.255490598 | 0.24688627 | 1.03485139 | 0.30073832 | 0.43207226 | LOC101928504 | 101928504 |
| ENSG000002 | 4.41475669 | -0.391687871 | 0.37859336 | -1.0345873 | 0.3008617 | 0.43223421 | NA | NA |
| ENSG000002 | 4.79986017 | -0.388544382 | 0.37562658 | -1.03439 | 0.30095387 | 0.43235131 | NA | NA |
| ENSG000002 | 2.99818896 | -0.388277612 | 0.37538904 | -1.0343339 | 0.3009801 | 0.43237367 | NA | NA |
| ENSG000002 | 2.73614677 | -0.38368552 | 0.37099081 | -1.0342184 | 0.30103408 | 0.43242058 | NA | NA |
| ENSG000002 | 5.59489362 | -0.382144386 | 0.36949859 | -1.0342242 | 0.30103136 | 0.43242058 | LINC00290 | 728081 |
| ENSG000002 | 11.1809861 | -0.340691362 | 0.32947132 | -1.0340547 | 0.30111059 | 0.43251517 | NA | NA |
| ENSG00000C | 116.555102 | 0.16315044 | 0.15779993 | 1.0339069 | 0.30117969 | 0.4325913 | SKAP2 | 8935 |
| ENSG000002 | 2.37923918 | -0.383488859 | 0.37091639 | -1.0338957 | 0.30118493 | 0.4325913 | NA | NA |
| ENSG000002 | 7.61208992 | -0.366733242 | 0.35471904 | -1.0338696 | 0.30119712 | 0.43259348 | NA | NA |
| ENSG000002 | 16.2995748 | 0.30892827 | 0.29885504 | 1.03370607 | 0.30127359 | 0.43268799 | NA | NA |
| ENSG000002 | 9.11812095 | -0.350478149 | 0.33906562 | -1.0336588 | 0.30129572 | 0.43270445 | NA | NA |
| ENSG000002 | 3.46738917 | -0.39120317 | 0.37854854 | -1.0334294 | 0.30140302 | 0.43284321 | NA | NA |
| ENSG000002 | 2.3919878 | -0.383645154 | 0.37127429 | -1.03332 | 0.30145416 | 0.43290133 | NA | NA |
| ENSG000002 | 5.59259722 | -0.381273993 | 0.36899941 | -1.0332645 | 0.30148013 | 0.43292329 | NA | NA |
| ENSG000002 | 13.7911649 | -0.327618847 | 0.31709918 | -1.0331747 | 0.30152216 | 0.43296831 | NA | NA |
| ENSG000001 | 117.483393 | 0.171867191 | 0.16638104 | 1.03297339 | 0.30161635 | 0.43305756 | TMEM8B | 51754 |
| ENSG000001 | 11.6427689 | -0.336416541 | 0.32567106 | -1.0329949 | 0.30160629 | 0.43305756 | CCDC37 | 348807 |
| ENSG000002 | 2.52567321 | -0.375812336 | 0.3638054 | -1.0330037 | 0.30160215 | 0.43305756 | NA | NA |
| ENSG000001 | 9.9064538 | -0.345739814 | 0.33474921 | -1.0328324 | 0.30168236 | 0.43307252 | CTF1 | 1489 |

| | | | | | | | | |
|---|---|---|---|---|---|---|---|---|
| ENSG000001 | 4.79685602 | 0.390162457 | 0.37774955 | 1.03286015 | 0.30166935 | 0.43307252 | NA | NA |
| ENSG000002 | 5.06113126 | -0.3852467 | 0.37298313 | -1.0328797 | 0.3016602 | 0.43307252 | NA | NA |
| ENSG000002 | 25.7742015 | 0.268863625 | 0.26029354 | 1.03292471 | 0.30163914 | 0.43307252 | NA | NA |
| ENSG000002 | 3.22498015 | 0.390528104 | 0.37812038 | 1.03281421 | 0.30169085 | 0.43307252 | NA | NA |
| ENSG000002 | 16.6032183 | 0.310335995 | 0.30047203 | 1.03282823 | 0.30168429 | 0.43307252 | TTC4 | 7268 |
| ENSG000002 | 29.1283659 | 0.266975994 | 0.2586131 | 1.03233748 | 0.30191405 | 0.43337758 | SNORD116-29 | 100033821 |
| ENSG000002 | 6.39985033 | -0.380195011 | 0.36830971 | -1.0322698 | 0.30194572 | 0.4334077 | NA | NA |
| ENSG000002 | 6.25833047 | -0.381723363 | 0.36980872 | -1.0322184 | 0.30196982 | 0.43342694 | LOC285627 | 285627 |
| ENSG000002 | 4.78190543 | -0.387689652 | 0.37561021 | -1.0321595 | 0.30199739 | 0.43345118 | RN7SL325P | 106481011 |
| ENSG000001 | 4.37041001 | -0.390302923 | 0.37815671 | -1.0321195 | 0.30201612 | 0.43346272 | NA | NA |
| ENSG000002 | 3.55330919 | -0.390181462 | 0.37804976 | -1.0320902 | 0.30202985 | 0.43346708 | NA | NA |
| ENSG000001 | 119.984489 | 0.168334439 | 0.16310839 | 1.03204035 | 0.30205322 | 0.43348528 | SETD9 | 133383 |
| ENSG000002 | 3.50506429 | -0.390365152 | 0.37827408 | -1.0319638 | 0.30208908 | 0.4335214 | NA | NA |
| ENSG000002 | 9.20432913 | -0.353767478 | 0.34282943 | -1.0319052 | 0.30211651 | 0.43354543 | NA | NA |
| ENSG000002 | 2.35692817 | -0.384117269 | 0.37228838 | -1.0317735 | 0.30217826 | 0.43361869 | NA | NA |
| ENSG000002 | 68.8841667 | 0.246697753 | 0.23912056 | 1.03168775 | 0.30221842 | 0.43366097 | NA | NA |
| ENSG000001 | 7.88287909 | -0.365003175 | 0.35380944 | -1.0316378 | 0.30224184 | 0.43367924 | LSMEM2 | 132228 |
| ENSG000002 | 2.71008445 | -0.38193576 | 0.37025105 | -1.0315589 | 0.30227881 | 0.43371693 | MYO16-AS1 | 100885782 |
| ENSG000001 | 45.065025 | 0.223277366 | 0.21645796 | 1.03150451 | 0.30230429 | 0.43373815 | TFR2 | 7036 |
| ENSG000002 | 4.21678192 | -0.389833826 | 0.37799734 | -1.0313137 | 0.30239374 | 0.43385114 | RNF224 | 643596 |
| ENSG000002 | 25.3176316 | -0.271269469 | 0.26304485 | -1.031267 | 0.30241563 | 0.43386719 | LOC100132077 | 100132077 |
| ENSG000002 | 3.50219563 | -0.390333864 | 0.37853704 | -1.0311643 | 0.3024638 | 0.43392094 | NA | NA |
| ENSG000002 | 11.4480955 | -0.35031575 | 0.33979569 | -1.03096 | 0.3025596 | 0.43402767 | NA | NA |
| ENSG000002 | 4.60970868 | -0.388735887 | 0.37705924 | -1.0309677 | 0.30255598 | 0.43402767 | NA | NA |
| ENSG000002 | 3.24467531 | -0.388941642 | 0.37727286 | -1.0309293 | 0.30257398 | 0.43403295 | NA | NA |
| ENSG000002 | 5.56075857 | -0.382535667 | 0.37113291 | -1.0307242 | 0.30267017 | 0.43415558 | NA | NA |
| ENSG000002 | 3.50928744 | -0.390084158 | 0.37856139 | -1.0304383 | 0.3028043 | 0.4343326 | NA | NA |
| ENSG000002 | 3.49338559 | -0.389388462 | 0.37795112 | -1.0302614 | 0.3028873 | 0.43443629 | NA | NA |
| ENSG000002 | 2.66761293 | -0.382264812 | 0.37106261 | -1.0301895 | 0.30292104 | 0.43445656 | NA | NA |

| | | | | | | | | |
|---|---|---|---|---|---|---|---|---|
| ENSG000002 | 4.78756965 | -0.386267536 | 0.37494944 | -1.0301857 | 0.30292286 | 0.43445656 | NA | NA |
| ENSG000001 | 119.558895 | -0.166690119 | 0.16182414 | -1.0300696 | 0.30297736 | 0.43451936 | MLKL | 197259 |
| ENSG000002 | 6.42082789 | -0.377198916 | 0.36621709 | -1.0299872 | 0.30301601 | 0.43455943 | NA | NA |
| ENSG000001 | 54.5404367 | -0.209423452 | 0.20334303 | -1.0299023 | 0.30305587 | 0.43460122 | ADGRE2 | 30817 |
| ENSG000002 | 2.36026125 | -0.382309199 | 0.37123668 | -1.029826 | 0.30309167 | 0.43463719 | NA | NA |
| ENSG000001 | 45.3263571 | 0.219929706 | 0.2136073 | 1.02959828 | 0.30319862 | 0.43477518 | NA | NA |
| ENSG000001 | 6.88024236 | -0.371887329 | 0.36120681 | -1.029569 | 0.30321238 | 0.43477954 | DOK7 | 285489 |
| ENSG000002 | 13.033054 | -0.327809076 | 0.31840778 | -1.029526 | 0.30323258 | 0.43479312 | NA | NA |
| ENSG000002 | 3.46913105 | -0.389479779 | 0.37833795 | -1.0294494 | 0.30326854 | 0.43482931 | NA | NA |
| ENSG000001 | 11.934461 | -0.332406225 | 0.32298855 | -1.0291579 | 0.30340547 | 0.43499488 | NA | NA |
| ENSG000002 | 2.36580651 | -0.382589657 | 0.37174858 | -1.0291624 | 0.30340338 | 0.43499488 | NA | NA |
| ENSG000001 | 2.71721548 | -0.381621044 | 0.37089113 | -1.0289301 | 0.30351253 | 0.43513299 | ZNF679 | 168417 |
| ENSG000002 | 10.2874301 | -0.350976295 | 0.34119547 | -1.0286663 | 0.3036365 | 0.43529533 | NA | NA |
| ENSG000002 | 9.86819692 | -0.343202749 | 0.33367618 | -1.0285503 | 0.30369102 | 0.4353581 | NA | NA |
| ENSG000002 | 3.28649503 | -0.387925342 | 0.37719652 | -1.0284436 | 0.30374121 | 0.43539927 | NA | NA |
| ENSG000002 | 3.75673934 | -0.389345647 | 0.37856956 | -1.0284653 | 0.30373102 | 0.43539927 | RNASE11 | 122651 |
| ENSG000002 | 5.91111952 | -0.381106153 | 0.3706065 | -1.028331 | 0.30379415 | 0.43545976 | FLJ38122 | 401289 |
| ENSG000002 | 3.71848246 | -0.389134618 | 0.37845013 | -1.0282322 | 0.3038406 | 0.43551094 | NA | NA |
| ENSG000002 | 6.49428079 | -0.378432677 | 0.3680535 | -1.0282002 | 0.30385568 | 0.43551716 | NA | NA |
| ENSG000001 | 10.8926021 | -0.338375033 | 0.32911185 | -1.028146 | 0.30388115 | 0.43553828 | LRRN4 | 164312 |
| ENSG000002 | 2.67345899 | -0.382423502 | 0.37200796 | -1.0279982 | 0.30395069 | 0.43557637 | NA | NA |
| ENSG000002 | 86.534676 | 0.182367643 | 0.17739558 | 1.02802809 | 0.30393662 | 0.43557637 | TRAM2-AS1 | 401264 |
| ENSG000002 | 23.0876688 | 0.277177224 | 0.26962293 | 1.02801798 | 0.30394137 | 0.43557637 | NA | NA |
| ENSG000002 | 5.3002773 | -0.383221085 | 0.37278049 | -1.0280074 | 0.30394636 | 0.43557637 | NA | NA |
| ENSG000001 | 17.9450403 | 0.299756699 | 0.2916077 | 1.02794508 | 0.30397566 | 0.43559676 | RAC2 | 5880 |
| ENSG000001 | 2.3662109 | -0.381231583 | 0.37088508 | -1.0278968 | 0.30399839 | 0.43561393 | OBP2B | 29989 |
| ENSG000002 | 13.0970763 | 0.322819116 | 0.31412492 | 1.02767751 | 0.30410155 | 0.43574567 | NA | NA |
| ENSG000002 | 28.05721 | 0.265891466 | 0.25873594 | 1.0276557 | 0.30411181 | 0.43574567 | NA | NA |
| ENSG000002 | 3.42580403 | -0.383749546 | 0.3734423 | -1.0276007 | 0.30413772 | 0.43576739 | NA | NA |

| | | | | | | | | |
|---|---|---|---|---|---|---|---|---|
| ENSG00000( | 53.5310049 | -0.210370014 | 0.20472451 | -1.0275761 | 0.30414928 | 0.43576856 | TMCC3 | 57458 |
| ENSG000002 | 3.61925919 | 0.388684481 | 0.3782809 | 1.02750226 | 0.30418402 | 0.43578992 | NA | NA |
| ENSG000002 | 5.31912048 | -0.38121147 | 0.37100919 | -1.0274987 | 0.30418568 | 0.43578992 | NA | NA |
| ENSG000002 | 3.91262399 | -0.386592091 | 0.37626732 | -1.02744 | 0.30421333 | 0.43581413 | NA | NA |
| ENSG000002 | 5.87798967 | -0.381415275 | 0.3712738 | -1.0273154 | 0.304272 | 0.43588277 | NA | NA |
| ENSG000002 | 3.0757331 | -0.383917952 | 0.37371825 | -1.0272925 | 0.30428276 | 0.4358828 | LOC101929341 | 101929341 |
| ENSG000002 | 25.2005653 | 0.263294218 | 0.25638249 | 1.02695865 | 0.30443994 | 0.43607714 | NA | NA |
| ENSG000002 | 4.00264522 | -0.388047178 | 0.37785975 | -1.0269609 | 0.30443888 | 0.43607714 | NA | NA |
| ENSG000001 | 16.9275599 | -0.303117331 | 0.29520749 | -1.0267942 | 0.30451739 | 0.43615727 | TNFSF9 | 8744 |
| ENSG000001 | 85.8719369 | 0.179288459 | 0.17460673 | 1.02681299 | 0.30450853 | 0.43615727 | PYGO1 | 26108 |
| ENSG000001 | 8.5988334 | -0.355579326 | 0.3463687 | -1.026592 | 0.30461263 | 0.436269 | C9orf9 | 11092 |
| ENSG000002 | 11.6864471 | -0.337199431 | 0.32846781 | -1.0265829 | 0.30461691 | 0.436269 | UBL5P4 | 106480800 |
| ENSG000001 | 73.1069798 | -0.186179806 | 0.18140455 | -1.0263238 | 0.30473899 | 0.43639759 | OXTR | 5021 |
| ENSG000002 | 2.36005273 | -0.381526692 | 0.37173391 | -1.0263435 | 0.30472969 | 0.43639759 | NA | NA |
| ENSG000002 | 20.2530775 | -0.299057647 | 0.29138159 | -1.0263437 | 0.30472962 | 0.43639759 | LINC00639 | 283547 |
| ENSG000002 | 14.3312696 | -0.312579647 | 0.30462207 | -1.0261228 | 0.30483371 | 0.43651783 | NA | NA |
| ENSG000001 | 3.46862306 | -0.388410049 | 0.37854852 | -1.0260509 | 0.30486759 | 0.43655092 | OR1K1 | 392392 |
| ENSG000001 | 13.3577477 | -0.321719927 | 0.31362522 | -1.0258101 | 0.3049811 | 0.43669804 | NHLRC1 | 378884 |
| ENSG000002 | 6.15908339 | -0.376632292 | 0.36716777 | -1.0257771 | 0.30499666 | 0.4367049 | NA | NA |
| ENSG000001 | 11.9319553 | 0.336013808 | 0.32762749 | 1.02559711 | 0.30508153 | 0.43681032 | ZDHHC19 | 131540 |
| ENSG000001 | 3.7201337 | -0.387985397 | 0.37831002 | -1.0255753 | 0.30509183 | 0.43681032 | INSM2 | 84684 |
| ENSG000002 | 3.42979728 | -0.388119346 | 0.37848646 | -1.0254511 | 0.3051504 | 0.43687876 | RNU6-3P | 106478919 |
| ENSG000002 | 4.77734587 | -0.385051679 | 0.37551943 | -1.0253842 | 0.30518196 | 0.43690851 | NA | NA |
| ENSG000002 | 28.3033435 | 0.270985351 | 0.26431381 | 1.02524098 | 0.3052495 | 0.43698977 | NA | NA |
| ENSG000002 | 2.40223643 | -0.379664424 | 0.37033688 | -1.0251866 | 0.30527513 | 0.43701104 | NA | NA |
| ENSG000001 | 21.565235 | -0.287311777 | 0.28029084 | -1.0250488 | 0.30534018 | 0.43708873 | TRIM17 | 51127 |
| ENSG000001 | 10.4650048 | -0.34442024 | 0.33601817 | -1.0250048 | 0.30536092 | 0.43710299 | OR51I2 | 390064 |
| ENSG000001 | 6.95419751 | -0.373385487 | 0.36431089 | -1.0249089 | 0.30540615 | 0.43713755 | CALHM1 | 255022 |
| ENSG000002 | 4.1859602 | -0.387899661 | 0.37847268 | -1.024908 | 0.30540662 | 0.43713755 | NA | NA |

| | | | | | | | | |
|---|---|---|---|---|---|---|---|---|
| ENSG000001 | 82.5305347 | 0.190613636 | 0.18599465 | 1.02483397 | 0.30544153 | 0.43715666 | CUX2 | 23316 |
| ENSG000001 | 13.2963128 | -0.327283107 | 0.31934965 | -1.0248425 | 0.30543749 | 0.43715666 | HHEX | 3087 |
| ENSG000002 | 8.6509037 | -0.354333076 | 0.34582355 | -1.0246066 | 0.30554887 | 0.43727941 | NA | NA |
| ENSG000002 | 5.09206854 | -0.382884141 | 0.37368457 | -1.0246185 | 0.30554321 | 0.43727941 | NA | NA |
| ENSG000002 | 2.36673959 | -0.380556625 | 0.37145438 | -1.0245044 | 0.30559712 | 0.43729519 | LOC101927532 | 101927532 |
| ENSG000002 | 4.77894395 | -0.385436145 | 0.37620773 | -1.0245301 | 0.30558495 | 0.43729519 | LINC01288 | 103689844 |
| ENSG000002 | 3.97204818 | -0.387051016 | 0.37777836 | -1.0245452 | 0.30557783 | 0.43729519 | NA | NA |
| ENSG000002 | 95.5160434 | 0.176952092 | 0.17272182 | 1.02449183 | 0.30560303 | 0.43729519 | NA | NA |
| ENSG000002 | 2.40517253 | -0.378326669 | 0.36930064 | -1.0244409 | 0.30562709 | 0.43729876 | NA | NA |
| ENSG000002 | 4.87211973 | -0.385456595 | 0.37625861 | -1.0244459 | 0.30562471 | 0.43729876 | NA | NA |
| ENSG000002 | 3.76690363 | -0.387776474 | 0.37857624 | -1.0243022 | 0.30569257 | 0.43737701 | NA | NA |
| ENSG000001 | 982.520136 | 0.261082467 | 0.25490651 | 1.02422831 | 0.30572745 | 0.43741149 | SMG7 | 9887 |
| ENSG000002 | 3.99049742 | -0.387284924 | 0.37813378 | -1.0242008 | 0.30574044 | 0.43741463 | NA | NA |
| ENSG000002 | 8.19452892 | -0.360553977 | 0.3520586 | -1.0241306 | 0.3057736 | 0.43744665 | NA | NA |
| ENSG000001 | 4.74602371 | -0.384820213 | 0.37577454 | -1.0240721 | 0.30580124 | 0.43747075 | BMP8A | 353500 |
| ENSG000002 | 3.07609782 | -0.382812883 | 0.37387443 | -1.0239076 | 0.30587891 | 0.43756643 | NA | NA |
| ENSG000002 | 4.37422026 | -0.387316559 | 0.37828236 | -1.0238822 | 0.30589094 | 0.4375682 | NA | NA |
| ENSG000002 | 7.14437623 | -0.371161987 | 0.36253387 | -1.0237995 | 0.30593 | 0.43759321 | NA | NA |
| ENSG000002 | 5.33934382 | -0.381641538 | 0.37276423 | -1.0238148 | 0.30592276 | 0.43759321 | NA | NA |
| ENSG000001 | 3.45445263 | -0.387496265 | 0.37851568 | -1.0237258 | 0.3059648 | 0.43762755 | PPY | 5539 |
| ENSG000002 | 4.57023797 | -0.385962865 | 0.37707357 | -1.0235744 | 0.30603633 | 0.43771442 | PABPN1L | 390748 |
| ENSG000001 | 26.1724489 | 0.275024633 | 0.26869918 | 1.02354103 | 0.30605211 | 0.43772155 | TP53TG1 | 11257 |
| ENSG000002 | 14.5647399 | -0.332850091 | 0.32522323 | -1.0234511 | 0.30609459 | 0.43774415 | NA | NA |
| ENSG000002 | 7.71446556 | -0.363898925 | 0.35556482 | -1.0234391 | 0.3061003 | 0.43774415 | NA | NA |
| ENSG000002 | 2.37909537 | -0.378828849 | 0.37014326 | -1.0234655 | 0.30608782 | 0.43774415 | NA | NA |
| ENSG000002 | 3.84466035 | -0.387216165 | 0.3783829 | -1.0233448 | 0.30614485 | 0.43777699 | NA | NA |
| ENSG000002 | 6.88128526 | -0.37163763 | 0.36315554 | -1.0233566 | 0.30613925 | 0.43777699 | NA | NA |
| ENSG000001 | 66.9123971 | 0.189873899 | 0.1855564 | 1.02326787 | 0.30618121 | 0.43781354 | DOK5 | 55816 |
| ENSG000001 | 13.3256423 | -0.329152083 | 0.32168162 | -1.0232232 | 0.30620235 | 0.43782833 | PBX4 | 80714 |

| | | | | | | | | |
|---|---|---|---|---|---|---|---|---|
| ENSG000002 | 4.63524918 | -0.386710327 | 0.37795656 | -1.0231608 | 0.30623184 | 0.43785506 | NA | NA |
| ENSG000001 | 352.090573 | 0.148027519 | 0.14469137 | 1.02305698 | 0.30628091 | 0.43790584 | CEP128 | 145508 |
| ENSG000001 | 2.38912328 | -0.378155773 | 0.36966424 | -1.0229709 | 0.30632159 | 0.43790584 | NA | NA |
| ENSG000002 | 4.88219992 | -0.384970831 | 0.37632254 | -1.0229811 | 0.3063168 | 0.43790584 | NA | NA |
| ENSG000002 | 5.82182893 | -0.381765816 | 0.37320137 | -1.0229486 | 0.30633215 | 0.43790584 | NA | NA |
| ENSG000002 | 5.09677211 | -0.382659539 | 0.37404489 | -1.0230311 | 0.30629316 | 0.43790584 | NA | NA |
| ENSG000002 | 3.74520509 | -0.387162395 | 0.37845909 | -1.0229967 | 0.30630941 | 0.43790584 | NA | NA |
| ENSG000002 | 4.54177258 | -0.385749868 | 0.37711252 | -1.0229039 | 0.30635328 | 0.43792061 | LOC102724392 | 102724392 |
| ENSG000002 | 3.20587786 | -0.386328505 | 0.37771784 | -1.0227966 | 0.30640405 | 0.43797774 | NA | NA |
| ENSG000002 | 3.62879479 | -0.385467092 | 0.37689459 | -1.0227451 | 0.30642838 | 0.43798165 | NA | NA |
| ENSG000002 | 14.2227669 | -0.314396204 | 0.30740132 | -1.0227549 | 0.30642375 | 0.43798165 | NA | NA |
| ENSG000002 | 3.20553817 | -0.386217505 | 0.37772232 | -1.0224906 | 0.30654877 | 0.43813828 | NA | NA |
| ENSG000002 | 6.94551505 | -0.3723764 | 0.36419834 | -1.0224549 | 0.30656562 | 0.43814692 | NA | NA |
| ENSG000002 | 116.995872 | -0.171954956 | 0.1681984 | -1.0223341 | 0.3066228 | 0.4382132 | LOC100507053 | 100507053 |
| ENSG000002 | 5.35025363 | -0.379906118 | 0.37163842 | -1.0222466 | 0.30666419 | 0.4382569 | NA | NA |
| ENSG000002 | 4.88139037 | -0.384813812 | 0.3764514 | -1.0222138 | 0.30667973 | 0.43826367 | NA | NA |
| ENSG000002 | 10.2708516 | 0.346745955 | 0.33925707 | 1.02207437 | 0.3067457 | 0.43834249 | NA | NA |
| ENSG000001 | 60.5277935 | 0.204020886 | 0.19966537 | 1.02181406 | 0.30686891 | 0.43850311 | SLC16A12 | 387700 |
| ENSG000001 | 60.8892173 | -0.212948662 | 0.20841581 | -1.0217491 | 0.30689967 | 0.4385316 | TDRD12 | 91646 |
| ENSG000002 | 3.83871015 | -0.386376933 | 0.37816214 | -1.021723 | 0.30691205 | 0.43853384 | NA | NA |
| ENSG000002 | 3.89030946 | 0.386673876 | 0.37847418 | 1.02166513 | 0.30693942 | 0.4385575 | NA | NA |
| ENSG000001 | 60.0745124 | 0.196378323 | 0.19222077 | 1.02162903 | 0.30695651 | 0.43856619 | Sep-04 | 5414 |
| ENSG000002 | 9.13495097 | -0.350938652 | 0.34351643 | -1.0216066 | 0.30696713 | 0.43856619 | NA | NA |
| ENSG000002 | 10.9542118 | 0.350683326 | 0.34327933 | 1.02156844 | 0.3069852 | 0.43857655 | NA | NA |
| ENSG000002 | 7.18510642 | -0.368994072 | 0.3612669 | -1.0213891 | 0.30707013 | 0.43866191 | NA | NA |
| ENSG000002 | 3.85659082 | -0.38589697 | 0.37782151 | -1.0213737 | 0.3070774 | 0.43866191 | LOC101927363 | 101927363 |
| ENSG000002 | 2.40656317 | -0.379263805 | 0.37131567 | -1.0214053 | 0.30706244 | 0.43866191 | NA | NA |
| ENSG000002 | 2.37016551 | -0.379733134 | 0.37181446 | -1.0212974 | 0.30711357 | 0.43869812 | NA | NA |
| ENSG000002 | 5.62026504 | -0.379827214 | 0.37195734 | -1.021158 | 0.30717958 | 0.43877697 | LOC100128233 | 100128233 |

| | | | | | | | | | |
|---|---|---|---|---|---|---|---|---|---|
| ENSG000001 | 10.9502284 | -0.341266751 | 0.33422099 | -1.0210812 | 0.30721599 | 0.43881351 | CDH15 | | 1013 |
| ENSG000002 | 5.2826947 | -0.381039018 | 0.37318197 | -1.0210542 | 0.30722875 | 0.43881629 | NA | NA | |
| ENSG000002 | 3.74992882 | -0.386443267 | 0.378489 | -1.0210158 | 0.30724693 | 0.4388268 | NA | NA | |
| ENSG000002 | 3.73370276 | -0.386472258 | 0.37855001 | -1.0209279 | 0.30728861 | 0.43887086 | NA | NA | |
| ENSG000002 | 3.97827858 | -0.385607976 | 0.37771342 | -1.0209009 | 0.30730139 | 0.43887366 | NA | NA | |
| ENSG000000 | 34.6319639 | -0.242672963 | 0.23772287 | -1.020823 | 0.30733832 | 0.43891094 | WSCD2 | | 9671 |
| ENSG000002 | 4.54541365 | -0.385455376 | 0.37760762 | -1.0207828 | 0.30735734 | 0.43892264 | NA | NA | |
| ENSG000001 | 2.39218368 | -0.377964042 | 0.37031232 | -1.0206629 | 0.30741418 | 0.4389729 | NA | NA | |
| ENSG000002 | 2.37090272 | -0.379504856 | 0.37181649 | -1.0206779 | 0.30740709 | 0.4389729 | NA | NA | |
| ENSG000002 | 5.5477171 | -0.378830502 | 0.37120568 | -1.0205407 | 0.3074721 | 0.43904015 | NA | NA | |
| ENSG000002 | 2.36782968 | -0.379404706 | 0.37180693 | -1.0204347 | 0.30752232 | 0.4390964 | NA | NA | |
| ENSG000001 | 8.85594171 | -0.36336441 | 0.35611081 | -1.0203689 | 0.30755352 | 0.43911002 | FBXL22 | | 283807 |
| ENSG000002 | 3.72072546 | -0.386312204 | 0.37859303 | -1.0203891 | 0.30754396 | 0.43911002 | NA | NA | |
| ENSG000002 | 3.99871819 | -0.385801375 | 0.37811523 | -1.0203275 | 0.30757316 | 0.4391226 | NA | NA | |
| ENSG000002 | 2.66017453 | 0.380645191 | 0.37309837 | 1.02022744 | 0.30762061 | 0.43917488 | NA | NA | |
| ENSG000000 | 55.6318683 | 0.23543642 | 0.23077449 | 1.02020125 | 0.30763302 | 0.43917714 | COL23A1 | | 91522 |
| ENSG000001 | 109.236296 | -0.19365362 | 0.1898245 | -1.0201719 | 0.30764693 | 0.43918154 | PHACTR1 | | 221692 |
| ENSG000002 | 5.0107585 | -0.381745228 | 0.37423485 | -1.0200686 | 0.30769591 | 0.439236 | LOC401127 | | 401127 |
| ENSG000002 | 2.36173622 | -0.379352431 | 0.37197717 | -1.0198272 | 0.30781042 | 0.43938399 | MIR511 | | 574445 |
| ENSG000002 | 53.2310487 | 0.211046828 | 0.20698364 | 1.01963046 | 0.30790375 | 0.43950175 | NA | NA | |
| ENSG000002 | 2.35297411 | -0.378536919 | 0.37142148 | -1.0191573 | 0.30812828 | 0.43980676 | NA | NA | |
| ENSG000002 | 8.64875766 | -0.351638792 | 0.34505581 | -1.019078 | 0.30816593 | 0.43984502 | NA | NA | |
| ENSG000002 | 5.39567345 | -0.380378315 | 0.3732845 | -1.0190038 | 0.30820117 | 0.43987983 | NA | NA | |
| ENSG000001 | 102.575518 | -0.173142611 | 0.16993254 | -1.0188902 | 0.30825507 | 0.43989979 | STXBP6 | | 29091 |
| ENSG000002 | 3.22162708 | -0.38460076 | 0.37748082 | -1.0188617 | 0.30826861 | 0.43989979 | NA | NA | |
| ENSG000002 | 426.818888 | 0.128017871 | 0.12563802 | 1.01894216 | 0.30823043 | 0.43989979 | CCDC169 | | 728591 |
| ENSG000002 | 6.19893444 | -0.378360827 | 0.37135645 | -1.0188616 | 0.30826867 | 0.43989979 | NA | NA | |
| ENSG000002 | 2.34275774 | -0.378147209 | 0.37114734 | -1.0188601 | 0.3082694 | 0.43989979 | NA | NA | |
| ENSG000000 | 27.5491631 | -0.256414964 | 0.25174682 | -1.018543 | 0.30841997 | 0.44008489 | NGEF | | 25791 |

| | | | | | | | | |
|---|---|---|---|---|---|---|---|---|
| ENSG000002 | 2.96950822 | -0.380908534 | 0.37397458 | -1.0185412 | 0.30842081 | 0.44008489 | NA | NA |
| ENSG000002 | 2.37992432 | -0.37659736 | 0.36976684 | -1.0184725 | 0.30845346 | 0.44011599 | NA | NA |
| ENSG000002 | 4.43204324 | -0.385401509 | 0.37845601 | -1.0183522 | 0.30851062 | 0.44018205 | NA | NA |
| ENSG000002 | 2.37992486 | -0.378011581 | 0.37124386 | -1.0182298 | 0.30856874 | 0.44024949 | NA | NA |
| ENSG000001 | 10.1955918 | -0.345637368 | 0.33948355 | -1.018127 | 0.3086176 | 0.44028822 | NKPD1 | 284353 |
| ENSG000002 | 21.179035 | 0.279135396 | 0.27416062 | 1.01814546 | 0.30860883 | 0.44028822 | RALY-AS1 | 101926888 |
| ENSG000001 | 60.6864029 | 0.203226881 | 0.19961799 | 1.01807898 | 0.30864042 | 0.44030529 | DNAJC28 | 54943 |
| ENSG000001 | 5.78751822 | -0.373995672 | 0.36739862 | -1.0179561 | 0.30869882 | 0.4403731 | T | 6862 |
| ENSG000002 | 2.94962443 | -0.381737721 | 0.37509835 | -1.0177004 | 0.30882038 | 0.44049205 | RNA5SP161 | 106480757 |
| ENSG000002 | 2.46022538 | -0.37123282 | 0.36476945 | -1.0177191 | 0.30881149 | 0.44049205 | NA | NA |
| ENSG000002 | 2.19913432 | -0.366541502 | 0.36016041 | -1.0177174 | 0.3088123 | 0.44049205 | NA | NA |
| ENSG000002 | 14.668657 | -0.311451428 | 0.30603784 | -1.0176893 | 0.30882565 | 0.44049205 | SP2-AS1 | 100506325 |
| ENSG000001 | 18.955381 | -0.293430172 | 0.2883687 | -1.0175521 | 0.30889086 | 0.44056957 | PIK3IP1 | 113791 |
| ENSG000002 | 4.53998605 | -0.383597449 | 0.37699891 | -1.0175028 | 0.3089143 | 0.4405875 | NA | NA |
| ENSG000002 | 3.85986636 | -0.384259222 | 0.37769424 | -1.0173817 | 0.30897187 | 0.44065411 | NA | NA |
| ENSG000001 | 40.997791 | 0.241042944 | 0.23696396 | 1.01721352 | 0.30905187 | 0.44069269 | PPM1J | 333926 |
| ENSG000002 | 6.06067479 | -0.373517884 | 0.36716602 | -1.0172997 | 0.30901088 | 0.44069269 | NA | NA |
| ENSG000002 | 3.25033267 | -0.383844792 | 0.37735037 | -1.0172106 | 0.30905326 | 0.44069269 | NA | NA |
| ENSG000002 | 2.36887955 | -0.377292346 | 0.37089602 | -1.0172456 | 0.3090366 | 0.44069269 | NA | NA |
| ENSG000002 | 3.75812202 | -0.385082625 | 0.3785524 | -1.0172505 | 0.30903427 | 0.44069269 | NA | NA |
| ENSG000002 | 3.76750513 | -0.384415786 | 0.3779228 | -1.0171807 | 0.30906747 | 0.44069745 | NA | NA |
| ENSG000002 | 6.37939197 | -0.375755773 | 0.3694816 | -1.016981 | 0.30916246 | 0.44081741 | LRRC73 | 221424 |
| ENSG000002 | 6.31471768 | -0.369431896 | 0.36328998 | -1.0169064 | 0.30919797 | 0.44085253 | NA | NA |
| ENSG000001 | 60.1290962 | 0.207473191 | 0.20404972 | 1.01677765 | 0.30925922 | 0.44092435 | SLC47A1 | 55244 |
| ENSG000002 | 3.49641483 | -0.384627769 | 0.37833293 | -1.0166384 | 0.3093255 | 0.44100336 | NA | NA |
| ENSG000002 | 9.11175839 | -0.348373567 | 0.34268576 | -1.0165977 | 0.30934483 | 0.4410154 | NA | NA |
| ENSG000001 | 2444.08478 | 0.144879206 | 0.14255606 | 1.01629637 | 0.30948828 | 0.4412044 | LAMC1 | 3915 |
| ENSG000002 | 2.35097851 | -0.37737417 | 0.37141838 | -1.0160353 | 0.3096126 | 0.44136611 | RN7SL41P | 106480933 |
| ENSG000002 | 3.18622722 | -0.383354519 | 0.37735209 | -1.0159067 | 0.30967381 | 0.44143786 | NA | NA |

| | | | | | | | | |
|---|---|---|---|---|---|---|---|---|
| ENSG000001 | 12.2653301 | -0.331463363 | 0.32628917 | -1.0158577 | 0.30969716 | 0.44145513 | PTGDR | 5729 |
| ENSG000002 | 2.40403147 | -0.376864238 | 0.37098941 | -1.0158356 | 0.3097077 | 0.44145513 | NA | NA |
| ENSG000002 | 4.25003295 | -0.382893516 | 0.37693681 | -1.0158029 | 0.30972324 | 0.44146176 | NA | NA |
| ENSG000002 | 11.0840936 | -0.343123646 | 0.33779664 | -1.0157699 | 0.30973899 | 0.4414687 | NA | NA |
| ENSG000002 | 2.37770114 | -0.377624491 | 0.371835 | -1.0155701 | 0.30983417 | 0.44158882 | NA | NA |
| ENSG000001 | 54.3938806 | 0.202906822 | 0.1998094 | 1.01550189 | 0.30986665 | 0.4416196 | CAPN13 | 92291 |
| ENSG000001 | 417.709997 | 0.18071964 | 0.17798358 | 1.01537255 | 0.30992827 | 0.4416919 | ZFP36L2 | 678 |
| ENSG000001 | 2.36046279 | 0.369193015 | 0.36370602 | 1.01508634 | 0.31006467 | 0.44187076 | NA | NA |
| ENSG000002 | 5.13662718 | -0.381317811 | 0.3756935 | -1.0149705 | 0.3101199 | 0.44191841 | NA | NA |
| ENSG000002 | 2.34725258 | -0.376292402 | 0.37073478 | -1.0149908 | 0.3101102 | 0.44191841 | NA | NA |
| ENSG000002 | 2.35669719 | 0.369720237 | 0.36435984 | 1.01471181 | 0.31024322 | 0.4420786 | NA | NA |
| ENSG000002 | 24.9522743 | 0.27895538 | 0.27494758 | 1.01457661 | 0.3103077 | 0.44215494 | KLHL7-AS1 | 100775104 |
| ENSG000002 | 3.22480317 | -0.382158486 | 0.37669585 | -1.0145014 | 0.31034354 | 0.44219048 | NA | NA |
| ENSG000001 | 7.82353734 | -0.357582544 | 0.35249059 | -1.0144457 | 0.31037015 | 0.44219731 | LOC105370792 | 105370792 |
| ENSG000002 | 3.7204139 | -0.383976154 | 0.37850306 | -1.0144598 | 0.31036338 | 0.44219731 | NA | NA |
| ENSG000001 | 3.27927848 | -0.381927264 | 0.37649901 | -1.0144177 | 0.31038348 | 0.44220077 | HOXB4 | 3214 |
| ENSG000002 | 2.34909969 | -0.377286897 | 0.37194659 | -1.0143577 | 0.31041209 | 0.442226 | NA | NA |
| ENSG000002 | 54.0559247 | 0.213375953 | 0.21038174 | 1.01423228 | 0.31047193 | 0.4422957 | PTOV1-AS1 | 100506033 |
| ENSG000002 | 2.61606388 | 0.379163365 | 0.37388804 | 1.01410937 | 0.31053057 | 0.4423637 | LOC101928602 | 101928602 |
| ENSG000001 | 46.6242974 | -0.21623363 | 0.21325254 | -1.0139791 | 0.3105927 | 0.44243667 | PLEKHG1 | 57480 |
| ENSG000002 | 5.38361051 | -0.378136029 | 0.37294563 | -1.0139173 | 0.31062222 | 0.44246317 | NA | NA |
| ENSG000002 | 3.18557399 | -0.381786996 | 0.37660257 | -1.0137663 | 0.31069428 | 0.44255027 | NA | NA |
| ENSG000002 | 2.3498369 | -0.377059806 | 0.37194867 | -1.0137415 | 0.31070611 | 0.44255157 | NA | NA |
| ENSG000002 | 16.4375839 | 0.310738837 | 0.30653486 | 1.01371451 | 0.310719 | 0.44255439 | NA | NA |
| ENSG000001 | 9.66142238 | 0.351574384 | 0.34683276 | 1.01367122 | 0.31073966 | 0.44256828 | NA | NA |
| ENSG000002 | 4.02145485 | -0.383128014 | 0.37809402 | -1.0133141 | 0.31091014 | 0.44279553 | NA | NA |
| ENSG000002 | 7.40592068 | -0.36452075 | 0.35974128 | -1.0132859 | 0.31092364 | 0.44279921 | NA | NA |
| ENSG000002 | 3.73672373 | -0.383514769 | 0.37851473 | -1.0132096 | 0.31096005 | 0.4428355 | NA | NA |
| ENSG000002 | 30.3314828 | -0.285820916 | 0.28211887 | -1.0131223 | 0.31100175 | 0.44284824 | NA | NA |

| | | | | | | | | |
|---|---|---|---|---|---|---|---|---|
| ENSG000002 | 3.22044527 | -0.382029605 | 0.37707303 | -1.0131449 | 0.31099097 | 0.44284824 | LOC102724020 | 102724020 |
| ENSG000002 | 2.37820967 | -0.375798115 | 0.37092262 | -1.0131442 | 0.31099127 | 0.44284824 | NA | NA |
| ENSG000001 | 27.5289618 | -0.259793938 | 0.25645953 | -1.0130017 | 0.31105936 | 0.44286806 | PMP22 | 5376 |
| ENSG000001 | 7.46870762 | -0.367871873 | 0.36315007 | -1.0130023 | 0.31105904 | 0.44286806 | CCDC116 | 164592 |
| ENSG000002 | 3.04755621 | -0.37853878 | 0.37367605 | -1.0130132 | 0.31105384 | 0.44286806 | NA | NA |
| ENSG000002 | 6.9659219 | 0.371949624 | 0.36715434 | 1.01306067 | 0.31103118 | 0.44286806 | NA | NA |
| ENSG000002 | 10.7406858 | 0.337098648 | 0.33278396 | 1.01296542 | 0.31107668 | 0.44287717 | NA | NA |
| ENSG000002 | 35.7655941 | 0.233656073 | 0.23067463 | 1.0129249 | 0.31109603 | 0.44288917 | TAPT1-AS1 | 202020 |
| ENSG000001 | 9.65506767 | -0.342601627 | 0.33831064 | -1.0126836 | 0.31121133 | 0.44303776 | TAS2R10 | 50839 |
| ENSG000002 | 6.1566834 | -0.372184002 | 0.36755332 | -1.0125987 | 0.3112519 | 0.44307997 | NA | NA |
| ENSG000002 | 8.58072144 | 0.35982859 | 0.35535981 | 1.01257537 | 0.31126303 | 0.44308025 | NA | NA |
| ENSG000001 | 29.8076257 | -0.244989501 | 0.24196596 | -1.0124957 | 0.31130108 | 0.44311886 | KLHL31 | 401265 |
| ENSG000002 | 15.1171311 | 0.326232472 | 0.32224152 | 1.01238496 | 0.31135403 | 0.44317751 | NA | NA |
| ENSG000002 | 7.32212544 | -0.365404913 | 0.36094229 | -1.0123638 | 0.31136414 | 0.44317751 | NA | NA |
| ENSG000002 | 2.63510881 | -0.379929163 | 0.37531747 | -1.0122874 | 0.31140064 | 0.44321391 | NA | NA |
| ENSG000002 | 2.95013297 | -0.378862277 | 0.37430154 | -1.0121847 | 0.31144977 | 0.44323716 | NA | NA |
| ENSG000002 | 6.79358149 | 0.366110088 | 0.36170187 | 1.01218743 | 0.31144844 | 0.44323716 | NA | NA |
| ENSG000002 | 3.7461106 | -0.383207578 | 0.37858604 | -1.0122074 | 0.31143892 | 0.44323716 | NA | NA |
| ENSG000002 | 64.6641338 | -0.196222644 | 0.19389918 | -1.0119829 | 0.31154625 | 0.443285 | C1RL-AS1 | 283314 |
| ENSG000002 | 20.2089808 | 0.286222181 | 0.28282826 | 1.01199995 | 0.31153808 | 0.443285 | SNORA2B | 677794 |
| ENSG000002 | 3.21635516 | -0.381682235 | 0.37717448 | -1.0119514 | 0.31156131 | 0.443285 | NA | NA |
| ENSG000002 | 2.40844199 | -0.374334441 | 0.36989059 | -1.012014 | 0.31153137 | 0.443285 | NA | NA |
| ENSG000002 | 2.9756204 | -0.377002201 | 0.3725383 | -1.0119824 | 0.31154647 | 0.443285 | NA | NA |
| ENSG000002 | 2.36853464 | -0.376258998 | 0.37181169 | -1.0119612 | 0.31155662 | 0.443285 | NA | NA |
| ENSG000002 | 7.72449673 | -0.367539875 | 0.36318583 | -1.0119885 | 0.31154357 | 0.443285 | NA | NA |
| ENSG000002 | 5.84151168 | -0.373306665 | 0.3689051 | -1.0119314 | 0.31157084 | 0.443285 | NA | NA |
| ENSG000002 | 3.02296557 | -0.376985765 | 0.37256289 | -1.0118715 | 0.31159951 | 0.44331024 | NA | NA |
| ENSG000002 | 5.85573176 | 0.380182254 | 0.37574257 | 1.01181577 | 0.31162615 | 0.44333259 | URAHP | 100130015 |
| ENSG000001 | 3.9792838 | -0.382426845 | 0.37797148 | -1.0117876 | 0.31163963 | 0.44333621 | RNU6-222P | 106479665 |

| | | | | | | | | |
|---|---|---|---|---|---|---|---|---|
| ENSG000002 | 4.53956129 | -0.382242242 | 0.37785751 | -1.0116042 | 0.31172733 | 0.44344542 | NA | NA |
| ENSG000000 | 8.81070503 | -0.349187948 | 0.34522308 | -1.0114849 | 0.31178439 | 0.44351102 | NA | NA |
| ENSG000002 | 5.59580689 | -0.37571762 | 0.37161455 | -1.0110412 | 0.31199671 | 0.44379749 | NA | NA |
| ENSG000001 | 12.5381179 | -0.32624649 | 0.32269118 | -1.0110177 | 0.31200796 | 0.44379792 | CSRP3 | 8048 |
| ENSG000002 | 5.37097921 | -0.377014618 | 0.37293226 | -1.0109466 | 0.31204197 | 0.44383073 | NA | NA |
| ENSG000001 | 4.60160001 | -0.381294385 | 0.37720087 | -1.0108524 | 0.3120871 | 0.44387935 | NA | NA |
| ENSG000002 | 6.06155635 | -0.370559935 | 0.36662682 | -1.0107278 | 0.3121467 | 0.44394855 | PCDHA7 | 56141 |
| ENSG000002 | 2.48725217 | -0.362369203 | 0.35853556 | -1.0106925 | 0.31216363 | 0.44395705 | NA | NA |
| ENSG000002 | 2.35758106 | -0.375397656 | 0.37143787 | -1.0106607 | 0.31217885 | 0.44396313 | NA | NA |
| ENSG000001 | 56.3107711 | 0.209332699 | 0.20716191 | 1.01047873 | 0.31226599 | 0.44407147 | CNKSR1 | 10256 |
| ENSG000001 | 4.99138881 | -0.378271584 | 0.37437821 | -1.0103996 | 0.3123039 | 0.44410981 | SERPINA6 | 866 |
| ENSG000001 | 4.87876738 | -0.381293785 | 0.37738736 | -1.0103512 | 0.31232704 | 0.44412715 | IL36A | 27179 |
| ENSG000001 | 109.279701 | 0.196221953 | 0.19422052 | 1.01030494 | 0.31234922 | 0.4441431 | BSN | 8927 |
| ENSG000002 | 18.6524037 | 0.294027315 | 0.29107165 | 1.01015444 | 0.31242131 | 0.44423003 | NA | NA |
| ENSG000002 | 2.34964103 | -0.375429051 | 0.37171472 | -1.0099924 | 0.31249892 | 0.44432482 | NA | NA |
| ENSG000002 | 19.6650484 | -0.300610113 | 0.29769656 | -1.009787 | 0.31259735 | 0.44444919 | LOC105372566 | 105372566 |
| ENSG000002 | 2.98587863 | -0.378673664 | 0.37501455 | -1.0097573 | 0.3126116 | 0.44445386 | NA | NA |
| ENSG000002 | 74.249003 | -0.1885908 | 0.18683756 | -1.0093838 | 0.31279061 | 0.44467719 | APOL6 | 80830 |
| ENSG000002 | 3.9651856 | -0.381239655 | 0.37768858 | -1.0094021 | 0.31278183 | 0.44467719 | NA | NA |
| ENSG000002 | 59.869203 | 0.224731906 | 0.2227072 | 1.00909135 | 0.31293082 | 0.44486092 | NA | NA |
| ENSG000002 | 2.34156352 | -0.372973675 | 0.3696637 | -1.008954 | 0.31299669 | 0.44493896 | NA | NA |
| ENSG000001 | 32.8435883 | -0.244236049 | 0.24210963 | -1.0087829 | 0.31307878 | 0.44504006 | MMP19 | 4327 |
| ENSG000002 | 3.5144721 | -0.381313051 | 0.3780728 | -1.0085704 | 0.31318069 | 0.44516931 | NA | NA |
| ENSG000002 | 4.04996135 | -0.381816212 | 0.37859258 | -1.0085148 | 0.3132074 | 0.44519167 | NA | NA |
| ENSG000002 | 3.24492308 | 0.381417849 | 0.37821803 | 1.00846024 | 0.31323357 | 0.44521326 | NA | NA |
| ENSG000002 | 3.99576248 | -0.381386746 | 0.3781976 | -1.0084325 | 0.31324689 | 0.44521659 | NA | NA |
| ENSG000002 | 31.3102571 | 0.24978666 | 0.2477249 | 1.00832277 | 0.31329954 | 0.44527581 | NA | NA |
| ENSG000001 | 17.0107448 | -0.299505778 | 0.29705436 | -1.0082524 | 0.3133333 | 0.44527698 | IGFBP7 | 3490 |
| ENSG000002 | 2.63010489 | -0.377871547 | 0.37477035 | -1.0082749 | 0.3133225 | 0.44527698 | NA | NA |

| | | | | | | | | |
|---|---|---|---|---|---|---|---|---|
| ENSG000002 | 23.3839973 | 0.266561043 | 0.26436838 | 1.00829397 | 0.31331336 | 0.44527698 | NA | NA |
| ENSG000002 | 4.94687421 | -0.381527359 | 0.37844614 | -1.0081418 | 0.31338641 | 0.44533685 | NA | NA |
| ENSG000002 | 3.95122369 | -0.381005999 | 0.37796574 | -1.0080437 | 0.31343346 | 0.4453725 | NA | NA |
| ENSG000002 | 9.70573108 | -0.345776035 | 0.34301006 | -1.0080638 | 0.31342381 | 0.4453725 | NA | NA |
| ENSG000001 | 22.0646478 | -0.28174031 | 0.27950847 | -1.0079849 | 0.31346173 | 0.44539706 | DYNAP | 284254 |
| ENSG000002 | 4.00383945 | -0.381110981 | 0.37811276 | -1.0079294 | 0.31348834 | 0.44541926 | NA | NA |
| ENSG000002 | 3.70541397 | -0.381549727 | 0.37857297 | -1.0078631 | 0.31352019 | 0.4454489 | NA | NA |
| ENSG000002 | 53.1950306 | 0.229551175 | 0.22776857 | 1.00782636 | 0.31353783 | 0.44545836 | NA | NA |
| ENSG000001 | 2.83277571 | -0.370000319 | 0.36715415 | -1.007752 | 0.31357355 | 0.44549351 | NA | NA |
| ENSG000002 | 4.06864589 | -0.381432809 | 0.37857205 | -1.0075567 | 0.31366732 | 0.44561111 | NA | NA |
| ENSG000002 | 3.24046176 | -0.380174098 | 0.37743108 | -1.0072676 | 0.3138062 | 0.44579278 | NA | NA |
| ENSG000002 | 2.41365389 | -0.370078494 | 0.36742525 | -1.0072212 | 0.3138285 | 0.44580885 | NA | NA |
| ENSG000002 | 2.36640732 | -0.373444052 | 0.37078867 | -1.0071614 | 0.3138572 | 0.445834 | NA | NA |
| ENSG000002 | 3.73946341 | -0.381265371 | 0.37857025 | -1.0071192 | 0.3138775 | 0.44584722 | NA | NA |
| ENSG000002 | 7.6926025 | -0.359973796 | 0.35748681 | -1.0069569 | 0.31395551 | 0.44592679 | NA | NA |
| ENSG000002 | 5.88784497 | -0.372859836 | 0.37028192 | -1.006962 | 0.31395302 | 0.44592679 | NA | NA |
| ENSG000002 | 4.20173598 | -0.379378421 | 0.37684218 | -1.0067302 | 0.31406443 | 0.44606587 | NA | NA |
| ENSG000002 | 2.34707652 | -0.373211928 | 0.37073719 | -1.0066752 | 0.31409089 | 0.44608476 | NA | NA |
| ENSG000002 | 51.6555983 | -0.20544575 | 0.20408718 | -1.0066568 | 0.31409973 | 0.44608476 | LOC100507487 | 100507487 |
| ENSG000002 | 12.0974552 | -0.324528313 | 0.32240863 | -1.0065745 | 0.31413929 | 0.44612531 | NA | NA |
| ENSG000002 | 13.3881218 | -0.322999639 | 0.32097575 | -1.0063054 | 0.31426867 | 0.44629343 | NA | NA |
| ENSG000001 | 16.5044318 | -0.311522198 | 0.30959181 | -1.0062353 | 0.31430241 | 0.44631008 | GIF | 2694 |
| ENSG000001 | 10.0977404 | -0.33932439 | 0.33721774 | -1.0062472 | 0.31429669 | 0.44631008 | NKAPL | 222698 |
| ENSG000002 | 6.52118044 | -0.363174867 | 0.36099116 | -1.0060492 | 0.3143919 | 0.4464059 | ZNF492 | 57615 |
| ENSG000002 | 20.2677577 | 0.286720414 | 0.2849925 | 1.006063 | 0.31438527 | 0.4464059 | PCDHGC4 | 56098 |
| ENSG000002 | 4.01143959 | -0.380502906 | 0.3782874 | -1.0058567 | 0.31448452 | 0.44652177 | NA | NA |
| ENSG000001 | 237.837842 | 0.139391766 | 0.13859825 | 1.0057253 | 0.31454773 | 0.44656553 | EXPH5 | 23086 |
| ENSG000002 | 4.66786882 | -0.380422192 | 0.37829061 | -1.0056348 | 0.31459129 | 0.44656553 | NA | NA |
| ENSG000002 | 3.21340686 | -0.37911332 | 0.37697775 | -1.005665 | 0.31457675 | 0.44656553 | NA | NA |

| | | | | | | | | |
|---|---|---|---|---|---|---|---|---|
| ENSG000002 | 3.77352524 | -0.380703965 | 0.37852351 | -1.0057604 | 0.31453083 | 0.44656553 | NA | NA |
| ENSG000002 | 3.78869456 | -0.380715459 | 0.37858253 | -1.005634 | 0.31459167 | 0.44656553 | NA | NA |
| ENSG000002 | 5.35221698 | -0.37461014 | 0.37252048 | -1.0056095 | 0.31460344 | 0.44656553 | NA | NA |
| ENSG000002 | 10.6207604 | 0.35186045 | 0.34989577 | 1.00561504 | 0.31460079 | 0.44656553 | NA | NA |
| ENSG000002 | 2.38384612 | -0.371516354 | 0.36941156 | -1.0056977 | 0.31456101 | 0.44656553 | NA | NA |
| ENSG000002 | 2.65999352 | -0.372881475 | 0.37082417 | -1.0055479 | 0.31463308 | 0.44659197 | NA | NA |
| ENSG000002 | 11.7133453 | -0.332369717 | 0.33054813 | -1.0055108 | 0.31465094 | 0.44660169 | NA | NA |
| ENSG000002 | 3.94982149 | -0.379711579 | 0.377653 | -1.005451 | 0.31467975 | 0.44662694 | NA | NA |
| ENSG000001 | 132.445308 | 0.165605699 | 0.16471527 | 1.00540586 | 0.31470146 | 0.44663998 | IRAK4 | 51135 |
| ENSG000002 | 3.88202397 | 0.380628729 | 0.3785896 | 1.00538612 | 0.31471096 | 0.44663998 | NA | NA |
| ENSG000002 | 5.16933496 | -0.379741517 | 0.37775851 | -1.0052494 | 0.31477677 | 0.44671775 | NA | NA |
| ENSG000002 | 5.59844328 | -0.372279722 | 0.37034432 | -1.005226 | 0.31478806 | 0.44671813 | NA | NA |
| ENSG000002 | 5.94010223 | 0.371852256 | 0.36993331 | 1.00518727 | 0.31480668 | 0.44672893 | NA | NA |
| ENSG000001 | 9.59967205 | -0.340606641 | 0.33889323 | -1.0050559 | 0.31486993 | 0.44680304 | CLDN2 | 9075 |
| ENSG000002 | 17.5652979 | -0.29715363 | 0.29566555 | -1.005033 | 0.31488097 | 0.44680304 | ATAD3C | 219293 |
| ENSG000002 | 9.73209981 | -0.345535084 | 0.34381253 | -1.0050101 | 0.31489196 | 0.44680304 | RC3H1-IT1 | 106481790 |
| ENSG000001 | 4.20654403 | -0.378542841 | 0.37672121 | -1.0048355 | 0.31497608 | 0.44685984 | HDAC10 | 83933 |
| ENSG000002 | 2.41140293 | -0.370189806 | 0.36838798 | -1.0048911 | 0.31494929 | 0.44685984 | NA | NA |
| ENSG000002 | 5.44710091 | 0.376885058 | 0.3750695 | 1.0048406 | 0.31497361 | 0.44685984 | NA | NA |
| ENSG000002 | 7.03448382 | -0.368380096 | 0.36660542 | -1.0048408 | 0.3149735 | 0.44685984 | NA | NA |
| ENSG000002 | 3.67750563 | -0.38030906 | 0.37852415 | -1.0047155 | 0.31503389 | 0.44690452 | NA | NA |
| ENSG000002 | 3.75092662 | -0.380373263 | 0.37859332 | -1.0047015 | 0.31504063 | 0.44690452 | NA | NA |
| ENSG000002 | 4.89385163 | -0.379003066 | 0.37721688 | -1.0047352 | 0.31502439 | 0.44690452 | NA | NA |
| ENSG000001 | 44.9071384 | -0.223587443 | 0.22258445 | -1.0045061 | 0.31513471 | 0.44697596 | ACTG2 | 72 |
| ENSG000002 | 3.93043807 | -0.379705015 | 0.37801058 | -1.0044825 | 0.3151461 | 0.44697596 | NA | NA |
| ENSG000002 | 13.368136 | -0.316178418 | 0.31475132 | -1.004534 | 0.31512127 | 0.44697596 | LOC102723895 | 102723895 |
| ENSG000002 | 31.3852224 | 0.250960548 | 0.24983542 | 1.00450349 | 0.31513599 | 0.44697596 | NA | NA |
| ENSG000002 | 3.00524852 | -0.376302574 | 0.37459779 | -1.004551 | 0.31511312 | 0.44697596 | NA | NA |
| ENSG000002 | 4.55251789 | -0.379795661 | 0.37814376 | -1.0043684 | 0.31520106 | 0.44703827 | NA | NA |

| | | | | | | | | |
|---|---|---|---|---|---|---|---|---|
| ENSG000002 | 4.15221212 | -0.380182087 | 0.37854134 | -1.0043344 | 0.31521746 | 0.44704589 | NA | NA |
| ENSG000002 | 6.05841228 | -0.367315787 | 0.3657466 | -1.0042904 | 0.31523868 | 0.44706035 | NA | NA |
| ENSG000002 | 2.17764429 | -0.361576807 | 0.36007548 | -1.0041695 | 0.31529693 | 0.44708041 | NA | NA |
| ENSG000002 | 3.48609487 | -0.379951773 | 0.37834975 | -1.0042342 | 0.31526572 | 0.44708041 | NA | NA |
| ENSG000002 | 17.2991687 | 0.29862744 | 0.29738465 | 1.00417907 | 0.31529231 | 0.44708041 | GHRLOS | 100126793 |
| ENSG000002 | 38.3197096 | 0.257973332 | 0.25690065 | 1.00417549 | 0.31529403 | 0.44708041 | RAB11B-AS1 | 100507567 |
| ENSG000002 | 3.71497223 | -0.380158774 | 0.37859127 | -1.0041404 | 0.31531097 | 0.44708469 | CCL3 | 6348 |
| ENSG000002 | 13.6874831 | -0.313465512 | 0.31221143 | -1.0040168 | 0.31537053 | 0.44715351 | LOC105371286 | 105371286 |
| ENSG000001 | 138.760722 | 0.152738797 | 0.15215139 | 1.00386068 | 0.31544577 | 0.44724455 | EPS8 | 2059 |
| ENSG000002 | 3.2396923 | -0.378513323 | 0.37707603 | -1.0038117 | 0.31546939 | 0.44726241 | LOC102723692 | 102723692 |
| ENSG000001 | 4.02780545 | -0.379979947 | 0.37858599 | -1.003682 | 0.31553191 | 0.44729795 | TAS1R3 | 83756 |
| ENSG000002 | 5.79471363 | -0.3715397 | 0.37018181 | -1.0036682 | 0.31553859 | 0.44729795 | LINC01317 | 104355287 |
| ENSG000002 | 37.345535 | -0.229528066 | 0.22868372 | -1.0036922 | 0.315527 | 0.44729795 | PCDHA9 | 9752 |
| ENSG000002 | 3.47137548 | -0.379744451 | 0.37833499 | -1.0037254 | 0.31551097 | 0.44729795 | NA | NA |
| ENSG00000C | 18.3923479 | 0.291543448 | 0.290547 | 1.00342954 | 0.31565366 | 0.44744543 | SUSD2 | 56241 |
| ENSG000002 | 35.3972978 | -0.23884064 | 0.2380448 | -1.0033432 | 0.31569529 | 0.44748881 | TTLL3 | 26140 |
| ENSG000002 | 4.22518969 | -0.378492443 | 0.37724364 | -1.0033103 | 0.31571116 | 0.44749565 | NA | NA |
| ENSG000002 | 19.5424859 | 0.293176332 | 0.29221635 | 1.00328518 | 0.31572328 | 0.4474972 | NA | NA |
| ENSG000002 | 5.80576004 | -0.369188693 | 0.36799768 | -1.0032365 | 0.31574677 | 0.44751485 | NA | NA |
| ENSG000002 | 5.08758504 | -0.377771218 | 0.37657275 | -1.0031826 | 0.31577278 | 0.44753606 | NA | NA |
| ENSG000002 | 6.57862119 | -0.366400712 | 0.36527788 | -1.0030739 | 0.3158252 | 0.44759472 | NA | NA |
| ENSG000002 | 1486.18848 | 0.126673975 | 0.12631288 | 1.0028587 | 0.31592904 | 0.44772624 | KCNQ1OT1 | 10984 |
| ENSG000001 | 219.447343 | 0.130487167 | 0.13013415 | 1.00271276 | 0.31599947 | 0.4478104 | ACP6 | 51205 |
| ENSG000002 | 2.33589461 | -0.371293579 | 0.37030923 | -1.0026582 | 0.31602581 | 0.44783207 | NA | NA |
| ENSG000002 | 4.25300185 | -0.378351553 | 0.37739359 | -1.0025384 | 0.31608365 | 0.44788273 | NA | NA |
| ENSG000002 | 3.97036579 | -0.378742976 | 0.37777926 | -1.002551 | 0.31607755 | 0.44788273 | NA | NA |
| ENSG000002 | 2.4053684 | -0.368245028 | 0.36733764 | -1.0024702 | 0.31611656 | 0.44791007 | NA | NA |
| ENSG000002 | 5.42261029 | -0.377694188 | 0.37677011 | -1.0024526 | 0.31612504 | 0.44791007 | NA | NA |
| ENSG000002 | 12.6120015 | -0.315190808 | 0.31445189 | -1.0023499 | 0.31617465 | 0.44794907 | NA | NA |

| | | | | | | | | |
|---|---|---|---|---|---|---|---|---|
| ENSG000002 | 3.51739666 | -0.379168994 | 0.37827539 | -1.0023623 | 0.31616864 | 0.44794907 | NA | NA |
| ENSG000002 | 3.26135893 | -0.377864916 | 0.37699728 | -1.0023014 | 0.31619803 | 0.44796655 | NA | NA |
| ENSG000002 | 7.08135253 | -0.357079597 | 0.35628325 | -1.0022351 | 0.31623004 | 0.44799623 | NA | NA |
| ENSG000001 | 2.98621002 | -0.37309915 | 0.37227808 | -1.0022055 | 0.31624433 | 0.44800084 | RBMXL2 | 27288 |
| ENSG000002 | 74.1957933 | 0.180686723 | 0.18033408 | 1.00195548 | 0.31636509 | 0.44815625 | NA | NA |
| ENSG000001 | 173.719159 | -0.153173183 | 0.15287783 | -1.0019319 | 0.31637646 | 0.4481567 | SMYD3 | 64754 |
| ENSG000002 | 4.03300973 | -0.379053846 | 0.3783388 | -1.00189 | 0.31639674 | 0.44816977 | NA | NA |
| ENSG000002 | 3.0415743 | -0.374317284 | 0.3736285 | -1.0018435 | 0.31641918 | 0.44818591 | NA | NA |
| ENSG000002 | 36.9870344 | -0.237406808 | 0.23697836 | -1.001808 | 0.31643635 | 0.44819457 | NA | NA |
| ENSG000002 | 9.84625119 | -0.335894168 | 0.33530008 | -1.0017718 | 0.31645381 | 0.44820364 | NA | NA |
| ENSG000001 | 97.8043002 | 0.172558198 | 0.17226821 | 1.00168333 | 0.31649656 | 0.44824854 | ATF7 | 11016 |
| ENSG000001 | 10.0311594 | -0.344991068 | 0.34448743 | -1.001462 | 0.3166035 | 0.44838434 | KCNK3 | 3777 |
| ENSG000002 | 11.9381539 | 0.331521178 | 0.33105352 | 1.00141264 | 0.31662736 | 0.4483868 | NA | NA |
| ENSG000002 | 2.64948776 | -0.37492126 | 0.37438673 | -1.0014277 | 0.31662006 | 0.4483868 | NA | NA |
| ENSG000001 | 57.1023218 | 0.214586833 | 0.21430996 | 1.00129194 | 0.31668569 | 0.44845374 | DGKQ | 1609 |
| ENSG000002 | 11.5420696 | 0.329060562 | 0.32864918 | 1.00125173 | 0.31670512 | 0.4484656 | NA | NA |
| ENSG000002 | 6.27726273 | -0.364497219 | 0.36411714 | -1.0010438 | 0.31680562 | 0.44859225 | SFTA1P | 207107 |
| ENSG000002 | 2.634508 | -0.374654583 | 0.37428431 | -1.0009893 | 0.31683199 | 0.44861391 | NA | NA |
| ENSG000002 | 3.46132273 | -0.378683085 | 0.37834756 | -1.0008868 | 0.31688153 | 0.4486684 | VCX3B | 425054 |
| ENSG000002 | 2.60699663 | -0.375148568 | 0.37492285 | -1.000602 | 0.31701924 | 0.44884771 | RN7SL466P | 106479393 |
| ENSG000002 | 3.46867568 | -0.378673553 | 0.37848326 | -1.0005028 | 0.31706725 | 0.44890001 | NA | NA |
| ENSG000001 | 19.7693623 | -0.278213208 | 0.27808173 | -1.0004728 | 0.31708176 | 0.44890488 | RAB26 | 25837 |
| ENSG000002 | 24.6477567 | 0.265445507 | 0.26534045 | 1.00039593 | 0.31711894 | 0.44894184 | C16orf96 | 342346 |
| ENSG000001 | 3.7470522 | -0.378658606 | 0.37852832 | -1.0003442 | 0.31714396 | 0.44896159 | NA | NA |
| ENSG000002 | 3.50430333 | -0.378385246 | 0.37827411 | -1.0002938 | 0.31716835 | 0.44898044 | LOC283214 | 283214 |
| ENSG000002 | 5.77649272 | -0.368716765 | 0.36861704 | -1.0002705 | 0.3171796 | 0.44898069 | RNU6-137P | 106481220 |
| ENSG000002 | 6.61430382 | -0.364933752 | 0.3649554 | -0.9999407 | 0.31733922 | 0.44919096 | NA | NA |
| ENSG000002 | 4.1479452 | -0.378320661 | 0.37841821 | -0.9997422 | 0.31743527 | 0.44930614 | NA | NA |
| ENSG000002 | 9.48330459 | -0.353202335 | 0.35330342 | -0.9997139 | 0.31744898 | 0.44930614 | NA | NA |

| | | | | | | | | |
|---|---|---|---|---|---|---|---|---|
| ENSG000002 | 18.4833816 | 0.285256484 | 0.28534098 | 0.99970389 | 0.31745383 | 0.44930614 | NA | NA |
| ENSG000001 | 5.65261684 | 0.372035771 | 0.37219942 | 0.99956032 | 0.31752334 | 0.44938883 | NA | NA |
| ENSG000002 | 6.57217831 | -0.361360259 | 0.36158587 | -0.9993761 | 0.31761256 | 0.44949941 | NA | NA |
| ENSG000002 | 5.00627609 | -0.375508259 | 0.37577608 | -0.9992873 | 0.31765554 | 0.44954456 | NA | NA |
| ENSG000002 | 4.69869256 | 0.374039434 | 0.37432874 | 0.99922714 | 0.31768467 | 0.4495701 | NA | NA |
| ENSG000002 | 3.9613759 | -0.377495138 | 0.37789181 | -0.9989503 | 0.31781877 | 0.44973997 | NA | NA |
| ENSG000002 | 2.94830622 | -0.373678009 | 0.37407695 | -0.9989335 | 0.3178269 | 0.44973997 | NA | NA |
| ENSG000002 | 8.82044084 | 0.348108698 | 0.34852112 | 0.99881665 | 0.31788352 | 0.4498044 | NA | NA |
| ENSG000001 | 57.5670053 | 0.210308929 | 0.21056437 | 0.99878688 | 0.31789794 | 0.44980912 | NA | NA |
| ENSG000001 | 34.16765 | -0.233136121 | 0.23345716 | -0.9986249 | 0.31797645 | 0.4498888 | ITGB4 | 3691 |
| ENSG000002 | 21.5796132 | 0.28409653 | 0.28448725 | 0.99862657 | 0.31797562 | 0.4498888 | NA | NA |
| ENSG000002 | 2.36386298 | -0.369210019 | 0.3697275 | -0.9986004 | 0.31798833 | 0.44988991 | RN7SL750P | 106481126 |
| ENSG000001 | 29.7243334 | 0.2563469 | 0.25673343 | 0.99849443 | 0.31803967 | 0.44994685 | RASGRP4 | 115727 |
| ENSG000001 | 10.5616021 | 0.333449249 | 0.33398005 | 0.99841068 | 0.31808026 | 0.44998857 | ICAM5 | 7087 |
| ENSG000001 | 12.8675595 | -0.317711233 | 0.31822665 | -0.9983803 | 0.31809496 | 0.44999368 | NDNF | 79625 |
| ENSG000002 | 4.60303357 | -0.377802911 | 0.37845323 | -0.9982816 | 0.31814281 | 0.45004567 | LOC102467147 | 102467147 |
| ENSG000002 | 3.71527358 | -0.377924241 | 0.37858489 | -0.998255 | 0.31815575 | 0.45004827 | FXYD1 | 5348 |
| ENSG000002 | 4.46716909 | -0.37451657 | 0.37522715 | -0.9981063 | 0.31822784 | 0.45013374 | NA | NA |
| ENSG000002 | 7.4801079 | -0.364900893 | 0.36560119 | -0.9980845 | 0.31823837 | 0.45013374 | NA | NA |
| ENSG000002 | 2.17682331 | -0.359924169 | 0.36063418 | -0.9980312 | 0.31826422 | 0.45015459 | NA | NA |
| ENSG000002 | 6.74172834 | -0.366649954 | 0.36739774 | -0.9979646 | 0.3182965 | 0.45018456 | NA | NA |
| ENSG000002 | 3.18415252 | -0.376216704 | 0.37706141 | -0.9977598 | 0.31839586 | 0.4503054 | NA | NA |
| ENSG000002 | 3.4378615 | -0.377111944 | 0.37796513 | -0.9977427 | 0.31840415 | 0.4503054 | NA | NA |
| ENSG000002 | 5.57730579 | -0.370399551 | 0.37131855 | -0.997525 | 0.31850972 | 0.45043899 | NA | NA |
| ENSG000001 | 7.37515499 | -0.359001148 | 0.35992287 | -0.9974391 | 0.31855141 | 0.45047748 | OR10R2 | 343406 |
| ENSG000002 | 5.07300608 | -0.373745547 | 0.37471112 | -0.9974232 | 0.31855915 | 0.45047748 | LOC100506682 | 100506682 |
| ENSG000002 | 2.39414682 | -0.367349122 | 0.36833429 | -0.9973253 | 0.31860662 | 0.45052889 | NA | NA |
| ENSG000002 | 2.34839473 | -0.369074673 | 0.37010577 | -0.997214 | 0.31866063 | 0.45058955 | NA | NA |
| ENSG000002 | 3.71823472 | -0.377509232 | 0.37859298 | -0.9971374 | 0.3186978 | 0.4506264 | NA | NA |

| | | | | | | | | |
|---|---|---|---|---|---|---|---|---|
| ENSG000001 | 162.684684 | 0.147786563 | 0.14822031 | 0.99707364 | 0.31872876 | 0.45065447 | DDX60 | 55601 |
| ENSG000001 | 10.1620309 | -0.333739333 | 0.3347647 | -0.996937 | 0.31879507 | 0.45070107 | TUBAL3 | 79861 |
| ENSG000002 | 4.03090121 | -0.377302985 | 0.37845206 | -0.9969638 | 0.3187821 | 0.45070107 | NA | NA |
| ENSG000002 | 11.3345322 | 0.337171785 | 0.33820511 | 0.99694467 | 0.31879137 | 0.45070107 | NDUFV2-AS1 | 101927275 |
| ENSG000002 | 2.60397566 | -0.373573305 | 0.37473691 | -0.9968949 | 0.31881553 | 0.45071429 | NA | NA |
| ENSG000002 | 11.1323897 | -0.331429246 | 0.33246947 | -0.9968712 | 0.31882702 | 0.45071482 | NA | NA |
| ENSG000002 | 10.6196519 | -0.329268886 | 0.33034016 | -0.9967571 | 0.31888244 | 0.45077745 | NA | NA |
| ENSG000002 | 15.3249338 | -0.307847524 | 0.30887274 | -0.9966808 | 0.31891949 | 0.45078551 | NA | NA |
| ENSG000002 | 97.1907924 | 0.171770815 | 0.17233848 | 0.99670608 | 0.3189072 | 0.45078551 | NA | NA |
| ENSG000002 | 2.40798498 | -0.366606317 | 0.36782874 | -0.9966766 | 0.31892149 | 0.45078551 | PRAMEF18 | 391003 |
| ENSG000002 | 17.2220227 | -0.296003491 | 0.29703671 | -0.9965216 | 0.31899678 | 0.45087621 | NA | NA |
| ENSG000002 | 11.8606468 | -0.320652839 | 0.32178383 | -0.9964853 | 0.31901443 | 0.45088544 | NA | NA |
| ENSG000002 | 6.35127189 | -0.363805201 | 0.36510682 | -0.996435 | 0.31903885 | 0.45090423 | NA | NA |
| ENSG000001 | 70.8817275 | 0.185660467 | 0.18633441 | 0.99638317 | 0.31906401 | 0.45092408 | IFIH1 | 64135 |
| ENSG000002 | 5.29160913 | -0.371802529 | 0.37316438 | -0.9963505 | 0.31907986 | 0.45093076 | NHEG1 | 100294720 |
| ENSG000002 | 2.66965991 | -0.372737737 | 0.37412766 | -0.9962849 | 0.31911174 | 0.45094439 | NA | NA |
| ENSG000002 | 2.39594131 | -0.366730389 | 0.36809202 | -0.9963009 | 0.31910399 | 0.45094439 | NA | NA |
| ENSG000002 | 7.86416184 | -0.353979363 | 0.35533796 | -0.9961766 | 0.31916435 | 0.45100302 | NA | NA |
| ENSG000001 | 385.381195 | 0.121680215 | 0.1221522 | 0.99613608 | 0.31918403 | 0.45101511 | MTMR10 | 54893 |
| ENSG000002 | 26.2401964 | 0.260434281 | 0.2614781 | 0.99600799 | 0.31924627 | 0.45108733 | NA | NA |
| ENSG000002 | 10.464859 | -0.334085351 | 0.33543992 | -0.9959618 | 0.31926869 | 0.4511033 | NA | NA |
| ENSG000002 | 2.36347864 | -0.368639538 | 0.37014933 | -0.9959211 | 0.31928847 | 0.45111553 | NA | NA |
| ENSG000002 | 2.38310946 | -0.368638627 | 0.37016051 | -0.9958886 | 0.31930428 | 0.45112104 | NA | NA |
| ENSG000002 | 5.8894887 | 0.369112006 | 0.37064376 | 0.99586731 | 0.31931462 | 0.45112104 | NA | NA |
| ENSG000002 | 11.1167163 | -0.325636555 | 0.32701018 | -0.9957995 | 0.31934759 | 0.45115191 | NA | NA |
| ENSG000002 | 3.49578951 | -0.37668071 | 0.37828826 | -0.9957505 | 0.3193714 | 0.45116982 | NA | NA |
| ENSG000000 | 15.6378447 | -0.311160706 | 0.31259186 | -0.9954217 | 0.31953123 | 0.45137988 | FKBP6 | 8468 |
| ENSG000002 | 3.17420392 | -0.37534709 | 0.37711214 | -0.9953196 | 0.31958087 | 0.45143427 | NA | NA |
| ENSG000002 | 3.99008038 | -0.376503523 | 0.37834193 | -0.9951409 | 0.31966774 | 0.45153978 | RNU7-90P | 106479060 |

| | | | | | | | | |
|---|---|---|---|---|---|---|---|---|
| ENSG000002 | 8.95719024 | -0.347252142 | 0.34895499 | -0.9951202 | 0.31967783 | 0.45153978 | LINC01097 | 285547 |
| ENSG000001 | 13.8412957 | 0.307336054 | 0.30887135 | 0.99502932 | 0.31972201 | 0.45158645 | RABGEF1 | 27342 |
| ENSG000002 | 27.1836241 | 0.258519125 | 0.25983243 | 0.99494555 | 0.31976275 | 0.45162827 | NA | NA |
| ENSG000002 | 4.47257741 | -0.373619676 | 0.37553071 | -0.9949111 | 0.3197795 | 0.4516362 | NA | NA |
| ENSG000002 | 4.54917271 | -0.37565121 | 0.37762391 | -0.994776 | 0.31984521 | 0.45171327 | NA | NA |
| ENSG000002 | 4.47898353 | -0.373944231 | 0.37602568 | -0.9944646 | 0.31999673 | 0.45191152 | NA | NA |
| ENSG000001 | 83.1779636 | 0.186750992 | 0.18783274 | 0.99424091 | 0.32010559 | 0.45204952 | FNDC4 | 64838 |
| ENSG000002 | 4.78890659 | -0.375073265 | 0.37726868 | -0.9941808 | 0.32013487 | 0.45207513 | NA | NA |
| ENSG000002 | 6.95671702 | -0.364809993 | 0.366956 | -0.9941519 | 0.32014894 | 0.45207924 | NA | NA |
| ENSG000001 | 35.396259 | 0.233084129 | 0.23447793 | 0.99405574 | 0.32019573 | 0.45209809 | PLSCR4 | 57088 |
| ENSG000002 | 4.9168431 | -0.375333336 | 0.37757607 | -0.9940602 | 0.32019357 | 0.45209809 | LINC01448 | 101928795 |
| ENSG000002 | 2.51840512 | -0.359199114 | 0.36133344 | -0.9940932 | 0.3201775 | 0.45209809 | NA | NA |
| ENSG000001 | 13.5607344 | -0.322714696 | 0.32476012 | -0.9937017 | 0.32036809 | 0.45232571 | PLEKHN1 | 84069 |
| ENSG000002 | 7.33529073 | -0.352261505 | 0.35453057 | -0.9935998 | 0.32041773 | 0.45238004 | NA | NA |
| ENSG000001 | 5.09433303 | -0.372528497 | 0.37495216 | -0.9935361 | 0.32044878 | 0.45240812 | HIGD1B | 51751 |
| ENSG000002 | 2.60044616 | -0.371798777 | 0.37424232 | -0.9934707 | 0.32048063 | 0.45243734 | TEN1 | 100134934 |
| ENSG000002 | 4.83462696 | -0.374701107 | 0.37720239 | -0.9933689 | 0.32053024 | 0.45249161 | NA | NA |
| ENSG000001 | 27.3766239 | -0.259879769 | 0.26162562 | -0.9933269 | 0.32055067 | 0.45250471 | NFE2 | 4778 |
| ENSG000001 | 45.3122544 | -0.214974862 | 0.2164429 | -0.9932174 | 0.320604 | 0.45256424 | LOX | 4015 |
| ENSG000001 | 4.46102356 | -0.372782492 | 0.37535031 | -0.9931589 | 0.32063253 | 0.45258143 | APOF | 319 |
| ENSG000002 | 2.34042136 | -0.364941657 | 0.36746 | -0.9931466 | 0.3206385 | 0.45258143 | NA | NA |
| ENSG000001 | 9.22505113 | -0.343777987 | 0.34621126 | -0.9929717 | 0.32072374 | 0.45268599 | POM121L9P | 29774 |
| ENSG000002 | 3.30133778 | 0.374967629 | 0.37763283 | 0.99294234 | 0.32073805 | 0.45269043 | NA | NA |
| ENSG000002 | 5.01088379 | -0.372202332 | 0.37494578 | -0.9926831 | 0.32086442 | 0.45285303 | NA | NA |
| ENSG000000 | 104.686679 | -0.174615789 | 0.17591032 | -0.992641 | 0.32088495 | 0.45285329 | THSD7A | 221981 |
| ENSG000002 | 3.68180405 | -0.375787033 | 0.37857452 | -0.9926369 | 0.32088695 | 0.45285329 | NA | NA |
| ENSG000002 | 5.03260696 | -0.373053217 | 0.37590765 | -0.9924066 | 0.32099924 | 0.452996 | NA | NA |
| ENSG000002 | 3.25716334 | -0.373513393 | 0.37641367 | -0.992295 | 0.32105365 | 0.45305702 | NA | NA |
| ENSG000002 | 12.7152745 | -0.32157303 | 0.32408381 | -0.9922527 | 0.32107429 | 0.45307037 | NA | NA |

| | | | | | | | | |
|---|---|---|---|---|---|---|---|---|
| ENSG000002 | 7.32971769 | -0.353780643 | 0.35656813 | -0.9921825 | 0.32110852 | 0.45310292 | NA | NA |
| ENSG000002 | 23.3061683 | 0.262749394 | 0.26486386 | 0.9920168 | 0.32118933 | 0.45320117 | NA | NA |
| ENSG000002 | 7.56420894 | 0.352481335 | 0.35534302 | 0.9919467 | 0.32122353 | 0.45323366 | NA | NA |
| ENSG000002 | 5.5515168 | -0.367987383 | 0.37100633 | -0.9918628 | 0.32126445 | 0.45327562 | MIR544B | 100422864 |
| ENSG000002 | 7.48696053 | -0.362262837 | 0.36524802 | -0.991827 | 0.32128194 | 0.45328452 | NA | NA |
| ENSG000002 | 2.40069729 | -0.365298319 | 0.36835333 | -0.9917063 | 0.32134082 | 0.45335183 | NA | NA |
| ENSG000002 | 2.3746281 | -0.367049817 | 0.37013703 | -0.9916593 | 0.32136377 | 0.45336843 | NA | NA |
| ENSG000001 | 38.6035744 | -0.23014779 | 0.23211202 | -0.9915376 | 0.32142315 | 0.45342884 | SLCO2A1 | 6578 |
| ENSG000002 | 2.61286142 | -0.37042513 | 0.37359374 | -0.9915186 | 0.32143243 | 0.45342884 | NA | NA |
| ENSG000002 | 5.08688835 | -0.372362432 | 0.37555359 | -0.9915028 | 0.32144013 | 0.45342884 | NA | NA |
| ENSG000002 | 2.38832191 | -0.364906409 | 0.36807743 | -0.9913849 | 0.32149766 | 0.45349422 | NA | NA |
| ENSG000002 | 3.94134013 | -0.374601082 | 0.37791481 | -0.9912316 | 0.32157253 | 0.45352983 | NA | NA |
| ENSG000002 | 3.97816201 | -0.374783222 | 0.37810324 | -0.9912193 | 0.32157851 | 0.45352983 | NA | NA |
| ENSG000002 | 17.4801015 | 0.289731665 | 0.29229844 | 0.99121864 | 0.32157883 | 0.45352983 | SNORD116-30 | 100873856 |
| ENSG000002 | 7.07711752 | -0.356783694 | 0.35994117 | -0.9912278 | 0.32157436 | 0.45352983 | LINC00515 | 282566 |
| ENSG000002 | 6.63189329 | -0.360161575 | 0.36335053 | -0.9912235 | 0.32157647 | 0.45352983 | NA | NA |
| ENSG000001 | 53.3033938 | 0.203443253 | 0.20527981 | 0.99105338 | 0.32165951 | 0.45358775 | LOC100049716 | 100049716 |
| ENSG000002 | 2.61863481 | -0.370188968 | 0.3735434 | -0.99102 | 0.32167583 | 0.45358775 | MIR610 | 693195 |
| ENSG000002 | 12.5176779 | -0.321126018 | 0.3240074 | -0.9911071 | 0.32163331 | 0.45358775 | NA | NA |
| ENSG000002 | 9.85772276 | 0.336802569 | 0.33984997 | 0.99103309 | 0.32166942 | 0.45358775 | NA | NA |
| ENSG000002 | 2.4014345 | -0.36505705 | 0.36835529 | -0.991046 | 0.32166311 | 0.45358775 | NA | NA |
| ENSG000002 | 4.52300317 | -0.374813513 | 0.37821915 | -0.9909956 | 0.32168773 | 0.45358877 | LINC00588 | 26138 |
| ENSG000001 | 150.966546 | 0.157232045 | 0.15867281 | 0.99091992 | 0.32172468 | 0.45360932 | ADCY5 | 111 |
| ENSG000002 | 7.3285718 | 0.359150376 | 0.3624335 | 0.99094144 | 0.32171418 | 0.45360932 | NA | NA |
| ENSG000002 | 5.17002369 | -0.372796873 | 0.37626699 | -0.9907775 | 0.32179423 | 0.4536916 | NA | NA |
| ENSG000001 | 3.18294494 | 0.373681667 | 0.37717853 | 0.9907289 | 0.32181798 | 0.45369353 | NA | NA |
| ENSG000002 | 3.72360372 | -0.375082475 | 0.37859106 | -0.9907325 | 0.32181621 | 0.45369353 | NA | NA |
| ENSG000001 | 136.272053 | 0.186224151 | 0.18801157 | 0.99049303 | 0.32193319 | 0.45384018 | NLGN2 | 57555 |
| ENSG000002 | 2.6242401 | -0.370980701 | 0.37455125 | -0.9904671 | 0.32194585 | 0.45384225 | NA | NA |

| | | | | | | | | |
|---|---|---|---|---|---|---|---|---|
| ENSG000002 | 2.37873064 | -0.364751979 | 0.36829007 | -0.9903932 | 0.32198197 | 0.45387739 | TBC1D3P7 | 106480773 |
| ENSG000002 | 12.5848045 | -0.329140106 | 0.33234477 | -0.9903574 | 0.32199946 | 0.45388627 | NA | NA |
| ENSG000001 | 579.613452 | 0.124041212 | 0.12527533 | 0.99014874 | 0.32210142 | 0.45401421 | ST6GAL2 | 84620 |
| ENSG000002 | 2.69243713 | -0.365491315 | 0.36913642 | -0.9901253 | 0.32211287 | 0.45401456 | NA | NA |
| ENSG000001 | 3.54290545 | -0.374191476 | 0.37795317 | -0.9900472 | 0.32215106 | 0.45404167 | OR6C74 | 254783 |
| ENSG000002 | 3.07731853 | -0.366781275 | 0.37047111 | -0.9900402 | 0.32215449 | 0.45404167 | NA | NA |
| ENSG000002 | 4.70319591 | 0.37198114 | 0.37574754 | 0.98997625 | 0.32218573 | 0.45406991 | ECSCR | 641700 |
| ENSG000002 | 3.4155618 | 0.369871792 | 0.37365152 | 0.98988436 | 0.32223064 | 0.45411743 | NA | NA |
| ENSG000002 | 24.7397316 | 0.262973019 | 0.26570825 | 0.98970589 | 0.32231789 | 0.4542246 | NA | NA |
| ENSG000002 | 3.6806619 | -0.374551652 | 0.37850975 | -0.9895429 | 0.32239757 | 0.45430531 | NA | NA |
| ENSG000002 | 3.85538645 | 0.374604452 | 0.37855826 | 0.98955561 | 0.32239138 | 0.45430531 | NA | NA |
| ENSG000002 | 4.38398971 | 0.374107159 | 0.37807081 | 0.98951612 | 0.32241069 | 0.45430801 | RN7SL535P | 106479417 |
| ENSG000002 | 17.5771712 | -0.286188203 | 0.28922868 | -0.9894876 | 0.32242462 | 0.45431185 | LOC100130476 | 100130476 |
| ENSG000001 | 41.6803001 | -0.215340901 | 0.21765815 | -0.9893537 | 0.3224901 | 0.45438832 | KLK13 | 26085 |
| ENSG000001 | 8.44506192 | -0.349022286 | 0.35280267 | -0.9892847 | 0.32252386 | 0.4544201 | CSN1S1 | 1446 |
| ENSG000002 | 3.69847806 | -0.374175397 | 0.37829486 | -0.9891104 | 0.32260911 | 0.45452442 | NA | NA |
| ENSG000001 | 31.3367388 | -0.247084451 | 0.24982839 | -0.9890167 | 0.32265497 | 0.45455745 | GMPR | 2766 |
| ENSG000002 | 2.40571386 | -0.363783999 | 0.36781809 | -0.9890324 | 0.3226473 | 0.45455745 | NA | NA |
| ENSG000002 | 2.72819021 | -0.36563976 | 0.36971771 | -0.9889701 | 0.32267778 | 0.45457379 | CIR1P3 | 106480783 |
| ENSG000002 | 5.33619012 | 0.367661631 | 0.37178397 | 0.988912 | 0.3227062 | 0.45459804 | NA | NA |
| ENSG000002 | 4.19111521 | -0.373013878 | 0.37725987 | -0.9887452 | 0.32278783 | 0.45469724 | NA | NA |
| ENSG000002 | 2.34343024 | -0.365513926 | 0.36970684 | -0.9886588 | 0.3228301 | 0.45474098 | NA | NA |
| ENSG000001 | 2.59609512 | -0.369748755 | 0.37403725 | -0.9885346 | 0.32289091 | 0.45479505 | HYPM | 25763 |
| ENSG000002 | 2.39326635 | -0.365041116 | 0.36926672 | -0.9885568 | 0.32288004 | 0.45479505 | NA | NA |
| ENSG000002 | 4.24806961 | -0.373533646 | 0.37787719 | -0.9885054 | 0.32290519 | 0.45479936 | NA | NA |
| ENSG000002 | 2.63644684 | -0.370181261 | 0.37451622 | -0.9884252 | 0.32294446 | 0.45483887 | NA | NA |
| ENSG000002 | 5.01927211 | -0.371309005 | 0.37567701 | -0.988373 | 0.32297002 | 0.45485908 | LINC01428 | 101929265 |
| ENSG000002 | 4.231676 | -0.372446582 | 0.37686985 | -0.9882631 | 0.3230238 | 0.45488741 | NA | NA |
| ENSG000002 | 19.8662928 | -0.283166159 | 0.28652642 | -0.9882724 | 0.32301925 | 0.45488741 | ALG1L2 | 644974 |

| | | | | | | | | |
|---|---|---|---|---|---|---|---|---|
| ENSG000002 | 6.43623054 | -0.366731868 | 0.37107349 | -0.9882998 | 0.32300583 | 0.45488741 | NA | NA |
| ENSG000002 | 7.65617951 | 0.352013636 | 0.35620615 | 0.98823008 | 0.32303998 | 0.4548944 | MIR8075 | 102465874 |
| ENSG000002 | 4.09915239 | 0.373024026 | 0.37757043 | 0.98795878 | 0.32317283 | 0.45506568 | NA | NA |
| ENSG000002 | 2.64661115 | -0.369221531 | 0.37374281 | -0.9879027 | 0.32320031 | 0.45508136 | NA | NA |
| ENSG000002 | 3.74605908 | -0.3740078 | 0.37859247 | -0.9878902 | 0.32320641 | 0.45508136 | NA | NA |
| ENSG000002 | 2.34883856 | -0.364821868 | 0.36933212 | -0.9877881 | 0.32325644 | 0.455136 | NA | NA |
| ENSG000002 | 4.0643418 | 0.373690414 | 0.37836698 | 0.98764013 | 0.32332893 | 0.45522225 | NA | NA |
| ENSG000002 | 3.41770264 | -0.373587654 | 0.37830013 | -0.987543 | 0.3233765 | 0.45527343 | NA | NA |
| ENSG000002 | 2.61438846 | -0.369861675 | 0.37453826 | -0.9875137 | 0.32339085 | 0.45527782 | NA | NA |
| ENSG000002 | 7.26852155 | -0.36252561 | 0.36714627 | -0.9874147 | 0.3234394 | 0.45533036 | NA | NA |
| ENSG000001 | 5.03945332 | -0.369848247 | 0.37460304 | -0.9873071 | 0.32349211 | 0.45538875 | TRIM49 | 57093 |
| ENSG000002 | 2.61456452 | -0.369288238 | 0.37407107 | -0.9872141 | 0.32353769 | 0.45542131 | NA | NA |
| ENSG000002 | 5.62558107 | -0.368981462 | 0.37375472 | -0.9872289 | 0.32353043 | 0.45542131 | NA | NA |
| ENSG000002 | 7.64933918 | -0.359888614 | 0.36456139 | -0.9871825 | 0.32355319 | 0.45542732 | NA | NA |
| ENSG000001 | 126.97458 | 0.155439343 | 0.15748176 | 0.98703075 | 0.32362756 | 0.45548457 | PRPF18 | 8559 |
| ENSG000001 | 96.206596 | -0.163177508 | 0.16532083 | -0.9870354 | 0.3236253 | 0.45548457 | MTSS1 | 9788 |
| ENSG000002 | 10.3923105 | 0.327097577 | 0.33139414 | 0.98703488 | 0.32362554 | 0.45548457 | SMIM1 | 388588 |
| ENSG000002 | 15.4778559 | -0.302154892 | 0.30615324 | -0.98694 | 0.32367203 | 0.45553134 | GTF3C2-AS1 | 100505624 |
| ENSG000001 | 2.38931174 | -0.362523785 | 0.36734525 | -0.9868748 | 0.323704 | 0.45554471 | NA | NA |
| ENSG000002 | 4.73393969 | -0.369016767 | 0.37392174 | -0.9868824 | 0.32370032 | 0.45554471 | NA | NA |
| ENSG000001 | 2.3703019 | -0.363730083 | 0.36858625 | -0.9868249 | 0.32372849 | 0.45556337 | CCL19 | 6363 |
| ENSG000002 | 13.6931168 | 0.311851128 | 0.31603854 | 0.9867503 | 0.32376507 | 0.45559903 | NA | NA |
| ENSG000002 | 3.7138108 | -0.373524187 | 0.37856428 | -0.9866863 | 0.32379644 | 0.45562737 | NA | NA |
| ENSG000002 | 2.32460079 | -0.363447706 | 0.36844645 | -0.9864329 | 0.32392072 | 0.45578643 | NA | NA |
| ENSG000002 | 14.1460701 | -0.308774998 | 0.31304629 | -0.9863557 | 0.32395859 | 0.4558239 | NA | NA |
| ENSG000002 | 3.95708902 | -0.373091143 | 0.3782996 | -0.9862319 | 0.32401932 | 0.45589353 | NA | NA |
| ENSG000002 | 3.9774576 | -0.372818908 | 0.37803609 | -0.9861993 | 0.32403535 | 0.45590026 | NA | NA |
| ENSG000001 | 4.44109936 | -0.370436265 | 0.37569437 | -0.9860043 | 0.324131 | 0.45601046 | LY6H | 4062 |
| ENSG000002 | 32.5363186 | 0.242029613 | 0.24546768 | 0.98599379 | 0.32413616 | 0.45601046 | NA | NA |

| | | | | | | | | | |
|---|---|---|---|---|---|---|---|---|---|
| ENSG000002 | 3.01556927 | -0.366861048 | 0.37210358 | -0.9859111 | 0.32417674 | 0.45605173 | NA | NA | |
| ENSG000001 | 7340.14298 | 0.127730527 | 0.12957537 | 0.98576238 | 0.32424973 | 0.45613858 | CCNG1 | | 900 |
| ENSG000002 | 16.0419289 | -0.298172717 | 0.30249013 | -0.9857271 | 0.32426705 | 0.45614712 | NA | NA | |
| ENSG000002 | 3.96328641 | -0.372964641 | 0.37840429 | -0.9856248 | 0.32431728 | 0.45620196 | NA | NA | |
| ENSG000001 | 19.1575309 | 0.283087887 | 0.28727138 | 0.98543713 | 0.3244094 | 0.45631571 | NA | NA | |
| ENSG000002 | 3.67917648 | -0.372990743 | 0.37854332 | -0.9853317 | 0.32446116 | 0.45637268 | NA | NA | |
| ENSG000002 | 2.3702691 | -0.363445105 | 0.36889865 | -0.9852167 | 0.32451765 | 0.45643631 | NA | NA | |
| ENSG000002 | 2.27292204 | 0.362735283 | 0.3682128 | 0.98512405 | 0.32456314 | 0.45648446 | NA | NA | |
| ENSG000002 | 56.1153984 | -0.199154516 | 0.20217181 | -0.9850756 | 0.32458693 | 0.45650209 | NA | NA | |
| ENSG000002 | 4.82889882 | -0.372193216 | 0.37786974 | -0.9849776 | 0.32463509 | 0.45655387 | NA | NA | |
| ENSG000002 | 4.24015736 | -0.371420857 | 0.37709431 | -0.9849548 | 0.32464627 | 0.45655387 | NA | NA | |
| ENSG000002 | 49.7897165 | 0.214893032 | 0.21818499 | 0.98491208 | 0.32466726 | 0.45656585 | NA | NA | |
| ENSG000002 | 3.86888353 | -0.37196217 | 0.37766812 | -0.9848916 | 0.32467731 | 0.45656585 | NA | NA | |
| ENSG000002 | 9.31564903 | -0.354068998 | 0.35953715 | -0.9847911 | 0.32472668 | 0.45660361 | NA | NA | |
| ENSG000002 | 8.29399008 | -0.354793681 | 0.3602686 | -0.9848032 | 0.32472073 | 0.45660361 | NA | NA | |
| ENSG000002 | 2.35055451 | -0.364091091 | 0.36973245 | -0.984742 | 0.3247508 | 0.45662169 | NA | NA | |
| ENSG000002 | 4.53439193 | -0.370890642 | 0.37668641 | -0.9846138 | 0.3248138 | 0.45667861 | NA | NA | |
| ENSG000002 | 5.78313548 | -0.362954776 | 0.36862538 | -0.9846169 | 0.32481228 | 0.45667861 | NA | NA | |
| ENSG000002 | 18.4034649 | 0.290713752 | 0.29534096 | 0.98433265 | 0.32495198 | 0.45685705 | NA | NA | |
| ENSG000002 | 2.58047043 | -0.368815866 | 0.37471034 | -0.9842692 | 0.32498315 | 0.45688502 | NA | NA | |
| ENSG000002 | 7.02407031 | 0.352493642 | 0.35817834 | 0.98412886 | 0.32505216 | 0.4569662 | NA | NA | |
| ENSG000002 | 14.9244106 | -0.302206994 | 0.30709477 | -0.9840838 | 0.3250743 | 0.45698148 | NA | NA | |
| ENSG000002 | 7.56932956 | -0.347225837 | 0.35286474 | -0.9840196 | 0.32510585 | 0.45701 | NA | NA | |
| ENSG000002 | 36.5971504 | 0.227584487 | 0.23128859 | 0.98398493 | 0.32512292 | 0.45701815 | NA | NA | |
| ENSG000002 | 2.35707197 | -0.362761281 | 0.36869557 | -0.9839046 | 0.3251624 | 0.4570578 | NA | NA | |
| ENSG000001 | 70.2159394 | 0.198143479 | 0.20141664 | 0.98374928 | 0.3252388 | 0.45714935 | ANKRD35 | | 148741 |
| ENSG000002 | 2.31508993 | -0.364225149 | 0.37025527 | -0.9837136 | 0.32525634 | 0.45715816 | NA | NA | |
| ENSG000002 | 4.61000869 | -0.371762462 | 0.37798188 | -0.9835457 | 0.32533892 | 0.45725838 | NA | NA | |
| ENSG000001 | 13.9929755 | -0.309604647 | 0.31480771 | -0.9834722 | 0.32537507 | 0.45729333 | C15orf26 | | 161502 |

| | | | | | | | | |
|---|---|---|---|---|---|---|---|---|
| ENSG000002 | 11.9722959 | -0.325112183 | 0.33059796 | -0.9834065 | 0.32540741 | 0.45732293 | MMP12 | 4321 |
| ENSG000002 | 4.72119107 | -0.368752563 | 0.3749835 | -0.9833834 | 0.32541876 | 0.45732303 | NA | NA |
| ENSG000002 | 3.99108614 | -0.372008437 | 0.37831903 | -0.9833194 | 0.32545027 | 0.45735147 | CACNA1C-AS4 | 100874234 |
| ENSG000002 | 5.62373092 | -0.365782725 | 0.37203791 | -0.9831867 | 0.32551555 | 0.45742736 | NA | NA |
| ENSG000001 | 28.3636847 | -0.245150409 | 0.24938169 | -0.9830329 | 0.32559125 | 0.45747692 | METTL21B | 25895 |
| ENSG000001 | 13.126282 | -0.31152721 | 0.31691093 | -0.9830119 | 0.3256016 | 0.45747692 | CD70 | 970 |
| ENSG000002 | 8.66739068 | -0.343373466 | 0.34931162 | -0.9830004 | 0.32560724 | 0.45747692 | NA | NA |
| ENSG000002 | 8.78472071 | -0.340718504 | 0.34658278 | -0.9830797 | 0.3255682 | 0.45747692 | LINC01609 | 101927487 |
| ENSG000002 | 2.32479667 | -0.36187155 | 0.3681277 | -0.9830055 | 0.32560473 | 0.45747692 | NA | NA |
| ENSG000002 | 29.5592442 | 0.247220538 | 0.25150355 | 0.98297038 | 0.32562202 | 0.45748184 | NA | NA |
| ENSG00000C | 29.7082629 | -0.23675462 | 0.24088195 | -0.9828658 | 0.32567351 | 0.45751725 | GPRC5A | 9052 |
| ENSG000001 | 224.633601 | 0.148296427 | 0.15088754 | 0.9828275 | 0.32569235 | 0.45751725 | IQCG | 84223 |
| ENSG000002 | 16.5616549 | -0.309805077 | 0.31520564 | -0.9828665 | 0.32567313 | 0.45751725 | DOCK4-AS1 | 100506413 |
| ENSG000002 | 17.7356844 | 0.293460912 | 0.29858748 | 0.98283059 | 0.32569082 | 0.45751725 | SNORA71B | 26776 |
| ENSG000002 | 7.56277588 | -0.352977949 | 0.35916351 | -0.9827779 | 0.32571678 | 0.45753572 | PP13 | 100129503 |
| ENSG000002 | 6.5479744 | -0.357626784 | 0.36392116 | -0.982704 | 0.32575314 | 0.45757094 | NA | NA |
| ENSG000001 | 261.777963 | 0.180510109 | 0.18371342 | 0.98256354 | 0.32582229 | 0.45763638 | USP4 | 7375 |
| ENSG000001 | 60.7796598 | 0.195991259 | 0.19946894 | 0.9825653 | 0.32582143 | 0.45763638 | ZNF341 | 84905 |
| ENSG000002 | 8.90386973 | -0.3439354 | 0.35005479 | -0.9825188 | 0.32584435 | 0.4576515 | NA | NA |
| ENSG000002 | 2.44379392 | -0.356873519 | 0.363296 | -0.9823216 | 0.32594142 | 0.45775613 | NA | NA |
| ENSG000002 | 4.05918033 | 0.37155497 | 0.37823381 | 0.98234203 | 0.32593137 | 0.45775613 | NA | NA |
| ENSG000002 | 4.82881558 | -0.370247689 | 0.37692651 | -0.9822808 | 0.32596152 | 0.4577685 | NA | NA |
| ENSG000002 | 3.68535426 | -0.371767717 | 0.37859006 | -0.9819796 | 0.3261099 | 0.45796103 | NA | NA |
| ENSG000002 | 4.49947665 | -0.369320576 | 0.37615062 | -0.9818423 | 0.32617756 | 0.45804018 | NA | NA |
| ENSG000002 | 2.18133797 | -0.353106075 | 0.35964988 | -0.9818051 | 0.32619589 | 0.45805005 | NA | NA |
| ENSG000002 | 4.26575048 | -0.370627117 | 0.37753539 | -0.9817017 | 0.32624685 | 0.45810575 | NA | NA |
| ENSG000002 | 3.24977206 | -0.36956336 | 0.37646753 | -0.9816607 | 0.32626705 | 0.45811826 | NA | NA |
| ENSG000002 | 3.69104353 | -0.371570507 | 0.3785563 | -0.9815462 | 0.32632345 | 0.45818159 | NA | NA |
| ENSG000001 | 4.77885995 | -0.369795172 | 0.37678441 | -0.9814503 | 0.32637073 | 0.45820037 | NA | NA |

| | | | | | | | | |
|---|---|---|---|---|---|---|---|---|
| ENSG000002 | 13.6116658 | -0.303500659 | 0.30922334 | -0.9814934 | 0.3263495 | 0.45820037 | TCTEX1D2 | 255758 |
| ENSG000002 | 7.77699864 | -0.344708369 | 0.35121797 | -0.9814656 | 0.32636317 | 0.45820037 | RN7SL239P | 106479309 |
| ENSG000001 | 4.54179117 | -0.36987011 | 0.37690549 | -0.9813338 | 0.32642814 | 0.4582651 | C1QL4 | 338761 |
| ENSG00000C | 165.702776 | 0.14094922 | 0.1436471 | 0.9812187 | 0.3264849 | 0.45827955 | C20orf194 | 25943 |
| ENSG000001 | 10.1621199 | -0.331799324 | 0.33812806 | -0.981283 | 0.3264532 | 0.45827955 | CST9 | 128822 |
| ENSG000001 | 1327.98902 | 0.136637495 | 0.13925899 | 0.98117544 | 0.32650624 | 0.45827955 | MDM4 | 4194 |
| ENSG000002 | 8.62017166 | -0.348718334 | 0.35539291 | -0.9812192 | 0.32648468 | 0.45827955 | NA | NA |
| ENSG000002 | 2.35238125 | -0.361726872 | 0.36864101 | -0.9812443 | 0.3264723 | 0.45827955 | LOC100506470 | 100506470 |
| ENSG000002 | 16.5301962 | -0.300141269 | 0.30589612 | -0.9811869 | 0.32650058 | 0.45827955 | NA | NA |
| ENSG000001 | 6.55622135 | -0.356020276 | 0.36292155 | -0.9809841 | 0.32660057 | 0.45836436 | NGB | 58157 |
| ENSG000002 | 3.9964877 | 0.371208971 | 0.37839023 | 0.98102155 | 0.32658212 | 0.45836436 | NA | NA |
| ENSG000002 | 2.43263291 | -0.360973366 | 0.36796793 | -0.9809914 | 0.326597 | 0.45836436 | RN7SL313P | 106481008 |
| ENSG000002 | 3.724974 | -0.371309436 | 0.37858772 | -0.9807752 | 0.32670363 | 0.45847725 | NA | NA |
| ENSG000002 | 2.5998533 | -0.367320636 | 0.37452019 | -0.9807766 | 0.32670293 | 0.45847725 | NA | NA |
| ENSG00000C | 37.4735979 | -0.224488421 | 0.22889676 | -0.9807409 | 0.32672051 | 0.45848508 | TIE1 | 7075 |
| ENSG000002 | 4.01194344 | -0.371044726 | 0.37834944 | -0.9806932 | 0.32674406 | 0.45850226 | NA | NA |
| ENSG000001 | 8.31769684 | -0.347979474 | 0.3548782 | -0.9805603 | 0.32680962 | 0.45854754 | ATP6V1B1 | 525 |
| ENSG000002 | 3.43454858 | -0.370865812 | 0.37821876 | -0.980559 | 0.32681025 | 0.45854754 | NA | NA |
| ENSG000002 | 2.66203996 | -0.366030228 | 0.37327141 | -0.9806007 | 0.32678967 | 0.45854754 | NA | NA |
| ENSG000002 | 3.14310248 | -0.368399227 | 0.37572115 | -0.9805124 | 0.32683328 | 0.45856398 | NA | NA |
| ENSG000002 | 2.34207314 | -0.362491739 | 0.3697072 | -0.9804833 | 0.3268476 | 0.45856821 | NA | NA |
| ENSG000001 | 14.0805528 | -0.301090455 | 0.30709698 | -0.9804409 | 0.32686851 | 0.45856916 | RPE65 | 6121 |
| ENSG000001 | 2.36565802 | -0.362503317 | 0.36973681 | -0.9804361 | 0.32687089 | 0.45856916 | OR13J1 | 392309 |
| ENSG000001 | 19.5303756 | -0.272703168 | 0.27819984 | -0.980242 | 0.32696668 | 0.45868768 | PCSK1N | 27344 |
| ENSG000002 | 2.33729735 | -0.361085498 | 0.36838698 | -0.9801798 | 0.32699735 | 0.45871483 | NA | NA |
| ENSG000002 | 4.19786679 | -0.369948735 | 0.37743988 | -0.9801527 | 0.32701073 | 0.45871772 | NA | NA |
| ENSG000002 | 6.3477181 | -0.357627191 | 0.3649377 | -0.9799678 | 0.32710202 | 0.45881712 | NA | NA |
| ENSG000002 | 3.81186643 | -0.370405211 | 0.37797864 | -0.9799633 | 0.32710421 | 0.45881712 | NA | NA |
| ENSG000002 | 2.59743315 | -0.366523566 | 0.37404039 | -0.9799037 | 0.32713366 | 0.45884255 | NA | NA |

| | | | | | | | | |
|---|---|---|---|---|---|---|---|---|
| ENSG000001 | 3.95209206 | -0.370611364 | 0.37832916 | -0.9796003 | 0.32728346 | 0.45896295 | DCD | 117159 |
| ENSG000001 | 11.2409458 | 0.31807583 | 0.32468678 | 0.97963899 | 0.32726435 | 0.45896295 | OXER1 | 165140 |
| ENSG000002 | 8.05865727 | -0.342112355 | 0.34920705 | -0.9796834 | 0.32724242 | 0.45896295 | LOC100507564 | 100507564 |
| ENSG000002 | 2.60132772 | -0.366878999 | 0.37452213 | -0.9795923 | 0.3272874 | 0.45896295 | RN7SL505P | 106481758 |
| ENSG000002 | 13.2261393 | 0.318132782 | 0.32473396 | 0.97967203 | 0.32724804 | 0.45896295 | NA | NA |
| ENSG000002 | 3.86672263 | -0.369516271 | 0.3772017 | -0.9796252 | 0.32727119 | 0.45896295 | NA | NA |
| ENSG000001 | 652.156489 | 0.153688168 | 0.15692027 | 0.97940292 | 0.32738094 | 0.4590683 | ABL2 | 27 |
| ENSG000002 | 8.21911214 | -0.348600612 | 0.35593489 | -0.9793943 | 0.32738517 | 0.4590683 | CBX3P2 | 645158 |
| ENSG000002 | 2.88344227 | -0.368643564 | 0.37643939 | -0.9792906 | 0.3274364 | 0.45911537 | NA | NA |
| ENSG000002 | 2.36895864 | -0.361077863 | 0.36871749 | -0.9792805 | 0.32744138 | 0.45911537 | NA | NA |
| ENSG000001 | 113.405256 | 0.15912095 | 0.16251582 | 0.97911054 | 0.32752536 | 0.45921724 | RALGDS | 5900 |
| ENSG000002 | 3.99994488 | -0.370639378 | 0.37855983 | -0.9790774 | 0.32754173 | 0.45922431 | NA | NA |
| ENSG000002 | 4.71118025 | 0.367950098 | 0.37587493 | 0.97891632 | 0.32762133 | 0.45932003 | RN7SL552P | 106481072 |
| ENSG00000C | 51.7448619 | -0.204593948 | 0.20901407 | -0.9788525 | 0.32765286 | 0.45934836 | LRRC23 | 10233 |
| ENSG000001 | 3.16749778 | -0.368978631 | 0.3769762 | -0.9787849 | 0.32768625 | 0.45936621 | PTGER1 | 5731 |
| ENSG000002 | 17.4789364 | -0.288583076 | 0.29484621 | -0.978758 | 0.32769958 | 0.45936621 | NA | NA |
| ENSG000002 | 10.9354763 | -0.325705972 | 0.33277186 | -0.9787666 | 0.32769533 | 0.45936621 | NA | NA |
| ENSG000001 | 120.583658 | -0.15088224 | 0.15418082 | -0.9786057 | 0.32777482 | 0.45943991 | MAP3K5 | 4217 |
| ENSG000002 | 2.67910682 | -0.364656149 | 0.37262624 | -0.978611 | 0.3277722 | 0.45943991 | NA | NA |
| ENSG000002 | 2.61635956 | -0.366504646 | 0.37454261 | -0.9785393 | 0.32780768 | 0.45945422 | NA | NA |
| ENSG000002 | 7.63877983 | 0.348671477 | 0.35631657 | 0.9785441 | 0.32780529 | 0.45945422 | NA | NA |
| ENSG000002 | 7.59586318 | -0.347715724 | 0.35535944 | -0.9784902 | 0.32783194 | 0.45946853 | NA | NA |
| ENSG000002 | 12.0803543 | 0.317078919 | 0.32405493 | 0.97847276 | 0.32784056 | 0.45946853 | LOC100506100 | 100506100 |
| ENSG000002 | 2.88913134 | -0.368529435 | 0.37668103 | -0.9783594 | 0.32789659 | 0.45949942 | NA | NA |
| ENSG000002 | 10.3683898 | -0.326929881 | 0.3341507 | -0.9783905 | 0.32788121 | 0.45949942 | NA | NA |
| ENSG000002 | 8.08063615 | -0.352136696 | 0.35992459 | -0.9783624 | 0.3278951 | 0.45949942 | NA | NA |
| ENSG000002 | 4.2049013 | -0.369139532 | 0.37733075 | -0.9782917 | 0.32793009 | 0.45953048 | NA | NA |
| ENSG000002 | 3.78179863 | -0.370232384 | 0.37847192 | -0.9782295 | 0.32796084 | 0.45955435 | NA | NA |
| ENSG000002 | 4.17714121 | -0.368989491 | 0.37720834 | -0.9782114 | 0.32796979 | 0.45955435 | NA | NA |

| | | | | | | | | |
|---|---|---|---|---|---|---|---|---|
| ENSG000002 | 8.36747239 | 0.344794476 | 0.35250288 | 0.97813236 | 0.32800887 | 0.45959071 | NA | NA |
| ENSG000002 | 14.2784552 | -0.313193894 | 0.32020213 | -0.9781131 | 0.3280184 | 0.45959071 | LOC100506834 | 100506834 |
| ENSG000002 | 3.92743029 | -0.36998942 | 0.37834466 | -0.9779163 | 0.32811571 | 0.45971116 | NA | NA |
| ENSG000001 | 4.88066024 | -0.369827676 | 0.37822749 | -0.9777916 | 0.32817738 | 0.45973403 | HTR6 | 3362 |
| ENSG000001 | 37.7224164 | -0.219705989 | 0.22469581 | -0.977793 | 0.32817673 | 0.45973403 | SERPINB8 | 5271 |
| ENSG000002 | 4.0722207 | 0.369308092 | 0.37768578 | 0.97781837 | 0.32816416 | 0.45973403 | NA | NA |
| ENSG000002 | 2.26837458 | 0.360788334 | 0.36897077 | 0.97782363 | 0.32816156 | 0.45973403 | MIR6859-1 | 102466751 |
| ENSG000001 | 3.9492928 | -0.369899649 | 0.37836819 | -0.9776183 | 0.32826316 | 0.45982243 | APOA1 | 335 |
| ENSG000002 | 48.3313934 | -0.20096435 | 0.20556129 | -0.9776371 | 0.32825382 | 0.45982243 | C6orf163 | 206412 |
| ENSG000002 | 3.73769724 | -0.370071377 | 0.37858128 | -0.9775216 | 0.328311 | 0.45987355 | LINC01502 | 100130954 |
| ENSG000002 | 4.185054 | -0.368825068 | 0.37733004 | -0.9774601 | 0.32834141 | 0.45990027 | NA | NA |
| ENSG000002 | 5.65546239 | -0.365432644 | 0.37404 | -0.9769881 | 0.32857503 | 0.46020426 | NA | NA |
| ENSG000002 | 3.25084066 | -0.36582629 | 0.37444766 | -0.9769758 | 0.32858114 | 0.46020426 | NA | NA |
| ENSG000002 | 4.53353849 | -0.36793268 | 0.37662615 | -0.9769175 | 0.32860999 | 0.46022877 | RN7SL20P | 106479232 |
| ENSG000001 | 84.1444777 | 0.178027828 | 0.1822621 | 0.97676824 | 0.3286839 | 0.46025517 | TBC1D19 | 55296 |
| ENSG000001 | 4.23981712 | -0.368641761 | 0.37741097 | -0.9767648 | 0.32868559 | 0.46025517 | MIRLET7D | 406886 |
| ENSG000002 | 30.073804 | 0.250866639 | 0.25683146 | 0.97677535 | 0.32868038 | 0.46025517 | NA | NA |
| ENSG000002 | 3.93698114 | -0.369688574 | 0.37846822 | -0.9768022 | 0.3286671 | 0.46025517 | NA | NA |
| ENSG000002 | 25.3918252 | 0.257809292 | 0.26393229 | 0.97680087 | 0.32866775 | 0.46025517 | NA | NA |
| ENSG000002 | 6.12146069 | -0.360373767 | 0.36895893 | -0.9767314 | 0.32870215 | 0.46026246 | NA | NA |
| ENSG000002 | 7.67793808 | 0.349214603 | 0.35757402 | 0.97662185 | 0.3287564 | 0.46031826 | SNORA37 | 677819 |
| ENSG000002 | 2.60457649 | -0.364313681 | 0.37304094 | -0.9766051 | 0.3287647 | 0.46031826 | MIR4276 | 100423042 |
| ENSG000002 | 3.07066446 | -0.36178102 | 0.37046739 | -0.976553 | 0.32879052 | 0.46033852 | NA | NA |
| ENSG000002 | 8.67680299 | 0.342184933 | 0.35041453 | 0.97651468 | 0.32880948 | 0.46034916 | NA | NA |
| ENSG000002 | 14.6920233 | -0.307517263 | 0.31496403 | -0.9763568 | 0.3288877 | 0.46044278 | NA | NA |
| ENSG000002 | 4.33302138 | -0.36959682 | 0.37856537 | -0.9763091 | 0.3289113 | 0.46045992 | NA | NA |
| ENSG000002 | 3.50007587 | 0.369539834 | 0.37855605 | 0.9761826 | 0.32897398 | 0.46051203 | NA | NA |
| ENSG000002 | 6.37129546 | -0.358536239 | 0.36729052 | -0.9761652 | 0.32898259 | 0.46051203 | NA | NA |
| ENSG000002 | 13.593776 | 0.307121605 | 0.31461114 | 0.9761943 | 0.32896819 | 0.46051203 | NA | NA |

| | | | | | | | | |
|---|---|---|---|---|---|---|---|---|
| ENSG000002 | 78.1822973 | 0.177838657 | 0.18219109 | 0.9761106 | 0.32900966 | 0.46053403 | ZNF702P | 79986 |
| ENSG000001 | 185.56041 | 0.142878204 | 0.14638324 | 0.97605573 | 0.32903685 | 0.46054814 | ADGRG6 | 57211 |
| ENSG000002 | 2.37315423 | -0.360168393 | 0.36902553 | -0.9759986 | 0.32906517 | 0.46054814 | NAALADL2-AS3 | 100862679 |
| ENSG000002 | 45.3803806 | -0.21858738 | 0.22395395 | -0.9760372 | 0.32904605 | 0.46054814 | NA | NA |
| ENSG000002 | 2.63927194 | -0.364836174 | 0.37379982 | -0.9760202 | 0.32905445 | 0.46054814 | NA | NA |
| ENSG000002 | 3.94046653 | -0.369235394 | 0.3783255 | -0.9759728 | 0.32907794 | 0.46055012 | NA | NA |
| ENSG000002 | 2.42034075 | -0.357068633 | 0.36589867 | -0.9758675 | 0.32913011 | 0.46060723 | NA | NA |
| ENSG000002 | 8.08024649 | 0.348263004 | 0.35691132 | 0.975769 | 0.32917895 | 0.46065968 | NA | NA |
| ENSG000002 | 2.88175933 | -0.367430886 | 0.37663586 | -0.97556 | 0.32928255 | 0.46078876 | NA | NA |
| ENSG000002 | 4.47234908 | -0.367096223 | 0.37633535 | -0.9754497 | 0.32933722 | 0.46084936 | LOC101927023 | 101927023 |
| ENSG000002 | 8.28037169 | -0.34026545 | 0.34886906 | -0.9753386 | 0.32939234 | 0.46091059 | NA | NA |
| ENSG000002 | 3.17999682 | -0.36618276 | 0.37548185 | -0.9752343 | 0.32944407 | 0.46096707 | NA | NA |
| ENSG000002 | 5.75619661 | -0.361765543 | 0.37100798 | -0.9750883 | 0.32951646 | 0.46105244 | NA | NA |
| ENSG000002 | 4.17569958 | -0.367493342 | 0.37707503 | -0.9745894 | 0.32976395 | 0.46138281 | NA | NA |
| ENSG000002 | 9.64803677 | -0.331271846 | 0.33992846 | -0.974534 | 0.32979146 | 0.46140538 | NA | NA |
| ENSG000002 | 4.27232001 | -0.36831034 | 0.3779585 | -0.974473 | 0.32982175 | 0.46143184 | LRRC24 | 441381 |
| ENSG000002 | 2.18865791 | -0.349366607 | 0.35859463 | -0.9742661 | 0.32992441 | 0.46155954 | NA | NA |
| ENSG000001 | 8.69709013 | 0.3475682 | 0.3567808 | 0.97417854 | 0.3299679 | 0.4616044 | DHDH | 27294 |
| ENSG000002 | 4.47702128 | -0.366110768 | 0.37582367 | -0.9741557 | 0.32997925 | 0.4616044 | NA | NA |
| ENSG000002 | 2.92581839 | -0.364164595 | 0.37385273 | -0.9740857 | 0.33001399 | 0.46163708 | NA | NA |
| ENSG000002 | 3.75804545 | -0.368702673 | 0.37853099 | -0.9740356 | 0.33003885 | 0.46165592 | NA | NA |
| ENSG000002 | 9.40542354 | -0.3352615 | 0.34420891 | -0.9740059 | 0.33005362 | 0.46166067 | LYZL6 | 57151 |
| ENSG000002 | 2.72023644 | -0.358376234 | 0.36795202 | -0.9739755 | 0.33006873 | 0.46166587 | OR1D5 | 8386 |
| ENSG000002 | 13.1952602 | 0.307211872 | 0.31544223 | 0.9739085 | 0.33010197 | 0.46168053 | NA | NA |
| ENSG000002 | 11.2041087 | 0.323814727 | 0.33248344 | 0.97392737 | 0.3300926 | 0.46168053 | NA | NA |
| ENSG000001 | 268.33256 | 0.145132452 | 0.14902779 | 0.97386167 | 0.33012523 | 0.46168427 | GORAB | 92344 |
| ENSG000001 | 784.852488 | 0.119480395 | 0.12269153 | 0.97382759 | 0.33014215 | 0.46168427 | RPS6KC1 | 26750 |
| ENSG000001 | 3.43623152 | -0.368028865 | 0.37793512 | -0.9737885 | 0.33016158 | 0.46168427 | NA | NA |
| ENSG000002 | 2.62404368 | -0.362558317 | 0.37231325 | -0.9737991 | 0.33015629 | 0.46168427 | NA | NA |

| | | | | | | | | |
|---|---|---|---|---|---|---|---|---|
| ENSG000002 | 22.1101217 | -0.267864283 | 0.27505794 | -0.9738467 | 0.33013264 | 0.46168427 | MIR2052HG | 441355 |
| ENSG000002 | 13.2044959 | 0.304031743 | 0.31223003 | 0.9737428 | 0.33018426 | 0.46170007 | NA | NA |
| ENSG000002 | 3.76787007 | -0.368593601 | 0.37854835 | -0.9737028 | 0.33020411 | 0.46171191 | NA | NA |
| ENSG000002 | 10.1598953 | 0.33126584 | 0.34022466 | 0.97366793 | 0.33022145 | 0.46172022 | NA | NA |
| ENSG000002 | 7.8056527 | -0.341983755 | 0.35125133 | -0.9736155 | 0.33024747 | 0.46172997 | NA | NA |
| ENSG000002 | 2.42764881 | -0.357697091 | 0.36739332 | -0.973608 | 0.33025119 | 0.46172997 | NA | NA |
| ENSG000002 | 14.0837299 | -0.29900383 | 0.30714865 | -0.9734825 | 0.33031357 | 0.46180126 | RABGAP1L-IT1 | 106480742 |
| ENSG000001 | 15.0482055 | -0.298731948 | 0.30688411 | -0.9734357 | 0.33033681 | 0.46180191 | POPDC2 | 64091 |
| ENSG000002 | 2.58231699 | -0.364092151 | 0.37401956 | -0.9734575 | 0.33032596 | 0.46180191 | NA | NA |
| ENSG000002 | 2.5952932 | -0.363682686 | 0.37367253 | -0.9732658 | 0.33042122 | 0.46190399 | MIR640 | 693225 |
| ENSG000002 | 19.7356115 | -0.271307466 | 0.27877641 | -0.9732081 | 0.33044988 | 0.46192812 | PAXIP1-AS2 | 100132707 |
| ENSG000002 | 3.56669885 | -0.36711678 | 0.37723318 | -0.9731826 | 0.33046254 | 0.46192991 | NA | NA |
| ENSG000002 | 3.3323394 | -0.364639337 | 0.37471575 | -0.9731092 | 0.33049904 | 0.461965 | NA | NA |
| ENSG000002 | 4.97498602 | -0.362502922 | 0.37254481 | -0.9730451 | 0.33053087 | 0.46199356 | NA | NA |
| ENSG000000 | 758.376991 | 0.118352982 | 0.12163478 | 0.97301925 | 0.33054373 | 0.46199562 | GPATCH2 | 55105 |
| ENSG000002 | 12.9908179 | -0.319754341 | 0.32866968 | -0.9728745 | 0.3306157 | 0.46208028 | NA | NA |
| ENSG000002 | 2.63275337 | -0.362804117 | 0.37293582 | -0.9728326 | 0.33063652 | 0.46209345 | RN7SL625P | 106479450 |
| ENSG000002 | 4.44559474 | -0.365502047 | 0.37573026 | -0.9727778 | 0.33066375 | 0.46211559 | NA | NA |
| ENSG000001 | 42.540547 | 0.212344813 | 0.21829321 | 0.97275044 | 0.33067735 | 0.46211866 | CLDN3 | 1365 |
| ENSG000002 | 2.31597095 | -0.359290833 | 0.36942228 | -0.9725749 | 0.33076462 | 0.4622247 | NA | NA |
| ENSG000002 | 5.47668566 | -0.361177051 | 0.37142511 | -0.9724088 | 0.33084721 | 0.46232417 | NA | NA |
| ENSG000002 | 5.79693233 | -0.360866893 | 0.37114733 | -0.9723009 | 0.33090086 | 0.46235134 | NA | NA |
| ENSG000002 | 4.03274832 | -0.368064096 | 0.3785416 | -0.9723214 | 0.33089069 | 0.46235134 | NA | NA |
| ENSG000002 | 4.28904719 | -0.367388564 | 0.37785087 | -0.972311 | 0.33089585 | 0.46235134 | NA | NA |
| ENSG000001 | 3.95190937 | -0.367799236 | 0.37832967 | -0.972166 | 0.33096798 | 0.4624292 | NA | NA |
| ENSG000002 | 6.5726631 | -0.35260319 | 0.36274517 | -0.972041 | 0.33103013 | 0.4625001 | ARID3C | 138715 |
| ENSG000001 | 182.814212 | 0.141963726 | 0.14605295 | 0.97200177 | 0.33104967 | 0.46251146 | C9orf3 | 84909 |
| ENSG000002 | 4.58265433 | -0.367499025 | 0.37812735 | -0.9718922 | 0.33110418 | 0.46257168 | LOC101927513 | 101927513 |
| ENSG000001 | 13.560621 | -0.302323067 | 0.31116276 | -0.9715914 | 0.33125385 | 0.46273565 | CCR10 | 2826 |

| | | | | | | | | |
|---|---|---|---|---|---|---|---|---|
| ENSG000002 | 2.59346645 | -0.363483214 | 0.37412163 | -0.9715643 | 0.33126736 | 0.46273565 | NA | NA |
| ENSG000002 | 2.62087367 | -0.362773215 | 0.37338133 | -0.9715891 | 0.33125503 | 0.46273565 | NA | NA |
| ENSG000002 | 6.82431624 | -0.352654002 | 0.36298391 | -0.9715417 | 0.33127861 | 0.46273565 | SMIM18 | 100507341 |
| ENSG000002 | 2.89223629 | -0.365483276 | 0.37618217 | -0.9715593 | 0.33126986 | 0.46273565 | NA | NA |
| ENSG000002 | 3.79236842 | -0.367722309 | 0.37851908 | -0.9714763 | 0.33131116 | 0.46276518 | NA | NA |
| ENSG000001 | 39.1611093 | 0.223269726 | 0.22984602 | 0.97138825 | 0.33135498 | 0.46281045 | ZNF404 | 342908 |
| ENSG000002 | 7.56478319 | -0.34282501 | 0.35295999 | -0.9712858 | 0.331406 | 0.46286577 | NA | NA |
| ENSG000002 | 7.56188008 | 0.347387965 | 0.35766914 | 0.97125506 | 0.33142128 | 0.46287117 | NA | NA |
| ENSG000002 | 7.34092435 | 0.356177656 | 0.36679357 | 0.97105751 | 0.33151964 | 0.46299259 | NA | NA |
| ENSG000002 | 12.299734 | -0.308735768 | 0.31798464 | -0.9709141 | 0.33159107 | 0.4630764 | NA | NA |
| ENSG000002 | 6.3280278 | -0.355484709 | 0.36617273 | -0.9708115 | 0.33164214 | 0.46313177 | NA | NA |
| ENSG000001 | 13.941988 | -0.310452709 | 0.31981512 | -0.9707255 | 0.33168497 | 0.46317563 | TRPV2 | 51393 |
| ENSG000002 | 2.67998784 | -0.361107804 | 0.37207332 | -0.9705286 | 0.33178307 | 0.46328072 | NA | NA |
| ENSG000002 | 10.7426084 | -0.327972073 | 0.33792414 | -0.9705494 | 0.33177271 | 0.46328072 | NA | NA |
| ENSG000002 | 6.5336164 | -0.352059414 | 0.36277616 | -0.9704591 | 0.33181772 | 0.46331315 | NA | NA |
| ENSG000001 | 19.072191 | -0.279612571 | 0.28813127 | -0.9704347 | 0.33182987 | 0.46331416 | SDSL | 113675 |
| ENSG000002 | 6.50265952 | -0.352797809 | 0.36358112 | -0.9703414 | 0.33187635 | 0.4633631 | BTNL2 | 56244 |
| ENSG000001 | 5.61346771 | -0.362689321 | 0.37378758 | -0.9703086 | 0.33189267 | 0.46336993 | RAPSN | 5913 |
| ENSG000002 | 4.20873584 | -0.366641182 | 0.37792682 | -0.970138 | 0.3319777 | 0.46347269 | LOC101929269 | 101929269 |
| ENSG000002 | 3.7323729 | -0.367132925 | 0.37854107 | -0.9698629 | 0.33211486 | 0.46364821 | P4HA2-AS1 | 100861518 |
| ENSG000001 | 11.0131215 | -0.327990546 | 0.33822795 | -0.9697322 | 0.33217998 | 0.46372317 | SDR16C5 | 195814 |
| ENSG000002 | 4.51318177 | -0.36601029 | 0.37746287 | -0.9696591 | 0.33221646 | 0.46375813 | NA | NA |
| ENSG000002 | 2.88978423 | -0.365201833 | 0.3766399 | -0.9696313 | 0.33223031 | 0.46376149 | NA | NA |
| ENSG000001 | 11.3644448 | -0.316724794 | 0.32671207 | -0.969431 | 0.33233021 | 0.46388497 | NA | NA |
| ENSG000001 | 196.033253 | 0.15935505 | 0.16439244 | 0.96935751 | 0.33236685 | 0.46390417 | NA | NA |
| ENSG000001 | 149.533937 | 0.147586721 | 0.15225008 | 0.96937041 | 0.33236041 | 0.46390417 | COL25A1 | 84570 |
| ENSG000002 | 3.18065494 | -0.363231769 | 0.37472466 | -0.9693298 | 0.33238068 | 0.46390751 | NA | NA |
| ENSG000002 | 3.95486976 | -0.366934009 | 0.37857578 | -0.9692485 | 0.33242121 | 0.46392376 | NA | NA |
| ENSG000002 | 2.85458078 | -0.365598688 | 0.37720233 | -0.9692376 | 0.33242664 | 0.46392376 | NA | NA |

| | | | | | | | | |
|---|---|---|---|---|---|---|---|---|
| ENSG000002 | 2.36161889 | -0.355974006 | 0.36726143 | -0.969266 | 0.3324125 | 0.46392376 | NA | NA |
| ENSG000002 | 6.09290609 | -0.357846983 | 0.3692569 | -0.9691003 | 0.33249513 | 0.46400338 | NA | NA |
| ENSG000002 | 2.28359433 | 0.358031211 | 0.36945984 | 0.96906666 | 0.33251193 | 0.46401085 | NA | NA |
| ENSG000002 | 6.13961014 | -0.362544962 | 0.3741371 | -0.9690163 | 0.33253704 | 0.46401395 | NA | NA |
| ENSG000002 | 4.73814344 | 0.36387647 | 0.37550676 | 0.96902774 | 0.33253135 | 0.46401395 | NA | NA |
| ENSG000001 | 13.2898841 | -0.302559749 | 0.3122843 | -0.96886 | 0.33261507 | 0.46410686 | SFTPD | 6441 |
| ENSG000001 | 58.8916269 | -0.211584191 | 0.21840493 | -0.9687702 | 0.33265986 | 0.46415339 | WIF1 | 11197 |
| ENSG000001 | 218.686697 | 0.152395805 | 0.15731572 | 0.96872584 | 0.33268199 | 0.4641683 | SOCS3 | 9021 |
| ENSG000002 | 5.26789702 | 0.362911748 | 0.37468733 | 0.96857225 | 0.33275866 | 0.46425929 | NA | NA |
| ENSG000002 | 5.07086558 | -0.364738766 | 0.37663313 | -0.9684192 | 0.33283504 | 0.46432305 | NA | NA |
| ENSG000002 | 6.06968573 | -0.358404178 | 0.37009477 | -0.9684119 | 0.33283871 | 0.46432305 | NA | NA |
| ENSG000002 | 2.85503082 | 0.36312628 | 0.37495668 | 0.96844863 | 0.33282036 | 0.46432305 | NA | NA |
| ENSG000002 | 2.56703721 | -0.361808001 | 0.37363393 | -0.9683489 | 0.33287015 | 0.46435094 | RN7SL242P | 106480496 |
| ENSG000002 | 4.89676391 | -0.366585918 | 0.37858416 | -0.9683076 | 0.33289078 | 0.46436374 | NA | NA |
| ENSG000001 | 2.87738106 | -0.364484228 | 0.37643417 | -0.9682549 | 0.33291709 | 0.46438079 | OR6M1 | 390261 |
| ENSG000002 | 2.8601255 | -0.364830243 | 0.3767984 | -0.9682372 | 0.3329259 | 0.46438079 | NA | NA |
| ENSG000002 | 3.49066088 | 0.366522323 | 0.37856929 | 0.96817765 | 0.33295566 | 0.46439034 | NA | NA |
| ENSG000002 | 4.96607316 | -0.366324351 | 0.37835932 | -0.9681917 | 0.33294865 | 0.46439034 | NA | NA |
| ENSG000001 | 146.939609 | 0.148701327 | 0.15360393 | 0.9680828 | 0.33300302 | 0.46440857 | CAP2 | 10486 |
| ENSG000002 | 2.34112741 | -0.357730405 | 0.3695334 | -0.9680597 | 0.33301453 | 0.46440857 | NA | NA |
| ENSG000002 | 20.794515 | 0.268561546 | 0.27742181 | 0.96806211 | 0.33301335 | 0.46440857 | NA | NA |
| ENSG000002 | 2.36774591 | -0.35691374 | 0.3686887 | -0.9680626 | 0.33301311 | 0.46440857 | NA | NA |
| ENSG000001 | 321.575441 | -0.121604528 | 0.12562386 | -0.968005 | 0.33304186 | 0.4644307 | CACNB4 | 785 |
| ENSG000002 | 2.86206848 | -0.363829882 | 0.37589039 | -0.9679148 | 0.33308691 | 0.46447756 | NA | NA |
| ENSG000001 | 2.88612301 | -0.363658029 | 0.37579456 | -0.9677044 | 0.33319204 | 0.46460818 | RNU6-862P | 106481920 |
| ENSG000002 | 4.73402956 | 0.365479887 | 0.37768836 | 0.9676758 | 0.33320631 | 0.4646121 | NA | NA |
| ENSG000001 | 11.6505964 | -0.318736562 | 0.32940662 | -0.9676083 | 0.33324005 | 0.46464318 | LDLRAD2 | 401944 |
| ENSG000002 | 5.23964879 | 0.363193294 | 0.37537415 | 0.9675501 | 0.33326911 | 0.46466771 | NA | NA |
| ENSG000002 | 23.9877703 | 0.267682268 | 0.27669298 | 0.96743426 | 0.33332699 | 0.46473243 | CEBPA | 1050 |

| | | | | | | | | |
|---|---|---|---|---|---|---|---|---|
| ENSG000002 | 4.22048557 | -0.365777691 | 0.37811651 | -0.9673677 | 0.33336026 | 0.46474686 | NA | NA |
| ENSG000002 | 9.75597995 | 0.338176464 | 0.34957709 | 0.96738737 | 0.33335042 | 0.46474686 | NA | NA |
| ENSG000001 | 7.79372958 | -0.345784267 | 0.35748133 | -0.9672792 | 0.33340445 | 0.46479249 | C17orf47 | 284083 |
| ENSG000001 | 10.6897823 | 0.316798298 | 0.32756424 | 0.96713333 | 0.33347738 | 0.46487818 | FIBIN | 387758 |
| ENSG000001 | 48.7134933 | 0.200601388 | 0.20745424 | 0.96696694 | 0.33356056 | 0.46496216 | PAQR9 | 344838 |
| ENSG000002 | 3.4189679 | -0.364670247 | 0.37712487 | -0.9669748 | 0.33355663 | 0.46496216 | NA | NA |
| ENSG000002 | 2.35687664 | -0.356751091 | 0.36897658 | -0.9668665 | 0.33361078 | 0.46501618 | LOC105370016 | 105370016 |
| ENSG000001 | 2.63126686 | -0.360925326 | 0.37330488 | -0.966838 | 0.33362504 | 0.46502008 | NA | NA |
| ENSG000002 | 7.33489877 | -0.346318427 | 0.35825348 | -0.9666855 | 0.3337013 | 0.46511038 | GPC6-AS1 | 100873972 |
| ENSG000002 | 2.6624243 | -0.359333013 | 0.37174655 | -0.9666075 | 0.33374028 | 0.46514872 | NA | NA |
| ENSG00000C | 301.766777 | -0.129321299 | 0.1338041 | -0.9664973 | 0.33379542 | 0.46520958 | SLC4A8 | 9498 |
| ENSG000002 | 7.37231421 | -0.353775081 | 0.36607216 | -0.966408 | 0.33384004 | 0.46525578 | NA | NA |
| ENSG000002 | 4.73784168 | -0.362467634 | 0.37509836 | -0.9663269 | 0.33388063 | 0.46528245 | NA | NA |
| ENSG000002 | 14.4980511 | -0.304018716 | 0.31461367 | -0.9663239 | 0.33388212 | 0.46528245 | NA | NA |
| ENSG000002 | 4.01060487 | -0.365760517 | 0.378536 | -0.9662503 | 0.33391896 | 0.46531779 | NA | NA |
| ENSG000002 | 4.55825489 | -0.36537315 | 0.3781741 | -0.9661506 | 0.33396881 | 0.46537127 | NA | NA |
| ENSG000002 | 2.65306879 | -0.358477508 | 0.37105822 | -0.966095 | 0.33399662 | 0.46539403 | NA | NA |
| ENSG000002 | 5.32634401 | -0.361341597 | 0.37407626 | -0.965957 | 0.33406568 | 0.46547425 | NA | NA |
| ENSG000001 | 35.4164518 | -0.235850659 | 0.24417362 | -0.9659138 | 0.33408734 | 0.46548026 | TRAM1L1 | 133022 |
| ENSG000001 | 41.4464444 | 0.218274355 | 0.22598492 | 0.96588019 | 0.33410414 | 0.46548026 | SHISA4 | 149345 |
| ENSG000002 | 2.56470138 | -0.36077459 | 0.37352696 | -0.9658596 | 0.33411445 | 0.46548026 | NA | NA |
| ENSG000002 | 4.25897057 | -0.365564126 | 0.37848693 | -0.9658567 | 0.3341159 | 0.46548026 | NA | NA |
| ENSG00000C | 4.4143252 | -0.363223591 | 0.37614524 | -0.9656472 | 0.33422075 | 0.46560749 | ST6GALNAC2 | 10610 |
| ENSG000001 | 182.131031 | 0.137376848 | 0.14228031 | 0.96553659 | 0.33427612 | 0.46560749 | COL9A1 | 1297 |
| ENSG000001 | 127.270324 | -0.150577747 | 0.15593922 | -0.9656182 | 0.33423526 | 0.46560749 | ASTN2 | 23245 |
| ENSG000002 | 5.80318368 | -0.358595773 | 0.37138256 | -0.9655698 | 0.33425951 | 0.46560749 | NA | NA |
| ENSG000002 | 2.84476139 | -0.363936585 | 0.37692275 | -0.9655469 | 0.33427096 | 0.46560749 | NA | NA |
| ENSG000002 | 8.80598957 | -0.336466438 | 0.34845456 | -0.9655963 | 0.33424622 | 0.46560749 | NA | NA |
| ENSG000002 | 2.38958029 | -0.352647446 | 0.36524665 | -0.9655049 | 0.33429196 | 0.46561357 | NA | NA |

| | | | | | | | | |
|---|---|---|---|---|---|---|---|---|
| ENSG000001 | 4.95620396 | -0.359642217 | 0.37264734 | -0.9651007 | 0.33449436 | 0.46587947 | ACPT | 93650 |
| ENSG000002 | 3.4910269 | -0.36431329 | 0.3775103 | -0.965042 | 0.33452379 | 0.46590446 | NA | NA |
| ENSG000002 | 4.45639212 | -0.363301676 | 0.37649125 | -0.9649671 | 0.33456128 | 0.46592597 | NA | NA |
| ENSG000002 | 2.61915185 | -0.359991987 | 0.37306213 | -0.9649652 | 0.33456222 | 0.46592597 | NA | NA |
| ENSG000002 | 3.96410293 | -0.365217943 | 0.37848803 | -0.9649392 | 0.33457525 | 0.46592812 | NA | NA |
| ENSG000002 | 2.37232507 | -0.354645219 | 0.36755239 | -0.9648834 | 0.33460319 | 0.46595103 | NA | NA |
| ENSG000002 | 2.89678375 | -0.362654659 | 0.37590995 | -0.9647381 | 0.334676 | 0.46603641 | NA | NA |
| ENSG000002 | 2.60209685 | -0.359082242 | 0.37228198 | -0.9645437 | 0.33477341 | 0.46615605 | PHBP5 | 106478939 |
| ENSG000002 | 4.93943711 | -0.359072305 | 0.37228653 | -0.9645052 | 0.33479269 | 0.46616689 | RPL23AP91 | 106480337 |
| ENSG000001 | 80.2220637 | 0.198287684 | 0.20559679 | 0.9644493 | 0.33482071 | 0.46617612 | MAFK | 7975 |
| ENSG000002 | 2.39682811 | -0.35228069 | 0.36526736 | -0.9644461 | 0.33482231 | 0.46617612 | NA | NA |
| ENSG000002 | 9.91110897 | 0.330707202 | 0.34291314 | 0.96440517 | 0.33484283 | 0.46618869 | MIR3143 | 100422934 |
| ENSG000002 | 5.27081544 | -0.362715524 | 0.3761176 | -0.9643673 | 0.3348618 | 0.46619909 | RNU6-1285P | 106481948 |
| ENSG000001 | 5.77052291 | -0.35826027 | 0.37152591 | -0.9642942 | 0.33489846 | 0.46621812 | EPO | 2056 |
| ENSG000002 | 3.68437573 | -0.365045152 | 0.37855331 | -0.9643164 | 0.33488734 | 0.46621812 | NA | NA |
| ENSG000001 | 10.896723 | -0.331274358 | 0.34357914 | -0.9641865 | 0.33495244 | 0.46627726 | DCST2 | 127579 |
| ENSG000001 | 2.58234979 | -0.360630065 | 0.37410982 | -0.9639685 | 0.33506174 | 0.4664134 | S100A12 | 6283 |
| ENSG000001 | 142.789453 | 0.150311881 | 0.15594723 | 0.96386377 | 0.33511423 | 0.46643865 | SLC44A5 | 204962 |
| ENSG000002 | 6.73401816 | -0.347072695 | 0.36009349 | -0.9638405 | 0.33512589 | 0.46643865 | NA | NA |
| ENSG000002 | 2.86300211 | -0.362917277 | 0.37652367 | -0.9638631 | 0.33511456 | 0.46643865 | LINC01525 | 104355292 |
| ENSG000002 | 5.07990366 | 0.362942354 | 0.37653193 | 0.96390857 | 0.33509177 | 0.46643865 | NA | NA |
| ENSG000001 | 14.3555589 | -0.306323876 | 0.31788736 | -0.9636239 | 0.3352345 | 0.46655778 | PTH1R | 5745 |
| ENSG000002 | 4.06394653 | -0.364820847 | 0.37858645 | -0.9636395 | 0.33522671 | 0.46655778 | MIR3193 | 100422904 |
| ENSG000002 | 5.85110349 | -0.357511828 | 0.37105035 | -0.963513 | 0.33529016 | 0.46661924 | NA | NA |
| ENSG000002 | 2.64009292 | -0.358711862 | 0.37232839 | -0.9634287 | 0.33533242 | 0.46666204 | NA | NA |
| ENSG000001 | 48.368037 | 0.210876164 | 0.21889631 | 0.96336097 | 0.3353664 | 0.46669331 | RGS19 | 10287 |
| ENSG000002 | 7.26078413 | -0.346787928 | 0.35998888 | -0.9633296 | 0.33538216 | 0.46669922 | NA | NA |
| ENSG000002 | 7.05098939 | -0.346423497 | 0.35962829 | -0.9632821 | 0.33540597 | 0.46671634 | SERTAD4-AS1 | 574036 |
| ENSG000002 | 14.0787193 | -0.300826409 | 0.31231799 | -0.9632055 | 0.3354444 | 0.46673778 | NA | NA |

| | | | | | | | | |
|---|---|---|---|---|---|---|---|---|
| ENSG000002 | 5.85002639 | -0.357686641 | 0.37134962 | -0.9632072 | 0.33544353 | 0.46673778 | NA | NA |
| ENSG000002 | 3.65060674 | -0.364524063 | 0.37847861 | -0.9631299 | 0.33548234 | 0.46677455 | NA | NA |
| ENSG000001 | 3.40828297 | -0.363979474 | 0.3779325 | -0.9630806 | 0.33550705 | 0.46679292 | CETN1 | 1068 |
| ENSG000001 | 4.24867152 | -0.363630806 | 0.37759905 | -0.9630077 | 0.33554364 | 0.46682781 | NA | NA |
| ENSG000002 | 4.98649781 | 0.359549199 | 0.3735034 | 0.9626397 | 0.33572837 | 0.46706879 | LOC375196 | 375196 |
| ENSG000002 | 11.696436 | -0.324438212 | 0.33704599 | -0.9625933 | 0.33575166 | 0.46708516 | NA | NA |
| ENSG000002 | 2.3385431 | -0.352060741 | 0.36577249 | -0.9625129 | 0.33579202 | 0.46712529 | RN7SKP154 | 106480881 |
| ENSG000002 | 2.84803031 | -0.362642576 | 0.37679386 | -0.9624429 | 0.33582717 | 0.46714283 | NA | NA |
| ENSG000002 | 4.46159192 | -0.362263184 | 0.37640005 | -0.9624419 | 0.33582767 | 0.46714283 | NA | NA |
| ENSG000002 | 2.858599 | -0.36236432 | 0.37651971 | -0.9624047 | 0.33584638 | 0.46715283 | NA | NA |
| ENSG000002 | 7.7386252 | -0.33891489 | 0.35216919 | -0.9623638 | 0.33586688 | 0.46716533 | NA | NA |
| ENSG000001 | 8.26977809 | -0.345113837 | 0.35862016 | -0.9623381 | 0.33587981 | 0.46716728 | FAM92B | 339145 |
| ENSG000002 | 3.96920492 | -0.364164375 | 0.37842546 | -0.9623147 | 0.33589156 | 0.4671676 | NA | NA |
| ENSG000002 | 8.39643859 | 0.338668785 | 0.35197786 | 0.96218775 | 0.3359553 | 0.46722778 | NA | NA |
| ENSG000002 | 19.0457863 | 0.275245479 | 0.28607047 | 0.96215969 | 0.33596939 | 0.46722778 | RPL13AP20 | 387841 |
| ENSG000002 | 6.03783016 | -0.354674709 | 0.36862266 | -0.962162 | 0.33596824 | 0.46722778 | NA | NA |
| ENSG000002 | 2.3855211 | -0.34942197 | 0.36321793 | -0.9620174 | 0.33604087 | 0.46731115 | NA | NA |
| ENSG000001 | 3.74491225 | -0.36411431 | 0.37852717 | -0.9619238 | 0.33608786 | 0.4673463 | TAS2R42 | 353164 |
| ENSG000002 | 3.44769364 | -0.363472171 | 0.37786065 | -0.9619212 | 0.33608919 | 0.4673463 | NA | NA |
| ENSG000002 | 7.31155682 | -0.343461379 | 0.35708338 | -0.9618521 | 0.33612393 | 0.46737857 | NA | NA |
| ENSG000002 | 4.98674784 | -0.360588212 | 0.3749503 | -0.961696 | 0.33620232 | 0.46747154 | NA | NA |
| ENSG000001 | 118.426981 | -0.148095829 | 0.15401166 | -0.9615884 | 0.3362564 | 0.46752951 | TMC5 | 79838 |
| ENSG000001 | 18.9963628 | 0.27673176 | 0.28779775 | 0.96154942 | 0.33627599 | 0.46752951 | PLAC1 | 10761 |
| ENSG000002 | 624.673498 | 0.117070779 | 0.12175288 | 0.96154424 | 0.33627859 | 0.46752951 | SFT2D2 | 375035 |
| ENSG000002 | 4.19057497 | -0.362683522 | 0.37722157 | -0.9614602 | 0.33632083 | 0.46757221 | NA | NA |
| ENSG000001 | 4.71512987 | -0.361697556 | 0.37626099 | -0.9612943 | 0.33640419 | 0.46767206 | GHRH | 2691 |
| ENSG000002 | 2.3862153 | -0.354007403 | 0.36831593 | -0.9611515 | 0.336476 | 0.46775586 | NA | NA |
| ENSG000002 | 2.58542228 | -0.359243052 | 0.37377269 | -0.9611271 | 0.33648827 | 0.46775688 | NA | NA |
| ENSG000002 | 18.6112865 | -0.279132027 | 0.29043202 | -0.9610925 | 0.33650567 | 0.46776503 | NA | NA |

| | | | | | | | | |
|---|---|---|---|---|---|---|---|---|
| ENSG000002 | 8.78309725 | -0.334793671 | 0.34836238 | -0.96105 | 0.33652702 | 0.46777868 | NA | NA |
| ENSG00000C | 29.0973894 | -0.255430127 | 0.26580907 | -0.9609534 | 0.33657561 | 0.46779811 | TP73 | 7161 |
| ENSG000002 | 6.59810643 | -0.350993516 | 0.36524923 | -0.9609699 | 0.3365673 | 0.46779811 | NA | NA |
| ENSG000002 | 21.4998403 | 0.262970228 | 0.27364299 | 0.9609975 | 0.33655342 | 0.46779811 | NA | NA |
| ENSG000002 | 5.49474209 | 0.359548014 | 0.37426255 | 0.96068391 | 0.33671112 | 0.46797041 | SNORA79 | 677845 |
| ENSG000002 | 2.70768845 | -0.355741051 | 0.37037877 | -0.9604791 | 0.33681417 | 0.46806826 | NA | NA |
| ENSG000002 | 2.92629225 | 0.361663485 | 0.37654645 | 0.9604751 | 0.33681616 | 0.46806826 | NA | NA |
| ENSG000002 | 13.032096 | -0.319080501 | 0.33220573 | -0.9604907 | 0.33680833 | 0.46806826 | LINC00554 | 100861542 |
| ENSG000002 | 3.2540301 | 0.362025769 | 0.37695827 | 0.96038687 | 0.33686055 | 0.4681103 | NA | NA |
| ENSG000002 | 6.74655662 | -0.347869738 | 0.36222505 | -0.9603691 | 0.33686949 | 0.4681103 | LOC101928844 | 101928844 |
| ENSG000002 | 2.64535744 | -0.355821445 | 0.37053929 | -0.9602799 | 0.33691435 | 0.46815659 | RNU6-988P | 106480003 |
| ENSG000001 | 4.73388817 | -0.359774487 | 0.37468065 | -0.9602163 | 0.33694634 | 0.468185 | AMELX | 265 |
| ENSG000002 | 6.8839249 | -0.351109219 | 0.36566884 | -0.9601836 | 0.33696282 | 0.46819186 | NA | NA |
| ENSG000002 | 9.6243732 | 0.33239062 | 0.34621849 | 0.96006027 | 0.33702488 | 0.46826205 | NA | NA |
| ENSG000002 | 113.81142 | -0.158415916 | 0.16508123 | -0.959624 | 0.33724447 | 0.46855108 | TEX41 | 401014 |
| ENSG00000C | 63.9246212 | 0.19106013 | 0.19913199 | 0.95946477 | 0.33732466 | 0.46864644 | LY75 | 4065 |
| ENSG000001 | 189.691362 | 0.143813236 | 0.14989587 | 0.95942091 | 0.33734675 | 0.46866107 | C1orf74 | 148304 |
| ENSG000002 | 6.88718913 | -0.348632735 | 0.36340661 | -0.9593461 | 0.33738439 | 0.46869731 | NA | NA |
| ENSG000002 | 10.2248208 | -0.332386858 | 0.34651529 | -0.9592271 | 0.33744436 | 0.46876455 | NA | NA |
| ENSG000001 | 125.363943 | 0.146232439 | 0.15245246 | 0.95920027 | 0.33745786 | 0.46876726 | SORCS1 | 114815 |
| ENSG000002 | 4.67601976 | -0.35960491 | 0.37493691 | -0.9591078 | 0.33750444 | 0.4688159 | NA | NA |
| ENSG000002 | 3.67706789 | -0.36272971 | 0.37822936 | -0.9590205 | 0.33754842 | 0.46884731 | NA | NA |
| ENSG000002 | 2.90611213 | 0.361234088 | 0.37667714 | 0.95900188 | 0.3375578 | 0.46884731 | NA | NA |
| ENSG000002 | 4.77552944 | -0.361394337 | 0.37684732 | -0.9589941 | 0.33756174 | 0.46884731 | NA | NA |
| ENSG000002 | 9.0399678 | -0.328816187 | 0.34289361 | -0.9589452 | 0.33758635 | 0.46886543 | NA | NA |
| ENSG000001 | 39.6617663 | 0.21898379 | 0.22838059 | 0.95885465 | 0.33763197 | 0.46889668 | KLF9 | 687 |
| ENSG000002 | 2.33355932 | -0.353318671 | 0.3684741 | -0.9588697 | 0.33762437 | 0.46889668 | NA | NA |
| ENSG000002 | 4.46352247 | -0.361642538 | 0.37719768 | -0.9587613 | 0.337679 | 0.46894593 | NA | NA |
| ENSG00000C | 15.5802358 | -0.29025243 | 0.30275166 | -0.9587146 | 0.33770255 | 0.46896257 | IL11 | 3589 |

| | | | | | | | | |
|---|---|---|---|---|---|---|---|---|
| ENSG000001 | 57.2229579 | -0.198904251 | 0.20751563 | -0.9585025 | 0.33780942 | 0.46907885 | IRF1 | 3659 |
| ENSG000002 | 5.00276593 | -0.357926901 | 0.37341909 | -0.9585126 | 0.33780435 | 0.46907885 | NA | NA |
| ENSG000002 | 2.84863113 | -0.361274339 | 0.37692606 | -0.9584754 | 0.33782311 | 0.4690818 | NA | NA |
| ENSG000002 | 4.80676728 | -0.360730184 | 0.37645444 | -0.9582307 | 0.33794646 | 0.46923701 | NA | NA |
| ENSG000002 | 2.6793027 | -0.355507477 | 0.37108042 | -0.9580335 | 0.33804586 | 0.46935701 | NA | NA |
| ENSG000002 | 18.4165611 | 0.271685924 | 0.28359305 | 0.95801334 | 0.33805603 | 0.46935701 | NA | NA |
| ENSG000002 | 2.32325863 | -0.353262846 | 0.36876369 | -0.9579654 | 0.33808022 | 0.46937451 | RN7SKP125 | 106479150 |
| ENSG00000C | 4.74902415 | -0.358993765 | 0.37477279 | -0.9578971 | 0.33811465 | 0.46938489 | HSPB6 | 126393 |
| ENSG000002 | 6.49282728 | -0.34947883 | 0.36483269 | -0.9579153 | 0.33810546 | 0.46938489 | MT1JP | 4498 |
| ENSG000002 | 3.71161236 | -0.362188004 | 0.3781135 | -0.9578817 | 0.33812242 | 0.46938489 | NA | NA |
| ENSG000002 | 26.2393068 | 0.24584301 | 0.25665895 | 0.9578587 | 0.33813402 | 0.46938493 | NA | NA |
| ENSG000001 | 3.96929665 | -0.362504522 | 0.37847596 | -0.9578007 | 0.33816329 | 0.46940949 | QRFP | 347148 |
| ENSG000002 | 12.7486243 | -0.305011555 | 0.31846392 | -0.9577586 | 0.3381845 | 0.46942287 | LOC645434 | 645434 |
| ENSG000002 | 15.7034784 | 0.296950028 | 0.31006049 | 0.95771643 | 0.33820577 | 0.46943632 | NA | NA |
| ENSG000002 | 4.18149966 | -0.361647448 | 0.37768951 | -0.9575258 | 0.33830193 | 0.46953765 | NA | NA |
| ENSG000002 | 6.36496102 | -0.352402178 | 0.368034 | -0.9575262 | 0.33830175 | 0.46953765 | NA | NA |
| ENSG000002 | 54.4153622 | 0.195648911 | 0.20433753 | 0.95747907 | 0.33832551 | 0.4695543 | C1orf210 | 149466 |
| ENSG000001 | 53.1956547 | -0.198021979 | 0.20683115 | -0.9574089 | 0.33836093 | 0.46957132 | RAMP2-AS1 | 100190938 |
| ENSG000002 | 2.87048568 | -0.359785468 | 0.37578715 | -0.9574182 | 0.33835621 | 0.46957132 | NA | NA |
| ENSG000002 | 4.70281451 | -0.358961265 | 0.37495403 | -0.9573474 | 0.33839194 | 0.4695963 | LOC101928259 | 101928259 |
| ENSG000002 | 6.13311052 | 0.351516523 | 0.36718532 | 0.95732729 | 0.33840209 | 0.4695963 | NA | NA |
| ENSG000002 | 2.30437994 | 0.353330241 | 0.36911705 | 0.95723089 | 0.33845073 | 0.46963838 | NA | NA |
| ENSG000002 | 4.96428093 | -0.356936307 | 0.37288797 | -0.9572213 | 0.33845557 | 0.46963838 | NA | NA |
| ENSG000002 | 8.35583512 | -0.337508351 | 0.35264577 | -0.9570747 | 0.33852954 | 0.46972494 | NA | NA |
| ENSG000001 | 5.58082135 | -0.358934948 | 0.37511974 | -0.9568543 | 0.33864079 | 0.46986322 | VGLL1 | 51442 |
| ENSG000002 | 63.5197092 | 0.181606573 | 0.18980631 | 0.95679944 | 0.33866849 | 0.46988559 | NEURL1B | 54492 |
| ENSG000001 | 10.9088253 | -0.323826065 | 0.33848639 | -0.9566886 | 0.33872447 | 0.46994717 | SAXO1 | 158297 |
| ENSG000002 | 2.68363447 | -0.355608529 | 0.37173768 | -0.9566115 | 0.3387634 | 0.46998511 | NA | NA |
| ENSG000002 | 8.28324363 | -0.334984932 | 0.35023368 | -0.9564612 | 0.33883927 | 0.47005477 | C9orf92 | 100129385 |

| | | | | | | | | |
|---|---|---|---|---|---|---|---|---|
| ENSG000002 | 4.73598214 | -0.359311966 | 0.37567645 | -0.95644 | 0.33885001 | 0.47005477 | NA | NA |
| ENSG000002 | 2.90535685 | -0.358919212 | 0.3752721 | -0.9564239 | 0.3388581 | 0.47005477 | NA | NA |
| ENSG000002 | 2.57579932 | -0.357067004 | 0.37333695 | -0.9564202 | 0.33885998 | 0.47005477 | NA | NA |
| ENSG000002 | 16.1651273 | -0.30609243 | 0.32007886 | -0.9563032 | 0.33891908 | 0.47012067 | NA | NA |
| ENSG000002 | 2.32164323 | 0.352413295 | 0.3685325 | 0.9562611 | 0.33894034 | 0.47013407 | NA | NA |
| ENSG000002 | 2.34133593 | -0.351525154 | 0.36762803 | -0.9561979 | 0.33897226 | 0.47016227 | NA | NA |
| ENSG000002 | 2.87043415 | -0.360000696 | 0.37652674 | -0.9561092 | 0.33901705 | 0.4702083 | NA | NA |
| ENSG000002 | 6.53464591 | -0.351356552 | 0.36753776 | -0.955974 | 0.33908535 | 0.47028696 | NA | NA |
| ENSG000002 | 2.56695289 | -0.356817683 | 0.37332416 | -0.9557851 | 0.3391808 | 0.47039363 | NA | NA |
| ENSG000002 | 6.60697268 | -0.349179902 | 0.36533658 | -0.9557759 | 0.33918546 | 0.47039363 | NA | NA |
| ENSG000002 | 9.77273669 | -0.320377705 | 0.33521656 | -0.9557335 | 0.33920688 | 0.47040725 | LOC100506122 | 100506122 |
| ENSG000002 | 4.46654378 | -0.360221127 | 0.3769523 | -0.9556146 | 0.33926697 | 0.47047449 | LOC101929011 | 101929011 |
| ENSG000001 | 4.97805906 | -0.357057363 | 0.37373842 | -0.955367 | 0.33939212 | 0.47059705 | ANKK1 | 255239 |
| ENSG000001 | 2.88066979 | -0.359389767 | 0.37617651 | -0.9553753 | 0.33938792 | 0.47059705 | RNU6-942P | 106481477 |
| ENSG000002 | 12.5729137 | -0.322484102 | 0.33755671 | -0.9553479 | 0.33940178 | 0.47059705 | NA | NA |
| ENSG000002 | 3.53195922 | 0.36156413 | 0.37844717 | 0.95538865 | 0.33938119 | 0.47059705 | NA | NA |
| ENSG000001 | 3.66702922 | -0.36163795 | 0.37858165 | -0.9552443 | 0.33945419 | 0.47063124 | NA | NA |
| ENSG000002 | 3.2142549 | 0.35805055 | 0.37482106 | 0.9552573 | 0.3394476 | 0.47063124 | RNU2-33P | 106480211 |
| ENSG000002 | 5.52449371 | -0.356725111 | 0.37344409 | -0.9552303 | 0.33946125 | 0.47063124 | LOC105375679 | 105375679 |
| ENSG000002 | 2.30367988 | -0.351999861 | 0.36852062 | -0.9551701 | 0.33949171 | 0.47064128 | LRRC3-AS1 | 100861510 |
| ENSG000002 | 6.66619068 | -0.352029992 | 0.36854431 | -0.9551904 | 0.33948141 | 0.47064128 | NA | NA |
| ENSG000002 | 2.38879853 | -0.349388816 | 0.36579981 | -0.9551367 | 0.33950858 | 0.47064859 | NA | NA |
| ENSG00000C | 148.505764 | 0.157631422 | 0.16508412 | 0.95485513 | 0.33965096 | 0.47081377 | ITGAE | 3682 |
| ENSG000001 | 21.3760008 | 0.273955798 | 0.28690205 | 0.95487571 | 0.33964055 | 0.47081377 | TRIM47 | 91107 |
| ENSG000001 | 34.3594756 | -0.226674638 | 0.23742029 | -0.95474 | 0.33970921 | 0.47086232 | TMPRSS13 | 84000 |
| ENSG000002 | 4.47383167 | -0.359616337 | 0.37666367 | -0.9547412 | 0.33970858 | 0.47086232 | NA | NA |
| ENSG000001 | 29.7878478 | 0.246058354 | 0.25773779 | 0.95468482 | 0.33973711 | 0.4708849 | ZBTB42 | 100128927 |
| ENSG000002 | 2.83995389 | -0.35976691 | 0.37692207 | -0.9544862 | 0.3398376 | 0.47099199 | NA | NA |
| ENSG000002 | 2.83995389 | -0.35976691 | 0.37692207 | -0.9544862 | 0.3398376 | 0.47099199 | NA | NA |

| | | | | | | | | |
|---|---|---|---|---|---|---|---|---|
| ENSG000001 | 109.53535 | 0.183252022 | 0.19200248 | 0.95442527 | 0.33986842 | 0.47100251 | TEF | 7008 |
| ENSG000002 | 2.35445649 | -0.350450919 | 0.36718274 | -0.9544319 | 0.33986507 | 0.47100251 | NA | NA |
| ENSG000002 | 3.79184007 | 0.361139627 | 0.37843377 | 0.95430074 | 0.33993143 | 0.47107373 | NA | NA |
| ENSG000001 | 25.0154484 | -0.252408337 | 0.2645121 | -0.9542412 | 0.33996157 | 0.47109623 | PON3 | 5446 |
| ENSG000002 | 2.34744956 | -0.350818699 | 0.36764865 | -0.9542227 | 0.3399709 | 0.47109623 | OR51B2 | 79345 |
| ENSG000002 | 7.73741707 | -0.349462239 | 0.36625916 | -0.9541393 | 0.34001316 | 0.47113868 | NA | NA |
| ENSG000002 | 2.83954895 | -0.35924727 | 0.37660576 | -0.9539081 | 0.34013019 | 0.47128474 | NA | NA |
| ENSG000002 | 11.8589434 | -0.311614413 | 0.3266932 | -0.9538442 | 0.34016251 | 0.47130857 | NA | NA |
| ENSG000002 | 4.98200249 | -0.358529895 | 0.3758852 | -0.9538282 | 0.34017063 | 0.47130857 | NA | NA |
| ENSG000001 | 183.407147 | 0.140641215 | 0.14751416 | 0.95340826 | 0.34038326 | 0.47153872 | MAML2 | 84441 |
| ENSG000002 | 7.30069168 | -0.34374255 | 0.3605377 | -0.9534164 | 0.34037914 | 0.47153872 | NA | NA |
| ENSG000002 | 14.2553556 | -0.294032362 | 0.30839676 | -0.9534223 | 0.34037612 | 0.47153872 | CT62 | 196993 |
| ENSG000002 | 2.88203953 | -0.35776359 | 0.375237 | -0.9534337 | 0.34037038 | 0.47153872 | NA | NA |
| ENSG000002 | 2.59908329 | -0.354610059 | 0.37201414 | -0.9532166 | 0.34048033 | 0.47165708 | LOC105372925 | 105372925 |
| ENSG000002 | 4.94909288 | -0.3556921 | 0.37320572 | -0.9530725 | 0.34055334 | 0.47170989 | NA | NA |
| ENSG000002 | 4.41358799 | -0.35901458 | 0.37668245 | -0.9530961 | 0.34054137 | 0.47170989 | NA | NA |
| ENSG000002 | 6.72033535 | -0.347011272 | 0.36409195 | -0.9530869 | 0.34054604 | 0.47170989 | NA | NA |
| ENSG000002 | 3.9974598 | -0.360780043 | 0.37859193 | -0.9529523 | 0.34061425 | 0.47177813 | NA | NA |
| ENSG000002 | 2.85451399 | 0.357876309 | 0.37556864 | 0.95289187 | 0.34064486 | 0.47180424 | NA | NA |
| ENSG000002 | 3.44819032 | -0.359637329 | 0.37742572 | -0.9528692 | 0.34065637 | 0.47180424 | NA | NA |
| ENSG000001 | 33.2532686 | 0.227272193 | 0.23852393 | 0.95282764 | 0.34067741 | 0.47181726 | IQSEC3 | 440073 |
| ENSG000002 | 3.13623887 | -0.360082179 | 0.37793167 | -0.9527706 | 0.34070632 | 0.4718412 | NA | NA |
| ENSG000002 | 3.48588542 | 0.36055488 | 0.37845566 | 0.95270044 | 0.34074187 | 0.47187431 | NA | NA |
| ENSG000002 | 6.02379701 | -0.350734826 | 0.36819942 | -0.9525676 | 0.34080922 | 0.47191923 | NA | NA |
| ENSG000002 | 4.97709351 | -0.356106344 | 0.37383 | -0.952589 | 0.34079835 | 0.47191923 | NA | NA |
| ENSG000002 | 2.84397156 | -0.357742622 | 0.37555069 | -0.9525814 | 0.34080218 | 0.47191923 | NA | NA |
| ENSG000002 | 2.93010838 | -0.357062889 | 0.37486394 | -0.9525133 | 0.34083673 | 0.4719412 | NA | NA |
| ENSG000002 | 4.14188569 | -0.359615129 | 0.37756337 | -0.952463 | 0.34086224 | 0.47196042 | NA | NA |
| ENSG000002 | 4.72264643 | -0.357393034 | 0.37532482 | -0.9522233 | 0.34098376 | 0.47211255 | RNU6-826P | 106479931 |

| | | | | | | | | | |
|---|---|---|---|---|---|---|---|---|---|
| ENSG000001 | 32.9294029 | -0.235689654 | 0.24757146 | -0.9520066 | 0.34109365 | 0.47224857 | JAG2 | | 3714 |
| ENSG000001 | 6.11685319 | -0.353675562 | 0.3715404 | -0.9519168 | 0.34113916 | 0.47226321 | SFTPC | | 6440 |
| ENSG000002 | 4.27661898 | -0.360332729 | 0.37852966 | -0.9519273 | 0.34113383 | 0.47226321 | NA | NA | |
| ENSG000002 | 2.90040554 | -0.35759894 | 0.37565243 | -0.951941 | 0.34112692 | 0.47226321 | NA | NA | |
| ENSG000001 | 13.4240394 | 0.296577544 | 0.3115942 | 0.951807 | 0.34119487 | 0.47232421 | HTR1B | | 3351 |
| ENSG000002 | 11.53585 | 0.329886381 | 0.34668143 | 0.95155481 | 0.34132281 | 0.4724852 | NA | NA | |
| ENSG000002 | 4.47577499 | -0.358737572 | 0.37703519 | -0.9514697 | 0.34136598 | 0.47251269 | NA | NA | |
| ENSG000002 | 3.10322635 | -0.359465623 | 0.37779322 | -0.9514878 | 0.34135683 | 0.47251269 | NA | NA | |
| ENSG000002 | 25.054814 | 0.258846818 | 0.27205643 | 0.95144534 | 0.34137835 | 0.47251369 | NA | NA | |
| ENSG000001 | 68.8815658 | -0.180163989 | 0.18938837 | -0.9512938 | 0.34145524 | 0.47260398 | MITF | | 4286 |
| ENSG000001 | 16.8778637 | -0.29039051 | 0.30532112 | -0.9510987 | 0.3415543 | 0.47268277 | GLRA2 | | 2742 |
| ENSG000002 | 2.59228542 | -0.354608675 | 0.37283701 | -0.9511091 | 0.34154899 | 0.47268277 | NA | NA | |
| ENSG000002 | 7.88912773 | -0.339907091 | 0.35736355 | -0.9511521 | 0.34152717 | 0.47268277 | NA | NA | |
| ENSG000002 | 10.3023866 | 0.329484343 | 0.34642822 | 0.9510898 | 0.34155879 | 0.47268277 | LOC399900 | | 399900 |
| ENSG000002 | 2.57664809 | -0.355207785 | 0.37349983 | -0.9510253 | 0.34159154 | 0.47271196 | MIR626 | | 693211 |
| ENSG000002 | 2.85065898 | -0.357707272 | 0.37621809 | -0.9507976 | 0.34170711 | 0.47285576 | NA | NA | |
| ENSG000002 | 6.5175911 | -0.34826648 | 0.36630725 | -0.9507496 | 0.34173149 | 0.47287336 | NA | NA | |
| ENSG000001 | 150.921604 | 0.149287563 | 0.15705298 | 0.95055545 | 0.3418301 | 0.47299366 | DFNA5 | | 1687 |
| ENSG000001 | 2.85711249 | -0.357839425 | 0.37647763 | -0.9504932 | 0.34186171 | 0.47302127 | IL10 | | 3586 |
| ENSG000001 | 107.759124 | 0.150849152 | 0.15871169 | 0.95046023 | 0.34187845 | 0.47302829 | FAM63A | | 55793 |
| ENSG000001 | 6.98563208 | -0.344224592 | 0.36219494 | -0.9503849 | 0.34191673 | 0.47306511 | CELA3A | | 10136 |
| ENSG000002 | 25.6123671 | -0.247636483 | 0.26062159 | -0.9501764 | 0.34202262 | 0.47319547 | NA | NA | |
| ENSG000002 | 6.46876578 | -0.347909593 | 0.36618977 | -0.9500801 | 0.34207158 | 0.47324706 | LOC101928521 | | 101928521 |
| ENSG000002 | 2.59807862 | -0.354596662 | 0.37325572 | -0.95001 | 0.34210717 | 0.47328016 | NA | NA | |
| ENSG000001 | 5.1784278 | -0.352748605 | 0.37133543 | -0.949946 | 0.34213969 | 0.473309 | NA | NA | |
| ENSG000001 | 65.5332043 | -0.180227256 | 0.18973239 | -0.9499024 | 0.34216185 | 0.4733235 | LEF1 | | 51176 |
| ENSG000001 | 7.0514464 | -0.342489762 | 0.36058383 | -0.9498201 | 0.34220368 | 0.47336523 | IFNK | | 56832 |
| ENSG000002 | 2.62392744 | -0.353964897 | 0.37267611 | -0.9497923 | 0.3422178 | 0.47336861 | NA | NA | |
| ENSG000002 | 4.23319443 | -0.359499209 | 0.37855388 | -0.9496646 | 0.34228272 | 0.47344226 | NA | NA | |

| | | | | | | | | |
|---|---|---|---|---|---|---|---|---|
| ENSG000002 | 25.2309872 | -0.25596433 | 0.26954878 | -0.949603 | 0.34231402 | 0.47346939 | NA | NA |
| ENSG000002 | 3.95005037 | -0.359306061 | 0.3784394 | -0.9494415 | 0.34239613 | 0.47356682 | NA | NA |
| ENSG000001 | 40.0801702 | 0.2232376 | 0.23513598 | 0.94939786 | 0.3424183 | 0.47358132 | CHSY3 | 337876 |
| ENSG000001 | 3.1357824 | -0.358870953 | 0.37806143 | -0.9492398 | 0.34249867 | 0.47367633 | NA | NA |
| ENSG000002 | 3.79647151 | -0.359014122 | 0.37823017 | -0.9491948 | 0.34252154 | 0.47369179 | NA | NA |
| ENSG00000C | 14.4257194 | -0.309831018 | 0.32643334 | -0.9491402 | 0.34254929 | 0.47371402 | EBF4 | 57593 |
| ENSG000002 | 3.11785379 | -0.358703064 | 0.37798813 | -0.9489797 | 0.34263093 | 0.473801 | NA | NA |
| ENSG000002 | 2.86002136 | -0.35679239 | 0.37597833 | -0.9489706 | 0.34263556 | 0.473801 | NA | NA |
| ENSG000002 | 5.28626029 | -0.356161776 | 0.37535701 | -0.9488614 | 0.34269111 | 0.47386167 | LINC01327 | 104310350 |
| ENSG000002 | 5.87998534 | -0.353391514 | 0.37245789 | -0.9488093 | 0.3427176 | 0.47388214 | FABP9 | 646480 |
| ENSG000002 | 6.29586872 | -0.350169739 | 0.36909483 | -0.9487257 | 0.34276015 | 0.47391002 | NA | NA |
| ENSG000002 | 12.0894684 | -0.309916585 | 0.32666684 | -0.9487237 | 0.34276114 | 0.47391002 | NA | NA |
| ENSG000002 | 3.73460882 | -0.358630447 | 0.37806812 | -0.9485869 | 0.34283078 | 0.47399014 | NA | NA |
| ENSG000002 | 5.79776545 | 0.34952102 | 0.36848074 | 0.94854625 | 0.34285144 | 0.47400255 | CAHM | 100526820 |
| ENSG000001 | 4.68755434 | -0.355939495 | 0.37536624 | -0.9482459 | 0.34300429 | 0.47418153 | NA | NA |
| ENSG000002 | 13.0361291 | 0.305684126 | 0.32236436 | 0.94825658 | 0.34299885 | 0.47418153 | NA | NA |
| ENSG000001 | 63.7392541 | 0.181596328 | 0.19151442 | 0.94821231 | 0.34302138 | 0.474189 | B3GAT2 | 135152 |
| ENSG000001 | 31.0377945 | 0.231299223 | 0.24395927 | 0.94810589 | 0.34307555 | 0.47419921 | RNF112 | 7732 |
| ENSG000002 | 4.69178826 | -0.356642016 | 0.37614058 | -0.9481615 | 0.34304724 | 0.47419921 | NA | NA |
| ENSG000002 | 10.8871336 | -0.316111439 | 0.33340775 | -0.9481226 | 0.34306702 | 0.47419921 | NA | NA |
| ENSG000002 | 2.62598253 | -0.352413794 | 0.37170046 | -0.9481123 | 0.34307227 | 0.47419921 | NA | NA |
| ENSG000002 | 2.63574243 | -0.353329298 | 0.37269142 | -0.9480478 | 0.3431051 | 0.4742239 | NA | NA |
| ENSG00000C | 159.260849 | 0.273122824 | 0.28812401 | 0.94793498 | 0.34316256 | 0.47425482 | SEMA3F | 6405 |
| ENSG000002 | 2.24634322 | 0.345281312 | 0.36423972 | 0.94795074 | 0.34315453 | 0.47425482 | NA | NA |
| ENSG000002 | 4.9767486 | -0.353884197 | 0.37331621 | -0.9479476 | 0.34315615 | 0.47425482 | NA | NA |
| ENSG000002 | 36.439838 | -0.225753322 | 0.23817603 | -0.9478423 | 0.34320973 | 0.47430385 | NA | NA |
| ENSG000002 | 3.74266119 | -0.358213901 | 0.37794585 | -0.9477916 | 0.34323556 | 0.47432338 | NA | NA |
| ENSG000002 | 4.7239203 | -0.355150731 | 0.37472883 | -0.9477539 | 0.34325473 | 0.47433371 | NA | NA |
| ENSG000002 | 6.17292818 | -0.353493691 | 0.37299572 | -0.9477151 | 0.34327449 | 0.47434485 | PRR29-AS1 | 400612 |

| | | | | | | | | |
|---|---|---|---|---|---|---|---|---|
| ENSG000002 | 2.30876758 | -0.346771856 | 0.36591906 | -0.9476737 | 0.34329561 | 0.47435787 | NA | NA |
| ENSG000002 | 46.1126055 | -0.206012411 | 0.21742961 | -0.9474901 | 0.34338908 | 0.47447086 | NA | NA |
| ENSG000001 | 95.60863 | -0.166919206 | 0.17617959 | -0.9474378 | 0.34341573 | 0.47447755 | ICA1L | 130026 |
| ENSG000002 | 2.87590664 | -0.354813734 | 0.3745031 | -0.9474254 | 0.34342207 | 0.47447755 | NA | NA |
| ENSG000002 | 4.99554992 | -0.354580577 | 0.37426296 | -0.9474103 | 0.34342975 | 0.47447755 | NA | NA |
| ENSG000002 | 2.83491717 | -0.356156383 | 0.37593479 | -0.9473887 | 0.34344073 | 0.47447755 | NA | NA |
| ENSG000001 | 17.3076883 | 0.275711565 | 0.29103378 | 0.94735246 | 0.34345921 | 0.47448691 | NBL1 | 4681 |
| ENSG000001 | 3.48195948 | 0.357953291 | 0.37786146 | 0.94731358 | 0.34347901 | 0.47449811 | MAGEB6 | 158809 |
| ENSG000002 | 7.8410193 | 0.335542187 | 0.35421969 | 0.94727141 | 0.34350049 | 0.47451162 | NA | NA |
| ENSG000002 | 4.46722968 | -0.356679825 | 0.37654719 | -0.9472381 | 0.34351749 | 0.47451893 | NA | NA |
| ENSG000001 | 2.35741688 | -0.345840956 | 0.36512867 | -0.9471756 | 0.34354932 | 0.47454673 | PLA2G1B | 5319 |
| ENSG000002 | 6.28106033 | -0.348155492 | 0.367643 | -0.9469934 | 0.34364214 | 0.47465878 | NA | NA |
| ENSG000002 | 3.11144712 | -0.357737412 | 0.37781616 | -0.9468558 | 0.34371228 | 0.47473949 | NA | NA |
| ENSG000002 | 4.95110809 | -0.353777883 | 0.37365152 | -0.9468124 | 0.34373439 | 0.47475386 | NA | NA |
| ENSG000001 | 20.289331 | -0.275227746 | 0.2907402 | -0.946645 | 0.34381972 | 0.47485554 | AOC3 | 8639 |
| ENSG000002 | 3.99336228 | -0.358275696 | 0.3785666 | -0.9464007 | 0.34394424 | 0.47497142 | NA | NA |
| ENSG000002 | 2.56902813 | -0.353329204 | 0.37332808 | -0.9464308 | 0.34392889 | 0.47497142 | NA | NA |
| ENSG000002 | 23.8426784 | -0.25815073 | 0.27275833 | -0.9464449 | 0.34392172 | 0.47497142 | NA | NA |
| ENSG000002 | 2.57155187 | -0.352625605 | 0.37260133 | -0.9463885 | 0.34395048 | 0.47497142 | NA | NA |
| ENSG000001 | 15.6088901 | -0.288592973 | 0.3049979 | -0.946213 | 0.34403996 | 0.47506353 | MMP13 | 4322 |
| ENSG000002 | 2.61188148 | 0.348987963 | 0.3688265 | 0.94621173 | 0.34404061 | 0.47506353 | NA | NA |
| ENSG000001 | 448.824577 | 0.118187995 | 0.12491662 | 0.94613503 | 0.34407972 | 0.47509602 | ELAVL2 | 1993 |
| ENSG000002 | 2.40994866 | -0.345227273 | 0.36489641 | -0.9460967 | 0.34409929 | 0.47509602 | RN7SL252P | 106481832 |
| ENSG000002 | 7.51174233 | -0.337164482 | 0.35636959 | -0.946109 | 0.34409298 | 0.47509602 | NA | NA |
| ENSG000002 | 4.99779425 | -0.354586545 | 0.37481848 | -0.946022 | 0.34413735 | 0.4751324 | PRPS1L1 | 221823 |
| ENSG000002 | 4.66978935 | -0.354980575 | 0.37532419 | -0.9457972 | 0.34425203 | 0.47527455 | NA | NA |
| ENSG000001 | 501.617278 | 0.127191935 | 0.13449105 | 0.94572785 | 0.34428742 | 0.47530722 | EXT1 | 2131 |
| ENSG00000C | 6.7518936 | -0.344576599 | 0.36440315 | -0.9455917 | 0.34435687 | 0.47537528 | SERPIND1 | 3053 |
| ENSG000002 | 23.711032 | 0.251654456 | 0.26613618 | 0.94558528 | 0.34436016 | 0.47537528 | NA | NA |

| | | | | | | | | |
|---|---|---|---|---|---|---|---|---|
| ENSG000001 | 55.3384711 | -0.209642745 | 0.22173009 | -0.9454862 | 0.34441071 | 0.47542887 | NA | NA |
| ENSG000002 | 2.84564186 | -0.354761503 | 0.37522523 | -0.9454628 | 0.34442265 | 0.47542918 | NA | NA |
| ENSG000002 | 4.90889334 | 0.353677728 | 0.37411754 | 0.94536528 | 0.34447242 | 0.47548169 | LOC102724334 | 102724334 |
| ENSG000002 | 4.5505432 | -0.3575656 | 0.37827413 | -0.9452552 | 0.34452859 | 0.47554303 | NA | NA |
| ENSG000002 | 5.18073138 | -0.350985069 | 0.37133997 | -0.9451853 | 0.3445643 | 0.47557613 | LINC01585 | 101929765 |
| ENSG000001 | 4.22129512 | -0.357784502 | 0.37855088 | -0.9451424 | 0.34458617 | 0.47559013 | XAGE2 | 9502 |
| ENSG000001 | 4.14936926 | -0.356876448 | 0.37765201 | -0.9449876 | 0.34466523 | 0.47567435 | CELP | 1057 |
| ENSG000002 | 6.26173086 | -0.346894743 | 0.36709334 | -0.9449769 | 0.34467065 | 0.47567435 | NA | NA |
| ENSG000001 | 11.6614681 | 0.309185251 | 0.32722165 | 0.94488019 | 0.34472005 | 0.47571015 | C20orf195 | 79025 |
| ENSG000002 | 5.38293333 | -0.35553271 | 0.37626513 | -0.9448994 | 0.34471022 | 0.47571015 | NA | NA |
| ENSG000001 | 33.6369449 | 0.239560529 | 0.25355778 | 0.94479659 | 0.34476273 | 0.47572048 | NA | NA |
| ENSG000002 | 3.1642395 | -0.356560046 | 0.37739186 | -0.9448006 | 0.3447607 | 0.47572048 | SNORD115-28 | 100036565 |
| ENSG000002 | 3.99229335 | -0.357691039 | 0.37857685 | -0.9448307 | 0.34474531 | 0.47572048 | NA | NA |
| ENSG000002 | 2.34645176 | -0.347383742 | 0.36772087 | -0.9446941 | 0.34481507 | 0.47577651 | NA | NA |
| ENSG000002 | 2138.43375 | -0.11181605 | 0.11838017 | -0.9445505 | 0.34488839 | 0.47586149 | ZNF850 | 342892 |
| ENSG000001 | 12.152216 | -0.308563575 | 0.32669072 | -0.9445128 | 0.34490766 | 0.47586938 | NA | NA |
| ENSG000002 | 2.89429171 | -0.354064106 | 0.37487198 | -0.9444934 | 0.34491758 | 0.47586938 | NA | NA |
| ENSG000002 | 2.81342714 | -0.355400637 | 0.37631087 | -0.9444336 | 0.3449481 | 0.4758953 | NA | NA |
| ENSG000002 | 4.43441283 | -0.355966078 | 0.37694008 | -0.9443572 | 0.34498715 | 0.47593148 | NA | NA |
| ENSG000002 | 2.32232554 | -0.347105839 | 0.3675659 | -0.9443363 | 0.3449978 | 0.47593148 | NA | NA |
| ENSG000002 | 3.11460504 | -0.356826712 | 0.37798832 | -0.9440152 | 0.34516188 | 0.47614164 | NA | NA |
| ENSG000002 | 3.50986102 | -0.355860339 | 0.37706306 | -0.9437688 | 0.34528783 | 0.47629917 | NA | NA |
| ENSG000002 | 3.08600359 | -0.356776507 | 0.37805768 | -0.9437092 | 0.34531827 | 0.47632496 | RNU6-1284P | 106480126 |
| ENSG000002 | 18.7866875 | 0.28209218 | 0.29893387 | 0.94366082 | 0.345343 | 0.47634287 | NA | NA |
| ENSG000001 | 113.906043 | 0.153028699 | 0.16217199 | 0.94361979 | 0.34536398 | 0.4763556 | CASP7 | 840 |
| ENSG000002 | 3.71724372 | 0.356957512 | 0.37830282 | 0.94357615 | 0.34538629 | 0.47637017 | NA | NA |
| ENSG000001 | 18.1024931 | -0.268962302 | 0.28507276 | -0.9434865 | 0.34543211 | 0.47641717 | KLK5 | 25818 |
| ENSG000002 | 2.613078 | -0.350872746 | 0.37195142 | -0.9433295 | 0.34551239 | 0.47651168 | NA | NA |
| ENSG000002 | 4.97745301 | -0.352860456 | 0.37409081 | -0.9432481 | 0.345554 | 0.47655185 | NA | NA |

| | | | | | | | | |
|---|---|---|---|---|---|---|---|---|
| ENSG000002 | 6.82504758 | -0.343578613 | 0.36425884 | -0.9432266 | 0.34556502 | 0.47655185 | NA | NA |
| ENSG000000 | 27.578987 | -0.241620829 | 0.25624018 | -0.9429467 | 0.34570818 | 0.47673306 | MAOB | 4129 |
| ENSG000002 | 5.73280669 | -0.346685036 | 0.36768228 | -0.942893 | 0.34573565 | 0.47673852 | NA | NA |
| ENSG000002 | 4.13006514 | 0.356976843 | 0.37859334 | 0.94290313 | 0.34573045 | 0.47673852 | NA | NA |
| ENSG000002 | 4.52659103 | -0.355854689 | 0.37749431 | -0.9426756 | 0.34584684 | 0.47687563 | NA | NA |
| ENSG000002 | 2.84866338 | -0.354361684 | 0.37597413 | -0.9425161 | 0.34592846 | 0.47697195 | NA | NA |
| ENSG000002 | 18.2055388 | 0.282060437 | 0.29935528 | 0.94222638 | 0.34607675 | 0.47716019 | NA | NA |
| ENSG000001 | 2.89869537 | -0.353898661 | 0.37564612 | -0.9421065 | 0.3461381 | 0.4772001 | LOC729173 | 729173 |
| ENSG000002 | 13.9796529 | -0.305192015 | 0.32393406 | -0.9421424 | 0.34611973 | 0.4772001 | LOC101927653 | 101927653 |
| ENSG000002 | 3.09087058 | -0.356045548 | 0.3779272 | -0.9421009 | 0.346141 | 0.4772001 | NA | NA |
| ENSG000002 | 2.84837522 | -0.354056925 | 0.37585318 | -0.9420086 | 0.34618824 | 0.47721655 | NA | NA |
| ENSG000002 | 4.97539884 | -0.355258326 | 0.37712627 | -0.9420143 | 0.34618534 | 0.47721655 | NA | NA |
| ENSG000002 | 3.94945731 | -0.356591381 | 0.37853117 | -0.9420397 | 0.34617233 | 0.47721655 | NA | NA |
| ENSG000002 | 4.68152036 | -0.353502431 | 0.37528234 | -0.9419639 | 0.3462111 | 0.47722062 | NA | NA |
| ENSG000002 | 6.66347048 | -0.346904493 | 0.36828066 | -0.9419569 | 0.34621473 | 0.47722062 | NA | NA |
| ENSG000002 | 3.11368479 | 0.356144826 | 0.37814982 | 0.94180879 | 0.34629054 | 0.4773089 | NA | NA |
| ENSG000002 | 9.96063777 | -0.315115339 | 0.33459637 | -0.9417775 | 0.34630655 | 0.47731474 | LOC105371083 | 105371083 |
| ENSG000001 | 2.84971325 | -0.354265949 | 0.37621644 | -0.9416546 | 0.34636949 | 0.47736304 | TREX2 | 11219 |
| ENSG000002 | 5.47918911 | -0.347876723 | 0.36942528 | -0.9416701 | 0.34636159 | 0.47736304 | PKNOX2-AS1 | 103695364 |
| ENSG000002 | 5.52242698 | -0.350966975 | 0.37271879 | -0.9416401 | 0.34637691 | 0.47736304 | PRSS2 | 5645 |
| ENSG000001 | 4.45791916 | -0.354780719 | 0.37681091 | -0.9415352 | 0.34643068 | 0.47742072 | RIPPLY1 | 92129 |
| ENSG000002 | 15.2750845 | -0.291837252 | 0.30996643 | -0.9415125 | 0.34644231 | 0.47742072 | NA | NA |
| ENSG000002 | 3.75789313 | -0.35571343 | 0.37784042 | -0.9414383 | 0.34648031 | 0.47745687 | LINC01563 | 101060544 |
| ENSG000002 | 3.71041305 | 0.356214186 | 0.37838632 | 0.94140344 | 0.34649815 | 0.47746522 | NA | NA |
| ENSG000002 | 7.89948901 | -0.335857904 | 0.35682176 | -0.9412484 | 0.34657759 | 0.47755846 | NA | NA |
| ENSG000001 | 29.2224828 | 0.236517788 | 0.25133224 | 0.94105631 | 0.346676 | 0.47764708 | NUDT17 | 200035 |
| ENSG000001 | 9.32770779 | -0.321556354 | 0.34169813 | -0.9410539 | 0.34667724 | 0.47764708 | IER5L | 389792 |
| ENSG000002 | 3.91123256 | -0.356201104 | 0.37850965 | -0.9410621 | 0.34667301 | 0.47764708 | NA | NA |
| ENSG000002 | 8.55870471 | 0.330723577 | 0.3514943 | 0.94090737 | 0.34675233 | 0.4777343 | FLJ30679 | 146512 |

| | | | | | | | | |
|---|---|---|---|---|---|---|---|---|
| ENSG000002 | 3.12710516 | -0.355614615 | 0.37796072 | -0.9408772 | 0.34676781 | 0.4777394 | NA | NA |
| ENSG000002 | 7.49851251 | -0.335005542 | 0.35607356 | -0.9408324 | 0.34679075 | 0.47775478 | NA | NA |
| ENSG000002 | 8.58553198 | -0.328940964 | 0.3496396 | -0.9408001 | 0.34680732 | 0.47776137 | NA | NA |
| ENSG000000 | 210.171963 | 0.151254294 | 0.16078608 | 0.9407176 | 0.3468496 | 0.47780338 | POMT2 | 29954 |
| ENSG000001 | 26.3256377 | -0.242964985 | 0.25828989 | -0.9406678 | 0.34687512 | 0.47782231 | ATP6V1C2 | 245973 |
| ENSG000002 | 2.90804302 | 0.353940823 | 0.37629508 | 0.94059382 | 0.34691305 | 0.47785832 | NA | NA |
| ENSG000002 | 6.60848588 | -0.3468081 | 0.36872856 | -0.9405512 | 0.34693489 | 0.47787217 | LINC01566 | 283914 |
| ENSG000002 | 2.33020663 | -0.345987369 | 0.36788802 | -0.9404693 | 0.34697691 | 0.47791382 | NA | NA |
| ENSG000002 | 5.24670848 | -0.351077081 | 0.37331234 | -0.9404379 | 0.34699297 | 0.47791971 | NA | NA |
| ENSG000001 | 7.78387691 | -0.332694662 | 0.35378522 | -0.940386 | 0.34701961 | 0.47794016 | ZNF467 | 168544 |
| ENSG000001 | 82.1123704 | -0.171238921 | 0.18213359 | -0.9401831 | 0.34712366 | 0.47806723 | DNAJC6 | 9829 |
| ENSG000001 | 66.10497 | 0.179571502 | 0.19101916 | 0.94007064 | 0.34718133 | 0.47809794 | GABRE | 2564 |
| ENSG000001 | 22.74079 | -0.250232258 | 0.26617863 | -0.9400915 | 0.34717064 | 0.47809794 | ISLR2 | 57611 |
| ENSG000002 | 3.99982865 | -0.355862466 | 0.37853979 | -0.9400926 | 0.34717005 | 0.47809794 | NA | NA |
| ENSG000001 | 66.7982667 | -0.180790476 | 0.19232233 | -0.9400389 | 0.34719759 | 0.4781041 | FHAD1 | 114827 |
| ENSG000001 | 49.4060751 | -0.194864724 | 0.20731092 | -0.9399636 | 0.34723623 | 0.47814099 | MUC3A | 4584 |
| ENSG000002 | 4.95848772 | -0.351834143 | 0.37431524 | -0.9399407 | 0.34724797 | 0.47814099 | LOC101927911 | 101927911 |
| ENSG000002 | 3.15015339 | -0.354695926 | 0.37738898 | -0.9398683 | 0.34728514 | 0.47817595 | NA | NA |
| ENSG000001 | 6.24258296 | -0.345563537 | 0.36770205 | -0.9397923 | 0.34732413 | 0.47821339 | CKM | 1158 |
| ENSG000002 | 3.57975938 | -0.35266118 | 0.37526942 | -0.9397546 | 0.34734343 | 0.47822373 | NA | NA |
| ENSG000002 | 11.8092877 | 0.303792875 | 0.32328223 | 0.93971411 | 0.34736422 | 0.47823612 | NA | NA |
| ENSG000001 | 78.5427784 | 0.173774445 | 0.18493792 | 0.93963663 | 0.34740398 | 0.47827462 | TK2 | 7084 |
| ENSG000002 | 3.0958866 | -0.354461781 | 0.37725926 | -0.9395708 | 0.34743776 | 0.47830488 | NA | NA |
| ENSG000001 | 22.8190923 | -0.25916473 | 0.27586104 | -0.9394756 | 0.3474866 | 0.47835587 | TLR7 | 51284 |
| ENSG000002 | 4.15794703 | -0.355260646 | 0.37821359 | -0.9393122 | 0.34757046 | 0.47845508 | NA | NA |
| ENSG000002 | 4.70417737 | -0.352694931 | 0.37552063 | -0.9392159 | 0.34761993 | 0.47850693 | NA | NA |
| ENSG000002 | 2.91741979 | -0.351598593 | 0.37436745 | -0.9391805 | 0.3476381 | 0.4785157 | NA | NA |
| ENSG000002 | 3.37970479 | -0.355408604 | 0.37846088 | -0.9390894 | 0.34768484 | 0.47854755 | NA | NA |
| ENSG000002 | 4.41030643 | -0.353953069 | 0.37691026 | -0.9390911 | 0.34768398 | 0.47854755 | NA | NA |

| | | | | | | | | |
|---|---|---|---|---|---|---|---|---|
| ENSG000002 | 4.74058245 | -0.353215219 | 0.37617788 | -0.938958 | 0.34775232 | 0.47862419 | NA | NA |
| ENSG000002 | 68.5611794 | -0.175264362 | 0.18669251 | -0.9387863 | 0.3478405 | 0.4787293 | NA | NA |
| ENSG000002 | 5.82336578 | 0.344867628 | 0.36743352 | 0.93858511 | 0.3479438 | 0.47885522 | LOC101559451 | 101559451 |
| ENSG000002 | 2.66390668 | -0.348871569 | 0.37174322 | -0.9384746 | 0.34800057 | 0.47891709 | NA | NA |
| ENSG000001 | 4.9364184 | -0.35526737 | 0.37858078 | -0.9384189 | 0.34802916 | 0.47894018 | RTP5 | 285093 |
| ENSG000002 | 3.21293721 | -0.350800553 | 0.37388026 | -0.9382698 | 0.34810578 | 0.47902937 | NA | NA |
| ENSG000002 | 12.2572232 | -0.30395837 | 0.32398836 | -0.9381768 | 0.34815355 | 0.4790626 | NA | NA |
| ENSG000002 | 4.97959296 | -0.350621177 | 0.37372015 | -0.9381918 | 0.34814585 | 0.4790626 | NA | NA |
| ENSG000001 | 27.3483921 | 0.241273097 | 0.25723002 | 0.93796632 | 0.34826172 | 0.47917891 | FN3K | 64122 |
| ENSG000002 | 22.7601931 | -0.257435024 | 0.2744606 | -0.9379672 | 0.34826129 | 0.47917891 | CCDC153 | 283152 |
| ENSG000002 | 14.8391892 | -0.29084011 | 0.31024015 | -0.9374677 | 0.34851805 | 0.47951533 | NA | NA |
| ENSG000001 | 45.9066201 | 0.217168375 | 0.23167381 | 0.93738853 | 0.34855874 | 0.47954395 | SLC44A3 | 126969 |
| ENSG000002 | 6.07395375 | -0.346770848 | 0.36993578 | -0.9373812 | 0.3485625 | 0.47954395 | NA | NA |
| ENSG000002 | 7.92902565 | -0.337513781 | 0.36009248 | -0.9372975 | 0.34860555 | 0.4795869 | NA | NA |
| ENSG000002 | 7.262248 | -0.336089565 | 0.35859573 | -0.9372381 | 0.34863612 | 0.47961269 | NA | NA |
| ENSG000002 | 17.1189516 | -0.28129065 | 0.30015059 | -0.9371651 | 0.34867365 | 0.47964805 | NA | NA |
| ENSG000002 | 10.3640243 | -0.31931892 | 0.34076236 | -0.9370722 | 0.34872143 | 0.47968839 | GGNBP1 | 449520 |
| ENSG000002 | 2.88243705 | -0.351636259 | 0.37525397 | -0.9370621 | 0.34872663 | 0.47968839 | NA | NA |
| ENSG000002 | 5.1845022 | -0.348143797 | 0.37165359 | -0.9367427 | 0.34889092 | 0.47989809 | NA | NA |
| ENSG000002 | 4.18938129 | -0.353789647 | 0.37771243 | -0.936664 | 0.34893141 | 0.47993751 | NA | NA |
| ENSG000002 | 2.38294638 | -0.342945118 | 0.3661482 | -0.9366293 | 0.3489493 | 0.47994583 | RN7SKP11 | 106479099 |
| ENSG000002 | 3.11697223 | -0.353178219 | 0.37709354 | -0.9365799 | 0.34897471 | 0.4799645 | NA | NA |
| ENSG000002 | 2.64422792 | -0.344723125 | 0.36812842 | -0.9364208 | 0.34905656 | 0.48004062 | RN7SKP53 | 106480849 |
| ENSG000002 | 22.0570104 | -0.26471472 | 0.28269306 | -0.9364033 | 0.34906557 | 0.48004062 | NA | NA |
| ENSG000002 | 3.11793866 | -0.353898088 | 0.37792963 | -0.9364127 | 0.34906077 | 0.48004062 | NA | NA |
| ENSG00000C | 37.3643307 | -0.231902507 | 0.24766933 | -0.9363392 | 0.34909855 | 0.48006971 | PTGS2 | 5743 |
| ENSG000001 | 203.913189 | 0.13542135 | 0.14467539 | 0.93603583 | 0.34925474 | 0.4802403 | ABCG2 | 9429 |
| ENSG000002 | 7.23175813 | -0.339251417 | 0.36244568 | -0.9360062 | 0.34926998 | 0.4802403 | NA | NA |
| ENSG000002 | 4.68151981 | -0.352170895 | 0.37624452 | -0.936016 | 0.34926495 | 0.4802403 | NA | NA |

| | | | | | | | | |
|---|---|---|---|---|---|---|---|---|
| ENSG000002 | 3.11061818 | -0.353360122 | 0.37750047 | -0.9360521 | 0.34924634 | 0.4802403 | NA | NA |
| ENSG000002 | 4.15396769 | -0.35397913 | 0.37821809 | -0.9359127 | 0.34931812 | 0.4802902 | NA | NA |
| ENSG000002 | 4.90953044 | -0.349828295 | 0.37384737 | -0.9357517 | 0.34940107 | 0.48038797 | NA | NA |
| ENSG000002 | 8.29133488 | -0.32657136 | 0.34901152 | -0.9357037 | 0.34942577 | 0.48040565 | NA | NA |
| ENSG000001 | 98.9964845 | 0.158184554 | 0.16908115 | 0.93555402 | 0.34950286 | 0.48049533 | DEPTOR | 64798 |
| ENSG000001 | 5.7477457 | -0.346682659 | 0.37063286 | -0.9353802 | 0.34959237 | 0.48060211 | OR6C2 | 341416 |
| ENSG000002 | 2.34383573 | -0.343915289 | 0.36771114 | -0.9352866 | 0.34964064 | 0.48065216 | NA | NA |
| ENSG000001 | 451.430609 | 0.117574464 | 0.12571831 | 0.93522146 | 0.34967418 | 0.48066569 | SHISA2 | 387914 |
| ENSG000002 | 3.17059409 | -0.352798281 | 0.37723368 | -0.9352248 | 0.34967248 | 0.48066569 | MTCO1P4 | 106480684 |
| ENSG000002 | 4.08448836 | 0.354029826 | 0.37857351 | 0.93516798 | 0.34970174 | 0.48068727 | NA | NA |
| ENSG000002 | 4.19878823 | -0.353764878 | 0.37831911 | -0.9350965 | 0.34973857 | 0.48072161 | NA | NA |
| ENSG000002 | 32.9374151 | 0.25278844 | 0.27035594 | 0.93502085 | 0.34977755 | 0.48075889 | NA | NA |
| ENSG000002 | 4.98576039 | 0.350507355 | 0.37488334 | 0.93497715 | 0.34980007 | 0.48077355 | LOC105375050 | 105375050 |
| ENSG000001 | 34.4590409 | 0.234991236 | 0.25134333 | 0.93494122 | 0.34981859 | 0.4807827 | GBGT1 | 26301 |
| ENSG000002 | 8.92436946 | -0.330157691 | 0.35319646 | -0.9347707 | 0.34990649 | 0.48088722 | NA | NA |
| ENSG000001 | 24.7075396 | -0.248232734 | 0.2656174 | -0.93455 | 0.35002027 | 0.48101097 | PTCH2 | 8643 |
| ENSG000002 | 2.8334629 | -0.351587636 | 0.37621008 | -0.9345514 | 0.35001956 | 0.48101097 | NA | NA |
| ENSG000001 | 24.2125977 | 0.24880367 | 0.26625471 | 0.93445735 | 0.35006803 | 0.48103728 | ABHD15 | 116236 |
| ENSG000002 | 4.21349978 | -0.353545464 | 0.37834855 | -0.9344438 | 0.350075 | 0.48103728 | OR5M10 | 390167 |
| ENSG000002 | 4.2147854 | -0.353598502 | 0.37840104 | -0.9344544 | 0.35006957 | 0.48103728 | NA | NA |
| ENSG000001 | 5.24256595 | -0.348870808 | 0.37335978 | -0.9344092 | 0.35009286 | 0.48104552 | LRIT1 | 26103 |
| ENSG000002 | 76.5791185 | 0.175803647 | 0.1881741 | 0.93426059 | 0.35016949 | 0.48113451 | LOC100506282 | 100506282 |
| ENSG000002 | 5.72762671 | -0.344161692 | 0.36840099 | -0.934204 | 0.35019866 | 0.48115829 | NA | NA |
| ENSG000002 | 79.2095929 | 0.176595888 | 0.18904499 | 0.9341474 | 0.35022786 | 0.48118211 | FBXO16 | 157574 |
| ENSG000002 | 5.66919768 | -0.342844194 | 0.36702944 | -0.9341054 | 0.35024951 | 0.48119555 | NA | NA |
| ENSG000002 | 9.89787825 | 0.325470954 | 0.34848598 | 0.9339571 | 0.35032602 | 0.48128436 | NA | NA |
| ENSG000002 | 6.64228693 | 0.339639665 | 0.36368033 | 0.93389616 | 0.35035746 | 0.48131125 | NA | NA |
| ENSG000002 | 2.47111425 | -0.334044715 | 0.35780031 | -0.9336066 | 0.35050688 | 0.4815002 | NA | NA |
| ENSG000002 | 2.38412632 | -0.337361566 | 0.36138272 | -0.9335299 | 0.35054646 | 0.48153826 | NA | NA |

| | | | | | | | | |
|---|---|---|---|---|---|---|---|---|
| ENSG000002 | 4.4448779 | -0.35195331 | 0.37702801 | -0.9334938 | 0.35056507 | 0.48154751 | NA | NA |
| ENSG000002 | 28.1720249 | 0.23394863 | 0.25063079 | 0.93343932 | 0.35059319 | 0.48156983 | NA | NA |
| ENSG000002 | 3.34708513 | -0.353332468 | 0.37854067 | -0.9334069 | 0.35060993 | 0.48157651 | NA | NA |
| ENSG000002 | 11.7612963 | 0.310254752 | 0.3324043 | 0.93336564 | 0.35063121 | 0.48158943 | NA | NA |
| ENSG000002 | 42.1334379 | 0.212971684 | 0.22819233 | 0.93329904 | 0.35066559 | 0.48160402 | NA | NA |
| ENSG000002 | 6.36210969 | -0.346316511 | 0.37106619 | -0.9333012 | 0.35066449 | 0.48160402 | NA | NA |
| ENSG000002 | 3.13393584 | -0.352474679 | 0.37770398 | -0.9332035 | 0.35071491 | 0.48165544 | NA | NA |
| ENSG000002 | 3.35872385 | -0.353219897 | 0.37855988 | -0.9330622 | 0.35078788 | 0.48173934 | NA | NA |
| ENSG000002 | 3.35020969 | -0.35312849 | 0.37851735 | -0.9329255 | 0.35085843 | 0.48175218 | NA | NA |
| ENSG000002 | 2.38246921 | -0.33800513 | 0.36231424 | -0.932906 | 0.35086851 | 0.48175218 | NA | NA |
| ENSG000002 | 3.18268928 | -0.351658776 | 0.37693461 | -0.9329437 | 0.35084904 | 0.48175218 | NA | NA |
| ENSG000002 | 2.39466441 | -0.341618757 | 0.36617671 | -0.9329342 | 0.35085397 | 0.48175218 | NA | NA |
| ENSG000002 | 2.38453236 | -0.342098342 | 0.36669022 | -0.9329355 | 0.35085325 | 0.48175218 | LINC00904 | 100652909 |
| ENSG000002 | 11.574031 | -0.309770059 | 0.33201701 | -0.9329945 | 0.35082279 | 0.48175218 | NA | NA |
| ENSG000002 | 2.84530959 | -0.349713045 | 0.37493722 | -0.9327243 | 0.35096235 | 0.48186471 | NA | NA |
| ENSG000002 | 7.98903499 | -0.327465254 | 0.35110068 | -0.9326819 | 0.35098422 | 0.48187843 | NA | NA |
| ENSG000002 | 7.80687395 | -0.332484736 | 0.35650591 | -0.9326205 | 0.35101592 | 0.48190563 | NA | NA |
| ENSG000002 | 3.10888372 | -0.352119396 | 0.37770423 | -0.9322623 | 0.35120101 | 0.48214341 | NA | NA |
| ENSG000001 | 35.1306082 | 0.221960646 | 0.23810161 | 0.93220978 | 0.35122812 | 0.48216431 | BOLA1 | 51027 |
| ENSG000002 | 91.5745128 | 0.180279729 | 0.19340287 | 0.9321461 | 0.35126102 | 0.48219315 | MIR4697HG | 283174 |
| ENSG000002 | 2.90919935 | -0.348855528 | 0.3743534 | -0.9318882 | 0.35139429 | 0.48235977 | NA | NA |
| ENSG000001 | 6.02756152 | -0.346089666 | 0.37151531 | -0.9315623 | 0.35156277 | 0.4825747 | ANKRD61 | 100310846 |
| ENSG000002 | 5.21676432 | -0.346817459 | 0.37234675 | -0.9314368 | 0.35162767 | 0.48263875 | NA | NA |
| ENSG000002 | 35.0824125 | 0.229835929 | 0.24675704 | 0.93142603 | 0.35163323 | 0.48263875 | NA | NA |
| ENSG000001 | 224.231632 | 0.127465512 | 0.13685733 | 0.93137512 | 0.35165956 | 0.48265855 | BBS2 | 583 |
| ENSG000002 | 60.1170587 | -0.185633291 | 0.19931942 | -0.9313357 | 0.35167995 | 0.4826702 | NR2F1-AS1 | 441094 |
| ENSG000002 | 4.68989071 | -0.349751062 | 0.37554946 | -0.9313049 | 0.35169586 | 0.48267569 | NA | NA |
| ENSG000002 | 5.88965668 | -0.349951123 | 0.37584828 | -0.9310968 | 0.35180351 | 0.48280709 | NA | NA |
| ENSG000002 | 2.30799867 | -0.341723448 | 0.36702301 | -0.9310682 | 0.35181829 | 0.48281104 | NA | NA |

| | | | | | | | | |
|---|---|---|---|---|---|---|---|---|
| ENSG000002 | 2.85688436 | -0.349921305 | 0.37594585 | -0.9307758 | 0.35196955 | 0.48300227 | NA | NA |
| ENSG000001 | 67.6480745 | -0.183761111 | 0.19743617 | -0.9307368 | 0.35198972 | 0.48301361 | VIPR1 | 7433 |
| ENSG000002 | 2.60326602 | -0.345461127 | 0.37118267 | -0.9307038 | 0.3520068 | 0.4830207 | NA | NA |
| ENSG000002 | 3.66161925 | -0.351577405 | 0.37777835 | -0.9306447 | 0.3520374 | 0.48304634 | NA | NA |
| ENSG000001 | 321.314174 | 0.173659178 | 0.18661999 | 0.93054972 | 0.35208653 | 0.48309742 | CMIP | 80790 |
| ENSG000002 | 4.22247759 | -0.351938073 | 0.37822745 | -0.9304932 | 0.35211577 | 0.48312119 | LOC105373519 | 105373519 |
| ENSG000001 | 4.92030274 | -0.347635401 | 0.37362077 | -0.9304499 | 0.3521382 | 0.48312622 | NA | NA |
| ENSG000002 | 10.6019496 | -0.31889879 | 0.34273974 | -0.9304401 | 0.35214327 | 0.48312622 | NA | NA |
| ENSG000002 | 3.72672288 | 0.352129618 | 0.37847488 | 0.93039098 | 0.35216869 | 0.48314475 | NA | NA |
| ENSG000002 | 52.5013866 | -0.192010053 | 0.20638777 | -0.9303364 | 0.35219695 | 0.4831565 | NA | NA |
| ENSG000002 | 3.98203154 | -0.352200706 | 0.37857676 | -0.9303284 | 0.35220109 | 0.4831565 | OR2L8 | 391190 |
| ENSG000002 | 5.18628405 | -0.345755941 | 0.37171965 | -0.9301524 | 0.35229216 | 0.48326509 | LINC00593 | 414926 |
| ENSG000002 | 6.12059784 | -0.347263937 | 0.37340025 | -0.9300046 | 0.35236872 | 0.48333741 | NA | NA |
| ENSG000002 | 4.201495 | -0.351973776 | 0.37845675 | -0.9300238 | 0.35235878 | 0.48333741 | NA | NA |
| ENSG000002 | 8.57418698 | -0.324591613 | 0.34904416 | -0.9299443 | 0.35239994 | 0.48336389 | LOC101926942 | 101926942 |
| ENSG000002 | 3.10566666 | -0.350983726 | 0.37749805 | -0.929763 | 0.3524938 | 0.48347627 | NA | NA |
| ENSG000002 | 3.36780603 | -0.351916709 | 0.37855983 | -0.9296198 | 0.35256798 | 0.48356167 | NA | NA |
| ENSG000002 | 2.3757708 | -0.340069489 | 0.36588275 | -0.9294494 | 0.35265626 | 0.48366639 | NA | NA |
| ENSG000002 | 3.09434856 | -0.351123294 | 0.37779371 | -0.9294048 | 0.35267933 | 0.48368167 | ARHGAP26-IT1 | 100874372 |
| ENSG000002 | 6.04897388 | -0.344020367 | 0.37019761 | -0.9292885 | 0.35273961 | 0.48374798 | NA | NA |
| ENSG000001 | 30.1279257 | 0.228333057 | 0.24573252 | 0.92919347 | 0.35278883 | 0.48379593 | ZNF653 | 115950 |
| ENSG000001 | 225.3226 | 0.140929814 | 0.15167199 | 0.92917492 | 0.35279844 | 0.48379593 | HPDL | 84842 |
| ENSG000002 | 3.41437643 | -0.350124408 | 0.37684542 | -0.9290929 | 0.35284093 | 0.48383784 | NA | NA |
| ENSG000001 | 43.5559176 | 0.209180804 | 0.22515169 | 0.92906612 | 0.35285482 | 0.48384052 | DES | 1674 |
| ENSG000001 | 10.2338552 | -0.309445113 | 0.3330967 | -0.9289948 | 0.35289177 | 0.48385846 | BPIFA4P | 317716 |
| ENSG000002 | 3.1251425 | -0.347444825 | 0.3739975 | -0.9290031 | 0.3528875 | 0.48385846 | NA | NA |
| ENSG000002 | 2.89217267 | -0.348745216 | 0.3755725 | -0.9285696 | 0.35311216 | 0.48414429 | NA | NA |
| ENSG000002 | 3.75814942 | 0.351298593 | 0.3783955 | 0.92838999 | 0.35320531 | 0.48425562 | NA | NA |
| ENSG000002 | 6.00613928 | 0.339106518 | 0.36527333 | 0.92836375 | 0.35321892 | 0.4842579 | NA | NA |

| | | | | | | | | |
|---|---|---|---|---|---|---|---|---|
| ENSG000001 | 5.2089681 | -0.345643357 | 0.37232798 | -0.9283303 | 0.35323624 | 0.48426529 | PIP | 5304 |
| ENSG000002 | 5.46307928 | -0.345125473 | 0.37177967 | -0.9283065 | 0.35324862 | 0.48426588 | NA | NA |
| ENSG000001 | 187.228041 | -0.140506363 | 0.15139604 | -0.9280716 | 0.35337044 | 0.48437152 | PDZD2 | 23037 |
| ENSG000001 | 9.50307738 | -0.314521846 | 0.33890039 | -0.9280658 | 0.35337346 | 0.48437152 | PRDM8 | 56978 |
| ENSG000001 | 5.47440067 | -0.344146842 | 0.37081641 | -0.9280788 | 0.35336671 | 0.48437152 | SLC2A7 | 155184 |
| ENSG000002 | 11.0030871 | -0.302601683 | 0.3260443 | -0.9280999 | 0.35335575 | 0.48437152 | CPS1-IT1 | 29034 |
| ENSG000002 | 5.94683754 | -0.339355268 | 0.365696 | -0.927971 | 0.35342264 | 0.48442256 | NA | NA |
| ENSG000002 | 2.84885926 | -0.348107045 | 0.37522644 | -0.9277252 | 0.35355011 | 0.48456225 | NA | NA |
| ENSG000002 | 4.18322203 | -0.350952334 | 0.37830148 | -0.9277054 | 0.3535604 | 0.48456225 | NA | NA |
| ENSG000002 | 11.233694 | -0.309620582 | 0.3337399 | -0.9277302 | 0.35354755 | 0.48456225 | NA | NA |
| ENSG000002 | 69.7601244 | 0.173718436 | 0.18728361 | 0.92756883 | 0.35363128 | 0.484643 | NA | NA |
| ENSG000002 | 9.87107106 | -0.319303217 | 0.34428505 | -0.9274385 | 0.3536989 | 0.48465249 | OR10C1 | 442194 |
| ENSG000002 | 15.885238 | 0.279991659 | 0.30187727 | 0.92750163 | 0.35366614 | 0.48465249 | LINC01503 | 100506119 |
| ENSG000002 | 7.58049255 | -0.333301574 | 0.35937277 | -0.9274536 | 0.35369106 | 0.48465249 | NA | NA |
| ENSG000002 | 3.15222863 | -0.349875566 | 0.37725796 | -0.9274173 | 0.35370991 | 0.48465249 | LOC101928850 | 101928850 |
| ENSG000002 | 2.65637106 | -0.342210299 | 0.36896317 | -0.9274918 | 0.35367127 | 0.48465249 | NA | NA |
| ENSG000002 | 4.92769893 | -0.348370653 | 0.37562607 | -0.92744 | 0.35369811 | 0.48465249 | NA | NA |
| ENSG000002 | 7.487611 | -0.331960551 | 0.35797898 | -0.9273186 | 0.35376116 | 0.48470634 | NA | NA |
| ENSG000002 | 3.37970534 | -0.350943381 | 0.37850493 | -0.9271831 | 0.35383147 | 0.48478629 | NA | NA |
| ENSG000002 | 3.15522889 | -0.348147192 | 0.37553776 | -0.9270631 | 0.35389378 | 0.48485529 | NA | NA |
| ENSG000002 | 3.38269351 | -0.350782584 | 0.3783903 | -0.9270391 | 0.35390625 | 0.48485599 | NA | NA |
| ENSG000002 | 5.96203747 | -0.340128669 | 0.36698188 | -0.9268269 | 0.35401643 | 0.48499055 | NA | NA |
| ENSG000002 | 18.3330214 | -0.266941553 | 0.28807154 | -0.9266502 | 0.35410817 | 0.48509984 | NA | NA |
| ENSG000001 | 249.775399 | 0.135367415 | 0.14612286 | 0.9263945 | 0.354241 | 0.48524902 | NOL4L | 140688 |
| ENSG000002 | 2.85498573 | -0.348147485 | 0.3758074 | -0.9263987 | 0.35423882 | 0.48524902 | NA | NA |
| ENSG000002 | 3.36285417 | -0.350616013 | 0.37849382 | -0.9263454 | 0.35426651 | 0.48526757 | NA | NA |
| ENSG000002 | 3.2384471 | -0.347315995 | 0.37495352 | -0.9262908 | 0.35429488 | 0.48526965 | NA | NA |
| ENSG000002 | 2.83614419 | -0.348158472 | 0.37585148 | -0.9263193 | 0.35428009 | 0.48526965 | NA | NA |
| ENSG000002 | 6.84398803 | -0.33905486 | 0.36604189 | -0.9262734 | 0.35430393 | 0.48526965 | NA | NA |

| | | | | | | | | |
|---|---|---|---|---|---|---|---|---|
| ENSG000001 | 5.46953314 | -0.344309056 | 0.37173349 | -0.9262255 | 0.35432878 | 0.4852873 | GPR183 | 1880 |
| ENSG000002 | 23.8050361 | 0.245429194 | 0.2650191 | 0.92608115 | 0.35440381 | 0.48537366 | NA | NA |
| ENSG000002 | 54.0477758 | 0.189093569 | 0.20420136 | 0.92601525 | 0.35443806 | 0.48540418 | NA | NA |
| ENSG000001 | 124.1961 | -0.148564042 | 0.16044691 | -0.9259389 | 0.35447771 | 0.48544209 | EHHADH | 1962 |
| ENSG000002 | 9.99846356 | -0.31167961 | 0.33662354 | -0.9258996 | 0.35449815 | 0.48544465 | RNU1-92P | 106481620 |
| ENSG000002 | 6.80460109 | -0.338953902 | 0.36608471 | -0.9258893 | 0.35450353 | 0.48544465 | NA | NA |
| ENSG000002 | 2.82862728 | -0.34698451 | 0.37484002 | -0.9256869 | 0.3546087 | 0.48557228 | NDUFB2-AS1 | 100134713 |
| ENSG000002 | 6.29975362 | -0.342426884 | 0.36997152 | -0.9255493 | 0.35468025 | 0.48565384 | LOC102724050 | 102724050 |
| ENSG000002 | 3.10481287 | -0.349613135 | 0.37775919 | -0.9254921 | 0.35471001 | 0.4856782 | NA | NA |
| ENSG000002 | 3.17829437 | -0.346504679 | 0.37441193 | -0.9254638 | 0.35472472 | 0.48568194 | TPTE | 7179 |
| ENSG000001 | 5.95959771 | -0.339353265 | 0.36671674 | -0.9253825 | 0.35476696 | 0.48572338 | C5orf38 | 153571 |
| ENSG000002 | 24.9734427 | 0.246337323 | 0.26622757 | 0.92528854 | 0.35481584 | 0.4857575 | NA | NA |
| ENSG000002 | 10.1145449 | 0.3104571 | 0.33551875 | 0.92530476 | 0.3548074 | 0.4857575 | NA | NA |
| ENSG000001 | 190.974844 | -0.150755458 | 0.16294437 | -0.9251959 | 0.35486404 | 0.48580708 | SULF2 | 55959 |
| ENSG000002 | 6.2195578 | -0.336656841 | 0.36399039 | -0.9249058 | 0.35501489 | 0.48599719 | NA | NA |
| ENSG000002 | 5.74542791 | -0.344302011 | 0.37230822 | -0.9247768 | 0.35508201 | 0.48607267 | LOC284379 | 284379 |
| ENSG000002 | 2.33635107 | -0.336413028 | 0.36379207 | -0.9247399 | 0.35510124 | 0.48608258 | NA | NA |
| ENSG000002 | 2.84011751 | -0.347656132 | 0.37599696 | -0.9246248 | 0.35516109 | 0.48614809 | LINC01591 | 286094 |
| ENSG000002 | 5.95403239 | -0.339384489 | 0.36709969 | -0.9245022 | 0.35522488 | 0.48620259 | RN7SL497P | 106481846 |
| ENSG000002 | 3.36029021 | -0.349965884 | 0.37854032 | -0.9245142 | 0.35521868 | 0.48620259 | NA | NA |
| ENSG000001 | 4.92291159 | -0.346753932 | 0.37510833 | -0.9244101 | 0.35527284 | 0.48625181 | CXCR3 | 2833 |
| ENSG000002 | 14.280017 | 0.289040677 | 0.3127244 | 0.92426647 | 0.35534759 | 0.48633772 | NA | NA |
| ENSG000001 | 40.7137529 | -0.212000035 | 0.22937866 | -0.9242361 | 0.35536342 | 0.48634296 | ZNF410 | 57862 |
| ENSG000001 | 12.360468 | -0.301270685 | 0.32597831 | -0.9242047 | 0.35537975 | 0.4863489 | CCK | 885 |
| ENSG000002 | 10.0745923 | -0.313065749 | 0.33884223 | -0.9239278 | 0.35552392 | 0.48652979 | NA | NA |
| ENSG000002 | 3.12482882 | -0.348503821 | 0.3772612 | -0.9237733 | 0.35560436 | 0.48662344 | RNU6-649P | 106479854 |
| ENSG000002 | 11.7254656 | 0.299547089 | 0.32433153 | 0.923583 | 0.35570347 | 0.48671119 | NA | NA |
| ENSG000002 | 3.95667177 | -0.349635166 | 0.37856469 | -0.923581 | 0.35570449 | 0.48671119 | NA | NA |
| ENSG000002 | 2.59051053 | -0.342801585 | 0.37115657 | -0.9236037 | 0.35569268 | 0.48671119 | NA | NA |

| | | | | | | | | |
|---|---|---|---|---|---|---|---|---|
| ENSG000002 | 2.62484147 | -0.343573384 | 0.37204844 | -0.9234641 | 0.35576539 | 0.48677809 | NA | NA |
| ENSG000002 | 25.9555581 | 0.236074175 | 0.25564846 | 0.9234328 | 0.35578171 | 0.48678399 | NA | NA |
| ENSG000002 | 3.17568179 | -0.347014553 | 0.3759071 | -0.9231391 | 0.3559347 | 0.48697689 | NA | NA |
| ENSG000001 | 11.8163352 | 0.300687365 | 0.3257431 | 0.9230813 | 0.35596484 | 0.4870017 | LRG1 | 116844 |
| ENSG000002 | 3.7584154 | -0.348662452 | 0.37775874 | -0.9229765 | 0.35601943 | 0.48705995 | NA | NA |
| ENSG000002 | 11.9385723 | -0.296063822 | 0.32077936 | -0.9229516 | 0.35603243 | 0.4870613 | NA | NA |
| ENSG000002 | 13.212756 | -0.28862284 | 0.31274436 | -0.9228715 | 0.3560742 | 0.48710201 | PRKAR2A-AS1 | 100506637 |
| ENSG000002 | 53.0922788 | -0.188338622 | 0.20409395 | -0.9228036 | 0.35610958 | 0.48713398 | NA | NA |
| ENSG000002 | 57.4904037 | 0.19664443 | 0.21310821 | 0.92274453 | 0.35614035 | 0.48715964 | NA | NA |
| ENSG000001 | 116.318 | -0.144032231 | 0.15611984 | -0.9225748 | 0.35622882 | 0.48726422 | NA | NA |
| ENSG000002 | 3.3770246 | -0.349153585 | 0.37848163 | -0.9225113 | 0.35626193 | 0.48729306 | OR52E6 | 390078 |
| ENSG000002 | 4.20681158 | -0.349176218 | 0.37854736 | -0.9224109 | 0.35631428 | 0.48734823 | NA | NA |
| ENSG000002 | 24.488976 | 0.244706051 | 0.26534108 | 0.92223206 | 0.35640755 | 0.48745935 | ABHD16A | 7920 |
| ENSG000001 | 9.77954068 | -0.314249591 | 0.34078146 | -0.922144 | 0.35645345 | 0.48750569 | NA | NA |
| ENSG000002 | 4.96001476 | -0.345303058 | 0.3745175 | -0.9219945 | 0.35653146 | 0.48759595 | NA | NA |
| ENSG000001 | 7.13429722 | -0.336123377 | 0.36460697 | -0.9218786 | 0.35659188 | 0.48764568 | AREG | 374 |
| ENSG000002 | 6.76467557 | -0.335180788 | 0.36357618 | -0.9218997 | 0.35658087 | 0.48764568 | NA | NA |
| ENSG000002 | 4.17825613 | -0.348772984 | 0.37834795 | -0.9218313 | 0.35661658 | 0.487663 | NA | NA |
| ENSG000002 | 11.0373968 | 0.306396531 | 0.33239967 | 0.92177145 | 0.3566478 | 0.48768926 | NA | NA |
| ENSG000002 | 25.4262236 | -0.242975578 | 0.26360722 | -0.9217334 | 0.35666765 | 0.48769996 | NA | NA |
| ENSG000002 | 5.65092416 | -0.339421381 | 0.36825401 | -0.9217045 | 0.35668273 | 0.48770413 | NA | NA |
| ENSG000002 | 43.9693781 | 0.214925444 | 0.23320384 | 0.9216205 | 0.35672656 | 0.48774761 | ZNF865 | 100507290 |
| ENSG000002 | 29.8955754 | -0.229592358 | 0.24913783 | -0.9215476 | 0.35676462 | 0.4877832 | LINC01140 | 339524 |
| ENSG000002 | 4.26099538 | -0.348682784 | 0.37837991 | -0.9215151 | 0.35678159 | 0.48778995 | NA | NA |
| ENSG000002 | 5.47964612 | -0.341463335 | 0.37065699 | -0.9212381 | 0.35692614 | 0.48797113 | NA | NA |
| ENSG000001 | 162.931174 | 0.137664423 | 0.14943856 | 0.92121082 | 0.35694037 | 0.48797414 | MERTK | 10461 |
| ENSG000002 | 3.12445633 | -0.347574196 | 0.37734068 | -0.9211151 | 0.35699033 | 0.48800954 | NA | NA |
| ENSG000002 | 5.49731326 | -0.342401104 | 0.37172317 | -0.9211185 | 0.35698854 | 0.48800954 | NA | NA |
| ENSG000001 | 10.7909254 | -0.310690302 | 0.33732309 | -0.9210467 | 0.35702607 | 0.48804194 | CYP4Z1 | 199974 |

| | | | | | | | | |
|---|---|---|---|---|---|---|---|---|
| ENSG000002 | 3.09898034 | -0.347899635 | 0.37779239 | -0.9208752 | 0.3571156 | 0.48813142 | NA | NA |
| ENSG000002 | 11.7917841 | -0.301644406 | 0.32756068 | -0.920881 | 0.35711256 | 0.48813142 | NA | NA |
| ENSG000002 | 9.04199131 | -0.3194534 | 0.34698012 | -0.9206677 | 0.35722393 | 0.48824658 | NA | NA |
| ENSG000002 | 2.59014656 | -0.342768699 | 0.37230026 | -0.9206781 | 0.35721851 | 0.48824658 | NA | NA |
| ENSG000002 | 2.89087431 | -0.344896828 | 0.37465855 | -0.9205631 | 0.35727859 | 0.48830482 | NA | NA |
| ENSG000002 | 7.34741851 | -0.334481578 | 0.3634039 | -0.9204127 | 0.35735712 | 0.48839569 | NA | NA |
| ENSG000002 | 24.9358826 | -0.237651744 | 0.25823005 | -0.9203102 | 0.3574107 | 0.48841964 | NA | NA |
| ENSG000002 | 3.35409206 | -0.348205648 | 0.37834506 | -0.9203388 | 0.35739572 | 0.48841964 | LOC102724321 | 102724321 |
| ENSG000002 | 172.010118 | 0.140778142 | 0.15296817 | 0.92031002 | 0.35741078 | 0.48841964 | ZNF879 | 345462 |
| ENSG000002 | 4.39963828 | -0.347663414 | 0.3778096 | -0.920208 | 0.35746407 | 0.48847601 | LINC01256 | 104355285 |
| ENSG000001 | 23.7271811 | 0.243841538 | 0.26501393 | 0.92010839 | 0.35751612 | 0.48851421 | C4orf19 | 55286 |
| ENSG000002 | 5.4448865 | -0.340190238 | 0.36971995 | -0.9201295 | 0.35750508 | 0.48851421 | NA | NA |
| ENSG000002 | 4.8217793 | -0.347721329 | 0.37796579 | -0.919981 | 0.35758271 | 0.48858874 | NA | NA |
| ENSG000002 | 2.52222104 | 0.342836683 | 0.37267129 | 0.9199439 | 0.35760207 | 0.48859873 | MIR559 | 693144 |
| ENSG000001 | 3.1103978 | -0.347114752 | 0.37734716 | -0.9198817 | 0.35763458 | 0.48861101 | SIX3 | 6496 |
| ENSG000002 | 3.72633981 | -0.347919925 | 0.37823052 | -0.9198621 | 0.35764482 | 0.48861101 | NA | NA |
| ENSG000002 | 20.9310786 | 0.253490453 | 0.27557578 | 0.91985753 | 0.35764721 | 0.48861101 | SLC10A5 | 347051 |
| ENSG000002 | 4.9691611 | -0.34488307 | 0.37501948 | -0.9196404 | 0.35776069 | 0.48874958 | NA | NA |
| ENSG000001 | 18.8331775 | -0.259093406 | 0.28182816 | -0.9193312 | 0.35792238 | 0.48889566 | MB | 4151 |
| ENSG000002 | 2.30461297 | -0.335811231 | 0.36529539 | -0.9192868 | 0.35794559 | 0.48889566 | LOC102724450 | 102724450 |
| ENSG000002 | 2.87741352 | -0.343506327 | 0.37365523 | -0.9193136 | 0.35793157 | 0.48889566 | NA | NA |
| ENSG000002 | 2.8365354 | -0.344248621 | 0.3744786 | -0.9192745 | 0.35795202 | 0.48889566 | NA | NA |
| ENSG000002 | 12.4445128 | 0.295051385 | 0.32092096 | 0.91938956 | 0.35789185 | 0.48889566 | NA | NA |
| ENSG000002 | 3.10720078 | -0.347117095 | 0.37756828 | -0.9193492 | 0.35791295 | 0.48889566 | NA | NA |
| ENSG000002 | 8.8667069 | -0.321731354 | 0.34994998 | -0.9193638 | 0.3579053 | 0.48889566 | NA | NA |
| ENSG000002 | 5.96542051 | -0.340836386 | 0.37087143 | -0.9190149 | 0.35808775 | 0.48906458 | NA | NA |
| ENSG000002 | 4.31110779 | 0.346990223 | 0.3775822 | 0.9189793 | 0.3581064 | 0.48907358 | NA | NA |
| ENSG000002 | 5.96727029 | -0.337537603 | 0.3673286 | -0.9188982 | 0.3581488 | 0.48909462 | NA | NA |
| ENSG000002 | 8.60748694 | 0.317341174 | 0.34535624 | 0.91888067 | 0.35815799 | 0.48909462 | NA | NA |

| | | | | | | | | |
|---|---|---|---|---|---|---|---|---|
| ENSG000002 | 7.57075789 | -0.334597924 | 0.3641328 | -0.9188898 | 0.35815321 | 0.48909462 | LOC101928144 | 101928144 |
| ENSG000002 | 3.33720102 | -0.347718823 | 0.37844773 | -0.9188028 | 0.35819874 | 0.4891338 | NA | NA |
| ENSG000002 | 2.86619182 | 0.34424407 | 0.37468174 | 0.91876393 | 0.35821907 | 0.48914508 | NA | NA |
| ENSG000002 | 3.35750947 | -0.347122879 | 0.37792713 | -0.9184916 | 0.35836158 | 0.4893232 | NA | NA |
| ENSG000002 | 43.2903508 | -0.196647328 | 0.21410443 | -0.9184645 | 0.35837572 | 0.48932603 | HS3ST5 | 222537 |
| ENSG000002 | 3.3816883 | -0.34743493 | 0.37828907 | -0.9184377 | 0.35838978 | 0.48932876 | TMF1P1 | 106480320 |
| ENSG000002 | 10.0206537 | -0.308306226 | 0.33577283 | -0.9181988 | 0.35851477 | 0.48948293 | NA | NA |
| ENSG000002 | 5.93417378 | -0.33626976 | 0.36624496 | -0.9181553 | 0.35853755 | 0.48949755 | LINC01489 | 101928340 |
| ENSG000002 | 3.60583138 | -0.347443117 | 0.37845553 | -0.9180553 | 0.3585899 | 0.48955253 | NA | NA |
| ENSG000002 | 6.351424 | -0.340350668 | 0.37076408 | -0.917971 | 0.35863405 | 0.48959633 | NA | NA |
| ENSG000001 | 105.760607 | 0.1478849 | 0.16110894 | 0.91791862 | 0.35866147 | 0.48961728 | NA | NA |
| ENSG000001 | 4.9227303 | -0.343657705 | 0.37444224 | -0.9177856 | 0.35873111 | 0.48969586 | KCNK7 | 10089 |
| ENSG000001 | 19.6643549 | -0.258330209 | 0.28156722 | -0.9174726 | 0.35889504 | 0.48990314 | ERICH6 | 131831 |
| ENSG000002 | 5.66922939 | -0.339618211 | 0.37018306 | -0.9174332 | 0.35891571 | 0.48991486 | NA | NA |
| ENSG000002 | 4.99216443 | -0.344286813 | 0.37532237 | -0.9173096 | 0.35898043 | 0.48998672 | NA | NA |
| ENSG000002 | 35.6331537 | 0.227248548 | 0.24775295 | 0.9172385 | 0.35901767 | 0.49000457 | NA | NA |
| ENSG000002 | 3.37426476 | -0.347183472 | 0.37850527 | -0.9172487 | 0.35901232 | 0.49000457 | NA | NA |
| ENSG000002 | 4.59229752 | 0.344227256 | 0.37530739 | 0.91718752 | 0.35904438 | 0.49002453 | NA | NA |
| ENSG000002 | 2.59199313 | -0.34113169 | 0.37194402 | -0.9171587 | 0.35905949 | 0.49002866 | LINC01107 | 151171 |
| ENSG000002 | 44.8895661 | 0.211281679 | 0.23037561 | 0.91711828 | 0.35908066 | 0.49004106 | NA | NA |
| ENSG000001 | 6.20408159 | -0.334772999 | 0.36505248 | -0.9170544 | 0.35911412 | 0.49007022 | DIO3 | 1735 |
| ENSG000002 | 15.3291848 | -0.29173216 | 0.31821084 | -0.9167889 | 0.35925328 | 0.49024363 | LOC100129461 | 100129461 |
| ENSG000002 | 4.34919691 | 0.346542179 | 0.37802919 | 0.91670746 | 0.35929596 | 0.49028537 | NA | NA |
| ENSG000002 | 5.12604093 | -0.341392916 | 0.37243388 | -0.9166538 | 0.3593241 | 0.49030728 | LINC00896 | 150197 |
| ENSG000002 | 10.2335233 | -0.305597332 | 0.33341376 | -0.9165709 | 0.35936756 | 0.49035008 | LINC00934 | 144742 |
| ENSG000002 | 4.59788492 | -0.346868873 | 0.37850017 | -0.9164299 | 0.35944145 | 0.4904344 | NA | NA |
| ENSG000002 | 3.08551487 | -0.346139629 | 0.37779191 | -0.9162177 | 0.35955273 | 0.49056972 | NA | NA |
| ENSG000002 | 2.60121005 | -0.337137368 | 0.36805502 | -0.9159972 | 0.35966837 | 0.49071099 | NA | NA |
| ENSG000001 | 71.5930892 | 0.169392671 | 0.18493245 | 0.9159705 | 0.35968237 | 0.49071358 | ACOT11 | 26027 |

| | | | | | | | | |
|---|---|---|---|---|---|---|---|---|
| ENSG000001 | 71.993349 | -0.166125178 | 0.18137526 | -0.9159197 | 0.359709 | 0.49072649 | GPNMB | 10457 |
| ENSG000002 | 5.18163341 | 0.340482831 | 0.37174417 | 0.91590631 | 0.35971604 | 0.49072649 | SNORA80B | 100302743 |
| ENSG000002 | 24.6033445 | 0.241484784 | 0.26373053 | 0.9156497 | 0.35985065 | 0.49089362 | NA | NA |
| ENSG000002 | 5.26115931 | -0.343207871 | 0.37490434 | -0.9154545 | 0.35995306 | 0.49101681 | NA | NA |
| ENSG000002 | 6.76600758 | -0.339884332 | 0.37130771 | -0.9153711 | 0.35999687 | 0.49105702 | NA | NA |
| ENSG000002 | 3.59669658 | -0.346545067 | 0.37859205 | -0.9153522 | 0.36000676 | 0.49105702 | RN7SL181P | 106480492 |
| ENSG000002 | 35.6173848 | 0.218175793 | 0.2383798 | 0.91524446 | 0.36006331 | 0.49111763 | NA | NA |
| ENSG000002 | 7.99317696 | -0.324206126 | 0.35427244 | -0.9151322 | 0.36012222 | 0.49118147 | NA | NA |
| ENSG000002 | 10.3392464 | -0.317060087 | 0.34648588 | -0.9150736 | 0.36015299 | 0.49120691 | NA | NA |
| ENSG000002 | 3.41123547 | -0.346126462 | 0.37826515 | -0.9150366 | 0.36017241 | 0.49120858 | NA | NA |
| ENSG000002 | 5.93992255 | -0.335189052 | 0.36631677 | -0.9150251 | 0.36017844 | 0.49120858 | NA | NA |
| ENSG000002 | 2.61129092 | -0.338973337 | 0.3704782 | -0.9149616 | 0.36021178 | 0.49123753 | SQSTM1P1 | 106480677 |
| ENSG000001 | 63.7342336 | 0.181633467 | 0.1985301 | 0.91489131 | 0.36024869 | 0.49126751 | PRSS16 | 10279 |
| ENSG000001 | 4.98502089 | -0.343787264 | 0.3757757 | -0.9148736 | 0.36025799 | 0.49126751 | SIX2 | 10736 |
| ENSG000002 | 4.97370856 | -0.342670844 | 0.37460054 | -0.9147634 | 0.36031587 | 0.49132992 | NA | NA |
| ENSG000002 | 3.65948406 | -0.346239518 | 0.378561 | -0.9146201 | 0.36039108 | 0.49141594 | NA | NA |
| ENSG000001 | 203.809078 | -0.13489578 | 0.14752428 | -0.9143971 | 0.36050822 | 0.4915553 | CNKSR3 | 154043 |
| ENSG000002 | 4.01237056 | -0.345153695 | 0.37747918 | -0.9143648 | 0.36052517 | 0.4915553 | NA | NA |
| ENSG000002 | 5.9984033 | 0.337553826 | 0.36917099 | 0.91435632 | 0.36052965 | 0.4915553 | NA | NA |
| ENSG000002 | 3.66943984 | -0.346108025 | 0.37854224 | -0.9143181 | 0.36054972 | 0.49156613 | NA | NA |
| ENSG000002 | 14.9633622 | -0.287935739 | 0.31494505 | -0.9142412 | 0.36059012 | 0.49160469 | NA | NA |
| ENSG000002 | 2.29512891 | 0.334534737 | 0.3659659 | 0.91411451 | 0.36065668 | 0.4916789 | PRELID3BP8 | 106480243 |
| ENSG000002 | 3.3488722 | -0.345848479 | 0.3784082 | -0.9139561 | 0.36073992 | 0.49177585 | NA | NA |
| ENSG000002 | 5.64888063 | -0.339133845 | 0.37108481 | -0.9138985 | 0.36077019 | 0.49180058 | NA | NA |
| ENSG000002 | 5.18392841 | -0.341054153 | 0.37321911 | -0.9138175 | 0.36081275 | 0.49184207 | NA | NA |
| ENSG000002 | 88.8335831 | 0.173671558 | 0.19006511 | 0.9137477 | 0.36084943 | 0.49187554 | DKFZP434I0714 | 54553 |
| ENSG000001 | 75.8701996 | 0.165019247 | 0.18060441 | 0.91370554 | 0.36087159 | 0.49188921 | ROR2 | 4920 |
| ENSG000001 | 66.0216998 | 0.177201009 | 0.19395342 | 0.91362665 | 0.36091306 | 0.4919292 | TIRAP | 114609 |
| ENSG00000C | 51.7390201 | -0.194662519 | 0.21309641 | -0.913495 | 0.36098224 | 0.49199041 | CDH7 | 1005 |

| | | | | | | | | |
|---|---|---|---|---|---|---|---|---|
| ENSG000002 | 7.70496175 | 0.325060648 | 0.3558363 | 0.91351177 | 0.36097345 | 0.49199041 | NA | NA |
| ENSG000002 | 6.29116072 | -0.3397516 | 0.3719675 | -0.9133905 | 0.36103718 | 0.49204875 | ORM1 | 5004 |
| ENSG000002 | 3.37477309 | -0.345482162 | 0.37827488 | -0.9133098 | 0.36107961 | 0.49209005 | NA | NA |
| ENSG000002 | 3.13950833 | -0.344437314 | 0.37726056 | -0.9129958 | 0.36124474 | 0.49229854 | NA | NA |
| ENSG000002 | 12.9381838 | -0.295661995 | 0.32387272 | -0.9128956 | 0.36129743 | 0.4923538 | NA | NA |
| ENSG000002 | 3.64672388 | -0.345537892 | 0.37857056 | -0.9127437 | 0.36137735 | 0.49243814 | NA | NA |
| ENSG000002 | 6.49989534 | -0.334597081 | 0.3665886 | -0.9127318 | 0.3613836 | 0.49243814 | NA | NA |
| ENSG000001 | 90.9831266 | 0.154528714 | 0.16931167 | 0.9126879 | 0.36140671 | 0.49245307 | ELMOD3 | 84173 |
| ENSG000001 | 219.465447 | 0.12360759 | 0.13544423 | 0.91260875 | 0.36144835 | 0.49249326 | NDUFAF6 | 137682 |
| ENSG000001 | 5.47198574 | -0.339141584 | 0.37165324 | -0.9125215 | 0.36149424 | 0.49252701 | MAGEC1 | 9947 |
| ENSG000001 | 7.31983919 | -0.333147927 | 0.36508741 | -0.9125155 | 0.36149741 | 0.49252701 | CCDC154 | 645811 |
| ENSG000002 | 5.22369913 | -0.34106834 | 0.37378265 | -0.9124777 | 0.3615173 | 0.49253756 | NA | NA |
| ENSG000001 | 6.36956951 | 0.334292107 | 0.36637407 | 0.91243387 | 0.36154037 | 0.49255244 | KLK4 | 9622 |
| ENSG000001 | 50.8191052 | -0.190905442 | 0.20923531 | -0.9123959 | 0.36156033 | 0.4925631 | C4orf32 | 132720 |
| ENSG000002 | 4.93366462 | -0.34196857 | 0.37481982 | -0.9123546 | 0.3615821 | 0.4925762 | ZNF32-AS2 | 414208 |
| ENSG000002 | 45.2691026 | 0.196225023 | 0.21510174 | 0.91224283 | 0.3616409 | 0.49263975 | NA | NA |
| ENSG000002 | 2.56998551 | -0.337010227 | 0.36944325 | -0.9122111 | 0.36165761 | 0.49264597 | NA | NA |
| ENSG000001 | 449.526226 | 0.140170972 | 0.15366939 | 0.91215936 | 0.36168483 | 0.4926665 | EIF4E3 | 317649 |
| ENSG000002 | 37.7444854 | 0.216650442 | 0.23753019 | 0.91209646 | 0.36171794 | 0.49268878 | LINC00960 | 401074 |
| ENSG000002 | 3.1779016 | -0.343703973 | 0.37683446 | -0.9120821 | 0.36172549 | 0.49268878 | NA | NA |
| ENSG000002 | 4.94527122 | -0.344438378 | 0.37766716 | -0.9120157 | 0.36176046 | 0.49270686 | NA | NA |
| ENSG000002 | 5.49746262 | -0.339078848 | 0.3717926 | -0.9120107 | 0.36176306 | 0.49270686 | LINC00269 | 100996279 |
| ENSG000002 | 36.1861908 | -0.233492194 | 0.25608519 | -0.9117755 | 0.36188692 | 0.49285899 | NA | NA |
| ENSG000001 | 2.31763428 | -0.330338161 | 0.36238971 | -0.911555 | 0.36200301 | 0.49300054 | CGB2 | 114336 |
| ENSG000002 | 4.65269133 | -0.342900349 | 0.37619343 | -0.9115001 | 0.36203193 | 0.49302337 | AADACL4 | 343066 |
| ENSG000002 | 12.2152477 | -0.290485721 | 0.31871125 | -0.9114385 | 0.36206435 | 0.49303441 | NA | NA |
| ENSG000002 | 3.11027829 | -0.343051984 | 0.37637729 | -0.9114577 | 0.36205425 | 0.49303441 | NA | NA |
| ENSG000002 | 3.07206184 | -0.343943852 | 0.37738007 | -0.9113991 | 0.36208513 | 0.49304615 | NA | NA |
| ENSG00000C | 89.6009117 | -0.159031519 | 0.17451749 | -0.9112641 | 0.36215626 | 0.49311791 | LRCH4 | 4034 |

| | | | | | | | | | |
|---|---|---|---|---|---|---|---|---|---|
| ENSG000001 | 22.5143666 | -0.247153379 | 0.27122371 | -0.9112529 | 0.36216216 | 0.49311791 | ADAP2 | | 55803 |
| ENSG000002 | 18.1500369 | -0.271192784 | 0.297634 | -0.911162 | 0.36221004 | 0.49314999 | RAD51AP2 | | 729475 |
| ENSG000002 | 3.61692876 | -0.344883234 | 0.37849975 | -0.9111848 | 0.36219799 | 0.49314999 | NA | NA | |
| ENSG000002 | 4.62778804 | -0.342931319 | 0.37638398 | -0.9111209 | 0.36223167 | 0.49316288 | NA | NA | |
| ENSG000002 | 6.65896074 | 0.330289754 | 0.36252245 | 0.91108772 | 0.36224915 | 0.49317013 | NA | NA | |
| ENSG000001 | 2.66215206 | -0.33586814 | 0.36865738 | -0.9110577 | 0.36226498 | 0.49317512 | IFNA21 | | 3452 |
| ENSG000001 | 64.9849556 | 0.176087722 | 0.19330809 | 0.91091751 | 0.36233884 | 0.4932375 | ZNF222 | | 7673 |
| ENSG000002 | 6.65889763 | -0.328147093 | 0.36023555 | -0.9109237 | 0.36233559 | 0.4932375 | NA | NA | |
| ENSG000002 | 10.2317553 | -0.305180086 | 0.33503084 | -0.9109015 | 0.36234729 | 0.4932375 | NA | NA | |
| ENSG000002 | 4.3070207 | -0.344789971 | 0.37854255 | -0.9108355 | 0.36238208 | 0.49326829 | NA | NA | |
| ENSG000001 | 3.42573495 | -0.342960074 | 0.37656988 | -0.9107475 | 0.36242843 | 0.49329827 | PTGDS | | 5730 |
| ENSG000002 | 6.20260772 | -0.332981202 | 0.36560973 | -0.9107558 | 0.36242403 | 0.49329827 | NA | NA | |
| ENSG000002 | 4.29752835 | -0.344654049 | 0.37845378 | -0.9106899 | 0.36245877 | 0.49332301 | NA | NA | |
| ENSG000002 | 3.57557869 | -0.344742165 | 0.37858526 | -0.9106064 | 0.36250279 | 0.49336057 | UGT1A8 | | 54576 |
| ENSG000002 | 14.7865816 | -0.286322813 | 0.31443611 | -0.9105914 | 0.3625107 | 0.49336057 | NA | NA | |
| ENSG000002 | 22.3203336 | 0.255530507 | 0.28070404 | 0.91032001 | 0.36265377 | 0.49352215 | EWSAT1 | | 283673 |
| ENSG000002 | 5.95739359 | -0.334408733 | 0.3673472 | -0.9103342 | 0.36264626 | 0.49352215 | NA | NA | |
| ENSG000002 | 5.17183845 | -0.340326413 | 0.3739635 | -0.9100525 | 0.36279482 | 0.49368097 | NA | NA | |
| ENSG000002 | 3.57964953 | -0.344517186 | 0.3785672 | -0.9100556 | 0.36279321 | 0.49368097 | NA | NA | |
| ENSG000002 | 5.87449304 | -0.340163536 | 0.37386131 | -0.9098656 | 0.3628934 | 0.49379613 | NA | NA | |
| ENSG000002 | 5.38307646 | -0.338128986 | 0.37163326 | -0.9098459 | 0.36290381 | 0.49379613 | NA | NA | |
| ENSG000002 | 3.96139462 | -0.343839067 | 0.3779354 | -0.9097826 | 0.36293715 | 0.49382493 | NA | NA | |
| ENSG000001 | 10.267101 | -0.319430262 | 0.35115196 | -0.9096639 | 0.36299979 | 0.49389358 | ANKRD1 | | 27063 |
| ENSG000001 | 7.43868057 | -0.335282146 | 0.36864816 | -0.9094909 | 0.36309105 | 0.49398461 | SASH3 | | 54440 |
| ENSG000002 | 8.93521376 | -0.312677942 | 0.34379232 | -0.9094966 | 0.36308805 | 0.49398461 | NA | NA | |
| ENSG000002 | 4.20190125 | -0.344058823 | 0.37835649 | -0.9093509 | 0.36316491 | 0.49403537 | TPRG1-AS1 | | 100874043 |
| ENSG000002 | 4.9228997 | -0.340596708 | 0.37454866 | -0.9093523 | 0.36316417 | 0.49403537 | NA | NA | |
| ENSG000002 | 6.41600495 | -0.329644599 | 0.36248701 | -0.909397 | 0.36314061 | 0.49403537 | BMS1P4 | | 729096 |
| ENSG000002 | 11.7220744 | -0.296233615 | 0.32585533 | -0.9090955 | 0.36329971 | 0.49420217 | NA | NA | |

| | | | | | | | | |
|---|---|---|---|---|---|---|---|---|
| ENSG000001 | 9.96191219 | -0.306040741 | 0.33670542 | -0.9089273 | 0.36338852 | 0.49430639 | CD300A | 11314 |
| ENSG000001 | 40.6706909 | 0.204196713 | 0.22466611 | 0.90888969 | 0.36340836 | 0.4943168 | KCTD13 | 253980 |
| ENSG000002 | 3.07854726 | -0.34291857 | 0.37734467 | -0.9087675 | 0.36347286 | 0.49438795 | NA | NA |
| ENSG000002 | 2.89914715 | -0.340125636 | 0.3743197 | -0.9086501 | 0.36353485 | 0.49445082 | NA | NA |
| ENSG000002 | 9.0946354 | -0.315044473 | 0.34672327 | -0.9086338 | 0.36354347 | 0.49445082 | NA | NA |
| ENSG000002 | 2.85887996 | -0.340402291 | 0.37464438 | -0.9086011 | 0.36356073 | 0.49445771 | NA | NA |
| ENSG000001 | 363.265203 | 0.110087587 | 0.12119298 | 0.90836607 | 0.36368484 | 0.49459416 | SYT11 | 23208 |
| ENSG000001 | 70.8073089 | 0.171316191 | 0.18859842 | 0.90836493 | 0.36368545 | 0.49459416 | NA | NA |
| ENSG000002 | 5.73061634 | -0.340568382 | 0.37498014 | -0.9082305 | 0.36375648 | 0.49467417 | NA | NA |
| ENSG000002 | 2.51992543 | 0.337212094 | 0.37140409 | 0.90793856 | 0.36391069 | 0.49486728 | NA | NA |
| ENSG000002 | 21.1822525 | -0.252432659 | 0.27803711 | -0.90791 | 0.36392579 | 0.49487123 | NA | NA |
| ENSG000002 | 5.44607167 | -0.338341469 | 0.37280531 | -0.9075554 | 0.36411317 | 0.49510942 | NA | NA |
| ENSG000002 | 2.86627954 | -0.340379581 | 0.37514351 | -0.9073317 | 0.36423144 | 0.49525363 | NA | NA |
| ENSG000002 | 3.3484237 | -0.343426669 | 0.37852225 | -0.9072826 | 0.36425735 | 0.49527226 | CACNA1G-AS1 | 253962 |
| ENSG000002 | 28.1743511 | 0.244932046 | 0.26997649 | 0.90723471 | 0.36428269 | 0.49529011 | NA | NA |
| ENSG000002 | 41.3439029 | 0.204628655 | 0.22557479 | 0.90714329 | 0.36433103 | 0.49533922 | LOC101930370 | 101930370 |
| ENSG000001 | 22.7062825 | 0.239081411 | 0.26358046 | 0.90705287 | 0.36437884 | 0.49538761 | LRRC3B | 116135 |
| ENSG000002 | 28.2177407 | -0.23173004 | 0.25549128 | -0.9069978 | 0.36440794 | 0.49541057 | NA | NA |
| ENSG000002 | 14.9215882 | 0.291796376 | 0.32172937 | 0.90696219 | 0.36442679 | 0.49541959 | SEC1P | 653677 |
| ENSG000002 | 3.32485734 | -0.342865534 | 0.37805803 | -0.9069125 | 0.36445309 | 0.49543873 | NA | NA |
| ENSG000002 | 4.75150324 | -0.342767568 | 0.3779688 | -0.9068674 | 0.36447695 | 0.49545455 | NA | NA |
| ENSG000001 | 10.9153513 | -0.299693762 | 0.33056745 | -0.906604 | 0.36461626 | 0.49554426 | AMDHD1 | 144193 |
| ENSG000001 | 19.3867423 | -0.254624334 | 0.28082138 | -0.9067128 | 0.3645587 | 0.49554426 | GPR39 | 2863 |
| ENSG000002 | 28.6521681 | -0.23363306 | 0.25768901 | -0.9066474 | 0.36459331 | 0.49554426 | ANAPC1P1 | 100286979 |
| ENSG000002 | 3.64835476 | -0.343248347 | 0.37858932 | -0.9066509 | 0.36459144 | 0.49554426 | NA | NA |
| ENSG000002 | 6.49861195 | -0.332370396 | 0.36660553 | -0.9066159 | 0.36460996 | 0.49554426 | RN7SL762P | 106479500 |
| ENSG000002 | 15.2678004 | 0.272817301 | 0.30089162 | 0.90669624 | 0.36456745 | 0.49554426 | NA | NA |
| ENSG000002 | 30.7881089 | 0.224614488 | 0.24776316 | 0.90656934 | 0.36463458 | 0.49555255 | NA | NA |
| ENSG000001 | 3.66184004 | -0.343129825 | 0.37859203 | -0.9063313 | 0.3647605 | 0.49569045 | NA | NA |

| | | | | | | | | |
|---|---|---|---|---|---|---|---|---|
| ENSG000002 | 4.80770112 | -0.342294172 | 0.37766845 | -0.9063351 | 0.36475851 | 0.49569045 | NA | NA |
| ENSG000001 | 85.2601291 | -0.168948741 | 0.1864258 | -0.9062519 | 0.36480253 | 0.49573095 | CPEB2 | 132864 |
| ENSG000002 | 5.28509705 | -0.339303399 | 0.37446212 | -0.9061087 | 0.3648783 | 0.4958173 | LOC101927866 | 101927866 |
| ENSG00000C | 48.2577309 | -0.190844101 | 0.21062868 | -0.9060689 | 0.36489936 | 0.49582931 | ACPP | 55 |
| ENSG000002 | 12.9472177 | -0.289336417 | 0.3193522 | -0.9060104 | 0.36493035 | 0.4958548 | NA | NA |
| ENSG000001 | 50.3198894 | -0.190167084 | 0.20991408 | -0.9059282 | 0.36497385 | 0.49589258 | CYP2J2 | 1573 |
| ENSG000002 | 10.9440971 | -0.298580342 | 0.32959102 | -0.9059117 | 0.36498262 | 0.49589258 | NA | NA |
| ENSG000002 | 2.83944502 | -0.339154294 | 0.37447843 | -0.9056711 | 0.36510996 | 0.49604899 | C10orf62 | 414157 |
| ENSG000002 | 3.74750849 | 0.342858115 | 0.37859087 | 0.90561643 | 0.36513891 | 0.49607169 | NA | NA |
| ENSG000001 | 103.96586 | 0.156840865 | 0.17321015 | 0.90549466 | 0.36520339 | 0.49614267 | DEPDC7 | 91614 |
| ENSG000001 | 8.28832795 | -0.320503909 | 0.35397345 | -0.9054462 | 0.36522906 | 0.49614431 | SOST | 50964 |
| ENSG000002 | 2.80380417 | -0.339605968 | 0.37506722 | -0.9054536 | 0.36522512 | 0.49614431 | NA | NA |
| ENSG000001 | 49.6874249 | 0.186290788 | 0.20577692 | 0.90530456 | 0.36530406 | 0.4962142 | ZC3HAV1L | 92092 |
| ENSG000002 | 3.64667181 | -0.342740568 | 0.37859218 | -0.9053028 | 0.36530498 | 0.4962142 | NA | NA |
| ENSG000002 | 2.86992562 | -0.338983658 | 0.37448959 | -0.9051885 | 0.36536556 | 0.49627985 | NA | NA |
| ENSG000001 | 22.0278263 | 0.246118159 | 0.27190945 | 0.90514751 | 0.36538725 | 0.49628274 | PRIMA1 | 145270 |
| ENSG000002 | 3.74784021 | 0.342555533 | 0.37846084 | 0.90512807 | 0.36539754 | 0.49628274 | RNU6-290P | 106481270 |
| ENSG000002 | 13.2761657 | 0.284639905 | 0.31447923 | 0.90511513 | 0.3654044 | 0.49628274 | WASH5P | 375690 |
| ENSG000002 | 5.1919533 | -0.338045396 | 0.37351655 | -0.9050346 | 0.36544707 | 0.49632407 | AIRN | 100271873 |
| ENSG000002 | 2.66127139 | -0.334253762 | 0.36935224 | -0.9049729 | 0.36547975 | 0.49633521 | NA | NA |
| ENSG000002 | 9.98508921 | -0.317709663 | 0.35106497 | -0.9049882 | 0.36547163 | 0.49633521 | NA | NA |
| ENSG000001 | 153.739089 | 0.134525657 | 0.1486727 | 0.90484437 | 0.36554784 | 0.49641105 | C11orf80 | 79703 |
| ENSG000001 | 5.43120513 | -0.336003189 | 0.37136366 | -0.9047821 | 0.36558085 | 0.49643925 | SPINT4 | 391253 |
| ENSG000002 | 9.66669713 | -0.305667066 | 0.33786641 | -0.904698 | 0.3656254 | 0.49648312 | NA | NA |
| ENSG000002 | 4.90825244 | -0.339133123 | 0.37494163 | -0.9044958 | 0.36573257 | 0.49661202 | NA | NA |
| ENSG000001 | 3.15442065 | -0.340316047 | 0.37630571 | -0.9043606 | 0.36580424 | 0.49669271 | KRTAP19-2 | 337969 |
| ENSG000001 | 81.4910769 | -0.172148195 | 0.19035973 | -0.904331 | 0.36581994 | 0.49669739 | PDE10A | 10846 |
| ENSG000001 | 85.1513094 | -0.152629515 | 0.16880273 | -0.9041887 | 0.36589538 | 0.49678319 | MASP1 | 5648 |
| ENSG000002 | 3.8702656 | -0.341948135 | 0.37830588 | -0.9038933 | 0.366052 | 0.4969792 | NA | NA |

| | | | | | | | | |
|---|---|---|---|---|---|---|---|---|
| ENSG000002 | 5.4562045 | -0.337076446 | 0.37294389 | -0.9038262 | 0.36608759 | 0.49701087 | NA | NA |
| ENSG000002 | 3.05324753 | -0.34123279 | 0.37759717 | -0.9036953 | 0.36615699 | 0.49708846 | NA | NA |
| ENSG000001 | 95.3295556 | 0.168250239 | 0.18618652 | 0.90366498 | 0.36617307 | 0.49709364 | GUSBP1 | 728411 |
| ENSG000002 | 6.65830422 | -0.326330551 | 0.36124293 | -0.9033549 | 0.36633759 | 0.49729824 | NA | NA |
| ENSG000002 | 51.4463183 | 0.186516347 | 0.20647536 | 0.90333465 | 0.36634831 | 0.49729824 | NA | NA |
| ENSG000002 | 168.839116 | 0.158808323 | 0.17584008 | 0.90314067 | 0.36645124 | 0.49742132 | PCDHA10 | 56139 |
| ENSG000001 | 8.60713431 | 0.312675348 | 0.34629782 | 0.90290879 | 0.36657431 | 0.49757171 | MAGEB3 | 4114 |
| ENSG000002 | 4.70107949 | -0.340487909 | 0.37711633 | -0.9028723 | 0.36659365 | 0.4975813 | SLCO1B7 | 338821 |
| ENSG000002 | 3.12906154 | 0.340928301 | 0.3776155 | 0.90284509 | 0.36660812 | 0.49758429 | RN7SL230P | 106479306 |
| ENSG000002 | 11.4292168 | -0.293700122 | 0.32534055 | -0.9027467 | 0.36666032 | 0.49763849 | NA | NA |
| ENSG000002 | 5.1658428 | -0.338010829 | 0.37446519 | -0.9026495 | 0.36671192 | 0.49769186 | NA | NA |
| ENSG000002 | 33.5394702 | -0.218133982 | 0.24167546 | -0.9025905 | 0.36674325 | 0.49771772 | NA | NA |
| ENSG000002 | 3.14854312 | -0.339564251 | 0.37623124 | -0.9025413 | 0.36676937 | 0.49773651 | NA | NA |
| ENSG000002 | 3.34814295 | -0.341461636 | 0.37836368 | -0.9024694 | 0.36680758 | 0.49777171 | NA | NA |
| ENSG000002 | 8.72537828 | -0.314220354 | 0.34819613 | -0.9024235 | 0.36683196 | 0.49778814 | NA | NA |
| ENSG000002 | 8.93896745 | -0.316847688 | 0.35112649 | -0.9023748 | 0.36685782 | 0.49780657 | NA | NA |
| ENSG000001 | 6.45537131 | -0.328847645 | 0.36448419 | -0.9022275 | 0.36693605 | 0.49789606 | LUZP4 | 51213 |
| ENSG000002 | 37.1749418 | 0.202487479 | 0.2244713 | 0.90206399 | 0.36702288 | 0.49799476 | NA | NA |
| ENSG000002 | 60.5117318 | 0.189236618 | 0.2097864 | 0.90204427 | 0.36703335 | 0.49799476 | NA | NA |
| ENSG000002 | 2.53396054 | -0.333012828 | 0.36921851 | -0.9019397 | 0.36708891 | 0.49805348 | NA | NA |
| ENSG000002 | 8.25363393 | -0.316771293 | 0.35124065 | -0.901864 | 0.36712912 | 0.49809137 | RP9P | 441212 |
| ENSG000001 | 18.9425226 | -0.257176026 | 0.28523097 | -0.9016413 | 0.36724744 | 0.49821856 | TMPRSS3 | 64699 |
| ENSG000002 | 8.94911763 | 0.318731135 | 0.35349848 | 0.90164783 | 0.36724397 | 0.49821856 | NA | NA |
| ENSG000002 | 13.8472408 | -0.27709677 | 0.30734079 | -0.9015945 | 0.3672723 | 0.49823562 | LOC100287042 | 100287042 |
| ENSG000002 | 2.87172011 | -0.33733616 | 0.37417311 | -0.901551 | 0.36729541 | 0.49824155 | USP17L4 | 645402 |
| ENSG000002 | 29.4408743 | -0.242927074 | 0.26945788 | -0.9015401 | 0.36730125 | 0.49824155 | NA | NA |
| ENSG000002 | 13.2780027 | 0.289860432 | 0.32153967 | 0.90147642 | 0.36733507 | 0.49827076 | LOC171391 | 171391 |
| ENSG000002 | 5.24871921 | -0.337047369 | 0.37398659 | -0.9012285 | 0.36746685 | 0.49843284 | NA | NA |
| ENSG000002 | 7.69930428 | -0.323345775 | 0.35880241 | -0.9011806 | 0.3674923 | 0.49845069 | NA | NA |

| | | | | | | | | |
|---|---|---|---|---|---|---|---|---|
| ENSG000001 | 3.59301859 | -0.340929999 | 0.37854994 | -0.9006209 | 0.3677899 | 0.49880429 | GML | 2765 |
| ENSG000002 | 21.7521032 | 0.254030146 | 0.28205771 | 0.90063181 | 0.36778412 | 0.49880429 | LOC101930114 | 101930114 |
| ENSG000002 | 4.22996115 | -0.340932156 | 0.37853711 | -0.9006572 | 0.36777064 | 0.49880429 | NA | NA |
| ENSG000002 | 9.48002088 | -0.309224451 | 0.34336301 | -0.9005759 | 0.36781385 | 0.49882009 | NA | NA |
| ENSG000002 | 6.14408173 | -0.328730002 | 0.36505913 | -0.9004843 | 0.36786259 | 0.49886951 | NA | NA |
| ENSG000002 | 7.15009171 | -0.320307141 | 0.35574996 | -0.9003716 | 0.36792255 | 0.49893414 | NA | NA |
| ENSG000002 | 8.10015911 | -0.32237386 | 0.35811698 | -0.9001915 | 0.36801835 | 0.49904736 | NA | NA |
| ENSG000002 | 8.51959488 | -0.316629532 | 0.35176933 | -0.9001056 | 0.36806408 | 0.49909267 | NA | NA |
| ENSG000001 | 87.043748 | -0.164910073 | 0.18323196 | -0.9000071 | 0.36811646 | 0.49914701 | HAPLN1 | 1404 |
| ENSG000001 | 5.40352379 | -0.334454581 | 0.37163796 | -0.8999473 | 0.3681483 | 0.4991568 | SCGB1C1 | 147199 |
| ENSG000002 | 3.83144833 | -0.340236758 | 0.37805377 | -0.8999692 | 0.36813663 | 0.4991568 | NA | NA |
| ENSG000002 | 3.36299798 | -0.340188358 | 0.37806653 | -0.8998108 | 0.36822094 | 0.4992386 | NA | NA |
| ENSG000002 | 4.66299944 | -0.338830463 | 0.37662 | -0.8996614 | 0.36830049 | 0.49932977 | NA | NA |
| ENSG000002 | 13.9111498 | -0.280075824 | 0.3113846 | -0.899453 | 0.36841141 | 0.49946344 | NA | NA |
| ENSG000002 | 3.89847529 | -0.340314842 | 0.37842554 | -0.8992914 | 0.36849746 | 0.49955436 | NA | NA |
| ENSG000002 | 7.96760355 | -0.319456895 | 0.35523598 | -0.8992808 | 0.36850311 | 0.49955436 | NA | NA |
| ENSG000002 | 6.89312069 | -0.32213716 | 0.35839028 | -0.8988446 | 0.36873545 | 0.49985262 | NA | NA |
| ENSG000002 | 2.91361606 | 0.334940763 | 0.37264912 | 0.89881001 | 0.36875387 | 0.49986087 | NA | NA |
| ENSG000000 | 74.3400314 | -0.162937186 | 0.18128846 | -0.8987731 | 0.36877356 | 0.49987085 | RAPGEF3 | 10411 |
| ENSG000002 | 7.1844205 | 0.327792233 | 0.36475695 | 0.89865933 | 0.36883414 | 0.49991956 | NA | NA |
| ENSG000002 | 3.86218918 | -0.339893962 | 0.3782157 | -0.8986776 | 0.36882444 | 0.49991956 | NA | NA |
| ENSG000000 | 5.26896328 | -0.337136038 | 0.3752196 | -0.8985033 | 0.3689173 | 0.5000153 | FGF22 | 27006 |
| ENSG000002 | 2.93367755 | -0.332447615 | 0.37001094 | -0.8984805 | 0.36892944 | 0.5000153 | NA | NA |
| ENSG000002 | 3.1389154 | -0.334309687 | 0.372116 | -0.8984018 | 0.36897138 | 0.50005542 | RNU6-1331P | 106480147 |
| ENSG000001 | 54.9961908 | -0.194646694 | 0.21668076 | -0.8983109 | 0.36901981 | 0.50010434 | FOXB1 | 27023 |
| ENSG000002 | 2.89373829 | -0.334601219 | 0.37259515 | -0.8980289 | 0.36917013 | 0.50028645 | NA | NA |
| ENSG000002 | 14.6023452 | -0.275676841 | 0.30698551 | -0.8980126 | 0.36917886 | 0.50028645 | DSG2-AS1 | 100652770 |
| ENSG000002 | 3.83617151 | -0.339587536 | 0.37817379 | -0.8979669 | 0.36920321 | 0.50030273 | MCIDAS | 345643 |
| ENSG000002 | 5.12609354 | -0.334737387 | 0.37281909 | -0.8978547 | 0.369263 | 0.50036704 | LOC101928716 | 101928716 |

| | | | | | | | | |
|---|---|---|---|---|---|---|---|---|
| ENSG000001 | 210.971479 | 0.122257389 | 0.13617411 | 0.89780201 | 0.36929111 | 0.50038841 | DPP10 | 57628 |
| ENSG000002 | 8.76421198 | -0.318179933 | 0.35445619 | -0.8976566 | 0.36936866 | 0.50047676 | NA | NA |
| ENSG000002 | 3.06509477 | -0.338541287 | 0.37722709 | -0.8974469 | 0.36948049 | 0.50059483 | NA | NA |
| ENSG000002 | 3.87493726 | -0.33962114 | 0.37842465 | -0.8974604 | 0.3694733 | 0.50059483 | NA | NA |
| ENSG000002 | 2.81345994 | -0.335997857 | 0.37446155 | -0.8972827 | 0.36956811 | 0.50069682 | NA | NA |
| ENSG000002 | 77.5682669 | 0.170384457 | 0.18998942 | 0.89681026 | 0.36982017 | 0.50102158 | NA | NA |
| ENSG000001 | 6.96915862 | -0.330193144 | 0.36829864 | -0.8965364 | 0.36996634 | 0.50120286 | CYP2D6 | 1565 |
| ENSG000002 | 3.67366456 | -0.338911548 | 0.37807156 | -0.8964217 | 0.37002758 | 0.50124514 | LINC01251 | 101929688 |
| ENSG000002 | 3.1633515 | 0.338845941 | 0.37800394 | 0.89640848 | 0.37003464 | 0.50124514 | NA | NA |
| ENSG000002 | 6.02883508 | 0.328102765 | 0.36601385 | 0.89642173 | 0.37002757 | 0.50124514 | NA | NA |
| ENSG000002 | 3.33927626 | -0.339007958 | 0.3782344 | -0.8962906 | 0.37009755 | 0.50131362 | NA | NA |
| ENSG000002 | 3.09317449 | -0.337952326 | 0.37710641 | -0.8961723 | 0.37016073 | 0.50138245 | NA | NA |
| ENSG000002 | 5.42524478 | 0.331854954 | 0.3703579 | 0.89603855 | 0.37023218 | 0.50146015 | NA | NA |
| ENSG000002 | 3.35190004 | -0.338548845 | 0.37783685 | -0.8960186 | 0.37024282 | 0.50146015 | NA | NA |
| ENSG000001 | 56.338159 | 0.192342128 | 0.21467361 | 0.89597473 | 0.37026626 | 0.50147514 | COX16 | 51241 |
| ENSG000002 | 3.15944538 | 0.337617165 | 0.37688461 | 0.89581044 | 0.37035402 | 0.50157724 | NA | NA |
| ENSG000001 | 5.30989628 | 0.337460779 | 0.37672756 | 0.89576875 | 0.37037628 | 0.50159064 | SIK1 | 150094 |
| ENSG000001 | 6.66060038 | -0.323350123 | 0.36101852 | -0.8956608 | 0.37043397 | 0.50165202 | GOLGA8M | 653720 |
| ENSG000002 | 8.98956452 | -0.310113601 | 0.34626129 | -0.8956057 | 0.37046337 | 0.50167508 | NA | NA |
| ENSG000002 | 19.8142942 | 0.256785047 | 0.28673562 | 0.89554638 | 0.37049508 | 0.50170127 | NA | NA |
| ENSG000002 | 2.6082705 | -0.330522383 | 0.36909083 | -0.8955042 | 0.37051763 | 0.50171504 | NA | NA |
| ENSG000002 | 6.42329242 | -0.325949027 | 0.36413368 | -0.8951356 | 0.37071459 | 0.50196499 | NA | NA |
| ENSG000002 | 11.2585388 | -0.294266302 | 0.32877422 | -0.8950407 | 0.37076535 | 0.50201695 | NA | NA |
| ENSG000002 | 3.68542488 | 0.338786126 | 0.37858145 | 0.89488306 | 0.3708496 | 0.50211427 | NA | NA |
| ENSG000001 | 45.9373027 | 0.191694017 | 0.21424859 | 0.89472709 | 0.370933 | 0.50221041 | NANOS1 | 340719 |
| ENSG000002 | 15.2854089 | 0.271673886 | 0.3036574 | 0.89467238 | 0.37096225 | 0.50223324 | NA | NA |
| ENSG000002 | 3.84707357 | -0.338274297 | 0.37814173 | -0.8945701 | 0.37101692 | 0.50229049 | NA | NA |
| ENSG000002 | 5.19120366 | -0.334756857 | 0.37424271 | -0.8944913 | 0.37105907 | 0.50233078 | NA | NA |
| ENSG000001 | 20.4519613 | -0.249686341 | 0.27919336 | -0.8943133 | 0.37115427 | 0.50239258 | TMEM51-AS1 | 200197 |

| | | | | | | | | |
|---|---|---|---|---|---|---|---|---|
| ENSG000001 | 31.1541277 | -0.219195831 | 0.2450982 | -0.8943184 | 0.37115156 | 0.50239258 | ZFP2 | 80108 |
| ENSG000002 | 13.1663785 | -0.279917345 | 0.3129963 | -0.8943152 | 0.37115327 | 0.50239258 | WWTR1-AS1 | 100128025 |
| ENSG000002 | 5.12437063 | -0.334467754 | 0.37397593 | -0.8943564 | 0.37113125 | 0.50239258 | NA | NA |
| ENSG000002 | 53.9800568 | 0.180954471 | 0.20235249 | 0.89425376 | 0.37118613 | 0.50241893 | HLA-DMB | 3109 |
| ENSG000001 | 14.4945358 | -0.282103685 | 0.31549472 | -0.8941629 | 0.37123472 | 0.5024437 | CRHR2 | 1395 |
| ENSG000001 | 119.490478 | 0.165112736 | 0.18465886 | 0.89415008 | 0.3712416 | 0.5024437 | TMPPE | 643853 |
| ENSG000002 | 3.65708461 | -0.338531554 | 0.37858902 | -0.8941927 | 0.37121877 | 0.5024437 | LINC00445 | 100507114 |
| ENSG000002 | 2.81622053 | -0.335379578 | 0.3751168 | -0.8940671 | 0.371286 | 0.50248702 | NA | NA |
| ENSG000001 | 39.5012261 | 0.217509364 | 0.24330402 | 0.89398178 | 0.37133164 | 0.50249848 | C11orf70 | 85016 |
| ENSG000002 | 86.7250395 | 0.163419625 | 0.18279657 | 0.89399722 | 0.37132338 | 0.50249848 | NA | NA |
| ENSG000002 | 3.37686042 | -0.337722987 | 0.37776085 | -0.8940127 | 0.37131511 | 0.50249848 | NA | NA |
| ENSG000002 | 9.45141968 | -0.320575359 | 0.3586669 | -0.8937969 | 0.37143057 | 0.50261558 | NA | NA |
| ENSG000002 | 3.73885955 | -0.337612456 | 0.37775476 | -0.8937345 | 0.37146398 | 0.50264401 | NA | NA |
| ENSG000002 | 2.55789883 | -0.331435629 | 0.37088839 | -0.8936263 | 0.37152186 | 0.50270556 | NA | NA |
| ENSG00000C | 56.405858 | 0.184538872 | 0.2065118 | 0.89359965 | 0.37153614 | 0.5027081 | OSBPL7 | 114881 |
| ENSG000002 | 7.16529219 | -0.318637043 | 0.3566123 | -0.8935111 | 0.37158354 | 0.50275547 | MIR5707 | 100847032 |
| ENSG000002 | 2.65870743 | -0.329790996 | 0.36914016 | -0.8934032 | 0.37164129 | 0.50281683 | NA | NA |
| ENSG000002 | 3.06235401 | -0.336046485 | 0.37615614 | -0.8933697 | 0.37165924 | 0.50282433 | NA | NA |
| ENSG000002 | 12.6232731 | -0.283185433 | 0.31701423 | -0.8932893 | 0.37170225 | 0.50286575 | NA | NA |
| ENSG000001 | 43.7159758 | -0.193626834 | 0.21676531 | -0.8932556 | 0.37172029 | 0.50287338 | RBKS | 64080 |
| ENSG000001 | 12.0287454 | -0.295389721 | 0.3307137 | -0.8931887 | 0.37175615 | 0.50289536 | FAM159A | 348378 |
| ENSG000002 | 3.0973095 | -0.335191097 | 0.37528339 | -0.8931679 | 0.37176724 | 0.50289536 | NA | NA |
| ENSG000002 | 22.0848941 | -0.243371989 | 0.27249885 | -0.893112 | 0.3717972 | 0.50289536 | DPY19L2P3 | 442524 |
| ENSG000002 | 11.1638013 | -0.294201612 | 0.32940407 | -0.8931329 | 0.37178601 | 0.50289536 | CACNA1C-AS1 | 100652846 |
| ENSG000002 | 3.69389565 | -0.337384524 | 0.37776505 | -0.8931068 | 0.37179999 | 0.50289536 | NA | NA |
| ENSG000002 | 5.39210711 | -0.331981951 | 0.37172437 | -0.8930863 | 0.37181095 | 0.50289536 | NA | NA |
| ENSG000002 | 3.63685243 | -0.337932894 | 0.37839816 | -0.8930617 | 0.37182415 | 0.50289644 | NA | NA |
| ENSG000002 | 4.72938967 | -0.337853334 | 0.37833164 | -0.8930084 | 0.37185267 | 0.50291825 | NA | NA |
| ENSG000001 | 11.6434366 | -0.288476767 | 0.32308709 | -0.8928762 | 0.37192349 | 0.50293576 | SLC6A2 | 6530 |

| | | | | | | | | |
|---|---|---|---|---|---|---|---|---|
| ENSG000001 | 3.82663287 | -0.337657891 | 0.37817206 | -0.8928684 | 0.37192764 | 0.50293576 | NA | NA |
| ENSG000002 | 12.457402 | -0.285974693 | 0.32026855 | -0.8929216 | 0.37189918 | 0.50293576 | NA | NA |
| ENSG000002 | 42.073792 | 0.202127392 | 0.22637871 | 0.89287278 | 0.3719253 | 0.50293576 | NA | NA |
| ENSG000002 | 3.42595465 | -0.337200644 | 0.37762089 | -0.8929608 | 0.37187817 | 0.50293576 | NA | NA |
| ENSG000000 | 53.8748423 | 0.178639631 | 0.20009496 | 0.89277429 | 0.37197806 | 0.50298717 | ZNF213 | 7760 |
| ENSG000001 | 573.431653 | 0.121747725 | 0.13639867 | 0.89258731 | 0.37207822 | 0.50310583 | CDH2 | 1000 |
| ENSG000002 | 5.78178514 | -0.334176264 | 0.37441434 | -0.8925306 | 0.37210859 | 0.50313012 | NA | NA |
| ENSG000002 | 19.5152968 | 0.256904998 | 0.28785738 | 0.8924732 | 0.37213935 | 0.50313816 | NA | NA |
| ENSG000002 | 2.82849177 | -0.334014947 | 0.37425216 | -0.8924863 | 0.37213231 | 0.50313816 | NA | NA |
| ENSG000002 | 15.4754044 | -0.277688208 | 0.3111565 | -0.8924391 | 0.37215765 | 0.50314512 | ABCC6P1 | 653190 |
| ENSG000002 | 2.93131504 | -0.330098289 | 0.36989231 | -0.8924173 | 0.37216932 | 0.50314512 | NA | NA |
| ENSG000001 | 104.093322 | -0.150628442 | 0.16883968 | -0.8921389 | 0.3723185 | 0.50333003 | PLXNC1 | 10154 |
| ENSG000001 | 65.6555531 | 0.16739414 | 0.18769358 | 0.89184799 | 0.37247442 | 0.50352402 | LRRC6 | 23639 |
| ENSG000001 | 136.319839 | 0.140138126 | 0.15717684 | 0.89159524 | 0.37260992 | 0.50368078 | AHR | 196 |
| ENSG000001 | 41.447794 | -0.195349902 | 0.21910967 | -0.8915622 | 0.37262764 | 0.50368078 | NA | NA |
| ENSG000001 | 45.3679458 | -0.196431873 | 0.22032014 | -0.8915748 | 0.3726209 | 0.50368078 | ZNF774 | 342132 |
| ENSG000001 | 186.783421 | 0.129481498 | 0.14523759 | 0.89151504 | 0.37265293 | 0.50368138 | MYB | 4602 |
| ENSG000001 | 20.4932072 | -0.24743637 | 0.27753957 | -0.8915355 | 0.37264197 | 0.50368138 | MIRLET7BHG | 400931 |
| ENSG000002 | 3.33239407 | -0.33734877 | 0.37844031 | -0.8914187 | 0.37270459 | 0.50373442 | NA | NA |
| ENSG000002 | 2.85882823 | -0.332828566 | 0.37339273 | -0.8913633 | 0.37273432 | 0.50375781 | NA | NA |
| ENSG000002 | 3.32579152 | -0.337259062 | 0.37838265 | -0.8913175 | 0.37275888 | 0.50377422 | NA | NA |
| ENSG000002 | 7.91727999 | -0.312031683 | 0.35015257 | -0.8911306 | 0.37285908 | 0.50389285 | NA | NA |
| ENSG000002 | 3.58903228 | -0.337323236 | 0.37856781 | -0.891051 | 0.3729018 | 0.50392811 | RNU6-1099P | 106481526 |
| ENSG000002 | 4.86612027 | -0.335194083 | 0.37618481 | -0.8910357 | 0.37291004 | 0.50392811 | NA | NA |
| ENSG000002 | 2.7959757 | -0.33314278 | 0.37398642 | -0.8907884 | 0.37304267 | 0.50408242 | NA | NA |
| ENSG000002 | 10.1811164 | -0.29975325 | 0.33650781 | -0.8907765 | 0.37304909 | 0.50408242 | NA | NA |
| ENSG000001 | 12914.5076 | 0.128471689 | 0.14425391 | 0.89059417 | 0.37314693 | 0.50419783 | ND1 | 4535 |
| ENSG000001 | 24.5707177 | 0.243147898 | 0.27303995 | 0.89052132 | 0.37318603 | 0.50421706 | ACADS | 35 |
| ENSG000001 | 25.5783127 | -0.225489498 | 0.2532105 | -0.8905219 | 0.37318572 | 0.50421706 | PHYHIPL | 84457 |

| | | | | | | | | |
|---|---|---|---|---|---|---|---|---|
| ENSG000002 | 2.9131217 | -0.332348844 | 0.37336222 | -0.8901512 | 0.37338468 | 0.50446866 | NA | NA |
| ENSG000002 | 4.1132428 | -0.336016545 | 0.37755763 | -0.8899742 | 0.37347975 | 0.50456978 | PSORS1C3 | 100130889 |
| ENSG000002 | 6.42461802 | -0.324067083 | 0.36413444 | -0.8899655 | 0.37348442 | 0.50456978 | NA | NA |
| ENSG000001 | 3.59352767 | -0.336890121 | 0.37856978 | -0.8899023 | 0.37351835 | 0.50459881 | NA | NA |
| ENSG000002 | 3.084465 | -0.335548746 | 0.37725479 | -0.8894486 | 0.37376204 | 0.5049112 | NA | NA |
| ENSG000002 | 3.57830737 | -0.336658172 | 0.37854194 | -0.889355 | 0.37381232 | 0.5049623 | NA | NA |
| ENSG000001 | 12.4600698 | -0.287159938 | 0.32292102 | -0.8892575 | 0.37386471 | 0.50499942 | OR51M1 | 390059 |
| ENSG000002 | 11.263965 | -0.294488836 | 0.33115885 | -0.8892676 | 0.37385928 | 0.50499942 | LOC100507506 | 100507506 |
| ENSG000002 | 3.04616837 | -0.33477807 | 0.37654402 | -0.8890808 | 0.37395964 | 0.50509401 | NA | NA |
| ENSG000002 | 3.09429649 | -0.334618505 | 0.37635862 | -0.8890948 | 0.37395211 | 0.50509401 | NA | NA |
| ENSG000002 | 3.55138887 | -0.336566658 | 0.37856645 | -0.8890557 | 0.37397315 | 0.50509543 | NA | NA |
| ENSG000002 | 37.9399065 | -0.21445617 | 0.24125112 | -0.8889334 | 0.3740389 | 0.50515058 | RIPPLY2 | 134701 |
| ENSG000002 | 84.7431715 | 0.165832518 | 0.18654837 | 0.88895182 | 0.37402897 | 0.50515058 | TMEM158 | 25907 |
| ENSG000002 | 4.13871969 | -0.335664065 | 0.37763401 | -0.8888608 | 0.37407789 | 0.50516256 | NA | NA |
| ENSG000002 | 9.15501789 | -0.30719407 | 0.34559487 | -0.8888849 | 0.37406493 | 0.50516256 | NA | NA |
| ENSG000002 | 2.32241041 | -0.322993566 | 0.36338475 | -0.8888473 | 0.37408514 | 0.50516256 | NA | NA |
| ENSG000001 | 11.3961408 | 0.289451098 | 0.32567554 | 0.88877137 | 0.37412597 | 0.50520087 | PRR7 | 80758 |
| ENSG000002 | 16.0801694 | -0.275903521 | 0.31052971 | -0.8884931 | 0.37427554 | 0.50538602 | NA | NA |
| ENSG00000C | 239.371013 | 0.115831305 | 0.13037543 | 0.88844426 | 0.37430183 | 0.50540345 | COL11A1 | 1301 |
| ENSG000002 | 25.5027992 | 0.232679471 | 0.26190526 | 0.88841085 | 0.37431979 | 0.50540345 | LINC00449 | 106478991 |
| ENSG000002 | 6.42501124 | -0.328250527 | 0.36948523 | -0.8883996 | 0.37432584 | 0.50540345 | NA | NA |
| ENSG000002 | 6.72755153 | -0.321884674 | 0.36234896 | -0.8883279 | 0.37436442 | 0.5054387 | NA | NA |
| ENSG000002 | 3.58466142 | -0.33623607 | 0.37859194 | -0.8881226 | 0.3744748 | 0.5055709 | NA | NA |
| ENSG000002 | 3.0962996 | -0.334593693 | 0.37679264 | -0.8880049 | 0.37453814 | 0.50563958 | NA | NA |
| ENSG000002 | 136.748513 | 0.135666634 | 0.15279201 | 0.88791707 | 0.37458536 | 0.5056865 | NA | NA |
| ENSG000002 | 201.211066 | -0.132683188 | 0.14943673 | -0.8878887 | 0.37460061 | 0.50569025 | EFCAB2 | 84288 |
| ENSG00000C | 132.672675 | -0.164503208 | 0.18530208 | -0.887757 | 0.3746715 | 0.50576911 | TENM1 | 10178 |
| ENSG000001 | 4.3908798 | 0.336014522 | 0.37852057 | 0.88770479 | 0.37469957 | 0.50579016 | SNORD9 | 692053 |
| ENSG000002 | 3.12893909 | -0.334149548 | 0.37643857 | -0.8876602 | 0.37472355 | 0.50580569 | NA | NA |

| | | | | | | | | |
|---|---|---|---|---|---|---|---|---|
| ENSG000002 | 7.60856476 | -0.32619106 | 0.36749924 | -0.8875966 | 0.37475781 | 0.50582797 | NA | NA |
| ENSG000002 | 2.54090331 | -0.328509073 | 0.37011638 | -0.8875832 | 0.374765 | 0.50582797 | NA | NA |
| ENSG000002 | 15.2832647 | -0.276984949 | 0.31207482 | -0.8875594 | 0.37477778 | 0.50582838 | NA | NA |
| ENSG000001 | 3.11722848 | -0.333517645 | 0.37588002 | -0.8872982 | 0.3749184 | 0.50595081 | OR5T3 | 390154 |
| ENSG000001 | 2.58638784 | -0.32494313 | 0.36620877 | -0.8873166 | 0.37490846 | 0.50595081 | RNU6-249P | 106479677 |
| ENSG000002 | 2.32615561 | -0.321988203 | 0.36286236 | -0.8873563 | 0.3748871 | 0.50595081 | NA | NA |
| ENSG000002 | 2.86849062 | -0.331797415 | 0.37393366 | -0.8873163 | 0.37490866 | 0.50595081 | NA | NA |
| ENSG000002 | 4.76140694 | -0.335273344 | 0.37789155 | -0.8872211 | 0.37495989 | 0.50598997 | CACTIN-AS1 | 404665 |
| ENSG000002 | 3.09228497 | -0.332216635 | 0.37452183 | -0.8870421 | 0.37505623 | 0.50610313 | RN7SL523P | 106479414 |
| ENSG000002 | 3.62486879 | -0.335770058 | 0.3785764 | -0.8869281 | 0.3751176 | 0.5061691 | PRAMEF6 | 440561 |
| ENSG000002 | 6.14120456 | -0.324970252 | 0.36650487 | -0.8866738 | 0.37525457 | 0.50633707 | NA | NA |
| ENSG000002 | 6.23630391 | 0.324541745 | 0.36606541 | 0.88656763 | 0.37531173 | 0.50639735 | NA | NA |
| ENSG000002 | 4.93577232 | -0.333220951 | 0.3759063 | -0.8864468 | 0.37537679 | 0.50646828 | NA | NA |
| ENSG000002 | 7.65361001 | -0.311869089 | 0.35186644 | -0.886328 | 0.37544079 | 0.50653778 | NA | NA |
| ENSG000002 | 3.60548646 | -0.335518788 | 0.37858064 | -0.8862545 | 0.37548041 | 0.50657438 | NA | NA |
| ENSG000001 | 1196.30451 | 0.273882776 | 0.30907397 | 0.88613991 | 0.37554214 | 0.50664081 | GPATCH4 | 54865 |
| ENSG000002 | 14.1221198 | -0.274971583 | 0.31034381 | -0.8860224 | 0.37560544 | 0.50669656 | IQCF3 | 401067 |
| ENSG000002 | 3.42193587 | -0.334632503 | 0.37768188 | -0.8860168 | 0.37560846 | 0.50669656 | NA | NA |
| ENSG000001 | 848.768533 | 0.10359828 | 0.11693435 | 0.88595252 | 0.37564311 | 0.50672645 | KIAA2018 | 205717 |
| ENSG000002 | 2.61347443 | -0.324716264 | 0.36655048 | -0.8858705 | 0.3756873 | 0.5067692 | NA | NA |
| ENSG000002 | 3.58678819 | -0.335324524 | 0.37857842 | -0.8857465 | 0.37575413 | 0.50684249 | NA | NA |
| ENSG000002 | 3.09647642 | -0.334110264 | 0.37725752 | -0.8856292 | 0.37581739 | 0.50684593 | LINC01117 | 102724224 |
| ENSG000002 | 8.05015866 | 0.320721641 | 0.3621118 | 0.88569784 | 0.37578037 | 0.50684593 | RNU1-13P | 26860 |
| ENSG000002 | 49.5879633 | 0.194907049 | 0.22006401 | 0.88568344 | 0.37578814 | 0.50684593 | RAD51-AS1 | 100505648 |
| ENSG000002 | 3.63019781 | -0.335151408 | 0.37842574 | -0.8856464 | 0.37580808 | 0.50684593 | NA | NA |
| ENSG000002 | 6.46650225 | -0.323039017 | 0.36475789 | -0.8856259 | 0.37581917 | 0.50684593 | NA | NA |
| ENSG000002 | 3.61122705 | -0.335094248 | 0.37859041 | -0.8851102 | 0.37609717 | 0.50720398 | NA | NA |
| ENSG000002 | 3.63625902 | -0.334767647 | 0.37824948 | -0.8850446 | 0.37613259 | 0.50723488 | NA | NA |
| ENSG000001 | 3.31657295 | -0.334781253 | 0.37833405 | -0.8848827 | 0.3762199 | 0.50733575 | NA | NA |

| | | | | | | | | |
|---|---|---|---|---|---|---|---|---|
| ENSG000001 | 10.2990531 | 0.298802301 | 0.33770768 | 0.8847957 | 0.37626683 | 0.50737971 | HIST1H2BA | 255626 |
| ENSG000002 | 7.18722693 | -0.316395651 | 0.35759978 | -0.8847759 | 0.37627753 | 0.50737971 | LINC01281 | 286442 |
| ENSG000001 | 69.043138 | 0.193783579 | 0.21904296 | 0.88468297 | 0.37632764 | 0.50743041 | NA | NA |
| ENSG000001 | 33.2628988 | 0.215537808 | 0.24367234 | 0.88453948 | 0.37640506 | 0.50751792 | HEY1 | 23462 |
| ENSG000001 | 7.24968573 | -0.322191119 | 0.36428976 | -0.8844364 | 0.37646069 | 0.50752543 | GSTA3 | 2940 |
| ENSG000002 | 5.70923729 | -0.327571701 | 0.37036813 | -0.884449 | 0.37645391 | 0.50752543 | NA | NA |
| ENSG000002 | 20.0090321 | 0.257502493 | 0.29114635 | 0.88444348 | 0.37645686 | 0.50752543 | NA | NA |
| ENSG000002 | 8.75127609 | -0.311667554 | 0.35236575 | -0.8845001 | 0.3764263 | 0.50752543 | NA | NA |
| ENSG000002 | 3.3539486 | -0.334136609 | 0.37784099 | -0.8843313 | 0.37651741 | 0.50758502 | NA | NA |
| ENSG000001 | 23.9090755 | 0.258703341 | 0.29256777 | 0.88425101 | 0.37656073 | 0.50760021 | KCNJ11 | 3767 |
| ENSG000002 | 5.95472549 | -0.326533941 | 0.36927868 | -0.884248 | 0.37656235 | 0.50760021 | NA | NA |
| ENSG000002 | 5.8900275 | -0.324511848 | 0.36699488 | -0.8842408 | 0.37656623 | 0.50760021 | NA | NA |
| ENSG000001 | 6.43203547 | -0.321509168 | 0.36376375 | -0.8838406 | 0.37678227 | 0.50785767 | PDE6H | 5149 |
| ENSG000002 | 3.11621574 | -0.331465521 | 0.37501999 | -0.8838609 | 0.37677128 | 0.50785767 | NA | NA |
| ENSG000002 | 3.10097126 | -0.333296157 | 0.37713467 | -0.883759 | 0.37682632 | 0.50790016 | NA | NA |
| ENSG000002 | 3.82917667 | -0.334541904 | 0.3785817 | -0.8836716 | 0.3768735 | 0.50794687 | ENPP7P14 | 106481720 |
| ENSG000002 | 6.58579386 | -0.329241853 | 0.37259576 | -0.8836436 | 0.37688864 | 0.50795039 | NA | NA |
| ENSG000002 | 3.10378785 | -0.331495978 | 0.37516777 | -0.883594 | 0.37691544 | 0.50796962 | NA | NA |
| ENSG000001 | 17.1268169 | -0.265198836 | 0.30018541 | -0.8834501 | 0.37699312 | 0.50805743 | BDKRB2 | 624 |
| ENSG000002 | 2.28124724 | -0.324534034 | 0.36743272 | -0.8832475 | 0.37710255 | 0.50818802 | NA | NA |
| ENSG000002 | 3.40093478 | -0.332930313 | 0.37705212 | -0.8829822 | 0.37724589 | 0.50836428 | NA | NA |
| ENSG000002 | 10.2353122 | 0.29962492 | 0.33936966 | 0.88288657 | 0.37729756 | 0.50840911 | FAM157C | 100996541 |
| ENSG000002 | 6.98848851 | -0.320805526 | 0.36336492 | -0.8828742 | 0.37730423 | 0.50840911 | NA | NA |
| ENSG000002 | 12.715099 | -0.28268532 | 0.32020626 | -0.8828226 | 0.37733213 | 0.50842816 | NA | NA |
| ENSG000002 | 10.8921114 | -0.293703375 | 0.33269463 | -0.8828017 | 0.37734344 | 0.50842816 | NA | NA |
| ENSG000002 | 3.37461709 | -0.333382524 | 0.37765925 | -0.8827601 | 0.37736589 | 0.50844151 | NA | NA |
| ENSG00000C | 40.4942403 | -0.202971481 | 0.22995792 | -0.8826462 | 0.37742746 | 0.50850758 | SEMA3G | 56920 |
| ENSG000001 | 109.757131 | -0.151093535 | 0.1711995 | -0.8825583 | 0.37747497 | 0.50855469 | ANKH | 56172 |
| ENSG000002 | 3.34756229 | -0.333791052 | 0.37822452 | -0.8825209 | 0.37749517 | 0.50856501 | NA | NA |

| | | | | | | | | |
|---|---|---|---|---|---|---|---|---|
| ENSG000002 | 4.08220194 | -0.333215613 | 0.37759116 | -0.8824773 | 0.37751877 | 0.5085799 | PSMB11 | 122706 |
| ENSG000001 | 5.68368056 | -0.32850383 | 0.37231372 | -0.8823307 | 0.37759799 | 0.50866972 | MC5R | 4161 |
| ENSG000002 | 5.12953797 | -0.330869481 | 0.37503908 | -0.8822267 | 0.37765424 | 0.50871171 | NA | NA |
| ENSG000002 | 5.86350236 | 0.328088904 | 0.3718829 | 0.88223712 | 0.37764859 | 0.50871171 | NA | NA |
| ENSG000001 | 139.878165 | 0.148476664 | 0.16830495 | 0.88218833 | 0.37767497 | 0.50872273 | HESX1 | 8820 |
| ENSG000002 | 2.31075905 | -0.320412492 | 0.36328747 | -0.8819806 | 0.37778731 | 0.50885714 | NA | NA |
| ENSG000001 | 124.305379 | 0.144204028 | 0.16352656 | 0.88183855 | 0.37786412 | 0.50892793 | CAMK1D | 57118 |
| ENSG000002 | 24.7417802 | 0.233940251 | 0.26528741 | 0.88183699 | 0.37786497 | 0.50892793 | NA | NA |
| ENSG000002 | 3.10706417 | -0.331738021 | 0.37621253 | -0.8817835 | 0.37789387 | 0.50894996 | NA | NA |
| ENSG000001 | 5.15714685 | -0.331150898 | 0.37557038 | -0.8817279 | 0.37792395 | 0.50895946 | SLC32A1 | 140679 |
| ENSG000002 | 2.23914841 | -0.326546393 | 0.37034985 | -0.8817241 | 0.37792602 | 0.50895946 | RN7SL675P | 106481107 |
| ENSG000002 | 6.10588978 | -0.322545464 | 0.36588054 | -0.8815595 | 0.37801507 | 0.50904011 | NA | NA |
| ENSG000002 | 13.3763508 | -0.276450958 | 0.31359945 | -0.8815416 | 0.37802476 | 0.50904011 | PCED1B-AS1 | 100233209 |
| ENSG000002 | 6.51676923 | -0.324710426 | 0.368326 | -0.8815843 | 0.37800164 | 0.50904011 | LOC339978 | 339978 |
| ENSG000002 | 2.8357398 | -0.329653832 | 0.37396045 | -0.8815206 | 0.37803612 | 0.50904011 | RN7SL117P | 106479264 |
| ENSG000002 | 3.12431287 | -0.331008383 | 0.37551348 | -0.881482 | 0.37805698 | 0.50905129 | NA | NA |
| ENSG000002 | 3.39488695 | 0.3336164 | 0.37853752 | 0.88132981 | 0.37813933 | 0.50914527 | NA | NA |
| ENSG000002 | 2.31572354 | -0.319730864 | 0.36281862 | -0.8812416 | 0.37818706 | 0.50918041 | NA | NA |
| ENSG000002 | 3.59680923 | -0.333618467 | 0.37858052 | -0.8812352 | 0.37819055 | 0.50918041 | NA | NA |
| ENSG000002 | 2.25725329 | -0.326398199 | 0.37041376 | -0.8811719 | 0.37822478 | 0.5092096 | NA | NA |
| ENSG000002 | 8.2850195 | -0.314168365 | 0.35655972 | -0.8811101 | 0.37825823 | 0.50923773 | NA | NA |
| ENSG000002 | 16.9818293 | -0.265211813 | 0.30102192 | -0.8810382 | 0.37829714 | 0.5092732 | NA | NA |
| ENSG000001 | 309.75406 | 0.111768104 | 0.12687623 | 0.88092236 | 0.37835984 | 0.50933487 | PCBP1-AS1 | 400960 |
| ENSG000002 | 6.9441294 | 0.317753402 | 0.36071157 | 0.88090716 | 0.37836807 | 0.50933487 | NA | NA |
| ENSG000002 | 4.10630799 | -0.332764091 | 0.37777291 | -0.8808575 | 0.37839496 | 0.50935416 | NA | NA |
| ENSG000001 | 10.8939949 | -0.310261992 | 0.35230551 | -0.8806618 | 0.37850092 | 0.50947987 | BPIFA2 | 140683 |
| ENSG000002 | 2.27722902 | -0.324548609 | 0.36856788 | -0.8805667 | 0.37855238 | 0.50948149 | C16orf90 | 646174 |
| ENSG000002 | 6.89832159 | -0.315780122 | 0.35859432 | -0.8806055 | 0.3785314 | 0.50948149 | NA | NA |
| ENSG000002 | 6.63727121 | -0.320136233 | 0.36355489 | -0.8805719 | 0.37854956 | 0.50948149 | PLBD1-AS1 | 101928290 |

| | | | | | | | | |
|---|---|---|---|---|---|---|---|---|
| ENSG000002 | 24.8839951 | 0.231849768 | 0.26328315 | 0.88060996 | 0.37852896 | 0.50948149 | NA | NA |
| ENSG000002 | 11.23977 | -0.290956514 | 0.33045624 | -0.8804691 | 0.37860523 | 0.50953099 | LEF1-AS1 | 641518 |
| ENSG000002 | 3.32580306 | -0.332348231 | 0.37747439 | -0.8804524 | 0.37861428 | 0.50953099 | NA | NA |
| ENSG000001 | 28.6960738 | 0.223176569 | 0.25350278 | 0.88037129 | 0.3786582 | 0.50957318 | ATP8B3 | 148229 |
| ENSG000002 | 4.98606468 | -0.331845375 | 0.37698013 | -0.8802729 | 0.37871151 | 0.50962801 | GZMAP1 | 106479035 |
| ENSG000002 | 5.64879664 | -0.327952545 | 0.37263763 | -0.8800844 | 0.37881361 | 0.50974849 | NA | NA |
| ENSG000001 | 20.1164254 | -0.25650054 | 0.29148492 | -0.8799788 | 0.37887081 | 0.50978213 | ARHGEF15 | 22899 |
| ENSG000002 | 14.3673561 | -0.281807802 | 0.32025601 | -0.8799454 | 0.3788889 | 0.50978213 | TTC34 | 100287898 |
| ENSG000002 | 3.59318744 | -0.33314701 | 0.37859271 | -0.8799615 | 0.37888016 | 0.50978213 | PLA2G4B | 100137049 |
| ENSG000002 | 2.91882341 | -0.327737978 | 0.37244519 | -0.879963 | 0.37887937 | 0.50978213 | NA | NA |
| ENSG000002 | 7.15122122 | -0.314129542 | 0.35703342 | -0.8798323 | 0.37895015 | 0.50984763 | FALEC | 100874054 |
| ENSG000002 | 12.843337 | -0.290975591 | 0.33072668 | -0.8798068 | 0.37896397 | 0.5098493 | NA | NA |
| ENSG000002 | 50.0854737 | 0.184883906 | 0.21017468 | 0.87966786 | 0.37903927 | 0.50993369 | NA | NA |
| ENSG000001 | 6.96753425 | -0.318754814 | 0.3623968 | -0.879574 | 0.37909012 | 0.50997756 | KIF2B | 84643 |
| ENSG000002 | 3.17636748 | -0.329344863 | 0.37445343 | -0.8795349 | 0.37911131 | 0.50997756 | ARHGEF7-AS2 | 100874238 |
| ENSG000002 | 12.2306024 | 0.284314162 | 0.32325639 | 0.87953144 | 0.37911319 | 0.50997756 | NA | NA |
| ENSG000002 | 8.98163659 | -0.307195925 | 0.34927884 | -0.8795149 | 0.37912218 | 0.50997756 | LOC100506286 | 100506286 |
| ENSG00000C | 16.1485829 | 0.26605991 | 0.30256926 | 0.87933557 | 0.37921936 | 0.50998939 | CD79B | 974 |
| ENSG000002 | 2.87324606 | -0.326404303 | 0.3712908 | -0.8791069 | 0.37934331 | 0.50998939 | NA | NA |
| ENSG000002 | 4.83278269 | 0.328362361 | 0.37343813 | 0.87929521 | 0.37924123 | 0.50998939 | NA | NA |
| ENSG000002 | 2.22705322 | -0.325893063 | 0.37067164 | -0.8791961 | 0.37929496 | 0.50998939 | NA | NA |
| ENSG000002 | 3.3836585 | -0.331516821 | 0.37698505 | -0.8793898 | 0.37918994 | 0.50998939 | NA | NA |
| ENSG000002 | 8.18512858 | -0.306118571 | 0.34817929 | -0.8791981 | 0.37929388 | 0.50998939 | NA | NA |
| ENSG000002 | 3.29950575 | -0.332245148 | 0.3778175 | -0.87938 | 0.37919528 | 0.50998939 | NA | NA |
| ENSG000002 | 2.32374756 | -0.315754355 | 0.35917064 | -0.8791207 | 0.37933582 | 0.50998939 | LOC101928973 | 101928973 |
| ENSG000002 | 2.32922093 | -0.315840067 | 0.35918925 | -0.8793138 | 0.37923115 | 0.50998939 | NA | NA |
| ENSG000002 | 4.34210961 | -0.330676207 | 0.37607927 | -0.8792726 | 0.37925348 | 0.50998939 | NA | NA |
| ENSG000002 | 2.22705322 | -0.325893063 | 0.37067164 | -0.8791961 | 0.37929496 | 0.50998939 | NA | NA |
| ENSG000002 | 2.22705322 | -0.325893063 | 0.37067164 | -0.8791961 | 0.37929496 | 0.50998939 | NA | NA |

| | | | | | | | | |
|---|---|---|---|---|---|---|---|---|
| ENSG000002 | 4.19352326 | -0.332840703 | 0.37859339 | -0.8791509 | 0.37931948 | 0.50998939 | NA | NA |
| ENSG000002 | 2.22705322 | -0.325893063 | 0.37067164 | -0.8791961 | 0.37929496 | 0.50998939 | NA | NA |
| ENSG000002 | 99.2021279 | 0.156910777 | 0.17842461 | 0.87942342 | 0.37917174 | 0.50998939 | NA | NA |
| ENSG000002 | 2.82717411 | -0.329371543 | 0.37451721 | -0.8794564 | 0.37915389 | 0.50998939 | GRAMD4P4 | 106480248 |
| ENSG000002 | 3.8323299 | -0.33245383 | 0.3781734 | -0.8791042 | 0.37934477 | 0.50998939 | NA | NA |
| ENSG000002 | 3.88033871 | -0.332596433 | 0.3784139 | -0.8789224 | 0.37944338 | 0.51010504 | NA | NA |
| ENSG000002 | 17.0448824 | 0.269327292 | 0.30650412 | 0.87870691 | 0.37956021 | 0.51023116 | RAET1K | 646024 |
| ENSG000002 | 3.3697295 | -0.331845767 | 0.37766407 | -0.8786798 | 0.37957494 | 0.51023116 | MSL3P1 | 151507 |
| ENSG000002 | 16.6036644 | -0.261238496 | 0.29730297 | -0.8786945 | 0.37956692 | 0.51023116 | NA | NA |
| ENSG000001 | 31.7500098 | -0.213412795 | 0.24291596 | -0.8785458 | 0.37964759 | 0.51029498 | GAS2 | 2620 |
| ENSG000002 | 8.01045812 | 0.308763504 | 0.35144251 | 0.8785605 | 0.37963962 | 0.51029498 | NA | NA |
| ENSG000002 | 3.61707257 | -0.332555113 | 0.37855341 | -0.8784893 | 0.37967826 | 0.51031929 | NA | NA |
| ENSG000002 | 4.41422858 | -0.33224397 | 0.37837859 | -0.8780729 | 0.37990419 | 0.51060603 | NA | NA |
| ENSG000002 | 2.26345089 | -0.324720952 | 0.36984532 | -0.8779912 | 0.37994848 | 0.51064864 | NA | NA |
| ENSG00000C | 21.3077552 | -0.241209476 | 0.27476443 | -0.8778774 | 0.38001027 | 0.51071476 | SMARCD3 | 6604 |
| ENSG000002 | 22.160436 | 0.241324645 | 0.27491998 | 0.8777996 | 0.38005248 | 0.51075456 | LINC00852 | 84657 |
| ENSG000001 | 48.8620406 | 0.194107619 | 0.22114031 | 0.87775773 | 0.38007521 | 0.51076818 | FAAH | 2166 |
| ENSG00000C | 96.2953752 | -0.153439065 | 0.17481706 | -0.8777122 | 0.38009993 | 0.51078448 | APOB | 338 |
| ENSG000002 | 5.22857965 | -0.329747466 | 0.37577632 | -0.87751 | 0.38020971 | 0.51091506 | NA | NA |
| ENSG000001 | 42.5036988 | 0.209769526 | 0.23908042 | 0.87740154 | 0.38026858 | 0.51096441 | SHROOM1 | 134549 |
| ENSG000002 | 2.24034209 | -0.323756418 | 0.36899695 | -0.8773959 | 0.38027163 | 0.51096441 | NA | NA |
| ENSG000001 | 487.839095 | 0.114980054 | 0.13105361 | 0.87735129 | 0.38029586 | 0.51098003 | CITED2 | 10370 |
| ENSG000002 | 2.22659621 | -0.324337518 | 0.36971614 | -0.8772609 | 0.38034493 | 0.51102904 | RN7SL464P | 106481049 |
| ENSG000002 | 10.6428923 | -0.291802088 | 0.33275835 | -0.8769189 | 0.38053068 | 0.51126167 | NA | NA |
| ENSG000001 | 508.616217 | 0.117513054 | 0.13403401 | 0.87674054 | 0.38062758 | 0.51132995 | ZBTB41 | 360023 |
| ENSG000002 | 2.27028156 | -0.32393266 | 0.36947718 | -0.8767325 | 0.38063194 | 0.51132995 | NUTF2P5 | 106481706 |
| ENSG000002 | 9.19157311 | 0.308209288 | 0.35153531 | 0.87675201 | 0.38062135 | 0.51132995 | NA | NA |
| ENSG000002 | 16.2438859 | -0.260400181 | 0.29701179 | -0.8767335 | 0.38063142 | 0.51132995 | NA | NA |
| ENSG000002 | 15.6339651 | 0.277714818 | 0.31677956 | 0.87668161 | 0.3806596 | 0.51135016 | NA | NA |

| | | | | | | | | |
|---|---|---|---|---|---|---|---|---|
| ENSG000001 | 16.7803966 | -0.259061612 | 0.29555845 | -0.8765157 | 0.38074976 | 0.51145434 | NA | NA |
| ENSG000001 | 305.99706 | 0.114887442 | 0.13108283 | 0.87644923 | 0.38078586 | 0.51146895 | INA | 9118 |
| ENSG000001 | 86.4783935 | 0.155807071 | 0.17776957 | 0.87645522 | 0.38078261 | 0.51146895 | ZMAT1 | 84460 |
| ENSG000002 | 3.62282525 | -0.331221876 | 0.37794462 | -0.8763768 | 0.38082525 | 0.51150491 | NA | NA |
| ENSG000002 | 2.24216883 | -0.322244614 | 0.36784807 | -0.8760264 | 0.38101569 | 0.51174375 | NA | NA |
| ENSG000001 | 168.683978 | 0.138943436 | 0.15862307 | 0.87593461 | 0.38106558 | 0.51179381 | TMEM106C | 79022 |
| ENSG000001 | 3.08832623 | -0.329199771 | 0.37586022 | -0.8758569 | 0.38110783 | 0.51181664 | DNASE1L2 | 1775 |
| ENSG000002 | 3.60141563 | -0.331581781 | 0.37857761 | -0.8758621 | 0.381105 | 0.51181664 | NA | NA |
| ENSG000002 | 29.6788883 | -0.217981079 | 0.24891245 | -0.8757339 | 0.38117469 | 0.51188948 | LOC101929705 | 101929705 |
| ENSG000001 | 18.3322817 | -0.256889252 | 0.29344632 | -0.8754216 | 0.38134454 | 0.51204975 | SPATA12 | 353324 |
| ENSG000001 | 4.37813514 | -0.329625204 | 0.37651199 | -0.8754707 | 0.38131786 | 0.51204975 | VN1R2 | 317701 |
| ENSG000002 | 3.84877612 | -0.331233405 | 0.37835305 | -0.8754612 | 0.38132302 | 0.51204975 | NA | NA |
| ENSG000002 | 7.16164666 | -0.312307811 | 0.35674626 | -0.875434 | 0.3813378 | 0.51204975 | NA | NA |
| ENSG000002 | 59.160277 | 0.18711267 | 0.21376183 | 0.87533248 | 0.38139303 | 0.51206516 | NA | NA |
| ENSG000002 | 3.82860397 | -0.331182999 | 0.37835238 | -0.8753295 | 0.38139464 | 0.51206516 | NA | NA |
| ENSG000002 | 15.1980402 | 0.272493752 | 0.31131197 | 0.87530766 | 0.38140653 | 0.51206516 | NA | NA |
| ENSG000002 | 5.9333856 | -0.323031978 | 0.36903893 | -0.8753331 | 0.38139271 | 0.51206516 | LOC102723385 | 102723385 |
| ENSG000002 | 4.49597547 | -0.331090497 | 0.37828736 | -0.8752354 | 0.38144583 | 0.51210097 | NA | NA |
| ENSG000002 | 2.24766202 | -0.322593354 | 0.368591 | -0.8752068 | 0.38146138 | 0.51210489 | NA | NA |
| ENSG000001 | 6.37659406 | -0.319623099 | 0.36529078 | -0.8749827 | 0.38158334 | 0.51223217 | FBP2 | 8789 |
| ENSG000002 | 3.30302261 | -0.330770479 | 0.37804946 | -0.8749397 | 0.38160671 | 0.51223217 | NA | NA |
| ENSG000002 | 3.88823876 | -0.331241939 | 0.3785834 | -0.874951 | 0.38160054 | 0.51223217 | NA | NA |
| ENSG000002 | 4.5800778 | 0.330062679 | 0.37721461 | 0.87499973 | 0.38157405 | 0.51223217 | NA | NA |
| ENSG000002 | 18.7641836 | 0.246848717 | 0.28221178 | 0.87469317 | 0.38174088 | 0.51236138 | NA | NA |
| ENSG000002 | 3.29334758 | -0.330639436 | 0.37800028 | -0.8747068 | 0.38173343 | 0.51236138 | NA | NA |
| ENSG000002 | 2.26454042 | -0.322355568 | 0.36852453 | -0.8747194 | 0.38172658 | 0.51236138 | NA | NA |
| ENSG000001 | 2.9175369 | -0.324593011 | 0.37112176 | -0.8746267 | 0.38177704 | 0.51239296 | NA | NA |
| ENSG000001 | 27.0352043 | -0.234079685 | 0.2676742 | -0.8744947 | 0.38184888 | 0.51247241 | LONRF3 | 79836 |
| ENSG000002 | 3.10521816 | -0.328838556 | 0.37605787 | -0.874436 | 0.38188084 | 0.51249834 | NA | NA |

| | | | | | | | | |
|---|---|---|---|---|---|---|---|---|
| ENSG000002 | 3.13737993 | -0.325672004 | 0.37251385 | -0.8742548 | 0.38197953 | 0.51261382 | NA | NA |
| ENSG000002 | 4.35975006 | -0.328921752 | 0.37631109 | -0.8740687 | 0.38208083 | 0.5127328 | NA | NA |
| ENSG000001 | 29.2498911 | 0.238424142 | 0.27288607 | 0.87371314 | 0.38227449 | 0.51296104 | KLHL17 | 339451 |
| ENSG000002 | 31.3993642 | 0.222054554 | 0.25415133 | 0.87370998 | 0.38227621 | 0.51296104 | LENG8-AS1 | 104355426 |
| ENSG000002 | 2.60895564 | -0.322104653 | 0.36871131 | -0.8735958 | 0.38233842 | 0.51302754 | NA | NA |
| ENSG000002 | 3.60827876 | -0.330643852 | 0.37854079 | -0.8734696 | 0.38240719 | 0.51310284 | NA | NA |
| ENSG000002 | 2.83678444 | -0.326522113 | 0.37392224 | -0.8732353 | 0.38253482 | 0.5132571 | NA | NA |
| ENSG00000C | 18.1218377 | -0.2677696 | 0.30670496 | -0.8730527 | 0.38263434 | 0.51333967 | DUSP13 | 51207 |
| ENSG000002 | 3.79183382 | -0.330374544 | 0.37840109 | -0.8730803 | 0.38261931 | 0.51333967 | NA | NA |
| ENSG000002 | 7.92530021 | -0.306177114 | 0.35069012 | -0.8730702 | 0.38262479 | 0.51333967 | LOC101927267 | 101927267 |
| ENSG000002 | 4.14008221 | -0.330123192 | 0.37813675 | -0.8730259 | 0.38264894 | 0.51334229 | NA | NA |
| ENSG000002 | 14.100169 | -0.278078253 | 0.31854802 | -0.8729555 | 0.38268733 | 0.5133768 | LOC729348 | 729348 |
| ENSG000002 | 7.06644667 | 0.316066833 | 0.36208695 | 0.87290313 | 0.38271588 | 0.5133867 | ARIH2OS | 646450 |
| ENSG000002 | 8.03948323 | 0.305280553 | 0.34973321 | 0.87289552 | 0.38272003 | 0.5133867 | NA | NA |
| ENSG000002 | 3.12175633 | -0.32729772 | 0.37500973 | -0.8727713 | 0.38278776 | 0.51346056 | NA | NA |
| ENSG000001 | 14.6979302 | -0.269992206 | 0.30938732 | -0.8726673 | 0.38284444 | 0.51346993 | NA | NA |
| ENSG000001 | 10.5232353 | 0.294349632 | 0.33729947 | 0.87266557 | 0.38284539 | 0.51346993 | CH17-340M24.3 | 158960 |
| ENSG000002 | 5.78797456 | -0.3267342 | 0.37440098 | -0.8726852 | 0.38283467 | 0.51346993 | NA | NA |
| ENSG000002 | 6.00770677 | -0.324600966 | 0.37194811 | -0.872705 | 0.3828239 | 0.51346993 | NA | NA |
| ENSG000002 | 4.10485428 | -0.329395677 | 0.37747264 | -0.8726346 | 0.38286228 | 0.51347561 | NA | NA |
| ENSG000002 | 3.80366842 | -0.3302441 | 0.37852144 | -0.8724581 | 0.38295852 | 0.51358768 | NA | NA |
| ENSG000002 | 5.91060329 | -0.323027115 | 0.37026781 | -0.8724148 | 0.38298213 | 0.51360236 | LOC101927964 | 101927964 |
| ENSG000002 | 134.178861 | 0.139442132 | 0.15985649 | 0.87229573 | 0.38304707 | 0.51367247 | METTL10 | 399818 |
| ENSG000002 | 6.71244702 | -0.317721764 | 0.36428757 | -0.872173 | 0.38311403 | 0.51374527 | LOC101929897 | 101929897 |
| ENSG000002 | 7.36114578 | 0.309216938 | 0.3545701 | 0.87208973 | 0.38315943 | 0.51377404 | HSBP1L1 | 440498 |
| ENSG000002 | 8.94782889 | -0.299931612 | 0.34392389 | -0.8720872 | 0.38316082 | 0.51377404 | NA | NA |
| ENSG000002 | 4.32014242 | -0.328759731 | 0.37699233 | -0.8720595 | 0.38317594 | 0.51377732 | NA | NA |
| ENSG000002 | 3.81294702 | -0.329866372 | 0.37829091 | -0.8719913 | 0.38321315 | 0.51381022 | NA | NA |
| ENSG000002 | 3.8457667 | -0.330071282 | 0.3785503 | -0.8719351 | 0.3832438 | 0.51382067 | TGFB2-AS1 | 728463 |

| | | | | | | | | |
|---|---|---|---|---|---|---|---|---|
| ENSG000002 | 7.14776672 | 0.310927075 | 0.35659616 | 0.87193052 | 0.38324628 | 0.51382067 | SCARNA23 | 677773 |
| ENSG000002 | 3.56604856 | -0.330074029 | 0.37859091 | -0.8718488 | 0.38329087 | 0.51386345 | RN7SL573P | 106481078 |
| ENSG000002 | 4.13965278 | -0.329581056 | 0.37809665 | -0.8716847 | 0.38338042 | 0.51396653 | NA | NA |
| ENSG000002 | 7.92195962 | -0.307155872 | 0.35242424 | -0.8715515 | 0.3834531 | 0.51404697 | NA | NA |
| ENSG000001 | 90.0539604 | -0.155257865 | 0.17814837 | -0.8715088 | 0.38347643 | 0.51406124 | ALDH1A3 | 220 |
| ENSG000002 | 3.19263863 | 0.328590968 | 0.37705988 | 0.87145567 | 0.3835054 | 0.51408309 | NA | NA |
| ENSG000002 | 2.25839489 | -0.321034506 | 0.36850414 | -0.8711829 | 0.3836543 | 0.51424892 | NA | NA |
| ENSG000002 | 3.83418169 | -0.329613419 | 0.37835171 | -0.8711826 | 0.38365447 | 0.51424892 | NA | NA |
| ENSG000002 | 2.60534126 | -0.31952079 | 0.36677657 | -0.8711592 | 0.38366722 | 0.51424901 | NA | NA |
| ENSG000001 | 497.973472 | 0.111759461 | 0.12830797 | 0.87102512 | 0.38374044 | 0.51433015 | TRIM5 | 85363 |
| ENSG000002 | 8.68400734 | -0.300623666 | 0.34517203 | -0.8709387 | 0.38378764 | 0.51437641 | LOC101928509 | 101928509 |
| ENSG000002 | 8.24438601 | -0.307541796 | 0.35321159 | -0.8707013 | 0.38391726 | 0.51453313 | NA | NA |
| ENSG000002 | 21.8592918 | 0.243034405 | 0.27920606 | 0.87044818 | 0.38405553 | 0.51470142 | NA | NA |
| ENSG000002 | 6.9880524 | -0.314080356 | 0.36086109 | -0.8703636 | 0.38410174 | 0.51474634 | NA | NA |
| ENSG000001 | 12.4004882 | -0.278930813 | 0.32054067 | -0.8701885 | 0.38419739 | 0.51485751 | LINC00917 | 732275 |
| ENSG000002 | 4.05763853 | -0.328857834 | 0.37795727 | -0.8700926 | 0.38424979 | 0.51491072 | NA | NA |
| ENSG000002 | 3.08551649 | 0.211621773 | 0.24323293 | 0.87003752 | 0.3842799 | 0.51491795 | NA | NA |
| ENSG000002 | 6.17360983 | -0.319330582 | 0.36703135 | -0.8700363 | 0.38428058 | 0.51491795 | NA | NA |
| ENSG000002 | 3.28085954 | -0.32839804 | 0.3774876 | -0.8699571 | 0.38432382 | 0.51495531 | NA | NA |
| ENSG000002 | 3.92607532 | 0.329340281 | 0.37857869 | 0.86993878 | 0.38433386 | 0.51495531 | NA | NA |
| ENSG000001 | 78.0692073 | 0.179035639 | 0.20581001 | 0.86990732 | 0.38435105 | 0.51496133 | CDC42BPG | 55561 |
| ENSG000001 | 209.448673 | 0.132899451 | 0.15279117 | 0.86981108 | 0.38440366 | 0.51501479 | ZNF485 | 220992 |
| ENSG000002 | 7.48543903 | -0.317350963 | 0.36489175 | -0.8697126 | 0.38445747 | 0.51505285 | NA | NA |
| ENSG000002 | 3.36834038 | 0.329176957 | 0.37848623 | 0.86971976 | 0.38445357 | 0.51505285 | NA | NA |
| ENSG000001 | 7.4388425 | -0.311033817 | 0.35765733 | -0.8696419 | 0.38449611 | 0.51507059 | NKX6-1 | 4825 |
| ENSG000002 | 9.67952759 | 0.295067879 | 0.3392933 | 0.86965429 | 0.38448936 | 0.51507059 | APOBEC3B-AS1 | 100874530 |
| ENSG000002 | 6.37870123 | -0.319821136 | 0.36777496 | -0.869611 | 0.38451305 | 0.51507626 | NA | NA |
| ENSG000001 | 5400.97586 | 0.111123126 | 0.12779437 | 0.86954637 | 0.38454836 | 0.51510517 | ND2 | 4536 |
| ENSG000002 | 32.0406743 | -0.217474335 | 0.25010705 | -0.869525 | 0.38456004 | 0.51510517 | NA | NA |

| | | | | | | | | |
|---|---|---|---|---|---|---|---|---|
| ENSG000002 | 4.15225705 | -0.3287508 | 0.37811133 | -0.869455 | 0.3845983 | 0.51513941 | NA | NA |
| ENSG000002 | 2.26248534 | -0.320945851 | 0.36921507 | -0.8692653 | 0.38470203 | 0.51525109 | NA | NA |
| ENSG000002 | 3.59082677 | -0.329061692 | 0.37855553 | -0.8692561 | 0.38470709 | 0.51525109 | NA | NA |
| ENSG000002 | 3.85769488 | -0.328879393 | 0.37842037 | -0.8690848 | 0.38480075 | 0.5153595 | NA | NA |
| ENSG000002 | 2.24712069 | -0.319674199 | 0.36786458 | -0.8689997 | 0.38484731 | 0.51538782 | NA | NA |
| ENSG000002 | 7.53092389 | -0.317931216 | 0.36585112 | -0.869018 | 0.38483727 | 0.51538782 | NA | NA |
| ENSG000001 | 5.97542026 | -0.321342821 | 0.3698061 | -0.8689495 | 0.38487476 | 0.51539055 | KRT28 | 162605 |
| ENSG000002 | 7.15578929 | -0.310349338 | 0.35714771 | -0.8689663 | 0.38486554 | 0.51539055 | NA | NA |
| ENSG000002 | 24.4481609 | 0.2397719 | 0.27606317 | 0.86853999 | 0.38509879 | 0.51567351 | NA | NA |
| ENSG000001 | 1318.27887 | -0.098356831 | 0.11325058 | -0.8684886 | 0.38512692 | 0.51569416 | SYT14 | 255928 |
| ENSG000002 | 9.97603876 | 0.294056533 | 0.33859995 | 0.86844826 | 0.38514899 | 0.51570667 | NA | NA |
| ENSG000001 | 34.2326014 | 0.227556631 | 0.26203859 | 0.86840885 | 0.38517055 | 0.51571852 | ERCC5 | 2073 |
| ENSG000002 | 2.22298238 | -0.32044408 | 0.36907694 | -0.8682311 | 0.38526783 | 0.51583173 | NA | NA |
| ENSG000002 | 7.38297729 | -0.309062763 | 0.3560614 | -0.8680041 | 0.38539208 | 0.51598105 | NA | NA |
| ENSG000001 | 32.2928416 | -0.204811917 | 0.23599036 | -0.8678826 | 0.38545863 | 0.51603812 | CNR1 | 1268 |
| ENSG000002 | 7.09994028 | 0.313647394 | 0.36139498 | 0.86787977 | 0.38546015 | 0.51603812 | NA | NA |
| ENSG000002 | 13.5408237 | -0.272061851 | 0.31350089 | -0.8678184 | 0.38549374 | 0.51606605 | RBPMS-AS1 | 100128750 |
| ENSG000000 | 60.5815637 | -0.180015062 | 0.2075145 | -0.8674818 | 0.38567806 | 0.51629575 | TLL2 | 7093 |
| ENSG000002 | 2.2708229 | -0.319809268 | 0.36867498 | -0.8674559 | 0.38569229 | 0.51629776 | NA | NA |
| ENSG000001 | 26.5948932 | -0.228117876 | 0.26299804 | -0.8673748 | 0.38573667 | 0.51634012 | PDZD7 | 79955 |
| ENSG000001 | 2032.72425 | 0.09610719 | 0.11081023 | 0.86731335 | 0.38577035 | 0.51635299 | ARMCX2 | 9823 |
| ENSG000002 | 5.74930434 | -0.326765141 | 0.37676679 | -0.8672875 | 0.38578448 | 0.51635299 | NA | NA |
| ENSG000002 | 5.65242014 | 0.319064807 | 0.36788692 | 0.86729045 | 0.38578289 | 0.51635299 | NA | NA |
| ENSG000002 | 6.52232188 | 0.315426428 | 0.36379614 | 0.86704171 | 0.38591916 | 0.5165162 | NA | NA |
| ENSG000002 | 2.20835549 | -0.320496795 | 0.36965567 | -0.8670144 | 0.38593411 | 0.51651916 | NA | NA |
| ENSG000002 | 3.59698528 | -0.328190742 | 0.37854112 | -0.8669883 | 0.3859484 | 0.51652124 | NA | NA |
| ENSG000002 | 2.76892363 | 0.326043094 | 0.37610521 | 0.86689331 | 0.38600047 | 0.51657388 | EIF4EBP3 | 8637 |
| ENSG000000 | 443.988136 | 0.235496664 | 0.27166862 | 0.86685265 | 0.38602275 | 0.51658665 | POLR2J | 5439 |
| ENSG000002 | 2.28351835 | -0.317886448 | 0.36672491 | -0.8668253 | 0.38603771 | 0.51658962 | NA | NA |

| | | | | | | | | |
|---|---|---|---|---|---|---|---|---|
| ENSG00000C | 172.12383 | 0.136545982 | 0.15758926 | 0.86646758 | 0.3862338 | 0.51675013 | ARAP2 | 116984 |
| ENSG000001 | 197.82818 | 0.126947505 | 0.14650564 | 0.86650253 | 0.38621464 | 0.51675013 | FBXO25 | 26260 |
| ENSG000001 | 55.6193535 | -0.176118033 | 0.20324269 | -0.8665406 | 0.3861938 | 0.51675013 | SCG2 | 7857 |
| ENSG000002 | 37.191859 | 0.196112945 | 0.22633631 | 0.866467 | 0.38623411 | 0.51675013 | PIGBOS1 | 101928527 |
| ENSG000002 | 4.07929362 | -0.326993084 | 0.37736241 | -0.8665227 | 0.3862036 | 0.51675013 | NA | NA |
| ENSG000002 | 11.0131472 | -0.294357998 | 0.33970229 | -0.8665176 | 0.3862064 | 0.51675013 | RGMB-AS1 | 503569 |
| ENSG000002 | 8.43665562 | -0.306837938 | 0.35418214 | -0.8663281 | 0.38631027 | 0.51683496 | NA | NA |
| ENSG000002 | 5.96930665 | -0.321450845 | 0.37107519 | -0.8662688 | 0.38634279 | 0.51686142 | NA | NA |
| ENSG000001 | 94.6029578 | -0.146391812 | 0.16901382 | -0.8661529 | 0.3864063 | 0.51692933 | SMTN | 6525 |
| ENSG000002 | 3.38257673 | -0.326501614 | 0.37698531 | -0.8660858 | 0.38644313 | 0.51696155 | RNU4-56P | 106480547 |
| ENSG000002 | 5.2102147 | 0.323650051 | 0.37372842 | 0.86600332 | 0.38648834 | 0.51700497 | NA | NA |
| ENSG000002 | 4.57104718 | 0.325978119 | 0.37645185 | 0.86592248 | 0.38653267 | 0.51704722 | NA | NA |
| ENSG000002 | 3.58616864 | -0.327779097 | 0.37857275 | -0.8658285 | 0.38658419 | 0.51709908 | NA | NA |
| ENSG000001 | 101.087416 | 0.194623742 | 0.22479153 | 0.86579659 | 0.38660172 | 0.51710546 | MSTO1 | 55154 |
| ENSG000002 | 2.26750191 | -0.319037262 | 0.36853397 | -0.8656929 | 0.38665857 | 0.51714739 | NA | NA |
| ENSG000002 | 2.24034263 | -0.320107822 | 0.3697681 | -0.8656988 | 0.38665533 | 0.51714739 | NA | NA |
| ENSG000002 | 4.60956536 | -0.324808956 | 0.37522231 | -0.865644 | 0.3866854 | 0.51716622 | NA | NA |
| ENSG000002 | 3.5740205 | -0.327485098 | 0.37834435 | -0.8655742 | 0.38672372 | 0.51720041 | NA | NA |
| ENSG000002 | 2.24117158 | -0.320036967 | 0.36977051 | -0.8655016 | 0.38676355 | 0.51722296 | NA | NA |
| ENSG000002 | 3.81557514 | -0.32764661 | 0.37856472 | -0.8654969 | 0.38676609 | 0.51722296 | NA | NA |
| ENSG000001 | 89.1498778 | 0.153989036 | 0.17792764 | 0.86545875 | 0.38678705 | 0.51723392 | SPRY3 | 10251 |
| ENSG000002 | 3.34995761 | -0.32709889 | 0.37798678 | -0.8653712 | 0.38683506 | 0.51728108 | NA | NA |
| ENSG000002 | 6.6384858 | -0.313866528 | 0.36274173 | -0.8652617 | 0.38689517 | 0.51733725 | NA | NA |
| ENSG000002 | 3.8306596 | -0.327422412 | 0.37841445 | -0.8652482 | 0.38690259 | 0.51733725 | NA | NA |
| ENSG000001 | 4.71562162 | -0.327120173 | 0.37809797 | -0.865173 | 0.38694382 | 0.51737532 | IQCF2 | 389123 |
| ENSG000001 | 18.3611587 | 0.258278901 | 0.29855079 | 0.86510874 | 0.38697911 | 0.51740545 | GNRH1 | 2796 |
| ENSG000001 | 2564.44626 | 0.10186239 | 0.11775886 | 0.86500832 | 0.38703422 | 0.51741089 | AHCTF1 | 25909 |
| ENSG000002 | 8.92451102 | -0.299721533 | 0.34647073 | -0.8650703 | 0.38700021 | 0.51741089 | NA | NA |
| ENSG000002 | 4.33858011 | -0.325587878 | 0.3763822 | -0.8650459 | 0.3870136 | 0.51741089 | LINGO1-AS2 | 101929478 |

| | | | | | | | | |
|---|---|---|---|---|---|---|---|---|
| ENSG000002 | 3.38755397 | -0.326377636 | 0.37731045 | -0.8650109 | 0.38703282 | 0.51741089 | NA | NA |
| ENSG000002 | 3.85686539 | -0.327434772 | 0.37858374 | -0.8648939 | 0.38709702 | 0.51746073 | NA | NA |
| ENSG000002 | 3.3222424 | -0.326968311 | 0.37804057 | -0.8649027 | 0.38709219 | 0.51746073 | NA | NA |
| ENSG000002 | 36.7457464 | 0.210917529 | 0.24389148 | 0.86480073 | 0.38714817 | 0.51751205 | NA | NA |
| ENSG000002 | 3.88461339 | -0.327342477 | 0.37855435 | -0.8647173 | 0.387194 | 0.51755625 | C2CD4D | 100191040 |
| ENSG000002 | 10.7109637 | -0.291440549 | 0.33709298 | -0.8645702 | 0.38727473 | 0.51764709 | NA | NA |
| ENSG000002 | 6.23205902 | -0.31785774 | 0.36766152 | -0.8645391 | 0.38729184 | 0.5176529 | NA | NA |
| ENSG000002 | 46.3134598 | 0.18874276 | 0.21843244 | 0.86407843 | 0.3875448 | 0.51795687 | NA | NA |
| ENSG000002 | 15.0508386 | -0.261337772 | 0.30243929 | -0.8640999 | 0.38753299 | 0.51795687 | NA | NA |
| ENSG000001 | 119.577302 | -0.137126765 | 0.15870972 | -0.8640099 | 0.38758246 | 0.51799012 | MCTP1 | 79772 |
| ENSG000002 | 2.75726509 | 0.325165296 | 0.37635924 | 0.86397585 | 0.38760116 | 0.51799804 | NA | NA |
| ENSG000002 | 5.9381041 | -0.318887362 | 0.36914447 | -0.8638552 | 0.38766745 | 0.51806956 | NA | NA |
| ENSG000002 | 2.22378376 | -0.317277587 | 0.36730768 | -0.8637924 | 0.38770193 | 0.51809857 | NA | NA |
| ENSG000002 | 12.9702224 | -0.283887823 | 0.32872352 | -0.8636067 | 0.387804 | 0.51821788 | NA | NA |
| ENSG000002 | 3.7998061 | -0.326914563 | 0.37857934 | -0.8635299 | 0.38784621 | 0.51825721 | NA | NA |
| ENSG000002 | 4.07914872 | -0.326602233 | 0.37824645 | -0.8634641 | 0.38788235 | 0.51828843 | NA | NA |
| ENSG000002 | 7.64263627 | -0.308426839 | 0.35722665 | -0.8633926 | 0.38792165 | 0.51832387 | LINC01210 | 100507274 |
| ENSG000002 | 5.87656182 | -0.318495585 | 0.36894551 | -0.8632591 | 0.387995 | 0.51840479 | NA | NA |
| ENSG000002 | 3.9288112 | -0.326634608 | 0.37841908 | -0.8631558 | 0.38805183 | 0.51846364 | NA | NA |
| ENSG000001 | 174.020721 | 0.123439568 | 0.14301807 | 0.86310468 | 0.38807992 | 0.51847727 | CENPP | 401541 |
| ENSG000002 | 5.4602382 | -0.323702872 | 0.37505082 | -0.8630907 | 0.38808761 | 0.51847727 | NA | NA |
| ENSG000002 | 3.82095176 | -0.326570425 | 0.3785485 | -0.8626911 | 0.38830733 | 0.51875372 | NA | NA |
| ENSG000001 | 7.70449872 | -0.305936903 | 0.35468745 | -0.8625535 | 0.38838301 | 0.51883773 | NBPF4 | 148545 |
| ENSG000002 | 3.84681353 | -0.326266391 | 0.37830548 | -0.8624416 | 0.38844454 | 0.51890284 | NA | NA |
| ENSG00000C | 10.6190027 | -0.287466737 | 0.33336689 | -0.8623134 | 0.38851507 | 0.51891991 | BARX2 | 8538 |
| ENSG00000C | 8.15047389 | -0.300455448 | 0.34841908 | -0.8623392 | 0.38850086 | 0.51891991 | TUBA3D | 113457 |
| ENSG00000C | 28.6197427 | -0.225827506 | 0.26186583 | -0.8623787 | 0.38847917 | 0.51891991 | MAPRE3 | 22924 |
| ENSG00000C | 6334.19935 | 0.119786818 | 0.1389173 | 0.8622887 | 0.38852867 | 0.51891991 | SEMA6A | 57556 |
| ENSG000001 | 37.6596041 | 0.21165345 | 0.24547818 | 0.86220882 | 0.38857262 | 0.51891991 | FUT2 | 2524 |

| | | | | | | | | |
|---|---|---|---|---|---|---|---|---|
| ENSG000002 | 9.70504449 | 0.293011784 | 0.33982004 | 0.86225576 | 0.38854679 | 0.51891991 | SNORD70 | 692110 |
| ENSG000002 | 3.6518848 | -0.326252494 | 0.37840161 | -0.8621858 | 0.38858529 | 0.51891991 | NA | NA |
| ENSG000002 | 82.8398473 | 0.152067002 | 0.17637228 | 0.86219333 | 0.38858114 | 0.51891991 | PCDHGB4 | 8641 |
| ENSG000002 | 3.34829895 | -0.325700948 | 0.37770326 | -0.8623197 | 0.38851163 | 0.51891991 | NA | NA |
| ENSG000002 | 2.25714118 | -0.316903052 | 0.36754876 | -0.8622068 | 0.38857371 | 0.51891991 | NA | NA |
| ENSG000002 | 4.09560346 | -0.325800263 | 0.37789453 | -0.862146 | 0.38860718 | 0.51893205 | NA | NA |
| ENSG000002 | 3.82816732 | -0.326153291 | 0.37832948 | -0.862088 | 0.38863911 | 0.51895761 | LINC00628 | 127841 |
| ENSG000002 | 4.09421281 | -0.32617898 | 0.37837074 | -0.8620619 | 0.38865348 | 0.51895971 | NA | NA |
| ENSG000001 | 3.55451343 | -0.326093955 | 0.37833205 | -0.8619253 | 0.38872865 | 0.51902589 | RN7SKP95 | 106480862 |
| ENSG000002 | 2.25858335 | -0.317624812 | 0.36850034 | -0.861939 | 0.38872109 | 0.51902589 | NA | NA |
| ENSG000002 | 7.38094093 | -0.308867525 | 0.3583743 | -0.8618574 | 0.38876602 | 0.5190587 | NA | NA |
| ENSG000001 | 141.306741 | 0.131824752 | 0.15295981 | 0.86182609 | 0.38878323 | 0.51906458 | TFCP2L1 | 29842 |
| ENSG000002 | 3.3078103 | -0.324436863 | 0.37646525 | -0.8617976 | 0.38879888 | 0.51906839 | LZTS1-AS1 | 100874051 |
| ENSG000002 | 5.70107655 | -0.320542674 | 0.37196193 | -0.861762 | 0.38881847 | 0.51907746 | NA | NA |
| ENSG000002 | 9.16174873 | -0.293950068 | 0.34114859 | -0.8616482 | 0.38888114 | 0.51914403 | NA | NA |
| ENSG000002 | 8.83534135 | -0.295945581 | 0.34358752 | -0.8613397 | 0.38905096 | 0.51935363 | LOC102724421 | 102724421 |
| ENSG000001 | 60.4773178 | -0.166282755 | 0.19308763 | -0.8611777 | 0.3891402 | 0.51945566 | NOVA1 | 4857 |
| ENSG000001 | 56.7978778 | -0.174170359 | 0.20225352 | -0.8611487 | 0.38915613 | 0.51945983 | SSPN | 8082 |
| ENSG000002 | 6.2313813 | -0.317397066 | 0.36859029 | -0.8611108 | 0.38917704 | 0.51947064 | NA | NA |
| ENSG000002 | 4.81418856 | 0.32196955 | 0.37405322 | 0.86075867 | 0.38937097 | 0.51971239 | NA | NA |
| ENSG000002 | 3.83460624 | -0.325652332 | 0.37837941 | -0.8606502 | 0.3894307 | 0.51974237 | NA | NA |
| ENSG000002 | 2.25872771 | -0.317264929 | 0.3686349 | -0.8606481 | 0.38943188 | 0.51974237 | NA | NA |
| ENSG000002 | 2.24153631 | -0.316993044 | 0.36830344 | -0.8606844 | 0.38941186 | 0.51974237 | NA | NA |
| ENSG000001 | 49.3871319 | 0.186433312 | 0.21664919 | 0.86053084 | 0.38949649 | 0.5198115 | CYP4X1 | 260293 |
| ENSG000001 | 63.4352804 | -0.16387896 | 0.19047391 | -0.8603749 | 0.38958244 | 0.51990909 | ZNF365 | 22891 |
| ENSG000002 | 2.22867145 | -0.317508044 | 0.36905763 | -0.8603211 | 0.38961209 | 0.51993155 | NA | NA |
| ENSG000001 | 25.9295455 | -0.221810594 | 0.25790133 | -0.8600599 | 0.38975602 | 0.52010651 | KCNJ8 | 3764 |
| ENSG000001 | 6.15786836 | -0.315873537 | 0.36729546 | -0.8599985 | 0.38978987 | 0.52013456 | WNT10A | 80326 |
| ENSG000002 | 2.22270218 | -0.317382309 | 0.36907884 | -0.8599309 | 0.38982711 | 0.52016714 | NA | NA |

| | | | | | | | | |
|---|---|---|---|---|---|---|---|---|
| ENSG000002 | 3.55900937 | -0.325462534 | 0.37858883 | -0.8596728 | 0.38996941 | 0.52033988 | XIAP-AS1 | 106480735 |
| ENSG000001 | 188.492076 | 0.120385593 | 0.14005149 | 0.85958096 | 0.39002007 | 0.52035402 | LCA5 | 167691 |
| ENSG000001 | 36.2953701 | 0.210428634 | 0.24480955 | 0.85956055 | 0.39003133 | 0.52035402 | CILP2 | 148113 |
| ENSG000002 | 10.5151433 | 0.288796775 | 0.33597528 | 0.85957744 | 0.39002201 | 0.52035402 | NA | NA |
| ENSG000002 | 6.29537755 | 0.313468891 | 0.36468376 | 0.85956362 | 0.39002964 | 0.52035402 | NA | NA |
| ENSG000002 | 4.54739725 | 0.323841297 | 0.37680952 | 0.85942971 | 0.39010349 | 0.52043316 | NA | NA |
| ENSG000002 | 2.50893998 | -0.32119938 | 0.37378675 | -0.8593118 | 0.39016849 | 0.52050276 | MIR4770 | 100616373 |
| ENSG000002 | 4.57927356 | -0.32237693 | 0.37532329 | -0.8589313 | 0.39037841 | 0.52076567 | RN7SL479P | 106479398 |
| ENSG000002 | 2.28425557 | -0.314251825 | 0.36590027 | -0.8588456 | 0.39042572 | 0.52081165 | NA | NA |
| ENSG000001 | 2.24404874 | -0.317001758 | 0.36915557 | -0.8587213 | 0.39049429 | 0.5208664 | NA | NA |
| ENSG000002 | 6.64142169 | -0.312213745 | 0.36358566 | -0.8587075 | 0.3905019 | 0.5208664 | NA | NA |
| ENSG000002 | 2.27147578 | -0.316584899 | 0.3686787 | -0.8587014 | 0.3905053 | 0.5208664 | NA | NA |
| ENSG000001 | 8.86169184 | -0.294859669 | 0.3435234 | -0.8583394 | 0.39070509 | 0.52111574 | NA | NA |
| ENSG000001 | 2.51494171 | -0.32056639 | 0.37350526 | -0.8582647 | 0.3907463 | 0.52115357 | HDGFL1 | 154150 |
| ENSG000002 | 12.9809321 | -0.282351571 | 0.32900562 | -0.8581968 | 0.39078381 | 0.52118238 | DENND6B | 414918 |
| ENSG000002 | 7.13210978 | 0.308366485 | 0.35932652 | 0.85817904 | 0.39079361 | 0.52118238 | RHOXF1P1 | 101928941 |
| ENSG000001 | 187.516671 | -0.120620835 | 0.14055931 | -0.858149 | 0.39081018 | 0.52118639 | SPINK5 | 11005 |
| ENSG000002 | 41.3305617 | 0.199846398 | 0.23288673 | 0.85812705 | 0.39082231 | 0.52118639 | LOC105274304 | 105274304 |
| ENSG000002 | 3.33585711 | 0.32434184 | 0.37801957 | 0.85800278 | 0.39089093 | 0.52124427 | NA | NA |
| ENSG000002 | 2.2639079 | -0.316442036 | 0.36881276 | -0.8580019 | 0.39089143 | 0.52124427 | NA | NA |
| ENSG000002 | 36.9069206 | 0.217748407 | 0.25380779 | 0.85792642 | 0.39093309 | 0.52128269 | GPRIN2 | 9721 |
| ENSG000001 | 2.21425308 | -0.316049036 | 0.3684729 | -0.8577267 | 0.3910434 | 0.52141264 | OR5M3 | 219482 |
| ENSG000000 | 30.3222006 | -0.221624873 | 0.25843711 | -0.8575582 | 0.39113645 | 0.52151955 | TNFRSF1B | 7133 |
| ENSG000001 | 10.9585883 | -0.288034899 | 0.33589703 | -0.8575095 | 0.39116336 | 0.52153829 | PMCHL2 | 5370 |
| ENSG000002 | 2.53051433 | -0.319057865 | 0.37209447 | -0.8574647 | 0.39118812 | 0.52155415 | NA | NA |
| ENSG000000 | 109.391588 | 0.139897995 | 0.16317519 | 0.85734846 | 0.39125233 | 0.52162261 | GABRP | 2568 |
| ENSG000002 | 4.60682513 | -0.32168323 | 0.37522409 | -0.8573096 | 0.39127378 | 0.52163405 | NA | NA |
| ENSG000002 | 19.2324274 | -0.243224314 | 0.28371544 | -0.8572826 | 0.39128871 | 0.52163681 | NA | NA |
| ENSG000002 | 3.70589042 | 0.324148765 | 0.37816971 | 0.85715158 | 0.39136112 | 0.52171619 | NA | NA |

| | | | | | | | | |
|---|---|---|---|---|---|---|---|---|
| ENSG000001 | 22.9829639 | -0.235368569 | 0.27460615 | -0.8571133 | 0.39138229 | 0.52172726 | SNX32 | 254122 |
| ENSG000001 | 67.7883256 | 0.17163 | 0.2002754 | 0.85696997 | 0.39146148 | 0.52179852 | MXRA8 | 54587 |
| ENSG000002 | 4.2934405 | -0.322633206 | 0.37647321 | -0.8569885 | 0.39145125 | 0.52179852 | NA | NA |
| ENSG000002 | 3.04401577 | -0.321744572 | 0.37546726 | -0.8569178 | 0.39149032 | 0.52180266 | NA | NA |
| ENSG000002 | 10.0204148 | -0.292894383 | 0.34179106 | -0.8569398 | 0.39147813 | 0.52180266 | NA | NA |
| ENSG000002 | 97.6374919 | 0.151102541 | 0.17633983 | 0.85688266 | 0.39150974 | 0.52181139 | NA | NA |
| ENSG000001 | 4.39339476 | -0.3236835 | 0.37784103 | -0.8566658 | 0.3916296 | 0.52195398 | NA | NA |
| ENSG000002 | 8.21189622 | 0.299132611 | 0.34925721 | 0.85648228 | 0.39173107 | 0.52207206 | NA | NA |
| ENSG000002 | 4.14085543 | -0.324180584 | 0.37853925 | -0.8563989 | 0.39177719 | 0.52211637 | RNU6-197P | 106479654 |
| ENSG000002 | 2.68338129 | -0.309976204 | 0.36198655 | -0.8563197 | 0.39182098 | 0.52215757 | DPYD-AS2 | 100873933 |
| ENSG000002 | 4.05854026 | -0.323496462 | 0.3778155 | -0.8562287 | 0.3918713 | 0.52220747 | NA | NA |
| ENSG000002 | 5.91829647 | -0.31738411 | 0.37070179 | -0.856171 | 0.39190322 | 0.52221568 | LOC101929427 | 101929427 |
| ENSG000002 | 3.5569862 | -0.324099142 | 0.37853553 | -0.8561921 | 0.39189152 | 0.52221568 | NA | NA |
| ENSG000002 | 4.89171981 | -0.323632032 | 0.37801761 | -0.8561295 | 0.39192615 | 0.52222907 | NA | NA |
| ENSG000001 | 158.15191 | 0.128034699 | 0.14956736 | 0.85603368 | 0.39197915 | 0.52226948 | TMEM241 | 85019 |
| ENSG000001 | 179.635622 | -0.126477691 | 0.14774946 | -0.8560281 | 0.39198223 | 0.52226948 | ZFP28 | 140612 |
| ENSG000002 | 7.31518578 | -0.314327112 | 0.36723149 | -0.8559373 | 0.39203248 | 0.52231927 | NA | NA |
| ENSG000002 | 3.84195286 | -0.324014394 | 0.37858297 | -0.855861 | 0.39207467 | 0.52232399 | NRG3-AS1 | 101929590 |
| ENSG000002 | 3.91602619 | -0.324004463 | 0.37856503 | -0.8558753 | 0.39206675 | 0.52232399 | NA | NA |
| ENSG000002 | 3.86380851 | -0.323992381 | 0.37854031 | -0.8558993 | 0.39205349 | 0.52232399 | NA | NA |
| ENSG000002 | 11.2284451 | -0.28356706 | 0.33135016 | -0.8557927 | 0.39211243 | 0.52235687 | CLYBL-AS2 | 100874063 |
| ENSG000002 | 6.27265705 | -0.318796332 | 0.3725258 | -0.8557698 | 0.39212512 | 0.52235687 | LOC101927735 | 101927735 |
| ENSG000002 | 3.4046605 | -0.322289629 | 0.37662263 | -0.8557362 | 0.3921437 | 0.52236447 | NA | NA |
| ENSG000002 | 8.93740365 | -0.295847009 | 0.34575511 | -0.8556548 | 0.39218877 | 0.52240734 | NA | NA |
| ENSG000001 | 118.777321 | -0.14231843 | 0.16634372 | -0.8555684 | 0.39223655 | 0.52243667 | ITIH5 | 80760 |
| ENSG000002 | 2.27094709 | -0.314144509 | 0.36716769 | -0.8555886 | 0.39222535 | 0.52243667 | NA | NA |
| ENSG000002 | 8.0677415 | -0.300535061 | 0.35127909 | -0.855545 | 0.39224952 | 0.52243676 | NA | NA |
| ENSG000002 | 4.09113976 | -0.323605133 | 0.37825472 | -0.8555217 | 0.39226238 | 0.52243676 | NA | NA |
| ENSG000002 | 2.62505641 | -0.311598671 | 0.36423058 | -0.8554984 | 0.39227531 | 0.52243681 | NA | NA |

| | | | | | | | | |
|---|---|---|---|---|---|---|---|---|
| ENSG000002 | 2.34861077 | -0.306280003 | 0.35804548 | -0.8554221 | 0.39231754 | 0.52247237 | NENFP1 | 106480294 |
| ENSG000002 | 3.54421274 | -0.323757094 | 0.37848463 | -0.8554035 | 0.39232778 | 0.52247237 | NA | NA |
| ENSG000002 | 2.17382624 | 0.314515951 | 0.3677487 | 0.85524694 | 0.39241445 | 0.52253965 | NA | NA |
| ENSG000002 | 17.9950081 | 0.264246694 | 0.30897286 | 0.85524243 | 0.39241695 | 0.52253965 | NA | NA |
| ENSG000002 | 6.92978037 | -0.309225938 | 0.36155325 | -0.8552708 | 0.39240125 | 0.52253965 | NA | NA |
| ENSG000002 | 2.28400816 | -0.314465583 | 0.36775155 | -0.8551033 | 0.39249394 | 0.522625 | NA | NA |
| ENSG000002 | 2.23522192 | -0.315563562 | 0.36908269 | -0.8549942 | 0.39255436 | 0.52267316 | RN7SKP85 | 106479132 |
| ENSG000002 | 23.1985837 | 0.227937438 | 0.26659835 | 0.85498441 | 0.39255978 | 0.52267316 | ST7-AS1 | 93653 |
| ENSG000002 | 19.8973779 | -0.239609696 | 0.28026333 | -0.8549449 | 0.39258167 | 0.52267316 | NA | NA |
| ENSG000002 | 4.32630865 | -0.321915701 | 0.37652955 | -0.8549547 | 0.39257624 | 0.52267316 | NA | NA |
| ENSG000001 | 79.7401699 | 0.167820528 | 0.19631015 | 0.85487442 | 0.39262067 | 0.52270772 | CAMTA2 | 23125 |
| ENSG000002 | 2.23360369 | -0.315420883 | 0.36897743 | -0.8548514 | 0.3926334 | 0.52270772 | NA | NA |
| ENSG000002 | 4.9014928 | -0.323350581 | 0.3783386 | -0.8546592 | 0.39273982 | 0.52283223 | NA | NA |
| ENSG000002 | 7.38415921 | -0.303649063 | 0.35531231 | -0.8545976 | 0.39277393 | 0.52286048 | NA | NA |
| ENSG000002 | 3.36856285 | -0.322083122 | 0.37690936 | -0.8545373 | 0.39280733 | 0.5228706 | NA | NA |
| ENSG000002 | 3.70844119 | 0.323355154 | 0.37839425 | 0.85454563 | 0.39280274 | 0.5228706 | RNU7-181P | 106479093 |
| ENSG000002 | 5.98458834 | 0.316849756 | 0.37080198 | 0.85449855 | 0.39282881 | 0.52288204 | NA | NA |
| ENSG000001 | 15.3438437 | -0.257769429 | 0.30168332 | -0.8544371 | 0.39286283 | 0.52291015 | CLCNKA | 1187 |
| ENSG000002 | 2.26706417 | -0.311788737 | 0.36495313 | -0.8543254 | 0.39292471 | 0.52294641 | NA | NA |
| ENSG000002 | 2.2206471 | -0.315825148 | 0.36970112 | -0.8542715 | 0.39295455 | 0.52294641 | NA | NA |
| ENSG000002 | 4.16803818 | -0.323351886 | 0.37851093 | -0.8542736 | 0.39295343 | 0.52294641 | NA | NA |
| ENSG000002 | 3.62382415 | -0.322572545 | 0.37759819 | -0.8542746 | 0.39295285 | 0.52294641 | NA | NA |
| ENSG000002 | 2.49857925 | -0.318650636 | 0.37300045 | -0.8542902 | 0.3929442 | 0.52294641 | NA | NA |
| ENSG000002 | 8.45433939 | -0.300092516 | 0.35130178 | -0.85423 | 0.39297757 | 0.52295989 | NA | NA |
| ENSG000002 | 14.7570779 | -0.258121737 | 0.3021829 | -0.8541904 | 0.3929995 | 0.5229719 | NA | NA |
| ENSG000002 | 35.5258722 | -0.211524814 | 0.24765136 | -0.8541234 | 0.39303664 | 0.52300415 | NA | NA |
| ENSG000002 | 3.79886147 | -0.32320279 | 0.37845066 | -0.8540157 | 0.39309632 | 0.52306641 | NA | NA |
| ENSG000001 | 43.517412 | -0.188538478 | 0.22079329 | -0.853914 | 0.39315264 | 0.52312419 | LCA5L | 150082 |
| ENSG000001 | 141.482486 | 0.14390036 | 0.16855177 | 0.85374574 | 0.39324588 | 0.52323109 | NPHP4 | 261734 |

| | | | | | | | | |
|---|---|---|---|---|---|---|---|---|
| ENSG000001 | 8.4280815 | -0.296855501 | 0.34774634 | -0.8536553 | 0.39329602 | 0.52326345 | APOA2 | 336 |
| ENSG000001 | 4.07610847 | -0.322668438 | 0.37797542 | -0.8536757 | 0.39328469 | 0.52326345 | KRT38 | 8687 |
| ENSG000002 | 3.8722491 | -0.323141574 | 0.37858871 | -0.8535425 | 0.3933585 | 0.523326 | NA | NA |
| ENSG000002 | 7.1841224 | -0.307614192 | 0.3604049 | -0.8535239 | 0.39336884 | 0.523326 | NA | NA |
| ENSG000001 | 7.94346959 | -0.302934571 | 0.35493279 | -0.8534984 | 0.39338296 | 0.52332762 | C11orf72 | 100505621 |
| ENSG000002 | 7.13262897 | -0.306776076 | 0.35944689 | -0.8534671 | 0.39340035 | 0.52333358 | NA | NA |
| ENSG000001 | 127.510654 | -0.13188005 | 0.15456523 | -0.8532323 | 0.39353047 | 0.52345517 | CCDC149 | 91050 |
| ENSG000002 | 2.26313823 | -0.314441522 | 0.36851975 | -0.8532555 | 0.3935176 | 0.52345517 | NA | NA |
| ENSG000002 | 4.08091185 | -0.322318851 | 0.37775854 | -0.8532404 | 0.393526 | 0.52345517 | NA | NA |
| ENSG000002 | 6.87916738 | -0.307049375 | 0.35989179 | -0.8531714 | 0.39356428 | 0.52348296 | NA | NA |
| ENSG000002 | 21.5659591 | 0.232453363 | 0.27250376 | 0.85302811 | 0.39364372 | 0.52357145 | NA | NA |
| ENSG000002 | 2.25932798 | -0.312740372 | 0.36663733 | -0.8529965 | 0.39366122 | 0.52357756 | RNU6-650P | 106481387 |
| ENSG000001 | 104.938263 | 0.302850133 | 0.3550932 | 0.85287504 | 0.3937286 | 0.52361006 | PALM3 | 342979 |
| ENSG000002 | 8.90864959 | -0.300652608 | 0.35251907 | -0.8528691 | 0.39373192 | 0.52361006 | NA | NA |
| ENSG000002 | 19.9599747 | 0.237429597 | 0.27839244 | 0.85285934 | 0.39373731 | 0.52361006 | NA | NA |
| ENSG000002 | 2.28406024 | -0.313475208 | 0.36753416 | -0.8529145 | 0.39370674 | 0.52361006 | NA | NA |
| ENSG000002 | 2.23141222 | -0.314784286 | 0.36911201 | -0.8528151 | 0.39376187 | 0.52362555 | NA | NA |
| ENSG000002 | 7.27697836 | -0.310392591 | 0.36398532 | -0.8527613 | 0.39379169 | 0.52364803 | LOC101928075 | 101928075 |
| ENSG000001 | 14.3100342 | -0.259875071 | 0.30477731 | -0.852672 | 0.39384123 | 0.52367957 | FGB | 2244 |
| ENSG000002 | 2.52222939 | -0.318496544 | 0.37351853 | -0.8526927 | 0.39382972 | 0.52367957 | NA | NA |
| ENSG000001 | 3.59618715 | -0.321791067 | 0.37743984 | -0.8525625 | 0.39390194 | 0.52374312 | SMIM2 | 79024 |
| ENSG000001 | 5.14434277 | -0.319538607 | 0.37481868 | -0.8525151 | 0.39392823 | 0.52375631 | KRT37 | 8688 |
| ENSG000002 | 5.21252658 | -0.322487939 | 0.37828582 | -0.8524981 | 0.39393769 | 0.52375631 | SAPCD1 | 401251 |
| ENSG000002 | 2.28016512 | -0.310262938 | 0.36396175 | -0.8524603 | 0.39395867 | 0.52376703 | KRTAP3-1 | 83896 |
| ENSG000002 | 4.79864084 | -0.31845278 | 0.37376567 | -0.8520119 | 0.39420751 | 0.52408068 | NA | NA |
| ENSG000002 | 6.14055332 | -0.313401982 | 0.36786941 | -0.8519381 | 0.39424842 | 0.52411789 | GOLGA8N | 643699 |
| ENSG000002 | 4.574871 | -0.319311633 | 0.37485331 | -0.8518309 | 0.39430795 | 0.52417983 | NA | NA |
| ENSG000001 | 35.4844544 | -0.200118746 | 0.23498206 | -0.8516341 | 0.39441719 | 0.52430786 | SIT1 | 27240 |
| ENSG000001 | 12.6958588 | 0.277472734 | 0.32584075 | 0.85155935 | 0.39445871 | 0.52434587 | MFSD6L | 162387 |

| | | | | | | | | |
|---|---|---|---|---|---|---|---|---|
| ENSG000002 | 2.49169651 | -0.317613419 | 0.37298914 | -0.8515353 | 0.39447207 | 0.52434644 | NA | NA |
| ENSG000002 | 9.73423051 | -0.293922195 | 0.34518741 | -0.8514858 | 0.39449952 | 0.52436575 | NA | NA |
| ENSG000002 | 10.1777467 | -0.285564709 | 0.33543809 | -0.8513187 | 0.39459235 | 0.52447194 | NA | NA |
| ENSG000001 | 59.9826837 | -0.166246075 | 0.19529229 | -0.851268 | 0.39462051 | 0.52449217 | MSRA | 4482 |
| ENSG000002 | 37.8776739 | 0.197974393 | 0.23257182 | 0.85123981 | 0.39463615 | 0.52449577 | NA | NA |
| ENSG000002 | 9.36902004 | -0.288794258 | 0.33927641 | -0.8512064 | 0.39465469 | 0.52450079 | NA | NA |
| ENSG000002 | 18.3289695 | -0.244419265 | 0.28715127 | -0.8511864 | 0.3946658 | 0.52450079 | NA | NA |
| ENSG000002 | 3.53527491 | -0.322129963 | 0.37855098 | -0.8509553 | 0.39479418 | 0.52465421 | NA | NA |
| ENSG000002 | 13.4786741 | 0.275017769 | 0.32320986 | 0.85089535 | 0.39482749 | 0.52468128 | NA | NA |
| ENSG000002 | 3.60953803 | -0.321970822 | 0.37842107 | -0.8508269 | 0.39486552 | 0.52471463 | NA | NA |
| ENSG000002 | 12.3501881 | 0.279101981 | 0.32805457 | 0.85077912 | 0.39489206 | 0.52473271 | NA | NA |
| ENSG000002 | 4.54154583 | -0.319298544 | 0.37532197 | -0.8507324 | 0.39491805 | 0.5247426 | LINC01431 | 100505683 |
| ENSG000002 | 2.26315087 | -0.312343498 | 0.36715922 | -0.8507031 | 0.39493428 | 0.5247426 | NA | NA |
| ENSG000002 | 4.71219649 | -0.321870082 | 0.37836094 | -0.8506958 | 0.39493833 | 0.5247426 | SSTR5-AS1 | 146336 |
| ENSG000001 | 3.38875506 | -0.320306837 | 0.37662935 | -0.8504564 | 0.39507139 | 0.5249022 | OR2D3 | 120775 |
| ENSG000002 | 3.53974959 | -0.321715482 | 0.37831695 | -0.8503861 | 0.39511046 | 0.5249369 | NA | NA |
| ENSG000001 | 423.419837 | 0.111257116 | 0.13083779 | 0.85034389 | 0.39513392 | 0.52495088 | DOCK5 | 80005 |
| ENSG000002 | 6.9293506 | -0.308544229 | 0.36286203 | -0.8503073 | 0.39515426 | 0.5249607 | NA | NA |
| ENSG000001 | 14.8076344 | -0.257790051 | 0.30318837 | -0.8502637 | 0.39517852 | 0.52497573 | ADORA2A | 135 |
| ENSG000002 | 4.8636077 | -0.318125229 | 0.37427089 | -0.8499866 | 0.39533255 | 0.52516315 | NA | NA |
| ENSG000002 | 2.23269022 | -0.313347333 | 0.36869802 | -0.8498753 | 0.39539443 | 0.52522815 | NA | NA |
| ENSG000002 | 4.81423383 | -0.317347033 | 0.3734638 | -0.8497397 | 0.3954698 | 0.52529385 | NA | NA |
| ENSG000002 | 7.18120669 | -0.306249386 | 0.36039898 | -0.849751 | 0.39546355 | 0.52529385 | NA | NA |
| ENSG000002 | 5.05629701 | -0.315953907 | 0.37186671 | -0.8496429 | 0.39552366 | 0.52534818 | FLJ33360 | 401172 |
| ENSG000002 | 4.29013933 | -0.320331771 | 0.37704376 | -0.8495878 | 0.39555431 | 0.52537168 | NA | NA |
| ENSG000001 | 25.4847543 | -0.225235373 | 0.26512545 | -0.8495426 | 0.39557942 | 0.52537308 | RIBC1 | 158787 |
| ENSG000001 | 5.8270753 | -0.315861798 | 0.37180364 | -0.8495393 | 0.39558128 | 0.52537308 | FOXE3 | 2301 |
| ENSG000002 | 3.81073539 | -0.321546353 | 0.37856319 | -0.8493862 | 0.39566642 | 0.52546895 | NA | NA |
| ENSG000001 | 43.5640007 | 0.187713336 | 0.22102274 | 0.84929423 | 0.39571759 | 0.5255197 | LMF1 | 64788 |

| | | | | | | | | |
|---|---|---|---|---|---|---|---|---|
| ENSG000002 | 3.35736187 | -0.320352158 | 0.37724709 | -0.8491839 | 0.39577896 | 0.52558399 | NA | NA |
| ENSG000002 | 2.23875143 | -0.311928695 | 0.36737002 | -0.8490859 | 0.39583351 | 0.52563922 | NA | NA |
| ENSG000002 | 4.61952957 | -0.320977686 | 0.37808236 | -0.8489623 | 0.39590225 | 0.52571328 | NA | NA |
| ENSG000001 | 21.1931153 | -0.237959372 | 0.2803514 | -0.8487897 | 0.39599833 | 0.52578921 | ZCCHC12 | 170261 |
| ENSG000002 | 4.33732695 | -0.320007608 | 0.37700631 | -0.8488123 | 0.39598572 | 0.52578921 | NA | NA |
| ENSG000002 | 2.25357997 | -0.312784302 | 0.36848664 | -0.8488349 | 0.39597319 | 0.52578921 | NA | NA |
| ENSG000001 | 6.61774184 | 0.31319753 | 0.36906579 | 0.84862248 | 0.39609139 | 0.52589556 | RAB43 | 339122 |
| ENSG000002 | 2.21724179 | -0.31257544 | 0.36835623 | -0.8485683 | 0.39612153 | 0.52591638 | NA | NA |
| ENSG000002 | 2.50916832 | -0.316337842 | 0.37279912 | -0.8485477 | 0.39613301 | 0.52591638 | NA | NA |
| ENSG000002 | 91.602241 | 0.153352909 | 0.18073211 | 0.84850948 | 0.3961543 | 0.52592741 | ALOX12P2 | 245 |
| ENSG000002 | 2.2144167 | -0.312514363 | 0.36834238 | -0.8484344 | 0.39619607 | 0.52596565 | NA | NA |
| ENSG000001 | 12.9369988 | -0.272222023 | 0.32089347 | -0.8483252 | 0.39625688 | 0.52602916 | PFN4 | 375189 |
| ENSG000002 | 2.24162859 | -0.313010192 | 0.36900718 | -0.8482496 | 0.39629898 | 0.52606782 | BRDTP1 | 643486 |
| ENSG000001 | 7965.81092 | 0.136985923 | 0.1615239 | 0.84808454 | 0.39639089 | 0.52617261 | PRRC2C | 23215 |
| ENSG000002 | 3.88340653 | -0.321047619 | 0.37858983 | -0.8480091 | 0.3964329 | 0.52621115 | NA | NA |
| ENSG000002 | 3.64347534 | -0.320572324 | 0.37806872 | -0.8479208 | 0.39648209 | 0.52624686 | NA | NA |
| ENSG000002 | 3.5957186 | -0.320086148 | 0.37749828 | -0.8479142 | 0.39648576 | 0.52624686 | NA | NA |
| ENSG000002 | 3.6205828 | -0.320584756 | 0.37812913 | -0.8478182 | 0.39653923 | 0.52628338 | HNRNPH3P1 | 106480233 |
| ENSG000002 | 3.90004111 | -0.320852451 | 0.37843934 | -0.8478306 | 0.39653232 | 0.52628338 | NA | NA |
| ENSG000002 | 15.125981 | 0.268143877 | 0.31629077 | 0.84777648 | 0.39656246 | 0.52629699 | LCAT | 3931 |
| ENSG000002 | 2.23481698 | -0.311482441 | 0.36746593 | -0.8476498 | 0.396633 | 0.52637338 | MIR4330 | 100422930 |
| ENSG000002 | 4.85862321 | 0.31679788 | 0.37378845 | 0.84753256 | 0.39669835 | 0.52644287 | MYCNUT | 103752554 |
| ENSG000002 | 15.2650439 | 0.262626984 | 0.30990447 | 0.84744497 | 0.39674715 | 0.5264904 | NA | NA |
| ENSG000002 | 27.3745698 | -0.210628127 | 0.24856193 | -0.8473869 | 0.39677949 | 0.52651609 | LINC01123 | 440894 |
| ENSG000002 | 6.89410585 | -0.310036891 | 0.36593443 | -0.8472471 | 0.3968574 | 0.52660224 | NA | NA |
| ENSG000002 | 6.90676666 | -0.311425009 | 0.36765094 | -0.8470671 | 0.39695773 | 0.52671813 | LOC283731 | 283731 |
| ENSG000002 | 2.25437612 | -0.310945867 | 0.36710143 | -0.8470298 | 0.39697849 | 0.52672844 | NA | NA |
| ENSG00000C | 50.7513773 | 0.17965896 | 0.21213711 | 0.84690021 | 0.39705075 | 0.52680708 | COL9A2 | 1298 |
| ENSG000002 | 2.53569452 | -0.314644078 | 0.37153726 | -0.8468709 | 0.39706711 | 0.52681156 | NA | NA |

| | | | | | | | | |
|---|---|---|---|---|---|---|---|---|
| ENSG000002 | 2.22133224 | -0.312541269 | 0.3690734 | -0.8468269 | 0.39709164 | 0.52682686 | RN7SL543P | 106480378 |
| ENSG000002 | 4.34650087 | -0.319618805 | 0.37744929 | -0.8467861 | 0.39711437 | 0.52683979 | NA | NA |
| ENSG000002 | 4.56215407 | -0.31871304 | 0.3763985 | -0.8467437 | 0.39713802 | 0.52685392 | NA | NA |
| ENSG000001 | 123.310766 | 0.131177663 | 0.15493569 | 0.84665881 | 0.39718533 | 0.52688932 | NCAM1 | 4684 |
| ENSG000002 | 2.22275426 | -0.311379969 | 0.36777921 | -0.8466492 | 0.39719069 | 0.52688932 | NA | NA |
| ENSG000001 | 15.64371 | -0.27791365 | 0.32828955 | -0.8465504 | 0.39724577 | 0.52691068 | RFPL2 | 10739 |
| ENSG000002 | 4.3376859 | -0.319520334 | 0.37743238 | -0.8465631 | 0.39723867 | 0.52691068 | RNU2-35P | 106480212 |
| ENSG000002 | 2.52242527 | -0.315806704 | 0.37304455 | -0.8465657 | 0.39723726 | 0.52691068 | NA | NA |
| ENSG000002 | 4.65801082 | -0.319547073 | 0.37749349 | -0.8464969 | 0.39727559 | 0.526933 | NA | NA |
| ENSG000002 | 11.0832027 | -0.287856426 | 0.34010208 | -0.8463824 | 0.39733944 | 0.52700044 | NA | NA |
| ENSG000002 | 5.86911848 | -0.313643254 | 0.37060906 | -0.8462914 | 0.39739022 | 0.52705055 | NA | NA |
| ENSG000002 | 8.82220152 | -0.291731943 | 0.3447526 | -0.8462066 | 0.39743748 | 0.52706152 | NA | NA |
| ENSG000002 | 9.48184829 | -0.297678971 | 0.35176144 | -0.8462524 | 0.39741195 | 0.52706152 | NA | NA |
| ENSG000002 | 3.38140057 | 0.320299428 | 0.37850295 | 0.84622704 | 0.3974261 | 0.52706152 | NA | NA |
| ENSG00000C | 72.9835936 | -0.151019373 | 0.17849069 | -0.846091 | 0.39750196 | 0.52709533 | CLEC2D | 29121 |
| ENSG000002 | 2.49357587 | -0.315864046 | 0.37330684 | -0.8461245 | 0.39748332 | 0.52709533 | NA | NA |
| ENSG000002 | 4.56428084 | -0.318786078 | 0.37677162 | -0.846099 | 0.39749753 | 0.52709533 | LOC105370458 | 105370458 |
| ENSG000002 | 10.5043816 | 0.292027899 | 0.3451809 | 0.84601408 | 0.39754489 | 0.52713501 | NA | NA |
| ENSG000001 | 20.9850181 | -0.244793888 | 0.28937529 | -0.8459391 | 0.39758669 | 0.5271732 | TDO2 | 6999 |
| ENSG000002 | 3.80554724 | -0.320183326 | 0.37850512 | -0.8459154 | 0.39759992 | 0.5271735 | NA | NA |
| ENSG000002 | 8.07806998 | -0.297444115 | 0.35168708 | -0.8457636 | 0.39768466 | 0.52726862 | EIF4A2P5 | 106480446 |
| ENSG000002 | 33.7380853 | -0.197765622 | 0.23388078 | -0.8455831 | 0.39778538 | 0.52738492 | NA | NA |
| ENSG000002 | 3.52143675 | -0.319942507 | 0.37840566 | -0.8455014 | 0.39783096 | 0.5274281 | NA | NA |
| ENSG000001 | 127.551875 | 0.127958826 | 0.15136116 | 0.84538745 | 0.39789455 | 0.52748746 | ZNF692 | 55657 |
| ENSG000002 | 3.81928843 | -0.320047036 | 0.37858608 | -0.8453745 | 0.39790175 | 0.52748746 | LINC01618 | 152578 |
| ENSG000002 | 2.21837598 | -0.311014322 | 0.36792381 | -0.8453226 | 0.39793073 | 0.52750863 | NA | NA |
| ENSG000002 | 2.25258218 | -0.310581407 | 0.36742269 | -0.8452973 | 0.39794487 | 0.52751013 | NA | NA |
| ENSG000002 | 2.7899889 | 0.318254601 | 0.37653304 | 0.84522357 | 0.39798603 | 0.52754744 | NA | NA |
| ENSG000002 | 17.8164918 | -0.248788728 | 0.29436451 | -0.8451723 | 0.39801464 | 0.52756813 | NA | NA |

| | | | | | | | | |
|---|---|---|---|---|---|---|---|---|
| ENSG000002 | 7.14235895 | 0.304859873 | 0.36072798 | 0.84512401 | 0.39804161 | 0.52758662 | NA | NA |
| ENSG000002 | 4.33215438 | -0.318857176 | 0.37736697 | -0.8449525 | 0.39813735 | 0.52769627 | LAT | 27040 |
| ENSG000002 | 7.48083192 | -0.303868417 | 0.35966951 | -0.8448545 | 0.39819206 | 0.52775155 | NA | NA |
| ENSG000002 | 3.42146468 | 0.319736573 | 0.37850033 | 0.84474583 | 0.39825277 | 0.52781475 | NA | NA |
| ENSG000001 | 156.047799 | 0.136545254 | 0.16164969 | 0.84469854 | 0.39827918 | 0.52781525 | CACNA2D3 | 55799 |
| ENSG000002 | 4.90525061 | -0.316550385 | 0.37474539 | -0.8447079 | 0.39827395 | 0.52781525 | NA | NA |
| ENSG000002 | 9.03479021 | -0.29038778 | 0.34380507 | -0.8446291 | 0.39831793 | 0.52784936 | NA | NA |
| ENSG000002 | 23.9958884 | 0.224376095 | 0.2656681 | 0.84457296 | 0.39834931 | 0.52787369 | NA | NA |
| ENSG000002 | 3.35248647 | 0.319347276 | 0.37822157 | 0.84433915 | 0.39847991 | 0.52801225 | NA | NA |
| ENSG000002 | 14.7272374 | 0.260877085 | 0.30896913 | 0.84434675 | 0.39847567 | 0.52801225 | NA | NA |
| ENSG000002 | 10.1962708 | 0.282564086 | 0.33479797 | 0.84398386 | 0.39867843 | 0.52825803 | NA | NA |
| ENSG000001 | 34.9179119 | -0.198752681 | 0.23552458 | -0.8438724 | 0.39874073 | 0.52827429 | IL17RE | 132014 |
| ENSG000002 | 3.28604007 | -0.318243688 | 0.3771256 | -0.8438666 | 0.39874397 | 0.52827429 | NA | NA |
| ENSG000002 | 8.90938855 | 0.292418925 | 0.34651088 | 0.84389537 | 0.39872787 | 0.52827429 | NA | NA |
| ENSG000002 | 17.2719169 | -0.251920746 | 0.29852548 | -0.8438836 | 0.39873447 | 0.52827429 | PRMT5-AS1 | 100505758 |
| ENSG000002 | 2.27284552 | -0.307979143 | 0.36498116 | -0.843822 | 0.39876886 | 0.52827429 | NA | NA |
| ENSG000002 | 17.4535899 | -0.247506586 | 0.29331183 | -0.8438343 | 0.398762 | 0.52827429 | NA | NA |
| ENSG000001 | 239.385925 | 0.253169343 | 0.30004567 | 0.84376935 | 0.3987983 | 0.52829604 | TYMS | 7298 |
| ENSG000001 | 24.2557366 | -0.230653288 | 0.27337933 | -0.8437115 | 0.39883063 | 0.52829941 | NA | NA |
| ENSG000002 | 2.2129024 | -0.309777598 | 0.3671725 | -0.8436841 | 0.39884595 | 0.52829941 | NA | NA |
| ENSG000002 | 2.21469744 | -0.311323309 | 0.36901008 | -0.8436716 | 0.39885297 | 0.52829941 | NA | NA |
| ENSG000002 | 5.48712196 | -0.318078891 | 0.37699998 | -0.8437106 | 0.39883112 | 0.52829941 | NA | NA |
| ENSG000001 | 150.896275 | 0.150372614 | 0.17824161 | 0.84364481 | 0.39886791 | 0.52830196 | SLC12A4 | 6560 |
| ENSG000002 | 3.52400126 | -0.319297656 | 0.37848636 | -0.8436173 | 0.39888327 | 0.52830504 | NA | NA |
| ENSG000001 | 6.5161513 | -0.311500193 | 0.36927113 | -0.8435542 | 0.39891859 | 0.52833457 | POTEI | 653269 |
| ENSG000002 | 2.22150851 | -0.310836319 | 0.36850217 | -0.8435129 | 0.39894168 | 0.52834789 | NA | NA |
| ENSG000002 | 7.9803734 | 0.298435967 | 0.35381696 | 0.84347559 | 0.39896251 | 0.52835823 | NA | NA |
| ENSG000002 | 2.74577111 | -0.317107699 | 0.37598612 | -0.8434027 | 0.39900328 | 0.52839402 | NA | NA |
| ENSG000002 | 35.668445 | -0.207093468 | 0.2455516 | -0.8433806 | 0.3990156 | 0.52839402 | NA | NA |

| | | | | | | | | |
|---|---|---|---|---|---|---|---|---|
| ENSG000001 | 17.7746976 | -0.250315175 | 0.29684169 | -0.8432615 | 0.39908221 | 0.5284477 | ADAD2 | 161931 |
| ENSG000002 | 3.85525382 | -0.319031774 | 0.37832635 | -0.8432714 | 0.3990767 | 0.5284477 | NA | NA |
| ENSG000001 | 159.89051 | -0.122465519 | 0.14525545 | -0.8431044 | 0.39917003 | 0.52854421 | CAV1 | 857 |
| ENSG000002 | 7.70808586 | -0.301250198 | 0.35731908 | -0.8430846 | 0.39918115 | 0.52854421 | NA | NA |
| ENSG000001 | 7.18233978 | -0.304976765 | 0.36178371 | -0.8429809 | 0.39923911 | 0.52859381 | CALML6 | 163688 |
| ENSG000002 | 4.04624865 | -0.318920775 | 0.37832949 | -0.842971 | 0.39924468 | 0.52859381 | NA | NA |
| ENSG000001 | 3.11406161 | 0.316134804 | 0.375057 | 0.84289803 | 0.39928547 | 0.5286133 | ACTL7A | 10881 |
| ENSG000002 | 4.58235456 | -0.316220823 | 0.3751566 | -0.8429035 | 0.3992824 | 0.5286133 | LOC102467655 | 102467655 |
| ENSG00000C | 24.8850334 | 0.224113046 | 0.26589402 | 0.84286606 | 0.39930336 | 0.5286197 | ACHE | 43 |
| ENSG000002 | 9.10998174 | -0.288747584 | 0.34258772 | -0.8428428 | 0.39931638 | 0.5286197 | NA | NA |
| ENSG000002 | 6.67098867 | -0.307409427 | 0.36476341 | -0.8427639 | 0.39936052 | 0.52866087 | NA | NA |
| ENSG000001 | 23.1906814 | 0.238570637 | 0.28312435 | 0.84263552 | 0.39943232 | 0.52873866 | TGFB1I1 | 7041 |
| ENSG000002 | 7.10295725 | -0.30270992 | 0.35927564 | -0.8425562 | 0.3994767 | 0.52878014 | NA | NA |
| ENSG000002 | 19.1789382 | 0.243761386 | 0.28934641 | 0.84245519 | 0.39953321 | 0.52883768 | NA | NA |
| ENSG000002 | 33.0681936 | -0.205939102 | 0.24449806 | -0.8422934 | 0.39962374 | 0.52894025 | KLLN | 100144748 |
| ENSG00000C | 54.9265669 | -0.169837069 | 0.2016743 | -0.8421354 | 0.39971217 | 0.52904002 | BCORL1 | 63035 |
| ENSG000002 | 9.41344305 | -0.287818652 | 0.34179072 | -0.8420903 | 0.39973741 | 0.52905616 | NA | NA |
| ENSG000002 | 5.91348155 | -0.312130865 | 0.37069692 | -0.8420109 | 0.39978183 | 0.52909768 | FOXP4-AS1 | 101060264 |
| ENSG000002 | 8.7752551 | 0.296897402 | 0.3526257 | 0.84196189 | 0.39980928 | 0.52911675 | NA | NA |
| ENSG000002 | 4.84092345 | -0.315374965 | 0.3746218 | -0.8418489 | 0.39987252 | 0.52918316 | NA | NA |
| ENSG000002 | 2.56423589 | 0.310749888 | 0.36918099 | 0.84172776 | 0.39994036 | 0.52925567 | NA | NA |
| ENSG000002 | 50.1055702 | -0.174170841 | 0.20699017 | -0.841445 | 0.40009869 | 0.52944792 | NA | NA |
| ENSG000002 | 19.6690979 | -0.238888986 | 0.28393324 | -0.8413562 | 0.40014842 | 0.52949645 | NA | NA |
| ENSG000002 | 2.48975298 | -0.313336821 | 0.3724298 | -0.8413312 | 0.4001624 | 0.52949767 | NA | NA |
| ENSG000002 | 5.35142056 | -0.313041129 | 0.3721467 | -0.8411767 | 0.40024896 | 0.52959489 | NA | NA |
| ENSG000002 | 7.93726833 | 0.307661655 | 0.36576164 | 0.84115342 | 0.40026199 | 0.52959489 | NA | NA |
| ENSG000002 | 4.34370803 | -0.317769464 | 0.37782614 | -0.8410468 | 0.40032173 | 0.52963936 | NA | NA |
| ENSG000002 | 4.86102447 | -0.314471871 | 0.37390236 | -0.8410535 | 0.40031799 | 0.52963936 | NA | NA |
| ENSG000002 | 6.40406241 | -0.308275748 | 0.36662637 | -0.8408444 | 0.40043508 | 0.52977205 | NA | NA |

| | | | | | | | | |
|---|---|---|---|---|---|---|---|---|
| ENSG000002 | 2.49433236 | -0.312820967 | 0.37204782 | -0.8408085 | 0.40045522 | 0.52978141 | NA | NA |
| ENSG000002 | 4.86979392 | -0.317480578 | 0.37760566 | -0.8407728 | 0.40047521 | 0.52978239 | NA | NA |
| ENSG000002 | 10.2401932 | -0.288634217 | 0.34330133 | -0.8407606 | 0.40048209 | 0.52978239 | LOC100128006 | 100128006 |
| ENSG000001 | 49.2877394 | 0.181020154 | 0.21532069 | 0.84070025 | 0.40051588 | 0.5298098 | OLFM3 | 118427 |
| ENSG000002 | 4.29633693 | -0.317331565 | 0.3774749 | -0.8406693 | 0.40053322 | 0.52981545 | NA | NA |
| ENSG000002 | 5.14487348 | -0.315332285 | 0.37512254 | -0.8406114 | 0.40056566 | 0.52984109 | NA | NA |
| ENSG000002 | 12.0022756 | -0.277929247 | 0.33065001 | -0.8405542 | 0.40059774 | 0.52986624 | NA | NA |
| ENSG000000 | 556.353038 | -0.110231108 | 0.13116524 | -0.8403988 | 0.40068484 | 0.52987734 | PKP2 | 5318 |
| ENSG000000 | 67.717009 | 0.170835184 | 0.20329551 | 0.84032937 | 0.40072374 | 0.52987734 | MAST3 | 23031 |
| ENSG000001 | 24.8924859 | 0.218866648 | 0.26041061 | 0.84046748 | 0.40064633 | 0.52987734 | FGF8 | 2253 |
| ENSG000002 | 6.26921532 | 0.307305507 | 0.3656837 | 0.84035878 | 0.40070725 | 0.52987734 | KIF25-AS1 | 100505879 |
| ENSG000002 | 9.82246431 | 0.293111771 | 0.34878946 | 0.84036877 | 0.40070165 | 0.52987734 | NA | NA |
| ENSG000002 | 2.28768148 | -0.306546526 | 0.36478437 | -0.8403499 | 0.40071221 | 0.52987734 | NA | NA |
| ENSG000002 | 21.5005998 | 0.231819535 | 0.27580737 | 0.84051246 | 0.40062112 | 0.52987734 | NA | NA |
| ENSG000002 | 4.55062059 | -0.315479581 | 0.37538617 | -0.8404134 | 0.40067662 | 0.52987734 | NA | NA |
| ENSG000002 | 2.4830708 | -0.311580594 | 0.37074884 | -0.8404088 | 0.4006792 | 0.52987734 | NA | NA |
| ENSG000002 | 2.24185672 | -0.308688829 | 0.36737823 | -0.840248 | 0.40076934 | 0.52992036 | NA | NA |
| ENSG000000 | 16.6709671 | -0.257263169 | 0.30619466 | -0.8401948 | 0.40079915 | 0.5299425 | CD22 | 933 |
| ENSG000001 | 41.8039617 | -0.188385144 | 0.22424462 | -0.8400877 | 0.40085922 | 0.52997008 | LRRTM1 | 347730 |
| ENSG000002 | 2.23997735 | -0.307675472 | 0.36623587 | -0.840102 | 0.40085122 | 0.52997008 | NA | NA |
| ENSG000002 | 3.87106394 | -0.318047899 | 0.37856915 | -0.8401316 | 0.40083461 | 0.52997008 | NA | NA |
| ENSG000002 | 2.24166084 | -0.308597757 | 0.36737857 | -0.8399993 | 0.40090875 | 0.53001829 | NA | NA |
| ENSG000002 | 4.3126143 | -0.317054776 | 0.37746093 | -0.8399671 | 0.40092681 | 0.53002487 | NA | NA |
| ENSG000002 | 4.07166102 | -0.31738097 | 0.37787889 | -0.8399013 | 0.40096373 | 0.53003913 | NA | NA |
| ENSG000002 | 15.7456357 | -0.253164951 | 0.30142039 | -0.8399065 | 0.4009608 | 0.53003913 | NA | NA |
| ENSG000001 | 18.8304488 | 0.245209197 | 0.2920345 | 0.83965833 | 0.40109999 | 0.53015012 | TMEM91 | 641649 |
| ENSG000001 | 55.1082842 | 0.171606812 | 0.20437547 | 0.83966443 | 0.40109657 | 0.53015012 | MYRIP | 25924 |
| ENSG000001 | 3.81127727 | -0.317830194 | 0.37851679 | -0.8396726 | 0.40109196 | 0.53015012 | OR4D5 | 219875 |
| ENSG000002 | 5.5489457 | -0.308980394 | 0.3679833 | -0.8396587 | 0.40109975 | 0.53015012 | NA | NA |

| | | | | | | | | |
|---|---|---|---|---|---|---|---|---|
| ENSG000002 | 2.20027907 | -0.309218157 | 0.36829065 | -0.8396036 | 0.40113068 | 0.5301734 | NA | NA |
| ENSG000002 | 8.1635415 | -0.298070987 | 0.35505857 | -0.8394981 | 0.40118987 | 0.53023436 | NA | NA |
| ENSG000002 | 4.07209712 | -0.317706753 | 0.37846442 | -0.8394627 | 0.40120973 | 0.53024332 | NA | NA |
| ENSG000001 | 21.7165365 | -0.231898895 | 0.27626225 | -0.8394158 | 0.40123602 | 0.53026078 | PNPLA7 | 375775 |
| ENSG000002 | 4.41770657 | -0.317734037 | 0.37859221 | -0.8392514 | 0.40132826 | 0.53033083 | NA | NA |
| ENSG000002 | 5.58418194 | -0.309339538 | 0.36858804 | -0.8392555 | 0.40132596 | 0.53033083 | LOC105375009 | 105375009 |
| ENSG000002 | 10.1199135 | -0.281609459 | 0.33553808 | -0.8392772 | 0.40131378 | 0.53033083 | NA | NA |
| ENSG000001 | 11.5187023 | -0.269866953 | 0.32163143 | -0.8390565 | 0.40143759 | 0.53045802 | P2RX3 | 5024 |
| ENSG000002 | 3.815844 | -0.317549951 | 0.37847733 | -0.8390197 | 0.40145823 | 0.530468 | RN7SL382P | 106481027 |
| ENSG000002 | 2.76065859 | -0.314318662 | 0.37465653 | -0.8389515 | 0.40149652 | 0.53050132 | NA | NA |
| ENSG000002 | 6.06100965 | -0.311588623 | 0.37142569 | -0.838899 | 0.40152602 | 0.530523 | NA | NA |
| ENSG000002 | 12.838501 | 0.269713752 | 0.32156261 | 0.83875968 | 0.40160418 | 0.53059831 | NA | NA |
| ENSG000002 | 2.23186869 | -0.30743489 | 0.36654921 | -0.8387275 | 0.40162226 | 0.53059831 | NA | NA |
| ENSG000002 | 5.31803143 | -0.311014062 | 0.37080904 | -0.8387445 | 0.40161267 | 0.53059831 | NA | NA |
| ENSG000002 | 4.27152592 | -0.316330565 | 0.37717207 | -0.8386903 | 0.40164311 | 0.53060856 | NA | NA |
| ENSG000001 | 143.32348 | -0.121846221 | 0.14530986 | -0.8385269 | 0.40173485 | 0.53071247 | SFMBT2 | 57713 |
| ENSG000002 | 4.14418117 | -0.317137277 | 0.37823145 | -0.8384741 | 0.40176448 | 0.53073432 | NA | NA |
| ENSG000001 | 4.36731849 | -0.31632788 | 0.37728818 | -0.8384251 | 0.401792 | 0.53075338 | NA | NA |
| ENSG000002 | 2.7317971 | -0.31556928 | 0.37647316 | -0.8382252 | 0.40190425 | 0.53086708 | HSPE1P25 | 106480260 |
| ENSG000002 | 2.46729762 | -0.313215659 | 0.37366422 | -0.8382276 | 0.40190289 | 0.53086708 | NA | NA |
| ENSG000002 | 2.5002294 | -0.309778774 | 0.36966058 | -0.8380087 | 0.40202583 | 0.53101037 | RNU6-681P | 106481911 |
| ENSG000001 | 11.7567467 | -0.269496226 | 0.32165179 | -0.8378509 | 0.40211447 | 0.53111016 | SEC16B | 89866 |
| ENSG000001 | 2.21416875 | -0.307684115 | 0.36727486 | -0.8377489 | 0.40217173 | 0.53116113 | RNU6-1273P | 106481947 |
| ENSG000002 | 4.08831631 | -0.316602724 | 0.37792683 | -0.8377355 | 0.40217927 | 0.53116113 | NA | NA |
| ENSG000002 | 3.31996497 | -0.314290116 | 0.37520445 | -0.8376503 | 0.40222714 | 0.53120706 | NA | NA |
| ENSG000002 | 14.6924093 | 0.258617782 | 0.30875111 | 0.83762542 | 0.40224111 | 0.53120821 | NA | NA |
| ENSG000002 | 4.34634497 | -0.315724084 | 0.37694694 | -0.8375823 | 0.40226534 | 0.5312229 | NA | NA |
| ENSG000002 | 4.37649241 | -0.316287518 | 0.37765624 | -0.8375011 | 0.40231096 | 0.53126076 | NA | NA |
| ENSG000002 | 2.23408032 | -0.308542305 | 0.36841548 | -0.8374846 | 0.4023202 | 0.53126076 | NA | NA |

| | | | | | | | | |
|---|---|---|---|---|---|---|---|---|
| ENSG000001 | 302.157759 | 0.10942487 | 0.13067516 | 0.83738082 | 0.40237855 | 0.53126665 | PYROXD1 | 79912 |
| ENSG000001 | 2.5065719 | -0.312184864 | 0.37279121 | -0.8374255 | 0.40235343 | 0.53126665 | NA | NA |
| ENSG000001 | 3.55580462 | -0.31648755 | 0.37794928 | -0.837381 | 0.40237844 | 0.53126665 | NA | NA |
| ENSG000002 | 4.57295972 | -0.314763369 | 0.37585798 | -0.8374529 | 0.40233802 | 0.53126665 | NA | NA |
| ENSG000002 | 6.84980891 | -0.304137155 | 0.3632095 | -0.8373601 | 0.40239017 | 0.53126665 | NA | NA |
| ENSG000002 | 4.30937839 | -0.315712082 | 0.37706924 | -0.8372788 | 0.40243587 | 0.53127509 | NA | NA |
| ENSG000002 | 4.80711479 | -0.313609824 | 0.37455829 | -0.8372791 | 0.40243571 | 0.53127509 | NA | NA |
| ENSG000002 | 2.49954535 | -0.312310433 | 0.37300528 | -0.8372815 | 0.40243433 | 0.53127509 | NA | NA |
| ENSG000001 | 21.1927282 | -0.239558372 | 0.28613066 | -0.8372342 | 0.40246094 | 0.53129089 | LOC284023 | 284023 |
| ENSG000002 | 11.503556 | -0.286190311 | 0.34183873 | -0.8372086 | 0.40247535 | 0.53129262 | NA | NA |
| ENSG000002 | 2.498547 | -0.311631238 | 0.37225107 | -0.8371534 | 0.40250636 | 0.53131626 | NA | NA |
| ENSG000002 | 3.86995403 | -0.316917867 | 0.37858337 | -0.8371151 | 0.40252785 | 0.53132733 | NA | NA |
| ENSG000002 | 2.2295334 | -0.308412508 | 0.36853265 | -0.8368662 | 0.4026678 | 0.53149476 | NA | NA |
| ENSG000001 | 35.2076555 | 0.197744829 | 0.23630023 | 0.83683723 | 0.40268406 | 0.53149893 | CRIP2 | 1397 |
| ENSG000002 | 5.70223066 | 0.308078189 | 0.36817418 | 0.83677293 | 0.40272022 | 0.53152934 | NA | NA |
| ENSG000002 | 8.09826697 | -0.296563739 | 0.3544363 | -0.8367194 | 0.4027503 | 0.53155174 | NA | NA |
| ENSG000001 | 35.1265807 | -0.19475561 | 0.23277896 | -0.8366547 | 0.40278668 | 0.53158247 | ADGRL4 | 64123 |
| ENSG00000C | 570.476338 | 0.115861693 | 0.13849497 | 0.83657691 | 0.40283043 | 0.5316229 | ARID4A | 5926 |
| ENSG000002 | 2.24802207 | -0.307442264 | 0.36751743 | -0.8365379 | 0.40285235 | 0.53163453 | NA | NA |
| ENSG000002 | 13.3711169 | -0.263012935 | 0.31442673 | -0.836484 | 0.40288266 | 0.53165723 | LOC100128494 | 100128494 |
| ENSG000002 | 2.50799425 | -0.311772799 | 0.37279385 | -0.8363142 | 0.40297816 | 0.53175321 | NA | NA |
| ENSG000002 | 25.1560029 | -0.217598139 | 0.26018898 | -0.8363081 | 0.40298162 | 0.53175321 | LOC101929140 | 101929140 |
| ENSG000002 | 2.46775408 | -0.311545857 | 0.37272078 | -0.8358693 | 0.40322844 | 0.53206157 | NA | NA |
| ENSG000002 | 3.06779044 | -0.313706803 | 0.3753214 | -0.8358351 | 0.40324767 | 0.53206964 | NA | NA |
| ENSG000002 | 3.38106088 | 0.316007795 | 0.37812093 | 0.83573208 | 0.40330564 | 0.53211153 | NA | NA |
| ENSG000002 | 2.75444889 | -0.314467701 | 0.37629914 | -0.8356854 | 0.40333191 | 0.53211153 | NA | NA |
| ENSG000002 | 4.59696109 | -0.314946429 | 0.37685252 | -0.8357286 | 0.4033076 | 0.53211153 | LOC101927495 | 101927495 |
| ENSG000002 | 2.75444889 | -0.314467701 | 0.37629914 | -0.8356854 | 0.40333191 | 0.53211153 | NA | NA |
| ENSG000001 | 4.88652063 | -0.313202614 | 0.37489933 | -0.8354312 | 0.40347495 | 0.53228292 | FMR1NB | 158521 |

| | | | | | | | | |
|---|---|---|---|---|---|---|---|---|
| ENSG000002 | 6.92441093 | -0.303297698 | 0.36311405 | -0.8352684 | 0.40356659 | 0.53238651 | NA | NA |
| ENSG000001 | 10.1321844 | -0.292027859 | 0.34969011 | -0.8351047 | 0.40365876 | 0.53246704 | CXCL13 | 10563 |
| ENSG000002 | 7.96939966 | -0.301619072 | 0.36118151 | -0.83509 | 0.40366703 | 0.53246704 | NA | NA |
| ENSG000002 | 2.30162323 | -0.303234529 | 0.36309687 | -0.835134 | 0.40364228 | 0.53246704 | NA | NA |
| ENSG000001 | 4.60692422 | 0.315449629 | 0.37777666 | 0.8350162 | 0.40370859 | 0.53250057 | ASCL2 | 430 |
| ENSG000002 | 4.82942112 | -0.313593555 | 0.37556195 | -0.8349982 | 0.40371872 | 0.53250057 | NA | NA |
| ENSG000002 | 8.82568705 | -0.297024259 | 0.35577041 | -0.8348762 | 0.40378739 | 0.53257383 | NA | NA |
| ENSG000001 | 16.3464409 | -0.253527375 | 0.3036848 | -0.8348372 | 0.40380937 | 0.53258549 | MAP1LC3C | 440738 |
| ENSG000002 | 7.12033581 | -0.302804734 | 0.36277769 | -0.834684 | 0.40389565 | 0.53268198 | NA | NA |
| ENSG000002 | 3.60754657 | -0.314802723 | 0.37716421 | -0.8346569 | 0.40391089 | 0.53268475 | RN7SL798P | 106480533 |
| ENSG000001 | 32.4275057 | -0.205351489 | 0.24607676 | -0.8345017 | 0.4039983 | 0.53275664 | TMEM106A | 113277 |
| ENSG000002 | 2.73413238 | -0.313117787 | 0.37522648 | -0.8344768 | 0.40401236 | 0.53275664 | NA | NA |
| ENSG000002 | 19.1634243 | -0.250963593 | 0.30074723 | -0.8344669 | 0.40401796 | 0.53275664 | NA | NA |
| ENSG000002 | 10.0846489 | -0.281679796 | 0.33753883 | -0.8345108 | 0.4039932 | 0.53275664 | NA | NA |
| ENSG000002 | 4.34885556 | 0.31590606 | 0.37859143 | 0.83442475 | 0.40404167 | 0.53277059 | RNU4ATAC18P | 106481866 |
| ENSG000002 | 2.54895147 | -0.309402782 | 0.37087396 | -0.8342532 | 0.40413832 | 0.5328807 | NA | NA |
| ENSG000002 | 4.10833226 | -0.3155609 | 0.37829037 | -0.8341764 | 0.40418159 | 0.53292042 | NA | NA |
| ENSG000002 | 5.31302751 | -0.309967324 | 0.37162237 | -0.8340922 | 0.40422902 | 0.53296563 | ATP11A-AS1 | 100874205 |
| ENSG000001 | 14.6825674 | -0.253881536 | 0.30442926 | -0.833959 | 0.40430406 | 0.53304724 | RASAL3 | 64926 |
| ENSG000002 | 2.7348696 | -0.312904788 | 0.37522698 | -0.833908 | 0.40433282 | 0.53306783 | LOC105379051 | 105379051 |
| ENSG000001 | 4.98644369 | -0.315596612 | 0.37847325 | -0.8338677 | 0.40435555 | 0.53308046 | TBX20 | 57057 |
| ENSG000001 | 146.943824 | -0.13024659 | 0.15621485 | -0.8337658 | 0.404413 | 0.53311561 | RASGEF1B | 153020 |
| ENSG000002 | 3.36819887 | -0.314584243 | 0.37731226 | -0.8337504 | 0.40442165 | 0.53311561 | NA | NA |
| ENSG000002 | 19.7325997 | -0.23469474 | 0.28148265 | -0.8337805 | 0.4044047 | 0.53311561 | LOC101928674 | 101928674 |
| ENSG000001 | 2.26035857 | -0.305639474 | 0.36663674 | -0.8336302 | 0.40448937 | 0.53317022 | NA | NA |
| ENSG000002 | 2.48656894 | -0.310740783 | 0.3727552 | -0.8336323 | 0.4044882 | 0.53317022 | NA | NA |
| ENSG000001 | 19.2186518 | -0.235828949 | 0.28291241 | -0.8335758 | 0.40452004 | 0.53317654 | PRSS36 | 146547 |
| ENSG000002 | 2.70928077 | 0.311318668 | 0.37349495 | 0.83352845 | 0.40454676 | 0.53317654 | NA | NA |
| ENSG000002 | 2.50865978 | -0.310447488 | 0.3724356 | -0.8335602 | 0.40452887 | 0.53317654 | NA | NA |

| | | | | | | | | |
|---|---|---|---|---|---|---|---|---|
| ENSG000002 | 25.1542253 | -0.215683316 | 0.25875687 | -0.8335366 | 0.40454217 | 0.53317654 | NA | NA |
| ENSG000002 | 4.5452955 | -0.314425555 | 0.37723819 | -0.8334934 | 0.4045665 | 0.53318522 | NA | NA |
| ENSG000002 | 2.20194882 | -0.304678855 | 0.36557053 | -0.8334338 | 0.40460009 | 0.53321217 | NA | NA |
| ENSG000002 | 2.18832915 | 0.304772525 | 0.36569797 | 0.83339955 | 0.40461943 | 0.53322032 | NA | NA |
| ENSG000002 | 10.4083072 | 0.279370942 | 0.33529678 | 0.83320497 | 0.40472914 | 0.53334757 | NA | NA |
| ENSG000001 | 18.9086086 | 0.243362874 | 0.29211813 | 0.83309745 | 0.40478977 | 0.53337632 | ARSA | 410 |
| ENSG000002 | 9.37640501 | 0.283675441 | 0.34051425 | 0.83307949 | 0.4047999 | 0.53337632 | NA | NA |
| ENSG000002 | 5.0377957 | -0.309964655 | 0.37206677 | -0.8330888 | 0.40479465 | 0.53337632 | NA | NA |
| ENSG000002 | 11.2410988 | -0.27785317 | 0.33352801 | -0.833073 | 0.40480357 | 0.53337632 | NA | NA |
| ENSG000002 | 5.08996224 | 0.309700605 | 0.37183937 | 0.83288815 | 0.40490781 | 0.53349634 | NA | NA |
| ENSG000002 | 5.4630512 | -0.312779292 | 0.37559812 | -0.8327499 | 0.40498581 | 0.53358176 | NA | NA |
| ENSG000002 | 8.68832002 | 0.297347584 | 0.35712298 | 0.83261958 | 0.40505931 | 0.53362046 | SNORD116-23 | 100033434 |
| ENSG000002 | 2.47530236 | -0.309833258 | 0.37210358 | -0.8326533 | 0.40504031 | 0.53362046 | NA | NA |
| ENSG000002 | 5.83153742 | -0.305254929 | 0.36662507 | -0.8326079 | 0.4050659 | 0.53362046 | NA | NA |
| ENSG000002 | 2.75018218 | -0.312922574 | 0.37583579 | -0.8326045 | 0.40506781 | 0.53362046 | NA | NA |
| ENSG000002 | 3.89115122 | -0.315156002 | 0.37857914 | -0.8324706 | 0.40514337 | 0.53368532 | NA | NA |
| ENSG000002 | 2.24879153 | -0.304913758 | 0.36626951 | -0.8324847 | 0.40513541 | 0.53368532 | NA | NA |
| ENSG000002 | 2.24406856 | -0.305831736 | 0.36739015 | -0.832444 | 0.40515836 | 0.53368773 | C17orf50 | 146853 |
| ENSG000002 | 7.94943045 | -0.300222212 | 0.36068064 | -0.8323768 | 0.40519627 | 0.53372033 | LOC101927877 | 101927877 |
| ENSG000002 | 4.83535171 | -0.311600765 | 0.37438882 | -0.8322918 | 0.40524423 | 0.53376616 | RNU1-59P | 106480167 |
| ENSG000002 | 5.54422252 | -0.306629258 | 0.36843986 | -0.8322369 | 0.40527522 | 0.53378618 | LINC00609 | 101101773 |
| ENSG000002 | 4.10125918 | -0.315071606 | 0.37859251 | -0.8322183 | 0.40528575 | 0.53378618 | NA | NA |
| ENSG000002 | 3.12332248 | -0.310733028 | 0.37342329 | -0.8321201 | 0.40534115 | 0.53382446 | NA | NA |
| ENSG000002 | 5.87463404 | -0.306692496 | 0.36856629 | -0.832123 | 0.4053395 | 0.53382446 | RN7SL849P | 106479531 |
| ENSG000002 | 2.47063125 | -0.309734008 | 0.37239617 | -0.8317325 | 0.40555994 | 0.53409525 | NA | NA |
| ENSG000002 | 62.7817392 | 0.179908066 | 0.21631221 | 0.83170556 | 0.40557516 | 0.53409795 | SNORD12B | 100113393 |
| ENSG000001 | 3.75697061 | -0.314844403 | 0.37859088 | -0.8316217 | 0.40562249 | 0.53413674 | PTH | 5741 |
| ENSG000002 | 2.76859916 | -0.311914172 | 0.37507413 | -0.8316067 | 0.40563096 | 0.53413674 | NA | NA |
| ENSG000001 | 13.3774253 | 0.272325118 | 0.32751548 | 0.83148778 | 0.40569813 | 0.53420784 | RNASE1 | 6035 |

| | | | | | | | | |
|---|---|---|---|---|---|---|---|---|
| ENSG000002 | 5.10197742 | -0.312150391 | 0.37547586 | -0.8313461 | 0.40577814 | 0.5342785 | NA | NA |
| ENSG000002 | 30.1345584 | 0.219692185 | 0.26425962 | 0.83134982 | 0.40577604 | 0.5342785 | NA | NA |
| ENSG000002 | 2.74121155 | -0.312514907 | 0.37593711 | -0.8312957 | 0.4058066 | 0.53429862 | NA | NA |
| ENSG000002 | 12.3242941 | -0.279672555 | 0.33646713 | -0.8312032 | 0.40585884 | 0.53435006 | NA | NA |
| ENSG000002 | 4.42078326 | -0.314321067 | 0.37816913 | -0.8311653 | 0.40588024 | 0.53436088 | NA | NA |
| ENSG000001 | 2.50578317 | -0.309554515 | 0.3724634 | -0.8311005 | 0.40591687 | 0.53439175 | GPR119 | 139760 |
| ENSG000002 | 4.12462945 | -0.314464878 | 0.37842519 | -0.830983 | 0.40598326 | 0.53446181 | LOC101928185 | 101928185 |
| ENSG000002 | 2.46041433 | -0.308950458 | 0.37182456 | -0.8309039 | 0.40602795 | 0.53450329 | NA | NA |
| ENSG000001 | 32.4038301 | 0.195941061 | 0.23582853 | 0.83086242 | 0.40605135 | 0.53451675 | PRRX2 | 51450 |
| ENSG000002 | 9.95242537 | -0.283849059 | 0.34172341 | -0.8306398 | 0.40617715 | 0.53466498 | NA | NA |
| ENSG000002 | 5.6665206 | -0.309401464 | 0.37254097 | -0.8305166 | 0.40624676 | 0.53473926 | NA | NA |
| ENSG000002 | 12.0652246 | -0.270906512 | 0.32622411 | -0.8304307 | 0.40629532 | 0.53478581 | NA | NA |
| ENSG000002 | 2.50250106 | -0.30862056 | 0.37168362 | -0.8303313 | 0.40635147 | 0.53484236 | TBC1D3B | 414059 |
| ENSG000002 | 3.65460497 | -0.313726755 | 0.37788161 | -0.830225 | 0.4064116 | 0.53484292 | NA | NA |
| ENSG000002 | 5.37935723 | -0.308706173 | 0.3718393 | -0.8302139 | 0.40641784 | 0.53484292 | LINC01564 | 101927171 |
| ENSG000002 | 7.1189478 | -0.299970074 | 0.36128177 | -0.830294 | 0.40637261 | 0.53484292 | NA | NA |
| ENSG000002 | 2.49666873 | -0.3094093 | 0.37267779 | -0.8302327 | 0.40640721 | 0.53484292 | NA | NA |
| ENSG000002 | 2.48904857 | -0.309185932 | 0.37239897 | -0.8302545 | 0.4063949 | 0.53484292 | OR10X1 | 128367 |
| ENSG000002 | 5.36972186 | -0.308806367 | 0.37197388 | -0.8301829 | 0.40643536 | 0.53484862 | NA | NA |
| ENSG000001 | 44.6926656 | 0.17546267 | 0.21138328 | 0.83006881 | 0.40649988 | 0.53488145 | FUK | 197258 |
| ENSG000002 | 2.18967961 | 0.303967668 | 0.3661828 | 0.83009815 | 0.40648329 | 0.53488145 | NA | NA |
| ENSG000002 | 2.72758246 | -0.311901851 | 0.37575379 | -0.8300697 | 0.40649935 | 0.53488145 | NA | NA |
| ENSG000001 | 212.619926 | -0.111110422 | 0.13388027 | -0.8299238 | 0.40658187 | 0.53497198 | SZT2 | 23334 |
| ENSG000002 | 2.50438042 | -0.309086818 | 0.37245781 | -0.8298573 | 0.40661948 | 0.53500411 | NA | NA |
| ENSG000002 | 2.52809472 | -0.308786665 | 0.37210885 | -0.8298289 | 0.40663555 | 0.53500789 | NA | NA |
| ENSG00000C | 70.2688347 | -0.155504533 | 0.1874396 | -0.8296247 | 0.40675098 | 0.53503824 | CALCR | 799 |
| ENSG000002 | 2.71195722 | -0.31164201 | 0.37562308 | -0.8296668 | 0.40672718 | 0.53503824 | HSFY1P1 | 27437 |
| ENSG000002 | 2.7509513 | -0.311460801 | 0.37541418 | -0.8296458 | 0.40673907 | 0.53503824 | NA | NA |
| ENSG000002 | 5.84765137 | -0.304517431 | 0.36701944 | -0.8297038 | 0.40670626 | 0.53503824 | NA | NA |

| | | | | | | | | |
|---|---|---|---|---|---|---|---|---|
| ENSG000002 | 4.32429235 | -0.31385117 | 0.37829697 | -0.8296423 | 0.40674105 | 0.53503824 | LOC100508631 | 100508631 |
| ENSG000002 | 2.46664418 | -0.30758225 | 0.3707145 | -0.8297012 | 0.40670777 | 0.53503824 | NA | NA |
| ENSG000002 | 2.49630346 | -0.308534441 | 0.37189584 | -0.829626 | 0.4067503 | 0.53503824 | NA | NA |
| ENSG000002 | 6.44171813 | -0.306529835 | 0.36953357 | -0.8295047 | 0.40681887 | 0.53511019 | NA | NA |
| ENSG000001 | 185.294326 | 0.119748709 | 0.14439305 | 0.82932463 | 0.40692074 | 0.53522682 | LRCH1 | 23143 |
| ENSG000002 | 2.21183292 | -0.303905543 | 0.36646391 | -0.8292919 | 0.40693927 | 0.53523383 | RNU6-309P | 106479704 |
| ENSG000002 | 15.5120691 | -0.251574801 | 0.30337564 | -0.8292518 | 0.40696192 | 0.53524627 | NA | NA |
| ENSG000002 | 6.10139005 | -0.306731157 | 0.36991278 | -0.8291986 | 0.40699204 | 0.53525116 | NA | NA |
| ENSG000002 | 3.81763774 | -0.31391241 | 0.37856881 | -0.8292083 | 0.40698654 | 0.53525116 | NA | NA |
| ENSG000001 | 4.33118938 | -0.31286162 | 0.37736076 | -0.8290783 | 0.4070601 | 0.5353233 | IL27 | 246778 |
| ENSG000001 | 41.6399551 | -0.186998069 | 0.22559604 | -0.8289067 | 0.40715721 | 0.53539927 | UBE2Q2P1 | 388165 |
| ENSG000002 | 8.6108658 | -0.29327027 | 0.35380392 | -0.8289062 | 0.40715747 | 0.53539927 | LOC105378231 | 105378231 |
| ENSG000002 | 4.57356054 | -0.311663751 | 0.37598868 | -0.8289179 | 0.40715086 | 0.53539927 | NA | NA |
| ENSG000002 | 2.48548682 | -0.308729063 | 0.37246994 | -0.8288697 | 0.40717812 | 0.53540905 | NA | NA |
| ENSG000002 | 2.7617881 | -0.310934847 | 0.37520061 | -0.8287163 | 0.40726498 | 0.5355059 | NA | NA |
| ENSG000001 | 19.9580025 | 0.232000932 | 0.27998301 | 0.82862505 | 0.40731661 | 0.5355231 | LGALS12 | 85329 |
| ENSG000002 | 2.22684416 | -0.302067932 | 0.36453859 | -0.8286309 | 0.40731331 | 0.5355231 | RNY4P37 | 106480314 |
| ENSG000002 | 9.85148536 | -0.278627609 | 0.3362537 | -0.8286232 | 0.40731767 | 0.5355231 | PWRN3 | 101928840 |
| ENSG000002 | 3.41031731 | -0.311517213 | 0.37597109 | -0.8285669 | 0.40734951 | 0.53554759 | NA | NA |
| ENSG000002 | 2.73575116 | -0.311493802 | 0.37597992 | -0.8284852 | 0.40739577 | 0.53557704 | NA | NA |
| ENSG000002 | 2.26835122 | -0.303093583 | 0.36584266 | -0.8284807 | 0.40739833 | 0.53557704 | NA | NA |
| ENSG000001 | 2.22118844 | -0.304273613 | 0.36730049 | -0.8284051 | 0.4074411 | 0.53561591 | RN7SKP72 | 106481748 |
| ENSG000002 | 3.98778405 | 0.313460989 | 0.37841113 | 0.82836091 | 0.40746614 | 0.53563146 | NA | NA |
| ENSG000002 | 5.06299239 | -0.308215239 | 0.3721275 | -0.8282517 | 0.40752796 | 0.535678 | NA | NA |
| ENSG000002 | 5.39094389 | -0.308726821 | 0.37274264 | -0.8282573 | 0.40752479 | 0.535678 | RNU6-84P | 106479603 |
| ENSG000002 | 11.8345595 | -0.267530985 | 0.32303134 | -0.8281889 | 0.4075635 | 0.53570736 | NA | NA |
| ENSG000002 | 2.79040219 | -0.309868887 | 0.37421901 | -0.8280415 | 0.40764698 | 0.53579971 | NA | NA |
| ENSG000001 | 58.9739904 | -0.167242223 | 0.20202649 | -0.8278232 | 0.40777062 | 0.53592279 | SLITRK3 | 22865 |
| ENSG000002 | 2.46164135 | -0.308935926 | 0.37320103 | -0.8278003 | 0.4077836 | 0.53592279 | NA | NA |

| | | | | | | | | |
|---|---|---|---|---|---|---|---|---|
| ENSG000002 | 2.51631199 | -0.307184514 | 0.37109311 | -0.8277829 | 0.40779349 | 0.53592279 | NA | NA |
| ENSG000002 | 2.23636408 | -0.30409465 | 0.36735944 | -0.827785 | 0.40779225 | 0.53592279 | NA | NA |
| ENSG00000C | 438.181561 | -0.098299383 | 0.11875441 | -0.8277535 | 0.4078101 | 0.53592725 | PPP1R12B | 4660 |
| ENSG000002 | 7.33162723 | -0.298073141 | 0.36011307 | -0.827721 | 0.40782853 | 0.53593411 | NA | NA |
| ENSG000001 | 2.45881626 | -0.308886287 | 0.37319297 | -0.8276852 | 0.40784883 | 0.53594341 | SCGB2A1 | 4246 |
| ENSG000002 | 7.59593028 | -0.297348669 | 0.35932511 | -0.8275199 | 0.40794246 | 0.53604908 | MIR550A3 | 100616354 |
| ENSG000002 | 32.0738899 | 0.196345229 | 0.23727811 | 0.82748984 | 0.40795948 | 0.53605407 | LINC01012 | 100507173 |
| ENSG000001 | 4.36468265 | -0.312777989 | 0.37800262 | -0.8274493 | 0.40798247 | 0.53606691 | NA | NA |
| ENSG000002 | 4.06424217 | -0.313077368 | 0.37838902 | -0.8273955 | 0.40801294 | 0.53608958 | NA | NA |
| ENSG000001 | 61.6802065 | 0.170026659 | 0.20557523 | 0.82707755 | 0.40819311 | 0.53628354 | BORCS7 | 119032 |
| ENSG000001 | 5.54251997 | -0.304779828 | 0.36850684 | -0.827067 | 0.40819911 | 0.53628354 | HSPE1P11 | 106481673 |
| ENSG000002 | 2.20733808 | -0.303063642 | 0.36643269 | -0.827065 | 0.40820023 | 0.53628354 | NA | NA |
| ENSG000002 | 2.72704167 | -0.311305816 | 0.37647105 | -0.8269051 | 0.40829087 | 0.53638524 | NA | NA |
| ENSG000002 | 3.52761508 | -0.312705393 | 0.37820909 | -0.8268056 | 0.40834726 | 0.53644194 | NA | NA |
| ENSG000002 | 4.07579637 | -0.31292676 | 0.37849872 | -0.8267578 | 0.40837437 | 0.53646017 | BOLA2P2 | 106480322 |
| ENSG000002 | 2.22391602 | -0.303399636 | 0.36698636 | -0.8267327 | 0.40838856 | 0.53646144 | NA | NA |
| ENSG000002 | 74.4169532 | 0.148758375 | 0.17994478 | 0.82668901 | 0.40841335 | 0.53647662 | BOLA3-AS1 | 100507171 |
| ENSG000001 | 22.4938828 | -0.230837746 | 0.27926018 | -0.8266046 | 0.40846121 | 0.53650473 | ASB9 | 140462 |
| ENSG000002 | 2.75999361 | -0.310117868 | 0.37516866 | -0.8266092 | 0.4084586 | 0.53650473 | LOC105375680 | 105375680 |
| ENSG000002 | 6.0810968 | -0.302535503 | 0.36607638 | -0.8264273 | 0.40856176 | 0.53660203 | NA | NA |
| ENSG000002 | 82.0603845 | -0.149411094 | 0.18078668 | -0.8264497 | 0.40854905 | 0.53660203 | NA | NA |
| ENSG000002 | 5.09580017 | -0.308665721 | 0.37352231 | -0.8263649 | 0.40859714 | 0.53663112 | NA | NA |
| ENSG000002 | 2.5091485 | -0.305828348 | 0.37012147 | -0.8262918 | 0.40863856 | 0.53666815 | NA | NA |
| ENSG000001 | 12.638244 | -0.26572738 | 0.32168885 | -0.8260385 | 0.40878224 | 0.5367873 | CRYGN | 155051 |
| ENSG000001 | 495.74451 | 0.098759826 | 0.11955486 | 0.8260628 | 0.40876847 | 0.5367873 | STXBP4 | 252983 |
| ENSG000002 | 17.9462948 | -0.239382043 | 0.28978048 | -0.8260806 | 0.40875834 | 0.5367873 | NA | NA |
| ENSG000002 | 5.58474996 | -0.305740692 | 0.37012592 | -0.8260451 | 0.40877853 | 0.5367873 | NA | NA |
| ENSG000002 | 11.4656561 | -0.278285288 | 0.33692633 | -0.8259529 | 0.40883079 | 0.53683367 | LOC284080 | 284080 |
| ENSG000002 | 8.8117846 | -0.287364165 | 0.34793904 | -0.8259038 | 0.40885869 | 0.53685291 | NA | NA |

| | | | | | | | | |
|---|---|---|---|---|---|---|---|---|
| ENSG000002 | 22.2490376 | -0.222706594 | 0.26966504 | -0.8258638 | 0.40888136 | 0.5368653 | LOC102724814 | 102724814 |
| ENSG000002 | 15.3072843 | -0.251418751 | 0.30446131 | -0.8257823 | 0.40892762 | 0.53690865 | NA | NA |
| ENSG000002 | 3.46800913 | -0.309355877 | 0.37466548 | -0.8256856 | 0.40898249 | 0.53692854 | HS6ST2-AS1 | 100874102 |
| ENSG000002 | 2.22247439 | -0.303834666 | 0.36797272 | -0.8256989 | 0.40897492 | 0.53692854 | NA | NA |
| ENSG000002 | 2.22668108 | -0.301934012 | 0.36567003 | -0.8257007 | 0.40897388 | 0.53692854 | NA | NA |
| ENSG000001 | 11.2424762 | -0.283569398 | 0.34348912 | -0.8255557 | 0.40905618 | 0.53700791 | PXT1 | 222659 |
| ENSG000002 | 4.45707313 | -0.31249557 | 0.37856614 | -0.8254715 | 0.40910394 | 0.53704749 | NA | NA |
| ENSG000002 | 2.74617605 | -0.309885541 | 0.37541139 | -0.8254559 | 0.40911282 | 0.53704749 | NA | NA |
| ENSG000002 | 8.69486508 | -0.289084892 | 0.35024329 | -0.8253831 | 0.40915413 | 0.53708433 | NA | NA |
| ENSG000002 | 5.83089717 | -0.304317786 | 0.36874511 | -0.8252795 | 0.40921292 | 0.53712673 | NA | NA |
| ENSG000002 | 3.06803839 | -0.309240083 | 0.37470687 | -0.8252853 | 0.40920962 | 0.53712673 | NA | NA |
| ENSG000001 | 4.80148575 | -0.308983603 | 0.37444037 | -0.8251877 | 0.40926501 | 0.53717771 | KCNK18 | 338567 |
| ENSG000002 | 2.47273819 | -0.306733433 | 0.3717471 | -0.8251132 | 0.40930733 | 0.53721588 | NA | NA |
| ENSG000002 | 3.82238596 | -0.312322582 | 0.37855546 | -0.8250378 | 0.40935011 | 0.53725462 | NA | NA |
| ENSG000002 | 2.49058716 | -0.306020732 | 0.37092973 | -0.82501 | 0.40936592 | 0.53725799 | NA | NA |
| ENSG000002 | 6.50398743 | 0.302697892 | 0.36696472 | 0.82486918 | 0.40944586 | 0.53731074 | GPR20 | 2843 |
| ENSG000002 | 2.53873565 | -0.306434633 | 0.3714925 | -0.8248743 | 0.40944294 | 0.53731074 | NA | NA |
| ENSG000002 | 4.97468904 | -0.31165819 | 0.37782022 | -0.8248849 | 0.40943695 | 0.53731074 | NA | NA |
| ENSG000001 | 29.9875168 | 0.203518414 | 0.24681037 | 0.82459425 | 0.40960199 | 0.53746344 | ADAMTS14 | 140766 |
| ENSG000002 | 2.45574377 | -0.307740921 | 0.37319011 | -0.8246224 | 0.40958601 | 0.53746344 | NA | NA |
| ENSG000002 | 4.59385689 | -0.310314692 | 0.37632307 | -0.8245965 | 0.40960069 | 0.53746344 | NA | NA |
| ENSG000002 | 98.590325 | -0.133114707 | 0.16144908 | -0.8244996 | 0.40965573 | 0.53751657 | STRADA | 92335 |
| ENSG000002 | 193.690071 | 0.161757243 | 0.19621319 | 0.82439537 | 0.40971495 | 0.53757687 | NA | NA |
| ENSG000002 | 44.7263695 | 0.178783802 | 0.21687429 | 0.82436604 | 0.40973161 | 0.53758134 | NA | NA |
| ENSG000002 | 10.3200354 | -0.274623402 | 0.33323047 | -0.8241245 | 0.4098688 | 0.53774395 | OR9A4 | 130075 |
| ENSG000002 | 2.19446526 | -0.299697003 | 0.36368446 | -0.8240578 | 0.40990674 | 0.53777632 | NA | NA |
| ENSG000001 | 58.2925966 | 0.166953531 | 0.20260834 | 0.82402102 | 0.40992761 | 0.5377863 | SGK3 | 23678 |
| ENSG000002 | 5.300396 | -0.306957117 | 0.37254008 | -0.8239573 | 0.40996382 | 0.5378164 | NA | NA |
| ENSG000002 | 3.82780225 | -0.311884012 | 0.37857629 | -0.823834 | 0.41003389 | 0.53789093 | NA | NA |

| | | | | | | | | |
|---|---|---|---|---|---|---|---|---|
| ENSG000002 | 3.33425361 | -0.310259569 | 0.37664571 | -0.8237438 | 0.41008514 | 0.53794076 | NA | NA |
| ENSG000002 | 2.51122374 | -0.305437347 | 0.37080664 | -0.8237106 | 0.41010404 | 0.53794814 | SLC9A9-AS2 | 106480356 |
| ENSG000002 | 2.74870775 | -0.309066188 | 0.37523794 | -0.8236539 | 0.41013625 | 0.53797299 | NA | NA |
| ENSG000001 | 2.23329159 | -0.30248605 | 0.36726009 | -0.823629 | 0.41015041 | 0.53797416 | FOXI1 | 2299 |
| ENSG000002 | 2.52461674 | -0.305405872 | 0.37082517 | -0.8235845 | 0.41017568 | 0.5379899 | NA | NA |
| ENSG000001 | 20.7405105 | -0.230601715 | 0.28000587 | -0.8235603 | 0.41018944 | 0.53799056 | TSHZ1 | 10194 |
| ENSG000002 | 2.21853961 | -0.301801762 | 0.3664901 | -0.8234923 | 0.41022812 | 0.53802388 | NA | NA |
| ENSG000002 | 6.31391026 | -0.298212015 | 0.36220049 | -0.8233341 | 0.41031803 | 0.53810719 | LOC105371795 | 105371795 |
| ENSG000002 | 13.0930215 | -0.267077854 | 0.32439425 | -0.8233125 | 0.41033028 | 0.53810719 | NA | NA |
| ENSG000002 | 6.04113793 | -0.299689381 | 0.3640053 | -0.8233105 | 0.41033145 | 0.53810719 | NA | NA |
| ENSG000002 | 5.0717864 | -0.306613138 | 0.37244391 | -0.8232465 | 0.41036784 | 0.53813751 | NA | NA |
| ENSG000002 | 2.99882829 | -0.310589733 | 0.37746178 | -0.8228376 | 0.41060037 | 0.53840761 | NA | NA |
| ENSG000002 | 6.35683293 | -0.30544919 | 0.37120727 | -0.8228535 | 0.41059133 | 0.53840761 | NA | NA |
| ENSG000002 | 2.72545515 | -0.308328088 | 0.37473297 | -0.8227941 | 0.41062508 | 0.53841983 | OSBPL9P5 | 106480232 |
| ENSG000002 | 2.20556319 | -0.301494659 | 0.36643656 | -0.8227745 | 0.41063624 | 0.53841983 | MIR3973 | 100616311 |
| ENSG000002 | 2.48978578 | -0.305896052 | 0.37182528 | -0.8226876 | 0.41068566 | 0.53846588 | NA | NA |
| ENSG000002 | 2.25836298 | -0.300927874 | 0.3657959 | -0.8226661 | 0.41069791 | 0.53846588 | NA | NA |
| ENSG000002 | 2.50538977 | -0.3023551 | 0.36755173 | -0.8226192 | 0.41072459 | 0.53848344 | NA | NA |
| ENSG000002 | 16.4302155 | -0.244086535 | 0.29682394 | -0.8223277 | 0.41089044 | 0.53868115 | NA | NA |
| ENSG000002 | 8.65769795 | -0.28710971 | 0.34915131 | -0.8223074 | 0.41090196 | 0.53868115 | NA | NA |
| ENSG000001 | 62.82456 | -0.160919032 | 0.19576275 | -0.8220105 | 0.41107093 | 0.53886782 | KIAA1462 | 57608 |
| ENSG000001 | 14.1555698 | -0.256657138 | 0.3122232 | -0.822031 | 0.41105928 | 0.53886782 | FGG | 2266 |
| ENSG000002 | 5.54457553 | -0.308265802 | 0.37503524 | -0.8219649 | 0.41109688 | 0.53888442 | NA | NA |
| ENSG000002 | 2.72630446 | -0.309058715 | 0.3761819 | -0.8215672 | 0.41132326 | 0.53916374 | NA | NA |
| ENSG000001 | 13.3459257 | -0.272583744 | 0.33180275 | -0.8215235 | 0.41134817 | 0.53917897 | ARL9 | 132946 |
| ENSG000002 | 2.4710558 | -0.305957255 | 0.37244162 | -0.8214905 | 0.41136692 | 0.5391803 | NA | NA |
| ENSG000002 | 2.5199536 | -0.305534655 | 0.37194258 | -0.8214565 | 0.41138628 | 0.5391803 | NA | NA |
| ENSG000002 | 10.0355319 | -0.275870816 | 0.33583332 | -0.8214516 | 0.41138908 | 0.5391803 | LINC00551 | 283483 |
| ENSG000002 | 2.24350796 | -0.301742645 | 0.36738759 | -0.8213196 | 0.41146422 | 0.53926135 | SERPINA2 | 390502 |

| | | | | | | | | |
|---|---|---|---|---|---|---|---|---|
| ENSG000002 | 2.74665267 | -0.308613516 | 0.37576656 | -0.8212905 | 0.41148081 | 0.53926566 | NA | NA |
| ENSG000002 | 24.3191954 | -0.227381038 | 0.27692736 | -0.8210855 | 0.41159757 | 0.53934897 | CERS6-AS1 | 100861402 |
| ENSG000002 | 2.55482634 | -0.29918444 | 0.36435671 | -0.8211306 | 0.41157189 | 0.53934897 | NA | NA |
| ENSG000002 | 5.30565584 | -0.305300801 | 0.3718203 | -0.8210977 | 0.4115906 | 0.53934897 | NA | NA |
| ENSG000002 | 7.0846648 | -0.295741111 | 0.36017442 | -0.8211053 | 0.41158632 | 0.53934897 | NA | NA |
| ENSG000002 | 7.08361982 | -0.297036051 | 0.36177112 | -0.8210607 | 0.41161169 | 0.53935004 | NA | NA |
| ENSG000001 | 23.463485 | -0.214371459 | 0.26112629 | -0.8209494 | 0.41167512 | 0.53939829 | SLC5A5 | 6528 |
| ENSG000002 | 8.34343117 | -0.287993701 | 0.35080059 | -0.8209613 | 0.41166832 | 0.53939829 | NA | NA |
| ENSG000002 | 4.07451748 | -0.310713071 | 0.37854235 | -0.8208145 | 0.41175191 | 0.53947996 | NA | NA |
| ENSG000002 | 10.8175961 | -0.270778849 | 0.32989898 | -0.8207932 | 0.41176405 | 0.53947996 | CDRT4 | 284040 |
| ENSG000002 | 14.7785766 | -0.252292987 | 0.3074061 | -0.8207156 | 0.41180828 | 0.53952047 | SERPINB4 | 6318 |
| ENSG000002 | 8.62071022 | -0.288231999 | 0.35134602 | -0.8203651 | 0.41200803 | 0.53966819 | CKLF | 51192 |
| ENSG000002 | 2.7722656 | -0.307972499 | 0.37538389 | -0.8204201 | 0.41197665 | 0.53966819 | NA | NA |
| ENSG000002 | 3.01894837 | -0.309569247 | 0.37736045 | -0.8203542 | 0.41201419 | 0.53966819 | NA | NA |
| ENSG000002 | 16.9941394 | -0.24232586 | 0.29537082 | -0.8204123 | 0.41198109 | 0.53966819 | NA | NA |
| ENSG000002 | 5.03442265 | -0.306763192 | 0.37388226 | -0.8204807 | 0.41194211 | 0.53966819 | NA | NA |
| ENSG000002 | 2.72777888 | -0.308643085 | 0.37618295 | -0.8204601 | 0.41195388 | 0.53966819 | NA | NA |
| ENSG000002 | 4.6264882 | -0.309572005 | 0.3773503 | -0.8203836 | 0.41199745 | 0.53966819 | NA | NA |
| ENSG000002 | 3.00451736 | -0.309721353 | 0.37759761 | -0.8202418 | 0.41207827 | 0.53973469 | NA | NA |
| ENSG000002 | 2.48594383 | -0.305296917 | 0.37242636 | -0.8197511 | 0.41235801 | 0.54008365 | NA | NA |
| ENSG000002 | 4.62772732 | -0.309538267 | 0.37761575 | -0.8197176 | 0.41237712 | 0.54009123 | NA | NA |
| ENSG000002 | 43.1955674 | 0.175943784 | 0.21464604 | 0.81969265 | 0.41239134 | 0.54009241 | SUCLG2-AS1 | 101927111 |
| ENSG000001 | 10.0195462 | -0.277361802 | 0.33845098 | -0.8195036 | 0.41249914 | 0.54018125 | TIMD4 | 91937 |
| ENSG000002 | 5.56587672 | -0.302279309 | 0.36885603 | -0.8195049 | 0.41249843 | 0.54018125 | NA | NA |
| ENSG000002 | 2.46992574 | -0.303463473 | 0.37029962 | -0.8195079 | 0.41249668 | 0.54018125 | NA | NA |
| ENSG000002 | 2.97115993 | -0.309306711 | 0.37749106 | -0.8193749 | 0.41257253 | 0.54024247 | NA | NA |
| ENSG000002 | 6.54263216 | -0.295653101 | 0.36082153 | -0.8193887 | 0.41256465 | 0.54024247 | BCAR4 | 400500 |
| ENSG000002 | 2.48548627 | -0.304334695 | 0.3714336 | -0.8193515 | 0.41258587 | 0.54024249 | NA | NA |
| ENSG000002 | 2.22674111 | -0.300019123 | 0.36618895 | -0.8193014 | 0.41261447 | 0.54024505 | RN7SL169P | 106480372 |

| | | | | | | | | |
|---|---|---|---|---|---|---|---|---|
| ENSG000002 | 3.63361671 | 0.310173884 | 0.37857384 | 0.81932201 | 0.41260272 | 0.54024505 | NA | NA |
| ENSG000002 | 5.53377802 | -0.301748166 | 0.36832463 | -0.8192451 | 0.41264657 | 0.54025219 | NA | NA |
| ENSG000002 | 2.79445204 | 0.307030399 | 0.37476897 | 0.81925245 | 0.41264239 | 0.54025219 | NA | NA |
| ENSG000001 | 190.818184 | -0.114132101 | 0.13933352 | -0.8191288 | 0.41271294 | 0.54032163 | SMYD2 | 56950 |
| ENSG000002 | 4.30072873 | -0.309303248 | 0.37768978 | -0.8189347 | 0.4128237 | 0.54044919 | NA | NA |
| ENSG000002 | 11.3730018 | -0.272310384 | 0.33258159 | -0.8187777 | 0.41291327 | 0.540549 | NA | NA |
| ENSG000001 | 2.47828345 | -0.304444058 | 0.37185662 | -0.8187136 | 0.41294986 | 0.54057946 | NA | NA |
| ENSG000002 | 2.19933334 | -0.300565865 | 0.36721444 | -0.8185023 | 0.41307043 | 0.54070238 | NA | NA |
| ENSG000002 | 2.53634775 | -0.303668624 | 0.37099467 | -0.8185256 | 0.41305715 | 0.54070238 | NA | NA |
| ENSG000002 | 3.86625534 | -0.309772161 | 0.37848887 | -0.8184446 | 0.41310338 | 0.54072805 | NA | NA |
| ENSG000001 | 11.3008543 | -0.268179897 | 0.32769429 | -0.8183844 | 0.41313772 | 0.54075555 | NA | NA |
| ENSG000002 | 6.27654616 | -0.298237781 | 0.36443741 | -0.8183512 | 0.41315669 | 0.54076293 | NA | NA |
| ENSG00000C | 98.7647916 | 0.159390677 | 0.19479717 | 0.81823916 | 0.41322064 | 0.54081381 | CLDN11 | 5010 |
| ENSG000002 | 2.49083565 | -0.304395638 | 0.37201431 | -0.8182364 | 0.41322224 | 0.54081381 | NA | NA |
| ENSG000002 | 2.18016765 | 0.297918092 | 0.36417414 | 0.81806494 | 0.41332011 | 0.54092444 | RNU7-125P | 106480818 |
| ENSG000002 | 2.52838702 | -0.300254773 | 0.36705571 | -0.8180087 | 0.41335221 | 0.54094899 | AP1S2P1 | 106480305 |
| ENSG000002 | 7.94328581 | -0.296366965 | 0.36235741 | -0.8178857 | 0.41342243 | 0.54102343 | NA | NA |
| ENSG000002 | 4.13044271 | -0.309527164 | 0.37853039 | -0.8177076 | 0.41352419 | 0.54113913 | NA | NA |
| ENSG000002 | 10.3200867 | -0.274029018 | 0.33525108 | -0.8173844 | 0.41370876 | 0.54134573 | LOC101929023 | 101929023 |
| ENSG000002 | 24.5743217 | -0.219617109 | 0.26868084 | -0.8173903 | 0.41370543 | 0.54134573 | LOC102724684 | 102724684 |
| ENSG000002 | 6.83144421 | -0.297660681 | 0.36420651 | -0.8172854 | 0.41376532 | 0.5413848 | C21orf62 | 56245 |
| ENSG000002 | 2.77246148 | -0.306632477 | 0.37517777 | -0.8172992 | 0.41375749 | 0.5413848 | NA | NA |
| ENSG000002 | 8.25862184 | 0.287673333 | 0.3520672 | 0.8170978 | 0.41387254 | 0.54150761 | NA | NA |
| ENSG000001 | 40.5213853 | 0.185593517 | 0.22714417 | 0.81707364 | 0.41388635 | 0.5415082 | KLF4 | 9314 |
| ENSG000002 | 2.7293775 | -0.307111574 | 0.37593045 | -0.8169372 | 0.41396432 | 0.54159274 | NA | NA |
| ENSG000002 | 5.58162595 | -0.301872492 | 0.3695775 | -0.8168043 | 0.41404027 | 0.54167464 | NA | NA |
| ENSG000002 | 32.0253299 | -0.198189596 | 0.24274248 | -0.8164603 | 0.41423693 | 0.54189695 | NA | NA |
| ENSG000002 | 23.2332939 | 0.218493725 | 0.26760521 | 0.81647783 | 0.4142269 | 0.54189695 | NA | NA |
| ENSG000001 | 5.39538741 | -0.304057878 | 0.37243963 | -0.8163951 | 0.41427422 | 0.54192825 | CTSG | 1511 |

| | | | | | | | | |
|---|---|---|---|---|---|---|---|---|
| ENSG000001 | 89.1758854 | 0.172556252 | 0.21137223 | 0.81636199 | 0.41429313 | 0.5419355 | AHDC1 | 27245 |
| ENSG000001 | 28.997182 | 0.206113234 | 0.25249074 | 0.81631999 | 0.41431715 | 0.54193798 | TMSB15B | 286527 |
| ENSG000001 | 5.09512169 | -0.306783886 | 0.3758385 | -0.8162652 | 0.41434848 | 0.54193798 | C7orf62 | 219557 |
| ENSG000001 | 19.824661 | 0.232180311 | 0.28444208 | 0.81626569 | 0.41434819 | 0.54193798 | MFSD7 | 84179 |
| ENSG000002 | 3.56832022 | -0.308811783 | 0.37830797 | -0.8162973 | 0.4143301 | 0.54193798 | NA | NA |
| ENSG000002 | 2.18792275 | -0.298342524 | 0.36550986 | -0.8162366 | 0.41436484 | 0.54194189 | NA | NA |
| ENSG000002 | 2.97637238 | -0.307876219 | 0.37723539 | -0.8161382 | 0.41442108 | 0.54199798 | RNU6-1124P | 106481534 |
| ENSG000001 | 96.7387158 | 0.140719049 | 0.17243944 | 0.8160491 | 0.41447205 | 0.54201639 | MTERF2 | 80298 |
| ENSG000002 | 2.43442946 | -0.303387157 | 0.37177154 | -0.816058 | 0.41446694 | 0.54201639 | NA | NA |
| ENSG000002 | 36.8914126 | -0.186677127 | 0.2287588 | -0.8160435 | 0.41447526 | 0.54201639 | NA | NA |
| ENSG000001 | 2.4940885 | 0.296854089 | 0.36381246 | 0.81595361 | 0.41452667 | 0.54203229 | OR52L1 | 338751 |
| ENSG000001 | 34.9858926 | 0.208337983 | 0.25532794 | 0.81596235 | 0.41452167 | 0.54203229 | NA | NA |
| ENSG000002 | 6.30572508 | 0.301763537 | 0.36982995 | 0.81595213 | 0.41452752 | 0.54203229 | NA | NA |
| ENSG000001 | 124.772649 | 0.165761699 | 0.20318397 | 0.81582075 | 0.41460266 | 0.54211307 | DIRAS1 | 148252 |
| ENSG000002 | 5.52371013 | -0.301989928 | 0.37018642 | -0.815778 | 0.41462713 | 0.54212758 | NA | NA |
| ENSG000002 | 10.0559654 | -0.273820843 | 0.33568299 | -0.8157126 | 0.41466453 | 0.54215878 | SNORD72 | 619564 |
| ENSG000002 | 25.5623036 | 0.208574931 | 0.25571328 | 0.81565935 | 0.41469499 | 0.54215878 | ENO1-IT1 | 106480431 |
| ENSG000002 | 5.32324477 | -0.30349684 | 0.37209529 | -0.8156428 | 0.41470447 | 0.54215878 | NA | NA |
| ENSG000002 | 2.74591546 | -0.30621516 | 0.37540938 | -0.8156833 | 0.41468129 | 0.54215878 | NA | NA |
| ENSG000002 | 2.74070301 | -0.306279485 | 0.37559859 | -0.8154436 | 0.41481841 | 0.54229025 | NA | NA |
| ENSG000002 | 2.26194456 | -0.296443707 | 0.36355774 | -0.8153965 | 0.4148454 | 0.54230805 | NA | NA |
| ENSG000002 | 4.52314093 | -0.307295839 | 0.37710762 | -0.8148757 | 0.41514346 | 0.5426802 | NA | NA |
| ENSG000002 | 4.86827991 | -0.306103788 | 0.37569193 | -0.8147734 | 0.41520203 | 0.5427227 | NA | NA |
| ENSG000002 | 3.79214289 | -0.308442293 | 0.37857349 | -0.8147488 | 0.41521612 | 0.5427227 | SPRR2F | 6705 |
| ENSG000002 | 17.9282798 | -0.236683357 | 0.2904947 | -0.8147596 | 0.4152099 | 0.5427227 | CEP83-AS1 | 144486 |
| ENSG000002 | 454.132183 | 0.11101101 | 0.13628539 | 0.81454814 | 0.41533101 | 0.54277269 | ERCC6 | 2074 |
| ENSG000002 | 6.07202313 | -0.297755551 | 0.3655539 | -0.8145325 | 0.41533993 | 0.54277269 | NA | NA |
| ENSG000002 | 5.54254013 | -0.300062094 | 0.36836039 | -0.8145884 | 0.41530796 | 0.54277269 | NA | NA |
| ENSG000002 | 2.7203548 | -0.30626402 | 0.37596845 | -0.8146003 | 0.41530112 | 0.54277269 | NA | NA |

| | | | | | | | | |
|---|---|---|---|---|---|---|---|---|
| ENSG000002 | 2.18267213 | 0.298050591 | 0.36587053 | 0.81463406 | 0.41528181 | 0.54277269 | NA | NA |
| ENSG000002 | 4.59020319 | -0.306525317 | 0.37632709 | -0.8145183 | 0.41534807 | 0.54277269 | NA | NA |
| ENSG000002 | 18.2464873 | -0.240154259 | 0.2948139 | -0.8145961 | 0.41530354 | 0.54277269 | NA | NA |
| ENSG000001 | 7.30242985 | -0.295774116 | 0.36315862 | -0.8144488 | 0.41538788 | 0.54278973 | SLC39A5 | 283375 |
| ENSG000002 | 4.84025094 | -0.306251269 | 0.37602056 | -0.8144535 | 0.41538519 | 0.54278973 | MIR4465 | 100616180 |
| ENSG000001 | 51.2564674 | 0.167000015 | 0.20510609 | 0.81421286 | 0.41552302 | 0.54294883 | C12orf60 | 144608 |
| ENSG000001 | 1016.83861 | 0.104678126 | 0.12858999 | 0.8140457 | 0.41561877 | 0.54300396 | ELK4 | 2005 |
| ENSG000001 | 43.466174 | 0.194158522 | 0.23849756 | 0.81409016 | 0.4155933 | 0.54300396 | NA | NA |
| ENSG000001 | 2.73406822 | -0.30580326 | 0.37565549 | -0.8140524 | 0.41561492 | 0.54300396 | RXFP3 | 51289 |
| ENSG000002 | 22.4663767 | 0.21618661 | 0.26555967 | 0.81407922 | 0.41559957 | 0.54300396 | CD302 | 9936 |
| ENSG000002 | 16.7060432 | -0.267750583 | 0.32897644 | -0.8138898 | 0.41570806 | 0.54310311 | NA | NA |
| ENSG000002 | 2.43735814 | -0.302166307 | 0.37133311 | -0.8137338 | 0.41579746 | 0.54320241 | NA | NA |
| ENSG000001 | 55.1342484 | -0.168651818 | 0.20729714 | -0.8135752 | 0.41588833 | 0.54330362 | NA | NA |
| ENSG000002 | 4.33555968 | -0.307304779 | 0.37778718 | -0.8134336 | 0.41596951 | 0.54339216 | NA | NA |
| ENSG00000C | 32.3724633 | -0.198884348 | 0.24454347 | -0.8132883 | 0.41605277 | 0.54348343 | USP2 | 9099 |
| ENSG00000C | 2.78232907 | 0.304860328 | 0.37491346 | 0.81314853 | 0.4161329 | 0.54353317 | NA | NA |
| ENSG000002 | 12.5931379 | -0.261455336 | 0.32151884 | -0.8131882 | 0.41611015 | 0.54353317 | NA | NA |
| ENSG000002 | 2.51573785 | -0.299522223 | 0.36836162 | -0.8131201 | 0.41614923 | 0.54353317 | NA | NA |
| ENSG000002 | 4.7743984 | -0.305437745 | 0.37564368 | -0.813105 | 0.41615787 | 0.54353317 | NA | NA |
| ENSG000002 | 16.2827881 | 0.246077418 | 0.30261655 | 0.81316576 | 0.41612303 | 0.54353317 | NA | NA |
| ENSG000002 | 2.78350735 | -0.305034102 | 0.37526268 | -0.8128549 | 0.41630127 | 0.54370295 | NA | NA |
| ENSG000001 | 9.8076119 | -0.276000641 | 0.33956525 | -0.8128059 | 0.41632935 | 0.54371226 | S100A2 | 6273 |
| ENSG000002 | 4.58372395 | -0.306950681 | 0.37765413 | -0.8127825 | 0.41634275 | 0.54371226 | NA | NA |
| ENSG000002 | 2.17879791 | 0.29326567 | 0.36082144 | 0.8127723 | 0.41634862 | 0.54371226 | NA | NA |
| ENSG000002 | 3.83884125 | -0.307647018 | 0.3785879 | -0.8126172 | 0.4164376 | 0.54381093 | NA | NA |
| ENSG000002 | 22.4551463 | -0.214895672 | 0.26446927 | -0.8125544 | 0.41647358 | 0.54384041 | FEZF1-AS1 | 154860 |
| ENSG000002 | 9.40430969 | -0.280925145 | 0.34574473 | -0.8125218 | 0.41649226 | 0.54384729 | NA | NA |
| ENSG000001 | 75.1651445 | 0.148984856 | 0.18339509 | 0.81237104 | 0.41657876 | 0.5439315 | CADM4 | 199731 |
| ENSG000002 | 2.22563141 | -0.297334495 | 0.36601202 | -0.8123626 | 0.41658358 | 0.5439315 | NA | NA |

| | | | | | | | | |
|---|---|---|---|---|---|---|---|---|
| ENSG000002 | 11.6325071 | -0.267402016 | 0.32924282 | -0.8121727 | 0.41669256 | 0.54405628 | NA | NA |
| ENSG000001 | 81.5637296 | 0.161118632 | 0.19839006 | 0.81213055 | 0.41671672 | 0.54406177 | ABHD3 | 171586 |
| ENSG000002 | 3.82743065 | -0.307444402 | 0.37859263 | -0.8120718 | 0.41675043 | 0.54406177 | NA | NA |
| ENSG000002 | 4.42762937 | 0.306092336 | 0.3769078 | 0.81211463 | 0.41672586 | 0.54406177 | NA | NA |
| ENSG000002 | 151.291629 | -0.121962896 | 0.15018382 | -0.8120908 | 0.41673953 | 0.54406177 | NBPF19 | 101060226 |
| ENSG000002 | 3.3521421 | 0.304995366 | 0.37563976 | 0.8119358 | 0.41682847 | 0.54414613 | ACY1 | 95 |
| ENSG000001 | 14.0030911 | -0.261086715 | 0.32162423 | -0.8117756 | 0.41692038 | 0.54421356 | SLC4A5 | 57835 |
| ENSG000001 | 40.0625458 | 0.188995059 | 0.23281639 | 0.81177729 | 0.41691943 | 0.54421356 | NA | NA |
| ENSG000002 | 2.50458949 | -0.301738085 | 0.37169495 | -0.8117896 | 0.41691238 | 0.54421356 | NA | NA |
| ENSG000002 | 2.4776906 | -0.301458241 | 0.37141726 | -0.8116431 | 0.41699647 | 0.54429536 | RNU1-49P | 106480162 |
| ENSG00000C | 394.531459 | 0.098583691 | 0.12147217 | 0.81157427 | 0.41703596 | 0.54431186 | TARBP1 | 6894 |
| ENSG000001 | 13.5311956 | -0.252650237 | 0.31130495 | -0.8115844 | 0.41703015 | 0.54431186 | CXCL14 | 9547 |
| ENSG000002 | 4.39976986 | -0.306899955 | 0.37818841 | -0.8115002 | 0.4170785 | 0.54434987 | LOC102724053 | 102724053 |
| ENSG000001 | 19.4514036 | -0.234888077 | 0.28946309 | -0.8114612 | 0.41710084 | 0.5443615 | FAM107A | 11170 |
| ENSG000002 | 8.89722146 | -0.284388278 | 0.3505513 | -0.8112601 | 0.41721632 | 0.5444947 | NA | NA |
| ENSG000002 | 4.51804059 | -0.305647754 | 0.37681427 | -0.8111364 | 0.41728737 | 0.54455236 | NA | NA |
| ENSG000002 | 2.23146484 | -0.297708357 | 0.36701837 | -0.8111538 | 0.41727734 | 0.54455236 | NA | NA |
| ENSG000002 | 3.57090115 | 0.306970296 | 0.37850632 | 0.81100441 | 0.41736314 | 0.54456362 | NA | NA |
| ENSG000002 | 5.40416141 | -0.30363871 | 0.37439292 | -0.8110162 | 0.41735639 | 0.54456362 | NA | NA |
| ENSG000002 | 8.81455653 | -0.285777455 | 0.35234632 | -0.8110698 | 0.41732559 | 0.54456362 | NA | NA |
| ENSG000002 | 2.70005845 | -0.304872628 | 0.37590479 | -0.8110368 | 0.41734452 | 0.54456362 | SLC25A30-AS1 | 100874259 |
| ENSG000002 | 4.65137636 | -0.306710114 | 0.37817816 | -0.8110202 | 0.41735409 | 0.54456362 | NA | NA |
| ENSG000002 | 4.21503219 | 0.306309206 | 0.37770233 | 0.81098044 | 0.4173769 | 0.54456405 | NA | NA |
| ENSG00000C | 83.3998764 | 0.159653799 | 0.19687478 | 0.81094083 | 0.41739965 | 0.54457621 | RAB27A | 5873 |
| ENSG000002 | 4.30344835 | -0.30637992 | 0.37782149 | -0.8109118 | 0.4174163 | 0.54458042 | NA | NA |
| ENSG000002 | 8.47723384 | 0.286182722 | 0.35302241 | 0.81066445 | 0.41755839 | 0.54474827 | NA | NA |
| ENSG000002 | 4.42804945 | -0.306885313 | 0.37858559 | -0.8106101 | 0.4175896 | 0.54477145 | NA | NA |
| ENSG000002 | 5.54469825 | -0.300672593 | 0.37094257 | -0.8105637 | 0.41761626 | 0.5447887 | NA | NA |
| ENSG000002 | 2.49812334 | -0.300201403 | 0.37039115 | -0.8104983 | 0.41765384 | 0.5448202 | LINC00552 | 100130386 |

| | | | | | | | | |
|---|---|---|---|---|---|---|---|---|
| ENSG000002 | 5.81760722 | -0.301214272 | 0.37169247 | -0.8103857 | 0.41771853 | 0.54488706 | NA | NA |
| ENSG000002 | 5.51151799 | -0.299941875 | 0.37016537 | -0.8102916 | 0.41777262 | 0.54494009 | LOC101928414 | 101928414 |
| ENSG000002 | 5.09342058 | -0.303245801 | 0.37425775 | -0.8102592 | 0.41779119 | 0.54494679 | NA | NA |
| ENSG000002 | 170.118442 | 0.131591098 | 0.1624207 | 0.81018674 | 0.41783286 | 0.5449836 | LOC105369306 | 105369306 |
| ENSG000002 | 2.46058352 | -0.29998529 | 0.37028058 | -0.8101567 | 0.41785012 | 0.54498859 | RNU6-1098P | 106481935 |
| ENSG000002 | 7.02285022 | 0.289762677 | 0.35768197 | 0.81011262 | 0.41787545 | 0.5450041 | NA | NA |
| ENSG000002 | 23.2404515 | 0.219187858 | 0.27057494 | 0.81008188 | 0.41789312 | 0.54500961 | NA | NA |
| ENSG000001 | 81.5395976 | 0.145836291 | 0.18003219 | 0.81005677 | 0.41790755 | 0.5450109 | DZIP1L | 199221 |
| ENSG000002 | 2.55473249 | -0.298058808 | 0.36797892 | -0.8099888 | 0.4179466 | 0.5450443 | NA | NA |
| ENSG000001 | 8.92491293 | -0.282926792 | 0.34936839 | -0.8098237 | 0.41804153 | 0.54514721 | LINC01555 | 439927 |
| ENSG000002 | 11.0583792 | 0.267398936 | 0.33020174 | 0.80980475 | 0.4180524 | 0.54514721 | NA | NA |
| ENSG00000C | 95.7813419 | 0.13913224 | 0.17183 | 0.80970868 | 0.41810763 | 0.5452017 | MZF1 | 7593 |
| ENSG000002 | 8.92395148 | -0.283253573 | 0.34983812 | -0.8096704 | 0.41812964 | 0.54521286 | NA | NA |
| ENSG000002 | 9.40913754 | 0.279938401 | 0.34580167 | 0.80953456 | 0.41820773 | 0.54529716 | ERVV-2 | 100271846 |
| ENSG000002 | 3.83557954 | -0.306387838 | 0.3784957 | -0.8094883 | 0.41823433 | 0.54531431 | NA | NA |
| ENSG000002 | 5.81749153 | -0.298303562 | 0.36856856 | -0.809357 | 0.4183098 | 0.54539518 | NA | NA |
| ENSG000001 | 33.6144376 | 0.193104129 | 0.23862783 | 0.8092272 | 0.41838447 | 0.545475 | USHBP1 | 83878 |
| ENSG000001 | 2.96521082 | -0.305636457 | 0.37780091 | -0.8089882 | 0.41852196 | 0.54563671 | OR13D1 | 286365 |
| ENSG000002 | 2.71241478 | -0.304093667 | 0.37591553 | -0.8089415 | 0.4185488 | 0.54565416 | NA | NA |
| ENSG000002 | 2.50228591 | -0.299796274 | 0.37078402 | -0.8085469 | 0.41877581 | 0.54593255 | NA | NA |
| ENSG000002 | 5.54903854 | -0.29886832 | 0.36971212 | -0.8083812 | 0.41887118 | 0.5459627 | NA | NA |
| ENSG000002 | 3.00448456 | -0.304487287 | 0.37666992 | -0.8083663 | 0.41887972 | 0.5459627 | NA | NA |
| ENSG000002 | 22.6639067 | -0.22856658 | 0.28271351 | -0.8084742 | 0.41881765 | 0.5459627 | NA | NA |
| ENSG000002 | 3.25170866 | -0.305586568 | 0.37800795 | -0.8084131 | 0.41885283 | 0.5459627 | NA | NA |
| ENSG000002 | 30.5960009 | 0.205995344 | 0.25482028 | 0.80839462 | 0.41886345 | 0.5459627 | NA | NA |
| ENSG000002 | 183.30992 | 0.112135361 | 0.1387136 | 0.80839488 | 0.4188633 | 0.5459627 | NA | NA |
| ENSG000002 | 8.87733638 | -0.286245639 | 0.35411708 | -0.8083362 | 0.41889709 | 0.54596779 | LL22NC03-63E9 | 648691 |
| ENSG000002 | 6.07449481 | -0.297684341 | 0.36832882 | -0.8082027 | 0.41897391 | 0.54604107 | NA | NA |
| ENSG000002 | 2.70120006 | -0.30323152 | 0.37519753 | -0.8081917 | 0.41898025 | 0.54604107 | NA | NA |

| | | | | | | | | |
|---|---|---|---|---|---|---|---|---|
| ENSG000002 | 2.96321991 | -0.304848394 | 0.37723206 | -0.808119 | 0.41902208 | 0.54607804 | NA | NA |
| ENSG000002 | 5.26007206 | -0.300793661 | 0.37223149 | -0.8080823 | 0.41904323 | 0.54608805 | NA | NA |
| ENSG000002 | 2.23999773 | -0.293997417 | 0.36385142 | -0.808015 | 0.41908193 | 0.54612093 | NA | NA |
| ENSG000002 | 2.48792678 | -0.30053554 | 0.37203395 | -0.8078175 | 0.41919564 | 0.54621646 | NA | NA |
| ENSG000002 | 5.33229992 | -0.300645607 | 0.37215443 | -0.8078517 | 0.41917593 | 0.54621646 | NA | NA |
| ENSG000002 | 2.17420933 | -0.295002627 | 0.36518199 | -0.8078236 | 0.41919215 | 0.54621646 | NA | NA |
| ENSG000002 | 2.46252237 | -0.300042972 | 0.37161122 | -0.807411 | 0.41942975 | 0.54648639 | NA | NA |
| ENSG000002 | 4.78623354 | -0.302795194 | 0.37501174 | -0.8074286 | 0.41941961 | 0.54648639 | NA | NA |
| ENSG000001 | 2.44436508 | -0.298898122 | 0.3702452 | -0.8072978 | 0.41949495 | 0.5465011 | GFRA4 | 64096 |
| ENSG000001 | 4.46370717 | 0.304248932 | 0.37686714 | 0.80731086 | 0.41948741 | 0.5465011 | NA | NA |
| ENSG000002 | 5.67052727 | -0.300679398 | 0.3724507 | -0.8072999 | 0.41949374 | 0.5465011 | NA | NA |
| ENSG000002 | 90.1975517 | 0.141679821 | 0.17549248 | 0.80732701 | 0.41947811 | 0.5465011 | NBPF15 | 284565 |
| ENSG000001 | 8.92911289 | 0.27641547 | 0.34241405 | 0.80725506 | 0.41951955 | 0.54651559 | TIGD3 | 220359 |
| ENSG000001 | 34.4619533 | 0.185558762 | 0.22992473 | 0.80704135 | 0.41964266 | 0.54662328 | LEAP2 | 116842 |
| ENSG000002 | 5.28629333 | -0.299718522 | 0.37136424 | -0.8070743 | 0.41962369 | 0.54662328 | NA | NA |
| ENSG000002 | 3.86181311 | -0.305503166 | 0.3785451 | -0.8070456 | 0.4196402 | 0.54662328 | NA | NA |
| ENSG000002 | 6.3001842 | -0.292936631 | 0.36302713 | -0.8069276 | 0.41970817 | 0.54669106 | NA | NA |
| ENSG000002 | 2.73099573 | -0.302840278 | 0.37540007 | -0.8067134 | 0.4198316 | 0.54683427 | NA | NA |
| ENSG000002 | 3.37787073 | 0.304747629 | 0.37778318 | 0.80667337 | 0.41985469 | 0.54684677 | NA | NA |
| ENSG000002 | 4.16914897 | -0.305358707 | 0.3785896 | -0.8065692 | 0.41991472 | 0.5468983 | MIR1295A | 100302178 |
| ENSG000002 | 8.38597942 | -0.288400921 | 0.35757052 | -0.8065568 | 0.4199219 | 0.5468983 | NA | NA |
| ENSG000002 | 5.79369814 | -0.297724713 | 0.36914069 | -0.8065345 | 0.41993471 | 0.5468983 | NA | NA |
| ENSG000002 | 12.387086 | -0.263907097 | 0.32723266 | -0.8064815 | 0.41996526 | 0.54692052 | MKRN2OS | 100129480 |
| ENSG000002 | 6.29465213 | -0.292612871 | 0.36298617 | -0.8061268 | 0.42016975 | 0.54716926 | NA | NA |
| ENSG000002 | 5.57177355 | -0.299757658 | 0.3719376 | -0.8059354 | 0.42028013 | 0.54729542 | NA | NA |
| ENSG000001 | 2.17642097 | -0.294291911 | 0.36519174 | -0.8058559 | 0.42032595 | 0.54733752 | MT1B | 4490 |
| ENSG000002 | 3.25176073 | -0.304464333 | 0.37792791 | -0.8056148 | 0.42046496 | 0.54750095 | NA | NA |
| ENSG000002 | 2.47626879 | -0.299990753 | 0.37240613 | -0.8055473 | 0.42050392 | 0.5475341 | NA | NA |
| ENSG000002 | 2.71136436 | -0.302228424 | 0.37520438 | -0.8055035 | 0.42052922 | 0.54754945 | MT1A | 4489 |

| | | | | | | | | |
|---|---|---|---|---|---|---|---|---|
| ENSG000002 | 3.00595953 | -0.303925708 | 0.37735163 | -0.8054178 | 0.42057864 | 0.54759623 | NA | NA |
| ENSG000002 | 3.00118428 | -0.304025252 | 0.37749679 | -0.8053718 | 0.42060518 | 0.5476132 | NA | NA |
| ENSG000002 | 2.51015402 | -0.298592543 | 0.37079978 | -0.8052662 | 0.42066606 | 0.54766269 | NA | NA |
| ENSG000002 | 2.73342797 | -0.301098147 | 0.37391463 | -0.8052591 | 0.4206702 | 0.54766269 | NA | NA |
| ENSG000002 | 8.89831483 | -0.287776611 | 0.35741059 | -0.8051709 | 0.42072105 | 0.5477113 | NA | NA |
| ENSG000002 | 2.31006492 | -0.29059618 | 0.36100859 | -0.8049564 | 0.42084483 | 0.54785486 | NA | NA |
| ENSG000002 | 9.81441031 | -0.272942478 | 0.33909008 | -0.8049262 | 0.42086229 | 0.54786 | NA | NA |
| ENSG000002 | 6.07611325 | -0.296340921 | 0.36825552 | -0.8047155 | 0.42098388 | 0.54800069 | NA | NA |
| ENSG000002 | 2.25936133 | -0.293637327 | 0.36491969 | -0.8046629 | 0.42101425 | 0.54802262 | RN7SKP227 | 106481818 |
| ENSG000001 | 52.4444926 | -0.171111196 | 0.21267477 | -0.8045674 | 0.42106934 | 0.54804183 | ALDH1L1 | 10840 |
| ENSG000001 | 425.684783 | 0.105841509 | 0.13155089 | 0.80456702 | 0.42106958 | 0.54804183 | PCLO | 27445 |
| ENSG000002 | 6.14199579 | 0.29304462 | 0.36422357 | 0.80457347 | 0.42106586 | 0.54804183 | NA | NA |
| ENSG000002 | 2.99330319 | -0.30350439 | 0.37723794 | -0.8045437 | 0.42108306 | 0.54804183 | NA | NA |
| ENSG000002 | 2.19714132 | -0.292587545 | 0.3636952 | -0.8044856 | 0.42111659 | 0.54806788 | NA | NA |
| ENSG000002 | 5.61124426 | -0.299815055 | 0.37270926 | -0.8044207 | 0.42115404 | 0.54809902 | NA | NA |
| ENSG000002 | 4.57279108 | -0.303107995 | 0.37690266 | -0.8042076 | 0.42127708 | 0.54824156 | NA | NA |
| ENSG000002 | 2.98536261 | -0.302877638 | 0.37664396 | -0.8041484 | 0.42131128 | 0.54826847 | NA | NA |
| ENSG000002 | 2.70863623 | -0.300689351 | 0.37394511 | -0.8041002 | 0.42133909 | 0.54826947 | NA | NA |
| ENSG000002 | 3.41793638 | -0.299126437 | 0.3719918 | -0.8041211 | 0.42132707 | 0.54826947 | NA | NA |
| ENSG000002 | 5.09967459 | -0.302013783 | 0.375644 | -0.8039894 | 0.42140311 | 0.54833518 | NA | NA |
| ENSG000001 | 13.0805983 | -0.256728902 | 0.31937265 | -0.8038537 | 0.42148145 | 0.54840192 | SPATA41 | 388182 |
| ENSG000002 | 2.71349167 | -0.301963299 | 0.37563866 | -0.8038664 | 0.42147414 | 0.54840192 | ABHD11-AS1 | 171022 |
| ENSG000002 | 15.167721 | -0.247150358 | 0.30747216 | -0.8038138 | 0.42150453 | 0.54841435 | NA | NA |
| ENSG000001 | 514.183706 | 0.108520622 | 0.13502336 | 0.80371736 | 0.42156022 | 0.54846922 | MME | 4311 |
| ENSG000002 | 2.47823159 | -0.298106972 | 0.37101674 | -0.8034866 | 0.42169355 | 0.54862508 | NA | NA |
| ENSG000002 | 4.75668541 | 0.303669762 | 0.37796028 | 0.80344359 | 0.42171839 | 0.5486398 | NA | NA |
| ENSG000002 | 8.44967547 | -0.288029597 | 0.35850894 | -0.8034098 | 0.42173791 | 0.54864759 | HTN3 | 3347 |
| ENSG000002 | 4.27599592 | -0.303886568 | 0.37826957 | -0.8033598 | 0.4217668 | 0.54866758 | NA | NA |
| ENSG000001 | 2.98727389 | -0.302042662 | 0.37601063 | -0.8032822 | 0.42181163 | 0.54870828 | GAST | 2520 |

| | | | | | | | | |
|---|---|---|---|---|---|---|---|---|
| ENSG00000C | 128.232692 | 0.135153471 | 0.16826477 | 0.80321906 | 0.42184813 | 0.54872057 | OPN3 | 23596 |
| ENSG000001 | 7.03673567 | -0.287778833 | 0.35828011 | -0.803223 | 0.42184584 | 0.54872057 | VSTM1 | 284415 |
| ENSG000002 | 70.1098594 | 0.146509153 | 0.18240828 | 0.80319356 | 0.42186287 | 0.54872213 | NA | NA |
| ENSG000002 | 2.70755466 | -0.301334998 | 0.37520371 | -0.8031237 | 0.42190323 | 0.54875703 | NA | NA |
| ENSG000001 | 323.34839 | 0.120899804 | 0.15054158 | 0.80309907 | 0.42191748 | 0.54875796 | SIPA1L2 | 57568 |
| ENSG00000C | 217.716067 | 0.111524854 | 0.13888423 | 0.80300591 | 0.42197132 | 0.54877518 | GRAMD1B | 57476 |
| ENSG000001 | 16.8253638 | -0.251341504 | 0.31299 | -0.8030337 | 0.42195528 | 0.54877518 | TMEM184A | 202915 |
| ENSG000002 | 2.49609494 | -0.29749379 | 0.3704671 | -0.8030235 | 0.42196115 | 0.54877518 | NA | NA |
| ENSG000001 | 117.430683 | 0.125910677 | 0.15681019 | 0.80294956 | 0.42200389 | 0.54879994 | EP400NL | 347918 |
| ENSG00000C | 2.97850024 | -0.303232252 | 0.37768512 | -0.8028705 | 0.42204957 | 0.54884175 | WNT11 | 7481 |
| ENSG000002 | 17.458237 | 0.235304718 | 0.29311304 | 0.80277806 | 0.42210303 | 0.54887605 | LOC100505715 | 100505715 |
| ENSG000002 | 3.0201337 | 0.303324833 | 0.3778334 | 0.80280048 | 0.42209006 | 0.54887605 | NA | NA |
| ENSG000002 | 2.97047534 | -0.303028078 | 0.37749055 | -0.8027435 | 0.42212302 | 0.54888445 | NA | NA |
| ENSG000001 | 2.72285927 | -0.301188124 | 0.37528498 | -0.8025584 | 0.42223 | 0.54897304 | LCN9 | 392399 |
| ENSG000002 | 2.75698114 | -0.301308405 | 0.37541926 | -0.8025918 | 0.42221073 | 0.54897304 | NA | NA |
| ENSG000002 | 2.21791952 | -0.292527835 | 0.36449551 | -0.8025554 | 0.42223176 | 0.54897304 | NA | NA |
| ENSG000002 | 31.046965 | -0.195230903 | 0.2432816 | -0.8024894 | 0.42226992 | 0.54900505 | NA | NA |
| ENSG000002 | 2.74140797 | -0.301152034 | 0.37530142 | -0.8024271 | 0.42230594 | 0.54903428 | NA | NA |
| ENSG000002 | 7.28244079 | -0.284301063 | 0.35442138 | -0.8021555 | 0.42246301 | 0.54920901 | NA | NA |
| ENSG000002 | 9.41475599 | 0.286921566 | 0.35769161 | 0.80214787 | 0.42246743 | 0.54920901 | NA | NA |
| ENSG000001 | 128.989812 | 0.125114417 | 0.15602523 | 0.8018858 | 0.42261902 | 0.54938527 | RWDD2B | 10069 |
| ENSG000002 | 11.1162874 | -0.271994065 | 0.33920112 | -0.8018666 | 0.4226301 | 0.54938527 | NA | NA |
| ENSG000001 | 4.28457425 | -0.303392167 | 0.37837867 | -0.8018215 | 0.42265619 | 0.54940157 | ASB10 | 136371 |
| ENSG000002 | 6.87580516 | -0.289631097 | 0.36124689 | -0.8017539 | 0.42269534 | 0.54943485 | NA | NA |
| ENSG000002 | 6.02141014 | -0.292715547 | 0.36513449 | -0.801665 | 0.42274676 | 0.54948407 | NA | NA |
| ENSG000002 | 3.98645375 | 0.303443599 | 0.37856785 | 0.80155671 | 0.42280943 | 0.5495479 | NA | NA |
| ENSG000001 | 5.49798118 | -0.296440002 | 0.36993411 | -0.8013319 | 0.42293954 | 0.54969843 | UGT2B28 | 54490 |
| ENSG000002 | 3.03481321 | 0.302537097 | 0.37755325 | 0.80130975 | 0.42295235 | 0.54969843 | NA | NA |
| ENSG000002 | 2.96308351 | -0.302246269 | 0.377232 | -0.8012212 | 0.42300361 | 0.54974744 | NA | NA |

| | | | | | | | | |
|---|---|---|---|---|---|---|---|---|
| ENSG000001 | 3.23485804 | -0.303028317 | 0.37823765 | -0.8011585 | 0.42303988 | 0.54976327 | NA | NA |
| ENSG000002 | 2.46793069 | -0.296998704 | 0.37071396 | -0.8011533 | 0.42304291 | 0.54976327 | NA | NA |
| ENSG000002 | 2.51317499 | -0.296727354 | 0.37038962 | -0.8011222 | 0.4230609 | 0.54976903 | NA | NA |
| ENSG000002 | 2.70279126 | -0.300555032 | 0.37519642 | -0.8010605 | 0.42309663 | 0.54978022 | NA | NA |
| ENSG000002 | 3.0215327 | -0.302195749 | 0.37724389 | -0.801062 | 0.42309575 | 0.54978022 | NA | NA |
| ENSG000002 | 2.77404028 | -0.299018092 | 0.37337064 | -0.8008613 | 0.42321195 | 0.54991246 | NA | NA |
| ENSG000002 | 2.76649256 | -0.299358723 | 0.3739059 | -0.8006258 | 0.42334829 | 0.55003673 | NA | NA |
| ENSG000002 | 6.7759423 | -0.288018809 | 0.35973568 | -0.8006401 | 0.42334004 | 0.55003673 | LOC101927870 | 101927870 |
| ENSG000002 | 58.3611142 | -0.162934258 | 0.20350715 | -0.8006316 | 0.42334494 | 0.55003673 | NA | NA |
| ENSG000002 | 3.64286809 | 0.303015208 | 0.37852373 | 0.80051839 | 0.42341051 | 0.55009995 | NA | NA |
| ENSG000001 | 4.48527657 | -0.301591387 | 0.37677913 | -0.8004461 | 0.42345238 | 0.55013671 | ITPKA | 3706 |
| ENSG000002 | 90.4653538 | 0.138180529 | 0.17264413 | 0.8003778 | 0.42349194 | 0.55017049 | NIFK-AS1 | 254128 |
| ENSG000002 | 2.76948073 | -0.299499062 | 0.37421546 | -0.8003386 | 0.42351467 | 0.5501824 | NA | NA |
| ENSG000001 | 125.34325 | 0.131568341 | 0.16444415 | 0.8000792 | 0.42366491 | 0.55033544 | FBXL2 | 25827 |
| ENSG000002 | 14.2062498 | -0.248540412 | 0.31065031 | -0.8000649 | 0.42367319 | 0.55033544 | NA | NA |
| ENSG000002 | 7.61617956 | -0.292025153 | 0.36499925 | -0.8000705 | 0.42366992 | 0.55033544 | NA | NA |
| ENSG000002 | 4.58942655 | -0.301805749 | 0.37730764 | -0.799893 | 0.42377279 | 0.55044717 | NA | NA |
| ENSG000002 | 2.52857135 | -0.296265222 | 0.37044266 | -0.79976 | 0.42384986 | 0.55052965 | NA | NA |
| ENSG000001 | 40.0060894 | 0.182793834 | 0.22859601 | 0.79963703 | 0.42392113 | 0.55060458 | MEDAG | 84935 |
| ENSG000001 | 50.3191313 | 0.162709824 | 0.20350744 | 0.79952763 | 0.42398453 | 0.5506693 | GIPC2 | 54810 |
| ENSG000002 | 2.99356432 | -0.301670866 | 0.37735042 | -0.7994449 | 0.42403248 | 0.55071394 | NA | NA |
| ENSG000002 | 2.75083561 | -0.300102394 | 0.37541534 | -0.7993877 | 0.42406566 | 0.55073939 | NA | NA |
| ENSG000002 | 11.6866582 | -0.265274002 | 0.33189956 | -0.7992599 | 0.42413974 | 0.55081795 | LOC101929551 | 101929551 |
| ENSG000002 | 167.284318 | 0.130776608 | 0.16363478 | 0.7991981 | 0.42417555 | 0.55084682 | NA | NA |
| ENSG000002 | 3.23252276 | -0.302393773 | 0.37848132 | -0.7989662 | 0.42431003 | 0.55100381 | PHBP17 | 106480420 |
| ENSG000002 | 5.27986739 | -0.29695674 | 0.37171834 | -0.7988757 | 0.42436251 | 0.55105431 | NA | NA |
| ENSG000001 | 5.56410651 | -0.297053043 | 0.37185233 | -0.7988468 | 0.42437925 | 0.5510584 | NA | NA |
| ENSG000002 | 4.3281682 | -0.302091868 | 0.37829982 | -0.7985515 | 0.42455055 | 0.55124553 | PRSS50 | 29122 |
| ENSG000002 | 2.73776637 | -0.298686231 | 0.37403253 | -0.7985568 | 0.42454743 | 0.55124553 | NA | NA |

| | | | | | | | | |
|---|---|---|---|---|---|---|---|---|
| ENSG000002 | 2.81673964 | 0.296699122 | 0.37156233 | 0.79851776 | 0.42457009 | 0.55125326 | NA | NA |
| ENSG000002 | 25.0415152 | -0.211302844 | 0.26472062 | -0.7982108 | 0.42474819 | 0.55146685 | NA | NA |
| ENSG000001 | 8.7473699 | -0.27327758 | 0.34246567 | -0.7979707 | 0.42488748 | 0.55153383 | KRT82 | 3888 |
| ENSG000002 | 2.22054296 | -0.290302077 | 0.36376892 | -0.7980398 | 0.42484739 | 0.55153383 | RN7SKP293 | 106480923 |
| ENSG000002 | 5.51120587 | -0.294864018 | 0.36946974 | -0.7980735 | 0.42482782 | 0.55153383 | NA | NA |
| ENSG000002 | 5.3525061 | 0.296151327 | 0.37111519 | 0.79800378 | 0.4248683 | 0.55153383 | NA | NA |
| ENSG000002 | 2.76847442 | -0.29598976 | 0.37096435 | -0.7978927 | 0.42493274 | 0.55153383 | RNU6-389P | 106481301 |
| ENSG000002 | 2.72291931 | -0.299722472 | 0.37558685 | -0.7980111 | 0.42486405 | 0.55153383 | NA | NA |
| ENSG000002 | 2.17715877 | 0.291553186 | 0.3653455 | 0.79802046 | 0.42485861 | 0.55153383 | NA | NA |
| ENSG000002 | 55.211445 | 0.167569859 | 0.21001272 | 0.7979034 | 0.42492655 | 0.55153383 | NA | NA |
| ENSG000002 | 2.97892853 | 0.30044009 | 0.37654444 | 0.79788747 | 0.4249358 | 0.55153383 | NA | NA |
| ENSG000002 | 28.6132394 | 0.20747172 | 0.26000645 | 0.79794837 | 0.42490045 | 0.55153383 | NA | NA |
| ENSG000002 | 4.32889298 | -0.301809399 | 0.37831458 | -0.7977736 | 0.42500186 | 0.55157346 | NA | NA |
| ENSG000002 | 4.96476585 | -0.301789574 | 0.37828304 | -0.7977877 | 0.42499367 | 0.55157346 | LOC101926940 | 101926940 |
| ENSG000002 | 5.13328341 | 0.298800259 | 0.37454692 | 0.79776455 | 0.42500714 | 0.55157346 | NA | NA |
| ENSG000002 | 5.85914161 | -0.294161403 | 0.36877044 | -0.7976816 | 0.42505527 | 0.55161828 | NA | NA |
| ENSG000002 | 2.96709019 | -0.301281437 | 0.37771637 | -0.7976393 | 0.42507982 | 0.55163248 | NA | NA |
| ENSG000002 | 5.06571723 | -0.299114606 | 0.37501498 | -0.7976071 | 0.42509854 | 0.55163912 | NA | NA |
| ENSG000002 | 4.77743177 | -0.300246658 | 0.37647175 | -0.7975277 | 0.42514461 | 0.55168125 | NA | NA |
| ENSG000002 | 8.9991768 | -0.278033584 | 0.34864834 | -0.7974614 | 0.42518313 | 0.55171358 | NA | NA |
| ENSG000002 | 14.5244651 | -0.248761054 | 0.31203908 | -0.7972112 | 0.42532838 | 0.55186674 | C22orf42 | 150297 |
| ENSG000002 | 2.7697339 | -0.298074999 | 0.37389271 | -0.7972207 | 0.42532288 | 0.55186674 | NA | NA |
| ENSG000001 | 2.98439782 | -0.300842121 | 0.37738934 | -0.7971664 | 0.42535437 | 0.5518828 | OR2Y1 | 134083 |
| ENSG000002 | 6.52880306 | -0.287488977 | 0.36074006 | -0.7969422 | 0.4254846 | 0.55203411 | NA | NA |
| ENSG000001 | 54.5827889 | -0.173849689 | 0.21815749 | -0.7968999 | 0.42550916 | 0.5520393 | POLN | 353497 |
| ENSG000001 | 25.1103434 | 0.218869381 | 0.27465499 | 0.79688843 | 0.42551583 | 0.5520393 | C7orf25 | 79020 |
| ENSG000002 | 2.47056654 | -0.295002011 | 0.37022389 | -0.7968206 | 0.42555525 | 0.55207278 | RN7SL734P | 106479490 |
| ENSG000001 | 223.880555 | 0.108359564 | 0.13599499 | 0.79679087 | 0.4255725 | 0.55207749 | ULK4 | 54986 |
| ENSG000001 | 6.58087729 | -0.289607793 | 0.36354925 | -0.7966123 | 0.42567625 | 0.55208024 | KIR3DL1 | 3811 |

| | | | | | | | | | |
|---|---|---|---|---|---|---|---|---|---|
| ENSG000001 | 2.94725031 | -0.300699112 | 0.37744998 | -0.7966595 | 0.42564881 | 0.55208024 | MIR135B | | 442891 |
| ENSG000002 | 2.48548736 | -0.295850782 | 0.37139202 | -0.7965997 | 0.42568353 | 0.55208024 | NA | NA | |
| ENSG000002 | 11.8567096 | -0.263928242 | 0.33129256 | -0.7966622 | 0.42564727 | 0.55208024 | NA | NA | |
| ENSG000002 | 2.98982576 | -0.300072775 | 0.37666767 | -0.7966513 | 0.42565359 | 0.55208024 | NA | NA | |
| ENSG000002 | 2.17970252 | -0.290704203 | 0.3648682 | -0.7967376 | 0.42560346 | 0.55208024 | NA | NA | |
| ENSG000002 | 2.21062715 | -0.291260539 | 0.36560348 | -0.7966569 | 0.42565032 | 0.55208024 | NA | NA | |
| ENSG000002 | 3.20941341 | -0.301119352 | 0.37798845 | -0.7966364 | 0.42566224 | 0.55208024 | CCL15 | | 6359 |
| ENSG000002 | 14.5041112 | -0.252129836 | 0.3165217 | -0.7965642 | 0.4257042 | 0.55208939 | NA | NA | |
| ENSG000001 | 15.469345 | 0.240747803 | 0.3022442 | 0.79653407 | 0.42572168 | 0.55209441 | MLLT4-AS1 | | 653483 |
| ENSG000001 | 5.330651 | 0.2963274 | 0.37206805 | 0.79643334 | 0.42578021 | 0.55215264 | NA | NA | |
| ENSG000002 | 25.5029692 | 0.216695432 | 0.2720907 | 0.7964088 | 0.42579446 | 0.55215348 | NA | NA | |
| ENSG000001 | 11.9555739 | 0.259993559 | 0.32650783 | 0.79628582 | 0.42586592 | 0.55218528 | CHST13 | | 166012 |
| ENSG000002 | 3.49061558 | -0.301479389 | 0.37859102 | -0.7963194 | 0.42584639 | 0.55218528 | NA | NA | |
| ENSG000002 | 4.51819605 | -0.300716205 | 0.37762926 | -0.7963265 | 0.42584226 | 0.55218528 | NA | NA | |
| ENSG000002 | 10.2668267 | 0.268800051 | 0.33757279 | 0.79627285 | 0.42587346 | 0.55218528 | NA | NA | |
| ENSG000002 | 10.0146748 | -0.270584335 | 0.33982562 | -0.7962447 | 0.42588984 | 0.55218887 | NA | NA | |
| ENSG000001 | 149.384998 | 0.125584633 | 0.15773494 | 0.7961751 | 0.42593027 | 0.55220764 | TRIM69 | | 140691 |
| ENSG000002 | 3.22358439 | -0.30126699 | 0.37839393 | -0.7961729 | 0.42593156 | 0.55220764 | NA | NA | |
| ENSG000001 | 28.2348072 | 0.197316319 | 0.24785596 | 0.79609268 | 0.42597817 | 0.55221669 | RAB32 | | 10981 |
| ENSG000001 | 60.5898495 | 0.155131615 | 0.1948653 | 0.79609668 | 0.42597585 | 0.55221669 | SLC25A20 | | 788 |
| ENSG000001 | 49.4480724 | -0.179776755 | 0.2258245 | -0.7960906 | 0.42597939 | 0.55221669 | KREMEN1 | | 83999 |
| ENSG000002 | 2.50032223 | -0.292538938 | 0.36753327 | -0.7959523 | 0.42605978 | 0.55230325 | NA | NA | |
| ENSG000002 | 25.0225154 | 0.209970608 | 0.26382268 | 0.79587778 | 0.42610308 | 0.55234171 | NA | NA | |
| ENSG000002 | 7.31161548 | -0.283569559 | 0.35631123 | -0.7958479 | 0.42612042 | 0.55234655 | NA | NA | |
| ENSG000002 | 116.456406 | 0.128599877 | 0.16161559 | 0.79571457 | 0.42619795 | 0.55242938 | EFCAB7 | | 84455 |
| ENSG000002 | 38.2346849 | 0.178195605 | 0.22396739 | 0.79563192 | 0.42624601 | 0.55246018 | NA | NA | |
| ENSG000002 | 2.77258602 | -0.296977391 | 0.37326216 | -0.7956268 | 0.42624896 | 0.55246018 | NA | NA | |
| ENSG000002 | 5.35361655 | 0.297092678 | 0.37345285 | 0.79552929 | 0.42630568 | 0.55248384 | NA | NA | |
| ENSG000002 | 15.9996771 | -0.250740041 | 0.31518807 | -0.7955252 | 0.42630809 | 0.55248384 | PCDHGA7 | | 56108 |

| | | | | | | | | |
|---|---|---|---|---|---|---|---|---|
| ENSG000002 | 4.52043323 | 0.29921696 | 0.37611236 | 0.79555204 | 0.42629245 | 0.55248384 | LINC01397 | 104355139 |
| ENSG000002 | 2.99125986 | -0.298480618 | 0.37521358 | -0.7954952 | 0.42632548 | 0.55248872 | NA | NA |
| ENSG000002 | 8.01263821 | -0.277932747 | 0.34940652 | -0.7954424 | 0.42635623 | 0.55251091 | NA | NA |
| ENSG000002 | 3.26872346 | -0.300082532 | 0.37726955 | -0.7954062 | 0.42637726 | 0.55252051 | NA | NA |
| ENSG000002 | 2.72986621 | -0.297811724 | 0.37448712 | -0.7952522 | 0.42646678 | 0.55260396 | NA | NA |
| ENSG000002 | 8.76087648 | -0.279815159 | 0.35185873 | -0.7952486 | 0.42646892 | 0.55260396 | NA | NA |
| ENSG000002 | 2.73862778 | -0.296036149 | 0.3722895 | -0.7951773 | 0.4265104 | 0.55264005 | NA | NA |
| ENSG000001 | 4.47755777 | 0.299899833 | 0.37715988 | 0.79515306 | 0.42652447 | 0.55264063 | VCX | 26609 |
| ENSG000001 | 27.0410639 | 0.205691003 | 0.25870426 | 0.79508161 | 0.42656603 | 0.55265916 | FAM98C | 147965 |
| ENSG000002 | 7.2807584 | -0.281811097 | 0.35443792 | -0.795093 | 0.42655943 | 0.55265916 | NA | NA |
| ENSG000002 | 2.79461683 | -0.29564274 | 0.37196473 | -0.7948139 | 0.42672176 | 0.55284327 | NA | NA |
| ENSG000002 | 8.03223569 | -0.280255077 | 0.35266344 | -0.7946814 | 0.42679885 | 0.55292547 | NA | NA |
| ENSG000001 | 14.9089158 | -0.249512754 | 0.31400718 | -0.7946084 | 0.42684131 | 0.55294515 | HTR1F | 3355 |
| ENSG000002 | 3.26309599 | -0.297901201 | 0.37489451 | -0.7946267 | 0.42683066 | 0.55294515 | NA | NA |
| ENSG000002 | 24.928761 | -0.211657866 | 0.26642258 | -0.7944442 | 0.42693689 | 0.55304585 | NA | NA |
| ENSG000002 | 15.8936243 | 0.236998884 | 0.29832646 | 0.79442798 | 0.42694632 | 0.55304585 | NA | NA |
| ENSG000001 | 5.86411687 | -0.295453667 | 0.37200659 | -0.7942162 | 0.42706957 | 0.55318783 | FOXD4L1 | 200350 |
| ENSG000002 | 9.10217417 | -0.274394855 | 0.34553801 | -0.794109 | 0.42713196 | 0.55325098 | IRGM | 345611 |
| ENSG000001 | 295.050065 | 0.101940554 | 0.12838903 | 0.79399735 | 0.42719697 | 0.55329983 | C1orf131 | 128061 |
| ENSG000002 | 9.80349453 | 0.271201952 | 0.34155696 | 0.79401676 | 0.42718568 | 0.55329983 | NA | NA |
| ENSG000002 | 18.9147289 | 0.228643724 | 0.28799943 | 0.79390338 | 0.42725168 | 0.55335302 | NA | NA |
| ENSG000002 | 5.32526073 | -0.295734056 | 0.37255044 | -0.7938094 | 0.42730638 | 0.55340618 | LOC102723505 | 102723505 |
| ENSG000001 | 80.7087283 | 0.139115483 | 0.1752569 | 0.79378033 | 0.42732332 | 0.55341045 | JAK3 | 3718 |
| ENSG000002 | 6.28530827 | -0.292227733 | 0.36816912 | -0.7937323 | 0.42735127 | 0.55342897 | NA | NA |
| ENSG000002 | 11.7154366 | -0.261185061 | 0.32911112 | -0.7936075 | 0.42742395 | 0.55350542 | NA | NA |
| ENSG000002 | 3.28218947 | -0.299711063 | 0.37767777 | -0.7935629 | 0.42744995 | 0.55352141 | NA | NA |
| ENSG000001 | 3.00847196 | -0.299308448 | 0.37724079 | -0.7934149 | 0.42753615 | 0.55361536 | DEFB4B | 100289462 |
| ENSG000002 | 14.9242257 | -0.239947486 | 0.30247063 | -0.7932919 | 0.4276078 | 0.55366284 | NA | NA |
| ENSG000002 | 3.23633192 | -0.299866223 | 0.37800682 | -0.7932826 | 0.42761321 | 0.55366284 | NA | NA |

| | | | | | | | | |
|---|---|---|---|---|---|---|---|---|
| ENSG000002 | 44.8340018 | 0.177121434 | 0.22327688 | 0.79328159 | 0.42761378 | 0.55366284 | NA | NA |
| ENSG000002 | 5.59966303 | -0.295586351 | 0.37267299 | -0.7931521 | 0.42768923 | 0.55374284 | DDX39B-AS1 | 106478957 |
| ENSG000002 | 3.22904478 | -0.29999349 | 0.37827886 | -0.7930485 | 0.42774956 | 0.55380327 | NA | NA |
| ENSG000002 | 2.23590761 | -0.289295438 | 0.36482147 | -0.7929781 | 0.42779058 | 0.55383871 | NA | NA |
| ENSG000002 | 4.62090461 | -0.299807858 | 0.37810639 | -0.7929193 | 0.42782483 | 0.55386536 | LINC01340 | 102546227 |
| ENSG000001 | 228.00609 | 0.108259093 | 0.13656931 | 0.79270441 | 0.42795006 | 0.55397443 | SAMD12 | 401474 |
| ENSG000002 | 2.77046588 | -0.296224523 | 0.37368455 | -0.7927128 | 0.42794517 | 0.55397443 | NA | NA |
| ENSG000002 | 4.47298611 | 0.299866713 | 0.37827658 | 0.79271817 | 0.42794204 | 0.55397443 | NA | NA |
| ENSG000002 | 7.51677954 | -0.283035094 | 0.3570618 | -0.7926782 | 0.42796535 | 0.55397653 | NA | NA |
| ENSG000001 | 2.71651264 | -0.297184998 | 0.37494964 | -0.7925998 | 0.42801104 | 0.55401799 | NA | NA |
| ENSG000002 | 2.97021475 | -0.299139809 | 0.37745356 | -0.7925208 | 0.42805704 | 0.55404217 | NA | NA |
| ENSG000002 | 34.002855 | -0.194859603 | 0.24587309 | -0.7925211 | 0.42805691 | 0.55404217 | NA | NA |
| ENSG000001 | 8.47809462 | -0.273838686 | 0.34559767 | -0.7923627 | 0.42814925 | 0.55410845 | NA | NA |
| ENSG000002 | 2.75183177 | -0.293928855 | 0.37094827 | -0.7923715 | 0.42814407 | 0.55410845 | RN7SL619P | 106481091 |
| ENSG000002 | 4.55555318 | -0.299153783 | 0.37754347 | -0.7923691 | 0.42814549 | 0.55410845 | NA | NA |
| ENSG00000C | 82.855258 | 0.140605725 | 0.17747295 | 0.79226565 | 0.4282058 | 0.55412858 | NOX4 | 50507 |
| ENSG000001 | 164.941565 | -0.122325593 | 0.15439647 | -0.7922823 | 0.42819608 | 0.55412858 | PDE4D | 5144 |
| ENSG000002 | 19.2955064 | 0.229250379 | 0.28935793 | 0.79227266 | 0.42820171 | 0.55412858 | NA | NA |
| ENSG000002 | 2.50795482 | -0.291241041 | 0.36761743 | -0.7922395 | 0.42822105 | 0.55413064 | NA | NA |
| ENSG000002 | 3.6386724 | 0.299896886 | 0.37859327 | 0.79213475 | 0.42828211 | 0.55417066 | NA | NA |
| ENSG000002 | 2.54286597 | -0.291514912 | 0.36801027 | -0.792138 | 0.42828022 | 0.55417066 | NA | NA |
| ENSG000002 | 10.7009296 | 0.261226459 | 0.32978304 | 0.79211611 | 0.42829297 | 0.55417066 | NA | NA |
| ENSG000002 | 3.38817405 | -0.296342837 | 0.37412958 | -0.7920861 | 0.42831048 | 0.55417562 | NA | NA |
| ENSG000002 | 5.35273444 | 0.294252391 | 0.37151837 | 0.79202648 | 0.42834523 | 0.55420291 | NA | NA |
| ENSG000002 | 4.88569943 | -0.298177921 | 0.37649196 | -0.7919901 | 0.42836643 | 0.55421265 | NA | NA |
| ENSG000002 | 8.03872756 | -0.277380142 | 0.35024769 | -0.7919542 | 0.42838736 | 0.55422205 | NA | NA |
| ENSG000002 | 2.78190873 | -0.296106719 | 0.37394446 | -0.7918468 | 0.42845002 | 0.55428543 | NA | NA |
| ENSG000001 | 68.4534948 | -0.149748448 | 0.18912572 | -0.7917931 | 0.4284813 | 0.55429719 | MYOCD | 93649 |
| ENSG000002 | 3.64527583 | 0.299702724 | 0.37851561 | 0.79178432 | 0.42848645 | 0.55429719 | NA | NA |

| | | | | | | | | |
|---|---|---|---|---|---|---|---|---|
| ENSG000002 | 3.0149417 | -0.296734223 | 0.37478914 | -0.7917365 | 0.42851436 | 0.55431562 | NA | NA |
| ENSG000002 | 4.20395594 | 0.299657972 | 0.37858602 | 0.79151885 | 0.42864128 | 0.5544621 | MIR4492 | 100616376 |
| ENSG000002 | 24.2943923 | 0.209361243 | 0.26451965 | 0.79147709 | 0.42866564 | 0.55447593 | RAD21-AS1 | 644660 |
| ENSG000002 | 2.97386028 | -0.297685819 | 0.37618914 | -0.7913195 | 0.42875755 | 0.55457713 | NA | NA |
| ENSG000002 | 2.75703321 | -0.296463631 | 0.37471525 | -0.7911705 | 0.42884453 | 0.55466854 | NA | NA |
| ENSG000002 | 60.6299023 | 0.15867554 | 0.20056277 | 0.79115152 | 0.42885558 | 0.55466854 | NA | NA |
| ENSG000002 | 2.21698175 | -0.289148259 | 0.36563128 | -0.7908193 | 0.42904947 | 0.55488391 | NA | NA |
| ENSG000002 | 11.1929714 | -0.269038947 | 0.34019569 | -0.7908359 | 0.42903979 | 0.55488391 | LOC101928441 | 101928441 |
| ENSG000002 | 2.50769423 | -0.291489801 | 0.36863665 | -0.7907239 | 0.42910514 | 0.55493821 | NA | NA |
| ENSG000002 | 2.96671695 | -0.297237377 | 0.37597673 | -0.7905739 | 0.4291927 | 0.55503375 | NA | NA |
| ENSG000001 | 3.46340423 | -0.299266544 | 0.37857201 | -0.7905142 | 0.42922754 | 0.5550611 | DLX2 | 1746 |
| ENSG000000 | 54.2992267 | 0.164109976 | 0.20762893 | 0.79040034 | 0.429294 | 0.55509597 | MVP | 9961 |
| ENSG000002 | 27.4749469 | -0.224824772 | 0.28444514 | -0.7903977 | 0.42929557 | 0.55509597 | NA | NA |
| ENSG000002 | 14.7824201 | -0.247421726 | 0.31302098 | -0.7904318 | 0.42927566 | 0.55509597 | LINC00989 | 100506035 |
| ENSG000001 | 159.018777 | -0.119322375 | 0.15097482 | -0.7903462 | 0.42932561 | 0.55509941 | NTRK3 | 4916 |
| ENSG000002 | 6.88735936 | -0.287449697 | 0.36369699 | -0.7903549 | 0.42932054 | 0.55509941 | NA | NA |
| ENSG000001 | 120.728873 | -0.129329541 | 0.16364872 | -0.7902875 | 0.42935987 | 0.55512601 | SETBP1 | 26040 |
| ENSG000002 | 24.24392 | 0.20743815 | 0.26254317 | 0.79011062 | 0.42946317 | 0.55521595 | APOM | 55937 |
| ENSG000002 | 3.03060637 | -0.296650902 | 0.37546087 | -0.790098 | 0.42947051 | 0.55521595 | NA | NA |
| ENSG000002 | 3.00501789 | 0.297999211 | 0.37714594 | 0.79014297 | 0.42944428 | 0.55521595 | NA | NA |
| ENSG000002 | 39.6693782 | -0.184050911 | 0.23300893 | -0.7898878 | 0.4295933 | 0.55533663 | LOC654841 | 654841 |
| ENSG000002 | 2.20346366 | -0.287305655 | 0.36371977 | -0.7899094 | 0.42958069 | 0.55533663 | NA | NA |
| ENSG000002 | 20.2455359 | -0.225249949 | 0.28517422 | -0.7898679 | 0.42960495 | 0.55533663 | NA | NA |
| ENSG000001 | 7.12560508 | -0.286428594 | 0.36266798 | -0.7897819 | 0.42965517 | 0.55537648 | NA | NA |
| ENSG000002 | 2.7466006 | -0.295127388 | 0.37368862 | -0.7897682 | 0.42966316 | 0.55537648 | NA | NA |
| ENSG000002 | 2.68665749 | -0.295292936 | 0.37392546 | -0.7897107 | 0.42969674 | 0.55540218 | NA | NA |
| ENSG000002 | 5.06958676 | -0.296448968 | 0.37541873 | -0.7896488 | 0.42973288 | 0.55543118 | RN7SL60P | 106480938 |
| ENSG000002 | 8.06477267 | -0.276867941 | 0.35068178 | -0.7895133 | 0.42981204 | 0.55551579 | NA | NA |
| ENSG000002 | 2.3848163 | 0.292717004 | 0.3707943 | 0.78943232 | 0.42985937 | 0.55555927 | NA | NA |

| | | | | | | | | |
|---|---|---|---|---|---|---|---|---|
| ENSG000002 | 2.71036711 | -0.295970083 | 0.37494507 | -0.7893692 | 0.42989626 | 0.55558425 | FAM99B | 100132464 |
| ENSG000002 | 14.7340483 | -0.240520743 | 0.30470644 | -0.7893523 | 0.42990611 | 0.55558425 | NA | NA |
| ENSG000002 | 3.23505392 | -0.298293951 | 0.37792874 | -0.7892862 | 0.42994473 | 0.55561646 | NA | NA |
| ENSG000002 | 2.98695382 | -0.296935491 | 0.37622617 | -0.7892473 | 0.42996748 | 0.55562815 | NA | NA |
| ENSG000002 | 2.7619647 | -0.294427708 | 0.37311217 | -0.7891131 | 0.43004589 | 0.55569406 | NA | NA |
| ENSG000002 | 7.85016858 | -0.279276781 | 0.35390708 | -0.7891246 | 0.43003919 | 0.55569406 | NA | NA |
| ENSG000001 | 16.4242725 | -0.236271146 | 0.29942988 | -0.78907 | 0.43007108 | 0.55570889 | B3GNT4 | 79369 |
| ENSG000002 | 6.01604672 | -0.289937572 | 0.36761072 | -0.7887082 | 0.4302826 | 0.55594678 | NA | NA |
| ENSG000002 | 3.0187525 | -0.296407074 | 0.37580779 | -0.7887199 | 0.43027575 | 0.55594678 | NA | NA |
| ENSG000001 | 25.0527052 | -0.202581325 | 0.25688271 | -0.7886141 | 0.4303376 | 0.55596274 | ALOX15 | 246 |
| ENSG000001 | 3.01094363 | -0.296953385 | 0.37656091 | -0.7885932 | 0.43034979 | 0.55596274 | LRRC70 | 100130733 |
| ENSG000002 | 3.01094363 | -0.296953385 | 0.37656091 | -0.7885932 | 0.43034979 | 0.55596274 | NA | NA |
| ENSG000002 | 5.0428723 | -0.295118241 | 0.3742005 | -0.7886634 | 0.43030876 | 0.55596274 | NA | NA |
| ENSG000002 | 3.0397536 | -0.296307899 | 0.37580229 | -0.7884675 | 0.43042329 | 0.55603997 | HSPE1P13 | 106480256 |
| ENSG000001 | 146.496377 | -0.127116435 | 0.16125779 | -0.7882809 | 0.43053241 | 0.55616322 | SYT16 | 83851 |
| ENSG000002 | 10.3867958 | 0.2673547 | 0.33917526 | 0.78824942 | 0.43055082 | 0.55616929 | NA | NA |
| ENSG000002 | 8.09365487 | -0.276202691 | 0.35041143 | -0.788224 | 0.43056571 | 0.5561708 | NA | NA |
| ENSG000002 | 2.96735078 | -0.297159912 | 0.37702582 | -0.7881686 | 0.4305981 | 0.55619492 | NA | NA |
| ENSG000002 | 2.97084783 | -0.296848954 | 0.37666475 | -0.7880986 | 0.43063905 | 0.55621239 | NA | NA |
| ENSG000002 | 7.54801792 | -0.278544746 | 0.35343621 | -0.7881047 | 0.43063545 | 0.55621239 | NA | NA |
| ENSG000002 | 10.4957202 | -0.261620737 | 0.3319907 | -0.7880363 | 0.43067545 | 0.55622396 | NA | NA |
| ENSG000002 | 8.27556576 | -0.275967164 | 0.35019323 | -0.7880425 | 0.43067182 | 0.55622396 | NA | NA |
| ENSG000001 | 13.57366 | -0.253005772 | 0.32116441 | -0.7877765 | 0.43082746 | 0.55640257 | ISLR | 3671 |
| ENSG000002 | 12.5754808 | 0.256341351 | 0.32541776 | 0.78773007 | 0.43085461 | 0.55641992 | SLC22A31 | 146429 |
| ENSG000001 | 84.0332099 | 0.150560222 | 0.1911871 | 0.78750199 | 0.43098806 | 0.55644215 | SEMA6C | 10500 |
| ENSG000001 | 219.916906 | -0.109262759 | 0.1387482 | -0.7874896 | 0.43099533 | 0.55644215 | FBN1 | 2200 |
| ENSG000001 | 334.411377 | 0.098293442 | 0.12480208 | 0.78759457 | 0.43093389 | 0.55644215 | DENND5B | 160518 |
| ENSG000001 | 15.045862 | 0.240221227 | 0.3050336 | 0.78752381 | 0.4309753 | 0.55644215 | GIPC3 | 126326 |
| ENSG000002 | 5.28243658 | -0.293907277 | 0.37320856 | -0.7875148 | 0.43098055 | 0.55644215 | EBLN1 | 340900 |

| | | | | | | | | |
|---|---|---|---|---|---|---|---|---|
| ENSG000002 | 2.47100428 | -0.292353743 | 0.37121971 | -0.7875491 | 0.43096051 | 0.55644215 | NA | NA |
| ENSG000002 | 3.25384806 | -0.297042665 | 0.37712428 | -0.7876519 | 0.43090033 | 0.55644215 | NA | NA |
| ENSG000002 | 2.56105462 | -0.288553939 | 0.36634681 | -0.7876524 | 0.43090007 | 0.55644215 | NA | NA |
| ENSG000002 | 7.8485826 | -0.279547607 | 0.35498532 | -0.7874906 | 0.43099472 | 0.55644215 | NA | NA |
| ENSG000002 | 2.23212982 | -0.284670459 | 0.3615072 | -0.7874545 | 0.43101588 | 0.55645096 | RN7SKP65 | 106479124 |
| ENSG000002 | 6.92986609 | -0.290260343 | 0.36862167 | -0.7874207 | 0.43103561 | 0.55645873 | NA | NA |
| ENSG000001 | 20.6831307 | -0.227103748 | 0.28851889 | -0.7871365 | 0.43120196 | 0.55665576 | SUMO4 | 387082 |
| ENSG000002 | 8.0228115 | -0.280940819 | 0.35696183 | -0.7870332 | 0.43126242 | 0.55671609 | NA | NA |
| ENSG00000C | 578.960529 | 0.10157043 | 0.12906103 | 0.78699534 | 0.4312846 | 0.55672008 | TMEM131 | 23505 |
| ENSG000002 | 3.49929336 | -0.297938687 | 0.37858433 | -0.786981 | 0.43129297 | 0.55672008 | NA | NA |
| ENSG000002 | 3.38506592 | 0.297165577 | 0.37771753 | 0.78674024 | 0.43143394 | 0.55688432 | NA | NA |
| ENSG000001 | 2.77283451 | -0.294215548 | 0.37399633 | -0.7866803 | 0.43146904 | 0.55691189 | GPR32 | 2854 |
| ENSG000002 | 19.7138279 | -0.227913944 | 0.28972862 | -0.7866463 | 0.43148896 | 0.55691988 | PKN2-AS1 | 101927891 |
| ENSG000002 | 2.74344324 | -0.29320198 | 0.37277253 | -0.786544 | 0.43154887 | 0.55697948 | NA | NA |
| ENSG000002 | 11.9759355 | -0.26329134 | 0.33477071 | -0.7864826 | 0.4315848 | 0.55700813 | NA | NA |
| ENSG000001 | 102.727553 | -0.130687307 | 0.16617704 | -0.7864342 | 0.43161314 | 0.55702697 | TBC1D30 | 23329 |
| ENSG000002 | 3.47985842 | -0.297680133 | 0.37858386 | -0.7862991 | 0.4316923 | 0.55711141 | LOC105369900 | 105369900 |
| ENSG000002 | 3.23554373 | -0.297429983 | 0.37828013 | -0.7862691 | 0.43170984 | 0.55711631 | NA | NA |
| ENSG000002 | 22.0760037 | 0.216502087 | 0.27538984 | 0.78616586 | 0.43177033 | 0.55714117 | RNU6-1016P | 106480630 |
| ENSG000002 | 5.28706246 | -0.294179848 | 0.37418033 | -0.786198 | 0.43175147 | 0.55714117 | NA | NA |
| ENSG000002 | 2.53887891 | -0.287091037 | 0.3651748 | -0.7861743 | 0.43176539 | 0.55714117 | NA | NA |
| ENSG000002 | 3.23255501 | -0.297252437 | 0.37813522 | -0.7861009 | 0.43180837 | 0.55717254 | NA | NA |
| ENSG000002 | 3.46269927 | -0.297585484 | 0.37859004 | -0.7860362 | 0.43184627 | 0.55720371 | NA | NA |
| ENSG000002 | 11.5091422 | -0.2562352 | 0.32602317 | -0.7859417 | 0.43190165 | 0.55725729 | NA | NA |
| ENSG000002 | 19.7541975 | -0.218733806 | 0.27831617 | -0.7859184 | 0.43191529 | 0.55725729 | EXTL3-AS1 | 101929402 |
| ENSG000001 | 169.808288 | 0.118570784 | 0.15088022 | 0.78586035 | 0.43194931 | 0.55728346 | PLEKHM3 | 389072 |
| ENSG000002 | 7.2888271 | -0.282016659 | 0.35889976 | -0.7857811 | 0.43199573 | 0.55732562 | NA | NA |
| ENSG000002 | 2.21842338 | -0.2858429 | 0.36378052 | -0.7857565 | 0.43201017 | 0.55732652 | NA | NA |
| ENSG000002 | 9.02093125 | 0.26925207 | 0.34270005 | 0.78567852 | 0.43205585 | 0.55735346 | NA | NA |

| | | | | | | | | |
|---|---|---|---|---|---|---|---|---|
| ENSG000002 | 5.38142062 | -0.294380488 | 0.37468532 | -0.7856739 | 0.43205854 | 0.55735346 | NA | NA |
| ENSG000002 | 2.94056344 | -0.296275035 | 0.37710982 | -0.7856466 | 0.43207457 | 0.55735641 | NA | NA |
| ENSG000002 | 2.43872897 | -0.291154795 | 0.37064969 | -0.7855255 | 0.43214551 | 0.55743018 | NA | NA |
| ENSG000002 | 8.0964147 | -0.276296089 | 0.35177639 | -0.785431 | 0.4322009 | 0.55746616 | NA | NA |
| ENSG000002 | 2.93976619 | -0.2958374 | 0.37665298 | -0.7854376 | 0.43219707 | 0.55746616 | NA | NA |
| ENSG000002 | 2.3914183 | 0.292059629 | 0.37187481 | 0.78537083 | 0.43223618 | 0.5574849 | NA | NA |
| ENSG000002 | 7.26965993 | -0.282396514 | 0.35957619 | -0.7853593 | 0.43224293 | 0.5574849 | NACA2 | 342538 |
| ENSG000002 | 2.302818 | -0.283252222 | 0.36068097 | -0.7853262 | 0.43226233 | 0.5574922 | NA | NA |
| ENSG000002 | 3.60573211 | 0.297100291 | 0.37839243 | 0.78516448 | 0.43235714 | 0.55759674 | LOC102724770 | 102724770 |
| ENSG000002 | 12.8716718 | 0.246436009 | 0.31392313 | 0.78502024 | 0.4324417 | 0.55768806 | NA | NA |
| ENSG000002 | 2.94045133 | -0.295657911 | 0.37665316 | -0.7849607 | 0.43247664 | 0.55769765 | NA | NA |
| ENSG000002 | 6.74324995 | -0.282886564 | 0.36038126 | -0.7849647 | 0.43247427 | 0.55769765 | PRRT3-AS1 | 100874032 |
| ENSG000002 | 9.01732957 | -0.271087897 | 0.3453684 | -0.7849238 | 0.43249822 | 0.55770774 | NA | NA |
| ENSG000001 | 265.66514 | 0.108409148 | 0.13812583 | 0.78485789 | 0.43253689 | 0.55773988 | CCSAP | 126731 |
| ENSG000002 | 2.76623197 | -0.293102792 | 0.37346854 | -0.7848125 | 0.43256349 | 0.55775643 | NA | NA |
| ENSG000002 | 2.97271033 | 0.295377868 | 0.37640857 | 0.78472674 | 0.4326138 | 0.55780357 | MIR298 | 100126296 |
| ENSG000001 | 105.491362 | 0.132097105 | 0.16834532 | 0.78467939 | 0.43264157 | 0.55782164 | CCDC122 | 160857 |
| ENSG000001 | 19.9576053 | -0.21974645 | 0.28014483 | -0.784403 | 0.43280367 | 0.55799155 | KCNAB3 | 9196 |
| ENSG000002 | 4.90010322 | -0.29615016 | 0.37754808 | -0.7844038 | 0.43280319 | 0.55799155 | LOC101927018 | 101927018 |
| ENSG000002 | 6.50626262 | -0.284004872 | 0.36207362 | -0.7843843 | 0.43281463 | 0.55799155 | LINC01533 | 100996665 |
| ENSG000001 | 15.345414 | -0.239084338 | 0.30482853 | -0.784324 | 0.43285001 | 0.55800167 | TBX6 | 6911 |
| ENSG000002 | 6.1246691 | 0.289361954 | 0.36892811 | 0.78433155 | 0.43284559 | 0.55800167 | NA | NA |
| ENSG000002 | 2.97634102 | -0.295483231 | 0.37680233 | -0.7841863 | 0.43293079 | 0.55807033 | NA | NA |
| ENSG000002 | 1773.12005 | 0.101145745 | 0.12898075 | 0.78419253 | 0.43292715 | 0.55807033 | ATP8 | 4509 |
| ENSG000002 | 273.419505 | 0.110613614 | 0.1410856 | 0.78401773 | 0.43302971 | 0.5581801 | ZNF670 | 93474 |
| ENSG00000C | 15.697999 | -0.255097163 | 0.3253963 | -0.7839584 | 0.43306452 | 0.55820451 | MS4A12 | 54860 |
| ENSG000002 | 3.99178708 | -0.296793692 | 0.37859306 | -0.7839385 | 0.43307617 | 0.55820451 | NA | NA |
| ENSG000001 | 5.75191938 | -0.289526163 | 0.36947631 | -0.7836122 | 0.43326767 | 0.55843173 | ZNF575 | 284346 |
| ENSG000002 | 8.47623562 | -0.270738966 | 0.34551046 | -0.7835912 | 0.43328 | 0.55843173 | NA | NA |

| | | | | | | | | |
|---|---|---|---|---|---|---|---|---|
| ENSG000002 | 12.6440199 | -0.245608027 | 0.31346266 | -0.783532 | 0.43331477 | 0.55845878 | SMTNL1 | 219537 |
| ENSG000002 | 6.73956369 | -0.284113834 | 0.36262724 | -0.7834873 | 0.43334101 | 0.55847486 | OR52I1 | 390037 |
| ENSG000000 | 8.95178394 | 0.271707179 | 0.34688096 | 0.78328651 | 0.43345888 | 0.55859567 | CCDC85A | 114800 |
| ENSG000001 | 78.2218783 | -0.139743336 | 0.17841308 | -0.7832572 | 0.43347608 | 0.55859567 | DISP1 | 84976 |
| ENSG000002 | 14.7576476 | -0.243778544 | 0.31123311 | -0.7832667 | 0.43347049 | 0.55859567 | LOC101929475 | 101929475 |
| ENSG000001 | 6.78181948 | -0.283484326 | 0.36201965 | -0.7830634 | 0.43358991 | 0.55867134 | DBH | 1621 |
| ENSG000002 | 3.04488118 | -0.292242882 | 0.37318718 | -0.7831 | 0.43356839 | 0.55867134 | RN7SL444P | 106479384 |
| ENSG000002 | 3.05242916 | 0.29475639 | 0.37639141 | 0.78311137 | 0.43356172 | 0.55867134 | NA | NA |
| ENSG000002 | 3.08959401 | -0.291683984 | 0.3724808 | -0.7830846 | 0.43357742 | 0.55867134 | NA | NA |
| ENSG000002 | 16.3971905 | 0.233102745 | 0.29772234 | 0.78295349 | 0.43365442 | 0.55870121 | LOC100507250 | 100507250 |
| ENSG000002 | 2.47481398 | -0.290173273 | 0.37059899 | -0.7829845 | 0.4336362 | 0.55870121 | NA | NA |
| ENSG000002 | 4.48630608 | -0.296171252 | 0.37826582 | -0.7829712 | 0.43364401 | 0.55870121 | NA | NA |
| ENSG000002 | 8.8725325 | -0.27454896 | 0.35068491 | -0.7828936 | 0.43368961 | 0.55872879 | HBD | 3045 |
| ENSG000002 | 3.44163346 | -0.296380974 | 0.37859203 | -0.7828505 | 0.43371488 | 0.55874359 | NA | NA |
| ENSG000001 | 11.6102626 | 0.260047626 | 0.33220263 | 0.78279821 | 0.43374562 | 0.55874769 | CHKB | 1120 |
| ENSG000002 | 13.1861262 | -0.246834063 | 0.31531339 | -0.7828214 | 0.43373202 | 0.55874769 | NA | NA |
| ENSG000002 | 2.94958614 | -0.294912013 | 0.37684362 | -0.7825846 | 0.43387109 | 0.55889157 | NA | NA |
| ENSG000002 | 24.103021 | -0.205490543 | 0.26262635 | -0.7824445 | 0.43395339 | 0.55897983 | ELFN1 | 392617 |
| ENSG000002 | 23.0134552 | 0.206598541 | 0.2640759 | 0.78234531 | 0.43401166 | 0.55903713 | NA | NA |
| ENSG000001 | 167.816831 | 0.120089447 | 0.15350803 | 0.78230075 | 0.43403784 | 0.55904019 | HEG1 | 57493 |
| ENSG000002 | 2.70247806 | -0.290926952 | 0.37188937 | -0.7822943 | 0.43404161 | 0.55904019 | NA | NA |
| ENSG000002 | 2.97511509 | -0.295217132 | 0.37738765 | -0.7822649 | 0.43405889 | 0.55904468 | NA | NA |
| ENSG000002 | 13.5678088 | 0.257781529 | 0.3295508 | 0.78222092 | 0.43408475 | 0.55905367 | NA | NA |
| ENSG000002 | 3.25711808 | -0.295757469 | 0.37810681 | -0.7822061 | 0.43409343 | 0.55905367 | NA | NA |
| ENSG000002 | 20.8956804 | 0.218207771 | 0.27903713 | 0.78200263 | 0.43421303 | 0.55918993 | NA | NA |
| ENSG000002 | 3.01194885 | -0.293314832 | 0.37513967 | -0.7818817 | 0.43428412 | 0.55926372 | PTTG2 | 10744 |
| ENSG000001 | 39.1106665 | -0.180493486 | 0.23086875 | -0.7818013 | 0.43433137 | 0.55928905 | CTHRC1 | 115908 |
| ENSG000002 | 2.70765934 | -0.293194072 | 0.37502117 | -0.7818067 | 0.43432817 | 0.55928905 | NA | NA |
| ENSG000002 | 2.72444526 | -0.291834091 | 0.37332938 | -0.7817067 | 0.43438694 | 0.55934284 | NA | NA |

| | | | | | | | | |
|---|---|---|---|---|---|---|---|---|
| ENSG000001 | 14.1422837 | 0.23972355 | 0.30671011 | 0.7815965 | 0.43445175 | 0.55940832 | FAM89B | 23625 |
| ENSG000002 | 3.27965777 | -0.294844345 | 0.37724464 | -0.7815733 | 0.43446538 | 0.55940832 | NA | NA |
| ENSG000002 | 6.0372437 | -0.287969858 | 0.36846807 | -0.7815327 | 0.43448923 | 0.55942126 | NA | NA |
| ENSG000002 | 3.23616829 | -0.294916858 | 0.37737113 | -0.7815035 | 0.43450642 | 0.55942563 | NA | NA |
| ENSG000002 | 6.98558114 | -0.282188791 | 0.36113466 | -0.7813949 | 0.43457025 | 0.55949006 | NA | NA |
| ENSG000002 | 13.1143312 | 0.255010322 | 0.32639203 | 0.78130069 | 0.43462566 | 0.55954363 | NA | NA |
| ENSG000002 | 2.75899601 | -0.291262211 | 0.37292615 | -0.7810185 | 0.43479163 | 0.55973953 | NA | NA |
| ENSG000001 | 121.994407 | 0.133948188 | 0.17151313 | 0.78097919 | 0.43481473 | 0.55974451 | TPCN2 | 219931 |
| ENSG000002 | 4.57473494 | -0.29481445 | 0.37750023 | -0.7809649 | 0.43482311 | 0.55974451 | NA | NA |
| ENSG000001 | 2.99071907 | -0.293158484 | 0.37539318 | -0.7809372 | 0.43483945 | 0.55974778 | NA | NA |
| ENSG000001 | 33.2673633 | -0.193822374 | 0.24824105 | -0.7807829 | 0.43493017 | 0.55984679 | ARL11 | 115761 |
| ENSG000002 | 27.7532694 | 0.200775789 | 0.25716128 | 0.78073879 | 0.43495614 | 0.55986244 | SNORD99 | 692212 |
| ENSG000002 | 5.2950086 | -0.292002887 | 0.37408008 | -0.7805892 | 0.43504412 | 0.55995791 | NA | NA |
| ENSG000002 | 53.0393296 | 0.178470867 | 0.22866483 | 0.78049112 | 0.43510185 | 0.56001445 | NA | NA |
| ENSG000002 | 6.73114649 | -0.283864996 | 0.36372866 | -0.7804307 | 0.43513743 | 0.56003799 | RNU6-898P | 106481462 |
| ENSG000002 | 6.99872326 | 0.281135217 | 0.36024982 | 0.78038962 | 0.43516158 | 0.56003799 | AKIRIN1P2 | 106480306 |
| ENSG000002 | 15.8000055 | -0.24165295 | 0.30965498 | -0.7803942 | 0.43515889 | 0.56003799 | L3MBTL4-AS1 | 101927150 |
| ENSG000002 | 2.75260287 | -0.291966558 | 0.37414998 | -0.7803463 | 0.43518706 | 0.5600523 | NA | NA |
| ENSG000002 | 20.2625567 | -0.216441487 | 0.27738229 | -0.7803003 | 0.43521413 | 0.5600523 | NA | NA |
| ENSG000002 | 4.28010011 | -0.295073939 | 0.37815092 | -0.7803073 | 0.43521 | 0.5600523 | NA | NA |
| ENSG000002 | 2.28629192 | -0.279506982 | 0.35824016 | -0.7802223 | 0.43526001 | 0.56009357 | NA | NA |
| ENSG000002 | 7.81615936 | -0.275124293 | 0.35265676 | -0.7801475 | 0.43530406 | 0.56013247 | NA | NA |
| ENSG000001 | 139.37479 | -0.125176595 | 0.16046835 | -0.7800703 | 0.43534949 | 0.56015538 | RBMS3 | 27303 |
| ENSG000002 | 6.04308833 | -0.289479833 | 0.37109202 | -0.7800756 | 0.43534637 | 0.56015538 | NA | NA |
| ENSG000002 | 16.6982126 | 0.237644471 | 0.30468788 | 0.77996037 | 0.4354142 | 0.56022088 | NA | NA |
| ENSG000002 | 56.737438 | 0.151635147 | 0.19442725 | 0.77990689 | 0.43544569 | 0.56024361 | SLC9A7P1 | 121456 |
| ENSG000002 | 9.31408062 | -0.26661128 | 0.34198068 | -0.7796092 | 0.43562094 | 0.56045131 | NA | NA |
| ENSG000002 | 11.0205443 | -0.25719513 | 0.33000726 | -0.779362 | 0.43576647 | 0.56062076 | NA | NA |
| ENSG000002 | 3.2078233 | -0.294761969 | 0.37822409 | -0.7793316 | 0.43578443 | 0.56062607 | NA | NA |

| | | | | | | | | |
|---|---|---|---|---|---|---|---|---|
| ENSG000002 | 9.80692207 | -0.266685635 | 0.34222043 | -0.7792803 | 0.43581462 | 0.56064713 | NA | NA |
| ENSG000002 | 3.23078775 | -0.294767039 | 0.37829976 | -0.7791891 | 0.43586835 | 0.56069845 | NA | NA |
| ENSG000002 | 16.9738219 | 0.229438189 | 0.29449976 | 0.77907768 | 0.43593396 | 0.56074728 | NA | NA |
| ENSG000002 | 3.72397723 | -0.29470407 | 0.37826827 | -0.7790875 | 0.43592819 | 0.56074728 | NA | NA |
| ENSG000001 | 2.68202625 | -0.291167203 | 0.37381464 | -0.778908 | 0.43603393 | 0.56085808 | A3GALT2 | 127550 |
| ENSG000002 | 5.82830783 | -0.290465292 | 0.37292481 | -0.7788843 | 0.43604785 | 0.5608582 | NA | NA |
| ENSG000002 | 2.93065972 | -0.293533925 | 0.37693392 | -0.7787411 | 0.43613226 | 0.56094897 | NA | NA |
| ENSG000002 | 8.36696934 | -0.278863605 | 0.35811273 | -0.7787034 | 0.43615445 | 0.56095972 | LINC00675 | 100289255 |
| ENSG000002 | 8.07905104 | 0.272157451 | 0.34956124 | 0.77856874 | 0.4362338 | 0.56104398 | NA | NA |
| ENSG000002 | 162.199297 | -0.111491992 | 0.14321423 | -0.778498 | 0.4362755 | 0.56107982 | NA | NA |
| ENSG000002 | 8.81674387 | -0.275853669 | 0.35437233 | -0.7784289 | 0.43631619 | 0.5610875 | LRRC72 | 100506049 |
| ENSG000002 | 3.263211 | -0.293160963 | 0.37661057 | -0.7784194 | 0.43632183 | 0.5610875 | NA | NA |
| ENSG000002 | 5.56322878 | -0.288684767 | 0.37086114 | -0.7784174 | 0.43632298 | 0.5610875 | NA | NA |
| ENSG000002 | 3.0133551 | 0.293245579 | 0.37674342 | 0.77836947 | 0.43635123 | 0.56108824 | NA | NA |
| ENSG000002 | 2.70234276 | -0.291909617 | 0.37501665 | -0.7783911 | 0.43633849 | 0.56108824 | NA | NA |
| ENSG000001 | 23.3623277 | 0.211020169 | 0.27112116 | 0.77832424 | 0.43637789 | 0.56110473 | STX1A | 6804 |
| ENSG000002 | 2.71045143 | -0.290813754 | 0.37366043 | -0.7782835 | 0.43640189 | 0.56110819 | NA | NA |
| ENSG000002 | 6.2670232 | -0.284221115 | 0.36519476 | -0.7782727 | 0.43640825 | 0.56110819 | ZNF709 | 163051 |
| ENSG000001 | 5.87936553 | -0.290696406 | 0.37355415 | -0.7781908 | 0.43645654 | 0.56113468 | KCNRG | 283518 |
| ENSG000002 | 9.76048797 | 0.268715705 | 0.34530103 | 0.77820708 | 0.43644694 | 0.56113468 | NA | NA |
| ENSG000001 | 6.08678751 | -0.286366414 | 0.368068 | -0.7780258 | 0.43655379 | 0.56118854 | RBP2 | 5948 |
| ENSG000001 | 17.4288314 | -0.228038018 | 0.2930929 | -0.7780401 | 0.43654539 | 0.56118854 | APLNR | 187 |
| ENSG000001 | 104.360072 | 0.12784555 | 0.16431418 | 0.77805549 | 0.4365363 | 0.56118854 | XPA | 7507 |
| ENSG000002 | 3.00637656 | -0.291624816 | 0.37481237 | -0.7780555 | 0.43653631 | 0.56118854 | NA | NA |
| ENSG000002 | 3.73726664 | -0.294238485 | 0.37825371 | -0.7778866 | 0.43663588 | 0.56127628 | NA | NA |
| ENSG000001 | 31.2812584 | 0.187703828 | 0.24134273 | 0.77774802 | 0.43671758 | 0.5613635 | NEIL1 | 79661 |
| ENSG000002 | 2.70594808 | -0.288631962 | 0.37125303 | -0.7774535 | 0.43689127 | 0.56156897 | NA | NA |
| ENSG000001 | 86.1394525 | -0.138045343 | 0.17757214 | -0.7774043 | 0.4369203 | 0.56157068 | LOC100506127 | 100506127 |
| ENSG000002 | 9.70913879 | -0.266587052 | 0.34291942 | -0.7774044 | 0.43692025 | 0.56157068 | NA | NA |

| | | | | | | | | |
|---|---|---|---|---|---|---|---|---|
| ENSG000001 | 52.6296092 | 0.161228117 | 0.20742005 | 0.77730248 | 0.43698033 | 0.56159487 | SLC30A3 | 7781 |
| ENSG000002 | 2.76384407 | -0.289877704 | 0.37292888 | -0.7773002 | 0.43698167 | 0.56159487 | LINC01014 | 100874330 |
| ENSG000002 | 3.27799464 | -0.293577736 | 0.37767549 | -0.777328 | 0.43696527 | 0.56159487 | NA | NA |
| ENSG000002 | 75.4309972 | 0.144557917 | 0.18597958 | 0.77727844 | 0.43699451 | 0.56159487 | LINC01138 | 388685 |
| ENSG000001 | 131.291446 | 0.122489642 | 0.15761508 | 0.7771442 | 0.4370737 | 0.56167884 | TRIM52 | 84851 |
| ENSG000002 | 2.94912967 | -0.292942819 | 0.37702034 | -0.7769947 | 0.43716188 | 0.56175655 | NA | NA |
| ENSG000002 | 3.22697805 | -0.29394371 | 0.37830071 | -0.7770107 | 0.43715244 | 0.56175655 | LOC103191607 | 103191607 |
| ENSG000002 | 5.57838983 | -0.289385128 | 0.37249388 | -0.7768856 | 0.43722627 | 0.56182149 | NA | NA |
| ENSG000002 | 18.0180002 | -0.225359031 | 0.29010657 | -0.7768146 | 0.43726814 | 0.56185261 | NA | NA |
| ENSG000002 | 9.81094731 | -0.267707669 | 0.34462989 | -0.7767976 | 0.4372782 | 0.56185261 | NA | NA |
| ENSG000002 | 12.051552 | -0.254440299 | 0.32756476 | -0.7767634 | 0.4372984 | 0.56186076 | NA | NA |
| ENSG000001 | 597.638104 | 0.097468604 | 0.12548735 | 0.77672054 | 0.43732367 | 0.56187542 | TUBB6 | 84617 |
| ENSG000002 | 10.1481093 | -0.273453286 | 0.3521186 | -0.7765943 | 0.43739818 | 0.56193917 | NA | NA |
| ENSG000002 | 14.9119892 | -0.23572459 | 0.30353822 | -0.7765895 | 0.437401 | 0.56193917 | LOC105372556 | 105372556 |
| ENSG000001 | 40.1790425 | -0.176785913 | 0.22771118 | -0.7763603 | 0.4375363 | 0.56204177 | FITM2 | 128486 |
| ENSG000002 | 25.0806497 | 0.205295476 | 0.26441189 | 0.77642302 | 0.43749926 | 0.56204177 | NA | NA |
| ENSG000002 | 2.96774308 | -0.292870814 | 0.37723268 | -0.7763665 | 0.43753262 | 0.56204177 | NA | NA |
| ENSG000002 | 5.73036877 | -0.288200843 | 0.37121787 | -0.7763658 | 0.43753305 | 0.56204177 | NA | NA |
| ENSG000002 | 10.9779905 | -0.286133444 | 0.36857121 | -0.7763315 | 0.43755327 | 0.56204576 | MIR3648-1 | 100500862 |
| ENSG000002 | 3.25324366 | -0.293597263 | 0.37822148 | -0.7762575 | 0.43759695 | 0.56207609 | NA | NA |
| ENSG000002 | 4.47262023 | -0.293223697 | 0.37774654 | -0.7762446 | 0.43760461 | 0.56207609 | NA | NA |
| ENSG000002 | 3.2227634 | -0.293504456 | 0.37817671 | -0.776104 | 0.43768758 | 0.56216486 | NA | NA |
| ENSG000002 | 3.22052344 | -0.292549351 | 0.37696408 | -0.7760669 | 0.43770947 | 0.56217518 | NA | NA |
| ENSG000001 | 3.00064349 | -0.292502055 | 0.37698745 | -0.7758934 | 0.43781196 | 0.56228899 | OR10A4 | 283297 |
| ENSG000002 | 4.80374423 | -0.292462718 | 0.37702103 | -0.7757199 | 0.43791438 | 0.56240272 | NA | NA |
| ENSG000002 | 10.7980084 | -0.257020772 | 0.33134459 | -0.7756903 | 0.43793189 | 0.5624074 | NA | NA |
| ENSG000002 | 7.369153 | -0.283793177 | 0.36595589 | -0.7754846 | 0.43805333 | 0.56254555 | NA | NA |
| ENSG000002 | 137.714555 | 0.120988125 | 0.15604184 | 0.77535696 | 0.43812876 | 0.56262459 | MCM3AP-AS1 | 114044 |
| ENSG000002 | 2.44970979 | -0.2868244 | 0.36995772 | -0.7752897 | 0.43816852 | 0.56264362 | NA | NA |

| | | | | | | | | |
|---|---|---|---|---|---|---|---|---|
| ENSG000002 | 2.38908302 | 0.287211313 | 0.37045906 | 0.7752849 | 0.43817133 | 0.56264362 | NA | NA |
| ENSG000002 | 2.45503935 | -0.286751984 | 0.36994298 | -0.7751248 | 0.43826591 | 0.56274725 | NA | NA |
| ENSG000001 | 22.4623019 | 0.209075089 | 0.26977887 | 0.7749869 | 0.4383474 | 0.56283407 | RAB36 | 9609 |
| ENSG000002 | 15.031287 | 0.250066787 | 0.32270629 | 0.77490521 | 0.43839567 | 0.56287822 | NA | NA |
| ENSG000001 | 124.894434 | 0.131157161 | 0.16930959 | 0.77465876 | 0.43854132 | 0.56304741 | SORBS3 | 10174 |
| ENSG000001 | 15.3059406 | -0.233894183 | 0.30207604 | -0.7742891 | 0.43875985 | 0.56331013 | WNT9A | 7483 |
| ENSG000001 | 58.6532897 | -0.15569218 | 0.2011112 | -0.7741597 | 0.43883637 | 0.56332643 | SLC22A3 | 6581 |
| ENSG000001 | 22.2013955 | -0.207281607 | 0.26775375 | -0.7741502 | 0.438842 | 0.56332643 | TSPAN10 | 83882 |
| ENSG000002 | 34.3546819 | 0.186439296 | 0.24081136 | 0.77421303 | 0.43880482 | 0.56332643 | LRRFIP1P1 | 101290506 |
| ENSG000002 | 5.76556142 | -0.285690779 | 0.36901643 | -0.7741953 | 0.43881532 | 0.56332643 | NA | NA |
| ENSG000002 | 3.21621292 | -0.29291633 | 0.37833095 | -0.7742331 | 0.43879298 | 0.56332643 | NA | NA |
| ENSG000002 | 7.08976357 | -0.280693507 | 0.36262335 | -0.7740635 | 0.43889322 | 0.56337435 | NA | NA |
| ENSG000002 | 12.252076 | -0.249333629 | 0.32216982 | -0.7739199 | 0.43897817 | 0.56346555 | NA | NA |
| ENSG000001 | 19.5062299 | -0.217481337 | 0.28104056 | -0.7738432 | 0.43902351 | 0.56350591 | PLEK2 | 26499 |
| ENSG000001 | 2.7300817 | -0.287565267 | 0.37162939 | -0.7737958 | 0.43905155 | 0.56350623 | RNU6-495P | 106481337 |
| ENSG000002 | 11.5890077 | -0.25605767 | 0.33090846 | -0.7738021 | 0.43904781 | 0.56350623 | LOC284648 | 284648 |
| ENSG000001 | 126.091028 | 0.125306752 | 0.1619692 | 0.77364555 | 0.43914043 | 0.56359819 | STAC | 6769 |
| ENSG000002 | 3.05638039 | -0.287341165 | 0.37142436 | -0.7736196 | 0.43915578 | 0.56359819 | NA | NA |
| ENSG000002 | 6.54001757 | -0.281046778 | 0.36329532 | -0.7736042 | 0.4391649 | 0.56359819 | NA | NA |
| ENSG000002 | 2.72683206 | -0.286641675 | 0.37054726 | -0.7735631 | 0.43918922 | 0.56360807 | NA | NA |
| ENSG000002 | 2.937086 | -0.291637451 | 0.3770146 | -0.7735442 | 0.43920039 | 0.56360807 | NA | NA |
| ENSG000002 | 10.5664574 | -0.264181665 | 0.34154481 | -0.7734905 | 0.43923216 | 0.563631 | TMEM75 | 641384 |
| ENSG000002 | 3.70165771 | -0.292692301 | 0.37845193 | -0.7733936 | 0.43928947 | 0.56368671 | NA | NA |
| ENSG000002 | 21.6504121 | -0.210872893 | 0.27267869 | -0.7733384 | 0.43932216 | 0.56371082 | NA | NA |
| ENSG000002 | 7.70935317 | -0.273982063 | 0.35436026 | -0.7731738 | 0.43941954 | 0.56381793 | NA | NA |
| ENSG000002 | 8.46019847 | -0.270193805 | 0.34956423 | -0.7729447 | 0.43955514 | 0.56397406 | NA | NA |
| ENSG000002 | 3.21928597 | -0.292328002 | 0.37830037 | -0.7727405 | 0.439676 | 0.56411129 | NA | NA |
| ENSG000002 | 12.3439306 | 0.253541729 | 0.32811987 | 0.77271069 | 0.43969362 | 0.56411605 | NA | NA |
| ENSG000001 | 71.394693 | 0.149244374 | 0.193174 | 0.77259039 | 0.43976484 | 0.56418957 | IL6R | 3570 |

| | | | | | | | | |
|---|---|---|---|---|---|---|---|---|
| ENSG000002 | 2.96065127 | -0.291122451 | 0.37684351 | -0.7725288 | 0.43980132 | 0.56420857 | NA | NA |
| ENSG000002 | 2.40296474 | 0.28757568 | 0.3722574 | 0.77251837 | 0.43980747 | 0.56420857 | MIR5009 | 100846993 |
| ENSG000001 | 38.7159934 | 0.173440618 | 0.22456013 | 0.77235712 | 0.43990294 | 0.56431225 | PLEKHF1 | 79156 |
| ENSG000002 | 7.5333393 | -0.274504203 | 0.35542876 | -0.7723185 | 0.43992582 | 0.56431225 | NA | NA |
| ENSG000002 | 2.20307191 | -0.279232955 | 0.3615549 | -0.7723114 | 0.43993004 | 0.56431225 | NA | NA |
| ENSG000002 | 111.914182 | 0.122381797 | 0.15846819 | 0.77227991 | 0.43994866 | 0.56431828 | LINC01128 | 643837 |
| ENSG000002 | 18.8404365 | 0.218324275 | 0.28274603 | 0.77215681 | 0.44002156 | 0.56439394 | NA | NA |
| ENSG000002 | 9.00802722 | -0.26613747 | 0.34468005 | -0.772129 | 0.44003801 | 0.56439719 | NA | NA |
| ENSG000002 | 2.48073138 | -0.285696633 | 0.37004782 | -0.7720533 | 0.44008288 | 0.56443688 | NA | NA |
| ENSG000001 | 2.75411663 | -0.286408676 | 0.37101314 | -0.7719637 | 0.44013593 | 0.56448706 | OR4L1 | 122742 |
| ENSG000002 | 6.49963304 | -0.280629859 | 0.36354848 | -0.7719187 | 0.4401626 | 0.56450341 | NA | NA |
| ENSG000002 | 5.12136315 | 0.290132365 | 0.37591632 | 0.77180039 | 0.44023266 | 0.56457541 | NA | NA |
| ENSG000001 | 4.28981348 | -0.292120935 | 0.37856381 | -0.7716557 | 0.44031835 | 0.564662 | GZMM | 3004 |
| ENSG000002 | 2.95450574 | -0.290861721 | 0.37693995 | -0.7716394 | 0.44032803 | 0.564662 | NA | NA |
| ENSG000002 | 2.23225357 | -0.279438066 | 0.36219313 | -0.7715168 | 0.44040069 | 0.56473733 | NA | NA |
| ENSG000002 | 3.9740129 | -0.291610187 | 0.3780025 | -0.7714504 | 0.44044001 | 0.56476988 | NA | NA |
| ENSG000002 | 3477.25808 | 0.110074975 | 0.14272627 | 0.77123135 | 0.44056981 | 0.56491847 | NA | NA |
| ENSG000002 | 3.48026281 | -0.291869656 | 0.37848011 | -0.7711625 | 0.44061063 | 0.56495294 | LOC100996679 | 100996679 |
| ENSG000001 | 4.05529946 | -0.291898059 | 0.37858685 | -0.7710201 | 0.44069504 | 0.5650433 | FIGLA | 344018 |
| ENSG000002 | 6.99421598 | 0.279914565 | 0.36312905 | 0.77084047 | 0.4408015 | 0.56516193 | NA | NA |
| ENSG000002 | 47.3398575 | -0.164138144 | 0.21299468 | -0.7706209 | 0.44093169 | 0.56531097 | DOC2B | 8447 |
| ENSG000001 | 67.744651 | 0.153431393 | 0.1991201 | 0.77054701 | 0.44097548 | 0.56531541 | CYP4V2 | 285440 |
| ENSG000002 | 3.832866 | -0.291163201 | 0.37786682 | -0.7705445 | 0.44097697 | 0.56531541 | NA | NA |
| ENSG000002 | 8.71276708 | -0.267315153 | 0.34691653 | -0.770546 | 0.44097607 | 0.56531541 | NA | NA |
| ENSG000002 | 3.52141646 | -0.291577382 | 0.37843805 | -0.7704759 | 0.44101766 | 0.56534971 | NA | NA |
| ENSG00000C | 356.024236 | 0.09509817 | 0.12344226 | 0.77038583 | 0.44107106 | 0.56537113 | ARHGAP28 | 79822 |
| ENSG000002 | 3.72314828 | -0.291537495 | 0.37842525 | -0.7703965 | 0.44106472 | 0.56537113 | NA | NA |
| ENSG000002 | 4.52453578 | 0.290113081 | 0.37658578 | 0.77037715 | 0.4410762 | 0.56537113 | NA | NA |
| ENSG000001 | 573.097285 | 0.283992878 | 0.36871349 | 0.77022643 | 0.44116559 | 0.56546783 | EIF3C | 8663 |

| | | | | | | | | |
|---|---|---|---|---|---|---|---|---|
| ENSG000001 | 19.0836967 | 0.226822654 | 0.29456288 | 0.77003134 | 0.4412813 | 0.5655973 | EXTL1 | 2134 |
| ENSG000002 | 7.07984757 | -0.279994645 | 0.36362511 | -0.7700091 | 0.44129449 | 0.5655973 | NA | NA |
| ENSG000001 | 47.9554679 | 0.162811986 | 0.21148371 | 0.76985591 | 0.44138537 | 0.5656959 | RNU5F-1 | 26828 |
| ENSG000002 | 5.26912288 | -0.287677629 | 0.37372252 | -0.7697626 | 0.44144071 | 0.56574894 | NA | NA |
| ENSG000002 | 8.1256921 | 0.268754768 | 0.34915812 | 0.76972223 | 0.44146468 | 0.56576178 | NA | NA |
| ENSG000002 | 15.3963573 | -0.232663256 | 0.3022918 | -0.7696645 | 0.44149895 | 0.56578297 | NA | NA |
| ENSG000002 | 2.70996161 | -0.286840664 | 0.37269104 | -0.7696473 | 0.44150912 | 0.56578297 | NA | NA |
| ENSG000002 | 7.43259569 | -0.281603842 | 0.3659258 | -0.7695654 | 0.44155773 | 0.5658095 | NA | NA |
| ENSG000002 | 3.50255165 | -0.29020552 | 0.37710064 | -0.7695705 | 0.44155472 | 0.5658095 | LOC105374344 | 105374344 |
| ENSG000001 | 114.157948 | 0.121847224 | 0.15835707 | 0.76944605 | 0.44162856 | 0.56585427 | TEC | 7006 |
| ENSG000002 | 3.84051632 | 0.29128357 | 0.37856246 | 0.76944652 | 0.44162828 | 0.56585427 | UBE2Q1-AS1 | 100874097 |
| ENSG000002 | 3.98570369 | -0.290827158 | 0.37797446 | -0.769436 | 0.44163453 | 0.56585427 | NA | NA |
| ENSG000001 | 179.085894 | 0.111319015 | 0.14468365 | 0.76939596 | 0.44165828 | 0.56586683 | MAMDC2 | 256691 |
| ENSG000002 | 3.20478272 | -0.290740751 | 0.37792977 | -0.7692984 | 0.44171621 | 0.56592317 | NA | NA |
| ENSG000001 | 15.2186052 | 0.238275737 | 0.30984346 | 0.76901974 | 0.44188159 | 0.56610979 | FAM53A | 152877 |
| ENSG000002 | 19.869329 | 0.215879393 | 0.28072527 | 0.76900592 | 0.4418898 | 0.56610979 | ACAD11 | 84129 |
| ENSG000002 | 11.1806872 | 0.251789517 | 0.32745331 | 0.76893259 | 0.44193333 | 0.56614768 | NA | NA |
| ENSG000002 | 2.52356797 | -0.283441069 | 0.36865858 | -0.7688444 | 0.44198571 | 0.5661969 | XXYLT1-AS1 | 100874528 |
| ENSG000001 | 30.6863254 | -0.18952999 | 0.24653888 | -0.7687631 | 0.44203396 | 0.56620504 | IL1RAPL1 | 11141 |
| ENSG000002 | 7.03234093 | 0.276478081 | 0.35963938 | 0.76876477 | 0.44203297 | 0.56620504 | NA | NA |
| ENSG000002 | 3.6975031 | -0.290784467 | 0.37824537 | -0.768772 | 0.44202865 | 0.56620504 | NA | NA |
| ENSG000001 | 656.152497 | 0.097753663 | 0.12716645 | 0.76870641 | 0.44206762 | 0.56623027 | AAK1 | 22848 |
| ENSG000002 | 2.2188085 | -0.278046075 | 0.36172436 | -0.7686684 | 0.44209021 | 0.56624131 | NA | NA |
| ENSG000002 | 3.68213844 | -0.290857286 | 0.37841846 | -0.7686128 | 0.44212319 | 0.56625081 | NA | NA |
| ENSG000002 | 127.153926 | 0.130901444 | 0.17030957 | 0.76860885 | 0.44212555 | 0.56625081 | NA | NA |
| ENSG000001 | 17.526148 | -0.226901656 | 0.29528223 | -0.768423 | 0.44223592 | 0.56637082 | TMEM176B | 28959 |
| ENSG000002 | 12.3830865 | -0.246239449 | 0.3204557 | -0.768404 | 0.44224719 | 0.56637082 | NA | NA |
| ENSG000002 | 3.73769085 | -0.290757011 | 0.37845735 | -0.7682689 | 0.44232745 | 0.56645572 | NA | NA |
| ENSG000002 | 3.49169302 | -0.290825587 | 0.37856406 | -0.7682335 | 0.44234848 | 0.56646477 | NA | NA |

| | | | | | | | | |
|---|---|---|---|---|---|---|---|---|
| ENSG000002 | 3.84955808 | 0.290637255 | 0.37836804 | 0.76813374 | 0.44240773 | 0.56650486 | NA | NA |
| ENSG000002 | 2.38844995 | 0.284018125 | 0.36974569 | 0.76814452 | 0.44240133 | 0.56650486 | NA | NA |
| ENSG000002 | 6.33414931 | -0.283448087 | 0.36906164 | -0.7680237 | 0.44247311 | 0.56657069 | NA | NA |
| ENSG000002 | 2.42915698 | -0.281188715 | 0.36613732 | -0.767987 | 0.44249489 | 0.56658068 | LOC102723896 | 102723896 |
| ENSG000002 | 6.37171472 | -0.282043339 | 0.36733236 | -0.7678151 | 0.44259704 | 0.56669358 | LOC101928924 | 101928924 |
| ENSG000002 | 9.9569936 | -0.257608499 | 0.33553 | -0.7677659 | 0.44262627 | 0.56671312 | NA | NA |
| ENSG000002 | 9.75775143 | -0.261055952 | 0.34015214 | -0.7674682 | 0.44280319 | 0.56692174 | NA | NA |
| ENSG000002 | 9.17259807 | 0.263485693 | 0.34334485 | 0.76740833 | 0.44283878 | 0.5669494 | NA | NA |
| ENSG000002 | 2.550314 | -0.278265297 | 0.36265476 | -0.7673008 | 0.44290268 | 0.56699541 | NA | NA |
| ENSG000002 | 3.21498645 | -0.28987968 | 0.37778123 | -0.7673216 | 0.44289035 | 0.56699541 | LINC00566 | 100861547 |
| ENSG000002 | 6.10999732 | 0.28507644 | 0.37158825 | 0.76718368 | 0.44297232 | 0.56706666 | NA | NA |
| ENSG000002 | 2.43457326 | -0.281381119 | 0.3668215 | -0.7670791 | 0.44303447 | 0.56712831 | LOC100508046 | 100508046 |
| ENSG000001 | 3.25754229 | -0.289599702 | 0.37756747 | -0.7670144 | 0.44307293 | 0.56714174 | S100G | 795 |
| ENSG000002 | 5.48537745 | -0.285323708 | 0.37199011 | -0.7670196 | 0.44306985 | 0.56714174 | NA | NA |
| ENSG000002 | 7.47241766 | -0.271756427 | 0.35432852 | -0.7669618 | 0.44310424 | 0.56716392 | NA | NA |
| ENSG000002 | 8.5689471 | -0.272841605 | 0.35578046 | -0.7668819 | 0.44315173 | 0.56720679 | NA | NA |
| ENSG000001 | 9.25427444 | -0.261436465 | 0.34093266 | -0.7668273 | 0.44318422 | 0.56723048 | CABP7 | 164633 |
| ENSG000002 | 9.93554902 | -0.257500245 | 0.33582156 | -0.766777 | 0.44321412 | 0.56725085 | NA | NA |
| ENSG000002 | 7.88448181 | 0.2721242 | 0.35491529 | 0.76673 | 0.44324206 | 0.5672687 | NA | NA |
| ENSG000001 | 8.96703042 | 0.267345019 | 0.34872192 | 0.76664242 | 0.44329415 | 0.56731746 | TCTE1 | 202500 |
| ENSG000002 | 32.9731949 | -0.179294622 | 0.23387986 | -0.7666099 | 0.44331347 | 0.56732428 | SRRM2-AS1 | 100128788 |
| ENSG000002 | 3.73942566 | -0.290200284 | 0.37858031 | -0.7665488 | 0.44334983 | 0.5673402 | NA | NA |
| ENSG000002 | 5.14550819 | -0.288921078 | 0.37691489 | -0.766542 | 0.44335389 | 0.5673402 | LINC01179 | 101928151 |
| ENSG000002 | 15.3874294 | 0.237419548 | 0.30976082 | 0.76646086 | 0.44340213 | 0.56738403 | NA | NA |
| ENSG000002 | 11.0624103 | -0.25684162 | 0.33512436 | -0.7664069 | 0.44343423 | 0.5674072 | NA | NA |
| ENSG000001 | 25.9976395 | -0.213656839 | 0.27880745 | -0.766324 | 0.44348355 | 0.5674524 | BEND2 | 139105 |
| ENSG000002 | 4.22669739 | -0.289346939 | 0.37759837 | -0.7662823 | 0.44350836 | 0.56746624 | NA | NA |
| ENSG000001 | 46.7655767 | -0.164216095 | 0.21433778 | -0.7661556 | 0.44358371 | 0.56754473 | C9orf43 | 257169 |
| ENSG000002 | 6.20138961 | 0.28017273 | 0.36571087 | 0.76610447 | 0.44361415 | 0.56754787 | NA | NA |

| | | | | | | | | |
|---|---|---|---|---|---|---|---|---|
| ENSG000002 | 3.03082285 | -0.286277274 | 0.37367812 | -0.7661066 | 0.44361288 | 0.56754787 | NA | NA |
| ENSG000001 | 42.8128513 | -0.164510729 | 0.2147756 | -0.7659656 | 0.44369675 | 0.56761535 | APOLD1 | 81575 |
| ENSG000001 | 9.0729376 | -0.272407949 | 0.35564938 | -0.7659452 | 0.44370888 | 0.56761535 | C15orf62 | 643338 |
| ENSG000002 | 3.42908181 | -0.289994808 | 0.37859328 | -0.7659798 | 0.44368832 | 0.56761535 | NA | NA |
| ENSG000002 | 3.0566559 | 0.28837457 | 0.37661332 | 0.76570464 | 0.44385206 | 0.5677806 | LOC152225 | 152225 |
| ENSG000002 | 9.01488602 | -0.263890072 | 0.34469733 | -0.7655704 | 0.44393196 | 0.56786489 | NA | NA |
| ENSG000001 | 25.2535678 | 0.199705386 | 0.26088922 | 0.76547965 | 0.44398598 | 0.56789817 | NCCRP1 | 342897 |
| ENSG000002 | 9.68159818 | -0.257955636 | 0.33698371 | -0.765484 | 0.44398339 | 0.56789817 | NA | NA |
| ENSG000002 | 3.27458988 | -0.289005638 | 0.37760047 | -0.7653741 | 0.44404878 | 0.56794632 | NA | NA |
| ENSG000002 | 2.18860843 | -0.277049859 | 0.36198191 | -0.7653693 | 0.44405164 | 0.56794632 | NA | NA |
| ENSG000001 | 5.2727954 | -0.286403804 | 0.37421812 | -0.7653392 | 0.44406958 | 0.56795135 | NA | NA |
| ENSG000001 | 44.7361953 | 0.163457692 | 0.21358694 | 0.76529816 | 0.44409402 | 0.56796469 | GFRA2 | 2675 |
| ENSG000002 | 2.91705024 | -0.28796112 | 0.37629118 | -0.7652614 | 0.44411589 | 0.56797476 | NA | NA |
| ENSG000002 | 2.46844032 | -0.283130132 | 0.37002332 | -0.7651683 | 0.44417131 | 0.5680098 | NA | NA |
| ENSG000002 | 9.18662595 | -0.261430217 | 0.34165474 | -0.7651883 | 0.44415942 | 0.5680098 | NA | NA |
| ENSG000002 | 2.40410635 | 0.284913069 | 0.3724529 | 0.76496403 | 0.44429296 | 0.56812954 | NA | NA |
| ENSG000002 | 2.5134954 | -0.281400533 | 0.36785443 | -0.764978 | 0.44428466 | 0.56812954 | NA | NA |
| ENSG000002 | 2.95305616 | -0.287464519 | 0.37582723 | -0.7648848 | 0.44434017 | 0.56817198 | NA | NA |
| ENSG000001 | 18.8744757 | 0.226067955 | 0.2956059 | 0.76476131 | 0.44441369 | 0.56824808 | CA3 | 761 |
| ENSG000002 | 6.23771016 | -0.280584897 | 0.3669095 | -0.7647251 | 0.44443526 | 0.56825775 | NA | NA |
| ENSG000002 | 10.2581983 | 0.256163777 | 0.33501945 | 0.76462361 | 0.44449571 | 0.56831711 | RN7SL674P | 106479468 |
| ENSG000002 | 4.8666065 | -0.289276937 | 0.37834056 | -0.764594 | 0.44451332 | 0.5683217 | NA | NA |
| ENSG000002 | 3.45982321 | -0.289387461 | 0.37857978 | -0.7644028 | 0.44462722 | 0.56841357 | NA | NA |
| ENSG000002 | 4.2086531 | -0.287932056 | 0.37667276 | -0.764409 | 0.44462354 | 0.56841357 | NA | NA |
| ENSG000002 | 2.70128438 | -0.284840856 | 0.37261591 | -0.7644356 | 0.44460772 | 0.56841357 | NA | NA |
| ENSG000001 | 6.80256686 | -0.279728455 | 0.36600663 | -0.7642716 | 0.44470544 | 0.56844188 | OR10H4 | 126541 |
| ENSG000001 | 3.23451368 | -0.289015876 | 0.3781342 | -0.7643209 | 0.44467602 | 0.56844188 | GM140 | 100287948 |
| ENSG000002 | 3.62780345 | 0.288567132 | 0.37755645 | 0.76430194 | 0.44468733 | 0.56844188 | NA | NA |
| ENSG000002 | 3.24626754 | -0.287768603 | 0.37651785 | -0.7642894 | 0.44469479 | 0.56844188 | NA | NA |

| | | | | | | | | |
|---|---|---|---|---|---|---|---|---|
| ENSG000002 | 11.8737117 | -0.246809156 | 0.32298107 | -0.7641598 | 0.44477201 | 0.56850905 | NA | NA |
| ENSG000002 | 3.70513569 | -0.289194653 | 0.3784867 | -0.7640814 | 0.44481873 | 0.56855085 | NA | NA |
| ENSG000002 | 7.48825829 | -0.273599909 | 0.35812974 | -0.7639687 | 0.4448859 | 0.56861717 | LOC100507175 | 100507175 |
| ENSG000002 | 3.70317146 | -0.289225391 | 0.37859339 | -0.7639473 | 0.44489866 | 0.56861717 | NA | NA |
| ENSG000001 | 291.663204 | 0.11180755 | 0.14636344 | 0.76390352 | 0.44492474 | 0.56861963 | JUND | 3727 |
| ENSG000002 | 2.96714171 | -0.287365842 | 0.37618402 | -0.763897 | 0.44492863 | 0.56861963 | RNU6-273P | 106480566 |
| ENSG000002 | 7.27521424 | -0.272274308 | 0.35648829 | -0.7637679 | 0.44500558 | 0.56870004 | NA | NA |
| ENSG000002 | 2.93409674 | -0.286497377 | 0.37513739 | -0.7637132 | 0.44503818 | 0.56870586 | NA | NA |
| ENSG000002 | 3.95366502 | -0.288455205 | 0.37769795 | -0.7637193 | 0.44503455 | 0.56870586 | NA | NA |
| ENSG000002 | 13.3767195 | 0.239760662 | 0.31395614 | 0.76367567 | 0.44506055 | 0.56871652 | NA | NA |
| ENSG000002 | 3.01214582 | -0.287272341 | 0.37619607 | -0.763624 | 0.44509135 | 0.56873797 | NA | NA |
| ENSG000002 | 6.12586988 | -0.284571871 | 0.37270942 | -0.7635221 | 0.4451521 | 0.56876182 | LINGO3 | 645191 |
| ENSG000002 | 6.19115428 | 0.28164353 | 0.36887222 | 0.76352599 | 0.44514977 | 0.56876182 | NA | NA |
| ENSG000002 | 6.38429527 | 0.277726148 | 0.36373281 | 0.7635444 | 0.4451388 | 0.56876182 | NA | NA |
| ENSG000002 | 6.7570562 | -0.276412604 | 0.36203807 | -0.7634904 | 0.44517097 | 0.56876801 | NA | NA |
| ENSG000002 | 2.46883152 | -0.280758054 | 0.36777511 | -0.763396 | 0.44522725 | 0.56880408 | NA | NA |
| ENSG000002 | 10.2538921 | -0.256312287 | 0.33575082 | -0.7634003 | 0.44522468 | 0.56880408 | NA | NA |
| ENSG000002 | 14.0358931 | -0.235095983 | 0.30800087 | -0.7632965 | 0.4452866 | 0.56884783 | NA | NA |
| ENSG000002 | 2.67006745 | -0.285271133 | 0.37373811 | -0.7632915 | 0.44528956 | 0.56884783 | NA | NA |
| ENSG000002 | 4.25518051 | 0.288681908 | 0.37827197 | 0.76315966 | 0.44536818 | 0.5689126 | NA | NA |
| ENSG000002 | 29.3488506 | 0.198189056 | 0.25969548 | 0.76315944 | 0.44536832 | 0.5689126 | NA | NA |
| ENSG000002 | 3.0510278 | -0.285627975 | 0.37430287 | -0.7630932 | 0.44540781 | 0.56894513 | LOC105373878 | 105373878 |
| ENSG000001 | 5.52179528 | -0.283601223 | 0.37168548 | -0.763014 | 0.44545504 | 0.56898754 | CD5 | 921 |
| ENSG000001 | 9.52711367 | -0.261300586 | 0.34248704 | -0.7629503 | 0.44549305 | 0.56901816 | MRGPRX1 | 259249 |
| ENSG000002 | 6.7303266 | -0.276260787 | 0.36211518 | -0.7629086 | 0.44551791 | 0.56903199 | NA | NA |
| ENSG000002 | 8.06001539 | -0.273388285 | 0.35841953 | -0.7627606 | 0.4456062 | 0.56912683 | LOC105372028 | 105372028 |
| ENSG000002 | 58.2362226 | 0.165109815 | 0.21649623 | 0.76264521 | 0.44567501 | 0.56919679 | NA | NA |
| ENSG000002 | 5.38011593 | -0.286119932 | 0.37520008 | -0.7625796 | 0.44571415 | 0.56922885 | NA | NA |
| ENSG000002 | 13.5623696 | 0.23639885 | 0.3100204 | 0.76252675 | 0.44574569 | 0.5692512 | NA | NA |

| | | | | | | | | |
|---|---|---|---|---|---|---|---|---|
| ENSG000001 | 124.337305 | -0.124192961 | 0.16288754 | -0.762446 | 0.44579384 | 0.56929476 | NPHP1 | 4867 |
| ENSG000002 | 3.23465007 | -0.288100707 | 0.37790686 | -0.762359 | 0.44584575 | 0.56934312 | NA | NA |
| ENSG000002 | 2.25970101 | -0.273972555 | 0.35940626 | -0.7622921 | 0.44588569 | 0.56934683 | NA | NA |
| ENSG000002 | 8.70880038 | -0.2660529 | 0.3490209 | -0.7622836 | 0.44589077 | 0.56934683 | PRRT4 | 401399 |
| ENSG000002 | 3.44781179 | -0.288577499 | 0.37854788 | -0.7623276 | 0.4458645 | 0.56934683 | NA | NA |
| ENSG000001 | 25.8300987 | -0.199639599 | 0.26192235 | -0.7622091 | 0.44593521 | 0.56934979 | ANKRD19P | 138649 |
| ENSG000002 | 2.23950791 | -0.272122429 | 0.35700406 | -0.762239 | 0.44591735 | 0.56934979 | NA | NA |
| ENSG000002 | 10.495937 | -0.254878898 | 0.33438478 | -0.7622324 | 0.44592133 | 0.56934979 | NA | NA |
| ENSG000001 | 407.015353 | 0.096139612 | 0.12614033 | 0.76216397 | 0.44596215 | 0.56935263 | SLC41A1 | 254428 |
| ENSG000002 | 3.04973295 | 0.287349355 | 0.37702515 | 0.76214904 | 0.44597106 | 0.56935263 | TCF4-AS2 | 106480339 |
| ENSG000002 | 10.0948161 | 0.259873722 | 0.34100236 | 0.76208774 | 0.44600764 | 0.56935263 | LOC101929704 | 101929704 |
| ENSG000002 | 3.45547162 | -0.288499123 | 0.37856264 | -0.7620908 | 0.44600579 | 0.56935263 | NA | NA |
| ENSG000002 | 40.6770595 | 0.168566175 | 0.22117743 | 0.76213101 | 0.44598182 | 0.56935263 | NA | NA |
| ENSG000001 | 2.94979465 | -0.285878455 | 0.37514832 | -0.7620411 | 0.44603545 | 0.56937021 | LCE5A | 254910 |
| ENSG000002 | 20.1893482 | 0.208970451 | 0.27428363 | 0.76187722 | 0.44613328 | 0.56947717 | ADAMTSL4-AS1 | 574406 |
| ENSG000002 | 5.83234631 | -0.284444972 | 0.37342383 | -0.7617215 | 0.44622622 | 0.56957787 | NA | NA |
| ENSG000001 | 23.963365 | 0.208833771 | 0.2742093 | 0.76158529 | 0.44630756 | 0.56966227 | NPY2R | 4887 |
| ENSG000002 | 3.27263808 | -0.286132132 | 0.37572821 | -0.7615402 | 0.44633448 | 0.56966227 | NA | NA |
| ENSG000002 | 9.3286428 | -0.264225582 | 0.3469556 | -0.7615545 | 0.44632597 | 0.56966227 | NA | NA |
| ENSG000002 | 3.47390931 | -0.288294542 | 0.37859316 | -0.761489 | 0.44636503 | 0.56968333 | NA | NA |
| ENSG000002 | 2.42726722 | 0.283202726 | 0.37198446 | 0.76132946 | 0.44646031 | 0.56978699 | NA | NA |
| ENSG000001 | 7.23764216 | -0.270968581 | 0.35607333 | -0.7609909 | 0.44666248 | 0.57002707 | CSF3 | 1440 |
| ENSG000002 | 3.86231969 | 0.288087976 | 0.37859284 | 0.76094406 | 0.44669048 | 0.57004487 | MIR301A | 407027 |
| ENSG000002 | 8.28019253 | -0.268870956 | 0.35338442 | -0.7608455 | 0.44674933 | 0.57010203 | NA | NA |
| ENSG000002 | 6.72789436 | -0.274302023 | 0.36068013 | -0.7605133 | 0.44694784 | 0.5703374 | HLA-DQA2 | 3118 |
| ENSG000002 | 40.1938792 | -0.172596335 | 0.22696344 | -0.7604587 | 0.44698042 | 0.57036102 | HMGN3-AS1 | 100288198 |
| ENSG000001 | 80.8170085 | 0.138930595 | 0.18273486 | 0.76028512 | 0.44708417 | 0.57044491 | DUSP19 | 142679 |
| ENSG000001 | 2.72007907 | -0.281660606 | 0.37047331 | -0.7602723 | 0.44709181 | 0.57044491 | NA | NA |
| ENSG000002 | 3.79652462 | -0.287814109 | 0.37858769 | -0.760231 | 0.4471165 | 0.57044491 | NA | NA |

| | | | | | | | | |
|---|---|---|---|---|---|---|---|---|
| ENSG000002 | 5.70028136 | -0.281627859 | 0.37043901 | -0.7602543 | 0.44710258 | 0.57044491 | LOC101928943 | 101928943 |
| ENSG000002 | 3.7565641 | -0.287812164 | 0.37853958 | -0.7603225 | 0.44706182 | 0.57044491 | GDF2 | 2658 |
| ENSG000002 | 7.20733632 | -0.274065992 | 0.36055034 | -0.7601324 | 0.44717546 | 0.57050218 | NA | NA |
| ENSG000002 | 3.96865719 | -0.287152509 | 0.37779313 | -0.7600787 | 0.44720755 | 0.57052518 | NA | NA |
| ENSG000002 | 2.76565042 | -0.28200576 | 0.37103447 | -0.7600527 | 0.44722306 | 0.57052702 | NA | NA |
| ENSG000002 | 3.48668875 | -0.287417051 | 0.37818438 | -0.7599919 | 0.44725945 | 0.57055549 | DHRSX-IT1 | 106478924 |
| ENSG000001 | 535.566895 | 0.096126986 | 0.12653116 | 0.75971 | 0.44742795 | 0.57073453 | DCAF6 | 55827 |
| ENSG000002 | 3.22145708 | -0.285840434 | 0.37624576 | -0.7597174 | 0.44742355 | 0.57073453 | NA | NA |
| ENSG000002 | 3.74957035 | -0.287592448 | 0.37859326 | -0.7596344 | 0.44747317 | 0.57077427 | NA | NA |
| ENSG000001 | 4.24647312 | -0.286491187 | 0.37723984 | -0.7594404 | 0.44758914 | 0.57088627 | MYF5 | 4617 |
| ENSG000001 | 29.6390381 | -0.188554862 | 0.24827423 | -0.7594621 | 0.4475762 | 0.57088627 | NOL3 | 8996 |
| ENSG000001 | 6.28332452 | -0.281491227 | 0.37069213 | -0.7593666 | 0.44763329 | 0.57090344 | GNRHR | 2798 |
| ENSG000001 | 507.979773 | -0.101668939 | 0.13389406 | -0.7593237 | 0.44765892 | 0.57090344 | CEP85L | 387119 |
| ENSG000002 | 3.44505937 | -0.287431494 | 0.37852455 | -0.759347 | 0.44764499 | 0.57090344 | NA | NA |
| ENSG000002 | 4.96722018 | -0.285067543 | 0.37540141 | -0.7593673 | 0.44763289 | 0.57090344 | NA | NA |
| ENSG000002 | 3.7032048 | -0.287332861 | 0.37845222 | -0.7592315 | 0.44771407 | 0.57095582 | NA | NA |
| ENSG000000 | 71.5107715 | 0.15007123 | 0.1976967 | 0.75909832 | 0.44779375 | 0.5710392 | YPEL3 | 83719 |
| ENSG000002 | 3.44376022 | -0.287154939 | 0.3782958 | -0.7590751 | 0.44780761 | 0.5710392 | NA | NA |
| ENSG000001 | 12.5047648 | -0.249462294 | 0.32867998 | -0.7589823 | 0.44786314 | 0.57107544 | C2orf66 | 401027 |
| ENSG000002 | 7.35114601 | -0.27397141 | 0.36097291 | -0.7589805 | 0.4478642 | 0.57107544 | NA | NA |
| ENSG000001 | 127.754689 | 0.119563394 | 0.15759594 | 0.75867054 | 0.44804967 | 0.57129397 | PHF11 | 51131 |
| ENSG000001 | 7.47944476 | -0.269792977 | 0.3556839 | -0.7585189 | 0.44814038 | 0.5713737 | C15orf43 | 145645 |
| ENSG000002 | 2.49632361 | -0.27842976 | 0.36706214 | -0.7585358 | 0.44813031 | 0.5713737 | NA | NA |
| ENSG000001 | 15.0750595 | -0.232617288 | 0.30669889 | -0.7584549 | 0.44817868 | 0.57138446 | CALML4 | 91860 |
| ENSG000001 | 14.1437534 | -0.238531454 | 0.31450514 | -0.7584342 | 0.44819109 | 0.57138446 | SLC5A2 | 6524 |
| ENSG000001 | 63.0978923 | 0.151356334 | 0.19956266 | 0.75844016 | 0.44818753 | 0.57138446 | ZNF713 | 349075 |
| ENSG000002 | 3.71495508 | -0.28701692 | 0.37851149 | -0.758278 | 0.4482846 | 0.5714857 | GS1-204I12.4 | 105371653 |
| ENSG000002 | 27.0574443 | 0.20288269 | 0.26759191 | 0.75817947 | 0.44834355 | 0.57154289 | NA | NA |
| ENSG000002 | 12.0314867 | -0.250465259 | 0.33039532 | -0.7580775 | 0.44840459 | 0.5715668 | GYPB | 2994 |

| | | | | | | | | |
|---|---|---|---|---|---|---|---|---|
| ENSG000002 | 21.8040061 | -0.204600111 | 0.26988593 | -0.7580985 | 0.44839204 | 0.5715668 | MAP3K14-AS1 | 100133991 |
| ENSG000002 | 3.70420205 | -0.286997261 | 0.37856529 | -0.7581183 | 0.44838019 | 0.5715668 | NA | NA |
| ENSG00000C | 11.9813743 | -0.25090688 | 0.33103052 | -0.7579569 | 0.44847676 | 0.57164083 | ST6GALNAC1 | 55808 |
| ENSG000002 | 3.46918558 | -0.286822177 | 0.37850544 | -0.7577756 | 0.44858535 | 0.57176126 | LOC100996385 | 100996385 |
| ENSG000002 | 3.22151766 | -0.286369802 | 0.37792786 | -0.7577367 | 0.44860863 | 0.57177297 | NA | NA |
| ENSG000001 | 866.75977 | 0.108340417 | 0.14299806 | 0.75763557 | 0.44866918 | 0.57183216 | AMOTL1 | 154810 |
| ENSG000002 | 3.17787119 | -0.286202904 | 0.37778273 | -0.757586 | 0.44869887 | 0.57183405 | NA | NA |
| ENSG000002 | 13.1906099 | -0.240151598 | 0.3169921 | -0.7575949 | 0.44869353 | 0.57183405 | LOC100128386 | 100128386 |
| ENSG000002 | 3.20676568 | -0.285674273 | 0.37720127 | -0.7573524 | 0.44883879 | 0.5719944 | NA | NA |
| ENSG000002 | 3.26703326 | 0.286228854 | 0.37798962 | 0.75723998 | 0.4489061 | 0.5720622 | RARA-AS1 | 101929693 |
| ENSG000002 | 23.569961 | -0.197870916 | 0.26132244 | -0.7571907 | 0.44893564 | 0.57208186 | NA | NA |
| ENSG000002 | 11.0111822 | -0.253982662 | 0.33548853 | -0.7570532 | 0.44901799 | 0.57216883 | LINC01446 | 401337 |
| ENSG000002 | 9.94174738 | -0.254113731 | 0.33567982 | -0.7570122 | 0.44904253 | 0.57218212 | NA | NA |
| ENSG000002 | 4.67474103 | 0.284115348 | 0.37538015 | 0.75687367 | 0.44912555 | 0.57226992 | NA | NA |
| ENSG000002 | 8.52603405 | -0.266812483 | 0.35262694 | -0.7566424 | 0.44926416 | 0.57242854 | NA | NA |
| ENSG000002 | 4.22382077 | -0.285544091 | 0.37739515 | -0.7566183 | 0.44927855 | 0.5724289 | LOC101927434 | 101927434 |
| ENSG000002 | 17.9600973 | -0.223719224 | 0.29578115 | -0.7563674 | 0.44942894 | 0.57260252 | TAS2R14 | 50840 |
| ENSG000002 | 29.6321118 | 0.191224632 | 0.25288826 | 0.75616254 | 0.44955175 | 0.57274099 | LOC100288069 | 100288069 |
| ENSG000001 | 83.9562033 | 0.13415502 | 0.17746126 | 0.75596791 | 0.44966843 | 0.57281664 | NA | NA |
| ENSG000002 | 5.46037664 | -0.282177679 | 0.37324599 | -0.7560099 | 0.44964323 | 0.57281664 | KCNQ1-AS1 | 338653 |
| ENSG000002 | 3.9642263 | -0.285943947 | 0.37825991 | -0.7559457 | 0.44968176 | 0.57281664 | NA | NA |
| ENSG000002 | 2.6962165 | -0.281892055 | 0.37287192 | -0.7560024 | 0.44964778 | 0.57281664 | NA | NA |
| ENSG000002 | 88.9300853 | 0.136680193 | 0.18080241 | 0.75596445 | 0.44967051 | 0.57281664 | USP27X | 389856 |
| ENSG00000C | 6.63978317 | -0.27894885 | 0.36905582 | -0.7558446 | 0.44974237 | 0.57283986 | P2RY10 | 27334 |
| ENSG000002 | 8.31556558 | 0.266246937 | 0.35224988 | 0.75584677 | 0.44974107 | 0.57283986 | NA | NA |
| ENSG000002 | 2.4115703 | 0.28082709 | 0.37153012 | 0.75586627 | 0.44972938 | 0.57283986 | NA | NA |
| ENSG000001 | 4.20298419 | -0.284593951 | 0.37656617 | -0.7557608 | 0.44979262 | 0.5728516 | NA | NA |
| ENSG000002 | 6.84236355 | -0.276710622 | 0.36613627 | -0.7557586 | 0.44979396 | 0.5728516 | NA | NA |
| ENSG000002 | 8.46615588 | -0.265218585 | 0.35092056 | -0.7557796 | 0.44978138 | 0.5728516 | NA | NA |

| | | | | | | | | |
|---|---|---|---|---|---|---|---|---|
| ENSG000002 | 4.23406995 | -0.285281618 | 0.37750334 | -0.7557062 | 0.44982536 | 0.5728736 | NA | NA |
| ENSG000002 | 2.73447371 | -0.281319395 | 0.37227515 | -0.755676 | 0.44984348 | 0.57287869 | NA | NA |
| ENSG000002 | 52.3593632 | 0.163435485 | 0.21632522 | 0.75550823 | 0.44994409 | 0.57298882 | NA | NA |
| ENSG000000 | 11.1471683 | -0.247023981 | 0.3269826 | -0.7554652 | 0.4499699 | 0.57300369 | SNAI2 | 6591 |
| ENSG000001 | 3.16706609 | -0.284648437 | 0.37681505 | -0.7554062 | 0.45000527 | 0.57303073 | NA | NA |
| ENSG000002 | 2.23976938 | -0.271729863 | 0.35973499 | -0.7553612 | 0.45003228 | 0.57304713 | NA | NA |
| ENSG000002 | 3.45564026 | -0.285715312 | 0.37830301 | -0.7552552 | 0.45009588 | 0.57311012 | NA | NA |
| ENSG000001 | 9.36929557 | -0.268759516 | 0.35587042 | -0.7552173 | 0.45011858 | 0.57312103 | CYP26C1 | 340665 |
| ENSG000002 | 11.3163168 | -0.251495213 | 0.33305452 | -0.7551172 | 0.45017864 | 0.5731795 | NA | NA |
| ENSG000001 | 11.4237837 | -0.245625474 | 0.32531685 | -0.7550346 | 0.45022823 | 0.57322464 | TMIGD1 | 388364 |
| ENSG000001 | 3.23642888 | -0.284135039 | 0.37634082 | -0.7549939 | 0.45025262 | 0.57323739 | C3orf22 | 152065 |
| ENSG000002 | 3.18305138 | -0.285228273 | 0.3778122 | -0.7549472 | 0.45028065 | 0.57323739 | NA | NA |
| ENSG000002 | 3.45586915 | -0.285763453 | 0.3785122 | -0.754965 | 0.45027 | 0.57323739 | NA | NA |
| ENSG000002 | 3.1732722 | -0.285152531 | 0.3777489 | -0.7548732 | 0.45032505 | 0.57325792 | NA | NA |
| ENSG000002 | 2.19620768 | -0.268937275 | 0.35626525 | -0.7548793 | 0.45032138 | 0.57325792 | NA | NA |
| ENSG000002 | 2.92028833 | -0.227863901 | 0.30190848 | -0.754745 | 0.45040202 | 0.57333791 | NA | NA |
| ENSG000002 | 3.22077249 | -0.284325619 | 0.37687277 | -0.754434 | 0.45058869 | 0.57355752 | NA | NA |
| ENSG000002 | 3.95714246 | -0.285302589 | 0.3782171 | -0.7543355 | 0.45064779 | 0.57361474 | NA | NA |
| ENSG000002 | 7.84930826 | -0.268383868 | 0.35582617 | -0.7542556 | 0.45069578 | 0.57365783 | NA | NA |
| ENSG000002 | 10.6935344 | -0.248048882 | 0.32889909 | -0.7541793 | 0.45074157 | 0.5736981 | LOC101928737 | 101928737 |
| ENSG000002 | 3.0080155 | -0.283389093 | 0.37579691 | -0.7541017 | 0.45078815 | 0.57373937 | NA | NA |
| ENSG000002 | 5.25848195 | -0.282526832 | 0.37469704 | -0.7540141 | 0.45084078 | 0.57378836 | NA | NA |
| ENSG000002 | 2.77941724 | -0.27972316 | 0.37101335 | -0.7539437 | 0.45088305 | 0.57382414 | NA | NA |
| ENSG000001 | 5.6770538 | -0.283022277 | 0.37544399 | -0.7538335 | 0.45094918 | 0.5738738 | LINC00521 | 256369 |
| ENSG000002 | 17.2800916 | -0.230320333 | 0.30553289 | -0.7538315 | 0.45095038 | 0.5738738 | NA | NA |
| ENSG000001 | 89.7262711 | 0.125291177 | 0.16622014 | 0.75376653 | 0.45098943 | 0.57388748 | RNF175 | 285533 |
| ENSG000002 | 4.0139087 | -0.284977284 | 0.37806797 | -0.7537726 | 0.45098576 | 0.57388748 | NA | NA |
| ENSG000001 | 33.0351184 | 0.184198478 | 0.24447384 | 0.75344861 | 0.45118038 | 0.57411245 | ETHE1 | 23474 |
| ENSG000002 | 2.405371 | -0.274522477 | 0.36449487 | -0.7531587 | 0.45135456 | 0.57430352 | NA | NA |

| | | | | | | | | |
|---|---|---|---|---|---|---|---|---|
| ENSG000002 | 3.45771516 | -0.285025188 | 0.37844335 | -0.7531515 | 0.45135886 | 0.57430352 | NA | NA |
| ENSG000001 | 199.332024 | 0.110400667 | 0.14659957 | 0.75307632 | 0.45140406 | 0.57432212 | KATNAL2 | 83473 |
| ENSG000001 | 71.8855056 | 0.13777734 | 0.18295751 | 0.75305649 | 0.45141597 | 0.57432212 | LRTOMT | 220074 |
| ENSG000002 | 8.31173645 | -0.272494771 | 0.36185102 | -0.7530579 | 0.45141512 | 0.57432212 | NA | NA |
| ENSG000001 | 2.94737395 | -0.281414047 | 0.37372766 | -0.7529923 | 0.45145454 | 0.57435318 | KAAG1 | 353219 |
| ENSG000002 | 8.69414192 | -0.272632425 | 0.36209152 | -0.7529379 | 0.45148724 | 0.57437675 | LINC01392 | 104355291 |
| ENSG000002 | 28.0283247 | 0.195329276 | 0.25943147 | 0.75291279 | 0.45150232 | 0.57437792 | NA | NA |
| ENSG000002 | 4.24106479 | -0.283686679 | 0.3768353 | -0.7528134 | 0.45156202 | 0.57443585 | NA | NA |
| ENSG000001 | 2.69663254 | -0.279228214 | 0.37094248 | -0.7527534 | 0.45159812 | 0.57445943 | NTN3 | 4917 |
| ENSG000002 | 2.44965772 | -0.276714236 | 0.36763679 | -0.7526837 | 0.45163998 | 0.57445943 | RNU6-550P | 106481355 |
| ENSG000002 | 3.24763947 | -0.284286484 | 0.37767877 | -0.7527203 | 0.45161799 | 0.57445943 | NA | NA |
| ENSG000002 | 3.72257503 | -0.284841439 | 0.37844397 | -0.7526648 | 0.45165139 | 0.57445943 | NA | NA |
| ENSG000002 | 8.80523364 | -0.262390158 | 0.34860208 | -0.7526925 | 0.45163474 | 0.57445943 | NA | NA |
| ENSG000001 | 235.777307 | -0.102795162 | 0.13662425 | -0.7523933 | 0.4518146 | 0.5745988 | NFKBIZ | 64332 |
| ENSG000002 | 12.6547126 | -0.240589694 | 0.31976629 | -0.7523923 | 0.45181518 | 0.5745988 | LOC101926966 | 101926966 |
| ENSG000002 | 2.20328795 | -0.272409271 | 0.36205947 | -0.7523882 | 0.45181765 | 0.5745988 | RN7SL451P | 106480377 |
| ENSG000002 | 4.47944192 | -0.283367822 | 0.37661948 | -0.7523982 | 0.45181163 | 0.5745988 | NA | NA |
| ENSG000001 | 120.030082 | 0.115180309 | 0.15309661 | 0.75233742 | 0.45184817 | 0.57461787 | ESYT3 | 83850 |
| ENSG000002 | 2.42997553 | 0.278575825 | 0.37029092 | 0.75231611 | 0.45186098 | 0.57461787 | NA | NA |
| ENSG000002 | 2.46579073 | -0.274957053 | 0.36553974 | -0.7521947 | 0.45193398 | 0.57469268 | NA | NA |
| ENSG000002 | 3.7245775 | -0.284672469 | 0.37858368 | -0.7519407 | 0.45208674 | 0.5748689 | NA | NA |
| ENSG000001 | 10.4897262 | 0.258636217 | 0.34404456 | 0.75175208 | 0.45220016 | 0.5749951 | CD8B | 926 |
| ENSG000002 | 2.42129775 | 0.279189643 | 0.37141986 | 0.75168205 | 0.45224229 | 0.57501927 | ZNF705D | 728957 |
| ENSG000002 | 10.6682136 | -0.250429481 | 0.33316265 | -0.7516733 | 0.45224753 | 0.57501927 | DNM1P35 | 100128285 |
| ENSG000002 | 19.6823389 | -0.20951449 | 0.27876008 | -0.7515943 | 0.45229507 | 0.57505265 | LINC01285 | 101928287 |
| ENSG000002 | 8.79891205 | -0.261282594 | 0.34764324 | -0.7515825 | 0.45230214 | 0.57505265 | NA | NA |
| ENSG000001 | 7.2141891 | 0.271350299 | 0.3610853 | 0.75148532 | 0.45236063 | 0.57510898 | PHOSPHO1 | 162466 |
| ENSG000002 | 2.65702599 | -0.279959537 | 0.37257178 | -0.7514244 | 0.45239727 | 0.57513753 | NA | NA |
| ENSG000001 | 10.7543624 | -0.250317116 | 0.33315273 | -0.7513585 | 0.45243696 | 0.57516995 | CLEC2A | 387836 |

| | | | | | | | | |
|---|---|---|---|---|---|---|---|---|
| ENSG000002 | 7.30825846 | -0.270783229 | 0.36041928 | -0.7513006 | 0.45247176 | 0.57519616 | NA | NA |
| ENSG000002 | 3.1538293 | -0.283423533 | 0.37730382 | -0.7511812 | 0.45254362 | 0.57526948 | NA | NA |
| ENSG000001 | 53.2062196 | 0.155364371 | 0.2068372 | 0.75114326 | 0.45256645 | 0.57528046 | ASGR1 | 432 |
| ENSG000002 | 4.30264797 | -0.284346161 | 0.37859339 | -0.7510595 | 0.45261685 | 0.57532649 | NA | NA |
| ENSG000002 | 5.39362049 | -0.281996967 | 0.37562189 | -0.7507469 | 0.452805 | 0.57554761 | NA | NA |
| ENSG000001 | 23.139822 | 0.205276121 | 0.27348406 | 0.75059628 | 0.45289566 | 0.57564481 | ACE | 1636 |
| ENSG000001 | 122.464753 | 0.120027234 | 0.15992616 | 0.75051656 | 0.45294366 | 0.57568777 | COL21A1 | 81578 |
| ENSG000002 | 12.2861002 | 0.24031105 | 0.32024196 | 0.75040463 | 0.45301104 | 0.57575537 | NA | NA |
| ENSG000002 | 19.3244995 | 0.208879865 | 0.27836536 | 0.75038024 | 0.45302573 | 0.57575599 | RBM14-RBM4 | 100526737 |
| ENSG000001 | 9.18215051 | -0.258543114 | 0.34459564 | -0.7502797 | 0.45308626 | 0.57581488 | SERPINA11 | 256394 |
| ENSG000002 | 7.01991296 | -0.271517943 | 0.36197854 | -0.750094 | 0.45319808 | 0.57593892 | LOC105376554 | 105376554 |
| ENSG000002 | 5.79692472 | -0.281763369 | 0.37573372 | -0.7499017 | 0.45331389 | 0.57606805 | LOC102724828 | 102724828 |
| ENSG000002 | 5.34187707 | 0.279405997 | 0.37260763 | 0.74986655 | 0.45333508 | 0.57607693 | NA | NA |
| ENSG000001 | 9.78342518 | -0.256849777 | 0.34253863 | -0.7498418 | 0.45335 | 0.57607783 | NAA11 | 84779 |
| ENSG000002 | 7.29474153 | 0.270816714 | 0.36118792 | 0.74979449 | 0.45337849 | 0.57609598 | NA | NA |
| ENSG000001 | 34.9997602 | 0.179408909 | 0.23941177 | 0.74937382 | 0.45363193 | 0.5763819 | NXPH2 | 11249 |
| ENSG000002 | 16.1596552 | -0.225779474 | 0.30128146 | -0.7493972 | 0.45361785 | 0.5763819 | NA | NA |
| ENSG000002 | 4.05742468 | 0.283409989 | 0.37836293 | 0.74904269 | 0.45383147 | 0.57661737 | NA | NA |
| ENSG000001 | 204.920378 | 0.140008465 | 0.18693857 | 0.7489544 | 0.45388469 | 0.57663079 | FLNC | 2318 |
| ENSG000001 | 226.586573 | 0.109037945 | 0.14558456 | 0.74896642 | 0.45387744 | 0.57663079 | SERPINF1 | 5176 |
| ENSG000002 | 12.2508649 | 0.242146827 | 0.32329567 | 0.74899495 | 0.45386025 | 0.57663079 | NA | NA |
| ENSG000002 | 18.9234729 | -0.213923412 | 0.28565653 | -0.7488833 | 0.45392753 | 0.57666715 | NA | NA |
| ENSG000002 | 7.74686162 | -0.265632478 | 0.35476284 | -0.7487607 | 0.45400144 | 0.57674297 | NA | NA |
| ENSG000002 | 7.01530622 | -0.270309555 | 0.36103585 | -0.7487056 | 0.45403468 | 0.57676713 | NA | NA |
| ENSG000002 | 13.3686522 | -0.236991375 | 0.31660457 | -0.7485406 | 0.45413415 | 0.57687541 | LOC101929759 | 101929759 |
| ENSG000002 | 3.17239009 | -0.281901767 | 0.37663481 | -0.7484751 | 0.45417365 | 0.57690753 | NA | NA |
| ENSG000002 | 13.4007142 | 0.236791932 | 0.31652124 | 0.74810755 | 0.45439529 | 0.5771359 | NA | NA |
| ENSG000002 | 68.9691155 | 0.170165212 | 0.2274685 | 0.74808254 | 0.45441037 | 0.5771359 | NA | NA |
| ENSG000002 | 2.65088046 | -0.278717141 | 0.37256551 | -0.7481024 | 0.45439841 | 0.5771359 | NA | NA |

| | | | | | | | | |
|---|---|---|---|---|---|---|---|---|
| ENSG000002 | 26.9529704 | -0.190073637 | 0.25407047 | -0.7481138 | 0.4543915 | 0.5771359 | NA | NA |
| ENSG000002 | 3.45480609 | -0.282879098 | 0.37816123 | -0.7480383 | 0.45443704 | 0.57715169 | NA | NA |
| ENSG000001 | 55.0553861 | 0.149501484 | 0.19986879 | 0.74799813 | 0.45446129 | 0.57716441 | SPOCK2 | 9806 |
| ENSG000001 | 96.9593684 | 0.151453909 | 0.20248932 | 0.74796 | 0.45448428 | 0.57717554 | SAC3D1 | 29901 |
| ENSG000002 | 7.31435659 | -0.270403175 | 0.36159057 | -0.7478159 | 0.45457121 | 0.57726786 | NA | NA |
| ENSG000001 | 24.8924885 | -0.199359119 | 0.26661984 | -0.747728 | 0.45462424 | 0.57731343 | NME5 | 8382 |
| ENSG000002 | 3.94708284 | -0.282385805 | 0.37766795 | -0.7477092 | 0.45463557 | 0.57731343 | NA | NA |
| ENSG000002 | 22.9556249 | -0.199725821 | 0.26712817 | -0.7476779 | 0.45465448 | 0.57731936 | BAZ2B | 29994 |
| ENSG000001 | 18.5900827 | -0.216961621 | 0.29020708 | -0.7476097 | 0.45469563 | 0.57735353 | NA | NA |
| ENSG000002 | 3.95644522 | -0.282419097 | 0.37777927 | -0.747577 | 0.45471534 | 0.57736048 | NA | NA |
| ENSG000002 | 3.44276297 | -0.282903623 | 0.37849801 | -0.7474375 | 0.45479949 | 0.57744925 | LINC01168 | 399829 |
| ENSG000002 | 12.4119257 | -0.240169299 | 0.32149536 | -0.7470382 | 0.45504053 | 0.5777372 | NA | NA |
| ENSG000001 | 3.68119326 | -0.282642213 | 0.37839662 | -0.746947 | 0.45509557 | 0.57778899 | NA | NA |
| ENSG000002 | 3.74683699 | -0.282547465 | 0.37831168 | -0.7468642 | 0.45514551 | 0.57783431 | NA | NA |
| ENSG000002 | 2.94592491 | -0.279775797 | 0.3746529 | -0.74676 | 0.45520844 | 0.57789611 | NA | NA |
| ENSG000002 | 11.248049 | -0.248177424 | 0.33235045 | -0.7467341 | 0.45522406 | 0.57789785 | NA | NA |
| ENSG000002 | 4.02120107 | -0.282596352 | 0.37845738 | -0.7467059 | 0.45524111 | 0.5779014 | NA | NA |
| ENSG000000 | 165.130746 | 0.106580437 | 0.14275358 | 0.74660432 | 0.45530244 | 0.57796116 | ATP8B1 | 5205 |
| ENSG000002 | 4.25944987 | -0.281740136 | 0.37738413 | -0.7465606 | 0.45532882 | 0.57797655 | VSIG8 | 391123 |
| ENSG000002 | 3.18521094 | -0.281866125 | 0.37758092 | -0.7465052 | 0.45536228 | 0.57800094 | NA | NA |
| ENSG000002 | 3.42272721 | -0.28242679 | 0.3784 | -0.746371 | 0.45544333 | 0.57808572 | NA | NA |
| ENSG000002 | 6.80943164 | -0.272749573 | 0.36544822 | -0.7463426 | 0.45546049 | 0.5780894 | NA | NA |
| ENSG000002 | 7.2543249 | 0.268064803 | 0.35935994 | 0.74595071 | 0.45569719 | 0.57837173 | NA | NA |
| ENSG000002 | 4.47997116 | -0.280234851 | 0.37569361 | -0.7459133 | 0.4557198 | 0.57838232 | NA | NA |
| ENSG000001 | 2.44512246 | -0.271411372 | 0.36391457 | -0.7458107 | 0.45578179 | 0.57844289 | VMO1 | 284013 |
| ENSG000002 | 4.72411546 | 0.279720563 | 0.37510593 | 0.74571084 | 0.45584211 | 0.57848324 | RN7SL128P | 106481826 |
| ENSG000002 | 3.20477063 | -0.281875356 | 0.37798772 | -0.7457262 | 0.45583281 | 0.57848324 | NA | NA |
| ENSG000002 | 4.58525929 | -0.282012472 | 0.37821429 | -0.745642 | 0.45588368 | 0.57851789 | NA | NA |
| ENSG000002 | 8.37926073 | 0.271895717 | 0.3648016 | 0.74532491 | 0.45607533 | 0.57874298 | MIR561 | 693146 |

| | | | | | | | | |
|---|---|---|---|---|---|---|---|---|
| ENSG000002 | 6.85356594 | 0.271510434 | 0.36430423 | 0.74528488 | 0.45609952 | 0.57875557 | NA | NA |
| ENSG000002 | 2.42125738 | -0.271948245 | 0.36497303 | -0.7451187 | 0.45619994 | 0.57886488 | SIGLEC14 | 100049587 |
| ENSG000001 | 7.75969956 | -0.267844985 | 0.3595183 | -0.7450107 | 0.45626524 | 0.57892962 | GPHA2 | 170589 |
| ENSG000001 | 3.75207014 | -0.282012501 | 0.37859272 | -0.7448968 | 0.45633412 | 0.57895082 | HIPK4 | 147746 |
| ENSG000002 | 10.0499554 | -0.253866271 | 0.34078012 | -0.7449562 | 0.45629818 | 0.57895082 | NA | NA |
| ENSG000002 | 3.24812818 | -0.280553879 | 0.3766387 | -0.7448886 | 0.45633906 | 0.57895082 | NA | NA |
| ENSG000002 | 7.82184842 | -0.26534775 | 0.35621053 | -0.7449183 | 0.45632109 | 0.57895082 | NA | NA |
| ENSG000001 | 191.200281 | 0.107695087 | 0.14459618 | 0.74479899 | 0.45639324 | 0.5789958 | GNPDA2 | 132789 |
| ENSG000002 | 4.20050455 | -0.280825726 | 0.37705724 | -0.7447827 | 0.45640307 | 0.5789958 | NA | NA |
| ENSG000002 | 2.41831809 | 0.272515999 | 0.36593305 | 0.74471547 | 0.45644375 | 0.57901118 | RNU6-7 | 101954275 |
| ENSG000002 | 68.1750049 | 0.142695389 | 0.19160511 | 0.74473687 | 0.45643081 | 0.57901118 | NA | NA |
| ENSG000001 | 46.1903954 | 0.163905591 | 0.22011461 | 0.74463749 | 0.4564909 | 0.57905288 | ERMN | 57471 |
| ENSG000001 | 2.24526999 | -0.266429728 | 0.35792656 | -0.7443698 | 0.45665278 | 0.5792401 | PRR23B | 389151 |
| ENSG00000C | 69.0446596 | 0.151692601 | 0.20384226 | 0.74416659 | 0.4567757 | 0.57937789 | IL20RA | 53832 |
| ENSG000002 | 2.72214964 | -0.276659449 | 0.37178869 | -0.7441309 | 0.45679729 | 0.57938716 | NA | NA |
| ENSG000002 | 18.2058983 | 0.21847049 | 0.29363914 | 0.74401011 | 0.45687036 | 0.57946172 | NA | NA |
| ENSG000001 | 74.8317013 | -0.140890428 | 0.18938724 | -0.7439278 | 0.45692016 | 0.57950675 | LRRC9 | 341883 |
| ENSG000002 | 4.7171472 | -0.279214927 | 0.3755264 | -0.7435294 | 0.45716121 | 0.57979434 | NA | NA |
| ENSG000001 | 9.45181342 | -0.254132614 | 0.341851 | -0.7434017 | 0.45723852 | 0.57987425 | BIK | 638 |
| ENSG000002 | 3.68723934 | -0.281390576 | 0.37854089 | -0.7433558 | 0.45726629 | 0.57989133 | NA | NA |
| ENSG000001 | 57.9255638 | -0.145348258 | 0.19555146 | -0.7432737 | 0.457316 | 0.57993086 | TMBIM4 | 51643 |
| ENSG000002 | 4.2369617 | -0.280370705 | 0.37721902 | -0.7432571 | 0.45732606 | 0.57993086 | NA | NA |
| ENSG000002 | 4.02456138 | -0.281365494 | 0.37858867 | -0.7431958 | 0.45736317 | 0.57995978 | NA | NA |
| ENSG000002 | 4.23155338 | -0.280235842 | 0.37708383 | -0.7431659 | 0.45738125 | 0.57996457 | NA | NA |
| ENSG000001 | 4.29548382 | -0.281263109 | 0.3785924 | -0.742918 | 0.45753133 | 0.58013674 | C17orf74 | 201243 |
| ENSG000002 | 4.97860142 | -0.277542591 | 0.37362663 | -0.7428341 | 0.45758215 | 0.58018303 | SLC23A3 | 151295 |
| ENSG000001 | 5.55732251 | 0.275377946 | 0.37073661 | 0.74278596 | 0.45761128 | 0.58019465 | HMX2 | 3167 |
| ENSG000002 | 5.56541898 | 0.274547911 | 0.36962626 | 0.74277167 | 0.45761993 | 0.58019465 | NA | NA |
| ENSG000002 | 3.02051828 | 0.279238554 | 0.37597298 | 0.7427091 | 0.45765782 | 0.58020641 | NA | NA |

| | | | | | | | | |
|---|---|---|---|---|---|---|---|---|
| ENSG000002 | 7.00462074 | -0.2676398 | 0.36035598 | -0.7427095 | 0.45765759 | 0.58020641 | NA | NA |
| ENSG000002 | 4.45525154 | -0.279676962 | 0.37658981 | -0.7426567 | 0.45768953 | 0.58022847 | NA | NA |
| ENSG000001 | 4.24501885 | -0.280181224 | 0.37730338 | -0.7425887 | 0.45773074 | 0.58026258 | OR6K6 | 128371 |
| ENSG000002 | 3.46091274 | -0.280788091 | 0.37814275 | -0.7425452 | 0.45775706 | 0.5802778 | NA | NA |
| ENSG000002 | 53.0115382 | 0.15158322 | 0.20421835 | 0.74226053 | 0.4579295 | 0.58047825 | NA | NA |
| ENSG000002 | 9.07213302 | -0.261899382 | 0.35285157 | -0.7422367 | 0.45794395 | 0.58047842 | NA | NA |
| ENSG000002 | 3.49013482 | -0.28081029 | 0.37834685 | -0.7422033 | 0.45796415 | 0.58048048 | NA | NA |
| ENSG000002 | 11.4337317 | -0.242910942 | 0.32729087 | -0.7421867 | 0.4579742 | 0.58048048 | NA | NA |
| ENSG000002 | 3.42715092 | -0.280898732 | 0.37850776 | -0.7421215 | 0.45801375 | 0.58051246 | NA | NA |
| ENSG000002 | 4.0860454 | 0.280566734 | 0.37814145 | 0.74196239 | 0.45811012 | 0.58061646 | GALNT4 | 8693 |
| ENSG000002 | 3.24662481 | 0.280472704 | 0.3780314 | 0.74192965 | 0.45812996 | 0.58062346 | NA | NA |
| ENSG000002 | 7.08479942 | -0.27115499 | 0.36551313 | -0.7418475 | 0.45817976 | 0.5806661 | LOC101928590 | 101928590 |
| ENSG000002 | 5.98553455 | -0.275296937 | 0.37110673 | -0.7418269 | 0.45819225 | 0.5806661 | LOC105369891 | 105369891 |
| ENSG000001 | 18.4024956 | -0.21409097 | 0.28865404 | -0.7416871 | 0.45827695 | 0.58075529 | CORO6 | 84940 |
| ENSG000001 | 33.16831 | 0.178778208 | 0.24109455 | 0.74152736 | 0.45837375 | 0.58084166 | POLR2J3 | 548644 |
| ENSG000002 | 3.44906549 | -0.280660675 | 0.37848125 | -0.7415445 | 0.45836338 | 0.58084166 | LINC00898 | 400932 |
| ENSG000001 | 8.66578907 | -0.257971793 | 0.34792957 | -0.7414483 | 0.45842166 | 0.58088422 | REP15 | 387849 |
| ENSG000002 | 3.01217807 | -0.277492811 | 0.37429955 | -0.7413656 | 0.4584718 | 0.58091145 | RN7SL681P | 106479471 |
| ENSG000002 | 3.48038711 | -0.280460931 | 0.37829656 | -0.7413785 | 0.45846398 | 0.58091145 | NA | NA |
| ENSG000002 | 5.28052572 | -0.278737396 | 0.37602808 | -0.7412675 | 0.45853126 | 0.58096863 | NA | NA |
| ENSG000002 | 2.90049191 | -0.277696622 | 0.37467515 | -0.7411664 | 0.45859257 | 0.58102815 | NA | NA |
| ENSG000002 | 3.4960118 | -0.280288947 | 0.37826695 | -0.7409819 | 0.45870444 | 0.58115174 | RNU6-1242P | 106480108 |
| ENSG000002 | 3.96620513 | -0.28007487 | 0.3780476 | -0.7408455 | 0.45878711 | 0.58123119 | LOC101928739 | 101928739 |
| ENSG000002 | 6.9964845 | -0.267631496 | 0.36125842 | -0.7408312 | 0.45879581 | 0.58123119 | NA | NA |
| ENSG000002 | 3.92531207 | -0.279901638 | 0.37784203 | -0.7407901 | 0.45882071 | 0.58124457 | NA | NA |
| ENSG000001 | 4.45433065 | -0.27915705 | 0.37690589 | -0.7406545 | 0.45890295 | 0.58133059 | STBD1 | 8987 |
| ENSG000002 | 3.22441389 | -0.27893947 | 0.37664515 | -0.7405896 | 0.45894234 | 0.58136233 | CYP21A2 | 1589 |
| ENSG000002 | 4.68595729 | -0.277219977 | 0.37433607 | -0.7405644 | 0.45895758 | 0.58136347 | NA | NA |
| ENSG000001 | 6.42858514 | -0.275404588 | 0.37193111 | -0.740472 | 0.45901362 | 0.58141629 | NA | NA |

| | | | | | | | | |
|---|---|---|---|---|---|---|---|---|
| ENSG000001 | 212.299873 | -0.109751656 | 0.1482467 | -0.7403312 | 0.45909907 | 0.58150637 | DNAH11 | 8701 |
| ENSG000001 | 6.80986883 | -0.270114361 | 0.36488063 | -0.7402814 | 0.45912924 | 0.58152642 | SSX1 | 6756 |
| ENSG000002 | 3.70453488 | -0.280146916 | 0.37852986 | -0.740092 | 0.45924418 | 0.58165384 | NA | NA |
| ENSG000001 | 435.51398 | -0.094169017 | 0.12725446 | -0.7400056 | 0.45929657 | 0.58168993 | CADM1 | 23705 |
| ENSG000002 | 3.92243546 | -0.279567071 | 0.3777945 | -0.7399977 | 0.45930137 | 0.58168993 | NA | NA |
| ENSG000002 | 3.18451917 | -0.279202718 | 0.37735197 | -0.7399 | 0.45936068 | 0.58172774 | NA | NA |
| ENSG000002 | 3.69602922 | -0.280039998 | 0.37848299 | -0.7399012 | 0.45935997 | 0.58172774 | NA | NA |
| ENSG000002 | 4.02361599 | -0.280009427 | 0.37845371 | -0.7398776 | 0.45937426 | 0.58172774 | NA | NA |
| ENSG000002 | 22.4514084 | -0.207725493 | 0.28078611 | -0.7397997 | 0.45942152 | 0.58176942 | NA | NA |
| ENSG000002 | 3.18937407 | -0.279232546 | 0.37760793 | -0.7394774 | 0.45961713 | 0.58198077 | NA | NA |
| ENSG000002 | 4.77730601 | -0.278399727 | 0.3764751 | -0.7394904 | 0.45960926 | 0.58198077 | NA | NA |
| ENSG000001 | 50.2397171 | -0.153337689 | 0.20750203 | -0.7389696 | 0.45992546 | 0.58235301 | LINC00476 | 100128782 |
| ENSG000002 | 4.47440574 | -0.278766208 | 0.37724869 | -0.7389455 | 0.45994012 | 0.58235338 | NA | NA |
| ENSG000001 | 3.94098829 | -0.279598783 | 0.37851152 | -0.7386797 | 0.4601015 | 0.58253953 | ENPP7 | 339221 |
| ENSG000001 | 131.378005 | 0.125696454 | 0.17018559 | 0.73858461 | 0.46015927 | 0.58255811 | SCN5A | 6331 |
| ENSG000002 | 4.48569268 | -0.277660361 | 0.37593046 | -0.738595 | 0.46015296 | 0.58255811 | NA | NA |
| ENSG000002 | 7.74995447 | -0.261643307 | 0.35423985 | -0.738605 | 0.46014691 | 0.58255811 | LOC100506557 | 100506557 |
| ENSG000001 | 67.9374394 | -0.141201725 | 0.19122024 | -0.7384246 | 0.46025648 | 0.58263996 | ABCB9 | 23457 |
| ENSG000002 | 10.0624629 | -0.250735317 | 0.33956239 | -0.7384072 | 0.46026704 | 0.58263996 | NA | NA |
| ENSG000002 | 4.24454277 | -0.278540663 | 0.37721283 | -0.7384178 | 0.46026061 | 0.58263996 | NA | NA |
| ENSG000002 | 7.18386112 | -0.271241171 | 0.36737094 | -0.7383305 | 0.46031363 | 0.58266257 | NA | NA |
| ENSG000002 | 30.9879742 | -0.17918609 | 0.24268835 | -0.7383382 | 0.46030894 | 0.58266257 | PCDHGA8 | 9708 |
| ENSG000002 | 6.44758367 | 0.268028951 | 0.36305513 | 0.73825964 | 0.46035669 | 0.58269888 | SNHG21 | 100505616 |
| ENSG000002 | 7.04682405 | -0.267637387 | 0.36257447 | -0.7381584 | 0.46041821 | 0.582755 | LOC101928107 | 101928107 |
| ENSG000002 | 3.45637245 | -0.279283519 | 0.37836151 | -0.7381394 | 0.46042977 | 0.582755 | NA | NA |
| ENSG000001 | 4.21243109 | -0.278288209 | 0.37706953 | -0.7380289 | 0.46049689 | 0.58282177 | TAAR3 | 9288 |
| ENSG000002 | 10.1763413 | -0.250763411 | 0.3398084 | -0.7379553 | 0.46054161 | 0.58286017 | GTF2IP14 | 106481735 |
| ENSG000002 | 9.28115028 | 0.251534858 | 0.34088411 | 0.73788965 | 0.46058151 | 0.58289247 | PSMB10 | 5699 |
| ENSG000002 | 4.73157353 | -0.276880844 | 0.37529874 | -0.7377612 | 0.46065959 | 0.58295261 | NA | NA |

| | | | | | | | | | |
|---|---|---|---|---|---|---|---|---|---|
| ENSG000002 | 3.45261984 | -0.279078213 | 0.37828777 | -0.7377405 | 0.46067215 | 0.58295261 | NA | NA | |
| ENSG000002 | 3.89088779 | 0.279281461 | 0.37855301 | 0.7377605 | 0.46066 | 0.58295261 | NA | NA | |
| ENSG000002 | 3.20276652 | -0.277943421 | 0.37680886 | -0.7376244 | 0.4607427 | 0.5830055 | NA | NA | |
| ENSG000002 | 62.1354756 | 0.157886221 | 0.21404606 | 0.7376273 | 0.46074096 | 0.5830055 | NA | NA | |
| ENSG000001 | 19.2006374 | -0.210935062 | 0.28600057 | -0.7375337 | 0.46079786 | 0.5830571 | TAPBPL | | 55080 |
| ENSG000002 | 3.24720547 | 0.278893969 | 0.37819959 | 0.73742536 | 0.46086372 | 0.58312224 | NA | NA | |
| ENSG000001 | 93.4281619 | 0.141109249 | 0.19138096 | 0.73732126 | 0.46092701 | 0.58315075 | FAM189B | | 10712 |
| ENSG000001 | 4.66355345 | -0.276104481 | 0.37448372 | -0.7372937 | 0.46094378 | 0.58315075 | GP1BA | | 2811 |
| ENSG000002 | 3.985275 | -0.278868966 | 0.37822387 | -0.7373119 | 0.46093269 | 0.58315075 | NA | NA | |
| ENSG000002 | 8.5308041 | -0.259042069 | 0.35133146 | -0.7373153 | 0.46093061 | 0.58315075 | NA | NA | |
| ENSG000002 | 4.19039974 | -0.278021599 | 0.37724013 | -0.7369884 | 0.4611294 | 0.58336739 | NA | NA | |
| ENSG000002 | 8.58888822 | -0.259384791 | 0.35198424 | -0.7369216 | 0.46117002 | 0.58338238 | NA | NA | |
| ENSG000002 | 4.08705529 | 0.278382564 | 0.37775417 | 0.73694107 | 0.46115819 | 0.58338238 | NA | NA | |
| ENSG000002 | 4.4662768 | -0.277547222 | 0.37668349 | -0.7368181 | 0.46123296 | 0.5834438 | NA | NA | |
| ENSG000002 | 44.6799647 | -0.162142269 | 0.22006949 | -0.7367776 | 0.46125761 | 0.58345678 | XKR5 | | 389610 |
| ENSG000002 | 3.76524188 | -0.278754392 | 0.37838701 | -0.7366912 | 0.46131014 | 0.58350502 | POTEM | | 641455 |
| ENSG000002 | 3.92815623 | -0.278665432 | 0.37829279 | -0.7366395 | 0.46134158 | 0.58352658 | NA | NA | |
| ENSG00000C | 12.4817782 | 0.236935059 | 0.32171449 | 0.73647618 | 0.46144096 | 0.58359767 | HCN2 | | 610 |
| ENSG000002 | 4.25230599 | -0.278336889 | 0.37791867 | -0.7364994 | 0.46142681 | 0.58359767 | NA | NA | |
| ENSG000002 | 4.18748314 | -0.278084471 | 0.37757314 | -0.7365049 | 0.46142348 | 0.58359767 | NA | NA | |
| ENSG000002 | 8.01317691 | -0.273316852 | 0.37113293 | -0.7364392 | 0.46146344 | 0.58360791 | NA | NA | |
| ENSG000002 | 4.68796648 | 0.278159771 | 0.377743 | 0.73637307 | 0.46150369 | 0.58364061 | NA | NA | |
| ENSG000001 | 17.4596461 | -0.21381744 | 0.29040858 | -0.7362642 | 0.46156993 | 0.58369351 | GNB3 | | 2784 |
| ENSG000002 | 15.191625 | 0.233939691 | 0.31774189 | 0.736257 | 0.46157431 | 0.58369351 | NA | NA | |
| ENSG000002 | 3.43107272 | -0.278624055 | 0.3785134 | -0.7361009 | 0.46166929 | 0.58379541 | NA | NA | |
| ENSG000001 | 28.94667 | 0.18469175 | 0.25091901 | 0.7360612 | 0.46169345 | 0.58380776 | NKX3-1 | | 4824 |
| ENSG000002 | 16.2477051 | -0.222846477 | 0.30278303 | -0.7359939 | 0.46173439 | 0.58384132 | NA | NA | |
| ENSG000002 | 3.47822037 | -0.277935193 | 0.37765653 | -0.735947 | 0.46176295 | 0.58385923 | NA | NA | |
| ENSG000002 | 3.6801753 | -0.278598247 | 0.37858781 | -0.7358881 | 0.46179882 | 0.58388638 | NA | NA | |

| | | | | | | | | |
|---|---|---|---|---|---|---|---|---|
| ENSG000002 | 5.47947088 | -0.272097881 | 0.36979328 | -0.7358108 | 0.46184586 | 0.58392765 | NA | NA |
| ENSG000001 | 5.88592427 | -0.274809132 | 0.37360278 | -0.735565 | 0.46199546 | 0.58409858 | TEX29 | 121793 |
| ENSG000002 | 11.6321134 | -0.238493974 | 0.32431351 | -0.7353809 | 0.46210753 | 0.58422206 | NA | NA |
| ENSG000002 | 3.28510981 | 0.278177457 | 0.37829983 | 0.73533594 | 0.46213493 | 0.58423848 | SNORD88A | 692202 |
| ENSG000002 | 8.6681772 | -0.264824263 | 0.3602527 | -0.7351069 | 0.46227436 | 0.58439654 | NA | NA |
| ENSG000002 | 3.44621317 | -0.277884795 | 0.37811713 | -0.7349172 | 0.4623899 | 0.58452438 | NA | NA |
| ENSG000002 | 2.21523938 | 0.265564606 | 0.3613973 | 0.73482731 | 0.46244467 | 0.58457539 | NA | NA |
| ENSG000002 | 3.44269184 | -0.277968908 | 0.3783133 | -0.7347585 | 0.46248659 | 0.58461016 | NA | NA |
| ENSG000002 | 5.18537629 | -0.273750151 | 0.3726115 | -0.7346798 | 0.46253451 | 0.5846525 | NA | NA |
| ENSG000002 | 5.24030338 | -0.273698938 | 0.37262156 | -0.7345226 | 0.46263032 | 0.58475538 | NA | NA |
| ENSG000002 | 4.13858872 | 0.27699427 | 0.37714817 | 0.73444415 | 0.46267809 | 0.58479753 | NA | NA |
| ENSG000002 | 5.62033508 | 0.274065722 | 0.37318906 | 0.73438842 | 0.46271204 | 0.58482222 | NA | NA |
| ENSG000002 | 3.93544391 | -0.277728405 | 0.37825047 | -0.7342447 | 0.46279961 | 0.58491467 | NA | NA |
| ENSG000002 | 64.6120156 | 0.147862745 | 0.20139353 | 0.73419808 | 0.46282802 | 0.58493234 | NA | NA |
| ENSG000001 | 9.45261613 | -0.252344658 | 0.34371606 | -0.734166 | 0.46284755 | 0.58493879 | PGPEP1L | 145814 |
| ENSG000001 | 26.5432592 | 0.200467309 | 0.27322738 | 0.73370138 | 0.46313076 | 0.58527846 | C20orf196 | 149840 |
| ENSG000002 | 4.10091251 | 0.277046746 | 0.37762902 | 0.73364793 | 0.46316334 | 0.5852907 | NA | NA |
| ENSG000002 | 21.7553912 | -0.200590882 | 0.27341999 | -0.7336365 | 0.46317031 | 0.5852907 | RBFADN | 100506070 |
| ENSG000002 | 18.5202827 | 0.213594093 | 0.29115305 | 0.73361447 | 0.46318374 | 0.5852907 | NA | NA |
| ENSG000002 | 10.624749 | -0.245431211 | 0.33459142 | -0.7335251 | 0.46323821 | 0.58534129 | NA | NA |
| ENSG000002 | 4.43019976 | -0.276083368 | 0.37645237 | -0.7333819 | 0.46332551 | 0.58543336 | NA | NA |
| ENSG00000C | 99.0594313 | 0.127048372 | 0.17325554 | 0.73330049 | 0.46337518 | 0.58547787 | ALG9 | 79796 |
| ENSG000002 | 3.18061141 | -0.276408409 | 0.3770036 | -0.7331718 | 0.46345366 | 0.58555879 | NA | NA |
| ENSG000002 | 5.48536792 | -0.272516945 | 0.37175765 | -0.73305 | 0.46352794 | 0.58560073 | RNU1-138P | 106480195 |
| ENSG000002 | 13.1279308 | -0.23619736 | 0.32220413 | -0.7330675 | 0.46351729 | 0.58560073 | HIF1A-AS1 | 100750246 |
| ENSG000002 | 9.2072501 | -0.253880924 | 0.34633681 | -0.7330463 | 0.46353018 | 0.58560073 | NA | NA |
| ENSG000002 | 4.04915074 | 0.276959493 | 0.37793001 | 0.73283277 | 0.46366043 | 0.58574704 | STEAP2-AS1 | 100874100 |
| ENSG000002 | 4.45526374 | -0.276459204 | 0.37727344 | -0.732782 | 0.46369137 | 0.58576788 | NA | NA |
| ENSG000001 | 49.9716754 | -0.159561349 | 0.21775996 | -0.7327396 | 0.46371725 | 0.58578232 | FSIP1 | 161835 |

| | | | | | | | | |
|---|---|---|---|---|---|---|---|---|
| ENSG000001 | 94.7761318 | -0.137332172 | 0.18743915 | -0.732676 | 0.46375606 | 0.58579672 | NOVA2 | 4858 |
| ENSG000002 | 3.68185879 | -0.277365606 | 0.37856641 | -0.7326736 | 0.46375755 | 0.58579672 | NA | NA |
| ENSG000002 | 8.73684938 | -0.255834979 | 0.34921076 | -0.7326091 | 0.46379689 | 0.58582817 | NA | NA |
| ENSG000002 | 5.2050399 | -0.272646356 | 0.37218331 | -0.7325593 | 0.46382724 | 0.58584826 | NA | NA |
| ENSG000002 | 2.66136663 | 0.274417985 | 0.37461884 | 0.73252585 | 0.46384767 | 0.58585582 | NA | NA |
| ENSG000001 | 5.69438743 | -0.269918166 | 0.36850948 | -0.7324592 | 0.46388833 | 0.58587068 | JCHAIN | 3512 |
| ENSG000002 | 79.7811587 | -0.129955181 | 0.17742149 | -0.7324658 | 0.46388428 | 0.58587068 | MLLT6 | 4302 |
| ENSG000002 | 10.9110675 | -0.243844242 | 0.33298951 | -0.7322881 | 0.46399276 | 0.58598432 | NA | NA |
| ENSG000002 | 18.9181099 | 0.208047825 | 0.28412321 | 0.73224508 | 0.46401899 | 0.58599919 | UBE2QL1 | 134111 |
| ENSG000001 | 21.4644468 | -0.200883194 | 0.27441139 | -0.7320512 | 0.46413731 | 0.58613036 | IDNK | 414328 |
| ENSG000002 | 3.22344909 | -0.2762092 | 0.37734717 | -0.7319763 | 0.46418301 | 0.58615157 | NA | NA |
| ENSG000002 | 4.96826881 | -0.272995628 | 0.37295627 | -0.7319776 | 0.46418225 | 0.58615157 | NA | NA |
| ENSG000002 | 28.0186187 | 0.180801995 | 0.24704355 | 0.73186286 | 0.46425228 | 0.58622078 | NA | NA |
| ENSG000002 | 4.22906602 | -0.276601729 | 0.37807033 | -0.7316145 | 0.46440391 | 0.58638709 | RBMS3-AS1 | 100873977 |
| ENSG000002 | 5.6224148 | 0.272451714 | 0.37240543 | 0.73159974 | 0.46441291 | 0.58638709 | CTAGE15 | 441294 |
| ENSG000002 | 3.24876864 | -0.27578923 | 0.37704232 | -0.7314543 | 0.46450171 | 0.58648095 | LINC00901 | 100506724 |
| ENSG000002 | 3.91418222 | -0.276522985 | 0.37812252 | -0.7313053 | 0.46459272 | 0.5865776 | NA | NA |
| ENSG000002 | 37.931761 | -0.171451209 | 0.23445963 | -0.7312611 | 0.46461967 | 0.58659337 | NA | NA |
| ENSG000002 | 2.66332907 | -0.271640902 | 0.37151035 | -0.7311799 | 0.46466927 | 0.58663773 | NA | NA |
| ENSG000002 | 21.3778392 | 0.209876191 | 0.28707475 | 0.73108551 | 0.46472692 | 0.58666996 | RN7SL396P | 106479368 |
| ENSG000002 | 16.079827 | 0.223350197 | 0.3055023 | 0.73109169 | 0.46472315 | 0.58666996 | NA | NA |
| ENSG000002 | 6.3119028 | -0.271407395 | 0.3712483 | -0.731067 | 0.46473821 | 0.58666996 | NA | NA |
| ENSG000002 | 4.02617906 | -0.276593377 | 0.37835734 | -0.7310374 | 0.4647563 | 0.58667453 | NA | NA |
| ENSG000001 | 6.95417664 | -0.265483231 | 0.36327229 | -0.7308106 | 0.46489487 | 0.58678589 | TAS1R1 | 80835 |
| ENSG000002 | 4.44127012 | -0.274931421 | 0.37620701 | -0.7307982 | 0.4649024 | 0.58678589 | NA | NA |
| ENSG000002 | 5.54589972 | 0.271575795 | 0.37159796 | 0.7308323 | 0.46488159 | 0.58678589 | NA | NA |
| ENSG000002 | 23.8738722 | 0.198166898 | 0.27115754 | 0.73081832 | 0.46489013 | 0.58678589 | NA | NA |
| ENSG000002 | 9.16510457 | -0.251666081 | 0.34438971 | -0.7307596 | 0.46492602 | 0.58679744 | NA | NA |
| ENSG000001 | 7.72113252 | 0.258306563 | 0.35352688 | 0.73065608 | 0.46498925 | 0.58684861 | JOSD2 | 126119 |

| | | | | | | | | |
|---|---|---|---|---|---|---|---|---|
| ENSG000002 | 5.46532061 | -0.269568923 | 0.36894609 | -0.7306458 | 0.46499551 | 0.58684861 | NA | NA |
| ENSG000001 | 317.588846 | 0.116632719 | 0.15963652 | 0.73061425 | 0.4650148 | 0.5868547 | OBSCN | 84033 |
| ENSG000001 | 298.240556 | 0.093568375 | 0.12812972 | 0.73026286 | 0.46522953 | 0.58709449 | VAMP4 | 8674 |
| ENSG000001 | 62.1198803 | -0.151144485 | 0.20697468 | -0.7302559 | 0.46523376 | 0.58709449 | FRMD3 | 257019 |
| ENSG000001 | 2.62534111 | 0.272528842 | 0.37325594 | 0.73013934 | 0.46530502 | 0.58714786 | VTRNA1-1 | 56664 |
| ENSG000002 | 3.70531893 | -0.276414259 | 0.37856998 | -0.7301537 | 0.46529626 | 0.58714786 | NA | NA |
| ENSG000002 | 6.88593475 | 0.261669279 | 0.35843376 | 0.73003525 | 0.46536864 | 0.58720987 | NA | NA |
| ENSG000002 | 6.47753807 | -0.267516688 | 0.3664915 | -0.7299397 | 0.46542706 | 0.58722472 | NA | NA |
| ENSG000002 | 21.5194384 | 0.201958852 | 0.2766776 | 0.7299429 | 0.46542509 | 0.58722472 | NA | NA |
| ENSG000002 | 4.66904664 | -0.273480959 | 0.37464887 | -0.7299661 | 0.46541093 | 0.58722472 | NA | NA |
| ENSG000002 | 4.46648642 | -0.274382215 | 0.3759066 | -0.7299212 | 0.46543833 | 0.58722472 | NA | NA |
| ENSG000002 | 7.67151091 | 0.26031566 | 0.35670513 | 0.72977828 | 0.46552572 | 0.58729843 | NA | NA |
| ENSG000002 | 3.93914968 | -0.276080182 | 0.37829715 | -0.7297972 | 0.46551418 | 0.58729843 | LINC00881 | 100498859 |
| ENSG000002 | 4.42122968 | -0.274132069 | 0.37568635 | -0.7296833 | 0.46558377 | 0.58731685 | RNU6-341P | 106481287 |
| ENSG000002 | 159.636904 | 0.106020023 | 0.14529033 | 0.72971148 | 0.46556656 | 0.58731685 | NA | NA |
| ENSG000002 | 28.3598206 | 0.181468555 | 0.24869312 | 0.72968867 | 0.46558051 | 0.58731685 | NA | NA |
| ENSG000001 | 74.1341482 | -0.13689005 | 0.18763057 | -0.7295722 | 0.46565172 | 0.58736603 | BEND7 | 222389 |
| ENSG000002 | 12.9849854 | 0.230742864 | 0.31626693 | 0.72958265 | 0.46564533 | 0.58736603 | NA | NA |
| ENSG000002 | 8.99227465 | -0.254388394 | 0.34870797 | -0.729517 | 0.46568546 | 0.58739032 | LOC102724604 | 102724604 |
| ENSG000002 | 4.44120596 | -0.274074266 | 0.37572189 | -0.7294605 | 0.46572004 | 0.58741566 | NA | NA |
| ENSG000002 | 3.223872 | -0.274122977 | 0.37581581 | -0.7294078 | 0.46575226 | 0.58743803 | LOC101927934 | 101927934 |
| ENSG000002 | 5.46486305 | -0.270671114 | 0.37115369 | -0.7292696 | 0.46583674 | 0.5875263 | NA | NA |
| ENSG000001 | 15.4950562 | -0.216704876 | 0.29717497 | -0.7292164 | 0.46586928 | 0.58752987 | CCDC87 | 55231 |
| ENSG000002 | 4.7068793 | -0.273946103 | 0.37568346 | -0.7291939 | 0.46588304 | 0.58752987 | NA | NA |
| ENSG000002 | 5.03164119 | -0.27402588 | 0.37577921 | -0.7292204 | 0.46586683 | 0.58752987 | LOC101927620 | 101927620 |
| ENSG000002 | 3.75231863 | -0.276056624 | 0.3785928 | -0.729165 | 0.46590073 | 0.58753392 | NA | NA |
| ENSG000002 | 5.26545985 | -0.271887305 | 0.37293055 | -0.7290561 | 0.46596733 | 0.58759962 | NA | NA |
| ENSG000002 | 15.5478825 | 0.221820344 | 0.30437614 | 0.72877047 | 0.46614208 | 0.58780171 | NA | NA |
| ENSG000002 | 3.71600496 | -0.275758398 | 0.37841512 | -0.7287193 | 0.46617339 | 0.58782291 | NA | NA |

| | | | | | | | | |
|---|---|---|---|---|---|---|---|---|
| ENSG000002 | 5.79584601 | -0.271638555 | 0.372784 | -0.7286755 | 0.46620021 | 0.58783845 | NA | NA |
| ENSG000001 | 801.942437 | 0.09648594 | 0.13242427 | 0.72861222 | 0.4662389 | 0.58786896 | INTS7 | 25896 |
| ENSG000001 | 44.8447675 | 0.154265132 | 0.21173659 | 0.72857096 | 0.46626415 | 0.58788252 | FOXJ1 | 2302 |
| ENSG000002 | 3.46356765 | -0.275152479 | 0.37767317 | -0.7285465 | 0.4662791 | 0.58788308 | NA | NA |
| ENSG000002 | 5.55224328 | -0.272417663 | 0.37394721 | -0.7284923 | 0.46631228 | 0.58790664 | NA | NA |
| ENSG000002 | 4.45100499 | -0.274386867 | 0.37673712 | -0.7283245 | 0.46641498 | 0.58799956 | NA | NA |
| ENSG000002 | 4.9161021 | -0.271439224 | 0.37268986 | -0.7283247 | 0.46641485 | 0.58799956 | LOC100129620 | 100129620 |
| ENSG000001 | 12.8640595 | -0.245485367 | 0.33708919 | -0.7282505 | 0.46646027 | 0.58803838 | AKAP4 | 8852 |
| ENSG000002 | 4.15756382 | -0.274800771 | 0.37736434 | -0.7282108 | 0.46648459 | 0.58805075 | NA | NA |
| ENSG000002 | 3.98217528 | -0.275502009 | 0.37834242 | -0.7281816 | 0.4665024 | 0.58805492 | NA | NA |
| ENSG000002 | 3.9415902 | -0.275190439 | 0.37799239 | -0.7280317 | 0.46659419 | 0.58815234 | NA | NA |
| ENSG000002 | 7.10377014 | -0.26709132 | 0.36693316 | -0.7279018 | 0.46667368 | 0.58823426 | NA | NA |
| ENSG000001 | 130.273002 | 0.116532154 | 0.16013812 | 0.72769776 | 0.46679862 | 0.58835517 | EXO5 | 64789 |
| ENSG000002 | 2.95515842 | -0.272591909 | 0.3745888 | -0.7277097 | 0.4667913 | 0.58835517 | NA | NA |
| ENSG000002 | 5.00289801 | 0.272144644 | 0.3740472 | 0.72756766 | 0.46687828 | 0.58843728 | NA | NA |
| ENSG000002 | 5.27268278 | 0.273804443 | 0.37649576 | 0.72724443 | 0.46707623 | 0.58866848 | NA | NA |
| ENSG000001 | 6.66250185 | -0.266782933 | 0.36687194 | -0.7271827 | 0.46711403 | 0.58869782 | NA | NA |
| ENSG000001 | 7.69553124 | -0.258215568 | 0.35511544 | -0.7271313 | 0.4671455 | 0.58870651 | FAM163A | 148753 |
| ENSG000002 | 2.62431553 | 0.27261971 | 0.37492875 | 0.72712406 | 0.46714996 | 0.58870651 | NA | NA |
| ENSG000002 | 3.26925678 | 0.274774912 | 0.37794731 | 0.7270191 | 0.46721426 | 0.58873808 | MTRNR2L1 | 100462977 |
| ENSG000002 | 5.72395476 | -0.268054675 | 0.36870733 | -0.7270121 | 0.46721857 | 0.58873808 | NA | NA |
| ENSG000002 | 22.9114792 | -0.207321661 | 0.28515095 | -0.7270593 | 0.46718961 | 0.58873808 | NA | NA |
| ENSG000002 | 5.46227928 | -0.269942118 | 0.37131676 | -0.7269861 | 0.46723448 | 0.58873984 | NA | NA |
| ENSG000002 | 5.21147466 | 0.274267535 | 0.37728805 | 0.72694467 | 0.46725985 | 0.58875351 | NA | NA |
| ENSG000002 | 2.91431798 | -0.272734829 | 0.37523403 | -0.7268393 | 0.46732442 | 0.58879828 | NA | NA |
| ENSG000002 | 6.80151177 | -0.265873081 | 0.36578218 | -0.7268618 | 0.46731064 | 0.58879828 | PHEX-AS1 | 100873942 |
| ENSG000002 | 6.94370792 | -0.271243712 | 0.37335362 | -0.7265062 | 0.46752852 | 0.58903713 | NA | NA |
| ENSG000002 | 35.9321069 | -0.171904935 | 0.23667761 | -0.7263253 | 0.46763937 | 0.58915849 | NA | NA |
| ENSG000001 | 231.443356 | 0.102313855 | 0.14087519 | 0.72627307 | 0.46767139 | 0.58917389 | TAF1A | 9015 |

| | | | | | | | | |
|---|---|---|---|---|---|---|---|---|
| ENSG000002 | 5.01212247 | -0.271978423 | 0.37449287 | -0.7262579 | 0.46768065 | 0.58917389 | NA | NA |
| ENSG000002 | 4.144535 | -0.274367166 | 0.37779965 | -0.726224 | 0.46770148 | 0.58918182 | TRIM80P | 106480328 |
| ENSG00000C | 1540.56653 | 0.086137224 | 0.11862286 | 0.72614352 | 0.46775079 | 0.58920733 | ADSS | 159 |
| ENSG000002 | 4.45384989 | -0.27293568 | 0.37586737 | -0.7261489 | 0.46774749 | 0.58920733 | NA | NA |
| ENSG000002 | 4.98035659 | -0.271751761 | 0.37430953 | -0.7260081 | 0.46783378 | 0.58929356 | NA | NA |
| ENSG000002 | 4.16007592 | -0.27403005 | 0.37756268 | -0.7257869 | 0.46796941 | 0.58944609 | NA | NA |
| ENSG000002 | 4.69009947 | -0.272635573 | 0.37565536 | -0.7257598 | 0.46798601 | 0.5894487 | NA | NA |
| ENSG000002 | 3.94617678 | -0.274558732 | 0.37838066 | -0.7256151 | 0.46807474 | 0.58954215 | LOC101927419 | 101927419 |
| ENSG000002 | 3.44593352 | -0.274352628 | 0.37811862 | -0.7255729 | 0.46810063 | 0.58955644 | NA | NA |
| ENSG000002 | 4.6958293 | -0.271449359 | 0.37422813 | -0.725358 | 0.46823244 | 0.58970414 | NA | NA |
| ENSG000002 | 4.75932621 | -0.272693022 | 0.37599596 | -0.7252552 | 0.46829548 | 0.58976521 | NA | NA |
| ENSG000001 | 9.28198445 | 0.248799861 | 0.34313933 | 0.72506949 | 0.4684094 | 0.58989036 | CLEC4G | 339390 |
| ENSG000002 | 4.16370936 | -0.27406637 | 0.37801373 | -0.725017 | 0.46844162 | 0.58991261 | NA | NA |
| ENSG000002 | 8.37389418 | 0.253538812 | 0.34980777 | 0.72479468 | 0.468578 | 0.59006604 | NA | NA |
| ENSG000001 | 11.84939 | 0.235122913 | 0.32442529 | 0.72473669 | 0.46861358 | 0.59007143 | CDKN2D | 1032 |
| ENSG000001 | 18.9673366 | 0.212637118 | 0.29341689 | 0.72469284 | 0.46864049 | 0.59007143 | DCAF12L2 | 340578 |
| ENSG000002 | 5.48945892 | -0.268636881 | 0.37065902 | -0.7247547 | 0.46860251 | 0.59007143 | NA | NA |
| ENSG000002 | 4.67405798 | -0.271875413 | 0.37515074 | -0.7247098 | 0.4686301 | 0.59007143 | NA | NA |
| ENSG000002 | 4.51026709 | -0.274085098 | 0.37824505 | -0.7246231 | 0.46868329 | 0.59008999 | CT83 | 203413 |
| ENSG000002 | 4.43495629 | -0.272358452 | 0.37586312 | -0.7246214 | 0.46868434 | 0.59008999 | NA | NA |
| ENSG000001 | 56.5837864 | 0.157774729 | 0.2177637 | 0.72452264 | 0.46874493 | 0.59014312 | TACR1 | 6869 |
| ENSG000002 | 3.51242656 | -0.273546612 | 0.37756335 | -0.7245052 | 0.46875564 | 0.59014312 | LINC01015 | 100507362 |
| ENSG000001 | 140.714238 | 0.109990332 | 0.15182543 | 0.72445264 | 0.46878789 | 0.5901654 | MTURN | 222166 |
| ENSG000002 | 3.04286542 | -0.265321322 | 0.36631653 | -0.7242953 | 0.46888449 | 0.59026869 | NA | NA |
| ENSG000002 | 3.77036347 | -0.274164263 | 0.37854528 | -0.7242575 | 0.46890765 | 0.59027952 | NA | NA |
| ENSG000002 | 23.7560074 | 0.190565879 | 0.26313559 | 0.72421171 | 0.46893577 | 0.59029659 | NA | NA |
| ENSG000002 | 3.49324011 | -0.273750763 | 0.37810362 | -0.7240099 | 0.46905966 | 0.59043422 | LOC646522 | 646522 |
| ENSG000002 | 25.0365862 | -0.195356902 | 0.26984605 | -0.7239569 | 0.46909221 | 0.59045686 | NA | NA |
| ENSG000001 | 83.8178514 | 0.134108421 | 0.18526303 | 0.72388118 | 0.46913868 | 0.5904669 | ZNF546 | 339327 |

| | | | | | | | | |
|---|---|---|---|---|---|---|---|---|
| ENSG000002 | 5.44916668 | -0.268269337 | 0.3706029 | -0.7238727 | 0.46914387 | 0.5904669 | LOC105369922 | 105369922 |
| ENSG000002 | 4.22981837 | -0.273343258 | 0.377591 | -0.7239136 | 0.46911878 | 0.5904669 | NA | NA |
| ENSG000002 | 6.92521334 | -0.258901999 | 0.35770543 | -0.7237855 | 0.46919744 | 0.59051599 | NA | NA |
| ENSG000002 | 3.66546443 | -0.273992935 | 0.37859123 | -0.7237171 | 0.46923945 | 0.59055054 | NA | NA |
| ENSG000002 | 5.83723533 | -0.270534584 | 0.37389753 | -0.7235528 | 0.46934036 | 0.5906592 | NA | NA |
| ENSG000002 | 6.7052849 | -0.261180703 | 0.36105086 | -0.7233903 | 0.46944011 | 0.59076641 | LOC105373383 | 105373383 |
| ENSG000002 | 5.01105278 | -0.27160901 | 0.37548939 | -0.7233467 | 0.46946691 | 0.5907818 | CYP2C58P | 106480430 |
| ENSG000002 | 2.68715562 | 0.271456029 | 0.37530747 | 0.72328971 | 0.46950191 | 0.59080751 | NA | NA |
| ENSG000002 | 5.96502035 | -0.265173582 | 0.36665592 | -0.7232219 | 0.46954358 | 0.5908416 | NA | NA |
| ENSG000002 | 2.64247193 | 0.271166884 | 0.37505937 | 0.72299722 | 0.46968159 | 0.59099693 | NA | NA |
| ENSG000001 | 161.90348 | 0.103908975 | 0.14372523 | 0.72296964 | 0.46969854 | 0.59099992 | ANTXR2 | 118429 |
| ENSG000002 | 2.64415487 | 0.271478046 | 0.37561835 | 0.72274969 | 0.46983368 | 0.59115162 | NA | NA |
| ENSG000002 | 4.76273684 | 0.271719811 | 0.37597723 | 0.72270283 | 0.46986247 | 0.59116951 | NA | NA |
| ENSG000001 | 230.987518 | 0.140391451 | 0.19427197 | 0.72265418 | 0.46989237 | 0.59117044 | TOB1 | 10140 |
| ENSG000002 | 12.7913548 | 0.232741424 | 0.32206234 | 0.72265955 | 0.46988907 | 0.59117044 | NA | NA |
| ENSG000002 | 4.18971514 | -0.272535867 | 0.37725963 | -0.7224093 | 0.47004287 | 0.59132484 | NA | NA |
| ENSG000002 | 9.28709327 | -0.251535203 | 0.34819041 | -0.722407 | 0.47004426 | 0.59132484 | NA | NA |
| ENSG000002 | 10.9795081 | -0.246678409 | 0.34161292 | -0.7220992 | 0.4702335 | 0.59154456 | STAU2-AS1 | 100128126 |
| ENSG000001 | 41.3128538 | -0.158614083 | 0.21966964 | -0.7220574 | 0.47025921 | 0.59155854 | ADAMTS5 | 11096 |
| ENSG000002 | 3.4785278 | -0.272397755 | 0.37734992 | -0.7218705 | 0.47037413 | 0.59168475 | NA | NA |
| ENSG000002 | 24.2402036 | 0.189076212 | 0.26198975 | 0.72169316 | 0.47048315 | 0.59180353 | NA | NA |
| ENSG000002 | 3.71801548 | -0.273199763 | 0.37859335 | -0.721618 | 0.47052939 | 0.59184334 | NA | NA |
| ENSG000002 | 4.95259618 | -0.270575762 | 0.37507434 | -0.7213924 | 0.47066814 | 0.59198114 | NA | NA |
| ENSG000002 | 5.91217646 | -0.263464361 | 0.36521051 | -0.7214041 | 0.47066092 | 0.59198114 | NA | NA |
| ENSG000002 | 6.67762973 | -0.259616148 | 0.35989726 | -0.7213618 | 0.47068692 | 0.59198639 | NA | NA |
| ENSG000001 | 263.451547 | 0.093889719 | 0.13016885 | 0.72129177 | 0.47073002 | 0.59199663 | ATP8A2 | 51761 |
| ENSG000001 | 11.1026821 | -0.239511212 | 0.33206532 | -0.7212774 | 0.47073885 | 0.59199663 | ABCG8 | 64241 |
| ENSG000001 | 78.8241338 | -0.133753907 | 0.18543555 | -0.7212959 | 0.47072747 | 0.59199663 | PIFO | 128344 |
| ENSG000002 | 11.8642462 | 0.234943955 | 0.32576766 | 0.72120098 | 0.47078587 | 0.59203739 | LOC105378663 | 105378663 |

| | | | | | | | | |
|---|---|---|---|---|---|---|---|---|
| ENSG000001 | 11.6710665 | 0.23613824 | 0.32752161 | 0.72098521 | 0.47091861 | 0.59218385 | ICAM4 | 3386 |
| ENSG000002 | 120.767531 | -0.117557561 | 0.16305603 | -0.7209642 | 0.47093154 | 0.59218385 | NA | NA |
| ENSG000002 | 7.46759851 | -0.259175502 | 0.35949904 | -0.7209352 | 0.4709494 | 0.59218794 | NA | NA |
| ENSG000002 | 6.16794158 | 0.267656039 | 0.37130385 | 0.72085447 | 0.47099906 | 0.59221776 | NA | NA |
| ENSG000002 | 5.3842578 | -0.272648781 | 0.37823277 | -0.7208492 | 0.47100232 | 0.59221776 | FLJ37505 | 400087 |
| ENSG000002 | 2.66487577 | 0.27049765 | 0.3752862 | 0.72077696 | 0.47104675 | 0.59225527 | NA | NA |
| ENSG000002 | 3.67092537 | -0.272782625 | 0.37854846 | -0.7206016 | 0.47115467 | 0.59237259 | NA | NA |
| ENSG000002 | 4.90044286 | 0.270142246 | 0.37490046 | 0.72057059 | 0.47117376 | 0.59237822 | NA | NA |
| ENSG000002 | 4.13515226 | 0.272644524 | 0.37845851 | 0.72040796 | 0.47127385 | 0.59248569 | NA | NA |
| ENSG000002 | 5.48035135 | -0.268482672 | 0.37280989 | -0.7201597 | 0.47142665 | 0.59265942 | EGFR-AS1 | 100507500 |
| ENSG000001 | 29.7241459 | -0.175375128 | 0.24356855 | -0.7200237 | 0.47151041 | 0.59274634 | CCDC159 | 126075 |
| ENSG000002 | 17.1018054 | -0.213728256 | 0.29687934 | -0.7199162 | 0.47157657 | 0.59281113 | NA | NA |
| ENSG000001 | 3.93490224 | -0.272524058 | 0.37858489 | -0.7198493 | 0.4716178 | 0.59284458 | OR1N1 | 138883 |
| ENSG000001 | 8.77589585 | -0.252438708 | 0.35077981 | -0.7196501 | 0.47174048 | 0.59297652 | MIR7-3HG | 284424 |
| ENSG000002 | 4.19659734 | -0.271994463 | 0.37797462 | -0.7196104 | 0.47176492 | 0.59297652 | MTCO2P4 | 106480296 |
| ENSG000002 | 147.327856 | -0.122682612 | 0.17048543 | -0.7196076 | 0.47176663 | 0.59297652 | NA | NA |
| ENSG000002 | 13.0096697 | -0.224965943 | 0.31265325 | -0.7195382 | 0.4718094 | 0.59298571 | NA | NA |
| ENSG000002 | 2.63729194 | 0.26955519 | 0.37460499 | 0.71957181 | 0.47178867 | 0.59298571 | NA | NA |
| ENSG000002 | 5.45698306 | -0.266143164 | 0.36988756 | -0.7195245 | 0.47181781 | 0.59298571 | NA | NA |
| ENSG000002 | 3.66853692 | -0.272386576 | 0.37858713 | -0.7194819 | 0.47184407 | 0.59300033 | NA | NA |
| ENSG000002 | 8.61852334 | -0.258947748 | 0.35996042 | -0.7193784 | 0.4719078 | 0.59306206 | NA | NA |
| ENSG000002 | 3.68111396 | -0.272254543 | 0.37850963 | -0.7192804 | 0.4719682 | 0.59311957 | NA | NA |
| ENSG000001 | 8.1949117 | -0.255816688 | 0.35571582 | -0.7191603 | 0.47204217 | 0.59317577 | SOWAHA | 134548 |
| ENSG000002 | 4.23864464 | -0.271791272 | 0.37791785 | -0.7191808 | 0.47202952 | 0.59317577 | NA | NA |
| ENSG000002 | 3.93520783 | -0.272187647 | 0.37859327 | -0.7189448 | 0.47217496 | 0.59330586 | NA | NA |
| ENSG000002 | 3.67766431 | -0.272177641 | 0.37857209 | -0.7189585 | 0.47216646 | 0.59330586 | BGLT3 | 103344929 |
| ENSG000002 | 3.93843318 | -0.271836436 | 0.37812064 | -0.7189146 | 0.47219355 | 0.59331084 | NA | NA |
| ENSG000002 | 3.49143264 | -0.271291987 | 0.37739938 | -0.7188459 | 0.47223589 | 0.59334286 | NA | NA |
| ENSG000002 | 2.73359248 | -0.265803519 | 0.36977463 | -0.7188257 | 0.4722483 | 0.59334286 | NA | NA |

| | | | | | | | | |
|---|---|---|---|---|---|---|---|---|
| ENSG000002 | 12.6953967 | 0.229260475 | 0.31899451 | 0.71869724 | 0.47232748 | 0.59342396 | NA | NA |
| ENSG000001 | 13.3804299 | 0.230868771 | 0.3212825 | 0.71858495 | 0.47239669 | 0.59349252 | AQP5 | 362 |
| ENSG000002 | 4.98387369 | -0.269797481 | 0.37549738 | -0.718507 | 0.47244476 | 0.59351614 | NA | NA |
| ENSG000002 | 4.03454996 | -0.272022205 | 0.3785928 | -0.7185087 | 0.47244371 | 0.59351614 | NA | NA |
| ENSG000002 | 2.73752693 | -0.265459909 | 0.36948473 | -0.7184598 | 0.47247385 | 0.59353429 | NA | NA |
| ENSG000002 | 17.5480155 | -0.21016183 | 0.29253099 | -0.7184259 | 0.47249474 | 0.59354215 | MIR325HG | 101928469 |
| ENSG000002 | 4.23931051 | -0.271585792 | 0.37804489 | -0.7183956 | 0.47251339 | 0.59354719 | NA | NA |
| ENSG000002 | 3.19800391 | -0.27031479 | 0.37630686 | -0.7183361 | 0.47255009 | 0.59357491 | NA | NA |
| ENSG000002 | 5.97071738 | -0.262766932 | 0.36587838 | -0.7181811 | 0.47264564 | 0.59367654 | NA | NA |
| ENSG000002 | 4.92753176 | -0.269123609 | 0.37479208 | -0.7180611 | 0.47271962 | 0.59373268 | LOC101929172 | 101929172 |
| ENSG000002 | 4.43634639 | -0.269899545 | 0.37586666 | -0.7180726 | 0.47271254 | 0.59373268 | NA | NA |
| ENSG00000C | 76.7002292 | 0.130174908 | 0.18132086 | 0.71792571 | 0.47280309 | 0.59380074 | SDK2 | 54549 |
| ENSG000001 | 88.3405728 | 0.149962558 | 0.20887742 | 0.71794527 | 0.47279104 | 0.59380074 | DPYSL4 | 10570 |
| ENSG000002 | 3.18206677 | -0.270644946 | 0.37704367 | -0.717808 | 0.4728757 | 0.59387353 | NA | NA |
| ENSG000001 | 199.388728 | 0.107895036 | 0.15033697 | 0.71768799 | 0.47294969 | 0.59394806 | ELF1 | 1997 |
| ENSG000002 | 10.3889081 | -0.244652113 | 0.34090594 | -0.7176528 | 0.47297137 | 0.59395689 | NA | NA |
| ENSG000001 | 4.69468606 | -0.270460984 | 0.37696486 | -0.71747 | 0.47308413 | 0.59404597 | SYT8 | 90019 |
| ENSG000001 | 41.1288256 | 0.166508366 | 0.23206656 | 0.71750263 | 0.47306401 | 0.59404597 | Sep-05 | 5413 |
| ENSG000002 | 11.8117656 | 0.231497674 | 0.32265987 | 0.71746658 | 0.47308625 | 0.59404597 | NA | NA |
| ENSG000002 | 4.42176251 | -0.270782742 | 0.37744186 | -0.7174158 | 0.47311759 | 0.59406692 | NA | NA |
| ENSG000002 | 4.71681079 | -0.26894435 | 0.37497472 | -0.7172333 | 0.47323016 | 0.59418988 | LINC00687 | 728450 |
| ENSG000002 | 13.1443426 | -0.22865036 | 0.3188441 | -0.7171228 | 0.47329836 | 0.59423871 | NA | NA |
| ENSG000002 | 4.97173131 | 0.267846623 | 0.37349112 | 0.71714321 | 0.47328574 | 0.59423871 | NA | NA |
| ENSG000001 | 288.94984 | 0.092353027 | 0.12881195 | 0.71696008 | 0.47339873 | 0.59433601 | DDX59 | 83479 |
| ENSG000002 | 15.8220653 | 0.2168834 | 0.30250856 | 0.71694964 | 0.47340517 | 0.59433601 | NA | NA |
| ENSG000002 | 6.28159081 | -0.266269477 | 0.37144545 | -0.7168468 | 0.47346864 | 0.59439729 | NA | NA |
| ENSG000002 | 30.3478061 | 0.175372238 | 0.24467064 | 0.71676862 | 0.47351687 | 0.59441829 | NA | NA |
| ENSG000002 | 5.42887088 | -0.264998951 | 0.3697238 | -0.7167484 | 0.47352934 | 0.59441829 | LOC101927588 | 101927588 |
| ENSG000002 | 4.94436386 | -0.268044766 | 0.37397027 | -0.7167542 | 0.47352576 | 0.59441829 | NA | NA |

| | | | | | | | | |
|---|---|---|---|---|---|---|---|---|
| ENSG000002 | 5.71550063 | -0.263969551 | 0.3684176 | -0.7164955 | 0.47368544 | 0.59459583 | NA | NA |
| ENSG000002 | 4.77641313 | -0.269992702 | 0.37700583 | -0.7161499 | 0.47389877 | 0.5948452 | NA | NA |
| ENSG000002 | 21.9876309 | 0.204710571 | 0.28585937 | 0.71612336 | 0.47391518 | 0.59484739 | NA | NA |
| ENSG000002 | 4.41188637 | -0.269926767 | 0.37695638 | -0.716069 | 0.47394876 | 0.59487112 | NA | NA |
| ENSG000002 | 9.2020901 | -0.250122882 | 0.34936548 | -0.7159347 | 0.47403165 | 0.59495674 | NA | NA |
| ENSG000002 | 4.90024077 | -0.268366657 | 0.37486078 | -0.7159102 | 0.4740468 | 0.59495735 | CISTR | 102216268 |
| ENSG000002 | 3.43986708 | -0.27071367 | 0.37816052 | -0.7158697 | 0.47407179 | 0.59497029 | NA | NA |
| ENSG000002 | 4.66072891 | -0.269016582 | 0.37583116 | -0.715791 | 0.47412038 | 0.59501286 | NA | NA |
| ENSG000002 | 10.1603801 | -0.241183347 | 0.33698258 | -0.7157146 | 0.47416761 | 0.59505371 | LNX1-AS1 | 100873939 |
| ENSG000001 | 5.70380297 | -0.263268112 | 0.36786726 | -0.7156606 | 0.47420093 | 0.59507712 | PABPC1L2B | 645974 |
| ENSG000001 | 184.399478 | -0.1090163 | 0.15233887 | -0.7156171 | 0.47422781 | 0.59509244 | SRGAP1 | 57522 |
| ENSG000001 | 5.19363007 | -0.266864302 | 0.37293859 | -0.7155717 | 0.47425585 | 0.5951092 | ODAM | 54959 |
| ENSG000002 | 5.04539536 | -0.269005152 | 0.37594567 | -0.7155426 | 0.47427381 | 0.59511332 | NA | NA |
| ENSG000001 | 79.4966993 | 0.142957761 | 0.19980958 | 0.71546999 | 0.47431868 | 0.59515121 | NA | NA |
| ENSG000001 | 66.1674419 | 0.14062738 | 0.19656674 | 0.71541796 | 0.47435082 | 0.59517312 | PURG | 29942 |
| ENSG000002 | 3.96350402 | -0.270807791 | 0.37857382 | -0.7153368 | 0.47440095 | 0.5952176 | LOC105375042 | 105375042 |
| ENSG000002 | 10.3608212 | 0.252345802 | 0.35279757 | 0.7152708 | 0.47444173 | 0.59525035 | NA | NA |
| ENSG000001 | 2.64879462 | 0.267329559 | 0.3737644 | 0.71523547 | 0.47446355 | 0.59525932 | NA | NA |
| ENSG000002 | 4.15793652 | -0.270218938 | 0.3778408 | -0.7151661 | 0.4745064 | 0.59529466 | NA | NA |
| ENSG000002 | 4.7060179 | -0.269212563 | 0.37646128 | -0.7151136 | 0.47453888 | 0.59531698 | NA | NA |
| ENSG000001 | 112.5172 | 0.136127012 | 0.19046743 | 0.71469969 | 0.47479463 | 0.5956194 | NA | NA |
| ENSG000002 | 3.46155822 | -0.269545846 | 0.37718988 | -0.7146158 | 0.47484649 | 0.5956476 | NA | NA |
| ENSG000002 | 3.89447121 | -0.270548591 | 0.37858429 | -0.7146324 | 0.47483623 | 0.5956476 | NA | NA |
| ENSG000001 | 3.00247024 | -0.26479018 | 0.37066192 | -0.7143711 | 0.47499771 | 0.59578258 | SPATA31D4 | 389761 |
| ENSG000002 | 18.2332643 | -0.205931887 | 0.28827046 | -0.7143704 | 0.47499817 | 0.59578258 | NA | NA |
| ENSG000002 | 11.8715555 | -0.233508801 | 0.32685913 | -0.7144019 | 0.47497867 | 0.59578258 | NA | NA |
| ENSG000002 | 4.41064083 | -0.268899425 | 0.37645393 | -0.7142957 | 0.47504435 | 0.59580364 | NA | NA |
| ENSG000002 | 23.214267 | 0.1911817 | 0.26764271 | 0.71431687 | 0.47503126 | 0.59580364 | C17orf96 | 100170841 |
| ENSG000002 | 3.44217066 | -0.27006846 | 0.37825217 | -0.7139905 | 0.47523305 | 0.59602188 | LINC00850 | 101241891 |

| | | | | | | | | |
|---|---|---|---|---|---|---|---|---|
| ENSG000002 | 3.49253812 | 0.270232046 | 0.37857139 | 0.71382056 | 0.47533814 | 0.59611681 | NA | NA |
| ENSG000002 | 3.96302849 | -0.270053082 | 0.37831903 | -0.7138237 | 0.47533622 | 0.59611681 | LINC01338 | 102546175 |
| ENSG000002 | 13.2153583 | 0.225179771 | 0.31549179 | 0.7137421 | 0.47538667 | 0.59615923 | NA | NA |
| ENSG000001 | 22.6962983 | -0.187801081 | 0.26317045 | -0.7136101 | 0.47546834 | 0.5962432 | CAMK2N1 | 55450 |
| ENSG000002 | 12.3101027 | -0.225734426 | 0.31636558 | -0.713524 | 0.47552158 | 0.59629153 | LOC105376671 | 105376671 |
| ENSG000001 | 20.0546482 | 0.201977242 | 0.2831128 | 0.71341614 | 0.47558829 | 0.59635674 | CBX8 | 57332 |
| ENSG000002 | 8.77005327 | 0.246466341 | 0.34549874 | 0.71336394 | 0.47562058 | 0.59637879 | NA | NA |
| ENSG00000C | 163.098629 | -0.111302357 | 0.15609584 | -0.7130386 | 0.4758219 | 0.59661277 | SOAT1 | 6646 |
| ENSG000002 | 7.67222946 | -0.252406874 | 0.35403372 | -0.7129459 | 0.47587926 | 0.59666625 | NA | NA |
| ENSG000002 | 5.92689977 | -0.262843502 | 0.36875763 | -0.7127812 | 0.47598117 | 0.59675713 | NA | NA |
| ENSG000002 | 4.22373789 | -0.269286391 | 0.37779094 | -0.7127921 | 0.47597443 | 0.59675713 | NA | NA |
| ENSG000001 | 73.1464526 | -0.1334817 | 0.18729914 | -0.7126658 | 0.47605257 | 0.59679128 | HCN1 | 348980 |
| ENSG000002 | 4.17487255 | -0.269126648 | 0.37762803 | -0.7126766 | 0.47604592 | 0.59679128 | NA | NA |
| ENSG000002 | 9.11532419 | -0.242083376 | 0.33968342 | -0.7126735 | 0.47604782 | 0.59679128 | NA | NA |
| ENSG000002 | 3.43831964 | -0.269146764 | 0.37767839 | -0.7126348 | 0.47607177 | 0.59679691 | NA | NA |
| ENSG000002 | 17.1248787 | -0.207886028 | 0.29176599 | -0.7125095 | 0.47614936 | 0.59687572 | LOC100507377 | 100507377 |
| ENSG000001 | 4.46378531 | -0.269021516 | 0.37767616 | -0.7123074 | 0.47627446 | 0.59697718 | GPT | 2875 |
| ENSG000002 | 6.50354244 | -0.263559833 | 0.36999689 | -0.7123299 | 0.47626054 | 0.59697718 | NA | NA |
| ENSG000002 | 15.0169432 | -0.215386898 | 0.30236462 | -0.7123416 | 0.47625326 | 0.59697718 | NA | NA |
| ENSG000001 | 104.141509 | -0.124206951 | 0.17441712 | -0.7121259 | 0.47638681 | 0.59709955 | TENM2 | 57451 |
| ENSG000001 | 110.399935 | 0.119377201 | 0.16766563 | 0.71199565 | 0.47646747 | 0.59718218 | XKR6 | 286046 |
| ENSG000001 | 20.8952172 | 0.194189746 | 0.2727499 | 0.71196999 | 0.47648336 | 0.59718364 | APCDD1L | 164284 |
| ENSG000001 | 372.238407 | 0.089964855 | 0.126377 | 0.7118768 | 0.47654107 | 0.5972006 | APAF1 | 317 |
| ENSG000002 | 5.70004458 | -0.262524531 | 0.36877261 | -0.7118873 | 0.47653457 | 0.5972006 | NA | NA |
| ENSG000002 | 7.98711549 | 0.252173214 | 0.35423696 | 0.71187719 | 0.47654083 | 0.5972006 | LOC102725254 | 102725254 |
| ENSG000002 | 2.68152489 | -0.263717763 | 0.37047398 | -0.7118388 | 0.47656459 | 0.59721162 | NA | NA |
| ENSG000002 | 5.43185959 | -0.263165969 | 0.36973287 | -0.7117733 | 0.4766052 | 0.59724405 | NA | NA |
| ENSG000002 | 5.15022491 | -0.265806701 | 0.37346275 | -0.7117355 | 0.47662858 | 0.59725489 | NA | NA |
| ENSG000002 | 13.2929154 | -0.221192218 | 0.3108488 | -0.711575 | 0.47672802 | 0.59736104 | NA | NA |

| | | | | | | | | |
|---|---|---|---|---|---|---|---|---|
| ENSG000001 | 36.7244417 | 0.164173073 | 0.23074642 | 0.71148699 | 0.47678251 | 0.59741086 | AMBP | 259 |
| ENSG000002 | 3.67857723 | -0.269325453 | 0.37858954 | -0.7113917 | 0.47684154 | 0.59746637 | CLDN34 | 100288814 |
| ENSG000002 | 7.68585911 | -0.251731863 | 0.3538837 | -0.7113407 | 0.47687317 | 0.59748752 | NA | NA |
| ENSG000002 | 20.0303719 | -0.198689045 | 0.27939912 | -0.7111298 | 0.4770038 | 0.59761907 | NA | NA |
| ENSG000002 | 4.24570454 | -0.268736643 | 0.37790425 | -0.7111236 | 0.47700763 | 0.59761907 | NA | NA |
| ENSG000001 | 253.312556 | 0.10604438 | 0.1491617 | 0.7109357 | 0.47712408 | 0.59774649 | SYT4 | 6860 |
| ENSG000002 | 3.39199955 | -0.268664134 | 0.37792563 | -0.7108915 | 0.47715145 | 0.59776231 | NA | NA |
| ENSG000002 | 3.15518052 | -0.267349103 | 0.37610533 | -0.7108357 | 0.47718605 | 0.59778719 | NA | NA |
| ENSG000002 | 5.15383461 | -0.264575611 | 0.37222363 | -0.7107975 | 0.47720976 | 0.59779843 | NA | NA |
| ENSG000002 | 6.43092715 | -0.25819701 | 0.36330677 | -0.7106859 | 0.47727889 | 0.59786655 | NA | NA |
| ENSG000001 | 44.0709945 | 0.153041402 | 0.21536311 | 0.71062031 | 0.47731955 | 0.59788169 | GEM | 2669 |
| ENSG000002 | 6.63974555 | -0.255412797 | 0.35942305 | -0.7106189 | 0.47732045 | 0.59788169 | PABPC5-AS1 | 102724167 |
| ENSG000002 | 3.98580861 | -0.268975714 | 0.37854537 | -0.7105508 | 0.47736265 | 0.5978976 | NA | NA |
| ENSG000002 | 5.43097858 | -0.262820778 | 0.36988194 | -0.7105531 | 0.47736119 | 0.5978976 | NA | NA |
| ENSG000002 | 5.21066426 | -0.265765609 | 0.37416374 | -0.7102923 | 0.47752292 | 0.59807987 | NA | NA |
| ENSG000001 | 24.5160435 | -0.202045302 | 0.28449082 | -0.7101997 | 0.47758029 | 0.59813324 | KCNJ1 | 3758 |
| ENSG000002 | 5.14111523 | 0.268023918 | 0.37742825 | 0.71013211 | 0.47762222 | 0.59815538 | NA | NA |
| ENSG000002 | 14.3815449 | -0.220325447 | 0.3102635 | -0.7101236 | 0.47762746 | 0.59815538 | LINC00987 | 100499405 |
| ENSG000002 | 4.96201836 | -0.266293314 | 0.37502831 | -0.7100619 | 0.47766575 | 0.59818486 | NA | NA |
| ENSG000001 | 6.52221544 | -0.261829573 | 0.36878404 | -0.7099808 | 0.47771606 | 0.59822939 | SLC16A11 | 162515 |
| ENSG000001 | 5.62906357 | -0.26133118 | 0.36810437 | -0.7099377 | 0.47774275 | 0.59824433 | FSCB | 84075 |
| ENSG000002 | 4.27021553 | -0.268472379 | 0.37823129 | -0.7098101 | 0.47782192 | 0.59832499 | NA | NA |
| ENSG000002 | 2.922147 | -0.265771466 | 0.37450024 | -0.7096697 | 0.47790899 | 0.59841555 | NA | NA |
| ENSG00000C | 25.7405713 | 0.181227839 | 0.25541819 | 0.70953379 | 0.47799329 | 0.59850262 | TP53INP2 | 58476 |
| ENSG000002 | 4.52626502 | -0.268524413 | 0.37850711 | -0.7094303 | 0.47805749 | 0.59856452 | NA | NA |
| ENSG000002 | 4.17397779 | -0.26815287 | 0.37804044 | -0.7093232 | 0.47812391 | 0.59862921 | LOC105372352 | 105372352 |
| ENSG000002 | 2.3861563 | -0.26329904 | 0.37123062 | -0.70926 | 0.47816312 | 0.59864622 | NA | NA |
| ENSG000002 | 8.08398392 | -0.258933105 | 0.36507823 | -0.7092537 | 0.47816703 | 0.59864622 | ESPNP | 284729 |
| ENSG000001 | 2.71012634 | -0.262187826 | 0.36978249 | -0.7090326 | 0.47830427 | 0.59879955 | FAM47A | 158724 |

| | | | | | | | | |
|---|---|---|---|---|---|---|---|---|
| ENSG000002 | 3.90218366 | -0.26832723 | 0.37857359 | -0.7087849 | 0.47845799 | 0.59896566 | NA | NA |
| ENSG000002 | 5.26265204 | -0.26684875 | 0.37649492 | -0.7087712 | 0.47846649 | 0.59896566 | TRPC7-AS2 | 106478968 |
| ENSG000002 | 10.4443786 | -0.239939338 | 0.3385468 | -0.7087331 | 0.4784901 | 0.59897672 | LHX1 | 3975 |
| ENSG000001 | 17.7139707 | -0.212024629 | 0.29926121 | -0.7084935 | 0.47863883 | 0.59912923 | DDX53 | 168400 |
| ENSG000002 | 4.90841798 | -0.26460467 | 0.37347732 | -0.7084893 | 0.47864148 | 0.59912923 | NA | NA |
| ENSG000002 | 6.07933818 | 0.260752796 | 0.36808898 | 0.70839609 | 0.47869932 | 0.59918314 | NA | NA |
| ENSG000002 | 34.3704322 | 0.167748736 | 0.23684765 | 0.70825586 | 0.47878638 | 0.59927361 | NA | NA |
| ENSG000002 | 49.2604445 | -0.155260731 | 0.21925371 | -0.7081327 | 0.47886283 | 0.5993508 | GJA5 | 2702 |
| ENSG000002 | 2.38874007 | -0.262690477 | 0.37098426 | -0.7080906 | 0.47888898 | 0.59936504 | NA | NA |
| ENSG000001 | 119.884493 | 0.110899142 | 0.15665753 | 0.70790816 | 0.47900229 | 0.59948834 | TPD52L1 | 7164 |
| ENSG000002 | 3.15225846 | -0.266152994 | 0.37601894 | -0.7078181 | 0.47905825 | 0.59953988 | NA | NA |
| ENSG000002 | 15.3471566 | 0.212676177 | 0.30047962 | 0.70778902 | 0.47907628 | 0.59954394 | NA | NA |
| ENSG000002 | 10.7669917 | 0.23442378 | 0.33124527 | 0.70770453 | 0.47912876 | 0.59959111 | LINC01003 | 100128822 |
| ENSG000002 | 6.69204865 | -0.254790403 | 0.36009148 | -0.7075713 | 0.47921151 | 0.59962064 | LOC105377888 | 105377888 |
| ENSG000002 | 27.7008913 | 0.18018736 | 0.25463468 | 0.70763087 | 0.47917452 | 0.59962064 | NA | NA |
| ENSG000002 | 7.71265308 | -0.251481972 | 0.35540875 | -0.7075852 | 0.47920288 | 0.59962064 | NA | NA |
| ENSG000002 | 2.68543535 | -0.208754952 | 0.29501869 | -0.7075991 | 0.47919426 | 0.59962064 | NA | NA |
| ENSG000002 | 3.70667069 | -0.267864188 | 0.37859336 | -0.7075248 | 0.47924042 | 0.59963831 | NA | NA |
| ENSG000002 | 4.48628086 | -0.267029822 | 0.37744206 | -0.7074723 | 0.47927299 | 0.59966057 | NA | NA |
| ENSG000002 | 5.92905988 | -0.260132634 | 0.36774549 | -0.7073714 | 0.47933571 | 0.59972054 | NA | NA |
| ENSG000002 | 4.43811744 | 0.267661915 | 0.3784802 | 0.70720189 | 0.47944102 | 0.59981529 | NA | NA |
| ENSG000002 | 18.2181003 | 0.211310674 | 0.29878892 | 0.70722393 | 0.47942733 | 0.59981529 | NA | NA |
| ENSG000002 | 6.19912718 | -0.257884345 | 0.36468742 | -0.707138 | 0.4794807 | 0.59984642 | NA | NA |
| ENSG000002 | 19.0653721 | -0.200426743 | 0.28344462 | -0.7071108 | 0.47949764 | 0.59984911 | RFX3-AS1 | 101929302 |
| ENSG000002 | 24.2116811 | 0.190439028 | 0.26939165 | 0.70692254 | 0.47961461 | 0.59995842 | ADAMTS19-AS1 | 103689846 |
| ENSG000002 | 2.36884867 | -0.263039156 | 0.37208018 | -0.7069421 | 0.47960244 | 0.59995842 | NA | NA |
| ENSG000001 | 14.9873982 | 0.220122961 | 0.31139258 | 0.70689854 | 0.47962953 | 0.59995857 | FOXL2 | 668 |
| ENSG000002 | 6.4352856 | -0.257902431 | 0.3648515 | -0.7068696 | 0.47964753 | 0.59996258 | NA | NA |
| ENSG000001 | 17.6378045 | -0.205672916 | 0.29098174 | -0.7068241 | 0.47967578 | 0.59997631 | ARHGAP36 | 158763 |

| | | | | | | | | |
|---|---|---|---|---|---|---|---|---|
| ENSG000002 | 17.8231138 | 0.210197974 | 0.29739204 | 0.70680431 | 0.4796881 | 0.59997631 | SNORA55 | 677834 |
| ENSG000002 | 47.6147117 | -0.163225307 | 0.23096649 | -0.7067056 | 0.47974946 | 0.60003456 | NA | NA |
| ENSG000002 | 13.298108 | -0.220210443 | 0.31165856 | -0.7065759 | 0.47983004 | 0.60011683 | NA | NA |
| ENSG000002 | 3.49017872 | -0.265991042 | 0.37654324 | -0.7064024 | 0.47993794 | 0.60023326 | NA | NA |
| ENSG000002 | 2.4012402 | -0.262064204 | 0.37101546 | -0.7063431 | 0.47997481 | 0.60026086 | NA | NA |
| ENSG000001 | 7.20016494 | -0.254959652 | 0.36100392 | -0.7062518 | 0.48003159 | 0.60028387 | SLC25A52 | 147407 |
| ENSG000001 | 10.321395 | 0.241667319 | 0.34217341 | 0.70627148 | 0.48001933 | 0.60028387 | FAM212A | 389119 |
| ENSG000002 | 4.07773203 | 0.267179937 | 0.37831213 | 0.70624206 | 0.48003762 | 0.60028387 | NA | NA |
| ENSG000002 | 18.2283149 | 0.206539009 | 0.29255265 | 0.7059892 | 0.48019486 | 0.60046198 | NA | NA |
| ENSG000001 | 4.40161847 | -0.265982823 | 0.37685517 | -0.7057959 | 0.48031509 | 0.6005938 | HSPB3 | 8988 |
| ENSG000002 | 3.90029254 | -0.267106354 | 0.37847711 | -0.7057398 | 0.48034998 | 0.60061891 | LOC105372440 | 105372440 |
| ENSG000001 | 10.8827027 | -0.235763502 | 0.33410138 | -0.7056646 | 0.48039677 | 0.6006589 | FADS6 | 283985 |
| ENSG00000C | 50.2644504 | -0.144742628 | 0.20513981 | -0.7055804 | 0.48044912 | 0.60070584 | CRLF1 | 9244 |
| ENSG000001 | 4.17828472 | -0.266602783 | 0.37796689 | -0.7053601 | 0.48058614 | 0.60084025 | CST2 | 1470 |
| ENSG000002 | 5.69892304 | -0.261123758 | 0.37020919 | -0.7053411 | 0.48059798 | 0.60084025 | NA | NA |
| ENSG000002 | 4.44645862 | -0.265966235 | 0.37708521 | -0.7053213 | 0.4806103 | 0.60084025 | NA | NA |
| ENSG000002 | 4.96511145 | -0.264865925 | 0.37552999 | -0.7053123 | 0.4806159 | 0.60084025 | NA | NA |
| ENSG000002 | 2.6980951 | -0.259560156 | 0.3680212 | -0.7052859 | 0.48063234 | 0.60084229 | NA | NA |
| ENSG000002 | 2.38735052 | -0.262175201 | 0.37178748 | -0.7051749 | 0.48070139 | 0.60091008 | C4B | 721 |
| ENSG000002 | 12.8196297 | -0.224118743 | 0.31786675 | -0.7050714 | 0.48076583 | 0.60097195 | MRPL30 | 51263 |
| ENSG000002 | 2.62949518 | 0.263977117 | 0.37441027 | 0.70504775 | 0.48078052 | 0.60097195 | NA | NA |
| ENSG000001 | 10.1552679 | 0.236486453 | 0.33544563 | 0.7049919 | 0.48081528 | 0.60099035 | NA | NA |
| ENSG000002 | 3.49587541 | -0.266006234 | 0.37734604 | -0.7049398 | 0.48084772 | 0.60099035 | NA | NA |
| ENSG000002 | 8.37222269 | -0.243554867 | 0.34550277 | -0.7049288 | 0.48085453 | 0.60099035 | NA | NA |
| ENSG000002 | 18.3933938 | -0.202360301 | 0.28704618 | -0.7049747 | 0.48082596 | 0.60099035 | NA | NA |
| ENSG000002 | 20.9791514 | -0.198420647 | 0.2814934 | -0.7048856 | 0.48088143 | 0.60100545 | NA | NA |
| ENSG000002 | 19.7569734 | -0.198579578 | 0.28173742 | -0.7048392 | 0.48091031 | 0.60102302 | NA | NA |
| ENSG000002 | 17.0076132 | -0.210538895 | 0.29873731 | -0.7047626 | 0.48095796 | 0.60104553 | NA | NA |
| ENSG000002 | 5.68434766 | -0.260670479 | 0.36986837 | -0.7047655 | 0.48095616 | 0.60104553 | NA | NA |

| | | | | | | | | |
|---|---|---|---|---|---|---|---|---|
| ENSG000002 | 4.38836207 | -0.265192189 | 0.37635054 | -0.7046414 | 0.4810334 | 0.60112128 | NA | NA |
| ENSG000002 | 3.44520372 | -0.266081827 | 0.37765502 | -0.7045632 | 0.48108211 | 0.60116362 | NA | NA |
| ENSG000002 | 19.7497136 | 0.214808161 | 0.30498981 | 0.70431258 | 0.48123813 | 0.60134006 | NA | NA |
| ENSG000002 | 4.29892003 | -0.266589465 | 0.37854956 | -0.7042393 | 0.48128379 | 0.60137857 | NA | NA |
| ENSG000001 | 17.1614339 | -0.205853017 | 0.29234729 | -0.7041386 | 0.48134645 | 0.60143834 | FUT3 | 2525 |
| ENSG000002 | 3.69292448 | -0.266516266 | 0.3785707 | -0.7040066 | 0.48142867 | 0.601504 | NA | NA |
| ENSG000002 | 3.93223558 | -0.266381347 | 0.37837391 | -0.7040162 | 0.48142271 | 0.601504 | NA | NA |
| ENSG000001 | 8.86468213 | -0.240653459 | 0.34184951 | -0.7039749 | 0.48144843 | 0.60151016 | HOXB1 | 3211 |
| ENSG000002 | 8.41625958 | -0.245299297 | 0.34847079 | -0.7039307 | 0.48147592 | 0.60152597 | NA | NA |
| ENSG000002 | 4.79555293 | 0.264759271 | 0.37618711 | 0.70379676 | 0.48155935 | 0.60161167 | MIR4735 | 100616363 |
| ENSG000002 | 3.22507942 | -0.263253885 | 0.3740745 | -0.7037472 | 0.48159022 | 0.6016317 | NA | NA |
| ENSG000002 | 86.3119325 | -0.126887874 | 0.18031814 | -0.7036889 | 0.48162654 | 0.60165854 | NA | NA |
| ENSG000001 | 51.1732655 | 0.151657223 | 0.21561824 | 0.7033599 | 0.48183149 | 0.60187748 | ST6GALNAC5 | 81849 |
| ENSG000002 | 35.9533918 | -0.166242759 | 0.23635258 | -0.7033677 | 0.48182665 | 0.60187748 | NA | NA |
| ENSG000002 | 4.94419522 | -0.263764515 | 0.37505418 | -0.7032704 | 0.48188724 | 0.60191458 | NA | NA |
| ENSG000002 | 13.4802694 | 0.219119192 | 0.31157433 | 0.70326459 | 0.48189088 | 0.60191458 | NA | NA |
| ENSG000002 | 5.6378978 | -0.259209299 | 0.36860897 | -0.7032094 | 0.48192525 | 0.60193897 | NA | NA |
| ENSG000002 | 2.66711986 | 0.262078314 | 0.37271047 | 0.70316865 | 0.48195066 | 0.60195217 | NA | NA |
| ENSG000002 | 13.3764167 | -0.220617213 | 0.31376498 | -0.7031289 | 0.48197545 | 0.6019646 | SATB2-AS1 | 150538 |
| ENSG000001 | 4.94488036 | -0.26362354 | 0.37505572 | -0.7028917 | 0.48212327 | 0.60211212 | GNAT1 | 2779 |
| ENSG000002 | 6.7192557 | -0.261102119 | 0.37146579 | -0.7028968 | 0.48212006 | 0.60211212 | NA | NA |
| ENSG000001 | 4.94245258 | -0.263147864 | 0.37442118 | -0.7028124 | 0.48217264 | 0.60215524 | TNFAIP8L2 | 79626 |
| ENSG000002 | 5.12972746 | -0.262738605 | 0.37395093 | -0.7026018 | 0.48230393 | 0.6022821 | NA | NA |
| ENSG000002 | 5.24850488 | -0.263056879 | 0.3744016 | -0.7026062 | 0.48230121 | 0.6022821 | NA | NA |
| ENSG000002 | 4.9982647 | -0.264660213 | 0.37673546 | -0.7025094 | 0.48236153 | 0.60233548 | NA | NA |
| ENSG000001 | 32.1943288 | 0.171234718 | 0.24376531 | 0.70245729 | 0.48239403 | 0.60235751 | LRRC8E | 80131 |
| ENSG000002 | 7.16385966 | -0.250193423 | 0.35624049 | -0.7023161 | 0.48248207 | 0.6024489 | NA | NA |
| ENSG000002 | 10.6649113 | -0.233075159 | 0.33190008 | -0.702245 | 0.48252641 | 0.60248571 | NA | NA |
| ENSG000001 | 65.7465992 | 0.135292141 | 0.19271423 | 0.70203502 | 0.48265732 | 0.60263062 | NUP62CL | 54830 |

| | | | | | | | | |
|---|---|---|---|---|---|---|---|---|
| ENSG000002 | 3.16918923 | -0.218829594 | 0.31172665 | -0.7019919 | 0.48268419 | 0.60264561 | NA | NA |
| ENSG000002 | 13.6718353 | -0.217457379 | 0.30979707 | -0.7019349 | 0.48271975 | 0.60267145 | NA | NA |
| ENSG000002 | 5.20765201 | -0.261620037 | 0.37273117 | -0.7019001 | 0.48274148 | 0.60268002 | NA | NA |
| ENSG000002 | 14.6648959 | -0.234251806 | 0.33384558 | -0.7016771 | 0.48288054 | 0.60283507 | NA | NA |
| ENSG000002 | 2.45419782 | 0.258950406 | 0.36924602 | 0.70129505 | 0.4831189 | 0.60311408 | NA | NA |
| ENSG000002 | 13.5794183 | -0.223199079 | 0.31829699 | -0.701229 | 0.48316011 | 0.60314695 | NA | NA |
| ENSG000002 | 14.0804659 | -0.22736344 | 0.32424857 | -0.7012011 | 0.48317755 | 0.60315015 | NA | NA |
| ENSG000002 | 2.674407 | 0.26283911 | 0.3749306 | 0.70103404 | 0.48328178 | 0.60326169 | NA | NA |
| ENSG000002 | 4.92769593 | -0.262619415 | 0.3746452 | -0.7009817 | 0.48331446 | 0.60328392 | NA | NA |
| ENSG000002 | 5.98173547 | -0.258716423 | 0.3691158 | -0.7009086 | 0.48336008 | 0.60330746 | NA | NA |
| ENSG000002 | 20.3274894 | 0.193608945 | 0.27622757 | 0.70090377 | 0.48336307 | 0.60330746 | LOC100287808 | 100287808 |
| ENSG000002 | 7.18005815 | -0.25007579 | 0.35682787 | -0.7008303 | 0.48340891 | 0.6033461 | NA | NA |
| ENSG000002 | 6.19329431 | -0.256543865 | 0.36614055 | -0.7006704 | 0.4835087 | 0.60345208 | NA | NA |
| ENSG000002 | 5023.91759 | 0.078581397 | 0.1121678 | 0.70057001 | 0.4835714 | 0.60351025 | CHML | 1122 |
| ENSG000002 | 3.71590136 | -0.265116163 | 0.37844106 | -0.7005481 | 0.48358508 | 0.60351025 | HPN-AS1 | 100128675 |
| ENSG000002 | 3.26252632 | 0.260763371 | 0.37224107 | 0.70052283 | 0.48360085 | 0.60351136 | NA | NA |
| ENSG000002 | 15.6457859 | 0.210010539 | 0.29980298 | 0.70049517 | 0.48361812 | 0.60351434 | NA | NA |
| ENSG000001 | 35.2491413 | -0.177584227 | 0.25353796 | -0.7004246 | 0.48366216 | 0.60355073 | ARC | 23237 |
| ENSG000002 | 2.36770706 | -0.260770107 | 0.3724033 | -0.7002358 | 0.48378008 | 0.6036793 | NA | NA |
| ENSG000002 | 6.18058958 | -0.257602403 | 0.36789721 | -0.7002021 | 0.48380109 | 0.60368694 | NA | NA |
| ENSG000002 | 2.65157895 | 0.26204901 | 0.37426546 | 0.70016884 | 0.48382187 | 0.60369429 | NA | NA |
| ENSG000002 | 6.92362323 | -0.251083709 | 0.35862644 | -0.700126 | 0.48384859 | 0.60370905 | NA | NA |
| ENSG000002 | 6.12734871 | -0.25622969 | 0.36608861 | -0.6999117 | 0.48398247 | 0.60385752 | DUXA | 503835 |
| ENSG00000C | 41.0351668 | 0.157331348 | 0.22487541 | 0.69963786 | 0.48415349 | 0.6040351 | ADAM11 | 4185 |
| ENSG000002 | 12.2946362 | -0.228694425 | 0.32687625 | -0.6996361 | 0.48415459 | 0.6040351 | NA | NA |
| ENSG000002 | 5.9172317 | -0.258506457 | 0.36950431 | -0.6996034 | 0.48417504 | 0.60404203 | NA | NA |
| ENSG000002 | 8.13734227 | -0.243277597 | 0.34776515 | -0.6995456 | 0.4842111 | 0.60406843 | NA | NA |
| ENSG000001 | 1230.85817 | 0.089263175 | 0.12764518 | 0.69930703 | 0.48436017 | 0.60423581 | UFM1 | 51569 |
| ENSG000002 | 12.1037209 | 0.228530529 | 0.32684427 | 0.69920311 | 0.48442511 | 0.60429823 | LOC105374428 | 105374428 |

| | | | | | | | | |
|---|---|---|---|---|---|---|---|---|
| ENSG000002 | 4.90503662 | -0.26251602 | 0.37546411 | -0.6991774 | 0.48444117 | 0.60429968 | NA | NA |
| ENSG000002 | 2.68456263 | -0.255114601 | 0.36493844 | -0.699062 | 0.48451327 | 0.60437103 | NA | NA |
| ENSG000002 | 5.45677454 | -0.260286302 | 0.37245026 | -0.6988485 | 0.48464672 | 0.60451889 | NA | NA |
| ENSG000002 | 9.40503494 | -0.237063185 | 0.33926329 | -0.6987587 | 0.48470283 | 0.60457028 | NA | NA |
| ENSG000002 | 3.93146557 | -0.264490182 | 0.37855622 | -0.6986814 | 0.48475117 | 0.60461198 | NA | NA |
| ENSG000002 | 13.949795 | -0.21784902 | 0.3118715 | -0.6985217 | 0.48485098 | 0.60471786 | LOC285593 | 285593 |
| ENSG000001 | 21.0108831 | 0.195843319 | 0.28038576 | 0.69847811 | 0.48487824 | 0.60473327 | IL2RB | 3560 |
| ENSG000002 | 2.39016243 | -0.258399584 | 0.36998503 | -0.6984055 | 0.48492364 | 0.60477129 | NA | NA |
| ENSG000002 | 6.16512189 | -0.256449288 | 0.3672606 | -0.6982761 | 0.48500457 | 0.60485362 | NA | NA |
| ENSG000002 | 10.8455457 | -0.22865119 | 0.32749307 | -0.6981863 | 0.48506067 | 0.60490498 | GLUD1P7 | 414212 |
| ENSG000001 | 9.16950214 | -0.24670354 | 0.35336497 | -0.6981551 | 0.48508023 | 0.60491077 | SLC38A8 | 146167 |
| ENSG000001 | 7.93889336 | -0.250599823 | 0.35898802 | -0.6980729 | 0.48513158 | 0.6049562 | IL17REL | 400935 |
| ENSG000001 | 104.445279 | 0.122712327 | 0.17584131 | 0.69785835 | 0.48526579 | 0.60506774 | CRTC2 | 200186 |
| ENSG000001 | 78.878602 | -0.124744668 | 0.17875349 | -0.6978586 | 0.4852656 | 0.60506774 | C6orf141 | 135398 |
| ENSG000002 | 6.94810809 | 0.258356537 | 0.37018911 | 0.69790421 | 0.4852371 | 0.60506774 | NA | NA |
| ENSG000002 | 6.94690013 | -0.251329619 | 0.36022248 | -0.6977067 | 0.48536067 | 0.60515893 | NA | NA |
| ENSG000002 | 4.92591463 | -0.26115345 | 0.3743096 | -0.6976937 | 0.48536877 | 0.60515893 | NA | NA |
| ENSG000002 | 8.9588341 | -0.243607344 | 0.3492753 | -0.6974651 | 0.48551176 | 0.60531859 | NA | NA |
| ENSG000001 | 86.2844246 | 0.132278312 | 0.18968583 | 0.69735472 | 0.48558083 | 0.6053861 | KLF5 | 688 |
| ENSG000002 | 3.66233156 | -0.263806587 | 0.3783295 | -0.6972932 | 0.48561932 | 0.60541547 | NA | NA |
| ENSG000002 | 17.204806 | -0.205040384 | 0.29407011 | -0.69725 | 0.48564636 | 0.60543057 | NA | NA |
| ENSG000002 | 11.0198006 | 0.238000332 | 0.34137574 | 0.69717999 | 0.48569016 | 0.60546656 | NA | NA |
| ENSG000002 | 2.45155669 | -0.252605582 | 0.36235795 | -0.6971162 | 0.4857301 | 0.60549774 | NA | NA |
| ENSG000002 | 2.67098164 | 0.260486375 | 0.37380159 | 0.69685733 | 0.48589209 | 0.60566242 | RNU6-757P | 106479901 |
| ENSG000002 | 3.94821235 | -0.263760911 | 0.37849303 | -0.6968713 | 0.48588337 | 0.60566242 | NA | NA |
| ENSG000002 | 27.0603371 | 0.18145634 | 0.26041049 | 0.69680886 | 0.48592242 | 0.60568162 | NA | NA |
| ENSG000001 | 16.2955787 | 0.207164048 | 0.29734462 | 0.69671362 | 0.48598203 | 0.60572825 | NRL | 4901 |
| ENSG000001 | 46.8889494 | 0.148361538 | 0.21294854 | 0.69670136 | 0.48598971 | 0.60572825 | NA | NA |
| ENSG000002 | 11.1094038 | -0.233177078 | 0.33471678 | -0.6966399 | 0.48602816 | 0.60575756 | NA | NA |

| | | | | | | | | |
|---|---|---|---|---|---|---|---|---|
| ENSG000002 | 7.1051829 | -0.247873467 | 0.35584999 | -0.6965673 | 0.48607363 | 0.60579561 | NA | NA |
| ENSG000001 | 5.0424865 | -0.262321005 | 0.37663343 | -0.6964889 | 0.4861227 | 0.60583815 | NEURL3 | 93082 |
| ENSG000001 | 25.9930597 | 0.181412281 | 0.26049486 | 0.69641404 | 0.48616957 | 0.60587794 | SLC35E4 | 339665 |
| ENSG000002 | 8.10730975 | -0.244827409 | 0.3516314 | -0.6962615 | 0.48626507 | 0.60597833 | COX5BP6 | 101234262 |
| ENSG000002 | 4.20943018 | -0.263522709 | 0.37857151 | -0.6960976 | 0.48636773 | 0.60608763 | LINC01386 | 104355294 |
| ENSG000002 | 5.75115101 | -0.260628019 | 0.37443137 | -0.6960635 | 0.48638905 | 0.60609558 | LIN37 | 55957 |
| ENSG000002 | 6.23335786 | -0.255098642 | 0.36652961 | -0.6959837 | 0.48643903 | 0.60613923 | NA | NA |
| ENSG000002 | 2.38900066 | -0.256194251 | 0.36813411 | -0.6959264 | 0.48647492 | 0.60616533 | NA | NA |
| ENSG000002 | 2.3578356 | -0.258460431 | 0.3714706 | -0.6957763 | 0.48656895 | 0.60626387 | NA | NA |
| ENSG000002 | 6.69094562 | -0.253669603 | 0.36460079 | -0.6957462 | 0.4865878 | 0.60626873 | NA | NA |
| ENSG000002 | 7.82242624 | 0.251115556 | 0.36094856 | 0.69571009 | 0.48661041 | 0.60627826 | NA | NA |
| ENSG000002 | 15.3643731 | -0.210296993 | 0.30230333 | -0.6956489 | 0.48664871 | 0.60630736 | NA | NA |
| ENSG000001 | 46.9357048 | -0.145713029 | 0.20953116 | -0.6954241 | 0.48678953 | 0.6063524 | ETNPPL | 64850 |
| ENSG000002 | 8.88617534 | 0.241651803 | 0.34747225 | 0.69545641 | 0.48676932 | 0.6063524 | NA | NA |
| ENSG000002 | 12.7522686 | -0.232379873 | 0.33410336 | -0.6955329 | 0.48672141 | 0.6063524 | NA | NA |
| ENSG000002 | 4.4192197 | -0.262541352 | 0.37745836 | -0.6955505 | 0.48671037 | 0.6063524 | NA | NA |
| ENSG000002 | 5.40610285 | -0.257971111 | 0.37091908 | -0.6954916 | 0.48674726 | 0.6063524 | LOC100507103 | 100507103 |
| ENSG000002 | 4.20167373 | -0.26323193 | 0.37848085 | -0.695496 | 0.4867445 | 0.6063524 | NA | NA |
| ENSG000002 | 2.61228234 | -0.260786717 | 0.37500271 | -0.6954262 | 0.48678824 | 0.6063524 | KCNJ18 | 100134444 |
| ENSG000002 | 3.18817984 | -0.26132965 | 0.37581127 | -0.6953747 | 0.48682051 | 0.60637235 | NA | NA |
| ENSG000002 | 9.72592798 | -0.237274978 | 0.34123095 | -0.6953501 | 0.48683592 | 0.60637293 | NA | NA |
| ENSG000002 | 2.33910561 | -0.258826798 | 0.37233377 | -0.6951473 | 0.48696302 | 0.60651261 | NA | NA |
| ENSG000001 | 45.5394605 | -0.150190273 | 0.21608454 | -0.6950533 | 0.4870219 | 0.60656732 | GABRB2 | 2561 |
| ENSG000001 | 357.879992 | 0.093858985 | 0.13504776 | 0.69500585 | 0.48705164 | 0.60658572 | ZNF695 | 57116 |
| ENSG000002 | 49.1613254 | 0.148858831 | 0.21423378 | 0.69484296 | 0.48715372 | 0.60669422 | SH3BP5-AS1 | 100505696 |
| ENSG000002 | 4.65281611 | -0.261367565 | 0.37619887 | -0.694759 | 0.48720633 | 0.60672247 | NA | NA |
| ENSG000002 | 4.33588519 | 0.262130358 | 0.37728997 | 0.6947716 | 0.48719845 | 0.60672247 | NA | NA |
| ENSG000002 | 3.24416144 | 0.259157175 | 0.37310432 | 0.69459709 | 0.48730783 | 0.60683024 | NA | NA |
| ENSG000002 | 2.91241934 | -0.258992622 | 0.37293072 | -0.6944792 | 0.48738176 | 0.60690366 | RNU6-665P | 106481392 |

| | | | | | | | | |
|---|---|---|---|---|---|---|---|---|
| ENSG000002 | 2.38960148 | -0.256306865 | 0.36908196 | -0.6944443 | 0.48740361 | 0.60691224 | NA | NA |
| ENSG000001 | 79.4728522 | 0.128455714 | 0.18499554 | 0.69437195 | 0.48744898 | 0.60695009 | DPY19L2 | 283417 |
| ENSG000002 | 15.7411802 | -0.207588168 | 0.29904627 | -0.6941674 | 0.48757725 | 0.60709116 | LOC100507537 | 100507537 |
| ENSG000002 | 6.06255208 | 0.25620207 | 0.36911377 | 0.69410054 | 0.48761916 | 0.6071247 | NA | NA |
| ENSG000001 | 544.149708 | -0.105610684 | 0.15216632 | -0.6940477 | 0.48765231 | 0.60714734 | HAS2 | 3037 |
| ENSG000002 | 10.1971177 | 0.22988797 | 0.33128149 | 0.69393545 | 0.48772269 | 0.60721632 | NA | NA |
| ENSG000002 | 5.86907984 | -0.254322422 | 0.36662648 | -0.6936826 | 0.48788126 | 0.60739509 | LINC01267 | 101927565 |
| ENSG000002 | 18.849844 | -0.196906019 | 0.28388704 | -0.6936069 | 0.48792874 | 0.60743556 | CRYGS | 1427 |
| ENSG000002 | 9.37764948 | 0.240467871 | 0.34673975 | 0.69351112 | 0.48798885 | 0.60749174 | NA | NA |
| ENSG000002 | 7.67662811 | -0.247350227 | 0.35685649 | -0.6931364 | 0.48822395 | 0.60775788 | NA | NA |
| ENSG000002 | 2.35803147 | -0.257257217 | 0.37115687 | -0.6931226 | 0.48823261 | 0.60775788 | NA | NA |
| ENSG000002 | 6.29083713 | 0.252611368 | 0.36450615 | 0.69302361 | 0.48829474 | 0.60781655 | NA | NA |
| ENSG000002 | 2.39483353 | -0.255444694 | 0.36868667 | -0.6928504 | 0.48840346 | 0.60793323 | RN7SL597P | 106479440 |
| ENSG000002 | 3.44223537 | -0.261764543 | 0.37783852 | -0.6927947 | 0.48843837 | 0.60795802 | NA | NA |
| ENSG000001 | 4.95352513 | -0.259804519 | 0.3750892 | -0.6926473 | 0.48853092 | 0.60805456 | H1FNT | 341567 |
| ENSG000002 | 4.67195083 | -0.260823554 | 0.37660941 | -0.6925572 | 0.48858748 | 0.60808763 | NA | NA |
| ENSG000002 | 3.90869013 | -0.262072313 | 0.37840916 | -0.6925633 | 0.48858362 | 0.60808763 | NHLRC4 | 283948 |
| ENSG000002 | 9.65043285 | -0.233404454 | 0.33703558 | -0.6925217 | 0.48860977 | 0.6080967 | NA | NA |
| ENSG000002 | 5.41755178 | -0.257741246 | 0.37237902 | -0.6921476 | 0.48884464 | 0.60837034 | NA | NA |
| ENSG000002 | 9.89793849 | 0.239524361 | 0.34607469 | 0.69211754 | 0.48886352 | 0.60837516 | NA | NA |
| ENSG000001 | 2.6061368 | -0.259522499 | 0.37499625 | -0.6920669 | 0.48889531 | 0.60839605 | NA | NA |
| ENSG000002 | 11.5255673 | -0.224474258 | 0.32437087 | -0.6920297 | 0.48891871 | 0.6084065 | NA | NA |
| ENSG000002 | 2.35582039 | -0.256833474 | 0.37115668 | -0.6919813 | 0.48894908 | 0.60842562 | NA | NA |
| ENSG000002 | 6.85887207 | -0.247932541 | 0.35835548 | -0.691862 | 0.48902399 | 0.60850016 | NA | NA |
| ENSG00000C | 416.646035 | 0.087508132 | 0.12649397 | 0.69179689 | 0.48906489 | 0.60853239 | TULP3 | 7289 |
| ENSG000002 | 7.85188582 | -0.243688713 | 0.35232054 | -0.6916676 | 0.48914608 | 0.60861073 | NA | NA |
| ENSG000002 | 324.472635 | 0.090318056 | 0.13058369 | 0.69164885 | 0.48915787 | 0.60861073 | PSMA3-AS1 | 379025 |
| ENSG000001 | 8.8788446 | -0.23885294 | 0.34538208 | -0.6915615 | 0.48921277 | 0.60866035 | ACRV1 | 56 |
| ENSG000002 | 7.71577518 | -0.250326535 | 0.36199852 | -0.6915126 | 0.48924348 | 0.60867989 | NA | NA |

| | | | | | | | | |
|---|---|---|---|---|---|---|---|---|
| ENSG000001 | 243.911607 | 0.106568795 | 0.15413363 | 0.69140522 | 0.48931093 | 0.60874513 | PRKAG2 | 51422 |
| ENSG000001 | 176.068639 | 0.111147477 | 0.16077423 | 0.69132643 | 0.48936043 | 0.60876935 | LRIG3 | 121227 |
| ENSG000002 | 5.45044055 | -0.257474134 | 0.37243229 | -0.6913314 | 0.48935731 | 0.60876935 | DCDC2B | 149069 |
| ENSG000002 | 5.69834238 | -0.256314208 | 0.37079579 | -0.6912544 | 0.48940571 | 0.60880701 | NA | NA |
| ENSG000001 | 100.85644 | 0.115048025 | 0.16644477 | 0.6912084 | 0.48943459 | 0.60882426 | FAM131A | 131408 |
| ENSG00000C | 99.8097697 | 0.119067448 | 0.17232459 | 0.69094868 | 0.48959779 | 0.60900859 | FGFR3 | 2261 |
| ENSG000002 | 2.38301909 | -0.255929312 | 0.37063069 | -0.6905238 | 0.48986484 | 0.60932208 | NA | NA |
| ENSG000001 | 47.5222755 | 0.152856657 | 0.22139516 | 0.69042457 | 0.48992723 | 0.60938099 | MMRN2 | 79812 |
| ENSG000001 | 338.746376 | 0.085833959 | 0.12432796 | 0.69038337 | 0.48995313 | 0.60939451 | DGKH | 160851 |
| ENSG000002 | 10.2743919 | -0.231414698 | 0.33520911 | -0.6903592 | 0.48996831 | 0.60939471 | NA | NA |
| ENSG000001 | 32.7410925 | 0.173148375 | 0.25091748 | 0.69006102 | 0.49015581 | 0.60960921 | SERTAD1 | 29950 |
| ENSG000001 | 252.701056 | 0.097437129 | 0.14121921 | 0.68997078 | 0.49021256 | 0.6096611 | FEZ1 | 9638 |
| ENSG000002 | 3.5221468 | -0.259755816 | 0.37651715 | -0.6898911 | 0.4902627 | 0.60970475 | NA | NA |
| ENSG000002 | 9.35915867 | -0.233172444 | 0.33801088 | -0.6898371 | 0.49029664 | 0.60972826 | RASGRF2-AS1 | 102524628 |
| ENSG000002 | 8.89590243 | 0.241623704 | 0.35032983 | 0.68970348 | 0.49038068 | 0.60979536 | NA | NA |
| ENSG000002 | 78.9801168 | 0.128370807 | 0.18612223 | 0.68971237 | 0.49037509 | 0.60979536 | GATS | 352954 |
| ENSG000001 | 119.971715 | 0.110400194 | 0.16008135 | 0.68965058 | 0.49041395 | 0.60979935 | DOPEY2 | 9980 |
| ENSG000002 | 26.476062 | -0.173811311 | 0.25202048 | -0.6896714 | 0.49040088 | 0.60979935 | NA | NA |
| ENSG000002 | 6.17871097 | -0.253777129 | 0.3680102 | -0.6895927 | 0.49045039 | 0.60982595 | NA | NA |
| ENSG000002 | 6.91226471 | -0.249181393 | 0.36146823 | -0.689359 | 0.49059735 | 0.60998636 | MACROD2-AS1 | 100379174 |
| ENSG000002 | 11.809326 | -0.222814084 | 0.32322825 | -0.6893398 | 0.49060948 | 0.60998636 | NA | NA |
| ENSG000001 | 64.323593 | -0.135156327 | 0.19611705 | -0.6891615 | 0.49072162 | 0.61010119 | YPEL2 | 388403 |
| ENSG000001 | 2.33241875 | -0.256354465 | 0.37198907 | -0.6891451 | 0.49073193 | 0.61010119 | NA | NA |
| ENSG000002 | 2.60018715 | -0.258351595 | 0.37498925 | -0.6889573 | 0.49085012 | 0.61019685 | NA | NA |
| ENSG000002 | 2.35706613 | -0.25595582 | 0.3715152 | -0.6889511 | 0.49085402 | 0.61019685 | MIR1302-8 | 100302223 |
| ENSG000002 | 3.31573443 | 0.259981678 | 0.37735673 | 0.68895467 | 0.49085179 | 0.61019685 | NA | NA |
| ENSG000002 | 4.88912357 | -0.259725329 | 0.37709274 | -0.6887572 | 0.49097609 | 0.61032989 | NA | NA |
| ENSG000002 | 14.2988296 | -0.220103512 | 0.31963065 | -0.6886183 | 0.4910635 | 0.61037069 | NA | NA |
| ENSG000002 | 26.0849353 | -0.182158367 | 0.26452307 | -0.6886294 | 0.4910565 | 0.61037069 | NA | NA |

| | | | | | | | | |
|---|---|---|---|---|---|---|---|---|
| ENSG000002 | 9.54665839 | -0.236262651 | 0.34306828 | -0.6886753 | 0.49102762 | 0.61037069 | RN7SL166P | 106479282 |
| ENSG000002 | 3.19852151 | -0.258728622 | 0.37572625 | -0.6886094 | 0.49106912 | 0.61037069 | NA | NA |
| ENSG000002 | 16.9227875 | 0.207822974 | 0.30188286 | 0.68842258 | 0.49118671 | 0.61049814 | NA | NA |
| ENSG000002 | 4.89475643 | -0.258116737 | 0.37501542 | -0.688283 | 0.49127458 | 0.61058864 | NA | NA |
| ENSG000002 | 8.42612407 | -0.24138311 | 0.35076969 | -0.6881527 | 0.49135662 | 0.61067189 | RN7SL558P | 106481074 |
| ENSG000002 | 10.0639705 | -0.229463387 | 0.33371157 | -0.6876099 | 0.49169846 | 0.611078 | NA | NA |
| ENSG000002 | 16.5312696 | -0.216211443 | 0.31447884 | -0.687523 | 0.49175319 | 0.6111273 | LOC255187 | 255187 |
| ENSG000002 | 5.2091989 | -0.256619411 | 0.37327783 | -0.6874756 | 0.49178306 | 0.61113115 | LOC284930 | 284930 |
| ENSG000002 | 3.49033626 | -0.259069299 | 0.37685748 | -0.6874463 | 0.4918015 | 0.61113115 | NA | NA |
| ENSG000002 | 5.66163834 | -0.254876774 | 0.37075087 | -0.687461 | 0.49179225 | 0.61113115 | NA | NA |
| ENSG000002 | 33.3450874 | -0.166086515 | 0.24172169 | -0.6870981 | 0.49202092 | 0.61138506 | NA | NA |
| ENSG000002 | 50.766193 | 0.159513346 | 0.23222863 | 0.68688061 | 0.49215797 | 0.61153662 | NA | NA |
| ENSG00000C | 27.9688677 | -0.172236264 | 0.25079108 | -0.6867719 | 0.49222649 | 0.61158429 | USH1C | 10083 |
| ENSG000002 | 7.20653207 | -0.248112407 | 0.36127043 | -0.6867775 | 0.49222294 | 0.61158429 | NA | NA |
| ENSG000002 | 3.72078905 | -0.259910646 | 0.37847616 | -0.6867292 | 0.49225336 | 0.61159894 | NA | NA |
| ENSG000002 | 7.17033392 | 0.247440533 | 0.36041625 | 0.686541 | 0.49237202 | 0.6117095 | NA | NA |
| ENSG000002 | 3.26824304 | -0.2556392 | 0.37235867 | -0.6865402 | 0.49237252 | 0.6117095 | NA | NA |
| ENSG000001 | 5.42022142 | -0.254868745 | 0.37130621 | -0.6864112 | 0.49245384 | 0.61175431 | APOBEC3H | 164668 |
| ENSG000001 | 42.288279 | -0.151406514 | 0.22056708 | -0.686442 | 0.49243441 | 0.61175431 | CSAD | 51380 |
| ENSG000002 | 3.66707491 | -0.259205467 | 0.37761512 | -0.6864277 | 0.49244346 | 0.61175431 | NA | NA |
| ENSG000002 | 5.13014504 | -0.256973446 | 0.37445047 | -0.6862682 | 0.49254401 | 0.61184757 | NA | NA |
| ENSG000001 | 8.08798801 | -0.238935021 | 0.34818356 | -0.6862329 | 0.49256625 | 0.61185645 | C1QL2 | 165257 |
| ENSG000002 | 8.02226542 | 0.242078411 | 0.35279759 | 0.68616798 | 0.49260719 | 0.61188857 | LOC101929095 | 101929095 |
| ENSG000002 | 6.15177539 | -0.252797232 | 0.36844224 | -0.6861245 | 0.49263463 | 0.6119039 | OR12D2 | 26529 |
| ENSG000001 | 41.3955052 | 0.15208849 | 0.22171094 | 0.68597649 | 0.49272793 | 0.61200105 | NA | NA |
| ENSG000002 | 6.67623407 | -0.251880197 | 0.36727517 | -0.6858079 | 0.49283428 | 0.6120926 | NA | NA |
| ENSG000002 | 2.68616992 | 0.25641521 | 0.3738829 | 0.6858169 | 0.49282858 | 0.6120926 | NA | NA |
| ENSG000002 | 12.8187558 | -0.220581614 | 0.32164703 | -0.6857878 | 0.49284692 | 0.6120926 | NA | NA |
| ENSG000002 | 8.95711146 | 0.238225219 | 0.34748143 | 0.68557684 | 0.49297999 | 0.61220161 | NA | NA |

| | | | | | | | | |
|---|---|---|---|---|---|---|---|---|
| ENSG000002 | 2.38220224 | -0.252543647 | 0.36834476 | -0.6856176 | 0.49295428 | 0.61220161 | NA | NA |
| ENSG000002 | 26.9325689 | 0.172622061 | 0.25178815 | 0.68558453 | 0.49297514 | 0.61220161 | NA | NA |
| ENSG000002 | 2.37579612 | -0.253714334 | 0.37010757 | -0.6855151 | 0.49301892 | 0.61223122 | UBE2FP3 | 106480360 |
| ENSG000000 | 106.465595 | -0.117030263 | 0.17074922 | -0.6853927 | 0.49309617 | 0.61228664 | CACNB1 | 782 |
| ENSG000002 | 34.8068455 | 0.166580966 | 0.24306867 | 0.6853247 | 0.49313905 | 0.61228664 | LINC01152 | 102606463 |
| ENSG000002 | 7.12567567 | -0.244192556 | 0.35628157 | -0.685392 | 0.49309656 | 0.61228664 | NA | NA |
| ENSG000002 | 4.44542747 | -0.25928912 | 0.37831925 | -0.6853712 | 0.49310971 | 0.61228664 | NA | NA |
| ENSG000002 | 2.36449021 | -0.254201683 | 0.37091896 | -0.6853294 | 0.49313606 | 0.61228664 | NA | NA |
| ENSG000001 | 2.64743425 | -0.256357432 | 0.3741727 | -0.6851313 | 0.49326107 | 0.61241939 | XCL2 | 6846 |
| ENSG000002 | 4.70712312 | -0.257994993 | 0.37660053 | -0.6850628 | 0.49330432 | 0.61245434 | NA | NA |
| ENSG000001 | 6.17021741 | -0.25284122 | 0.36913291 | -0.6849598 | 0.49336926 | 0.61247196 | BANF2 | 140836 |
| ENSG000001 | 94.0727377 | 0.114051507 | 0.16650402 | 0.6849775 | 0.49335812 | 0.61247196 | PPP1R1A | 5502 |
| ENSG000001 | 7.86923342 | -0.240897433 | 0.35169766 | -0.684956 | 0.49337167 | 0.61247196 | NA | NA |
| ENSG000002 | 2.37708208 | -0.252809635 | 0.36909506 | -0.6849445 | 0.49337893 | 0.61247196 | ATP2B2-IT2 | 100874325 |
| ENSG000002 | 2.63257868 | -0.256054678 | 0.37386099 | -0.6848927 | 0.4934116 | 0.61249377 | NA | NA |
| ENSG000002 | 3.18960219 | -0.256918629 | 0.37519122 | -0.6847672 | 0.49349085 | 0.61257339 | NA | NA |
| ENSG000002 | 2.34881291 | -0.25227889 | 0.36856047 | -0.684498 | 0.49366078 | 0.61276557 | MIR6870 | 102465525 |
| ENSG000001 | 11.045226 | -0.228070336 | 0.33323873 | -0.6844053 | 0.4937193 | 0.61280744 | HPD | 3242 |
| ENSG000002 | 5.79981073 | -0.256462042 | 0.37472721 | -0.6843966 | 0.49372474 | 0.61280744 | NA | NA |
| ENSG000002 | 5.25082011 | -0.257662585 | 0.37659154 | -0.6841964 | 0.49385113 | 0.6129268 | NA | NA |
| ENSG000002 | 12.1834967 | 0.226647304 | 0.33125842 | 0.68420089 | 0.49384832 | 0.6129268 | MTHFS | 10588 |
| ENSG000002 | 4.68887105 | 0.25830345 | 0.37760674 | 0.68405413 | 0.49394098 | 0.61300079 | NA | NA |
| ENSG000002 | 5.38991643 | -0.254232202 | 0.37164437 | -0.6840739 | 0.49392852 | 0.61300079 | NA | NA |
| ENSG000002 | 7.25643197 | -0.250261414 | 0.36586703 | -0.6840229 | 0.49396073 | 0.61300654 | KBTBD11-OT1 | 104266957 |
| ENSG000001 | 8.22459044 | 0.238178605 | 0.34828845 | 0.68385444 | 0.49406708 | 0.61308224 | NA | NA |
| ENSG000002 | 8.3545696 | -0.238413608 | 0.34862865 | -0.6838612 | 0.49406281 | 0.61308224 | LOC101929526 | 101929526 |
| ENSG000002 | 31.3396113 | 0.176334941 | 0.25783905 | 0.68389542 | 0.49404121 | 0.61308224 | PARD6G-AS1 | 100130522 |
| ENSG000001 | 122.081916 | -0.107026791 | 0.15651657 | -0.6838049 | 0.49409839 | 0.61308356 | EPB41L4A | 64097 |
| ENSG000002 | 4.52795602 | -0.25855927 | 0.378109 | -0.683822 | 0.49408755 | 0.61308356 | RN7SL775P | 106481859 |

| | | | | | | | | |
|---|---|---|---|---|---|---|---|---|
| ENSG000001 | 24.8239827 | -0.179842733 | 0.26301254 | -0.6837801 | 0.49411406 | 0.61308424 | EVA1A | 84141 |
| ENSG000001 | 21.0608863 | 0.195286157 | 0.28566289 | 0.68362453 | 0.49421229 | 0.61318736 | CYGB | 114757 |
| ENSG00000C | 113.961291 | -0.113842259 | 0.16657354 | -0.6834354 | 0.49433174 | 0.6132622 | FOXJ2 | 55810 |
| ENSG000002 | 8.51227908 | 0.250513541 | 0.36651452 | 0.68350237 | 0.49428945 | 0.6132622 | NA | NA |
| ENSG000002 | 3.18997433 | -0.256227226 | 0.37491184 | -0.6834333 | 0.4943331 | 0.6132622 | NA | NA |
| ENSG000002 | 2.35396118 | -0.253916346 | 0.37150745 | -0.6834758 | 0.49430624 | 0.6132622 | MIR548BB | 103504735 |
| ENSG000002 | 3.55717325 | 0.258492096 | 0.37824412 | 0.68340017 | 0.49435401 | 0.61326938 | NA | NA |
| ENSG000002 | 3.93539294 | -0.258654958 | 0.37859171 | -0.6832029 | 0.49447863 | 0.61340521 | NA | NA |
| ENSG000002 | 2.63480045 | 0.253323679 | 0.37081568 | 0.68315255 | 0.49451045 | 0.61342592 | NA | NA |
| ENSG000002 | 2.38775654 | 0.248997868 | 0.36452868 | 0.68306797 | 0.49456389 | 0.61347345 | NA | NA |
| ENSG000002 | 2.3428638 | -0.253483997 | 0.37112144 | -0.6830217 | 0.49459313 | 0.61347673 | NA | NA |
| ENSG000002 | 9.34348659 | -0.23324819 | 0.34149746 | -0.6830159 | 0.4945968 | 0.61347673 | NA | NA |
| ENSG000002 | 5.19256811 | -0.256302693 | 0.37526959 | -0.6829828 | 0.49461769 | 0.61348387 | NA | NA |
| ENSG000002 | 4.02368104 | -0.258208246 | 0.37811128 | -0.6828896 | 0.49467664 | 0.61353822 | NA | NA |
| ENSG000002 | 33.7651236 | 0.16432133 | 0.24064771 | 0.68282941 | 0.49471465 | 0.6135666 | DFNB59 | 494513 |
| ENSG000002 | 2.61549973 | -0.256299211 | 0.3753697 | -0.6827914 | 0.49473865 | 0.6135776 | NA | NA |
| ENSG000002 | 13.2277174 | 0.215746074 | 0.31600384 | 0.68273244 | 0.49477592 | 0.61360507 | NA | NA |
| ENSG000002 | 2.59404162 | -0.255703816 | 0.37472624 | -0.682375 | 0.49500186 | 0.6138665 | NA | NA |
| ENSG000002 | 2.69398575 | 0.255139433 | 0.37393697 | 0.68230598 | 0.49504549 | 0.61390183 | RNU6-240P | 106479673 |
| ENSG000002 | 3.74605577 | -0.257680591 | 0.3776788 | -0.6822744 | 0.49506543 | 0.61390778 | NA | NA |
| ENSG000002 | 4.94030761 | -0.256182951 | 0.37552859 | -0.6821929 | 0.49511696 | 0.6139529 | NA | NA |
| ENSG000001 | 21.0022176 | 0.192941501 | 0.28284906 | 0.68213591 | 0.49515302 | 0.61397884 | ORAI3 | 93129 |
| ENSG000001 | 85.7974515 | 0.123121012 | 0.18051214 | 0.682065 | 0.49519785 | 0.61399688 | TIMP3 | 7078 |
| ENSG000001 | 10.8896684 | -0.225959181 | 0.33127598 | -0.6820874 | 0.49518367 | 0.61399688 | TCAF2 | 285966 |
| ENSG000002 | 14.6368412 | 0.223885567 | 0.32827191 | 0.68201257 | 0.495231 | 0.61401921 | NA | NA |
| ENSG000002 | 2.63047154 | -0.254450104 | 0.3731671 | -0.6818664 | 0.49532344 | 0.61411504 | NA | NA |
| ENSG000002 | 2.17912084 | -0.246945199 | 0.36228449 | -0.6816334 | 0.49547081 | 0.61427898 | NA | NA |
| ENSG000002 | 24.8568406 | -0.177067081 | 0.25985747 | -0.6814008 | 0.49561793 | 0.61444259 | NA | NA |
| ENSG000002 | 7.16465893 | -0.249250999 | 0.36584474 | -0.6813027 | 0.49567999 | 0.61450075 | DPYD-AS1 | 100873932 |

| | | | | | | | | |
|---|---|---|---|---|---|---|---|---|
| ENSG000002 | 5.44104646 | -0.252972842 | 0.3713254 | -0.68127 | 0.49570068 | 0.61450761 | NA | NA |
| ENSG000002 | 5.50551613 | -0.255119935 | 0.37464299 | -0.6809681 | 0.49589166 | 0.61472557 | NA | NA |
| ENSG000002 | 9.80267327 | -0.230138694 | 0.3380054 | -0.6808728 | 0.49595196 | 0.61478152 | NA | NA |
| ENSG000002 | 6.39312206 | -0.248295762 | 0.36470813 | -0.6808068 | 0.49599377 | 0.61481455 | NA | NA |
| ENSG000001 | 124.859996 | -0.102850274 | 0.15112241 | -0.6805759 | 0.49613987 | 0.61497686 | PARD3B | 117583 |
| ENSG000001 | 12.5842376 | -0.223356622 | 0.32820015 | -0.68055 | 0.49615625 | 0.61497837 | C1orf228 | 339541 |
| ENSG000002 | 4.10696293 | -0.257583685 | 0.37857563 | -0.6804022 | 0.49624984 | 0.61507556 | RN7SL414P | 106481036 |
| ENSG000002 | 5.13147456 | -0.255772591 | 0.37597158 | -0.6802977 | 0.496316 | 0.61510116 | NA | NA |
| ENSG000002 | 2.39170952 | -0.251711141 | 0.36998988 | -0.680319 | 0.4963025 | 0.61510116 | NA | NA |
| ENSG000002 | 6.40507399 | -0.247425531 | 0.36369561 | -0.6803094 | 0.4963086 | 0.61510116 | NA | NA |
| ENSG000002 | 8.88233969 | -0.238672151 | 0.35086995 | -0.6802297 | 0.49635905 | 0.61511692 | NA | NA |
| ENSG000002 | 2.63702201 | -0.254070059 | 0.37349519 | -0.6802499 | 0.49634626 | 0.61511692 | NA | NA |
| ENSG000002 | 14.3240193 | 0.214095835 | 0.31476849 | 0.6801692 | 0.49639733 | 0.61514556 | NA | NA |
| ENSG000002 | 3.91706735 | -0.257444985 | 0.3785758 | -0.6800355 | 0.49648198 | 0.61520021 | IGSF23 | 147710 |
| ENSG000002 | 3.65753103 | -0.257131283 | 0.37809691 | -0.6800671 | 0.49646195 | 0.61520021 | NA | NA |
| ENSG000002 | 2.36191385 | -0.251171291 | 0.36935452 | -0.6800277 | 0.49648695 | 0.61520021 | NA | NA |
| ENSG000002 | 15.197583 | 0.207133365 | 0.30462448 | 0.67996296 | 0.49652792 | 0.61523218 | NA | NA |
| ENSG000001 | 30.096931 | 0.165000848 | 0.24270391 | 0.6798442 | 0.49660312 | 0.61530656 | POLR3GL | 84265 |
| ENSG000002 | 5.63839314 | -0.252513094 | 0.37147253 | -0.6797625 | 0.49665486 | 0.61535187 | NA | NA |
| ENSG000001 | 17.3235976 | 0.202453771 | 0.29785616 | 0.67970316 | 0.49669244 | 0.61536446 | KCNA3 | 3738 |
| ENSG000002 | 2.36284694 | -0.251042513 | 0.36935573 | -0.6796768 | 0.49670915 | 0.61536446 | LINC01445 | 102723605 |
| ENSG000002 | 2.8637409 | -0.256138379 | 0.37685445 | -0.6796746 | 0.49671055 | 0.61536446 | NA | NA |
| ENSG000002 | 4.32677109 | 0.257107569 | 0.37831913 | 0.67960499 | 0.49675461 | 0.61540024 | NA | NA |
| ENSG000002 | 2.38705713 | -0.250495002 | 0.36866114 | -0.6794722 | 0.4968387 | 0.61548562 | NA | NA |
| ENSG000002 | 8.52846944 | -0.235403837 | 0.34654796 | -0.6792821 | 0.49695911 | 0.61558456 | C6orf25 | 80739 |
| ENSG000002 | 17.2876264 | -0.198879394 | 0.2927704 | -0.6793016 | 0.49694679 | 0.61558456 | NA | NA |
| ENSG000002 | 16.9678433 | 0.197836809 | 0.29124733 | 0.67927424 | 0.49696411 | 0.61558456 | NA | NA |
| ENSG000002 | 5.3665988 | -0.253871873 | 0.37376421 | -0.67923 | 0.49699213 | 0.61560046 | NA | NA |
| ENSG000002 | 8.92533171 | 0.235979135 | 0.34745485 | 0.6791649 | 0.49703339 | 0.61563276 | NA | NA |

| | | | | | | | | |
|---|---|---|---|---|---|---|---|---|
| ENSG000001 | 2.41586764 | -0.250710711 | 0.3691925 | -0.6790786 | 0.49708809 | 0.6156441 | OR5M8 | 219484 |
| ENSG000002 | 2.59918935 | -0.254625611 | 0.37493711 | -0.6791155 | 0.49706466 | 0.6156441 | LINC01063 | 101929769 |
| ENSG000002 | 12.0045227 | -0.218424405 | 0.3216399 | -0.6790961 | 0.49707696 | 0.6156441 | NA | NA |
| ENSG000002 | 5.59230029 | -0.252775945 | 0.37243353 | -0.6787143 | 0.49731893 | 0.61591119 | NA | NA |
| ENSG000001 | 10.314536 | -0.234389142 | 0.34543677 | -0.6785298 | 0.49743584 | 0.61603008 | NA | NA |
| ENSG000002 | 5.66304218 | -0.250380824 | 0.36902607 | -0.6784909 | 0.49746051 | 0.61603008 | NA | NA |
| ENSG000002 | 5.70763253 | -0.253655161 | 0.37384353 | -0.6785062 | 0.49745079 | 0.61603008 | NA | NA |
| ENSG000000 | 31.8492259 | 0.172575932 | 0.25440945 | 0.67833933 | 0.49755657 | 0.61611278 | DERL3 | 91319 |
| ENSG000001 | 10.5130771 | -0.226628162 | 0.33409348 | -0.6783376 | 0.49755768 | 0.61611278 | HS3ST2 | 9956 |
| ENSG000001 | 11.6910667 | -0.22743796 | 0.33533988 | -0.6782312 | 0.49762513 | 0.61615867 | SPEF1 | 25876 |
| ENSG000002 | 6.68454904 | -0.251304084 | 0.37052628 | -0.6782355 | 0.49762237 | 0.61615867 | NA | NA |
| ENSG000002 | 3.72188155 | 0.256704723 | 0.37852284 | 0.67817498 | 0.49766075 | 0.61618397 | NA | NA |
| ENSG000002 | 3.70176201 | 0.256724562 | 0.37856918 | 0.67814438 | 0.49768015 | 0.61618918 | NA | NA |
| ENSG000001 | 39.6250023 | -0.153686448 | 0.22670243 | -0.6779215 | 0.49782147 | 0.61634533 | RNF32 | 140545 |
| ENSG000001 | 39.7135967 | 0.152966151 | 0.22566216 | 0.67785468 | 0.49786384 | 0.61636751 | ZNF862 | 643641 |
| ENSG000001 | 105.937538 | 0.117519688 | 0.1733847 | 0.67779735 | 0.49790019 | 0.61636751 | RSPH3 | 83861 |
| ENSG000002 | 9.08531867 | 0.239233145 | 0.35293952 | 0.67783042 | 0.49787922 | 0.61636751 | NA | NA |
| ENSG000002 | 7.08843532 | -0.243483375 | 0.35922304 | -0.6778056 | 0.49789499 | 0.61636751 | NA | NA |
| ENSG000002 | 2.63349195 | -0.252322935 | 0.37238958 | -0.6775779 | 0.49803938 | 0.61652101 | NA | NA |
| ENSG000002 | 2.34773133 | -0.252061305 | 0.3720284 | -0.6775324 | 0.4980682 | 0.61653786 | NA | NA |
| ENSG000001 | 47.4449889 | 0.144375281 | 0.21312587 | 0.67741792 | 0.49814083 | 0.61660894 | MBD6 | 114785 |
| ENSG000002 | 4.67525276 | -0.255594309 | 0.37736168 | -0.6773192 | 0.49820345 | 0.61666763 | NA | NA |
| ENSG000001 | 3.73497047 | -0.255200533 | 0.37680684 | -0.6772715 | 0.49823371 | 0.61668625 | C19orf80 | 55908 |
| ENSG000002 | 6.01334465 | 0.249414978 | 0.36836761 | 0.67708173 | 0.4983541 | 0.61679762 | MIR5687 | 100847019 |
| ENSG000002 | 6.26085712 | 0.246455256 | 0.36398643 | 0.67710011 | 0.49834244 | 0.61679762 | NA | NA |
| ENSG000002 | 2.86166315 | 0.254905007 | 0.37651781 | 0.67700651 | 0.49840183 | 0.61683786 | NA | NA |
| ENSG000002 | 4.75064952 | -0.255512907 | 0.37758225 | -0.6767079 | 0.49859127 | 0.61705349 | NA | NA |
| ENSG000001 | 38.7239828 | -0.167625092 | 0.24780196 | -0.6764478 | 0.49875637 | 0.61718309 | KEL | 3792 |
| ENSG000002 | 4.46077285 | -0.255784983 | 0.37809803 | -0.6765044 | 0.49872045 | 0.61718309 | NA | NA |

| | | | | | | | | |
|---|---|---|---|---|---|---|---|---|
| ENSG000002 | 5.37039542 | 0.251787435 | 0.37222048 | 0.67644703 | 0.49875687 | 0.61718309 | NA | NA |
| ENSG000002 | 9.91341356 | -0.228957979 | 0.33844989 | -0.67649 | 0.49872957 | 0.61718309 | NA | NA |
| ENSG000002 | 47.0999226 | 0.143336117 | 0.21191473 | 0.67638583 | 0.49879571 | 0.61721232 | CD27-AS1 | 678655 |
| ENSG000001 | 24.9558448 | -0.175287099 | 0.25925908 | -0.6761079 | 0.49897217 | 0.61741183 | CBR3 | 874 |
| ENSG000001 | 20.2301473 | -0.186167901 | 0.27537594 | -0.67605 | 0.49900891 | 0.61741961 | GGACT | 87769 |
| ENSG000002 | 15.9122324 | -0.201220501 | 0.29763652 | -0.6760612 | 0.49900179 | 0.61741961 | SNORA19 | 641451 |
| ENSG000002 | 8.40907333 | -0.242892991 | 0.35933256 | -0.675956 | 0.49906859 | 0.61747462 | CASC18 | 101929110 |
| ENSG000002 | 2.86576407 | -0.254865245 | 0.37710248 | -0.6758514 | 0.49913499 | 0.61753792 | NA | NA |
| ENSG000001 | 224.330249 | 0.095324443 | 0.14105436 | 0.67579932 | 0.49916806 | 0.61754116 | RPGR | 6103 |
| ENSG000002 | 4.48787982 | -0.255585802 | 0.37819009 | -0.6758131 | 0.49915934 | 0.61754116 | NA | NA |
| ENSG000002 | 2.853016 | -0.254754808 | 0.37702097 | -0.6757046 | 0.49922821 | 0.61759673 | NA | NA |
| ENSG000001 | 7.32139695 | -0.243463167 | 0.36033234 | -0.6756628 | 0.49925475 | 0.61761072 | TEX38 | 374973 |
| ENSG000001 | 362.495301 | 0.088839238 | 0.1315435 | 0.67536014 | 0.49944698 | 0.61775429 | ZNF808 | 388558 |
| ENSG000002 | 6.38563926 | -0.246625848 | 0.36516198 | -0.6753875 | 0.49942958 | 0.61775429 | NA | NA |
| ENSG000002 | 81.0658502 | 0.121103266 | 0.17931241 | 0.67537582 | 0.49943702 | 0.61775429 | PTPRG-AS1 | 100506994 |
| ENSG000002 | 2.59446617 | -0.253248905 | 0.37498129 | -0.6753641 | 0.49944446 | 0.61775429 | NA | NA |
| ENSG000002 | 33.1770114 | 0.1644362 | 0.24345921 | 0.67541581 | 0.49941162 | 0.61775429 | NA | NA |
| ENSG000001 | 14.2812585 | -0.214948981 | 0.31832265 | -0.6752551 | 0.49951373 | 0.61781801 | OTOP1 | 133060 |
| ENSG000001 | 22.2810068 | -0.188468801 | 0.27914894 | -0.675155 | 0.4995773 | 0.61787779 | GJC3 | 349149 |
| ENSG000002 | 2.19494141 | -0.241576147 | 0.35782348 | -0.6751266 | 0.49959533 | 0.61788125 | NA | NA |
| ENSG000002 | 7.6360549 | -0.241790903 | 0.3581862 | -0.6750425 | 0.49964878 | 0.6179285 | NA | NA |
| ENSG000002 | 25.4829203 | 0.172813132 | 0.25605842 | 0.67489728 | 0.49974103 | 0.6180049 | HSD17B8 | 7923 |
| ENSG000002 | 8.34408577 | -0.235053328 | 0.34827086 | -0.6749153 | 0.49972959 | 0.6180049 | NA | NA |
| ENSG000002 | 2.60059154 | -0.251553843 | 0.37277056 | -0.6748222 | 0.49978875 | 0.61804506 | MED15P8 | 106480229 |
| ENSG000002 | 13.7341272 | 0.207580767 | 0.30762645 | 0.67478191 | 0.49981434 | 0.61805785 | LOC102724623 | 102724623 |
| ENSG000002 | 2.37507907 | -0.249240316 | 0.36938554 | -0.674743 | 0.49983904 | 0.61806956 | NA | NA |
| ENSG000002 | 8.79559346 | -0.231607084 | 0.34346769 | -0.6743199 | 0.50010799 | 0.61834555 | NA | NA |
| ENSG000002 | 8.14783918 | -0.237245111 | 0.35181605 | -0.6743442 | 0.50009251 | 0.61834555 | NA | NA |
| ENSG000002 | 2.35634908 | -0.24980188 | 0.37043699 | -0.6743438 | 0.50009278 | 0.61834555 | NA | NA |

| | | | | | | | | |
|---|---|---|---|---|---|---|---|---|
| ENSG000001 | 15.0375178 | -0.214153244 | 0.31772148 | -0.6740282 | 0.50029337 | 0.6185559 | SHBG | 6462 |
| ENSG00000C | 12.3801984 | 0.217275888 | 0.32243358 | 0.67386247 | 0.50039875 | 0.61864847 | MRPS24 | 64951 |
| ENSG000001 | 26.7062862 | -0.176197755 | 0.26146882 | -0.6738767 | 0.50038968 | 0.61864847 | PROK2 | 60675 |
| ENSG000002 | 2.37633277 | -0.250115463 | 0.37120565 | -0.6737922 | 0.50044345 | 0.61868487 | NA | NA |
| ENSG000002 | 8.82319762 | 0.23585455 | 0.3501027 | 0.67367247 | 0.50051957 | 0.61876011 | POLR2J2 | 246721 |
| ENSG000002 | 2.34938202 | -0.250232321 | 0.37149906 | -0.6735746 | 0.50058183 | 0.61881821 | RN7SL568P | 106479429 |
| ENSG000002 | 2.63537132 | -0.251186053 | 0.37296278 | -0.6734883 | 0.50063672 | 0.6188672 | NA | NA |
| ENSG000002 | 4.39084116 | -0.25471239 | 0.37822334 | -0.6734444 | 0.50066461 | 0.61888281 | LOC102724763 | 102724763 |
| ENSG000002 | 5.42754136 | -0.25098712 | 0.3727329 | -0.6733699 | 0.50071198 | 0.6189225 | LOC105374436 | 105374436 |
| ENSG000002 | 2.34547927 | -0.248151264 | 0.36854219 | -0.673332 | 0.50073608 | 0.61893342 | RNU4-60P | 106479585 |
| ENSG000002 | 2.3427274 | -0.249733042 | 0.37090846 | -0.673301 | 0.50075581 | 0.61893894 | RN7SL201P | 106481830 |
| ENSG000001 | 2.57915413 | -0.252677368 | 0.37532649 | -0.6732202 | 0.50080722 | 0.61898362 | OR8K1 | 390157 |
| ENSG000002 | 9.59181686 | 0.226619062 | 0.33666263 | 0.673134 | 0.50086205 | 0.61903252 | NA | NA |
| ENSG000001 | 7.17658902 | -0.242045045 | 0.35965411 | -0.672994 | 0.50095113 | 0.61912375 | C20orf173 | 140873 |
| ENSG000001 | 100.457851 | -0.113434744 | 0.16855943 | -0.6729659 | 0.500969 | 0.61912697 | DYRK3 | 8444 |
| ENSG000001 | 44.281815 | 0.144741243 | 0.21509872 | 0.67290611 | 0.50100702 | 0.61915509 | ACRC | 93953 |
| ENSG000002 | 3.88566586 | -0.254691095 | 0.37858803 | -0.6727394 | 0.50111308 | 0.61926728 | NA | NA |
| ENSG00000C | 43.5125354 | -0.151053242 | 0.2245748 | -0.6726188 | 0.50118982 | 0.61934058 | SYN1 | 6853 |
| ENSG000002 | 2.38034358 | -0.247947937 | 0.36864198 | -0.6725982 | 0.50120294 | 0.61934058 | NA | NA |
| ENSG000002 | 6.63598324 | -0.243203072 | 0.36167593 | -0.6724337 | 0.50130766 | 0.6194511 | LOC101928424 | 101928424 |
| ENSG000002 | 2.37334405 | -0.248905562 | 0.37020958 | -0.6723369 | 0.50136924 | 0.61950832 | RN7SL296P | 106481003 |
| ENSG000002 | 10.2917498 | -0.238480215 | 0.35482068 | -0.6721148 | 0.50151064 | 0.61966416 | NA | NA |
| ENSG000001 | 67.3956037 | -0.134452581 | 0.20005192 | -0.6720884 | 0.50152739 | 0.61966597 | SUGCT | 79783 |
| ENSG000001 | 10.1170885 | -0.227244078 | 0.33820015 | -0.6719219 | 0.50163344 | 0.61967944 | NEUROD1 | 4760 |
| ENSG000001 | 26.8297625 | 0.172531406 | 0.25676396 | 0.67194558 | 0.50161834 | 0.61967944 | SFN | 2810 |
| ENSG000002 | 9.31289757 | 0.229184494 | 0.34109165 | 0.6719147 | 0.501638 | 0.61967944 | MTCP1 | 4515 |
| ENSG000002 | 2.88821999 | -0.253189011 | 0.37682359 | -0.6719033 | 0.50164526 | 0.61967944 | NA | NA |
| ENSG000002 | 2.38916463 | -0.247196988 | 0.36789384 | -0.6719248 | 0.50163158 | 0.61967944 | NA | NA |
| ENSG000002 | 7.11541836 | 0.242129339 | 0.36036213 | 0.67190561 | 0.50164379 | 0.61967944 | LOC101928530 | 101928530 |

| | | | | | | | | |
|---|---|---|---|---|---|---|---|---|
| ENSG000002 | 5.67784568 | -0.249109744 | 0.37072347 | -0.6719557 | 0.50161191 | 0.61967944 | LOC100507534 | 100507534 |
| ENSG000002 | 6.43221995 | -0.247877918 | 0.36896907 | -0.6718122 | 0.50170327 | 0.61969662 | NA | NA |
| ENSG000002 | 2.89795919 | -0.251957337 | 0.37504286 | -0.6718094 | 0.50170501 | 0.61969662 | NA | NA |
| ENSG000002 | 2.66405675 | -0.248395193 | 0.36972835 | -0.6718316 | 0.5016909 | 0.61969662 | PKD1P6 | 353511 |
| ENSG000002 | 6.87481683 | -0.244204744 | 0.36352347 | -0.6717716 | 0.50172911 | 0.61970751 | NA | NA |
| ENSG000001 | 6.65351954 | -0.243025066 | 0.36179125 | -0.6717273 | 0.50175731 | 0.61972346 | C7orf72 | 100130988 |
| ENSG000002 | 6.29360851 | 0.244377549 | 0.3638782 | 0.67159161 | 0.50184372 | 0.61981131 | NA | NA |
| ENSG000001 | 20.7579856 | -0.193995632 | 0.28902838 | -0.6711992 | 0.50209361 | 0.62010105 | CCDC17 | 149483 |
| ENSG000002 | 5.87448153 | -0.248026415 | 0.36962855 | -0.6710153 | 0.50221078 | 0.62022687 | NA | NA |
| ENSG000002 | 2.38580397 | -0.246811553 | 0.3678829 | -0.6708971 | 0.50228611 | 0.62030101 | NA | NA |
| ENSG000002 | 5.64566266 | -0.24797335 | 0.36965578 | -0.6708223 | 0.50233373 | 0.62034093 | NA | NA |
| ENSG000002 | 13.9033453 | -0.209679596 | 0.31259578 | -0.6707691 | 0.50236762 | 0.62036388 | LOC440028 | 440028 |
| ENSG000001 | 175.07758 | 0.107039561 | 0.15960947 | 0.67063413 | 0.50245363 | 0.62045121 | REL | 5966 |
| ENSG000002 | 4.41324421 | -0.253584797 | 0.37817773 | -0.670544 | 0.50251108 | 0.62046547 | NA | NA |
| ENSG000002 | 5.15756034 | -0.251757498 | 0.37545059 | -0.6705476 | 0.50250876 | 0.62046547 | NA | NA |
| ENSG000002 | 5.24641491 | 0.249311208 | 0.37178378 | 0.67058118 | 0.50248737 | 0.62046547 | NA | NA |
| ENSG000002 | 2.9026926 | 0.252482082 | 0.37658334 | 0.67045474 | 0.50256795 | 0.62051679 | RN7SKP248 | 106480476 |
| ENSG000002 | 8.74588038 | 0.24369104 | 0.36351721 | 0.67037002 | 0.50262194 | 0.62054566 | NA | NA |
| ENSG000002 | 2.59848439 | -0.250376654 | 0.37348969 | -0.670371 | 0.50262132 | 0.62054566 | NA | NA |
| ENSG000002 | 12.9657851 | -0.21378092 | 0.31899719 | -0.6701655 | 0.50275228 | 0.62068769 | NA | NA |
| ENSG00000C | 198.739952 | 0.098930688 | 0.14765053 | 0.67003275 | 0.50283691 | 0.62074911 | MTMR7 | 9108 |
| ENSG000002 | 2.33933429 | -0.248505679 | 0.37089546 | -0.6700154 | 0.50284796 | 0.62074911 | NA | NA |
| ENSG000002 | 2.5778161 | -0.250894084 | 0.37443487 | -0.6700607 | 0.50281911 | 0.62074911 | RASSF1-AS1 | 102060282 |
| ENSG000002 | 2.32021201 | -0.247749076 | 0.36979087 | -0.6699708 | 0.50287643 | 0.62076536 | RN7SL135P | 106480958 |
| ENSG000001 | 54.598319 | 0.139944777 | 0.20894244 | 0.6697767 | 0.50300015 | 0.62089918 | CCDC88B | 283234 |
| ENSG000002 | 2.33799626 | -0.247666134 | 0.36983773 | -0.6696616 | 0.50307352 | 0.62097084 | NA | NA |
| ENSG000002 | 2.59256719 | -0.25008456 | 0.37350175 | -0.6695673 | 0.50313366 | 0.62102618 | NA | NA |
| ENSG000002 | 8.1678159 | -0.236558704 | 0.35339993 | -0.6693796 | 0.50325336 | 0.62115502 | NA | NA |
| ENSG000001 | 27.2536253 | -0.171314564 | 0.25596367 | -0.6692925 | 0.50330891 | 0.62120467 | SEC14L2 | 23541 |

| | | | | | | | | |
|---|---|---|---|---|---|---|---|---|
| ENSG000002 | 2.57910152 | -0.249922406 | 0.37346612 | -0.669197 | 0.50336982 | 0.62126094 | NA | NA |
| ENSG000002 | 3.49434525 | 0.253143469 | 0.37830278 | 0.66915572 | 0.50339615 | 0.62127453 | NA | NA |
| ENSG000002 | 11.5679867 | -0.220693042 | 0.32983584 | -0.6690996 | 0.50343191 | 0.62129976 | NA | NA |
| ENSG000001 | 11.390429 | -0.226958146 | 0.33923148 | -0.6690362 | 0.50347237 | 0.62131187 | CD8A | 925 |
| ENSG000002 | 3.90251159 | -0.253270883 | 0.37854823 | -0.6690584 | 0.5034582 | 0.62131187 | TAS2R41 | 259287 |
| ENSG000002 | 8.61920711 | -0.231519164 | 0.34607253 | -0.6689903 | 0.50350167 | 0.62132912 | NA | NA |
| ENSG000001 | 57.3303346 | -0.13271263 | 0.19839118 | -0.6689442 | 0.50353108 | 0.6213465 | XDH | 7498 |
| ENSG000002 | 4.24501994 | -0.253128741 | 0.37850084 | -0.6687667 | 0.50364434 | 0.62146735 | NA | NA |
| ENSG00000C | 57.1731891 | 0.134537329 | 0.20118936 | 0.66870997 | 0.50368051 | 0.62147416 | FKBP7 | 51661 |
| ENSG000002 | 17.1828729 | 0.203957977 | 0.30499523 | 0.66872514 | 0.50367083 | 0.62147416 | NA | NA |
| ENSG000002 | 2.62306761 | -0.250063141 | 0.3740552 | -0.6685194 | 0.50380213 | 0.6215864 | NA | NA |
| ENSG000002 | 7.88144016 | -0.237133295 | 0.35470809 | -0.6685308 | 0.5037948 | 0.6215864 | NA | NA |
| ENSG000002 | 4.55425576 | 0.252609701 | 0.37796645 | 0.66833896 | 0.50391725 | 0.62170952 | NA | NA |
| ENSG000002 | 3.11112863 | -0.25244694 | 0.37783131 | -0.6681472 | 0.50403962 | 0.62184157 | NA | NA |
| ENSG000001 | 14.7594428 | -0.210954757 | 0.31581774 | -0.6679636 | 0.50415682 | 0.62196724 | SCOC-AS1 | 100129858 |
| ENSG000002 | 4.43198271 | -0.252315399 | 0.37780502 | -0.6678455 | 0.50423219 | 0.62202238 | NA | NA |
| ENSG000002 | 2.89988015 | 0.251723117 | 0.37690937 | 0.66786113 | 0.50422224 | 0.62202238 | NA | NA |
| ENSG000002 | 27.0620181 | 0.169002934 | 0.25316309 | 0.66756547 | 0.50441101 | 0.62222404 | TNKS2-AS1 | 100507633 |
| ENSG000002 | 2.61439038 | -0.249817783 | 0.37433252 | -0.6673686 | 0.5045367 | 0.62236015 | NA | NA |
| ENSG000002 | 2.38401689 | -0.245985093 | 0.36864703 | -0.6672645 | 0.50460317 | 0.62242322 | NA | NA |
| ENSG000002 | 11.3919956 | 0.218729934 | 0.32786609 | 0.66713192 | 0.50468787 | 0.62250876 | NA | NA |
| ENSG000002 | 5.44334665 | 0.246741948 | 0.36996016 | 0.66694194 | 0.50480922 | 0.62263949 | NA | NA |
| ENSG000002 | 4.42191592 | -0.251680159 | 0.37741775 | -0.6668477 | 0.50486942 | 0.6226948 | NA | NA |
| ENSG000002 | 3.1746295 | -0.248866287 | 0.37325342 | -0.6667488 | 0.50493258 | 0.62275376 | NA | NA |
| ENSG000002 | 2.64346756 | -0.247782476 | 0.37170264 | -0.6666148 | 0.50501822 | 0.6228215 | NA | NA |
| ENSG000002 | 3.12362821 | -0.251324554 | 0.37701157 | -0.6666229 | 0.50501302 | 0.6228215 | NA | NA |
| ENSG000002 | 11.1180674 | 0.221707469 | 0.33264849 | 0.66649173 | 0.50509685 | 0.62289953 | NA | NA |
| ENSG000002 | 3.11255098 | -0.251766586 | 0.37780067 | -0.6664006 | 0.50515509 | 0.62295241 | LOC105372429 | 105372429 |
| ENSG000001 | 7.72298951 | -0.239022932 | 0.3587466 | -0.6662723 | 0.50523705 | 0.62301183 | VPS37D | 155382 |

| | | | | | | | | |
|---|---|---|---|---|---|---|---|---|
| ENSG000002 | 3.76094014 | 0.252093329 | 0.3783744 | 0.66625366 | 0.50524898 | 0.62301183 | NA | NA |
| ENSG000002 | 2.3948539 | -0.245774232 | 0.36889021 | -0.6662531 | 0.50524936 | 0.62301183 | NA | NA |
| ENSG000002 | 9.56346187 | -0.227856008 | 0.34217814 | -0.6658988 | 0.50547579 | 0.62327208 | LINC01142 | 284688 |
| ENSG000002 | 3.75956985 | 0.251866577 | 0.37837397 | 0.66565513 | 0.50563156 | 0.62344519 | MIR186 | 406962 |
| ENSG000000 | 9.51503208 | -0.229018427 | 0.34406692 | -0.6656218 | 0.5056529 | 0.62345254 | RUNX3 | 864 |
| ENSG000002 | 11.8547929 | -0.221105331 | 0.33230366 | -0.6653713 | 0.50581301 | 0.62358409 | VAC14-AS1 | 100130894 |
| ENSG000002 | 4.16514022 | -0.251822708 | 0.3784903 | -0.6653346 | 0.50583648 | 0.62358409 | NA | NA |
| ENSG000002 | 2.3453955 | -0.24675241 | 0.37086757 | -0.6653383 | 0.50583412 | 0.62358409 | NA | NA |
| ENSG000002 | 6.90450204 | -0.240423298 | 0.36132605 | -0.6653915 | 0.50580009 | 0.62358409 | LOC641367 | 641367 |
| ENSG000002 | 2.8950721 | 0.250872841 | 0.37704357 | 0.6653683 | 0.50581495 | 0.62358409 | NA | NA |
| ENSG000000 | 74.5399445 | 0.145282825 | 0.21838936 | 0.6652468 | 0.50589265 | 0.62361548 | DTNBP1 | 84062 |
| ENSG000002 | 2.58433378 | -0.24845104 | 0.37347202 | -0.6652467 | 0.5058927 | 0.62361548 | NA | NA |
| ENSG000001 | 53.9355444 | -0.137661964 | 0.20695379 | -0.6651821 | 0.505934 | 0.62364744 | KCTD21 | 283219 |
| ENSG000001 | 52.03113 | 0.140754199 | 0.2116171 | 0.66513623 | 0.50596336 | 0.62366467 | GAS2L1 | 10634 |
| ENSG000001 | 2.60821259 | -0.249378231 | 0.37499785 | -0.6650124 | 0.50604254 | 0.62372435 | LOC102724659 | 102724659 |
| ENSG000002 | 16.4078403 | -0.210097204 | 0.31592963 | -0.6650127 | 0.50604239 | 0.62372435 | NA | NA |
| ENSG000002 | 3.95448297 | -0.251691663 | 0.37857775 | -0.6648348 | 0.50615616 | 0.62384543 | NA | NA |
| ENSG000002 | 2.36304336 | -0.245913795 | 0.36990854 | -0.6647962 | 0.50618085 | 0.62385689 | NA | NA |
| ENSG000002 | 3.67544058 | -0.251357871 | 0.37813778 | -0.6647256 | 0.50622601 | 0.62387463 | NA | NA |
| ENSG000002 | 2.32027149 | -0.246001144 | 0.37006746 | -0.6647467 | 0.50621249 | 0.62387463 | NA | NA |
| ENSG000002 | 2.34227094 | -0.246645919 | 0.37111877 | -0.664601 | 0.50630571 | 0.62395389 | NA | NA |
| ENSG000001 | 18.5658845 | -0.192549925 | 0.28984103 | -0.6643294 | 0.50647949 | 0.62414909 | BRINP2 | 57795 |
| ENSG000001 | 87.884566 | -0.121505357 | 0.18293231 | -0.6642094 | 0.50655631 | 0.62419404 | CNIH3 | 149111 |
| ENSG000001 | 6.94713587 | -0.242405764 | 0.36495885 | -0.6642003 | 0.50656215 | 0.62419404 | C19orf45 | 374877 |
| ENSG000002 | 23.1618167 | -0.178261446 | 0.2683752 | -0.6642247 | 0.50654648 | 0.62419404 | NA | NA |
| ENSG000002 | 2.34132466 | -0.245716492 | 0.3700134 | -0.6640746 | 0.50664258 | 0.62427418 | NA | NA |
| ENSG000002 | 2.86736214 | -0.249035324 | 0.37502967 | -0.6640417 | 0.50666365 | 0.62428117 | NA | NA |
| ENSG000002 | 55.6321423 | -0.144682146 | 0.21788981 | -0.6640152 | 0.50668058 | 0.62428306 | LOC101928978 | 101928978 |
| ENSG000002 | 2.34392163 | -0.246111724 | 0.37086911 | -0.663608 | 0.50694121 | 0.62449034 | RNU6-302P | 106479701 |

| | | | | | | | | |
|---|---|---|---|---|---|---|---|---|
| ENSG000002 | 4.50666177 | -0.251222708 | 0.37856501 | -0.6636184 | 0.50693457 | 0.62449034 | LOC105377763 | 105377763 |
| ENSG000002 | 2.40872376 | -0.243079354 | 0.3662484 | -0.6637008 | 0.50688183 | 0.62449034 | NA | NA |
| ENSG000002 | 3.35707474 | -0.251245481 | 0.37856767 | -0.6636739 | 0.50689905 | 0.62449034 | NA | NA |
| ENSG000002 | 3.15291535 | -0.250539634 | 0.37753387 | -0.6636216 | 0.50693252 | 0.62449034 | NA | NA |
| ENSG000002 | 10.7721289 | -0.2228069 | 0.33573236 | -0.6636444 | 0.50691793 | 0.62449034 | NA | NA |
| ENSG000002 | 2.85394908 | -0.24972122 | 0.37632981 | -0.6635701 | 0.50696549 | 0.62450088 | LL22NC03-75H1 | 101927722 |
| ENSG000002 | 2.28015574 | 0.245158546 | 0.36946698 | 0.66354657 | 0.50698057 | 0.62450088 | NA | NA |
| ENSG000002 | 2.31918195 | -0.246179571 | 0.37105565 | -0.6634573 | 0.50703773 | 0.62453335 | NA | NA |
| ENSG000002 | 2.58122883 | -0.248000994 | 0.37379265 | -0.6634721 | 0.50702824 | 0.62453335 | NA | NA |
| ENSG000001 | 11.1976823 | 0.223156688 | 0.33640606 | 0.66335515 | 0.50710313 | 0.62459493 | SPATA31E1 | 286234 |
| ENSG000001 | 1090.76563 | 0.06801553 | 0.10254796 | 0.66325578 | 0.50716676 | 0.6246215 | DIEXF | 27042 |
| ENSG000002 | 2.35419007 | -0.246155718 | 0.37114951 | -0.6632252 | 0.50718631 | 0.6246215 | NA | NA |
| ENSG000002 | 2.84410954 | -0.248312668 | 0.37440035 | -0.6632277 | 0.50718476 | 0.6246215 | NA | NA |
| ENSG000002 | 2.38256263 | -0.244526866 | 0.3686686 | -0.6632701 | 0.50715757 | 0.6246215 | NA | NA |
| ENSG000001 | 6.14884701 | -0.243565772 | 0.36737735 | -0.6629853 | 0.50733999 | 0.62479017 | NA | NA |
| ENSG000001 | 59.9638931 | 0.131492351 | 0.19834033 | 0.66296325 | 0.50735409 | 0.62479017 | SP140L | 93349 |
| ENSG000002 | 4.9284411 | -0.249192958 | 0.37593765 | -0.6628571 | 0.50742205 | 0.62485489 | NA | NA |
| ENSG000002 | 2.84197068 | -0.249862385 | 0.37697375 | -0.6628111 | 0.50745155 | 0.62486888 | NA | NA |
| ENSG000002 | 2.59054436 | -0.247384193 | 0.37324599 | -0.6627913 | 0.50746423 | 0.62486888 | NA | NA |
| ENSG000002 | 3.71511504 | 0.250757929 | 0.37858769 | 0.66235099 | 0.5077463 | 0.62518237 | NA | NA |
| ENSG000002 | 2.334951 | -0.245787121 | 0.3710993 | -0.6623217 | 0.50776507 | 0.62518237 | NA | NA |
| ENSG000002 | 2.37997906 | -0.244665931 | 0.36939774 | -0.6623374 | 0.50775503 | 0.62518237 | NA | NA |
| ENSG000002 | 2.41918863 | -0.242991425 | 0.36690409 | -0.662275 | 0.50779497 | 0.6252002 | RNU6-758P | 106481419 |
| ENSG000002 | 74.4536552 | 0.12732133 | 0.1922793 | 0.66216869 | 0.50786312 | 0.62526512 | NA | NA |
| ENSG000001 | 2.63024231 | -0.244405817 | 0.36916616 | -0.6620483 | 0.50794029 | 0.62532401 | OR52B6 | 340980 |
| ENSG000002 | 10.0180148 | -0.22551061 | 0.34062684 | -0.6620459 | 0.50794179 | 0.62532401 | PGCP1 | 441897 |
| ENSG000002 | 13.081866 | -0.219837521 | 0.33218931 | -0.6617838 | 0.50810976 | 0.6255118 | NA | NA |
| ENSG000002 | 10.9609352 | -0.227013775 | 0.34309513 | -0.6616642 | 0.50818644 | 0.62556934 | NA | NA |
| ENSG000002 | 7.88504211 | -0.234423401 | 0.35429436 | -0.6616628 | 0.50818735 | 0.62556934 | NA | NA |

| | | | | | | | | |
|---|---|---|---|---|---|---|---|---|
| ENSG000002 | 12.0980177 | -0.218944776 | 0.33094346 | -0.6615776 | 0.50824197 | 0.62561759 | ST3GAL6-AS1 | 100874207 |
| ENSG000002 | 6.36321036 | -0.242113309 | 0.36606923 | -0.6613867 | 0.50836437 | 0.62574926 | NA | NA |
| ENSG000002 | 2.85435403 | -0.249096451 | 0.37666856 | -0.6613147 | 0.50841053 | 0.62578708 | NA | NA |
| ENSG000001 | 2.86829578 | -0.24874612 | 0.37618674 | -0.6612304 | 0.50846455 | 0.62583457 | IFNL2 | 282616 |
| ENSG000002 | 2.387723 | -0.243239131 | 0.36788319 | -0.6611858 | 0.50849318 | 0.62585082 | MIR5706 | 100847085 |
| ENSG000002 | 2.36053147 | -0.244958618 | 0.37052937 | -0.6611045 | 0.50854532 | 0.625896 | NA | NA |
| ENSG000001 | 9.88462324 | -0.22697645 | 0.34341906 | -0.6609314 | 0.50865628 | 0.62597557 | LIM2 | 3982 |
| ENSG000002 | 10.5365404 | -0.219365157 | 0.33189615 | -0.6609452 | 0.50864748 | 0.62597557 | NA | NA |
| ENSG000002 | 3.37289531 | -0.250015782 | 0.37827396 | -0.6609384 | 0.50865182 | 0.62597557 | NA | NA |
| ENSG000002 | 3.1004558 | -0.24949747 | 0.37780022 | -0.6603953 | 0.50900021 | 0.62636081 | NA | NA |
| ENSG000002 | 4.50527974 | -0.250002544 | 0.37855673 | -0.6604097 | 0.50899094 | 0.62636081 | NA | NA |
| ENSG000002 | 8.61431454 | -0.231102191 | 0.35002016 | -0.660254 | 0.50909087 | 0.62645336 | NA | NA |
| ENSG000002 | 41.8367794 | 0.146015656 | 0.22120803 | 0.66008299 | 0.50920058 | 0.62656934 | FAM83H-AS1 | 100128338 |
| ENSG000002 | 4.1464744 | -0.24984495 | 0.37858373 | -0.6599463 | 0.50928826 | 0.62663921 | TTLL13P | 440307 |
| ENSG000002 | 11.3021176 | -0.215884821 | 0.32712028 | -0.6599555 | 0.5092824 | 0.62663921 | NA | NA |
| ENSG000002 | 2.64223445 | -0.244742493 | 0.37088524 | -0.6598874 | 0.50932609 | 0.62666674 | NA | NA |
| ENSG000002 | 8.45968959 | -0.236490731 | 0.35855097 | -0.6595735 | 0.50952754 | 0.62689557 | NA | NA |
| ENSG000002 | 2.55710261 | -0.246281772 | 0.37344985 | -0.6594775 | 0.50958919 | 0.6269524 | NA | NA |
| ENSG000002 | 2.9332659 | 0.246707195 | 0.37413902 | 0.65939981 | 0.50963907 | 0.62699475 | RN7SL519P | 106479412 |
| ENSG000002 | 13.7529069 | -0.208742581 | 0.31663935 | -0.659244 | 0.50973912 | 0.62709881 | NA | NA |
| ENSG000002 | 2.62448963 | -0.24516933 | 0.37192455 | -0.659191 | 0.5097731 | 0.62710793 | NA | NA |
| ENSG000002 | 11.6336267 | 0.219976298 | 0.33372207 | 0.65916017 | 0.50979292 | 0.62710793 | NA | NA |
| ENSG000002 | 3.12826796 | -0.248858272 | 0.37753421 | -0.6591675 | 0.50978823 | 0.62710793 | NA | NA |
| ENSG000001 | 215.384287 | -0.097912629 | 0.14856498 | -0.6590559 | 0.50985985 | 0.62717123 | CARD8 | 22900 |
| ENSG000002 | 2.81965912 | -0.248430377 | 0.37696557 | -0.6590267 | 0.50987862 | 0.6271753 | NA | NA |
| ENSG000002 | 7.04591191 | 0.240679077 | 0.365262 | 0.65892174 | 0.50994602 | 0.62723918 | LOC101929549 | 101929549 |
| ENSG000001 | 2.8373585 | -0.247966944 | 0.37637342 | -0.6588322 | 0.5100035 | 0.62725279 | HOXD1 | 3231 |
| ENSG000002 | 16.1206624 | -0.20088341 | 0.3049045 | -0.6588404 | 0.50999825 | 0.62725279 | TOB1-AS1 | 400604 |
| ENSG000002 | 3.16309947 | -0.248410257 | 0.37702655 | -0.6588668 | 0.50998134 | 0.62725279 | NA | NA |

| | | | | | | | | |
|---|---|---|---|---|---|---|---|---|
| ENSG000002 | 5.39269457 | 0.245288497 | 0.37237068 | 0.65872129 | 0.51007476 | 0.62732141 | NA | NA |
| ENSG000001 | 12.0387614 | -0.213489878 | 0.32413135 | -0.6586524 | 0.51011904 | 0.62735684 | TSKS | 60385 |
| ENSG000002 | 2.58652614 | -0.246290184 | 0.37400818 | -0.6585155 | 0.51020694 | 0.62744591 | NA | NA |
| ENSG000002 | 24.2365552 | 0.185556641 | 0.28183413 | 0.65838954 | 0.51028785 | 0.62752639 | NA | NA |
| ENSG000001 | 39.9351906 | 0.146648412 | 0.22281461 | 0.65816336 | 0.51043316 | 0.62761562 | CDK5RAP3 | 80279 |
| ENSG000001 | 9.44312899 | -0.234118212 | 0.3557084 | -0.6581745 | 0.51042599 | 0.62761562 | ECM1 | 1893 |
| ENSG000001 | 11.053288 | -0.217553818 | 0.33054829 | -0.6581605 | 0.51043502 | 0.62761562 | GPR22 | 2845 |
| ENSG000002 | 2.5639859 | -0.246601157 | 0.37468488 | -0.6581562 | 0.5104378 | 0.62761562 | NA | NA |
| ENSG000002 | 7.4016426 | -0.240540076 | 0.36543774 | -0.6582245 | 0.51039388 | 0.62761562 | NA | NA |
| ENSG000002 | 3.96120622 | -0.248360015 | 0.37738735 | -0.6581037 | 0.5104715 | 0.62763803 | NA | NA |
| ENSG000002 | 4.62365026 | 0.24735362 | 0.3759444 | 0.65795266 | 0.51056855 | 0.62772361 | NA | NA |
| ENSG000002 | 31.8433873 | -0.166958313 | 0.25375786 | -0.6579434 | 0.51057449 | 0.62772361 | ADAM20P1 | 317760 |
| ENSG000002 | 48.634083 | 0.143928498 | 0.21876189 | 0.6579231 | 0.51058755 | 0.62772361 | RDM1 | 201299 |
| ENSG000002 | 2.59634499 | -0.245121525 | 0.37259457 | -0.6578773 | 0.51061696 | 0.62774074 | NA | NA |
| ENSG000002 | 2.38069645 | -0.241998785 | 0.36786866 | -0.6578402 | 0.51064083 | 0.62775106 | NA | NA |
| ENSG000002 | 2.57906981 | -0.246399633 | 0.37470491 | -0.6575831 | 0.51080603 | 0.6279351 | NA | NA |
| ENSG000001 | 3.10405001 | -0.248214283 | 0.3775014 | -0.6575188 | 0.51084736 | 0.62796688 | CECR5-AS1 | 100130717 |
| ENSG000002 | 3.73511834 | 0.248847605 | 0.37857534 | 0.6573265 | 0.510971 | 0.62808079 | NA | NA |
| ENSG000002 | 18.8012949 | -0.189293968 | 0.28796733 | -0.6573453 | 0.51095892 | 0.62808079 | NA | NA |
| ENSG000001 | 8.86261788 | -0.234234831 | 0.35638659 | -0.6572493 | 0.51102065 | 0.62811846 | NA | NA |
| ENSG000002 | 14.0989457 | 0.206822136 | 0.31468731 | 0.65723063 | 0.51103263 | 0.62811846 | NA | NA |
| ENSG000002 | 3.39352447 | -0.248600872 | 0.37834466 | -0.6570751 | 0.5111326 | 0.62818572 | AGAP2-AS1 | 100130776 |
| ENSG000002 | 12.2030837 | -0.213027483 | 0.32420661 | -0.6570732 | 0.51113383 | 0.62818572 | NA | NA |
| ENSG000002 | 12.0356613 | -0.22934482 | 0.34902136 | -0.6571083 | 0.51111131 | 0.62818572 | MAPT-AS1 | 100128977 |
| ENSG000001 | 27.2495857 | -0.189152486 | 0.28792083 | -0.6569601 | 0.5112066 | 0.62823708 | CAGE1 | 285782 |
| ENSG000002 | 6.28248682 | 0.242298871 | 0.36881205 | 0.65697113 | 0.51119948 | 0.62823708 | NA | NA |
| ENSG000001 | 29.6554072 | 0.171566974 | 0.26121287 | 0.65680905 | 0.5113037 | 0.62831886 | NPAS1 | 4861 |
| ENSG000002 | 2.35596474 | -0.242581738 | 0.36933411 | -0.6568084 | 0.51130414 | 0.62831886 | NA | NA |
| ENSG000002 | 2.28536764 | 0.244021672 | 0.37156056 | 0.65674805 | 0.51134293 | 0.62833027 | LOC102724719 | 102724719 |

| | | | | | | | | |
|---|---|---|---|---|---|---|---|---|
| ENSG000002 | 7.06967291 | 0.240140891 | 0.36566224 | 0.65672871 | 0.51135537 | 0.62833027 | NA | NA |
| ENSG000002 | 13.1803192 | -0.209107752 | 0.31841154 | -0.6567217 | 0.5113599 | 0.62833027 | NA | NA |
| ENSG000002 | 7.10697794 | -0.235850451 | 0.35925431 | -0.6565 | 0.51150247 | 0.6284864 | NA | NA |
| ENSG000002 | 8.31892701 | -0.232303432 | 0.35391492 | -0.6563821 | 0.51157828 | 0.6285605 | RN7SKP247 | 106479205 |
| ENSG000002 | 2.88939351 | -0.244920783 | 0.3731768 | -0.656313 | 0.51162276 | 0.62859611 | NA | NA |
| ENSG000002 | 2.86886937 | -0.246343686 | 0.37535973 | -0.6562869 | 0.51163952 | 0.62859765 | NA | NA |
| ENSG000002 | 39.7537199 | 0.144551967 | 0.22028997 | 0.65618951 | 0.51170219 | 0.62865561 | NA | NA |
| ENSG000002 | 9.05797174 | 0.241990186 | 0.36891978 | 0.65594256 | 0.51186108 | 0.62883176 | NA | NA |
| ENSG000002 | 2.64705906 | 0.241651008 | 0.36846331 | 0.65583465 | 0.51193052 | 0.62889801 | NA | NA |
| ENSG000002 | 4.53863521 | 0.24745677 | 0.37735856 | 0.65576032 | 0.51197835 | 0.62893772 | NA | NA |
| ENSG000002 | 5.50154788 | 0.242961591 | 0.37052241 | 0.65572711 | 0.51199972 | 0.62894492 | RN7SL752P | 106481127 |
| ENSG000002 | 2.35721104 | -0.242761446 | 0.37029537 | -0.6555887 | 0.51208882 | 0.62903531 | SNORA77 | 677843 |
| ENSG000002 | 2.59546452 | -0.245238053 | 0.37410868 | -0.6555262 | 0.512129 | 0.62906562 | NA | NA |
| ENSG000002 | 2.60881232 | -0.243379471 | 0.37130078 | -0.6554779 | 0.51216009 | 0.62908474 | NA | NA |
| ENSG000002 | 2.33022782 | -0.242971013 | 0.37083079 | -0.6552072 | 0.51233434 | 0.62927971 | LOC101927123 | 101927123 |
| ENSG000002 | 3.09183804 | -0.247639667 | 0.37799143 | -0.6551463 | 0.51237359 | 0.62930887 | NA | NA |
| ENSG000001 | 4.63004319 | 0.245454604 | 0.37495983 | 0.65461574 | 0.51271519 | 0.62967439 | GPR139 | 124274 |
| ENSG000002 | 2.57622545 | -0.244896327 | 0.37408418 | -0.6546557 | 0.51268948 | 0.62967439 | NA | NA |
| ENSG000002 | 2.87017514 | -0.246354881 | 0.37633742 | -0.6546117 | 0.51271778 | 0.62967439 | NA | NA |
| ENSG000002 | 23.406038 | 0.178456903 | 0.27264195 | 0.65454676 | 0.51275961 | 0.6297067 | GOLGA6L10 | 647042 |
| ENSG000002 | 2.92500933 | -0.237272451 | 0.36257703 | -0.6544056 | 0.51285051 | 0.62979548 | NA | NA |
| ENSG000002 | 27.8068659 | -0.166890408 | 0.25503348 | -0.6543863 | 0.51286297 | 0.62979548 | NA | NA |
| ENSG000002 | 2.8671021 | -0.246241408 | 0.37633457 | -0.6543151 | 0.51290879 | 0.62983267 | NA | NA |
| ENSG000001 | 3.13565979 | -0.246931379 | 0.3775335 | -0.6540648 | 0.51307005 | 0.63001162 | NA | NA |
| ENSG000002 | 15.0194025 | 0.197427638 | 0.30186942 | 0.65401669 | 0.51310105 | 0.63003002 | TDRG1 | 732253 |
| ENSG000002 | 4.67744533 | -0.246700469 | 0.37723561 | -0.6539692 | 0.51313165 | 0.63003002 | NA | NA |
| ENSG000002 | 4.79492885 | 0.244621429 | 0.37405291 | 0.65397547 | 0.51312761 | 0.63003002 | NA | NA |
| ENSG000002 | 4.62804039 | 0.245826293 | 0.37592133 | 0.65393015 | 0.51315681 | 0.63004184 | NA | NA |
| ENSG000002 | 2.84336078 | -0.246457255 | 0.37692703 | -0.6538593 | 0.51320244 | 0.63005971 | NA | NA |

| | | | | | | | | |
|---|---|---|---|---|---|---|---|---|
| ENSG000002 | 26.7859747 | 0.16753771 | 0.25622419 | 0.65387156 | 0.51319456 | 0.63005971 | NA | NA |
| ENSG000002 | 12.6931323 | 0.210440587 | 0.32191153 | 0.65372181 | 0.51329105 | 0.63014941 | NA | NA |
| ENSG000002 | 6.8709033 | -0.237453966 | 0.36326007 | -0.6536748 | 0.51332132 | 0.63016019 | NA | NA |
| ENSG000002 | 9.29041552 | 0.227691523 | 0.34833329 | 0.65365996 | 0.51333091 | 0.63016019 | NA | NA |
| ENSG000002 | 2.83001875 | -0.245541127 | 0.37565777 | -0.6536298 | 0.51335031 | 0.63016494 | LOC154449 | 154449 |
| ENSG000002 | 3.99746336 | -0.247236038 | 0.37829186 | -0.653559 | 0.51339598 | 0.63020192 | NA | NA |
| ENSG000001 | 569.751205 | 0.086703811 | 0.13269699 | 0.65339699 | 0.51350038 | 0.63030013 | MPDU1 | 9526 |
| ENSG000001 | 27.9252507 | 0.167463272 | 0.25630044 | 0.6533866 | 0.51350707 | 0.63030013 | EFNA1 | 1942 |
| ENSG000001 | 3.08716693 | -0.246995094 | 0.37806361 | -0.6533162 | 0.51355243 | 0.63033673 | OR4F5 | 79501 |
| ENSG000001 | 11.9496218 | 0.213146803 | 0.32634917 | 0.65312501 | 0.51367569 | 0.63043076 | RBBP8NL | 140893 |
| ENSG000002 | 8.39458273 | -0.228592393 | 0.34997866 | -0.653161 | 0.51365252 | 0.63043076 | NA | NA |
| ENSG000002 | 15.7053027 | 0.198586085 | 0.30404917 | 0.65313805 | 0.51366728 | 0.63043076 | NA | NA |
| ENSG000002 | 3.13274319 | -0.246140346 | 0.3770097 | -0.6528754 | 0.51383663 | 0.63059011 | NA | NA |
| ENSG000002 | 3.59102925 | -0.24716608 | 0.37857948 | -0.6528777 | 0.51383516 | 0.63059011 | NA | NA |
| ENSG000002 | 2.63431404 | -0.242147945 | 0.37093913 | -0.652797 | 0.51388717 | 0.63063305 | RNU6-1089P | 106481523 |
| ENSG000001 | 37.1092883 | -0.151008373 | 0.23141469 | -0.6525445 | 0.51405001 | 0.63076395 | TMEM169 | 92691 |
| ENSG000002 | 18.6513474 | 0.197864394 | 0.30321826 | 0.65254774 | 0.51404788 | 0.63076395 | NA | NA |
| ENSG000002 | 2.27807995 | 0.242445756 | 0.37154441 | 0.65253507 | 0.51405605 | 0.63076395 | NA | NA |
| ENSG000002 | 2.83818058 | -0.245808849 | 0.37665912 | -0.6526029 | 0.51401229 | 0.63076395 | NA | NA |
| ENSG000001 | 33.3242013 | 0.154543763 | 0.23689547 | 0.65237112 | 0.51416179 | 0.6308746 | SLC29A4 | 222962 |
| ENSG000001 | 21.1098376 | 0.184900181 | 0.2834817 | 0.65224733 | 0.51424163 | 0.6309153 | SHB | 6461 |
| ENSG000002 | 3.08252773 | -0.246572011 | 0.37801833 | -0.6522753 | 0.51422361 | 0.6309153 | NA | NA |
| ENSG000002 | 2.56654986 | -0.243582628 | 0.37344695 | -0.652255 | 0.51423671 | 0.6309153 | NA | NA |
| ENSG000002 | 4.6548553 | 0.245340214 | 0.37616815 | 0.65220889 | 0.51426643 | 0.63092663 | NA | NA |
| ENSG000002 | 3.6021787 | -0.246870545 | 0.37856998 | -0.6521134 | 0.51432804 | 0.63098313 | NA | NA |
| ENSG000002 | 6.21408723 | -0.241488954 | 0.37038357 | -0.6519969 | 0.51440315 | 0.63105618 | NA | NA |
| ENSG000002 | 35.0852598 | -0.153033022 | 0.23473115 | -0.6519502 | 0.51443329 | 0.63107407 | EIF1B-AS1 | 440952 |
| ENSG000002 | 2.84023621 | -0.245753619 | 0.37697438 | -0.6519107 | 0.5144588 | 0.63108627 | NA | NA |
| ENSG000002 | 3.09827696 | -0.246277892 | 0.37779695 | -0.651879 | 0.51447924 | 0.63109226 | NA | NA |

| | | | | | | | | |
|---|---|---|---|---|---|---|---|---|
| ENSG000002 | 3.32257571 | -0.246614513 | 0.37852112 | -0.6515211 | 0.51471013 | 0.63135638 | NA | NA |
| ENSG000001 | 287.585214 | -0.087127548 | 0.13374071 | -0.6514661 | 0.51474562 | 0.63136747 | STK17A | 9263 |
| ENSG000002 | 2.59479037 | -0.23946103 | 0.36757659 | -0.6514589 | 0.51475031 | 0.63136747 | NA | NA |
| ENSG000002 | 2.6200789 | -0.242084151 | 0.3716193 | -0.6514305 | 0.51476862 | 0.63137082 | NA | NA |
| ENSG000002 | 3.37838904 | -0.246556601 | 0.3785172 | -0.6513749 | 0.51480452 | 0.63139576 | NA | NA |
| ENSG000002 | 20.8539258 | 0.179940991 | 0.27626145 | 0.65134311 | 0.51482503 | 0.63140182 | NA | NA |
| ENSG000002 | 17.5522085 | -0.188525939 | 0.28955879 | -0.65108 | 0.51499484 | 0.63159098 | NA | NA |
| ENSG000002 | 2.35880801 | -0.236572446 | 0.36344636 | -0.6509143 | 0.51510179 | 0.63170304 | HLA-DQB1-AS1 | 106480429 |
| ENSG000002 | 18.5586222 | 0.190115211 | 0.29213053 | 0.65078858 | 0.51518297 | 0.63178349 | GRID2IP | 392862 |
| ENSG000001 | 6.65845822 | -0.238466705 | 0.36649895 | -0.6506614 | 0.51526511 | 0.63186511 | HOXC9 | 3225 |
| ENSG000002 | 3.47087648 | -0.243570301 | 0.37439941 | -0.6505627 | 0.5153288 | 0.6319241 | NA | NA |
| ENSG000002 | 3.64560603 | -0.245777944 | 0.37783496 | -0.6504902 | 0.51537562 | 0.6319624 | NA | NA |
| ENSG000002 | 2.88361195 | -0.241012189 | 0.37068119 | -0.6501873 | 0.51557126 | 0.63216474 | NA | NA |
| ENSG000002 | 5.40641083 | -0.242860015 | 0.37352366 | -0.6501864 | 0.51557181 | 0.63216474 | NA | NA |
| ENSG000001 | 200.405565 | 0.093172993 | 0.14333517 | 0.65003582 | 0.51566908 | 0.6322649 | CMC1 | 152100 |
| ENSG000001 | 210.636914 | 0.102137196 | 0.15715076 | 0.64993128 | 0.51573661 | 0.63232858 | SUSD5 | 26032 |
| ENSG000002 | 4.66286038 | 0.244455188 | 0.37617883 | 0.64983771 | 0.51579706 | 0.63238357 | NA | NA |
| ENSG000002 | 2.64859622 | -0.241900771 | 0.37236193 | -0.6496388 | 0.51592553 | 0.63252196 | NA | NA |
| ENSG000002 | 2.59443337 | -0.241276704 | 0.37148216 | -0.6494974 | 0.51601691 | 0.63261487 | NA | NA |
| ENSG000002 | 11.0506268 | -0.214458269 | 0.33020404 | -0.649472 | 0.51603336 | 0.6326159 | NA | NA |
| ENSG000001 | 4.20415446 | -0.245754227 | 0.37859334 | -0.6491245 | 0.51625788 | 0.63287201 | NA | NA |
| ENSG000001 | 19.2766316 | -0.190086424 | 0.29285551 | -0.6490792 | 0.51628718 | 0.6328888 | RAI1 | 10743 |
| ENSG000002 | 10.6579045 | 0.218437944 | 0.33656568 | 0.64902026 | 0.51632528 | 0.63291637 | RTEL1 | 51750 |
| ENSG000002 | 2.5962855 | -0.24178801 | 0.37257762 | -0.6489601 | 0.51636417 | 0.63294491 | MIR1269A | 100302177 |
| ENSG000002 | 2.55143371 | -0.24250414 | 0.37374946 | -0.6488414 | 0.51644087 | 0.63300206 | NA | NA |
| ENSG000002 | 3.14463041 | -0.244675043 | 0.37711035 | -0.6488155 | 0.51645763 | 0.63300206 | NA | NA |
| ENSG000002 | 3.87465101 | -0.245488197 | 0.37835212 | -0.6488353 | 0.51644486 | 0.63300206 | NA | NA |
| ENSG000001 | 20.9564664 | 0.179577584 | 0.27679627 | 0.6487717 | 0.51648595 | 0.63301765 | MAMDC4 | 158056 |
| ENSG000002 | 7.87335241 | -0.230079736 | 0.35474073 | -0.6485856 | 0.51660625 | 0.63314595 | NA | NA |

| | | | | | | | | |
|---|---|---|---|---|---|---|---|---|
| ENSG000002 | 2.87789206 | -0.243648637 | 0.37567882 | -0.6485557 | 0.5166256 | 0.63315053 | NA | NA |
| ENSG000002 | 2.59938577 | -0.242481439 | 0.37391076 | -0.6485008 | 0.51666107 | 0.63317486 | NA | NA |
| ENSG000002 | 3.07633013 | -0.245086658 | 0.37799031 | -0.648394 | 0.51673013 | 0.63324036 | NA | NA |
| ENSG000002 | 6.5789368 | -0.236665899 | 0.36505967 | -0.6482937 | 0.51679498 | 0.6333007 | NA | NA |
| ENSG000002 | 17.0704881 | -0.190802358 | 0.29443001 | -0.6480398 | 0.51695923 | 0.6334642 | LINC01347 | 731275 |
| ENSG000002 | 3.86215089 | -0.245215339 | 0.37839589 | -0.6480391 | 0.51695965 | 0.6334642 | NA | NA |
| ENSG000002 | 3.106517 | -0.244571128 | 0.37742616 | -0.6479973 | 0.51698671 | 0.63347821 | NA | NA |
| ENSG000001 | 3.35325239 | -0.245196456 | 0.37851827 | -0.6477797 | 0.51712744 | 0.6336315 | TEX43 | 389320 |
| ENSG000001 | 2.4033479 | -0.235700553 | 0.36402739 | -0.6474803 | 0.51732116 | 0.63381142 | CLDN24 | 100132463 |
| ENSG000002 | 64.654967 | 0.12888434 | 0.19905512 | 0.64748067 | 0.5173209 | 0.63381142 | NA | NA |
| ENSG000002 | 15.871013 | -0.19586454 | 0.30249834 | -0.6474896 | 0.5173151 | 0.63381142 | NA | NA |
| ENSG000002 | 3.14294692 | -0.24374651 | 0.37648336 | -0.6474297 | 0.51735388 | 0.63383236 | NA | NA |
| ENSG000002 | 4.11378572 | -0.244505092 | 0.37769851 | -0.6473552 | 0.51740209 | 0.63386951 | NA | NA |
| ENSG000002 | 12.4696016 | -0.210769722 | 0.3255963 | -0.6473345 | 0.51741547 | 0.63386951 | NA | NA |
| ENSG000001 | 85.9152054 | -0.116413138 | 0.17986705 | -0.6472177 | 0.51749106 | 0.63394297 | CACNB2 | 783 |
| ENSG000002 | 3.83648609 | -0.244698591 | 0.37816412 | -0.6470698 | 0.51758675 | 0.63404104 | UBXN10-AS1 | 101928017 |
| ENSG000002 | 9.01702802 | -0.224652404 | 0.34721271 | -0.6470166 | 0.51762117 | 0.63406405 | NA | NA |
| ENSG000002 | 7.3584691 | -0.232753427 | 0.35977388 | -0.6469436 | 0.51766842 | 0.63410277 | NA | NA |
| ENSG000002 | 2.85198594 | -0.243398538 | 0.37627539 | -0.6468628 | 0.51772077 | 0.63413642 | OR5H6 | 79295 |
| ENSG000002 | 2.86004309 | -0.243332934 | 0.37617972 | -0.6468529 | 0.51772717 | 0.63413642 | NA | NA |
| ENSG000002 | 2.33022672 | -0.237363972 | 0.36701704 | -0.6467383 | 0.51780134 | 0.63416053 | NA | NA |
| ENSG000002 | 3.35186284 | -0.244842009 | 0.37858054 | -0.6467369 | 0.51780223 | 0.63416053 | NA | NA |
| ENSG000002 | 2.67961782 | -0.239633894 | 0.37053398 | -0.6467258 | 0.51780941 | 0.63416053 | NA | NA |
| ENSG000002 | 2.81939798 | -0.24299135 | 0.37569549 | -0.6467774 | 0.51777602 | 0.63416053 | NA | NA |
| ENSG000001 | 13.1639143 | -0.2023541 | 0.31294941 | -0.6466032 | 0.51788876 | 0.63420026 | USP50 | 373509 |
| ENSG000002 | 33.9818278 | -0.15592312 | 0.24113536 | -0.6466207 | 0.51787744 | 0.63420026 | NA | NA |
| ENSG000002 | 3.53099087 | 0.244489385 | 0.3781007 | 0.64662506 | 0.51787464 | 0.63420026 | NA | NA |
| ENSG000002 | 10.362782 | -0.220637759 | 0.34125653 | -0.6465452 | 0.51792636 | 0.63422715 | NA | NA |
| ENSG000002 | 24.3000037 | -0.170098605 | 0.26313606 | -0.6464283 | 0.518002 | 0.63428146 | NA | NA |

| | | | | | | | | |
|---|---|---|---|---|---|---|---|---|
| ENSG000002 | 3.72903643 | 0.244643742 | 0.3784522 | 0.64643235 | 0.5179994 | 0.63428146 | NA | NA |
| ENSG000002 | 3.67660968 | 0.244698478 | 0.37856889 | 0.64637767 | 0.5180348 | 0.63430247 | NA | NA |
| ENSG000002 | 3.33783622 | -0.244607007 | 0.37856222 | -0.6461474 | 0.51818388 | 0.63446585 | NA | NA |
| ENSG000002 | 160.90985 | 0.094909079 | 0.14692327 | 0.64597716 | 0.51829415 | 0.6345817 | NUTM2B-AS1 | 101060691 |
| ENSG000002 | 2.81763595 | -0.242616123 | 0.37559999 | -0.6459428 | 0.51831637 | 0.63458975 | NA | NA |
| ENSG000002 | 32.3168351 | -0.154115574 | 0.23861715 | -0.6458696 | 0.51836378 | 0.63462864 | NA | NA |
| ENSG000002 | 70.304901 | -0.123148325 | 0.19068695 | -0.6458141 | 0.51839973 | 0.63465349 | PCDHA11 | 56138 |
| ENSG000002 | 2.82648925 | -0.24188072 | 0.37460399 | -0.6456971 | 0.51847552 | 0.63472711 | NA | NA |
| ENSG000002 | 3.44415019 | 0.242078699 | 0.37494568 | 0.64563673 | 0.51851465 | 0.63475585 | NA | NA |
| ENSG000002 | 3.75370398 | 0.244279988 | 0.3784169 | 0.64553139 | 0.51858288 | 0.63480106 | NA | NA |
| ENSG000002 | 10.1264662 | 0.220959541 | 0.34229036 | 0.64553247 | 0.51858218 | 0.63480106 | NA | NA |
| ENSG000002 | 5.47527117 | -0.242667303 | 0.37597314 | -0.6454379 | 0.51864347 | 0.63485606 | LOC102723924 | 102723924 |
| ENSG000002 | 6.40106862 | -0.237289274 | 0.36770013 | -0.6453337 | 0.51871098 | 0.63491953 | NA | NA |
| ENSG000002 | 2.56480744 | -0.240645997 | 0.37294717 | -0.6452549 | 0.518762 | 0.63496281 | NA | NA |
| ENSG000001 | 19.2212209 | -0.185832804 | 0.28801412 | -0.6452211 | 0.51878388 | 0.63497043 | ISM1 | 140862 |
| ENSG000002 | 2.632463 | -0.240581833 | 0.37304262 | -0.6449178 | 0.51898045 | 0.63519184 | NA | NA |
| ENSG000002 | 7.64677677 | -0.230883425 | 0.35802417 | -0.6448822 | 0.51900351 | 0.6352009 | NA | NA |
| ENSG000002 | 17.2788394 | 0.196280433 | 0.30438997 | 0.64483213 | 0.51903598 | 0.63522146 | LOC440600 | 440600 |
| ENSG000001 | 75.5671538 | 0.117370946 | 0.18204433 | 0.64473825 | 0.51909682 | 0.63527675 | MYH2 | 4620 |
| ENSG000002 | 2.83175485 | -0.242891841 | 0.37679761 | -0.6446215 | 0.5191725 | 0.63529267 | YWHAQP9 | 101290508 |
| ENSG000002 | 12.9418908 | 0.205454414 | 0.31871044 | 0.64464288 | 0.51915864 | 0.63529267 | NA | NA |
| ENSG000002 | 2.57596954 | -0.239950553 | 0.37219406 | -0.644692 | 0.51912677 | 0.63529267 | NA | NA |
| ENSG000002 | 5.45467535 | -0.241650708 | 0.37485205 | -0.6446562 | 0.51914999 | 0.63529267 | NA | NA |
| ENSG000001 | 3.10037202 | -0.243681962 | 0.37806395 | -0.6445522 | 0.5192174 | 0.63532844 | NA | NA |
| ENSG00000C | 155.64266 | 0.095698962 | 0.14849522 | 0.64445819 | 0.51927836 | 0.63538387 | TGFBR3 | 7049 |
| ENSG000001 | 9.23175477 | -0.224890111 | 0.34900011 | -0.6443841 | 0.51932641 | 0.63541944 | HPCA | 3208 |
| ENSG000002 | 7.83161396 | -0.232032354 | 0.36009459 | -0.644365 | 0.51933877 | 0.63541944 | SUMO2P18 | 106481679 |
| ENSG000002 | 2.32878565 | -0.237409653 | 0.36849282 | -0.6442721 | 0.51939899 | 0.63547394 | NA | NA |
| ENSG000002 | 134.548177 | 0.106672166 | 0.1655998 | 0.64415635 | 0.51947405 | 0.6355466 | NA | NA |

| | | | | | | | | |
|---|---|---|---|---|---|---|---|---|
| ENSG000002 | 2.86123732 | -0.242528735 | 0.37653204 | -0.6441118 | 0.51950293 | 0.63556276 | NA | NA |
| ENSG000002 | 3.14230667 | -0.242248868 | 0.37611346 | -0.6440846 | 0.51952061 | 0.63556521 | NA | NA |
| ENSG000002 | 3.5924516 | -0.243739952 | 0.37858863 | -0.6438121 | 0.51969727 | 0.63576216 | HID1-AS1 | 102723641 |
| ENSG000002 | 2.85463477 | -0.242182129 | 0.37622087 | -0.6437233 | 0.51975489 | 0.63579429 | NA | NA |
| ENSG000002 | 2.59866011 | -0.23664814 | 0.36761438 | -0.6437402 | 0.51974394 | 0.63579429 | NA | NA |
| ENSG000002 | 5.64192408 | -0.239648261 | 0.37232123 | -0.6436599 | 0.51979597 | 0.63581544 | NA | NA |
| ENSG000002 | 7.83017453 | -0.231690113 | 0.35996904 | -0.643639 | 0.51980955 | 0.63581544 | KRT18P55 | 284085 |
| ENSG000002 | 3.07154169 | -0.243076001 | 0.37766765 | -0.6436241 | 0.51981922 | 0.63581544 | OR51G1 | 79324 |
| ENSG000002 | 2.61803198 | -0.24033737 | 0.3735757 | -0.6433431 | 0.52000146 | 0.63601916 | NA | NA |
| ENSG000001 | 4.93191908 | -0.242330561 | 0.3767339 | -0.6432406 | 0.52006796 | 0.63608131 | HTR5A | 3361 |
| ENSG000002 | 2.81140589 | -0.241648534 | 0.37577308 | -0.6430704 | 0.52017844 | 0.63619724 | NA | NA |
| ENSG000002 | 2.88199013 | -0.241498662 | 0.3756247 | -0.6429254 | 0.52027251 | 0.6362931 | NA | NA |
| ENSG000002 | 10.4141219 | 0.216056608 | 0.33608942 | 0.64285454 | 0.52031849 | 0.63633015 | LOC101928489 | 101928489 |
| ENSG000002 | 2.59441464 | -0.240253829 | 0.37380741 | -0.6427209 | 0.52040523 | 0.63641703 | NA | NA |
| ENSG00000C | 256.501019 | 0.084140246 | 0.1309177 | 0.64269571 | 0.52042157 | 0.63641782 | SCYL3 | 57147 |
| ENSG000002 | 14.5022969 | 0.205138392 | 0.31929125 | 0.64248047 | 0.52056126 | 0.63656946 | NA | NA |
| ENSG000002 | 2.82173436 | -0.241860231 | 0.37651581 | -0.6423641 | 0.52063681 | 0.63662345 | DUXAP11 | 106481741 |
| ENSG000002 | 2.6506839 | -0.238225352 | 0.37085561 | -0.6423669 | 0.52063502 | 0.63662345 | NA | NA |
| ENSG000001 | 70.8089564 | 0.121508665 | 0.18917304 | 0.64231491 | 0.52066874 | 0.63664329 | MDH1B | 130752 |
| ENSG000002 | 4.38977489 | -0.242327716 | 0.37730762 | -0.642255 | 0.52070761 | 0.63667162 | NA | NA |
| ENSG000001 | 44.6929826 | -0.137857325 | 0.21466629 | -0.6421936 | 0.52074746 | 0.63670114 | LPXN | 9404 |
| ENSG000001 | 94.7504638 | 0.1076391 | 0.16764717 | 0.64205735 | 0.52083594 | 0.63679013 | SLC1A6 | 6511 |
| ENSG000002 | 2.81300451 | -0.241135317 | 0.37559761 | -0.6420044 | 0.52087033 | 0.63681297 | NA | NA |
| ENSG000002 | 2.59798826 | -0.238553252 | 0.37168117 | -0.6418223 | 0.52098859 | 0.63693836 | NA | NA |
| ENSG000002 | 54.0618094 | 0.1317955 | 0.20536923 | 0.64174902 | 0.52103615 | 0.63696641 | NA | NA |
| ENSG000002 | 10.8871032 | 0.213051403 | 0.33199097 | 0.64173855 | 0.52104295 | 0.63696641 | NA | NA |
| ENSG000002 | 4.95860174 | -0.24228168 | 0.37769354 | -0.6414769 | 0.52121289 | 0.63715495 | RD3L | 647286 |
| ENSG000001 | 29.934238 | 0.161650327 | 0.25201363 | 0.64143486 | 0.52124019 | 0.63716911 | PAQR6 | 79957 |
| ENSG000001 | 15.2390976 | -0.193587239 | 0.30184365 | -0.6413494 | 0.52129571 | 0.63721777 | TEPP | 374739 |

| | | | | | | | | |
|---|---|---|---|---|---|---|---|---|
| ENSG000001 | 16.8274482 | -0.192483467 | 0.30015525 | -0.6412797 | 0.52134099 | 0.63725391 | GDF6 | 392255 |
| ENSG000002 | 2.31678153 | 0.237163608 | 0.36987432 | 0.64120052 | 0.52139241 | 0.63729755 | NA | NA |
| ENSG000002 | 3.73207701 | 0.242700148 | 0.37853261 | 0.64116047 | 0.52141843 | 0.63731014 | NA | NA |
| ENSG000001 | 8.55126668 | -0.223308755 | 0.34831134 | -0.6411182 | 0.52144586 | 0.63732447 | CA9 | 768 |
| ENSG000002 | 3.35254798 | -0.242647401 | 0.37854234 | -0.6410047 | 0.52151966 | 0.63739545 | NA | NA |
| ENSG000002 | 2.83214661 | -0.241360619 | 0.37656512 | -0.6409532 | 0.52155309 | 0.63741709 | NA | NA |
| ENSG000001 | 23.2257957 | 0.17578802 | 0.27429184 | 0.64087951 | 0.52160097 | 0.6374564 | ESPNL | 339768 |
| ENSG000001 | 13.2831522 | -0.205206909 | 0.32021909 | -0.6408328 | 0.52163129 | 0.63747425 | LRRN4CL | 221091 |
| ENSG000002 | 6.59890986 | 0.232639708 | 0.36308917 | 0.64072334 | 0.52170245 | 0.63754199 | NA | NA |
| ENSG000002 | 39.1077954 | 0.15274916 | 0.23843097 | 0.64064312 | 0.52175458 | 0.63758649 | CLDN23 | 137075 |
| ENSG000002 | 82.2454008 | 0.11299964 | 0.17640248 | 0.64057853 | 0.52179655 | 0.63759935 | NA | NA |
| ENSG000002 | 16.4520137 | -0.19418765 | 0.3031423 | -0.6405825 | 0.52179398 | 0.63759935 | LOC100130111 | 100130111 |
| ENSG00000C | 160.744101 | -0.101640502 | 0.15867842 | -0.6405439 | 0.52181903 | 0.63760556 | ACACB | 32 |
| ENSG000002 | 18.6249474 | -0.186140451 | 0.29060728 | -0.6405223 | 0.52183308 | 0.63760556 | NA | NA |
| ENSG000001 | 215.205525 | 0.089283324 | 0.13940568 | 0.64045686 | 0.52187563 | 0.63763833 | AFF2 | 2334 |
| ENSG000002 | 5.63797535 | 0.23821115 | 0.37201953 | 0.64031894 | 0.52196527 | 0.63772864 | NA | NA |
| ENSG000002 | 3.1311972 | -0.241042958 | 0.3764841 | -0.6402474 | 0.52201179 | 0.63776626 | NA | NA |
| ENSG000002 | 2.63932559 | -0.236728865 | 0.36977635 | -0.6401947 | 0.52204602 | 0.63778887 | NA | NA |
| ENSG000002 | 3.07229155 | -0.24133112 | 0.37708881 | -0.6399848 | 0.52218246 | 0.63793634 | NA | NA |
| ENSG000002 | 2.56298756 | -0.2381607 | 0.37217619 | -0.6399139 | 0.5222286 | 0.63797349 | NA | NA |
| ENSG000002 | 2.33908634 | -0.234942917 | 0.36724844 | -0.6397384 | 0.52234271 | 0.63809367 | NA | NA |
| ENSG000002 | 5.32871462 | -0.240307386 | 0.37565129 | -0.6397087 | 0.52236202 | 0.63809803 | NA | NA |
| ENSG000001 | 2.63983446 | -0.237910216 | 0.37200228 | -0.6395397 | 0.52247191 | 0.63814485 | NA | NA |
| ENSG000001 | 3.59406948 | -0.242119582 | 0.37858318 | -0.6395413 | 0.52247085 | 0.63814485 | NA | NA |
| ENSG000002 | 2.90868498 | -0.238392039 | 0.37273236 | -0.6395797 | 0.52244591 | 0.63814485 | LOC400558 | 400558 |
| ENSG000002 | 3.36676993 | -0.241950579 | 0.37827795 | -0.6396106 | 0.5224258 | 0.63814485 | NA | NA |
| ENSG000002 | 8.36924938 | -0.234166834 | 0.3661553 | -0.6395287 | 0.52247903 | 0.63814485 | NA | NA |
| ENSG00000C | 155.409658 | 0.106892314 | 0.16715065 | 0.63949686 | 0.52249976 | 0.63815095 | ITGA3 | 3675 |
| ENSG000001 | 56.3474215 | 0.124802685 | 0.19519911 | 0.63936094 | 0.52258815 | 0.63823968 | HTR7P1 | 93164 |

| | | | | | | | | |
|---|---|---|---|---|---|---|---|---|
| ENSG000002 | 10.4955571 | -0.215570788 | 0.3372256 | -0.639248 | 0.52266162 | 0.63831018 | NA | NA |
| ENSG000002 | 10.9003611 | -0.220578898 | 0.34513285 | -0.639113 | 0.52274941 | 0.63839817 | NA | NA |
| ENSG000002 | 21.7875747 | 0.190569638 | 0.29821569 | 0.63903291 | 0.52280152 | 0.63844259 | INE1 | 8552 |
| ENSG000002 | 2.5997833 | -0.238412397 | 0.37315026 | -0.6389179 | 0.52287631 | 0.63849546 | NA | NA |
| ENSG000002 | 3.14841003 | 0.241754057 | 0.37836824 | 0.63893855 | 0.52286291 | 0.63849546 | NA | NA |
| ENSG000002 | 2.36781916 | -0.235379362 | 0.36859882 | -0.6385787 | 0.52309703 | 0.6387302 | RNU6-1187P | 106481554 |
| ENSG000002 | 2.28587617 | 0.236689739 | 0.37066759 | 0.63854987 | 0.5231158 | 0.6387302 | NA | NA |
| ENSG000002 | 2.30903759 | 0.23488882 | 0.36784681 | 0.63855065 | 0.52311529 | 0.6387302 | NA | NA |
| ENSG000001 | 32.4313316 | -0.156057104 | 0.24441166 | -0.6385011 | 0.52314756 | 0.63873597 | RTN1 | 6252 |
| ENSG000002 | 36.7555306 | 0.148847427 | 0.2331226 | 0.6384942 | 0.52315203 | 0.63873597 | NA | NA |
| ENSG000002 | 2.3413167 | -0.231326963 | 0.36232737 | -0.6384474 | 0.5231825 | 0.63874957 | NA | NA |
| ENSG000002 | 6.39665287 | -0.235531568 | 0.36892385 | -0.6384287 | 0.52319467 | 0.63874957 | RN7SKP96 | 106479137 |
| ENSG000002 | 2.9073758 | 0.240326281 | 0.37652495 | 0.63827451 | 0.523295 | 0.63885283 | NA | NA |
| ENSG000002 | 2.32697797 | -0.233879779 | 0.3666287 | -0.63792 | 0.52352576 | 0.63911531 | NA | NA |
| ENSG000002 | 3.83762824 | -0.24120887 | 0.37822637 | -0.6377368 | 0.52364504 | 0.63924167 | NA | NA |
| ENSG000002 | 4.57711536 | -0.239365207 | 0.37537411 | -0.6376711 | 0.52368783 | 0.63927466 | RNU1-65P | 106481614 |
| ENSG000002 | 3.07039954 | -0.240363967 | 0.37697891 | -0.6376059 | 0.52373027 | 0.63930724 | NA | NA |
| ENSG000001 | 159.079335 | -0.090437375 | 0.1418574 | -0.6375231 | 0.52378416 | 0.63933452 | ITGA9 | 3680 |
| ENSG000002 | 3.10233515 | -0.240857705 | 0.37779643 | -0.637533 | 0.5237777 | 0.63933452 | NA | NA |
| ENSG000002 | 6.50303354 | -0.236175872 | 0.37048407 | -0.6374792 | 0.52381278 | 0.63935022 | NA | NA |
| ENSG000001 | 6.66894314 | -0.234597799 | 0.36809242 | -0.637334 | 0.52390734 | 0.63941305 | SSX3 | 10214 |
| ENSG000002 | 2.86074805 | -0.239657885 | 0.37606329 | -0.6372807 | 0.523942 | 0.63941305 | TOLLIP-AS1 | 255512 |
| ENSG000002 | 4.35431086 | -0.239754763 | 0.37621631 | -0.637279 | 0.52394311 | 0.63941305 | NA | NA |
| ENSG000002 | 3.640945 | -0.241276755 | 0.37856056 | -0.6373531 | 0.52389486 | 0.63941305 | LOC645967 | 645967 |
| ENSG000002 | 11.5276537 | -0.207309837 | 0.3252912 | -0.6373054 | 0.52392594 | 0.63941305 | NA | NA |
| ENSG00000C | 21.7504043 | -0.181398716 | 0.28474123 | -0.6370652 | 0.5240824 | 0.6395638 | RIMBP2 | 23504 |
| ENSG000001 | 26.8714147 | 0.165126914 | 0.25925619 | 0.63692563 | 0.52417329 | 0.63965546 | RASSF10 | 644943 |
| ENSG000002 | 2.61871712 | -0.237564786 | 0.37302262 | -0.6368643 | 0.52421324 | 0.63968497 | NA | NA |
| ENSG000002 | 30.4026008 | -0.164820671 | 0.25885287 | -0.6367349 | 0.52429751 | 0.63976854 | RGAG4 | 340526 |

| | | | | | | | | |
|---|---|---|---|---|---|---|---|---|
| ENSG000001 | 177.822637 | 0.109009509 | 0.17122366 | 0.63664981 | 0.52435297 | 0.63981697 | MEF2D | 4209 |
| ENSG000001 | 11.1839247 | -0.206182111 | 0.32396289 | -0.6364374 | 0.52449134 | 0.6399473 | P2RY2 | 5029 |
| ENSG000002 | 27.9029694 | 0.158505905 | 0.24904678 | 0.63645034 | 0.52448294 | 0.6399473 | PSMG3-AS1 | 114796 |
| ENSG000002 | 12.5388767 | -0.204418247 | 0.32121869 | -0.6363834 | 0.52452655 | 0.63995175 | ST7-OT4 | 338069 |
| ENSG000002 | 3.54868193 | -0.240935784 | 0.37859339 | -0.6363972 | 0.52451754 | 0.63995175 | NA | NA |
| ENSG000002 | 2.25176226 | 0.234395011 | 0.36833961 | 0.63635569 | 0.52454461 | 0.63995453 | NA | NA |
| ENSG000002 | 2.87153691 | -0.236919831 | 0.37236951 | -0.6362493 | 0.52461396 | 0.64001987 | NA | NA |
| ENSG000001 | 180.032773 | 0.094942287 | 0.14923615 | 0.63618828 | 0.52465371 | 0.64004912 | DCBLD1 | 285761 |
| ENSG000001 | 11.23117 | -0.22227964 | 0.34944798 | -0.6360879 | 0.52471912 | 0.64007114 | SLA2 | 84174 |
| ENSG000002 | 2.34150093 | -0.232834334 | 0.36603822 | -0.636093 | 0.52471584 | 0.64007114 | NA | NA |
| ENSG000002 | 2.60405743 | -0.237017657 | 0.3726005 | -0.6361174 | 0.52469992 | 0.64007114 | NA | NA |
| ENSG000002 | 2.86128939 | -0.239165029 | 0.37603258 | -0.6360221 | 0.52476202 | 0.64008885 | NA | NA |
| ENSG000002 | 4.12751178 | -0.2407054 | 0.37845737 | -0.6360172 | 0.5247652 | 0.64008885 | NA | NA |
| ENSG000001 | 2786.89475 | 0.072014411 | 0.11323586 | 0.63596824 | 0.52479712 | 0.64010853 | PBX1 | 5087 |
| ENSG000002 | 2.30030774 | 0.234821544 | 0.36926954 | 0.63590824 | 0.52483623 | 0.64011772 | NA | NA |
| ENSG000002 | 4.80554907 | 0.237752733 | 0.37387873 | 0.63590868 | 0.52483594 | 0.64011772 | NA | NA |
| ENSG000002 | 2.58802725 | -0.23617957 | 0.37148567 | -0.6357703 | 0.52492613 | 0.64018885 | NA | NA |
| ENSG000002 | 5.85946408 | -0.237240335 | 0.37314286 | -0.6357896 | 0.52491359 | 0.64018885 | NA | NA |
| ENSG000002 | 3.38285163 | -0.240519699 | 0.37832995 | -0.6357406 | 0.52494553 | 0.64019326 | RNU2-44P | 106481648 |
| ENSG000002 | 3.85286008 | -0.240573505 | 0.3785545 | -0.6355056 | 0.52509872 | 0.64036083 | NA | NA |
| ENSG000002 | 5.63695774 | 0.236068009 | 0.37149005 | 0.63546253 | 0.52512679 | 0.6403758 | NA | NA |
| ENSG000002 | 2.61721492 | -0.235581869 | 0.37080893 | -0.6353188 | 0.52522054 | 0.64047086 | NA | NA |
| ENSG000002 | 3.36566079 | -0.240319428 | 0.37833308 | -0.635206 | 0.52529409 | 0.6405413 | NA | NA |
| ENSG000001 | 29.2281443 | -0.155877731 | 0.24541429 | -0.6351616 | 0.52532303 | 0.64055731 | FAM188B | 84182 |
| ENSG000001 | 47.9004355 | 0.134909791 | 0.21245497 | 0.63500417 | 0.52542569 | 0.64066324 | LRRC17 | 10234 |
| ENSG000002 | 2.81558883 | -0.238901676 | 0.3762599 | -0.6349379 | 0.52546891 | 0.64069667 | NA | NA |
| ENSG000002 | 3.17019128 | -0.238568533 | 0.3757803 | -0.6348617 | 0.52551859 | 0.64073798 | NA | NA |
| ENSG000002 | 16.066678 | -0.201046584 | 0.31671727 | -0.6347825 | 0.52557027 | 0.64078172 | NA | NA |
| ENSG000002 | 2.59084493 | -0.236324377 | 0.37232053 | -0.6347337 | 0.52560213 | 0.6408013 | NA | NA |

| | | | | | | | | |
|---|---|---|---|---|---|---|---|---|
| ENSG000002 | 3.05629437 | -0.239337176 | 0.37713488 | -0.6346196 | 0.52567656 | 0.64085205 | NA | NA |
| ENSG000002 | 4.34049276 | -0.239250343 | 0.37700776 | -0.6346032 | 0.52568722 | 0.64085205 | NA | NA |
| ENSG000002 | 2.83429756 | -0.235835679 | 0.37163052 | -0.6345972 | 0.52569117 | 0.64085205 | NA | NA |
| ENSG000002 | 10.8842191 | 0.21284401 | 0.33543611 | 0.63452922 | 0.52573551 | 0.64088683 | NA | NA |
| ENSG000002 | 2.35858022 | -0.231246958 | 0.36454923 | -0.6343367 | 0.52586111 | 0.64102068 | NA | NA |
| ENSG000001 | 58.782393 | -0.123725687 | 0.19506388 | -0.6342829 | 0.52589621 | 0.64104419 | KIAA1407 | 57577 |
| ENSG000001 | 58.3560879 | -0.129819913 | 0.20469158 | -0.634222 | 0.52593593 | 0.64107334 | NUTM1 | 256646 |
| ENSG000002 | 5.13750083 | -0.238197183 | 0.37566031 | -0.634076 | 0.52603123 | 0.64117023 | NA | NA |
| ENSG000000 | 19.2108626 | -0.181489543 | 0.28626651 | -0.633988 | 0.52608863 | 0.64122092 | OPRK1 | 4986 |
| ENSG000001 | 13.8965275 | -0.195305806 | 0.30807932 | -0.6339465 | 0.52611573 | 0.64123451 | NHLH1 | 4807 |
| ENSG000002 | 2.8424473 | -0.238235834 | 0.37582661 | -0.6338983 | 0.52614722 | 0.64123451 | NA | NA |
| ENSG000002 | 3.18692566 | -0.237858189 | 0.3752284 | -0.6339024 | 0.5261445 | 0.64123451 | NA | NA |
| ENSG000002 | 2.92769015 | -0.235018829 | 0.37080029 | -0.6338151 | 0.52620149 | 0.64128137 | NA | NA |
| ENSG000001 | 121.398991 | 0.098726453 | 0.15578302 | 0.63374336 | 0.52624832 | 0.64131918 | ATP7A | 538 |
| ENSG000001 | 98.9460906 | 0.104319398 | 0.16462297 | 0.63368677 | 0.52628526 | 0.64132224 | ZNF230 | 7773 |
| ENSG000002 | 3.38152157 | -0.23972302 | 0.37831082 | -0.6336668 | 0.52629827 | 0.64132224 | NA | NA |
| ENSG000002 | 5.09267644 | -0.236572046 | 0.37332056 | -0.6336968 | 0.52627871 | 0.64132224 | NA | NA |
| ENSG000002 | 3.56848956 | -0.239852603 | 0.37855046 | -0.633608 | 0.52633667 | 0.64134975 | NA | NA |
| ENSG000002 | 3.33491939 | -0.239725858 | 0.378401 | -0.6335233 | 0.52639196 | 0.64139784 | NA | NA |
| ENSG000002 | 2.82605314 | -0.238378461 | 0.37631369 | -0.6334568 | 0.52643539 | 0.64140193 | NA | NA |
| ENSG000002 | 2.67929883 | 0.23490599 | 0.3708352 | 0.63345116 | 0.52643907 | 0.64140193 | NA | NA |
| ENSG000002 | 2.8224195 | -0.238166486 | 0.37598576 | -0.6334455 | 0.52644276 | 0.64140193 | NA | NA |
| ENSG000002 | 41.16899 | 0.14380839 | 0.22705993 | 0.63334993 | 0.52650515 | 0.64145449 | NA | NA |
| ENSG000002 | 7.18457171 | -0.233600123 | 0.36885779 | -0.6333067 | 0.52653336 | 0.64145449 | NA | NA |
| ENSG000002 | 2.57178267 | -0.236716735 | 0.37377621 | -0.6333114 | 0.52653031 | 0.64145449 | NA | NA |
| ENSG000002 | 3.08660599 | -0.238773829 | 0.37710471 | -0.6331765 | 0.52661841 | 0.64153883 | NA | NA |
| ENSG000001 | 72.1676573 | -0.123396937 | 0.19495166 | -0.6329617 | 0.52675864 | 0.64163701 | PLA2G7 | 7941 |
| ENSG000002 | 6.73104666 | 0.231349877 | 0.36550697 | 0.63295613 | 0.52676229 | 0.64163701 | MSMP | 692094 |
| ENSG000002 | 2.32937906 | -0.232388059 | 0.36714395 | -0.6329617 | 0.52675866 | 0.64163701 | NA | NA |

| | | | | | | | | |
|---|---|---|---|---|---|---|---|---|
| ENSG000002 | 10.5372155 | -0.211393015 | 0.3339428 | -0.6330216 | 0.52671952 | 0.64163701 | LOC101927481 | 101927481 |
| ENSG000002 | 17.9742558 | 0.189075875 | 0.29873809 | 0.63291518 | 0.52678903 | 0.64165031 | PET100 | 100131801 |
| ENSG000002 | 25.6108532 | 0.163146232 | 0.25780447 | 0.63282935 | 0.52684509 | 0.64169931 | NA | NA |
| ENSG000001 | 16.393832 | -0.197406492 | 0.31198418 | -0.6327452 | 0.52690005 | 0.64174698 | BSND | 7809 |
| ENSG000002 | 4.12798894 | -0.239069727 | 0.37787001 | -0.6326772 | 0.52694448 | 0.64178182 | NA | NA |
| ENSG000002 | 3.13330468 | -0.238495429 | 0.37700397 | -0.6326072 | 0.52699018 | 0.6418182 | NA | NA |
| ENSG000002 | 3.11044383 | -0.238139417 | 0.37661249 | -0.6323195 | 0.52717813 | 0.64198926 | NA | NA |
| ENSG000002 | 28.0661506 | -0.173436064 | 0.27427984 | -0.6323325 | 0.52716961 | 0.64198926 | DLGAP4-AS1 | 101926987 |
| ENSG000002 | 4.1389494 | -0.239311798 | 0.37845084 | -0.6323458 | 0.52716094 | 0.64198926 | NA | NA |
| ENSG000002 | 2.88784784 | -0.236490584 | 0.37407636 | -0.6321987 | 0.52725706 | 0.64206609 | NA | NA |
| ENSG000001 | 27.3990319 | 0.159557634 | 0.25239709 | 0.63216908 | 0.5272764 | 0.64207036 | SLC31A2 | 1318 |
| ENSG000002 | 15.8205992 | -0.188615983 | 0.29838697 | -0.6321187 | 0.52730931 | 0.64209116 | NA | NA |
| ENSG000002 | 3.3538775 | -0.23846106 | 0.37728928 | -0.6320377 | 0.52736222 | 0.64213631 | NA | NA |
| ENSG000002 | 3.30911058 | -0.239195143 | 0.37847869 | -0.631991 | 0.52739273 | 0.64215418 | NA | NA |
| ENSG000002 | 3.85325293 | -0.239038322 | 0.37840015 | -0.6317078 | 0.52757784 | 0.6423273 | NA | NA |
| ENSG000002 | 3.32826532 | -0.239119764 | 0.37854259 | -0.6316852 | 0.52759258 | 0.6423273 | NA | NA |
| ENSG000002 | 3.08051286 | -0.238616477 | 0.37773875 | -0.6316971 | 0.52758481 | 0.6423273 | NA | NA |
| ENSG000002 | 5.71922515 | 0.23371915 | 0.36999816 | 0.63167652 | 0.52759827 | 0.6423273 | NA | NA |
| ENSG000002 | 2.83309288 | -0.237798859 | 0.3765192 | -0.6315717 | 0.52766681 | 0.64239145 | RN7SL604P | 106479442 |
| ENSG000002 | 13.3944164 | -0.19880381 | 0.31479286 | -0.6315385 | 0.52768849 | 0.64239855 | NA | NA |
| ENSG000002 | 5.13437319 | -0.237917389 | 0.37679275 | -0.6314277 | 0.52776089 | 0.64246741 | NA | NA |
| ENSG000001 | 70.9276977 | 0.124231651 | 0.19677202 | 0.63134815 | 0.52781291 | 0.64251144 | WDSUB1 | 151525 |
| ENSG000002 | 3.60416962 | -0.238892826 | 0.37859336 | -0.6310011 | 0.52803981 | 0.64276836 | NA | NA |
| ENSG000002 | 3.09250302 | -0.236977096 | 0.37558941 | -0.6309472 | 0.52807504 | 0.64279194 | NA | NA |
| ENSG000002 | 4.08436308 | -0.238502552 | 0.37803964 | -0.630893 | 0.52811051 | 0.64281583 | NA | NA |
| ENSG000001 | 2.60457895 | -0.234219301 | 0.37128859 | -0.6308282 | 0.52815289 | 0.64282846 | TMED6 | 146456 |
| ENSG000002 | 3.61544884 | -0.238823674 | 0.37858759 | -0.630828 | 0.52815296 | 0.64282846 | NA | NA |
| ENSG000002 | 3.08553606 | -0.238146888 | 0.37752891 | -0.6308044 | 0.52816845 | 0.64282846 | NA | NA |
| ENSG000002 | 2.56080131 | -0.234280928 | 0.37142382 | -0.6307644 | 0.52819458 | 0.64284097 | RN7SKP110 | 106480867 |

| | | | | | | | | |
|---|---|---|---|---|---|---|---|---|
| ENSG000002 | 2.27582431 | 0.229260041 | 0.36352693 | 0.63065491 | 0.52826619 | 0.64290289 | NA | NA |
| ENSG000002 | 7.77361821 | -0.225169528 | 0.35705029 | -0.6306381 | 0.52827716 | 0.64290289 | NA | NA |
| ENSG000002 | 2.40021049 | -0.228649808 | 0.36261093 | -0.6305651 | 0.5283249 | 0.6429417 | NA | NA |
| ENSG000002 | 4.84479999 | -0.235596838 | 0.37369716 | -0.6304486 | 0.52840114 | 0.64296945 | NA | NA |
| ENSG000002 | 5.95174542 | 0.231626594 | 0.36739125 | 0.63046301 | 0.5283917 | 0.64296945 | NA | NA |
| ENSG000002 | 32.9115724 | 0.150763152 | 0.23912732 | 0.63047231 | 0.52838562 | 0.64296945 | TAS2R31 | 259290 |
| ENSG000002 | 2.82034481 | -0.237272628 | 0.37636437 | -0.6304333 | 0.52841114 | 0.64296945 | NA | NA |
| ENSG000002 | 3.36171448 | -0.238435804 | 0.37831285 | -0.6302609 | 0.52852389 | 0.64308736 | NA | NA |
| ENSG000002 | 2.66390771 | -0.230882803 | 0.36636339 | -0.6302016 | 0.52856268 | 0.64311525 | NA | NA |
| ENSG000002 | 4.85450783 | -0.235626789 | 0.37396177 | -0.6300826 | 0.52864057 | 0.64313284 | NA | NA |
| ENSG000002 | 2.19181685 | -0.220114587 | 0.34933373 | -0.6300983 | 0.52863028 | 0.64313284 | NA | NA |
| ENSG000002 | 4.3688262 | -0.237784021 | 0.37737025 | -0.630108 | 0.52862391 | 0.64313284 | NA | NA |
| ENSG000002 | 8.94560926 | 0.216604044 | 0.34375189 | 0.63011739 | 0.52861778 | 0.64313284 | NA | NA |
| ENSG000002 | 3.61137821 | -0.238520042 | 0.37858251 | -0.6300345 | 0.52867201 | 0.6431518 | RNU6-935P | 106480403 |
| ENSG000002 | 2.58563935 | -0.233647792 | 0.37091036 | -0.6299306 | 0.52873998 | 0.64321518 | NA | NA |
| ENSG000001 | 3.37634138 | -0.238140018 | 0.37811432 | -0.6298096 | 0.52881919 | 0.64327294 | GPR45 | 11250 |
| ENSG000002 | 15.9410061 | 0.195651758 | 0.31064326 | 0.6298278 | 0.52880726 | 0.64327294 | NA | NA |
| ENSG000001 | 13.3323206 | -0.198915266 | 0.31588922 | -0.6296994 | 0.52889125 | 0.6433413 | GFRA3 | 2676 |
| ENSG000001 | 14.1278644 | -0.196449978 | 0.31203184 | -0.6295831 | 0.52896738 | 0.64335913 | CES4A | 283848 |
| ENSG000002 | 3.12806161 | 0.237952294 | 0.37795398 | 0.62958007 | 0.52896937 | 0.64335913 | NA | NA |
| ENSG000002 | 2.27233881 | 0.232659968 | 0.3695139 | 0.62963794 | 0.5289315 | 0.64335913 | NA | NA |
| ENSG000002 | 3.0769632 | -0.237499674 | 0.37722573 | -0.6295956 | 0.52895918 | 0.64335913 | CTD-2194D22.4 | 101929081 |
| ENSG000002 | 3.0594465 | -0.237495054 | 0.37730121 | -0.6294574 | 0.52904962 | 0.64343744 | NA | NA |
| ENSG000002 | 2.35049116 | -0.229646721 | 0.36487017 | -0.629393 | 0.52909182 | 0.64346947 | LOC101928031 | 101928031 |
| ENSG000002 | 6.66873122 | -0.23367437 | 0.37132192 | -0.629304 | 0.52915005 | 0.64350839 | NA | NA |
| ENSG000002 | 2.59973013 | -0.232602933 | 0.36962429 | -0.6292956 | 0.52915556 | 0.64350839 | NA | NA |
| ENSG000001 | 4.5965307 | -0.236742299 | 0.37625055 | -0.6292145 | 0.52920864 | 0.64351504 | NA | NA |
| ENSG000002 | 14.6705265 | -0.195348075 | 0.31045467 | -0.6292322 | 0.52919706 | 0.64351504 | NA | NA |
| ENSG000002 | 5.10460543 | -0.236560865 | 0.37594681 | -0.6292403 | 0.52919178 | 0.64351504 | NA | NA |

| | | | | | | | | |
|---|---|---|---|---|---|---|---|---|
| ENSG000002 | 5.14122136 | -0.235939765 | 0.37507492 | -0.629047 | 0.52931826 | 0.64362904 | NA | NA |
| ENSG000002 | 4.82816119 | 0.235353145 | 0.37421623 | 0.62892287 | 0.52939955 | 0.64370858 | NA | NA |
| ENSG000002 | 3.31822521 | -0.238002252 | 0.37845206 | -0.6288835 | 0.52942534 | 0.64372063 | NA | NA |
| ENSG000002 | 3.58462257 | -0.23804503 | 0.37854343 | -0.6288447 | 0.52945073 | 0.64373221 | NA | NA |
| ENSG000001 | 3.58710331 | -0.237983801 | 0.37856932 | -0.62864 | 0.52958479 | 0.6438759 | NA | NA |
| ENSG000002 | 2.56621018 | -0.233161699 | 0.37102601 | -0.6284241 | 0.52972613 | 0.64398982 | NA | NA |
| ENSG000002 | 3.04453886 | -0.236986664 | 0.37710195 | -0.6284419 | 0.5297145 | 0.64398982 | NA | NA |
| ENSG000002 | 4.83784457 | -0.235533163 | 0.37479238 | -0.6284364 | 0.52971812 | 0.64398982 | NA | NA |
| ENSG000002 | 9.44442785 | -0.211498616 | 0.33659821 | -0.6283415 | 0.52978027 | 0.64403632 | LINC01569 | 100507501 |
| ENSG000002 | 4.60326943 | -0.236680577 | 0.37674511 | -0.6282247 | 0.52985676 | 0.64411 | NA | NA |
| ENSG00000C | 68.7343168 | -0.119645229 | 0.19047342 | -0.6281466 | 0.52990791 | 0.64414841 | AFP | 174 |
| ENSG000002 | 2.26561208 | 0.232087324 | 0.36949052 | 0.62812795 | 0.52992013 | 0.64414841 | NA | NA |
| ENSG000002 | 2.81523541 | -0.235539071 | 0.37501407 | -0.6280806 | 0.52995112 | 0.64416678 | NA | NA |
| ENSG000002 | 9.52153534 | -0.212748856 | 0.33879455 | -0.6279583 | 0.53003124 | 0.64422555 | NA | NA |
| ENSG000002 | 3.30953479 | -0.237607913 | 0.37836802 | -0.627981 | 0.53001641 | 0.64422555 | NA | NA |
| ENSG000002 | 3.57455579 | -0.237595848 | 0.37851898 | -0.6276986 | 0.53020138 | 0.64441302 | NA | NA |
| ENSG000001 | 3.0807644 | -0.234503111 | 0.37360642 | -0.6276742 | 0.53021739 | 0.64441316 | NA | NA |
| ENSG000002 | 3.06933006 | -0.236364405 | 0.37676998 | -0.627344 | 0.53043374 | 0.64465679 | NA | NA |
| ENSG000002 | 4.84004401 | -0.234893881 | 0.37445262 | -0.6272993 | 0.53046303 | 0.64466802 | NA | NA |
| ENSG000002 | 5.94256861 | -0.233810217 | 0.37273575 | -0.6272814 | 0.53047477 | 0.64466802 | NA | NA |
| ENSG000002 | 2.64854306 | -0.228133752 | 0.36376466 | -0.6271465 | 0.53056318 | 0.64475613 | NA | NA |
| ENSG000001 | 27.5469807 | -0.155755777 | 0.24837993 | -0.6270868 | 0.53060233 | 0.64477345 | NA | NA |
| ENSG000002 | 8.31400505 | -0.221729478 | 0.35359251 | -0.6270763 | 0.53060923 | 0.64477345 | NA | NA |
| ENSG000002 | 4.09030064 | -0.236626876 | 0.37736586 | -0.627049 | 0.53062713 | 0.64477588 | NA | NA |
| ENSG00000C | 46.0287496 | 0.135371934 | 0.21594994 | 0.62686721 | 0.53074628 | 0.64488614 | OTUB2 | 78990 |
| ENSG000001 | 5.07303352 | -0.232986732 | 0.37167147 | -0.626862 | 0.53074967 | 0.64488614 | SLC25A47 | 283600 |
| ENSG000001 | 63.8256604 | -0.119147057 | 0.19007868 | -0.6268302 | 0.53077054 | 0.64489218 | GRAMD1C | 54762 |
| ENSG000002 | 3.58395008 | -0.236805869 | 0.37792737 | -0.6265909 | 0.53092741 | 0.64506344 | NA | NA |
| ENSG000001 | 92.804258 | 0.117322449 | 0.18729095 | 0.62641812 | 0.53104072 | 0.64518178 | SLC45A3 | 85414 |

| | | | | | | | | |
|---|---|---|---|---|---|---|---|---|
| ENSG000001 | 2.56179389 | -0.231684351 | 0.36987325 | -0.6263885 | 0.53106014 | 0.64518605 | NEUROG3 | 50674 |
| ENSG000002 | 17.5309946 | 0.191788442 | 0.30623303 | 0.62628268 | 0.53112954 | 0.6452317 | NA | NA |
| ENSG000002 | 2.81064906 | -0.234349036 | 0.37418923 | -0.6262848 | 0.53112814 | 0.6452317 | NA | NA |
| ENSG000002 | 44.299896 | 0.136835182 | 0.21854367 | 0.62612283 | 0.53123438 | 0.64532546 | NA | NA |
| ENSG000002 | 4.07495581 | -0.236384041 | 0.37754005 | -0.6261165 | 0.53123855 | 0.64532546 | NA | NA |
| ENSG000001 | 14.6097486 | 0.198847044 | 0.31761863 | 0.62605599 | 0.53127822 | 0.64533973 | PRR16 | 51334 |
| ENSG000002 | 2.84982672 | -0.234386972 | 0.37439017 | -0.62605 | 0.53128213 | 0.64533973 | NA | NA |
| ENSG000002 | 3.073826 | -0.236389725 | 0.37760578 | -0.6260225 | 0.53130017 | 0.64534231 | NA | NA |
| ENSG000002 | 18.139311 | -0.179621802 | 0.28694919 | -0.6259708 | 0.53133412 | 0.64536422 | NA | NA |
| ENSG000002 | 4.59654976 | -0.236135675 | 0.37727398 | -0.6258997 | 0.53138072 | 0.6454015 | NAV2-AS4 | 399876 |
| ENSG000002 | 2.3248512 | -0.229202188 | 0.36634181 | -0.6256512 | 0.53154376 | 0.64556084 | NA | NA |
| ENSG000002 | 3.82304078 | -0.236598334 | 0.37816271 | -0.6256522 | 0.53154309 | 0.64556084 | NA | NA |
| ENSG000002 | 3.81536875 | -0.236549304 | 0.37810857 | -0.6256121 | 0.53156938 | 0.64557263 | LOC100128568 | 100128568 |
| ENSG000002 | 3.79963196 | -0.236634289 | 0.37829778 | -0.6255239 | 0.53162729 | 0.64562361 | NA | NA |
| ENSG000002 | 3.58243131 | -0.236756228 | 0.37857647 | -0.6253855 | 0.53171809 | 0.64571455 | NA | NA |
| ENSG000002 | 2.33037163 | -0.229622138 | 0.36721846 | -0.6253012 | 0.53177341 | 0.64576239 | NA | NA |
| ENSG000002 | 4.55949975 | -0.235265598 | 0.37628304 | -0.6252357 | 0.53181636 | 0.64578282 | NA | NA |
| ENSG000002 | 6.33355529 | -0.227820097 | 0.36439396 | -0.6252027 | 0.53183801 | 0.64578282 | NA | NA |
| ENSG000002 | 5.50170334 | 0.232583752 | 0.37200949 | 0.6252092 | 0.53183377 | 0.64578282 | EGFL8 | 80864 |
| ENSG000001 | 54.6328868 | -0.12543788 | 0.20067776 | -0.6250712 | 0.53192435 | 0.64586832 | ACCS | 84680 |
| ENSG000002 | 3.08153048 | -0.235839849 | 0.37738097 | -0.6249384 | 0.53201151 | 0.64595481 | RN7SL42P | 106479240 |
| ENSG000001 | 14.7457735 | 0.196390743 | 0.31434841 | 0.62475501 | 0.53213186 | 0.64608159 | CHST1 | 8534 |
| ENSG000001 | 48.3254716 | -0.137385318 | 0.21994506 | -0.6246347 | 0.53221085 | 0.6461388 | EXD3 | 54932 |
| ENSG000002 | 3.59504856 | -0.236484358 | 0.37859084 | -0.6246436 | 0.53220497 | 0.6461388 | MIR4696 | 100616402 |
| ENSG000002 | 10.9688095 | 0.210059444 | 0.3363067 | 0.62460677 | 0.53222917 | 0.64614171 | NA | NA |
| ENSG000002 | 5.76786487 | -0.234478031 | 0.37543047 | -0.6245578 | 0.53226132 | 0.64616139 | NA | NA |
| ENSG000002 | 2.30608058 | 0.231354431 | 0.37047978 | 0.62447249 | 0.53231733 | 0.64621003 | NA | NA |
| ENSG000002 | 3.04532869 | -0.235869808 | 0.37772615 | -0.6244466 | 0.53233433 | 0.64621133 | RNU6-1246P | 106480110 |
| ENSG000002 | 6.32224573 | -0.22986153 | 0.36823579 | -0.6242238 | 0.53248064 | 0.64636959 | NA | NA |

| | | | | | | | | |
|---|---|---|---|---|---|---|---|---|
| ENSG000001 | 144.642339 | 0.091985858 | 0.14742358 | 0.6239562 | 0.53265635 | 0.64648611 | TBC1D12 | 23232 |
| ENSG000002 | 2.64589622 | -0.231550046 | 0.37107916 | -0.623991 | 0.53263352 | 0.64648611 | NA | NA |
| ENSG000002 | 3.82258546 | 0.236232498 | 0.3785539 | 0.62403928 | 0.53260179 | 0.64648611 | NA | NA |
| ENSG000002 | 2.36456931 | -0.225352172 | 0.3611535 | -0.6239789 | 0.53264143 | 0.64648611 | NA | NA |
| ENSG000002 | 17.4682617 | -0.178885988 | 0.2866657 | -0.624023 | 0.53261248 | 0.64648611 | NA | NA |
| ENSG000002 | 4.3428812 | -0.235099209 | 0.37682937 | -0.6238877 | 0.53270133 | 0.64652136 | NA | NA |
| ENSG000001 | 29.737604 | 0.153270487 | 0.24570975 | 0.62378676 | 0.53276764 | 0.64654107 | DUSP28 | 285193 |
| ENSG000002 | 3.33934289 | -0.235771425 | 0.37796933 | -0.6237845 | 0.53276909 | 0.64654107 | NA | NA |
| ENSG000002 | 4.32402593 | -0.235313764 | 0.37724693 | -0.6237659 | 0.53278135 | 0.64654107 | NA | NA |
| ENSG000002 | 8.54318073 | -0.215439521 | 0.34535894 | -0.6238134 | 0.53275016 | 0.64654107 | NA | NA |
| ENSG000002 | 3.85742791 | -0.236055339 | 0.37845426 | -0.6237355 | 0.53280133 | 0.64654598 | LOC100192426 | 100192426 |
| ENSG000001 | 92.4467991 | 0.113263336 | 0.18162233 | 0.6236201 | 0.53287711 | 0.64658051 | SLC2A11 | 66035 |
| ENSG000002 | 4.07530868 | -0.235325814 | 0.37735492 | -0.6236193 | 0.53287763 | 0.64658051 | NA | NA |
| ENSG000002 | 3.55707951 | -0.236094533 | 0.37857853 | -0.6236342 | 0.53286782 | 0.64658051 | RN7SL40P | 106480483 |
| ENSG000001 | 26.8952199 | -0.164079506 | 0.26321413 | -0.6233689 | 0.53304212 | 0.64671497 | NA | NA |
| ENSG000002 | 3.38857316 | -0.235686979 | 0.37810993 | -0.6233292 | 0.53306818 | 0.64671497 | RNU4ATAC16P | 106481160 |
| ENSG000002 | 3.6280798 | -0.235943311 | 0.37851136 | -0.6233454 | 0.53305758 | 0.64671497 | NA | NA |
| ENSG000002 | 2.30281962 | 0.2299539 | 0.36888633 | 0.62337333 | 0.53303922 | 0.64671497 | NA | NA |
| ENSG000002 | 15.6232385 | -0.196770155 | 0.31565414 | -0.6233726 | 0.53303967 | 0.64671497 | NA | NA |
| ENSG000001 | 290.839516 | -0.11756408 | 0.18864457 | -0.6232041 | 0.53315038 | 0.64675664 | NOTCH1 | 4851 |
| ENSG000002 | 3.16642317 | 0.235229802 | 0.37743225 | 0.62323715 | 0.53312869 | 0.64675664 | MAGEA3 | 4102 |
| ENSG000002 | 4.08168234 | -0.23543533 | 0.37777494 | -0.6232159 | 0.53314268 | 0.64675664 | NA | NA |
| ENSG000002 | 2.56063322 | -0.232065154 | 0.37252644 | -0.6229495 | 0.53331771 | 0.64694028 | NA | NA |
| ENSG000002 | 293.236474 | 0.08127617 | 0.13051801 | 0.62271998 | 0.53346855 | 0.6471039 | SHISA9 | 729993 |
| ENSG000002 | 5.68395577 | 0.230508315 | 0.37017864 | 0.6226948 | 0.5334851 | 0.64710417 | NA | NA |
| ENSG000002 | 9.94768576 | -0.206787722 | 0.33209785 | -0.6226711 | 0.5335007 | 0.64710417 | NA | NA |
| ENSG000001 | 5.20794106 | -0.234553868 | 0.37676191 | -0.622552 | 0.53357899 | 0.64714106 | AZU1 | 566 |
| ENSG000002 | 5.56851862 | -0.230538146 | 0.3703073 | -0.622559 | 0.53357436 | 0.64714106 | NA | NA |
| ENSG000002 | 4.59307342 | -0.233997674 | 0.37586308 | -0.6225609 | 0.53357308 | 0.64714106 | NA | NA |

| | | | | | | | | |
|---|---|---|---|---|---|---|---|---|
| ENSG000002 | 13.3409365 | 0.193934613 | 0.31153453 | 0.62251403 | 0.53360392 | 0.64715195 | NA | NA |
| ENSG000002 | 8.57773094 | -0.217256321 | 0.34904653 | -0.622428 | 0.53366049 | 0.6472012 | NA | NA |
| ENSG000002 | 4.87619318 | -0.234391786 | 0.37664331 | -0.6223177 | 0.53373301 | 0.64726979 | NA | NA |
| ENSG000002 | 2.94534018 | 0.233421926 | 0.37530885 | 0.62194623 | 0.53397723 | 0.64754659 | NA | NA |
| ENSG000002 | 2.25570836 | 0.228823722 | 0.36795187 | 0.62188493 | 0.53401754 | 0.64757611 | RNU6-271P | 106481264 |
| ENSG000002 | 24.9071282 | 0.17657694 | 0.2840136 | 0.62172002 | 0.53412599 | 0.64768825 | LOC102467213 | 102467213 |
| ENSG000002 | 2.29071647 | 0.229295046 | 0.36884429 | 0.62165812 | 0.5341667 | 0.64771825 | NA | NA |
| ENSG000002 | 3.85797645 | -0.235165307 | 0.37858984 | -0.6211612 | 0.53449359 | 0.64809525 | NA | NA |
| ENSG000002 | 3.16715983 | 0.234757023 | 0.37800465 | 0.62104269 | 0.53457153 | 0.64813752 | LINC00845 | 100507058 |
| ENSG000002 | 4.79765719 | 0.23454799 | 0.37765384 | 0.62106608 | 0.53455614 | 0.64813752 | NA | NA |
| ENSG000002 | 4.57415443 | -0.233539319 | 0.37606309 | -0.621011 | 0.53459238 | 0.64813752 | NA | NA |
| ENSG000002 | 3.33567665 | -0.234758732 | 0.3780237 | -0.6210159 | 0.53458916 | 0.64813752 | NA | NA |
| ENSG000002 | 4.78349068 | 0.233188134 | 0.37554008 | 0.62094073 | 0.53463862 | 0.6481742 | NA | NA |
| ENSG000002 | 2.59506753 | -0.231317075 | 0.37254163 | -0.620916 | 0.53465487 | 0.64817452 | ZNF341-AS1 | 101929746 |
| ENSG000001 | 5.92506192 | 0.227747335 | 0.36696326 | 0.62062707 | 0.53484502 | 0.64825103 | NA | NA |
| ENSG000001 | 67.0737372 | 0.118583899 | 0.19106239 | 0.62065536 | 0.53482641 | 0.64825103 | ELOVL2 | 54898 |
| ENSG000002 | 11.3536768 | -0.202636445 | 0.32649126 | -0.6206489 | 0.53483065 | 0.64825103 | LOC285696 | 285696 |
| ENSG000002 | 2.27788353 | 0.229640341 | 0.37001418 | 0.62062579 | 0.53484587 | 0.64825103 | NA | NA |
| ENSG000002 | 7.740532 | -0.218108717 | 0.3513859 | -0.6207099 | 0.5347905 | 0.64825103 | NA | NA |
| ENSG000002 | 3.9819375 | 0.23471469 | 0.3781567 | 0.62068103 | 0.53480951 | 0.64825103 | NA | NA |
| ENSG000002 | 2.28044846 | -0.226355541 | 0.36470222 | -0.6206585 | 0.53482432 | 0.64825103 | NA | NA |
| ENSG000002 | 13.3039786 | 0.195380401 | 0.31479312 | 0.62066288 | 0.53482146 | 0.64825103 | NA | NA |
| ENSG000001 | 2.84726331 | -0.233299814 | 0.3759934 | -0.6204891 | 0.53493581 | 0.64832129 | NA | NA |
| ENSG000002 | 3.0538095 | -0.234099246 | 0.37726873 | -0.6205106 | 0.53492167 | 0.64832129 | NA | NA |
| ENSG000002 | 3.07180337 | -0.234342771 | 0.37772762 | -0.6204015 | 0.53499351 | 0.64836742 | NA | NA |
| ENSG000002 | 28.2676674 | 0.153670659 | 0.247703 | 0.62038271 | 0.53500585 | 0.64836742 | NA | NA |
| ENSG000002 | 19.8092456 | 0.171930216 | 0.2772069 | 0.62022343 | 0.53511069 | 0.6484598 | FBXL19-AS1 | 283932 |
| ENSG000002 | 16.9231536 | -0.185383241 | 0.29889998 | -0.6202183 | 0.53511406 | 0.6484598 | NA | NA |
| ENSG000002 | 4.02368275 | -0.23427751 | 0.37780227 | -0.6201061 | 0.53518791 | 0.64852991 | NA | NA |

| | | | | | | | | |
|---|---|---|---|---|---|---|---|---|
| ENSG000002 | 5.17503323 | -0.233281764 | 0.37621424 | -0.6200769 | 0.53520716 | 0.64853385 | LOC102724691 | 102724691 |
| ENSG000002 | 4.82092261 | -0.231593027 | 0.3735124 | -0.6200411 | 0.53523075 | 0.64854306 | NA | NA |
| ENSG000002 | 3.61926651 | -0.234691881 | 0.37856156 | -0.619957 | 0.53528611 | 0.64857139 | LOC105374825 | 105374825 |
| ENSG000002 | 2.81279653 | -0.23309084 | 0.37597725 | -0.61996 | 0.53528415 | 0.64857139 | NA | NA |
| ENSG000001 | 4.80722826 | -0.231975926 | 0.37424886 | -0.619844 | 0.53536047 | 0.64862437 | DRD5 | 1816 |
| ENSG000002 | 4.05402662 | -0.234316177 | 0.37802567 | -0.619842 | 0.53536183 | 0.64862437 | NA | NA |
| ENSG000002 | 3.82777639 | -0.234564246 | 0.37856943 | -0.6196069 | 0.53551659 | 0.64877311 | NA | NA |
| ENSG000002 | 6.81038287 | -0.224073262 | 0.36162991 | -0.6196204 | 0.53550775 | 0.64877311 | NA | NA |
| ENSG000002 | 5.40492899 | -0.232641664 | 0.37551435 | -0.619528 | 0.53556857 | 0.64881669 | NA | NA |
| ENSG000001 | 8.72246296 | -0.214878812 | 0.34689358 | -0.6194373 | 0.53562833 | 0.6488697 | THBD | 7056 |
| ENSG000002 | 4.10760828 | -0.234335737 | 0.37839383 | -0.6192906 | 0.53572495 | 0.64896736 | NA | NA |
| ENSG000001 | 52.5558563 | 0.127572819 | 0.2060643 | 0.61909229 | 0.53585556 | 0.64906741 | IFI6 | 2537 |
| ENSG000002 | 3.13378151 | -0.233451802 | 0.37707852 | -0.6191066 | 0.53584613 | 0.64906741 | NA | NA |
| ENSG000002 | 4.34244435 | -0.233119009 | 0.37653471 | -0.6191169 | 0.53583934 | 0.64906741 | NA | NA |
| ENSG000002 | 4.27468431 | -0.233434842 | 0.37710439 | -0.6190192 | 0.53590374 | 0.64910638 | NA | NA |
| ENSG00000C | 214.001712 | 0.08236716 | 0.13307767 | 0.61894052 | 0.53595554 | 0.64912489 | TRPM3 | 80036 |
| ENSG000001 | 4.97956351 | -0.234038427 | 0.37811728 | -0.6189572 | 0.53594453 | 0.64912489 | OCSTAMP | 128506 |
| ENSG000002 | 3.5823994 | -0.234312128 | 0.37858038 | -0.6189231 | 0.53596704 | 0.64912489 | NA | NA |
| ENSG000001 | 70.3488357 | -0.127852564 | 0.2066273 | -0.6187593 | 0.53607494 | 0.64912594 | BBC3 | 27113 |
| ENSG000002 | 2.54719945 | -0.22873868 | 0.36961612 | -0.6188547 | 0.53601207 | 0.64912594 | NA | NA |
| ENSG000002 | 97.3583916 | -0.109662133 | 0.17721926 | -0.6187935 | 0.53605239 | 0.64912594 | PLXNA4 | 91584 |
| ENSG000002 | 17.7456372 | 0.186089881 | 0.30075219 | 0.61874821 | 0.53608225 | 0.64912594 | NA | NA |
| ENSG000002 | 3.53686749 | -0.234291567 | 0.37858972 | -0.6188535 | 0.53601289 | 0.64912594 | NA | NA |
| ENSG000002 | 3.12254881 | 0.233764175 | 0.3778145 | 0.61872738 | 0.53609597 | 0.64912594 | LINC00051 | 619434 |
| ENSG000002 | 2.17880818 | -0.218486732 | 0.35308712 | -0.6187899 | 0.5360548 | 0.64912594 | NA | NA |
| ENSG000002 | 11.177312 | 0.202900301 | 0.32790784 | 0.61877233 | 0.53606635 | 0.64912594 | NA | NA |
| ENSG000002 | 2.8021677 | -0.231286884 | 0.37386604 | -0.6186357 | 0.53615637 | 0.64917968 | NA | NA |
| ENSG000002 | 3.07453096 | -0.233571423 | 0.37760501 | -0.6185602 | 0.53620614 | 0.64922056 | NA | NA |
| ENSG000002 | 34.6606315 | 0.14279079 | 0.23085389 | 0.61853317 | 0.53622394 | 0.64922273 | NA | NA |

| | | | | | | | | |
|---|---|---|---|---|---|---|---|---|
| ENSG000002 | 4.07624177 | -0.233528594 | 0.37758342 | -0.6184821 | 0.53625758 | 0.64922805 | NA | NA |
| ENSG000002 | 7.91790576 | 0.220415682 | 0.35638408 | 0.61847791 | 0.53626036 | 0.64922805 | HRAT92 | 441307 |
| ENSG000002 | 4.33864564 | -0.233956049 | 0.3782931 | -0.6184518 | 0.53627756 | 0.6492295 | NA | NA |
| ENSG000002 | 3.79794901 | -0.233942545 | 0.37829389 | -0.6184148 | 0.53630193 | 0.64923962 | NA | NA |
| ENSG000001 | 13.2113582 | -0.197909077 | 0.3200971 | -0.6182783 | 0.53639193 | 0.64927104 | C6orf58 | 352999 |
| ENSG000002 | 2.29902941 | -0.226141351 | 0.36575377 | -0.6182885 | 0.53638518 | 0.64927104 | NA | NA |
| ENSG000002 | 22.7176071 | 0.166560278 | 0.26937599 | 0.61831895 | 0.53636511 | 0.64927104 | AGPAT4-IT1 | 79992 |
| ENSG000002 | 3.156174 | 0.233524777 | 0.37768433 | 0.61830677 | 0.53637314 | 0.64927104 | NA | NA |
| ENSG000002 | 2.33693353 | 0.228893159 | 0.37023479 | 0.61823785 | 0.53641857 | 0.6492839 | MIR548AM | 100616428 |
| ENSG000002 | 4.82959205 | 0.232899995 | 0.37673075 | 0.61821339 | 0.53643469 | 0.64928403 | NA | NA |
| ENSG000002 | 2.95532876 | 0.231501191 | 0.37455641 | 0.61806763 | 0.53653076 | 0.64934217 | NA | NA |
| ENSG000002 | 9.03168379 | -0.215023827 | 0.34788496 | -0.6180889 | 0.53651675 | 0.64934217 | NA | NA |
| ENSG000002 | 29.1223872 | 0.160961257 | 0.26041545 | 0.61809412 | 0.5365133 | 0.64934217 | NA | NA |
| ENSG000002 | 6.58258609 | -0.224994502 | 0.36407489 | -0.6179896 | 0.53658218 | 0.64936894 | NA | NA |
| ENSG000002 | 5.02439632 | -0.230331632 | 0.37272831 | -0.6179612 | 0.53660092 | 0.64936894 | NA | NA |
| ENSG000002 | 2.58521589 | -0.230235068 | 0.37256729 | -0.6179691 | 0.53659572 | 0.64936894 | NA | NA |
| ENSG000002 | 3.57079347 | -0.233847274 | 0.37854408 | -0.6177544 | 0.53673724 | 0.64949515 | NA | NA |
| ENSG000002 | 3.93240585 | 0.232908099 | 0.37702028 | 0.61776014 | 0.53673346 | 0.64949515 | NA | NA |
| ENSG000001 | 1153.62125 | 0.07936066 | 0.12850517 | 0.61756781 | 0.53686027 | 0.64962463 | LITAF | 9516 |
| ENSG000002 | 6.14315907 | -0.230102333 | 0.37268591 | -0.6174162 | 0.53696022 | 0.6497068 | NA | NA |
| ENSG000002 | 83.3165017 | 0.109505298 | 0.17735945 | 0.61742016 | 0.53695763 | 0.6497068 | NA | NA |
| ENSG000002 | 3.13318232 | 0.233106981 | 0.37761348 | 0.61731636 | 0.53702608 | 0.6497671 | NA | NA |
| ENSG000001 | 4.30172701 | -0.232804694 | 0.37716218 | -0.6172535 | 0.5370675 | 0.64978064 | FAM170B | 170370 |
| ENSG000002 | 3.31759248 | -0.233594353 | 0.37844318 | -0.6172508 | 0.53706931 | 0.64978064 | PRAMEF19 | 645414 |
| ENSG000001 | 18.1668978 | -0.18094281 | 0.29323292 | -0.6170617 | 0.53719402 | 0.64985378 | STMN2 | 11075 |
| ENSG000001 | 89.7813191 | -0.115773064 | 0.18761082 | -0.6170916 | 0.5371743 | 0.64985378 | HIF3A | 64344 |
| ENSG000002 | 2.88474997 | -0.229822855 | 0.37245131 | -0.6170548 | 0.53719861 | 0.64985378 | NA | NA |
| ENSG000002 | 154.990006 | 0.091398074 | 0.14811045 | 0.61709403 | 0.5371727 | 0.64985378 | NA | NA |
| ENSG000002 | 4.05788949 | -0.232955435 | 0.37753844 | -0.6170377 | 0.53720989 | 0.64985378 | NA | NA |

| | | | | | | | | |
|---|---|---|---|---|---|---|---|---|
| ENSG000002 | 5.56752136 | -0.227318735 | 0.36848079 | -0.616908 | 0.53729543 | 0.64993786 | LOC101929076 | 101929076 |
| ENSG000002 | 36.0585023 | 0.140648804 | 0.22800457 | 0.61686836 | 0.53732156 | 0.64995008 | NA | NA |
| ENSG000002 | 4.28689215 | -0.23205999 | 0.3762064 | -0.6168422 | 0.5373388 | 0.64995155 | NA | NA |
| ENSG000002 | 3.84334956 | -0.23341663 | 0.37858194 | -0.6165551 | 0.53752821 | 0.65016126 | LOC105376199 | 105376199 |
| ENSG000002 | 8.76911089 | -0.211169763 | 0.34253636 | -0.6164886 | 0.53757209 | 0.65019495 | C1QTNF9 | 338872 |
| ENSG000002 | 8.27736505 | -0.219886827 | 0.35671863 | -0.6164153 | 0.53762045 | 0.65019525 | NA | NA |
| ENSG000002 | 2.59783259 | -0.228670435 | 0.37096088 | -0.6164274 | 0.5376125 | 0.65019525 | NA | NA |
| ENSG000002 | 12.4766028 | -0.201220127 | 0.32641121 | -0.6164621 | 0.5375896 | 0.65019525 | NA | NA |
| ENSG000002 | 3.83810541 | -0.233057558 | 0.3781681 | -0.6162803 | 0.53770952 | 0.65026419 | NA | NA |
| ENSG000002 | 3.83078547 | -0.233031793 | 0.37811481 | -0.616299 | 0.53769718 | 0.65026419 | NA | NA |
| ENSG000001 | 3.8799103 | -0.233104876 | 0.37847767 | -0.6159013 | 0.53795968 | 0.65054731 | DEFB123 | 245936 |
| ENSG000001 | 6.84126175 | 0.223191033 | 0.36248575 | 0.61572361 | 0.53807695 | 0.65066973 | C16orf86 | 388284 |
| ENSG000002 | 3.31773629 | -0.232970487 | 0.37841081 | -0.6156549 | 0.53812228 | 0.65070513 | LINC00424 | 100874182 |
| ENSG000002 | 4.05448364 | -0.232656324 | 0.37793958 | -0.6155913 | 0.53816429 | 0.65073652 | NA | NA |
| ENSG000002 | 2.56100515 | -0.228262191 | 0.37082862 | -0.6155463 | 0.53819399 | 0.65075303 | NA | NA |
| ENSG000002 | 4.34438787 | -0.23190605 | 0.37692007 | -0.6152659 | 0.53837916 | 0.6509187 | NA | NA |
| ENSG000002 | 6.54041035 | -0.222116743 | 0.36098596 | -0.6153058 | 0.53835281 | 0.6509187 | GSTA1 | 2938 |
| ENSG000002 | 4.07260071 | -0.232631538 | 0.37808677 | -0.6152861 | 0.53836579 | 0.6509187 | NA | NA |
| ENSG000002 | 8.54418786 | -0.218486872 | 0.35521332 | -0.6150864 | 0.53849768 | 0.65104258 | NA | NA |
| ENSG000002 | 6.77499052 | -0.220607677 | 0.35879465 | -0.6148578 | 0.53864865 | 0.6512057 | NDE1P2 | 106480247 |
| ENSG000002 | 4.84898735 | 0.231607899 | 0.37675896 | 0.6147376 | 0.53872802 | 0.65128224 | NUCB1-AS1 | 100874085 |
| ENSG000002 | 21.138525 | -0.170343122 | 0.2771378 | -0.6146513 | 0.538785 | 0.65131229 | NA | NA |
| ENSG000002 | 4.57281749 | -0.230430546 | 0.37488279 | -0.6146736 | 0.53877031 | 0.65131229 | NA | NA |
| ENSG000002 | 3.57358577 | -0.232680946 | 0.37859125 | -0.6145967 | 0.53882108 | 0.65133648 | NA | NA |
| ENSG000002 | 3.80955604 | -0.232357831 | 0.37809321 | -0.6145517 | 0.53885081 | 0.651353 | NA | NA |
| ENSG000002 | 4.09634224 | -0.232197182 | 0.37785162 | -0.6145195 | 0.5388721 | 0.65135932 | NA | NA |
| ENSG000002 | 6.08043394 | -0.224310134 | 0.3650935 | -0.6143909 | 0.53895703 | 0.65141118 | NA | NA |
| ENSG000002 | 6.30611974 | -0.226368494 | 0.36846392 | -0.6143573 | 0.53897926 | 0.65141118 | NA | NA |
| ENSG000002 | 4.5773644 | -0.230426408 | 0.37503646 | -0.6144107 | 0.53894397 | 0.65141118 | NA | NA |

| | | | | | | | | |
|---|---|---|---|---|---|---|---|---|
| ENSG000002 | 2.82238725 | -0.230474164 | 0.37513929 | -0.6143696 | 0.53897114 | 0.65141118 | NA | NA |
| ENSG000001 | 53.2656962 | -0.129423168 | 0.21073917 | -0.6141391 | 0.5391234 | 0.65156597 | SLC4A3 | 6508 |
| ENSG000002 | 17.9869812 | 0.177910455 | 0.28977138 | 0.61396835 | 0.53923624 | 0.65168293 | LINC01465 | 283416 |
| ENSG000002 | 3.53724017 | -0.232408336 | 0.37857841 | -0.6138975 | 0.53928306 | 0.65169384 | NA | NA |
| ENSG000002 | 4.58314521 | -0.230595602 | 0.37563523 | -0.6138817 | 0.53929349 | 0.65169384 | NA | NA |
| ENSG000002 | 3.79319358 | -0.232153754 | 0.37815467 | -0.6139122 | 0.53927336 | 0.65169384 | NA | NA |
| ENSG000002 | 9.05402616 | 0.211858326 | 0.34517453 | 0.6137716 | 0.53936626 | 0.65176236 | NA | NA |
| ENSG000002 | 4.83528858 | -0.230010776 | 0.37489251 | -0.6135379 | 0.5395207 | 0.65192956 | RNU6-118P | 106479618 |
| ENSG000002 | 5.07486081 | -0.228496451 | 0.3724494 | -0.6134966 | 0.53954802 | 0.65194313 | LOC105377391 | 105377391 |
| ENSG000001 | 10.5514691 | 0.211105743 | 0.34426886 | 0.61320023 | 0.53974395 | 0.65208382 | TAS2R3 | 50831 |
| ENSG000001 | 24.0255722 | -0.162234804 | 0.26453613 | -0.6132803 | 0.53969101 | 0.65208382 | SAXO2 | 283726 |
| ENSG000002 | 4.530417 | -0.230782224 | 0.37633861 | -0.6132303 | 0.5397241 | 0.65208382 | NA | NA |
| ENSG000002 | 13.8585186 | -0.191907169 | 0.31296058 | -0.6131992 | 0.53974465 | 0.65208382 | DKFZP434L187 | 26082 |
| ENSG000002 | 4.29720026 | -0.231215095 | 0.37707876 | -0.6131745 | 0.53976093 | 0.65208382 | NA | NA |
| ENSG000002 | 3.31070809 | -0.231240483 | 0.37711873 | -0.6131769 | 0.53975939 | 0.65208382 | NA | NA |
| ENSG00000C | 57.6432454 | 0.132929241 | 0.21683803 | 0.61303471 | 0.53985339 | 0.65217609 | GUCY1B3 | 2983 |
| ENSG000002 | 5.98649186 | 0.224896369 | 0.3669082 | 0.61294997 | 0.53990942 | 0.65222435 | MAN1B1-AS1 | 100289341 |
| ENSG000001 | 35.2246303 | 0.140390591 | 0.22905598 | 0.61290951 | 0.53993617 | 0.65223483 | KLK10 | 5655 |
| ENSG000002 | 4.81942759 | -0.230028001 | 0.37531804 | -0.6128882 | 0.53995026 | 0.65223483 | NA | NA |
| ENSG000002 | 3.59927039 | -0.231957182 | 0.3785629 | -0.6127309 | 0.5400543 | 0.65232163 | NA | NA |
| ENSG000002 | 12.1377757 | -0.198418154 | 0.32382208 | -0.6127382 | 0.54004947 | 0.65232163 | NA | NA |
| ENSG000002 | 4.58363359 | -0.231149628 | 0.37729278 | -0.6126532 | 0.54010568 | 0.65236426 | RN7SL200P | 106480976 |
| ENSG000002 | 2.60072325 | -0.225951646 | 0.36884967 | -0.6125846 | 0.54015101 | 0.65238929 | NA | NA |
| ENSG000002 | 11.3060307 | -0.203618309 | 0.33239833 | -0.6125732 | 0.54015858 | 0.65238929 | NA | NA |
| ENSG000001 | 3.09053336 | -0.230871215 | 0.37696585 | -0.612446 | 0.54024273 | 0.6524715 | NA | NA |
| ENSG000001 | 4.77651987 | -0.228632624 | 0.37339278 | -0.6123113 | 0.54033181 | 0.65255964 | OR1F1 | 4992 |
| ENSG000001 | 60.8177447 | 0.135638285 | 0.2215471 | 0.61223229 | 0.54038408 | 0.6525902 | EGR1 | 1958 |
| ENSG000001 | 78.1642719 | -0.107410307 | 0.17544271 | -0.6122244 | 0.5403893 | 0.6525902 | LACC1 | 144811 |
| ENSG000002 | 17.305684 | 0.179650444 | 0.29349461 | 0.61210816 | 0.5404662 | 0.65266363 | NA | NA |

| | | | | | | | | |
|---|---|---|---|---|---|---|---|---|
| ENSG000002 | 8.10008108 | -0.219634799 | 0.35888148 | -0.6119981 | 0.54053898 | 0.65267378 | NA | NA |
| ENSG000002 | 3.09192346 | -0.230951066 | 0.37734972 | -0.6120346 | 0.5405149 | 0.65267378 | NA | NA |
| ENSG000002 | 3.80618243 | -0.231653166 | 0.37848404 | -0.6120553 | 0.54050117 | 0.65267378 | NA | NA |
| ENSG000002 | 4.22634578 | 0.231219732 | 0.37779833 | 0.61201894 | 0.54052522 | 0.65267378 | NA | NA |
| ENSG000002 | 2.27605678 | 0.224905912 | 0.36757576 | 0.61186274 | 0.54062857 | 0.65271442 | NA | NA |
| ENSG000002 | 9.92160252 | 0.206677305 | 0.33776564 | 0.61189559 | 0.54060684 | 0.65271442 | NA | NA |
| ENSG000002 | 14.1178365 | -0.193523421 | 0.31625515 | -0.6119218 | 0.5405895 | 0.65271442 | NA | NA |
| ENSG000002 | 5.31285391 | -0.226518325 | 0.37021874 | -0.61185 | 0.54063702 | 0.65271442 | NA | NA |
| ENSG000001 | 116.274645 | -0.100255201 | 0.16388375 | -0.6117458 | 0.54070593 | 0.65277818 | DSC3 | 1825 |
| ENSG00000C | 118.228757 | 0.101183011 | 0.16541261 | 0.61170069 | 0.5407358 | 0.65279481 | SMAP2 | 64744 |
| ENSG000001 | 120.954116 | -0.098386191 | 0.16086254 | -0.6116165 | 0.54079149 | 0.6528426 | AMPD3 | 272 |
| ENSG000002 | 7.0601792 | -0.218526257 | 0.35731485 | -0.611579 | 0.54081633 | 0.65285315 | NA | NA |
| ENSG000002 | 4.06011267 | -0.231265761 | 0.37816347 | -0.6115497 | 0.54083572 | 0.65285712 | BUD31P1 | 106480331 |
| ENSG000002 | 6.38742511 | -0.224102032 | 0.3665513 | -0.6113797 | 0.54094823 | 0.6529735 | NA | NA |
| ENSG00000C | 6.55332618 | -0.223143221 | 0.36505866 | -0.6112531 | 0.54103207 | 0.65303865 | TBXA2R | 6915 |
| ENSG000001 | 39.3630383 | 0.134895814 | 0.22069741 | 0.61122518 | 0.54105052 | 0.65303865 | HYAL1 | 3373 |
| ENSG000001 | 694.437304 | 0.075582172 | 0.12365467 | 0.61123591 | 0.54104341 | 0.65303865 | PRDM5 | 11107 |
| ENSG00000C | 287.599945 | -0.08076732 | 0.13215358 | -0.6111625 | 0.54109198 | 0.65306497 | ZNF37A | 7587 |
| ENSG000002 | 4.14330153 | -0.23117412 | 0.37827989 | -0.6111192 | 0.54112064 | 0.65306497 | NA | NA |
| ENSG000002 | 4.30968774 | -0.23021171 | 0.37669468 | -0.6111361 | 0.5411095 | 0.65306497 | NA | NA |
| ENSG000002 | 6.52792341 | -0.220519239 | 0.36090865 | -0.6110112 | 0.54119214 | 0.65313183 | NA | NA |
| ENSG000002 | 3.81166319 | -0.23118258 | 0.37841249 | -0.6109275 | 0.54124761 | 0.65317932 | NA | NA |
| ENSG000002 | 6.97333729 | 0.220115868 | 0.36035928 | 0.61082337 | 0.54131652 | 0.65324305 | NA | NA |
| ENSG000002 | 8.37320737 | -0.215604621 | 0.35299516 | -0.6107863 | 0.54134104 | 0.6532532 | NA | NA |
| ENSG000002 | 8.97835275 | -0.209767447 | 0.3434587 | -0.6107501 | 0.54136501 | 0.65326268 | NA | NA |
| ENSG000002 | 7.92364679 | 0.217337892 | 0.35594131 | 0.61060036 | 0.54146418 | 0.65336291 | NA | NA |
| ENSG000002 | 7.9878365 | -0.212965003 | 0.34879823 | -0.6105679 | 0.54148568 | 0.65336941 | NA | NA |
| ENSG000002 | 3.36305251 | -0.230755423 | 0.3779614 | -0.6105264 | 0.54151314 | 0.6533831 | NA | NA |
| ENSG000001 | 47.5472795 | -0.128202574 | 0.21001028 | -0.6104586 | 0.54155808 | 0.65341789 | AGAP2 | 116986 |

| | | | | | | | | |
|---|---|---|---|---|---|---|---|---|
| ENSG000002 | 5.75765413 | -0.22390638 | 0.36680015 | -0.6104315 | 0.54157598 | 0.65342004 | NA | NA |
| ENSG000001 | 10.9885356 | -0.200586593 | 0.32865771 | -0.6103207 | 0.54164941 | 0.65348537 | C7orf55 | 154791 |
| ENSG000001 | 8.04245419 | -0.217671046 | 0.35666172 | -0.6103011 | 0.54166236 | 0.65348537 | VN1R1 | 57191 |
| ENSG000001 | 4.87052537 | -0.228546173 | 0.37452319 | -0.6102324 | 0.5417079 | 0.65352087 | RHOXF1 | 158800 |
| ENSG000002 | 3.32195094 | -0.230683619 | 0.37805243 | -0.6101895 | 0.54173629 | 0.65353568 | NA | NA |
| ENSG000001 | 61.8353207 | 0.121083103 | 0.19845763 | 0.61012066 | 0.54178188 | 0.65357123 | GSTM3 | 2947 |
| ENSG000002 | 9.48933227 | -0.209122889 | 0.34279301 | -0.6100559 | 0.5418248 | 0.65360357 | NA | NA |
| ENSG000002 | 3.08873309 | -0.229273088 | 0.37585941 | -0.6099969 | 0.54186384 | 0.65363121 | NA | NA |
| ENSG000002 | 3.56487607 | -0.230831056 | 0.37859262 | -0.6097083 | 0.54205507 | 0.65384244 | NA | NA |
| ENSG000002 | 3.64458661 | -0.230622175 | 0.37832861 | -0.6095816 | 0.54213898 | 0.65390475 | NA | NA |
| ENSG000002 | 4.34042914 | -0.229849856 | 0.37705815 | -0.6095873 | 0.54213523 | 0.65390475 | NA | NA |
| ENSG000000 | 6.66707675 | -0.225951909 | 0.37075326 | -0.6094401 | 0.54223276 | 0.65397896 | RGS11 | 8786 |
| ENSG000002 | 5.03182781 | -0.227491992 | 0.37327088 | -0.6094555 | 0.54222256 | 0.65397896 | NA | NA |
| ENSG000001 | 87.5091655 | 0.110236387 | 0.18089939 | 0.60937954 | 0.5422729 | 0.65398847 | DUSP4 | 1846 |
| ENSG000002 | 4.27548155 | -0.229853519 | 0.37718097 | -0.6093985 | 0.54226032 | 0.65398847 | NA | NA |
| ENSG000002 | 5.03270102 | 0.228651423 | 0.37531963 | 0.60921786 | 0.54238004 | 0.65409823 | ARHGEF9-IT1 | 100874355 |
| ENSG000002 | 4.3170592 | -0.2300531 | 0.37766258 | -0.6091498 | 0.54242513 | 0.65413315 | LOC105373518 | 105373518 |
| ENSG000002 | 8.18670441 | 0.213426454 | 0.35042577 | 0.60904898 | 0.54249198 | 0.65419431 | NA | NA |
| ENSG000001 | 8.60331917 | -0.214544236 | 0.35234007 | -0.6089124 | 0.54258251 | 0.65428403 | GPR35 | 2859 |
| ENSG000002 | 3.54526507 | -0.23050521 | 0.3785788 | -0.6088698 | 0.54261072 | 0.6542986 | NA | NA |
| ENSG000002 | 2.56264843 | -0.224562638 | 0.36887269 | -0.6087809 | 0.54266969 | 0.65435024 | NA | NA |
| ENSG000002 | 3.1708654 | 0.230177785 | 0.37813626 | 0.6087165 | 0.54271237 | 0.65438224 | NA | NA |
| ENSG000002 | 2.30066049 | -0.221370068 | 0.36368437 | -0.6086873 | 0.54273171 | 0.65438611 | NA | NA |
| ENSG000002 | 6.98900066 | -0.218600378 | 0.35931765 | -0.6083764 | 0.54293786 | 0.6546152 | NA | NA |
| ENSG000002 | 50.3737813 | -0.134641293 | 0.22134506 | -0.6082869 | 0.54299723 | 0.6546505 | FRRS1L | 23732 |
| ENSG000002 | 4.31984353 | -0.229858249 | 0.3778801 | -0.6082836 | 0.54299943 | 0.6546505 | NA | NA |
| ENSG000002 | 4.52232053 | -0.228594977 | 0.37586666 | -0.608181 | 0.5430674 | 0.65471299 | NA | NA |
| ENSG000002 | 3.85362596 | -0.230185506 | 0.37857025 | -0.6080391 | 0.54316156 | 0.65480704 | NA | NA |
| ENSG000002 | 6.78390907 | -0.219196089 | 0.3605372 | -0.6079708 | 0.54320684 | 0.65484216 | LOC101927337 | 101927337 |

| | | | | | | | | |
|---|---|---|---|---|---|---|---|---|
| ENSG000002 | 3.62198613 | -0.229873352 | 0.3781314 | -0.6079192 | 0.54324104 | 0.65485693 | PRB1 | 5542 |
| ENSG000002 | 7.6785995 | -0.221321153 | 0.36410048 | -0.6078573 | 0.54328209 | 0.65485693 | NA | NA |
| ENSG000002 | 5.73924018 | 0.226394751 | 0.37244865 | 0.60785494 | 0.54328368 | 0.65485693 | NA | NA |
| ENSG000002 | 3.37613523 | 0.230098006 | 0.37853811 | 0.60785956 | 0.54328062 | 0.65485693 | NA | NA |
| ENSG000002 | 21.8776892 | -0.180238507 | 0.29654215 | -0.6078006 | 0.5433197 | 0.65488088 | NA | NA |
| ENSG000001 | 91.819137 | -0.116533489 | 0.1917665 | -0.6076843 | 0.54339689 | 0.65488156 | CLIP3 | 25999 |
| ENSG000001 | 48.3636761 | 0.133378022 | 0.21948353 | 0.60769035 | 0.54339286 | 0.65488156 | SLC6A11 | 6538 |
| ENSG000001 | 78.9112605 | -0.112824138 | 0.18564562 | -0.6077393 | 0.54336039 | 0.65488156 | NTNG1 | 22854 |
| ENSG000002 | 5.92101557 | 0.224009075 | 0.36863117 | 0.60767806 | 0.54340102 | 0.65488156 | NA | NA |
| ENSG000002 | 7.96227509 | -0.213181865 | 0.35075973 | -0.6077718 | 0.54333884 | 0.65488156 | NA | NA |
| ENSG000002 | 5.06093888 | -0.225922133 | 0.3718389 | -0.6075807 | 0.54346561 | 0.65493994 | NA | NA |
| ENSG000001 | 21.0774951 | -0.168645191 | 0.27766306 | -0.6073735 | 0.54360306 | 0.65507648 | NA | NA |
| ENSG000002 | 3.09215179 | -0.228968675 | 0.37700331 | -0.6073386 | 0.54362621 | 0.65507648 | LINC00574 | 80069 |
| ENSG000002 | 29.9092513 | 0.153011893 | 0.25193908 | 0.60733687 | 0.54362737 | 0.65507648 | NA | NA |
| ENSG000001 | 57.5948194 | -0.120706824 | 0.19876851 | -0.6072734 | 0.54366951 | 0.65508148 | FMNL1 | 752 |
| ENSG000001 | 3.84500967 | 0.229723469 | 0.37832582 | 0.60721066 | 0.54371112 | 0.65508148 | RNU1-72P | 106481955 |
| ENSG000002 | 4.30322159 | -0.229195884 | 0.3774325 | -0.60725 | 0.54368501 | 0.65508148 | NA | NA |
| ENSG000002 | 4.3467963 | -0.229705575 | 0.37829746 | -0.6072089 | 0.5437123 | 0.65508148 | NA | NA |
| ENSG000002 | 2.88355629 | -0.226156973 | 0.37244342 | -0.607225 | 0.54370157 | 0.65508148 | NA | NA |
| ENSG000001 | 32.6035153 | -0.143757759 | 0.23676314 | -0.6071797 | 0.54373169 | 0.65508538 | ISG20 | 3669 |
| ENSG000002 | 12.3472919 | -0.194993836 | 0.32118743 | -0.6071029 | 0.5437826 | 0.65510779 | NA | NA |
| ENSG000002 | 52.0639565 | 0.127011472 | 0.20920846 | 0.60710487 | 0.54378132 | 0.65510779 | NA | NA |
| ENSG000002 | 4.12219574 | -0.229817523 | 0.37858091 | -0.60705 | 0.54381776 | 0.65513069 | NA | NA |
| ENSG000001 | 12.36151 | -0.202148444 | 0.333184 | -0.6067171 | 0.54403864 | 0.65537731 | SOSTDC1 | 25928 |
| ENSG000001 | 44.920427 | 0.132449183 | 0.21843106 | 0.60636606 | 0.5442717 | 0.65558016 | CFAP58 | 159686 |
| ENSG000002 | 4.26347014 | -0.228915659 | 0.37747515 | -0.606439 | 0.54422326 | 0.65558016 | NA | NA |
| ENSG000002 | 2.32945815 | -0.221244497 | 0.36484769 | -0.6064024 | 0.5442476 | 0.65558016 | NA | NA |
| ENSG000002 | 6.18024696 | 0.222003572 | 0.36611018 | 0.6063846 | 0.54425939 | 0.65558016 | PTGES2-AS1 | 389791 |
| ENSG000002 | 15.1224154 | 0.185093443 | 0.30536523 | 0.60613791 | 0.54442318 | 0.65574276 | NA | NA |

| | | | | | | | | |
|---|---|---|---|---|---|---|---|---|
| ENSG000002 | 2.62674202 | -0.224508057 | 0.37040564 | -0.606114 | 0.54443903 | 0.65574276 | OR8B4 | 283162 |
| ENSG000001 | 36.0170785 | 0.138491917 | 0.22854652 | 0.60596818 | 0.54453588 | 0.65583993 | TNFAIP8L1 | 126282 |
| ENSG000002 | 2.31176964 | 0.222586605 | 0.36736368 | 0.60590258 | 0.54457944 | 0.6558587 | NA | NA |
| ENSG000002 | 15.0135117 | -0.182812983 | 0.30172531 | -0.6058921 | 0.5445864 | 0.6558587 | NA | NA |
| ENSG000002 | 6.54348284 | -0.219952757 | 0.36303524 | -0.6058716 | 0.54459999 | 0.6558587 | NA | NA |
| ENSG000001 | 4.55008117 | -0.227391359 | 0.37534392 | -0.6058213 | 0.54463339 | 0.65587945 | CCL20 | 6364 |
| ENSG000002 | 4.04496216 | 0.229290729 | 0.3785879 | 0.60564727 | 0.54474901 | 0.65597717 | LINC01201 | 104266960 |
| ENSG000002 | 2.56350983 | -0.224282833 | 0.37031741 | -0.6056503 | 0.54474702 | 0.65597717 | NA | NA |
| ENSG000002 | 3.69301125 | 0.228999409 | 0.37812011 | 0.6056261 | 0.54476306 | 0.65597717 | LOC101927059 | 101927059 |
| ENSG000001 | 7.78144542 | -0.2135965 | 0.35272274 | -0.6055649 | 0.54480374 | 0.65598849 | RPRML | 388394 |
| ENSG000002 | 12.9494427 | 0.20375881 | 0.33650524 | 0.60551452 | 0.54483718 | 0.65598849 | NA | NA |
| ENSG000002 | 3.16022339 | -0.227577147 | 0.37583837 | -0.6055187 | 0.54483442 | 0.65598849 | NA | NA |
| ENSG000002 | 4.30675164 | -0.227990807 | 0.37649922 | -0.6055545 | 0.5448106 | 0.65598849 | NA | NA |
| ENSG000002 | 5.29928877 | -0.225569391 | 0.37261502 | -0.6053685 | 0.54493418 | 0.65606633 | NA | NA |
| ENSG000002 | 3.34531236 | -0.228813673 | 0.37796141 | -0.605389 | 0.54492057 | 0.65606633 | NA | NA |
| ENSG000001 | 3.03818536 | -0.228461035 | 0.37741801 | -0.6053263 | 0.54496223 | 0.65607695 | SERF1A | 8293 |
| ENSG000001 | 110.942516 | 0.103795494 | 0.17147924 | 0.60529481 | 0.54498313 | 0.65607695 | ZNF583 | 147949 |
| ENSG000002 | 2.85423058 | -0.226677143 | 0.37449831 | -0.6052821 | 0.54499154 | 0.65607695 | NA | NA |
| ENSG000002 | 5.32850289 | -0.225511083 | 0.37264448 | -0.6051642 | 0.54506993 | 0.65610264 | NA | NA |
| ENSG000002 | 2.54029154 | 0.225839709 | 0.37315769 | 0.60521253 | 0.54503779 | 0.65610264 | LOC101929073 | 101929073 |
| ENSG000002 | 9.92549045 | -0.205503098 | 0.33957166 | -0.6051833 | 0.54505721 | 0.65610264 | HNRNPA1P72 | 106480236 |
| ENSG000002 | 16.1166308 | 0.182522906 | 0.30161467 | 0.60515261 | 0.5450776 | 0.65610264 | NA | NA |
| ENSG000002 | 4.60385835 | -0.227608578 | 0.37615215 | -0.6050971 | 0.54511448 | 0.65612755 | NA | NA |
| ENSG000002 | 2.79834591 | -0.227014289 | 0.37527031 | -0.6049354 | 0.54522193 | 0.65623741 | NA | NA |
| ENSG000002 | 3.58638571 | -0.228944279 | 0.37848004 | -0.6049045 | 0.54524245 | 0.65624263 | NA | NA |
| ENSG000002 | 3.3844508 | -0.228556128 | 0.37786325 | -0.6048647 | 0.54526893 | 0.65625502 | NA | NA |
| ENSG000002 | 5.07966852 | -0.226109754 | 0.37386902 | -0.6047833 | 0.54532296 | 0.65630057 | NA | NA |
| ENSG000002 | 3.75143231 | 0.228909508 | 0.37854532 | 0.60470833 | 0.54537281 | 0.65634109 | SNORA80A | 677846 |
| ENSG000002 | 3.64621748 | -0.22850985 | 0.37792633 | -0.6046412 | 0.54541739 | 0.65637526 | NA | NA |

| | | | | | | | | |
|---|---|---|---|---|---|---|---|---|
| ENSG000002 | 3.66245485 | -0.228209572 | 0.37746022 | -0.6045924 | 0.54544987 | 0.65639486 | NA | NA |
| ENSG000001 | 297.940273 | 0.076838908 | 0.12712781 | 0.60442252 | 0.54556277 | 0.65651124 | SYNPO2 | 171024 |
| ENSG000001 | 177.535685 | -0.086987191 | 0.14393093 | -0.6043676 | 0.54559929 | 0.6565357 | FAM105A | 54491 |
| ENSG000002 | 3.8157726 | -0.228711738 | 0.37847544 | -0.6042974 | 0.54564591 | 0.65657232 | NA | NA |
| ENSG000002 | 2.60219822 | -0.223814744 | 0.37040909 | -0.6042366 | 0.54568632 | 0.65660147 | RNF225 | 646862 |
| ENSG000002 | 3.16456203 | -0.226025732 | 0.37423301 | -0.6039706 | 0.5458632 | 0.6567948 | NA | NA |
| ENSG000002 | 3.94421087 | 0.228413668 | 0.37833617 | 0.60373204 | 0.54602181 | 0.65696616 | NA | NA |
| ENSG000002 | 6.06411615 | -0.221912106 | 0.36764721 | -0.6036007 | 0.54610916 | 0.65703358 | NA | NA |
| ENSG000002 | 6.17888757 | -0.226475417 | 0.37520838 | -0.603599 | 0.54611025 | 0.65703358 | NA | NA |
| ENSG000002 | 2.80614192 | -0.225952436 | 0.37436304 | -0.603565 | 0.54613287 | 0.6570413 | NA | NA |
| ENSG000002 | 4.14503097 | -0.228433477 | 0.37851332 | -0.6035018 | 0.54617491 | 0.65707238 | NA | NA |
| ENSG000002 | 3.06801294 | -0.227159617 | 0.37647928 | -0.6033788 | 0.54625672 | 0.65715131 | NA | NA |
| ENSG000002 | 15.8477998 | 0.179515289 | 0.29756182 | 0.60328737 | 0.54631753 | 0.65720497 | B3GALT4 | 8705 |
| ENSG000001 | 3.79967274 | 0.228252768 | 0.37851762 | 0.60301755 | 0.54649701 | 0.65740137 | NA | NA |
| ENSG000002 | 4.31697543 | -0.227270155 | 0.37693906 | -0.6029361 | 0.54655122 | 0.65744708 | PYY2 | 23615 |
| ENSG000001 | 68.7331977 | 0.119487465 | 0.19818688 | 0.602903 | 0.54657322 | 0.6574533 | LTBR | 4055 |
| ENSG000002 | 3.52963983 | -0.228236409 | 0.37859243 | -0.6028552 | 0.54660503 | 0.6574533 | CELSR3-AS1 | 102724368 |
| ENSG000002 | 3.30911112 | -0.228017176 | 0.37822017 | -0.6028689 | 0.54659591 | 0.6574533 | NA | NA |
| ENSG000002 | 7.30544017 | -0.214972768 | 0.35664962 | -0.6027562 | 0.54667089 | 0.65751301 | NA | NA |
| ENSG000002 | 2.52660983 | 0.22581362 | 0.37466642 | 0.60270579 | 0.54670443 | 0.65751435 | NA | NA |
| ENSG000002 | 58.3003628 | -0.120090837 | 0.19925193 | -0.6027085 | 0.5467026 | 0.65751435 | NA | NA |
| ENSG000002 | 11.6985202 | -0.197678634 | 0.32801718 | -0.6026472 | 0.54674342 | 0.65754174 | CLDN9 | 9080 |
| ENSG000002 | 8.14915991 | 0.213943851 | 0.35515043 | 0.60240347 | 0.5469056 | 0.65771728 | NA | NA |
| ENSG000002 | 4.38318086 | -0.227777782 | 0.3782012 | -0.6022661 | 0.54699699 | 0.65778817 | NA | NA |
| ENSG000002 | 3.08185055 | -0.226377525 | 0.37586768 | -0.6022798 | 0.54698788 | 0.65778817 | NA | NA |
| ENSG000001 | 114.487283 | -0.099205386 | 0.16476162 | -0.6021147 | 0.54709781 | 0.6578899 | RGL1 | 23179 |
| ENSG000001 | 12.454725 | -0.197532856 | 0.32810677 | -0.6020383 | 0.54714862 | 0.65791279 | BTNL8 | 79908 |
| ENSG000002 | 6.51383731 | -0.218872863 | 0.36355364 | -0.6020373 | 0.54714929 | 0.65791279 | NA | NA |
| ENSG000002 | 10.3919573 | -0.20116084 | 0.33430841 | -0.6017224 | 0.54735897 | 0.6581454 | NA | NA |

| | | | | | | | | |
|---|---|---|---|---|---|---|---|---|
| ENSG000002 | 8.9554211 | -0.20612447 | 0.34259421 | -0.6016578 | 0.54740196 | 0.65817758 | NA | NA |
| ENSG000001 | 4.55996527 | -0.226187738 | 0.37597897 | -0.6015968 | 0.54744257 | 0.65820688 | FAM209A | 200232 |
| ENSG000002 | 4.12166715 | -0.227708176 | 0.37853727 | -0.6015476 | 0.54747534 | 0.65822677 | NA | NA |
| ENSG000001 | 21.9403832 | -0.160683255 | 0.2671818 | -0.6014005 | 0.54757329 | 0.65832502 | ABTB1 | 80325 |
| ENSG000002 | 2.52492689 | 0.224696349 | 0.37367068 | 0.60132185 | 0.54762564 | 0.6583294 | NA | NA |
| ENSG000002 | 4.77805323 | -0.225560983 | 0.37509591 | -0.6013422 | 0.54761212 | 0.6583294 | NA | NA |
| ENSG000002 | 6.04765752 | -0.220777625 | 0.36715201 | -0.6013248 | 0.54762364 | 0.6583294 | NA | NA |
| ENSG000002 | 4.9740177 | -0.227602572 | 0.37857292 | -0.601212 | 0.54769881 | 0.65839784 | NA | NA |
| ENSG000002 | 4.3084971 | -0.227467154 | 0.37857434 | -0.600852 | 0.54793856 | 0.65866652 | NA | NA |
| ENSG000001 | 59.6458064 | -0.114652173 | 0.19092385 | -0.6005126 | 0.54816468 | 0.65891856 | ANKDD1A | 348094 |
| ENSG000002 | 2.63426251 | -0.220058932 | 0.36649056 | -0.6004491 | 0.54820697 | 0.65891856 | NA | NA |
| ENSG000002 | 3.80051352 | -0.227256782 | 0.37848393 | -0.6004397 | 0.54821322 | 0.65891856 | NA | NA |
| ENSG000002 | 4.05903165 | -0.226928067 | 0.37792772 | -0.6004536 | 0.54820396 | 0.65891856 | NA | NA |
| ENSG000002 | 3.3515201 | 0.227145054 | 0.37839191 | 0.60029046 | 0.54831268 | 0.65901857 | NA | NA |
| ENSG000002 | 30.625253 | 0.149741176 | 0.24947068 | 0.60023558 | 0.54834925 | 0.65904298 | NA | NA |
| ENSG000001 | 140.567679 | 0.091459938 | 0.15243783 | 0.5999819 | 0.5485183 | 0.65922663 | SPOCK1 | 6695 |
| ENSG000002 | 29.4972633 | 0.155385634 | 0.25900729 | 0.59992764 | 0.54855446 | 0.65923358 | C17orf67 | 339210 |
| ENSG000002 | 2.51212674 | 0.224033562 | 0.37343795 | 0.59992179 | 0.54855836 | 0.65923358 | NA | NA |
| ENSG000002 | 3.86517295 | -0.227117154 | 0.37859166 | -0.5999 | 0.54857285 | 0.65923358 | NA | NA |
| ENSG000002 | 2.28861771 | -0.219292689 | 0.36561627 | -0.5997892 | 0.54864673 | 0.65929058 | NA | NA |
| ENSG000002 | 5.0837067 | -0.225061144 | 0.37523944 | -0.5997801 | 0.5486528 | 0.65929058 | NA | NA |
| ENSG000002 | 4.53018942 | -0.225431332 | 0.37587778 | -0.5997464 | 0.54867528 | 0.65929806 | NA | NA |
| ENSG000002 | 61.059387 | -0.119085589 | 0.19860693 | -0.5996044 | 0.54876991 | 0.65939222 | NA | NA |
| ENSG000002 | 3.06229196 | -0.22604765 | 0.37701262 | -0.5995758 | 0.54878897 | 0.65939558 | NA | NA |
| ENSG000002 | 15.2855116 | -0.184815688 | 0.30829745 | -0.599472 | 0.54885818 | 0.6594592 | ECEL1P2 | 347694 |
| ENSG000001 | 7.09837679 | 0.215541336 | 0.35962552 | 0.59934939 | 0.54893992 | 0.65953787 | MYL7 | 58498 |
| ENSG000002 | 3.34788428 | -0.226411002 | 0.3777768 | -0.5993248 | 0.54895632 | 0.65953803 | NA | NA |
| ENSG000002 | 18.6920172 | -0.171604358 | 0.28642338 | -0.5991283 | 0.54908732 | 0.65967588 | NA | NA |
| ENSG000001 | 32.4019354 | 0.15447886 | 0.25787714 | 0.59904053 | 0.54914586 | 0.65970711 | ARHGAP25 | 9938 |

| | | | | | | | | |
|---|---|---|---|---|---|---|---|---|
| ENSG000002 | 7.98846882 | -0.212243216 | 0.35429951 | -0.5990503 | 0.54913936 | 0.65970711 | NA | NA |
| ENSG000002 | 7.23021204 | -0.21228795 | 0.3545132 | -0.5988154 | 0.54929602 | 0.65986796 | NA | NA |
| ENSG000002 | 32.1837482 | 0.147656228 | 0.2466616 | 0.59861862 | 0.54942724 | 0.66000604 | NA | NA |
| ENSG000001 | 60.2520935 | -0.118117671 | 0.19739109 | -0.5983941 | 0.54957697 | 0.66016636 | BCL2L14 | 79370 |
| ENSG000002 | 4.05929169 | -0.226483445 | 0.37853954 | -0.5983086 | 0.54963407 | 0.66021538 | NA | NA |
| ENSG000002 | 4.30974036 | -0.22530183 | 0.37661173 | -0.5982337 | 0.54968401 | 0.66025581 | NA | NA |
| ENSG000002 | 2.58293213 | -0.221659213 | 0.3705524 | -0.5981859 | 0.54971591 | 0.66027458 | NA | NA |
| ENSG000001 | 33.639856 | 0.141049096 | 0.23585439 | 0.59803464 | 0.54981682 | 0.66035666 | ADORA2B | 136 |
| ENSG000002 | 7.23293753 | 0.212929895 | 0.35604463 | 0.5980427 | 0.54981144 | 0.66035666 | NA | NA |
| ENSG000002 | 5.66006141 | -0.224493342 | 0.37541676 | -0.5979843 | 0.54985038 | 0.66035785 | LINC01269 | 103695436 |
| ENSG000002 | 6.58409625 | -0.216866312 | 0.36264752 | -0.5980085 | 0.54983424 | 0.66035785 | NA | NA |
| ENSG000002 | 6.36887173 | -0.219175278 | 0.36654473 | -0.5979496 | 0.54987356 | 0.66036613 | NA | NA |
| ENSG000001 | 260.482302 | 0.082988898 | 0.13880709 | 0.59787217 | 0.54992523 | 0.66039023 | ZNF793 | 390927 |
| ENSG000002 | 30.4404583 | 0.145355265 | 0.24312157 | 0.59787072 | 0.5499262 | 0.66039023 | NA | NA |
| ENSG000002 | 2.9014144 | 0.221264146 | 0.37016002 | 0.59775268 | 0.55000497 | 0.660459 | NA | NA |
| ENSG000002 | 3.81442357 | -0.226225876 | 0.37849834 | -0.5976932 | 0.55004468 | 0.660459 | NA | NA |
| ENSG000002 | 2.24247102 | 0.217652823 | 0.36415837 | 0.59768728 | 0.55004862 | 0.660459 | NA | NA |
| ENSG000002 | 2.49922222 | 0.222295064 | 0.37190355 | 0.59772234 | 0.55002521 | 0.660459 | NA | NA |
| ENSG000002 | 3.739533 | 0.226225088 | 0.37853434 | 0.59763425 | 0.550084 | 0.66048193 | NA | NA |
| ENSG000001 | 166.597975 | 0.084219537 | 0.1409518 | 0.59750594 | 0.55016964 | 0.66049297 | ZNF382 | 84911 |
| ENSG000002 | 6.79164114 | -0.216078051 | 0.36158538 | -0.5975851 | 0.55011679 | 0.66049297 | LOC100506731 | 100506731 |
| ENSG000002 | 2.81695856 | -0.222915645 | 0.37306064 | -0.5975319 | 0.55015229 | 0.66049297 | NA | NA |
| ENSG000002 | 2.81026483 | -0.223680343 | 0.37436137 | -0.5974985 | 0.55017464 | 0.66049297 | LOC105377658 | 105377658 |
| ENSG000002 | 6.06429276 | -0.220513811 | 0.36904492 | -0.5975257 | 0.55015648 | 0.66049297 | NA | NA |
| ENSG000002 | 7.33093935 | -0.217775538 | 0.36450284 | -0.5974591 | 0.55020091 | 0.66050496 | NA | NA |
| ENSG000001 | 9.22498455 | -0.205516282 | 0.34406405 | -0.5973198 | 0.55029387 | 0.66057744 | IDI2 | 91734 |
| ENSG000002 | 34.7665968 | -0.137549117 | 0.23027012 | -0.5973381 | 0.55028166 | 0.66057744 | NA | NA |
| ENSG000001 | 52.9989563 | 0.13245563 | 0.22178949 | 0.59721329 | 0.55036499 | 0.66064326 | TMEM27 | 57393 |
| ENSG000002 | 4.84547699 | 0.225394721 | 0.37743972 | 0.59716746 | 0.55039558 | 0.66066043 | SNORA30 | 677813 |

| | | | | | | | | |
|---|---|---|---|---|---|---|---|---|
| ENSG000001 | 12.0307937 | -0.196377201 | 0.32889272 | -0.5970859 | 0.55045004 | 0.66068669 | PRSS53 | 339105 |
| ENSG000002 | 5.34880755 | -0.222864291 | 0.37324298 | -0.5971024 | 0.55043899 | 0.66068669 | NA | NA |
| ENSG000001 | 41.5975057 | 0.133095734 | 0.22292774 | 0.59703531 | 0.5504838 | 0.660699 | NKAIN4 | 128414 |
| ENSG000002 | 5.74854745 | -0.219658821 | 0.36792435 | -0.5970217 | 0.55049289 | 0.660699 | NA | NA |
| ENSG000002 | 3.03087786 | -0.224629017 | 0.37628735 | -0.5969614 | 0.55053316 | 0.66072778 | NA | NA |
| ENSG000002 | 6.26586749 | -0.220582898 | 0.36956622 | -0.5968698 | 0.55059434 | 0.66078068 | ERI3-IT1 | 100874278 |
| ENSG000002 | 7.57650417 | -0.214018663 | 0.35858238 | -0.5968466 | 0.55060983 | 0.66078068 | NA | NA |
| ENSG000002 | 4.81366717 | -0.224026797 | 0.37543439 | -0.5967136 | 0.55069863 | 0.66086769 | THOC7-AS1 | 100874039 |
| ENSG000002 | 3.5223846 | -0.22579912 | 0.37849874 | -0.5965651 | 0.55079781 | 0.66096715 | LINC00708 | 100507143 |
| ENSG000002 | 6.49953712 | -0.217150734 | 0.36403864 | -0.5965046 | 0.55083816 | 0.66097646 | NA | NA |
| ENSG000002 | 15.4813771 | -0.177341554 | 0.29729604 | -0.596515 | 0.55083122 | 0.66097646 | LOC101927765 | 101927765 |
| ENSG000001 | 8.97377393 | -0.2051028 | 0.34394798 | -0.5963192 | 0.55096197 | 0.6610859 | MMP27 | 64066 |
| ENSG000001 | 54.2007906 | 0.12252255 | 0.20545983 | 0.59633337 | 0.55095255 | 0.6610859 | N6AMT2 | 221143 |
| ENSG000002 | 4.85769276 | -0.223992666 | 0.37565343 | -0.5962748 | 0.55099168 | 0.66110198 | NA | NA |
| ENSG000002 | 8.82050697 | 0.208773563 | 0.35021737 | 0.59612567 | 0.55109128 | 0.66116831 | NA | NA |
| ENSG000002 | 4.23283512 | 0.224660272 | 0.37685674 | 0.59614238 | 0.55108011 | 0.66116831 | NA | NA |
| ENSG000002 | 11.2721353 | -0.202058828 | 0.33895732 | -0.5961188 | 0.55109588 | 0.66116831 | NA | NA |
| ENSG000002 | 2.25551248 | 0.21690573 | 0.36389272 | 0.59607055 | 0.5511281 | 0.66118741 | RN7SL564P | 106479428 |
| ENSG000001 | 87.0116703 | -0.10347576 | 0.17361888 | -0.5959937 | 0.55117942 | 0.66122941 | SLC22A5 | 6584 |
| ENSG000001 | 22.0056382 | -0.159795873 | 0.26813394 | -0.5959554 | 0.55120502 | 0.66124056 | KCNIP3 | 30818 |
| ENSG000002 | 40.5023719 | -0.149643419 | 0.25116869 | -0.5957885 | 0.55131652 | 0.66135476 | NA | NA |
| ENSG000002 | 37.6243693 | 0.139459081 | 0.23411669 | 0.59568193 | 0.55138773 | 0.66142062 | KIF4B | 285643 |
| ENSG000002 | 8.63353573 | -0.214087163 | 0.35954155 | -0.5954448 | 0.55154616 | 0.66159109 | LOC100506801 | 100506801 |
| ENSG000001 | 23.5507707 | 0.16759604 | 0.28150073 | 0.59536627 | 0.55159867 | 0.6616345 | NA | NA |
| ENSG000002 | 8.10444978 | 0.210770423 | 0.35407682 | 0.5952675 | 0.55166468 | 0.66169411 | NA | NA |
| ENSG000001 | 128.332604 | 0.107512628 | 0.18064676 | 0.59515393 | 0.55174058 | 0.66171315 | GPRIN1 | 114787 |
| ENSG000002 | 4.59562893 | 0.225085346 | 0.37818205 | 0.59517723 | 0.55172501 | 0.66171315 | NA | NA |
| ENSG000002 | 5.55877941 | -0.220255746 | 0.37008686 | -0.5951461 | 0.55174583 | 0.66171315 | ARHGDIG | 398 |
| ENSG000002 | 3.13571981 | -0.22341685 | 0.37537252 | -0.595187 | 0.55171847 | 0.66171315 | NA | NA |

| | | | | | | | | |
|---|---|---|---|---|---|---|---|---|
| ENSG000002 | 4.58821384 | -0.223914931 | 0.3763366 | -0.5949858 | 0.55185297 | 0.66182207 | NA | NA |
| ENSG000002 | 4.22719867 | 0.224556417 | 0.3774727 | 0.59489447 | 0.55191402 | 0.66187571 | NA | NA |
| ENSG000002 | 2.31638869 | 0.219401992 | 0.36884002 | 0.59484323 | 0.55194827 | 0.66189721 | SNORA38B | 100124536 |
| ENSG000001 | 8.36807562 | 0.20851963 | 0.35058427 | 0.59477749 | 0.55199222 | 0.66193034 | NA | NA |
| ENSG000002 | 2.31105401 | -0.216668403 | 0.36433778 | -0.594691 | 0.55205003 | 0.66198009 | NA | NA |
| ENSG000002 | 5.48769426 | -0.219467562 | 0.36912461 | -0.5945623 | 0.55213611 | 0.66206189 | NA | NA |
| ENSG000002 | 7.00481024 | -0.213002812 | 0.3582648 | -0.5945402 | 0.55215089 | 0.66206189 | NA | NA |
| ENSG000002 | 3.31067673 | -0.224266698 | 0.37727441 | -0.5944392 | 0.5522184 | 0.66212325 | NA | NA |
| ENSG000002 | 3.84524929 | -0.224978636 | 0.37851621 | -0.5943699 | 0.55226475 | 0.66213967 | NA | NA |
| ENSG000002 | 3.83280864 | -0.22494539 | 0.37844875 | -0.594388 | 0.55225265 | 0.66213967 | CXADRP3 | 440224 |
| ENSG000002 | 7.01832115 | -0.213289596 | 0.35886794 | -0.5943401 | 0.55228469 | 0.66214397 | NA | NA |
| ENSG000002 | 5.06404418 | -0.222176416 | 0.3738357 | -0.5943157 | 0.552301 | 0.66214397 | NA | NA |
| ENSG000002 | 3.38277974 | -0.223423808 | 0.37597541 | -0.5942511 | 0.55234417 | 0.66215658 | NA | NA |
| ENSG000002 | 5.09993572 | 0.221668601 | 0.37301794 | 0.59425722 | 0.55234009 | 0.66215658 | NA | NA |
| ENSG000002 | 5.06504163 | -0.222429485 | 0.37432855 | -0.5942092 | 0.55237217 | 0.66217057 | NA | NA |
| ENSG000002 | 4.76451587 | -0.223204197 | 0.37580626 | -0.5939342 | 0.55255612 | 0.6623715 | NA | NA |
| ENSG000002 | 12.878095 | -0.192462563 | 0.32413576 | -0.5937715 | 0.55266498 | 0.66248241 | NA | NA |
| ENSG000002 | 4.38553629 | -0.224407324 | 0.37803605 | -0.5936135 | 0.55277062 | 0.66258946 | NA | NA |
| ENSG000002 | 9.70465758 | -0.199912157 | 0.33680628 | -0.5935523 | 0.55281157 | 0.66261896 | LINC00870 | 201617 |
| ENSG00000C | 188.387968 | 0.081582645 | 0.13745715 | 0.59351328 | 0.5528377 | 0.66263069 | NA | NA |
| ENSG000002 | 3.15559196 | -0.221832473 | 0.3737938 | -0.5934621 | 0.55287191 | 0.66265211 | NA | NA |
| ENSG000002 | 7.45849676 | -0.210472789 | 0.35477287 | -0.5932607 | 0.55300672 | 0.6627941 | NA | NA |
| ENSG000001 | 45.4300917 | -0.125136024 | 0.21096433 | -0.593162 | 0.55307274 | 0.66280848 | PCDH17 | 27253 |
| ENSG000001 | 52.3679686 | -0.122777069 | 0.20699334 | -0.593145 | 0.5530841 | 0.66280848 | ANKMY1 | 51281 |
| ENSG000001 | 6.07229805 | -0.219077239 | 0.36933545 | -0.593166 | 0.55307005 | 0.66280848 | LCN15 | 389812 |
| ENSG000002 | 46.0810038 | -0.129975445 | 0.21910261 | -0.5932172 | 0.55303579 | 0.66280848 | NA | NA |
| ENSG000002 | 4.13157794 | -0.224404983 | 0.37835019 | -0.5931145 | 0.55310453 | 0.66281338 | NA | NA |
| ENSG000001 | 36.624097 | 0.139199806 | 0.23472064 | 0.59304458 | 0.55315132 | 0.66283027 | HIST1H1T | 3010 |
| ENSG000002 | 4.24429633 | -0.224009754 | 0.37772499 | -0.5930499 | 0.55314778 | 0.66283027 | NA | NA |

| | | | | | | | | |
|---|---|---|---|---|---|---|---|---|
| ENSG000002 | 5.51613173 | -0.219552933 | 0.37026652 | -0.5929592 | 0.55320847 | 0.66287188 | NA | NA |
| ENSG000002 | 4.30310502 | -0.223857824 | 0.37755186 | -0.5929194 | 0.55323509 | 0.66287188 | LOC101928896 | 101928896 |
| ENSG000002 | 16.1396796 | -0.176429263 | 0.29755917 | -0.5929216 | 0.55323361 | 0.66287188 | BCDIN3D-AS1 | 100286844 |
| ENSG000002 | 3.54213996 | -0.224436055 | 0.37854636 | -0.5928892 | 0.5532553 | 0.66287651 | NA | NA |
| ENSG000002 | 4.05208309 | -0.224301199 | 0.37835723 | -0.5928292 | 0.55329549 | 0.66290464 | NA | NA |
| ENSG000002 | 3.66954644 | -0.223838573 | 0.37759206 | -0.5928053 | 0.55331147 | 0.66290464 | NA | NA |
| ENSG000002 | 9.53818018 | -0.205875101 | 0.34734662 | -0.592708 | 0.55337662 | 0.66296311 | NA | NA |
| ENSG000002 | 9.22700682 | -0.203832606 | 0.34415407 | -0.5922714 | 0.55366888 | 0.66329365 | NA | NA |
| ENSG000001 | 5.61661271 | -0.222271899 | 0.37531387 | -0.5922294 | 0.553697 | 0.66330774 | TMEM105 | 284186 |
| ENSG000002 | 3.55172381 | -0.22416454 | 0.37857335 | -0.5921297 | 0.55376371 | 0.66336806 | NA | NA |
| ENSG000002 | 3.57320044 | -0.223872952 | 0.37814465 | -0.5920299 | 0.55383055 | 0.66341934 | NA | NA |
| ENSG000002 | 15.5323072 | -0.184618268 | 0.31184626 | -0.5920169 | 0.55383924 | 0.66341934 | NA | NA |
| ENSG000002 | 5.50798264 | -0.219398366 | 0.37068229 | -0.5918771 | 0.55393287 | 0.6634923 | NA | NA |
| ENSG000002 | 5.5196814 | -0.21952108 | 0.37088443 | -0.5918854 | 0.55392733 | 0.6634923 | NA | NA |
| ENSG000002 | 25.0719706 | -0.162335807 | 0.27431622 | -0.5917835 | 0.55399559 | 0.66352509 | RASAL2-AS1 | 100302401 |
| ENSG000002 | 3.57335764 | -0.223942493 | 0.37843277 | -0.591763 | 0.55400933 | 0.66352509 | CYCSP52 | 360155 |
| ENSG000002 | 4.27660534 | 0.22317933 | 0.37711901 | 0.59180079 | 0.55398399 | 0.66352509 | NA | NA |
| ENSG000001 | 6.70749958 | -0.214315975 | 0.3622113 | -0.5916877 | 0.55405973 | 0.66354947 | HCRTR1 | 3061 |
| ENSG000002 | 8.4723116 | -0.205073934 | 0.34659386 | -0.5916837 | 0.55406242 | 0.66354947 | NA | NA |
| ENSG000002 | 5.12476713 | 0.220577213 | 0.37283064 | 0.59162845 | 0.55409942 | 0.66357419 | NA | NA |
| ENSG000002 | 3.84342941 | -0.223961736 | 0.37857769 | -0.5915873 | 0.55412701 | 0.66358763 | CELA3B | 23436 |
| ENSG000002 | 3.97999363 | 0.223945414 | 0.37857762 | 0.59154424 | 0.55415582 | 0.66360253 | NA | NA |
| ENSG000002 | 3.94204544 | -0.22323047 | 0.37742215 | -0.591461 | 0.5542116 | 0.66364974 | NA | NA |
| ENSG000002 | 4.7898495 | -0.221774192 | 0.37502299 | -0.5913616 | 0.55427817 | 0.66366078 | NA | NA |
| ENSG000002 | 3.60372635 | -0.22345718 | 0.37787669 | -0.5913495 | 0.55428629 | 0.66366078 | NA | NA |
| ENSG000002 | 18.8637066 | 0.169442773 | 0.28651561 | 0.59139107 | 0.55425842 | 0.66366078 | NA | NA |
| ENSG000002 | 2.57485551 | -0.217952918 | 0.36854508 | -0.5913874 | 0.55426088 | 0.66366078 | MIR466 | 100423038 |
| ENSG000002 | 4.3039794 | -0.223241827 | 0.37758004 | -0.5912437 | 0.55435714 | 0.66372601 | NA | NA |
| ENSG000002 | 6.32080144 | -0.218774671 | 0.37004643 | -0.5912087 | 0.55438059 | 0.66372813 | NA | NA |

| | | | | | | | | |
|---|---|---|---|---|---|---|---|---|
| ENSG000002 | 6.29892042 | -0.216506005 | 0.36623669 | -0.5911642 | 0.55441044 | 0.66372813 | NA | NA |
| ENSG000002 | 9.49334163 | -0.208437206 | 0.35258429 | -0.5911699 | 0.55440662 | 0.66372813 | NA | NA |
| ENSG000002 | 3.32300101 | -0.223303235 | 0.37774803 | -0.5911433 | 0.55442438 | 0.66372813 | NA | NA |
| ENSG000002 | 21.677356 | 0.163167051 | 0.27604907 | 0.59107989 | 0.5544669 | 0.66375944 | NA | NA |
| ENSG000001 | 7.51454619 | -0.209914278 | 0.3552548 | -0.5908837 | 0.55459833 | 0.66389025 | GALR2 | 8811 |
| ENSG000002 | 7.28576425 | -0.21113978 | 0.35733837 | -0.5908679 | 0.55460892 | 0.66389025 | NA | NA |
| ENSG000001 | 3.55069245 | -0.22338942 | 0.37814856 | -0.5907451 | 0.5546912 | 0.66391036 | PROK1 | 84432 |
| ENSG000002 | 4.59518908 | -0.222363429 | 0.37636665 | -0.590816 | 0.55464373 | 0.66391036 | NA | NA |
| ENSG000002 | 6.06119598 | -0.216483845 | 0.36644742 | -0.5907637 | 0.55467874 | 0.66391036 | NA | NA |
| ENSG000002 | 3.07758418 | -0.221845248 | 0.37553068 | -0.5907513 | 0.55468705 | 0.66391036 | NA | NA |
| ENSG000002 | 6.51960622 | -0.213316108 | 0.36118279 | -0.5906043 | 0.55478558 | 0.66396453 | NA | NA |
| ENSG000002 | 3.31986543 | 0.222887268 | 0.37738229 | 0.59061401 | 0.55477907 | 0.66396453 | LINC00659 | 100652730 |
| ENSG000002 | 4.04857395 | -0.223222464 | 0.37792607 | -0.5906511 | 0.55475419 | 0.66396453 | NA | NA |
| ENSG000002 | 3.12040757 | -0.221507882 | 0.37513008 | -0.5904829 | 0.55486697 | 0.66398355 | NA | NA |
| ENSG000002 | 5.72071568 | -0.217565182 | 0.36844797 | -0.5904909 | 0.55486157 | 0.66398355 | NA | NA |
| ENSG000002 | 3.6520826 | -0.222738616 | 0.3772064 | -0.5904953 | 0.55485862 | 0.66398355 | LOC105371242 | 105371242 |
| ENSG000002 | 33.1428496 | -0.145518384 | 0.24643344 | -0.5904977 | 0.55485702 | 0.66398355 | NA | NA |
| ENSG000002 | 4.03608702 | -0.223010649 | 0.37771506 | -0.5904203 | 0.55490889 | 0.66401411 | FRG2MP | 106481739 |
| ENSG000001 | 122.049323 | -0.094236618 | 0.15962562 | -0.5903602 | 0.55494917 | 0.66403298 | PRSS23 | 11098 |
| ENSG000001 | 40.9741152 | 0.13383792 | 0.22671024 | 0.59034794 | 0.55495741 | 0.66403298 | GPR135 | 64582 |
| ENSG000002 | 4.64120896 | -0.223266847 | 0.37825954 | -0.5902478 | 0.55502456 | 0.66409374 | NA | NA |
| ENSG000002 | 6.71675703 | -0.213736044 | 0.36215883 | -0.5901721 | 0.55507526 | 0.66413481 | NA | NA |
| ENSG000001 | 17.5719944 | -0.175979155 | 0.29824179 | -0.5900553 | 0.55515356 | 0.6642089 | TM4SF1 | 4071 |
| ENSG000002 | 4.59586103 | -0.222271131 | 0.37683389 | -0.5898385 | 0.55529894 | 0.66436323 | NA | NA |
| ENSG000001 | 15.8262446 | 0.178091316 | 0.30205243 | 0.58960399 | 0.55545618 | 0.66453174 | LRRC37A | 9884 |
| ENSG000002 | 8.13894769 | 0.206974158 | 0.35108165 | 0.58953283 | 0.55550389 | 0.66456922 | NA | NA |
| ENSG000001 | 39.6249004 | 0.142753605 | 0.24222603 | 0.58934048 | 0.5556329 | 0.66467328 | NA | NA |
| ENSG000002 | 5.76105944 | -0.217531595 | 0.36910382 | -0.5893507 | 0.55562604 | 0.66467328 | NA | NA |
| ENSG000002 | 5.05175559 | -0.222449065 | 0.37746107 | -0.5893298 | 0.55564005 | 0.66467328 | NA | NA |

| | | | | | | | | |
|---|---|---|---|---|---|---|---|---|
| ENSG000002 | 5.56069899 | -0.217912899 | 0.36981085 | -0.589255 | 0.55569022 | 0.66471369 | NA | NA |
| ENSG000002 | 5.54942913 | -0.220406387 | 0.37407171 | -0.5892089 | 0.55572114 | 0.66473107 | NA | NA |
| ENSG000002 | 3.35985656 | 0.223036289 | 0.37856671 | 0.58915981 | 0.55575408 | 0.6647422 | NA | NA |
| ENSG000002 | 4.8084751 | -0.221035977 | 0.37518021 | -0.5891462 | 0.55576323 | 0.6647422 | NA | NA |
| ENSG000002 | 11.2181724 | -0.194240502 | 0.32973003 | -0.5890895 | 0.55580124 | 0.66476805 | NA | NA |
| ENSG000002 | 4.17031177 | -0.222965633 | 0.37854666 | -0.5890044 | 0.55585835 | 0.66481675 | NA | NA |
| ENSG000002 | 24.2228094 | -0.16020053 | 0.27201701 | -0.5889357 | 0.55590441 | 0.66485223 | NA | NA |
| ENSG000002 | 7.37764189 | -0.213494978 | 0.36279408 | -0.5884743 | 0.55621401 | 0.66520288 | NA | NA |
| ENSG000002 | 2.54367049 | -0.217335994 | 0.36958925 | -0.5880474 | 0.55650049 | 0.66552587 | NA | NA |
| ENSG000001 | 5.56145547 | -0.218071988 | 0.37087418 | -0.5879945 | 0.55653597 | 0.66554867 | NA | NA |
| ENSG000002 | 3.53243213 | -0.222346286 | 0.37826267 | -0.5878092 | 0.55666035 | 0.66567779 | NA | NA |
| ENSG000002 | 7.49922031 | 0.208579023 | 0.3548664 | 0.58776774 | 0.5566882 | 0.66569146 | NA | NA |
| ENSG000001 | 5.05620709 | -0.221462479 | 0.37683494 | -0.5876909 | 0.55673976 | 0.66573349 | IRX6 | 79190 |
| ENSG000002 | 2.56084827 | 0.219558967 | 0.37362106 | 0.58765147 | 0.55676626 | 0.66574554 | NA | NA |
| ENSG000002 | 8.23272065 | -0.204954577 | 0.34880787 | -0.5875859 | 0.5568103 | 0.66577857 | SMLR1 | 100507203 |
| ENSG000002 | 6.78832448 | -0.214575965 | 0.36520212 | -0.587554 | 0.5568317 | 0.66578453 | NA | NA |
| ENSG000002 | 2.59167442 | -0.216718792 | 0.36908109 | -0.5871848 | 0.55707963 | 0.66606133 | NA | NA |
| ENSG000002 | 140.354433 | -0.110364984 | 0.18799531 | -0.5870624 | 0.55716178 | 0.66613991 | ERVMER61-1 | 339476 |
| ENSG000002 | 2.62473691 | -0.17148343 | 0.29211998 | -0.5870308 | 0.557183 | 0.66614565 | LOC105374631 | 105374631 |
| ENSG000002 | 8.17839327 | -0.20570355 | 0.35046369 | -0.5869468 | 0.55723942 | 0.66619346 | NA | NA |
| ENSG000002 | 14.6834025 | -0.184541927 | 0.31443182 | -0.586906 | 0.55726683 | 0.66619543 | NA | NA |
| ENSG000002 | 4.82207552 | -0.221308386 | 0.37708316 | -0.5868954 | 0.55727393 | 0.66619543 | LOC101927081 | 101927081 |
| ENSG000002 | 6.09118642 | -0.216461808 | 0.36886337 | -0.5868347 | 0.55731476 | 0.6662246 | NA | NA |
| ENSG000002 | 4.03933612 | -0.221815858 | 0.37808333 | -0.5866851 | 0.55741521 | 0.66632503 | NA | NA |
| ENSG000002 | 10.1566625 | 0.201176055 | 0.34294457 | 0.58661391 | 0.55746304 | 0.66636257 | NA | NA |
| ENSG000002 | 2.84621289 | -0.218312942 | 0.37225404 | -0.5864622 | 0.55756492 | 0.66646471 | NA | NA |
| ENSG000002 | 7.27677422 | -0.209564309 | 0.3573747 | -0.5863994 | 0.55760715 | 0.66649553 | NA | NA |
| ENSG000001 | 22.0344004 | -0.163127688 | 0.27825412 | -0.5862544 | 0.5577046 | 0.66659237 | EIF4E1B | 253314 |
| ENSG000001 | 16.1835608 | -0.173839071 | 0.29661671 | -0.5860731 | 0.5578264 | 0.66671829 | KCNQ4 | 9132 |

| | | | | | | | | |
|---|---|---|---|---|---|---|---|---|
| ENSG000001 | 50.2885582 | 0.121315938 | 0.2070169 | 0.58601948 | 0.55786243 | 0.66672205 | CSF1 | 1435 |
| ENSG000002 | 3.38752492 | 0.221862727 | 0.37859078 | 0.58602253 | 0.55786037 | 0.66672205 | NA | NA |
| ENSG000002 | 10.1893334 | -0.196066222 | 0.33474872 | -0.5857116 | 0.55806931 | 0.66691458 | MIR573 | 693158 |
| ENSG000002 | 2.82637355 | -0.218659416 | 0.3733416 | -0.5856819 | 0.5580893 | 0.66691458 | NA | NA |
| ENSG000002 | 9.21313785 | -0.200056682 | 0.34156443 | -0.585707 | 0.55807243 | 0.66691458 | NA | NA |
| ENSG000002 | 4.03304623 | -0.22163569 | 0.37841391 | -0.5856965 | 0.55807951 | 0.66691458 | NA | NA |
| ENSG000002 | 5.54454506 | -0.217420666 | 0.37127067 | -0.5856123 | 0.55813609 | 0.66695084 | NA | NA |
| ENSG000002 | 3.59147471 | -0.221632471 | 0.37848669 | -0.5855753 | 0.55816092 | 0.66696086 | MIR3145 | 100423001 |
| ENSG000002 | 24.1445832 | 0.154893639 | 0.26456856 | 0.58545746 | 0.55824016 | 0.66703588 | NA | NA |
| ENSG000002 | 2.33870914 | -0.145048237 | 0.24790827 | -0.5850883 | 0.55848832 | 0.66731274 | NA | NA |
| ENSG000002 | 4.06163673 | 0.221271059 | 0.37857449 | 0.58448486 | 0.55889415 | 0.66777797 | NA | NA |
| ENSG000002 | 3.35669846 | -0.220548861 | 0.37738156 | -0.5844187 | 0.55893862 | 0.66781143 | NA | NA |
| ENSG000002 | 5.28372377 | -0.21841931 | 0.37376995 | -0.5843683 | 0.55897255 | 0.66783229 | NA | NA |
| ENSG000001 | 3.36548166 | 0.220427249 | 0.37729457 | 0.58423117 | 0.55906479 | 0.66792282 | GNRH2 | 2797 |
| ENSG000001 | 45.5976524 | 0.125547747 | 0.21492913 | 0.58413556 | 0.55912911 | 0.66796795 | ANXA9 | 8416 |
| ENSG000002 | 4.0975478 | -0.221051638 | 0.37843201 | -0.5841251 | 0.55913615 | 0.66796795 | SORCS3-AS1 | 100505890 |
| ENSG000002 | 6.23792778 | -0.213716573 | 0.36589833 | -0.5840873 | 0.55916157 | 0.66796795 | CD81-AS1 | 101927682 |
| ENSG000002 | 5.78863083 | -0.216240577 | 0.37022609 | -0.5840771 | 0.55916845 | 0.66796795 | NA | NA |
| ENSG000002 | 12.4690675 | 0.189799 | 0.32502132 | 0.58395862 | 0.55924815 | 0.66801795 | LOC100134368 | 100134368 |
| ENSG000002 | 5.58885607 | -0.21751332 | 0.37249169 | -0.5839414 | 0.55925974 | 0.66801795 | NA | NA |
| ENSG000002 | 5.0389915 | -0.218252682 | 0.37375528 | -0.5839454 | 0.55925704 | 0.66801795 | NA | NA |
| ENSG000002 | 6.52445394 | -0.212177621 | 0.36337909 | -0.5839016 | 0.55928654 | 0.66803029 | NA | NA |
| ENSG000001 | 6.03389203 | -0.215494211 | 0.36912521 | -0.5837971 | 0.55935686 | 0.66809461 | CCDC89 | 220388 |
| ENSG000002 | 5.28757324 | -0.217278674 | 0.37230551 | -0.5836032 | 0.55948733 | 0.66823076 | NA | NA |
| ENSG000002 | 26.1847387 | 0.154962084 | 0.26559806 | 0.58344585 | 0.5595932 | 0.66833752 | NA | NA |
| ENSG000001 | 39.6035812 | -0.130704537 | 0.22404794 | -0.5833775 | 0.55963918 | 0.66837275 | FNDC7 | 163479 |
| ENSG000002 | 16.1318964 | -0.173518398 | 0.29746033 | -0.5833329 | 0.55966922 | 0.66838894 | LMF1 | 64788 |
| ENSG000002 | 4.73062234 | -0.219739108 | 0.37671757 | -0.5832993 | 0.55969181 | 0.66839624 | NA | NA |
| ENSG000001 | 11.9108252 | 0.186856788 | 0.32037267 | 0.58324821 | 0.55972622 | 0.66841765 | CLCNKB | 1188 |

| | | | | | | | | |
|---|---|---|---|---|---|---|---|---|
| ENSG000002 | 6.73496639 | -0.211081402 | 0.36198282 | -0.5831255 | 0.55980884 | 0.66849662 | NA | NA |
| ENSG000002 | 25.2503307 | -0.151549098 | 0.26005765 | -0.5827519 | 0.56006031 | 0.66877722 | PCDHGA5 | 56110 |
| ENSG000002 | 6.28791432 | -0.213078092 | 0.36577858 | -0.582533 | 0.56020772 | 0.66891385 | SBK3 | 100130827 |
| ENSG000002 | 3.13917056 | -0.21847654 | 0.37504379 | -0.5825361 | 0.56020564 | 0.66891385 | NA | NA |
| ENSG000002 | 5.24383658 | -0.216259411 | 0.37135118 | -0.5823582 | 0.56032542 | 0.66903469 | NA | NA |
| ENSG000002 | 16.9035445 | 0.172482635 | 0.29629677 | 0.58212795 | 0.5604805 | 0.66920015 | NA | NA |
| ENSG000002 | 8.04639758 | -0.205396452 | 0.35288761 | -0.582045 | 0.5605364 | 0.66924719 | NA | NA |
| ENSG000001 | 5.00968544 | -0.217817187 | 0.37429579 | -0.5819387 | 0.56060801 | 0.66931298 | ALPP | 250 |
| ENSG000002 | 4.75664807 | -0.218338531 | 0.37521684 | -0.5818996 | 0.56063431 | 0.66932467 | NA | NA |
| ENSG000002 | 3.34103246 | -0.218670415 | 0.37592615 | -0.5816845 | 0.56077921 | 0.66947796 | NA | NA |
| ENSG000002 | 4.52424011 | -0.219280751 | 0.37702855 | -0.5816025 | 0.56083445 | 0.66950448 | NA | NA |
| ENSG000002 | 4.27162069 | 0.219259793 | 0.37699064 | 0.5816054 | 0.5608325 | 0.66950448 | MIR7161 | 102465694 |
| ENSG000002 | 3.32829235 | -0.218939042 | 0.37647745 | -0.5815462 | 0.56087236 | 0.66951032 | NA | NA |
| ENSG000002 | 22.645472 | -0.164226886 | 0.28239372 | -0.5815529 | 0.56086787 | 0.66951032 | NA | NA |
| ENSG000001 | 9.69494212 | -0.198216142 | 0.34094662 | -0.58137 | 0.5609911 | 0.66963235 | C1orf145 | 574407 |
| ENSG000001 | 143.525702 | 0.089367723 | 0.15372823 | 0.58133579 | 0.56101416 | 0.66964016 | METTL25 | 84190 |
| ENSG000002 | 25.0653841 | 0.15749326 | 0.27097598 | 0.58120745 | 0.56110065 | 0.66972368 | SUB1P3 | 106480442 |
| ENSG000001 | 38.1436343 | 0.145100416 | 0.24981016 | 0.58084273 | 0.56134645 | 0.66999735 | CNIH2 | 254263 |
| ENSG000001 | 146.30899 | 0.09177196 | 0.1580942 | 0.58048911 | 0.56158483 | 0.67020296 | NDUFV2 | 4729 |
| ENSG000002 | 3.37508319 | -0.218819961 | 0.37692912 | -0.5805334 | 0.56155494 | 0.67020296 | NA | NA |
| ENSG000002 | 2.57482282 | 0.216420991 | 0.37282481 | 0.58048977 | 0.56158438 | 0.67020296 | NA | NA |
| ENSG000002 | 40.8057333 | 0.133600948 | 0.23014537 | 0.58050678 | 0.56157291 | 0.67020296 | NA | NA |
| ENSG000002 | 8.37176112 | 0.205399126 | 0.35390265 | 0.58038312 | 0.56165628 | 0.67026851 | NA | NA |
| ENSG000002 | 9.43820627 | -0.198439661 | 0.34194955 | -0.5803185 | 0.56169983 | 0.67030075 | NA | NA |
| ENSG000002 | 25.6611003 | 0.147854311 | 0.25479943 | 0.58027725 | 0.56172767 | 0.67031425 | NA | NA |
| ENSG000001 | 71.9758982 | -0.113915448 | 0.19647072 | -0.5798088 | 0.56204358 | 0.67067149 | STEAP1B | 256227 |
| ENSG000001 | 224.9958 | 0.080490856 | 0.13885802 | 0.57966298 | 0.56214191 | 0.67074935 | IFT172 | 26160 |
| ENSG000002 | 4.24020658 | 0.218449486 | 0.37685302 | 0.5796676 | 0.56213879 | 0.67074935 | NA | NA |
| ENSG000002 | 11.6303968 | -0.191117412 | 0.32972391 | -0.5796286 | 0.56216509 | 0.67075727 | NA | NA |

| | | | | | | | | |
|---|---|---|---|---|---|---|---|---|
| ENSG000001 | 84.6453128 | -0.101143836 | 0.17451414 | -0.5795739 | 0.56220202 | 0.67077755 | PDE2A | 5138 |
| ENSG000002 | 7.25956897 | 0.209937874 | 0.36224017 | 0.57955437 | 0.56221517 | 0.67077755 | NA | NA |
| ENSG000002 | 3.37005945 | -0.217840354 | 0.37598332 | -0.5793883 | 0.56232717 | 0.67089144 | NA | NA |
| ENSG000002 | 2.64993459 | -0.212113791 | 0.36629247 | -0.5790831 | 0.56253308 | 0.67111735 | NA | NA |
| ENSG000002 | 4.78198801 | -0.217835486 | 0.37621435 | -0.5790196 | 0.56257594 | 0.67114874 | NA | NA |
| ENSG000002 | 7.796527 | -0.209134749 | 0.3612886 | -0.5788579 | 0.56268509 | 0.67125921 | NA | NA |
| ENSG000002 | 5.34969949 | 0.214735147 | 0.37097993 | 0.57883225 | 0.56270237 | 0.67126008 | NA | NA |
| ENSG000001 | 7.42912619 | -0.206263327 | 0.35646763 | -0.5786313 | 0.56283799 | 0.67140211 | SNX15 | 29907 |
| ENSG000002 | 3.79976422 | -0.219043262 | 0.37858829 | -0.5785791 | 0.56287323 | 0.67141895 | NA | NA |
| ENSG000002 | 3.36703739 | -0.215489456 | 0.37245743 | -0.5785613 | 0.56288522 | 0.67141895 | NA | NA |
| ENSG000002 | 5.26051834 | -0.215463192 | 0.3725604 | -0.5783309 | 0.56304074 | 0.67158469 | NA | NA |
| ENSG000002 | 6.99402554 | -0.21074863 | 0.36446454 | -0.5782418 | 0.56310088 | 0.67163668 | NA | NA |
| ENSG000002 | 3.04074987 | -0.217675408 | 0.37647088 | -0.5781999 | 0.56312919 | 0.67165069 | IFT74-AS1 | 101929602 |
| ENSG000002 | 5.51888002 | -0.214044816 | 0.37021602 | -0.578162 | 0.56315478 | 0.67166146 | NA | NA |
| ENSG000002 | 8.74011335 | -0.200274337 | 0.34644616 | -0.5780821 | 0.56320867 | 0.67170597 | AP4B1-AS1 | 100287722 |
| ENSG000002 | 5.05328191 | 0.216771382 | 0.37508027 | 0.57793331 | 0.56330915 | 0.67180605 | NA | NA |
| ENSG000002 | 23.8032648 | -0.152148189 | 0.26336193 | -0.5777152 | 0.56345642 | 0.67196193 | ADNP-AS1 | 101927631 |
| ENSG000002 | 7.04483152 | 0.2074475 | 0.35924858 | 0.57744835 | 0.56363662 | 0.67215706 | NA | NA |
| ENSG000002 | 11.895792 | -0.193004709 | 0.33429148 | -0.5773546 | 0.56369996 | 0.67220509 | NA | NA |
| ENSG000002 | 4.76332274 | -0.217272817 | 0.3763345 | -0.5773396 | 0.56371005 | 0.67220509 | LOC285629 | 285629 |
| ENSG000002 | 4.18069564 | 0.218343503 | 0.3782681 | 0.57721891 | 0.56379158 | 0.67228254 | NA | NA |
| ENSG00000C | 16.3402684 | -0.175147313 | 0.30350886 | -0.5770748 | 0.56388893 | 0.67235908 | GIPR | 2696 |
| ENSG000002 | 3.41298916 | 0.218417385 | 0.37848386 | 0.57708508 | 0.56388198 | 0.67235908 | NA | NA |
| ENSG00000C | 5.24881273 | -0.215790479 | 0.37401911 | -0.5769504 | 0.56397296 | 0.6724395 | FOXC1 | 2296 |
| ENSG000002 | 3.1828366 | 0.217335979 | 0.3767893 | 0.57681038 | 0.56406756 | 0.67253252 | NA | NA |
| ENSG000001 | 192.558272 | 0.08493666 | 0.14728297 | 0.5766903 | 0.56414869 | 0.67260947 | ABCB10 | 23456 |
| ENSG000002 | 42.7777151 | -0.125973185 | 0.21845841 | -0.5766461 | 0.56417856 | 0.67262531 | TAS2R30 | 259293 |
| ENSG000001 | 22.454885 | -0.162820942 | 0.28241992 | -0.5765207 | 0.56426325 | 0.67266695 | HSH2D | 84941 |
| ENSG000002 | 5.35127454 | -0.216024783 | 0.37469798 | -0.5765304 | 0.56425672 | 0.67266695 | NA | NA |

| | | | | | | | | |
|---|---|---|---|---|---|---|---|---|
| ENSG000002 | 18.1758287 | 0.166790316 | 0.28929939 | 0.57653186 | 0.56425574 | 0.67266695 | NA | NA |
| ENSG000001 | 11.6299803 | 0.186694531 | 0.32389747 | 0.57640009 | 0.56434479 | 0.67274438 | NA | NA |
| ENSG000002 | 9.17921262 | -0.197465009 | 0.34277349 | -0.5760802 | 0.56456099 | 0.67298233 | LOC101929124 | 101929124 |
| ENSG000002 | 2.48952648 | 0.214373197 | 0.37215743 | 0.57602825 | 0.56459608 | 0.67300438 | NA | NA |
| ENSG000001 | 9.48203269 | -0.198868481 | 0.34528502 | -0.5759546 | 0.5646459 | 0.67300627 | ALG1L | 200810 |
| ENSG000002 | 8.58888955 | 0.198532944 | 0.34469626 | 0.57596489 | 0.56463891 | 0.67300627 | NA | NA |
| ENSG000002 | 3.93561597 | 0.218049071 | 0.37858881 | 0.57595223 | 0.56464747 | 0.67300627 | NA | NA |
| ENSG000001 | 112.175123 | -0.098065982 | 0.17030569 | -0.5758233 | 0.56473463 | 0.6730735 | SP110 | 3431 |
| ENSG000002 | 4.23126876 | 0.217477038 | 0.37768253 | 0.57581968 | 0.56473707 | 0.6730735 | NA | NA |
| ENSG000002 | 2.2813821 | -0.207744858 | 0.36082219 | -0.5757541 | 0.56478139 | 0.67310655 | NA | NA |
| ENSG000002 | 3.4138426 | 0.217953361 | 0.3785779 | 0.57571602 | 0.56480715 | 0.67311746 | NA | NA |
| ENSG000002 | 5.55039088 | -0.213152983 | 0.37035138 | -0.5755426 | 0.56492441 | 0.67323742 | RN7SL351P | 106479351 |
| ENSG000001 | 68.3308946 | -0.107938792 | 0.18762484 | -0.5752905 | 0.56509483 | 0.67339515 | NCALD | 83988 |
| ENSG000002 | 5.57926033 | -0.214239055 | 0.37240956 | -0.5752781 | 0.56510325 | 0.67339515 | NA | NA |
| ENSG000002 | 3.31966793 | -0.217212834 | 0.37758209 | -0.5752731 | 0.56510658 | 0.67339515 | NA | NA |
| ENSG000001 | 4.26807818 | -0.216948297 | 0.37728887 | -0.5750191 | 0.5652784 | 0.67355383 | OR2AP1 | 121129 |
| ENSG000002 | 5.11223701 | -0.216528573 | 0.37656743 | -0.5750061 | 0.56528716 | 0.67355383 | NA | NA |
| ENSG000002 | 4.3810092 | -0.217553173 | 0.37835167 | -0.5750026 | 0.56528957 | 0.67355383 | NA | NA |
| ENSG000002 | 3.08852533 | -0.215941895 | 0.37566107 | -0.5748317 | 0.56540512 | 0.67367171 | NA | NA |
| ENSG000002 | 5.49904461 | -0.21447761 | 0.37324209 | -0.574634 | 0.56553882 | 0.67381122 | NA | NA |
| ENSG000002 | 3.31552683 | 0.216599925 | 0.37708551 | 0.57440532 | 0.56569355 | 0.67397577 | LOC100507661 | 100507661 |
| ENSG000000 | 101.16847 | -0.105658591 | 0.18402671 | -0.5741481 | 0.56586757 | 0.67416329 | CHGB | 1114 |
| ENSG000002 | 4.84673973 | -0.216060781 | 0.37642771 | -0.5739768 | 0.56598348 | 0.67428157 | NA | NA |
| ENSG000002 | 9.20593418 | -0.200675999 | 0.34968836 | -0.573871 | 0.56605512 | 0.67434711 | NA | NA |
| ENSG000002 | 34.203306 | 0.133056268 | 0.23187826 | 0.57381951 | 0.56608994 | 0.67436878 | LOC100131496 | 100131496 |
| ENSG000001 | 3.78714034 | -0.21722372 | 0.37858134 | -0.5737835 | 0.56611433 | 0.67437802 | FOXF1 | 2294 |
| ENSG000002 | 12.656219 | -0.189058684 | 0.3295403 | -0.5737043 | 0.56616793 | 0.67442206 | NA | NA |
| ENSG000001 | 9.6312805 | -0.196047403 | 0.34192604 | -0.573362 | 0.5663996 | 0.67465839 | CEP295NL | 100653515 |
| ENSG000002 | 18.9700411 | 0.176462855 | 0.30776224 | 0.57337396 | 0.56639152 | 0.67465839 | NA | NA |

| | | | | | | | | |
|---|---|---|---|---|---|---|---|---|
| ENSG000001 | 5.03481652 | -0.215055138 | 0.3751391 | -0.5732677 | 0.56646343 | 0.6747146 | OR4P4 | 81300 |
| ENSG000002 | 4.12544012 | -0.216986585 | 0.37856872 | -0.5731762 | 0.56652539 | 0.67476858 | NA | NA |
| ENSG000002 | 2.53368845 | 0.213092937 | 0.37198249 | 0.57285745 | 0.56674122 | 0.67500582 | NA | NA |
| ENSG000002 | 5.52950133 | -0.21192974 | 0.36998692 | -0.5728033 | 0.56677787 | 0.67500982 | NA | NA |
| ENSG000002 | 7.66056143 | -0.205306422 | 0.35841178 | -0.5728228 | 0.56676471 | 0.67500982 | NA | NA |
| ENSG000002 | 3.83486373 | -0.216751233 | 0.37848609 | -0.5726795 | 0.56686171 | 0.67507056 | NA | NA |
| ENSG000002 | 41.5277767 | 0.135998009 | 0.23747692 | 0.57267884 | 0.56686217 | 0.67507056 | RNF139-AS1 | 101927612 |
| ENSG000002 | 2.5609326 | 0.21349167 | 0.3728201 | 0.57263991 | 0.56688853 | 0.67508213 | RNU6ATAC11P | 106481162 |
| ENSG000002 | 2.48893362 | 0.211401039 | 0.36920826 | 0.57257939 | 0.56692952 | 0.67511112 | NA | NA |
| ENSG000002 | 5.35257236 | -0.215020742 | 0.37569324 | -0.5723306 | 0.56709802 | 0.67529194 | LOC102724714 | 102724714 |
| ENSG000002 | 5.38305766 | 0.213676338 | 0.37355959 | 0.57200068 | 0.56732151 | 0.67549857 | RN7SL233P | 106480985 |
| ENSG000002 | 7.30403241 | -0.20688064 | 0.3616679 | -0.5720183 | 0.56730961 | 0.67549857 | NA | NA |
| ENSG000002 | 2.34283057 | 0.210211603 | 0.3674994 | 0.57200529 | 0.56731839 | 0.67549857 | NA | NA |
| ENSG000002 | 2.59232731 | -0.210590487 | 0.36819915 | -0.5719472 | 0.56735772 | 0.67552184 | RNU6-1275P | 106480122 |
| ENSG000002 | 55.902258 | 0.122424529 | 0.2140823 | 0.57185733 | 0.56741864 | 0.67557454 | SYT3 | 84258 |
| ENSG000001 | 57.283696 | 0.120608328 | 0.21094162 | 0.57176166 | 0.56748346 | 0.67563188 | PGBD4 | 161779 |
| ENSG000002 | 11.5532682 | -0.184709532 | 0.32307205 | -0.5717286 | 0.56750586 | 0.67563871 | NA | NA |
| ENSG000002 | 3.25419199 | -0.215734186 | 0.37736678 | -0.571683 | 0.56753673 | 0.67565563 | NA | NA |
| ENSG000001 | 204.01466 | -0.081942406 | 0.14336042 | -0.5715832 | 0.56760439 | 0.67571634 | STARD9 | 57519 |
| ENSG000001 | 27.0131551 | 0.148336182 | 0.25961744 | 0.57136448 | 0.5677526 | 0.67587293 | NA | NA |
| ENSG000001 | 11.3702467 | -0.187251141 | 0.32779658 | -0.5712419 | 0.56783568 | 0.675952 | LHFPL3 | 375612 |
| ENSG000002 | 2.30304927 | -0.20738886 | 0.36321013 | -0.5709887 | 0.56800733 | 0.67613648 | NA | NA |
| ENSG000002 | 9.48894263 | 0.195168018 | 0.34187557 | 0.57087442 | 0.56808477 | 0.67620881 | NA | NA |
| ENSG000002 | 24.2811295 | 0.15229545 | 0.26682988 | 0.57075861 | 0.56816329 | 0.67628242 | NA | NA |
| ENSG000002 | 5.07413625 | -0.215065253 | 0.37692471 | -0.5705788 | 0.56828519 | 0.67640766 | LOC101928668 | 101928668 |
| ENSG000001 | 6.09267458 | -0.210228115 | 0.36847256 | -0.5705394 | 0.5683119 | 0.6764196 | FUT7 | 2529 |
| ENSG000001 | 86.4879449 | -0.096111224 | 0.16846873 | -0.5704989 | 0.56833937 | 0.67643245 | KIAA0513 | 9764 |
| ENSG000002 | 3.58052275 | 0.215937795 | 0.3785816 | 0.5703864 | 0.56841565 | 0.67650338 | NA | NA |
| ENSG000002 | 13.4798338 | -0.175994998 | 0.30859774 | -0.5703055 | 0.56847049 | 0.67654879 | NA | NA |

| | | | | | | | | |
|---|---|---|---|---|---|---|---|---|
| ENSG000002 | 48.216143 | -0.126406079 | 0.22166811 | -0.5702493 | 0.56850864 | 0.67657433 | NA | NA |
| ENSG000002 | 18.8510649 | 0.164520765 | 0.28856627 | 0.57013165 | 0.56858841 | 0.67663576 | NA | NA |
| ENSG000002 | 189.881748 | -0.087949167 | 0.15426324 | -0.570124 | 0.56859363 | 0.67663576 | ZNF229 | 7772 |
| ENSG000001 | 16.4393934 | -0.170270666 | 0.29869957 | -0.5700399 | 0.56865065 | 0.67668377 | CDH20 | 28316 |
| ENSG000001 | 14.2386121 | -0.178659363 | 0.31345617 | -0.569966 | 0.56870075 | 0.67672353 | AGR2 | 10551 |
| ENSG000002 | 3.18085453 | 0.213390656 | 0.37442385 | 0.56991736 | 0.56873375 | 0.67673544 | LOC105377849 | 105377849 |
| ENSG000002 | 8.52116897 | -0.201566278 | 0.35368584 | -0.569902 | 0.56874414 | 0.67673544 | NA | NA |
| ENSG000002 | 6.74394518 | -0.206815395 | 0.36306878 | -0.5696314 | 0.5689277 | 0.67693398 | NA | NA |
| ENSG000002 | 12.2949238 | 0.185522173 | 0.32572199 | 0.56957214 | 0.56896793 | 0.67696199 | NA | NA |
| ENSG000002 | 11.0180337 | 0.195335161 | 0.34300639 | 0.56947966 | 0.56903067 | 0.67701678 | NA | NA |
| ENSG000002 | 29.0544121 | 0.140112529 | 0.2460479 | 0.56945224 | 0.56904928 | 0.67701905 | NA | NA |
| ENSG000002 | 28.7457 | 0.144503495 | 0.25380136 | 0.56935666 | 0.56911412 | 0.67705647 | DICER1-AS1 | 400242 |
| ENSG000002 | 3.0569941 | -0.212982409 | 0.37407042 | -0.5693645 | 0.56910882 | 0.67705647 | NA | NA |
| ENSG000001 | 6.24980508 | 0.208591637 | 0.36640884 | 0.56928658 | 0.56916167 | 0.67707331 | ULBP2 | 80328 |
| ENSG000002 | 12.5779721 | -0.179827996 | 0.31587269 | -0.5693053 | 0.56914897 | 0.67707331 | NA | NA |
| ENSG000001 | 11.1096142 | -0.186468751 | 0.32757286 | -0.5692436 | 0.56919084 | 0.67708524 | S100B | 6285 |
| ENSG000001 | 151.87091 | -0.094184218 | 0.16546114 | -0.5692226 | 0.56920509 | 0.67708524 | CTNNA3 | 29119 |
| ENSG000001 | 29.5921947 | -0.144029827 | 0.25304861 | -0.5691785 | 0.56923501 | 0.67710096 | PCDH8 | 5100 |
| ENSG000002 | 7.70499253 | 0.20425888 | 0.35891048 | 0.56910814 | 0.56928275 | 0.67713789 | NA | NA |
| ENSG000002 | 28.1213896 | 0.143439381 | 0.25207971 | 0.5690239 | 0.56933992 | 0.67718603 | SCAMP1-AS1 | 728769 |
| ENSG000001 | 6.4857199 | -0.209455058 | 0.36813865 | -0.568957 | 0.56938533 | 0.67722017 | NA | NA |
| ENSG000002 | 42.9645159 | 0.127352435 | 0.22385267 | 0.56891185 | 0.56941596 | 0.67723674 | NA | NA |
| ENSG000002 | 5.96690952 | -0.210357087 | 0.36979921 | -0.5688414 | 0.5694638 | 0.67727377 | NA | NA |
| ENSG000002 | 11.095544 | 0.191429761 | 0.33654793 | 0.56880386 | 0.56948925 | 0.67728418 | NA | NA |
| ENSG000001 | 248.071646 | 0.084587078 | 0.14880793 | 0.56843124 | 0.56974218 | 0.67756156 | ZNF654 | 55279 |
| ENSG000002 | 15.8497761 | -0.170808711 | 0.3005021 | -0.568411 | 0.5697559 | 0.67756156 | NA | NA |
| ENSG000002 | 4.58186776 | -0.21417526 | 0.37688159 | -0.5682826 | 0.56984307 | 0.67764535 | NA | NA |
| ENSG000002 | 7.32392907 | -0.2077612 | 0.36562051 | -0.5682427 | 0.56987015 | 0.67765768 | NA | NA |
| ENSG000001 | 9.78001741 | -0.194655709 | 0.34262759 | -0.5681262 | 0.56994928 | 0.67771202 | C10orf54 | 64115 |

| | | | | | | | | |
|---|---|---|---|---|---|---|---|---|
| ENSG000002 | 6.14830283 | 0.208916103 | 0.3677133 | 0.56814944 | 0.5699335 | 0.67771202 | NA | NA |
| ENSG000002 | 8.80614398 | 0.203159849 | 0.35765067 | 0.5680399 | 0.57000788 | 0.67776183 | NA | NA |
| ENSG000002 | 4.47512628 | -0.214438583 | 0.37757133 | -0.5679419 | 0.57007441 | 0.67782107 | NA | NA |
| ENSG000002 | 6.02265782 | -0.211145888 | 0.37179694 | -0.5679065 | 0.57009848 | 0.67782981 | NA | NA |
| ENSG000001 | 2.5635374 | -0.20837235 | 0.36700385 | -0.5677661 | 0.57019379 | 0.67792325 | LYZL2 | 119180 |
| ENSG000001 | 5.50412808 | -0.21250983 | 0.3743506 | -0.5676759 | 0.57025503 | 0.67797619 | HUS1B | 135458 |
| ENSG000002 | 4.19646448 | 0.214600307 | 0.37838524 | 0.56714767 | 0.57061386 | 0.67838291 | ULK4P2 | 100288380 |
| ENSG000002 | 2.9556488 | 0.208186176 | 0.36710902 | 0.56709632 | 0.57064874 | 0.67840449 | NA | NA |
| ENSG000002 | 3.60371588 | 0.214618891 | 0.37858075 | 0.56690387 | 0.5707795 | 0.67854004 | NA | NA |
| ENSG000002 | 5.83470521 | -0.210230438 | 0.3711896 | -0.5663694 | 0.57114268 | 0.67895189 | NA | NA |
| ENSG000002 | 4.58423474 | -0.214118241 | 0.37807149 | -0.5663433 | 0.57116046 | 0.67895312 | NA | NA |
| ENSG000002 | 27.5956925 | 0.14427801 | 0.25478113 | 0.56628217 | 0.57120198 | 0.67898257 | LOC101928103 | 101928103 |
| ENSG000002 | 5.80963139 | -0.211762164 | 0.37399989 | -0.5662092 | 0.5712516 | 0.67902165 | NA | NA |
| ENSG000001 | 36.7999697 | -0.136604605 | 0.24127987 | -0.5661666 | 0.57128052 | 0.67903612 | GUCY1B2 | 2974 |
| ENSG000001 | 168.528211 | 0.080941308 | 0.14303201 | 0.56589645 | 0.57146417 | 0.67923451 | LMLN | 89782 |
| ENSG000002 | 11.4131461 | -0.190611704 | 0.33708206 | -0.5654757 | 0.57175026 | 0.67955463 | NA | NA |
| ENSG000001 | 3.15637348 | -0.210332788 | 0.37199276 | -0.5654217 | 0.57178696 | 0.67957832 | NA | NA |
| ENSG000002 | 5.80826298 | -0.209423748 | 0.37046186 | -0.5653045 | 0.57186668 | 0.67965093 | NA | NA |
| ENSG000002 | 8.84529529 | 0.194652266 | 0.34434506 | 0.56528259 | 0.57188157 | 0.67965093 | NA | NA |
| ENSG000001 | 38.4174739 | 0.129991791 | 0.23001079 | 0.56515519 | 0.57196821 | 0.67973399 | IGSF5 | 150084 |
| ENSG000002 | 3.54489979 | -0.213312635 | 0.37754714 | -0.564996 | 0.57207646 | 0.6798427 | NA | NA |
| ENSG000002 | 13.4985513 | -0.174855353 | 0.30967171 | -0.5646475 | 0.57231355 | 0.68010453 | NA | NA |
| ENSG000000 | 31.7663655 | -0.136662847 | 0.24205049 | -0.5646047 | 0.57234266 | 0.68011919 | ARHGAP6 | 395 |
| ENSG000001 | 3.5941632 | -0.213419541 | 0.37802238 | -0.5645685 | 0.57236727 | 0.6801285 | BATF | 10538 |
| ENSG000002 | 6.42765864 | 0.20517982 | 0.36344903 | 0.56453534 | 0.57238985 | 0.68013541 | NA | NA |
| ENSG000002 | 9.72489981 | 0.189191591 | 0.33520902 | 0.56439887 | 0.57248271 | 0.68022581 | NA | NA |
| ENSG000002 | 3.37655869 | 0.213308364 | 0.37806104 | 0.56421674 | 0.57260664 | 0.68035313 | RN7SL618P | 106479447 |
| ENSG000001 | 215.421223 | 0.080689116 | 0.14308192 | 0.56393648 | 0.57279736 | 0.6805598 | SWT1 | 54823 |
| ENSG000001 | 4.9782261 | -0.212536403 | 0.37696275 | -0.5638128 | 0.57288157 | 0.68063991 | UPK3A | 7380 |

| | | | | | | | | |
|---|---|---|---|---|---|---|---|---|
| ENSG000002 | 5.80319182 | -0.20980655 | 0.37218273 | -0.5637192 | 0.57294525 | 0.68069562 | NA | NA |
| ENSG000002 | 6.07033022 | -0.207633495 | 0.36860071 | -0.5633019 | 0.5732293 | 0.68099319 | NA | NA |
| ENSG000002 | 4.10925483 | -0.213253894 | 0.37856961 | -0.5633149 | 0.57322049 | 0.68099319 | NA | NA |
| ENSG000001 | 24.5985895 | 0.146103539 | 0.2594725 | 0.56307909 | 0.57338103 | 0.68115349 | FAM86B3P | 286042 |
| ENSG000002 | 3.58003349 | 0.213128036 | 0.37858791 | 0.56295521 | 0.57346538 | 0.68123373 | NA | NA |
| ENSG000002 | 16.9037718 | -0.167337812 | 0.29726567 | -0.5629234 | 0.57348702 | 0.68123949 | NA | NA |
| ENSG000002 | 11.64608 | -0.182657463 | 0.32449961 | -0.5628896 | 0.57351005 | 0.68124688 | NA | NA |
| ENSG000001 | 28.4947827 | -0.143402208 | 0.25485422 | -0.5626833 | 0.57365057 | 0.68139384 | ZNF385A | 25946 |
| ENSG000001 | 15.2499029 | -0.173533845 | 0.3085654 | -0.5623892 | 0.57385088 | 0.68161181 | HDC | 3067 |
| ENSG000001 | 510.950747 | 0.064409563 | 0.11456995 | 0.56218551 | 0.57398964 | 0.68172076 | MARK1 | 4139 |
| ENSG000001 | 27.0650328 | -0.1407353 | 0.25031984 | -0.5622219 | 0.57396483 | 0.68172076 | PTGES | 9536 |
| ENSG000002 | 5.70272181 | -0.209711777 | 0.37303282 | -0.5621805 | 0.57399304 | 0.68172076 | NA | NA |
| ENSG000002 | 20.2709806 | 0.158171516 | 0.28148736 | 0.56191339 | 0.57417503 | 0.68191693 | NA | NA |
| ENSG000002 | 10.7609896 | 0.188758257 | 0.33599747 | 0.56178475 | 0.57426268 | 0.68200106 | NA | NA |
| ENSG000002 | 14.8693436 | -0.173275723 | 0.30857412 | -0.5615368 | 0.57443165 | 0.68217932 | LINC00310 | 114036 |
| ENSG000002 | 5.46215812 | -0.21014011 | 0.37423767 | -0.5615151 | 0.57444643 | 0.68217932 | NA | NA |
| ENSG000001 | 144.392705 | 0.105435468 | 0.18779313 | 0.56144476 | 0.57449438 | 0.68221628 | TENM4 | 26011 |
| ENSG000002 | 3.73139116 | -0.212229839 | 0.3781787 | -0.5611893 | 0.5746685 | 0.68240306 | NA | NA |
| ENSG000001 | 208.609393 | 0.083432657 | 0.14870799 | 0.56105026 | 0.57476328 | 0.68249563 | UVSSA | 57654 |
| ENSG00000C | 17.1655032 | -0.166312253 | 0.2965455 | -0.5608322 | 0.57491196 | 0.68261562 | SOX30 | 11063 |
| ENSG000001 | 60.0587799 | 0.110234667 | 0.19655701 | 0.56082796 | 0.57491483 | 0.68261562 | SLCO4A1 | 28231 |
| ENSG000002 | 40.3915293 | -0.128941317 | 0.22990129 | -0.5608551 | 0.57489631 | 0.68261562 | SEC24B-AS1 | 100533182 |
| ENSG000002 | 6.43844337 | 0.203873111 | 0.36365142 | 0.56062784 | 0.57505127 | 0.68275763 | NA | NA |
| ENSG000002 | 6.39023594 | 0.204723064 | 0.36519447 | 0.56058642 | 0.57507951 | 0.68277117 | NA | NA |
| ENSG000002 | 4.30848685 | -0.211893913 | 0.37816219 | -0.5603255 | 0.57525745 | 0.68296243 | RN7SL605P | 106481087 |
| ENSG000001 | 9.68697029 | -0.194418961 | 0.34708238 | -0.5601522 | 0.57537562 | 0.68308273 | CYP1A1 | 1543 |
| ENSG000002 | 14.1808144 | -0.179810665 | 0.32108358 | -0.560012 | 0.57547124 | 0.68317624 | LOC101928266 | 101928266 |
| ENSG000002 | 38.3302737 | 0.128241909 | 0.22902228 | 0.55995385 | 0.57551091 | 0.68320335 | NA | NA |
| ENSG000001 | 4.26944883 | 0.211733055 | 0.37815856 | 0.55990549 | 0.5755439 | 0.68320572 | FAM163B | 642968 |

| | | | | | | | | |
|---|---|---|---|---|---|---|---|---|
| ENSG000002 | 11.344083 | -0.186973578 | 0.33395888 | -0.5598701 | 0.57556808 | 0.68320572 | NA | NA |
| ENSG000002 | 4.58323063 | -0.211393805 | 0.37757461 | -0.5598729 | 0.57556611 | 0.68320572 | A2ML1-AS2 | 106478979 |
| ENSG000002 | 11.028817 | 0.183118671 | 0.327084 | 0.55985213 | 0.5755803 | 0.68320572 | NA | NA |
| ENSG000002 | 3.53216467 | -0.211593252 | 0.3779859 | -0.5597914 | 0.57562174 | 0.6832349 | NA | NA |
| ENSG000001 | 7.45954043 | 0.200901758 | 0.35893361 | 0.55971843 | 0.57567151 | 0.68327397 | NA | NA |
| ENSG000002 | 4.30018931 | -0.211797879 | 0.37846184 | -0.5596281 | 0.57573314 | 0.68332712 | NA | NA |
| ENSG000002 | 3.28930424 | -0.21080916 | 0.37671779 | -0.5595944 | 0.57575614 | 0.68333442 | NA | NA |
| ENSG000002 | 10.5973891 | 0.186706591 | 0.33367892 | 0.55953967 | 0.57579347 | 0.68335873 | NA | NA |
| ENSG000002 | 4.56680804 | -0.211818409 | 0.37859337 | -0.5594879 | 0.57582879 | 0.68338064 | NA | NA |
| ENSG000001 | 44.6548158 | 0.122680018 | 0.21934586 | 0.55929945 | 0.57595737 | 0.68349323 | RPH3AL | 9501 |
| ENSG000002 | 3.35328647 | 0.211082122 | 0.37740222 | 0.55930281 | 0.57595508 | 0.68349323 | RBM5-AS1 | 100775107 |
| ENSG000002 | 12.4856305 | -0.176520918 | 0.31564249 | -0.5592432 | 0.57599575 | 0.68351877 | NA | NA |
| ENSG000002 | 7.15898308 | -0.200700822 | 0.35890123 | -0.5592091 | 0.57601905 | 0.68352642 | NA | NA |
| ENSG000002 | 5.04483791 | -0.210233825 | 0.37611841 | -0.5589565 | 0.57619142 | 0.68371096 | LOC105371485 | 105371485 |
| ENSG000001 | 61.0600349 | 0.11266798 | 0.20160893 | 0.5588442 | 0.57626805 | 0.68378188 | SEMA7A | 8482 |
| ENSG000002 | 15.6484118 | 0.180126251 | 0.32234815 | 0.55879412 | 0.57630224 | 0.68380243 | LOC105378074 | 105378074 |
| ENSG000001 | 15.3828342 | 0.17021574 | 0.30473179 | 0.5585756 | 0.5764514 | 0.6839594 | GOLGA6D | 653643 |
| ENSG000002 | 4.52472938 | -0.211043168 | 0.37808288 | -0.5581929 | 0.57671268 | 0.6842494 | STPG2-AS1 | 101410545 |
| ENSG000001 | 8.39733024 | -0.19656695 | 0.35222919 | -0.5580655 | 0.57679967 | 0.68433259 | FITM1 | 161247 |
| ENSG000001 | 70.4894327 | 0.101825034 | 0.18247515 | 0.55802138 | 0.57682978 | 0.68434829 | ZNF565 | 147929 |
| ENSG000002 | 6.69112841 | 0.200784679 | 0.35988511 | 0.55791327 | 0.57690361 | 0.68441586 | NA | NA |
| ENSG000001 | 3.58926775 | -0.210081034 | 0.37662597 | -0.5577975 | 0.57698265 | 0.68448067 | SNORD1B | 677849 |
| ENSG000002 | 2.74650636 | 0.208568913 | 0.37392428 | 0.55778383 | 0.57699201 | 0.68448067 | NA | NA |
| ENSG000002 | 2.55439063 | 0.204744268 | 0.36716423 | 0.55763676 | 0.57709245 | 0.6845798 | NA | NA |
| ENSG000001 | 799.048505 | 0.066489121 | 0.11929201 | 0.5573644 | 0.57727849 | 0.68478046 | ABCA1 | 19 |
| ENSG000002 | 4.34973945 | -0.210938917 | 0.37858617 | -0.5571754 | 0.57740757 | 0.68491354 | NA | NA |
| ENSG000002 | 8.37681628 | 0.194847946 | 0.34981935 | 0.55699591 | 0.57753022 | 0.68503899 | NA | NA |
| ENSG000002 | 6.25129648 | -0.204279169 | 0.36684067 | -0.5568607 | 0.57762258 | 0.6851285 | NA | NA |
| ENSG000002 | 4.17510388 | 0.210750282 | 0.37852642 | 0.55676506 | 0.57768796 | 0.68516597 | NA | NA |

| | | | | | | | | |
|---|---|---|---|---|---|---|---|---|
| ENSG000002 | 6.02982837 | -0.206943614 | 0.37168906 | -0.5567654 | 0.57768772 | 0.68516597 | NA | NA |
| ENSG000001 | 66.4943723 | 0.105382921 | 0.18931062 | 0.55666672 | 0.57775516 | 0.68522563 | TUBB4A | 10382 |
| ENSG000001 | 15.4045533 | -0.168589376 | 0.30287834 | -0.5566241 | 0.5777843 | 0.68524014 | POTEF | 728378 |
| ENSG000002 | 12.3291821 | -0.179602155 | 0.32276734 | -0.5564446 | 0.57790693 | 0.68536554 | IFITM10 | 402778 |
| ENSG000002 | 3.75097101 | -0.2105829 | 0.37846659 | -0.5564108 | 0.57793005 | 0.68537292 | NA | NA |
| ENSG000002 | 7.26038672 | -0.202439595 | 0.36389986 | -0.5563058 | 0.57800183 | 0.685438 | LINC01399 | 104310353 |
| ENSG000002 | 3.74998431 | -0.209790011 | 0.37748985 | -0.5557501 | 0.57838173 | 0.68586845 | NA | NA |
| ENSG000002 | 7.34988434 | 0.185627727 | 0.33410314 | 0.55560007 | 0.57848428 | 0.68597 | NA | NA |
| ENSG000001 | 12.2988904 | -0.178160924 | 0.32072726 | -0.5554904 | 0.57855925 | 0.68603884 | CD164L2 | 388611 |
| ENSG000002 | 10.6538806 | -0.183723425 | 0.33081455 | -0.5553668 | 0.57864378 | 0.68611901 | LINC00327 | 100506697 |
| ENSG000001 | 97.6330743 | 0.093110948 | 0.16768529 | 0.555272 | 0.57870863 | 0.68617583 | CDHR1 | 92211 |
| ENSG000002 | 6.65704205 | -0.20426502 | 0.36810457 | -0.5549103 | 0.57895601 | 0.68644314 | GOLGA6L17P | 642402 |
| ENSG000002 | 25.220969 | -0.14442302 | 0.26027189 | -0.5548929 | 0.57896793 | 0.68644314 | MANEA-AS1 | 101927288 |
| ENSG000002 | 3.3564835 | 0.209444922 | 0.37757158 | 0.5547158 | 0.57908907 | 0.6865667 | NA | NA |
| ENSG000002 | 2.58463391 | 0.206389682 | 0.3722217 | 0.55448053 | 0.57925003 | 0.68673746 | NA | NA |
| ENSG000002 | 27.978924 | 0.138491315 | 0.24986472 | 0.55426519 | 0.57939737 | 0.68689205 | NA | NA |
| ENSG000001 | 611.013297 | 0.071725785 | 0.12941406 | 0.55423488 | 0.57941811 | 0.68689656 | E2F7 | 144455 |
| ENSG000002 | 2.778245 | 0.208281595 | 0.37583244 | 0.55418738 | 0.57945061 | 0.68691501 | NA | NA |
| ENSG000002 | 28.1362145 | 0.139729203 | 0.25218071 | 0.55408364 | 0.57952161 | 0.68697909 | NA | NA |
| ENSG000002 | 3.00064609 | -0.207167521 | 0.37410676 | -0.5537658 | 0.57973915 | 0.68721688 | NA | NA |
| ENSG000002 | 3.20859443 | 0.206278847 | 0.37267967 | 0.55350175 | 0.57991988 | 0.68741103 | RNU6-107P | 106479614 |
| ENSG000002 | 5.34669592 | -0.207476492 | 0.3749374 | -0.553363 | 0.58001486 | 0.68748341 | NA | NA |
| ENSG000002 | 6.77004751 | -0.202376333 | 0.36570726 | -0.5533834 | 0.58000089 | 0.68748341 | NA | NA |
| ENSG000001 | 209.951499 | 0.099836007 | 0.1804826 | 0.55316138 | 0.58015291 | 0.68762095 | CYR61 | 3491 |
| ENSG000002 | 6.18126622 | -0.203441327 | 0.36779089 | -0.553144 | 0.58016481 | 0.68762095 | NA | NA |
| ENSG000000 | 344.064751 | 0.089087269 | 0.16116208 | 0.55278059 | 0.58041366 | 0.68787568 | NUAK1 | 9891 |
| ENSG000002 | 4.5319848 | -0.20860323 | 0.37735763 | -0.5527998 | 0.5804005 | 0.68787568 | NA | NA |
| ENSG000001 | 6.24228565 | -0.203848371 | 0.3688307 | -0.5526882 | 0.58047694 | 0.68793057 | EDN3 | 1908 |
| ENSG000001 | 11.326674 | -0.180102762 | 0.32591799 | -0.5526015 | 0.58053633 | 0.68798085 | OR9Q1 | 219956 |

| | | | | | | | | | |
|---|---|---|---|---|---|---|---|---|---|
| ENSG000002 | 7.22702297 | -0.198626593 | 0.35948785 | -0.5525266 | 0.58058763 | 0.68798504 | LINC01007 | | 100506527 |
| ENSG000002 | 2.57477173 | -0.20277606 | 0.36700088 | -0.552522 | 0.58059076 | 0.68798504 | NA | NA | |
| ENSG000002 | 10.0699336 | -0.185380139 | 0.33549848 | -0.5525513 | 0.58057067 | 0.68798504 | ACTG1P17 | | 283693 |
| ENSG000002 | 10.0604274 | -0.184929195 | 0.33481391 | -0.5523343 | 0.58071935 | 0.68811731 | NA | NA | |
| ENSG000001 | 24.5250043 | -0.146450361 | 0.2652069 | -0.5522117 | 0.5808033 | 0.68819667 | TMEM155 | | 132332 |
| ENSG000002 | 10.3418012 | -0.182587131 | 0.33069557 | -0.5521306 | 0.5808589 | 0.68824244 | NA | NA | |
| ENSG000001 | 21.0747007 | -0.154680987 | 0.28017073 | -0.5520955 | 0.58088295 | 0.68825083 | DUSP18 | | 150290 |
| ENSG000002 | 5.23502295 | -0.206288852 | 0.37368515 | -0.5520392 | 0.58092149 | 0.68827638 | NA | NA | |
| ENSG000002 | 4.30366787 | 0.208843968 | 0.3783707 | 0.55195597 | 0.58097852 | 0.68832383 | NA | NA | |
| ENSG000002 | 66.7143234 | 0.105408159 | 0.191026 | 0.55180006 | 0.58108534 | 0.68843029 | NA | NA | |
| ENSG000002 | 5.46640031 | -0.205668856 | 0.37276379 | -0.5517404 | 0.5811262 | 0.68845858 | NA | NA | |
| ENSG000001 | 93.6905934 | 0.094020181 | 0.17041916 | 0.5516996 | 0.58115418 | 0.68847161 | BAIAP2 | | 10458 |
| ENSG000000 | 647.464753 | -0.070550616 | 0.12791018 | -0.5515637 | 0.58124729 | 0.6885618 | CYFIP2 | | 26999 |
| ENSG000001 | 8.43852641 | 0.193265186 | 0.35059878 | 0.55124318 | 0.58146698 | 0.68880193 | NA | NA | |
| ENSG000002 | 3.00294194 | -0.205252664 | 0.37243721 | -0.5511068 | 0.58156049 | 0.68889258 | NA | NA | |
| ENSG000001 | 7.72950042 | 0.194217654 | 0.35256382 | 0.55087234 | 0.58172119 | 0.6890628 | VSTM2L | | 128434 |
| ENSG000002 | 11.5523238 | -0.178506937 | 0.32434429 | -0.5503625 | 0.58207076 | 0.68945674 | NA | NA | |
| ENSG000001 | 98.0915653 | 0.095681083 | 0.1738665 | 0.5503135 | 0.58210436 | 0.68945756 | FAM189A2 | | 9413 |
| ENSG000001 | 60.420551 | 0.108814223 | 0.19773191 | 0.55031191 | 0.58210546 | 0.68945756 | MAPK4 | | 5596 |
| ENSG000001 | 173.549781 | -0.079964068 | 0.14539822 | -0.5499659 | 0.58234273 | 0.68971845 | PPARGC1A | | 10891 |
| ENSG000002 | 4.71669089 | 0.206536349 | 0.37556819 | 0.54993035 | 0.58236714 | 0.68972721 | LINC01137 | | 728431 |
| ENSG000002 | 18.9390539 | -0.161401318 | 0.29355239 | -0.5498212 | 0.58244204 | 0.68979577 | NA | NA | |
| ENSG000001 | 57.476885 | -0.111912039 | 0.20359651 | -0.5496756 | 0.58254187 | 0.68989385 | ZNF23 | | 7571 |
| ENSG000001 | 6.97837556 | -0.200785912 | 0.36531013 | -0.5496314 | 0.58257223 | 0.68990965 | NA | NA | |
| ENSG000001 | 105.372719 | 0.090950526 | 0.16549773 | 0.54955753 | 0.58262289 | 0.68994951 | ZNF287 | | 57336 |
| ENSG000001 | 25.4342569 | 0.153239705 | 0.27898919 | 0.54926753 | 0.58282187 | 0.69014482 | WFIKKN1 | | 117166 |
| ENSG000002 | 6.04632756 | -0.204331592 | 0.3720045 | -0.5492718 | 0.58281891 | 0.69014482 | NA | NA | |
| ENSG000001 | 8.64017771 | -0.195071251 | 0.35516519 | -0.5492409 | 0.58284014 | 0.69014631 | COMMD3 | | 23412 |
| ENSG000001 | 10.1186603 | 0.188530879 | 0.34348965 | 0.54886916 | 0.58309524 | 0.69042822 | VSTM2B | | 342865 |

| | | | | | | | | |
|---|---|---|---|---|---|---|---|---|
| ENSG00000( | 39.3540874 | 0.121916916 | 0.22234479 | 0.54832369 | 0.58346966 | 0.69084876 | GLTSCR1 | 29998 |
| ENSG000001 | 89.5166371 | -0.097535232 | 0.17788594 | -0.5483021 | 0.58348448 | 0.69084876 | COL7A1 | 1294 |
| ENSG000001 | 9.76292401 | 0.186835818 | 0.34087076 | 0.54811336 | 0.58361407 | 0.69097571 | AMH | 268 |
| ENSG000001 | 20.1800424 | -0.156615981 | 0.28574537 | -0.5480963 | 0.58362578 | 0.69097571 | IGF2BP2-AS1 | 646600 |
| ENSG000001 | 91.2251572 | 0.102911245 | 0.18779866 | 0.54798711 | 0.58370075 | 0.6910443 | NAT1 | 9 |
| ENSG000002 | 6.70317153 | 0.196716816 | 0.35900366 | 0.54795211 | 0.58372479 | 0.69105257 | NA | NA |
| ENSG000002 | 20.4499891 | -0.159145244 | 0.29055166 | -0.5477348 | 0.58387403 | 0.69120908 | NA | NA |
| ENSG000001 | 4.7655151 | -0.206407089 | 0.37686637 | -0.547693 | 0.58390269 | 0.69122283 | NTSR2 | 23620 |
| ENSG000001 | 50.4234467 | 0.120984081 | 0.22092232 | 0.54763176 | 0.58394477 | 0.69125247 | BCDIN3D | 144233 |
| ENSG000002 | 7.53818968 | 0.198841147 | 0.36316189 | 0.54752757 | 0.58401634 | 0.69130292 | NA | NA |
| ENSG000002 | 2.53072186 | -0.20079363 | 0.36673292 | -0.5475201 | 0.58402149 | 0.69130292 | NA | NA |
| ENSG000002 | 6.96050459 | -0.200512439 | 0.36629066 | -0.5474135 | 0.58409471 | 0.69136941 | NA | NA |
| ENSG000002 | 144.975941 | 0.085869566 | 0.15690164 | 0.54728276 | 0.58418448 | 0.69145549 | ZBTB8B | 728116 |
| ENSG000002 | 126.517307 | 0.096466799 | 0.17628464 | 0.54722181 | 0.58422635 | 0.69148486 | NA | NA |
| ENSG000002 | 6.88451985 | -0.198855829 | 0.36341249 | -0.5471904 | 0.58424792 | 0.69149021 | LOC102724699 | 102724699 |
| ENSG000001 | 16.9794436 | 0.16476063 | 0.30113806 | 0.54712656 | 0.58429179 | 0.69152194 | OLIG3 | 167826 |
| ENSG000002 | 5.61560923 | 0.20303475 | 0.37121257 | 0.54695009 | 0.58441302 | 0.69164524 | NA | NA |
| ENSG000001 | 1314.58436 | -0.073679782 | 0.13472118 | -0.5469057 | 0.58444353 | 0.69165286 | AEN | 64782 |
| ENSG000001 | 21.8497448 | -0.151301681 | 0.2766578 | -0.5468911 | 0.58445357 | 0.69165286 | FAM181B | 220382 |
| ENSG000002 | 10.6816149 | 0.182390313 | 0.33352099 | 0.54686306 | 0.58447281 | 0.69165544 | NA | NA |
| ENSG000002 | 2.5726553 | 0.203443879 | 0.37206547 | 0.54679592 | 0.58451895 | 0.69168985 | NA | NA |
| ENSG000002 | 3.74518999 | -0.206821594 | 0.37834237 | -0.5466519 | 0.58461788 | 0.6917835 | NA | NA |
| ENSG000002 | 4.46209403 | 0.206394233 | 0.377575 | 0.54663109 | 0.58463221 | 0.6917835 | NA | NA |
| ENSG000002 | 2.52953886 | 0.201276614 | 0.36851769 | 0.54617898 | 0.58494292 | 0.69213096 | NA | NA |
| ENSG000002 | 19.9563057 | -0.154850555 | 0.28368977 | -0.5458447 | 0.58517271 | 0.69238266 | NA | NA |
| ENSG000002 | 4.29226931 | -0.206345506 | 0.3781353 | -0.5456923 | 0.5852775 | 0.69248644 | NA | NA |
| ENSG000001 | 40.0963674 | -0.12209295 | 0.22376402 | -0.5456326 | 0.5853185 | 0.69251474 | LEMD1 | 93273 |
| ENSG000002 | 3.63777716 | 0.206446899 | 0.37858801 | 0.54530755 | 0.58554201 | 0.69275899 | NA | NA |
| ENSG000001 | 50.9767445 | 0.116042241 | 0.21282386 | 0.54525016 | 0.58558148 | 0.69278546 | POMZP3 | 22932 |

| | | | | | | | | |
|---|---|---|---|---|---|---|---|---|
| ENSG000002 | 57.5291902 | 0.110806432 | 0.20334881 | 0.54490818 | 0.58581667 | 0.69304349 | NA | NA |
| ENSG000002 | 4.29735796 | 0.205865309 | 0.37794839 | 0.54469159 | 0.58596565 | 0.69317929 | CLRN2 | 645104 |
| ENSG000002 | 61.4432832 | 0.107842882 | 0.19798872 | 0.54469205 | 0.58596533 | 0.69317929 | NA | NA |
| ENSG000002 | 3.31150609 | -0.203837056 | 0.37431051 | -0.5445667 | 0.58605153 | 0.69326067 | NA | NA |
| ENSG000002 | 8.26372983 | -0.194770526 | 0.35800011 | -0.5440516 | 0.58640598 | 0.69365972 | NA | NA |
| ENSG000001 | 5.057189 | -0.204647853 | 0.37619584 | -0.5439929 | 0.58644638 | 0.69368727 | IL25 | 64806 |
| ENSG000001 | 16.7954779 | -0.157706371 | 0.29005635 | -0.5437094 | 0.58664144 | 0.69389776 | SLC45A1 | 50651 |
| ENSG000001 | 2.80415796 | 0.204297077 | 0.37582355 | 0.54359838 | 0.58671787 | 0.69396793 | MIR103A2 | 406896 |
| ENSG000001 | 9.58151221 | -0.185369308 | 0.34106576 | -0.5435002 | 0.58678545 | 0.69402762 | NPPC | 4880 |
| ENSG000002 | 6.40715435 | 0.199682003 | 0.36742646 | 0.54346115 | 0.58681234 | 0.69403917 | NA | NA |
| ENSG000002 | 5.8091607 | 0.201010701 | 0.36996744 | 0.54331997 | 0.58690952 | 0.69413386 | ACTN1-AS1 | 161159 |
| ENSG000000 | 118.228248 | 0.089723434 | 0.16515439 | 0.54327005 | 0.58694388 | 0.69415426 | MPPED2 | 744 |
| ENSG000001 | 31.0570173 | -0.136752158 | 0.25175993 | -0.5431848 | 0.5870026 | 0.69420346 | FRS3 | 10817 |
| ENSG000001 | 45.1366728 | 0.118910445 | 0.21894086 | 0.54311674 | 0.58704943 | 0.6942386 | PLA2G15 | 23659 |
| ENSG000001 | 52.6929743 | -0.123378299 | 0.22722108 | -0.5429879 | 0.58713812 | 0.69432324 | FOXP4 | 116113 |
| ENSG000002 | 10.76427 | -0.180286376 | 0.33213695 | -0.5428073 | 0.58726246 | 0.69445002 | NA | NA |
| ENSG000002 | 6.26756669 | -0.202464403 | 0.37318016 | -0.542538 | 0.58744796 | 0.69464912 | NA | NA |
| ENSG000002 | 8.83350371 | 0.186297767 | 0.34342345 | 0.54247247 | 0.58749307 | 0.6946822 | ZGLP1 | 100125288 |
| ENSG000001 | 228.136097 | 0.072097721 | 0.13293435 | 0.54235582 | 0.58757341 | 0.69475694 | RAB30 | 27314 |
| ENSG000002 | 7.97444459 | -0.193382544 | 0.35669988 | -0.5421436 | 0.58771961 | 0.69490955 | NAPA-AS1 | 100505681 |
| ENSG000002 | 12.7807948 | -0.173187569 | 0.31946532 | -0.542117 | 0.58773794 | 0.69491096 | NA | NA |
| ENSG000001 | 6.67127688 | 0.197024696 | 0.36352218 | 0.5419881 | 0.5878267 | 0.69499565 | AQP8 | 343 |
| ENSG000001 | 49.9337761 | 0.116041828 | 0.21412946 | 0.5419237 | 0.58787107 | 0.69502783 | KLHL29 | 114818 |
| ENSG000002 | 6.05636124 | -0.202254338 | 0.37324746 | -0.5418773 | 0.58790301 | 0.69504534 | NA | NA |
| ENSG000001 | 41.113309 | -0.120155139 | 0.22184671 | -0.5416134 | 0.58808489 | 0.69524009 | CACNB3 | 784 |
| ENSG000002 | 8.5576402 | -0.193390304 | 0.35722205 | -0.5413728 | 0.58825064 | 0.69541577 | NA | NA |
| ENSG000000 | 74.8248662 | 0.100471104 | 0.18559584 | 0.54134351 | 0.58827083 | 0.69541937 | GPR137B | 7107 |
| ENSG000002 | 2.23946298 | -0.19995157 | 0.36950614 | -0.5411319 | 0.58841665 | 0.69557147 | NA | NA |
| ENSG000001 | 17.5490958 | -0.158024044 | 0.29209328 | -0.5410054 | 0.58850385 | 0.69565427 | F12 | 2161 |

| | | | | | | | | |
|---|---|---|---|---|---|---|---|---|
| ENSG000002 | 2.2466658 | -0.199279169 | 0.36845775 | -0.5408467 | 0.58861323 | 0.69576328 | NA | NA |
| ENSG000002 | 2.53733419 | 0.200584004 | 0.37097077 | 0.5407003 | 0.58871417 | 0.69586232 | NA | NA |
| ENSG000001 | 80.5173511 | 0.098315196 | 0.18186333 | 0.54059934 | 0.58878377 | 0.69590401 | CCDC57 | 284001 |
| ENSG000002 | 2.53456339 | 0.196549588 | 0.36357403 | 0.54060403 | 0.58878054 | 0.69590401 | NA | NA |
| ENSG000001 | 6.61775535 | 0.204242071 | 0.37790468 | 0.54045923 | 0.58888037 | 0.6959979 | FDCSP | 260436 |
| ENSG000001 | 27.1462515 | 0.138588503 | 0.25647986 | 0.54034849 | 0.58895673 | 0.69606786 | MMP9 | 4318 |
| ENSG000002 | 2.49678634 | -0.201204421 | 0.37245818 | -0.5402067 | 0.58905446 | 0.69616308 | NA | NA |
| ENSG000002 | 35.7004233 | 0.128275962 | 0.23748195 | 0.54015035 | 0.58909335 | 0.69617902 | SNORD20 | 6082 |
| ENSG000002 | 11.7370155 | -0.175694427 | 0.32528884 | -0.5401182 | 0.58911551 | 0.69617902 | NA | NA |
| ENSG000002 | 4.07292154 | -0.20424666 | 0.37815578 | -0.5401125 | 0.58911946 | 0.69617902 | RN7SL130P | 106479269 |
| ENSG000002 | 6.05050387 | -0.200711286 | 0.37170745 | -0.5399711 | 0.58921695 | 0.69625365 | NA | NA |
| ENSG000002 | 2.81757135 | 0.20311 | 0.37613824 | 0.53998764 | 0.58920555 | 0.69625365 | NA | NA |
| ENSG000002 | 2.49747148 | -0.201023234 | 0.3724594 | -0.5397185 | 0.58939117 | 0.69641893 | NA | NA |
| ENSG000002 | 3.59749757 | 0.204323738 | 0.37856297 | 0.53973514 | 0.5893797 | 0.69641893 | NA | NA |
| ENSG000002 | 3.33069396 | 0.203085351 | 0.37660866 | 0.53924769 | 0.58971596 | 0.6967824 | NA | NA |
| ENSG000001 | 10.1126225 | -0.179347622 | 0.33264014 | -0.5391641 | 0.58977362 | 0.69683023 | GPR146 | 115330 |
| ENSG000001 | 10.5818342 | -0.181166819 | 0.3360529 | -0.5391021 | 0.58981641 | 0.69686048 | NA | NA |
| ENSG000002 | 2.77628187 | 0.201492994 | 0.37412953 | 0.5385648 | 0.59018718 | 0.69727822 | NA | NA |
| ENSG000002 | 2.26871732 | -0.199001841 | 0.36960402 | -0.538419 | 0.59028778 | 0.69737676 | NA | NA |
| ENSG000002 | 7.40545989 | -0.193039208 | 0.3585572 | -0.5383777 | 0.59031631 | 0.69739014 | NA | NA |
| ENSG000002 | 3.60372742 | 0.203737857 | 0.37859158 | 0.53814683 | 0.59047568 | 0.6975581 | NA | NA |
| ENSG000002 | 2.29282338 | -0.196594922 | 0.36537035 | -0.5380703 | 0.59052853 | 0.69760022 | NA | NA |
| ENSG000001 | 4.89195102 | -0.203668504 | 0.3785712 | -0.5379926 | 0.59058215 | 0.69764323 | TMEM204 | 79652 |
| ENSG000002 | 2.79799261 | 0.201969219 | 0.37557533 | 0.53775954 | 0.59074307 | 0.69781299 | NA | NA |
| ENSG000001 | 19.8368401 | -0.149872572 | 0.27875761 | -0.5376448 | 0.59082232 | 0.69785546 | PADI3 | 51702 |
| ENSG000002 | 10.8340656 | -0.176607607 | 0.32849119 | -0.5376327 | 0.59083065 | 0.69785546 | CCDC183-AS1 | 100131193 |
| ENSG000002 | 3.33980673 | -0.201012501 | 0.37388342 | -0.5376342 | 0.59082964 | 0.69785546 | NA | NA |
| ENSG000001 | 91.8730151 | 0.100766898 | 0.18744634 | 0.5375773 | 0.59086891 | 0.69788032 | MAML3 | 55534 |
| ENSG00000C | 15.3122918 | 0.163771138 | 0.30466177 | 0.53755067 | 0.5908873 | 0.69788171 | DPEP1 | 1800 |

| | | | | | | | | |
|---|---|---|---|---|---|---|---|---|
| ENSG000001 | 205.445546 | -0.071175762 | 0.13259226 | -0.5368018 | 0.59140454 | 0.69845255 | KIF13B | 23303 |
| ENSG000002 | 5.28601972 | -0.20200508 | 0.37631277 | -0.536801 | 0.59140506 | 0.69845255 | NA | NA |
| ENSG000002 | 10.1025425 | -0.178436131 | 0.33245034 | -0.5367302 | 0.591454 | 0.69849 | LOC100505915 | 100505915 |
| ENSG000002 | 38.5664947 | 0.130513244 | 0.24329097 | 0.53644919 | 0.59164815 | 0.69869893 | NA | NA |
| ENSG000002 | 2.26356904 | -0.19537897 | 0.36427382 | -0.5363519 | 0.59171534 | 0.69875793 | NA | NA |
| ENSG000002 | 2.23712715 | -0.197664222 | 0.36880963 | -0.5359519 | 0.59199179 | 0.69906404 | NA | NA |
| ENSG000001 | 28.5743401 | -0.144547599 | 0.269773 | -0.535812 | 0.59208852 | 0.6991579 | RAB40B | 10966 |
| ENSG000002 | 2.22916229 | -0.197921138 | 0.36947289 | -0.5356851 | 0.59217618 | 0.69924105 | NA | NA |
| ENSG000002 | 14.1393408 | -0.163702808 | 0.30563769 | -0.5356107 | 0.59222765 | 0.69926111 | NA | NA |
| ENSG000002 | 30.4673322 | -0.133776983 | 0.24976067 | -0.5356207 | 0.59222072 | 0.69926111 | PTGES3L | 100885848 |
| ENSG000002 | 2.24675067 | -0.197909796 | 0.36953038 | -0.5355711 | 0.592255 | 0.69927304 | NA | NA |
| ENSG000002 | 7.29681435 | 0.193772287 | 0.36182872 | 0.53553595 | 0.59227931 | 0.69928138 | NA | NA |
| ENSG000002 | 14.3274436 | -0.172735874 | 0.32275868 | -0.5351858 | 0.59252136 | 0.6995314 | NA | NA |
| ENSG000002 | 10.8097877 | -0.176796629 | 0.33035001 | -0.5351797 | 0.59252558 | 0.6995314 | NA | NA |
| ENSG000002 | 9.55027021 | -0.180296888 | 0.33699746 | -0.5350096 | 0.59264319 | 0.69964989 | NA | NA |
| ENSG000002 | 3.6440348 | 0.202408029 | 0.37836548 | 0.53495374 | 0.59268184 | 0.69967515 | NA | NA |
| ENSG000002 | 4.71969 | -0.201690911 | 0.37710987 | -0.5348333 | 0.59276514 | 0.69975312 | NA | NA |
| ENSG000002 | 12.6503977 | -0.171410524 | 0.32052753 | -0.5347763 | 0.59280455 | 0.69977927 | C19orf38 | 255809 |
| ENSG000001 | 68.5287268 | 0.105778029 | 0.19786814 | 0.53458849 | 0.59293445 | 0.69991223 | SRRM3 | 222183 |
| ENSG000002 | 10.7953566 | -0.177268249 | 0.33165302 | -0.5344991 | 0.59299625 | 0.69996481 | PRR29 | 92340 |
| ENSG000002 | 2.24267983 | -0.196750285 | 0.36826205 | -0.5342671 | 0.59315678 | 0.70013391 | NA | NA |
| ENSG000001 | 34.5776408 | 0.123852975 | 0.23185429 | 0.53418452 | 0.59321388 | 0.70018093 | NA | NA |
| ENSG000001 | 5.57230901 | 0.19890012 | 0.37253385 | 0.53391154 | 0.59340274 | 0.70038346 | FAM166A | 401565 |
| ENSG000001 | 16.1118274 | 0.164513678 | 0.30815313 | 0.5338699 | 0.59343155 | 0.70039708 | DKKL1 | 27120 |
| ENSG000002 | 3.99246638 | -0.201977871 | 0.37839853 | -0.5337702 | 0.59350051 | 0.70045808 | NA | NA |
| ENSG000002 | 11.3431241 | -0.181777404 | 0.34073743 | -0.5334823 | 0.59369973 | 0.70065242 | LRRC10B | 390205 |
| ENSG000002 | 12.3900721 | 0.171325505 | 0.32113438 | 0.53350098 | 0.59368683 | 0.70065242 | NA | NA |
| ENSG000002 | 3.98434457 | 0.201570955 | 0.37811739 | 0.53309093 | 0.59397063 | 0.70095173 | LOC100996288 | 100996288 |
| ENSG000002 | 7.6376673 | -0.188584066 | 0.35382837 | -0.5329818 | 0.59404621 | 0.70102051 | NA | NA |

| | | | | | | | | |
|---|---|---|---|---|---|---|---|---|
| ENSG000002 | 9.62896401 | -0.18066199 | 0.33934206 | -0.532389 | 0.59445658 | 0.70146396 | CDIPT-AS1 | 440356 |
| ENSG000002 | 2.7727199 | 0.197892402 | 0.37168997 | 0.53241254 | 0.5944403 | 0.70146396 | NA | NA |
| ENSG000001 | 42.3539183 | -0.117721114 | 0.22114812 | -0.532318 | 0.5945058 | 0.70150162 | RUNDC3B | 154661 |
| ENSG000002 | 11.9834196 | 0.177487208 | 0.33349674 | 0.53220073 | 0.59458698 | 0.70157693 | NA | NA |
| ENSG000002 | 2.75077661 | -0.199606615 | 0.37507644 | -0.5321758 | 0.59460422 | 0.70157693 | NA | NA |
| ENSG000002 | 2.20510776 | -0.196539881 | 0.36938466 | -0.5320738 | 0.59467491 | 0.70163992 | LOC101927727 | 101927727 |
| ENSG000002 | 2.46531872 | 0.151616835 | 0.28500892 | 0.53197224 | 0.59474522 | 0.70170246 | NA | NA |
| ENSG000002 | 45.639933 | 0.116798012 | 0.21967454 | 0.53168662 | 0.59494306 | 0.70191545 | NA | NA |
| ENSG000002 | 5.48025863 | -0.19869054 | 0.37377562 | -0.531577 | 0.59501897 | 0.70198458 | NA | NA |
| ENSG000001 | 33.7263962 | -0.137871593 | 0.25938108 | -0.5315407 | 0.59504417 | 0.70199389 | APOL4 | 80832 |
| ENSG000001 | 80.5530946 | -0.101665254 | 0.19127979 | -0.5315002 | 0.59507218 | 0.70200651 | C16orf89 | 146556 |
| ENSG000002 | 8.88066161 | -0.183272886 | 0.34492755 | -0.5313373 | 0.59518503 | 0.70211922 | NA | NA |
| ENSG000002 | 7.62583696 | 0.190040289 | 0.35768343 | 0.53130862 | 0.59520493 | 0.70212227 | NA | NA |
| ENSG000001 | 86.0811713 | 0.101751071 | 0.19152073 | 0.53127968 | 0.59522498 | 0.7021255 | SMAGP | 57228 |
| ENSG000002 | 3.28169812 | -0.199004159 | 0.37475653 | -0.5310225 | 0.59540317 | 0.70226482 | NA | NA |
| ENSG000002 | 5.45806853 | -0.198857682 | 0.37446185 | -0.5310492 | 0.59538465 | 0.70226482 | FAM83C-AS1 | 140846 |
| ENSG000002 | 3.87882348 | 0.200645877 | 0.37780519 | 0.5310829 | 0.59536133 | 0.70226482 | MIR3685 | 100500802 |
| ENSG000002 | 6.6978681 | 0.195179854 | 0.36756394 | 0.53100926 | 0.59541236 | 0.70226482 | NA | NA |
| ENSG000002 | 13.4502718 | -0.17016336 | 0.3205314 | -0.5308789 | 0.59550269 | 0.70235093 | NA | NA |
| ENSG000002 | 2.49479129 | -0.198534429 | 0.37402063 | -0.5308114 | 0.59554945 | 0.70236522 | NA | NA |
| ENSG000002 | 14.9851998 | 0.168066834 | 0.31661398 | 0.53082569 | 0.59553958 | 0.70236522 | NA | NA |
| ENSG000002 | 2.25975933 | -0.194689429 | 0.36709172 | -0.5303564 | 0.59586488 | 0.70269875 | NA | NA |
| ENSG000002 | 5.78411706 | -0.199097502 | 0.37542303 | -0.5303284 | 0.59588424 | 0.70269875 | NA | NA |
| ENSG000002 | 15.2780832 | 0.160892814 | 0.30337725 | 0.53033908 | 0.59587686 | 0.70269875 | NA | NA |
| ENSG000001 | 214.521011 | 0.071292498 | 0.13445262 | 0.53024254 | 0.59594378 | 0.70274853 | DLG2 | 1740 |
| ENSG000001 | 362.022244 | -0.065903674 | 0.12438367 | -0.5298419 | 0.59622158 | 0.70305567 | KIAA1217 | 56243 |
| ENSG00000C | 121.488253 | 0.081628308 | 0.15407468 | 0.52979702 | 0.59625267 | 0.70306674 | COL16A1 | 1307 |
| ENSG000001 | 21.5892293 | -0.151843872 | 0.28661776 | -0.5297783 | 0.59626564 | 0.70306674 | SLC10A4 | 201780 |
| ENSG000002 | 33.1902564 | -0.12994348 | 0.24529157 | -0.5297511 | 0.59628451 | 0.70306853 | NA | NA |

| | | | | | | | | |
|---|---|---|---|---|---|---|---|---|
| ENSG000002 | 35.3676839 | -0.125844713 | 0.2376884 | -0.5294525 | 0.5964916 | 0.70329227 | NA | NA |
| ENSG000002 | 7.21461568 | -0.191365433 | 0.3614683 | -0.5294114 | 0.59652011 | 0.70330543 | NA | NA |
| ENSG000002 | 10.1553094 | 0.178352491 | 0.33701852 | 0.52920679 | 0.59666201 | 0.70345227 | NA | NA |
| ENSG000002 | 26.3819737 | 0.148430313 | 0.28059734 | 0.52897976 | 0.59681949 | 0.70361749 | NA | NA |
| ENSG000002 | 5.60800082 | 0.195458881 | 0.36965048 | 0.52876674 | 0.59696727 | 0.70375079 | NA | NA |
| ENSG000002 | 2.45844514 | -0.197088801 | 0.37272111 | -0.5287836 | 0.59695559 | 0.70375079 | NA | NA |
| ENSG000002 | 2.47800428 | -0.196778645 | 0.37218397 | -0.5287134 | 0.59700429 | 0.70375352 | NA | NA |
| ENSG000002 | 2.24225473 | -0.19268049 | 0.36442958 | -0.528718 | 0.59700106 | 0.70375352 | NA | NA |
| ENSG000001 | 133.519639 | 0.08248999 | 0.15602812 | 0.52868669 | 0.59702281 | 0.70375489 | EGFR | 1956 |
| ENSG000002 | 6.87101234 | 0.190927189 | 0.36125783 | 0.52850672 | 0.59714768 | 0.70388163 | NA | NA |
| ENSG000001 | 40.9151023 | 0.121026151 | 0.22912698 | 0.52820558 | 0.59735666 | 0.70410749 | BCAN | 63827 |
| ENSG000002 | 3.80955637 | -0.199441684 | 0.37761446 | -0.5281622 | 0.59738677 | 0.70412251 | LOC102724927 | 102724927 |
| ENSG000002 | 14.5163811 | 0.16755037 | 0.31724996 | 0.52813362 | 0.5974066 | 0.70412542 | LOC105377348 | 105377348 |
| ENSG000001 | 2.50971102 | -0.195369738 | 0.3700042 | -0.5280203 | 0.59748523 | 0.7041407 | RNU4-82P | 106481198 |
| ENSG000002 | 2.2242625 | -0.195293021 | 0.36984431 | -0.5280412 | 0.59747076 | 0.7041407 | NA | NA |
| ENSG000002 | 19.3828044 | -0.155116228 | 0.29373264 | -0.5280864 | 0.59743934 | 0.7041407 | NA | NA |
| ENSG000002 | 14.3337175 | 0.165988687 | 0.31436367 | 0.52801486 | 0.59748902 | 0.7041407 | NA | NA |
| ENSG000001 | 1077.91349 | 0.063994358 | 0.1212073 | 0.52797446 | 0.59751706 | 0.70415328 | ZBTB37 | 84614 |
| ENSG000002 | 2.51530781 | -0.194054718 | 0.36776102 | -0.5276653 | 0.59773168 | 0.70438573 | NA | NA |
| ENSG000002 | 6.21392842 | -0.194300616 | 0.36826346 | -0.5276131 | 0.59776792 | 0.70440797 | NA | NA |
| ENSG000002 | 47.394969 | 0.111994878 | 0.21235259 | 0.52740059 | 0.59791544 | 0.70456132 | PCDHA12 | 56137 |
| ENSG000002 | 9.92879089 | -0.180173393 | 0.34166397 | -0.5273409 | 0.59795686 | 0.70457033 | LOC100996342 | 100996342 |
| ENSG000002 | 64.3506081 | 0.099489237 | 0.18866258 | 0.52733953 | 0.59795783 | 0.70457033 | NA | NA |
| ENSG000001 | 241.276664 | 0.067637593 | 0.12829324 | 0.52721088 | 0.59804715 | 0.7046367 | NA | NA |
| ENSG000002 | 17.7631501 | 0.151906553 | 0.28813381 | 0.52720835 | 0.59804891 | 0.7046367 | NA | NA |
| ENSG000002 | 6.42412086 | 0.192472328 | 0.36516232 | 0.52708704 | 0.59813314 | 0.70471547 | NA | NA |
| ENSG000002 | 30.4747904 | 0.129401972 | 0.2455836 | 0.52691618 | 0.5982518 | 0.70483479 | NA | NA |
| ENSG000001 | 7.76083378 | 0.186751292 | 0.35448501 | 0.52682423 | 0.59831565 | 0.70487148 | NA | NA |
| ENSG000002 | 2.21146236 | -0.194611858 | 0.36940774 | -0.5268213 | 0.5983177 | 0.70487148 | NA | NA |

| | | | | | | | | |
|---|---|---|---|---|---|---|---|---|
| ENSG000002 | 2.46647004 | -0.19661526 | 0.37322994 | -0.5267939 | 0.59833671 | 0.70487339 | NA | NA |
| ENSG000002 | 2.24833665 | -0.194641548 | 0.36953386 | -0.5267218 | 0.59838678 | 0.7049119 | RNU6-25P | 106480704 |
| ENSG000002 | 9.93838115 | 0.178576451 | 0.33905946 | 0.52668182 | 0.59841457 | 0.70492415 | NA | NA |
| ENSG000001 | 62.4756207 | 0.105144752 | 0.19967681 | 0.52657468 | 0.59848898 | 0.7049895 | PHC2 | 1912 |
| ENSG000001 | 10.1235428 | -0.17987888 | 0.34162064 | -0.5265457 | 0.59850911 | 0.7049895 | RSPO3 | 84870 |
| ENSG000002 | 10.0002955 | -0.178844103 | 0.33966758 | -0.5265269 | 0.5985222 | 0.7049895 | NA | NA |
| ENSG000002 | 2.23536044 | -0.195002964 | 0.37047453 | -0.52636 | 0.59863812 | 0.70509986 | NA | NA |
| ENSG000002 | 2.23272405 | -0.193416292 | 0.36747271 | -0.5263419 | 0.59865066 | 0.70509986 | NA | NA |
| ENSG000001 | 13.0489549 | 0.170468163 | 0.32394822 | 0.52622041 | 0.59873507 | 0.7051788 | NPM2 | 10361 |
| ENSG000001 | 78.8487805 | -0.098496141 | 0.18719577 | -0.5261665 | 0.59877253 | 0.70520243 | ZNF799 | 90576 |
| ENSG000002 | 3.9627509 | -0.199163937 | 0.37858715 | -0.5260716 | 0.59883848 | 0.70525962 | RN7SL851P | 106480537 |
| ENSG00000C | 204.046926 | 0.069277203 | 0.13169523 | 0.52604187 | 0.59885912 | 0.70526344 | DOCK3 | 1795 |
| ENSG000002 | 2.23911753 | -0.192842198 | 0.36668052 | -0.5259134 | 0.59894838 | 0.70534808 | NA | NA |
| ENSG000002 | 2.27433416 | -0.191714244 | 0.36455395 | -0.5258872 | 0.59896661 | 0.70534906 | NA | NA |
| ENSG000002 | 2.25021546 | -0.193332371 | 0.36775586 | -0.5257085 | 0.59909078 | 0.70547458 | RN7SL817P | 106481144 |
| ENSG000002 | 66.502059 | 0.105140485 | 0.20000712 | 0.5256837 | 0.59910799 | 0.70547458 | NA | NA |
| ENSG000001 | 4.56388743 | -0.198998837 | 0.37858102 | -0.525644 | 0.59913559 | 0.70548659 | IHH | 3549 |
| ENSG00000C | 7.66379762 | -0.185113006 | 0.35225364 | -0.5255105 | 0.59922832 | 0.7055753 | EPYC | 1833 |
| ENSG000002 | 2.98174295 | -0.198170709 | 0.37713854 | -0.5254587 | 0.59926439 | 0.70559728 | NA | NA |
| ENSG000002 | 7.45824542 | 0.188697062 | 0.35915871 | 0.52538629 | 0.59931468 | 0.70561552 | NA | NA |
| ENSG000002 | 2.49752409 | -0.196009971 | 0.37306614 | -0.5254027 | 0.59930326 | 0.70561552 | MIR2355 | 100423036 |
| ENSG000002 | 30.9245003 | 0.130675641 | 0.24875789 | 0.52531254 | 0.59936594 | 0.70565538 | NA | NA |
| ENSG000002 | 2.73509985 | -0.196638441 | 0.37445469 | -0.5251328 | 0.59949087 | 0.70578198 | NA | NA |
| ENSG000002 | 7.9520119 | 0.195490906 | 0.372334 | 0.52504177 | 0.59955415 | 0.70583598 | NA | NA |
| ENSG000002 | 2.49914232 | -0.195888937 | 0.37311536 | -0.525009 | 0.59957691 | 0.70584228 | NA | NA |
| ENSG000001 | 46.4335492 | -0.109322728 | 0.2082624 | -0.5249278 | 0.59963336 | 0.70588825 | SHISA6 | 388336 |
| ENSG000002 | 2.99617397 | -0.197643032 | 0.37654422 | -0.5248866 | 0.599662 | 0.70590058 | RN7SL57P | 106480484 |
| ENSG000002 | 2.2379244 | -0.193592502 | 0.36885122 | -0.5248526 | 0.5996857 | 0.70590058 | NA | NA |
| ENSG000002 | 15.2543119 | -0.159964436 | 0.30478842 | -0.5248376 | 0.59969606 | 0.70590058 | NA | NA |

| | | | | | | | | |
|---|---|---|---|---|---|---|---|---|
| ENSG000002 | 6.87216798 | -0.196370052 | 0.37428324 | -0.5246563 | 0.59982215 | 0.70602851 | NA | NA |
| ENSG000002 | 2.53717086 | -0.19420427 | 0.37024056 | -0.5245354 | 0.59990622 | 0.70610696 | NA | NA |
| ENSG000002 | 2.21586546 | -0.193354922 | 0.36873196 | -0.524378 | 0.60001567 | 0.70621529 | NA | NA |
| ENSG000002 | 3.87988077 | 0.198193378 | 0.3780231 | 0.52428907 | 0.6000775 | 0.70626756 | OR2W1 | 26692 |
| ENSG000002 | 7.13803227 | -0.187673323 | 0.35798736 | -0.5242457 | 0.60010768 | 0.70628258 | NA | NA |
| ENSG000002 | 31.0636145 | 0.127561054 | 0.24333825 | 0.52421292 | 0.60013045 | 0.70628889 | SARNP | 84324 |
| ENSG000001 | 6.05196851 | 0.191380964 | 0.36515117 | 0.52411433 | 0.60019902 | 0.70632859 | MYF6 | 4618 |
| ENSG000001 | 6.94973654 | -0.19139962 | 0.36516962 | -0.5241389 | 0.6001819 | 0.70632859 | CIDEA | 1149 |
| ENSG000002 | 2.20496395 | -0.192898914 | 0.36812192 | -0.5240082 | 0.60027282 | 0.70639494 | NA | NA |
| ENSG000001 | 12.1852989 | -0.170611575 | 0.32566898 | -0.5238803 | 0.60036177 | 0.70647911 | NA | NA |
| ENSG000002 | 3.85847506 | 0.19811152 | 0.37827716 | 0.52372055 | 0.60047292 | 0.7065894 | RN7SL472P | 106479395 |
| ENSG000002 | 2.8058409 | 0.19624122 | 0.37478534 | 0.52360965 | 0.60055007 | 0.70665968 | NA | NA |
| ENSG000001 | 27.7935396 | -0.134685658 | 0.2572597 | -0.5235397 | 0.60059875 | 0.70667905 | LSMEM1 | 286006 |
| ENSG000002 | 2.46687498 | -0.195710956 | 0.3738253 | -0.5235359 | 0.60060139 | 0.70667905 | NA | NA |
| ENSG000002 | 12.4106181 | -0.178177768 | 0.34044911 | -0.5233609 | 0.6007231 | 0.70680175 | NA | NA |
| ENSG000002 | 2.21959139 | -0.193466226 | 0.36982873 | -0.5231238 | 0.60088808 | 0.70695989 | NA | NA |
| ENSG000002 | 2.74326113 | -0.196529413 | 0.37568873 | -0.5231177 | 0.60089237 | 0.70695989 | NA | NA |
| ENSG000002 | 2.75621773 | -0.196007075 | 0.37480872 | -0.5229523 | 0.60100746 | 0.70701324 | NA | NA |
| ENSG000002 | 17.9869058 | -0.160405378 | 0.30672871 | -0.5229552 | 0.60100542 | 0.70701324 | NA | NA |
| ENSG000002 | 3.44219555 | 0.197679577 | 0.37800005 | 0.52296178 | 0.60100085 | 0.70701324 | NA | NA |
| ENSG000002 | 2.2053689 | -0.193177186 | 0.36938663 | -0.5229675 | 0.6009969 | 0.70701324 | NA | NA |
| ENSG00000C | 71.4588883 | 0.097501889 | 0.18647194 | 0.52287701 | 0.60105985 | 0.70705436 | PALM | 5064 |
| ENSG000001 | 105.383801 | 0.09610568 | 0.18384318 | 0.52275901 | 0.60114197 | 0.70713045 | HHLA3 | 11147 |
| ENSG000002 | 2.72681491 | -0.19670584 | 0.37635999 | -0.5226534 | 0.60121545 | 0.70719637 | NA | NA |
| ENSG000002 | 2.7713537 | -0.195865092 | 0.37482048 | -0.5225571 | 0.60128252 | 0.70725475 | NA | NA |
| ENSG000002 | 2.23917014 | -0.19306674 | 0.36950096 | -0.5225067 | 0.60131756 | 0.70725494 | NA | NA |
| ENSG000002 | 2.23917014 | -0.19306674 | 0.36950096 | -0.5225067 | 0.60131756 | 0.70725494 | NA | NA |
| ENSG000002 | 23.6424525 | 0.142703196 | 0.27319905 | 0.52234148 | 0.60143259 | 0.70736972 | NA | NA |
| ENSG000001 | 10.6610647 | -0.176107436 | 0.33719635 | -0.5222697 | 0.60148257 | 0.70740798 | KRT26 | 353288 |

| | | | | | | | | |
|---|---|---|---|---|---|---|---|---|
| ENSG000002 | 42.4081386 | 0.115890102 | 0.22194212 | 0.52216363 | 0.60155641 | 0.7074743 | NA | NA |
| ENSG000001 | 15.2350083 | -0.158296451 | 0.30319743 | -0.5220903 | 0.60160743 | 0.7074834 | SOX10 | 6663 |
| ENSG000002 | 4.90524431 | 0.195574111 | 0.37457765 | 0.522119 | 0.60158748 | 0.7074834 | CCDC183 | 84960 |
| ENSG000002 | 6.16408491 | -0.192545097 | 0.3688057 | -0.5220773 | 0.60161648 | 0.7074834 | NA | NA |
| ENSG000002 | 3.24310523 | -0.197359056 | 0.37807919 | -0.5220045 | 0.60166717 | 0.70752248 | NA | NA |
| ENSG000002 | 2.77507913 | 0.196234918 | 0.37597042 | 0.52194243 | 0.60171041 | 0.70755282 | RNU6-536P | 106481350 |
| ENSG000002 | 2.71338115 | -0.195348765 | 0.37433683 | -0.5218529 | 0.60177274 | 0.7076056 | NA | NA |
| ENSG000002 | 2.50256769 | -0.193646396 | 0.37115259 | -0.5217434 | 0.60184904 | 0.70761323 | NA | NA |
| ENSG000002 | 2.20391408 | -0.192739042 | 0.36937814 | -0.5217933 | 0.60181425 | 0.70761323 | NA | NA |
| ENSG000002 | 2.25855302 | -0.19110049 | 0.36625775 | -0.521765 | 0.60183393 | 0.70761323 | NA | NA |
| ENSG000002 | 9.01594539 | 0.181081356 | 0.347068 | 0.52174605 | 0.60184716 | 0.70761323 | NA | NA |
| ENSG000002 | 2.45353321 | 0.188933439 | 0.36221706 | 0.52160282 | 0.6019469 | 0.70768726 | NA | NA |
| ENSG000002 | 9.08676254 | 0.179817913 | 0.34473219 | 0.52161625 | 0.60193755 | 0.70768726 | NA | NA |
| ENSG000002 | 2.46599397 | -0.194698164 | 0.37353305 | -0.5212341 | 0.6022037 | 0.70789112 | RNU6-727P | 106481409 |
| ENSG000002 | 10.2168351 | 0.180504532 | 0.34624092 | 0.5213264 | 0.60213941 | 0.70789112 | TAX1BP3 | 30851 |
| ENSG000002 | 3.44598185 | 0.195601572 | 0.37526637 | 0.52123395 | 0.60220381 | 0.70789112 | NA | NA |
| ENSG000002 | 2.46078152 | -0.194920084 | 0.37396281 | -0.5212285 | 0.60220759 | 0.70789112 | RNU6-976P | 106481488 |
| ENSG000002 | 4.97449141 | 0.195776097 | 0.3755531 | 0.5213007 | 0.60215731 | 0.70789112 | NA | NA |
| ENSG000002 | 12.9540293 | -0.170517381 | 0.32720862 | -0.5211274 | 0.60227802 | 0.70795339 | NA | NA |
| ENSG000001 | 6.14438992 | -0.190241416 | 0.36513516 | -0.5210164 | 0.60235533 | 0.70802374 | NA | NA |
| ENSG000002 | 2.21968313 | -0.192461365 | 0.36943501 | -0.5209614 | 0.60239369 | 0.70804831 | NA | NA |
| ENSG000001 | 69.9384303 | -0.094968831 | 0.18231522 | -0.5209046 | 0.60243326 | 0.70807429 | SGSM1 | 129049 |
| ENSG000002 | 2.22519559 | -0.192052922 | 0.36876546 | -0.5207996 | 0.60250635 | 0.70811915 | NA | NA |
| ENSG000002 | 7.90469108 | -0.188228503 | 0.3614099 | -0.5208172 | 0.6024941 | 0.70811915 | NA | NA |
| ENSG000001 | 77.2659691 | -0.102144604 | 0.19616295 | -0.520713 | 0.60256669 | 0.70814437 | L3MBTL4 | 91133 |
| ENSG000001 | 4.21462394 | -0.196400171 | 0.37718949 | -0.5206936 | 0.6025802 | 0.70814437 | IFNA2 | 3440 |
| ENSG000002 | 2.73753927 | -0.194456674 | 0.37345631 | -0.5206946 | 0.60257956 | 0.70814437 | RN7SL452P | 106479387 |
| ENSG000002 | 3.00875374 | -0.196333737 | 0.37718192 | -0.520528 | 0.60269564 | 0.70821846 | NA | NA |
| ENSG000002 | 2.96598187 | -0.19643671 | 0.37737027 | -0.520541 | 0.60268654 | 0.70821846 | LOC101929926 | 101929926 |

| | | | | | | | | |
|---|---|---|---|---|---|---|---|---|
| ENSG000002 | 2.46390608 | -0.194434548 | 0.37352628 | -0.5205378 | 0.60268879 | 0.70821846 | NA | NA |
| ENSG000002 | 2.77320087 | 0.195055867 | 0.37477182 | 0.52046568 | 0.60273904 | 0.70824894 | NA | NA |
| ENSG000001 | 45.4871514 | 0.112912572 | 0.21695825 | 0.52043457 | 0.60276072 | 0.70825389 | LRRN3 | 54674 |
| ENSG000002 | 2.46701138 | -0.193820519 | 0.37250413 | -0.5203178 | 0.60284211 | 0.70828851 | NA | NA |
| ENSG000002 | 36.1524492 | 0.132746977 | 0.25512708 | 0.52031709 | 0.60284259 | 0.70828851 | NA | NA |
| ENSG000002 | 18.4098108 | -0.151943477 | 0.29200732 | -0.5203413 | 0.60282569 | 0.70828851 | NA | NA |
| ENSG000002 | 21.7224322 | 0.141510681 | 0.27200014 | 0.52025958 | 0.60288266 | 0.70831508 | NA | NA |
| ENSG000002 | 2.80387755 | 0.195002337 | 0.37483649 | 0.52023307 | 0.60290114 | 0.70831626 | NA | NA |
| ENSG000002 | 19.0785497 | 0.151554047 | 0.29135036 | 0.52017799 | 0.60293952 | 0.70834083 | NA | NA |
| ENSG000002 | 2.45298475 | -0.193522496 | 0.37208923 | -0.520097 | 0.60299597 | 0.70838662 | NA | NA |
| ENSG000002 | 2.95823772 | -0.196381554 | 0.37760799 | -0.5200673 | 0.60301669 | 0.70839045 | NA | NA |
| ENSG000001 | 7.90276924 | -0.182483478 | 0.35102823 | -0.5198541 | 0.60316525 | 0.70848286 | BRS3 | 680 |
| ENSG000001 | 805.02205 | 0.078787025 | 0.15153708 | 0.51991911 | 0.60311996 | 0.70848286 | SDC2 | 6383 |
| ENSG000002 | 2.2697746 | -0.19026139 | 0.36596361 | -0.5198915 | 0.60313917 | 0.70848286 | NA | NA |
| ENSG000002 | 5.92571464 | -0.191559068 | 0.36848018 | -0.5198626 | 0.60315934 | 0.70848286 | NA | NA |
| ENSG000002 | 2.51334521 | -0.193618521 | 0.37248431 | -0.5198032 | 0.60320078 | 0.70850246 | NA | NA |
| ENSG000002 | 3.02014009 | 0.196153771 | 0.3773966 | 0.51975501 | 0.60323434 | 0.70850246 | NA | NA |
| ENSG000002 | 35.8698045 | 0.124489498 | 0.23950468 | 0.51977898 | 0.60321763 | 0.70850246 | TRABD2B | 388630 |
| ENSG000002 | 2.24987469 | -0.185759068 | 0.3574163 | -0.5197275 | 0.60325354 | 0.70850448 | NA | NA |
| ENSG000002 | 2.72331677 | -0.193836934 | 0.37298678 | -0.5196885 | 0.60328071 | 0.70851588 | NA | NA |
| ENSG000001 | 9.15932673 | 0.182446582 | 0.35124311 | 0.51943107 | 0.60346017 | 0.70868326 | WDR38 | 401551 |
| ENSG000002 | 8.04626682 | -0.181818571 | 0.35006594 | -0.5193838 | 0.60349314 | 0.70868326 | NA | NA |
| ENSG000002 | 99.8767193 | -0.088823556 | 0.17101046 | -0.5194042 | 0.60347889 | 0.70868326 | NA | NA |
| ENSG000002 | 2.22707496 | -0.192211286 | 0.370073 | -0.5193875 | 0.60349055 | 0.70868326 | NA | NA |
| ENSG000002 | 2.77121736 | 0.195010982 | 0.37549063 | 0.51934979 | 0.60351684 | 0.70869056 | HIST2H3DP1 | 106479023 |
| ENSG000002 | 2.46508049 | -0.193979719 | 0.37352916 | -0.5193161 | 0.60354031 | 0.7086976 | NA | NA |
| ENSG000002 | 2.22246801 | -0.191484094 | 0.36875501 | -0.5192718 | 0.60357119 | 0.70871334 | NA | NA |
| ENSG000002 | 2.49786433 | -0.193428839 | 0.37256282 | -0.5191845 | 0.60363209 | 0.70876433 | NA | NA |
| ENSG000002 | 2.48537684 | -0.193614748 | 0.37309265 | -0.5189455 | 0.60379875 | 0.70893949 | NA | NA |

| | | | | | | | | |
|---|---|---|---|---|---|---|---|---|
| ENSG000002 | 130.432157 | -0.077393895 | 0.14916021 | -0.5188642 | 0.60385544 | 0.7089402 | TRHDE-AS1 | 283392 |
| ENSG000002 | 3.96128656 | -0.196013043 | 0.37780601 | -0.5188193 | 0.60388677 | 0.7089402 | NA | NA |
| ENSG000002 | 46.7902098 | -0.111939791 | 0.21575089 | -0.5188381 | 0.60387361 | 0.7089402 | IQCH-AS1 | 100506686 |
| ENSG000002 | 27.9203877 | -0.132600978 | 0.2555418 | -0.5189013 | 0.60382955 | 0.7089402 | NA | NA |
| ENSG000002 | 75.4603504 | -0.093011601 | 0.17925189 | -0.5188877 | 0.60383906 | 0.7089402 | NA | NA |
| ENSG000002 | 4.96345438 | -0.19404424 | 0.37405234 | -0.5187623 | 0.60392653 | 0.70894583 | NA | NA |
| ENSG000002 | 2.73941918 | -0.195143719 | 0.37616956 | -0.5187653 | 0.60392441 | 0.70894583 | NA | NA |
| ENSG000002 | 11.4813686 | 0.177303195 | 0.34179947 | 0.51873456 | 0.60394586 | 0.708948 | NA | NA |
| ENSG000002 | 7.1488622 | -0.184580362 | 0.35598784 | -0.5185019 | 0.60410815 | 0.70911798 | NA | NA |
| ENSG000002 | 3.91499302 | 0.196015128 | 0.37810656 | 0.5184124 | 0.60417057 | 0.70917071 | NA | NA |
| ENSG000002 | 14.9070224 | 0.158075766 | 0.30519024 | 0.51795813 | 0.60448749 | 0.70952217 | NA | NA |
| ENSG000002 | 2.26402605 | -0.18949487 | 0.36593768 | -0.5178337 | 0.60457429 | 0.70958298 | NA | NA |
| ENSG000002 | 4.82367776 | -0.19600347 | 0.37849351 | -0.5178516 | 0.60456182 | 0.70958298 | NA | NA |
| ENSG000002 | 7.13731467 | -0.186022719 | 0.35928304 | -0.5177609 | 0.60462509 | 0.70961289 | NA | NA |
| ENSG000002 | 27.583043 | -0.130369835 | 0.25180218 | -0.517747 | 0.60463477 | 0.70961289 | NA | NA |
| ENSG000001 | 33.2618811 | 0.121448551 | 0.23460105 | 0.51768119 | 0.60468073 | 0.70964629 | SMTNL2 | 342527 |
| ENSG000002 | 6.22297039 | -0.19242277 | 0.37176705 | -0.5175896 | 0.60474462 | 0.70970073 | WWTR1-IT1 | 106478977 |
| ENSG000001 | 53.4601431 | 0.108387209 | 0.2094364 | 0.51751849 | 0.60479427 | 0.7097088 | S100A4 | 6275 |
| ENSG000002 | 2.24527625 | -0.189530154 | 0.36621373 | -0.5175397 | 0.60477944 | 0.7097088 | RNU6-1165P | 106480079 |
| ENSG000002 | 3.01100584 | 0.195304027 | 0.37739577 | 0.51750454 | 0.604804 | 0.7097088 | NA | NA |
| ENSG000002 | 40.2882485 | 0.120448829 | 0.23277203 | 0.51745405 | 0.60483924 | 0.70972961 | MSH5 | 4439 |
| ENSG000002 | 10.6295345 | -0.174434028 | 0.33727868 | -0.5171807 | 0.60503001 | 0.70991239 | LINC00612 | 253128 |
| ENSG000002 | 2.4888937 | -0.192797128 | 0.37277565 | -0.5171935 | 0.60502112 | 0.70991239 | RN7SL863P | 106480538 |
| ENSG000001 | 35.5852114 | 0.123615233 | 0.23909395 | 0.51701531 | 0.60514547 | 0.70994515 | OR2B6 | 26212 |
| ENSG000002 | 2.98618627 | -0.194682014 | 0.37654209 | -0.5170259 | 0.60513808 | 0.70994515 | NA | NA |
| ENSG000002 | 8.80372352 | -0.177237507 | 0.34277867 | -0.5170611 | 0.60511352 | 0.70994515 | NA | NA |
| ENSG000002 | 4.96371531 | -0.193373672 | 0.37398128 | -0.5170678 | 0.60510884 | 0.70994515 | NA | NA |
| ENSG000002 | 17.175506 | -0.155062196 | 0.29991073 | -0.5170278 | 0.60513672 | 0.70994515 | FLJ38576 | 651430 |
| ENSG000002 | 2.83410986 | 0.193727641 | 0.37483298 | 0.51683724 | 0.60526978 | 0.71007045 | NA | NA |

| | | | | | | | | |
|---|---|---|---|---|---|---|---|---|
| ENSG000002 | 8.14853721 | -0.184036477 | 0.3561141 | -0.5167908 | 0.60530223 | 0.71008797 | NA | NA |
| ENSG000002 | 4.45897143 | -0.194299823 | 0.37601333 | -0.5167365 | 0.60534009 | 0.71011184 | NA | NA |
| ENSG000002 | 5.8421768 | 0.1901564 | 0.36801639 | 0.51670633 | 0.60536118 | 0.71011603 | NA | NA |
| ENSG000001 | 3.00616942 | -0.193920369 | 0.37540666 | -0.5165608 | 0.60546278 | 0.71019412 | NA | NA |
| ENSG000002 | 8.17271592 | -0.18208293 | 0.35247725 | -0.5165807 | 0.60544892 | 0.71019412 | NA | NA |
| ENSG000002 | 2.55936585 | -0.190021305 | 0.3679302 | -0.5164602 | 0.60553303 | 0.71025598 | NA | NA |
| ENSG000002 | 2.50919005 | -0.192233241 | 0.3722395 | -0.5164235 | 0.60555863 | 0.71026547 | NA | NA |
| ENSG000002 | 2.45045305 | -0.191893136 | 0.37170811 | -0.5162468 | 0.60568202 | 0.71038965 | NA | NA |
| ENSG000001 | 79.0534126 | 0.100452476 | 0.19462441 | 0.51613503 | 0.60576011 | 0.71046068 | ZNF454 | 285676 |
| ENSG000002 | 2.46806921 | -0.192989356 | 0.37397495 | -0.5160489 | 0.60582027 | 0.71051069 | NA | NA |
| ENSG000002 | 4.95599097 | -0.19243565 | 0.37294004 | -0.5159962 | 0.60585705 | 0.71053328 | MIR548AQ | 100847078 |
| ENSG000002 | 7.60604653 | -0.18123994 | 0.35133757 | -0.515857 | 0.6059543 | 0.71062678 | NA | NA |
| ENSG000001 | 13.8944371 | 0.164788817 | 0.31956064 | 0.51567308 | 0.60608276 | 0.71063357 | PTGER4 | 5734 |
| ENSG000001 | 8.19592136 | -0.185021837 | 0.35872817 | -0.5157717 | 0.60601388 | 0.71063357 | CTSL3P | 392360 |
| ENSG000002 | 2.69613822 | -0.193605821 | 0.37541725 | -0.5157084 | 0.60605811 | 0.71063357 | NA | NA |
| ENSG000002 | 11.37124 | 0.169147318 | 0.3280114 | 0.51567512 | 0.60608133 | 0.71063357 | NA | NA |
| ENSG000002 | 12.7411437 | 0.166113796 | 0.32206987 | 0.51576944 | 0.60601545 | 0.71063357 | HHIP-AS1 | 646576 |
| ENSG000002 | 3.02833329 | 0.194722356 | 0.37758102 | 0.51571013 | 0.60605688 | 0.71063357 | NA | NA |
| ENSG000002 | 2.4505646 | -0.192219222 | 0.37265649 | -0.5158081 | 0.60598847 | 0.71063357 | NA | NA |
| ENSG000001 | 36.9487628 | -0.120677527 | 0.23405848 | -0.5155871 | 0.60614283 | 0.71064236 | NXPH3 | 11248 |
| ENSG000002 | 2.70810922 | -0.193356886 | 0.37499237 | -0.5156288 | 0.60611366 | 0.71064236 | NA | NA |
| ENSG000002 | 2.20437109 | -0.190658872 | 0.3697745 | -0.5156085 | 0.60612788 | 0.71064236 | NA | NA |
| ENSG000001 | 12.6566809 | -0.168278137 | 0.32651005 | -0.5153842 | 0.60628454 | 0.71078795 | REM2 | 161253 |
| ENSG000002 | 2.18529474 | 0.188567144 | 0.36589765 | 0.51535489 | 0.60630505 | 0.71079144 | NA | NA |
| ENSG000002 | 7.44106175 | -0.186639827 | 0.36222169 | -0.5152641 | 0.60636849 | 0.71084527 | NA | NA |
| ENSG000001 | 2.50671042 | -0.191758006 | 0.3722332 | -0.5151556 | 0.60644431 | 0.7109136 | LRRC52 | 440699 |
| ENSG000001 | 91.2716872 | 0.0918401 | 0.17834752 | 0.51495026 | 0.60658778 | 0.71094495 | ERG | 2078 |
| ENSG000002 | 2.48642616 | -0.190919542 | 0.3707899 | -0.5148995 | 0.60662323 | 0.71094495 | RNU6-466P | 106481770 |
| ENSG000002 | 2.23738307 | -0.188976278 | 0.36700646 | -0.5149127 | 0.60661404 | 0.71094495 | NA | NA |

| | | | | | | | | |
|---|---|---|---|---|---|---|---|---|
| ENSG000002 | 2.71784428 | -0.193618644 | 0.37600312 | -0.5149389 | 0.6065957 | 0.71094495 | NA | NA |
| ENSG000002 | 3.46386261 | -0.194971455 | 0.37858405 | -0.5150018 | 0.60655179 | 0.71094495 | NA | NA |
| ENSG000002 | 6.64236316 | -0.186695459 | 0.36256203 | -0.5149338 | 0.60659925 | 0.71094495 | NA | NA |
| ENSG000002 | 8.09236515 | -0.182136557 | 0.35366109 | -0.5150031 | 0.60655085 | 0.71094495 | NA | NA |
| ENSG000002 | 2.23818498 | -0.189630074 | 0.36823921 | -0.5149644 | 0.60657792 | 0.71094495 | NA | NA |
| ENSG000002 | 2.19489194 | -0.190174405 | 0.3693485 | -0.5148915 | 0.60662884 | 0.71094495 | NA | NA |
| ENSG000002 | 3.71716775 | -0.194829107 | 0.37843206 | -0.5148325 | 0.6066701 | 0.71097276 | NA | NA |
| ENSG000002 | 3.00727877 | -0.193035449 | 0.37497672 | -0.5147932 | 0.60669757 | 0.7109844 | NA | NA |
| ENSG000002 | 2.72590143 | -0.193437989 | 0.37583724 | -0.5146855 | 0.60677279 | 0.71104355 | NA | NA |
| ENSG000002 | 3.02392197 | -0.193616251 | 0.37619439 | -0.5146708 | 0.60678311 | 0.71104355 | NA | NA |
| ENSG000002 | 6.91557653 | 0.186207963 | 0.36183642 | 0.51461919 | 0.60681916 | 0.71106524 | NA | NA |
| ENSG000002 | 5.93687783 | -0.188233207 | 0.36581736 | -0.5145552 | 0.60686391 | 0.71109713 | NA | NA |
| ENSG000002 | 5.96878045 | -0.189119429 | 0.36759479 | -0.514478 | 0.60691785 | 0.71113978 | NA | NA |
| ENSG000002 | 11.522574 | 0.167071829 | 0.3248398 | 0.51432069 | 0.6070278 | 0.71124807 | NA | NA |
| ENSG000002 | 2.45998428 | -0.191143926 | 0.37171629 | -0.5142199 | 0.60709823 | 0.71131004 | LOC101927557 | 101927557 |
| ENSG000002 | 2.71503183 | -0.192832221 | 0.37504457 | -0.5141582 | 0.60714142 | 0.71134009 | NA | NA |
| ENSG000001 | 3.00038862 | -0.192489633 | 0.37441208 | -0.5141117 | 0.6071739 | 0.71135495 | IGFL1 | 374918 |
| ENSG000002 | 2.22449118 | -0.189930235 | 0.36944951 | -0.5140898 | 0.60718919 | 0.71135495 | NA | NA |
| ENSG000002 | 7.13672676 | 0.185116131 | 0.36010906 | 0.51405574 | 0.60721302 | 0.71135528 | LOC101926908 | 101926908 |
| ENSG000002 | 2.97309329 | -0.194001363 | 0.37740575 | -0.5140392 | 0.60722456 | 0.71135528 | NA | NA |
| ENSG000002 | 6.48334372 | -0.187642896 | 0.36517915 | -0.5138379 | 0.60736531 | 0.71149962 | SACS-AS1 | 100506680 |
| ENSG000002 | 2.7563819 | -0.192621475 | 0.3748971 | -0.5137983 | 0.60739305 | 0.71150287 | NA | NA |
| ENSG000002 | 2.82582438 | 0.192149163 | 0.37398839 | 0.51378377 | 0.60740318 | 0.71150287 | NA | NA |
| ENSG000001 | 4.19216857 | -0.193414743 | 0.37651899 | -0.5136919 | 0.60746744 | 0.71154435 | LENEP | 55891 |
| ENSG000002 | 32.5343961 | 0.122626565 | 0.23872034 | 0.51368293 | 0.60747369 | 0.71154435 | DBIL5P | 100131454 |
| ENSG000001 | 40.0626621 | 0.115309105 | 0.22453975 | 0.51353537 | 0.60757688 | 0.71161855 | KCTD8 | 386617 |
| ENSG000002 | 2.49098159 | -0.191152858 | 0.3722082 | -0.5135643 | 0.60755661 | 0.71161855 | NA | NA |
| ENSG000002 | 2.52334066 | -0.190543845 | 0.3710565 | -0.5135171 | 0.60758968 | 0.71161855 | NA | NA |
| ENSG000002 | 2.93692341 | -0.193406079 | 0.37667631 | -0.5134543 | 0.60763355 | 0.71164938 | NA | NA |

| | | | | | | | | |
|---|---|---|---|---|---|---|---|---|
| ENSG000001 | 2.95797713 | -0.193929868 | 0.37773144 | -0.5134067 | 0.60766683 | 0.71165754 | IGF2 | 3481 |
| ENSG000002 | 2.50837299 | -0.189325125 | 0.3687895 | -0.5133691 | 0.60769317 | 0.71165754 | NA | NA |
| ENSG000002 | 12.8874893 | -0.161407359 | 0.31439923 | -0.5133834 | 0.60768313 | 0.71165754 | GAS6-AS2 | 100506394 |
| ENSG000001 | 15.8460969 | -0.154872038 | 0.30170099 | -0.5133296 | 0.60772081 | 0.71166936 | OTX1 | 5013 |
| ENSG000002 | 4.25293322 | -0.194001934 | 0.37803614 | -0.5131836 | 0.60782293 | 0.71174783 | LOC100128714 | 100128714 |
| ENSG000002 | 3.25253232 | -0.193845398 | 0.37773064 | -0.5131842 | 0.60782247 | 0.71174783 | NA | NA |
| ENSG000002 | 2.2871024 | -0.186482555 | 0.36341634 | -0.5131375 | 0.60785514 | 0.711765 | RNU11-2P | 106479561 |
| ENSG000002 | 2.48580084 | -0.190406844 | 0.37115122 | -0.5130169 | 0.60793951 | 0.71184324 | NA | NA |
| ENSG000002 | 2.23155019 | -0.187385314 | 0.36529694 | -0.5129671 | 0.60797433 | 0.71186346 | NA | NA |
| ENSG000002 | 8.64352282 | -0.177958602 | 0.34696322 | -0.5129034 | 0.60801892 | 0.71189511 | NA | NA |
| ENSG000002 | 2.51440249 | -0.190334111 | 0.37117104 | -0.5127935 | 0.60809575 | 0.7119645 | NA | NA |
| ENSG000002 | 6.05335064 | 0.1882418 | 0.36713933 | 0.51272578 | 0.60814315 | 0.71199945 | NA | NA |
| ENSG000001 | 14.6448088 | 0.156630169 | 0.30550246 | 0.51269692 | 0.60816334 | 0.71200253 | KIAA1683 | 80726 |
| ENSG000002 | 4.44625526 | -0.193102194 | 0.37677193 | -0.5125175 | 0.6082889 | 0.71212897 | NA | NA |
| ENSG000001 | 23.9272184 | -0.139191358 | 0.27164084 | -0.5124095 | 0.60836442 | 0.71219682 | CLIC2 | 1193 |
| ENSG000002 | 4.19532539 | -0.193602973 | 0.37795292 | -0.512241 | 0.60848236 | 0.71229377 | NA | NA |
| ENSG000002 | 6.64819658 | -0.18431236 | 0.35979884 | -0.512265 | 0.60846554 | 0.71229377 | NA | NA |
| ENSG000001 | 63.0645302 | 0.101291121 | 0.1977628 | 0.51218491 | 0.6085216 | 0.71230592 | C3orf70 | 285382 |
| ENSG000002 | 3.7011185 | -0.193782884 | 0.37835218 | -0.5121759 | 0.60852788 | 0.71230592 | NA | NA |
| ENSG000002 | 5.75627575 | -0.190642582 | 0.37233182 | -0.5120233 | 0.60863468 | 0.71234694 | NA | NA |
| ENSG000002 | 6.63024393 | 0.187229468 | 0.36566145 | 0.51202955 | 0.60863033 | 0.71234694 | NA | NA |
| ENSG000002 | 2.18900644 | -0.188012454 | 0.36721156 | -0.5120004 | 0.60865075 | 0.71234694 | NA | NA |
| ENSG000002 | 30.325612 | 0.139825419 | 0.27305332 | 0.51208101 | 0.60859431 | 0.71234694 | NA | NA |
| ENSG000002 | 7.68768774 | -0.18354061 | 0.35845662 | -0.5120302 | 0.60862988 | 0.71234694 | NA | NA |
| ENSG000002 | 4.71034567 | -0.191761796 | 0.37458144 | -0.5119362 | 0.60869567 | 0.71237895 | NA | NA |
| ENSG000002 | 8.78047126 | -0.176749931 | 0.3452923 | -0.511885 | 0.60873153 | 0.71240035 | NA | NA |
| ENSG000001 | 6.62022024 | -0.184419625 | 0.36037827 | -0.511739 | 0.60883367 | 0.71241925 | TMIGD2 | 126259 |
| ENSG000002 | 3.97996447 | -0.193620586 | 0.37839713 | -0.5116862 | 0.60887066 | 0.71241925 | NA | NA |
| ENSG000002 | 23.7608674 | 0.141478854 | 0.27648446 | 0.51170636 | 0.60885654 | 0.71241925 | ZBED5-AS1 | 729013 |

| | | | | | | | | |
|---|---|---|---|---|---|---|---|---|
| ENSG000002 | 2.74980365 | -0.190212519 | 0.37170927 | -0.5117239 | 0.60884425 | 0.71241925 | NA | NA |
| ENSG000002 | 4.19546974 | -0.192984166 | 0.3771373 | -0.511708 | 0.6088554 | 0.71241925 | NA | NA |
| ENSG000002 | 2.66658219 | -0.146090741 | 0.28545301 | -0.5117856 | 0.60880107 | 0.71241925 | NA | NA |
| ENSG000002 | 10.7685178 | -0.170089055 | 0.3323118 | -0.5118357 | 0.60876598 | 0.71241925 | NA | NA |
| ENSG000002 | 40.6609616 | -0.113350972 | 0.22153551 | -0.5116605 | 0.60888864 | 0.71241973 | NA | NA |
| ENSG000002 | 2.71924703 | -0.192156523 | 0.37560527 | -0.5115917 | 0.60893682 | 0.71244959 | NA | NA |
| ENSG000002 | 6.60730297 | -0.184682449 | 0.36100839 | -0.5115738 | 0.60894929 | 0.71244959 | NA | NA |
| ENSG000002 | 3.73484045 | -0.193640818 | 0.37857839 | -0.5114946 | 0.60900474 | 0.71247397 | RNU6-853P | 106479943 |
| ENSG000002 | 3.48321323 | -0.193646716 | 0.37859049 | -0.5114939 | 0.60900527 | 0.71247397 | NA | NA |
| ENSG000001 | 4.68703793 | 0.192497575 | 0.37651086 | 0.51126699 | 0.60916411 | 0.71253996 | NA | NA |
| ENSG000001 | 31.7429035 | 0.122024889 | 0.23872042 | 0.51116233 | 0.60923739 | 0.71253996 | IZUMO1 | 284359 |
| ENSG000002 | 2.72347332 | -0.191056541 | 0.37365286 | -0.511321 | 0.60912632 | 0.71253996 | NA | NA |
| ENSG000002 | 2.51059259 | -0.18937763 | 0.37047122 | -0.5111804 | 0.60922473 | 0.71253996 | PRR9 | 574414 |
| ENSG000002 | 3.23636684 | -0.193487728 | 0.37835868 | -0.511387 | 0.60908007 | 0.71253996 | NA | NA |
| ENSG000002 | 2.19804875 | -0.187574561 | 0.36685094 | -0.51131 | 0.60913399 | 0.71253996 | NA | NA |
| ENSG000002 | 6.92649325 | -0.185563151 | 0.36299087 | -0.5112061 | 0.60920674 | 0.71253996 | NA | NA |
| ENSG000002 | 4.53251661 | -0.193266609 | 0.37808582 | -0.5111713 | 0.60923113 | 0.71253996 | RN7SL124P | 106480955 |
| ENSG000002 | 2.23018025 | -0.187892082 | 0.36746054 | -0.5113259 | 0.60912289 | 0.71253996 | HMGN1P12 | 106480725 |
| ENSG000002 | 2.20177468 | -0.188289179 | 0.36828972 | -0.5112529 | 0.609174 | 0.71253996 | NA | NA |
| ENSG000001 | 22.9584845 | -0.136513438 | 0.26711655 | -0.5110632 | 0.6093068 | 0.71260059 | STAC3 | 246329 |
| ENSG000002 | 5.42504215 | -0.188725931 | 0.36941262 | -0.5108811 | 0.6094343 | 0.71268804 | NA | NA |
| ENSG000002 | 2.23068878 | -0.188210364 | 0.36839147 | -0.5108977 | 0.60942268 | 0.71268804 | NA | NA |
| ENSG000002 | 5.1998492 | -0.190648938 | 0.37314594 | -0.5109233 | 0.60940479 | 0.71268804 | NA | NA |
| ENSG000002 | 5.91209296 | -0.188677903 | 0.36937723 | -0.5108 | 0.60949108 | 0.71273388 | OLIG2 | 10215 |
| ENSG000002 | 3.45896282 | -0.193373045 | 0.37859319 | -0.5107674 | 0.60951396 | 0.71274009 | NA | NA |
| ENSG000002 | 3.22065728 | -0.19305691 | 0.37805171 | -0.5106627 | 0.60958726 | 0.71280524 | RN7SL815P | 106479519 |
| ENSG000002 | 8.9143767 | -0.179325293 | 0.35121326 | -0.5105881 | 0.60963954 | 0.71284582 | NA | NA |
| ENSG000002 | 2.22229971 | -0.188585913 | 0.36944393 | -0.5104588 | 0.60973006 | 0.7129311 | NA | NA |
| ENSG000002 | 7.12827291 | -0.182803277 | 0.35813469 | -0.5104316 | 0.6097491 | 0.71293281 | NA | NA |

| | | | | | | | | |
|---|---|---|---|---|---|---|---|---|
| ENSG000002 | 2.71421035 | 0.190100351 | 0.37248303 | 0.51035977 | 0.60979944 | 0.71297111 | NA | NA |
| ENSG000001 | 2.48987168 | -0.189577494 | 0.37163134 | -0.5101225 | 0.60996567 | 0.71312434 | NA | NA |
| ENSG000002 | 2.76048499 | -0.190950165 | 0.37431162 | -0.5101369 | 0.60995557 | 0.71312434 | NA | NA |
| ENSG000002 | 3.67168279 | 0.193024528 | 0.37848466 | 0.509993 | 0.61005636 | 0.71320981 | NA | NA |
| ENSG000002 | 2.96461192 | -0.192457013 | 0.37740299 | -0.5099509 | 0.61008583 | 0.7132237 | NA | NA |
| ENSG000002 | 2.96508821 | -0.192346188 | 0.37725078 | -0.5098629 | 0.6101475 | 0.71325467 | IFITM4P | 340198 |
| ENSG000002 | 3.58281151 | -0.191287633 | 0.37515875 | -0.5098845 | 0.61013238 | 0.71325467 | NA | NA |
| ENSG000002 | 2.72429087 | 0.189019106 | 0.37077829 | 0.50979011 | 0.61019852 | 0.71326548 | NA | NA |
| ENSG000002 | 2.45671013 | -0.189317764 | 0.37134169 | -0.5098209 | 0.61017693 | 0.71326548 | NA | NA |
| ENSG000002 | 2.24046078 | -0.186149311 | 0.36516017 | -0.5097744 | 0.61020952 | 0.71326548 | NA | NA |
| ENSG000001 | 43.5253828 | -0.115631073 | 0.22685048 | -0.5097237 | 0.61024502 | 0.71327267 | FAM150B | 285016 |
| ENSG000002 | 124.119466 | -0.078919619 | 0.15483074 | -0.5097154 | 0.61025084 | 0.71327267 | NA | NA |
| ENSG000001 | 33.4767877 | -0.124080697 | 0.24347648 | -0.5096209 | 0.6103171 | 0.71332442 | MSS51 | 118490 |
| ENSG000002 | 5.72617593 | -0.187922379 | 0.36876301 | -0.509602 | 0.6103303 | 0.71332442 | RNU6-1045P | 106480025 |
| ENSG000002 | 7.60364544 | -0.180390658 | 0.35403434 | -0.5095287 | 0.61038168 | 0.71334335 | NA | NA |
| ENSG000002 | 3.71107428 | -0.1928486 | 0.37847991 | -0.5095346 | 0.61037757 | 0.71334335 | NA | NA |
| ENSG000001 | 38.3469062 | -0.113592246 | 0.22298122 | -0.5094252 | 0.61045423 | 0.71339087 | NA | NA |
| ENSG000002 | 3.72568191 | -0.192781885 | 0.37843372 | -0.5094205 | 0.61045753 | 0.71339087 | NA | NA |
| ENSG000002 | 2.2310411 | -0.186488335 | 0.3661541 | -0.5093165 | 0.61053037 | 0.71343488 | NA | NA |
| ENSG000002 | 2.44587388 | -0.189530949 | 0.37212016 | -0.5093273 | 0.61052284 | 0.71343488 | RPL23AP90 | 106480690 |
| ENSG000002 | 2.44282045 | -0.189685731 | 0.37245284 | -0.5092879 | 0.61055043 | 0.71343775 | NA | NA |
| ENSG000001 | 4.69274933 | -0.191250153 | 0.37565316 | -0.5091136 | 0.61067257 | 0.71347821 | KCNA10 | 3744 |
| ENSG000002 | 4.69571878 | -0.190551347 | 0.37420951 | -0.5092103 | 0.6106048 | 0.71347821 | NA | NA |
| ENSG000002 | 2.47233592 | -0.19031926 | 0.37383196 | -0.5091038 | 0.61067949 | 0.71347821 | NA | NA |
| ENSG000002 | 7.12797245 | -0.182136315 | 0.35774549 | -0.5091226 | 0.61066629 | 0.71347821 | LURAP1L-AS1 | 101929467 |
| ENSG000002 | 31.036804 | 0.173012675 | 0.33981153 | 0.50914303 | 0.61065197 | 0.71347821 | NA | NA |
| ENSG000002 | 2.49417031 | -0.187847728 | 0.36898879 | -0.5090879 | 0.61069062 | 0.71347821 | NA | NA |
| ENSG000002 | 7.87471989 | -0.181323608 | 0.35620141 | -0.509048 | 0.61071859 | 0.71349034 | NA | NA |
| ENSG000002 | 3.88526186 | 0.192374407 | 0.37802007 | 0.50889998 | 0.61082233 | 0.71357416 | NA | NA |

| | | | | | | | | |
|---|---|---|---|---|---|---|---|---|
| ENSG000002 | 2.48334102 | -0.189243668 | 0.37187144 | -0.5088954 | 0.61082554 | 0.71357416 | NA | NA |
| ENSG000002 | 3.48656647 | -0.192598446 | 0.37848514 | -0.5088666 | 0.61084573 | 0.7135772 | LOC105375732 | 105375732 |
| ENSG000001 | 7.91121869 | -0.178855558 | 0.35151094 | -0.5088193 | 0.61087887 | 0.71359535 | NA | NA |
| ENSG000002 | 2.19039654 | -0.187346492 | 0.36824542 | -0.5087544 | 0.61092436 | 0.71362793 | NA | NA |
| ENSG000001 | 7.91464351 | -0.180993858 | 0.3558073 | -0.5086851 | 0.610973 | 0.7136394 | OASL | 8638 |
| ENSG000002 | 2.23197475 | -0.18651982 | 0.36667016 | -0.5086856 | 0.61097264 | 0.7136394 | NA | NA |
| ENSG000002 | 2.51935436 | -0.187633644 | 0.3688746 | -0.5086651 | 0.61098697 | 0.7136394 | NA | NA |
| ENSG000002 | 2.73543266 | -0.190927208 | 0.37544292 | -0.5085386 | 0.61107569 | 0.71370191 | NA | NA |
| ENSG000002 | 4.97939011 | 0.189741332 | 0.37310871 | 0.50854168 | 0.61107352 | 0.71370191 | NA | NA |
| ENSG000002 | 2.95567376 | -0.191911436 | 0.37740159 | -0.5085072 | 0.61109767 | 0.71370702 | NA | NA |
| ENSG000001 | 3.95679171 | -0.192278489 | 0.37818533 | -0.508424 | 0.61115604 | 0.71371353 | ARL14 | 80117 |
| ENSG000002 | 2.51559596 | -0.186984219 | 0.36775791 | -0.5084438 | 0.61114216 | 0.71371353 | NA | NA |
| ENSG000002 | 34.6805788 | 0.121409894 | 0.23878512 | 0.50844832 | 0.61113897 | 0.71371353 | NA | NA |
| ENSG000002 | 3.20315322 | -0.192273466 | 0.37823482 | -0.5083442 | 0.611212 | 0.71375832 | NA | NA |
| ENSG000002 | 6.38221471 | -0.184820743 | 0.3636389 | -0.5082535 | 0.61127558 | 0.71378182 | NA | NA |
| ENSG000002 | 2.97451529 | -0.191734116 | 0.37725101 | -0.5082402 | 0.61128492 | 0.71378182 | NA | NA |
| ENSG000002 | 4.76225397 | 0.190875706 | 0.37554367 | 0.50826501 | 0.61126751 | 0.71378182 | RBM12B-AS1 | 55472 |
| ENSG000002 | 2.23476703 | -0.186752588 | 0.36747565 | -0.508204 | 0.6113103 | 0.7137909 | NA | NA |
| ENSG000001 | 15.0931078 | -0.155654641 | 0.30638423 | -0.5080374 | 0.61142713 | 0.71386291 | MAMSTR | 284358 |
| ENSG000002 | 2.71884263 | -0.191091777 | 0.37615336 | -0.5080156 | 0.6114424 | 0.71386291 | NA | NA |
| ENSG000002 | 77.1349752 | 0.089384161 | 0.17592638 | 0.50807709 | 0.61139928 | 0.71386291 | SOCS2-AS1 | 144481 |
| ENSG000002 | 2.25590474 | -0.186459946 | 0.3670001 | -0.5080651 | 0.61140768 | 0.71386291 | NA | NA |
| ENSG00000C | 7.65366611 | -0.179348123 | 0.35309133 | -0.5079369 | 0.61149758 | 0.71388624 | MADCAM1 | 8174 |
| ENSG000002 | 3.22873369 | -0.192026107 | 0.37804855 | -0.5079403 | 0.61149519 | 0.71388624 | NA | NA |
| ENSG000002 | 5.63192194 | -0.187609553 | 0.36943716 | -0.5078253 | 0.61157584 | 0.71395704 | NA | NA |
| ENSG000002 | 21.7131176 | 0.145404176 | 0.2863992 | 0.50769756 | 0.61166546 | 0.71399146 | NA | NA |
| ENSG000002 | 6.85739628 | -0.184503538 | 0.3634228 | -0.5076829 | 0.61167575 | 0.71399146 | ENO1-AS1 | 100505975 |
| ENSG000002 | 20.8869531 | -0.146367699 | 0.28827633 | -0.507734 | 0.61163989 | 0.71399146 | PTGES3P4 | 106480775 |
| ENSG000002 | 3.22808135 | -0.192098695 | 0.37835812 | -0.5077166 | 0.61165211 | 0.71399146 | RNU6-1281P | 106480125 |

| | | | | | | | | |
|---|---|---|---|---|---|---|---|---|
| ENSG000001 | 6.02788675 | 0.186418412 | 0.36722881 | 0.50763558 | 0.61170893 | 0.71400964 | CLDND2 | 125875 |
| ENSG000002 | 2.20604139 | -0.187151167 | 0.3686942 | -0.5076054 | 0.6117301 | 0.7140138 | RN7SL842P | 106479529 |
| ENSG000001 | 67.113683 | 0.096308536 | 0.1898379 | 0.50731985 | 0.61193041 | 0.71415235 | CPAMD8 | 27151 |
| ENSG000001 | 21.1239369 | 0.13742151 | 0.27084797 | 0.50737507 | 0.61189167 | 0.71415235 | KCNH6 | 81033 |
| ENSG000002 | 2.45656687 | -0.189476947 | 0.37351142 | -0.5072856 | 0.61195448 | 0.71415235 | NA | NA |
| ENSG000002 | 2.47483984 | -0.188282321 | 0.37115201 | -0.5072917 | 0.61195019 | 0.71415235 | NA | NA |
| ENSG000002 | 4.74314353 | 0.190777504 | 0.37606475 | 0.50729962 | 0.6119446 | 0.71415235 | PCBP2P1 | 101060045 |
| ENSG000002 | 2.19966719 | -0.186358791 | 0.36734385 | -0.5073143 | 0.6119343 | 0.71415235 | NA | NA |
| ENSG000001 | 28.4336517 | -0.128868207 | 0.25406355 | -0.5072282 | 0.61199468 | 0.71417872 | MFNG | 4242 |
| ENSG000002 | 2.2286215 | -0.186594674 | 0.36792485 | -0.5071543 | 0.61204655 | 0.71421783 | NA | NA |
| ENSG000002 | 3.44245712 | -0.191994973 | 0.37859104 | -0.5071303 | 0.61206342 | 0.71421783 | NA | NA |
| ENSG000002 | 8.59060725 | -0.175345636 | 0.3458299 | -0.5070286 | 0.61213478 | 0.71428055 | NA | NA |
| ENSG000002 | 2.69408314 | -0.190433583 | 0.37564861 | -0.5069461 | 0.61219267 | 0.71432754 | NA | NA |
| ENSG000002 | 4.1819522 | -0.191166083 | 0.37715076 | -0.5068691 | 0.61224664 | 0.71436649 | NA | NA |
| ENSG000002 | 4.17896403 | -0.190832679 | 0.37650849 | -0.5068483 | 0.61226128 | 0.71436649 | NA | NA |
| ENSG000001 | 66.1262079 | -0.095672297 | 0.18879657 | -0.5067481 | 0.6123316 | 0.714381 | METTL20 | 254013 |
| ENSG000002 | 3.01450752 | -0.190438966 | 0.37581931 | -0.5067301 | 0.61234419 | 0.714381 | NA | NA |
| ENSG000002 | 3.99245196 | -0.191856273 | 0.37856102 | -0.5068041 | 0.61229229 | 0.714381 | NA | NA |
| ENSG000002 | 2.97479605 | -0.190953479 | 0.37683439 | -0.5067305 | 0.61234393 | 0.714381 | NA | NA |
| ENSG000002 | 3.23801753 | -0.191600607 | 0.37817182 | -0.5066496 | 0.61240069 | 0.71442636 | NA | NA |
| ENSG000001 | 2.18700722 | -0.127207911 | 0.25112089 | -0.5065604 | 0.61246327 | 0.71445826 | CCL24 | 6369 |
| ENSG000002 | 2.75530425 | -0.18897165 | 0.37303714 | -0.506576 | 0.61245238 | 0.71445826 | NA | NA |
| ENSG000001 | 30.1772354 | -0.125405461 | 0.24761055 | -0.5064625 | 0.612532 | 0.71451789 | LRRC55 | 219527 |
| ENSG000002 | 4.19857524 | -0.190813401 | 0.37678727 | -0.5064221 | 0.61256036 | 0.71453041 | NA | NA |
| ENSG000002 | 3.97751207 | -0.19166829 | 0.3785088 | -0.5063774 | 0.61259176 | 0.71454649 | NA | NA |
| ENSG00000C | 25.4681635 | -0.14264742 | 0.28176439 | -0.5062649 | 0.6126707 | 0.71458004 | MLXIPL | 51085 |
| ENSG000002 | 5.6973056 | -0.187372942 | 0.37012966 | -0.506236 | 0.61269102 | 0.71458004 | CXorf65 | 158830 |
| ENSG000002 | 2.7390539 | -0.189367457 | 0.3740468 | -0.5062667 | 0.6126694 | 0.71458004 | NA | NA |
| ENSG000002 | 3.46340615 | -0.191653683 | 0.37858386 | -0.5062384 | 0.61268931 | 0.71458004 | NA | NA |

| | | | | | | | | |
|---|---|---|---|---|---|---|---|---|
| ENSG000002 | 6.71570016 | -0.183963014 | 0.36351887 | -0.5060618 | 0.61281327 | 0.71469238 | LOC100130172 | 100130172 |
| ENSG000002 | 3.46927148 | -0.19158662 | 0.37859338 | -0.5060485 | 0.61282258 | 0.71469238 | NA | NA |
| ENSG000002 | 5.23586307 | -0.188384772 | 0.372358 | -0.5059238 | 0.61291014 | 0.71477393 | LOC101928847 | 101928847 |
| ENSG000002 | 6.19505084 | -0.184470175 | 0.36464601 | -0.5058884 | 0.61293502 | 0.71478239 | NA | NA |
| ENSG000001 | 104.3665 | 0.087514114 | 0.17302496 | 0.50578894 | 0.61300482 | 0.71484323 | EDA | 1896 |
| ENSG000001 | 3.68420742 | 0.191458995 | 0.37856417 | 0.50575044 | 0.61303186 | 0.7148542 | C19orf71 | 100128569 |
| ENSG000001 | 322.9467 | 0.063922074 | 0.12640775 | 0.50568159 | 0.6130802 | 0.71489001 | LAMA2 | 3908 |
| ENSG000002 | 4.70484561 | -0.189542133 | 0.3749094 | -0.5055678 | 0.61316007 | 0.71494475 | NA | NA |
| ENSG000002 | 4.95232562 | -0.188247619 | 0.37235133 | -0.5055645 | 0.6131624 | 0.71494475 | NA | NA |
| ENSG000002 | 6.10576853 | -0.184669248 | 0.36536824 | -0.5054332 | 0.61325459 | 0.71503168 | NA | NA |
| ENSG000002 | 3.03757998 | 0.189291589 | 0.37455195 | 0.5053814 | 0.61329098 | 0.71505355 | NA | NA |
| ENSG000002 | 2.2193112 | -0.186147507 | 0.36835424 | -0.5053492 | 0.61331362 | 0.71505939 | NA | NA |
| ENSG000002 | 2.70837777 | -0.188891434 | 0.37380881 | -0.5053156 | 0.61333717 | 0.71506628 | LARP1BP2 | 106479034 |
| ENSG000002 | 4.69115867 | -0.189538204 | 0.37518409 | -0.5051872 | 0.61342735 | 0.71515086 | LOC340581 | 340581 |
| ENSG000002 | 2.25451443 | -0.18355045 | 0.36340496 | -0.5050852 | 0.61349903 | 0.71521386 | NA | NA |
| ENSG000002 | 2.68432378 | -0.189881176 | 0.37597715 | -0.5050338 | 0.61353508 | 0.71523534 | NA | NA |
| ENSG000002 | 2.24407048 | -0.185951504 | 0.36826053 | -0.5049455 | 0.6135971 | 0.71528707 | RNU6-739P | 106481413 |
| ENSG000002 | 2.97118221 | -0.190671751 | 0.37773325 | -0.5047788 | 0.61371418 | 0.71538242 | COX6CP14 | 106481683 |
| ENSG000002 | 3.69539752 | -0.190930211 | 0.37823092 | -0.504798 | 0.61370072 | 0.71538242 | NA | NA |
| ENSG000002 | 2.78621694 | 0.188485904 | 0.37344866 | 0.50471704 | 0.61375759 | 0.71541246 | BNIP3P11 | 106480286 |
| ENSG000002 | 2.23342955 | -0.186053453 | 0.36866777 | -0.5046643 | 0.61379465 | 0.7154351 | NA | NA |
| ENSG000002 | 2.18493615 | -0.185739645 | 0.36822521 | -0.5044186 | 0.61396725 | 0.71561571 | NA | NA |
| ENSG000001 | 112.203072 | 0.085085787 | 0.16871018 | 0.50433107 | 0.61402875 | 0.71564624 | ZNF333 | 84449 |
| ENSG000002 | 3.89243242 | 0.190742477 | 0.37820813 | 0.50433204 | 0.61402807 | 0.71564624 | NA | NA |
| ENSG000002 | 5.80808141 | 0.185352765 | 0.36776527 | 0.50399747 | 0.61426315 | 0.71578547 | FOXB2 | 442425 |
| ENSG000002 | 72.8061609 | 0.097773429 | 0.19394525 | 0.50412903 | 0.61417071 | 0.71578547 | NA | NA |
| ENSG000002 | 2.71967158 | -0.189450331 | 0.37583198 | -0.5040825 | 0.61420339 | 0.71578547 | NA | NA |
| ENSG000002 | 6.12864147 | -0.184728748 | 0.36648989 | -0.5040487 | 0.61422717 | 0.71578547 | NA | NA |
| ENSG000002 | 5.91202193 | -0.18440286 | 0.36584729 | -0.5040433 | 0.61423098 | 0.71578547 | NA | NA |

| | | | | | | | | |
|---|---|---|---|---|---|---|---|---|
| ENSG000002 | 2.19724558 | 0.186027192 | 0.36910662 | 0.5039931 | 0.61426622 | 0.71578547 | NA | NA |
| ENSG000002 | 2.2714777 | -0.184173133 | 0.3654336 | -0.5039852 | 0.61427177 | 0.71578547 | PPIAL4A | 653505 |
| ENSG000002 | 3.95272087 | -0.190514748 | 0.37805143 | -0.5039387 | 0.61430445 | 0.71580299 | CEMP1 | 752014 |
| ENSG000001 | 5.89226861 | 0.18586622 | 0.36885587 | 0.50389931 | 0.61433214 | 0.71581468 | SPINK2 | 6691 |
| ENSG000002 | 4.4229459 | -0.189754432 | 0.37659161 | -0.5038732 | 0.61435046 | 0.71581546 | NA | NA |
| ENSG000001 | 39.4988387 | 0.112553563 | 0.22341644 | 0.50378372 | 0.61441337 | 0.71585802 | IFIT2 | 3433 |
| ENSG000001 | 696.019339 | 0.065154821 | 0.12936645 | 0.50364544 | 0.61451056 | 0.71585802 | LIN9 | 286826 |
| ENSG000001 | 709.541366 | 0.073296101 | 0.14551113 | 0.50371473 | 0.61446186 | 0.71585802 | FAM3C | 10447 |
| ENSG000002 | 3.85233694 | 0.190662751 | 0.37851607 | 0.50371112 | 0.6144644 | 0.71585802 | NA | NA |
| ENSG000002 | 2.94335023 | -0.190193795 | 0.37760208 | -0.5036884 | 0.61448034 | 0.71585802 | NA | NA |
| ENSG000002 | 2.70083004 | -0.189410463 | 0.37599087 | -0.5037635 | 0.61442761 | 0.71585802 | NA | NA |
| ENSG000002 | 3.80152957 | -0.190220052 | 0.377686 | -0.503646 | 0.61451015 | 0.71585802 | NA | NA |
| ENSG000002 | 3.46981282 | -0.190651546 | 0.37859063 | -0.5035823 | 0.61455493 | 0.71588914 | NA | NA |
| ENSG000002 | 5.44015885 | -0.185985536 | 0.3694043 | -0.5034742 | 0.61463091 | 0.71595709 | LOC102546228 | 102546228 |
| ENSG000001 | 3920.8866 | -0.061272111 | 0.12173064 | -0.5033417 | 0.61472405 | 0.71602444 | ASCC3 | 10973 |
| ENSG000002 | 2.43980003 | -0.187296857 | 0.37210435 | -0.503345 | 0.61472174 | 0.71602444 | RNU1-67P | 106480169 |
| ENSG000002 | 2.20216518 | 0.184465021 | 0.36653103 | 0.5032726 | 0.61477263 | 0.71604001 | NA | NA |
| ENSG000002 | 2.94654692 | -0.189313701 | 0.37616543 | -0.5032725 | 0.61477273 | 0.71604001 | NA | NA |
| ENSG000002 | 13.3666769 | -0.159925346 | 0.31781275 | -0.5032062 | 0.6148193 | 0.71607369 | NA | NA |
| ENSG000002 | 2.48785647 | -0.186543976 | 0.37074572 | -0.5031588 | 0.61485262 | 0.71609192 | NA | NA |
| ENSG000001 | 2.1757691 | -0.183583423 | 0.3650596 | -0.5028862 | 0.61504431 | 0.71626325 | TPSAB1 | 7177 |
| ENSG000001 | 30.6075893 | 0.121336831 | 0.24135898 | 0.50272349 | 0.61515869 | 0.71626325 | NA | NA |
| ENSG000002 | 3.45404377 | -0.190399 | 0.37858553 | -0.502922 | 0.6150191 | 0.71626325 | NA | NA |
| ENSG000002 | 2.20972735 | -0.182833467 | 0.36365314 | -0.5027688 | 0.6151268 | 0.71626325 | NA | NA |
| ENSG000002 | 3.4640915 | -0.190367864 | 0.37859077 | -0.5028328 | 0.61508182 | 0.71626325 | NA | NA |
| ENSG000002 | 2.48008007 | -0.18728789 | 0.37249473 | -0.5027934 | 0.61510954 | 0.71626325 | NA | NA |
| ENSG000002 | 2.74901491 | -0.187547755 | 0.37303857 | -0.502757 | 0.61513512 | 0.71626325 | LOC105375744 | 105375744 |
| ENSG000002 | 2.20718355 | -0.185709158 | 0.36938872 | -0.5027472 | 0.615142 | 0.71626325 | LOC105369920 | 105369920 |
| ENSG000002 | 2.18916861 | 0.185892725 | 0.36969848 | 0.50282254 | 0.61508905 | 0.71626325 | NA | NA |

| | | | | | | | | |
|---|---|---|---|---|---|---|---|---|
| ENSG000002 | 2.17933141 | -0.183439716 | 0.36507577 | -0.5024703 | 0.61533677 | 0.71641177 | NA | NA |
| ENSG000002 | 5.39872447 | -0.186855778 | 0.37184928 | -0.5025041 | 0.61531299 | 0.71641177 | NA | NA |
| ENSG000002 | 33.755265 | 0.122846035 | 0.24448591 | 0.50246673 | 0.61533925 | 0.71641177 | NA | NA |
| ENSG000002 | 3.4176133 | -0.190207102 | 0.3785806 | -0.5024217 | 0.61537093 | 0.71642809 | NA | NA |
| ENSG000002 | 2.95851846 | -0.189435122 | 0.37713145 | -0.5023053 | 0.61545278 | 0.71648224 | NA | NA |
| ENSG000002 | 3.21057785 | -0.190109202 | 0.37845513 | -0.5023296 | 0.61543572 | 0.71648224 | NA | NA |
| ENSG000002 | 2.27440638 | -0.183062959 | 0.36448881 | -0.5022458 | 0.61549466 | 0.71650394 | RN7SL398P | 106481032 |
| ENSG000002 | 3.86244087 | 0.189987838 | 0.3782896 | 0.50222854 | 0.61550677 | 0.71650394 | NA | NA |
| ENSG000001 | 2.22462757 | -0.183855703 | 0.36611585 | -0.502179 | 0.61554159 | 0.71652391 | NA | NA |
| ENSG000002 | 2.47830485 | -0.186346813 | 0.37109792 | -0.50215 | 0.61556203 | 0.71652713 | MIR576 | 693161 |
| ENSG000001 | 11.443397 | 0.167406963 | 0.33349047 | 0.50198425 | 0.6156786 | 0.71664226 | PLA2G2A | 5320 |
| ENSG000001 | 29.2208371 | 0.128892359 | 0.25682503 | 0.50186837 | 0.61576012 | 0.71671644 | UNC5A | 90249 |
| ENSG000002 | 4.21804985 | -0.189664443 | 0.37793551 | -0.5018434 | 0.61577768 | 0.71671644 | NA | NA |
| ENSG000002 | 2.73754036 | -0.188405057 | 0.37544593 | -0.5018168 | 0.61579643 | 0.71671769 | MIR3163 | 100423029 |
| ENSG000002 | 2.24544042 | -0.183324915 | 0.36534705 | -0.5017829 | 0.61582022 | 0.7167248 | NA | NA |
| ENSG000002 | 4.97772949 | -0.187773009 | 0.37430403 | -0.5016591 | 0.61590736 | 0.71680566 | NA | NA |
| ENSG000002 | 263.661534 | 0.189616132 | 0.37806083 | 0.50154927 | 0.61598461 | 0.71687499 | GTF2IP1 | 2970 |
| ENSG000001 | 6.9151636 | -0.180031247 | 0.35922609 | -0.5011642 | 0.61625558 | 0.71702571 | FAM132A | 388581 |
| ENSG000001 | 2.17447701 | 0.183301906 | 0.3656472 | 0.50130811 | 0.6161543 | 0.71702571 | NA | NA |
| ENSG000002 | 4.74228129 | -0.188222429 | 0.37554036 | -0.5012043 | 0.61622737 | 0.71702571 | NA | NA |
| ENSG000002 | 3.70643542 | -0.18971761 | 0.37843206 | -0.5013254 | 0.61614212 | 0.71702571 | NA | NA |
| ENSG000002 | 3.71424373 | -0.189763781 | 0.37859292 | -0.5012344 | 0.61620619 | 0.71702571 | NA | NA |
| ENSG000002 | 3.73071754 | -0.18968934 | 0.37847907 | -0.5011885 | 0.6162385 | 0.71702571 | NA | NA |
| ENSG000002 | 3.50877444 | -0.189583913 | 0.37823379 | -0.5012347 | 0.61620593 | 0.71702571 | NA | NA |
| ENSG000002 | 3.00394624 | -0.188783263 | 0.37659832 | -0.5012855 | 0.61617023 | 0.71702571 | NA | NA |
| ENSG000002 | 5.2191155 | -0.187014798 | 0.37320713 | -0.5011019 | 0.61629942 | 0.71705614 | NA | NA |
| ENSG000002 | 4.88925631 | -0.188119744 | 0.37544693 | -0.5010555 | 0.61633208 | 0.71707356 | NA | NA |
| ENSG000002 | 2.26987929 | -0.183511457 | 0.36628695 | -0.5010046 | 0.61636786 | 0.71709463 | NA | NA |
| ENSG000002 | 6.927348 | -0.181912352 | 0.36320523 | -0.5008528 | 0.61647473 | 0.71715723 | NA | NA |

| | | | | | | | | |
|---|---|---|---|---|---|---|---|---|
| ENSG000002 | 2.71434758 | -0.187939305 | 0.37522601 | -0.5008696 | 0.61646289 | 0.71715723 | NA | NA |
| ENSG000002 | 4.23031368 | -0.189426259 | 0.37819449 | -0.50087 | 0.61646265 | 0.71715723 | MIR3976 | 100616244 |
| ENSG000002 | 2.17903857 | -0.184395465 | 0.36820199 | -0.5007998 | 0.61651206 | 0.71718008 | NA | NA |
| ENSG000002 | 8.88199697 | 0.177610435 | 0.35468627 | 0.50075362 | 0.61654453 | 0.71719632 | C6orf226 | 441150 |
| ENSG000002 | 2.20082715 | 0.184134183 | 0.36773172 | 0.50072967 | 0.61656139 | 0.71719632 | CYB561D2 | 11068 |
| ENSG000002 | 2.96568902 | -0.188817693 | 0.37713288 | -0.5006662 | 0.61660605 | 0.71722769 | NA | NA |
| ENSG000001 | 2.18867935 | 0.184969139 | 0.36947524 | 0.50062661 | 0.61663393 | 0.71723955 | OR4X2 | 119764 |
| ENSG000001 | 51.4281091 | -0.110056531 | 0.2198502 | -0.5005978 | 0.6166542 | 0.71724256 | NA | NA |
| ENSG000002 | 6.01032606 | -0.186063986 | 0.37179217 | -0.5004516 | 0.61675713 | 0.7173417 | RNU6-60P | 106480750 |
| ENSG000002 | 7.07043729 | -0.179693896 | 0.35912047 | -0.5003722 | 0.61681303 | 0.71738615 | NA | NA |
| ENSG000002 | 5.15645222 | -0.185976663 | 0.37175379 | -0.5002684 | 0.61688612 | 0.71745057 | NA | NA |
| ENSG000002 | 3.4567391 | -0.189323211 | 0.37859285 | -0.5000707 | 0.61702526 | 0.71759182 | NA | NA |
| ENSG000002 | 3.94808256 | -0.188835735 | 0.37764371 | -0.5000368 | 0.6170492 | 0.71759907 | RNU1-57P | 106480166 |
| ENSG000002 | 2.17385658 | 0.183719592 | 0.36746124 | 0.49996998 | 0.61709621 | 0.71761259 | NA | NA |
| ENSG000002 | 2.51679073 | -0.185576242 | 0.37116969 | -0.4999768 | 0.61709139 | 0.71761259 | NA | NA |
| ENSG00000C | 189.454668 | 0.082571949 | 0.16518987 | 0.49986084 | 0.61717307 | 0.71763337 | TFAP2C | 7022 |
| ENSG000002 | 3.89119887 | 0.18916033 | 0.37843068 | 0.49985463 | 0.61717744 | 0.71763337 | NA | NA |
| ENSG000002 | 4.14010144 | 0.188475434 | 0.37704495 | 0.49987523 | 0.61716293 | 0.71763337 | NA | NA |
| ENSG000002 | 3.17460385 | -0.188994287 | 0.37810649 | -0.4998441 | 0.61718488 | 0.71763337 | NA | NA |
| ENSG000002 | 5.9838762 | 0.184948062 | 0.37004712 | 0.49979597 | 0.61721875 | 0.71765218 | RN7SL192P | 106480974 |
| ENSG000001 | 13.9606073 | 0.156389314 | 0.31294045 | 0.49974145 | 0.61725714 | 0.71767625 | LRRC32 | 2615 |
| ENSG000002 | 5.4062929 | -0.185170526 | 0.37062178 | -0.4996213 | 0.61734177 | 0.71775406 | NA | NA |
| ENSG000001 | 91.3944806 | -0.093917174 | 0.18804066 | -0.4994514 | 0.61746141 | 0.71787258 | S100A6 | 6277 |
| ENSG000002 | 2.4673516 | -0.185843327 | 0.37215513 | -0.4993706 | 0.61751833 | 0.71791817 | NA | NA |
| ENSG000002 | 2.99973159 | -0.188318803 | 0.37713775 | -0.4993369 | 0.61754205 | 0.71792517 | NA | NA |
| ENSG000002 | 4.22387531 | -0.188408478 | 0.3774409 | -0.4991735 | 0.61765719 | 0.71803844 | NA | NA |
| ENSG000002 | 3.67709677 | -0.188849818 | 0.37838521 | -0.4990941 | 0.61771311 | 0.71806228 | SCARNA11 | 677780 |
| ENSG000002 | 3.2428526 | -0.188593141 | 0.37785479 | -0.4991154 | 0.6176981 | 0.71806228 | NA | NA |
| ENSG000002 | 4.77186573 | 0.188766225 | 0.37825264 | 0.499048 | 0.61774557 | 0.71807942 | NA | NA |

| | | | | | | | | |
|---|---|---|---|---|---|---|---|---|
| ENSG000001 | 15.6068818 | -0.155729475 | 0.31211153 | -0.4989546 | 0.61781139 | 0.71809418 | NPIPB15 | 440348 |
| ENSG000002 | 8.68639384 | -0.175952547 | 0.35263247 | -0.4989686 | 0.61780147 | 0.71809418 | NA | NA |
| ENSG000002 | 7.21761798 | 0.186128215 | 0.37301232 | 0.49898678 | 0.6177887 | 0.71809418 | NA | NA |
| ENSG000002 | 3.6917363 | -0.18889071 | 0.37859237 | -0.498929 | 0.6178294 | 0.71809453 | NA | NA |
| ENSG000002 | 3.45105003 | -0.188839529 | 0.37854742 | -0.498853 | 0.61788292 | 0.71813548 | NA | NA |
| ENSG000002 | 8.11349837 | -0.175731396 | 0.35228805 | -0.4988287 | 0.61790006 | 0.71813548 | NA | NA |
| ENSG000002 | 2.22918211 | -0.181705471 | 0.3643348 | -0.4987321 | 0.61796811 | 0.71819399 | OR5R1 | 219479 |
| ENSG000001 | 110.290224 | 0.081166172 | 0.16277172 | 0.49865034 | 0.61802573 | 0.71819921 | CYP1B1 | 1545 |
| ENSG000002 | 2.28636573 | -0.181813 | 0.36459082 | -0.4986768 | 0.61800706 | 0.71819921 | NA | NA |
| ENSG000002 | 3.68782963 | -0.188673016 | 0.37835063 | -0.4986724 | 0.61801019 | 0.71819921 | NA | NA |
| ENSG000002 | 4.72561859 | -0.187036018 | 0.3751261 | -0.4985951 | 0.61806469 | 0.7182239 | SOGA3 | 387104 |
| ENSG000002 | 4.69344243 | -0.18741501 | 0.37591763 | -0.4985534 | 0.61809404 | 0.71823743 | NA | NA |
| ENSG000002 | 13.6437526 | -0.154778826 | 0.31052949 | -0.4984352 | 0.61817734 | 0.71827432 | NA | NA |
| ENSG000002 | 2.18220802 | -0.181237923 | 0.36361547 | -0.4984329 | 0.61817893 | 0.71827432 | NA | NA |
| ENSG000002 | 2.44094218 | -0.185748482 | 0.37263772 | -0.4984693 | 0.61815327 | 0.71827432 | NA | NA |
| ENSG000002 | 24.4000232 | -0.131605243 | 0.26412411 | -0.4982705 | 0.61829342 | 0.71834173 | CCNI2 | 645121 |
| ENSG000002 | 2.46234768 | -0.184922355 | 0.37110489 | -0.4983021 | 0.61827111 | 0.71834173 | NA | NA |
| ENSG000002 | 2.29416195 | -0.181081638 | 0.36341795 | -0.4982738 | 0.61829108 | 0.71834173 | NA | NA |
| ENSG000002 | 12.1494882 | -0.159277007 | 0.31967283 | -0.4982501 | 0.61830781 | 0.71834173 | NA | NA |
| ENSG000001 | 28.0614744 | 0.129210518 | 0.25936459 | 0.49818103 | 0.61835645 | 0.71837767 | MRPS28 | 28957 |
| ENSG000002 | 3.18903541 | -0.18845775 | 0.37835906 | -0.4980923 | 0.61841896 | 0.71842971 | NA | NA |
| ENSG000002 | 2.4581851 | -0.185483053 | 0.37247995 | -0.4979679 | 0.61850668 | 0.71844928 | NA | NA |
| ENSG000002 | 3.42503738 | -0.188498669 | 0.37850876 | -0.4980035 | 0.61848161 | 0.71844928 | NA | NA |
| ENSG000002 | 3.44869906 | -0.188103967 | 0.37773934 | -0.4979729 | 0.61850312 | 0.71844928 | MIR5690 | 100847048 |
| ENSG000002 | 2.50477953 | -0.184833618 | 0.37114582 | -0.4980081 | 0.61847835 | 0.71844928 | LINC01443 | 400644 |
| ENSG000002 | 8.92512674 | 0.174356398 | 0.35016416 | 0.4979276 | 0.61853507 | 0.71846168 | NA | NA |
| ENSG000002 | 9.29361956 | -0.171349174 | 0.34417482 | -0.4978551 | 0.61858621 | 0.71850049 | HIST2H2BC | 337873 |
| ENSG000001 | 35.1963753 | -0.116268631 | 0.23357587 | -0.4977767 | 0.61864142 | 0.71853518 | GCK | 2645 |
| ENSG000001 | 81.132846 | 0.088906391 | 0.1786121 | 0.49776242 | 0.61865151 | 0.71853518 | NIPAL1 | 152519 |

| | | | | | | | | |
|---|---|---|---|---|---|---|---|---|
| ENSG000001 | 4.42510601 | -0.187025093 | 0.37581625 | -0.4976504 | 0.6187305 | 0.7185446 | DMRTC1B | 728656 |
| ENSG000002 | 3.20668327 | -0.188167439 | 0.37810589 | -0.497658 | 0.61872509 | 0.7185446 | NA | NA |
| ENSG000002 | 3.89804273 | 0.188383961 | 0.37853812 | 0.4976618 | 0.61872244 | 0.7185446 | NA | NA |
| ENSG000002 | 7.17664675 | 0.179219473 | 0.36012405 | 0.49766038 | 0.61872344 | 0.7185446 | NA | NA |
| ENSG000002 | 4.92323959 | -0.186384967 | 0.37464731 | -0.4974945 | 0.61884039 | 0.71861246 | AWAT1 | 158833 |
| ENSG000002 | 18.8296965 | -0.142329005 | 0.28607244 | -0.4975278 | 0.61881687 | 0.71861246 | NA | NA |
| ENSG000002 | 3.69822151 | -0.188340669 | 0.37858024 | -0.4974921 | 0.61884209 | 0.71861246 | RN7SL600P | 106481086 |
| ENSG000001 | 42.9097149 | 0.110535746 | 0.22222019 | 0.49741541 | 0.61889614 | 0.71865464 | AKR1C3 | 8644 |
| ENSG000002 | 15.0513737 | 0.167664472 | 0.3371672 | 0.49727398 | 0.61899586 | 0.71874985 | NA | NA |
| ENSG000002 | 4.12465336 | 0.187898914 | 0.37788167 | 0.49724273 | 0.61901789 | 0.71875485 | NA | NA |
| ENSG000001 | 14.8193388 | 0.156822143 | 0.31541801 | 0.49718829 | 0.61905628 | 0.71877884 | NUTM2F | 54754 |
| ENSG000002 | 4.43592321 | -0.187014252 | 0.37622859 | -0.4970761 | 0.61913537 | 0.71885009 | NA | NA |
| ENSG000002 | 2.72974414 | -0.186610887 | 0.37543828 | -0.4970481 | 0.61915513 | 0.71885245 | NA | NA |
| ENSG000002 | 3.22214434 | -0.1877055 | 0.37769708 | -0.4969737 | 0.61920762 | 0.71887224 | NA | NA |
| ENSG000002 | 4.20344278 | -0.187623185 | 0.37752516 | -0.4969819 | 0.61920179 | 0.71887224 | GDF10 | 2662 |
| ENSG000002 | 3.01274025 | -0.187026805 | 0.37635054 | -0.4969484 | 0.61922543 | 0.71887233 | NA | NA |
| ENSG000002 | 2.72488402 | -0.186231845 | 0.37478733 | -0.4969001 | 0.61925953 | 0.71889134 | RNU2-31P | 106480210 |
| ENSG000001 | 32.3659614 | -0.123375803 | 0.24834635 | -0.4967893 | 0.61933767 | 0.7189368 | CELF4 | 56853 |
| ENSG000001 | 54.4328859 | 0.10559652 | 0.21256487 | 0.49677316 | 0.61934903 | 0.7189368 | CCDC121 | 79635 |
| ENSG000002 | 4.43087961 | -0.186578213 | 0.37558337 | -0.4967691 | 0.61935188 | 0.7189368 | NA | NA |
| ENSG000002 | 2.94982052 | -0.186272453 | 0.37506274 | -0.4966435 | 0.61944051 | 0.7190059 | NA | NA |
| ENSG000002 | 3.91774556 | -0.187702901 | 0.37796165 | -0.4966189 | 0.61945781 | 0.7190059 | NA | NA |
| ENSG000002 | 2.68909903 | -0.186095204 | 0.37475987 | -0.4965718 | 0.61949101 | 0.7190059 | NA | NA |
| ENSG000002 | 19.165358 | -0.15450975 | 0.3111609 | -0.496559 | 0.61950006 | 0.7190059 | NA | NA |
| ENSG000002 | 3.72582571 | -0.187997324 | 0.37859286 | -0.4965686 | 0.61949331 | 0.7190059 | LINC00251 | 552859 |
| ENSG000002 | 2.91466392 | -0.187080365 | 0.37681804 | -0.496474 | 0.61956001 | 0.7190549 | RNU6-253P | 106480697 |
| ENSG000002 | 3.95569445 | -0.187932296 | 0.37859289 | -0.4963968 | 0.61961449 | 0.71909754 | MIR5586 | 100847088 |
| ENSG000002 | 2.46718819 | -0.184364385 | 0.37147351 | -0.4963056 | 0.61967881 | 0.7191387 | NA | NA |
| ENSG000002 | 4.93456511 | -0.1862203 | 0.37522005 | -0.4962962 | 0.61968542 | 0.7191387 | NA | NA |

| | | | | | | | | |
|---|---|---|---|---|---|---|---|---|
| ENSG000001 | 164.910217 | -0.073610061 | 0.14836106 | -0.4961549 | 0.61978516 | 0.71918191 | ANXA4 | 307 |
| ENSG000002 | 2.22911077 | -0.181661882 | 0.36614831 | -0.4961429 | 0.6197936 | 0.71918191 | NA | NA |
| ENSG000002 | 11.5739162 | 0.161048222 | 0.32457782 | 0.49617754 | 0.61976916 | 0.71918191 | NA | NA |
| ENSG000002 | 5.71349954 | -0.183993228 | 0.37082463 | -0.4961732 | 0.61977225 | 0.71918191 | NA | NA |
| ENSG000002 | 2.4790547 | -0.18282345 | 0.36853686 | -0.4960791 | 0.61983858 | 0.71919294 | NA | NA |
| ENSG000002 | 2.47524479 | -0.183930813 | 0.37076178 | -0.4960889 | 0.61983168 | 0.71919294 | NA | NA |
| ENSG000002 | 4.66944052 | -0.185748643 | 0.37448616 | -0.4960094 | 0.61988781 | 0.71920097 | NA | NA |
| ENSG000002 | 6.70060382 | -0.180325189 | 0.36361595 | -0.4959221 | 0.61994936 | 0.71920097 | NA | NA |
| ENSG000002 | 6.4006524 | -0.179922811 | 0.36280501 | -0.4959215 | 0.61994978 | 0.71920097 | NA | NA |
| ENSG000002 | 4.39653648 | -0.186931929 | 0.37689066 | -0.4959845 | 0.61990534 | 0.71920097 | NA | NA |
| ENSG000002 | 3.23670707 | -0.187385127 | 0.37785552 | -0.4959174 | 0.61995269 | 0.71920097 | NA | NA |
| ENSG000002 | 2.70063361 | -0.185973882 | 0.37502797 | -0.4958934 | 0.61996965 | 0.71920097 | NA | NA |
| ENSG000002 | 6.13283138 | -0.180841832 | 0.36457202 | -0.4960387 | 0.61986711 | 0.71920097 | NA | NA |
| ENSG000001 | 62.9859081 | 0.097352818 | 0.19634513 | 0.49582497 | 0.6200179 | 0.71923638 | MPZ | 4359 |
| ENSG000000 | 9.6402483 | -0.174692852 | 0.35240157 | -0.495721 | 0.62009128 | 0.71925977 | APOH | 350 |
| ENSG000001 | 4.42708842 | -0.186778229 | 0.37674284 | -0.4957711 | 0.62005588 | 0.71925977 | ACTA1 | 58 |
| ENSG000002 | 2.75332847 | -0.184921619 | 0.37303314 | -0.4957244 | 0.62008889 | 0.71925977 | NA | NA |
| ENSG000000 | 103.99653 | 0.085621735 | 0.17273723 | 0.4956762 | 0.62012288 | 0.71926855 | NA | NA |
| ENSG000002 | 2.45241117 | -0.183111111 | 0.36942887 | -0.49566 | 0.62013432 | 0.71926855 | NA | NA |
| ENSG000001 | 4.20537345 | -0.187076014 | 0.37756561 | -0.4954795 | 0.6202617 | 0.71936435 | SAA1 | 6288 |
| ENSG000002 | 2.46610552 | -0.183877493 | 0.37111126 | -0.4954781 | 0.62026269 | 0.71936435 | NA | NA |
| ENSG000002 | 4.1512436 | -0.186796106 | 0.37700978 | -0.4954675 | 0.62027014 | 0.71936435 | LOC100506700 | 100506700 |
| ENSG000002 | 7.17503185 | -0.178261424 | 0.35995664 | -0.4952303 | 0.62043759 | 0.71953797 | NA | NA |
| ENSG000002 | 7.42969964 | -0.177287259 | 0.35824611 | -0.4948756 | 0.62068794 | 0.71980772 | NA | NA |
| ENSG000002 | 2.94460339 | -0.186492114 | 0.37690957 | -0.4947927 | 0.62074644 | 0.71984197 | NA | NA |
| ENSG000002 | 5.14208395 | 0.185674119 | 0.37526339 | 0.49478347 | 0.62075298 | 0.71984197 | NA | NA |
| ENSG000001 | 4.74022587 | -0.186078603 | 0.37626867 | -0.4945365 | 0.62092731 | 0.72001554 | KRTAP4-1 | 85285 |
| ENSG000002 | 4.18377208 | -0.186460948 | 0.37705352 | -0.4945212 | 0.62093816 | 0.72001554 | NA | NA |
| ENSG000001 | 23.8039966 | 0.130168217 | 0.26326135 | 0.49444484 | 0.62099205 | 0.72003862 | CCDC74B | 91409 |

| | | | | | | | | |
|---|---|---|---|---|---|---|---|---|
| ENSG000002 | 19.1337293 | 0.146091764 | 0.29548257 | 0.49441753 | 0.62101133 | 0.72003862 | NA | NA |
| ENSG000002 | 3.03143804 | 0.186759322 | 0.37772369 | 0.4944337 | 0.62099992 | 0.72003862 | NA | NA |
| ENSG000002 | 3.49614923 | -0.186417789 | 0.37712651 | -0.494311 | 0.62108654 | 0.72010523 | NA | NA |
| ENSG000002 | 4.41573044 | -0.186045042 | 0.37640199 | -0.4942722 | 0.62111395 | 0.72011642 | LOC101928471 | 101928471 |
| ENSG000001 | 10.1308722 | -0.169287495 | 0.34251931 | -0.4942422 | 0.62113514 | 0.7201204 | C1orf127 | 148345 |
| ENSG000001 | 2.52409769 | -0.182531904 | 0.36935654 | -0.4941889 | 0.62117277 | 0.72014344 | OR52D1 | 390066 |
| ENSG000002 | 2.48064102 | -0.183695453 | 0.37175761 | -0.4941269 | 0.62121653 | 0.72017358 | NA | NA |
| ENSG000002 | 4.20550985 | -0.186703032 | 0.37795097 | -0.4939874 | 0.62131505 | 0.72022601 | NA | NA |
| ENSG000002 | 2.43920717 | -0.184346974 | 0.37317507 | -0.493996 | 0.62130904 | 0.72022601 | NA | NA |
| ENSG000002 | 3.85786294 | 0.186839511 | 0.37822347 | 0.49399238 | 0.62131156 | 0.72022601 | NA | NA |
| ENSG000002 | 2.24383384 | -0.178620948 | 0.36171699 | -0.4938141 | 0.62143748 | 0.72034734 | NA | NA |
| ENSG000002 | 7.45968304 | 0.177153991 | 0.35882258 | 0.49370914 | 0.62151161 | 0.72040571 | SNORD71 | 692111 |
| ENSG000002 | 3.47106597 | -0.186833077 | 0.37844018 | -0.4936925 | 0.62152336 | 0.72040571 | NA | NA |
| ENSG000002 | 55.6330102 | 0.106439663 | 0.21563174 | 0.49361779 | 0.62157613 | 0.72044629 | NA | NA |
| ENSG000002 | 2.4462384 | -0.182553149 | 0.36985517 | -0.4935801 | 0.62160277 | 0.72045656 | NA | NA |
| ENSG000001 | 3.94276564 | -0.186644189 | 0.37825345 | -0.4934368 | 0.62170395 | 0.72051206 | RTP3 | 83597 |
| ENSG000002 | 7.603138 | -0.174416679 | 0.35347185 | -0.4934387 | 0.62170267 | 0.72051206 | NA | NA |
| ENSG000002 | 3.4771005 | -0.186798696 | 0.37854958 | -0.493459 | 0.62168831 | 0.72051206 | NA | NA |
| ENSG000002 | 2.91569397 | -0.185025647 | 0.37505426 | -0.4933303 | 0.62177919 | 0.72056463 | C1GALT1P3 | 106481730 |
| ENSG000002 | 3.20368191 | -0.186487837 | 0.37802432 | -0.4933223 | 0.62178485 | 0.72056463 | NA | NA |
| ENSG000002 | 2.22254765 | -0.180834203 | 0.36660918 | -0.4932615 | 0.62182781 | 0.72059382 | NA | NA |
| ENSG000002 | 2.940421 | -0.185271885 | 0.3757266 | -0.4931029 | 0.62193987 | 0.72068859 | NA | NA |
| ENSG000002 | 4.66898296 | -0.185748188 | 0.37669822 | -0.4930955 | 0.62194513 | 0.72068859 | NA | NA |
| ENSG000002 | 2.50917078 | -0.182657913 | 0.3704942 | -0.4930115 | 0.62200444 | 0.72073671 | NA | NA |
| ENSG000002 | 7.12016734 | 0.177386547 | 0.3598294 | 0.49297402 | 0.62203095 | 0.72074683 | NA | NA |
| ENSG000002 | 3.71665976 | -0.186519477 | 0.37839248 | -0.492926 | 0.62206491 | 0.72075369 | NA | NA |
| ENSG000002 | 2.22864111 | -0.180336506 | 0.36585696 | -0.4929153 | 0.62207241 | 0.72075369 | OR4C16 | 219428 |
| ENSG000002 | 2.75202345 | -0.184163132 | 0.37367133 | -0.4928479 | 0.62212006 | 0.7207677 | NA | NA |
| ENSG000002 | 4.11478156 | 0.186023309 | 0.37742716 | 0.49287208 | 0.62210297 | 0.7207677 | NA | NA |

| | | | | | | | | |
|---|---|---|---|---|---|---|---|---|
| ENSG000002 | 4.73422448 | -0.184920985 | 0.37526696 | -0.4927718 | 0.62217382 | 0.7208094 | NA | NA |
| ENSG000001 | 46.8316983 | -0.107727194 | 0.21864113 | -0.4927124 | 0.62221583 | 0.72083747 | FAM110B | 90362 |
| ENSG000001 | 50.9164739 | 0.100713478 | 0.20444076 | 0.49262916 | 0.62227464 | 0.72088501 | LPPR5 | 163404 |
| ENSG000001 | 15.247198 | -0.149409207 | 0.30333136 | -0.492561 | 0.62232279 | 0.72088917 | PHF24 | 23349 |
| ENSG000002 | 18.1273114 | -0.140021443 | 0.28427944 | -0.4925486 | 0.62233157 | 0.72088917 | LINC00115 | 79854 |
| ENSG000002 | 25.4089024 | 0.125938692 | 0.25567446 | 0.4925744 | 0.62231334 | 0.72088917 | NA | NA |
| ENSG000002 | 3.23716354 | -0.185710232 | 0.37708807 | -0.492485 | 0.62237649 | 0.72090591 | NA | NA |
| ENSG000002 | 3.17785349 | -0.186241984 | 0.37817331 | -0.4924779 | 0.62238157 | 0.72090591 | RN7SL727P | 106481121 |
| ENSG000002 | 2.92623041 | -0.184567812 | 0.37490054 | -0.4923114 | 0.62249922 | 0.7209804 | NA | NA |
| ENSG000002 | 3.91190017 | 0.186276036 | 0.37835725 | 0.49232844 | 0.62248718 | 0.7209804 | NA | NA |
| ENSG000002 | 5.24794833 | 0.183031404 | 0.37175468 | 0.49234458 | 0.62247577 | 0.7209804 | NA | NA |
| ENSG000002 | 3.91523367 | -0.186235856 | 0.37832672 | -0.492262 | 0.62253416 | 0.72100028 | NA | NA |
| ENSG000001 | 75.7979465 | 0.090237162 | 0.18339559 | 0.49203562 | 0.62269416 | 0.72116499 | GLIPR2 | 152007 |
| ENSG000001 | 14.4214729 | 0.152174962 | 0.30931039 | 0.49198142 | 0.62273248 | 0.72116817 | UCMA | 221044 |
| ENSG000002 | 2.71333017 | -0.184517917 | 0.37504138 | -0.4919935 | 0.62272395 | 0.72116817 | NA | NA |
| ENSG000002 | 4.67361574 | -0.185133914 | 0.376359 | -0.4919078 | 0.62278455 | 0.72120788 | LOC101928137 | 101928137 |
| ENSG000002 | 3.4818433 | -0.185974929 | 0.37809196 | -0.4918775 | 0.62280594 | 0.72121205 | NA | NA |
| ENSG000002 | 2.75445081 | -0.183278631 | 0.37266148 | -0.49181 | 0.62285369 | 0.72124125 | NA | NA |
| ENSG000002 | 5.22392626 | 0.183065501 | 0.37224208 | 0.49179153 | 0.62286673 | 0.72124125 | NA | NA |
| ENSG000002 | 2.28149819 | -0.178799344 | 0.3636357 | -0.4916991 | 0.62293208 | 0.72129072 | NA | NA |
| ENSG000002 | 24.1392778 | 0.129204854 | 0.26280888 | 0.49163047 | 0.6229806 | 0.72129072 | NA | NA |
| ENSG000002 | 11.3624983 | 0.164041735 | 0.3336622 | 0.49164015 | 0.62297376 | 0.72129072 | WWC2-AS2 | 152641 |
| ENSG000002 | 3.44323399 | -0.186016233 | 0.37835704 | -0.4916421 | 0.6229724 | 0.72129072 | NA | NA |
| ENSG000002 | 3.71708845 | -0.185897855 | 0.37814583 | -0.4916036 | 0.62299959 | 0.72129212 | NA | NA |
| ENSG000002 | 5.50057859 | -0.183121551 | 0.37253333 | -0.4915575 | 0.6230322 | 0.72130928 | NA | NA |
| ENSG000001 | 167.758839 | 0.072549472 | 0.14760901 | 0.4914976 | 0.62307455 | 0.72133771 | ARNTL | 406 |
| ENSG000002 | 6.565954 | -0.177963974 | 0.3621373 | -0.4914268 | 0.62312461 | 0.72137218 | NA | NA |
| ENSG000002 | 7.37419179 | -0.178112664 | 0.36247242 | -0.4913827 | 0.62315582 | 0.72137218 | NA | NA |
| ENSG000002 | 24.5830213 | 0.137932627 | 0.28070459 | 0.49138003 | 0.62315769 | 0.72137218 | NA | NA |

| | | | | | | | | |
|---|---|---|---|---|---|---|---|---|
| ENSG000002 | 4.43305845 | -0.185068574 | 0.37665939 | -0.491342 | 0.62318459 | 0.72138272 | LINC01324 | 104355289 |
| ENSG000002 | 3.97062987 | -0.185922952 | 0.37845486 | -0.4912685 | 0.62323655 | 0.72142229 | NA | NA |
| ENSG000002 | 6.62395358 | -0.178193219 | 0.36279665 | -0.4911656 | 0.62330935 | 0.72148596 | NA | NA |
| ENSG000002 | 5.19100549 | 0.182100116 | 0.37077065 | 0.49113951 | 0.62332778 | 0.72148669 | CATIP-AS1 | 101928513 |
| ENSG000001 | 38.1517335 | -0.111102585 | 0.22622919 | -0.4911063 | 0.62335124 | 0.72149124 | SMAD9 | 4093 |
| ENSG000002 | 3.41379096 | -0.185893151 | 0.37857543 | -0.4910333 | 0.62340288 | 0.72149124 | NA | NA |
| ENSG000002 | 2.94707616 | -0.185277428 | 0.37728912 | -0.4910755 | 0.62337308 | 0.72149124 | RNU6-542P | 106480588 |
| ENSG000002 | 3.9511228 | -0.185574035 | 0.37791709 | -0.4910443 | 0.62339511 | 0.72149124 | C7orf77 | 154872 |
| ENSG000002 | 3.22718052 | -0.1851169 | 0.37724227 | -0.4907109 | 0.62363092 | 0.72173457 | HSPE1P6 | 106480254 |
| ENSG000002 | 14.0976511 | -0.155797052 | 0.31751523 | -0.4906758 | 0.62365574 | 0.72174269 | NA | NA |
| ENSG000002 | 4.52273487 | -0.185722689 | 0.37856585 | -0.4905955 | 0.6237126 | 0.72178789 | LOC101929331 | 101929331 |
| ENSG000001 | 209.721609 | -0.071768329 | 0.14630922 | -0.490525 | 0.62376245 | 0.72182499 | DLGAP1 | 9229 |
| ENSG000002 | 3.1644718 | -0.184956177 | 0.37710186 | -0.4904674 | 0.62380317 | 0.72185151 | NA | NA |
| ENSG000002 | 4.15187634 | -0.185319354 | 0.37789523 | -0.4903988 | 0.62385175 | 0.72188713 | NA | NA |
| ENSG000002 | 3.406171 | -0.18563675 | 0.37858437 | -0.4903445 | 0.62389017 | 0.72191098 | NA | NA |
| ENSG000002 | 4.16783419 | -0.185130756 | 0.37764298 | -0.4902269 | 0.62397334 | 0.72198661 | NA | NA |
| ENSG000002 | 4.97780251 | -0.183565294 | 0.37450342 | -0.4901565 | 0.62402314 | 0.72200304 | NA | NA |
| ENSG000002 | 4.17127936 | -0.184814858 | 0.37704401 | -0.4901679 | 0.62401511 | 0.72200304 | NA | NA |
| ENSG000002 | 2.43553331 | -0.182084392 | 0.37166911 | -0.4899099 | 0.62419763 | 0.72218431 | NA | NA |
| ENSG000002 | 2.74228316 | -0.18214834 | 0.37182724 | -0.4898736 | 0.62422336 | 0.72219348 | NA | NA |
| ENSG000001 | 4.17429979 | -0.185020952 | 0.3779377 | -0.4895541 | 0.62444946 | 0.72241351 | RNU6-228P | 106479668 |
| ENSG000002 | 2.21234392 | -0.17943639 | 0.36658456 | -0.4894816 | 0.62450079 | 0.72241351 | NA | NA |
| ENSG000002 | 3.16893494 | -0.185146006 | 0.37818748 | -0.4895614 | 0.62444428 | 0.72241351 | NA | NA |
| ENSG000002 | 5.61408483 | -0.181844596 | 0.37150642 | -0.489479 | 0.62450261 | 0.72241351 | NA | NA |
| ENSG000002 | 6.10180441 | 0.179766791 | 0.36723396 | 0.4895157 | 0.62447664 | 0.72241351 | NA | NA |
| ENSG000002 | 5.68228261 | -0.180191472 | 0.36827244 | -0.4892885 | 0.62463746 | 0.72254888 | NA | NA |
| ENSG000001 | 41.0676169 | -0.105399895 | 0.21551075 | -0.4890702 | 0.62479197 | 0.7226724 | ADARB2 | 105 |
| ENSG000002 | 4.99592032 | 0.1823281 | 0.37279677 | 0.48908176 | 0.62478381 | 0.7226724 | NA | NA |
| ENSG000002 | 22.1139217 | 0.133553449 | 0.27308073 | 0.48906215 | 0.6247977 | 0.7226724 | GVQW2 | 100507462 |

| | | | | | | | | |
|---|---|---|---|---|---|---|---|---|
| ENSG00000( | 2.96965573 | -0.183554137 | 0.37536082 | -0.4890072 | 0.62483661 | 0.72267618 | ITIH4 | 3700 |
| ENSG000001 | 5.15011115 | -0.181754603 | 0.37166422 | -0.4890291 | 0.62482113 | 0.72267618 | ANKRD23 | 200539 |
| ENSG000002 | 2.46658214 | -0.18175959 | 0.3717252 | -0.4889623 | 0.62486842 | 0.72269235 | NA | NA |
| ENSG000001 | 76.3178778 | 0.09295269 | 0.19017843 | 0.48876567 | 0.62500761 | 0.72269894 | SPAG1 | 6674 |
| ENSG000002 | 3.26097844 | 0.184363598 | 0.37721003 | 0.48875582 | 0.62501458 | 0.72269894 | NA | NA |
| ENSG000002 | 4.43037127 | 0.184984339 | 0.37847108 | 0.48876744 | 0.62500635 | 0.72269894 | NA | NA |
| ENSG000002 | 5.65077175 | -0.180133059 | 0.36857334 | -0.4887306 | 0.62503246 | 0.72269894 | YWHAQP6 | 101290504 |
| ENSG000002 | 2.20669428 | -0.179373536 | 0.36702802 | -0.4887189 | 0.62504071 | 0.72269894 | NA | NA |
| ENSG000002 | 10.9238425 | -0.15934239 | 0.32598671 | -0.4888003 | 0.62498312 | 0.72269894 | PPT2-EGFL8 | 100532746 |
| ENSG000002 | 3.18350976 | -0.184691272 | 0.37786101 | -0.488781 | 0.62499677 | 0.72269894 | NA | NA |
| ENSG000002 | 2.20379551 | 0.1804106 | 0.36914731 | 0.48872251 | 0.62503816 | 0.72269894 | NA | NA |
| ENSG000002 | 3.23786909 | 0.184913914 | 0.37829674 | 0.48880652 | 0.62497868 | 0.72269894 | NA | NA |
| ENSG000002 | 2.72536064 | -0.183264008 | 0.37500117 | -0.4887025 | 0.62505233 | 0.72269894 | NA | NA |
| ENSG000002 | 6.96114423 | -0.178322415 | 0.36493382 | -0.4886432 | 0.62509435 | 0.72272691 | RNF208 | 727800 |
| ENSG000002 | 4.42424461 | -0.183852337 | 0.37648673 | -0.4883368 | 0.62531127 | 0.7229571 | LOC440300 | 440300 |
| ENSG000001 | 4.08811045 | 0.18432511 | 0.37753962 | 0.48822719 | 0.62538892 | 0.72298587 | ARL6IP4 | 51329 |
| ENSG000002 | 41.375838 | -0.107529535 | 0.22023157 | -0.4882567 | 0.62536804 | 0.72298587 | LOC643623 | 643623 |
| ENSG000002 | 4.21041726 | -0.184599786 | 0.378103 | -0.4882262 | 0.62538964 | 0.72298587 | NA | NA |
| ENSG000002 | 111.987163 | 0.077437127 | 0.15863927 | 0.48813339 | 0.62545536 | 0.72304123 | NA | NA |
| ENSG000002 | 2.20997474 | -0.175506076 | 0.35959622 | -0.4880643 | 0.6255043 | 0.72307719 | NA | NA |
| ENSG000001 | 7.80298013 | 0.176567999 | 0.36181807 | 0.48800216 | 0.62554831 | 0.72307899 | FAM110D | 79927 |
| ENSG000002 | 5.59127887 | 0.182580165 | 0.37417705 | 0.48795126 | 0.62558436 | 0.72307899 | NA | NA |
| ENSG000002 | 4.50379948 | -0.183961769 | 0.3770201 | -0.4879362 | 0.62559501 | 0.72307899 | NA | NA |
| ENSG000002 | 4.4769204 | -0.183789088 | 0.37662851 | -0.4879851 | 0.62556042 | 0.72307899 | NA | NA |
| ENSG000002 | 6.13674547 | -0.180313224 | 0.36947763 | -0.488022 | 0.62553424 | 0.72307899 | NA | NA |
| ENSG000001 | 56.3401889 | 0.10016497 | 0.20530731 | 0.48787825 | 0.62563608 | 0.72308929 | MAP3K6 | 9064 |
| ENSG000002 | 2.48096109 | -0.180726177 | 0.3704367 | -0.4878733 | 0.62563959 | 0.72308929 | NA | NA |
| ENSG000002 | 2.46362567 | -0.180124215 | 0.36923074 | -0.4878365 | 0.62566569 | 0.72309885 | NA | NA |
| ENSG000002 | 2.96415546 | -0.183673581 | 0.37652989 | -0.4878061 | 0.62568719 | 0.72310309 | NA | NA |

| | | | | | | | | |
|---|---|---|---|---|---|---|---|---|
| ENSG00000C | 7.10159527 | -0.17591777 | 0.36068641 | -0.4877305 | 0.62574073 | 0.72314435 | THPO | 7066 |
| ENSG000002 | 11.1628599 | -0.161202483 | 0.33067026 | -0.4875022 | 0.62590248 | 0.72327787 | NA | NA |
| ENSG000002 | 3.88996565 | -0.184433945 | 0.37833231 | -0.4874919 | 0.62590977 | 0.72327787 | NA | NA |
| ENSG000002 | 3.24540915 | -0.183880846 | 0.37716735 | -0.4875312 | 0.62588195 | 0.72327787 | NA | NA |
| ENSG000001 | 31.7649309 | 0.120439475 | 0.24716904 | 0.48727574 | 0.62606294 | 0.72336266 | ZNF157 | 7712 |
| ENSG000001 | 70.0854075 | 0.094344993 | 0.19362955 | 0.4872448 | 0.62608486 | 0.72336266 | ZNF311 | 282890 |
| ENSG000002 | 4.71347078 | -0.183164544 | 0.37594414 | -0.4872121 | 0.62610801 | 0.72336266 | NA | NA |
| ENSG000002 | 2.95403103 | -0.1835241 | 0.37667528 | -0.487221 | 0.62610174 | 0.72336266 | NA | NA |
| ENSG000002 | 12.5880184 | -0.156657369 | 0.3215305 | -0.487224 | 0.62609962 | 0.72336266 | NA | NA |
| ENSG000002 | 2.6953691 | -0.182510952 | 0.37451368 | -0.4873279 | 0.626026 | 0.72336266 | NA | NA |
| ENSG000002 | 6.31699614 | 0.176815908 | 0.36282998 | 0.48732442 | 0.62602845 | 0.72336266 | NA | NA |
| ENSG000002 | 2.43618511 | -0.178293805 | 0.36601241 | -0.487125 | 0.62616973 | 0.72341336 | NA | NA |
| ENSG000001 | 93.4169271 | 0.080138201 | 0.16452871 | 0.48707732 | 0.62620354 | 0.72343181 | TCEANC | 170082 |
| ENSG000002 | 2.20784908 | -0.178530332 | 0.36655497 | -0.4870493 | 0.62622341 | 0.72343415 | NA | NA |
| ENSG000002 | 6.44954944 | -0.179580879 | 0.36876144 | -0.4869839 | 0.62626975 | 0.72346708 | NA | NA |
| ENSG000002 | 3.49446707 | -0.18339002 | 0.37663071 | -0.4869226 | 0.62631316 | 0.7234966 | NA | NA |
| ENSG000002 | 19.2587098 | 0.139924386 | 0.28744668 | 0.48678379 | 0.62641156 | 0.72358966 | MAP2K1P1 | 29778 |
| ENSG000001 | 3.95030106 | -0.1841754 | 0.37845094 | -0.4866559 | 0.62650217 | 0.72366909 | KISS1R | 84634 |
| ENSG000002 | 5.39356299 | -0.182087726 | 0.37417612 | -0.4866364 | 0.62651601 | 0.72366909 | LOC101927424 | 101927424 |
| ENSG000002 | 2.92177534 | -0.183238345 | 0.37661591 | -0.486539 | 0.62658503 | 0.72370891 | NA | NA |
| ENSG000002 | 4.42067533 | -0.18357633 | 0.37731184 | -0.4865374 | 0.62658618 | 0.72370891 | NA | NA |
| ENSG000002 | 18.7835023 | 0.138101105 | 0.2839869 | 0.48629393 | 0.62675878 | 0.72388765 | NA | NA |
| ENSG000001 | 10.2484198 | -0.164941429 | 0.33928839 | -0.4861393 | 0.62686839 | 0.7239613 | PAGE4 | 9506 |
| ENSG000001 | 2.26194558 | -0.173423002 | 0.35673097 | -0.4861451 | 0.62686431 | 0.7239613 | KRTAP19-6 | 337973 |
| ENSG000002 | 6.46886139 | -0.179083301 | 0.36846956 | -0.4860193 | 0.62695351 | 0.7239613 | NA | NA |
| ENSG000002 | 3.45356247 | -0.183960253 | 0.37847997 | -0.4860502 | 0.6269316 | 0.7239613 | NA | NA |
| ENSG000002 | 2.45412635 | -0.178662529 | 0.36759693 | -0.4860283 | 0.62694707 | 0.7239613 | NA | NA |
| ENSG000002 | 4.16889202 | -0.183321853 | 0.37712055 | -0.4861094 | 0.62688959 | 0.7239613 | NA | NA |
| ENSG000002 | 3.92049342 | 0.183997333 | 0.37859337 | 0.48600252 | 0.62696538 | 0.7239613 | NA | NA |

| | | | | | | | | |
|---|---|---|---|---|---|---|---|---|
| ENSG000002 | 13.5845453 | -0.153774173 | 0.31635921 | -0.4860746 | 0.62691428 | 0.7239613 | NA | NA |
| ENSG000002 | 312.497595 | -0.063061101 | 0.12977199 | -0.4859377 | 0.62701135 | 0.72397679 | MBD5 | 55777 |
| ENSG000002 | 4.94668897 | -0.181840795 | 0.37420942 | -0.4859332 | 0.6270145 | 0.72397679 | NA | NA |
| ENSG000002 | 4.17097193 | -0.183479513 | 0.37764043 | -0.4858577 | 0.62706806 | 0.72401802 | NA | NA |
| ENSG000002 | 2.45571343 | -0.180878822 | 0.3723725 | -0.485747 | 0.62714656 | 0.72408804 | NA | NA |
| ENSG000002 | 18.0117825 | 0.142320013 | 0.29301874 | 0.48570277 | 0.62717792 | 0.72410363 | NKILA | 105416157 |
| ENSG000001 | 3.27073725 | 0.183688092 | 0.37830717 | 0.48555277 | 0.62728429 | 0.7241852 | NA | NA |
| ENSG000002 | 4.16902786 | -0.183367465 | 0.37764159 | -0.4855595 | 0.62727951 | 0.7241852 | NA | NA |
| ENSG000002 | 4.91780732 | -0.181055505 | 0.37311 | -0.4852604 | 0.62749164 | 0.72436272 | NA | NA |
| ENSG000002 | 7.18968413 | -0.176034615 | 0.36275101 | -0.4852767 | 0.62748008 | 0.72436272 | NA | NA |
| ENSG000002 | 2.93097485 | -0.183101811 | 0.37731753 | -0.4852725 | 0.62748308 | 0.72436272 | NA | NA |
| ENSG000002 | 2.19963473 | -0.176381149 | 0.36360669 | -0.4850877 | 0.62761411 | 0.72438028 | NA | NA |
| ENSG000002 | 4.64946926 | -0.182875136 | 0.37699199 | -0.4850902 | 0.62761233 | 0.72438028 | NA | NA |
| ENSG000002 | 2.22363774 | -0.178011277 | 0.36687452 | -0.4852103 | 0.62752721 | 0.72438028 | NA | NA |
| ENSG000002 | 7.30503663 | -0.177293403 | 0.36550151 | -0.4850689 | 0.6276275 | 0.72438028 | LOC101928694 | 101928694 |
| ENSG000002 | 5.91427323 | -0.179058413 | 0.36905504 | -0.4851808 | 0.62754811 | 0.72438028 | NA | NA |
| ENSG000002 | 3.21504044 | -0.183398812 | 0.37809304 | -0.4850627 | 0.6276319 | 0.72438028 | MIR4292 | 100422860 |
| ENSG000002 | 5.21457075 | 0.179995837 | 0.37100097 | 0.48516272 | 0.62756093 | 0.72438028 | NA | NA |
| ENSG000002 | 3.19946727 | -0.183009539 | 0.37731669 | -0.485029 | 0.62765577 | 0.72438721 | NA | NA |
| ENSG000002 | 4.17018983 | -0.18315339 | 0.37763846 | -0.4849967 | 0.62767872 | 0.72439308 | LOC101927348 | 101927348 |
| ENSG000002 | 2.76523575 | -0.180411142 | 0.37201881 | -0.4849517 | 0.62771063 | 0.72440929 | NA | NA |
| ENSG000002 | 2.94084576 | -0.182471081 | 0.37634306 | -0.4848531 | 0.62778059 | 0.72446941 | NA | NA |
| ENSG000001 | 12.276467 | -0.157652362 | 0.3251907 | -0.4847997 | 0.62781842 | 0.72449245 | SPATA22 | 84690 |
| ENSG000002 | 62.7545856 | 0.096652386 | 0.19939781 | 0.48472139 | 0.62787399 | 0.72453596 | NA | NA |
| ENSG000002 | 2.70290528 | -0.181707344 | 0.37503148 | -0.4845122 | 0.62802238 | 0.72468657 | NA | NA |
| ENSG00000C | 4.96105548 | -0.181059891 | 0.37378052 | -0.4844016 | 0.62810087 | 0.72471527 | TYMP | 1890 |
| ENSG000002 | 2.19013651 | -0.177530323 | 0.36648425 | -0.4844146 | 0.62809165 | 0.72471527 | NA | NA |
| ENSG000002 | 2.66325797 | -0.181654161 | 0.37499572 | -0.4844166 | 0.62809022 | 0.72471527 | NA | NA |
| ENSG000002 | 5.18098499 | 0.179986935 | 0.37167279 | 0.4842618 | 0.62820008 | 0.72477859 | NA | NA |

| | | | | | | | | |
|---|---|---|---|---|---|---|---|---|
| ENSG000002 | 2.44780532 | -0.179097483 | 0.3698653 | -0.4842235 | 0.62822724 | 0.72477859 | NA | NA |
| ENSG000002 | 4.43947232 | -0.182469209 | 0.3768165 | -0.4842389 | 0.62821633 | 0.72477859 | LOC101929288 | 101929288 |
| ENSG000002 | 14.2173604 | -0.149879086 | 0.30949801 | -0.4842651 | 0.62819774 | 0.72477859 | NA | NA |
| ENSG000001 | 3.17125095 | -0.182778303 | 0.3775942 | -0.4840602 | 0.62834318 | 0.72489173 | OR52K2 | 119774 |
| ENSG000002 | 5.8209131 | -0.179087295 | 0.37008741 | -0.4839054 | 0.62845301 | 0.72496525 | NA | NA |
| ENSG000002 | 2.21571463 | 0.178328854 | 0.36850228 | 0.48392877 | 0.62843642 | 0.72496525 | RN7SL47P | 106479242 |
| ENSG000002 | 2.19503574 | -0.176038791 | 0.3637956 | -0.4838948 | 0.62846054 | 0.72496525 | NA | NA |
| ENSG000002 | 5.742558 | -0.180198029 | 0.37244273 | -0.4838275 | 0.62850831 | 0.72499973 | NA | NA |
| ENSG000002 | 2.45332498 | -0.178721709 | 0.3694254 | -0.483783 | 0.62853991 | 0.7250061 | NA | NA |
| ENSG000002 | 2.96856598 | -0.181528177 | 0.3752566 | -0.4837441 | 0.62856747 | 0.7250061 | NA | NA |
| ENSG000002 | 3.44072265 | -0.183150039 | 0.37859082 | -0.4837678 | 0.62855069 | 0.7250061 | NA | NA |
| ENSG000002 | 3.95244047 | -0.183077113 | 0.37855657 | -0.4836189 | 0.62865636 | 0.725088 | NA | NA |
| ENSG000002 | 3.06229099 | 0.182425922 | 0.37726758 | 0.48354519 | 0.62870868 | 0.72512773 | NA | NA |
| ENSG000002 | 2.47161832 | -0.179211453 | 0.37074973 | -0.4833758 | 0.62882891 | 0.72524577 | NA | NA |
| ENSG000002 | 5.70485456 | -0.180279418 | 0.37299906 | -0.4833241 | 0.62886566 | 0.72526753 | NA | NA |
| ENSG000001 | 4.421825 | -0.181723292 | 0.37621753 | -0.4830272 | 0.62907644 | 0.72542808 | TAAR2 | 9287 |
| ENSG000002 | 4.17811059 | -0.182359433 | 0.37750024 | -0.483071 | 0.62904529 | 0.72542808 | NA | NA |
| ENSG000002 | 3.26900279 | 0.182436913 | 0.37768072 | 0.48304534 | 0.62906354 | 0.72542808 | RNU6-667P | 106479862 |
| ENSG000002 | 2.67059718 | -0.18038359 | 0.37338881 | -0.4830985 | 0.62902577 | 0.72542808 | NA | NA |
| ENSG000002 | 17.0124187 | 0.142888576 | 0.29585689 | 0.48296518 | 0.62912046 | 0.72545822 | NA | NA |
| ENSG000002 | 6.44393359 | -0.176811315 | 0.36612405 | -0.4829273 | 0.62914733 | 0.72546857 | NA | NA |
| ENSG000001 | 36.1364547 | 0.111799876 | 0.23155965 | 0.48281243 | 0.62922892 | 0.72554203 | SPSB1 | 80176 |
| ENSG000002 | 3.94794617 | -0.182559836 | 0.37815156 | -0.482769 | 0.62925979 | 0.72555699 | LINC01276 | 103106903 |
| ENSG000001 | 14.97377 | -0.159298452 | 0.33005774 | -0.4826381 | 0.62935269 | 0.72564348 | GGN | 199720 |
| ENSG000001 | 7.5674357 | -0.175893309 | 0.36456276 | -0.4824774 | 0.62946682 | 0.72569857 | B3GNT8 | 374907 |
| ENSG000001 | 3.24518844 | -0.181768146 | 0.37673705 | -0.4824801 | 0.6294649 | 0.72569857 | NA | NA |
| ENSG000002 | 5.23255088 | 0.179070563 | 0.37112509 | 0.48250729 | 0.62944562 | 0.72569857 | NA | NA |
| ENSG000002 | 3.17953588 | -0.181921651 | 0.37708278 | -0.4824449 | 0.62948996 | 0.72569857 | NA | NA |
| ENSG000002 | 3.67003776 | -0.182618974 | 0.37852651 | -0.482447 | 0.62948845 | 0.72569857 | NA | NA |

| | | | | | | | | |
|---|---|---|---|---|---|---|---|---|
| ENSG000001 | 10234.372 | -0.05957764 | 0.12351129 | -0.482366 | 0.629546 | 0.72570129 | ND5 | 4540 |
| ENSG000002 | 3.47095497 | -0.18253271 | 0.37840685 | -0.4823716 | 0.62954201 | 0.72570129 | NA | NA |
| ENSG000002 | 4.18911624 | -0.182066785 | 0.37740779 | -0.482414 | 0.62951191 | 0.72570129 | OOSP1P2 | 106480252 |
| ENSG000001 | 10.53746 | -0.169348143 | 0.35112377 | -0.4823033 | 0.62959052 | 0.72573198 | DYDC1 | 143241 |
| ENSG000002 | 3.14842255 | -0.18169605 | 0.37695411 | -0.4820111 | 0.62979809 | 0.72595061 | NA | NA |
| ENSG000002 | 22.5036351 | 0.129707896 | 0.26911817 | 0.48197376 | 0.62982458 | 0.72596051 | NA | NA |
| ENSG000001 | 53.4178865 | 0.09568496 | 0.19858248 | 0.48183989 | 0.62991969 | 0.72596696 | NA | NA |
| ENSG000001 | 4.67479787 | -0.180545247 | 0.3746483 | -0.481906 | 0.62987276 | 0.72596696 | DNAJB8 | 165721 |
| ENSG000002 | 5.61521568 | -0.178066062 | 0.36952222 | -0.4818819 | 0.62988986 | 0.72596696 | NA | NA |
| ENSG000002 | 6.39958272 | -0.176981169 | 0.36727053 | -0.4818823 | 0.62988956 | 0.72596696 | NA | NA |
| ENSG000002 | 2.4120796 | -0.178165573 | 0.36975562 | -0.4818468 | 0.62991476 | 0.72596696 | NA | NA |
| ENSG000002 | 11.6215916 | -0.154625247 | 0.32093136 | -0.4818016 | 0.62994688 | 0.72597766 | LOC105376736 | 105376736 |
| ENSG000002 | 2.22159495 | -0.175278983 | 0.36389126 | -0.4816796 | 0.63003359 | 0.72605696 | NA | NA |
| ENSG000002 | 2.17983994 | -0.174362064 | 0.36205751 | -0.4815867 | 0.63009961 | 0.7261124 | RN7SKP134 | 106479154 |
| ENSG000002 | 3.16798203 | -0.181779393 | 0.37748153 | -0.4815584 | 0.6301197 | 0.72611492 | NA | NA |
| ENSG000002 | 2.26963113 | -0.173829419 | 0.36110479 | -0.4813822 | 0.63024489 | 0.72623854 | NA | NA |
| ENSG000002 | 3.02739911 | 0.181687106 | 0.377528 | 0.48125466 | 0.63033552 | 0.72632234 | CHKB-CPT1B | 386593 |
| ENSG000002 | 10.0536903 | 0.162617731 | 0.33793637 | 0.48120814 | 0.63036858 | 0.72633979 | NA | NA |
| ENSG000002 | 8.07047422 | -0.170111468 | 0.35355244 | -0.4811492 | 0.63041047 | 0.72636742 | NA | NA |
| ENSG000002 | 64.7272911 | 0.091860228 | 0.19096201 | 0.48103929 | 0.63048858 | 0.72643678 | TTC23L | 153657 |
| ENSG000002 | 14.5766505 | -0.147718092 | 0.30712909 | -0.4809642 | 0.63054196 | 0.72644149 | NA | NA |
| ENSG000002 | 2.90839419 | -0.180866713 | 0.37602626 | -0.4809949 | 0.63052013 | 0.72644149 | NA | NA |
| ENSG000002 | 2.95007697 | -0.180926443 | 0.37617937 | -0.4809579 | 0.63054641 | 0.72644149 | NA | NA |
| ENSG000002 | 16.1337197 | 0.143276735 | 0.29801276 | 0.48077383 | 0.63067725 | 0.72657158 | NA | NA |
| ENSG000001 | 77.5435748 | 0.099113673 | 0.20616727 | 0.48074398 | 0.63069847 | 0.72657539 | ARHGEF17 | 9828 |
| ENSG000002 | 2.18934722 | -0.175651324 | 0.36563805 | -0.4803967 | 0.63094532 | 0.72683911 | NA | NA |
| ENSG000002 | 6.40950625 | -0.175386736 | 0.36515788 | -0.4803038 | 0.63101135 | 0.72689453 | NA | NA |
| ENSG000001 | 148.538733 | -0.070888227 | 0.147606 | -0.480253 | 0.6310475 | 0.72691552 | CMYA5 | 202333 |
| ENSG000002 | 3.19223244 | -0.181118732 | 0.37718199 | -0.4801892 | 0.63109284 | 0.7269471 | NA | NA |

| | | | | | | | | |
|---|---|---|---|---|---|---|---|---|
| ENSG000002 | 2.83151345 | 0.17806345 | 0.37092446 | 0.48005314 | 0.63118961 | 0.72703792 | NA | NA |
| ENSG000002 | 3.25001575 | -0.181077648 | 0.3773047 | -0.4799242 | 0.63128131 | 0.72712289 | NA | NA |
| ENSG000002 | 9.77402252 | 0.16465354 | 0.34311194 | 0.47988286 | 0.63131069 | 0.72713607 | NA | NA |
| ENSG000002 | 3.700402 | -0.181630024 | 0.37857886 | -0.479768 | 0.63139237 | 0.72718884 | NA | NA |
| ENSG000002 | 2.19600184 | -0.17583967 | 0.36650623 | -0.4797727 | 0.63138906 | 0.72718884 | NA | NA |
| ENSG000002 | 2.53244768 | -0.175593746 | 0.3660404 | -0.4797114 | 0.63143263 | 0.72721456 | NA | NA |
| ENSG000001 | 7.16210672 | -0.173703402 | 0.36214978 | -0.4796452 | 0.63147971 | 0.72724812 | SPNS1 | 83985 |
| ENSG000002 | 2.18888787 | 0.176330169 | 0.36771283 | 0.47953228 | 0.63156001 | 0.72731995 | NA | NA |
| ENSG000001 | 2.23530892 | -0.175562271 | 0.3661641 | -0.4794634 | 0.63160903 | 0.72733508 | UCN2 | 90226 |
| ENSG000002 | 4.48733211 | -0.18144923 | 0.37843402 | -0.4794739 | 0.63160154 | 0.72733508 | NA | NA |
| ENSG000002 | 3.88666448 | -0.181376777 | 0.37837228 | -0.4793606 | 0.63168209 | 0.7273366 | NA | NA |
| ENSG000002 | 2.93360297 | -0.179861486 | 0.37520986 | -0.4793624 | 0.63168086 | 0.7273366 | NA | NA |
| ENSG000002 | 5.62669795 | -0.176643644 | 0.36849255 | -0.4793683 | 0.63167664 | 0.7273366 | NA | NA |
| ENSG000002 | 5.18628176 | 0.177938996 | 0.37117197 | 0.47939772 | 0.63165572 | 0.7273366 | LOC100288846 | 100288846 |
| ENSG000002 | 11.6759303 | -0.154987961 | 0.32336568 | -0.4792963 | 0.63172788 | 0.7273443 | NA | NA |
| ENSG000002 | 2.46422704 | -0.177185392 | 0.36969417 | -0.4792756 | 0.63174259 | 0.7273443 | NA | NA |
| ENSG000002 | 10.0046662 | -0.161552304 | 0.33707591 | -0.4792757 | 0.63174248 | 0.7273443 | NA | NA |
| ENSG000002 | 12.6824625 | 0.156167901 | 0.32585944 | 0.47924928 | 0.6317613 | 0.7273452 | NA | NA |
| ENSG000002 | 2.67479256 | -0.178652204 | 0.37296944 | -0.4789996 | 0.63193893 | 0.72752904 | RN7SKP253 | 106479208 |
| ENSG000002 | 4.65252986 | -0.179871259 | 0.37553925 | -0.4789679 | 0.63196144 | 0.72753431 | NA | NA |
| ENSG000002 | 3.19189993 | -0.181016071 | 0.37800155 | -0.4788765 | 0.63202646 | 0.72758851 | NA | NA |
| ENSG000002 | 7.36221887 | 0.171676711 | 0.35851782 | 0.47885126 | 0.63204445 | 0.72758855 | NA | NA |
| ENSG000001 | 4.74469621 | -0.180438339 | 0.37695095 | -0.4786786 | 0.63216732 | 0.7276274 | TMEM121 | 80757 |
| ENSG000002 | 2.44558551 | -0.175029887 | 0.3655844 | -0.4787674 | 0.63210412 | 0.7276274 | LRRC3C | 100505591 |
| ENSG000002 | 8.87306271 | -0.169061676 | 0.35314746 | -0.4787283 | 0.63213194 | 0.7276274 | NA | NA |
| ENSG000002 | 3.89144007 | -0.181190599 | 0.3785125 | -0.4786912 | 0.63215833 | 0.7276274 | NA | NA |
| ENSG000002 | 7.87518073 | -0.170164253 | 0.35548813 | -0.4786777 | 0.63216791 | 0.7276274 | NA | NA |
| ENSG000002 | 2.418154 | -0.177659588 | 0.37125745 | -0.4785347 | 0.63226966 | 0.72772385 | NA | NA |
| ENSG000002 | 3.46567727 | -0.180924366 | 0.37814711 | -0.4784497 | 0.63233018 | 0.72777286 | NA | NA |

| | | | | | | | | |
|---|---|---|---|---|---|---|---|---|
| ENSG000002 | 19.0663609 | 0.13612742 | 0.28455372 | 0.47838918 | 0.63237324 | 0.72780176 | NA | NA |
| ENSG000002 | 8.41720363 | -0.169345459 | 0.35404233 | -0.4783198 | 0.6324226 | 0.72782049 | NA | NA |
| ENSG000002 | 2.17907137 | -0.175271471 | 0.36643458 | -0.4783159 | 0.63242541 | 0.72782049 | NA | NA |
| ENSG000002 | 4.87611234 | -0.178692045 | 0.37362099 | -0.4782709 | 0.63245741 | 0.72783666 | NA | NA |
| ENSG000002 | 3.28062135 | 0.18089733 | 0.37825308 | 0.47824417 | 0.63247643 | 0.72783789 | NA | NA |
| ENSG000002 | 5.76594226 | 0.17822212 | 0.37268576 | 0.47821016 | 0.63250063 | 0.72784509 | NA | NA |
| ENSG000002 | 3.40680407 | -0.180907686 | 0.37841492 | -0.478067 | 0.63260252 | 0.72794167 | NA | NA |
| ENSG000002 | 2.95192304 | 0.179043552 | 0.37456359 | 0.47800575 | 0.63264611 | 0.72797118 | NA | NA |
| ENSG000002 | 3.27040443 | 0.180821708 | 0.37835938 | 0.47790994 | 0.6327143 | 0.72802413 | NA | NA |
| ENSG000002 | 11.2154901 | 0.157044457 | 0.32862006 | 0.47789066 | 0.63272803 | 0.72802413 | NA | NA |
| ENSG00000C | 50.0601245 | -0.09931966 | 0.20786528 | -0.4778078 | 0.63278698 | 0.72805382 | SLC17A6 | 57084 |
| ENSG000001 | 138.780332 | 0.077488387 | 0.16217611 | 0.47780396 | 0.63278974 | 0.72805382 | NEDD9 | 4739 |
| ENSG000002 | 4.02541708 | -0.180563539 | 0.37800248 | -0.4776782 | 0.63287927 | 0.72813617 | NA | NA |
| ENSG000001 | 67.7151835 | 0.09214511 | 0.19292236 | 0.47762794 | 0.63291504 | 0.72814725 | PBLD | 64081 |
| ENSG000002 | 2.95251728 | -0.179957175 | 0.37682335 | -0.4775638 | 0.63296073 | 0.72814725 | NA | NA |
| ENSG000002 | 3.02687793 | 0.179544496 | 0.37594261 | 0.47758485 | 0.63294572 | 0.72814725 | RN7SL525P | 106480516 |
| ENSG000002 | 2.19422406 | 0.17568289 | 0.36784731 | 0.47759732 | 0.63293684 | 0.72814725 | GAGE12B | 729428 |
| ENSG000002 | 2.96142698 | -0.17864131 | 0.37408775 | -0.4775385 | 0.6329787 | 0.72814726 | NA | NA |
| ENSG000001 | 365.916288 | 0.064086051 | 0.13420845 | 0.47751131 | 0.63299807 | 0.72814889 | CHST9 | 83539 |
| ENSG000001 | 3.80862117 | -0.179193005 | 0.37537053 | -0.4773763 | 0.63309417 | 0.72818705 | NBPF6 | 653149 |
| ENSG000002 | 5.4230473 | -0.177421139 | 0.37166859 | -0.4773638 | 0.63310307 | 0.72818705 | NA | NA |
| ENSG000002 | 3.43340251 | -0.18052019 | 0.37813366 | -0.4773978 | 0.63307886 | 0.72818705 | NA | NA |
| ENSG000002 | 3.28255169 | 0.180716777 | 0.37851326 | 0.47743843 | 0.63304995 | 0.72818705 | NA | NA |
| ENSG000002 | 4.928416 | -0.178360367 | 0.37367392 | -0.4773155 | 0.63313745 | 0.72820594 | MIR2113 | 100302164 |
| ENSG000002 | 20.0654546 | 0.137122163 | 0.28738182 | 0.47714279 | 0.63326044 | 0.72832674 | NA | NA |
| ENSG000002 | 4.13867234 | -0.180051805 | 0.37743937 | -0.477035 | 0.63333717 | 0.72839433 | NA | NA |
| ENSG000002 | 2.1973757 | -0.169710224 | 0.35587811 | -0.4768774 | 0.63344943 | 0.72848212 | NA | NA |
| ENSG000002 | 2.23939903 | -0.173922306 | 0.36469336 | -0.4769001 | 0.63343325 | 0.72848212 | MIR3921 | 100500859 |
| ENSG000001 | 6.12968701 | -0.175761794 | 0.36862017 | -0.47681 | 0.6334974 | 0.7285109 | GOLGA8F | 100132565 |

| | | | | | | | | |
|---|---|---|---|---|---|---|---|---|
| ENSG000001 | 138.855841 | 0.074042606 | 0.15529582 | 0.47678428 | 0.63351574 | 0.7285109 | ANKRD36B | 57730 |
| ENSG000002 | 2.91167575 | -0.179311382 | 0.3760989 | -0.4767666 | 0.63352835 | 0.7285109 | RN7SL551P | 106479422 |
| ENSG000001 | 2.45404313 | -0.176882439 | 0.37104566 | -0.4767134 | 0.63356622 | 0.72851312 | OR5AP2 | 338675 |
| ENSG000001 | 140.574398 | -0.07453016 | 0.15634068 | -0.4767164 | 0.63356412 | 0.72851312 | NLGN3 | 54413 |
| ENSG000001 | 26.8880338 | 0.124719983 | 0.26168209 | 0.47660879 | 0.63364072 | 0.72857813 | C10orf10 | 11067 |
| ENSG000002 | 3.70926793 | -0.18040708 | 0.37857603 | -0.4765412 | 0.63368886 | 0.72861282 | NA | NA |
| ENSG000001 | 2.72520833 | -0.176037064 | 0.36943895 | -0.4764984 | 0.63371936 | 0.72862723 | CST5 | 1473 |
| ENSG000002 | 26.9483578 | 0.122078036 | 0.25623906 | 0.47642243 | 0.63377346 | 0.72866877 | NA | NA |
| ENSG000002 | 8.00622417 | -0.167912895 | 0.35248922 | -0.4763632 | 0.63381564 | 0.72867595 | NA | NA |
| ENSG000002 | 8.26333979 | -0.168099199 | 0.35286263 | -0.4763871 | 0.63379864 | 0.72867595 | NA | NA |
| ENSG000002 | 7.36317721 | 0.170826889 | 0.35864784 | 0.47630815 | 0.63385486 | 0.72870038 | NA | NA |
| ENSG000002 | 5.34791594 | -0.176998336 | 0.37162593 | -0.4762809 | 0.63387427 | 0.72870203 | NA | NA |
| ENSG000002 | 3.01479568 | -0.178302254 | 0.37440106 | -0.4762333 | 0.63390817 | 0.72870546 | NA | NA |
| ENSG000002 | 7.1052553 | -0.175564315 | 0.36865736 | -0.4762263 | 0.63391319 | 0.72870546 | NA | NA |
| ENSG000002 | 2.69752866 | -0.177807406 | 0.37340654 | -0.4761765 | 0.63394867 | 0.72872558 | NA | NA |
| ENSG000002 | 7.295865 | -0.173107025 | 0.36357419 | -0.4761257 | 0.63398481 | 0.72874647 | NA | NA |
| ENSG000002 | 3.41636811 | -0.180096231 | 0.37829324 | -0.4760757 | 0.63402043 | 0.72876675 | NA | NA |
| ENSG000002 | 2.4557655 | -0.176644135 | 0.37108445 | -0.4760214 | 0.63405914 | 0.72879059 | MTRNR2L7 | 100288485 |
| ENSG000002 | 2.69198361 | -0.17759213 | 0.37316368 | -0.4759095 | 0.63413888 | 0.72886159 | NA | NA |
| ENSG000002 | 4.18126795 | -0.179605289 | 0.37757238 | -0.4756844 | 0.63429924 | 0.72902304 | NA | NA |
| ENSG000002 | 4.44180869 | -0.179057408 | 0.37643845 | -0.4756618 | 0.63431531 | 0.72902304 | NA | NA |
| ENSG000002 | 34.5100968 | -0.113514642 | 0.23866187 | -0.4756296 | 0.63433831 | 0.72902881 | NA | NA |
| ENSG000001 | 173.405041 | 0.076507626 | 0.16086758 | 0.4755938 | 0.63436379 | 0.72903744 | PHF2 | 5253 |
| ENSG000002 | 4.4052146 | -0.179207541 | 0.37688048 | -0.4755023 | 0.63442899 | 0.7290917 | NA | NA |
| ENSG000002 | 3.22703174 | 0.179839682 | 0.37825734 | 0.47544268 | 0.63447148 | 0.72911987 | RNU6-195P | 106479653 |
| ENSG000002 | 6.3248678 | -0.174534281 | 0.3671686 | -0.4753519 | 0.6345362 | 0.72917358 | NA | NA |
| ENSG000002 | 6.14285242 | -0.173902073 | 0.36593063 | -0.4752324 | 0.63462137 | 0.72923657 | NA | NA |
| ENSG000002 | 2.71944324 | -0.177240935 | 0.37296255 | -0.4752245 | 0.63462698 | 0.72923657 | NA | NA |
| ENSG000002 | 4.17195219 | 0.179577658 | 0.37794611 | 0.47514091 | 0.63468654 | 0.72928435 | NA | NA |

| | | | | | | | | |
|---|---|---|---|---|---|---|---|---|
| ENSG000002 | 2.71113815 | -0.176206751 | 0.37090223 | -0.475076 | 0.6347328 | 0.72930625 | NA | NA |
| ENSG000002 | 3.74192778 | -0.179830896 | 0.37856071 | -0.4750385 | 0.63475956 | 0.72930625 | NA | NA |
| ENSG000002 | 11.8082517 | -0.154649709 | 0.32554044 | -0.4750553 | 0.63474756 | 0.72930625 | NA | NA |
| ENSG000002 | 4.16855853 | 0.17919836 | 0.37731968 | 0.4749245 | 0.63484078 | 0.72937891 | NA | NA |
| ENSG000001 | 35.5916844 | -0.109796354 | 0.23120902 | -0.4748792 | 0.63487309 | 0.72939536 | ZNF396 | 252884 |
| ENSG000001 | 20.154879 | 0.136275859 | 0.28708125 | 0.47469439 | 0.63500482 | 0.72952603 | TMC4 | 147798 |
| ENSG000001 | 201.020961 | 0.073298595 | 0.15448331 | 0.47447583 | 0.63516063 | 0.72965389 | ZNF568 | 374900 |
| ENSG000002 | 5.24334084 | 0.176352086 | 0.37174742 | 0.47438685 | 0.63522407 | 0.72965389 | PSMB8-AS1 | 100507463 |
| ENSG000002 | 3.67558247 | -0.179543414 | 0.37844224 | -0.4744275 | 0.63519507 | 0.72965389 | NA | NA |
| ENSG000002 | 2.53013908 | -0.173143727 | 0.3649294 | -0.4744581 | 0.63517324 | 0.72965389 | NA | NA |
| ENSG000002 | 5.65283947 | -0.176234808 | 0.37143647 | -0.4744682 | 0.63516604 | 0.72965389 | NA | NA |
| ENSG000002 | 4.90375976 | -0.178180166 | 0.37559583 | -0.4743933 | 0.63521943 | 0.72965389 | NA | NA |
| ENSG000002 | 2.97399467 | -0.178507731 | 0.37634113 | -0.4743243 | 0.63526869 | 0.72968447 | NA | NA |
| ENSG000001 | 259.282845 | 0.069812294 | 0.14722578 | 0.47418525 | 0.63536781 | 0.72976684 | NA | NA |
| ENSG000002 | 3.96144276 | -0.179508786 | 0.37859208 | -0.4741483 | 0.63539416 | 0.72976684 | NA | NA |
| ENSG000002 | 2.97948752 | -0.178194474 | 0.37582038 | -0.474148 | 0.63539438 | 0.72976684 | NA | NA |
| ENSG000002 | 3.02178968 | 0.177923408 | 0.37539427 | 0.4739641 | 0.63552551 | 0.72983475 | NA | NA |
| ENSG000002 | 5.11908955 | -0.176482256 | 0.37233456 | -0.4739884 | 0.63550819 | 0.72983475 | NA | NA |
| ENSG000002 | 3.64841155 | -0.179448842 | 0.37857543 | -0.4740108 | 0.6354922 | 0.72983475 | NA | NA |
| ENSG000002 | 2.44985278 | -0.173711069 | 0.36647455 | -0.4740058 | 0.63549576 | 0.72983475 | NA | NA |
| ENSG000002 | 12.956169 | 0.152092171 | 0.32092054 | 0.4739247 | 0.63555361 | 0.72984635 | NA | NA |
| ENSG000001 | 85.4261092 | 0.088503006 | 0.18682122 | 0.47373102 | 0.63569173 | 0.7299111 | SEC14L5 | 9717 |
| ENSG000001 | 5.44129795 | 0.175466553 | 0.37039602 | 0.47372688 | 0.63569468 | 0.7299111 | NGF | 4803 |
| ENSG000002 | 4.20061857 | -0.178794565 | 0.37742713 | -0.4737194 | 0.63569999 | 0.7299111 | NA | NA |
| ENSG000002 | 2.19188878 | 0.17193358 | 0.36289788 | 0.47377952 | 0.63565714 | 0.7299111 | NA | NA |
| ENSG000002 | 3.69129443 | -0.179332448 | 0.37854176 | -0.4737455 | 0.63568141 | 0.7299111 | NA | NA |
| ENSG000002 | 11.1745718 | -0.154595678 | 0.32636639 | -0.4736875 | 0.63572277 | 0.72991659 | NA | NA |
| ENSG000002 | 19.5263639 | -0.135288012 | 0.28563958 | -0.4736319 | 0.63576244 | 0.72994147 | NA | NA |
| ENSG000001 | 11.0784973 | 0.166516042 | 0.35163389 | 0.47354947 | 0.63582122 | 0.72994443 | NA | NA |

| | | | | | | | | |
|---|---|---|---|---|---|---|---|---|
| ENSG000002 | 2.20374344 | 0.173582075 | 0.36657247 | 0.47352731 | 0.63583702 | 0.72994443 | RNU7-174P | 106480829 |
| ENSG000002 | 3.68565777 | -0.179289632 | 0.37858807 | -0.4735744 | 0.63580341 | 0.72994443 | NA | NA |
| ENSG000002 | 10.1067257 | 0.171957982 | 0.36314072 | 0.47352988 | 0.63583519 | 0.72994443 | NA | NA |
| ENSG000001 | 5.17027954 | -0.179007258 | 0.37811081 | -0.4734254 | 0.63590972 | 0.72996589 | ZP1 | 22917 |
| ENSG000002 | 2.42283083 | 0.175770855 | 0.37126502 | 0.47343769 | 0.63590094 | 0.72996589 | NA | NA |
| ENSG000002 | 3.21008913 | -0.179002574 | 0.37807413 | -0.4734589 | 0.6358858 | 0.72996589 | NA | NA |
| ENSG000002 | 3.42062218 | -0.179191136 | 0.37852963 | -0.4733873 | 0.63593686 | 0.72997638 | CSAG4 | 100130935 |
| ENSG000001 | 3.6641806 | -0.17917669 | 0.37857628 | -0.4732908 | 0.63600569 | 0.73003473 | FOXD4 | 2298 |
| ENSG000002 | 3.63690867 | -0.179146518 | 0.37854823 | -0.4732462 | 0.63603753 | 0.7300506 | NA | NA |
| ENSG000002 | 33.284195 | -0.111339599 | 0.23528882 | -0.4732039 | 0.63606768 | 0.73006455 | NA | NA |
| ENSG000001 | 6.37778045 | -0.172513734 | 0.36473369 | -0.4729855 | 0.63622355 | 0.73014012 | GJC2 | 57165 |
| ENSG000002 | 5.87099282 | -0.174767708 | 0.36946868 | -0.4730244 | 0.63619576 | 0.73014012 | NA | NA |
| ENSG000002 | 2.69223135 | -0.175963906 | 0.37200628 | -0.4730133 | 0.63620371 | 0.73014012 | NA | NA |
| ENSG000002 | 4.17064684 | -0.178432343 | 0.37724466 | -0.4729884 | 0.63622147 | 0.73014012 | NA | NA |
| ENSG000002 | 3.21188362 | -0.178823196 | 0.37799899 | -0.4730785 | 0.63615718 | 0.73014012 | NA | NA |
| ENSG000002 | 3.96373428 | -0.179051753 | 0.37859314 | -0.4729398 | 0.63625615 | 0.73015687 | LOC101927815 | 101927815 |
| ENSG000001 | 51.7286818 | -0.099540672 | 0.21056662 | -0.4727277 | 0.63640747 | 0.73027716 | CBX7 | 23492 |
| ENSG000002 | 2.96116227 | -0.178066707 | 0.37667184 | -0.472737 | 0.63640084 | 0.73027716 | BCKDHA | 593 |
| ENSG000002 | 2.70293754 | -0.176232645 | 0.37280782 | -0.4727171 | 0.636415 | 0.73027716 | NA | NA |
| ENSG000001 | 100.099411 | 0.080325418 | 0.16995255 | 0.47263438 | 0.63647405 | 0.73030359 | SLC1A2 | 6506 |
| ENSG000002 | 4.95966593 | -0.176915047 | 0.37429924 | -0.4726567 | 0.63645812 | 0.73030359 | NA | NA |
| ENSG000002 | 6.5770723 | -0.171044839 | 0.36199642 | -0.4725042 | 0.63656692 | 0.73036882 | NA | NA |
| ENSG000002 | 3.04350042 | 0.178371916 | 0.37749884 | 0.47250984 | 0.63656292 | 0.73036882 | NA | NA |
| ENSG000002 | 3.21876637 | -0.178502024 | 0.37785183 | -0.4724128 | 0.6366322 | 0.73042305 | NA | NA |
| ENSG000001 | 51.8171377 | 0.098011249 | 0.20750361 | 0.47233515 | 0.63668759 | 0.73046593 | PROSER2 | 254427 |
| ENSG000002 | 12.8145717 | 0.149154772 | 0.31592236 | 0.47212478 | 0.63683773 | 0.73061752 | NA | NA |
| ENSG000002 | 18.9783478 | -0.138613045 | 0.29361544 | -0.4720904 | 0.63686223 | 0.73062496 | NA | NA |
| ENSG000001 | 116.445322 | 0.077926437 | 0.16515081 | 0.47185015 | 0.63703375 | 0.73073903 | DAAM2 | 23500 |
| ENSG000002 | 2.41436391 | -0.175080157 | 0.37102295 | -0.471885 | 0.63700889 | 0.73073903 | NA | NA |

| | | | | | | | | |
|---|---|---|---|---|---|---|---|---|
| ENSG000002 | 3.69036168 | -0.178639792 | 0.37859259 | -0.4718523 | 0.6370322 | 0.73073903 | NA | NA |
| ENSG000002 | 4.94533903 | 0.177014184 | 0.37513775 | 0.4718645 | 0.63702351 | 0.73073903 | NA | NA |
| ENSG000001 | 226.100107 | 0.07014772 | 0.14868013 | 0.47180293 | 0.63706746 | 0.73075703 | CSMD2 | 114784 |
| ENSG000001 | 97.6076129 | 0.081614346 | 0.17302541 | 0.47168996 | 0.63714811 | 0.73082887 | FAM46A | 55603 |
| ENSG000002 | 4.14235785 | -0.178154496 | 0.3777436 | -0.4716281 | 0.63719227 | 0.73085885 | NA | NA |
| ENSG000002 | 2.46223633 | -0.174114097 | 0.36922471 | -0.4715668 | 0.63723606 | 0.73086194 | NA | NA |
| ENSG000002 | 3.21900768 | -0.178185878 | 0.37785266 | -0.471575 | 0.63723016 | 0.73086194 | NA | NA |
| ENSG000002 | 2.21822942 | -0.172220541 | 0.36522333 | -0.4715486 | 0.63724904 | 0.73086194 | NA | NA |
| ENSG000002 | 2.7146682 | -0.176201617 | 0.37372333 | -0.4714761 | 0.63730079 | 0.73090062 | NA | NA |
| ENSG000002 | 21.9162639 | 0.143917268 | 0.30527386 | 0.47143659 | 0.63732899 | 0.7309123 | NA | NA |
| ENSG000001 | 2.65799291 | -0.175142903 | 0.37154495 | -0.4713909 | 0.63736163 | 0.73092905 | IFNA6 | 3443 |
| ENSG000002 | 4.91651614 | -0.176884685 | 0.37527733 | -0.4713439 | 0.63739516 | 0.73094684 | NA | NA |
| ENSG000001 | 26.5920091 | -0.119871926 | 0.25433735 | -0.4713107 | 0.63741884 | 0.73095333 | UNC5CL | 222643 |
| ENSG000002 | 3.95421545 | -0.178368853 | 0.37849456 | -0.4712587 | 0.63745601 | 0.73097527 | NA | NA |
| ENSG000002 | 15.1533671 | 0.143505557 | 0.30457073 | 0.47117317 | 0.63751707 | 0.73102463 | NA | NA |
| ENSG000002 | 3.18220508 | -0.178054063 | 0.37791946 | -0.4711429 | 0.6375387 | 0.73102875 | NA | NA |
| ENSG000001 | 5.39734208 | -0.175220173 | 0.37205846 | -0.470948 | 0.6376779 | 0.73116769 | CD2 | 914 |
| ENSG000002 | 18.8188999 | -0.139022734 | 0.29526031 | -0.470848 | 0.63774927 | 0.73122884 | ZNF252P-AS1 | 286103 |
| ENSG000001 | 4.17199676 | -0.178110876 | 0.37829739 | -0.4708224 | 0.63776758 | 0.73122917 | KLK14 | 43847 |
| ENSG000002 | 3.93128249 | 0.178152276 | 0.37842852 | 0.47076863 | 0.63780597 | 0.7312525 | NA | NA |
| ENSG000001 | 6.62234058 | -0.172778206 | 0.36707525 | -0.4706888 | 0.63786299 | 0.73127548 | NA | NA |
| ENSG000002 | 9.81453524 | 0.162018053 | 0.34423658 | 0.47065902 | 0.63788425 | 0.73127548 | NA | NA |
| ENSG000002 | 11.1892836 | -0.156448718 | 0.33241726 | -0.4706396 | 0.63789814 | 0.73127548 | LINC00298 | 339788 |
| ENSG000002 | 41.0494065 | 0.110163701 | 0.23407125 | 0.47064174 | 0.63789659 | 0.73127548 | NA | NA |
| ENSG000002 | 6.64958702 | -0.172009492 | 0.3656072 | -0.4704762 | 0.63801483 | 0.73134223 | NA | NA |
| ENSG000002 | 3.91377995 | -0.177853601 | 0.37800545 | -0.4705054 | 0.63799398 | 0.73134223 | NA | NA |
| ENSG000002 | 2.9650882 | -0.176519591 | 0.37520556 | -0.470461 | 0.63802567 | 0.73134223 | CAB39P1 | 106481732 |
| ENSG000002 | 7.20651734 | 0.168964265 | 0.35914918 | 0.47045705 | 0.63802851 | 0.73134223 | NA | NA |
| ENSG000002 | 4.90209831 | -0.175923358 | 0.37406967 | -0.4702957 | 0.63814377 | 0.73145367 | LOC105369893 | 105369893 |

| | | | | | | | | |
|---|---|---|---|---|---|---|---|---|
| ENSG00000C | 50.2202002 | 0.104205331 | 0.22161423 | 0.47021047 | 0.63820465 | 0.73150278 | MYBPC2 | 4606 |
| ENSG000002 | 3.26855787 | 0.177672818 | 0.37799572 | 0.47003923 | 0.63832699 | 0.73162232 | MIR5702 | 100847053 |
| ENSG000002 | 3.18963623 | -0.177369748 | 0.37747592 | -0.4698836 | 0.63843817 | 0.73172907 | NA | NA |
| ENSG000002 | 2.58219424 | -0.16901297 | 0.35976237 | -0.4697906 | 0.63850465 | 0.73178458 | NA | NA |
| ENSG000002 | 2.4408254 | -0.173819686 | 0.37002887 | -0.4697463 | 0.63853629 | 0.73180017 | NA | NA |
| ENSG000002 | 2.19625987 | 0.172205266 | 0.3666212 | 0.46970897 | 0.63856296 | 0.73181004 | NA | NA |
| ENSG000001 | 11.1858332 | -0.15339715 | 0.32661441 | -0.4696582 | 0.6385992 | 0.7318309 | ELAVL3 | 1995 |
| ENSG000001 | 2.72509972 | -0.174873067 | 0.37237188 | -0.4696194 | 0.63862695 | 0.73184201 | PPP1R27 | 116729 |
| ENSG000001 | 76.7004524 | -0.086033208 | 0.1832742 | -0.4694234 | 0.63876699 | 0.73196479 | DISC1 | 27185 |
| ENSG000002 | 28.1514517 | 0.129160516 | 0.27514976 | 0.46941897 | 0.63877019 | 0.73196479 | LOC105370040 | 105370040 |
| ENSG000002 | 5.18110156 | 0.175445436 | 0.37378359 | 0.46937705 | 0.63880015 | 0.73197844 | NA | NA |
| ENSG000001 | 200.871038 | -0.067801623 | 0.1445038 | -0.469203 | 0.63892451 | 0.73203819 | EPM2A | 7957 |
| ENSG000002 | 6.32706779 | -0.171405509 | 0.3652838 | -0.4692393 | 0.63889861 | 0.73203819 | LOC105372290 | 105372290 |
| ENSG000002 | 2.69877496 | -0.175318725 | 0.37364428 | -0.4692129 | 0.63891745 | 0.73203819 | NA | NA |
| ENSG000002 | 5.07891355 | -0.176013698 | 0.37510755 | -0.4692353 | 0.63890147 | 0.73203819 | NA | NA |
| ENSG000002 | 10.2133845 | 0.161025452 | 0.34332758 | 0.46901402 | 0.63905961 | 0.73211806 | NA | NA |
| ENSG000002 | 2.93893972 | -0.175520278 | 0.37419702 | -0.4690585 | 0.63902785 | 0.73211806 | NA | NA |
| ENSG000002 | 3.72739208 | -0.177372999 | 0.37819042 | -0.4690045 | 0.63906643 | 0.73211806 | NA | NA |
| ENSG000002 | 2.40561961 | 0.174366583 | 0.37176256 | 0.46902675 | 0.63905052 | 0.73211806 | NA | NA |
| ENSG000002 | 33.1089376 | 0.118217355 | 0.25208825 | 0.46895226 | 0.63910376 | 0.73214014 | NA | NA |
| ENSG000001 | 2.671198 | -0.174712994 | 0.37258777 | -0.4689177 | 0.63912847 | 0.73214777 | NA | NA |
| ENSG000002 | 2.4155754 | 0.17445329 | 0.37241859 | 0.46843336 | 0.63947472 | 0.73252371 | NA | NA |
| ENSG000002 | 16.3366455 | -0.137664579 | 0.29391875 | -0.4683763 | 0.63951551 | 0.73254974 | LOC442497 | 442497 |
| ENSG000002 | 3.1581183 | -0.176739766 | 0.37755441 | -0.4681173 | 0.63970068 | 0.73274116 | OR6V1 | 346517 |
| ENSG000001 | 15.9757112 | -0.14782797 | 0.31585465 | -0.4680253 | 0.63976649 | 0.73279583 | AMT | 275 |
| ENSG000002 | 2.40886276 | -0.171964366 | 0.3674978 | -0.4679331 | 0.63983247 | 0.7328507 | RNU6-979P | 106479999 |
| ENSG000002 | 7.63219669 | -0.168678218 | 0.36054515 | -0.4678421 | 0.63989748 | 0.73290446 | NA | NA |
| ENSG000002 | 2.65686449 | -0.174758402 | 0.37357537 | -0.4677996 | 0.63992792 | 0.73291863 | NA | NA |
| ENSG000002 | 3.27585741 | 0.176957362 | 0.3783609 | 0.46769463 | 0.64000298 | 0.73298246 | NA | NA |

| | | | | | | | | |
|---|---|---|---|---|---|---|---|---|
| ENSG000002 | 2.42258234 | 0.174557679 | 0.3732488 | 0.46767111 | 0.64001981 | 0.73298246 | NA | NA |
| ENSG000002 | 3.68124746 | -0.177007664 | 0.37857633 | -0.4675614 | 0.64009831 | 0.73303279 | NA | NA |
| ENSG000002 | 3.00453914 | 0.17562144 | 0.37561333 | 0.46755912 | 0.6400999 | 0.73303279 | NA | NA |
| ENSG000002 | 4.10629168 | 0.176591448 | 0.3778287 | 0.46738495 | 0.64022449 | 0.73311879 | RN7SL608P | 106480521 |
| ENSG000002 | 3.4584422 | -0.176751818 | 0.37817719 | -0.4673783 | 0.64022923 | 0.73311879 | NA | NA |
| ENSG000002 | 14.5180227 | 0.145176006 | 0.31061388 | 0.46738415 | 0.64022506 | 0.73311879 | MAP2K4P1 | 139201 |
| ENSG000002 | 4.69778715 | -0.176028299 | 0.37677862 | -0.4671929 | 0.64036187 | 0.73324996 | NA | NA |
| ENSG000001 | 5.8856012 | -0.172476093 | 0.36922708 | -0.4671274 | 0.64040872 | 0.73328041 | NA | NA |
| ENSG000002 | 23.0088494 | 0.131577497 | 0.28170233 | 0.4670799 | 0.64044271 | 0.73328041 | NA | NA |
| ENSG000002 | 5.98197678 | 0.17414968 | 0.37283617 | 0.46709438 | 0.64043236 | 0.73328041 | NA | NA |
| ENSG000002 | 3.18687674 | -0.176316563 | 0.37763593 | -0.4668956 | 0.64057456 | 0.73341067 | NA | NA |
| ENSG000001 | 39.5001069 | 0.103247401 | 0.22118117 | 0.46680014 | 0.64064287 | 0.73342676 | BHMT | 635 |
| ENSG000001 | 3.67217303 | -0.176726493 | 0.37857541 | -0.4668198 | 0.64062882 | 0.73342676 | CNPY1 | 285888 |
| ENSG000002 | 3.20124772 | -0.175767619 | 0.37652225 | -0.4668187 | 0.64062961 | 0.73342676 | NA | NA |
| ENSG000002 | 3.48216446 | -0.17655575 | 0.3782818 | -0.4667307 | 0.64069253 | 0.73346289 | NA | NA |
| ENSG000001 | 25.5031673 | 0.12180353 | 0.26099382 | 0.46669124 | 0.6407208 | 0.73346861 | NA | NA |
| ENSG000002 | 4.43512183 | -0.175911473 | 0.37694786 | -0.4666732 | 0.64073369 | 0.73346861 | NA | NA |
| ENSG000002 | 4.94651889 | -0.176403904 | 0.37805155 | -0.4666134 | 0.64077652 | 0.73349693 | NA | NA |
| ENSG000002 | 2.49226754 | -0.170624285 | 0.36582828 | -0.4664054 | 0.64092535 | 0.73360516 | NA | NA |
| ENSG000002 | 5.41917268 | -0.173751046 | 0.37250058 | -0.466445 | 0.64089698 | 0.73360516 | NA | NA |
| ENSG000002 | 11.2314553 | 0.153037124 | 0.3281179 | 0.46640894 | 0.64092281 | 0.73360516 | NA | NA |
| ENSG000002 | 4.60347641 | -0.175279912 | 0.37585053 | -0.4663554 | 0.64096116 | 0.73362544 | NA | NA |
| ENSG000002 | 5.15780267 | -0.174936746 | 0.37518751 | -0.4662648 | 0.64102594 | 0.73367889 | NA | NA |
| ENSG000002 | 8.66200509 | 0.162872462 | 0.34936484 | 0.46619592 | 0.64107527 | 0.73371463 | NA | NA |
| ENSG000001 | 3.50656335 | -0.17580514 | 0.37717274 | -0.4661131 | 0.64113458 | 0.7337618 | IFNA16 | 3449 |
| ENSG000002 | 3.14475721 | -0.175700121 | 0.37717215 | -0.4658354 | 0.64133336 | 0.73396859 | NA | NA |
| ENSG000002 | 14.5492902 | -0.145284741 | 0.31200056 | -0.4656554 | 0.64146219 | 0.73408427 | LINC01535 | 101927667 |
| ENSG000002 | 5.11553988 | -0.17427148 | 0.37425939 | -0.4656436 | 0.64147065 | 0.73408427 | NA | NA |
| ENSG000001 | 85.0553688 | 0.082756741 | 0.17780714 | 0.46542979 | 0.64162371 | 0.73423871 | EDNRA | 1909 |

| | | | | | | | | |
|---|---|---|---|---|---|---|---|---|
| ENSG000002 | 3.39688827 | -0.176057555 | 0.37844104 | -0.4652179 | 0.6417754 | 0.73439158 | NA | NA |
| ENSG000001 | 26.1156228 | 0.12104033 | 0.26027731 | 0.46504373 | 0.64190014 | 0.73450962 | IL1A | 3552 |
| ENSG000002 | 2.18058512 | -0.167884007 | 0.3610228 | -0.4650233 | 0.64191478 | 0.73450962 | LOC100996630 | 100996630 |
| ENSG000001 | 45.1247291 | -0.099976263 | 0.21503368 | -0.464933 | 0.64197941 | 0.73456284 | OSBP2 | 23762 |
| ENSG000002 | 3.6973612 | -0.175927768 | 0.37858351 | -0.4647 | 0.64214629 | 0.73473305 | NA | NA |
| ENSG000002 | 2.89696398 | -0.173992913 | 0.37457365 | -0.4645092 | 0.64228295 | 0.73486869 | NA | NA |
| ENSG000002 | 2.48608234 | -0.171497727 | 0.36927019 | -0.4644234 | 0.64234441 | 0.7348768 | RNU6-697P | 106479874 |
| ENSG000002 | 3.28400684 | 0.175380096 | 0.37760991 | 0.4644478 | 0.64232695 | 0.7348768 | NA | NA |
| ENSG000002 | 3.29635794 | 0.175346443 | 0.37752755 | 0.46446 | 0.64231821 | 0.7348768 | NA | NA |
| ENSG000002 | 3.43559399 | -0.174609603 | 0.37606097 | -0.464312 | 0.64242426 | 0.73494742 | NA | NA |
| ENSG000001 | 29.3837058 | -0.116949627 | 0.25194718 | -0.4641831 | 0.64251655 | 0.73497006 | HSD11B1L | 374875 |
| ENSG000002 | 4.63828865 | -0.17457477 | 0.3760864 | -0.4641879 | 0.6425131 | 0.73497006 | LOC101928280 | 101928280 |
| ENSG000002 | 10.877373 | -0.152676775 | 0.32890485 | -0.4641974 | 0.64250633 | 0.73497006 | NA | NA |
| ENSG000002 | 8.67115198 | 0.163123898 | 0.35141036 | 0.46419775 | 0.64250607 | 0.73497006 | NA | NA |
| ENSG000001 | 10.3867914 | -0.154901923 | 0.33379469 | -0.4640635 | 0.64260227 | 0.73499648 | TREML2 | 79865 |
| ENSG000002 | 8.93043935 | 0.159839889 | 0.34443848 | 0.46405933 | 0.64260524 | 0.73499648 | NA | NA |
| ENSG000002 | 40.4818218 | 0.103461604 | 0.22292868 | 0.46410181 | 0.6425748 | 0.73499648 | NA | NA |
| ENSG000002 | 2.20166819 | 0.170108412 | 0.3665737 | 0.46404969 | 0.64261215 | 0.73499648 | NA | NA |
| ENSG000002 | 7.53054971 | 0.168330663 | 0.36297152 | 0.46375722 | 0.64282169 | 0.73519468 | IBA57-AS1 | 574432 |
| ENSG000002 | 4.40394901 | -0.174721004 | 0.37674358 | -0.4637664 | 0.64281514 | 0.73519468 | NA | NA |
| ENSG000002 | 4.16234154 | -0.175236394 | 0.37788902 | -0.4637245 | 0.64284515 | 0.73520077 | NA | NA |
| ENSG000001 | 57.2748787 | 0.098909012 | 0.21332768 | 0.46364827 | 0.64289977 | 0.73522176 | ELF3 | 1999 |
| ENSG000002 | 10.144687 | 0.154222464 | 0.33261365 | 0.46366848 | 0.64288528 | 0.73522176 | NA | NA |
| ENSG000001 | 3.69588699 | -0.175445161 | 0.37842668 | -0.4636173 | 0.64292195 | 0.7352264 | OR2A20P | 401428 |
| ENSG000002 | 3.13974532 | -0.174159621 | 0.37572545 | -0.4635289 | 0.64298527 | 0.73525734 | NA | NA |
| ENSG000002 | 5.57974334 | -0.17225006 | 0.37160152 | -0.4635343 | 0.64298142 | 0.73525734 | RNU6-1201P | 106481942 |
| ENSG000002 | 20.0350508 | -0.129367508 | 0.27921395 | -0.4633275 | 0.64312962 | 0.73540166 | SNORD53 | 26796 |
| ENSG000001 | 62.5172885 | 0.090769634 | 0.1960269 | 0.46304683 | 0.6433308 | 0.73554874 | GPAT3 | 84803 |
| ENSG000002 | 3.28670685 | 0.175113012 | 0.37816702 | 0.46305733 | 0.64332328 | 0.73554874 | NA | NA |

| | | | | | | | | |
|---|---|---|---|---|---|---|---|---|
| ENSG000002 | 6.94321709 | -0.171200793 | 0.3696787 | -0.463107 | 0.64328767 | 0.73554874 | NA | NA |
| ENSG000002 | 6.27485256 | -0.173643947 | 0.37497133 | -0.4630859 | 0.64330278 | 0.73554874 | NA | NA |
| ENSG000002 | 3.73538995 | -0.175142649 | 0.37834292 | -0.4629204 | 0.64342144 | 0.73563163 | NA | NA |
| ENSG000002 | 2.21085452 | 0.169273626 | 0.36585687 | 0.46267719 | 0.64359577 | 0.73580125 | TMEM88B | 643965 |
| ENSG000002 | 8.76698891 | -0.163627333 | 0.35366433 | -0.4626628 | 0.64360608 | 0.73580125 | NA | NA |
| ENSG000002 | 3.46880183 | -0.174437783 | 0.37711811 | -0.4625548 | 0.64368353 | 0.73585125 | NA | NA |
| ENSG000002 | 2.71875865 | -0.172824546 | 0.37363336 | -0.4625512 | 0.64368612 | 0.73585125 | NA | NA |
| ENSG000002 | 3.18521286 | -0.174317542 | 0.376932 | -0.4624642 | 0.6437485 | 0.73590182 | NA | NA |
| ENSG000002 | 24.0188491 | -0.122155191 | 0.26416661 | -0.4624172 | 0.64378215 | 0.73591955 | NA | NA |
| ENSG000002 | 9.93473075 | -0.154195185 | 0.33360099 | -0.4622144 | 0.64392757 | 0.73606502 | NA | NA |
| ENSG000002 | 3.4756983 | -0.174802693 | 0.37823461 | -0.4621541 | 0.64397078 | 0.73609367 | NA | NA |
| ENSG000002 | 5.4773435 | 0.172064307 | 0.37256034 | 0.46184279 | 0.64419407 | 0.73632814 | NA | NA |
| ENSG000002 | 3.28314455 | 0.174788904 | 0.37850372 | 0.46178913 | 0.64423255 | 0.73635137 | NA | NA |
| ENSG000002 | 5.25510248 | -0.173808118 | 0.37646137 | -0.4616891 | 0.64430428 | 0.7364126 | NA | NA |
| ENSG000002 | 2.39779059 | 0.171416483 | 0.37137023 | 0.46157841 | 0.64438368 | 0.7364826 | SNORD115-36 | 100033810 |
| ENSG000002 | 5.09749559 | -0.17240191 | 0.37362492 | -0.4614304 | 0.64448982 | 0.73658315 | NA | NA |
| ENSG000002 | 3.69792813 | -0.173942433 | 0.3770515 | -0.4613228 | 0.64456707 | 0.73665067 | NA | NA |
| ENSG000002 | 3.68145632 | -0.174510059 | 0.3784706 | -0.4610928 | 0.64473205 | 0.73681845 | NA | NA |
| ENSG000002 | 5.20585671 | 0.173878771 | 0.37730998 | 0.46083798 | 0.64491485 | 0.7370066 | NA | NA |
| ENSG000002 | 4.99811902 | -0.17333025 | 0.37620907 | -0.4607285 | 0.64499338 | 0.73707557 | NA | NA |
| ENSG000002 | 12.2898474 | 0.148277924 | 0.32189622 | 0.46063892 | 0.64505769 | 0.73712828 | NA | NA |
| ENSG000001 | 12.9469914 | -0.150288569 | 0.32642211 | -0.4604117 | 0.64522071 | 0.7372462 | CHRNE | 1145 |
| ENSG000001 | 49.5468678 | -0.100184172 | 0.21764251 | -0.4603153 | 0.64528994 | 0.7372462 | CAMK1 | 8536 |
| ENSG000001 | 13.0157408 | -0.149021004 | 0.32378852 | -0.4602418 | 0.64534268 | 0.7372462 | FERMT3 | 83706 |
| ENSG000001 | 108.076612 | 0.080568951 | 0.17498384 | 0.46043652 | 0.64520292 | 0.7372462 | VENTX | 27287 |
| ENSG000001 | 2.73389462 | -0.171893303 | 0.37343009 | -0.4603092 | 0.6452943 | 0.7372462 | CDCP2 | 200008 |
| ENSG000002 | 13.6123692 | -0.144281997 | 0.31341268 | -0.4603579 | 0.64525936 | 0.7372462 | NA | NA |
| ENSG000002 | 79.4083126 | -0.082911517 | 0.18010992 | -0.4603384 | 0.64527332 | 0.7372462 | NA | NA |
| ENSG000002 | 3.42131886 | -0.173329141 | 0.37659621 | -0.4602519 | 0.64533539 | 0.7372462 | NA | NA |

| | | | | | | | | |
|---|---|---|---|---|---|---|---|---|
| ENSG000002 | 4.12615957 | -0.173987715 | 0.3779525 | -0.4603428 | 0.64527018 | 0.7372462 | NA | NA |
| ENSG000002 | 20.5165742 | 0.128338717 | 0.27883257 | 0.46027162 | 0.64532127 | 0.7372462 | GET4 | 51608 |
| ENSG000002 | 10.9348644 | -0.151814235 | 0.32994089 | -0.4601256 | 0.6454261 | 0.73732072 | NA | NA |
| ENSG000001 | 160.228024 | 0.073643587 | 0.16008779 | 0.46002001 | 0.64550186 | 0.73738649 | KDM1B | 221656 |
| ENSG000001 | 11.0658817 | 0.156824539 | 0.34097153 | 0.45993441 | 0.6455633 | 0.73740507 | PDE6G | 5148 |
| ENSG000002 | 23.3152549 | -0.125921678 | 0.2737676 | -0.4599583 | 0.64554616 | 0.73740507 | ABCA9-AS1 | 104355297 |
| ENSG000002 | 2.93401433 | -0.171582234 | 0.37306864 | -0.4599214 | 0.64557267 | 0.73740507 | NA | NA |
| ENSG000002 | 3.4370243 | -0.173946509 | 0.3782433 | -0.45988 | 0.64560239 | 0.73741825 | NA | NA |
| ENSG000002 | 2.46725932 | -0.167875865 | 0.36509842 | -0.4598099 | 0.64565268 | 0.73745492 | NA | NA |
| ENSG000001 | 12.1814365 | 0.151308348 | 0.3291582 | 0.45968275 | 0.64574395 | 0.73750829 | MAGEB17 | 645864 |
| ENSG000002 | 5.85842874 | -0.169977388 | 0.36978231 | -0.4596688 | 0.64575398 | 0.73750829 | NA | NA |
| ENSG000002 | 4.10351932 | -0.174031559 | 0.37858173 | -0.4596935 | 0.64573624 | 0.73750829 | ASCL5 | 647219 |
| ENSG000002 | 5.2825308 | 0.170653735 | 0.37132911 | 0.45957543 | 0.645821 | 0.73754632 | NA | NA |
| ENSG000002 | 10.3870464 | -0.153361636 | 0.33370554 | -0.4595717 | 0.64582365 | 0.73754632 | NA | NA |
| ENSG000002 | 19.8133254 | 0.131393984 | 0.28608923 | 0.45927623 | 0.64603581 | 0.73776784 | PLS3-AS1 | 101927352 |
| ENSG000002 | 4.21968878 | -0.173717765 | 0.37833593 | -0.4591628 | 0.6461173 | 0.73779856 | CADM2-AS2 | 100874037 |
| ENSG000002 | 4.64766289 | -0.173427335 | 0.37769109 | -0.4591777 | 0.64610654 | 0.73779856 | NA | NA |
| ENSG000002 | 2.41805503 | 0.170432676 | 0.3711514 | 0.45919988 | 0.64609064 | 0.73779856 | NA | NA |
| ENSG000002 | 3.49189156 | -0.173571589 | 0.37805895 | -0.4591125 | 0.64615338 | 0.73781899 | NA | NA |
| ENSG000002 | 7.92472734 | 0.160723862 | 0.3500967 | 0.4590842 | 0.6461737 | 0.73782142 | NA | NA |
| ENSG000002 | 4.63745447 | -0.172334993 | 0.37542435 | -0.4590405 | 0.64620506 | 0.73783645 | NA | NA |
| ENSG000002 | 2.9275897 | -0.172699656 | 0.37634163 | -0.4588907 | 0.64631269 | 0.73793856 | NA | NA |
| ENSG000001 | 125.354003 | 0.083359026 | 0.18168604 | 0.45880808 | 0.64637199 | 0.73798549 | IER3 | 8870 |
| ENSG000002 | 2.73547342 | -0.170677577 | 0.37203642 | -0.4587658 | 0.64640237 | 0.7379994 | NA | NA |
| ENSG000001 | 54.0839613 | -0.099488792 | 0.21691559 | -0.4586521 | 0.64648401 | 0.73807182 | PROS1 | 5627 |
| ENSG000001 | 10.24169 | -0.152679181 | 0.33292387 | -0.4586009 | 0.6465208 | 0.73809304 | MGP | 4256 |
| ENSG000001 | 22.7559189 | -0.121293283 | 0.26452635 | -0.4585301 | 0.64657167 | 0.73813034 | CNTFR | 1271 |
| ENSG000002 | 5.4139657 | -0.171565035 | 0.37419349 | -0.4584928 | 0.6465984 | 0.73814008 | IFNA13 | 3447 |
| ENSG000001 | 45.202395 | -0.097784968 | 0.21332767 | -0.4583792 | 0.64668003 | 0.73820383 | NA | NA |

| | | | | | | | | |
|---|---|---|---|---|---|---|---|---|
| ENSG000002 | 3.67994943 | -0.173449658 | 0.37843089 | -0.4583391 | 0.64670886 | 0.73820383 | NA | NA |
| ENSG000002 | 18.9869089 | -0.128560716 | 0.28047915 | -0.458361 | 0.64669308 | 0.73820383 | NA | NA |
| ENSG000002 | 33.4891499 | 0.111677622 | 0.24371265 | 0.45823481 | 0.64678375 | 0.73826853 | PALM2 | 114299 |
| ENSG000002 | 4.16046196 | 0.173092853 | 0.37793268 | 0.45799917 | 0.64695304 | 0.73844098 | NA | NA |
| ENSG000001 | 32.0386198 | 0.108584497 | 0.23712301 | 0.45792475 | 0.6470065 | 0.73848122 | KCNIP2 | 30819 |
| ENSG00000C | 8.69261027 | -0.157813076 | 0.34465326 | -0.4578894 | 0.6470319 | 0.73848942 | P2RX6 | 9127 |
| ENSG000002 | 5.31366505 | 0.171228927 | 0.37408104 | 0.45773217 | 0.64714487 | 0.73859757 | NA | NA |
| ENSG000001 | 6.43186211 | 0.169635107 | 0.37076894 | 0.45752243 | 0.64729558 | 0.73871721 | NTSR1 | 4923 |
| ENSG000001 | 343.093799 | 0.058370967 | 0.12758396 | 0.45751023 | 0.64730435 | 0.73871721 | PHF21A | 51317 |
| ENSG000002 | 6.05640152 | -0.169816153 | 0.3711531 | -0.4575367 | 0.64728535 | 0.73871721 | NA | NA |
| ENSG000002 | 5.35443471 | -0.169795871 | 0.37129776 | -0.4573038 | 0.64745271 | 0.73886573 | NA | NA |
| ENSG000002 | 2.94246454 | -0.171716608 | 0.37552721 | -0.4572681 | 0.64747837 | 0.73887422 | NA | NA |
| ENSG000002 | 5.69757559 | -0.170891999 | 0.37376754 | -0.4572147 | 0.64751677 | 0.73889724 | NA | NA |
| ENSG000001 | 6.85601739 | -0.166258042 | 0.36367095 | -0.4571661 | 0.64755164 | 0.7389047 | NA | NA |
| ENSG000001 | 4.18496163 | -0.172757655 | 0.37789746 | -0.4571549 | 0.64755975 | 0.7389047 | OR52H1 | 390067 |
| ENSG000002 | 4.98084165 | -0.172797084 | 0.37816973 | -0.45693 | 0.64772138 | 0.73906834 | NA | NA |
| ENSG000002 | 40.7420058 | -0.102918362 | 0.22530431 | -0.4567971 | 0.64781686 | 0.73915649 | ZNF192P1 | 651302 |
| ENSG000002 | 18.9467524 | 0.129008419 | 0.28256219 | 0.45656647 | 0.64798269 | 0.73932489 | NA | NA |
| ENSG000001 | 71.1530256 | -0.085405059 | 0.18707982 | -0.4565167 | 0.64801849 | 0.73934493 | OSMR | 9180 |
| ENSG000001 | 59.7390346 | -0.089645271 | 0.19640954 | -0.4564201 | 0.64808789 | 0.73940331 | STK32B | 55351 |
| ENSG000002 | 7.64334801 | 0.162293319 | 0.35567258 | 0.45629978 | 0.64817443 | 0.73946947 | C4orf48 | 401115 |
| ENSG000002 | 3.28983883 | 0.172613667 | 0.37829918 | 0.45628876 | 0.64818234 | 0.73946947 | NA | NA |
| ENSG000001 | 9.48808342 | -0.154907722 | 0.33956149 | -0.4561993 | 0.64824665 | 0.73952203 | CHRND | 1144 |
| ENSG000002 | 4.19288183 | -0.17256115 | 0.37832896 | -0.456114 | 0.64830797 | 0.73955037 | LTB | 4050 |
| ENSG000002 | 2.86999811 | 0.169743682 | 0.37214211 | 0.45612598 | 0.64829939 | 0.73955037 | NA | NA |
| ENSG000002 | 3.9505501 | -0.172603218 | 0.37845519 | -0.4560731 | 0.64833744 | 0.73956318 | NA | NA |
| ENSG000002 | 2.39659692 | 0.169343738 | 0.37137941 | 0.4559858 | 0.64840019 | 0.73959315 | NA | NA |
| ENSG000002 | 3.88928737 | -0.172399577 | 0.3780612 | -0.4560097 | 0.648383 | 0.73959315 | NA | NA |
| ENSG000002 | 2.91935628 | -0.171540229 | 0.37640488 | -0.4557333 | 0.64858179 | 0.73977949 | RN7SL377P | 106479360 |

| | | | | | | | | |
|---|---|---|---|---|---|---|---|---|
| ENSG000002 | 9.96523608 | -0.155065632 | 0.34029407 | -0.4556813 | 0.64861919 | 0.73980134 | NA | NA |
| ENSG000002 | 2.51010441 | -0.167508899 | 0.36762686 | -0.4556492 | 0.64864223 | 0.7398068 | LINC01056 | 100144597 |
| ENSG000002 | 4.92687597 | -0.171891943 | 0.37734798 | -0.4555263 | 0.64873063 | 0.73988681 | NA | NA |
| ENSG000002 | 6.64980817 | -0.168009764 | 0.36887293 | -0.4554679 | 0.64877268 | 0.73991397 | NA | NA |
| ENSG000002 | 7.43476247 | -0.166398493 | 0.36543731 | -0.4553407 | 0.64886413 | 0.73993505 | NA | NA |
| ENSG000002 | 26.3331836 | -0.121004865 | 0.26574218 | -0.4553469 | 0.64885973 | 0.73993505 | NA | NA |
| ENSG000002 | 5.6838385 | 0.168884357 | 0.37083822 | 0.45541249 | 0.64881251 | 0.73993505 | UTAT33 | 101927492 |
| ENSG000002 | 16.5624384 | 0.134065802 | 0.29442966 | 0.45534069 | 0.64886416 | 0.73993505 | NA | NA |
| ENSG000002 | 4.67164085 | -0.171505296 | 0.37671392 | -0.4552667 | 0.64891739 | 0.73997495 | NA | NA |
| ENSG000002 | 6.55440882 | 0.165040069 | 0.36261478 | 0.45513883 | 0.64900936 | 0.74005901 | NA | NA |
| ENSG000002 | 2.43706542 | 0.169655658 | 0.37280609 | 0.45507749 | 0.64905349 | 0.74008853 | RNU6-444P | 106479762 |
| ENSG000002 | 2.20539702 | -0.165286467 | 0.36331949 | -0.4549342 | 0.64915657 | 0.74018524 | NA | NA |
| ENSG000002 | 4.91231413 | -0.170841573 | 0.37574028 | -0.45468 | 0.64933951 | 0.74037302 | NA | NA |
| ENSG000002 | 9.88877397 | 0.157570663 | 0.34662315 | 0.45458782 | 0.6494058 | 0.74041101 | NA | NA |
| ENSG000002 | 4.38149212 | 0.170908971 | 0.37596878 | 0.45458289 | 0.64940935 | 0.74041101 | NA | NA |
| ENSG00000C | 26.0482917 | 0.134336726 | 0.2957505 | 0.45422316 | 0.64966822 | 0.74053251 | PRICKLE3 | 4007 |
| ENSG000001 | 3.26107376 | 0.171816195 | 0.37817325 | 0.45433196 | 0.64958991 | 0.74053251 | NA | NA |
| ENSG000002 | 2.38529046 | 0.168083885 | 0.37000102 | 0.45427952 | 0.64962765 | 0.74053251 | RNU6-831P | 106481443 |
| ENSG000002 | 2.44321096 | 0.169555088 | 0.37329964 | 0.45420641 | 0.64968027 | 0.74053251 | LOC105373185 | 105373185 |
| ENSG000002 | 6.10976389 | -0.167246854 | 0.36810283 | -0.4543482 | 0.64957823 | 0.74053251 | NA | NA |
| ENSG000002 | 11.2724519 | 0.150883027 | 0.33217821 | 0.45422314 | 0.64966823 | 0.74053251 | NA | NA |
| ENSG000002 | 15.3332087 | 0.141162124 | 0.31067033 | 0.45437916 | 0.64955595 | 0.74053251 | KRTAP5-AS1 | 338651 |
| ENSG000002 | 6.7664446 | 0.166544343 | 0.36665136 | 0.45423081 | 0.64966271 | 0.74053251 | NA | NA |
| ENSG000002 | 4.21175528 | -0.171960009 | 0.37858137 | -0.4542221 | 0.64966901 | 0.74053251 | NA | NA |
| ENSG000002 | 3.69119029 | -0.171616419 | 0.37793862 | -0.4540854 | 0.64976734 | 0.74061094 | NA | NA |
| ENSG000002 | 33.0738525 | 0.10874212 | 0.23949295 | 0.45405145 | 0.64979179 | 0.74061799 | NA | NA |
| ENSG000002 | 5.18143405 | 0.169864893 | 0.37413298 | 0.45402277 | 0.64981244 | 0.74062071 | RNU6-26P | 100873741 |
| ENSG000002 | 2.39736604 | 0.168725321 | 0.37180513 | 0.45380041 | 0.64997249 | 0.74078231 | NA | NA |
| ENSG000002 | 10.2456121 | 0.157412525 | 0.34691171 | 0.45375385 | 0.650006 | 0.74079968 | LOC101928725 | 101928725 |

| | | | | | | | | | |
|---|---|---|---|---|---|---|---|---|---|
| ENSG000002 | 6.55703222 | 0.166521358 | 0.36707534 | 0.45364354 | 0.65008541 | 0.74086936 | NA | NA | |
| ENSG000001 | 81.4986566 | 0.085504782 | 0.18863248 | 0.45328769 | 0.6503416 | 0.74114049 | RFTN1 | | 23180 |
| ENSG000002 | 5.09583226 | -0.16935103 | 0.37364963 | -0.4532348 | 0.65037965 | 0.74114854 | NA | NA | |
| ENSG000002 | 3.46143092 | -0.171249692 | 0.37784521 | -0.4532271 | 0.65038522 | 0.74114854 | NA | NA | |
| ENSG000002 | 10.3091881 | 0.154569704 | 0.3411145 | 0.45313144 | 0.6504541 | 0.74120621 | NA | NA | |
| ENSG000001 | 94.6669092 | 0.088249567 | 0.19483027 | 0.45295613 | 0.65058033 | 0.74132346 | MURC | | 347273 |
| ENSG000002 | 2.40323172 | 0.168267204 | 0.37150182 | 0.45293776 | 0.65059356 | 0.74132346 | NA | NA | |
| ENSG000001 | 14.521058 | -0.142816691 | 0.31538564 | -0.4528319 | 0.65066977 | 0.74136863 | ZP3 | | 7784 |
| ENSG000002 | 5.86357005 | -0.167347383 | 0.36953889 | -0.4528546 | 0.65065345 | 0.74136863 | FLJ35934 | | 400579 |
| ENSG000002 | 28.6931737 | 0.114399614 | 0.25268716 | 0.4527322 | 0.65074159 | 0.74142963 | NA | NA | |
| ENSG000002 | 10.2546303 | -0.157615871 | 0.34817905 | -0.4526863 | 0.65077467 | 0.7414465 | NA | NA | |
| ENSG000002 | 19.9513386 | 0.126000006 | 0.2783726 | 0.45263077 | 0.65081464 | 0.7414712 | NA | NA | |
| ENSG000002 | 14.7303497 | -0.136784236 | 0.3024309 | -0.4522826 | 0.6510654 | 0.74173606 | NA | NA | |
| ENSG000001 | 8.33840734 | -0.160759343 | 0.35554606 | -0.4521477 | 0.65116256 | 0.74180506 | SQRDL | | 58472 |
| ENSG000002 | 2.39570326 | 0.166776676 | 0.36884519 | 0.45215901 | 0.65115444 | 0.74180506 | NA | NA | |
| ENSG000001 | 46.7291164 | -0.098895007 | 0.21875785 | -0.4520752 | 0.65121479 | 0.74184372 | ANKRD29 | | 147463 |
| ENSG000002 | 23.0764658 | 0.124511168 | 0.27545165 | 0.45202549 | 0.65125062 | 0.74186369 | NA | NA | |
| ENSG000002 | 4.36188091 | 0.170223293 | 0.37661175 | 0.45198614 | 0.65127897 | 0.74187515 | NA | NA | |
| ENSG000001 | 8.93971779 | -0.154821616 | 0.34257792 | -0.4519311 | 0.65131862 | 0.74189948 | SYCE2 | | 256126 |
| ENSG000002 | 4.17252611 | 0.170980422 | 0.37840508 | 0.45184495 | 0.65138069 | 0.74194933 | LINC00492 | | 100861468 |
| ENSG000002 | 6.85950836 | -0.165033279 | 0.36530309 | -0.4517708 | 0.65143409 | 0.74196848 | NA | NA | |
| ENSG000002 | 6.51902767 | 0.163575959 | 0.36205697 | 0.45179619 | 0.65141581 | 0.74196848 | NA | NA | |
| ENSG000002 | 2.50011617 | -0.164729212 | 0.36469646 | -0.4516885 | 0.65149337 | 0.74201516 | NA | NA | |
| ENSG000002 | 2.92270918 | -0.169624582 | 0.37560864 | -0.4515993 | 0.65155771 | 0.74203605 | NA | NA | |
| ENSG000002 | 17.4136578 | 0.135351912 | 0.29971262 | 0.45160564 | 0.6515531 | 0.74203605 | NA | NA | |
| ENSG000002 | 5.39254811 | -0.168402276 | 0.37291223 | -0.4515869 | 0.65156661 | 0.74203605 | NA | NA | |
| ENSG000002 | 11.325451 | -0.147101408 | 0.32576572 | -0.4515558 | 0.651589 | 0.74204071 | LINC00936 | | 338758 |
| ENSG000002 | 3.72494579 | -0.170860591 | 0.37846056 | -0.4514621 | 0.65165655 | 0.74209679 | MIR183 | | 406959 |
| ENSG000002 | 12.7170165 | -0.146595822 | 0.32495147 | -0.4511314 | 0.65189482 | 0.74234729 | NA | NA | |

| | | | | | | | | |
|---|---|---|---|---|---|---|---|---|
| ENSG000002 | 8.5033389 | -0.15637989 | 0.34666604 | -0.4510966 | 0.65191989 | 0.74235499 | NA | NA |
| ENSG000002 | 5.36540779 | -0.168604185 | 0.37382114 | -0.451029 | 0.65196862 | 0.74238964 | NA | NA |
| ENSG000002 | 4.88933725 | -0.169156183 | 0.3752619 | -0.4507683 | 0.65215652 | 0.74258274 | NA | NA |
| ENSG000001 | 44.4495939 | 0.096696591 | 0.21462628 | 0.45053471 | 0.65232493 | 0.74271194 | ADAMTS15 | 170689 |
| ENSG000002 | 3.14827406 | -0.169888463 | 0.3770684 | -0.4505508 | 0.65231335 | 0.74271194 | NA | NA |
| ENSG000002 | 4.14851189 | -0.170230692 | 0.37780833 | -0.4505742 | 0.65229646 | 0.74271194 | NA | NA |
| ENSG000002 | 17.5199871 | -0.130445924 | 0.28959937 | -0.4504358 | 0.65239623 | 0.74277226 | NA | NA |
| ENSG000001 | 15.6767962 | -0.135237925 | 0.30030661 | -0.4503328 | 0.65247047 | 0.74283593 | NA | NA |
| ENSG000002 | 2.21751202 | -0.163267 | 0.36261936 | -0.4502435 | 0.65253489 | 0.74288841 | OR51F1 | 256892 |
| ENSG000001 | 53.5015852 | 0.093008621 | 0.20659808 | 0.45019111 | 0.65257264 | 0.74291053 | MCOLN2 | 255231 |
| ENSG00000C | 135.492307 | 0.068266452 | 0.15165289 | 0.45014937 | 0.65260274 | 0.74292394 | JAK2 | 3717 |
| ENSG000002 | 11.1107935 | -0.148602959 | 0.3302173 | -0.4500157 | 0.65269914 | 0.74299197 | NA | NA |
| ENSG000002 | 4.34666027 | 0.169439055 | 0.37651617 | 0.45001801 | 0.65269746 | 0.74299197 | NA | NA |
| ENSG000002 | 9.21035182 | -0.152970567 | 0.33999297 | -0.4499227 | 0.65276615 | 0.74304739 | NA | NA |
| ENSG000002 | 3.28191807 | 0.170150327 | 0.37822071 | 0.44987047 | 0.65280384 | 0.74306943 | NA | NA |
| ENSG000002 | 29.7912388 | 0.113510697 | 0.25242395 | 0.44968276 | 0.65293921 | 0.74320265 | PCOLCE-AS1 | 100129845 |
| ENSG000001 | 22.3109955 | 0.125687705 | 0.2795741 | 0.44956848 | 0.65302162 | 0.74323387 | GJA3 | 2700 |
| ENSG000001 | 120.369755 | -0.070525264 | 0.15685642 | -0.4496167 | 0.65298685 | 0.74323387 | AJAP1 | 55966 |
| ENSG000002 | 3.67959732 | -0.17018026 | 0.37852345 | -0.4495897 | 0.65300628 | 0.74323387 | NA | NA |
| ENSG000002 | 10.6773411 | 0.152596347 | 0.33946313 | 0.4495226 | 0.65305471 | 0.74325068 | NA | NA |
| ENSG000001 | 17.5530375 | -0.135797743 | 0.30218675 | -0.4493835 | 0.65315503 | 0.74332312 | RNF133 | 168433 |
| ENSG000002 | 4.21083549 | -0.170019203 | 0.37832003 | -0.4494058 | 0.65313897 | 0.74332312 | RNU6-611P | 106479836 |
| ENSG000002 | 4.67627284 | -0.169028649 | 0.37633934 | -0.4491389 | 0.65333144 | 0.74348215 | NA | NA |
| ENSG000002 | 16.5274397 | 0.138292305 | 0.30789666 | 0.44915169 | 0.65332224 | 0.74348215 | NA | NA |
| ENSG000001 | 79.171194 | 0.085885587 | 0.19127267 | 0.44902173 | 0.65341598 | 0.7435575 | RET | 5979 |
| ENSG000002 | 10.8892865 | -0.149509683 | 0.33304033 | -0.4489237 | 0.65348673 | 0.74361714 | LOC105378732 | 105378732 |
| ENSG000002 | 10.347797 | 0.150397364 | 0.33505743 | 0.4488704 | 0.65352515 | 0.74363999 | NA | NA |
| ENSG00000C | 2.43559389 | -0.165971806 | 0.37001319 | -0.4485565 | 0.65375165 | 0.74387494 | PRKY | 5616 |
| ENSG000002 | 3.6877779 | -0.169646844 | 0.3782257 | -0.4485334 | 0.65376832 | 0.74387494 | NA | NA |

| | | | | | | | | |
|---|---|---|---|---|---|---|---|---|
| ENSG000002 | 3.05593673 | 0.167262247 | 0.37299599 | 0.44842907 | 0.65384356 | 0.74393968 | NA | NA |
| ENSG000002 | 2.4628225 | 0.166580395 | 0.37151363 | 0.448383 | 0.65387681 | 0.74395663 | NA | NA |
| ENSG000001 | 41.2432904 | 0.104769095 | 0.23377191 | 0.44816802 | 0.65403194 | 0.74411226 | TXNRD2 | 10587 |
| ENSG000002 | 5.55348548 | -0.167070985 | 0.37280817 | -0.448142 | 0.65405074 | 0.74411278 | NA | NA |
| ENSG000001 | 6.31225224 | 0.165835065 | 0.37015518 | 0.44801497 | 0.65414239 | 0.74418212 | SOX7 | 83595 |
| ENSG000002 | 9.27192502 | -0.154694876 | 0.34529594 | -0.4480066 | 0.6541484 | 0.74418212 | NA | NA |
| ENSG000001 | 92.2430359 | -0.07731608 | 0.17259477 | -0.447963 | 0.65417986 | 0.74419704 | KCNQ3 | 3786 |
| ENSG000002 | 6.55797002 | 0.162771967 | 0.36338537 | 0.44793209 | 0.65420221 | 0.74420158 | NA | NA |
| ENSG000002 | 76.9618704 | -0.08033398 | 0.17938604 | -0.4478274 | 0.65427778 | 0.74426667 | NA | NA |
| ENSG000002 | 10.2450027 | -0.15135826 | 0.3380101 | -0.4477921 | 0.65430323 | 0.74427474 | NA | NA |
| ENSG000002 | 10.1668717 | -0.149170646 | 0.33320774 | -0.4476806 | 0.65438371 | 0.74434541 | NA | NA |
| ENSG00000C | 14.0240892 | -0.138479693 | 0.30948364 | -0.447454 | 0.65454729 | 0.7445106 | MAMLD1 | 10046 |
| ENSG000002 | 15.2118176 | -0.14078119 | 0.31465875 | -0.4474091 | 0.6545797 | 0.74452657 | NA | NA |
| ENSG000001 | 4.34309852 | 0.168621281 | 0.37691841 | 0.44736812 | 0.65460929 | 0.74453934 | PABPC1L2A | 340529 |
| ENSG000001 | 152.11591 | -0.07172842 | 0.16042769 | -0.4471075 | 0.65479746 | 0.74473247 | SATB2 | 23314 |
| ENSG000002 | 9.03562303 | -0.153605871 | 0.34360173 | -0.4470463 | 0.65484165 | 0.74476185 | NA | NA |
| ENSG000001 | 44.3582837 | 0.100985978 | 0.22594259 | 0.44695415 | 0.65490816 | 0.74481661 | DOC2A | 8448 |
| ENSG000002 | 7.97511118 | -0.159847113 | 0.35777327 | -0.4467833 | 0.65503154 | 0.74493603 | NA | NA |
| ENSG000002 | 10.1513407 | 0.150206802 | 0.33624831 | 0.44671393 | 0.65508162 | 0.74497209 | NA | NA |
| ENSG000001 | 45.9007827 | 0.095056627 | 0.21287607 | 0.44653504 | 0.65521081 | 0.74507721 | SYTL1 | 84958 |
| ENSG000002 | 4.60260214 | 0.167491554 | 0.37507769 | 0.44655163 | 0.65519883 | 0.74507721 | RN7SL45P | 106481975 |
| ENSG000002 | 3.53891059 | 0.169021947 | 0.37858969 | 0.44645153 | 0.65527112 | 0.7451249 | PANDAR | 101154753 |
| ENSG000002 | 4.17003471 | -0.168774203 | 0.37807288 | -0.4464065 | 0.65530362 | 0.74514096 | NA | NA |
| ENSG00000C | 60.610179 | 0.097998304 | 0.21957981 | 0.44629924 | 0.6553811 | 0.74517555 | RELT | 84957 |
| ENSG000002 | 3.44347507 | -0.16840358 | 0.3773858 | -0.4462372 | 0.65542592 | 0.74517555 | NA | NA |
| ENSG000002 | 5.19696127 | -0.167769044 | 0.37593174 | -0.4462753 | 0.65539841 | 0.74517555 | NA | NA |
| ENSG000002 | 25.1265894 | 0.115075904 | 0.25786499 | 0.44626415 | 0.65540645 | 0.74517555 | NA | NA |
| ENSG000002 | 8.93900129 | -0.151899931 | 0.34038994 | -0.4462527 | 0.65541471 | 0.74517555 | NA | NA |
| ENSG000002 | 10.5449886 | 0.147971834 | 0.33166179 | 0.44615279 | 0.65548688 | 0.74522397 | NA | NA |

| | | | | | | | | |
|---|---|---|---|---|---|---|---|---|
| ENSG000002 | 5.21619345 | 0.166806661 | 0.3739881 | 0.44602131 | 0.65558186 | 0.74531105 | NA | NA |
| ENSG000001 | 121.248369 | -0.071389044 | 0.16012605 | -0.4458303 | 0.65571984 | 0.74544702 | FAM227A | 646851 |
| ENSG000001 | 150.204921 | -0.069938825 | 0.15688387 | -0.4458 | 0.65574174 | 0.74545102 | SLIT2 | 9353 |
| ENSG00000C | 39.7459041 | 0.103321408 | 0.23178973 | 0.4457549 | 0.65577431 | 0.74546714 | UNC13D | 201294 |
| ENSG000001 | 6.54991431 | 0.163521762 | 0.36689303 | 0.44569329 | 0.65581881 | 0.74547594 | NA | NA |
| ENSG000002 | 4.37465766 | 0.168674964 | 0.37845054 | 0.44569883 | 0.65581481 | 0.74547594 | NA | NA |
| ENSG000001 | 17.5617547 | -0.141502062 | 0.31751458 | -0.4456553 | 0.65584624 | 0.74548622 | AKAP14 | 158798 |
| ENSG000002 | 9.89398204 | -0.148910469 | 0.33420919 | -0.4455607 | 0.65591463 | 0.74554306 | NA | NA |
| ENSG000002 | 5.85701075 | -0.167021363 | 0.37503376 | -0.4453502 | 0.65606669 | 0.74566969 | NA | NA |
| ENSG000002 | 38.4556211 | -0.101165322 | 0.22716928 | -0.4453301 | 0.65608121 | 0.74566969 | NA | NA |
| ENSG000002 | 4.13182009 | 0.168337098 | 0.37797946 | 0.44536044 | 0.65605929 | 0.74566969 | NA | NA |
| ENSG000002 | 18.2812664 | -0.128159263 | 0.28795093 | -0.4450733 | 0.65626681 | 0.74585973 | NA | NA |
| ENSG000002 | 3.51036067 | 0.168428282 | 0.37852214 | 0.44496283 | 0.65634662 | 0.74592953 | NA | NA |
| ENSG000002 | 3.19435974 | -0.167384771 | 0.37624209 | -0.4448858 | 0.65640227 | 0.74597187 | NA | NA |
| ENSG000001 | 9.89100825 | 0.150314761 | 0.3378962 | 0.44485484 | 0.65642466 | 0.74597641 | LCN2 | 3934 |
| ENSG000002 | 2.40593173 | 0.165606648 | 0.3724205 | 0.44467651 | 0.65655355 | 0.74610198 | NA | NA |
| ENSG000002 | 6.39374516 | -0.16379775 | 0.36868929 | -0.4442704 | 0.65684708 | 0.74639371 | NA | NA |
| ENSG000002 | 5.48393015 | 0.164433276 | 0.37010905 | 0.44428331 | 0.65683776 | 0.74639371 | NA | NA |
| ENSG000001 | 98.3636619 | 0.076338996 | 0.17184049 | 0.44424336 | 0.65686664 | 0.74639503 | PPP1R1B | 84152 |
| ENSG000002 | 8.12014499 | 0.155636293 | 0.35038906 | 0.44418137 | 0.65691146 | 0.74642503 | NA | NA |
| ENSG000002 | 4.3871886 | 0.167314955 | 0.37672154 | 0.44413429 | 0.6569455 | 0.74644279 | NA | NA |
| ENSG000002 | 42.2382137 | -0.100112339 | 0.22542924 | -0.4440965 | 0.65697281 | 0.74645292 | APOBEC3G | 60489 |
| ENSG000001 | 19.063006 | 0.127160043 | 0.28650582 | 0.44383057 | 0.65716508 | 0.74662953 | PRR19 | 284338 |
| ENSG000002 | 3.50490028 | 0.168031178 | 0.37858193 | 0.44384363 | 0.65715564 | 0.74662953 | NA | NA |
| ENSG000002 | 3.30851784 | 0.167408142 | 0.37723013 | 0.44378253 | 0.65719982 | 0.74664808 | TEX13A | 56157 |
| ENSG000002 | 2.21205163 | -0.160719016 | 0.3621878 | -0.443745 | 0.65722698 | 0.74665801 | NA | NA |
| ENSG000001 | 66.3316269 | 0.082929751 | 0.18697986 | 0.44352236 | 0.65738795 | 0.74681996 | TDRD7 | 23424 |
| ENSG000002 | 7.01037794 | -0.161605002 | 0.36439024 | -0.4434943 | 0.65740823 | 0.74682208 | NA | NA |
| ENSG000002 | 3.52862175 | 0.167842477 | 0.37857151 | 0.44335739 | 0.65750725 | 0.74691365 | NA | NA |

| | | | | | | | | |
|---|---|---|---|---|---|---|---|---|
| ENSG000002 | 8.12780521 | -0.164724273 | 0.37156351 | -0.4433274 | 0.65752896 | 0.74691739 | NA | NA |
| ENSG000001 | 34.1227989 | -0.1032151 | 0.23291476 | -0.4431454 | 0.65766059 | 0.74704598 | CYP11A1 | 1583 |
| ENSG000002 | 3.86102039 | -0.167720472 | 0.3785063 | -0.4431114 | 0.65768513 | 0.74705294 | NA | NA |
| ENSG000002 | 2.67397157 | -0.164642311 | 0.37158118 | -0.4430857 | 0.65770374 | 0.74705315 | RNU6-1209P | 106480095 |
| ENSG000002 | 5.46207506 | 0.163909741 | 0.36996427 | 0.44304208 | 0.6577353 | 0.74706806 | MGC15885 | 197003 |
| ENSG000002 | 7.74270578 | -0.157343287 | 0.35517011 | -0.4430082 | 0.65775977 | 0.74707494 | NA | NA |
| ENSG000002 | 4.68288726 | -0.16705894 | 0.37753198 | -0.4425028 | 0.65812544 | 0.74746932 | NA | NA |
| ENSG000002 | 3.15155617 | -0.166763701 | 0.37702096 | -0.4423194 | 0.65825806 | 0.74759901 | NA | NA |
| ENSG000001 | 27.9532783 | -0.110637615 | 0.25018945 | -0.4422154 | 0.65833338 | 0.74766361 | CACNG6 | 59285 |
| ENSG000002 | 8.52964814 | 0.156026184 | 0.35286979 | 0.44216362 | 0.65837081 | 0.74766424 | HMGN2P15 | 100288165 |
| ENSG000002 | 12.2938656 | 0.144240395 | 0.32621244 | 0.44216706 | 0.65836832 | 0.74766424 | CNTF | 1270 |
| ENSG000000 | 158.110473 | 0.062248844 | 0.14080395 | 0.44209586 | 0.65841984 | 0.74767805 | PDE8A | 5151 |
| ENSG000002 | 2.21308168 | -0.159848005 | 0.36155723 | -0.4421098 | 0.65840973 | 0.74767805 | RN7SKP16 | 106480837 |
| ENSG000002 | 19.9170338 | 0.123078802 | 0.27842466 | 0.44205424 | 0.65844996 | 0.7476913 | NA | NA |
| ENSG000002 | 6.59454395 | 0.160151772 | 0.36242345 | 0.44189131 | 0.65856786 | 0.74780425 | NA | NA |
| ENSG000002 | 5.65000317 | -0.164172123 | 0.37171623 | -0.4416598 | 0.65873539 | 0.74797354 | NA | NA |
| ENSG000002 | 3.18103863 | -0.166317094 | 0.37661388 | -0.4416117 | 0.65877022 | 0.74799214 | NA | NA |
| ENSG000002 | 3.02236411 | -0.164172392 | 0.37185315 | -0.4414979 | 0.65885257 | 0.74806469 | BIRC6-AS2 | 103752586 |
| ENSG000002 | 2.96272843 | -0.164929133 | 0.37364429 | -0.4414068 | 0.65891852 | 0.74811863 | LOC105373343 | 105373343 |
| ENSG000002 | 3.29576887 | 0.166759914 | 0.37791412 | 0.44126405 | 0.65902185 | 0.748215 | NA | NA |
| ENSG000002 | 5.41219265 | 0.164889457 | 0.37376743 | 0.44115523 | 0.65910062 | 0.74828348 | NA | NA |
| ENSG000000 | 96.944001 | 0.078754877 | 0.17855091 | 0.44107798 | 0.65915654 | 0.74830507 | BTN3A1 | 11119 |
| ENSG000002 | 8.97578429 | -0.151316309 | 0.34305648 | -0.4410828 | 0.65915306 | 0.74830507 | NA | NA |
| ENSG000001 | 8.07692376 | -0.155520583 | 0.35271945 | -0.4409186 | 0.6592719 | 0.74839412 | TWIST1 | 7291 |
| ENSG000002 | 7.53109241 | 0.158993929 | 0.36057824 | 0.44094155 | 0.65925531 | 0.74839412 | NA | NA |
| ENSG000002 | 2.94198825 | -0.164911805 | 0.37404425 | -0.4408885 | 0.65929373 | 0.74839795 | GRIFIN | 402635 |
| ENSG000002 | 5.47457519 | 0.163242087 | 0.37034207 | 0.44078732 | 0.65936698 | 0.7484392 | NA | NA |
| ENSG000002 | 3.37590678 | -0.166453234 | 0.37761264 | -0.4408042 | 0.65935476 | 0.7484392 | PRELID3BP2 | 106481667 |
| ENSG000002 | 16.5944977 | 0.131934156 | 0.29933299 | 0.4407605 | 0.65938639 | 0.74844028 | CCDC177 | 56936 |

| | | | | | | | | |
|---|---|---|---|---|---|---|---|---|
| ENSG000002 | 11.748779 | 0.144956765 | 0.32900716 | 0.44058848 | 0.65951094 | 0.7485607 | NA | NA |
| ENSG000002 | 8.78736858 | 0.152182397 | 0.34565613 | 0.44027108 | 0.65974078 | 0.74880062 | NA | NA |
| ENSG000001 | 45.7746714 | 0.097461392 | 0.22141549 | 0.44017423 | 0.65981092 | 0.74885927 | STARD5 | 80765 |
| ENSG000001 | 31.9459632 | -0.108657954 | 0.24688252 | -0.4401201 | 0.65985014 | 0.74888281 | CKMT2 | 1160 |
| ENSG000002 | 7.58941252 | -0.160581182 | 0.36502256 | -0.4399212 | 0.65999414 | 0.74902529 | LINC01176 | 100506516 |
| ENSG000002 | 26.8200936 | 0.113849117 | 0.25885547 | 0.43981732 | 0.66006942 | 0.74906879 | SRP14-AS1 | 100131089 |
| ENSG000002 | 2.39734622 | 0.163081521 | 0.37078045 | 0.43983312 | 0.66005798 | 0.74906879 | NA | NA |
| ENSG000000 | 40.7989166 | -0.107065992 | 0.24353538 | -0.4396322 | 0.66020353 | 0.74916702 | SOX8 | 30812 |
| ENSG000002 | 97.8730735 | 0.073065301 | 0.16618374 | 0.43966575 | 0.66017921 | 0.74916702 | NA | NA |
| ENSG000002 | 13.5257965 | -0.138881453 | 0.31591156 | -0.4396213 | 0.6602114 | 0.74916702 | NA | NA |
| ENSG000002 | 53.3589688 | 0.095500144 | 0.21724978 | 0.43958684 | 0.66023637 | 0.74917439 | NA | NA |
| ENSG000001 | 6.57513384 | 0.15977315 | 0.36356957 | 0.43945689 | 0.66033051 | 0.74926025 | CLCF1 | 23529 |
| ENSG000002 | 816.232439 | -0.047052194 | 0.10710287 | -0.4393178 | 0.66043131 | 0.74935365 | LINC00707 | 100507127 |
| ENSG000002 | 2.44319114 | 0.163500754 | 0.37226503 | 0.43920525 | 0.66051282 | 0.74940798 | NA | NA |
| ENSG000002 | 3.91029066 | -0.166258855 | 0.37854875 | -0.4392006 | 0.66051615 | 0.74940798 | NA | NA |
| ENSG000002 | 10.5690748 | 0.145516143 | 0.33134206 | 0.43917197 | 0.66053694 | 0.74941059 | NA | NA |
| ENSG000002 | 16.6089706 | -0.130452109 | 0.29706962 | -0.4391298 | 0.66056752 | 0.74942432 | PCDHGB3 | 56102 |
| ENSG000001 | 18.780036 | 0.133170951 | 0.30327882 | 0.43910403 | 0.66058616 | 0.7494245 | PGAM2 | 5224 |
| ENSG000002 | 17.3451355 | -0.127265284 | 0.28985619 | -0.4390635 | 0.6606155 | 0.74943682 | NA | NA |
| ENSG000002 | 48.826778 | 0.096168007 | 0.21904633 | 0.43903044 | 0.66063948 | 0.74944306 | NA | NA |
| ENSG000001 | 119.256269 | 0.068259033 | 0.15549268 | 0.43898552 | 0.66067203 | 0.74945902 | PUS10 | 150962 |
| ENSG000002 | 2.41964514 | 0.163538347 | 0.37273216 | 0.43875567 | 0.66083858 | 0.74962698 | NA | NA |
| ENSG000002 | 2.4380106 | 0.163236287 | 0.37225231 | 0.4385098 | 0.66101676 | 0.74980813 | NA | NA |
| ENSG000002 | 17.4857099 | -0.1268366 | 0.28926423 | -0.4384801 | 0.66103831 | 0.74981159 | LINC00958 | 100506305 |
| ENSG000002 | 2.45898 | 0.16266455 | 0.3711327 | 0.43829215 | 0.66117452 | 0.74990316 | NA | NA |
| ENSG000002 | 7.84665442 | -0.155889704 | 0.35565884 | -0.4383125 | 0.66115979 | 0.74990316 | NA | NA |
| ENSG000002 | 8.67588439 | -0.151092203 | 0.34472249 | -0.438301 | 0.66116812 | 0.74990316 | NA | NA |
| ENSG000002 | 2.42562292 | 0.161220272 | 0.36788836 | 0.4382315 | 0.66121847 | 0.74993204 | NA | NA |
| ENSG000002 | 4.40064261 | -0.16553161 | 0.37785775 | -0.4380792 | 0.66132889 | 0.7500363 | THCAT158 | 102724508 |

| | | | | | | | | |
|---|---|---|---|---|---|---|---|---|
| ENSG000002 | 16.5506153 | -0.127460328 | 0.29096998 | -0.4380532 | 0.66134773 | 0.75003668 | LINC00926 | 283663 |
| ENSG000001 | 78.0771894 | 0.083619351 | 0.1909268 | 0.4379655 | 0.66141129 | 0.75008779 | ALG14 | 199857 |
| ENSG000002 | 25.8422747 | -0.112906439 | 0.25793468 | -0.4377327 | 0.66158008 | 0.75025823 | ARRDC3-AS1 | 100129716 |
| ENSG000002 | 3.09256349 | 0.162136669 | 0.37043108 | 0.43769727 | 0.66160575 | 0.75026636 | NA | NA |
| ENSG000001 | 4.38216428 | 0.165056217 | 0.37718669 | 0.4375982 | 0.66167758 | 0.75032683 | NA | NA |
| ENSG000001 | 181.055725 | -0.068372001 | 0.15630546 | -0.4374255 | 0.66180276 | 0.75038631 | NCOA6 | 23054 |
| ENSG000002 | 7.56826238 | -0.1588472 | 0.36314258 | -0.4374238 | 0.66180404 | 0.75038631 | LINC01608 | 101927459 |
| ENSG000002 | 2.96779761 | -0.163996846 | 0.37488145 | -0.4374632 | 0.66177544 | 0.75038631 | NA | NA |
| ENSG000002 | 13.4862415 | -0.136970634 | 0.31310756 | -0.4374555 | 0.66178102 | 0.75038631 | NA | NA |
| ENSG000001 | 132.339141 | -0.066536068 | 0.15214126 | -0.4373309 | 0.66187142 | 0.75044172 | EIF5A2 | 56648 |
| ENSG000002 | 4.71252594 | -0.165309534 | 0.37802473 | -0.4372982 | 0.6618951 | 0.75044758 | LOC101927908 | 101927908 |
| ENSG000002 | 13.2145858 | -0.142440346 | 0.32594847 | -0.4370027 | 0.66210942 | 0.75066959 | LOC101929988 | 101929988 |
| ENSG000002 | 26.2454788 | 0.113190846 | 0.25920906 | 0.43667782 | 0.66234502 | 0.7509157 | NA | NA |
| ENSG000002 | 8.94323575 | -0.148977414 | 0.3413273 | -0.436465 | 0.6624994 | 0.75106973 | LOC105370809 | 105370809 |
| ENSG000002 | 8.66391764 | -0.151478014 | 0.34716802 | -0.4363248 | 0.6626011 | 0.75116402 | HCG23 | 414764 |
| ENSG000002 | 6.52496488 | 0.160945451 | 0.36912621 | 0.4360174 | 0.66282411 | 0.75139583 | NA | NA |
| ENSG000002 | 5.88885658 | -0.162604655 | 0.37297136 | -0.4359709 | 0.66285786 | 0.75141309 | NA | NA |
| ENSG000001 | 177.875623 | 0.064900185 | 0.1488996 | 0.4358654 | 0.66293439 | 0.75146742 | B3GLCT | 145173 |
| ENSG000002 | 14.100622 | 0.136228899 | 0.31255645 | 0.43585374 | 0.66294285 | 0.75146742 | NA | NA |
| ENSG000001 | 68.518959 | 0.084567077 | 0.19408062 | 0.4357317 | 0.6630314 | 0.75154679 | UGT3A1 | 133688 |
| ENSG000001 | 100.172665 | 0.076330447 | 0.17523543 | 0.43558798 | 0.66313569 | 0.75164399 | CRADD | 8738 |
| ENSG000002 | 5.92720084 | 0.160026101 | 0.36741882 | 0.43554139 | 0.6631695 | 0.75166131 | NA | NA |
| ENSG000002 | 20.5722859 | -0.119776183 | 0.27504262 | -0.4354823 | 0.66321241 | 0.75168893 | NA | NA |
| ENSG000001 | 820.036397 | -0.052487496 | 0.12058033 | -0.4352907 | 0.66335143 | 0.75182548 | AKT3 | 10000 |
| ENSG000001 | 29.8244958 | 0.110365047 | 0.25375311 | 0.43493081 | 0.66361265 | 0.75209961 | TPGS1 | 91978 |
| ENSG000002 | 3.8923626 | -0.164638084 | 0.37858208 | -0.4348808 | 0.66364894 | 0.75209961 | NA | NA |
| ENSG000002 | 8.16407056 | 0.151341329 | 0.34800223 | 0.4348861 | 0.66364511 | 0.75209961 | TGFBR3L | 100507588 |
| ENSG000002 | 22.596026 | -0.120434391 | 0.27697298 | -0.4348236 | 0.66369047 | 0.75212566 | AARSD1 | 80755 |
| ENSG000002 | 11.3184652 | -0.149881268 | 0.34472399 | -0.4347863 | 0.66371755 | 0.75213533 | DEPDC1-AS1 | 101927220 |

| | | | | | | | | |
|---|---|---|---|---|---|---|---|---|
| ENSG000002 | 4.20254945 | -0.164411241 | 0.37857799 | -0.4342863 | 0.66408054 | 0.75252563 | LINC01574 | 102577424 |
| ENSG000001 | 6.59195059 | -0.157339133 | 0.36258379 | -0.4339387 | 0.66433296 | 0.75279064 | MLNR | 2862 |
| ENSG000002 | 3.39985772 | -0.163990276 | 0.37794012 | -0.4339054 | 0.6643571 | 0.75279696 | NA | NA |
| ENSG000002 | 11.9266966 | 0.13955032 | 0.32178063 | 0.43368155 | 0.66451971 | 0.75293775 | NA | NA |
| ENSG000002 | 6.61578618 | -0.160072882 | 0.36912269 | -0.4336577 | 0.66453706 | 0.75293775 | NA | NA |
| ENSG000002 | 29.9707919 | 0.108937064 | 0.25119143 | 0.43368146 | 0.66451977 | 0.75293775 | NA | NA |
| ENSG000002 | 12.0763097 | 0.140594466 | 0.32425649 | 0.43359029 | 0.66458599 | 0.75297215 | NA | NA |
| ENSG000002 | 45.7467576 | -0.092558219 | 0.21352239 | -0.4334825 | 0.66466428 | 0.75303982 | LOC105378405 | 105378405 |
| ENSG000002 | 7.54647487 | -0.153129322 | 0.35337128 | -0.4333383 | 0.66476899 | 0.75311805 | NA | NA |
| ENSG000002 | 20.3044976 | -0.121502805 | 0.28038917 | -0.4333363 | 0.66477047 | 0.75311805 | NA | NA |
| ENSG000001 | 200.535312 | 0.060124817 | 0.13878106 | 0.43323504 | 0.66484403 | 0.75316067 | CAMK4 | 814 |
| ENSG000002 | 43.6363992 | -0.0943376 | 0.21775239 | -0.4332334 | 0.66484524 | 0.75316067 | PCDHGB1 | 56104 |
| ENSG000002 | 2.93020572 | -0.162246018 | 0.37456503 | -0.4331585 | 0.66489964 | 0.75320125 | NA | NA |
| ENSG000002 | 59.10809 | 0.087585559 | 0.20224476 | 0.43306713 | 0.664966 | 0.75325538 | NA | NA |
| ENSG000002 | 2.4503945 | 0.159848548 | 0.36920739 | 0.43295057 | 0.66505068 | 0.75333026 | NA | NA |
| ENSG000001 | 395.379475 | 0.05531032 | 0.12778705 | 0.43283196 | 0.66513686 | 0.75340611 | TLE4 | 7091 |
| ENSG000002 | 9.87647833 | -0.147178822 | 0.34005627 | -0.4328073 | 0.6651548 | 0.75340611 | ZNF436-AS1 | 148898 |
| ENSG000002 | 2.4129062 | 0.160647565 | 0.37127798 | 0.43268811 | 0.66524137 | 0.75348313 | NA | NA |
| ENSG000002 | 253.573127 | 0.06582657 | 0.15219368 | 0.43251842 | 0.66536467 | 0.75358593 | LINC01356 | 100996702 |
| ENSG000002 | 6.05998848 | 0.160772473 | 0.37171791 | 0.43251204 | 0.6653693 | 0.75358593 | NA | NA |
| ENSG000002 | 5.7026002 | 0.159270439 | 0.36835957 | 0.43237764 | 0.66546697 | 0.7536755 | RN7SL67P | 106479248 |
| ENSG00000C | 48.053755 | -0.092111336 | 0.21306651 | -0.4323126 | 0.66551423 | 0.75370797 | SEMA3C | 10512 |
| ENSG000002 | 2.66768685 | 0.162002372 | 0.3748269 | 0.43220583 | 0.66559182 | 0.75377479 | NA | NA |
| ENSG000002 | 5.70736359 | 0.159922328 | 0.3700631 | 0.43214881 | 0.66563326 | 0.75380067 | NA | NA |
| ENSG000002 | 3.15299813 | -0.162688136 | 0.3764897 | -0.4321184 | 0.66565534 | 0.75380463 | NA | NA |
| ENSG000001 | 14.0616295 | -0.13400744 | 0.31031737 | -0.43184 | 0.66585774 | 0.75401276 | SPINK1 | 6690 |
| ENSG000001 | 109.876183 | 0.069623987 | 0.16124215 | 0.43179768 | 0.66588847 | 0.75402651 | WNT5A | 7474 |
| ENSG000002 | 7.94143157 | -0.151869648 | 0.35175723 | -0.4317456 | 0.6659263 | 0.75404829 | NA | NA |
| ENSG000001 | 84.6345151 | 0.077019214 | 0.17842661 | 0.43165767 | 0.66599024 | 0.75409963 | SYT17 | 51760 |

| | | | | | | | | |
|---|---|---|---|---|---|---|---|---|
| ENSG000002 | 2.91164459 | -0.161548168 | 0.37437485 | -0.4315145 | 0.66609433 | 0.75419644 | NA | NA |
| ENSG000001 | 25.7904456 | -0.112893292 | 0.26174937 | -0.431303 | 0.66624806 | 0.75434943 | PRSS27 | 83886 |
| ENSG000000 | 7.79987997 | -0.158864789 | 0.36886459 | -0.4306859 | 0.66669675 | 0.75483638 | ADRB1 | 153 |
| ENSG000002 | 7.46851862 | -0.151867348 | 0.3526424 | -0.4306554 | 0.66671897 | 0.75484046 | NA | NA |
| ENSG000001 | 12.2675101 | -0.137043265 | 0.3183699 | -0.430453 | 0.66686618 | 0.75495526 | ROM1 | 6094 |
| ENSG000002 | 11.7949167 | -0.139654359 | 0.32444623 | -0.4304392 | 0.66687622 | 0.75495526 | NA | NA |
| ENSG000002 | 2.63385315 | 0.160849961 | 0.37368613 | 0.43044135 | 0.66687462 | 0.75495526 | NA | NA |
| ENSG000000 | 98.0380524 | 0.087207985 | 0.20265154 | 0.43033468 | 0.66695221 | 0.75502021 | ANKRD24 | 170961 |
| ENSG000002 | 15.0076917 | -0.133073564 | 0.30925138 | -0.4303087 | 0.66697108 | 0.7550205 | LOC729739 | 729739 |
| ENSG000002 | 4.61980564 | 0.161141939 | 0.3745807 | 0.43019285 | 0.66705536 | 0.75509483 | NA | NA |
| ENSG000002 | 20.9723883 | -0.121367473 | 0.28230793 | -0.4299117 | 0.66725989 | 0.75530527 | WDR86-AS1 | 100131176 |
| ENSG000001 | 25.7685756 | 0.110632528 | 0.25742041 | 0.42977372 | 0.66736025 | 0.75539778 | IRAK2 | 3656 |
| ENSG000002 | 3.54065356 | 0.162159684 | 0.37759347 | 0.42945574 | 0.6675916 | 0.75563855 | NA | NA |
| ENSG000002 | 3.85930465 | -0.162481966 | 0.37857419 | -0.4291945 | 0.66778168 | 0.75581421 | UPK1A-AS1 | 100862728 |
| ENSG000002 | 4.20085466 | -0.162480825 | 0.37857443 | -0.4291912 | 0.66778407 | 0.75581421 | NA | NA |
| ENSG000001 | 60.906441 | -0.087931704 | 0.20497646 | -0.4289844 | 0.66793458 | 0.75596347 | ARID5B | 84159 |
| ENSG000001 | 83.2965223 | -0.081150719 | 0.18918966 | -0.4289384 | 0.66796802 | 0.75598022 | F8 | 2157 |
| ENSG000001 | 77.1211272 | -0.076075511 | 0.17741587 | -0.4287977 | 0.66807048 | 0.75607508 | FBXL13 | 222235 |
| ENSG000002 | 2.65761484 | 0.159157887 | 0.37131721 | 0.42863052 | 0.66819213 | 0.75619165 | RNU6-658P | 106479858 |
| ENSG000002 | 17.7531811 | 0.123615434 | 0.28853077 | 0.42843068 | 0.66833759 | 0.75633516 | NA | NA |
| ENSG000002 | 14.3708512 | 0.13159602 | 0.30723074 | 0.42832959 | 0.66841118 | 0.75639732 | NA | NA |
| ENSG000002 | 3.47716024 | 0.161994567 | 0.37822716 | 0.42829967 | 0.66843296 | 0.75640087 | NA | NA |
| ENSG000002 | 2.65946608 | 0.160389076 | 0.37482323 | 0.42790591 | 0.66871963 | 0.75670414 | NA | NA |
| ENSG000002 | 2.19315802 | -0.153662602 | 0.3591656 | -0.4278322 | 0.66877331 | 0.75671099 | NA | NA |
| ENSG000002 | 11.5868672 | 0.140749952 | 0.32897759 | 0.42784055 | 0.66876721 | 0.75671099 | RN7SL809P | 106479516 |
| ENSG000002 | 7.27779893 | -0.158570764 | 0.37064771 | -0.4278207 | 0.66878167 | 0.75671099 | NA | NA |
| ENSG000002 | 3.24000795 | 0.161111264 | 0.37665492 | 0.42774236 | 0.66883871 | 0.75675442 | LOC100507406 | 100507406 |
| ENSG000002 | 6.80945146 | -0.154552653 | 0.36135123 | -0.4277076 | 0.66886405 | 0.75676198 | NA | NA |
| ENSG000001 | 32.7044865 | 0.104889878 | 0.2453538 | 0.4275046 | 0.66901184 | 0.75690807 | TMEM215 | 401498 |

| | | | | | | | | | |
|---|---|---|---|---|---|---|---|---|---|
| ENSG000002 | 3.94206895 | -0.161689419 | 0.37824262 | -0.4274754 | 0.66903309 | 0.756911 | NA | NA | |
| ENSG000002 | 7.02027288 | -0.152461896 | 0.35677733 | -0.4273307 | 0.66913849 | 0.75700913 | NA | NA | |
| ENSG000001 | 102.29283 | 0.078457942 | 0.18368743 | 0.42712745 | 0.6692865 | 0.75715545 | ITPKB | | 3707 |
| ENSG000002 | 2.70372703 | -0.156353876 | 0.36614092 | -0.427032 | 0.66935605 | 0.757213 | NA | NA | |
| ENSG000002 | 2.40792244 | 0.156816323 | 0.36725812 | 0.42699212 | 0.66938507 | 0.7572247 | NA | NA | |
| ENSG000002 | 7.25127888 | -0.151617325 | 0.3551756 | -0.4268799 | 0.66946681 | 0.75729604 | NA | NA | |
| ENSG000002 | 11.2185601 | 0.144869106 | 0.33943304 | 0.4267973 | 0.66952697 | 0.75732185 | NA | NA | |
| ENSG000002 | 2.19833801 | -0.153019429 | 0.35852265 | -0.4268055 | 0.66952102 | 0.75732185 | NA | NA | |
| ENSG000002 | 5.71251655 | 0.157573287 | 0.36972369 | 0.42619203 | 0.66996793 | 0.75779949 | NA | NA | |
| ENSG000002 | 8.03026483 | -0.150158264 | 0.35254349 | -0.4259283 | 0.67016007 | 0.75799567 | LOC102723617 | | 102723617 |
| ENSG000002 | 34.644708 | 0.097664479 | 0.22931417 | 0.42589815 | 0.67018206 | 0.75799941 | NA | NA | |
| ENSG000002 | 82.8375197 | -0.075918428 | 0.17827554 | -0.4258488 | 0.67021801 | 0.75801893 | CPEB1 | | 64506 |
| ENSG000002 | 14.4428464 | 0.1334631 | 0.31343004 | 0.42581465 | 0.67024291 | 0.75802595 | NA | NA | |
| ENSG000001 | 33.6659228 | -0.115880368 | 0.27260785 | -0.4250808 | 0.67077776 | 0.75859303 | DCAF4L1 | | 285429 |
| ENSG000002 | 4.61009814 | 0.160050853 | 0.37652347 | 0.42507537 | 0.67078174 | 0.75859303 | NA | NA | |
| ENSG000002 | 3.50472367 | 0.160881209 | 0.37854787 | 0.42499568 | 0.67083983 | 0.75863757 | NA | NA | |
| ENSG000002 | 2.66122811 | 0.158962956 | 0.37434786 | 0.42463968 | 0.67109936 | 0.75890634 | SHISA8 | | 440829 |
| ENSG000002 | 4.83612079 | 0.1593718 | 0.37532953 | 0.42461833 | 0.67111493 | 0.75890634 | NA | NA | |
| ENSG00000C | 146.455244 | 0.064492942 | 0.15190186 | 0.42456979 | 0.67115032 | 0.7589252 | KIFAP3 | | 22920 |
| ENSG000002 | 7.48687646 | -0.15103029 | 0.35581948 | -0.4244576 | 0.6712321 | 0.75899651 | NA | NA | |
| ENSG000002 | 7.29555743 | -0.151857723 | 0.35787745 | -0.4243288 | 0.67132601 | 0.75908154 | INE2 | | 8551 |
| ENSG000002 | 163.619898 | 0.062372958 | 0.14708875 | 0.42404982 | 0.67152948 | 0.75929043 | NA | NA | |
| ENSG000001 | 225.430798 | 0.057584404 | 0.13582222 | 0.42396896 | 0.67158845 | 0.75933593 | TTC7B | | 145567 |
| ENSG000001 | 30.3592849 | -0.105554538 | 0.24901207 | -0.4238933 | 0.67164367 | 0.75937719 | ADPRHL1 | | 113622 |
| ENSG000002 | 8.35542901 | 0.147311828 | 0.34765811 | 0.42372613 | 0.67176555 | 0.75947265 | SNORA7B | | 677797 |
| ENSG000002 | 5.91027666 | 0.156110764 | 0.36841571 | 0.42373537 | 0.67175882 | 0.75947265 | NA | NA | |
| ENSG000002 | 2.21711662 | 0.153557221 | 0.36243665 | 0.42368017 | 0.67179908 | 0.75948937 | NA | NA | |
| ENSG000002 | 19.3431022 | 0.119670202 | 0.28251336 | 0.4235913 | 0.6718639 | 0.75954149 | NA | NA | |
| ENSG000002 | 2.45081905 | 0.154504585 | 0.36484918 | 0.42347521 | 0.67194858 | 0.75959486 | NA | NA | |

| | | | | | | | | |
|---|---|---|---|---|---|---|---|---|
| ENSG000002 | 74.4522715 | 0.078280206 | 0.18484656 | 0.42348749 | 0.67193962 | 0.75959486 | PWAR1 | 145624 |
| ENSG000002 | 24.9993399 | -0.118873928 | 0.28076832 | -0.423388 | 0.67201222 | 0.75964563 | NA | NA |
| ENSG000002 | 3.75151943 | 0.160263381 | 0.37858954 | 0.42331699 | 0.672064 | 0.75968298 | NA | NA |
| ENSG000001 | 13.8797584 | 0.132958274 | 0.31419291 | 0.42317402 | 0.6721683 | 0.7597797 | PNMT | 5409 |
| ENSG000001 | 22.0949856 | -0.122372622 | 0.28934871 | -0.4229244 | 0.67235044 | 0.75996439 | MST1R | 4486 |
| ENSG000001 | 3.22085459 | -0.158519706 | 0.37490245 | -0.4228292 | 0.67241987 | 0.76000051 | NA | NA |
| ENSG000002 | 4.58354436 | 0.158988528 | 0.37598982 | 0.42285328 | 0.67240231 | 0.76000051 | NA | NA |
| ENSG000001 | 42.221154 | -0.097081515 | 0.22962957 | -0.4227745 | 0.67245983 | 0.76002449 | PCDHB14 | 56122 |
| ENSG000001 | 300.946235 | 0.158733498 | 0.37561426 | 0.42259711 | 0.67258924 | 0.76014956 | HMGN5 | 79366 |
| ENSG000001 | 47.5958469 | 0.094154857 | 0.22283093 | 0.42253944 | 0.67263132 | 0.76017594 | TRPM4 | 54795 |
| ENSG000002 | 4.11535612 | -0.159863898 | 0.37857936 | -0.4222731 | 0.67282565 | 0.76037437 | NA | NA |
| ENSG000001 | 8.98151333 | -0.144855713 | 0.34312564 | -0.4221652 | 0.67290443 | 0.7604422 | GIP | 2695 |
| ENSG000001 | 81.1736704 | 0.079936369 | 0.18942685 | 0.42199071 | 0.6730318 | 0.76054375 | NA | NA |
| ENSG000002 | 8.78921535 | 0.149336864 | 0.35388476 | 0.42199293 | 0.67303018 | 0.76054375 | NA | NA |
| ENSG000002 | 12.1936112 | 0.1345609 | 0.31899338 | 0.42182976 | 0.67314928 | 0.76065531 | NA | NA |
| ENSG000002 | 3.30640925 | 0.159353281 | 0.37794107 | 0.42163526 | 0.67329126 | 0.76077322 | NA | NA |
| ENSG000002 | 2.42954886 | 0.156362385 | 0.37086885 | 0.421611 | 0.67330898 | 0.76077322 | NA | NA |
| ENSG000002 | 2.49302257 | 0.15550776 | 0.36884292 | 0.42160972 | 0.67330991 | 0.76077322 | NA | NA |
| ENSG000002 | 9.15498285 | -0.14454661 | 0.34288117 | -0.4215647 | 0.67334275 | 0.76078913 | NA | NA |
| ENSG000001 | 7.47417489 | -0.150114478 | 0.35616684 | -0.4214724 | 0.67341019 | 0.76084413 | CYSRT1 | 375791 |
| ENSG000002 | 2.38256289 | 0.154313167 | 0.36617388 | 0.42142046 | 0.67344808 | 0.76086574 | NA | NA |
| ENSG000001 | 13.447369 | 0.131839214 | 0.31295861 | 0.42126725 | 0.67355994 | 0.76097092 | GPS2 | 2874 |
| ENSG000002 | 6.14435625 | 0.154121051 | 0.36597213 | 0.42112783 | 0.67366174 | 0.76106472 | JMJD7 | 100137047 |
| ENSG000001 | 63.5641094 | 0.084314089 | 0.20022693 | 0.42109264 | 0.67368743 | 0.76107255 | TTYH2 | 94015 |
| ENSG000002 | 7.02575367 | -0.151634729 | 0.36022995 | -0.4209387 | 0.67379984 | 0.76117833 | LOC339568 | 339568 |
| ENSG000002 | 7.86192734 | -0.151019874 | 0.35896329 | -0.4207112 | 0.67396599 | 0.76134481 | NA | NA |
| ENSG000001 | 9.25036671 | 0.145143294 | 0.34504249 | 0.42065339 | 0.6740082 | 0.76137129 | LMNTD2 | 256329 |
| ENSG000001 | 474.435418 | -0.050648929 | 0.120413 | -0.4206268 | 0.67402765 | 0.76137205 | SORBS2 | 8470 |
| ENSG000001 | 16.5454097 | -0.125005987 | 0.29733836 | -0.4204166 | 0.67418114 | 0.76152421 | IL15RA | 3601 |

| | | | | | | | | |
|---|---|---|---|---|---|---|---|---|
| ENSG000002 | 6.80438337 | -0.151161548 | 0.35958305 | -0.4203801 | 0.67420783 | 0.76153315 | NA | NA |
| ENSG000001 | 325.879701 | -0.053775213 | 0.12800805 | -0.4200924 | 0.67441793 | 0.76174076 | MIER2 | 54531 |
| ENSG000002 | 6.81281374 | 0.150894996 | 0.35920794 | 0.42007701 | 0.6744292 | 0.76174076 | NA | NA |
| ENSG000002 | 8.56095455 | 0.147674249 | 0.35168055 | 0.41991019 | 0.67455106 | 0.76184 | NA | NA |
| ENSG000002 | 2.43833156 | 0.153817998 | 0.36631592 | 0.4199053 | 0.67455464 | 0.76184 | RN7SL336P | 106479346 |
| ENSG000001 | 7.74595051 | -0.147919633 | 0.35232948 | -0.4198333 | 0.67460726 | 0.76187821 | DDN | 23109 |
| ENSG000002 | 4.34854748 | -0.158874246 | 0.37858916 | -0.4196482 | 0.6747425 | 0.76200972 | NA | NA |
| ENSG000002 | 13.4551412 | -0.131220957 | 0.31272155 | -0.4196096 | 0.67477069 | 0.76202034 | NA | NA |
| ENSG000002 | 2.63291951 | 0.157088296 | 0.37472349 | 0.41921123 | 0.67506176 | 0.76232782 | NA | NA |
| ENSG000002 | 20.5529464 | 0.116879457 | 0.27897265 | 0.41896385 | 0.67524255 | 0.76251074 | TUBB3 | 10381 |
| ENSG000002 | 11.9153431 | 0.136966917 | 0.32711505 | 0.41871176 | 0.6754268 | 0.76269756 | LOC100507670 | 100507670 |
| ENSG000002 | 7.20873569 | -0.149712308 | 0.35760579 | -0.4186518 | 0.67547062 | 0.76272581 | NA | NA |
| ENSG000001 | 20.825045 | -0.12764982 | 0.30501332 | -0.4185057 | 0.67557741 | 0.76281542 | OR6A2 | 8590 |
| ENSG000002 | 80.832435 | -0.075534408 | 0.18049198 | -0.4184918 | 0.67558761 | 0.76281542 | LPAL2 | 80350 |
| ENSG000001 | 7.48784963 | -0.148848565 | 0.35573124 | -0.4184298 | 0.67563288 | 0.76284529 | NA | NA |
| ENSG000002 | 2.42165214 | -0.153317125 | 0.36648031 | -0.4183502 | 0.67569106 | 0.76288975 | NA | NA |
| ENSG000002 | 78.7389415 | -0.076231257 | 0.18225028 | -0.4182779 | 0.67574397 | 0.76292824 | ZNF337-AS1 | 102724826 |
| ENSG000002 | 17.8089781 | 0.124541059 | 0.29786082 | 0.41811829 | 0.67586063 | 0.76303871 | NA | NA |
| ENSG000000 | 7.66974196 | 0.148773262 | 0.35594491 | 0.4179671 | 0.67597117 | 0.76314226 | CMTM1 | 113540 |
| ENSG000001 | 52.2083314 | 0.098503925 | 0.23570432 | 0.41791311 | 0.67601065 | 0.76314434 | ATP1B2 | 482 |
| ENSG000002 | 122.417454 | 0.067647873 | 0.16186932 | 0.41791658 | 0.67600811 | 0.76314434 | GOLGA2P5 | 55592 |
| ENSG000002 | 2.40171653 | 0.154309392 | 0.36935097 | 0.41778526 | 0.67610413 | 0.76322862 | NA | NA |
| ENSG000001 | 4.45418538 | -0.158044666 | 0.37838033 | -0.4176873 | 0.67617574 | 0.76328761 | IL3 | 3562 |
| ENSG000002 | 2.48524562 | 0.154147228 | 0.36907143 | 0.41766232 | 0.67619403 | 0.76328761 | NA | NA |
| ENSG000002 | 11.0417596 | 0.138567793 | 0.33205987 | 0.41729761 | 0.67646073 | 0.76356741 | NA | NA |
| ENSG000002 | 81.1026857 | 0.074510272 | 0.17858044 | 0.41723646 | 0.67650546 | 0.76359664 | CCPG1 | 9236 |
| ENSG000001 | 20.1191363 | -0.117123157 | 0.28082215 | -0.4170724 | 0.67662548 | 0.76371085 | C1QTNF2 | 114898 |
| ENSG000001 | 5.34418899 | -0.155091942 | 0.37196538 | -0.4169526 | 0.67671306 | 0.76378845 | GLOD5 | 392465 |
| ENSG000002 | 13.6379326 | -0.132373016 | 0.31769412 | -0.4166681 | 0.67692116 | 0.76400206 | NA | NA |

| | | | | | | | | |
|---|---|---|---|---|---|---|---|---|
| ENSG000002 | 2.64438163 | 0.156393706 | 0.37545564 | 0.41654377 | 0.67701215 | 0.7640716 | MIR3679 | 100500878 |
| ENSG000002 | 7.4251735 | -0.147989508 | 0.3552893 | -0.4165324 | 0.67702046 | 0.7640716 | NA | NA |
| ENSG000001 | 6.58428574 | -0.151511109 | 0.36389743 | -0.4163566 | 0.67714905 | 0.76419546 | UGT1A6 | 54578 |
| ENSG000001 | 49.9541577 | -0.094805972 | 0.22790035 | -0.4159975 | 0.67741185 | 0.76447076 | GRM2 | 2912 |
| ENSG000001 | 11.9443167 | 0.13441489 | 0.32334849 | 0.41569666 | 0.67763198 | 0.7646979 | LRFN2 | 57497 |
| ENSG000001 | 82.6059909 | -0.073431286 | 0.17673488 | -0.4154884 | 0.67778443 | 0.76484865 | PCSK6 | 5046 |
| ENSG000002 | 4.82644521 | 0.155668975 | 0.37469978 | 0.41544988 | 0.6778126 | 0.76485916 | NA | NA |
| ENSG000002 | 8.26726013 | -0.146228631 | 0.35215167 | -0.4152433 | 0.67796378 | 0.76498718 | NA | NA |
| ENSG000002 | 3.12778239 | -0.155455162 | 0.37435157 | -0.4152651 | 0.67794781 | 0.76498718 | NA | NA |
| ENSG000002 | 7.0836912 | -0.152180836 | 0.36651708 | -0.415208 | 0.67798961 | 0.76499504 | CTD-2270F17.1 | 101928033 |
| ENSG000001 | 24.2903877 | 0.11173291 | 0.26917206 | 0.41509847 | 0.67806982 | 0.76506425 | KLC3 | 147700 |
| ENSG000002 | 6.99482879 | -0.148473988 | 0.35793865 | -0.4148029 | 0.6782862 | 0.7652871 | NA | NA |
| ENSG000002 | 274.126697 | -0.057866596 | 0.13952853 | -0.4147295 | 0.67833994 | 0.76532644 | NBPF1 | 55672 |
| ENSG000000 | 6.80944623 | -0.15012644 | 0.36205372 | -0.4146524 | 0.67839639 | 0.76534754 | FLT3LG | 2323 |
| ENSG000001 | 741.914811 | 0.04742863 | 0.11437496 | 0.41467668 | 0.67837861 | 0.76534754 | NEB | 4703 |
| ENSG000002 | 8.42789208 | 0.14388693 | 0.34703793 | 0.41461442 | 0.67842419 | 0.76535761 | NA | NA |
| ENSG000002 | 4.3616223 | -0.156856893 | 0.37850812 | -0.4144083 | 0.67857513 | 0.7654853 | NA | NA |
| ENSG000002 | 4.41442184 | -0.156714798 | 0.37814972 | -0.4144253 | 0.67856269 | 0.7654853 | RN7SL219P | 106479302 |
| ENSG000002 | 6.49084164 | -0.151054629 | 0.36465765 | -0.4142368 | 0.67870067 | 0.76560562 | NA | NA |
| ENSG000001 | 24.0155352 | -0.111031231 | 0.26816154 | -0.4140461 | 0.67884037 | 0.7657419 | HSF2BP | 11077 |
| ENSG000002 | 3.17850747 | -0.155525569 | 0.37582267 | -0.413827 | 0.67900081 | 0.76590158 | NA | NA |
| ENSG000002 | 5.31017922 | -0.153537689 | 0.37114115 | -0.4136908 | 0.67910055 | 0.76599278 | NA | NA |
| ENSG000002 | 5.5739294 | -0.152382312 | 0.36844064 | -0.4135871 | 0.67917649 | 0.76605712 | NA | NA |
| ENSG000000 | 38.7111496 | 0.101698264 | 0.24596126 | 0.41347269 | 0.67926033 | 0.76613038 | SLC9A3R2 | 9351 |
| ENSG000002 | 10.8548625 | -0.138254134 | 0.33474608 | -0.413012 | 0.67959785 | 0.76648104 | NA | NA |
| ENSG000002 | 7.27075404 | -0.14832047 | 0.35913233 | -0.4129967 | 0.67960903 | 0.76648104 | NA | NA |
| ENSG000002 | 28.6984142 | -0.103678314 | 0.25112282 | -0.412859 | 0.67970993 | 0.76657351 | LINC00886 | 730091 |
| ENSG000002 | 2.67128052 | 0.154894713 | 0.37524002 | 0.41278836 | 0.67976168 | 0.76661056 | MIR3138 | 100423011 |
| ENSG000001 | 5.32531465 | -0.15329161 | 0.37184046 | -0.412251 | 0.68015545 | 0.76701679 | SPRR2D | 6703 |

| | | | | | | | | |
|---|---|---|---|---|---|---|---|---|
| ENSG000002 | 6.74352647 | -0.150409076 | 0.36485344 | -0.4122452 | 0.68015972 | 0.76701679 | NA | NA |
| ENSG000002 | 4.87938138 | 0.154335134 | 0.37447626 | 0.41213596 | 0.68023977 | 0.76708573 | NA | NA |
| ENSG000002 | 26.672624 | 0.111107919 | 0.26962981 | 0.41207579 | 0.68028387 | 0.76711413 | NA | NA |
| ENSG000002 | 2.90343102 | -0.152973577 | 0.37125577 | -0.4120436 | 0.68030744 | 0.76711938 | NA | NA |
| ENSG000001 | 19.1192835 | 0.118401756 | 0.28754925 | 0.41176166 | 0.68051413 | 0.7673311 | CFP | 5199 |
| ENSG000002 | 6.42353815 | 0.151831446 | 0.3688058 | 0.411684 | 0.68057105 | 0.76737395 | NA | NA |
| ENSG000002 | 2.43016231 | 0.150708329 | 0.36620027 | 0.4115462 | 0.68067207 | 0.76744518 | NA | NA |
| ENSG000002 | 6.46411139 | -0.150341666 | 0.36529851 | -0.4115584 | 0.68066314 | 0.76744518 | NA | NA |
| ENSG000002 | 4.66143556 | 0.154588723 | 0.37577797 | 0.41138314 | 0.68079161 | 0.76755862 | NA | NA |
| ENSG000002 | 6.99645499 | -0.146866427 | 0.35713584 | -0.4112341 | 0.6809009 | 0.76765099 | NA | NA |
| ENSG000002 | 8.25999086 | -0.145126225 | 0.35291647 | -0.4112198 | 0.6809114 | 0.76765099 | NA | NA |
| ENSG000001 | 94.0135589 | 0.073038568 | 0.17764511 | 0.41114878 | 0.68096344 | 0.76768831 | SNTG2 | 54221 |
| ENSG000001 | 7.08494244 | 0.148919014 | 0.36223054 | 0.41111667 | 0.68098699 | 0.76769351 | LRRC29 | 26231 |
| ENSG000002 | 5.32687339 | -0.153245305 | 0.37295054 | -0.4108998 | 0.68114601 | 0.76785143 | NA | NA |
| ENSG000001 | 31.037793 | -0.102505302 | 0.24967348 | -0.4105574 | 0.68139709 | 0.76811312 | HPCAL4 | 51440 |
| ENSG000002 | 5.918882 | 0.152444237 | 0.37151416 | 0.41033224 | 0.68156225 | 0.76827794 | NA | NA |
| ENSG000002 | 10.2853511 | -0.14021354 | 0.34183896 | -0.4101742 | 0.68167813 | 0.76838721 | CTB-178M22.2 | 101927862 |
| ENSG000002 | 2.65232165 | 0.153961141 | 0.37539803 | 0.41012773 | 0.68171225 | 0.76840431 | NA | NA |
| ENSG000002 | 2.68329173 | 0.153742579 | 0.37525391 | 0.4097028 | 0.68202397 | 0.76873431 | NA | NA |
| ENSG000002 | 3.77322957 | -0.153575245 | 0.37497868 | -0.4095573 | 0.68213074 | 0.768816 | NA | NA |
| ENSG000002 | 11.922298 | 0.133771022 | 0.32664803 | 0.40952649 | 0.68215333 | 0.768816 | NA | NA |
| ENSG000002 | 12.7672784 | -0.129808176 | 0.31696787 | -0.409531 | 0.68215 | 0.768816 | NA | NA |
| ENSG000002 | 6.17944814 | 0.149042901 | 0.36398537 | 0.40947497 | 0.68219113 | 0.76883723 | NA | NA |
| ENSG000002 | 8.47450486 | -0.144350157 | 0.3529516 | -0.40898 | 0.68255432 | 0.76922517 | RN7SL832P | 106479524 |
| ENSG000002 | 20.1178928 | -0.11793474 | 0.2885016 | -0.4087837 | 0.68269843 | 0.76936621 | NA | NA |
| ENSG000002 | 5.54618936 | -0.150697325 | 0.36891681 | -0.4084859 | 0.68291696 | 0.76959108 | NA | NA |
| ENSG000002 | 24.823493 | 0.108930062 | 0.26668727 | 0.40845617 | 0.6829388 | 0.76959431 | NA | NA |
| ENSG000001 | 145.144945 | 0.059793327 | 0.14647551 | 0.40821381 | 0.68311671 | 0.76977341 | KIAA0825 | 285600 |
| ENSG00000C | 123.572449 | 0.064699944 | 0.15853191 | 0.40811938 | 0.68318603 | 0.76983013 | GABRA3 | 2556 |

| | | | | | | | | |
|---|---|---|---|---|---|---|---|---|
| ENSG000002 | 7.00981814 | 0.148256461 | 0.36344504 | 0.40791989 | 0.68333249 | 0.76997377 | NA | NA |
| ENSG000002 | 2.66777063 | 0.152901775 | 0.37492342 | 0.40782135 | 0.68340484 | 0.77003389 | NA | NA |
| ENSG000000 | 354.981118 | 0.055643282 | 0.13661047 | 0.40731345 | 0.68377779 | 0.7704327 | RFX3 | 5991 |
| ENSG000002 | 5.76901402 | -0.149324866 | 0.36687056 | -0.4070233 | 0.68399087 | 0.77062997 | NA | NA |
| ENSG000002 | 12.5155087 | 0.133845188 | 0.32883073 | 0.4070337 | 0.68398324 | 0.77062997 | NA | NA |
| ENSG000001 | 4.36441671 | -0.153690027 | 0.37785752 | -0.4067407 | 0.68419845 | 0.77082101 | NA | NA |
| ENSG000002 | 11.6009627 | -0.132638856 | 0.32608224 | -0.406765 | 0.68418056 | 0.77082101 | NA | NA |
| ENSG000002 | 6.76616617 | -0.147270485 | 0.36213469 | -0.4066732 | 0.68424801 | 0.77085542 | NA | NA |
| ENSG000002 | 5.76049245 | -0.149991551 | 0.36899739 | -0.406484 | 0.68438698 | 0.77099056 | NA | NA |
| ENSG000001 | 731.722415 | -0.106845255 | 0.26292745 | -0.4063678 | 0.68447236 | 0.77106533 | NA | NA |
| ENSG000002 | 3.76336578 | 0.153695824 | 0.37834626 | 0.40623058 | 0.68457318 | 0.77115749 | NA | NA |
| ENSG000002 | 15.148468 | 0.123697639 | 0.30457225 | 0.40613562 | 0.68464296 | 0.77121466 | NA | NA |
| ENSG000002 | 7.30552301 | 0.144468015 | 0.35595117 | 0.4058647 | 0.68484201 | 0.77141746 | NA | NA |
| ENSG000001 | 5.93448688 | 0.149654079 | 0.36879987 | 0.40578669 | 0.68489934 | 0.77141775 | LOC102723701 | 102723701 |
| ENSG000001 | 5.05752643 | -0.152056993 | 0.37469476 | -0.4058156 | 0.68487806 | 0.77141775 | FAM180A | 389558 |
| ENSG000002 | 8.78880248 | 0.142285654 | 0.35062141 | 0.40580995 | 0.68488225 | 0.77141775 | NA | NA |
| ENSG000001 | 590.217597 | 0.058307827 | 0.14376002 | 0.4055914 | 0.68504285 | 0.77152937 | XPC | 7508 |
| ENSG000002 | 10.299564 | -0.134898148 | 0.33261032 | -0.4055742 | 0.68505553 | 0.77152937 | NA | NA |
| ENSG000002 | 6.79682345 | -0.146597846 | 0.36145216 | -0.4055802 | 0.68505107 | 0.77152937 | NA | NA |
| ENSG000002 | 91.297121 | -0.074094583 | 0.18276395 | -0.4054114 | 0.68517515 | 0.77164267 | LINC00941 | 100287314 |
| ENSG000002 | 8.51355052 | 0.144462078 | 0.35638707 | 0.40535163 | 0.68521906 | 0.77167069 | NA | NA |
| ENSG000002 | 2.38309092 | 0.148480228 | 0.36635501 | 0.40529056 | 0.68526394 | 0.7716998 | NA | NA |
| ENSG000002 | 6.54190651 | -0.147247343 | 0.36352393 | -0.4050554 | 0.68543676 | 0.77187298 | NA | NA |
| ENSG000002 | 14.8074158 | -0.125550892 | 0.30999752 | -0.4050061 | 0.68547301 | 0.77189237 | NA | NA |
| ENSG000002 | 5.66395755 | 0.150365043 | 0.37134378 | 0.4049214 | 0.68553529 | 0.77194106 | NA | NA |
| ENSG000001 | 5.05726605 | -0.150947701 | 0.37304311 | -0.4046388 | 0.68574306 | 0.77215358 | STX11 | 8676 |
| ENSG000001 | 6.76471177 | -0.145797463 | 0.36084875 | -0.4040404 | 0.68618303 | 0.77262753 | LOC100996455 | 100996455 |
| ENSG000001 | 52.1967598 | -0.087348468 | 0.21623577 | -0.4039501 | 0.68624939 | 0.7726808 | PROM2 | 150696 |
| ENSG000002 | 11.3206868 | 0.13388582 | 0.33151666 | 0.4038585 | 0.68631678 | 0.77273522 | NA | NA |

| | | | | | | | | |
|---|---|---|---|---|---|---|---|---|
| ENSG000001 | 13.4891676 | 0.126252553 | 0.31267376 | 0.40378365 | 0.68637182 | 0.77277573 | ZNF20 | 7568 |
| ENSG000002 | 8.85999438 | 0.142266232 | 0.35236982 | 0.40374126 | 0.686403 | 0.77278938 | NA | NA |
| ENSG000002 | 4.64393538 | -0.152584053 | 0.37795476 | -0.4037098 | 0.6864261 | 0.77279394 | NA | NA |
| ENSG000002 | 23.1656082 | 0.113027324 | 0.28023801 | 0.40332618 | 0.68670829 | 0.77309017 | NA | NA |
| ENSG000001 | 16.2222843 | 0.123105167 | 0.3052496 | 0.40329346 | 0.68673235 | 0.77309579 | THSD1 | 55901 |
| ENSG000002 | 11.7812966 | -0.130449326 | 0.323546 | -0.4031863 | 0.68681115 | 0.77316304 | NA | NA |
| ENSG000001 | 85.9607521 | 0.074231779 | 0.18427154 | 0.40283908 | 0.68706661 | 0.77342914 | RGS10 | 6001 |
| ENSG000002 | 2.64160453 | -0.146403615 | 0.36367518 | -0.402567 | 0.68726682 | 0.77362265 | RNU1-91P | 106480179 |
| ENSG000002 | 4.80964999 | -0.150322312 | 0.37344592 | -0.4025277 | 0.68729574 | 0.77362265 | NA | NA |
| ENSG000002 | 142.698691 | -0.060428133 | 0.15011753 | -0.4025388 | 0.68728754 | 0.77362265 | NA | NA |
| ENSG000002 | 11.7982269 | -0.130924839 | 0.3252968 | -0.4024781 | 0.68733219 | 0.77364221 | NA | NA |
| ENSG000001 | 74.7831262 | 0.072928633 | 0.18125251 | 0.4023593 | 0.68741962 | 0.77371913 | ATAT1 | 79969 |
| ENSG000001 | 5.53202548 | 0.149786565 | 0.3724268 | 0.40219062 | 0.68754375 | 0.77383737 | RAB24 | 53917 |
| ENSG000002 | 8.58610296 | 0.140053596 | 0.34832754 | 0.40207442 | 0.68762925 | 0.77391213 | NA | NA |
| ENSG000002 | 6.15358692 | 0.146905202 | 0.36545076 | 0.40198358 | 0.68769611 | 0.77394441 | NA | NA |
| ENSG000002 | 2.67214172 | 0.150674893 | 0.37480491 | 0.40200886 | 0.68767751 | 0.77394441 | NA | NA |
| ENSG000001 | 6.02730347 | -0.147357235 | 0.36663805 | -0.4019147 | 0.68774678 | 0.77397995 | ZNF763 | 284390 |
| ENSG000002 | 2.68090404 | 0.149858596 | 0.37321938 | 0.40152951 | 0.68803031 | 0.77425606 | NA | NA |
| ENSG000002 | 3.72841859 | 0.151492244 | 0.37728377 | 0.40153396 | 0.68802704 | 0.77425606 | NA | NA |
| ENSG000002 | 4.63008616 | 0.151160591 | 0.37672097 | 0.40125345 | 0.68823353 | 0.77446326 | NA | NA |
| ENSG000001 | 6.24885048 | -0.146240087 | 0.36462107 | -0.4010742 | 0.68836553 | 0.7745903 | IZUMO1R | 390243 |
| ENSG00000C | 123.611041 | -0.067263215 | 0.16785214 | -0.400729 | 0.68861967 | 0.77485478 | PER3 | 8863 |
| ENSG000002 | 5.75468672 | 0.147641875 | 0.36862539 | 0.4005201 | 0.68877348 | 0.77500634 | NA | NA |
| ENSG000002 | 17.6912992 | 0.119365363 | 0.29809744 | 0.40042398 | 0.68884427 | 0.77506448 | NA | NA |
| ENSG000002 | 15.9742062 | -0.118971956 | 0.29728519 | -0.4001947 | 0.68901313 | 0.77523296 | TWIST2 | 117581 |
| ENSG000001 | 29.3425694 | 0.101706807 | 0.25417257 | 0.40014863 | 0.68904705 | 0.77524962 | LRRN2 | 10446 |
| ENSG000001 | 85.5919012 | -0.073765783 | 0.18445813 | -0.3999053 | 0.68922626 | 0.77542974 | ZNF441 | 126068 |
| ENSG000002 | 27.0783892 | 0.111104573 | 0.27786922 | 0.39984484 | 0.6892708 | 0.77545834 | NA | NA |
| ENSG000001 | 108.779872 | 0.064584693 | 0.16154696 | 0.39978898 | 0.68931195 | 0.77548311 | RNF182 | 221687 |

| | | | | | | | | |
|---|---|---|---|---|---|---|---|---|
| ENSG000001 | 18.46801 | 0.116368442 | 0.29116095 | 0.3996705 | 0.68939922 | 0.77555124 | NA | NA |
| ENSG000002 | 6.94286592 | -0.143320958 | 0.35861184 | -0.3996548 | 0.68941076 | 0.77555124 | NA | NA |
| ENSG000002 | 2.71400087 | 0.148511309 | 0.37177487 | 0.3994657 | 0.68955009 | 0.77568647 | NA | NA |
| ENSG000001 | 42.6771844 | -0.09067399 | 0.22702656 | -0.3993981 | 0.68959986 | 0.77572093 | APCDD1 | 147495 |
| ENSG000002 | 6.85969559 | -0.145900948 | 0.36533128 | -0.3993662 | 0.68962343 | 0.77572593 | NA | NA |
| ENSG000002 | 2.6854504 | 0.149987537 | 0.37566935 | 0.39925412 | 0.68970597 | 0.77579725 | NA | NA |
| ENSG00000C | 52.4249814 | 0.085635381 | 0.21453562 | 0.39916627 | 0.6897707 | 0.77584854 | NGFR | 4804 |
| ENSG000001 | 13.8120034 | 0.125539986 | 0.31463491 | 0.39900209 | 0.68989166 | 0.77596231 | TSGA10IP | 254187 |
| ENSG000002 | 4.33327888 | -0.150263057 | 0.37662081 | -0.398977 | 0.68991012 | 0.77596231 | ARHGEF3-AS1 | 100874200 |
| ENSG000001 | 36.5023352 | -0.093482145 | 0.23440004 | -0.3988145 | 0.69002986 | 0.77597129 | FLI1 | 2313 |
| ENSG000001 | 46.3435264 | 0.085108186 | 0.21340513 | 0.3988104 | 0.69003291 | 0.77597129 | DEPDC4 | 120863 |
| ENSG000002 | 47.407355 | 0.087766218 | 0.22003762 | 0.39886914 | 0.68998963 | 0.77597129 | NA | NA |
| ENSG000002 | 5.55255038 | -0.148050672 | 0.37118134 | -0.3988635 | 0.68999381 | 0.77597129 | ZNF32-AS1 | 414197 |
| ENSG000002 | 2.46072277 | 0.14348533 | 0.35977796 | 0.39881634 | 0.69002854 | 0.77597129 | NA | NA |
| ENSG000002 | 8.53556668 | 0.140207501 | 0.35156203 | 0.39881298 | 0.69003101 | 0.77597129 | NA | NA |
| ENSG000002 | 2.70808292 | 0.149497661 | 0.37496561 | 0.398697 | 0.69011648 | 0.77604374 | NA | NA |
| ENSG000001 | 160.851168 | 0.058293539 | 0.14629129 | 0.39847581 | 0.69027949 | 0.776184 | PLCG2 | 5336 |
| ENSG000002 | 18.5128759 | 0.11389693 | 0.28582221 | 0.39848873 | 0.69026997 | 0.776184 | NA | NA |
| ENSG000001 | 138.069951 | -0.059425562 | 0.14914787 | -0.3984339 | 0.6903104 | 0.77619723 | MCTP2 | 55784 |
| ENSG000001 | 223.240033 | -0.056399085 | 0.14156195 | -0.3984057 | 0.69033117 | 0.77619907 | SEPSECS | 51091 |
| ENSG000001 | 35.5712378 | -0.092492988 | 0.23221413 | -0.398309 | 0.6904024 | 0.77623611 | IL21R | 50615 |
| ENSG000002 | 9.34967863 | -0.136392034 | 0.34240967 | -0.39833 | 0.69038697 | 0.77623611 | NA | NA |
| ENSG000002 | 3.70628774 | 0.150283791 | 0.37736022 | 0.39825022 | 0.69044575 | 0.77626333 | RNU6-431P | 106481314 |
| ENSG000001 | 94.7264091 | 0.067612304 | 0.16992969 | 0.397884 | 0.6907157 | 0.7765453 | ECHDC2 | 55268 |
| ENSG000002 | 4.87969349 | 0.148618605 | 0.37362415 | 0.3977757 | 0.69079553 | 0.77661352 | NA | NA |
| ENSG000002 | 62.0474526 | 0.077124908 | 0.19393622 | 0.39768181 | 0.69086475 | 0.7766698 | NA | NA |
| ENSG000002 | 32.8421962 | 0.09426599 | 0.23733276 | 0.39718912 | 0.69122801 | 0.77705663 | NA | NA |
| ENSG000001 | 46.95311 | 0.084037488 | 0.21172083 | 0.39692594 | 0.69142208 | 0.77725325 | NDST3 | 9348 |
| ENSG000001 | 5.50596935 | -0.146023322 | 0.36796381 | -0.3968415 | 0.69148432 | 0.77730168 | NT5C1B | 93034 |

| | | | | | | | | |
|---|---|---|---|---|---|---|---|---|
| ENSG000002 | 2.67779854 | 0.148324376 | 0.37429754 | 0.39627398 | 0.69190292 | 0.77775067 | NA | NA |
| ENSG000002 | 14.2496007 | 0.121602707 | 0.30693758 | 0.39618058 | 0.69197182 | 0.77780655 | SCARNA18 | 677765 |
| ENSG000002 | 4.37750556 | -0.149465154 | 0.3773331 | -0.3961093 | 0.69202439 | 0.77784408 | NA | NA |
| ENSG000000 | 72.744116 | 0.070298726 | 0.1774976 | 0.39605452 | 0.69206481 | 0.77785963 | FAM214B | 80256 |
| ENSG000002 | 6.7742758 | -0.142937196 | 0.36091737 | -0.3960386 | 0.69207658 | 0.77785963 | MIR659 | 724029 |
| ENSG000002 | 4.79600825 | -0.147859281 | 0.37351935 | -0.3958544 | 0.69221244 | 0.77799076 | NA | NA |
| ENSG000002 | 4.31191718 | -0.149754586 | 0.37843885 | -0.3957167 | 0.69231401 | 0.77808335 | NA | NA |
| ENSG000001 | 33.4143303 | -0.093687588 | 0.23692155 | -0.3954372 | 0.69252027 | 0.7782936 | SLC25A18 | 83733 |
| ENSG000001 | 20.1070186 | -0.114194022 | 0.28882561 | -0.3953736 | 0.69256719 | 0.778312 | RGS20 | 8601 |
| ENSG000002 | 5.06076444 | 0.147356032 | 0.37271076 | 0.39536297 | 0.69257503 | 0.778312 | ATP5L2 | 267020 |
| ENSG000001 | 48.3785099 | -0.08454594 | 0.21387303 | -0.395309 | 0.69261484 | 0.77833517 | FHIT | 2272 |
| ENSG000002 | 6.72135814 | -0.143753822 | 0.3638122 | -0.3951319 | 0.69274551 | 0.77846043 | NA | NA |
| ENSG000002 | 2.65045114 | 0.144723323 | 0.36641348 | 0.3949727 | 0.69286303 | 0.77857092 | NA | NA |
| ENSG000002 | 46.6467139 | 0.085928135 | 0.21758903 | 0.39491023 | 0.69290914 | 0.77857958 | PPP1R3E | 90673 |
| ENSG000002 | 8.6970311 | 0.136433941 | 0.34547799 | 0.39491355 | 0.69290669 | 0.77857958 | NA | NA |
| ENSG000001 | 27.0023494 | 0.100347161 | 0.25423736 | 0.39469872 | 0.69306524 | 0.77871183 | IGFBP6 | 3489 |
| ENSG000001 | 10.3072752 | -0.133649601 | 0.33859606 | -0.3947169 | 0.6930518 | 0.77871183 | NA | NA |
| ENSG000001 | 27.4096155 | 0.103988355 | 0.26356211 | 0.39454972 | 0.69317523 | 0.77879225 | APLN | 8862 |
| ENSG000002 | 11.5449445 | 0.127054785 | 0.3220087 | 0.39456941 | 0.69316069 | 0.77879225 | NA | NA |
| ENSG000002 | 7.74548274 | -0.139778631 | 0.35449814 | -0.3943 | 0.69335954 | 0.77894038 | NA | NA |
| ENSG000002 | 7.52413065 | -0.143572519 | 0.36411757 | -0.3943026 | 0.69335762 | 0.77894038 | NA | NA |
| ENSG000002 | 3.78926751 | -0.146873993 | 0.37249958 | -0.394293 | 0.6933647 | 0.77894038 | NA | NA |
| ENSG000002 | 4.32363609 | -0.148391874 | 0.37659689 | -0.3940337 | 0.69355615 | 0.77913387 | NA | NA |
| ENSG000002 | 6.92862306 | -0.144632871 | 0.36713147 | -0.3939539 | 0.69361508 | 0.77915874 | NA | NA |
| ENSG000002 | 38.2010965 | -0.093002827 | 0.23607674 | -0.3939517 | 0.69361672 | 0.77915874 | NA | NA |
| ENSG000002 | 8.90280893 | 0.13459413 | 0.34170616 | 0.39388851 | 0.69366335 | 0.77918954 | NA | NA |
| ENSG000001 | 13.7665469 | 0.123059036 | 0.31255694 | 0.39371717 | 0.69378986 | 0.77931007 | SULT1A2 | 6799 |
| ENSG000001 | 42.0629933 | -0.087598577 | 0.2225678 | -0.3935815 | 0.69389001 | 0.77937938 | SPDYA | 245711 |
| ENSG000002 | 3.98380419 | 0.149001405 | 0.3785727 | 0.3935873 | 0.69388576 | 0.77937938 | AOAH-IT1 | 100874264 |

| | | | | | | | | |
|---|---|---|---|---|---|---|---|---|
| ENSG000001 | 6.50394248 | -0.143830065 | 0.36552672 | -0.3934871 | 0.69395972 | 0.77943609 | RNASE6 | 6039 |
| ENSG000001 | 4.85544228 | -0.148347126 | 0.37705298 | -0.3934384 | 0.6939957 | 0.77945492 | TUBA3E | 112714 |
| ENSG000002 | 5.08189008 | 0.147071715 | 0.37398193 | 0.39325888 | 0.69412829 | 0.779528 | NA | NA |
| ENSG000002 | 3.75655418 | 0.148843429 | 0.37842821 | 0.39332012 | 0.69408306 | 0.779528 | NA | NA |
| ENSG000002 | 8.61052181 | 0.136178721 | 0.34626841 | 0.39327503 | 0.69411636 | 0.779528 | NA | NA |
| ENSG000002 | 5.77261576 | -0.145763021 | 0.37066608 | -0.3932462 | 0.69413766 | 0.779528 | NA | NA |
| ENSG000002 | 21.8860478 | 0.108978543 | 0.27717101 | 0.3931816 | 0.69418536 | 0.77955997 | NA | NA |
| ENSG000002 | 28.1926532 | 0.098292124 | 0.25003126 | 0.39311935 | 0.69423134 | 0.77959001 | LOC283856 | 283856 |
| ENSG000002 | 4.6393243 | -0.148046461 | 0.37683589 | -0.3928672 | 0.69441757 | 0.77977755 | NA | NA |
| ENSG000002 | 5.0465406 | -0.147667569 | 0.37603099 | -0.3927005 | 0.69454068 | 0.7798726 | NA | NA |
| ENSG000002 | 56.1311256 | 0.082220024 | 0.20935935 | 0.392722 | 0.69452482 | 0.7798726 | NA | NA |
| ENSG000002 | 5.0402188 | -0.1467223 | 0.37370418 | -0.3926162 | 0.694603 | 0.77989939 | LOC101928271 | 101928271 |
| ENSG000002 | 5.37593536 | -0.146916996 | 0.37418689 | -0.39263 | 0.69459279 | 0.77989939 | NA | NA |
| ENSG000002 | 3.34102015 | 0.146422083 | 0.37319905 | 0.39234313 | 0.6948047 | 0.78010425 | NA | NA |
| ENSG000002 | 2.87736128 | -0.144511344 | 0.36853432 | -0.3921245 | 0.69496621 | 0.78026399 | NA | NA |
| ENSG000002 | 5.5534972 | -0.144567571 | 0.36873875 | -0.3920596 | 0.69501417 | 0.78029623 | LOC101927124 | 101927124 |
| ENSG000001 | 10.2978887 | -0.130843639 | 0.33376434 | -0.392024 | 0.69504046 | 0.78030415 | ZSCAN1 | 284312 |
| ENSG000002 | 7.41037089 | -0.140029928 | 0.35732844 | -0.3918802 | 0.69514676 | 0.78038028 | IFIT1B | 439996 |
| ENSG000002 | 13.8654257 | -0.123923486 | 0.31622461 | -0.3918844 | 0.69514364 | 0.78038028 | NA | NA |
| ENSG000002 | 5.52212219 | -0.145116802 | 0.37046377 | -0.3917166 | 0.69526764 | 0.78049436 | OR2A14 | 135941 |
| ENSG000002 | 5.56227858 | -0.144694997 | 0.36950953 | -0.3915866 | 0.69536366 | 0.78058055 | NA | NA |
| ENSG000002 | 59.5009465 | 0.12711493 | 0.32494023 | 0.3911948 | 0.69565325 | 0.78088401 | NA | NA |
| ENSG000001 | 11.4256151 | -0.129006123 | 0.32982362 | -0.3911367 | 0.69569619 | 0.7809106 | MUC1 | 4582 |
| ENSG000001 | 3.98208049 | 0.147847252 | 0.37813523 | 0.39099042 | 0.69580432 | 0.78096886 | NA | NA |
| ENSG000002 | 39.4389531 | -0.088439877 | 0.22619069 | -0.390997 | 0.69579947 | 0.78096886 | NKAPP1 | 158801 |
| ENSG000002 | 2.90019919 | 0.147597525 | 0.37749856 | 0.39098831 | 0.69580587 | 0.78096886 | NA | NA |
| ENSG000001 | 6.24404099 | -0.145691375 | 0.37282337 | -0.3907785 | 0.69596094 | 0.78112128 | MS4A15 | 219995 |
| ENSG000002 | 5.052503 | -0.146634057 | 0.37528333 | -0.3907289 | 0.6959976 | 0.78114082 | NA | NA |
| ENSG000002 | 6.46215587 | -0.143819847 | 0.36819847 | -0.3906041 | 0.69608987 | 0.78120112 | NA | NA |

| | | | | | | | | |
|---|---|---|---|---|---|---|---|---|
| ENSG000002 | 2.86679259 | -0.144414343 | 0.36971598 | -0.3906089 | 0.69608637 | 0.78120112 | NA | NA |
| ENSG000002 | 4.82143307 | -0.146324253 | 0.37468926 | -0.3905216 | 0.69615088 | 0.78124798 | NA | NA |
| ENSG000001 | 12.1450857 | -0.128628869 | 0.32964085 | -0.3902091 | 0.69638192 | 0.78148562 | RLN1 | 6013 |
| ENSG000002 | 5.04926186 | -0.144871668 | 0.3714149 | -0.3900535 | 0.69649701 | 0.78159316 | NA | NA |
| ENSG000002 | 5.02399349 | -0.145171808 | 0.37231266 | -0.3899191 | 0.69659639 | 0.78168304 | NA | NA |
| ENSG000001 | 2.87868237 | -0.146583742 | 0.37596556 | -0.3898861 | 0.69662078 | 0.78168879 | NA | NA |
| ENSG000002 | 3.82447625 | -0.147561637 | 0.37858682 | -0.3897696 | 0.69670692 | 0.78176381 | NA | NA |
| ENSG000001 | 112.701136 | -0.062233877 | 0.15971845 | -0.3896474 | 0.69679731 | 0.78182197 | CUL9 | 23113 |
| ENSG000002 | 5.85566733 | -0.144726283 | 0.37142666 | -0.3896497 | 0.6967956 | 0.78182197 | LOC105379175 | 105379175 |
| ENSG000002 | 5.5165058 | -0.144813398 | 0.37169931 | -0.3895982 | 0.69683365 | 0.78184111 | NA | NA |
| ENSG000002 | 8.01208972 | -0.140310301 | 0.36023927 | -0.389492 | 0.69691225 | 0.78190767 | NA | NA |
| ENSG000002 | 6.17621093 | 0.142139291 | 0.3650323 | 0.38938826 | 0.69698896 | 0.78197209 | NA | NA |
| ENSG000002 | 14.5759085 | 0.124981667 | 0.32100125 | 0.38934947 | 0.69701765 | 0.78198264 | NA | NA |
| ENSG000002 | 15.5032166 | -0.117399828 | 0.30155524 | -0.3893145 | 0.69704351 | 0.78199002 | NA | NA |
| ENSG000002 | 4.84545505 | 0.145981782 | 0.37504032 | 0.3892429 | 0.69709647 | 0.7820278 | NA | NA |
| ENSG000001 | 19.0578918 | -0.110376663 | 0.28370256 | -0.3890577 | 0.69723347 | 0.78212174 | EMX1 | 2016 |
| ENSG000001 | 5.49575243 | -0.144056566 | 0.37027637 | -0.3890515 | 0.69723807 | 0.78212174 | FOXE1 | 2304 |
| ENSG000002 | 6.8267553 | -0.141951976 | 0.36484112 | -0.3890789 | 0.69721779 | 0.78212174 | NA | NA |
| ENSG000002 | 5.99035238 | -0.142394844 | 0.36611107 | -0.3889389 | 0.69732133 | 0.7821935 | NA | NA |
| ENSG000002 | 5.04463012 | 0.145007767 | 0.3728592 | 0.38890757 | 0.69734452 | 0.78219788 | LINC01023 | 100652853 |
| ENSG000002 | 10.4940349 | -0.130553169 | 0.33575815 | -0.388831 | 0.69740118 | 0.78221816 | NA | NA |
| ENSG000002 | 47.0580376 | 0.084300627 | 0.21679699 | 0.38884594 | 0.69739012 | 0.78221816 | LOC101927204 | 101927204 |
| ENSG000002 | 4.08301453 | -0.146805784 | 0.37764506 | -0.3887401 | 0.6974684 | 0.78227192 | NA | NA |
| ENSG000001 | 12.0132842 | 0.129693159 | 0.33366826 | 0.38868893 | 0.69750627 | 0.78227396 | MAPK15 | 225689 |
| ENSG000002 | 4.15518923 | -0.147133182 | 0.37854043 | -0.3886855 | 0.6975088 | 0.78227396 | NA | NA |
| ENSG000002 | 4.80144937 | -0.145297917 | 0.37390815 | -0.3885925 | 0.69757759 | 0.78232934 | NA | NA |
| ENSG000002 | 4.01365028 | 0.146749894 | 0.37766985 | 0.38856661 | 0.69759677 | 0.78232934 | NA | NA |
| ENSG000001 | 6.77770813 | -0.140862053 | 0.36270304 | -0.3883675 | 0.69774413 | 0.78245846 | CCL22 | 6367 |
| ENSG000002 | 114.166501 | 0.067452556 | 0.17368616 | 0.38835884 | 0.6977505 | 0.78245846 | NA | NA |

| | | | | | | | | |
|---|---|---|---|---|---|---|---|---|
| ENSG000001 | 99.4436542 | 0.066510988 | 0.17130125 | 0.38826914 | 0.69781687 | 0.78251125 | C1orf94 | 84970 |
| ENSG000001 | 144.627364 | 0.058846487 | 0.15168076 | 0.38796276 | 0.69804359 | 0.7827222 | DYSF | 8291 |
| ENSG000002 | 4.53923778 | -0.146266838 | 0.37700464 | -0.3879709 | 0.69803755 | 0.7827222 | NA | NA |
| ENSG000001 | 14.2770091 | -0.119411781 | 0.30791721 | -0.3878048 | 0.69816048 | 0.78280997 | PDXDC2P | 283970 |
| ENSG000002 | 4.8519328 | -0.145672617 | 0.37561518 | -0.3878241 | 0.69814621 | 0.78280997 | RN7SL385P | 106481840 |
| ENSG000002 | 7.55101948 | 0.138680955 | 0.35790597 | 0.38747874 | 0.69840182 | 0.78305893 | NA | NA |
| ENSG000002 | 4.56360029 | -0.145331045 | 0.37515486 | -0.3873895 | 0.69846789 | 0.78311136 | NA | NA |
| ENSG000001 | 17.8657722 | 0.116740951 | 0.30142364 | 0.38729859 | 0.69853517 | 0.78314552 | B9D2 | 80776 |
| ENSG000001 | 5.2809775 | -0.144021706 | 0.37186945 | -0.387291 | 0.69854076 | 0.78314552 | NA | NA |
| ENSG000002 | 14.0711383 | 0.118661847 | 0.30640596 | 0.38727004 | 0.6985563 | 0.78314552 | NA | NA |
| ENSG000002 | 2.66191271 | 0.14366778 | 0.37106015 | 0.38718191 | 0.69862154 | 0.783197 | NA | NA |
| ENSG000002 | 5.78015139 | -0.142877602 | 0.3690633 | -0.3871358 | 0.6986557 | 0.78320268 | C1orf53 | 388722 |
| ENSG000002 | 3.94598308 | 0.146501912 | 0.37843774 | 0.38712289 | 0.69866523 | 0.78320268 | NA | NA |
| ENSG000001 | 47.0056619 | -0.0817694 | 0.2112811 | -0.3870171 | 0.69874354 | 0.78326881 | MPP3 | 4356 |
| ENSG000002 | 4.81126876 | -0.144553856 | 0.37376411 | -0.3867516 | 0.69894013 | 0.78344587 | NA | NA |
| ENSG000002 | 10.0024226 | 0.132223783 | 0.34186883 | 0.38676759 | 0.69892828 | 0.78344587 | NA | NA |
| ENSG000001 | 22.4394477 | -0.106302136 | 0.27491165 | -0.3866775 | 0.69899501 | 0.78348572 | PGF | 5228 |
| ENSG000002 | 6.20277688 | -0.141712877 | 0.36652484 | -0.3866392 | 0.69902332 | 0.7834958 | NA | NA |
| ENSG000002 | 4.33471387 | -0.145862949 | 0.37732466 | -0.3865715 | 0.69907348 | 0.78353037 | NA | NA |
| ENSG000001 | 25.5203198 | -0.104977983 | 0.27160259 | -0.3865132 | 0.69911663 | 0.78355708 | ADAM8 | 101 |
| ENSG000002 | 3.96965193 | 0.146130954 | 0.37813624 | 0.38645054 | 0.69916303 | 0.78358742 | NA | NA |
| ENSG000002 | 4.75313224 | -0.144707738 | 0.37450266 | -0.3863998 | 0.69920063 | 0.7836079 | NA | NA |
| ENSG000002 | 5.23645798 | -0.143356831 | 0.37108378 | -0.3863193 | 0.6992602 | 0.783653 | NA | NA |
| ENSG000001 | 2.87828462 | 0.145414313 | 0.37658833 | 0.386136 | 0.69939595 | 0.78378348 | GALR3 | 8484 |
| ENSG000002 | 9.81640448 | 0.13143608 | 0.34042266 | 0.38609674 | 0.69942502 | 0.7837944 | NA | NA |
| ENSG000002 | 4.58728178 | -0.145469576 | 0.37687562 | -0.3859883 | 0.69950534 | 0.78386274 | NA | NA |
| ENSG000002 | 5.06363971 | -0.144891865 | 0.37551147 | -0.385852 | 0.69960625 | 0.78395416 | NA | NA |
| ENSG000002 | 14.5839275 | 0.121800475 | 0.31580356 | 0.3856843 | 0.69973049 | 0.78407171 | NA | NA |
| ENSG000002 | 15.3085035 | -0.118200801 | 0.30658793 | -0.3855364 | 0.69984005 | 0.78417281 | NA | NA |

| | | | | | | | | |
|---|---|---|---|---|---|---|---|---|
| ENSG000001 | 56.3080157 | 0.080117099 | 0.20784217 | 0.38547085 | 0.6998886 | 0.78420554 | KBTBD3 | 143879 |
| ENSG000002 | 6.4773776 | -0.139867211 | 0.36289104 | -0.3854248 | 0.6999227 | 0.78422002 | NA | NA |
| ENSG000002 | 4.16218402 | 0.145748362 | 0.37819871 | 0.38537509 | 0.69995954 | 0.78422002 | NA | NA |
| ENSG000002 | 2.9040089 | 0.145401798 | 0.37728648 | 0.38538831 | 0.69994974 | 0.78422002 | NA | NA |
| ENSG000002 | 3.7558103 | 0.145599089 | 0.37785633 | 0.38532923 | 0.69999351 | 0.78423641 | NA | NA |
| ENSG000001 | 6.29686746 | -0.142200903 | 0.36908949 | -0.3852749 | 0.70003378 | 0.78425986 | ATOH7 | 220202 |
| ENSG000001 | 12.4916546 | -0.124903889 | 0.32434975 | -0.3850901 | 0.70017064 | 0.78439151 | MYCT1 | 80177 |
| ENSG000002 | 2.90239067 | 0.145228239 | 0.37728688 | 0.38492788 | 0.70029085 | 0.78450451 | NA | NA |
| ENSG000002 | 7.80043595 | 0.135003183 | 0.351054 | 0.38456529 | 0.70055951 | 0.7847838 | NA | NA |
| ENSG000002 | 18.785377 | -0.114191171 | 0.29702882 | -0.3844448 | 0.70064883 | 0.78486218 | NA | NA |
| ENSG000002 | 2.64665875 | -0.142923001 | 0.37196914 | -0.3842335 | 0.7008054 | 0.78499735 | NA | NA |
| ENSG000002 | 5.99205416 | 0.141329153 | 0.36782466 | 0.38422969 | 0.70080822 | 0.78499735 | NA | NA |
| ENSG000002 | 4.55596719 | 0.145267738 | 0.37826798 | 0.38403393 | 0.7009533 | 0.78513817 | NA | NA |
| ENSG000002 | 3.80110741 | -0.145378357 | 0.37858436 | -0.3840052 | 0.70097461 | 0.78514035 | NA | NA |
| ENSG000002 | 39.3183884 | 0.085893139 | 0.22378649 | 0.38381735 | 0.70111382 | 0.78524617 | LYRM9 | 201229 |
| ENSG000002 | 3.10429987 | -0.14509327 | 0.3780446 | -0.3837993 | 0.70112717 | 0.78524617 | NA | NA |
| ENSG000002 | 4.84527853 | -0.144059563 | 0.37531084 | -0.3838407 | 0.70109655 | 0.78524617 | NA | NA |
| ENSG000001 | 27.1166139 | 0.100237423 | 0.26121079 | 0.38374152 | 0.70117004 | 0.78527249 | FBXO4 | 26272 |
| ENSG000001 | 220.878153 | 0.053026966 | 0.13836022 | 0.38325297 | 0.70153221 | 0.78563089 | CCDC171 | 203238 |
| ENSG000002 | 4.08283827 | -0.1448232 | 0.37789823 | -0.3832333 | 0.70154676 | 0.78563089 | NA | NA |
| ENSG000002 | 10.5572367 | -0.126966061 | 0.33130389 | -0.3832314 | 0.70154818 | 0.78563089 | NA | NA |
| ENSG000002 | 4.00062146 | 0.144624295 | 0.37746705 | 0.38314416 | 0.70161288 | 0.78568165 | NA | NA |
| ENSG000002 | 11.4407497 | -0.128129693 | 0.33447203 | -0.3830804 | 0.70166012 | 0.78571285 | NA | NA |
| ENSG000001 | 88.747724 | -0.067120362 | 0.17525792 | -0.3829805 | 0.70173424 | 0.78576317 | BICC1 | 80114 |
| ENSG000001 | 543.219669 | 0.047728118 | 0.12462705 | 0.38296758 | 0.7017438 | 0.78576317 | ZNF677 | 342926 |
| ENSG000002 | 3.84631925 | -0.144897775 | 0.37842723 | -0.3828947 | 0.70179785 | 0.78580198 | LINC01432 | 100270679 |
| ENSG000002 | 9.72022204 | -0.134016133 | 0.35022066 | -0.382662 | 0.70197041 | 0.78597349 | NA | NA |
| ENSG000001 | 173.936317 | -0.05769802 | 0.15082557 | -0.382548 | 0.70205494 | 0.78600303 | TTLL7 | 79739 |
| ENSG000002 | 31.5606752 | 0.092385949 | 0.24149079 | 0.3825651 | 0.70204225 | 0.78600303 | DDX11-AS1 | 100506660 |

| | | | | | | | | | |
|---|---|---|---|---|---|---|---|---|---|
| ENSG000002 | 15.4356235 | 0.116705028 | 0.3050501 | 0.38257659 | 0.70203373 | 0.78600303 | LINC01144 | | 400752 |
| ENSG000000 | 76.9561951 | 0.069228603 | 0.18101108 | 0.38245506 | 0.70212385 | 0.78605848 | CA12 | | 771 |
| ENSG000002 | 4.93856798 | 0.144331054 | 0.37752478 | 0.38230883 | 0.70223231 | 0.7861582 | SNORA46 | | 677827 |
| ENSG000001 | 302.803527 | -0.053786159 | 0.14075829 | -0.3821172 | 0.70237447 | 0.78616542 | RNF144B | | 255488 |
| ENSG000001 | 8.95815447 | -0.13323512 | 0.34860232 | -0.382198 | 0.70231449 | 0.78616542 | PPP1R32 | | 220004 |
| ENSG000002 | 2.39188893 | -0.136366027 | 0.35683659 | -0.3821526 | 0.70234818 | 0.78616542 | NA | NA | |
| ENSG000002 | 4.12221436 | -0.144595472 | 0.3783888 | -0.3821347 | 0.70236149 | 0.78616542 | NA | NA | |
| ENSG000002 | 5.24108966 | -0.142782891 | 0.37362635 | -0.3821542 | 0.70234697 | 0.78616542 | NA | NA | |
| ENSG000002 | 5.30576457 | -0.141681788 | 0.37072713 | -0.3821727 | 0.70233327 | 0.78616542 | NA | NA | |
| ENSG000002 | 3.74732296 | 0.144658883 | 0.37842573 | 0.38226492 | 0.70226487 | 0.78616542 | MIR3162 | | 100422880 |
| ENSG000001 | 27.4719694 | -0.10396582 | 0.27218064 | -0.3819736 | 0.70248094 | 0.7862629 | FNDC5 | | 252995 |
| ENSG000001 | 255.113739 | 0.048956534 | 0.12819235 | 0.38189903 | 0.70253626 | 0.78626698 | AKAP5 | | 9495 |
| ENSG000002 | 6.97715698 | -0.137596853 | 0.36030469 | -0.3818903 | 0.70254276 | 0.78626698 | NA | NA | |
| ENSG000002 | 4.32943726 | -0.143799188 | 0.37653451 | -0.3819017 | 0.70253424 | 0.78626698 | NA | NA | |
| ENSG000002 | 6.24876636 | -0.140182617 | 0.36718734 | -0.3817741 | 0.70262893 | 0.78634171 | NA | NA | |
| ENSG000002 | 3.82572309 | -0.144305148 | 0.37815281 | -0.3816054 | 0.70275409 | 0.78646008 | NA | NA | |
| ENSG000002 | 3.82076455 | 0.144427226 | 0.3785688 | 0.38150853 | 0.70282594 | 0.78651879 | NA | NA | |
| ENSG000002 | 3.14425929 | -0.14369052 | 0.37673585 | -0.3814092 | 0.70289964 | 0.78657955 | NA | NA | |
| ENSG000001 | 60.462846 | -0.073229008 | 0.1920376 | -0.3813264 | 0.70296107 | 0.78662658 | FAM198A | | 729085 |
| ENSG000001 | 195.787743 | 0.052682849 | 0.13816633 | 0.3813002 | 0.7029805 | 0.78662663 | FCHO2 | | 115548 |
| ENSG000002 | 14.3312584 | -0.117052606 | 0.30701192 | -0.381264 | 0.70300733 | 0.78663494 | NA | NA | |
| ENSG000002 | 5.48762394 | -0.141866273 | 0.37213374 | -0.3812239 | 0.70303712 | 0.78664657 | NA | NA | |
| ENSG000002 | 4.52367278 | -0.143503691 | 0.37653861 | -0.3811128 | 0.70311954 | 0.78671708 | NA | NA | |
| ENSG000002 | 6.2726248 | -0.139613387 | 0.36641086 | -0.3810296 | 0.70318128 | 0.78676445 | NA | NA | |
| ENSG000002 | 4.99744747 | -0.142019937 | 0.37294274 | -0.380809 | 0.70334501 | 0.78692593 | NA | NA | |
| ENSG000002 | 2.91701756 | 0.143774573 | 0.37763706 | 0.38072157 | 0.70340986 | 0.78697678 | NA | NA | |
| ENSG000002 | 4.80777171 | -0.14226599 | 0.37376067 | -0.3806339 | 0.70347491 | 0.78702784 | NA | NA | |
| ENSG000000 | 668.621395 | -0.053421249 | 0.14038826 | -0.3805251 | 0.7035557 | 0.78709067 | TM7SF3 | | 51768 |
| ENSG000002 | 11.1245725 | -0.128905912 | 0.33883928 | -0.3804338 | 0.70362343 | 0.78709067 | NA | NA | |

| | | | | | | | | |
|---|---|---|---|---|---|---|---|---|
| ENSG000002 | 3.40064956 | -0.143188464 | 0.37632566 | -0.3804908 | 0.7035811 | 0.78709067 | NA | NA |
| ENSG000002 | 5.47095327 | -0.141661158 | 0.37237362 | -0.3804275 | 0.70362812 | 0.78709067 | NA | NA |
| ENSG000002 | 74.7141568 | 0.071082197 | 0.18682981 | 0.38046497 | 0.70360029 | 0.78709067 | NA | NA |
| ENSG000001 | 56.1873383 | 0.076018359 | 0.19995273 | 0.38018165 | 0.70381058 | 0.78727305 | IRF4 | 3662 |
| ENSG000002 | 5.04290063 | -0.141915501 | 0.37359395 | -0.3798656 | 0.70404516 | 0.78751373 | NA | NA |
| ENSG000002 | 6.90961353 | -0.137179666 | 0.36121159 | -0.3797765 | 0.70411135 | 0.78756604 | NA | NA |
| ENSG00000C | 173.118859 | 0.066587659 | 0.1753736 | 0.37969033 | 0.7041753 | 0.78760538 | GNAO1 | 2775 |
| ENSG000002 | 5.07036734 | -0.142791736 | 0.37608763 | -0.3796768 | 0.70418536 | 0.78760538 | NA | NA |
| ENSG000001 | 4.51785951 | -0.142520768 | 0.37572611 | -0.3793209 | 0.70444958 | 0.78783048 | FAM87B | 400728 |
| ENSG000001 | 252.578135 | 0.056948755 | 0.15014132 | 0.37930103 | 0.70446434 | 0.78783048 | ZNF69 | 7620 |
| ENSG000002 | 4.03061376 | 0.143616592 | 0.37856001 | 0.37937602 | 0.70440865 | 0.78783048 | NA | NA |
| ENSG000002 | 3.34475279 | -0.14349121 | 0.37827854 | -0.3793269 | 0.70444516 | 0.78783048 | SLC25A36P1 | 106480240 |
| ENSG000001 | 33.5534266 | 0.098228438 | 0.25902093 | 0.37922973 | 0.70451728 | 0.78786796 | BHLHE41 | 79365 |
| ENSG00000C | 56.4070416 | 0.074951042 | 0.19767599 | 0.37916108 | 0.70456825 | 0.7878815 | KLHL22 | 84861 |
| ENSG000002 | 7.73333632 | 0.135211463 | 0.35660552 | 0.37916257 | 0.70456715 | 0.7878815 | NA | NA |
| ENSG000002 | 3.06881514 | -0.142891109 | 0.37691751 | -0.3791045 | 0.7046103 | 0.7879068 | NA | NA |
| ENSG000001 | 5.74928734 | -0.139075314 | 0.36712955 | -0.3788181 | 0.70482296 | 0.78812287 | TMEM72 | 643236 |
| ENSG000002 | 52.8983228 | -0.078630586 | 0.20789736 | -0.3782183 | 0.70526844 | 0.7885775 | ATP10A | 57194 |
| ENSG000002 | 4.51282368 | -0.142197817 | 0.37594873 | -0.3782373 | 0.70525435 | 0.7885775 | NA | NA |
| ENSG000002 | 4.53867215 | -0.142458016 | 0.37669248 | -0.3781812 | 0.70529599 | 0.78858657 | HLA-DRB6 | 3128 |
| ENSG000002 | 4.53204258 | -0.142380766 | 0.37655501 | -0.3781141 | 0.70534582 | 0.78862053 | NA | NA |
| ENSG000002 | 4.03148089 | -0.14280742 | 0.37777871 | -0.3780187 | 0.70541669 | 0.78867804 | NA | NA |
| ENSG000002 | 4.63755704 | -0.142880502 | 0.37802813 | -0.3779626 | 0.70545836 | 0.78870288 | NA | NA |
| ENSG000002 | 3.1355498 | -0.142670018 | 0.37753801 | -0.3778958 | 0.70550802 | 0.78873665 | NA | NA |
| ENSG000002 | 3.10250538 | -0.142691395 | 0.37773831 | -0.377752 | 0.70561485 | 0.78883354 | LOC101929464 | 101929464 |
| ENSG000002 | 8.65045149 | -0.130900169 | 0.34654726 | -0.3777267 | 0.70563359 | 0.78883354 | NA | NA |
| ENSG000002 | 20.9864486 | 0.110935207 | 0.29375706 | 0.37764269 | 0.70569604 | 0.78886452 | NA | NA |
| ENSG000002 | 5.04413319 | -0.141332899 | 0.37425588 | -0.3776371 | 0.70570021 | 0.78886452 | NA | NA |
| ENSG000002 | 4.77299228 | -0.141422339 | 0.37459623 | -0.3775327 | 0.70577774 | 0.78892944 | NA | NA |

| | | | | | | | | |
|---|---|---|---|---|---|---|---|---|
| ENSG000002 | 5.29794057 | -0.141506638 | 0.37489217 | -0.3774596 | 0.7058321 | 0.78896845 | NA | NA |
| ENSG000002 | 2.91824404 | 0.142358704 | 0.37729488 | 0.37731417 | 0.70594014 | 0.78906747 | NA | NA |
| ENSG000002 | 3.61040105 | -0.142674535 | 0.37827926 | -0.3771672 | 0.70604934 | 0.78916478 | FAM106A | 80039 |
| ENSG000002 | 57.1867185 | 0.077451458 | 0.20536275 | 0.37714464 | 0.70606612 | 0.78916478 | LOC100507283 | 100507283 |
| ENSG000002 | 2.63133991 | -0.109510346 | 0.29040721 | -0.3770924 | 0.70610496 | 0.78918643 | NA | NA |
| ENSG000002 | 10.0541692 | 0.127677266 | 0.33863239 | 0.37703796 | 0.70614539 | 0.78920987 | NA | NA |
| ENSG000001 | 3.87527357 | -0.142677129 | 0.37853782 | -0.3769164 | 0.7062357 | 0.78928905 | OR4K14 | 122740 |
| ENSG000002 | 3.6310432 | -0.142669055 | 0.37855192 | -0.3768811 | 0.70626198 | 0.78929667 | RN7SL127P | 106480956 |
| ENSG00000C | 121.541048 | -0.059074694 | 0.15678589 | -0.3767858 | 0.70633281 | 0.78935407 | CP | 1356 |
| ENSG000002 | 3.61849046 | -0.142293558 | 0.3777104 | -0.3767266 | 0.70637679 | 0.78938146 | NA | NA |
| ENSG000002 | 3.45207705 | -0.140695269 | 0.37356134 | -0.3766323 | 0.70644688 | 0.78943803 | NA | NA |
| ENSG000002 | 10.0312306 | -0.125636498 | 0.33379119 | -0.3763925 | 0.70662513 | 0.78959371 | NA | NA |
| ENSG000002 | 3.31962136 | -0.142394292 | 0.37830237 | -0.3764034 | 0.70661703 | 0.78959371 | NA | NA |
| ENSG000001 | 20.7071325 | 0.105674338 | 0.28085278 | 0.37626239 | 0.70672184 | 0.78968001 | ASIC2 | 40 |
| ENSG000002 | 3.14641907 | -0.141795807 | 0.37688668 | -0.3762293 | 0.70674645 | 0.78968575 | NA | NA |
| ENSG000001 | 116.369017 | 0.061364972 | 0.16320122 | 0.37600806 | 0.70691091 | 0.78984774 | C4orf33 | 132321 |
| ENSG000002 | 12.6648458 | 0.119146299 | 0.31690455 | 0.37596904 | 0.70693991 | 0.78985839 | NA | NA |
| ENSG00000C | 98.10631 | -0.063123395 | 0.16805045 | -0.3756217 | 0.70719815 | 0.79012515 | ERO1B | 56605 |
| ENSG000002 | 219.606561 | 0.058995107 | 0.15709968 | 0.37552659 | 0.70726887 | 0.79017218 | NA | NA |
| ENSG000002 | 12.1571185 | 0.123473629 | 0.32881347 | 0.37551268 | 0.70727922 | 0.79017218 | NA | NA |
| ENSG000002 | 4.07724148 | -0.141845101 | 0.37780213 | -0.3754481 | 0.70732722 | 0.79020404 | NA | NA |
| ENSG000002 | 3.0964714 | -0.141974684 | 0.37819741 | -0.3753984 | 0.70736423 | 0.79022361 | NA | NA |
| ENSG000001 | 5.96788187 | -0.138842784 | 0.36995982 | -0.3752915 | 0.70744367 | 0.79028865 | TNFSF13 | 8741 |
| ENSG000002 | 13.2703457 | 0.118336346 | 0.31533852 | 0.37526765 | 0.70746143 | 0.79028865 | NA | NA |
| ENSG000001 | 4.10976996 | -0.141744733 | 0.37782309 | -0.3751617 | 0.70754025 | 0.7903378 | TTC36 | 143941 |
| ENSG000002 | 6.03649809 | -0.138315679 | 0.36868837 | -0.3751561 | 0.7075444 | 0.7903378 | NA | NA |
| ENSG000001 | 13.6039618 | -0.119484028 | 0.31852983 | -0.375111 | 0.70757795 | 0.7903535 | OR56A3 | 390083 |
| ENSG000002 | 3.97522945 | 0.141767055 | 0.37803985 | 0.37500558 | 0.70765632 | 0.79041196 | NA | NA |
| ENSG000002 | 5.31743554 | -0.139716628 | 0.37258944 | -0.3749882 | 0.70766927 | 0.79041196 | NA | NA |

| | | | | | | | | |
|---|---|---|---|---|---|---|---|---|
| ENSG000002 | 13.6605396 | -0.119806125 | 0.31954077 | -0.3749322 | 0.70771089 | 0.79042305 | NA | NA |
| ENSG000002 | 4.78061169 | -0.140780192 | 0.37551781 | -0.3748962 | 0.70773768 | 0.79042305 | PEX5L-AS2 | 101928790 |
| ENSG000002 | 3.81598334 | -0.141771809 | 0.37815043 | -0.3749085 | 0.70772852 | 0.79042305 | NA | NA |
| ENSG000002 | 5.4880609 | -0.138646979 | 0.36988092 | -0.3748422 | 0.70777783 | 0.79044612 | NA | NA |
| ENSG000002 | 3.11346583 | -0.140332455 | 0.3744679 | -0.3747516 | 0.7078452 | 0.79048897 | NA | NA |
| ENSG000002 | 18.7899621 | -0.107872737 | 0.28787084 | -0.3747262 | 0.70786413 | 0.79048897 | NA | NA |
| ENSG000002 | 21.7410889 | -0.105082684 | 0.28043587 | -0.374712 | 0.70787467 | 0.79048897 | NA | NA |
| ENSG000002 | 8.82355682 | -0.127981092 | 0.34160237 | -0.3746493 | 0.70792129 | 0.79051926 | NA | NA |
| ENSG000002 | 3.55983351 | -0.14178069 | 0.37850899 | -0.3745768 | 0.70797522 | 0.79053594 | NA | NA |
| ENSG000002 | 4.01875263 | -0.141465378 | 0.37764272 | -0.3746011 | 0.70795715 | 0.79053594 | NA | NA |
| ENSG000002 | 3.59719706 | -0.141706236 | 0.37851254 | -0.3743766 | 0.70812415 | 0.79068046 | NA | NA |
| ENSG000001 | 71.9023104 | -0.0681335 | 0.18205763 | -0.3742414 | 0.70822474 | 0.79077101 | ANKRD18A | 253650 |
| ENSG00000C | 17.1951604 | -0.11016744 | 0.29442726 | -0.3741754 | 0.70827382 | 0.79080404 | ZMYND12 | 84217 |
| ENSG000002 | 2.61732064 | -0.140120614 | 0.37461675 | -0.3740372 | 0.70837661 | 0.79089703 | HTR3E-AS1 | 106478970 |
| ENSG000001 | 3.88933244 | -0.141583654 | 0.37858925 | -0.373977 | 0.70842146 | 0.79092532 | GRP | 2922 |
| ENSG000002 | 33.3262027 | -0.096587156 | 0.25833962 | -0.3738767 | 0.70849607 | 0.79098685 | NA | NA |
| ENSG000002 | 3.39735881 | -0.139675405 | 0.37362457 | -0.3738389 | 0.70852416 | 0.79099643 | MIR625 | 693210 |
| ENSG000001 | 4.07609988 | -0.14098033 | 0.37736867 | -0.3735878 | 0.70871102 | 0.79118325 | DEFB108B | 245911 |
| ENSG000002 | 5.57348557 | -0.139654854 | 0.37400527 | -0.3734034 | 0.7088482 | 0.79131462 | NA | NA |
| ENSG000002 | 3.09715654 | -0.1411575 | 0.37811146 | -0.3733225 | 0.70890846 | 0.7913601 | NA | NA |
| ENSG000002 | 4.34767853 | -0.140634796 | 0.37682345 | -0.3732114 | 0.70899114 | 0.7914306 | NA | NA |
| ENSG000002 | 6.63658018 | 0.134962234 | 0.36186 | 0.37296809 | 0.70917219 | 0.79160283 | NA | NA |
| ENSG000002 | 2.84339461 | -0.140608122 | 0.3770144 | -0.3729516 | 0.70918447 | 0.79160283 | RN7SL847P | 106481152 |
| ENSG000002 | 5.28262276 | -0.139782173 | 0.37489719 | -0.3728547 | 0.7092566 | 0.79166156 | NA | NA |
| ENSG000001 | 3.59310606 | -0.141129563 | 0.37857481 | -0.3727917 | 0.70930345 | 0.79169206 | INS-IGF2 | 723961 |
| ENSG000002 | 3.80665266 | -0.141053382 | 0.37843957 | -0.3727237 | 0.70935412 | 0.79172322 | PSORS1C2 | 170680 |
| ENSG000002 | 2.60074967 | -0.139061713 | 0.37311793 | -0.3727018 | 0.70937042 | 0.79172322 | NA | NA |
| ENSG000002 | 37.3258778 | 0.084507018 | 0.22676125 | 0.37266958 | 0.70939438 | 0.79172818 | MYCNOS | 10408 |
| ENSG000002 | 2.64070964 | -0.139270411 | 0.37400959 | -0.3723712 | 0.70961647 | 0.79195425 | NA | NA |

| | | | | | | | | | |
|---|---|---|---|---|---|---|---|---|---|
| ENSG000001 | 21.8892913 | -0.102607649 | 0.2756831 | -0.3721942 | 0.70974827 | 0.79205975 | TAS2R4 | | 50832 |
| ENSG000002 | 4.00681927 | 0.140768719 | 0.37821556 | 0.37219177 | 0.70975007 | 0.79205975 | APOBEC3D | | 140564 |
| ENSG000001 | 11.4022151 | 0.122047332 | 0.32795891 | 0.37214214 | 0.70978702 | 0.79207919 | VMAC | | 400673 |
| ENSG000002 | 4.80816849 | -0.140285379 | 0.37701822 | -0.3720918 | 0.70982452 | 0.79208177 | HSPB9 | | 94086 |
| ENSG000002 | 12.4018521 | 0.120603166 | 0.32412664 | 0.37208656 | 0.7098284 | 0.79208177 | NA | NA | |
| ENSG000001 | 112.176114 | -0.060643885 | 0.16302712 | -0.3719865 | 0.70990292 | 0.79209953 | COLGALT2 | | 23127 |
| ENSG000002 | 6.94601915 | -0.13428418 | 0.36098326 | -0.3719956 | 0.70989614 | 0.79209953 | NA | NA | |
| ENSG000002 | 15.0081282 | 0.114724402 | 0.30837137 | 0.37203324 | 0.7098681 | 0.79209953 | NA | NA | |
| ENSG000002 | 3.58385558 | -0.140708696 | 0.37857822 | -0.3716767 | 0.71013359 | 0.79233511 | NA | NA | |
| ENSG000002 | 6.0599505 | -0.139120725 | 0.37435185 | -0.3716309 | 0.71016766 | 0.79235132 | NA | NA | |
| ENSG000001 | 2.36278903 | -0.137792596 | 0.37097201 | -0.3714366 | 0.71031234 | 0.79243782 | MIR26A1 | | 407015 |
| ENSG000002 | 2.86585052 | -0.139907174 | 0.37663285 | -0.3714683 | 0.71028874 | 0.79243782 | NA | NA | |
| ENSG000002 | 2.36907892 | -0.137710929 | 0.37073218 | -0.3714566 | 0.71029745 | 0.79243782 | NA | NA | |
| ENSG000002 | 14.5707755 | 0.115459595 | 0.31085838 | 0.37142185 | 0.71032336 | 0.79243782 | LOC283887 | | 283887 |
| ENSG000002 | 3.08907957 | -0.140227824 | 0.3776382 | -0.3713285 | 0.71039288 | 0.79249358 | NA | NA | |
| ENSG000002 | 12.9424016 | 0.117446996 | 0.31639692 | 0.37120145 | 0.71048749 | 0.79257732 | NA | NA | |
| ENSG000002 | 2.60610647 | -0.139258832 | 0.3752414 | -0.371118 | 0.71054966 | 0.79258054 | RN7SKP58 | | 106480850 |
| ENSG000002 | 21.4686617 | 0.101506927 | 0.27349644 | 0.37114533 | 0.71052929 | 0.79258054 | NA | NA | |
| ENSG000002 | 3.07678796 | -0.139751049 | 0.37659345 | -0.3710926 | 0.71056855 | 0.79258054 | NA | NA | |
| ENSG000002 | 23.7254439 | 0.102771388 | 0.27694179 | 0.37109383 | 0.71056765 | 0.79258054 | NA | NA | |
| ENSG000002 | 3.82825514 | -0.140306118 | 0.37820495 | -0.3709791 | 0.71065312 | 0.79265306 | FANK1-AS1 | | 101410540 |
| ENSG000002 | 3.80104941 | 0.140321299 | 0.37828348 | 0.37094218 | 0.7106806 | 0.79266191 | NA | NA | |
| ENSG000002 | 5.72668706 | -0.136682435 | 0.36885341 | -0.3705603 | 0.71096505 | 0.79295736 | NA | NA | |
| ENSG000002 | 6.68686741 | -0.135999454 | 0.36707458 | -0.3704954 | 0.71101338 | 0.79296765 | CA3-AS1 | | 100996348 |
| ENSG000002 | 9.11417092 | 0.1271114 | 0.34307171 | 0.37050971 | 0.71100274 | 0.79296765 | JMJD1C-AS1 | | 84989 |
| ENSG000002 | 25.4464484 | -0.096626052 | 0.26087595 | -0.3703908 | 0.71109132 | 0.79303276 | NA | NA | |
| ENSG000001 | 4.27068782 | -0.13935203 | 0.3766715 | -0.3699564 | 0.71141498 | 0.793163 | KRT33B | | 3884 |
| ENSG000001 | 8.89792409 | -0.128926515 | 0.34836985 | -0.3700852 | 0.71131904 | 0.793163 | NA | NA | |
| ENSG000002 | 2.85275698 | -0.139200646 | 0.37622319 | -0.3699949 | 0.71138632 | 0.793163 | RNU1-105P | | 106480185 |

| | | | | | | | | |
|---|---|---|---|---|---|---|---|---|
| ENSG000002 | 4.10164058 | -0.140094958 | 0.37858938 | -0.3700446 | 0.71134925 | 0.793163 | NA | NA |
| ENSG000002 | 720.712177 | 0.044372638 | 0.11985941 | 0.37020572 | 0.71122922 | 0.793163 | NA | NA |
| ENSG000002 | 2.335205 | -0.137005036 | 0.37028834 | -0.3699955 | 0.71138581 | 0.793163 | NA | NA |
| ENSG000002 | 4.0639124 | -0.139651818 | 0.37736343 | -0.3700725 | 0.71132849 | 0.793163 | NA | NA |
| ENSG000002 | 4.55902037 | -0.139909745 | 0.37798904 | -0.3701423 | 0.71127644 | 0.793163 | LOC401463 | 401463 |
| ENSG000002 | 3.62734505 | -0.140105823 | 0.37857589 | -0.3700865 | 0.71131805 | 0.793163 | NA | NA |
| ENSG000002 | 3.95732045 | 0.139964308 | 0.37833937 | 0.36994381 | 0.71142436 | 0.793163 | TMEM56-RWDI | 100527978 |
| ENSG000002 | 3.34850356 | -0.139900084 | 0.3781911 | -0.369919 | 0.71144282 | 0.793163 | NA | NA |
| ENSG000002 | 12.9247035 | -0.117193545 | 0.31663362 | -0.3701235 | 0.71129047 | 0.793163 | NA | NA |
| ENSG000002 | 2.9286965 | 0.139158055 | 0.3762623 | 0.3698432 | 0.71149932 | 0.79318238 | RNU6-786P | 106481429 |
| ENSG000002 | 3.80567469 | -0.139985154 | 0.37848657 | -0.369855 | 0.71149053 | 0.79318238 | NA | NA |
| ENSG000002 | 2.58484423 | -0.138251817 | 0.37385247 | -0.3698031 | 0.71152917 | 0.79319385 | NA | NA |
| ENSG000002 | 3.06195255 | -0.139701962 | 0.37796621 | -0.3696149 | 0.71166941 | 0.79330799 | NA | NA |
| ENSG000002 | 2.87595011 | -0.138959916 | 0.37596033 | -0.3696132 | 0.71167069 | 0.79330799 | NA | NA |
| ENSG000001 | 98.6295166 | -0.064934672 | 0.17571814 | -0.3695388 | 0.71172615 | 0.79332686 | SDK1 | 221935 |
| ENSG000002 | 3.55767429 | -0.139894204 | 0.37859202 | -0.3695118 | 0.71174631 | 0.79332686 | NA | NA |
| ENSG000002 | 3.8404024 | -0.139884529 | 0.37856155 | -0.3695159 | 0.71174319 | 0.79332686 | NA | NA |
| ENSG000002 | 12.3655283 | -0.117717212 | 0.31866046 | -0.3694127 | 0.71182015 | 0.79338736 | NA | NA |
| ENSG000002 | 2.35862646 | -0.13631412 | 0.36908819 | -0.3693267 | 0.71188423 | 0.79343698 | NA | NA |
| ENSG000002 | 5.09307111 | 0.13816354 | 0.37413132 | 0.36929156 | 0.71191041 | 0.79344435 | NUDT18 | 79873 |
| ENSG000002 | 3.82639559 | -0.139680864 | 0.37833862 | -0.3691954 | 0.71198212 | 0.79350245 | NA | NA |
| ENSG000002 | 26.7713407 | 0.097845664 | 0.2650448 | 0.36916651 | 0.71200361 | 0.7935046 | PCDHGB2 | 56103 |
| ENSG000001 | 145.996609 | 0.057439827 | 0.15561234 | 0.36912127 | 0.71203733 | 0.79352037 | SYP | 6855 |
| ENSG000001 | 33.3489467 | -0.08744522 | 0.23693611 | -0.3690667 | 0.71207804 | 0.79352134 | CCDC180 | 100499483 |
| ENSG000002 | 2.35211619 | -0.137356463 | 0.37219802 | -0.3690414 | 0.7120969 | 0.79352134 | NA | NA |
| ENSG000002 | 18.7326314 | 0.106518253 | 0.28860057 | 0.36908539 | 0.71206408 | 0.79352134 | NA | NA |
| ENSG000001 | 83.8649652 | 0.070601127 | 0.19137689 | 0.36891146 | 0.71219372 | 0.7935638 | OPRD1 | 4985 |
| ENSG000002 | 3.8370814 | -0.139630746 | 0.3784749 | -0.36893 | 0.7121799 | 0.7935638 | NA | NA |
| ENSG000002 | 2.83958436 | -0.138129607 | 0.37441606 | -0.3689201 | 0.71218729 | 0.7935638 | NA | NA |

| | | | | | | | | |
|---|---|---|---|---|---|---|---|---|
| ENSG000001 | 174.709055 | 0.053872118 | 0.14610286 | 0.36872733 | 0.71233098 | 0.79369494 | SYNJ1 | 8867 |
| ENSG000002 | 3.77512254 | -0.139466289 | 0.37832165 | -0.3686447 | 0.71239254 | 0.79374172 | NA | NA |
| ENSG000002 | 6.47472966 | -0.134477167 | 0.3648258 | -0.3686065 | 0.71242103 | 0.79375166 | NA | NA |
| ENSG000002 | 2.33961607 | -0.137032922 | 0.37183177 | -0.3685347 | 0.71247455 | 0.79378948 | NA | NA |
| ENSG000002 | 2.35986035 | -0.13665239 | 0.37096643 | -0.3683686 | 0.7125984 | 0.79390564 | NA | NA |
| ENSG000002 | 2.83533725 | -0.13850789 | 0.37610123 | -0.3682729 | 0.71266976 | 0.79396333 | NA | NA |
| ENSG000002 | 31.9273545 | -0.090610157 | 0.24608452 | -0.3682075 | 0.71271854 | 0.79399587 | NA | NA |
| ENSG000002 | 2.60931536 | -0.135917056 | 0.36918849 | -0.3681508 | 0.71276076 | 0.79402109 | RNU6-510P | 106481342 |
| ENSG000002 | 2.6032414 | -0.135691259 | 0.36865184 | -0.3680743 | 0.71281785 | 0.79405574 | NA | NA |
| ENSG000002 | 2.3457616 | -0.136519206 | 0.37092763 | -0.3680481 | 0.71283737 | 0.79405574 | NA | NA |
| ENSG000002 | 17.5731684 | 0.110412842 | 0.30001016 | 0.36803034 | 0.71285061 | 0.79405574 | FAM72C | 554282 |
| ENSG000002 | 28.3030162 | 0.09373353 | 0.25473179 | 0.36796951 | 0.71289597 | 0.79408445 | RNF144A-AS1 | 386597 |
| ENSG000002 | 2.85810637 | -0.13860273 | 0.3767617 | -0.367879 | 0.71296343 | 0.79413778 | NA | NA |
| ENSG000001 | 44.4641219 | -0.084152324 | 0.22888802 | -0.3676572 | 0.71312888 | 0.79429008 | TSACC | 128229 |
| ENSG000002 | 14.5647562 | 0.116290784 | 0.3163143 | 0.36764315 | 0.71313934 | 0.79429008 | NA | NA |
| ENSG000001 | 17.6123922 | -0.106851098 | 0.29067171 | -0.3676006 | 0.71317106 | 0.79430359 | CCDC65 | 85478 |
| ENSG000002 | 7.72455352 | -0.131443792 | 0.3579051 | -0.3672588 | 0.713426 | 0.79456571 | CYP2A6 | 1548 |
| ENSG000002 | 9.94433772 | -0.131147192 | 0.35730518 | -0.3670453 | 0.71358522 | 0.79461237 | NA | NA |
| ENSG000002 | 3.08506135 | -0.138773654 | 0.37811057 | -0.3670187 | 0.71360506 | 0.79461237 | NA | NA |
| ENSG000002 | 4.26178191 | -0.13850922 | 0.3773455 | -0.3670621 | 0.71357272 | 0.79461237 | RN7SL328P | 106481837 |
| ENSG000002 | 31.7783627 | -0.090081781 | 0.24537333 | -0.3671213 | 0.71352853 | 0.79461237 | LOC101927550 | 101927550 |
| ENSG000002 | 2.34907507 | -0.134484739 | 0.36640395 | -0.3670395 | 0.71358952 | 0.79461237 | KIRREL3-AS2 | 100874251 |
| ENSG000002 | 2.90194762 | -0.136067276 | 0.37069114 | -0.3670637 | 0.71357147 | 0.79461237 | NA | NA |
| ENSG000002 | 4.00432162 | -0.13863726 | 0.37773013 | -0.3670273 | 0.71359867 | 0.79461237 | NA | NA |
| ENSG00000C | 103.327265 | -0.064312465 | 0.17529969 | -0.3668715 | 0.71371485 | 0.79463105 | ARHGAP31 | 57514 |
| ENSG000001 | 13.7489948 | -0.115294457 | 0.3142357 | -0.3669044 | 0.71369034 | 0.79463105 | NXPH4 | 11247 |
| ENSG000002 | 16.4651359 | -0.108803021 | 0.29657519 | -0.3668649 | 0.71371981 | 0.79463105 | NA | NA |
| ENSG000002 | 3.79301889 | -0.13871844 | 0.37804032 | -0.3669409 | 0.71366313 | 0.79463105 | NA | NA |
| ENSG000002 | 9.41941738 | -0.127911586 | 0.34864747 | -0.3668794 | 0.71370895 | 0.79463105 | ANP32AP1 | 723972 |

| | | | | | | | | |
|---|---|---|---|---|---|---|---|---|
| ENSG000002 | 2.35308174 | -0.136139559 | 0.37131751 | -0.3666392 | 0.71388815 | 0.79479665 | NA | NA |
| ENSG000002 | 2.61853393 | -0.136284227 | 0.37176147 | -0.3665905 | 0.71392449 | 0.79481528 | NA | NA |
| ENSG000002 | 3.12785968 | 0.137930274 | 0.3763813 | 0.3664642 | 0.71401872 | 0.79489837 | NA | NA |
| ENSG000002 | 2.35752451 | -0.135847879 | 0.37073758 | -0.366426 | 0.71404724 | 0.7949083 | RN7SKP226 | 106480903 |
| ENSG000002 | 4.75591298 | -0.136888171 | 0.37365618 | -0.3663479 | 0.71410547 | 0.79492947 | RN7SKP30 | 106479108 |
| ENSG000002 | 13.1249433 | -0.11727108 | 0.32009109 | -0.3663678 | 0.71409061 | 0.79492947 | LOC101243545 | 101243545 |
| ENSG000000 | 108.345654 | 0.05857281 | 0.15990378 | 0.36630035 | 0.71414097 | 0.79494717 | PRKCH | 5583 |
| ENSG000002 | 78.1564907 | 0.071354007 | 0.19484327 | 0.36621233 | 0.71420664 | 0.79499845 | AMZ2P1 | 201283 |
| ENSG000002 | 4.06309032 | -0.138487285 | 0.37820795 | -0.366167 | 0.71424044 | 0.79501424 | NA | NA |
| ENSG000000 | 35.7674713 | -0.089642956 | 0.24497592 | -0.3659256 | 0.71442061 | 0.79516583 | EHD2 | 30846 |
| ENSG000000 | 103.904288 | -0.063575792 | 0.17373672 | -0.3659318 | 0.71441597 | 0.79516583 | PTPN3 | 5774 |
| ENSG000002 | 5.99544196 | -0.135272413 | 0.36969201 | -0.3659057 | 0.71443545 | 0.79516583 | GLT6D1 | 360203 |
| ENSG000002 | 2.61114265 | -0.136848373 | 0.37417704 | -0.3657316 | 0.71456535 | 0.79528858 | NA | NA |
| ENSG000001 | 10.1014423 | -0.124967462 | 0.34178754 | -0.3656291 | 0.71464188 | 0.7953301 | MESP2 | 145873 |
| ENSG000002 | 5.08381948 | -0.138191864 | 0.37793694 | -0.3656479 | 0.7146278 | 0.7953301 | NA | NA |
| ENSG000002 | 2.36540561 | -0.135960465 | 0.37189413 | -0.3655892 | 0.71467166 | 0.79534141 | NA | NA |
| ENSG000002 | 21.7677665 | -0.100971198 | 0.27623222 | -0.3655301 | 0.71471572 | 0.79536862 | NA | NA |
| ENSG000002 | 47.7276631 | -0.076151482 | 0.20835069 | -0.3654967 | 0.71474069 | 0.79537459 | C8orf44 | 56260 |
| ENSG000002 | 3.81615995 | -0.138154846 | 0.37820139 | -0.3652944 | 0.71489166 | 0.79552076 | NA | NA |
| ENSG000002 | 5.01700695 | -0.137294519 | 0.37596767 | -0.3651764 | 0.71497973 | 0.79559693 | NA | NA |
| ENSG000002 | 30.8953283 | 0.098126181 | 0.26873627 | 0.36513932 | 0.71500741 | 0.79560589 | NA | NA |
| ENSG000001 | 31.4036595 | -0.088941284 | 0.24365597 | -0.3650281 | 0.7150904 | 0.7956716 | KDM6B | 23135 |
| ENSG000002 | 3.57055875 | -0.138186835 | 0.37858614 | -0.3650076 | 0.7151057 | 0.7956716 | NA | NA |
| ENSG000002 | 14.5782054 | 0.110707627 | 0.30336238 | 0.36493526 | 0.71515973 | 0.79570988 | TINCR | 257000 |
| ENSG000002 | 3.61993283 | -0.138107608 | 0.37847793 | -0.3649027 | 0.71518406 | 0.79571511 | NA | NA |
| ENSG000002 | 5.04374994 | -0.13608669 | 0.37299487 | -0.3648487 | 0.71522436 | 0.79573812 | NA | NA |
| ENSG000002 | 8.50597679 | 0.128276302 | 0.35186474 | 0.3645614 | 0.71543884 | 0.79594616 | NA | NA |
| ENSG000002 | 2.91636654 | -0.136495747 | 0.37442706 | -0.3645456 | 0.71545061 | 0.79594616 | NA | NA |
| ENSG000002 | 5.08302284 | 0.135845219 | 0.37270479 | 0.36448477 | 0.71549604 | 0.79597487 | NA | NA |

| | | | | | | | | |
|---|---|---|---|---|---|---|---|---|
| ENSG000002 | 3.04806232 | -0.13729328 | 0.37674238 | -0.3644222 | 0.71554277 | 0.79599298 | NA | NA |
| ENSG000002 | 3.10033318 | -0.136810222 | 0.37544394 | -0.3643959 | 0.71556242 | 0.79599298 | NA | NA |
| ENSG000002 | 3.58149273 | -0.137879677 | 0.37839104 | -0.3643841 | 0.71557121 | 0.79599298 | NA | NA |
| ENSG000001 | 13.204346 | -0.118355203 | 0.32487841 | -0.3643062 | 0.7156294 | 0.79601404 | BBS5 | 129880 |
| ENSG000002 | 46.3193627 | 0.078985199 | 0.21680298 | 0.36431786 | 0.71562066 | 0.79601404 | NA | NA |
| ENSG000002 | 3.0744725 | -0.137724246 | 0.37810919 | -0.3642446 | 0.71567534 | 0.7960433 | NA | NA |
| ENSG000002 | 8.65702285 | 0.126040058 | 0.34608346 | 0.36418977 | 0.7157163 | 0.79606703 | NA | NA |
| ENSG000002 | 3.0619203 | -0.137640823 | 0.37803958 | -0.364091 | 0.71579003 | 0.79611827 | NA | NA |
| ENSG000002 | 22.7969446 | -0.097832948 | 0.2687161 | -0.3640755 | 0.71580163 | 0.79611827 | NA | NA |
| ENSG000002 | 23.3524686 | 0.098676042 | 0.27108097 | 0.36400947 | 0.71585093 | 0.79612943 | NA | NA |
| ENSG000002 | 4.04698956 | -0.137550258 | 0.3778732 | -0.3640117 | 0.71584929 | 0.79612943 | NA | NA |
| ENSG000002 | 18.2426678 | -0.104619193 | 0.28746621 | -0.3639356 | 0.71590608 | 0.79616893 | NA | NA |
| ENSG000002 | 3.96655908 | 0.137676503 | 0.37834242 | 0.36389391 | 0.71593723 | 0.79618173 | NA | NA |
| ENSG000002 | 5.98021347 | -0.135023753 | 0.3711489 | -0.3637994 | 0.7160078 | 0.79621981 | NA | NA |
| ENSG000002 | 4.80406038 | -0.136765078 | 0.37593947 | -0.3637955 | 0.71601074 | 0.79621981 | NA | NA |
| ENSG000002 | 2.85844605 | -0.136857725 | 0.37622221 | -0.3637683 | 0.71603101 | 0.79622052 | NA | NA |
| ENSG000002 | 3.75194744 | 0.137547999 | 0.37826518 | 0.36362849 | 0.71613545 | 0.79631481 | NA | NA |
| ENSG000002 | 6.4242838 | 0.132364623 | 0.36405399 | 0.36358515 | 0.71616782 | 0.79632897 | NA | NA |
| ENSG000002 | 4.03749003 | -0.137563743 | 0.37838277 | -0.3635571 | 0.71618877 | 0.79633043 | NA | NA |
| ENSG000001 | 9.06354118 | -0.124142451 | 0.34165712 | -0.3633539 | 0.71634053 | 0.79647733 | GPR151 | 134391 |
| ENSG000002 | 2.3297446 | -0.134313231 | 0.36971307 | -0.3632905 | 0.71638792 | 0.79650818 | RNU6ATAC24P | 106479554 |
| ENSG000002 | 2.68456973 | 0.134633042 | 0.37072649 | 0.36316003 | 0.71648534 | 0.79659466 | NA | NA |
| ENSG000001 | 10.2647484 | -0.121134202 | 0.33364897 | -0.3630588 | 0.71656094 | 0.79664679 | CAPS | 828 |
| ENSG000001 | 3.31249785 | -0.137401396 | 0.37846968 | -0.3630447 | 0.71657152 | 0.79664679 | NA | NA |
| ENSG000002 | 6.99122437 | -0.132085401 | 0.3638913 | -0.3629804 | 0.71661955 | 0.79667834 | NA | NA |
| ENSG000002 | 39.2143892 | -0.090406234 | 0.24913157 | -0.3628855 | 0.71669043 | 0.7967353 | NA | NA |
| ENSG000002 | 4.04463908 | 0.137075502 | 0.37791599 | 0.36271421 | 0.71681838 | 0.7968557 | NA | NA |
| ENSG000002 | 3.84110735 | -0.137218365 | 0.37847984 | -0.3625513 | 0.71694008 | 0.79696914 | NA | NA |
| ENSG000002 | 2.61094677 | -0.135456135 | 0.37366704 | -0.3625049 | 0.71697476 | 0.79698584 | NA | NA |

| | | | | | | | | |
|---|---|---|---|---|---|---|---|---|
| ENSG000002 | 2.61099884 | -0.135430329 | 0.37366692 | -0.362436 | 0.71702627 | 0.79702125 | LOC100996694 | 100996694 |
| ENSG000002 | 16.0985849 | 0.106940183 | 0.2951639 | 0.3623078 | 0.71712203 | 0.79710522 | NA | NA |
| ENSG000002 | 5.28326181 | -0.13548541 | 0.3739775 | -0.3622822 | 0.71714112 | 0.79710522 | NA | NA |
| ENSG000002 | 3.08079464 | -0.136672725 | 0.37742275 | -0.3621211 | 0.71726157 | 0.79721724 | NA | NA |
| ENSG000002 | 2.80371538 | -0.13625328 | 0.37640733 | -0.3619836 | 0.71736429 | 0.79726871 | NA | NA |
| ENSG000002 | 2.38000025 | -0.134291141 | 0.37101341 | -0.3619576 | 0.71738368 | 0.79726871 | NA | NA |
| ENSG000002 | 2.34901091 | -0.133471851 | 0.36875378 | -0.3619538 | 0.71738652 | 0.79726871 | RN7SL793P | 106481137 |
| ENSG000002 | 4.32104763 | -0.136817158 | 0.37793979 | -0.3620078 | 0.71734619 | 0.79726871 | NA | NA |
| ENSG000002 | 4.11243805 | -0.136856255 | 0.37825457 | -0.3618099 | 0.71749408 | 0.7973664 | NA | NA |
| ENSG000002 | 2.34542933 | -0.134320186 | 0.37129477 | -0.3617616 | 0.7175302 | 0.79738468 | NA | NA |
| ENSG000002 | 2.88309399 | -0.135864695 | 0.37559899 | -0.3617281 | 0.71755525 | 0.79739067 | NA | NA |
| ENSG000002 | 3.07711282 | -0.13578724 | 0.37543335 | -0.3616813 | 0.71759017 | 0.79740762 | RN7SKP229 | 106480904 |
| ENSG000002 | 2.93876851 | 0.136324103 | 0.37702654 | 0.36157694 | 0.7176682 | 0.79747248 | NA | NA |
| ENSG000002 | 10.4086456 | 0.120750873 | 0.33404822 | 0.36147737 | 0.71774262 | 0.79751147 | NA | NA |
| ENSG000002 | 2.36005622 | -0.133491904 | 0.36928105 | -0.3614913 | 0.71773217 | 0.79751147 | NA | NA |
| ENSG000002 | 2.8339477 | -0.1352165 | 0.37421684 | -0.361332 | 0.71785131 | 0.79757233 | DLEU1-AS1 | 103689915 |
| ENSG000002 | 3.5846369 | -0.136650034 | 0.37821887 | -0.3612988 | 0.71787607 | 0.79757233 | NA | NA |
| ENSG000002 | 4.07060723 | -0.136429311 | 0.37760373 | -0.3613029 | 0.71787305 | 0.79757233 | NA | NA |
| ENSG000002 | 4.52485491 | -0.135721255 | 0.3755985 | -0.3613466 | 0.71784034 | 0.79757233 | NA | NA |
| ENSG000002 | 3.06221259 | -0.135769348 | 0.37597381 | -0.3611138 | 0.71801435 | 0.79765239 | NA | NA |
| ENSG000002 | 2.91944512 | 0.135744848 | 0.37590136 | 0.36111827 | 0.71801104 | 0.79765239 | NA | NA |
| ENSG000002 | 3.77049075 | -0.136715291 | 0.37859027 | -0.3611168 | 0.71801216 | 0.79765239 | NA | NA |
| ENSG000002 | 8.86069387 | 0.124630601 | 0.34514422 | 0.36109717 | 0.71802681 | 0.79765239 | NA | NA |
| ENSG000002 | 4.73735165 | -0.135362785 | 0.37489682 | -0.3610668 | 0.71804953 | 0.79765578 | NA | NA |
| ENSG000001 | 4.80726318 | -0.135546686 | 0.37546223 | -0.3610128 | 0.71808985 | 0.79765686 | NA | NA |
| ENSG000002 | 19.2392093 | -0.100464079 | 0.27826787 | -0.3610337 | 0.71807425 | 0.79765686 | BZRAP1-AS1 | 100506779 |
| ENSG000001 | 262.402141 | -0.050280778 | 0.13936606 | -0.3607821 | 0.71826236 | 0.79769069 | DENND4A | 10260 |
| ENSG000001 | 7.63064031 | -0.129567927 | 0.3591511 | -0.3607616 | 0.71827767 | 0.79769069 | NA | NA |
| ENSG000002 | 3.52519046 | -0.136601271 | 0.37858457 | -0.360821 | 0.71823325 | 0.79769069 | NA | NA |

| | | | | | | | | |
|---|---|---|---|---|---|---|---|---|
| ENSG000002 | 4.78608059 | -0.135330382 | 0.37499523 | -0.3608856 | 0.71818496 | 0.79769069 | LOC100131635 | 100131635 |
| ENSG000002 | 2.63089821 | -0.134565217 | 0.37298557 | -0.3607786 | 0.71826495 | 0.79769069 | NA | NA |
| ENSG000002 | 3.11158811 | -0.136397899 | 0.3779389 | -0.3608993 | 0.7181747 | 0.79769069 | NA | NA |
| ENSG000002 | 3.37784962 | -0.136295889 | 0.37770014 | -0.3608574 | 0.71820605 | 0.79769069 | NA | NA |
| ENSG000002 | 3.3724658 | -0.136049044 | 0.37704578 | -0.360829 | 0.71822729 | 0.79769069 | NA | NA |
| ENSG000001 | 2.84836608 | -0.135041228 | 0.37441258 | -0.3606749 | 0.71834247 | 0.79769712 | HES4 | 57801 |
| ENSG000002 | 2.84836608 | -0.135041228 | 0.37441258 | -0.3606749 | 0.71834247 | 0.79769712 | IQCF5 | 389124 |
| ENSG000002 | 2.85490859 | -0.134719689 | 0.37346676 | -0.3607274 | 0.71830325 | 0.79769712 | NA | NA |
| ENSG000001 | 726.377026 | -0.042221904 | 0.11707292 | -0.3606462 | 0.71836394 | 0.79769911 | NAV1 | 89796 |
| ENSG000002 | 2.81976463 | -0.135334599 | 0.375316 | -0.3605884 | 0.71840716 | 0.79772526 | NA | NA |
| ENSG000002 | 3.12182375 | -0.135414837 | 0.37557616 | -0.3605523 | 0.71843418 | 0.79773341 | CHRM3-AS1 | 100873984 |
| ENSG000002 | 2.87619619 | 0.135325428 | 0.37548365 | 0.36040298 | 0.7185458 | 0.79781366 | NA | NA |
| ENSG000002 | 2.85631929 | -0.135124013 | 0.37491625 | -0.3604112 | 0.71853965 | 0.79781366 | NA | NA |
| ENSG000001 | 190.21982 | 0.05450373 | 0.15125811 | 0.36033592 | 0.71859594 | 0.79784749 | TTC13 | 79573 |
| ENSG000002 | 4.2489493 | -0.136120442 | 0.37778833 | -0.3603088 | 0.71861626 | 0.7978482 | NA | NA |
| ENSG000002 | 9.166917 | 0.124795897 | 0.34639144 | 0.36027419 | 0.7186421 | 0.79785505 | OCLM | 10896 |
| ENSG000002 | 4.73838877 | -0.135269468 | 0.37572567 | -0.3600219 | 0.71883076 | 0.79798999 | NA | NA |
| ENSG000002 | 2.35515698 | -0.133875683 | 0.3718703 | -0.3600064 | 0.71884235 | 0.79798999 | NA | NA |
| ENSG000002 | 6.91498971 | 0.128836742 | 0.35786298 | 0.36001696 | 0.71883445 | 0.79798999 | NA | NA |
| ENSG000002 | 10.7417598 | -0.119772866 | 0.33263589 | -0.360072 | 0.7187933 | 0.79798999 | NA | NA |
| ENSG000002 | 6.69684446 | 0.132116278 | 0.36712733 | 0.359865 | 0.71894809 | 0.79808552 | NA | NA |
| ENSG000002 | 26.1560995 | -0.093598461 | 0.26027021 | -0.3596203 | 0.71913108 | 0.79826679 | RAP2C-AS1 | 101928578 |
| ENSG000002 | 7.19863948 | -0.129973058 | 0.36161397 | -0.3594249 | 0.71927726 | 0.79838474 | NA | NA |
| ENSG000002 | 4.63301102 | -0.135670105 | 0.37745184 | -0.3594369 | 0.7192683 | 0.79838474 | NA | NA |
| ENSG000002 | 3.30630025 | -0.136016744 | 0.37845578 | -0.3593993 | 0.71929639 | 0.79838474 | NA | NA |
| ENSG000002 | 3.53906108 | -0.136032565 | 0.37858496 | -0.3593185 | 0.71935687 | 0.79843001 | NA | NA |
| ENSG000002 | 4.56046363 | -0.134988923 | 0.37574586 | -0.3592559 | 0.71940365 | 0.79846008 | NA | NA |
| ENSG000002 | 13.7918402 | -0.112177474 | 0.31228227 | -0.3592182 | 0.71943186 | 0.79846954 | NA | NA |
| ENSG000002 | 2.83147548 | -0.134767323 | 0.375196 | -0.3591918 | 0.71945162 | 0.79846961 | NA | NA |

| | | | | | | | | |
|---|---|---|---|---|---|---|---|---|
| ENSG000001 | 70.8156444 | -0.068899037 | 0.19187045 | -0.3590915 | 0.71952668 | 0.79849992 | HACD4 | 401494 |
| ENSG000002 | 4.51830412 | -0.13577469 | 0.3781221 | -0.3590763 | 0.719538 | 0.79849992 | NA | NA |
| ENSG000002 | 5.26102934 | -0.133883112 | 0.37285138 | -0.359079 | 0.71953597 | 0.79849992 | NA | NA |
| ENSG000002 | 3.32790162 | -0.135667527 | 0.37795928 | -0.3589475 | 0.71963439 | 0.79857125 | NA | NA |
| ENSG000002 | 2.81595492 | -0.135376546 | 0.37715885 | -0.3589377 | 0.71964166 | 0.79857125 | NA | NA |
| ENSG000001 | 2.93327532 | 0.135373988 | 0.37726965 | 0.35882554 | 0.7197256 | 0.79864254 | NA | NA |
| ENSG000001 | 45.0728793 | -0.07644579 | 0.2131139 | -0.3587086 | 0.71981309 | 0.79871776 | PRKCG | 5582 |
| ENSG000002 | 7.53038958 | -0.130187287 | 0.36302606 | -0.3586169 | 0.71988169 | 0.79875891 | STK4-AS1 | 100505826 |
| ENSG000002 | 10.5034755 | -0.12307661 | 0.34320808 | -0.3586064 | 0.71988957 | 0.79875891 | NA | NA |
| ENSG000001 | 2115.70257 | 0.044495695 | 0.12416972 | 0.35834577 | 0.72008457 | 0.7987715 | CAMSAP2 | 23271 |
| ENSG000002 | 2.83746456 | -0.134708951 | 0.3759376 | -0.358328 | 0.7200979 | 0.7987715 | RNU6-813P | 106481438 |
| ENSG000002 | 2.3626728 | -0.132966659 | 0.37097061 | -0.3584291 | 0.72002223 | 0.7987715 | NA | NA |
| ENSG000002 | 4.30811608 | -0.135215125 | 0.3773161 | -0.3583603 | 0.72007367 | 0.7987715 | NA | NA |
| ENSG000002 | 5.31988313 | -0.134032445 | 0.3739161 | -0.3584559 | 0.72000214 | 0.7987715 | NA | NA |
| ENSG000002 | 2.80317405 | -0.134682383 | 0.37585797 | -0.3583332 | 0.72009399 | 0.7987715 | NA | NA |
| ENSG000002 | 3.30265396 | -0.135708988 | 0.37852815 | -0.3585176 | 0.71995603 | 0.7987715 | NA | NA |
| ENSG000002 | 2.58635145 | -0.134004622 | 0.37394071 | -0.3583579 | 0.72007546 | 0.7987715 | NA | NA |
| ENSG000002 | 5.13179644 | 0.133481867 | 0.37235362 | 0.35848145 | 0.71998305 | 0.7987715 | NA | NA |
| ENSG000002 | 4.27304371 | -0.135475931 | 0.37785232 | -0.358542 | 0.71993774 | 0.7987715 | NA | NA |
| ENSG000002 | 16.6294873 | -0.106522 | 0.29738996 | -0.3581896 | 0.7202014 | 0.79886447 | LINC00683 | 400660 |
| ENSG000002 | 25.0498491 | -0.093673321 | 0.26159682 | -0.3580828 | 0.72028135 | 0.79888759 | MFSD2B | 388931 |
| ENSG000002 | 3.84789914 | -0.135535755 | 0.37847723 | -0.3581081 | 0.72026243 | 0.79888759 | NA | NA |
| ENSG000002 | 2.35419144 | -0.133173968 | 0.37186535 | -0.3581242 | 0.72025039 | 0.79888759 | NA | NA |
| ENSG000002 | 2.31856235 | -0.130355538 | 0.3640698 | -0.358051 | 0.72030518 | 0.79889217 | NA | NA |
| ENSG000002 | 2.81834282 | -0.134954886 | 0.3770012 | -0.3579694 | 0.72036623 | 0.79893802 | NA | NA |
| ENSG000001 | 104.976928 | -0.062812598 | 0.17550331 | -0.3578998 | 0.72041831 | 0.79897393 | NUAK2 | 81788 |
| ENSG000001 | 4.55314369 | -0.134397345 | 0.3757057 | -0.3577197 | 0.72055306 | 0.79903815 | GPR34 | 2857 |
| ENSG000002 | 337.340807 | -0.049081164 | 0.13719104 | -0.3577578 | 0.72052459 | 0.79903815 | ZNF891 | 101060200 |
| ENSG000002 | 4.25731263 | 0.135081551 | 0.37762116 | 0.35771711 | 0.72055503 | 0.79903815 | SNORD111 | 692214 |

| | | | | | | | | |
|---|---|---|---|---|---|---|---|---|
| ENSG000002 | 3.5333073 | -0.135433159 | 0.37857634 | -0.3577433 | 0.7205354 | 0.79903815 | XXYLT1-AS2 | 101410543 |
| ENSG000002 | 3.0433388 | -0.13474767 | 0.37673977 | -0.3576678 | 0.72059196 | 0.79905725 | NA | NA |
| ENSG000000 | 299.338515 | -0.044303867 | 0.1238923 | -0.3575999 | 0.72064279 | 0.79909176 | EFHC1 | 114327 |
| ENSG000002 | 3.31327472 | -0.135261977 | 0.37827879 | -0.3575722 | 0.72066349 | 0.79909287 | NA | NA |
| ENSG000002 | 2.34688305 | -0.127671273 | 0.3570846 | -0.3575379 | 0.72068916 | 0.79909948 | NA | NA |
| ENSG000002 | 2.61168398 | -0.133267573 | 0.37282761 | -0.3574509 | 0.72075426 | 0.79914981 | NA | NA |
| ENSG000001 | 4.79222633 | -0.134132245 | 0.37527808 | -0.3574209 | 0.7207767 | 0.79915284 | IL1F10 | 84639 |
| ENSG000002 | 2.55632766 | -0.13381361 | 0.37453903 | -0.3572755 | 0.72088559 | 0.79925171 | NA | NA |
| ENSG000002 | 2.55606652 | -0.133234277 | 0.37304904 | -0.3571495 | 0.72097989 | 0.79933442 | NA | NA |
| ENSG000001 | 163.47306 | 0.055612162 | 0.15577706 | 0.35699841 | 0.72109299 | 0.79943795 | SYTL2 | 54843 |
| ENSG000002 | 9.66186077 | -0.124336381 | 0.34832869 | -0.3569513 | 0.72112826 | 0.79945519 | NA | NA |
| ENSG000000 | 3.09972105 | -0.13470256 | 0.37759995 | -0.3567335 | 0.7212913 | 0.79957134 | DEFB127 | 140850 |
| ENSG000002 | 3.13523714 | -0.134221631 | 0.37628081 | -0.356706 | 0.7213119 | 0.79957134 | NA | NA |
| ENSG000002 | 2.56747657 | -0.132507824 | 0.37141315 | -0.3567666 | 0.72126651 | 0.79957134 | NA | NA |
| ENSG000002 | 7.92842766 | -0.126517309 | 0.35466424 | -0.3567242 | 0.72129831 | 0.79957134 | NA | NA |
| ENSG000001 | 15.7025252 | -0.108597214 | 0.30455623 | -0.3565752 | 0.72140981 | 0.79964779 | BEST4 | 266675 |
| ENSG000001 | 42.6860115 | -0.081733556 | 0.22922726 | -0.3565612 | 0.7214203 | 0.79964779 | AKR1C1 | 1645 |
| ENSG000002 | 5.50437162 | -0.133055824 | 0.37335989 | -0.3563742 | 0.72156036 | 0.79978117 | NCR3 | 259197 |
| ENSG000002 | 30.8989451 | -0.089883286 | 0.25229946 | -0.3562564 | 0.72164859 | 0.79985282 | FTCDNL1 | 348751 |
| ENSG000002 | 2.32564206 | -0.132741876 | 0.37262428 | -0.3562352 | 0.72166445 | 0.79985282 | NA | NA |
| ENSG000000 | 42.7259929 | 0.078017155 | 0.21903457 | 0.3561865 | 0.7217009 | 0.79987136 | FMO4 | 2329 |
| ENSG000002 | 2.83450885 | -0.134159298 | 0.37670656 | -0.3561374 | 0.72173767 | 0.79988215 | NA | NA |
| ENSG000002 | 3.51364059 | 0.13434099 | 0.37723431 | 0.35612081 | 0.72175009 | 0.79988215 | MIR3942 | 100500904 |
| ENSG000002 | 3.29627921 | -0.134240847 | 0.37701678 | -0.3560607 | 0.72179513 | 0.79991021 | RN7SKP56 | 106481807 |
| ENSG000002 | 4.10695283 | -0.134700854 | 0.37838385 | -0.35599 | 0.72184808 | 0.79994702 | NA | NA |
| ENSG000001 | 11.4321567 | -0.116077626 | 0.32624203 | -0.3558022 | 0.72198872 | 0.79997424 | PVALB | 5816 |
| ENSG000001 | 4.74344532 | -0.133579255 | 0.37541657 | -0.3558161 | 0.7219783 | 0.79997424 | FLJ13224 | 79857 |
| ENSG000002 | 3.13067627 | 0.134512529 | 0.37798196 | 0.35587023 | 0.72193775 | 0.79997424 | MIR130A | 406919 |
| ENSG000002 | 2.82073017 | -0.133945683 | 0.37641128 | -0.3558493 | 0.72195344 | 0.79997424 | NA | NA |

| | | | | | | | | | |
|---|---|---|---|---|---|---|---|---|---|
| ENSG000002 | 3.53189738 | -0.134621617 | 0.3783641 | -0.3557991 | 0.721991 | 0.79997424 | NA | NA | |
| ENSG000002 | 13.2303396 | -0.115667037 | 0.32508692 | -0.3558034 | 0.72198779 | 0.79997424 | NA | NA | |
| ENSG000002 | 18.4478585 | -0.102234979 | 0.28737676 | -0.3557524 | 0.72202598 | 0.79999113 | C10orf105 | | 414152 |
| ENSG000001 | 34.9590785 | -0.084919416 | 0.23879128 | -0.3556219 | 0.72212372 | 0.80007756 | DNER | | 92737 |
| ENSG00000C | 74.4661001 | -0.065120429 | 0.18314945 | -0.355559 | 0.72217088 | 0.80008609 | KIF9 | | 64147 |
| ENSG000002 | 5.24615398 | 0.133886204 | 0.3765461 | 0.35556391 | 0.72216718 | 0.80008609 | NA | NA | |
| ENSG000002 | 2.36808112 | -0.13129086 | 0.36930033 | -0.3555124 | 0.72220573 | 0.80010284 | NA | NA | |
| ENSG000002 | 19.6801284 | -0.099872852 | 0.28106177 | -0.3553413 | 0.72233393 | 0.80022301 | NA | NA | |
| ENSG000002 | 3.10632359 | -0.13417507 | 0.37779162 | -0.3551563 | 0.72247251 | 0.80035467 | NA | NA | |
| ENSG00000C | 132.762554 | -0.054851733 | 0.15450612 | -0.3550133 | 0.72257961 | 0.80042308 | MXD1 | | 4084 |
| ENSG000002 | 2.85085835 | -0.132834845 | 0.3742159 | -0.3549685 | 0.72261321 | 0.80042308 | NA | NA | |
| ENSG000002 | 8.07823627 | -0.123497985 | 0.34790649 | -0.3549746 | 0.72260858 | 0.80042308 | NA | NA | |
| ENSG000002 | 4.8102519 | -0.13331404 | 0.37555737 | -0.3549765 | 0.7226072 | 0.80042308 | NA | NA | |
| ENSG000001 | 2.84760925 | -0.133256121 | 0.37546958 | -0.3549052 | 0.72266059 | 0.80045368 | MAGEA6 | | 4105 |
| ENSG000001 | 42.3507352 | 0.080953475 | 0.22815536 | 0.35481732 | 0.72272645 | 0.80047162 | PYROXD2 | | 84795 |
| ENSG000002 | 3.09863117 | -0.133834928 | 0.37718209 | -0.3548284 | 0.72271813 | 0.80047162 | NA | NA | |
| ENSG000002 | 5.37973401 | 0.131445378 | 0.3704726 | 0.35480458 | 0.722736 | 0.80047162 | NA | NA | |
| ENSG000002 | 5.16392827 | 0.132763957 | 0.3742932 | 0.35470577 | 0.72281003 | 0.80053175 | NA | NA | |
| ENSG000002 | 2.58094259 | -0.131274615 | 0.37015048 | -0.354652 | 0.7228503 | 0.80055448 | NA | NA | |
| ENSG000001 | 39.2118224 | -0.078441069 | 0.22121878 | -0.3545859 | 0.72289981 | 0.80055994 | IFIT3 | | 3437 |
| ENSG000001 | 15.0803518 | 0.112138049 | 0.31628086 | 0.35455212 | 0.72292515 | 0.80055994 | PRICKLE4 | | 29964 |
| ENSG000002 | 2.31742129 | -0.131618375 | 0.371221 | -0.3545553 | 0.72292277 | 0.80055994 | NA | NA | |
| ENSG000002 | 2.92293475 | 0.132977018 | 0.37509697 | 0.35451371 | 0.72295393 | 0.80055994 | NA | NA | |
| ENSG000002 | 2.84222054 | -0.13338043 | 0.37621129 | -0.354536 | 0.72293726 | 0.80055994 | NA | NA | |
| ENSG000002 | 8.66196154 | -0.123724027 | 0.3491901 | -0.3543171 | 0.72310124 | 0.8007012 | NA | NA | |
| ENSG000001 | 57.4387125 | -0.072959499 | 0.20599042 | -0.3541888 | 0.7231974 | 0.8007858 | ABCG1 | | 9619 |
| ENSG000001 | 31.5057119 | 0.08650093 | 0.2443452 | 0.35401117 | 0.72333051 | 0.80091133 | CA4 | | 762 |
| ENSG000002 | 15.0876038 | 0.108192469 | 0.30568381 | 0.35393589 | 0.72338693 | 0.80095193 | NA | NA | |
| ENSG000002 | 22.5103414 | -0.101611545 | 0.28723723 | -0.3537548 | 0.72352266 | 0.80108033 | C1orf220 | | 400798 |

| | | | | | | | | |
|---|---|---|---|---|---|---|---|---|
| ENSG000001 | 33.7817999 | -0.084972954 | 0.24023718 | -0.3537044 | 0.7235604 | 0.80110025 | ALOX12 | 239 |
| ENSG000002 | 2.55106314 | -0.132389754 | 0.37453154 | -0.3534809 | 0.72372793 | 0.80126386 | NA | NA |
| ENSG000001 | 3.5567291 | -0.133775015 | 0.37853547 | -0.3534015 | 0.72378743 | 0.80130786 | NUTM2A | 728118 |
| ENSG000002 | 17.7901123 | -0.108778118 | 0.30783489 | -0.3533651 | 0.72381472 | 0.80131619 | NA | NA |
| ENSG000001 | 9.39312954 | 0.122582836 | 0.34701633 | 0.35324804 | 0.7239025 | 0.80131959 | NOBOX | 135935 |
| ENSG000002 | 3.51893811 | 0.13280571 | 0.37595181 | 0.35325195 | 0.72389957 | 0.80131959 | NA | NA |
| ENSG000002 | 2.3106818 | -0.129587652 | 0.3667975 | -0.3532948 | 0.72386744 | 0.80131959 | NA | NA |
| ENSG000002 | 2.36930759 | -0.130382353 | 0.36908317 | -0.3532601 | 0.72389346 | 0.80131959 | NA | NA |
| ENSG000002 | 3.16666368 | -0.132440636 | 0.37494245 | -0.3532292 | 0.72391659 | 0.80131959 | NA | NA |
| ENSG000002 | 5.7981334 | -0.131316501 | 0.37185409 | -0.3531399 | 0.72398359 | 0.80137188 | NA | NA |
| ENSG000002 | 2.6033985 | -0.131628963 | 0.37281302 | -0.3530697 | 0.72403622 | 0.80140825 | NA | NA |
| ENSG000002 | 2.33088676 | -0.130388583 | 0.36945073 | -0.3529255 | 0.7241443 | 0.801506 | NA | NA |
| ENSG000001 | 160.909098 | -0.051230523 | 0.14520681 | -0.3528107 | 0.72423033 | 0.8015078 | KIAA1328 | 57536 |
| ENSG000002 | 2.5736549 | -0.131308989 | 0.37217188 | -0.3528181 | 0.72422479 | 0.8015078 | NA | NA |
| ENSG000002 | 3.28315954 | -0.133407426 | 0.37817405 | -0.3527673 | 0.72426293 | 0.8015078 | NA | NA |
| ENSG000002 | 6.70508068 | 0.126616954 | 0.35892705 | 0.35276514 | 0.72426452 | 0.8015078 | NA | NA |
| ENSG000002 | 2.36153119 | -0.131215705 | 0.37188181 | -0.3528425 | 0.72420652 | 0.8015078 | NA | NA |
| ENSG000002 | 2.92677692 | 0.133032845 | 0.37706733 | 0.35280926 | 0.72423144 | 0.8015078 | NA | NA |
| ENSG000001 | 2.33196386 | -0.13070105 | 0.37066922 | -0.3526083 | 0.7243821 | 0.80160756 | MRGPRD | 116512 |
| ENSG000002 | 9.79344664 | -0.119241946 | 0.33818658 | -0.3525922 | 0.72439419 | 0.80160756 | NA | NA |
| ENSG000001 | 9.24849286 | -0.121656895 | 0.3450691 | -0.3525581 | 0.72441978 | 0.80161399 | TMEM151A | 256472 |
| ENSG000002 | 3.05489354 | -0.132870074 | 0.37698599 | -0.3524536 | 0.7244981 | 0.80167053 | SSX9 | 280660 |
| ENSG000002 | 2.36619489 | -0.130084337 | 0.36909934 | -0.3524372 | 0.72451041 | 0.80167053 | NA | NA |
| ENSG000001 | 2.37840163 | -0.130029547 | 0.36912856 | -0.3522609 | 0.72464263 | 0.80178962 | NA | NA |
| ENSG000002 | 2.5777337 | -0.13202974 | 0.37482793 | -0.3522409 | 0.72465758 | 0.80178962 | MIR6076 | 102464828 |
| ENSG000002 | 2.38549344 | -0.130402987 | 0.37041405 | -0.3520465 | 0.72480336 | 0.80184152 | NA | NA |
| ENSG000002 | 3.32732096 | -0.132802905 | 0.37719536 | -0.3520799 | 0.72477834 | 0.80184152 | NA | NA |
| ENSG000002 | 4.02405737 | -0.133309715 | 0.37859052 | -0.3521211 | 0.72474743 | 0.80184152 | LOC101929441 | 101929441 |
| ENSG000002 | 2.59292174 | -0.131974396 | 0.37484665 | -0.3520757 | 0.72478149 | 0.80184152 | NA | NA |

| | | | | | | | | |
|---|---|---|---|---|---|---|---|---|
| ENSG000002 | 2.38549344 | -0.130402987 | 0.37041405 | -0.3520465 | 0.72480336 | 0.80184152 | NA | NA |
| ENSG000002 | 2.84597893 | -0.132531364 | 0.37652389 | -0.3519866 | 0.72484831 | 0.80186938 | RNU6-151P | 106479633 |
| ENSG000002 | 3.02799486 | -0.132806273 | 0.37734243 | -0.3519516 | 0.72487456 | 0.80187654 | NA | NA |
| ENSG000002 | 3.98463873 | -0.133134578 | 0.37841422 | -0.3518223 | 0.7249715 | 0.80193128 | NA | NA |
| ENSG000002 | 3.05649127 | -0.131823356 | 0.37470413 | -0.3518065 | 0.72498337 | 0.80193128 | NA | NA |
| ENSG000002 | 6.90085154 | 0.126309179 | 0.35900156 | 0.35183462 | 0.72496229 | 0.80193128 | NA | NA |
| ENSG000002 | 8.24449896 | 0.122047466 | 0.34696724 | 0.35175502 | 0.725022 | 0.80195213 | NA | NA |
| ENSG000002 | 2.57847091 | -0.131834591 | 0.37482862 | -0.3517196 | 0.72504852 | 0.80195959 | NA | NA |
| ENSG000002 | 2.34453567 | -0.130555978 | 0.37124277 | -0.3516728 | 0.72508367 | 0.8019766 | NA | NA |
| ENSG000002 | 2.81447309 | -0.132016972 | 0.37544341 | -0.3516295 | 0.72511615 | 0.80199064 | RNU6-36P | 100873757 |
| ENSG000002 | 2.79876352 | -0.132244391 | 0.37623542 | -0.3514937 | 0.72521798 | 0.80208139 | NA | NA |
| ENSG000001 | 70.938723 | -0.06970564 | 0.19834659 | -0.3514335 | 0.72526314 | 0.80210946 | HSPA12A | 259217 |
| ENSG000002 | 3.53205414 | -0.133034671 | 0.37859271 | -0.3513926 | 0.72529385 | 0.80212155 | NA | NA |
| ENSG000002 | 4.28689461 | -0.132443348 | 0.37700781 | -0.3513013 | 0.72536229 | 0.80215349 | NA | NA |
| ENSG000002 | 3.80859173 | -0.132979011 | 0.37853253 | -0.3513014 | 0.72536224 | 0.80215349 | NA | NA |
| ENSG000001 | 41.0440744 | 0.079266116 | 0.22565684 | 0.3512684 | 0.725387 | 0.80215894 | PRELID3A | 10650 |
| ENSG000002 | 3.55441254 | -0.132954452 | 0.37857045 | -0.3512014 | 0.7254373 | 0.80219268 | THRIL | 102659353 |
| ENSG000002 | 12.0684922 | 0.11513345 | 0.32789032 | 0.35113403 | 0.7254878 | 0.80222665 | NA | NA |
| ENSG000002 | 2.60217257 | -0.13080714 | 0.37271916 | -0.3509536 | 0.72562314 | 0.80234368 | RN7SL634P | 106481096 |
| ENSG000002 | 4.06035606 | 0.132799783 | 0.37841143 | 0.3509402 | 0.72563321 | 0.80234368 | LOC101928449 | 101928449 |
| ENSG000002 | 2.38351687 | -0.098248404 | 0.27999199 | -0.3508972 | 0.72566548 | 0.80235749 | NA | NA |
| ENSG000001 | 3.06806638 | -0.132621864 | 0.37803787 | -0.3508163 | 0.72572614 | 0.80240268 | GSX2 | 170825 |
| ENSG000002 | 3.14450178 | 0.131706838 | 0.3755604 | 0.35069415 | 0.72581781 | 0.80248216 | NA | NA |
| ENSG000001 | 4.02578028 | -0.132332442 | 0.37747754 | -0.3505704 | 0.72591068 | 0.80254271 | PMCH | 5367 |
| ENSG000002 | 2.59441568 | -0.129983426 | 0.37077906 | -0.3505684 | 0.72591216 | 0.80254271 | NA | NA |
| ENSG000002 | 4.63626012 | -0.132622539 | 0.37838267 | -0.3504985 | 0.72596465 | 0.80257886 | NA | NA |
| ENSG000002 | 3.56496252 | -0.132668126 | 0.37859339 | -0.3504238 | 0.72602068 | 0.80261892 | NA | NA |
| ENSG000002 | 4.0301908 | -0.132448085 | 0.37801971 | -0.3503735 | 0.72605842 | 0.80263876 | NA | NA |
| ENSG000002 | 2.37954378 | -0.129879946 | 0.3707876 | -0.3502813 | 0.72612763 | 0.80269339 | NA | NA |

| | | | | | | | | |
|---|---|---|---|---|---|---|---|---|
| ENSG000002 | 3.31671304 | -0.132476276 | 0.37827879 | -0.350208 | 0.72618257 | 0.80270566 | NA | NA |
| ENSG000002 | 3.25947715 | -0.132401728 | 0.3780883 | -0.3501873 | 0.72619812 | 0.80270566 | NA | NA |
| ENSG000002 | 36.6532515 | -0.080050948 | 0.22856679 | -0.35023 | 0.7261661 | 0.80270566 | TMCC1-AS1 | 100507032 |
| ENSG000002 | 3.3647891 | -0.132310768 | 0.37786329 | -0.3501551 | 0.72622228 | 0.80271049 | NA | NA |
| ENSG000002 | 2.84910349 | -0.131613891 | 0.37594112 | -0.3500918 | 0.72626983 | 0.80271928 | NA | NA |
| ENSG000002 | 2.34317803 | -0.128941213 | 0.3682831 | -0.3501144 | 0.72625285 | 0.80271928 | NA | NA |
| ENSG000002 | 9.84651385 | -0.119953115 | 0.34273752 | -0.3499854 | 0.72634968 | 0.80274627 | NA | NA |
| ENSG000002 | 3.0895892 | -0.13224112 | 0.37785326 | -0.3499801 | 0.72635363 | 0.80274627 | NA | NA |
| ENSG000002 | 3.87741297 | -0.13215014 | 0.37758772 | -0.3499853 | 0.72634975 | 0.80274627 | LINC01607 | 101927067 |
| ENSG000002 | 3.61322704 | -0.132381788 | 0.37829415 | -0.349944 | 0.72638069 | 0.80275429 | NA | NA |
| ENSG000002 | 2.61867828 | -0.130182384 | 0.37214385 | -0.3498174 | 0.72647576 | 0.8028156 | NA | NA |
| ENSG000002 | 2.364264 | -0.129685219 | 0.37070877 | -0.3498305 | 0.72646594 | 0.8028156 | NA | NA |
| ENSG000002 | 2.33560994 | -0.126622958 | 0.36201549 | -0.3497722 | 0.72650966 | 0.80283119 | NA | NA |
| ENSG000002 | 5.91910394 | -0.129946985 | 0.37160996 | -0.3496865 | 0.72657399 | 0.80283664 | NA | NA |
| ENSG000002 | 3.38891531 | -0.132094961 | 0.37773099 | -0.3497064 | 0.72655902 | 0.80283664 | NA | NA |
| ENSG000002 | 2.81544605 | -0.131444666 | 0.37586216 | -0.3497151 | 0.72655253 | 0.80283664 | NA | NA |
| ENSG000002 | 2.57374739 | -0.130463335 | 0.37311935 | -0.3496558 | 0.72659705 | 0.80284024 | NA | NA |
| ENSG000002 | 4.78298107 | -0.131253875 | 0.37552981 | -0.3495165 | 0.72670157 | 0.80292098 | NA | NA |
| ENSG000002 | 3.11887579 | -0.131922303 | 0.37745397 | -0.3495057 | 0.72670972 | 0.80292098 | NA | NA |
| ENSG000002 | 10.4224109 | 0.118638118 | 0.33947334 | 0.34947698 | 0.72673125 | 0.80292289 | NA | NA |
| ENSG000001 | 51.7888331 | -0.069675515 | 0.19949117 | -0.3492662 | 0.72688951 | 0.80296648 | CCDC173 | 129881 |
| ENSG000001 | 19.1904759 | -0.098588999 | 0.28226823 | -0.3492742 | 0.7268835 | 0.80296648 | SH3RF3 | 344558 |
| ENSG000002 | 2.62243667 | -0.130350219 | 0.37313842 | -0.3493348 | 0.72683801 | 0.80296648 | NA | NA |
| ENSG000002 | 3.32610767 | -0.132173765 | 0.37836885 | -0.3493252 | 0.7268452 | 0.80296648 | LINC01221 | 104266961 |
| ENSG000002 | 3.84857632 | -0.132212169 | 0.37849535 | -0.3493099 | 0.72685668 | 0.80296648 | NA | NA |
| ENSG000002 | 7.41963278 | -0.125403704 | 0.35894772 | -0.3493648 | 0.72681545 | 0.80296648 | NA | NA |
| ENSG000002 | 2.36386702 | -0.129556894 | 0.37097192 | -0.3492364 | 0.72691185 | 0.80296928 | NA | NA |
| ENSG000001 | 46.2253904 | -0.073525962 | 0.21059126 | -0.3491406 | 0.72698375 | 0.80302683 | NA | NA |
| ENSG000002 | 2.37411564 | -0.129051152 | 0.3697047 | -0.3490655 | 0.72704014 | 0.80306725 | NA | NA |

| | | | | | | | | |
|---|---|---|---|---|---|---|---|---|
| ENSG000002 | 2.34047802 | -0.129505137 | 0.37128066 | -0.3488066 | 0.72723452 | 0.8032017 | NA | NA |
| ENSG000002 | 8.82975029 | -0.120180884 | 0.34454206 | -0.3488134 | 0.72722941 | 0.8032017 | NA | NA |
| ENSG000002 | 2.5900374 | -0.129445076 | 0.37105961 | -0.3488525 | 0.72720004 | 0.8032017 | NA | NA |
| ENSG000002 | 64.7561202 | -0.066545404 | 0.19078503 | -0.3487978 | 0.7272411 | 0.8032017 | NA | NA |
| ENSG000001 | 41.2592217 | -0.076665425 | 0.21982699 | -0.3487535 | 0.7272744 | 0.80320545 | LPPR1 | 54886 |
| ENSG000002 | 2.84071441 | -0.131322895 | 0.37656332 | -0.3487405 | 0.7272841 | 0.80320545 | NA | NA |
| ENSG000001 | 39.2121952 | 0.077995689 | 0.22378618 | 0.34852773 | 0.72744389 | 0.80325518 | PRR14 | 78994 |
| ENSG000001 | 25.8110079 | -0.088774731 | 0.25474152 | -0.3484894 | 0.72747264 | 0.80325518 | PHLDB3 | 653583 |
| ENSG000001 | 14.5151119 | 0.109620969 | 0.3145687 | 0.34848022 | 0.72747957 | 0.80325518 | SOWAHB | 345079 |
| ENSG000002 | 3.09088702 | -0.131446146 | 0.37704056 | -0.348626 | 0.7273701 | 0.80325518 | NA | NA |
| ENSG000002 | 7.76583657 | 0.123992476 | 0.35578844 | 0.34850058 | 0.72746428 | 0.80325518 | SSR4P1 | 728039 |
| ENSG000002 | 2.56528564 | -0.130351811 | 0.37406947 | -0.3484695 | 0.7274876 | 0.80325518 | NA | NA |
| ENSG000002 | 4.24406902 | -0.131944173 | 0.37858853 | -0.348516 | 0.72745268 | 0.80325518 | LOC101928134 | 101928134 |
| ENSG000002 | 2.55459264 | -0.130553916 | 0.37453284 | -0.348578 | 0.72740613 | 0.80325518 | NA | NA |
| ENSG000001 | 4.80725795 | -0.131284543 | 0.3767824 | -0.348436 | 0.72751279 | 0.80326112 | ACR | 49 |
| ENSG000002 | 3.07919657 | -0.131610587 | 0.37776496 | -0.3483928 | 0.72754522 | 0.80327505 | NA | NA |
| ENSG000001 | 33.4461817 | -0.094514414 | 0.2713143 | -0.3483577 | 0.7275716 | 0.80328231 | CASKIN1 | 57524 |
| ENSG000002 | 3.53567593 | -0.131820283 | 0.37858533 | -0.3481917 | 0.72769619 | 0.80337612 | NA | NA |
| ENSG000002 | 13.5462872 | 0.10902281 | 0.31308949 | 0.34821612 | 0.72767788 | 0.80337612 | NA | NA |
| ENSG000001 | 18.6816114 | 0.101909081 | 0.29275083 | 0.3481086 | 0.72775863 | 0.8034013 | NA | NA |
| ENSG000002 | 3.78558795 | -0.131715916 | 0.37837563 | -0.3481089 | 0.72775842 | 0.8034013 | NA | NA |
| ENSG000001 | 26.2711468 | -0.089664639 | 0.25765342 | -0.3480048 | 0.72783654 | 0.8034217 | CD53 | 963 |
| ENSG000002 | 2.58746135 | -0.13045377 | 0.37484024 | -0.348025 | 0.72782142 | 0.8034217 | NA | NA |
| ENSG000002 | 4.48702079 | 0.130995996 | 0.37641003 | 0.34801409 | 0.7278296 | 0.8034217 | NA | NA |
| ENSG000001 | 40.9390653 | 0.079044173 | 0.22722031 | 0.34787459 | 0.72793437 | 0.80343909 | NA | NA |
| ENSG000001 | 5.32427833 | 0.1301787 | 0.37419119 | 0.34789355 | 0.72792013 | 0.80343909 | NA | NA |
| ENSG000002 | 6.27435939 | 0.129208563 | 0.37135145 | 0.34794146 | 0.72788415 | 0.80343909 | NA | NA |
| ENSG000002 | 21.7373103 | -0.09506442 | 0.27333989 | -0.3477883 | 0.72799917 | 0.80343909 | LINC01534 | 101927621 |
| ENSG000002 | 3.78327218 | -0.131676032 | 0.37858557 | -0.3478105 | 0.72798251 | 0.80343909 | LOC100505887 | 100505887 |

| | | | | | | | | |
|---|---|---|---|---|---|---|---|---|
| ENSG000002 | 6.51734323 | 0.12656798 | 0.36391759 | 0.34779297 | 0.72799567 | 0.80343909 | LINC01336 | 104326191 |
| ENSG000002 | 7.12492999 | -0.126813726 | 0.3646453 | -0.3477728 | 0.7280108 | 0.80343909 | NA | NA |
| ENSG000002 | 5.35400863 | 0.128694923 | 0.37002349 | 0.34780203 | 0.72798886 | 0.80343909 | NA | NA |
| ENSG000002 | 9.45016071 | 0.118207404 | 0.34000425 | 0.34766449 | 0.72809217 | 0.80350702 | NA | NA |
| ENSG000002 | 8.95135748 | -0.122414105 | 0.35217662 | -0.3475929 | 0.72814591 | 0.80354447 | GNAT3 | 346562 |
| ENSG000002 | 2.33526558 | -0.129219807 | 0.37181792 | -0.3475352 | 0.72818929 | 0.80357047 | NA | NA |
| ENSG000001 | 2.85147102 | -0.129926398 | 0.37388261 | -0.3475059 | 0.72821131 | 0.8035729 | CTRB2 | 440387 |
| ENSG000002 | 6.23567392 | -0.128396104 | 0.36953991 | -0.3474485 | 0.72825437 | 0.80359854 | NA | NA |
| ENSG000002 | 3.07082621 | -0.130997187 | 0.37718631 | -0.347301 | 0.72836519 | 0.80369896 | MIR1302-3 | 100302128 |
| ENSG000002 | 4.80494751 | -0.130729104 | 0.37653606 | -0.3471888 | 0.72844948 | 0.8037701 | NA | NA |
| ENSG000002 | 2.60020943 | -0.130114625 | 0.37485398 | -0.3471075 | 0.72851057 | 0.80381563 | NA | NA |
| ENSG000001 | 63.1812396 | 0.067855043 | 0.19555179 | 0.34699269 | 0.72859681 | 0.8038233 | SLC25A27 | 9481 |
| ENSG000002 | 2.57543033 | -0.129737659 | 0.37383327 | -0.3470468 | 0.72855612 | 0.8038233 | NA | NA |
| ENSG000002 | 2.30313353 | -0.127191075 | 0.36649962 | -0.3470429 | 0.72855912 | 0.8038233 | NA | NA |
| ENSG000002 | 3.5651903 | -0.131342099 | 0.37850227 | -0.3470048 | 0.72858773 | 0.8038233 | WBP1LP5 | 106480780 |
| ENSG000002 | 2.33222479 | -0.1284123 | 0.37013736 | -0.3469315 | 0.7286428 | 0.80385217 | NA | NA |
| ENSG000001 | 2.33204873 | -0.128938491 | 0.37180822 | -0.3467876 | 0.72875087 | 0.80393157 | NA | NA |
| ENSG000001 | 22.8007144 | -0.09489714 | 0.27365667 | -0.3467744 | 0.72876077 | 0.80393157 | HTR1A | 3350 |
| ENSG000002 | 3.11462889 | -0.130860382 | 0.37738406 | -0.3467565 | 0.72877425 | 0.80393157 | NA | NA |
| ENSG000002 | 2.61250049 | -0.129535216 | 0.37366309 | -0.3466631 | 0.72884441 | 0.80396523 | NA | NA |
| ENSG000002 | 2.63823741 | -0.128443885 | 0.37049374 | -0.346683 | 0.72882948 | 0.80396523 | NA | NA |
| ENSG000002 | 4.10993413 | -0.131086556 | 0.37825446 | -0.3465565 | 0.7289245 | 0.8040317 | NA | NA |
| ENSG000001 | 2.85230107 | -0.130446363 | 0.3766207 | -0.34636 | 0.72907215 | 0.80413147 | TNP1 | 7141 |
| ENSG000001 | 6.21224539 | -0.127577463 | 0.36832873 | -0.3463685 | 0.72906581 | 0.80413147 | NRN1 | 51299 |
| ENSG000002 | 2.57232538 | -0.129715099 | 0.37455638 | -0.3463166 | 0.72910478 | 0.80413147 | NA | NA |
| ENSG000002 | 2.82005801 | -0.13005902 | 0.37556291 | -0.3463042 | 0.7291141 | 0.80413147 | NA | NA |
| ENSG000002 | 4.55960222 | -0.130550872 | 0.37694234 | -0.3463418 | 0.72908589 | 0.80413147 | NA | NA |
| ENSG000002 | 7.59014148 | 0.123452379 | 0.3565911 | 0.34620152 | 0.72919127 | 0.80419471 | RNU6-762P | 106479903 |
| ENSG000002 | 3.05546211 | -0.130689094 | 0.37766047 | -0.3460492 | 0.72930576 | 0.80423348 | NA | NA |

| | | | | | | | | |
|---|---|---|---|---|---|---|---|---|
| ENSG000002 | 3.3614158 | -0.130870808 | 0.37818507 | -0.3460496 | 0.72930541 | 0.80423348 | NA | NA |
| ENSG000002 | 2.32939969 | -0.128061596 | 0.36998879 | -0.3461229 | 0.72925035 | 0.80423348 | NA | NA |
| ENSG000002 | 3.18114022 | 0.13085166 | 0.37812956 | 0.34604981 | 0.72930529 | 0.80423348 | LINC01243 | 101929620 |
| ENSG000002 | 3.31585905 | -0.130924756 | 0.37841539 | -0.3459816 | 0.72935655 | 0.80426762 | NA | NA |
| ENSG000002 | 8.55353244 | -0.119480706 | 0.34544277 | -0.345877 | 0.72943516 | 0.80433244 | NA | NA |
| ENSG000002 | 3.07019314 | -0.130381983 | 0.37701164 | -0.3458301 | 0.72947038 | 0.80434567 | NA | NA |
| ENSG000002 | 7.97196397 | -0.123311418 | 0.35658901 | -0.3458082 | 0.72948683 | 0.80434567 | NA | NA |
| ENSG000002 | 2.36950347 | -0.126877099 | 0.36703543 | -0.3456808 | 0.72958262 | 0.80442941 | NA | NA |
| ENSG000002 | 2.68835948 | 0.127054069 | 0.36764919 | 0.34558507 | 0.72965457 | 0.80448687 | NA | NA |
| ENSG000002 | 2.83355072 | -0.129378085 | 0.37443089 | -0.3455326 | 0.729694 | 0.80449039 | NA | NA |
| ENSG000002 | 2.60988447 | -0.128574462 | 0.37213851 | -0.3455016 | 0.72971728 | 0.80449039 | NA | NA |
| ENSG000002 | 17.6688302 | -0.104412385 | 0.30220134 | -0.345506 | 0.72971398 | 0.80449039 | NA | NA |
| ENSG000002 | 2.82921208 | -0.130124324 | 0.37670227 | -0.3454302 | 0.72977101 | 0.80452775 | DEFB110 | 245913 |
| ENSG000002 | 2.56876307 | -0.129301268 | 0.37435538 | -0.3453971 | 0.72979585 | 0.80453326 | NA | NA |
| ENSG000002 | 2.53717291 | -0.128188312 | 0.37116664 | -0.3453659 | 0.72981929 | 0.80453723 | NA | NA |
| ENSG000002 | 8.1137805 | -0.122640641 | 0.35528303 | -0.3451914 | 0.7299505 | 0.80466 | NA | NA |
| ENSG000001 | 233.425565 | 0.046551869 | 0.13495639 | 0.34494008 | 0.73013942 | 0.80479358 | TET2 | 54790 |
| ENSG000002 | 8.85161858 | -0.120685936 | 0.34989078 | -0.3449246 | 0.73015106 | 0.80479358 | TUSC8 | 400128 |
| ENSG000002 | 2.57539787 | -0.128951688 | 0.37382944 | -0.3449479 | 0.73013352 | 0.80479358 | NA | NA |
| ENSG000002 | 3.55725028 | -0.130577709 | 0.37855208 | -0.3449399 | 0.73013957 | 0.80479358 | NA | NA |
| ENSG000002 | 17.4018654 | 0.102130124 | 0.29618388 | 0.34481999 | 0.7302297 | 0.80485838 | NA | NA |
| ENSG000002 | 5.53140105 | 0.128062322 | 0.37157412 | 0.34464812 | 0.73035892 | 0.80497893 | NA | NA |
| ENSG000002 | 2.5650696 | -0.12835542 | 0.37251857 | -0.3445611 | 0.73042433 | 0.80502914 | NA | NA |
| ENSG000002 | 2.60880703 | -0.126453309 | 0.36714032 | -0.3444277 | 0.73052463 | 0.8051178 | NA | NA |
| ENSG000002 | 3.06177594 | -0.129444075 | 0.37598291 | -0.3442818 | 0.73063437 | 0.80520435 | RNU6-817P | 106479927 |
| ENSG000002 | 7.13619948 | -0.122899193 | 0.3569844 | -0.3442705 | 0.73064288 | 0.80520435 | NA | NA |
| ENSG000001 | 48.5680402 | -0.070788489 | 0.20567154 | -0.3441822 | 0.73070924 | 0.80523487 | KCNE3 | 10008 |
| ENSG000001 | 49.6511911 | 0.076638259 | 0.22269495 | 0.34414009 | 0.73074093 | 0.80523487 | LUZP2 | 338645 |
| ENSG000002 | 2.31047383 | -0.127935932 | 0.37175049 | -0.3441446 | 0.73073753 | 0.80523487 | RNU6-10P | 100873745 |

| | | | | | | | | |
|---|---|---|---|---|---|---|---|---|
| ENSG000002 | 2.29965665 | -0.127154649 | 0.36949808 | -0.344128 | 0.73075 | 0.80523487 | NA | NA |
| ENSG000001 | 25.2225553 | 0.093064029 | 0.27048141 | 0.34406812 | 0.73079506 | 0.80526264 | MORN3 | 283385 |
| ENSG000002 | 3.57414884 | -0.130187911 | 0.37857537 | -0.343889 | 0.73092976 | 0.80538866 | RNU6-936P | 106481924 |
| ENSG000002 | 3.14027997 | 0.129744436 | 0.37731408 | 0.34386323 | 0.73094915 | 0.80538866 | NA | NA |
| ENSG000002 | 2.32044226 | -0.126740641 | 0.36866499 | -0.3437827 | 0.73100972 | 0.80543086 | NA | NA |
| ENSG000002 | 2.59919147 | -0.128010835 | 0.37238488 | -0.3437595 | 0.73102717 | 0.80543086 | NA | NA |
| ENSG000002 | 2.33953229 | -0.127906476 | 0.3721615 | -0.3436854 | 0.73108289 | 0.80547036 | RN7SKP50 | 106480459 |
| ENSG000001 | 182.724056 | 0.053688759 | 0.15636934 | 0.34334581 | 0.73133832 | 0.80571169 | HYLS1 | 219844 |
| ENSG000002 | 4.23506489 | 0.129639228 | 0.3776102 | 0.34331495 | 0.73136153 | 0.80571169 | NA | NA |
| ENSG000002 | 2.33317715 | -0.125772101 | 0.36634156 | -0.3433192 | 0.73135832 | 0.80571169 | NA | NA |
| ENSG000002 | 2.58834237 | -0.128419246 | 0.37414559 | -0.3432334 | 0.73142291 | 0.80575742 | NA | NA |
| ENSG000002 | 21.5995428 | 0.093144778 | 0.27141564 | 0.34318132 | 0.73146206 | 0.80577865 | NA | NA |
| ENSG000002 | 2.94762202 | 0.128980357 | 0.37591931 | 0.3431065 | 0.73151834 | 0.80580476 | NA | NA |
| ENSG000002 | 3.0732273 | -0.129503539 | 0.37745461 | -0.343097 | 0.7315255 | 0.80580476 | NA | NA |
| ENSG000001 | 2.56637572 | -0.128461574 | 0.37454826 | -0.3429774 | 0.73161549 | 0.80588199 | C11orf42 | 160298 |
| ENSG000002 | 2.79548197 | -0.128356588 | 0.37430642 | -0.3429185 | 0.73165979 | 0.8059089 | NA | NA |
| ENSG000002 | 5.56753095 | 0.126326985 | 0.36858548 | 0.34273457 | 0.73179815 | 0.80603941 | NA | NA |
| ENSG000002 | 3.16844422 | 0.129485713 | 0.37790382 | 0.34264198 | 0.73186781 | 0.80609424 | NA | NA |
| ENSG000002 | 63.7751441 | 0.066660373 | 0.1945792 | 0.34258735 | 0.73190892 | 0.80609573 | NA | NA |
| ENSG000002 | 3.28042638 | -0.129454448 | 0.37785112 | -0.342607 | 0.73189412 | 0.80609573 | NA | NA |
| ENSG000001 | 52.9181796 | -0.072708858 | 0.21226966 | -0.3425306 | 0.73195161 | 0.80612085 | SMIM10 | 644538 |
| ENSG000002 | 75.3530426 | 0.067312361 | 0.19657608 | 0.34242397 | 0.73203186 | 0.80618733 | NA | NA |
| ENSG000002 | 3.186581 | 0.129500248 | 0.37824126 | 0.34237473 | 0.7320689 | 0.80620624 | NA | NA |
| ENSG000001 | 12.6819676 | -0.11077085 | 0.32361802 | -0.3422889 | 0.7321335 | 0.80625548 | OPRL1 | 4987 |
| ENSG000002 | 2.33134377 | -0.126289955 | 0.36900221 | -0.3422471 | 0.73216491 | 0.80626817 | NA | NA |
| ENSG000002 | 3.15262276 | 0.129211268 | 0.37764811 | 0.34214726 | 0.73224007 | 0.80632905 | NA | NA |
| ENSG000002 | 2.35020512 | -0.126619744 | 0.37018126 | -0.342048 | 0.7323148 | 0.80638944 | NA | NA |
| ENSG000001 | 12.8598836 | -0.109225852 | 0.31957242 | -0.3417875 | 0.73251084 | 0.8065834 | SMPD3 | 55512 |
| ENSG000002 | 2.63210908 | 0.125341344 | 0.36676156 | 0.34175158 | 0.73253785 | 0.80659124 | NA | NA |

| | | | | | | | | |
|---|---|---|---|---|---|---|---|---|
| ENSG000002 | 4.24215905 | -0.129276287 | 0.37831827 | -0.341713 | 0.73256686 | 0.80660128 | NA | NA |
| ENSG000002 | 2.58977681 | -0.126992385 | 0.37171397 | -0.3416401 | 0.73262178 | 0.80661796 | NA | NA |
| ENSG000002 | 36.2998368 | 0.080651538 | 0.23606612 | 0.34164808 | 0.73261575 | 0.80661796 | NA | NA |
| ENSG000002 | 9.07970197 | -0.11853784 | 0.34701933 | -0.3415886 | 0.73266053 | 0.80663872 | NA | NA |
| ENSG000002 | 2.60912799 | -0.127464513 | 0.37334563 | -0.3414116 | 0.73279378 | 0.80674704 | NA | NA |
| ENSG000002 | 9.28936803 | 0.118771207 | 0.34797009 | 0.34132591 | 0.73285825 | 0.80674704 | CYP4F26P | 106478958 |
| ENSG000002 | 3.28927336 | -0.129177219 | 0.37847635 | -0.3413086 | 0.7328713 | 0.80674704 | NA | NA |
| ENSG000002 | 5.60401293 | 0.125593618 | 0.36794584 | 0.34133724 | 0.73284971 | 0.80674704 | NA | NA |
| ENSG000002 | 2.60382326 | -0.12640517 | 0.37034813 | -0.3413145 | 0.73286686 | 0.80674704 | NA | NA |
| ENSG000002 | 3.09814225 | -0.128662718 | 0.37697916 | -0.3412993 | 0.73287829 | 0.80674704 | LOC641367 | 641367 |
| ENSG000002 | 3.07091108 | -0.128838793 | 0.37753068 | -0.3412671 | 0.73290253 | 0.80675182 | BOLA2P3 | 106480323 |
| ENSG000002 | 6.66504081 | -0.12326934 | 0.361249 | -0.341231 | 0.73292972 | 0.80675986 | NA | NA |
| ENSG000002 | 3.78476621 | -0.129130325 | 0.37852664 | -0.3411393 | 0.73299869 | 0.80681388 | NA | NA |
| ENSG000002 | 2.84615553 | -0.128378447 | 0.3764213 | -0.3410499 | 0.73306602 | 0.80684418 | NA | NA |
| ENSG000002 | 2.58616299 | -0.126907383 | 0.37210402 | -0.3410535 | 0.7330633 | 0.80684418 | NA | NA |
| ENSG000001 | 14475.8857 | 0.044526127 | 0.13063632 | 0.34084033 | 0.73322379 | 0.80697723 | ATP6 | 4508 |
| ENSG000002 | 2.34263703 | -0.125843801 | 0.36922049 | -0.3408365 | 0.73322671 | 0.80697723 | NA | NA |
| ENSG000002 | 3.35966152 | -0.128752688 | 0.37809203 | -0.3405327 | 0.73345542 | 0.80720705 | NA | NA |
| ENSG000001 | 251.896994 | 0.049250386 | 0.14471014 | 0.34033818 | 0.73360187 | 0.8073463 | NA | NA |
| ENSG000001 | 2.34246097 | -0.126193092 | 0.37090917 | -0.3402264 | 0.73368604 | 0.80737429 | OR8I2 | 120586 |
| ENSG000002 | 2.33108318 | -0.126187744 | 0.37088076 | -0.340238 | 0.73367727 | 0.80737429 | NA | NA |
| ENSG000002 | 9.04750513 | -0.119308135 | 0.35067412 | -0.3402251 | 0.73368703 | 0.80737429 | NA | NA |
| ENSG000001 | 109.167219 | 0.060247639 | 0.17711211 | 0.34016669 | 0.733731 | 0.80739596 | PARP9 | 83666 |
| ENSG000002 | 10.6221146 | 0.115266917 | 0.33887477 | 0.34014606 | 0.73374654 | 0.80739596 | NA | NA |
| ENSG000001 | 3.49929699 | -0.128503733 | 0.37790163 | -0.3400455 | 0.73382229 | 0.8074355 | NA | NA |
| ENSG000002 | 2.61014484 | -0.126175887 | 0.37104704 | -0.3400536 | 0.73381615 | 0.8074355 | NA | NA |
| ENSG000002 | 42.8367593 | -0.074227187 | 0.21835228 | -0.3399423 | 0.73389995 | 0.80749903 | NA | NA |
| ENSG000002 | 2.34300973 | -0.125430896 | 0.36903718 | -0.3398869 | 0.73394172 | 0.80752308 | NA | NA |
| ENSG000002 | 2.54795785 | -0.126924312 | 0.37357773 | -0.3397534 | 0.73404223 | 0.80760141 | LTC4S | 4056 |

| | | | | | | | | |
|---|---|---|---|---|---|---|---|---|
| ENSG000002 | 2.83033442 | -0.127811708 | 0.37620508 | -0.3397395 | 0.73405274 | 0.80760141 | NA | NA |
| ENSG000002 | 3.53871617 | -0.128600375 | 0.37858545 | -0.3396865 | 0.73409261 | 0.80762336 | POU5F1P2 | 100129512 |
| ENSG000001 | 11.5002889 | -0.111802534 | 0.32916501 | -0.339655 | 0.73411637 | 0.80762759 | FZD9 | 8326 |
| ENSG000002 | 12.0297202 | 0.114065652 | 0.33589851 | 0.33958368 | 0.73417007 | 0.80766476 | LVCAT5 | 105375475 |
| ENSG000001 | 2.91791067 | 0.127720683 | 0.37627502 | 0.33943439 | 0.73428252 | 0.80776654 | PAEP | 5047 |
| ENSG000002 | 2.31691275 | -0.125639004 | 0.37023092 | -0.3393531 | 0.73434376 | 0.807812 | NA | NA |
| ENSG000002 | 2.3405497 | -0.125437202 | 0.369733 | -0.3392643 | 0.73441064 | 0.80786366 | NA | NA |
| ENSG000002 | 2.84115161 | -0.127268082 | 0.37545516 | -0.3389701 | 0.73463223 | 0.80808549 | RN7SL152P | 106480962 |
| ENSG000001 | 2.8349334 | -0.127390179 | 0.3758637 | -0.3389265 | 0.73466508 | 0.8080997 | HIST1H2AA | 221613 |
| ENSG000001 | 74.0018784 | -0.061406868 | 0.18123344 | -0.3388275 | 0.7347397 | 0.80813809 | AGPAT3 | 56894 |
| ENSG000002 | 54.1913205 | -0.075568749 | 0.22303028 | -0.3388273 | 0.73473983 | 0.80813809 | NA | NA |
| ENSG000001 | 33.0800768 | -0.082674404 | 0.24405176 | -0.3387577 | 0.7347923 | 0.80815858 | DACT3 | 147906 |
| ENSG000002 | 2.65070563 | -0.125387827 | 0.37014894 | -0.3387497 | 0.73479833 | 0.80815858 | NA | NA |
| ENSG000001 | 2.96463715 | 0.127019642 | 0.37510177 | 0.33862715 | 0.73489063 | 0.80819434 | FGFBP2 | 83888 |
| ENSG000001 | 21.5538418 | -0.098744862 | 0.2915939 | -0.3386383 | 0.73488223 | 0.80819434 | FBXO43 | 286151 |
| ENSG000001 | 128.510209 | 0.052769727 | 0.15582216 | 0.33865355 | 0.73487073 | 0.80819434 | ZNF79 | 7633 |
| ENSG000002 | 2.31417198 | -0.124555631 | 0.36816667 | -0.3383132 | 0.7351272 | 0.80843259 | RNU1-109P | 106481625 |
| ENSG000001 | 56.413828 | 0.068443606 | 0.20236321 | 0.33822159 | 0.73519621 | 0.80846597 | RCN3 | 57333 |
| ENSG000002 | 23.8653616 | -0.090783169 | 0.26841458 | -0.33822 | 0.73519743 | 0.80846597 | NA | NA |
| ENSG000002 | 52.1022079 | 0.068323918 | 0.2020559 | 0.33814364 | 0.73525494 | 0.80848564 | SBF2-AS1 | 283104 |
| ENSG000002 | 2.59469697 | -0.126501448 | 0.37410602 | -0.3381433 | 0.73525519 | 0.80848564 | NA | NA |
| ENSG000002 | 3.26298738 | -0.127512948 | 0.3772002 | -0.3380511 | 0.73532467 | 0.80849626 | NA | NA |
| ENSG000002 | 3.10931231 | -0.127422697 | 0.37690617 | -0.3380754 | 0.73530638 | 0.80849626 | NA | NA |
| ENSG000002 | 102.412614 | 0.059402222 | 0.17569237 | 0.33810359 | 0.73528513 | 0.80849626 | NA | NA |
| ENSG000002 | 5.21074116 | 0.127527266 | 0.37735617 | 0.33794933 | 0.73540137 | 0.80855867 | NA | NA |
| ENSG000002 | 3.82642893 | -0.127911139 | 0.37853306 | -0.3379127 | 0.73542895 | 0.80856708 | NA | NA |
| ENSG000002 | 8.81537178 | -0.117556022 | 0.34809946 | -0.3377081 | 0.73558314 | 0.80867245 | NA | NA |
| ENSG000002 | 21.6726625 | 0.100078012 | 0.29634641 | 0.33770618 | 0.73558462 | 0.80867245 | NA | NA |
| ENSG000002 | 14.9721139 | -0.103669447 | 0.30697182 | -0.3377165 | 0.73557684 | 0.80867245 | NA | NA |

| | | | | | | | | |
|---|---|---|---|---|---|---|---|---|
| ENSG000002 | 3.82130823 | -0.12781668 | 0.3785718 | -0.3376286 | 0.73564306 | 0.80871477 | GACAT1 | 104326057 |
| ENSG000002 | 8.47202774 | 0.116774747 | 0.34594829 | 0.33754971 | 0.73570254 | 0.80873631 | NA | NA |
| ENSG000002 | 4.39449585 | -0.127640222 | 0.37813427 | -0.3375526 | 0.73570033 | 0.80873631 | NA | NA |
| ENSG000002 | 4.0863815 | -0.127760257 | 0.37858739 | -0.3374657 | 0.73576587 | 0.808784 | CCDC144A | 9720 |
| ENSG000002 | 2.59260908 | -0.126235545 | 0.37410004 | -0.3374379 | 0.73578679 | 0.80878507 | NA | NA |
| ENSG000001 | 23.3580692 | -0.089492519 | 0.26527227 | -0.337361 | 0.73584478 | 0.80882689 | TACSTD2 | 4070 |
| ENSG000002 | 4.30071979 | -0.127366593 | 0.37772484 | -0.3371941 | 0.73597057 | 0.80892131 | NA | NA |
| ENSG000002 | 24.0000284 | 0.0909487 | 0.26971651 | 0.33720108 | 0.73596532 | 0.80892131 | NA | NA |
| ENSG000002 | 3.16162564 | 0.127399526 | 0.37804429 | 0.3369963 | 0.73611969 | 0.80903511 | NA | NA |
| ENSG000002 | 3.30381593 | -0.127485336 | 0.37832018 | -0.3369774 | 0.73613397 | 0.80903511 | LOC100130548 | 100130548 |
| ENSG000002 | 2.80266606 | -0.126905714 | 0.3765417 | -0.3370296 | 0.73609456 | 0.80903511 | NA | NA |
| ENSG000002 | 8.5389256 | -0.11825604 | 0.35099745 | -0.3369142 | 0.73618155 | 0.80904962 | NA | NA |
| ENSG000002 | 4.54273199 | -0.126873924 | 0.37658451 | -0.3369069 | 0.73618707 | 0.80904962 | MT1L | 4500 |
| ENSG000002 | 11.6024509 | -0.108410292 | 0.32196878 | -0.3367106 | 0.73633509 | 0.80917931 | NA | NA |
| ENSG000002 | 7.462085 | -0.121784897 | 0.36170426 | -0.3366974 | 0.73634499 | 0.80917931 | NA | NA |
| ENSG000002 | 2.54983721 | -0.126052208 | 0.37452467 | -0.3365658 | 0.73644422 | 0.80926642 | NA | NA |
| ENSG000002 | 2.29684474 | -0.12446319 | 0.36989879 | -0.336479 | 0.73650965 | 0.80931639 | NA | NA |
| ENSG000002 | 11.6426392 | -0.109494523 | 0.32546785 | -0.3364219 | 0.73655271 | 0.80934178 | NA | NA |
| ENSG000002 | 4.29900672 | 0.126691974 | 0.37684073 | 0.33619501 | 0.73672381 | 0.80950785 | LOC105369332 | 105369332 |
| ENSG000002 | 2.31496923 | -0.124367132 | 0.37022593 | -0.3359223 | 0.73692949 | 0.80971191 | NA | NA |
| ENSG000001 | 3.0687 | -0.126668721 | 0.37717884 | -0.335832 | 0.73699756 | 0.80975921 | OR13C4 | 138804 |
| ENSG000002 | 3.992725 | 0.127129367 | 0.3785926 | 0.33579464 | 0.73702573 | 0.80975921 | NA | NA |
| ENSG000002 | 3.0448408 | -0.126106373 | 0.37555608 | -0.3357857 | 0.73703245 | 0.80975921 | NA | NA |
| ENSG000002 | 12.2904421 | 0.109863485 | 0.32724463 | 0.3357228 | 0.73707991 | 0.80978942 | NA | NA |
| ENSG000001 | 54.2194842 | 0.066567483 | 0.19830904 | 0.33567548 | 0.7371156 | 0.80980669 | NA | NA |
| ENSG000001 | 4.49915047 | -0.126838575 | 0.37805691 | -0.3355013 | 0.73724699 | 0.80988521 | CABS1 | 85438 |
| ENSG000002 | 2.37341752 | -0.122470151 | 0.36501749 | -0.3355186 | 0.73723393 | 0.80988521 | NA | NA |
| ENSG000002 | 2.55092675 | -0.125060379 | 0.37274012 | -0.3355163 | 0.73723566 | 0.80988521 | NA | NA |
| ENSG000002 | 2.86683567 | -0.125307617 | 0.373634 | -0.3353753 | 0.73734199 | 0.80996763 | NA | NA |

| | | | | | | | | |
|---|---|---|---|---|---|---|---|---|
| ENSG000002 | 3.28239062 | -0.12688336 | 0.37838316 | -0.3353304 | 0.7373759 | 0.80998293 | NA | NA |
| ENSG000002 | 5.63014304 | 0.124702636 | 0.37195058 | 0.33526668 | 0.73742392 | 0.81001375 | NA | NA |
| ENSG000002 | 3.08070976 | -0.125526188 | 0.37458775 | -0.3351049 | 0.73754596 | 0.81012585 | NA | NA |
| ENSG000002 | 2.56042607 | -0.125192629 | 0.37381049 | -0.3349094 | 0.73769347 | 0.81026593 | NA | NA |
| ENSG000002 | 16.3320422 | 0.099141432 | 0.29611014 | 0.33481269 | 0.73776639 | 0.81032407 | NA | NA |
| ENSG000002 | 2.81546621 | -0.125983399 | 0.37640406 | -0.3347026 | 0.73784948 | 0.81039338 | NA | NA |
| ENSG000001 | 29.0987709 | -0.083035128 | 0.24812261 | -0.3346536 | 0.7378864 | 0.8104119 | ADAMTS13 | 11093 |
| ENSG000002 | 4.9370357 | -0.126096588 | 0.37682704 | -0.3346272 | 0.7379063 | 0.8104119 | NA | NA |
| ENSG000002 | 3.33468077 | -0.12550647 | 0.37526422 | -0.3344483 | 0.73804132 | 0.81047237 | NA | NA |
| ENSG000002 | 3.16443754 | 0.126532007 | 0.37831651 | 0.33446071 | 0.73803194 | 0.81047237 | NA | NA |
| ENSG000002 | 7.16488191 | 0.119629258 | 0.35765062 | 0.33448637 | 0.73801257 | 0.81047237 | GOLGA8H | 728498 |
| ENSG000002 | 2.80510603 | -0.125845831 | 0.37619301 | -0.3345246 | 0.7379837 | 0.81047237 | LOC101928982 | 101928982 |
| ENSG000001 | 7.10117315 | -0.118948101 | 0.35576893 | -0.3343409 | 0.73812234 | 0.8105394 | NMBR | 4829 |
| ENSG000002 | 2.32692061 | -0.123063171 | 0.36820842 | -0.3342215 | 0.73821242 | 0.81061636 | NA | NA |
| ENSG000002 | 22.6892117 | 0.088790662 | 0.26580103 | 0.33404935 | 0.73834232 | 0.81073705 | NA | NA |
| ENSG000002 | 3.10042359 | 0.125806658 | 0.37664801 | 0.33401652 | 0.7383671 | 0.81074231 | NA | NA |
| ENSG000002 | 29.3564125 | 0.081875708 | 0.24534296 | 0.33371941 | 0.7385913 | 0.81096653 | FAM212B-AS1 | 100506343 |
| ENSG000002 | 8.56203997 | -0.116308441 | 0.34857797 | -0.3336655 | 0.73863199 | 0.81098924 | LINC00605 | 100131366 |
| ENSG000002 | 2.87873554 | -0.124237218 | 0.37253369 | -0.3334926 | 0.7387625 | 0.81111058 | NA | NA |
| ENSG000001 | 7.86456636 | -0.117914248 | 0.35363642 | -0.3334336 | 0.73880704 | 0.81111556 | GPR82 | 27197 |
| ENSG000002 | 3.48558391 | -0.126182257 | 0.3784164 | -0.3334482 | 0.738796 | 0.81111556 | NA | NA |
| ENSG000002 | 8.48516791 | 0.115709912 | 0.347118 | 0.33334461 | 0.73887417 | 0.81114668 | NA | NA |
| ENSG000002 | 7.12559853 | -0.119168759 | 0.35749592 | -0.333343 | 0.73887539 | 0.81114668 | NA | NA |
| ENSG000002 | 2.37316991 | -0.122744786 | 0.36833329 | -0.3332438 | 0.73895026 | 0.81120691 | NA | NA |
| ENSG000002 | 2.31918387 | -0.123549418 | 0.37084727 | -0.3331545 | 0.7390177 | 0.81125898 | NA | NA |
| ENSG000002 | 2.58288198 | -0.124259303 | 0.37312335 | -0.3330247 | 0.73911561 | 0.81131695 | NA | NA |
| ENSG000002 | 6.65257348 | -0.120540528 | 0.36197815 | -0.333005 | 0.73913052 | 0.81131695 | NA | NA |
| ENSG000002 | 8.07835968 | -0.118342187 | 0.35537617 | -0.3330054 | 0.7391302 | 0.81131695 | NA | NA |
| ENSG000002 | 4.07466543 | -0.125989737 | 0.37841955 | -0.3329367 | 0.7391821 | 0.81132964 | OOEP | 441161 |

| | | | | | | | | |
|---|---|---|---|---|---|---|---|---|
| ENSG000002 | 2.32069075 | -0.123388849 | 0.3705927 | -0.33295 | 0.73917201 | 0.81132964 | NA | NA |
| ENSG000002 | 2.27854389 | 0.119559471 | 0.3594771 | 0.33259273 | 0.73944173 | 0.81158591 | NA | NA |
| ENSG000002 | 11.2869153 | 0.110415789 | 0.33200333 | 0.33257434 | 0.73945561 | 0.81158591 | LOC101927811 | 101927811 |
| ENSG000002 | 16.2373777 | 0.103573218 | 0.31147721 | 0.33252262 | 0.73949466 | 0.8116068 | NA | NA |
| ENSG000002 | 7.82757495 | -0.116713254 | 0.35106018 | -0.3324594 | 0.7395424 | 0.81161526 | NA | NA |
| ENSG000002 | 2.28147203 | 0.122573832 | 0.36867232 | 0.33247366 | 0.73953163 | 0.81161526 | EPB41L4A-AS2 | 54508 |
| ENSG000001 | 49.3199595 | 0.071497208 | 0.21518769 | 0.3322551 | 0.73969664 | 0.81176256 | HAP1 | 9001 |
| ENSG000002 | 15.2858627 | 0.099983704 | 0.3010191 | 0.3321507 | 0.73977547 | 0.8118271 | NA | NA |
| ENSG000002 | 10.6688524 | 0.110589813 | 0.3330047 | 0.33209685 | 0.73981613 | 0.81184975 | NA | NA |
| ENSG000002 | 3.33171296 | -0.12555865 | 0.37827421 | -0.331925 | 0.73994592 | 0.81197021 | NA | NA |
| ENSG000002 | 2.59304628 | -0.123768096 | 0.3732519 | -0.331594 | 0.74019584 | 0.81222247 | RN7SL473P | 106479396 |
| ENSG000002 | 21.672013 | 0.091422945 | 0.27584848 | 0.3314245 | 0.74032386 | 0.81234097 | RNF39 | 80352 |
| ENSG000001 | 8.07040312 | -0.115972806 | 0.35000583 | -0.3313454 | 0.74038364 | 0.81235078 | GMFG | 9535 |
| ENSG000002 | 9.06664755 | 0.116001395 | 0.35010507 | 0.33133309 | 0.7403929 | 0.81235078 | NA | NA |
| ENSG000002 | 2.35672369 | -0.122627457 | 0.37006262 | -0.3313695 | 0.74036542 | 0.81235078 | NA | NA |
| ENSG000001 | 98.9279668 | 0.05997564 | 0.18103638 | 0.33129054 | 0.74042504 | 0.81236406 | MORN1 | 79906 |
| ENSG000002 | 8.56138139 | 0.114583066 | 0.3459223 | 0.33123931 | 0.74046373 | 0.81238453 | NA | NA |
| ENSG000002 | 2.59970089 | -0.12380741 | 0.37394809 | -0.3310818 | 0.74058269 | 0.81249307 | NA | NA |
| ENSG000002 | 4.23668601 | -0.125004798 | 0.3775997 | -0.3310511 | 0.74060589 | 0.81249653 | NA | NA |
| ENSG000002 | 8.01065958 | 0.117010237 | 0.35354612 | 0.33096173 | 0.74067339 | 0.81254861 | NA | NA |
| ENSG000002 | 2.29746462 | -0.121113959 | 0.36622374 | -0.3307103 | 0.74086333 | 0.81273499 | NA | NA |
| ENSG000002 | 5.83408114 | 0.122653049 | 0.37098341 | 0.330616 | 0.74093456 | 0.81277105 | NA | NA |
| ENSG000002 | 2.86501634 | -0.123277902 | 0.37287595 | -0.3306137 | 0.74093629 | 0.81277105 | NA | NA |
| ENSG000001 | 42.3015392 | -0.074014049 | 0.22391554 | -0.3305445 | 0.74098858 | 0.81280643 | OLFML2B | 25903 |
| ENSG000002 | 23.1439557 | 0.086421986 | 0.26151736 | 0.33046367 | 0.74104964 | 0.81285142 | NA | NA |
| ENSG000001 | 10.9907692 | -0.110300178 | 0.33388695 | -0.3303519 | 0.7411341 | 0.81292207 | NA | NA |
| ENSG000001 | 269.341489 | 0.044987673 | 0.13619924 | 0.33030781 | 0.74116739 | 0.81293659 | PARM1 | 25849 |
| ENSG000002 | 2.28266571 | 0.122487672 | 0.37095839 | 0.33019248 | 0.74125453 | 0.81300755 | NA | NA |
| ENSG000002 | 5.35867233 | 0.122766504 | 0.37182915 | 0.33016912 | 0.74127218 | 0.81300755 | NA | NA |

| | | | | | | | | |
|---|---|---|---|---|---|---|---|---|
| ENSG000001 | 257.978226 | -0.043264445 | 0.13111009 | -0.3299856 | 0.74141082 | 0.81313762 | WDPCP | 51057 |
| ENSG000002 | 9.36081816 | 0.11243996 | 0.3407913 | 0.32993789 | 0.74144689 | 0.81315519 | NA | NA |
| ENSG000002 | 3.24920925 | -0.124607251 | 0.37772819 | -0.329886 | 0.74148608 | 0.8131578 | HCG14 | 414760 |
| ENSG000002 | 2.32564261 | -0.122463382 | 0.37123429 | -0.3298817 | 0.74148939 | 0.8131578 | NA | NA |
| ENSG000002 | 2.76742927 | -0.123432824 | 0.3743588 | -0.329718 | 0.74161307 | 0.81326836 | NA | NA |
| ENSG000002 | 3.33376784 | -0.124315891 | 0.37706313 | -0.3296952 | 0.74163031 | 0.81326836 | COX6CP16 | 106481965 |
| ENSG000002 | 6.09578874 | 0.124324279 | 0.37725112 | 0.32955311 | 0.74173766 | 0.81336409 | NA | NA |
| ENSG000002 | 2.3652373 | -0.120725186 | 0.36642943 | -0.3294637 | 0.74180525 | 0.8134162 | NA | NA |
| ENSG000001 | 2.3731503 | -0.120756095 | 0.36670148 | -0.3293035 | 0.74192627 | 0.81352691 | PCP2 | 126006 |
| ENSG000001 | 2.36329846 | -0.118705412 | 0.36059293 | -0.3291951 | 0.74200826 | 0.81359481 | NA | NA |
| ENSG000001 | 3.03590318 | -0.124081651 | 0.37700313 | -0.3291263 | 0.74206021 | 0.81362978 | NA | NA |
| ENSG000002 | 2.96164809 | 0.123618591 | 0.37616016 | 0.32863286 | 0.74243321 | 0.81399473 | SNORD116-5 | 100033417 |
| ENSG000002 | 11.9709629 | -0.109774358 | 0.33402318 | -0.3286429 | 0.74242559 | 0.81399473 | NA | NA |
| ENSG000002 | 7.8664324 | -0.116191191 | 0.35361072 | -0.328585 | 0.74246936 | 0.81401236 | NA | NA |
| ENSG000002 | 13.8976655 | -0.106070186 | 0.3228404 | -0.328553 | 0.74249357 | 0.81401689 | RNVU1-17 | 101954269 |
| ENSG000002 | 8.54046529 | -0.115645644 | 0.35209202 | -0.3284529 | 0.74256926 | 0.81407786 | NA | NA |
| ENSG000001 | 86.3968837 | 0.058582293 | 0.17841568 | 0.32834722 | 0.74264914 | 0.81414343 | LINC00174 | 285908 |
| ENSG000001 | 191.986554 | -0.053048745 | 0.16165206 | -0.3281662 | 0.74278599 | 0.81423346 | PLEKHA6 | 22874 |
| ENSG000002 | 4.51075415 | 0.123589659 | 0.3765833 | 0.32818678 | 0.74277045 | 0.81423346 | MIR4766 | 100616283 |
| ENSG000002 | 6.88191692 | -0.11757142 | 0.35827586 | -0.3281589 | 0.74279151 | 0.81423346 | NA | NA |
| ENSG000001 | 2.57977669 | -0.122551096 | 0.37361752 | -0.3280122 | 0.74290246 | 0.81433307 | OR6C76 | 390326 |
| ENSG000001 | 47.2336708 | -0.072065105 | 0.21973309 | -0.3279666 | 0.74293695 | 0.81434887 | LRRC3 | 81543 |
| ENSG000002 | 2.54911906 | -0.121081908 | 0.36952226 | -0.3276715 | 0.74316007 | 0.81457141 | NA | NA |
| ENSG000000 | 59.5212355 | 0.068092108 | 0.20790248 | 0.32751946 | 0.74327502 | 0.81461741 | CD4 | 920 |
| ENSG000001 | 14.9713969 | 0.105552298 | 0.32221189 | 0.3275866 | 0.74322426 | 0.81461741 | TMEM132E | 124842 |
| ENSG000002 | 2.85949626 | -0.121293271 | 0.37035014 | -0.3275097 | 0.74328239 | 0.81461741 | ARMS2 | 387715 |
| ENSG000002 | 2.57087056 | -0.121644754 | 0.37140489 | -0.327526 | 0.74327009 | 0.81461741 | LOC100506125 | 100506125 |
| ENSG000002 | 7.11310582 | -0.117329885 | 0.35829662 | -0.3274658 | 0.74331562 | 0.81463181 | NA | NA |
| ENSG000001 | 17.3962705 | -0.09597608 | 0.29314195 | -0.3274048 | 0.74336174 | 0.81466034 | NA | NA |

| | | | | | | | | |
|---|---|---|---|---|---|---|---|---|
| ENSG000001 | 18.9850462 | 0.094607701 | 0.28923419 | 0.32709722 | 0.74359435 | 0.81487122 | BBS1 | 582 |
| ENSG000002 | 12.5757311 | -0.106810797 | 0.32652424 | -0.3271144 | 0.74358133 | 0.81487122 | NA | NA |
| ENSG000001 | 65.2460551 | -0.063039944 | 0.19278764 | -0.3269916 | 0.74367422 | 0.81493672 | ADCY7 | 113 |
| ENSG000002 | 3.1301172 | -0.122053884 | 0.37334174 | -0.3269227 | 0.74372636 | 0.81497183 | NA | NA |
| ENSG000002 | 3.16255928 | 0.12322613 | 0.37698468 | 0.32687305 | 0.74376391 | 0.81499095 | KLHL6-AS1 | 100874019 |
| ENSG000002 | 6.06521109 | -0.123101177 | 0.37664159 | -0.326839 | 0.74378963 | 0.81499711 | NA | NA |
| ENSG000001 | 32.2564608 | -0.079495067 | 0.24328592 | -0.3267557 | 0.74385265 | 0.81501317 | SERTM1 | 400120 |
| ENSG000002 | 3.3691277 | -0.123164791 | 0.37689695 | -0.3267864 | 0.74382946 | 0.81501317 | NA | NA |
| ENSG000002 | 8.53945475 | 0.113822564 | 0.34835827 | 0.32673995 | 0.74386458 | 0.81501317 | NA | NA |
| ENSG000002 | 3.57457339 | -0.123619197 | 0.37839882 | -0.3266902 | 0.74390219 | 0.81502317 | LOC100190940 | 100190940 |
| ENSG000002 | 3.19429269 | 0.123296347 | 0.37742846 | 0.32667474 | 0.7439139 | 0.81502317 | NA | NA |
| ENSG000002 | 7.38921287 | -0.115944774 | 0.3552505 | -0.3263747 | 0.74414088 | 0.81524982 | RN7SL81P | 106479253 |
| ENSG000002 | 4263.6843 | 0.045251185 | 0.13877912 | 0.32606624 | 0.74437424 | 0.81548345 | MTATP6P1 | 106480796 |
| ENSG000002 | 7.90290588 | -0.117838609 | 0.36158194 | -0.3258974 | 0.74450199 | 0.81560137 | NA | NA |
| ENSG000002 | 3.04078314 | -0.122631278 | 0.37649586 | -0.3257175 | 0.74463813 | 0.81572847 | NA | NA |
| ENSG000001 | 57.0467104 | -0.064860022 | 0.19919285 | -0.3256142 | 0.74471627 | 0.81576647 | MAP3K15 | 389840 |
| ENSG000002 | 2.54818652 | -0.121454581 | 0.37298876 | -0.3256253 | 0.74470786 | 0.81576647 | NA | NA |
| ENSG000002 | 9.59719412 | 0.112312819 | 0.34494971 | 0.32559187 | 0.74473317 | 0.81576647 | NA | NA |
| ENSG000002 | 2.84962033 | -0.1218228 | 0.3742047 | -0.3255512 | 0.74476392 | 0.81577812 | NA | NA |
| ENSG00000C | 11.8704623 | -0.108287762 | 0.332863 | -0.3253223 | 0.74493714 | 0.81594582 | TEKT2 | 27285 |
| ENSG000002 | 11.5773523 | -0.110105909 | 0.33864789 | -0.3251339 | 0.74507973 | 0.81607996 | LOC100129434 | 100129434 |
| ENSG000002 | 3.09004622 | -0.122253711 | 0.37628395 | -0.3248975 | 0.74525865 | 0.81625388 | NA | NA |
| ENSG000002 | 2.37889886 | -0.118513411 | 0.36484138 | -0.3248354 | 0.74530562 | 0.81628327 | NA | NA |
| ENSG000002 | 2.28966861 | -0.119373305 | 0.36762591 | -0.3247141 | 0.74539748 | 0.81636184 | NA | NA |
| ENSG000002 | 50.80959 | -0.070365443 | 0.21675391 | -0.3246329 | 0.74545895 | 0.81640711 | LINC01011 | 401232 |
| ENSG000002 | 2.31964088 | -0.120346184 | 0.37084699 | -0.3245171 | 0.74554659 | 0.81643693 | NA | NA |
| ENSG000002 | 6.69587371 | -0.118982796 | 0.36658961 | -0.3245667 | 0.745509 | 0.81643693 | NA | NA |
| ENSG000002 | 2.31964088 | -0.120346184 | 0.37084699 | -0.3245171 | 0.74554659 | 0.81643693 | NA | NA |
| ENSG000002 | 4.22259647 | 0.122487821 | 0.37748746 | 0.32448183 | 0.74557328 | 0.81644412 | NA | NA |

| | | | | | | | | |
|---|---|---|---|---|---|---|---|---|
| ENSG000002 | 3.28509783 | -0.122302325 | 0.37696767 | -0.3244372 | 0.74560708 | 0.81645908 | NA | NA |
| ENSG000000 | 25.1574425 | -0.084418215 | 0.26036794 | -0.3242266 | 0.74576648 | 0.81661158 | PCDHB4 | 56131 |
| ENSG000001 | 41.6314482 | -0.077050985 | 0.23766998 | -0.3241932 | 0.74579179 | 0.81661724 | VILL | 50853 |
| ENSG000002 | 2.82191254 | -0.120925604 | 0.37330102 | -0.3239359 | 0.74598656 | 0.81680845 | NA | NA |
| ENSG000002 | 3.20215397 | 0.122160142 | 0.37735847 | 0.3237244 | 0.7461467 | 0.81696173 | NA | NA |
| ENSG000001 | 7.16385077 | 0.11559707 | 0.35724261 | 0.32358141 | 0.74625496 | 0.81705821 | FBXO6 | 26270 |
| ENSG000001 | 55.656026 | 0.077018003 | 0.23807515 | 0.32350291 | 0.74631441 | 0.81710124 | TLCD2 | 727910 |
| ENSG000002 | 7.17701209 | 0.11785095 | 0.36438427 | 0.32342491 | 0.74637347 | 0.81714383 | LOC101928161 | 101928161 |
| ENSG000002 | 3.75724186 | -0.12238572 | 0.37859137 | -0.323266 | 0.7464938 | 0.81725351 | NA | NA |
| ENSG000002 | 4.24752282 | -0.122219623 | 0.3782112 | -0.3231518 | 0.7465803 | 0.81732614 | NA | NA |
| ENSG000000 | 292.103324 | 0.042620388 | 0.13195976 | 0.32298019 | 0.74671024 | 0.81744633 | TRAF5 | 7188 |
| ENSG000002 | 6.67816247 | -0.116441525 | 0.36058475 | -0.3229242 | 0.74675268 | 0.81747072 | NA | NA |
| ENSG000001 | 22.9826309 | -0.088083807 | 0.27290174 | -0.3227675 | 0.74687134 | 0.81757855 | C1QL3 | 389941 |
| ENSG000002 | 164.323591 | -0.048748117 | 0.1510619 | -0.3227029 | 0.74692023 | 0.81761 | NA | NA |
| ENSG000001 | 21.0451472 | 0.089790275 | 0.2784113 | 0.32250944 | 0.74706678 | 0.81772627 | TSNAXIP1 | 55815 |
| ENSG000001 | 3.79897706 | -0.122098087 | 0.37857743 | -0.3225181 | 0.7470602 | 0.81772627 | CSTL1 | 128817 |
| ENSG000002 | 6.83112756 | -0.115717952 | 0.35891253 | -0.3224127 | 0.74714008 | 0.81778442 | NA | NA |
| ENSG000002 | 6.9340638 | -0.116429656 | 0.36122391 | -0.3223199 | 0.74721036 | 0.81783927 | NA | NA |
| ENSG000001 | 48.6583496 | -0.069501814 | 0.21570564 | -0.3222068 | 0.74729607 | 0.81791101 | C6orf223 | 221416 |
| ENSG000001 | 19.1473862 | -0.092999554 | 0.28870188 | -0.3221301 | 0.74735417 | 0.81793045 | CARD9 | 64170 |
| ENSG000002 | 4.50257271 | 0.120987451 | 0.37558516 | 0.32213054 | 0.7473538 | 0.81793045 | NA | NA |
| ENSG000002 | 2.54735758 | -0.119870147 | 0.37230266 | -0.3219696 | 0.74747571 | 0.81802078 | NA | NA |
| ENSG000002 | 2.79713321 | -0.120290595 | 0.37361059 | -0.3219678 | 0.74747706 | 0.81802078 | NA | NA |
| ENSG000002 | 18.2058789 | 0.092319327 | 0.286936 | 0.32174188 | 0.74764825 | 0.81818605 | NA | NA |
| ENSG000001 | 41.9331752 | -0.073672497 | 0.22915423 | -0.3214974 | 0.74783345 | 0.81836664 | SOCS1 | 8651 |
| ENSG000002 | 2.8689239 | -0.11888001 | 0.36993895 | -0.3213503 | 0.7479449 | 0.81846652 | NA | NA |
| ENSG000002 | 2.33881435 | -0.116853318 | 0.36370197 | -0.3212887 | 0.74799165 | 0.81849558 | NA | NA |
| ENSG000001 | 15.335423 | -0.100042975 | 0.31153561 | -0.3211285 | 0.74811298 | 0.81850811 | TNFRSF13C | 115650 |
| ENSG000001 | 32.4683318 | -0.077819991 | 0.24224503 | -0.3212449 | 0.74802477 | 0.81850811 | GAB3 | 139716 |

| | | | | | | | | |
|---|---|---|---|---|---|---|---|---|
| ENSG000002 | 673.820189 | 0.03945447 | 0.12285099 | 0.32115713 | 0.74809132 | 0.81850811 | COX20 | 116228 |
| ENSG000002 | 6.5644928 | -0.116711524 | 0.36340949 | -0.3211571 | 0.74809137 | 0.81850811 | NA | NA |
| ENSG000002 | 2.28147149 | 0.119156736 | 0.37096112 | 0.32121084 | 0.74805061 | 0.81850811 | NA | NA |
| ENSG000002 | 14.7930553 | -0.09991699 | 0.31115764 | -0.3211137 | 0.74812421 | 0.81850811 | LOC441242 | 441242 |
| ENSG000001 | 51.0298077 | -0.072656338 | 0.22649559 | -0.3207848 | 0.7483735 | 0.81875877 | SCUBE1 | 80274 |
| ENSG000002 | 2.30654398 | 0.118437044 | 0.369352 | 0.32066171 | 0.74846676 | 0.81883871 | NA | NA |
| ENSG000000 | 24.1170881 | 0.085615587 | 0.26708298 | 0.320558 | 0.74854537 | 0.81890159 | CYP26A1 | 1592 |
| ENSG000001 | 18.8890979 | 0.091047967 | 0.28406398 | 0.32051922 | 0.74857476 | 0.81890159 | C6orf52 | 347744 |
| ENSG000002 | 7.63128456 | -0.113361298 | 0.35369483 | -0.3205059 | 0.74858482 | 0.81890159 | LOC101927769 | 101927769 |
| ENSG000001 | 7.62510602 | -0.11420878 | 0.35643431 | -0.3204203 | 0.74864976 | 0.81895054 | NA | NA |
| ENSG000002 | 2.60715688 | -0.118690763 | 0.37053291 | -0.3203245 | 0.74872237 | 0.81900787 | NA | NA |
| ENSG000002 | 7.15643242 | -0.114881246 | 0.35868638 | -0.3202833 | 0.74875361 | 0.81901995 | SMKR1 | 100287482 |
| ENSG000002 | 3.98177087 | 0.121233845 | 0.37859319 | 0.32022194 | 0.74880009 | 0.8190487 | NA | NA |
| ENSG000002 | 3.05326322 | -0.120461941 | 0.37627584 | -0.3201426 | 0.7488602 | 0.81909235 | NA | NA |
| ENSG000002 | 3.05210201 | -0.12012222 | 0.37526754 | -0.3200975 | 0.74889439 | 0.81910765 | NA | NA |
| ENSG000002 | 6.06016754 | 0.116890458 | 0.3652563 | 0.32002311 | 0.74895081 | 0.81914727 | NA | NA |
| ENSG000002 | 11.6031969 | -0.103676795 | 0.32399924 | -0.3199909 | 0.74897526 | 0.81915192 | NA | NA |
| ENSG000001 | 24.346166 | 0.086059279 | 0.26900624 | 0.31991555 | 0.74903235 | 0.81919226 | LYSMD2 | 256586 |
| ENSG000002 | 9.86473693 | 0.108272864 | 0.33855308 | 0.3198106 | 0.74911191 | 0.81923509 | LOC100133985 | 100133985 |
| ENSG000002 | 2.54275859 | -0.119067765 | 0.3723009 | -0.3198159 | 0.74910786 | 0.81923509 | NA | NA |
| ENSG000002 | 3.09949346 | -0.120230017 | 0.3763763 | -0.319441 | 0.74939213 | 0.81949734 | NA | NA |
| ENSG000002 | 3.27214124 | -0.119997089 | 0.37561915 | -0.3194648 | 0.74937409 | 0.81949734 | NA | NA |
| ENSG000002 | 6.59343448 | -0.115466034 | 0.36158396 | -0.319334 | 0.74947329 | 0.81956399 | NA | NA |
| ENSG000002 | 3.28509893 | -0.120696873 | 0.37804905 | -0.3192625 | 0.74952749 | 0.81960116 | NA | NA |
| ENSG000002 | 2.30550271 | -0.117252322 | 0.36733985 | -0.319193 | 0.74958014 | 0.81963662 | UBTFL1 | 642623 |
| ENSG000001 | 23.4462834 | 0.084378126 | 0.2647087 | 0.31875842 | 0.74990971 | 0.81997489 | C16orf93 | 90835 |
| ENSG000002 | 3.74892119 | -0.119952215 | 0.37638507 | -0.3186955 | 0.74995745 | 0.81998657 | NA | NA |
| ENSG000002 | 3.42719047 | 0.120627078 | 0.37851587 | 0.31868433 | 0.7499659 | 0.81998657 | NA | NA |
| ENSG000002 | 6.62166378 | -0.115571509 | 0.36267475 | -0.3186643 | 0.74998106 | 0.81998657 | LBX2-AS1 | 151534 |

| | | | | | | | | |
|---|---|---|---|---|---|---|---|---|
| ENSG000002 | 363.243781 | 0.042517388 | 0.13346829 | 0.31855798 | 0.75006172 | 0.82005265 | NA | NA |
| ENSG000002 | 2.87524624 | -0.118523806 | 0.37213127 | -0.3185 | 0.7501057 | 0.82007862 | NA | NA |
| ENSG000002 | 2.30092299 | -0.11601888 | 0.36443351 | -0.318354 | 0.7502164 | 0.82017754 | NA | NA |
| ENSG000001 | 157.542971 | 0.046149593 | 0.14497807 | 0.3183212 | 0.75024131 | 0.82018266 | RORB | 6096 |
| ENSG000001 | 71.7350619 | -0.062562793 | 0.19661447 | -0.3182003 | 0.75033297 | 0.82022619 | LPPR4 | 9890 |
| ENSG000001 | 70.2880332 | 0.058505782 | 0.18387134 | 0.31818869 | 0.75034181 | 0.82022619 | BANK1 | 55024 |
| ENSG000002 | 7.34388479 | -0.11274501 | 0.35429594 | -0.3182227 | 0.75031602 | 0.82022619 | MAGEB1 | 4112 |
| ENSG000002 | 3.48790656 | -0.119843492 | 0.3767173 | -0.3181258 | 0.75038952 | 0.82025623 | NA | NA |
| ENSG000002 | 4.94977202 | -0.119733577 | 0.37648497 | -0.3180302 | 0.75046206 | 0.82031341 | ZNF649-AS1 | 101928571 |
| ENSG000002 | 6.6711943 | -0.115951861 | 0.36464969 | -0.3179815 | 0.75049896 | 0.82033164 | NA | NA |
| ENSG000002 | 13.2288983 | -0.099045721 | 0.31159912 | -0.3178627 | 0.75058912 | 0.82038596 | NA | NA |
| ENSG000002 | 2.6550046 | -0.115613136 | 0.36370082 | -0.3178798 | 0.75057613 | 0.82038596 | NA | NA |
| ENSG000002 | 11.9060877 | 0.103589884 | 0.32596184 | 0.31779758 | 0.75063849 | 0.82039569 | NA | NA |
| ENSG000002 | 2.34878366 | -0.117037337 | 0.36826333 | -0.3178088 | 0.75062996 | 0.82039569 | NA | NA |
| ENSG000002 | 3.52007085 | -0.120007941 | 0.37773956 | -0.3177002 | 0.75071235 | 0.8204543 | NA | NA |
| ENSG000002 | 2.31616345 | -0.116901167 | 0.36817102 | -0.3175187 | 0.75085009 | 0.82058272 | NA | NA |
| ENSG000002 | 9.75851195 | -0.108935977 | 0.34323712 | -0.3173782 | 0.75095666 | 0.82067707 | NA | NA |
| ENSG000001 | 95.5086233 | 0.058674553 | 0.18495824 | 0.31723135 | 0.75106806 | 0.82075458 | TTYH1 | 57348 |
| ENSG000002 | 3.33513591 | 0.119901252 | 0.3779503 | 0.31724079 | 0.7510609 | 0.82075458 | NA | NA |
| ENSG000002 | 12.411651 | 0.103126966 | 0.32513011 | 0.31718676 | 0.75110189 | 0.82076943 | NA | NA |
| ENSG000002 | 2.7749977 | -0.118859778 | 0.37480392 | -0.3171252 | 0.75114858 | 0.82079833 | NA | NA |
| ENSG000002 | 4.20291702 | -0.119983564 | 0.37851868 | -0.3169819 | 0.75125735 | 0.82089506 | NA | NA |
| ENSG000001 | 33.321971 | -0.074745664 | 0.23585758 | -0.3169102 | 0.75131177 | 0.82093241 | JAKMIP1 | 152789 |
| ENSG000002 | 2.379584 | -0.114864509 | 0.36251623 | -0.3168534 | 0.75135482 | 0.8209474 | NA | NA |
| ENSG000002 | 2.30662795 | 0.117156098 | 0.36976572 | 0.31683872 | 0.75136598 | 0.8209474 | NA | NA |
| ENSG000002 | 16.2117271 | 0.094482229 | 0.29836533 | 0.31666625 | 0.75149686 | 0.82106828 | NA | NA |
| ENSG000002 | 2.2801012 | 0.117141591 | 0.36996663 | 0.31662745 | 0.7515263 | 0.82107833 | NA | NA |
| ENSG000002 | 6.1670957 | -0.116395338 | 0.36773949 | -0.3165157 | 0.75161108 | 0.8211327 | NA | NA |
| ENSG000002 | 7.10723406 | -0.11339552 | 0.35827005 | -0.3165085 | 0.75161657 | 0.8211327 | NA | NA |

| | | | | | | | | |
|---|---|---|---|---|---|---|---|---|
| ENSG000001 | 55.9111425 | 0.065215873 | 0.20616735 | 0.31632494 | 0.75175588 | 0.82124065 | PHYHD1 | 254295 |
| ENSG000002 | 9.94479236 | 0.106133516 | 0.33551272 | 0.31633232 | 0.75175028 | 0.82124065 | DNLZ | 728489 |
| ENSG000002 | 5.21494455 | -0.119029116 | 0.37644882 | -0.3161894 | 0.75185877 | 0.82133092 | NA | NA |
| ENSG000002 | 3.78391276 | -0.119637965 | 0.37850491 | -0.3160803 | 0.75194152 | 0.82137953 | NA | NA |
| ENSG000002 | 17.6562969 | 0.092573358 | 0.29288196 | 0.31607737 | 0.75194378 | 0.82137953 | NA | NA |
| ENSG000002 | 3.15214593 | 0.119375283 | 0.3777924 | 0.31598117 | 0.7520168 | 0.82143661 | NA | NA |
| ENSG000002 | 2.61877111 | -0.116715303 | 0.36943256 | -0.3159313 | 0.75205466 | 0.82143661 | NA | NA |
| ENSG000002 | 7.38137199 | -0.112603462 | 0.35642076 | -0.3159285 | 0.75205681 | 0.82143661 | NA | NA |
| ENSG000002 | 2.95720512 | 0.118427384 | 0.37494137 | 0.31585574 | 0.75211201 | 0.82147477 | NA | NA |
| ENSG000002 | 4.30986702 | -0.119506421 | 0.37856502 | -0.3156827 | 0.75224337 | 0.82159613 | NA | NA |
| ENSG000002 | 2.55675275 | -0.117384355 | 0.37231277 | -0.3152843 | 0.75254584 | 0.82190434 | NA | NA |
| ENSG000001 | 8.50336807 | 0.109807799 | 0.34844189 | 0.31513949 | 0.75265575 | 0.82200224 | CEL | 1056 |
| ENSG000002 | 2.32262513 | 0.117102245 | 0.37163525 | 0.31509994 | 0.75268577 | 0.82201289 | NA | NA |
| ENSG000002 | 187.597173 | -0.044379393 | 0.1410074 | -0.3147309 | 0.75296595 | 0.82229673 | PVT1 | 5820 |
| ENSG000002 | 17.511964 | 0.09153422 | 0.29086413 | 0.31469752 | 0.75299133 | 0.8223023 | PITRM1-AS1 | 100507034 |
| ENSG000002 | 7.85188113 | 0.112662573 | 0.35810416 | 0.31460839 | 0.75305901 | 0.82230978 | NA | NA |
| ENSG000002 | 3.06933307 | -0.117858834 | 0.37458854 | -0.3146355 | 0.75303846 | 0.82230978 | NA | NA |
| ENSG000002 | 3.50317992 | -0.118599998 | 0.3769496 | -0.3146309 | 0.75304191 | 0.82230978 | NA | NA |
| ENSG000002 | 6.27246482 | 0.117116157 | 0.3724016 | 0.31448887 | 0.75314977 | 0.82238674 | NA | NA |
| ENSG000002 | 29.7535384 | 0.079369439 | 0.25239856 | 0.31446075 | 0.75317113 | 0.82238791 | NA | NA |
| ENSG000002 | 10.6600308 | -0.103072008 | 0.32804152 | -0.3142042 | 0.75336599 | 0.82257854 | NA | NA |
| ENSG000001 | 11.53485 | 0.109494588 | 0.34858858 | 0.31410837 | 0.75343874 | 0.82263582 | PLIN4 | 729359 |
| ENSG00000C | 16.2624027 | -0.095959858 | 0.30565464 | -0.3139486 | 0.75356005 | 0.82273048 | IL32 | 9235 |
| ENSG000001 | 203.471912 | -0.04391198 | 0.13987345 | -0.3139408 | 0.75356602 | 0.82273048 | ARV1 | 64801 |
| ENSG000002 | 3.08327537 | -0.117969713 | 0.37583499 | -0.313887 | 0.75360688 | 0.82275294 | NA | NA |
| ENSG000002 | 3.55733047 | -0.118775434 | 0.37845584 | -0.3138423 | 0.75364085 | 0.82276788 | TUBB8P2 | 106478982 |
| ENSG000001 | 685.698866 | -0.038757089 | 0.12354152 | -0.3137171 | 0.7537359 | 0.82282734 | PDE4DIP | 9659 |
| ENSG000002 | 5.86602068 | 0.116191512 | 0.37033906 | 0.3137436 | 0.75371578 | 0.82282734 | NA | NA |
| ENSG000002 | 3.95671033 | -0.118759325 | 0.37858865 | -0.3136896 | 0.75375679 | 0.822828 | NA | NA |

| | | | | | | | | |
|---|---|---|---|---|---|---|---|---|
| ENSG000002 | 6.86501388 | -0.112499426 | 0.35867504 | -0.3136528 | 0.75378476 | 0.82283638 | NA | NA |
| ENSG000001 | 91.6643022 | -0.060121059 | 0.19175628 | -0.3135285 | 0.75387917 | 0.82288551 | NA | NA |
| ENSG000002 | 2.30540519 | -0.114575275 | 0.3654529 | -0.3135158 | 0.75388878 | 0.82288551 | NA | NA |
| ENSG000002 | 24.6022249 | -0.086415788 | 0.27563667 | -0.3135134 | 0.75389065 | 0.82288551 | EID3 | 493861 |
| ENSG000002 | 17.6492555 | -0.093673806 | 0.29886094 | -0.3134361 | 0.75394937 | 0.82292745 | NA | NA |
| ENSG000002 | 6.3548215 | -0.113867516 | 0.36332953 | -0.3134001 | 0.7539767 | 0.82293513 | NA | NA |
| ENSG000002 | 7.22840635 | -0.113769751 | 0.36310722 | -0.3133227 | 0.75403547 | 0.82297713 | LINC00443 | 100874173 |
| ENSG000002 | 4.46178359 | 0.11777558 | 0.37607521 | 0.31317028 | 0.7541513 | 0.8230814 | NA | NA |
| ENSG000001 | 36.6003029 | -0.073068624 | 0.2335314 | -0.3128857 | 0.75436754 | 0.82329524 | GHRHR | 2692 |
| ENSG000002 | 22.7601315 | 0.084895739 | 0.27143091 | 0.31277108 | 0.75445459 | 0.82336809 | LOC105378421 | 105378421 |
| ENSG000002 | 2.27817066 | 0.11464439 | 0.3665813 | 0.31273932 | 0.75447872 | 0.82337226 | CTAGE6 | 340307 |
| ENSG000001 | 16.0728716 | -0.097123614 | 0.31060761 | -0.3126891 | 0.75451688 | 0.82339175 | GPR141 | 353345 |
| ENSG000002 | 14.1928706 | -0.108094802 | 0.34586258 | -0.3125368 | 0.75463256 | 0.82349583 | NA | NA |
| ENSG00000C | 43.1916468 | 0.069878404 | 0.22367317 | 0.31241299 | 0.75472668 | 0.82357637 | DGAT2 | 84649 |
| ENSG000001 | 10.4662837 | -0.103158186 | 0.33023203 | -0.3123809 | 0.75475104 | 0.8235808 | FGF18 | 8817 |
| ENSG000002 | 5.92455589 | -0.114179228 | 0.36561002 | -0.3122979 | 0.75481416 | 0.82362751 | LOC105375968 | 105375968 |
| ENSG000002 | 2.32962892 | -0.114003777 | 0.36508002 | -0.3122707 | 0.75483484 | 0.82362791 | NA | NA |
| ENSG000001 | 247.227239 | 0.042021667 | 0.13462433 | 0.31214021 | 0.75493397 | 0.82364968 | NMNAT2 | 23057 |
| ENSG000001 | 3.23597246 | -0.117847536 | 0.37754096 | -0.312145 | 0.7549303 | 0.82364968 | NA | NA |
| ENSG000001 | 2.29664309 | -0.113319614 | 0.36299742 | -0.3121775 | 0.75490566 | 0.82364968 | OR6B3 | 150681 |
| ENSG000002 | 3.05172374 | -0.116925293 | 0.37459549 | -0.3121375 | 0.75493604 | 0.82364968 | NA | NA |
| ENSG000001 | 36.2757147 | -0.074411382 | 0.23843649 | -0.3120805 | 0.75497934 | 0.82367476 | TLR2 | 7097 |
| ENSG000002 | 2.56675951 | -0.113942731 | 0.36524466 | -0.3119628 | 0.75506883 | 0.82375023 | NA | NA |
| ENSG000002 | 10.3023495 | 0.10565572 | 0.33897248 | 0.3116941 | 0.75527301 | 0.82393843 | NA | NA |
| ENSG000002 | 7.87425249 | 0.111786834 | 0.35865634 | 0.3116823 | 0.75528198 | 0.82393843 | SCARNA8 | 677776 |
| ENSG000002 | 10.928224 | 0.103307209 | 0.33149443 | 0.31164086 | 0.75531348 | 0.82395063 | NA | NA |
| ENSG000002 | 2.85414818 | -0.116604252 | 0.37419926 | -0.3116101 | 0.75533689 | 0.82395401 | NA | NA |
| ENSG000002 | 3.55685364 | -0.117613957 | 0.37761606 | -0.3114644 | 0.75544761 | 0.82405261 | NA | NA |
| ENSG000001 | 12.9980075 | -0.100070718 | 0.32133661 | -0.3114202 | 0.75548119 | 0.82406708 | ATRIP | 84126 |

| | | | | | | | | |
|---|---|---|---|---|---|---|---|---|
| ENSG000001 | 381.601662 | -0.048281386 | 0.15508342 | -0.3113253 | 0.75555337 | 0.82412365 | FOXO4 | 4303 |
| ENSG000002 | 5.96453513 | -0.114333452 | 0.3674542 | -0.3111502 | 0.75568643 | 0.82424661 | NA | NA |
| ENSG000002 | 7.38902963 | -0.111820172 | 0.35942506 | -0.3111084 | 0.75571818 | 0.82425907 | NA | NA |
| ENSG000000 | 103.566727 | 0.058467142 | 0.1879672 | 0.3110497 | 0.75576284 | 0.82428561 | PPP2R2C | 5522 |
| ENSG000002 | 4.77378772 | 0.116897349 | 0.37587391 | 0.3110015 | 0.75579948 | 0.8243034 | NA | NA |
| ENSG000002 | 3.40443474 | 0.117338593 | 0.37748118 | 0.31084621 | 0.75591754 | 0.82436565 | NA | NA |
| ENSG000002 | 3.05167256 | -0.117034988 | 0.37644283 | -0.3108971 | 0.75587884 | 0.82436565 | NA | NA |
| ENSG000002 | 7.59574089 | -0.110850022 | 0.35658273 | -0.3108676 | 0.75590127 | 0.82436565 | NA | NA |
| ENSG000002 | 2.31300162 | 0.115400269 | 0.37141503 | 0.31070436 | 0.75602538 | 0.82446108 | NA | NA |
| ENSG000002 | 3.41396522 | 0.117546723 | 0.37857384 | 0.3104988 | 0.75618167 | 0.82460934 | RNU6-525P | 106479799 |
| ENSG000001 | 28.1792419 | 0.081603594 | 0.2630679 | 0.31019974 | 0.75640907 | 0.82481668 | MAFA | 389692 |
| ENSG000002 | 18.6768475 | 0.09041515 | 0.29147819 | 0.31019525 | 0.75641249 | 0.82481668 | NA | NA |
| ENSG000002 | 13.1270351 | -0.099284548 | 0.32010438 | -0.310163 | 0.75643697 | 0.8248212 | NA | NA |
| ENSG000001 | 183.160579 | 0.04516821 | 0.14568259 | 0.31004535 | 0.75652647 | 0.82489661 | ITGA2 | 3673 |
| ENSG000002 | 8.14840272 | -0.112679389 | 0.36351453 | -0.3099722 | 0.75658212 | 0.82491293 | LINC00703 | 100507059 |
| ENSG000002 | 2.29390341 | -0.113700476 | 0.36679127 | -0.3099869 | 0.75657094 | 0.82491293 | NA | NA |
| ENSG000002 | 9.58341415 | 0.105830789 | 0.34154194 | 0.30986177 | 0.75666608 | 0.82498229 | C17orf49 | 124944 |
| ENSG000002 | 3.06928042 | -0.116319639 | 0.37565222 | -0.3096471 | 0.7568293 | 0.82513806 | NA | NA |
| ENSG000002 | 5.59656218 | 0.115650834 | 0.37362641 | 0.30953603 | 0.75691381 | 0.82520801 | NA | NA |
| ENSG000002 | 11.8783067 | 0.099156745 | 0.32039469 | 0.30948311 | 0.75695406 | 0.8252297 | PLIN5 | 440503 |
| ENSG000002 | 6.13619884 | -0.112462158 | 0.36342239 | -0.309453 | 0.75697694 | 0.82523247 | NA | NA |
| ENSG000001 | 42.0573261 | 0.072699155 | 0.23511923 | 0.30920123 | 0.75716846 | 0.82541906 | NA | NA |
| ENSG000002 | 31.9237229 | 0.074165931 | 0.23991048 | 0.30914001 | 0.75721502 | 0.82544763 | NA | NA |
| ENSG000000 | 26.9804808 | 0.077632497 | 0.25131369 | 0.30890675 | 0.75739246 | 0.82561886 | FAM65C | 140876 |
| ENSG000002 | 3.30142838 | -0.116063103 | 0.37582264 | -0.3088241 | 0.75745531 | 0.82566518 | NA | NA |
| ENSG000002 | 18.9439936 | 0.090214031 | 0.29237214 | 0.30855892 | 0.75765707 | 0.82586291 | NA | NA |
| ENSG000001 | 24.9540128 | 0.081091106 | 0.26285221 | 0.30850457 | 0.75769843 | 0.82588579 | PCSK4 | 54760 |
| ENSG000001 | 39.0520384 | -0.070389446 | 0.2282053 | -0.3084479 | 0.75774154 | 0.82591057 | SYPL2 | 284612 |
| ENSG000002 | 4.24991937 | 0.116593785 | 0.37806055 | 0.30839976 | 0.75777816 | 0.8259283 | NA | NA |

| | | | | | | | | |
|---|---|---|---|---|---|---|---|---|
| ENSG000001 | 7.19390697 | -0.111612636 | 0.36213444 | -0.3082077 | 0.75792426 | 0.82606533 | NA | NA |
| ENSG000002 | 4.01658928 | -0.11663597 | 0.37859337 | -0.3080772 | 0.75802362 | 0.82613109 | NA | NA |
| ENSG000002 | 2.29008924 | 0.114280884 | 0.37095163 | 0.3080749 | 0.75802534 | 0.82613109 | NA | NA |
| ENSG000001 | 130.945924 | -0.045915875 | 0.14908198 | -0.3079908 | 0.75808935 | 0.82615644 | RAD51B | 5890 |
| ENSG000002 | 5.90799756 | -0.113091936 | 0.3671733 | -0.308007 | 0.75807703 | 0.82615644 | NA | NA |
| ENSG000001 | 95.8957641 | 0.05370634 | 0.17441285 | 0.30792651 | 0.75813825 | 0.82618753 | GPRIN3 | 285513 |
| ENSG000002 | 6.67512825 | 0.112564065 | 0.36568746 | 0.30781495 | 0.75822315 | 0.82625784 | CSE1L-AS1 | 102723483 |
| ENSG000002 | 6.61561458 | 0.113781277 | 0.36973147 | 0.30774031 | 0.75827994 | 0.82627864 | NA | NA |
| ENSG000002 | 5.19075005 | 0.116420554 | 0.37831269 | 0.30773631 | 0.75828298 | 0.82627864 | LOC101928600 | 101928600 |
| ENSG000001 | 45.5032699 | 0.065130995 | 0.21171751 | 0.3076316 | 0.75836267 | 0.82632787 | RFESD | 317671 |
| ENSG000002 | 7.47160005 | 0.110223675 | 0.35830719 | 0.30762339 | 0.75836892 | 0.82632787 | NA | NA |
| ENSG000002 | 34.5384693 | -0.070899578 | 0.23056059 | -0.3075095 | 0.75845557 | 0.82640009 | LOC102723582 | 102723582 |
| ENSG000001 | 119.821477 | -0.049692441 | 0.16161698 | -0.3074704 | 0.75848533 | 0.8264103 | ATL1 | 51062 |
| ENSG000002 | 14.5867387 | 0.093713648 | 0.30495808 | 0.3073001 | 0.75861496 | 0.82652933 | NA | NA |
| ENSG000001 | 71.5972945 | -0.059634819 | 0.19428454 | -0.3069458 | 0.75888465 | 0.82677777 | MYO7A | 4647 |
| ENSG000001 | 52.7787127 | 0.065801742 | 0.21439978 | 0.30691143 | 0.75891079 | 0.82677777 | INPP1 | 3628 |
| ENSG000002 | 4.54226413 | 0.11545649 | 0.37621044 | 0.30689337 | 0.75892453 | 0.82677777 | LOC399886 | 399886 |
| ENSG000002 | 2.81288985 | -0.113743426 | 0.37056864 | -0.3069429 | 0.75888681 | 0.82677777 | NA | NA |
| ENSG000002 | 2.30982498 | 0.113179025 | 0.36937208 | 0.30640926 | 0.75929306 | 0.82715702 | NA | NA |
| ENSG000001 | 30.113218 | -0.076134433 | 0.2486045 | -0.3062472 | 0.75941643 | 0.8272692 | BTNL9 | 153579 |
| ENSG000001 | 33.2318236 | -0.071223012 | 0.23272454 | -0.30604 | 0.75957421 | 0.82741884 | NUPR1 | 26471 |
| ENSG000001 | 38.7094077 | -0.069817671 | 0.22860863 | -0.3054026 | 0.76005953 | 0.8278808 | TRAPPC5 | 126003 |
| ENSG000002 | 4.76805844 | 0.114757157 | 0.37574678 | 0.30541088 | 0.76005323 | 0.8278808 | RNU1-27P | 26865 |
| ENSG000002 | 7.88897284 | -0.109855212 | 0.3596996 | -0.3054082 | 0.76005527 | 0.8278808 | NA | NA |
| ENSG000001 | 23.4304515 | -0.082615534 | 0.2706568 | -0.3052409 | 0.76018266 | 0.82799267 | ATP2A1 | 487 |
| ENSG000002 | 13.4186381 | 0.096340904 | 0.31566146 | 0.30520325 | 0.76021136 | 0.82800168 | NA | NA |
| ENSG000001 | 15.1217288 | -0.095179514 | 0.31229026 | -0.304779 | 0.76053447 | 0.82833137 | C6orf118 | 168090 |
| ENSG000002 | 7.32502343 | -0.108715404 | 0.35674834 | -0.3047398 | 0.76056437 | 0.82834168 | NA | NA |
| ENSG000001 | 28.0007079 | -0.075862651 | 0.24917049 | -0.3044608 | 0.76077684 | 0.82855083 | ADHFE1 | 137872 |

| | | | | | | | | |
|---|---|---|---|---|---|---|---|---|
| ENSG000002 | 4.74939296 | 0.114233981 | 0.37550854 | 0.30421141 | 0.76096683 | 0.82873549 | NA | NA |
| ENSG000002 | 3.04496609 | -0.114252416 | 0.37570989 | -0.3040974 | 0.76105366 | 0.82880779 | LOC105377682 | 105377682 |
| ENSG00000C | 13.5336451 | 0.095257663 | 0.31333879 | 0.30400852 | 0.7611214 | 0.82881799 | PRSS22 | 64063 |
| ENSG000001 | 74.9019389 | -0.056079943 | 0.18448695 | -0.3039778 | 0.76114478 | 0.82881799 | PTK2B | 2185 |
| ENSG000002 | 10.01379 | -0.103320442 | 0.33987142 | -0.3039986 | 0.76112895 | 0.82881799 | TMEM191A | 84222 |
| ENSG000002 | 10.5245767 | -0.102079182 | 0.3357561 | -0.3040278 | 0.76110672 | 0.82881799 | NA | NA |
| ENSG000002 | 4.57186158 | -0.114975134 | 0.37853479 | -0.3037373 | 0.76132804 | 0.82895076 | NA | NA |
| ENSG000002 | 2.83427583 | -0.112997571 | 0.37201878 | -0.3037416 | 0.76132478 | 0.82895076 | NA | NA |
| ENSG000002 | 5.44670239 | -0.113109342 | 0.37233832 | -0.3037811 | 0.76129467 | 0.82895076 | NA | NA |
| ENSG000002 | 2.29992127 | 0.11150464 | 0.36716602 | 0.30368998 | 0.76136409 | 0.82896776 | NA | NA |
| ENSG000002 | 15.4985087 | -0.093230094 | 0.30703119 | -0.3036502 | 0.76139437 | 0.82897847 | NA | NA |
| ENSG000002 | 24.3240089 | -0.080394724 | 0.26494626 | -0.3034379 | 0.7615562 | 0.8291324 | NA | NA |
| ENSG000001 | 11.4052723 | -0.100838094 | 0.33241125 | -0.3033534 | 0.76162053 | 0.82918018 | NUDT8 | 254552 |
| ENSG000002 | 25.9002837 | -0.078515694 | 0.25913483 | -0.3029917 | 0.76189621 | 0.82945804 | NA | NA |
| ENSG000001 | 87.3224431 | -0.0523211 | 0.1727243 | -0.3029168 | 0.76195323 | 0.82949785 | LRGUK | 136332 |
| ENSG000002 | 8.2774791 | -0.106914481 | 0.35312474 | -0.3027669 | 0.76206751 | 0.82959999 | NA | NA |
| ENSG000002 | 2.3057512 | -0.111440266 | 0.36812967 | -0.3027201 | 0.76210315 | 0.82961651 | NA | NA |
| ENSG000001 | 2.31473663 | 0.11243222 | 0.37162418 | 0.3025428 | 0.76223831 | 0.82972887 | C20orf85 | 128602 |
| ENSG000002 | 5.85048719 | 0.111283871 | 0.36787549 | 0.30250418 | 0.76226775 | 0.82972887 | PTPRD-AS2 | 101929428 |
| ENSG000002 | 5.50514849 | -0.114389629 | 0.37813824 | -0.3025074 | 0.76226526 | 0.82972887 | NA | NA |
| ENSG000002 | 6.40638793 | -0.110060108 | 0.36392622 | -0.3024242 | 0.76232868 | 0.82975921 | NA | NA |
| ENSG000002 | 6.92468377 | -0.109939524 | 0.36353989 | -0.3024139 | 0.76233654 | 0.82975921 | LOC101928333 | 101928333 |
| ENSG000002 | 4.97104868 | -0.113495496 | 0.37559346 | -0.3021764 | 0.76251756 | 0.82992091 | NA | NA |
| ENSG000002 | 6.35234927 | -0.110374215 | 0.36527756 | -0.3021653 | 0.76252603 | 0.82992091 | NA | NA |
| ENSG000002 | 6.36546242 | -0.110081074 | 0.36452446 | -0.3019854 | 0.76266318 | 0.83004789 | NA | NA |
| ENSG000001 | 47.913151 | -0.064370591 | 0.21320557 | -0.301918 | 0.76271461 | 0.83006619 | CHP2 | 63928 |
| ENSG000002 | 6.87035673 | -0.110650594 | 0.36650232 | -0.3019097 | 0.76272093 | 0.83006619 | NA | NA |
| ENSG000002 | 22.2671559 | 0.082935438 | 0.27490536 | 0.30168724 | 0.7628905 | 0.83022845 | NA | NA |
| ENSG000002 | 4.95201377 | 0.112737626 | 0.37403208 | 0.30141165 | 0.76310061 | 0.83043483 | RNU6-33P | 100873748 |

| | | | | | | | | |
|---|---|---|---|---|---|---|---|---|
| ENSG000002 | 42.5818651 | -0.065850023 | 0.21857902 | -0.3012642 | 0.76321307 | 0.83049271 | NA | NA |
| ENSG000002 | 5.9787975 | -0.112113562 | 0.37210918 | -0.3012921 | 0.76319175 | 0.83049271 | NA | NA |
| ENSG000002 | 4.28706743 | 0.114052164 | 0.37858218 | 0.30126131 | 0.76321524 | 0.83049271 | NA | NA |
| ENSG000002 | 5.53637665 | 0.112587752 | 0.37389482 | 0.30112145 | 0.76332188 | 0.83058646 | NA | NA |
| ENSG000001 | 97.4277196 | -0.051335953 | 0.17061086 | -0.300895 | 0.76349457 | 0.83065694 | SNCA | 6622 |
| ENSG000002 | 3.63363362 | 0.113945177 | 0.37856579 | 0.30099174 | 0.76342079 | 0.83065694 | NA | NA |
| ENSG000002 | 2.32908332 | 0.111834172 | 0.37166445 | 0.30090091 | 0.76349005 | 0.83065694 | NA | NA |
| ENSG000002 | 9.51495589 | -0.103515367 | 0.34404736 | -0.3008753 | 0.76350956 | 0.83065694 | NA | NA |
| ENSG000002 | 5.69377856 | -0.11072443 | 0.36799165 | -0.3008884 | 0.76349957 | 0.83065694 | NA | NA |
| ENSG000002 | 4.76003355 | 0.113117262 | 0.37586481 | 0.30095199 | 0.7634511 | 0.83065694 | NA | NA |
| ENSG000001 | 11.2857726 | -0.100061446 | 0.33292983 | -0.3005482 | 0.76375907 | 0.8308842 | GUCY2D | 3000 |
| ENSG000002 | 12.750037 | 0.096416774 | 0.32080359 | 0.30054768 | 0.76375943 | 0.8308842 | NA | NA |
| ENSG000002 | 3.00242234 | -0.11259506 | 0.3746811 | -0.300509 | 0.7637889 | 0.83089397 | NA | NA |
| ENSG000001 | 25.7261262 | -0.078415116 | 0.26108628 | -0.3003418 | 0.76391647 | 0.83101046 | ACOX2 | 8309 |
| ENSG000002 | 33.1474001 | -0.073374299 | 0.24438557 | -0.3002399 | 0.76399417 | 0.8310683 | NA | NA |
| ENSG000002 | 7.37735327 | 0.107698539 | 0.35873407 | 0.30021832 | 0.76401063 | 0.8310683 | NA | NA |
| ENSG000002 | 11.4346713 | 0.097703622 | 0.3254711 | 0.30019139 | 0.76403117 | 0.83106835 | NA | NA |
| ENSG000001 | 9.23796913 | -0.102046535 | 0.34009718 | -0.3000511 | 0.76413818 | 0.83116245 | CMTM5 | 116173 |
| ENSG000002 | 21.8195292 | -0.085685604 | 0.28594668 | -0.2996559 | 0.76443966 | 0.83146807 | NA | NA |
| ENSG000001 | 82.8159924 | 0.055944711 | 0.18676449 | 0.29954682 | 0.76452285 | 0.83150351 | PLA2G4C | 8605 |
| ENSG000001 | 6.3490034 | 0.110809767 | 0.36994234 | 0.29953253 | 0.76453375 | 0.83150351 | NA | NA |
| ENSG000002 | 5.90064537 | -0.110150399 | 0.36771041 | -0.2995575 | 0.76451473 | 0.83150351 | C1orf185 | 284546 |
| ENSG000001 | 25.4176805 | 0.079386533 | 0.26517738 | 0.29937144 | 0.76465665 | 0.83161487 | ZGPAT | 84619 |
| ENSG000001 | 124.80939 | 0.046924116 | 0.15687769 | 0.29911274 | 0.76485403 | 0.83180318 | CR2 | 1380 |
| ENSG000001 | 6.3958515 | -0.109115653 | 0.3648246 | -0.2990907 | 0.76487083 | 0.83180318 | MRAP | 56246 |
| ENSG000001 | 62.6494381 | -0.06181346 | 0.20677503 | -0.2989406 | 0.76498534 | 0.8319016 | EYA1 | 2138 |
| ENSG000001 | 10.0336109 | -0.10123553 | 0.33867287 | -0.2989183 | 0.76500236 | 0.8319016 | PCDHB9 | 56127 |
| ENSG000002 | 2.31351015 | 0.1108918 | 0.37106629 | 0.29884633 | 0.7650573 | 0.83192338 | NA | NA |
| ENSG000002 | 9.91984943 | 0.101433403 | 0.3394257 | 0.29883831 | 0.76506342 | 0.83192338 | NA | NA |

| | | | | | | | | |
|---|---|---|---|---|---|---|---|---|
| ENSG000002 | 2.33166744 | 0.109978266 | 0.36806656 | 0.29879994 | 0.7650927 | 0.83193291 | NA | NA |
| ENSG000002 | 287.658847 | 0.039617436 | 0.13269376 | 0.29856291 | 0.76527357 | 0.83210727 | CASC15 | 401237 |
| ENSG000002 | 3.65112469 | 0.113020102 | 0.37858916 | 0.29852969 | 0.76529892 | 0.83211252 | NA | NA |
| ENSG000002 | 6.10947676 | -0.108562938 | 0.36371987 | -0.2984795 | 0.76533719 | 0.83213182 | LOC100240735 | 100240735 |
| ENSG000002 | 5.68251178 | -0.1115357 | 0.37374219 | -0.2984295 | 0.76537537 | 0.83215102 | LINC01570 | 101926950 |
| ENSG000002 | 19.055104 | -0.085682503 | 0.28737181 | -0.298159 | 0.76558178 | 0.83230849 | NA | NA |
| ENSG000002 | 2.27248159 | 0.108454205 | 0.36371785 | 0.29818225 | 0.76556407 | 0.83230849 | NA | NA |
| ENSG000002 | 4.74776498 | -0.112741067 | 0.37807009 | -0.2982015 | 0.76554938 | 0.83230849 | NA | NA |
| ENSG000002 | 3.57902714 | 0.112467088 | 0.37758748 | 0.29785703 | 0.76581228 | 0.83253676 | NA | NA |
| ENSG000002 | 98.5624156 | -0.052669248 | 0.17688628 | -0.2977577 | 0.76588812 | 0.83259689 | NA | NA |
| ENSG000001 | 87.2215291 | 0.057401645 | 0.19282515 | 0.29768754 | 0.76594166 | 0.83263277 | RBFOX3 | 146713 |
| ENSG000002 | 5.85283711 | -0.109467985 | 0.36816386 | -0.297335 | 0.7662108 | 0.83290302 | NA | NA |
| ENSG000002 | 5.79484249 | 0.110103068 | 0.37036446 | 0.29728303 | 0.76625044 | 0.83292378 | NA | NA |
| ENSG000001 | 60.3852798 | -0.06432485 | 0.21645383 | -0.2971758 | 0.76633227 | 0.8329904 | PLA2G4A | 5321 |
| ENSG000002 | 10.1731632 | -0.099725 | 0.33565212 | -0.2971082 | 0.7663839 | 0.83302419 | NA | NA |
| ENSG000002 | 3.18156891 | 0.11168185 | 0.37594804 | 0.29706725 | 0.76641517 | 0.83303585 | NA | NA |
| ENSG000002 | 2.28869914 | 0.109993559 | 0.37036374 | 0.29698793 | 0.76647573 | 0.83307934 | NA | NA |
| ENSG000002 | 3.45290344 | 0.111397248 | 0.37519132 | 0.29690785 | 0.76653686 | 0.83312346 | NA | NA |
| ENSG000002 | 361.092865 | 0.043286655 | 0.14593108 | 0.29662395 | 0.76675362 | 0.83333672 | SRGAP2 | 23380 |
| ENSG000002 | 4.68639976 | -0.112245825 | 0.37858871 | -0.2964849 | 0.76685982 | 0.83342979 | TPRKBP2 | 106480804 |
| ENSG000002 | 69.6257492 | -0.054835092 | 0.18501601 | -0.2963803 | 0.7669397 | 0.83347194 | SPIRE2 | 84501 |
| ENSG000002 | 5.97967886 | -0.109957527 | 0.37096808 | -0.296407 | 0.7669193 | 0.83347194 | NA | NA |
| ENSG000002 | 2.34066997 | 0.109678303 | 0.37015364 | 0.29630481 | 0.76699731 | 0.83349999 | MIR577 | 693162 |
| ENSG000002 | 3.04954177 | -0.086750812 | 0.29284083 | -0.2962388 | 0.76704773 | 0.83349999 | NA | NA |
| ENSG000002 | 9.25207674 | 0.10172604 | 0.34336616 | 0.29626112 | 0.76703068 | 0.83349999 | NA | NA |
| ENSG000002 | 2.32652712 | 0.10864277 | 0.3666747 | 0.29629198 | 0.76700711 | 0.83349999 | NA | NA |
| ENSG000002 | 2.5482947 | 0.110882013 | 0.37439805 | 0.29616076 | 0.76710731 | 0.83354239 | NA | NA |
| ENSG000002 | 21.338739 | -0.086973811 | 0.29377318 | -0.2960577 | 0.76718602 | 0.83360558 | NA | NA |
| ENSG000002 | 5.63671947 | -0.108595864 | 0.36688474 | -0.2959945 | 0.76723428 | 0.83363569 | NA | NA |

| | | | | | | | | |
|---|---|---|---|---|---|---|---|---|
| ENSG000001 | 10.0102298 | 0.101495997 | 0.34305926 | 0.29585558 | 0.76734038 | 0.83366161 | NA | NA |
| ENSG000002 | 12.9888512 | -0.093590605 | 0.31633047 | -0.2958634 | 0.76733441 | 0.83366161 | NA | NA |
| ENSG000002 | 12.3764039 | 0.094523231 | 0.31945907 | 0.29588526 | 0.76731771 | 0.83366161 | NA | NA |
| ENSG000002 | 3.67648947 | 0.111957308 | 0.37841155 | 0.29586123 | 0.76733606 | 0.83366161 | NA | NA |
| ENSG000002 | 32.0170295 | -0.073073881 | 0.24703209 | -0.2958072 | 0.76737729 | 0.83367938 | NA | NA |
| ENSG000002 | 5.8537702 | -0.109018884 | 0.36858179 | -0.2957794 | 0.76739859 | 0.83368018 | NA | NA |
| ENSG000001 | 47.1819814 | 0.077510103 | 0.26217986 | 0.29563714 | 0.76750721 | 0.83377585 | C19orf68 | 374920 |
| ENSG000002 | 2.30827789 | 0.109656744 | 0.3710549 | 0.29552701 | 0.76759132 | 0.83384488 | NA | NA |
| ENSG000001 | 19.8568669 | 0.083876908 | 0.2839887 | 0.29535297 | 0.76772425 | 0.83396695 | HTRA3 | 94031 |
| ENSG000002 | 6.25812158 | 0.108809393 | 0.3684439 | 0.29532147 | 0.76774832 | 0.83397075 | NA | NA |
| ENSG000001 | 101.055481 | -0.048471312 | 0.1642243 | -0.2951531 | 0.76787693 | 0.83408811 | VIL1 | 7429 |
| ENSG000002 | 7.04763097 | -0.106486312 | 0.36125845 | -0.2947649 | 0.76817348 | 0.83438788 | NA | NA |
| ENSG000002 | 5.37576104 | -0.109701896 | 0.37223647 | -0.2947102 | 0.76821525 | 0.8344109 | NA | NA |
| ENSG000002 | 34.1240358 | 0.072979477 | 0.24769729 | 0.29463171 | 0.76827523 | 0.8344537 | USP46-AS1 | 643783 |
| ENSG000002 | 4.96900155 | -0.110090863 | 0.3737226 | -0.2945791 | 0.76831544 | 0.83447502 | NA | NA |
| ENSG000001 | 9.44122669 | -0.100989725 | 0.34296439 | -0.2944613 | 0.76840546 | 0.83455045 | KLK7 | 5650 |
| ENSG000002 | 4.55496048 | 0.111077502 | 0.37735102 | 0.29436121 | 0.76848191 | 0.83460742 | NA | NA |
| ENSG000002 | 28.1427381 | 0.077007325 | 0.26162825 | 0.29433872 | 0.76849909 | 0.83460742 | NA | NA |
| ENSG000002 | 36.9769041 | 0.068364539 | 0.2323486 | 0.29423262 | 0.76858015 | 0.83467311 | PSMB9 | 5698 |
| ENSG000002 | 5.88601848 | -0.108119861 | 0.36765025 | -0.2940835 | 0.76869412 | 0.83475939 | NA | NA |
| ENSG000002 | 29.6908415 | 0.074197225 | 0.25230735 | 0.29407477 | 0.76870077 | 0.83475939 | NA | NA |
| ENSG000002 | 3.39706254 | 0.11096492 | 0.37740966 | 0.29401717 | 0.76874479 | 0.83478483 | NA | NA |
| ENSG000001 | 41.7916028 | -0.066592555 | 0.22651925 | -0.2939819 | 0.76877175 | 0.83479175 | LRRC27 | 80313 |
| ENSG000002 | 27.2095283 | -0.075143888 | 0.25584121 | -0.293713 | 0.76897722 | 0.83499251 | NA | NA |
| ENSG000002 | 3.65201835 | 0.111113737 | 0.37858596 | 0.29349672 | 0.76914251 | 0.83514962 | NA | NA |
| ENSG000002 | 10.4479637 | -0.098777971 | 0.33675201 | -0.2933256 | 0.76927332 | 0.83524693 | NA | NA |
| ENSG000002 | 27.3461304 | 0.075592187 | 0.25769597 | 0.29333864 | 0.76926332 | 0.83524693 | NA | NA |
| ENSG000001 | 158.293378 | 0.048966494 | 0.16699211 | 0.29322639 | 0.76934912 | 0.83530686 | EXOG | 9941 |
| ENSG000002 | 8.7844659 | 0.10112154 | 0.34497878 | 0.29312394 | 0.76942742 | 0.83536951 | NA | NA |

| | | | | | | | | |
|---|---|---|---|---|---|---|---|---|
| ENSG000002 | 6.38033857 | -0.106487908 | 0.3637741 | -0.2927309 | 0.76972787 | 0.83567334 | LINC01030 | 101937451 |
| ENSG000002 | 2.26004563 | 0.10728003 | 0.3665372 | 0.29268524 | 0.76976275 | 0.83568883 | NA | NA |
| ENSG000001 | 13.5834357 | 0.091261969 | 0.31187811 | 0.29262063 | 0.76981215 | 0.83571573 | NA | NA |
| ENSG000002 | 26.9299707 | 0.077726296 | 0.2656411 | 0.29259891 | 0.76982875 | 0.83571573 | NA | NA |
| ENSG000001 | 29.8327848 | -0.073574404 | 0.25147599 | -0.2925703 | 0.76985062 | 0.83571711 | SERPINF2 | 5345 |
| ENSG000002 | 4.04207825 | -0.110368155 | 0.37727808 | -0.2925379 | 0.76987535 | 0.83572158 | NA | NA |
| ENSG000002 | 13.4404388 | -0.090570579 | 0.309857 | -0.292298 | 0.77005879 | 0.83587596 | NA | NA |
| ENSG000002 | 3.98233156 | -0.110645399 | 0.37852771 | -0.2923046 | 0.77005373 | 0.83587596 | MIR4677 | 100616343 |
| ENSG000002 | 3.43380511 | 0.110539182 | 0.37831598 | 0.29218745 | 0.7701433 | 0.83594532 | NA | NA |
| ENSG000002 | 5.49545437 | -0.110143877 | 0.37726755 | -0.2919516 | 0.77032361 | 0.83611864 | DNAJB5-AS1 | 101926900 |
| ENSG000001 | 18.1481095 | -0.085801071 | 0.29396154 | -0.2918786 | 0.77037948 | 0.83615184 | IQCD | 115811 |
| ENSG000002 | 11.7148623 | 0.094660881 | 0.32434897 | 0.29184887 | 0.77040218 | 0.83615184 | FAM229A | 100128071 |
| ENSG000002 | 23.1800379 | -0.079803402 | 0.27345785 | -0.2918307 | 0.77041605 | 0.83615184 | NA | NA |
| ENSG000002 | 21.5850388 | -0.0809308 | 0.277393 | -0.291755 | 0.77047394 | 0.83619229 | NA | NA |
| ENSG000002 | 3.64262296 | 0.110365097 | 0.3783858 | 0.29167346 | 0.7705363 | 0.83623759 | NA | NA |
| ENSG000002 | 9.23678323 | 0.098590081 | 0.33820084 | 0.29151341 | 0.77065869 | 0.83634803 | NA | NA |
| ENSG000002 | 2.28158861 | 0.104629344 | 0.35940455 | 0.29111859 | 0.77096063 | 0.83665331 | NA | NA |
| ENSG000002 | 10.6024826 | 0.096521529 | 0.33166349 | 0.29102248 | 0.77103413 | 0.83671069 | NA | NA |
| ENSG000002 | 5.22102127 | -0.108496134 | 0.37305559 | -0.290831 | 0.77118056 | 0.8368472 | NA | NA |
| ENSG000002 | 24.3690655 | 0.077281206 | 0.26579247 | 0.2907577 | 0.77123664 | 0.83688566 | NA | NA |
| ENSG000002 | 2.5512271 | -0.106076975 | 0.36520684 | -0.2904572 | 0.77146645 | 0.83711263 | TTLL10-AS1 | 100506376 |
| ENSG000001 | 14.7590994 | 0.091253062 | 0.31421768 | 0.29041352 | 0.7714999 | 0.83712653 | RHBDL1 | 9028 |
| ENSG000001 | 104.248539 | 0.047163697 | 0.16245009 | 0.29032732 | 0.77156584 | 0.83717567 | QPCT | 25797 |
| ENSG000001 | 21.9733344 | -0.079883156 | 0.27548165 | -0.2899763 | 0.77183435 | 0.83743367 | NA | NA |
| ENSG000002 | 12.2263773 | -0.095609741 | 0.32973197 | -0.289962 | 0.77184531 | 0.83743367 | NA | NA |
| ENSG000002 | 5.94554978 | -0.107265129 | 0.36996202 | -0.2899355 | 0.77186557 | 0.83743367 | NA | NA |
| ENSG000002 | 10.4499608 | 0.096289258 | 0.33251091 | 0.28958225 | 0.77213585 | 0.83770449 | LINC00601 | 101101772 |
| ENSG000002 | 21.9782515 | 0.080313148 | 0.27769141 | 0.28921726 | 0.77241512 | 0.83797092 | COLCA2 | 120376 |
| ENSG000002 | 2.30005712 | 0.107199109 | 0.3706653 | 0.2892073 | 0.77242275 | 0.83797092 | NA | NA |

| | | | | | | | | |
|---|---|---|---|---|---|---|---|---|
| ENSG000002 | 2.32006096 | 0.104737398 | 0.36256121 | 0.28888197 | 0.7726717 | 0.83821857 | MIR3920 | 100500823 |
| ENSG000001 | 62.7825256 | -0.058300313 | 0.20189119 | -0.288771 | 0.77275665 | 0.83824562 | CER1 | 9350 |
| ENSG000002 | 14.8301864 | 0.088925728 | 0.30794832 | 0.28876835 | 0.77275865 | 0.83824562 | NA | NA |
| ENSG000002 | 9.72234564 | -0.098930222 | 0.34255216 | -0.2888034 | 0.77273185 | 0.83824562 | NA | NA |
| ENSG000001 | 6.26661049 | 0.106605496 | 0.36932446 | 0.28864997 | 0.77284925 | 0.83827956 | Mar-11 | 441061 |
| ENSG000002 | 26.873195 | -0.073890686 | 0.25599029 | -0.2886464 | 0.77285196 | 0.83827956 | ERICH6-AS1 | 101928085 |
| ENSG000002 | 3.40146454 | 0.109241447 | 0.37844057 | 0.28866209 | 0.77283997 | 0.83827956 | NA | NA |
| ENSG000002 | 2.2901534 | 0.106530442 | 0.3693248 | 0.28844649 | 0.77300499 | 0.83842312 | NA | NA |
| ENSG00000C | 21.5670117 | 0.083618341 | 0.29029552 | 0.28804558 | 0.77331185 | 0.83873352 | F7 | 2155 |
| ENSG000002 | 3.39888132 | 0.108907712 | 0.37825095 | 0.28792449 | 0.77340454 | 0.83881161 | LOC105372582 | 105372582 |
| ENSG00000C | 30.8266381 | 0.073790645 | 0.25640754 | 0.28778657 | 0.77351012 | 0.83890368 | SCARF1 | 8578 |
| ENSG000001 | 5.18435062 | -0.107178858 | 0.37290976 | -0.2874123 | 0.77379663 | 0.83913556 | CYLC1 | 1538 |
| ENSG000001 | 6.54706948 | -0.104184686 | 0.36250864 | -0.2873992 | 0.77380669 | 0.83913556 | NA | NA |
| ENSG000002 | 8.68336162 | 0.100240467 | 0.3487522 | 0.28742605 | 0.77378612 | 0.83913556 | NA | NA |
| ENSG000002 | 6.3015793 | -0.106078751 | 0.36907056 | -0.2874213 | 0.77378973 | 0.83913556 | NA | NA |
| ENSG000001 | 36.2663088 | -0.066216192 | 0.2306183 | -0.2871246 | 0.7740169 | 0.83932637 | SLC16A3 | 9123 |
| ENSG000002 | 6.54521517 | -0.105159306 | 0.3662616 | -0.2871153 | 0.77402404 | 0.83932637 | NA | NA |
| ENSG000002 | 3.49455787 | 0.108058598 | 0.37688608 | 0.28671422 | 0.77433115 | 0.83963694 | NA | NA |
| ENSG000002 | 2.56682413 | 0.105763843 | 0.36921045 | 0.28645951 | 0.7745262 | 0.83982598 | NA | NA |
| ENSG000002 | 211.96153 | -0.043155176 | 0.15071914 | -0.2863284 | 0.77462657 | 0.83989253 | ANKRD18B | 441459 |
| ENSG000002 | 7.48890679 | 0.103071889 | 0.35998181 | 0.28632527 | 0.774629 | 0.83989253 | NA | NA |
| ENSG000002 | 5.63100317 | -0.106148879 | 0.37076759 | -0.2862949 | 0.77465225 | 0.83989528 | NA | NA |
| ENSG000002 | 7.67002964 | 0.100452142 | 0.35114316 | 0.28607176 | 0.77482316 | 0.84003566 | NA | NA |
| ENSG000002 | 4.77090205 | 0.107689946 | 0.37641646 | 0.28609255 | 0.77480724 | 0.84003566 | NA | NA |
| ENSG000002 | 4.96501524 | -0.107076813 | 0.37440355 | -0.285993 | 0.77488345 | 0.84007857 | NA | NA |
| ENSG000002 | 18.2776588 | 0.082615247 | 0.28898473 | 0.28588101 | 0.77496926 | 0.84014914 | NA | NA |
| ENSG000002 | 6.28661813 | 0.104154658 | 0.36440385 | 0.28582205 | 0.77501442 | 0.84017563 | NA | NA |
| ENSG000001 | 64.77463 | -0.05573526 | 0.19506756 | -0.2857229 | 0.7750904 | 0.84023554 | SYNM | 23336 |
| ENSG000001 | 46.1365675 | -0.061083348 | 0.21383545 | -0.2856559 | 0.77514172 | 0.84026871 | RGPD3 | 653489 |

| | | | | | | | | |
|---|---|---|---|---|---|---|---|---|
| ENSG000002 | 15.9850626 | -0.088338639 | 0.30929435 | -0.2856135 | 0.77517417 | 0.84028143 | NA | NA |
| ENSG000001 | 2.52245309 | 0.106194927 | 0.37186503 | 0.28557384 | 0.77520454 | 0.84029188 | BIRC7 | 79444 |
| ENSG000002 | 3.63515922 | 0.108100628 | 0.37859301 | 0.28553255 | 0.77523617 | 0.84030371 | NA | NA |
| ENSG000002 | 2.55378735 | 0.106970101 | 0.37467931 | 0.28549775 | 0.77526283 | 0.84031014 | NA | NA |
| ENSG000002 | 18.9127931 | -0.085535638 | 0.29967509 | -0.2854279 | 0.77531632 | 0.84034566 | NA | NA |
| ENSG000001 | 19.6939201 | 0.082920538 | 0.29085304 | 0.28509428 | 0.77557191 | 0.84060022 | NECAB2 | 54550 |
| ENSG000002 | 10.1503206 | 0.096077581 | 0.33706034 | 0.28504565 | 0.77560917 | 0.84061813 | NA | NA |
| ENSG000002 | 2.27477666 | 0.104694777 | 0.36734567 | 0.28500343 | 0.77564151 | 0.84063072 | NA | NA |
| ENSG000002 | 5.36018897 | -0.10570537 | 0.37095672 | -0.2849534 | 0.77567986 | 0.84064981 | NA | NA |
| ENSG000001 | 9.40103834 | 0.100894206 | 0.35418517 | 0.28486287 | 0.7757492 | 0.84070249 | MYLPF | 29895 |
| ENSG000001 | 35.5123928 | 0.066705976 | 0.23430424 | 0.28469812 | 0.77587543 | 0.84079435 | CNRIP1 | 25927 |
| ENSG000001 | 13.1845043 | -0.090644564 | 0.31837165 | -0.284713 | 0.77586399 | 0.84079435 | PPP1R14C | 81706 |
| ENSG000001 | 5.61094312 | -0.105013916 | 0.36899014 | -0.2845982 | 0.77595202 | 0.84083241 | CD79A | 973 |
| ENSG000002 | 6.06930538 | 0.104026343 | 0.36549858 | 0.28461491 | 0.77593919 | 0.84083241 | NA | NA |
| ENSG000001 | 98.3302228 | 0.050300829 | 0.1768548 | 0.2844188 | 0.77608946 | 0.84095886 | IL1RAP | 3556 |
| ENSG000002 | 7.94697282 | 0.09985574 | 0.35125006 | 0.28428675 | 0.77619064 | 0.84104603 | LOC100506368 | 100506368 |
| ENSG000002 | 4.72460013 | 0.106568083 | 0.37493846 | 0.2842282 | 0.77623551 | 0.84107217 | MIR616 | 693201 |
| ENSG00000C | 359.265018 | 0.038812965 | 0.13663285 | 0.2840676 | 0.77635858 | 0.84118304 | ADAM22 | 53616 |
| ENSG000002 | 11.7501626 | -0.092686863 | 0.32635504 | -0.2840062 | 0.77640561 | 0.84121153 | RFPL3S | 10737 |
| ENSG000002 | 3.65139771 | 0.107474914 | 0.37853179 | 0.28392572 | 0.7764673 | 0.84123342 | NA | NA |
| ENSG000002 | 7.16970042 | 0.101948108 | 0.35904028 | 0.28394616 | 0.77645164 | 0.84123342 | NA | NA |
| ENSG000002 | 16.8226552 | 0.08608905 | 0.30325681 | 0.28388167 | 0.77650106 | 0.84124752 | NA | NA |
| ENSG000001 | 44.2597997 | -0.061997146 | 0.21847595 | -0.283771 | 0.77658585 | 0.84124956 | CRB2 | 286204 |
| ENSG000002 | 2.51784615 | 0.105969119 | 0.3733947 | 0.28379921 | 0.77656426 | 0.84124956 | NA | NA |
| ENSG000002 | 32.9078858 | 0.070162856 | 0.2471958 | 0.28383515 | 0.77653672 | 0.84124956 | NA | NA |
| ENSG000002 | 16.5783345 | -0.085456418 | 0.30114578 | -0.2837709 | 0.77658593 | 0.84124956 | NA | NA |
| ENSG000002 | 5.41518056 | -0.105419651 | 0.37160993 | -0.2836836 | 0.77665285 | 0.84129958 | NA | NA |
| ENSG000002 | 2.33275697 | 0.104011199 | 0.36670268 | 0.28363905 | 0.77668701 | 0.84131411 | NA | NA |
| ENSG00000C | 25.9100437 | 0.075821531 | 0.26748649 | 0.28345929 | 0.77682478 | 0.84144087 | SH2D3C | 10044 |

| | | | | | | | | |
|---|---|---|---|---|---|---|---|---|
| ENSG000001 | 126.474754 | 0.044818618 | 0.15814937 | 0.28339422 | 0.77687466 | 0.84147241 | SLC10A7 | 84068 |
| ENSG000002 | 9.38383609 | -0.096500915 | 0.34064462 | -0.2832891 | 0.77695522 | 0.84149224 | NA | NA |
| ENSG000002 | 14.9678327 | -0.089675394 | 0.31652039 | -0.2833163 | 0.77693436 | 0.84149224 | LOC643770 | 643770 |
| ENSG000002 | 17.8993333 | 0.083676602 | 0.29537253 | 0.28329175 | 0.7769532 | 0.84149224 | NA | NA |
| ENSG000002 | 35.4665538 | -0.067524301 | 0.23842274 | -0.2832125 | 0.77701395 | 0.84153337 | LINC00894 | 100272228 |
| ENSG000002 | 6.08610448 | 0.103491226 | 0.36552512 | 0.28313027 | 0.77707698 | 0.84157916 | NA | NA |
| ENSG000001 | 15.7082256 | 0.08788374 | 0.31065743 | 0.28289598 | 0.77725657 | 0.84175118 | KCNK15 | 60598 |
| ENSG000002 | 5.66804075 | -0.106148301 | 0.37536712 | -0.2827853 | 0.77734143 | 0.8418206 | NA | NA |
| ENSG000002 | 46.0231167 | -0.065935699 | 0.23322283 | -0.2827155 | 0.77739496 | 0.84185609 | NA | NA |
| ENSG000002 | 3.87533228 | 0.10683415 | 0.37795962 | 0.28266022 | 0.77743731 | 0.84187947 | LOC387720 | 387720 |
| ENSG000002 | 2.52711679 | 0.105713103 | 0.37418147 | 0.28251828 | 0.77754613 | 0.84197483 | NA | NA |
| ENSG000002 | 3.46801075 | 0.106878289 | 0.37840113 | 0.28244706 | 0.77760073 | 0.84201146 | NA | NA |
| ENSG000002 | 2.3076895 | -0.102079596 | 0.36185949 | -0.2820973 | 0.77786888 | 0.84227933 | NA | NA |
| ENSG000002 | 4.68933852 | -0.105978702 | 0.37578421 | -0.2820201 | 0.7779281 | 0.84232096 | NA | NA |
| ENSG000002 | 20.8496338 | -0.07831602 | 0.27803849 | -0.2816733 | 0.77819402 | 0.8425639 | NA | NA |
| ENSG000002 | 5.70040868 | -0.104304176 | 0.37028986 | -0.2816825 | 0.77818697 | 0.8425639 | REV3L-IT1 | 106478974 |
| ENSG000001 | 62.1882101 | 0.056209406 | 0.19958229 | 0.28163523 | 0.77822322 | 0.84256453 | IGSF10 | 285313 |
| ENSG000002 | 5.43346092 | -0.104593791 | 0.37140259 | -0.2816184 | 0.77823616 | 0.84256453 | NA | NA |
| ENSG000001 | 15.2885295 | -0.084864276 | 0.30156758 | -0.2814105 | 0.77839558 | 0.8427127 | TMEM171 | 134285 |
| ENSG000002 | 2.33106683 | 0.102210779 | 0.36324083 | 0.28138571 | 0.77841458 | 0.8427127 | NA | NA |
| ENSG000002 | 12.2798165 | -0.089569441 | 0.31863213 | -0.2811061 | 0.77862901 | 0.84285584 | NA | NA |
| ENSG000002 | 11.0696746 | -0.095315019 | 0.3390262 | -0.2811435 | 0.77860032 | 0.84285584 | NA | NA |
| ENSG000002 | 14.4579403 | -0.089216348 | 0.31735114 | -0.2811282 | 0.77861209 | 0.84285584 | NA | NA |
| ENSG000002 | 4.72300359 | -0.105819682 | 0.37644197 | -0.2811049 | 0.77862994 | 0.84285584 | NA | NA |
| ENSG000002 | 4.71615344 | -0.106016475 | 0.3772656 | -0.2810128 | 0.77870056 | 0.84290978 | NA | NA |
| ENSG000002 | 5.35241236 | -0.104144552 | 0.37067842 | -0.2809566 | 0.77874368 | 0.84293396 | NA | NA |
| ENSG000002 | 5.01856467 | -0.106158977 | 0.37821845 | -0.2806816 | 0.77895459 | 0.84313975 | LOC105373495 | 105373495 |
| ENSG000002 | 2.57280604 | 0.104162275 | 0.37115441 | 0.28064404 | 0.77898343 | 0.84314846 | NA | NA |
| ENSG000002 | 47.3447673 | -0.059161822 | 0.21085394 | -0.280582 | 0.77903101 | 0.84317746 | NA | NA |

| | | | | | | | | |
|---|---|---|---|---|---|---|---|---|
| ENSG000002 | 3.58498508 | -0.104665616 | 0.37309389 | -0.2805343 | 0.77906765 | 0.84319461 | NA | NA |
| ENSG000002 | 2.29237747 | -0.10207081 | 0.36393746 | -0.2804625 | 0.77912269 | 0.84323167 | NA | NA |
| ENSG000001 | 14.2572799 | 0.089138138 | 0.31801045 | 0.2802994 | 0.77924781 | 0.84334458 | PQLC2L | 152078 |
| ENSG000001 | 14.0399297 | 0.088812637 | 0.31728688 | 0.27991273 | 0.77954446 | 0.84364153 | S100A1 | 6271 |
| ENSG000002 | 55.7994815 | -0.056007274 | 0.20010636 | -0.2798875 | 0.7795638 | 0.84364153 | NA | NA |
| ENSG000002 | 3.2155863 | 0.105145815 | 0.37570996 | 0.27985901 | 0.77958568 | 0.84364269 | NA | NA |
| ENSG000002 | 20.6447158 | 0.079995911 | 0.28603278 | 0.27967393 | 0.77972768 | 0.84377384 | TCF24 | 100129654 |
| ENSG000000 | 43.4440222 | -0.060924868 | 0.21804371 | -0.2794158 | 0.77992571 | 0.84396562 | KCNAB2 | 8514 |
| ENSG000002 | 5.15080972 | -0.103915685 | 0.37195416 | -0.2793777 | 0.77995501 | 0.8439748 | NA | NA |
| ENSG000002 | 3.8873633 | 0.105707157 | 0.37844166 | 0.2793222 | 0.77999757 | 0.84399833 | NA | NA |
| ENSG000001 | 87.9800541 | 0.049886881 | 0.17867382 | 0.27920644 | 0.7800864 | 0.84407193 | SYN2 | 6854 |
| ENSG000002 | 4.94837349 | -0.104612501 | 0.37475993 | -0.2791454 | 0.78013326 | 0.84410011 | NA | NA |
| ENSG000002 | 2.33543771 | 0.102056292 | 0.36619386 | 0.27869471 | 0.78047912 | 0.84445179 | NA | NA |
| ENSG000002 | 5.14217181 | -0.104075096 | 0.37359658 | -0.2785761 | 0.78057012 | 0.84452772 | NA | NA |
| ENSG000002 | 5.64289836 | -0.102945004 | 0.37007508 | -0.2781733 | 0.78087933 | 0.84483973 | NA | NA |
| ENSG000002 | 8.53702542 | 0.098490553 | 0.35423273 | 0.27803911 | 0.78098234 | 0.84492863 | NA | NA |
| ENSG000002 | 4.65347642 | -0.104814222 | 0.37706329 | -0.2779751 | 0.78103144 | 0.8449592 | SNORA80E | 677823 |
| ENSG000002 | 18.7911825 | -0.078810008 | 0.28383377 | -0.2776625 | 0.78127141 | 0.84519627 | CDC37L1-AS1 | 101929351 |
| ENSG000002 | 4.43039117 | -0.104677445 | 0.3770668 | -0.2776098 | 0.78131189 | 0.84521752 | NA | NA |
| ENSG000002 | 28.4862865 | 0.07190069 | 0.25906107 | 0.27754339 | 0.78136289 | 0.84525014 | HLA-F-AS1 | 285830 |
| ENSG000002 | 3.87796846 | 0.104993689 | 0.37858675 | 0.27733059 | 0.78152627 | 0.84540432 | NA | NA |
| ENSG000002 | 7.92291794 | 0.097753329 | 0.35260357 | 0.27723295 | 0.78160124 | 0.84544031 | NA | NA |
| ENSG000002 | 9.32405013 | 0.095564457 | 0.34468183 | 0.27725412 | 0.78158498 | 0.84544031 | NA | NA |
| ENSG000002 | 96.4880707 | -0.053655287 | 0.19359322 | -0.2771548 | 0.78166126 | 0.84548268 | NA | NA |
| ENSG000002 | 25.9520114 | 0.07231906 | 0.26109764 | 0.2769809 | 0.78179477 | 0.84560454 | NA | NA |
| ENSG000002 | 9.63976555 | -0.093415808 | 0.33737099 | -0.2768934 | 0.78186194 | 0.84565464 | SNORD63 | 26785 |
| ENSG000002 | 4.98518164 | 0.103642183 | 0.37437289 | 0.27684212 | 0.78190134 | 0.84567469 | NA | NA |
| ENSG000002 | 15.7501793 | 0.082918334 | 0.29963662 | 0.27672964 | 0.78198771 | 0.84574555 | NA | NA |
| ENSG000002 | 5.21960722 | -0.103102207 | 0.37268693 | -0.2766456 | 0.78205223 | 0.84579277 | LOC102724539 | 102724539 |

| | | | | | | | | |
|---|---|---|---|---|---|---|---|---|
| ENSG000001 | 297.497901 | -0.037612314 | 0.13598943 | -0.2765826 | 0.7821006 | 0.84582253 | DDX25 | 29118 |
| ENSG000001 | 73.2887334 | 0.051128714 | 0.1849145 | 0.27649921 | 0.78216466 | 0.84585321 | SIAE | 54414 |
| ENSG000002 | 47.2464964 | -0.063331835 | 0.22906424 | -0.2764807 | 0.7821789 | 0.84585321 | NA | NA |
| ENSG000002 | 5.14310579 | -0.10265731 | 0.37132225 | -0.2764642 | 0.78219155 | 0.84585321 | NA | NA |
| ENSG000002 | 8.47628549 | -0.097321909 | 0.35208971 | -0.2764123 | 0.78223145 | 0.84586147 | NA | NA |
| ENSG000002 | 4.48514743 | -0.10411836 | 0.37671806 | -0.2763827 | 0.78225413 | 0.84586147 | LOC101927230 | 101927230 |
| ENSG000002 | 7.46424406 | 0.098938653 | 0.35798987 | 0.27637277 | 0.78226177 | 0.84586147 | NA | NA |
| ENSG000002 | 4.98728304 | -0.103891547 | 0.37605569 | -0.2762664 | 0.78234347 | 0.84592726 | NA | NA |
| ENSG000002 | 5.55444652 | -0.104457084 | 0.37824348 | -0.2761636 | 0.78242241 | 0.84598517 | NA | NA |
| ENSG000002 | 7.9872905 | -0.097612252 | 0.3534853 | -0.2761423 | 0.78243875 | 0.84598517 | NA | NA |
| ENSG000002 | 10.1573924 | 0.092601704 | 0.33566277 | 0.2758772 | 0.78264239 | 0.84618278 | NA | NA |
| ENSG000002 | 10.4345211 | -0.093613258 | 0.33973389 | -0.2755488 | 0.78289466 | 0.84643297 | NA | NA |
| ENSG000001 | 11.5488294 | -0.088633172 | 0.3217043 | -0.2755113 | 0.78292344 | 0.84644151 | ARMC12 | 221481 |
| ENSG000001 | 26.3530101 | 0.070432159 | 0.2556911 | 0.27545799 | 0.78296439 | 0.84646323 | RAI2 | 10742 |
| ENSG000002 | 6.34463111 | 0.101680033 | 0.36921692 | 0.27539375 | 0.78301374 | 0.84649401 | NA | NA |
| ENSG000002 | 5.0132864 | 0.102499673 | 0.37224455 | 0.27535574 | 0.78304294 | 0.84650301 | NA | NA |
| ENSG000002 | 4.94040641 | 0.103354079 | 0.37554951 | 0.2752076 | 0.78315675 | 0.84660347 | NA | NA |
| ENSG000002 | 6.85747336 | -0.100731037 | 0.3660992 | -0.2751468 | 0.78320343 | 0.84663136 | NA | NA |
| ENSG000001 | 4.67511135 | -0.1036026 | 0.37668902 | -0.2750348 | 0.78328948 | 0.8466599 | NA | NA |
| ENSG000002 | 7.80298219 | -0.099942787 | 0.36338743 | -0.2750309 | 0.78329247 | 0.8466599 | KANSL1-AS1 | 644246 |
| ENSG000002 | 5.17064055 | -0.103991364 | 0.37809472 | -0.2750405 | 0.78328512 | 0.8466599 | NA | NA |
| ENSG000002 | 12.8151486 | -0.087702898 | 0.31893616 | -0.2749857 | 0.78332719 | 0.84667487 | GAPLINC | 100505592 |
| ENSG00000C | 62.2057223 | -0.058497649 | 0.21284899 | -0.2748317 | 0.78344554 | 0.84674296 | MOXD1 | 26002 |
| ENSG000002 | 4.64977141 | -0.102983811 | 0.37472804 | -0.2748228 | 0.78345237 | 0.84674296 | MIR221 | 407006 |
| ENSG000002 | 2.31141509 | 0.10151912 | 0.36939926 | 0.27482221 | 0.78345283 | 0.84674296 | NA | NA |
| ENSG00000C | 62.4700899 | 0.054316507 | 0.197754 | 0.27466704 | 0.78357205 | 0.84678154 | XYLB | 9942 |
| ENSG000001 | 8.55822033 | 0.098415001 | 0.35820424 | 0.2747455 | 0.78351177 | 0.84678154 | MGARP | 84709 |
| ENSG000001 | 16.9375339 | -0.080643939 | 0.29358128 | -0.2746903 | 0.78355416 | 0.84678154 | FOXI2 | 399823 |
| ENSG000002 | 2.35621034 | 0.101000011 | 0.36769623 | 0.27468329 | 0.78355957 | 0.84678154 | NA | NA |

| | | | | | | | | |
|---|---|---|---|---|---|---|---|---|
| ENSG000002 | 4.87991621 | -0.102348519 | 0.37284749 | -0.2745051 | 0.78369652 | 0.84689347 | NA | NA |
| ENSG000002 | 12.7413188 | -0.090782808 | 0.33088183 | -0.2743663 | 0.78380317 | 0.84698615 | LOC102725045 | 102725045 |
| ENSG000002 | 4.65957063 | -0.102824756 | 0.3750784 | -0.274142 | 0.78397547 | 0.84714977 | NA | NA |
| ENSG000002 | 6.09934202 | 0.101218199 | 0.3693119 | 0.2740724 | 0.78402899 | 0.84718502 | NA | NA |
| ENSG000001 | 39.5586895 | -0.062709332 | 0.22891028 | -0.2739472 | 0.78412519 | 0.84726035 | ADAMTSL4 | 54507 |
| ENSG000002 | 11.3359462 | 0.09306946 | 0.3397597 | 0.2739273 | 0.78414049 | 0.84726035 | NA | NA |
| ENSG000002 | 8.16179574 | 0.09882019 | 0.36118669 | 0.27359865 | 0.78439307 | 0.8474881 | NA | NA |
| ENSG000002 | 4.19718651 | -0.103393853 | 0.37787806 | -0.273617 | 0.78437899 | 0.8474881 | WFDC6 | 140870 |
| ENSG000002 | 4.95499564 | -0.102556789 | 0.37490989 | -0.2735505 | 0.78443008 | 0.8475055 | NA | NA |
| ENSG000002 | 13.3605199 | -0.08556327 | 0.31304462 | -0.2733261 | 0.78460254 | 0.84766925 | NA | NA |
| ENSG000002 | 4.41548408 | -0.10258265 | 0.37574468 | -0.2730116 | 0.78484431 | 0.84790787 | NA | NA |
| ENSG000002 | 2.30460403 | 0.100232529 | 0.36720096 | 0.27296369 | 0.78488113 | 0.84792505 | NA | NA |
| ENSG000002 | 2.55287353 | 0.102045454 | 0.37422291 | 0.27268628 | 0.78509438 | 0.84811024 | NA | NA |
| ENSG000002 | 4.65447531 | -0.102051332 | 0.3742343 | -0.2726937 | 0.78508868 | 0.84811024 | NA | NA |
| ENSG000002 | 12.1929496 | -0.086430115 | 0.31705552 | -0.2726025 | 0.78515882 | 0.84815726 | NA | NA |
| ENSG000001 | 63.5860076 | -0.053701318 | 0.19713775 | -0.272405 | 0.78531059 | 0.84828733 | SUFU | 51684 |
| ENSG000002 | 11.795128 | -0.087263054 | 0.32035905 | -0.2723914 | 0.78532107 | 0.84828733 | NA | NA |
| ENSG000001 | 7.90978477 | 0.095425892 | 0.350409 | 0.27232717 | 0.78537047 | 0.84831809 | CFD | 1675 |
| ENSG00000C | 56.2259536 | -0.053523027 | 0.1965705 | -0.2722841 | 0.78540356 | 0.84833124 | ADAMTS2 | 9509 |
| ENSG000002 | 18.4607528 | 0.079793252 | 0.29316441 | 0.27217919 | 0.78548424 | 0.84839034 | ATE1-AS1 | 100130887 |
| ENSG000002 | 6.21077626 | -0.0931346 | 0.34220716 | -0.2721585 | 0.78550012 | 0.84839034 | NA | NA |
| ENSG000002 | 12.2650756 | 0.089465703 | 0.32877219 | 0.27212066 | 0.78552925 | 0.8483992 | MROH6 | 642475 |
| ENSG000001 | 8.95430162 | 0.095805762 | 0.35212197 | 0.27208118 | 0.7855596 | 0.84840939 | PDCD1 | 5133 |
| ENSG000002 | 5.43226759 | -0.101318344 | 0.37256371 | -0.271949 | 0.7856612 | 0.84849652 | NA | NA |
| ENSG000001 | 92.4999952 | -0.046606046 | 0.17141546 | -0.2718894 | 0.78570707 | 0.84852346 | MAATS1 | 89876 |
| ENSG000001 | 136.162338 | -0.041422666 | 0.15238263 | -0.2718333 | 0.78575023 | 0.84854747 | ZNF559 | 84527 |
| ENSG000002 | 8.87518479 | -0.093240185 | 0.34317576 | -0.2716981 | 0.7858542 | 0.84863715 | HCG2040054 | 644093 |
| ENSG000001 | 1177.13357 | 0.035389957 | 0.13033538 | 0.27152993 | 0.78598348 | 0.84875416 | VSNL1 | 7447 |
| ENSG000002 | 3.61212254 | 0.102707101 | 0.37841721 | 0.27141234 | 0.78607391 | 0.8488292 | LOC101928167 | 101928167 |

| | | | | | | | | |
|---|---|---|---|---|---|---|---|---|
| ENSG000002 | 2.59206438 | 0.101102281 | 0.37257121 | 0.27136365 | 0.78611136 | 0.84884704 | NA | NA |
| ENSG000002 | 4.41013992 | -0.102232801 | 0.37710825 | -0.2710967 | 0.78631666 | 0.84904612 | NA | NA |
| ENSG000002 | 4.22731466 | -0.102371802 | 0.37815708 | -0.2707124 | 0.78661226 | 0.84934268 | NA | NA |
| ENSG000002 | 2.33400272 | 0.099011293 | 0.36592046 | 0.27058146 | 0.78671295 | 0.84941428 | NA | NA |
| ENSG000002 | 6.59050454 | 0.097526323 | 0.3604454 | 0.2705717 | 0.78672047 | 0.84941428 | NA | NA |
| ENSG000002 | 3.67123705 | 0.102405017 | 0.37856913 | 0.27050546 | 0.78677142 | 0.84944668 | LOC101929180 | 101929180 |
| ENSG000002 | 3.51170416 | -0.102093192 | 0.37773657 | -0.2702762 | 0.7869478 | 0.84961449 | LOC105369984 | 105369984 |
| ENSG000002 | 5.58539411 | -0.100430108 | 0.37176214 | -0.2701461 | 0.78704782 | 0.84969986 | NA | NA |
| ENSG000002 | 4.66451487 | -0.101721482 | 0.37670978 | -0.2700261 | 0.78714015 | 0.84977691 | NA | NA |
| ENSG000002 | 5.215013 | 0.100512123 | 0.37229817 | 0.26997748 | 0.78717758 | 0.84979469 | NA | NA |
| ENSG000001 | 25.0570611 | -0.073938697 | 0.27409147 | -0.2697592 | 0.78734551 | 0.84995335 | ADGRE5 | 976 |
| ENSG000002 | 2.53505682 | 0.100343707 | 0.37249377 | 0.26938358 | 0.78763452 | 0.85024271 | NA | NA |
| ENSG000001 | 4.66563219 | -0.101228687 | 0.375833 | -0.2693449 | 0.78766431 | 0.85025224 | NA | NA |
| ENSG000002 | 5.7296434 | -0.100528114 | 0.37356091 | -0.2691077 | 0.78784681 | 0.85041209 | NA | NA |
| ENSG000002 | 3.91993165 | 0.101847881 | 0.37847891 | 0.26909791 | 0.78785434 | 0.85041209 | NA | NA |
| ENSG000002 | 4.87292933 | -0.100509279 | 0.37358817 | -0.2690376 | 0.78790072 | 0.85043951 | NA | NA |
| ENSG000002 | 6.5669593 | 0.0984378 | 0.3659338 | 0.2690044 | 0.7879263 | 0.85044449 | NA | NA |
| ENSG000002 | 6.76357076 | 0.097092125 | 0.36122616 | 0.26878487 | 0.78809524 | 0.85060419 | NA | NA |
| ENSG000002 | 4.69038426 | -0.100550156 | 0.37414968 | -0.2687431 | 0.78812736 | 0.85061622 | NA | NA |
| ENSG000001 | 23.2183581 | -0.070907227 | 0.26418468 | -0.2684002 | 0.78839127 | 0.85086756 | ZNF524 | 147807 |
| ENSG000002 | 15.8490151 | -0.081280178 | 0.30286453 | -0.2683714 | 0.78841345 | 0.85086756 | IRF9 | 10379 |
| ENSG000002 | 7.90918451 | 0.094196953 | 0.3510113 | 0.26835875 | 0.78842319 | 0.85086756 | NA | NA |
| ENSG000001 | 61.903696 | -0.054469685 | 0.2031082 | -0.2681806 | 0.78856029 | 0.85095975 | ARMC2 | 84071 |
| ENSG000001 | 42.6483338 | 0.061193687 | 0.22824308 | 0.26810752 | 0.78861655 | 0.85095975 | SLITRK6 | 84189 |
| ENSG000002 | 6.58830511 | 0.096468539 | 0.3598442 | 0.26808418 | 0.78863452 | 0.85095975 | NA | NA |
| ENSG000002 | 4.92869777 | -0.100289953 | 0.37402007 | -0.2681406 | 0.78859112 | 0.85095975 | NA | NA |
| ENSG000002 | 4.21058747 | -0.101297139 | 0.37771691 | -0.2681827 | 0.78855869 | 0.85095975 | NA | NA |
| ENSG000002 | 2.53512187 | 0.099532439 | 0.37121985 | 0.26812262 | 0.78860493 | 0.85095975 | NA | NA |
| ENSG000002 | 2.28276158 | 0.097744961 | 0.36470299 | 0.26801251 | 0.78868969 | 0.85099664 | MIR4705 | 100616239 |

| | | | | | | | | |
|---|---|---|---|---|---|---|---|---|
| ENSG000001 | 5.18071586 | 0.100067016 | 0.37344694 | 0.26795511 | 0.78873387 | 0.85102167 | IGFLR1 | 79713 |
| ENSG000001 | 166.774092 | 0.047105436 | 0.17595969 | 0.26770583 | 0.78892576 | 0.85120606 | LINGO1 | 84894 |
| ENSG000002 | 5.44457129 | -0.10006136 | 0.37381801 | -0.267674 | 0.78895027 | 0.85120985 | NA | NA |
| ENSG000001 | 42.0597611 | -0.059843994 | 0.22381571 | -0.2673807 | 0.78917608 | 0.85143084 | PSMD9 | 5715 |
| ENSG000002 | 41.4848401 | -0.057558149 | 0.21533714 | -0.2672932 | 0.78924343 | 0.85145935 | NA | NA |
| ENSG000002 | 4.80968762 | 0.100457428 | 0.37587268 | 0.26726451 | 0.78926551 | 0.85145935 | NA | NA |
| ENSG000002 | 26.0572091 | -0.067262226 | 0.25165992 | -0.2672743 | 0.78925799 | 0.85145935 | GHET1 | 102723099 |
| ENSG000001 | 45.0332545 | 0.057536647 | 0.21536459 | 0.26715927 | 0.78934653 | 0.85150146 | DYDC2 | 84332 |
| ENSG000002 | 4.96803711 | -0.100428495 | 0.37588595 | -0.2671781 | 0.78933203 | 0.85150146 | NA | NA |
| ENSG000002 | 4.69958322 | -0.10027223 | 0.37542895 | -0.2670871 | 0.7894021 | 0.85153874 | STX16-NPEPL1 | 100534593 |
| ENSG000001 | 4.1069515 | 0.100819528 | 0.37753423 | 0.26704738 | 0.78943268 | 0.85154908 | GOLGA6C | 653641 |
| ENSG000002 | 20.6435105 | 0.072993032 | 0.27339368 | 0.26698873 | 0.78947784 | 0.85157078 | NA | NA |
| ENSG000002 | 5.83847942 | -0.098053034 | 0.36728563 | -0.2669667 | 0.7894948 | 0.85157078 | NA | NA |
| ENSG000002 | 17.4179283 | -0.083484287 | 0.31289041 | -0.2668164 | 0.78961054 | 0.85167297 | LOC101929333 | 101929333 |
| ENSG000001 | 23.7237769 | -0.069774497 | 0.26157809 | -0.2667444 | 0.78966595 | 0.85171008 | LGI4 | 163175 |
| ENSG000002 | 2.75933684 | 0.100041007 | 0.37523195 | 0.26661111 | 0.7897686 | 0.85179814 | NA | NA |
| ENSG000001 | 11.3058652 | -0.086947893 | 0.32623828 | -0.2665165 | 0.78984143 | 0.85185404 | CXCL1 | 2919 |
| ENSG000002 | 2.55538542 | 0.099804946 | 0.37469581 | 0.26636259 | 0.78995997 | 0.85195923 | NA | NA |
| ENSG000002 | 5.14942813 | -0.098986828 | 0.37173002 | -0.2662869 | 0.79001827 | 0.85199944 | NA | NA |
| ENSG000002 | 38.5931854 | -0.064689249 | 0.24307228 | -0.2661317 | 0.79013775 | 0.85210563 | NA | NA |
| ENSG000001 | 49.0403685 | 0.057316564 | 0.2154884 | 0.26598445 | 0.79025118 | 0.85220529 | CDADC1 | 81602 |
| ENSG000002 | 6.5125607 | 0.097241223 | 0.36575057 | 0.26586759 | 0.79034118 | 0.85227969 | MIR4296 | 100423041 |
| ENSG000000 | 13.5323371 | 0.084091636 | 0.3165645 | 0.26563824 | 0.79051783 | 0.85236374 | SPAG4 | 6676 |
| ENSG000001 | 30.9561268 | -0.066471974 | 0.25023198 | -0.2656414 | 0.7905154 | 0.85236374 | TMEM17 | 200728 |
| ENSG000002 | 4.73729889 | -0.100326941 | 0.37758842 | -0.2657045 | 0.7904668 | 0.85236374 | NA | NA |
| ENSG000002 | 5.55718466 | -0.099216383 | 0.37351355 | -0.2656299 | 0.79052422 | 0.85236374 | NA | NA |
| ENSG000002 | 3.95417705 | -0.100549479 | 0.37839005 | -0.2657297 | 0.79044738 | 0.85236374 | NA | NA |
| ENSG000001 | 5.12950392 | -0.098659186 | 0.37167163 | -0.2654472 | 0.790665 | 0.85245287 | DAND5 | 199699 |
| ENSG000002 | 2.49955085 | -0.064054062 | 0.24131209 | -0.2654407 | 0.79066995 | 0.85245287 | SUMO2P13 | 106480263 |

| | | | | | | | | |
|---|---|---|---|---|---|---|---|---|
| ENSG000002 | 2.56135489 | 0.099225166 | 0.37379098 | 0.26545629 | 0.79065798 | 0.85245287 | RN7SL49P | 106480935 |
| ENSG000002 | 3.9910045 | -0.100424371 | 0.37851302 | -0.2653129 | 0.79076846 | 0.85253641 | NA | NA |
| ENSG000001 | 4.41158188 | -0.100149589 | 0.37755423 | -0.2652588 | 0.79081009 | 0.85255862 | NA | NA |
| ENSG000002 | 4.92449556 | -0.099548896 | 0.37535741 | -0.265211 | 0.79084696 | 0.8525757 | NA | NA |
| ENSG000002 | 26.0445327 | 0.072966671 | 0.2751568 | 0.26518215 | 0.79086915 | 0.85257696 | YJEFN3 | 374887 |
| ENSG000001 | 4.94213044 | -0.099381415 | 0.37492893 | -0.2650673 | 0.79095758 | 0.85264963 | SBK2 | 646643 |
| ENSG000001 | 106.282606 | -0.04484989 | 0.16972906 | -0.264244 | 0.79159189 | 0.85331073 | LRRC34 | 151827 |
| ENSG000002 | 5.39186434 | -0.098261328 | 0.37190453 | -0.2642112 | 0.79161721 | 0.85331533 | CT45A10 | 102723631 |
| ENSG000002 | 4.39427446 | -0.099108778 | 0.3758555 | -0.2636885 | 0.79201995 | 0.85372676 | NA | NA |
| ENSG000002 | 16.4446966 | 0.082305718 | 0.31219556 | 0.26363513 | 0.79206108 | 0.85374841 | NA | NA |
| ENSG000001 | 5.27196904 | 0.098811436 | 0.37484431 | 0.26360661 | 0.79208307 | 0.85374941 | TSHB | 7252 |
| ENSG000002 | 4.42993525 | -0.098957437 | 0.37582787 | -0.2633052 | 0.79231534 | 0.85397708 | LINC01072 | 104355137 |
| ENSG000002 | 41.0057288 | 0.061610849 | 0.2340526 | 0.26323505 | 0.79236941 | 0.85401266 | LOC100133091 | 100133091 |
| ENSG000001 | 11.2974722 | -0.088526224 | 0.33644693 | -0.2631209 | 0.79245739 | 0.85408477 | NA | NA |
| ENSG000002 | 6.57220413 | 0.096548367 | 0.36712879 | 0.26298229 | 0.79256423 | 0.85417722 | NA | NA |
| ENSG000002 | 4.42601344 | -0.099039686 | 0.37681218 | -0.2628357 | 0.79267723 | 0.8542763 | NA | NA |
| ENSG000001 | 53.3771978 | 0.056767018 | 0.21600634 | 0.26280256 | 0.79270276 | 0.85428112 | NUDT13 | 25961 |
| ENSG000002 | 4.94605802 | -0.098262967 | 0.37406743 | -0.2626878 | 0.79279118 | 0.8543537 | NA | NA |
| ENSG000002 | 4.21060764 | -0.099242646 | 0.37788092 | -0.2626294 | 0.79283623 | 0.85437953 | NA | NA |
| ENSG000002 | 4.41957452 | -0.098853426 | 0.37653358 | -0.2625355 | 0.79290863 | 0.85443485 | HSPE1P26 | 106481677 |
| ENSG000001 | 13.5987095 | -0.082420647 | 0.31418686 | -0.26233 | 0.79306701 | 0.85458282 | HLA-DRB5 | 3127 |
| ENSG000002 | 3.73976163 | -0.099110789 | 0.37836599 | -0.2619442 | 0.79336444 | 0.85488059 | LOC102546298 | 102546298 |
| ENSG000002 | 25.5722773 | -0.069088798 | 0.26387038 | -0.2618285 | 0.79345363 | 0.85495399 | RORA-AS1 | 101928784 |
| ENSG000002 | 2.58221219 | 0.097628762 | 0.37351157 | 0.26138083 | 0.79379884 | 0.85530322 | NA | NA |
| ENSG000002 | 6.36949344 | 0.095416081 | 0.36544103 | 0.26109844 | 0.7940166 | 0.85551512 | NA | NA |
| ENSG000002 | 2.56027779 | 0.097592334 | 0.37395241 | 0.26097528 | 0.79411157 | 0.85559472 | NA | NA |
| ENSG000002 | 4.78644222 | 0.098242525 | 0.37652326 | 0.2609202 | 0.79415404 | 0.85561774 | NA | NA |
| ENSG000001 | 4.45307651 | -0.098004123 | 0.37585651 | -0.2607488 | 0.79428625 | 0.85572999 | NA | NA |
| ENSG000002 | 4.97998346 | -0.097766978 | 0.37497348 | -0.2607304 | 0.79430043 | 0.85572999 | NA | NA |

| | | | | | | | | |
|---|---|---|---|---|---|---|---|---|
| ENSG000001 | 9.62000497 | -0.089401749 | 0.34292989 | -0.2606998 | 0.79432403 | 0.85573268 | CPZ | 8532 |
| ENSG000002 | 5.17352154 | -0.097107581 | 0.37265215 | -0.2605851 | 0.79441252 | 0.85580527 | NA | NA |
| ENSG000001 | 10.5936324 | -0.086984202 | 0.33404412 | -0.2603973 | 0.79455729 | 0.85593849 | CST1 | 1469 |
| ENSG000002 | 5.11528122 | -0.097369211 | 0.37399707 | -0.2603475 | 0.79459572 | 0.85595715 | NA | NA |
| ENSG000002 | 13.8205118 | 0.080786181 | 0.31058042 | 0.26011357 | 0.79477617 | 0.8560962 | LOC100268168 | 100268168 |
| ENSG000002 | 8.65972029 | -0.091164864 | 0.35068642 | -0.2599612 | 0.7948937 | 0.8560962 | NA | NA |
| ENSG000002 | 15.650942 | -0.083197815 | 0.31994689 | -0.2600363 | 0.79483575 | 0.8560962 | NA | NA |
| ENSG000002 | 20.6950617 | -0.071677162 | 0.27572151 | -0.2599622 | 0.79489295 | 0.8560962 | UBR5-AS1 | 101927221 |
| ENSG000002 | 9.41587762 | -0.090579872 | 0.34840237 | -0.2599864 | 0.79487428 | 0.8560962 | NA | NA |
| ENSG000002 | 2.52766499 | 0.096519286 | 0.37121583 | 0.26000854 | 0.79485719 | 0.8560962 | RNU6-881P | 106479956 |
| ENSG000002 | 5.24513342 | 0.096430352 | 0.370883 | 0.26000208 | 0.79486217 | 0.8560962 | NA | NA |
| ENSG000002 | 4.12521204 | 0.098272329 | 0.37795495 | 0.26001069 | 0.79485553 | 0.8560962 | NA | NA |
| ENSG000002 | 16.45227 | -0.077723494 | 0.29904676 | -0.2599042 | 0.79493771 | 0.85612087 | NA | NA |
| ENSG000002 | 3.45125196 | -0.098269854 | 0.37818992 | -0.2598426 | 0.79498519 | 0.85612652 | NA | NA |
| ENSG000002 | 3.50942094 | -0.098335963 | 0.37842517 | -0.2598558 | 0.79497504 | 0.85612652 | NA | NA |
| ENSG000001 | 3.73920102 | -0.098313857 | 0.37859236 | -0.2596826 | 0.7951086 | 0.85623669 | NA | NA |
| ENSG000002 | 133.450027 | 0.042623047 | 0.16418879 | 0.25959779 | 0.79517404 | 0.85628442 | NA | NA |
| ENSG000001 | 61.2121417 | -0.051893866 | 0.20018548 | -0.2592289 | 0.79545862 | 0.85656812 | TNK1 | 8711 |
| ENSG000001 | 5.53915222 | -0.0966812 | 0.37319523 | -0.2590633 | 0.79558638 | 0.85668295 | H2BFWT | 158983 |
| ENSG000002 | 14.9533738 | -0.082064981 | 0.31685269 | -0.2590004 | 0.79563491 | 0.85671246 | NA | NA |
| ENSG000002 | 4.71505888 | -0.097587326 | 0.37690403 | -0.2589182 | 0.79569833 | 0.85673525 | NA | NA |
| ENSG000002 | 5.57660696 | -0.096797047 | 0.3738224 | -0.2589386 | 0.79568262 | 0.85673525 | NA | NA |
| ENSG000001 | 11.2867458 | 0.085545389 | 0.33043836 | 0.25888456 | 0.79572431 | 0.85674048 | FAM26F | 441168 |
| ENSG000002 | 4.16800461 | -0.097574335 | 0.37735289 | -0.2585758 | 0.79596254 | 0.85697421 | RNU6-323P | 106481281 |
| ENSG000002 | 3.4682673 | -0.097839057 | 0.37856662 | -0.2584461 | 0.79606266 | 0.85705926 | NA | NA |
| ENSG000001 | 5.26426401 | 0.09655807 | 0.37388468 | 0.25825629 | 0.79620912 | 0.85719418 | PRR18 | 285800 |
| ENSG000002 | 11.114931 | 0.085014995 | 0.32931846 | 0.25815436 | 0.79628778 | 0.85725611 | NA | NA |
| ENSG000002 | 10.0743647 | -0.087273508 | 0.3382681 | -0.258001 | 0.79640614 | 0.857313 | SNORD62B | 692093 |
| ENSG000002 | 8.39688189 | 0.089039674 | 0.34510597 | 0.25800676 | 0.79640169 | 0.857313 | BRWD1-IT2 | 103091865 |

| | | | | | | | | |
|---|---|---|---|---|---|---|---|---|
| ENSG000002 | 13.6358066 | 0.080681241 | 0.31268708 | 0.25802551 | 0.79638722 | 0.857313 | NA | NA |
| ENSG000002 | 3.47200532 | -0.097600198 | 0.37833009 | -0.2579763 | 0.7964252 | 0.857313 | NA | NA |
| ENSG000002 | 4.9762353 | 0.096474549 | 0.37402368 | 0.25793701 | 0.79645552 | 0.85732289 | RNU6-925P | 106479975 |
| ENSG000001 | 36.2758724 | -0.060021199 | 0.23273605 | -0.2578939 | 0.79648882 | 0.85733598 | FSTL5 | 56884 |
| ENSG000002 | 7.44755025 | -0.092895937 | 0.36059455 | -0.2576188 | 0.79670112 | 0.85754173 | NA | NA |
| ENSG000002 | 5.60858006 | -0.096587635 | 0.37517066 | -0.2574499 | 0.79683152 | 0.85761232 | NA | NA |
| ENSG000002 | 4.95880589 | -0.096732159 | 0.37576944 | -0.2574242 | 0.7968513 | 0.85761232 | NA | NA |
| ENSG000002 | 3.92496674 | 0.097307456 | 0.37797338 | 0.25744526 | 0.79683506 | 0.85761232 | RPL29P2 | 118432 |
| ENSG000002 | 44.3956727 | -0.058442965 | 0.22701995 | -0.2574354 | 0.7968427 | 0.85761232 | ZSCAN16-AS1 | 100129195 |
| ENSG000001 | 4.40414592 | -0.096761753 | 0.37649354 | -0.2570077 | 0.7971728 | 0.85793557 | NA | NA |
| ENSG000001 | 39.7155826 | 0.057666037 | 0.22440263 | 0.25697577 | 0.79719748 | 0.85793936 | MFI2 | 4241 |
| ENSG000002 | 5.25584736 | 0.095935226 | 0.37346575 | 0.25687824 | 0.79727277 | 0.85799761 | NA | NA |
| ENSG000002 | 3.67572731 | -0.097199107 | 0.3784763 | -0.2568169 | 0.79732013 | 0.85802581 | NA | NA |
| ENSG000001 | 588.342861 | -0.03717063 | 0.14482967 | -0.2566507 | 0.79744847 | 0.85814115 | ID3 | 3399 |
| ENSG000002 | 8.62640158 | 0.089465871 | 0.34864679 | 0.25660891 | 0.7974807 | 0.85815306 | NA | NA |
| ENSG000002 | 5.2334947 | 0.095290543 | 0.37140288 | 0.25656921 | 0.79751135 | 0.85816326 | NA | NA |
| ENSG000002 | 50.8270625 | -0.05193885 | 0.20247521 | -0.2565195 | 0.79754969 | 0.85818175 | NA | NA |
| ENSG000002 | 14.2414289 | -0.079024229 | 0.308126 | -0.2564673 | 0.79759006 | 0.85820242 | NA | NA |
| ENSG000002 | 19.7209711 | -0.070866818 | 0.27656414 | -0.2562401 | 0.79776546 | 0.85836837 | NA | NA |
| ENSG000001 | 3.92278331 | -0.096764242 | 0.37804921 | -0.2559567 | 0.79798424 | 0.85855821 | DEFB124 | 245937 |
| ENSG000002 | 7.68166982 | -0.090037099 | 0.35173648 | -0.2559788 | 0.79796717 | 0.85855821 | NA | NA |
| ENSG000002 | 3.69497434 | -0.096887683 | 0.37859015 | -0.2559171 | 0.79801487 | 0.85856838 | COX6CP17 | 106480269 |
| ENSG000002 | 4.66204374 | -0.095993006 | 0.37532153 | -0.2557621 | 0.79813458 | 0.85867439 | NA | NA |
| ENSG000002 | 16.5579453 | 0.078969744 | 0.30880375 | 0.25572793 | 0.79816093 | 0.85867996 | GPR162 | 27239 |
| ENSG000001 | 38.2927891 | 0.062969071 | 0.24632872 | 0.25563024 | 0.79823636 | 0.85873834 | TULP1 | 7287 |
| ENSG000002 | 2.76909675 | 0.095404846 | 0.37328939 | 0.25557878 | 0.79827611 | 0.85875831 | NA | NA |
| ENSG000002 | 48.7256149 | 0.052570273 | 0.20576491 | 0.25548706 | 0.79834694 | 0.85881172 | CTSO | 1519 |
| ENSG000002 | 2.79420801 | 0.096023842 | 0.37591586 | 0.25543972 | 0.79838349 | 0.85882826 | NA | NA |
| ENSG000002 | 3.64701333 | 0.096580111 | 0.37816251 | 0.25539314 | 0.79841947 | 0.85884418 | NA | NA |

| | | | | | | | | |
|---|---|---|---|---|---|---|---|---|
| ENSG000001 | 21.6352585 | 0.069443393 | 0.27205718 | 0.25525293 | 0.79852776 | 0.85893788 | TAGLN3 | 29114 |
| ENSG000002 | 4.17432675 | -0.096228622 | 0.37710728 | -0.2551757 | 0.79858738 | 0.85897923 | NA | NA |
| ENSG000002 | 9.71048535 | 0.085748125 | 0.33623587 | 0.25502373 | 0.79870477 | 0.85908271 | NA | NA |
| ENSG000002 | 4.17728246 | -0.096326599 | 0.37798584 | -0.2548418 | 0.79884528 | 0.85921104 | NA | NA |
| ENSG000002 | 2.78435671 | 0.095513935 | 0.37532101 | 0.25448598 | 0.79912013 | 0.85948388 | NA | NA |
| ENSG000001 | 38.7479812 | -0.057018926 | 0.2241091 | -0.2544249 | 0.79916735 | 0.85951186 | FZD1 | 8321 |
| ENSG000002 | 11.7214351 | 0.083619584 | 0.3287008 | 0.25439422 | 0.79919101 | 0.85951452 | NA | NA |
| ENSG000001 | 58.9286861 | -0.054448206 | 0.21407609 | -0.2543404 | 0.79923256 | 0.8595364 | BTN2A3P | 54718 |
| ENSG000002 | 5.17090585 | -0.094831725 | 0.37310357 | -0.25417 | 0.79936424 | 0.85965522 | NA | NA |
| ENSG000002 | 6.80707875 | 0.091449951 | 0.36009782 | 0.25395863 | 0.79952752 | 0.85980801 | NA | NA |
| ENSG000002 | 4.68031224 | -0.09515614 | 0.37476028 | -0.253912 | 0.79956353 | 0.85982081 | NA | NA |
| ENSG000002 | 5.45414804 | 0.094479059 | 0.37212839 | 0.25388834 | 0.79958182 | 0.85982081 | MIR5188 | 100847004 |
| ENSG000002 | 4.70012435 | -0.095530627 | 0.37631167 | -0.2538604 | 0.79960342 | 0.85982122 | NA | NA |
| ENSG000001 | 11.7403802 | -0.082402475 | 0.32467048 | -0.2538034 | 0.79964744 | 0.85984576 | C16orf71 | 146562 |
| ENSG000001 | 117.903996 | 0.04382908 | 0.17279472 | 0.25364826 | 0.79976731 | 0.85990625 | BTBD9 | 114781 |
| ENSG000002 | 2.55262538 | 0.094072564 | 0.3708635 | 0.25365819 | 0.79975964 | 0.85990625 | NA | NA |
| ENSG000002 | 32.3963756 | -0.066736219 | 0.26306024 | -0.2536918 | 0.79973369 | 0.85990625 | MAP1LC3B2 | 643246 |
| ENSG000001 | 15.502164 | -0.077501654 | 0.30577771 | -0.2534575 | 0.7999147 | 0.86004192 | BCL2L15 | 440603 |
| ENSG000002 | 2.78799776 | 0.094138378 | 0.37167842 | 0.25327911 | 0.80005254 | 0.86016731 | NA | NA |
| ENSG000002 | 5.33229337 | -0.093879302 | 0.37096963 | -0.2530647 | 0.80021825 | 0.86032266 | NA | NA |
| ENSG00000C | 254.802967 | 0.032343577 | 0.12785201 | 0.25297669 | 0.80028623 | 0.86035013 | RORA | 6095 |
| ENSG000002 | 3.46696893 | -0.095463808 | 0.37732504 | -0.2530015 | 0.80026704 | 0.86035013 | LOC100130587 | 100130587 |
| ENSG000002 | 4.37743627 | -0.095361717 | 0.37701704 | -0.2529374 | 0.80031657 | 0.86035994 | TPBGL | 100507050 |
| ENSG000002 | 7.83172942 | -0.09114107 | 0.36060258 | -0.2527466 | 0.80046405 | 0.86047286 | EGFLAM-AS4 | 100852408 |
| ENSG000002 | 5.12432428 | -0.09407678 | 0.37219091 | -0.2527649 | 0.80044993 | 0.86047286 | NA | NA |
| ENSG000002 | 6.54071488 | -0.093592643 | 0.37052636 | -0.2525938 | 0.80058216 | 0.86057701 | RNU6-808P | 106481436 |
| ENSG000002 | 2.79292205 | 0.095156847 | 0.37678627 | 0.2525486 | 0.80061705 | 0.8605917 | NA | NA |
| ENSG000002 | 4.18624065 | -0.095268245 | 0.37726966 | -0.2525203 | 0.80063893 | 0.86059241 | TAS2R40 | 259286 |
| ENSG000002 | 18.0906858 | -0.07453248 | 0.29528661 | -0.2524072 | 0.8007263 | 0.86066351 | NA | NA |

| | | | | | | | | |
|---|---|---|---|---|---|---|---|---|
| ENSG000002 | 3.22483308 | -0.094921087 | 0.37618532 | -0.2523253 | 0.80078961 | 0.86070874 | NA | NA |
| ENSG000002 | 2.78453298 | 0.094693852 | 0.37533844 | 0.25228925 | 0.8008175 | 0.8607159 | NA | NA |
| ENSG000002 | 6.62748242 | 0.091410953 | 0.36242509 | 0.25222027 | 0.80087081 | 0.86075039 | NA | NA |
| ENSG000001 | 5.39710402 | -0.094637348 | 0.37526085 | -0.2521908 | 0.80089356 | 0.86075203 | INHBC | 3626 |
| ENSG000002 | 3.20620006 | -0.09518571 | 0.37768131 | -0.2520265 | 0.80102055 | 0.8608657 | NA | NA |
| ENSG000002 | 3.97691316 | -0.095276214 | 0.37855866 | -0.2516815 | 0.80128725 | 0.8611295 | NA | NA |
| ENSG000002 | 5.38850294 | -0.094901529 | 0.37739651 | -0.2514637 | 0.80145561 | 0.86127395 | NA | NA |
| ENSG000002 | 12.8873569 | 0.078707756 | 0.31301221 | 0.25145267 | 0.80146415 | 0.86127395 | NA | NA |
| ENSG000002 | 5.42512945 | 0.09399413 | 0.37386222 | 0.25141382 | 0.80149418 | 0.8612834 | NA | NA |
| ENSG000002 | 4.42825231 | -0.094648039 | 0.37668318 | -0.251267 | 0.80160772 | 0.86138258 | NA | NA |
| ENSG000002 | 14.6515296 | -0.07764136 | 0.30948534 | -0.2508725 | 0.80191269 | 0.86168746 | NA | NA |
| ENSG000002 | 3.88888118 | 0.094826559 | 0.37843766 | 0.25057379 | 0.80214365 | 0.8619128 | SNRPGP18 | 106480680 |
| ENSG000002 | 25.9114307 | 0.063825216 | 0.25477977 | 0.25051132 | 0.80219195 | 0.86194186 | NA | NA |
| ENSG000002 | 2.76830692 | 0.094043039 | 0.37629865 | 0.24991596 | 0.80265234 | 0.86241369 | NA | NA |
| ENSG000002 | 8.20487662 | 0.087418273 | 0.35003874 | 0.24973885 | 0.80278931 | 0.86251558 | NA | NA |
| ENSG000002 | 11.0991956 | 0.083028965 | 0.33246383 | 0.24973834 | 0.80278971 | 0.86251558 | NA | NA |
| ENSG000002 | 9.64838385 | -0.086091182 | 0.34493958 | -0.2495834 | 0.80290956 | 0.86262149 | NA | NA |
| ENSG000002 | 6.78825019 | 0.090862335 | 0.36416982 | 0.2495054 | 0.80296987 | 0.86266343 | NA | NA |
| ENSG000002 | 11.294469 | 0.082421333 | 0.33067486 | 0.24925189 | 0.80316594 | 0.86285122 | NA | NA |
| ENSG000002 | 4.6978808 | -0.093848631 | 0.37662763 | -0.2491815 | 0.8032204 | 0.86288687 | NA | NA |
| ENSG000001 | 35.1638985 | -0.060980844 | 0.24490849 | -0.2489944 | 0.80336511 | 0.8629966 | BMP1 | 649 |
| ENSG000002 | 7.03857422 | 0.088668324 | 0.35607106 | 0.24901862 | 0.80334638 | 0.8629966 | NA | NA |
| ENSG000001 | 4.38184679 | -0.093860164 | 0.37702646 | -0.2489485 | 0.80340063 | 0.8630119 | ZNF890P | 645700 |
| ENSG000002 | 5.16544546 | -0.092970924 | 0.37359168 | -0.2488571 | 0.80347136 | 0.863065 | GAS5-AS1 | 100506046 |
| ENSG000002 | 2.97303428 | -0.093047195 | 0.37411302 | -0.2487141 | 0.80358192 | 0.8631609 | NA | NA |
| ENSG000002 | 8.53397894 | -0.087784712 | 0.35313344 | -0.2485879 | 0.80367954 | 0.86323447 | LOC101927708 | 101927708 |
| ENSG000002 | 7.28509435 | -0.089423316 | 0.35975023 | -0.2485706 | 0.80369298 | 0.86323447 | NA | NA |
| ENSG000002 | 3.933849 | -0.093885867 | 0.37780034 | -0.2485066 | 0.80374249 | 0.86326478 | NA | NA |
| ENSG000002 | 4.15682956 | -0.094055985 | 0.3785904 | -0.2484373 | 0.80379606 | 0.86329945 | NA | NA |

| | | | | | | | | |
|---|---|---|---|---|---|---|---|---|
| ENSG000001 | 62.5359193 | 0.049780049 | 0.20054431 | 0.24822469 | 0.80396056 | 0.86345326 | TMC7 | 79905 |
| ENSG000002 | 3.90587266 | -0.093786941 | 0.37807167 | -0.2480666 | 0.8040829 | 0.86356178 | MIR4502 | 100616227 |
| ENSG000002 | 5.28130295 | 0.092442743 | 0.3729865 | 0.24784474 | 0.80425454 | 0.86372324 | NA | NA |
| ENSG000002 | 3.96188243 | -0.093658437 | 0.37813301 | -0.2476865 | 0.80437699 | 0.86382057 | NA | NA |
| ENSG000002 | 2.61179957 | 0.09164951 | 0.37004306 | 0.24767255 | 0.80438777 | 0.86382057 | NA | NA |
| ENSG000002 | 3.89446262 | -0.093720318 | 0.37849308 | -0.2476143 | 0.80443281 | 0.86382318 | NA | NA |
| ENSG000002 | 2.59141094 | 0.092418464 | 0.3732142 | 0.24762848 | 0.80442188 | 0.86382318 | NA | NA |
| ENSG000002 | 3.99171742 | -0.093724583 | 0.37858191 | -0.2475675 | 0.80446905 | 0.86383099 | NA | NA |
| ENSG000002 | 5.11276987 | -0.092224183 | 0.37254786 | -0.2475499 | 0.80448268 | 0.86383099 | NA | NA |
| ENSG000002 | 2.57360687 | 0.091741388 | 0.37066208 | 0.24750681 | 0.80451602 | 0.86384391 | NA | NA |
| ENSG000002 | 8.24790401 | -0.087152035 | 0.35219446 | -0.2474543 | 0.80455665 | 0.86386466 | LOC101927497 | 101927497 |
| ENSG000001 | 23.2154578 | 0.070530644 | 0.28505941 | 0.24742437 | 0.80457982 | 0.86386666 | C2CD4C | 126567 |
| ENSG000002 | 4.26440542 | -0.093661323 | 0.37858778 | -0.2473966 | 0.80460132 | 0.86386688 | NA | NA |
| ENSG000000 | 5.30062146 | 0.092367 | 0.37376521 | 0.24712573 | 0.80481092 | 0.86406081 | SLC7A4 | 6545 |
| ENSG000002 | 20.8098438 | 0.0740038 | 0.29947946 | 0.2471081 | 0.80482456 | 0.86406081 | NA | NA |
| ENSG000002 | 4.97600436 | -0.093287988 | 0.37775994 | -0.2469505 | 0.80494657 | 0.86416892 | NA | NA |
| ENSG000002 | 4.41229535 | -0.092857741 | 0.37618637 | -0.2468397 | 0.80503227 | 0.86423804 | NA | NA |
| ENSG000002 | 3.65688632 | -0.093405579 | 0.3785527 | -0.2467439 | 0.80510641 | 0.8642725 | NA | NA |
| ENSG000002 | 3.67353638 | -0.0934032 | 0.37854422 | -0.2467432 | 0.805107 | 0.8642725 | NA | NA |
| ENSG000002 | 4.98516571 | 0.092517376 | 0.37517563 | 0.24659751 | 0.80521973 | 0.86435941 | NA | NA |
| ENSG000002 | 6.53857726 | 0.090291675 | 0.36617081 | 0.24658349 | 0.80523058 | 0.86435941 | NA | NA |
| ENSG000002 | 135.596095 | 0.038724425 | 0.15706598 | 0.24654878 | 0.80525745 | 0.86436536 | NA | NA |
| ENSG000002 | 12.4548469 | -0.083504379 | 0.33875287 | -0.2465053 | 0.8052911 | 0.86437861 | NOTO | 344022 |
| ENSG000002 | 35.1216437 | 0.058716924 | 0.23824809 | 0.24645286 | 0.80533169 | 0.86439929 | NA | NA |
| ENSG000002 | 4.4340463 | -0.092971902 | 0.37741832 | -0.2463365 | 0.80542177 | 0.8644731 | NA | NA |
| ENSG000001 | 14.4413158 | -0.077219383 | 0.31353581 | -0.2462857 | 0.80546109 | 0.86449242 | LINC00176 | 284739 |
| ENSG000002 | 3.90582114 | -0.093049724 | 0.37803639 | -0.2461396 | 0.80557417 | 0.86459091 | NA | NA |
| ENSG000002 | 25.9694629 | 0.065453577 | 0.26629911 | 0.2457897 | 0.80584503 | 0.86485872 | LOC102724190 | 102724190 |
| ENSG000002 | 3.24495371 | -0.092845462 | 0.37780706 | -0.2457484 | 0.80587704 | 0.86487018 | NA | NA |

| | | | | | | | | |
|---|---|---|---|---|---|---|---|---|
| ENSG000002 | 2.34581716 | 0.089578268 | 0.36461274 | 0.24568058 | 0.80592951 | 0.8649036 | NA | NA |
| ENSG000001 | 7.42460706 | -0.087042198 | 0.35444885 | -0.2455705 | 0.80601469 | 0.86494792 | CYP1A2 | 1544 |
| ENSG000001 | 4.9954347 | 0.092686027 | 0.37747128 | 0.24554458 | 0.80603479 | 0.86494792 | NDST2 | 8509 |
| ENSG000002 | 4.14243515 | 0.092637148 | 0.37726431 | 0.24554973 | 0.80603081 | 0.86494792 | NA | NA |
| ENSG000002 | 4.68605249 | -0.092585995 | 0.37727394 | -0.2454079 | 0.80614063 | 0.8650386 | HIST1H3PS1 | 100289545 |
| ENSG000001 | 4.05437463 | 0.092643316 | 0.37777339 | 0.24523515 | 0.80627436 | 0.86515921 | CSN2 | 1447 |
| ENSG000002 | 17.1580581 | -0.073850517 | 0.301207 | -0.2451819 | 0.80631556 | 0.86518052 | NA | NA |
| ENSG000002 | 4.62004099 | -0.092399059 | 0.37697615 | -0.2451058 | 0.80637448 | 0.86522084 | NA | NA |
| ENSG000002 | 4.13489547 | -0.092547168 | 0.37772513 | -0.2450119 | 0.80644718 | 0.86526442 | CPT1B | 1375 |
| ENSG000002 | 2.82317452 | 0.091806789 | 0.37472421 | 0.24499828 | 0.80645776 | 0.86526442 | NA | NA |
| ENSG000001 | 9.97885523 | 0.082847032 | 0.33829117 | 0.2448986 | 0.80653495 | 0.86530145 | NA | NA |
| ENSG000001 | 34.4919883 | -0.058134785 | 0.23736129 | -0.2449211 | 0.80651754 | 0.86530145 | JAKMIP3 | 282973 |
| ENSG000002 | 2.58259676 | 0.091489084 | 0.37374139 | 0.24479249 | 0.80661711 | 0.86536234 | NA | NA |
| ENSG000002 | 3.47025767 | -0.092273384 | 0.37697969 | -0.2447702 | 0.80663439 | 0.86536234 | NA | NA |
| ENSG000002 | 7.54422604 | -0.087377144 | 0.35730969 | -0.2445418 | 0.80681125 | 0.86550629 | NA | NA |
| ENSG000002 | 3.90324399 | -0.092486003 | 0.37816696 | -0.2445639 | 0.80679408 | 0.86550629 | NA | NA |
| ENSG000002 | 3.88361317 | -0.092422911 | 0.37800332 | -0.2445029 | 0.80684135 | 0.86551568 | NA | NA |
| ENSG000002 | 3.70401685 | -0.092524678 | 0.37847188 | -0.2444691 | 0.80686753 | 0.86552087 | NA | NA |
| ENSG000002 | 2.78962067 | 0.091512437 | 0.37460135 | 0.24429286 | 0.80700401 | 0.86564437 | NA | NA |
| ENSG000002 | 3.17926891 | -0.092075083 | 0.37746244 | -0.2439318 | 0.80728366 | 0.86592143 | NA | NA |
| ENSG000002 | 3.68745907 | -0.092274837 | 0.37841745 | -0.243844 | 0.80735162 | 0.86597142 | RN7SL788P | 106481135 |
| ENSG000002 | 3.67097243 | -0.09225534 | 0.37847782 | -0.2437536 | 0.80742164 | 0.86602362 | NA | NA |
| ENSG000001 | 303.916713 | -0.036385567 | 0.14930445 | -0.2437005 | 0.8074628 | 0.86604486 | FRMD4A | 55691 |
| ENSG000001 | 21.1339199 | 0.067819422 | 0.27840171 | 0.24360275 | 0.8075385 | 0.86610314 | ZNF223 | 7766 |
| ENSG000002 | 4.87409164 | -0.091205947 | 0.37446343 | -0.2435644 | 0.80756823 | 0.86611212 | NA | NA |
| ENSG000002 | 4.12733391 | -0.092133937 | 0.37835301 | -0.2435132 | 0.80760789 | 0.86613175 | NA | NA |
| ENSG000002 | 4.65738005 | -0.091524786 | 0.37591405 | -0.2434726 | 0.80763928 | 0.8661425 | NA | NA |
| ENSG000001 | 70.7966191 | 0.046685853 | 0.19189078 | 0.24329388 | 0.80777774 | 0.86626809 | EDARADD | 128178 |
| ENSG000001 | 65.9157835 | 0.047304564 | 0.19448167 | 0.24323405 | 0.80782409 | 0.86627197 | TLR3 | 7098 |

| | | | | | | | | |
|---|---|---|---|---|---|---|---|---|
| ENSG000002 | 3.45799885 | -0.092052716 | 0.37842945 | -0.2432493 | 0.80781224 | 0.86627197 | NA | NA |
| ENSG000002 | 3.22910034 | -0.091957395 | 0.3782248 | -0.2431289 | 0.80790551 | 0.86632178 | NA | NA |
| ENSG000002 | 2.61990879 | 0.089831405 | 0.36949572 | 0.24311893 | 0.80791327 | 0.86632178 | NA | NA |
| ENSG000002 | 3.22263419 | -0.09148526 | 0.37660086 | -0.2429237 | 0.80806454 | 0.86641526 | NA | NA |
| ENSG000002 | 2.79978498 | 0.091572364 | 0.37691763 | 0.2429506 | 0.80804367 | 0.86641526 | NA | NA |
| ENSG000002 | 3.92332409 | -0.091964213 | 0.3785639 | -0.2429292 | 0.80806027 | 0.86641526 | NA | NA |
| ENSG000002 | 3.93929012 | -0.091644892 | 0.37787139 | -0.2425293 | 0.80837004 | 0.8667199 | NA | NA |
| ENSG000000 | 33.9281809 | -0.056825283 | 0.23435073 | -0.2424796 | 0.80840853 | 0.86673825 | BAIAP3 | 8938 |
| ENSG000001 | 50.8828999 | -0.053585394 | 0.22102082 | -0.242445 | 0.80843537 | 0.86674411 | WLS | 79971 |
| ENSG000002 | 3.02129994 | 0.091373239 | 0.37705276 | 0.24233542 | 0.80852027 | 0.86678931 | SPRR2A | 6700 |
| ENSG000002 | 3.92278386 | -0.091589698 | 0.37794496 | -0.2423361 | 0.80851976 | 0.86678931 | NA | NA |
| ENSG000001 | 19.8832624 | 0.068303091 | 0.28195036 | 0.24225218 | 0.80858476 | 0.86681261 | ADCK5 | 203054 |
| ENSG000002 | 72.6030997 | 0.045735832 | 0.18878648 | 0.24226223 | 0.80857698 | 0.86681261 | NA | NA |
| ENSG000002 | 10.7422279 | -0.080387854 | 0.33216509 | -0.2420117 | 0.80877106 | 0.86696649 | NA | NA |
| ENSG000002 | 5.29247721 | -0.090547014 | 0.37410526 | -0.2420362 | 0.80875212 | 0.86696649 | NA | NA |
| ENSG000002 | 14.6812019 | 0.073665216 | 0.30448684 | 0.24193235 | 0.80883259 | 0.86700952 | NA | NA |
| ENSG000002 | 3.45973441 | -0.091526615 | 0.37859199 | -0.2417553 | 0.80896979 | 0.86711075 | NA | NA |
| ENSG000002 | 83.8785037 | -0.041994672 | 0.17369825 | -0.241768 | 0.80895996 | 0.86711075 | NA | NA |
| ENSG000001 | 56.0930533 | -0.050757737 | 0.21004955 | -0.2416465 | 0.80905409 | 0.86715527 | PRR5L | 79899 |
| ENSG000002 | 3.72811583 | -0.091353883 | 0.37803259 | -0.2416561 | 0.80904664 | 0.86715527 | NA | NA |
| ENSG000002 | 4.64212008 | -0.090522249 | 0.37526823 | -0.2412201 | 0.80938452 | 0.86741772 | NA | NA |
| ENSG000002 | 4.1639936 | -0.091207836 | 0.37798481 | -0.2413003 | 0.8093224 | 0.86741772 | NA | NA |
| ENSG000002 | 3.67424134 | -0.091311512 | 0.37850618 | -0.2412418 | 0.80936772 | 0.86741772 | ALG1L10P | 106479037 |
| ENSG000002 | 2.7902215 | 0.090593251 | 0.37553152 | 0.24124007 | 0.80936905 | 0.86741772 | NA | NA |
| ENSG000001 | 3.42706213 | -0.091285622 | 0.37859166 | -0.2411189 | 0.80946293 | 0.86747883 | TNFRSF18 | 8784 |
| ENSG000002 | 4.17953114 | 0.091228576 | 0.37847246 | 0.24104416 | 0.80952089 | 0.86751801 | NA | NA |
| ENSG000002 | 14.8923949 | 0.075103426 | 0.3117132 | 0.24093759 | 0.80960349 | 0.8675836 | NA | NA |
| ENSG000001 | 50.8212667 | -0.050086491 | 0.20799486 | -0.2408064 | 0.80970518 | 0.86764673 | BMPR1B | 658 |
| ENSG000002 | 3.69641705 | -0.091093679 | 0.37832937 | -0.2407788 | 0.80972659 | 0.86764673 | NA | NA |

| | | | | | | | | |
|---|---|---|---|---|---|---|---|---|
| ENSG000002 | 4.11706319 | 0.090931685 | 0.37762639 | 0.24079801 | 0.80971167 | 0.86764673 | NA | NA |
| ENSG000002 | 4.46551117 | -0.091035214 | 0.37858023 | -0.2404648 | 0.80996996 | 0.86788457 | NA | NA |
| ENSG000002 | 2.57318231 | 0.089321123 | 0.3715762 | 0.2403844 | 0.81003227 | 0.86792841 | NA | NA |
| ENSG000002 | 26.793087 | -0.062410113 | 0.25973432 | -0.2402844 | 0.81010976 | 0.86798851 | NA | NA |
| ENSG000002 | 10.7441738 | -0.079249933 | 0.32997762 | -0.2401676 | 0.81020033 | 0.86806261 | ZNF346-IT1 | 106480741 |
| ENSG000001 | 157.293159 | 0.038652784 | 0.16118018 | 0.23981102 | 0.81047676 | 0.86824408 | MANSC1 | 54682 |
| ENSG000002 | 2.842421 | 0.089299018 | 0.37230115 | 0.23985695 | 0.81044115 | 0.86824408 | NA | NA |
| ENSG000002 | 3.47580348 | -0.090807764 | 0.37858706 | -0.2398597 | 0.81043905 | 0.86824408 | NA | NA |
| ENSG000002 | 4.67040908 | -0.090250768 | 0.37633463 | -0.2398152 | 0.81047352 | 0.86824408 | RNU6-856P | 106481451 |
| ENSG000002 | 9.30930956 | 0.083379431 | 0.34757583 | 0.23988846 | 0.81041673 | 0.86824408 | NA | NA |
| ENSG000001 | 55.2005099 | -0.049749008 | 0.20785672 | -0.2393428 | 0.81083978 | 0.8685707 | ANO1 | 55107 |
| ENSG000001 | 177.433621 | 0.035327227 | 0.14760584 | 0.23933489 | 0.81084591 | 0.8685707 | RAPGEF6 | 51735 |
| ENSG000002 | 4.67277716 | -0.089890033 | 0.37555226 | -0.2393543 | 0.81083089 | 0.8685707 | NA | NA |
| ENSG000001 | 21.5597848 | 0.065933803 | 0.27552121 | 0.23930573 | 0.81086853 | 0.86857198 | KLK6 | 5653 |
| ENSG000002 | 3.93202204 | -0.090442542 | 0.37831856 | -0.2390645 | 0.81105556 | 0.86874938 | NA | NA |
| ENSG000002 | 11.6424262 | 0.078470541 | 0.32840479 | 0.23894457 | 0.81114857 | 0.8688031 | NA | NA |
| ENSG000002 | 4.40093737 | -0.089964438 | 0.37648372 | -0.2389597 | 0.81113683 | 0.8688031 | NA | NA |
| ENSG000002 | 3.4354193 | -0.090391729 | 0.37854047 | -0.2387901 | 0.81126832 | 0.86886252 | NA | NA |
| ENSG000002 | 2.82440011 | 0.089884326 | 0.37632874 | 0.23884523 | 0.81122559 | 0.86886252 | NA | NA |
| ENSG000002 | 3.92517559 | 0.090388474 | 0.37848437 | 0.23881693 | 0.81124754 | 0.86886252 | GACAT2 | 100287082 |
| ENSG000002 | 34.3031501 | 0.058366826 | 0.24447259 | 0.23874589 | 0.81130263 | 0.86887632 | DGCR5 | 26220 |
| ENSG000001 | 190.813854 | -0.038869538 | 0.16285554 | -0.2386749 | 0.81135765 | 0.86891229 | NOS1AP | 9722 |
| ENSG000002 | 3.69623327 | -0.090333196 | 0.37853002 | -0.2386421 | 0.81138312 | 0.86891663 | NA | NA |
| ENSG000001 | 67.190485 | 0.046103689 | 0.19323905 | 0.2385837 | 0.8114284 | 0.86891922 | BOC | 91653 |
| ENSG000001 | 4.06246369 | 0.09033025 | 0.37859287 | 0.23859469 | 0.81141988 | 0.86891922 | OR5B12 | 390191 |
| ENSG000002 | 23.2462663 | 0.063874768 | 0.26824786 | 0.23811846 | 0.81178922 | 0.86928264 | NA | NA |
| ENSG000001 | 30.6289831 | -0.058768853 | 0.24701938 | -0.2379119 | 0.81194941 | 0.86941599 | ITGB1BP2 | 26548 |
| ENSG000001 | 4.16359379 | 0.090051037 | 0.37852057 | 0.23790263 | 0.81195662 | 0.86941599 | OR7E12P | 10821 |
| ENSG000002 | 2.94626013 | -0.089268275 | 0.37544197 | -0.2377685 | 0.81206065 | 0.86950442 | NA | NA |

| | | | | | | | | |
|---|---|---|---|---|---|---|---|---|
| ENSG000002 | 2.38026467 | 0.085457949 | 0.35966826 | 0.23760215 | 0.81218969 | 0.86957864 | NA | NA |
| ENSG000002 | 10.7707775 | -0.078524748 | 0.33049665 | -0.2375962 | 0.8121943 | 0.86957864 | NA | NA |
| ENSG000002 | 5.25965618 | 0.088445907 | 0.37222661 | 0.23761307 | 0.81218121 | 0.86957864 | NA | NA |
| ENSG000001 | 22.2088049 | -0.067368957 | 0.28359057 | -0.2375571 | 0.81222461 | 0.86958814 | RBM34 | 23029 |
| ENSG000002 | 16.405282 | -0.070166845 | 0.29540755 | -0.2375256 | 0.81224909 | 0.86959138 | NA | NA |
| ENSG000002 | 31.3563705 | -0.05665676 | 0.23865098 | -0.2374043 | 0.81234318 | 0.86966916 | LOC100507291 | 100507291 |
| ENSG000001 | 3.49152045 | -0.089809176 | 0.37853678 | -0.2372535 | 0.81246013 | 0.86970252 | HRASLS2 | 54979 |
| ENSG000001 | 57.5847829 | -0.04821219 | 0.20314902 | -0.2373243 | 0.81240525 | 0.86970252 | CPNE9 | 151835 |
| ENSG000002 | 3.43374103 | -0.089612533 | 0.3776658 | -0.23728 | 0.81243959 | 0.86970252 | NA | NA |
| ENSG000002 | 7.20106479 | -0.084200446 | 0.35481537 | -0.2373078 | 0.81241803 | 0.86970252 | NA | NA |
| ENSG000001 | 75.1606127 | -0.04242005 | 0.17889315 | -0.2371251 | 0.81255977 | 0.86976326 | ST3GAL3 | 6487 |
| ENSG000002 | 3.87326454 | -0.089709266 | 0.37829819 | -0.237139 | 0.81254893 | 0.86976326 | NA | NA |
| ENSG000002 | 3.87251873 | 0.089724534 | 0.37846768 | 0.23707318 | 0.81260002 | 0.86978219 | NA | NA |
| ENSG000002 | 13.3338407 | -0.073412279 | 0.30969507 | -0.237047 | 0.81262035 | 0.86978219 | NA | NA |
| ENSG000002 | 14.572737 | 0.074913531 | 0.31625677 | 0.23687566 | 0.81275325 | 0.86990147 | NA | NA |
| ENSG000002 | 59.2435566 | 0.052083778 | 0.21991685 | 0.23683396 | 0.81278561 | 0.86991314 | NA | NA |
| ENSG000001 | 115.123844 | -0.037732104 | 0.15945772 | -0.2366276 | 0.81294567 | 0.87005197 | GUCY1A2 | 2977 |
| ENSG000002 | 3.90136517 | -0.089487378 | 0.37820391 | -0.2366115 | 0.81295823 | 0.87005197 | NA | NA |
| ENSG000002 | 3.93666145 | -0.08944353 | 0.37818235 | -0.236509 | 0.81303772 | 0.87011408 | NA | NA |
| ENSG000002 | 7.21879753 | -0.08499628 | 0.35949742 | -0.2364309 | 0.81309836 | 0.87015601 | NA | NA |
| ENSG000002 | 23.3353742 | -0.062435703 | 0.26414639 | -0.2363678 | 0.81314727 | 0.87018539 | LINC01270 | 284751 |
| ENSG000002 | 2.7196969 | -0.087954675 | 0.37232779 | -0.2362291 | 0.81325486 | 0.87027756 | NA | NA |
| ENSG000002 | 4.42589797 | -0.088871417 | 0.37665853 | -0.2359469 | 0.81347387 | 0.87048895 | RN7SL35P | 106480931 |
| ENSG000002 | 15.5147071 | 0.074774084 | 0.3170871 | 0.23581559 | 0.81357577 | 0.87056648 | NA | NA |
| ENSG000002 | 8.58262285 | 0.081466261 | 0.34549143 | 0.23579821 | 0.81358925 | 0.87056648 | NA | NA |
| ENSG000002 | 2.77385163 | 0.08857525 | 0.37578627 | 0.23570646 | 0.81366046 | 0.8706197 | NA | NA |
| ENSG000002 | 6.10010338 | -0.087475714 | 0.37130858 | -0.2355876 | 0.81375266 | 0.87069538 | NA | NA |
| ENSG000001 | 66.731604 | 0.044648023 | 0.18962826 | 0.23545026 | 0.81385928 | 0.87077789 | GPR156 | 165829 |
| ENSG000001 | 139.412191 | -0.036146758 | 0.15354164 | -0.2354199 | 0.81388283 | 0.87077789 | DLEU1 | 10301 |

| | | | | | | | | |
|---|---|---|---|---|---|---|---|---|
| ENSG000002 | 5.7495605 | 0.087698104 | 0.37254095 | 0.23540527 | 0.81389419 | 0.87077789 | NA | NA |
| ENSG000002 | 3.06245358 | 0.088743437 | 0.37710154 | 0.23533035 | 0.81395234 | 0.87079415 | NA | NA |
| ENSG000002 | 3.43454569 | -0.089095838 | 0.37858785 | -0.2353373 | 0.81394696 | 0.87079415 | NA | NA |
| ENSG000002 | 3.68744643 | -0.089022417 | 0.3784495 | -0.2352293 | 0.81403075 | 0.87085506 | NA | NA |
| ENSG000002 | 3.70233446 | -0.088979773 | 0.3784515 | -0.2351154 | 0.81411917 | 0.8709037 | DEFB134 | 613211 |
| ENSG000002 | 2.76330244 | 0.088171891 | 0.37499086 | 0.23513078 | 0.81410723 | 0.8709037 | NA | NA |
| ENSG000002 | 8.61623764 | -0.083512348 | 0.35540022 | -0.2349811 | 0.81422337 | 0.87099219 | DND1 | 373863 |
| ENSG000002 | 39.8606302 | -0.055367943 | 0.23569102 | -0.2349175 | 0.81427278 | 0.87102206 | LINC01060 | 401164 |
| ENSG000002 | 4.32229226 | 0.088645886 | 0.37747151 | 0.23484126 | 0.81433194 | 0.87106237 | NA | NA |
| ENSG000001 | 77.6149189 | 0.04399517 | 0.18739607 | 0.23477104 | 0.81438644 | 0.87108007 | LOC100379224 | 100379224 |
| ENSG000002 | 4.46329804 | -0.077371442 | 0.32960917 | -0.2347369 | 0.81441293 | 0.87108007 | NA | NA |
| ENSG000002 | 3.67873639 | -0.088872572 | 0.37859008 | -0.2347462 | 0.81440575 | 0.87108007 | NA | NA |
| ENSG000002 | 17.8813631 | 0.067430816 | 0.28733094 | 0.23467997 | 0.81445713 | 0.87110437 | LOC100130950 | 100130950 |
| ENSG000002 | 9.77657455 | 0.079734256 | 0.33979819 | 0.23465179 | 0.81447901 | 0.87110479 | NA | NA |
| ENSG000002 | 63.234718 | 0.045304693 | 0.1932337 | 0.23445545 | 0.81463141 | 0.87124482 | NA | NA |
| ENSG000002 | 5.31452893 | -0.087603928 | 0.37378895 | -0.2343674 | 0.8146998 | 0.87129498 | NA | NA |
| ENSG000002 | 2.72391209 | -0.087769018 | 0.37467667 | -0.2342527 | 0.81478881 | 0.87136719 | NA | NA |
| ENSG000002 | 18.6929961 | -0.068957913 | 0.29462704 | -0.2340515 | 0.81494497 | 0.8715112 | NA | NA |
| ENSG000001 | 96.652749 | -0.039538866 | 0.16922496 | -0.2336468 | 0.81525921 | 0.87182427 | CADM2 | 253559 |
| ENSG000002 | 7.93887928 | 0.082409497 | 0.352953 | 0.23348575 | 0.81538423 | 0.87193497 | NA | NA |
| ENSG000002 | 17.8928122 | 0.066497399 | 0.28484539 | 0.23345085 | 0.81541133 | 0.87194096 | NA | NA |
| ENSG000002 | 3.18229043 | -0.088186029 | 0.37813895 | -0.2332106 | 0.81559784 | 0.8720944 | NA | NA |
| ENSG000002 | 2.80918015 | 0.087207934 | 0.37390771 | 0.23323385 | 0.81557982 | 0.8720944 | NA | NA |
| ENSG000002 | 17.4306712 | -0.069349785 | 0.2974329 | -0.2331611 | 0.8156363 | 0.87209589 | NA | NA |
| ENSG000002 | 3.92515912 | -0.088269558 | 0.37858996 | -0.2331535 | 0.81564225 | 0.87209589 | NA | NA |
| ENSG000002 | 3.62997096 | 0.088017267 | 0.37756085 | 0.23312075 | 0.81566765 | 0.87210005 | NA | NA |
| ENSG000002 | 10.3939849 | 0.078386586 | 0.33639246 | 0.23302123 | 0.81574492 | 0.87214073 | NA | NA |
| ENSG000002 | 2.52941209 | 0.086171497 | 0.36985276 | 0.23298865 | 0.81577022 | 0.87214073 | NA | NA |
| ENSG000002 | 4.39138237 | 0.08810823 | 0.37813513 | 0.23300726 | 0.81575577 | 0.87214073 | NA | NA |

| | | | | | | | | |
|---|---|---|---|---|---|---|---|---|
| ENSG000002 | 3.66292825 | -0.088177956 | 0.3785832 | -0.2329157 | 0.8158269 | 0.87217834 | NA | NA |
| ENSG000002 | 6.85242688 | 0.083610751 | 0.3591136 | 0.23282535 | 0.81589703 | 0.87218433 | NA | NA |
| ENSG000002 | 3.69624079 | -0.08814486 | 0.37856682 | -0.2328383 | 0.81588696 | 0.87218433 | LINC01186 | 101927574 |
| ENSG000002 | 11.1654261 | 0.07633313 | 0.32779969 | 0.23286517 | 0.81586611 | 0.87218433 | NA | NA |
| ENSG000002 | 16.8440221 | -0.068563606 | 0.29460482 | -0.2327308 | 0.81597047 | 0.87223985 | LOC101928812 | 101928812 |
| ENSG000002 | 3.72792791 | -0.088058695 | 0.37858476 | -0.2325997 | 0.81607227 | 0.87231281 | NA | NA |
| ENSG000002 | 5.49264569 | 0.085982153 | 0.36967662 | 0.23258747 | 0.81608175 | 0.87231281 | NA | NA |
| ENSG000001 | 2.74881194 | 0.086925242 | 0.37387856 | 0.23249593 | 0.81615285 | 0.87236581 | UBL4B | 164153 |
| ENSG000001 | 54.25975 | 0.052770244 | 0.22710231 | 0.23236331 | 0.81625584 | 0.87242991 | NA | NA |
| ENSG000002 | 3.21103568 | -0.087657181 | 0.37723655 | -0.2323666 | 0.81625328 | 0.87242991 | NA | NA |
| ENSG000001 | 26.5167448 | 0.059920396 | 0.25800937 | 0.23224116 | 0.81635071 | 0.87245674 | CALB2 | 794 |
| ENSG000002 | 64.2330667 | -0.043681536 | 0.18805962 | -0.2322749 | 0.81632448 | 0.87245674 | TRAF3IP2-AS1 | 643749 |
| ENSG000002 | 2.76797409 | 0.085978319 | 0.37024482 | 0.23222018 | 0.816367 | 0.87245674 | COPS8P3 | 106479058 |
| ENSG000002 | 4.13869408 | -0.08758009 | 0.37708106 | -0.232258 | 0.81633764 | 0.87245674 | NA | NA |
| ENSG000002 | 6.8182229 | -0.083975711 | 0.36177184 | -0.2321234 | 0.81644217 | 0.87251407 | NA | NA |
| ENSG000002 | 7.01558961 | -0.085096875 | 0.36671609 | -0.2320511 | 0.81649832 | 0.87255108 | NA | NA |
| ENSG000002 | 3.6778361 | -0.087789628 | 0.37841761 | -0.2319914 | 0.8165447 | 0.87257112 | NA | NA |
| ENSG000002 | 5.73908081 | 0.085659454 | 0.36926706 | 0.23197155 | 0.81656011 | 0.87257112 | NA | NA |
| ENSG000002 | 10.7947876 | -0.076339567 | 0.32920088 | -0.2318936 | 0.81662068 | 0.87261285 | NA | NA |
| ENSG000002 | 4.61768784 | -0.087435405 | 0.37723021 | -0.2317826 | 0.81670686 | 0.87268195 | NA | NA |
| ENSG000001 | 4.37710725 | 0.087091149 | 0.37591373 | 0.23167855 | 0.81678769 | 0.87274532 | KRTDAP | 388533 |
| ENSG000002 | 9.72046522 | 0.079158641 | 0.34191167 | 0.23151781 | 0.81691255 | 0.87285574 | SNORD82 | 25826 |
| ENSG000002 | 3.20980921 | -0.087239397 | 0.37687855 | -0.2314788 | 0.81694285 | 0.87286511 | NA | NA |
| ENSG000002 | 2.99121139 | -0.086707351 | 0.37477814 | -0.2313565 | 0.81703787 | 0.87289764 | NA | NA |
| ENSG000002 | 3.93346411 | -0.087478924 | 0.378093 | -0.2313688 | 0.8170283 | 0.87289764 | NA | NA |
| ENSG000002 | 3.03644855 | 0.087185868 | 0.37682859 | 0.23136744 | 0.81702936 | 0.87289764 | NA | NA |
| ENSG000002 | 3.68432187 | -0.087530483 | 0.37849838 | -0.2312572 | 0.81711498 | 0.87295702 | NA | NA |
| ENSG000002 | 4.43206997 | -0.087398286 | 0.37825036 | -0.2310594 | 0.81726869 | 0.87309823 | NA | NA |
| ENSG000001 | 121.843035 | 0.039016005 | 0.16906554 | 0.23077444 | 0.81749004 | 0.87326788 | CTNND2 | 1501 |

| | | | | | | | | |
|---|---|---|---|---|---|---|---|---|
| ENSG000002 | 2.75353095 | -0.0846655 | 0.36687975 | -0.2307718 | 0.81749209 | 0.87326788 | NA | NA |
| ENSG000002 | 5.26709245 | 0.086543589 | 0.37493798 | 0.23082108 | 0.8174538 | 0.87326788 | NA | NA |
| ENSG000002 | 12.9606536 | 0.072731156 | 0.3156851 | 0.23039148 | 0.81778758 | 0.87356052 | NA | NA |
| ENSG000001 | 9.58124884 | 0.079981952 | 0.34726594 | 0.23031902 | 0.81784388 | 0.87359764 | CCR8 | 1237 |
| ENSG000001 | 3.89134155 | 0.087072649 | 0.37839888 | 0.23010811 | 0.81800776 | 0.87374968 | NA | NA |
| ENSG000002 | 3.73042737 | -0.087065643 | 0.37852968 | -0.2300101 | 0.81808394 | 0.87380802 | NA | NA |
| ENSG000002 | 5.70602385 | 0.084914678 | 0.36933375 | 0.22991313 | 0.81815927 | 0.87386548 | NA | NA |
| ENSG000002 | 4.41735538 | 0.086604271 | 0.37693906 | 0.22975669 | 0.81828084 | 0.8739723 | NA | NA |
| ENSG000002 | 4.40860242 | -0.086732324 | 0.37755878 | -0.2297187 | 0.81831034 | 0.87398079 | NA | NA |
| ENSG000001 | 228.083075 | -0.032064694 | 0.13976846 | -0.2294129 | 0.81854797 | 0.87418858 | ZNF283 | 284349 |
| ENSG000002 | 9.29822621 | -0.079578591 | 0.34687932 | -0.2294129 | 0.81854801 | 0.87418858 | LINC00857 | 439990 |
| ENSG000002 | 6.93218671 | -0.082529495 | 0.36024231 | -0.2290944 | 0.81879555 | 0.87442991 | NA | NA |
| ENSG000002 | 2.99301907 | -0.086102121 | 0.3759671 | -0.229015 | 0.81885723 | 0.87447275 | NA | NA |
| ENSG000002 | 3.45278696 | -0.086690196 | 0.37858726 | -0.2289834 | 0.81888182 | 0.87447599 | NA | NA |
| ENSG000001 | 21.826504 | 0.063657356 | 0.2782748 | 0.22875717 | 0.81905766 | 0.8746177 | CYP7B1 | 9420 |
| ENSG000002 | 5.96225855 | 0.084299417 | 0.36847296 | 0.22878047 | 0.81903955 | 0.8746177 | NA | NA |
| ENSG000002 | 3.23075158 | -0.086515126 | 0.37828056 | -0.2287062 | 0.81909724 | 0.87463694 | NA | NA |
| ENSG000002 | 5.72124394 | 0.083731041 | 0.36623963 | 0.22862365 | 0.81916144 | 0.87468246 | NA | NA |
| ENSG000002 | 2.95211337 | -0.086150806 | 0.37694087 | -0.2285526 | 0.81921669 | 0.87471841 | NA | NA |
| ENSG000002 | 2.99973027 | 0.085989822 | 0.3765778 | 0.22834544 | 0.8193777 | 0.87484427 | NA | NA |
| ENSG000002 | 27.5902327 | -0.058724653 | 0.257157 | -0.2283611 | 0.81936553 | 0.87484427 | NA | NA |
| ENSG000001 | 6.74812016 | -0.082729159 | 0.36273088 | -0.2280731 | 0.81958941 | 0.8749338 | KRT36 | 8689 |
| ENSG000002 | 2.51454026 | -0.084264688 | 0.36936737 | -0.2281325 | 0.81954326 | 0.8749338 | TMEM240 | 339453 |
| ENSG000002 | 11.4742322 | 0.076809275 | 0.33659486 | 0.22819503 | 0.81949462 | 0.8749338 | ZBED6 | 100381270 |
| ENSG000002 | 3.42657341 | -0.086341659 | 0.3785706 | -0.2280728 | 0.81958964 | 0.8749338 | NA | NA |
| ENSG000002 | 5.43589871 | 0.084754401 | 0.37161404 | 0.22807104 | 0.81959101 | 0.8749338 | NA | NA |
| ENSG000002 | 2.77869901 | 0.085700181 | 0.3757008 | 0.22810753 | 0.81956265 | 0.8749338 | NA | NA |
| ENSG000002 | 2.95669254 | -0.085389051 | 0.37449589 | -0.2280107 | 0.81963796 | 0.87494671 | NA | NA |
| ENSG000002 | 5.94743001 | 0.083539226 | 0.36640017 | 0.22799997 | 0.81964627 | 0.87494671 | NA | NA |

| | | | | | | | | |
|---|---|---|---|---|---|---|---|---|
| ENSG000002 | 3.43491097 | -0.086283355 | 0.37858779 | -0.2279084 | 0.81971742 | 0.87499963 | NA | NA |
| ENSG000002 | 4.63631479 | -0.085752391 | 0.37667904 | -0.2276537 | 0.81991544 | 0.87518797 | LINC01429 | 101927678 |
| ENSG000002 | 9.55271869 | -0.076723927 | 0.33721596 | -0.2275216 | 0.82001815 | 0.87523276 | NA | NA |
| ENSG000002 | 6.20544049 | 0.084115888 | 0.36971337 | 0.22751649 | 0.82002215 | 0.87523276 | NA | NA |
| ENSG000002 | 38.9496335 | 0.055791888 | 0.24516724 | 0.22756665 | 0.81998315 | 0.87523276 | NA | NA |
| ENSG000002 | 2.76657136 | 0.084916631 | 0.37330652 | 0.2274716 | 0.82005705 | 0.87524697 | LOC100240734 | 100240734 |
| ENSG000002 | 3.70096417 | -0.086068993 | 0.37857633 | -0.2273491 | 0.82015229 | 0.87532558 | NA | NA |
| ENSG000002 | 12.1207317 | -0.074914551 | 0.32968108 | -0.2272334 | 0.82024226 | 0.87539857 | NA | NA |
| ENSG000001 | 11.5472068 | 0.075465619 | 0.33229208 | 0.22710628 | 0.8203411 | 0.87546084 | CCR7 | 1236 |
| ENSG000001 | 92.0229579 | 0.037623865 | 0.16568912 | 0.22707505 | 0.82036538 | 0.87546084 | SYK | 6850 |
| ENSG000002 | 12.316394 | 0.07298871 | 0.32142213 | 0.22708054 | 0.82036112 | 0.87546084 | OLA1P2 | 106479059 |
| ENSG000002 | 3.73458253 | -0.085942943 | 0.37859327 | -0.227006 | 0.82041908 | 0.87549511 | NA | NA |
| ENSG000002 | 2.48881543 | -0.084064605 | 0.37057433 | -0.2268495 | 0.82054076 | 0.87559932 | NA | NA |
| ENSG000002 | 2.48593881 | -0.084391951 | 0.37205775 | -0.2268249 | 0.82055992 | 0.87559932 | NA | NA |
| ENSG000002 | 3.68286705 | -0.085815866 | 0.37858446 | -0.2266756 | 0.82067599 | 0.87570013 | NA | NA |
| ENSG000002 | 6.4532437 | -0.083042114 | 0.36641741 | -0.2266326 | 0.82070947 | 0.87571282 | NA | NA |
| ENSG000002 | 5.98380841 | 0.082939987 | 0.36641153 | 0.22635747 | 0.8209234 | 0.87589499 | NA | NA |
| ENSG000002 | 7.52599229 | -0.08136674 | 0.35943161 | -0.2263761 | 0.82090889 | 0.87589499 | NA | NA |
| ENSG000001 | 21.8697078 | -0.061989148 | 0.27394705 | -0.2262815 | 0.82098249 | 0.87593499 | CDNF | 441549 |
| ENSG000002 | 3.1980793 | -0.085540298 | 0.37813747 | -0.2262148 | 0.82103435 | 0.87596727 | NA | NA |
| ENSG000001 | 12.5980382 | -0.071069922 | 0.31423054 | -0.2261713 | 0.82106821 | 0.87598035 | PTX3 | 5806 |
| ENSG000002 | 3.46736178 | -0.085413604 | 0.37804939 | -0.2259324 | 0.821254 | 0.87615551 | NA | NA |
| ENSG000002 | 3.20269833 | -0.085282825 | 0.37760706 | -0.2258507 | 0.82131753 | 0.87620024 | NA | NA |
| ENSG000002 | 3.66249902 | -0.085469735 | 0.37857344 | -0.2257679 | 0.82138194 | 0.87624589 | NA | NA |
| ENSG000001 | 3.45507072 | -0.085399214 | 0.3784813 | -0.2256365 | 0.82148411 | 0.87626269 | LAIR2 | 3904 |
| ENSG000002 | 2.8087228 | 0.084952771 | 0.37647524 | 0.22565301 | 0.82147131 | 0.87626269 | NA | NA |
| ENSG000002 | 2.8087228 | 0.084952771 | 0.37647524 | 0.22565301 | 0.82147131 | 0.87626269 | NA | NA |
| ENSG000002 | 4.1535821 | -0.085269045 | 0.37779079 | -0.2257044 | 0.82143133 | 0.87626269 | LOC101928775 | 101928775 |
| ENSG000002 | 31.9866523 | 0.055087178 | 0.24417239 | 0.22560773 | 0.82150653 | 0.87626355 | SLC44A4 | 80736 |

| | | | | | | | | |
|---|---|---|---|---|---|---|---|---|
| ENSG000001 | 3.2338048 | -0.085032976 | 0.37699504 | -0.2255546 | 0.82154783 | 0.87628455 | RXFP4 | 339403 |
| ENSG000002 | 18.9659499 | -0.063987342 | 0.28379052 | -0.2254739 | 0.82161066 | 0.87630547 | NA | NA |
| ENSG000002 | 3.654559 | -0.085337096 | 0.37844454 | -0.2254943 | 0.82159474 | 0.87630547 | LOC101929384 | 101929384 |
| ENSG000002 | 3.45131308 | -0.085328246 | 0.37855575 | -0.2254047 | 0.82166446 | 0.87633979 | NA | NA |
| ENSG000002 | 43.8626332 | 0.048679909 | 0.21602468 | 0.2253442 | 0.82171152 | 0.87636694 | NA | NA |
| ENSG000001 | 61.1359582 | -0.043706119 | 0.19398576 | -0.2253058 | 0.82174138 | 0.87637574 | SSTR2 | 6752 |
| ENSG000002 | 13.1721994 | 0.070806915 | 0.31431717 | 0.22527218 | 0.82176754 | 0.87638059 | NA | NA |
| ENSG000002 | 7.06566844 | 0.082554341 | 0.36655844 | 0.22521468 | 0.82181227 | 0.87640524 | NA | NA |
| ENSG000001 | 5.66055142 | 0.083678616 | 0.37163169 | 0.22516545 | 0.82185057 | 0.87642304 | NA | NA |
| ENSG000001 | 59.9198813 | -0.043575128 | 0.1936402 | -0.2250314 | 0.82195484 | 0.87651118 | ALS2CL | 259173 |
| ENSG000002 | 3.66719552 | -0.085166859 | 0.37859112 | -0.2249574 | 0.82201245 | 0.87654956 | NA | NA |
| ENSG000001 | 71.089491 | -0.041240784 | 0.18335609 | -0.2249218 | 0.8220401 | 0.876556 | KIAA1644 | 85352 |
| ENSG000002 | 3.17144099 | -0.08508174 | 0.37837966 | -0.2248581 | 0.82208965 | 0.8765745 | NA | NA |
| ENSG000002 | 2.96448931 | -0.084730813 | 0.37684277 | -0.2248439 | 0.82210068 | 0.8765745 | NA | NA |
| ENSG000002 | 13.0864269 | -0.073015439 | 0.32495364 | -0.2246949 | 0.8222166 | 0.87667504 | NA | NA |
| ENSG000002 | 6.09767295 | -0.083159795 | 0.3702475 | -0.224606 | 0.82228583 | 0.87672581 | NA | NA |
| ENSG000002 | 4.36072554 | 0.084540957 | 0.37663052 | 0.22446656 | 0.82239429 | 0.87680322 | NA | NA |
| ENSG000002 | 3.70748219 | -0.084977793 | 0.37859266 | -0.2244571 | 0.82240168 | 0.87680322 | NA | NA |
| ENSG000002 | 3.18621168 | -0.084722515 | 0.37758068 | -0.2243825 | 0.82245965 | 0.87684197 | NA | NA |
| ENSG000001 | 14.6892057 | 0.069203489 | 0.3085714 | 0.22427059 | 0.82254676 | 0.87691179 | VEGFC | 7424 |
| ENSG000002 | 3.21919717 | -0.084804507 | 0.37827941 | -0.2241848 | 0.82261348 | 0.87695986 | NA | NA |
| ENSG000002 | 3.19088335 | -0.084560085 | 0.37729188 | -0.2241238 | 0.82266098 | 0.87698745 | LOC105370906 | 105370906 |
| ENSG000002 | 5.69499893 | 0.083629677 | 0.37351321 | 0.22390019 | 0.82283497 | 0.87714987 | NA | NA |
| ENSG000002 | 3.19130049 | -0.084551061 | 0.3776823 | -0.2238682 | 0.82285985 | 0.87715333 | NA | NA |
| ENSG000002 | 2.94765078 | -0.084507166 | 0.37768125 | -0.2237526 | 0.82294981 | 0.87722616 | NA | NA |
| ENSG000002 | 3.22059971 | -0.084484014 | 0.37763627 | -0.223718 | 0.82297677 | 0.87723185 | NA | NA |
| ENSG000001 | 97.2606849 | -0.036776054 | 0.16450573 | -0.2235548 | 0.8231037 | 0.87733731 | PIK3AP1 | 118788 |
| ENSG000002 | 2.79718856 | 0.083598049 | 0.37399652 | 0.22352627 | 0.82312594 | 0.87733731 | NA | NA |
| ENSG000002 | 4.39156235 | -0.084282383 | 0.3770899 | -0.2235074 | 0.82314062 | 0.87733731 | NA | NA |

| | | | | | | | | |
|---|---|---|---|---|---|---|---|---|
| ENSG000001 | 46.2311128 | -0.051414788 | 0.23018993 | -0.2233581 | 0.8232568 | 0.87743808 | ZNF436 | 80818 |
| ENSG000001 | 51.0119455 | 0.04537318 | 0.2033042 | 0.22317876 | 0.82339638 | 0.87756378 | RASAL1 | 8437 |
| ENSG000001 | 28.6459638 | -0.060059034 | 0.26921474 | -0.2230897 | 0.8234657 | 0.87761459 | POU3F1 | 5453 |
| ENSG000001 | 3.64202607 | -0.084402598 | 0.3784744 | -0.2230074 | 0.82352973 | 0.8776367 | TUBA3C | 7278 |
| ENSG000002 | 406.890069 | 0.07558551 | 0.33893355 | 0.22300982 | 0.82352786 | 0.8776367 | NA | NA |
| ENSG000002 | 15.7237994 | -0.066197257 | 0.29693849 | -0.2229326 | 0.823588 | 0.87767573 | LINC00535 | 642924 |
| ENSG000001 | 3.38757879 | -0.084128257 | 0.37796275 | -0.2225835 | 0.82385971 | 0.87793558 | NA | NA |
| ENSG000002 | 4.14694731 | -0.084097298 | 0.37785729 | -0.2225637 | 0.82387513 | 0.87793558 | NA | NA |
| ENSG000002 | 2.96352397 | -0.083989876 | 0.37755502 | -0.2224573 | 0.8239579 | 0.87800071 | NA | NA |
| ENSG000002 | 3.04316712 | 0.083924293 | 0.37746843 | 0.2223346 | 0.82405342 | 0.87807942 | NA | NA |
| ENSG000002 | 6.51059833 | -0.081109668 | 0.36485558 | -0.2223062 | 0.8240755 | 0.87807987 | NA | NA |
| ENSG000002 | 3.88136962 | -0.084004015 | 0.3784575 | -0.2219642 | 0.82434176 | 0.87831742 | NA | NA |
| ENSG000002 | 9.92709296 | -0.077644298 | 0.34977118 | -0.221986 | 0.8243248 | 0.87831742 | NA | NA |
| ENSG000002 | 4.60439235 | -0.083579079 | 0.3767191 | -0.2218605 | 0.8244225 | 0.87837714 | NA | NA |
| ENSG000002 | 21.707194 | -0.061251102 | 0.27610917 | -0.2218365 | 0.82444114 | 0.87837714 | NA | NA |
| ENSG00000C | 6.78953768 | -0.080124073 | 0.36166098 | -0.2215447 | 0.82466833 | 0.87859612 | LINC01587 | 10141 |
| ENSG000001 | 86.7962379 | -0.038043583 | 0.17179011 | -0.2214539 | 0.82473906 | 0.87864839 | SLC22A15 | 55356 |
| ENSG000002 | 4.17741438 | -0.08378692 | 0.37842922 | -0.2214071 | 0.82477546 | 0.87866408 | NA | NA |
| ENSG000002 | 8.68184059 | 0.07618704 | 0.34417774 | 0.22135958 | 0.82481246 | 0.87868042 | RPP21 | 79897 |
| ENSG000002 | 3.04074697 | 0.083641903 | 0.37806427 | 0.22123726 | 0.8249077 | 0.87873909 | NA | NA |
| ENSG000002 | 6.93517576 | -0.079216398 | 0.35806744 | -0.2212332 | 0.82491088 | 0.87873909 | NA | NA |
| ENSG000002 | 67.6394855 | -0.045394244 | 0.20528343 | -0.2211296 | 0.82499152 | 0.87880191 | NA | NA |
| ENSG000002 | 3.8691827 | -0.083611414 | 0.37815826 | -0.2211016 | 0.82501329 | 0.87880201 | NA | NA |
| ENSG000001 | 152.221523 | 0.0356582 | 0.1613157 | 0.22104606 | 0.82505657 | 0.87882503 | DPPA2 | 151871 |
| ENSG000002 | 3.6636721 | -0.083544362 | 0.37814828 | -0.2209302 | 0.82514681 | 0.87889807 | RN7SL254P | 106480991 |
| ENSG000001 | 89.7058223 | -0.039798409 | 0.180262 | -0.2207809 | 0.82526303 | 0.87890792 | KIAA1549L | 25758 |
| ENSG000002 | 3.97512773 | -0.08354499 | 0.37845757 | -0.2207513 | 0.8252861 | 0.87890792 | NA | NA |
| ENSG000002 | 3.91919019 | -0.083491969 | 0.37817746 | -0.2207746 | 0.82526796 | 0.87890792 | NA | NA |
| ENSG000002 | 3.65470246 | -0.083527568 | 0.37827645 | -0.2208109 | 0.82523967 | 0.87890792 | NA | NA |

| | | | | | | | | |
|---|---|---|---|---|---|---|---|---|
| ENSG000002 | 6.47279129 | -0.079684821 | 0.360967 | -0.2207538 | 0.82528417 | 0.87890792 | NA | NA |
| ENSG000002 | 2.84497666 | 0.082910858 | 0.37537466 | 0.22087495 | 0.8251898 | 0.87890792 | NA | NA |
| ENSG000002 | 2.45733737 | -0.082116502 | 0.37224496 | -0.220598 | 0.82540544 | 0.87901192 | NA | NA |
| ENSG000002 | 2.94907313 | -0.083181585 | 0.37733999 | -0.220442 | 0.82552693 | 0.87909513 | HMX1 | 3166 |
| ENSG000002 | 4.12952736 | -0.083389049 | 0.37827585 | -0.2204451 | 0.82552454 | 0.87909513 | NA | NA |
| ENSG000002 | 3.23960542 | -0.083260221 | 0.37797751 | -0.2202782 | 0.82565447 | 0.87920785 | NA | NA |
| ENSG000002 | 2.98205285 | -0.082594103 | 0.37543874 | -0.2199936 | 0.82587617 | 0.87942085 | NA | NA |
| ENSG000001 | 56.9285914 | -0.044631146 | 0.20294772 | -0.2199145 | 0.82593775 | 0.87945894 | CCT6B | 10693 |
| ENSG000002 | 2.97701322 | 0.082593889 | 0.37561129 | 0.21989192 | 0.82595533 | 0.87945894 | NA | NA |
| ENSG000002 | 2.70865954 | -0.082307836 | 0.37474752 | -0.2196354 | 0.82615509 | 0.87964855 | NA | NA |
| ENSG000002 | 6.97798009 | -0.078261989 | 0.35649705 | -0.2195305 | 0.82623679 | 0.87971245 | NA | NA |
| ENSG000002 | 4.89814559 | -0.082669837 | 0.37729163 | -0.2191139 | 0.82656133 | 0.88003488 | NA | NA |
| ENSG000001 | 4.19751073 | 0.082764564 | 0.37800696 | 0.21894984 | 0.82668912 | 0.88005474 | CLPS | 1208 |
| ENSG000001 | 4.41589282 | 0.08242431 | 0.37627465 | 0.21905358 | 0.82660831 | 0.88005474 | NPB | 256933 |
| ENSG000002 | 2.73429264 | -0.081758007 | 0.37353101 | -0.2188788 | 0.82674449 | 0.88005474 | RNU6-1029P | 106480019 |
| ENSG000002 | 2.48400792 | -0.08155712 | 0.37263312 | -0.2188671 | 0.8267536 | 0.88005474 | NA | NA |
| ENSG000002 | 7.05138592 | 0.078248243 | 0.35733742 | 0.21897579 | 0.82666891 | 0.88005474 | NA | NA |
| ENSG000002 | 2.70242207 | -0.081905546 | 0.37413733 | -0.2189184 | 0.82671361 | 0.88005474 | NA | NA |
| ENSG000002 | 2.46829095 | -0.081591307 | 0.37260337 | -0.2189763 | 0.82666851 | 0.88005474 | NA | NA |
| ENSG000002 | 2.98066329 | -0.082610879 | 0.37719721 | -0.2190124 | 0.82664036 | 0.88005474 | NA | NA |
| ENSG000002 | 80.9861072 | 0.039689454 | 0.18140393 | 0.21879049 | 0.82681326 | 0.88009514 | KC6 | 641516 |
| ENSG000002 | 2.727181 | -0.081890005 | 0.37437011 | -0.2187408 | 0.82685199 | 0.88011327 | LINC00851 | 440757 |
| ENSG000001 | 2.6954222 | -0.081783227 | 0.37399554 | -0.2186743 | 0.82690374 | 0.88014524 | P2RY4 | 5030 |
| ENSG000002 | 9.33192383 | -0.074674396 | 0.34157468 | -0.2186181 | 0.82694758 | 0.8801688 | FRMD6-AS1 | 145438 |
| ENSG000002 | 3.06673239 | 0.082543018 | 0.37771077 | 0.21853499 | 0.8270123 | 0.88021459 | NA | NA |
| ENSG000001 | 12.7954677 | 0.070069844 | 0.32097227 | 0.21830498 | 0.8271915 | 0.8803822 | SLC35D3 | 340146 |
| ENSG000001 | 18.9396924 | 0.062538749 | 0.28667433 | 0.21815259 | 0.82731022 | 0.88038563 | RIBC2 | 26150 |
| ENSG000001 | 48.0588763 | 0.045840451 | 0.21006962 | 0.21821552 | 0.82726119 | 0.88038563 | ADGRG5 | 221188 |
| ENSG000002 | 4.192363 | 0.082435708 | 0.3776834 | 0.2182667 | 0.82722132 | 0.88038563 | NA | NA |

| | | | | | | | | | |
|---|---|---|---|---|---|---|---|---|---|
| ENSG000002 | 2.82081799 | 0.082123692 | 0.37648336 | 0.21813365 | 0.82732498 | 0.88038563 | NA | NA | |
| ENSG000002 | 7.26786005 | -0.078236638 | 0.35864427 | -0.2181455 | 0.82731574 | 0.88038563 | NA | NA | |
| ENSG000002 | 4.40909866 | -0.082405246 | 0.37772754 | -0.2181605 | 0.82730402 | 0.88038563 | NA | NA | |
| ENSG000002 | 13.4163262 | -0.070209477 | 0.32197974 | -0.2180556 | 0.82738581 | 0.88042726 | NA | NA | |
| ENSG000002 | 2.80106298 | 0.081803236 | 0.37544544 | 0.21788316 | 0.82752015 | 0.88054711 | NA | NA | |
| ENSG000002 | 3.19652034 | -0.082027175 | 0.37716596 | -0.217483 | 0.82783197 | 0.8808558 | NA | NA | |
| ENSG000001 | 81.6096987 | -0.038430722 | 0.17673536 | -0.2174478 | 0.82785935 | 0.88086182 | TPK1 | | 27010 |
| ENSG000002 | 6.24647601 | -0.078956092 | 0.3631867 | -0.2173981 | 0.82789813 | 0.88087996 | RN7SL217P | | 106479301 |
| ENSG000001 | 11.4053495 | 0.07109373 | 0.32722684 | 0.21726131 | 0.82800471 | 0.88097025 | FAM167B | | 84734 |
| ENSG000002 | 3.17209387 | -0.082122711 | 0.37833195 | -0.2170652 | 0.82815751 | 0.8811097 | NA | NA | |
| ENSG000002 | 3.71089513 | 0.08204766 | 0.37825189 | 0.21691275 | 0.82827634 | 0.88121301 | NA | NA | |
| ENSG000002 | 6.66951342 | -0.078812036 | 0.36339005 | -0.21688 | 0.82830186 | 0.88121704 | NA | NA | |
| ENSG000002 | 9.10812796 | 0.074866772 | 0.34525146 | 0.21684708 | 0.82832752 | 0.88122122 | NA | NA | |
| ENSG000001 | 7.31052722 | -0.078376996 | 0.36180122 | -0.21663 | 0.82849671 | 0.88136277 | LINC01620 | | 140834 |
| ENSG000002 | 9.07829886 | -0.074168464 | 0.34238881 | -0.2166206 | 0.82850404 | 0.88136277 | NA | NA | |
| ENSG000002 | 7.59062581 | -0.078735972 | 0.36355552 | -0.2165721 | 0.82854185 | 0.88137987 | NA | NA | |
| ENSG000001 | 9.5576457 | -0.07307818 | 0.33754983 | -0.216496 | 0.82860116 | 0.88138029 | POMC | | 5443 |
| ENSG000001 | 137.472105 | 0.03301037 | 0.15248136 | 0.2164879 | 0.82860746 | 0.88138029 | DAB2 | | 1601 |
| ENSG000002 | 10.7610583 | -0.072763449 | 0.33610465 | -0.2164905 | 0.82860546 | 0.88138029 | LOC105377466 | | 105377466 |
| ENSG000001 | 6.94365712 | -0.077321811 | 0.35728268 | -0.2164163 | 0.82866323 | 0.8814165 | SLC25A34 | | 284723 |
| ENSG000002 | 3.06279794 | 0.081549039 | 0.37706083 | 0.21627555 | 0.82877297 | 0.8815101 | NA | NA | |
| ENSG000001 | 17.7662448 | -0.062772878 | 0.29042694 | -0.21614 | 0.82887862 | 0.88159935 | NTNG2 | | 84628 |
| ENSG000001 | 59.999868 | 0.041253553 | 0.1909976 | 0.2159899 | 0.82899562 | 0.88170067 | CALCB | | 797 |
| ENSG000002 | 19.4571928 | -0.063687013 | 0.29495849 | -0.2159186 | 0.82905123 | 0.88171819 | NA | NA | |
| ENSG000002 | 4.34066199 | -0.081556901 | 0.37773042 | -0.215913 | 0.82905558 | 0.88171819 | NA | NA | |
| ENSG000002 | 2.25386312 | -0.078503532 | 0.36377209 | -0.2158042 | 0.8291404 | 0.88178527 | NA | NA | |
| ENSG000002 | 6.64661681 | 0.079149439 | 0.36695348 | 0.21569339 | 0.82922676 | 0.88185398 | NA | NA | |
| ENSG000002 | 2.71656024 | -0.081010718 | 0.37573855 | -0.215604 | 0.82929647 | 0.88190499 | NA | NA | |
| ENSG000001 | 32.9191774 | -0.053779211 | 0.24971225 | -0.2153647 | 0.82948297 | 0.88205705 | GTF2IRD2B | | 389524 |

| | | | | | | | | |
|---|---|---|---|---|---|---|---|---|
| ENSG000002 | 7.0012768 | -0.077555718 | 0.36007353 | -0.2153886 | 0.82946439 | 0.88205705 | NA | NA |
| ENSG000002 | 2.69949304 | -0.079231488 | 0.36827316 | -0.2151433 | 0.82965563 | 0.88220494 | NA | NA |
| ENSG000002 | 5.91987126 | 0.079857526 | 0.37120498 | 0.21513053 | 0.82966555 | 0.88220494 | NA | NA |
| ENSG000002 | 6.6792218 | -0.077379443 | 0.35973865 | -0.2150991 | 0.82969009 | 0.8822079 | NA | NA |
| ENSG000002 | 2.7247614 | -0.08091314 | 0.37627915 | -0.2150349 | 0.82974013 | 0.88223798 | NA | NA |
| ENSG000002 | 2.96247409 | -0.081198424 | 0.37768203 | -0.2149915 | 0.82977395 | 0.88225081 | NA | NA |
| ENSG000001 | 226.043974 | 0.027940919 | 0.1300346 | 0.21487296 | 0.82986637 | 0.88232593 | TBC1D32 | 221322 |
| ENSG000002 | 3.03521446 | 0.080747905 | 0.37605097 | 0.21472596 | 0.82998098 | 0.88242465 | NA | NA |
| ENSG000002 | 3.16515907 | -0.080810956 | 0.37665633 | -0.2145483 | 0.83011954 | 0.88254883 | NA | NA |
| ENSG000002 | 114.091396 | 0.035090772 | 0.16358178 | 0.21451516 | 0.83014535 | 0.88255313 | SDHAP2 | 727956 |
| ENSG000002 | 8.40241832 | 0.076866964 | 0.35842322 | 0.21445867 | 0.8301894 | 0.88257682 | RN7SL93P | 106480947 |
| ENSG000002 | 2.9478144 | -0.080900207 | 0.37730105 | -0.2144182 | 0.83022096 | 0.88258724 | LOC105371782 | 105371782 |
| ENSG000001 | 2.47626377 | -0.079953616 | 0.37294523 | -0.2143843 | 0.83024735 | 0.88259216 | FATE1 | 89885 |
| ENSG000001 | 34.975941 | 0.050093672 | 0.23390425 | 0.21416316 | 0.83041983 | 0.88268295 | DTX1 | 1840 |
| ENSG000001 | 3.44757472 | -0.081035224 | 0.37836639 | -0.2141713 | 0.83041347 | 0.88268295 | PRR32 | 100130613 |
| ENSG000002 | 3.05426451 | 0.080963444 | 0.37799487 | 0.21419191 | 0.8303974 | 0.88268295 | NA | NA |
| ENSG000002 | 3.94252733 | -0.081062045 | 0.37841599 | -0.2142141 | 0.83038009 | 0.88268295 | NA | NA |
| ENSG000002 | 3.40980656 | -0.081048341 | 0.37854938 | -0.2141024 | 0.83046718 | 0.88271015 | NA | NA |
| ENSG000001 | 2.69692942 | -0.080470042 | 0.37594802 | -0.2140457 | 0.83051145 | 0.88273406 | AVPR2 | 554 |
| ENSG000002 | 3.21314393 | -0.080860762 | 0.37787333 | -0.2139891 | 0.83055558 | 0.88274248 | NA | NA |
| ENSG000002 | 4.17778023 | -0.081000713 | 0.37854396 | -0.2139797 | 0.83056291 | 0.88274248 | NA | NA |
| ENSG000002 | 4.09209931 | -0.080925202 | 0.37855406 | -0.2137745 | 0.83072292 | 0.88288038 | RNU6-48P | 100887744 |
| ENSG000002 | 3.08710042 | 0.080672462 | 0.37747282 | 0.21371728 | 0.83076754 | 0.88288038 | RNU6-130P | 106481218 |
| ENSG000002 | 3.96179844 | -0.080898029 | 0.37849243 | -0.2137375 | 0.83075176 | 0.88288038 | PLCH1-AS1 | 100874035 |
| ENSG000002 | 4.39727886 | 0.08040013 | 0.37632444 | 0.21364578 | 0.83082329 | 0.88288038 | NA | NA |
| ENSG000002 | 3.03049039 | 0.080678437 | 0.37761364 | 0.2136534 | 0.83081735 | 0.88288038 | NA | NA |
| ENSG000002 | 2.96549506 | -0.080646378 | 0.37745348 | -0.2136591 | 0.83081289 | 0.88288038 | NA | NA |
| ENSG000002 | 7.52365538 | 0.07587842 | 0.35521813 | 0.21361077 | 0.8308506 | 0.88288626 | NA | NA |
| ENSG000001 | 131.640713 | 0.032874637 | 0.15408569 | 0.21335296 | 0.83105167 | 0.88307678 | HFM1 | 164045 |

| | | | | | | | | |
|---|---|---|---|---|---|---|---|---|
| ENSG000002 | 4.1644458 | 0.080521341 | 0.37759857 | 0.21324589 | 0.83113518 | 0.88312583 | LMO7DN-IT1 | 104326189 |
| ENSG000002 | 3.48152534 | -0.080593099 | 0.37794916 | -0.2132379 | 0.83114138 | 0.88312583 | NA | NA |
| ENSG000002 | 3.69522392 | -0.08065019 | 0.3785265 | -0.2130635 | 0.83127742 | 0.88324723 | MIR4449 | 100616436 |
| ENSG000002 | 4.16837296 | -0.080525561 | 0.37807219 | -0.2129899 | 0.83133483 | 0.88328509 | NA | NA |
| ENSG000002 | 7.57620642 | 0.07525031 | 0.35339971 | 0.21293257 | 0.83137955 | 0.88330946 | NA | NA |
| ENSG000002 | 3.17987082 | -0.080544472 | 0.37837869 | -0.2128674 | 0.83143042 | 0.88334036 | NA | NA |
| ENSG000002 | 14.7429247 | -0.066259318 | 0.31183775 | -0.2124801 | 0.8317325 | 0.88362469 | NA | NA |
| ENSG000002 | 2.75364326 | -0.079261927 | 0.37310181 | -0.2124405 | 0.83176341 | 0.88362469 | NA | NA |
| ENSG000002 | 9.49108687 | -0.073880459 | 0.34772792 | -0.2124663 | 0.83174328 | 0.88362469 | NA | NA |
| ENSG000002 | 4.1230099 | -0.080282451 | 0.37818148 | -0.2122855 | 0.83188429 | 0.88371671 | NA | NA |
| ENSG000002 | 6.25604711 | -0.078256358 | 0.36865804 | -0.2122736 | 0.83189361 | 0.88371671 | NA | NA |
| ENSG000002 | 2.69459359 | -0.079732989 | 0.37572086 | -0.2122134 | 0.83194058 | 0.88374211 | NA | NA |
| ENSG000002 | 6.94712191 | -0.076304105 | 0.35960775 | -0.212187 | 0.83196112 | 0.88374211 | NA | NA |
| ENSG000002 | 12.9967076 | -0.068698046 | 0.32383567 | -0.2121386 | 0.83199891 | 0.88375911 | GBP1P1 | 400759 |
| ENSG000001 | 2.91762698 | -0.079878842 | 0.37663879 | -0.2120834 | 0.83204196 | 0.88378168 | OR5B21 | 219968 |
| ENSG000002 | 10.2799157 | 0.070557646 | 0.33274944 | 0.21204437 | 0.83207242 | 0.88379089 | NA | NA |
| ENSG000002 | 7.45439118 | -0.075639351 | 0.35685933 | -0.2119584 | 0.83213945 | 0.88383894 | NA | NA |
| ENSG000002 | 3.07054209 | 0.079913928 | 0.37718873 | 0.21186722 | 0.83221063 | 0.88388428 | NA | NA |
| ENSG000002 | 2.93410078 | -0.079870934 | 0.37705561 | -0.211828 | 0.83224123 | 0.88388428 | NA | NA |
| ENSG000002 | 7.32582998 | 0.07593464 | 0.35848679 | 0.21181991 | 0.83224753 | 0.88388428 | NA | NA |
| ENSG000002 | 2.84883898 | 0.079058231 | 0.37334963 | 0.21175388 | 0.83229905 | 0.8838927 | NA | NA |
| ENSG000002 | 2.9609598 | -0.079673854 | 0.37622541 | -0.2117716 | 0.83228523 | 0.8838927 | NA | NA |
| ENSG000002 | 2.70372784 | -0.079573291 | 0.37626235 | -0.2114835 | 0.83250998 | 0.8840704 | NA | NA |
| ENSG000002 | 7.1085241 | 0.07594429 | 0.35905933 | 0.21150903 | 0.83249009 | 0.8840704 | NA | NA |
| ENSG000002 | 2.75107909 | -0.079054462 | 0.37388658 | -0.2114397 | 0.83254419 | 0.88408357 | NA | NA |
| ENSG000002 | 2.2095447 | -0.077248378 | 0.36550054 | -0.2113496 | 0.83261451 | 0.8841351 | NA | NA |
| ENSG000002 | 7.51056184 | 0.075257574 | 0.35627813 | 0.21123265 | 0.83270574 | 0.88420881 | NA | NA |
| ENSG000001 | 785.519559 | -0.029069891 | 0.1376543 | -0.2111804 | 0.8327465 | 0.88421815 | PHLDA1 | 22822 |
| ENSG000002 | 7.49773107 | -0.075921657 | 0.3595363 | -0.2111655 | 0.83275814 | 0.88421815 | LINC00488 | 677779 |

| | | | | | | | | |
|---|---|---|---|---|---|---|---|---|
| ENSG000002 | 3.46969816 | -0.07977784 | 0.37810679 | -0.2109929 | 0.83289283 | 0.88433801 | NA | NA |
| ENSG000002 | 5.99656116 | 0.077520667 | 0.36761215 | 0.21087624 | 0.83298385 | 0.88441149 | NA | NA |
| ENSG000002 | 2.67305116 | -0.07911482 | 0.37553862 | -0.2106703 | 0.83314458 | 0.88455899 | NA | NA |
| ENSG000002 | 3.22067956 | -0.07956999 | 0.37795559 | -0.2105274 | 0.83325611 | 0.88465423 | NA | NA |
| ENSG000001 | 4.17520363 | -0.079622061 | 0.37839575 | -0.2104201 | 0.83333983 | 0.88471995 | GATA3-AS1 | 399717 |
| ENSG000002 | 2.21788967 | -0.077084641 | 0.36666067 | -0.2102343 | 0.83348483 | 0.8848044 | NA | NA |
| ENSG000002 | 2.6737363 | -0.078952332 | 0.3755391 | -0.2102373 | 0.83348246 | 0.8848044 | NA | NA |
| ENSG000002 | 3.0653619 | 0.079360398 | 0.37739103 | 0.21028692 | 0.83344374 | 0.8848044 | NA | NA |
| ENSG000002 | 2.22678817 | -0.07684572 | 0.36564541 | -0.2101646 | 0.83353921 | 0.88483897 | NA | NA |
| ENSG000002 | 2.26162709 | -0.075861211 | 0.36142933 | -0.2098922 | 0.83375178 | 0.88504145 | NA | NA |
| ENSG000002 | 3.1465134 | -0.079358641 | 0.37820096 | -0.2098319 | 0.83379885 | 0.88506824 | NA | NA |
| ENSG000001 | 2.91699425 | -0.078762221 | 0.37553711 | -0.2097322 | 0.8338767 | 0.88512771 | RNU6-79P | 100873779 |
| ENSG000002 | 31.8708752 | 0.052074365 | 0.24842092 | 0.2096215 | 0.8339631 | 0.88519625 | NA | NA |
| ENSG000002 | 3.25019428 | -0.078718721 | 0.37568486 | -0.2095339 | 0.83403146 | 0.88524564 | NA | NA |
| ENSG000001 | 34.307454 | -0.049050375 | 0.23435868 | -0.2092962 | 0.83421704 | 0.88537765 | KCNK12 | 56660 |
| ENSG000002 | 46.915767 | 0.044157469 | 0.21098632 | 0.20929068 | 0.83422133 | 0.88537765 | LINC00632 | 286411 |
| ENSG000002 | 3.65854551 | -0.079245721 | 0.37858346 | -0.2093217 | 0.83419714 | 0.88537765 | NA | NA |
| ENSG000001 | 12.4367238 | 0.06717876 | 0.32111987 | 0.2092015 | 0.83429094 | 0.88538996 | KCNK17 | 89822 |
| ENSG000002 | 2.48687245 | -0.077949707 | 0.37263417 | -0.2091856 | 0.83430334 | 0.88538996 | RNU6-1081P | 106480636 |
| ENSG000002 | 2.92186178 | -0.079017042 | 0.37767315 | -0.2092207 | 0.83427595 | 0.88538996 | NA | NA |
| ENSG000002 | 2.43676082 | -0.077922633 | 0.37254335 | -0.2091639 | 0.83432027 | 0.88538996 | NA | NA |
| ENSG000001 | 2.49942465 | -0.077991436 | 0.37298439 | -0.2091011 | 0.83436934 | 0.88541886 | GPR148 | 344561 |
| ENSG000002 | 2.24910769 | -0.076881697 | 0.3679844 | -0.2089265 | 0.83450561 | 0.8855403 | NA | NA |
| ENSG000002 | 3.49124059 | -0.078908789 | 0.37782481 | -0.2088502 | 0.83456518 | 0.88555716 | NA | NA |
| ENSG000002 | 16.7294501 | -0.061340715 | 0.2936811 | -0.2088684 | 0.83455094 | 0.88555716 | DNAJC9-AS1 | 414245 |
| ENSG000002 | 8.87505162 | 0.072002537 | 0.34484478 | 0.20879694 | 0.83460676 | 0.88557811 | NA | NA |
| ENSG000002 | 6.04345136 | -0.07690813 | 0.36838888 | -0.2087689 | 0.83462867 | 0.88557819 | NA | NA |
| ENSG000002 | 3.02956506 | 0.078491264 | 0.37607596 | 0.2087112 | 0.8346737 | 0.88560279 | NA | NA |
| ENSG000001 | 46.1284229 | -0.044951228 | 0.21546977 | -0.2086196 | 0.83474517 | 0.88561134 | NA | NA |

| | | | | | | | | |
|---|---|---|---|---|---|---|---|---|
| ENSG000001 | 77.6841948 | 0.037874262 | 0.18154621 | 0.20862051 | 0.8347445 | 0.88561134 | PPP1R1C | 151242 |
| ENSG000002 | 15.198731 | 0.063226588 | 0.30308787 | 0.20860811 | 0.83475418 | 0.88561134 | NA | NA |
| ENSG000002 | 2.68869601 | -0.078360384 | 0.37566885 | -0.208589 | 0.83476911 | 0.88561134 | NA | NA |
| ENSG00000C | 214.027545 | -0.030687625 | 0.14715326 | -0.2085419 | 0.83480585 | 0.88562714 | RBMS2 | 5939 |
| ENSG000001 | 9.52311401 | -0.072200015 | 0.34641914 | -0.2084181 | 0.83490253 | 0.88570654 | GYPE | 2996 |
| ENSG000002 | 14.4997838 | -0.063354575 | 0.30416409 | -0.2082908 | 0.83500193 | 0.88578882 | LINC00562 | 100861549 |
| ENSG000002 | 2.70367577 | -0.07816775 | 0.37549306 | -0.2081736 | 0.8350934 | 0.88583951 | NA | NA |
| ENSG000002 | 18.21212 | 0.059905843 | 0.28773113 | 0.20820077 | 0.83507221 | 0.88583951 | NA | NA |
| ENSG000002 | 2.49183694 | -0.077122538 | 0.37057316 | -0.2081169 | 0.83513769 | 0.88586331 | NA | NA |
| ENSG000002 | 2.21014552 | -0.076623079 | 0.3685725 | -0.2078915 | 0.83531371 | 0.88602684 | NA | NA |
| ENSG000002 | 13.2049357 | 0.065491998 | 0.31541628 | 0.20763671 | 0.83551264 | 0.88621467 | NA | NA |
| ENSG000002 | 45.1785813 | 0.070555233 | 0.33986652 | 0.20759689 | 0.83554373 | 0.88622447 | NA | NA |
| ENSG000002 | 7.47492554 | 0.074721325 | 0.36026776 | 0.20740497 | 0.8356936 | 0.88636024 | NA | NA |
| ENSG000001 | 58.6752779 | 0.040545664 | 0.19558728 | 0.20730215 | 0.8357739 | 0.88642223 | C1orf56 | 54964 |
| ENSG000002 | 6.72553637 | -0.07482961 | 0.36102439 | -0.2072702 | 0.83579882 | 0.88642547 | NA | NA |
| ENSG000001 | 66.3401532 | 0.042749108 | 0.20634488 | 0.2071731 | 0.83587468 | 0.88648274 | PPP3CC | 5533 |
| ENSG000002 | 72.9799959 | -0.040427802 | 0.19518737 | -0.207123 | 0.83591377 | 0.88650101 | NA | NA |
| ENSG000002 | 4.61035756 | 0.078149393 | 0.37738518 | 0.20708125 | 0.83594641 | 0.88651244 | NA | NA |
| ENSG000002 | 2.83323435 | 0.076924381 | 0.37177313 | 0.20691216 | 0.83607846 | 0.8866293 | NA | NA |
| ENSG000002 | 2.59730798 | 0.076263337 | 0.36867539 | 0.20685768 | 0.83612101 | 0.88665123 | NA | NA |
| ENSG000002 | 3.22634881 | -0.078029097 | 0.37737641 | -0.2067673 | 0.8361916 | 0.8867029 | RHOXF2B | 727940 |
| ENSG000002 | 32.7217114 | 0.049778795 | 0.24080767 | 0.20671599 | 0.83623167 | 0.8867222 | CCDC163P | 126661 |
| ENSG000002 | 5.76653596 | -0.075844712 | 0.36730318 | -0.2064908 | 0.83640758 | 0.88688554 | NA | NA |
| ENSG00000C | 55.9918154 | -0.040974993 | 0.19853788 | -0.2063838 | 0.83649116 | 0.88695097 | PTPRN | 5798 |
| ENSG000002 | 2.25858684 | -0.075777978 | 0.36735432 | -0.2062804 | 0.83657192 | 0.88701342 | NA | NA |
| ENSG000001 | 4.59629827 | 0.077719338 | 0.37685411 | 0.2062319 | 0.83660977 | 0.88703035 | UPK2 | 7379 |
| ENSG000001 | 2.97263894 | -0.077730917 | 0.37706401 | -0.2061478 | 0.83667546 | 0.88707599 | CRYGA | 1418 |
| ENSG000002 | 2.458271 | -0.077056952 | 0.37384368 | -0.2061208 | 0.83669656 | 0.88707599 | NA | NA |
| ENSG000001 | 66.9185174 | 0.038304284 | 0.18592312 | 0.20602217 | 0.8367736 | 0.88710515 | INPP5J | 27124 |

| | | | | | | | | |
|---|---|---|---|---|---|---|---|---|
| ENSG000002 | 3.0775694 | 0.077287203 | 0.37515858 | 0.20601209 | 0.83678147 | 0.88710515 | NA | NA |
| ENSG000002 | 2.74035418 | -0.077183894 | 0.37467628 | -0.2060015 | 0.8367897 | 0.88710515 | NA | NA |
| ENSG000002 | 2.24350294 | -0.075649324 | 0.36729898 | -0.2059612 | 0.83682125 | 0.88711541 | NA | NA |
| ENSG000002 | 4.18477006 | 0.077896266 | 0.37847047 | 0.20581861 | 0.8369326 | 0.88721026 | NA | NA |
| ENSG000002 | 2.25937613 | -0.075546952 | 0.36735641 | -0.2056503 | 0.83706409 | 0.88732645 | NA | NA |
| ENSG000001 | 310.734486 | -0.02619848 | 0.12744208 | -0.2055717 | 0.83712552 | 0.88733429 | FLRT2 | 23768 |
| ENSG000002 | 20.5880166 | 0.058082124 | 0.2825985 | 0.20552877 | 0.83715902 | 0.88733429 | VSTM5 | 387804 |
| ENSG000002 | 2.45523021 | -0.076257669 | 0.3709159 | -0.2055929 | 0.83710894 | 0.88733429 | NA | NA |
| ENSG000002 | 2.96224596 | -0.077563654 | 0.377381 | -0.2055314 | 0.83715695 | 0.88733429 | NA | NA |
| ENSG000002 | 23.727268 | 0.05450807 | 0.26525026 | 0.20549676 | 0.83718403 | 0.88733761 | NA | NA |
| ENSG000002 | 2.76870768 | -0.076451947 | 0.3724088 | -0.2052904 | 0.83734525 | 0.88747284 | NA | NA |
| ENSG000002 | 2.69997021 | -0.077080004 | 0.37549188 | -0.2052774 | 0.83735539 | 0.88747284 | NA | NA |
| ENSG000001 | 17.2778033 | -0.059070738 | 0.28780257 | -0.2052474 | 0.83737881 | 0.88747447 | KIF12 | 113220 |
| ENSG000002 | 2.48445812 | 0.058303669 | 0.28417434 | 0.20516866 | 0.83744036 | 0.8875165 | NA | NA |
| ENSG000002 | 3.4688045 | -0.077516705 | 0.37828414 | -0.2049166 | 0.83763727 | 0.88770199 | NA | NA |
| ENSG000001 | 30.2465743 | -0.053828753 | 0.2628581 | -0.2047826 | 0.83774202 | 0.88778979 | NKAIN2 | 154215 |
| ENSG000001 | 180.012186 | 0.034546824 | 0.16882763 | 0.20462779 | 0.83786294 | 0.88789474 | PURA | 5813 |
| ENSG000001 | 153.28371 | 0.029553105 | 0.14455043 | 0.20444841 | 0.8380031 | 0.88797366 | AP3B2 | 8120 |
| ENSG000002 | 3.39981887 | -0.07739559 | 0.3784848 | -0.204488 | 0.83797219 | 0.88797366 | NA | NA |
| ENSG000002 | 3.67008727 | -0.077410751 | 0.37859146 | -0.2044704 | 0.83798591 | 0.88797366 | LOC102723833 | 102723833 |
| ENSG000002 | 2.48896033 | -0.075637 | 0.37027779 | -0.204271 | 0.83814176 | 0.88809737 | ADARB2-AS1 | 642394 |
| ENSG000001 | 17.1439728 | 0.059444787 | 0.29108269 | 0.20421959 | 0.83818191 | 0.88811671 | GNGT2 | 2793 |
| ENSG000002 | 13.7447216 | -0.063429305 | 0.31094068 | -0.2039917 | 0.83836002 | 0.88828222 | NA | NA |
| ENSG000002 | 3.44512768 | -0.077090764 | 0.37805529 | -0.203914 | 0.83842071 | 0.88832331 | TCL1B | 9623 |
| ENSG000001 | 35.1485062 | 0.049772729 | 0.24428562 | 0.20374809 | 0.83855036 | 0.88843747 | PRR4 | 11272 |
| ENSG000002 | 4.12721932 | -0.077052672 | 0.37837428 | -0.2036414 | 0.83863373 | 0.88850258 | NCOA7-AS1 | 104355145 |
| ENSG000002 | 21.6260967 | -0.058063158 | 0.28531707 | -0.203504 | 0.83874114 | 0.88856995 | SNORD69 | 692109 |
| ENSG000002 | 8.9945282 | 0.071198489 | 0.34981497 | 0.20353185 | 0.83871935 | 0.88856995 | NA | NA |
| ENSG000002 | 3.48414934 | -0.07652547 | 0.37668809 | -0.2031534 | 0.83901513 | 0.88883699 | NA | NA |

| | | | | | | | | |
|---|---|---|---|---|---|---|---|---|
| ENSG000002 | 4.66115379 | 0.076122579 | 0.37483839 | 0.20308106 | 0.83907167 | 0.88885046 | RN7SL65P | 106479247 |
| ENSG000002 | 3.26385238 | 0.076652395 | 0.37741725 | 0.20309722 | 0.83905904 | 0.88885046 | NA | NA |
| ENSG000001 | 19.977455 | 0.057017368 | 0.28095765 | 0.20293936 | 0.83918243 | 0.88894456 | AMER3 | 205147 |
| ENSG000002 | 31.3984052 | 0.049022439 | 0.24161387 | 0.2028958 | 0.83921648 | 0.88895741 | UBAC2-AS1 | 100289373 |
| ENSG000001 | 9.44018197 | 0.071547332 | 0.35271125 | 0.20284959 | 0.83925259 | 0.88897244 | NA | NA |
| ENSG000002 | 2.43684459 | -0.073758265 | 0.36380638 | -0.2027404 | 0.83933791 | 0.88901637 | NA | NA |
| ENSG000002 | 2.46959673 | -0.075547586 | 0.37259946 | -0.2027582 | 0.83932405 | 0.88901637 | NA | NA |
| ENSG000001 | 3.45620023 | -0.076598293 | 0.37803279 | -0.2026234 | 0.83942939 | 0.88902038 | SOX1 | 6656 |
| ENSG000002 | 3.18037991 | -0.07665708 | 0.37819892 | -0.2026898 | 0.83937746 | 0.88902038 | CACYBPP3 | 106480809 |
| ENSG000002 | 2.66095597 | -0.076045887 | 0.37528372 | -0.2026357 | 0.83941977 | 0.88902038 | NA | NA |
| ENSG000002 | 47.6576379 | -0.04343223 | 0.21434911 | -0.2026238 | 0.83942909 | 0.88902038 | TMEM150C | 441027 |
| ENSG000002 | 2.69089544 | -0.07611021 | 0.37571715 | -0.2025732 | 0.83946867 | 0.88903876 | NA | NA |
| ENSG000001 | 113.574758 | 0.034402772 | 0.17000014 | 0.20236908 | 0.8396282 | 0.88916127 | Mar-02 | 54996 |
| ENSG000002 | 4.20297078 | 0.076539157 | 0.37820077 | 0.20237705 | 0.83962197 | 0.88916127 | NA | NA |
| ENSG000002 | 3.08291803 | 0.076353377 | 0.37747801 | 0.20227238 | 0.83970379 | 0.88921809 | NA | NA |
| ENSG000002 | 32.9793093 | 0.04776236 | 0.23617078 | 0.20223653 | 0.83973182 | 0.88922455 | LOC103611081 | 103611081 |
| ENSG000002 | 2.25312645 | -0.074275115 | 0.36756848 | -0.2020715 | 0.83986083 | 0.88933794 | LINC01058 | 103724387 |
| ENSG000002 | 2.43741371 | -0.07531239 | 0.37287244 | -0.201979 | 0.83993317 | 0.88939132 | NA | NA |
| ENSG000002 | 2.89187024 | -0.075972582 | 0.3762118 | -0.201941 | 0.83996286 | 0.88939953 | NA | NA |
| ENSG000002 | 7.29515104 | -0.074072381 | 0.36697462 | -0.2018461 | 0.84003707 | 0.88945489 | NA | NA |
| ENSG000001 | 4.32585293 | 0.076161397 | 0.37776593 | 0.20161002 | 0.84022161 | 0.88961952 | NA | NA |
| ENSG000002 | 10.2068372 | 0.06974406 | 0.34596801 | 0.20159106 | 0.84023643 | 0.88961952 | NA | NA |
| ENSG000002 | 2.97622052 | -0.075985053 | 0.37706183 | -0.2015188 | 0.84029292 | 0.88963287 | NA | NA |
| ENSG000002 | 2.99785777 | 0.075541434 | 0.37481704 | 0.20154216 | 0.84027467 | 0.88963287 | NA | NA |
| ENSG000002 | 2.94539459 | -0.075724108 | 0.3759376 | -0.2014273 | 0.84036445 | 0.88968537 | NA | NA |
| ENSG000001 | 29.1800361 | 0.049213247 | 0.24453279 | 0.20125418 | 0.84049983 | 0.88978223 | CDK20 | 23552 |
| ENSG000002 | 16.5099936 | 0.061046879 | 0.30330198 | 0.20127425 | 0.84048413 | 0.88978223 | DGUOK-AS1 | 100874048 |
| ENSG000002 | 6.02374397 | -0.073870327 | 0.36736481 | -0.2010817 | 0.84063472 | 0.8899018 | NA | NA |
| ENSG000001 | 63.012433 | -0.040333308 | 0.20070172 | -0.2009615 | 0.84072872 | 0.88997808 | TMEM44 | 93109 |

| | | | | | | | | |
|---|---|---|---|---|---|---|---|---|
| ENSG000002 | 4.66883322 | 0.075525158 | 0.3761262 | 0.20079739 | 0.84085701 | 0.89009065 | NA | NA |
| ENSG000002 | 3.28067095 | 0.075871401 | 0.37806008 | 0.2006861 | 0.84094403 | 0.89015953 | NA | NA |
| ENSG000002 | 3.17872922 | -0.075852776 | 0.37813277 | -0.2005983 | 0.84101272 | 0.890209 | NA | NA |
| ENSG000000 | 486.599973 | 0.030469942 | 0.1519556 | 0.20051871 | 0.84107492 | 0.89022837 | RUNX1T1 | 862 |
| ENSG000002 | 2.96210216 | -0.075534727 | 0.37664485 | -0.2005463 | 0.84105336 | 0.89022837 | NA | NA |
| ENSG000001 | 2.69003404 | -0.075316708 | 0.37566757 | -0.2004876 | 0.84109922 | 0.89023085 | NA | NA |
| ENSG000002 | 3.59214366 | -0.075724606 | 0.37853955 | -0.2000441 | 0.84144609 | 0.89057473 | LOC101927795 | 101927795 |
| ENSG000002 | 6.69794684 | 0.073589798 | 0.36799254 | 0.19997633 | 0.84149909 | 0.89059878 | NA | NA |
| ENSG000002 | 3.14147701 | -0.075277027 | 0.37646251 | -0.1999589 | 0.84151273 | 0.89059878 | RN7SL430P | 106481040 |
| ENSG000001 | 2.9430069 | -0.075299604 | 0.37678434 | -0.199848 | 0.84159944 | 0.8906673 | PSCA | 8000 |
| ENSG000002 | 3.07763335 | 0.07534494 | 0.37724235 | 0.19972556 | 0.84169522 | 0.89074542 | NA | NA |
| ENSG000001 | 13.3108866 | -0.062979365 | 0.31541531 | -0.1996712 | 0.84173771 | 0.89075413 | LIN7B | 64130 |
| ENSG000002 | 2.45298688 | -0.074319134 | 0.37223057 | -0.1996589 | 0.84174738 | 0.89075413 | NA | NA |
| ENSG000001 | 3.92063257 | -0.075562249 | 0.37857614 | -0.1995959 | 0.84179663 | 0.89075975 | NPIPB6 | 728741 |
| ENSG000002 | 2.93808784 | -0.075338535 | 0.37744555 | -0.1996011 | 0.84179259 | 0.89075975 | NA | NA |
| ENSG000002 | 4.59060179 | 0.07504889 | 0.37615536 | 0.19951568 | 0.84185938 | 0.89077966 | NA | NA |
| ENSG000002 | 2.68895714 | -0.074922011 | 0.37547911 | -0.1995371 | 0.84184263 | 0.89077966 | NA | NA |
| ENSG000002 | 2.46248565 | -0.074466687 | 0.373398 | -0.1994298 | 0.84192656 | 0.8908275 | NA | NA |
| ENSG000002 | 6.59823762 | 0.073248462 | 0.36743677 | 0.19934984 | 0.84198909 | 0.89087042 | NA | NA |
| ENSG000002 | 2.91535118 | -0.074969363 | 0.37616607 | -0.1992986 | 0.84202919 | 0.8908896 | NA | NA |
| ENSG000002 | 3.04961268 | 0.074978339 | 0.37665347 | 0.19906451 | 0.84221228 | 0.89105386 | NA | NA |
| ENSG000002 | 5.78232984 | -0.073759698 | 0.37056996 | -0.1990439 | 0.84222839 | 0.89105386 | NA | NA |
| ENSG000002 | 2.88611646 | -0.07482737 | 0.37620873 | -0.1988985 | 0.8423421 | 0.89115092 | NA | NA |
| ENSG000002 | 2.23874751 | -0.073609141 | 0.37033047 | -0.1987661 | 0.84244572 | 0.89123729 | NA | NA |
| ENSG000002 | 4.67209551 | 0.07484793 | 0.37663226 | 0.19872947 | 0.84247437 | 0.89124434 | NA | NA |
| ENSG000001 | 233.477984 | 0.027330198 | 0.13759249 | 0.19863146 | 0.84255104 | 0.8913022 | TAPT1 | 202018 |
| ENSG000002 | 3.41955407 | -0.074815785 | 0.3767302 | -0.1985925 | 0.84258154 | 0.89131121 | NA | NA |
| ENSG000001 | 3.27621612 | 0.074989652 | 0.37780479 | 0.19848783 | 0.84266341 | 0.89135252 | PRR34 | 55267 |
| ENSG000002 | 17.2915423 | 0.057494467 | 0.28966456 | 0.19848637 | 0.84266455 | 0.89135252 | NA | NA |

| | | | | | | | | |
|---|---|---|---|---|---|---|---|---|
| ENSG000002 | 9.01968821 | -0.068893454 | 0.34721902 | -0.198415 | 0.8427204 | 0.89138834 | ADAT3 | 113179 |
| ENSG000002 | 2.44591991 | -0.073359738 | 0.36984263 | -0.1983539 | 0.84276816 | 0.89141561 | NA | NA |
| ENSG000002 | 5.05371776 | -0.074610262 | 0.37628324 | -0.1982822 | 0.84282429 | 0.89142848 | NA | NA |
| ENSG000002 | 6.47989083 | -0.072209714 | 0.3641263 | -0.1983095 | 0.8428029 | 0.89142848 | NA | NA |
| ENSG000002 | 2.49035043 | -0.073641206 | 0.37147846 | -0.1982382 | 0.84285873 | 0.89144166 | NA | NA |
| ENSG000001 | 31.8604261 | 0.047471295 | 0.239565 | 0.19815622 | 0.84292284 | 0.89147461 | NKD2 | 85409 |
| ENSG000001 | 178.507818 | 0.029210908 | 0.14748675 | 0.19805784 | 0.84299981 | 0.89147461 | SCN9A | 6335 |
| ENSG000002 | 2.46433221 | -0.073755136 | 0.37235222 | -0.1980789 | 0.8429833 | 0.89147461 | NA | NA |
| ENSG000002 | 3.02797874 | 0.074245317 | 0.37480718 | 0.19808937 | 0.84297514 | 0.89147461 | NA | NA |
| ENSG000002 | 4.83520485 | 0.074030783 | 0.37370421 | 0.19809994 | 0.84296687 | 0.89147461 | NA | NA |
| ENSG000001 | 68.4357547 | -0.037747159 | 0.19070605 | -0.1979337 | 0.84309692 | 0.89154387 | TGFA | 7039 |
| ENSG000002 | 5.80410858 | -0.073031055 | 0.36907184 | -0.1978776 | 0.84314082 | 0.89154387 | NA | NA |
| ENSG000002 | 8.0032177 | 0.070469295 | 0.3561347 | 0.19787259 | 0.84314475 | 0.89154387 | NA | NA |
| ENSG000002 | 34.818057 | 0.046536694 | 0.23519805 | 0.19786173 | 0.84315325 | 0.89154387 | STAM-AS1 | 102723166 |
| ENSG000002 | 2.42574775 | -0.073185071 | 0.37031655 | -0.1976284 | 0.8433358 | 0.89171115 | NA | NA |
| ENSG000002 | 3.17730686 | -0.074649605 | 0.37777504 | -0.1976033 | 0.84335543 | 0.89171115 | MIR378J | 102465136 |
| ENSG000002 | 6.05345477 | -0.072509019 | 0.36702221 | -0.1975603 | 0.8433891 | 0.89172349 | NA | NA |
| ENSG000002 | 2.92624473 | -0.073854225 | 0.37390808 | -0.1975197 | 0.84342084 | 0.89173381 | NA | NA |
| ENSG000002 | 2.94656865 | -0.074174931 | 0.37582647 | -0.1973648 | 0.84354204 | 0.8918387 | NA | NA |
| ENSG000002 | 9.15521006 | -0.069888061 | 0.35465381 | -0.1970599 | 0.84378063 | 0.89206769 | NA | NA |
| ENSG000002 | 3.44267507 | -0.074130311 | 0.37647745 | -0.1969051 | 0.84390184 | 0.89217258 | NA | NA |
| ENSG000002 | 2.70158844 | -0.073456188 | 0.37326837 | -0.1967919 | 0.8439904 | 0.89224294 | NA | NA |
| ENSG000002 | 3.15339669 | -0.074388773 | 0.37822167 | -0.1966804 | 0.84407768 | 0.89231195 | NA | NA |
| ENSG000002 | 5.32821355 | -0.072889363 | 0.37093316 | -0.1965027 | 0.84421673 | 0.89243568 | NA | NA |
| ENSG000002 | 2.71294696 | -0.073859068 | 0.37595475 | -0.1964573 | 0.84425223 | 0.89244994 | C12orf71 | 728858 |
| ENSG000002 | 16.9482683 | -0.060467287 | 0.30808057 | -0.196271 | 0.84439805 | 0.89253428 | CFL1P1 | 142913 |
| ENSG000002 | 4.37479606 | -0.074304179 | 0.37849527 | -0.1963147 | 0.84436387 | 0.89253428 | NA | NA |
| ENSG000002 | 2.21051059 | -0.071404919 | 0.36379059 | -0.1962803 | 0.84439079 | 0.89253428 | MIR4790 | 100616334 |
| ENSG000002 | 2.90307177 | -0.073616665 | 0.37540664 | -0.1960985 | 0.8445331 | 0.89265377 | CCDC188 | 388849 |

| | | | | | | | | | |
|---|---|---|---|---|---|---|---|---|---|
| ENSG000002 | 2.92852326 | -0.073231284 | 0.37350561 | -0.1960648 | 0.84455949 | 0.89265839 | NA | NA | |
| ENSG000002 | 3.22611635 | 0.073964076 | 0.37738591 | 0.19599056 | 0.84461756 | 0.8926965 | NA | NA | |
| ENSG000002 | 59.0483913 | 0.038065795 | 0.19425176 | 0.19596114 | 0.84464058 | 0.89269757 | NA | NA | |
| ENSG000002 | 5.54868311 | -0.072779072 | 0.37160556 | -0.1958503 | 0.84472731 | 0.89274834 | TIFAB | | 497189 |
| ENSG000002 | 4.45066521 | -0.074102277 | 0.37837495 | -0.1958435 | 0.84473266 | 0.89274834 | NA | NA | |
| ENSG000001 | 6.00597157 | -0.071545429 | 0.36549959 | -0.1957469 | 0.84480824 | 0.89280495 | KCNF1 | | 3754 |
| ENSG000001 | 28.7387195 | 0.050481679 | 0.25793472 | 0.19571495 | 0.84483328 | 0.89280815 | CISH | | 1154 |
| ENSG00000C | 84.7541653 | 0.034251785 | 0.17509359 | 0.19561986 | 0.84490771 | 0.89286354 | BZRAP1 | | 9256 |
| ENSG000002 | 3.69993412 | -0.073754351 | 0.37719972 | -0.1955313 | 0.84497704 | 0.89291353 | NA | NA | |
| ENSG000001 | 16.8002393 | -0.056984506 | 0.29172918 | -0.1953336 | 0.84513181 | 0.89302922 | JPH2 | | 57158 |
| ENSG000002 | 2.91030738 | -0.073481389 | 0.37623522 | -0.195307 | 0.84515258 | 0.89302922 | NA | NA | |
| ENSG000002 | 2.45750154 | -0.072772742 | 0.37257558 | -0.1953234 | 0.84513975 | 0.89302922 | NA | NA | |
| ENSG000002 | 3.18405356 | -0.073734486 | 0.37767625 | -0.195232 | 0.84521133 | 0.89306803 | NA | NA | |
| ENSG000002 | 7.25429706 | -0.070869496 | 0.36337041 | -0.1950338 | 0.84536652 | 0.89320873 | NA | NA | |
| ENSG000001 | 38.5481509 | 0.044871041 | 0.23020123 | 0.19492094 | 0.84545484 | 0.89327877 | AGMO | | 392636 |
| ENSG000001 | 11.4004405 | 0.063901529 | 0.3279052 | 0.19487806 | 0.84548841 | 0.89329097 | ZNF280A | | 129025 |
| ENSG000002 | 2.90746323 | -0.073306158 | 0.37631919 | -0.1947978 | 0.84555123 | 0.89333406 | RN7SKP97 | | 106480463 |
| ENSG000002 | 4.82567451 | 0.073526081 | 0.37778662 | 0.1946233 | 0.84568786 | 0.89345513 | NA | NA | |
| ENSG000002 | 2.45836274 | -0.071727483 | 0.36902038 | -0.1943727 | 0.84588408 | 0.89363915 | NA | NA | |
| ENSG000002 | 2.66250252 | -0.072355457 | 0.37232367 | -0.1943348 | 0.84591372 | 0.89364718 | NA | NA | |
| ENSG000001 | 40.8406869 | -0.042750352 | 0.22005716 | -0.1942693 | 0.84596502 | 0.89367532 | VSTM4 | | 196740 |
| ENSG000002 | 14.1300295 | 0.061227218 | 0.31522695 | 0.19423218 | 0.84599408 | 0.89367532 | NA | NA | |
| ENSG000002 | 3.0646566 | 0.07335721 | 0.37774353 | 0.19419846 | 0.84602049 | 0.89367532 | NA | NA | |
| ENSG000002 | 3.46292373 | 0.073346089 | 0.37770616 | 0.19418822 | 0.8460285 | 0.89367532 | NA | NA | |
| ENSG000002 | 5.96525065 | 0.071343628 | 0.36765578 | 0.19405007 | 0.84613668 | 0.89376631 | NA | NA | |
| ENSG000002 | 5.9926303 | -0.071735523 | 0.36974675 | -0.1940126 | 0.84616603 | 0.89377402 | NA | NA | |
| ENSG000002 | 2.67324758 | -0.072864212 | 0.37569841 | -0.1939434 | 0.84622023 | 0.89380799 | NA | NA | |
| ENSG000002 | 4.4019184 | 0.073339202 | 0.37831111 | 0.1938595 | 0.8462859 | 0.89385407 | NA | NA | |
| ENSG000002 | 2.44691303 | -0.072166208 | 0.37231985 | -0.1938285 | 0.84631015 | 0.89385641 | NA | NA | |

| | | | | | | | | |
|---|---|---|---|---|---|---|---|---|
| ENSG000002 | 2.43855586 | -0.072256857 | 0.37304802 | -0.1936932 | 0.84641613 | 0.89392178 | NA | NA |
| ENSG000002 | 3.37404252 | -0.07326182 | 0.37820033 | -0.1937117 | 0.84640165 | 0.89392178 | NA | NA |
| ENSG000002 | 2.45907292 | -0.071772119 | 0.37155637 | -0.1931662 | 0.84682884 | 0.89433435 | NA | NA |
| ENSG000002 | 6.05450868 | 0.071146886 | 0.36846328 | 0.19309084 | 0.84688782 | 0.89437336 | NA | NA |
| ENSG000001 | 67.1659447 | -0.037541491 | 0.1945427 | -0.192973 | 0.8469801 | 0.89444751 | FAM84A | 151354 |
| ENSG000002 | 2.46974576 | -0.07088585 | 0.3675689 | -0.1928505 | 0.84707605 | 0.89452554 | MIR4762 | 100616253 |
| ENSG000002 | 5.27352255 | -0.071504944 | 0.37113524 | -0.1926655 | 0.84722097 | 0.89465528 | NA | NA |
| ENSG000002 | 10.6971905 | -0.063621555 | 0.33087017 | -0.1922856 | 0.84751853 | 0.8949462 | NA | NA |
| ENSG00000C | 180.845773 | -0.029321993 | 0.15259047 | -0.1921614 | 0.8476158 | 0.89502561 | ATP9A | 10079 |
| ENSG00000C | 573.678059 | -0.022474738 | 0.11699908 | -0.1920933 | 0.84766912 | 0.8950586 | IFT74 | 80173 |
| ENSG000001 | 2.66457035 | -0.0714469 | 0.37216392 | -0.191977 | 0.84776025 | 0.89513151 | NA | NA |
| ENSG00000C | 160.232418 | -0.028832742 | 0.15034794 | -0.1917734 | 0.84791967 | 0.89527653 | CCDC80 | 151887 |
| ENSG000001 | 13.8984576 | -0.059661662 | 0.31148411 | -0.19154 | 0.84810256 | 0.8952797 | GJD2 | 57369 |
| ENSG000002 | 5.10925066 | -0.071843912 | 0.37487734 | -0.1916465 | 0.84801915 | 0.8952797 | NA | NA |
| ENSG000002 | 2.67017509 | -0.071993237 | 0.37587503 | -0.191535 | 0.84810643 | 0.8952797 | FRG2 | 448831 |
| ENSG000002 | 2.43368832 | -0.071446702 | 0.37303441 | -0.1915284 | 0.8481116 | 0.8952797 | NA | NA |
| ENSG000002 | 3.06716904 | 0.07208537 | 0.37602358 | 0.19170439 | 0.84797377 | 0.8952797 | NA | NA |
| ENSG000002 | 2.22029772 | -0.070528349 | 0.36816937 | -0.191565 | 0.84808299 | 0.8952797 | RN7SL783P | 106479505 |
| ENSG000002 | 6.43731146 | 0.070029824 | 0.36569202 | 0.19149946 | 0.8481343 | 0.8952797 | NDUFA7 | 4701 |
| ENSG000002 | 4.92194302 | 0.072035077 | 0.37596234 | 0.19160184 | 0.8480541 | 0.8952797 | NA | NA |
| ENSG000002 | 15.6799882 | 0.057649975 | 0.30077475 | 0.19167159 | 0.84799946 | 0.8952797 | NA | NA |
| ENSG000002 | 3.59320095 | -0.072360676 | 0.37788664 | -0.1914878 | 0.84814345 | 0.8952797 | NA | NA |
| ENSG000002 | 2.69886827 | -0.071580039 | 0.37418618 | -0.1912953 | 0.84829428 | 0.89541561 | NA | NA |
| ENSG000002 | 3.21311202 | -0.072142349 | 0.37737486 | -0.1911689 | 0.84839324 | 0.89549676 | NA | NA |
| ENSG000001 | 80.2576612 | 0.034483007 | 0.18049792 | 0.19104379 | 0.84849128 | 0.89557693 | PHYKPL | 85007 |
| ENSG000001 | 56.6664057 | 0.044975242 | 0.23571846 | 0.19080068 | 0.84868175 | 0.89575465 | TGFB1 | 7040 |
| ENSG000002 | 5.86296936 | -0.071627771 | 0.37554806 | -0.1907286 | 0.8487382 | 0.8957676 | MIR4799 | 100616246 |
| ENSG000002 | 20.1737028 | -0.059581127 | 0.31237748 | -0.1907344 | 0.8487337 | 0.8957676 | NA | NA |
| ENSG000002 | 4.14848033 | 0.072132678 | 0.37829882 | 0.19067646 | 0.84877909 | 0.89578703 | NA | NA |

| | | | | | | | | |
|---|---|---|---|---|---|---|---|---|
| ENSG000002 | 50.1805364 | 0.041262318 | 0.2164311 | 0.19064875 | 0.84880079 | 0.89578703 | NA | NA |
| ENSG000002 | 3.15999923 | -0.072034432 | 0.37806189 | -0.1905361 | 0.84888907 | 0.89585688 | NA | NA |
| ENSG000002 | 21.6079616 | -0.052346579 | 0.2749778 | -0.1903666 | 0.8490219 | 0.89597374 | NA | NA |
| ENSG000002 | 4.64299684 | 0.071463102 | 0.37546132 | 0.19033413 | 0.84904731 | 0.89597724 | NA | NA |
| ENSG000002 | 4.40838834 | 0.071676731 | 0.37685831 | 0.19019543 | 0.84915599 | 0.89606861 | NA | NA |
| ENSG000002 | 3.30503794 | 0.071966886 | 0.37856388 | 0.190105 | 0.84922686 | 0.89612007 | NA | NA |
| ENSG000001 | 52.8617538 | -0.039932845 | 0.21013556 | -0.1900337 | 0.84928269 | 0.89613236 | CSPG5 | 10675 |
| ENSG000002 | 4.57575309 | 0.071483106 | 0.37612527 | 0.19005133 | 0.84926891 | 0.89613236 | NA | NA |
| ENSG000002 | 6.82505603 | -0.0679863 | 0.35784557 | -0.1899878 | 0.84931867 | 0.896147 | NA | NA |
| ENSG000002 | 3.25680568 | 0.071720611 | 0.37769932 | 0.18988811 | 0.84939681 | 0.89620613 | SNORD62A | 26786 |
| ENSG000002 | 2.48607575 | -0.06981628 | 0.36777391 | -0.1898348 | 0.8494386 | 0.8962269 | MIR1205 | 100302161 |
| ENSG000002 | 2.47850264 | -0.070171219 | 0.36972875 | -0.1897911 | 0.84947285 | 0.89623971 | NA | NA |
| ENSG000002 | 4.59026863 | 0.071492251 | 0.37690428 | 0.18968278 | 0.84955772 | 0.89630594 | NA | NA |
| ENSG000002 | 2.71647646 | -0.071205936 | 0.37549551 | -0.1896319 | 0.84959757 | 0.89632466 | NDUFS5P6 | 106480785 |
| ENSG000002 | 2.72698045 | -0.070324198 | 0.37113089 | -0.1894862 | 0.84971174 | 0.89642179 | NA | NA |
| ENSG000002 | 2.21825528 | -0.069744941 | 0.36816052 | -0.1894417 | 0.84974667 | 0.89643532 | NA | NA |
| ENSG000002 | 4.85313307 | -0.071237779 | 0.37610334 | -0.1894101 | 0.84977139 | 0.89643808 | NA | NA |
| ENSG000001 | 4.67936204 | 0.071257308 | 0.37632901 | 0.18934843 | 0.84981974 | 0.89646576 | FAM43B | 163933 |
| ENSG000002 | 2.66857647 | -0.071122354 | 0.37569391 | -0.1893093 | 0.8498504 | 0.89647478 | NA | NA |
| ENSG000002 | 3.29359543 | 0.071465625 | 0.37777198 | 0.18917662 | 0.84995439 | 0.89655838 | NA | NA |
| ENSG000002 | 2.23763815 | -0.069365883 | 0.36672077 | -0.1891518 | 0.84997387 | 0.89655838 | NA | NA |
| ENSG000002 | 2.18410858 | -0.069565019 | 0.36784192 | -0.1891166 | 0.85000142 | 0.89656412 | NA | NA |
| ENSG000002 | 2.90806384 | -0.070969862 | 0.3755995 | -0.1889509 | 0.8501313 | 0.89667779 | NA | NA |
| ENSG000001 | 80.5924971 | 0.034090626 | 0.18044756 | 0.18892261 | 0.85015347 | 0.89667785 | STAT5A | 6776 |
| ENSG000001 | 31.1190177 | -0.04618258 | 0.24455285 | -0.188845 | 0.85021431 | 0.89669538 | CNNM1 | 26507 |
| ENSG000002 | 5.93793909 | -0.069427908 | 0.36763001 | -0.1888527 | 0.85020829 | 0.89669538 | NA | NA |
| ENSG000002 | 6.17535468 | 0.068991167 | 0.36539713 | 0.18881146 | 0.85024059 | 0.89669977 | NA | NA |
| ENSG000002 | 152.367446 | 0.030483929 | 0.16148439 | 0.18877322 | 0.85027056 | 0.89670806 | NA | NA |
| ENSG000002 | 2.45773021 | -0.070512546 | 0.37383591 | -0.188619 | 0.85039146 | 0.89681223 | NA | NA |

| | | | | | | | | | |
|---|---|---|---|---|---|---|---|---|---|
| ENSG000002 | 72.0977795 | 0.036889321 | 0.1957338 | 0.18846679 | 0.85051074 | 0.89691471 | NA | NA | |
| ENSG000002 | 2.19071112 | -0.069679024 | 0.36978721 | -0.1884301 | 0.85053954 | 0.89692175 | NA | NA | |
| ENSG000001 | 33.7259957 | -0.047083996 | 0.25007226 | -0.1882816 | 0.85065594 | 0.89697923 | KLHDC8A | | 55220 |
| ENSG000002 | 6.54034336 | -0.068769046 | 0.36516624 | -0.1883226 | 0.85062379 | 0.89697923 | NA | NA | |
| ENSG000002 | 2.20386359 | -0.068592068 | 0.36431686 | -0.1882759 | 0.85066041 | 0.89697923 | NA | NA | |
| ENSG000001 | 2.43832752 | -0.069913205 | 0.37189507 | -0.1879917 | 0.85088312 | 0.89719074 | NA | NA | |
| ENSG000002 | 5.51913614 | -0.069218606 | 0.36836286 | -0.1879088 | 0.85094818 | 0.89723601 | NA | NA | |
| ENSG000002 | 6.00162904 | 0.069395963 | 0.36943963 | 0.18784115 | 0.85100118 | 0.89726856 | NA | NA | |
| ENSG000001 | 12.5549739 | 0.061809197 | 0.32912549 | 0.18779827 | 0.85103479 | 0.8972729 | NR5A1 | | 2516 |
| ENSG000002 | 2.24242137 | -0.069278058 | 0.36893312 | -0.1877794 | 0.85104955 | 0.8972729 | NA | NA | |
| ENSG000002 | 14.1968781 | -0.057751632 | 0.3078149 | -0.187618 | 0.85117608 | 0.89735964 | NA | NA | |
| ENSG000002 | 5.52175197 | -0.069951654 | 0.37282322 | -0.1876269 | 0.85116916 | 0.89735964 | NA | NA | |
| ENSG000001 | 93.74771 | 0.031530576 | 0.16808975 | 0.18758179 | 0.8512045 | 0.89736627 | LCTL | | 197021 |
| ENSG000002 | 2.73549461 | -0.069789002 | 0.3726263 | -0.1872895 | 0.85143364 | 0.8975845 | GPC5-AS1 | | 100873969 |
| ENSG000002 | 2.47751803 | -0.069244929 | 0.3698093 | -0.187245 | 0.85146857 | 0.89759799 | NA | NA | |
| ENSG000002 | 2.96265125 | -0.070132436 | 0.3748339 | -0.1871027 | 0.85158011 | 0.89769224 | NA | NA | |
| ENSG000002 | 4.64542386 | 0.070225362 | 0.37544726 | 0.18704454 | 0.85162571 | 0.89771697 | NA | NA | |
| ENSG000001 | 8.1633772 | -0.065278655 | 0.34927792 | -0.186896 | 0.85174215 | 0.89781638 | NA | NA | |
| ENSG000002 | 3.53273068 | 0.070584005 | 0.37858156 | 0.18644332 | 0.85209712 | 0.8981672 | NA | NA | |
| ENSG000002 | 2.9306241 | -0.070012009 | 0.37563313 | -0.186384 | 0.85214363 | 0.89819288 | NA | NA | |
| ENSG000002 | 2.48910468 | -0.068992996 | 0.37061829 | -0.1861565 | 0.85232205 | 0.89835759 | MAGEA9B | | 728269 |
| ENSG000002 | 5.74752612 | -0.06833489 | 0.36717116 | -0.1861118 | 0.85235708 | 0.89837116 | NA | NA | |
| ENSG00000C | 2.20326332 | -0.068685799 | 0.36920428 | -0.1860374 | 0.85241544 | 0.89840932 | NA | NA | |
| ENSG000002 | 20.0322262 | -0.052197637 | 0.2808764 | -0.1858385 | 0.85257145 | 0.89855039 | NA | NA | |
| ENSG000002 | 2.24306629 | -0.068104372 | 0.36679591 | -0.1856737 | 0.85270062 | 0.89866317 | NA | NA | |
| ENSG000002 | 2.4898419 | -0.068794087 | 0.3706193 | -0.1856193 | 0.85274333 | 0.89868123 | NA | NA | |
| ENSG000002 | 3.5231672 | 0.07021623 | 0.37832962 | 0.18559538 | 0.85276208 | 0.89868123 | NA | NA | |
| ENSG000001 | 58.3301155 | 0.041465905 | 0.22357558 | 0.18546706 | 0.85286271 | 0.89876215 | CRTC1 | | 23373 |
| ENSG000002 | 6.18369203 | 0.067454387 | 0.36381213 | 0.18540995 | 0.85290751 | 0.89876215 | NA | NA | |

| | | | | | | | | |
|---|---|---|---|---|---|---|---|---|
| ENSG000002 | 3.33464514 | 0.069988285 | 0.37753054 | 0.18538443 | 0.85292752 | 0.89876215 | NA | NA |
| ENSG000002 | 3.42817292 | -0.070069536 | 0.37794624 | -0.1853955 | 0.85291883 | 0.89876215 | NA | NA |
| ENSG000002 | 2.6768295 | -0.069323396 | 0.374162 | -0.1852764 | 0.85301223 | 0.89882806 | NA | NA |
| ENSG000002 | 2.47170477 | -0.069145084 | 0.37340985 | -0.1851721 | 0.85309405 | 0.89889092 | NA | NA |
| ENSG000000 | 98.0038391 | 0.033249131 | 0.17971578 | 0.18500952 | 0.85322157 | 0.89900192 | ZFHX4 | 79776 |
| ENSG000002 | 5.05181943 | -0.068918523 | 0.37270308 | -0.1849154 | 0.85329542 | 0.89903302 | NA | NA |
| ENSG000002 | 2.23797804 | -0.067922226 | 0.36726975 | -0.1849383 | 0.85327747 | 0.89903302 | NA | NA |
| ENSG000002 | 2.20186058 | -0.068233595 | 0.36919499 | -0.1848172 | 0.85337239 | 0.89906739 | NA | NA |
| ENSG000002 | 4.33678218 | -0.069962992 | 0.3785134 | -0.1848362 | 0.85335749 | 0.89906739 | NA | NA |
| ENSG000002 | 2.2188561 | -0.068334996 | 0.36988166 | -0.1847483 | 0.85342649 | 0.89910102 | NA | NA |
| ENSG000002 | 7.91642612 | -0.06515483 | 0.35304305 | -0.1845521 | 0.85358037 | 0.89919306 | NA | NA |
| ENSG000002 | 4.40286325 | 0.069671703 | 0.37749307 | 0.18456419 | 0.85357088 | 0.89919306 | ENPP7P12 | 106480230 |
| ENSG000002 | 2.19736573 | -0.068325666 | 0.3701879 | -0.1845702 | 0.85356613 | 0.89919306 | NA | NA |
| ENSG000002 | 27.4877108 | 0.050033381 | 0.27119708 | 0.18449086 | 0.8536284 | 0.89922029 | NA | NA |
| ENSG000002 | 13.4818142 | -0.056841749 | 0.30817826 | -0.1844444 | 0.85366485 | 0.89923534 | NA | NA |
| ENSG000001 | 141.938394 | -0.028566779 | 0.15498566 | -0.1843189 | 0.85376332 | 0.8993157 | ITGB8 | 3696 |
| ENSG000002 | 3.12561678 | -0.069420167 | 0.37675402 | -0.1842586 | 0.85381059 | 0.89934213 | NA | NA |
| ENSG000002 | 2.47535084 | -0.067999161 | 0.36919171 | -0.1841839 | 0.85386921 | 0.89935715 | NA | NA |
| ENSG000002 | 2.24598312 | -0.067347543 | 0.36562569 | -0.1841981 | 0.85385808 | 0.89935715 | NA | NA |
| ENSG000002 | 3.14437399 | -0.069447325 | 0.37731809 | -0.1840551 | 0.85397022 | 0.89944018 | NA | NA |
| ENSG000002 | 4.68485545 | 0.06933884 | 0.37684834 | 0.18399667 | 0.85401607 | 0.89945854 | NA | NA |
| ENSG000002 | 2.41604788 | -0.068157246 | 0.37046746 | -0.1839763 | 0.85403202 | 0.89945854 | NA | NA |
| ENSG000002 | 5.12806555 | 0.068765162 | 0.37399304 | 0.18386749 | 0.85411741 | 0.89952511 | NA | NA |
| ENSG000002 | 5.96225475 | -0.067762833 | 0.36862936 | -0.1838238 | 0.85415172 | 0.89953789 | NA | NA |
| ENSG000002 | 5.06666281 | -0.069039857 | 0.37575632 | -0.1837357 | 0.85422079 | 0.89958726 | NA | NA |
| ENSG000002 | 2.48517023 | -0.068049085 | 0.37060394 | -0.1836167 | 0.85431413 | 0.8996622 | LOC105376992 | 105376992 |
| ENSG000002 | 2.93508669 | -0.06904009 | 0.37607152 | -0.1835823 | 0.85434112 | 0.89966725 | NA | NA |
| ENSG000002 | 8.02496036 | 0.064500598 | 0.35146045 | 0.18352164 | 0.85438874 | 0.89969404 | NA | NA |
| ENSG000002 | 5.71238901 | 0.068263695 | 0.37206155 | 0.1834742 | 0.85442596 | 0.89970987 | NA | NA |

| | | | | | | | | |
|---|---|---|---|---|---|---|---|---|
| ENSG000002 | 4.99151094 | -0.069385837 | 0.37825884 | -0.1834348 | 0.85445687 | 0.89971905 | NA | NA |
| ENSG000002 | 36.0673358 | 0.042722456 | 0.23300983 | 0.18335044 | 0.85452306 | 0.89974202 | NA | NA |
| ENSG000002 | 2.18817941 | -0.067489533 | 0.36804653 | -0.1833723 | 0.85450592 | 0.89974202 | NA | NA |
| ENSG000002 | 19.6823815 | 0.051816207 | 0.28269548 | 0.18329337 | 0.85456784 | 0.89976581 | ARHGAP31-AS1 | 100874246 |
| ENSG000002 | 6.52827127 | 0.067956174 | 0.37093291 | 0.18320341 | 0.85463842 | 0.89981676 | NA | NA |
| ENSG000002 | 2.19571504 | -0.067794916 | 0.37017939 | -0.1831407 | 0.85468761 | 0.89984519 | NA | NA |
| ENSG000002 | 2.46450882 | -0.067851958 | 0.37076314 | -0.1830062 | 0.85479314 | 0.89993293 | NA | NA |
| ENSG000002 | 2.22510577 | -0.067178423 | 0.36721974 | -0.1829379 | 0.85484671 | 0.89996597 | RNA5SP302 | 106479004 |
| ENSG000001 | 134.786498 | -0.02810556 | 0.15370811 | -0.1828502 | 0.85491556 | 0.89999489 | PTPRM | 5797 |
| ENSG000001 | 2.23072316 | -0.067503751 | 0.36929718 | -0.1827898 | 0.85496297 | 0.89999489 | RNU6-836P | 106481918 |
| ENSG000002 | 2.65128093 | -0.068338923 | 0.37375425 | -0.1828445 | 0.85492 | 0.89999489 | NA | NA |
| ENSG000002 | 2.21583513 | -0.06749692 | 0.36924319 | -0.182798 | 0.85495651 | 0.89999489 | NA | NA |
| ENSG000002 | 6.68493562 | 0.066537711 | 0.36414902 | 0.1827211 | 0.85501686 | 0.90002567 | NA | NA |
| ENSG000002 | 3.26362316 | 0.069092104 | 0.37818081 | 0.18269595 | 0.85503659 | 0.90002567 | RN7SL354P | 106481838 |
| ENSG000001 | 94.7725202 | 0.030707544 | 0.168113 | 0.18266015 | 0.85506469 | 0.90003188 | RNF180 | 285671 |
| ENSG000002 | 2.21134029 | -0.06759885 | 0.37023445 | -0.1825839 | 0.85512452 | 0.90007149 | EIF1AX-AS1 | 100874078 |
| ENSG000001 | 2.66343615 | -0.068310695 | 0.3744064 | -0.1824507 | 0.85522907 | 0.90015818 | OR5B17 | 219965 |
| ENSG000002 | 2.67458561 | -0.068321784 | 0.37488974 | -0.182245 | 0.85539045 | 0.90025951 | NA | NA |
| ENSG000002 | 3.15460335 | -0.068395817 | 0.37525054 | -0.1822671 | 0.85537314 | 0.90025951 | NA | NA |
| ENSG000002 | 2.23146037 | -0.067302265 | 0.36929938 | -0.1822431 | 0.85539195 | 0.90025951 | NA | NA |
| ENSG000001 | 3.50244575 | 0.068944915 | 0.37856371 | 0.18212235 | 0.85548671 | 0.90033587 | LCN12 | 286256 |
| ENSG000002 | 6.04045448 | 0.067193793 | 0.36926343 | 0.18196709 | 0.85560855 | 0.90044073 | NTF4 | 4909 |
| ENSG000002 | 2.44166115 | -0.067606199 | 0.37162934 | -0.1819184 | 0.8556468 | 0.90045761 | NA | NA |
| ENSG000002 | 2.92640924 | -0.068241778 | 0.37559564 | -0.1816895 | 0.85582642 | 0.90062327 | RNU4-59P | 106481764 |
| ENSG000002 | 2.23413588 | -0.066847753 | 0.36821459 | -0.1815456 | 0.85593931 | 0.90069728 | NA | NA |
| ENSG000002 | 2.6521625 | -0.067941683 | 0.37424511 | -0.1815433 | 0.85594117 | 0.90069728 | NA | NA |
| ENSG000002 | 3.34112295 | 0.068666372 | 0.37830024 | 0.18151289 | 0.85596501 | 0.900699 | NA | NA |
| ENSG000002 | 2.21529435 | -0.067057991 | 0.36964538 | -0.1814117 | 0.85604445 | 0.90075921 | NA | NA |
| ENSG000002 | 2.91059575 | -0.067959573 | 0.37475878 | -0.1813422 | 0.856099 | 0.90079324 | NA | NA |

| | | | | | | | | |
|---|---|---|---|---|---|---|---|---|
| ENSG000002 | 2.70745432 | -0.0679989 | 0.37510624 | -0.181279 | 0.85614856 | 0.90082201 | NA | NA |
| ENSG000002 | 9.92831265 | 0.061169503 | 0.33778986 | 0.18108745 | 0.85629894 | 0.90095686 | NA | NA |
| ENSG000002 | 4.8761579 | -0.068118412 | 0.3764682 | -0.1809407 | 0.85641414 | 0.90105469 | NA | NA |
| ENSG000001 | 103.907358 | -0.033489374 | 0.18533726 | -0.1806942 | 0.85660759 | 0.90119484 | TMEM62 | 80021 |
| ENSG000002 | 3.4625033 | -0.067902612 | 0.37572684 | -0.1807233 | 0.85658473 | 0.90119484 | NA | NA |
| ENSG000002 | 2.45832362 | -0.067133497 | 0.37154778 | -0.180686 | 0.85661402 | 0.90119484 | NA | NA |
| ENSG000002 | 2.24343843 | -0.065380326 | 0.3620397 | -0.1805888 | 0.85669032 | 0.90125173 | NA | NA |
| ENSG000002 | 2.22124345 | -0.066056875 | 0.36587327 | -0.1805458 | 0.85672412 | 0.9012639 | NA | NA |
| ENSG000002 | 27.4315682 | -0.04602897 | 0.25505392 | -0.1804676 | 0.85678549 | 0.9012817 | NA | NA |
| ENSG000002 | 2.44475401 | -0.067410984 | 0.37352314 | -0.1804734 | 0.85678095 | 0.9012817 | NA | NA |
| ENSG00000C | 87.5528968 | -0.032932602 | 0.18271634 | -0.180239 | 0.85696498 | 0.90142786 | TCF7 | 6932 |
| ENSG000001 | 2.2096896 | -0.066655933 | 0.36984717 | -0.1802256 | 0.85697545 | 0.90142786 | MOS | 4342 |
| ENSG000002 | 2.92251522 | -0.067936371 | 0.37704627 | -0.1801805 | 0.85701089 | 0.90142786 | NA | NA |
| ENSG000002 | 17.6911235 | 0.053041571 | 0.29438537 | 0.18017733 | 0.85701335 | 0.90142786 | NA | NA |
| ENSG000001 | 12.0137645 | 0.061130667 | 0.33966009 | 0.17997601 | 0.85717141 | 0.90154734 | IRX3 | 79191 |
| ENSG000002 | 5.95047864 | -0.066317045 | 0.36842467 | -0.1800016 | 0.85715128 | 0.90154734 | RIMBP3 | 85376 |
| ENSG000002 | 2.19896435 | -0.066088787 | 0.36736091 | -0.1799015 | 0.85722988 | 0.90158546 | NA | NA |
| ENSG000002 | 5.35532813 | -0.067414962 | 0.37579038 | -0.1793951 | 0.85762746 | 0.90196676 | NA | NA |
| ENSG000002 | 3.10393143 | 0.067273932 | 0.3750294 | 0.17938309 | 0.85763691 | 0.90196676 | RN7SL672P | 106481106 |
| ENSG000002 | 7.76080603 | 0.063656622 | 0.35518666 | 0.17922019 | 0.85776481 | 0.90203109 | CYP2D7 | 1564 |
| ENSG000002 | 9.82235622 | 0.061598006 | 0.34369617 | 0.17922227 | 0.85776318 | 0.90203109 | NA | NA |
| ENSG000002 | 7.42186582 | -0.063907439 | 0.35647555 | -0.1792758 | 0.85772115 | 0.90203109 | NA | NA |
| ENSG000002 | 4.8295676 | -0.067712695 | 0.37845336 | -0.1789195 | 0.85800089 | 0.90225596 | NA | NA |
| ENSG000001 | 86.5565754 | 0.031813035 | 0.17795515 | 0.17876996 | 0.85811833 | 0.90235004 | RASGRP1 | 10125 |
| ENSG000002 | 2.9427595 | -0.066515102 | 0.37211474 | -0.1787489 | 0.85813487 | 0.90235004 | NA | NA |
| ENSG000002 | 5.26187734 | 0.067197018 | 0.37615812 | 0.17864035 | 0.85822011 | 0.90241627 | NA | NA |
| ENSG000002 | 2.19478196 | -0.065725811 | 0.36806908 | -0.1785692 | 0.85827596 | 0.90242894 | NA | NA |
| ENSG000002 | 3.61736017 | -0.067462082 | 0.37785418 | -0.17854 | 0.85829892 | 0.90242894 | NA | NA |
| ENSG000002 | 12.6213655 | -0.058982963 | 0.33034797 | -0.178548 | 0.85829265 | 0.90242894 | TCP10L | 140290 |

| | | | | | | | | | |
|---|---|---|---|---|---|---|---|---|---|
| ENSG000002 | 3.40486301 | -0.067196378 | 0.37657225 | -0.1784422 | 0.85837571 | 0.90248629 | NA | NA | |
| ENSG000001 | 36.3168373 | 0.043740297 | 0.24519491 | 0.17838991 | 0.85841677 | 0.90250606 | EMP3 | | 2014 |
| ENSG000001 | 54.3097244 | -0.035534362 | 0.19922749 | -0.1783607 | 0.85843968 | 0.90250674 | HCLS1 | | 3059 |
| ENSG000002 | 3.3003737 | 0.067400072 | 0.37821029 | 0.17820793 | 0.85855969 | 0.9026095 | NA | NA | |
| ENSG000002 | 2.47344719 | -0.065882579 | 0.37022946 | -0.1779507 | 0.85876172 | 0.9027985 | NA | NA | |
| ENSG000001 | 3.51362801 | 0.067266259 | 0.37847142 | 0.17773141 | 0.85893392 | 0.90295612 | NA | NA | |
| ENSG000002 | 10.5187413 | 0.059455383 | 0.33492009 | 0.1775211 | 0.85909909 | 0.90310635 | NA | NA | |
| ENSG000002 | 5.04391962 | -0.06620819 | 0.37353006 | -0.17725 | 0.85931205 | 0.90328716 | NA | NA | |
| ENSG000002 | 2.22115968 | -0.065521205 | 0.36966382 | -0.1772454 | 0.85931565 | 0.90328716 | NA | NA | |
| ENSG000002 | 2.18009036 | -0.065337083 | 0.36870544 | -0.1772067 | 0.85934601 | 0.90329566 | NA | NA | |
| ENSG000002 | 7.57047684 | 0.063106687 | 0.35633552 | 0.17709906 | 0.85943058 | 0.90336114 | NA | NA | |
| ENSG000002 | 2.21091518 | -0.063544081 | 0.359021 | -0.1769927 | 0.85951416 | 0.90342557 | NA | NA | |
| ENSG000002 | 5.14986858 | 0.066064313 | 0.37336808 | 0.17694151 | 0.85955433 | 0.90344438 | OR2A7 | | 401427 |
| ENSG000001 | 10.2126267 | -0.059448126 | 0.33606096 | -0.1768969 | 0.85958941 | 0.90345784 | MS4A6E | | 245802 |
| ENSG000001 | 98.4236821 | 0.031204601 | 0.17656523 | 0.17673129 | 0.85971946 | 0.9035711 | ZNF624 | | 57547 |
| ENSG000001 | 16.3493508 | 0.052871559 | 0.29927169 | 0.17666742 | 0.85976963 | 0.90360041 | CLIC6 | | 54102 |
| ENSG000002 | 5.48236623 | -0.066231531 | 0.37530605 | -0.1764734 | 0.85992205 | 0.90373718 | NA | NA | |
| ENSG000002 | 17.349234 | -0.052417528 | 0.29714665 | -0.1764029 | 0.85997743 | 0.90377196 | NA | NA | |
| ENSG000001 | 356.0711 | 0.029584716 | 0.1678075 | 0.17630151 | 0.86005707 | 0.90383223 | SYT2 | | 127833 |
| ENSG000001 | 72.9509848 | 0.031388121 | 0.1781288 | 0.17621025 | 0.86012877 | 0.9038759 | TVP23C | | 201158 |
| ENSG000002 | 2.45279797 | -0.064771827 | 0.36762096 | -0.1761919 | 0.8601432 | 0.9038759 | NA | NA | |
| ENSG000002 | 19.5140411 | 0.050423903 | 0.28670636 | 0.17587298 | 0.86039373 | 0.90411574 | C8orf34-AS1 | | 286189 |
| ENSG000002 | 6.51037458 | 0.063607527 | 0.36182585 | 0.17579597 | 0.86045423 | 0.90413071 | NA | NA | |
| ENSG000002 | 5.45795938 | -0.065354269 | 0.37175844 | -0.1757977 | 0.8604529 | 0.90413071 | NA | NA | |
| ENSG000002 | 2.17427921 | 0.064629924 | 0.3676966 | 0.17576971 | 0.86047487 | 0.90413071 | NA | NA | |
| ENSG000002 | 2.43501316 | -0.064324905 | 0.36624817 | -0.175632 | 0.86058305 | 0.90419752 | NA | NA | |
| ENSG000002 | 3.50632072 | 0.06649941 | 0.37859292 | 0.17564885 | 0.86056982 | 0.90419752 | NA | NA | |
| ENSG000002 | 4.89602157 | 0.065498339 | 0.37315651 | 0.17552511 | 0.86066704 | 0.90426234 | NA | NA | |
| ENSG000002 | 2.93057237 | -0.065951064 | 0.37592612 | -0.1754362 | 0.86073687 | 0.90429219 | NA | NA | |

| | | | | | | | | |
|---|---|---|---|---|---|---|---|---|
| ENSG000002 | 11.3594956 | -0.057965461 | 0.33041522 | -0.1754322 | 0.86074006 | 0.90429219 | NA | NA |
| ENSG000002 | 5.78837428 | -0.064929182 | 0.37018541 | -0.1753964 | 0.86076818 | 0.90429831 | NA | NA |
| ENSG000001 | 33.4977041 | -0.042346928 | 0.24158736 | -0.1752862 | 0.86085476 | 0.90435256 | WNT5B | 81029 |
| ENSG000001 | 24.1734341 | -0.046433319 | 0.2649261 | -0.1752689 | 0.86086831 | 0.90435256 | TFEB | 7942 |
| ENSG000002 | 12.4983794 | -0.057189846 | 0.3263413 | -0.1752455 | 0.86088673 | 0.90435256 | NA | NA |
| ENSG000001 | 23.5983342 | 0.048121814 | 0.27522965 | 0.17484241 | 0.86120347 | 0.90461168 | INHBB | 3625 |
| ENSG000002 | 2.41613275 | -0.064912547 | 0.37131818 | -0.1748165 | 0.86122382 | 0.90461168 | NA | NA |
| ENSG000002 | 5.28107744 | -0.065649482 | 0.37559138 | -0.1747896 | 0.86124493 | 0.90461168 | NA | NA |
| ENSG000002 | 3.0843723 | 0.065722033 | 0.37598992 | 0.17479733 | 0.86123889 | 0.90461168 | NA | NA |
| ENSG000002 | 5.04545804 | 0.065719369 | 0.3757822 | 0.17488686 | 0.86116853 | 0.90461168 | NA | NA |
| ENSG000001 | 4.64250723 | 0.066009949 | 0.37828449 | 0.17449816 | 0.86147397 | 0.90475852 | GPR62 | 118442 |
| ENSG000002 | 22.9592531 | -0.046752692 | 0.26789956 | -0.1745157 | 0.86146016 | 0.90475852 | NA | NA |
| ENSG000002 | 6.2250738 | 0.063603674 | 0.36437456 | 0.17455575 | 0.86142872 | 0.90475852 | NA | NA |
| ENSG000002 | 5.4991602 | -0.064568292 | 0.36993075 | -0.1745416 | 0.86143987 | 0.90475852 | NA | NA |
| ENSG000001 | 24.6813595 | -0.047477826 | 0.27217851 | -0.1744364 | 0.86152255 | 0.9047861 | STAC2 | 342667 |
| ENSG000002 | 3.49590792 | 0.06601054 | 0.37856389 | 0.17437094 | 0.86157395 | 0.90479322 | NA | NA |
| ENSG000002 | 5.29078617 | -0.064962319 | 0.37253497 | -0.1743791 | 0.86156752 | 0.90479322 | NA | NA |
| ENSG000002 | 2.18413086 | 0.064441915 | 0.36966715 | 0.17432416 | 0.86161071 | 0.90480839 | NA | NA |
| ENSG000002 | 55.8188124 | 0.035457827 | 0.20362839 | 0.17413007 | 0.86176324 | 0.90494513 | PTCHD2 | 57540 |
| ENSG000002 | 3.06054589 | 0.065477061 | 0.37624083 | 0.17402965 | 0.86184216 | 0.9049577 | NA | NA |
| ENSG000002 | 2.67390081 | -0.06468333 | 0.37157756 | -0.1740776 | 0.86180448 | 0.9049577 | NA | NA |
| ENSG000002 | 6.72726278 | 0.062722218 | 0.36038959 | 0.17404004 | 0.86183399 | 0.9049577 | NA | NA |
| ENSG000002 | 8.35004563 | 0.062183949 | 0.35753361 | 0.17392477 | 0.86192459 | 0.90502082 | NA | NA |
| ENSG000002 | 8.55941743 | 0.062117218 | 0.3572085 | 0.17389625 | 0.861947 | 0.90502092 | NA | NA |
| ENSG000002 | 14.4697411 | -0.05412801 | 0.31142343 | -0.1738084 | 0.86201604 | 0.90506997 | NA | NA |
| ENSG000002 | 7.49745918 | -0.063836639 | 0.36742375 | -0.1737412 | 0.86206886 | 0.90507456 | NA | NA |
| ENSG000002 | 2.1993889 | -0.064222276 | 0.36958597 | -0.1737682 | 0.86204766 | 0.90507456 | NA | NA |
| ENSG000002 | 4.88204702 | 0.064873336 | 0.37344125 | 0.17371765 | 0.86208737 | 0.90507456 | NA | NA |
| ENSG000002 | 8.56674467 | -0.060089445 | 0.34604072 | -0.1736485 | 0.86214173 | 0.9051082 | NA | NA |

| | | | | | | | | |
|---|---|---|---|---|---|---|---|---|
| ENSG000001 | 4.63693454 | 0.065075652 | 0.37510314 | 0.17348736 | 0.86226837 | 0.90521771 | SNORA60 | 677837 |
| ENSG000001 | 36.1911419 | 0.040458787 | 0.23344079 | 0.17331498 | 0.86240385 | 0.90525609 | ELF4 | 2000 |
| ENSG000001 | 94.8113555 | -0.031175156 | 0.17987127 | -0.1733193 | 0.86240048 | 0.90525609 | ZNF440 | 126070 |
| ENSG000002 | 32.8064464 | 0.044146087 | 0.25467998 | 0.17333945 | 0.86238462 | 0.90525609 | NA | NA |
| ENSG000002 | 5.57942003 | -0.064374641 | 0.37139242 | -0.1733332 | 0.86238952 | 0.90525609 | NA | NA |
| ENSG000002 | 2.21123614 | -0.06339919 | 0.36583739 | -0.1732988 | 0.86241655 | 0.90525609 | NA | NA |
| ENSG000001 | 159.300809 | 0.025736797 | 0.14854183 | 0.17326296 | 0.86244473 | 0.90526225 | FAM110C | 642273 |
| ENSG000002 | 2.43151646 | -0.063733832 | 0.36814365 | -0.1731222 | 0.86255539 | 0.90530809 | PCGEM1 | 64002 |
| ENSG000002 | 9.95859669 | 0.058595068 | 0.33835883 | 0.17317434 | 0.86251439 | 0.90530809 | NA | NA |
| ENSG000002 | 15.7600146 | -0.051284228 | 0.29618373 | -0.1731501 | 0.86253348 | 0.90530809 | NA | NA |
| ENSG000001 | 13.0253207 | 0.05397617 | 0.31187467 | 0.17307007 | 0.86259635 | 0.90531856 | SGCG | 6445 |
| ENSG000002 | 5.47310233 | -0.064698978 | 0.37386865 | -0.1730527 | 0.86261001 | 0.90531856 | NA | NA |
| ENSG000001 | 5.24132709 | -0.06473863 | 0.37432646 | -0.172947 | 0.8626931 | 0.90536911 | ASB12 | 142689 |
| ENSG000002 | 2.67509469 | -0.06478323 | 0.37461113 | -0.1729346 | 0.86270283 | 0.90536911 | NA | NA |
| ENSG000002 | 2.19919303 | -0.063678997 | 0.36918534 | -0.1724852 | 0.86305612 | 0.90571643 | NA | NA |
| ENSG000002 | 2.20825461 | -0.062411489 | 0.36191067 | -0.17245 | 0.86308378 | 0.90572202 | NA | NA |
| ENSG000002 | 62.1390902 | 0.033932584 | 0.19725811 | 0.17202124 | 0.86342083 | 0.90605228 | ZNF790-AS1 | 284408 |
| ENSG000002 | 3.2346889 | 0.064803697 | 0.37697857 | 0.17190287 | 0.86351389 | 0.90612648 | RNU6-1143P | 106480069 |
| ENSG000002 | 4.85870953 | 0.06423329 | 0.37423342 | 0.17163964 | 0.86372084 | 0.90632019 | NA | NA |
| ENSG000002 | 2.41424074 | -0.063544044 | 0.37028642 | -0.1716078 | 0.86374586 | 0.90632299 | RNU6-1111P | 106481530 |
| ENSG000002 | 2.69065467 | -0.063786592 | 0.37179651 | -0.1715632 | 0.86378095 | 0.90633637 | NA | NA |
| ENSG000002 | 2.46686948 | -0.062663055 | 0.36545399 | -0.1714663 | 0.8638571 | 0.90639281 | NA | NA |
| ENSG00000C | 231.197551 | 0.023479677 | 0.13702577 | 0.17135227 | 0.86394678 | 0.90644 | IKZF2 | 22807 |
| ENSG000002 | 10.4636688 | -0.05739715 | 0.33493985 | -0.1713655 | 0.86393635 | 0.90644 | NA | NA |
| ENSG000002 | 2.42064741 | -0.063586455 | 0.37184976 | -0.1710004 | 0.86422346 | 0.90670682 | RN7SL147P | 106479274 |
| ENSG000001 | 2.20177681 | -0.062830503 | 0.3679017 | -0.1707807 | 0.86439622 | 0.90679423 | RNU6-1079P | 106480040 |
| ENSG000002 | 8.01438786 | -0.059916148 | 0.35078119 | -0.1708078 | 0.86437493 | 0.90679423 | NA | NA |
| ENSG000002 | 10.7483841 | 0.058264375 | 0.34104476 | 0.17084085 | 0.86434891 | 0.90679423 | NA | NA |
| ENSG000002 | 2.65597241 | -0.063487734 | 0.37174522 | -0.1707829 | 0.86439446 | 0.90679423 | NA | NA |

| | | | | | | | | |
|---|---|---|---|---|---|---|---|---|
| ENSG000001 | 36.2653165 | 0.040803019 | 0.23938716 | 0.17044782 | 0.86465797 | 0.90699843 | TMEM81 | 388730 |
| ENSG000002 | 2.19827955 | -0.062935535 | 0.36917858 | -0.1704745 | 0.86463698 | 0.90699843 | GTF3AP2 | 106480682 |
| ENSG000002 | 10.3975399 | -0.056927485 | 0.33390781 | -0.1704886 | 0.86462587 | 0.90699843 | NA | NA |
| ENSG000000 | 2.61479895 | -0.063138102 | 0.37078364 | -0.1702829 | 0.86478768 | 0.90706411 | PAGE1 | 8712 |
| ENSG000002 | 20.2376762 | -0.050881351 | 0.29877757 | -0.1702984 | 0.86477545 | 0.90706411 | NA | NA |
| ENSG000002 | 3.33414792 | 0.064447699 | 0.3783914 | 0.1703202 | 0.86475833 | 0.90706411 | NA | NA |
| ENSG000002 | 5.75502264 | -0.063206191 | 0.37161673 | -0.1700843 | 0.8649438 | 0.90720439 | NA | NA |
| ENSG000002 | 2.41771872 | -0.06329448 | 0.37248934 | -0.1699229 | 0.86507075 | 0.90731407 | NA | NA |
| ENSG000001 | 2.71400479 | -0.063414434 | 0.37326454 | -0.1698914 | 0.86509555 | 0.90731661 | NA | NA |
| ENSG000002 | 9.63052349 | 0.05719177 | 0.33670458 | 0.16985742 | 0.86512227 | 0.90732117 | NA | NA |
| ENSG000002 | 6.18579863 | 0.062241335 | 0.36682793 | 0.16967447 | 0.86526615 | 0.90744861 | RPS6KA2-IT1 | 100874353 |
| ENSG000002 | 2.45107615 | -0.063157862 | 0.37231514 | -0.1696355 | 0.86529681 | 0.90745729 | NA | NA |
| ENSG000002 | 5.74001827 | -0.062811212 | 0.37051993 | -0.1695218 | 0.86538621 | 0.90752758 | NA | NA |
| ENSG000002 | 3.3212957 | 0.064121989 | 0.37834587 | 0.16947982 | 0.86541925 | 0.90753875 | NA | NA |
| ENSG000001 | 3.51934899 | 0.064106915 | 0.37859337 | 0.16932921 | 0.8655377 | 0.90763006 | LMO1 | 4004 |
| ENSG000002 | 3.3385909 | 0.064022945 | 0.37813547 | 0.16931219 | 0.86555109 | 0.90763006 | NA | NA |
| ENSG000000 | 3.5471289 | 0.064068506 | 0.37859062 | 0.16922898 | 0.86561653 | 0.90767522 | NA | NA |
| ENSG000002 | 5.04483309 | -0.063081106 | 0.37284701 | -0.1691876 | 0.86564905 | 0.90768584 | NA | NA |
| ENSG000002 | 4.80736835 | -0.063051051 | 0.37349027 | -0.1688158 | 0.86594155 | 0.90794559 | NA | NA |
| ENSG000002 | 2.1775264 | -0.062320878 | 0.36910396 | -0.1688437 | 0.86591958 | 0.90794559 | NA | NA |
| ENSG000002 | 4.41900911 | 0.063643373 | 0.37724021 | 0.16870782 | 0.86602647 | 0.90801115 | NA | NA |
| ENSG000002 | 20.3459521 | 0.04743779 | 0.28127834 | 0.1686507 | 0.8660714 | 0.90803478 | NA | NA |
| ENSG000002 | 2.18239338 | -0.061774532 | 0.36649667 | -0.1685541 | 0.86614735 | 0.90809094 | NA | NA |
| ENSG000002 | 2.47378001 | -0.062485491 | 0.3708988 | -0.1684705 | 0.86621318 | 0.908113 | NA | NA |
| ENSG000002 | 2.17673711 | -0.062113525 | 0.3686878 | -0.1684719 | 0.86621207 | 0.908113 | NA | NA |
| ENSG000001 | 22.6386837 | -0.045718619 | 0.27169815 | -0.1682699 | 0.86637096 | 0.90815165 | RAB9B | 51209 |
| ENSG000002 | 2.18524021 | 0.061877422 | 0.36745815 | 0.16839311 | 0.86627403 | 0.90815165 | NA | NA |
| ENSG000002 | 2.44159699 | -0.062483627 | 0.37136757 | -0.1682528 | 0.86638442 | 0.90815165 | NA | NA |
| ENSG000002 | 4.98340237 | -0.06282253 | 0.37330871 | -0.1682857 | 0.8663585 | 0.90815165 | NA | NA |

| | | | | | | | | |
|---|---|---|---|---|---|---|---|---|
| ENSG000002 | 15.2055829 | 0.051406601 | 0.30540165 | 0.16832457 | 0.86632795 | 0.90815165 | NA | NA |
| ENSG000002 | 14.8594095 | -0.051672318 | 0.30701815 | -0.1683038 | 0.86634429 | 0.90815165 | NA | NA |
| ENSG000000 | 70.2411326 | -0.031222682 | 0.18590769 | -0.1679472 | 0.86662479 | 0.90838013 | SCML1 | 6322 |
| ENSG000002 | 14.5934777 | 0.063366206 | 0.37736924 | 0.16791566 | 0.86664963 | 0.90838268 | NA | NA |
| ENSG000002 | 5.24379965 | -0.06222662 | 0.3707569 | -0.1678367 | 0.86671174 | 0.9084243 | NA | NA |
| ENSG000002 | 2.86078448 | -0.062794681 | 0.37442554 | -0.1677094 | 0.8668119 | 0.9085058 | NA | NA |
| ENSG000002 | 5.56989352 | 0.062720106 | 0.37459344 | 0.16743514 | 0.86702768 | 0.90870847 | NA | NA |
| ENSG000000 | 111.074259 | -0.027536849 | 0.16478079 | -0.167112 | 0.86728191 | 0.90895143 | SLC4A4 | 8671 |
| ENSG000002 | 2.46732127 | -0.061457988 | 0.36819374 | -0.1669175 | 0.86743494 | 0.90908531 | NA | NA |
| ENSG000002 | 6.93024776 | 0.059657914 | 0.35746272 | 0.16689269 | 0.86745449 | 0.90908531 | NA | NA |
| ENSG000001 | 53.5253824 | -0.036241777 | 0.21733534 | -0.1667551 | 0.86756275 | 0.90917527 | CREB3L1 | 90993 |
| ENSG000001 | 2.20437301 | -0.060888866 | 0.36545229 | -0.1666124 | 0.86767507 | 0.90921158 | OR10AD1 | 121275 |
| ENSG000001 | 383.238388 | 0.024109838 | 0.14469588 | 0.16662422 | 0.86766574 | 0.90921158 | ZNF470 | 388566 |
| ENSG000002 | 3.07060571 | 0.062758122 | 0.37682618 | 0.16654395 | 0.8677289 | 0.90921158 | NA | NA |
| ENSG000002 | 17.552721 | -0.04915478 | 0.29490734 | -0.1666787 | 0.86762285 | 0.90921158 | MIR124-2HG | 100130155 |
| ENSG000002 | 2.74525451 | -0.061533872 | 0.36948381 | -0.1665401 | 0.86773193 | 0.90921158 | RN7SL644P | 106481853 |
| ENSG000002 | 2.4227549 | -0.061777068 | 0.3708311 | -0.1665909 | 0.86769199 | 0.90921158 | NA | NA |
| ENSG000002 | 8.02757662 | 0.06036916 | 0.36255946 | 0.1665083 | 0.86775695 | 0.90921431 | NA | NA |
| ENSG000002 | 2.2037096 | 0.061129082 | 0.36725688 | 0.16644775 | 0.8678046 | 0.90924074 | NA | NA |
| ENSG000002 | 3.56164424 | 0.062952711 | 0.37857823 | 0.16628719 | 0.86793095 | 0.90934962 | NA | NA |
| ENSG000002 | 6.75839593 | 0.060163005 | 0.36197155 | 0.16620921 | 0.86799231 | 0.90939042 | HM13-IT1 | 106480432 |
| ENSG000001 | 158.712559 | 0.023912053 | 0.14405647 | 0.16599082 | 0.86816417 | 0.90949998 | FREM1 | 158326 |
| ENSG000002 | 3.30639503 | 0.062521961 | 0.37665197 | 0.16599399 | 0.86816168 | 0.90949998 | LINC00456 | 103625683 |
| ENSG000002 | 2.20240988 | -0.060734762 | 0.3658023 | -0.1660317 | 0.86813204 | 0.90949998 | LOC102723640 | 102723640 |
| ENSG000002 | 87.0796861 | -0.034895142 | 0.21029592 | -0.1659335 | 0.86820927 | 0.9095181 | FOXI3 | 344167 |
| ENSG000002 | 5.50077843 | -0.061906926 | 0.37313144 | -0.1659118 | 0.86822632 | 0.9095181 | NA | NA |
| ENSG000002 | 6.39938566 | 0.060451562 | 0.36454778 | 0.16582617 | 0.86829375 | 0.90956524 | NA | NA |
| ENSG000002 | 28.7523505 | 0.062736404 | 0.37848722 | 0.16575567 | 0.86834923 | 0.90959986 | NA | NA |
| ENSG000002 | 5.45630401 | -0.061286055 | 0.37000214 | -0.165637 | 0.86844261 | 0.90967417 | C20orf202 | 400831 |

| | | | | | | | | |
|---|---|---|---|---|---|---|---|---|
| ENSG000002 | 157.335226 | 0.0255384 | 0.15422713 | 0.16558954 | 0.86847998 | 0.90968982 | MAP10 | 54627 |
| ENSG000001 | 2.91138558 | -0.062111824 | 0.37534761 | -0.1654781 | 0.86856765 | 0.90973612 | NA | NA |
| ENSG000002 | 2.19417646 | -0.060873441 | 0.3678679 | -0.1654764 | 0.86856905 | 0.90973612 | NA | NA |
| ENSG000002 | 2.67283133 | -0.061354491 | 0.37106364 | -0.1653476 | 0.86867038 | 0.90981875 | CLDN25 | 644672 |
| ENSG000002 | 2.21196071 | -0.060720179 | 0.36737292 | -0.1652821 | 0.86872192 | 0.90982574 | MIR548F2 | 100313771 |
| ENSG000002 | 10.1848661 | -0.055770551 | 0.3373694 | -0.16531 | 0.86869995 | 0.90982574 | NA | NA |
| ENSG000002 | 8.53971342 | 0.057506085 | 0.34825508 | 0.16512633 | 0.86884455 | 0.90990716 | NA | NA |
| ENSG000002 | 5.1391624 | 0.061459438 | 0.37216802 | 0.16513895 | 0.86883461 | 0.90990716 | NA | NA |
| ENSG000002 | 3.35871098 | 0.062398166 | 0.3779775 | 0.16508434 | 0.8688776 | 0.90991827 | NA | NA |
| ENSG000002 | 2.43604431 | -0.061211621 | 0.37085685 | -0.1650546 | 0.86890102 | 0.90991931 | NA | NA |
| ENSG000002 | 2.68989819 | -0.06147999 | 0.37299076 | -0.1648298 | 0.86907795 | 0.91005759 | NA | NA |
| ENSG000002 | 6.01744254 | -0.06086004 | 0.36921419 | -0.1648367 | 0.86907254 | 0.91005759 | LOC102724889 | 102724889 |
| ENSG000002 | 2.22835507 | -0.060132611 | 0.36500986 | -0.1647424 | 0.86914672 | 0.91007018 | NA | NA |
| ENSG000002 | 12.9793216 | -0.051767628 | 0.31416301 | -0.1647795 | 0.86911753 | 0.91007018 | NA | NA |
| ENSG000002 | 2.64666986 | -0.061408114 | 0.37278271 | -0.164729 | 0.86915731 | 0.91007018 | NA | NA |
| ENSG000002 | 3.24755376 | 0.062076116 | 0.37722581 | 0.16455956 | 0.86929066 | 0.91018631 | MAGEA12 | 4111 |
| ENSG000002 | 2.20581518 | -0.060661842 | 0.36878994 | -0.1644889 | 0.8693463 | 0.91022107 | NA | NA |
| ENSG000002 | 2.19680513 | -0.060046493 | 0.36577332 | -0.1641631 | 0.86960273 | 0.91046604 | ZFAT-AS1 | 594840 |
| ENSG000002 | 2.2162618 | 0.059930185 | 0.36541079 | 0.1640077 | 0.86972508 | 0.91057063 | NA | NA |
| ENSG000002 | 11.6537314 | 0.05307706 | 0.32379369 | 0.16392247 | 0.86979218 | 0.91061737 | NA | NA |
| ENSG000002 | 296.582909 | -0.024154502 | 0.147433 | -0.1638337 | 0.86986202 | 0.91066698 | ZNF737 | 100129842 |
| ENSG000002 | 2.19587867 | -0.059640381 | 0.36464896 | -0.1635556 | 0.870081 | 0.91087271 | NA | NA |
| ENSG000002 | 2.70571931 | -0.060868193 | 0.372633 | -0.1633462 | 0.87024584 | 0.91099825 | NA | NA |
| ENSG000002 | 2.1973354 | 0.06036593 | 0.36949562 | 0.16337387 | 0.87022407 | 0.91099825 | NA | NA |
| ENSG000001 | 62.9089179 | -0.033195684 | 0.20335581 | -0.1632394 | 0.87032993 | 0.9110538 | HNF4G | 3174 |
| ENSG000001 | 2.18936291 | 0.059375917 | 0.36383815 | 0.16319321 | 0.87036631 | 0.9110538 | NA | NA |
| ENSG000002 | 3.27601881 | 0.06155133 | 0.37710487 | 0.16322072 | 0.87034465 | 0.9110538 | NA | NA |
| ENSG000002 | 8.88902368 | -0.057981834 | 0.35550881 | -0.1630954 | 0.87044336 | 0.91108742 | NA | NA |
| ENSG000002 | 3.76787887 | 0.061725671 | 0.37843854 | 0.1631062 | 0.87043482 | 0.91108742 | NA | NA |

| | | | | | | | | |
|---|---|---|---|---|---|---|---|---|
| ENSG000001 | 17.356854 | -0.050405181 | 0.30918604 | -0.1630254 | 0.87049842 | 0.91112154 | CD248 | 57124 |
| ENSG000002 | 97.2241574 | 0.027766482 | 0.17040334 | 0.16294565 | 0.87056123 | 0.91116375 | N4BP2L2-IT2 | 116828 |
| ENSG000001 | 30.9870855 | 0.042081336 | 0.25830934 | 0.16291063 | 0.8705888 | 0.9111691 | LMO2 | 4005 |
| ENSG000002 | 6.67978645 | 0.058718331 | 0.36061832 | 0.16282681 | 0.8706548 | 0.91121465 | NA | NA |
| ENSG000001 | 5.55472058 | -0.060804423 | 0.37366226 | -0.1627256 | 0.87073447 | 0.91127452 | SULT1E1 | 6783 |
| ENSG000002 | 2.21335027 | -0.05844716 | 0.3596095 | -0.1625295 | 0.87088888 | 0.91136555 | NA | NA |
| ENSG000002 | 4.12795964 | -0.061531729 | 0.3785021 | -0.1625664 | 0.87085983 | 0.91136555 | NA | NA |
| ENSG000002 | 5.24642887 | -0.0606063 | 0.37282875 | -0.162558 | 0.87086645 | 0.91136555 | NA | NA |
| ENSG000002 | 6.73263089 | 0.058308103 | 0.35883796 | 0.16249146 | 0.87091885 | 0.9113734 | NA | NA |
| ENSG000002 | 2.45195716 | -0.060069462 | 0.36983709 | -0.1624214 | 0.87097401 | 0.91139603 | NA | NA |
| ENSG000002 | 7.98413181 | 0.057341936 | 0.35307572 | 0.16240691 | 0.87098543 | 0.91139603 | NA | NA |
| ENSG000002 | 5.52605964 | -0.060016735 | 0.36980435 | -0.1622932 | 0.87107496 | 0.91146619 | NEGR1-IT1 | 100852409 |
| ENSG000001 | 63.4847153 | 0.030911197 | 0.19059107 | 0.16218597 | 0.87115941 | 0.91152038 | TIFA | 92610 |
| ENSG000002 | 2.63585266 | -0.060597439 | 0.37366532 | -0.1621704 | 0.8711717 | 0.91152038 | NOP56P3 | 106481967 |
| ENSG000002 | 72.230407 | -0.029563804 | 0.18238058 | -0.1620995 | 0.87122749 | 0.91155523 | NA | NA |
| ENSG000002 | 2.19120756 | -0.05937188 | 0.36652812 | -0.1619845 | 0.87131805 | 0.91162645 | NA | NA |
| ENSG000002 | 5.1283656 | 0.060505126 | 0.37379261 | 0.16186817 | 0.87140967 | 0.91169879 | NA | NA |
| ENSG000002 | 2.42651308 | -0.059649383 | 0.36893433 | -0.1616802 | 0.87155768 | 0.91183012 | MIR3142 | 100422938 |
| ENSG000000 | 59.7332865 | -0.033690256 | 0.2085532 | -0.1615427 | 0.87166595 | 0.91191987 | NA | NA |
| ENSG000002 | 3.81344358 | 0.061104075 | 0.37856605 | 0.16140928 | 0.87177106 | 0.9120063 | NA | NA |
| ENSG000002 | 10.1751395 | -0.053552824 | 0.33206986 | -0.1612698 | 0.87188095 | 0.9120742 | NA | NA |
| ENSG000002 | 2.18643231 | -0.05942455 | 0.3684644 | -0.1612762 | 0.87187584 | 0.9120742 | NA | NA |
| ENSG000001 | 4.77088637 | -0.060478546 | 0.37531736 | -0.1611398 | 0.87198333 | 0.91215777 | UPK1A | 11045 |
| ENSG000002 | 24.4281284 | 0.043733604 | 0.27145815 | 0.16110625 | 0.87200972 | 0.91216185 | NA | NA |
| ENSG000002 | 2.22158423 | -0.0591463 | 0.36740989 | -0.1609818 | 0.87210775 | 0.91224085 | NA | NA |
| ENSG000002 | 2.21025267 | 0.058846264 | 0.36574178 | 0.16089566 | 0.87217558 | 0.91228828 | MIR593 | 693178 |
| ENSG000000 | 97.2017104 | 0.029825435 | 0.18540534 | 0.1608661 | 0.87219886 | 0.9122891 | ST3GAL6 | 10402 |
| ENSG000002 | 11.922107 | 0.053576662 | 0.33345459 | 0.16067154 | 0.87235211 | 0.91242585 | NA | NA |
| ENSG000001 | 558.78125 | -0.020886834 | 0.1300852 | -0.1605627 | 0.87243782 | 0.91249196 | RCOR3 | 55758 |

| | | | | | | | | |
|---|---|---|---|---|---|---|---|---|
| ENSG000002 | 9.94631293 | -0.053448078 | 0.33299489 | -0.1605072 | 0.87248155 | 0.91251417 | NA | NA |
| ENSG000001 | 15.2053355 | 0.049497859 | 0.30865339 | 0.16036713 | 0.87259188 | 0.91260602 | GLYATL2 | 219970 |
| ENSG000002 | 3.07144706 | 0.060210686 | 0.37564412 | 0.16028651 | 0.87265538 | 0.9126489 | NA | NA |
| ENSG000001 | 5.00131171 | -0.059751455 | 0.37303041 | -0.1601785 | 0.87274046 | 0.91271433 | GOLGA8DP | 100132979 |
| ENSG000002 | 4.66318143 | 0.060537132 | 0.37818444 | 0.16007304 | 0.87282354 | 0.91275414 | NA | NA |
| ENSG000002 | 2.67859228 | -0.059845201 | 0.37384319 | -0.160081 | 0.87281725 | 0.91275414 | NA | NA |
| ENSG000002 | 2.44561576 | -0.059280748 | 0.37051736 | -0.1599945 | 0.87288539 | 0.91279528 | RN7SL689P | 106481111 |
| ENSG000002 | 4.75866643 | -0.059769851 | 0.37390265 | -0.159854 | 0.87299605 | 0.91288746 | NA | NA |
| ENSG000002 | 2.17729806 | -0.058521338 | 0.36620745 | -0.1598038 | 0.87303564 | 0.91290532 | MIR545 | 664614 |
| ENSG000002 | 5.76205694 | -0.059202417 | 0.37136427 | -0.1594187 | 0.87333899 | 0.91319898 | NA | NA |
| ENSG000001 | 42.7296773 | 0.036785208 | 0.23093472 | 0.15928834 | 0.87344171 | 0.91328005 | RBP7 | 116362 |
| ENSG000002 | 2.21153671 | -0.058697069 | 0.36855495 | -0.1592627 | 0.87346188 | 0.91328005 | NA | NA |
| ENSG000002 | 2.21387254 | -0.058619529 | 0.36813327 | -0.1592345 | 0.8734841 | 0.91328005 | NA | NA |
| ENSG000001 | 2.18669482 | 0.058281474 | 0.3661581 | 0.15917025 | 0.87353475 | 0.91330946 | RNU6-661P | 106479859 |
| ENSG000001 | 8.52806674 | 0.055242039 | 0.34745896 | 0.15898867 | 0.8736778 | 0.91343548 | GCM2 | 9247 |
| ENSG000001 | 5.72764244 | -0.05867943 | 0.36938743 | -0.1588561 | 0.87378228 | 0.91352116 | NOXA1 | 10811 |
| ENSG000002 | 3.73234261 | 0.060103668 | 0.37851309 | 0.15878887 | 0.87383521 | 0.91355295 | NA | NA |
| ENSG000002 | 4.12119145 | -0.059939376 | 0.37787456 | -0.1586224 | 0.87396636 | 0.9136665 | CTD-2201E9.1 | 101929338 |
| ENSG000001 | 8.26950101 | -0.055356068 | 0.34935295 | -0.1584531 | 0.87409975 | 0.91375884 | GNMT | 27232 |
| ENSG000002 | 24.4507401 | 0.041874673 | 0.26426916 | 0.15845463 | 0.87409857 | 0.91375884 | LOC400685 | 400685 |
| ENSG000002 | 6.53774159 | 0.057768847 | 0.36479372 | 0.15836031 | 0.87417289 | 0.91381173 | NA | NA |
| ENSG000002 | 6.99860748 | -0.058078671 | 0.3673482 | -0.1581025 | 0.87437603 | 0.91400052 | NA | NA |
| ENSG000001 | 27.2941202 | -0.040632669 | 0.25724589 | -0.1579526 | 0.87449411 | 0.91405048 | SLC35F3 | 148641 |
| ENSG000002 | 3.76373635 | 0.059708685 | 0.37835082 | 0.15781302 | 0.87460414 | 0.91405048 | NA | NA |
| ENSG000002 | 372.116154 | -0.050326668 | 0.31889062 | -0.157818 | 0.87460024 | 0.91405048 | GTF2IP4 | 100093631 |
| ENSG000002 | 8.75128391 | 0.054300286 | 0.34403032 | 0.15783576 | 0.87458622 | 0.91405048 | NA | NA |
| ENSG000002 | 14.5368276 | -0.048839867 | 0.3094101 | -0.1578483 | 0.87457632 | 0.91405048 | NA | NA |
| ENSG000002 | 45.7479983 | 0.034459137 | 0.21815471 | 0.15795734 | 0.87449042 | 0.91405048 | KCTD21-AS1 | 100289388 |
| ENSG000002 | 5.20475371 | -0.058620631 | 0.37115794 | -0.1579399 | 0.87450419 | 0.91405048 | LOC105371919 | 105371919 |

| | | | | | | | | |
|---|---|---|---|---|---|---|---|---|
| ENSG000002 | 6.37929041 | 0.058397574 | 0.3699714 | 0.15784348 | 0.87458014 | 0.91405048 | NA | NA |
| ENSG000000 | 11.1076519 | -0.054026202 | 0.34248445 | -0.1577479 | 0.87465545 | 0.91408055 | EFNA2 | 1943 |
| ENSG000002 | 3.76215789 | 0.059639464 | 0.37857777 | 0.15753557 | 0.87482278 | 0.91423185 | NA | NA |
| ENSG000002 | 3.51164417 | 0.059544429 | 0.37839145 | 0.15736198 | 0.87495957 | 0.91430412 | SP9 | 100131390 |
| ENSG000002 | 2.42345986 | -0.058328097 | 0.37054164 | -0.1574131 | 0.87491931 | 0.91430412 | NA | NA |
| ENSG000002 | 11.4670428 | 0.052680551 | 0.33474634 | 0.15737454 | 0.87494968 | 0.91430412 | SYNPR-AS1 | 100874016 |
| ENSG000000 | 40.2590772 | -0.037630881 | 0.23930785 | -0.1572488 | 0.87504874 | 0.91435017 | TBC1D2 | 55357 |
| ENSG000002 | 7.41719525 | -0.055897571 | 0.35544226 | -0.157262 | 0.87503835 | 0.91435017 | NA | NA |
| ENSG000002 | 2.18028678 | -0.057914902 | 0.36844087 | -0.1571891 | 0.87509579 | 0.91437577 | NA | NA |
| ENSG000002 | 2.1810559 | -0.057504314 | 0.36622092 | -0.1570208 | 0.87522843 | 0.9144838 | NA | NA |
| ENSG000002 | 2.19599737 | 0.058195584 | 0.37067081 | 0.15700072 | 0.87524428 | 0.9144838 | NA | NA |
| ENSG000002 | 2.43451837 | -0.057796798 | 0.36839058 | -0.15689 | 0.87533154 | 0.91455141 | NA | NA |
| ENSG000002 | 2.20630602 | 0.057792156 | 0.36846712 | 0.15684481 | 0.87536715 | 0.91455746 | NA | NA |
| ENSG000002 | 2.39676969 | -0.057761161 | 0.36831505 | -0.1568254 | 0.87538244 | 0.91455746 | NA | NA |
| ENSG000002 | 3.56596303 | 0.059305995 | 0.37855306 | 0.15666495 | 0.87550891 | 0.91466603 | NA | NA |
| ENSG000002 | 18.2809197 | -0.045094601 | 0.28820413 | -0.1564676 | 0.87566448 | 0.91478175 | NA | NA |
| ENSG000002 | 3.80803471 | 0.059194114 | 0.37831651 | 0.15646717 | 0.8756648 | 0.91478175 | MIR3682 | 100500850 |
| ENSG000002 | 4.5809619 | -0.058973791 | 0.37700314 | -0.1564279 | 0.87569579 | 0.91479056 | LOC101928700 | 101928700 |
| ENSG000001 | 475.97502 | -0.020750159 | 0.13270942 | -0.1563578 | 0.87575097 | 0.91481589 | NRP2 | 8828 |
| ENSG000002 | 4.54312925 | -0.058740351 | 0.37572221 | -0.1563398 | 0.87576515 | 0.91481589 | NA | NA |
| ENSG000002 | 5.23017077 | -0.058436613 | 0.37387408 | -0.1563003 | 0.87579636 | 0.91482492 | NA | NA |
| ENSG000002 | 2.19231028 | -0.056675334 | 0.3628853 | -0.1561797 | 0.87589134 | 0.91490057 | NA | NA |
| ENSG000002 | 4.09705482 | -0.059113819 | 0.3785904 | -0.1561419 | 0.87592118 | 0.91490817 | NA | NA |
| ENSG000002 | 2.19747179 | 0.057693149 | 0.36972743 | 0.15604238 | 0.87599962 | 0.91496653 | NA | NA |
| ENSG000001 | 47.6530765 | 0.032832597 | 0.21055544 | 0.15593326 | 0.87608563 | 0.91500923 | UBXN10 | 127733 |
| ENSG000002 | 22.9751533 | 0.042627428 | 0.27332995 | 0.15595593 | 0.87606776 | 0.91500923 | JHDM1D-AS1 | 100134229 |
| ENSG000002 | 5.26955715 | -0.058462433 | 0.37526346 | -0.1557904 | 0.87619826 | 0.9151033 | NA | NA |
| ENSG000002 | 5.46810691 | -0.057613166 | 0.37049239 | -0.1555043 | 0.87642375 | 0.91523276 | PRH1 | 5554 |
| ENSG000002 | 11.538037 | -0.051316815 | 0.33003299 | -0.15549 | 0.87643507 | 0.91523276 | NA | NA |

| | | | | | | | | |
|---|---|---|---|---|---|---|---|---|
| ENSG000002 | 5.10716729 | 0.058163767 | 0.37390493 | 0.15555764 | 0.87638172 | 0.91523276 | RNU6-126P | 106479622 |
| ENSG000002 | 2.66388052 | -0.057827471 | 0.37176688 | -0.1555477 | 0.87638958 | 0.91523276 | NA | NA |
| ENSG000002 | 20.7578819 | -0.043865504 | 0.28200919 | -0.1555464 | 0.87639061 | 0.91523276 | NA | NA |
| ENSG000002 | 24.4001611 | -0.040876177 | 0.26335146 | -0.1552153 | 0.87665158 | 0.91543379 | AGAP9 | 642517 |
| ENSG000002 | 7.12953996 | 0.055862118 | 0.36002947 | 0.15515985 | 0.8766953 | 0.91543379 | NA | NA |
| ENSG000002 | 2.21736375 | 0.05683441 | 0.36627935 | 0.15516684 | 0.87668979 | 0.91543379 | RNU6-1267P | 106481580 |
| ENSG000002 | 12.0334223 | -0.050572361 | 0.32631657 | -0.1549794 | 0.87683752 | 0.91555872 | METTL11B | 149281 |
| ENSG000002 | 4.79560598 | -0.05791527 | 0.37381853 | -0.1549288 | 0.87687742 | 0.91556747 | NA | NA |
| ENSG000002 | 5.1497835 | 0.057795412 | 0.37308657 | 0.15491153 | 0.87689106 | 0.91556747 | RHOA-IT1 | 106480739 |
| ENSG000002 | 5.52119211 | -0.057591605 | 0.37208474 | -0.1547809 | 0.87699406 | 0.91565144 | NA | NA |
| ENSG000002 | 4.92665086 | 0.058038643 | 0.37528643 | 0.15465159 | 0.877096 | 0.91573429 | LOC100996263 | 100996263 |
| ENSG000002 | 2.2036041 | -0.056415544 | 0.36491792 | -0.1545979 | 0.87713833 | 0.9157549 | NA | NA |
| ENSG000002 | 2.91856892 | -0.057496682 | 0.37223089 | -0.1544651 | 0.87724303 | 0.91581706 | NA | NA |
| ENSG000002 | 9.03684339 | -0.053740943 | 0.34787859 | -0.1544819 | 0.87722979 | 0.91581706 | NA | NA |
| ENSG000002 | 8.45153697 | -0.054205208 | 0.35101565 | -0.1544239 | 0.8772755 | 0.91582739 | LINC00638 | 196872 |
| ENSG000002 | 3.75920163 | 0.0584209 | 0.37848127 | 0.15435612 | 0.87732895 | 0.9158596 | NA | NA |
| ENSG000002 | 2.19713794 | -0.056490179 | 0.36660732 | -0.1540891 | 0.87753951 | 0.91605583 | NA | NA |
| ENSG000002 | 15.5061222 | -0.048883083 | 0.31744758 | -0.1539879 | 0.87761928 | 0.91611552 | GTF2IRD1P1 | 729156 |
| ENSG000001 | 33.0204047 | 0.03766615 | 0.24477788 | 0.15387889 | 0.87770523 | 0.91618165 | ADGRB1 | 575 |
| ENSG00000C | 219.832838 | 0.022087759 | 0.14370846 | 0.15369839 | 0.87784755 | 0.91630663 | MYO3A | 53904 |
| ENSG000001 | 51.3310062 | -0.032706281 | 0.21297713 | -0.1535671 | 0.87795108 | 0.91636751 | SERPINE1 | 5054 |
| ENSG000002 | 14.3705702 | -0.048798219 | 0.31776234 | -0.1535683 | 0.87795014 | 0.91636751 | NA | NA |
| ENSG000002 | 3.78631601 | 0.058103269 | 0.37856246 | 0.15348397 | 0.87801663 | 0.91641234 | NA | NA |
| ENSG000001 | 5.13657862 | 0.05721164 | 0.37287367 | 0.15343438 | 0.87805574 | 0.91642957 | NA | NA |
| ENSG000001 | 50.8743682 | 0.031500568 | 0.20539052 | 0.15336914 | 0.87810718 | 0.91645967 | CDH26 | 60437 |
| ENSG000002 | 3.11358665 | 0.057291154 | 0.37390274 | 0.15322475 | 0.87822104 | 0.91655492 | RNU6-519P | 106479796 |
| ENSG000001 | 2.93333275 | -0.057082049 | 0.37262683 | -0.1531882 | 0.87824983 | 0.91656138 | UBQLNL | 143630 |
| ENSG000002 | 5.30271668 | 0.057499683 | 0.37544433 | 0.15315102 | 0.87827918 | 0.91656842 | NA | NA |
| ENSG000002 | 5.46171343 | -0.056705432 | 0.37033453 | -0.1531195 | 0.87830405 | 0.91657079 | MIR3936 | 100500865 |

| | | | | | | | | |
|---|---|---|---|---|---|---|---|---|
| ENSG000002 | 2.41606859 | -0.056280891 | 0.36786868 | -0.1529918 | 0.87840474 | 0.91665227 | NA | NA |
| ENSG000001 | 25.8703582 | 0.039159091 | 0.25608383 | 0.15291512 | 0.87846521 | 0.91669178 | SLC37A2 | 219855 |
| ENSG000002 | 2.92989519 | -0.056947821 | 0.37253484 | -0.1528658 | 0.87850414 | 0.91670882 | NA | NA |
| ENSG000002 | 5.04375152 | -0.057074163 | 0.37372566 | -0.1527167 | 0.87862166 | 0.91680786 | NA | NA |
| ENSG000002 | 5.11211218 | 0.056981443 | 0.37338169 | 0.1526091 | 0.87870655 | 0.91687284 | NA | NA |
| ENSG000002 | 14.9962558 | -0.046599064 | 0.30543286 | -0.1525673 | 0.87873952 | 0.91688365 | NA | NA |
| ENSG000002 | 2.20225343 | -0.055560977 | 0.36451339 | -0.1524251 | 0.87885169 | 0.91697709 | NA | NA |
| ENSG000002 | 5.28325871 | -0.056936657 | 0.37375346 | -0.1523375 | 0.87892077 | 0.91702558 | NA | NA |
| ENSG000002 | 5.30785003 | 0.056528527 | 0.37130283 | 0.15224373 | 0.87899471 | 0.91707913 | NA | NA |
| ENSG000002 | 27.9967391 | -0.038444249 | 0.25261034 | -0.1521879 | 0.8790387 | 0.91710143 | NA | NA |
| ENSG000002 | 9.44678095 | -0.051315701 | 0.33746637 | -0.1520617 | 0.87913829 | 0.91718174 | NA | NA |
| ENSG000002 | 4.5277976 | -0.057089379 | 0.37563742 | -0.15198 | 0.8792027 | 0.91722534 | NA | NA |
| ENSG000001 | 51.5636195 | 0.031211324 | 0.20550836 | 0.15187374 | 0.87928652 | 0.91728918 | DENND2D | 79961 |
| ENSG000002 | 10.0994058 | 0.051495778 | 0.33937291 | 0.15173803 | 0.87939356 | 0.91735364 | NA | NA |
| ENSG000002 | 9.73465733 | -0.051580167 | 0.33989088 | -0.1517551 | 0.87938011 | 0.91735364 | NPM1P5 | 4874 |
| ENSG000002 | 5.91626832 | -0.056278921 | 0.37123782 | -0.151598 | 0.87950399 | 0.91744524 | NA | NA |
| ENSG000002 | 7.17350516 | 0.053824433 | 0.35520404 | 0.15153103 | 0.87955684 | 0.91745727 | NA | NA |
| ENSG000002 | 4.78764091 | -0.056858897 | 0.37524176 | -0.151526 | 0.87956077 | 0.91745727 | NA | NA |
| ENSG000001 | 47.2093775 | -0.03162031 | 0.20969263 | -0.1507936 | 0.88013852 | 0.91801268 | CHRNA7 | 1139 |
| ENSG000002 | 2.22456656 | 0.055464094 | 0.36780132 | 0.15079906 | 0.88013423 | 0.91801268 | NA | NA |
| ENSG000002 | 2.20205858 | 0.055036108 | 0.36519035 | 0.15070526 | 0.88020822 | 0.91806177 | NA | NA |
| ENSG000001 | 181.396095 | -0.022497485 | 0.14937442 | -0.1506114 | 0.8802823 | 0.91809181 | GPR161 | 23432 |
| ENSG000002 | 7.30074125 | -0.053410932 | 0.35462221 | -0.1506136 | 0.88028052 | 0.91809181 | LOC101929697 | 101929697 |
| ENSG000002 | 23.4117335 | 0.042144079 | 0.28001115 | 0.15050858 | 0.88036339 | 0.91813808 | NA | NA |
| ENSG000002 | 9.62426967 | -0.052107685 | 0.34623571 | -0.1504977 | 0.88037195 | 0.91813808 | NA | NA |
| ENSG000001 | 5.17125426 | 0.056205088 | 0.37370619 | 0.15039913 | 0.88044973 | 0.91817196 | LYZL4 | 131375 |
| ENSG000002 | 2.63713121 | -0.055711627 | 0.37035622 | -0.1504271 | 0.88042763 | 0.91817196 | MIR4418 | 100616433 |
| ENSG000002 | 27.1137124 | -0.037836444 | 0.25171968 | -0.1503118 | 0.88051861 | 0.91822018 | PCDHGA4 | 56111 |
| ENSG000002 | 3.74024242 | 0.0568747 | 0.37857945 | 0.15023187 | 0.88058168 | 0.91826234 | RN7SKP269 | 106480915 |

| | | | | | | | | |
|---|---|---|---|---|---|---|---|---|
| ENSG000002 | 22.5864938 | -0.040606759 | 0.27100638 | -0.1498369 | 0.88089328 | 0.91856047 | NA | NA |
| ENSG000002 | 24.9928888 | 0.039119169 | 0.26112159 | 0.14981208 | 0.88091288 | 0.91856047 | NA | NA |
| ENSG000002 | 15.5556232 | -0.044664828 | 0.29824252 | -0.1497601 | 0.88095389 | 0.91857962 | LOC101926911 | 101926911 |
| ENSG000002 | 11.8856233 | 0.04816706 | 0.32209771 | 0.14954176 | 0.88112616 | 0.91873561 | NA | NA |
| ENSG000002 | 10.1035414 | 0.049563198 | 0.33167135 | 0.14943467 | 0.88121066 | 0.9188001 | NA | NA |
| ENSG000001 | 49.0737752 | 0.031714603 | 0.2123597 | 0.14934379 | 0.88128236 | 0.91882257 | SPX | 80763 |
| ENSG000001 | 550.924791 | 0.023348545 | 0.15642061 | 0.1492677 | 0.8813424 | 0.91882257 | RGS5 | 8490 |
| ENSG000002 | 7.40463021 | -0.054947977 | 0.36789261 | -0.1493587 | 0.88127057 | 0.91882257 | LOC101928336 | 101928336 |
| ENSG000002 | 4.04817552 | 0.056515101 | 0.37855004 | 0.14929361 | 0.88132196 | 0.91882257 | LOC645485 | 645485 |
| ENSG000002 | 10.5684456 | 0.049799811 | 0.3336363 | 0.14926377 | 0.8813455 | 0.91882257 | NA | NA |
| ENSG000002 | 2.2378039 | 0.054609828 | 0.36636245 | 0.14905956 | 0.88150663 | 0.91896693 | NA | NA |
| ENSG000001 | 68.2251086 | 0.028841507 | 0.19373745 | 0.14886904 | 0.88165697 | 0.9190764 | NIPAL2 | 79815 |
| ENSG000001 | 38.8811102 | 0.032870014 | 0.22078965 | 0.14887479 | 0.88165243 | 0.9190764 | SAMD14 | 201191 |
| ENSG000002 | 5.36888265 | 0.055227189 | 0.37114937 | 0.14880044 | 0.8817111 | 0.91908557 | NA | NA |
| ENSG000002 | 9.68374031 | 0.050561321 | 0.3397417 | 0.14882283 | 0.88169344 | 0.91908557 | NA | NA |
| ENSG000002 | 6.91363209 | -0.054007714 | 0.36350566 | -0.1485746 | 0.8818893 | 0.9192477 | NA | NA |
| ENSG000002 | 10.1445137 | 0.049541717 | 0.33357559 | 0.14851721 | 0.8819346 | 0.91927129 | NA | NA |
| ENSG000002 | 5.28529428 | -0.055329167 | 0.37277735 | -0.1484242 | 0.88200803 | 0.91932419 | NA | NA |
| ENSG000002 | 5.36217596 | 0.05485385 | 0.36980698 | 0.14833103 | 0.88208153 | 0.91937717 | MIR1302-6 | 100302140 |
| ENSG000002 | 12.4286209 | -0.046811652 | 0.31597086 | -0.1481518 | 0.88222297 | 0.91950095 | NA | NA |
| ENSG000002 | 15.5810582 | 0.044347517 | 0.29996747 | 0.14784109 | 0.88246818 | 0.91973289 | NA | NA |
| ENSG000002 | 2.4104007 | 0.054666216 | 0.37001357 | 0.14774111 | 0.88254708 | 0.91979148 | NA | NA |
| ENSG000002 | 2.44449534 | 0.054761357 | 0.37106776 | 0.14757778 | 0.88267599 | 0.91986276 | NIFKP9 | 106480805 |
| ENSG000002 | 2.20439441 | 0.053695619 | 0.36386976 | 0.14756824 | 0.88268352 | 0.91986276 | ZBTB20-AS3 | 104413890 |
| ENSG000002 | 11.9717607 | 0.047348788 | 0.32078463 | 0.14760304 | 0.88265605 | 0.91986276 | NA | NA |
| ENSG000002 | 7.19930946 | -0.053489093 | 0.36258303 | -0.1475223 | 0.88271976 | 0.91987687 | NA | NA |
| ENSG000002 | 4.52492078 | -0.055664727 | 0.37746278 | -0.1474708 | 0.88276045 | 0.91989564 | NA | NA |
| ENSG000002 | 9.17598406 | -0.050778867 | 0.34441928 | -0.1474333 | 0.88279003 | 0.91990283 | SND1-IT1 | 27099 |
| ENSG000001 | 14.7147212 | -0.045013996 | 0.30554278 | -0.1473247 | 0.88287574 | 0.91995332 | LRRC56 | 115399 |

| | | | | | | | | |
|---|---|---|---|---|---|---|---|---|
| ENSG000002 | 2.20246249 | -0.05400117 | 0.36657085 | -0.1473144 | 0.88288386 | 0.91995332 | NA | NA |
| ENSG000002 | 5.13317277 | 0.05485185 | 0.37247757 | 0.14726216 | 0.8829251 | 0.91996344 | NA | NA |
| ENSG000002 | 11.8753496 | 0.049116166 | 0.3335685 | 0.14724462 | 0.88293895 | 0.91996344 | NA | NA |
| ENSG000002 | 2.41713964 | 0.054649713 | 0.37186808 | 0.14695995 | 0.88316363 | 0.92015027 | NA | NA |
| ENSG000002 | 20.6845833 | 0.04128239 | 0.28088681 | 0.14697162 | 0.88315442 | 0.92015027 | NA | NA |
| ENSG000002 | 4.94926985 | 0.054949995 | 0.37447225 | 0.14673983 | 0.88333738 | 0.92030764 | NA | NA |
| ENSG000001 | 5.35462804 | 0.054492202 | 0.37171799 | 0.14659555 | 0.88345127 | 0.92033249 | TRH | 7200 |
| ENSG000002 | 5.38250511 | 0.054647057 | 0.37277673 | 0.1465946 | 0.88345201 | 0.92033249 | NA | NA |
| ENSG000002 | 4.01431444 | 0.055505089 | 0.37858527 | 0.14661185 | 0.8834384 | 0.92033249 | NA | NA |
| ENSG000002 | 4.52904335 | -0.055123368 | 0.37596416 | -0.1466187 | 0.88343301 | 0.92033249 | NA | NA |
| ENSG000002 | 6.96307662 | -0.054439589 | 0.37151046 | -0.1465358 | 0.88349842 | 0.92035719 | LINC01103 | 101927360 |
| ENSG000002 | 2.22275225 | 0.054015363 | 0.36878356 | 0.14646901 | 0.88355115 | 0.92038848 | NA | NA |
| ENSG000002 | 5.35406709 | 0.054212845 | 0.37092557 | 0.14615559 | 0.88379857 | 0.92062256 | NA | NA |
| ENSG000001 | 5.62556196 | 0.053957772 | 0.3692777 | 0.14611706 | 0.88382898 | 0.92063059 | MTNR1A | 4543 |
| ENSG000002 | 8.78633114 | 0.049923279 | 0.34569684 | 0.14441346 | 0.88517398 | 0.92200792 | NA | NA |
| ENSG000002 | 4.00336066 | 0.054604602 | 0.37822273 | 0.14437155 | 0.88520708 | 0.92201871 | NA | NA |
| ENSG000002 | 10.3281687 | 0.048528464 | 0.33641999 | 0.14424964 | 0.88530334 | 0.92209529 | MLIP-IT1 | 100874282 |
| ENSG000001 | 62.975987 | 0.028167421 | 0.19548341 | 0.1440911 | 0.88542853 | 0.92220199 | GALNT16 | 57452 |
| ENSG000002 | 2.38518338 | -0.052913273 | 0.36745281 | -0.1440002 | 0.88550032 | 0.92222939 | NA | NA |
| ENSG000002 | 12.1404168 | 0.048221801 | 0.3348559 | 0.14400762 | 0.88549445 | 0.92222939 | NA | NA |
| ENSG000002 | 37.2371797 | -0.032953872 | 0.22892845 | -0.1439483 | 0.88554126 | 0.92224834 | NA | NA |
| ENSG000002 | 4.51445568 | -0.054232704 | 0.3772991 | -0.1437393 | 0.88570635 | 0.92239658 | NA | NA |
| ENSG000002 | 5.26226457 | -0.05362419 | 0.37338141 | -0.1436177 | 0.88580234 | 0.92247286 | NA | NA |
| ENSG000002 | 18.2779549 | 0.041642098 | 0.29008358 | 0.14355207 | 0.88585419 | 0.92250317 | C5orf58 | 133874 |
| ENSG000002 | 4.49307024 | -0.053855455 | 0.37525847 | -0.1435156 | 0.88588297 | 0.92250944 | NA | NA |
| ENSG000002 | 17.5200598 | 0.043216222 | 0.30152453 | 0.14332573 | 0.88603294 | 0.92259563 | NA | NA |
| ENSG000002 | 2.23139723 | 0.052433664 | 0.36583906 | 0.1433244 | 0.88603398 | 0.92259563 | NA | NA |
| ENSG000002 | 5.03885173 | -0.053694855 | 0.37450046 | -0.1433773 | 0.88599223 | 0.92259563 | RN7SL577P | 106481080 |
| ENSG000002 | 2.2313634 | -0.05128924 | 0.35833653 | -0.1431315 | 0.88618634 | 0.92273058 | NA | NA |

| | | | | | | | | |
|---|---|---|---|---|---|---|---|---|
| ENSG000001 | 19.4133615 | -0.040352329 | 0.28210472 | -0.1430402 | 0.88625839 | 0.92275821 | ENG | 2022 |
| ENSG000002 | 7.39160976 | -0.051610727 | 0.36074474 | -0.1430672 | 0.88623714 | 0.92275821 | NA | NA |
| ENSG000001 | 2.17608477 | -0.051695835 | 0.36172763 | -0.1429137 | 0.88635834 | 0.92278977 | C14orf180 | 400258 |
| ENSG000002 | 2.17374997 | 0.052385415 | 0.36662225 | 0.14288662 | 0.88637973 | 0.92278977 | NA | NA |
| ENSG000002 | 2.44825352 | 0.052548782 | 0.36760771 | 0.14294798 | 0.88633126 | 0.92278977 | NA | NA |
| ENSG000002 | 4.08271565 | -0.054104632 | 0.37859206 | -0.1429101 | 0.88636118 | 0.92278977 | LOC101927131 | 101927131 |
| ENSG000002 | 6.91113713 | 0.051681289 | 0.36221044 | 0.14268305 | 0.88654051 | 0.92293346 | NA | NA |
| ENSG000002 | 15.6404842 | 0.042703157 | 0.29940268 | 0.14262784 | 0.88658411 | 0.92295516 | NA | NA |
| ENSG00000C | 84.8783945 | -0.025381823 | 0.17799804 | -0.1425961 | 0.88660919 | 0.92295757 | RAPGEF4 | 11069 |
| ENSG000001 | 35.5981834 | 0.032776024 | 0.23012803 | 0.14242517 | 0.88674418 | 0.9230744 | ZCCHC18 | 644353 |
| ENSG000002 | 13.1959773 | 0.045547894 | 0.31993966 | 0.14236402 | 0.88679248 | 0.92310099 | NA | NA |
| ENSG000002 | 5.16666692 | 0.053362836 | 0.37511093 | 0.14225881 | 0.88687558 | 0.92316378 | NA | NA |
| ENSG000002 | 6.69006699 | 0.05165767 | 0.36322721 | 0.14221861 | 0.88690733 | 0.92317314 | NA | NA |
| ENSG000002 | 38.7517721 | 0.034725869 | 0.24440215 | 0.14208496 | 0.8870129 | 0.92325933 | LRRC69 | 100130742 |
| ENSG000002 | 3.59704411 | 0.053689825 | 0.37807342 | 0.14200899 | 0.8870729 | 0.92328993 | NA | NA |
| ENSG000002 | 3.73367954 | 0.053740936 | 0.37848374 | 0.14199008 | 0.88708784 | 0.92328993 | NA | NA |
| ENSG000002 | 10.061876 | -0.048908225 | 0.34493438 | -0.1417899 | 0.88724593 | 0.92343077 | NA | NA |
| ENSG000001 | 2.42734415 | 0.05157458 | 0.36416854 | 0.14162283 | 0.88737793 | 0.92349705 | NA | NA |
| ENSG000002 | 7.29883053 | -0.050307299 | 0.35507839 | -0.1416794 | 0.88733323 | 0.92349705 | NA | NA |
| ENSG000002 | 3.55784663 | 0.053600249 | 0.37844838 | 0.1416316 | 0.88737101 | 0.92349705 | NA | NA |
| ENSG00000C | 54.7561344 | -0.029000814 | 0.20503858 | -0.1414408 | 0.88752175 | 0.92359268 | PSD | 5662 |
| ENSG000002 | 9.23132829 | -0.048239766 | 0.34103876 | -0.1414495 | 0.88751484 | 0.92359268 | NA | NA |
| ENSG000002 | 7.88892302 | -0.050197377 | 0.35495245 | -0.14142 | 0.88753815 | 0.92359268 | LOC100996419 | 100996419 |
| ENSG000002 | 22.8970446 | -0.037879172 | 0.26840843 | -0.1411251 | 0.8877711 | 0.92381138 | NA | NA |
| ENSG000002 | 2.66615742 | -0.05221907 | 0.37038619 | -0.1409855 | 0.88788142 | 0.92390248 | NA | NA |
| ENSG000002 | 2.65085225 | -0.052018441 | 0.36905222 | -0.1409514 | 0.88790831 | 0.92390675 | NA | NA |
| ENSG000002 | 3.77644421 | 0.053285388 | 0.37852839 | 0.14076986 | 0.88805175 | 0.92401974 | NA | NA |
| ENSG000002 | 10.5804125 | 0.050075983 | 0.35576374 | 0.14075629 | 0.88806248 | 0.92401974 | NA | NA |
| ENSG000002 | 2.42918276 | 0.052472374 | 0.37295283 | 0.1406944 | 0.88811137 | 0.92404691 | VAV3-AS1 | 100873946 |

| | | | | | | | | |
|---|---|---|---|---|---|---|---|---|
| ENSG000002 | 19.3356905 | -0.04057667 | 0.28872618 | -0.1405369 | 0.88823583 | 0.92415269 | BTBD19 | 149478 |
| ENSG000001 | 21.1063989 | -0.038298043 | 0.27266264 | -0.1404594 | 0.88829699 | 0.92419261 | GPX2 | 2877 |
| ENSG000001 | 2.41432719 | 0.052428476 | 0.37353809 | 0.14035644 | 0.88837837 | 0.92421365 | WFDC5 | 149708 |
| ENSG000002 | 2.64100252 | 0.051865768 | 0.36955298 | 0.14034731 | 0.88838558 | 0.92421365 | NA | NA |
| ENSG000002 | 17.6684347 | 0.040582124 | 0.2890383 | 0.14040397 | 0.88834082 | 0.92421365 | NA | NA |
| ENSG000002 | 8.94607511 | -0.048437501 | 0.34599634 | -0.1399943 | 0.88866453 | 0.92448012 | NA | NA |
| ENSG000002 | 8.05746847 | -0.049656554 | 0.35494002 | -0.1399013 | 0.88873801 | 0.92453285 | NA | NA |
| ENSG000002 | 9.64585319 | -0.047315971 | 0.33841982 | -0.1398144 | 0.88880662 | 0.92458051 | CTXN2 | 399697 |
| ENSG000001 | 5.18820334 | -0.052401618 | 0.37510726 | -0.1396977 | 0.88889885 | 0.92461317 | SGSM3 | 27352 |
| ENSG000002 | 4.31453154 | -0.052755731 | 0.37764611 | -0.1396962 | 0.88890002 | 0.92461317 | NA | NA |
| ENSG000002 | 2.21376201 | 0.050749431 | 0.3633053 | 0.13968811 | 0.88890642 | 0.92461317 | NA | NA |
| ENSG000002 | 21.5376547 | -0.038948832 | 0.27907471 | -0.1395642 | 0.88900435 | 0.92469131 | MAGEL2 | 54551 |
| ENSG000002 | 3.83992714 | -0.052759352 | 0.37844107 | -0.1394123 | 0.88912433 | 0.92479238 | NA | NA |
| ENSG000002 | 6.05085098 | 0.051790812 | 0.37185474 | 0.139277 | 0.88923127 | 0.92487989 | LOC101927809 | 101927809 |
| ENSG000002 | 38.4397042 | 0.032319512 | 0.23220764 | 0.13918367 | 0.88930501 | 0.92493287 | DENND1C | 79958 |
| ENSG000002 | 3.34520533 | 0.052511678 | 0.37772498 | 0.13902093 | 0.88943361 | 0.92504289 | RNU1-124P | 106481629 |
| ENSG000002 | 2.42082559 | 0.051392242 | 0.37004237 | 0.13888205 | 0.88954336 | 0.92510959 | NA | NA |
| ENSG000002 | 7.43588167 | -0.050251179 | 0.36182634 | -0.138882 | 0.88954337 | 0.92510959 | NA | NA |
| ENSG000002 | 5.32941119 | 0.051707234 | 0.37258763 | 0.13877872 | 0.88962501 | 0.92517077 | NA | NA |
| ENSG000002 | 4.81430426 | -0.05193795 | 0.37562914 | -0.1382692 | 0.89002765 | 0.92556576 | RN7SL141P | 106480959 |
| ENSG000002 | 9.20450925 | -0.047469742 | 0.3439524 | -0.1380125 | 0.89023051 | 0.92575298 | NA | NA |
| ENSG000001 | 60.1272327 | -0.027849873 | 0.20215356 | -0.1377659 | 0.89042541 | 0.92583478 | NPHP3 | 27031 |
| ENSG000002 | 23.207523 | 0.036809445 | 0.26723924 | 0.13773967 | 0.89044616 | 0.92583478 | NA | NA |
| ENSG000002 | 10.6813311 | 0.046501924 | 0.33749806 | 0.13778427 | 0.89041092 | 0.92583478 | NA | NA |
| ENSG000002 | 4.06007127 | -0.052153551 | 0.37831141 | -0.1378588 | 0.89035203 | 0.92583478 | NA | NA |
| ENSG000002 | 4.73015782 | -0.051605879 | 0.37454901 | -0.1377814 | 0.8904132 | 0.92583478 | LOC100506082 | 100506082 |
| ENSG000002 | 5.62169941 | 0.051293576 | 0.37237765 | 0.13774612 | 0.89044107 | 0.92583478 | NA | NA |
| ENSG000001 | 28.7028223 | 0.034002381 | 0.24712134 | 0.13759387 | 0.8905614 | 0.92593086 | TDH | 157739 |
| ENSG000001 | 6.84929828 | -0.049160638 | 0.35763442 | -0.1374606 | 0.89066674 | 0.92601664 | NA | NA |

| | | | | | | | | |
|---|---|---|---|---|---|---|---|---|
| ENSG000002 | 6.864531 | -0.049715068 | 0.36203646 | -0.1373206 | 0.89077738 | 0.92610792 | GAS2L1P2 | 340508 |
| ENSG000001 | 6.85485075 | -0.049679057 | 0.36193972 | -0.1372578 | 0.89082701 | 0.92613578 | MS4A5 | 64232 |
| ENSG000002 | 10.428553 | 0.047040628 | 0.34309625 | 0.13710621 | 0.89094684 | 0.9262188 | RN7SL381P | 106479362 |
| ENSG000002 | 7.69071258 | 0.049431454 | 0.360553 | 0.13709899 | 0.89095255 | 0.9262188 | NA | NA |
| ENSG000001 | 2.42670971 | -0.050584888 | 0.36907254 | -0.1370595 | 0.89098379 | 0.92622753 | NA | NA |
| ENSG000002 | 2.44721516 | 0.050355866 | 0.36790333 | 0.13687255 | 0.89113154 | 0.92634262 | NA | NA |
| ENSG000002 | 4.22871723 | 0.05168429 | 0.37763906 | 0.13686161 | 0.89114018 | 0.92634262 | NA | NA |
| ENSG000001 | 103.008442 | -0.022877675 | 0.16720635 | -0.136823 | 0.8911707 | 0.92635059 | DLC1 | 10395 |
| ENSG000001 | 2.428478 | 0.050882059 | 0.37210318 | 0.1367418 | 0.89123489 | 0.92639357 | RNU6-208P | 106479658 |
| ENSG000001 | 138.631721 | -0.02154653 | 0.157655 | -0.1366689 | 0.89129254 | 0.92642975 | PLEKHH2 | 130271 |
| ENSG000002 | 2.20923416 | 0.050520388 | 0.3697826 | 0.13662186 | 0.89132969 | 0.92644462 | NA | NA |
| ENSG000002 | 4.06087319 | -0.051554139 | 0.37753599 | -0.1365542 | 0.89138315 | 0.92647644 | KCND3-AS1 | 100873995 |
| ENSG000002 | 4.76851917 | -0.050989977 | 0.37377521 | -0.1364188 | 0.89149019 | 0.92656394 | NA | NA |
| ENSG000002 | 5.35962011 | 0.051506104 | 0.3776841 | 0.1363735 | 0.89152602 | 0.92656709 | P3H2-AS1 | 101929152 |
| ENSG000002 | 27.4593675 | -0.036032683 | 0.26425217 | -0.1363572 | 0.89153892 | 0.92656709 | NA | NA |
| ENSG000001 | 4.52474472 | -0.05119073 | 0.37564813 | -0.1362731 | 0.8916054 | 0.92661243 | XAGE3 | 170626 |
| ENSG000001 | 28.2120257 | -0.033656872 | 0.24728809 | -0.1361039 | 0.89173915 | 0.92670394 | F13A1 | 2162 |
| ENSG000001 | 80.4642891 | -0.02572668 | 0.18900253 | -0.1361182 | 0.89172786 | 0.92670394 | NPR2 | 4882 |
| ENSG000002 | 7.19235914 | 0.048871251 | 0.35921022 | 0.13605195 | 0.89178021 | 0.92672286 | NA | NA |
| ENSG000002 | 3.60705441 | -0.051358804 | 0.37765371 | -0.1359944 | 0.89182568 | 0.92674636 | NA | NA |
| ENSG000001 | 132.92755 | 0.020473717 | 0.15068651 | 0.13586961 | 0.89192436 | 0.92677765 | PDLIM3 | 27295 |
| ENSG000002 | 11.9765005 | -0.043200281 | 0.31785959 | -0.13591 | 0.89189247 | 0.92677765 | NA | NA |
| ENSG000002 | 20.393989 | -0.037835014 | 0.27845466 | -0.135875 | 0.89192013 | 0.92677765 | NA | NA |
| ENSG000002 | 6.64941083 | -0.049659833 | 0.36592077 | -0.135712 | 0.89204897 | 0.92678878 | NA | NA |
| ENSG000002 | 3.80837908 | 0.051370051 | 0.37841134 | 0.13575188 | 0.89201743 | 0.92678878 | NA | NA |
| ENSG000002 | 8.55222466 | 0.047016904 | 0.34631208 | 0.13576455 | 0.89200742 | 0.92678878 | NA | NA |
| ENSG000002 | 5.48781039 | -0.050773369 | 0.3741272 | -0.1357115 | 0.89204934 | 0.92678878 | NA | NA |
| ENSG000002 | 5.34799792 | 0.050432092 | 0.37144808 | 0.13577158 | 0.89200186 | 0.92678878 | NA | NA |
| ENSG000002 | 2.43766412 | 0.050592002 | 0.37297331 | 0.1356451 | 0.89210185 | 0.92681959 | NA | NA |

| | | | | | | | | |
|---|---|---|---|---|---|---|---|---|
| ENSG000002 | 4.94458544 | -0.050900451 | 0.37538027 | -0.135597 | 0.89213985 | 0.92683532 | NA | NA |
| ENSG000002 | 4.26446255 | 0.051094175 | 0.37742965 | 0.13537403 | 0.89231615 | 0.92697618 | NA | NA |
| ENSG000002 | 4.09066886 | -0.05108722 | 0.37747656 | -0.1353388 | 0.89234402 | 0.92697618 | GATSL3 | 652968 |
| ENSG000002 | 13.9227645 | -0.042125333 | 0.31124534 | -0.1353445 | 0.89233953 | 0.92697618 | CRTC3-AS1 | 101926895 |
| ENSG000002 | 9.21187354 | 0.046810421 | 0.34636965 | 0.13514585 | 0.89249656 | 0.92711089 | NA | NA |
| ENSG000001 | 4.46221729 | -0.051004269 | 0.37748739 | -0.1351152 | 0.89252082 | 0.92711235 | GDPD3 | 79153 |
| ENSG000002 | 10.4601034 | 0.045031375 | 0.33341921 | 0.13505933 | 0.89256496 | 0.92713445 | NA | NA |
| ENSG000002 | 29.1174133 | -0.034058064 | 0.25243707 | -0.1349171 | 0.89267746 | 0.92722755 | NA | NA |
| ENSG000002 | 29.2478237 | 0.034093225 | 0.25279086 | 0.13486732 | 0.89271678 | 0.92724465 | SLFNL1-AS1 | 100507178 |
| ENSG000002 | 4.28722791 | -0.050723616 | 0.37629926 | -0.134796 | 0.89277321 | 0.9272795 | NA | NA |
| ENSG000002 | 28.49139 | -0.033879219 | 0.25186783 | -0.1345119 | 0.8929978 | 0.92748902 | LIFR-AS1 | 100506495 |
| ENSG000002 | 2.43307699 | 0.049997101 | 0.37221629 | 0.13432271 | 0.89314739 | 0.92761546 | NA | NA |
| ENSG000002 | 10.5802649 | -0.04476099 | 0.33329087 | -0.1343001 | 0.89316529 | 0.92761546 | NA | NA |
| ENSG000001 | 11.2812148 | 0.04582948 | 0.34192414 | 0.13403406 | 0.89337564 | 0.92781016 | MALL | 7851 |
| ENSG000002 | 4.11853365 | -0.050689521 | 0.37842833 | -0.1339475 | 0.89344411 | 0.9278435 | LOC392196 | 392196 |
| ENSG000002 | 12.4503506 | 0.042466814 | 0.31706892 | 0.13393559 | 0.89345351 | 0.9278435 | NA | NA |
| ENSG000002 | 7.18309533 | 0.047872852 | 0.35752407 | 0.13390106 | 0.89348081 | 0.9278481 | NA | NA |
| ENSG000001 | 15.2818528 | -0.040568843 | 0.30318687 | -0.133808 | 0.89355436 | 0.92788604 | ANGPTL5 | 253935 |
| ENSG000002 | 4.52958 | -0.050190783 | 0.37518378 | -0.1337765 | 0.89357929 | 0.92788604 | NA | NA |
| ENSG000002 | 2.45084939 | 0.049690425 | 0.37146708 | 0.13376805 | 0.89358599 | 0.92788604 | NA | NA |
| ENSG000002 | 8.85412309 | 0.046672727 | 0.34904725 | 0.13371464 | 0.89362823 | 0.92790613 | SYCE1L | 100130958 |
| ENSG000002 | 6.43981828 | -0.048879465 | 0.36568664 | -0.1336649 | 0.89366757 | 0.92790916 | NA | NA |
| ENSG000002 | 11.2854107 | -0.044763627 | 0.33499655 | -0.1336241 | 0.89369979 | 0.92790916 | NA | NA |
| ENSG000002 | 5.37804839 | 0.049973699 | 0.37392243 | 0.13364724 | 0.89368153 | 0.92790916 | NA | NA |
| ENSG000002 | 6.37519111 | -0.049225211 | 0.36867834 | -0.133518 | 0.89378369 | 0.92797252 | NA | NA |
| ENSG000002 | 6.94028335 | -0.048111024 | 0.36111166 | -0.1332303 | 0.89401123 | 0.92818499 | GAGE10 | 102724473 |
| ENSG000001 | 41.8899281 | 0.029617528 | 0.222705 | 0.13298996 | 0.89420132 | 0.92834368 | NAAA | 27163 |
| ENSG000002 | 4.05429081 | -0.050289736 | 0.37817757 | -0.1329792 | 0.89420986 | 0.92834368 | LOC100505622 | 100505622 |
| ENSG000002 | 137.907094 | -0.024511095 | 0.18438537 | -0.1329341 | 0.89424554 | 0.92835695 | EPPK1 | 83481 |

| | | | | | | | | |
|---|---|---|---|---|---|---|---|---|
| ENSG000002 | 4.25220018 | -0.050032056 | 0.37685948 | -0.1327605 | 0.89438279 | 0.92847567 | NA | NA |
| ENSG000001 | 59.7506665 | -0.025791587 | 0.19431808 | -0.1327287 | 0.89440794 | 0.92847801 | SLC13A4 | 26266 |
| ENSG000001 | 27.0367525 | -0.048296087 | 0.36422409 | -0.1325999 | 0.89450979 | 0.92855997 | NA | NA |
| ENSG000002 | 3.82802804 | -0.050149431 | 0.37854051 | -0.132481 | 0.89460384 | 0.92863382 | NA | NA |
| ENSG000002 | 5.19327264 | 0.04961232 | 0.37549568 | 0.13212488 | 0.89488552 | 0.92890244 | NA | NA |
| ENSG000002 | 6.69884528 | -0.048061817 | 0.36414381 | -0.1319858 | 0.89499551 | 0.92899283 | NA | NA |
| ENSG000001 | 114.487511 | -0.020849408 | 0.15815346 | -0.1318302 | 0.89511857 | 0.92902544 | PRLR | 5618 |
| ENSG000002 | 9.48842495 | -0.045118985 | 0.34218003 | -0.1318574 | 0.89509705 | 0.92902544 | NA | NA |
| ENSG000002 | 2.1801757 | 0.048480006 | 0.36764588 | 0.13186604 | 0.89509025 | 0.92902544 | NA | NA |
| ENSG000002 | 4.2535308 | -0.049676989 | 0.37682418 | -0.1318307 | 0.89511822 | 0.92902544 | NA | NA |
| ENSG000001 | 22.4591985 | 0.035646014 | 0.27062094 | 0.13171935 | 0.89520628 | 0.9290927 | PAK6 | 56924 |
| ENSG000002 | 4.29739276 | -0.049524966 | 0.37616436 | -0.1316578 | 0.89525498 | 0.92911946 | NA | NA |
| ENSG000002 | 8.97768365 | -0.046417894 | 0.35266427 | -0.1316206 | 0.89528437 | 0.92912618 | NA | NA |
| ENSG000002 | 4.49170792 | -0.049570597 | 0.37675743 | -0.1315717 | 0.89532311 | 0.92914261 | NA | NA |
| ENSG000001 | 105.557986 | 0.020840669 | 0.15901075 | 0.13106453 | 0.89572426 | 0.92945588 | SERTAD2 | 9792 |
| ENSG000002 | 17.2772385 | 0.039500346 | 0.30119809 | 0.13114408 | 0.89566133 | 0.92945588 | LOC101928358 | 101928358 |
| ENSG000002 | 11.926796 | 0.042244861 | 0.32232541 | 0.13106277 | 0.89572565 | 0.92945588 | NA | NA |
| ENSG000002 | 34.3006579 | -0.031432852 | 0.23986275 | -0.1310452 | 0.89573958 | 0.92945588 | NA | NA |
| ENSG000002 | 32.8597123 | -0.030635928 | 0.23375901 | -0.1310577 | 0.89572963 | 0.92945588 | NA | NA |
| ENSG000002 | 2.25177791 | 0.0480321 | 0.36676381 | 0.13096194 | 0.89580542 | 0.92950041 | NA | NA |
| ENSG000001 | 216.13638 | 0.019007208 | 0.14528317 | 0.1308287 | 0.89591082 | 0.92958599 | METTL18 | 92342 |
| ENSG000002 | 19.9973735 | 0.036107701 | 0.27615527 | 0.13075145 | 0.89597193 | 0.92962561 | NA | NA |
| ENSG000001 | 46.8770013 | 0.02910637 | 0.22277895 | 0.13065135 | 0.89605112 | 0.92968399 | LGALS3 | 3958 |
| ENSG000002 | 5.84881118 | 0.048378958 | 0.37040773 | 0.13061001 | 0.89608382 | 0.92968871 | NA | NA |
| ENSG000002 | 14.5815604 | -0.040544908 | 0.31048044 | -0.1305876 | 0.89610152 | 0.92968871 | LOC101928063 | 101928063 |
| ENSG00000C | 643.358466 | 0.016368081 | 0.12546521 | 0.13045913 | 0.89620319 | 0.9297537 | REV3L | 5980 |
| ENSG00000C | 18.2836598 | -0.039162858 | 0.30021244 | -0.1304505 | 0.89621003 | 0.9297537 | KCNQ1 | 3784 |
| ENSG000002 | 11.0350417 | -0.043681156 | 0.33492349 | -0.1304213 | 0.89623312 | 0.92975387 | NA | NA |
| ENSG000002 | 7.82173873 | -0.04681443 | 0.35906943 | -0.1303771 | 0.89626809 | 0.92976636 | NA | NA |

| | | | | | | | | |
|---|---|---|---|---|---|---|---|---|
| ENSG00000( | 368.022694 | -0.019090342 | 0.14665462 | -0.1301721 | 0.89643025 | 0.92982909 | TIGAR | 57103 |
| ENSG000001 | 96.486342 | -0.024294426 | 0.18651656 | -0.1302535 | 0.89636591 | 0.92982909 | CPEB3 | 22849 |
| ENSG000002 | 43.5459415 | 0.029961963 | 0.23007468 | 0.13022712 | 0.89638674 | 0.92982909 | NA | NA |
| ENSG000002 | 7.42033229 | 0.046215663 | 0.35506666 | 0.13016052 | 0.89643943 | 0.92982909 | NA | NA |
| ENSG000002 | 4.27727267 | -0.048956858 | 0.37614055 | -0.1301558 | 0.8964432 | 0.92982909 | NA | NA |
| ENSG000001 | 4.53235303 | -0.048963528 | 0.37633246 | -0.1301071 | 0.89648169 | 0.92984523 | NA | NA |
| ENSG000001 | 2.39283661 | 0.047939548 | 0.36923318 | 0.12983543 | 0.89669663 | 0.93000505 | RPS4XP21 | 126235 |
| ENSG000001 | 8.69657722 | 0.045554517 | 0.35096905 | 0.12979639 | 0.89672752 | 0.93000505 | LINC00173 | 100287569 |
| ENSG000002 | 2.45844973 | 0.048348146 | 0.3722726 | 0.12987296 | 0.89666693 | 0.93000505 | NA | NA |
| ENSG000002 | 8.65776131 | -0.045551588 | 0.35088082 | -0.1298207 | 0.8967083 | 0.93000505 | NA | NA |
| ENSG000001 | 61.326402 | -0.02682227 | 0.20693478 | -0.129617 | 0.89686944 | 0.93010466 | SUSD4 | 55061 |
| ENSG000002 | 2.21324029 | 0.047529409 | 0.36662337 | 0.12964097 | 0.89685049 | 0.93010466 | NA | NA |
| ENSG000002 | 12.8594002 | 0.042109229 | 0.32506893 | 0.12953939 | 0.89693086 | 0.93014458 | LINC01134 | 100133612 |
| ENSG000002 | 4.02302339 | 0.048861422 | 0.37739887 | 0.12946892 | 0.89698662 | 0.93017861 | NA | NA |
| ENSG000002 | 10.5228205 | -0.043804885 | 0.33859919 | -0.1293709 | 0.89706417 | 0.93023524 | NA | NA |
| ENSG000002 | 18.1962771 | 0.0388319 | 0.30055153 | 0.12920214 | 0.8971977 | 0.93034992 | NA | NA |
| ENSG00000( | 15.5019442 | 0.039451962 | 0.30548747 | 0.12914429 | 0.89724348 | 0.93037359 | MASP2 | 10747 |
| ENSG000002 | 10.3355644 | 0.042811289 | 0.33158294 | 0.12911185 | 0.89726914 | 0.93037642 | NA | NA |
| ENSG000002 | 32.6433837 | -0.03106194 | 0.24069303 | -0.1290521 | 0.89731642 | 0.93040166 | LINC00865 | 643529 |
| ENSG000001 | 7.91222628 | 0.046144085 | 0.35788639 | 0.12893501 | 0.89740908 | 0.93045222 | TMEM221 | 100130519 |
| ENSG000002 | 2.45846934 | 0.047767091 | 0.37048146 | 0.12893247 | 0.89741108 | 0.93045222 | NA | NA |
| ENSG000002 | 2.40630922 | -0.047309295 | 0.36749303 | -0.1287352 | 0.89756717 | 0.93059026 | NA | NA |
| ENSG000002 | 2.43922252 | 0.047734218 | 0.3712954 | 0.1285613 | 0.89770479 | 0.93070915 | NA | NA |
| ENSG000002 | 6.43335954 | -0.046638133 | 0.36424645 | -0.12804 | 0.89811728 | 0.93108919 | NA | NA |
| ENSG000002 | 5.363983 | 0.048135422 | 0.37587797 | 0.1280613 | 0.89810046 | 0.93108919 | NA | NA |
| ENSG000002 | 4.05273453 | 0.048295983 | 0.37819003 | 0.12770295 | 0.89838405 | 0.93131814 | NA | NA |
| ENSG000002 | 28.3045625 | 0.032370619 | 0.25344208 | 0.12772393 | 0.89836745 | 0.93131814 | NA | NA |
| ENSG000002 | 4.30651683 | 0.048293258 | 0.37838477 | 0.12763003 | 0.89844177 | 0.93133035 | CELA2B | 51032 |
| ENSG000002 | 7.26424141 | 0.04587968 | 0.35945652 | 0.12763624 | 0.89843685 | 0.93133035 | NA | NA |

| | | | | | | | | |
|---|---|---|---|---|---|---|---|---|
| ENSG000002 | 7.08322589 | -0.046765644 | 0.36706431 | -0.1274045 | 0.89862026 | 0.93149157 | NA | NA |
| ENSG000002 | 7.10822909 | -0.045652242 | 0.35857607 | -0.1273154 | 0.8986908 | 0.93152793 | NA | NA |
| ENSG000002 | 2.45533703 | 0.047106746 | 0.37012337 | 0.12727309 | 0.89872426 | 0.93152793 | NA | NA |
| ENSG000002 | 2.17378223 | 0.046082456 | 0.36205695 | 0.12727958 | 0.89871912 | 0.93152793 | MIR4728 | 100616132 |
| ENSG000001 | 21.1038165 | -0.035962183 | 0.28341826 | -0.1268873 | 0.89902958 | 0.93172562 | TNFRSF11B | 4982 |
| ENSG000002 | 4.22284449 | 0.048010791 | 0.37803507 | 0.12700089 | 0.89893969 | 0.93172562 | NA | NA |
| ENSG000002 | 3.76064048 | -0.048005921 | 0.37842271 | -0.1268579 | 0.89905284 | 0.93172562 | NA | NA |
| ENSG000002 | 2.20428231 | 0.046711297 | 0.36804782 | 0.12691638 | 0.89900658 | 0.93172562 | NA | NA |
| ENSG000002 | 7.39578563 | -0.046057206 | 0.36285424 | -0.1269303 | 0.89899555 | 0.93172562 | NA | NA |
| ENSG000002 | 5.17737167 | 0.047355425 | 0.37326467 | 0.12686822 | 0.8990447 | 0.93172562 | NA | NA |
| ENSG000002 | 7.77633098 | -0.046896164 | 0.36995844 | -0.1267606 | 0.89912986 | 0.93178162 | NA | NA |
| ENSG000002 | 2.42357801 | 0.047220209 | 0.3727283 | 0.12668801 | 0.89918733 | 0.93181737 | NA | NA |
| ENSG000001 | 127.846132 | -0.019734413 | 0.15583738 | -0.1266347 | 0.89922957 | 0.93183732 | FRMD4B | 23150 |
| ENSG000002 | 6.14583558 | -0.046160847 | 0.36591316 | -0.1261525 | 0.89961124 | 0.93216803 | NA | NA |
| ENSG000002 | 6.39923822 | -0.045841497 | 0.36325055 | -0.126198 | 0.89957518 | 0.93216803 | NA | NA |
| ENSG000002 | 11.0891727 | 0.041403336 | 0.32822187 | 0.12614435 | 0.89961766 | 0.93216803 | NA | NA |
| ENSG000002 | 4.06849582 | 0.047624836 | 0.37779227 | 0.12606091 | 0.89968371 | 0.93218883 | NA | NA |
| ENSG000002 | 4.36541283 | -0.047677507 | 0.37818408 | -0.1260696 | 0.89967685 | 0.93218883 | LOC100505978 | 100505978 |
| ENSG000001 | 21.4338676 | -0.034605697 | 0.27463244 | -0.1260073 | 0.89972613 | 0.93220896 | C2orf40 | 84417 |
| ENSG000001 | 75.967574 | 0.023569631 | 0.18709441 | 0.1259772 | 0.89974997 | 0.93220984 | ZNF665 | 79788 |
| ENSG000001 | 17.6645054 | 0.037520487 | 0.29801944 | 0.12589946 | 0.89981151 | 0.93224978 | RASD1 | 51655 |
| ENSG000001 | 7.24138034 | 0.04478004 | 0.35586558 | 0.12583414 | 0.89986321 | 0.93225571 | OSCAR | 126014 |
| ENSG000002 | 7.26950571 | 0.044809347 | 0.35609525 | 0.12583528 | 0.89986231 | 0.93225571 | NA | NA |
| ENSG000002 | 2.21265211 | 0.046003616 | 0.36577189 | 0.12577133 | 0.89991294 | 0.93228341 | NA | NA |
| ENSG000001 | 114.310841 | -0.047207333 | 0.37553096 | -0.1257082 | 0.89996288 | 0.93229403 | NA | NA |
| ENSG000001 | 45.9376116 | 0.028479767 | 0.22656882 | 0.12570029 | 0.89996917 | 0.93229403 | HYKK | 123688 |
| ENSG000002 | 2.44936755 | 0.046640944 | 0.37146633 | 0.12555901 | 0.90008101 | 0.93238607 | NA | NA |
| ENSG000001 | 4.01548563 | -0.047346967 | 0.37797963 | -0.1252633 | 0.90031513 | 0.93248636 | PLVAP | 83483 |
| ENSG000002 | 4.69376015 | -0.047094344 | 0.37558416 | -0.1253896 | 0.90021513 | 0.93248636 | NA | NA |

| | | | | | | | | |
|---|---|---|---|---|---|---|---|---|
| ENSG000002 | 26.5836325 | 0.034086457 | 0.27209026 | 0.12527628 | 0.90030483 | 0.93248636 | NA | NA |
| ENSG000002 | 2.19808471 | 0.045694635 | 0.36484469 | 0.12524407 | 0.90033033 | 0.93248636 | NA | NA |
| ENSG000002 | 20.2916689 | 0.035225162 | 0.28102239 | 0.12534646 | 0.90024927 | 0.93248636 | NA | NA |
| ENSG000002 | 7.40975553 | 0.04443003 | 0.35462924 | 0.12528586 | 0.90029724 | 0.93248636 | NA | NA |
| ENSG000002 | 5.549176 | 0.046530872 | 0.37155329 | 0.12523337 | 0.9003388 | 0.93248636 | NA | NA |
| ENSG000002 | 3.81933796 | -0.046972091 | 0.37546431 | -0.125104 | 0.90044121 | 0.93256861 | VENTXP1 | 139538 |
| ENSG000002 | 3.76085884 | 0.046890178 | 0.37520801 | 0.12497115 | 0.90054639 | 0.93265372 | RNU6-32P | 106480708 |
| ENSG000002 | 3.56933365 | -0.047049385 | 0.37676999 | -0.1248756 | 0.90062202 | 0.93270823 | NA | NA |
| ENSG000002 | 3.76393636 | 0.04723892 | 0.37847138 | 0.12481504 | 0.90066998 | 0.93273408 | NA | NA |
| ENSG000000 | 75.8535568 | 0.023593502 | 0.18922334 | 0.12468601 | 0.90077214 | 0.93276841 | PLAUR | 5329 |
| ENSG000002 | 4.7245925 | -0.046889284 | 0.37590365 | -0.1247375 | 0.90073136 | 0.93276841 | NA | NA |
| ENSG000002 | 5.59149162 | 0.046043745 | 0.36927184 | 0.12468794 | 0.90077061 | 0.93276841 | NA | NA |
| ENSG000002 | 48.2214559 | 0.027941709 | 0.22439292 | 0.12452135 | 0.9009025 | 0.93287958 | CCT6P3 | 643180 |
| ENSG000001 | 42.2247858 | 0.027344365 | 0.21978801 | 0.12441245 | 0.90098872 | 0.93294504 | EPAS1 | 2034 |
| ENSG000002 | 4.27513518 | 0.046869664 | 0.37747151 | 0.12416742 | 0.90118272 | 0.93312209 | NA | NA |
| ENSG000002 | 2.48183873 | 0.046067297 | 0.37116235 | 0.1241163 | 0.90122319 | 0.93314017 | NA | NA |
| ENSG000002 | 3.50224126 | 0.046610275 | 0.37681506 | 0.12369536 | 0.90155649 | 0.93346143 | NA | NA |
| ENSG000002 | 4.06487987 | -0.046609758 | 0.37721384 | -0.1235632 | 0.90166111 | 0.93354593 | NA | NA |
| ENSG000002 | 6.93400919 | -0.045556158 | 0.3689888 | -0.1234622 | 0.90174113 | 0.93360494 | NA | NA |
| ENSG000001 | 170.059744 | -0.020386404 | 0.16524797 | -0.1233686 | 0.90181525 | 0.93365784 | NACC2 | 138151 |
| ENSG000002 | 4.27558973 | -0.04639406 | 0.37617409 | -0.1233314 | 0.90184471 | 0.9336645 | LOC101929460 | 101929460 |
| ENSG000001 | 2.20565259 | 0.045095452 | 0.36576063 | 0.12329226 | 0.90187567 | 0.93367272 | RNU4-44P | 106479578 |
| ENSG000002 | 4.19282755 | 0.046566057 | 0.37807697 | 0.12316555 | 0.90197601 | 0.93375275 | NA | NA |
| ENSG000001 | 237.670988 | -0.016633902 | 0.13523799 | -0.1229973 | 0.90210927 | 0.93386687 | ID4 | 3400 |
| ENSG000002 | 6.74613492 | 0.045634307 | 0.37126484 | 0.12291578 | 0.90217379 | 0.93390982 | NA | NA |
| ENSG000002 | 5.48422937 | -0.046083948 | 0.37503548 | -0.1228789 | 0.90220299 | 0.93391621 | NA | NA |
| ENSG000001 | 31.6346846 | 0.030093734 | 0.2449725 | 0.12284536 | 0.90222955 | 0.93391986 | TMEM86A | 144110 |
| ENSG000002 | 3.81054937 | -0.046454721 | 0.37846605 | -0.1227448 | 0.90230923 | 0.93397849 | NA | NA |
| ENSG000001 | 19.7265512 | 0.034779695 | 0.28390461 | 0.12250486 | 0.9024992 | 0.93412962 | ACTL8 | 81569 |

| | | | | | | | | |
|---|---|---|---|---|---|---|---|---|
| ENSG000002 | 4.05188385 | -0.046237534 | 0.37753212 | -0.1224731 | 0.90252434 | 0.93412962 | TCEB3B | 51224 |
| ENSG000002 | 6.61020379 | -0.044357 | 0.36211576 | -0.122494 | 0.90250782 | 0.93412962 | NA | NA |
| ENSG000002 | 20.0214419 | 0.034684183 | 0.283392 | 0.12238942 | 0.90259062 | 0.93417438 | NA | NA |
| ENSG000002 | 7.66582856 | 0.043676007 | 0.35709501 | 0.12230921 | 0.90265414 | 0.93421628 | MIR3939 | 100500857 |
| ENSG000002 | 3.82046049 | -0.045992256 | 0.3782833 | -0.1215815 | 0.90323045 | 0.93478889 | NA | NA |
| ENSG000002 | 21.1974579 | -0.033630076 | 0.27706839 | -0.1213782 | 0.90339144 | 0.93493164 | NA | NA |
| ENSG00000C | 29.586691 | 0.029510096 | 0.24319798 | 0.12134187 | 0.90342026 | 0.9349376 | GADD45B | 4616 |
| ENSG000001 | 47.2374541 | -0.025580301 | 0.21112586 | -0.1211614 | 0.90356321 | 0.93504438 | FRMPD3 | 84443 |
| ENSG000002 | 73.2803021 | -0.022862683 | 0.18870859 | -0.1211534 | 0.90356955 | 0.93504438 | NA | NA |
| ENSG00000C | 148.751223 | 0.01767139 | 0.14595342 | 0.12107555 | 0.9036312 | 0.93507981 | CD44 | 960 |
| ENSG000002 | 2.45405882 | 0.044621558 | 0.36861505 | 0.12105192 | 0.90364992 | 0.93507981 | NA | NA |
| ENSG000002 | 65.129756 | 0.025410347 | 0.21019118 | 0.1208916 | 0.9037769 | 0.93515315 | NA | NA |
| ENSG000002 | 2.67915481 | 0.04482994 | 0.37087822 | 0.1208751 | 0.90378997 | 0.93515315 | NA | NA |
| ENSG000002 | 7.2486082 | 0.043523604 | 0.35996402 | 0.12091099 | 0.90376154 | 0.93515315 | NA | NA |
| ENSG000002 | 44.817475 | 0.026006826 | 0.21529157 | 0.12079816 | 0.9038509 | 0.93519234 | NA | NA |
| ENSG000002 | 4.02418214 | -0.045677802 | 0.37855777 | -0.1206627 | 0.90395821 | 0.93527949 | NA | NA |
| ENSG000002 | 2.44401817 | 0.044920958 | 0.37248722 | 0.12059731 | 0.90401 | 0.93530922 | NA | NA |
| ENSG000002 | 20.2683116 | -0.034004102 | 0.28219184 | -0.1204999 | 0.90408712 | 0.93536514 | NA | NA |
| ENSG000002 | 4.26662102 | 0.045388051 | 0.37722585 | 0.12032063 | 0.90422917 | 0.93548824 | NA | NA |
| ENSG000002 | 2.47170634 | 0.044789395 | 0.37254035 | 0.12022696 | 0.90430336 | 0.93554112 | NA | NA |
| ENSG000001 | 2.23638135 | 0.043680878 | 0.36350151 | 0.12016698 | 0.90435088 | 0.93554255 | HOXA6 | 3203 |
| ENSG000002 | 2.45075711 | 0.044925278 | 0.37377009 | 0.12019495 | 0.90432872 | 0.93554255 | NA | NA |
| ENSG000001 | 240.470443 | 0.018293932 | 0.15228565 | 0.12012906 | 0.90438091 | 0.93554848 | FAM210B | 116151 |
| ENSG000002 | 3.79987811 | 0.045469394 | 0.37859143 | 0.12010149 | 0.90440275 | 0.93554848 | NA | NA |
| ENSG000002 | 20.1553337 | -0.03368126 | 0.28152393 | -0.1196391 | 0.90476907 | 0.93590354 | NA | NA |
| ENSG000002 | 3.78516477 | -0.045221191 | 0.37821497 | -0.1195648 | 0.90482792 | 0.93591666 | NA | NA |
| ENSG000002 | 4.02357658 | 0.045269364 | 0.37857583 | 0.11957806 | 0.9048174 | 0.93591666 | LOC105372419 | 105372419 |
| ENSG000001 | 29.9252624 | 0.029837613 | 0.24965412 | 0.1195158 | 0.90486672 | 0.93591669 | GLP1R | 2740 |
| ENSG000002 | 2.4838864 | 0.044274542 | 0.37047818 | 0.11950648 | 0.90487411 | 0.93591669 | RN7SL101P | 106479260 |

| | | | | | | | | |
|---|---|---|---|---|---|---|---|---|
| ENSG000002 | 2.64621442 | 0.044316927 | 0.37168935 | 0.11923109 | 0.90509227 | 0.93611847 | KLRC2 | 3822 |
| ENSG000002 | 11.0215387 | -0.03899455 | 0.32728093 | -0.119147 | 0.90515888 | 0.93616348 | NA | NA |
| ENSG000002 | 11.2624297 | -0.038831341 | 0.32616313 | -0.119055 | 0.90523181 | 0.93621503 | LOC100132215 | 100132215 |
| ENSG000001 | 5.0369091 | -0.044840661 | 0.37718042 | -0.1188839 | 0.90536737 | 0.93630748 | ZNF645 | 158506 |
| ENSG000002 | 3.54978736 | -0.045004969 | 0.37855832 | -0.1188852 | 0.90536633 | 0.93630748 | NA | NA |
| ENSG000002 | 3.82421868 | -0.044951159 | 0.37856858 | -0.1187398 | 0.9054815 | 0.93640162 | NA | NA |
| ENSG000001 | 15.1417273 | -0.038351166 | 0.32312955 | -0.1186867 | 0.90552361 | 0.93642129 | F10 | 2159 |
| ENSG000002 | 2.44588957 | 0.044061104 | 0.37189815 | 0.11847626 | 0.9056903 | 0.93656365 | RNU6-1309P | 106480410 |
| ENSG000002 | 4.27055547 | 0.044682689 | 0.37721362 | 0.1184546 | 0.90570746 | 0.93656365 | NA | NA |
| ENSG000001 | 221.43851 | 0.017281234 | 0.14601268 | 0.11835434 | 0.9057869 | 0.93659802 | VEPH1 | 79674 |
| ENSG000002 | 3.82316105 | -0.044763038 | 0.37818028 | -0.1183643 | 0.90577902 | 0.93659802 | NA | NA |
| ENSG000002 | 2.6810066 | 0.043930609 | 0.37130279 | 0.11831478 | 0.90581825 | 0.93660655 | NA | NA |
| ENSG000002 | 10.5724168 | -0.040190303 | 0.34026559 | -0.1181145 | 0.90597693 | 0.93674674 | NA | NA |
| ENSG000002 | 3.7976649 | -0.044661267 | 0.37835233 | -0.1180415 | 0.9060348 | 0.93678269 | NA | NA |
| ENSG000002 | 3.57121302 | -0.044537585 | 0.3774005 | -0.1180115 | 0.90605858 | 0.93678339 | NA | NA |
| ENSG000002 | 11.0841029 | 0.038622653 | 0.32777437 | 0.11783305 | 0.90619994 | 0.93688202 | NA | NA |
| ENSG000002 | 29.2182614 | 0.029496725 | 0.25032705 | 0.11783275 | 0.90620018 | 0.93688202 | NA | NA |
| ENSG000001 | 56.1507837 | -0.02472069 | 0.20987032 | -0.1177903 | 0.90623381 | 0.9368929 | FAM46C | 54855 |
| ENSG000002 | 4.21508547 | -0.044433848 | 0.37734746 | -0.1177531 | 0.90626326 | 0.93689946 | GATA6-AS1 | 100128893 |
| ENSG000002 | 3.59136748 | 0.044412322 | 0.37726711 | 0.11772116 | 0.9062886 | 0.93690178 | NA | NA |
| ENSG000002 | 2.45544803 | 0.043685857 | 0.37133857 | 0.11764428 | 0.90634952 | 0.93691698 | NA | NA |
| ENSG000002 | 2.69046071 | 0.044102976 | 0.37480342 | 0.11766962 | 0.90632944 | 0.93691698 | NA | NA |
| ENSG000002 | 58.5739328 | -0.022541023 | 0.19172226 | -0.1175712 | 0.9064074 | 0.93692904 | NA | NA |
| ENSG000002 | 2.66266119 | 0.044145513 | 0.37547293 | 0.11757309 | 0.90640593 | 0.93692904 | MIR3125 | 100422986 |
| ENSG000002 | 4.04067125 | 0.044437344 | 0.37819208 | 0.1174994 | 0.90646432 | 0.93694011 | NA | NA |
| ENSG000002 | 2.46578894 | 0.043841897 | 0.37303858 | 0.11752644 | 0.90644289 | 0.93694011 | NA | NA |
| ENSG000000 | 26.6319153 | 0.03038695 | 0.25889938 | 0.11736973 | 0.90656707 | 0.93697467 | EPN3 | 55040 |
| ENSG000002 | 4.07801307 | -0.044413085 | 0.37824997 | -0.1174173 | 0.90652939 | 0.93697467 | LOC101927686 | 101927686 |
| ENSG000002 | 2.89914795 | 0.044062259 | 0.3753426 | 0.11739211 | 0.90654934 | 0.93697467 | NA | NA |

| | | | | | | | | |
|---|---|---|---|---|---|---|---|---|
| ENSG000001 | 60.3879294 | -0.023195472 | 0.19779025 | -0.1172731 | 0.90664365 | 0.93702994 | RASIP1 | 54922 |
| ENSG00000C | 172.866477 | 0.017287928 | 0.14759245 | 0.11713288 | 0.90675476 | 0.93712088 | MAP2 | 4133 |
| ENSG000002 | 4.03155661 | 0.04426429 | 0.37816374 | 0.11705059 | 0.90681996 | 0.93716439 | NA | NA |
| ENSG000001 | 89.9386008 | -0.020896447 | 0.17862747 | -0.1169834 | 0.90687322 | 0.93717166 | ZNF699 | 374879 |
| ENSG000002 | 4.06005255 | -0.044210954 | 0.37785186 | -0.117006 | 0.90685526 | 0.93717166 | NA | NA |
| ENSG000002 | 2.44540086 | 0.043185939 | 0.36959554 | 0.11684648 | 0.90698171 | 0.937236 | LOC101929626 | 101929626 |
| ENSG000002 | 6.38828464 | -0.04255963 | 0.36417026 | -0.1168674 | 0.90696514 | 0.937236 | NA | NA |
| ENSG000002 | 2.44953063 | 0.043552548 | 0.37283043 | 0.11681597 | 0.90700589 | 0.93723711 | MAGI2-AS2 | 100874021 |
| ENSG000002 | 16.1010928 | 0.035193485 | 0.30155791 | 0.11670556 | 0.90709338 | 0.93730363 | NA | NA |
| ENSG000002 | 2.24525969 | 0.042689854 | 0.36591424 | 0.11666628 | 0.90712451 | 0.93731191 | RNU6-638P | 106481383 |
| ENSG000002 | 5.83568211 | 0.043015865 | 0.3689136 | 0.11660146 | 0.90717588 | 0.9373411 | NA | NA |
| ENSG000002 | 20.3868949 | -0.032233867 | 0.27685977 | -0.1164267 | 0.90731438 | 0.93746033 | LINC01018 | 255167 |
| ENSG000002 | 2.68753725 | 0.043344465 | 0.37255929 | 0.11634246 | 0.90738113 | 0.93750541 | NA | NA |
| ENSG000002 | 4.251029 | 0.043988957 | 0.37853884 | 0.11620725 | 0.90748829 | 0.93752058 | NA | NA |
| ENSG000002 | 3.11448537 | -0.043405739 | 0.37329733 | -0.1162766 | 0.90743334 | 0.93752058 | MIR1256 | 100302155 |
| ENSG000002 | 2.68044545 | 0.043673055 | 0.37565246 | 0.1162592 | 0.90744712 | 0.93752058 | NA | NA |
| ENSG000002 | 3.80613362 | -0.043930726 | 0.37801774 | -0.1162134 | 0.90748342 | 0.93752058 | NA | NA |
| ENSG000002 | 9.77096443 | 0.03952611 | 0.34048993 | 0.11608599 | 0.9075844 | 0.93759598 | NA | NA |
| ENSG000002 | 7.46051209 | 0.040907219 | 0.35267913 | 0.11598991 | 0.90766054 | 0.93762687 | NA | NA |
| ENSG000002 | 4.04809375 | -0.043848885 | 0.37802432 | -0.1159949 | 0.9076566 | 0.93762687 | NA | NA |
| ENSG000002 | 2.43449879 | 0.043082447 | 0.37187889 | 0.11585075 | 0.90777083 | 0.93771348 | NA | NA |
| ENSG000002 | 4.51879561 | -0.043628738 | 0.37667559 | -0.1158258 | 0.90779063 | 0.93771348 | NA | NA |
| ENSG000002 | 6.35229889 | -0.042186795 | 0.3643191 | -0.1157963 | 0.907814 | 0.93771373 | NA | NA |
| ENSG000002 | 36.4967269 | 0.026602842 | 0.22988484 | 0.11572247 | 0.90787249 | 0.93775026 | LINC00577 | 100113403 |
| ENSG000002 | 2.45437093 | 0.04313533 | 0.37301478 | 0.11563973 | 0.90793807 | 0.93779411 | RN7SL670P | 106481105 |
| ENSG000002 | 8.76553438 | -0.040575152 | 0.35097196 | -0.115608 | 0.90796326 | 0.93779624 | NA | NA |
| ENSG000002 | 39.4695013 | -0.026274064 | 0.22736134 | -0.1155608 | 0.90800061 | 0.93781093 | NA | NA |
| ENSG000002 | 9.90492378 | 0.039357197 | 0.34089136 | 0.11545378 | 0.90808545 | 0.93787467 | ORM2 | 5005 |
| ENSG000002 | 7.07883162 | 0.041921006 | 0.36330908 | 0.11538662 | 0.90813868 | 0.93790576 | NA | NA |

| | | | | | | | | |
|---|---|---|---|---|---|---|---|---|
| ENSG000002 | 6.32665658 | 0.042235816 | 0.3662238 | 0.11532788 | 0.90818524 | 0.93792995 | NA | NA |
| ENSG000001 | 98.1977943 | 0.019359203 | 0.16805914 | 0.1151928 | 0.9082923 | 0.93798101 | MRO | 83876 |
| ENSG000001 | 64.3149039 | 0.023364328 | 0.20285416 | 0.11517796 | 0.90830407 | 0.93798101 | EFCAB10 | 100130771 |
| ENSG000002 | 3.34420561 | -0.04357178 | 0.37816657 | -0.1152185 | 0.90827195 | 0.93798101 | NA | NA |
| ENSG000001 | 3.37075356 | 0.043298388 | 0.3761759 | 0.11510144 | 0.90836472 | 0.93801523 | ATP5EP2 | 432369 |
| ENSG000002 | 8.1656544 | -0.040926523 | 0.35564227 | -0.1150778 | 0.90838347 | 0.93801523 | NA | NA |
| ENSG000002 | 2.46340104 | 0.042711178 | 0.37135621 | 0.11501404 | 0.90843399 | 0.93804351 | NA | NA |
| ENSG000002 | 9.74386191 | 0.038958404 | 0.33905144 | 0.11490411 | 0.90852113 | 0.9381096 | NA | NA |
| ENSG000001 | 4.68108397 | -0.043258973 | 0.37687572 | -0.1147831 | 0.90861702 | 0.93817669 | NOXO1 | 124056 |
| ENSG000002 | 2.65030487 | 0.042948906 | 0.37424137 | 0.11476258 | 0.90863331 | 0.93817669 | MIR1278 | 100302163 |
| ENSG000002 | 5.66735944 | 0.042380529 | 0.36937889 | 0.11473457 | 0.90865551 | 0.93817669 | NA | NA |
| ENSG000002 | 14.0615324 | 0.037880366 | 0.33117915 | 0.11438028 | 0.90893634 | 0.93839497 | NA | NA |
| ENSG000002 | 23.1627843 | -0.030950658 | 0.2705539 | -0.1143974 | 0.90892279 | 0.93839497 | CECR3 | 27442 |
| ENSG000002 | 4.03408695 | -0.043184603 | 0.37744186 | -0.1144139 | 0.90890968 | 0.93839497 | NA | NA |
| ENSG000001 | 48.5678974 | -0.02415987 | 0.2117674 | -0.1140868 | 0.90916897 | 0.93861124 | PXMP4 | 11264 |
| ENSG000002 | 2.62756135 | 0.042167927 | 0.37000647 | 0.11396538 | 0.90926524 | 0.93863894 | NA | NA |
| ENSG000002 | 8.4632458 | -0.039665351 | 0.34800235 | -0.1139801 | 0.90925355 | 0.93863894 | NA | NA |
| ENSG000002 | 11.053306 | -0.037044449 | 0.32492606 | -0.1140089 | 0.90923077 | 0.93863894 | NA | NA |
| ENSG000002 | 3.28510354 | -0.042864774 | 0.37627044 | -0.1139201 | 0.90930112 | 0.93865208 | NA | NA |
| ENSG000001 | 4.09184381 | -0.042977201 | 0.37847884 | -0.1135525 | 0.90959258 | 0.93892349 | OR2K2 | 26248 |
| ENSG000002 | 2.24779139 | 0.041442865 | 0.36503877 | 0.11353004 | 0.90961035 | 0.93892349 | NA | NA |
| ENSG000001 | 48.3520284 | 0.023895621 | 0.21066489 | 0.11342954 | 0.90969002 | 0.93896545 | MBNL2 | 10150 |
| ENSG000001 | 9.3634653 | -0.038740003 | 0.34156133 | -0.1134203 | 0.90969731 | 0.93896545 | NA | NA |
| ENSG000002 | 18.0922684 | -0.032750388 | 0.28890823 | -0.1133591 | 0.90974583 | 0.93899163 | NA | NA |
| ENSG000002 | 6.33716312 | -0.041138528 | 0.36309879 | -0.1132984 | 0.90979395 | 0.9390174 | NA | NA |
| ENSG000001 | 6.04309822 | 0.041468409 | 0.36659145 | 0.11311886 | 0.90993632 | 0.93908714 | RBP4 | 5950 |
| ENSG000001 | 13.3951485 | 0.035478757 | 0.31357912 | 0.11314132 | 0.90991851 | 0.93908714 | FOXN4 | 121643 |
| ENSG000002 | 3.79455939 | -0.042827018 | 0.37858108 | -0.1131251 | 0.90993139 | 0.93908714 | NA | NA |
| ENSG000002 | 10.571747 | -0.037543772 | 0.33222009 | -0.1130087 | 0.91002363 | 0.93908714 | NA | NA |

| | | | | | | | | |
|---|---|---|---|---|---|---|---|---|
| ENSG000002 | 8.39692165 | -0.039610057 | 0.35035155 | -0.113058 | 0.90998455 | 0.93908714 | NA | NA |
| ENSG000002 | 8.02912344 | -0.039939902 | 0.35335998 | -0.1130289 | 0.91000761 | 0.93908714 | NA | NA |
| ENSG000002 | 2.24710625 | 0.041273783 | 0.36503654 | 0.11306754 | 0.90997701 | 0.93908714 | NA | NA |
| ENSG000002 | 11.9164911 | 0.038526503 | 0.34109057 | 0.11295095 | 0.91006945 | 0.93911052 | NA | NA |
| ENSG000002 | 3.5901917 | -0.042745048 | 0.3785932 | -0.112905 | 0.91010591 | 0.93912425 | NA | NA |
| ENSG000002 | 2.89482917 | 0.042425072 | 0.3758703 | 0.11287157 | 0.91013237 | 0.93912766 | NA | NA |
| ENSG000002 | 7.63903893 | 0.039523215 | 0.35246436 | 0.11213393 | 0.91071721 | 0.93970722 | NA | NA |
| ENSG000002 | 34.1038696 | 0.02702335 | 0.24114739 | 0.11206155 | 0.91077461 | 0.93974253 | NA | NA |
| ENSG000002 | 6.59791194 | -0.040625016 | 0.36312913 | -0.1118748 | 0.91092264 | 0.93987135 | NA | NA |
| ENSG000002 | 4.0084282 | 0.04227205 | 0.3781888 | 0.11177499 | 0.91100182 | 0.93991373 | NA | NA |
| ENSG000002 | 2.67568947 | 0.042044904 | 0.37619163 | 0.11176459 | 0.91101006 | 0.93991373 | NA | NA |
| ENSG000002 | 3.81984198 | 0.042237956 | 0.37826432 | 0.11166254 | 0.91109098 | 0.93997329 | NA | NA |
| ENSG000001 | 102.171706 | -0.020269901 | 0.18192113 | -0.1114214 | 0.91128222 | 0.94006599 | FAAH2 | 158584 |
| ENSG000002 | 2.68101346 | 0.041727828 | 0.37456622 | 0.11140307 | 0.91129672 | 0.94006599 | NA | NA |
| ENSG000002 | 6.22535112 | -0.04119811 | 0.36977578 | -0.1114138 | 0.91128825 | 0.94006599 | NA | NA |
| ENSG000002 | 15.7151558 | -0.034002062 | 0.30514783 | -0.1114282 | 0.91127683 | 0.94006599 | NA | NA |
| ENSG000002 | 5.18871666 | 0.041816397 | 0.37536134 | 0.11140305 | 0.91129674 | 0.94006599 | NA | NA |
| ENSG000002 | 10.880838 | 0.037094736 | 0.33352398 | 0.1112206 | 0.91144142 | 0.94017747 | NA | NA |
| ENSG000002 | 5.63306872 | 0.04113135 | 0.36985866 | 0.11120829 | 0.91145118 | 0.94017747 | MIR210HG | 100506211 |
| ENSG000002 | 10.4615169 | 0.037411591 | 0.33676968 | 0.11108954 | 0.91154534 | 0.94020285 | NA | NA |
| ENSG000002 | 18.1366495 | 0.032002239 | 0.28804562 | 0.11110129 | 0.91153603 | 0.94020285 | NA | NA |
| ENSG000002 | 21.7980001 | 0.029923388 | 0.26922868 | 0.11114487 | 0.91150146 | 0.94020285 | LOC105372233 | 105372233 |
| ENSG000002 | 3.8370114 | -0.042043319 | 0.37857441 | -0.111057 | 0.91157119 | 0.94020559 | NA | NA |
| ENSG000002 | 16.3822547 | -0.032699374 | 0.29470404 | -0.1109567 | 0.91165072 | 0.94026371 | NA | NA |
| ENSG000001 | 301.716087 | 0.016458462 | 0.14843792 | 0.11087775 | 0.91171329 | 0.94030433 | TANC1 | 85461 |
| ENSG000001 | 27.8752734 | 0.028423387 | 0.25700893 | 0.11059299 | 0.9119391 | 0.94036981 | STARD8 | 9754 |
| ENSG000001 | 32.9717656 | 0.027315446 | 0.24679784 | 0.11067944 | 0.91187055 | 0.94036981 | TMEM79 | 84283 |
| ENSG000001 | 35.6701684 | -0.026702767 | 0.2414293 | -0.1106028 | 0.91193129 | 0.94036981 | TVP23A | 780776 |
| ENSG000002 | 4.26403725 | 0.041677008 | 0.37681444 | 0.11060353 | 0.91193075 | 0.94036981 | NA | NA |

| | | | | | | | | |
|---|---|---|---|---|---|---|---|---|
| ENSG000002 | 3.823025 | -0.041924235 | 0.37856446 | -0.1107453 | 0.91181832 | 0.94036981 | NA | NA |
| ENSG000002 | 5.39613813 | 0.041494457 | 0.37478701 | 0.11071477 | 0.91184254 | 0.94036981 | RN7SL559P | 106481849 |
| ENSG000002 | 11.0421434 | -0.03614681 | 0.32681892 | -0.1106019 | 0.911932 | 0.94036981 | NA | NA |
| ENSG000002 | 3.35780802 | -0.04175947 | 0.37789667 | -0.110505 | 0.91200889 | 0.94041785 | NA | NA |
| ENSG000002 | 3.40856014 | -0.041026938 | 0.37157212 | -0.1104145 | 0.91208068 | 0.94046796 | NA | NA |
| ENSG000002 | 2.46137787 | 0.0412029 | 0.37334241 | 0.11036223 | 0.91212211 | 0.94048399 | NA | NA |
| ENSG000002 | 12.5299633 | -0.036573553 | 0.33147322 | -0.1103364 | 0.91214261 | 0.94048399 | NA | NA |
| ENSG000002 | 29.9901554 | 0.027799887 | 0.25208112 | 0.11028151 | 0.91218612 | 0.94050494 | RNU4ATAC | 100151683 |
| ENSG000002 | 2.42475153 | 0.040996787 | 0.37199564 | 0.11020771 | 0.91224465 | 0.94054137 | NA | NA |
| ENSG000002 | 3.79394693 | -0.041698621 | 0.37850485 | -0.1101667 | 0.91227719 | 0.94055101 | RNU6-1138P | 106480641 |
| ENSG000002 | 222.724622 | -0.017771547 | 0.16136306 | -0.1101339 | 0.91230316 | 0.94055387 | NA | NA |
| ENSG000002 | 3.29030389 | -0.041631156 | 0.37817615 | -0.110084 | 0.91234273 | 0.94057075 | NA | NA |
| ENSG000001 | 29.5737462 | -0.026854446 | 0.24407774 | -0.1100242 | 0.91239022 | 0.9405958 | CCDC151 | 115948 |
| ENSG000002 | 7.93723139 | 0.038614663 | 0.35181076 | 0.10975976 | 0.91259991 | 0.94078805 | NA | NA |
| ENSG000002 | 4.73645017 | 0.041280781 | 0.37720873 | 0.1094375 | 0.91285549 | 0.94097977 | NA | NA |
| ENSG000002 | 2.91631953 | 0.040952877 | 0.37418867 | 0.10944446 | 0.91284997 | 0.94097977 | LOC105372107 | 105372107 |
| ENSG000002 | 7.56554008 | -0.039667203 | 0.36235597 | -0.1094703 | 0.91282951 | 0.94097977 | NA | NA |
| ENSG000002 | 3.79143483 | -0.041234502 | 0.3785881 | -0.1089165 | 0.9132687 | 0.94138177 | NA | NA |
| ENSG000002 | 6.12494658 | -0.039664094 | 0.36549164 | -0.1085226 | 0.91358117 | 0.94167993 | RNU6-171P | 106481230 |
| ENSG000002 | 8.74172869 | -0.037391173 | 0.34517823 | -0.1083243 | 0.91373849 | 0.94181814 | NA | NA |
| ENSG000001 | 161.503766 | 0.015900197 | 0.14701965 | 0.10815015 | 0.91387659 | 0.94193596 | SUSD1 | 64420 |
| ENSG000002 | 15.2226553 | -0.032732602 | 0.30273882 | -0.1081216 | 0.91389924 | 0.94193596 | MC1R | 4157 |
| ENSG000002 | 17.9976378 | 0.031163774 | 0.28833656 | 0.10808124 | 0.91393125 | 0.941945 | SNORD100 | 594838 |
| ENSG000002 | 5.91609838 | 0.039710282 | 0.36759925 | 0.10802601 | 0.91397506 | 0.94196621 | NA | NA |
| ENSG000002 | 7.24437319 | 0.039079064 | 0.36266827 | 0.1077543 | 0.9141906 | 0.94216441 | NA | NA |
| ENSG000001 | 3.33515101 | -0.040686649 | 0.37827711 | -0.1075578 | 0.91434648 | 0.94230112 | EN1 | 2019 |
| ENSG000000 | 7.45140486 | 0.038228294 | 0.35566735 | 0.10748328 | 0.91440559 | 0.94233808 | ENTPD2 | 954 |
| ENSG000002 | 3.73948327 | -0.040632133 | 0.37857082 | -0.1073303 | 0.91452692 | 0.94243916 | NA | NA |
| ENSG000001 | 6.30798313 | 0.039100697 | 0.36457337 | 0.10725056 | 0.91459021 | 0.94245648 | CCL25 | 6370 |

| | | | | | | | | |
|---|---|---|---|---|---|---|---|---|
| ENSG000002 | 5.64079326 | 0.039606411 | 0.36919172 | 0.10727871 | 0.91456788 | 0.94245648 | NA | NA |
| ENSG000001 | 17.3418765 | 0.031263358 | 0.29168889 | 0.10718049 | 0.91464579 | 0.94248981 | NLRP10 | 338322 |
| ENSG000002 | 4.50947458 | 0.04049634 | 0.37803695 | 0.1071227 | 0.91469164 | 0.9425131 | LINC00613 | 100507528 |
| ENSG000002 | 2.70875371 | 0.039805572 | 0.3718543 | 0.10704615 | 0.91475236 | 0.94255172 | MIR4254 | 100423028 |
| ENSG00000C | 49.8845469 | -0.022380699 | 0.20932083 | -0.1069206 | 0.91485201 | 0.94260649 | DAZL | 1618 |
| ENSG000002 | 3.78393884 | -0.04042548 | 0.37801144 | -0.1069425 | 0.91483461 | 0.94260649 | NA | NA |
| ENSG000001 | 899.777934 | 0.01304825 | 0.12209845 | 0.10686662 | 0.91489479 | 0.94261832 | ZNF829 | 374899 |
| ENSG000002 | 3.10171767 | -0.04027241 | 0.37691493 | -0.1068475 | 0.91490998 | 0.94261832 | NA | NA |
| ENSG000002 | 3.77357846 | -0.04043454 | 0.37858989 | -0.106803 | 0.91494526 | 0.94263072 | NA | NA |
| ENSG000002 | 2.69296485 | 0.040106144 | 0.37584583 | 0.10670903 | 0.91501982 | 0.94268359 | NA | NA |
| ENSG000002 | 6.88745962 | -0.038514981 | 0.36126581 | -0.1066112 | 0.91509743 | 0.94272313 | NA | NA |
| ENSG000002 | 2.67905535 | 0.03995918 | 0.37484442 | 0.10660204 | 0.9151047 | 0.94272313 | NA | NA |
| ENSG000002 | 3.34892968 | -0.040325978 | 0.3784387 | -0.1065588 | 0.915139 | 0.94273452 | NAV2-AS2 | 100874014 |
| ENSG000002 | 18.3852842 | 0.031033265 | 0.29211022 | 0.1062382 | 0.91539336 | 0.9429726 | NA | NA |
| ENSG000002 | 2.47569266 | 0.039228326 | 0.36943345 | 0.10618509 | 0.9154355 | 0.94297604 | NA | NA |
| ENSG000002 | 18.869604 | -0.031984158 | 0.30123896 | -0.1061754 | 0.91544321 | 0.94297604 | NA | NA |
| ENSG000002 | 2.49254382 | 0.0393253 | 0.37056473 | 0.10612262 | 0.91548506 | 0.94299519 | NA | NA |
| ENSG000002 | 16.9617164 | 0.03114649 | 0.29362626 | 0.10607529 | 0.91552262 | 0.94300992 | NA | NA |
| ENSG000002 | 5.90306744 | -0.038712116 | 0.36579886 | -0.105829 | 0.91571805 | 0.94316331 | NA | NA |
| ENSG000002 | 12.9255552 | -0.033867796 | 0.31999698 | -0.1058379 | 0.915711 | 0.94316331 | NA | NA |
| ENSG000001 | 35.6606267 | 0.024154596 | 0.22860961 | 0.10565871 | 0.91585314 | 0.94327849 | RASSF6 | 166824 |
| ENSG000002 | 6.4265179 | -0.039173834 | 0.371645 | -0.1054066 | 0.91605318 | 0.94346056 | LINC01029 | 101927715 |
| ENSG000001 | 54.3833766 | -0.020871587 | 0.19831673 | -0.1052437 | 0.91618243 | 0.94349782 | CHRNB4 | 1143 |
| ENSG000001 | 10.7320643 | -0.034759833 | 0.33027543 | -0.105245 | 0.91618141 | 0.94349782 | ZACN | 353174 |
| ENSG000002 | 2.45558443 | 0.038807268 | 0.36862549 | 0.1052756 | 0.91615713 | 0.94349782 | NA | NA |
| ENSG000002 | 2.45173508 | 0.038554426 | 0.36615905 | 0.10529421 | 0.91614236 | 0.94349782 | NA | NA |
| ENSG000001 | 107.943677 | -0.0195132 | 0.1859125 | -0.1049591 | 0.9164083 | 0.94370646 | FCHSD2 | 9873 |
| ENSG000002 | 3.78158982 | -0.039702737 | 0.37851049 | -0.104892 | 0.91646148 | 0.94373725 | NA | NA |
| ENSG000002 | 7.86040236 | 0.037556222 | 0.35825595 | 0.1048307 | 0.91651015 | 0.94376341 | NA | NA |

| | | | | | | | | |
|---|---|---|---|---|---|---|---|---|
| ENSG000002 | 2.6700004 | 0.039363123 | 0.37565247 | 0.10478601 | 0.91654561 | 0.94377596 | NA | NA |
| ENSG000000 | 229.187566 | -0.014530816 | 0.13895789 | -0.1045699 | 0.91671708 | 0.94390459 | SDCCAG8 | 10806 |
| ENSG000002 | 3.2722913 | -0.039456108 | 0.37728012 | -0.1045804 | 0.91670876 | 0.94390459 | NA | NA |
| ENSG000002 | 6.35415864 | -0.037762503 | 0.36168898 | -0.104406 | 0.91684716 | 0.94401455 | NA | NA |
| ENSG000002 | 2.69789743 | 0.039012197 | 0.37406839 | 0.10429162 | 0.91693793 | 0.94408404 | NA | NA |
| ENSG000002 | 10.3120797 | -0.035419756 | 0.34017876 | -0.104121 | 0.91707332 | 0.94419946 | ZNF350-AS1 | 101669766 |
| ENSG000002 | 3.31699406 | -0.039214664 | 0.37728353 | -0.1039395 | 0.91721735 | 0.94429981 | NA | NA |
| ENSG000002 | 8.67304329 | -0.03559952 | 0.34245375 | -0.1039542 | 0.91720566 | 0.94429981 | NA | NA |
| ENSG000001 | 4.45692264 | 0.039026299 | 0.37597832 | 0.10379934 | 0.91732859 | 0.94439036 | FAM90A27P | 646508 |
| ENSG000002 | 3.75589724 | -0.039193466 | 0.37785681 | -0.1037257 | 0.91738702 | 0.94440623 | NA | NA |
| ENSG000002 | 6.03798007 | -0.038405428 | 0.37027553 | -0.1037212 | 0.91739059 | 0.94440623 | NA | NA |
| ENSG000002 | 4.49799814 | -0.039191444 | 0.37804398 | -0.103669 | 0.91743202 | 0.94442491 | NA | NA |
| ENSG000001 | 38.7785966 | -0.023963777 | 0.23182169 | -0.1033716 | 0.91766806 | 0.94458783 | NFIC | 4782 |
| ENSG000002 | 3.53185964 | -0.039145759 | 0.37848898 | -0.1034264 | 0.91762455 | 0.94458783 | NA | NA |
| ENSG000002 | 23.3851776 | 0.028012735 | 0.27104156 | 0.10335218 | 0.91768347 | 0.94458783 | NA | NA |
| ENSG000002 | 23.4690718 | 0.027601182 | 0.26695325 | 0.10339332 | 0.91765081 | 0.94458783 | LOC102723373 | 102723373 |
| ENSG000002 | 4.02899108 | -0.039041864 | 0.37852017 | -0.1031434 | 0.91784915 | 0.94472764 | NA | NA |
| ENSG000002 | 2.9108382 | 0.03879306 | 0.37618489 | 0.10312232 | 0.91786589 | 0.94472764 | RRM1-AS1 | 106478966 |
| ENSG000002 | 3.97824638 | -0.039018665 | 0.37851346 | -0.103084 | 0.91789634 | 0.944735 | NA | NA |
| ENSG000002 | 9.29881176 | 0.035009662 | 0.34070996 | 0.10275503 | 0.9181574 | 0.94497971 | PRR13P2 | 106480303 |
| ENSG000002 | 3.35874908 | -0.038810558 | 0.37826693 | -0.102601 | 0.91827966 | 0.94508156 | NA | NA |
| ENSG000002 | 2.6893312 | 0.038542427 | 0.37600091 | 0.10250621 | 0.91835489 | 0.94512413 | NA | NA |
| ENSG000002 | 6.30668286 | -0.037259488 | 0.36354217 | -0.1024901 | 0.91836764 | 0.94512413 | NA | NA |
| ENSG000002 | 4.44363323 | 0.03849356 | 0.37641634 | 0.10226325 | 0.91854772 | 0.94528547 | NA | NA |
| ENSG000002 | 3.3303361 | -0.038652883 | 0.37809975 | -0.1022293 | 0.91857465 | 0.94528919 | NA | NA |
| ENSG000002 | 31.2562682 | -0.025924516 | 0.25366498 | -0.1021998 | 0.91859807 | 0.9452893 | NA | NA |
| ENSG000001 | 151.087248 | 0.015993504 | 0.15710208 | 0.10180326 | 0.91891284 | 0.94558922 | ZNF600 | 162966 |
| ENSG000002 | 4.6749398 | 0.038310116 | 0.3767306 | 0.10169101 | 0.91900194 | 0.94565691 | NA | NA |
| ENSG000002 | 4.52462265 | 0.038174135 | 0.37597952 | 0.10153249 | 0.91912777 | 0.94573984 | FMR1-IT1 | 106478973 |

| | | | | | | | | |
|---|---|---|---|---|---|---|---|---|
| ENSG000002 | 3.60594147 | -0.038428126 | 0.37848765 | -0.1015307 | 0.91912917 | 0.94573984 | NA | NA |
| ENSG000001 | 24.0784254 | -0.027774349 | 0.27388659 | -0.1014082 | 0.91922642 | 0.9458159 | SMOC2 | 64094 |
| ENSG000002 | 10.4787121 | -0.034935375 | 0.344771 | -0.1013292 | 0.91928912 | 0.94585642 | LOC728752 | 728752 |
| ENSG000001 | 4.75681607 | -0.038315738 | 0.37831925 | -0.1012788 | 0.91932911 | 0.94586761 | C11orf96 | 387763 |
| ENSG000002 | 13.1148269 | -0.032294452 | 0.31893631 | -0.1012567 | 0.91934665 | 0.94586761 | NA | NA |
| ENSG000002 | 3.5728965 | -0.038312656 | 0.37854192 | -0.1012111 | 0.91938286 | 0.94588087 | KRTAP3-2 | 83897 |
| ENSG000002 | 2.9603049 | 0.037871884 | 0.3744165 | 0.10114908 | 0.91943212 | 0.94590755 | NA | NA |
| ENSG000002 | 5.77621497 | 0.037400422 | 0.37042095 | 0.10096735 | 0.91957638 | 0.94600819 | NA | NA |
| ENSG000002 | 3.37010808 | 0.037598794 | 0.37238671 | 0.10096707 | 0.9195766 | 0.94600819 | NA | NA |
| ENSG000001 | 8.59583634 | -0.035064022 | 0.34739511 | -0.1009341 | 0.91960275 | 0.94601109 | PHLDA2 | 7262 |
| ENSG000002 | 4.20071312 | 0.038134829 | 0.37851896 | 0.10074747 | 0.91975093 | 0.94613952 | NA | NA |
| ENSG000001 | 5.87519214 | 0.036981699 | 0.36740177 | 0.10065738 | 0.91982245 | 0.94618909 | RAMP1 | 10267 |
| ENSG000001 | 8.19261367 | -0.035958152 | 0.35789131 | -0.1004723 | 0.9199694 | 0.94628942 | LHX2 | 9355 |
| ENSG000002 | 13.379296 | -0.031582901 | 0.31442562 | -0.1004463 | 0.91998999 | 0.94628942 | ZNF564 | 163050 |
| ENSG000002 | 6.54171748 | 0.036622109 | 0.36448486 | 0.10047635 | 0.91996616 | 0.94628942 | NA | NA |
| ENSG000002 | 4.2284275 | -0.037925537 | 0.37768717 | -0.1004152 | 0.92001469 | 0.94629083 | NA | NA |
| ENSG000001 | 12.9638679 | 0.032087229 | 0.31987044 | 0.1003132 | 0.92009567 | 0.94635012 | RTBDN | 83546 |
| ENSG000001 | 6.11846991 | 0.036766312 | 0.36718355 | 0.10013061 | 0.92024064 | 0.9464272 | CD7 | 924 |
| ENSG000001 | 3.7915662 | -0.037913681 | 0.37845388 | -0.1001805 | 0.92020107 | 0.9464272 | OR5T1 | 390155 |
| ENSG000002 | 23.0375657 | -0.028410247 | 0.28368827 | -0.100146 | 0.92022841 | 0.9464272 | NA | NA |
| ENSG000002 | 3.53524011 | -0.037888625 | 0.37859337 | -0.1000774 | 0.92028291 | 0.94644667 | NA | NA |
| ENSG000002 | 5.75008937 | -0.037312179 | 0.37354956 | -0.0998855 | 0.92043524 | 0.94657933 | NA | NA |
| ENSG000002 | 7.02788625 | -0.036086743 | 0.3615898 | -0.0998002 | 0.92050293 | 0.94660093 | NA | NA |
| ENSG000002 | 20.8127099 | 0.027192445 | 0.27244379 | 0.09980937 | 0.92049567 | 0.94660093 | NA | NA |
| ENSG00000C | 72.0205311 | 0.019106211 | 0.19154477 | 0.09974802 | 0.92054438 | 0.94661954 | SLC4A11 | 83959 |
| ENSG00000C | 9.64299225 | -0.034293668 | 0.34480807 | -0.0994573 | 0.92077522 | 0.94679244 | CXCL2 | 2920 |
| ENSG000001 | 75.1805494 | -0.018203781 | 0.1830482 | -0.099448 | 0.92078256 | 0.94679244 | HES1 | 3280 |
| ENSG000002 | 2.437636 | 0.036853264 | 0.37039776 | 0.09949645 | 0.9207441 | 0.94679244 | NA | NA |
| ENSG000001 | 51.8066266 | 0.020355929 | 0.20493983 | 0.09932637 | 0.92087914 | 0.94686775 | SYCP2L | 221711 |

| | | | | | | | | |
|---|---|---|---|---|---|---|---|---|
| ENSG000001 | 6.10510916 | 0.036180937 | 0.36453388 | 0.0992526 | 0.92093771 | 0.94690395 | CCDC42B | 387885 |
| ENSG000002 | 12.0377792 | 0.032737304 | 0.33007357 | 0.09918184 | 0.92099389 | 0.94693771 | NA | NA |
| ENSG000002 | 12.7209675 | 0.031514061 | 0.31791284 | 0.09912799 | 0.92103665 | 0.94695766 | NA | NA |
| ENSG000001 | 27.6452401 | -0.025246548 | 0.25476946 | -0.0990957 | 0.92106232 | 0.94696004 | ST8SIA6 | 338596 |
| ENSG000002 | 12.8105563 | -0.031349561 | 0.3168533 | -0.0989403 | 0.92118567 | 0.94705206 | NA | NA |
| ENSG000002 | 3.77222799 | -0.037447225 | 0.37854497 | -0.0989241 | 0.92119853 | 0.94705206 | NA | NA |
| ENSG000002 | 2.24634922 | 0.035756943 | 0.36200012 | 0.09877605 | 0.92131608 | 0.94710088 | NA | NA |
| ENSG000002 | 2.69191498 | 0.037040376 | 0.3748773 | 0.09880667 | 0.92129178 | 0.94710088 | NA | NA |
| ENSG000002 | 6.08407505 | 0.036146913 | 0.36584157 | 0.09880483 | 0.92129324 | 0.94710088 | NA | NA |
| ENSG000001 | 5.85422172 | -0.036007409 | 0.36537186 | -0.09855 | 0.92149555 | 0.94720739 | FAM131C | 348487 |
| ENSG000002 | 2.42935051 | 0.036160151 | 0.36668892 | 0.09861261 | 0.92144586 | 0.94720739 | NA | NA |
| ENSG000002 | 6.84063707 | -0.035937395 | 0.36485229 | -0.0984985 | 0.92153648 | 0.94720739 | LINC00441 | 100862704 |
| ENSG000002 | 2.92137554 | 0.036734582 | 0.37278464 | 0.09854103 | 0.92150269 | 0.94720739 | NA | NA |
| ENSG000002 | 2.94695566 | 0.037085003 | 0.37646421 | 0.09850871 | 0.92152836 | 0.94720739 | NA | NA |
| ENSG000002 | 2.69250783 | 0.036967554 | 0.37566958 | 0.09840444 | 0.92161115 | 0.94724446 | NA | NA |
| ENSG000002 | 6.59857491 | 0.03617422 | 0.36764581 | 0.09839421 | 0.92161927 | 0.94724446 | NA | NA |
| ENSG000001 | 15.3915458 | -0.030217166 | 0.30786523 | -0.0981506 | 0.92181268 | 0.94740415 | C22orf15 | 150248 |
| ENSG000001 | 17.9536722 | -0.028007203 | 0.28570159 | -0.0980296 | 0.92190882 | 0.94740415 | SPHK1 | 8877 |
| ENSG000002 | 3.40738628 | -0.036776659 | 0.37518777 | -0.098022 | 0.92191482 | 0.94740415 | NA | NA |
| ENSG000002 | 5.18878722 | -0.036708 | 0.37435179 | -0.0980575 | 0.92188664 | 0.94740415 | NA | NA |
| ENSG000002 | 83.5573993 | -0.016919494 | 0.17254449 | -0.0980587 | 0.92188566 | 0.94740415 | NA | NA |
| ENSG000002 | 5.87860302 | 0.036347585 | 0.37073998 | 0.09804064 | 0.92190002 | 0.94740415 | NA | NA |
| ENSG000001 | 38.5806523 | 0.022793262 | 0.23278033 | 0.09791747 | 0.92199782 | 0.94745212 | OSCP1 | 127700 |
| ENSG000001 | 45.8808248 | 0.020872568 | 0.21364498 | 0.09769745 | 0.92217254 | 0.94745212 | ZC2HC1C | 79696 |
| ENSG000002 | 32.1279804 | 0.023931494 | 0.2447785 | 0.09776796 | 0.92211655 | 0.94745212 | DDX47 | 51202 |
| ENSG000002 | 3.28136607 | -0.036954611 | 0.37836588 | -0.097669 | 0.92219514 | 0.94745212 | NA | NA |
| ENSG000002 | 2.67884628 | 0.036621224 | 0.37456869 | 0.09776905 | 0.92211569 | 0.94745212 | NA | NA |
| ENSG000002 | 13.2508835 | 0.030884604 | 0.31597145 | 0.09774492 | 0.92213485 | 0.94745212 | NA | NA |
| ENSG000002 | 3.55641006 | -0.037047224 | 0.3785696 | -0.0978611 | 0.92204262 | 0.94745212 | NA | NA |

| | | | | | | | | |
|---|---|---|---|---|---|---|---|---|
| ENSG000002 | 8.69166192 | -0.034061659 | 0.34817219 | -0.0978299 | 0.92206735 | 0.94745212 | NA | NA |
| ENSG000002 | 2.68420308 | 0.036653506 | 0.37525476 | 0.09767633 | 0.92218932 | 0.94745212 | NA | NA |
| ENSG000002 | 2.88121914 | 0.036798243 | 0.37638007 | 0.09776884 | 0.92211585 | 0.94745212 | NA | NA |
| ENSG000002 | 7.28645804 | -0.035540499 | 0.36428365 | -0.0975627 | 0.92227954 | 0.94751482 | NA | NA |
| ENSG000001 | 10.9504219 | -0.032130425 | 0.32977436 | -0.0974315 | 0.9223837 | 0.947575 | ISM2 | 145501 |
| ENSG000002 | 6.98076536 | 0.035616806 | 0.36559882 | 0.09742046 | 0.9223925 | 0.947575 | NA | NA |
| ENSG000002 | 5.91339591 | -0.035704977 | 0.36657835 | -0.0974007 | 0.92240821 | 0.947575 | NA | NA |
| ENSG000002 | 4.07969133 | -0.036805641 | 0.37804224 | -0.0973585 | 0.92244167 | 0.94758536 | NA | NA |
| ENSG000002 | 15.1960688 | -0.030291582 | 0.31196735 | -0.0970986 | 0.92264812 | 0.94777343 | NA | NA |
| ENSG000002 | 2.51473881 | 0.035814227 | 0.36951047 | 0.09692344 | 0.92278719 | 0.94789228 | LOC105378328 | 105378328 |
| ENSG000002 | 158.086939 | -0.014373667 | 0.14841323 | -0.096849 | 0.92284634 | 0.94792903 | NA | NA |
| ENSG000002 | 3.05227854 | -0.036473927 | 0.37696665 | -0.0967564 | 0.92291987 | 0.94798054 | NA | NA |
| ENSG000002 | 15.5019847 | 0.029121372 | 0.30114505 | 0.09670215 | 0.92296294 | 0.94800076 | DDX50P2 | 100421328 |
| ENSG000002 | 104.085527 | -0.021545982 | 0.22295459 | -0.0966384 | 0.92301354 | 0.94802872 | NA | NA |
| ENSG000001 | 22.9803219 | -0.026611397 | 0.27557218 | -0.0965678 | 0.92306964 | 0.94806233 | RASL11A | 387496 |
| ENSG00000C | 79.0283448 | 0.018977166 | 0.19663433 | 0.09650994 | 0.92311558 | 0.94807101 | ACSM3 | 6296 |
| ENSG000002 | 14.961069 | 0.02980896 | 0.30890669 | 0.09649827 | 0.92312485 | 0.94807101 | NA | NA |
| ENSG000002 | 2.66880618 | 0.036220489 | 0.37565451 | 0.09641968 | 0.92318726 | 0.94810816 | NA | NA |
| ENSG000002 | 2.90826184 | 0.036249133 | 0.37616725 | 0.0963644 | 0.92323116 | 0.94810816 | NA | NA |
| ENSG000002 | 8.01841914 | -0.034553425 | 0.35847907 | -0.096389 | 0.92321166 | 0.94810816 | NA | NA |
| ENSG000001 | 3.98049844 | -0.0364457 | 0.3784466 | -0.0963034 | 0.9232796 | 0.94813389 | FOXS1 | 2307 |
| ENSG000002 | 4.24433376 | 0.036290038 | 0.37807875 | 0.09598539 | 0.92353217 | 0.94836925 | ZNF197-AS1 | 100874094 |
| ENSG000002 | 3.30008396 | -0.036293567 | 0.37836476 | -0.0959222 | 0.92358239 | 0.9483968 | NA | NA |
| ENSG000002 | 8.91227061 | -0.032755308 | 0.34258735 | -0.0956116 | 0.92382909 | 0.9486261 | NA | NA |
| ENSG000002 | 19.3892348 | 0.026708552 | 0.27963353 | 0.09551269 | 0.92390761 | 0.94868271 | NA | NA |
| ENSG00000C | 5.36986483 | -0.035818576 | 0.37527816 | -0.0954454 | 0.92396105 | 0.94871356 | DRD4 | 1815 |
| ENSG000002 | 597.429206 | 0.01318979 | 0.13865202 | 0.09512872 | 0.92421258 | 0.9489478 | NA | NA |
| ENSG000002 | 3.08335276 | -0.035884129 | 0.37763816 | -0.0950225 | 0.92429694 | 0.94901039 | NA | NA |
| ENSG000001 | 3.75339332 | -0.035915954 | 0.37858951 | -0.0948678 | 0.92441983 | 0.94911253 | NA | NA |

| | | | | | | | | |
|---|---|---|---|---|---|---|---|---|
| ENSG000002 | 2.9425529 | 0.035435129 | 0.37386285 | 0.09478109 | 0.9244887 | 0.94915921 | NA | NA |
| ENSG000002 | 6.58112603 | -0.034375234 | 0.36397957 | -0.0944428 | 0.92475745 | 0.94941109 | NA | NA |
| ENSG000001 | 20.3625372 | -0.026314546 | 0.27908792 | -0.0942877 | 0.92488065 | 0.94951354 | DRD3 | 1814 |
| ENSG000001 | 8.94086451 | -0.032751872 | 0.34800331 | -0.0941137 | 0.92501885 | 0.9496225 | TAS2R5 | 54429 |
| ENSG000002 | 3.26701937 | -0.035534394 | 0.37764344 | -0.0940951 | 0.92503362 | 0.9496225 | NA | NA |
| ENSG000001 | 508.072101 | 0.013421578 | 0.14272203 | 0.09403999 | 0.92507739 | 0.94964339 | BCL11A | 53335 |
| ENSG000001 | 50.6126726 | -0.020231173 | 0.2153256 | -0.0939562 | 0.92514396 | 0.94968769 | ZNF169 | 169841 |
| ENSG000002 | 2.48200649 | 0.034933823 | 0.37192873 | 0.09392612 | 0.92516784 | 0.94968816 | NA | NA |
| ENSG000001 | 19.9755418 | 0.026787407 | 0.2860961 | 0.0936308 | 0.92540244 | 0.9498226 | MCHR1 | 2847 |
| ENSG000002 | 3.60406266 | -0.035452363 | 0.37849985 | -0.0936655 | 0.92537491 | 0.9498226 | NA | NA |
| ENSG000002 | 2.68941553 | 0.035170162 | 0.37530759 | 0.09371023 | 0.92533934 | 0.9498226 | NA | NA |
| ENSG000002 | 7.71210516 | -0.033826719 | 0.36145703 | -0.0935843 | 0.92543935 | 0.9498226 | NA | NA |
| ENSG000002 | 46.7748727 | 0.019999952 | 0.21366609 | 0.09360377 | 0.92542391 | 0.9498226 | NA | NA |
| ENSG000002 | 4.52620145 | 0.0352455 | 0.37652799 | 0.09360659 | 0.92542168 | 0.9498226 | NA | NA |
| ENSG000002 | 11.2765367 | -0.030477314 | 0.32586728 | -0.0935268 | 0.92548508 | 0.9498455 | NA | NA |
| ENSG000002 | 2.90922739 | 0.035082424 | 0.37536031 | 0.09346333 | 0.92553548 | 0.94986702 | NA | NA |
| ENSG000002 | 10.2398064 | 0.03147227 | 0.33681288 | 0.09344141 | 0.92555289 | 0.94986702 | NA | NA |
| ENSG000002 | 70.8639325 | -0.019163986 | 0.20572569 | -0.0931531 | 0.92578193 | 0.95007803 | NA | NA |
| ENSG000002 | 6.34576372 | 0.033949973 | 0.36499239 | 0.09301556 | 0.9258912 | 0.95016611 | NA | NA |
| ENSG000002 | 4.00626259 | -0.035154665 | 0.3785718 | -0.0928613 | 0.92601376 | 0.95026784 | NA | NA |
| ENSG000002 | 3.60440234 | -0.03511224 | 0.3783326 | -0.0928079 | 0.92605621 | 0.95028735 | NA | NA |
| ENSG000002 | 7.76185494 | 0.03350081 | 0.36143477 | 0.0926884 | 0.92615111 | 0.9503566 | NA | NA |
| ENSG000002 | 2.88613819 | 0.03485904 | 0.37626708 | 0.09264441 | 0.92618606 | 0.9503566 | NA | NA |
| ENSG000002 | 2.83362866 | -0.034790378 | 0.37556645 | -0.0926344 | 0.926194 | 0.9503566 | NA | NA |
| ENSG000002 | 2.44657416 | 0.034054292 | 0.367818 | 0.09258463 | 0.92623356 | 0.95037314 | NA | NA |
| ENSG000002 | 3.53910209 | -0.034947334 | 0.37797816 | -0.0924586 | 0.92633368 | 0.95045182 | NA | NA |
| ENSG000002 | 2.67470376 | 0.034689378 | 0.37549428 | 0.09238324 | 0.92639356 | 0.9504706 | NA | NA |
| ENSG000002 | 17.8271815 | -0.026784865 | 0.28995303 | -0.0923766 | 0.92639886 | 0.9504706 | NA | NA |
| ENSG000002 | 3.07929456 | -0.034628048 | 0.37564577 | -0.0921827 | 0.92655287 | 0.95060456 | MIR6723 | 102465432 |

| | | | | | | | | |
|---|---|---|---|---|---|---|---|---|
| ENSG000001 | 92.3737688 | -0.017744793 | 0.19288221 | -0.0919981 | 0.92669956 | 0.95073101 | ABTB2 | 25841 |
| ENSG000002 | 6.10270749 | -0.034085702 | 0.37074738 | -0.0919378 | 0.92674745 | 0.95075609 | NA | NA |
| ENSG000001 | 3.5266546 | -0.034724474 | 0.3782285 | -0.0918082 | 0.92685044 | 0.95083769 | BIRC8 | 112401 |
| ENSG000002 | 2.84485024 | -0.034177686 | 0.37243041 | -0.0917693 | 0.92688132 | 0.95084531 | NA | NA |
| ENSG000002 | 11.7880997 | 0.030298573 | 0.33104789 | 0.09152323 | 0.92707685 | 0.95102184 | NA | NA |
| ENSG000002 | 7.99873601 | -0.03229053 | 0.35305504 | -0.0914603 | 0.92712683 | 0.95104905 | NA | NA |
| ENSG000002 | 6.12119032 | 0.033271149 | 0.36400886 | 0.09140203 | 0.92717315 | 0.95107251 | NA | NA |
| ENSG000002 | 9.98145693 | 0.030524244 | 0.33434224 | 0.0912964 | 0.92725708 | 0.95113454 | NA | NA |
| ENSG00000C | 157.661077 | -0.013690906 | 0.1502556 | -0.0911174 | 0.92739927 | 0.95125634 | DYRK4 | 8798 |
| ENSG000002 | 2.93379079 | 0.034134183 | 0.3751785 | 0.09098118 | 0.92750754 | 0.95134333 | RN7SL530P | 106481067 |
| ENSG000002 | 6.09468895 | 0.033212906 | 0.36558292 | 0.09084917 | 0.92761244 | 0.95142685 | NA | NA |
| ENSG000002 | 3.0748714 | -0.034291929 | 0.37773145 | -0.0907839 | 0.92766432 | 0.95145599 | NA | NA |
| ENSG000001 | 25.8995927 | -0.024811682 | 0.27362558 | -0.0906775 | 0.92774885 | 0.95151863 | NA | NA |
| ENSG000002 | 13.2922005 | -0.02797419 | 0.30914004 | -0.0904903 | 0.92789756 | 0.95164709 | NA | NA |
| ENSG000001 | 10.2509777 | -0.029982887 | 0.33216538 | -0.0902649 | 0.92807668 | 0.95180672 | GPR182 | 11318 |
| ENSG000002 | 17.0215239 | -0.026279769 | 0.29226927 | -0.0899163 | 0.92835373 | 0.95206677 | NA | NA |
| ENSG000001 | 4.10413591 | 0.033935851 | 0.37837636 | 0.08968808 | 0.92853509 | 0.95220459 | FGF6 | 2251 |
| ENSG000002 | 9.17119347 | -0.031097255 | 0.34664927 | -0.0897081 | 0.92851916 | 0.95220459 | NA | NA |
| ENSG000002 | 9.51249421 | 0.030433802 | 0.33959012 | 0.08961922 | 0.92858981 | 0.95223663 | NA | NA |
| ENSG000001 | 4.49022745 | 0.033598245 | 0.37560256 | 0.08945159 | 0.92872303 | 0.952301 | TMEM61 | 199964 |
| ENSG000001 | 5.90036846 | 0.033240576 | 0.3715727 | 0.08945914 | 0.92871702 | 0.952301 | ALPPL2 | 251 |
| ENSG000002 | 3.77474784 | -0.033866572 | 0.3784968 | -0.0894765 | 0.92870322 | 0.952301 | NA | NA |
| ENSG000002 | 3.0825833 | -0.033709601 | 0.37790737 | -0.0892007 | 0.92892241 | 0.95244622 | NA | NA |
| ENSG000002 | 2.48055188 | 0.032569555 | 0.36519216 | 0.0891847 | 0.92893512 | 0.95244622 | NA | NA |
| ENSG000002 | 3.06749939 | -0.033725904 | 0.37808276 | -0.0892024 | 0.92892103 | 0.95244622 | NA | NA |
| ENSG000001 | 300.112159 | -0.01252816 | 0.14065109 | -0.0890726 | 0.92902421 | 0.95250744 | GATAD2B | 57459 |
| ENSG000002 | 19.0419432 | -0.025108772 | 0.28196117 | -0.0890505 | 0.92904181 | 0.95250744 | NA | NA |
| ENSG000002 | 3.03701858 | -0.033483519 | 0.37688713 | -0.0888423 | 0.92920724 | 0.95265297 | NA | NA |
| ENSG00000C | 186.167011 | 0.011890222 | 0.13403918 | 0.08870706 | 0.92931472 | 0.95271498 | MAST4 | 375449 |

| | | | | | | | | |
|---|---|---|---|---|---|---|---|---|
| ENSG000002 | 3.78619068 | -0.033564991 | 0.37835203 | -0.0887137 | 0.92930948 | 0.95271498 | NA | NA |
| ENSG000001 | 6.04847876 | 0.032471637 | 0.3665825 | 0.08857934 | 0.92941622 | 0.95279079 | EBI3 | 10148 |
| ENSG000002 | 13.6172772 | -0.028744282 | 0.32459282 | -0.0885549 | 0.92943566 | 0.95279079 | NA | NA |
| ENSG000002 | 4.32320294 | 0.033324976 | 0.37664966 | 0.08847738 | 0.92949726 | 0.95282986 | NA | NA |
| ENSG000001 | 68.6185462 | -0.017771797 | 0.20118421 | -0.0883359 | 0.92960967 | 0.95289691 | NCAN | 1463 |
| ENSG000002 | 2.46265587 | 0.032810976 | 0.37137304 | 0.08835045 | 0.92959814 | 0.95289691 | NA | NA |
| ENSG000002 | 4.70866937 | 0.033150764 | 0.37558282 | 0.08826486 | 0.92966617 | 0.95290665 | RNU6-652P | 106481388 |
| ENSG000002 | 3.75886745 | -0.033420681 | 0.37859293 | -0.088276 | 0.92965729 | 0.95290665 | NA | NA |
| ENSG000002 | 3.09675372 | -0.033344449 | 0.37801322 | -0.0882097 | 0.92970997 | 0.95292746 | NA | NA |
| ENSG000002 | 3.0922066 | -0.033136048 | 0.37629396 | -0.0880589 | 0.92982983 | 0.95302622 | NA | NA |
| ENSG000001 | 4.47747937 | 0.033109252 | 0.3762394 | 0.08800049 | 0.92987629 | 0.95304975 | CDK5R2 | 8941 |
| ENSG000002 | 8.18291095 | -0.031513486 | 0.35872365 | -0.0878489 | 0.92999675 | 0.95310602 | NA | NA |
| ENSG000002 | 4.52813234 | 0.033026431 | 0.3759724 | 0.0878427 | 0.9300017 | 0.95310602 | NA | NA |
| ENSG000002 | 3.07243143 | -0.033081782 | 0.37641629 | -0.0878862 | 0.92996716 | 0.95310602 | NA | NA |
| ENSG000001 | 11.1341933 | 0.028910155 | 0.32945079 | 0.08775257 | 0.93007333 | 0.95315534 | B3GNT3 | 10331 |
| ENSG000002 | 12.5471257 | -0.028139116 | 0.32091685 | -0.0876835 | 0.93012823 | 0.95318751 | NA | NA |
| ENSG000002 | 3.24988801 | -0.033061866 | 0.37810596 | -0.0874407 | 0.93032119 | 0.95336116 | NA | NA |
| ENSG000002 | 17.8092666 | -0.026283882 | 0.30115753 | -0.0872762 | 0.93045198 | 0.9534711 | AGER | 177 |
| ENSG000002 | 2.72188671 | 0.032383729 | 0.37214951 | 0.08701806 | 0.93065716 | 0.95365725 | LOC105369507 | 105369507 |
| ENSG000002 | 2.64041325 | 0.032321641 | 0.37163382 | 0.08697174 | 0.93069397 | 0.95367088 | NA | NA |
| ENSG000002 | 5.00932664 | 0.032501944 | 0.37418146 | 0.08686145 | 0.93078164 | 0.95373661 | NA | NA |
| ENSG000002 | 6.07417031 | -0.031691717 | 0.36536288 | -0.0867404 | 0.93087787 | 0.95381111 | NA | NA |
| ENSG000002 | 2.68249998 | 0.032475727 | 0.37526001 | 0.08654193 | 0.93103562 | 0.95394865 | HPVC1 | 3262 |
| ENSG000001 | 104.721324 | 0.014423574 | 0.16687937 | 0.08643114 | 0.93112369 | 0.95396422 | NMI | 9111 |
| ENSG000002 | 6.09247051 | -0.031782182 | 0.36795649 | -0.0863748 | 0.93116845 | 0.95396422 | NA | NA |
| ENSG000002 | 4.74504777 | 0.032477703 | 0.37576464 | 0.08643097 | 0.93112383 | 0.95396422 | NA | NA |
| ENSG000002 | 2.90778443 | 0.031928027 | 0.36951917 | 0.08640425 | 0.93114507 | 0.95396422 | RNU2-34P | 106481960 |
| ENSG000002 | 31.3368035 | 0.020744836 | 0.23984509 | 0.08649264 | 0.9310748 | 0.95396422 | NA | NA |
| ENSG000001 | 4.70907411 | 0.03263088 | 0.37796885 | 0.08633219 | 0.93120235 | 0.95397485 | TFF3 | 7033 |

| | | | | | | | | | |
|---|---|---|---|---|---|---|---|---|---|
| ENSG000001 | 6.32478169 | 0.03124278 | 0.36248079 | 0.08619155 | 0.93131415 | 0.95404118 | OR52N2 | | 390077 |
| ENSG000001 | 45.0448122 | 0.019224432 | 0.22301897 | 0.08620088 | 0.93130673 | 0.95404118 | ANKRD45 | | 339416 |
| ENSG000002 | 3.29859689 | -0.032512032 | 0.3778438 | -0.0860462 | 0.93142967 | 0.95413542 | NA | NA | |
| ENSG000002 | 3.49274898 | -0.032471242 | 0.37858708 | -0.0857695 | 0.93164962 | 0.9542643 | NA | NA | |
| ENSG000002 | 3.07761162 | -0.032352519 | 0.37713099 | -0.0857859 | 0.93163662 | 0.9542643 | NA | NA | |
| ENSG000002 | 4.33101898 | 0.032414978 | 0.37772835 | 0.08581558 | 0.93161302 | 0.9542643 | CTAGE8 | | 100142659 |
| ENSG000002 | 2.83686532 | -0.032169083 | 0.37497165 | -0.0857907 | 0.93163279 | 0.9542643 | NA | NA | |
| ENSG000002 | 14.3807064 | -0.02637025 | 0.30955637 | -0.0851872 | 0.93211254 | 0.95471435 | NA | NA | |
| ENSG00000C | 5.19252534 | -0.031808206 | 0.37432879 | -0.084974 | 0.93228208 | 0.95474566 | STON1-GTF2A1 | | 286749 |
| ENSG000001 | 73.8772719 | 0.015959759 | 0.1876405 | 0.08505498 | 0.93221768 | 0.95474566 | MOB3C | | 148932 |
| ENSG000002 | 8.48059584 | -0.029746693 | 0.35013403 | -0.084958 | 0.93229477 | 0.95474566 | NA | NA | |
| ENSG000002 | 13.6045956 | -0.026965235 | 0.31695032 | -0.0850772 | 0.93220004 | 0.95474566 | NA | NA | |
| ENSG000002 | 3.51566135 | -0.032217721 | 0.37858918 | -0.0850994 | 0.93218235 | 0.95474566 | NA | NA | |
| ENSG000002 | 8.82391148 | 0.0301571 | 0.35501188 | 0.08494673 | 0.93230374 | 0.95474566 | NA | NA | |
| ENSG000002 | 12.0659507 | 0.02713323 | 0.31943442 | 0.08494147 | 0.93230792 | 0.95474566 | NA | NA | |
| ENSG000002 | 2.93699968 | 0.032010084 | 0.37741468 | 0.08481409 | 0.93240919 | 0.95482525 | RN7SL50P | | 106479243 |
| ENSG00000C | 22.1036345 | -0.022626227 | 0.26771009 | -0.0845176 | 0.93264487 | 0.95499425 | TTC22 | | 55001 |
| ENSG000002 | 3.27801338 | -0.031907402 | 0.37751517 | -0.0845195 | 0.93264338 | 0.95499425 | NA | NA | |
| ENSG000002 | 9.55385911 | 0.028940745 | 0.34221345 | 0.08456928 | 0.93260382 | 0.95499425 | GTSE1-AS1 | | 150384 |
| ENSG000001 | 3.79304117 | -0.03195188 | 0.37843184 | -0.0844323 | 0.93271271 | 0.95501548 | THRSP | | 7069 |
| ENSG000002 | 32.1997664 | -0.021119241 | 0.25011622 | -0.0844377 | 0.93270842 | 0.95501548 | NA | NA | |
| ENSG000001 | 51.4160355 | 0.01899494 | 0.225377 | 0.08428074 | 0.93283323 | 0.95511476 | MEGF6 | | 1953 |
| ENSG000001 | 29.4511452 | 0.020918977 | 0.24831339 | 0.08424426 | 0.93286223 | 0.95512034 | DNAAF3 | | 352909 |
| ENSG000002 | 3.08021501 | -0.031713728 | 0.37667352 | -0.0841942 | 0.93290203 | 0.95513697 | MIR4753 | | 100616224 |
| ENSG000001 | 99.8239546 | 0.014650082 | 0.17418501 | 0.08410644 | 0.9329718 | 0.95518429 | TMTC2 | | 160335 |
| ENSG000002 | 3.58761622 | -0.031610355 | 0.37712235 | -0.0838199 | 0.93319963 | 0.95539343 | NA | NA | |
| ENSG000002 | 4.74858891 | 0.031467907 | 0.37602766 | 0.08368509 | 0.93330681 | 0.95547904 | NA | NA | |
| ENSG000002 | 4.75652949 | 0.031321291 | 0.37444385 | 0.0836475 | 0.9333367 | 0.95548551 | NA | NA | |
| ENSG000002 | 2.70355315 | 0.031353269 | 0.37506614 | 0.08359398 | 0.93337926 | 0.95550495 | NA | NA | |

| | | | | | | | | |
|---|---|---|---|---|---|---|---|---|
| ENSG000002 | 4.28223225 | -0.031604078 | 0.37853032 | -0.0834915 | 0.93346071 | 0.95556421 | LOC105376557 | 105376557 |
| ENSG000001 | 3.81539295 | -0.031529548 | 0.37859337 | -0.0832808 | 0.93362829 | 0.95566339 | MESP1 | 55897 |
| ENSG000001 | 19.7447444 | 0.024979478 | 0.29987183 | 0.08330051 | 0.93361259 | 0.95566339 | UFSP1 | 402682 |
| ENSG000002 | 8.6418809 | 0.029009066 | 0.34819134 | 0.08331358 | 0.9336022 | 0.95566339 | NA | NA |
| ENSG000002 | 6.58078908 | -0.030512037 | 0.36652915 | -0.0832459 | 0.93365604 | 0.95566767 | NA | NA |
| ENSG000002 | 70.3676993 | -0.015762292 | 0.18957186 | -0.0831468 | 0.93373482 | 0.95572418 | NA | NA |
| ENSG000002 | 17.4494254 | 0.024232913 | 0.29177532 | 0.08305333 | 0.93380913 | 0.955752 | FXYD7 | 53822 |
| ENSG000002 | 5.65068902 | 0.030834855 | 0.37116172 | 0.0830766 | 0.93379063 | 0.955752 | NA | NA |
| ENSG000002 | 3.34118608 | -0.031354768 | 0.37785796 | -0.0829803 | 0.9338672 | 0.95578731 | NA | NA |
| ENSG000002 | 3.10183012 | -0.031005048 | 0.37481585 | -0.0827208 | 0.93407359 | 0.95591196 | NA | NA |
| ENSG000002 | 3.10654588 | -0.031205624 | 0.37729624 | -0.0827085 | 0.93408329 | 0.95591196 | NA | NA |
| ENSG000002 | 3.5332249 | -0.031307476 | 0.37841596 | -0.082733 | 0.93406387 | 0.95591196 | NA | NA |
| ENSG000002 | 3.5486218 | -0.031335834 | 0.37857865 | -0.0827723 | 0.93403258 | 0.95591196 | NA | NA |
| ENSG000002 | 3.30925101 | -0.031209101 | 0.37816658 | -0.0825274 | 0.93422734 | 0.95598523 | NA | NA |
| ENSG000002 | 3.08568859 | -0.031182292 | 0.3779045 | -0.0825137 | 0.93423824 | 0.95598523 | NA | NA |
| ENSG000002 | 2.95858199 | 0.030906926 | 0.37442807 | 0.08254436 | 0.93421384 | 0.95598523 | NA | NA |
| ENSG000002 | 7.90318499 | 0.029056876 | 0.35220493 | 0.08249991 | 0.93424919 | 0.95598523 | NA | NA |
| ENSG000001 | 4.26664722 | -0.031211625 | 0.37849405 | -0.0824627 | 0.93427881 | 0.95599142 | NOG | 9241 |
| ENSG000002 | 2.67196354 | 0.030691937 | 0.37252696 | 0.0823885 | 0.93433778 | 0.95602764 | NA | NA |
| ENSG000002 | 3.72348719 | -0.031172302 | 0.37856466 | -0.0823434 | 0.93437364 | 0.95604021 | NA | NA |
| ENSG000002 | 3.31531222 | -0.031139247 | 0.37854846 | -0.0822596 | 0.93444028 | 0.95608426 | NA | NA |
| ENSG000002 | 6.37863134 | 0.0297715 | 0.36215892 | 0.08220562 | 0.9344832 | 0.95610406 | NA | NA |
| ENSG000002 | 3.14635908 | 0.030979501 | 0.37714253 | 0.08214269 | 0.93453325 | 0.95610702 | NA | NA |
| ENSG000002 | 9.34599702 | 0.028828846 | 0.35085545 | 0.0821673 | 0.93451368 | 0.95610702 | PRAF2 | 11230 |
| ENSG000002 | 2.92372953 | 0.030545051 | 0.3734399 | 0.08179375 | 0.93481072 | 0.95636677 | LOC105371660 | 105371660 |
| ENSG000001 | 14.4471287 | 0.025665465 | 0.31462433 | 0.08157495 | 0.93498472 | 0.95650789 | TTPA | 7274 |
| ENSG000002 | 2.95385084 | 0.030728708 | 0.37675749 | 0.08156098 | 0.93499583 | 0.95650789 | NA | NA |
| ENSG000002 | 6.638306 | 0.029719983 | 0.36471032 | 0.08148928 | 0.93505285 | 0.95654208 | NA | NA |
| ENSG000002 | 2.91413903 | 0.030562073 | 0.37521859 | 0.08145138 | 0.93508299 | 0.95654878 | NA | NA |

| | | | | | | | | |
|---|---|---|---|---|---|---|---|---|
| ENSG000001 | 91.4030696 | -0.014333839 | 0.17622336 | -0.081339 | 0.93517233 | 0.95655104 | SIL1 | 64374 |
| ENSG000001 | 2.91008117 | 0.030615035 | 0.3763956 | 0.08133739 | 0.93517364 | 0.95655104 | TBC1D3P2 | 440452 |
| ENSG000002 | 4.07368735 | 0.03078083 | 0.37845986 | 0.08133182 | 0.93517807 | 0.95655104 | LOC101927482 | 101927482 |
| ENSG000002 | 3.18690039 | 0.030584773 | 0.37605791 | 0.08132995 | 0.93517955 | 0.95655104 | NA | NA |
| ENSG000001 | 123.367469 | 0.013365098 | 0.16461731 | 0.08118889 | 0.93529173 | 0.95664165 | PPFIBP2 | 8495 |
| ENSG000002 | 2.88996054 | 0.030542983 | 0.37648615 | 0.08112645 | 0.93534139 | 0.95666832 | NA | NA |
| ENSG000002 | 10.4671193 | 0.026840538 | 0.33150298 | 0.08096621 | 0.93546883 | 0.95677453 | NA | NA |
| ENSG000002 | 11.3936857 | -0.026958167 | 0.33350854 | -0.080832 | 0.93557555 | 0.95685954 | NA | NA |
| ENSG000002 | 6.30199296 | 0.029367057 | 0.36349662 | 0.08079045 | 0.9356086 | 0.95686921 | NA | NA |
| ENSG00000C | 154.985955 | 0.011838244 | 0.14662323 | 0.08073921 | 0.93564935 | 0.95688676 | NRCAM | 4897 |
| ENSG000002 | 3.10804754 | -0.030180705 | 0.37582254 | -0.0803057 | 0.9359941 | 0.95721519 | NA | NA |
| ENSG000002 | 3.29106923 | -0.030239225 | 0.37750915 | -0.080102 | 0.93615616 | 0.95733264 | NA | NA |
| ENSG000002 | 16.2173804 | 0.023954647 | 0.29894457 | 0.08013073 | 0.93613328 | 0.95733264 | LRRC2-AS1 | 83598 |
| ENSG000002 | 3.0851478 | -0.030242725 | 0.37773048 | -0.0800643 | 0.93618612 | 0.95733913 | NA | NA |
| ENSG000002 | 3.10922195 | -0.030100449 | 0.37638913 | -0.0799716 | 0.93625983 | 0.95734208 | NA | NA |
| ENSG000002 | 10.0577799 | -0.026991905 | 0.33734537 | -0.0800127 | 0.93622718 | 0.95734208 | TUBB8 | 347688 |
| ENSG000002 | 16.5160748 | -0.023343268 | 0.29183684 | -0.0799874 | 0.93624728 | 0.95734208 | NA | NA |
| ENSG000002 | 8.43465397 | -0.028045362 | 0.35082528 | -0.0799411 | 0.93628409 | 0.95734275 | LOC100996246 | 100996246 |
| ENSG000002 | 13.8122396 | 0.025392061 | 0.31794156 | 0.07986393 | 0.93634548 | 0.95738138 | FLJ42393 | 401105 |
| ENSG000002 | 12.598172 | -0.025699156 | 0.3221766 | -0.0797673 | 0.93642233 | 0.95743582 | RPEL1 | 729020 |
| ENSG000001 | 27.1945582 | 0.019930691 | 0.25024904 | 0.07964342 | 0.93652086 | 0.95751241 | PARP10 | 84875 |
| ENSG00000C | 8.53044633 | -0.028032662 | 0.35283411 | -0.07945 | 0.93667472 | 0.95762936 | SLC7A9 | 11136 |
| ENSG000002 | 6.09494349 | -0.029025444 | 0.36537466 | -0.0794402 | 0.93668248 | 0.95762936 | MIR1254-1 | 100302273 |
| ENSG000002 | 11.4728364 | 0.027351255 | 0.3447924 | 0.07932673 | 0.93677274 | 0.95769751 | NA | NA |
| ENSG000001 | 15.6652806 | -0.024045579 | 0.30355962 | -0.079212 | 0.93686396 | 0.95776661 | MSC | 9242 |
| ENSG000002 | 10.2654415 | -0.026620328 | 0.33620976 | -0.0791777 | 0.93689125 | 0.95777037 | LOC101927905 | 101927905 |
| ENSG000001 | 6.35250156 | 0.028576945 | 0.36341227 | 0.07863506 | 0.9373229 | 0.95813205 | NA | NA |
| ENSG000001 | 64.9546684 | 0.015940776 | 0.20275255 | 0.07862183 | 0.93733342 | 0.95813205 | SULT1C2 | 6819 |
| ENSG000002 | 7.21246128 | -0.028574025 | 0.36347194 | -0.0786141 | 0.93733956 | 0.95813205 | NA | NA |

| | | | | | | | | | |
|---|---|---|---|---|---|---|---|---|---|
| ENSG000002 | 4.69408345 | -0.029740192 | 0.37807069 | -0.078663 | 0.93730064 | 0.95813205 | NA | NA | |
| ENSG000001 | 11.2542324 | -0.025941421 | 0.33071405 | -0.0784406 | 0.93747755 | 0.95824894 | IRX4 | | 50805 |
| ENSG000001 | 33.7173793 | -0.018903308 | 0.24180306 | -0.0781765 | 0.93768769 | 0.95840288 | LHX6 | | 26468 |
| ENSG000002 | 3.05201109 | -0.029507931 | 0.37752187 | -0.0781622 | 0.93769905 | 0.95840288 | NA | NA | |
| ENSG000002 | 8.47722255 | -0.027434292 | 0.35095309 | -0.0781708 | 0.93769217 | 0.95840288 | NA | NA | |
| ENSG000002 | 29.9889385 | 0.019989838 | 0.25638441 | 0.07796823 | 0.93785333 | 0.9585364 | NA | NA | |
| ENSG000002 | 4.55062679 | 0.029351226 | 0.37700021 | 0.07785467 | 0.93794366 | 0.95860457 | SNORA2A | | 677793 |
| ENSG000001 | 36.5312276 | -0.018835244 | 0.24205999 | -0.0778123 | 0.93797736 | 0.95861485 | KRTCAP2 | | 200185 |
| ENSG000002 | 25.9575967 | 0.020265027 | 0.26063747 | 0.07775178 | 0.9380255 | 0.95863989 | NA | NA | |
| ENSG000002 | 3.01843797 | -0.028946525 | 0.37259552 | -0.0776889 | 0.93807555 | 0.95866687 | NA | NA | |
| ENSG000002 | 6.85334784 | 0.028219917 | 0.36351938 | 0.07762975 | 0.93812258 | 0.95867588 | NA | NA | |
| ENSG000002 | 4.75290736 | 0.029203663 | 0.37624691 | 0.07761835 | 0.93813165 | 0.95867588 | NA | NA | |
| ENSG000002 | 10.4625237 | -0.026189208 | 0.33852839 | -0.0773619 | 0.93833563 | 0.95886017 | NA | NA | |
| ENSG000002 | 3.50367031 | -0.029238217 | 0.37858238 | -0.0772308 | 0.93843995 | 0.9589426 | NA | NA | |
| ENSG000002 | 2.80783966 | -0.029019499 | 0.37608553 | -0.077162 | 0.9384947 | 0.95897438 | NA | NA | |
| ENSG000001 | 6.13770901 | 0.028347574 | 0.36780233 | 0.07707285 | 0.93856559 | 0.9589985 | TREH | | 11181 |
| ENSG000002 | 3.23707522 | -0.029102876 | 0.37749501 | -0.0770947 | 0.93854818 | 0.9589985 | NA | NA | |
| ENSG000001 | 3.96989031 | -0.02910931 | 0.37843326 | -0.0769206 | 0.93868672 | 0.95907393 | A1BG | | 1 |
| ENSG000001 | 718.369073 | -0.009809431 | 0.12749823 | -0.0769378 | 0.93867304 | 0.95907393 | IRF2BP2 | | 359948 |
| ENSG000001 | 14.6404705 | 0.023488463 | 0.30649125 | 0.07663665 | 0.9389126 | 0.95925638 | KRT24 | | 192666 |
| ENSG000002 | 2.91847025 | 0.028924175 | 0.3774122 | 0.07663816 | 0.9389114 | 0.95925638 | MIR3174 | | 100422841 |
| ENSG000002 | 18.0156468 | 0.022387655 | 0.29245737 | 0.07655015 | 0.93898142 | 0.95930252 | NA | NA | |
| ENSG000002 | 2.89375118 | 0.028617419 | 0.37482579 | 0.07634859 | 0.93914177 | 0.95942244 | NA | NA | |
| ENSG000002 | 4.95830098 | 0.028530491 | 0.373714 | 0.07634312 | 0.93914612 | 0.95942244 | NA | NA | |
| ENSG000001 | 33.0995658 | 0.018240748 | 0.23946209 | 0.07617384 | 0.93928079 | 0.95953584 | C14orf79 | | 122616 |
| ENSG000002 | 4.77187399 | 0.028496237 | 0.37430245 | 0.07613158 | 0.93931441 | 0.95954602 | NA | NA | |
| ENSG000002 | 3.04620524 | -0.028708053 | 0.37781715 | -0.075984 | 0.93943184 | 0.9596418 | NA | NA | |
| ENSG000002 | 2.94855353 | 0.02849116 | 0.3768972 | 0.07559398 | 0.93974212 | 0.95993458 | NA | NA | |
| ENSG000002 | 2.97502821 | 0.028286905 | 0.37591758 | 0.07524763 | 0.94001769 | 0.96019188 | NA | NA | |

| | | | | | | | | |
|---|---|---|---|---|---|---|---|---|
| ENSG000002 | 9.26100082 | -0.02633746 | 0.35051073 | -0.0751402 | 0.94010313 | 0.96025496 | NA | NA |
| ENSG000002 | 21.8262947 | 0.021821063 | 0.29094254 | 0.07500128 | 0.94021369 | 0.9603282 | GS1-24F4.2 | 100652791 |
| ENSG000002 | 5.17579836 | -0.028011235 | 0.37352996 | -0.0749906 | 0.94022219 | 0.9603282 | NA | NA |
| ENSG000001 | 10.5263566 | 0.024904638 | 0.33243133 | 0.07491664 | 0.94028104 | 0.96036411 | TSSK3 | 81629 |
| ENSG000002 | 6.8173466 | 0.026869164 | 0.35937808 | 0.07476573 | 0.94040111 | 0.96044503 | NA | NA |
| ENSG000002 | 7.45224419 | -0.026576944 | 0.35550858 | -0.0747575 | 0.94040763 | 0.96044503 | NA | NA |
| ENSG000002 | 274.31947 | -0.010171456 | 0.13616492 | -0.0746995 | 0.94045378 | 0.96046797 | ITGA1 | 3672 |
| ENSG000002 | 3.16631554 | 0.02815069 | 0.37793349 | 0.07448583 | 0.94062382 | 0.96056905 | NA | NA |
| ENSG000002 | 32.7779017 | -0.0183829 | 0.24670538 | -0.0745136 | 0.94060174 | 0.96056905 | HEXA-AS1 | 80072 |
| ENSG000002 | 3.28400987 | -0.02817061 | 0.37804118 | -0.0745173 | 0.94059877 | 0.96056905 | NA | NA |
| ENSG000002 | 2.84725829 | -0.027932097 | 0.37557758 | -0.074371 | 0.94071515 | 0.96062706 | NA | NA |
| ENSG000002 | 2.82884097 | -0.027983633 | 0.37650307 | -0.0743251 | 0.94075169 | 0.96062706 | NA | NA |
| ENSG000002 | 2.59064403 | -0.027313855 | 0.36740953 | -0.0743417 | 0.94073848 | 0.96062706 | NA | NA |
| ENSG000002 | 3.04116906 | -0.02801847 | 0.37772585 | -0.0741767 | 0.94086976 | 0.96072343 | NA | NA |
| ENSG000001 | 114.119641 | -0.012919632 | 0.17431479 | -0.0741167 | 0.94091755 | 0.96074804 | ACSBG1 | 23205 |
| ENSG000002 | 6.79811939 | 0.026749935 | 0.36152786 | 0.07399135 | 0.94101727 | 0.96077728 | NA | NA |
| ENSG000002 | 3.09171789 | -0.02794199 | 0.37762338 | -0.0739943 | 0.9410149 | 0.96077728 | NA | NA |
| ENSG000002 | 2.90606186 | 0.027876838 | 0.37670112 | 0.07400254 | 0.94100837 | 0.96077728 | NA | NA |
| ENSG00000C | 25.066634 | 0.019247593 | 0.26056188 | 0.07386956 | 0.94111417 | 0.96082703 | NTN1 | 9423 |
| ENSG000002 | 3.13734881 | 0.027920289 | 0.37811474 | 0.07384078 | 0.94113707 | 0.96082703 | NA | NA |
| ENSG000002 | 10.503795 | -0.024485891 | 0.33149722 | -0.0738645 | 0.94111817 | 0.96082703 | NA | NA |
| ENSG00000C | 247.790093 | -0.009935367 | 0.13463829 | -0.073793 | 0.94117508 | 0.96084163 | NNAT | 4826 |
| ENSG000001 | 65.5036944 | -0.015915089 | 0.21607689 | -0.0736548 | 0.9412851 | 0.96086573 | LDOC1 | 23641 |
| ENSG000001 | 4.37289001 | 0.027830103 | 0.37788009 | 0.07364798 | 0.9412905 | 0.96086573 | NA | NA |
| ENSG000002 | 4.09180443 | 0.027894784 | 0.37856975 | 0.07368466 | 0.9412613 | 0.96086573 | NA | NA |
| ENSG000002 | 6.56744589 | 0.026608844 | 0.36131595 | 0.07364425 | 0.94129346 | 0.96086573 | LOC105372367 | 105372367 |
| ENSG000002 | 11.7417652 | 0.024414975 | 0.33217118 | 0.07350118 | 0.9414073 | 0.96095775 | NA | NA |
| ENSG000002 | 4.96356256 | -0.027530371 | 0.37471353 | -0.0734704 | 0.94143176 | 0.96095852 | NA | NA |
| ENSG000002 | 5.15832711 | -0.027230064 | 0.37090314 | -0.0734156 | 0.94147543 | 0.96097891 | MYZAP | 100820829 |

| | | | | | | | | |
|---|---|---|---|---|---|---|---|---|
| ENSG00000C | 38.6285889 | 0.01756685 | 0.23961463 | 0.07331293 | 0.9415571 | 0.96103332 | CBARP | 255057 |
| ENSG000002 | 3.37406425 | 0.027728091 | 0.37833895 | 0.07328902 | 0.94157613 | 0.96103332 | NA | NA |
| ENSG000001 | 32.0165578 | 0.019428893 | 0.26521081 | 0.0732583 | 0.94160058 | 0.96103408 | PDGFRL | 5157 |
| ENSG000002 | 5.88179916 | 0.027042621 | 0.36937711 | 0.07321142 | 0.94163788 | 0.96104797 | NA | NA |
| ENSG000002 | 6.35363087 | 0.02687196 | 0.36745879 | 0.07312918 | 0.94170332 | 0.96108606 | NA | NA |
| ENSG000002 | 3.75971655 | -0.027672777 | 0.37853499 | -0.0731049 | 0.94172261 | 0.96108606 | NA | NA |
| ENSG000001 | 27.7064251 | -0.019525158 | 0.268432 | -0.0727378 | 0.94201475 | 0.96129322 | SLCO1B1 | 10599 |
| ENSG000001 | 88.0577904 | 0.013166941 | 0.18100023 | 0.07274544 | 0.94200869 | 0.96129322 | PRRG4 | 79056 |
| ENSG000001 | 10.211046 | 0.024968327 | 0.3429537 | 0.07280378 | 0.94196226 | 0.96129322 | GH2 | 2689 |
| ENSG000002 | 3.13592646 | 0.027491115 | 0.37798504 | 0.07273069 | 0.94202042 | 0.96129322 | NA | NA |
| ENSG000002 | 27.1820856 | -0.018183975 | 0.25023598 | -0.0726673 | 0.94207086 | 0.9613205 | NA | NA |
| ENSG000002 | 3.29633429 | -0.027469269 | 0.37852602 | -0.072569 | 0.94214907 | 0.96137611 | NA | NA |
| ENSG000001 | 3.17383135 | 0.027397806 | 0.37824116 | 0.07243475 | 0.94225593 | 0.96138277 | NEU2 | 4759 |
| ENSG000001 | 19.5062824 | 0.020597568 | 0.28419233 | 0.07247756 | 0.94222186 | 0.96138277 | ZDHHC1 | 29800 |
| ENSG000002 | 6.92078117 | 0.026587616 | 0.367172 | 0.07241188 | 0.94227413 | 0.96138277 | LOC105374546 | 105374546 |
| ENSG000002 | 2.96387875 | 0.027209164 | 0.37526943 | 0.07250568 | 0.94219949 | 0.96138277 | NA | NA |
| ENSG000002 | 7.96081924 | -0.025492728 | 0.35183525 | -0.0724564 | 0.94223868 | 0.96138277 | NA | NA |
| ENSG000002 | 8.00776767 | 0.026303792 | 0.36397087 | 0.07226895 | 0.94238787 | 0.96147463 | NA | NA |
| ENSG000002 | 2.92709542 | 0.02720221 | 0.37751764 | 0.07205547 | 0.94255777 | 0.96162377 | NA | NA |
| ENSG000002 | 3.31967122 | -0.02716719 | 0.37806745 | -0.0718581 | 0.94271487 | 0.96175986 | NA | NA |
| ENSG000002 | 45.1468641 | 0.015612828 | 0.21750964 | 0.07177993 | 0.94277704 | 0.96179908 | NA | NA |
| ENSG000002 | 2.89331378 | 0.02695032 | 0.37564485 | 0.07174415 | 0.94280552 | 0.96180394 | NA | NA |
| ENSG000001 | 175.646053 | 0.009802213 | 0.13727178 | 0.07140734 | 0.94307357 | 0.96200478 | CDH11 | 1009 |
| ENSG000002 | 6.61320683 | 0.025915492 | 0.36290703 | 0.07141083 | 0.94307079 | 0.96200478 | NA | NA |
| ENSG000002 | 3.52576149 | -0.027040062 | 0.3785792 | -0.0714251 | 0.94305942 | 0.96200478 | NA | NA |
| ENSG000001 | 12.8019548 | 0.022819356 | 0.31986808 | 0.0713399 | 0.94312724 | 0.96203534 | NA | NA |
| ENSG000002 | 2.92778056 | 0.026893686 | 0.37714411 | 0.07130878 | 0.94315201 | 0.9620364 | NA | NA |
| ENSG000002 | 3.74593788 | -0.026968471 | 0.37857917 | -0.071236 | 0.94320992 | 0.96207127 | NA | NA |
| ENSG000002 | 32.3032479 | 0.017411642 | 0.24498404 | 0.07107256 | 0.94334001 | 0.96211002 | NA | NA |

| | | | | | | | | |
|---|---|---|---|---|---|---|---|---|
| ENSG000002 | 31.8140191 | 0.018968319 | 0.26676071 | 0.07110612 | 0.9433133 | 0.96211002 | NA | NA |
| ENSG000002 | 13.8301865 | 0.022950502 | 0.32253485 | 0.07115666 | 0.94327308 | 0.96211002 | LOC101929241 | 101929241 |
| ENSG000002 | 3.07393363 | -0.026777809 | 0.37678587 | -0.071069 | 0.94334281 | 0.96211002 | NA | NA |
| ENSG000002 | 3.29345712 | -0.026794275 | 0.37836159 | -0.0708166 | 0.94354373 | 0.96229074 | NA | NA |
| ENSG000002 | 32.5426046 | -0.016844861 | 0.23824472 | -0.070704 | 0.94363332 | 0.96235789 | HLA-F | 3134 |
| ENSG000002 | 8.22152945 | 0.02446654 | 0.34806569 | 0.07029288 | 0.94396055 | 0.96266741 | NA | NA |
| ENSG000002 | 3.42415627 | 0.026517979 | 0.37831468 | 0.07009503 | 0.94411802 | 0.96280379 | LOC101928891 | 101928891 |
| ENSG000002 | 2.94428682 | 0.026328145 | 0.37655398 | 0.06991865 | 0.94425841 | 0.9629176 | RNU6-674P | 106481773 |
| ENSG000002 | 2.9429686 | 0.026339806 | 0.37684738 | 0.06989515 | 0.94427711 | 0.9629176 | ASH1L-IT1 | 106478976 |
| ENSG000002 | 20.0889389 | 0.019368514 | 0.27766744 | 0.06975436 | 0.94438918 | 0.96300766 | NA | NA |
| ENSG000002 | 6.58892328 | 0.025191337 | 0.36296534 | 0.06940425 | 0.94466785 | 0.9632676 | LOC105378272 | 105378272 |
| ENSG000002 | 2.8227351 | -0.025960167 | 0.37451989 | -0.0693159 | 0.9447382 | 0.96331512 | NA | NA |
| ENSG000002 | 7.25120096 | 0.024963237 | 0.36080661 | 0.06918731 | 0.94484053 | 0.96339523 | NA | NA |
| ENSG000002 | 14.3861759 | -0.020970218 | 0.30345465 | -0.069105 | 0.94490608 | 0.96343784 | LOC101926943 | 101926943 |
| ENSG000002 | 5.42762125 | 0.025710938 | 0.37348447 | 0.06884072 | 0.94511641 | 0.96362807 | NA | NA |
| ENSG000002 | 3.16019648 | 0.02592645 | 0.3773974 | 0.06869801 | 0.94523 | 0.96371966 | NA | NA |
| ENSG000002 | 4.81541498 | 0.025624271 | 0.37406575 | 0.06850205 | 0.94538599 | 0.96383835 | TMEM238 | 388564 |
| ENSG000002 | 6.34121021 | -0.025280877 | 0.36910683 | -0.068492 | 0.94539396 | 0.96383835 | NA | NA |
| ENSG000002 | 14.1996773 | -0.021022735 | 0.30712457 | -0.0684502 | 0.94542727 | 0.96384808 | TAS2R20 | 259295 |
| ENSG000002 | 14.5146688 | 0.02223518 | 0.32574716 | 0.06825902 | 0.94557945 | 0.96397899 | DPH6-AS1 | 100507466 |
| ENSG000001 | 3.13524167 | 0.025684686 | 0.37683041 | 0.0681598 | 0.94565843 | 0.96403527 | S100P | 6286 |
| ENSG000001 | 11.8183778 | 0.02225654 | 0.32702839 | 0.06805691 | 0.94574034 | 0.96409453 | NA | NA |
| ENSG000002 | 2.85093215 | -0.025500822 | 0.37615384 | -0.0677936 | 0.94594994 | 0.96428396 | NA | NA |
| ENSG000002 | 3.09148976 | -0.025588648 | 0.37790277 | -0.0677123 | 0.9460147 | 0.96432573 | NA | NA |
| ENSG000001 | 2.96335703 | 0.025047578 | 0.37022427 | 0.06765515 | 0.94606015 | 0.96434783 | IFNA8 | 3445 |
| ENSG000002 | 5.20552006 | 0.025314143 | 0.37458775 | 0.06757867 | 0.94612103 | 0.96438564 | NA | NA |
| ENSG000001 | 7.88271156 | -0.024291159 | 0.36056181 | -0.0673703 | 0.94628691 | 0.96449298 | NA | NA |
| ENSG000002 | 2.8327556 | -0.02520032 | 0.37445175 | -0.0672992 | 0.94634348 | 0.96449298 | NA | NA |
| ENSG000002 | 6.08045356 | -0.025048843 | 0.37212221 | -0.0673135 | 0.94633214 | 0.96449298 | NA | NA |

| | | | | | | | | |
|---|---|---|---|---|---|---|---|---|
| ENSG000002 | 10.5300232 | 0.023619144 | 0.35096873 | 0.067297 | 0.94634526 | 0.96449298 | NA | NA |
| ENSG000002 | 6.58275814 | 0.0247549 | 0.36732662 | 0.06739207 | 0.94626958 | 0.96449298 | NA | NA |
| ENSG000001 | 5.59444596 | -0.024700956 | 0.36817969 | -0.0670894 | 0.94651053 | 0.96463717 | MRGPRX2 | 117194 |
| ENSG000001 | 12.7637346 | -0.021344309 | 0.31836405 | -0.0670437 | 0.9465469 | 0.96465 | RBP5 | 83758 |
| ENSG000002 | 2.5609084 | -0.025028136 | 0.37351572 | -0.0670069 | 0.9465762 | 0.96465561 | NA | NA |
| ENSG000002 | 13.0441381 | 0.02124903 | 0.31734046 | 0.06695972 | 0.94661377 | 0.96466965 | NA | NA |
| ENSG000002 | 33.0587424 | -0.015980478 | 0.23993936 | -0.0666022 | 0.94689843 | 0.9649355 | LOC101928238 | 101928238 |
| ENSG000002 | 10.999466 | -0.022079905 | 0.33209244 | -0.0664872 | 0.94698993 | 0.96500449 | RN7SL751P | 106479495 |
| ENSG000002 | 3.51907462 | -0.025135563 | 0.3784452 | -0.066418 | 0.94704506 | 0.96500451 | ZFX-AS1 | 100873922 |
| ENSG000002 | 3.54793721 | -0.025156724 | 0.37854244 | -0.0664568 | 0.94701414 | 0.96500451 | NA | NA |
| ENSG000002 | 6.77720668 | -0.024148909 | 0.36370198 | -0.0663975 | 0.94706135 | 0.96500451 | NA | NA |
| ENSG000002 | 2.55690227 | -0.024616013 | 0.37160501 | -0.0662424 | 0.94718483 | 0.96505759 | NA | NA |
| ENSG000002 | 6.83982179 | 0.023783203 | 0.35890376 | 0.06626624 | 0.94716586 | 0.96505759 | NA | NA |
| ENSG000002 | 2.79927398 | -0.024971381 | 0.37676702 | -0.066278 | 0.94715646 | 0.96505759 | NA | NA |
| ENSG000001 | 23.4710034 | 0.017581793 | 0.26576521 | 0.06615536 | 0.94725414 | 0.96510395 | SLC52A3 | 113278 |
| ENSG000001 | 58.6793721 | -0.013274901 | 0.20151168 | -0.0658766 | 0.94747608 | 0.96523369 | MEMO1 | 51072 |
| ENSG000002 | 13.952323 | 0.020944768 | 0.31774867 | 0.06591615 | 0.94744459 | 0.96523369 | NA | NA |
| ENSG000002 | 3.28539976 | -0.024763158 | 0.37590657 | -0.0658758 | 0.94747668 | 0.96523369 | NA | NA |
| ENSG000002 | 2.93489164 | 0.024858522 | 0.37714819 | 0.06591182 | 0.94744803 | 0.96523369 | NA | NA |
| ENSG000001 | 133.709469 | -0.011688816 | 0.17759777 | -0.0658162 | 0.94752413 | 0.96524855 | RBM43 | 375287 |
| ENSG000002 | 4.88837933 | -0.024636007 | 0.37442039 | -0.0657977 | 0.94753887 | 0.96524855 | CFB | 629 |
| ENSG000001 | 43.3451206 | -0.014217856 | 0.21653518 | -0.0656607 | 0.94764795 | 0.96533517 | PLP2 | 5355 |
| ENSG000002 | 8.66336983 | 0.022473037 | 0.34243153 | 0.06562783 | 0.94767413 | 0.96533517 | NA | NA |
| ENSG000002 | 12.0553126 | -0.02103174 | 0.32059988 | -0.0656012 | 0.94769532 | 0.96533517 | LINC01273 | 101927541 |
| ENSG000002 | 116.291475 | 0.010719455 | 0.16434705 | 0.06522451 | 0.94799525 | 0.96561642 | NA | NA |
| ENSG000002 | 51.1925499 | 0.014012488 | 0.21539795 | 0.06505395 | 0.94813105 | 0.96573048 | NA | NA |
| ENSG000001 | 35.0626448 | 0.015506994 | 0.23859061 | 0.06499415 | 0.94817866 | 0.96575472 | ERICH5 | 203111 |
| ENSG000002 | 48.2267519 | -0.013782182 | 0.21219622 | -0.0649502 | 0.94821367 | 0.96576613 | NA | NA |
| ENSG000002 | 3.37160389 | 0.024535966 | 0.37824714 | 0.06486755 | 0.94827946 | 0.96580722 | NA | NA |

| | | | | | | | | |
|---|---|---|---|---|---|---|---|---|
| ENSG000002 | 4.54728599 | 0.024349537 | 0.37553451 | 0.06483968 | 0.94830165 | 0.96580722 | LOC284454 | 284454 |
| ENSG000002 | 5.45384721 | 0.02404359 | 0.37165159 | 0.0646939 | 0.94841772 | 0.96590117 | NA | NA |
| ENSG000001 | 27.712715 | 0.01608501 | 0.24899336 | 0.06460015 | 0.94849236 | 0.96593802 | CRYL1 | 51084 |
| ENSG000002 | 2.98857821 | 0.024202469 | 0.37471719 | 0.06458863 | 0.94850154 | 0.96593802 | NA | NA |
| ENSG000002 | 2.82926552 | -0.0242619 | 0.37635418 | -0.0644656 | 0.94859949 | 0.96601351 | NA | NA |
| ENSG000001 | 33.4215386 | -0.015385165 | 0.23934222 | -0.064281 | 0.94874646 | 0.96613891 | RASSF9 | 9182 |
| ENSG000002 | 4874.60513 | -0.011145289 | 0.17382234 | -0.0641189 | 0.94887559 | 0.96624615 | NA | NA |
| ENSG000002 | 2.85987053 | -0.024050138 | 0.37564141 | -0.0640242 | 0.94895096 | 0.96629863 | NA | NA |
| ENSG000002 | 3.12437206 | 0.024123527 | 0.37796065 | 0.0638255 | 0.94910918 | 0.96642887 | NA | NA |
| ENSG000002 | 2.84025156 | -0.023961924 | 0.3758697 | -0.0637506 | 0.9491688 | 0.96642887 | NA | NA |
| ENSG000002 | 14.4090389 | 0.019313471 | 0.30298563 | 0.06374385 | 0.94917419 | 0.96642887 | NA | NA |
| ENSG000002 | 2.95334142 | 0.024072537 | 0.37742459 | 0.06378105 | 0.94914457 | 0.96642887 | NA | NA |
| ENSG000002 | 7.45573492 | -0.022314261 | 0.35359059 | -0.0631076 | 0.9496808 | 0.96692041 | NA | NA |
| ENSG000002 | 2.86055567 | -0.023645799 | 0.37506441 | -0.0630446 | 0.94973096 | 0.9669472 | NA | NA |
| ENSG000002 | 4.76401739 | 0.023539433 | 0.37460714 | 0.06283765 | 0.94989578 | 0.96709073 | NA | NA |
| ENSG000002 | 3.97319944 | -0.023763454 | 0.37846838 | -0.0627885 | 0.94993494 | 0.96710632 | NA | NA |
| ENSG000002 | 2.83194681 | -0.023449795 | 0.37465217 | -0.0625908 | 0.95009232 | 0.9672144 | LOC101060385 | 101060385 |
| ENSG000002 | 2.7816856 | -0.023404982 | 0.37408886 | -0.0625653 | 0.95011266 | 0.9672144 | NA | NA |
| ENSG000002 | 5.16496245 | -0.023236926 | 0.37134888 | -0.0625744 | 0.95010543 | 0.9672144 | NA | NA |
| ENSG000001 | 30.800206 | -0.015192276 | 0.24349088 | -0.0623936 | 0.95024938 | 0.9673293 | FAM222A | 84915 |
| ENSG000002 | 3.42107004 | 0.023516114 | 0.37785925 | 0.06223512 | 0.9503756 | 0.9674335 | NA | NA |
| ENSG000000 | 12.305116 | -0.019646138 | 0.31632612 | -0.0621072 | 0.95047744 | 0.96744003 | TBX5 | 6910 |
| ENSG000001 | 31.058969 | -0.015306298 | 0.24612775 | -0.0621884 | 0.95041278 | 0.96744003 | MICALL2 | 79778 |
| ENSG000001 | 39.7729381 | 0.014118903 | 0.2271498 | 0.06215679 | 0.95043797 | 0.96744003 | NA | NA |
| ENSG000002 | 3.3904388 | 0.023476795 | 0.37783533 | 0.06213499 | 0.95045533 | 0.96744003 | NA | NA |
| ENSG000002 | 18.7031858 | 0.017865185 | 0.28779578 | 0.06207591 | 0.95050238 | 0.96744113 | NA | NA |
| ENSG000002 | 5.20626992 | 0.023049069 | 0.37265766 | 0.06185052 | 0.95068187 | 0.96759953 | NA | NA |
| ENSG000002 | 2.83732288 | -0.023261545 | 0.37665322 | -0.0617585 | 0.95075514 | 0.96764982 | NA | NA |
| ENSG000001 | 98.2162853 | 0.010390783 | 0.16835584 | 0.06171917 | 0.95078647 | 0.96765742 | EML2 | 24139 |

| | | | | | | | | |
|---|---|---|---|---|---|---|---|---|
| ENSG000001 | 15.2851758 | 0.019039156 | 0.30942556 | 0.06153065 | 0.9509366 | 0.9677842 | GABARAP | 11337 |
| ENSG000002 | 15.7744576 | 0.019030681 | 0.30942783 | 0.06150281 | 0.95095878 | 0.9677842 | NA | NA |
| ENSG000002 | 3.15952398 | 0.023191487 | 0.37813414 | 0.06133138 | 0.9510953 | 0.96789885 | NA | NA |
| ENSG000002 | 3.17276132 | 0.023129908 | 0.37778316 | 0.06122535 | 0.95117974 | 0.96796049 | NA | NA |
| ENSG000002 | 5.42330431 | -0.02274618 | 0.37209417 | -0.0611302 | 0.95125554 | 0.96801334 | NA | NA |
| ENSG000002 | 4.8030713 | 0.022813097 | 0.37402938 | 0.06099279 | 0.95136495 | 0.96808179 | NA | NA |
| ENSG000002 | 5.37885896 | 0.022818931 | 0.37416825 | 0.06098575 | 0.95137056 | 0.96808179 | LOC285804 | 285804 |
| ENSG000002 | 3.06060455 | -0.02302011 | 0.37800512 | -0.0608989 | 0.95143969 | 0.96810356 | GH1 | 2688 |
| ENSG000002 | 4.79196072 | 0.022913019 | 0.37620659 | 0.06090542 | 0.95143454 | 0.96810356 | NA | NA |
| ENSG000002 | 2.83865294 | -0.022753821 | 0.3743063 | -0.0607893 | 0.95152701 | 0.9681681 | NA | NA |
| ENSG000002 | 22.3462564 | 0.017198848 | 0.28308099 | 0.06075593 | 0.95155359 | 0.96817086 | NA | NA |
| ENSG000002 | 2.80705093 | -0.022858107 | 0.37648799 | -0.060714 | 0.95158695 | 0.96818051 | NA | NA |
| ENSG000001 | 54.8465142 | -0.011941485 | 0.19682667 | -0.0606701 | 0.95162198 | 0.96819186 | MPZL2 | 10205 |
| ENSG000002 | 3.29020802 | -0.022913345 | 0.37786325 | -0.0606393 | 0.95164651 | 0.96819253 | NA | NA |
| ENSG000002 | 65.4029497 | -0.011595558 | 0.19139826 | -0.0605834 | 0.95169099 | 0.96821349 | ROCK1P1 | 727758 |
| ENSG000002 | 9.75804938 | -0.020219514 | 0.33397244 | -0.0605425 | 0.9517236 | 0.96822237 | NA | NA |
| ENSG000002 | 5.05572443 | 0.02255176 | 0.37383611 | 0.06032526 | 0.95189658 | 0.96834977 | NA | NA |
| ENSG000002 | 2.81344496 | -0.022641163 | 0.37523223 | -0.0603391 | 0.95188559 | 0.96834977 | RN7SL720P | 106480528 |
| ENSG000001 | 2.92823702 | 0.022704094 | 0.37655486 | 0.06029425 | 0.95192128 | 0.96835059 | NA | NA |
| ENSG000001 | 107.114169 | -0.011539666 | 0.19162938 | -0.0602187 | 0.95198148 | 0.96838754 | MAPKBP1 | 23005 |
| ENSG000002 | 2.62557579 | -0.0223674 | 0.37204864 | -0.0601196 | 0.95206041 | 0.96844354 | RN7SKP105 | 106479141 |
| ENSG000002 | 3.0269397 | -0.022453542 | 0.37500343 | -0.0598756 | 0.95225474 | 0.96860856 | NA | NA |
| ENSG000002 | 2.82852886 | -0.022381497 | 0.37411051 | -0.0598259 | 0.9522943 | 0.96860856 | NA | NA |
| ENSG000002 | 6.83976175 | 0.02150664 | 0.35939536 | 0.05984117 | 0.95228213 | 0.96860856 | NA | NA |
| ENSG000002 | 2.36958937 | -0.021718401 | 0.36335311 | -0.0597722 | 0.9523371 | 0.9686278 | NA | NA |
| ENSG000001 | 77.7582193 | 0.010758532 | 0.18035412 | 0.05965227 | 0.95243259 | 0.96868611 | KCNK10 | 54207 |
| ENSG000002 | 50.3016152 | 0.013255012 | 0.22224964 | 0.05964019 | 0.95244221 | 0.96868611 | NA | NA |
| ENSG000002 | 13.2683852 | -0.018656366 | 0.3139229 | -0.0594298 | 0.9526098 | 0.96883226 | NA | NA |
| ENSG000002 | 5.05150944 | 0.022133961 | 0.37282045 | 0.05936895 | 0.95265824 | 0.96885723 | NA | NA |

| | | | | | | | | |
|---|---|---|---|---|---|---|---|---|
| ENSG000002 | 2.50227038 | 0.021499123 | 0.36337779 | 0.05916466 | 0.95282096 | 0.96899841 | NA | NA |
| ENSG000001 | 40.1682088 | -0.01292654 | 0.21883216 | -0.0590706 | 0.9528959 | 0.96904458 | TMEM26 | 219623 |
| ENSG000002 | 12.7864004 | 0.018518043 | 0.31362769 | 0.05904467 | 0.95291653 | 0.96904458 | UNQ6494 | 100129066 |
| ENSG000002 | 5.18107323 | 0.022017713 | 0.37306997 | 0.05901765 | 0.95293805 | 0.96904458 | UXT-AS1 | 100133957 |
| ENSG000002 | 9.79248425 | -0.019789129 | 0.33562816 | -0.0589615 | 0.9529828 | 0.96906579 | NA | NA |
| ENSG000001 | 8.92314995 | -0.02088108 | 0.35447832 | -0.0589065 | 0.95302657 | 0.969086 | MDS2 | 259283 |
| ENSG000002 | 6.86450918 | 0.021255435 | 0.36129796 | 0.05883076 | 0.95308691 | 0.96912305 | MIR3976HG | 645355 |
| ENSG000002 | 7.64097565 | -0.020713084 | 0.35269132 | -0.0587286 | 0.95316824 | 0.96918145 | NA | NA |
| ENSG000002 | 6.8317691 | 0.021375174 | 0.36494486 | 0.05857097 | 0.95329383 | 0.96928485 | NA | NA |
| ENSG000002 | 2.56298364 | -0.021865196 | 0.37427277 | -0.0584205 | 0.9534137 | 0.96938242 | NA | NA |
| ENSG000002 | 2.80289577 | -0.021816355 | 0.37578443 | -0.0580555 | 0.95370442 | 0.96965369 | NA | NA |
| ENSG000002 | 3.01228742 | -0.021841735 | 0.37664777 | -0.0579898 | 0.95375674 | 0.96968258 | NA | NA |
| ENSG000002 | 16.338897 | 0.017678179 | 0.30624988 | 0.05772469 | 0.95396793 | 0.96987298 | NA | NA |
| ENSG000002 | 11.923297 | -0.018747221 | 0.32513687 | -0.0576595 | 0.95401987 | 0.96990147 | NA | NA |
| ENSG000001 | 71.4062681 | 0.010831305 | 0.18816828 | 0.0575618 | 0.95409768 | 0.96995625 | ASB3 | 51130 |
| ENSG000002 | 4.95043048 | -0.021629973 | 0.37646047 | -0.0574562 | 0.95418183 | 0.96996885 | NA | NA |
| ENSG000002 | 43.8605054 | 0.012585211 | 0.21900714 | 0.05746484 | 0.95417491 | 0.96996885 | NA | NA |
| ENSG000002 | 3.43414376 | 0.021746057 | 0.37837993 | 0.05747149 | 0.95416962 | 0.96996885 | NA | NA |
| ENSG000001 | 9.05008125 | 0.019842952 | 0.34553875 | 0.05742613 | 0.95420575 | 0.96996885 | CHRNA10 | 57053 |
| ENSG000002 | 27.0690424 | -0.014737969 | 0.25733264 | -0.0572721 | 0.95432848 | 0.97004497 | NA | NA |
| ENSG000002 | 2.64122064 | -0.021238717 | 0.37070119 | -0.0572934 | 0.95431151 | 0.97004497 | NA | NA |
| ENSG000002 | 2.56045193 | -0.02133987 | 0.37397987 | -0.0570615 | 0.95449617 | 0.9701911 | NA | NA |
| ENSG000002 | 2.98447491 | 0.021148038 | 0.37154042 | 0.05691989 | 0.95460901 | 0.97028148 | NA | NA |
| ENSG000002 | 2.86604886 | -0.021333324 | 0.3752307 | -0.0568539 | 0.95466159 | 0.9703106 | NA | NA |
| ENSG000002 | 3.18458851 | 0.021454114 | 0.37778167 | 0.05678972 | 0.95471271 | 0.97033824 | NA | NA |
| ENSG000002 | 14.9318877 | -0.018050567 | 0.31848787 | -0.0566758 | 0.95480342 | 0.97040283 | NA | NA |
| ENSG000002 | 2.94459892 | 0.021302727 | 0.37604204 | 0.05664985 | 0.95482412 | 0.97040283 | LOC105369723 | 105369723 |
| ENSG000002 | 3.18394358 | 0.021267344 | 0.37664483 | 0.05646525 | 0.95497118 | 0.97047932 | NA | NA |
| ENSG000002 | 2.84222266 | -0.021285859 | 0.37669386 | -0.0565071 | 0.95493788 | 0.97047932 | NA | NA |

| | | | | | | | | |
|---|---|---|---|---|---|---|---|---|
| ENSG000002 | 2.54138913 | -0.021038597 | 0.37256375 | -0.0564698 | 0.95496756 | 0.97047932 | NA | NA |
| ENSG000002 | 6.78062518 | -0.020331749 | 0.36100281 | -0.0563202 | 0.95508673 | 0.97057242 | DGCR9 | 25787 |
| ENSG000001 | 3.09171843 | -0.021227785 | 0.37729787 | -0.0562627 | 0.95513256 | 0.97059467 | FAM228A | 653140 |
| ENSG000002 | 6.16942727 | 0.020631087 | 0.36808743 | 0.05604942 | 0.95530244 | 0.9707328 | NA | NA |
| ENSG000002 | 3.04629752 | -0.021089361 | 0.37638108 | -0.0560319 | 0.95531636 | 0.9707328 | NA | NA |
| ENSG00000C | 21.1065811 | 0.01607539 | 0.28735959 | 0.05594172 | 0.95538824 | 0.97077128 | GNA15 | 2769 |
| ENSG000002 | 9.80070447 | -0.01901531 | 0.34020152 | -0.0558943 | 0.95542605 | 0.97077128 | FAM230B | 642633 |
| ENSG000002 | 3.43061391 | 0.021165532 | 0.37852673 | 0.05591556 | 0.95540908 | 0.97077128 | NA | NA |
| ENSG000002 | 2.57368927 | -0.020814767 | 0.37352562 | -0.0557251 | 0.95556077 | 0.97088384 | RNU6-377P | 106479733 |
| ENSG000002 | 2.5711848 | -0.020843779 | 0.37447974 | -0.0556606 | 0.95561216 | 0.97091172 | NA | NA |
| ENSG000002 | 3.20134271 | 0.020968189 | 0.37728311 | 0.0555768 | 0.95567894 | 0.97094459 | NA | NA |
| ENSG000002 | 4.44559378 | -0.020951173 | 0.37709148 | -0.0555599 | 0.95569239 | 0.97094459 | NA | NA |
| ENSG000002 | 2.78439914 | -0.020808012 | 0.37493204 | -0.0554981 | 0.95574165 | 0.97097031 | NA | NA |
| ENSG000002 | 3.2698596 | -0.020883326 | 0.37755348 | -0.0553122 | 0.95588972 | 0.9710964 | NA | NA |
| ENSG000001 | 3.56486026 | -0.020877301 | 0.37780393 | -0.0552596 | 0.95593163 | 0.97110219 | ACTL10 | 170487 |
| ENSG000002 | 4.18032896 | -0.020904551 | 0.37839744 | -0.055245 | 0.95594331 | 0.97110219 | NA | NA |
| ENSG000002 | 7.19161946 | -0.019683041 | 0.35655786 | -0.0552029 | 0.95597679 | 0.97111187 | NA | NA |
| ENSG000002 | 2.79592129 | -0.020747695 | 0.37633919 | -0.0551303 | 0.95603465 | 0.97114632 | NA | NA |
| ENSG000002 | 2.81009172 | -0.020772764 | 0.37710772 | -0.0550844 | 0.9560712 | 0.97115912 | NA | NA |
| ENSG000002 | 2.80960245 | -0.020735647 | 0.37668041 | -0.0550484 | 0.95609992 | 0.97116397 | NA | NA |
| ENSG000001 | 68.7381555 | 0.010623771 | 0.19340781 | 0.05492938 | 0.95619472 | 0.97121161 | SPRYD3 | 84926 |
| ENSG000002 | 5.15898054 | -0.020398439 | 0.37119608 | -0.0549533 | 0.95617569 | 0.97121161 | NA | NA |
| ENSG000001 | 19.9084004 | 0.015353254 | 0.28016333 | 0.05480108 | 0.95629694 | 0.9712911 | C19orf57 | 79173 |
| ENSG00000C | 67.0204386 | 0.010101862 | 0.18466205 | 0.05470459 | 0.95637381 | 0.97134092 | ZNF343 | 79175 |
| ENSG000001 | 11.8409643 | -0.017553663 | 0.32116752 | -0.0546558 | 0.95641269 | 0.97134092 | OSGIN1 | 29948 |
| ENSG000001 | 107.692905 | -0.008710995 | 0.15939808 | -0.0546493 | 0.95641786 | 0.97134092 | FAM49A | 81553 |
| ENSG000001 | 86.64669 | 0.010096364 | 0.18524533 | 0.05450266 | 0.95653469 | 0.97143525 | CCDC40 | 55036 |
| ENSG000002 | 27.4701349 | -0.013853314 | 0.25466694 | -0.0543978 | 0.95661825 | 0.97149579 | LOC105371814 | 105371814 |
| ENSG000002 | 2.9573798 | 0.020455194 | 0.37672185 | 0.05429787 | 0.95669785 | 0.97155229 | NA | NA |

| | | | | | | | | |
|---|---|---|---|---|---|---|---|---|
| ENSG000002 | 3.41966033 | 0.020502082 | 0.37852812 | 0.05416264 | 0.95680559 | 0.97163737 | NA | NA |
| ENSG000002 | 5.35193078 | -0.020203361 | 0.37359311 | -0.0540785 | 0.95687261 | 0.97168109 | NA | NA |
| ENSG000002 | 5.28634607 | 0.020204699 | 0.37440176 | 0.05396529 | 0.95696282 | 0.97172403 | NA | NA |
| ENSG000002 | 2.34158078 | -0.019933935 | 0.3693564 | -0.0539694 | 0.95695956 | 0.97172403 | NA | NA |
| ENSG000002 | 2.76373772 | -0.020205209 | 0.37475775 | -0.0539154 | 0.95700258 | 0.97174007 | LOC440602 | 440602 |
| ENSG000002 | 170.224459 | -0.008359264 | 0.15528317 | -0.0538324 | 0.95706871 | 0.97178288 | NBPF11 | 200030 |
| ENSG000001 | 7.40152054 | -0.018996226 | 0.35395239 | -0.0536689 | 0.95719898 | 0.9718665 | FLRT1 | 23769 |
| ENSG000002 | 3.74542955 | -0.020307924 | 0.37828039 | -0.0536848 | 0.95718626 | 0.9718665 | NA | NA |
| ENSG000001 | 68.3187912 | 0.010075733 | 0.18832955 | 0.05350054 | 0.9573331 | 0.97197833 | TIGD6 | 81789 |
| ENSG000002 | 14.2851234 | 0.016658065 | 0.3125746 | 0.05329309 | 0.95749839 | 0.97212181 | NA | NA |
| ENSG000002 | 2.96631783 | 0.019883953 | 0.37341343 | 0.05324916 | 0.95753338 | 0.972133 | NA | NA |
| ENSG000002 | 8.0630084 | -0.018895765 | 0.35554281 | -0.0531462 | 0.95761539 | 0.97219191 | NA | NA |
| ENSG000002 | 2.5656842 | -0.019792216 | 0.37281193 | -0.053089 | 0.95766099 | 0.97221387 | NA | NA |
| ENSG000002 | 2.82866525 | -0.019846288 | 0.37509985 | -0.0529093 | 0.95780414 | 0.97233485 | NA | NA |
| ENSG000002 | 6.55674268 | -0.019297653 | 0.36579999 | -0.0527547 | 0.95792739 | 0.9723626 | NA | NA |
| ENSG000002 | 2.60991829 | -0.019644149 | 0.37228696 | -0.0527661 | 0.95791823 | 0.9723626 | RN7SL549P | 106481071 |
| ENSG000002 | 4.53393709 | 0.019936865 | 0.37747323 | 0.05281663 | 0.95787801 | 0.9723626 | NA | NA |
| ENSG000002 | 12.8068603 | 0.016647466 | 0.31545428 | 0.05277299 | 0.95791278 | 0.9723626 | NA | NA |
| ENSG000002 | 3.18877035 | 0.019910746 | 0.37811583 | 0.05265779 | 0.95800457 | 0.9724166 | NA | NA |
| ENSG000001 | 56.8907027 | 0.010224261 | 0.19487555 | 0.05246559 | 0.95815771 | 0.972501 | PADI2 | 11240 |
| ENSG000001 | 79.1583621 | 0.009084905 | 0.17302788 | 0.05250544 | 0.95812596 | 0.972501 | ZSCAN5A | 79149 |
| ENSG000002 | 3.43744458 | 0.019852918 | 0.37841649 | 0.05246314 | 0.95815966 | 0.972501 | NA | NA |
| ENSG000001 | 125.403418 | -0.008262545 | 0.15777683 | -0.0523686 | 0.95823503 | 0.97255315 | SLC9B2 | 133308 |
| ENSG000002 | 4.7743255 | 0.019598316 | 0.37464721 | 0.05231139 | 0.95828058 | 0.97256707 | NA | NA |
| ENSG000002 | 145.849452 | -0.007917523 | 0.15141231 | -0.0522911 | 0.95829671 | 0.97256707 | NA | NA |
| ENSG000001 | 61.5468639 | -0.009905464 | 0.18974678 | -0.0522036 | 0.95836647 | 0.97261353 | ZNF599 | 148103 |
| ENSG000002 | 7.00004473 | -0.018738668 | 0.35957036 | -0.0521141 | 0.95843782 | 0.97266159 | NA | NA |
| ENSG000002 | 2.51927813 | -0.0188711 | 0.36398142 | -0.0518463 | 0.95865114 | 0.97285374 | NA | NA |
| ENSG000002 | 5.26201079 | 0.019209192 | 0.37138803 | 0.0517227 | 0.95874965 | 0.97292935 | NA | NA |

| | | | | | | | | |
|---|---|---|---|---|---|---|---|---|
| ENSG000002 | 3.39412396 | 0.01949812 | 0.37852974 | 0.05151014 | 0.95891902 | 0.97307688 | SCARNA20 | 677681 |
| ENSG000001 | 59.1104497 | -0.01059808 | 0.20749738 | -0.0510757 | 0.95926517 | 0.97335507 | VAV1 | 7409 |
| ENSG000002 | 5.43406454 | -0.019195946 | 0.3754736 | -0.0511246 | 0.95922621 | 0.97335507 | NA | NA |
| ENSG000002 | 12.8304504 | 0.016014886 | 0.31350747 | 0.05108295 | 0.95925942 | 0.97335507 | NA | NA |
| ENSG000002 | 22.7925354 | 0.01400033 | 0.27533164 | 0.05084897 | 0.95944587 | 0.97351406 | NA | NA |
| ENSG000002 | 3.42254935 | 0.01914945 | 0.37801264 | 0.05065823 | 0.95959786 | 0.97363475 | SUCLA2-AS1 | 101929410 |
| ENSG000002 | 2.78684652 | -0.019032371 | 0.37584086 | -0.0506394 | 0.95961283 | 0.97363475 | NA | NA |
| ENSG000002 | 2.33870417 | -0.018721025 | 0.37018499 | -0.0505721 | 0.95966651 | 0.97366485 | NA | NA |
| ENSG000002 | 2.80487927 | -0.019023169 | 0.37667757 | -0.0505025 | 0.95972194 | 0.97369672 | NA | NA |
| ENSG000001 | 34.691684 | 0.01187313 | 0.23563252 | 0.05038833 | 0.95981293 | 0.97376468 | TM6SF1 | 53346 |
| ENSG00000C | 141.814948 | -0.008950899 | 0.17829777 | -0.050202 | 0.95996144 | 0.97379657 | IGSF9 | 57549 |
| ENSG000001 | 193.626138 | 0.007253971 | 0.14428471 | 0.0502754 | 0.95990293 | 0.97379657 | CCDC181 | 57821 |
| ENSG000001 | 93.5380567 | -0.008495409 | 0.16923728 | -0.0501982 | 0.95996444 | 0.97379657 | COL5A1 | 1289 |
| ENSG000002 | 6.95093787 | -0.018026115 | 0.35839741 | -0.0502964 | 0.95988616 | 0.97379657 | NA | NA |
| ENSG000002 | 2.78925403 | -0.018903178 | 0.37633181 | -0.0502301 | 0.95993904 | 0.97379657 | NA | NA |
| ENSG000001 | 170.210632 | 0.007435579 | 0.1483282 | 0.05012923 | 0.96001941 | 0.97382797 | ELMO1 | 9844 |
| ENSG000002 | 2.55874938 | -0.018692105 | 0.37350446 | -0.0500452 | 0.96008637 | 0.97387154 | NA | NA |
| ENSG000002 | 7.43178064 | -0.017947473 | 0.35913999 | -0.0499735 | 0.96014353 | 0.97389402 | LOC105377443 | 105377443 |
| ENSG000002 | 6.81369181 | 0.01802809 | 0.36087142 | 0.0499571 | 0.96015657 | 0.97389402 | NA | NA |
| ENSG000002 | 2.57520357 | -0.018650453 | 0.37383607 | -0.0498894 | 0.96021053 | 0.97392253 | NA | NA |
| ENSG000002 | 3.15656772 | 0.018779589 | 0.37663457 | 0.04986156 | 0.96023271 | 0.97392253 | NA | NA |
| ENSG000002 | 3.20555702 | 0.01878148 | 0.37778651 | 0.04971453 | 0.96034988 | 0.974017 | MIR1305 | 100302270 |
| ENSG000002 | 3.1890183 | 0.018690332 | 0.37696349 | 0.04958128 | 0.96045606 | 0.97410034 | NA | NA |
| ENSG000002 | 12.5310068 | -0.016208898 | 0.327297 | -0.0495235 | 0.9605021 | 0.97412266 | NA | NA |
| ENSG000002 | 5.62168563 | -0.018366103 | 0.37114864 | -0.0494845 | 0.96053319 | 0.97412984 | NA | NA |
| ENSG000002 | 5.16984468 | 0.018578495 | 0.37580611 | 0.04943638 | 0.96057153 | 0.97414436 | NA | NA |
| ENSG000002 | 19.1349556 | 0.014484624 | 0.29360616 | 0.04933351 | 0.96065351 | 0.97417877 | NA | NA |
| ENSG000002 | 3.03614641 | -0.018578224 | 0.37638342 | -0.0493598 | 0.96063253 | 0.97417877 | NA | NA |
| ENSG000001 | 48.1589881 | -0.01031452 | 0.20965816 | -0.0491968 | 0.96076243 | 0.97426485 | WFDC1 | 58189 |

| | | | | | | | | | |
|---|---|---|---|---|---|---|---|---|---|
| ENSG000000 | 136.296303 | 0.007656345 | 0.15667352 | 0.04886815 | 0.96102437 | 0.97445524 | PPP1R13B | | 23368 |
| ENSG000001 | 20.451356 | -0.013419807 | 0.27476722 | -0.0488406 | 0.96104629 | 0.97445524 | BATF2 | | 116071 |
| ENSG000002 | 5.30837725 | -0.018402739 | 0.37610341 | -0.04893 | 0.96097508 | 0.97445524 | NA | NA | |
| ENSG000002 | 2.38026275 | -0.018073751 | 0.36960729 | -0.0488999 | 0.96099909 | 0.97445524 | NA | NA | |
| ENSG000002 | 2.55729457 | -0.018203529 | 0.37317696 | -0.0487799 | 0.96109471 | 0.97447996 | NA | NA | |
| ENSG000002 | 5.35165776 | -0.018082351 | 0.37264427 | -0.0485244 | 0.96129829 | 0.97463764 | NA | NA | |
| ENSG000002 | 7.46158874 | -0.017460226 | 0.35961056 | -0.0485532 | 0.96127541 | 0.97463764 | NA | NA | |
| ENSG000002 | 2.36517885 | -0.017862774 | 0.370253 | -0.0482448 | 0.96152116 | 0.97479049 | MIR30C1 | | 407031 |
| ENSG000002 | 3.91176913 | 0.018267096 | 0.37841662 | 0.04827245 | 0.96149911 | 0.97479049 | NA | NA | |
| ENSG000002 | 21.5584696 | 0.013495906 | 0.27966416 | 0.04825755 | 0.96151099 | 0.97479049 | NA | NA | |
| ENSG000001 | 48.2086232 | -0.010197861 | 0.21156438 | -0.0482022 | 0.96155512 | 0.97479129 | SPARCL1 | | 8404 |
| ENSG000002 | 2.6210683 | -0.017974206 | 0.37303687 | -0.0481835 | 0.96157003 | 0.97479129 | NA | NA | |
| ENSG000002 | 10.8949457 | -0.015770766 | 0.32767019 | -0.04813 | 0.96161264 | 0.97481011 | NA | NA | |
| ENSG000002 | 2.57810035 | -0.017986068 | 0.37404777 | -0.0480849 | 0.96164854 | 0.97482214 | NA | NA | |
| ENSG000002 | 2.59109637 | -0.017927774 | 0.37406147 | -0.0479273 | 0.96177414 | 0.97492509 | NA | NA | |
| ENSG000002 | 72.2657641 | 0.009439461 | 0.19772138 | 0.04774122 | 0.96192248 | 0.97505108 | PCDHGB7 | | 56099 |
| ENSG000001 | 10.1962961 | -0.015944058 | 0.33518162 | -0.0475684 | 0.96206021 | 0.97516631 | DKK1 | | 22943 |
| ENSG000001 | 11.4499813 | 0.015718282 | 0.33128784 | 0.047446 | 0.96215777 | 0.97524083 | CYP2A13 | | 1553 |
| ENSG000002 | 5.24437014 | 0.017661085 | 0.37367681 | 0.04726299 | 0.96230362 | 0.97536428 | NA | NA | |
| ENSG000001 | 3.43474403 | 0.017846517 | 0.37785917 | 0.0472306 | 0.96232944 | 0.97536607 | MSX1 | | 4487 |
| ENSG000002 | 4.88478532 | -0.017662836 | 0.37426825 | -0.047193 | 0.96235942 | 0.97537208 | NA | NA | |
| ENSG000002 | 5.20138824 | -0.01764024 | 0.37428294 | -0.0471308 | 0.96240901 | 0.97539796 | NA | NA | |
| ENSG000001 | 5.32440644 | -0.017478302 | 0.37225247 | -0.0469528 | 0.96255083 | 0.97551732 | KCNJ4 | | 3761 |
| ENSG000002 | 4.65289788 | -0.017510903 | 0.37578104 | -0.0465987 | 0.96283308 | 0.97575459 | NA | NA | |
| ENSG000002 | 7.63622043 | -0.016553843 | 0.35520523 | -0.0466036 | 0.96282916 | 0.97575459 | NA | NA | |
| ENSG000002 | 4.8902138 | -0.017393967 | 0.37462821 | -0.0464299 | 0.96296757 | 0.9758617 | NA | NA | |
| ENSG000002 | 22.7709279 | 0.012355081 | 0.26633004 | 0.04639011 | 0.96299932 | 0.9758617 | A2M-AS1 | | 144571 |
| ENSG000002 | 2.55724305 | -0.017322911 | 0.37402178 | -0.0463152 | 0.96305899 | 0.9758617 | NA | NA | |
| ENSG000002 | 3.45266488 | 0.017536135 | 0.37858458 | 0.04632026 | 0.96305499 | 0.9758617 | NA | NA | |

| | | | | | | | | |
|---|---|---|---|---|---|---|---|---|
| ENSG000002 | 2.77910237 | -0.017423891 | 0.3763253 | -0.0463001 | 0.96307108 | 0.9758617 | MIR4652 | 100616206 |
| ENSG000002 | 4.87357617 | -0.017334372 | 0.37451446 | -0.0462849 | 0.96308316 | 0.9758617 | NA | NA |
| ENSG000001 | 55.4811626 | 0.009099543 | 0.19684357 | 0.04622728 | 0.9631291 | 0.97587723 | HBEGF | 1839 |
| ENSG000002 | 7.81202707 | 0.016390785 | 0.35473825 | 0.04620529 | 0.96314662 | 0.97587723 | NA | NA |
| ENSG000002 | 5.2876037 | 0.017191659 | 0.37303922 | 0.04608539 | 0.96324219 | 0.97594968 | SNORD67 | 692108 |
| ENSG000002 | 7.65607251 | -0.016361136 | 0.35558245 | -0.0460122 | 0.96330051 | 0.97598439 | RN7SL74P | 106480942 |
| ENSG000002 | 3.16657503 | 0.017356484 | 0.37778568 | 0.04594267 | 0.96335594 | 0.97599177 | NA | NA |
| ENSG000002 | 19.5947589 | 0.01344038 | 0.29252223 | 0.04594653 | 0.96335287 | 0.97599177 | IKBKG | 8517 |
| ENSG000001 | 4.96857499 | -0.017308355 | 0.37744659 | -0.0458564 | 0.96342468 | 0.97601682 | SYNGR4 | 23546 |
| ENSG000001 | 3.32519721 | -0.017129658 | 0.37359184 | -0.0458513 | 0.9634288 | 0.97601682 | UCN3 | 114131 |
| ENSG000001 | 2.59787518 | -0.017071437 | 0.37355444 | -0.0457 | 0.96354937 | 0.97609207 | GPR150 | 285601 |
| ENSG000002 | 3.42954444 | 0.017209145 | 0.37658689 | 0.04569767 | 0.96355122 | 0.97609207 | OR52B4 | 143496 |
| ENSG000001 | 61.5644434 | -0.008566864 | 0.1884518 | -0.0454592 | 0.96374131 | 0.97617109 | NA | NA |
| ENSG000001 | 2.37368003 | -0.01682327 | 0.37023862 | -0.045439 | 0.9637574 | 0.97617109 | NA | NA |
| ENSG000002 | 3.42060606 | 0.017195456 | 0.37837523 | 0.04544551 | 0.9637522 | 0.97617109 | NA | NA |
| ENSG000002 | 29.8291071 | -0.0113922 | 0.25082676 | -0.0454186 | 0.96377366 | 0.97617109 | NA | NA |
| ENSG000002 | 2.36898876 | -0.016646067 | 0.36601302 | -0.0454794 | 0.96372516 | 0.97617109 | NA | NA |
| ENSG000002 | 2.55891335 | -0.017009376 | 0.37381296 | -0.0455024 | 0.96370689 | 0.97617109 | NA | NA |
| ENSG000001 | 17773.4491 | 0.005931808 | 0.1308058 | 0.04534821 | 0.96382976 | 0.97620353 | ND4 | 4538 |
| ENSG000001 | 27.9214772 | 0.011274023 | 0.24969698 | 0.04515082 | 0.96398709 | 0.9763385 | RHOV | 171177 |
| ENSG000001 | 231.69775 | -0.005942018 | 0.1318617 | -0.0450625 | 0.96405749 | 0.97638541 | KATNAL1 | 84056 |
| ENSG000002 | 4.13396092 | 0.016955702 | 0.37847037 | 0.04480061 | 0.96426624 | 0.97657244 | RN7SL745P | 106481858 |
| ENSG000002 | 2.54215914 | -0.016651598 | 0.37219586 | -0.0447388 | 0.9643155 | 0.97659794 | NA | NA |
| ENSG000002 | 2.81051627 | -0.016815378 | 0.37663438 | -0.0446464 | 0.96438914 | 0.97664812 | NA | NA |
| ENSG000002 | 2.58114058 | -0.016700883 | 0.37475039 | -0.0445654 | 0.96445376 | 0.97668917 | NA | NA |
| ENSG000001 | 75.3641847 | -0.008089824 | 0.18179774 | -0.044499 | 0.96450662 | 0.97669392 | SMAD6 | 4091 |
| ENSG000002 | 2.81987865 | -0.016595075 | 0.37289988 | -0.0445028 | 0.96450365 | 0.97669392 | NA | NA |
| ENSG000002 | 20.2236461 | 0.012868425 | 0.2898974 | 0.04438959 | 0.96459386 | 0.97675787 | NA | NA |
| ENSG000002 | 2.33882094 | -0.016384288 | 0.36948839 | -0.0443432 | 0.96463086 | 0.97677094 | NA | NA |

| | | | | | | | | |
|---|---|---|---|---|---|---|---|---|
| ENSG000002 | 3.02147408 | -0.016700688 | 0.37715449 | -0.0442808 | 0.96468061 | 0.97679693 | NA | NA |
| ENSG000002 | 4.43978106 | -0.016660958 | 0.37704262 | -0.0441885 | 0.96475413 | 0.97684698 | MIR1265 | 100302116 |
| ENSG000002 | 5.17287208 | 0.016545558 | 0.37595284 | 0.04400966 | 0.9648967 | 0.97696694 | NA | NA |
| ENSG000002 | 4.83992591 | -0.016456759 | 0.37459447 | -0.0439322 | 0.96495845 | 0.97700507 | NA | NA |
| ENSG000002 | 5.00130532 | 0.016344694 | 0.37278133 | 0.04384526 | 0.96502775 | 0.97705084 | NA | NA |
| ENSG000002 | 2.31192205 | -0.016117596 | 0.36795824 | -0.0438028 | 0.96506161 | 0.97706072 | OR13C5 | 138799 |
| ENSG000002 | 3.67975681 | 0.016455993 | 0.37848382 | 0.04347872 | 0.96531992 | 0.97729785 | NA | NA |
| ENSG000002 | 3.62685255 | 0.016397176 | 0.37856866 | 0.04331361 | 0.96545154 | 0.9774067 | NA | NA |
| ENSG000001 | 60.1407819 | -0.008531636 | 0.19753575 | -0.0431903 | 0.96554981 | 0.97748178 | RNFT1 | 51136 |
| ENSG000002 | 2.35262686 | -0.015740325 | 0.36597067 | -0.0430098 | 0.96569372 | 0.97757867 | NA | NA |
| ENSG000002 | 29.4107537 | 0.011407719 | 0.26505656 | 0.04303881 | 0.9656706 | 0.97757867 | LOC100129520 | 100129520 |
| ENSG000002 | 5.7261427 | 0.015974379 | 0.37180611 | 0.04296427 | 0.96573001 | 0.977591 | NA | NA |
| ENSG000002 | 5.52110971 | 0.01586988 | 0.37060462 | 0.04282159 | 0.96584375 | 0.97768173 | NA | NA |
| ENSG000002 | 2.82213538 | -0.016041428 | 0.37578656 | -0.0426876 | 0.96595056 | 0.97776544 | EIF1P5 | 106481690 |
| ENSG000002 | 5.30425445 | -0.015844519 | 0.37166373 | -0.0426313 | 0.96599542 | 0.97778644 | NA | NA |
| ENSG000001 | 3.41337045 | 0.016126142 | 0.37858586 | 0.04259573 | 0.9660238 | 0.97779076 | NA | NA |
| ENSG000001 | 3.41680801 | 0.016085515 | 0.37859287 | 0.04248763 | 0.96610997 | 0.97785357 | NA | NA |
| ENSG000002 | 2.36939349 | -0.01539469 | 0.3640188 | -0.0422909 | 0.96626678 | 0.97798788 | NA | NA |
| ENSG000002 | 42.0702363 | 0.009207994 | 0.21802735 | 0.04223321 | 0.96631279 | 0.97801003 | NA | NA |
| ENSG000002 | 3.39384322 | 0.015927938 | 0.37847736 | 0.04208426 | 0.96643153 | 0.97808139 | NA | NA |
| ENSG000002 | 5.13148879 | -0.015649263 | 0.37163659 | -0.042109 | 0.96641177 | 0.97808139 | NA | NA |
| ENSG000001 | 2.50885379 | -0.015496231 | 0.36907239 | -0.041987 | 0.96650909 | 0.97813547 | OR1A2 | 26189 |
| ENSG000002 | 20.4586099 | 0.011647988 | 0.27802923 | 0.04189483 | 0.96658254 | 0.97816098 | NA | NA |
| ENSG000002 | 4.66364373 | -0.015816099 | 0.37744338 | -0.0419032 | 0.96657584 | 0.97816098 | SNRPGP16 | 106480270 |
| ENSG000001 | 700.948435 | -0.004808742 | 0.11488478 | -0.0418571 | 0.96661263 | 0.97816414 | DNAH14 | 127602 |
| ENSG000002 | 2500.56038 | 0.005094288 | 0.12178439 | 0.04183039 | 0.96663391 | 0.97816414 | ND4L | 4539 |
| ENSG000002 | 8.99648279 | 0.014362884 | 0.34456036 | 0.04168467 | 0.96675008 | 0.97825728 | NA | NA |
| ENSG000002 | 2.56224697 | -0.01562704 | 0.37534751 | -0.0416335 | 0.96679085 | 0.97827413 | NA | NA |
| ENSG000001 | 11.6714237 | 0.01355959 | 0.32636233 | 0.04154766 | 0.9668593 | 0.97831898 | MAP1LC3A | 84557 |

| | | | | | | | | |
|---|---|---|---|---|---|---|---|---|
| ENSG000002 | 3.65214076 | 0.015672295 | 0.37852477 | 0.04140362 | 0.96697413 | 0.97841076 | NA | NA |
| ENSG000002 | 2.53640537 | -0.015390501 | 0.37218475 | -0.0413518 | 0.96701546 | 0.97842816 | NA | NA |
| ENSG000002 | 11.5326145 | -0.01361534 | 0.33004108 | -0.0412535 | 0.96709383 | 0.97845601 | NA | NA |
| ENSG000002 | 28.1382361 | 0.010418143 | 0.25270535 | 0.04122644 | 0.96711537 | 0.97845601 | ARHGAP8 | 23779 |
| ENSG000002 | 2.32309211 | -0.015096217 | 0.36603739 | -0.0412423 | 0.96710275 | 0.97845601 | NA | NA |
| ENSG000002 | 4.63555103 | -0.01540054 | 0.37571604 | -0.0409898 | 0.967304 | 0.97859802 | RN7SL434P | 106480509 |
| ENSG000002 | 5.38093249 | -0.015264931 | 0.37237642 | -0.0409933 | 0.96730125 | 0.97859802 | NA | NA |
| ENSG000002 | 4.90630848 | -0.015272362 | 0.37360179 | -0.0408787 | 0.96739258 | 0.97866322 | NA | NA |
| ENSG000002 | 41.256968 | -0.009616259 | 0.23561324 | -0.0408137 | 0.96744438 | 0.97869121 | HCP5 | 10866 |
| ENSG000001 | 5.1449024 | -0.015206105 | 0.37370696 | -0.0406899 | 0.9675431 | 0.97874224 | SERHL | 94009 |
| ENSG000002 | 2.56492249 | -0.015124746 | 0.37156129 | -0.0407059 | 0.96753034 | 0.97874224 | NA | NA |
| ENSG000002 | 5.46343661 | 0.015153884 | 0.3728774 | 0.04064039 | 0.96758258 | 0.97875777 | LY6G5B | 58496 |
| ENSG000001 | 7.69886499 | -0.014371924 | 0.35510621 | -0.0404722 | 0.96771668 | 0.978869 | RPL3L | 6123 |
| ENSG000002 | 5.06864081 | 0.015111004 | 0.37381861 | 0.04042336 | 0.96775561 | 0.97888395 | NA | NA |
| ENSG000001 | 175.405808 | -0.005694724 | 0.14136611 | -0.0402835 | 0.9678671 | 0.9789723 | RAD51C | 5889 |
| ENSG000002 | 37.9844623 | 0.009364028 | 0.23341417 | 0.04011765 | 0.96799933 | 0.97908163 | LOC339803 | 339803 |
| ENSG000002 | 4.89179315 | -0.014978796 | 0.37409774 | -0.0400398 | 0.9680614 | 0.97911999 | RN7SL19P | 106480370 |
| ENSG000002 | 2.79226759 | -0.01496404 | 0.37440127 | -0.0399679 | 0.9681187 | 0.97915353 | RNU6-1237P | 106480651 |
| ENSG000002 | 2.60336837 | -0.014862313 | 0.37431395 | -0.0397055 | 0.96832794 | 0.97934072 | NA | NA |
| ENSG000002 | 5.26657014 | 0.014582226 | 0.37170404 | 0.03923074 | 0.96870642 | 0.97962578 | SRRM5 | 100170229 |
| ENSG000002 | 4.68506163 | -0.014808188 | 0.37727999 | -0.0392499 | 0.96869118 | 0.97962578 | NA | NA |
| ENSG000002 | 8.77145326 | 0.013707028 | 0.34934514 | 0.03923635 | 0.96870195 | 0.97962578 | LOC100289473 | 100289473 |
| ENSG000002 | 39.5324903 | -0.009226923 | 0.23510242 | -0.0392464 | 0.96869394 | 0.97962578 | NA | NA |
| ENSG000002 | 3.65637523 | 0.014835422 | 0.3785579 | 0.03918931 | 0.96873946 | 0.97963476 | NA | NA |
| ENSG000002 | 5.11630074 | -0.01455815 | 0.37416514 | -0.0389084 | 0.96896346 | 0.97981241 | NA | NA |
| ENSG000002 | 2.58017559 | -0.014534938 | 0.37352857 | -0.0389125 | 0.96896013 | 0.97981241 | NA | NA |
| ENSG000001 | 13.0601699 | 0.012263328 | 0.31548473 | 0.03887139 | 0.96899293 | 0.97981778 | NA | NA |
| ENSG000001 | 5.47226344 | 0.014347824 | 0.37011974 | 0.03876536 | 0.96907747 | 0.97985574 | TUBA3FP | 113691 |
| ENSG000002 | 4.64963693 | -0.014548787 | 0.37532006 | -0.0387637 | 0.9690788 | 0.97985574 | NA | NA |

| | | | | | | | | |
|---|---|---|---|---|---|---|---|---|
| ENSG000001 | 21.3854356 | -0.010711446 | 0.27982034 | -0.0382797 | 0.96946465 | 0.98009925 | NAGS | 162417 |
| ENSG000001 | 45.2643655 | 0.008169991 | 0.21280293 | 0.03839228 | 0.96937491 | 0.98009925 | FRZB | 2487 |
| ENSG000002 | 3.39355018 | 0.014509489 | 0.37828735 | 0.03835573 | 0.96940405 | 0.98009925 | NA | NA |
| ENSG000002 | 2.32014282 | -0.014054898 | 0.36673552 | -0.0383243 | 0.96942908 | 0.98009925 | GLIS3-AS1 | 84850 |
| ENSG000002 | 43.481802 | 0.008583123 | 0.22404341 | 0.03831009 | 0.96944044 | 0.98009925 | NA | NA |
| ENSG000002 | 4.21544952 | -0.014474612 | 0.37806059 | -0.0382865 | 0.96945927 | 0.98009925 | NA | NA |
| ENSG000002 | 2.7912772 | -0.014272897 | 0.37367468 | -0.0381961 | 0.96953137 | 0.98011647 | ASPDH | 554235 |
| ENSG000002 | 4.69056244 | -0.014299113 | 0.37464182 | -0.0381674 | 0.96955419 | 0.98011647 | NA | NA |
| ENSG000002 | 4.86494598 | -0.014288313 | 0.37377135 | -0.0382274 | 0.96950636 | 0.98011647 | NA | NA |
| ENSG000002 | 5.1683559 | -0.014349254 | 0.37705287 | -0.0380563 | 0.96964276 | 0.98018157 | NA | NA |
| ENSG000002 | 3.64427208 | 0.014279552 | 0.37838381 | 0.03773828 | 0.96989636 | 0.98041348 | ARL14EPL | 644100 |
| ENSG000002 | 3.45218826 | 0.014275711 | 0.37859157 | 0.03770742 | 0.96992096 | 0.98041391 | NA | NA |
| ENSG000001 | 2.56742717 | -0.014069488 | 0.3744647 | -0.0375723 | 0.97002872 | 0.98049839 | NANOS3 | 342977 |
| ENSG000002 | 12.7415985 | 0.012176406 | 0.32572085 | 0.03738295 | 0.97017967 | 0.98062653 | BSN-AS2 | 100132677 |
| ENSG000001 | 292.815167 | 0.005441289 | 0.14627789 | 0.0371983 | 0.9703269 | 0.9807509 | PIK3C2B | 5287 |
| ENSG000001 | 8.63573888 | -0.013197881 | 0.35537748 | -0.0371376 | 0.97037526 | 0.98077533 | NA | NA |
| ENSG000002 | 6.82035233 | -0.013441775 | 0.36261567 | -0.0370689 | 0.97043005 | 0.98078182 | HTR2A-AS1 | 100874082 |
| ENSG000002 | 2.94093771 | 0.01388283 | 0.37443936 | 0.03707631 | 0.97042416 | 0.98078182 | NA | NA |
| ENSG000001 | 29.5615379 | -0.009357166 | 0.2545333 | -0.0367621 | 0.97067473 | 0.98098021 | CPO | 130749 |
| ENSG000001 | 279.092966 | 0.005118847 | 0.13921851 | 0.03676844 | 0.97066964 | 0.98098021 | RAPH1 | 65059 |
| ENSG000002 | 19.0448753 | 0.010331183 | 0.28367772 | 0.03641873 | 0.97094848 | 0.98123241 | NOL12 | 79159 |
| ENSG000001 | 26.9396776 | 0.009463976 | 0.26039276 | 0.03634501 | 0.97100726 | 0.98125563 | EVA1C | 59271 |
| ENSG000002 | 21.1664225 | -0.010248442 | 0.28209916 | -0.0363292 | 0.97101985 | 0.98125563 | NA | NA |
| ENSG000001 | 18.7566781 | 0.010611814 | 0.29436887 | 0.03604938 | 0.97124299 | 0.98139852 | LYPD3 | 27076 |
| ENSG000001 | 57.1327576 | 0.007116824 | 0.19736645 | 0.03605893 | 0.97123537 | 0.98139852 | FAM71F1 | 84691 |
| ENSG000001 | 50.5197255 | 0.007893931 | 0.2190904 | 0.03603048 | 0.97125806 | 0.98139852 | PIDD1 | 55367 |
| ENSG000002 | 3.2069918 | 0.013605315 | 0.3775107 | 0.03603955 | 0.97125082 | 0.98139852 | NA | NA |
| ENSG000002 | 2.33122115 | -0.013285817 | 0.3700141 | -0.0359062 | 0.97135712 | 0.9814497 | NA | NA |
| ENSG000002 | 2.57032394 | -0.013455747 | 0.37473549 | -0.0359073 | 0.97135626 | 0.9814497 | NA | NA |

| | | | | | | | | |
|---|---|---|---|---|---|---|---|---|
| ENSG000002 | 7.18976135 | -0.012750715 | 0.35635635 | -0.0357808 | 0.97145714 | 0.98152631 | NA | NA |
| ENSG000001 | 36.5343484 | 0.008600698 | 0.24077687 | 0.03572061 | 0.97150513 | 0.98155034 | ANO9 | 338440 |
| ENSG000001 | 4.63523782 | -0.01335735 | 0.37631034 | -0.0354956 | 0.97168458 | 0.98170719 | LBX2 | 85474 |
| ENSG000001 | 10.6747446 | 0.012075392 | 0.34178534 | 0.03533034 | 0.97181633 | 0.98181584 | RASGEF1C | 255426 |
| ENSG000002 | 6.50391805 | -0.012850823 | 0.36484312 | -0.0352229 | 0.97190202 | 0.98185349 | HAUS7 | 55559 |
| ENSG000002 | 5.04367246 | 0.013191937 | 0.37443415 | 0.03523166 | 0.97189502 | 0.98185349 | RNU6-583P | 106479824 |
| ENSG000001 | 3.96085787 | -0.013193429 | 0.37847082 | -0.0348598 | 0.97219151 | 0.98204808 | KRT85 | 3891 |
| ENSG000002 | 26.5013802 | -0.009176399 | 0.26279093 | -0.034919 | 0.97214432 | 0.98204808 | SEC14L6 | 730005 |
| ENSG000002 | 2.58598885 | -0.013009033 | 0.37314664 | -0.0348631 | 0.97218894 | 0.98204808 | NA | NA |
| ENSG000002 | 4.37919213 | -0.013177519 | 0.37709948 | -0.0349444 | 0.97212407 | 0.98204808 | NA | NA |
| ENSG000002 | 4.39976902 | -0.013075168 | 0.37618685 | -0.0347571 | 0.97227342 | 0.98209308 | NA | NA |
| ENSG000002 | 16.1560866 | 0.010567439 | 0.30442449 | 0.03471284 | 0.97230872 | 0.98209308 | NA | NA |
| ENSG000002 | 3.20594159 | 0.013131885 | 0.37806795 | 0.03473419 | 0.9722917 | 0.98209308 | NA | NA |
| ENSG000002 | 2.77668277 | -0.013003731 | 0.37583114 | -0.0345999 | 0.97239876 | 0.98215956 | NA | NA |
| ENSG000001 | 8.42161595 | -0.012078129 | 0.35090362 | -0.0344201 | 0.97254218 | 0.98226003 | OR10AG1 | 282770 |
| ENSG000001 | 15.7968255 | -0.010694858 | 0.31076853 | -0.0344142 | 0.97254684 | 0.98226003 | GPR1 | 2825 |
| ENSG000002 | 5.31224733 | -0.012927577 | 0.37597587 | -0.0343841 | 0.97257089 | 0.98226003 | NA | NA |
| ENSG000001 | 22.044713 | -0.009310692 | 0.27324176 | -0.0340749 | 0.97281741 | 0.98248452 | GPR157 | 80045 |
| ENSG000001 | 13.8588202 | -0.010666058 | 0.31454749 | -0.0339092 | 0.97294955 | 0.98259351 | DUSP8 | 1850 |
| ENSG000002 | 2.53457862 | -0.012620603 | 0.37441989 | -0.0337071 | 0.97311073 | 0.98273181 | NA | NA |
| ENSG000001 | 47.4105801 | -0.006973277 | 0.20819952 | -0.0334932 | 0.97328125 | 0.98283518 | TOM1L2 | 146691 |
| ENSG000001 | 50.1894283 | 0.007119853 | 0.21244785 | 0.03351342 | 0.97326517 | 0.98283518 | PIPSL | 266971 |
| ENSG000002 | 26.8249919 | 0.008529274 | 0.25469983 | 0.03348755 | 0.97328579 | 0.98283518 | ANKHD1-EIF4EE | 404734 |
| ENSG000002 | 2.53711032 | -0.012451297 | 0.37251482 | -0.033425 | 0.9733357 | 0.9828611 | NA | NA |
| ENSG000001 | 24.0801198 | 0.009358508 | 0.2812482 | 0.03327491 | 0.97345536 | 0.98286922 | ZFPM1 | 161882 |
| ENSG000001 | 2.34295161 | -0.012272797 | 0.36811422 | -0.0333396 | 0.97340374 | 0.98286922 | OR6T1 | 219874 |
| ENSG000002 | 24.9855241 | 0.008585444 | 0.25810868 | 0.0332629 | 0.97346494 | 0.98286922 | CORO7 | 79585 |
| ENSG000002 | 5.1130522 | -0.012414242 | 0.37207828 | -0.0333646 | 0.97338384 | 0.98286922 | NA | NA |
| ENSG000002 | 9.29235254 | -0.011293953 | 0.33939828 | -0.0332764 | 0.97345417 | 0.98286922 | NA | NA |

| | | | | | | | | |
|---|---|---|---|---|---|---|---|---|
| ENSG000002 | 2.32467809 | -0.012315734 | 0.371129 | -0.0331845 | 0.97352745 | 0.98290787 | NA | NA |
| ENSG000002 | 2.29616097 | -0.012247812 | 0.36948727 | -0.0331481 | 0.97355646 | 0.98291269 | NA | NA |
| ENSG000002 | 3.70091436 | 0.012492973 | 0.37854803 | 0.03300235 | 0.97367272 | 0.98300559 | KRTAP21-3 | 100288323 |
| ENSG000001 | 12.658891 | 0.010859569 | 0.32949898 | 0.03295782 | 0.97370822 | 0.98301696 | C2orf78 | 388960 |
| ENSG000002 | 4.15601134 | 0.012435264 | 0.37844144 | 0.03285915 | 0.97378691 | 0.98307192 | NA | NA |
| ENSG000002 | 2.31183828 | -0.012005435 | 0.36803023 | -0.0326208 | 0.97397699 | 0.98323934 | NA | NA |
| ENSG000002 | 3.6696046 | 0.012236698 | 0.37838803 | 0.03233902 | 0.97420169 | 0.98344169 | NA | NA |
| ENSG000002 | 9.32698206 | 0.011360341 | 0.35164016 | 0.03230672 | 0.97422745 | 0.98344321 | FAR1-IT1 | 106478978 |
| ENSG000002 | 7.22878823 | -0.011586193 | 0.35956955 | -0.0322224 | 0.97429469 | 0.98348661 | RN7SL145P | 106481753 |
| ENSG000002 | 6.47625004 | -0.011662429 | 0.36316775 | -0.0321131 | 0.97438189 | 0.98355014 | MMP28 | 79148 |
| ENSG000002 | 2.32214638 | -0.011892013 | 0.37167821 | -0.0319955 | 0.97447568 | 0.98362033 | RN7SL711P | 106479482 |
| ENSG000001 | 210.71557 | 0.005100606 | 0.159963 | 0.03188616 | 0.97456284 | 0.98368382 | NA | NA |
| ENSG000002 | 4.4325532 | -0.01200066 | 0.37850481 | -0.0317054 | 0.97470696 | 0.98380481 | NA | NA |
| ENSG000002 | 6.1148144 | -0.011715683 | 0.37239369 | -0.0314605 | 0.97490231 | 0.98397749 | NA | NA |
| ENSG000002 | 10.4842392 | -0.010543073 | 0.33595241 | -0.0313826 | 0.97496439 | 0.98401565 | NA | NA |
| ENSG000002 | 3.22091429 | 0.011698857 | 0.37385939 | 0.03129213 | 0.97503657 | 0.98406401 | NA | NA |
| ENSG000002 | 2.30711455 | -0.01154495 | 0.37070414 | -0.0311433 | 0.97515526 | 0.98415931 | NA | NA |
| ENSG000002 | 2.3424629 | -0.011482163 | 0.36910001 | -0.0311085 | 0.97518298 | 0.98416279 | NA | NA |
| ENSG000002 | 3.15540575 | 0.011606648 | 0.37403799 | 0.03103067 | 0.97524508 | 0.98420097 | NA | NA |
| ENSG00000C | 64.9645645 | -0.006127001 | 0.19873734 | -0.0308296 | 0.9754054 | 0.98432722 | TLE2 | 7089 |
| ENSG000001 | 7.3198905 | 0.010917636 | 0.35431998 | 0.03081293 | 0.97541873 | 0.98432722 | LOC113230 | 113230 |
| ENSG000002 | 6.28091747 | -0.011254797 | 0.36635548 | -0.030721 | 0.97549206 | 0.98437672 | NA | NA |
| ENSG000002 | 3.46946399 | 0.011602816 | 0.37847081 | 0.03065709 | 0.97554301 | 0.98440364 | LOC105370014 | 105370014 |
| ENSG000001 | 473.628119 | 0.003534398 | 0.11569233 | 0.03054998 | 0.97562843 | 0.98446534 | NRXN1 | 9378 |
| ENSG000001 | 52.4590279 | 0.006428412 | 0.2109498 | 0.03047366 | 0.9756893 | 0.98450226 | APLP1 | 333 |
| ENSG000002 | 3.04762889 | -0.011307273 | 0.37534281 | -0.0301252 | 0.97596721 | 0.98475818 | NA | NA |
| ENSG000002 | 2.84877403 | -0.011123159 | 0.37034854 | -0.0300343 | 0.9760397 | 0.98480681 | NA | NA |
| ENSG000002 | 13.6930239 | 0.009425189 | 0.31458903 | 0.02996032 | 0.9760987 | 0.98484184 | NA | NA |
| ENSG000002 | 2.30873278 | -0.010917671 | 0.36720478 | -0.0297318 | 0.97628092 | 0.98500119 | NA | NA |

| | | | | | | | | |
|---|---|---|---|---|---|---|---|---|
| ENSG000001 | 17.6395565 | 0.008632783 | 0.29099057 | 0.02966688 | 0.97633272 | 0.98502894 | SNAI3 | 333929 |
| ENSG000001 | 33.7939047 | -0.007349503 | 0.24945128 | -0.0294627 | 0.97649558 | 0.98516252 | TNNC2 | 7125 |
| ENSG000002 | 4.13720206 | 0.011114078 | 0.3782997 | 0.02937903 | 0.9765623 | 0.98516252 | NA | NA |
| ENSG000002 | 3.45401501 | 0.011131097 | 0.37857385 | 0.02940271 | 0.97654341 | 0.98516252 | MDC1-AS1 | 106478956 |
| ENSG000002 | 4.68420064 | -0.011087154 | 0.37719408 | -0.0293938 | 0.97655055 | 0.98516252 | NA | NA |
| ENSG000002 | 2.55768742 | -0.010989619 | 0.37471743 | -0.0293277 | 0.9766032 | 0.98517927 | NA | NA |
| ENSG000002 | 2.31858463 | -0.010880329 | 0.37201043 | -0.0292474 | 0.9766673 | 0.98519491 | NA | NA |
| ENSG000002 | 2.57189031 | -0.010939227 | 0.37398536 | -0.0292504 | 0.97666487 | 0.98519491 | NA | NA |
| ENSG000002 | 2.30583655 | -0.010808153 | 0.37008538 | -0.0292045 | 0.9767015 | 0.98520491 | NA | NA |
| ENSG000000 | 211.991842 | -0.004249535 | 0.14604461 | -0.0290975 | 0.97678682 | 0.98521745 | LAMC2 | 3918 |
| ENSG000001 | 13.7539529 | 0.008993248 | 0.30900138 | 0.02910423 | 0.97678146 | 0.98521745 | PTGER2 | 5732 |
| ENSG000002 | 7.30355528 | 0.010396017 | 0.35687339 | 0.02913083 | 0.97676025 | 0.98521745 | SNORA20 | 677806 |
| ENSG000001 | 9.39583815 | 0.009780521 | 0.33684598 | 0.02903559 | 0.97683621 | 0.98524276 | FAM19A3 | 284467 |
| ENSG000002 | 2.57370964 | -0.010753651 | 0.37241593 | -0.0288754 | 0.97696398 | 0.98534712 | LOC101928108 | 101928108 |
| ENSG000002 | 6.12067716 | -0.010800454 | 0.37487313 | -0.028811 | 0.97701536 | 0.98537444 | NA | NA |
| ENSG000000 | 106.458975 | 0.005027755 | 0.17489617 | 0.02874709 | 0.9770663 | 0.9854013 | CASP10 | 843 |
| ENSG000002 | 2.30350072 | -0.010623623 | 0.37162927 | -0.0285866 | 0.97719429 | 0.98550587 | NA | NA |
| ENSG000002 | 2.60358406 | -0.01049125 | 0.36794846 | -0.0285128 | 0.97725314 | 0.98551546 | NA | NA |
| ENSG000002 | 14.5267297 | -0.008661828 | 0.30410218 | -0.0284833 | 0.9772767 | 0.98551546 | LOC283683 | 283683 |
| ENSG000002 | 10.4565315 | -0.009559894 | 0.33523309 | -0.0285172 | 0.97724969 | 0.98551546 | NA | NA |
| ENSG000000 | 36.7658279 | 0.006773722 | 0.24007732 | 0.02821475 | 0.97749087 | 0.9856379 | ARHGAP33 | 115703 |
| ENSG000002 | 6.54848497 | -0.010349668 | 0.36671971 | -0.0282223 | 0.97748486 | 0.9856379 | ABCC6P2 | 730013 |
| ENSG000002 | 2.8272512 | -0.010497891 | 0.3723029 | -0.0281972 | 0.97750489 | 0.9856379 | NA | NA |
| ENSG000002 | 2.76990417 | -0.010532196 | 0.37376486 | -0.0281787 | 0.97751965 | 0.9856379 | NA | NA |
| ENSG000002 | 3.65663526 | 0.010710844 | 0.37857639 | 0.02829243 | 0.97742892 | 0.9856379 | NA | NA |
| ENSG000002 | 34.3908931 | -0.006794676 | 0.24417931 | -0.0278266 | 0.97780046 | 0.98589653 | NA | NA |
| ENSG000002 | 10.8270977 | -0.009160768 | 0.33205768 | -0.0275879 | 0.97799085 | 0.98606397 | NUTM2D | 728130 |
| ENSG000001 | 2.56154256 | -0.010167049 | 0.37013897 | -0.0274682 | 0.97808631 | 0.9861357 | NA | NA |
| ENSG000002 | 2.36341213 | -0.010053464 | 0.36737076 | -0.027366 | 0.97816783 | 0.98619337 | NA | NA |

| | | | | | | | | |
|---|---|---|---|---|---|---|---|---|
| ENSG000002 | 6.28631213 | -0.010175257 | 0.37357785 | -0.0272373 | 0.97827045 | 0.98627232 | NA | NA |
| ENSG000002 | 10.8827937 | -0.008927252 | 0.32879211 | -0.0271517 | 0.97833877 | 0.98631667 | LINC01592 | 100505718 |
| ENSG000001 | 3.89866729 | 0.010205869 | 0.37789928 | 0.02700685 | 0.97845427 | 0.98639675 | RNF151 | 146310 |
| ENSG000002 | 8.23304131 | 0.009474276 | 0.3510152 | 0.02699107 | 0.97846686 | 0.98639675 | NA | NA |
| ENSG000002 | 4.63188513 | -0.010130794 | 0.37641554 | -0.0269139 | 0.97852844 | 0.98643432 | NA | NA |
| ENSG000002 | 4.39597913 | -0.010098653 | 0.37569928 | -0.0268796 | 0.97855575 | 0.98643732 | NA | NA |
| ENSG000002 | 3.88976913 | 0.010139274 | 0.37840527 | 0.02679475 | 0.97862344 | 0.98645652 | NA | NA |
| ENSG000002 | 131.138221 | 0.004054523 | 0.15116126 | 0.0268225 | 0.97860131 | 0.98645652 | TYW1B | 441250 |
| ENSG000002 | 3.44274081 | 0.010105741 | 0.37849905 | 0.02669951 | 0.9786994 | 0.98650856 | NA | NA |
| ENSG000002 | 5.53696218 | 0.00981296 | 0.37078647 | 0.02646526 | 0.97888624 | 0.98667237 | NA | NA |
| ENSG000002 | 9.34577282 | 0.009216857 | 0.34967024 | 0.02635871 | 0.97897123 | 0.9867335 | NA | NA |
| ENSG000002 | 14.5838559 | -0.008033068 | 0.3058144 | -0.0262678 | 0.97904375 | 0.98678207 | NA | NA |
| ENSG000002 | 26.1296279 | 0.006807603 | 0.25980101 | 0.02620314 | 0.97909531 | 0.98680951 | NA | NA |
| ENSG00000C | 201.088292 | 0.003643944 | 0.13944204 | 0.02613232 | 0.9791518 | 0.98683737 | ZFAT | 57623 |
| ENSG000001 | 42.3574953 | 0.006542511 | 0.2508924 | 0.02607696 | 0.97919596 | 0.98683737 | CYP27C1 | 339761 |
| ENSG000002 | 3.63505296 | 0.009879728 | 0.37848737 | 0.02610319 | 0.97917503 | 0.98683737 | NA | NA |
| ENSG000002 | 12.2886902 | 0.00865193 | 0.3328868 | 0.02599061 | 0.97926483 | 0.98688225 | NA | NA |
| ENSG000002 | 11.7443033 | 0.008309327 | 0.32078284 | 0.02590328 | 0.97933449 | 0.98692792 | LOC101929066 | 101929066 |
| ENSG000002 | 49.2437709 | -0.005504294 | 0.21339981 | -0.0257933 | 0.97942217 | 0.98699176 | NA | NA |
| ENSG000002 | 3.98850194 | -0.009744579 | 0.378593 | -0.0257389 | 0.97946557 | 0.98701097 | NA | NA |
| ENSG000002 | 4.69906396 | -0.009646751 | 0.37603013 | -0.0256542 | 0.97953316 | 0.98705455 | CLDN10-AS1 | 100874194 |
| ENSG000002 | 2.53435049 | -0.009560504 | 0.37397924 | -0.0255643 | 0.97960489 | 0.98709964 | LOC101927159 | 101927159 |
| ENSG000002 | 12.7305576 | -0.00831737 | 0.32569809 | -0.0255371 | 0.97962659 | 0.98709964 | NA | NA |
| ENSG000001 | 15.6778142 | 0.007547856 | 0.29907833 | 0.02523705 | 0.97986588 | 0.98731622 | ZFP92 | 139735 |
| ENSG000001 | 298.245249 | -0.003450256 | 0.13818514 | -0.0249684 | 0.9800802 | 0.98750084 | NME7 | 29922 |
| ENSG000002 | 11.0382235 | 0.008173472 | 0.32764289 | 0.02494628 | 0.98009781 | 0.98750084 | NA | NA |
| ENSG000002 | 37.5162113 | 0.005978408 | 0.24019333 | 0.02488998 | 0.98014272 | 0.98752155 | LOC102724885 | 102724885 |
| ENSG000002 | 4.41938077 | -0.009338737 | 0.37586197 | -0.0248462 | 0.98017765 | 0.98753221 | NA | NA |
| ENSG000001 | 31.3312709 | -0.006309284 | 0.25470991 | -0.0247705 | 0.98023805 | 0.98754724 | SCHIP1 | 29970 |

| | | | | | | | | |
|---|---|---|---|---|---|---|---|---|
| ENSG000002 | 2.55222703 | -0.009273567 | 0.37444106 | -0.0247664 | 0.98024127 | 0.98754724 | LOC105371426 | 105371426 |
| ENSG000002 | 6.25523251 | -0.009046102 | 0.3661573 | -0.0247055 | 0.98028986 | 0.98757165 | NA | NA |
| ENSG000001 | 17.9841747 | -0.007082784 | 0.2888821 | -0.0245179 | 0.9804395 | 0.98767333 | LOC648987 | 648987 |
| ENSG000002 | 5.27432638 | 0.009178734 | 0.37391986 | 0.02454733 | 0.98041603 | 0.98767333 | NA | NA |
| ENSG000001 | 4.46445622 | -0.0091006 | 0.37660151 | -0.0241651 | 0.98072094 | 0.9878036 | DNMT3L | 29947 |
| ENSG000001 | 26.766069 | -0.006161213 | 0.25520717 | -0.024142 | 0.98073934 | 0.9878036 | HCN3 | 57657 |
| ENSG000001 | 3.67450439 | 0.009159076 | 0.3784581 | 0.02420103 | 0.98069226 | 0.9878036 | ALPI | 248 |
| ENSG000001 | 21.0042796 | -0.00699444 | 0.28755449 | -0.0243239 | 0.98059427 | 0.9878036 | LINC00910 | 100130581 |
| ENSG000002 | 3.39796537 | 0.009141247 | 0.37783928 | 0.02419348 | 0.98069828 | 0.9878036 | NA | NA |
| ENSG000002 | 4.38997741 | -0.0090754 | 0.37589812 | -0.0241432 | 0.98073835 | 0.9878036 | NA | NA |
| ENSG000002 | 3.70282564 | 0.009178375 | 0.37853424 | 0.02424715 | 0.98065547 | 0.9878036 | NA | NA |
| ENSG000002 | 30.4492607 | -0.005781409 | 0.24082093 | -0.0240071 | 0.98084696 | 0.98788746 | HMSD | 284293 |
| ENSG000001 | 11.8264236 | 0.007836434 | 0.32854802 | 0.02385171 | 0.98097089 | 0.98796245 | SERP2 | 387923 |
| ENSG000002 | 5.51245064 | 0.008839862 | 0.37155445 | 0.02379157 | 0.98101887 | 0.98796245 | NA | NA |
| ENSG000002 | 36.3444491 | 0.005538779 | 0.23260298 | 0.02381216 | 0.98100244 | 0.98796245 | NA | NA |
| ENSG000002 | 12.0599621 | -0.007771675 | 0.32544724 | -0.02388 | 0.98094834 | 0.98796245 | NA | NA |
| ENSG000002 | 16.6123009 | -0.006929661 | 0.29270861 | -0.0236743 | 0.98111243 | 0.98803214 | NA | NA |
| ENSG000002 | 3.44805739 | 0.008912715 | 0.37854237 | 0.02354483 | 0.98121568 | 0.98811158 | NA | NA |
| ENSG000002 | 5.22100488 | 0.008790761 | 0.37405953 | 0.02350097 | 0.98125067 | 0.98812227 | MS4A18 | 728588 |
| ENSG000002 | 4.38681263 | -0.008811286 | 0.37602011 | -0.023433 | 0.98130487 | 0.98812802 | NA | NA |
| ENSG000002 | 2.55208322 | -0.008751761 | 0.37348479 | -0.0234327 | 0.98130511 | 0.98812802 | NA | NA |
| ENSG000001 | 25.3941464 | -0.006315895 | 0.27104557 | -0.023302 | 0.9814094 | 0.9882085 | TPPP | 11076 |
| ENSG000001 | 11.0745622 | 0.007585402 | 0.33112807 | 0.02290776 | 0.98172385 | 0.98845149 | PPAPDC3 | 84814 |
| ENSG000002 | 4.65555488 | -0.008593812 | 0.37443638 | -0.0229513 | 0.9816891 | 0.98845149 | NA | NA |
| ENSG000002 | 13.798825 | 0.007146076 | 0.31189064 | 0.02291212 | 0.98172037 | 0.98845149 | NA | NA |
| ENSG000002 | 8.03481118 | 0.008010707 | 0.35048557 | 0.02285602 | 0.98176512 | 0.9884685 | NA | NA |
| ENSG000002 | 3.43582582 | 0.008620512 | 0.37847605 | 0.0227769 | 0.98182823 | 0.9885075 | NA | NA |
| ENSG000001 | 41.5822874 | 0.005145514 | 0.22627478 | 0.02274011 | 0.98185758 | 0.98851251 | SLC16A6 | 9120 |
| ENSG000002 | 15.5897969 | 0.006804674 | 0.30214465 | 0.02252125 | 0.98203216 | 0.98863918 | NA | NA |

| | | | | | | | | |
|---|---|---|---|---|---|---|---|---|
| ENSG000002 | 4.41299392 | -0.008477586 | 0.37607945 | -0.022542 | 0.9820156 | 0.98863918 | NA | NA |
| ENSG000000 | 18.5346679 | -0.006764791 | 0.30269071 | -0.0223489 | 0.98216968 | 0.98875308 | VPS9D1 | 9605 |
| ENSG000001 | 203.343381 | 0.003172328 | 0.14250707 | 0.02226085 | 0.98223988 | 0.9887992 | TIAM2 | 26230 |
| ENSG000002 | 5.98733938 | -0.008159479 | 0.36705937 | -0.0222293 | 0.98226503 | 0.98879997 | NA | NA |
| ENSG000001 | 2.35264701 | -0.008246186 | 0.37175073 | -0.022182 | 0.98230275 | 0.9888134 | NA | NA |
| ENSG000000 | 11.217799 | -0.007448501 | 0.33673797 | -0.0221196 | 0.98235257 | 0.988839 | NA | NA |
| ENSG000002 | 2.56614951 | -0.008219738 | 0.37329538 | -0.0220194 | 0.98243249 | 0.98889489 | NA | NA |
| ENSG000001 | 21.4189159 | -0.005877868 | 0.27114512 | -0.0216779 | 0.98270486 | 0.98889904 | MIXL1 | 83881 |
| ENSG000002 | 3.44110994 | 0.008246318 | 0.37856264 | 0.02178323 | 0.98262087 | 0.98889904 | NA | NA |
| ENSG000002 | 3.68924806 | 0.008209441 | 0.37846075 | 0.02169166 | 0.98269392 | 0.98889904 | LOC101929565 | 101929565 |
| ENSG000002 | 4.38469328 | -0.008271729 | 0.37757763 | -0.0219074 | 0.98252185 | 0.98889904 | NA | NA |
| ENSG000002 | 3.67811822 | 0.008271212 | 0.37849273 | 0.02185303 | 0.98256519 | 0.98889904 | NA | NA |
| ENSG000002 | 4.68412441 | -0.008184888 | 0.37755066 | -0.0216789 | 0.98270408 | 0.98889904 | NA | NA |
| ENSG000002 | 283.991211 | -0.003181433 | 0.14646532 | -0.0217214 | 0.98267019 | 0.98889904 | NA | NA |
| ENSG000002 | 6.20718349 | -0.007930461 | 0.36467676 | -0.0217465 | 0.98265013 | 0.98889904 | SNORD92 | 692209 |
| ENSG000002 | 2.52815289 | -0.008110175 | 0.37392241 | -0.0216895 | 0.98269567 | 0.98889904 | NA | NA |
| ENSG000002 | 4.20677227 | -0.008206764 | 0.37810826 | -0.0217048 | 0.98268343 | 0.98889904 | NA | NA |
| ENSG000002 | 5.07629954 | 0.00825234 | 0.37651439 | 0.02191773 | 0.98251358 | 0.98889904 | NA | NA |
| ENSG000001 | 46.232502 | -0.004865477 | 0.22534318 | -0.0215914 | 0.98277388 | 0.98891942 | MGAT5B | 146664 |
| ENSG000002 | 10.0010363 | 0.007471904 | 0.34599454 | 0.02159544 | 0.98277067 | 0.98891942 | HM13-AS1 | 100874042 |
| ENSG000002 | 4.41397245 | -0.008073922 | 0.37593436 | -0.0214769 | 0.98286519 | 0.98898676 | LOC101929516 | 101929516 |
| ENSG000002 | 11.7485251 | 0.007028309 | 0.32830528 | 0.02140785 | 0.98292031 | 0.98901768 | LINCR-0001 | 101929191 |
| ENSG000002 | 4.4315556 | -0.008009007 | 0.37610356 | -0.0212947 | 0.98301058 | 0.98905943 | NA | NA |
| ENSG000002 | 29.3190061 | -0.005305806 | 0.24901491 | -0.0213072 | 0.98300061 | 0.98905943 | NA | NA |
| ENSG000001 | 25.9255895 | -0.005604224 | 0.26372489 | -0.0212503 | 0.98304602 | 0.98906176 | FGF16 | 8823 |
| ENSG000001 | 2.32198994 | -0.00785784 | 0.37011808 | -0.0212306 | 0.98306168 | 0.98906176 | RNU6-170P | 106479641 |
| ENSG000002 | 2.32466545 | -0.007694742 | 0.36334131 | -0.0211777 | 0.98310388 | 0.98907968 | NA | NA |
| ENSG000002 | 5.55604481 | 0.007829435 | 0.37040489 | 0.0211375 | 0.98313597 | 0.98908742 | NA | NA |
| ENSG000001 | 26.8250991 | -0.005242524 | 0.25101262 | -0.0208855 | 0.98333699 | 0.98926512 | PCDHB10 | 56126 |

| | | | | | | | | |
|---|---|---|---|---|---|---|---|---|
| ENSG000002 | 3.88600354 | 0.007871252 | 0.37798908 | 0.02082402 | 0.98338604 | 0.98928992 | NA | NA |
| ENSG000001 | 567.21141 | 0.002616179 | 0.12582273 | 0.02079257 | 0.98341112 | 0.98929061 | C1orf21 | 81563 |
| ENSG000001 | 42.2261625 | -0.004703538 | 0.22657339 | -0.0207594 | 0.98343755 | 0.98929265 | NA | NA |
| ENSG000001 | 55.1739869 | -0.00415766 | 0.20093982 | -0.0206911 | 0.98349209 | 0.98932298 | ZNF563 | 147837 |
| ENSG000002 | 3.90172051 | 0.007808208 | 0.37826842 | 0.02064198 | 0.98353125 | 0.98933783 | NA | NA |
| ENSG000002 | 3.65383522 | 0.007764176 | 0.37852467 | 0.02051168 | 0.9836352 | 0.98941784 | EIF1P6 | 106480277 |
| ENSG000001 | 32.0628283 | -0.004883133 | 0.23857443 | -0.020468 | 0.98367007 | 0.98942837 | RAB3A | 5864 |
| ENSG000001 | 3.45619385 | 0.007701322 | 0.37847353 | 0.02034838 | 0.98376547 | 0.98943361 | CLEC4A | 50856 |
| ENSG000001 | 120.668232 | -0.003130414 | 0.154038 | -0.0203223 | 0.98378623 | 0.98943361 | ARHGEF3 | 50650 |
| ENSG000002 | 2.33893339 | -0.007513401 | 0.37052133 | -0.0202779 | 0.98382167 | 0.98943361 | NA | NA |
| ENSG000002 | 2.35094425 | -0.007561836 | 0.37055172 | -0.020407 | 0.98371873 | 0.98943361 | NA | NA |
| ENSG000002 | 5.4693587 | 0.007503009 | 0.36971393 | 0.02029409 | 0.98380877 | 0.98943361 | NA | NA |
| ENSG000002 | 4.43875135 | -0.007679699 | 0.37824543 | -0.0203035 | 0.98380128 | 0.98943361 | NA | NA |
| ENSG00000C | 46.6613687 | -0.004442328 | 0.22008865 | -0.0201843 | 0.98389638 | 0.98944607 | ATP2C2 | 9914 |
| ENSG000002 | 2.30605334 | 0.007365894 | 0.36517919 | 0.02017063 | 0.98390726 | 0.98944607 | NA | NA |
| ENSG000002 | 7.12916112 | -0.007364 | 0.36496485 | -0.0201773 | 0.98390195 | 0.98944607 | NA | NA |
| ENSG000002 | 7.51929336 | 0.00711834 | 0.35477377 | 0.02006445 | 0.98399196 | 0.98950671 | NA | NA |
| ENSG000002 | 2.5660057 | -0.007421521 | 0.37127376 | -0.0199894 | 0.98405187 | 0.98951787 | NA | NA |
| ENSG000002 | 4.16808528 | -0.007557334 | 0.37793303 | -0.0199965 | 0.98404617 | 0.98951787 | NA | NA |
| ENSG00000C | 15.4155249 | 0.005961713 | 0.29883341 | 0.01994996 | 0.98408329 | 0.98952494 | NA | NA |
| ENSG000001 | 43.5402986 | -0.004388661 | 0.22057128 | -0.0198968 | 0.98412571 | 0.98954305 | CR1L | 1379 |
| ENSG000001 | 141.865744 | 0.003077616 | 0.15536603 | 0.01980881 | 0.98419589 | 0.98958908 | SKIDA1 | 387640 |
| ENSG000001 | 523.326769 | -0.00246175 | 0.1253527 | -0.0196386 | 0.98433168 | 0.98970107 | CRABP2 | 1382 |
| ENSG000002 | 11.1521616 | -0.006580945 | 0.33621816 | -0.0195734 | 0.98438365 | 0.98972879 | LOC101929231 | 101929231 |
| ENSG000002 | 30.6720091 | -0.005008099 | 0.25904391 | -0.019333 | 0.98457545 | 0.98989708 | NA | NA |
| ENSG000002 | 2.3206078 | -0.007121962 | 0.3692825 | -0.0192859 | 0.984613 | 0.98991029 | NA | NA |
| ENSG000002 | 7.46288696 | 0.006983706 | 0.36351219 | 0.01921175 | 0.98467218 | 0.98994525 | NA | NA |
| ENSG000001 | 10.429705 | -0.006264514 | 0.33217802 | -0.0188589 | 0.98495366 | 0.99020369 | ACOT1 | 641371 |
| ENSG000001 | 81.5072941 | -0.003348129 | 0.17812563 | -0.0187964 | 0.98500349 | 0.99022923 | LOC101926980 | 101926980 |

| | | | | | | | | | |
|---|---|---|---|---|---|---|---|---|---|
| ENSG000002 | 3.65299421 | 0.007084486 | 0.37857638 | 0.01871349 | 0.98506967 | 0.99027121 | NA | NA | |
| ENSG000001 | 4.47473242 | -0.007046263 | 0.37792904 | -0.0186444 | 0.98512478 | 0.99030206 | DEFB126 | | 81623 |
| ENSG000002 | 3.89157527 | 0.007043317 | 0.37846762 | 0.01861009 | 0.98515215 | 0.99030503 | NA | NA | |
| ENSG000002 | 5.98789639 | 0.006742884 | 0.36585465 | 0.0184305 | 0.98529542 | 0.99039995 | NA | NA | |
| ENSG000002 | 2.33409309 | -0.006819784 | 0.3695841 | -0.0184526 | 0.9852778 | 0.99039995 | NA | NA | |
| ENSG000001 | 10.7999999 | 0.006127707 | 0.33339091 | 0.01837995 | 0.98533575 | 0.99041593 | NA | NA | |
| ENSG000001 | 13.7950143 | 0.005654937 | 0.30929082 | 0.01828356 | 0.98541264 | 0.99046867 | PNMA5 | | 114824 |
| ENSG000002 | 4.41251819 | -0.006846865 | 0.37632712 | -0.0181939 | 0.98548415 | 0.990516 | NA | NA | |
| ENSG000002 | 3.91640361 | -0.006779958 | 0.37844112 | -0.0179155 | 0.98570627 | 0.99069014 | NA | NA | |
| ENSG000002 | 8.12161167 | -0.006320368 | 0.35232826 | -0.0179389 | 0.98568763 | 0.99069014 | NA | NA | |
| ENSG000001 | 2.33370133 | -0.006583246 | 0.36958532 | -0.0178125 | 0.98578842 | 0.99074814 | C11orf40 | | 143501 |
| ENSG000002 | 10.0745942 | 0.00595246 | 0.33578382 | 0.01772706 | 0.98585659 | 0.99079211 | NA | NA | |
| ENSG000002 | 236.537883 | -0.002499029 | 0.14153383 | -0.0176568 | 0.98591268 | 0.99082391 | ZNF432 | | 9668 |
| ENSG000002 | 5.24747922 | 0.006539319 | 0.3721863 | 0.01757002 | 0.98598188 | 0.9908689 | NA | NA | |
| ENSG000002 | 5.02398882 | 0.00658854 | 0.37597959 | 0.01752367 | 0.98601885 | 0.99088151 | NA | NA | |
| ENSG000001 | 77.4324344 | 0.003123319 | 0.17910239 | 0.01743874 | 0.98608661 | 0.99090793 | PDGFRB | | 5159 |
| ENSG000001 | 38.6381689 | -0.003978489 | 0.22866439 | -0.0173988 | 0.98611846 | 0.99090793 | FAM57B | | 83723 |
| ENSG000001 | 16.8693169 | 0.005196021 | 0.29838026 | 0.01741409 | 0.98610627 | 0.99090793 | NA | NA | |
| ENSG000000 | 63.7369171 | -0.003365258 | 0.19429071 | -0.0173207 | 0.98618074 | 0.99094596 | RNASET2 | | 8635 |
| ENSG000002 | 3.68384716 | 0.006542082 | 0.37857528 | 0.0172808 | 0.98621261 | 0.99095343 | NA | NA | |
| ENSG000001 | 45.5085916 | -0.003777123 | 0.21905842 | -0.0172425 | 0.98624313 | 0.99095954 | GRK4 | | 2868 |
| ENSG000002 | 3.88672059 | 0.006474498 | 0.3779542 | 0.01713038 | 0.98633261 | 0.99102489 | NA | NA | |
| ENSG000002 | 2.5294711 | -0.006306802 | 0.36931525 | -0.017077 | 0.98637518 | 0.99104311 | NA | NA | |
| ENSG000001 | 55.9947357 | 0.003531878 | 0.2072236 | 0.0170438 | 0.98640167 | 0.99104517 | THBS3 | | 7059 |
| ENSG000002 | 5.80871686 | 0.006233839 | 0.37204543 | 0.01675559 | 0.9866316 | 0.99125162 | NA | NA | |
| ENSG000000 | 30.5813317 | -0.004051957 | 0.24577779 | -0.0164863 | 0.98684646 | 0.99144293 | CACNG4 | | 27092 |
| ENSG000001 | 4.659137 | -0.006125184 | 0.37537314 | -0.0163176 | 0.98698103 | 0.99155355 | TEX12 | | 56158 |
| ENSG000001 | 10.3952027 | 0.005673533 | 0.3495282 | 0.01623197 | 0.98704933 | 0.99157304 | GJB5 | | 2709 |
| ENSG000002 | 2.34589598 | 0.005815791 | 0.35808773 | 0.01624125 | 0.98704193 | 0.99157304 | NA | NA | |

| | | | | | | | | |
|---|---|---|---|---|---|---|---|---|
| ENSG000001 | 22.7970748 | -0.004764512 | 0.29495626 | -0.0161533 | 0.98711211 | 0.99161154 | PRR36 | 80164 |
| ENSG000002 | 4.18212489 | -0.006031903 | 0.37749351 | -0.0159788 | 0.98725128 | 0.99172678 | NA | NA |
| ENSG000002 | 43.1321385 | 0.00371255 | 0.23400327 | 0.01586538 | 0.98734179 | 0.99179313 | TMEM179 | 388021 |
| ENSG000002 | 9.72874697 | -0.005499112 | 0.34980598 | -0.0157205 | 0.9874574 | 0.99188469 | NA | NA |
| ENSG000002 | 15.1180523 | -0.004815451 | 0.30813187 | -0.0156279 | 0.98753126 | 0.99192019 | NA | NA |
| ENSG000002 | 14.7338988 | -0.004798067 | 0.30727606 | -0.0156148 | 0.98754166 | 0.99192019 | LOC653160 | 653160 |
| ENSG000002 | 6.30972296 | -0.005656579 | 0.36554863 | -0.0154742 | 0.98765385 | 0.99200831 | LOC105379194 | 105379194 |
| ENSG000002 | 2.29518299 | -0.005686644 | 0.36976309 | -0.0153792 | 0.98772969 | 0.99203566 | NA | NA |
| ENSG000002 | 2.28421068 | 0.005632461 | 0.36624933 | 0.01537876 | 0.98773001 | 0.99203566 | NA | NA |
| ENSG000001 | 31.116686 | -0.003745813 | 0.24585129 | -0.0152361 | 0.98784383 | 0.99205169 | SOD3 | 6649 |
| ENSG000001 | 25.9188392 | -0.003964085 | 0.25953617 | -0.0152737 | 0.9878138 | 0.99205169 | PNCK | 139728 |
| ENSG000002 | 27.4280401 | 0.003875751 | 0.25290957 | 0.01532465 | 0.98777318 | 0.99205169 | NA | NA |
| ENSG000002 | 4.41580637 | -0.005740254 | 0.37644926 | -0.0152484 | 0.987834 | 0.99205169 | NA | NA |
| ENSG000001 | 122.280144 | 0.002470631 | 0.16359026 | 0.01510256 | 0.98795036 | 0.99211797 | WDR17 | 116966 |
| ENSG000002 | 4.69080352 | -0.005702829 | 0.37787037 | -0.015092 | 0.98795876 | 0.99211797 | NA | NA |
| ENSG000002 | 12.1001418 | 0.00481757 | 0.32157325 | 0.01498125 | 0.98804714 | 0.99218215 | SUGT1P3 | 283507 |
| ENSG000002 | 4.84906125 | -0.005596017 | 0.37447418 | -0.0149437 | 0.98807712 | 0.99218769 | NA | NA |
| ENSG000002 | 3.87763841 | 0.005560241 | 0.37843586 | 0.01469269 | 0.98827735 | 0.99236418 | NA | NA |
| ENSG000002 | 3.23197563 | -0.005396922 | 0.37004697 | -0.0145844 | 0.98836373 | 0.99242633 | NA | NA |
| ENSG000002 | 2.29655327 | -0.005348485 | 0.36976666 | -0.0144645 | 0.98845941 | 0.99248365 | NA | NA |
| ENSG000002 | 20.851615 | -0.004146211 | 0.28690501 | -0.0144515 | 0.98846976 | 0.99248365 | NA | NA |
| ENSG000001 | 2.30076901 | 0.005302724 | 0.36862709 | 0.01438506 | 0.98852277 | 0.9924902 | C7orf61 | 402573 |
| ENSG000002 | 2.30896166 | -0.005349751 | 0.37197576 | -0.014382 | 0.98852523 | 0.9924902 | MIR548H3 | 100302287 |
| ENSG000001 | 24.6265901 | 0.003710832 | 0.26014355 | 0.01426456 | 0.98861892 | 0.992541 | FAM3B | 54097 |
| ENSG000002 | 2.31134956 | -0.005270637 | 0.36968247 | -0.0142572 | 0.98862479 | 0.992541 | NA | NA |
| ENSG000002 | 23.5026754 | 0.003922746 | 0.27734921 | 0.01414371 | 0.98871533 | 0.9925745 | NA | NA |
| ENSG000002 | 4.11868005 | 0.005328094 | 0.37725453 | 0.01412334 | 0.98873158 | 0.9925745 | NA | NA |
| ENSG000002 | 2.34055183 | -0.005252156 | 0.37116068 | -0.0141506 | 0.98870981 | 0.9925745 | NA | NA |
| ENSG000001 | 198.307445 | -0.001947734 | 0.13883678 | -0.014029 | 0.98880688 | 0.99262552 | CLN5 | 1203 |

| | | | | | | | | |
|---|---|---|---|---|---|---|---|---|
| ENSG000001 | 23.2349593 | 0.003641257 | 0.26205581 | 0.01389497 | 0.98891378 | 0.99268225 | BATF3 | 55509 |
| ENSG000002 | 3.4195673 | 0.005225884 | 0.37711494 | 0.01385754 | 0.98894364 | 0.99268225 | NA | NA |
| ENSG000002 | 19.662433 | 0.003990716 | 0.28749579 | 0.01388095 | 0.98892496 | 0.99268225 | NA | NA |
| ENSG000002 | 15.1924339 | -0.004182734 | 0.30232153 | -0.0138354 | 0.98896132 | 0.99268225 | TNFSF12 | 8742 |
| ENSG000002 | 7.11187832 | -0.004994779 | 0.36260974 | -0.0137745 | 0.98900986 | 0.99270641 | NA | NA |
| ENSG000002 | 13.1060985 | 0.004291955 | 0.31562226 | 0.01359839 | 0.98915039 | 0.99282288 | LY6G5C | 80741 |
| ENSG000001 | 2.36501109 | -0.004896942 | 0.36139657 | -0.0135501 | 0.98918895 | 0.99283702 | NA | NA |
| ENSG000002 | 4.17023408 | 0.005085489 | 0.37859216 | 0.01343263 | 0.98928263 | 0.99288255 | NA | NA |
| ENSG000002 | 10.3744728 | -0.004534072 | 0.33756211 | -0.0134318 | 0.98928328 | 0.99288255 | NA | NA |
| ENSG000001 | 198.754225 | 0.002470789 | 0.18492455 | 0.01336106 | 0.98933973 | 0.9929132 | EPHA2 | 1969 |
| ENSG000002 | 2.31281158 | -0.004912761 | 0.36848973 | -0.0133322 | 0.9893628 | 0.9929132 | NA | NA |
| ENSG000002 | 2.55010026 | -0.004926165 | 0.37237745 | -0.013229 | 0.98944513 | 0.99297125 | MIR585 | 693170 |
| ENSG000001 | 30.8355675 | 0.003149629 | 0.24022681 | 0.01311106 | 0.98953918 | 0.99304107 | CYP39A1 | 51302 |
| ENSG000002 | 2.32151386 | -0.00483635 | 0.37073414 | -0.0130453 | 0.98959163 | 0.99306912 | NA | NA |
| ENSG000001 | 4.37141662 | -0.004869439 | 0.37666718 | -0.0129277 | 0.98968548 | 0.99312454 | GOLGA8G | 283768 |
| ENSG000002 | 3.86693277 | 0.004870737 | 0.37804493 | 0.01288402 | 0.98972033 | 0.99312454 | NA | NA |
| ENSG000002 | 6.48689969 | -0.004683954 | 0.36279953 | -0.0129106 | 0.98969913 | 0.99312454 | NA | NA |
| ENSG000002 | 8.04219563 | 0.004512395 | 0.35222392 | 0.01281115 | 0.98977846 | 0.9931583 | NA | NA |
| ENSG000002 | 8.70297278 | 0.004517784 | 0.35577872 | 0.0126983 | 0.9898685 | 0.99322407 | NA | NA |
| ENSG000002 | 16.1785647 | 0.003834355 | 0.30318095 | 0.01264708 | 0.98990936 | 0.99324049 | NA | NA |
| ENSG000002 | 3.70503638 | 0.004752091 | 0.37846192 | 0.01255632 | 0.98998177 | 0.99326705 | TAS2R38 | 5726 |
| ENSG000002 | 3.98922672 | -0.004752287 | 0.37859305 | -0.0125525 | 0.98998482 | 0.99326705 | NA | NA |
| ENSG000002 | 2.35637294 | -0.004508902 | 0.36554934 | -0.0123346 | 0.99015867 | 0.9934169 | NA | NA |
| ENSG000001 | 4076.74429 | -0.00129652 | 0.10611114 | -0.0122185 | 0.99025128 | 0.99348524 | RABGAP1L | 9910 |
| ENSG000002 | 9.4372039 | -0.004168471 | 0.34444859 | -0.0121019 | 0.99034434 | 0.99355402 | FAM225B | 100128385 |
| ENSG000002 | 2.29006282 | 0.004340825 | 0.3615028 | 0.01200772 | 0.99041945 | 0.99360479 | NA | NA |
| ENSG000001 | 12.8766084 | -0.003794954 | 0.31703915 | -0.01197 | 0.99044956 | 0.99361041 | OGFRP1 | 388906 |
| ENSG000001 | 3.9049294 | 0.004493706 | 0.37802091 | 0.01188745 | 0.99051541 | 0.99363614 | C19orf67 | 646457 |
| ENSG000002 | 11.6013382 | 0.003998471 | 0.33667329 | 0.01187641 | 0.99052422 | 0.99363614 | LOC155060 | 155060 |

| | | | | | | | | |
|---|---|---|---|---|---|---|---|---|
| ENSG000002 | 4.86054376 | -0.004363836 | 0.3755687 | -0.0116193 | 0.99072937 | 0.99381736 | NA | NA |
| ENSG000002 | 2.52806891 | -0.004256408 | 0.36738981 | -0.0115855 | 0.99075629 | 0.99381977 | NA | NA |
| ENSG000002 | 5.98479523 | -0.004223873 | 0.36735484 | -0.0114981 | 0.99082607 | 0.99382923 | NA | NA |
| ENSG000002 | 5.75266802 | 0.00420659 | 0.36637755 | 0.01148157 | 0.99083923 | 0.99382923 | NA | NA |
| ENSG000002 | 11.1627262 | -0.00391296 | 0.34015506 | -0.0115035 | 0.99082177 | 0.99382923 | NA | NA |
| ENSG000002 | 36.5059398 | 0.002603391 | 0.22867553 | 0.01138465 | 0.99091656 | 0.9938822 | NA | NA |
| ENSG000001 | 13.1421159 | -0.00363288 | 0.32114047 | -0.0113124 | 0.99097418 | 0.99391077 | NPW | 283869 |
| ENSG000002 | 3.45759638 | 0.004261123 | 0.3775081 | 0.0112875 | 0.99099407 | 0.99391077 | NA | NA |
| ENSG000002 | 3.87089206 | 0.004160158 | 0.3781122 | 0.01100244 | 0.9912215 | 0.99411429 | NA | NA |
| ENSG000002 | 7.74387329 | 0.003914132 | 0.35725657 | 0.01095608 | 0.99125849 | 0.99412038 | NA | NA |
| ENSG000002 | 6.4323762 | -0.004040025 | 0.36951297 | -0.0109334 | 0.9912766 | 0.99412038 | NA | NA |
| ENSG000001 | 61.9421522 | 0.002145955 | 0.20104697 | 0.0106739 | 0.99148362 | 0.99421193 | MN1 | 4330 |
| ENSG000002 | 2.29026393 | -0.003963012 | 0.37123112 | -0.0106753 | 0.99148249 | 0.99421193 | NA | NA |
| ENSG000002 | 2.35219055 | -0.003987555 | 0.36962527 | -0.0107881 | 0.99139251 | 0.99421193 | NA | NA |
| ENSG000002 | 2.33558015 | -0.003927468 | 0.36807816 | -0.0106702 | 0.99148657 | 0.99421193 | NA | NA |
| ENSG000002 | 25.3225743 | -0.002785729 | 0.2611954 | -0.0106653 | 0.99149048 | 0.99421193 | NA | NA |
| ENSG000001 | 177.983823 | 0.001497924 | 0.14276536 | 0.01049221 | 0.99162858 | 0.99432456 | AOX1 | 316 |
| ENSG000002 | 16.2985245 | 0.003126853 | 0.29884689 | 0.01046306 | 0.99165184 | 0.99432456 | NA | NA |
| ENSG000002 | 6.53872671 | -0.003782838 | 0.36502423 | -0.0103633 | 0.99173147 | 0.99436006 | NA | NA |
| ENSG000002 | 6.43995337 | 0.00380108 | 0.3669983 | 0.01035721 | 0.99173629 | 0.99436006 | NA | NA |
| ENSG00000C | 66.0406762 | -0.002064661 | 0.20311609 | -0.0101649 | 0.9918897 | 0.99448929 | GSDMB | 55876 |
| ENSG00000C | 85.0221823 | 0.001875747 | 0.18592881 | 0.01008852 | 0.99195066 | 0.99452582 | LPCAT2 | 54947 |
| ENSG000002 | 6.68347137 | -0.003607758 | 0.3623486 | -0.0099566 | 0.99205592 | 0.99458315 | NA | NA |
| ENSG000002 | 8.34013129 | -0.003439478 | 0.34548991 | -0.0099554 | 0.9920569 | 0.99458315 | CACNA1C-AS2 | 100874235 |
| ENSG000002 | 3.69282909 | 0.003721951 | 0.37857086 | 0.00983158 | 0.99215566 | 0.99465757 | NA | NA |
| ENSG000002 | 5.32366024 | 0.003651577 | 0.37577693 | 0.00971741 | 0.99224675 | 0.99469971 | NA | NA |
| ENSG000002 | 3.91565376 | -0.003682333 | 0.37806686 | -0.0097399 | 0.99222881 | 0.99469971 | NA | NA |
| ENSG000002 | 5.97324118 | -0.00352489 | 0.3675327 | -0.0095907 | 0.99234786 | 0.99477648 | NA | NA |
| ENSG000001 | 22.5905018 | -0.002447583 | 0.26677794 | -0.0091746 | 0.99267983 | 0.99508466 | GRK5 | 2869 |

| | | | | | | | | |
|---|---|---|---|---|---|---|---|---|
| ENSG000002 | 3.66981258 | 0.003436036 | 0.37841712 | 0.00908002 | 0.99275529 | 0.9951357 | NA | NA |
| ENSG000002 | 3.89269727 | 0.003335277 | 0.37855043 | 0.00881065 | 0.99297021 | 0.99532653 | NA | NA |
| ENSG000002 | 3.95419137 | -0.003122502 | 0.37814021 | -0.0082575 | 0.99341152 | 0.99574428 | NA | NA |
| ENSG000002 | 3.44360222 | 0.003095801 | 0.37757901 | 0.00819908 | 0.99345815 | 0.99576641 | NA | NA |
| ENSG000002 | 21.6416544 | 0.002182413 | 0.26882926 | 0.00811821 | 0.99352268 | 0.99580647 | NA | NA |
| ENSG000002 | 2.33691076 | -0.002920199 | 0.36450138 | -0.0080115 | 0.99360783 | 0.9958672 | NA | NA |
| ENSG000001 | 2.51342848 | -0.002900065 | 0.36594288 | -0.0079249 | 0.9936769 | 0.99590398 | OR2C1 | 4993 |
| ENSG000002 | 3.94099479 | 0.002987544 | 0.37798223 | 0.00790393 | 0.99369364 | 0.99590398 | OR2A2 | 442361 |
| ENSG000002 | 18.3613632 | -0.002243228 | 0.28565567 | -0.0078529 | 0.99373435 | 0.99592016 | NA | NA |
| ENSG000002 | 10.6113594 | -0.002612672 | 0.33512542 | -0.0077961 | 0.99377967 | 0.99592555 | NA | NA |
| ENSG000002 | 5.7506443 | -0.00286336 | 0.36926639 | -0.0077542 | 0.99381312 | 0.99592555 | NA | NA |
| ENSG000002 | 2.33182252 | -0.002860029 | 0.36885418 | -0.0077538 | 0.99381341 | 0.99592555 | NA | NA |
| ENSG000002 | 66.6168376 | -0.00157055 | 0.20896774 | -0.0075158 | 0.99400335 | 0.99609129 | NA | NA |
| ENSG000002 | 9.44834182 | 0.002533007 | 0.33884183 | 0.00747549 | 0.99403548 | 0.99609887 | NA | NA |
| ENSG000002 | 2.30285579 | -0.002673941 | 0.36339118 | -0.0073583 | 0.99412898 | 0.99616794 | NA | NA |
| ENSG000002 | 4.11655274 | 0.002716834 | 0.37730322 | 0.00720066 | 0.99425475 | 0.99626935 | NA | NA |
| ENSG000002 | 20.8349163 | -0.001960501 | 0.27507206 | -0.0071272 | 0.99431334 | 0.99630345 | LPP-AS2 | 339929 |
| ENSG00000C | 42.4653671 | -0.001550757 | 0.22573253 | -0.0068699 | 0.99451867 | 0.99648455 | SEMA4G | 57715 |
| ENSG000002 | 4.16486933 | -0.002548895 | 0.37693868 | -0.0067621 | 0.99460467 | 0.99654611 | NA | NA |
| ENSG000001 | 70.0301826 | 0.001277834 | 0.19182697 | 0.00666139 | 0.99468502 | 0.99660199 | LRRIQ3 | 127255 |
| ENSG000001 | 55.5131605 | 0.001333365 | 0.20301778 | 0.00656772 | 0.99475975 | 0.99662762 | SLC8A2 | 6543 |
| ENSG000002 | 5.52860871 | -0.002452323 | 0.37209344 | -0.0065906 | 0.99474149 | 0.99662762 | NA | NA |
| ENSG000001 | 132.17783 | 0.000970792 | 0.1500132 | 0.00647138 | 0.99483663 | 0.99668001 | PPP1R17 | 10842 |
| ENSG000002 | 4.3726505 | -0.002414942 | 0.3768998 | -0.0064074 | 0.99488768 | 0.99670653 | NA | NA |
| ENSG000002 | 2.54323713 | -0.002350685 | 0.37102384 | -0.0063357 | 0.9949449 | 0.99673923 | NA | NA |
| ENSG000002 | 5.54277446 | 0.002344322 | 0.3744853 | 0.00626012 | 0.99500518 | 0.996775 | NA | NA |
| ENSG000002 | 6.19271335 | 0.002256154 | 0.36639356 | 0.00615773 | 0.99508687 | 0.99683221 | NA | NA |
| ENSG000001 | 26.1551649 | 0.001592596 | 0.26224626 | 0.0060729 | 0.99515455 | 0.99687538 | WASH2P | 375260 |
| ENSG000002 | 30.0956892 | -0.001491966 | 0.24915914 | -0.005988 | 0.99522229 | 0.99691861 | PCDHGB8P | 56120 |

| | | | | | | | | |
|---|---|---|---|---|---|---|---|---|
| ENSG000002 | 4.35425224 | -0.00221596 | 0.37736707 | -0.0058722 | 0.99531472 | 0.99698657 | NA | NA |
| ENSG000002 | 8.24515522 | 0.002017587 | 0.34931434 | 0.00577585 | 0.99539156 | 0.99703891 | NA | NA |
| ENSG000002 | 6.25419037 | -0.002070988 | 0.36305619 | -0.0057043 | 0.99544864 | 0.99707145 | NA | NA |
| ENSG000001 | 14.1079038 | 0.001750739 | 0.31321851 | 0.00558951 | 0.99554024 | 0.99712826 | SLC22A4 | 6583 |
| ENSG000002 | 10.7158552 | -0.001865436 | 0.33481175 | -0.0055716 | 0.99555453 | 0.99712826 | NA | NA |
| ENSG000002 | 3.93856668 | -0.002069369 | 0.37778092 | -0.0054777 | 0.99562945 | 0.99717867 | NA | NA |
| ENSG000002 | 2.32612081 | -0.001976865 | 0.37111725 | -0.0053268 | 0.99574985 | 0.99722659 | NA | NA |
| ENSG000002 | 12.4193723 | -0.001694421 | 0.31818571 | -0.0053253 | 0.99575108 | 0.99722659 | NA | NA |
| ENSG000002 | 2.32612081 | -0.001976865 | 0.37111725 | -0.0053268 | 0.99574985 | 0.99722659 | NA | NA |
| ENSG000001 | 422.782548 | 0.000651356 | 0.1274783 | 0.00510954 | 0.99592319 | 0.99724957 | NBEAL1 | 65065 |
| ENSG000001 | 24.4101561 | -0.001390061 | 0.26733983 | -0.0051996 | 0.99585134 | 0.99724957 | KCNB2 | 9312 |
| ENSG000002 | 6.02709933 | 0.001863274 | 0.36583122 | 0.00509326 | 0.99593618 | 0.99724957 | NA | NA |
| ENSG000002 | 26.6640412 | -0.001390648 | 0.2644033 | -0.0052596 | 0.99580349 | 0.99724957 | NA | NA |
| ENSG000002 | 17.7490298 | 0.001454634 | 0.28701126 | 0.00506821 | 0.99595617 | 0.99724957 | NA | NA |
| ENSG000002 | 26.5272565 | 0.001319325 | 0.25616761 | 0.00515024 | 0.99589072 | 0.99724957 | NA | NA |
| ENSG000002 | 4.34759783 | -0.001905491 | 0.37733095 | -0.0050499 | 0.99597076 | 0.99724957 | NA | NA |
| ENSG000002 | 35.7283935 | -0.001238745 | 0.2436893 | -0.0050833 | 0.99594413 | 0.99724957 | NA | NA |
| ENSG000001 | 6.24627647 | -0.001887676 | 0.37617287 | -0.0050181 | 0.99599615 | 0.99725036 | MSLN | 10232 |
| ENSG000002 | 21.4237117 | -0.00138555 | 0.27825199 | -0.0049795 | 0.99602697 | 0.9972566 | NA | NA |
| ENSG000002 | 2.27171001 | -0.001738697 | 0.36468791 | -0.0047676 | 0.996196 | 0.99737659 | NA | NA |
| ENSG000002 | 2.28233132 | 0.001762551 | 0.36743912 | 0.00479685 | 0.99617268 | 0.99737659 | RN7SKP114 | 106480868 |
| ENSG000002 | 4.15744525 | -0.001784169 | 0.37683414 | -0.0047346 | 0.99622233 | 0.99737832 | HSPE1P27 | 106480261 |
| ENSG000001 | 97.1640335 | -0.000759852 | 0.16866127 | -0.0045052 | 0.99640539 | 0.99753697 | IKZF4 | 64375 |
| ENSG000002 | 4.44938062 | -0.001688125 | 0.3782543 | -0.0044629 | 0.9964391 | 0.99754609 | NA | NA |
| ENSG000002 | 7.61246971 | -0.001572803 | 0.35666066 | -0.0044098 | 0.9964815 | 0.99756391 | NA | NA |
| ENSG000002 | 6.18927613 | 0.001589254 | 0.36447796 | 0.00436036 | 0.99652095 | 0.99757878 | NA | NA |
| ENSG000002 | 7.85457386 | 0.001454296 | 0.34985505 | 0.00415685 | 0.99668332 | 0.99771669 | NA | NA |
| ENSG000002 | 2.32601667 | -0.001517428 | 0.36852166 | -0.0041176 | 0.99671463 | 0.9977234 | NA | NA |
| ENSG000002 | 2.33862094 | -0.00141492 | 0.36958671 | -0.0038284 | 0.9969454 | 0.99792977 | NA | NA |

| | | | | | | | | |
|---|---|---|---|---|---|---|---|---|
| ENSG000002 | 6.43594646 | 0.001334693 | 0.36309135 | 0.00367591 | 0.99706705 | 0.99802691 | MT1M | 4499 |
| ENSG000002 | 4.41240195 | 0.001364992 | 0.37692669 | 0.00362137 | 0.99711057 | 0.99804583 | LOC101927447 | 101927447 |
| ENSG000001 | 43.2476541 | 0.000776264 | 0.22814489 | 0.00340251 | 0.9972852 | 0.99812207 | GBAP1 | 2630 |
| ENSG000002 | 7.60803401 | 0.001210942 | 0.35499452 | 0.00341116 | 0.9972783 | 0.99812207 | NA | NA |
| ENSG000002 | 23.1946852 | 0.000909604 | 0.26697314 | 0.0034071 | 0.99728153 | 0.99812207 | NA | NA |
| ENSG000002 | 28.6254777 | 0.000863508 | 0.25146188 | 0.00343395 | 0.99726011 | 0.99812207 | NA | NA |
| ENSG000002 | 25.0895557 | -0.000898213 | 0.27247527 | -0.0032965 | 0.99736978 | 0.99818209 | NA | NA |
| ENSG000002 | 16.4681259 | 0.000948625 | 0.29738826 | 0.00318985 | 0.99745487 | 0.99824261 | NA | NA |
| ENSG000001 | 21.3537902 | 0.000837242 | 0.27634834 | 0.00302966 | 0.99758268 | 0.99832125 | SCNN1G | 6340 |
| ENSG000002 | 2.28167843 | -0.001114529 | 0.36568293 | -0.0030478 | 0.99756821 | 0.99832125 | NA | NA |
| ENSG000002 | 4.17246444 | 0.001117399 | 0.37842667 | 0.00295275 | 0.99764405 | 0.99835803 | NA | NA |
| ENSG000002 | 16.6058562 | 0.000844412 | 0.29227979 | 0.00288905 | 0.99769487 | 0.99838425 | NA | NA |
| ENSG000002 | 2.31969487 | 0.000899737 | 0.36890918 | 0.00243891 | 0.99805403 | 0.99869867 | NA | NA |
| ENSG000002 | 3.95923572 | 0.000921048 | 0.37848367 | 0.00243352 | 0.99805833 | 0.99869867 | NA | NA |
| ENSG000002 | 22.0732184 | 0.000635443 | 0.26762556 | 0.00237437 | 0.99810553 | 0.99872125 | NFAM1 | 150372 |
| ENSG000002 | 24.9366848 | -0.000591185 | 0.25734332 | -0.0022973 | 0.99816705 | 0.99875816 | ATP1A1-AS1 | 84852 |
| ENSG000002 | 4.1528066 | -0.00084725 | 0.37771327 | -0.0022431 | 0.99821026 | 0.99877676 | NA | NA |
| ENSG000002 | 15.9125365 | 0.000631377 | 0.30231398 | 0.00208848 | 0.99833364 | 0.99887555 | NA | NA |
| ENSG000002 | 14.5402448 | 0.000623521 | 0.30605698 | 0.00203727 | 0.99837449 | 0.99889179 | NA | NA |
| ENSG000002 | 2.30457837 | -0.00069987 | 0.36718046 | -0.0019061 | 0.99847918 | 0.99897188 | NA | NA |
| ENSG000002 | 7.36496988 | 0.000659023 | 0.36130265 | 0.00182402 | 0.99854464 | 0.99901273 | NA | NA |
| ENSG000001 | 2.30933414 | -0.000612133 | 0.36799385 | -0.0016634 | 0.99867277 | 0.99906697 | CRHR1 | 1394 |
| ENSG000001 | 85.1745583 | -0.000300103 | 0.18031293 | -0.0016643 | 0.99867205 | 0.99906697 | KAZN | 23254 |
| ENSG000002 | 2.28880933 | -0.000625734 | 0.36712894 | -0.0017044 | 0.99864009 | 0.99906697 | NA | NA |
| ENSG000002 | 54.4605085 | 0.000341057 | 0.2096965 | 0.00162643 | 0.9987023 | 0.99907186 | NA | NA |
| ENSG000002 | 5.57565605 | 0.000581366 | 0.3768366 | 0.00154275 | 0.99876906 | 0.999114 | NA | NA |
| ENSG000001 | 12.0367139 | -0.000479213 | 0.32552026 | -0.0014721 | 0.9988254 | 0.99914571 | ASTL | 431705 |
| ENSG000002 | 3.95633929 | -0.000526093 | 0.37855083 | -0.0013898 | 0.99889114 | 0.99918682 | LINC01479 | 101927922 |
| ENSG000002 | 2.26013599 | -0.000462575 | 0.3650436 | -0.0012672 | 0.99898894 | 0.99926 | NA | NA |

| | | | | | | | | |
|---|---|---|---|---|---|---|---|---|
| ENSG000002 | 4.61384039 | 0.000402367 | 0.37760114 | 0.00106559 | 0.99914978 | 0.99935474 | RN7SL623P | 106480522 |
| ENSG000002 | 4.19554622 | -0.000404989 | 0.37773944 | -0.0010721 | 0.99914456 | 0.99935474 | NA | NA |
| ENSG000002 | 4.35906084 | 0.000398315 | 0.37726074 | 0.00105581 | 0.99915759 | 0.99935474 | C9orf173-AS1 | 100129722 |
| ENSG000001 | 4.39074796 | -0.000370699 | 0.37674771 | -0.0009839 | 0.99921493 | 0.99938744 | NA | NA |
| ENSG000002 | 2.32959858 | 0.000264782 | 0.36228172 | 0.00073087 | 0.99941685 | 0.99956474 | NA | NA |
| ENSG000002 | 76.4438476 | 0.000127137 | 0.18478636 | 0.00068802 | 0.99945104 | 0.99957429 | NA | NA |
| ENSG000001 | 77.7818695 | 0.000112028 | 0.17832464 | 0.00062822 | 0.99949875 | 0.99959735 | NR3C2 | 4306 |
| ENSG000001 | 8.69532598 | 0.000166414 | 0.34884245 | 0.00047705 | 0.99961937 | 0.9996753 | NA | NA |
| ENSG000002 | 6.51362286 | 0.000170822 | 0.36442739 | 0.00046874 | 0.999626 | 0.9996753 | IFI30 | 10437 |
| ENSG000002 | 16.063579 | 1.56E-06 | 0.30182319 | 5.17E-06 | 0.99999588 | 0.9999977 | PCDHGA12 | 26025 |
| ENSG000002 | 14.5124834 | 8.92E-07 | 0.30922118 | 2.88E-06 | 0.9999977 | 0.9999977 | NA | NA |
| ENSG000000 | 0 | NA | NA | NA | NA | NA | NA | NA |
| ENSG000000 | 0 | NA | NA | NA | NA | NA | KDM5D | 8284 |
| ENSG000000 | 39.6514099 | 0.354101793 | 0.37485691 | 0.94463189 | NA | NA | NA | NA |
| ENSG000000 | 2.00256932 | 0.356954428 | 0.356681 | 1.00076657 | 0.31693967 | NA | GUCA2B | 2981 |
| ENSG000000 | 1.37774598 | -0.247461006 | 0.332322 | -0.7446423 | 0.45648801 | NA | GSC2 | 2928 |
| ENSG000000 | 1.3272688 | -0.35479644 | 0.32493874 | -1.0918872 | 0.2748827 | NA | NA | NA |
| ENSG000000 | 0.12685397 | -0.090215768 | 0.12901919 | -0.699243 | 0.48440017 | NA | DDX3Y | 8653 |
| ENSG000000 | 0.13280363 | -0.093394593 | 0.13115644 | -0.7120855 | 0.47641185 | NA | ZFY | 7544 |
| ENSG000000 | 0 | NA | NA | NA | NA | NA | KIAA1107 | 23285 |
| ENSG000000 | 1.71503947 | 0.069941559 | 0.34312849 | 0.20383489 | 0.83848252 | NA | SCT | 6343 |
| ENSG000000 | 0.52028267 | -0.202047067 | 0.25039858 | -0.8069018 | 0.41972305 | NA | PRLH | 51052 |
| ENSG000000 | 0.12685397 | -0.090215768 | 0.12901919 | -0.699243 | 0.48440017 | NA | CFHR2 | 3080 |
| ENSG000000 | 1.92947663 | 0.203581355 | 0.3638436 | 0.55952985 | 0.57580016 | NA | HBQ1 | 3049 |
| ENSG000000 | 1.17823651 | -0.398539152 | 0.313524 | -1.2711599 | 0.20367176 | NA | MT3 | 4504 |
| ENSG000000 | 2.17235667 | -0.653103702 | 0.36561162 | -1.7863319 | 0.07404553 | NA | POU4F3 | 5459 |
| ENSG000000 | 0.11934008 | 0.083564698 | 0.12760091 | 0.65489106 | 0.5125379 | NA | TBL1Y | 90665 |
| ENSG000000 | 1.17630158 | -0.776782023 | 0.3188478 | -2.4362157 | 0.01484183 | NA | TPSD1 | 23430 |
| ENSG000000 | 0 | NA | NA | NA | NA | NA | AMELY | 266 |

| | | | | | | | | | |
|---|---|---|---|---|---|---|---|---|---|
| ENSG000000 | 1.7112976 | -0.70048047 | 0.34840342 | -2.0105442 | 0.04437362 | NA | | IGF2-AS | 51214 |
| ENSG000000 | 1.22243045 | 0.091661752 | 0.32393886 | 0.28296004 | 0.77720747 | NA | | GSTT2 | 2953 |
| ENSG000000 | 1.57453136 | 0.143252573 | 0.34458869 | 0.41572048 | 0.67761455 | NA | | OSM | 5008 |
| ENSG000001 | 0 | NA | NA | NA | NA | NA | | NA | NA |
| ENSG000001 | 1.12148459 | -0.265820875 | 0.31957363 | -0.8317985 | 0.40552268 | NA | | GGTLC2 | 91227 |
| ENSG000001 | 0.12685397 | -0.090215768 | 0.12901919 | -0.699243 | 0.48440017 | NA | | AVP | 551 |
| ENSG000001 | 0.38064837 | -0.102706251 | 0.21886737 | -0.4692625 | 0.63888201 | NA | | OXT | 5020 |
| ENSG000001 | 1.47250393 | -0.177553828 | 0.3426021 | -0.5182509 | 0.60428326 | NA | | OPN1LW | 5956 |
| ENSG000001 | 2.10381754 | -0.510662194 | 0.36605969 | -1.3950244 | 0.16300848 | NA | | RAB40AL | 282808 |
| ENSG000001 | 1.88067967 | -0.530361781 | 0.35898443 | -1.4773949 | 0.1395698 | NA | | MT4 | 84560 |
| ENSG000001 | 0.11131518 | 0.081951103 | 0.12663804 | 0.64712866 | 0.51754868 | NA | | NA | NA |
| ENSG000001 | 0 | NA | NA | NA | NA | NA | | CORO7-PAM16 | 100529144 |
| ENSG000001 | 1.08553342 | -0.095142482 | 0.30677394 | -0.3101387 | 0.75645546 | NA | | RRN3P2 | 653390 |
| ENSG000001 | 0.9898443 | -0.039993873 | 0.31291302 | -0.1278115 | 0.89829817 | NA | | LHB | 3972 |
| ENSG000001 | 0.39479706 | -0.300008631 | 0.22317059 | -1.3443018 | 0.17885081 | NA | | CGB | 1082 |
| ENSG000001 | 1.06492152 | 0.345299456 | 0.31973731 | 1.07994735 | 0.28016563 | NA | | RETN | 56729 |
| ENSG000001 | 1.6995046 | -0.053337647 | 0.35359838 | -0.1508425 | 0.8801 | NA | | OVOL3 | 728361 |
| ENSG000001 | 0.70611255 | 0.126099889 | 0.27476576 | 0.45893596 | 0.64628015 | NA | | NKG7 | 4818 |
| ENSG000001 | 1.35716943 | -0.242082124 | 0.33186485 | -0.72946 | 0.46572034 | NA | | ZNRF4 | 148066 |
| ENSG000001 | 1.26630356 | -0.614239306 | 0.31999369 | -1.9195357 | 0.05491658 | NA | | SIGLEC5 | 8778 |
| ENSG000001 | 1.29966743 | 0.447505719 | 0.32747129 | 1.36654946 | 0.17176657 | NA | | PIK3R2 | 5296 |
| ENSG000001 | 2.12470867 | -0.389668185 | 0.36714254 | -1.061354 | 0.28852907 | NA | | NA | NA |
| ENSG000001 | 0 | NA | NA | NA | NA | NA | | NA | NA |
| ENSG000001 | 1.2308063 | -0.043187953 | 0.33003757 | -0.1308577 | 0.89588789 | NA | | PITX3 | 5309 |
| ENSG000001 | 1.23134709 | -0.049422726 | 0.32417164 | -0.1524585 | 0.87882531 | NA | | NA | NA |
| ENSG000001 | 0 | NA | NA | NA | NA | NA | | PTGES3L-AARSI | 100885850 |
| ENSG000001 | 0.90161618 | -0.276593441 | 0.30057786 | -0.9202056 | 0.35746531 | NA | | APOC3 | 345 |
| ENSG000001 | 0 | NA | NA | NA | NA | NA | | NA | NA |
| ENSG000001 | 0 | NA | NA | NA | NA | NA | | USP9Y | 8287 |

| | | | | | | | | | |
|---|---|---|---|---|---|---|---|---|---|
| ENSG000001 | 1.71881366 | -0.197675249 | 0.35732976 | -0.5532012 | 0.58012565 | NA | NA | NA | |
| ENSG000001 | 1.98924373 | 0.666041886 | 0.36225642 | 1.83859234 | 0.06597517 | NA | INO80B | | 83444 |
| ENSG000001 | 1.35762836 | -0.086739535 | 0.31671479 | -0.2738727 | 0.78418245 | NA | WNT6 | | 7475 |
| ENSG000001 | 1.06694524 | 0.351442479 | 0.31748731 | 1.1069497 | 0.26831568 | NA | VAX2 | | 25806 |
| ENSG000001 | 0.51413714 | -0.199727707 | 0.24976394 | -0.7996659 | 0.42390439 | NA | NA | NA | |
| ENSG000001 | 0.84160602 | 0.189944396 | 0.28885129 | 0.65758542 | 0.51080457 | NA | NA | NA | |
| ENSG000001 | 1.77668251 | -0.762884323 | 0.3532344 | -2.1597113 | 0.03079503 | NA | HOXB8 | | 3218 |
| ENSG000001 | 1.71895453 | -0.441183091 | 0.35005239 | -1.2603345 | 0.20754873 | NA | NPPB | | 4879 |
| ENSG000001 | 1.69874915 | 0.067552593 | 0.35058303 | 0.19268643 | 0.84720455 | NA | PRB2 | | 653247 |
| ENSG000001 | 1.49550008 | -0.191781812 | 0.34496035 | -0.5559532 | 0.57824284 | NA | NA | NA | |
| ENSG000001 | 2.04564246 | -0.876413344 | 0.35958278 | -2.4373062 | 0.01479714 | NA | OBP2A | | 29991 |
| ENSG000001 | 1.05195366 | -0.505828659 | 0.30598087 | -1.6531382 | 0.09830275 | NA | PRM2 | | 5620 |
| ENSG000001 | 0.63296622 | -0.299126688 | 0.27328461 | -1.0945611 | 0.27370899 | NA | NA | NA | |
| ENSG000001 | 2.12721809 | -0.494816815 | 0.36173888 | -1.367884 | 0.1713484 | NA | ESX1 | | 80712 |
| ENSG000001 | 1.05889644 | -0.50235593 | 0.30635604 | -1.6397781 | 0.10105131 | NA | MAGEA9 | | 4108 |
| ENSG000001 | 1.60530564 | -0.370244404 | 0.34418369 | -1.0757175 | 0.28205364 | NA | GTSF1L | | 149699 |
| ENSG000001 | 0 | NA | NA | NA | NA | NA | TMEM189-UBE2 | | 387522 |
| ENSG000001 | 1.02414946 | -0.501551323 | 0.30565594 | -1.6409016 | 0.10081784 | NA | IRGC | | 56269 |
| ENSG000001 | 1.37565347 | -0.710687365 | 0.33258243 | -2.1368758 | 0.03260809 | NA | SCGB1D2 | | 10647 |
| ENSG000001 | 1.9943503 | -0.431545349 | 0.3584994 | -1.2037547 | 0.22868439 | NA | WNT1 | | 7471 |
| ENSG000001 | 1.0187584 | -0.046501627 | 0.30759758 | -0.1511768 | 0.87983623 | NA | NA | NA | |
| ENSG000001 | 1.6701445 | -0.71318402 | 0.35069358 | -2.0336387 | 0.04198804 | NA | BARHL1 | | 56751 |
| ENSG000001 | 0.89113496 | 0.030496727 | 0.28181048 | 0.10821715 | 0.91382344 | NA | NPBWR2 | | 2832 |
| ENSG000001 | 1.65337711 | -0.68734296 | 0.34615865 | -1.9856299 | 0.04707444 | NA | BHLHE23 | | 128408 |
| ENSG000001 | 0 | NA | NA | NA | NA | NA | NA | NA | |
| ENSG000001 | 2.02984576 | -0.580353189 | 0.36113434 | -1.6070285 | 0.10804811 | NA | NKX2-2 | | 4821 |
| ENSG000001 | 1.73099764 | -0.467167572 | 0.3549239 | -1.3162472 | 0.1880911 | NA | C20orf187 | | 728434 |
| ENSG000001 | 0 | NA | NA | NA | NA | NA | NA | NA | |
| ENSG000001 | 0.86455511 | -0.103797748 | 0.29753822 | -0.3488552 | 0.72719804 | NA | SLURP1 | | 57152 |

| | | | | | | | | | |
|---|---|---|---|---|---|---|---|---|---|
| ENSG000001 | 1.63590798 | -0.556254453 | 0.35080597 | -1.585647 | 0.1128194 | NA | CTAG2 | | 30848 |
| ENSG000001 | 0.97866101 | -0.180785309 | 0.30829057 | -0.5864121 | 0.55759865 | NA | OR7C2 | | 26658 |
| ENSG000001 | 0.85704121 | 0.053051046 | 0.29513872 | 0.17974953 | 0.85734921 | NA | GNG13 | | 51764 |
| ENSG000001 | 1.02732801 | -0.35149678 | 0.31009385 | -1.1335174 | 0.25699704 | NA | IGLL1 | | 3543 |
| ENSG000001 | 0.75873415 | -0.034571091 | 0.2860415 | -0.1208604 | 0.90380161 | NA | MYOD1 | | 4654 |
| ENSG000001 | 0.85732354 | 0.199144281 | 0.28973083 | 0.68734238 | 0.491867 | NA | CD68 | | 968 |
| ENSG000001 | 0 | NA | NA | NA | NA | NA | NA | NA | |
| ENSG000001 | 0 | NA | NA | NA | NA | NA | RPS4Y1 | | 6192 |
| ENSG000001 | 0 | NA | NA | NA | NA | NA | NA | NA | |
| ENSG000001 | 0 | NA | NA | NA | NA | NA | VCY1B | | 353513 |
| ENSG000001 | 0 | NA | NA | NA | NA | NA | VCY | | 9084 |
| ENSG000001 | 0 | NA | NA | NA | NA | NA | CDY2B | | 203611 |
| ENSG000001 | 0 | NA | NA | NA | NA | NA | GDF1 | | 2657 |
| ENSG000001 | 0.6181343 | 0.056012223 | 0.26406445 | 0.21211573 | 0.83201675 | NA | FUT5 | | 2527 |
| ENSG000001 | 0.85772389 | -0.116826005 | 0.3022732 | -0.3864914 | 0.69913274 | NA | SCO2 | | 9997 |
| ENSG000001 | 0.97470641 | -0.190151385 | 0.30809904 | -0.6171762 | 0.53711853 | NA | OR11H1 | | 81061 |
| ENSG000001 | 2.06649546 | 0.13041324 | 0.36605528 | 0.35626652 | 0.72164098 | NA | TNNI2 | | 7136 |
| ENSG000001 | 1.67871718 | 0.317142803 | 0.3416667 | 0.92822274 | 0.35329204 | NA | H19 | | 283120 |
| ENSG000001 | 0.87645442 | -0.100062288 | 0.29039953 | -0.3445677 | 0.73041942 | NA | HBZ | | 3050 |
| ENSG000001 | 0 | NA | NA | NA | NA | NA | TXLNGY | | 246126 |
| ENSG000001 | 0 | NA | NA | NA | NA | NA | NA | NA | |
| ENSG000001 | 0 | NA | NA | NA | NA | NA | NA | NA | |
| ENSG000001 | 1.03578818 | -0.535870025 | 0.31393025 | -1.7069716 | 0.08782733 | NA | GJA9 | | 81025 |
| ENSG000001 | 0.37330862 | -0.105391168 | 0.21773449 | -0.4840352 | 0.62836086 | NA | NA | NA | |
| ENSG000001 | 0 | NA | NA | NA | NA | NA | NA | NA | |
| ENSG000001 | 0 | NA | NA | NA | NA | NA | NA | NA | |
| ENSG000001 | 1.07289243 | 0.188082351 | 0.32081141 | 0.58627076 | 0.55769358 | NA | NR0B2 | | 8431 |
| ENSG000001 | 1.34368414 | -0.249519063 | 0.33430322 | -0.7463855 | 0.4554346 | NA | NA | NA | |
| ENSG000001 | 0 | NA | NA | NA | NA | NA | SLX1A | | 548593 |

| | | | | | | | | | |
|---|---|---|---|---|---|---|---|---|---|
| ENSG000001 | 0.25260017 | -0.015978705 | 0.18049464 | -0.0885273 | 0.92945758 | NA | FTHL17 | | 53940 |
| ENSG000001 | 1.71662219 | -0.193919323 | 0.35626518 | -0.5443118 | 0.58622696 | NA | SCP2D1 | | 140856 |
| ENSG000001 | 1.24693795 | -0.185607464 | 0.32357432 | -0.5736162 | 0.56622756 | NA | SSTR4 | | 6754 |
| ENSG000001 | 0.23868015 | 0.154625155 | 0.16997278 | 0.90970538 | 0.36297791 | NA | LINC01260 | | 79015 |
| ENSG000001 | 1.08863974 | 0.028312312 | 0.3149176 | 0.08990387 | 0.9283636 | NA | NA | NA | |
| ENSG000001 | 1.91112381 | 0.17822773 | 0.35474686 | 0.5024082 | 0.61538041 | NA | NA | NA | |
| ENSG000001 | 0.11934008 | 0.083564698 | 0.12760091 | 0.65489106 | 0.5125379 | NA | GSTM1 | | 2944 |
| ENSG000001 | 1.84654866 | -0.249825715 | 0.3562877 | -0.701191 | 0.48318385 | NA | TAAR5 | | 9038 |
| ENSG000001 | 2.1265577 | -0.234924416 | 0.36201762 | -0.6489309 | 0.51638305 | NA | CHST5 | | 23563 |
| ENSG000001 | 1.62607998 | -0.257820688 | 0.35088435 | -0.734774 | 0.46247716 | NA | NA | NA | |
| ENSG000001 | 0.48416733 | -0.018553864 | 0.24528998 | -0.0756405 | 0.93970509 | NA | NKX2-8 | | 26257 |
| ENSG000001 | 0.9878256 | -0.340395897 | 0.30848007 | -1.1034616 | 0.26982677 | NA | CFC1 | | 55997 |
| ENSG000001 | 0.95785132 | 0.111017849 | 0.30669792 | 0.36197784 | 0.71736859 | NA | OR1J1 | | 347168 |
| ENSG000001 | 1.42377793 | 0.090910943 | 0.33998812 | 0.26739447 | 0.78916546 | NA | OR1L4 | | 254973 |
| ENSG000001 | 0 | NA | NA | NA | NA | NA | NA | NA | |
| ENSG000001 | 0 | NA | NA | NA | NA | NA | NPFF | | 8620 |
| ENSG000001 | 0.75478244 | -0.522418532 | 0.27835884 | -1.8767808 | 0.06054814 | NA | CELF6 | | 60677 |
| ENSG000001 | 2.15848818 | -0.521234833 | 0.36643369 | -1.4224534 | 0.15489468 | NA | LMAN1L | | 79748 |
| ENSG000001 | 0.82155648 | 0.046914577 | 0.28690296 | 0.16352071 | 0.87010847 | NA | CTRL | | 1506 |
| ENSG000001 | 1.50380484 | -0.201076246 | 0.34631354 | -0.5806191 | 0.56149722 | NA | NA | NA | |
| ENSG000001 | 1.75814585 | -0.329753512 | 0.35447197 | -0.9302668 | 0.35223295 | NA | PTH2 | | 113091 |
| ENSG000001 | 0 | NA | NA | NA | NA | NA | NA | NA | |
| ENSG000001 | 2.13617607 | -0.503101565 | 0.36361508 | -1.3836103 | 0.16647782 | NA | S100A7 | | 6278 |
| ENSG000001 | 0 | NA | NA | NA | NA | NA | NA | NA | |
| ENSG000001 | 0 | NA | NA | NA | NA | NA | NICN1 | | 84276 |
| ENSG000001 | 1.76149326 | -0.18540446 | 0.34663215 | -0.5348738 | 0.59273712 | NA | TM4SF19 | | 116211 |
| ENSG000001 | 0.25993993 | -0.013144386 | 0.18211683 | -0.0721756 | 0.94246219 | NA | NA | NA | |
| ENSG000001 | 1.28474015 | -0.635699112 | 0.32342489 | -1.9655232 | 0.04935372 | NA | FERD3L | | 222894 |
| ENSG000001 | 1.52877127 | -0.324311664 | 0.34108981 | -0.9508102 | 0.34170075 | NA | LOC541473 | | 541473 |

| | | | | | | | | |
|---|---|---|---|---|---|---|---|---|
| ENSG000001 | 0 | NA | NA | NA | NA | NA | TTTY7 | 246122 |
| ENSG000001 | 0 | NA | NA | NA | NA | NA | TTTY7B | 100101120 |
| ENSG000001 | 1.70452254 | 0.070355556 | 0.35468925 | 0.1983583 | 0.84276474 | NA | OR5C1 | 392391 |
| ENSG000001 | 1.12882434 | -0.263175918 | 0.3198263 | -0.8228714 | 0.41058112 | NA | ADIRF | 10974 |
| ENSG000001 | 1.02077966 | -0.191281245 | 0.30841641 | -0.6202045 | 0.53512316 | NA | SAA4 | 6291 |
| ENSG000001 | 1.11806664 | -0.279759399 | 0.32553234 | -0.8593905 | 0.39012512 | NA | OR5F1 | 338674 |
| ENSG000001 | 1.86304908 | -0.130340629 | 0.35642556 | -0.3656882 | 0.71459774 | NA | GGTLC1 | 92086 |
| ENSG000001 | 0 | NA | NA | NA | NA | NA | OR5M9 | 390162 |
| ENSG000001 | 0.39462238 | -0.104329077 | 0.21264929 | -0.4906157 | 0.6236983 | NA | NA | NA |
| ENSG000001 | 0 | NA | NA | NA | NA | NA | KCNA6 | 3742 |
| ENSG000001 | 0.97453226 | -0.037252082 | 0.31645905 | -0.1177153 | 0.90629322 | NA | MSGN1 | 343930 |
| ENSG000001 | 1.61589003 | -0.121530648 | 0.34743769 | -0.3497912 | 0.7264954 | NA | NA | NA |
| ENSG000001 | 0 | NA | NA | NA | NA | NA | CFC1B | 653275 |
| ENSG000001 | 0.98144884 | 0.104238647 | 0.30775483 | 0.33870678 | 0.73483063 | NA | NA | NA |
| ENSG000001 | 14.8041215 | 0.029608665 | 0.26868318 | 0.11019918 | NA | NA | MAGED4 | 728239 |
| ENSG000001 | 0 | NA | NA | NA | NA | NA | TMSB4Y | 9087 |
| ENSG000001 | 0.72441173 | -0.021440533 | 0.27745778 | -0.0772749 | 0.93840483 | NA | NA | NA |
| ENSG000001 | 1.84740398 | -0.569339582 | 0.34535696 | -1.6485539 | 0.09923905 | NA | CLDN17 | 26285 |
| ENSG000001 | 0.11131518 | 0.081951103 | 0.12663804 | 0.64712866 | 0.51754868 | NA | NA | NA |
| ENSG000001 | 0.85960326 | -0.112061982 | 0.30237674 | -0.3706038 | 0.71093262 | NA | COX6A2 | 1339 |
| ENSG000001 | 1.97920848 | -0.328863974 | 0.36430996 | -0.9027038 | 0.36668314 | NA | NA | NA |
| ENSG000001 | 0.84878552 | -0.106052095 | 0.28885264 | -0.3671495 | 0.71350752 | NA | PRAC1 | 84366 |
| ENSG000001 | 1.98239307 | -0.32260484 | 0.36352149 | -0.8874437 | 0.37484009 | NA | LCE2B | 26239 |
| ENSG000001 | 2.02435665 | 0.120842647 | 0.35668615 | 0.33879265 | 0.73476593 | NA | AGRP | 181 |
| ENSG000001 | 0.11934008 | 0.083564698 | 0.12760091 | 0.65489106 | 0.5125379 | NA | NA | NA |
| ENSG000001 | 1.73696849 | -0.334594517 | 0.3559282 | -0.9400618 | 0.34718584 | NA | TFF2 | 7032 |
| ENSG000001 | 1.03510304 | -0.536100349 | 0.31389681 | -1.7078872 | 0.08765727 | NA | TFF1 | 7031 |
| ENSG000001 | 0.37304803 | -0.090842893 | 0.20939682 | -0.4338313 | 0.66441096 | NA | NA | NA |
| ENSG000001 | 0.84813476 | 0.043577475 | 0.3014628 | 0.14455341 | 0.88506349 | NA | NA | NA |

| | | | | | | | | |
|---|---|---|---|---|---|---|---|---|
| ENSG000001 | 1.42443971 | -0.697268301 | 0.33084331 | -2.1075484 | 0.03507007 | NA | LCN1 | 3933 |
| ENSG000001 | 1.85867418 | -0.376876663 | 0.35385153 | -1.06507 | 0.28684424 | NA | TMEM190 | 147744 |
| ENSG000001 | 0.62570027 | -0.108408956 | 0.26508355 | -0.4089615 | 0.68256794 | NA | SCGB3A1 | 92304 |
| ENSG000001 | 0.79602198 | -0.363127991 | 0.28386389 | -1.2792328 | 0.2008151 | NA | HCRT | 3060 |
| ENSG000001 | 0.99212456 | -0.346343867 | 0.31027043 | -1.1162645 | 0.26430893 | NA | OR7G1 | 125962 |
| ENSG000001 | 0 | NA | NA | NA | NA | NA | NA | NA |
| ENSG000001 | 1.46132009 | -0.295730285 | 0.33471496 | -0.8835287 | 0.37695066 | NA | HS3ST6 | 64711 |
| ENSG000001 | 1.17047221 | -0.167069354 | 0.21772035 | -0.7673575 | 0.44286897 | NA | NA | NA |
| ENSG000001 | 1.85339915 | -0.23691102 | 0.35225893 | -0.672548 | 0.50123492 | NA | OR2M5 | 127059 |
| ENSG000001 | 1.41761921 | 0.081920671 | 0.33103814 | 0.24746596 | 0.80454763 | NA | NA | NA |
| ENSG000001 | 1.87239261 | -0.612547666 | 0.35057874 | -1.747247 | 0.0805945 | NA | TEKT4 | 150483 |
| ENSG000001 | 0 | NA | NA | NA | NA | NA | TMOD4 | 29765 |
| ENSG000001 | 1.87620052 | -0.261618739 | 0.36109273 | -0.7245195 | 0.46874683 | NA | LCE3D | 84648 |
| ENSG000001 | 1.62824972 | -0.406146372 | 0.35094166 | -1.1573045 | 0.24714797 | NA | S100A9 | 6280 |
| ENSG000001 | 1.71095039 | 0.201562322 | 0.3538161 | 0.56968104 | 0.56889405 | NA | NA | NA |
| ENSG000001 | 1.46558234 | -0.049215847 | 0.34531977 | -0.1425225 | 0.88666728 | NA | UCN | 7349 |
| ENSG000001 | 1.23336387 | 0.081140129 | 0.32258418 | 0.25153164 | 0.8014031 | NA | CLEC3B | 7123 |
| ENSG000001 | 2.14158357 | -0.627182233 | 0.36301805 | -1.7276888 | 0.08404403 | NA | CAMP | 820 |
| ENSG000001 | 1.28912556 | -0.515686166 | 0.33274927 | -1.549774 | 0.12119577 | NA | CSF2 | 1437 |
| ENSG000001 | 1.13329085 | -0.530087503 | 0.30863378 | -1.7175291 | 0.08588254 | NA | TLX3 | 30012 |
| ENSG000001 | 0.49512483 | -0.344267812 | 0.23617548 | -1.457678 | 0.14492931 | NA | NA | NA |
| ENSG000001 | 1.8919523 | -0.639444675 | 0.35470844 | -1.8027332 | 0.07143013 | NA | DEFA6 | 1671 |
| ENSG000001 | 2.02276446 | -0.708266458 | 0.35852606 | -1.9754951 | 0.04821199 | NA | DEFB1 | 1672 |
| ENSG000001 | 1.40163442 | -0.418362943 | 0.33660919 | -1.2428744 | 0.21391413 | NA | NCF1C | 654817 |
| ENSG000001 | 0.2596576 | -0.193002035 | 0.1837913 | -1.0501152 | 0.29366517 | NA | RPL10L | 140801 |
| ENSG000001 | 0.34344486 | 0.238503239 | 0.20662376 | 1.15428757 | 0.24838229 | NA | PRAP1 | 118471 |
| ENSG000001 | 0.23202362 | -0.008302493 | 0.17586207 | -0.0472103 | 0.96234565 | NA | OPN1MW2 | 728458 |
| ENSG000001 | 1.12566698 | -0.285529985 | 0.32708227 | -0.8729608 | 0.38268447 | NA | POTED | 317754 |
| ENSG000001 | 1.8746402 | -0.389105732 | 0.35917869 | -1.0833208 | 0.27866607 | NA | OR10A5 | 144124 |

| | | | | | | | | | |
|---|---|---|---|---|---|---|---|---|---|
| ENSG000001 | 1.93965792 | -0.504264579 | 0.34863392 | -1.4464014 | 0.14806466 | NA | | C16orf46 | 123775 |
| ENSG000001 | 1.46945207 | -0.044282 | 0.34212985 | -0.1294304 | 0.8970171 | NA | | NA | NA |
| ENSG000001 | 1.88775437 | -0.257199955 | 0.3587034 | -0.7170268 | 0.47335755 | NA | | SAA3P | 6290 |
| ENSG000001 | 1.23168327 | -0.315834546 | 0.32054695 | -0.9852989 | 0.32447729 | NA | | MIR202HG | 101927671 |
| ENSG000001 | 1.0898015 | 0.035516904 | 0.3212039 | 0.11057432 | 0.91195391 | NA | | CCDC103 | 388389 |
| ENSG000001 | 1.24210816 | -0.330667888 | 0.32465358 | -1.0185253 | 0.30842837 | NA | | NA | NA |
| ENSG000001 | 1.35298861 | -0.107811048 | 0.32953526 | -0.3271609 | 0.7435462 | NA | | GNG8 | 94235 |
| ENSG000001 | 0.3593886 | 0.071604659 | 0.20679693 | 0.34625591 | 0.7291504 | NA | | C19orf33 | 64073 |
| ENSG000001 | 1.17991541 | -0.721654832 | 0.31085787 | -2.3214945 | 0.02026017 | NA | | LY6D | 8581 |
| ENSG000001 | 0.13280363 | -0.093394593 | 0.13115644 | -0.7120855 | 0.47641185 | NA | | NA | NA |
| ENSG000001 | 0.35801832 | 0.071082974 | 0.20657063 | 0.34410977 | 0.73076373 | NA | | NA | NA |
| ENSG000001 | 1.638209 | -0.128826997 | 0.34870069 | -0.3694486 | 0.71179335 | NA | | OR5I1 | 10798 |
| ENSG000001 | 1.62593955 | -0.530379706 | 0.34717646 | -1.5276949 | 0.12658831 | NA | | PRR25 | 388199 |
| ENSG000001 | 1.51249314 | -0.349363781 | 0.34704373 | -1.0066852 | 0.31408608 | NA | | NA | NA |
| ENSG000001 | 2.09476059 | -0.127957776 | 0.36820861 | -0.3475143 | 0.72820494 | NA | | PTF1A | 256297 |
| ENSG000001 | 2.10564429 | -0.509746415 | 0.36538496 | -1.3950941 | 0.16298745 | NA | | SCGB1D1 | 10648 |
| ENSG000001 | 0 | NA | NA | NA | NA | NA | | TSPY2 | 64591 |
| ENSG000001 | 1.73870507 | -0.198363809 | 0.35544356 | -0.558074 | 0.57679386 | NA | | CTRB1 | 1504 |
| ENSG000001 | 1.33226771 | -0.117697624 | 0.33558524 | -0.3507235 | 0.7257958 | NA | | NA | NA |
| ENSG000001 | 1.1088698 | -0.103242421 | 0.31341425 | -0.329412 | 0.74184428 | NA | | SPRR1B | 6699 |
| ENSG000001 | 1.24376174 | -0.75926109 | 0.31575072 | -2.4046219 | 0.01618921 | NA | | SPRR1A | 6698 |
| ENSG000001 | 1.19795829 | -0.027411642 | 0.31605263 | -0.0867313 | 0.93088513 | NA | | CRCT1 | 54544 |
| ENSG000001 | 1.97670243 | -0.454105217 | 0.36315878 | -1.2504316 | 0.21114193 | NA | | VPREB1 | 7441 |
| ENSG000001 | 0.88815263 | -0.103100371 | 0.29106497 | -0.3542177 | 0.72317573 | NA | | CLIC3 | 9022 |
| ENSG000001 | 0.3854038 | -0.110128691 | 0.21952326 | -0.5016721 | 0.61589821 | NA | | NA | NA |
| ENSG000001 | 1.62431932 | -0.12367196 | 0.34755777 | -0.3558314 | 0.72196685 | NA | | GP9 | 2815 |
| ENSG000001 | 1.01521163 | -0.523036317 | 0.31168178 | -1.67811 | 0.09332562 | NA | | ACTRT2 | 140625 |
| ENSG000001 | 0 | NA | NA | NA | NA | NA | | NA | NA |
| ENSG000001 | 0 | NA | NA | NA | NA | NA | | PRY | 9081 |

| | | | | | | | | |
|---|---|---|---|---|---|---|---|---|
| ENSG000001 | 0 | NA | NA | NA | NA | NA | NA | NA |
| ENSG000001 | 0 | NA | NA | NA | NA | NA | PRY2 | 442862 |
| ENSG000001 | 0 | NA | NA | NA | NA | NA | NA | NA |
| ENSG000001 | 0 | NA | NA | NA | NA | NA | NA | NA |
| ENSG000001 | 0 | NA | NA | NA | NA | NA | NA | NA |
| ENSG000001 | 1.14410412 | -0.258048109 | 0.31817605 | -0.811023 | 0.41735244 | NA | GPR25 | 2848 |
| ENSG000001 | 0 | NA | NA | NA | NA | NA | NA | NA |
| ENSG000001 | 0 | NA | NA | NA | NA | NA | HOXD4 | 3233 |
| ENSG000001 | 1.59937325 | 0.01768925 | 0.34914786 | 0.05066407 | 0.95959321 | NA | HSPB2 | 3316 |
| ENSG000001 | 2.14636586 | 0.297205268 | 0.36518671 | 0.8138447 | 0.41573392 | NA | COX7B2 | 170712 |
| ENSG000001 | 0.85681254 | 0.050296224 | 0.30189457 | 0.16660195 | 0.86768326 | NA | RASA4B | 100271927 |
| ENSG000001 | 0.83902224 | 0.203987147 | 0.29630441 | 0.68843777 | 0.49117714 | NA | HSFX1 | 100506164 |
| ENSG000001 | 2.15640242 | -0.370869569 | 0.3636641 | -1.0198135 | 0.3078169 | NA | RLN3 | 117579 |
| ENSG000001 | 1.26920136 | -0.491955005 | 0.32713564 | -1.5038258 | 0.13262623 | NA | KRTAP4-4 | 84616 |
| ENSG000001 | 1.96463366 | -0.300031542 | 0.35744025 | -0.8393894 | 0.40125086 | NA | OR1L6 | 392390 |
| ENSG000001 | 0 | NA | NA | NA | NA | NA | NA | NA |
| ENSG000001 | 0.12070844 | -0.088503695 | 0.12783931 | -0.6923042 | 0.48874631 | NA | SPACA5 | 389852 |
| ENSG000001 | 0 | NA | NA | NA | NA | NA | NA | NA |
| ENSG000001 | 0.74365079 | -0.022786451 | 0.27880612 | -0.0817287 | 0.93486249 | NA | LKAAEAR1 | 198437 |
| ENSG000001 | 1.12594718 | -0.265952722 | 0.3175336 | -0.8375577 | 0.40227914 | NA | DEFB4A | 1673 |
| ENSG000001 | 0.23936338 | -0.005479057 | 0.17759977 | -0.0308506 | 0.9753887 | NA | NXNL1 | 115861 |
| ENSG000001 | 1.56876282 | -0.954515901 | 0.33785461 | -2.8252268 | 0.00472472 | NA | NA | NA |
| ENSG000001 | 0.25260017 | -0.015978705 | 0.18049464 | -0.0885273 | 0.92945758 | NA | LCE1D | 353134 |
| ENSG000001 | 1.60988988 | 0.006181251 | 0.3450083 | 0.01791624 | 0.98570567 | NA | NA | NA |
| ENSG000001 | 0.99277745 | -0.337160118 | 0.30872563 | -1.0921028 | 0.27478793 | NA | NA | NA |
| ENSG000001 | 0 | NA | NA | NA | NA | NA | NA | NA |
| ENSG000001 | 0 | NA | NA | NA | NA | NA | CDY1 | 9085 |
| ENSG000001 | 0 | NA | NA | NA | NA | NA | NA | NA |
| ENSG000001 | 0 | NA | NA | NA | NA | NA | NA | NA |

| | | | | | | | | | |
|---|---|---|---|---|---|---|---|---|---|
| ENSG000001 | 0 | NA | NA | NA | NA | NA | NA | NA | |
| ENSG000001 | 0 | NA | NA | NA | NA | NA | CDY1B | | 253175 |
| ENSG000001 | 1.76523362 | -0.615846091 | 0.35398102 | -1.7397715 | 0.08189915 | NA | INSL5 | | 10022 |
| ENSG000001 | 1.75268197 | -0.601426072 | 0.35277354 | -1.7048503 | 0.08822235 | NA | OR5AR1 | | 219493 |
| ENSG000001 | 0 | NA | NA | NA | NA | NA | HSFY1 | | 86614 |
| ENSG000001 | 0.8776481 | -0.117739236 | 0.30325589 | -0.3882505 | 0.6978307 | NA | NA | NA | |
| ENSG000001 | 1.32013827 | 0.30688053 | 0.33353467 | 0.92008586 | 0.35752789 | NA | HOXC5 | | 3222 |
| ENSG000001 | 0.3863157 | -0.274134926 | 0.21457207 | -1.277589 | 0.20139437 | NA | NA | NA | |
| ENSG000001 | 1.11229167 | -0.404529268 | 0.31628561 | -1.2789999 | 0.20089709 | NA | NA | NA | |
| ENSG000001 | 0 | NA | NA | NA | NA | NA | NA | NA | |
| ENSG000001 | 0 | NA | NA | NA | NA | NA | NA | NA | |
| ENSG000001 | 1.97607477 | -0.060306948 | 0.35700912 | -0.1689227 | 0.86585744 | NA | RNASE8 | | 122665 |
| ENSG000001 | 2.10091016 | 0.121280702 | 0.36167337 | 0.33533213 | 0.73737456 | NA | NA | NA | |
| ENSG000001 | 0 | NA | NA | NA | NA | NA | NA | NA | |
| ENSG000001 | 1.03510304 | -0.536100349 | 0.31389681 | -1.7078872 | 0.08765727 | NA | NA | NA | |
| ENSG000001 | 0.35346928 | -0.088570748 | 0.20618147 | -0.4295767 | 0.66750362 | NA | NA | NA | |
| ENSG000001 | 1.72862956 | -0.445543091 | 0.35132644 | -1.2681741 | 0.20473578 | NA | NA | NA | |
| ENSG000001 | 1.73762138 | -0.336453312 | 0.35737597 | -0.9414548 | 0.34647186 | NA | NA | NA | |
| ENSG000001 | 0.6275579 | -0.281336606 | 0.26701873 | -1.0536213 | 0.29205636 | NA | NA | NA | |
| ENSG000001 | 1.59573961 | 0.016185629 | 0.35325089 | 0.04581907 | 0.96345446 | NA | NA | NA | |
| ENSG000001 | 0.62041457 | -0.27371262 | 0.26650407 | -1.0270486 | 0.3043976 | NA | AMY1B | | 277 |
| ENSG000001 | 1.49669822 | -0.449249377 | 0.33824881 | -1.3281625 | 0.18412443 | NA | OR10P1 | | 121130 |
| ENSG000001 | 1.48315448 | -0.333572098 | 0.34240819 | -0.9741943 | 0.32996009 | NA | NA | NA | |
| ENSG000001 | 1.37274205 | -0.257040375 | 0.33493962 | -0.767423 | 0.44283005 | NA | GPR152 | | 390212 |
| ENSG000001 | 0.12070844 | -0.088503695 | 0.12783931 | -0.6923042 | 0.48874631 | NA | NA | NA | |
| ENSG000001 | 2.14309182 | -0.510211513 | 0.36560687 | -1.3955195 | 0.16285924 | NA | NA | NA | |
| ENSG000001 | 1.9358594 | -0.537048111 | 0.35943402 | -1.4941494 | 0.13513659 | NA | NA | NA | |
| ENSG000001 | 0.73681957 | -0.033780691 | 0.28998188 | -0.1164924 | 0.90726229 | NA | NA | NA | |
| ENSG000001 | 1.84372177 | 0.128868657 | 0.35838578 | 0.35958083 | 0.71916062 | NA | ASCL3 | | 56676 |

| | | | | | | | | | |
|---|---|---|---|---|---|---|---|---|---|
| ENSG000001 | 0.11131518 | 0.081951103 | 0.12663804 | 0.64712866 | 0.51754868 | NA | AMIGO3 | | 386724 |
| ENSG000001 | 1.50053826 | -0.617336966 | 0.34012137 | -1.815049 | 0.06951638 | NA | NA | NA | |
| ENSG000001 | 0.50497064 | -0.183718113 | 0.24165995 | -0.760234 | 0.44711472 | NA | NA | NA | |
| ENSG000001 | 0.12070844 | -0.088503695 | 0.12783931 | -0.6923042 | 0.48874631 | NA | NA | NA | |
| ENSG000001 | 1.02255084 | -0.536784374 | 0.31334684 | -1.7130678 | 0.08670007 | NA | NA | NA | |
| ENSG000001 | 0.12070844 | -0.088503695 | 0.12783931 | -0.6923042 | 0.48874631 | NA | NA | NA | |
| ENSG000001 | 1.52615311 | -0.501169091 | 0.34804844 | -1.4399406 | 0.1498842 | NA | OR51B6 | | 390058 |
| ENSG000001 | 0.75537934 | -0.187046077 | 0.28061071 | -0.6665679 | 0.50504821 | NA | NA | NA | |
| ENSG000001 | 1.15008603 | -0.257736615 | 0.31587482 | -0.8159454 | 0.41453134 | NA | NA | NA | |
| ENSG000001 | 1.60613012 | -0.128055524 | 0.35358351 | -0.3621649 | 0.71722884 | NA | NA | NA | |
| ENSG000001 | 0 | NA | NA | NA | NA | NA | TGIF2LY | | 90655 |
| ENSG000001 | 0.78754062 | -0.365675631 | 0.28493535 | -1.2833635 | 0.19936476 | NA | FOXC2 | | 2303 |
| ENSG000001 | 0 | NA | NA | NA | NA | NA | NA | NA | |
| ENSG000001 | 1.84500478 | -0.113705991 | 0.35309617 | -0.3220256 | 0.74743333 | NA | NA | NA | |
| ENSG000001 | 1.24435864 | -0.464330056 | 0.32286007 | -1.4381774 | 0.15038372 | NA | NA | NA | |
| ENSG000001 | 1.38768215 | -0.262193137 | 0.33745926 | -0.7769623 | 0.43718099 | NA | DEFB104A | | 140596 |
| ENSG000001 | 0.12070844 | -0.088503695 | 0.12783931 | -0.6923042 | 0.48874631 | NA | DEFB103A | | 414325 |
| ENSG000001 | 1.55790303 | 0.288883467 | 0.34659103 | 0.83349955 | 0.40456305 | NA | NA | NA | |
| ENSG000001 | 1.29696742 | 0.484450625 | 0.33447069 | 1.44840982 | 0.14750247 | NA | NA | NA | |
| ENSG000001 | 0.90112692 | -0.259399981 | 0.29324956 | -0.8845708 | 0.37638819 | NA | DEFB104B | | 503618 |
| ENSG000001 | 0.96451232 | -0.025804315 | 0.30694787 | -0.0840674 | 0.93300283 | NA | OR2T35 | | 403244 |
| ENSG000001 | 1.62955337 | -0.508951541 | 0.34303584 | -1.4836687 | 0.13789683 | NA | OR2M7 | | 391196 |
| ENSG000001 | 0.12685397 | -0.090215768 | 0.12901919 | -0.699243 | 0.48440017 | NA | SPACA4 | | 171169 |
| ENSG000001 | 0.48573157 | -0.181932918 | 0.23941992 | -0.7598905 | 0.44732006 | NA | DEFB103B | | 55894 |
| ENSG000001 | 1.24088278 | -0.318319133 | 0.32382206 | -0.9830063 | 0.32560433 | NA | NA | NA | |
| ENSG000001 | 0.12070844 | -0.088503695 | 0.12783931 | -0.6923042 | 0.48874631 | NA | NA | NA | |
| ENSG000001 | 1.95473736 | -0.312030718 | 0.36006966 | -0.8665843 | 0.38616981 | NA | NA | NA | |
| ENSG000001 | 1.76320397 | -0.186140565 | 0.35282078 | -0.5275782 | 0.5977921 | NA | NA | NA | |
| ENSG000001 | 2.13984644 | -0.765049766 | 0.36237553 | -2.111207 | 0.03475452 | NA | NA | NA | |

| | | | | | | | | | |
|---|---|---|---|---|---|---|---|---|---|
| ENSG000001 | 1.20094254 | 0.263948939 | 0.33158271 | 0.79602745 | 0.42601608 | NA | OR2G3 | | 81469 |
| ENSG000001 | 1.82700212 | -0.752450496 | 0.35138595 | -2.141379 | 0.03224349 | NA | OR2G2 | | 81470 |
| ENSG000001 | 1.11897854 | -0.414906694 | 0.31891638 | -1.300989 | 0.19326223 | NA | VCX2 | | 51480 |
| ENSG000001 | 2.16872791 | 0.175099706 | 0.36738552 | 0.47661026 | 0.63363968 | NA | FAM187B | | 148109 |
| ENSG000001 | 1.6718508 | 0.078441653 | 0.35319196 | 0.22209354 | 0.82424107 | NA | NA | NA | |
| ENSG000001 | 0.96928948 | 0.113164225 | 0.29784144 | 0.37994789 | 0.7039841 | NA | C2orf57 | | 165100 |
| ENSG000001 | 1.80254668 | -0.467784272 | 0.35357946 | -1.3229962 | 0.18583665 | NA | DEFB114 | | 245928 |
| ENSG000001 | 0.61818637 | 0.059474041 | 0.27041681 | 0.21993471 | 0.82592201 | NA | OR4F4 | | 26682 |
| ENSG000001 | 0.24756242 | -0.187034346 | 0.18119817 | -1.0322088 | 0.30197431 | NA | NA | NA | |
| ENSG000001 | 2.13594594 | -0.130918652 | 0.36266239 | -0.3609932 | 0.71810454 | NA | NA | NA | |
| ENSG000001 | 0.73969619 | -0.030242014 | 0.26941072 | -0.1122525 | 0.91062324 | NA | ULK4P3 | | 89837 |
| ENSG000001 | 1.43851088 | 0.238090717 | 0.34570167 | 0.68871728 | 0.4910012 | NA | FAM27L | | 284123 |
| ENSG000001 | 1.88948938 | -0.26275847 | 0.35930704 | -0.7312923 | 0.46460063 | NA | PRM3 | | 58531 |
| ENSG000001 | 1.94860909 | 0.062422978 | 0.36427246 | 0.17136343 | 0.86393801 | NA | SPAG11A | | 653423 |
| ENSG000001 | 0.72241029 | 0.133267595 | 0.27759156 | 0.48008518 | 0.63116682 | NA | CALML5 | | 51806 |
| ENSG000001 | 2.13873921 | -0.620988362 | 0.36283139 | -1.7115067 | 0.08698761 | NA | CCDC185 | | 164127 |
| ENSG000001 | 1.23533147 | -0.202069 | 0.33412968 | -0.6047622 | 0.54533701 | NA | LINC00314 | | 246705 |
| ENSG000001 | 0.70341063 | -0.018084693 | 0.26668241 | -0.0678136 | 0.94593402 | NA | NA | NA | |
| ENSG000001 | 1.82182162 | -0.229650583 | 0.34669389 | -0.6624016 | 0.50771389 | NA | NA | NA | |
| ENSG000001 | 1.92527638 | -0.647572464 | 0.35457787 | -1.8263194 | 0.06780216 | NA | OTOS | | 150677 |
| ENSG000001 | 0.35733509 | 0.244646831 | 0.20894085 | 1.17089037 | 0.24164285 | NA | NA | NA | |
| ENSG000001 | 1.33374426 | 0.026937582 | 0.33875353 | 0.07951971 | 0.93661926 | NA | LGALS7B | | 653499 |
| ENSG000001 | 2.02418022 | -0.838146936 | 0.35688725 | -2.3484922 | 0.01884959 | NA | TAS1R2 | | 80834 |
| ENSG000001 | 0 | NA | NA | NA | NA | NA | NA | NA | |
| ENSG000001 | 1.63512604 | 0.492562299 | 0.34876407 | 1.41230804 | 0.15785928 | NA | NA | NA | |
| ENSG000001 | 1.53783013 | -0.616819047 | 0.34086926 | -1.8095473 | 0.07036602 | NA | NA | NA | |
| ENSG000001 | 1.60434402 | -0.62871754 | 0.33891299 | -1.8551002 | 0.06358192 | NA | ZCCHC5 | | 203430 |
| ENSG000001 | 0.63921993 | 0.0498278 | 0.25845554 | 0.19279061 | 0.84712296 | NA | NA | NA | |
| ENSG000001 | 1.36975388 | -0.240005816 | 0.330003 | -0.7272837 | 0.46705216 | NA | NA | NA | |

| | | | | | | | | |
|---|---|---|---|---|---|---|---|---|
| ENSG000001 | 1.52701397 | -0.466749454 | 0.33877389 | -1.377761 | 0.16827709 | NA | NA | NA |
| ENSG000001 | 0 | NA | NA | NA | NA | NA | HNRNPCL4 | 101060301 |
| ENSG000001 | 0.37304803 | -0.090842893 | 0.20939682 | -0.4338313 | 0.66441096 | NA | SYCN | 342898 |
| ENSG000001 | 0 | NA | NA | NA | NA | NA | NA | NA |
| ENSG000001 | 2.04851261 | -0.451464478 | 0.36192847 | -1.2473859 | 0.21225603 | NA | PIK3CD-AS1 | 644997 |
| ENSG000001 | 1.92785977 | 0.306739045 | 0.35723087 | 0.85865771 | 0.39052939 | NA | OR10A7 | 121364 |
| ENSG000001 | 2.13506885 | -0.372549078 | 0.36263634 | -1.0273352 | 0.30426265 | NA | OR1E1 | 8387 |
| ENSG000001 | 1.84571971 | -0.260483773 | 0.35992259 | -0.7237216 | 0.46923664 | NA | NA | NA |
| ENSG000001 | 0.78731386 | -0.195162287 | 0.28101843 | -0.6944822 | 0.48737989 | NA | NA | NA |
| ENSG000001 | 1.63908865 | -0.264851 | 0.35216759 | -0.7520596 | 0.45201524 | NA | FAM71E2 | 284418 |
| ENSG000001 | 1.84586407 | -0.251770009 | 0.35732169 | -0.7046032 | 0.4810572 | NA | NKX2-6 | 137814 |
| ENSG000001 | 1.43956659 | 0.090397944 | 0.33883842 | 0.26678776 | 0.78963258 | NA | NA | NA |
| ENSG000001 | 2.01948056 | -0.324170358 | 0.36110384 | -0.8977206 | 0.36933452 | NA | NA | NA |
| ENSG000001 | 2.14192151 | 0.186504949 | 0.36779429 | 0.50709039 | 0.6120914 | NA | TMEM86B | 255043 |
| ENSG000001 | 1.77457536 | -0.732438217 | 0.35016925 | -2.0916691 | 0.03646812 | NA | NA | NA |
| ENSG000001 | 1.73625301 | -0.195816973 | 0.35694028 | -0.5485987 | 0.58328088 | NA | NA | NA |
| ENSG000001 | 1.98551938 | 0.790248942 | 0.35932891 | 2.1992356 | 0.02786117 | NA | NA | NA |
| ENSG000001 | 0.83504177 | 0.050250725 | 0.29558221 | 0.17000592 | 0.86500548 | NA | KRTAP9-7 | 100505724 |
| ENSG000001 | 1.51978082 | -0.346912549 | 0.34716896 | -0.9992614 | 0.31766807 | NA | NA | NA |
| ENSG000001 | 1.90719324 | -0.51192199 | 0.35597956 | -1.4380657 | 0.15041542 | NA | NA | NA |
| ENSG000001 | 1.31243187 | 0.165961012 | 0.33498821 | 0.49542344 | 0.62030125 | NA | OR10Q1 | 219960 |
| ENSG000001 | 0.88612946 | -0.120234611 | 0.30365646 | -0.395956 | 0.69213746 | NA | OR2A4 | 79541 |
| ENSG000001 | 1.45829465 | -0.050770594 | 0.3439483 | -0.1476111 | 0.88264968 | NA | NA | NA |
| ENSG000001 | 1.64819904 | -0.558988467 | 0.35095727 | -1.5927537 | 0.11121547 | NA | NA | NA |
| ENSG000001 | 1.54673359 | -0.757336684 | 0.33618725 | -2.2527228 | 0.02427663 | NA | NA | NA |
| ENSG000001 | 1.86240066 | -0.398572514 | 0.36141505 | -1.1028111 | 0.27010923 | NA | MAP6D1 | 79929 |
| ENSG000001 | 1.08790095 | -0.115040445 | 0.32144923 | -0.3578806 | 0.72043266 | NA | CSNK1G2-AS1 | 255193 |
| ENSG000001 | 1.79642259 | 0.129621909 | 0.35496177 | 0.36517147 | 0.71498342 | NA | DEFB112 | 245915 |
| ENSG000001 | 0 | NA | NA | NA | NA | NA | TTTY11 | 83866 |

| | | | | | | | | | |
|---|---|---|---|---|---|---|---|---|---|
| ENSG000001 | 2.07600495 | 0.001008534 | 0.36456278 | 0.00276642 | 0.99779272 | NA | NA | NA | |
| ENSG000001 | 1.00627483 | -0.356078214 | 0.31079499 | -1.1457013 | 0.25191877 | NA | NA | NA | |
| ENSG000001 | 1.26712666 | -0.511089193 | 0.33221499 | -1.538429 | 0.12394375 | NA | SPEM1 | | 374768 |
| ENSG000001 | 2.0707721 | -0.451018927 | 0.360498 | -1.2510997 | 0.21089812 | NA | NA | NA | |
| ENSG000001 | 0.85551123 | -0.243943637 | 0.28822537 | -0.8463642 | 0.39734961 | NA | NA | NA | |
| ENSG000001 | 0.23202362 | -0.008302493 | 0.17586207 | -0.0472103 | 0.96234565 | NA | NA | NA | |
| ENSG000001 | 0.85058056 | -0.103122652 | 0.29515749 | -0.3493818 | 0.7268027 | NA | NA | NA | |
| ENSG000001 | 1.06788693 | 0.037171252 | 0.31892133 | 0.11655304 | 0.90721425 | NA | NA | NA | |
| ENSG000001 | 0.86302513 | -0.41454039 | 0.29130684 | -1.4230369 | 0.15472545 | NA | OR8K5 | | 219453 |
| ENSG000001 | 0.99969403 | -0.200271688 | 0.31387515 | -0.6380616 | 0.52343357 | NA | OR8H3 | | 390152 |
| ENSG000001 | 2.16071348 | -0.387526557 | 0.36453134 | -1.0630816 | 0.28774495 | NA | NA | NA | |
| ENSG000001 | 0.60162804 | 0.055843814 | 0.26262531 | 0.21263683 | 0.83161024 | NA | ACTL9 | | 284382 |
| ENSG000001 | 0.75401097 | -0.030402718 | 0.27950383 | -0.1087739 | 0.91338184 | NA | NA | NA | |
| ENSG000001 | 1.64359264 | -0.541839615 | 0.34877615 | -1.5535455 | 0.12029287 | NA | FTMT | | 94033 |
| ENSG000001 | 1.36910312 | -0.112545917 | 0.33492786 | -0.3360303 | 0.736848 | NA | OR4C6 | | 219432 |
| ENSG000001 | 1.56272709 | 0.01542393 | 0.34932614 | 0.04415338 | 0.96478214 | NA | OR4C15 | | 81309 |
| ENSG000001 | 1.0851623 | 0.035481716 | 0.32104825 | 0.11051833 | 0.91199831 | NA | NA | NA | |
| ENSG000001 | 1.63501197 | -0.125062893 | 0.34865202 | -0.3587041 | 0.71981649 | NA | NA | NA | |
| ENSG000001 | 1.61182365 | -0.35822613 | 0.34199733 | -1.047453 | 0.2948907 | NA | OR4A15 | | 81328 |
| ENSG000001 | 1.71891918 | -0.063921618 | 0.35044614 | -0.1824007 | 0.85526829 | NA | NEUROG1 | | 4762 |
| ENSG000001 | 0.62618953 | -0.114303343 | 0.27141404 | -0.4211401 | 0.67365276 | NA | NA | NA | |
| ENSG000001 | 1.24820272 | -0.319428029 | 0.3240314 | -0.9857934 | 0.32423449 | NA | NA | NA | |
| ENSG000001 | 1.27429365 | -0.19122064 | 0.3287161 | -0.5817197 | 0.56075548 | NA | NA | NA | |
| ENSG000001 | 0.7214772 | 0.14040389 | 0.28371685 | 0.49487328 | 0.62068957 | NA | OR6B2 | | 389090 |
| ENSG000001 | 1.10312125 | -0.112535241 | 0.31575395 | -0.3564017 | 0.72153976 | NA | MRGPRG | | 386746 |
| ENSG000001 | 0.26813897 | -0.197007041 | 0.18550319 | -1.0620143 | 0.2882292 | NA | MBD3L3 | | 653657 |
| ENSG000001 | 0.62889593 | -0.276614073 | 0.26719437 | -1.0352541 | 0.30055024 | NA | GLTPD2 | | 388323 |
| ENSG000001 | 2.01188469 | -0.609029536 | 0.36470253 | -1.6699351 | 0.09493221 | NA | PRAMEF8 | | 391002 |
| ENSG000001 | 0.23890637 | -0.010420674 | 0.17747379 | -0.0587167 | 0.95317776 | NA | NA | NA | |

| | | | | | | | | |
|---|---|---|---|---|---|---|---|---|
| ENSG000001 | 1.72638602 | -0.453428168 | 0.35319528 | -1.2837889 | 0.19921584 | NA | NA | NA |
| ENSG000001 | 1.61552283 | -0.271292275 | 0.35185546 | -0.7710333 | 0.4406872 | NA | LOC339059 | 339059 |
| ENSG000001 | 2.07392916 | -0.010197968 | 0.36105838 | -0.0282447 | 0.97746703 | NA | NA | NA |
| ENSG000001 | 1.43663653 | -0.032135787 | 0.33558361 | -0.0957609 | 0.92371048 | NA | IFNL1 | 282618 |
| ENSG000001 | 0 | NA | NA | NA | NA | NA | CDY2A | 9426 |
| ENSG000001 | 1.15461726 | -0.614542622 | 0.32342171 | -1.9001279 | 0.05741633 | NA | KCNK4 | 50801 |
| ENSG000001 | 1.95173949 | -0.058669219 | 0.34960162 | -0.1678174 | 0.86672697 | NA | NA | NA |
| ENSG000001 | 1.51778276 | -0.440955885 | 0.33413075 | -1.3197106 | 0.18693166 | NA | KRTAP11-1 | 337880 |
| ENSG000001 | 0.85940684 | -0.110668361 | 0.29120881 | -0.3800309 | 0.70392244 | NA | NA | NA |
| ENSG000001 | 0 | NA | NA | NA | NA | NA | CMC4 | 100272147 |
| ENSG000001 | 0.63270563 | -0.25685676 | 0.259029 | -0.9916139 | 0.3213859 | NA | LOC401589 | 401589 |
| ENSG000001 | 0.76628085 | -0.185682035 | 0.28133178 | -0.6600109 | 0.50924684 | NA | KRTAP13-2 | 337959 |
| ENSG000001 | 1.86154274 | -0.129302521 | 0.36151407 | -0.3576694 | 0.72059074 | NA | NA | NA |
| ENSG000001 | 2.12154973 | -0.495177019 | 0.36290479 | -1.3644819 | 0.17241595 | NA | ODF3L1 | 161753 |
| ENSG000001 | 0 | NA | NA | NA | NA | NA | NA | NA |
| ENSG000001 | 1.9756829 | -0.552914954 | 0.35693412 | -1.5490673 | 0.12136555 | NA | OR1G1 | 8390 |
| ENSG000001 | 1.63627407 | -0.414455152 | 0.35166136 | -1.1785632 | 0.23857214 | NA | NKX2-5 | 1482 |
| ENSG000001 | 0 | NA | NA | NA | NA | NA | PRORY | 100533178 |
| ENSG000001 | 1.33230154 | 0.013027052 | 0.33306516 | 0.03911262 | 0.9688006 | NA | GJD3 | 125111 |
| ENSG000001 | 1.76983822 | -0.197849741 | 0.35403998 | -0.5588345 | 0.5762747 | NA | NA | NA |
| ENSG000001 | 0.49082195 | -0.014647497 | 0.23849846 | -0.0614155 | 0.95102832 | NA | RIMBP3C | 150221 |
| ENSG000001 | 1.57402428 | -0.215984998 | 0.33194375 | -0.6506675 | 0.51526117 | NA | ABHD16B | 140701 |
| ENSG000001 | 1.69908251 | 0.083495253 | 0.35459646 | 0.23546555 | 0.81384741 | NA | OR52E8 | 390079 |
| ENSG000001 | 0.93204246 | -0.430357711 | 0.29493135 | -1.4591793 | 0.14451577 | NA | OR5P2 | 120065 |
| ENSG000001 | 0.47667325 | 0.138618663 | 0.22766399 | 0.6088739 | 0.54260802 | NA | NA | NA |
| ENSG000001 | 1.54897904 | -0.642473395 | 0.34397877 | -1.8677705 | 0.06179407 | NA | OR2T34 | 127068 |
| ENSG000001 | 0.56681244 | 0.393392237 | 0.25469451 | 1.54456504 | 0.12245151 | NA | BOLA2 | 552900 |
| ENSG000001 | 0 | NA | NA | NA | NA | NA | NA | NA |
| ENSG000001 | 0.48646878 | -0.181653651 | 0.2395107 | -0.7584365 | 0.44818973 | NA | TMEM89 | 440955 |

| | | | | | | | | |
|---|---|---|---|---|---|---|---|---|
| ENSG000001 | 1.73739462 | -0.187984426 | 0.35101847 | -0.53554 | 0.59227652 | NA | URAD | 646625 |
| ENSG000001 | 0.51550551 | -0.37860525 | 0.24578489 | -1.5403927 | 0.12346466 | NA | KLRC4 | 8302 |
| ENSG000001 | 0.36553413 | 0.069401489 | 0.20780452 | 0.33397488 | 0.73839851 | NA | NA | NA |
| ENSG000001 | 1.51123475 | -0.299341337 | 0.33406718 | -0.8960513 | 0.37022535 | NA | NA | NA |
| ENSG000001 | 1.46598536 | -0.170401643 | 0.3359405 | -0.5072376 | 0.61198813 | NA | NA | NA |
| ENSG000001 | 0.91627341 | -0.460143215 | 0.30152955 | -1.5260303 | 0.12700233 | NA | NA | NA |
| ENSG000001 | 0 | NA | NA | NA | NA | NA | NA | NA |
| ENSG000001 | 0 | NA | NA | NA | NA | NA | BPY2 | 9083 |
| ENSG000001 | 0.79556497 | -0.367634006 | 0.28377548 | -1.2955101 | 0.19514432 | NA | NA | NA |
| ENSG000001 | 0 | NA | NA | NA | NA | NA | BPY2B | 442867 |
| ENSG000001 | 0.11865494 | 0.083440866 | 0.12751478 | 0.65436232 | 0.5128784 | NA | UTY | 7404 |
| ENSG000001 | 0.72563629 | -0.175145842 | 0.27692276 | -0.6324718 | 0.52707862 | NA | NA | NA |
| ENSG000001 | 1.98666226 | -0.175352257 | 0.35809697 | -0.4896781 | 0.62436168 | NA | NA | NA |
| ENSG000001 | 1.39838779 | -0.25127298 | 0.33388438 | -0.7525748 | 0.45170544 | NA | NA | NA |
| ENSG000001 | 2.0313991 | 0.392065591 | 0.36467495 | 1.07510974 | 0.2823256 | NA | NA | NA |
| ENSG000001 | 1.75694826 | -0.064396278 | 0.34953772 | -0.1842327 | 0.8538309 | NA | KRTAP20-2 | 337976 |
| ENSG000001 | 0 | NA | NA | NA | NA | NA | CTAG1B | 1485 |
| ENSG000001 | 2.00379562 | -0.519050048 | 0.352756 | -1.4714138 | 0.14117924 | NA | NA | NA |
| ENSG000001 | 0.7710561 | -0.188233817 | 0.28164389 | -0.6683398 | 0.50391673 | NA | NA | NA |
| ENSG000001 | 2.11036589 | -0.608369398 | 0.36068938 | -1.6866851 | 0.09166392 | NA | LOC100288570 | 100288570 |
| ENSG000001 | 1.47464278 | -0.18810567 | 0.34284914 | -0.5486543 | 0.58324268 | NA | NA | NA |
| ENSG000001 | 1.84526619 | -0.005052953 | 0.35417219 | -0.0142669 | 0.98861702 | NA | OR10J5 | 127385 |
| ENSG000001 | 1.52117092 | -0.342983509 | 0.34720467 | -0.9878424 | 0.32322982 | NA | OLIG1 | 116448 |
| ENSG000001 | 1.8836618 | -0.399631953 | 0.35922903 | -1.1124712 | 0.2659356 | NA | NA | NA |
| ENSG000001 | 1.58426894 | -0.467128571 | 0.33913144 | -1.3774263 | 0.16838048 | NA | NA | NA |
| ENSG000001 | 2.13604023 | -0.467574609 | 0.35832176 | -1.3049015 | 0.19192638 | NA | MRGPRE | 116534 |
| ENSG000001 | 1.5085082 | -0.198620045 | 0.34205448 | -0.5806679 | 0.56146432 | NA | KRTAP19-1 | 337882 |
| ENSG000001 | 1.13875137 | 0.027452037 | 0.31663529 | 0.08669923 | 0.93091059 | NA | NA | NA |
| ENSG000001 | 2.14536294 | -0.481505949 | 0.3617991 | -1.3308655 | 0.18323327 | NA | NA | NA |

| | | | | | | | | |
|---|---|---|---|---|---|---|---|---|
| ENSG000001 | 1.73363753 | -0.178564455 | 0.35097254 | -0.5087704 | 0.61091314 | NA | NA | NA |
| ENSG000001 | 1.56500868 | -0.639101352 | 0.34310673 | -1.8626896 | 0.06250594 | NA | NA | NA |
| ENSG000001 | 0.49782676 | -0.186543761 | 0.24081583 | -0.7746325 | 0.43855686 | NA | LRRC26 | 389816 |
| ENSG000001 | 0.4933859 | -0.021085811 | 0.24629532 | -0.0856119 | 0.93177494 | NA | KRTAP6-1 | 337966 |
| ENSG000001 | 2.14562544 | -0.372242559 | 0.3645039 | -1.0212307 | 0.30714516 | NA | NA | NA |
| ENSG000001 | 0.12070844 | -0.088503695 | 0.12783931 | -0.6923042 | 0.48874631 | NA | NA | NA |
| ENSG000001 | 2.10041686 | -0.124976991 | 0.36561541 | -0.3418264 | 0.73248156 | NA | NA | NA |
| ENSG000001 | 0.84970146 | 0.05060011 | 0.2947463 | 0.17167344 | 0.86369427 | NA | NA | NA |
| ENSG000001 | 0 | NA | NA | NA | NA | NA | SRY | 6736 |
| ENSG000001 | 0.85800868 | 0.195090245 | 0.28727085 | 0.67911605 | 0.49706433 | NA | NA | NA |
| ENSG000001 | 0 | NA | NA | NA | NA | NA | NA | NA |
| ENSG000001 | 0 | NA | NA | NA | NA | NA | TTTY13 | 83868 |
| ENSG000001 | 1.86800094 | -0.134965072 | 0.36156658 | -0.3732786 | 0.70894109 | NA | LRRC14B | 389257 |
| ENSG000001 | 1.86900512 | -0.245442368 | 0.35751661 | -0.6865202 | 0.49238514 | NA | AQP12B | 653437 |
| ENSG000001 | 0.9617216 | 0.12384308 | 0.30683493 | 0.40361467 | 0.68649609 | NA | NA | NA |
| ENSG000001 | 0.75711827 | -0.523135555 | 0.27850213 | -1.8783898 | 0.06032786 | NA | LINC00313 | 114038 |
| ENSG000001 | 0 | NA | NA | NA | NA | NA | NA | NA |
| ENSG000001 | 1.1141969 | -0.257546194 | 0.31462896 | -0.8185712 | 0.41303112 | NA | NA | NA |
| ENSG000001 | 0.50750235 | -0.197774682 | 0.24907427 | -0.794039 | 0.42717272 | NA | NUPR1L | 389493 |
| ENSG000001 | 0.90802231 | -0.279957619 | 0.29926735 | -0.9354766 | 0.34954272 | NA | NA | NA |
| ENSG000001 | 0.87710676 | -0.118147501 | 0.30323109 | -0.3896286 | 0.69681121 | NA | NA | NA |
| ENSG000001 | 1.64142448 | -0.249522423 | 0.34632918 | -0.7204776 | 0.47123098 | NA | NA | NA |
| ENSG000001 | 1.97285792 | -0.065724134 | 0.3649785 | -0.1800767 | 0.85709233 | NA | SPATA8 | 145946 |
| ENSG000001 | 1.38789067 | -0.24826172 | 0.3295875 | -0.7532498 | 0.45129982 | NA | NA | NA |
| ENSG000001 | 0.8814961 | -0.275762091 | 0.2995484 | -0.9205928 | 0.35726308 | NA | NA | NA |
| ENSG000001 | 0 | NA | NA | NA | NA | NA | NA | NA |
| ENSG000001 | 1.92852573 | -0.175009944 | 0.35587597 | -0.4917723 | 0.62288032 | NA | TMEM210 | 100505993 |
| ENSG000001 | 1.22718451 | -0.048179678 | 0.33351147 | -0.1444618 | 0.88513579 | NA | ATP6V0C | 527 |
| ENSG000001 | 0 | NA | NA | NA | NA | NA | BPY2C | 442868 |

| | | | | | | | | | |
|---|---|---|---|---|---|---|---|---|---|
| ENSG000001 | 1.00943219 | -0.339564349 | 0.30698159 | -1.1061391 | 0.2686663 | NA | OR4C46 | | 119749 |
| ENSG000001 | 1.09134009 | 0.02790342 | 0.3149945 | 0.08858383 | 0.92941266 | NA | KRTAP5-5 | | 439915 |
| ENSG000001 | 0.52147635 | -0.19458195 | 0.24341208 | -0.7993932 | 0.42406247 | NA | LCE3A | | 353142 |
| ENSG000001 | 1.5760699 | -0.322844329 | 0.33765775 | -0.9561289 | 0.33900713 | NA | LCE3E | | 353145 |
| ENSG000001 | 1.51117684 | -0.199808227 | 0.3452114 | -0.5787996 | 0.56272439 | NA | TAL2 | | 6887 |
| ENSG000001 | 0.7585039 | -0.199073479 | 0.28684023 | -0.6940222 | 0.48766831 | NA | OR5D14 | | 219436 |
| ENSG000001 | 0.38270188 | -0.272446727 | 0.2140009 | -1.2731102 | 0.20297894 | NA | OR5D18 | | 219438 |
| ENSG000001 | 1.50693578 | -0.314899178 | 0.33947318 | -0.9276114 | 0.3536092 | NA | NA | NA | |
| ENSG000001 | 0.6530863 | -0.285158107 | 0.26914708 | -1.059488 | 0.28937758 | NA | LOC440895 | | 440895 |
| ENSG000001 | 0.37304803 | -0.090842893 | 0.20939682 | -0.4338313 | 0.66441096 | NA | NA | NA | |
| ENSG000001 | 0.76215602 | -0.362469896 | 0.28326072 | -1.2796335 | 0.20067407 | NA | DEFB105A | | 245908 |
| ENSG000001 | 1.88602183 | -0.110094051 | 0.34940148 | -0.3150933 | 0.75269085 | NA | DEFB107A | | 245910 |
| ENSG000001 | 0.51459361 | -0.192534203 | 0.24265108 | -0.7934611 | 0.42750921 | NA | DEFB106A | | 245909 |
| ENSG000001 | 0.50548109 | -0.025402968 | 0.24760631 | -0.1025942 | 0.91828506 | NA | DEFB105B | | 504180 |
| ENSG000001 | 0.23816916 | -0.010712105 | 0.1772998 | -0.060418 | 0.9518227 | NA | NA | NA | |
| ENSG000001 | 1.55500625 | 0.304212603 | 0.35178908 | 0.86475851 | 0.38717135 | NA | NA | NA | |
| ENSG000001 | 1.97856006 | -0.582151034 | 0.36130927 | -1.6112264 | 0.10713038 | NA | NA | NA | |
| ENSG000001 | 0.23936338 | -0.005479057 | 0.17759977 | -0.0308506 | 0.9753887 | NA | LCE1A | | 353131 |
| ENSG000001 | 1.3391893 | -0.257194319 | 0.33769001 | -0.7616284 | 0.44628179 | NA | KRTAP17-1 | | 83902 |
| ENSG000001 | 1.92054928 | -0.390561405 | 0.35856027 | -1.089249 | 0.27604411 | NA | NA | NA | |
| ENSG000001 | 1.55489645 | -0.621501489 | 0.3411506 | -1.8217805 | 0.0684883 | NA | FGF3 | | 2248 |
| ENSG000001 | 1.5266939 | -0.43166562 | 0.33274563 | -1.2972841 | 0.19453345 | NA | KRTAP22-1 | | 337979 |
| ENSG000001 | 0.64813445 | -0.288472774 | 0.26867189 | -1.0736991 | 0.28295756 | NA | KRTAP6-2 | | 337967 |
| ENSG000001 | 1.09273466 | -0.243785855 | 0.31473352 | -0.7745786 | 0.43858869 | NA | NA | NA | |
| ENSG000001 | 1.29182749 | -0.350914226 | 0.33012851 | -1.0629625 | 0.28779896 | NA | OR13C8 | | 138802 |
| ENSG000001 | 1.36417245 | 0.019184852 | 0.33939761 | 0.05652619 | 0.95492264 | NA | KRTAP19-4 | | 337971 |
| ENSG000001 | 0.35073255 | 0.241740392 | 0.2078501 | 1.16305162 | 0.24480856 | NA | KRTAP15-1 | | 254950 |
| ENSG000001 | 1.58973384 | -0.256543424 | 0.34854184 | -0.7360477 | 0.46170166 | NA | KRTAP23-1 | | 337963 |
| ENSG000001 | 1.74490851 | -0.314124437 | 0.3493437 | -0.8991845 | 0.3685544 | NA | NA | NA | |

| | | | | | | | | | |
|---|---|---|---|---|---|---|---|---|---|
| ENSG000001 | 1.6745511 | -0.411525035 | 0.35152826 | -1.1706741 | 0.2417298 | NA | KRTAP21-2 | | 337978 |
| ENSG000001 | 1.60211348 | 0.012836246 | 0.34987241 | 0.03668836 | 0.97073349 | NA | DEFB106B | | 503841 |
| ENSG000001 | 1.41959494 | -0.419263729 | 0.33698939 | -1.2441452 | 0.21344614 | NA | LCE4A | | 199834 |
| ENSG000001 | 0.61927241 | -0.278564577 | 0.26634985 | -1.0458597 | 0.29562581 | NA | LCE2A | | 353139 |
| ENSG000001 | 1.71142756 | 0.206441291 | 0.35046665 | 0.5890469 | 0.55582981 | NA | KRTAP12-1 | | 353332 |
| ENSG000001 | 1.37798483 | -0.426722426 | 0.33955458 | -1.2567123 | 0.20885782 | NA | LCE2C | | 353140 |
| ENSG000001 | 2.01180254 | 0.006286622 | 0.35843688 | 0.01753899 | 0.98600663 | NA | LOC730098 | | 730098 |
| ENSG000001 | 0 | NA | NA | NA | NA | NA | DAZ3 | | 57054 |
| ENSG000001 | 1.33565533 | 0.017803369 | 0.33313315 | 0.0534422 | 0.95737958 | NA | LCE2D | | 353141 |
| ENSG000001 | 0.46227394 | 0.141974152 | 0.23581243 | 0.60206391 | 0.5471316 | NA | LCE3B | | 353143 |
| ENSG000001 | 0.85681042 | -0.103241883 | 0.28933509 | -0.3568246 | 0.72122311 | NA | KRTAP9-8 | | 83901 |
| ENSG000001 | 0.67756292 | -0.43472079 | 0.25931557 | -1.6764161 | 0.09365672 | NA | NA | NA | |
| ENSG000001 | 1.94431494 | -0.164211172 | 0.35518572 | -0.4623248 | 0.64384841 | NA | NA | NA | |
| ENSG000001 | 2.13352971 | -0.359547003 | 0.36051313 | -0.9973201 | 0.31860914 | NA | FOXD4L3 | | 286380 |
| ENSG000001 | 0 | NA | NA | NA | NA | NA | NA | NA | |
| ENSG000001 | 1.07816388 | 0.021826176 | 0.20325886 | 0.10738118 | 0.91448659 | NA | C5orf52 | | 100190949 |
| ENSG000001 | 0.3934287 | -0.107186984 | 0.22072612 | -0.4856108 | 0.62724315 | NA | HAPLN4 | | 404037 |
| ENSG000001 | 1.63715563 | -0.381815899 | 0.34585394 | -1.1039802 | 0.26960173 | NA | CXorf67 | | 340602 |
| ENSG000001 | 1.43829502 | -0.918725396 | 0.33479517 | -2.7441417 | 0.00606693 | NA | SLC18A3 | | 6572 |
| ENSG000001 | 1.40180911 | -0.582783045 | 0.33866 | -1.72085 | 0.08527805 | NA | NA | NA | |
| ENSG000001 | 0.47144099 | 0.149959821 | 0.2369265 | 0.63293816 | 0.52677403 | NA | AMY1C | | 278 |
| ENSG000001 | 1.34643814 | -0.114409297 | 0.33364265 | -0.3429097 | 0.73166641 | NA | KRTAP10-8 | | 386681 |
| ENSG000001 | 1.90229754 | 0.200098068 | 0.36050178 | 0.55505431 | 0.57885751 | NA | HELT | | 391723 |
| ENSG000001 | 0.73887308 | -0.197609267 | 0.2855683 | -0.691986 | 0.48894611 | NA | PLSCR3 | | 57048 |
| ENSG000001 | 1.18375114 | 0.267101794 | 0.33241774 | 0.80351245 | 0.42167861 | NA | ZCCHC13 | | 389874 |
| ENSG000001 | 1.20253767 | -0.034017046 | 0.32776517 | -0.1037848 | 0.91734012 | NA | NA | NA | |
| ENSG000001 | 0.13280363 | -0.093394593 | 0.13115644 | -0.7120855 | 0.47641185 | NA | PRSS45 | | 377047 |
| ENSG000001 | 0.11934008 | 0.083564698 | 0.12760091 | 0.65489106 | 0.5125379 | NA | FAM25A | | 643161 |
| ENSG000001 | 0 | NA | NA | NA | NA | NA | DAZ1 | | 1617 |

| | | | | | | | | |
|---|---|---|---|---|---|---|---|---|
| ENSG000001 | 0.98731706 | -0.318643437 | 0.30305984 | -1.0514209 | 0.29306534 | NA | KRTAP10-6 | 386674 |
| ENSG000001 | 0 | NA | NA | NA | NA | NA | NA | NA |
| ENSG000001 | 1.69627186 | -0.310669463 | 0.34994005 | -0.8877791 | 0.37465959 | NA | OR7D1P | 26656 |
| ENSG000001 | 1.12640419 | -0.276028344 | 0.32347082 | -0.8533331 | 0.39347463 | NA | NA | NA |
| ENSG000001 | 1.75052296 | -0.577693145 | 0.34902225 | -1.6551757 | 0.09788887 | NA | C10orf99 | 387695 |
| ENSG000001 | 1.26507507 | -0.191864637 | 0.32997141 | -0.5814584 | 0.56093157 | NA | TCTEX1D4 | 343521 |
| ENSG000001 | 0 | NA | NA | NA | NA | NA | NA | NA |
| ENSG000001 | 0.48416733 | -0.018553864 | 0.24528998 | -0.0756405 | 0.93970509 | NA | NA | NA |
| ENSG000001 | 2.08189982 | -0.602299974 | 0.36253717 | -1.6613468 | 0.09664381 | NA | MAGEB5 | 347541 |
| ENSG000001 | 1.97864596 | -0.436629817 | 0.35836342 | -1.2183995 | 0.22307219 | NA | NANOS2 | 339345 |
| ENSG000001 | 1.24267123 | -0.205275703 | 0.33726848 | -0.6086418 | 0.54276187 | NA | NA | NA |
| ENSG000001 | 1.39586 | -0.568695747 | 0.33793579 | -1.6828515 | 0.09240384 | NA | NA | NA |
| ENSG000001 | 1.20116876 | 0.105261357 | 0.33139086 | 0.31763506 | 0.75076178 | NA | HBA2 | 3040 |
| ENSG000001 | 0.87855945 | -0.279392772 | 0.29937887 | -0.9332415 | 0.35069532 | NA | HMX3 | 340784 |
| ENSG000001 | 0.72421585 | -0.021287011 | 0.27744113 | -0.0767262 | 0.93884134 | NA | ZNF177 | 7730 |
| ENSG000001 | 1.00925751 | -0.200650023 | 0.312748 | -0.6415709 | 0.52115182 | NA | NA | NA |
| ENSG000001 | 0 | NA | NA | NA | NA | NA | NA | NA |
| ENSG000001 | 0.11934008 | 0.083564698 | 0.12760091 | 0.65489106 | 0.5125379 | NA | LINC00319 | 284836 |
| ENSG000001 | 1.09714126 | 0.037831979 | 0.32144521 | 0.1176934 | 0.9063106 | NA | NA | NA |
| ENSG000001 | 1.9979985 | -0.324028203 | 0.36334748 | -0.891786 | 0.37250764 | NA | OR56A5 | 390084 |
| ENSG000001 | 1.9525398 | -0.671354059 | 0.357367 | -1.8786123 | 0.06029745 | NA | KRTAP24-1 | 643803 |
| ENSG000001 | 1.16076279 | -0.608304719 | 0.3222027 | -1.8879566 | 0.05903178 | NA | BSX | 390259 |
| ENSG000001 | 1.84162509 | -0.44721324 | 0.34872083 | -1.2824391 | 0.19968864 | NA | VHLL | 391104 |
| ENSG000001 | 2.01840036 | -0.195970972 | 0.36063615 | -0.5434036 | 0.58685197 | NA | CGB5 | 93659 |
| ENSG000001 | 1.91730602 | -0.509723866 | 0.35614536 | -1.4312242 | 0.15236597 | NA | FAM25G | 100133093 |
| ENSG000001 | 1.03304953 | -0.354196327 | 0.31193091 | -1.1354961 | 0.25616753 | NA | NA | NA |
| ENSG000001 | 0.63330253 | 0.047732352 | 0.25793088 | 0.1850587 | 0.853183 | NA | GRAPL | 400581 |
| ENSG000001 | 0.98306761 | 0.112072446 | 0.30614197 | 0.36607998 | 0.71430539 | NA | NA | NA |
| ENSG000001 | 0.74823601 | -0.179761241 | 0.28015682 | -0.6416451 | 0.52110366 | NA | KRTAP10-12 | 386685 |

| | | | | | | | | |
|---|---|---|---|---|---|---|---|---|
| ENSG000001 | 1.32942389 | -0.105010903 | 0.32984147 | -0.3183678 | 0.75020596 | NA | OR14I1 | 401994 |
| ENSG000001 | 0 | NA | NA | NA | NA | NA | BTBD8 | 284697 |
| ENSG000001 | 2.1173042 | -0.376311237 | 0.36486228 | -1.0313788 | 0.3023632 | NA | SPANXN3 | 139067 |
| ENSG000001 | 2.06749054 | -0.435542482 | 0.35834827 | -1.2154167 | 0.22420719 | NA | SPANXN4 | 441525 |
| ENSG000001 | 0.51322524 | -0.027081322 | 0.24101349 | -0.1123643 | 0.91053453 | NA | NA | NA |
| ENSG000001 | 1.878049 | -0.122382185 | 0.35717045 | -0.3426436 | 0.73186656 | NA | MMP23B | 8510 |
| ENSG000001 | 0.9948826 | 0.112032491 | 0.30668173 | 0.3653054 | 0.71488345 | NA | RNU1-75P | 106480172 |
| ENSG000001 | 0.34407602 | 0.071624546 | 0.21281613 | 0.33655599 | 0.73645163 | NA | NA | NA |
| ENSG000001 | 0.83646412 | 0.050725812 | 0.29565982 | 0.17156816 | 0.86377704 | NA | NA | NA |
| ENSG000001 | 0.60150597 | 0.230273195 | 0.26371223 | 0.87319879 | 0.38255474 | NA | NA | NA |
| ENSG000001 | 1.75615995 | -0.598196182 | 0.35289045 | -1.6951328 | 0.09005023 | NA | NA | NA |
| ENSG000001 | 1.78376386 | -0.096727884 | 0.34441343 | -0.2808482 | 0.77882685 | NA | OR5K4 | 403278 |
| ENSG000001 | 1.93467181 | -0.181606498 | 0.36198678 | -0.5016937 | 0.61588298 | NA | KRTAP4-3 | 85290 |
| ENSG000001 | 1.72387762 | -0.181013955 | 0.35281862 | -0.513051 | 0.60791563 | NA | HRCT1 | 646962 |
| ENSG000001 | 0.5964156 | 0.058115253 | 0.26216088 | 0.22167783 | 0.82456469 | NA | KRTAP5-3 | 387266 |
| ENSG000001 | 1.92365657 | -0.832766135 | 0.35845191 | -2.3232297 | 0.02016682 | NA | OR10J3 | 441911 |
| ENSG000001 | 0.12685397 | -0.090215768 | 0.12901919 | -0.699243 | 0.48440017 | NA | NA | NA |
| ENSG000001 | 1.36297877 | 0.024373954 | 0.34224315 | 0.07121824 | 0.94322407 | NA | NA | NA |
| ENSG000001 | 0.78141628 | -0.193205819 | 0.28064196 | -0.6884424 | 0.49117424 | NA | NA | NA |
| ENSG000001 | 2.17348956 | -0.854403868 | 0.35579043 | -2.4014245 | 0.01633138 | NA | CGB7 | 94027 |
| ENSG000001 | 1.56270122 | -0.095861714 | 0.33925203 | -0.2825678 | 0.77750813 | NA | PRSS29P | 123787 |
| ENSG000001 | 1.45095677 | -0.368929855 | 0.3246363 | -1.1364406 | 0.25577223 | NA | NA | NA |
| ENSG000001 | 1.56062598 | -0.105860256 | 0.34450487 | -0.3072823 | 0.75862849 | NA | SPANXD | 64648 |
| ENSG000001 | 0.51433302 | -0.186673425 | 0.24269587 | -0.7691661 | 0.44179474 | NA | PFN3 | 345456 |
| ENSG000001 | 2.13310413 | -0.50715811 | 0.36556447 | -1.3873288 | 0.16534154 | NA | OR5AC2 | 81050 |
| ENSG000001 | 1.5496102 | -0.808979811 | 0.3427739 | -2.3600975 | 0.01827013 | NA | NA | NA |
| ENSG000001 | 1.87837664 | -0.502253439 | 0.3551342 | -1.4142638 | 0.15728447 | NA | LCE1B | 353132 |
| ENSG000001 | 1.10935906 | -0.105371602 | 0.31595365 | -0.3335034 | 0.73875436 | NA | SPINK14 | 408187 |
| ENSG000001 | 0.76078766 | -0.199688475 | 0.28697083 | -0.6958494 | 0.48652316 | NA | SPRR2B | 6701 |

| | | | | | | | | |
|---|---|---|---|---|---|---|---|---|
| ENSG000001 | 0 | NA | NA | NA | NA | NA | ZNF709 | 163051 |
| ENSG000001 | 0.64676608 | -0.114974633 | 0.26681508 | -0.430915 | 0.66653014 | NA | HIST1H2AD | 3013 |
| ENSG000001 | 2.0227534 | -0.565697402 | 0.35837532 | -1.5785055 | 0.11444954 | NA | OR2T4 | 127074 |
| ENSG000001 | 1.84461428 | -0.507548691 | 0.35476449 | -1.4306637 | 0.15252661 | NA | NA | NA |
| ENSG000001 | 1.40177877 | -0.417958899 | 0.33875905 | -1.233794 | 0.21727966 | NA | NA | NA |
| ENSG000001 | 0 | NA | NA | NA | NA | NA | NA | NA |
| ENSG000001 | 2.16848171 | -0.651727013 | 0.36510476 | -1.7850411 | 0.07425465 | NA | NA | NA |
| ENSG000001 | 1.13839175 | -0.566917627 | 0.31494538 | -1.8000506 | 0.07185264 | NA | LCE1C | 353133 |
| ENSG000001 | 0 | NA | NA | NA | NA | NA | NA | NA |
| ENSG000001 | 1.15136953 | -0.411416243 | 0.31513186 | -1.3055368 | 0.19171012 | NA | IFNL3 | 282617 |
| ENSG000001 | 1.34696221 | -0.511364303 | 0.3262526 | -1.5673877 | 0.11702411 | NA | NA | NA |
| ENSG000001 | 1.86478992 | -0.222491962 | 0.34710699 | -0.6409896 | 0.52152946 | NA | NA | NA |
| ENSG000001 | 1.29843613 | 0.026882929 | 0.33073441 | 0.08128253 | 0.93521727 | NA | OR1J2 | 26740 |
| ENSG000001 | 1.11436691 | -0.34927398 | 0.301767 | -1.1574293 | 0.24709699 | NA | TPSB2 | 64499 |
| ENSG000001 | 1.96744556 | -0.297607607 | 0.35559035 | -0.8369395 | 0.40262658 | NA | GUCA2A | 2980 |
| ENSG000001 | 1.41051025 | 0.236964065 | 0.3413822 | 0.69413129 | 0.48759988 | NA | LYPD2 | 137797 |
| ENSG000001 | 0 | NA | NA | NA | NA | NA | HIST1H3D | 8351 |
| ENSG000001 | 1.79771395 | -0.93371913 | 0.35259101 | -2.6481649 | 0.008093 | NA | GGT3P | 2679 |
| ENSG000001 | 1.25322199 | -0.041993678 | 0.32056166 | -0.1310003 | 0.89577506 | NA | NA | NA |
| ENSG000001 | 1.67119574 | -0.517854984 | 0.34368545 | -1.5067702 | 0.13186958 | NA | OR11L1 | 391189 |
| ENSG000001 | 0.66293603 | -0.489755159 | 0.27215878 | -1.7995199 | 0.07193647 | NA | NA | NA |
| ENSG000001 | 0.62812434 | -0.408089682 | 0.2528562 | -1.61392 | 0.10654477 | NA | NA | NA |
| ENSG000001 | 1.09309684 | 0.168727515 | 0.31294013 | 0.53916868 | 0.58977047 | NA | NA | NA |
| ENSG000001 | 1.0247963 | -0.3508469 | 0.30999594 | -1.131779 | 0.25772735 | NA | C14orf178 | 283579 |
| ENSG000001 | 1.13722083 | -0.265912275 | 0.31792134 | -0.836409 | 0.40292485 | NA | SCGB1D4 | 404552 |
| ENSG000001 | 1.7033854 | -0.187760037 | 0.35661139 | -0.5265116 | 0.59853279 | NA | OR52M1 | 119772 |
| ENSG000001 | 1.71758327 | 0.062859084 | 0.35107067 | 0.17904966 | 0.85789871 | NA | NA | NA |
| ENSG000001 | 1.45656019 | -0.045383323 | 0.3439152 | -0.1319608 | 0.8950153 | NA | OR5M12P | 81188 |
| ENSG000001 | 1.63285667 | -0.40253458 | 0.35101724 | -1.1467658 | 0.25147841 | NA | OR1S2 | 219958 |

| | | | | | | | | |
|---|---|---|---|---|---|---|---|---|
| ENSG000001 | 2.10938803 | -0.11773185 | 0.36489089 | -0.3226495 | 0.74696073 | NA | HES5 | 388585 |
| ENSG000001 | 1.14868934 | -0.409600281 | 0.31754893 | -1.2898808 | 0.19709204 | NA | NA | NA |
| ENSG000001 | 1.48249107 | -0.198401287 | 0.34716207 | -0.5714947 | 0.56766435 | NA | NA | NA |
| ENSG000001 | 0.27408862 | -0.199826037 | 0.18669465 | -1.0703362 | 0.28446801 | NA | SPANXA1 | 30014 |
| ENSG000001 | 0.61335714 | -0.10408451 | 0.2640241 | -0.3942235 | 0.69341602 | NA | KRTAP9-9 | 81870 |
| ENSG000001 | 1.34166309 | -0.110506425 | 0.3343234 | -0.3305375 | 0.74099386 | NA | KRTAP4-6 | 81871 |
| ENSG000001 | 1.09296044 | 0.193163571 | 0.32526029 | 0.59387381 | 0.55259652 | NA | DEFB107B | 503614 |
| ENSG000001 | 1.98544493 | -0.456868866 | 0.36401277 | -1.2550902 | 0.20944602 | NA | NA | NA |
| ENSG000001 | 0 | NA | NA | NA | NA | NA | NA | NA |
| ENSG000001 | 1.3657155 | -0.276605243 | 0.34297312 | -0.8064925 | 0.41995894 | NA | KRTAP4-5 | 85289 |
| ENSG000001 | 1.4238214 | 0.38593557 | 0.34084093 | 1.13230405 | 0.25750663 | NA | KRTAP13-1 | 140258 |
| ENSG000001 | 2.09936307 | 0.117626341 | 0.36380904 | 0.32331891 | 0.74645374 | NA | NA | NA |
| ENSG000001 | 2.03477533 | -0.736333721 | 0.36115326 | -2.03884 | 0.041466 | NA | PRR31 | 101928638 |
| ENSG000001 | 1.01296617 | -0.644917897 | 0.29961632 | -2.1524792 | 0.03135963 | NA | NA | NA |
| ENSG000001 | 0.24756242 | -0.187034346 | 0.18119817 | -1.0322088 | 0.30197431 | NA | SPANXC | 64663 |
| ENSG000001 | 1.52820662 | -0.650044064 | 0.34550795 | -1.8814156 | 0.0599154 | NA | OR2M2 | 391194 |
| ENSG000001 | 0 | NA | NA | NA | NA | NA | EIF1AY | 9086 |
| ENSG000001 | 0.90138751 | -0.278391458 | 0.29894121 | -0.9312582 | 0.35172002 | NA | NA | NA |
| ENSG000001 | 1.64828549 | -0.415292708 | 0.35240599 | -1.1784496 | 0.23861741 | NA | OR4D12P | 282754 |
| ENSG000001 | 1.46442504 | -0.309896251 | 0.33869621 | -0.9149682 | 0.36020834 | NA | C1orf68 | 100129271 |
| ENSG000001 | 0 | NA | NA | NA | NA | NA | NA | NA |
| ENSG000001 | 1.52401474 | -0.317170991 | 0.33725967 | -0.9404356 | 0.34699418 | NA | NA | NA |
| ENSG000001 | 0.38884741 | -0.29745092 | 0.22235466 | -1.3377319 | 0.18098386 | NA | NA | NA |
| ENSG000001 | 0 | NA | NA | NA | NA | NA | NA | NA |
| ENSG000001 | 1.5763033 | -0.106360134 | 0.34542798 | -0.3079083 | 0.75815213 | NA | MIRLET7A2 | 406882 |
| ENSG000001 | 0.58799035 | 0.401250203 | 0.25669282 | 1.56315322 | 0.11801656 | NA | MIR429 | 554210 |
| ENSG000001 | 0.37125511 | 0.067313264 | 0.2170755 | 0.31009148 | 0.75649139 | NA | NA | NA |
| ENSG000001 | 0.89410229 | -0.104970982 | 0.29138816 | -0.3602445 | 0.7186643 | NA | MIR449A | 554213 |
| ENSG000001 | 2.0888368 | -0.009336907 | 0.36574401 | -0.0255285 | 0.97963339 | NA | MIR345 | 442910 |

| | | | | | | | | | |
|---|---|---|---|---|---|---|---|---|---|
| ENSG000001 | 1.27762673 | -0.192968955 | 0.32606246 | -0.5918159 | 0.55397386 | NA | MIR16-2 | | 406951 |
| ENSG000001 | 1.10306058 | 0.179115613 | 0.31326539 | 0.57176955 | 0.56747811 | NA | NA | NA | |
| ENSG000001 | 2.05466891 | -0.118007333 | 0.35705755 | -0.3304995 | 0.74102259 | NA | MIR107 | | 406901 |
| ENSG000001 | 1.47566232 | -0.053992454 | 0.34916796 | -0.1546318 | 0.87711164 | NA | MIR448 | | 554212 |
| ENSG000001 | 0.51433302 | -0.186673425 | 0.24269587 | -0.7691661 | 0.44179474 | NA | NA | NA | |
| ENSG000001 | 0.24411881 | -0.012946855 | 0.17864669 | -0.0724718 | 0.94222641 | NA | MIR412 | | 574433 |
| ENSG000001 | 0.99391514 | -0.03501723 | 0.30655428 | -0.1142285 | 0.90905668 | NA | NA | NA | |
| ENSG000001 | 1.00635056 | -0.03588081 | 0.30120837 | -0.1191229 | 0.905178 | NA | MIR1-1 | | 406904 |
| ENSG000001 | 0.24550891 | -0.007886114 | 0.17900103 | -0.0440563 | 0.96485956 | NA | NA | NA | |
| ENSG000001 | 1.13574539 | -0.43525825 | 0.32353461 | -1.3453221 | 0.17852125 | NA | NA | NA | |
| ENSG000001 | 0 | NA | NA | NA | NA | NA | MIR369 | | 442914 |
| ENSG000001 | 1.89122831 | -0.525863424 | 0.3580415 | -1.468722 | 0.14190822 | NA | MIRLET7C | | 406885 |
| ENSG000001 | 1.40612981 | -0.418696198 | 0.33884716 | -1.2356491 | 0.21658902 | NA | MIR371A | | 442916 |
| ENSG000001 | 1.57963425 | -0.252533317 | 0.34748917 | -0.7267372 | 0.46738694 | NA | MIR425 | | 494337 |
| ENSG000001 | 1.34926323 | -0.122357603 | 0.34061647 | -0.359224 | 0.71942754 | NA | MIR103A1 | | 406895 |
| ENSG000001 | 0.12685397 | -0.090215768 | 0.12901919 | -0.699243 | 0.48440017 | NA | MIR219A1 | | 407002 |
| ENSG000001 | 0.11934008 | 0.083564698 | 0.12760091 | 0.65489106 | 0.5125379 | NA | MIR210 | | 406992 |
| ENSG000001 | 0.12070844 | -0.088503695 | 0.12783931 | -0.6923042 | 0.48874631 | NA | NA | NA | |
| ENSG000001 | 0.77632645 | -0.361571052 | 0.28424434 | -1.2720431 | 0.2033578 | NA | NA | NA | |
| ENSG000001 | 1.23905795 | -0.180522776 | 0.32930107 | -0.5481998 | 0.58355472 | NA | NA | NA | |
| ENSG000001 | 0.39479706 | -0.300008631 | 0.22317059 | -1.3443018 | 0.17885081 | NA | NA | NA | |
| ENSG000001 | 0.96033095 | 0.108661823 | 0.30514451 | 0.35609955 | 0.72176602 | NA | MIR129-2 | | 406918 |
| ENSG000001 | 0.73998086 | -0.354752955 | 0.28183273 | -1.2587358 | 0.20812577 | NA | MIR133B | | 442890 |
| ENSG000001 | 0.59027007 | 0.063652523 | 0.26808709 | 0.23743226 | 0.81232147 | NA | MIR342 | | 442909 |
| ENSG000001 | 1.86223001 | 0.006812587 | 0.35790151 | 0.01903481 | 0.98481334 | NA | NA | NA | |
| ENSG000001 | 1.00037971 | -0.179662206 | 0.30756199 | -0.5841496 | 0.55911969 | NA | NA | NA | |
| ENSG000001 | 0.41537361 | -0.286583533 | 0.21871501 | -1.3103057 | 0.19009243 | NA | MIR326 | | 442900 |
| ENSG000001 | 0.11131518 | 0.081951103 | 0.12663804 | 0.64712866 | 0.51754868 | NA | NA | NA | |
| ENSG000001 | 1.37735526 | -0.126156488 | 0.33283524 | -0.3790358 | 0.70466124 | NA | NA | NA | |

| | | | | | | | | |
|---|---|---|---|---|---|---|---|---|
| ENSG000001 | 0.67141739 | -0.461190807 | 0.26562301 | -1.7362608 | 0.08251773 | NA | MIR372 | 442917 |
| ENSG000001 | 0.48573157 | -0.181932918 | 0.23941992 | -0.7598905 | 0.44732006 | NA | MIR346 | 442911 |
| ENSG000001 | 0.11865494 | 0.083440866 | 0.12751478 | 0.65436232 | 0.5128784 | NA | NA | NA |
| ENSG000001 | 0.84699315 | 0.046452071 | 0.29623822 | 0.15680648 | 0.87539737 | NA | MIR411 | 693121 |
| ENSG000001 | 0.47408893 | 0.142247222 | 0.23727093 | 0.59951389 | 0.54883025 | NA | NA | NA |
| ENSG000001 | 1.59039328 | -0.632413724 | 0.34231631 | -1.8474542 | 0.06468134 | NA | MIR148B | 442892 |
| ENSG000001 | 0.11131518 | 0.081951103 | 0.12663804 | 0.64712866 | 0.51754868 | NA | MIR383 | 494332 |
| ENSG000001 | 0.49819341 | -0.026649049 | 0.23931956 | -0.1113534 | 0.9113361 | NA | NA | NA |
| ENSG000001 | 1.13702496 | -0.279999728 | 0.32379574 | -0.8647419 | 0.38718049 | NA | MIR450A1 | 554214 |
| ENSG000001 | 1.11374201 | -0.12138282 | 0.32078272 | -0.3783958 | 0.70513662 | NA | MIR101-1 | 406893 |
| ENSG000001 | 1.03029107 | -0.194319334 | 0.3088027 | -0.6292669 | 0.52917431 | NA | MIR373 | 442918 |
| ENSG000001 | 0.49699973 | -0.021154807 | 0.23920592 | -0.0884376 | 0.92952885 | NA | MIR337 | 442905 |
| ENSG000001 | 0.24756242 | -0.187034346 | 0.18119817 | -1.0322088 | 0.30197431 | NA | MIR30D | 407033 |
| ENSG000001 | 0.3934287 | -0.107186984 | 0.22072612 | -0.4856108 | 0.62724315 | NA | MIR422A | 494334 |
| ENSG000001 | 1.25016394 | -0.472607307 | 0.32432071 | -1.4572221 | 0.14505508 | NA | NA | NA |
| ENSG000001 | 0.47207214 | -0.012756135 | 0.23624418 | -0.0539956 | 0.95693871 | NA | MIR96 | 407053 |
| ENSG000001 | 0.54319505 | -0.072630779 | 0.15595508 | -0.465716 | 0.64141878 | NA | MIR126 | 406913 |
| ENSG000001 | 1.7710647 | -0.204025858 | 0.35727931 | -0.5710542 | 0.56796289 | NA | MIRLET7A1 | 406881 |
| ENSG000001 | 1.25621263 | 0.096665511 | 0.32321988 | 0.29907043 | 0.7648863 | NA | MIR331 | 442903 |
| ENSG000001 | 1.02319981 | -0.187512967 | 0.30852835 | -0.6077658 | 0.54334284 | NA | MIR31 | 407035 |
| ENSG000001 | 0.80925878 | -0.344388394 | 0.27575198 | -1.2489064 | 0.21169933 | NA | NA | NA |
| ENSG000001 | 1.2310865 | -0.046903466 | 0.33004793 | -0.1421111 | 0.88699227 | NA | NA | NA |
| ENSG000001 | 0.59146429 | 0.065095808 | 0.26170762 | 0.24873486 | 0.80356588 | NA | NA | NA |
| ENSG000001 | 2.04091226 | -0.436263813 | 0.3593753 | -1.2139505 | 0.22476664 | NA | NA | NA |
| ENSG000001 | 1.27093636 | -0.505739972 | 0.33096118 | -1.5280945 | 0.12648906 | NA | NA | NA |
| ENSG000001 | 1.56206314 | 0.142070138 | 0.34515842 | 0.41160849 | 0.68062641 | NA | RNU105C | 26766 |
| ENSG000001 | 0.90705484 | -0.464035783 | 0.30257984 | -1.5335978 | 0.12512862 | NA | RNU6-1123P | 106480060 |
| ENSG000001 | 1.61924876 | -0.264773171 | 0.35078669 | -0.7547982 | 0.45037006 | NA | RN7SKP184 | 106480470 |
| ENSG000001 | 0.49036493 | -0.020766486 | 0.24598337 | -0.0844223 | 0.93272066 | NA | RNU6-500P | 106479788 |

| | | | | | | | | |
|---|---|---|---|---|---|---|---|---|
| ENSG000001 | 0 | NA | NA | NA | NA | NA | NA | NA |
| ENSG000001 | 0.61605906 | 0.051157992 | 0.2638843 | 0.19386523 | 0.84628141 | NA | NA | NA |
| ENSG000001 | 0.49105007 | -0.020523257 | 0.24605963 | -0.0834077 | 0.9335274 | NA | NA | NA |
| ENSG000001 | 0.7265173 | -0.177755649 | 0.27865224 | -0.6379121 | 0.52353088 | NA | NA | NA |
| ENSG000001 | 1.18747954 | 0.380170738 | 0.31412379 | 1.21025769 | 0.22618003 | NA | NA | NA |
| ENSG000001 | 0.61864338 | 0.060567647 | 0.26410487 | 0.22933181 | 0.81861104 | NA | NA | NA |
| ENSG000001 | 0.58075474 | 0.403955941 | 0.25749097 | 1.56881594 | 0.11669083 | NA | RNU6-834P | 106479935 |
| ENSG000001 | 0.62545232 | -0.114578523 | 0.27135287 | -0.4222492 | 0.67284316 | NA | NA | NA |
| ENSG000001 | 0.84107575 | 0.044472277 | 0.29591903 | 0.15028529 | 0.88053954 | NA | NA | NA |
| ENSG000001 | 0.61258555 | -0.25554054 | 0.25718763 | -0.9935958 | 0.32041971 | NA | RNU6-896P | 106481922 |
| ENSG000001 | 1.24637096 | -0.054514582 | 0.32460283 | -0.1679424 | 0.86662859 | NA | RNU6-28P | 100873756 |
| ENSG000001 | 0.64420212 | -0.102988282 | 0.25928072 | -0.3972076 | 0.69121435 | NA | RNU6-664P | 106479861 |
| ENSG000001 | 0.60236525 | 0.059515457 | 0.26911653 | 0.22115125 | 0.82497467 | NA | NA | NA |
| ENSG000001 | 0 | NA | NA | NA | NA | NA | RNA5S12 | 100169763 |
| ENSG000001 | 0.99294768 | -0.182417453 | 0.30147696 | -0.6050792 | 0.54512634 | NA | RNU6-349P | 106479721 |
| ENSG000001 | 0 | NA | NA | NA | NA | NA | NA | NA |
| ENSG000001 | 1.85181427 | -0.238484388 | 0.35499422 | -0.671798 | 0.50171233 | NA | NA | NA |
| ENSG000001 | 0.23868015 | 0.154625155 | 0.16997278 | 0.90970538 | 0.36297791 | NA | NA | NA |
| ENSG000001 | 0.2596576 | -0.193002035 | 0.1837913 | -1.0501152 | 0.29366517 | NA | RNU1-58P | 106481612 |
| ENSG000001 | 1.03083633 | -0.502952708 | 0.30594337 | -1.6439405 | 0.10018849 | NA | NA | NA |
| ENSG000001 | 1.25681992 | -0.357482642 | 0.33407357 | -1.0700716 | 0.28458708 | NA | RNU6-1094P | 106480047 |
| ENSG000001 | 0.24485602 | -0.012655878 | 0.17881722 | -0.0707755 | 0.94357643 | NA | RNU6-502P | 106481899 |
| ENSG000001 | 0.62545232 | -0.114578523 | 0.27135287 | -0.4222492 | 0.67284316 | NA | NA | NA |
| ENSG000001 | 1.39468751 | -0.411835239 | 0.33864396 | -1.2161305 | 0.22393523 | NA | NA | NA |
| ENSG000001 | 0 | NA | NA | NA | NA | NA | NA | NA |
| ENSG000001 | 0.25260017 | -0.015978705 | 0.18049464 | -0.0885273 | 0.92945758 | NA | RNU6-1312P | 106481594 |
| ENSG000001 | 0.60717723 | -0.256800472 | 0.2582777 | -0.9942805 | 0.32008634 | NA | NA | NA |
| ENSG000001 | 0.75213216 | -0.027056401 | 0.28563056 | -0.0947252 | 0.92453313 | NA | RNU6-858P | 106479945 |
| ENSG000001 | 1.26296992 | -0.650114613 | 0.32694875 | -1.9884297 | 0.04676418 | NA | SNORD115-34 | 100033808 |

| | | | | | | | | |
|---|---|---|---|---|---|---|---|---|
| ENSG000001 | 0.24411881 | -0.012946855 | 0.17864669 | -0.0724718 | 0.94222641 | NA | NA | NA |
| ENSG000001 | 1.23057604 | -0.201957268 | 0.33699085 | -0.599296 | 0.54897552 | NA | RN7SKP25 | 106480364 |
| ENSG000001 | 1.12217131 | -0.11910319 | 0.31641857 | -0.3764102 | 0.70661193 | NA | NA | NA |
| ENSG000001 | 0.75133883 | -0.359102617 | 0.28257827 | -1.2708076 | 0.2037971 | NA | NA | NA |
| ENSG000001 | 1.08686698 | 0.195202704 | 0.32505488 | 0.6005223 | 0.5481582 | NA | RNU4-39P | 106479575 |
| ENSG000001 | 0.35664995 | 0.244349217 | 0.20882949 | 1.17008962 | 0.2419649 | NA | RNU6-1298P | 106481590 |
| ENSG000001 | 0.62478892 | 0.058386395 | 0.26463671 | 0.22062848 | 0.82538172 | NA | RNU6-230P | 106479669 |
| ENSG000001 | 0.73064371 | 0.138033838 | 0.28433319 | 0.48546509 | 0.62734647 | NA | NA | NA |
| ENSG000001 | 0 | NA | NA | NA | NA | NA | RNA5S11 | 100169762 |
| ENSG000001 | 0.37125511 | 0.067313264 | 0.2170755 | 0.31009148 | 0.75649139 | NA | RNU6-204P | 106481241 |
| ENSG000001 | 0 | NA | NA | NA | NA | NA | RNA5S3 | 100169754 |
| ENSG000001 | 1.10226726 | -0.105488567 | 0.31318323 | -0.336827 | 0.73624732 | NA | RNU6-766P | 106481422 |
| ENSG000001 | 0.39388516 | -0.104610737 | 0.21253504 | -0.4922047 | 0.62257467 | NA | NA | NA |
| ENSG000001 | 0 | NA | NA | NA | NA | NA | RNA5S1 | 100169751 |
| ENSG000001 | 0.74703594 | -0.027676606 | 0.26993737 | -0.1025297 | 0.91833622 | NA | NA | NA |
| ENSG000001 | 0.81551701 | 0.20201966 | 0.28897753 | 0.69908431 | 0.48449934 | NA | RNU6-1115P | 106480639 |
| ENSG000001 | 0.99260276 | -0.190905426 | 0.3121099 | -0.6116609 | 0.54076213 | NA | RNU1-150P | 106480413 |
| ENSG000001 | 1.83854604 | -0.109124236 | 0.35311677 | -0.3090316 | 0.7572975 | NA | NA | NA |
| ENSG000001 | 0 | NA | NA | NA | NA | NA | NA | NA |
| ENSG000001 | 1.27506511 | -0.661844382 | 0.32866954 | -2.0137077 | 0.04404023 | NA | NA | NA |
| ENSG000001 | 1.46791817 | -0.051632502 | 0.34904661 | -0.1479244 | 0.88240245 | NA | RNU6-1084P | 106480042 |
| ENSG000001 | 0.73664489 | 0.119060265 | 0.27862315 | 0.42731649 | 0.66914883 | NA | NA | NA |
| ENSG000001 | 1.73605713 | -0.201059748 | 0.35930595 | -0.5595781 | 0.57576723 | NA | NA | NA |
| ENSG000001 | 0.11934008 | 0.083564698 | 0.12760091 | 0.65489106 | 0.5125379 | NA | NA | NA |
| ENSG000001 | 2.00317711 | -0.44205624 | 0.36164231 | -1.2223576 | 0.22157241 | NA | RNU6-369P | 106479730 |
| ENSG000001 | 0.23070733 | 0.167416683 | 0.17664244 | 0.94777156 | 0.34324576 | NA | SNORD114-7 | 767583 |
| ENSG000001 | 0.37304803 | -0.090842893 | 0.20939682 | -0.4338313 | 0.66441096 | NA | NA | NA |
| ENSG000001 | 1.11949112 | -0.114871106 | 0.32248769 | -0.3562031 | 0.72168849 | NA | RNU6-600P | 106479831 |
| ENSG000001 | 0.35022155 | 0.06940789 | 0.21379411 | 0.32464827 | 0.74544729 | NA | NA | NA |

| | | | | | | | | |
|---|---|---|---|---|---|---|---|---|
| ENSG000001 | 0 | NA | NA | NA | NA | NA | RNA5S5 | 100169756 |
| ENSG000001 | 0.90958655 | -0.438566765 | 0.29615636 | -1.4808622 | 0.13864329 | NA | NA | NA |
| ENSG000001 | 1.10445873 | -0.117549567 | 0.31580194 | -0.3722256 | 0.70972489 | NA | NA | NA |
| ENSG000001 | 0.51931521 | -0.354214731 | 0.23897171 | -1.4822455 | 0.13827499 | NA | NA | NA |
| ENSG000001 | 1.59391286 | 0.019901618 | 0.35090852 | 0.05671455 | 0.95477258 | NA | NA | NA |
| ENSG000001 | 1.24907577 | -0.324112181 | 0.3240456 | -1.0002055 | 0.31721108 | NA | NA | NA |
| ENSG000001 | 0.76940933 | -0.516275266 | 0.27702168 | -1.8636638 | 0.06236894 | NA | NA | NA |
| ENSG000001 | 0.28256998 | -0.18483211 | 0.18021686 | -1.0256094 | 0.30507573 | NA | NA | NA |
| ENSG000001 | 2.07162091 | -0.004770809 | 0.36365788 | -0.0131189 | 0.98953289 | NA | NA | NA |
| ENSG000001 | 0.35750923 | 0.072271886 | 0.21496454 | 0.33620376 | 0.73671722 | NA | RNA5SP370 | 106480765 |
| ENSG000001 | 0.49082386 | 0.135175125 | 0.22941993 | 0.58920394 | 0.55572448 | NA | RN7SKP170 | 106479171 |
| ENSG000001 | 0.49219223 | -0.014160136 | 0.23865874 | -0.0593321 | 0.95268755 | NA | NA | NA |
| ENSG000001 | 0.35682409 | 0.072008558 | 0.21485623 | 0.33514763 | 0.73751372 | NA | NA | NA |
| ENSG000001 | 0.84223772 | 0.049919309 | 0.28809738 | 0.17327235 | 0.86243736 | NA | RNU1-108P | 106481957 |
| ENSG000001 | 0.84241378 | 0.050867159 | 0.30116699 | 0.16890018 | 0.86587516 | NA | NA | NA |
| ENSG000001 | 0.75867859 | -0.356397589 | 0.28312748 | -1.2587884 | 0.20810676 | NA | NA | NA |
| ENSG000001 | 0.40327843 | -0.281391716 | 0.21700406 | -1.2967118 | 0.19473038 | NA | NA | NA |
| ENSG000001 | 2.11101836 | -0.126712802 | 0.36348528 | -0.348605 | 0.72738584 | NA | RNU6-543P | 106479807 |
| ENSG000001 | 0.37928001 | 0.070449581 | 0.21829793 | 0.32272217 | 0.74690566 | NA | RNU6-845P | 106479940 |
| ENSG000001 | 1.21369922 | -0.047190354 | 0.32958242 | -0.1431823 | 0.88614624 | NA | NA | NA |
| ENSG000001 | 0.14128499 | -0.097772014 | 0.13403235 | -0.7294658 | 0.4657168 | NA | SNORD115-12 | 100033449 |
| ENSG000001 | 0.73664489 | 0.119060265 | 0.27862315 | 0.42731649 | 0.66914883 | NA | NA | NA |
| ENSG000001 | 1.90875785 | 0.317932115 | 0.35768467 | 0.88886146 | 0.37407754 | NA | NA | NA |
| ENSG000001 | 0.63865529 | -0.262882814 | 0.2611518 | -1.0066284 | 0.31411339 | NA | RNU4-35P | 106481182 |
| ENSG000001 | 1.86464558 | -0.249131479 | 0.35192177 | -0.7079172 | 0.47899667 | NA | NA | NA |
| ENSG000001 | 0.49168123 | -0.197384925 | 0.247356 | -0.7979791 | 0.4248826 | NA | RNU6-1216P | 106480098 |
| ENSG000001 | 0.38614101 | -0.1098476 | 0.21963552 | -0.5001359 | 0.61697942 | NA | NA | NA |
| ENSG000001 | 0.11205239 | 0.082118251 | 0.12671962 | 0.64803108 | 0.51696485 | NA | NA | NA |
| ENSG000001 | 0.75637659 | -0.193693286 | 0.28066413 | -0.6901248 | 0.49011569 | NA | RNU6-556P | 106481357 |

| | | | | | | | | |
|---|---|---|---|---|---|---|---|---|
| ENSG000001 | 0.85723517 | -0.086923536 | 0.28025668 | -0.3101569 | 0.75644166 | NA | RNU1-62P | 106481613 |
| ENSG000001 | 0.50107449 | -0.372905725 | 0.24426561 | -1.5266403 | 0.12685049 | NA | NA | NA |
| ENSG000001 | 0.56539008 | 0.39817126 | 0.25602908 | 1.55517983 | 0.11990325 | NA | NA | NA |
| ENSG000001 | 0.50630812 | -0.189615696 | 0.24175631 | -0.7843257 | 0.432849 | NA | NA | NA |
| ENSG000001 | 0 | NA | NA | NA | NA | NA | NA | NA |
| ENSG000001 | 1.21503725 | -0.04601322 | 0.33625988 | -0.1368383 | 0.89115863 | NA | RN7SKP104 | 106480865 |
| ENSG000001 | 0.48472985 | 0.140480862 | 0.23855057 | 0.58889343 | 0.55593276 | NA | SNORA31 | 677814 |
| ENSG000001 | 0.47523053 | 0.12360509 | 0.22747273 | 0.54338422 | 0.58686529 | NA | RNA5SP389 | 106480766 |
| ENSG000001 | 0.8359875 | 0.042370493 | 0.28775603 | 0.1472445 | 0.88293904 | NA | RNU6-633P | 106481381 |
| ENSG000001 | 0 | NA | NA | NA | NA | NA | RNU4-15P | 106479565 |
| ENSG000001 | 0.38884741 | -0.29745092 | 0.22235466 | -1.3377319 | 0.18098386 | NA | NA | NA |
| ENSG000001 | 0.50428741 | -0.01691703 | 0.23137633 | -0.0731148 | 0.94171478 | NA | NA | NA |
| ENSG000001 | 1.13611595 | -0.5889264 | 0.32009134 | -1.8398698 | 0.06578735 | NA | SNORD115-25 | 100033801 |
| ENSG000001 | 0.35802023 | 0.220517756 | 0.19914056 | 1.1073473 | 0.2681438 | NA | NA | NA |
| ENSG000001 | 0.97471088 | -0.450815093 | 0.29492584 | -1.528571 | 0.12637082 | NA | RNU6-1313P | 106480139 |
| ENSG000001 | 0.61950267 | -0.112567707 | 0.27088102 | -0.4155614 | 0.67773094 | NA | RNU6-247P | 106479676 |
| ENSG000001 | 1.67321465 | -0.255970488 | 0.34257643 | -0.7471923 | 0.45494746 | NA | NA | NA |
| ENSG000001 | 1.10867392 | -0.112034002 | 0.32214076 | -0.3477797 | 0.72800567 | NA | NA | NA |
| ENSG000001 | 1.65451926 | -0.70157855 | 0.34876258 | -2.0116222 | 0.04425978 | NA | RNU6-235P | 106479671 |
| ENSG000001 | 1.44779808 | -0.050092704 | 0.34764866 | -0.14409 | 0.88542938 | NA | NA | NA |
| ENSG000001 | 1.01486226 | -0.193402712 | 0.30816768 | -0.6275892 | 0.53027307 | NA | NA | NA |
| ENSG000001 | 0.72304144 | -0.021905802 | 0.27736023 | -0.0789796 | 0.93704884 | NA | RNU6-386P | 106481300 |
| ENSG000001 | 1.81915925 | 0.140629442 | 0.36045165 | 0.39014786 | 0.69642721 | NA | RNA5SP226 | 106479000 |
| ENSG000001 | 1.16130413 | -0.607940092 | 0.32222668 | -1.8866845 | 0.05920277 | NA | NA | NA |
| ENSG000001 | 0.3593886 | 0.071604659 | 0.20679693 | 0.34625591 | 0.7291504 | NA | RNU6-462P | 106479770 |
| ENSG000001 | 1.88331297 | -0.131990757 | 0.35722428 | -0.3694899 | 0.71176262 | NA | NA | NA |
| ENSG000001 | 0.60856498 | 0.209703707 | 0.25574566 | 0.81996975 | 0.41223335 | NA | NA | NA |
| ENSG000001 | 0.97418343 | 0.27333574 | 0.30819636 | 0.88688827 | 0.37513906 | NA | RNU6-315P | 106479707 |
| ENSG000001 | 0.47092999 | -0.01913341 | 0.24378131 | -0.078486 | 0.9374415 | NA | NA | NA |

| | | | | | | | | |
|---|---|---|---|---|---|---|---|---|
| ENSG000001 | 0.25214371 | -0.009862892 | 0.18046657 | -0.0546522 | 0.95641556 | NA | NA | NA |
| ENSG000001 | 0 | NA | NA | NA | NA | NA | NA | NA |
| ENSG000001 | 0.26062507 | -0.012888319 | 0.18226503 | -0.070712 | 0.94362699 | NA | NA | NA |
| ENSG000001 | 1.41534996 | -0.270555967 | 0.33675068 | -0.8034311 | 0.42172561 | NA | RNU6-37P | 106478920 |
| ENSG000001 | 1.22990644 | -0.543241002 | 0.30825228 | -1.762326 | 0.07801424 | NA | NA | NA |
| ENSG000001 | 0.73699426 | -0.182203658 | 0.27937391 | -0.6521857 | 0.51428141 | NA | NA | NA |
| ENSG000001 | 0.47616417 | 0.157689149 | 0.24487722 | 0.64395189 | 0.51960663 | NA | NA | NA |
| ENSG000001 | 1.51795353 | -0.35112914 | 0.34712483 | -1.0115357 | 0.31176012 | NA | NA | NA |
| ENSG000001 | 0 | NA | NA | NA | NA | NA | RNA5SP174 | 106478998 |
| ENSG000001 | 0 | NA | NA | NA | NA | NA | SNORD18C | 595100 |
| ENSG000001 | 0.62271367 | 0.053535606 | 0.26445898 | 0.20243444 | 0.83957711 | NA | SNORD114-1 | 767577 |
| ENSG000001 | 0.74387701 | -0.194370845 | 0.28592582 | -0.6797947 | 0.49663448 | NA | RNU6-822P | 106479930 |
| ENSG000001 | 0.36277375 | 0.069994797 | 0.21578753 | 0.32436906 | 0.74565864 | NA | NA | NA |
| ENSG000001 | 0.11205239 | 0.082118251 | 0.12671962 | 0.64803108 | 0.51696485 | NA | NA | NA |
| ENSG000001 | 1.77559647 | -0.460571415 | 0.35372876 | -1.3020468 | 0.19290037 | NA | NA | NA |
| ENSG000001 | 0.75982019 | -0.379788636 | 0.28855699 | -1.3161651 | 0.18811864 | NA | NA | NA |
| ENSG000001 | 1.02602882 | -0.46014062 | 0.29742969 | -1.5470568 | 0.12184955 | NA | SNORD114-14 | 767592 |
| ENSG000001 | 0.73545121 | 0.138033976 | 0.28989858 | 0.47614575 | 0.63397055 | NA | RNU6-1301P | 106481591 |
| ENSG000001 | 2.13332799 | -0.261126564 | 0.36682131 | -0.7118631 | 0.47654954 | NA | NA | NA |
| ENSG000001 | 0.25333738 | -0.015686291 | 0.1806605 | -0.0868275 | 0.93080867 | NA | RNU6-859P | 106481452 |
| ENSG000001 | 1.26143155 | -0.323765509 | 0.3214056 | -1.0073425 | 0.31377023 | NA | RNU6-562P | 106479816 |
| ENSG000001 | 1.43979547 | 0.101579583 | 0.34349588 | 0.29572286 | 0.76744174 | NA | RNU6-951P | 106481480 |
| ENSG000001 | 0.99761878 | -0.204823835 | 0.31377682 | -0.6527692 | 0.5139051 | NA | NA | NA |
| ENSG000001 | 0.25993993 | -0.013144386 | 0.18211683 | -0.0721756 | 0.94246219 | NA | NA | NA |
| ENSG000001 | 0.8847611 | 0.040212685 | 0.29823024 | 0.13483772 | 0.89274018 | NA | NA | NA |
| ENSG000001 | 1.36682273 | -0.395625145 | 0.33210897 | -1.191251 | 0.23355505 | NA | RN7SKP20 | 106479104 |
| ENSG000001 | 1.14400054 | -0.090378212 | 0.20820484 | -0.4340831 | 0.66422807 | NA | RNU6-989P | 106481492 |
| ENSG000001 | 0.78588993 | -0.359597889 | 0.28326202 | -1.2694886 | 0.20426687 | NA | RNU6-1010P | 106481929 |
| ENSG000001 | 0.3588776 | -0.099902505 | 0.21555794 | -0.4634601 | 0.6430346 | NA | NA | NA |

| | | | | | | | | | |
|---|---|---|---|---|---|---|---|---|---|
| ENSG000001 | 0 | NA | NA | NA | NA | NA | | SNORD33 | 26818 |
| ENSG000001 | 1.80906358 | -0.113333047 | 0.35285414 | -0.3211895 | 0.74806678 | NA | | RNU4-79P | 106481197 |
| ENSG000001 | 0.13280363 | -0.093394593 | 0.13115644 | -0.7120855 | 0.47641185 | NA | | NA | NA |
| ENSG000001 | 1.13135605 | -0.282292155 | 0.32727603 | -0.8625506 | 0.38838458 | NA | | NA | NA |
| ENSG000001 | 0.64235948 | -0.482579697 | 0.27062934 | -1.7831759 | 0.07455768 | NA | | RN7SKP294 | 106479226 |
| ENSG000001 | 1.67153013 | -0.398170682 | 0.34835477 | -1.1430034 | 0.25303719 | NA | | RNU4-90P | 106479600 |
| ENSG000001 | 0 | NA | NA | NA | NA | NA | | RNU6-1330P | 106480411 |
| ENSG000001 | 1.45139465 | -0.429172408 | 0.33279727 | -1.2895911 | 0.19719265 | NA | | RNU6-1272P | 106480121 |
| ENSG000001 | 0.87224315 | -0.418038832 | 0.29179694 | -1.4326361 | 0.15196186 | NA | | RNU1-35P | 106480157 |
| ENSG000001 | 1.38817087 | -0.265266482 | 0.3352626 | -0.79122 | 0.42881561 | NA | | NA | NA |
| ENSG000001 | 0.77174124 | -0.198363094 | 0.28768104 | -0.6895244 | 0.49049333 | NA | | NA | NA |
| ENSG000001 | 1.86589399 | -0.119857566 | 0.35601816 | -0.3366614 | 0.73637218 | NA | | NA | NA |
| ENSG000001 | 1.14784057 | -0.444178472 | 0.32529103 | -1.3654802 | 0.17210216 | NA | | RNU4-21P | 106481178 |
| ENSG000001 | 0 | NA | NA | NA | NA | NA | | NA | NA |
| ENSG000001 | 0.53969588 | -0.357270382 | 0.23978341 | -1.4899712 | 0.13623181 | NA | | NA | NA |
| ENSG000001 | 0.60698135 | -0.252864344 | 0.25666105 | -0.9852073 | 0.32452224 | NA | | NA | NA |
| ENSG000001 | 2.07649814 | -0.24000276 | 0.36302454 | -0.6611199 | 0.5085354 | NA | | RN7SKP185 | 106479178 |
| ENSG000001 | 0.70420183 | 0.127448135 | 0.26885587 | 0.47403887 | 0.63547219 | NA | | RNU1-38P | 106481608 |
| ENSG000001 | 0.11131518 | 0.081951103 | 0.12663804 | 0.64712866 | 0.51754868 | NA | | RNU6-1128P | 106480062 |
| ENSG000001 | 1.74232528 | -0.315638883 | 0.35245522 | -0.8955432 | 0.37049677 | NA | | RNU6-446P | 106481319 |
| ENSG000001 | 0.4693335 | 0.149163697 | 0.23666968 | 0.63026112 | 0.52852375 | NA | | NA | NA |
| ENSG000001 | 1.43683841 | -0.421051964 | 0.33811026 | -1.2453096 | 0.21301796 | NA | | RNU6-223P | 106481248 |
| ENSG000001 | 1.47044987 | -0.050846142 | 0.34661416 | -0.1466938 | 0.88337373 | NA | | NA | NA |
| ENSG000001 | 0.11934008 | 0.083564698 | 0.12760091 | 0.65489106 | 0.5125379 | NA | | SNORD115-29 | 100033803 |
| ENSG000001 | 1.09543658 | -0.115418275 | 0.3241919 | -0.3560184 | 0.7218268 | NA | | NA | NA |
| ENSG000001 | 1.85941543 | -0.128958161 | 0.35868076 | -0.3595346 | 0.71919516 | NA | | RNU4-23P | 106479568 |
| ENSG000001 | 0.25333738 | -0.015686291 | 0.1806605 | -0.0868275 | 0.93080867 | NA | | NA | NA |
| ENSG000001 | 0.46415331 | 0.147192669 | 0.23603211 | 0.6236129 | 0.53288183 | NA | | NA | NA |
| ENSG000001 | 0.12685397 | -0.090215768 | 0.12901919 | -0.699243 | 0.48440017 | NA | | SNORD115-2 | 100033437 |

| | | | | | | | | | |
|---|---|---|---|---|---|---|---|---|---|
| ENSG000001 | 1.09301251 | 0.193189919 | 0.32526205 | 0.5939516 | 0.55254448 | NA | NA | NA | |
| ENSG000001 | 1.00427532 | -0.044259671 | 0.31348448 | -0.1411862 | 0.88772288 | NA | NA | NA | |
| ENSG000001 | 1.34303283 | -0.120420964 | 0.33582675 | -0.3585806 | 0.71990885 | NA | NA | NA | |
| ENSG000001 | 1.46285608 | -0.112428045 | 0.19942965 | -0.5637479 | 0.57292572 | NA | RN7SKP74 | | 106479127 |
| ENSG000001 | 0.12070844 | -0.088503695 | 0.12783931 | -0.6923042 | 0.48874631 | NA | RNU6-655P | | 106479856 |
| ENSG000001 | 0.40689225 | -0.283036689 | 0.21754773 | -1.3010326 | 0.19324729 | NA | NA | NA | |
| ENSG000001 | 1.55862008 | 0.283617749 | 0.34553113 | 0.82081679 | 0.41175064 | NA | NA | NA | |
| ENSG000001 | 0 | NA | NA | NA | NA | NA | RNU6-227P | | 106481883 |
| ENSG000001 | 1.50261563 | -0.447020553 | 0.33837492 | -1.3210806 | 0.18647449 | NA | NA | NA | |
| ENSG000001 | 0 | NA | NA | NA | NA | NA | SNORD36A | | 26815 |
| ENSG000001 | 0.52165104 | -0.403921832 | 0.25252134 | -1.5995552 | 0.10969729 | NA | NA | NA | |
| ENSG000001 | 1.39238202 | -0.576337849 | 0.33852701 | -1.7024871 | 0.0886641 | NA | NA | NA | |
| ENSG000001 | 1.08291876 | 0.0372422 | 0.3209687 | 0.11603063 | 0.90762827 | NA | NA | NA | |
| ENSG000001 | 0.42385498 | -0.262217565 | 0.21037102 | -1.2464529 | 0.21259817 | NA | NA | NA | |
| ENSG000001 | 0.70080974 | 0.261528332 | 0.26671791 | 0.98054283 | 0.32681824 | NA | RNU6-363P | | 106479727 |
| ENSG000001 | 0.11934008 | 0.083564698 | 0.12760091 | 0.65489106 | 0.5125379 | NA | RNU6-78P | | 100873778 |
| ENSG000001 | 1.66880647 | -0.716954736 | 0.3506257 | -2.0447866 | 0.0408759 | NA | NA | NA | |
| ENSG000001 | 0.36351096 | 0.070290861 | 0.21590418 | 0.32556508 | 0.74475344 | NA | RNU1-76P | | 106481617 |
| ENSG000001 | 0.72423759 | 0.136226018 | 0.27602641 | 0.4935253 | 0.62164147 | NA | NA | NA | |
| ENSG000001 | 1.86129076 | -0.390011751 | 0.35805626 | -1.0892471 | 0.27604493 | NA | NA | NA | |
| ENSG000001 | 0.75808126 | -0.038810283 | 0.28599502 | -0.1357027 | 0.89205635 | NA | SNORD115-9 | | 100033446 |
| ENSG000001 | 0.26813897 | -0.197007041 | 0.18550319 | -1.0620143 | 0.2882292 | NA | NA | NA | |
| ENSG000001 | 0.23276084 | -0.008011382 | 0.17603982 | -0.0455089 | 0.96370166 | NA | RNU6-1287P | | 106480127 |
| ENSG000001 | 0 | NA | NA | NA | NA | NA | SNORA52 | | 619565 |
| ENSG000001 | 2.10592269 | -0.124894073 | 0.36488158 | -0.3422866 | 0.73213522 | NA | NA | NA | |
| ENSG000001 | 1.80826123 | 0.393806277 | 0.35411384 | 1.11208948 | 0.26609968 | NA | NA | NA | |
| ENSG000001 | 0.24550891 | -0.007886114 | 0.17900103 | -0.0440563 | 0.96485956 | NA | RNU6-451P | | 106479765 |
| ENSG000001 | 1.00601425 | -0.360230169 | 0.31375627 | -1.148121 | 0.25091861 | NA | RNU6-1233P | | 106481569 |
| ENSG000001 | 0.91320037 | -0.446160783 | 0.29797291 | -1.4973199 | 0.13431003 | NA | RNU6-924P | | 106480621 |

| | | | | | | | | | |
|---|---|---|---|---|---|---|---|---|---|
| ENSG000001 | 0.24550891 | -0.007886114 | 0.17900103 | -0.0440563 | 0.96485956 | NA | NA | NA | |
| ENSG000001 | 0.23936338 | -0.005479057 | 0.17759977 | -0.0308506 | 0.9753887 | NA | SNORD114-5 | | 767581 |
| ENSG000001 | 0.61446491 | -0.254170746 | 0.25900567 | -0.9813328 | 0.32642867 | NA | NA | NA | |
| ENSG000001 | 0.35346928 | -0.088570748 | 0.20618147 | -0.4295767 | 0.66750362 | NA | RNU6-1159P | | 106481939 |
| ENSG000001 | 1.13016183 | -0.273706642 | 0.32211704 | -0.8497118 | 0.39548535 | NA | NA | NA | |
| ENSG000001 | 1.3051169 | -0.329716441 | 0.32394702 | -1.0178098 | 0.30876836 | NA | RNU1-134P | | 106480194 |
| ENSG000001 | 1.40085946 | -0.253913928 | 0.33281959 | -0.7629176 | 0.44551253 | NA | NA | NA | |
| ENSG000001 | 0.89318635 | -0.270387539 | 0.29448624 | -0.918167 | 0.35853146 | NA | RNU6-428P | | 106481894 |
| ENSG000001 | 0.47753253 | -0.016714525 | 0.24454657 | -0.068349 | 0.94550778 | NA | RNU6-1260P | | 106481578 |
| ENSG000001 | 0 | NA | NA | NA | NA | NA | NA | NA | |
| ENSG000001 | 0.48441528 | -0.017227682 | 0.23774503 | -0.0724628 | 0.94223357 | NA | RNU6-199P | | 106481881 |
| ENSG000001 | 0.64460494 | -0.261028787 | 0.26005827 | -1.0037319 | 0.31550784 | NA | RNU6-1290P | | 106480129 |
| ENSG000001 | 1.39164481 | -0.576618076 | 0.33850737 | -1.7034137 | 0.08849068 | NA | RN7SKP291 | | 106480922 |
| ENSG000001 | 0.23936338 | -0.005479057 | 0.17759977 | -0.0308506 | 0.9753887 | NA | NA | NA | |
| ENSG000001 | 0.23890637 | -0.010420674 | 0.17747379 | -0.0587167 | 0.95317776 | NA | SNORD115-21 | | 100033603 |
| ENSG000001 | 0.96103837 | 0.259364832 | 0.29780785 | 0.87091336 | 0.38380146 | NA | RNU1-47P | | 106480161 |
| ENSG000001 | 0.47758652 | 0.147793567 | 0.23768536 | 0.6218034 | 0.53407115 | NA | NA | NA | |
| ENSG000001 | 0.25370795 | -0.171331747 | 0.17411721 | -0.9840024 | 0.32511435 | NA | NA | NA | |
| ENSG000001 | 0.13280363 | -0.093394593 | 0.13115644 | -0.7120855 | 0.47641185 | NA | NA | NA | |
| ENSG000001 | 0.25260017 | -0.015978705 | 0.18049464 | -0.0885273 | 0.92945758 | NA | NA | NA | |
| ENSG000001 | 0.63370343 | -0.298851213 | 0.27334848 | -1.0932975 | 0.2742632 | NA | NA | NA | |
| ENSG000001 | 0.59485136 | 0.227885061 | 0.26312451 | 0.86607312 | 0.38645007 | NA | NA | NA | |
| ENSG000001 | 1.81928187 | 0.010612986 | 0.36100517 | 0.02939843 | 0.97654682 | NA | RNU6-490P | | 106479783 |
| ENSG000001 | 0.72235822 | 0.133238945 | 0.27758738 | 0.47998921 | 0.63123507 | NA | NA | NA | |
| ENSG000001 | 1.50989884 | -0.196696202 | 0.34759736 | -0.5658737 | 0.57147965 | NA | NA | NA | |
| ENSG000001 | 0.72967624 | -0.02677813 | 0.28955569 | -0.0924801 | 0.92631662 | NA | RNU6-582P | | 106481365 |
| ENSG000001 | 1.32642624 | 0.17527743 | 0.3386323 | 0.51760399 | 0.6047346 | NA | NA | NA | |
| ENSG000001 | 0.46748638 | 0.139734475 | 0.23646399 | 0.59093341 | 0.55456504 | NA | NA | NA | |
| ENSG000001 | 0 | NA | NA | NA | NA | NA | NA | NA | |

| | | | | | | | | |
|---|---|---|---|---|---|---|---|---|
| ENSG000001 | 0.65493287 | -0.26892161 | 0.26251527 | -1.0244037 | 0.30564465 | NA | NA | NA |
| ENSG000001 | 0.95365841 | 0.247986291 | 0.29746881 | 0.83365476 | 0.40447556 | NA | NA | NA |
| ENSG000001 | 1.31732973 | 0.025102068 | 0.33118394 | 0.07579494 | 0.93958224 | NA | RNU6-888P | 106479959 |
| ENSG000001 | 1.87662665 | -0.128000246 | 0.36075816 | -0.354809 | 0.72273268 | NA | NA | NA |
| ENSG000001 | 0.49970612 | -0.194507174 | 0.24827631 | -0.7834303 | 0.43337449 | NA | RN7SKP90 | 106481809 |
| ENSG000001 | 0.86748772 | -0.38803067 | 0.28521484 | -1.3604855 | 0.17367633 | NA | NA | NA |
| ENSG000001 | 1.26082818 | -0.190887737 | 0.32986196 | -0.5786898 | 0.56279853 | NA | RNU6-330P | 106481283 |
| ENSG000001 | 1.33699199 | -0.10121427 | 0.32750148 | -0.3090498 | 0.75728364 | NA | NA | NA |
| ENSG000001 | 1.09271292 | -0.403776013 | 0.31556307 | -1.2795414 | 0.20070646 | NA | NA | NA |
| ENSG000001 | 2.01497712 | -0.066878519 | 0.36261448 | -0.1844342 | 0.85367283 | NA | NA | NA |
| ENSG000001 | 1.63861437 | -0.660950786 | 0.34324123 | -1.925616 | 0.05415235 | NA | NA | NA |
| ENSG000001 | 1.13980707 | -0.102437236 | 0.30966021 | -0.3308053 | 0.74079156 | NA | NA | NA |
| ENSG000001 | 0.96121252 | 0.104196805 | 0.29038685 | 0.35882067 | 0.71972925 | NA | RNU6-492P | 106481335 |
| ENSG000001 | 0.24550891 | -0.007886114 | 0.17900103 | -0.0440563 | 0.96485956 | NA | NA | NA |
| ENSG000001 | 0 | NA | NA | NA | NA | NA | RNA5S10 | 100169761 |
| ENSG000001 | 0.48353618 | 0.145822039 | 0.2384055 | 0.61165553 | 0.54076569 | NA | NA | NA |
| ENSG000001 | 0.87970161 | -0.274795643 | 0.2994662 | -0.9176182 | 0.35881877 | NA | NA | NA |
| ENSG000001 | 0 | NA | NA | NA | NA | NA | NA | NA |
| ENSG000001 | 0.63352874 | -0.1085548 | 0.25832188 | -0.4202308 | 0.67431688 | NA | SNORD114-16 | 767594 |
| ENSG000001 | 0.35210091 | 0.074791332 | 0.21412433 | 0.34928928 | 0.72687215 | NA | RNU6-161P | 106480387 |
| ENSG000001 | 1.70591318 | 0.084543533 | 0.35774871 | 0.236321 | 0.8131836 | NA | RNU6-1252P | 106480113 |
| ENSG000001 | 0 | NA | NA | NA | NA | NA | NA | NA |
| ENSG000001 | 1.10389676 | -0.262060301 | 0.32129252 | -0.8156439 | 0.41470381 | NA | NA | NA |
| ENSG000001 | 0.90298455 | -0.457507079 | 0.30248048 | -1.5125177 | 0.13040221 | NA | NA | NA |
| ENSG000001 | 0.89956268 | -0.114598785 | 0.29931506 | -0.3828701 | 0.70181608 | NA | NA | NA |
| ENSG000001 | 1.15375989 | -0.259698665 | 0.31847076 | -0.8154553 | 0.41481175 | NA | NA | NA |
| ENSG000001 | 0.88310849 | -0.114847963 | 0.298483 | -0.3847722 | 0.70040619 | NA | NA | NA |
| ENSG000001 | 0.96306823 | -0.179450239 | 0.30593533 | -0.5865627 | 0.55749747 | NA | RN7SKP75 | 106480856 |
| ENSG000001 | 0.75982019 | -0.379788636 | 0.28855699 | -1.3161651 | 0.18811864 | NA | SNORD114-19 | 767597 |

| | | | | | | | | |
|---|---|---|---|---|---|---|---|---|
| ENSG000001 | 0.64602887 | -0.11525331 | 0.26675303 | -0.43206 | 0.66569782 | NA | RNU6-653P | 106480397 |
| ENSG000001 | 0.2596576 | -0.193002035 | 0.1837913 | -1.0501152 | 0.29366517 | NA | NA | NA |
| ENSG000001 | 1.08115907 | -0.147776524 | 0.2109005 | -0.7006931 | 0.48349456 | NA | RNA5SP443 | 106481745 |
| ENSG000001 | 0.89748744 | -0.1192697 | 0.29920235 | -0.3986255 | 0.69016914 | NA | NA | NA |
| ENSG000001 | 1.64959731 | -0.260994837 | 0.34504906 | -0.7563992 | 0.44940988 | NA | SNORD115-14 | 100033451 |
| ENSG000001 | 0.4988463 | -0.023573523 | 0.24689417 | -0.0954803 | 0.92393336 | NA | RNU6-605P | 106481372 |
| ENSG000001 | 0.63504146 | -0.278824355 | 0.26770076 | -1.0415524 | 0.29761927 | NA | NA | NA |
| ENSG000001 | 0.35022155 | 0.06940789 | 0.21379411 | 0.32464827 | 0.74544729 | NA | SNORD115-19 | 100033458 |
| ENSG000001 | 0.77658704 | -0.341413701 | 0.27605383 | -1.236765 | 0.21617435 | NA | SNORD115-3 | 100033440 |
| ENSG000001 | 1.24862089 | -0.204586271 | 0.33622263 | -0.6084845 | 0.54286616 | NA | RNU6-797P | 106479918 |
| ENSG000001 | 0.51413714 | -0.199727707 | 0.24976394 | -0.7996659 | 0.42390439 | NA | RN7SKP243 | 106481819 |
| ENSG000001 | 0.87243903 | -0.103976508 | 0.16036052 | -0.6483922 | 0.51673132 | NA | NA | NA |
| ENSG000001 | 0.9164481 | -0.645992679 | 0.30062045 | -2.1488647 | 0.03164513 | NA | NA | NA |
| ENSG000002 | 1.62328714 | -0.233034888 | 0.339621 | -0.6861616 | 0.49261121 | NA | RNU6-986P | 106481491 |
| ENSG000002 | 0.34407602 | 0.071624546 | 0.21281613 | 0.33655599 | 0.73645163 | NA | NA | NA |
| ENSG000002 | 1.59250048 | -0.123223125 | 0.34217562 | -0.3601166 | 0.71875993 | NA | NA | NA |
| ENSG000002 | 0.23276084 | -0.008011382 | 0.17603982 | -0.0455089 | 0.96370166 | NA | RNU6-623P | 106481378 |
| ENSG000002 | 1.09762861 | -0.106657454 | 0.31555812 | -0.3379962 | 0.73536603 | NA | NA | NA |
| ENSG000002 | 0.47958604 | -0.1795445 | 0.23869266 | -0.7521995 | 0.4519311 | NA | NA | NA |
| ENSG000002 | 1.28707206 | -0.360660004 | 0.33479808 | -1.0772463 | 0.28137023 | NA | NA | NA |
| ENSG000002 | 2.01464315 | -0.716637964 | 0.36014696 | -1.9898487 | 0.0466076 | NA | RNU6-642P | 106481384 |
| ENSG000002 | 1.02274672 | -0.497155251 | 0.30480652 | -1.6310519 | 0.10287937 | NA | RNU6-403P | 106480578 |
| ENSG000002 | 0.5301324 | -0.3843196 | 0.24728844 | -1.5541349 | 0.12015224 | NA | RNU4-73P | 106481195 |
| ENSG000002 | 0.87708503 | -0.279954352 | 0.29929796 | -0.9353701 | 0.34959762 | NA | NA | NA |
| ENSG000002 | 0.39246123 | -0.276864021 | 0.21549122 | -1.2848042 | 0.19886073 | NA | NA | NA |
| ENSG000002 | 0.61950267 | -0.112567707 | 0.27088102 | -0.4155614 | 0.67773094 | NA | NA | NA |
| ENSG000002 | 2.09384869 | -0.002954672 | 0.36807567 | -0.0080273 | 0.99359517 | NA | RN7SKP115 | 106479146 |
| ENSG000002 | 0.38651158 | -0.274258716 | 0.21461355 | -1.2779189 | 0.20127801 | NA | RNU6-812P | 106479926 |
| ENSG000002 | 1.13323487 | -0.277572658 | 0.32368507 | -0.8575393 | 0.39114692 | NA | NA | NA |

| | | | | | | | | |
|---|---|---|---|---|---|---|---|---|
| ENSG000002 | 0.50135681 | -0.182211159 | 0.24123516 | -0.7553259 | 0.45005347 | NA | NA | NA |
| ENSG000002 | 1.36269645 | -0.120290223 | 0.33961273 | -0.3541982 | 0.72319034 | NA | NA | NA |
| ENSG000002 | 0 | NA | NA | NA | NA | NA | RN7SKP63 | 106479123 |
| ENSG000002 | 0.97615262 | 0.124394399 | 0.31229827 | 0.3983192 | 0.69039491 | NA | NA | NA |
| ENSG000002 | 0.81221529 | 0.20432271 | 0.29316762 | 0.69694842 | 0.48583508 | NA | NA | NA |
| ENSG000002 | 0.52760069 | -0.383393289 | 0.24704506 | -1.5519164 | 0.12068222 | NA | RNU4-58P | 106481190 |
| ENSG000002 | 1.75243885 | -0.163069156 | 0.34120904 | -0.4779157 | 0.63271021 | NA | NA | NA |
| ENSG000002 | 0.11865494 | 0.083440866 | 0.12751478 | 0.65436232 | 0.5128784 | NA | NA | NA |
| ENSG000002 | 0.49264869 | -0.021364872 | 0.24621241 | -0.0867741 | 0.93085104 | NA | NA | NA |
| ENSG000002 | 1.48826604 | -0.06001222 | 0.34570689 | -0.1735928 | 0.86218551 | NA | RNU4-80P | 106480386 |
| ENSG000002 | 1.37237149 | -0.123219709 | 0.33646177 | -0.366222 | 0.71419941 | NA | NA | NA |
| ENSG000002 | 1.73901748 | -0.659562124 | 0.34137171 | -1.9320937 | 0.05334795 | NA | NA | NA |
| ENSG000002 | 1.50352497 | -0.179301736 | 0.33977173 | -0.5277123 | 0.59769901 | NA | NA | NA |
| ENSG000002 | 0 | NA | NA | NA | NA | NA | NA | NA |
| ENSG000002 | 0.89950657 | -0.435154337 | 0.29566513 | -1.471781 | 0.14108001 | NA | NA | NA |
| ENSG000002 | 0.74313979 | -0.194647915 | 0.28587591 | -0.6808825 | 0.49594582 | NA | SNORD114-31 | 767612 |
| ENSG000002 | 1.78772391 | -0.457481153 | 0.35198931 | -1.2997019 | 0.19370317 | NA | RN7SKP163 | 106479168 |
| ENSG000002 | 0.87738909 | 0.041650852 | 0.29784854 | 0.13983903 | 0.88878717 | NA | RNU6-181P | 106480559 |
| ENSG000002 | 0.67736705 | -0.463272559 | 0.26608476 | -1.7410714 | 0.08167107 | NA | RNU6-1167P | 106480407 |
| ENSG000002 | 0.94100262 | 0.276036131 | 0.30502224 | 0.9049705 | 0.36548102 | NA | RNU6-508P | 106480585 |
| ENSG000002 | 1.86026338 | -0.225376549 | 0.3505416 | -0.6429381 | 0.52026427 | NA | RNU6-251P | 106481257 |
| ENSG000002 | 0.74826634 | -0.352264801 | 0.2807694 | -1.254641 | 0.20960911 | NA | RNU6-984P | 106481927 |
| ENSG000002 | 2.13241844 | -0.529789951 | 0.36759078 | -1.4412493 | 0.14951427 | NA | RNU6-1249P | 106480652 |
| ENSG000002 | 0.13280363 | -0.093394593 | 0.13115644 | -0.7120855 | 0.47641185 | NA | RNU6-774P | 106481425 |
| ENSG000002 | 0.25351207 | -0.189965547 | 0.1824784 | -1.0410303 | 0.29786148 | NA | NA | NA |
| ENSG000002 | 1.4687452 | -0.197338802 | 0.3483005 | -0.5665763 | 0.5710021 | NA | NA | NA |
| ENSG000002 | 0 | NA | NA | NA | NA | NA | NA | NA |
| ENSG000002 | 0.98283948 | 0.122754107 | 0.31257693 | 0.39271647 | 0.69452891 | NA | RN7SKP77 | 106479128 |
| ENSG000002 | 0.66936389 | -0.269876139 | 0.26211055 | -1.0296272 | 0.30318506 | NA | RNU6-1021P | 106481930 |

| | | | | | | | | |
|---|---|---|---|---|---|---|---|---|
| ENSG000002 | 1.45046776 | 0.089269944 | 0.34371618 | 0.25971994 | 0.79507981 | NA | NA | NA |
| ENSG000002 | 0.14128499 | -0.097772014 | 0.13403235 | -0.7294658 | 0.4657168 | NA | RNY1P10 | 106480308 |
| ENSG000002 | 0.39499294 | -0.277908454 | 0.21584201 | -1.287555 | 0.19790091 | NA | RNU6-778P | 106479910 |
| ENSG000002 | 1.12555935 | -0.568173249 | 0.31531052 | -1.8019483 | 0.07155355 | NA | NA | NA |
| ENSG000002 | 0.5301324 | -0.3843196 | 0.24728844 | -1.5541349 | 0.12015224 | NA | RNU6-544P | 106481353 |
| ENSG000002 | 0.12070844 | -0.088503695 | 0.12783931 | -0.6923042 | 0.48874631 | NA | NA | NA |
| ENSG000002 | 0.39462238 | -0.104329077 | 0.21264929 | -0.4906157 | 0.6236983 | NA | RN7SKP231 | 106479198 |
| ENSG000002 | 0.99707745 | -0.172027674 | 0.29293416 | -0.5872571 | 0.55703103 | NA | RNU6-216P | 106479662 |
| ENSG000002 | 0.35664995 | 0.244349217 | 0.20882949 | 1.17008962 | 0.2419649 | NA | RN7SKP145 | 106480878 |
| ENSG000002 | 0.12070844 | -0.088503695 | 0.12783931 | -0.6923042 | 0.48874631 | NA | NA | NA |
| ENSG000002 | 1.00450207 | -0.204287946 | 0.3180715 | -0.6422705 | 0.52069756 | NA | SNORD115-18 | 100033456 |
| ENSG000002 | 1.11100975 | -0.116740449 | 0.32604992 | -0.3580447 | 0.72030985 | NA | NA | NA |
| ENSG000002 | 2.01692244 | -0.474618005 | 0.36540463 | -1.2988834 | 0.19398395 | NA | RNA5SP350 | 106481794 |
| ENSG000002 | 0.513452 | -0.199963756 | 0.2496885 | -0.8008529 | 0.42321682 | NA | NA | NA |
| ENSG000002 | 0 | NA | NA | NA | NA | NA | NA | NA |
| ENSG000002 | 0 | NA | NA | NA | NA | NA | NA | NA |
| ENSG000002 | 1.59568808 | 0.022821496 | 0.3420755 | 0.06671479 | 0.94680876 | NA | RNU6-238P | 106481253 |
| ENSG000002 | 1.09963004 | -0.259399898 | 0.31885155 | -0.8135444 | 0.415906 | NA | RNU6-832P | 106479934 |
| ENSG000002 | 1.56725414 | -0.466392076 | 0.33951673 | -1.373694 | 0.16953671 | NA | NA | NA |
| ENSG000002 | 0.8417807 | 0.048619213 | 0.29595319 | 0.16428008 | 0.86951067 | NA | NA | NA |
| ENSG000002 | 0.76325988 | -0.182583956 | 0.27949837 | -0.6532559 | 0.51359132 | NA | RN7SKP211 | 106479189 |
| ENSG000002 | 0.60970501 | 0.062117743 | 0.26972855 | 0.23029725 | 0.8178608 | NA | NA | NA |
| ENSG000002 | 1.12091347 | -0.108006689 | 0.31635556 | -0.3414092 | 0.73279557 | NA | NA | NA |
| ENSG000002 | 0.23816916 | -0.010712105 | 0.1772998 | -0.060418 | 0.9518227 | NA | NA | NA |
| ENSG000002 | 1.22713244 | -0.049503608 | 0.33655116 | -0.1470909 | 0.88306029 | NA | NA | NA |
| ENSG000002 | 0.72972831 | -0.025248506 | 0.28417722 | -0.0888477 | 0.92920291 | NA | NA | NA |
| ENSG000002 | 0.38133351 | -0.102466351 | 0.2189713 | -0.4679442 | 0.63982449 | NA | NA | NA |
| ENSG000002 | 2.09203638 | -0.374786752 | 0.3638698 | -1.0300023 | 0.3030089 | NA | RNU6-992P | 106481493 |
| ENSG000002 | 0.98159319 | 0.111594572 | 0.30607873 | 0.36459434 | 0.71541424 | NA | SNORD113-6 | 767566 |

| | | | | | | | | | |
|---|---|---|---|---|---|---|---|---|---|
| ENSG000002 | 0.35273207 | -0.08884607 | 0.20605566 | -0.4311751 | 0.66634105 | NA | | RNU6-745P | 106481415 |
| ENSG000002 | 0.35961481 | -0.099625193 | 0.21567743 | -0.4619176 | 0.64414043 | NA | | RNU6-839P | 106481446 |
| ENSG000002 | 0.84648854 | -0.247366387 | 0.2902103 | -0.8523694 | 0.39400908 | NA | | RNU6-179P | 106479646 |
| ENSG000002 | 1.00498027 | -0.038668406 | 0.30871656 | -0.1252554 | 0.90032139 | NA | | RNU6-626P | 106481379 |
| ENSG000002 | 1.1304447 | -0.113660466 | 0.32284156 | -0.3520627 | 0.72479125 | NA | | NA | NA |
| ENSG000002 | 0.25260017 | -0.015978705 | 0.18049464 | -0.0885273 | 0.92945758 | NA | | NA | NA |
| ENSG000002 | 0.37330862 | -0.105391168 | 0.21773449 | -0.4840352 | 0.62836086 | NA | | RNU1-23P | 106478980 |
| ENSG000002 | 0.520565 | -0.024378065 | 0.24183609 | -0.1008041 | 0.91970599 | NA | | NA | NA |
| ENSG000002 | 0.26560726 | -0.177019426 | 0.17672274 | -1.0016788 | 0.31649875 | NA | | NA | NA |
| ENSG000002 | 1.29076523 | 0.156208116 | 0.326292 | 0.47873719 | 0.6321256 | NA | | NA | NA |
| ENSG000002 | 1.39312272 | -0.275911813 | 0.34075575 | -0.8097055 | 0.41810944 | NA | | RNA5SP263 | 106479002 |
| ENSG000002 | 1.14368149 | -0.122054925 | 0.32325304 | -0.3775832 | 0.70574023 | NA | | RNU6-254P | 106479679 |
| ENSG000002 | 1.27919097 | -0.335388722 | 0.3257012 | -1.0297436 | 0.30313039 | NA | | NA | NA |
| ENSG000002 | 0.75458656 | -0.522322277 | 0.27834054 | -1.8765584 | 0.06057864 | NA | | RNU6-1147P | 106480071 |
| ENSG000002 | 1.52099623 | -0.19083606 | 0.34150944 | -0.5588017 | 0.57629706 | NA | | NA | NA |
| ENSG000002 | 0.48945358 | 0.14806159 | 0.23910487 | 0.61923286 | 0.53576297 | NA | | RNU6-529P | 106481348 |
| ENSG000002 | 1.39169879 | -0.409765381 | 0.3372028 | -1.2151897 | 0.22429373 | NA | | NA | NA |
| ENSG000002 | 1.00971452 | -0.196447329 | 0.31277776 | -0.6280732 | 0.52995598 | NA | | RNU6-97P | 106480551 |
| ENSG000002 | 0 | NA | NA | NA | NA | NA | | SNORD35A | 26816 |
| ENSG000002 | 0.49076933 | -0.025124039 | 0.23846109 | -0.1053591 | 0.91609089 | NA | | NA | NA |
| ENSG000002 | 0.63370343 | -0.298851213 | 0.27334848 | -1.0932975 | 0.2742632 | NA | | NA | NA |
| ENSG000002 | 0.40122492 | -0.101896709 | 0.21365861 | -0.4769136 | 0.63342361 | NA | | RNU6-66P | 100873770 |
| ENSG000002 | 0.50818749 | -0.197539377 | 0.24915052 | -0.7928516 | 0.42786432 | NA | | RNU6-838P | 106479937 |
| ENSG000002 | 0 | NA | NA | NA | NA | NA | | RNU4-32P | 106479572 |
| ENSG000002 | 0.23868015 | 0.154625155 | 0.16997278 | 0.90970538 | 0.36297791 | NA | | RNA5SP199 | 106480438 |
| ENSG000002 | 1.61956197 | -0.236048144 | 0.34092914 | -0.6923672 | 0.48870678 | NA | | NA | NA |
| ENSG000002 | 0 | NA | NA | NA | NA | NA | | SNORD114-10 | 767588 |
| ENSG000002 | 0.85382829 | -0.24463488 | 0.29064849 | -0.8416864 | 0.39996351 | NA | | RNU6-624P | 106479842 |
| ENSG000002 | 0.7164389 | -0.025867019 | 0.28328556 | -0.0913108 | 0.92724567 | NA | | NA | NA |

| | | | | | | | | |
|---|---|---|---|---|---|---|---|---|
| ENSG000002 | 0.25333738 | -0.015686291 | 0.1806605 | -0.0868275 | 0.93080867 | NA | NA | NA |
| ENSG000002 | 0.60561299 | -0.101020012 | 0.26336431 | -0.3835752 | 0.70129334 | NA | NA | NA |
| ENSG000002 | 0.76289324 | -0.362184658 | 0.28331563 | -1.2783787 | 0.20111594 | NA | NA | NA |
| ENSG000002 | 0.35067856 | 0.074230421 | 0.21389444 | 0.34704231 | 0.72855953 | NA | NA | NA |
| ENSG000002 | 1.18193624 | -0.264706739 | 0.31394795 | -0.8431549 | 0.39914184 | NA | NA | NA |
| ENSG000002 | 0.46478446 | -0.015417091 | 0.23534337 | -0.0655089 | 0.9477688 | NA | RNU6-527P | 106480587 |
| ENSG000002 | 1.04239018 | -0.535722969 | 0.31261827 | -1.713665 | 0.08659028 | NA | RNU1-83P | 106480176 |
| ENSG000002 | 1.14097902 | -0.276474224 | 0.32245111 | -0.8574144 | 0.39121591 | NA | NA | NA |
| ENSG000002 | 0.61084662 | 0.056873999 | 0.26981345 | 0.21079009 | 0.83305108 | NA | NA | NA |
| ENSG000002 | 0.48279897 | 0.145542372 | 0.23831717 | 0.6107087 | 0.54139244 | NA | RNU6-1255P | 106481946 |
| ENSG000002 | 0.97379505 | -0.036042115 | 0.31226337 | -0.1154222 | 0.90811051 | NA | NA | NA |
| ENSG000002 | 1.21208074 | -0.566062503 | 0.3145352 | -1.7996794 | 0.07191128 | NA | NA | NA |
| ENSG000002 | 1.18691843 | -0.40486134 | 0.3120343 | -1.2974899 | 0.19446269 | NA | NA | NA |
| ENSG000002 | 1.14807083 | -0.263168712 | 0.31829328 | -0.826812 | 0.40834361 | NA | NA | NA |
| ENSG000002 | 0.76073209 | -0.532127553 | 0.28034426 | -1.8981218 | 0.05768004 | NA | NA | NA |
| ENSG000002 | 0 | NA | NA | NA | NA | NA | NA | NA |
| ENSG000002 | 0.14128499 | -0.097772014 | 0.13403235 | -0.7294658 | 0.4657168 | NA | NA | NA |
| ENSG000002 | 0.12685397 | -0.090215768 | 0.12901919 | -0.699243 | 0.48440017 | NA | NA | NA |
| ENSG000002 | 1.62474408 | -0.114960838 | 0.34623505 | -0.3320312 | 0.7398657 | NA | NA | NA |
| ENSG000002 | 0 | NA | NA | NA | NA | NA | NA | NA |
| ENSG000002 | 0.23890637 | -0.010420674 | 0.17747379 | -0.0587167 | 0.95317776 | NA | NA | NA |
| ENSG000002 | 0 | NA | NA | NA | NA | NA | RNA5SP333 | 106480763 |
| ENSG000002 | 0 | NA | NA | NA | NA | NA | RNA5S8 | 100169759 |
| ENSG000002 | 1.35260078 | -0.402288501 | 0.33560232 | -1.198706 | 0.2306423 | NA | NA | NA |
| ENSG000002 | 0.40689225 | -0.283036689 | 0.21754773 | -1.3010326 | 0.19324729 | NA | RNU6-485P | 106481333 |
| ENSG000002 | 0.88017981 | -0.118396126 | 0.30337318 | -0.3902656 | 0.69634014 | NA | NA | NA |
| ENSG000002 | 1.15261774 | -0.268049598 | 0.32007412 | -0.837461 | 0.40233347 | NA | NA | NA |
| ENSG000002 | 0.99647718 | -0.207034513 | 0.31775868 | -0.6515464 | 0.51469386 | NA | RNU6-833P | 106481444 |
| ENSG000002 | 1.12424305 | -0.416784188 | 0.31908831 | -1.3061719 | 0.19149411 | NA | RNU6-792P | 106481431 |

| | | | | | | | | |
|---|---|---|---|---|---|---|---|---|
| ENSG000002 | 1.49413427 | -0.461023938 | 0.34171791 | -1.349136 | 0.17729328 | NA | NA | NA |
| ENSG000002 | 1.62725193 | -0.411784317 | 0.35155337 | -1.171328 | 0.24146695 | NA | RNU6-511P | 106479793 |
| ENSG000002 | 1.14391217 | -0.568291986 | 0.3160277 | -1.7982347 | 0.07213982 | NA | SNORD113-8 | 767568 |
| ENSG000002 | 1.37888758 | -0.242311002 | 0.32802642 | -0.7386936 | 0.4600931 | NA | RNU6-666P | 106480600 |
| ENSG000002 | 0 | NA | NA | NA | NA | NA | RNA5S17 | 100169768 |
| ENSG000002 | 0 | NA | NA | NA | NA | NA | NA | NA |
| ENSG000002 | 1.87002411 | -0.129918486 | 0.36070843 | -0.3601759 | 0.71871559 | NA | NA | NA |
| ENSG000002 | 0.86954123 | -0.540317036 | 0.28124427 | -1.9211664 | 0.05471073 | NA | NA | NA |
| ENSG000002 | 0 | NA | NA | NA | NA | NA | NA | NA |
| ENSG000002 | 0.73681957 | -0.033780691 | 0.28998188 | -0.1164924 | 0.90726229 | NA | RN7SKP45 | 106479115 |
| ENSG000002 | 0 | NA | NA | NA | NA | NA | RNA5S4 | 100169755 |
| ENSG000002 | 0.84702987 | -0.241400035 | 0.28775544 | -0.8389069 | 0.40152153 | NA | NA | NA |
| ENSG000002 | 0.51208364 | -0.022981898 | 0.24832117 | -0.0925491 | 0.92626179 | NA | RNU6-618P | 106481376 |
| ENSG000002 | 1.03120689 | -0.710702297 | 0.31018902 | -2.291191 | 0.02195237 | NA | RNU6-509P | 106479792 |
| ENSG000002 | 1.61854243 | -0.403690611 | 0.35082077 | -1.1507033 | 0.24985433 | NA | NA | NA |
| ENSG000002 | 0.47744217 | 0.124442156 | 0.22775401 | 0.54638843 | 0.58479896 | NA | NA | NA |
| ENSG000002 | 1.13500817 | -0.435538509 | 0.32350753 | -1.346301 | 0.17820547 | NA | RNU6-698P | 106481401 |
| ENSG000002 | 1.08790095 | -0.115040445 | 0.32144923 | -0.3578806 | 0.72043266 | NA | SNORD115-24 | 100036563 |
| ENSG000002 | 0.42385498 | -0.262217565 | 0.21037102 | -1.2464529 | 0.21259817 | NA | SNORD114-23 | 767603 |
| ENSG000002 | 0.26062507 | -0.012888319 | 0.18226503 | -0.070712 | 0.94362699 | NA | RNU6-1169P | 106480080 |
| ENSG000002 | 0.7802226 | -0.200980091 | 0.2881869 | -0.697395 | 0.48555566 | NA | NA | NA |
| ENSG000002 | 0 | NA | NA | NA | NA | NA | SNORD114-26 | 767606 |
| ENSG000002 | 0 | NA | NA | NA | NA | NA | NA | NA |
| ENSG000002 | 0.84943449 | 0.167194785 | 0.27964629 | 0.59787951 | 0.54992034 | NA | NA | NA |
| ENSG000002 | 0.26813897 | -0.197007041 | 0.18550319 | -1.0620143 | 0.2882292 | NA | NA | NA |
| ENSG000002 | 0.11205239 | 0.082118251 | 0.12671962 | 0.64803108 | 0.51696485 | NA | NA | NA |
| ENSG000002 | 1.60256782 | -0.125616747 | 0.34590791 | -0.3631508 | 0.71649221 | NA | RNU6-1155P | 106481544 |
| ENSG000002 | 0.39499294 | -0.277908454 | 0.21584201 | -1.287555 | 0.19790091 | NA | NA | NA |
| ENSG000002 | 0.3412853 | 0.237533224 | 0.20625411 | 1.15165331 | 0.24946357 | NA | RNU4-81P | 106479595 |

| | | | | | | | | |
|---|---|---|---|---|---|---|---|---|
| ENSG000002 | 1.24719853 | -0.192800664 | 0.32950263 | -0.5851263 | 0.55846278 | NA | NA | NA |
| ENSG000002 | 0.66156766 | -0.287991838 | 0.26979811 | -1.0674346 | 0.28577564 | NA | NA | NA |
| ENSG000002 | 1.35362086 | -0.226829437 | 0.32742486 | -0.6927679 | 0.48845521 | NA | NA | NA |
| ENSG000002 | 1.01201597 | -0.366610227 | 0.31542515 | -1.1622733 | 0.24512447 | NA | NA | NA |
| ENSG000002 | 0.4846238 | -0.022877807 | 0.23773721 | -0.0962315 | 0.92333672 | NA | RNU6-1066P | 106480035 |
| ENSG000002 | 0.76440148 | -0.201033375 | 0.28720316 | -0.6999692 | 0.48394652 | NA | RNU6-1339P | 106481603 |
| ENSG000002 | 0.14128499 | -0.097772014 | 0.13403235 | -0.7294658 | 0.4657168 | NA | RNU6-1085P | 106480043 |
| ENSG000002 | 1.47011356 | -0.335929248 | 0.34518797 | -0.9731777 | 0.33046497 | NA | NA | NA |
| ENSG000002 | 1.2509779 | -0.052455609 | 0.32909155 | -0.1593952 | 0.87335753 | NA | RNU6-1112P | 106480055 |
| ENSG000002 | 0.50067359 | -0.016990261 | 0.23962179 | -0.0709045 | 0.94347377 | NA | RNU6-1097P | 106480048 |
| ENSG000002 | 0 | NA | NA | NA | NA | NA | SNORD118 | 727676 |
| ENSG000002 | 1.10967619 | -0.388182503 | 0.31332704 | -1.2389052 | 0.2153806 | NA | RNA5SP403 | 106479012 |
| ENSG000002 | 0 | NA | NA | NA | NA | NA | NA | NA |
| ENSG000002 | 0.37285215 | -0.099309896 | 0.2177345 | -0.4561055 | 0.64831414 | NA | RNU6-1125P | 106480061 |
| ENSG000002 | 0.6606002 | -0.463974368 | 0.2663045 | -1.7422701 | 0.08146119 | NA | NA | NA |
| ENSG000002 | 2.0999274 | -1.016342633 | 0.35676789 | -2.8487503 | 0.00438913 | NA | RN7SKP162 | 106480884 |
| ENSG000002 | 0.6530863 | -0.285158107 | 0.26914708 | -1.059488 | 0.28937758 | NA | SNORD115-41 | 100033815 |
| ENSG000002 | 0.62087103 | -0.279525375 | 0.26646004 | -1.049033 | 0.29416293 | NA | SNORD114-28 | 767609 |
| ENSG000002 | 0.73223883 | -0.161122453 | 0.27005503 | -0.5966282 | 0.55075562 | NA | RNU6-1017P | 106480014 |
| ENSG000002 | 0.66156766 | -0.287991838 | 0.26979811 | -1.0674346 | 0.28577564 | NA | RNU6-1244P | 106481945 |
| ENSG000002 | 0.90779205 | -0.463756956 | 0.30262298 | -1.5324578 | 0.12540949 | NA | NA | NA |
| ENSG000002 | 0.38019136 | -0.10756138 | 0.21876967 | -0.491665 | 0.62295621 | NA | SNORD115-11 | 100033448 |
| ENSG000002 | 0.50497064 | -0.183718113 | 0.24165995 | -0.760234 | 0.44711472 | NA | NA | NA |
| ENSG000002 | 0.59815061 | 0.048708909 | 0.25313511 | 0.19242257 | 0.84741122 | NA | NA | NA |
| ENSG000002 | 1.24743113 | -0.499312218 | 0.33037796 | -1.5113363 | 0.13070278 | NA | RNU6-1205P | 106480093 |
| ENSG000002 | 0.11934008 | 0.083564698 | 0.12760091 | 0.65489106 | 0.5125379 | NA | SNORD115-5 | 100033442 |
| ENSG000002 | 0.65473699 | -0.286059797 | 0.2692552 | -1.0624114 | 0.28804896 | NA | NA | NA |
| ENSG000002 | 0.39499294 | -0.277908454 | 0.21584201 | -1.287555 | 0.19790091 | NA | NA | NA |
| ENSG000002 | 0 | NA | NA | NA | NA | NA | NA | NA |

| | | | | | | | | | |
|---|---|---|---|---|---|---|---|---|---|
| ENSG000002 | 0.64850501 | -0.484783006 | 0.2711005 | -1.788204 | 0.07374309 | NA | | RNU6-1214P | 106481781 |
| ENSG000002 | 0.25214371 | -0.009862892 | 0.18046657 | -0.0546522 | 0.95641556 | NA | | NA | NA |
| ENSG000002 | 0.94419636 | -0.023163351 | 0.30189025 | -0.0767277 | 0.93884015 | NA | | RNU6-957P | 106479989 |
| ENSG000002 | 0.92825262 | 0.112337809 | 0.29587259 | 0.37968305 | 0.7041807 | NA | | NA | NA |
| ENSG000002 | 0.64195858 | -0.101091878 | 0.25907734 | -0.3901996 | 0.69638895 | NA | | NA | NA |
| ENSG000002 | 0 | NA | NA | NA | NA | NA | | NA | NA |
| ENSG000002 | 0.51322524 | -0.027081322 | 0.24101349 | -0.1123643 | 0.91053453 | NA | | NA | NA |
| ENSG000002 | 0.64608094 | -0.115215656 | 0.26675803 | -0.4319107 | 0.6658063 | NA | | NA | NA |
| ENSG000002 | 0.88052864 | -0.456101693 | 0.30132726 | -1.5136423 | 0.13011659 | NA | | NA | NA |
| ENSG000002 | 0.25260017 | -0.015978705 | 0.18049464 | -0.0885273 | 0.92945758 | NA | | NA | NA |
| ENSG000002 | 0.48624257 | -0.012072666 | 0.23796621 | -0.0507327 | 0.95953853 | NA | | NA | NA |
| ENSG000002 | 1.48895118 | -0.061053652 | 0.34830502 | -0.1752879 | 0.86085343 | NA | | NA | NA |
| ENSG000002 | 0.85380656 | -0.412184383 | 0.29077846 | -1.4175203 | 0.15633085 | NA | | RNU6-1020P | 106481503 |
| ENSG000002 | 0.76650706 | -0.381422914 | 0.28896362 | -1.3199686 | 0.1868455 | NA | | RNU6-103P | 106480552 |
| ENSG000002 | 0.24411881 | -0.012946855 | 0.17864669 | -0.0724718 | 0.94222641 | NA | | RNA5SP429 | 106480767 |
| ENSG000002 | 0.77888457 | -0.19524685 | 0.28212634 | -0.6920547 | 0.48890301 | NA | | RNU6-288P | 106479694 |
| ENSG000002 | 2.02064837 | -0.464079981 | 0.36424183 | -1.2740985 | 0.20262849 | NA | | SNORD115-39 | 100033813 |
| ENSG000002 | 0.50548109 | -0.025402968 | 0.24760631 | -0.1025942 | 0.91828506 | NA | | RNU6-829P | 106479933 |
| ENSG000002 | 0.74140533 | -0.17512453 | 0.27799747 | -0.6299501 | 0.52872725 | NA | | NA | NA |
| ENSG000002 | 1.97949774 | -0.276134742 | 0.35449727 | -0.7789474 | 0.43601068 | NA | | RNU6-414P | 106479749 |
| ENSG000002 | 0.3433388 | 0.065348438 | 0.20407693 | 0.32021472 | 0.74880557 | NA | | NA | NA |
| ENSG000002 | 2.16415317 | -0.239784361 | 0.35766159 | -0.6704224 | 0.50258853 | NA | | NA | NA |
| ENSG000002 | 0 | NA | NA | NA | NA | NA | | SNORD115-33 | 100033807 |
| ENSG000002 | 0.52779657 | -0.383508725 | 0.24707466 | -1.5521977 | 0.12061492 | NA | | RNU6-824P | 106480612 |
| ENSG000002 | 1.89705939 | -0.129170409 | 0.35838709 | -0.3604215 | 0.71853196 | NA | | RNU1-87P | 106480178 |
| ENSG000002 | 0.87736578 | -0.270351876 | 0.29776284 | -0.9079437 | 0.36390799 | NA | | NA | NA |
| ENSG000002 | 1.38028683 | -0.573449315 | 0.33828623 | -1.6951601 | 0.09004506 | NA | | RNA5SP50 | 106478992 |
| ENSG000002 | 0.35690842 | 0.061733055 | 0.20636749 | 0.29914137 | 0.76483218 | NA | | NA | NA |
| ENSG000002 | 0.13280363 | -0.093394593 | 0.13115644 | -0.7120855 | 0.47641185 | NA | | SNORD114-11 | 767589 |

| | | | | | | | | | |
|---|---|---|---|---|---|---|---|---|---|
| ENSG000002 | 0.73957996 | -0.027414423 | 0.28481756 | -0.0962526 | 0.92331998 | NA | | NA | NA |
| ENSG000002 | 1.00546954 | -0.039858873 | 0.31353044 | -0.1271292 | 0.89883815 | NA | | SNORD114-25 | 767605 |
| ENSG000002 | 0.35750923 | 0.072271886 | 0.21496454 | 0.33620376 | 0.73671722 | NA | | NA | NA |
| ENSG000002 | 1.08567777 | -0.086031192 | 0.30676317 | -0.2804482 | 0.77913363 | NA | | NA | NA |
| ENSG000002 | 0.22410479 | 0.147643983 | 0.16638082 | 0.88738586 | 0.37487119 | NA | | NA | NA |
| ENSG000002 | 0.24756242 | -0.187034346 | 0.18119817 | -1.0322088 | 0.30197431 | NA | | NA | NA |
| ENSG000002 | 0.9704099 | -0.027130345 | 0.30553049 | -0.0887975 | 0.92924284 | NA | | NA | NA |
| ENSG000002 | 0 | NA | NA | NA | NA | NA | | RNU6-1059P | 106480031 |
| ENSG000002 | 0 | NA | NA | NA | NA | NA | | SNORD18A | 595098 |
| ENSG000002 | 0 | NA | NA | NA | NA | NA | | RNA5S6 | 100169757 |
| ENSG000002 | 1.12080294 | 0.021549692 | 0.31048766 | 0.06940595 | 0.94466649 | NA | | NA | NA |
| ENSG000002 | 0.72082377 | 0.119191211 | 0.27750365 | 0.42951222 | 0.6675505 | NA | | RNY1P11 | 106481710 |
| ENSG000002 | 0.90707658 | -0.278767286 | 0.30084136 | -0.9266255 | 0.35412099 | NA | | NA | NA |
| ENSG000002 | 0.36538978 | 0.059033792 | 0.20773944 | 0.28417229 | 0.77627835 | NA | | SNORD113-7 | 767567 |
| ENSG000002 | 0.37441639 | -0.268856313 | 0.21277471 | -1.2635727 | 0.20638344 | NA | | NA | NA |
| ENSG000002 | 1.00288172 | -0.325669509 | 0.30114663 | -1.0814317 | 0.27950513 | NA | | SNORD114-27 | 767608 |
| ENSG000002 | 0.47761686 | -0.01948631 | 0.23692179 | -0.0822479 | 0.93444962 | NA | | NA | NA |
| ENSG000002 | 0.7651387 | -0.200755189 | 0.28725161 | -0.6988827 | 0.48462532 | NA | | SNORD115-37 | 100033811 |
| ENSG000002 | 0.63233507 | -0.116461163 | 0.27190168 | -0.4283209 | 0.66841754 | NA | | RNU6-1210P | 106481562 |
| ENSG000002 | 1.10277634 | -0.108641013 | 0.32042825 | -0.3390494 | 0.7345725 | NA | | NA | NA |
| ENSG000002 | 0 | NA | NA | NA | NA | NA | | RNU6-226P | 106481249 |
| ENSG000002 | 1.51892099 | -0.206002123 | 0.34734228 | -0.593081 | 0.55312692 | NA | | NA | NA |
| ENSG000002 | 1.09480543 | 0.036457866 | 0.31510314 | 0.11570137 | 0.90788922 | NA | | NA | NA |
| ENSG000002 | 1.05464959 | 0.033033915 | 0.30521948 | 0.10823004 | 0.91381322 | NA | | RNU4-92P | 106479601 |
| ENSG000002 | 0 | NA | NA | NA | NA | NA | | SNORA5B | 677795 |
| ENSG000002 | 1.21194977 | 0.389231557 | 0.32305591 | 1.20484272 | 0.22826402 | NA | | SNORD116-10 | 100033422 |
| ENSG000002 | 0.53374623 | -0.385819457 | 0.24767828 | -1.5577444 | 0.11929385 | NA | | NA | NA |
| ENSG000002 | 0.96512174 | -0.315400769 | 0.29939956 | -1.0534443 | 0.2921374 | NA | | RNU6-1188P | 106480408 |
| ENSG000002 | 0.36576034 | -0.0934779 | 0.20820828 | -0.4489634 | 0.65345805 | NA | | RNU6-912P | 106479970 |

| | | | | | | | | | |
|---|---|---|---|---|---|---|---|---|---|
| ENSG000002 | 0.22410479 | 0.147643983 | 0.16638082 | 0.88738586 | 0.37487119 | NA | | RN7SKP174 | 106479172 |
| ENSG000002 | 1.31838207 | 0.172685696 | 0.33509846 | 0.51532824 | 0.60632367 | NA | | RN7SKP160 | 106479166 |
| ENSG000002 | 0.3854038 | -0.110128691 | 0.21952326 | -0.5016721 | 0.61589821 | NA | | SNORD115-4 | 100033441 |
| ENSG000002 | 1.0761077 | 0.036127328 | 0.30693382 | 0.11770396 | 0.90630223 | NA | | RNU6-263P | 106479683 |
| ENSG000002 | 0.70369083 | -0.024418318 | 0.27598945 | -0.0884756 | 0.92949872 | NA | | NA | NA |
| ENSG000002 | 0.24141689 | -0.165257896 | 0.17127401 | -0.9648743 | 0.33460776 | NA | | NA | NA |
| ENSG000002 | 1.11083507 | 0.034634703 | 0.32575494 | 0.10632134 | 0.9153274 | NA | | RNA5SP335 | 106479007 |
| ENSG000002 | 1.12535432 | -0.267689318 | 0.32198277 | -0.8313778 | 0.40576025 | NA | | NA | NA |
| ENSG000002 | 0.24619405 | -0.007630671 | 0.17915639 | -0.0425922 | 0.96602658 | NA | | NA | NA |
| ENSG000002 | 1.49871109 | -0.053439787 | 0.34392619 | -0.1553816 | 0.87652052 | NA | | RNU6-873P | 106479951 |
| ENSG000002 | 0.60282172 | 0.050569991 | 0.26272347 | 0.19248372 | 0.84736332 | NA | | RN7SKP93 | 106479136 |
| ENSG000002 | 1.00513323 | -0.365079386 | 0.31516187 | -1.1583869 | 0.24670617 | NA | | NA | NA |
| ENSG000002 | 1.34285924 | 0.032335182 | 0.33405455 | 0.09679611 | 0.92288831 | NA | | NA | NA |
| ENSG000002 | 0.50684946 | -0.202346264 | 0.24895102 | -0.8127955 | 0.41633533 | NA | | RNU6-682P | 106479867 |
| ENSG000002 | 1.67292185 | 0.205960141 | 0.35072056 | 0.58724856 | 0.55703678 | NA | | NA | NA |
| ENSG000002 | 1.00023536 | -0.191785415 | 0.30918425 | -0.6202949 | 0.53506365 | NA | | RN7SKP103 | 106479140 |
| ENSG000002 | 0.51931521 | -0.354214731 | 0.23897171 | -1.4822455 | 0.13827499 | NA | | NA | NA |
| ENSG000002 | 0.35004741 | 0.261422124 | 0.21550076 | 1.21309144 | 0.22509486 | NA | | RNU6-931P | 106479978 |
| ENSG000002 | 0.64255536 | -0.451090545 | 0.26335299 | -1.7128742 | 0.08673568 | NA | | SNORD115-8 | 100033445 |
| ENSG000002 | 0.38036605 | -0.293832733 | 0.22119083 | -1.3284128 | 0.18404178 | NA | | RNU6-194P | 106479652 |
| ENSG000002 | 0.36416385 | 0.068880976 | 0.20758164 | 0.33182596 | 0.74002068 | NA | | NA | NA |
| ENSG000002 | 0.54456342 | -0.359011254 | 0.24026449 | -1.4942335 | 0.1351146 | NA | | NA | NA |
| ENSG000002 | 0.75962432 | -0.361712227 | 0.28310455 | -1.277663 | 0.20136826 | NA | | NA | NA |
| ENSG000002 | 1.35042712 | 0.031468497 | 0.33705378 | 0.09336343 | 0.92561484 | NA | | RNU1-31P | 106481606 |
| ENSG000002 | 1.01298344 | -0.206632615 | 0.31837313 | -0.6490266 | 0.51632121 | NA | | NA | NA |
| ENSG000002 | 0.3575613 | 0.072305152 | 0.21497462 | 0.33634273 | 0.73661243 | NA | | NA | NA |
| ENSG000002 | 1.1992682 | -0.043133742 | 0.32918469 | -0.131032 | 0.89574996 | NA | | NA | NA |
| ENSG000002 | 1.46728859 | 0.224273603 | 0.34250601 | 0.65480195 | 0.51259527 | NA | | NA | NA |
| ENSG000002 | 1.02892662 | -0.35232479 | 0.31014919 | -1.1359849 | 0.25596289 | NA | | RNU6-236P | 106481252 |

| | | | | | | | | | |
|---|---|---|---|---|---|---|---|---|---|
| ENSG000002 | 0.77362006 | -0.203375959 | 0.28776237 | -0.7067497 | 0.47972205 | NA | | SNORD115-16 | 100033454 |
| ENSG000002 | 0.97063611 | -0.191821921 | 0.31273107 | -0.6133766 | 0.53962736 | NA | | RNU6-884P | 106479957 |
| ENSG000002 | 1.10556651 | -0.23847821 | 0.30696414 | -0.7768927 | 0.43722206 | NA | | RN7SKP113 | 106480465 |
| ENSG000002 | 0.26062507 | -0.012888319 | 0.18226503 | -0.070712 | 0.94362699 | NA | | RNU6-961P | 106481483 |
| ENSG000002 | 0.89319039 | 0.031156117 | 0.28192378 | 0.11051256 | 0.91200289 | NA | | NA | NA |
| ENSG000002 | 0.38614101 | -0.1098476 | 0.21963552 | -0.5001359 | 0.61697942 | NA | | RNU6-478P | 106479778 |
| ENSG000002 | 2.10947518 | -0.363102827 | 0.36140285 | -1.0047038 | 0.31503948 | NA | | RNU6-105P | 106481210 |
| ENSG000002 | 1.41198468 | 0.23274845 | 0.33926518 | 0.68603695 | 0.49268981 | NA | | RN7SKP180 | 106480888 |
| ENSG000002 | 1.31962782 | 0.173599782 | 0.3384934 | 0.51286016 | 0.60804914 | NA | | NA | NA |
| ENSG000002 | 0 | NA | NA | NA | NA | NA | | RNU6-631P | 106479845 |
| ENSG000002 | 1.83132282 | -0.576446827 | 0.34852131 | -1.6539787 | 0.09813184 | NA | | RN7SKP250 | 106479206 |
| ENSG000002 | 0.47670496 | 0.137439173 | 0.23758963 | 0.57847295 | 0.56294486 | NA | | RNU6-753P | 106479899 |
| ENSG000002 | 0.63296622 | -0.299126688 | 0.27328461 | -1.0945611 | 0.27370899 | NA | | RNU6-770P | 106481423 |
| ENSG000002 | 1.46259967 | -0.195705526 | 0.34820558 | -0.5620402 | 0.57408865 | NA | | RNU1-130P | 106480192 |
| ENSG000002 | 0 | NA | NA | NA | NA | NA | | NA | NA |
| ENSG000002 | 1.07768513 | -0.097936383 | 0.3122934 | -0.3136038 | 0.753822 | NA | | RNU1-32P | 106480155 |
| ENSG000002 | 0.99030131 | -0.03729337 | 0.31705927 | -0.1176227 | 0.90636661 | NA | | SNORD115-6 | 100033443 |
| ENSG000002 | 1.54152331 | -0.102520241 | 0.33891904 | -0.3024918 | 0.76227716 | NA | | RNU6-595P | 106479829 |
| ENSG000002 | 0.92970663 | -0.439225344 | 0.29724343 | -1.4776621 | 0.13949825 | NA | | RNU6-1091P | 106480637 |
| ENSG000002 | 0 | NA | NA | NA | NA | NA | | RNU6-899P | 106479964 |
| ENSG000002 | 1.68212111 | -0.248323603 | 0.34425704 | -0.721332 | 0.4707053 | NA | | RNU6-1204P | 106481560 |
| ENSG000002 | 0.51459361 | -0.192534203 | 0.24265108 | -0.7934611 | 0.42750921 | NA | | NA | NA |
| ENSG000002 | 0.26813897 | -0.197007041 | 0.18550319 | -1.0620143 | 0.2882292 | NA | | SNORD114-2 | 767578 |
| ENSG000002 | 0.97498928 | -0.0299656 | 0.30740826 | -0.0974782 | 0.92234666 | NA | | RNU6-324P | 106479710 |
| ENSG000002 | 0 | NA | NA | NA | NA | NA | | NA | NA |
| ENSG000002 | 0.47821768 | -0.016471461 | 0.24462483 | -0.0673336 | 0.94631616 | NA | | NA | NA |
| ENSG000002 | 1.39201145 | -0.419962981 | 0.33854026 | -1.2405112 | 0.21478638 | NA | | NA | NA |
| ENSG000002 | 1.62537311 | -0.42166247 | 0.35465937 | -1.1889224 | 0.2344702 | NA | | NA | NA |
| ENSG000002 | 0.64506141 | -0.288091388 | 0.2684363 | -1.0732207 | 0.28317211 | NA | | NA | NA |

| | | | | | | | | |
|---|---|---|---|---|---|---|---|---|
| ENSG000002 | 0.49401706 | -0.184896882 | 0.240355 | -0.7692658 | 0.44173553 | NA | NA | NA |
| ENSG000002 | 1.6002108 | -0.267987074 | 0.35166054 | -0.7620618 | 0.44602311 | NA | NA | NA |
| ENSG000002 | 0.6164817 | -0.105910369 | 0.26430836 | -0.4007076 | 0.68863541 | NA | NA | NA |
| ENSG000002 | 0.14128499 | -0.097772014 | 0.13403235 | -0.7294658 | 0.4657168 | NA | NA | NA |
| ENSG000002 | 0.87970161 | -0.274795643 | 0.2994662 | -0.9176182 | 0.35881877 | NA | RN7SKP36 | 106480458 |
| ENSG000002 | 1.1248077 | -0.110431972 | 0.31649034 | -0.3489268 | 0.72714424 | NA | RNU6-457P | 106479768 |
| ENSG000002 | 1.90477129 | -0.116408525 | 0.34915888 | -0.333397 | 0.73883462 | NA | RNU6-810P | 106481437 |
| ENSG000002 | 0.49168123 | -0.197384925 | 0.247356 | -0.7979791 | 0.4248826 | NA | NA | NA |
| ENSG000002 | 1.06686951 | 0.03892517 | 0.31238871 | 0.12460492 | 0.90083633 | NA | NA | NA |
| ENSG000002 | 1.47634397 | -0.322446747 | 0.34231053 | -0.9419714 | 0.34620726 | NA | RNU6-340P | 106479717 |
| ENSG000002 | 1.17410777 | -0.275953692 | 0.32123378 | -0.8590432 | 0.39031669 | NA | RNU6-560P | 106480590 |
| ENSG000002 | 1.39417651 | -0.573197502 | 0.33784913 | -1.6966079 | 0.08977082 | NA | NA | NA |
| ENSG000002 | 0.36345889 | 0.070257596 | 0.21589421 | 0.32542602 | 0.74485866 | NA | NA | NA |
| ENSG000002 | 1.72117123 | -0.058193482 | 0.35892343 | -0.1621334 | 0.87120079 | NA | RNU1-146P | 106480198 |
| ENSG000002 | 0.74177143 | -0.029328749 | 0.28496727 | -0.1029197 | 0.91802669 | NA | RNU6-598P | 106479830 |
| ENSG000002 | 0.50405874 | -0.025925695 | 0.24745105 | -0.104771 | 0.91655752 | NA | RNU4-13P | 106481763 |
| ENSG000002 | 1.66701547 | -0.39798553 | 0.34757175 | -1.1450457 | 0.25219023 | NA | RNA5SP99 | 106478993 |
| ENSG000002 | 0.73687356 | 0.124652094 | 0.27862992 | 0.44737512 | 0.65460424 | NA | NA | NA |
| ENSG000002 | 0.11934008 | 0.083564698 | 0.12760091 | 0.65489106 | 0.5125379 | NA | RNU6-944P | 106479983 |
| ENSG000002 | 0.3588776 | -0.099902505 | 0.21555794 | -0.4634601 | 0.6430346 | NA | NA | NA |
| ENSG000002 | 1.23820809 | -0.172009711 | 0.31531371 | -0.5455193 | 0.58539642 | NA | RNU6-1243P | 106481573 |
| ENSG000002 | 0.99312237 | -0.330085154 | 0.30326887 | -1.0884241 | 0.27640792 | NA | RNU6-627P | 106480596 |
| ENSG000002 | 1.59317811 | 0.138521671 | 0.33002626 | 0.41972924 | 0.67468326 | NA | RNU1-42P | 106481609 |
| ENSG000002 | 0.35802023 | 0.220517756 | 0.19914056 | 1.1073473 | 0.2681438 | NA | RNU6-854P | 106481450 |
| ENSG000002 | 1.60741259 | -0.387998596 | 0.34674364 | -1.1189783 | 0.26314941 | NA | RNA5SP435 | 106479013 |
| ENSG000002 | 0.76881256 | -0.186406664 | 0.28149285 | -0.6622075 | 0.50783827 | NA | RNU6-553P | 106480589 |
| ENSG000002 | 0.60191037 | 0.224859928 | 0.26377907 | 0.85245553 | 0.39396131 | NA | RNU6-960P | 106479991 |
| ENSG000002 | 0.50016259 | -0.200428691 | 0.24824437 | -0.8073846 | 0.41944491 | NA | RNU6-694P | 106479872 |
| ENSG000002 | 0.81983524 | 0.192181941 | 0.28753909 | 0.66836804 | 0.50389869 | NA | RNU6-501P | 106481339 |

| | | | | | | | | | |
|---|---|---|---|---|---|---|---|---|---|
| ENSG000002 | 0.74410726 | -0.031277165 | 0.29043452 | -0.1076909 | 0.91424086 | NA | | SNORD115-32 | 100033806 |
| ENSG000002 | 1.01063034 | -0.690960342 | 0.30717159 | -2.2494279 | 0.02448528 | NA | | RNU6-801P | 106479920 |
| ENSG000002 | 0.49970612 | -0.194507174 | 0.24827631 | -0.7834303 | 0.43337449 | NA | | RNU6-289P | 106479695 |
| ENSG000002 | 1.75231141 | -0.462777103 | 0.35345903 | -1.3092808 | 0.19043926 | NA | | RN7SKP87 | 106479133 |
| ENSG000002 | 0.74755087 | -0.190398856 | 0.2861764 | -0.6653199 | 0.50584591 | NA | | NA | NA |
| ENSG000002 | 1.84961935 | 0.126206205 | 0.35783955 | 0.35268936 | 0.72432133 | NA | | RNU4-87P | 106479598 |
| ENSG000002 | 0.81270238 | -0.53011708 | 0.27974244 | -1.8950184 | 0.05808996 | NA | | RNU1-7P | 106481952 |
| ENSG000002 | 0.80382012 | -0.054907665 | 0.15978555 | -0.3436335 | 0.73112194 | NA | | NA | NA |
| ENSG000002 | 0.85112135 | -0.11399948 | 0.29680331 | -0.384091 | 0.70091101 | NA | | NA | NA |
| ENSG000002 | 0 | NA | NA | NA | NA | NA | | SNORA45A | 619562 |
| ENSG000002 | 1.07440515 | 0.042057775 | 0.3160678 | 0.13306567 | 0.89414144 | NA | | RNU6-819P | 106479928 |
| ENSG000002 | 0.76881256 | -0.186406664 | 0.28149285 | -0.6622075 | 0.50783827 | NA | | SNORD115-30 | 100033804 |
| ENSG000002 | 1.7820222 | -0.473949757 | 0.35545303 | -1.3333682 | 0.18241101 | NA | | NA | NA |
| ENSG000002 | 0.23799501 | 0.170958857 | 0.17834799 | 0.95856902 | 0.33777591 | NA | | NA | NA |
| ENSG000002 | 0.88035449 | -0.282626859 | 0.30431514 | -0.9287309 | 0.35302858 | NA | | NA | NA |
| ENSG000002 | 0.66908156 | -0.466940971 | 0.26696073 | -1.7490999 | 0.08027376 | NA | | NA | NA |
| ENSG000002 | 1.15985089 | -0.407388318 | 0.31369388 | -1.2986811 | 0.19405339 | NA | | RNU6-270P | 106479686 |
| ENSG000002 | 0.8310558 | 0.050746957 | 0.28920356 | 0.17547141 | 0.86070923 | NA | | NA | NA |
| ENSG000002 | 2.14693535 | -0.38396465 | 0.36501864 | -1.0519042 | 0.29284349 | NA | | NA | NA |
| ENSG000002 | 0.9828172 | -0.045861653 | 0.3077518 | -0.1490216 | 0.88153662 | NA | | RN7SKP224 | 106480902 |
| ENSG000002 | 1.22966257 | -0.178942633 | 0.32475786 | -0.5510032 | 0.58163145 | NA | | NA | NA |
| ENSG000002 | 0.63603926 | -0.266971526 | 0.26084558 | -1.0234849 | 0.30607861 | NA | | NA | NA |
| ENSG000002 | 0.98508887 | -0.033772222 | 0.31272058 | -0.1079949 | 0.91399975 | NA | | NA | NA |
| ENSG000002 | 0.3863157 | -0.274134926 | 0.21457207 | -1.277589 | 0.20139437 | NA | | RNU6-280P | 106481886 |
| ENSG000002 | 0.51959753 | -0.202283781 | 0.25032396 | -0.808088 | 0.41903995 | NA | | RNU6-374P | 106481296 |
| ENSG000002 | 1.11169489 | -0.110683234 | 0.32072136 | -0.3451072 | 0.73001382 | NA | | RNU6-743P | 106480605 |
| ENSG000002 | 0.11865494 | 0.083440866 | 0.12751478 | 0.65436232 | 0.5128784 | NA | | RNY3P11 | 106481713 |
| ENSG000002 | 0.26813897 | -0.197007041 | 0.18550319 | -1.0620143 | 0.2882292 | NA | | NA | NA |
| ENSG000002 | 0.38884741 | -0.29745092 | 0.22235466 | -1.3377319 | 0.18098386 | NA | | NA | NA |

| | | | | | | | | |
|---|---|---|---|---|---|---|---|---|
| ENSG000002 | 0.38019136 | -0.10756138 | 0.21876967 | -0.491665 | 0.62295621 | NA | NA | NA |
| ENSG000002 | 1.20544594 | -0.588493051 | 0.31880259 | -1.8459481 | 0.06489974 | NA | RNU6-704P | 106479877 |
| ENSG000002 | 1.01884664 | -0.349130761 | 0.30975853 | -1.1271062 | 0.25969758 | NA | RNU1-96P | 106480181 |
| ENSG000002 | 0.83213895 | 0.182958625 | 0.28831404 | 0.63458105 | 0.52570169 | NA | NA | NA |
| ENSG000002 | 1.58203942 | 0.137597545 | 0.34330847 | 0.40079857 | 0.68856843 | NA | NA | NA |
| ENSG000002 | 0.71171572 | -0.021607924 | 0.27657551 | -0.0781267 | 0.93772729 | NA | NA | NA |
| ENSG000002 | 1.10052641 | 0.031328069 | 0.32156879 | 0.0974226 | 0.92239079 | NA | NA | NA |
| ENSG000002 | 0.63939408 | -0.113476381 | 0.26621037 | -0.4262658 | 0.66991416 | NA | NA | NA |
| ENSG000002 | 0.5983253 | -0.096573125 | 0.25511974 | -0.3785404 | 0.7050292 | NA | RNU6-268P | 106479685 |
| ENSG000002 | 1.69737783 | -0.049217188 | 0.35356871 | -0.1392012 | 0.88929116 | NA | NA | NA |
| ENSG000002 | 1.21668636 | -0.182966787 | 0.32869298 | -0.5566495 | 0.57776691 | NA | RNU6-668P | 106481393 |
| ENSG000002 | 1.67318691 | -0.048742721 | 0.34986091 | -0.1393203 | 0.88919706 | NA | NA | NA |
| ENSG000002 | 0.23139247 | 0.167748926 | 0.17680352 | 0.94878726 | 0.34272882 | NA | RNU6-461P | 106481897 |
| ENSG000002 | 0.50630812 | -0.189615696 | 0.24175631 | -0.7843257 | 0.432849 | NA | RN7SKP280 | 106479220 |
| ENSG000002 | 1.21324722 | -0.314700808 | 0.32458706 | -0.9695421 | 0.3322748 | NA | RNU4-84P | 106481873 |
| ENSG000002 | 0.73480279 | -0.177544794 | 0.27752528 | -0.6397428 | 0.52233984 | NA | NA | NA |
| ENSG000002 | 1.35906144 | -0.235483489 | 0.32891274 | -0.7159452 | 0.4740252 | NA | NA | NA |
| ENSG000002 | 1.04223939 | 0.514342331 | 0.31600612 | 1.62763408 | 0.1036025 | NA | NA | NA |
| ENSG000002 | 1.68707502 | -0.18137829 | 0.35314091 | -0.5136145 | 0.60752154 | NA | RNU4-51P | 106479581 |
| ENSG000002 | 2.10466239 | -0.246019324 | 0.36433298 | -0.6752596 | 0.49951088 | NA | RNU6-188P | 106479649 |
| ENSG000002 | 1.36989131 | -0.122894546 | 0.32830201 | -0.3743338 | 0.70815597 | NA | RN7SKP214 | 106480473 |
| ENSG000002 | 0 | NA | NA | NA | NA | NA | NA | NA |
| ENSG000002 | 1.71086449 | 0.076358086 | 0.35473827 | 0.2152519 | 0.82957093 | NA | RNU6-173P | 106479643 |
| ENSG000002 | 1.02278109 | -0.350503169 | 0.30990351 | -1.1310074 | 0.25805198 | NA | NA | NA |
| ENSG000002 | 0.90707658 | -0.278767286 | 0.30084136 | -0.9266255 | 0.35412099 | NA | NA | NA |
| ENSG000002 | 0 | NA | NA | NA | NA | NA | RNU6-266P | 106479684 |
| ENSG000002 | 0.24485602 | -0.012655878 | 0.17881722 | -0.0707755 | 0.94357643 | NA | NA | NA |
| ENSG000002 | 1.7672028 | -0.746183295 | 0.35073904 | -2.12746 | 0.03338188 | NA | RNU6-366P | 106481294 |
| ENSG000002 | 1.34794639 | 0.014035775 | 0.33341562 | 0.04209693 | 0.96642143 | NA | NA | NA |

| | | | | | | | | | |
|---|---|---|---|---|---|---|---|---|---|
| ENSG000002 | 1.34647698 | -0.246025463 | 0.33437183 | -0.7357841 | 0.46186213 | NA | | RNU6-11P | 106478915 |
| ENSG000002 | 0.88414246 | -0.430982109 | 0.29486454 | -1.4616274 | 0.14384333 | NA | | NA | NA |
| ENSG000002 | 1.72624571 | -0.178478772 | 0.3495983 | -0.5105253 | 0.6096835 | NA | | RNU6-647P | 106480598 |
| ENSG000002 | 0.65698638 | -0.462594106 | 0.26600012 | -1.7390748 | 0.0820216 | NA | | NA | NA |
| ENSG000002 | 0.87878971 | -0.122783908 | 0.30329377 | -0.4048349 | 0.68559886 | NA | | NA | NA |
| ENSG000002 | 0.37945415 | -0.107841678 | 0.21865587 | -0.4932027 | 0.62186934 | NA | | NA | NA |
| ENSG000002 | 0.73699426 | -0.182203658 | 0.27937391 | -0.6521857 | 0.51428141 | NA | | NA | NA |
| ENSG000002 | 1.12247379 | 0.034055144 | 0.32076758 | 0.10616766 | 0.91544933 | NA | | NA | NA |
| ENSG000002 | 0.12070844 | -0.088503695 | 0.12783931 | -0.6923042 | 0.48874631 | NA | | RNU6-777P | 106481426 |
| ENSG000002 | 1.37305526 | -0.266740823 | 0.34037924 | -0.7836577 | 0.43324097 | NA | | RNU6-353P | 106479723 |
| ENSG000002 | 1.32942389 | -0.105010903 | 0.32984147 | -0.3183678 | 0.75020596 | NA | | RN7SKP128 | 106479151 |
| ENSG000002 | 1.59383634 | -0.777584475 | 0.33841438 | -2.2977289 | 0.02157723 | NA | | NA | NA |
| ENSG000002 | 0.23202362 | -0.008302493 | 0.17586207 | -0.0472103 | 0.96234565 | NA | | SNORD115-42 | 100033816 |
| ENSG000002 | 0.3513637 | 0.074494292 | 0.21400438 | 0.34809704 | 0.72776731 | NA | | RNA5SP175 | 106480758 |
| ENSG000002 | 1.37805864 | -0.264224578 | 0.33725118 | -0.7834653 | 0.43335395 | NA | | NA | NA |
| ENSG000002 | 1.1080766 | 0.172116177 | 0.31599767 | 0.54467546 | 0.58597675 | NA | | RNU6-371P | 106479731 |
| ENSG000002 | 1.09682668 | -0.09957035 | 0.31298698 | -0.3181294 | 0.75038681 | NA | | NA | NA |
| ENSG000002 | 0.3867939 | -0.105148852 | 0.21977268 | -0.4784437 | 0.63233444 | NA | | NA | NA |
| ENSG000002 | 0.36351096 | 0.070290861 | 0.21590418 | 0.32556508 | 0.74475344 | NA | | NA | NA |
| ENSG000002 | 0.91897589 | -0.264865153 | 0.2941721 | -0.9003748 | 0.36792081 | NA | | RNU6-454P | 106480582 |
| ENSG000002 | 0 | NA | NA | NA | NA | NA | | RNU6-1229P | 106481568 |
| ENSG000002 | 0.6039295 | -0.097743897 | 0.2540715 | -0.3847102 | 0.70045213 | NA | | NA | NA |
| ENSG000002 | 1.22892982 | -0.175153154 | 0.2222731 | -0.7880088 | 0.43069158 | NA | | RNU1-132P | 106481958 |
| ENSG000002 | 0.86802459 | -0.107225507 | 0.28995809 | -0.3697966 | 0.71153407 | NA | | NA | NA |
| ENSG000002 | 0.60310247 | 0.05638274 | 0.26275708 | 0.21458124 | 0.83009382 | NA | | RNU6-1322P | 106480143 |
| ENSG000002 | 1.26236464 | -0.312734268 | 0.31758613 | -0.9847227 | 0.32476031 | NA | | RNU6-115P | 106479617 |
| ENSG000002 | 1.66916669 | 0.351359911 | 0.35041732 | 1.00268991 | 0.3160105 | NA | | RNU6-670P | 106479863 |
| ENSG000002 | 0.70100617 | 0.294547947 | 0.27599089 | 1.06723793 | 0.28586441 | NA | | RNU4-68P | 106479589 |
| ENSG000002 | 1.0199888 | -0.349635103 | 0.31142893 | -1.1226802 | 0.26157333 | NA | | NA | NA |

| | | | | | | | | |
|---|---|---|---|---|---|---|---|---|
| ENSG000002 | 0.37199232 | 0.06760863 | 0.21718977 | 0.31128828 | 0.75558148 | NA | NA | NA |
| ENSG000002 | 0.63387812 | -0.454271587 | 0.26413408 | -1.7198522 | 0.08545931 | NA | RNU6-879P | 106481456 |
| ENSG000002 | 1.26009097 | -0.191147347 | 0.32984252 | -0.5795109 | 0.56224447 | NA | NA | NA |
| ENSG000002 | 1.88440293 | -0.615737616 | 0.35001239 | -1.7591881 | 0.07854557 | NA | NA | NA |
| ENSG000002 | 1.53756316 | -0.4818045 | 0.34372647 | -1.401709 | 0.16100215 | NA | NA | NA |
| ENSG000002 | 0.74621284 | -0.194950297 | 0.28606354 | -0.681493 | 0.49555961 | NA | NA | NA |
| ENSG000002 | 0.59553458 | 0.061438566 | 0.26853563 | 0.22879111 | 0.81903127 | NA | NA | NA |
| ENSG000002 | 0.89088936 | -0.427376756 | 0.2952839 | -1.4473419 | 0.14780119 | NA | NA | NA |
| ENSG000002 | 0.35824644 | 0.072568381 | 0.21508272 | 0.33739754 | 0.73581724 | NA | RNU4-40P | 106479576 |
| ENSG000002 | 2.14477264 | -0.634636415 | 0.36300064 | -1.7483066 | 0.08041095 | NA | RNU6-1305P | 106481592 |
| ENSG000002 | 1.74066876 | -0.194026402 | 0.35373291 | -0.548511 | 0.58334106 | NA | NA | NA |
| ENSG000002 | 1.58517194 | 0.145698767 | 0.34684695 | 0.42006645 | 0.67443691 | NA | RNU4-50P | 106481871 |
| ENSG000002 | 0.37216701 | -0.099548723 | 0.21762847 | -0.4574251 | 0.64736554 | NA | NA | NA |
| ENSG000002 | 0.89954094 | -0.281346535 | 0.30044017 | -0.9364478 | 0.34904268 | NA | SNORD114-9 | 767585 |
| ENSG000002 | 0.40942396 | -0.284058841 | 0.21788566 | -1.303706 | 0.19233384 | NA | RNU6-978P | 106479998 |
| ENSG000002 | 0.84434466 | 0.043418337 | 0.29446077 | 0.14745033 | 0.88277658 | NA | NA | NA |
| ENSG000002 | 1.54507897 | -0.460861017 | 0.3391295 | -1.3589529 | 0.17416151 | NA | RNU6-654P | 106481909 |
| ENSG000002 | 0.13280363 | -0.093394593 | 0.13115644 | -0.7120855 | 0.47641185 | NA | NA | NA |
| ENSG000002 | 1.31992457 | -0.112565686 | 0.33383114 | -0.3371935 | 0.73597105 | NA | SNORD114-13 | 767591 |
| ENSG000002 | 0.35073255 | 0.241740392 | 0.2078501 | 1.16305162 | 0.24480856 | NA | NA | NA |
| ENSG000002 | 1.09181883 | 0.191195975 | 0.32144555 | 0.59480051 | 0.55197683 | NA | RNU6-1075P | 106480406 |
| ENSG000002 | 0.22263036 | 0.146922267 | 0.16600367 | 0.88505432 | 0.37612734 | NA | SNORD114-6 | 767582 |
| ENSG000002 | 0.48487229 | -0.012559814 | 0.237804 | -0.0528158 | 0.95787865 | NA | RNU6-755P | 106481915 |
| ENSG000002 | 0.63435632 | -0.27906164 | 0.26763939 | -1.0426778 | 0.29709757 | NA | RNU1-112P | 106480187 |
| ENSG000002 | 0.49810908 | -0.023852705 | 0.24681213 | -0.0966432 | 0.92300978 | NA | RNU6-776P | 106480608 |
| ENSG000002 | 0.47212421 | -0.012721785 | 0.23625154 | -0.0538485 | 0.95705589 | NA | NA | NA |
| ENSG000002 | 0.24619405 | -0.007630671 | 0.17915639 | -0.0425922 | 0.96602658 | NA | NA | NA |
| ENSG000002 | 0 | NA | NA | NA | NA | NA | NA | NA |
| ENSG000002 | 0.2596576 | -0.193002035 | 0.1837913 | -1.0501152 | 0.29366517 | NA | NA | NA |

| | | | | | | | | |
|---|---|---|---|---|---|---|---|---|
| ENSG000002 | 1.20632913 | 0.103222932 | 0.33153287 | 0.31135052 | 0.75553416 | NA | RN7SKP171 | 106480887 |
| ENSG000002 | 2.14867037 | -0.389240674 | 0.36534847 | -1.0653956 | 0.28669693 | NA | RN7SKP76 | 106481808 |
| ENSG000002 | 1.27360851 | -0.20053095 | 0.33375589 | -0.6008312 | 0.54795243 | NA | RNU1-34P | 106481607 |
| ENSG000002 | 1.02709988 | -0.352484354 | 0.31169902 | -1.1308484 | 0.25811889 | NA | RNU6-153P | 106479634 |
| ENSG000002 | 0.97825819 | -0.032021801 | 0.31244243 | -0.1024886 | 0.91836883 | NA | RNU6-1019P | 106480015 |
| ENSG000002 | 1.36270092 | -0.35183413 | 0.32392598 | -1.086156 | 0.27741 | NA | RNU4-41P | 106481184 |
| ENSG000002 | 0.97665903 | -0.036084117 | 0.29806623 | -0.1210607 | 0.90364293 | NA | RNU6-1164P | 106480644 |
| ENSG000002 | 0.73787528 | -0.192474738 | 0.28554293 | -0.6740658 | 0.50026945 | NA | SNORD115-27 | 100036564 |
| ENSG000002 | 0.51316968 | -0.377777671 | 0.24556403 | -1.538408 | 0.12394889 | NA | RNA5SP130 | 106478995 |
| ENSG000002 | 1.18211093 | -0.423351136 | 0.31860864 | -1.3287497 | 0.18393057 | NA | RNU6-512P | 106480394 |
| ENSG000002 | 0.26199343 | -0.194018184 | 0.18422801 | -1.0531416 | 0.29227609 | NA | NA | NA |
| ENSG000002 | 1.00774926 | -0.355511756 | 0.31086197 | -1.1436322 | 0.25277621 | NA | NA | NA |
| ENSG000002 | 1.12607138 | -0.251295745 | 0.31041593 | -0.8095453 | 0.41820157 | NA | NA | NA |
| ENSG000002 | 2.15860582 | -0.15838851 | 0.35877929 | -0.441465 | 0.65887637 | NA | SNORD114-30 | 767611 |
| ENSG000002 | 0.87457898 | -0.427920645 | 0.29435302 | -1.4537668 | 0.14601097 | NA | RNU6-361P | 106480574 |
| ENSG000002 | 0.25260017 | -0.015978705 | 0.18049464 | -0.0885273 | 0.92945758 | NA | SNORD115-22 | 100033799 |
| ENSG000002 | 0.4889793 | -0.336999675 | 0.23406338 | -1.4397796 | 0.14992977 | NA | NA | NA |
| ENSG000002 | 0.13280363 | -0.093394593 | 0.13115644 | -0.7120855 | 0.47641185 | NA | SNORD32B | 692092 |
| ENSG000002 | 0.73798346 | 0.133274977 | 0.27868582 | 0.47822663 | 0.63248891 | NA | SNORD115-23 | 100033800 |
| ENSG000002 | 0.64645151 | -0.283338099 | 0.26861796 | -1.0547995 | 0.29151702 | NA | NA | NA |
| ENSG000002 | 0.23139247 | 0.167748926 | 0.17680352 | 0.94878726 | 0.34272882 | NA | NA | NA |
| ENSG000002 | 0.88269033 | -0.253490119 | 0.29224055 | -0.8674023 | 0.38572163 | NA | RNU6-421P | 106481311 |
| ENSG000002 | 1.10465516 | -0.113219034 | 0.32048724 | -0.3532716 | 0.72388485 | NA | RNU4-38P | 106479574 |
| ENSG000002 | 0.48536101 | -0.024029524 | 0.24539296 | -0.0979226 | 0.92199373 | NA | NA | NA |
| ENSG000002 | 1.24859055 | -0.043155302 | 0.32901933 | -0.1311634 | 0.89564603 | NA | RNU105B | 26767 |
| ENSG000002 | 0.73701545 | -0.03254472 | 0.28464749 | -0.1143334 | 0.90897349 | NA | NA | NA |
| ENSG000002 | 1.00720792 | -0.350938794 | 0.30924522 | -1.1348237 | 0.25644921 | NA | NA | NA |
| ENSG000002 | 0 | NA | NA | NA | NA | NA | RNA5S14 | 100169765 |
| ENSG000002 | 1.21574412 | 0.109561395 | 0.3317607 | 0.33024223 | 0.74121694 | NA | NA | NA |

| | | | | | | | | |
|---|---|---|---|---|---|---|---|---|
| ENSG000002 | 0.39479706 | -0.300008631 | 0.22317059 | -1.3443018 | 0.17885081 | NA | RN7SKP193 | 106479181 |
| ENSG000002 | 0.83919638 | 0.037335998 | 0.28969729 | 0.12887935 | 0.89745312 | NA | NA | NA |
| ENSG000002 | 0.77521868 | -0.191196496 | 0.28023911 | -0.682262 | 0.49507329 | NA | NA | NA |
| ENSG000002 | 1.13955509 | -0.436452319 | 0.32365488 | -1.3485115 | 0.17749393 | NA | RN7SKP37 | 106480844 |
| ENSG000002 | 1.46151205 | -0.04178823 | 0.34216753 | -0.122128 | 0.90279766 | NA | NA | NA |
| ENSG000002 | 0.61611113 | 0.054662593 | 0.27024684 | 0.20226913 | 0.83970633 | NA | NA | NA |
| ENSG000002 | 1.12364573 | -0.124826171 | 0.32261416 | -0.3869209 | 0.69881474 | NA | NA | NA |
| ENSG000002 | 1.47469485 | -0.201041312 | 0.34945221 | -0.5753042 | 0.5650856 | NA | NA | NA |
| ENSG000002 | 1.01677587 | -0.697407278 | 0.30816713 | -2.2630813 | 0.02363068 | NA | RNU6-1251P | 106480112 |
| ENSG000002 | 0.37404583 | -0.105111695 | 0.21784993 | -0.4824959 | 0.62945372 | NA | RNU4-76P | 106480549 |
| ENSG000002 | 0.6256482 | -0.108446506 | 0.2650784 | -0.4091111 | 0.68245816 | NA | RNU6-558P | 106481358 |
| ENSG000002 | 0 | NA | NA | NA | NA | NA | NA | NA |
| ENSG000002 | 1.13363981 | -0.273516128 | 0.32371431 | -0.8449306 | 0.39814958 | NA | RNU6-1163P | 106481547 |
| ENSG000002 | 0 | NA | NA | NA | NA | NA | NA | NA |
| ENSG000002 | 1.19707673 | -0.040198335 | 0.32609146 | -0.1232732 | 0.90189077 | NA | RNU6-1141P | 106481540 |
| ENSG000002 | 0.52398687 | -0.355918066 | 0.23944695 | -1.4864172 | 0.1371688 | NA | RNU6-1078P | 106481519 |
| ENSG000002 | 0.50667477 | -0.027805704 | 0.23162709 | -0.1200451 | 0.9044474 | NA | NA | NA |
| ENSG000002 | 0.87375195 | -0.2849797 | 0.30397068 | -0.9375236 | 0.34848927 | NA | RN7SKP257 | 106479210 |
| ENSG000002 | 0.81500601 | 0.045368592 | 0.28825076 | 0.15739279 | 0.87493529 | NA | NA | NA |
| ENSG000002 | 0 | NA | NA | NA | NA | NA | NA | NA |
| ENSG000002 | 0 | NA | NA | NA | NA | NA | NA | NA |
| ENSG000002 | 1.00006459 | -0.336771554 | 0.30659048 | -1.098441 | 0.27201197 | NA | NA | NA |
| ENSG000002 | 0.37404583 | -0.105111695 | 0.21784993 | -0.4824959 | 0.62945372 | NA | NA | NA |
| ENSG000002 | 0.75916785 | -0.330053211 | 0.2749051 | -1.2006078 | 0.22990337 | NA | NA | NA |
| ENSG000002 | 1.63173259 | 0.461691749 | 0.3430313 | 1.34591726 | 0.1783292 | NA | NA | NA |
| ENSG000002 | 1.54713406 | -0.488319962 | 0.34385328 | -1.4201405 | 0.15556679 | NA | NA | NA |
| ENSG000002 | 1.48753172 | -0.477514147 | 0.34452816 | -1.3859946 | 0.16574857 | NA | NA | NA |
| ENSG000002 | 1.50139967 | -0.618366063 | 0.34012753 | -1.8180418 | 0.06905775 | NA | NA | NA |
| ENSG000002 | 0.35733509 | 0.244646831 | 0.20894085 | 1.17089037 | 0.24164285 | NA | RN7SKP246 | 106479204 |

| | | | | | | | | |
|---|---|---|---|---|---|---|---|---|
| ENSG000002 | 0.59892612 | -0.099249839 | 0.26277623 | -0.3776972 | 0.70565557 | NA | NA | NA |
| ENSG000002 | 0.64469531 | -0.422865637 | 0.25651704 | -1.6484894 | 0.09925228 | NA | NA | NA |
| ENSG000002 | 0.25370795 | -0.171331747 | 0.17411721 | -0.9840024 | 0.32511435 | NA | RNU6-335P | 106479715 |
| ENSG000002 | 0.89753951 | -0.113520912 | 0.29331672 | -0.387025 | 0.69873768 | NA | RNU4-24P | 106481179 |
| ENSG000002 | 1.59742008 | -0.126194835 | 0.35003038 | -0.3605254 | 0.71845428 | NA | RNU4-67P | 106481193 |
| ENSG000002 | 0.49168123 | -0.197384925 | 0.247356 | -0.7979791 | 0.4248826 | NA | NA | NA |
| ENSG000002 | 0.23936338 | -0.005479057 | 0.17759977 | -0.0308506 | 0.9753887 | NA | RNU6-646P | 106481386 |
| ENSG000002 | 1.25115259 | -0.20743628 | 0.33745935 | -0.6147 | 0.53875283 | NA | NA | NA |
| ENSG000002 | 1.65425633 | -0.265356359 | 0.3482963 | -0.7618696 | 0.44613785 | NA | RNU6-1082P | 106480041 |
| ENSG000002 | 1.14343397 | -0.768426485 | 0.31780149 | -2.4179449 | 0.01560844 | NA | NA | NA |
| ENSG000002 | 0.36391535 | 0.058449071 | 0.20749591 | 0.28168782 | 0.77818289 | NA | NA | NA |
| ENSG000002 | 1.35632224 | -0.128283524 | 0.3434903 | -0.3734706 | 0.70879824 | NA | NA | NA |
| ENSG000002 | 0.5008998 | -0.200150804 | 0.24832882 | -0.805991 | 0.42024801 | NA | RNU6-1317P | 106481596 |
| ENSG000002 | 2.0743273 | -0.118791121 | 0.36081094 | -0.3292337 | 0.74197906 | NA | NA | NA |
| ENSG000002 | 1.74129323 | 0.070015679 | 0.34649531 | 0.20206819 | 0.83986342 | NA | NA | NA |
| ENSG000002 | 0.24141689 | -0.165257896 | 0.17127401 | -0.9648743 | 0.33460776 | NA | NA | NA |
| ENSG000002 | 0.84471523 | -0.099319924 | 0.28861894 | -0.3441213 | 0.73075507 | NA | NA | NA |
| ENSG000002 | 0.25214371 | -0.009862892 | 0.18046657 | -0.0546522 | 0.95641556 | NA | NA | NA |
| ENSG000002 | 0.88667417 | -0.451945682 | 0.30012996 | -1.5058333 | 0.13210997 | NA | NA | NA |
| ENSG000002 | 0.39246123 | -0.276864021 | 0.21549122 | -1.2848042 | 0.19886073 | NA | RNU6-164P | 106479639 |
| ENSG000002 | 0.62775378 | -0.28146231 | 0.26704095 | -1.0540043 | 0.29188095 | NA | NA | NA |
| ENSG000002 | 0.23202362 | -0.008302493 | 0.17586207 | -0.0472103 | 0.96234565 | NA | NA | NA |
| ENSG000002 | 0 | NA | NA | NA | NA | NA | SNORD45B | 26804 |
| ENSG000002 | 0.36416385 | 0.068880976 | 0.20758164 | 0.33182596 | 0.74002068 | NA | NA | NA |
| ENSG000002 | 1.58818942 | -0.11353394 | 0.34568389 | -0.3284328 | 0.74258442 | NA | RNU1-141P | 106481633 |
| ENSG000002 | 1.76142132 | -0.613069793 | 0.33433919 | -1.8336761 | 0.06670208 | NA | RN7SKP275 | 106480917 |
| ENSG000002 | 1.57584965 | -0.327284691 | 0.33037449 | -0.9906476 | 0.32185769 | NA | NA | NA |
| ENSG000002 | 1.10690274 | 0.188111887 | 0.32046629 | 0.5869943 | 0.55720753 | NA | RNU6-312P | 106479706 |
| ENSG000002 | 1.5335267 | -0.344209703 | 0.34553097 | -0.9961761 | 0.31916457 | NA | NA | NA |

| | | | | | | | | |
|---|---|---|---|---|---|---|---|---|
| ENSG000002 | 0.38056192 | -0.244105065 | 0.20391883 | -1.1970698 | 0.23127937 | NA | NA | NA |
| ENSG000002 | 2.09911512 | 0.120458964 | 0.36545075 | 0.32961751 | 0.74168899 | NA | RN7SKP217 | 106479191 |
| ENSG000002 | 0.38884741 | -0.29745092 | 0.22235466 | -1.3377319 | 0.18098386 | NA | NA | NA |
| ENSG000002 | 0.58087681 | 0.226918272 | 0.26184027 | 0.86662863 | 0.38614552 | NA | NA | NA |
| ENSG000002 | 0 | NA | NA | NA | NA | NA | NA | NA |
| ENSG000002 | 1.34194329 | -0.121160917 | 0.34046782 | -0.355866 | 0.72194093 | NA | NA | NA |
| ENSG000002 | 0.50067359 | -0.016990261 | 0.23962179 | -0.0709045 | 0.94347377 | NA | RNU6-645P | 106479852 |
| ENSG000002 | 0.98896775 | -0.328473032 | 0.30614961 | -1.0729167 | 0.28330849 | NA | NA | NA |
| ENSG000002 | 0.11205239 | 0.082118251 | 0.12671962 | 0.64803108 | 0.51696485 | NA | RNU6-450P | 106481321 |
| ENSG000002 | 1.80692609 | 0.017885238 | 0.35632327 | 0.05019385 | 0.95996791 | NA | NA | NA |
| ENSG000002 | 0.8814961 | -0.275762091 | 0.2995484 | -0.9205928 | 0.35726308 | NA | NA | NA |
| ENSG000002 | 0.75253306 | -0.354317709 | 0.28272428 | -1.253227 | 0.21012309 | NA | NA | NA |
| ENSG000002 | 1.23192322 | -0.434493095 | 0.31582632 | -1.3757343 | 0.16890392 | NA | RN7SKP237 | 106479200 |
| ENSG000002 | 0.60507165 | -0.101443751 | 0.26332021 | -0.3852486 | 0.70005322 | NA | NA | NA |
| ENSG000002 | 0.89634583 | -0.106757902 | 0.29151321 | -0.3662198 | 0.71420108 | NA | NA | NA |
| ENSG000002 | 1.30580204 | -0.314750997 | 0.31886781 | -0.9870893 | 0.32359887 | NA | SNORA16B | 692157 |
| ENSG000002 | 1.51042237 | -0.623614869 | 0.34090852 | -1.8292734 | 0.06735867 | NA | RNU4-85P | 106479597 |
| ENSG000002 | 0.37859486 | 0.064147216 | 0.20986618 | 0.30565771 | 0.75986527 | NA | NA | NA |
| ENSG000002 | 1.91595072 | -0.932826225 | 0.35140075 | -2.6545937 | 0.0079404 | NA | NA | NA |
| ENSG000002 | 0.81556908 | 0.199131673 | 0.28725247 | 0.69322876 | 0.488166 | NA | NA | NA |
| ENSG000002 | 1.10964084 | 0.027618367 | 0.31734393 | 0.08702976 | 0.93064785 | NA | RNU6-726P | 106480604 |
| ENSG000002 | 0.50722002 | -0.375409304 | 0.24493426 | -1.5326941 | 0.12535123 | NA | NA | NA |
| ENSG000002 | 1.4488509 | 0.217256288 | 0.3408022 | 0.63748499 | 0.52380898 | NA | NA | NA |
| ENSG000002 | 0.46956163 | 0.155192681 | 0.24412227 | 0.63571702 | 0.52496088 | NA | SNORD114-15 | 767593 |
| ENSG000002 | 0.82694879 | 0.370612883 | 0.29155396 | 1.27116397 | 0.20367032 | NA | RNU6-973P | 106479996 |
| ENSG000002 | 0.61147777 | -0.109180311 | 0.26384761 | -0.4138006 | 0.67902011 | NA | NA | NA |
| ENSG000002 | 0.76033119 | -0.194152683 | 0.28698065 | -0.6765358 | 0.49870053 | NA | NA | NA |
| ENSG000002 | 0.38064837 | -0.102706251 | 0.21886737 | -0.4692625 | 0.63888201 | NA | NA | NA |
| ENSG000002 | 0.11205239 | 0.082118251 | 0.12671962 | 0.64803108 | 0.51696485 | NA | SNORD114-17 | 767595 |

| | | | | | | | | | |
|---|---|---|---|---|---|---|---|---|---|
| ENSG000002 | 0.86981963 | -0.10589843 | 0.29781108 | -0.3555893 | 0.72214817 | NA | | RNU6-343P | 106479719 |
| ENSG000002 | 1.98688263 | -0.18299948 | 0.35924864 | -0.5093951 | 0.61047534 | NA | | RN7SKP78 | 106479129 |
| ENSG000002 | 1.1045491 | -0.256637025 | 0.31680796 | -0.8100713 | 0.41789922 | NA | | RN7SKP191 | 106480471 |
| ENSG000002 | 0 | NA | NA | NA | NA | NA | | NA | NA |
| ENSG000002 | 0 | NA | NA | NA | NA | NA | | RNA5S2 | 100169753 |
| ENSG000002 | 0.14128499 | -0.097772014 | 0.13403235 | -0.7294658 | 0.4657168 | NA | | RNU6-99P | 106479610 |
| ENSG000002 | 0.11934008 | 0.083564698 | 0.12760091 | 0.65489106 | 0.5125379 | NA | | NA | NA |
| ENSG000002 | 0.58207103 | 0.244437201 | 0.26804193 | 0.91193641 | 0.36180219 | NA | | NA | NA |
| ENSG000002 | 1.88956036 | -0.951472172 | 0.35290341 | -2.6961263 | 0.00701511 | NA | | NA | NA |
| ENSG000002 | 1.63494106 | -0.684920014 | 0.34740656 | -1.971523 | 0.04866409 | NA | | RNU6-219P | 106479664 |
| ENSG000002 | 1.54123894 | -0.310806114 | 0.33757022 | -0.9207155 | 0.35719901 | NA | | RN7SKP124 | 106480871 |
| ENSG000002 | 0.39499294 | -0.277908454 | 0.21584201 | -1.287555 | 0.19790091 | NA | | NA | NA |
| ENSG000002 | 0.11205239 | 0.082118251 | 0.12671962 | 0.64803108 | 0.51696485 | NA | | RNU6-740P | 106479893 |
| ENSG000002 | 0.70809743 | 0.287649333 | 0.27661527 | 1.03988958 | 0.2983912 | NA | | RNU4-16P | 106479566 |
| ENSG000002 | 0.48441528 | -0.017227682 | 0.23774503 | -0.0724628 | 0.94223357 | NA | | RNU4-42P | 106479577 |
| ENSG000002 | 0.12685397 | -0.090215768 | 0.12901919 | -0.699243 | 0.48440017 | NA | | RNU4-28P | 106479570 |
| ENSG000002 | 1.46552835 | -0.186173051 | 0.34268795 | -0.5432728 | 0.58694197 | NA | | NA | NA |
| ENSG000002 | 1.45712663 | -0.162770016 | 0.33355189 | -0.4879901 | 0.62555686 | NA | | RN7SKP15 | 106480456 |
| ENSG000002 | 1.22399523 | -0.039896728 | 0.32835984 | -0.1215031 | 0.90329259 | NA | | RNU6-200P | 106479655 |
| ENSG000002 | 0.37330862 | -0.105391168 | 0.21773449 | -0.4840352 | 0.62836086 | NA | | NA | NA |
| ENSG000002 | 0.34407602 | 0.071624546 | 0.21281613 | 0.33655599 | 0.73645163 | NA | | NA | NA |
| ENSG000002 | 1.5156177 | -0.336915967 | 0.34296816 | -0.9823535 | 0.32592574 | NA | | NA | NA |
| ENSG000002 | 0.99761878 | -0.204823835 | 0.31377682 | -0.6527692 | 0.5139051 | NA | | RNU6-621P | 106481907 |
| ENSG000002 | 1.87149818 | -0.111917912 | 0.34802497 | -0.3215801 | 0.74777081 | NA | | RNU6-923P | 106481471 |
| ENSG000002 | 1.70261381 | -0.309344689 | 0.35166587 | -0.8796551 | 0.37904616 | NA | | NA | NA |
| ENSG000002 | 0.3842622 | -0.095651135 | 0.21113931 | -0.4530238 | 0.65053159 | NA | | NA | NA |
| ENSG000002 | 0.6567905 | -0.46248946 | 0.26597757 | -1.7388288 | 0.08206488 | NA | | NA | NA |
| ENSG000002 | 0.51570138 | -0.378721611 | 0.24581506 | -1.540677 | 0.12339542 | NA | | NA | NA |
| ENSG000002 | 0.34931019 | 0.26109409 | 0.21538338 | 1.21222954 | 0.22542453 | NA | | SNORD115-48 | 100033822 |

| | | | | | | | | |
|---|---|---|---|---|---|---|---|---|
| ENSG000002 | 0 | NA | NA | NA | NA | NA | NA | NA |
| ENSG000002 | 0.75799536 | -0.171531325 | 0.27367317 | -0.6267744 | 0.53080715 | NA | NA | NA |
| ENSG000002 | 0.76380067 | -0.180699362 | 0.27231745 | -0.6635614 | 0.50697104 | NA | RN7SKP28 | 106479107 |
| ENSG000002 | 1.52083014 | -0.336253059 | 0.34231113 | -0.9823025 | 0.32595087 | NA | RNU6-865P | 106479948 |
| ENSG000002 | 0.33541997 | 0.234895243 | 0.20524277 | 1.14447511 | 0.25242665 | NA | NA | NA |
| ENSG000002 | 1.08818328 | 0.036463773 | 0.32504328 | 0.11218128 | 0.91067967 | NA | RNU4-91P | 106481201 |
| ENSG000002 | 1.70278458 | -0.180610792 | 0.35148777 | -0.5138466 | 0.60735925 | NA | RNY4P34 | 106481712 |
| ENSG000002 | 0.13280363 | -0.093394593 | 0.13115644 | -0.7120855 | 0.47641185 | NA | RNU6-283P | 106479692 |
| ENSG000002 | 0.35095876 | 0.063746151 | 0.20537869 | 0.31038348 | 0.75626936 | NA | NA | NA |
| ENSG000002 | 0.87124356 | 0.043728526 | 0.29752478 | 0.1469744 | 0.88315223 | NA | NA | NA |
| ENSG000002 | 1.95900305 | -0.428695001 | 0.35914298 | -1.1936611 | 0.23261057 | NA | NA | NA |
| ENSG000002 | 0.72372659 | -0.023335709 | 0.28377892 | -0.082232 | 0.93446222 | NA | NA | NA |
| ENSG000002 | 0.64007922 | -0.113235616 | 0.26626803 | -0.4252693 | 0.67064038 | NA | NA | NA |
| ENSG000002 | 1.84661337 | -0.256959095 | 0.35732498 | -0.7191188 | 0.47206775 | NA | NA | NA |
| ENSG000002 | 1.50543571 | -0.195792119 | 0.34124351 | -0.5737607 | 0.56612974 | NA | NA | NA |
| ENSG000002 | 0 | NA | NA | NA | NA | NA | SNORD113-4 | 767564 |
| ENSG000002 | 0 | NA | NA | NA | NA | NA | SNORD32A | 26819 |
| ENSG000002 | 0.92259363 | -0.647830102 | 0.30090942 | -2.1529074 | 0.03132596 | NA | NA | NA |
| ENSG000002 | 1.22374729 | -0.041358701 | 0.33208023 | -0.1245443 | 0.90088433 | NA | SNORD115-15 | 100033453 |
| ENSG000002 | 0.25370795 | -0.171331747 | 0.17411721 | -0.9840024 | 0.32511435 | NA | NA | NA |
| ENSG000002 | 0.99038563 | -0.038010516 | 0.30807592 | -0.1233804 | 0.90180591 | NA | NA | NA |
| ENSG000002 | 1.12268273 | -0.585761118 | 0.31880745 | -1.8373508 | 0.06615813 | NA | RN7SKP239 | 106480907 |
| ENSG000002 | 1.11021643 | -0.400031377 | 0.31365869 | -1.2753716 | 0.20217775 | NA | RNU6-1107P | 106481529 |
| ENSG000002 | 0.63640983 | -0.455149466 | 0.26433333 | -1.7218769 | 0.08509182 | NA | SNORD114-29 | 767610 |
| ENSG000002 | 1.36861385 | -0.119928995 | 0.34100495 | -0.3516928 | 0.72506864 | NA | NA | NA |
| ENSG000002 | 0.11131518 | 0.081951103 | 0.12663804 | 0.64712866 | 0.51754868 | NA | NA | NA |
| ENSG000002 | 0.27408862 | -0.199826037 | 0.18669465 | -1.0703362 | 0.28446801 | NA | SNORD113-3 | 767563 |
| ENSG000002 | 1.40416221 | -0.129915774 | 0.33490328 | -0.3879203 | 0.69807503 | NA | NA | NA |
| ENSG000002 | 0.60333272 | 0.225368532 | 0.26390364 | 0.85398039 | 0.39311586 | NA | NA | NA |

| | | | | | | | | | |
|---|---|---|---|---|---|---|---|---|---|
| ENSG000002 | 0.94547593 | 0.111575953 | 0.29850566 | 0.3737817 | 0.70856673 | NA | | RNU6-818P | 106481439 |
| ENSG000002 | 0.50544884 | -0.019721866 | 0.2401671 | -0.0821173 | 0.93455346 | NA | | NA | NA |
| ENSG000002 | 0.65268349 | -0.105755209 | 0.26000422 | -0.4067442 | 0.68419588 | NA | | RNU6-686P | 106481397 |
| ENSG000002 | 0.26813897 | -0.197007041 | 0.18550319 | -1.0620143 | 0.2882292 | NA | | NA | NA |
| ENSG000002 | 0.71561187 | 0.138425705 | 0.28332471 | 0.48857619 | 0.62514177 | NA | | RNU6-1303P | 106481949 |
| ENSG000002 | 1.79469255 | -0.317689645 | 0.34988424 | -0.907985 | 0.36388614 | NA | | RN7SKP133 | 106481812 |
| ENSG000002 | 0.26813897 | -0.197007041 | 0.18550319 | -1.0620143 | 0.2882292 | NA | | NA | NA |
| ENSG000002 | 0.59824098 | -0.099488676 | 0.26271446 | -0.3786951 | 0.7049143 | NA | | NA | NA |
| ENSG000002 | 0.4716672 | -0.017393026 | 0.23619084 | -0.0736397 | 0.94129707 | NA | | RNU6-304P | 106481275 |
| ENSG000002 | 1.13704669 | -0.12054989 | 0.32304537 | -0.3731671 | 0.70902411 | NA | | NA | NA |
| ENSG000002 | 1.57593398 | -0.362877622 | 0.34006766 | -1.0670748 | 0.28593807 | NA | | NA | NA |
| ENSG000002 | 0.77613058 | -0.361458377 | 0.28422771 | -1.2717211 | 0.20347223 | NA | | NA | NA |
| ENSG000002 | 1.39449109 | -0.432696765 | 0.34154512 | -1.2668802 | 0.20519812 | NA | | NA | NA |
| ENSG000002 | 1.01298344 | -0.206632615 | 0.31837313 | -0.6490266 | 0.51632121 | NA | | RNU1-133P | 106481631 |
| ENSG000002 | 0.24550891 | -0.007886114 | 0.17900103 | -0.0440563 | 0.96485956 | NA | | NA | NA |
| ENSG000002 | 0.36484899 | 0.075158314 | 0.21612264 | 0.34775772 | 0.72802215 | NA | | RNU6-563P | 106481359 |
| ENSG000002 | 0.2596576 | -0.193002035 | 0.1837913 | -1.0501152 | 0.29366517 | NA | | RNU6-34P | 100873749 |
| ENSG000002 | 0.62243135 | -0.107944104 | 0.26481366 | -0.4076229 | 0.68355057 | NA | | RNU6-828P | 106479932 |
| ENSG000002 | 0.47075585 | 0.160115757 | 0.24423976 | 0.65556794 | 0.51210216 | NA | | NA | NA |
| ENSG000002 | 0.76559571 | -0.196311958 | 0.28730218 | -0.6832943 | 0.49442087 | NA | | RNU6-119P | 106481214 |
| ENSG000002 | 0 | NA | NA | NA | NA | NA | | SNORD52 | 26797 |
| ENSG000002 | 0.75341407 | -0.349034172 | 0.2811917 | -1.2412677 | 0.21450687 | NA | | NA | NA |
| ENSG000002 | 0.70773078 | 0.128255771 | 0.27653919 | 0.46378876 | 0.6427991 | NA | | RN7SKP55 | 106479119 |
| ENSG000002 | 0.64235948 | -0.482579697 | 0.27062934 | -1.7831759 | 0.07455768 | NA | | RNU6-336P | 106481285 |
| ENSG000002 | 0.520565 | -0.024378065 | 0.24183609 | -0.1008041 | 0.91970599 | NA | | NA | NA |
| ENSG000002 | 0 | NA | NA | NA | NA | NA | | RNU6-384P | 106481299 |
| ENSG000002 | 0.46552167 | -0.013602584 | 0.2265546 | -0.0600411 | 0.95212291 | NA | | NA | NA |
| ENSG000002 | 0.88602183 | -0.42604663 | 0.29504313 | -1.4440147 | 0.14873483 | NA | | RNU6-1325P | 106480145 |
| ENSG000002 | 0.38884741 | -0.29745092 | 0.22235466 | -1.3377319 | 0.18098386 | NA | | NA | NA |

| | | | | | | | | | |
|---|---|---|---|---|---|---|---|---|---|
| ENSG000002 | 1.24537708 | -0.328556548 | 0.32392634 | -1.014294 | 0.31044248 | NA | | RNU6-275P | 106481265 |
| ENSG000002 | 0.11865494 | 0.083440866 | 0.12751478 | 0.65436232 | 0.5128784 | NA | | NA | NA |
| ENSG000002 | 1.25297989 | -0.188458472 | 0.32966021 | -0.5716749 | 0.56754223 | NA | | NA | NA |
| ENSG000002 | 0 | NA | NA | NA | NA | NA | | RNU6-1071P | 106481517 |
| ENSG000002 | 1.91297927 | -0.52949348 | 0.35822127 | -1.4781185 | 0.13937606 | NA | | NA | NA |
| ENSG000002 | 0.47758652 | 0.147793567 | 0.23768536 | 0.6218034 | 0.53407115 | NA | | NA | NA |
| ENSG000002 | 0.83575724 | -0.109698178 | 0.29597774 | -0.3706298 | 0.71091327 | NA | | RN7SKP130 | 106480873 |
| ENSG000002 | 0.9779238 | -0.181055242 | 0.30825798 | -0.5873497 | 0.55696885 | NA | | NA | NA |
| ENSG000002 | 0.61290067 | -0.091310841 | 0.25649069 | -0.3560006 | 0.72184011 | NA | | RNU6-1180P | 106481552 |
| ENSG000002 | 0.47761686 | -0.01948631 | 0.23692179 | -0.0822479 | 0.93444962 | NA | | NA | NA |
| ENSG000002 | 0.77288285 | -0.203654568 | 0.28771447 | -0.7078357 | 0.47904731 | NA | | NA | NA |
| ENSG000002 | 0.70084804 | 0.12831363 | 0.2743736 | 0.46766027 | 0.64002756 | NA | | NA | NA |
| ENSG000002 | 1.1319872 | -0.440911523 | 0.32481316 | -1.3574312 | 0.17464423 | NA | | NA | NA |
| ENSG000002 | 1.62514689 | -0.25796247 | 0.35087271 | -0.7352024 | 0.46221621 | NA | | NA | NA |
| ENSG000002 | 1.09460901 | 0.028966755 | 0.31985179 | 0.09056305 | 0.92783979 | NA | | NA | NA |
| ENSG000002 | 0.23890637 | -0.010420674 | 0.17747379 | -0.0587167 | 0.95317776 | NA | | RNU6-915P | 106481923 |
| ENSG000002 | 0.12070844 | -0.088503695 | 0.12783931 | -0.6923042 | 0.48874631 | NA | | RNU6-526P | 106481347 |
| ENSG000002 | 1.95314176 | -0.153666918 | 0.34853251 | -0.4408969 | 0.65928761 | NA | | NA | NA |
| ENSG000002 | 0.7721977 | -0.203893177 | 0.28767039 | -0.7087736 | 0.47846499 | NA | | NA | NA |
| ENSG000002 | 0.74438746 | -0.033988507 | 0.28512681 | -0.1192049 | 0.90511304 | NA | | NA | NA |
| ENSG000002 | 1.25371518 | -0.341250739 | 0.33063059 | -1.0321209 | 0.30201549 | NA | | NA | NA |
| ENSG000002 | 0.59237565 | -0.09452067 | 0.25455693 | -0.3713145 | 0.71040332 | NA | | RNA5SP149 | 106480437 |
| ENSG000002 | 0 | NA | NA | NA | NA | NA | | SNORD48 | 26801 |
| ENSG000002 | 0.47280936 | -0.012478229 | 0.23633508 | -0.0527989 | 0.95789215 | NA | | RNU6-1131P | 106480064 |
| ENSG000002 | 0.47596775 | 0.137153722 | 0.23749987 | 0.57748967 | 0.56360872 | NA | | RNU6-867P | 106479949 |
| ENSG000002 | 0.4842194 | -0.018518118 | 0.24529708 | -0.0754926 | 0.93982277 | NA | | NA | NA |
| ENSG000002 | 0.76033119 | -0.194152683 | 0.28698065 | -0.6765358 | 0.49870053 | NA | | NA | NA |
| ENSG000002 | 0.5681808 | 0.209349632 | 0.25180656 | 0.83139071 | 0.40575295 | NA | | SNORD115-1 | 338433 |
| ENSG000002 | 1.38660705 | -0.689001796 | 0.32996275 | -2.0881199 | 0.03678702 | NA | | SNORD114-3 | 767579 |

| | | | | | | | | |
|---|---|---|---|---|---|---|---|---|
| ENSG000002 | 0.46136259 | 0.353033165 | 0.24467579 | 1.44286102 | 0.14905963 | NA | NA | NA |
| ENSG000002 | 0.26813897 | -0.197007041 | 0.18550319 | -1.0620143 | 0.2882292 | NA | NA | NA |
| ENSG000002 | 0.86438042 | 0.046609194 | 0.29716529 | 0.15684602 | 0.8753662 | NA | NA | NA |
| ENSG000002 | 1.15461726 | -0.614542622 | 0.32342171 | -1.9001279 | 0.05741633 | NA | RNU6-702P | 106481912 |
| ENSG000002 | 0.36621735 | -0.097238239 | 0.21673053 | -0.4486596 | 0.65367721 | NA | NA | NA |
| ENSG000002 | 0.40689225 | -0.283036689 | 0.21754773 | -1.3010326 | 0.19324729 | NA | NA | NA |
| ENSG000002 | 0.50516651 | -0.183853222 | 0.24168957 | -0.7606999 | 0.44683635 | NA | RNA5SP298 | 106480761 |
| ENSG000002 | 1.86871696 | -0.246456893 | 0.35509209 | -0.6940647 | 0.48764162 | NA | NA | NA |
| ENSG000002 | 1.0966046 | 0.023518608 | 0.30685176 | 0.07664485 | 0.93890608 | NA | NA | NA |
| ENSG000002 | 0.24756242 | -0.187034346 | 0.18119817 | -1.0322088 | 0.30197431 | NA | NA | NA |
| ENSG000002 | 1.0138736 | -0.498211149 | 0.30523646 | -1.6322137 | 0.10263447 | NA | RNU6-294P | 106479697 |
| ENSG000002 | 1.7089019 | 0.084003197 | 0.35359408 | 0.23756958 | 0.81221494 | NA | NA | NA |
| ENSG000002 | 0.73197769 | -0.177345773 | 0.27736038 | -0.6394056 | 0.52255912 | NA | NA | NA |
| ENSG000002 | 1.385064 | -0.418353415 | 0.33950217 | -1.2322555 | 0.21785366 | NA | NA | NA |
| ENSG000002 | 0 | NA | NA | NA | NA | NA | NA | NA |
| ENSG000002 | 0.84172863 | 0.052461019 | 0.30554709 | 0.17169536 | 0.86367703 | NA | NA | NA |
| ENSG000002 | 0.35346928 | -0.088570748 | 0.20618147 | -0.4295767 | 0.66750362 | NA | RNU6-967P | 106479993 |
| ENSG000002 | 1.13523046 | -0.270004135 | 0.32001209 | -0.843731 | 0.39881972 | NA | RN7SKP153 | 106481813 |
| ENSG000002 | 1.0634471 | 0.337967293 | 0.31747537 | 1.0645465 | 0.2870812 | NA | NA | NA |
| ENSG000002 | 0.25370795 | -0.171331747 | 0.17411721 | -0.9840024 | 0.32511435 | NA | SNORD114-24 | 767604 |
| ENSG000002 | 1.15805694 | -0.428595256 | 0.32209403 | -1.3306526 | 0.18330335 | NA | RNY1P13 | 106480310 |
| ENSG000002 | 0.87169844 | -0.120477128 | 0.30728461 | -0.3920702 | 0.69500637 | NA | RN7SKP48 | 106479116 |
| ENSG000002 | 0.86934088 | -0.273305014 | 0.29732012 | -0.9192281 | 0.35797627 | NA | SNORD115-38 | 100033812 |
| ENSG000002 | 0.23070733 | 0.167416683 | 0.17664244 | 0.94777156 | 0.34324576 | NA | RNU6-590P | 106480593 |
| ENSG000002 | 2.12621637 | -0.261464586 | 0.36529633 | -0.7157602 | 0.47413941 | NA | RNU1-140P | 106480669 |
| ENSG000002 | 1.48538942 | -0.049385827 | 0.34119637 | -0.1447431 | 0.88491369 | NA | RN7SKP189 | 106479180 |
| ENSG000002 | 1.13520405 | -0.419702156 | 0.31944658 | -1.3138415 | 0.18889956 | NA | NA | NA |
| ENSG000002 | 0.37285215 | -0.099309896 | 0.2177345 | -0.4561055 | 0.64831414 | NA | NA | NA |
| ENSG000002 | 0.3433388 | 0.065348438 | 0.20407693 | 0.32021472 | 0.74880557 | NA | RNU6-1022P | 106480016 |

| | | | | | | | | | |
|---|---|---|---|---|---|---|---|---|---|
| ENSG000002 | 0.46267888 | 0.146619746 | 0.23584798 | 0.62167056 | 0.53415852 | NA | | RNA5SP442 | 106480768 |
| ENSG000002 | 0.27408862 | -0.199826037 | 0.18669465 | -1.0703362 | 0.28446801 | NA | | RNU1-43P | 106480159 |
| ENSG000002 | 0.24485602 | -0.012655878 | 0.17881722 | -0.0707755 | 0.94357643 | NA | | NA | NA |
| ENSG000002 | 0 | NA | NA | NA | NA | NA | | RNA5S15 | 100169766 |
| ENSG000002 | 1.38506345 | -0.389736396 | 0.32961011 | -1.1824164 | 0.23704052 | NA | | RNA5SP172 | 106478997 |
| ENSG000002 | 0.40327843 | -0.281391716 | 0.21700406 | -1.2967118 | 0.19473038 | NA | | NA | NA |
| ENSG000002 | 1.63048467 | -0.125413035 | 0.35046586 | -0.3578467 | 0.72045807 | NA | | NA | NA |
| ENSG000002 | 1.10788115 | -0.407411504 | 0.31609669 | -1.2888825 | 0.19743892 | NA | | NA | NA |
| ENSG000002 | 1.54149063 | 0.445314583 | 0.34793977 | 1.27986112 | 0.20059398 | NA | | NA | NA |
| ENSG000002 | 0.57125542 | 0.368737025 | 0.24765945 | 1.48888735 | 0.13651704 | NA | | SNORD115-10 | 100033447 |
| ENSG000002 | 0.38270188 | -0.272446727 | 0.2140009 | -1.2731102 | 0.20297894 | NA | | NA | NA |
| ENSG000002 | 0.90298455 | -0.457507079 | 0.30248048 | -1.5125177 | 0.13040221 | NA | | SNORD113-9 | 767569 |
| ENSG000002 | 1.27311924 | -0.184355333 | 0.32430083 | -0.5684701 | 0.56971579 | NA | | RNU6-673P | 106481394 |
| ENSG000002 | 1.35165058 | -0.117156649 | 0.33454627 | -0.3501957 | 0.72619187 | NA | | RNY3P1 | 106478984 |
| ENSG000002 | 2.13338645 | -0.376364653 | 0.3657873 | -1.0289167 | 0.30351883 | NA | | NA | NA |
| ENSG000002 | 0 | NA | NA | NA | NA | NA | | NA | NA |
| ENSG000002 | 1.08428713 | -0.112189908 | 0.31980027 | -0.3508124 | 0.72572908 | NA | | NA | NA |
| ENSG000002 | 0 | NA | NA | NA | NA | NA | | NA | NA |
| ENSG000002 | 0.37996515 | 0.064665424 | 0.21008146 | 0.3078112 | 0.758226 | NA | | NA | NA |
| ENSG000002 | 1.92157218 | -0.518810563 | 0.35677883 | -1.4541518 | 0.14590424 | NA | | RN7SKP22 | 106481805 |
| ENSG000002 | 0.97481404 | 0.105928641 | 0.30746122 | 0.34452684 | 0.73045012 | NA | | NA | NA |
| ENSG000002 | 0.23890637 | -0.010420674 | 0.17747379 | -0.0587167 | 0.95317776 | NA | | SNORD115-20 | 100033460 |
| ENSG000002 | 1.24500706 | -0.205716836 | 0.33731973 | -0.6098571 | 0.54195646 | NA | | NA | NA |
| ENSG000002 | 0.98690977 | 0.11631077 | 0.31276288 | 0.37188163 | 0.70998099 | NA | | NA | NA |
| ENSG000002 | 0.84676639 | 0.21049294 | 0.30167962 | 0.6977367 | 0.48534187 | NA | | NA | NA |
| ENSG000002 | 1.84121818 | 0.138188676 | 0.35561889 | 0.38858644 | 0.6975821 | NA | | NA | NA |
| ENSG000002 | 2.13338591 | -0.391404285 | 0.367181 | -1.065971 | 0.28643677 | NA | | NA | NA |
| ENSG000002 | 0.63159785 | -0.116736614 | 0.27184111 | -0.4294296 | 0.66761063 | NA | | SNORD115-35 | 100033809 |
| ENSG000002 | 0.12070844 | -0.088503695 | 0.12783931 | -0.6923042 | 0.48874631 | NA | | NA | NA |

| | | | | | | | | |
|---|---|---|---|---|---|---|---|---|
| ENSG000002 | 0.87335105 | 0.041351406 | 0.28988809 | 0.1426461 | 0.88656969 | NA | NA | NA |
| ENSG000002 | 1.36220773 | -0.101905445 | 0.33060776 | -0.3082367 | 0.75790223 | NA | RNU6-619P | 106479840 |
| ENSG000002 | 0.90015946 | -0.421486757 | 0.29333848 | -1.4368614 | 0.15075737 | NA | RNU6-806P | 106481435 |
| ENSG000002 | 0.25351207 | -0.189965547 | 0.1824784 | -1.0410303 | 0.29786148 | NA | NA | NA |
| ENSG000002 | 0 | NA | NA | NA | NA | NA | NA | NA |
| ENSG000002 | 0.60558265 | 0.061136384 | 0.2538335 | 0.24085231 | 0.80966959 | NA | NA | NA |
| ENSG000002 | 1.9156752 | -0.046850168 | 0.34777501 | -0.134714 | 0.89283799 | NA | RNU6-934P | 106481475 |
| ENSG000002 | 0 | NA | NA | NA | NA | NA | NA | NA |
| ENSG000002 | 0.73861249 | -0.192197989 | 0.28559321 | -0.6729781 | 0.5009612 | NA | RNU6-1134P | 106480065 |
| ENSG000002 | 0.36007182 | -0.08619198 | 0.20729252 | -0.4157988 | 0.67755724 | NA | NA | NA |
| ENSG000002 | 1.73173039 | -0.18768226 | 0.34947418 | -0.5370418 | 0.59123871 | NA | RNU6-580P | 106480395 |
| ENSG000002 | 0 | NA | NA | NA | NA | NA | SNORD38A | 94162 |
| ENSG000002 | 0.35346928 | -0.088570748 | 0.20618147 | -0.4295767 | 0.66750362 | NA | RNU6-399P | 106481892 |
| ENSG000002 | 0.11934008 | 0.083564698 | 0.12760091 | 0.65489106 | 0.5125379 | NA | NA | NA |
| ENSG000002 | 0.96314395 | 0.122644775 | 0.3052351 | 0.40180429 | 0.68782806 | NA | RNU6-215P | 106481245 |
| ENSG000002 | 1.79631525 | -0.584903409 | 0.35065194 | -1.6680456 | 0.09530668 | NA | NA | NA |
| ENSG000002 | 0.23890637 | -0.010420674 | 0.17747379 | -0.0587167 | 0.95317776 | NA | NA | NA |
| ENSG000002 | 1.065185 | -0.091363262 | 0.30414736 | -0.3003914 | 0.7638786 | NA | NA | NA |
| ENSG000002 | 1.23789406 | -0.352848947 | 0.33361871 | -1.0576414 | 0.29021898 | NA | NA | NA |
| ENSG000002 | 1.27938347 | -0.053150098 | 0.32298242 | -0.1645603 | 0.86929005 | NA | SNORD114-20 | 767598 |
| ENSG000002 | 0.520565 | -0.024378065 | 0.24183609 | -0.1008041 | 0.91970599 | NA | RNU6-391P | 106479739 |
| ENSG000002 | 0 | NA | NA | NA | NA | NA | RNU6-382P | 106479735 |
| ENSG000002 | 0.73197769 | -0.177345773 | 0.27736038 | -0.6394056 | 0.52255912 | NA | NA | NA |
| ENSG000002 | 0 | NA | NA | NA | NA | NA | NA | NA |
| ENSG000002 | 1.62190777 | -0.378964637 | 0.34216492 | -1.1075497 | 0.26805635 | NA | NA | NA |
| ENSG000002 | 1.28038519 | -0.359394614 | 0.33464267 | -1.0739653 | 0.28283824 | NA | NA | NA |
| ENSG000002 | 1.61433363 | -0.522221724 | 0.34608025 | -1.5089614 | 0.13130866 | NA | NA | NA |
| ENSG000002 | 0.51894464 | -0.193743146 | 0.24314987 | -0.7968055 | 0.42556401 | NA | RNU4-66P | 106479588 |
| ENSG000002 | 0.50107449 | -0.372905725 | 0.24426561 | -1.5266403 | 0.12685049 | NA | RNU6-1175P | 106480082 |

| | | | | | | | | | |
|---|---|---|---|---|---|---|---|---|---|
| ENSG000002 | 0.77265259 | -0.365549345 | 0.28395139 | -1.2873659 | 0.19796678 | NA | | RNU6-715P | 106481405 |
| ENSG000002 | 1.555357 | 0.147367006 | 0.34427625 | 0.42804872 | 0.66861565 | NA | | NA | NA |
| ENSG000002 | 0.60438651 | -0.101682894 | 0.2632591 | -0.3862465 | 0.69931415 | NA | | NA | NA |
| ENSG000002 | 1.15299745 | -0.71148901 | 0.30941994 | -2.2994284 | 0.02148062 | NA | | RNU6-1280P | 106481584 |
| ENSG000002 | 0.61204084 | 0.061633846 | 0.26990716 | 0.22835202 | 0.81937259 | NA | | RNU6-1306P | 106480136 |
| ENSG000002 | 1.13690289 | -0.114057358 | 0.31690175 | -0.3599139 | 0.7189115 | NA | | NA | NA |
| ENSG000002 | 0 | NA | NA | NA | NA | NA | | NA | NA |
| ENSG000002 | 0.25145857 | -0.010118565 | 0.18031426 | -0.0561163 | 0.95524918 | NA | | RNU6-1172P | 106481549 |
| ENSG000002 | 1.01133432 | -0.045176087 | 0.30728854 | -0.1470152 | 0.88312002 | NA | | RNU6-234P | 106479670 |
| ENSG000002 | 1.13711375 | -0.589529324 | 0.31927247 | -1.8464772 | 0.06482295 | NA | | NA | NA |
| ENSG000002 | 0.11934008 | 0.083564698 | 0.12760091 | 0.65489106 | 0.5125379 | NA | | RNU4-12P | 106479564 |
| ENSG000002 | 1.76739693 | 0.129974041 | 0.35145039 | 0.36982188 | 0.71151521 | NA | | VTRNA1-2 | 56663 |
| ENSG000002 | 0.51336555 | -0.346899948 | 0.23688898 | -1.4643988 | 0.143085 | NA | | RNU6-690P | 106481398 |
| ENSG000002 | 0.58826863 | 0.215244732 | 0.25535246 | 0.8429319 | 0.39926653 | NA | | NA | NA |
| ENSG000002 | 1.17829021 | 0.245772589 | 0.32593704 | 0.75404929 | 0.45081963 | NA | | RNU1-100P | 106480183 |
| ENSG000002 | 0.70938393 | 0.278216169 | 0.26727878 | 1.04092128 | 0.29791208 | NA | | NA | NA |
| ENSG000002 | 0.25333738 | -0.015686291 | 0.1806605 | -0.0868275 | 0.93080867 | NA | | SNORD114-18 | 767596 |
| ENSG000002 | 0 | NA | NA | NA | NA | NA | | NA | NA |
| ENSG000002 | 1.32652712 | -0.110430861 | 0.33398098 | -0.3306502 | 0.74090876 | NA | | NA | NA |
| ENSG000002 | 0.75918904 | -0.198835545 | 0.28688515 | -0.6930841 | 0.48825676 | NA | | NA | NA |
| ENSG000002 | 1.82735774 | -0.003580302 | 0.35151379 | -0.0101854 | 0.99187338 | NA | | RNU6-407P | 106479745 |
| ENSG000002 | 0.796903 | -0.342445501 | 0.27749094 | -1.2340781 | 0.2171738 | NA | | NA | NA |
| ENSG000002 | 0.3561712 | 0.067399341 | 0.21474131 | 0.31386296 | 0.75362513 | NA | | RNU4-11P | 106481175 |
| ENSG000002 | 1.48928983 | -0.152176778 | 0.32476534 | -0.4685746 | 0.63937376 | NA | | RNU6-742P | 106481414 |
| ENSG000002 | 0.24141689 | -0.165257896 | 0.17127401 | -0.9648743 | 0.33460776 | NA | | NA | NA |
| ENSG000002 | 0.40191006 | -0.101653297 | 0.21376195 | -0.4755444 | 0.63439899 | NA | | NA | NA |
| ENSG000002 | 1.00262718 | -0.490562014 | 0.30482559 | -1.6093203 | 0.10754632 | NA | | RNU1-114P | 106481626 |
| ENSG000002 | 1.83929982 | -0.350580025 | 0.35018044 | -1.0011411 | 0.3167586 | NA | | NA | NA |
| ENSG000002 | 0.27408862 | -0.199826037 | 0.18669465 | -1.0703362 | 0.28446801 | NA | | NA | NA |

| | | | | | | | | |
|---|---|---|---|---|---|---|---|---|
| ENSG000002 | 0.62160824 | -0.2792447 | 0.26652831 | -1.0477112 | 0.29477168 | NA | NA | NA |
| ENSG000002 | 1.15091306 | -0.418132637 | 0.3183879 | -1.3132806 | 0.18908843 | NA | NA | NA |
| ENSG000002 | 0.11131518 | 0.081951103 | 0.12663804 | 0.64712866 | 0.51754868 | NA | NA | NA |
| ENSG000002 | 1.11097538 | -0.258164454 | 0.31924678 | -0.8086674 | 0.41870651 | NA | NA | NA |
| ENSG000002 | 0.89039618 | -0.112302745 | 0.29885955 | -0.375771 | 0.70708717 | NA | RNU6-497P | 106479786 |
| ENSG000002 | 0.90591323 | -0.45283099 | 0.30113132 | -1.5037658 | 0.13264168 | NA | RNA5SP355 | 106479009 |
| ENSG000002 | 0.14128499 | -0.097772014 | 0.13403235 | -0.7294658 | 0.4657168 | NA | SNORD115-31 | 100033805 |
| ENSG000002 | 1.32194136 | 0.019398996 | 0.33130398 | 0.05855346 | 0.95330778 | NA | NA | NA |
| ENSG000002 | 1.03012085 | -0.347757024 | 0.31023074 | -1.1209625 | 0.26230383 | NA | SNORD113-1 | 767561 |
| ENSG000002 | 0.97727249 | -0.037548228 | 0.29990922 | -0.1251986 | 0.90036629 | NA | NA | NA |
| ENSG000002 | 1.11678472 | 0.032855708 | 0.32594084 | 0.10080267 | 0.9197071 | NA | NA | NA |
| ENSG000002 | 0 | NA | NA | NA | NA | NA | RN7SK | 125050 |
| ENSG000002 | 0.9283686 | -0.4437535 | 0.2971005 | -1.4936141 | 0.1352765 | NA | RNU1-115P | 106480188 |
| ENSG000002 | 0.87018573 | 0.032252572 | 0.2896955 | 0.11133267 | 0.91135255 | NA | RNU6-1034P | 106480632 |
| ENSG000002 | 1.37160045 | -0.260994636 | 0.33836573 | -0.7713388 | 0.44050611 | NA | RNU6-169P | 106481229 |
| ENSG000002 | 1.11060481 | -0.120723725 | 0.32603316 | -0.3702805 | 0.71117349 | NA | NA | NA |
| ENSG000002 | 1.48187039 | -0.669412488 | 0.325806 | -2.0546353 | 0.03991425 | NA | RNU1-51P | 106480163 |
| ENSG000002 | 0 | NA | NA | NA | NA | NA | NA | NA |
| ENSG000002 | 1.66868969 | -0.66124392 | 0.34353439 | -1.924826 | 0.05425113 | NA | RN7SKP47 | 106481974 |
| ENSG000002 | 1.59220764 | -0.125896316 | 0.35111775 | -0.3585587 | 0.71992527 | NA | NA | NA |
| ENSG000002 | 1.49990923 | -0.338246279 | 0.34687877 | -0.9751138 | 0.32950381 | NA | NA | NA |
| ENSG000002 | 1.51230173 | -0.581637334 | 0.33461574 | -1.7382247 | 0.08217124 | NA | RNU6-282P | 106479691 |
| ENSG000002 | 0.50613398 | -0.020909936 | 0.24769218 | -0.084419 | 0.93272327 | NA | NA | NA |
| ENSG000002 | 0.3854038 | -0.110128691 | 0.21952326 | -0.5016721 | 0.61589821 | NA | RNU6-53P | 100873762 |
| ENSG000002 | 0.36502313 | -0.093755175 | 0.2080862 | -0.4505593 | 0.6523072 | NA | RNU6-396P | 106481303 |
| ENSG000002 | 1.79685762 | -0.45183983 | 0.34948842 | -1.2928606 | 0.19605926 | NA | RNU6-737P | 106481914 |
| ENSG000002 | 1.39613573 | -0.253902403 | 0.32977486 | -0.7699265 | 0.44134349 | NA | RNU6-1276P | 106481583 |
| ENSG000002 | 1.49641301 | -0.201151165 | 0.34497036 | -0.583097 | 0.55982803 | NA | RNU6-728P | 106479888 |
| ENSG000002 | 0.76169956 | -0.374892198 | 0.28875268 | -1.2983159 | 0.19417881 | NA | NA | NA |

| | | | | | | | | |
|---|---|---|---|---|---|---|---|---|
| ENSG000002 | 0.12070844 | -0.088503695 | 0.12783931 | -0.6923042 | 0.48874631 | NA | NA | NA |
| ENSG000002 | 0 | NA | NA | NA | NA | NA | SNORD14C | 85389 |
| ENSG000002 | 1.10443755 | -0.260315399 | 0.31679411 | -0.8217179 | 0.41123745 | NA | NA | NA |
| ENSG000002 | 0.73775266 | -0.030682722 | 0.27839171 | -0.1102142 | 0.91223949 | NA | NA | NA |
| ENSG000002 | 0 | NA | NA | NA | NA | NA | RNA5S16 | 100169767 |
| ENSG000002 | 0.36576034 | -0.0934779 | 0.20820828 | -0.4489634 | 0.65345805 | NA | RNU6-1318P | 106480141 |
| ENSG000002 | 1.86354226 | -0.406713404 | 0.36222754 | -1.1228119 | 0.26151737 | NA | RN7SKP69 | 106480461 |
| ENSG000002 | 0.97126727 | -0.324678945 | 0.30226839 | -1.0741412 | 0.28275939 | NA | SNORD115-44 | 100033818 |
| ENSG000002 | 1.02255084 | -0.536784374 | 0.31334684 | -1.7130678 | 0.08670007 | NA | NA | NA |
| ENSG000002 | 0.97888722 | -0.325982549 | 0.30262632 | -1.0771784 | 0.28140055 | NA | RNU6-596P | 106481369 |
| ENSG000002 | 0.87745615 | -0.395586073 | 0.28572351 | -1.3845066 | 0.1662034 | NA | NA | NA |
| ENSG000002 | 0.89353572 | -0.602673612 | 0.2930423 | -2.0566096 | 0.03972379 | NA | NA | NA |
| ENSG000002 | 0.90053014 | 0.033569332 | 0.2823269 | 0.11890235 | 0.90535272 | NA | SNORD114-12 | 767590 |
| ENSG000002 | 0.52165104 | -0.403921832 | 0.25252134 | -1.5995552 | 0.10969729 | NA | NA | NA |
| ENSG000002 | 0.86438042 | 0.046609194 | 0.29716529 | 0.15684602 | 0.8753662 | NA | NA | NA |
| ENSG000002 | 0 | NA | NA | NA | NA | NA | NA | NA |
| ENSG000002 | 0.60644002 | -0.257083821 | 0.25820471 | -0.9956589 | 0.31941589 | NA | NA | NA |
| ENSG000002 | 0.71831827 | -0.020963193 | 0.28341454 | -0.0739665 | 0.94103701 | NA | NA | NA |
| ENSG000002 | 0.73701545 | -0.03254472 | 0.28464749 | -0.1143334 | 0.90897349 | NA | NA | NA |
| ENSG000002 | 0.47687965 | -0.021216903 | 0.24445659 | -0.0867921 | 0.93083676 | NA | RNU6-117P | 106481213 |
| ENSG000002 | 0 | NA | NA | NA | NA | NA | NA | NA |
| ENSG000002 | 0.62061044 | -0.252611122 | 0.25798806 | -0.9791582 | 0.32750182 | NA | SNORD114-22 | 767600 |
| ENSG000002 | 0.38019136 | -0.10756138 | 0.21876967 | -0.491665 | 0.62295621 | NA | RNU6-1335P | 106480149 |
| ENSG000002 | 1.79658922 | 0.262446184 | 0.34892821 | 0.75214951 | 0.45196115 | NA | RNY3P12 | 106480315 |
| ENSG000002 | 1.60098686 | -0.379312983 | 0.34573357 | -1.0971251 | 0.2725867 | NA | RNU6-487P | 106480393 |
| ENSG000002 | 0.52147635 | -0.19458195 | 0.24341208 | -0.7993932 | 0.42406247 | NA | RNU6-1130P | 106480063 |
| ENSG000002 | 0.60466671 | -0.033939957 | 0.1573742 | -0.215664 | 0.82924963 | NA | NA | NA |
| ENSG000002 | 1.5567232 | -0.632328189 | 0.34217467 | -1.847969 | 0.06460683 | NA | RNU6-996P | 106481928 |
| ENSG000002 | 1.62926871 | -0.274907764 | 0.35202699 | -0.7809281 | 0.4348448 | NA | NA | NA |

| | | | | | | | | |
|---|---|---|---|---|---|---|---|---|
| ENSG000002 | 0.89158985 | -0.117378003 | 0.29890796 | -0.3926895 | 0.69454886 | NA | NA | NA |
| ENSG000002 | 0.68351258 | -0.46544448 | 0.26656372 | -1.7460909 | 0.08079516 | NA | RNU1-64P | 106480168 |
| ENSG000002 | 1.49755516 | -0.191355342 | 0.34186808 | -0.5597345 | 0.57566057 | NA | RNU1-84P | 106481618 |
| ENSG000002 | 0.12685397 | -0.090215768 | 0.12901919 | -0.699243 | 0.48440017 | NA | NA | NA |
| ENSG000002 | 0.37441639 | -0.268856313 | 0.21277471 | -1.2635727 | 0.20638344 | NA | NA | NA |
| ENSG000002 | 0.22263036 | 0.146922267 | 0.16600367 | 0.88505432 | 0.37612734 | NA | RNU6-609P | 106481906 |
| ENSG000002 | 1.13747985 | -0.440959123 | 0.32354223 | -1.3629106 | 0.1729107 | NA | NA | NA |
| ENSG000002 | 1.36415071 | -0.125681144 | 0.33628771 | -0.373731 | 0.70860446 | NA | NA | NA |
| ENSG000002 | 0 | NA | NA | NA | NA | NA | NA | NA |
| ENSG000002 | 0.89355691 | -0.424018087 | 0.29292988 | -1.4475071 | 0.14775494 | NA | RNU6-359P | 106479725 |
| ENSG000002 | 0.50516651 | -0.183853222 | 0.24168957 | -0.7606999 | 0.44683635 | NA | RNU4-45P | 106481186 |
| ENSG000002 | 0.64030597 | -0.296505743 | 0.27391229 | -1.0824843 | 0.27903741 | NA | RNU6-1051P | 106481932 |
| ENSG000002 | 1.0081542 | -0.346299707 | 0.30931545 | -1.1195681 | 0.26289786 | NA | RN7SKP208 | 106481817 |
| ENSG000002 | 1.87708099 | -0.245856923 | 0.35273786 | -0.6969961 | 0.48580521 | NA | NA | NA |
| ENSG000002 | 0.6530863 | -0.285158107 | 0.26914708 | -1.059488 | 0.28937758 | NA | RNU6-326P | 106481282 |
| ENSG000002 | 1.25132728 | -0.356280157 | 0.33523239 | -1.0627856 | 0.2878792 | NA | RN7SKP204 | 106480896 |
| ENSG000002 | 1.83566708 | 0.259979717 | 0.35447667 | 0.7334184 | 0.46330328 | NA | RNY3 | 6085 |
| ENSG000002 | 0.84155395 | 0.212634111 | 0.30139838 | 0.70549188 | 0.48050418 | NA | NA | NA |
| ENSG000002 | 1.30949734 | 0.324994517 | 0.33859381 | 0.95983598 | 0.33713777 | NA | NA | NA |
| ENSG000002 | 0.71459391 | 0.123448103 | 0.27538578 | 0.44827334 | 0.65395594 | NA | RN7SKP249 | 106480910 |
| ENSG000002 | 0 | NA | NA | NA | NA | NA | NA | NA |
| ENSG000002 | 1.41071268 | -0.123426758 | 0.33087405 | -0.3730325 | 0.70912429 | NA | NA | NA |
| ENSG000002 | 0.61950267 | -0.112567707 | 0.27088102 | -0.4155614 | 0.67773094 | NA | SNORD115-43 | 100033817 |
| ENSG000002 | 0.13280363 | -0.093394593 | 0.13115644 | -0.7120855 | 0.47641185 | NA | NA | NA |
| ENSG000002 | 0.87761776 | 0.046456494 | 0.29623136 | 0.15682503 | 0.87538274 | NA | RNU1-55P | 106481954 |
| ENSG000002 | 1.11609958 | 0.032649418 | 0.32592019 | 0.10017611 | 0.92020451 | NA | NA | NA |
| ENSG000002 | 1.38441515 | -0.115467494 | 0.3344695 | -0.3452258 | 0.72992464 | NA | NA | NA |
| ENSG000002 | 0 | NA | NA | NA | NA | NA | NA | NA |
| ENSG000002 | 0.62459249 | 0.048506127 | 0.26461252 | 0.18331002 | 0.85455477 | NA | NA | NA |

| | | | | | | | | |
|---|---|---|---|---|---|---|---|---|
| ENSG000002 | 0.89725718 | -0.276865406 | 0.29875216 | -0.9267394 | 0.35406184 | NA | NA | NA |
| ENSG000002 | 0.26199343 | -0.194018184 | 0.18422801 | -1.0531416 | 0.29227609 | NA | NA | NA |
| ENSG000002 | 0.74772501 | -0.357780902 | 0.28233051 | -1.2672414 | 0.205069 | NA | RNU6-61P | 106480435 |
| ENSG000002 | 1.54331388 | 0.16019835 | 0.35021354 | 0.45743048 | 0.64736166 | NA | NA | NA |
| ENSG000002 | 0.65174827 | -0.268696557 | 0.26059936 | -1.0310714 | 0.30250733 | NA | NA | NA |
| ENSG000002 | 0 | NA | NA | NA | NA | NA | RN7SKP187 | 106480890 |
| ENSG000002 | 0.53608206 | -0.386623811 | 0.24788931 | -1.5596631 | 0.11883952 | NA | NA | NA |
| ENSG000002 | 0.51550551 | -0.37860525 | 0.24578489 | -1.5403927 | 0.12346466 | NA | NA | NA |
| ENSG000002 | 0.59755775 | 0.062883211 | 0.26225914 | 0.2397751 | 0.81050461 | NA | RNY3P13 | 106481714 |
| ENSG000002 | 1.50352251 | -0.347719516 | 0.34690448 | -1.0023495 | 0.31617485 | NA | NA | NA |
| ENSG000002 | 0.47621624 | 0.147284149 | 0.23752141 | 0.62008787 | 0.53519994 | NA | NA | NA |
| ENSG000002 | 0.35664995 | 0.244349217 | 0.20882949 | 1.17008962 | 0.2419649 | NA | NA | NA |
| ENSG000002 | 1.07537049 | 0.037947705 | 0.3143994 | 0.12069904 | 0.90392942 | NA | RNU6-827P | 106481442 |
| ENSG000002 | 0.47482614 | 0.153008257 | 0.24473789 | 0.62519236 | 0.53184482 | NA | RNU6-744P | 106479895 |
| ENSG000002 | 0.59072708 | 0.064826349 | 0.26164045 | 0.24776883 | 0.80431327 | NA | RNU4-48P | 106479580 |
| ENSG000002 | 1.70036388 | -0.188687846 | 0.35329177 | -0.534085 | 0.5932827 | NA | NA | NA |
| ENSG000002 | 1.17547613 | -0.438122439 | 0.32256215 | -1.3582574 | 0.17438201 | NA | RNU6-438P | 106479760 |
| ENSG000002 | 0.23139247 | 0.167748926 | 0.17680352 | 0.94878726 | 0.34272882 | NA | RNU6-54P | 106478986 |
| ENSG000002 | 0.513452 | -0.199963756 | 0.2496885 | -0.8008529 | 0.42321682 | NA | NA | NA |
| ENSG000002 | 0.23065526 | 0.167384503 | 0.17662711 | 0.94767164 | 0.34329664 | NA | NA | NA |
| ENSG000002 | 0.35733509 | 0.244646831 | 0.20894085 | 1.17089037 | 0.24164285 | NA | NA | NA |
| ENSG000002 | 0 | NA | NA | NA | NA | NA | NA | NA |
| ENSG000002 | 0 | NA | NA | NA | NA | NA | NA | NA |
| ENSG000002 | 1.1717502 | -0.410650333 | 0.31411072 | -1.3073426 | 0.19109637 | NA | NA | NA |
| ENSG000002 | 1.92327711 | 0.072484954 | 0.36378308 | 0.19925323 | 0.84206466 | NA | NA | NA |
| ENSG000002 | 0.72695258 | -0.317778071 | 0.270123 | -1.1764199 | 0.23942713 | NA | RNU4-17P | 106481177 |
| ENSG000002 | 0.52554719 | -0.191309836 | 0.24390646 | -0.7843574 | 0.43283043 | NA | NA | NA |
| ENSG000002 | 1.79907299 | 0.27248894 | 0.35860226 | 0.75986398 | 0.44733589 | NA | NA | NA |
| ENSG000002 | 0.99809753 | -0.038514945 | 0.31323918 | -0.122957 | 0.90214116 | NA | RNU4-20P | 106479567 |

| | | | | | | | | |
|---|---|---|---|---|---|---|---|---|
| ENSG000002 | 1.37800657 | -0.279862056 | 0.34319938 | -0.8154503 | 0.41481458 | NA | NA | NA |
| ENSG000002 | 1.26669139 | -0.355480447 | 0.33432958 | -1.0632635 | 0.28766246 | NA | RN7SKP81 | 106480858 |
| ENSG000002 | 1.02730682 | -0.526470818 | 0.31214921 | -1.6865999 | 0.09168032 | NA | NA | NA |
| ENSG000002 | 0 | NA | NA | NA | NA | NA | NA | NA |
| ENSG000002 | 0.34197044 | 0.237837536 | 0.20637034 | 1.15247924 | 0.2491242 | NA | RNU6-81P | 100873780 |
| ENSG000002 | 2.02374319 | 0.127808917 | 0.36146352 | 0.35358732 | 0.72364818 | NA | NA | NA |
| ENSG000002 | 0.87321062 | -0.269491476 | 0.29754442 | -0.9057185 | 0.36508489 | NA | RNU6-499P | 106479787 |
| ENSG000002 | 1.3293734 | 0.022435368 | 0.32954706 | 0.06807941 | 0.94572242 | NA | RNU6-597P | 106481905 |
| ENSG000002 | 1.08888611 | -0.106844667 | 0.3135368 | -0.3407723 | 0.73327498 | NA | RNU6-716P | 106479883 |
| ENSG000002 | 0.60338479 | 0.225396395 | 0.26390915 | 0.85406814 | 0.39306724 | NA | NA | NA |
| ENSG000002 | 0.50565578 | -0.183478728 | 0.24174047 | -0.7589905 | 0.44785822 | NA | NA | NA |
| ENSG000002 | 0 | NA | NA | NA | NA | NA | SNORD34 | 26817 |
| ENSG000002 | 0.48962772 | -0.021045505 | 0.24589998 | -0.0855856 | 0.93179582 | NA | NA | NA |
| ENSG000002 | 1.88678691 | -0.377786857 | 0.35411726 | -1.0668411 | 0.28604359 | NA | RNU6-805P | 106479922 |
| ENSG000002 | 1.32744294 | -0.185440714 | 0.21974547 | -0.8438887 | 0.39873162 | NA | NA | NA |
| ENSG000002 | 0 | NA | NA | NA | NA | NA | RNA5S7 | 100169758 |
| ENSG000002 | 1.10119429 | -0.406058575 | 0.31585998 | -1.2855651 | 0.19859487 | NA | NA | NA |
| ENSG000002 | 0 | NA | NA | NA | NA | NA | RNA5S13 | 100169764 |
| ENSG000002 | 0 | NA | NA | NA | NA | NA | SNORD18B | 595099 |
| ENSG000002 | 0.26199343 | -0.194018184 | 0.18422801 | -1.0531416 | 0.29227609 | NA | RNU6-395P | 106479740 |
| ENSG000002 | 1.01660118 | -0.506439653 | 0.30700104 | -1.649635 | 0.09901762 | NA | NA | NA |
| ENSG000002 | 1.3076789 | 0.296183428 | 0.32290171 | 0.91725567 | 0.35900868 | NA | RNU6-1329P | 106481600 |
| ENSG000002 | 0.12070844 | -0.088503695 | 0.12783931 | -0.6923042 | 0.48874631 | NA | NA | NA |
| ENSG000002 | 0.63621395 | -0.448723671 | 0.2628158 | -1.7073694 | 0.08775341 | NA | NA | NA |
| ENSG000002 | 1.3437166 | -0.250597488 | 0.33430294 | -0.749612 | 0.45348844 | NA | NA | NA |
| ENSG000002 | 1.79236064 | -0.61038773 | 0.35318892 | -1.7282188 | 0.08394901 | NA | NA | NA |
| ENSG000002 | 0.52554719 | -0.191309836 | 0.24390646 | -0.7843574 | 0.43283043 | NA | NA | NA |
| ENSG000002 | 0.97729073 | -0.181250952 | 0.30823001 | -0.588038 | 0.55650681 | NA | NA | NA |
| ENSG000002 | 1.2426757 | -0.465418585 | 0.32415015 | -1.4358117 | 0.15105592 | NA | NA | NA |

| | | | | | | | | | |
|---|---|---|---|---|---|---|---|---|---|
| ENSG000002 | 1.92914922 | -0.837220926 | 0.35844955 | -2.335673 | 0.01950829 | NA | NA | NA | |
| ENSG000002 | 1.83675368 | 0.009559668 | 0.35405481 | 0.02700053 | 0.97845931 | NA | NA | NA | |
| ENSG000002 | 1.73124516 | 0.070901248 | 0.35497014 | 0.19973862 | 0.841685 | NA | NA | NA | |
| ENSG000002 | 2.16372628 | -0.640133521 | 0.36430427 | -1.7571398 | 0.07889399 | NA | NA | NA | |
| ENSG000002 | 1.74899972 | -0.282039291 | 0.34368631 | -0.82063 | 0.41185706 | NA | NA | NA | |
| ENSG000002 | 1.72424215 | -0.183901692 | 0.35086348 | -0.5241403 | 0.60018096 | NA | NA | NA | |
| ENSG000002 | 0.47212421 | -0.012721785 | 0.23625154 | -0.0538485 | 0.95705589 | NA | NA | NA | |
| ENSG000002 | 1.34687788 | -0.521593387 | 0.32914965 | -1.5846694 | 0.11304146 | NA | NA | NA | |
| ENSG000002 | 0.63621395 | -0.448723671 | 0.2628158 | -1.7073694 | 0.08775341 | NA | NA | NA | |
| ENSG000002 | 1.62326698 | -0.267301552 | 0.34966857 | -0.7644426 | 0.44460353 | NA | NA | NA | |
| ENSG000002 | 2.07242066 | -0.560465592 | 0.35605525 | -1.5740973 | 0.11546496 | NA | NA | NA | |
| ENSG000002 | 1.11484979 | -0.269878318 | 0.32162998 | -0.8390957 | 0.40141563 | NA | NA | NA | |
| ENSG000002 | 1.88499646 | -0.124596258 | 0.35472445 | -0.351248 | 0.72540229 | NA | NA | NA | |
| ENSG000002 | 1.92724866 | -0.966292918 | 0.35400742 | -2.7295838 | 0.00634143 | NA | NA | NA | |
| ENSG000002 | 1.71872989 | -0.175007166 | 0.34826215 | -0.5025156 | 0.61530487 | NA | NA | NA | |
| ENSG000002 | 0.93611275 | -0.437104539 | 0.29505376 | -1.4814403 | 0.13848927 | NA | NA | NA | |
| ENSG000002 | 1.08818328 | 0.036463773 | 0.32504328 | 0.11218128 | 0.91067967 | NA | NA | NA | |
| ENSG000002 | 1.90281889 | -0.811991578 | 0.35607864 | -2.2803715 | 0.02258567 | NA | NA | NA | |
| ENSG000002 | 0.6181343 | 0.056012223 | 0.26406445 | 0.21211573 | 0.83201675 | NA | NA | NA | |
| ENSG000002 | 1.84486681 | -0.239665451 | 0.35731184 | -0.6707459 | 0.50238242 | NA | NA | NA | |
| ENSG000002 | 0 | NA | NA | NA | NA | NA | GP1BB | | 2812 |
| ENSG000002 | 0.12070844 | -0.088503695 | 0.12783931 | -0.6923042 | 0.48874631 | NA | OR2T5 | | 401993 |
| ENSG000002 | 0.77018092 | -0.359515013 | 0.28385235 | -1.2665564 | 0.20531394 | NA | NA | NA | |
| ENSG000002 | 0.49196355 | -0.021608171 | 0.24613638 | -0.0877894 | 0.93004404 | NA | LOR | | 4014 |
| ENSG000002 | 2.16054169 | -0.6481677 | 0.36486472 | -1.7764603 | 0.07565708 | NA | LELP1 | | 149018 |
| ENSG000002 | 0 | NA | NA | NA | NA | NA | NA | NA | |
| ENSG000002 | 0.49307133 | -0.179259067 | 0.24031383 | -0.7459374 | 0.45570524 | NA | SOX18 | | 54345 |
| ENSG000002 | 1.74334482 | -0.196836095 | 0.35849574 | -0.5490612 | 0.58296346 | NA | SPATA42 | | 642864 |
| ENSG000002 | 1.97624946 | -0.183261204 | 0.36073329 | -0.5080241 | 0.61143644 | NA | SPANXN1 | | 494118 |

| | | | | | | | | | |
|---|---|---|---|---|---|---|---|---|---|
| ENSG000002 | 0.39246123 | -0.276864021 | 0.21549122 | -1.2848042 | 0.19886073 | NA | SPANXA2 | | 728712 |
| ENSG000002 | 0 | NA | NA | NA | NA | NA | NA | NA | |
| ENSG000002 | 0.77925514 | -0.363111002 | 0.28443192 | -1.2766183 | 0.20173703 | NA | NA | NA | |
| ENSG000002 | 1.24375727 | -0.485606071 | 0.3264512 | -1.4875304 | 0.13687479 | NA | NA | NA | |
| ENSG000002 | 1.2705658 | -0.361363611 | 0.33439162 | -1.0806599 | 0.27984844 | NA | NA | NA | |
| ENSG000002 | 1.32113999 | 0.02693777 | 0.33977886 | 0.0792803 | 0.93680967 | NA | NA | NA | |
| ENSG000002 | 0.87898613 | -0.113387039 | 0.30333236 | -0.3738046 | 0.70854968 | NA | NA | NA | |
| ENSG000002 | 0 | NA | NA | NA | NA | NA | NA | NA | |
| ENSG000002 | 1.42101784 | -0.398587945 | 0.3341756 | -1.19275 | 0.2329673 | NA | CYP21A1P | | 1590 |
| ENSG000002 | 0 | NA | NA | NA | NA | NA | NA | NA | |
| ENSG000002 | 0.24756242 | -0.187034346 | 0.18119817 | -1.0322088 | 0.30197431 | NA | NA | NA | |
| ENSG000002 | 0 | NA | NA | NA | NA | NA | NA | NA | |
| ENSG000002 | 1.0870634 | 0.194805741 | 0.31976166 | 0.60922169 | 0.5423775 | NA | LY6G6F | | 259215 |
| ENSG000002 | 2.00911516 | -0.309773209 | 0.35878055 | -0.8634058 | 0.38791439 | NA | NA | NA | |
| ENSG000002 | 0.38064837 | -0.102706251 | 0.21886737 | -0.4692625 | 0.63888201 | NA | TRIM64 | | 120146 |
| ENSG000002 | 0.25260017 | -0.015978705 | 0.18049464 | -0.0885273 | 0.92945758 | NA | NA | NA | |
| ENSG000002 | 0.60968328 | -0.108157951 | 0.26370283 | -0.4101509 | 0.68169526 | NA | PRAMEF7 | | 441871 |
| ENSG000002 | 1.54720786 | -0.328995455 | 0.34139737 | -0.9636731 | 0.33520986 | NA | MCCD1 | | 401250 |
| ENSG000002 | 1.41982362 | -0.417025197 | 0.33776472 | -1.2346618 | 0.21695639 | NA | CFTRP3 | | 106481718 |
| ENSG000002 | 0.13280363 | -0.093394593 | 0.13115644 | -0.7120855 | 0.47641185 | NA | NA | NA | |
| ENSG000002 | 2.12400179 | -0.491238729 | 0.3613362 | -1.359506 | 0.17398631 | NA | KRTAP5-11 | | 440051 |
| ENSG000002 | 1.78293568 | -0.451787083 | 0.35081577 | -1.2878186 | 0.1978091 | NA | KRTAP5-10 | | 387273 |
| ENSG000002 | 1.64438541 | -0.266747717 | 0.35111001 | -0.7597269 | 0.44741785 | NA | CST9LP2 | | 106480788 |
| ENSG000002 | 1.80312946 | -0.410941506 | 0.33659716 | -1.2208704 | 0.22213509 | NA | NA | NA | |
| ENSG000002 | 0.53627793 | -0.360851627 | 0.24080453 | -1.4985251 | 0.13399688 | NA | NA | NA | |
| ENSG000002 | 0 | NA | NA | NA | NA | NA | NA | NA | |
| ENSG000002 | 0 | NA | NA | NA | NA | NA | NA | NA | |
| ENSG000002 | 0.3854038 | -0.110128691 | 0.21952326 | -0.5016721 | 0.61589821 | NA | NA | NA | |
| ENSG000002 | 0.97655352 | -0.181520872 | 0.30819736 | -0.5889761 | 0.55587733 | NA | FOXD4L5 | | 653427 |

| | | | | | | | | |
|---|---|---|---|---|---|---|---|---|
| ENSG000002 | 0 | NA | NA | NA | NA | NA | NA | NA |
| ENSG000002 | 0 | NA | NA | NA | NA | NA | NA | NA |
| ENSG000002 | 0.24485602 | -0.012655878 | 0.17881722 | -0.0707755 | 0.94357643 | NA | NA | NA |
| ENSG000002 | 0 | NA | NA | NA | NA | NA | LOC100240735 | 100240735 |
| ENSG000002 | 0.11131518 | 0.081951103 | 0.12663804 | 0.64712866 | 0.51754868 | NA | NA | NA |
| ENSG000002 | 0.24004852 | -0.005223787 | 0.17775844 | -0.029387 | 0.97655594 | NA | NA | NA |
| ENSG000002 | 2.00061206 | -0.449159544 | 0.36065854 | -1.2453872 | 0.21298945 | NA | NAT8B | 51471 |
| ENSG000002 | 1.5213668 | -0.322265821 | 0.34096263 | -0.9451646 | 0.34457483 | NA | KRTAP9-3 | 83900 |
| ENSG000002 | 1.43718724 | -0.748210531 | 0.33724444 | -2.2186001 | 0.02651395 | NA | KRTAP4-8 | 728224 |
| ENSG000002 | 0.36576034 | -0.0934779 | 0.20820828 | -0.4489634 | 0.65345805 | NA | KRTAP1-4 | 728255 |
| ENSG000002 | 0.14128499 | -0.097772014 | 0.13403235 | -0.7294658 | 0.4657168 | NA | PRR20B | 729233 |
| ENSG000002 | 0 | NA | NA | NA | NA | NA | PRR20A | 122183 |
| ENSG000002 | 1.0755949 | 0.336371533 | 0.31578919 | 1.06517746 | 0.28679563 | NA | PRSS3P4 | 106480774 |
| ENSG000002 | 1.05041507 | -0.532808254 | 0.31301235 | -1.7021956 | 0.0887187 | NA | NA | NA |
| ENSG000002 | 1.48926968 | -0.189354006 | 0.34172182 | -0.5541174 | 0.5794985 | NA | NA | NA |
| ENSG000002 | 2.12063782 | -0.371485153 | 0.36437504 | -1.0195132 | 0.3079594 | NA | OR5D16 | 390144 |
| ENSG000002 | 0.64602887 | -0.11525331 | 0.26675303 | -0.43206 | 0.66569782 | NA | OR5L2 | 26338 |
| ENSG000002 | 0 | NA | NA | NA | NA | NA | LGALS7 | 3963 |
| ENSG000002 | 1.09083313 | 0.159369539 | 0.30718598 | 0.51880473 | 0.60389692 | NA | DKFZp779M065 | 374387 |
| ENSG000002 | 1.52974053 | -0.607551053 | 0.33948964 | -1.7896012 | 0.07351805 | NA | C11orf91 | 100131378 |
| ENSG000002 | 0.11131518 | 0.081951103 | 0.12663804 | 0.64712866 | 0.51754868 | NA | NA | NA |
| ENSG000002 | 1.53909924 | 0.15335665 | 0.34711184 | 0.4418076 | 0.65862844 | NA | NA | NA |
| ENSG000002 | 0 | NA | NA | NA | NA | NA | SAP25 | 100316904 |
| ENSG000002 | 1.01976012 | -0.369210986 | 0.31570354 | -1.1694864 | 0.24220773 | NA | OR6C68 | 403284 |
| ENSG000002 | 1.03558316 | -0.074533892 | 0.20908263 | -0.3564806 | 0.72148071 | NA | NA | NA |
| ENSG000002 | 1.18583686 | -0.389876056 | 0.31067936 | -1.2549146 | 0.20950978 | NA | NA | NA |
| ENSG000002 | 0.9818194 | -0.042650985 | 0.31258931 | -0.1364442 | 0.89147017 | NA | NA | NA |
| ENSG000002 | 1.510332 | -0.492966009 | 0.34602836 | -1.4246405 | 0.15426115 | NA | KRTAP12-3 | 386683 |
| ENSG000002 | 1.72366911 | -0.167878591 | 0.34832204 | -0.4819637 | 0.62983171 | NA | NA | NA |

| | | | | | | | | | |
|---|---|---|---|---|---|---|---|---|---|
| ENSG000002 | 1.87070733 | -0.267239613 | 0.36188282 | -0.73847 | 0.46022888 | NA | KRTAP10-2 | | 386679 |
| ENSG000002 | 0 | NA | NA | NA | NA | NA | NA | NA | |
| ENSG000002 | 1.42195255 | 0.226558042 | 0.34025702 | 0.66584384 | 0.50551092 | NA | OR52J3 | | 119679 |
| ENSG000002 | 1.76323568 | -0.199916223 | 0.35397198 | -0.5647798 | 0.57222355 | NA | OR51A2 | | 401667 |
| ENSG000002 | 1.56664944 | 0.026759104 | 0.34935903 | 0.07659485 | 0.93894585 | NA | C2CD4B | | 388125 |
| ENSG000002 | 1.78920225 | -0.750392745 | 0.3522043 | -2.130561 | 0.03312533 | NA | NA | NA | |
| ENSG000002 | 167.053987 | 0.121860043 | 0.30974775 | 0.39341705 | NA | NA | EIF3CL | | 728689 |
| ENSG000002 | 1.97738757 | -0.444287565 | 0.36145947 | -1.229149 | 0.21901593 | NA | NA | NA | |
| ENSG000002 | 1.21668581 | -0.191952121 | 0.32868203 | -0.5840055 | 0.5592166 | NA | NA | NA | |
| ENSG000002 | 0.23202362 | -0.008302493 | 0.17586207 | -0.0472103 | 0.96234565 | NA | NA | NA | |
| ENSG000002 | 1.362186 | -0.255854817 | 0.33692645 | -0.7593788 | 0.44762597 | NA | NA | NA | |
| ENSG000002 | 0.23202362 | -0.008302493 | 0.17586207 | -0.0472103 | 0.96234565 | NA | MBD3L4 | | 653656 |
| ENSG000002 | 2.08048198 | -0.377219243 | 0.36468415 | -1.0343725 | 0.30096207 | NA | NA | NA | |
| ENSG000002 | 1.06643424 | 0.194797787 | 0.3205561 | 0.60768704 | 0.54339506 | NA | NA | NA | |
| ENSG000002 | 1.41800079 | -0.735574162 | 0.33637908 | -2.1867417 | 0.02876138 | NA | KLRC3 | | 3823 |
| ENSG000002 | 0.49031286 | -0.020802257 | 0.24597635 | -0.0845702 | 0.93260313 | NA | KRTAP5-6 | | 440023 |
| ENSG000002 | 1.18471861 | 0.422394406 | 0.32970366 | 1.28113353 | 0.20014677 | NA | KRTAP5-2 | | 440021 |
| ENSG000002 | 1.38597748 | -0.414937428 | 0.33843986 | -1.22603 | 0.22018738 | NA | KRTAP5-1 | | 387264 |
| ENSG000002 | 1.33098524 | 0.153362082 | 0.33399846 | 0.45917003 | 0.64611207 | NA | DEFB135 | | 613209 |
| ENSG000002 | 0.73178182 | -0.190358911 | 0.28514777 | -0.6675799 | 0.50440181 | NA | DEFB136 | | 613210 |
| ENSG000002 | 0.40942396 | -0.284058841 | 0.21788566 | -1.303706 | 0.19233384 | NA | BHLHA9 | | 727857 |
| ENSG000002 | 0 | NA | NA | NA | NA | NA | DAZ4 | | 57135 |
| ENSG000002 | 0 | NA | NA | NA | NA | NA | NA | NA | |
| ENSG000002 | 0 | NA | NA | NA | NA | NA | NA | NA | |
| ENSG000002 | 0.49076933 | -0.025124039 | 0.23846109 | -0.1053591 | 0.91609089 | NA | USP17L6P | | 391622 |
| ENSG000002 | 1.65146778 | 0.498315835 | 0.35226298 | 1.41461313 | 0.15718196 | NA | NA | NA | |
| ENSG000002 | 0.12685397 | -0.090215768 | 0.12901919 | -0.699243 | 0.48440017 | NA | DEFB109P1B | | 641517 |
| ENSG000002 | 0.75367466 | -0.377752651 | 0.28818257 | -1.3108102 | 0.1899219 | NA | IFITM5 | | 387733 |
| ENSG000002 | 1.17157998 | -0.576043938 | 0.31605524 | -1.8226052 | 0.0683632 | NA | DEFB109P1B | | 641517 |

| | | | | | | | | |
|---|---|---|---|---|---|---|---|---|
| ENSG000002 | 1.00260152 | -0.321579681 | 0.3011477 | -1.067847 | 0.28558952 | NA | NA | NA |
| ENSG000002 | 1.51268956 | -0.345060282 | 0.34821429 | -0.9909423 | 0.32171374 | NA | NA | NA |
| ENSG000002 | 1.72079909 | -0.053716511 | 0.35383022 | -0.1518144 | 0.87933335 | NA | NA | NA |
| ENSG000002 | 2.07521295 | -0.589068246 | 0.36077505 | -1.6327854 | 0.10251413 | NA | NA | NA |
| ENSG000002 | 1.29086002 | -0.494105352 | 0.32693305 | -1.511335 | 0.13070313 | NA | KRTAP19-8 | 728299 |
| ENSG000002 | 2.11291699 | -0.123241424 | 0.36491116 | -0.3377299 | 0.73556671 | NA | KRTAP20-3 | 337985 |
| ENSG000002 | 0.93092421 | 0.453634956 | 0.30756779 | 1.47491049 | 0.14023661 | NA | KRTAP20-4 | 100151643 |
| ENSG000002 | 1.49321844 | -0.048396607 | 0.34453173 | -0.1404707 | 0.88828811 | NA | KRTAP22-2 | 100288287 |
| ENSG000002 | 1.98751919 | 0.058088481 | 0.36146847 | 0.16070138 | 0.8723286 | NA | KRTAP27-1 | 643812 |
| ENSG000002 | 0 | NA | NA | NA | NA | NA | GYG2P1 | 352887 |
| ENSG000002 | 1.26347454 | -0.344014384 | 0.33087791 | -1.0397019 | 0.29847843 | NA | NA | NA |
| ENSG000002 | 1.49851968 | -0.301037468 | 0.33679293 | -0.8938355 | 0.37140992 | NA | HBA1 | 3039 |
| ENSG000002 | 0.88855353 | -0.453244102 | 0.30180501 | -1.5017779 | 0.13315447 | NA | HBM | 3042 |
| ENSG000002 | 0.23065526 | 0.167384503 | 0.17662711 | 0.94767164 | 0.34329664 | NA | NA | NA |
| ENSG000002 | 0.60375535 | 0.060734453 | 0.26281561 | 0.23109149 | 0.81724372 | NA | PRR23A | 729627 |
| ENSG000002 | 0.87238359 | -0.115669067 | 0.30300345 | -0.3817417 | 0.70265293 | NA | NA | NA |
| ENSG000002 | 1.90412019 | -0.489058843 | 0.35182603 | -1.3900587 | 0.16451105 | NA | LRRC30 | 339291 |
| ENSG000002 | 0 | NA | NA | NA | NA | NA | NA | NA |
| ENSG000002 | 2.14610832 | -0.260966445 | 0.36611155 | -0.7128058 | 0.47596592 | NA | OR5K3 | 403277 |
| ENSG000002 | 0.71786125 | -0.023725508 | 0.27701985 | -0.0856455 | 0.93174822 | NA | NA | NA |
| ENSG000002 | 0.76747453 | -0.201328414 | 0.28738472 | -0.7005536 | 0.48358162 | NA | RNU6-1292P | 106480656 |
| ENSG000002 | 2.09156897 | 0.250364448 | 0.36472854 | 0.68644051 | 0.49243536 | NA | RNU6-46P | 100873760 |
| ENSG000002 | 0.47753253 | -0.016714525 | 0.24454657 | -0.068349 | 0.94550778 | NA | RNU6-132P | 106481219 |
| ENSG000002 | 1.40272101 | -0.730849657 | 0.33526437 | -2.1799204 | 0.02926336 | NA | NA | NA |
| ENSG000002 | 0.73659282 | 0.11903157 | 0.27861905 | 0.42721979 | 0.66921925 | NA | RNU6-47P | 106480710 |
| ENSG000002 | 0.47753253 | -0.016714525 | 0.24454657 | -0.068349 | 0.94550778 | NA | RNU6-877P | 106480617 |
| ENSG000002 | 0 | NA | NA | NA | NA | NA | SNORA57 | 692158 |
| ENSG000002 | 1.27853863 | -0.347525264 | 0.33125322 | -1.0491227 | 0.29412165 | NA | RNU6-1286P | 106481586 |
| ENSG000002 | 0.52876404 | -0.191903645 | 0.2442426 | -0.7857092 | 0.4320379 | NA | RNU6-841P | 106480699 |

| | | | | | | | | |
|---|---|---|---|---|---|---|---|---|
| ENSG000002 | 0 | NA | NA | NA | NA | NA | SNORD58A | 26791 |
| ENSG000002 | 0.47735839 | 0.152206097 | 0.23764319 | 0.64048164 | 0.52185952 | NA | RNU6-425P | 106481312 |
| ENSG000002 | 2.13782339 | -0.247072629 | 0.36260478 | -0.6813827 | 0.49562936 | NA | RNU6-946P | 106481478 |
| ENSG000002 | 0.38614101 | -0.1098476 | 0.21963552 | -0.5001359 | 0.61697942 | NA | RNU6-1296P | 106480132 |
| ENSG000002 | 0 | NA | NA | NA | NA | NA | SNORD24 | 26820 |
| ENSG000002 | 0.59090122 | -0.095071922 | 0.25441426 | -0.3736894 | 0.70863539 | NA | RNU6-1336P | 106480661 |
| ENSG000002 | 0.3867939 | -0.105148852 | 0.21977268 | -0.4784437 | 0.63233444 | NA | RNU6-477P | 106481330 |
| ENSG000002 | 1.45425819 | 0.084221553 | 0.34198209 | 0.24627474 | 0.80546956 | NA | RNU6-338P | 106481286 |
| ENSG000002 | 0.57953878 | 0.207961418 | 0.25452889 | 0.81704446 | 0.41390302 | NA | RNU6-741P | 106479894 |
| ENSG000002 | 0.61818637 | 0.059474041 | 0.27041681 | 0.21993471 | 0.82592201 | NA | NA | NA |
| ENSG000002 | 0.4009426 | -0.302672773 | 0.2240145 | -1.3511303 | 0.17665371 | NA | RNU6-1264P | 106480117 |
| ENSG000002 | 0 | NA | NA | NA | NA | NA | SNORD45C | 692085 |
| ENSG000002 | 0.26560726 | -0.177019426 | 0.17672274 | -1.0016788 | 0.31649875 | NA | SNORA69 | 26779 |
| ENSG000002 | 0 | NA | NA | NA | NA | NA | RNU1-39P | 106481953 |
| ENSG000002 | 1.13639827 | -0.415231483 | 0.31952479 | -1.2995282 | 0.19376272 | NA | RNU6-969P | 106479994 |
| ENSG000002 | 1.28363237 | -0.492074457 | 0.32674128 | -1.5060064 | 0.13206553 | NA | RNU1-63P | 106480664 |
| ENSG000002 | 0 | NA | NA | NA | NA | NA | SNORD60 | 26788 |
| ENSG000002 | 0.74593051 | -0.356732965 | 0.28222667 | -1.2639945 | 0.206232 | NA | RNU6-657P | 106481389 |
| ENSG000002 | 1.76293217 | -0.47981441 | 0.35625349 | -1.3468343 | 0.17803362 | NA | RNU6-1062P | 106480634 |
| ENSG000002 | 0.63182461 | -0.278261844 | 0.26744298 | -1.040453 | 0.29812951 | NA | NA | NA |
| ENSG000002 | 0.23936338 | -0.005479057 | 0.17759977 | -0.0308506 | 0.9753887 | NA | NA | NA |
| ENSG000002 | 1.89954605 | -0.518039147 | 0.35593966 | -1.4554128 | 0.14555501 | NA | RNU6-672P | 106481910 |
| ENSG000002 | 0 | NA | NA | NA | NA | NA | NA | NA |
| ENSG000002 | 0.49970612 | -0.194507174 | 0.24827631 | -0.7834303 | 0.43337449 | NA | NA | NA |
| ENSG000002 | 0.51413714 | -0.199727707 | 0.24976394 | -0.7996659 | 0.42390439 | NA | RNU6-449P | 106481896 |
| ENSG000002 | 0.76698526 | -0.191380706 | 0.28135307 | -0.6802154 | 0.49636805 | NA | RNU6-1279P | 106480655 |
| ENSG000002 | 0.26062507 | -0.012888319 | 0.18226503 | -0.070712 | 0.94362699 | NA | NA | NA |
| ENSG000002 | 0.35961481 | -0.099625193 | 0.21567743 | -0.4619176 | 0.64414043 | NA | NA | NA |
| ENSG000002 | 1.99554005 | -0.200222558 | 0.36455545 | -0.5492239 | 0.58285182 | NA | NA | NA |

| | | | | | | | | | |
|---|---|---|---|---|---|---|---|---|---|
| ENSG000002 | 1.00142903 | -0.185902062 | 0.30759884 | -0.6043653 | 0.5456008 | NA | | SNORA70G | 100379132 |
| ENSG000002 | 1.51686983 | -0.405450365 | 0.32780755 | -1.2368549 | 0.21614096 | NA | | NA | NA |
| ENSG000002 | 0.89820738 | -0.590908788 | 0.29086846 | -2.0315327 | 0.04220098 | NA | | RNU6-608P | 106481373 |
| ENSG000002 | 1.89539144 | -0.534414373 | 0.36004626 | -1.4842936 | 0.13773104 | NA | | RNU6-1039P | 106481931 |
| ENSG000002 | 1.33369164 | 0.017533882 | 0.33535292 | 0.05228486 | 0.95830171 | NA | | NA | NA |
| ENSG000002 | 0.13280363 | -0.093394593 | 0.13115644 | -0.7120855 | 0.47641185 | NA | | NA | NA |
| ENSG000002 | 0.42385498 | -0.262217565 | 0.21037102 | -1.2464529 | 0.21259817 | NA | | NA | NA |
| ENSG000002 | 1.20954461 | -0.036847327 | 0.32796246 | -0.1123523 | 0.9105441 | NA | | NA | NA |
| ENSG000002 | 0.74316153 | -0.023172289 | 0.27877499 | -0.0831218 | 0.93375466 | NA | | RNU6-573P | 106480591 |
| ENSG000002 | 1.01036528 | -0.325356495 | 0.30231847 | -1.0762045 | 0.28183581 | NA | | NA | NA |
| ENSG000002 | 1.47067663 | -0.187977133 | 0.34578481 | -0.5436246 | 0.58669982 | NA | | RNU6-471P | 106479774 |
| ENSG000002 | 0.12070844 | -0.088503695 | 0.12783931 | -0.6923042 | 0.48874631 | NA | | NA | NA |
| ENSG000002 | 0.4869258 | -0.176879301 | 0.23959723 | -0.738236 | 0.46037105 | NA | | RNU6-945P | 106479984 |
| ENSG000002 | 1.09831375 | -0.11120947 | 0.32027537 | -0.3472308 | 0.72841794 | NA | | NA | NA |
| ENSG000002 | 0.24619405 | -0.007630671 | 0.17915639 | -0.0425922 | 0.96602658 | NA | | NA | NA |
| ENSG000002 | 1.38581832 | -0.694644516 | 0.33090848 | -2.0992043 | 0.03579889 | NA | | RNU6-144P | 106481222 |
| ENSG000002 | 1.27967674 | -0.667342165 | 0.32869822 | -2.0302579 | 0.04233033 | NA | | NA | NA |
| ENSG000002 | 0 | NA | NA | NA | NA | NA | | SNORD21 | 6083 |
| ENSG000002 | 1.47373186 | -0.57845782 | 0.33465496 | -1.7285201 | 0.08389502 | NA | | RNU6-730P | 106479889 |
| ENSG000002 | 0.50702415 | -0.349150675 | 0.23755647 | -1.4697586 | 0.14162714 | NA | | RNU6-157P | 106479636 |
| ENSG000002 | 0.63279153 | -0.115881167 | 0.26564848 | -0.43622 | 0.66267715 | NA | | RNU6-1189P | 106480646 |
| ENSG000002 | 0.26813897 | -0.197007041 | 0.18550319 | -1.0620143 | 0.2882292 | NA | | RNU6-1036P | 106480021 |
| ENSG000002 | 1.91550863 | -0.131995061 | 0.35385464 | -0.3730206 | 0.70913313 | NA | | NA | NA |
| ENSG000002 | 1.37435657 | 0.022023965 | 0.33398162 | 0.06594364 | 0.94742269 | NA | | NA | NA |
| ENSG000002 | 0 | NA | NA | NA | NA | NA | | SNORA56 | 677835 |
| ENSG000002 | 1.16494126 | -0.268874891 | 0.31884064 | -0.8432893 | 0.39906669 | NA | | RNU6-442P | 106479761 |
| ENSG000002 | 0.49671898 | -0.027206884 | 0.23914704 | -0.1137663 | 0.90942302 | NA | | NA | NA |
| ENSG000002 | 0.3588776 | -0.099902505 | 0.21555794 | -0.4634601 | 0.6430346 | NA | | NA | NA |
| ENSG000002 | 0.62339882 | 0.057228974 | 0.27083985 | 0.2113019 | 0.8326517 | NA | | RNU6-723P | 106479886 |

| | | | | | | | | |
|---|---|---|---|---|---|---|---|---|
| ENSG000002 | 0.70072597 | 0.318240822 | 0.28328746 | 1.1233848 | 0.2612741 | NA | RNU6-1040P | 106480023 |
| ENSG000002 | 1.11331121 | -0.264197415 | 0.32159812 | -0.8215142 | 0.41135346 | NA | NA | NA |
| ENSG000002 | 0.11865494 | 0.083440866 | 0.12751478 | 0.65436232 | 0.5128784 | NA | RNU6-128P | 106481217 |
| ENSG000002 | 1.27153326 | -0.204836522 | 0.33369673 | -0.6138404 | 0.53932082 | NA | NA | NA |
| ENSG000002 | 0 | NA | NA | NA | NA | NA | NA | NA |
| ENSG000002 | 0.74603815 | -0.033182053 | 0.27895729 | -0.1189503 | 0.90531473 | NA | RNU6-1227P | 106481567 |
| ENSG000002 | 1.45659402 | 0.088352743 | 0.33916362 | 0.26050183 | 0.79447671 | NA | RNU6-1080P | 106481520 |
| ENSG000002 | 0.4933859 | -0.021085811 | 0.24629532 | -0.0856119 | 0.93177494 | NA | RNU6-147P | 106480556 |
| ENSG000002 | 1.22692604 | 0.10732395 | 0.32996694 | 0.32525667 | 0.74498682 | NA | NA | NA |
| ENSG000002 | 0.36212533 | -0.235926492 | 0.20089643 | -1.1743688 | 0.24024735 | NA | NA | NA |
| ENSG000002 | 1.73913018 | -0.190327084 | 0.35436295 | -0.5370965 | 0.59120099 | NA | NA | NA |
| ENSG000002 | 1.09042873 | 0.176989402 | 0.31531523 | 0.5613094 | 0.57458663 | NA | RNU6-133P | 106481877 |
| ENSG000002 | 0.98506768 | -0.166200817 | 0.30110626 | -0.5519673 | 0.58097074 | NA | NA | NA |
| ENSG000002 | 0.38056192 | -0.244105065 | 0.20391883 | -1.1970698 | 0.23127937 | NA | RNU6-546P | 106479809 |
| ENSG000002 | 2.1332738 | -0.383883881 | 0.36493422 | -1.0519263 | 0.29283339 | NA | NA | NA |
| ENSG000002 | 1.10758596 | 0.036273691 | 0.31795669 | 0.11408375 | 0.90917141 | NA | RNU6-1074P | 106480635 |
| ENSG000002 | 1.95261914 | -0.171767391 | 0.35518817 | -0.4835955 | 0.62867299 | NA | RNU6-1056P | 106481514 |
| ENSG000002 | 1.00015104 | -0.196087072 | 0.31390581 | -0.6246685 | 0.53218865 | NA | RNU6-756P | 106479900 |
| ENSG000002 | 0.11934008 | 0.083564698 | 0.12760091 | 0.65489106 | 0.5125379 | NA | RNU6-1027P | 106481505 |
| ENSG000002 | 0.51433302 | -0.186673425 | 0.24269587 | -0.7691661 | 0.44179474 | NA | RNU6-259P | 106481260 |
| ENSG000002 | 1.8681772 | -0.129663327 | 0.35709863 | -0.3631023 | 0.71652848 | NA | NA | NA |
| ENSG000002 | 0.35870346 | 0.077368606 | 0.21515516 | 0.35959447 | 0.71915042 | NA | RNU6-1126P | 106481535 |
| ENSG000002 | 0.81614816 | 0.049257558 | 0.28656509 | 0.1718896 | 0.86352432 | NA | NA | NA |
| ENSG000002 | 0.65524745 | -0.109156678 | 0.2585811 | -0.4221371 | 0.67292494 | NA | NA | NA |
| ENSG000002 | 0.11131518 | 0.081951103 | 0.12663804 | 0.64712866 | 0.51754868 | NA | RNU6-641P | 106481908 |
| ENSG000002 | 0.98608667 | -0.036898187 | 0.30788773 | -0.119843 | 0.90460752 | NA | NA | NA |
| ENSG000002 | 2.11864848 | -0.26136848 | 0.36552883 | -0.715042 | 0.47458306 | NA | NA | NA |
| ENSG000002 | 1.48793219 | -0.192974398 | 0.34606391 | -0.5576265 | 0.57709948 | NA | NA | NA |
| ENSG000002 | 1.29349991 | -0.1959662 | 0.32843024 | -0.5966753 | 0.55072422 | NA | NA | NA |

| | | | | | | | | |
|---|---|---|---|---|---|---|---|---|
| ENSG000002 | 0.36345889 | 0.070257596 | 0.21589421 | 0.32542602 | 0.74485866 | NA | RNU6-484P | 106479781 |
| ENSG000002 | 0.98457787 | -0.19372119 | 0.31177452 | -0.6213503 | 0.53436917 | NA | RNU6-620P | 106481377 |
| ENSG000002 | 0.60717723 | -0.256800472 | 0.2582777 | -0.9942805 | 0.32008634 | NA | RNU6-643P | 106479851 |
| ENSG000002 | 1.76981704 | -0.335114904 | 0.35626683 | -0.940629 | 0.34689503 | NA | RNU6-142P | 106479628 |
| ENSG000002 | 0.3867939 | -0.105148852 | 0.21977268 | -0.4784437 | 0.63233444 | NA | RNU6-245P | 106481255 |
| ENSG000002 | 0 | NA | NA | NA | NA | NA | RNU6-1083P | 106481521 |
| ENSG000002 | 1.00495854 | -0.209381385 | 0.3180633 | -0.658301 | 0.51034474 | NA | NA | NA |
| ENSG000002 | 1.49005896 | -0.202292366 | 0.34862612 | -0.5802559 | 0.56174205 | NA | NA | NA |
| ENSG000002 | 0.73055726 | -0.021363932 | 0.26874046 | -0.0794965 | 0.9366377 | NA | RNU6-317P | 106479708 |
| ENSG000002 | 0.60905212 | 0.050569842 | 0.25573684 | 0.19774172 | 0.84324715 | NA | RNU6-787P | 106479914 |
| ENSG000002 | 0 | NA | NA | NA | NA | NA | SNORA6 | 574040 |
| ENSG000002 | 1.78382831 | -0.53749458 | 0.34287296 | -1.5676202 | 0.1169698 | NA | RNU6-418P | 106481310 |
| ENSG000002 | 1.12942517 | -0.245986523 | 0.31054022 | -0.7921245 | 0.42828807 | NA | RNU6-152P | 106481223 |
| ENSG000002 | 0.81887182 | 0.385445377 | 0.29640387 | 1.30040601 | 0.19346185 | NA | RNU6-203P | 106479657 |
| ENSG000002 | 0.96226294 | 0.124254798 | 0.3068576 | 0.40492659 | 0.68553148 | NA | RNU6-435P | 106479758 |
| ENSG000002 | 1.27830837 | -0.503237477 | 0.33041853 | -1.5230304 | 0.12775109 | NA | RNU6-949P | 106481925 |
| ENSG000002 | 0.37945415 | -0.107841678 | 0.21865587 | -0.4932027 | 0.62186934 | NA | RNU6-116P | 106480553 |
| ENSG000002 | 0.99275572 | -0.500108375 | 0.30598225 | -1.6344359 | 0.10216735 | NA | NA | NA |
| ENSG000002 | 2.07971094 | -0.47201483 | 0.35864926 | -1.3160904 | 0.18814371 | NA | RNU6-44P | 106478922 |
| ENSG000002 | 0.69819371 | 0.293816096 | 0.27742409 | 1.05908644 | 0.28956041 | NA | RNU6-211P | 106479660 |
| ENSG000002 | 0.57928031 | 0.403399703 | 0.25735148 | 1.56750488 | 0.11699673 | NA | SNORD37 | 26812 |
| ENSG000002 | 1.31501407 | 0.025568338 | 0.31859023 | 0.08025462 | 0.93603475 | NA | NA | NA |
| ENSG000002 | 0.3934287 | -0.107186984 | 0.22072612 | -0.4856108 | 0.62724315 | NA | RNU6-637P | 106479848 |
| ENSG000002 | 2.1467502 | 0.290745177 | 0.36433474 | 0.79801662 | 0.42486085 | NA | RNU6-213P | 106481244 |
| ENSG000002 | 0.25333738 | -0.015686291 | 0.1806605 | -0.0868275 | 0.93080867 | NA | NA | NA |
| ENSG000002 | 1.04309705 | -0.311271508 | 0.29732247 | -1.0469155 | 0.29513855 | NA | RNU6-224P | 106479666 |
| ENSG000002 | 2.11252469 | -0.135170662 | 0.3676266 | -0.3676847 | 0.71310838 | NA | NA | NA |
| ENSG000002 | 2.05414985 | 0.005765247 | 0.36154097 | 0.01594632 | 0.98727722 | NA | RNU6-701P | 106481402 |
| ENSG000002 | 1.24088066 | -0.484277 | 0.32637477 | -1.4838065 | 0.13786027 | NA | RNU6-76P | 100873777 |

| | | | | | | | | | |
|---|---|---|---|---|---|---|---|---|---|
| ENSG000002 | 0.23890637 | -0.010420674 | 0.17747379 | -0.0587167 | 0.95317776 | NA | | RNU6-629P | 106479844 |
| ENSG000002 | 1.1897008 | 0.257097027 | 0.32917631 | 0.78103138 | 0.43478404 | NA | | NA | NA |
| ENSG000002 | 1.49783399 | -0.342397862 | 0.3468306 | -0.9872193 | 0.32353515 | NA | | NA | NA |
| ENSG000002 | 1.61603541 | 0.00494624 | 0.34818521 | 0.01420577 | 0.98866582 | NA | | NA | NA |
| ENSG000002 | 0.25260017 | -0.015978705 | 0.18049464 | -0.0885273 | 0.92945758 | NA | | RNU6-811P | 106479925 |
| ENSG000002 | 0.11865494 | 0.083440866 | 0.12751478 | 0.65436232 | 0.5128784 | NA | | NA | NA |
| ENSG000002 | 0 | NA | NA | NA | NA | NA | | SNORA32 | 692063 |
| ENSG000002 | 0.63389931 | -0.262485172 | 0.25907614 | -1.0131584 | 0.3109845 | NA | | NA | NA |
| ENSG000002 | 0.4009426 | -0.302672773 | 0.2240145 | -1.3511303 | 0.17665371 | NA | | RNU6-639P | 106479849 |
| ENSG000002 | 1.50853048 | -0.051268352 | 0.34299176 | -0.149474 | 0.88117962 | NA | | RNU6-926P | 106481472 |
| ENSG000002 | 0.74157948 | -0.350585661 | 0.28031838 | -1.2506695 | 0.21105507 | NA | | RNU6-968P | 106481485 |
| ENSG000002 | 0.54250991 | -0.072841599 | 0.15594206 | -0.4671068 | 0.64042346 | NA | | RNU6-1293P | 106480130 |
| ENSG000002 | 0.12685397 | -0.090215768 | 0.12901919 | -0.699243 | 0.48440017 | NA | | NA | NA |
| ENSG000002 | 1.19415055 | -0.762704184 | 0.31679854 | -2.4075369 | 0.01606054 | NA | | NA | NA |
| ENSG000002 | 0.39388516 | -0.104610737 | 0.21253504 | -0.4922047 | 0.62257467 | NA | | RNU6-815P | 106480611 |
| ENSG000002 | 0 | NA | NA | NA | NA | NA | | SNORA10 | 574042 |
| ENSG000002 | 1.69160424 | -0.049468716 | 0.35080664 | -0.1410142 | 0.88785873 | NA | | RNU6-38P | 106480709 |
| ENSG000002 | 0.35961481 | -0.099625193 | 0.21567743 | -0.4619176 | 0.64414043 | NA | | NA | NA |
| ENSG000002 | 0.3854038 | -0.110128691 | 0.21952326 | -0.5016721 | 0.61589821 | NA | | NA | NA |
| ENSG000002 | 1.50283929 | -0.19459516 | 0.34333177 | -0.5667846 | 0.57086053 | NA | | RNU6-483P | 106479780 |
| ENSG000002 | 0.3854038 | -0.110128691 | 0.21952326 | -0.5016721 | 0.61589821 | NA | | NA | NA |
| ENSG000002 | 1.61282145 | -0.397472265 | 0.35076595 | -1.1331552 | 0.25714908 | NA | | RNU6-849P | 106481449 |
| ENSG000002 | 0.71482259 | 0.130920764 | 0.2770477 | 0.47255676 | 0.63652944 | NA | | RNU6-260P | 106479682 |
| ENSG000002 | 0.6044405 | 0.057164517 | 0.25530853 | 0.22390367 | 0.82283226 | NA | | NA | NA |
| ENSG000002 | 0.72907542 | -0.027873462 | 0.27778572 | -0.1003416 | 0.92007313 | NA | | NA | NA |
| ENSG000002 | 0.40191006 | -0.101653297 | 0.21376195 | -0.4755444 | 0.63439899 | NA | | RNU6-974P | 106481487 |
| ENSG000002 | 0.90064872 | -0.456924471 | 0.30237983 | -1.5110944 | 0.1307644 | NA | | RNU6-1032P | 106481506 |
| ENSG000002 | 1.57551189 | -0.244958908 | 0.34669723 | -0.7065499 | 0.47984621 | NA | | NA | NA |
| ENSG000002 | 0.84711522 | -0.106440978 | 0.28873962 | -0.36864 | 0.71239607 | NA | | RNU6-20P | 106480706 |

| | | | | | | | | |
|---|---|---|---|---|---|---|---|---|
| ENSG000002 | 0 | NA | NA | NA | NA | NA | SNORA1 | 677792 |
| ENSG000002 | 1.73381167 | -0.323932494 | 0.3542217 | -0.9144908 | 0.360459 | NA | RNU1-74P | 106480171 |
| ENSG000002 | 1.19036204 | -0.603323933 | 0.32092623 | -1.8799458 | 0.06011546 | NA | RNU6-746P | 106479896 |
| ENSG000002 | 0 | NA | NA | NA | NA | NA | NA | NA |
| ENSG000002 | 0.47477407 | 0.142502628 | 0.23735343 | 0.60038158 | 0.54825196 | NA | RNU6-409P | 106481307 |
| ENSG000002 | 0.68351258 | -0.46544448 | 0.26656372 | -1.7460909 | 0.08079516 | NA | RNU6-413P | 106479748 |
| ENSG000002 | 0.12070844 | -0.088503695 | 0.12783931 | -0.6923042 | 0.48874631 | NA | RNU6-206P | 106481882 |
| ENSG000002 | 0.47070378 | 0.149674697 | 0.23683599 | 0.63197616 | 0.52740246 | NA | NA | NA |
| ENSG000002 | 0.11934008 | 0.083564698 | 0.12760091 | 0.65489106 | 0.5125379 | NA | RNU6-209P | 106481242 |
| ENSG000002 | 0.7888781 | -0.345530318 | 0.27689228 | -1.2478871 | 0.21207239 | NA | NA | NA |
| ENSG000002 | 0.85840903 | -0.116597839 | 0.30230797 | -0.3856922 | 0.69972461 | NA | NA | NA |
| ENSG000002 | 2.09808103 | -0.118561938 | 0.36181737 | -0.3276845 | 0.74315024 | NA | RNU6-890P | 106480618 |
| ENSG000002 | 0.88653386 | -0.113194521 | 0.29096748 | -0.3890281 | 0.69725537 | NA | NA | NA |
| ENSG000002 | 0.83765834 | 0.055260734 | 0.2957186 | 0.18686932 | 0.85176309 | NA | NA | NA |
| ENSG000002 | 0.71489639 | 0.303414967 | 0.2785771 | 1.08915976 | 0.27608344 | NA | RNU6-895P | 106479962 |
| ENSG000002 | 1.25420335 | -0.314489147 | 0.31823375 | -0.9882332 | 0.32303847 | NA | NA | NA |
| ENSG000002 | 0.22410479 | 0.147643983 | 0.16638082 | 0.88738586 | 0.37487119 | NA | NA | NA |
| ENSG000002 | 1.28107033 | -0.348105534 | 0.33131486 | -1.0506789 | 0.29340608 | NA | RNU6-571P | 106479819 |
| ENSG000002 | 0.48761094 | -0.176642206 | 0.23968055 | -0.7369902 | 0.46112833 | NA | RNU6-214P | 106479661 |
| ENSG000002 | 1.48884355 | -0.344233728 | 0.34667693 | -0.9929525 | 0.32073309 | NA | NA | NA |
| ENSG000002 | 1.24605884 | -0.039221063 | 0.32035643 | -0.1224295 | 0.90255891 | NA | RNU6-767P | 106481916 |
| ENSG000002 | 0.11934008 | 0.083564698 | 0.12760091 | 0.65489106 | 0.5125379 | NA | NA | NA |
| ENSG000002 | 0 | NA | NA | NA | NA | NA | NA | NA |
| ENSG000002 | 1.2516096 | -0.197896186 | 0.33454004 | -0.5915471 | 0.55415392 | NA | RNU6-207P | 106481766 |
| ENSG000002 | 0.73889482 | -0.029099289 | 0.29011813 | -0.1003015 | 0.92010495 | NA | NA | NA |
| ENSG000002 | 0.23890637 | -0.010420674 | 0.17747379 | -0.0587167 | 0.95317776 | NA | RNU6-187P | 106481880 |
| ENSG000002 | 2.16230555 | -0.064018365 | 0.36641965 | -0.1747132 | 0.86130496 | NA | RNU6-356P | 106479724 |
| ENSG000002 | 0.37267746 | 0.067870636 | 0.21729405 | 0.31234465 | 0.75477861 | NA | SNORA70F | 100337591 |
| ENSG000002 | 0.82761219 | 0.208279728 | 0.29564517 | 0.70449225 | 0.48112628 | NA | RNU6-398P | 106479742 |

| | | | | | | | | | |
|---|---|---|---|---|---|---|---|---|---|
| ENSG000002 | 0.76394502 | -0.170709309 | 0.27234044 | -0.6268232 | 0.53077513 | NA | | RNU6-533P | 106481901 |
| ENSG000002 | 0.7958473 | -0.182163495 | 0.27199033 | -0.6697425 | 0.50302193 | NA | | RNU6-761P | 106480607 |
| ENSG000002 | 0.85183737 | -0.238226003 | 0.28355105 | -0.8401521 | 0.40082313 | NA | | RNU6-1103P | 106480051 |
| ENSG000002 | 0.36277375 | 0.069994797 | 0.21578753 | 0.32436906 | 0.74565864 | NA | | NA | NA |
| ENSG000002 | 0.25351207 | -0.189965547 | 0.1824784 | -1.0410303 | 0.29786148 | NA | | RNU6-1310P | 106480658 |
| ENSG000002 | 0.7381576 | -0.029362007 | 0.29007186 | -0.1012232 | 0.91937327 | NA | | RNU6-190P | 106481236 |
| ENSG000002 | 0.38747904 | -0.104908247 | 0.21987514 | -0.4771265 | 0.63327207 | NA | | NA | NA |
| ENSG000002 | 0.11131518 | 0.081951103 | 0.12663804 | 0.64712866 | 0.51754868 | NA | | NA | NA |
| ENSG000002 | 0.37404583 | -0.105111695 | 0.21784993 | -0.4824959 | 0.62945372 | NA | | RNU6-1008P | 106481498 |
| ENSG000002 | 1.11583899 | 0.036134933 | 0.32206294 | 0.11219836 | 0.91066613 | NA | | RNU6-1200P | 106481559 |
| ENSG000002 | 0.2596576 | -0.193002035 | 0.1837913 | -1.0501152 | 0.29366517 | NA | | RNU6-217P | 106481246 |
| ENSG000002 | 1.44201648 | -0.558260247 | 0.33408786 | -1.6709983 | 0.09472201 | NA | | RNU6-42P | 100302742 |
| ENSG000002 | 0.36345889 | 0.070257596 | 0.21589421 | 0.32542602 | 0.74485866 | NA | | NA | NA |
| ENSG000002 | 0.73543394 | -0.320531528 | 0.27073292 | -1.18394 | 0.23643682 | NA | | NA | NA |
| ENSG000002 | 1.23823375 | -0.313402246 | 0.32074378 | -0.9771109 | 0.32851425 | NA | | NA | NA |
| ENSG000002 | 0.59687261 | 0.065989748 | 0.26864914 | 0.24563543 | 0.80596446 | NA | | NA | NA |
| ENSG000002 | 1.13941478 | -0.123173018 | 0.32161128 | -0.3829872 | 0.70172923 | NA | | NA | NA |
| ENSG000002 | 0.25214371 | -0.009862892 | 0.18046657 | -0.0546522 | 0.95641556 | NA | | RNU6-458P | 106481324 |
| ENSG000002 | 0.63114139 | -0.097702348 | 0.25814354 | -0.3784807 | 0.70507353 | NA | | RNU6-1013P | 106480012 |
| ENSG000002 | 1.14778054 | -0.147512562 | 0.20735851 | -0.711389 | 0.47684322 | NA | | RNU1-136P | 106481632 |
| ENSG000002 | 0 | NA | NA | NA | NA | NA | | NA | NA |
| ENSG000002 | 0 | NA | NA | NA | NA | NA | | SNORA29 | 677812 |
| ENSG000002 | 1.03695152 | -0.333146353 | 0.30524194 | -1.0914174 | 0.27508927 | NA | | NA | NA |
| ENSG000002 | 0 | NA | NA | NA | NA | NA | | NA | NA |
| ENSG000002 | 0.86572706 | -0.287838831 | 0.30354833 | -0.9482471 | 0.34300366 | NA | | NA | NA |
| ENSG000002 | 0.4009426 | -0.302672773 | 0.2240145 | -1.3511303 | 0.17665371 | NA | | RNU6-439P | 106481317 |
| ENSG000002 | 0.81036655 | -0.540716938 | 0.28190939 | -1.9180522 | 0.05510439 | NA | | RNU1-52P | 106480164 |
| ENSG000002 | 1.25528134 | -0.32833966 | 0.32668095 | -1.0050775 | 0.31485955 | NA | | NA | NA |
| ENSG000002 | 1.859677 | -0.658200269 | 0.35656215 | -1.8459622 | 0.06489769 | NA | | RNU6-80P | 106478990 |

| | | | | | | | | | |
|---|---|---|---|---|---|---|---|---|---|
| ENSG000002 | 1.05752807 | -0.312875709 | 0.29708136 | -1.0531651 | 0.29226535 | NA | | RNU6-432P | 106479757 |
| ENSG000002 | 0.90753304 | -0.266763647 | 0.29355604 | -0.9087316 | 0.36349183 | NA | | RNU6-689P | 106479870 |
| ENSG000002 | 0.87010195 | 0.048098246 | 0.29583509 | 0.16258466 | 0.87084547 | NA | | NA | NA |
| ENSG000002 | 1.08712992 | -0.233085789 | 0.30625434 | -0.7610857 | 0.44660589 | NA | | RNU6-1024P | 106481504 |
| ENSG000002 | 0 | NA | NA | NA | NA | NA | | NA | NA |
| ENSG000002 | 0.48536101 | -0.024029524 | 0.24539296 | -0.0979226 | 0.92199373 | NA | | NA | NA |
| ENSG000002 | 0.63828472 | -0.112219173 | 0.26613064 | -0.4216695 | 0.67326627 | NA | | RNU6-1042P | 106480024 |
| ENSG000002 | 0 | NA | NA | NA | NA | NA | | RNU6-514P | 106480586 |
| ENSG000002 | 0.98466219 | -0.189310584 | 0.30686736 | -0.6169134 | 0.53729185 | NA | | RNU4-52P | 106481188 |
| ENSG000002 | 0.96198006 | -0.036154158 | 0.31177589 | -0.115962 | 0.90768265 | NA | | SNORA70B | 100124537 |
| ENSG000002 | 1.18005742 | -0.266759845 | 0.31684873 | -0.8419155 | 0.39983528 | NA | | RNU4-31P | 106480546 |
| ENSG000002 | 0.47075585 | 0.160115757 | 0.24423976 | 0.65556794 | 0.51210216 | NA | | RNU6-281P | 106481267 |
| ENSG000002 | 0.36277375 | 0.069994797 | 0.21578753 | 0.32436906 | 0.74565864 | NA | | SNORD15A | 6079 |
| ENSG000002 | 1.08769593 | 0.178511065 | 0.31523925 | 0.56627169 | 0.5712091 | NA | | RNU6-708P | 106480602 |
| ENSG000002 | 0.48211382 | 0.155752635 | 0.24556107 | 0.63427249 | 0.525903 | NA | | NA | NA |
| ENSG000002 | 0.12070844 | -0.088503695 | 0.12783931 | -0.6923042 | 0.48874631 | NA | | SNORA36A | 677817 |
| ENSG000002 | 0.37945415 | -0.107841678 | 0.21865587 | -0.4932027 | 0.62186934 | NA | | RNU6-1324P | 106480144 |
| ENSG000002 | 1.57157685 | -0.775020691 | 0.3388747 | -2.2870421 | 0.02219337 | NA | | NA | NA |
| ENSG000002 | 0.46131052 | 0.308572634 | 0.23046384 | 1.33891996 | 0.18059673 | NA | | NA | NA |
| ENSG000002 | 0 | NA | NA | NA | NA | NA | | NA | NA |
| ENSG000002 | 1.60701169 | -0.11126047 | 0.34082886 | -0.3264409 | 0.74409082 | NA | | NA | NA |
| ENSG000002 | 0.23868015 | 0.154625155 | 0.16997278 | 0.90970538 | 0.36297791 | NA | | NA | NA |
| ENSG000002 | 0.36351096 | 0.070290861 | 0.21590418 | 0.32556508 | 0.74475344 | NA | | NA | NA |
| ENSG000002 | 0.89079899 | -0.240517948 | 0.28586709 | -0.8413629 | 0.40014469 | NA | | RNU6-793P | 106479917 |
| ENSG000002 | 0.47551128 | 0.153262041 | 0.24481524 | 0.62603146 | 0.53129431 | NA | | NA | NA |
| ENSG000002 | 0.34197044 | 0.237837536 | 0.20637034 | 1.15247924 | 0.2491242 | NA | | RNU6-74P | 106480752 |
| ENSG000002 | 0.8943285 | -0.291158821 | 0.30492371 | -0.9548579 | 0.33964956 | NA | | RNU6-675P | 106479864 |
| ENSG000002 | 1.01566363 | -0.19029424 | 0.30246948 | -0.6291353 | 0.52926045 | NA | | NA | NA |
| ENSG000002 | 0.37267746 | 0.067870636 | 0.21729405 | 0.31234465 | 0.75477861 | NA | | NA | NA |

| | | | | | | | | |
|---|---|---|---|---|---|---|---|---|
| ENSG000002 | 0.85960326 | -0.112061982 | 0.30237674 | -0.3706038 | 0.71093262 | NA | RNU6-1316P | 106480140 |
| ENSG000002 | 0.45481211 | 0.301948681 | 0.2282923 | 1.32264067 | 0.1859549 | NA | RNU6-474P | 106481329 |
| ENSG000002 | 0.3593886 | 0.071604659 | 0.20679693 | 0.34625591 | 0.7291504 | NA | RNU6-17P | 106480705 |
| ENSG000002 | 1.10693755 | -0.225780473 | 0.30700426 | -0.7354311 | 0.46207701 | NA | RNU6-1145P | 106481541 |
| ENSG000002 | 0.53627793 | -0.360851627 | 0.24080453 | -1.4985251 | 0.13399688 | NA | RNU6-1144P | 106481938 |
| ENSG000002 | 0.99138735 | -0.346626079 | 0.31023635 | -1.1172968 | 0.26386744 | NA | RNU6-13P | 106480423 |
| ENSG000002 | 0.96109905 | -0.034595124 | 0.30680131 | -0.1127607 | 0.91022029 | NA | NA | NA |
| ENSG000002 | 0.23276084 | -0.008011382 | 0.17603982 | -0.0455089 | 0.96370166 | NA | NA | NA |
| ENSG000002 | 0.61836106 | -0.107182493 | 0.27082484 | -0.3957632 | 0.69227976 | NA | NA | NA |
| ENSG000002 | 0 | NA | NA | NA | NA | NA | SNORD61 | 26787 |
| ENSG000002 | 1.24790541 | -0.043973975 | 0.33048596 | -0.1330585 | 0.89414709 | NA | RNU6-662P | 106479860 |
| ENSG000002 | 0.62041457 | -0.27371262 | 0.26650407 | -1.0270486 | 0.3043976 | NA | RNU6-40P | 106480703 |
| ENSG000002 | 2.14690344 | -0.337939108 | 0.3564723 | -0.9480094 | 0.34312465 | NA | NA | NA |
| ENSG000002 | 0.60150597 | 0.230273195 | 0.26371223 | 0.87319879 | 0.38255474 | NA | RNU6-49P | 106478923 |
| ENSG000002 | 0.99809541 | -0.188630508 | 0.30744749 | -0.6135373 | 0.53952112 | NA | RNU6-198P | 106481239 |
| ENSG000002 | 0.49196355 | -0.021608171 | 0.24613638 | -0.0877894 | 0.93004404 | NA | NA | NA |
| ENSG000002 | 0 | NA | NA | NA | NA | NA | NA | NA |
| ENSG000002 | 1.45277113 | 0.086286582 | 0.33976788 | 0.25395744 | 0.79952844 | NA | RNU6-574P | 106479820 |
| ENSG000002 | 0.76121029 | -0.351991847 | 0.28168635 | -1.2495879 | 0.21145014 | NA | NA | NA |
| ENSG000002 | 0.92222307 | -0.44186707 | 0.29679498 | -1.4887956 | 0.13654119 | NA | RNU6-842P | 106481447 |
| ENSG000002 | 1.03877243 | -0.196725068 | 0.30913895 | -0.6363645 | 0.52453885 | NA | RNU6-524P | 106481346 |
| ENSG000002 | 1.33397239 | 0.024028053 | 0.34167488 | 0.07032432 | 0.94393553 | NA | NA | NA |
| ENSG000002 | 1.15461333 | -0.253198734 | 0.31306354 | -0.8087775 | 0.41864317 | NA | RNU6-622P | 106479841 |
| ENSG000002 | 0.88356742 | 0.042138785 | 0.2904594 | 0.14507633 | 0.8846506 | NA | RNU6-820P | 106479929 |
| ENSG000002 | 0.36827086 | -0.266056975 | 0.21181163 | -1.2561018 | 0.20907905 | NA | NA | NA |
| ENSG000002 | 0.49150709 | -0.015856992 | 0.24611816 | -0.0644284 | 0.94862914 | NA | RNU6-301P | 106481274 |
| ENSG000002 | 0.98865062 | -0.031116886 | 0.30041571 | -0.1035794 | 0.91750312 | NA | RNU6-891P | 106479960 |
| ENSG000002 | 1.11393844 | -0.112377039 | 0.3207957 | -0.3503072 | 0.72610817 | NA | NA | NA |
| ENSG000002 | 0.68098087 | -0.43598206 | 0.25960849 | -1.6793829 | 0.09307744 | NA | RNU6-1295P | 106481589 |

| | | | | | | | | |
|---|---|---|---|---|---|---|---|---|
| ENSG000002 | 0 | NA | NA | NA | NA | NA | NA | NA |
| ENSG000002 | 1.85569205 | -0.510712685 | 0.35554169 | -1.4364355 | 0.15087847 | NA | RNU6-72P | 100873775 |
| ENSG000002 | 0.6039295 | -0.097743897 | 0.2540715 | -0.3847102 | 0.70045213 | NA | RNU6-804P | 106479921 |
| ENSG000002 | 1.36810285 | -0.262625323 | 0.33704128 | -0.7792082 | 0.43585709 | NA | RNU6-167P | 106481228 |
| ENSG000002 | 0.7872583 | -0.560227253 | 0.2858936 | -1.9595656 | 0.05004659 | NA | NA | NA |
| ENSG000002 | 0.9882478 | 0.120464272 | 0.31280431 | 0.38511065 | 0.70015544 | NA | NA | NA |
| ENSG000002 | 2.09855014 | -0.135255139 | 0.36713012 | -0.368412 | 0.71256605 | NA | RNU6-975P | 106479997 |
| ENSG000002 | 0.12685397 | -0.090215768 | 0.12901919 | -0.699243 | 0.48440017 | NA | NA | NA |
| ENSG000002 | 0.23276084 | -0.008011382 | 0.17603982 | -0.0455089 | 0.96370166 | NA | RNU6-472P | 106481328 |
| ENSG000002 | 0.79579523 | -0.182201703 | 0.271987 | -0.6698912 | 0.50292714 | NA | NA | NA |
| ENSG000002 | 1.97212126 | -0.055937303 | 0.36195719 | -0.1545412 | 0.87718302 | NA | RNU6-43P | 106478921 |
| ENSG000002 | 0.5301324 | -0.3843196 | 0.24728844 | -1.5541349 | 0.12015224 | NA | NA | NA |
| ENSG000002 | 2.07891715 | -0.70645914 | 0.35745493 | -1.9763586 | 0.04811417 | NA | RNU6-154P | 106481878 |
| ENSG000002 | 0.36827086 | -0.266056975 | 0.21181163 | -1.2561018 | 0.20907905 | NA | RNY3P14 | 106480316 |
| ENSG000002 | 0.60965294 | 0.058699225 | 0.2633339 | 0.22290797 | 0.82360713 | NA | RNU6-339P | 106481888 |
| ENSG000002 | 1.40655462 | 0.084134625 | 0.32774393 | 0.25670842 | 0.79740387 | NA | RNU6-196P | 106481238 |
| ENSG000002 | 0.24411881 | -0.012946855 | 0.17864669 | -0.0724718 | 0.94222641 | NA | RNU6-1226P | 106480102 |
| ENSG000002 | 0.34936227 | 0.241139221 | 0.20762307 | 1.16142789 | 0.24546793 | NA | RNU6-532P | 106481349 |
| ENSG000002 | 1.10875928 | 0.015767976 | 0.3082019 | 0.05116119 | 0.95919708 | NA | RNU6-886P | 106479958 |
| ENSG000002 | 0 | NA | NA | NA | NA | NA | NA | NA |
| ENSG000002 | 0.49902098 | -0.194741582 | 0.24819886 | -0.7846192 | 0.43267688 | NA | NA | NA |
| ENSG000002 | 0 | NA | NA | NA | NA | NA | SNORA27 | 619499 |
| ENSG000002 | 1.33186681 | 0.171804368 | 0.34003583 | 0.50525372 | 0.61338064 | NA | RNU6-378P | 106481297 |
| ENSG000002 | 0.13280363 | -0.093394593 | 0.13115644 | -0.7120855 | 0.47641185 | NA | RNU6-937P | 106479980 |
| ENSG000002 | 0.99560255 | -0.313468749 | 0.30084953 | -1.0419453 | 0.29743705 | NA | RNU1-8P | 106481604 |
| ENSG000002 | 1.36232397 | -0.122887266 | 0.337674 | -0.3639228 | 0.71591566 | NA | RNU6-784P | 106481428 |
| ENSG000002 | 0.90816262 | -0.635180171 | 0.29877972 | -2.1259146 | 0.03351037 | NA | RNU6-480P | 106480583 |
| ENSG000002 | 0.63911175 | -0.286025981 | 0.26796114 | -1.0674159 | 0.28578407 | NA | NA | NA |
| ENSG000002 | 0.22997012 | 0.167051788 | 0.17646558 | 0.94665368 | 0.34381528 | NA | NA | NA |

| | | | | | | | | |
|---|---|---|---|---|---|---|---|---|
| ENSG000002 | 0.75982019 | -0.379788636 | 0.28855699 | -1.3161651 | 0.18811864 | NA | NA | NA |
| ENSG000002 | 0 | NA | NA | NA | NA | NA | NA | NA |
| ENSG000002 | 1.77664716 | -0.325746963 | 0.35089008 | -0.9283448 | 0.35322876 | NA | RNU6-1207P | 106480648 |
| ENSG000002 | 0.37199232 | 0.06760863 | 0.21718977 | 0.31128828 | 0.75558148 | NA | RNU6-39P | 100873758 |
| ENSG000002 | 0.98475256 | -0.340128925 | 0.30835223 | -1.1030532 | 0.27000405 | NA | NA | NA |
| ENSG000002 | 0.11131518 | 0.081951103 | 0.12663804 | 0.64712866 | 0.51754868 | NA | NA | NA |
| ENSG000002 | 0.55070895 | -0.361402396 | 0.24091882 | -1.5001003 | 0.13358842 | NA | RNU6-1113P | 106480056 |
| ENSG000002 | 0.66184999 | -0.106729201 | 0.25913995 | -0.4118593 | 0.68044255 | NA | RNU6-1119P | 106481533 |
| ENSG000002 | 1.32615438 | -0.658162744 | 0.32733981 | -2.0106407 | 0.04436342 | NA | NA | NA |
| ENSG000002 | 1.71246365 | 0.083139837 | 0.35410593 | 0.23478804 | 0.81437325 | NA | RNU6-616P | 106479838 |
| ENSG000002 | 1.00971844 | -0.504920681 | 0.30670098 | -1.6462963 | 0.09970277 | NA | RNU6-22P | 106478917 |
| ENSG000002 | 0.38651158 | -0.274258716 | 0.21461355 | -1.2779189 | 0.20127801 | NA | NA | NA |
| ENSG000002 | 0.86893998 | 0.04033367 | 0.29741211 | 0.13561543 | 0.89212531 | NA | NA | NA |
| ENSG000002 | 1.58228193 | -0.765426173 | 0.33683727 | -2.2723916 | 0.02306287 | NA | RNU6-242P | 106481254 |
| ENSG000002 | 1.02100779 | -0.186034549 | 0.30271766 | -0.614548 | 0.53885323 | NA | RNU6-517P | 106479795 |
| ENSG000002 | 1.73036649 | -0.316079292 | 0.35231972 | -0.8971377 | 0.36964546 | NA | NA | NA |
| ENSG000002 | 1.88503071 | -0.550925022 | 0.3620825 | -1.5215456 | 0.12812299 | NA | NA | NA |
| ENSG000002 | 0.37928001 | 0.070449581 | 0.21829793 | 0.32272217 | 0.74690566 | NA | NA | NA |
| ENSG000002 | 0.59507757 | 0.053522528 | 0.26203995 | 0.20425332 | 0.83815555 | NA | RNU6-424P | 106479753 |
| ENSG000002 | 0.26560726 | -0.177019426 | 0.17672274 | -1.0016788 | 0.31649875 | NA | RNU6-614P | 106481375 |
| ENSG000002 | 0.36690249 | -0.096999144 | 0.21683809 | -0.4473344 | 0.65463361 | NA | NA | NA |
| ENSG000002 | 1.11783851 | -0.270043428 | 0.3232201 | -0.8354785 | 0.40344837 | NA | RNU6-166P | 106481879 |
| ENSG000002 | 0.64127344 | -0.108386707 | 0.26638026 | -0.4068872 | 0.68409087 | NA | NA | NA |
| ENSG000002 | 0.60680666 | -0.099379886 | 0.25590479 | -0.3883471 | 0.69775918 | NA | NA | NA |
| ENSG000002 | 0.96061382 | 0.276962861 | 0.30756026 | 0.90051576 | 0.36784584 | NA | RNU6-434P | 106480581 |
| ENSG000002 | 0.24550891 | -0.007886114 | 0.17900103 | -0.0440563 | 0.96485956 | NA | NA | NA |
| ENSG000002 | 0 | NA | NA | NA | NA | NA | SNORA25 | 684959 |
| ENSG000002 | 0.36416385 | 0.068880976 | 0.20758164 | 0.33182596 | 0.74002068 | NA | RNU6-16P | 100873753 |
| ENSG000002 | 0.8752602 | -0.104672008 | 0.29033686 | -0.3605192 | 0.71845891 | NA | RNU6-348P | 106481289 |

| | | | | | | | | | |
|---|---|---|---|---|---|---|---|---|---|
| ENSG000002 | 2.1441172 | -0.478425471 | 0.36080492 | -1.3259949 | 0.1848414 | NA | | RNU6-364P | 106481293 |
| ENSG000002 | 0.83577898 | 0.052536527 | 0.30082402 | 0.17464206 | 0.8613609 | NA | | RNU6-31P | 100873740 |
| ENSG000002 | 0 | NA | NA | NA | NA | NA | | SNORD14D | 85390 |
| ENSG000002 | 1.78688582 | -0.194830288 | 0.35315033 | -0.5516922 | 0.58115924 | NA | | NA | NA |
| ENSG000002 | 1.64179292 | -0.532293506 | 0.34727685 | -1.5327642 | 0.12533397 | NA | | RNU6-1230P | 106481944 |
| ENSG000002 | 1.01867408 | -0.042709894 | 0.30758828 | -0.1388541 | 0.88956544 | NA | | RNU6-591P | 106479827 |
| ENSG000002 | 1.11453321 | 0.02425141 | 0.30839891 | 0.0786365 | 0.93732176 | NA | | NA | NA |
| ENSG000002 | 0.23202362 | -0.008302493 | 0.17586207 | -0.0472103 | 0.96234565 | NA | | RNU6-163P | 106479638 |
| ENSG000002 | 0.99414582 | -0.488149888 | 0.30445976 | -1.6033314 | 0.10886151 | NA | | NA | NA |
| ENSG000002 | 1.86561461 | -0.006898336 | 0.3562554 | -0.0193635 | 0.98455116 | NA | | RNU6-729P | 106481410 |
| ENSG000002 | 1.99301281 | -0.453440054 | 0.36334082 | -1.2479744 | 0.21204042 | NA | | NA | NA |
| ENSG000002 | 1.13098548 | -0.12236573 | 0.32285096 | -0.3790162 | 0.70467586 | NA | | NA | NA |
| ENSG000002 | 0.50722002 | -0.375409304 | 0.24493426 | -1.5326941 | 0.12535123 | NA | | NA | NA |
| ENSG000002 | 1.92216504 | -0.506542338 | 0.35317469 | -1.4342544 | 0.15149969 | NA | | NA | NA |
| ENSG000002 | 0.35022155 | 0.06940789 | 0.21379411 | 0.32464827 | 0.74544729 | NA | | RNU6-106P | 106479613 |
| ENSG000002 | 0.73129447 | -0.021100375 | 0.26879465 | -0.0785 | 0.93743034 | NA | | RNU6-452P | 106481322 |
| ENSG000002 | 0.93844858 | -0.403733255 | 0.28607426 | -1.4112883 | 0.15815963 | NA | | RNU6-869P | 106480616 |
| ENSG000002 | 1.27235872 | -0.505217311 | 0.3310053 | -1.5263119 | 0.12693221 | NA | | NA | NA |
| ENSG000002 | 0.80925878 | -0.344388394 | 0.27575198 | -1.2489064 | 0.21169933 | NA | | NA | NA |
| ENSG000002 | 1.11169489 | -0.110683234 | 0.32072136 | -0.3451072 | 0.73001382 | NA | | RNU6-1297P | 106480657 |
| ENSG000002 | 0 | NA | NA | NA | NA | NA | | SNORA18 | 677805 |
| ENSG000002 | 0.2596576 | -0.193002035 | 0.1837913 | -1.0501152 | 0.29366517 | NA | | NA | NA |
| ENSG000002 | 1.60607613 | -0.255659069 | 0.34786645 | -0.7349345 | 0.46237937 | NA | | NA | NA |
| ENSG000002 | 0.50204195 | -0.181972198 | 0.24131623 | -0.7540819 | 0.45080006 | NA | | NA | NA |
| ENSG000002 | 0.11131518 | 0.081951103 | 0.12663804 | 0.64712866 | 0.51754868 | NA | | RNU6-465P | 106481326 |
| ENSG000002 | 0.22410479 | 0.147643983 | 0.16638082 | 0.88738586 | 0.37487119 | NA | | RNU6-185P | 106481235 |
| ENSG000002 | 1.32711329 | 0.293886295 | 0.32675529 | 0.89940791 | 0.36843543 | NA | | NA | NA |
| ENSG000002 | 0.36212533 | -0.235926492 | 0.20089643 | -1.1743688 | 0.24024735 | NA | | RNU1-46P | 106480663 |
| ENSG000002 | 0.61033616 | -0.109831062 | 0.27015137 | -0.4065538 | 0.68433575 | NA | | RNY1P14 | 106480311 |

| | | | | | | | | |
|---|---|---|---|---|---|---|---|---|
| ENSG000002 | 0 | NA | NA | NA | NA | NA | RNU6-505P | 106479790 |
| ENSG000002 | 0.49082195 | -0.014647497 | 0.23849846 | -0.0614155 | 0.95102832 | NA | NA | NA |
| ENSG000002 | 0.63347722 | -0.105383092 | 0.26574722 | -0.3965539 | 0.69169647 | NA | RNU6-929P | 106481473 |
| ENSG000002 | 1.00587393 | -0.041191161 | 0.30119101 | -0.1367609 | 0.89121977 | NA | RNU6-1289P | 106481587 |
| ENSG000002 | 0.82106172 | 0.188709219 | 0.28513822 | 0.66181665 | 0.50808874 | NA | NA | NA |
| ENSG000002 | 1.08800859 | 0.183931037 | 0.32134943 | 0.57237082 | 0.56707078 | NA | RNU6-905P | 106479967 |
| ENSG000002 | 0.3867939 | -0.105148852 | 0.21977268 | -0.4784437 | 0.63233444 | NA | NA | NA |
| ENSG000002 | 0 | NA | NA | NA | NA | NA | SNORA70 | 26778 |
| ENSG000002 | 0 | NA | NA | NA | NA | NA | SNORA68 | 26780 |
| ENSG000002 | 0.85243819 | -0.263033797 | 0.29646769 | -0.8872258 | 0.37495733 | NA | RNU6-1340P | 106480151 |
| ENSG000002 | 0.37928001 | 0.070449581 | 0.21829793 | 0.32272217 | 0.74690566 | NA | SNORA15 | 677803 |
| ENSG000002 | 1.28977791 | -0.48678606 | 0.32536114 | -1.4961408 | 0.13461698 | NA | RNU6-24P | 106480707 |
| ENSG000002 | 0.66751732 | -0.272810278 | 0.26356336 | -1.0350842 | 0.30062957 | NA | RNU6-1215P | 106480097 |
| ENSG000002 | 0.86346715 | 0.046464513 | 0.28678161 | 0.16202054 | 0.87128968 | NA | NA | NA |
| ENSG000002 | 0.50428741 | -0.01691703 | 0.23137633 | -0.0731148 | 0.94171478 | NA | RNU6-1162P | 106480078 |
| ENSG000002 | 0.94063848 | -0.603768454 | 0.29306415 | -2.0601922 | 0.03938017 | NA | NA | NA |
| ENSG000002 | 0.49819341 | -0.026649049 | 0.23931956 | -0.1113534 | 0.9113361 | NA | NA | NA |
| ENSG000002 | 0.49375647 | -0.17902118 | 0.24039617 | -0.7446923 | 0.45645774 | NA | RNU6-1122P | 106481936 |
| ENSG000002 | 0.48627291 | -0.18151782 | 0.23948009 | -0.7579662 | 0.44847119 | NA | RNU6-411P | 106479747 |
| ENSG000002 | 0.61924208 | -0.030634071 | 0.15758473 | -0.1943975 | 0.84586468 | NA | RNU6-843P | 106480614 |
| ENSG000002 | 1.87825615 | -0.241248607 | 0.35517682 | -0.6792352 | 0.49698883 | NA | RNU6-1157P | 106481545 |
| ENSG000002 | 0.78090774 | -0.190404338 | 0.28228031 | -0.6745222 | 0.49997937 | NA | NA | NA |
| ENSG000002 | 0.64469531 | -0.422865637 | 0.25651704 | -1.6484894 | 0.09925228 | NA | NA | NA |
| ENSG000002 | 1.82367121 | -0.112238607 | 0.35299326 | -0.3179625 | 0.75051341 | NA | RNU6-809P | 106479924 |
| ENSG000002 | 0.84434466 | 0.043418337 | 0.29446077 | 0.14745033 | 0.88277658 | NA | NA | NA |
| ENSG000002 | 1.38384859 | -0.544624832 | 0.33283798 | -1.6363061 | 0.10177553 | NA | RNU6-1026P | 106480631 |
| ENSG000002 | 1.11376375 | 0.033097757 | 0.32584955 | 0.10157374 | 0.91909502 | NA | NA | NA |
| ENSG000002 | 0.76083973 | -0.199648527 | 0.28697475 | -0.6957007 | 0.4866163 | NA | NA | NA |
| ENSG000002 | 0.35750923 | 0.072271886 | 0.21496454 | 0.33620376 | 0.73671722 | NA | RNU6-1195P | 106480647 |

| | | | | | | | | | |
|---|---|---|---|---|---|---|---|---|---|
| ENSG000002 | 2.0157853 | -0.712863062 | 0.36018534 | -1.9791562 | 0.04779842 | NA | | NA | NA |
| ENSG000002 | 0.61818637 | 0.059474041 | 0.27041681 | 0.21993471 | 0.82592201 | NA | | RNU6-45P | 100873759 |
| ENSG000002 | 0.4682236 | 0.140020865 | 0.23655532 | 0.59191594 | 0.55390688 | NA | | RNU1-148P | 106481635 |
| ENSG000002 | 0.24756242 | -0.187034346 | 0.18119817 | -1.0322088 | 0.30197431 | NA | | RNU6-800P | 106481433 |
| ENSG000002 | 0.87141612 | -0.272396215 | 0.29905577 | -0.9108542 | 0.36237218 | NA | | RNU6-71P | 100873774 |
| ENSG000002 | 1.15756768 | -0.397158554 | 0.31367529 | -1.2661455 | 0.20546101 | NA | | RNU6-393P | 106481302 |
| ENSG000002 | 2.01077299 | -0.18856675 | 0.35931956 | -0.5247884 | 0.59973027 | NA | | RNU6-1035P | 106481507 |
| ENSG000002 | 0.11205239 | 0.082118251 | 0.12671962 | 0.64803108 | 0.51696485 | NA | | NA | NA |
| ENSG000002 | 0.23890637 | -0.010420674 | 0.17747379 | -0.0587167 | 0.95317776 | NA | | RNU6-790P | 106481430 |
| ENSG000002 | 1.4143825 | -0.430099914 | 0.34002971 | -1.2648892 | 0.20591104 | NA | | NA | NA |
| ENSG000002 | 1.50975237 | -0.302991382 | 0.33074088 | -0.916099 | 0.35961498 | NA | | NA | NA |
| ENSG000002 | 0.35801832 | 0.071082974 | 0.20657063 | 0.34410977 | 0.73076373 | NA | | RNU6-102P | 106481209 |
| ENSG000002 | 1.62093562 | -0.508874233 | 0.3424103 | -1.4861534 | 0.13723855 | NA | | NA | NA |
| ENSG000002 | 0.95261901 | -0.40225911 | 0.28686768 | -1.4022462 | 0.16084175 | NA | | NA | NA |
| ENSG000002 | 0.11865494 | 0.083440866 | 0.12751478 | 0.65436232 | 0.5128784 | NA | | SNORA70E | 100379250 |
| ENSG000002 | 0.94529912 | 0.10342672 | 0.29851332 | 0.34647271 | 0.72898749 | NA | | NA | NA |
| ENSG000002 | 0.49699973 | -0.021154807 | 0.23920592 | -0.0884376 | 0.92952885 | NA | | RNU6-692P | 106479871 |
| ENSG000002 | 0 | NA | NA | NA | NA | NA | | RNU6-900P | 106481463 |
| ENSG000002 | 0.13280363 | -0.093394593 | 0.13115644 | -0.7120855 | 0.47641185 | NA | | NA | NA |
| ENSG000002 | 0.3934287 | -0.107186984 | 0.22072612 | -0.4856108 | 0.62724315 | NA | | NA | NA |
| ENSG000002 | 0.75332583 | -0.032209697 | 0.28569609 | -0.1127411 | 0.9102358 | NA | | RNU6-110P | 106479615 |
| ENSG000002 | 1.00583956 | -0.191334055 | 0.30777044 | -0.6216778 | 0.53415374 | NA | | NA | NA |
| ENSG000002 | 0 | NA | NA | NA | NA | NA | | SNORD45A | 26805 |
| ENSG000002 | 1.59938769 | -0.390833328 | 0.3466171 | -1.127565 | 0.25950365 | NA | | RNU6-1065P | 106480034 |
| ENSG000002 | 0 | NA | NA | NA | NA | NA | | NA | NA |
| ENSG000002 | 0.26199343 | -0.194018184 | 0.18422801 | -1.0531416 | 0.29227609 | NA | | NA | NA |
| ENSG000002 | 0.90614294 | -0.271362536 | 0.29346145 | -0.9246957 | 0.35512423 | NA | | NA | NA |
| ENSG000002 | 2.01264875 | 0.358097092 | 0.35818235 | 0.99976197 | 0.31742572 | NA | | RNU6-1005P | 106481497 |
| ENSG000002 | 1.38509837 | -0.273265904 | 0.34184847 | -0.7993773 | 0.42407168 | NA | | NA | NA |

| | | | | | | | | |
|---|---|---|---|---|---|---|---|---|
| ENSG000002 | 0.87115711 | -0.109553848 | 0.29624207 | -0.3698119 | 0.71152263 | NA | RNU6-342P | 106479718 |
| ENSG000002 | 0.26560726 | -0.177019426 | 0.17672274 | -1.0016788 | 0.31649875 | NA | NA | NA |
| ENSG000002 | 0.74498828 | -0.028170431 | 0.27889408 | -0.1010076 | 0.9195444 | NA | RNU6-1117P | 106481532 |
| ENSG000002 | 0.706623 | 0.287142598 | 0.27650879 | 1.0384574 | 0.29905716 | NA | RNU6-880P | 106479955 |
| ENSG000002 | 0 | NA | NA | NA | NA | NA | RNU6-18P | 106478916 |
| ENSG000002 | 0.71621269 | 0.14258294 | 0.28335356 | 0.50319798 | 0.61482509 | NA | NA | NA |
| ENSG000002 | 0.73770114 | -0.022487212 | 0.2847055 | -0.0789841 | 0.93704526 | NA | RNU6-738P | 106479892 |
| ENSG000002 | 1.28021105 | -0.19181929 | 0.33035959 | -0.5806379 | 0.56148455 | NA | RNU6-15P | 100302741 |
| ENSG000002 | 0.38614101 | -0.1098476 | 0.21963552 | -0.5001359 | 0.61697942 | NA | NA | NA |
| ENSG000002 | 0.72876434 | 0.133181361 | 0.28422461 | 0.46857787 | 0.6393714 | NA | RNU6-1241P | 106481572 |
| ENSG000002 | 0.24485602 | -0.012655878 | 0.17881722 | -0.0707755 | 0.94357643 | NA | SNORA70C | 100124538 |
| ENSG000002 | 0.62160824 | -0.2792447 | 0.26652831 | -1.0477112 | 0.29477168 | NA | RN7SKP62 | 106480460 |
| ENSG000002 | 0.49765262 | -0.018104365 | 0.24679454 | -0.073358 | 0.9415212 | NA | RNU6-601P | 106479832 |
| ENSG000002 | 0.84383421 | -0.106858859 | 0.29642339 | -0.360494 | 0.71847772 | NA | NA | NA |
| ENSG000002 | 0.46273095 | 0.157073337 | 0.24331041 | 0.64556769 | 0.51855937 | NA | NA | NA |
| ENSG000002 | 1.30307396 | -0.099495138 | 0.32985632 | -0.3016317 | 0.7629328 | NA | RNU6-441P | 106480392 |
| ENSG000002 | 0.39479706 | -0.300008631 | 0.22317059 | -1.3443018 | 0.17885081 | NA | RNU6-1160P | 106481546 |
| ENSG000002 | 0 | NA | NA | NA | NA | NA | NA | NA |
| ENSG000002 | 1.62741163 | -0.125222682 | 0.35042397 | -0.3573462 | 0.72083263 | NA | NA | NA |
| ENSG000002 | 0.63091271 | -0.116973765 | 0.27178557 | -0.4303899 | 0.66691204 | NA | NA | NA |
| ENSG000002 | 0.72884867 | 0.129253689 | 0.28424157 | 0.45473183 | 0.64930218 | NA | NA | NA |
| ENSG000002 | 1.67566046 | -0.386654006 | 0.34520111 | -1.1200833 | 0.26267826 | NA | NA | NA |
| ENSG000002 | 0.47092999 | -0.01913341 | 0.24378131 | -0.078486 | 0.9374415 | NA | RNU6-1274P | 106481582 |
| ENSG000002 | 0 | NA | NA | NA | NA | NA | RNU6-1235P | 106480701 |
| ENSG000002 | 2.03410332 | -0.278799949 | 0.35164423 | -0.7928467 | 0.42786717 | NA | NA | NA |
| ENSG000002 | 0.7536964 | -0.192464368 | 0.28656601 | -0.6716232 | 0.50182363 | NA | RNU6-30P | 100873747 |
| ENSG000002 | 0.98224204 | -0.180910714 | 0.30431097 | -0.5944929 | 0.55218248 | NA | NA | NA |
| ENSG000002 | 1.19423449 | 0.106919908 | 0.32137285 | 0.33269739 | 0.73936272 | NA | NA | NA |
| ENSG000002 | 0 | NA | NA | NA | NA | NA | NA | NA |

| | | | | | | | | | |
|---|---|---|---|---|---|---|---|---|---|
| ENSG000002 | 0.7283922 | 0.119158341 | 0.27073941 | 0.44012189 | 0.65984883 | NA | | RNU6-851P | 106479942 |
| ENSG000002 | 0.99121266 | -0.205080843 | 0.31756778 | -0.645786 | 0.51841792 | NA | | RNU6-140P | 106481221 |
| ENSG000002 | 0.22997012 | 0.167051788 | 0.17646558 | 0.94665368 | 0.34381528 | NA | | SNORD7 | 692076 |
| ENSG000002 | 0.59141222 | 0.061254288 | 0.25406951 | 0.24109263 | 0.80948332 | NA | | RNU6-83P | 100873782 |
| ENSG000002 | 0.25351207 | -0.189965547 | 0.1824784 | -1.0410303 | 0.29786148 | NA | | NA | NA |
| ENSG000002 | 0.97590063 | -0.193816421 | 0.31294085 | -0.6193388 | 0.53569316 | NA | | NA | NA |
| ENSG000002 | 0.12685397 | -0.090215768 | 0.12901919 | -0.699243 | 0.48440017 | NA | | NA | NA |
| ENSG000002 | 1.49208021 | -0.324669709 | 0.34183286 | -0.9497908 | 0.34221855 | NA | | NA | NA |
| ENSG000002 | 0 | NA | NA | NA | NA | NA | | SNORA8 | 654320 |
| ENSG000002 | 0.34999534 | 0.241411141 | 0.20772605 | 1.16216113 | 0.24517002 | NA | | NA | NA |
| ENSG000002 | 0.91356702 | -0.257298229 | 0.28712292 | -0.8961257 | 0.37018563 | NA | | RNU6-1152P | 106480073 |
| ENSG000002 | 0.47417772 | -0.160107059 | 0.22811633 | -0.7018658 | 0.48276283 | NA | | RNU6-145P | 106479630 |
| ENSG000002 | 0 | NA | NA | NA | NA | NA | | RNU6-878P | 106479954 |
| ENSG000002 | 1.13207153 | -0.423120938 | 0.31930944 | -1.3251125 | 0.18513383 | NA | | RNU4-70P | 106480548 |
| ENSG000002 | 1.11097538 | -0.258164454 | 0.31924678 | -0.8086674 | 0.41870651 | NA | | RNU6-1127P | 106480640 |
| ENSG000002 | 0.99354849 | -0.198387564 | 0.31363452 | -0.6325438 | 0.52703161 | NA | | RNU6-429P | 106479755 |
| ENSG000002 | 0.4009426 | -0.302672773 | 0.2240145 | -1.3511303 | 0.17665371 | NA | | NA | NA |
| ENSG000002 | 1.1567784 | -0.421135158 | 0.31773373 | -1.3254342 | 0.18502717 | NA | | RNU6-1184P | 106481553 |
| ENSG000002 | 1.20265974 | -0.184062079 | 0.3283039 | -0.5606454 | 0.57503928 | NA | | NA | NA |
| ENSG000002 | 0.90551233 | -0.110719234 | 0.29373998 | -0.3769294 | 0.70622607 | NA | | NA | NA |
| ENSG000002 | 0.83421419 | 0.200993891 | 0.29605006 | 0.67891859 | 0.49718945 | NA | | RNU6-160P | 106479637 |
| ENSG000002 | 1.27654111 | -0.489270034 | 0.32739408 | -1.4944376 | 0.13506128 | NA | | RNU1-89P | 106480412 |
| ENSG000002 | 0.74112301 | -0.344869609 | 0.28036958 | -1.2300536 | 0.21867704 | NA | | RNU6-901P | 106479965 |
| ENSG000002 | 1.38560479 | -0.402977302 | 0.33335507 | -1.2088531 | 0.2267193 | NA | | NA | NA |
| ENSG000002 | 2.14770639 | -0.262361547 | 0.36401466 | -0.7207445 | 0.47106676 | NA | | NA | NA |
| ENSG000002 | 1.20738292 | -0.176227633 | 0.32242496 | -0.5465695 | 0.58467456 | NA | | RNU6-636P | 106480597 |
| ENSG000002 | 1.67993769 | -0.552008823 | 0.34888028 | -1.58223 | 0.11359708 | NA | | NA | NA |
| ENSG000002 | 1.41503756 | 0.237128904 | 0.34253965 | 0.69226702 | 0.48876965 | NA | | RNU6-1263P | 106480653 |
| ENSG000002 | 0 | NA | NA | NA | NA | NA | | NA | NA |

| | | | | | | | | |
|---|---|---|---|---|---|---|---|---|
| ENSG000002 | 1.36357555 | -0.271424155 | 0.34113909 | -0.7956407 | 0.42624089 | NA | RNU6-680P | 106480601 |
| ENSG000002 | 1.59306692 | -0.254904666 | 0.34560669 | -0.7375571 | 0.46078364 | NA | RNU6-12P | 106481973 |
| ENSG000002 | 1.39081723 | -0.424164077 | 0.33849111 | -1.2531026 | 0.21016838 | NA | RNU6-296P | 106479698 |
| ENSG000002 | 0.7868574 | -0.180771671 | 0.27568303 | -0.6557229 | 0.51200244 | NA | NA | NA |
| ENSG000002 | 1.55948383 | -0.114412453 | 0.34829882 | -0.3284894 | 0.74254169 | NA | RNU6-225P | 106479667 |
| ENSG000002 | 0.61750123 | 0.055806918 | 0.26401083 | 0.21138117 | 0.83258985 | NA | RNU6-1222P | 106480100 |
| ENSG000002 | 0.95688789 | 0.256211939 | 0.30021216 | 0.85343626 | 0.39341742 | NA | RNU6-306P | 106479703 |
| ENSG000002 | 1.05751176 | 0.435215038 | 0.2990819 | 1.45517009 | 0.14562218 | NA | RNU6-540P | 106479806 |
| ENSG000002 | 0.3934287 | -0.107186984 | 0.22072612 | -0.4856108 | 0.62724315 | NA | RNU6-1257P | 106481577 |
| ENSG000002 | 0.52165104 | -0.403921832 | 0.25252134 | -1.5995552 | 0.10969729 | NA | NA | NA |
| ENSG000002 | 1.89359852 | -0.386768066 | 0.35774077 | -1.0811406 | 0.27963458 | NA | RNU6-14P | 100873746 |
| ENSG000002 | 1.65249855 | -0.512948051 | 0.34421753 | -1.4901857 | 0.1361754 | NA | RNU6-178P | 106479645 |
| ENSG000002 | 0.50016259 | -0.200428691 | 0.24824437 | -0.8073846 | 0.41944491 | NA | RNU6-422P | 106480580 |
| ENSG000002 | 0.8261595 | 0.365072387 | 0.28985807 | 1.25948669 | 0.20785459 | NA | RNU6-939P | 106481476 |
| ENSG000002 | 1.02027112 | -0.196917746 | 0.31469423 | -0.6257431 | 0.53148346 | NA | NA | NA |
| ENSG000002 | 1.33785127 | -0.261215775 | 0.33765189 | -0.7736245 | 0.43915288 | NA | RNU6-297P | 106479699 |
| ENSG000002 | 0.11865494 | 0.083440866 | 0.12751478 | 0.65436232 | 0.5128784 | NA | NA | NA |
| ENSG000002 | 1.50903901 | -0.056754322 | 0.34480022 | -0.1646006 | 0.86925836 | NA | NA | NA |
| ENSG000002 | 0.27408862 | -0.199826037 | 0.18669465 | -1.0703362 | 0.28446801 | NA | NA | NA |
| ENSG000002 | 1.10391794 | -0.117748111 | 0.32445569 | -0.3629097 | 0.71667236 | NA | RNU6-748P | 106481416 |
| ENSG000002 | 1.68343336 | -0.715150439 | 0.35017885 | -2.0422434 | 0.0411274 | NA | NA | NA |
| ENSG000002 | 0.24619405 | -0.007630671 | 0.17915639 | -0.0425922 | 0.96602658 | NA | RNU6-950P | 106479986 |
| ENSG000002 | 0 | NA | NA | NA | NA | NA | NA | NA |
| ENSG000002 | 0.73095424 | -0.032435186 | 0.2762491 | -0.1174128 | 0.90653292 | NA | NA | NA |
| ENSG000002 | 0.98236521 | -0.330952361 | 0.30584019 | -1.0821088 | 0.27920419 | NA | RNU6-310P | 106481277 |
| ENSG000002 | 0.51931521 | -0.354214731 | 0.23897171 | -1.4822455 | 0.13827499 | NA | NA | NA |
| ENSG000002 | 0.94297431 | -0.608628191 | 0.29393761 | -2.0706033 | 0.03839588 | NA | NA | NA |
| ENSG000002 | 0.6439198 | -0.261315757 | 0.26001163 | -1.0050157 | 0.31488931 | NA | NA | NA |
| ENSG000002 | 0.73659282 | 0.11903157 | 0.27861905 | 0.42721979 | 0.66921925 | NA | RNU6-136P | 106479626 |

| | | | | | | | | | |
|---|---|---|---|---|---|---|---|---|---|
| ENSG000002 | 0 | NA | NA | NA | NA | NA | | RNU6-1060P | 106480032 |
| ENSG000002 | 0 | NA | NA | NA | NA | NA | | NA | NA |
| ENSG000002 | 0.11934008 | 0.083564698 | 0.12760091 | 0.65489106 | 0.5125379 | NA | | RNU6-1179P | 106480084 |
| ENSG000002 | 1.8305188 | 0.244488776 | 0.35251169 | 0.69356218 | 0.48795682 | NA | | RNU6-1011P | 106480011 |
| ENSG000002 | 0.24756242 | -0.187034346 | 0.18119817 | -1.0322088 | 0.30197431 | NA | | NA | NA |
| ENSG000002 | 0.78130864 | -0.565696235 | 0.28699154 | -1.9711251 | 0.04870956 | NA | | RNU6-959P | 106479990 |
| ENSG000002 | 1.47697512 | -0.465843676 | 0.34136952 | -1.3646317 | 0.17236883 | NA | | NA | NA |
| ENSG000002 | 0.96408968 | 0.117920628 | 0.3052888 | 0.38625926 | 0.69930467 | NA | | NA | NA |
| ENSG000002 | 0 | NA | NA | NA | NA | NA | | SNORA64 | 26784 |
| ENSG000002 | 0 | NA | NA | NA | NA | NA | | SNORA41 | 619569 |
| ENSG000002 | 1.95020396 | -0.675945973 | 0.35798623 | -1.8881898 | 0.05900047 | NA | | NA | NA |
| ENSG000002 | 1.61483824 | -0.269636956 | 0.35405537 | -0.7615672 | 0.44631836 | NA | | NA | NA |
| ENSG000002 | 0.60762977 | 0.053874087 | 0.26315632 | 0.20472276 | 0.83778874 | NA | | NA | NA |
| ENSG000002 | 1.10391849 | -0.101185902 | 0.31746157 | -0.3187343 | 0.74992797 | NA | | RNU6-455P | 106479767 |
| ENSG000002 | 0.26813897 | -0.197007041 | 0.18550319 | -1.0620143 | 0.2882292 | NA | | RNU6-768P | 106481775 |
| ENSG000002 | 1.19978367 | -0.157221704 | 0.31831139 | -0.4939242 | 0.6213597 | NA | | RNU6-1151P | 106481543 |
| ENSG000002 | 0.36556447 | -0.101943371 | 0.21659044 | -0.4706735 | 0.63787395 | NA | | NA | NA |
| ENSG000002 | 0.60350741 | 0.060879422 | 0.26278835 | 0.23166713 | 0.81679657 | NA | | NA | NA |
| ENSG000002 | 0 | NA | NA | NA | NA | NA | | NA | NA |
| ENSG000002 | 0.51459361 | -0.192534203 | 0.24265108 | -0.7934611 | 0.42750921 | NA | | RNU6-95P | 106479608 |
| ENSG000002 | 0.12070844 | -0.088503695 | 0.12783931 | -0.6923042 | 0.48874631 | NA | | NA | NA |
| ENSG000002 | 0.38884741 | -0.29745092 | 0.22235466 | -1.3377319 | 0.18098386 | NA | | RNU6-906P | 106481465 |
| ENSG000002 | 0 | NA | NA | NA | NA | NA | | NA | NA |
| ENSG000002 | 0 | NA | NA | NA | NA | NA | | RNU6-1072P | 106480037 |
| ENSG000002 | 1.61905289 | -0.271170812 | 0.35310407 | -0.7679629 | 0.44250926 | NA | | RNU6-114P | 106481765 |
| ENSG000002 | 1.79437798 | -0.47588868 | 0.35455276 | -1.3422225 | 0.17952387 | NA | | NA | NA |
| ENSG000002 | 1.4709351 | -0.311058832 | 0.33625669 | -0.9250636 | 0.35493281 | NA | | NA | NA |
| ENSG000002 | 0.42385498 | -0.262217565 | 0.21037102 | -1.2464529 | 0.21259817 | NA | | RNU6-541P | 106481352 |
| ENSG000002 | 0.25333738 | -0.015686291 | 0.1806605 | -0.0868275 | 0.93080867 | NA | | RNU6-417P | 106479750 |

| | | | | | | | | | |
|---|---|---|---|---|---|---|---|---|---|
| ENSG000002 | 0.88581489 | -0.272748145 | 0.29980043 | -0.9097657 | 0.3629461 | NA | | SNORD56B | 319139 |
| ENSG000002 | 0.95813866 | -0.003584877 | 0.20190319 | -0.0177554 | 0.98583397 | NA | | RNU6-520P | 106481345 |
| ENSG000002 | 0.22336757 | 0.163792188 | 0.17487321 | 0.93663397 | 0.34894687 | NA | | RNU6-19P | 100873754 |
| ENSG000002 | 1.69432441 | -0.053116967 | 0.35059657 | -0.1515045 | 0.87957774 | NA | | NA | NA |
| ENSG000002 | 0 | NA | NA | NA | NA | NA | | RNU6-291P | 106479696 |
| ENSG000002 | 0.47667517 | 0.31075097 | 0.23106181 | 1.34488244 | 0.1786632 | NA | | RNU6-606P | 106479834 |
| ENSG000002 | 0.9572029 | -0.181518609 | 0.30566846 | -0.5938415 | 0.55261815 | NA | | RNU6-535P | 106479803 |
| ENSG000002 | 0.12685397 | -0.090215768 | 0.12901919 | -0.699243 | 0.48440017 | NA | | RNU6-1104P | 106481528 |
| ENSG000002 | 0.25260017 | -0.015978705 | 0.18049464 | -0.0885273 | 0.92945758 | NA | | RNU6-331P | 106479713 |
| ENSG000002 | 0.26560726 | -0.177019426 | 0.17672274 | -1.0016788 | 0.31649875 | NA | | RNU6-997P | 106480007 |
| ENSG000002 | 0.71029337 | -0.023758903 | 0.2828661 | -0.0839935 | 0.93306163 | NA | | RNU6-476P | 106479777 |
| ENSG000002 | 0.47280936 | -0.012478229 | 0.23633508 | -0.0527989 | 0.95789215 | NA | | NA | NA |
| ENSG000002 | 0.72995644 | -0.027793814 | 0.27785056 | -0.1000315 | 0.9203193 | NA | | RNU6-1311P | 106480138 |
| ENSG000002 | 1.10623204 | -0.261221652 | 0.31685444 | -0.8244216 | 0.40970003 | NA | | RNU6-376P | 106479732 |
| ENSG000002 | 0.64833032 | -0.271392816 | 0.26190928 | -1.0362092 | 0.30010454 | NA | | NA | NA |
| ENSG000002 | 0.99832566 | -0.040897058 | 0.30841276 | -0.1326049 | 0.89450582 | NA | | NA | NA |
| ENSG000002 | 1.75614496 | -0.189670467 | 0.35562248 | -0.5333478 | 0.59379285 | NA | | RNU6-41P | 106478912 |
| ENSG000002 | 1.12879793 | -0.385193702 | 0.31179537 | -1.2354055 | 0.21667964 | NA | | NA | NA |
| ENSG000002 | 0 | NA | NA | NA | NA | NA | | RNU6-286P | 106480567 |
| ENSG000002 | 0.89683902 | -0.429202453 | 0.29559086 | -1.4520153 | 0.14649734 | NA | | NA | NA |
| ENSG000002 | 0.634074 | -0.448061355 | 0.26266228 | -1.7058458 | 0.08803679 | NA | | NA | NA |
| ENSG000002 | 2.00398296 | -0.708964498 | 0.35896451 | -1.9750267 | 0.04826511 | NA | | RNU6-1067P | 106481516 |
| ENSG000002 | 1.37317733 | -0.41576081 | 0.33815795 | -1.229487 | 0.21888926 | NA | | NA | NA |
| ENSG000002 | 0.92222307 | -0.44186707 | 0.29679498 | -1.4887956 | 0.13654119 | NA | | RNU6-1044P | 106480633 |
| ENSG000002 | 1.71342708 | -0.055958964 | 0.35884586 | -0.1559415 | 0.87607913 | NA | | NA | NA |
| ENSG000002 | 0.7366151 | 0.27950748 | 0.27181582 | 1.02829733 | 0.30380999 | NA | | RNU6-713P | 106479882 |
| ENSG000002 | 0.62906615 | -0.101175312 | 0.25634904 | -0.394678 | 0.69308058 | NA | | NA | NA |
| ENSG000002 | 1.14362538 | -0.420876861 | 0.31811118 | -1.3230496 | 0.18581889 | NA | | NA | NA |
| ENSG000002 | 0 | NA | NA | NA | NA | NA | | SNORA7A | 619563 |

| | | | | | | | | | |
|---|---|---|---|---|---|---|---|---|---|
| ENSG000002 | 1.95506783 | 0.059543345 | 0.35863129 | 0.16602942 | 0.8681338 | NA | NA | NA | |
| ENSG000002 | 1.28084008 | -0.513734283 | 0.33255513 | -1.5448094 | 0.12239237 | NA | NA | NA | |
| ENSG000002 | 0 | NA | NA | NA | NA | NA | SNORD102 | | 26771 |
| ENSG000002 | 0 | NA | NA | NA | NA | NA | RNVU1-14 | | 101954266 |
| ENSG000002 | 0.4682236 | 0.140020865 | 0.23655532 | 0.59191594 | 0.55390688 | NA | NA | NA | |
| ENSG000002 | 0.40942396 | -0.284058841 | 0.21788566 | -1.303706 | 0.19233384 | NA | NA | NA | |
| ENSG000002 | 1.72619069 | -0.395980536 | 0.34238238 | -1.1565447 | 0.24745841 | NA | RNU6-1031P | | 106480020 |
| ENSG000002 | 0.85951681 | -0.245407694 | 0.28844915 | -0.8507832 | 0.39488978 | NA | RNU6-1105P | | 106480052 |
| ENSG000002 | 1.89917161 | 0.0764444 | 0.35313691 | 0.21647241 | 0.82861953 | NA | RNU6-9 | | 101954271 |
| ENSG000002 | 0.23890637 | -0.010420674 | 0.17747379 | -0.0587167 | 0.95317776 | NA | RNU6-771P | | 106479907 |
| ENSG000002 | 0.61699269 | 0.06131205 | 0.26397339 | 0.23226602 | 0.8163314 | NA | NA | NA | |
| ENSG000002 | 0 | NA | NA | NA | NA | NA | RNU1-3 | | 26869 |
| ENSG000002 | 0 | NA | NA | NA | NA | NA | RNU6-555P | | 106479813 |
| ENSG000002 | 0.34931019 | 0.26109409 | 0.21538338 | 1.21222954 | 0.22542453 | NA | NA | NA | |
| ENSG000002 | 0.68678367 | 0.298085096 | 0.27650287 | 1.07805424 | 0.28100955 | NA | NA | NA | |
| ENSG000002 | 0.38019136 | -0.10756138 | 0.21876967 | -0.491665 | 0.62295621 | NA | NA | NA | |
| ENSG000002 | 1.23241869 | 0.103640945 | 0.33220778 | 0.31197627 | 0.75505856 | NA | MIR521-2 | | 574481 |
| ENSG000002 | 1.41866028 | -0.546194618 | 0.33232099 | -1.6435754 | 0.10026393 | NA | MIR99B | | 407056 |
| ENSG000002 | 0.59829496 | 0.063152461 | 0.26232549 | 0.24074084 | 0.80975598 | NA | MIR608 | | 693193 |
| ENSG000002 | 0 | NA | NA | NA | NA | NA | MIR647 | | 693232 |
| ENSG000002 | 0.73704825 | -0.025096307 | 0.27834669 | -0.090162 | 0.92815845 | NA | MIR578 | | 693163 |
| ENSG000002 | 1.86612266 | -0.115821083 | 0.35643011 | -0.3249475 | 0.74522078 | NA | MIR635 | | 693220 |
| ENSG000002 | 0.47761686 | -0.01948631 | 0.23692179 | -0.0822479 | 0.93444962 | NA | NA | NA | |
| ENSG000002 | 0.23816916 | -0.010712105 | 0.1772998 | -0.060418 | 0.9518227 | NA | MIR23B | | 407011 |
| ENSG000002 | 0.12685397 | -0.090215768 | 0.12901919 | -0.699243 | 0.48440017 | NA | MIR203A | | 406986 |
| ENSG000002 | 0.77379475 | -0.378828349 | 0.28947278 | -1.3086838 | 0.19064148 | NA | NA | NA | |
| ENSG000002 | 0 | NA | NA | NA | NA | NA | MIR615 | | 693200 |
| ENSG000002 | 0.6545623 | -0.10940044 | 0.2585229 | -0.423175 | 0.67216755 | NA | MIR661 | | 724031 |
| ENSG000002 | 1.58905116 | -0.103432884 | 0.34125114 | -0.303099 | 0.76181441 | NA | MIR649 | | 693234 |

| | | | | | | | | |
|---|---|---|---|---|---|---|---|---|
| ENSG000002 | 0.66654986 | -0.459691143 | 0.26528715 | -1.7328059 | 0.08313017 | NA | MIR587 | 693172 |
| ENSG000002 | 0.52876404 | -0.191903645 | 0.2442426 | -0.7857092 | 0.4320379 | NA | MIR662 | 724032 |
| ENSG000002 | 0.36416385 | 0.068880976 | 0.20758164 | 0.33182596 | 0.74002068 | NA | MIR526B | 574468 |
| ENSG000002 | 0.26062507 | -0.012888319 | 0.18226503 | -0.070712 | 0.94362699 | NA | MIR199B | 406978 |
| ENSG000002 | 1.26259936 | -0.509082513 | 0.33211466 | -1.5328517 | 0.12531241 | NA | MIR30B | 407030 |
| ENSG000002 | 1.77132763 | -0.597278778 | 0.3523416 | -1.6951696 | 0.09004325 | NA | MIR606 | 693191 |
| ENSG000002 | 0.26813897 | -0.197007041 | 0.18550319 | -1.0620143 | 0.2882292 | NA | MIR181D | 574457 |
| ENSG000002 | 0.63893761 | -0.11408674 | 0.27243711 | -0.4187636 | 0.67538892 | NA | MIR508 | 574513 |
| ENSG000002 | 0.46346816 | 0.146936161 | 0.23594779 | 0.62274862 | 0.53344973 | NA | MIR215 | 406997 |
| ENSG000002 | 0.8678023 | -0.271126522 | 0.29886542 | -0.907186 | 0.36430845 | NA | MIR520A | 574467 |
| ENSG000002 | 1.48802099 | -0.457533742 | 0.34038476 | -1.3441664 | 0.17889459 | NA | MIR181A2 | 406954 |
| ENSG000002 | 0.38747904 | -0.104908247 | 0.21987514 | -0.4771265 | 0.63327207 | NA | MIR490 | 574443 |
| ENSG000002 | 0 | NA | NA | NA | NA | NA | MIR124-3 | 406909 |
| ENSG000002 | 1.66948272 | 0.083009335 | 0.35426809 | 0.2343122 | 0.81474261 | NA | MIR520E | 574461 |
| ENSG000002 | 1.00720792 | -0.350938794 | 0.30924522 | -1.1348237 | 0.25644921 | NA | NA | NA |
| ENSG000002 | 2.06159959 | -0.112088654 | 0.36320573 | -0.3086093 | 0.75761877 | NA | NA | NA |
| ENSG000002 | 1.10332117 | 0.185906891 | 0.32035547 | 0.5803144 | 0.56170262 | NA | MIR191 | 406966 |
| ENSG000002 | 1.13611306 | -0.12191497 | 0.31519416 | -0.3867932 | 0.69890928 | NA | MIR554 | 693139 |
| ENSG000002 | 1.10363562 | -0.260803268 | 0.31898894 | -0.8175934 | 0.41358937 | NA | MIR200A | 406983 |
| ENSG000002 | 0.50360227 | -0.018733935 | 0.23996924 | -0.0780681 | 0.93777391 | NA | MIR127 | 406914 |
| ENSG000002 | 0.36690249 | -0.096999144 | 0.21683809 | -0.4473344 | 0.65463361 | NA | MIR149 | 406941 |
| ENSG000002 | 1.23108513 | -0.180577358 | 0.32758196 | -0.5512433 | 0.58146691 | NA | MIR604 | 693189 |
| ENSG000002 | 0.24619405 | -0.007630671 | 0.17915639 | -0.0425922 | 0.96602658 | NA | MIR181C | 406957 |
| ENSG000002 | 0 | NA | NA | NA | NA | NA | MIR193A | 406968 |
| ENSG000002 | 0.7356259 | -0.02722741 | 0.28456398 | -0.0956812 | 0.92377381 | NA | NA | NA |
| ENSG000002 | 0.6103579 | 0.059265293 | 0.25585983 | 0.23163188 | 0.81682395 | NA | NA | NA |
| ENSG000002 | 1.25246889 | -0.350158116 | 0.33199199 | -1.0547186 | 0.29155405 | NA | MIR3074 | 100422842 |
| ENSG000002 | 1.61512248 | 0.014613906 | 0.35185742 | 0.0415336 | 0.96687051 | NA | MIR516A2 | 574499 |
| ENSG000002 | 0.71368044 | 0.133536879 | 0.28320862 | 0.47151417 | 0.6372736 | NA | MIR619 | 693204 |

| | | | | | | | | |
|---|---|---|---|---|---|---|---|---|
| ENSG000002 | 0.58826863 | 0.215244732 | 0.25535246 | 0.8429319 | 0.39926653 | NA | MIR194-1 | 406969 |
| ENSG000002 | 0.22263036 | 0.146922267 | 0.16600367 | 0.88505432 | 0.37612734 | NA | MIR128-2 | 406916 |
| ENSG000002 | 1.60939904 | -0.128356998 | 0.35362446 | -0.3629755 | 0.71662322 | NA | MIR562 | 693147 |
| ENSG000002 | 0.96724458 | -0.038194427 | 0.31199407 | -0.1224204 | 0.90256611 | NA | MIR581 | 693166 |
| ENSG000002 | 0.65493287 | -0.26892161 | 0.26251527 | -1.0244037 | 0.30564465 | NA | MIR651 | 723779 |
| ENSG000002 | 1.99933249 | -0.592513463 | 0.36231834 | -1.6353394 | 0.10197792 | NA | MIR526A1 | 574475 |
| ENSG000002 | 2.04646825 | -0.542484259 | 0.35484536 | -1.5287906 | 0.12631636 | NA | MIR7-3 | 407045 |
| ENSG000002 | 0.3867939 | -0.105148852 | 0.21977268 | -0.4784437 | 0.63233444 | NA | MIR641 | 693226 |
| ENSG000002 | 1.24799131 | 0.091184099 | 0.32898078 | 0.27717151 | 0.78164842 | NA | MIR588 | 693173 |
| ENSG000002 | 0.35067856 | 0.074230421 | 0.21389444 | 0.34704231 | 0.72855953 | NA | MIR521-1 | 574494 |
| ENSG000002 | 0.11205239 | 0.082118251 | 0.12671962 | 0.64803108 | 0.51696485 | NA | MIR499A | 574501 |
| ENSG000002 | 1.08997619 | -0.110626046 | 0.32152214 | -0.3440698 | 0.73079382 | NA | MIR595 | 693180 |
| ENSG000002 | 0.60644002 | -0.257083821 | 0.25820471 | -0.9956589 | 0.31941589 | NA | MIR99A | 407055 |
| ENSG000002 | 0.13280363 | -0.093394593 | 0.13115644 | -0.7120855 | 0.47641185 | NA | MIR193B | 574455 |
| ENSG000002 | 1.10286224 | 0.030862346 | 0.32164622 | 0.09595122 | 0.92355932 | NA | MIR510 | 574515 |
| ENSG000002 | 1.70752949 | -0.050999229 | 0.35368283 | -0.1441948 | 0.88534664 | NA | MIR571 | 693156 |
| ENSG000002 | 1.7439715 | -0.064683431 | 0.35344068 | -0.1830107 | 0.85478961 | NA | MIR512-2 | 574459 |
| ENSG000002 | 0.48032325 | -0.179266136 | 0.23878452 | -0.7507444 | 0.45280651 | NA | MIR512-1 | 574458 |
| ENSG000002 | 0.97723866 | -0.189527453 | 0.31301 | -0.6054997 | 0.54484704 | NA | MIR655 | 724025 |
| ENSG000002 | 0.77362006 | -0.203375959 | 0.28776237 | -0.7067497 | 0.47972205 | NA | MIR153-1 | 406944 |
| ENSG000002 | 0.62550439 | -0.114540246 | 0.27135801 | -0.4221001 | 0.67295194 | NA | NA | NA |
| ENSG000002 | 1.00564368 | -0.209156113 | 0.31808973 | -0.6575381 | 0.51083499 | NA | MIR570 | 693155 |
| ENSG000002 | 0.47707552 | -0.019925931 | 0.23686191 | -0.0841247 | 0.93295731 | NA | MIR28 | 407020 |
| ENSG000002 | 1.74229248 | -0.319399337 | 0.35211101 | -0.9070984 | 0.36435477 | NA | MIR558 | 693143 |
| ENSG000002 | 1.45123777 | 0.092848732 | 0.34189893 | 0.27156778 | 0.78595438 | NA | NA | NA |
| ENSG000002 | 0.50585166 | -0.183613822 | 0.24177005 | -0.7594564 | 0.44757957 | NA | MIR489 | 574442 |
| ENSG000002 | 2.10567463 | -0.630027731 | 0.36249967 | -1.7380091 | 0.0822092 | NA | NA | NA |
| ENSG000002 | 1.58643906 | -0.322369356 | 0.33145411 | -0.9725912 | 0.33075651 | NA | MIR548A2 | 693126 |
| ENSG000002 | 0.9622326 | 0.263755889 | 0.29782064 | 0.88561991 | 0.37582238 | NA | NA | NA |

| | | | | | | | | | |
|---|---|---|---|---|---|---|---|---|---|
| ENSG000002 | 0.23065526 | 0.167384503 | 0.17662711 | 0.94767164 | 0.34329664 | NA | MIR602 | | 693187 |
| ENSG000002 | 1.51860888 | -0.181814818 | 0.34146128 | -0.532461 | 0.59440674 | NA | MIR184 | | 406960 |
| ENSG000002 | 2.07628864 | 0.241207754 | 0.36387421 | 0.66288774 | 0.50740246 | NA | MIR32 | | 407036 |
| ENSG000002 | 1.15788618 | -0.576254973 | 0.31642482 | -1.8211434 | 0.06858506 | NA | MIR518A2 | | 574491 |
| ENSG000002 | 2.10073198 | 0.003091722 | 0.36552533 | 0.0084583 | 0.99325134 | NA | MIR597 | | 693182 |
| ENSG000002 | 1.09441313 | 0.029121997 | 0.31984474 | 0.09105042 | 0.92745253 | NA | MIR211 | | 406993 |
| ENSG000002 | 1.36503036 | -0.269439487 | 0.3402089 | -0.7919825 | 0.42837089 | NA | MIR7-2 | | 407044 |
| ENSG000002 | 0.86247987 | -0.11365785 | 0.30252319 | -0.3756996 | 0.70714021 | NA | NA | NA | |
| ENSG000002 | 0 | NA | NA | NA | NA | NA | MIR129-1 | | 406917 |
| ENSG000002 | 2.01692299 | -0.455224533 | 0.36349078 | -1.2523689 | 0.21043549 | NA | MIR518F | | 574472 |
| ENSG000002 | 0.24411881 | -0.012946855 | 0.17864669 | -0.0724718 | 0.94222641 | NA | MIR141 | | 406933 |
| ENSG000002 | 0.86339122 | -0.275317148 | 0.2986111 | -0.9219924 | 0.35653256 | NA | MIR525 | | 574470 |
| ENSG000002 | 1.80191928 | 0.381147742 | 0.3524413 | 1.08145027 | 0.27949687 | NA | MIR627 | | 693212 |
| ENSG000002 | 0.50016259 | -0.200428691 | 0.24824437 | -0.8073846 | 0.41944491 | NA | NA | NA | |
| ENSG000002 | 1.3798642 | -0.429160126 | 0.34127177 | -1.2575319 | 0.20856111 | NA | NA | NA | |
| ENSG000002 | 0 | NA | NA | NA | NA | NA | MIR572 | | 693157 |
| ENSG000002 | 2.00254699 | -0.199367058 | 0.36329333 | -0.5487771 | 0.58315843 | NA | MIR551B | | 693136 |
| ENSG000002 | 1.43225875 | 0.075036198 | 0.33387551 | 0.22474304 | 0.82217918 | NA | MIR623 | | 693208 |
| ENSG000002 | 0.64534373 | -0.115494329 | 0.26669592 | -0.4330562 | 0.66497392 | NA | MIR520B | | 574473 |
| ENSG000002 | 1.91168791 | 0.458979855 | 0.36074452 | 1.27231275 | 0.20326202 | NA | MIR222 | | 407007 |
| ENSG000002 | 1.08957338 | 0.040533153 | 0.32508124 | 0.12468623 | 0.90077196 | NA | NA | NA | |
| ENSG000002 | 0.26560726 | -0.177019426 | 0.17672274 | -1.0016788 | 0.31649875 | NA | MIR449B | | 693123 |
| ENSG000002 | 1.60866182 | -0.128586002 | 0.35361548 | -0.3636323 | 0.71613262 | NA | MIR200B | | 406984 |
| ENSG000002 | 0.49307133 | -0.179259067 | 0.24031383 | -0.7459374 | 0.45570524 | NA | MIR506 | | 574511 |
| ENSG000002 | 2.06900096 | 0.239539862 | 0.36384054 | 0.65836496 | 0.51030365 | NA | MIR218-1 | | 407000 |
| ENSG000002 | 1.89516076 | -0.124685127 | 0.35556386 | -0.3506687 | 0.72583689 | NA | MIR517A | | 574479 |
| ENSG000002 | 0.73636311 | -0.02696217 | 0.28461296 | -0.0947328 | 0.9245271 | NA | MIR520D | | 574482 |
| ENSG000002 | 0.38651158 | -0.274258716 | 0.21461355 | -1.2779189 | 0.20127801 | NA | MIR657 | | 724027 |
| ENSG000002 | 0.11934008 | 0.083564698 | 0.12760091 | 0.65489106 | 0.5125379 | NA | MIR181B2 | | 406956 |

| | | | | | | | | |
|---|---|---|---|---|---|---|---|---|
| ENSG000002 | 0.38270188 | -0.272446727 | 0.2140009 | -1.2731102 | 0.20297894 | NA | MIR520C | 574476 |
| ENSG000002 | 1.21458024 | -0.048578891 | 0.33319255 | -0.1457982 | 0.88408066 | NA | MIR218-2 | 407001 |
| ENSG000002 | 0.11934008 | 0.083564698 | 0.12760091 | 0.65489106 | 0.5125379 | NA | NA | NA |
| ENSG000002 | 0.73516888 | -0.031630387 | 0.28453243 | -0.1111662 | 0.91148456 | NA | NA | NA |
| ENSG000002 | 1.84709403 | 0.006747101 | 0.35298377 | 0.01911448 | 0.98474978 | NA | MIR10B | 406903 |
| ENSG000002 | 0.52760069 | -0.383393289 | 0.24704506 | -1.5519164 | 0.12068222 | NA | MIR575 | 693160 |
| ENSG000002 | 0 | NA | NA | NA | NA | NA | NA | NA |
| ENSG000002 | 1.75582809 | 0.536787887 | 0.3527364 | 1.52178194 | 0.12806373 | NA | MIR553 | 693138 |
| ENSG000002 | 0.61491745 | 0.056458936 | 0.26379319 | 0.21402727 | 0.83052579 | NA | MIR199A1 | 406976 |
| ENSG000002 | 0.36345889 | 0.070257596 | 0.21589421 | 0.32542602 | 0.74485866 | NA | NA | NA |
| ENSG000002 | 1.32475594 | 0.167415204 | 0.33380341 | 0.50153832 | 0.61599232 | NA | MIR450A2 | 574505 |
| ENSG000002 | 1.32091377 | 0.177390384 | 0.33850573 | 0.52403952 | 0.60025105 | NA | MIR580 | 693165 |
| ENSG000002 | 1.31866385 | 0.312966069 | 0.33566815 | 0.93236749 | 0.35114664 | NA | MIR93 | 407050 |
| ENSG000002 | 0.64813445 | -0.288472774 | 0.26867189 | -1.0736991 | 0.28295756 | NA | MIR532 | 693124 |
| ENSG000002 | 1.02337395 | -0.351378104 | 0.3099324 | -1.133725 | 0.25690992 | NA | MIR181A1 | 406995 |
| ENSG000002 | 0.25370795 | -0.171331747 | 0.17411721 | -0.9840024 | 0.32511435 | NA | NA | NA |
| ENSG000002 | 0.35022155 | 0.06940789 | 0.21379411 | 0.32464827 | 0.74544729 | NA | MIR329-2 | 574409 |
| ENSG000002 | 0.59479737 | 0.057784305 | 0.26201081 | 0.22054169 | 0.82544931 | NA | MIR617 | 693202 |
| ENSG000002 | 1.0138736 | -0.498211149 | 0.30523646 | -1.6322137 | 0.10263447 | NA | MIR516A1 | 574498 |
| ENSG000002 | 0.58867549 | 0.367778179 | 0.24725052 | 1.48747184 | 0.13689023 | NA | NA | NA |
| ENSG000002 | 1.58971265 | -0.384433244 | 0.34648668 | -1.1095181 | 0.26720674 | NA | MIR586 | 693171 |
| ENSG000002 | 0.4680172 | 0.307231342 | 0.22996067 | 1.33601689 | 0.18154375 | NA | MIR642A | 693227 |
| ENSG000002 | 0.86304239 | 0.044424632 | 0.30221493 | 0.14699681 | 0.88313454 | NA | MIR548A1 | 693125 |
| ENSG000002 | 0 | NA | NA | NA | NA | NA | MIR551A | 693135 |
| ENSG000002 | 0.57267777 | 0.42409321 | 0.26296456 | 1.61273905 | 0.10680121 | NA | NA | NA |
| ENSG000002 | 0.72500459 | -0.030476671 | 0.27582839 | -0.1104914 | 0.91201965 | NA | MIR15B | 406949 |
| ENSG000002 | 2.03970173 | 0.239646925 | 0.36120804 | 0.66345956 | 0.50703627 | NA | MIR648 | 693233 |
| ENSG000002 | 0.12685397 | -0.090215768 | 0.12901919 | -0.699243 | 0.48440017 | NA | NA | NA |
| ENSG000002 | 0.84935263 | 0.355549886 | 0.28886249 | 1.23086209 | 0.21837445 | NA | MIR500A | 574502 |

| | | | | | | | | | |
|---|---|---|---|---|---|---|---|---|---|
| ENSG000002 | 0.59342901 | 0.227374733 | 0.26299808 | 0.86454904 | 0.38728635 | NA | MIR519C | | 574466 |
| ENSG000002 | 0.63233507 | -0.116461163 | 0.27190168 | -0.4283209 | 0.66841754 | NA | MIR26A2 | | 407016 |
| ENSG000002 | 0.76949578 | -0.359754798 | 0.28380236 | -1.2676244 | 0.20493211 | NA | MIR187 | | 406963 |
| ENSG000002 | 0.60644002 | -0.257083821 | 0.25820471 | -0.9956589 | 0.31941589 | NA | MIR216A | | 406998 |
| ENSG000002 | 0.35543399 | 0.061147018 | 0.20611885 | 0.29665903 | 0.76672684 | NA | MIR520G | | 574484 |
| ENSG000002 | 0.3588776 | -0.099902505 | 0.21555794 | -0.4634601 | 0.6430346 | NA | NA | NA | |
| ENSG000002 | 0.63603926 | -0.266971526 | 0.26084558 | -1.0234849 | 0.30607861 | NA | MIR646 | | 693231 |
| ENSG000002 | 0.74638752 | -0.326267471 | 0.27400982 | -1.1907145 | 0.23376568 | NA | MIR518A1 | | 574488 |
| ENSG000002 | 0.25370795 | -0.171331747 | 0.17411721 | -0.9840024 | 0.32511435 | NA | MIR599 | | 693184 |
| ENSG000002 | 0.35073255 | 0.241740392 | 0.2078501 | 1.16305162 | 0.24480856 | NA | NA | NA | |
| ENSG000002 | 0.95790977 | -0.026156233 | 0.30081211 | -0.0869521 | 0.93070962 | NA | MIR95 | | 407052 |
| ENSG000002 | 0 | NA | NA | NA | NA | NA | MIR27A | | 407018 |
| ENSG000002 | 0.89227499 | -0.122214336 | 0.30394926 | -0.4020879 | 0.6876193 | NA | MIR519E | | 574463 |
| ENSG000002 | 0 | NA | NA | NA | NA | NA | NA | NA | |
| ENSG000002 | 1.41746108 | -0.040317943 | 0.32260186 | -0.1249774 | 0.90054144 | NA | MIR605 | | 693190 |
| ENSG000002 | 1.45801391 | -0.056242421 | 0.34520176 | -0.1629262 | 0.87057652 | NA | MIR147A | | 406939 |
| ENSG000002 | 1.28482822 | 0.144120588 | 0.31908499 | 0.45166833 | 0.65150793 | NA | MIR563 | | 693148 |
| ENSG000002 | 1.87898447 | 0.461484657 | 0.35690518 | 1.29301754 | 0.19600499 | NA | MIR105-2 | | 406898 |
| ENSG000002 | 0.69617693 | 0.132831392 | 0.27567226 | 0.48184534 | 0.62991582 | NA | MIR519B | | 574469 |
| ENSG000002 | 0.12685397 | -0.090215768 | 0.12901919 | -0.699243 | 0.48440017 | NA | MIR596 | | 693181 |
| ENSG000002 | 0.96881286 | 0.122086005 | 0.31200215 | 0.3912986 | 0.69557653 | NA | MIR30A | | 407029 |
| ENSG000002 | 0.97544362 | -0.189881147 | 0.30813177 | -0.6162336 | 0.53774036 | NA | MIR517C | | 574492 |
| ENSG000002 | 0.3701135 | 0.066858436 | 0.20853829 | 0.32060509 | 0.74850968 | NA | MIR33B | | 693120 |
| ENSG000002 | 0.59667619 | 0.052746379 | 0.26217118 | 0.20119061 | 0.84054954 | NA | MIR520H | | 574493 |
| ENSG000002 | 1.09638231 | -0.121548462 | 0.32558861 | -0.3733191 | 0.70891093 | NA | MIR518B | | 574474 |
| ENSG000002 | 2.0259972 | -0.430676383 | 0.35859335 | -1.2010161 | 0.22974496 | NA | MIR125B2 | | 406912 |
| ENSG000002 | 0.79746769 | 0.530002434 | 0.28491603 | 1.86020574 | 0.06285642 | NA | NA | NA | |
| ENSG000002 | 1.52587079 | -0.637487515 | 0.34361166 | -1.8552558 | 0.06355971 | NA | MIR514A2 | | 574517 |
| ENSG000002 | 1.55120332 | -0.594448246 | 0.33684811 | -1.7647368 | 0.07760801 | NA | MIR514A3 | | 574518 |

| | | | | | | | | |
|---|---|---|---|---|---|---|---|---|
| ENSG000002 | 0.98366651 | -0.034243466 | 0.31266311 | -0.1095219 | 0.91278853 | NA | MIR514A1 | 574516 |
| ENSG000002 | 0.92353047 | 0.265080526 | 0.30180748 | 0.87830998 | 0.37977552 | NA | MIR498 | 574460 |
| ENSG000002 | 0.14128499 | -0.097772014 | 0.13403235 | -0.7294658 | 0.4657168 | NA | MIR513B | 100313822 |
| ENSG000002 | 1.212053 | -0.327441866 | 0.32728785 | -1.0004706 | 0.31708283 | NA | MIR513A1 | 574509 |
| ENSG000002 | 0.60470109 | 0.059032226 | 0.26929751 | 0.21920821 | 0.82648786 | NA | MIR566 | 693151 |
| ENSG000002 | 0.50341032 | -0.342798738 | 0.23572799 | -1.4542131 | 0.14588723 | NA | MIR196A2 | 406973 |
| ENSG000002 | 0.70938393 | 0.278216169 | 0.26727878 | 1.04092128 | 0.29791208 | NA | MIR516B2 | 574485 |
| ENSG000002 | 0.46933541 | 0.307746265 | 0.23012247 | 1.33731513 | 0.18111979 | NA | MIR135A1 | 406925 |
| ENSG000002 | 0.2596576 | -0.193002035 | 0.1837913 | -1.0501152 | 0.29366517 | NA | MIR33A | 407039 |
| ENSG000002 | 1.03355999 | -0.202825642 | 0.31368466 | -0.6465909 | 0.51789675 | NA | NA | NA |
| ENSG000002 | 0.27408862 | -0.199826037 | 0.18669465 | -1.0703362 | 0.28446801 | NA | MIR654 | 724024 |
| ENSG000002 | 0.7717195 | -0.360573024 | 0.28228954 | -1.2773163 | 0.20149061 | NA | MIR567 | 693152 |
| ENSG000002 | 0.6331621 | -0.262817417 | 0.25902408 | -1.0146447 | 0.31027521 | NA | MIR552 | 693137 |
| ENSG000002 | 0.52193336 | -0.189821458 | 0.24349877 | -0.7795582 | 0.43565098 | NA | MIR136 | 406927 |
| ENSG000002 | 0.67503122 | -0.469043005 | 0.26742254 | -1.7539397 | 0.07944084 | NA | MIR634 | 693219 |
| ENSG000002 | 0.34948434 | 0.063159058 | 0.2051263 | 0.30790326 | 0.75815594 | NA | MIR574 | 693159 |
| ENSG000002 | 0.89244968 | -0.286343813 | 0.30487766 | -0.9392089 | 0.34762349 | NA | MIR516B1 | 574490 |
| ENSG000002 | 1.24492061 | -0.347767768 | 0.33178274 | -1.0481792 | 0.29455605 | NA | NA | NA |
| ENSG000002 | 0.12685397 | -0.090215768 | 0.12901919 | -0.699243 | 0.48440017 | NA | NA | NA |
| ENSG000002 | 0.47958604 | -0.1795445 | 0.23869266 | -0.7521995 | 0.4519311 | NA | MIR624 | 693209 |
| ENSG000002 | 0.8497318 | -0.108769682 | 0.2967386 | -0.3665505 | 0.71395434 | NA | MIR579 | 693164 |
| ENSG000002 | 0.49763088 | -0.186408494 | 0.24078583 | -0.7741672 | 0.43883193 | NA | MIR105-1 | 406897 |
| ENSG000002 | 0.39841089 | -0.251751984 | 0.20668213 | -1.2180636 | 0.2231998 | NA | MIR656 | 724026 |
| ENSG000002 | 1.27941773 | -0.514249845 | 0.33251177 | -1.5465613 | 0.12196906 | NA | MIR153-2 | 406945 |
| ENSG000002 | 0.60624414 | -0.253150918 | 0.25658854 | -0.9866026 | 0.3238375 | NA | MIR496 | 574454 |
| ENSG000002 | 1.7280878 | 0.066175292 | 0.35319669 | 0.18736102 | 0.85137758 | NA | MIR569 | 693154 |
| ENSG000002 | 1.97701308 | -0.063490769 | 0.36110566 | -0.1758232 | 0.86043281 | NA | MIR629 | 693214 |
| ENSG000002 | 1.87542893 | -0.380999917 | 0.35718606 | -1.0666707 | 0.28612055 | NA | MIR620 | 693205 |
| ENSG000002 | 1.50420765 | -0.347497732 | 0.34691644 | -1.0016756 | 0.3165003 | NA | MIR507 | 574512 |

| | | | | | | | | | |
|---|---|---|---|---|---|---|---|---|---|
| ENSG000002 | 0.25145857 | -0.010118565 | 0.18031426 | -0.0561163 | 0.95524918 | NA | NA | NA | |
| ENSG000002 | 1.62417888 | -0.416308141 | 0.3525836 | -1.180736 | 0.23770763 | NA | MIR125B1 | | 406911 |
| ENSG000002 | 0 | NA | NA | NA | NA | NA | MIR638 | | 693223 |
| ENSG000002 | 1.76481256 | -0.317120458 | 0.34749026 | -0.9126024 | 0.36145168 | NA | MIR589 | | 693174 |
| ENSG000002 | 1.92487765 | 0.193639626 | 0.35908996 | 0.53925102 | 0.58971366 | NA | MIR557 | | 693142 |
| ENSG000002 | 1.24918341 | -0.051604364 | 0.32904311 | -0.1568316 | 0.87537755 | NA | MIR181B1 | | 406955 |
| ENSG000002 | 2.17356374 | 0.18744079 | 0.36614077 | 0.51193641 | 0.60869552 | NA | MIR607 | | 693192 |
| ENSG000002 | 0.83651619 | 0.050756481 | 0.29566295 | 0.17167008 | 0.86369691 | NA | MIR154 | | 406946 |
| ENSG000002 | 1.22135747 | -0.187884833 | 0.33029671 | -0.5688365 | 0.56946709 | NA | MIR527 | | 574497 |
| ENSG000002 | 0 | NA | NA | NA | NA | NA | MIR23A | | 407010 |
| ENSG000002 | 1.39748198 | 0.220401906 | 0.33108866 | 0.66568848 | 0.50561024 | NA | MIR519D | | 574480 |
| ENSG000002 | 0.92384342 | -0.260230075 | 0.2918757 | -0.8915784 | 0.37261895 | NA | MIR613 | | 693198 |
| ENSG000002 | 0.73952597 | -0.193293011 | 0.28563587 | -0.6767113 | 0.49858917 | NA | MIR513A2 | | 574510 |
| ENSG000002 | 1.15786499 | -0.772067853 | 0.31826127 | -2.4258932 | 0.01527076 | NA | MIR518E | | 574487 |
| ENSG000002 | 0.25370795 | -0.171331747 | 0.17411721 | -0.9840024 | 0.32511435 | NA | MIR493 | | 574450 |
| ENSG000002 | 1.33753861 | -0.249738849 | 0.33494629 | -0.7456087 | 0.45590386 | NA | MIR182 | | 406958 |
| ENSG000002 | 1.11729505 | -0.370086191 | 0.30481136 | -1.2141483 | 0.2246911 | NA | MIR519A1 | | 574496 |
| ENSG000002 | 0.40327843 | -0.281391716 | 0.21700406 | -1.2967118 | 0.19473038 | NA | MIR134 | | 406924 |
| ENSG000002 | 0.47895489 | -0.016192723 | 0.24471014 | -0.066171 | 0.94724166 | NA | MIR100 | | 406892 |
| ENSG000002 | 1.1839814 | 0.416910738 | 0.3283158 | 1.26984672 | 0.20413923 | NA | MIR325 | | 442899 |
| ENSG000002 | 0.26560726 | -0.177019426 | 0.17672274 | -1.0016788 | 0.31649875 | NA | MIR509-1 | | 574514 |
| ENSG000002 | 0.72537515 | -0.168191868 | 0.26954542 | -0.6239834 | 0.53263849 | NA | MIR431 | | 574038 |
| ENSG000002 | 1.41651198 | 0.241893121 | 0.34448504 | 0.70218759 | 0.48256218 | NA | MIR643 | | 693228 |
| ENSG000002 | 0.98283948 | 0.122754107 | 0.31257693 | 0.39271647 | 0.69452891 | NA | MIR549A | | 693132 |
| ENSG000002 | 0.25333738 | -0.015686291 | 0.1806605 | -0.0868275 | 0.93080867 | NA | MIR323B | | 574410 |
| ENSG000002 | 0.25145857 | -0.010118565 | 0.18031426 | -0.0561163 | 0.95524918 | NA | MIR503 | | 574506 |
| ENSG000002 | 0.12070844 | -0.088503695 | 0.12783931 | -0.6923042 | 0.48874631 | NA | MIR16-1 | | 406950 |
| ENSG000002 | 0 | NA | NA | NA | NA | NA | MIR125A | | 406910 |
| ENSG000002 | 0.7731096 | -0.379060994 | 0.28942548 | -1.3097015 | 0.19029682 | NA | MIRLET7F2 | | 406889 |

| | | | | | | | | | |
|---|---|---|---|---|---|---|---|---|---|
| ENSG000002 | 0.73906896 | -0.197744251 | 0.28558485 | -0.6924186 | 0.48867451 | NA | NA | NA | |
| ENSG000002 | 0.8531754 | -0.262752222 | 0.29650959 | -0.8861508 | 0.37553624 | NA | MIR653 | | 724023 |
| ENSG000002 | 0.23276084 | -0.008011382 | 0.17603982 | -0.0455089 | 0.96370166 | NA | MIR362 | | 574030 |
| ENSG000002 | 1.27748739 | -0.474239409 | 0.32510827 | -1.4587122 | 0.14464433 | NA | MIR140 | | 406932 |
| ENSG000002 | 1.59551148 | 0.019100521 | 0.34977119 | 0.05460862 | 0.95645027 | NA | MIR645 | | 693230 |
| ENSG000002 | 1.52845806 | -0.328344696 | 0.34107626 | -0.9626724 | 0.33571196 | NA | MIR618 | | 693203 |
| ENSG000002 | 0 | NA | NA | NA | NA | NA | MIR199A2 | | 406977 |
| ENSG000002 | 1.30481191 | 0.153599372 | 0.32530579 | 0.4721692 | 0.63680602 | NA | MIR591 | | 693176 |
| ENSG000002 | 0.38270188 | -0.272446727 | 0.2140009 | -1.2731102 | 0.20297894 | NA | NA | NA | |
| ENSG000002 | 1.08331966 | -0.254808717 | 0.31602308 | -0.8062978 | 0.42007115 | NA | MIR548A3 | | 693127 |
| ENSG000002 | 0.23816916 | -0.010712105 | 0.1772998 | -0.060418 | 0.9518227 | NA | MIR106B | | 406900 |
| ENSG000002 | 1.08567276 | 0.191027833 | 0.32503696 | 0.58771111 | 0.55672622 | NA | MIR320A | | 407037 |
| ENSG000002 | 0 | NA | NA | NA | NA | NA | SNORD73A | | 8944 |
| ENSG000002 | 1.69124957 | 0.325027145 | 0.34550407 | 0.9407332 | 0.3468416 | NA | SNORA35 | | 677816 |
| ENSG000002 | 0.66202413 | -0.272682834 | 0.26145371 | -1.0429488 | 0.296972 | NA | SNORD96B | | 692226 |
| ENSG000002 | 0 | NA | NA | NA | NA | NA | NA | NA | |
| ENSG000002 | 0 | NA | NA | NA | NA | NA | SNORD83B | | 116938 |
| ENSG000002 | 0 | NA | NA | NA | NA | NA | SNORA48 | | 652965 |
| ENSG000002 | 0.35346928 | -0.088570748 | 0.20618147 | -0.4295767 | 0.66750362 | NA | SNORD105 | | 692229 |
| ENSG000002 | 0.83695147 | -0.103511903 | 0.29440911 | -0.3515921 | 0.72514423 | NA | SNORD41 | | 26810 |
| ENSG000002 | 1.183577 | 0.421534339 | 0.32825309 | 1.28417479 | 0.19908081 | NA | NA | NA | |
| ENSG000002 | 1.78569214 | -0.192711927 | 0.35420204 | -0.5440734 | 0.58639093 | NA | NA | NA | |
| ENSG000002 | 0 | NA | NA | NA | NA | NA | NA | NA | |
| ENSG000002 | 0.34999534 | 0.241411141 | 0.20772605 | 1.16216113 | 0.24517002 | NA | NA | NA | |
| ENSG000002 | 0 | NA | NA | NA | NA | NA | NA | NA | |
| ENSG000002 | 0 | NA | NA | NA | NA | NA | NA | NA | |
| ENSG000002 | 0 | NA | NA | NA | NA | NA | NA | NA | |
| ENSG000002 | 0.11865494 | 0.083440866 | 0.12751478 | 0.65436232 | 0.5128784 | NA | NA | NA | |
| ENSG000002 | 0.3575613 | 0.072305152 | 0.21497462 | 0.33634273 | 0.73661243 | NA | NA | NA | |

| | | | | | | | | |
|---|---|---|---|---|---|---|---|---|
| ENSG000002 | 0 | NA | NA | NA | NA | NA | NA | NA |
| ENSG000002 | 0.12070844 | -0.088503695 | 0.12783931 | -0.6923042 | 0.48874631 | NA | NA | NA |
| ENSG000002 | 0.75873415 | -0.034571091 | 0.2860415 | -0.1208604 | 0.90380161 | NA | NA | NA |
| ENSG000002 | 0.60419063 | -0.101535721 | 0.26323679 | -0.3857201 | 0.69970398 | NA | NA | NA |
| ENSG000002 | 1.64048927 | -0.410673344 | 0.35173247 | -1.167573 | 0.24297903 | NA | MIR196A1 | 406972 |
| ENSG000002 | 0 | NA | NA | NA | NA | NA | SNORA40 | 677822 |
| ENSG000002 | 0.95542959 | -0.033551797 | 0.29624078 | -0.1132585 | 0.90982559 | NA | NA | NA |
| ENSG000002 | 0.60396387 | 0.05533717 | 0.26282077 | 0.21055098 | 0.83323767 | NA | RNU6ATAC26P | 106480542 |
| ENSG000002 | 0.81551701 | 0.20201966 | 0.28897753 | 0.69908431 | 0.48449934 | NA | MIR190A | 406965 |
| ENSG000002 | 1.77765698 | 0.381281383 | 0.34759129 | 1.09692445 | 0.27267442 | NA | MIR320D1 | 100313896 |
| ENSG000002 | 1.01207154 | -0.046453419 | 0.31227181 | -0.1487596 | 0.88174336 | NA | NA | NA |
| ENSG000002 | 0.51180131 | -0.185828522 | 0.2424294 | -0.7665263 | 0.44336319 | NA | MIR216B | 100126319 |
| ENSG000002 | 1.09657874 | -0.108425755 | 0.32174398 | -0.3369939 | 0.73612151 | NA | MIR526A2 | 574486 |
| ENSG000002 | 0.49902098 | -0.194741582 | 0.24819886 | -0.7846192 | 0.43267688 | NA | MIR501 | 574503 |
| ENSG000002 | 1.22306215 | -0.043529992 | 0.33645225 | -0.1293794 | 0.89705744 | NA | MIR320B1 | 100302117 |
| ENSG000002 | 1.11778589 | -0.249078355 | 0.31011034 | -0.8031927 | 0.42186338 | NA | NA | NA |
| ENSG000002 | 0.97811576 | 0.103538885 | 0.3001056 | 0.34500818 | 0.73008822 | NA | MIR670 | 100313777 |
| ENSG000002 | 2.13441404 | -0.516058195 | 0.36551815 | -1.4118538 | 0.15799301 | NA | MIR770 | 768222 |
| ENSG000002 | 0.47624849 | 0.143072764 | 0.23753303 | 0.60232788 | 0.5469559 | NA | NA | NA |
| ENSG000002 | 1.64002174 | -0.248936109 | 0.3463064 | -0.718832 | 0.47224446 | NA | MIR766 | 768218 |
| ENSG000002 | 0.61053204 | -0.095342013 | 0.25468495 | -0.3743528 | 0.70814188 | NA | MIR759 | 100313778 |
| ENSG000002 | 0.64460494 | -0.261028787 | 0.26005827 | -1.0037319 | 0.31550784 | NA | MIR769 | 768217 |
| ENSG000002 | 1.36502431 | -0.109972556 | 0.32512217 | -0.3382499 | 0.73517484 | NA | MIR765 | 768220 |
| ENSG000002 | 0.74017673 | -0.321754383 | 0.2710983 | -1.186855 | 0.23528481 | NA | MIR758 | 768212 |
| ENSG000002 | 1.00592601 | -0.044485265 | 0.31202772 | -0.1425683 | 0.88663113 | NA | MIR767 | 768215 |
| ENSG000002 | 1.12310494 | -0.120294979 | 0.32641626 | -0.3685324 | 0.71247626 | NA | MIR802 | 768219 |
| ENSG000002 | 0.22997012 | 0.167051788 | 0.17646558 | 0.94665368 | 0.34381528 | NA | MIR762 | 100313837 |
| ENSG000002 | 1.36980753 | -0.127713282 | 0.34374615 | -0.3715337 | 0.71024005 | NA | NA | NA |
| ENSG000002 | 1.24757157 | -0.176311843 | 0.32359085 | -0.5448604 | 0.58584953 | NA | NA | NA |

| | | | | | | | | |
|---|---|---|---|---|---|---|---|---|
| ENSG000002 | 0.65445467 | -0.486865861 | 0.27154464 | -1.7929496 | 0.07298098 | NA | NA | NA |
| ENSG000002 | 0 | NA | NA | NA | NA | NA | NA | NA |
| ENSG000002 | 0.35067856 | 0.074230421 | 0.21389444 | 0.34704231 | 0.72855953 | NA | NA | NA |
| ENSG000002 | 1.54977901 | 0.031382194 | 0.33912254 | 0.09253939 | 0.92626949 | NA | NA | NA |
| ENSG000002 | 0.59760982 | 0.066257965 | 0.26871197 | 0.24657616 | 0.80523626 | NA | NA | NA |
| ENSG000002 | 1.85320573 | -0.103843752 | 0.34987723 | -0.2968005 | 0.76661879 | NA | NA | NA |
| ENSG000002 | 1.02824148 | -0.352569214 | 0.31011878 | -1.1368845 | 0.25558659 | NA | NA | NA |
| ENSG000002 | 1.15688838 | -0.551936344 | 0.31271549 | -1.7649792 | 0.07756725 | NA | NA | NA |
| ENSG000002 | 0.78136421 | -0.193245175 | 0.28063824 | -0.6885918 | 0.49108021 | NA | NA | NA |
| ENSG000002 | 1.31314789 | 0.027743513 | 0.33080013 | 0.0838679 | 0.93316146 | NA | NA | NA |
| ENSG000002 | 1.36973372 | -0.266937291 | 0.33706855 | -0.7919377 | 0.42839701 | NA | NA | NA |
| ENSG000002 | 1.01958598 | -0.197147843 | 0.31466692 | -0.6265287 | 0.53096824 | NA | NA | NA |
| ENSG000002 | 0.75190136 | -0.191187334 | 0.28037014 | -0.6819105 | 0.49529557 | NA | NA | NA |
| ENSG000002 | 1.45229697 | 0.219585364 | 0.3359576 | 0.65361036 | 0.51336287 | NA | NA | NA |
| ENSG000002 | 1.2200779 | -0.31153904 | 0.32322724 | -0.9638391 | 0.33512661 | NA | NA | NA |
| ENSG000002 | 1.39836997 | -0.712620308 | 0.33261404 | -2.1424841 | 0.03215454 | NA | NA | NA |
| ENSG000002 | 1.50019347 | -0.046146349 | 0.33923624 | -0.1360301 | 0.89179746 | NA | NA | NA |
| ENSG000002 | 1.13295467 | -0.283033201 | 0.32731671 | -0.8647075 | 0.38719937 | NA | NA | NA |
| ENSG000002 | 0 | NA | NA | NA | NA | NA | NA | NA |
| ENSG000002 | 1.04065125 | -0.197389758 | 0.30672345 | -0.6435431 | 0.51987177 | NA | NA | NA |
| ENSG000002 | 0.86344734 | 0.04862751 | 0.30222795 | 0.1608968 | 0.87217468 | NA | NA | NA |
| ENSG000002 | 0.25214371 | -0.009862892 | 0.18046657 | -0.0546522 | 0.95641556 | NA | NA | NA |
| ENSG000002 | 0.88974721 | -0.438397978 | 0.29679137 | -1.4771251 | 0.13964211 | NA | NA | NA |
| ENSG000002 | 0.76152487 | -0.199410494 | 0.28701953 | -0.6947628 | 0.48720396 | NA | NA | NA |
| ENSG000002 | 1.10804435 | 0.178525633 | 0.3176327 | 0.56205054 | 0.57408158 | NA | NA | NA |
| ENSG000002 | 0.76485795 | -0.196355366 | 0.28121884 | -0.6982298 | 0.48503351 | NA | NA | NA |
| ENSG000002 | 0.76306345 | -0.192579149 | 0.27944362 | -0.6891521 | 0.49072757 | NA | NA | NA |
| ENSG000002 | 0.79075747 | -0.360972421 | 0.28354666 | -1.2730618 | 0.20299612 | NA | NA | NA |
| ENSG000002 | 1.09657819 | -0.11594456 | 0.32021555 | -0.3620829 | 0.71729011 | NA | NA | NA |

| | | | | | | | | |
|---|---|---|---|---|---|---|---|---|
| ENSG000002 | 1.40903208 | -0.538820278 | 0.33213997 | -1.6222687 | 0.10474583 | NA | NA | NA |
| ENSG000002 | 0.87392664 | -0.422259382 | 0.29440295 | -1.4342906 | 0.15148935 | NA | NA | NA |
| ENSG000002 | 1.0081542 | -0.346299707 | 0.30931545 | -1.1195681 | 0.26289786 | NA | NA | NA |
| ENSG000002 | 1.51721686 | -0.342937442 | 0.34714166 | -0.9878891 | 0.32320699 | NA | NA | NA |
| ENSG000002 | 1.46019733 | 0.237308618 | 0.34527842 | 0.68729641 | 0.49189596 | NA | NA | NA |
| ENSG000002 | 1.13881788 | -0.436732811 | 0.32362791 | -1.3494906 | 0.17717945 | NA | NA | NA |
| ENSG000002 | 0.25214371 | -0.009862892 | 0.18046657 | -0.0546522 | 0.95641556 | NA | NA | NA |
| ENSG000002 | 0.36416385 | 0.068880976 | 0.20758164 | 0.33182596 | 0.74002068 | NA | NA | NA |
| ENSG000002 | 1.28089427 | -0.326776517 | 0.32575194 | -1.0031453 | 0.31579078 | NA | NA | NA |
| ENSG000002 | 0.76078766 | -0.199688475 | 0.28697083 | -0.6958494 | 0.48652316 | NA | NA | NA |
| ENSG000002 | 1.40386316 | -0.683857485 | 0.32906025 | -2.0782136 | 0.03768969 | NA | NA | NA |
| ENSG000002 | 1.24506105 | -0.045484982 | 0.32892924 | -0.138282 | 0.89001758 | NA | NA | NA |
| ENSG000002 | 1.26370479 | -0.201252851 | 0.33482765 | -0.601064 | 0.54779736 | NA | NA | NA |
| ENSG000002 | 1.47833434 | -0.32310618 | 0.34226571 | -0.9440215 | 0.34515867 | NA | NA | NA |
| ENSG000002 | 1.33742808 | -0.111060971 | 0.32752539 | -0.3390912 | 0.73454105 | NA | NA | NA |
| ENSG000002 | 0 | NA | NA | NA | NA | NA | NA | NA |
| ENSG000002 | 0 | NA | NA | NA | NA | NA | NA | NA |
| ENSG000002 | 0.12070844 | -0.088503695 | 0.12783931 | -0.6923042 | 0.48874631 | NA | NA | NA |
| ENSG000002 | 0.14128499 | -0.097772014 | 0.13403235 | -0.7294658 | 0.4657168 | NA | NA | NA |
| ENSG000002 | 0.64118699 | -0.259786178 | 0.25976884 | -1.0000667 | 0.31727822 | NA | NA | NA |
| ENSG000002 | 0.25370795 | -0.171331747 | 0.17411721 | -0.9840024 | 0.32511435 | NA | NA | NA |
| ENSG000002 | 1.49232153 | -0.312992751 | 0.33821959 | -0.9254128 | 0.35475121 | NA | NA | NA |
| ENSG000002 | 1.60098576 | -0.371644146 | 0.34101959 | -1.0898029 | 0.27579996 | NA | NA | NA |
| ENSG000002 | 0.74017673 | -0.321754383 | 0.2710983 | -1.186855 | 0.23528481 | NA | NA | NA |
| ENSG000002 | 0.58475569 | -0.031541521 | 0.15708832 | -0.2007885 | 0.84086399 | NA | NA | NA |
| ENSG000002 | 0.91140588 | -0.4294125 | 0.2938468 | -1.4613482 | 0.1439199 | NA | NA | NA |
| ENSG000002 | 0.63965309 | -0.285618162 | 0.26800573 | -1.0657166 | 0.28655176 | NA | NA | NA |
| ENSG000002 | 1.6850019 | -0.046685651 | 0.34653791 | -0.1347202 | 0.89283311 | NA | NA | NA |
| ENSG000002 | 0.2596576 | -0.193002035 | 0.1837913 | -1.0501152 | 0.29366517 | NA | NA | NA |

| | | | | | | | | |
|---|---|---|---|---|---|---|---|---|
| ENSG000002 | 1.00376432 | -0.198698202 | 0.309319 | -0.6423731 | 0.52063097 | NA | NA | NA |
| ENSG000002 | 1.21302047 | -0.177846276 | 0.32629465 | -0.5450481 | 0.58572044 | NA | NA | NA |
| ENSG000002 | 1.21967061 | 0.104974983 | 0.33321107 | 0.3150405 | 0.75273091 | NA | NA | NA |
| ENSG000002 | 1.46285008 | -0.036103254 | 0.33855177 | -0.1066403 | 0.91507437 | NA | NA | NA |
| ENSG000002 | 1.6023416 | 0.009586288 | 0.35104141 | 0.02730814 | 0.97821396 | NA | NA | NA |
| ENSG000002 | 2.00878186 | -0.442619653 | 0.36168237 | -1.22378 | 0.22103522 | NA | NA | NA |
| ENSG000002 | 1.33343105 | 0.02163561 | 0.33307459 | 0.06495725 | 0.94820804 | NA | NA | NA |
| ENSG000002 | 1.49234326 | -0.18689102 | 0.34491331 | -0.5418493 | 0.58792235 | NA | NA | NA |
| ENSG000002 | 0.72244254 | 0.129271463 | 0.27760219 | 0.46567163 | 0.64145056 | NA | NA | NA |
| ENSG000002 | 0.76215602 | -0.362469896 | 0.28326072 | -1.2796335 | 0.20067407 | NA | NA | NA |
| ENSG000002 | 0.64506141 | -0.288091388 | 0.2684363 | -1.0732207 | 0.28317211 | NA | NA | NA |
| ENSG000002 | 0.4682236 | 0.140020865 | 0.23655532 | 0.59191594 | 0.55390688 | NA | NA | NA |
| ENSG000002 | 1.58148328 | -0.112576534 | 0.34980914 | -0.3218227 | 0.74758699 | NA | NA | NA |
| ENSG000002 | 0.74908945 | -0.183477852 | 0.27852971 | -0.6587371 | 0.51006461 | NA | NA | NA |
| ENSG000002 | 0.99781521 | -0.202823463 | 0.31782692 | -0.638157 | 0.52337151 | NA | NA | NA |
| ENSG000002 | 1.58216596 | -0.260411381 | 0.35029385 | -0.7434084 | 0.45723448 | NA | NA | NA |
| ENSG000002 | 0.96716217 | 0.110385505 | 0.29957659 | 0.36847173 | 0.71252152 | NA | NA | NA |
| ENSG000002 | 0.89495965 | -0.460377077 | 0.30201036 | -1.5243751 | 0.12741503 | NA | NA | NA |
| ENSG000002 | 1.15548601 | 0.585137796 | 0.3249288 | 1.80081848 | 0.07173149 | NA | NA | NA |
| ENSG000002 | 1.35794047 | -0.119356884 | 0.33615055 | -0.3550697 | 0.72253735 | NA | NA | NA |
| ENSG000002 | 1.98481323 | -0.328183689 | 0.36404274 | -0.9014977 | 0.36732376 | NA | NA | NA |
| ENSG000002 | 1.39238202 | -0.576337849 | 0.33852701 | -1.7024871 | 0.0886641 | NA | NA | NA |
| ENSG000002 | 0.26560726 | -0.177019426 | 0.17672274 | -1.0016788 | 0.31649875 | NA | NA | NA |
| ENSG000002 | 0.25333738 | -0.015686291 | 0.1806605 | -0.0868275 | 0.93080867 | NA | NA | NA |
| ENSG000002 | 0.11865494 | 0.083440866 | 0.12751478 | 0.65436232 | 0.5128784 | NA | NA | NA |
| ENSG000002 | 0.22997012 | 0.167051788 | 0.17646558 | 0.94665368 | 0.34381528 | NA | NA | NA |
| ENSG000002 | 0.11934008 | 0.083564698 | 0.12760091 | 0.65489106 | 0.5125379 | NA | NA | NA |
| ENSG000002 | 0 | NA | NA | NA | NA | NA | NA | NA |
| ENSG000002 | 0.35004741 | 0.261422124 | 0.21550076 | 1.21309144 | 0.22509486 | NA | NA | NA |

| | | | | | | | | |
|---|---|---|---|---|---|---|---|---|
| ENSG000002 | 0.23936338 | -0.005479057 | 0.17759977 | -0.0308506 | 0.9753887 | NA | NA | NA |
| ENSG000002 | 2.00946508 | -0.580413564 | 0.36154686 | -1.6053619 | 0.10841418 | NA | NA | NA |
| ENSG000002 | 0.92307183 | -0.408029112 | 0.28913245 | -1.4112187 | 0.15818015 | NA | NA | NA |
| ENSG000002 | 1.10440721 | -0.118910289 | 0.32584268 | -0.3649316 | 0.71516247 | NA | NA | NA |
| ENSG000002 | 0.8261595 | 0.365072387 | 0.28985807 | 1.25948669 | 0.20785459 | NA | NA | NA |
| ENSG000002 | 0.99767332 | -0.035058095 | 0.30673943 | -0.1142928 | 0.90900573 | NA | NA | NA |
| ENSG000002 | 1.95433351 | -0.298234935 | 0.3596753 | -0.8291782 | 0.40700356 | NA | NA | NA |
| ENSG000002 | 1.70637156 | 0.207611361 | 0.35201119 | 0.58978625 | 0.55533396 | NA | NA | NA |
| ENSG000002 | 0.87531619 | -0.427631553 | 0.29439996 | -1.452553 | 0.1463479 | NA | NA | NA |
| ENSG000002 | 1.29299138 | -0.184864206 | 0.32481114 | -0.5691437 | 0.56925864 | NA | NA | NA |
| ENSG000002 | 2.01780209 | -0.576212928 | 0.36065623 | -1.5976791 | 0.11011441 | NA | NA | NA |
| ENSG000002 | 1.94722189 | -0.303508785 | 0.35950149 | -0.844249 | 0.39853026 | NA | NA | NA |
| ENSG000002 | 1.74980301 | -0.202578625 | 0.35855453 | -0.5649869 | 0.57208265 | NA | NA | NA |
| ENSG000002 | 1.38392589 | -0.116671939 | 0.33524779 | -0.348017 | 0.7278274 | NA | NA | NA |
| ENSG000002 | 1.91725365 | -0.057276771 | 0.36013574 | -0.1590422 | 0.87363565 | NA | NA | NA |
| ENSG000002 | 0.61630755 | 0.061075249 | 0.26391444 | 0.23142064 | 0.81698803 | NA | NA | NA |
| ENSG000002 | 1.36522815 | -0.115399079 | 0.33484564 | -0.3446337 | 0.73036979 | NA | NA | NA |
| ENSG000002 | 1.99130524 | -0.197341862 | 0.3614176 | -0.5460217 | 0.585051 | NA | NA | NA |
| ENSG000002 | 0.24141689 | -0.165257896 | 0.17127401 | -0.9648743 | 0.33460776 | NA | NA | NA |
| ENSG000002 | 0.74484447 | -0.03101441 | 0.29048039 | -0.1067694 | 0.91497194 | NA | NA | NA |
| ENSG000002 | 0.88630415 | -0.272387831 | 0.299825 | -0.9084894 | 0.36361972 | NA | NA | NA |
| ENSG000002 | 0.69338621 | 0.315777391 | 0.28279036 | 1.11664835 | 0.26414471 | NA | NA | NA |
| ENSG000002 | 2.09760675 | -0.492708915 | 0.36296752 | -1.3574463 | 0.17463945 | NA | NA | NA |
| ENSG000002 | 1.17075295 | -0.419773892 | 0.318252 | -1.3189984 | 0.18716964 | NA | NA | NA |
| ENSG000002 | 2.1166907 | -0.422424526 | 0.35179455 | -1.2007705 | 0.22984023 | NA | NA | NA |
| ENSG000002 | 1.08391601 | 0.026940877 | 0.30636788 | 0.08793636 | 0.92992725 | NA | NA | NA |
| ENSG000002 | 1.00450207 | -0.204287946 | 0.3180715 | -0.6422705 | 0.52069756 | NA | NA | NA |
| ENSG000002 | 0.70072597 | 0.318240822 | 0.28328746 | 1.1233848 | 0.2612741 | NA | NA | NA |
| ENSG000002 | 1.10102014 | -0.255084409 | 0.31891345 | -0.7998547 | 0.423795 | NA | NA | NA |

| | | | | | | | | |
|---|---|---|---|---|---|---|---|---|
| ENSG000002 | 0.4974562 | -0.028348769 | 0.24672499 | -0.1149003 | 0.90852417 | NA | NA | NA |
| ENSG000002 | 0.65542213 | -0.28582049 | 0.26931472 | -1.061288 | 0.28855903 | NA | NA | NA |
| ENSG000002 | 1.60737641 | -0.119583418 | 0.35015989 | -0.3415109 | 0.732719 | NA | NA | NA |
| ENSG000002 | 0.59100728 | 0.060563842 | 0.26166992 | 0.23145129 | 0.81696422 | NA | NA | NA |
| ENSG000002 | 0.72372659 | -0.023335709 | 0.28377892 | -0.082232 | 0.93446222 | NA | NA | NA |
| ENSG000002 | 1.92803037 | -0.524530514 | 0.35681557 | -1.4700326 | 0.14155293 | NA | NA | NA |
| ENSG000002 | 0.11131518 | 0.081951103 | 0.12663804 | 0.64712866 | 0.51754868 | NA | NA | NA |
| ENSG000002 | 0.65542213 | -0.28582049 | 0.26931472 | -1.061288 | 0.28855903 | NA | NA | NA |
| ENSG000002 | 1.82876048 | 0.003265043 | 0.3596596 | 0.00907815 | 0.99275679 | NA | NA | NA |
| ENSG000002 | 1.4797355 | -0.445309204 | 0.33792563 | -1.3177728 | 0.18757972 | NA | NA | NA |
| ENSG000002 | 0.49879422 | -0.023609326 | 0.24688725 | -0.095628 | 0.92381605 | NA | NA | NA |
| ENSG000002 | 0.98524182 | -0.339777001 | 0.30837415 | -1.1018336 | 0.27053402 | NA | NA | NA |
| ENSG000002 | 0.96833466 | -0.030372942 | 0.30130547 | -0.1008045 | 0.91970567 | NA | NA | NA |
| ENSG000002 | 1.85239281 | -0.37967581 | 0.35590977 | -1.0667755 | 0.28607324 | NA | NA | NA |
| ENSG000002 | 1.26442474 | -0.64331086 | 0.32575398 | -1.9748365 | 0.0482867 | NA | NA | NA |
| ENSG000002 | 1.50008337 | -0.494128146 | 0.34651689 | -1.4259857 | 0.15387246 | NA | NA | NA |
| ENSG000002 | 0.35870346 | 0.077368606 | 0.21515516 | 0.35959447 | 0.71915042 | NA | NA | NA |
| ENSG000002 | 0.52125014 | -0.010203417 | 0.15567753 | -0.065542 | 0.94774247 | NA | NA | NA |
| ENSG000002 | 0.76550926 | -0.332240521 | 0.27535282 | -1.2065993 | 0.2275865 | NA | NA | NA |
| ENSG000002 | 0.38056192 | -0.244105065 | 0.20391883 | -1.1970698 | 0.23127937 | NA | NA | NA |
| ENSG000002 | 1.43413202 | -0.256442304 | 0.32927417 | -0.7788109 | 0.43609113 | NA | NA | NA |
| ENSG000002 | 0.67004903 | -0.269626612 | 0.26217122 | -1.0284371 | 0.30374426 | NA | NA | NA |
| ENSG000002 | 1.83413902 | 0.133692421 | 0.35662235 | 0.37488515 | 0.70774589 | NA | NA | NA |
| ENSG000002 | 1.37527898 | -0.266706439 | 0.33842831 | -0.7880737 | 0.43065362 | NA | NA | NA |
| ENSG000002 | 1.9769888 | -0.05454575 | 0.35885149 | -0.1520009 | 0.87918622 | NA | NA | NA |
| ENSG000002 | 0.83765834 | 0.055260734 | 0.2957186 | 0.18686932 | 0.85176309 | NA | NA | NA |
| ENSG000002 | 1.57257928 | 0.020510362 | 0.339137 | 0.0604781 | 0.95177486 | NA | NA | NA |
| ENSG000002 | 0.73036138 | -0.025051354 | 0.28421872 | -0.0881411 | 0.92976452 | NA | NA | NA |
| ENSG000002 | 1.24381338 | -0.195477696 | 0.33435427 | -0.5846424 | 0.55878817 | NA | NA | NA |

| | | | | | | | | |
|---|---|---|---|---|---|---|---|---|
| ENSG000002 | 0.97486857 | 0.269601735 | 0.30661793 | 0.87927582 | 0.37925174 | NA | NA | NA |
| ENSG000002 | 0.47804353 | 0.152460376 | 0.23772494 | 0.64133101 | 0.52130765 | NA | NA | NA |
| ENSG000002 | 0.72170533 | 0.13625606 | 0.28374667 | 0.48020321 | 0.6310829 | NA | NA | NA |
| ENSG000002 | 1.47244939 | -0.324086001 | 0.3414809 | -0.9490604 | 0.34258989 | NA | NA | NA |
| ENSG000002 | 1.40601111 | -0.522709551 | 0.32893378 | -1.5891027 | 0.11203721 | NA | NA | NA |
| ENSG000002 | 1.50010511 | -0.321865877 | 0.34198136 | -0.9411796 | 0.34661283 | NA | NA | NA |
| ENSG000002 | 0.12685397 | -0.090215768 | 0.12901919 | -0.699243 | 0.48440017 | NA | NA | NA |
| ENSG000002 | 1.5753372 | -0.120647423 | 0.35089346 | -0.3438292 | 0.73097473 | NA | NA | NA |
| ENSG000002 | 0.49168123 | -0.197384925 | 0.247356 | -0.7979791 | 0.4248826 | NA | NA | NA |
| ENSG000002 | 1.3816765 | 0.021216393 | 0.3363836 | 0.06307202 | 0.94970916 | NA | NA | NA |
| ENSG000002 | 1.42414632 | -0.731835289 | 0.33578438 | -2.1794798 | 0.02929604 | NA | NA | NA |
| ENSG000002 | 0.87215491 | -0.121172621 | 0.30297358 | -0.3999445 | 0.68919738 | NA | NA | NA |
| ENSG000002 | 0.53627793 | -0.360851627 | 0.24080453 | -1.4985251 | 0.13399688 | NA | NA | NA |
| ENSG000002 | 0.37285215 | -0.099309896 | 0.2177345 | -0.4561055 | 0.64831414 | NA | NA | NA |
| ENSG000002 | 0 | NA | NA | NA | NA | NA | NA | NA |
| ENSG000002 | 0.14128499 | -0.097772014 | 0.13403235 | -0.7294658 | 0.4657168 | NA | NA | NA |
| ENSG000002 | 0.12070844 | -0.088503695 | 0.12783931 | -0.6923042 | 0.48874631 | NA | NA | NA |
| ENSG000002 | 0 | NA | NA | NA | NA | NA | NA | NA |
| ENSG000002 | 0.11934008 | 0.083564698 | 0.12760091 | 0.65489106 | 0.5125379 | NA | NA | NA |
| ENSG000002 | 0.11131518 | 0.081951103 | 0.12663804 | 0.64712866 | 0.51754868 | NA | NA | NA |
| ENSG000002 | 0 | NA | NA | NA | NA | NA | NA | NA |
| ENSG000002 | 0.22410479 | 0.147643983 | 0.16638082 | 0.88738586 | 0.37487119 | NA | NA | NA |
| ENSG000002 | 0 | NA | NA | NA | NA | NA | NA | NA |
| ENSG000002 | 0 | NA | NA | NA | NA | NA | NA | NA |
| ENSG000002 | 0.35346928 | -0.088570748 | 0.20618147 | -0.4295767 | 0.66750362 | NA | NA | NA |
| ENSG000002 | 0.37236288 | -0.09108266 | 0.20928647 | -0.4352057 | 0.66341314 | NA | NA | NA |
| ENSG000002 | 0.11205239 | 0.082118251 | 0.12671962 | 0.64803108 | 0.51696485 | NA | NA | NA |
| ENSG000002 | 0.11934008 | 0.083564698 | 0.12760091 | 0.65489106 | 0.5125379 | NA | NA | NA |
| ENSG000002 | 0 | NA | NA | NA | NA | NA | NA | NA |

| | | | | | | | | |
|---|---|---|---|---|---|---|---|---|
| ENSG000002 | 0 | NA | NA | NA | NA | NA | NA | NA |
| ENSG000002 | 0.13280363 | -0.093394593 | 0.13115644 | -0.7120855 | 0.47641185 | NA | NA | NA |
| ENSG000002 | 0.96375337 | -0.179216179 | 0.30596636 | -0.5857382 | 0.55805147 | NA | NA | NA |
| ENSG000002 | 0.33468275 | 0.234558662 | 0.20511328 | 1.14355667 | 0.25280754 | NA | NA | NA |
| ENSG000002 | 0.33541997 | 0.234895243 | 0.20524277 | 1.14447511 | 0.25242665 | NA | NA | NA |
| ENSG000002 | 0.86228792 | -0.414834899 | 0.2912593 | -1.4242803 | 0.15436534 | NA | NA | NA |
| ENSG000002 | 1.46777829 | -0.297529909 | 0.33323232 | -0.8928603 | 0.37193199 | NA | NA | NA |
| ENSG000002 | 0.35004741 | 0.261422124 | 0.21550076 | 1.21309144 | 0.22509486 | NA | NA | NA |
| ENSG000002 | 1.03735242 | -0.712412845 | 0.31042809 | -2.2949368 | 0.02173675 | NA | NA | NA |
| ENSG000002 | 2.05517173 | -0.230628319 | 0.36259424 | -0.6360507 | 0.52474338 | NA | NA | NA |
| ENSG000002 | 0.90064872 | -0.456924471 | 0.30237983 | -1.5110944 | 0.1307644 | NA | NA | NA |
| ENSG000002 | 1.64658081 | -0.555712174 | 0.35029907 | -1.5863935 | 0.11265007 | NA | NA | NA |
| ENSG000002 | 0.38064837 | -0.102706251 | 0.21886737 | -0.4692625 | 0.63888201 | NA | NA | NA |
| ENSG000002 | 1.11761017 | -0.254407609 | 0.31727531 | -0.8018513 | 0.422639 | NA | NA | NA |
| ENSG000002 | 0.73812323 | -0.16407654 | 0.26803682 | -0.6121418 | 0.54044394 | NA | NA | NA |
| ENSG000002 | 0.23890637 | -0.010420674 | 0.17747379 | -0.0587167 | 0.95317776 | NA | NA | NA |
| ENSG000002 | 0.37831254 | -0.102004459 | 0.21855316 | -0.4667261 | 0.64069587 | NA | NA | NA |
| ENSG000002 | 0.97555125 | 0.104456027 | 0.30583208 | 0.34154699 | 0.73269184 | NA | NA | NA |
| ENSG000002 | 0.66751732 | -0.272810278 | 0.26356336 | -1.0350842 | 0.30062957 | NA | NA | NA |
| ENSG000002 | 1.01958598 | -0.197147843 | 0.31466692 | -0.6265287 | 0.53096824 | NA | NA | NA |
| ENSG000002 | 0.99258158 | -0.337050613 | 0.30871569 | -1.0917832 | 0.27492839 | NA | NA | NA |
| ENSG000002 | 0.5854929 | -0.091197421 | 0.25231745 | -0.3614392 | 0.71777113 | NA | NA | NA |
| ENSG000002 | 1.5828833 | -0.745280393 | 0.33485883 | -2.2256555 | 0.02603727 | NA | NA | NA |
| ENSG000002 | 1.59682565 | -0.262188315 | 0.35162541 | -0.7456467 | 0.45588089 | NA | NA | NA |
| ENSG000002 | 0.87440484 | -0.268724769 | 0.29922967 | -0.8980552 | 0.36915611 | NA | NA | NA |
| ENSG000002 | 0.42385498 | -0.262217565 | 0.21037102 | -1.2464529 | 0.21259817 | NA | NA | NA |
| ENSG000002 | 0.58087681 | 0.226918272 | 0.26184027 | 0.86662863 | 0.38614552 | NA | NA | NA |
| ENSG000002 | 0.84864521 | 0.185926723 | 0.29101789 | 0.63888417 | 0.52289829 | NA | NA | NA |
| ENSG000002 | 0 | NA | NA | NA | NA | NA | NA | NA |

| | | | | | | | | |
|---|---|---|---|---|---|---|---|---|
| ENSG000002 | 0.24141689 | -0.165257896 | 0.17127401 | -0.9648743 | 0.33460776 | NA | NA | NA |
| ENSG000002 | 0.11205239 | 0.082118251 | 0.12671962 | 0.64803108 | 0.51696485 | NA | NA | NA |
| ENSG000002 | 0.26813897 | -0.197007041 | 0.18550319 | -1.0620143 | 0.2882292 | NA | NA | NA |
| ENSG000002 | 1.70708458 | -0.064328393 | 0.34897049 | -0.1843376 | 0.8537486 | NA | NA | NA |
| ENSG000002 | 0.24756242 | -0.187034346 | 0.18119817 | -1.0322088 | 0.30197431 | NA | NA | NA |
| ENSG000002 | 1.67609023 | -0.362780694 | 0.33997864 | -1.0670691 | 0.28594064 | NA | NA | NA |
| ENSG000002 | 0.49224375 | -0.024566489 | 0.23863564 | -0.1029456 | 0.91800615 | NA | NA | NA |
| ENSG000002 | 1.37567074 | -0.266930403 | 0.33843612 | -0.7887173 | 0.43027728 | NA | MIR509-2 | 100126301 |
| ENSG000002 | 0.88855353 | -0.453244102 | 0.30180501 | -1.5017779 | 0.13315447 | NA | MIR509-3 | 100126337 |
| ENSG000002 | 1.96867361 | -0.185492775 | 0.36029205 | -0.51484 | 0.6066648 | NA | NA | NA |
| ENSG000002 | 1.20785562 | 0.101665842 | 0.32947806 | 0.30856635 | 0.75765142 | NA | NA | NA |
| ENSG000002 | 0.35073255 | 0.241740392 | 0.2078501 | 1.16305162 | 0.24480856 | NA | NA | NA |
| ENSG000002 | 0.72238856 | -0.029280768 | 0.28909343 | -0.1012848 | 0.91932439 | NA | MIR320C2 | 100302195 |
| ENSG000002 | 0.61679627 | 0.054898003 | 0.27030252 | 0.20309838 | 0.83905814 | NA | MIR764 | 100313838 |
| ENSG000002 | 1.32916488 | 0.029321414 | 0.33994199 | 0.08625417 | 0.93126437 | NA | MIR301B | 100126318 |
| ENSG000002 | 0.2596576 | -0.193002035 | 0.1837913 | -1.0501152 | 0.29366517 | NA | RNU6-1168P | 106481548 |
| ENSG000002 | 1.35893882 | -0.115090545 | 0.33617163 | -0.3423565 | 0.73208261 | NA | NA | NA |
| ENSG000002 | 1.21321187 | 0.099326632 | 0.32950394 | 0.30144293 | 0.76307677 | NA | RNU6-696P | 106479873 |
| ENSG000002 | 1.08026444 | 0.190857655 | 0.3234979 | 0.58998113 | 0.5552033 | NA | NA | NA |
| ENSG000002 | 1.1907326 | -0.785634655 | 0.32002203 | -2.4549393 | 0.01409084 | NA | RNU6-1320P | 106480659 |
| ENSG000002 | 1.39603414 | -0.739041643 | 0.33642356 | -2.1967595 | 0.02803763 | NA | NA | NA |
| ENSG000002 | 1.84444418 | -0.116613203 | 0.35845259 | -0.3253239 | 0.74493593 | NA | RNU6-910P | 106481777 |
| ENSG000002 | 1.12355983 | -0.256093723 | 0.3174782 | -0.8066498 | 0.41986828 | NA | RNU6-1106P | 106480053 |
| ENSG000002 | 1.46259967 | -0.195705526 | 0.34820558 | -0.5620402 | 0.57408865 | NA | NA | NA |
| ENSG000002 | 0.90614294 | -0.271362536 | 0.29346145 | -0.9246957 | 0.35512423 | NA | RNU1-82P | 106480175 |
| ENSG000002 | 0.12685397 | -0.090215768 | 0.12901919 | -0.699243 | 0.48440017 | NA | RNA5SP377 | 106479011 |
| ENSG000002 | 0.25214371 | -0.009862892 | 0.18046657 | -0.0546522 | 0.95641556 | NA | RNU6-576P | 106479821 |
| ENSG000002 | 1.08311518 | 0.044344248 | 0.31468935 | 0.14091436 | 0.8879376 | NA | RNU6-1319P | 106480142 |
| ENSG000002 | 0.34344486 | 0.238503239 | 0.20662376 | 1.15428757 | 0.24838229 | NA | SNORD66 | 692107 |

| | | | | | | | | |
|---|---|---|---|---|---|---|---|---|
| ENSG000002 | 0.52876404 | -0.191903645 | 0.2442426 | -0.7857092 | 0.4320379 | NA | NA | NA |
| ENSG000002 | 1.11330886 | 0.173688474 | 0.31617693 | 0.54933949 | 0.58277249 | NA | NA | NA |
| ENSG000002 | 0.72828614 | -0.028172726 | 0.27772921 | -0.1014396 | 0.91920154 | NA | NA | NA |
| ENSG000002 | 0.87889968 | -0.231115046 | 0.28252001 | -0.8180484 | 0.41332955 | NA | RNU6-763P | 106481421 |
| ENSG000002 | 0.49512483 | -0.344267812 | 0.23617548 | -1.457678 | 0.14492931 | NA | NA | NA |
| ENSG000002 | 0.520565 | -0.024378065 | 0.24183609 | -0.1008041 | 0.91970599 | NA | RNU1-77P | 106480173 |
| ENSG000002 | 0 | NA | NA | NA | NA | NA | NA | NA |
| ENSG000002 | 0.61158383 | 0.053697626 | 0.26349596 | 0.20378918 | 0.83851825 | NA | NA | NA |
| ENSG000002 | 0.28256998 | -0.18483211 | 0.18021686 | -1.0256094 | 0.30507573 | NA | NA | NA |
| ENSG000002 | 0.47614243 | -0.020043108 | 0.23674202 | -0.0846622 | 0.93252992 | NA | NA | NA |
| ENSG000002 | 1.86586737 | -0.217527123 | 0.34544442 | -0.6297022 | 0.52888942 | NA | NA | NA |
| ENSG000002 | 1.48109237 | 0.083652159 | 0.33726542 | 0.24803064 | 0.8041107 | NA | RNU6-440P | 106481895 |
| ENSG000002 | 1.17120942 | -0.395728007 | 0.31013339 | -1.2759929 | 0.20195802 | NA | RNU6-258P | 106481259 |
| ENSG000002 | 0.5008998 | -0.200150804 | 0.24832882 | -0.805991 | 0.42024801 | NA | NA | NA |
| ENSG000002 | 0.26062507 | -0.012888319 | 0.18226503 | -0.070712 | 0.94362699 | NA | RNU6-328P | 106479711 |
| ENSG000002 | 1.41557834 | 0.219063468 | 0.3379559 | 0.64820134 | 0.51685473 | NA | RNU6-298P | 106481273 |
| ENSG000002 | 0.12070844 | -0.088503695 | 0.12783931 | -0.6923042 | 0.48874631 | NA | NA | NA |
| ENSG000002 | 1.60140502 | -0.260926289 | 0.3505536 | -0.7443264 | 0.45667905 | NA | RNU6-127P | 106480554 |
| ENSG000002 | 0.9572029 | -0.181518609 | 0.30566846 | -0.5938415 | 0.55261815 | NA | NA | NA |
| ENSG000002 | 1.13286822 | -0.426550225 | 0.3212771 | -1.3276708 | 0.18428688 | NA | NA | NA |
| ENSG000002 | 0.76468381 | -0.03650702 | 0.28641547 | -0.1274618 | 0.89857494 | NA | NA | NA |
| ENSG000002 | 1.21984475 | -0.050486312 | 0.33332732 | -0.1514617 | 0.87961155 | NA | RNU6-1321P | 106481597 |
| ENSG000002 | 1.48767106 | -0.18727469 | 0.34092715 | -0.54931 | 0.58279273 | NA | RNU6-913P | 106481467 |
| ENSG000002 | 0 | NA | NA | NA | NA | NA | RNU6-816P | 106480401 |
| ENSG000002 | 1.23291187 | -0.194262013 | 0.33275774 | -0.5837941 | 0.55935883 | NA | RNU6-604P | 106479833 |
| ENSG000002 | 0.26813897 | -0.197007041 | 0.18550319 | -1.0620143 | 0.2882292 | NA | RNU6-220P | 106481247 |
| ENSG000002 | 1.79477028 | -0.464417974 | 0.35392264 | -1.3122019 | 0.18945201 | NA | RNU6-907P | 106480620 |
| ENSG000002 | 0.11934008 | 0.083564698 | 0.12760091 | 0.65489106 | 0.5125379 | NA | NA | NA |
| ENSG000002 | 0 | NA | NA | NA | NA | NA | NA | NA |

| | | | | | | | | | |
|---|---|---|---|---|---|---|---|---|---|
| ENSG000002 | 0.13280363 | -0.093394593 | 0.13115644 | -0.7120855 | 0.47641185 | NA | | RNU6-747P | 106480399 |
| ENSG000002 | 0.11934008 | 0.083564698 | 0.12760091 | 0.65489106 | 0.5125379 | NA | | NA | NA |
| ENSG000002 | 0 | NA | NA | NA | NA | NA | | NA | NA |
| ENSG000002 | 0.11865494 | 0.083440866 | 0.12751478 | 0.65436232 | 0.5128784 | NA | | RNU6-930P | 106480622 |
| ENSG000002 | 0.64741897 | -0.110552013 | 0.26688537 | -0.4142303 | 0.67870543 | NA | | NA | NA |
| ENSG000002 | 1.27887302 | -0.195926161 | 0.33032017 | -0.5931402 | 0.55308735 | NA | | NA | NA |
| ENSG000002 | 1.70685837 | 0.066694972 | 0.35093888 | 0.19004726 | 0.8492721 | NA | | RNU6-360P | 106479726 |
| ENSG000002 | 0.38494734 | -0.095409837 | 0.21124663 | -0.4516514 | 0.65152014 | NA | | RNU6-278P | 106481767 |
| ENSG000002 | 1.56804559 | 0.14407473 | 0.34448318 | 0.41823445 | 0.67577571 | NA | | RNU6-750P | 106479897 |
| ENSG000002 | 1.13127118 | -0.246256639 | 0.30973008 | -0.7950685 | 0.42657364 | NA | | NA | NA |
| ENSG000002 | 0.14128499 | -0.097772014 | 0.13403235 | -0.7294658 | 0.4657168 | NA | | RNA5SP376 | 106479010 |
| ENSG000002 | 1.06537853 | 0.353826399 | 0.32115177 | 1.10174201 | 0.27057385 | NA | | RNU6-1176P | 106481551 |
| ENSG000002 | 1.33693779 | -0.243942345 | 0.33087681 | -0.7372603 | 0.46096405 | NA | | NA | NA |
| ENSG000002 | 0.51316968 | -0.377777671 | 0.24556403 | -1.538408 | 0.12394889 | NA | | RNU6-308P | 106480568 |
| ENSG000002 | 0.98842195 | -0.040465456 | 0.31285588 | -0.1293422 | 0.89708691 | NA | | RNU6-724P | 106479887 |
| ENSG000002 | 1.62318108 | -0.383808774 | 0.34577726 | -1.1099885 | 0.26700399 | NA | | NA | NA |
| ENSG000002 | 0.6558786 | -0.270452548 | 0.26091264 | -1.0365636 | 0.29993928 | NA | | NA | NA |
| ENSG000002 | 1.74508375 | -0.462482681 | 0.3550996 | -1.3024027 | 0.19277876 | NA | | NA | NA |
| ENSG000002 | 0.36482725 | -0.102221582 | 0.21647267 | -0.4722147 | 0.63677353 | NA | | RNU6-788P | 106480609 |
| ENSG000002 | 0.38019136 | -0.10756138 | 0.21876967 | -0.491665 | 0.62295621 | NA | | NA | NA |
| ENSG000002 | 0.25333738 | -0.015686291 | 0.1806605 | -0.0868275 | 0.93080867 | NA | | NA | NA |
| ENSG000002 | 0.89469906 | -0.449032547 | 0.30060706 | -1.4937525 | 0.13524032 | NA | | NA | NA |
| ENSG000002 | 0.24756242 | -0.187034346 | 0.18119817 | -1.0322088 | 0.30197431 | NA | | NA | NA |
| ENSG000002 | 1.13153019 | -0.439364526 | 0.32336125 | -1.358742 | 0.17422835 | NA | | NA | NA |
| ENSG000002 | 0.49970612 | -0.194507174 | 0.24827631 | -0.7834303 | 0.43337449 | NA | | NA | NA |
| ENSG000002 | 1.5208886 | -0.486676546 | 0.34542716 | -1.4089122 | 0.15886112 | NA | | RNU6-578P | 106481904 |
| ENSG000002 | 1.75656978 | 0.261354977 | 0.34915824 | 0.74852874 | 0.4541413 | NA | | SNORD89 | 692205 |
| ENSG000002 | 0 | NA | NA | NA | NA | NA | | NA | NA |
| ENSG000002 | 0.85767182 | -0.111725312 | 0.29715703 | -0.3759807 | 0.70693124 | NA | | RNU6-239P | 106480563 |

| | | | | | | | | |
|---|---|---|---|---|---|---|---|---|
| ENSG000002 | 0 | NA | NA | NA | NA | NA | NA | NA |
| ENSG000002 | 0.87782224 | -0.29156826 | 0.30412205 | -0.9587212 | 0.3376992 | NA | NA | NA |
| ENSG000002 | 0.14128499 | -0.097772014 | 0.13403235 | -0.7294658 | 0.4657168 | NA | RNU6-821P | 106481440 |
| ENSG000002 | 0.49031286 | -0.020802257 | 0.24597635 | -0.0845702 | 0.93260313 | NA | RNU6-1009P | 106481499 |
| ENSG000002 | 1.27585789 | -0.328889922 | 0.325628 | -1.0100173 | 0.31248699 | NA | NA | NA |
| ENSG000002 | 0.12070844 | -0.088503695 | 0.12783931 | -0.6923042 | 0.48874631 | NA | RNU6-1154P | 106480074 |
| ENSG000002 | 0 | NA | NA | NA | NA | NA | SNORD12 | 692057 |
| ENSG000002 | 1.98191287 | -0.052353691 | 0.34700944 | -0.1508711 | 0.88007741 | NA | RNU6-679P | 106479866 |
| ENSG000002 | 1.61919332 | 0.01573338 | 0.34291034 | 0.04588191 | 0.96340438 | NA | NA | NA |
| ENSG000002 | 1.00450557 | 0.090528942 | 0.29247378 | 0.3095284 | 0.75691961 | NA | NA | NA |
| ENSG000002 | 0.37194025 | 0.067575365 | 0.21717998 | 0.31114914 | 0.75568724 | NA | NA | NA |
| ENSG000002 | 0.23890637 | -0.010420674 | 0.17747379 | -0.0587167 | 0.95317776 | NA | NA | NA |
| ENSG000002 | 0.8783029 | 0.043318042 | 0.2875928 | 0.15062283 | 0.88027325 | NA | RNU6-1307P | 106481593 |
| ENSG000002 | 1.52322443 | -0.480975637 | 0.34421448 | -1.3973138 | 0.16231923 | NA | RNU6-1228P | 106480103 |
| ENSG000002 | 1.03823866 | -0.253440997 | 0.2066667 | -1.2263272 | 0.22007557 | NA | NA | NA |
| ENSG000002 | 0.83489988 | 0.207754519 | 0.29444023 | 0.70559147 | 0.48044223 | NA | RNU6-129P | 106479623 |
| ENSG000002 | 0.48536101 | -0.024029524 | 0.24539296 | -0.0979226 | 0.92199373 | NA | NA | NA |
| ENSG000002 | 0.9134827 | -0.282124842 | 0.29953083 | -0.9418892 | 0.34624939 | NA | RNU6-882P | 106481457 |
| ENSG000002 | 1.52179761 | -0.203484562 | 0.34537897 | -0.5891632 | 0.55575183 | NA | RNU6-316P | 106481279 |
| ENSG000002 | 0 | NA | NA | NA | NA | NA | NA | NA |
| ENSG000002 | 0.84992767 | -0.114089031 | 0.30189177 | -0.3779137 | 0.70549472 | NA | RNA5SP297 | 106479003 |
| ENSG000002 | 1.90017775 | -0.634757563 | 0.35327955 | -1.7967572 | 0.07237418 | NA | RNU6-780P | 106481427 |
| ENSG000002 | 0.69470251 | 0.122385865 | 0.2663967 | 0.45941209 | 0.64593827 | NA | NA | NA |
| ENSG000002 | 0.38036605 | -0.293832733 | 0.22119083 | -1.3284128 | 0.18404178 | NA | NA | NA |
| ENSG000002 | 1.80980471 | -0.331531666 | 0.34624942 | -0.9574938 | 0.33831809 | NA | NA | NA |
| ENSG000002 | 0.14128499 | -0.097772014 | 0.13403235 | -0.7294658 | 0.4657168 | NA | RNU6-823P | 106481441 |
| ENSG000002 | 0.12070844 | -0.088503695 | 0.12783931 | -0.6923042 | 0.48874631 | NA | NA | NA |
| ENSG000002 | 1.04409485 | -0.348707367 | 0.30837037 | -1.130807 | 0.25813634 | NA | RNU6-1300P | 106480134 |
| ENSG000002 | 1.32136675 | -0.110136035 | 0.33533907 | -0.3284319 | 0.74258516 | NA | RNU6-1177P | 106480645 |

| | | | | | | | | | |
|---|---|---|---|---|---|---|---|---|---|
| ENSG000002 | 0.50341032 | -0.342798738 | 0.23572799 | -1.4542131 | 0.14588723 | NA | NA | NA | |
| ENSG000002 | 0.79075747 | -0.360972421 | 0.28354666 | -1.2730618 | 0.20299612 | NA | RNU6-1000P | | 106481496 |
| ENSG000002 | 0.73889482 | -0.029099289 | 0.29011813 | -0.1003015 | 0.92010495 | NA | RNU6-482P | | 106481332 |
| ENSG000002 | 0.90454487 | -0.274182372 | 0.29914056 | -0.916567 | 0.35936957 | NA | NA | NA | |
| ENSG000002 | 1.28949613 | -0.678631562 | 0.33112094 | -2.0494976 | 0.04041349 | NA | RNU6-401P | | 106479743 |
| ENSG000002 | 0.24485602 | -0.012655878 | 0.17881722 | -0.0707755 | 0.94357643 | NA | NA | NA | |
| ENSG000002 | 1.31316908 | 0.166174963 | 0.33500439 | 0.49603816 | 0.61986749 | NA | NA | NA | |
| ENSG000002 | 0.63990507 | 0.050063835 | 0.25851563 | 0.19365883 | 0.84644303 | NA | RNU6-269P | | 106481263 |
| ENSG000002 | 2.12724092 | -0.366535431 | 0.36406373 | -1.0067892 | 0.31403608 | NA | NA | NA | |
| ENSG000002 | 1.0431835 | -0.179124424 | 0.30100312 | -0.5950916 | 0.55178225 | NA | RNU6-159P | | 106480557 |
| ENSG000002 | 1.36907223 | 0.019460374 | 0.32961145 | 0.05904034 | 0.95291997 | NA | NA | NA | |
| ENSG000002 | 0.51208364 | -0.022981898 | 0.24832117 | -0.0925491 | 0.92626179 | NA | SNORD113-2 | | 767562 |
| ENSG000002 | 0.61927241 | -0.278564577 | 0.26634985 | -1.0458597 | 0.29562581 | NA | NA | NA | |
| ENSG000002 | 1.13685082 | -0.120415313 | 0.32303868 | -0.3727582 | 0.70932842 | NA | RNU6-796P | | 106481432 |
| ENSG000002 | 1.22348423 | -0.185891099 | 0.32657983 | -0.5692057 | 0.56921656 | NA | NA | NA | |
| ENSG000002 | 0.23890637 | -0.010420674 | 0.17747379 | -0.0587167 | 0.95317776 | NA | NA | NA | |
| ENSG000002 | 0.12685397 | -0.090215768 | 0.12901919 | -0.699243 | 0.48440017 | NA | NA | NA | |
| ENSG000002 | 1.28679131 | -0.347951506 | 0.32923718 | -1.0568415 | 0.29058394 | NA | NA | NA | |
| ENSG000002 | 0.63352874 | -0.1085548 | 0.25832188 | -0.4202308 | 0.67431688 | NA | NA | NA | |
| ENSG000002 | 0.86757608 | -0.09663148 | 0.28990707 | -0.3333188 | 0.73889364 | NA | RNU6-1248P | | 106480111 |
| ENSG000002 | 0.37899768 | -0.10176489 | 0.21865765 | -0.4654074 | 0.64163973 | NA | NA | NA | |
| ENSG000002 | 0.11205239 | 0.082118251 | 0.12671962 | 0.64803108 | 0.51696485 | NA | RNU6-663P | | 106481391 |
| ENSG000002 | 0.513452 | -0.199963756 | 0.2496885 | -0.8008529 | 0.42321682 | NA | NA | NA | |
| ENSG000002 | 1.47601115 | -0.327558211 | 0.34229374 | -0.9569506 | 0.33859217 | NA | RNU6-932P | | 106481474 |
| ENSG000002 | 0.62684242 | -0.109923117 | 0.27148862 | -0.4048903 | 0.68555813 | NA | NA | NA | |
| ENSG000002 | 0.47667517 | 0.31075097 | 0.23106181 | 1.34488244 | 0.1786632 | NA | RNU11-3P | | 106479562 |
| ENSG000002 | 0.40942396 | -0.284058841 | 0.21788566 | -1.303706 | 0.19233384 | NA | NA | NA | |
| ENSG000002 | 0.13280363 | -0.093394593 | 0.13115644 | -0.7120855 | 0.47641185 | NA | NA | NA | |
| ENSG000002 | 0.61611113 | 0.054662593 | 0.27024684 | 0.20226913 | 0.83970633 | NA | RNY4P36 | | 106480313 |

| | | | | | | | | |
|---|---|---|---|---|---|---|---|---|
| ENSG000002 | 0.25993993 | -0.013144386 | 0.18211683 | -0.0721756 | 0.94246219 | NA | NA | NA |
| ENSG000002 | 0.62496306 | -0.108686629 | 0.26501936 | -0.4101083 | 0.68172654 | NA | NA | NA |
| ENSG000002 | 0.71559461 | -0.093823799 | 0.1588217 | -0.5907492 | 0.55468845 | NA | RNU1-119P | 106480668 |
| ENSG000002 | 1.64010818 | -0.117602353 | 0.34567119 | -0.3402145 | 0.733695 | NA | NA | NA |
| ENSG000002 | 0.5301324 | -0.3843196 | 0.24728844 | -1.5541349 | 0.12015224 | NA | NA | NA |
| ENSG000002 | 0.73292343 | -0.195671801 | 0.28518655 | -0.6861186 | 0.49263832 | NA | RNU11-6P | 106480544 |
| ENSG000002 | 1.34923098 | -0.122258384 | 0.34335912 | -0.3560656 | 0.72179141 | NA | NA | NA |
| ENSG000002 | 0.51894464 | -0.193743146 | 0.24314987 | -0.7968055 | 0.42556401 | NA | NA | NA |
| ENSG000002 | 1.59696754 | -0.398267299 | 0.34938673 | -1.1399039 | 0.25432633 | NA | NA | NA |
| ENSG000002 | 0.90798793 | -0.437671584 | 0.29609134 | -1.4781641 | 0.13936386 | NA | NA | NA |
| ENSG000002 | 0 | NA | NA | NA | NA | NA | RNU6-1269P | 106480119 |
| ENSG000002 | 0.98480654 | -0.179926523 | 0.306894 | -0.5862823 | 0.55768582 | NA | RNU6-243P | 106479675 |
| ENSG000002 | 1.57143654 | -0.46457584 | 0.33891022 | -1.3707932 | 0.17043942 | NA | NA | NA |
| ENSG000002 | 2.13715323 | -0.612474124 | 0.36124694 | -1.6954445 | 0.08999114 | NA | RNU6-131P | 106479624 |
| ENSG000002 | 0.81221529 | 0.20432271 | 0.29316762 | 0.69694842 | 0.48583508 | NA | SNORD90 | 692206 |
| ENSG000002 | 0.59892612 | -0.099249839 | 0.26277623 | -0.3776972 | 0.70565557 | NA | NA | NA |
| ENSG000002 | 0.39499294 | -0.277908454 | 0.21584201 | -1.287555 | 0.19790091 | NA | RNU6-241P | 106479674 |
| ENSG000002 | 1.74396333 | -0.30600074 | 0.35074583 | -0.8724287 | 0.38297453 | NA | RNU11-4P | 106481174 |
| ENSG000002 | 0.75637659 | -0.193693286 | 0.28066413 | -0.6901248 | 0.49011569 | NA | NA | NA |
| ENSG000002 | 1.33237589 | 0.178284715 | 0.34160402 | 0.52190462 | 0.60173674 | NA | NA | NA |
| ENSG000002 | 0.39274356 | -0.107428268 | 0.22062504 | -0.4869269 | 0.62631013 | NA | RNU6-695P | 106481400 |
| ENSG000002 | 0.72309543 | 0.133495179 | 0.27763978 | 0.48082152 | 0.63064336 | NA | RNU6-460P | 106479769 |
| ENSG000002 | 0.98594678 | -0.319137342 | 0.3029928 | -1.0532836 | 0.29221104 | NA | NA | NA |
| ENSG000002 | 2.13307325 | -0.36453513 | 0.36262237 | -1.0052748 | 0.31476455 | NA | RNU6-952P | 106479987 |
| ENSG000002 | 1.36591138 | -0.269331361 | 0.34022741 | -0.7916216 | 0.42858136 | NA | RNU6-754P | 106481418 |
| ENSG000002 | 0.52147635 | -0.19458195 | 0.24341208 | -0.7993932 | 0.42406247 | NA | RNU6-1158P | 106480076 |
| ENSG000002 | 0.58895377 | 0.229836184 | 0.2625711 | 0.87532932 | 0.38139475 | NA | RNU1-101P | 106480667 |
| ENSG000002 | 2.11167768 | -0.989096633 | 0.35436706 | -2.7911642 | 0.00525188 | NA | RNU6-400P | 106481304 |
| ENSG000002 | 0.51111617 | -0.186068642 | 0.2423498 | -0.7677689 | 0.44262448 | NA | NA | NA |

| | | | | | | | | | |
|---|---|---|---|---|---|---|---|---|---|
| ENSG000002 | 0.46341609 | 0.157328408 | 0.24338973 | 0.64640528 | 0.51801692 | NA | | RNU6-530P | 106479801 |
| ENSG000002 | 1.25434825 | -0.341056435 | 0.33064861 | -1.031477 | 0.30231718 | NA | | RNU6-1197P | 106481558 |
| ENSG000002 | 1.10093327 | 0.174786793 | 0.31322142 | 0.55802951 | 0.57682423 | NA | | RNU6-1109P | 106480054 |
| ENSG000002 | 1.90474905 | 0.181003686 | 0.3579473 | 0.50567133 | 0.6130874 | NA | | NA | NA |
| ENSG000002 | 0.72096812 | 0.1287583 | 0.27749713 | 0.46399867 | 0.6426487 | NA | | RNU6-332P | 106481284 |
| ENSG000002 | 0.23202362 | -0.008302493 | 0.17586207 | -0.0472103 | 0.96234565 | NA | | RNU6-1093P | 106481524 |
| ENSG000002 | 1.08197303 | 0.041394262 | 0.31940169 | 0.12959938 | 0.89688339 | NA | | NA | NA |
| ENSG000002 | 0 | NA | NA | NA | NA | NA | | SNORD86 | 692201 |
| ENSG000002 | 1.59702153 | -0.257345393 | 0.34983497 | -0.7356194 | 0.46196235 | NA | | NA | NA |
| ENSG000002 | 0.96294753 | 0.120250787 | 0.31176357 | 0.38571148 | 0.69971036 | NA | | NA | NA |
| ENSG000002 | 0 | NA | NA | NA | NA | NA | | RNU6-972P | 106479995 |
| ENSG000002 | 0.62541795 | -0.259640357 | 0.25836648 | -1.0049305 | 0.31493031 | NA | | NA | NA |
| ENSG000002 | 0 | NA | NA | NA | NA | NA | | RNU6-1268P | 106480654 |
| ENSG000002 | 1.02502655 | -0.195047533 | 0.31020996 | -0.6287597 | 0.52950636 | NA | | NA | NA |
| ENSG000002 | 0.23202362 | -0.008302493 | 0.17586207 | -0.0472103 | 0.96234565 | NA | | RNU11-5P | 106479563 |
| ENSG000002 | 1.34189122 | -0.124546688 | 0.343216 | -0.3628814 | 0.71669351 | NA | | RNU6-1250P | 106481575 |
| ENSG000002 | 1.59633246 | 0.014126755 | 0.34616168 | 0.0408097 | 0.9674476 | NA | | RNU6-918P | 106481469 |
| ENSG000002 | 1.85602994 | -0.12083015 | 0.35246841 | -0.3428113 | 0.73174043 | NA | | NA | NA |
| ENSG000002 | 0.90027816 | -0.281064798 | 0.30047954 | -0.9353875 | 0.34958865 | NA | | NA | NA |
| ENSG000002 | 0.12070844 | -0.088503695 | 0.12783931 | -0.6923042 | 0.48874631 | NA | | NA | NA |
| ENSG000002 | 1.50446824 | -0.34267273 | 0.34410213 | -0.995846 | 0.31932498 | NA | | NA | NA |
| ENSG000002 | 0.36345889 | 0.070257596 | 0.21589421 | 0.32542602 | 0.74485866 | NA | | NA | NA |
| ENSG000002 | 1.16882694 | 0.521692163 | 0.31455195 | 1.65852464 | 0.09721162 | NA | | NA | NA |
| ENSG000002 | 1.52889334 | -0.477860623 | 0.34225863 | -1.3961975 | 0.16265504 | NA | | NA | NA |
| ENSG000002 | 1.45627999 | -0.041476136 | 0.34320022 | -0.1208511 | 0.90380895 | NA | | NA | NA |
| ENSG000002 | 0.6102458 | 0.044525027 | 0.25426768 | 0.17511084 | 0.86099254 | NA | | NA | NA |
| ENSG000002 | 1.27239309 | -0.333373724 | 0.32714039 | -1.019054 | 0.30817732 | NA | | RNU6-337P | 106479716 |
| ENSG000002 | 0.37422052 | -0.268731395 | 0.21273223 | -1.2632378 | 0.20650373 | NA | | RNU6-995P | 106481494 |
| ENSG000002 | 0.48032325 | -0.179266136 | 0.23878452 | -0.7507444 | 0.45280651 | NA | | NA | NA |

| | | | | | | | | |
|---|---|---|---|---|---|---|---|---|
| ENSG000002 | 0.5008998 | -0.200150804 | 0.24832882 | -0.805991 | 0.42024801 | NA | RNU1-78P | 106480665 |
| ENSG000002 | 1.45958508 | -0.270163764 | 0.32691256 | -0.8264099 | 0.40857164 | NA | NA | NA |
| ENSG000002 | 0.36827086 | -0.266056975 | 0.21181163 | -1.2561018 | 0.20907905 | NA | NA | NA |
| ENSG000002 | 1.51182973 | -0.198498224 | 0.34644358 | -0.5729597 | 0.56667195 | NA | RNU6-192P | 106481237 |
| ENSG000002 | 1.24147701 | -0.19493898 | 0.32934838 | -0.5918929 | 0.55392229 | NA | NA | NA |
| ENSG000002 | 0.11131518 | 0.081951103 | 0.12663804 | 0.64712866 | 0.51754868 | NA | NA | NA |
| ENSG000002 | 1.7813411 | -0.181899829 | 0.34899985 | -0.5212032 | 0.60222525 | NA | NA | NA |
| ENSG000002 | 0.72037177 | -0.172939881 | 0.27655343 | -0.6253398 | 0.53174804 | NA | RNU6-630P | 106481380 |
| ENSG000002 | 0.61335714 | -0.10408451 | 0.2640241 | -0.3942235 | 0.69341602 | NA | RNU6-381P | 106480576 |
| ENSG000002 | 0.90844494 | -0.459619446 | 0.30273017 | -1.5182479 | 0.12895191 | NA | RNU6-1326P | 106481951 |
| ENSG000002 | 0.24756242 | -0.187034346 | 0.18119817 | -1.0322088 | 0.30197431 | NA | NA | NA |
| ENSG000002 | 1.82821675 | -0.226900949 | 0.35227589 | -0.6441001 | 0.51951051 | NA | NA | NA |
| ENSG000002 | 0.37831254 | -0.102004459 | 0.21855316 | -0.4667261 | 0.64069587 | NA | RNU6-1254P | 106481576 |
| ENSG000002 | 1.37001447 | -0.270087418 | 0.34030908 | -0.7936533 | 0.4273973 | NA | NA | NA |
| ENSG000002 | 0.72400679 | -0.023630267 | 0.26825607 | -0.0880885 | 0.92980636 | NA | RNU6-903P | 106481464 |
| ENSG000002 | 1.40634925 | 0.35348949 | 0.33228684 | 1.06380827 | 0.28741556 | NA | NA | NA |
| ENSG000002 | 2.08058569 | 0.115784972 | 0.36460277 | 0.31756471 | 0.75081515 | NA | NA | NA |
| ENSG000002 | 0.23202362 | -0.008302493 | 0.17586207 | -0.0472103 | 0.96234565 | NA | NA | NA |
| ENSG000002 | 0.86992726 | 0.189740827 | 0.29047191 | 0.65321574 | 0.5136172 | NA | NA | NA |
| ENSG000002 | 0.46956163 | 0.155192681 | 0.24412227 | 0.63571702 | 0.52496088 | NA | RNU6-587P | 106481366 |
| ENSG000002 | 0.93087023 | 0.272945625 | 0.30456466 | 0.89618285 | 0.37015511 | NA | NA | NA |
| ENSG000002 | 1.12452729 | -0.114036451 | 0.32113983 | -0.3550991 | 0.72251538 | NA | NA | NA |
| ENSG000002 | 0.83512609 | 0.048464843 | 0.30079428 | 0.16112289 | 0.87199661 | NA | NA | NA |
| ENSG000002 | 1.10002234 | -0.228799984 | 0.30943992 | -0.7394004 | 0.45966393 | NA | NA | NA |
| ENSG000002 | 0.24141689 | -0.165257896 | 0.17127401 | -0.9648743 | 0.33460776 | NA | NA | NA |
| ENSG000002 | 1.39306874 | -0.421064208 | 0.33855629 | -1.2437052 | 0.21360809 | NA | NA | NA |
| ENSG000002 | 0.61121718 | -0.095100843 | 0.25474802 | -0.3733134 | 0.70891523 | NA | RNU6-876P | 106481455 |
| ENSG000002 | 1.87353187 | -0.260033445 | 0.35917881 | -0.7239666 | 0.46908627 | NA | NA | NA |
| ENSG000002 | 0.4983681 | -0.186127524 | 0.2408746 | -0.7727154 | 0.4396908 | NA | RNU6-279P | 106479690 |

| | | | | | | | | |
|---|---|---|---|---|---|---|---|---|
| ENSG000002 | 1.27992873 | -0.347943608 | 0.33271397 | -1.045774 | 0.2956654 | NA | NA | NA |
| ENSG000002 | 0.85866962 | -0.11093676 | 0.2911669 | -0.3810075 | 0.70319771 | NA | RNU1-56P | 106480165 |
| ENSG000002 | 0.85858372 | -0.26887183 | 0.29839338 | -0.901065 | 0.36755376 | NA | NA | NA |
| ENSG000002 | 0.26813897 | -0.197007041 | 0.18550319 | -1.0620143 | 0.2882292 | NA | RNU1-139P | 106480196 |
| ENSG000002 | 0.50750235 | -0.197774682 | 0.24907427 | -0.794039 | 0.42717272 | NA | NA | NA |
| ENSG000002 | 1.25246889 | -0.350158116 | 0.33199199 | -1.0547186 | 0.29155405 | NA | NA | NA |
| ENSG000002 | 0 | NA | NA | NA | NA | NA | RNU6-1239P | 106481571 |
| ENSG000002 | 0.66088252 | -0.288231726 | 0.26973913 | -1.0685573 | 0.28526918 | NA | NA | NA |
| ENSG000002 | 1.56738666 | 0.026687647 | 0.34646664 | 0.07702804 | 0.93860124 | NA | NA | NA |
| ENSG000002 | 0.12685397 | -0.090215768 | 0.12901919 | -0.699243 | 0.48440017 | NA | RNU6-493P | 106481336 |
| ENSG000002 | 0.86552066 | -0.097333404 | 0.28979247 | -0.3358728 | 0.7369668 | NA | NA | NA |
| ENSG000002 | 1.12993561 | -0.111392587 | 0.3166667 | -0.351766 | 0.72501374 | NA | NA | NA |
| ENSG000002 | 0.97370916 | -0.181124969 | 0.30640194 | -0.5911352 | 0.55442985 | NA | NA | NA |
| ENSG000002 | 0.25993993 | -0.013144386 | 0.18211683 | -0.0721756 | 0.94246219 | NA | NA | NA |
| ENSG000002 | 1.29590225 | -0.684848914 | 0.33117534 | -2.0679345 | 0.03864618 | NA | KRTAP16-1 | 100505753 |
| ENSG000002 | 0.90642369 | -0.282996904 | 0.30077944 | -0.9408785 | 0.34676713 | NA | KRTAP29-1 | 100533177 |
| ENSG000002 | 1.21394908 | 0.098133254 | 0.32803809 | 0.29915201 | 0.76482406 | NA | KRTAP9-6 | 100507608 |
| ENSG000002 | 0.86295807 | 0.04613514 | 0.29709074 | 0.15528973 | 0.87659292 | NA | NA | NA |
| ENSG000002 | 1.61148343 | -0.396510634 | 0.34959656 | -1.1341949 | 0.25671281 | NA | NA | NA |
| ENSG000002 | 1.21890306 | 0.253244834 | 0.32855806 | 0.77077651 | 0.44083941 | NA | NA | NA |
| ENSG000002 | 1.00257064 | -0.193701868 | 0.30928282 | -0.6262937 | 0.53112235 | NA | KRTAP2-3 | 730755 |
| ENSG000002 | 0.26199343 | -0.194018184 | 0.18422801 | -1.0531416 | 0.29227609 | NA | KRTAP2-1 | 81872 |
| ENSG000002 | 1.96665841 | -0.1966569 | 0.3651481 | -0.5385675 | 0.59018532 | NA | NA | NA |
| ENSG000002 | 0 | NA | NA | NA | NA | NA | NA | NA |
| ENSG000002 | 0 | NA | NA | NA | NA | NA | NA | NA |
| ENSG000002 | 0.11865494 | 0.083440866 | 0.12751478 | 0.65436232 | 0.5128784 | NA | NA | NA |
| ENSG000002 | 0.50611225 | -0.202624884 | 0.24886748 | -0.8141879 | 0.41553732 | NA | KRTAP12-4 | 386684 |
| ENSG000002 | 0.47070378 | 0.149674697 | 0.23683599 | 0.63197616 | 0.52740246 | NA | KRTAP10-3 | 386682 |
| ENSG000002 | 1.34809287 | 0.167712111 | 0.33579001 | 0.49945533 | 0.61745865 | NA | KRTAP6-3 | 337968 |

| | | | | | | | | | |
|---|---|---|---|---|---|---|---|---|---|
| ENSG000002 | 1.65093185 | -0.566255054 | 0.35203515 | -1.6085185 | 0.10772166 | NA | NA | NA | |
| ENSG000002 | 0.61446491 | -0.254170746 | 0.25900567 | -0.9813328 | 0.32642867 | NA | NA | NA | |
| ENSG000002 | 1.66576726 | -0.530223682 | 0.3443451 | -1.5398032 | 0.12360833 | NA | NA | NA | |
| ENSG000002 | 0.73820967 | -0.027876881 | 0.28472784 | -0.0979071 | 0.92200605 | NA | NA | NA | |
| ENSG000002 | 0 | NA | NA | NA | NA | NA | KLK9 | | 284366 |
| ENSG000002 | 1.57624459 | -0.474415156 | 0.34189035 | -1.3876237 | 0.16525166 | NA | NA | NA | |
| ENSG000002 | 1.76372549 | -0.192784184 | 0.35461801 | -0.543639 | 0.58668991 | NA | NA | NA | |
| ENSG000002 | 0 | NA | NA | NA | NA | NA | SPHAR | | 10638 |
| ENSG000002 | 1.96639782 | -0.191395925 | 0.36467836 | -0.5248349 | 0.59969798 | NA | CGB8 | | 94115 |
| ENSG000002 | 1.29447832 | -0.83958934 | 0.32629903 | -2.5730672 | 0.01008016 | NA | NA | NA | |
| ENSG000002 | 1.46567271 | -0.18705364 | 0.34376618 | -0.5441304 | 0.58635172 | NA | NA | NA | |
| ENSG000002 | 0.47687965 | -0.021216903 | 0.24445659 | -0.0867921 | 0.93083676 | NA | NA | NA | |
| ENSG000002 | 1.40183085 | -0.430126919 | 0.34170917 | -1.2587515 | 0.20812011 | NA | NA | NA | |
| ENSG000002 | 1.63015082 | -0.247684662 | 0.34820594 | -0.7113166 | 0.47688808 | NA | NA | NA | |
| ENSG000002 | 1.88314742 | -0.251068282 | 0.35527247 | -0.7066922 | 0.47975778 | NA | NA | NA | |
| ENSG000002 | 1.64498219 | -0.568596558 | 0.35256221 | -1.6127552 | 0.10679769 | NA | NA | NA | |
| ENSG000002 | 2.12256871 | -0.393148111 | 0.36780556 | -1.068902 | 0.28511382 | NA | NA | NA | |
| ENSG000002 | 1.59410819 | 0.010651286 | 0.34476727 | 0.03089413 | 0.97535397 | NA | NA | NA | |
| ENSG000002 | 0 | NA | NA | NA | NA | NA | NA | NA | |
| ENSG000002 | 0.90816262 | -0.635180171 | 0.29877972 | -2.1259146 | 0.03351037 | NA | NA | NA | |
| ENSG000002 | 0.61409435 | -0.103807472 | 0.26408954 | -0.3930768 | 0.69426276 | NA | NA | NA | |
| ENSG000002 | 1.36343174 | -0.263970666 | 0.33834154 | -0.7801899 | 0.43527908 | NA | NA | NA | |
| ENSG000002 | 2.15445172 | -0.379532487 | 0.364193 | -1.0421191 | 0.29735645 | NA | NA | NA | |
| ENSG000002 | 1.06387863 | -0.719575284 | 0.31142394 | -2.3105972 | 0.02085511 | NA | NA | NA | |
| ENSG000002 | 0.36827086 | -0.266056975 | 0.21181163 | -1.2561018 | 0.20907905 | NA | NA | NA | |
| ENSG000002 | 1.73869882 | -0.281229858 | 0.34316954 | -0.8195071 | 0.41249717 | NA | CRIP1 | | 1396 |
| ENSG000002 | 0 | NA | NA | NA | NA | NA | NA | NA | |
| ENSG000002 | 1.48100064 | 0.089187182 | 0.34353754 | 0.25961408 | 0.79516147 | NA | NA | NA | |
| ENSG000002 | 1.59508672 | 0.011635934 | 0.34690222 | 0.0335424 | 0.97324205 | NA | NA | NA | |

| | | | | | | | | | |
|---|---|---|---|---|---|---|---|---|---|
| ENSG000002 | 0.77018092 | -0.359515013 | 0.28385235 | -1.2665564 | 0.20531394 | NA | NA | NA | |
| ENSG000002 | 2.00539032 | -0.325179428 | 0.36423437 | -0.8927752 | 0.37197755 | NA | NA | NA | |
| ENSG000002 | 1.70603121 | -0.30293894 | 0.34679606 | -0.8735363 | 0.38237085 | NA | NA | NA | |
| ENSG000002 | 0.35733509 | 0.244646831 | 0.20894085 | 1.17089037 | 0.24164285 | NA | NA | NA | |
| ENSG000002 | 1.8756052 | -0.381434244 | 0.3570276 | -1.0683607 | 0.28535786 | NA | NA | NA | |
| ENSG000002 | 2.10355653 | -0.004477907 | 0.36231655 | -0.0123591 | 0.99013911 | NA | NA | NA | |
| ENSG000002 | 1.80857177 | 0.263835113 | 0.35768295 | 0.73762283 | 0.46074368 | NA | NA | NA | |
| ENSG000002 | 0.83572691 | 0.048149579 | 0.28948181 | 0.16633024 | 0.86789707 | NA | NA | NA | |
| ENSG000002 | 0.84263999 | -0.111449283 | 0.2963531 | -0.3760692 | 0.70686543 | NA | NA | NA | |
| ENSG000002 | 0.61221498 | -0.115173106 | 0.27027142 | -0.4261387 | 0.6700068 | NA | NA | NA | |
| ENSG000002 | 0.66683218 | -0.273056421 | 0.26350214 | -1.0362588 | 0.30008141 | NA | NA | NA | |
| ENSG000002 | 0.77521868 | -0.191196496 | 0.28023911 | -0.682262 | 0.49507329 | NA | NA | NA | |
| ENSG000002 | 1.56394038 | 0.010139056 | 0.33515519 | 0.03025182 | 0.97586622 | NA | NA | NA | |
| ENSG000002 | 1.99608139 | -0.192549435 | 0.36144678 | -0.5327186 | 0.59422837 | NA | NA | NA | |
| ENSG000002 | 0.92763139 | -0.444046912 | 0.29705627 | -1.4948242 | 0.13496032 | NA | NA | NA | |
| ENSG000002 | 1.24653513 | -0.041113343 | 0.32205889 | -0.1276578 | 0.89841975 | NA | NA | NA | |
| ENSG000002 | 1.22756146 | -0.100713652 | 0.20736149 | -0.4856912 | 0.62718613 | NA | NA | NA | |
| ENSG000002 | 1.99782381 | -0.195440019 | 0.36239937 | -0.5392946 | 0.5896836 | NA | CSH2 | | 1443 |
| ENSG000002 | 1.19033387 | 0.240870155 | 0.32190739 | 0.74825916 | 0.45430385 | NA | NA | NA | |
| ENSG000002 | 1.71186229 | 0.075192157 | 0.35783206 | 0.21013254 | 0.83356423 | NA | NA | NA | |
| ENSG000002 | 1.09415466 | 0.197336826 | 0.32527781 | 0.60667165 | 0.54406884 | NA | NA | NA | |
| ENSG000002 | 0 | NA | NA | NA | NA | NA | NA | NA | |
| ENSG000002 | 0.60698135 | -0.252864344 | 0.25666105 | -0.9852073 | 0.32452224 | NA | NA | NA | |
| ENSG000002 | 1.50614067 | -0.192033064 | 0.34125537 | -0.5627254 | 0.57362189 | NA | NA | NA | |
| ENSG000002 | 0.13280363 | -0.093394593 | 0.13115644 | -0.7120855 | 0.47641185 | NA | NA | NA | |
| ENSG000002 | 1.11647014 | -0.118804332 | 0.32621597 | -0.3641892 | 0.71571673 | NA | NA | NA | |
| ENSG000002 | 0 | NA | NA | NA | NA | NA | NA | NA | |
| ENSG000002 | 1.38487162 | -0.121160834 | 0.33527117 | -0.3613816 | 0.71781419 | NA | NA | NA | |
| ENSG000002 | 0.84497582 | -0.111985652 | 0.29647309 | -0.3777262 | 0.705634 | NA | NA | NA | |

| | | | | | | | | | |
|---|---|---|---|---|---|---|---|---|---|
| ENSG000002 | 0.84952486 | 0.047897142 | 0.30152928 | 0.1588474 | 0.87378911 | NA | NA | NA | |
| ENSG000002 | 0.99349293 | -0.499820888 | 0.30602254 | -1.6332813 | 0.10240985 | NA | NA | NA | |
| ENSG000002 | 1.89963587 | -0.683615639 | 0.35864144 | -1.9061256 | 0.05663391 | NA | NA | NA | |
| ENSG000002 | 0 | NA | NA | NA | NA | NA | NA | NA | |
| ENSG000002 | 1.82126832 | 0.271181437 | 0.35776267 | 0.75799254 | 0.44845545 | NA | NA | NA | |
| ENSG000002 | 0.72008656 | 0.117012389 | 0.27576463 | 0.42431979 | 0.67133261 | NA | NA | NA | |
| ENSG000002 | 1.25593891 | -0.339557165 | 0.32845548 | -1.0337997 | 0.30122983 | NA | NA | NA | |
| ENSG000002 | 1.49637918 | -0.344394437 | 0.34753258 | -0.9909702 | 0.32170012 | NA | NA | NA | |
| ENSG000002 | 2.14915103 | -0.131285413 | 0.36538608 | -0.359306 | 0.71936619 | NA | NA | NA | |
| ENSG000002 | 0 | NA | NA | NA | NA | NA | NA | NA | |
| ENSG000002 | 0 | NA | NA | NA | NA | NA | NA | NA | |
| ENSG000002 | 2.17289904 | -0.490915408 | 0.35985044 | -1.3642207 | 0.17249813 | NA | NA | NA | |
| ENSG000002 | 1.72618508 | -0.678443036 | 0.34434694 | -1.970231 | 0.0488119 | NA | NA | NA | |
| ENSG000002 | 0.62980336 | -0.100896909 | 0.25641496 | -0.3934907 | 0.69395707 | NA | NA | NA | |
| ENSG000002 | 1.60135645 | 0.007622996 | 0.33962285 | 0.02244547 | 0.98209261 | NA | NA | NA | |
| ENSG000002 | 1.54007663 | -0.328162071 | 0.33860086 | -0.9691708 | 0.33245997 | NA | NA | NA | |
| ENSG000002 | 1.3775501 | -0.267721831 | 0.34046787 | -0.7863351 | 0.43167121 | NA | NA | NA | |
| ENSG000002 | 0.78130864 | -0.565696235 | 0.28699154 | -1.9711251 | 0.04870956 | NA | NA | NA | |
| ENSG000002 | 0 | NA | NA | NA | NA | NA | PTPRCAP | | 5790 |
| ENSG000002 | 1.87096634 | -0.39505294 | 0.35970992 | -1.0982542 | 0.2720935 | NA | NA | NA | |
| ENSG000002 | 1.35225195 | -0.117574996 | 0.33940064 | -0.3464195 | 0.72902748 | NA | NA | NA | |
| ENSG000002 | 0.11934008 | 0.083564698 | 0.12760091 | 0.65489106 | 0.5125379 | NA | PVRIG | | 79037 |
| ENSG000002 | 1.68115581 | 0.211751244 | 0.35080263 | 0.60361931 | 0.54609677 | NA | NA | NA | |
| ENSG000002 | 0.66313191 | -0.429680649 | 0.25813351 | -1.6645675 | 0.09599908 | NA | KRTAP2-4 | | 85294 |
| ENSG000002 | 1.91796107 | 0.076645619 | 0.35645666 | 0.21502087 | 0.82975105 | NA | NA | NA | |
| ENSG000002 | 0.77632645 | -0.361571052 | 0.28424434 | -1.2720431 | 0.2033578 | NA | NA | NA | |
| ENSG000002 | 0.37945415 | -0.107841678 | 0.21865587 | -0.4932027 | 0.62186934 | NA | NA | NA | |
| ENSG000002 | 1.1098676 | -0.124685886 | 0.32928509 | -0.3786563 | 0.70494308 | NA | NA | NA | |
| ENSG000002 | 2.02164851 | -0.852229305 | 0.35831388 | -2.3784434 | 0.0173859 | NA | NA | NA | |

| | | | | | | | | | |
|---|---|---|---|---|---|---|---|---|---|
| ENSG000002 | 0 | NA | NA | NA | NA | NA | NA | NA | |
| ENSG000002 | 1.49971281 | -0.342555996 | 0.34566573 | -0.9910036 | 0.3216838 | NA | NA | NA | |
| ENSG000002 | 0.75982019 | -0.379788636 | 0.28855699 | -1.3161651 | 0.18811864 | NA | NA | NA | |
| ENSG000002 | 1.51818378 | -0.206246766 | 0.34733075 | -0.5938051 | 0.55264249 | NA | NA | NA | |
| ENSG000002 | 0.47690138 | 0.157973135 | 0.24496158 | 0.64488945 | 0.51899883 | NA | NA | NA | |
| ENSG000002 | 0.12070844 | -0.088503695 | 0.12783931 | -0.6923042 | 0.48874631 | NA | NA | NA | |
| ENSG000002 | 1.02460042 | -0.343357349 | 0.3076009 | -1.116243 | 0.26431813 | NA | NA | NA | |
| ENSG000002 | 1.51046137 | -0.055466927 | 0.34303139 | -0.1616964 | 0.87154497 | NA | NA | NA | |
| ENSG000002 | 1.62824972 | -0.406146372 | 0.35094166 | -1.1573045 | 0.24714797 | NA | NA | NA | |
| ENSG000002 | 1.8370438 | -0.61475026 | 0.35362146 | -1.7384416 | 0.08213303 | NA | NA | NA | |
| ENSG000002 | 1.45573653 | -0.178427213 | 0.34035124 | -0.5242443 | 0.6001086 | NA | NA | NA | |
| ENSG000002 | 1.4035981 | -0.399599586 | 0.33406895 | -1.196159 | 0.2316345 | NA | NA | NA | |
| ENSG000002 | 2.00463083 | -0.453105667 | 0.36063073 | -1.2564255 | 0.20896172 | NA | NA | NA | |
| ENSG000002 | 1.00495854 | -0.209381385 | 0.3180633 | -0.658301 | 0.51034474 | NA | NA | NA | |
| ENSG000002 | 0.58682646 | 0.225002094 | 0.26240694 | 0.85745481 | 0.39119358 | NA | NA | NA | |
| ENSG000002 | 0.85268614 | -0.249316733 | 0.29056575 | -0.858039 | 0.39087096 | NA | NA | NA | |
| ENSG000002 | 0.87078709 | 0.048970072 | 0.29749719 | 0.16460684 | 0.86925345 | NA | NA | NA | |
| ENSG000002 | 1.90264632 | -0.507877801 | 0.35534848 | -1.4292387 | 0.15293563 | NA | KRT8P41 | | 283102 |
| ENSG000002 | 0.38270188 | -0.272446727 | 0.2140009 | -1.2731102 | 0.20297894 | NA | NA | NA | |
| ENSG000002 | 0.61955474 | -0.11252945 | 0.27088619 | -0.4154123 | 0.67784012 | NA | NA | NA | |
| ENSG000002 | 1.23808994 | -0.352957627 | 0.33362414 | -1.0579499 | 0.29007829 | NA | NA | NA | |
| ENSG000002 | 2.09693008 | -0.690589281 | 0.35642163 | -1.9375628 | 0.05267659 | NA | NA | NA | |
| ENSG000002 | 1.37071888 | -0.27495688 | 0.34030983 | -0.8079604 | 0.41911335 | NA | NA | NA | |
| ENSG000002 | 0 | NA | NA | NA | NA | NA | CPLX3 | | 594855 |
| ENSG000002 | 0.88012774 | -0.113321964 | 0.2983266 | -0.3798587 | 0.70405029 | NA | NA | NA | |
| ENSG000002 | 0 | NA | NA | NA | NA | NA | ZBTB9 | | 221504 |
| ENSG000002 | 2.05358249 | -0.836163664 | 0.35539911 | -2.3527455 | 0.01863538 | NA | NA | NA | |
| ENSG000002 | 0 | NA | NA | NA | NA | NA | NA | NA | |
| ENSG000002 | 0.7536964 | -0.192464368 | 0.28656601 | -0.6716232 | 0.50182363 | NA | NA | NA | |

| | | | | | | | | |
|---|---|---|---|---|---|---|---|---|
| ENSG000002 | 1.61089002 | -0.393618657 | 0.34879503 | -1.1285099 | 0.25910463 | NA | NA | NA |
| ENSG000002 | 1.65969745 | -0.26511029 | 0.34947757 | -0.7585903 | 0.44809768 | NA | NA | NA |
| ENSG000002 | 1.00746906 | -0.346543074 | 0.30928431 | -1.1204677 | 0.26251452 | NA | NA | NA |
| ENSG000002 | 1.79679562 | 0.143713023 | 0.35624553 | 0.40341004 | 0.6866466 | NA | NA | NA |
| ENSG000002 | 1.38079494 | 0.012557135 | 0.3363838 | 0.03732978 | 0.97022206 | NA | NA | NA |
| ENSG000002 | 0.91527561 | -0.434940576 | 0.2965341 | -1.4667472 | 0.14244483 | NA | NA | NA |
| ENSG000002 | 0.86569088 | 0.038333103 | 0.2803667 | 0.13672488 | 0.89124827 | NA | NA | NA |
| ENSG000002 | 0.38056192 | -0.244105065 | 0.20391883 | -1.1970698 | 0.23127937 | NA | NA | NA |
| ENSG000002 | 0.23139247 | 0.167748926 | 0.17680352 | 0.94878726 | 0.34272882 | NA | NA | NA |
| ENSG000002 | 0.77632645 | -0.361571052 | 0.28424434 | -1.2720431 | 0.2033578 | NA | NA | NA |
| ENSG000002 | 1.63866252 | -0.406523517 | 0.35172408 | -1.1558023 | 0.24776201 | NA | NA | NA |
| ENSG000002 | 1.13451891 | -0.419945847 | 0.31942037 | -1.3147122 | 0.18860665 | NA | NA | NA |
| ENSG000002 | 0.99400551 | -0.183277502 | 0.30727742 | -0.5964561 | 0.55087055 | NA | NA | NA |
| ENSG000002 | 0 | NA | NA | NA | NA | NA | TREX1 | 11277 |
| ENSG000002 | 1.86402489 | -0.723725674 | 0.34723915 | -2.0842283 | 0.0371394 | NA | NA | NA |
| ENSG000002 | 1.96552264 | -0.297200762 | 0.35788545 | -0.8304355 | 0.40629263 | NA | NA | NA |
| ENSG000002 | 1.59924683 | -0.129510459 | 0.34938684 | -0.3706793 | 0.71087643 | NA | NA | NA |
| ENSG000002 | 1.72080246 | -0.299668416 | 0.34751347 | -0.8623217 | 0.3885105 | NA | NA | NA |
| ENSG000002 | 0.86074541 | -0.094852058 | 0.28952449 | -0.3276132 | 0.7432041 | NA | NA | NA |
| ENSG000002 | 1.96322369 | -0.78003268 | 0.35174822 | -2.2175881 | 0.02658293 | NA | NA | NA |
| ENSG000002 | 0.96289354 | -0.037113289 | 0.31181034 | -0.1190252 | 0.90525539 | NA | NA | NA |
| ENSG000002 | 0.93497115 | -0.441253409 | 0.29749306 | -1.4832393 | 0.13801084 | NA | NA | NA |
| ENSG000002 | 1.13844731 | -0.288686568 | 0.32745486 | -0.8816072 | 0.37798925 | NA | NA | NA |
| ENSG000002 | 1.18534764 | 0.101952732 | 0.32722326 | 0.31156933 | 0.75536784 | NA | NA | NA |
| ENSG000002 | 1.04201961 | -0.35345347 | 0.30825225 | -1.1466371 | 0.25153163 | NA | NA | NA |
| ENSG000002 | 2.00147299 | -0.570151651 | 0.36031833 | -1.5823554 | 0.11356846 | NA | NA | NA |
| ENSG000002 | 1.91414473 | -0.253015659 | 0.35686945 | -0.7089866 | 0.47833279 | NA | NA | NA |
| ENSG000002 | 1.37848723 | 0.023827031 | 0.3196548 | 0.07453988 | 0.94058081 | NA | NA | NA |
| ENSG000002 | 1.0354388 | -0.186040249 | 0.30068253 | -0.6187265 | 0.53609655 | NA | NA | NA |

| | | | | | | | | |
|---|---|---|---|---|---|---|---|---|
| ENSG000002 | 1.38627032 | -0.403923104 | 0.33336697 | -1.211647 | 0.22564752 | NA | NA | NA |
| ENSG000002 | 1.39280815 | -0.420628827 | 0.33964832 | -1.2384246 | 0.21555867 | NA | NA | NA |
| ENSG000002 | 1.84842802 | -0.251725288 | 0.35838013 | -0.7023974 | 0.48243134 | NA | EIF1P7 | 106481691 |
| ENSG000002 | 0.8937316 | -0.589694461 | 0.29065376 | -2.0288554 | 0.04247302 | NA | NA | NA |
| ENSG000002 | 1.5359725 | -0.480062151 | 0.34444221 | -1.3937379 | 0.16339678 | NA | NA | NA |
| ENSG000002 | 1.03698186 | -0.533697484 | 0.31242205 | -1.708258 | 0.08758847 | NA | NA | NA |
| ENSG000002 | 1.38854198 | -0.410266197 | 0.33852124 | -1.2119363 | 0.22553675 | NA | NA | NA |
| ENSG000002 | 1.44667958 | 0.212103227 | 0.33299016 | 0.63696545 | 0.52414735 | NA | NA | NA |
| ENSG000002 | 0.5726257 | 0.369251316 | 0.24779848 | 1.49012745 | 0.13619073 | NA | NA | NA |
| ENSG000002 | 1.84800593 | -0.108907378 | 0.35191522 | -0.3094705 | 0.75696365 | NA | KLRK1 | 22914 |
| ENSG000002 | 1.74308231 | -0.318496028 | 0.35432681 | -0.8988765 | 0.36871847 | NA | NA | NA |
| ENSG000002 | 1.30753263 | 0.169795121 | 0.33339327 | 0.50929379 | 0.61054631 | NA | NA | NA |
| ENSG000002 | 0.97544362 | -0.189881147 | 0.30813177 | -0.6162336 | 0.53774036 | NA | NA | NA |
| ENSG000002 | 0.6009429 | 0.055606074 | 0.26256469 | 0.21178047 | 0.83227831 | NA | NA | NA |
| ENSG000002 | 0.50611225 | -0.202624884 | 0.24886748 | -0.8141879 | 0.41553732 | NA | NA | NA |
| ENSG000002 | 1.18448836 | 0.264173058 | 0.3311343 | 0.79778223 | 0.42499687 | NA | NA | NA |
| ENSG000002 | 0.76417123 | -0.337057922 | 0.27522973 | -1.2246421 | 0.22071008 | NA | NA | NA |
| ENSG000002 | 1.53119245 | -0.185275878 | 0.33335111 | -0.555798 | 0.57834896 | NA | NA | NA |
| ENSG000002 | 0.83231364 | 0.038511274 | 0.28751249 | 0.13394644 | 0.89344493 | NA | NA | NA |
| ENSG000002 | 1.60815083 | -0.262292919 | 0.34879168 | -0.7520045 | 0.45204834 | NA | NA | NA |
| ENSG000002 | 1.86675519 | -0.130959443 | 0.36155442 | -0.3622123 | 0.71719342 | NA | NA | NA |
| ENSG000002 | 0.90064872 | -0.456924471 | 0.30237983 | -1.5110944 | 0.1307644 | NA | NA | NA |
| ENSG000002 | 1.61734821 | -0.415075988 | 0.35249936 | -1.1775227 | 0.23898696 | NA | NA | NA |
| ENSG000002 | 0.7461825 | -0.025114767 | 0.28524947 | -0.0880449 | 0.92984097 | NA | NA | NA |
| ENSG000002 | 1.54320408 | -0.794162793 | 0.34164186 | -2.3245477 | 0.02009616 | NA | NA | NA |
| ENSG000002 | 1.21417797 | 0.109330819 | 0.33172076 | 0.32958691 | 0.74171212 | NA | MDP1 | 145553 |
| ENSG000002 | 0 | NA | NA | NA | NA | NA | CCL27 | 10850 |
| ENSG000002 | 1.93560098 | 0.072937486 | 0.36306634 | 0.200893 | 0.84078224 | NA | NA | NA |
| ENSG000002 | 1.47629189 | -0.322484251 | 0.34230978 | -0.9420831 | 0.34615011 | NA | NA | NA |

| | | | | | | | | |
|---|---|---|---|---|---|---|---|---|
| ENSG000002 | 1.23464688 | -0.184415723 | 0.32917871 | -0.5602298 | 0.5753227 | NA | NA | NA |
| ENSG000002 | 1.43188656 | -0.367999697 | 0.3224512 | -1.141257 | 0.25376299 | NA | NA | NA |
| ENSG000002 | 1.25132728 | -0.356280157 | 0.33523239 | -1.0627856 | 0.2878792 | NA | NA | NA |
| ENSG000002 | 1.64270247 | -0.265877467 | 0.35221007 | -0.7548832 | 0.45031907 | NA | NA | NA |
| ENSG000002 | 1.6375542 | -0.264655659 | 0.34757437 | -0.7614361 | 0.44639666 | NA | NA | NA |
| ENSG000002 | 1.36198097 | 0.026397348 | 0.33219288 | 0.07946392 | 0.93666363 | NA | NA | NA |
| ENSG000002 | 1.12352758 | -0.252266516 | 0.31748643 | -0.7945742 | 0.42686125 | NA | NA | NA |
| ENSG000002 | 1.25636504 | -0.203697155 | 0.33464405 | -0.608698 | 0.54272466 | NA | NA | NA |
| ENSG000002 | 1.9450498 | 0.204096698 | 0.36424675 | 0.56032538 | 0.57525752 | NA | IFNA7 | 3444 |
| ENSG000002 | 1.60476622 | -0.24985492 | 0.34943724 | -0.7150209 | 0.47459615 | NA | NA | NA |
| ENSG000002 | 1.71852329 | -0.046864524 | 0.35041639 | -0.1337395 | 0.89360854 | NA | NA | NA |
| ENSG000002 | 1.91123842 | -0.367768886 | 0.35137027 | -1.0466705 | 0.29525159 | NA | NA | NA |
| ENSG000002 | 2.09402283 | -0.134260285 | 0.3675429 | -0.3652915 | 0.71489385 | NA | NA | NA |
| ENSG000002 | 1.13801416 | 0.027212205 | 0.31661103 | 0.08594838 | 0.93150745 | NA | NA | NA |
| ENSG000002 | 1.51972684 | -0.485763825 | 0.34412322 | -1.4115985 | 0.15806821 | NA | NA | NA |
| ENSG000002 | 1.50394864 | -0.189253131 | 0.33981798 | -0.556925 | 0.57757867 | NA | NA | NA |
| ENSG000002 | 1.93728392 | 0.067824541 | 0.36125717 | 0.18774587 | 0.85107587 | NA | CYP4F30P | 100132708 |
| ENSG000002 | 1.70722676 | -0.650217081 | 0.34053217 | -1.9094146 | 0.05620863 | NA | NA | NA |
| ENSG000002 | 1.56001465 | 0.027561531 | 0.34280727 | 0.08039949 | 0.93591953 | NA | NA | NA |
| ENSG000002 | 1.88298167 | -0.643377867 | 0.35533669 | -1.8106148 | 0.0702005 | NA | NA | NA |
| ENSG000002 | 2.05624879 | -0.437899428 | 0.35821693 | -1.222442 | 0.22154052 | NA | NA | NA |
| ENSG000002 | 0.11131518 | 0.081951103 | 0.12663804 | 0.64712866 | 0.51754868 | NA | NA | NA |
| ENSG000002 | 1.45730733 | -0.738800499 | 0.33645198 | -2.1958572 | 0.02810217 | NA | NA | NA |
| ENSG000002 | 0.62476718 | -0.114815422 | 0.27129677 | -0.4232097 | 0.67214228 | NA | NA | NA |
| ENSG000002 | 1.62816383 | -0.546616398 | 0.34940374 | -1.5644263 | 0.11771749 | NA | NA | NA |
| ENSG000002 | 0 | NA | NA | NA | NA | NA | NA | NA |
| ENSG000002 | 0.12070844 | -0.088503695 | 0.12783931 | -0.6923042 | 0.48874631 | NA | NA | NA |
| ENSG000002 | 2.15732484 | -0.644927106 | 0.3644864 | -1.7694133 | 0.07682492 | NA | NA | NA |
| ENSG000002 | 2.00710752 | -0.739397865 | 0.36231293 | -2.0407714 | 0.04127355 | NA | NA | NA |

| | | | | | | | | |
|---|---|---|---|---|---|---|---|---|
| ENSG000002 | 1.00105847 | -0.04335497 | 0.31183898 | -0.13903 | 0.88942645 | NA | NA | NA |
| ENSG000002 | 0.85484336 | 0.203775775 | 0.29716883 | 0.68572391 | 0.49288723 | NA | NA | NA |
| ENSG000002 | 1.49666205 | -0.180907533 | 0.34323033 | -0.5270733 | 0.59814271 | NA | NA | NA |
| ENSG000002 | 0.76949578 | -0.359754798 | 0.28380236 | -1.2676244 | 0.20493211 | NA | NA | NA |
| ENSG000002 | 1.83464406 | -0.123005737 | 0.35847198 | -0.3431391 | 0.73149385 | NA | NA | NA |
| ENSG000002 | 2.08508446 | -0.13113846 | 0.3647827 | -0.3594975 | 0.71922296 | NA | NA | NA |
| ENSG000002 | 0.36212533 | -0.235926492 | 0.20089643 | -1.1743688 | 0.24024735 | NA | NA | NA |
| ENSG000002 | 1.38666262 | -0.426962511 | 0.34142534 | -1.2505296 | 0.21110614 | NA | NA | NA |
| ENSG000002 | 0 | NA | NA | NA | NA | NA | NA | NA |
| ENSG000002 | 1.65616995 | -0.701792877 | 0.34872185 | -2.0124717 | 0.04417023 | NA | NA | NA |
| ENSG000002 | 0 | NA | NA | NA | NA | NA | NA | NA |
| ENSG000002 | 0.76647672 | -0.185811905 | 0.28134795 | -0.6604345 | 0.50897501 | NA | NA | NA |
| ENSG000002 | 1.88362797 | -0.51093905 | 0.35576129 | -1.4361851 | 0.15094969 | NA | NA | NA |
| ENSG000002 | 1.60704147 | -0.266226853 | 0.35062098 | -0.759301 | 0.44767253 | NA | NA | NA |
| ENSG000002 | 1.9846168 | -0.32173659 | 0.36218201 | -0.8883285 | 0.37436409 | NA | NA | NA |
| ENSG000002 | 0.9576012 | -0.608655462 | 0.29388709 | -2.071052 | 0.03835393 | NA | NA | NA |
| ENSG000002 | 1.2666702 | -0.499623751 | 0.33090623 | -1.509865 | 0.13107786 | NA | NA | NA |
| ENSG000002 | 1.87867824 | -0.389352289 | 0.35880851 | -1.0851256 | 0.27786605 | NA | NA | NA |
| ENSG000002 | 1.86322569 | -0.128912196 | 0.35871462 | -0.3593726 | 0.71931639 | NA | NA | NA |
| ENSG000002 | 1.12177083 | -0.42067392 | 0.31896151 | -1.3188861 | 0.18720718 | NA | AS3MT | 57412 |
| ENSG000002 | 0 | NA | NA | NA | NA | NA | CCDC103 | 388389 |
| ENSG000002 | 2.046662 | -0.734192235 | 0.36126043 | -2.0323074 | 0.04212255 | NA | NA | NA |
| ENSG000002 | 1.81672626 | -0.76435673 | 0.35301445 | -2.1652279 | 0.03037024 | NA | NA | NA |
| ENSG000002 | 0.24756242 | -0.187034346 | 0.18119817 | -1.0322088 | 0.30197431 | NA | KRTAP2-2 | 728279 |
| ENSG000002 | 1.75452619 | -0.198055419 | 0.35560674 | -0.5569507 | 0.57756112 | NA | NA | NA |
| ENSG000002 | 1.69281828 | -0.042792917 | 0.35167626 | -0.1216827 | 0.90315031 | NA | NA | NA |
| ENSG000002 | 1.06263988 | 0.480649502 | 0.30995126 | 1.55072609 | 0.12096734 | NA | NA | NA |
| ENSG000002 | 0.60424462 | 0.061148521 | 0.26285402 | 0.23263301 | 0.81604639 | NA | NA | NA |
| ENSG000002 | 1.25161152 | -0.042498023 | 0.32679361 | -0.1300455 | 0.89653047 | NA | NA | NA |

| | | | | | | | | |
|---|---|---|---|---|---|---|---|---|
| ENSG000002 | 1.93532257 | -0.290810521 | 0.35653692 | -0.8156533 | 0.41469845 | NA | NA | NA |
| ENSG000002 | 1.20517026 | -0.30442644 | 0.31977154 | -0.9520123 | 0.34109075 | NA | NA | NA |
| ENSG000002 | 0.4680172 | 0.307231342 | 0.22996067 | 1.33601689 | 0.18154375 | NA | NA | NA |
| ENSG000002 | 0.96977629 | -0.038831764 | 0.31209637 | -0.1244223 | 0.90098088 | NA | NA | NA |
| ENSG000002 | 0.99698818 | -0.037407537 | 0.31319711 | -0.1194377 | 0.90492861 | NA | NA | NA |
| ENSG000002 | 1.39124391 | -0.274903686 | 0.34196402 | -0.8038965 | 0.42145673 | NA | NA | NA |
| ENSG000002 | 0.76678939 | -0.201566736 | 0.28734031 | -0.7014913 | 0.48299644 | NA | NA | NA |
| ENSG000002 | 1.77490921 | -0.61782184 | 0.35473822 | -1.7416275 | 0.08157364 | NA | DEFB113 | 245927 |
| ENSG000002 | 1.49388577 | -0.475938928 | 0.34298326 | -1.3876448 | 0.16524525 | NA | NA | NA |
| ENSG000002 | 0.37216701 | -0.099548723 | 0.21762847 | -0.4574251 | 0.64736554 | NA | NA | NA |
| ENSG000002 | 1.63940322 | -0.124881658 | 0.34871461 | -0.35812 | 0.72025355 | NA | NA | NA |
| ENSG000002 | 2.05876983 | -0.685925654 | 0.35460957 | -1.9343123 | 0.05307474 | NA | NA | NA |
| ENSG000002 | 0.35346928 | -0.088570748 | 0.20618147 | -0.4295767 | 0.66750362 | NA | NA | NA |
| ENSG000002 | 0.52165104 | -0.403921832 | 0.25252134 | -1.5995552 | 0.10969729 | NA | NA | NA |
| ENSG000002 | 0 | NA | NA | NA | NA | NA | NA | NA |
| ENSG000002 | 0 | NA | NA | NA | NA | NA | TOMM6 | 100188893 |
| ENSG000002 | 0.36416385 | 0.068880976 | 0.20758164 | 0.33182596 | 0.74002068 | NA | NA | NA |
| ENSG000002 | 1.77161098 | -0.299864993 | 0.34246742 | -0.8756015 | 0.3812467 | NA | NA | NA |
| ENSG000002 | 0.23065526 | 0.167384503 | 0.17662711 | 0.94767164 | 0.34329664 | NA | NA | NA |
| ENSG000002 | 1.84261551 | -0.218794758 | 0.3491444 | -0.6266598 | 0.53088226 | NA | NA | NA |
| ENSG000002 | 1.60737833 | 0.016447105 | 0.34804149 | 0.04725616 | 0.96230907 | NA | NA | NA |
| ENSG000002 | 1.76858899 | -0.457444148 | 0.3517687 | -1.3004118 | 0.19345988 | NA | NA | NA |
| ENSG000002 | 1.50093873 | -0.340898416 | 0.34500565 | -0.9880952 | 0.32310604 | NA | NA | NA |
| ENSG000002 | 1.84398525 | -0.259487431 | 0.36117076 | -0.7184619 | 0.47247253 | NA | NA | NA |
| ENSG000002 | 1.83253848 | 0.006622049 | 0.36196108 | 0.01829492 | 0.98540358 | NA | NA | NA |
| ENSG000002 | 1.10642399 | 0.030809945 | 0.325626 | 0.09461758 | 0.92461858 | NA | NA | NA |
| ENSG000002 | 1.69839874 | 0.217153588 | 0.35679707 | 0.60861932 | 0.5427768 | NA | NA | NA |
| ENSG000002 | 0.99789953 | -0.19936691 | 0.31380972 | -0.6353115 | 0.5252253 | NA | NA | NA |
| ENSG000002 | 0.24004852 | -0.005223787 | 0.17775844 | -0.029387 | 0.97655594 | NA | LOC105377213 | 105377213 |

| | | | | | | | | | |
|---|---|---|---|---|---|---|---|---|---|
| ENSG000002 | 0.87966935 | -0.27079777 | 0.29949138 | -0.9041922 | 0.3658935 | NA | NA | NA | |
| ENSG000002 | 1.88494426 | -0.66283079 | 0.35678071 | -1.8578101 | 0.06319599 | NA | NA | NA | |
| ENSG000002 | 1.54568005 | 0.015431271 | 0.34261248 | 0.04504001 | 0.96407541 | NA | NA | NA | |
| ENSG000002 | 1.72619014 | -0.456755529 | 0.35387146 | -1.2907386 | 0.19679435 | NA | GSG1L2 | | 644070 |
| ENSG000002 | 1.99025435 | -0.322029407 | 0.36329932 | -0.8864025 | 0.3754007 | NA | NA | NA | |
| ENSG000002 | 1.53640724 | -0.626143317 | 0.34074241 | -1.8375855 | 0.06612351 | NA | NA | NA | |
| ENSG000002 | 1.11806664 | -0.279759399 | 0.32553234 | -0.8593905 | 0.39012512 | NA | NA | NA | |
| ENSG000002 | 0.75053511 | 0.123730904 | 0.27954741 | 0.44261152 | 0.65804675 | NA | NA | NA | |
| ENSG000002 | 1.38575072 | -0.263110833 | 0.33598763 | -0.7830968 | 0.43357028 | NA | NA | NA | |
| ENSG000002 | 0.61909773 | -0.103740153 | 0.25703858 | -0.4035976 | 0.68650866 | NA | NA | NA | |
| ENSG000002 | 1.36048133 | -0.408181557 | 0.33575112 | -1.2157266 | 0.2240891 | NA | NA | NA | |
| ENSG000002 | 1.03698186 | -0.533697484 | 0.31242205 | -1.708258 | 0.08758847 | NA | NA | NA | |
| ENSG000002 | 1.86291166 | -0.248774801 | 0.35909748 | -0.6927779 | 0.48844894 | NA | NA | NA | |
| ENSG000002 | 1.58574945 | -0.117502226 | 0.34919434 | -0.3364952 | 0.73649745 | NA | NA | NA | |
| ENSG000002 | 1.49066892 | -0.469465887 | 0.34160698 | -1.3742866 | 0.16935273 | NA | NA | NA | |
| ENSG000002 | 1.99802883 | -0.457693556 | 0.36249055 | -1.2626358 | 0.20672009 | NA | NA | NA | |
| ENSG000002 | 0.24619405 | -0.007630671 | 0.17915639 | -0.0425922 | 0.96602658 | NA | NA | NA | |
| ENSG000002 | 0.90895539 | -0.266226439 | 0.29363402 | -0.9066607 | 0.36458623 | NA | NA | NA | |
| ENSG000002 | 1.93288406 | 0.318144824 | 0.35675751 | 0.89176771 | 0.37251745 | NA | NA | NA | |
| ENSG000002 | 2.17326803 | -0.805625629 | 0.36555229 | -2.2038588 | 0.02753427 | NA | FAM74A1 | | 401507 |
| ENSG000002 | 0 | NA | NA | NA | NA | NA | CXorf49B | | 100132994 |
| ENSG000002 | 0 | NA | NA | NA | NA | NA | CXorf49 | | 100130361 |
| ENSG000002 | 1.60130191 | -0.056155387 | 0.24630918 | -0.2279874 | 0.81965604 | NA | NA | NA | |
| ENSG000002 | 0.83231811 | -0.227557769 | 0.27972229 | -0.8135132 | 0.4159239 | NA | NA | NA | |
| ENSG000002 | 0.72165134 | -0.028079792 | 0.28363433 | -0.099 | 0.92113829 | NA | NA | NA | |
| ENSG000002 | 1.15142406 | -0.243039792 | 0.31132902 | -0.7806525 | 0.43500688 | NA | NA | NA | |
| ENSG000002 | 2.05639494 | -0.837309408 | 0.35606355 | -2.3515729 | 0.01869422 | NA | NA | NA | |
| ENSG000002 | 0.64235948 | -0.482579697 | 0.27062934 | -1.7831759 | 0.07455768 | NA | NA | NA | |
| ENSG000002 | 0 | NA | NA | NA | NA | NA | NA | NA | |

| | | | | | | | | |
|---|---|---|---|---|---|---|---|---|
| ENSG000002 | 1.85123813 | 0.135823231 | 0.35883789 | 0.37850861 | 0.7050528 | NA | NA | NA |
| ENSG000002 | 2.10002511 | -0.12823194 | 0.36756588 | -0.3488679 | 0.72718848 | NA | NA | NA |
| ENSG000002 | 0.49793494 | 0.145453577 | 0.24009604 | 0.60581415 | 0.54463818 | NA | NA | NA |
| ENSG000002 | 2.03920367 | -0.333936754 | 0.36385958 | -0.9177627 | 0.35874311 | NA | NA | NA |
| ENSG000002 | 0.72167308 | 0.133011302 | 0.2775391 | 0.47925248 | 0.63175902 | NA | NA | NA |
| ENSG000002 | 0.87958291 | -0.415226017 | 0.29226182 | -1.420733 | 0.15539439 | NA | NA | NA |
| ENSG000002 | 0.3842622 | -0.095651135 | 0.21113931 | -0.4530238 | 0.65053159 | NA | NA | NA |
| ENSG000002 | 2.07475674 | -0.122611406 | 0.36233234 | -0.3383949 | 0.73506564 | NA | NA | NA |
| ENSG000002 | 1.36166248 | 0.164335397 | 0.33610037 | 0.48894739 | 0.62487895 | NA | NA | NA |
| ENSG000002 | 1.7231171 | -0.429900616 | 0.34824672 | -1.2344714 | 0.2170273 | NA | NA | NA |
| ENSG000002 | 1.00271363 | -0.297903796 | 0.29455401 | -1.0113724 | 0.31183823 | NA | NA | NA |
| ENSG000002 | 1.71741147 | -0.187824682 | 0.35348205 | -0.5313556 | 0.59517236 | NA | NA | NA |
| ENSG000002 | 1.35611372 | -0.118847398 | 0.33947874 | -0.3500879 | 0.72627274 | NA | NA | NA |
| ENSG000002 | 0.24756242 | -0.187034346 | 0.18119817 | -1.0322088 | 0.30197431 | NA | NA | NA |
| ENSG000002 | 1.06763038 | 0.329909831 | 0.31010757 | 1.06385608 | 0.2873939 | NA | NA | NA |
| ENSG000002 | 1.61571926 | -0.264885012 | 0.3524726 | -0.7515053 | 0.45234863 | NA | NA | NA |
| ENSG000002 | 1.72088891 | -0.195898316 | 0.35827802 | -0.5467774 | 0.58453169 | NA | NA | NA |
| ENSG000002 | 2.02979369 | -0.580393533 | 0.36113409 | -1.6071414 | 0.10802336 | NA | LL22NC01-81G9 | 100506241 |
| ENSG000002 | 1.18224077 | -0.011317376 | 0.19743789 | -0.0573212 | 0.95428934 | NA | NA | NA |
| ENSG000002 | 1.36325706 | -0.121177515 | 0.33626537 | -0.3603628 | 0.71857587 | NA | NA | NA |
| ENSG000002 | 1.83007677 | -0.119404313 | 0.35306022 | -0.3381982 | 0.73521387 | NA | KRTAP10-4 | 386672 |
| ENSG000002 | 1.39582912 | -0.396863815 | 0.33361459 | -1.1895877 | 0.23420849 | NA | KRTAP10-1 | 386677 |
| ENSG000002 | 1.48129685 | -0.197738169 | 0.34472175 | -0.5736168 | 0.56622717 | NA | NA | NA |
| ENSG000002 | 1.28949613 | -0.678631562 | 0.33112094 | -2.0494976 | 0.04041349 | NA | NA | NA |
| ENSG000002 | 2.03313748 | 0.124163322 | 0.36193435 | 0.34305481 | 0.73155722 | NA | NA | NA |
| ENSG000002 | 0 | NA | NA | NA | NA | NA | RPL17-C18orf32 | 100526842 |
| ENSG000002 | 1.15435667 | -0.589639773 | 0.31991462 | -1.843116 | 0.06531207 | NA | NA | NA |
| ENSG000002 | 1.64944908 | 0.197291317 | 0.34264287 | 0.57579286 | 0.5647552 | NA | NA | NA |
| ENSG000002 | 0 | NA | NA | NA | NA | NA | NA | NA |

| | | | | | | | | |
|---|---|---|---|---|---|---|---|---|
| ENSG000002 | 0 | NA | NA | NA | NA | NA | NA | NA |
| ENSG000002 | 1.47955847 | 0.088286092 | 0.34666077 | 0.25467575 | 0.79897355 | NA | LINC00189 | 193629 |
| ENSG000002 | 0 | NA | NA | NA | NA | NA | NA | NA |
| ENSG000002 | 0 | NA | NA | NA | NA | NA | NA | NA |
| ENSG000002 | 1.87825348 | -0.400611176 | 0.35975795 | -1.1135575 | 0.26546905 | NA | NA | NA |
| ENSG000002 | 1.84751263 | -0.379008357 | 0.35694004 | -1.0618264 | 0.2883145 | NA | NA | NA |
| ENSG000002 | 1.54074775 | -0.449745058 | 0.33754 | -1.33242 | 0.1827222 | NA | NA | NA |
| ENSG000002 | 0 | NA | NA | NA | NA | NA | TTTY5 | 83863 |
| ENSG000002 | 0 | NA | NA | NA | NA | NA | GAB4 | 128954 |
| ENSG000002 | 1.25997886 | -0.19203964 | 0.32756309 | -0.5862676 | 0.55769568 | NA | NA | NA |
| ENSG000002 | 0 | NA | NA | NA | NA | NA | BCORP1 | 286554 |
| ENSG000002 | 0 | NA | NA | NA | NA | NA | NA | NA |
| ENSG000002 | 0 | NA | NA | NA | NA | NA | NA | NA |
| ENSG000002 | 0 | NA | NA | NA | NA | NA | NA | NA |
| ENSG000002 | 0.63256532 | 0.047465397 | 0.25786486 | 0.18407083 | 0.85395789 | NA | NA | NA |
| ENSG000002 | 0 | NA | NA | NA | NA | NA | RSC1A1 | 6248 |
| ENSG000002 | 0.38747904 | -0.104908247 | 0.21987514 | -0.4771265 | 0.63327207 | NA | NA | NA |
| ENSG000002 | 1.70056631 | -0.543278271 | 0.3467803 | -1.5666354 | 0.11719995 | NA | NA | NA |
| ENSG000002 | 0.97894334 | -0.031802496 | 0.31247024 | -0.1017777 | 0.91893314 | NA | LRRC37A14P | 106479025 |
| ENSG000002 | 1.37314061 | -0.121047845 | 0.33250224 | -0.3640512 | 0.71581976 | NA | NA | NA |
| ENSG000002 | 1.43268384 | 0.094405035 | 0.3415562 | 0.27639679 | 0.78224333 | NA | NA | NA |
| ENSG000002 | 0 | NA | NA | NA | NA | NA | NA | NA |
| ENSG000002 | 1.69694447 | 0.205337319 | 0.34776679 | 0.59044545 | 0.55489205 | NA | NA | NA |
| ENSG000002 | 1.08519903 | -0.255410762 | 0.31836767 | -0.8022509 | 0.42240782 | NA | NA | NA |
| ENSG000002 | 0.39499294 | -0.277908454 | 0.21584201 | -1.287555 | 0.19790091 | NA | NA | NA |
| ENSG000002 | 1.89185554 | -0.375195791 | 0.35492157 | -1.0571231 | 0.29045541 | NA | NA | NA |
| ENSG000002 | 1.99756377 | -0.179142217 | 0.36048463 | -0.4969483 | 0.61922552 | NA | NA | NA |
| ENSG000002 | 0.74296511 | -0.035856325 | 0.29036133 | -0.1234886 | 0.90172017 | NA | MMP23A | 8511 |
| ENSG000002 | 2.16905045 | 0.293788535 | 0.36388842 | 0.80735885 | 0.41945977 | NA | MIR942 | 100126331 |

| | | | | | | | | |
|---|---|---|---|---|---|---|---|---|
| ENSG000002 | 0.61016148 | 0.056638442 | 0.2697572 | 0.20996082 | 0.83369825 | NA | NA | NA |
| ENSG000002 | 1.75287785 | -0.585519923 | 0.35026886 | -1.67163 | 0.09459731 | NA | MIR892A | 100126342 |
| ENSG000002 | 1.53138729 | -0.332138478 | 0.34458391 | -0.9638827 | 0.33510472 | NA | MIR921 | 100126349 |
| ENSG000002 | 0.33468275 | 0.234558662 | 0.20511328 | 1.14355667 | 0.25280754 | NA | MIR300 | 100126297 |
| ENSG000002 | 0.37330862 | -0.105391168 | 0.21773449 | -0.4840352 | 0.62836086 | NA | MIR297 | 100126354 |
| ENSG000002 | 1.60249456 | -0.265334711 | 0.35056365 | -0.7568803 | 0.44912159 | NA | MIR876 | 100126310 |
| ENSG000002 | 1.18895307 | 0.372690382 | 0.31422071 | 1.18607835 | 0.23559136 | NA | MIR933 | 100126350 |
| ENSG000002 | 0.13280363 | -0.093394593 | 0.13115644 | -0.7120855 | 0.47641185 | NA | MIR208B | 100126336 |
| ENSG000002 | 0.48761094 | -0.176642206 | 0.23968055 | -0.7369902 | 0.46112833 | NA | NA | NA |
| ENSG000002 | 0.13280363 | -0.093394593 | 0.13115644 | -0.7120855 | 0.47641185 | NA | MIR888 | 100126306 |
| ENSG000002 | 0.59573046 | 0.057877409 | 0.26209971 | 0.2208221 | 0.82523095 | NA | NA | NA |
| ENSG000002 | 0.98101219 | 0.111438579 | 0.30605418 | 0.3641139 | 0.71577296 | NA | MIR938 | 100126327 |
| ENSG000002 | 0 | NA | NA | NA | NA | NA | MIR891A | 100126341 |
| ENSG000002 | 0.61818637 | 0.059474041 | 0.27041681 | 0.21993471 | 0.82592201 | NA | MIR934 | 100126324 |
| ENSG000002 | 0.64694077 | -0.282974687 | 0.26865662 | -1.0532951 | 0.29220579 | NA | MIR891B | 100126304 |
| ENSG000002 | 0.51413714 | -0.199727707 | 0.24976394 | -0.7996659 | 0.42390439 | NA | MIR875 | 100126309 |
| ENSG000002 | 2.04816186 | -0.2780054 | 0.34884812 | -0.7969239 | 0.42549524 | NA | MIR890 | 100126303 |
| ENSG000002 | 0.44747236 | 0.298956135 | 0.22733644 | 1.31503835 | 0.18849701 | NA | MIR892B | 100126307 |
| ENSG000002 | 0.5837196 | 0.05342512 | 0.25176116 | 0.21220557 | 0.83194666 | NA | NA | NA |
| ENSG000002 | 0 | NA | NA | NA | NA | NA | NA | NA |
| ENSG000002 | 0.11205239 | 0.082118251 | 0.12671962 | 0.64803108 | 0.51696485 | NA | NA | NA |
| ENSG000002 | 0 | NA | NA | NA | NA | NA | NA | NA |
| ENSG000002 | 0.11865494 | 0.083440866 | 0.12751478 | 0.65436232 | 0.5128784 | NA | MIR513C | 100302114 |
| ENSG000002 | 0.39246123 | -0.276864021 | 0.21549122 | -1.2848042 | 0.19886073 | NA | MIR541 | 100126308 |
| ENSG000002 | 0.64099112 | -0.296274405 | 0.27396996 | -1.081412 | 0.27951389 | NA | MIR8485 | 103504737 |
| ENSG000002 | 0.958593 | -0.167779531 | 0.29986034 | -0.5595256 | 0.57580308 | NA | MIR920 | 100126320 |
| ENSG000002 | 0.23799501 | 0.170958857 | 0.17834799 | 0.95856902 | 0.33777591 | NA | NA | NA |
| ENSG000002 | 1.70979207 | -0.176383809 | 0.35231447 | -0.5006431 | 0.61662232 | NA | NA | NA |
| ENSG000002 | 0.7863464 | -0.370445737 | 0.28479156 | -1.3007609 | 0.1933403 | NA | NA | NA |

| | | | | | | | | | |
|---|---|---|---|---|---|---|---|---|---|
| ENSG000002 | 0.91320037 | -0.446160783 | 0.29797291 | -1.4973199 | 0.13431003 | NA | NA | NA | |
| ENSG000002 | 0.59548251 | 0.058022206 | 0.26207208 | 0.22139789 | 0.82478264 | NA | NA | NA | |
| ENSG000002 | 0.74296511 | -0.035856325 | 0.29036133 | -0.1234886 | 0.90172017 | NA | NA | NA | |
| ENSG000002 | 1.66871768 | -0.387157801 | 0.3443513 | -1.1243106 | 0.2608813 | NA | NA | NA | |
| ENSG000002 | 1.26072054 | -0.504639735 | 0.33211407 | -1.5194771 | 0.12864244 | NA | NA | NA | |
| ENSG000002 | 1.10067076 | 0.038310684 | 0.31530818 | 0.12150235 | 0.90329316 | NA | NA | NA | |
| ENSG000002 | 0.85066488 | -0.108643662 | 0.29679137 | -0.3660607 | 0.71431977 | NA | NA | NA | |
| ENSG000002 | 1.59178746 | 0.142019407 | 0.34218234 | 0.41504014 | 0.67811252 | NA | NA | NA | |
| ENSG000002 | 1.65121609 | -0.264928346 | 0.35003845 | -0.756855 | 0.44913675 | NA | NA | NA | |
| ENSG000002 | 0.28256998 | -0.18483211 | 0.18021686 | -1.0256094 | 0.30507573 | NA | NA | NA | |
| ENSG000002 | 1.72390988 | -0.189441704 | 0.35529257 | -0.5331992 | 0.59389571 | NA | NA | NA | |
| ENSG000002 | 1.76340879 | -0.448660892 | 0.35102337 | -1.2781511 | 0.20119617 | NA | NA | NA | |
| ENSG000002 | 0 | NA | NA | NA | NA | NA | NA | NA | |
| ENSG000002 | 0.24619405 | -0.007630671 | 0.17915639 | -0.0425922 | 0.96602658 | NA | NA | NA | |
| ENSG000002 | 0.88305642 | -0.119986999 | 0.30351552 | -0.3953241 | 0.69260372 | NA | NA | NA | |
| ENSG000002 | 1.17026586 | 0.255585071 | 0.32714755 | 0.78125321 | 0.43465359 | NA | NA | NA | |
| ENSG000002 | 1.94722189 | -0.303508785 | 0.35950149 | -0.844249 | 0.39853026 | NA | NA | NA | |
| ENSG000002 | 0.49763088 | -0.186408494 | 0.24078583 | -0.7741672 | 0.43883193 | NA | NA | NA | |
| ENSG000002 | 1.63439526 | -0.393769795 | 0.34579349 | -1.1387426 | 0.25481051 | NA | NA | NA | |
| ENSG000002 | 1.96829837 | -0.056952173 | 0.36000065 | -0.1582002 | 0.87429905 | NA | NA | NA | |
| ENSG000002 | 1.47941995 | -0.048761577 | 0.33568294 | -0.1452608 | 0.88450495 | NA | NA | NA | |
| ENSG000002 | 0 | NA | NA | NA | NA | NA | PRRC2CP1 | | 360018 |
| ENSG000002 | 1.91827106 | -0.059949491 | 0.35962823 | -0.1666985 | 0.86760728 | NA | NA | NA | |
| ENSG000002 | 0 | NA | NA | NA | NA | NA | NA | NA | |
| ENSG000002 | 0.34862505 | 0.240809381 | 0.20749856 | 1.16053522 | 0.24583096 | NA | NA | NA | |
| ENSG000002 | 0 | NA | NA | NA | NA | NA | NA | NA | |
| ENSG000002 | 1.46040372 | 0.08512774 | 0.34514759 | 0.24664156 | 0.80518564 | NA | NA | NA | |
| ENSG000002 | 1.97034795 | 0.066073893 | 0.36090124 | 0.18308026 | 0.85473504 | NA | NA | NA | |
| ENSG000002 | 1.63769505 | -0.538562931 | 0.3483566 | -1.5460104 | 0.12210205 | NA | NA | NA | |

| | | | | | | | | |
|---|---|---|---|---|---|---|---|---|
| ENSG000002 | 0.88379363 | -0.119722389 | 0.30355205 | -0.3944048 | 0.69328219 | NA | NA | NA |
| ENSG000002 | 2.06795807 | -0.591323997 | 0.36135495 | -1.6364076 | 0.1017543 | NA | NA | NA |
| ENSG000002 | 1.71393753 | 0.078048732 | 0.35690658 | 0.21868112 | 0.82689846 | NA | NA | NA |
| ENSG000002 | 0.49270076 | -0.021329091 | 0.24621941 | -0.0866264 | 0.93096851 | NA | NA | NA |
| ENSG000002 | 1.9895904 | -0.195675827 | 0.36321164 | -0.5387378 | 0.59006781 | NA | RPL10P1 | 100288252 |
| ENSG000002 | 1.90654307 | 0.068894082 | 0.35562466 | 0.19372696 | 0.84638968 | NA | NA | NA |
| ENSG000002 | 0.83031859 | 0.052807184 | 0.29531712 | 0.17881518 | 0.85808283 | NA | NA | NA |
| ENSG000002 | 1.51040738 | -0.196599152 | 0.34520124 | -0.5695204 | 0.56900302 | NA | NA | NA |
| ENSG000002 | 1.65407194 | -0.939486871 | 0.33566162 | -2.7989106 | 0.00512753 | NA | NA | NA |
| ENSG000002 | 0.37945415 | -0.107841678 | 0.21865587 | -0.4932027 | 0.62186934 | NA | NA | NA |
| ENSG000002 | 0 | NA | NA | NA | NA | NA | NA | NA |
| ENSG000002 | 1.61355523 | -0.003549288 | 0.3436669 | -0.0103277 | 0.99175984 | NA | NA | NA |
| ENSG000002 | 1.40962707 | -0.397439135 | 0.33307023 | -1.1932593 | 0.23276786 | NA | TXNP7 | 106480275 |
| ENSG000002 | 1.36819164 | -0.559353207 | 0.33601039 | -1.6646902 | 0.09597458 | NA | NA | NA |
| ENSG000002 | 2.10088886 | -0.502681519 | 0.36504715 | -1.3770318 | 0.16850243 | NA | NA | NA |
| ENSG000002 | 1.59576994 | -0.124286851 | 0.35001039 | -0.3550947 | 0.72251862 | NA | NA | NA |
| ENSG000002 | 1.62440564 | -0.557971626 | 0.35127873 | -1.5884014 | 0.1121956 | NA | NA | NA |
| ENSG000002 | 2.04159711 | -0.002159836 | 0.3638436 | -0.0059362 | 0.99526365 | NA | NA | NA |
| ENSG000002 | 1.8170601 | -0.579316473 | 0.34834498 | -1.6630539 | 0.09630165 | NA | NA | NA |
| ENSG000002 | 1.12310494 | -0.120294979 | 0.32641626 | -0.3685324 | 0.71247626 | NA | NA | NA |
| ENSG000002 | 2.02759719 | -0.300481998 | 0.35504659 | -0.8463171 | 0.39737588 | NA | NA | NA |
| ENSG000002 | 0.76198188 | -0.18457699 | 0.28103812 | -0.6567685 | 0.51132977 | NA | NA | NA |
| ENSG000002 | 0.7523063 | -0.206231678 | 0.2915808 | -0.7072883 | 0.47938735 | NA | NA | NA |
| ENSG000002 | 1.5430082 | -0.808703482 | 0.3434393 | -2.3547203 | 0.01853665 | NA | NA | NA |
| ENSG000002 | 1.26991738 | -0.664859047 | 0.32844574 | -2.0242584 | 0.04294357 | NA | NA | NA |
| ENSG000002 | 0 | NA | NA | NA | NA | NA | NA | NA |
| ENSG000002 | 0.25333738 | -0.015686291 | 0.1806605 | -0.0868275 | 0.93080867 | NA | NA | NA |
| ENSG000002 | 1.11784043 | -0.112555685 | 0.32092276 | -0.3507252 | 0.72579455 | NA | NA | NA |
| ENSG000002 | 1.20797824 | -0.040377975 | 0.3294209 | -0.1225726 | 0.90244556 | NA | NA | NA |

| | | | | | | | | |
|---|---|---|---|---|---|---|---|---|
| ENSG000002 | 1.0247963 | -0.3508469 | 0.30999594 | -1.131779 | 0.25772735 | NA | NA | NA |
| ENSG000002 | 1.97030231 | -0.820886164 | 0.35632108 | -2.3037822 | 0.02123487 | NA | NA | NA |
| ENSG000002 | 1.23113857 | -0.046281183 | 0.32855899 | -0.1408611 | 0.88797967 | NA | NA | NA |
| ENSG000002 | 1.13030226 | 0.024574705 | 0.31082357 | 0.0790632 | 0.93698236 | NA | NA | NA |
| ENSG000002 | 1.42619591 | -0.575905519 | 0.33852949 | -1.7011975 | 0.08890591 | NA | NA | NA |
| ENSG000002 | 0 | NA | NA | NA | NA | NA | NA | NA |
| ENSG000002 | 1.417714 | -0.557207997 | 0.3335337 | -1.6706198 | 0.0947968 | NA | NA | NA |
| ENSG000002 | 1.02553509 | -0.20565425 | 0.3133652 | -0.6562766 | 0.51164616 | NA | NA | NA |
| ENSG000002 | 2.00303668 | -0.713779486 | 0.35903748 | -1.9880361 | 0.04680769 | NA | NA | NA |
| ENSG000002 | 1.56611253 | -0.461684873 | 0.33951594 | -1.3598327 | 0.17388288 | NA | NA | NA |
| ENSG000002 | 1.42648378 | -0.046653129 | 0.33377665 | -0.1397735 | 0.88883895 | NA | NA | NA |
| ENSG000002 | 1.77385542 | -0.323641726 | 0.35210096 | -0.9191731 | 0.35800505 | NA | NA | NA |
| ENSG000002 | 1.75131518 | -0.341987035 | 0.35805237 | -0.9551313 | 0.33951129 | NA | NA | NA |
| ENSG000002 | 1.74792769 | 0.064919865 | 0.34935998 | 0.18582514 | 0.85258189 | NA | NA | NA |
| ENSG000002 | 2.03172402 | -0.601911013 | 0.36337964 | -1.6564247 | 0.09763584 | NA | NA | NA |
| ENSG000002 | 1.81438546 | -0.236419411 | 0.35214808 | -0.6713636 | 0.50198891 | NA | NA | NA |
| ENSG000002 | 2.08986664 | -0.237093253 | 0.36371188 | -0.6518711 | 0.51448435 | NA | NA | NA |
| ENSG000002 | 1.79292371 | -0.475486588 | 0.35617019 | -1.3349983 | 0.18187691 | NA | NA | NA |
| ENSG000002 | 0.61887151 | 0.056280962 | 0.26412849 | 0.21308175 | 0.8312632 | NA | NA | NA |
| ENSG000002 | 1.55272888 | 0.160009918 | 0.34631488 | 0.46203593 | 0.64405556 | NA | NA | NA |
| ENSG000002 | 2.13882702 | -0.376864094 | 0.36498797 | -1.0325384 | 0.30181997 | NA | NA | NA |
| ENSG000002 | 1.1160652 | -0.112312741 | 0.31620668 | -0.3551878 | 0.72244894 | NA | NA | NA |
| ENSG000002 | 1.18173436 | 0.112098435 | 0.32709193 | 0.34271233 | 0.73181489 | NA | FCF1P10 | 106479046 |
| ENSG000002 | 0.37097279 | -0.10468565 | 0.21741418 | -0.4815033 | 0.63015882 | NA | NA | NA |
| ENSG000002 | 0.22997012 | 0.167051788 | 0.17646558 | 0.94665368 | 0.34381528 | NA | NA | NA |
| ENSG000002 | 1.75428059 | -0.623821723 | 0.3559947 | -1.7523343 | 0.07971634 | NA | NA | NA |
| ENSG000002 | 1.34555712 | -0.116968496 | 0.3358784 | -0.3482466 | 0.72765502 | NA | NA | NA |
| ENSG000002 | 0.6567905 | -0.46248946 | 0.26597757 | -1.7388288 | 0.08206488 | NA | NA | NA |
| ENSG000002 | 1.37228367 | -0.402748108 | 0.33604493 | -1.1984948 | 0.23072444 | NA | NA | NA |

| | | | | | | | | | |
|---|---|---|---|---|---|---|---|---|---|
| ENSG000002 | 2.0007873 | -0.593018287 | 0.36232793 | -1.6366894 | 0.10169538 | NA | NA | NA | |
| ENSG000002 | 0.24756242 | -0.187034346 | 0.18119817 | -1.0322088 | 0.30197431 | NA | NA | NA | |
| ENSG000002 | 1.76483327 | -0.328810584 | 0.35622746 | -0.9230355 | 0.35598872 | NA | NA | NA | |
| ENSG000002 | 0.3934287 | -0.107186984 | 0.22072612 | -0.4856108 | 0.62724315 | NA | NA | NA | |
| ENSG000002 | 0.85300071 | -0.109179228 | 0.29690999 | -0.3677183 | 0.71308332 | NA | NA | NA | |
| ENSG000002 | 0.54222759 | -0.38902668 | 0.24851205 | -1.5654238 | 0.11748356 | NA | NA | NA | |
| ENSG000002 | 1.12988108 | -0.2525582 | 0.31055504 | -0.8132478 | 0.41607601 | NA | FAM138C | | 654835 |
| ENSG000002 | 1.41487176 | -0.417183304 | 0.3368995 | -1.238302 | 0.21560412 | NA | NA | NA | |
| ENSG000002 | 0.97618487 | 0.119043716 | 0.31079741 | 0.38302673 | 0.70169995 | NA | SUMO2P12 | | 106480262 |
| ENSG000002 | 1.29387048 | -0.329874019 | 0.32609465 | -1.0115898 | 0.31173423 | NA | NA | NA | |
| ENSG000002 | 2.11130056 | -0.5090502 | 0.36508431 | -1.394336 | 0.16321616 | NA | NA | NA | |
| ENSG000002 | 2.0453255 | 0.128901972 | 0.3615902 | 0.35648636 | 0.72147637 | NA | SOD1P1 | | 106479055 |
| ENSG000002 | 0.37899768 | -0.10176489 | 0.21865765 | -0.4654074 | 0.64163973 | NA | NA | NA | |
| ENSG000002 | 2.11066549 | -0.128664622 | 0.36255057 | -0.3548874 | 0.72267396 | NA | NA | NA | |
| ENSG000002 | 1.77164275 | -0.409228728 | 0.34383378 | -1.1901935 | 0.23397034 | NA | NA | NA | |
| ENSG000002 | 1.95273784 | -0.064910589 | 0.36485865 | -0.1779061 | 0.8587967 | NA | NA | NA | |
| ENSG000002 | 0.38036605 | -0.293832733 | 0.22119083 | -1.3284128 | 0.18404178 | NA | NA | NA | |
| ENSG000002 | 0.61932853 | 0.060804402 | 0.26416359 | 0.23017707 | 0.81795418 | NA | NA | NA | |
| ENSG000002 | 0.88650003 | -0.284549442 | 0.30460367 | -0.9341629 | 0.35021988 | NA | NA | NA | |
| ENSG000002 | 1.86001276 | -0.389490113 | 0.35805001 | -1.0878093 | 0.27667932 | NA | NA | NA | |
| ENSG000002 | 0.75776669 | -0.19935134 | 0.28679134 | -0.6951093 | 0.48698678 | NA | NA | NA | |
| ENSG000002 | 2.08190591 | -0.251278943 | 0.36367767 | -0.6909386 | 0.48960413 | NA | NA | NA | |
| ENSG000002 | 1.37525257 | -0.391710721 | 0.3331579 | -1.175751 | 0.2396944 | NA | NA | NA | |
| ENSG000002 | 0.85951681 | -0.245407694 | 0.28844915 | -0.8507832 | 0.39488978 | NA | NA | NA | |
| ENSG000002 | 1.70987976 | -0.441871596 | 0.35039196 | -1.261078 | 0.20728074 | NA | NA | NA | |
| ENSG000002 | 1.88926317 | -0.127836695 | 0.35892133 | -0.3561691 | 0.72171391 | NA | NA | NA | |
| ENSG000002 | 0.26813897 | -0.197007041 | 0.18550319 | -1.0620143 | 0.2882292 | NA | NA | NA | |
| ENSG000002 | 0.98446631 | -0.179925643 | 0.30106788 | -0.5976248 | 0.55009028 | NA | NA | NA | |
| ENSG000002 | 1.56098599 | -0.332461558 | 0.34041524 | -0.9766353 | 0.32874971 | NA | NA | NA | |

| | | | | | | | | |
|---|---|---|---|---|---|---|---|---|
| ENSG000002 | 1.23746091 | -0.040450793 | 0.32872303 | -0.1230543 | 0.90206407 | NA | NA | NA |
| ENSG000002 | 1.56728502 | -0.607581748 | 0.33870237 | -1.7938515 | 0.07283688 | NA | NA | NA |
| ENSG000002 | 1.34322925 | -0.111836712 | 0.33582708 | -0.3330187 | 0.73912014 | NA | NA | NA |
| ENSG000002 | 0.38064837 | -0.102706251 | 0.21886737 | -0.4692625 | 0.63888201 | NA | NA | NA |
| ENSG000002 | 0.75637659 | -0.193693286 | 0.28066413 | -0.6901248 | 0.49011569 | NA | NA | NA |
| ENSG000002 | 1.65887187 | -0.547884439 | 0.34860587 | -1.5716444 | 0.11603305 | NA | NA | NA |
| ENSG000002 | 1.88989433 | -0.267721455 | 0.36202688 | -0.7395071 | 0.45959912 | NA | NA | NA |
| ENSG000002 | 0.59687261 | 0.065989748 | 0.26864914 | 0.24563543 | 0.80596446 | NA | NA | NA |
| ENSG000002 | 2.07938384 | -0.108510374 | 0.36081701 | -0.3007352 | 0.76361643 | NA | NA | NA |
| ENSG000002 | 2.12527119 | -0.251958529 | 0.36390141 | -0.6923813 | 0.48869791 | NA | NA | NA |
| ENSG000002 | 0.46079952 | 0.128217767 | 0.22562762 | 0.56827158 | 0.56985057 | NA | NA | NA |
| ENSG000002 | 2.10538528 | -0.358953893 | 0.36140284 | -0.9932238 | 0.32060091 | NA | NA | NA |
| ENSG000002 | 1.88352149 | -0.134931113 | 0.35618902 | -0.3788188 | 0.70482239 | NA | NA | NA |
| ENSG000002 | 1.87153291 | 0.001225172 | 0.35238408 | 0.00347681 | 0.99722591 | NA | NA | NA |
| ENSG000002 | 1.90040048 | 0.319157733 | 0.35822222 | 0.89094901 | 0.37295652 | NA | NA | NA |
| ENSG000002 | 1.31710445 | -0.593647818 | 0.31645461 | -1.8759335 | 0.06066441 | NA | NA | NA |
| ENSG000002 | 0 | NA | NA | NA | NA | NA | NA | NA |
| ENSG000002 | 0.91800842 | -0.426865864 | 0.29424669 | -1.4507074 | 0.14686134 | NA | NA | NA |
| ENSG000002 | 0.98620929 | -0.180604596 | 0.30694877 | -0.5883868 | 0.55627272 | NA | NA | NA |
| ENSG000002 | 2.15927069 | 0.310256826 | 0.36673003 | 0.84600878 | 0.39754784 | NA | NA | NA |
| ENSG000002 | 0.61053204 | -0.095342013 | 0.25468495 | -0.3743528 | 0.70814188 | NA | NA | NA |
| ENSG000002 | 1.20592419 | 0.095733894 | 0.32782219 | 0.29202994 | 0.77026373 | NA | NA | NA |
| ENSG000002 | 2.12929388 | -0.50496084 | 0.36418628 | -1.3865455 | 0.1655804 | NA | NA | NA |
| ENSG000002 | 1.37066681 | -0.274993272 | 0.34030881 | -0.8080698 | 0.4190504 | NA | NA | NA |
| ENSG000002 | 0.61881753 | -0.112804359 | 0.27082437 | -0.4165222 | 0.67702794 | NA | NA | NA |
| ENSG000002 | 0.87576874 | -0.122499967 | 0.30315437 | -0.4040845 | 0.68615062 | NA | NA | NA |
| ENSG000002 | 2.1476205 | -0.376580978 | 0.36206046 | -1.0401052 | 0.29829101 | NA | NA | NA |
| ENSG000002 | 1.02237615 | -0.338677438 | 0.3045704 | -1.1119841 | 0.26614499 | NA | NA | NA |
| ENSG000002 | 0.87494563 | -0.264608026 | 0.29350413 | -0.9015479 | 0.36729708 | NA | NA | NA |

| | | | | | | | | |
|---|---|---|---|---|---|---|---|---|
| ENSG000002 | 0.61818637 | 0.059474041 | 0.27041681 | 0.21993471 | 0.82592201 | NA | NA | NA |
| ENSG000002 | 0.75982019 | -0.379788636 | 0.28855699 | -1.3161651 | 0.18811864 | NA | NA | NA |
| ENSG000002 | 1.25371518 | -0.341250739 | 0.33063059 | -1.0321209 | 0.30201549 | NA | NA | NA |
| ENSG000002 | 1.90472624 | -0.535228262 | 0.3591718 | -1.4901734 | 0.13617865 | NA | NA | NA |
| ENSG000002 | 1.3537602 | 0.025713061 | 0.33917137 | 0.07581141 | 0.93956914 | NA | NA | NA |
| ENSG000002 | 0.52554719 | -0.191309836 | 0.24390646 | -0.7843574 | 0.43283043 | NA | NA | NA |
| ENSG000002 | 1.7231944 | -0.049008563 | 0.35047191 | -0.1398359 | 0.88878963 | NA | NA | NA |
| ENSG000002 | 1.41634618 | -0.383897681 | 0.33037237 | -1.1620151 | 0.24522933 | NA | NA | NA |
| ENSG000002 | 2.16595158 | -0.516595807 | 0.36486811 | -1.4158426 | 0.1568216 | NA | NA | NA |
| ENSG000002 | 0.73095424 | -0.032435186 | 0.2762491 | -0.1174128 | 0.90653292 | NA | NA | NA |
| ENSG000002 | 1.05551154 | 0.042784266 | 0.3111423 | 0.13750707 | 0.89063 | NA | FCF1P5 | 106479045 |
| ENSG000002 | 2.07202235 | -0.245850078 | 0.36502389 | -0.6735178 | 0.50061794 | NA | NA | NA |
| ENSG000002 | 1.70697046 | 0.078867266 | 0.35216618 | 0.22394901 | 0.82279698 | NA | NA | NA |
| ENSG000002 | 1.26746297 | -0.192304165 | 0.33003219 | -0.5826831 | 0.56010668 | NA | NA | NA |
| ENSG000002 | 1.4984888 | -0.190947452 | 0.34623706 | -0.5514934 | 0.58129549 | NA | NA | NA |
| ENSG000002 | 0.40942396 | -0.284058841 | 0.21788566 | -1.303706 | 0.19233384 | NA | NA | NA |
| ENSG000002 | 1.90323718 | -0.127676486 | 0.35361764 | -0.361058 | 0.71805607 | NA | NA | NA |
| ENSG000002 | 1.50352251 | -0.347719516 | 0.34690448 | -1.0023495 | 0.31617485 | NA | NA | NA |
| ENSG000002 | 0.91655574 | -0.263011952 | 0.29148811 | -0.9023076 | 0.36689347 | NA | NA | NA |
| ENSG000002 | 1.27112832 | -0.19245513 | 0.32424488 | -0.5935487 | 0.55281399 | NA | NA | NA |
| ENSG000002 | 1.7594244 | -0.326850253 | 0.35383441 | -0.9237379 | 0.3556228 | NA | NA | NA |
| ENSG000002 | 1.4573505 | 0.084808883 | 0.34062757 | 0.24897833 | 0.80337755 | NA | NA | NA |
| ENSG000002 | 1.0869037 | -0.094690096 | 0.3068257 | -0.308612 | 0.75761669 | NA | NA | NA |
| ENSG000002 | 0.75133883 | -0.359102617 | 0.28257827 | -1.2708076 | 0.2037971 | NA | NA | NA |
| ENSG000002 | 0.3854038 | -0.110128691 | 0.21952326 | -0.5016721 | 0.61589821 | NA | NA | NA |
| ENSG000002 | 0.74863691 | -0.520297052 | 0.27793971 | -1.8719781 | 0.06120964 | NA | NA | NA |
| ENSG000002 | 0.48436321 | -0.017262119 | 0.23773787 | -0.0726099 | 0.94211656 | NA | NA | NA |
| ENSG000002 | 2.17294953 | -0.608837194 | 0.36002059 | -1.6911177 | 0.09081432 | NA | NA | NA |
| ENSG000002 | 2.0011856 | -0.965057223 | 0.35386762 | -2.7271702 | 0.00638801 | NA | NA | NA |

| | | | | | | | | |
|---|---|---|---|---|---|---|---|---|
| ENSG000002 | 1.60644512 | -0.515791605 | 0.34347427 | -1.5016892 | 0.13317739 | NA | NA | NA |
| ENSG000002 | 1.40063662 | -0.42922408 | 0.34045438 | -1.2607389 | 0.20740293 | NA | NA | NA |
| ENSG000002 | 0.75042693 | -0.191749474 | 0.2802669 | -0.6841674 | 0.49386947 | NA | NA | NA |
| ENSG000002 | 1.45702058 | -0.291491181 | 0.33299352 | -0.8753659 | 0.38137483 | NA | NA | NA |
| ENSG000002 | 1.0124749 | -0.183380921 | 0.3080917 | -0.5952154 | 0.55169951 | NA | IGBP1-AS2 | 106478965 |
| ENSG000002 | 1.89919668 | -0.246523964 | 0.34972498 | -0.7049081 | 0.48086744 | NA | NA | NA |
| ENSG000002 | 0 | NA | NA | NA | NA | NA | NA | NA |
| ENSG000002 | 0 | NA | NA | NA | NA | NA | SNORD88C | 692204 |
| ENSG000002 | 0.64218479 | -0.28641133 | 0.26819919 | -1.0679053 | 0.28556324 | NA | RNU6ATAC29P | 106481168 |
| ENSG000002 | 0.77265259 | -0.365549345 | 0.28395139 | -1.2873659 | 0.19796678 | NA | MIR1323 | 100302255 |
| ENSG000002 | 0.49261432 | -0.167130763 | 0.23040378 | -0.725382 | 0.46821769 | NA | RNU6ATAC19P | 106481165 |
| ENSG000002 | 0.37079864 | 0.073139042 | 0.21703538 | 0.33699134 | 0.73612343 | NA | MIR1250 | 100302229 |
| ENSG000002 | 0 | NA | NA | NA | NA | NA | MIR1231 | 100302158 |
| ENSG000002 | 0.60168011 | 0.059279687 | 0.26905947 | 0.22032188 | 0.82562048 | NA | NA | NA |
| ENSG000002 | 0.72152927 | 0.131317808 | 0.27584275 | 0.47606039 | 0.63403135 | NA | MIR1272 | 100302184 |
| ENSG000002 | 0.49856397 | -0.186262775 | 0.24090455 | -0.7731808 | 0.43941538 | NA | NA | NA |
| ENSG000002 | 1.51900923 | -0.494935448 | 0.34616123 | -1.429783 | 0.1527793 | NA | RNU6ATAC7P | 106479546 |
| ENSG000002 | 1.29843003 | -0.348523892 | 0.33030916 | -1.0551445 | 0.29135924 | NA | NA | NA |
| ENSG000002 | 1.9595994 | -0.194410364 | 0.36510493 | -0.5324781 | 0.59439494 | NA | NA | NA |
| ENSG000002 | 1.35151316 | -0.23887896 | 0.33123075 | -0.7211859 | 0.47079512 | NA | NA | NA |
| ENSG000002 | 1.99201459 | 0.049164719 | 0.34871538 | 0.1409881 | 0.88787934 | NA | NA | NA |
| ENSG000002 | 0.50955585 | -0.376243729 | 0.24515812 | -1.5346982 | 0.12485797 | NA | RNU6ATAC38P | 106479559 |
| ENSG000002 | 0.63982777 | -0.42118588 | 0.25611206 | -1.6445375 | 0.10006523 | NA | RNU6ATAC6P | 106479545 |
| ENSG000002 | 0.37945415 | -0.107841678 | 0.21865587 | -0.4932027 | 0.62186934 | NA | NA | NA |
| ENSG000002 | 0 | NA | NA | NA | NA | NA | MIR1233-2 | 100422845 |
| ENSG000002 | 0.3863157 | -0.274134926 | 0.21457207 | -1.277589 | 0.20139437 | NA | MIR320D2 | 100302169 |
| ENSG000002 | 2.08743618 | 0.117295038 | 0.36463183 | 0.32168075 | 0.74769457 | NA | NA | NA |
| ENSG000002 | 0.63026429 | -0.446551733 | 0.2623193 | -1.7023213 | 0.08869516 | NA | MIR1302-5 | 100302146 |
| ENSG000002 | 0.37945415 | -0.107841678 | 0.21865587 | -0.4932027 | 0.62186934 | NA | NA | NA |

| | | | | | | | | |
|---|---|---|---|---|---|---|---|---|
| ENSG000002 | 2.07502697 | 0.0081748 | 0.36048056 | 0.02267751 | 0.98190752 | NA | SNORA11B | 100124539 |
| ENSG000002 | 0.37216701 | -0.099548723 | 0.21762847 | -0.4574251 | 0.64736554 | NA | RNU6ATAC30P | 106481868 |
| ENSG000002 | 0 | NA | NA | NA | NA | NA | SNORD110 | 692213 |
| ENSG000002 | 0 | NA | NA | NA | NA | NA | MIR1224 | 100187716 |
| ENSG000002 | 0 | NA | NA | NA | NA | NA | NA | NA |
| ENSG000002 | 0.52623233 | -0.191068033 | 0.24398404 | -0.7831169 | 0.43355844 | NA | NA | NA |
| ENSG000002 | 2.12989023 | -0.258336492 | 0.36531682 | -0.7071574 | 0.47946867 | NA | NA | NA |
| ENSG000002 | 1.17237143 | 0.100353203 | 0.32071935 | 0.31290036 | 0.75435636 | NA | MIR1207 | 100302175 |
| ENSG000002 | 2.03687448 | 0.007311317 | 0.36434776 | 0.02006686 | 0.98399003 | NA | MIR548M | 100313772 |
| ENSG000002 | 0 | NA | NA | NA | NA | NA | MIR1184-2 | 100422985 |
| ENSG000002 | 0.6275579 | -0.281336606 | 0.26701873 | -1.0536213 | 0.29205636 | NA | MIR1262 | 100302279 |
| ENSG000002 | 0.36827086 | -0.266056975 | 0.21181163 | -1.2561018 | 0.20907905 | NA | RNU6ATAC15P | 106481867 |
| ENSG000002 | 1.71245961 | -0.187230818 | 0.3551678 | -0.5271616 | 0.59808138 | NA | MIR548E | 100313921 |
| ENSG000002 | 1.62824972 | -0.406146372 | 0.35094166 | -1.1573045 | 0.24714797 | NA | MIR548L | 100302275 |
| ENSG000002 | 0.35682409 | 0.072008558 | 0.21485623 | 0.33514763 | 0.73751372 | NA | MIR1285-2 | 100302268 |
| ENSG000002 | 0 | NA | NA | NA | NA | NA | MIR1258 | 100302172 |
| ENSG000002 | 0.25260017 | -0.015978705 | 0.18049464 | -0.0885273 | 0.92945758 | NA | NA | NA |
| ENSG000002 | 0.65473699 | -0.286059797 | 0.2692552 | -1.0624114 | 0.28804896 | NA | MIR1263 | 100302148 |
| ENSG000002 | 2.01307891 | -0.594060208 | 0.36329217 | -1.6352134 | 0.10200433 | NA | RNU6ATAC37P | 106479558 |
| ENSG000002 | 1.0965399 | 0.005519065 | 0.19854379 | 0.02779772 | 0.97782348 | NA | MIR1208 | 100302281 |
| ENSG000002 | 0.66253513 | -0.036612812 | 0.15813333 | -0.2315313 | 0.81690208 | NA | MIR548P | 100302288 |
| ENSG000002 | 0.56391566 | 0.358682873 | 0.24477034 | 1.46538538 | 0.1428158 | NA | MIR1284 | 100302112 |
| ENSG000002 | 0 | NA | NA | NA | NA | NA | MIR1237 | 100302280 |
| ENSG000002 | 1.40061543 | -0.578228492 | 0.33868716 | -1.7072643 | 0.08777295 | NA | MIR548I2 | 100302277 |
| ENSG000002 | 0.12070844 | -0.088503695 | 0.12783931 | -0.6923042 | 0.48874631 | NA | MIR1289-2 | 100302134 |
| ENSG000002 | 0.74131889 | -0.324084021 | 0.27363922 | -1.1843479 | 0.23627539 | NA | MIR663B | 100313824 |
| ENSG000002 | 1.98282677 | -0.571614673 | 0.36023842 | -1.5867676 | 0.11256528 | NA | MIR548T | 100422849 |
| ENSG000002 | 0.61335714 | -0.10408451 | 0.2640241 | -0.3942235 | 0.69341602 | NA | NA | NA |
| ENSG000002 | 0.38036605 | -0.293832733 | 0.22119083 | -1.3284128 | 0.18404178 | NA | MIR548I3 | 100302186 |

| | | | | | | | | | |
|---|---|---|---|---|---|---|---|---|---|
| ENSG000002 | 0.98896328 | -0.041455006 | 0.31700684 | -0.1307701 | 0.8959572 | NA | | MIR1200 | 100302113 |
| ENSG000002 | 1.2466041 | -0.338625114 | 0.33045422 | -1.0247263 | 0.30549237 | NA | | MIR548Q | 100313841 |
| ENSG000002 | 1.58358379 | -0.450957931 | 0.33535084 | -1.3447347 | 0.17871091 | NA | | NA | NA |
| ENSG000002 | 0.73041345 | -0.026515459 | 0.28960252 | -0.0915581 | 0.92704914 | NA | | RNU6ATAC18P | 106479551 |
| ENSG000002 | 0.66313191 | -0.429680649 | 0.25813351 | -1.6645675 | 0.09599908 | NA | | NA | NA |
| ENSG000002 | 0.11131518 | 0.081951103 | 0.12663804 | 0.64712866 | 0.51754868 | NA | | MIR548F5 | 100302239 |
| ENSG000002 | 0.77953746 | -0.190889057 | 0.28218837 | -0.6764597 | 0.49874883 | NA | | MIR1288 | 100302124 |
| ENSG000002 | 0.40191006 | -0.101653297 | 0.21376195 | -0.4755444 | 0.63439899 | NA | | RNU6ATAC20P | 106480541 |
| ENSG000002 | 0 | NA | NA | NA | NA | NA | | MIR1228 | 100302201 |
| ENSG000002 | 0.11205239 | 0.082118251 | 0.12671962 | 0.64803108 | 0.51696485 | NA | | MIR548G | 100313938 |
| ENSG000002 | 1.85232864 | -0.368919577 | 0.35633538 | -1.0353156 | 0.30052155 | NA | | RNU6ATAC23P | 106481166 |
| ENSG000002 | 0 | NA | NA | NA | NA | NA | | SNORD88B | 692203 |
| ENSG000002 | 0 | NA | NA | NA | NA | NA | | MIR1229 | 100302156 |
| ENSG000002 | 0 | NA | NA | NA | NA | NA | | NA | NA |
| ENSG000002 | 0.50107449 | -0.372905725 | 0.24426561 | -1.5266403 | 0.12685049 | NA | | NA | NA |
| ENSG000002 | 0 | NA | NA | NA | NA | NA | | MIR1238 | 100302226 |
| ENSG000002 | 0 | NA | NA | NA | NA | NA | | MIR1227 | 100302283 |
| ENSG000002 | 0 | NA | NA | NA | NA | NA | | SNORA81 | 677847 |
| ENSG000002 | 1.6231216 | -0.391835812 | 0.34696243 | -1.1293321 | 0.25875777 | NA | | MIR1283-1 | 100302265 |
| ENSG000002 | 1.42052858 | -0.418575515 | 0.33914767 | -1.2341984 | 0.21712898 | NA | | MIR1294 | 100302181 |
| ENSG000002 | 1.48189664 | -0.309467697 | 0.33802287 | -0.9155229 | 0.35991716 | NA | | MIR548F3 | 100302159 |
| ENSG000002 | 0.28256998 | -0.18483211 | 0.18021686 | -1.0256094 | 0.30507573 | NA | | MIR548F4 | 100313895 |
| ENSG000002 | 0.26813897 | -0.197007041 | 0.18550319 | -1.0620143 | 0.2882292 | NA | | NA | NA |
| ENSG000002 | 0.96266733 | 0.119136402 | 0.30688904 | 0.38820676 | 0.69786303 | NA | | NA | NA |
| ENSG000002 | 0 | NA | NA | NA | NA | NA | | MIR1202 | 100302259 |
| ENSG000002 | 0.46478446 | -0.015417091 | 0.23534337 | -0.0655089 | 0.9477688 | NA | | SNORA11C | 100124540 |
| ENSG000002 | 1.37625037 | -0.402162916 | 0.33469775 | -1.2015704 | 0.22953001 | NA | | NA | NA |
| ENSG000002 | 0.99033165 | -0.197978355 | 0.31350967 | -0.6314904 | 0.5277199 | NA | | MIR1271 | 100302203 |
| ENSG000002 | 0.76940933 | -0.516275266 | 0.27702168 | -1.8636638 | 0.06236894 | NA | | MIR548AJ2 | 100616252 |

| | | | | | | | | | |
|---|---|---|---|---|---|---|---|---|---|
| ENSG000002 | 0 | NA | NA | NA | NA | NA | | SNORA11D | 100124541 |
| ENSG000002 | 1.5143725 | -0.320068258 | 0.33944796 | -0.9429082 | 0.34572787 | NA | | MIR1827 | 100302217 |
| ENSG000002 | 2.10904917 | -0.002992691 | 0.36035506 | -0.0083048 | 0.99337377 | NA | | MIR1251 | 100302289 |
| ENSG000002 | 1.55586608 | 0.160257498 | 0.35038926 | 0.45736989 | 0.6474052 | NA | | SNORA34 | 677815 |
| ENSG000002 | 1.49737944 | -0.177633615 | 0.33966529 | -0.5229666 | 0.60099747 | NA | | MIR548I4 | 100302191 |
| ENSG000002 | 1.04221604 | -0.365573339 | 0.31513426 | -1.1600558 | 0.24602607 | NA | | NA | NA |
| ENSG000002 | 0.96838673 | -0.033819419 | 0.31203989 | -0.1083817 | 0.9136929 | NA | | NA | NA |
| ENSG000002 | 0.37928001 | 0.070449581 | 0.21829793 | 0.32272217 | 0.74690566 | NA | | NA | NA |
| ENSG000002 | 1.22821848 | -0.331385651 | 0.32589828 | -1.0168377 | 0.30923066 | NA | | RNU6ATAC40P | 106479560 |
| ENSG000002 | 0.59621972 | 0.061674489 | 0.26859331 | 0.22962035 | 0.81838679 | NA | | MIR548O | 100313829 |
| ENSG000002 | 0.88035449 | -0.282626859 | 0.30431514 | -0.9287309 | 0.35302858 | NA | | RNU6ATAC16P | 106479550 |
| ENSG000002 | 0.23070733 | 0.167416683 | 0.17664244 | 0.94777156 | 0.34324576 | NA | | MIR1285-1 | 100302218 |
| ENSG000002 | 1.06621446 | -0.706075938 | 0.30933542 | -2.2825577 | 0.02245644 | NA | | MIR1185-1 | 100302157 |
| ENSG000002 | 0.24141689 | -0.165257896 | 0.17127401 | -0.9648743 | 0.33460776 | NA | | RNU6ATAC41P | 106481869 |
| ENSG000002 | 0.11865494 | 0.083440866 | 0.12751478 | 0.65436232 | 0.5128784 | NA | | MIR1184-1 | 100302111 |
| ENSG000002 | 0.78842164 | -0.365546386 | 0.28500054 | -1.2826165 | 0.19962647 | NA | | MIR548H1 | 100313830 |
| ENSG000002 | 0.26062507 | -0.012888319 | 0.18226503 | -0.070712 | 0.94362699 | NA | | MIR1180 | 100302256 |
| ENSG000002 | 1.60777923 | -0.248369173 | 0.34716104 | -0.7154293 | 0.47434382 | NA | | MIR1255B2 | 100313835 |
| ENSG000002 | 0.95629155 | -0.029485772 | 0.30999769 | -0.0951161 | 0.92422261 | NA | | MIR1283-2 | 100302205 |
| ENSG000002 | 1.66108755 | -0.261224941 | 0.34949928 | -0.7474263 | 0.4548063 | NA | | MIR1303 | 100302284 |
| ENSG000002 | 1.84278902 | 0.123375668 | 0.35100181 | 0.35149582 | 0.72521641 | NA | | RNU6ATAC10P | 106480540 |
| ENSG000002 | 1.04005435 | -0.513708114 | 0.3079622 | -1.6680882 | 0.09529821 | NA | | RNU6ATAC42P | 106481173 |
| ENSG000002 | 0.60373362 | -0.112425234 | 0.26959077 | -0.4170218 | 0.67666245 | NA | | RNU6ATAC33P | 106481170 |
| ENSG000002 | 1.05923088 | 0.182746517 | 0.31652139 | 0.57735914 | 0.56369687 | NA | | RNU6ATAC21P | 106479552 |
| ENSG000002 | 0 | NA | NA | NA | NA | NA | | MIR1226 | 100302232 |
| ENSG000002 | 0.74911118 | -0.025129523 | 0.27918326 | -0.0900109 | 0.92827859 | NA | | MIR1261 | 100302228 |
| ENSG000002 | 0.11934008 | 0.083564698 | 0.12760091 | 0.65489106 | 0.5125379 | NA | | MIR548F1 | 100302192 |
| ENSG000002 | 0.82877608 | 0.362390266 | 0.28752859 | 1.26036253 | 0.20753861 | NA | | NA | NA |
| ENSG000002 | 0.23139247 | 0.167748926 | 0.17680352 | 0.94878726 | 0.34272882 | NA | | MIR1184-3 | 100422977 |

| | | | | | | | | |
|---|---|---|---|---|---|---|---|---|
| ENSG000002 | 1.32584389 | 0.027491631 | 0.32943385 | 0.08345114 | 0.93349282 | NA | MIR1293 | 100302220 |
| ENSG000002 | 0.23936338 | -0.005479057 | 0.17759977 | -0.0308506 | 0.9753887 | NA | MIR1185-2 | 100302209 |
| ENSG000002 | 0.74635664 | -0.195125005 | 0.28607594 | -0.6820742 | 0.49519206 | NA | MIR548H4 | 100313884 |
| ENSG000002 | 2.0872804 | -0.378887118 | 0.36545975 | -1.036741 | 0.29985656 | NA | MIR1302-4 | 100302130 |
| ENSG000002 | 0.26813897 | -0.197007041 | 0.18550319 | -1.0620143 | 0.2882292 | NA | MIR1179 | 100302235 |
| ENSG000002 | 0.39274356 | -0.107428268 | 0.22062504 | -0.4869269 | 0.62631013 | NA | MIR1276 | 100302121 |
| ENSG000002 | 0.76687763 | -0.526528684 | 0.2791665 | -1.886074 | 0.05928498 | NA | NA | NA |
| ENSG000002 | 0.74640871 | -0.195085107 | 0.28607993 | -0.6819252 | 0.49528626 | NA | MIR1268A | 100302233 |
| ENSG000002 | 0 | NA | NA | NA | NA | NA | MIR1233-1 | 100302160 |
| ENSG000002 | 1.19923787 | 0.110863539 | 0.33266551 | 0.33325829 | 0.73893932 | NA | MIR1267 | 100302286 |
| ENSG000002 | 0 | NA | NA | NA | NA | NA | MIR1225 | 100188847 |
| ENSG000002 | 0.85587891 | 0.035512315 | 0.28889024 | 0.12292667 | 0.90216517 | NA | MIR1290 | 100302276 |
| ENSG000002 | 0.59687261 | 0.065989748 | 0.26864914 | 0.24563543 | 0.80596446 | NA | MIR548N | 100302152 |
| ENSG000002 | 0.98318777 | -0.200674067 | 0.3132137 | -0.6406938 | 0.52172166 | NA | NA | NA |
| ENSG000002 | 1.64484188 | -0.269557438 | 0.35045677 | -0.7691603 | 0.44179814 | NA | MIR1275 | 100302123 |
| ENSG000002 | 0.24004852 | -0.005223787 | 0.17775844 | -0.029387 | 0.97655594 | NA | MIR302E | 100313774 |
| ENSG000002 | 0 | NA | NA | NA | NA | NA | SNORA11E | 101340250 |
| ENSG000002 | 1.66277037 | -0.847347122 | 0.3453377 | -2.4536769 | 0.0141404 | NA | MIR1298 | 100302153 |
| ENSG000002 | 1.56329655 | 0.027783434 | 0.33511208 | 0.08290789 | 0.93392478 | NA | NA | NA |
| ENSG000002 | 1.75655171 | -0.556136407 | 0.34603237 | -1.6071803 | 0.10801482 | NA | NA | NA |
| ENSG000002 | 1.16866911 | -0.108149451 | 0.30795218 | -0.3511891 | 0.72544649 | NA | MIR548I1 | 100302204 |
| ENSG000002 | 0.25145857 | -0.010118565 | 0.18031426 | -0.0561163 | 0.95524918 | NA | MIR1203 | 100302211 |
| ENSG000002 | 1.88719185 | -0.404697817 | 0.36108039 | -1.120797 | 0.26237428 | NA | MIR1197 | 100302250 |
| ENSG000002 | 0 | NA | NA | NA | NA | NA | NA | NA |
| ENSG000002 | 0.62138203 | -0.094107809 | 0.25725582 | -0.3658141 | 0.71450379 | NA | MIR1273A | 100302165 |
| ENSG000002 | 0.39462238 | -0.104329077 | 0.21264929 | -0.4906157 | 0.6236983 | NA | MIR1260A | 100302236 |
| ENSG000002 | 1.41229891 | 0.364651594 | 0.33638726 | 1.08402321 | 0.27835451 | NA | MIR548J | 100313914 |
| ENSG000002 | 1.18966855 | 0.252172659 | 0.32545169 | 0.774839 | 0.4384348 | NA | MIR1289-1 | 100302125 |
| ENSG000002 | 0.13280363 | -0.093394593 | 0.13115644 | -0.7120855 | 0.47641185 | NA | NA | NA |

| | | | | | | | | |
|---|---|---|---|---|---|---|---|---|
| ENSG000002 | 1.33001833 | 0.022685525 | 0.33525789 | 0.0676659 | 0.9460516 | NA | MIR1252 | 100302136 |
| ENSG000002 | 0.61332276 | -0.255253496 | 0.25725954 | -0.9922023 | 0.32109886 | NA | MIR548H2 | 100313773 |
| ENSG000002 | 0 | NA | NA | NA | NA | NA | SNORD23 | 692091 |
| ENSG000002 | 0.88855353 | -0.453244102 | 0.30180501 | -1.5017779 | 0.13315447 | NA | RNU6ATAC34P | 106479557 |
| ENSG000002 | 0.49970612 | -0.194507174 | 0.24827631 | -0.7834303 | 0.43337449 | NA | NA | NA |
| ENSG000002 | 0.76371634 | -0.201271588 | 0.28715856 | -0.7009075 | 0.48336075 | NA | KRTAP10-9 | 386676 |
| ENSG000002 | 1.00357291 | -0.492055205 | 0.30397229 | -1.6187502 | 0.10550102 | NA | OR4A5 | 81318 |
| ENSG000002 | 2.06694843 | -0.116772722 | 0.36323978 | -0.3214756 | 0.74785001 | NA | KRTAP1-5 | 83895 |
| ENSG000002 | 0 | NA | NA | NA | NA | NA | NA | NA |
| ENSG000002 | 0.26062507 | -0.012888319 | 0.18226503 | -0.070712 | 0.94362699 | NA | KRTAP10-10 | 353333 |
| ENSG000002 | 1.24437591 | -0.047748512 | 0.33393885 | -0.1429858 | 0.8863014 | NA | KRTAP12-2 | 353323 |
| ENSG000002 | 0 | NA | NA | NA | NA | NA | ZNF816-ZNF32 | 100529240 |
| ENSG000002 | 1.83419163 | 0.141527541 | 0.35659798 | 0.39688262 | 0.69145402 | NA | NA | NA |
| ENSG000002 | 1.00460971 | 0.098741016 | 0.30126824 | 0.32775116 | 0.74309982 | NA | KRTAP1-3 | 81850 |
| ENSG000002 | 0.83922024 | 0.360156003 | 0.29071922 | 1.23884485 | 0.21540296 | NA | NA | NA |
| ENSG000002 | 1.7245931 | -0.333791893 | 0.35725078 | -0.934335 | 0.35013113 | NA | POM121L12 | 285877 |
| ENSG000002 | 1.25046745 | -0.202045457 | 0.33450343 | -0.6040161 | 0.54583294 | NA | OR4C12 | 283093 |
| ENSG000002 | 1.10955302 | -0.247681823 | 0.31448246 | -0.7875855 | 0.4309392 | NA | PRR21 | 643905 |
| ENSG000002 | 2.05307058 | -0.574526366 | 0.35989982 | -1.5963508 | 0.11041049 | NA | TMEM14EP | 645843 |
| ENSG000002 | 0.9952835 | -0.192292992 | 0.31221661 | -0.6158961 | 0.53796309 | NA | LOC101928881 | 101928881 |
| ENSG000002 | 0 | NA | NA | NA | NA | NA | NA | NA |
| ENSG000002 | 2.07395783 | 0.009873139 | 0.36399636 | 0.02712428 | 0.97836061 | NA | NA | NA |
| ENSG000002 | 0.87096069 | -0.114966285 | 0.29185658 | -0.3939136 | 0.6936448 | NA | NA | NA |
| ENSG000002 | 1.49236019 | 0.048502491 | 0.19080465 | 0.25419973 | 0.79934126 | NA | NA | NA |
| ENSG000002 | 1.23011903 | -0.197756493 | 0.33268161 | -0.5944317 | 0.55222341 | NA | RNU4-62P | 106479586 |
| ENSG000002 | 0.50451363 | -0.188485218 | 0.24157334 | -0.7802401 | 0.43524954 | NA | RNU4-86P | 106481199 |
| ENSG000002 | 1.66353961 | -0.259737569 | 0.34662702 | -0.7493287 | 0.45365912 | NA | RN7SKP285 | 106480920 |
| ENSG000002 | 0.86757608 | -0.09663148 | 0.28990707 | -0.3333188 | 0.73889364 | NA | MIR1915 | 100302129 |
| ENSG000002 | 0.27408862 | -0.199826037 | 0.18669465 | -1.0703362 | 0.28446801 | NA | NA | NA |

| | | | | | | | | | |
|---|---|---|---|---|---|---|---|---|---|
| ENSG000002 | 0.50630812 | -0.189615696 | 0.24175631 | -0.7843257 | 0.432849 | NA | | RNU6-721P | 106481774 |
| ENSG000002 | 0.75258862 | -0.032475079 | 0.28564831 | -0.113689 | 0.90948431 | NA | | RNU6-908P | 106479968 |
| ENSG000002 | 0.65542213 | -0.28582049 | 0.26931472 | -1.061288 | 0.28855903 | NA | | RNU2-65P | 106480226 |
| ENSG000002 | 0.53627793 | -0.360851627 | 0.24080453 | -1.4985251 | 0.13399688 | NA | | NA | NA |
| ENSG000002 | 1.41597954 | -0.588443959 | 0.34023424 | -1.729526 | 0.083715 | NA | | RN7SKP32 | 106479109 |
| ENSG000002 | 1.98748983 | -0.273912487 | 0.35318445 | -0.7755508 | 0.43801425 | NA | | RN7SKP232 | 106480905 |
| ENSG000002 | 2.1223097 | -0.255766133 | 0.366001 | -0.6988127 | 0.48466911 | NA | | RN7SKP117 | 106479147 |
| ENSG000002 | 0.57683685 | 0.056087234 | 0.2526609 | 0.22198621 | 0.82432462 | NA | | RNA5SP317 | 106479006 |
| ENSG000002 | 1.46595236 | -0.174422296 | 0.33373903 | -0.5226308 | 0.60123121 | NA | | RN7SKP148 | 106480467 |
| ENSG000002 | 0.12685397 | -0.090215768 | 0.12901919 | -0.699243 | 0.48440017 | NA | | RNU6-985P | 106480001 |
| ENSG000002 | 0.14128499 | -0.097772014 | 0.13403235 | -0.7294658 | 0.4657168 | NA | | NA | NA |
| ENSG000002 | 1.3851827 | -0.255932083 | 0.33361049 | -0.7671584 | 0.44298735 | NA | | RNU6-648P | 106479853 |
| ENSG000002 | 0.5226185 | -0.189580046 | 0.24357686 | -0.7783172 | 0.43638207 | NA | | NA | NA |
| ENSG000002 | 0.38056192 | -0.244105065 | 0.20391883 | -1.1970698 | 0.23127937 | NA | | RNU2-9P | 106481636 |
| ENSG000002 | 1.30355658 | -0.492008822 | 0.32730262 | -1.503223 | 0.13278155 | NA | | NA | NA |
| ENSG000002 | 0.64255536 | -0.451090545 | 0.26335299 | -1.7128742 | 0.08673568 | NA | | RNU6-380P | 106481298 |
| ENSG000002 | 1.38426028 | 0.019336641 | 0.33083148 | 0.05844861 | 0.95339129 | NA | | NA | NA |
| ENSG000002 | 1.73180969 | -0.179157423 | 0.34842784 | -0.514188 | 0.60712053 | NA | | RNU4-37P | 106481183 |
| ENSG000002 | 0.36484899 | 0.075158314 | 0.21612264 | 0.34775772 | 0.72802215 | NA | | NA | NA |
| ENSG000002 | 1.51890373 | -0.592722088 | 0.33551204 | -1.7666194 | 0.07729199 | NA | | RN7SKP218 | 106480900 |
| ENSG000002 | 0 | NA | NA | NA | NA | NA | | NA | NA |
| ENSG000002 | 1.46862258 | -0.052205315 | 0.34090645 | -0.1531368 | 0.87829041 | NA | | RN7SKP151 | 106479162 |
| ENSG000002 | 0.70869879 | 0.301373543 | 0.27649914 | 1.08996196 | 0.2757299 | NA | | RNU6-257P | 106479681 |
| ENSG000002 | 0.74689798 | -0.194712996 | 0.28610925 | -0.6805547 | 0.4961533 | NA | | RN7SKP146 | 106479160 |
| ENSG000002 | 1.01928192 | -0.529368 | 0.31174608 | -1.6980743 | 0.08949375 | NA | | RNU4-30P | 106479571 |
| ENSG000002 | 0.48536101 | -0.024029524 | 0.24539296 | -0.0979226 | 0.92199373 | NA | | NA | NA |
| ENSG000002 | 1.92007108 | -0.485464438 | 0.35093048 | -1.3833636 | 0.16655342 | NA | | RN7SKP26 | 106479106 |
| ENSG000002 | 0.9908421 | -0.044700626 | 0.31294954 | -0.1428365 | 0.88641929 | NA | | NA | NA |
| ENSG000002 | 0 | NA | NA | NA | NA | NA | | NA | NA |

| | | | | | | | | | |
|---|---|---|---|---|---|---|---|---|---|
| ENSG000002 | 0.89079899 | -0.240517948 | 0.28586709 | -0.8413629 | 0.40014469 | NA | | RNU4-26P | 106479569 |
| ENSG000002 | 0.86572706 | -0.287838831 | 0.30354833 | -0.9482471 | 0.34300366 | NA | | NA | NA |
| ENSG000002 | 0.51911933 | -0.349228947 | 0.23754574 | -1.4701545 | 0.14151991 | NA | | NA | NA |
| ENSG000002 | 0.2596576 | -0.193002035 | 0.1837913 | -1.0501152 | 0.29366517 | NA | | NA | NA |
| ENSG000002 | 1.05760404 | 0.518576503 | 0.31655472 | 1.63818914 | 0.10138224 | NA | | NA | NA |
| ENSG000002 | 2.08219167 | -0.454795823 | 0.35841816 | -1.2688973 | 0.2044777 | NA | | NA | NA |
| ENSG000002 | 1.32147684 | 0.308031071 | 0.32953606 | 0.93474164 | 0.34992146 | NA | | RNU6-588P | 106479826 |
| ENSG000002 | 1.03968378 | -0.360260375 | 0.3105369 | -1.160121 | 0.24599955 | NA | | RN7SKP129 | 106479152 |
| ENSG000002 | 1.39271991 | -0.126615619 | 0.33819803 | -0.3743831 | 0.70811933 | NA | | RNU2-17P | 106480204 |
| ENSG000002 | 1.60439511 | -0.124627354 | 0.34858861 | -0.3575199 | 0.72070266 | NA | | NA | NA |
| ENSG000002 | 0 | NA | NA | NA | NA | NA | | RNU6-447P | 106481320 |
| ENSG000002 | 0.84938242 | 0.183162426 | 0.28933211 | 0.63305253 | 0.52669934 | NA | | RNU2-54P | 106481651 |
| ENSG000002 | 0.39246123 | -0.276864021 | 0.21549122 | -1.2848042 | 0.19886073 | NA | | NA | NA |
| ENSG000002 | 1.6573597 | -0.415381076 | 0.35193231 | -1.1802868 | 0.23788615 | NA | | NA | NA |
| ENSG000002 | 0.61750123 | 0.055806918 | 0.26401083 | 0.21138117 | 0.83258985 | NA | | RNU2-43P | 106480216 |
| ENSG000002 | 0.77476221 | -0.188343058 | 0.28188047 | -0.6681664 | 0.50402738 | NA | | RN7SKP156 | 106480882 |
| ENSG000002 | 0.84858964 | -0.113323305 | 0.2966718 | -0.3819821 | 0.70247467 | NA | | NA | NA |
| ENSG000002 | 0.37285215 | -0.099309896 | 0.2177345 | -0.4561055 | 0.64831414 | NA | | MIR1471 | 100302126 |
| ENSG000002 | 0.25214371 | -0.009862892 | 0.18046657 | -0.0546522 | 0.95641556 | NA | | NA | NA |
| ENSG000002 | 0 | NA | NA | NA | NA | NA | | RNU6-262P | 106481261 |
| ENSG000002 | 1.97585367 | -0.444815548 | 0.36087539 | -1.2326015 | 0.21772448 | NA | | NA | NA |
| ENSG000002 | 1.40506521 | 0.534425461 | 0.34010268 | 1.57136503 | 0.11609788 | NA | | RNU6-101P | 106479611 |
| ENSG000002 | 0.11205239 | 0.082118251 | 0.12671962 | 0.64803108 | 0.51696485 | NA | | RN7SKP199 | 106479184 |
| ENSG000002 | 1.5867711 | 0.152134612 | 0.34610989 | 0.43955581 | 0.66025885 | NA | | RNU6-892P | 106481460 |
| ENSG000002 | 0.22336757 | 0.163792188 | 0.17487321 | 0.93663397 | 0.34894687 | NA | | NA | NA |
| ENSG000002 | 0.99394994 | -0.48444416 | 0.30363703 | -1.5954713 | 0.11060689 | NA | | RN7SKP111 | 106479144 |
| ENSG000002 | 1.46584293 | -0.05368833 | 0.3453306 | -0.1554694 | 0.8764513 | NA | | RNU6-584P | 106479825 |
| ENSG000002 | 0.48555743 | -0.013775713 | 0.2454629 | -0.0561214 | 0.95524512 | NA | | NA | NA |
| ENSG000002 | 1.81824577 | 0.133494369 | 0.35542038 | 0.3755957 | 0.70721749 | NA | | RNU6-1043P | 106481510 |

| | | | | | | | | | |
|---|---|---|---|---|---|---|---|---|---|
| ENSG000002 | 0.11934008 | 0.083564698 | 0.12760091 | 0.65489106 | 0.5125379 | NA | | RNU2-36P | 106481645 |
| ENSG000002 | 0.58608925 | 0.207285028 | 0.2535965 | 0.81738127 | 0.41371058 | NA | | RNU6-1156P | 106480075 |
| ENSG000002 | 1.15842696 | -0.564260926 | 0.31397543 | -1.7971499 | 0.07231182 | NA | | RNU2-21P | 106480206 |
| ENSG000002 | 0.96468646 | -0.179071102 | 0.30600965 | -0.5851812 | 0.55842586 | NA | | NA | NA |
| ENSG000002 | 1.56042292 | -0.468441669 | 0.33807974 | -1.3855952 | 0.16587054 | NA | | NA | NA |
| ENSG000002 | 2.14628693 | -0.659529429 | 0.36624463 | -1.8007894 | 0.07173609 | NA | | NA | NA |
| ENSG000002 | 1.06948554 | 0.035065284 | 0.3142081 | 0.11159892 | 0.91114143 | NA | | NA | NA |
| ENSG000002 | 0 | NA | NA | NA | NA | NA | | RN7SKP267 | 106480478 |
| ENSG000002 | 0.24550891 | -0.007886114 | 0.17900103 | -0.0440563 | 0.96485956 | NA | | RNU6-1118P | 106480058 |
| ENSG000002 | 0.65542213 | -0.28582049 | 0.26931472 | -1.061288 | 0.28855903 | NA | | NA | NA |
| ENSG000002 | 0.77288285 | -0.203654568 | 0.28771447 | -0.7078357 | 0.47904731 | NA | | RNU6-1050P | 106481512 |
| ENSG000002 | 1.52708565 | -0.428053175 | 0.32981251 | -1.2978682 | 0.19433263 | NA | | MIR1912 | 100302144 |
| ENSG000002 | 1.86732218 | -0.243502056 | 0.35576595 | -0.6844445 | 0.49369451 | NA | | RNU6-855P | 106479944 |
| ENSG000002 | 0 | NA | NA | NA | NA | NA | | NA | NA |
| ENSG000002 | 0 | NA | NA | NA | NA | NA | | RNU6-1076P | 106480038 |
| ENSG000002 | 0.87893002 | -0.428855974 | 0.2945988 | -1.4557288 | 0.1454676 | NA | | NA | NA |
| ENSG000002 | 0.87733544 | -0.0935169 | 0.28328198 | -0.3301195 | 0.74130969 | NA | | RN7SKP225 | 106479195 |
| ENSG000002 | 0 | NA | NA | NA | NA | NA | | RNU6-174P | 106481231 |
| ENSG000002 | 1.67743381 | -0.049714792 | 0.34991059 | -0.1420786 | 0.88701796 | NA | | RN7SKP139 | 106480876 |
| ENSG000002 | 1.08777833 | 0.032037272 | 0.32365735 | 0.09898515 | 0.92115006 | NA | | RNU6-613P | 106479837 |
| ENSG000002 | 1.63000996 | 0.005615137 | 0.34836854 | 0.01611838 | 0.98713995 | NA | | NA | NA |
| ENSG000002 | 0.36553413 | 0.069401489 | 0.20780452 | 0.33397488 | 0.73839851 | NA | | RNY1P15 | 106481711 |
| ENSG000002 | 0 | NA | NA | NA | NA | NA | | RN7SKP221 | 106480474 |
| ENSG000002 | 0.24141689 | -0.165257896 | 0.17127401 | -0.9648743 | 0.33460776 | NA | | RNU6-710P | 106479880 |
| ENSG000002 | 0.60219111 | 0.230513192 | 0.26377177 | 0.87391153 | 0.38216644 | NA | | RNU6-355P | 106481889 |
| ENSG000002 | 1.12191072 | -0.115808006 | 0.31808296 | -0.3640811 | 0.71579742 | NA | | RNU6-1186P | 106480086 |
| ENSG000002 | 1.63151809 | -0.398849153 | 0.34706673 | -1.1492002 | 0.25047344 | NA | | RNU4-34P | 106480385 |
| ENSG000002 | 1.35518255 | 0.025669261 | 0.33716403 | 0.07613286 | 0.9393134 | NA | | SNORA36B | 677818 |
| ENSG000002 | 2.02844247 | -0.567064553 | 0.35803335 | -1.5838317 | 0.11323203 | NA | | RN7SKP202 | 106480472 |

| | | | | | | | | |
|---|---|---|---|---|---|---|---|---|
| ENSG000002 | 1.65622394 | -0.54175765 | 0.34860755 | -1.5540617 | 0.12016969 | NA | RN7SKP127 | 106480466 |
| ENSG000002 | 1.14758156 | -0.274129086 | 0.32267169 | -0.8495604 | 0.39556955 | NA | RN7SKP152 | 106479163 |
| ENSG000002 | 1.44273859 | -0.300860304 | 0.33431535 | -0.8999297 | 0.36815767 | NA | NA | NA |
| ENSG000002 | 1.21947866 | -0.176971665 | 0.32191945 | -0.549739 | 0.58249843 | NA | NA | NA |
| ENSG000002 | 0.62408396 | 0.054009456 | 0.26457561 | 0.20413618 | 0.83824708 | NA | RN7SKP86 | 106480859 |
| ENSG000002 | 0.95790977 | -0.026156233 | 0.30081211 | -0.0869521 | 0.93070962 | NA | RNU2-28P | 106480670 |
| ENSG000002 | 1.0989449 | -0.259635601 | 0.31882725 | -0.8143457 | 0.41544693 | NA | NA | NA |
| ENSG000002 | 0.23065526 | 0.167384503 | 0.17662711 | 0.94767164 | 0.34329664 | NA | RNU6-554P | 106479812 |
| ENSG000002 | 0.53608206 | -0.386623811 | 0.24788931 | -1.5596631 | 0.11883952 | NA | RNU6-791P | 106479916 |
| ENSG000002 | 0.6261592 | 0.054226177 | 0.2557073 | 0.21206346 | 0.83205752 | NA | RN7SKP281 | 106480369 |
| ENSG000002 | 1.11564257 | 0.028662032 | 0.32591296 | 0.08794382 | 0.92992132 | NA | RNU4-65P | 106481192 |
| ENSG000002 | 0.9510791 | -0.025685198 | 0.3046664 | -0.084306 | 0.93281316 | NA | NA | NA |
| ENSG000002 | 2.09493686 | -0.121452098 | 0.36394544 | -0.3337096 | 0.73859868 | NA | RNU2-59P | 106480223 |
| ENSG000002 | 1.13710928 | -0.269584175 | 0.31790711 | -0.8479967 | 0.39643982 | NA | NA | NA |
| ENSG000002 | 1.16041396 | -0.277276481 | 0.32306597 | -0.8582658 | 0.3907457 | NA | NA | NA |
| ENSG000002 | 1.63064112 | -0.124007786 | 0.34555134 | -0.3588694 | 0.71969283 | NA | NA | NA |
| ENSG000002 | 1.16274979 | -0.272266724 | 0.32088754 | -0.8484802 | 0.39617059 | NA | NA | NA |
| ENSG000002 | 0.96826411 | 0.111433667 | 0.30549214 | 0.3647677 | 0.71528482 | NA | RNU2-50P | 106481650 |
| ENSG000002 | 0.4009426 | -0.302672773 | 0.2240145 | -1.3511303 | 0.17665371 | NA | NA | NA |
| ENSG000002 | 1.87980057 | -0.376220326 | 0.35593785 | -1.0569832 | 0.29051926 | NA | NA | NA |
| ENSG000002 | 1.11655784 | -0.148460626 | 0.20910371 | -0.7099856 | 0.47771305 | NA | RNU6-955P | 106480624 |
| ENSG000002 | 1.67916665 | -0.682409476 | 0.34513516 | -1.9772239 | 0.04801633 | NA | NA | NA |
| ENSG000002 | 0.49224375 | -0.024566489 | 0.23863564 | -0.1029456 | 0.91800615 | NA | RNU6-141P | 106480555 |
| ENSG000002 | 1.38757397 | -0.575608538 | 0.33767266 | -1.7046347 | 0.08826257 | NA | NA | NA |
| ENSG000002 | 0.64013129 | -0.113197991 | 0.26627307 | -0.4251199 | 0.67074925 | NA | RN7SKP278 | 106480918 |
| ENSG000002 | 0.24550891 | -0.007886114 | 0.17900103 | -0.0440563 | 0.96485956 | NA | RNU6-1077P | 106480039 |
| ENSG000002 | 1.30696956 | 0.025029102 | 0.3332487 | 0.07510638 | 0.94013008 | NA | RNU6-994P | 106480006 |
| ENSG000002 | 1.05988376 | 0.190362626 | 0.31882915 | 0.59706782 | 0.5504621 | NA | RNU2-26P | 106480414 |
| ENSG000002 | 0.76678939 | -0.201566736 | 0.28734031 | -0.7014913 | 0.48299644 | NA | RN7SKP150 | 106480880 |

| | | | | | | | | | |
|---|---|---|---|---|---|---|---|---|---|
| ENSG000002 | 1.31604569 | 0.15853353 | 0.331112 | 0.47879126 | 0.63208714 | NA | | RNU6-1140P | 106480068 |
| ENSG000002 | 0.49031286 | -0.020802257 | 0.24597635 | -0.0845702 | 0.93260313 | NA | | RN7SKP143 | 106479158 |
| ENSG000002 | 0.50565578 | -0.183478728 | 0.24174047 | -0.7589905 | 0.44785822 | NA | | RN7SKP59 | 106479121 |
| ENSG000002 | 0.88837884 | -0.27742775 | 0.29989645 | -0.9250785 | 0.35492508 | NA | | NA | NA |
| ENSG000002 | 0.86339122 | -0.275317148 | 0.2986111 | -0.9219924 | 0.35653256 | NA | | RNU6-121P | 106481215 |
| ENSG000002 | 0.64813445 | -0.288472774 | 0.26867189 | -1.0736991 | 0.28295756 | NA | | NA | NA |
| ENSG000002 | 1.72896924 | -0.406025346 | 0.34232309 | -1.1860881 | 0.23558751 | NA | | RN7SKP271 | 106480916 |
| ENSG000002 | 1.76739531 | -0.479266953 | 0.35689687 | -1.3428724 | 0.1793133 | NA | | NA | NA |
| ENSG000002 | 0.97883281 | 0.106929823 | 0.30597012 | 0.349478 | 0.72673049 | NA | | NA | NA |
| ENSG000002 | 0.62001367 | 0.056407614 | 0.25515095 | 0.22107546 | 0.82503368 | NA | | NA | NA |
| ENSG000002 | 0.12685397 | -0.090215768 | 0.12901919 | -0.699243 | 0.48440017 | NA | | RNU2-23P | 106481641 |
| ENSG000002 | 0.60299641 | -0.1063938 | 0.26311939 | -0.4043556 | 0.68595125 | NA | | RNU6-77P | 106481792 |
| ENSG000002 | 1.40832521 | -0.741865774 | 0.3366745 | -2.2035105 | 0.02755879 | NA | | RNU6-285P | 106479693 |
| ENSG000002 | 0.50955585 | -0.376243729 | 0.24515812 | -1.5346982 | 0.12485797 | NA | | NA | NA |
| ENSG000002 | 0.26813897 | -0.197007041 | 0.18550319 | -1.0620143 | 0.2882292 | NA | | RNU4-25P | 106481870 |
| ENSG000002 | 0.12070844 | -0.088503695 | 0.12783931 | -0.6923042 | 0.48874631 | NA | | NA | NA |
| ENSG000002 | 0.12070844 | -0.088503695 | 0.12783931 | -0.6923042 | 0.48874631 | NA | | NA | NA |
| ENSG000002 | 0.77925514 | -0.363111002 | 0.28443192 | -1.2766183 | 0.20173703 | NA | | NA | NA |
| ENSG000002 | 1.18677212 | 0.258729444 | 0.32908448 | 0.7862098 | 0.43174458 | NA | | RN7SKP223 | 106479194 |
| ENSG000002 | 0.64739724 | -0.288756313 | 0.26860627 | -1.075017 | 0.28236711 | NA | | RN7SKP109 | 106479143 |
| ENSG000002 | 0.61633981 | 0.053145758 | 0.25640938 | 0.20726917 | 0.83579965 | NA | | MIR1468 | 100302115 |
| ENSG000002 | 1.97875594 | -0.578339689 | 0.36076177 | -1.6031069 | 0.10891105 | NA | | RNU6-705P | 106481403 |
| ENSG000002 | 1.00421975 | -0.344920059 | 0.30916743 | -1.1156416 | 0.26457556 | NA | | RNU2-39P | 106480214 |
| ENSG000002 | 0.87357726 | -0.11559895 | 0.29798362 | -0.3879373 | 0.69806246 | NA | | RN7SKP220 | 106480901 |
| ENSG000002 | 1.58662867 | 0.284333841 | 0.34728946 | 0.81872292 | 0.41294451 | NA | | RNU6-735P | 106481412 |
| ENSG000002 | 0.89199425 | -0.115430831 | 0.29126591 | -0.3963074 | 0.69187828 | NA | | NA | NA |
| ENSG000002 | 1.18341458 | -0.565547003 | 0.31490977 | -1.7959018 | 0.07251014 | NA | | NA | NA |
| ENSG000002 | 0.78159097 | -0.381686531 | 0.28991249 | -1.3165577 | 0.18798691 | NA | | RNU6-1064P | 106481515 |
| ENSG000002 | 0.58884772 | 0.059786227 | 0.26147338 | 0.22865129 | 0.81913996 | NA | | RNU4-43P | 106481185 |

| | | | | | | | | | |
|---|---|---|---|---|---|---|---|---|---|
| ENSG000002 | 1.14436663 | -0.115515271 | 0.31715264 | -0.3642261 | 0.71568918 | NA | | RNU6-1025P | 106480018 |
| ENSG000002 | 0.61272598 | 0.058460947 | 0.26359129 | 0.22178634 | 0.82448022 | NA | | RN7SKP61 | 106479122 |
| ENSG000002 | 1.55705976 | 0.159810265 | 0.35271001 | 0.45309252 | 0.65048212 | NA | | NA | NA |
| ENSG000002 | 0.3854038 | -0.110128691 | 0.21952326 | -0.5016721 | 0.61589821 | NA | | NA | NA |
| ENSG000002 | 0.61881753 | -0.112804359 | 0.27082437 | -0.4165222 | 0.67702794 | NA | | RNU2-47P | 106480672 |
| ENSG000002 | 0.59027007 | 0.063652523 | 0.26808709 | 0.23743226 | 0.81232147 | NA | | RNU2-66P | 106481655 |
| ENSG000002 | 0.52779657 | -0.383508725 | 0.24707466 | -1.5521977 | 0.12061492 | NA | | RN7SKP222 | 106479193 |
| ENSG000002 | 0.38056192 | -0.244105065 | 0.20391883 | -1.1970698 | 0.23127937 | NA | | NA | NA |
| ENSG000002 | 0.72261881 | 0.135321717 | 0.28380279 | 0.47681602 | 0.63349314 | NA | | RN7SKP159 | 106480468 |
| ENSG000002 | 1.1248077 | -0.110431972 | 0.31649034 | -0.3489268 | 0.72714424 | NA | | RNU6-887P | 106481459 |
| ENSG000002 | 0.85345773 | -0.103254778 | 0.29530461 | -0.3496552 | 0.72659751 | NA | | RN7SKP235 | 106479199 |
| ENSG000002 | 0.73773147 | -0.181923771 | 0.2794266 | -0.651061 | 0.51500709 | NA | | RNU2-49P | 106480219 |
| ENSG000002 | 1.64435104 | -0.411681338 | 0.35177783 | -1.1702879 | 0.24188512 | NA | | NA | NA |
| ENSG000002 | 0.12685397 | -0.090215768 | 0.12901919 | -0.699243 | 0.48440017 | NA | | NA | NA |
| ENSG000002 | 0.23799501 | 0.170958857 | 0.17834799 | 0.95856902 | 0.33777591 | NA | | RNU6-628P | 106480396 |
| ENSG000002 | 0.46938748 | 0.335416007 | 0.2391134 | 1.40274868 | 0.16069179 | NA | | NA | NA |
| ENSG000002 | 0.8522635 | -0.114605707 | 0.30200126 | -0.3794875 | 0.70432587 | NA | | RNU4-69P | 106479590 |
| ENSG000002 | 0.74268725 | -0.479320364 | 0.26912835 | -1.7810103 | 0.07491077 | NA | | RNU6-402P | 106481305 |
| ENSG000002 | 0.38270188 | -0.272446727 | 0.2140009 | -1.2731102 | 0.20297894 | NA | | NA | NA |
| ENSG000002 | 0.50722002 | -0.375409304 | 0.24493426 | -1.5326941 | 0.12535123 | NA | | NA | NA |
| ENSG000002 | 0.72492081 | -0.016982134 | 0.27749364 | -0.0611983 | 0.9512013 | NA | | NA | NA |
| ENSG000002 | 0.24756242 | -0.187034346 | 0.18119817 | -1.0322088 | 0.30197431 | NA | | RN7SKP27 | 106480841 |
| ENSG000002 | 0.87440484 | -0.268724769 | 0.29922967 | -0.8980552 | 0.36915611 | NA | | RN7SKP254 | 106481820 |
| ENSG000002 | 0 | NA | NA | NA | NA | NA | | RNU6-1100P | 106480049 |
| ENSG000002 | 0 | NA | NA | NA | NA | NA | | NA | NA |
| ENSG000002 | 1.35467155 | -0.12434192 | 0.34072502 | -0.3649333 | 0.71516117 | NA | | RNU2-48P | 106481649 |
| ENSG000002 | 0.96723406 | 0.047453621 | 0.20346905 | 0.23322279 | 0.81558841 | NA | | RNU2-37P | 106480213 |
| ENSG000002 | 1.04947309 | 0.333002763 | 0.31232421 | 1.06620863 | 0.28632934 | NA | | RNU2-42P | 106481647 |
| ENSG000002 | 0 | NA | NA | NA | NA | NA | | RN7SKP131 | 106479153 |

| | | | | | | | | | |
|---|---|---|---|---|---|---|---|---|---|
| ENSG000002 | 0.85129604 | -0.271505613 | 0.29798313 | -0.9111443 | 0.36221937 | NA | | RNU6-1203P | 106480092 |
| ENSG000002 | 0 | NA | NA | NA | NA | NA | | NA | NA |
| ENSG000002 | 1.35009026 | -0.269708286 | 0.33990144 | -0.7934897 | 0.42749258 | NA | | RNU2-51P | 106480415 |
| ENSG000002 | 0.86669452 | -0.113860766 | 0.29762461 | -0.382565 | 0.70204231 | NA | | RNU2-16P | 106481639 |
| ENSG000002 | 0.38019136 | -0.10756138 | 0.21876967 | -0.491665 | 0.62295621 | NA | | NA | NA |
| ENSG000002 | 1.00675146 | -0.359950375 | 0.31378855 | -1.1471112 | 0.25133565 | NA | | RNU2-70P | 106480227 |
| ENSG000002 | 1.20791757 | 0.075301205 | 0.19400588 | 0.38813878 | 0.69791333 | NA | | RNU6-248P | 106481256 |
| ENSG000002 | 1.388912 | -0.539098482 | 0.33166551 | -1.6254282 | 0.10407134 | NA | | NA | NA |
| ENSG000002 | 1.10582765 | -0.266648132 | 0.32133778 | -0.8298064 | 0.40664827 | NA | | RNU2-8P | 106480200 |
| ENSG000002 | 0.48624257 | -0.012072666 | 0.23796621 | -0.0507327 | 0.95953853 | NA | | RNU4-55P | 106481189 |
| ENSG000002 | 1.10332117 | 0.185906891 | 0.32035547 | 0.5803144 | 0.56170262 | NA | | RN7SKP123 | 106479149 |
| ENSG000002 | 0.47417772 | -0.160107059 | 0.22811633 | -0.7018658 | 0.48276283 | NA | | RNU6-394P | 106480698 |
| ENSG000002 | 0.57478335 | 0.229074688 | 0.26124934 | 0.87684314 | 0.38057184 | NA | | NA | NA |
| ENSG000002 | 0.72509495 | -0.178274683 | 0.2785491 | -0.6400117 | 0.52216499 | NA | | RN7SKP213 | 106480899 |
| ENSG000002 | 1.80872255 | -0.610332956 | 0.35225729 | -1.732634 | 0.08316074 | NA | | NA | NA |
| ENSG000002 | 0.87141612 | -0.272396215 | 0.29905577 | -0.9108542 | 0.36237218 | NA | | RN7SKP119 | 106481749 |
| ENSG000002 | 0.95960806 | 0.245966518 | 0.29776913 | 0.82603095 | 0.40878653 | NA | | RNU4-72P | 106479591 |
| ENSG000002 | 0.36690249 | -0.096999144 | 0.21683809 | -0.4473344 | 0.65463361 | NA | | RNU6-733P | 106479891 |
| ENSG000002 | 0.62541795 | -0.259640357 | 0.25836648 | -1.0049305 | 0.31493031 | NA | | RN7SKP120 | 106480870 |
| ENSG000002 | 2.05648301 | -0.126356659 | 0.36319289 | -0.3479051 | 0.72791145 | NA | | NA | NA |
| ENSG000002 | 0.96038549 | 0.268436707 | 0.30352712 | 0.88439118 | 0.37648508 | NA | | NA | NA |
| ENSG000002 | 1.15282276 | -0.593764928 | 0.31977783 | -1.8568045 | 0.06333897 | NA | | RN7SKP182 | 106479177 |
| ENSG000002 | 1.85469034 | -0.267546328 | 0.36175933 | -0.7395699 | 0.45956099 | NA | | RN7SKP39 | 106480845 |
| ENSG000002 | 1.60015927 | -0.243876457 | 0.34571167 | -0.7054331 | 0.48054075 | NA | | RN7SKP89 | 106479134 |
| ENSG000002 | 2.01453509 | -0.453673886 | 0.3620668 | -1.2530116 | 0.2102015 | NA | | RN7SKP51 | 106479117 |
| ENSG000002 | 1.57105605 | 0.308168708 | 0.35198576 | 0.87551471 | 0.38129391 | NA | | RN7SKP38 | 106479112 |
| ENSG000002 | 1.17547613 | -0.438122439 | 0.32256215 | -1.3582574 | 0.17438201 | NA | | MIR1911 | 100302222 |
| ENSG000002 | 0.24141689 | -0.165257896 | 0.17127401 | -0.9648743 | 0.33460776 | NA | | NA | NA |
| ENSG000002 | 0 | NA | NA | NA | NA | NA | | NA | NA |

| | | | | | | | | |
|---|---|---|---|---|---|---|---|---|
| ENSG000002 | 0.89318847 | -0.11082335 | 0.29132878 | -0.3804065 | 0.70364372 | NA | NA | NA |
| ENSG000002 | 0.12685397 | -0.090215768 | 0.12901919 | -0.699243 | 0.48440017 | NA | RNU6-1190P | 106480087 |
| ENSG000002 | 0.6606002 | -0.463974368 | 0.2663045 | -1.7422701 | 0.08146119 | NA | RN7SKP166 | 106481814 |
| ENSG000002 | 0 | NA | NA | NA | NA | NA | NA | NA |
| ENSG000002 | 1.66384764 | 0.217624339 | 0.35431261 | 0.61421561 | 0.53907286 | NA | RN7SKP206 | 106479187 |
| ENSG000002 | 0.38494734 | -0.095409837 | 0.21124663 | -0.4516514 | 0.65152014 | NA | RNU4-53P | 106479582 |
| ENSG000002 | 2.0692633 | -0.577531917 | 0.35921986 | -1.6077394 | 0.10789227 | NA | RNU6-684P | 106481396 |
| ENSG000002 | 1.11761175 | -0.119416656 | 0.32242399 | -0.3703715 | 0.71110571 | NA | RN7SKP106 | 106479142 |
| ENSG000002 | 0.62431017 | -0.119347761 | 0.27124333 | -0.4400026 | 0.65993525 | NA | RNU2-40P | 106480671 |
| ENSG000002 | 1.7857503 | 0.144816914 | 0.35721851 | 0.40540148 | 0.68518243 | NA | RNU6-233P | 106480562 |
| ENSG000002 | 0.64030597 | -0.296505743 | 0.27391229 | -1.0824843 | 0.27903741 | NA | NA | NA |
| ENSG000002 | 1.40502651 | -0.544472366 | 0.33334893 | -1.6333407 | 0.10239737 | NA | RN7SKP212 | 106481750 |
| ENSG000002 | 0.72992419 | -0.021564412 | 0.26869413 | -0.0802564 | 0.93603337 | NA | RN7SKP91 | 106480861 |
| ENSG000002 | 0.60327873 | 0.055099477 | 0.26276039 | 0.20969476 | 0.83390592 | NA | RNU2-38P | 106481646 |
| ENSG000002 | 1.78245748 | -0.618992203 | 0.35470959 | -1.7450676 | 0.08097311 | NA | RNU4-14P | 106481176 |
| ENSG000002 | 0.14128499 | -0.097772014 | 0.13403235 | -0.7294658 | 0.4657168 | NA | RNU6-67P | 100873771 |
| ENSG000002 | 0.11205239 | 0.082118251 | 0.12671962 | 0.64803108 | 0.51696485 | NA | RNU6-860P | 106479946 |
| ENSG000002 | 0.63142371 | 0.056618775 | 0.26519971 | 0.21349486 | 0.830941 | NA | RNU4-83P | 106479596 |
| ENSG000002 | 0.23816916 | -0.010712105 | 0.1772998 | -0.060418 | 0.9518227 | NA | RNU6-383P | 106479736 |
| ENSG000002 | 1.89498752 | -0.363781032 | 0.35072 | -1.0372406 | 0.29962373 | NA | RNU2-62P | 106480224 |
| ENSG000002 | 1.51243055 | -0.186636481 | 0.34212282 | -0.5455248 | 0.58539263 | NA | NA | NA |
| ENSG000002 | 0.24141689 | -0.165257896 | 0.17127401 | -0.9648743 | 0.33460776 | NA | RNU4-47P | 106479579 |
| ENSG000002 | 1.09589359 | -0.108646251 | 0.32172105 | -0.3377033 | 0.73558681 | NA | RNU2-68P | 106481962 |
| ENSG000002 | 0.14128499 | -0.097772014 | 0.13403235 | -0.7294658 | 0.4657168 | NA | RNU4-27P | 106481180 |
| ENSG000002 | 0.3575613 | 0.072305152 | 0.21497462 | 0.33634273 | 0.73661243 | NA | RN7SKP286 | 106481822 |
| ENSG000002 | 1.23667865 | -0.185041599 | 0.2236618 | -0.8273277 | 0.40805136 | NA | MIR1537 | 100302139 |
| ENSG000002 | 0.74558505 | -0.096038636 | 0.15912748 | -0.6035327 | 0.54615438 | NA | NA | NA |
| ENSG000002 | 0 | NA | NA | NA | NA | NA | RNA5SP100 | 106480755 |
| ENSG000002 | 1.22833113 | -0.301123454 | 0.32046828 | -0.9396358 | 0.34740443 | NA | NA | NA |

| | | | | | | | | | |
|---|---|---|---|---|---|---|---|---|---|
| ENSG000002 | 0.75170548 | -0.191057177 | 0.28035362 | -0.6814864 | 0.49556378 | NA | | RN7SKP168 | 106479170 |
| ENSG000002 | 2.12971652 | -0.637869762 | 0.36411734 | -1.7518247 | 0.07980395 | NA | | RN7SKP42 | 106480846 |
| ENSG000002 | 0 | NA | NA | NA | NA | NA | | NA | NA |
| ENSG000002 | 0.12685397 | -0.090215768 | 0.12901919 | -0.699243 | 0.48440017 | NA | | RNA5SP59 | 106480357 |
| ENSG000002 | 0.79368615 | -0.367708356 | 0.2853167 | -1.2887726 | 0.19747714 | NA | | RNU6-920P | 106481470 |
| ENSG000002 | 1.15442138 | -0.57583688 | 0.31630001 | -1.8205402 | 0.06867678 | NA | | RN7SKP136 | 106479155 |
| ENSG000002 | 1.00601425 | -0.360230169 | 0.31375627 | -1.148121 | 0.25091861 | NA | | RNU6-1086P | 106481934 |
| ENSG000002 | 0 | NA | NA | NA | NA | NA | | RNU6-565P | 106481360 |
| ENSG000002 | 1.40929267 | -0.580237942 | 0.33885573 | -1.7123451 | 0.0868331 | NA | | RNU6-1328P | 106480146 |
| ENSG000002 | 1.11703267 | 0.029979084 | 0.31759487 | 0.09439411 | 0.9247961 | NA | | RN7SKP33 | 106480843 |
| ENSG000002 | 0.7585039 | -0.199073479 | 0.28684023 | -0.6940222 | 0.48766831 | NA | | RN7SKP261 | 106480477 |
| ENSG000002 | 0.37422052 | -0.268731395 | 0.21273223 | -1.2632378 | 0.20650373 | NA | | NA | NA |
| ENSG000002 | 1.18302283 | -0.578110385 | 0.31731702 | -1.8218701 | 0.0684747 | NA | | RN7SKP29 | 106480457 |
| ENSG000002 | 0.64810411 | -0.103253154 | 0.25961327 | -0.3977191 | 0.69083726 | NA | | RNU6-1068P | 106481933 |
| ENSG000002 | 1.72736883 | -0.058574263 | 0.35714767 | -0.1640057 | 0.86972664 | NA | | RNU6-1133P | 106481537 |
| ENSG000002 | 0.49475427 | -0.18461639 | 0.24044438 | -0.7678133 | 0.44259811 | NA | | RNU6-193P | 106480560 |
| ENSG000002 | 0.55665861 | -0.363599063 | 0.24151673 | -1.5054819 | 0.13220023 | NA | | RNU6-564P | 106479817 |
| ENSG000002 | 0.59072708 | 0.064826349 | 0.26164045 | 0.24776883 | 0.80431327 | NA | | NA | NA |
| ENSG000002 | 0.87027801 | 0.044007719 | 0.29585006 | 0.14875007 | 0.88175085 | NA | | NA | NA |
| ENSG000002 | 0.51894464 | -0.193743146 | 0.24314987 | -0.7968055 | 0.42556401 | NA | | RNU2-41P | 106480215 |
| ENSG000002 | 0.87852912 | -0.112522425 | 0.29825115 | -0.3772741 | 0.70596994 | NA | | RNU6-1148P | 106480642 |
| ENSG000002 | 0.13280363 | -0.093394593 | 0.13115644 | -0.7120855 | 0.47641185 | NA | | RN7SKP205 | 106479186 |
| ENSG000002 | 0.11205239 | 0.082118251 | 0.12671962 | 0.64803108 | 0.51696485 | NA | | RNU6-172P | 106479642 |
| ENSG000002 | 1.29472976 | -0.483483679 | 0.32553199 | -1.485211 | 0.13748795 | NA | | RN7SKP284 | 106479222 |
| ENSG000002 | 0.95817036 | -0.032501573 | 0.30498796 | -0.1065667 | 0.9151327 | NA | | NA | NA |
| ENSG000002 | 1.00285296 | -0.044731359 | 0.31342837 | -0.1427164 | 0.88651419 | NA | | RN7SKP262 | 106479212 |
| ENSG000002 | 1.41506819 | -0.416783209 | 0.33904439 | -1.229288 | 0.21896384 | NA | | NA | NA |
| ENSG000002 | 1.35649638 | -0.267867943 | 0.33678021 | -0.7953791 | 0.42639301 | NA | | NA | NA |
| ENSG000002 | 1.13349546 | -0.268419773 | 0.31778498 | -0.8446585 | 0.39830155 | NA | | RNU6-272P | 106479687 |

| | | | | | | | | |
|---|---|---|---|---|---|---|---|---|
| ENSG000002 | 1.30281047 | 0.320582442 | 0.33647419 | 0.95276979 | 0.34070673 | NA | NA | NA |
| ENSG000002 | 0.2596576 | -0.193002035 | 0.1837913 | -1.0501152 | 0.29366517 | NA | RNU6-651P | 106479855 |
| ENSG000002 | 2.00658345 | -0.317543226 | 0.36102057 | -0.879571 | 0.37909176 | NA | RNU2-25P | 106480208 |
| ENSG000002 | 1.27876484 | -0.518119334 | 0.33245216 | -1.5584778 | 0.11912002 | NA | RN7SKP228 | 106479196 |
| ENSG000002 | 0.98412086 | -0.176386317 | 0.29842657 | -0.5910543 | 0.55448401 | NA | RN7SKP94 | 106480365 |
| ENSG000002 | 0.78067907 | -0.193490064 | 0.28059267 | -0.6895763 | 0.49046066 | NA | RN7SKP167 | 106479169 |
| ENSG000002 | 0.96308996 | -0.026279449 | 0.30688538 | -0.0856328 | 0.93175834 | NA | RN7SKP216 | 106479190 |
| ENSG000002 | 0.38651158 | -0.274258716 | 0.21461355 | -1.2779189 | 0.20127801 | NA | NA | NA |
| ENSG000002 | 0.66040432 | -0.457482581 | 0.26479424 | -1.7276908 | 0.08404367 | NA | RNU2-14P | 106481638 |
| ENSG000002 | 0.87048303 | -0.254947844 | 0.29156045 | -0.8744253 | 0.38188666 | NA | RN7SKP68 | 106480854 |
| ENSG000002 | 0.3513637 | 0.074494292 | 0.21400438 | 0.34809704 | 0.72776731 | NA | NA | NA |
| ENSG000002 | 1.07782702 | -0.230958799 | 0.30769401 | -0.7506119 | 0.45288623 | NA | NA | NA |
| ENSG000002 | 0.38747904 | -0.104908247 | 0.21987514 | -0.4771265 | 0.63327207 | NA | RN7SKP259 | 106481751 |
| ENSG000002 | 1.99548606 | -0.331544727 | 0.36416525 | -0.9104238 | 0.36259903 | NA | RNU2-61P | 106480673 |
| ENSG000002 | 0.3588776 | -0.099902505 | 0.21555794 | -0.4634601 | 0.6430346 | NA | NA | NA |
| ENSG000002 | 1.13303899 | -0.267079798 | 0.31941963 | -0.8361408 | 0.40307567 | NA | RN7SKP73 | 106480855 |
| ENSG000002 | 2.09837387 | -0.135050008 | 0.36756055 | -0.3674225 | 0.71330391 | NA | RN7SKP264 | 106479213 |
| ENSG000002 | 0.85595517 | 0.337381311 | 0.28272854 | 1.19330474 | 0.23275005 | NA | RNU6-1135P | 106481538 |
| ENSG000002 | 0.72216234 | 0.140629872 | 0.28376217 | 0.49559063 | 0.62018326 | NA | NA | NA |
| ENSG000002 | 0.59214943 | 0.061522215 | 0.25414046 | 0.24207958 | 0.8087185 | NA | RNU6-55P | 100873763 |
| ENSG000002 | 1.24412392 | -0.329721345 | 0.3247143 | -1.0154199 | 0.30990573 | NA | NA | NA |
| ENSG000002 | 0.85017562 | -0.092873595 | 0.27904922 | -0.3328215 | 0.73926899 | NA | RNU6-284P | 106481268 |
| ENSG000002 | 0.97501046 | 0.120063134 | 0.31226553 | 0.38449051 | 0.70061493 | NA | RN7SKP268 | 106479215 |
| ENSG000002 | 0.77330548 | -0.35598868 | 0.28246534 | -1.2602915 | 0.20756421 | NA | RN7SKP144 | 106479159 |
| ENSG000002 | 0.38270188 | -0.272446727 | 0.2140009 | -1.2731102 | 0.20297894 | NA | RN7SKP161 | 106479167 |
| ENSG000002 | 0 | NA | NA | NA | NA | NA | NA | NA |
| ENSG000002 | 1.86173725 | -0.242654574 | 0.356832 | -0.6800247 | 0.49648882 | NA | RNU6-847P | 106481448 |
| ENSG000002 | 1.22529034 | -0.313426374 | 0.32337776 | -0.9692268 | 0.33243207 | NA | RN7SKP169 | 106480886 |
| ENSG000002 | 0 | NA | NA | NA | NA | NA | NA | NA |

| | | | | | | | | |
|---|---|---|---|---|---|---|---|---|
| ENSG000002 | 0 | NA | NA | NA | NA | NA | RNU6-1245P | 106480109 |
| ENSG000002 | 1.35973159 | 0.165102953 | 0.33382114 | 0.49458507 | 0.62089304 | NA | RNU6-325P | 106480390 |
| ENSG000002 | 1.59448982 | -0.251240692 | 0.34770331 | -0.722572 | 0.46994285 | NA | RN7SKP126 | 106480872 |
| ENSG000002 | 1.97185821 | -0.186967718 | 0.3622292 | -0.5161586 | 0.60574364 | NA | NA | NA |
| ENSG000002 | 0.73701791 | 0.121533762 | 0.26717046 | 0.45489222 | 0.64918678 | NA | RNU2-55P | 106481961 |
| ENSG000002 | 1.15173672 | -0.274900133 | 0.32280229 | -0.8516053 | 0.39443321 | NA | NA | NA |
| ENSG000002 | 1.0654306 | 0.331238536 | 0.3137935 | 1.05559399 | 0.29115375 | NA | NA | NA |
| ENSG000002 | 0.9134827 | -0.282124842 | 0.29953083 | -0.9418892 | 0.34624939 | NA | RMRPP5 | 106481874 |
| ENSG000002 | 1.93597312 | 0.069972262 | 0.36066221 | 0.19401052 | 0.84616765 | NA | NA | NA |
| ENSG000002 | 1.69801193 | 0.06283812 | 0.34459966 | 0.18235108 | 0.85530721 | NA | NA | NA |
| ENSG000002 | 0.84036027 | 0.217450247 | 0.30131285 | 0.72167598 | 0.47049371 | NA | RNU2-72P | 106480702 |
| ENSG000002 | 0 | NA | NA | NA | NA | NA | MIR1538 | 100302119 |
| ENSG000002 | 0.59755775 | 0.062883211 | 0.26225914 | 0.2397751 | 0.81050461 | NA | NA | NA |
| ENSG000002 | 1.20311228 | 0.098676064 | 0.32621897 | 0.30248414 | 0.76228302 | NA | RN7SKP296 | 106481823 |
| ENSG000002 | 1.03831989 | -0.50780213 | 0.30797528 | -1.6488405 | 0.09918031 | NA | RN7SKP102 | 106481810 |
| ENSG000002 | 1.24770899 | -0.054038667 | 0.33402409 | -0.1617807 | 0.87147852 | NA | RN7SKP181 | 106479176 |
| ENSG000002 | 0.62041457 | -0.27371262 | 0.26650407 | -1.0270486 | 0.3043976 | NA | RNU2-32P | 106481644 |
| ENSG000002 | 0.23202362 | -0.008302493 | 0.17586207 | -0.0472103 | 0.96234565 | NA | RN7SKP263 | 106480913 |
| ENSG000002 | 0.40327843 | -0.281391716 | 0.21700406 | -1.2967118 | 0.19473038 | NA | RN7SKP140 | 106480366 |
| ENSG000002 | 0.60168011 | 0.059279687 | 0.26905947 | 0.22032188 | 0.82562048 | NA | RN7SKP207 | 106480897 |
| ENSG000002 | 1.63327751 | -0.107729648 | 0.33791366 | -0.3188082 | 0.74987196 | NA | NA | NA |
| ENSG000002 | 0.74709385 | -0.19484779 | 0.28612562 | -0.6809869 | 0.4958798 | NA | RN7SKP252 | 106479207 |
| ENSG000002 | 1.57530937 | -0.635919595 | 0.34247688 | -1.8568249 | 0.06333607 | NA | RN7SKP112 | 106479145 |
| ENSG000002 | 0.75184983 | -0.181113777 | 0.28040834 | -0.645893 | 0.51834867 | NA | RNU4-88P | 106479599 |
| ENSG000002 | 0.23890637 | -0.010420674 | 0.17747379 | -0.0587167 | 0.95317776 | NA | RNU2-18P | 106480205 |
| ENSG000002 | 0.58412454 | 0.062473285 | 0.26103841 | 0.23932603 | 0.81085278 | NA | NA | NA |
| ENSG000002 | 1.78063265 | -0.472763324 | 0.35605439 | -1.327784 | 0.18424947 | NA | NA | NA |
| ENSG000002 | 0.7802226 | -0.200980091 | 0.2881869 | -0.697395 | 0.48555566 | NA | RN7SKP142 | 106480877 |
| ENSG000002 | 1.15472736 | -0.103702717 | 0.30746576 | -0.3372822 | 0.7359042 | NA | NA | NA |

| | | | | | | | | | |
|---|---|---|---|---|---|---|---|---|---|
| ENSG000002 | 1.2144061 | 0.110025405 | 0.33609244 | 0.3273665 | 0.7433907 | NA | | RN7SKP17 | 106479102 |
| ENSG000002 | 0 | NA | NA | NA | NA | NA | | RNU4-61P | 106481191 |
| ENSG000002 | 1.45776379 | -0.73526632 | 0.33519743 | -2.1935321 | 0.02826906 | NA | | NA | NA |
| ENSG000002 | 1.01810943 | -0.349417427 | 0.30972533 | -1.1281526 | 0.25925549 | NA | | RNU6-373P | 106480391 |
| ENSG000002 | 0.69159117 | 0.269756142 | 0.26835544 | 1.00521959 | 0.31479112 | NA | | RNU6-1199P | 106480091 |
| ENSG000002 | 0.99573997 | -0.200130962 | 0.3137209 | -0.6379268 | 0.52352135 | NA | | NA | NA |
| ENSG000002 | 1.29660913 | -0.489383409 | 0.32796158 | -1.4921974 | 0.13564741 | NA | | RNU6-177P | 106481232 |
| ENSG000002 | 1.10207138 | -0.114235881 | 0.32192216 | -0.3548556 | 0.72269777 | NA | | RN7SKP100 | 106479139 |
| ENSG000002 | 1.33464081 | -0.3557986 | 0.32513615 | -1.0943065 | 0.27382058 | NA | | RNU6-293P | 106481271 |
| ENSG000002 | 0.48348219 | -0.018797026 | 0.24521266 | -0.076656 | 0.9388972 | NA | | RNU6-1001P | 106480628 |
| ENSG000002 | 1.43393951 | -0.540355442 | 0.32976174 | -1.6386238 | 0.10129163 | NA | | RN7SKP297 | 106479227 |
| ENSG000002 | 2.05570963 | 0.112442504 | 0.36304233 | 0.30972285 | 0.75677173 | NA | | RNA5SP221 | 106480759 |
| ENSG000002 | 0.4009426 | -0.302672773 | 0.2240145 | -1.3511303 | 0.17665371 | NA | | RNU6-1217P | 106481564 |
| ENSG000002 | 0.3433388 | 0.065348438 | 0.20407693 | 0.32021472 | 0.74880557 | NA | | RNU4-74P | 106479592 |
| ENSG000002 | 0.60102723 | 0.047720838 | 0.25498529 | 0.18715134 | 0.85154198 | NA | | RNU6-607P | 106481772 |
| ENSG000002 | 0.23139247 | 0.167748926 | 0.17680352 | 0.94878726 | 0.34272882 | NA | | RNU6-938P | 106479981 |
| ENSG000002 | 0.70090011 | 0.137394752 | 0.28231267 | 0.48667582 | 0.62648808 | NA | | NA | NA |
| ENSG000002 | 0.52779657 | -0.383508725 | 0.24707466 | -1.5521977 | 0.12061492 | NA | | RN7SKP192 | 106480892 |
| ENSG000002 | 1.09819505 | -0.231627166 | 0.30849124 | -0.7508387 | 0.45274972 | NA | | RNU6-1054P | 106480029 |
| ENSG000002 | 0.49953144 | -0.023330163 | 0.24696918 | -0.0944659 | 0.92473908 | NA | | RN7SKP270 | 106479216 |
| ENSG000002 | 0.65542213 | -0.28582049 | 0.26931472 | -1.061288 | 0.28855903 | NA | | RNU4-63P | 106479587 |
| ENSG000002 | 0.59507757 | 0.053522528 | 0.26203995 | 0.20425332 | 0.83815555 | NA | | NA | NA |
| ENSG000002 | 0.37330862 | -0.105391168 | 0.21773449 | -0.4840352 | 0.62836086 | NA | | RNU6-785P | 106479913 |
| ENSG000002 | 1.28006861 | -0.05034067 | 0.31578802 | -0.1594129 | 0.87334361 | NA | | RNU6-575P | 106481363 |
| ENSG000002 | 0.38884741 | -0.29745092 | 0.22235466 | -1.3377319 | 0.18098386 | NA | | NA | NA |
| ENSG000002 | 1.43575497 | -0.041916135 | 0.33805861 | -0.1239907 | 0.90132261 | NA | | RN7SKP194 | 106480893 |
| ENSG000002 | 0.75099 | -0.033089548 | 0.29085543 | -0.1137663 | 0.90942305 | NA | | NA | NA |
| ENSG000002 | 1.12589566 | -0.270975439 | 0.32347778 | -0.8376941 | 0.40220251 | NA | | RNU6-387P | 106481891 |
| ENSG000002 | 0.23276084 | -0.008011382 | 0.17603982 | -0.0455089 | 0.96370166 | NA | | RNU6-592P | 106481368 |

| | | | | | | | | |
|---|---|---|---|---|---|---|---|---|
| ENSG000002 | 0.51570138 | -0.378721611 | 0.24581506 | -1.540677 | 0.12339542 | NA | NA | NA |
| ENSG000002 | 0.74268725 | -0.479320364 | 0.26912835 | -1.7810103 | 0.07491077 | NA | RNA5SP498 | 106480770 |
| ENSG000002 | 1.0310322 | -0.510514485 | 0.30759561 | -1.6596937 | 0.0969761 | NA | RNU6-57P | 100873765 |
| ENSG000002 | 0.11865494 | 0.083440866 | 0.12751478 | 0.65436232 | 0.5128784 | NA | RNU6-954P | 106481481 |
| ENSG000002 | 1.52317044 | -0.637554371 | 0.34355766 | -1.855742 | 0.06349035 | NA | NA | NA |
| ENSG000002 | 1.48884355 | -0.344233728 | 0.34667693 | -0.9929525 | 0.32073309 | NA | NA | NA |
| ENSG000002 | 0.75830802 | -0.198939032 | 0.28682413 | -0.6935924 | 0.48793787 | NA | NA | NA |
| ENSG000002 | 0.51459361 | -0.192534203 | 0.24265108 | -0.7934611 | 0.42750921 | NA | RNY3P8 | 106478985 |
| ENSG000002 | 1.430491 | -0.029006884 | 0.33245644 | -0.0872502 | 0.93047265 | NA | NA | NA |
| ENSG000002 | 1.62312318 | -0.273359138 | 0.35195139 | -0.7766957 | 0.43733831 | NA | NA | NA |
| ENSG000002 | 0.87394783 | -0.26920868 | 0.29758521 | -0.904644 | 0.36565401 | NA | RN7SKP101 | 106480864 |
| ENSG000002 | 0.25351207 | -0.189965547 | 0.1824784 | -1.0410303 | 0.29786148 | NA | RNU6-722P | 106481407 |
| ENSG000002 | 0.23936338 | -0.005479057 | 0.17759977 | -0.0308506 | 0.9753887 | NA | RNU6-516P | 106479794 |
| ENSG000002 | 0.90779205 | -0.463756956 | 0.30262298 | -1.5324578 | 0.12540949 | NA | RN7SKP273 | 106481821 |
| ENSG000002 | 0.37945415 | -0.107841678 | 0.21865587 | -0.4932027 | 0.62186934 | NA | RNU2-71P | 106480228 |
| ENSG000002 | 0.11865494 | 0.083440866 | 0.12751478 | 0.65436232 | 0.5128784 | NA | RNU6-1052P | 106480028 |
| ENSG000002 | 1.53184376 | -0.346444454 | 0.34736143 | -0.9973602 | 0.31858972 | NA | NA | NA |
| ENSG000002 | 0.49671898 | -0.027206884 | 0.23914704 | -0.1137663 | 0.90942302 | NA | KLF2P3 | 106481672 |
| ENSG000002 | 0.88874941 | -0.417513114 | 0.29274791 | -1.4261865 | 0.15381452 | NA | NA | NA |
| ENSG000002 | 0 | NA | NA | NA | NA | NA | NA | NA |
| ENSG000002 | 2.12397592 | -0.617773564 | 0.36219383 | -1.7056436 | 0.08807447 | NA | NA | NA |
| ENSG000002 | 0.53374623 | -0.385819457 | 0.24767828 | -1.5577444 | 0.11929385 | NA | NA | NA |
| ENSG000002 | 0.11865494 | 0.083440866 | 0.12751478 | 0.65436232 | 0.5128784 | NA | NA | NA |
| ENSG000002 | 1.07372443 | -0.474243424 | 0.29954887 | -1.5831922 | 0.11337767 | NA | NA | NA |
| ENSG000002 | 1.38664088 | -0.569535981 | 0.33650383 | -1.6925096 | 0.09054884 | NA | SDCBP2P1 | 106480801 |
| ENSG000002 | 0.12070844 | -0.088503695 | 0.12783931 | -0.6923042 | 0.48874631 | NA | NA | NA |
| ENSG000002 | 0.11865494 | 0.083440866 | 0.12751478 | 0.65436232 | 0.5128784 | NA | RBM22P7 | 106478953 |
| ENSG000002 | 0.27408862 | -0.199826037 | 0.18669465 | -1.0703362 | 0.28446801 | NA | NA | NA |
| ENSG000002 | 2.07187987 | -0.583574515 | 0.36077531 | -1.6175567 | 0.10575817 | NA | NA | NA |

| | | | | | | | | |
|---|---|---|---|---|---|---|---|---|
| ENSG000002 | 0 | NA | NA | NA | NA | NA | NA | NA |
| ENSG000002 | 0 | NA | NA | NA | NA | NA | USP9YP18 | 106480424 |
| ENSG000002 | 1.85451565 | -0.135532228 | 0.36146508 | -0.3749525 | 0.70769581 | NA | NA | NA |
| ENSG000002 | 1.86157308 | -0.268768803 | 0.36181031 | -0.7428445 | 0.45757582 | NA | NA | NA |
| ENSG000002 | 2.06698226 | -0.003577671 | 0.3630829 | -0.0098536 | 0.9921381 | NA | NA | NA |
| ENSG000002 | 0 | NA | NA | NA | NA | NA | NA | NA |
| ENSG000002 | 1.33189714 | 0.019209306 | 0.33872838 | 0.05671006 | 0.95477616 | NA | NA | NA |
| ENSG000002 | 0.87827925 | -0.275315466 | 0.29938851 | -0.9195926 | 0.35778568 | NA | NA | NA |
| ENSG000002 | 0.98415523 | -0.043136961 | 0.31268046 | -0.1379586 | 0.89027313 | NA | NA | NA |
| ENSG000002 | 0.64986946 | -0.085197438 | 0.15786165 | -0.5396969 | 0.58940609 | NA | NA | NA |
| ENSG000002 | 0.38747904 | -0.104908247 | 0.21987514 | -0.4771265 | 0.63327207 | NA | NA | NA |
| ENSG000002 | 1.4917767 | -0.603320914 | 0.33858212 | -1.7819042 | 0.07476485 | NA | NA | NA |
| ENSG000002 | 1.83386367 | -0.909165794 | 0.35034862 | -2.5950317 | 0.00945822 | NA | NA | NA |
| ENSG000002 | 1.85975217 | -0.375163143 | 0.35575322 | -1.0545601 | 0.29162657 | NA | NA | NA |
| ENSG000002 | 0.3513637 | 0.074494292 | 0.21400438 | 0.34809704 | 0.72776731 | NA | NA | NA |
| ENSG000002 | 0.71698024 | -0.023804784 | 0.27695399 | -0.0859521 | 0.93150447 | NA | NA | NA |
| ENSG000002 | 1.99257519 | 0.061367542 | 0.36256907 | 0.16925752 | 0.86559409 | NA | NA | NA |
| ENSG000002 | 0 | NA | NA | NA | NA | NA | NA | NA |
| ENSG000002 | 0.11205239 | 0.082118251 | 0.12671962 | 0.64803108 | 0.51696485 | NA | FAM90A7P | 441317 |
| ENSG000002 | 0.38064837 | -0.102706251 | 0.21886737 | -0.4692625 | 0.63888201 | NA | NA | NA |
| ENSG000002 | 1.24702619 | -0.482311223 | 0.32573958 | -1.4806651 | 0.13869584 | NA | NA | NA |
| ENSG000002 | 2.06838015 | -0.725576814 | 0.35948684 | -2.0183682 | 0.04355293 | NA | NA | NA |
| ENSG000002 | 1.24751256 | -0.056713994 | 0.31762646 | -0.1785556 | 0.85828663 | NA | NA | NA |
| ENSG000002 | 2.04051996 | -0.462337469 | 0.36274916 | -1.2745377 | 0.20247293 | NA | NA | NA |
| ENSG000002 | 1.46421397 | 0.088081567 | 0.34074112 | 0.25849996 | 0.79602108 | NA | NA | NA |
| ENSG000002 | 2.03615845 | 0.125279149 | 0.36289473 | 0.34522173 | 0.72992768 | NA | NA | NA |
| ENSG000002 | 0 | NA | NA | NA | NA | NA | NA | NA |
| ENSG000002 | 1.27244909 | -0.632577694 | 0.32307695 | -1.9579784 | 0.05023255 | NA | NA | NA |
| ENSG000002 | 0.88741138 | -0.451663868 | 0.3001743 | -1.504672 | 0.13240842 | NA | NA | NA |

| | | | | | | | | |
|---|---|---|---|---|---|---|---|---|
| ENSG000002 | 1.13565659 | -0.124737422 | 0.32299795 | -0.3861864 | 0.69935861 | NA | NA | NA |
| ENSG000002 | 0.60717723 | -0.256800472 | 0.2582777 | -0.9942805 | 0.32008634 | NA | NA | NA |
| ENSG000002 | 0 | NA | NA | NA | NA | NA | NA | NA |
| ENSG000002 | 0 | NA | NA | NA | NA | NA | NA | NA |
| ENSG000002 | 0.64850501 | -0.484783006 | 0.2711005 | -1.788204 | 0.07374309 | NA | NA | NA |
| ENSG000002 | 1.11838176 | -0.107372848 | 0.31626856 | -0.339499 | 0.73423388 | NA | NA | NA |
| ENSG000002 | 1.58219876 | -0.242763888 | 0.34544703 | -0.7027529 | 0.48220978 | NA | NA | NA |
| ENSG000002 | 2.13256088 | -0.621537152 | 0.36240286 | -1.7150448 | 0.08633701 | NA | NA | NA |
| ENSG000002 | 0.82750614 | 0.044980518 | 0.28725749 | 0.15658606 | 0.87557109 | NA | LINC01044 | 104266956 |
| ENSG000002 | 1.37371866 | -0.407035138 | 0.33605448 | -1.2112177 | 0.22581198 | NA | NA | NA |
| ENSG000002 | 0 | NA | NA | NA | NA | NA | NA | NA |
| ENSG000002 | 0 | NA | NA | NA | NA | NA | RBMY2EP | 159125 |
| ENSG000002 | 0.3575613 | 0.072305152 | 0.21497462 | 0.33634273 | 0.73661243 | NA | NA | NA |
| ENSG000002 | 0 | NA | NA | NA | NA | NA | NA | NA |
| ENSG000002 | 0.6606002 | -0.463974368 | 0.2663045 | -1.7422701 | 0.08146119 | NA | NA | NA |
| ENSG000002 | 1.91045361 | -0.668762236 | 0.35766968 | -1.8697761 | 0.06151491 | NA | NA | NA |
| ENSG000002 | 0 | NA | NA | NA | NA | NA | NA | NA |
| ENSG000002 | 0.50630812 | -0.189615696 | 0.24175631 | -0.7843257 | 0.432849 | NA | KRT8P51 | 106480676 |
| ENSG000002 | 2.14261657 | -0.375255094 | 0.36261939 | -1.0348456 | 0.300741 | NA | NA | NA |
| ENSG000002 | 0.47551128 | 0.153262041 | 0.24481524 | 0.62603146 | 0.53129431 | NA | NA | NA |
| ENSG000002 | 1.17268384 | -0.425206713 | 0.32068845 | -1.3259184 | 0.18486672 | NA | NA | NA |
| ENSG000002 | 0.90344102 | -0.436658607 | 0.29584239 | -1.4759839 | 0.13994822 | NA | NA | NA |
| ENSG000002 | 2.06194505 | -0.104311918 | 0.35821266 | -0.2912011 | 0.77089753 | NA | NA | NA |
| ENSG000002 | 0 | NA | NA | NA | NA | NA | NA | NA |
| ENSG000002 | 0.78961532 | -0.345233677 | 0.27694769 | -1.2465664 | 0.21255654 | NA | RAET1E-AS1 | 100652739 |
| ENSG000002 | 0.75093793 | -0.03172043 | 0.28555189 | -0.1110846 | 0.91154923 | NA | NA | NA |
| ENSG000002 | 1.75772027 | -0.464792063 | 0.35522836 | -1.3084317 | 0.19072692 | NA | NA | NA |
| ENSG000002 | 0.36690249 | -0.096999144 | 0.21683809 | -0.4473344 | 0.65463361 | NA | NA | NA |
| ENSG000002 | 1.19118072 | 0.095947538 | 0.31402809 | 0.30553808 | 0.75995637 | NA | NA | NA |

| | | | | | | | | |
|---|---|---|---|---|---|---|---|---|
| ENSG000002 | 1.51838212 | -0.048298686 | 0.33251093 | -0.1452544 | 0.88450999 | NA | NA | NA |
| ENSG000002 | 1.75860287 | -0.332180565 | 0.35615846 | -0.9326763 | 0.35098711 | NA | LOC105376310 | 105376310 |
| ENSG000002 | 1.46575648 | -0.187243339 | 0.34130199 | -0.5486148 | 0.58326979 | NA | NA | NA |
| ENSG000002 | 0.25333738 | -0.015686291 | 0.1806605 | -0.0868275 | 0.93080867 | NA | NA | NA |
| ENSG000002 | 1.10295048 | -0.261039145 | 0.31896476 | -0.8183949 | 0.41313171 | NA | NA | NA |
| ENSG000002 | 0 | NA | NA | NA | NA | NA | NA | NA |
| ENSG000002 | 1.59847634 | -0.264530919 | 0.3522544 | -0.7509655 | 0.45267339 | NA | TCEB1P30 | 106481670 |
| ENSG000002 | 1.16227159 | -0.426492051 | 0.32032078 | -1.331453 | 0.18304001 | NA | NA | NA |
| ENSG000002 | 1.30352625 | -0.324081329 | 0.32390447 | -1.000546 | 0.31704634 | NA | NA | NA |
| ENSG000002 | 0.49082386 | 0.135175125 | 0.22941993 | 0.58920394 | 0.55572448 | NA | NA | NA |
| ENSG000002 | 0.85838785 | -0.251108848 | 0.29088494 | -0.8632583 | 0.38799544 | NA | NA | NA |
| ENSG000002 | 1.45738275 | 0.085366683 | 0.3450935 | 0.24737262 | 0.80461986 | NA | NA | NA |
| ENSG000002 | 1.86860017 | -0.248990287 | 0.35751132 | -0.6964543 | 0.48614438 | NA | NA | NA |
| ENSG000002 | 1.09262413 | -0.111048627 | 0.31538162 | -0.3521088 | 0.72475671 | NA | BOLA3P4 | 106480324 |
| ENSG000002 | 1.06578135 | 0.190961914 | 0.32055074 | 0.59573069 | 0.55135516 | NA | NA | NA |
| ENSG000002 | 0 | NA | NA | NA | NA | NA | NA | NA |
| ENSG000002 | 0 | NA | NA | NA | NA | NA | IGKV1OR2-2 | 106481689 |
| ENSG000002 | 1.70038562 | -0.051519665 | 0.35066252 | -0.1469209 | 0.88319443 | NA | LOC102723377 | 102723377 |
| ENSG000002 | 0.73124048 | -0.17762673 | 0.27730728 | -0.6405412 | 0.52182083 | NA | NA | NA |
| ENSG000002 | 1.98669651 | -0.557260545 | 0.35866294 | -1.5537166 | 0.12025204 | NA | NA | NA |
| ENSG000002 | 1.79941808 | -0.753095998 | 0.35179251 | -2.1407391 | 0.03229508 | NA | LINC01342 | 254099 |
| ENSG000002 | 1.65758996 | -0.268946398 | 0.35062034 | -0.7670587 | 0.4430466 | NA | NA | NA |
| ENSG000002 | 1.77710707 | -0.742528176 | 0.35140024 | -2.1130554 | 0.03459603 | NA | NA | NA |
| ENSG000002 | 0.77953746 | -0.190889057 | 0.28218837 | -0.6764597 | 0.49874883 | NA | NA | NA |
| ENSG000002 | 1.06834394 | 0.041230317 | 0.31893038 | 0.12927686 | 0.89713858 | NA | NA | NA |
| ENSG000002 | 0.49475427 | -0.18461639 | 0.24044438 | -0.7678133 | 0.44259811 | NA | NA | NA |
| ENSG000002 | 0 | NA | NA | NA | NA | NA | BRD2-IT1 | 106478972 |
| ENSG000002 | 1.72399228 | -0.325830568 | 0.35411002 | -0.9201394 | 0.35749993 | NA | NA | NA |
| ENSG000002 | 1.13149985 | -0.253030021 | 0.3152611 | -0.8026046 | 0.42220329 | NA | NA | NA |

| | | | | | | | | |
|---|---|---|---|---|---|---|---|---|
| ENSG000002 | 0.97539312 | -0.041442074 | 0.29717562 | -0.1394531 | 0.88909208 | NA | NA | NA |
| ENSG000002 | 0 | NA | NA | NA | NA | NA | NA | NA |
| ENSG000002 | 1.6603463 | -0.547320936 | 0.34863048 | -1.5699171 | 0.11643441 | NA | NA | NA |
| ENSG000002 | 1.86556254 | -0.007459966 | 0.3511529 | -0.0212442 | 0.98305085 | NA | NA | NA |
| ENSG000002 | 0.35824644 | 0.072568381 | 0.21508272 | 0.33739754 | 0.73581724 | NA | NA | NA |
| ENSG000002 | 0.48216589 | 0.145317976 | 0.23824352 | 0.60995564 | 0.54189119 | NA | NA | NA |
| ENSG000002 | 0 | NA | NA | NA | NA | NA | NA | NA |
| ENSG000002 | 0.9779238 | -0.181055242 | 0.30825798 | -0.5873497 | 0.55696885 | NA | NA | NA |
| ENSG000002 | 2.06737243 | -0.115746833 | 0.360319 | -0.3212343 | 0.74803282 | NA | NA | NA |
| ENSG000002 | 0.11934008 | 0.083564698 | 0.12760091 | 0.65489106 | 0.5125379 | NA | NA | NA |
| ENSG000002 | 0 | NA | NA | NA | NA | NA | NA | NA |
| ENSG000002 | 0.40942396 | -0.284058841 | 0.21788566 | -1.303706 | 0.19233384 | NA | NA | NA |
| ENSG000002 | 2.07050339 | -1.088880081 | 0.3479678 | -3.1292553 | 0.0017525 | NA | NA | NA |
| ENSG000002 | 0 | NA | NA | NA | NA | NA | NA | NA |
| ENSG000002 | 1.63957186 | -0.04861865 | 0.24366022 | -0.1995346 | 0.84184456 | NA | NA | NA |
| ENSG000002 | 0.35346928 | -0.088570748 | 0.20618147 | -0.4295767 | 0.66750362 | NA | NA | NA |
| ENSG000002 | 0.37330862 | -0.105391168 | 0.21773449 | -0.4840352 | 0.62836086 | NA | NA | NA |
| ENSG000002 | 0.63939408 | -0.113476381 | 0.26621037 | -0.4262658 | 0.66991416 | NA | NA | NA |
| ENSG000002 | 0.86823757 | -0.416677564 | 0.29158263 | -1.4290205 | 0.15299834 | NA | NA | NA |
| ENSG000002 | 1.70266141 | -0.052332649 | 0.34304018 | -0.1525554 | 0.87874886 | NA | NA | NA |
| ENSG000002 | 0.12685397 | -0.090215768 | 0.12901919 | -0.699243 | 0.48440017 | NA | C1QTNF5 | 114902 |
| ENSG000002 | 0 | NA | NA | NA | NA | NA | NA | NA |
| ENSG000002 | 0 | NA | NA | NA | NA | NA | NA | NA |
| ENSG000002 | 0.99030131 | -0.03729337 | 0.31705927 | -0.1176227 | 0.90636661 | NA | NA | NA |
| ENSG000002 | 0 | NA | NA | NA | NA | NA | NA | NA |
| ENSG000002 | 0.74498828 | -0.028170431 | 0.27889408 | -0.1010076 | 0.9195444 | NA | NA | NA |
| ENSG000002 | 1.57285889 | -0.447612078 | 0.33516222 | -1.3355088 | 0.18170989 | NA | NA | NA |
| ENSG000002 | 1.93605532 | -0.058321174 | 0.35670721 | -0.1634987 | 0.87012578 | NA | NA | NA |
| ENSG000002 | 1.29106037 | -0.829442333 | 0.32502034 | -2.5519705 | 0.01071156 | NA | BNIP3P42 | 106481697 |

| | | | | | | | | |
|---|---|---|---|---|---|---|---|---|
| ENSG000002 | 0 | NA | NA | NA | NA | NA | NA | NA |
| ENSG000002 | 0.86225311 | 0.041993224 | 0.29705792 | 0.14136375 | 0.88758259 | NA | NA | NA |
| ENSG000002 | 0 | NA | NA | NA | NA | NA | NA | NA |
| ENSG000002 | 2.09999422 | -0.004531838 | 0.36743223 | -0.0123338 | 0.9901593 | NA | SMCR2 | 105371564 |
| ENSG000002 | 1.84769996 | -0.463075436 | 0.3466759 | -1.3357589 | 0.18162807 | NA | NA | NA |
| ENSG000002 | 1.29564166 | -0.666808983 | 0.329186 | -2.0256299 | 0.04280273 | NA | NA | NA |
| ENSG000002 | 0.37831254 | -0.102004459 | 0.21855316 | -0.4667261 | 0.64069587 | NA | NA | NA |
| ENSG000002 | 1.97994782 | -0.193922937 | 0.36446317 | -0.5320783 | 0.59467178 | NA | NA | NA |
| ENSG000002 | 2.04791124 | -0.44624473 | 0.35873264 | -1.2439479 | 0.21351872 | NA | LINC01441 | 101927796 |
| ENSG000002 | 1.13423659 | -0.58937406 | 0.31917699 | -1.8465431 | 0.06481339 | NA | NA | NA |
| ENSG000002 | 1.79773731 | -0.116888699 | 0.35461223 | -0.329624 | 0.74168409 | NA | NA | NA |
| ENSG000002 | 0 | NA | NA | NA | NA | NA | NA | NA |
| ENSG000002 | 0 | NA | NA | NA | NA | NA | NA | NA |
| ENSG000002 | 0.23202362 | -0.008302493 | 0.17586207 | -0.0472103 | 0.96234565 | NA | NA | NA |
| ENSG000002 | 0.7872583 | -0.560227253 | 0.2858936 | -1.9595656 | 0.05004659 | NA | NA | NA |
| ENSG000002 | 1.23649482 | -0.046741475 | 0.32431292 | -0.1441246 | 0.88540207 | NA | NA | NA |
| ENSG000002 | 0 | NA | NA | NA | NA | NA | NA | NA |
| ENSG000002 | 1.74013135 | -0.442103369 | 0.3484469 | -1.2687826 | 0.20451861 | NA | NA | NA |
| ENSG000002 | 0.24619405 | -0.007630671 | 0.17915639 | -0.0425922 | 0.96602658 | NA | NA | NA |
| ENSG000002 | 0.63210694 | -0.105863447 | 0.26563058 | -0.3985364 | 0.69023486 | NA | NA | NA |
| ENSG000002 | 1.67321916 | -0.052984007 | 0.35146845 | -0.1507504 | 0.88017262 | NA | NA | NA |
| ENSG000002 | 1.10119036 | -0.112716949 | 0.32037233 | -0.3518311 | 0.72496493 | NA | NA | NA |
| ENSG000002 | 0 | NA | NA | NA | NA | NA | TTTY22 | 252954 |
| ENSG000002 | 1.83827962 | 0.008879726 | 0.35661865 | 0.02489978 | 0.9801349 | NA | NA | NA |
| ENSG000002 | 1.89407851 | -0.650065846 | 0.35544385 | -1.8288848 | 0.06741687 | NA | NA | NA |
| ENSG000002 | 1.16195701 | -0.611962471 | 0.32373155 | -1.8903393 | 0.0587126 | NA | NA | NA |
| ENSG000002 | 0.99238515 | -0.346602368 | 0.30863497 | -1.1230171 | 0.26143021 | NA | NA | NA |
| ENSG000002 | 0 | NA | NA | NA | NA | NA | CT47A10 | 728036 |
| ENSG000002 | 2.10499195 | -0.47767259 | 0.36104412 | -1.3230311 | 0.18582505 | NA | NA | NA |

| | | | | | | | | |
|---|---|---|---|---|---|---|---|---|
| ENSG000002 | 1.95638255 | -0.182484308 | 0.36020841 | -0.5066076 | 0.6124302 | NA | NA | NA |
| ENSG000002 | 1.3035496 | 0.449081495 | 0.32891812 | 1.36532914 | 0.17214962 | NA | NA | NA |
| ENSG000002 | 2.09080573 | -0.537504802 | 0.35316955 | -1.5219455 | 0.12802274 | NA | LINC00633 | 100129515 |
| ENSG000002 | 1.80808517 | 0.405781971 | 0.35602293 | 1.13976358 | 0.25438481 | NA | NA | NA |
| ENSG000002 | 1.82312638 | -0.373504014 | 0.35672753 | -1.0470288 | 0.29508628 | NA | NA | NA |
| ENSG000002 | 0.62821079 | -0.27685072 | 0.26713242 | -1.0363801 | 0.30002486 | NA | NA | NA |
| ENSG000002 | 1.28698719 | -0.315791477 | 0.31832182 | -0.992051 | 0.32117265 | NA | NA | NA |
| ENSG000002 | 2.09761939 | -0.506987469 | 0.36452413 | -1.3908201 | 0.16428 | NA | NA | NA |
| ENSG000002 | 1.39836997 | -0.712620308 | 0.33261404 | -2.1424841 | 0.03215454 | NA | NA | NA |
| ENSG000002 | 0 | NA | NA | NA | NA | NA | NA | NA |
| ENSG000002 | 0.11934008 | 0.083564698 | 0.12760091 | 0.65489106 | 0.5125379 | NA | NA | NA |
| ENSG000002 | 1.59637076 | -0.125315202 | 0.35346452 | -0.354534 | 0.7229387 | NA | NA | NA |
| ENSG000002 | 1.91944704 | 0.059802855 | 0.35004304 | 0.17084429 | 0.8643462 | NA | NA | NA |
| ENSG000002 | 0 | NA | NA | NA | NA | NA | NA | NA |
| ENSG000002 | 0 | NA | NA | NA | NA | NA | HSFY6P | 106480722 |
| ENSG000002 | 0.89615388 | -0.429446277 | 0.29554868 | -1.4530475 | 0.14621056 | NA | NA | NA |
| ENSG000002 | 1.23515679 | -0.053033599 | 0.33016043 | -0.1606298 | 0.87238499 | NA | NA | NA |
| ENSG000002 | 1.46541212 | -0.17785971 | 0.34129828 | -0.5211269 | 0.60227838 | NA | NA | NA |
| ENSG000002 | 1.02943762 | -0.176037254 | 0.30126548 | -0.584326 | 0.55900099 | NA | NA | NA |
| ENSG000002 | 0.88718517 | -0.272269348 | 0.29987432 | -0.9079449 | 0.36390735 | NA | NA | NA |
| ENSG000002 | 0 | NA | NA | NA | NA | NA | NA | NA |
| ENSG000002 | 1.20504764 | -0.180552496 | 0.32606759 | -0.5537272 | 0.57976555 | NA | NA | NA |
| ENSG000002 | 1.9648916 | 0.293295969 | 0.35086575 | 0.83592077 | 0.40319948 | NA | NA | NA |
| ENSG000002 | 1.24019956 | -0.1852684 | 0.32932573 | -0.5625689 | 0.5737285 | NA | NA | NA |
| ENSG000002 | 1.28249077 | -0.514441873 | 0.33258783 | -1.546785 | 0.12191509 | NA | NA | NA |
| ENSG000002 | 0.6265601 | -0.275936484 | 0.2670182 | -1.0333995 | 0.30141697 | NA | NA | NA |
| ENSG000002 | 0 | NA | NA | NA | NA | NA | NA | NA |
| ENSG000002 | 0.49168123 | -0.197384925 | 0.247356 | -0.7979791 | 0.4248826 | NA | LINC00403 | 100505996 |
| ENSG000002 | 1.23236661 | 0.101453381 | 0.33001379 | 0.30742164 | 0.75852246 | NA | NA | NA |

| | | | | | | | | |
|---|---|---|---|---|---|---|---|---|
| ENSG000002 | 0.8737197 | -0.255299395 | 0.29348454 | -0.8698904 | 0.38436029 | NA | NA | NA |
| ENSG000002 | 1.96277952 | 0.06181281 | 0.3632677 | 0.17015774 | 0.86488609 | NA | VWC2L-IT1 | 100885780 |
| ENSG000002 | 1.8325404 | 0.140685389 | 0.36055971 | 0.3901861 | 0.69639894 | NA | NA | NA |
| ENSG000002 | 0.99121266 | -0.205080843 | 0.31756778 | -0.645786 | 0.51841792 | NA | NA | NA |
| ENSG000002 | 0.49512483 | -0.344267812 | 0.23617548 | -1.457678 | 0.14492931 | NA | NA | NA |
| ENSG000002 | 0.77860225 | -0.362322636 | 0.28273614 | -1.2814868 | 0.20002273 | NA | NA | NA |
| ENSG000002 | 1.72618725 | -0.06421755 | 0.35325469 | -0.1817882 | 0.85574891 | NA | NA | NA |
| ENSG000002 | 2.13919312 | -0.23986608 | 0.35986376 | -0.6665469 | 0.50506158 | NA | NA | NA |
| ENSG000002 | 1.41461117 | -0.420707232 | 0.33901265 | -1.240978 | 0.21461389 | NA | NA | NA |
| ENSG000002 | 1.78609249 | -0.479369945 | 0.35547314 | -1.3485405 | 0.17748459 | NA | NA | NA |
| ENSG000002 | 1.92367541 | -0.253287534 | 0.35442124 | -0.7146511 | 0.47482465 | NA | NA | NA |
| ENSG000002 | 1.1373057 | -0.280478515 | 0.32612245 | -0.8600405 | 0.38976672 | NA | NA | NA |
| ENSG000002 | 1.86994681 | -0.531550253 | 0.35888142 | -1.4811306 | 0.13857177 | NA | NA | NA |
| ENSG000002 | 1.27153272 | -0.196168992 | 0.32342423 | -0.6065377 | 0.54415775 | NA | NA | NA |
| ENSG000002 | 0.36538978 | 0.059033792 | 0.20773944 | 0.28417229 | 0.77627835 | NA | NA | NA |
| ENSG000002 | 1.04742793 | -0.339245138 | 0.30303175 | -1.1195036 | 0.26292536 | NA | NA | NA |
| ENSG000002 | 0 | NA | NA | NA | NA | NA | NA | NA |
| ENSG000002 | 1.76367096 | -0.335775992 | 0.35452236 | -0.9471222 | 0.34357652 | NA | NA | NA |
| ENSG000002 | 1.00558812 | -0.503374107 | 0.30654321 | -1.6420984 | 0.10056962 | NA | NA | NA |
| ENSG000002 | 1.31111612 | 0.333444614 | 0.33856756 | 0.98486877 | 0.32468854 | NA | NA | NA |
| ENSG000002 | 1.12282262 | -0.256369936 | 0.31745204 | -0.8075864 | 0.41932875 | NA | NA | NA |
| ENSG000002 | 1.86433559 | -0.124874245 | 0.36066368 | -0.3462346 | 0.72916641 | NA | NA | NA |
| ENSG000002 | 0.61238967 | -0.276614426 | 0.26576012 | -1.0408425 | 0.29794866 | NA | NA | NA |
| ENSG000002 | 0.51364788 | -0.186913858 | 0.24261662 | -0.7704083 | 0.44105773 | NA | NA | NA |
| ENSG000002 | 0.53627793 | -0.360851627 | 0.24080453 | -1.4985251 | 0.13399688 | NA | NA | NA |
| ENSG000002 | 1.6618462 | 0.354234762 | 0.35289319 | 1.00380165 | 0.31547423 | NA | NA | NA |
| ENSG000002 | 1.50648792 | -0.62331103 | 0.3409861 | -1.8279661 | 0.06755464 | NA | NA | NA |
| ENSG000002 | 0 | NA | NA | NA | NA | NA | NA | NA |
| ENSG000002 | 0.46273095 | 0.157073337 | 0.24331041 | 0.64556769 | 0.51855937 | NA | NA | NA |

| | | | | | | | | |
|---|---|---|---|---|---|---|---|---|
| ENSG000002 | 0 | NA | NA | NA | NA | NA | NA | NA |
| ENSG000002 | 0.3575613 | 0.072305152 | 0.21497462 | 0.33634273 | 0.73661243 | NA | NA | NA |
| ENSG000002 | 0.77476221 | -0.188343058 | 0.28188047 | -0.6681664 | 0.50402738 | NA | NA | NA |
| ENSG000002 | 1.70942041 | -0.058923033 | 0.33696533 | -0.1748638 | 0.86118667 | NA | NA | NA |
| ENSG000002 | 0.99203969 | -0.323421637 | 0.3006454 | -1.0757578 | 0.28203558 | NA | NA | NA |
| ENSG000002 | 1.89265224 | -0.397789286 | 0.35930928 | -1.1070944 | 0.26825312 | NA | NA | NA |
| ENSG000002 | 0 | NA | NA | NA | NA | NA | NA | NA |
| ENSG000002 | 1.35691956 | -0.388038162 | 0.332737 | -1.1662008 | 0.24353326 | NA | NA | NA |
| ENSG000002 | 2.1090556 | -0.860685402 | 0.35793905 | -2.4045586 | 0.01619201 | NA | NA | NA |
| ENSG000002 | 0.23065526 | 0.167384503 | 0.17662711 | 0.94767164 | 0.34329664 | NA | ATP5J2LP | 101241877 |
| ENSG000002 | 1.56429537 | 0.153234616 | 0.34748143 | 0.44098648 | 0.65922278 | NA | NA | NA |
| ENSG000002 | 0.97407972 | -0.314513624 | 0.30067042 | -1.0460411 | 0.29554204 | NA | NA | NA |
| ENSG000002 | 1.53064953 | -0.336653813 | 0.34322641 | -0.9808505 | 0.32666645 | NA | NA | NA |
| ENSG000002 | 1.62926926 | -0.272994301 | 0.35422527 | -0.7706799 | 0.44089669 | NA | NA | NA |
| ENSG000002 | 0 | NA | NA | NA | NA | NA | NA | NA |
| ENSG000002 | 0.85131777 | -0.102855071 | 0.29519776 | -0.3484277 | 0.72751902 | NA | NA | NA |
| ENSG000002 | 2.08218666 | -0.254045434 | 0.36589718 | -0.6943083 | 0.48748891 | NA | NA | NA |
| ENSG000002 | 0.74026318 | -0.19301616 | 0.28568604 | -0.6756234 | 0.49927981 | NA | NA | NA |
| ENSG000002 | 0.39479706 | -0.300008631 | 0.22317059 | -1.3443018 | 0.17885081 | NA | NA | NA |
| ENSG000002 | 1.64953707 | -0.510386537 | 0.34287264 | -1.4885601 | 0.13660324 | NA | NA | NA |
| ENSG000002 | 1.49615297 | -0.188109813 | 0.3449735 | -0.5452877 | 0.58555564 | NA | NA | NA |
| ENSG000002 | 2.13967187 | -0.126356988 | 0.36170952 | -0.3493328 | 0.7268395 | NA | NA | NA |
| ENSG000002 | 1.01013907 | -0.160240613 | 0.29351615 | -0.5459346 | 0.58511092 | NA | NA | NA |
| ENSG000002 | 0.36621735 | -0.097238239 | 0.21673053 | -0.4486596 | 0.65367721 | NA | NA | NA |
| ENSG000002 | 1.73850294 | -0.308980889 | 0.35207977 | -0.8775877 | 0.38016749 | NA | NA | NA |
| ENSG000002 | 0.76536895 | -0.034725567 | 0.28026919 | -0.1239008 | 0.90139385 | NA | NA | NA |
| ENSG000002 | 1.67219801 | -0.867127591 | 0.34801465 | -2.4916411 | 0.01271545 | NA | NA | NA |
| ENSG000002 | 0 | NA | NA | NA | NA | NA | NA | NA |
| ENSG000002 | 0.38651158 | -0.274258716 | 0.21461355 | -1.2779189 | 0.20127801 | NA | NA | NA |

| | | | | | | | | |
|---|---|---|---|---|---|---|---|---|
| ENSG000002 | 0 | NA | NA | NA | NA | NA | NA | NA |
| ENSG000002 | 1.13987124 | -0.121108491 | 0.31447087 | -0.3851183 | 0.70014976 | NA | NA | NA |
| ENSG000002 | 0.37304803 | -0.090842893 | 0.20939682 | -0.4338313 | 0.66441096 | NA | NA | NA |
| ENSG000002 | 1.48070297 | -0.311800701 | 0.34025469 | -0.9163744 | 0.35947054 | NA | NA | NA |
| ENSG000002 | 0 | NA | NA | NA | NA | NA | NA | NA |
| ENSG000002 | 0.9608773 | -0.059240678 | 0.20215151 | -0.2930509 | 0.76948326 | NA | NA | NA |
| ENSG000002 | 1.34419268 | -0.254404909 | 0.33430786 | -0.7609899 | 0.4466631 | NA | NA | NA |
| ENSG000002 | 0.98819616 | -0.449758006 | 0.2956505 | -1.5212489 | 0.12819738 | NA | NA | NA |
| ENSG000002 | 1.56226791 | -0.595738192 | 0.33521193 | -1.7771986 | 0.07553556 | NA | NA | NA |
| ENSG000002 | 1.69354014 | -0.264668857 | 0.34051071 | -0.7772703 | 0.43699932 | NA | NA | NA |
| ENSG000002 | 0.76656263 | -0.039500232 | 0.27864491 | -0.1417583 | 0.88727092 | NA | NA | NA |
| ENSG000002 | 1.49302841 | -0.176784503 | 0.33959181 | -0.5205794 | 0.6026598 | NA | NA | NA |
| ENSG000002 | 1.34178414 | -0.362162506 | 0.32530627 | -1.113297 | 0.26558088 | NA | NA | NA |
| ENSG000002 | 2.05096096 | -0.707553683 | 0.35812964 | -1.9756915 | 0.04818973 | NA | NA | NA |
| ENSG000002 | 0.25370795 | -0.171331747 | 0.17411721 | -0.9840024 | 0.32511435 | NA | NA | NA |
| ENSG000002 | 1.53486473 | -0.340269823 | 0.34556479 | -0.9846773 | 0.32478258 | NA | NA | NA |
| ENSG000002 | 1.72239771 | -0.055042127 | 0.3549597 | -0.1550658 | 0.87676941 | NA | NA | NA |
| ENSG000002 | 0 | NA | NA | NA | NA | NA | NA | NA |
| ENSG000002 | 0 | NA | NA | NA | NA | NA | NA | NA |
| ENSG000002 | 0.50204195 | -0.181972198 | 0.24131623 | -0.7540819 | 0.45080006 | NA | NA | NA |
| ENSG000002 | 0.76830156 | -0.364532338 | 0.28365742 | -1.2851147 | 0.19875219 | NA | NA | NA |
| ENSG000002 | 1.62502428 | -0.125032218 | 0.35154488 | -0.355665 | 0.72209144 | NA | NA | NA |
| ENSG000002 | 1.271555 | -0.049522394 | 0.32962218 | -0.1502399 | 0.88057537 | NA | NA | NA |
| ENSG000002 | 1.26301965 | -0.1925383 | 0.32991709 | -0.583596 | 0.55949214 | NA | NA | NA |
| ENSG000002 | 0.38019136 | -0.10756138 | 0.21876967 | -0.491665 | 0.62295621 | NA | NA | NA |
| ENSG000002 | 2.03773681 | -0.684684993 | 0.35541072 | -1.9264613 | 0.0540468 | NA | NA | NA |
| ENSG000002 | 0.87878971 | -0.122783908 | 0.30329377 | -0.4048349 | 0.68559886 | NA | NA | NA |
| ENSG000002 | 1.43663152 | 0.2276317 | 0.34297973 | 0.6636885 | 0.5068897 | NA | NA | NA |
| ENSG000002 | 0.74268725 | -0.479320364 | 0.26912835 | -1.7810103 | 0.07491077 | NA | NA | NA |

| | | | | | | | | |
|---|---|---|---|---|---|---|---|---|
| ENSG000002 | 1.36365987 | -0.260601918 | 0.33551163 | -0.7767299 | 0.43731814 | NA | NA | NA |
| ENSG000002 | 0 | NA | NA | NA | NA | NA | NA | NA |
| ENSG000002 | 0.12070844 | -0.088503695 | 0.12783931 | -0.6923042 | 0.48874631 | NA | NA | NA |
| ENSG000002 | 1.83262672 | -0.245948922 | 0.3572376 | -0.6884743 | 0.49115412 | NA | NA | NA |
| ENSG000002 | 1.74144155 | -1.010614395 | 0.34263729 | -2.9495166 | 0.00318271 | NA | NA | NA |
| ENSG000002 | 1.85859095 | -0.390870493 | 0.36000011 | -1.085751 | 0.27758916 | NA | NA | NA |
| ENSG000002 | 0.62431017 | -0.119347761 | 0.27124333 | -0.4400026 | 0.65993525 | NA | NA | NA |
| ENSG000002 | 1.71016208 | -0.326983077 | 0.35615335 | -0.9180963 | 0.35856845 | NA | NA | NA |
| ENSG000002 | 0 | NA | NA | NA | NA | NA | NA | NA |
| ENSG000002 | 2.09331625 | -0.693124889 | 0.35680541 | -1.9425852 | 0.0520663 | NA | NA | NA |
| ENSG000002 | 1.63898047 | -0.549933347 | 0.34953821 | -1.5733139 | 0.11564614 | NA | NA | NA |
| ENSG000002 | 1.21091297 | -0.171024181 | 0.32167988 | -0.5316596 | 0.59496181 | NA | NA | NA |
| ENSG000002 | 0.72540953 | -0.028108977 | 0.28387695 | -0.0990182 | 0.92112384 | NA | NA | NA |
| ENSG000002 | 1.42210388 | -0.731005464 | 0.3357513 | -2.177223 | 0.02946393 | NA | NA | NA |
| ENSG000002 | 1.92843319 | -0.675949492 | 0.35778454 | -1.8892641 | 0.05885645 | NA | LINC00885 | 401109 |
| ENSG000002 | 0.2596576 | -0.193002035 | 0.1837913 | -1.0501152 | 0.29366517 | NA | NA | NA |
| ENSG000002 | 0 | NA | NA | NA | NA | NA | NA | NA |
| ENSG000002 | 1.6299544 | -0.272785719 | 0.35423406 | -0.770072 | 0.4412572 | NA | NA | NA |
| ENSG000002 | 1.40003984 | -0.122333927 | 0.33331021 | -0.3670272 | 0.7135987 | NA | NA | NA |
| ENSG000002 | 2.11022366 | -0.511329651 | 0.36540483 | -1.3993511 | 0.16170773 | NA | NA | NA |
| ENSG000002 | 0.92102939 | -0.436637211 | 0.29681748 | -1.471063 | 0.14127408 | NA | NA | NA |
| ENSG000002 | 1.14012165 | 0.024948849 | 0.30659338 | 0.08137439 | 0.93514422 | NA | NA | NA |
| ENSG000002 | 0.37928001 | 0.070449581 | 0.21829793 | 0.32272217 | 0.74690566 | NA | PDIA3P2 | 106481687 |
| ENSG000002 | 0.60375535 | 0.060734453 | 0.26281561 | 0.23109149 | 0.81724372 | NA | NA | NA |
| ENSG000002 | 0 | NA | NA | NA | NA | NA | NA | NA |
| ENSG000002 | 0 | NA | NA | NA | NA | NA | NA | NA |
| ENSG000002 | 1.86479205 | -0.125477882 | 0.35812558 | -0.350374 | 0.72605805 | NA | NA | NA |
| ENSG000002 | 1.77556401 | -0.450290854 | 0.35186739 | -1.2797175 | 0.20064451 | NA | NA | NA |
| ENSG000002 | 0 | NA | NA | NA | NA | NA | NA | NA |

| | | | | | | | | |
|---|---|---|---|---|---|---|---|---|
| ENSG000002 | 0.35824644 | 0.072568381 | 0.21508272 | 0.33739754 | 0.73581724 | NA | NA | NA |
| ENSG000002 | 0 | NA | NA | NA | NA | NA | NA | NA |
| ENSG000002 | 0.22336757 | 0.163792188 | 0.17487321 | 0.93663397 | 0.34894687 | NA | NA | NA |
| ENSG000002 | 0.58608925 | 0.207285028 | 0.2535965 | 0.81738127 | 0.41371058 | NA | NA | NA |
| ENSG000002 | 0.50377641 | -0.188767022 | 0.24148558 | -0.7816906 | 0.4343964 | NA | NA | NA |
| ENSG000002 | 2.16202173 | -0.24032873 | 0.35991776 | -0.6677324 | 0.50430446 | NA | NA | NA |
| ENSG000002 | 1.38643586 | -0.253341452 | 0.33202357 | -0.7630225 | 0.44544998 | NA | NA | NA |
| ENSG000002 | 0.87554252 | 0.04253079 | 0.29775404 | 0.14283866 | 0.8864176 | NA | NA | NA |
| ENSG000002 | 1.5571491 | -0.090583241 | 0.3360715 | -0.2695356 | 0.78751753 | NA | NA | NA |
| ENSG000002 | 2.0239278 | -0.583482204 | 0.36107176 | -1.615973 | 0.10610016 | NA | NA | NA |
| ENSG000002 | 2.04962698 | -0.438332406 | 0.35875085 | -1.2218296 | 0.22177208 | NA | LINC01167 | 103695432 |
| ENSG000002 | 0.2596576 | -0.193002035 | 0.1837913 | -1.0501152 | 0.29366517 | NA | NA | NA |
| ENSG000002 | 1.15216285 | -0.12281731 | 0.32201543 | -0.3814019 | 0.70290502 | NA | NA | NA |
| ENSG000002 | 2.16105426 | -0.385335132 | 0.36582448 | -1.0533334 | 0.29218822 | NA | NA | NA |
| ENSG000002 | 0.48032325 | -0.179266136 | 0.23878452 | -0.7507444 | 0.45280651 | NA | NA | NA |
| ENSG000002 | 1.9494564 | 0.440927674 | 0.35763627 | 1.23289416 | 0.21761525 | NA | NA | NA |
| ENSG000002 | 0.72976056 | -0.027640562 | 0.27783407 | -0.0994859 | 0.92075251 | NA | NA | NA |
| ENSG000002 | 1.65454045 | -0.484307529 | 0.3364311 | -1.4395444 | 0.14999633 | NA | IQCF4 | 100506840 |
| ENSG000002 | 1.11009436 | -0.256908689 | 0.31863869 | -0.8062696 | 0.42008741 | NA | NA | NA |
| ENSG000002 | 0.9908421 | -0.044700626 | 0.31294954 | -0.1428365 | 0.88641929 | NA | NA | NA |
| ENSG000002 | 1.89125784 | 0.565673847 | 0.35168224 | 1.60847998 | 0.1077301 | NA | LINC00853 | 100874253 |
| ENSG000002 | 1.45935194 | -0.053201038 | 0.34671931 | -0.1534412 | 0.87805033 | NA | NA | NA |
| ENSG000002 | 1.50008392 | -0.468680459 | 0.34313908 | -1.3658615 | 0.17198243 | NA | NA | NA |
| ENSG000002 | 0.7442814 | -0.19995417 | 0.28591302 | -0.6993531 | 0.48433137 | NA | NA | NA |
| ENSG000002 | 0 | NA | NA | NA | NA | NA | NA | NA |
| ENSG000002 | 1.88996758 | -0.125121649 | 0.35351137 | -0.3539395 | 0.7233842 | NA | NA | NA |
| ENSG000002 | 0 | NA | NA | NA | NA | NA | NA | NA |
| ENSG000002 | 1.32040277 | 0.026419059 | 0.33847199 | 0.0780539 | 0.93778518 | NA | NA | NA |
| ENSG000002 | 1.5037745 | -0.053404778 | 0.34152445 | -0.1563718 | 0.87574 | NA | NA | NA |

| | | | | | | | | |
|---|---|---|---|---|---|---|---|---|
| ENSG000002 | 0 | NA | NA | NA | NA | NA | NA | NA |
| ENSG000002 | 0 | NA | NA | NA | NA | NA | USP9YP25 | 106480715 |
| ENSG000002 | 0 | NA | NA | NA | NA | NA | NA | NA |
| ENSG000002 | 1.13972978 | -0.566869226 | 0.31589767 | -1.7944711 | 0.07273801 | NA | NA | NA |
| ENSG000002 | 2.17157419 | 0.296878334 | 0.36590501 | 0.81135356 | 0.41716266 | NA | NA | NA |
| ENSG000002 | 0.79716359 | -0.340510436 | 0.27496505 | -1.2383771 | 0.21557626 | NA | NA | NA |
| ENSG000002 | 0.76033119 | -0.194152683 | 0.28698065 | -0.6765358 | 0.49870053 | NA | GAGE12H | 729442 |
| ENSG000002 | 1.84523738 | -0.330653089 | 0.34795263 | -0.9502819 | 0.34196903 | NA | NA | NA |
| ENSG000002 | 0.35870346 | 0.077368606 | 0.21515516 | 0.35959447 | 0.71915042 | NA | APOC2 | 344 |
| ENSG000002 | 0 | NA | NA | NA | NA | NA | NA | NA |
| ENSG000002 | 0.75304351 | -0.205958918 | 0.29162776 | -0.7062391 | 0.48003948 | NA | NA | NA |
| ENSG000002 | 0.11934008 | 0.083564698 | 0.12760091 | 0.65489106 | 0.5125379 | NA | NA | NA |
| ENSG000002 | 1.22695829 | 0.107113873 | 0.33338994 | 0.32128706 | 0.74799286 | NA | NA | NA |
| ENSG000002 | 0.76169956 | -0.374892198 | 0.28875268 | -1.2983159 | 0.19417881 | NA | NA | NA |
| ENSG000002 | 0.62682069 | -0.281617951 | 0.26695108 | -1.0549422 | 0.29145177 | NA | NA | NA |
| ENSG000002 | 0.35824644 | 0.072568381 | 0.21508272 | 0.33739754 | 0.73581724 | NA | NA | NA |
| ENSG000002 | 1.09000653 | -0.25629258 | 0.31628782 | -0.8103144 | 0.4177595 | NA | NA | NA |
| ENSG000002 | 1.72487734 | -0.05221675 | 0.34915703 | -0.1495509 | 0.88111895 | NA | NA | NA |
| ENSG000002 | 0 | NA | NA | NA | NA | NA | NA | NA |
| ENSG000002 | 0.63504146 | -0.278824355 | 0.26770076 | -1.0415524 | 0.29761927 | NA | NA | NA |
| ENSG000002 | 0 | NA | NA | NA | NA | NA | NA | NA |
| ENSG000002 | 1.64042264 | -0.532777645 | 0.34725311 | -1.5342631 | 0.12496493 | NA | NA | NA |
| ENSG000002 | 0.93204246 | -0.430357711 | 0.29493135 | -1.4591793 | 0.14451577 | NA | NA | NA |
| ENSG000002 | 1.54579215 | 0.026569277 | 0.33869131 | 0.07844688 | 0.93747258 | NA | NA | NA |
| ENSG000002 | 1.94798296 | -0.050523751 | 0.36269538 | -0.1393008 | 0.88921247 | NA | NA | NA |
| ENSG000002 | 1.30774059 | 0.158387617 | 0.32986362 | 0.48016092 | 0.63111298 | NA | NA | NA |
| ENSG000002 | 0 | NA | NA | NA | NA | NA | NA | NA |
| ENSG000002 | 1.78128662 | 0.130527155 | 0.35508229 | 0.36759692 | 0.71317381 | NA | LINC01526 | 101928770 |
| ENSG000002 | 2.03929054 | -0.699561784 | 0.35823826 | -1.9527836 | 0.05084525 | NA | NA | NA |

| | | | | | | | | |
|---|---|---|---|---|---|---|---|---|
| ENSG000002 | 1.93100144 | 0.062971221 | 0.35862201 | 0.17559218 | 0.86061435 | NA | NA | NA |
| ENSG000002 | 0.75871242 | -0.194288255 | 0.2808106 | -0.6918836 | 0.48901042 | NA | NA | NA |
| ENSG000002 | 1.83422977 | -0.759636095 | 0.35215886 | -2.1570836 | 0.03099915 | NA | NA | NA |
| ENSG000002 | 1.76873279 | -0.454247385 | 0.35107422 | -1.2938785 | 0.19570739 | NA | NA | NA |
| ENSG000002 | 0.22263036 | 0.146922267 | 0.16600367 | 0.88505432 | 0.37612734 | NA | NA | NA |
| ENSG000002 | 1.84293538 | -0.243731518 | 0.35729369 | -0.6821602 | 0.49513769 | NA | NA | NA |
| ENSG000002 | 0.86046309 | -0.246260481 | 0.29102415 | -0.8461857 | 0.39744914 | NA | NA | NA |
| ENSG000002 | 1.94719406 | -0.819883267 | 0.35645389 | -2.3001103 | 0.02144197 | NA | NA | NA |
| ENSG000002 | 1.38032066 | -0.416483401 | 0.3369171 | -1.2361599 | 0.21639914 | NA | NA | NA |
| ENSG000002 | 1.64819512 | -0.266089729 | 0.35050104 | -0.7591696 | 0.44775112 | NA | NA | NA |
| ENSG000002 | 1.23784199 | -0.34175653 | 0.33020316 | -1.0349887 | 0.3006742 | NA | NA | NA |
| ENSG000002 | 1.21201628 | -0.042906594 | 0.32953263 | -0.1302044 | 0.89640472 | NA | NA | NA |
| ENSG000002 | 1.66129257 | -0.561241171 | 0.35176165 | -1.5955155 | 0.110597 | NA | NA | NA |
| ENSG000002 | 1.56633624 | 0.022045958 | 0.34165203 | 0.06452752 | 0.9485502 | NA | NA | NA |
| ENSG000002 | 2.07284951 | 0.120798481 | 0.36366788 | 0.33216704 | 0.73976313 | NA | NA | NA |
| ENSG000002 | 1.20842206 | -0.03358472 | 0.31037221 | -0.1082079 | 0.91383079 | NA | NA | NA |
| ENSG000002 | 0.47781273 | -0.018045496 | 0.22812977 | -0.0791019 | 0.93695158 | NA | NA | NA |
| ENSG000002 | 0.5935178 | -0.088291727 | 0.25308915 | -0.3488562 | 0.72719725 | NA | NA | NA |
| ENSG000002 | 1.12589566 | -0.270975439 | 0.32347778 | -0.8376941 | 0.40220251 | NA | DPY19L2P5 | 106480421 |
| ENSG000002 | 0.87901647 | -0.275034858 | 0.29942893 | -0.9185314 | 0.35834075 | NA | NA | NA |
| ENSG000002 | 1.37849583 | -0.275466927 | 0.34171187 | -0.806138 | 0.42016328 | NA | NA | NA |
| ENSG000002 | 0.47280936 | -0.012478229 | 0.23633508 | -0.0527989 | 0.95789215 | NA | NA | NA |
| ENSG000002 | 1.50842285 | -0.329738961 | 0.34286281 | -0.9617227 | 0.3361889 | NA | NA | NA |
| ENSG000002 | 0 | NA | NA | NA | NA | NA | NA | NA |
| ENSG000002 | 0.11934008 | 0.083564698 | 0.12760091 | 0.65489106 | 0.5125379 | NA | NA | NA |
| ENSG000002 | 0.8602884 | -0.106640097 | 0.29730239 | -0.3586924 | 0.71982524 | NA | NA | NA |
| ENSG000002 | 1.24050562 | -0.052863681 | 0.31464785 | -0.168009 | 0.86657617 | NA | NA | NA |
| ENSG000002 | 0.87210284 | -0.116133774 | 0.29790572 | -0.389834 | 0.69665931 | NA | NA | NA |
| ENSG000002 | 1.49555215 | -0.19413231 | 0.34618578 | -0.5607749 | 0.57495098 | NA | NA | NA |

| | | | | | | | | |
|---|---|---|---|---|---|---|---|---|
| ENSG000002 | 0.98867044 | -0.02671387 | 0.29859721 | -0.0894646 | 0.92871271 | NA | NA | NA |
| ENSG000002 | 2.00648041 | -0.306329375 | 0.36043026 | -0.8498992 | 0.39538115 | NA | NA | NA |
| ENSG000002 | 0.38133351 | -0.102466351 | 0.2189713 | -0.4679442 | 0.63982449 | NA | NA | NA |
| ENSG000002 | 1.97605358 | -0.186646577 | 0.36225603 | -0.5152339 | 0.60638961 | NA | NA | NA |
| ENSG000002 | 1.61368128 | -0.470179115 | 0.33672878 | -1.396314 | 0.16261995 | NA | NA | NA |
| ENSG000002 | 1.62286259 | -0.275193704 | 0.35414273 | -0.77707 | 0.43711745 | NA | NA | NA |
| ENSG000002 | 1.95279237 | 0.072668445 | 0.36262634 | 0.20039484 | 0.8411718 | NA | NA | NA |
| ENSG000002 | 0.82032451 | 0.203072529 | 0.29363926 | 0.69157145 | 0.4892065 | NA | NA | NA |
| ENSG000002 | 0.37899768 | -0.10176489 | 0.21865765 | -0.4654074 | 0.64163973 | NA | NA | NA |
| ENSG000002 | 0.83902224 | 0.203987147 | 0.29630441 | 0.68843777 | 0.49117714 | NA | NA | NA |
| ENSG000002 | 1.76270596 | -0.317922564 | 0.3519804 | -0.9032394 | 0.36639885 | NA | NA | NA |
| ENSG000002 | 0.98187394 | 0.108706565 | 0.29758993 | 0.3652898 | 0.71489509 | NA | NA | NA |
| ENSG000002 | 0 | NA | NA | NA | NA | NA | NA | NA |
| ENSG000002 | 0.13280363 | -0.093394593 | 0.13115644 | -0.7120855 | 0.47641185 | NA | NA | NA |
| ENSG000002 | 1.51328634 | -0.627917275 | 0.34227032 | -1.8345654 | 0.06657011 | NA | NA | NA |
| ENSG000002 | 1.09712007 | -0.101581159 | 0.31553898 | -0.321929 | 0.74750646 | NA | NA | NA |
| ENSG000002 | 1.88457774 | -0.249383959 | 0.35639757 | -0.6997353 | 0.48409263 | NA | NA | NA |
| ENSG000002 | 1.20572026 | 0.390737729 | 0.31986706 | 1.22156287 | 0.22187298 | NA | NA | NA |
| ENSG000002 | 1.23040135 | -0.048563667 | 0.33359206 | -0.145578 | 0.88425453 | NA | NA | NA |
| ENSG000002 | 0.11205239 | 0.082118251 | 0.12671962 | 0.64803108 | 0.51696485 | NA | NA | NA |
| ENSG000002 | 1.91010657 | -0.789480693 | 0.35401238 | -2.2300935 | 0.02574124 | NA | NA | NA |
| ENSG000002 | 1.52771736 | -0.645979563 | 0.34483151 | -1.8733194 | 0.06102429 | NA | NA | NA |
| ENSG000002 | 0 | NA | NA | NA | NA | NA | NA | NA |
| ENSG000002 | 0.87096123 | -0.120737462 | 0.30724927 | -0.3929626 | 0.69434713 | NA | NA | NA |
| ENSG000002 | 0.58150796 | 0.05262168 | 0.25154459 | 0.20919424 | 0.83429661 | NA | NA | NA |
| ENSG000002 | 1.91314235 | -0.497574857 | 0.3516491 | -1.4149755 | 0.15707568 | NA | NA | NA |
| ENSG000002 | 1.0869568 | 0.032733848 | 0.31484866 | 0.10396693 | 0.91719559 | NA | NA | NA |
| ENSG000002 | 0 | NA | NA | NA | NA | NA | NA | NA |
| ENSG000002 | 0.12685397 | -0.090215768 | 0.12901919 | -0.699243 | 0.48440017 | NA | NA | NA |

| | | | | | | | | |
|---|---|---|---|---|---|---|---|---|
| ENSG000002 | 1.88214963 | -0.270733173 | 0.36111456 | -0.7497155 | 0.45342608 | NA | NA | NA |
| ENSG000002 | 0.50016259 | -0.200428691 | 0.24824437 | -0.8073846 | 0.41944491 | NA | NA | NA |
| ENSG000002 | 1.35737795 | -0.258948028 | 0.33681651 | -0.7688104 | 0.44200588 | NA | NA | NA |
| ENSG000002 | 0.59490343 | 0.227912985 | 0.26313009 | 0.86616087 | 0.38640195 | NA | NA | NA |
| ENSG000002 | 1.87132776 | -0.216812742 | 0.34440324 | -0.6295317 | 0.52900105 | NA | NA | NA |
| ENSG000002 | 1.74932614 | -0.635183172 | 0.33819442 | -1.8781598 | 0.06035931 | NA | DNAJC19P4 | 106480691 |
| ENSG000002 | 1.76819511 | -0.552664021 | 0.34432491 | -1.6050655 | 0.1084794 | NA | NA | NA |
| ENSG000002 | 0 | NA | NA | NA | NA | NA | NA | NA |
| ENSG000002 | 1.45787649 | -0.184735712 | 0.34363269 | -0.5375965 | 0.59085568 | NA | USP17L3 | 645836 |
| ENSG000002 | 0.39499294 | -0.277908454 | 0.21584201 | -1.287555 | 0.19790091 | NA | LOC105377138 | 105377138 |
| ENSG000002 | 1.83815766 | -0.368051707 | 0.35367609 | -1.0406463 | 0.29803975 | NA | NA | NA |
| ENSG000002 | 0 | NA | NA | NA | NA | NA | NA | NA |
| ENSG000002 | 1.74040817 | -0.189640492 | 0.35372694 | -0.5361211 | 0.59187483 | NA | NA | NA |
| ENSG000002 | 1.52683983 | -0.343028305 | 0.34542648 | -0.9930573 | 0.32068201 | NA | NA | NA |
| ENSG000002 | 1.81603222 | 0.006291748 | 0.35912523 | 0.01751965 | 0.98602206 | NA | NA | NA |
| ENSG000002 | 1.12512565 | -0.262441568 | 0.31751435 | -0.8265503 | 0.40849202 | NA | NA | NA |
| ENSG000002 | 1.79246883 | -0.333563712 | 0.35588984 | -0.9372668 | 0.34862136 | NA | NA | NA |
| ENSG000002 | 1.02278109 | -0.350503169 | 0.30990351 | -1.1310074 | 0.25805198 | NA | NA | NA |
| ENSG000002 | 0.99030131 | -0.03729337 | 0.31705927 | -0.1176227 | 0.90636661 | NA | NA | NA |
| ENSG000002 | 0.59078107 | 0.217502869 | 0.25401457 | 0.85626139 | 0.39185321 | NA | NA | NA |
| ENSG000002 | 0 | NA | NA | NA | NA | NA | NA | NA |
| ENSG000002 | 0 | NA | NA | NA | NA | NA | IGKV1OR9-2 | 106480274 |
| ENSG000002 | 0.99818185 | -0.042262952 | 0.31324248 | -0.1349209 | 0.89267443 | NA | NA | NA |
| ENSG000002 | 1.6598681 | -0.695800072 | 0.34732523 | -2.0033099 | 0.04514403 | NA | NA | NA |
| ENSG000002 | 0.11131518 | 0.081951103 | 0.12663804 | 0.64712866 | 0.51754868 | NA | NA | NA |
| ENSG000002 | 0.11865494 | 0.083440866 | 0.12751478 | 0.65436232 | 0.5128784 | NA | NA | NA |
| ENSG000002 | 1.57886725 | -0.108104495 | 0.34354551 | -0.314673 | 0.75300995 | NA | NA | NA |
| ENSG000002 | 0.37928001 | 0.070449581 | 0.21829793 | 0.32272217 | 0.74690566 | NA | NA | NA |
| ENSG000002 | 0 | NA | NA | NA | NA | NA | NA | NA |

| | | | | | | | | |
|---|---|---|---|---|---|---|---|---|
| ENSG000002 | 0.64099112 | -0.296274405 | 0.27396996 | -1.081412 | 0.27951389 | NA | NA | NA |
| ENSG000002 | 1.89135485 | -0.938243224 | 0.35157424 | -2.6686916 | 0.00761473 | NA | NA | NA |
| ENSG000002 | 1.82932747 | -0.122833877 | 0.35942696 | -0.3417492 | 0.73253964 | NA | NA | NA |
| ENSG000002 | 1.75168429 | -0.051969405 | 0.34468625 | -0.1507731 | 0.88015473 | NA | NA | NA |
| ENSG000002 | 1.73435638 | -0.56575311 | 0.34651719 | -1.6326841 | 0.10253545 | NA | NA | NA |
| ENSG000002 | 1.27515101 | -0.51710732 | 0.33236375 | -1.5558475 | 0.11974435 | NA | NA | NA |
| ENSG000002 | 0 | NA | NA | NA | NA | NA | MTND1P12 | 106480744 |
| ENSG000002 | 0.51336555 | -0.346899948 | 0.23688898 | -1.4643988 | 0.143085 | NA | NA | NA |
| ENSG000002 | 1.01884664 | -0.349130761 | 0.30975853 | -1.1271062 | 0.25969758 | NA | NA | NA |
| ENSG000002 | 0.98347009 | -0.043356565 | 0.31265282 | -0.1386732 | 0.8897084 | NA | NA | NA |
| ENSG000002 | 0 | NA | NA | NA | NA | NA | NA | NA |
| ENSG000002 | 1.14252207 | -0.603485769 | 0.32163895 | -1.8762832 | 0.0606164 | NA | NA | NA |
| ENSG000002 | 0 | NA | NA | NA | NA | NA | NA | NA |
| ENSG000002 | 1.36067328 | -0.127819868 | 0.34243366 | -0.373269 | 0.70894825 | NA | NA | NA |
| ENSG000002 | 0.51459361 | -0.192534203 | 0.24265108 | -0.7934611 | 0.42750921 | NA | NA | NA |
| ENSG000002 | 0.75025279 | -0.033352346 | 0.29080996 | -0.1146878 | 0.90869261 | NA | NA | NA |
| ENSG000002 | 0.9726529 | -0.038824562 | 0.30731214 | -0.1263359 | 0.89946602 | NA | NA | NA |
| ENSG000002 | 0.69085396 | 0.269492921 | 0.26829421 | 1.00446789 | 0.31515314 | NA | NA | NA |
| ENSG000002 | 2.10771314 | -0.389895067 | 0.36688732 | -1.0627107 | 0.28791318 | NA | NA | NA |
| ENSG000002 | 1.8863299 | -0.351849606 | 0.3461265 | -1.0165347 | 0.30937483 | NA | NA | NA |
| ENSG000002 | 0.87073447 | 0.03701048 | 0.29150635 | 0.12696286 | 0.89896979 | NA | NA | NA |
| ENSG000002 | 1.3435186 | -0.383889833 | 0.32994502 | -1.1634964 | 0.24462818 | NA | NA | NA |
| ENSG000002 | 0.51316968 | -0.377777671 | 0.24556403 | -1.538408 | 0.12394889 | NA | LOC100240728 | 100240728 |
| ENSG000002 | 0 | NA | NA | NA | NA | NA | NA | NA |
| ENSG000002 | 2.06024817 | -1.028373482 | 0.35798782 | -2.8726494 | 0.00407046 | NA | KCNMA1-AS2 | 101929310 |
| ENSG000002 | 0 | NA | NA | NA | NA | NA | NA | NA |
| ENSG000002 | 0.88764163 | -0.277708849 | 0.29985649 | -0.9261392 | 0.35437365 | NA | NA | NA |
| ENSG000002 | 0.14128499 | -0.097772014 | 0.13403235 | -0.7294658 | 0.4657168 | NA | NA | NA |
| ENSG000002 | 0 | NA | NA | NA | NA | NA | NA | NA |

| | | | | | | | | |
|---|---|---|---|---|---|---|---|---|
| ENSG000002 | 0 | NA | NA | NA | NA | NA | TTTY16 | 252948 |
| ENSG000002 | 1.95997345 | -0.057073063 | 0.35688804 | -0.1599187 | 0.87294514 | NA | NA | NA |
| ENSG000002 | 0.24141689 | -0.165257896 | 0.17127401 | -0.9648743 | 0.33460776 | NA | NA | NA |
| ENSG000002 | 1.83613696 | -0.248758739 | 0.35827824 | -0.6943172 | 0.48748332 | NA | NA | NA |
| ENSG000002 | 1.52253874 | -0.464719669 | 0.33868585 | -1.372126 | 0.1700242 | NA | NA | NA |
| ENSG000002 | 1.56940502 | 0.274787026 | 0.34239715 | 0.80253889 | 0.4222413 | NA | NA | NA |
| ENSG000002 | 1.96664823 | -0.044081546 | 0.35333575 | -0.1247582 | 0.90071495 | NA | NA | NA |
| ENSG000002 | 2.11312785 | -0.480803833 | 0.36127534 | -1.3308515 | 0.18323787 | NA | NA | NA |
| ENSG000002 | 1.00971844 | -0.504920681 | 0.30670098 | -1.6462963 | 0.09970277 | NA | NA | NA |
| ENSG000002 | 0.62545232 | -0.114578523 | 0.27135287 | -0.4222492 | 0.67284316 | NA | NA | NA |
| ENSG000002 | 1.13758748 | -0.127700509 | 0.32155037 | -0.39714 | 0.69126424 | NA | NA | NA |
| ENSG000002 | 0 | NA | NA | NA | NA | NA | NA | NA |
| ENSG000002 | 1.97059802 | 0.190797738 | 0.35902573 | 0.53143193 | 0.5951195 | NA | NA | NA |
| ENSG000002 | 1.41292823 | -0.393548917 | 0.33316071 | -1.1812585 | 0.23750005 | NA | TRIM53AP | 642569 |
| ENSG000002 | 1.5451927 | 0.149991621 | 0.34623806 | 0.43320373 | 0.66486677 | NA | NA | NA |
| ENSG000002 | 0 | NA | NA | NA | NA | NA | NA | NA |
| ENSG000002 | 1.58673053 | -0.490100772 | 0.3405741 | -1.439043 | 0.15013835 | NA | NA | NA |
| ENSG000002 | 0 | NA | NA | NA | NA | NA | NA | NA |
| ENSG000002 | 0.82806866 | 0.203127889 | 0.29569413 | 0.68695272 | 0.49211252 | NA | NA | NA |
| ENSG000002 | 1.58706629 | -0.240695455 | 0.34429124 | -0.6991042 | 0.48448691 | NA | NA | NA |
| ENSG000002 | 1.33383599 | 0.025659238 | 0.33386331 | 0.07685552 | 0.93873848 | NA | NA | NA |
| ENSG000002 | 0 | NA | NA | NA | NA | NA | NA | NA |
| ENSG000002 | 1.03120242 | -0.357910397 | 0.31020733 | -1.153778 | 0.24859119 | NA | NA | NA |
| ENSG000002 | 1.52055229 | -0.793167814 | 0.34106687 | -2.3255493 | 0.02004261 | NA | NA | NA |
| ENSG000002 | 0.47614243 | -0.020043108 | 0.23674202 | -0.0846622 | 0.93252992 | NA | NA | NA |
| ENSG000002 | 1.64270193 | -0.271024188 | 0.35108155 | -0.7719693 | 0.4401326 | NA | NA | NA |
| ENSG000002 | 0.73889482 | -0.029099289 | 0.29011813 | -0.1003015 | 0.92010495 | NA | NA | NA |
| ENSG000002 | 1.46142772 | -0.037641071 | 0.3407535 | -0.1104642 | 0.91204122 | NA | NA | NA |
| ENSG000002 | 1.98985098 | -0.201763309 | 0.36403926 | -0.554235 | 0.57941801 | NA | NA | NA |

| | | | | | | | | |
|---|---|---|---|---|---|---|---|---|
| ENSG000002 | 1.11536237 | 0.031101987 | 0.32205545 | 0.09657339 | 0.92306519 | NA | NA | NA |
| ENSG000002 | 1.21423141 | 0.247065898 | 0.32846404 | 0.75218554 | 0.45193949 | NA | NA | NA |
| ENSG000002 | 0.98022291 | 0.11794959 | 0.31248556 | 0.37745612 | 0.70583466 | NA | NA | NA |
| ENSG000002 | 0 | NA | NA | NA | NA | NA | NA | NA |
| ENSG000002 | 2.1723845 | -0.172009628 | 0.36556044 | -0.4705368 | 0.63797157 | NA | NA | NA |
| ENSG000002 | 1.53421184 | -0.334092353 | 0.34254753 | -0.9753168 | 0.32940315 | NA | NA | NA |
| ENSG000002 | 1.00350427 | -0.171931705 | 0.30194223 | -0.5694192 | 0.56907169 | NA | NA | NA |
| ENSG000002 | 0.73775266 | -0.030682722 | 0.27839171 | -0.1102142 | 0.91223949 | NA | NA | NA |
| ENSG000002 | 1.27216442 | -0.33871914 | 0.32711516 | -1.0354737 | 0.30044775 | NA | NA | NA |
| ENSG000002 | 2.11986679 | -0.510523617 | 0.36579024 | -1.3956732 | 0.16281294 | NA | NA | NA |
| ENSG000002 | 1.49492257 | 0.083210856 | 0.34572452 | 0.24068543 | 0.80979893 | NA | NA | NA |
| ENSG000002 | 2.01088455 | -0.201072637 | 0.36333771 | -0.5534043 | 0.57998663 | NA | NA | NA |
| ENSG000002 | 1.01173365 | -0.497723116 | 0.30515291 | -1.6310613 | 0.10287738 | NA | NA | NA |
| ENSG000002 | 0 | NA | NA | NA | NA | NA | NA | NA |
| ENSG000002 | 2.16742655 | -0.519393647 | 0.36648514 | -1.4172297 | 0.15641577 | NA | NA | NA |
| ENSG000002 | 0.73861249 | -0.192197989 | 0.28559321 | -0.6729781 | 0.5009612 | NA | NA | NA |
| ENSG000002 | 0 | NA | NA | NA | NA | NA | NA | NA |
| ENSG000002 | 1.6308335 | -0.414050251 | 0.3526794 | -1.1740131 | 0.24038976 | NA | LOC105373559 | 105373559 |
| ENSG000002 | 1.36353938 | 0.016453594 | 0.32869368 | 0.05005753 | 0.96007654 | NA | NA | NA |
| ENSG000002 | 0.3588776 | -0.099902505 | 0.21555794 | -0.4634601 | 0.6430346 | NA | NA | NA |
| ENSG000002 | 1.71428794 | -0.061106862 | 0.35312118 | -0.1730479 | 0.86261381 | NA | NA | NA |
| ENSG000002 | 1.47036343 | -0.17748746 | 0.33791951 | -0.5252359 | 0.59941921 | NA | NA | NA |
| ENSG000002 | 1.9870672 | -0.165756458 | 0.35505837 | -0.4668428 | 0.64061233 | NA | NA | NA |
| ENSG000002 | 1.7733699 | 0.390926934 | 0.35331454 | 1.10645584 | 0.26852927 | NA | NA | NA |
| ENSG000002 | 1.81335808 | 0.124738078 | 0.35468706 | 0.35168488 | 0.7250746 | NA | NA | NA |
| ENSG000002 | 1.60081162 | -0.261317564 | 0.3505449 | -0.745461 | 0.45599306 | NA | NA | NA |
| ENSG000002 | 1.09299078 | 0.029167801 | 0.32132133 | 0.09077455 | 0.92767173 | NA | LOC100129216 | 100129216 |
| ENSG000002 | 0 | NA | NA | NA | NA | NA | NA | NA |
| ENSG000002 | 0.8488199 | 0.043797333 | 0.30149722 | 0.14526612 | 0.88450076 | NA | NA | NA |

| | | | | | | | | |
|---|---|---|---|---|---|---|---|---|
| ENSG000002 | 0 | NA | NA | NA | NA | NA | NA | NA |
| ENSG000002 | 1.71416778 | 0.207548269 | 0.3531823 | 0.58765196 | 0.55676593 | NA | NA | NA |
| ENSG000002 | 0.12685397 | -0.090215768 | 0.12901919 | -0.699243 | 0.48440017 | NA | NA | NA |
| ENSG000002 | 1.73238774 | -0.463339481 | 0.35495519 | -1.3053464 | 0.19177492 | NA | NA | NA |
| ENSG000002 | 1.27480218 | -0.194837179 | 0.32794948 | -0.5941073 | 0.55244035 | NA | TRMT2B-AS1 | 106481789 |
| ENSG000002 | 0 | NA | NA | NA | NA | NA | NA | NA |
| ENSG000002 | 1.91659914 | -0.684531259 | 0.35954875 | -1.9038621 | 0.05692814 | NA | NA | NA |
| ENSG000002 | 0.63296622 | -0.299126688 | 0.27328461 | -1.0945611 | 0.27370899 | NA | NA | NA |
| ENSG000002 | 2.03599129 | -0.594445783 | 0.36292543 | -1.6379282 | 0.10143668 | NA | NA | NA |
| ENSG000002 | 0.48279897 | 0.145542372 | 0.23831717 | 0.6107087 | 0.54139244 | NA | NA | NA |
| ENSG000002 | 0.4855357 | -0.181797071 | 0.23938927 | -0.7594203 | 0.44760118 | NA | NA | NA |
| ENSG000002 | 1.35492997 | -0.821603875 | 0.32378469 | -2.5375007 | 0.01116472 | NA | NA | NA |
| ENSG000002 | 2.07769126 | -0.254103148 | 0.36580533 | -0.6946404 | 0.48728071 | NA | NA | NA |
| ENSG000002 | 0 | NA | NA | NA | NA | NA | NA | NA |
| ENSG000002 | 0 | NA | NA | NA | NA | NA | NA | NA |
| ENSG000002 | 1.53494016 | 0.451788776 | 0.34958914 | 1.29234212 | 0.19623869 | NA | NA | NA |
| ENSG000002 | 0 | NA | NA | NA | NA | NA | NA | NA |
| ENSG000002 | 0 | NA | NA | NA | NA | NA | NA | NA |
| ENSG000002 | 1.92327494 | -0.483285223 | 0.35019368 | -1.3800512 | 0.16757087 | NA | FRG2B | 441581 |
| ENSG000002 | 1.96395436 | -0.442357019 | 0.36079491 | -1.2260623 | 0.22017525 | NA | NA | NA |
| ENSG000002 | 0.42385498 | -0.262217565 | 0.21037102 | -1.2464529 | 0.21259817 | NA | NA | NA |
| ENSG000002 | 0.57330892 | 0.211197273 | 0.25231239 | 0.83704676 | 0.40256628 | NA | NA | NA |
| ENSG000002 | 1.98775234 | -0.154349369 | 0.34881137 | -0.442501 | 0.65812673 | NA | NA | NA |
| ENSG000002 | 1.14018624 | -0.602964954 | 0.32157057 | -1.8750626 | 0.06078411 | NA | NA | NA |
| ENSG000002 | 1.35330127 | -0.124759761 | 0.34069718 | -0.3661896 | 0.71422361 | NA | NA | NA |
| ENSG000002 | 2.13509863 | -0.512887793 | 0.36388762 | -1.4094676 | 0.15869696 | NA | NA | NA |
| ENSG000002 | 1.06043502 | -0.511783882 | 0.30711936 | -1.6664006 | 0.09563365 | NA | NA | NA |
| ENSG000002 | 1.25454413 | -0.328619244 | 0.32665941 | -1.0059996 | 0.31441575 | NA | NA | NA |
| ENSG000002 | 1.34828717 | 0.029351419 | 0.32489822 | 0.09034035 | 0.92801675 | NA | LOC101927362 | 101927362 |

| | | | | | | | | |
|---|---|---|---|---|---|---|---|---|
| ENSG000002 | 0.70118031 | 0.116096956 | 0.26691515 | 0.43495829 | 0.66359271 | NA | NA | NA |
| ENSG000002 | 2.14837944 | -0.252105317 | 0.3645451 | -0.6915614 | 0.48921281 | NA | NA | NA |
| ENSG000002 | 0.63921993 | 0.0498278 | 0.25845554 | 0.19279061 | 0.84712296 | NA | NA | NA |
| ENSG000002 | 1.46240324 | -0.190772563 | 0.34124538 | -0.559048 | 0.57612893 | NA | NA | NA |
| ENSG000002 | 1.27164324 | -0.321201663 | 0.32466844 | -0.9893221 | 0.32250557 | NA | NA | NA |
| ENSG000002 | 2.01315621 | -0.178366726 | 0.35828291 | -0.4978377 | 0.61859846 | NA | NA | NA |
| ENSG000002 | 1.7081843 | 0.080514097 | 0.34811119 | 0.23128845 | 0.81709072 | NA | NA | NA |
| ENSG000002 | 0 | NA | NA | NA | NA | NA | NA | NA |
| ENSG000002 | 0.64144758 | -0.286694252 | 0.26813299 | -1.0692241 | 0.28496871 | NA | NA | NA |
| ENSG000002 | 1.81285342 | -0.586814459 | 0.34978312 | -1.6776523 | 0.09341499 | NA | NA | NA |
| ENSG000002 | 1.89581757 | -0.370683167 | 0.35491443 | -1.0444297 | 0.29628664 | NA | NA | NA |
| ENSG000002 | 1.51254913 | -0.628204347 | 0.34225307 | -1.8354966 | 0.06643214 | NA | NA | NA |
| ENSG000002 | 0.60492784 | -0.101259155 | 0.26330325 | -0.3845724 | 0.70055427 | NA | NA | NA |
| ENSG000002 | 0.77613058 | -0.361458377 | 0.28422771 | -1.2717211 | 0.20347223 | NA | NA | NA |
| ENSG000002 | 1.969645 | -0.326664791 | 0.36425514 | -0.8968022 | 0.3698245 | NA | NA | NA |
| ENSG000002 | 0.60942426 | 0.05292158 | 0.26330637 | 0.20098861 | 0.84070748 | NA | NA | NA |
| ENSG000002 | 2.1067159 | -0.370014399 | 0.36395257 | -1.0166555 | 0.30931732 | NA | NA | NA |
| ENSG000002 | 0.13280363 | -0.093394593 | 0.13115644 | -0.7120855 | 0.47641185 | NA | NA | NA |
| ENSG000002 | 0 | NA | NA | NA | NA | NA | NA | NA |
| ENSG000002 | 0.47826975 | -0.016435741 | 0.244632 | -0.0671856 | 0.94643397 | NA | NA | NA |
| ENSG000002 | 0.54222759 | -0.38902668 | 0.24851205 | -1.5654238 | 0.11748356 | NA | NA | NA |
| ENSG000002 | 0.23139247 | 0.167748926 | 0.17680352 | 0.94878726 | 0.34272882 | NA | NA | NA |
| ENSG000002 | 1.66958198 | -0.774188249 | 0.33629796 | -2.3020903 | 0.02133008 | NA | NA | NA |
| ENSG000002 | 0 | NA | NA | NA | NA | NA | CT47A6 | 728062 |
| ENSG000002 | 0 | NA | NA | NA | NA | NA | NA | NA |
| ENSG000002 | 0.78708361 | -0.370158269 | 0.28484481 | -1.2995086 | 0.19376945 | NA | NA | NA |
| ENSG000002 | 1.87506049 | -0.12745805 | 0.36020781 | -0.3538459 | 0.72345439 | NA | NA | NA |
| ENSG000002 | 1.55470057 | -0.634523718 | 0.34322646 | -1.8487028 | 0.06450074 | NA | NA | NA |
| ENSG000002 | 1.98945008 | 0.052216219 | 0.35905015 | 0.14542876 | 0.88437236 | NA | NA | NA |

| | | | | | | | | |
|---|---|---|---|---|---|---|---|---|
| ENSG000002 | 0 | NA | NA | NA | NA | NA | NA | NA |
| ENSG000002 | 0.48573157 | -0.181932918 | 0.23941992 | -0.7598905 | 0.44732006 | NA | NA | NA |
| ENSG000002 | 1.60246073 | -0.397751808 | 0.35013266 | -1.1360031 | 0.25595525 | NA | NA | NA |
| ENSG000002 | 0 | NA | NA | NA | NA | NA | NA | NA |
| ENSG000002 | 0.49036493 | -0.020766486 | 0.24598337 | -0.0844223 | 0.93272066 | NA | NA | NA |
| ENSG000002 | 0.11934008 | 0.083564698 | 0.12760091 | 0.65489106 | 0.5125379 | NA | NA | NA |
| ENSG000002 | 0 | NA | NA | NA | NA | NA | NA | NA |
| ENSG000002 | 1.09125576 | 0.031599837 | 0.31500431 | 0.10031557 | 0.9200938 | NA | NA | NA |
| ENSG000002 | 1.75661195 | -0.33942832 | 0.35612368 | -0.9531192 | 0.34052967 | NA | NA | NA |
| ENSG000002 | 0 | NA | NA | NA | NA | NA | NA | NA |
| ENSG000002 | 1.73119063 | -0.068262646 | 0.35059443 | -0.1947054 | 0.84562355 | NA | NA | NA |
| ENSG000002 | 1.83891269 | 0.009293091 | 0.36116059 | 0.02573119 | 0.97947175 | NA | NA | NA |
| ENSG000002 | 0.62478892 | 0.058386395 | 0.26463671 | 0.22062848 | 0.82538172 | NA | NA | NA |
| ENSG000002 | 0.24485602 | -0.012655878 | 0.17881722 | -0.0707755 | 0.94357643 | NA | NA | NA |
| ENSG000002 | 1.19166998 | 0.107653642 | 0.32738762 | 0.32882624 | 0.74228703 | NA | FCF1P4 | 106480799 |
| ENSG000002 | 0 | NA | NA | NA | NA | NA | NA | NA |
| ENSG000002 | 1.35976585 | -0.267680392 | 0.34010603 | -0.7870498 | 0.4312527 | NA | NA | NA |
| ENSG000002 | 0.37422052 | -0.268731395 | 0.21273223 | -1.2632378 | 0.20650373 | NA | NA | NA |
| ENSG000002 | 0 | NA | NA | NA | NA | NA | NA | NA |
| ENSG000002 | 1.34275105 | -0.24575029 | 0.3326445 | -0.7387776 | 0.46004208 | NA | NA | NA |
| ENSG000002 | 1.48271975 | -0.198534033 | 0.34957872 | -0.5679237 | 0.57008679 | NA | NA | NA |
| ENSG000002 | 1.2570158 | -0.339529029 | 0.32848607 | -1.0336177 | 0.3013149 | NA | NA | NA |
| ENSG000002 | 1.24973216 | -0.046153606 | 0.32468674 | -0.1421481 | 0.88696302 | NA | NA | NA |
| ENSG000002 | 0.9155362 | -0.460427183 | 0.30148657 | -1.5271897 | 0.12671384 | NA | NA | NA |
| ENSG000002 | 1.39898751 | -0.241051542 | 0.32306223 | -0.7461459 | 0.45557931 | NA | NA | NA |
| ENSG000002 | 1.60892084 | -0.26518727 | 0.35177887 | -0.7538465 | 0.4509414 | NA | NA | NA |
| ENSG000002 | 0.12070844 | -0.088503695 | 0.12783931 | -0.6923042 | 0.48874631 | NA | NA | NA |
| ENSG000002 | 1.84555195 | -0.234535506 | 0.35558768 | -0.6595715 | 0.50952884 | NA | NA | NA |
| ENSG000002 | 0.12685397 | -0.090215768 | 0.12901919 | -0.699243 | 0.48440017 | NA | NA | NA |

| | | | | | | | | | |
|---|---|---|---|---|---|---|---|---|---|
| ENSG000002 | 1.96643165 | -0.067520531 | 0.36367502 | -0.1856617 | 0.85271005 | NA | NA | NA | |
| ENSG000002 | 1.29495652 | -0.667043108 | 0.32916087 | -2.0264957 | 0.04271401 | NA | NA | NA | |
| ENSG000002 | 1.0643207 | 0.312452913 | 0.31009849 | 1.00759249 | 0.31365013 | NA | NA | NA | |
| ENSG000002 | 1.92365603 | -0.835076649 | 0.35780015 | -2.3339192 | 0.01959995 | NA | NA | NA | |
| ENSG000002 | 1.97688061 | -0.302464472 | 0.35861652 | -0.8434204 | 0.39899339 | NA | NA | NA | |
| ENSG000002 | 0.14128499 | -0.097772014 | 0.13403235 | -0.7294658 | 0.4657168 | NA | LOC105378933 | | 105378933 |
| ENSG000002 | 1.85237624 | -0.127551388 | 0.3614418 | -0.3528961 | 0.72416636 | NA | NA | NA | |
| ENSG000002 | 1.62082619 | -0.378242753 | 0.34442524 | -1.0981854 | 0.27212356 | NA | NA | NA | |
| ENSG000002 | 1.51851997 | -0.45725139 | 0.33802257 | -1.3527244 | 0.17614368 | NA | NA | NA | |
| ENSG000002 | 1.77311821 | -0.336451574 | 0.35568969 | -0.9459132 | 0.34419286 | NA | NA | NA | |
| ENSG000002 | 1.35965821 | -0.538693234 | 0.33229031 | -1.6211524 | 0.10498496 | NA | NA | NA | |
| ENSG000002 | 1.01640531 | -0.527971857 | 0.31163794 | -1.6941835 | 0.09023042 | NA | C8orf59P1 | | 106481719 |
| ENSG000002 | 1.10257992 | -0.123362737 | 0.32578092 | -0.3786678 | 0.70493458 | NA | NA | NA | |
| ENSG000002 | 1.11878266 | -0.408794676 | 0.31731931 | -1.2882755 | 0.19765009 | NA | NA | NA | |
| ENSG000002 | 1.53321212 | -0.490682288 | 0.34561151 | -1.419751 | 0.15568018 | NA | NA | NA | |
| ENSG000002 | 1.7365136 | -0.203374061 | 0.35842691 | -0.5674073 | 0.57043746 | NA | NA | NA | |
| ENSG000002 | 0 | NA | NA | NA | NA | NA | NA | NA | |
| ENSG000002 | 1.8813957 | -0.658293844 | 0.3567798 | -1.8450984 | 0.06502322 | NA | NA | NA | |
| ENSG000002 | 0.90913008 | -0.459386821 | 0.30276954 | -1.5172821 | 0.12919546 | NA | NA | NA | |
| ENSG000002 | 0.11934008 | 0.083564698 | 0.12760091 | 0.65489106 | 0.5125379 | NA | NA | NA | |
| ENSG000002 | 2.11545921 | -0.256928291 | 0.36574825 | -0.7024731 | 0.4823842 | NA | NA | NA | |
| ENSG000002 | 1.63200948 | -0.252691217 | 0.34480225 | -0.7328584 | 0.46364481 | NA | NA | NA | |
| ENSG000002 | 2.14030536 | -0.574515197 | 0.35655535 | -1.6112932 | 0.10711584 | NA | LINC00381 | | 100874151 |
| ENSG000002 | 1.71616573 | -0.187230785 | 0.35520962 | -0.5270994 | 0.59812455 | NA | NA | NA | |
| ENSG000002 | 1.78426766 | -0.324836846 | 0.35290743 | -0.9204591 | 0.35733292 | NA | NA | NA | |
| ENSG000002 | 1.71039076 | -0.301104035 | 0.35011465 | -0.8600155 | 0.38978049 | NA | NA | NA | |
| ENSG000002 | 1.73369352 | -0.402014313 | 0.3433441 | -1.1708788 | 0.24164751 | NA | NA | NA | |
| ENSG000002 | 1.93216342 | 0.068388162 | 0.35659121 | 0.19178308 | 0.84791212 | NA | NA | NA | |
| ENSG000002 | 1.81970003 | 0.131288917 | 0.35516656 | 0.3696545 | 0.71163994 | NA | NA | NA | |

| | | | | | | | | |
|---|---|---|---|---|---|---|---|---|
| ENSG000002 | 1.75680837 | -0.342427484 | 0.3589983 | -0.9538415 | 0.34016386 | NA | NA | NA |
| ENSG000002 | 0 | NA | NA | NA | NA | NA | NA | NA |
| ENSG000002 | 0.72921923 | -0.029686978 | 0.28414149 | -0.1044796 | 0.91678879 | NA | NA | NA |
| ENSG000002 | 1.8763574 | -0.811509506 | 0.35579304 | -2.280847 | 0.0225575 | NA | NA | NA |
| ENSG000002 | 1.2116521 | -0.035556644 | 0.32649664 | -0.1089036 | 0.91327899 | NA | NA | NA |
| ENSG000002 | 0.51459361 | -0.192534203 | 0.24265108 | -0.7934611 | 0.42750921 | NA | NA | NA |
| ENSG000002 | 1.63171451 | -0.407477718 | 0.35121816 | -1.1601841 | 0.24597386 | NA | NA | NA |
| ENSG000002 | 0.97271136 | -0.166893475 | 0.30054029 | -0.5553115 | 0.57868163 | NA | NA | NA |
| ENSG000002 | 1.6920607 | -0.055146213 | 0.35286304 | -0.1562822 | 0.87581058 | NA | NA | NA |
| ENSG000002 | 1.62030502 | -0.37344692 | 0.34562796 | -1.0804882 | 0.27992486 | NA | NA | NA |
| ENSG000002 | 1.45209705 | -0.048121965 | 0.34200903 | -0.1407038 | 0.88810395 | NA | NA | NA |
| ENSG000002 | 1.12763012 | -0.256728942 | 0.31512023 | -0.8147016 | 0.41524316 | NA | NA | NA |
| ENSG000002 | 1.97149816 | -0.195511658 | 0.36128673 | -0.5411537 | 0.58840164 | NA | NA | NA |
| ENSG000002 | 1.6578707 | -0.265894833 | 0.35128158 | -0.7569279 | 0.44909305 | NA | NA | NA |
| ENSG000002 | 1.65910055 | -0.542923514 | 0.34864133 | -1.5572552 | 0.11940992 | NA | NA | NA |
| ENSG000002 | 0.99786728 | -0.195540788 | 0.31382159 | -0.6230954 | 0.53322184 | NA | NA | NA |
| ENSG000002 | 0.12685397 | -0.090215768 | 0.12901919 | -0.699243 | 0.48440017 | NA | NA | NA |
| ENSG000002 | 1.78890886 | -0.706286771 | 0.34649626 | -2.0383677 | 0.04151318 | NA | NA | NA |
| ENSG000002 | 1.58198969 | -0.232510418 | 0.33949947 | -0.6848624 | 0.49343076 | NA | NA | NA |
| ENSG000002 | 1.48622752 | -0.315604775 | 0.33655454 | -0.9377523 | 0.34837174 | NA | NA | NA |
| ENSG000002 | 0 | NA | NA | NA | NA | NA | NA | NA |
| ENSG000002 | 1.72167819 | -0.188970241 | 0.35526639 | -0.5319114 | 0.59478737 | NA | NA | NA |
| ENSG000002 | 0 | NA | NA | NA | NA | NA | NA | NA |
| ENSG000002 | 0 | NA | NA | NA | NA | NA | NA | NA |
| ENSG000002 | 2.11758707 | -0.243883835 | 0.36525447 | -0.6677094 | 0.50431912 | NA | NA | NA |
| ENSG000002 | 1.09885845 | -0.408520357 | 0.31659154 | -1.2903704 | 0.19692208 | NA | PARD3-AS1 | 100505601 |
| ENSG000002 | 1.31137905 | -0.09229363 | 0.32159349 | -0.2869885 | 0.77412113 | NA | NA | NA |
| ENSG000002 | 2.06420381 | -0.870266589 | 0.35878652 | -2.4255833 | 0.01528381 | NA | NA | NA |
| ENSG000002 | 1.67034038 | -0.692217704 | 0.34787907 | -1.9898228 | 0.04661046 | NA | NA | NA |

| | | | | | | | | |
|---|---|---|---|---|---|---|---|---|
| ENSG000002 | 2.10457147 | -0.129689296 | 0.36344742 | -0.356831 | 0.72121835 | NA | NA | NA |
| ENSG000002 | 0.63387812 | -0.454271587 | 0.26413408 | -1.7198522 | 0.08545931 | NA | NA | NA |
| ENSG000002 | 0 | NA | NA | NA | NA | NA | NA | NA |
| ENSG000002 | 1.3511098 | -0.123136859 | 0.34065379 | -0.3614722 | 0.71774652 | NA | NA | NA |
| ENSG000002 | 1.36841743 | -0.131590655 | 0.34371724 | -0.3828457 | 0.7018342 | NA | NA | NA |
| ENSG000002 | 0.6181343 | 0.056012223 | 0.26406445 | 0.21211573 | 0.83201675 | NA | NA | NA |
| ENSG000002 | 0 | NA | NA | NA | NA | NA | NA | NA |
| ENSG000002 | 0.11205239 | 0.082118251 | 0.12671962 | 0.64803108 | 0.51696485 | NA | MRPL23-AS1 | 100133545 |
| ENSG000002 | 0.91724088 | -0.262759522 | 0.29152474 | -0.9013284 | 0.36741375 | NA | NA | NA |
| ENSG000002 | 1.75447701 | -0.589511486 | 0.35218713 | -1.6738587 | 0.09415837 | NA | NA | NA |
| ENSG000002 | 0.77544544 | -0.361698684 | 0.2841781 | -1.2727887 | 0.20309302 | NA | NA | NA |
| ENSG000002 | 1.49927753 | -0.188398043 | 0.34048215 | -0.5533272 | 0.58003936 | NA | NA | NA |
| ENSG000002 | 1.6255478 | -0.550370248 | 0.35070315 | -1.5693336 | 0.11657022 | NA | NA | NA |
| ENSG000002 | 0.3588776 | -0.099902505 | 0.21555794 | -0.4634601 | 0.6430346 | NA | NA | NA |
| ENSG000002 | 2.02192637 | -0.46171979 | 0.36376696 | -1.2692736 | 0.20434351 | NA | NA | NA |
| ENSG000002 | 1.40498978 | -0.275436387 | 0.34099026 | -0.8077544 | 0.41923198 | NA | NA | NA |
| ENSG000002 | 0.85084115 | -0.109944538 | 0.29679278 | -0.3704421 | 0.71105312 | NA | NA | NA |
| ENSG000002 | 1.26467013 | -0.191343396 | 0.32736865 | -0.5844891 | 0.55889132 | NA | NA | NA |
| ENSG000002 | 0 | NA | NA | NA | NA | NA | NA | NA |
| ENSG000002 | 1.29339228 | -0.512825987 | 0.33289561 | -1.5405009 | 0.1234383 | NA | NA | NA |
| ENSG000002 | 1.73874179 | -0.474517143 | 0.35601811 | -1.3328455 | 0.18258251 | NA | NA | NA |
| ENSG000002 | 0 | NA | NA | NA | NA | NA | NA | NA |
| ENSG000002 | 2.0591557 | 0.128615188 | 0.36602331 | 0.35138523 | 0.72529936 | NA | NA | NA |
| ENSG000002 | 2.02719672 | -0.593906948 | 0.36250873 | -1.6383246 | 0.10135401 | NA | NA | NA |
| ENSG000002 | 0.86133772 | -0.113024524 | 0.29735512 | -0.3800995 | 0.70387157 | NA | NA | NA |
| ENSG000002 | 2.06235281 | -0.333240858 | 0.35595396 | -0.9361909 | 0.3491749 | NA | NA | NA |
| ENSG000002 | 0 | NA | NA | NA | NA | NA | NA | NA |
| ENSG000002 | 1.01045565 | -0.504631893 | 0.30674057 | -1.6451423 | 0.09994046 | NA | NA | NA |
| ENSG000002 | 1.51707384 | -0.220314438 | 0.21157969 | -1.0412835 | 0.297744 | NA | NA | NA |

| | | | | | | | | |
|---|---|---|---|---|---|---|---|---|
| ENSG000002 | 2.15271267 | -0.656891767 | 0.3654839 | -1.7973207 | 0.07228473 | NA | NA | NA |
| ENSG000002 | 1.76648396 | -0.329423082 | 0.35624463 | -0.9247103 | 0.35511661 | NA | NA | NA |
| ENSG000002 | 1.21637528 | -0.039133079 | 0.32964778 | -0.1187118 | 0.90550369 | NA | NA | NA |
| ENSG000002 | 0.62468073 | -0.25993165 | 0.25829707 | -1.0063283 | 0.31425767 | NA | NA | NA |
| ENSG000002 | 1.19679653 | -0.036135982 | 0.32528729 | -0.1110894 | 0.91154543 | NA | NA | NA |
| ENSG000002 | 1.41157294 | -0.913019348 | 0.33424483 | -2.7315886 | 0.00630298 | NA | NA | NA |
| ENSG000002 | 0 | NA | NA | NA | NA | NA | NA | NA |
| ENSG000002 | 1.33308277 | 0.305903792 | 0.33067811 | 0.92508026 | 0.35492416 | NA | NA | NA |
| ENSG000002 | 1.25436807 | -0.305216047 | 0.32005921 | -0.9536237 | 0.34027415 | NA | NA | NA |
| ENSG000002 | 0.76696407 | -0.358999402 | 0.28364792 | -1.2656514 | 0.2056379 | NA | NA | NA |
| ENSG000002 | 1.47722264 | 0.087570618 | 0.34541932 | 0.25351974 | 0.79986661 | NA | NA | NA |
| ENSG000002 | 1.98472678 | -0.438209639 | 0.35931743 | -1.2195613 | 0.22263122 | NA | NA | NA |
| ENSG000002 | 0 | NA | NA | NA | NA | NA | NA | NA |
| ENSG000002 | 0.8662935 | -0.382379665 | 0.2821115 | -1.3554203 | 0.17528367 | NA | NA | NA |
| ENSG000002 | 0.11131518 | 0.081951103 | 0.12663804 | 0.64712866 | 0.51754868 | NA | NA | NA |
| ENSG000002 | 1.97050194 | -0.071567867 | 0.36198984 | -0.1977068 | 0.84327444 | NA | NA | NA |
| ENSG000002 | 1.76959317 | -0.57938653 | 0.35035402 | -1.6537174 | 0.09818495 | NA | NA | NA |
| ENSG000002 | 0.36351096 | 0.070290861 | 0.21590418 | 0.32556508 | 0.74475344 | NA | NA | NA |
| ENSG000002 | 0.87050422 | -0.120430766 | 0.30290137 | -0.3975907 | 0.69093192 | NA | NA | NA |
| ENSG000002 | 2.01897595 | -0.606733848 | 0.36378114 | -1.6678541 | 0.0953447 | NA | NA | NA |
| ENSG000002 | 1.75416201 | -0.186651527 | 0.35273183 | -0.5291599 | 0.59669456 | NA | NA | NA |
| ENSG000002 | 1.50196176 | 0.066298084 | 0.33358018 | 0.19874707 | 0.8424606 | NA | NA | NA |
| ENSG000002 | 0 | NA | NA | NA | NA | NA | NA | NA |
| ENSG000002 | 0 | NA | NA | NA | NA | NA | NA | NA |
| ENSG000002 | 0.23070733 | 0.167416683 | 0.17664244 | 0.94777156 | 0.34324576 | NA | NA | NA |
| ENSG000002 | 2.16976184 | -0.519632494 | 0.3653667 | -1.4222218 | 0.15496187 | NA | NA | NA |
| ENSG000002 | 0 | NA | NA | NA | NA | NA | SNORD57 | 26792 |
| ENSG000002 | 2.14955585 | -0.657133216 | 0.36598158 | -1.7955363 | 0.07256829 | NA | NA | NA |
| ENSG000002 | 1.49508 | -0.474251271 | 0.34354437 | -1.3804659 | 0.16744326 | NA | NA | NA |

| | | | | | | | | |
|---|---|---|---|---|---|---|---|---|
| ENSG000002 | 2.00536859 | -0.464503601 | 0.36414385 | -1.2756047 | 0.20209529 | NA | NA | NA |
| ENSG000002 | 2.07651823 | -0.120657551 | 0.35528265 | -0.33961 | 0.73415023 | NA | NA | NA |
| ENSG000002 | 0.13280363 | -0.093394593 | 0.13115644 | -0.7120855 | 0.47641185 | NA | NA | NA |
| ENSG000002 | 0.98965289 | -0.311626105 | 0.30054797 | -1.0368598 | 0.29980121 | NA | NA | NA |
| ENSG000002 | 1.1008449 | -0.105972886 | 0.31313268 | -0.3384281 | 0.73504063 | NA | NA | NA |
| ENSG000002 | 0.90112692 | -0.259399981 | 0.29324956 | -0.8845708 | 0.37638819 | NA | NA | NA |
| ENSG000002 | 0.78364447 | -0.5663609 | 0.28711516 | -1.9725914 | 0.04854213 | NA | NA | NA |
| ENSG000002 | 0 | NA | NA | NA | NA | NA | CT47A9 | 728042 |
| ENSG000002 | 0 | NA | NA | NA | NA | NA | NA | NA |
| ENSG000002 | 0.7888781 | -0.345530318 | 0.27689228 | -1.2478871 | 0.21207239 | NA | NA | NA |
| ENSG000002 | 0.85109156 | 0.052630826 | 0.28861122 | 0.1823589 | 0.85530107 | NA | NA | NA |
| ENSG000002 | 1.27768126 | -0.012985106 | 0.19189327 | -0.0676684 | 0.94604962 | NA | NA | NA |
| ENSG000002 | 1.08040824 | 0.178971696 | 0.31661589 | 0.56526442 | 0.57189392 | NA | NA | NA |
| ENSG000002 | 1.25354241 | -0.045522183 | 0.3231058 | -0.1408894 | 0.88795731 | NA | NA | NA |
| ENSG000002 | 1.70708959 | -0.305416145 | 0.35032081 | -0.8718184 | 0.38330743 | NA | LOC101929574 | 101929574 |
| ENSG000002 | 1.4923736 | -0.334237201 | 0.34488269 | -0.969133 | 0.33247883 | NA | NA | NA |
| ENSG000002 | 0.99058151 | -0.038157036 | 0.30808504 | -0.1238523 | 0.90143224 | NA | NA | NA |
| ENSG000002 | 0.99881301 | -0.200372067 | 0.31383792 | -0.6384572 | 0.52317613 | NA | NA | NA |
| ENSG000002 | 1.67976964 | -0.050675767 | 0.35086017 | -0.1444329 | 0.88515861 | NA | NA | NA |
| ENSG000002 | 0 | NA | NA | NA | NA | NA | CT47A12 | 100507170 |
| ENSG000002 | 1.78492213 | -0.179048519 | 0.3440493 | -0.5204153 | 0.60277415 | NA | NA | NA |
| ENSG000002 | 1.18207655 | -0.582709231 | 0.31722838 | -1.8368761 | 0.06622819 | NA | NA | NA |
| ENSG000002 | 1.34275105 | -0.24575029 | 0.3326445 | -0.7387776 | 0.46004208 | NA | NA | NA |
| ENSG000002 | 0.87169844 | -0.120477128 | 0.30728461 | -0.3920702 | 0.69500637 | NA | NA | NA |
| ENSG000002 | 1.35705945 | -0.128046002 | 0.34350451 | -0.3727637 | 0.70932434 | NA | LOC101928525 | 101928525 |
| ENSG000002 | 1.41344783 | -0.576857365 | 0.33823811 | -1.7054771 | 0.08810548 | NA | NA | NA |
| ENSG000002 | 0.46273095 | 0.157073337 | 0.24331041 | 0.64556769 | 0.51855937 | NA | NA | NA |
| ENSG000002 | 0 | NA | NA | NA | NA | NA | PABPC1L2B-AS1 | 101928345 |
| ENSG000002 | 1.59887464 | -0.625196766 | 0.34100999 | -1.8333679 | 0.06674788 | NA | TMEM256P2 | 106479054 |

| | | | | | | | | |
|---|---|---|---|---|---|---|---|---|
| ENSG000002 | 0.87650595 | -0.117161493 | 0.29813851 | -0.3929767 | 0.69433668 | NA | NA | NA |
| ENSG000002 | 1.7068028 | -0.191147079 | 0.35401314 | -0.5399435 | 0.58923603 | NA | NA | NA |
| ENSG000002 | 0.50704533 | -0.189333637 | 0.24184367 | -0.7828761 | 0.43369986 | NA | NA | NA |
| ENSG000002 | 2.00311006 | -0.068064088 | 0.36389762 | -0.1870419 | 0.85162781 | NA | NA | NA |
| ENSG000002 | 0.37996515 | 0.064665424 | 0.21008146 | 0.3078112 | 0.758226 | NA | NA | NA |
| ENSG000002 | 0.90725072 | -0.437963416 | 0.29604611 | -1.4793757 | 0.13903994 | NA | NA | NA |
| ENSG000002 | 0.49076933 | -0.025124039 | 0.23846109 | -0.1053591 | 0.91609089 | NA | NA | NA |
| ENSG000002 | 0.11205239 | 0.082118251 | 0.12671962 | 0.64803108 | 0.51696485 | NA | NA | NA |
| ENSG000002 | 1.72572862 | -0.693363702 | 0.34721034 | -1.9969558 | 0.04582998 | NA | NA | NA |
| ENSG000002 | 0.34999534 | 0.241411141 | 0.20772605 | 1.16216113 | 0.24517002 | NA | NA | NA |
| ENSG000002 | 1.73028664 | -0.533961081 | 0.34346788 | -1.5546172 | 0.12003726 | NA | NA | NA |
| ENSG000002 | 2.03072677 | -0.586715537 | 0.36204082 | -1.6205784 | 0.10510808 | NA | NA | NA |
| ENSG000002 | 1.317468 | -0.309448353 | 0.31777533 | -0.973796 | 0.33015783 | NA | MED15P3 | 106481657 |
| ENSG000002 | 1.72775438 | -0.593892191 | 0.35207516 | -1.6868335 | 0.09163537 | NA | NA | NA |
| ENSG000002 | 1.78205253 | -0.585696651 | 0.3501579 | -1.6726644 | 0.09439339 | NA | NA | NA |
| ENSG000002 | 0 | NA | NA | NA | NA | NA | NA | NA |
| ENSG000002 | 0.78090774 | -0.190404338 | 0.28228031 | -0.6745222 | 0.49997937 | NA | NA | NA |
| ENSG000002 | 1.34479012 | 0.027476789 | 0.33153354 | 0.08287786 | 0.93394866 | NA | NA | NA |
| ENSG000002 | 0 | NA | NA | NA | NA | NA | NA | NA |
| ENSG000002 | 0.36345889 | 0.070257596 | 0.21589421 | 0.32542602 | 0.74485866 | NA | NA | NA |
| ENSG000002 | 1.1603967 | -0.732688452 | 0.31257823 | -2.3440162 | 0.01907734 | NA | NA | NA |
| ENSG000002 | 1.22599028 | -0.050917913 | 0.32991524 | -0.1543363 | 0.87734454 | NA | NA | NA |
| ENSG000002 | 2.0748152 | -0.247941906 | 0.3665191 | -0.6764774 | 0.49873756 | NA | NA | NA |
| ENSG000002 | 0 | NA | NA | NA | NA | NA | NA | NA |
| ENSG000002 | 1.69675661 | -0.534586106 | 0.34529151 | -1.5482168 | 0.12157011 | NA | NA | NA |
| ENSG000002 | 2.02002773 | -0.421018823 | 0.35613409 | -1.1821918 | 0.23712959 | NA | NA | NA |
| ENSG000002 | 0.96243708 | -0.033512237 | 0.31598804 | -0.1060554 | 0.9155384 | NA | NA | NA |
| ENSG000002 | 0.75227192 | -0.325771172 | 0.27195766 | -1.1978746 | 0.23096584 | NA | NA | NA |
| ENSG000002 | 0.34862505 | 0.240809381 | 0.20749856 | 1.16053522 | 0.24583096 | NA | NA | NA |

| | | | | | | | | | |
|---|---|---|---|---|---|---|---|---|---|
| ENSG000002 | 0.78039674 | -0.368519854 | 0.28442264 | -1.295677 | 0.1950868 | NA | NA | NA | |
| ENSG000002 | 0.71212066 | -0.015062431 | 0.26735149 | -0.0563394 | 0.95507141 | NA | NA | NA | |
| ENSG000002 | 0.64850501 | -0.484783006 | 0.2711005 | -1.788204 | 0.07374309 | NA | NA | NA | |
| ENSG000002 | 1.50122553 | -0.486858888 | 0.3459139 | -1.4074568 | 0.15929197 | NA | NA | NA | |
| ENSG000002 | 0.37765965 | -0.098061994 | 0.21009083 | -0.46676 | 0.64067161 | NA | NA | NA | |
| ENSG000002 | 2.06032547 | -0.592854336 | 0.36221567 | -1.636744 | 0.10168397 | NA | NA | NA | |
| ENSG000002 | 0 | NA | NA | NA | NA | NA | NA | NA | |
| ENSG000002 | 0.36576034 | -0.0934779 | 0.20820828 | -0.4489634 | 0.65345805 | NA | NA | NA | |
| ENSG000002 | 2.12785496 | -0.250396703 | 0.36530186 | -0.6854515 | 0.49305907 | NA | NA | NA | |
| ENSG000002 | 0 | NA | NA | NA | NA | NA | NA | NA | |
| ENSG000002 | 0 | NA | NA | NA | NA | NA | NA | NA | |
| ENSG000002 | 1.58971669 | -0.101654086 | 0.33885146 | -0.299996 | 0.7641802 | NA | NA | NA | |
| ENSG000002 | 0 | NA | NA | NA | NA | NA | CT47A11 | | 255313 |
| ENSG000002 | 0.7632638 | -0.52523984 | 0.27891552 | -1.8831503 | 0.05968001 | NA | NA | NA | |
| ENSG000002 | 0 | NA | NA | NA | NA | NA | NA | NA | |
| ENSG000002 | 1.07861822 | -0.094901702 | 0.30924484 | -0.3068821 | 0.75893312 | NA | NA | NA | |
| ENSG000002 | 0.86265401 | -0.262022423 | 0.2928341 | -0.8947811 | 0.37090411 | NA | NA | NA | |
| ENSG000002 | 2.13096417 | -0.488140227 | 0.36271069 | -1.3458115 | 0.17836331 | NA | NA | NA | |
| ENSG000002 | 1.89981736 | 0.185856938 | 0.35719572 | 0.52032241 | 0.60283888 | NA | NA | NA | |
| ENSG000002 | 1.67723576 | -0.713921383 | 0.35010244 | -2.0391785 | 0.04143221 | NA | NA | NA | |
| ENSG000002 | 1.52559951 | 0.599837504 | 0.34580866 | 1.73459366 | 0.0828128 | NA | NA | NA | |
| ENSG000002 | 0.96630097 | 0.117434284 | 0.30291998 | 0.38767428 | 0.69825709 | NA | NA | NA | |
| ENSG000002 | 0.94089656 | 0.113931567 | 0.2982781 | 0.38196423 | 0.70248789 | NA | NA | NA | |
| ENSG000002 | 0 | NA | NA | NA | NA | NA | NA | NA | |
| ENSG000002 | 1.90497556 | -0.381968362 | 0.3542918 | -1.078118 | 0.28098111 | NA | NA | NA | |
| ENSG000002 | 0.65084084 | -0.46034757 | 0.26550072 | -1.7338844 | 0.08293859 | NA | NA | NA | |
| ENSG000002 | 0.87420896 | -0.26470766 | 0.29762086 | -0.8894123 | 0.37378152 | NA | NA | NA | |
| ENSG000002 | 1.94360142 | -0.662495038 | 0.35581397 | -1.8619141 | 0.0626152 | NA | NA | NA | |
| ENSG000002 | 0.50593756 | -0.029751655 | 0.24018594 | -0.1238693 | 0.90141879 | NA | NA | NA | |

| | | | | | | | | |
|---|---|---|---|---|---|---|---|---|
| ENSG000002 | 0.77242638 | -0.187749988 | 0.28173694 | -0.6664017 | 0.50515435 | NA | NA | NA |
| ENSG000002 | 0.83546049 | 0.192120541 | 0.28847857 | 0.66597855 | 0.50542481 | NA | NA | NA |
| ENSG000002 | 0.72967624 | -0.02677813 | 0.28955569 | -0.0924801 | 0.92631662 | NA | NA | NA |
| ENSG000002 | 0.24756242 | -0.187034346 | 0.18119817 | -1.0322088 | 0.30197431 | NA | NA | NA |
| ENSG000002 | 0.37996515 | 0.064665424 | 0.21008146 | 0.3078112 | 0.758226 | NA | NA | NA |
| ENSG000002 | 2.13322428 | -0.779420997 | 0.36340806 | -2.1447543 | 0.0319725 | NA | NA | NA |
| ENSG000002 | 1.48781012 | -0.044788431 | 0.34123406 | -0.1312543 | 0.89557416 | NA | NA | NA |
| ENSG000002 | 1.00262326 | -0.181379752 | 0.30765704 | -0.5895518 | 0.5554912 | NA | NA | NA |
| ENSG000002 | 1.88943944 | -0.124858238 | 0.35598216 | -0.350743 | 0.72578119 | NA | NA | NA |
| ENSG000002 | 0.76482412 | -0.332489359 | 0.27530009 | -1.2077343 | 0.22714948 | NA | NA | NA |
| ENSG000002 | 1.07719586 | -0.105604382 | 0.31956186 | -0.3304662 | 0.74104776 | NA | NA | NA |
| ENSG000002 | 0.73613635 | 0.131710401 | 0.28471888 | 0.46259806 | 0.6436525 | NA | NA | NA |
| ENSG000002 | 1.73713925 | -0.1821707 | 0.35002193 | -0.5204551 | 0.60274641 | NA | NA | NA |
| ENSG000002 | 2.00633496 | -0.311193262 | 0.35700433 | -0.8716792 | 0.38338339 | NA | NA | NA |
| ENSG000002 | 1.72691557 | 0.685115476 | 0.35083481 | 1.95281496 | 0.05084152 | NA | NA | NA |
| ENSG000002 | 0.50127037 | -0.342056569 | 0.23551462 | -1.4523793 | 0.14639616 | NA | NA | NA |
| ENSG000002 | 1.59147289 | 0.0194779 | 0.34470725 | 0.05650563 | 0.95493901 | NA | NA | NA |
| ENSG000002 | 0.82087134 | 0.04669114 | 0.28686246 | 0.1627649 | 0.87070354 | NA | NA | NA |
| ENSG000002 | 1.26023319 | -0.312788086 | 0.32270027 | -0.9692836 | 0.3324037 | NA | NA | NA |
| ENSG000002 | 1.67002852 | -0.15984021 | 0.34611327 | -0.4618147 | 0.64421418 | NA | NA | NA |
| ENSG000002 | 1.32711138 | 0.177557207 | 0.33991241 | 0.52236165 | 0.60141855 | NA | NA | NA |
| ENSG000002 | 0.23276084 | -0.008011382 | 0.17603982 | -0.0455089 | 0.96370166 | NA | NA | NA |
| ENSG000002 | 0.6988466 | 0.297978543 | 0.27743105 | 1.07406342 | 0.28279426 | NA | PIGQP1 | 106480812 |
| ENSG000002 | 0 | NA | NA | NA | NA | NA | NA | NA |
| ENSG000002 | 1.59196215 | 0.018046526 | 0.33063512 | 0.05458139 | 0.95647196 | NA | CRIP1P3 | 106479040 |
| ENSG000002 | 1.86542108 | -0.398200974 | 0.35965754 | -1.107167 | 0.26822172 | NA | NA | NA |
| ENSG000002 | 0.12685397 | -0.090215768 | 0.12901919 | -0.699243 | 0.48440017 | NA | NA | NA |
| ENSG000002 | 0.36212533 | -0.235926492 | 0.20089643 | -1.1743688 | 0.24024735 | NA | NA | NA |
| ENSG000002 | 0.8943285 | -0.291158821 | 0.30492371 | -0.9548579 | 0.33964956 | NA | NA | NA |

| | | | | | | | | |
|---|---|---|---|---|---|---|---|---|
| ENSG000002 | 1.11060694 | 0.035289427 | 0.31568464 | 0.11178696 | 0.91099232 | NA | USP17L5 | 728386 |
| ENSG000002 | 1.54417154 | -0.618293628 | 0.34097777 | -1.813296 | 0.06978618 | NA | NA | NA |
| ENSG000002 | 0.8747492 | -0.246588133 | 0.28492303 | -0.8654552 | 0.38678897 | NA | NA | NA |
| ENSG000002 | 0.65084084 | -0.46034757 | 0.26550072 | -1.7338844 | 0.08293859 | NA | NA | NA |
| ENSG000002 | 1.64824931 | -0.119103759 | 0.34437969 | -0.3458501 | 0.72945535 | NA | NA | NA |
| ENSG000002 | 0.83094974 | -0.09579464 | 0.28779869 | -0.3328529 | 0.73924529 | NA | NA | NA |
| ENSG000002 | 1.52573151 | -0.087357829 | 0.24101787 | -0.3624537 | 0.71701299 | NA | NA | NA |
| ENSG000002 | 0 | NA | NA | NA | NA | NA | NA | NA |
| ENSG000002 | 1.56218473 | -0.102027329 | 0.33841273 | -0.3014879 | 0.76304251 | NA | NA | NA |
| ENSG000002 | 1.93082773 | -0.295868944 | 0.3582136 | -0.8259568 | 0.40882862 | NA | NA | NA |
| ENSG000002 | 1.5912123 | 0.016472586 | 0.34104933 | 0.04829972 | 0.96147738 | NA | NA | NA |
| ENSG000002 | 1.74490906 | -0.334119393 | 0.3574547 | -0.9347181 | 0.34993358 | NA | NA | NA |
| ENSG000002 | 0.52760069 | -0.383393289 | 0.24704506 | -1.5519164 | 0.12068222 | NA | NA | NA |
| ENSG000002 | 0.94510771 | -0.1645116 | 0.29658991 | -0.554677 | 0.57911562 | NA | MGAT3-AS1 | 104502417 |
| ENSG000002 | 0 | NA | NA | NA | NA | NA | NA | NA |
| ENSG000002 | 0.8776481 | -0.117739236 | 0.30325589 | -0.3882505 | 0.6978307 | NA | C6orf47-AS1 | 106480731 |
| ENSG000002 | 1.38527252 | -0.424164219 | 0.33972496 | -1.2485518 | 0.21182904 | NA | NA | NA |
| ENSG000002 | 0 | NA | NA | NA | NA | NA | NA | NA |
| ENSG000002 | 1.11655447 | -0.122432317 | 0.32621359 | -0.3753134 | 0.70742743 | NA | NA | NA |
| ENSG000002 | 1.97968826 | -0.066170954 | 0.35581434 | -0.1859705 | 0.85246793 | NA | NA | NA |
| ENSG000002 | 0.63398576 | -0.111029822 | 0.26576185 | -0.4177794 | 0.67610842 | NA | NA | NA |
| ENSG000002 | 1.32726433 | -0.115725993 | 0.33887988 | -0.3414956 | 0.73273051 | NA | NA | NA |
| ENSG000002 | 0.5074159 | -0.309594122 | 0.22571446 | -1.3716185 | 0.17018225 | NA | NA | NA |
| ENSG000002 | 0.8687113 | 0.034869 | 0.29740388 | 0.11724461 | 0.90666622 | NA | NA | NA |
| ENSG000002 | 1.50550951 | -0.056494452 | 0.33934499 | -0.1664809 | 0.86777853 | NA | SPATA13-AS1 | 100874231 |
| ENSG000002 | 1.24793766 | -0.046390719 | 0.32749586 | -0.1416528 | 0.88735423 | NA | NA | NA |
| ENSG000002 | 1.98546666 | -0.30804851 | 0.36183421 | -0.8513526 | 0.39457349 | NA | NA | NA |
| ENSG000002 | 1.62434093 | -0.498280622 | 0.34117112 | -1.4605006 | 0.14415254 | NA | NA | NA |
| ENSG000002 | 1.47725936 | -0.186930663 | 0.34466173 | -0.5423598 | 0.58757067 | NA | NA | NA |

| | | | | | | | | |
|---|---|---|---|---|---|---|---|---|
| ENSG000002 | 0 | NA | NA | NA | NA | NA | NA | NA |
| ENSG000002 | 1.828414 | -0.118187498 | 0.35673943 | -0.3312992 | 0.74041847 | NA | NA | NA |
| ENSG000002 | 2.10794182 | -0.386460148 | 0.36706469 | -1.0528393 | 0.29241464 | NA | NA | NA |
| ENSG000002 | 0.13280363 | -0.093394593 | 0.13115644 | -0.7120855 | 0.47641185 | NA | NA | NA |
| ENSG000002 | 0.76129866 | -0.025684083 | 0.27094315 | -0.0947951 | 0.92447758 | NA | NA | NA |
| ENSG000002 | 1.64175892 | 0.351564229 | 0.35005999 | 1.0042971 | 0.31523543 | NA | NA | NA |
| ENSG000002 | 0.25260017 | -0.015978705 | 0.18049464 | -0.0885273 | 0.92945758 | NA | LOC102724762 | 102724762 |
| ENSG000002 | 0.91869356 | -0.426614088 | 0.29428785 | -1.449649 | 0.14715642 | NA | NA | NA |
| ENSG000002 | 2.10921514 | -0.317910192 | 0.35305616 | -0.9004522 | 0.36787966 | NA | NA | NA |
| ENSG000002 | 1.99915545 | -0.074344958 | 0.36215673 | -0.2052839 | 0.83735029 | NA | NA | NA |
| ENSG000002 | 1.38307967 | -0.533382574 | 0.33156984 | -1.6086583 | 0.10769108 | NA | NA | NA |
| ENSG000002 | 0.22997012 | 0.167051788 | 0.17646558 | 0.94665368 | 0.34381528 | NA | NA | NA |
| ENSG000002 | 0 | NA | NA | NA | NA | NA | NA | NA |
| ENSG000002 | 1.49274362 | -0.483453536 | 0.34365595 | -1.4067952 | 0.15948813 | NA | NA | NA |
| ENSG000002 | 1.93881837 | 0.077688695 | 0.35968145 | 0.21599305 | 0.82899316 | NA | NA | NA |
| ENSG000002 | 1.7285392 | -0.313698692 | 0.35032618 | -0.8954475 | 0.37054793 | NA | NA | NA |
| ENSG000002 | 0.13280363 | -0.093394593 | 0.13115644 | -0.7120855 | 0.47641185 | NA | NA | NA |
| ENSG000002 | 1.11713355 | -0.274062992 | 0.32318008 | -0.8480195 | 0.39642713 | NA | NA | NA |
| ENSG000002 | 0 | NA | NA | NA | NA | NA | NA | NA |
| ENSG000002 | 1.35091729 | -0.359572568 | 0.32218561 | -1.1160417 | 0.26440429 | NA | NA | NA |
| ENSG000002 | 1.92538406 | 0.061182664 | 0.35839735 | 0.17071182 | 0.86445037 | NA | NA | NA |
| ENSG000002 | 1.88234925 | 0.459798436 | 0.35985398 | 1.27773616 | 0.20134247 | NA | NA | NA |
| ENSG000002 | 0.85029432 | 0.041498221 | 0.29478589 | 0.14077411 | 0.8880484 | NA | NA | NA |
| ENSG000002 | 0.25993993 | -0.013144386 | 0.18211683 | -0.0721756 | 0.94246219 | NA | NA | NA |
| ENSG000002 | 0.85112135 | -0.11399948 | 0.29680331 | -0.384091 | 0.70091101 | NA | NA | NA |
| ENSG000002 | 1.36591193 | -0.25223585 | 0.3370141 | -0.748443 | 0.45419298 | NA | NA | NA |
| ENSG000002 | 0.84172863 | 0.052461019 | 0.30554709 | 0.17169536 | 0.86367703 | NA | NA | NA |
| ENSG000002 | 1.0015194 | -0.344814197 | 0.30904115 | -1.115755 | 0.26452704 | NA | NA | NA |
| ENSG000002 | 1.38327108 | -0.27754501 | 0.34179991 | -0.8120102 | 0.41678578 | NA | NA | NA |

| | | | | | | | | | |
|---|---|---|---|---|---|---|---|---|---|
| ENSG000002 | 1.08297138 | 0.044525411 | 0.31468424 | 0.14149234 | 0.88748101 | NA | NA | NA | |
| ENSG000002 | 1.60350412 | 0.02327779 | 0.33824763 | 0.06881878 | 0.94513387 | NA | NA | NA | |
| ENSG000002 | 1.34345601 | -0.247694157 | 0.33508295 | -0.7392025 | 0.45978404 | NA | NA | NA | |
| ENSG000002 | 0.97296748 | 0.106783203 | 0.30737842 | 0.34739981 | 0.72829098 | NA | NA | NA | |
| ENSG000002 | 0.14128499 | -0.097772014 | 0.13403235 | -0.7294658 | 0.4657168 | NA | NA | NA | |
| ENSG000002 | 0.50955585 | -0.376243729 | 0.24515812 | -1.5346982 | 0.12485797 | NA | NA | NA | |
| ENSG000002 | 1.82249233 | 0.13595475 | 0.36101688 | 0.37658835 | 0.70647954 | NA | NA | NA | |
| ENSG000002 | 0.61546271 | -0.255876753 | 0.25745011 | -0.9938887 | 0.32027706 | NA | NA | NA | |
| ENSG000002 | 1.35022823 | -0.106452386 | 0.32353536 | -0.3290286 | 0.74213407 | NA | NA | NA | |
| ENSG000002 | 1.97146782 | -0.067324267 | 0.36252253 | -0.1857106 | 0.85267173 | NA | NA | NA | |
| ENSG000002 | 0.62023988 | -0.112292793 | 0.27094277 | -0.4144521 | 0.67854306 | NA | NA | NA | |
| ENSG000002 | 0 | NA | NA | NA | NA | NA | NA | NA | |
| ENSG000002 | 0.85932093 | -0.268592362 | 0.29843482 | -0.9000034 | 0.36811843 | NA | NA | NA | |
| ENSG000002 | 0.27408862 | -0.199826037 | 0.18669465 | -1.0703362 | 0.28446801 | NA | NA | NA | |
| ENSG000002 | 1.479158 | -0.184318522 | 0.34293011 | -0.5374813 | 0.5909352 | NA | NA | NA | |
| ENSG000002 | 2.1723606 | -0.936499771 | 0.36330986 | -2.5776889 | 0.00994635 | NA | NA | NA | |
| ENSG000002 | 1.82121276 | 0.006699777 | 0.35959508 | 0.01863145 | 0.98513511 | NA | NA | NA | |
| ENSG000002 | 0 | NA | NA | NA | NA | NA | NA | NA | |
| ENSG000002 | 0.92375698 | -0.407772392 | 0.28917518 | -1.4101224 | 0.15850356 | NA | NA | NA | |
| ENSG000002 | 1.05792897 | -0.708645707 | 0.30976443 | -2.2876923 | 0.02215544 | NA | NA | NA | |
| ENSG000002 | 0 | NA | NA | NA | NA | NA | NA | NA | |
| ENSG000002 | 0 | NA | NA | NA | NA | NA | NA | NA | |
| ENSG000002 | 1.92187045 | 0.318508452 | 0.35980182 | 0.88523303 | 0.37603096 | NA | KGFLP1 | | 387628 |
| ENSG000002 | 1.07600164 | -0.105155063 | 0.31477936 | -0.3340596 | 0.73833459 | NA | NA | NA | |
| ENSG000002 | 1.88013229 | -0.36834929 | 0.35289303 | -1.0437987 | 0.29657853 | NA | NA | NA | |
| ENSG000002 | 1.87518256 | -0.263551013 | 0.36016112 | -0.7317586 | 0.46431589 | NA | NA | NA | |
| ENSG000002 | 1.85670712 | -0.132122021 | 0.35805826 | -0.3689959 | 0.7121308 | NA | NA | NA | |
| ENSG000002 | 2.16808136 | -0.354321552 | 0.36067665 | -0.9823801 | 0.32591264 | NA | NA | NA | |
| ENSG000002 | 1.2239735 | -0.198445151 | 0.33385168 | -0.5944111 | 0.55223717 | NA | NA | NA | |

| | | | | | | | | |
|---|---|---|---|---|---|---|---|---|
| ENSG000002 | 1.89674836 | 0.451040414 | 0.35755331 | 1.2614634 | 0.20714194 | NA | NA | NA |
| ENSG000002 | 1.10867392 | -0.112034002 | 0.32214076 | -0.3477797 | 0.72800567 | NA | NCAM1-AS1 | 100288346 |
| ENSG000002 | 0 | NA | NA | NA | NA | NA | NA | NA |
| ENSG000002 | 2.1227434 | -0.516199673 | 0.36645195 | -1.4086422 | 0.158941 | NA | NA | NA |
| ENSG000002 | 0 | NA | NA | NA | NA | NA | NA | NA |
| ENSG000002 | 1.50147305 | 0.080978723 | 0.34142706 | 0.23717723 | 0.8125193 | NA | LOC102724009 | 102724009 |
| ENSG000002 | 1.09524016 | -0.121692347 | 0.32168899 | -0.3782919 | 0.70521374 | NA | NA | NA |
| ENSG000002 | 1.34763181 | -0.117417662 | 0.33217192 | -0.3534846 | 0.72372517 | NA | NA | NA |
| ENSG000002 | 2.08938652 | -0.488600849 | 0.36310198 | -1.3456298 | 0.17842193 | NA | NA | NA |
| ENSG000002 | 1.46512745 | 0.094588918 | 0.34886949 | 0.27112981 | 0.7862912 | NA | NA | NA |
| ENSG000002 | 1.6162066 | -0.406026611 | 0.35142742 | -1.1553641 | 0.24794137 | NA | NA | NA |
| ENSG000002 | 1.67293233 | -0.411098349 | 0.34928662 | -1.1769656 | 0.23920922 | NA | NA | NA |
| ENSG000002 | 1.6662138 | 0.067981049 | 0.34251788 | 0.19847445 | 0.84267387 | NA | NA | NA |
| ENSG000002 | 0.75936373 | -0.356158746 | 0.28317817 | -1.2577196 | 0.20849317 | NA | NA | NA |
| ENSG000002 | 1.08602159 | -0.113284216 | 0.31514512 | -0.3594668 | 0.71924589 | NA | NA | NA |
| ENSG000002 | 1.38144625 | -0.117671857 | 0.336649 | -0.3495387 | 0.72668491 | NA | NA | NA |
| ENSG000002 | 1.58845584 | -0.237957104 | 0.3443067 | -0.6911196 | 0.48949039 | NA | NA | NA |
| ENSG000002 | 0.75304351 | -0.205958918 | 0.29162776 | -0.7062391 | 0.48003948 | NA | NA | NA |
| ENSG000002 | 0.34999534 | 0.241411141 | 0.20772605 | 1.16216113 | 0.24517002 | NA | NA | NA |
| ENSG000002 | 1.23276943 | -0.038820424 | 0.31714557 | -0.1224057 | 0.90257773 | NA | NA | NA |
| ENSG000002 | 1.97940648 | -0.190244891 | 0.36282306 | -0.5243462 | 0.60003777 | NA | NA | NA |
| ENSG000002 | 0.13280363 | -0.093394593 | 0.13115644 | -0.7120855 | 0.47641185 | NA | USP17L12 | 100287205 |
| ENSG000002 | 0.59815061 | 0.048708909 | 0.25313511 | 0.19242257 | 0.84741122 | NA | NA | NA |
| ENSG000002 | 0.34270765 | 0.258172734 | 0.21433185 | 1.20454674 | 0.22837833 | NA | NA | NA |
| ENSG000002 | 1.46963536 | -0.452859584 | 0.33771611 | -1.3409475 | 0.17993749 | NA | NA | NA |
| ENSG000002 | 0.40191006 | -0.101653297 | 0.21376195 | -0.4755444 | 0.63439899 | NA | NA | NA |
| ENSG000002 | 1.40609948 | -0.232006765 | 0.32501375 | -0.7138368 | 0.47532812 | NA | NA | NA |
| ENSG000002 | 1.72755851 | -0.596333436 | 0.35246196 | -1.6919086 | 0.09066339 | NA | NA | NA |
| ENSG000002 | 0.91009755 | -0.261525473 | 0.29370035 | -0.89045 | 0.37322432 | NA | NA | NA |

| | | | | | | | | |
|---|---|---|---|---|---|---|---|---|
| ENSG000002 | 2.11614244 | -0.39483305 | 0.36752528 | -1.0743017 | 0.28268746 | NA | NA | NA |
| ENSG000002 | 1.35358594 | -0.381816805 | 0.33020243 | -1.1563113 | 0.24755384 | NA | NA | NA |
| ENSG000002 | 1.12407186 | 0.022789121 | 0.31614842 | 0.07208361 | 0.94253537 | NA | NA | NA |
| ENSG000002 | 2.00765506 | -0.176056348 | 0.35337754 | -0.4982103 | 0.61833579 | NA | FRMPD3-AS1 | 100874122 |
| ENSG000002 | 2.06459623 | -0.288846766 | 0.35183908 | -0.8209627 | 0.41166751 | NA | NA | NA |
| ENSG000002 | 0.37441639 | -0.268856313 | 0.21277471 | -1.2635727 | 0.20638344 | NA | NA | NA |
| ENSG000002 | 0.46341609 | 0.157328408 | 0.24338973 | 0.64640528 | 0.51801692 | NA | NA | NA |
| ENSG000002 | 1.61552283 | -0.271292275 | 0.35185546 | -0.7710333 | 0.4406872 | NA | NA | NA |
| ENSG000002 | 0 | NA | NA | NA | NA | NA | NA | NA |
| ENSG000002 | 0 | NA | NA | NA | NA | NA | NA | NA |
| ENSG000002 | 0 | NA | NA | NA | NA | NA | USP9YP21 | 106480355 |
| ENSG000002 | 1.23336676 | -0.339632216 | 0.33009147 | -1.0289034 | 0.30352509 | NA | NA | NA |
| ENSG000002 | 1.54456797 | -0.614646489 | 0.33872497 | -1.8145886 | 0.06958715 | NA | NA | NA |
| ENSG000002 | 1.68844078 | -0.389245989 | 0.34537329 | -1.1270298 | 0.25972988 | NA | NA | NA |
| ENSG000002 | 1.73633733 | -0.186963772 | 0.35100216 | -0.532657 | 0.59427101 | NA | NA | NA |
| ENSG000002 | 1.51297368 | -0.543669253 | 0.32842547 | -1.6553809 | 0.09784725 | NA | NA | NA |
| ENSG000002 | 0.87275415 | -0.250217546 | 0.29171115 | -0.8577579 | 0.39102615 | NA | NA | NA |
| ENSG000002 | 1.20288552 | 0.230141324 | 0.32149634 | 0.71584431 | 0.47408749 | NA | NA | NA |
| ENSG000002 | 1.23786373 | -0.193856899 | 0.33421021 | -0.5800448 | 0.56188439 | NA | NA | NA |
| ENSG000002 | 1.75589702 | -0.189590255 | 0.35562113 | -0.5331243 | 0.59394756 | NA | NA | NA |
| ENSG000002 | 1.13267234 | -0.440678296 | 0.32483798 | -1.3566095 | 0.17490532 | NA | NA | NA |
| ENSG000002 | 1.98623715 | -0.190249352 | 0.36231993 | -0.5250866 | 0.59952296 | NA | NA | NA |
| ENSG000002 | 1.45707265 | -0.29146283 | 0.3329972 | -0.8752711 | 0.38142641 | NA | NA | NA |
| ENSG000002 | 1.14029442 | -0.271156243 | 0.32393658 | -0.8370658 | 0.40255557 | NA | NA | NA |
| ENSG000002 | 0.47821768 | -0.016471461 | 0.24462483 | -0.0673336 | 0.94631616 | NA | NA | NA |
| ENSG000002 | 1.79338073 | -0.465810191 | 0.3550494 | -1.3119588 | 0.18953403 | NA | NA | NA |
| ENSG000002 | 0.44599794 | 0.298341626 | 0.22713929 | 1.31347433 | 0.18902317 | NA | NA | NA |
| ENSG000002 | 1.79571209 | -0.190686559 | 0.34964504 | -0.5453718 | 0.58549781 | NA | NA | NA |
| ENSG000002 | 1.37869171 | -0.27229286 | 0.3404782 | -0.7997366 | 0.42386345 | NA | NA | NA |

| | | | | | | | | |
|---|---|---|---|---|---|---|---|---|
| ENSG000002 | 1.91858346 | -0.512251246 | 0.35504123 | -1.4427937 | 0.1490786 | NA | NA | NA |
| ENSG000002 | 1.08909463 | -0.119821021 | 0.32148453 | -0.3727116 | 0.70936308 | NA | NA | NA |
| ENSG000002 | 1.84867597 | -0.248531438 | 0.35725778 | -0.6956642 | 0.48663916 | NA | NA | NA |
| ENSG000002 | 1.95610819 | -0.294014883 | 0.35737202 | -0.8227138 | 0.41067078 | NA | NA | NA |
| ENSG000002 | 1.62152511 | -0.259516539 | 0.350161 | -0.7411349 | 0.45861165 | NA | NA | NA |
| ENSG000002 | 0.99989045 | -0.180022187 | 0.3075418 | -0.5853584 | 0.55830674 | NA | NA | NA |
| ENSG000002 | 0.50818749 | -0.197539377 | 0.24915052 | -0.7928516 | 0.42786432 | NA | NA | NA |
| ENSG000002 | 1.05997855 | -0.50274004 | 0.30639411 | -1.640828 | 0.10083312 | NA | NA | NA |
| ENSG000002 | 2.13666342 | -0.657578696 | 0.36572244 | -1.7980267 | 0.07217278 | NA | NA | NA |
| ENSG000002 | 1.93324124 | -0.680156797 | 0.35877896 | -1.8957544 | 0.05799253 | NA | NA | NA |
| ENSG000002 | 2.08318228 | -0.888241584 | 0.36116706 | -2.4593649 | 0.01391831 | NA | NA | NA |
| ENSG000002 | 1.97755937 | -0.198011461 | 0.36313592 | -0.5452819 | 0.58555962 | NA | NA | NA |
| ENSG000002 | 1.00450153 | -0.198427005 | 0.30935055 | -0.6414309 | 0.52124276 | NA | NA | NA |
| ENSG000002 | 0.24550891 | -0.007886114 | 0.17900103 | -0.0440563 | 0.96485956 | NA | NA | NA |
| ENSG000002 | 2.01834534 | -0.449776209 | 0.36264368 | -1.2402704 | 0.21487541 | NA | NA | NA |
| ENSG000002 | 0.85767182 | -0.111725312 | 0.29715703 | -0.3759807 | 0.70693124 | NA | NA | NA |
| ENSG000002 | 0 | NA | NA | NA | NA | NA | NA | NA |
| ENSG000002 | 0.86023441 | -0.265815184 | 0.29686837 | -0.8953975 | 0.37057466 | NA | NA | NA |
| ENSG000002 | 1.16013164 | -0.426075742 | 0.32025662 | -1.3304198 | 0.18338001 | NA | NA | NA |
| ENSG000002 | 1.4152944 | -0.588676637 | 0.34021666 | -1.7302992 | 0.08357682 | NA | NA | NA |
| ENSG000002 | 0 | NA | NA | NA | NA | NA | NA | NA |
| ENSG000002 | 0 | NA | NA | NA | NA | NA | NA | NA |
| ENSG000002 | 0.14128499 | -0.097772014 | 0.13403235 | -0.7294658 | 0.4657168 | NA | NA | NA |
| ENSG000002 | 1.54468008 | -0.610012013 | 0.33877114 | -1.8006611 | 0.07175632 | NA | NA | NA |
| ENSG000002 | 1.235894 | -0.05222314 | 0.32869152 | -0.1588819 | 0.87376191 | NA | NA | NA |
| ENSG000002 | 1.00546954 | -0.039858873 | 0.31353044 | -0.1271292 | 0.89883815 | NA | NA | NA |
| ENSG000002 | 0.12070844 | -0.088503695 | 0.12783931 | -0.6923042 | 0.48874631 | NA | NA | NA |
| ENSG000002 | 1.01064761 | -0.198945547 | 0.31431679 | -0.632946 | 0.52676892 | NA | NA | NA |
| ENSG000002 | 0 | NA | NA | NA | NA | NA | NA | NA |

| | | | | | | | | |
|---|---|---|---|---|---|---|---|---|
| ENSG000002 | 0 | NA | NA | NA | NA | NA | NA | NA |
| ENSG000002 | 0 | NA | NA | NA | NA | NA | NA | NA |
| ENSG000002 | 0.11205239 | 0.082118251 | 0.12671962 | 0.64803108 | 0.51696485 | NA | NA | NA |
| ENSG000002 | 1.38176475 | -0.25494589 | 0.33353562 | -0.7643738 | 0.44464451 | NA | NA | NA |
| ENSG000002 | 0.11934008 | 0.083564698 | 0.12760091 | 0.65489106 | 0.5125379 | NA | NA | NA |
| ENSG000002 | 1.23643925 | -0.334106796 | 0.32794682 | -1.0187835 | 0.30830577 | NA | NA | NA |
| ENSG000002 | 1.15355295 | -0.108535401 | 0.31199783 | -0.3478723 | 0.72793609 | NA | NA | NA |
| ENSG000002 | 1.89188725 | -0.364381404 | 0.34908187 | -1.0438279 | 0.29656501 | NA | NA | NA |
| ENSG000002 | 1.71962468 | -0.054229536 | 0.35381693 | -0.15327 | 0.87818532 | NA | NA | NA |
| ENSG000002 | 0.35543399 | 0.061147018 | 0.20611885 | 0.29665903 | 0.76672684 | NA | NA | NA |
| ENSG000002 | 0 | NA | NA | NA | NA | NA | NA | NA |
| ENSG000002 | 0.11131518 | 0.081951103 | 0.12663804 | 0.64712866 | 0.51754868 | NA | NA | NA |
| ENSG000002 | 2.09023431 | -0.003024136 | 0.35893736 | -0.0084252 | 0.99327771 | NA | NA | NA |
| ENSG000002 | 0 | NA | NA | NA | NA | NA | NA | NA |
| ENSG000002 | 0.84108022 | -0.239563598 | 0.28742516 | -0.8334816 | 0.40457314 | NA | NA | NA |
| ENSG000002 | 1.85264322 | -0.243284741 | 0.3567581 | -0.6819319 | 0.49528199 | NA | NA | NA |
| ENSG000002 | 1.36013194 | -0.124803194 | 0.33955955 | -0.3675443 | 0.71321302 | NA | NA | NA |
| ENSG000002 | 0.99184382 | -0.351960866 | 0.3102115 | -1.1345836 | 0.25654984 | NA | FRMD8P1 | 83957 |
| ENSG000002 | 2.07543116 | -0.004764787 | 0.35951378 | -0.0132534 | 0.98942561 | NA | NA | NA |
| ENSG000002 | 0 | NA | NA | NA | NA | NA | NA | NA |
| ENSG000002 | 0.5834394 | 0.062235112 | 0.26097587 | 0.23847075 | 0.811516 | NA | NA | NA |
| ENSG000002 | 0.70983636 | -0.026536747 | 0.2764396 | -0.0959947 | 0.92352475 | NA | NA | NA |
| ENSG000002 | 1.38263993 | -0.123830374 | 0.33809744 | -0.3662565 | 0.71417371 | NA | NA | NA |
| ENSG000002 | 2.08258943 | -0.840928571 | 0.35666193 | -2.357775 | 0.01838483 | NA | NA | NA |
| ENSG000002 | 1.10880977 | -0.110672854 | 0.30767267 | -0.3597097 | 0.71906421 | NA | NA | NA |
| ENSG000002 | 1.0018855 | -0.193935658 | 0.30925346 | -0.6271091 | 0.53058772 | NA | NA | NA |
| ENSG000002 | 1.90295643 | -0.137213861 | 0.35743536 | -0.3838844 | 0.70106413 | NA | NA | NA |
| ENSG000002 | 1.38874701 | -0.706088395 | 0.33249283 | -2.1236199 | 0.03370194 | NA | NA | NA |
| ENSG000002 | 1.25954221 | -0.052052953 | 0.19633047 | -0.2651293 | 0.79090988 | NA | NA | NA |

| | | | | | | | | | |
|---|---|---|---|---|---|---|---|---|---|
| ENSG000002 | 0 | NA | NA | NA | NA | NA | NA | NA | |
| ENSG000002 | 0.5889017 | 0.229808217 | 0.26256547 | 0.87524159 | 0.38144247 | NA | NA | NA | |
| ENSG000002 | 1.40179859 | -0.372166623 | 0.32758681 | -1.1360855 | 0.25592079 | NA | NA | NA | |
| ENSG000002 | 0.37692244 | -0.098341116 | 0.20997214 | -0.4683532 | 0.63953205 | NA | NA | NA | |
| ENSG000002 | 1.14353714 | -0.124807375 | 0.31714394 | -0.3935354 | 0.69392407 | NA | NA | NA | |
| ENSG000002 | 0.71766538 | -0.02357197 | 0.27700307 | -0.0850964 | 0.93218474 | NA | NA | NA | |
| ENSG000002 | 0.62197881 | -0.414424795 | 0.25447679 | -1.6285367 | 0.10341113 | NA | NA | NA | |
| ENSG000002 | 1.99047004 | -0.309520056 | 0.36091915 | -0.8575883 | 0.39111982 | NA | NA | NA | |
| ENSG000002 | 1.44043497 | -0.901672555 | 0.33285334 | -2.7089185 | 0.00675029 | NA | NA | NA | |
| ENSG000002 | 1.66917703 | -0.866955169 | 0.34796773 | -2.4914815 | 0.01272116 | NA | NA | NA | |
| ENSG000002 | 0.61102131 | -0.109822872 | 0.27020887 | -0.406437 | 0.68442157 | NA | NA | NA | |
| ENSG000002 | 1.21412377 | -0.042244342 | 0.32958914 | -0.1281727 | 0.89801227 | NA | NA | NA | |
| ENSG000002 | 1.70585865 | -0.060657596 | 0.35585759 | -0.1704547 | 0.86465256 | NA | NA | NA | |
| ENSG000002 | 1.548811 | -0.101909566 | 0.34059826 | -0.2992075 | 0.7647817 | NA | NA | NA | |
| ENSG000002 | 1.94032361 | 0.060885971 | 0.36012176 | 0.16907051 | 0.86574118 | NA | NA | NA | |
| ENSG000002 | 0.88632534 | -0.11528753 | 0.29864358 | -0.3860372 | 0.69946912 | NA | NA | NA | |
| ENSG000002 | 1.49717442 | 0.079222991 | 0.33038301 | 0.23979136 | 0.81049201 | NA | NA | NA | |
| ENSG000002 | 1.00357291 | -0.492055205 | 0.30397229 | -1.6187502 | 0.10550102 | NA | NA | NA | |
| ENSG000002 | 1.89667336 | -0.817031791 | 0.35668534 | -2.2906234 | 0.0219852 | NA | IFNA14 | | 3448 |
| ENSG000002 | 1.10944147 | -0.257146232 | 0.31698087 | -0.8112358 | 0.41723027 | NA | NA | NA | |
| ENSG000002 | 1.7622515 | -0.760156315 | 0.35308213 | -2.1529164 | 0.03132525 | NA | COX6CP15 | | 106480268 |
| ENSG000002 | 1.65454099 | -0.516327886 | 0.34472741 | -1.497786 | 0.13418886 | NA | NA | NA | |
| ENSG000002 | 1.52476301 | -0.496208817 | 0.34624859 | -1.4330999 | 0.15182929 | NA | NA | NA | |
| ENSG000002 | 1.46308229 | -0.548344973 | 0.33177279 | -1.6527726 | 0.09837715 | NA | NA | NA | |
| ENSG000002 | 0.96789747 | -0.034189063 | 0.31202114 | -0.1095729 | 0.9127481 | NA | NA | NA | |
| ENSG000002 | 1.30704583 | 0.312563146 | 0.32913986 | 0.94963626 | 0.34229711 | NA | NA | NA | |
| ENSG000002 | 0.25214371 | -0.009862892 | 0.18046657 | -0.0546522 | 0.95641556 | NA | NA | NA | |
| ENSG000002 | 1.81945384 | -0.756338608 | 0.35201096 | -2.1486223 | 0.03166435 | NA | NA | NA | |
| ENSG000002 | 0.59892612 | -0.099249839 | 0.26277623 | -0.3776972 | 0.70565557 | NA | NA | NA | |

| | | | | | | | | |
|---|---|---|---|---|---|---|---|---|
| ENSG000002 | 0 | NA | NA | NA | NA | NA | NA | NA |
| ENSG000002 | 2.05583672 | -0.232590826 | 0.36037641 | -0.6454108 | 0.51866101 | NA | NA | NA |
| ENSG000002 | 2.162356 | -0.618854269 | 0.3615906 | -1.7114778 | 0.08699295 | NA | NA | NA |
| ENSG000002 | 1.08886842 | 0.03543426 | 0.3211724 | 0.11032785 | 0.91214937 | NA | NA | NA |
| ENSG000002 | 2.12923597 | -0.377236281 | 0.36441866 | -1.0351728 | 0.30058822 | NA | LOC102724188 | 102724188 |
| ENSG000002 | 1.38546448 | -0.127084196 | 0.33300937 | -0.3816235 | 0.70274066 | NA | NA | NA |
| ENSG000002 | 1.03211432 | -0.521938666 | 0.31001467 | -1.6835935 | 0.09226026 | NA | NA | NA |
| ENSG000002 | 0.98357615 | 0.106999703 | 0.30617926 | 0.3494675 | 0.72673836 | NA | NA | NA |
| ENSG000002 | 1.15953486 | -0.112296443 | 0.31036235 | -0.3618237 | 0.71748381 | NA | NA | NA |
| ENSG000002 | 1.51994445 | -0.322768867 | 0.34093674 | -0.9467119 | 0.34378561 | NA | NA | NA |
| ENSG000002 | 0.36576034 | -0.0934779 | 0.20820828 | -0.4489634 | 0.65345805 | NA | NA | NA |
| ENSG000002 | 0.7461825 | -0.025114767 | 0.28524947 | -0.0880449 | 0.92984097 | NA | NA | NA |
| ENSG000002 | 0 | NA | NA | NA | NA | NA | TCEB1P14 | 100462876 |
| ENSG000002 | 2.05575082 | -0.333606977 | 0.35750218 | -0.9331607 | 0.35073702 | NA | NA | NA |
| ENSG000002 | 1.71579619 | 0.065202637 | 0.34913642 | 0.18675404 | 0.85185348 | NA | NA | NA |
| ENSG000002 | 0 | NA | NA | NA | NA | NA | NA | NA |
| ENSG000002 | 0.12070844 | -0.088503695 | 0.12783931 | -0.6923042 | 0.48874631 | NA | NA | NA |
| ENSG000002 | 1.37279412 | -0.257004092 | 0.33494059 | -0.7673125 | 0.44289575 | NA | NA | NA |
| ENSG000002 | 0.96258088 | -0.028120982 | 0.29923895 | -0.093975 | 0.92512901 | NA | NA | NA |
| ENSG000002 | 0 | NA | NA | NA | NA | NA | NA | NA |
| ENSG000002 | 1.11871953 | -0.260732043 | 0.31949692 | -0.8160706 | 0.41445975 | NA | NA | NA |
| ENSG000002 | 0 | NA | NA | NA | NA | NA | NA | NA |
| ENSG000002 | 0.8565842 | 0.046409282 | 0.28894415 | 0.16061679 | 0.87239523 | NA | NA | NA |
| ENSG000002 | 1.69356733 | -0.511258202 | 0.34191426 | -1.4952819 | 0.13484089 | NA | NA | NA |
| ENSG000002 | 0.94644127 | 0.120225999 | 0.31106787 | 0.38649443 | 0.69913052 | NA | NA | NA |
| ENSG000002 | 1.31422942 | -0.488715442 | 0.32603475 | -1.4989673 | 0.13388211 | NA | NA | NA |
| ENSG000002 | 1.60363007 | -0.602383505 | 0.33629155 | -1.7912537 | 0.07325259 | NA | NA | NA |
| ENSG000002 | 2.01473097 | -0.456765551 | 0.36260452 | -1.2596797 | 0.20778493 | NA | BMS1P17 | 101101776 |
| ENSG000002 | 0 | NA | NA | NA | NA | NA | NA | NA |

| | | | | | | | | |
|---|---|---|---|---|---|---|---|---|
| ENSG000002 | 1.5320507 | -0.480140938 | 0.34364059 | -1.3972183 | 0.16234792 | NA | NA | NA |
| ENSG000002 | 1.32893675 | 0.032400727 | 0.33731989 | 0.09605341 | 0.92347815 | NA | NA | NA |
| ENSG000002 | 0.4842194 | -0.018518118 | 0.24529708 | -0.0754926 | 0.93982277 | NA | GUSBP11 | 91316 |
| ENSG000002 | 0.75070768 | -0.185851474 | 0.28031532 | -0.6630086 | 0.50732504 | NA | NA | NA |
| ENSG000002 | 1.85476606 | 0.005824819 | 0.35783759 | 0.01627783 | 0.98701275 | NA | NA | NA |
| ENSG000002 | 0.78805107 | -0.194877761 | 0.2810673 | -0.6933491 | 0.4880905 | NA | NA | NA |
| ENSG000002 | 1.17766603 | -0.557826762 | 0.3134051 | -1.7798905 | 0.07509388 | NA | NA | NA |
| ENSG000002 | 1.67979835 | -0.84292955 | 0.34519882 | -2.4418669 | 0.01461153 | NA | NA | NA |
| ENSG000002 | 1.8484261 | -0.388791246 | 0.35895271 | -1.0831267 | 0.27875221 | NA | NA | NA |
| ENSG000002 | 1.38420417 | -0.250846814 | 0.33113331 | -0.7575403 | 0.44872626 | NA | NA | NA |
| ENSG000002 | 1.62849767 | -0.401408617 | 0.34981349 | -1.1474932 | 0.25117783 | NA | NA | NA |
| ENSG000002 | 1.698377 | 0.069966573 | 0.3520921 | 0.19871668 | 0.84248438 | NA | NA | NA |
| ENSG000002 | 1.84640485 | -0.265816689 | 0.36169509 | -0.7349193 | 0.46238867 | NA | NA | NA |
| ENSG000002 | 1.4564566 | -0.027993694 | 0.32341547 | -0.0865564 | 0.93102409 | NA | NA | NA |
| ENSG000002 | 1.59155922 | -0.396526642 | 0.34931617 | -1.1351511 | 0.256312 | NA | NA | NA |
| ENSG000002 | 2.02715008 | 0.247365278 | 0.36110578 | 0.68502165 | 0.49333026 | NA | NA | NA |
| ENSG000002 | 1.20253767 | -0.034017046 | 0.32776517 | -0.1037848 | 0.91734012 | NA | NA | NA |
| ENSG000002 | 1.38481605 | -0.413916498 | 0.33842138 | -1.2230802 | 0.2212994 | NA | NA | NA |
| ENSG000002 | 0 | NA | NA | NA | NA | NA | NA | NA |
| ENSG000002 | 0 | NA | NA | NA | NA | NA | NA | NA |
| ENSG000002 | 1.00220454 | -0.34457114 | 0.30907251 | -1.1148554 | 0.26491241 | NA | NA | NA |
| ENSG000002 | 1.60089594 | -0.267774634 | 0.35166997 | -0.7614373 | 0.44639591 | NA | LOC101927534 | 101927534 |
| ENSG000002 | 1.38988414 | -0.401899387 | 0.3326274 | -1.208257 | 0.22694843 | NA | NA | NA |
| ENSG000002 | 0.63270563 | -0.25685676 | 0.259029 | -0.9916139 | 0.3213859 | NA | HSPC324 | 101928612 |
| ENSG000002 | 1.26593244 | -0.505393704 | 0.33003692 | -1.5313247 | 0.12568915 | NA | NA | NA |
| ENSG000002 | 2.14072277 | -0.019285466 | 0.36087762 | -0.0534405 | 0.95738097 | NA | NA | NA |
| ENSG000002 | 0 | NA | NA | NA | NA | NA | NA | NA |
| ENSG000002 | 2.02150428 | -0.312435463 | 0.36112321 | -0.8651769 | 0.3869417 | NA | NA | NA |
| ENSG000002 | 0.85823489 | 0.050766245 | 0.30196548 | 0.16811937 | 0.86648937 | NA | ARL5AP3 | 106480778 |

| | | | | | | | | | |
|---|---|---|---|---|---|---|---|---|---|
| ENSG000002 | 1.16155557 | -0.265847841 | 0.3162311 | -0.8406758 | 0.40052957 | NA | NA | NA | |
| ENSG000002 | 1.80496683 | -0.436944608 | 0.34784368 | -1.2561522 | 0.2090608 | NA | NA | NA | |
| ENSG000002 | 1.75575714 | -0.453782978 | 0.35311134 | -1.2850989 | 0.19875774 | NA | NA | NA | |
| ENSG000002 | 1.2643682 | -0.336304039 | 0.32690827 | -1.0287413 | 0.30360125 | NA | NA | NA | |
| ENSG000002 | 0.58412454 | 0.062473285 | 0.26103841 | 0.23932603 | 0.81085278 | NA | NA | NA | |
| ENSG000002 | 2.16902305 | -0.649632707 | 0.36490596 | -1.7802743 | 0.07503108 | NA | NA | NA | |
| ENSG000002 | 0 | NA | NA | NA | NA | NA | NA | NA | |
| ENSG000002 | 1.85978251 | -0.495307654 | 0.35216248 | -1.4064748 | 0.15958319 | NA | NA | NA | |
| ENSG000002 | 0.95418597 | 0.121684757 | 0.30650365 | 0.39700916 | 0.69136071 | NA | NA | NA | |
| ENSG000002 | 0 | NA | NA | NA | NA | NA | NA | NA | |
| ENSG000002 | 0.93722053 | -0.60694943 | 0.29365529 | -2.0668772 | 0.03874572 | NA | NA | NA | |
| ENSG000002 | 0.98480654 | -0.179926523 | 0.306894 | -0.5862823 | 0.55768582 | NA | NA | NA | |
| ENSG000002 | 2.16727522 | 0.312152947 | 0.36909771 | 0.845719 | 0.39770952 | NA | NA | NA | |
| ENSG000002 | 0.65465054 | -0.455468986 | 0.2643416 | -1.7230318 | 0.08488278 | NA | NA | NA | |
| ENSG000002 | 0 | NA | NA | NA | NA | NA | CT47A7 | | 653282 |
| ENSG000002 | 0 | NA | NA | NA | NA | NA | NA | NA | |
| ENSG000002 | 0.64235948 | -0.482579697 | 0.27062934 | -1.7831759 | 0.07455768 | NA | NA | NA | |
| ENSG000002 | 0.88009336 | -0.251804905 | 0.28961173 | -0.8694569 | 0.38459729 | NA | NA | NA | |
| ENSG000002 | 1.8591545 | -0.123121504 | 0.35323559 | -0.3485535 | 0.72742453 | NA | NA | NA | |
| ENSG000002 | 1.17556045 | -0.407455977 | 0.31337997 | -1.3001979 | 0.19353316 | NA | NA | NA | |
| ENSG000002 | 1.89717797 | -0.527077147 | 0.35808985 | -1.4719131 | 0.14104434 | NA | NA | NA | |
| ENSG000002 | 1.4643386 | -0.437263084 | 0.33524287 | -1.3043173 | 0.19212542 | NA | NA | NA | |
| ENSG000002 | 1.42766548 | 0.536085499 | 0.33980687 | 1.5776182 | 0.11465335 | NA | NA | NA | |
| ENSG000002 | 1.05568568 | -0.0946104 | 0.30377862 | -0.3114452 | 0.75546218 | NA | NA | NA | |
| ENSG000002 | 0 | NA | NA | NA | NA | NA | HSFY7P | | 106478935 |
| ENSG000002 | 1.13269353 | -0.257982953 | 0.31529159 | -0.8182361 | 0.41322239 | NA | NA | NA | |
| ENSG000002 | 0 | NA | NA | NA | NA | NA | NA | NA | |
| ENSG000002 | 2.03451474 | -0.751614973 | 0.36329791 | -2.0688668 | 0.03855859 | NA | NA | NA | |
| ENSG000002 | 0.76189543 | -0.351748879 | 0.28173689 | -1.2485013 | 0.21184752 | NA | NA | NA | |

| | | | | | | | | |
|---|---|---|---|---|---|---|---|---|
| ENSG000002 | 1.38743366 | -0.265532018 | 0.33524646 | -0.7920502 | 0.42833142 | NA | NA | NA |
| ENSG000002 | 1.74514846 | -0.443541941 | 0.35036472 | -1.2659435 | 0.20553331 | NA | NA | NA |
| ENSG000002 | 1.3332252 | 0.15687055 | 0.31891243 | 0.49189223 | 0.62279553 | NA | NA | NA |
| ENSG000002 | 1.64660254 | -0.408741557 | 0.35181949 | -1.1617934 | 0.24531941 | NA | NA | NA |
| ENSG000002 | 2.09712855 | -0.604865274 | 0.35990379 | -1.6806305 | 0.09283471 | NA | NA | NA |
| ENSG000002 | 1.9639584 | -0.180258283 | 0.36065132 | -0.4998132 | 0.61720662 | NA | NA | NA |
| ENSG000002 | 0.88749783 | -0.277546523 | 0.29984721 | -0.9256265 | 0.35464012 | NA | NA | NA |
| ENSG000002 | 0.12685397 | -0.090215768 | 0.12901919 | -0.699243 | 0.48440017 | NA | NA | NA |
| ENSG000002 | 0.73406558 | -0.177820193 | 0.27747212 | -0.6408579 | 0.52161503 | NA | NA | NA |
| ENSG000002 | 1.2791354 | -0.659029751 | 0.32727673 | -2.0136774 | 0.04404341 | NA | NA | NA |
| ENSG000002 | 0.26813897 | -0.197007041 | 0.18550319 | -1.0620143 | 0.2882292 | NA | NA | NA |
| ENSG000002 | 1.2390574 | -0.198464816 | 0.33422895 | -0.593799 | 0.55264657 | NA | LOC105376527 | 105376527 |
| ENSG000002 | 0.79437129 | -0.367466481 | 0.28536516 | -1.2877062 | 0.19784826 | NA | NA | NA |
| ENSG000002 | 1.98923737 | -0.83654405 | 0.35656313 | -2.3461317 | 0.01896941 | NA | NA | NA |
| ENSG000002 | 1.12577696 | -0.39284248 | 0.31389786 | -1.2514978 | 0.21075291 | NA | NA | NA |
| ENSG000002 | 1.89131668 | -0.258799685 | 0.3593248 | -0.7202389 | 0.47137794 | NA | NA | NA |
| ENSG000002 | 1.02232463 | -0.350176688 | 0.31151676 | -1.1241022 | 0.26096966 | NA | NA | NA |
| ENSG000002 | 0.89207857 | -0.119364867 | 0.29126901 | -0.4098097 | 0.68194555 | NA | NA | NA |
| ENSG000002 | 0.23202362 | -0.008302493 | 0.17586207 | -0.0472103 | 0.96234565 | NA | NA | NA |
| ENSG000002 | 1.2366695 | -0.189059345 | 0.32922736 | -0.5742516 | 0.56579754 | NA | NA | NA |
| ENSG000002 | 1.11148583 | -0.10221147 | 0.3077569 | -0.3321176 | 0.73980049 | NA | NA | NA |
| ENSG000002 | 1.36250313 | -0.512562455 | 0.32792745 | -1.5630361 | 0.11804409 | NA | NA | NA |
| ENSG000002 | 1.06697557 | 0.195204077 | 0.32057318 | 0.60892205 | 0.54257611 | NA | NA | NA |
| ENSG000002 | 0.63205432 | -0.109111113 | 0.258189 | -0.4226017 | 0.67258588 | NA | NA | NA |
| ENSG000002 | 2.02408997 | -0.183081907 | 0.35963759 | -0.5090733 | 0.61070082 | NA | NA | NA |
| ENSG000002 | 2.15546912 | -0.341127474 | 0.35606847 | -0.958039 | 0.3380431 | NA | NA | NA |
| ENSG000002 | 1.99615002 | -0.460269376 | 0.36360734 | -1.2658418 | 0.20556972 | NA | NA | NA |
| ENSG000002 | 1.46276934 | -0.057028355 | 0.33451477 | -0.1704808 | 0.86463201 | NA | NA | NA |
| ENSG000002 | 0.471387 | -0.012999703 | 0.2361605 | -0.0550461 | 0.95610178 | NA | NA | NA |

| | | | | | | | | |
|---|---|---|---|---|---|---|---|---|
| ENSG000002 | 0 | NA | NA | NA | NA | NA | NA | NA |
| ENSG000002 | 1.30793093 | -0.187483777 | 0.32137227 | -0.583385 | 0.55963412 | NA | NA | NA |
| ENSG000002 | 0.5821231 | 0.217379449 | 0.25471767 | 0.85341332 | 0.39343014 | NA | NA | NA |
| ENSG000002 | 1.11421864 | -0.111453877 | 0.31614263 | -0.352543 | 0.72443107 | NA | NA | NA |
| ENSG000002 | 1.42568095 | 0.351684295 | 0.33052212 | 1.06402649 | 0.2873167 | NA | NA | NA |
| ENSG000002 | 1.0307137 | -0.332427312 | 0.30497862 | -1.090002 | 0.27571226 | NA | NA | NA |
| ENSG000002 | 1.71195266 | -0.0544045 | 0.35591181 | -0.1528595 | 0.87850908 | NA | NA | NA |
| ENSG000002 | 0.11865494 | 0.083440866 | 0.12751478 | 0.65436232 | 0.5128784 | NA | NA | NA |
| ENSG000002 | 0 | NA | NA | NA | NA | NA | NLGN4Y-AS1 | 100874056 |
| ENSG000002 | 1.48707236 | -0.057494952 | 0.3493439 | -0.1645798 | 0.86927473 | NA | NA | NA |
| ENSG000002 | 0.97478425 | 0.273331145 | 0.30658847 | 0.89152453 | 0.37264784 | NA | NA | NA |
| ENSG000002 | 0.885162 | -0.288936305 | 0.30450304 | -0.9488782 | 0.34268255 | NA | NA | NA |
| ENSG000002 | 0.87363325 | -0.379412923 | 0.28258355 | -1.3426575 | 0.1793829 | NA | LAMA5-AS1 | 101928158 |
| ENSG000002 | 0.90064872 | -0.456924471 | 0.30237983 | -1.5110944 | 0.1307644 | NA | NA | NA |
| ENSG000002 | 1.42164845 | -0.566129195 | 0.33506933 | -1.6895882 | 0.09110676 | NA | NA | NA |
| ENSG000002 | 1.83962817 | -0.127898249 | 0.36134415 | -0.3539514 | 0.72337534 | NA | NA | NA |
| ENSG000002 | 1.75365101 | -0.328907058 | 0.35442798 | -0.9279941 | 0.35341065 | NA | NA | NA |
| ENSG000002 | 0.25260017 | -0.015978705 | 0.18049464 | -0.0885273 | 0.92945758 | NA | NA | NA |
| ENSG000002 | 0.14128499 | -0.097772014 | 0.13403235 | -0.7294658 | 0.4657168 | NA | NA | NA |
| ENSG000002 | 0.64488726 | -0.109815036 | 0.26668753 | -0.4117742 | 0.68050496 | NA | NA | NA |
| ENSG000002 | 1.1448196 | -0.438281139 | 0.32380937 | -1.3535159 | 0.17589087 | NA | NA | NA |
| ENSG000002 | 0 | NA | NA | NA | NA | NA | NA | NA |
| ENSG000002 | 0.12070844 | -0.088503695 | 0.12783931 | -0.6923042 | 0.48874631 | NA | NA | NA |
| ENSG000002 | 0.50613398 | -0.020909936 | 0.24769218 | -0.084419 | 0.93272327 | NA | NA | NA |
| ENSG000002 | 0 | NA | NA | NA | NA | NA | USP17L30 | 728419 |
| ENSG000002 | 1.97926405 | -0.071289722 | 0.36502349 | -0.1953017 | 0.84515674 | NA | NA | NA |
| ENSG000002 | 1.97223695 | -0.072093829 | 0.35700921 | -0.2019383 | 0.83996497 | NA | NA | NA |
| ENSG000002 | 0.23065526 | 0.167384503 | 0.17662711 | 0.94767164 | 0.34329664 | NA | NA | NA |
| ENSG000002 | 1.87295726 | -0.385245603 | 0.35917108 | -1.0725964 | 0.28345225 | NA | NA | NA |

| | | | | | | | | |
|---|---|---|---|---|---|---|---|---|
| ENSG000002 | 0 | NA | NA | NA | NA | NA | NA | NA |
| ENSG000002 | 1.72094447 | 0.072579799 | 0.35375451 | 0.20516996 | 0.83743934 | NA | NA | NA |
| ENSG000002 | 0.7226728 | 0.292544338 | 0.27764568 | 1.05366068 | 0.2920383 | NA | NA | NA |
| ENSG000002 | 0.85670436 | -0.256192558 | 0.29077187 | -0.8810775 | 0.37827586 | NA | NA | NA |
| ENSG000002 | 0.79670713 | -0.362882081 | 0.28391224 | -1.2781488 | 0.20119698 | NA | NA | NA |
| ENSG000002 | 0.88086495 | -0.113054583 | 0.29836514 | -0.3789135 | 0.70475209 | NA | LOC389199 | 389199 |
| ENSG000002 | 0 | NA | NA | NA | NA | NA | TSPY3 | 728137 |
| ENSG000002 | 0.40327843 | -0.281391716 | 0.21700406 | -1.2967118 | 0.19473038 | NA | NA | NA |
| ENSG000002 | 1.9430628 | -0.056257753 | 0.36138192 | -0.155674 | 0.87629003 | NA | NA | NA |
| ENSG000002 | 0 | NA | NA | NA | NA | NA | NA | NA |
| ENSG000002 | 0.11934008 | 0.083564698 | 0.12760091 | 0.65489106 | 0.5125379 | NA | NA | NA |
| ENSG000002 | 1.01064761 | -0.198945547 | 0.31431679 | -0.632946 | 0.52676892 | NA | NA | NA |
| ENSG000002 | 1.58887647 | 0.017228668 | 0.34326211 | 0.050191 | 0.95997018 | NA | NA | NA |
| ENSG000002 | 1.49100469 | -0.201581821 | 0.34610755 | -0.5824254 | 0.56028019 | NA | NA | NA |
| ENSG000002 | 1.76429193 | -0.323028591 | 0.35454637 | -0.9111039 | 0.36224061 | NA | OR2A20P | 401428 |
| ENSG000002 | 0.25351207 | -0.189965547 | 0.1824784 | -1.0410303 | 0.29786148 | NA | NA | NA |
| ENSG000002 | 0.61605906 | 0.051157992 | 0.2638843 | 0.19386523 | 0.84628141 | NA | NA | NA |
| ENSG000002 | 1.34717693 | 0.016937297 | 0.33160838 | 0.0510762 | 0.9592648 | NA | NA | NA |
| ENSG000002 | 1.62986795 | -0.400952835 | 0.34983402 | -1.146123 | 0.25174424 | NA | NA | NA |
| ENSG000002 | 1.89188937 | -0.250935465 | 0.35290186 | -0.711063 | 0.47704518 | NA | NA | NA |
| ENSG000002 | 1.07694128 | -0.50735963 | 0.30706395 | -1.652293 | 0.09847483 | NA | NA | NA |
| ENSG000002 | 1.50195827 | -0.205311559 | 0.34880431 | -0.5886153 | 0.55611936 | NA | NA | NA |
| ENSG000002 | 0.34936227 | 0.241139221 | 0.20762307 | 1.16142789 | 0.24546793 | NA | NA | NA |
| ENSG000002 | 1.57616423 | -0.242993074 | 0.34399293 | -0.7063897 | 0.47994581 | NA | NA | NA |
| ENSG000002 | 0 | NA | NA | NA | NA | NA | NA | NA |
| ENSG000002 | 1.13130398 | -0.273022612 | 0.32364448 | -0.843588 | 0.39889966 | NA | NA | NA |
| ENSG000002 | 0.74997047 | -0.186132153 | 0.28026364 | -0.6641324 | 0.5066056 | NA | NA | NA |
| ENSG000002 | 1.86675911 | -0.377482991 | 0.35581681 | -1.0608914 | 0.28873926 | NA | NA | NA |
| ENSG000002 | 1.65397401 | -0.410461402 | 0.3501272 | -1.1723208 | 0.24106828 | NA | LINC01038 | 102724076 |

| | | | | | | | | |
|---|---|---|---|---|---|---|---|---|
| ENSG000002 | 0.25993993 | -0.013144386 | 0.18211683 | -0.0721756 | 0.94246219 | NA | NA | NA |
| ENSG000002 | 1.83297163 | -0.258139962 | 0.35884227 | -0.7193689 | 0.47191367 | NA | NA | NA |
| ENSG000002 | 2.01922602 | -0.439481913 | 0.35972584 | -1.2217135 | 0.22181599 | NA | NA | NA |
| ENSG000002 | 1.63362422 | -0.552084266 | 0.35013938 | -1.5767557 | 0.11485175 | NA | NA | NA |
| ENSG000002 | 2.15060763 | -0.507597025 | 0.36515154 | -1.3900997 | 0.16449862 | NA | NA | NA |
| ENSG000002 | 1.79959042 | -0.459725588 | 0.3509847 | -1.3098166 | 0.19025788 | NA | NA | NA |
| ENSG000002 | 1.97331494 | -0.062996704 | 0.36571544 | -0.1722561 | 0.86323622 | NA | NA | NA |
| ENSG000002 | 1.35472575 | 0.024077925 | 0.33580345 | 0.07170243 | 0.94283872 | NA | NA | NA |
| ENSG000002 | 2.09744525 | -0.384343219 | 0.36698377 | -1.047303 | 0.29495983 | NA | NA | NA |
| ENSG000002 | 1.9676476 | 0.067675019 | 0.3623998 | 0.18674132 | 0.85186346 | NA | NA | NA |
| ENSG000002 | 1.57186134 | 0.017264429 | 0.35294588 | 0.04891523 | 0.96098685 | NA | NA | NA |
| ENSG000002 | 1.54672912 | -0.47048938 | 0.33911847 | -1.3873894 | 0.16532305 | NA | NA | NA |
| ENSG000002 | 1.09506602 | 0.035890705 | 0.3285751 | 0.10923136 | 0.91301899 | NA | NA | NA |
| ENSG000002 | 1.49120111 | -0.200569359 | 0.34970457 | -0.5735394 | 0.56627952 | NA | NA | NA |
| ENSG000002 | 1.28911859 | 0.43537804 | 0.32393356 | 1.34403501 | 0.17893707 | NA | NA | NA |
| ENSG000002 | 1.65243814 | 0.215609397 | 0.34765456 | 0.62018285 | 0.53513741 | NA | NA | NA |
| ENSG000002 | 1.67500702 | -0.359408879 | 0.33392244 | -1.0763244 | 0.28178219 | NA | NA | NA |
| ENSG000002 | 0.8586919 | 0.050622524 | 0.29079913 | 0.17408073 | 0.86180202 | NA | NA | NA |
| ENSG000002 | 1.6395218 | -0.567013042 | 0.35250079 | -1.608544 | 0.10771608 | NA | NA | NA |
| ENSG000002 | 0.36482725 | -0.102221582 | 0.21647267 | -0.4722147 | 0.63677353 | NA | NA | NA |
| ENSG000002 | 1.60242647 | 0.013764275 | 0.34700122 | 0.03966636 | 0.96835912 | NA | NA | NA |
| ENSG000002 | 1.00023536 | -0.191785415 | 0.30918425 | -0.6202949 | 0.53506365 | NA | NA | NA |
| ENSG000002 | 1.83943676 | -0.366869348 | 0.35556659 | -1.031788 | 0.30217144 | NA | AGBL5-IT1 | 106481744 |
| ENSG000002 | 0 | NA | NA | NA | NA | NA | NA | NA |
| ENSG000002 | 1.88912328 | -0.370677525 | 0.35603783 | -1.0411184 | 0.29782061 | NA | NA | NA |
| ENSG000002 | 0 | NA | NA | NA | NA | NA | NA | NA |
| ENSG000002 | 1.24235665 | -0.319018776 | 0.32386488 | -0.9850367 | 0.32460606 | NA | NA | NA |
| ENSG000002 | 0.75734405 | -0.037539772 | 0.27972812 | -0.1342009 | 0.8932437 | NA | NA | NA |
| ENSG000002 | 0.76576985 | -0.381700686 | 0.28891137 | -1.3211688 | 0.18644509 | NA | NA | NA |

| | | | | | | | | | |
|---|---|---|---|---|---|---|---|---|---|
| ENSG000002 | 0.14128499 | -0.097772014 | 0.13403235 | -0.7294658 | 0.4657168 | NA | NA | NA | |
| ENSG000002 | 0 | NA | NA | NA | NA | NA | NA | NA | |
| ENSG000002 | 1.49937896 | -0.466284626 | 0.34178641 | -1.3642574 | 0.17248658 | NA | NA | NA | |
| ENSG000002 | 0 | NA | NA | NA | NA | NA | NA | NA | |
| ENSG000002 | 1.09231202 | -0.109500448 | 0.32007435 | -0.3421094 | 0.73226856 | NA | NA | NA | |
| ENSG000002 | 2.01046425 | -0.442133926 | 0.36068918 | -1.2258031 | 0.22027277 | NA | NA | NA | |
| ENSG000002 | 1.26763712 | -0.360055083 | 0.33432432 | -1.0769635 | 0.28149658 | NA | NA | NA | |
| ENSG000002 | 0.83623791 | 0.197487193 | 0.28602229 | 0.69046085 | 0.48990442 | NA | NA | NA | |
| ENSG000002 | 1.76853692 | -0.47512972 | 0.35530444 | -1.3372468 | 0.18114209 | NA | NA | NA | |
| ENSG000002 | 1.76343265 | -0.179830477 | 0.35203082 | -0.5108373 | 0.60946498 | NA | PGA4 | | 643847 |
| ENSG000002 | 0 | NA | NA | NA | NA | NA | NA | NA | |
| ENSG000002 | 2.14557253 | 0.048497624 | 0.36099203 | 0.13434542 | 0.89312944 | NA | NA | NA | |
| ENSG000002 | 1.36328931 | -0.122304576 | 0.3340225 | -0.3661567 | 0.71424815 | NA | NA | NA | |
| ENSG000002 | 1.87167642 | 0.457138625 | 0.35847407 | 1.27523483 | 0.20222614 | NA | NA | NA | |
| ENSG000002 | 1.25517081 | -0.191866236 | 0.32547379 | -0.5894983 | 0.55552707 | NA | NA | NA | |
| ENSG000002 | 1.0989449 | -0.259635601 | 0.31882725 | -0.8143457 | 0.41544693 | NA | NA | NA | |
| ENSG000002 | 0 | NA | NA | NA | NA | NA | NA | NA | |
| ENSG000002 | 1.9332994 | -0.298543885 | 0.35763983 | -0.8347613 | 0.40385213 | NA | NA | NA | |
| ENSG000002 | 1.2395684 | -0.039786161 | 0.32877853 | -0.121012 | 0.9036815 | NA | NA | NA | |
| ENSG000002 | 1.56873069 | -0.314205397 | 0.33671181 | -0.9331582 | 0.35073828 | NA | NA | NA | |
| ENSG000002 | 2.04683129 | 0.122963848 | 0.36385448 | 0.33794788 | 0.73540247 | NA | NA | NA | |
| ENSG000002 | 0.26199343 | -0.194018184 | 0.18422801 | -1.0531416 | 0.29227609 | NA | NA | NA | |
| ENSG000002 | 0.35141577 | 0.074527565 | 0.21401461 | 0.34823588 | 0.72766304 | NA | KRT18P53 | | 106478925 |
| ENSG000002 | 0 | NA | NA | NA | NA | NA | NA | NA | |
| ENSG000002 | 0 | NA | NA | NA | NA | NA | NA | NA | |
| ENSG000002 | 1.96899658 | -0.546833378 | 0.35622923 | -1.5350604 | 0.124769 | NA | NA | NA | |
| ENSG000002 | 0 | NA | NA | NA | NA | NA | TTTY10 | | 246119 |
| ENSG000002 | 0.14128499 | -0.097772014 | 0.13403235 | -0.7294658 | 0.4657168 | NA | NA | NA | |
| ENSG000002 | 1.46954657 | -0.17481226 | 0.33269594 | -0.5254415 | 0.5992763 | NA | NA | NA | |

| | | | | | | | | |
|---|---|---|---|---|---|---|---|---|
| ENSG000002 | 1.02466046 | -0.305793162 | 0.29647229 | -1.0314393 | 0.30233487 | NA | NA | NA |
| ENSG000002 | 0 | NA | NA | NA | NA | NA | USP9YP31 | 106478930 |
| ENSG000002 | 1.20489076 | 0.395074319 | 0.32358849 | 1.22091584 | 0.22211789 | NA | NA | NA |
| ENSG000002 | 1.22249908 | -0.19000916 | 0.32884347 | -0.5778104 | 0.56339216 | NA | NA | NA |
| ENSG000002 | 1.70084509 | 0.086274766 | 0.3475332 | 0.24824899 | 0.80394176 | NA | NA | NA |
| ENSG000002 | 0.60580886 | -0.101167169 | 0.26338657 | -0.3841015 | 0.70090324 | NA | NA | NA |
| ENSG000002 | 1.11055274 | -0.12075786 | 0.32603143 | -0.3703872 | 0.71109403 | NA | NA | NA |
| ENSG000002 | 0 | NA | NA | NA | NA | NA | NA | NA |
| ENSG000002 | 0 | NA | NA | NA | NA | NA | NA | NA |
| ENSG000002 | 0 | NA | NA | NA | NA | NA | NA | NA |
| ENSG000002 | 1.57493281 | -0.11094729 | 0.3468449 | -0.3198758 | 0.74906248 | NA | NA | NA |
| ENSG000002 | 0 | NA | NA | NA | NA | NA | NA | NA |
| ENSG000002 | 0.72193154 | -0.030606144 | 0.27727904 | -0.1103803 | 0.91210778 | NA | NA | NA |
| ENSG000002 | 0.88132196 | -0.102972083 | 0.29244783 | -0.3521041 | 0.72476018 | NA | LOC101927827 | 101927827 |
| ENSG000002 | 1.10252784 | -0.123396877 | 0.32577918 | -0.3787746 | 0.70485525 | NA | NA | NA |
| ENSG000002 | 1.94483882 | 0.060353305 | 0.35477995 | 0.17011476 | 0.86491989 | NA | NA | NA |
| ENSG000002 | 0.88874941 | -0.417513114 | 0.29274791 | -1.4261865 | 0.15381452 | NA | LOC102031319 | 102031319 |
| ENSG000002 | 1.22632714 | 0.242255647 | 0.32464311 | 0.74622142 | 0.45553367 | NA | GK-IT1 | 106480740 |
| ENSG000002 | 1.38287018 | 0.017593814 | 0.33977587 | 0.05178065 | 0.95870348 | NA | NA | NA |
| ENSG000002 | 0 | NA | NA | NA | NA | NA | NA | NA |
| ENSG000002 | 1.54678523 | -0.199276238 | 0.34267496 | -0.5815314 | 0.56088239 | NA | NA | NA |
| ENSG000002 | 0 | NA | NA | NA | NA | NA | NA | NA |
| ENSG000002 | 1.47782188 | -0.049209377 | 0.34106693 | -0.1442807 | 0.88527881 | NA | PIN1P1 | 5301 |
| ENSG000002 | 0.48627291 | -0.18151782 | 0.23948009 | -0.7579662 | 0.44847119 | NA | NA | NA |
| ENSG000002 | 1.2381783 | -0.043858898 | 0.32874547 | -0.1334129 | 0.89386681 | NA | NA | NA |
| ENSG000002 | 2.08741926 | -0.219983877 | 0.35676075 | -0.6166146 | 0.53748899 | NA | NA | NA |
| ENSG000002 | 0.9850368 | -0.031630121 | 0.30619917 | -0.1032992 | 0.91772553 | NA | NA | NA |
| ENSG000002 | 0.52526486 | -0.351685369 | 0.238233 | -1.4762244 | 0.13988366 | NA | NA | NA |
| ENSG000002 | 1.61071588 | -0.254535866 | 0.35068627 | -0.7258222 | 0.4679478 | NA | NA | NA |

| | | | | | | | | |
|---|---|---|---|---|---|---|---|---|
| ENSG000002 | 0.85152629 | -0.109713239 | 0.29682987 | -0.3696166 | 0.7116682 | NA | NA | NA |
| ENSG000002 | 2.03271136 | -0.002164622 | 0.35717441 | -0.0060604 | 0.99516453 | NA | NA | NA |
| ENSG000002 | 0.85863579 | -0.268830487 | 0.29839657 | -0.9009168 | 0.36763255 | NA | NA | NA |
| ENSG000002 | 1.96477746 | -0.313420021 | 0.36111945 | -0.8679123 | 0.38544233 | NA | NA | NA |
| ENSG000002 | 0.35346928 | -0.088570748 | 0.20618147 | -0.4295767 | 0.66750362 | NA | NA | NA |
| ENSG000002 | 1.119435 | -0.436060276 | 0.32298771 | -1.3500832 | 0.1769893 | NA | NA | NA |
| ENSG000002 | 1.71539818 | -0.045228163 | 0.34780567 | -0.1300386 | 0.89653589 | NA | NA | NA |
| ENSG000002 | 1.62405627 | -0.267016002 | 0.34967906 | -0.7636031 | 0.44510383 | NA | NA | NA |
| ENSG000002 | 1.99020036 | -0.466462996 | 0.36480775 | -1.2786543 | 0.20101882 | NA | NA | NA |
| ENSG000002 | 0 | NA | NA | NA | NA | NA | NA | NA |
| ENSG000002 | 0.63965309 | -0.285618162 | 0.26800573 | -1.0657166 | 0.28655176 | NA | NA | NA |
| ENSG000002 | 1.54035737 | -0.345528747 | 0.34563663 | -0.9996879 | 0.31746158 | NA | NA | NA |
| ENSG000002 | 0 | NA | NA | NA | NA | NA | NA | NA |
| ENSG000002 | 1.12561491 | -0.27633688 | 0.32344409 | -0.8543575 | 0.39290694 | NA | NA | NA |
| ENSG000002 | 0.98956355 | -0.041520291 | 0.30047041 | -0.1381843 | 0.89009477 | NA | NA | NA |
| ENSG000002 | 1.6692565 | 0.223403388 | 0.35434811 | 0.63046304 | 0.52839168 | NA | NA | NA |
| ENSG000002 | 0.62197881 | -0.414424795 | 0.25447679 | -1.6285367 | 0.10341113 | NA | NA | NA |
| ENSG000002 | 1.70614152 | 0.079466684 | 0.3529217 | 0.22516803 | 0.82184856 | NA | NA | NA |
| ENSG000002 | 0.75776669 | -0.19935134 | 0.28679134 | -0.6951093 | 0.48698678 | NA | NA | NA |
| ENSG000002 | 0 | NA | NA | NA | NA | NA | NA | NA |
| ENSG000002 | 1.7364277 | -0.332277384 | 0.35737299 | -0.9297775 | 0.35248628 | NA | NA | NA |
| ENSG000002 | 0.83897464 | -0.092974912 | 0.28827292 | -0.3225239 | 0.74705582 | NA | NA | NA |
| ENSG000002 | 0 | NA | NA | NA | NA | NA | NA | NA |
| ENSG000002 | 0.48991005 | 0.142435647 | 0.23916196 | 0.59556147 | 0.55146823 | NA | NA | NA |
| ENSG000002 | 0.98406933 | -0.180198027 | 0.30686151 | -0.5872292 | 0.55704981 | NA | KSR1P1 | 106479050 |
| ENSG000002 | 1.13414835 | -0.274933331 | 0.32223582 | -0.8532054 | 0.39354543 | NA | ALG13-AS1 | 106478971 |
| ENSG000002 | 2.07244053 | -0.126872349 | 0.36169785 | -0.3507689 | 0.72576175 | NA | NA | NA |
| ENSG000002 | 1.2596345 | -0.188819205 | 0.33130071 | -0.569933 | 0.56872315 | NA | NA | NA |
| ENSG000002 | 0.11934008 | 0.083564698 | 0.12760091 | 0.65489106 | 0.5125379 | NA | NA | NA |

| | | | | | | | | |
|---|---|---|---|---|---|---|---|---|
| ENSG000002 | 0.97752098 | -0.032273661 | 0.31241232 | -0.1033047 | 0.91772114 | NA | NA | NA |
| ENSG000002 | 0.76129866 | -0.025684083 | 0.27094315 | -0.0947951 | 0.92447758 | NA | NA | NA |
| ENSG000002 | 1.14538425 | -0.114761673 | 0.31549737 | -0.3637484 | 0.71604587 | NA | NA | NA |
| ENSG000002 | 0.91573208 | -0.410925219 | 0.28867193 | -1.4235025 | 0.15459053 | NA | NA | NA |
| ENSG000002 | 2.13676893 | -0.490915012 | 0.36033717 | -1.3623768 | 0.17307899 | NA | NA | NA |
| ENSG000002 | 1.81996509 | -0.122936897 | 0.3606639 | -0.3408628 | 0.73320689 | NA | NA | NA |
| ENSG000002 | 0.72426738 | -0.032746305 | 0.28379902 | -0.1153855 | 0.90813953 | NA | UBE2DNL | 100131816 |
| ENSG000002 | 0 | NA | NA | NA | NA | NA | TSPY8 | 728403 |
| ENSG000002 | 0 | NA | NA | NA | NA | NA | NA | NA |
| ENSG000002 | 0 | NA | NA | NA | NA | NA | USP9YP17 | 106480714 |
| ENSG000002 | 1.43629979 | 0.226178021 | 0.33679323 | 0.67156345 | 0.50186165 | NA | NA | NA |
| ENSG000002 | 0.36553413 | 0.069401489 | 0.20780452 | 0.33397488 | 0.73839851 | NA | NA | NA |
| ENSG000002 | 0.86339122 | -0.275317148 | 0.2986111 | -0.9219924 | 0.35653256 | NA | NA | NA |
| ENSG000002 | 1.94246194 | -0.515730246 | 0.35637576 | -1.447153 | 0.14785407 | NA | NA | NA |
| ENSG000002 | 1.70187268 | -0.058059438 | 0.35531846 | -0.1634011 | 0.87020261 | NA | PRAMEF25 | 441873 |
| ENSG000002 | 0 | NA | NA | NA | NA | NA | USP17L26 | 728379 |
| ENSG000002 | 0 | NA | NA | NA | NA | NA | NA | NA |
| ENSG000002 | 1.55604469 | -0.218516308 | 0.33692114 | -0.6485681 | 0.51661758 | NA | NA | NA |
| ENSG000002 | 1.23954667 | -0.189155412 | 0.32930355 | -0.5744105 | 0.56569006 | NA | NA | NA |
| ENSG000002 | 0.73402886 | 0.123685362 | 0.27843246 | 0.44422034 | 0.65688329 | NA | NA | NA |
| ENSG000002 | 0.53517016 | -0.075146746 | 0.15579947 | -0.4823299 | 0.6295716 | NA | MTATP8P2 | 106479042 |
| ENSG000002 | 0.7710561 | -0.188233817 | 0.28164389 | -0.6683398 | 0.50391673 | NA | NA | NA |
| ENSG000002 | 1.94284302 | -0.83460773 | 0.35800213 | -2.3312926 | 0.01973794 | NA | NA | NA |
| ENSG000002 | 0 | NA | NA | NA | NA | NA | BSNDP4 | 106481726 |
| ENSG000002 | 1.72962928 | -0.328900438 | 0.35586179 | -0.9242364 | 0.35536323 | NA | NA | NA |
| ENSG000002 | 1.63632827 | -0.258683389 | 0.34828631 | -0.742732 | 0.45764395 | NA | NA | NA |
| ENSG000002 | 1.50763007 | -0.633395953 | 0.34331455 | -1.8449435 | 0.06504576 | NA | LINC01264 | 104266963 |
| ENSG000002 | 1.99742333 | -0.429240092 | 0.35942555 | -1.1942392 | 0.2323844 | NA | NA | NA |
| ENSG000002 | 1.25072996 | -0.052346994 | 0.32908534 | -0.1590681 | 0.87361519 | NA | PRAC2 | 360205 |

| | | | | | | | | |
|---|---|---|---|---|---|---|---|---|
| ENSG000002 | 2.07098838 | 0.002418892 | 0.36728544 | 0.00658586 | 0.99474528 | NA | NA | NA |
| ENSG000002 | 1.10073335 | -0.116805618 | 0.32035519 | -0.3646129 | 0.71540042 | NA | CYP2C56P | 106478959 |
| ENSG000002 | 0 | NA | NA | NA | NA | NA | LINC00280 | 100873964 |
| ENSG000002 | 0.24756242 | -0.187034346 | 0.18119817 | -1.0322088 | 0.30197431 | NA | NA | NA |
| ENSG000002 | 1.99438075 | -0.064025093 | 0.36116 | -0.1772763 | 0.8592914 | NA | NA | NA |
| ENSG000002 | 1.6693929 | 0.218809284 | 0.35373588 | 0.61856684 | 0.53620175 | NA | NA | NA |
| ENSG000002 | 0.96340454 | 0.124375372 | 0.31176932 | 0.39893397 | 0.68994186 | NA | NA | NA |
| ENSG000002 | 1.09106381 | -0.241433991 | 0.30910225 | -0.7810813 | 0.43475467 | NA | BUD31P2 | 106480332 |
| ENSG000002 | 0.4009426 | -0.302672773 | 0.2240145 | -1.3511303 | 0.17665371 | NA | PRR20C | 729240 |
| ENSG000002 | 1.94730779 | -0.189731152 | 0.36112404 | -0.5253905 | 0.59931173 | NA | NA | NA |
| ENSG000002 | 1.64096747 | -0.26403205 | 0.3504134 | -0.7534873 | 0.45115713 | NA | NA | NA |
| ENSG000002 | 0.75962432 | -0.361712227 | 0.28310455 | -1.277663 | 0.20136826 | NA | NA | NA |
| ENSG000002 | 0.8868146 | -0.114915491 | 0.29866695 | -0.3847613 | 0.70041426 | NA | NA | NA |
| ENSG000002 | 0 | NA | NA | NA | NA | NA | SNORD56 | 26793 |
| ENSG000002 | 0.98438199 | -0.203442562 | 0.31731215 | -0.6411433 | 0.52142958 | NA | NA | NA |
| ENSG000002 | 1.01064761 | -0.198945547 | 0.31431679 | -0.632946 | 0.52676892 | NA | NA | NA |
| ENSG000002 | 1.01068144 | -0.049223259 | 0.30726426 | -0.1601985 | 0.87272475 | NA | NA | NA |
| ENSG000002 | 1.57763685 | 0.158240708 | 0.35069773 | 0.4512168 | 0.65183331 | NA | NA | NA |
| ENSG000002 | 1.76630927 | -0.194230748 | 0.35510609 | -0.5469654 | 0.58440249 | NA | NA | NA |
| ENSG000002 | 0 | NA | NA | NA | NA | NA | USP9YP36 | 106478932 |
| ENSG000002 | 0.37441639 | -0.268856313 | 0.21277471 | -1.2635727 | 0.20638344 | NA | NA | NA |
| ENSG000002 | 1.72166049 | -0.055262708 | 0.35495237 | -0.1556905 | 0.87627699 | NA | NA | NA |
| ENSG000002 | 1.00968014 | -0.325606339 | 0.30228547 | -1.0771485 | 0.28141393 | NA | NA | NA |
| ENSG000002 | 0 | NA | NA | NA | NA | NA | NA | NA |
| ENSG000002 | 0.36538978 | 0.059033792 | 0.20773944 | 0.28417229 | 0.77627835 | NA | NA | NA |
| ENSG000002 | 0.9663377 | -0.156888774 | 0.29759573 | -0.5271876 | 0.59806333 | NA | NA | NA |
| ENSG000002 | 0.76673731 | -0.201606703 | 0.28733642 | -0.7016399 | 0.48290375 | NA | NA | NA |
| ENSG000002 | 0.13280363 | -0.093394593 | 0.13115644 | -0.7120855 | 0.47641185 | NA | NA | NA |
| ENSG000002 | 1.11843329 | -0.121270036 | 0.32093578 | -0.3778639 | 0.70553173 | NA | NA | NA |

| | | | | | | | | |
|---|---|---|---|---|---|---|---|---|
| ENSG000002 | 0.77427295 | -0.18871474 | 0.2818501 | -0.6695571 | 0.50314016 | NA | NA | NA |
| ENSG000002 | 0 | NA | NA | NA | NA | NA | NA | NA |
| ENSG000002 | 0.36553413 | 0.069401489 | 0.20780452 | 0.33397488 | 0.73839851 | NA | NA | NA |
| ENSG000002 | 1.91135853 | 0.076512025 | 0.35884639 | 0.21321665 | 0.83115798 | NA | NA | NA |
| ENSG000002 | 0.6929292 | 0.29595641 | 0.27700004 | 1.06843454 | 0.28532454 | NA | NA | NA |
| ENSG000002 | 1.74511758 | -0.32064344 | 0.35248558 | -0.909664 | 0.36299973 | NA | NA | NA |
| ENSG000002 | 1.50640359 | -0.633993594 | 0.34328477 | -1.8468445 | 0.06476968 | NA | NA | NA |
| ENSG000002 | 1.90219046 | -0.037922238 | 0.34474832 | -0.1099998 | 0.91240955 | NA | NA | NA |
| ENSG000002 | 1.02672931 | -0.193044961 | 0.30865692 | -0.6254354 | 0.53168534 | NA | NA | NA |
| ENSG000002 | 1.04525632 | 0.166025785 | 0.30534572 | 0.54373051 | 0.58662694 | NA | NA | NA |
| ENSG000002 | 0 | NA | NA | NA | NA | NA | NA | NA |
| ENSG000002 | 1.95507018 | -0.313232743 | 0.36162439 | -0.8661826 | 0.38639006 | NA | NA | NA |
| ENSG000002 | 1.39008057 | -0.419986964 | 0.33959601 | -1.2367252 | 0.2161891 | NA | NA | NA |
| ENSG000002 | 1.40544412 | -0.408387336 | 0.33378739 | -1.2234954 | 0.22114264 | NA | NA | NA |
| ENSG000002 | 1.90327198 | -0.507633053 | 0.35605389 | -1.4257197 | 0.15394926 | NA | LINC00399 | 104326054 |
| ENSG000002 | 1.41253648 | -0.393362048 | 0.33315258 | -1.1807264 | 0.23771143 | NA | NA | NA |
| ENSG000002 | 2.03650391 | 0.12656913 | 0.36345214 | 0.34824154 | 0.7276588 | NA | NA | NA |
| ENSG000002 | 1.90893234 | 0.192475313 | 0.35774249 | 0.53802754 | 0.59055804 | NA | NA | NA |
| ENSG000002 | 2.0764915 | -0.588443337 | 0.36078687 | -1.6309999 | 0.10289034 | NA | NA | NA |
| ENSG000002 | 1.90517602 | -0.11658467 | 0.35037685 | -0.3327408 | 0.73932996 | NA | NA | NA |
| ENSG000002 | 0 | NA | NA | NA | NA | NA | NA | NA |
| ENSG000002 | 1.57886725 | -0.108104495 | 0.34354551 | -0.314673 | 0.75300995 | NA | NA | NA |
| ENSG000002 | 0.37236288 | -0.09108266 | 0.20928647 | -0.4352057 | 0.66341314 | NA | NA | NA |
| ENSG000002 | 2.05833013 | -0.12441231 | 0.36619959 | -0.3397391 | 0.73405303 | NA | NA | NA |
| ENSG000002 | 1.39691124 | -0.408565256 | 0.33655877 | -1.2139492 | 0.22476711 | NA | NA | NA |
| ENSG000002 | 0.47596775 | 0.137153722 | 0.23749987 | 0.57748967 | 0.56360872 | NA | NA | NA |
| ENSG000002 | 2.10675207 | -0.626768769 | 0.3631338 | -1.7259995 | 0.08434749 | NA | NA | NA |
| ENSG000002 | 0.38651158 | -0.274258716 | 0.21461355 | -1.2779189 | 0.20127801 | NA | NA | NA |
| ENSG000002 | 2.04145168 | -0.580218313 | 0.36081207 | -1.6080901 | 0.10781545 | NA | NA | NA |

| | | | | | | | | |
|---|---|---|---|---|---|---|---|---|
| ENSG000002 | 0.72967624 | -0.02677813 | 0.28955569 | -0.0924801 | 0.92631662 | NA | NA | NA |
| ENSG000002 | 1.49760566 | -0.33167325 | 0.34401295 | -0.9641301 | 0.33498069 | NA | LOC101927762 | 101927762 |
| ENSG000002 | 1.69172549 | -0.293157768 | 0.34726251 | -0.8441964 | 0.39855965 | NA | OGFR-AS1 | 101409261 |
| ENSG000002 | 0.97200448 | -0.340708546 | 0.30777906 | -1.1069907 | 0.26829797 | NA | NA | NA |
| ENSG000002 | 2.01898042 | -0.855632009 | 0.35824843 | -2.3883761 | 0.01692301 | NA | NA | NA |
| ENSG000002 | 1.42799649 | -0.152813071 | 0.32493357 | -0.4702902 | 0.63814771 | NA | NA | NA |
| ENSG000002 | 0.2596576 | -0.193002035 | 0.1837913 | -1.0501152 | 0.29366517 | NA | NA | NA |
| ENSG000002 | 1.61942345 | -0.415954211 | 0.35364115 | -1.1762042 | 0.23951328 | NA | NA | NA |
| ENSG000002 | 1.72182199 | -0.188191978 | 0.3548309 | -0.5303709 | 0.59585481 | NA | NA | NA |
| ENSG000002 | 1.19407749 | 0.102827219 | 0.32137759 | 0.31995765 | 0.74900043 | NA | NA | NA |
| ENSG000002 | 0.85615965 | 0.048127972 | 0.30623801 | 0.15715871 | 0.87511976 | NA | NA | NA |
| ENSG000002 | 0.78255843 | -0.188463567 | 0.28072977 | -0.6713344 | 0.5020075 | NA | NA | NA |
| ENSG000002 | 1.11516884 | -0.421140184 | 0.32071865 | -1.3131141 | 0.1891445 | NA | NA | NA |
| ENSG000002 | 1.01315758 | -0.371579399 | 0.31542085 | -1.1780432 | 0.23877937 | NA | NA | NA |
| ENSG000002 | 2.01376405 | -0.59383716 | 0.36329814 | -1.6345725 | 0.10213869 | NA | NA | NA |
| ENSG000002 | 1.00515688 | -0.030150895 | 0.29848467 | -0.1010132 | 0.91953997 | NA | NA | NA |
| ENSG000002 | 0.13280363 | -0.093394593 | 0.13115644 | -0.7120855 | 0.47641185 | NA | NA | NA |
| ENSG000002 | 1.63604786 | -0.25759715 | 0.34828131 | -0.7396238 | 0.45952829 | NA | NA | NA |
| ENSG000002 | 1.22923856 | -0.172191786 | 0.32305089 | -0.5330175 | 0.59402145 | NA | NA | NA |
| ENSG000002 | 0 | NA | NA | NA | NA | NA | NA | NA |
| ENSG000002 | 0.74231669 | -0.350298432 | 0.28037475 | -1.2493937 | 0.21152112 | NA | NA | NA |
| ENSG000002 | 0.73617116 | -0.320233323 | 0.27079112 | -1.1825843 | 0.23697394 | NA | NA | NA |
| ENSG000002 | 0.11934008 | 0.083564698 | 0.12760091 | 0.65489106 | 0.5125379 | NA | NA | NA |
| ENSG000002 | 0 | NA | NA | NA | NA | NA | NA | NA |
| ENSG000002 | 1.3823959 | -0.412106326 | 0.33698594 | -1.2229185 | 0.2213605 | NA | NA | NA |
| ENSG000002 | 1.19696458 | -0.600879249 | 0.32120275 | -1.8707164 | 0.0613844 | NA | NA | NA |
| ENSG000002 | 1.36345293 | -0.107705315 | 0.32295088 | -0.3335037 | 0.73875409 | NA | NA | NA |
| ENSG000002 | 1.48103626 | -0.187324824 | 0.3415791 | -0.5484083 | 0.58341155 | NA | NA | NA |
| ENSG000002 | 0.35824644 | 0.072568381 | 0.21508272 | 0.33739754 | 0.73581724 | NA | NA | NA |

| | | | | | | | | |
|---|---|---|---|---|---|---|---|---|
| ENSG000002 | 0.39479706 | -0.300008631 | 0.22317059 | -1.3443018 | 0.17885081 | NA | NA | NA |
| ENSG000002 | 0.76670294 | -0.358458639 | 0.28198122 | -1.2712146 | 0.20365232 | NA | NA | NA |
| ENSG000002 | 0.97769512 | -0.186564347 | 0.30823367 | -0.6052692 | 0.54500015 | NA | NA | NA |
| ENSG000002 | 1.22223849 | -0.173647622 | 0.32201569 | -0.5392521 | 0.58971295 | NA | NA | NA |
| ENSG000002 | 1.58694967 | -0.448921249 | 0.33472007 | -1.3411841 | 0.17986068 | NA | NA | NA |
| ENSG000002 | 2.0314308 | 0.35909409 | 0.35987304 | 0.99783549 | 0.31835914 | NA | NHS-AS1 | 100873920 |
| ENSG000002 | 0 | NA | NA | NA | NA | NA | NA | NA |
| ENSG000002 | 0.83082959 | 0.188233236 | 0.27848724 | 0.67591333 | 0.49909567 | NA | NA | NA |
| ENSG000002 | 0 | NA | NA | NA | NA | NA | NA | NA |
| ENSG000002 | 0.11131518 | 0.081951103 | 0.12663804 | 0.64712866 | 0.51754868 | NA | POTEB2 | 100287399 |
| ENSG000002 | 1.71386181 | -0.195820495 | 0.35518574 | -0.5513186 | 0.58141531 | NA | NA | NA |
| ENSG000002 | 2.06691809 | 0.007827413 | 0.3641542 | 0.02149477 | 0.98285097 | NA | NA | NA |
| ENSG000002 | 1.24518175 | -0.350323405 | 0.3338189 | -1.0494415 | 0.29397497 | NA | LINC00364 | 100874145 |
| ENSG000002 | 1.59852977 | -0.10945476 | 0.33983929 | -0.322078 | 0.7473936 | NA | NA | NA |
| ENSG000002 | 0.13280363 | -0.093394593 | 0.13115644 | -0.7120855 | 0.47641185 | NA | NA | NA |
| ENSG000002 | 0.39274356 | -0.107428268 | 0.22062504 | -0.4869269 | 0.62631013 | NA | NA | NA |
| ENSG000002 | 0.12070844 | -0.088503695 | 0.12783931 | -0.6923042 | 0.48874631 | NA | NA | NA |
| ENSG000002 | 2.02887323 | 0.242886013 | 0.35813029 | 0.67820572 | 0.49764127 | NA | NA | NA |
| ENSG000002 | 0 | NA | NA | NA | NA | NA | NA | NA |
| ENSG000002 | 0 | NA | NA | NA | NA | NA | NA | NA |
| ENSG000002 | 0.26813897 | -0.197007041 | 0.18550319 | -1.0620143 | 0.2882292 | NA | NA | NA |
| ENSG000002 | 2.14335476 | -0.925021887 | 0.36257343 | -2.5512677 | 0.01073318 | NA | NA | NA |
| ENSG000002 | 1.13089904 | -0.262818004 | 0.31771408 | -0.8272155 | 0.40811495 | NA | NA | NA |
| ENSG000002 | 0.91049845 | -0.644235083 | 0.3003423 | -2.1450028 | 0.03195262 | NA | TCERG1L-AS1 | 101927489 |
| ENSG000002 | 0.58728293 | 0.202071621 | 0.25373362 | 0.79639279 | 0.42580377 | NA | NA | NA |
| ENSG000002 | 2.00628441 | -0.82723996 | 0.35504524 | -2.3299565 | 0.01980845 | NA | NA | NA |
| ENSG000002 | 0 | NA | NA | NA | NA | NA | NA | NA |
| ENSG000002 | 1.55813971 | -0.464678403 | 0.33935577 | -1.3692957 | 0.17090685 | NA | NA | NA |
| ENSG000002 | 0.74590018 | -0.179161319 | 0.28000754 | -0.6398446 | 0.52227362 | NA | NA | NA |

| | | | | | | | | |
|---|---|---|---|---|---|---|---|---|
| ENSG000002 | 0.58133382 | 0.231430989 | 0.26184763 | 0.88383838 | 0.37678347 | NA | NA | NA |
| ENSG000002 | 1.46374182 | -0.182844471 | 0.34373485 | -0.5319346 | 0.59477127 | NA | LOC101929721 | 101929721 |
| ENSG000002 | 1.73321435 | -0.058822108 | 0.35507518 | -0.165661 | 0.86842374 | NA | NA | NA |
| ENSG000002 | 2.15102972 | -0.640221831 | 0.3639697 | -1.7589976 | 0.07857792 | NA | NA | NA |
| ENSG000002 | 1.57738781 | 0.128735245 | 0.34200866 | 0.37640931 | 0.70661262 | NA | NA | NA |
| ENSG000002 | 1.64970729 | -0.381939512 | 0.34406307 | -1.1100858 | 0.26696207 | NA | NA | NA |
| ENSG000002 | 0.50722002 | -0.375409304 | 0.24493426 | -1.5326941 | 0.12535123 | NA | NA | NA |
| ENSG000002 | 0.35801832 | 0.071082974 | 0.20657063 | 0.34410977 | 0.73076373 | NA | NA | NA |
| ENSG000002 | 0.48416733 | -0.018553864 | 0.24528998 | -0.0756405 | 0.93970509 | NA | NA | NA |
| ENSG000002 | 1.94198374 | -0.662469618 | 0.35688136 | -1.8562741 | 0.0634145 | NA | NA | NA |
| ENSG000002 | 1.87589646 | -0.526911234 | 0.35797366 | -1.4719274 | 0.14104048 | NA | NA | NA |
| ENSG000002 | 2.01970456 | -0.301528547 | 0.35847016 | -0.8411538 | 0.40026177 | NA | LINC01433 | 728228 |
| ENSG000002 | 0.24619405 | -0.007630671 | 0.17915639 | -0.0425922 | 0.96602658 | NA | OR4F3 | 26683 |
| ENSG000002 | 1.09345014 | -0.384935017 | 0.31040807 | -1.2400935 | 0.21494082 | NA | NA | NA |
| ENSG000002 | 0.38884741 | -0.29745092 | 0.22235466 | -1.3377319 | 0.18098386 | NA | NA | NA |
| ENSG000002 | 1.12632179 | -0.11014525 | 0.31654154 | -0.3479646 | 0.72786677 | NA | NA | NA |
| ENSG000002 | 1.50517375 | -0.325890021 | 0.34281443 | -0.950631 | 0.34179173 | NA | NA | NA |
| ENSG000002 | 1.60999985 | -0.114274411 | 0.34526743 | -0.3309736 | 0.74066441 | NA | NA | NA |
| ENSG000002 | 1.42195255 | 0.226558042 | 0.34025702 | 0.66584384 | 0.50551092 | NA | LOC105375556 | 105375556 |
| ENSG000002 | 0.6606002 | -0.463974368 | 0.2663045 | -1.7422701 | 0.08146119 | NA | NA | NA |
| ENSG000002 | 1.83297163 | -0.258139962 | 0.35884227 | -0.7193689 | 0.47191367 | NA | NA | NA |
| ENSG000002 | 0.12070844 | -0.088503695 | 0.12783931 | -0.6923042 | 0.48874631 | NA | NA | NA |
| ENSG000002 | 1.14781024 | -0.258028638 | 0.31827659 | -0.8107057 | 0.41753471 | NA | NA | NA |
| ENSG000002 | 1.39554096 | -0.409043483 | 0.3365263 | -1.2154874 | 0.22418024 | NA | NA | NA |
| ENSG000002 | 0.52526486 | -0.351685369 | 0.238233 | -1.4762244 | 0.13988366 | NA | NA | NA |
| ENSG000002 | 1.96182095 | -0.819429499 | 0.35625099 | -2.3001466 | 0.02143992 | NA | NA | NA |
| ENSG000002 | 1.7284028 | -0.334856443 | 0.35728567 | -0.9372233 | 0.34864372 | NA | EIF2B5-AS1 | 100874079 |
| ENSG000002 | 1.34578737 | 0.024364904 | 0.34305628 | 0.07102305 | 0.94337941 | NA | NA | NA |
| ENSG000002 | 1.00197429 | -0.523960978 | 0.31107783 | -1.6843404 | 0.09211589 | NA | NA | NA |

| | | | | | | | | |
|---|---|---|---|---|---|---|---|---|
| ENSG000002 | 1.56750426 | 0.146825203 | 0.34657854 | 0.42364193 | 0.67182697 | NA | NA | NA |
| ENSG000002 | 1.73019337 | -0.056371714 | 0.35126183 | -0.1604835 | 0.87250025 | NA | NA | NA |
| ENSG000002 | 2.1076595 | -0.483507195 | 0.36124319 | -1.3384534 | 0.18074867 | NA | NA | NA |
| ENSG000002 | 1.23470479 | -0.31794512 | 0.32448757 | -0.9798376 | 0.32716629 | NA | NA | NA |
| ENSG000002 | 0.71712404 | -0.025635761 | 0.28333197 | -0.0904796 | 0.92790611 | NA | NA | NA |
| ENSG000002 | 0.26199343 | -0.194018184 | 0.18422801 | -1.0531416 | 0.29227609 | NA | NA | NA |
| ENSG000002 | 1.96372471 | 0.043154349 | 0.3544892 | 0.12173671 | 0.90310754 | NA | NA | NA |
| ENSG000002 | 1.1055711 | 0.041842929 | 0.31295229 | 0.13370386 | 0.89363675 | NA | NA | NA |
| ENSG000002 | 0.88154817 | -0.275720673 | 0.29955151 | -0.920445 | 0.35734028 | NA | NA | NA |
| ENSG000002 | 0.96335247 | 0.111957535 | 0.29936595 | 0.37398219 | 0.70841756 | NA | NA | NA |
| ENSG000002 | 0.8794426 | -0.118660638 | 0.30333647 | -0.3911849 | 0.69566059 | NA | NA | NA |
| ENSG000002 | 0.62821079 | -0.27685072 | 0.26713242 | -1.0363801 | 0.30002486 | NA | NA | NA |
| ENSG000002 | 2.0783392 | -0.103740306 | 0.35490917 | -0.292301 | 0.77005649 | NA | NA | NA |
| ENSG000002 | 2.07620028 | -0.006544233 | 0.36732447 | -0.0178159 | 0.98578568 | NA | NA | NA |
| ENSG000002 | 1.06006893 | -0.665566918 | 0.30288553 | -2.1974206 | 0.02799042 | NA | NA | NA |
| ENSG000002 | 1.09899889 | -0.106208641 | 0.31560672 | -0.3365221 | 0.73647717 | NA | NA | NA |
| ENSG000002 | 1.62776292 | -0.251079657 | 0.34685626 | -0.7238724 | 0.4691441 | NA | NA | NA |
| ENSG000002 | 1.71681469 | 0.079762737 | 0.35118316 | 0.22712575 | 0.82032596 | NA | NA | NA |
| ENSG000002 | 0.50585166 | -0.183613822 | 0.24177005 | -0.7594564 | 0.44757957 | NA | NA | NA |
| ENSG000002 | 1.38964921 | -0.538798843 | 0.33168606 | -1.6244241 | 0.10428532 | NA | NA | NA |
| ENSG000002 | 0 | NA | NA | NA | NA | NA | NA | NA |
| ENSG000002 | 0.98347009 | -0.043356565 | 0.31265282 | -0.1386732 | 0.8897084 | NA | NA | NA |
| ENSG000002 | 1.47097339 | -0.448712147 | 0.3377601 | -1.3284936 | 0.18401508 | NA | NA | NA |
| ENSG000002 | 1.66897201 | -0.527411156 | 0.34641092 | -1.5225015 | 0.12788346 | NA | NA | NA |
| ENSG000002 | 1.61726231 | -0.527260917 | 0.34683346 | -1.5202135 | 0.12845732 | NA | NA | NA |
| ENSG000002 | 1.60848714 | 0.007072456 | 0.34709991 | 0.02037585 | 0.98374355 | NA | NA | NA |
| ENSG000002 | 1.48207133 | -0.456125821 | 0.34027888 | -1.3404471 | 0.18010005 | NA | NA | NA |
| ENSG000002 | 1.28021105 | -0.19181929 | 0.33035959 | -0.5806379 | 0.56148455 | NA | NA | NA |
| ENSG000002 | 1.23831862 | -0.323953456 | 0.32621429 | -0.9930695 | 0.3206761 | NA | NA | NA |

| | | | | | | | | | |
|---|---|---|---|---|---|---|---|---|---|
| ENSG000002 | 0.97755324 | -0.036057819 | 0.31241351 | -0.115417 | 0.90811463 | NA | NA | NA | |
| ENSG000002 | 0.62061044 | -0.252611122 | 0.25798806 | -0.9791582 | 0.32750182 | NA | NA | NA | |
| ENSG000002 | 0 | NA | NA | NA | NA | NA | CT47A8 | | 728049 |
| ENSG000002 | 1.90312954 | -0.37603415 | 0.35502305 | -1.0591824 | 0.28951674 | NA | NA | NA | |
| ENSG000002 | 1.28277309 | -0.328875741 | 0.32495727 | -1.0120584 | 0.31151012 | NA | NA | NA | |
| ENSG000002 | 1.35156681 | -0.117785364 | 0.33938648 | -0.3470538 | 0.72855091 | NA | NA | NA | |
| ENSG000002 | 1.21728663 | -0.188097604 | 0.32870337 | -0.5722412 | 0.56715859 | NA | NA | NA | |
| ENSG000002 | 1.50420765 | -0.347497732 | 0.34691644 | -1.0016756 | 0.3165003 | NA | NA | NA | |
| ENSG000002 | 1.79264139 | -0.601192792 | 0.35257194 | -1.7051635 | 0.08816394 | NA | NA | NA | |
| ENSG000002 | 0 | NA | NA | NA | NA | NA | NA | NA | |
| ENSG000002 | 1.79810396 | 0.004461278 | 0.35521118 | 0.01255951 | 0.98997923 | NA | NA | NA | |
| ENSG000002 | 1.60899814 | 0.140939533 | 0.34509908 | 0.40840309 | 0.68297776 | NA | NA | NA | |
| ENSG000002 | 1.55518039 | 0.149130909 | 0.3485812 | 0.42782259 | 0.66878029 | NA | NA | NA | |
| ENSG000002 | 1.57965811 | 0.022292425 | 0.34764855 | 0.06412345 | 0.94887193 | NA | NA | NA | |
| ENSG000002 | 1.06972957 | 0.334694147 | 0.31557988 | 1.06056873 | 0.28888594 | NA | NA | NA | |
| ENSG000002 | 0.67503122 | -0.469043005 | 0.26742254 | -1.7539397 | 0.07944084 | NA | NA | NA | |
| ENSG000002 | 0.62504738 | -0.112837574 | 0.26501179 | -0.4257832 | 0.67026581 | NA | NA | NA | |
| ENSG000002 | 0 | NA | NA | NA | NA | NA | NA | NA | |
| ENSG000002 | 1.65981657 | -0.669949151 | 0.34495253 | -1.9421488 | 0.05211909 | NA | NA | NA | |
| ENSG000002 | 0.37267746 | 0.067870636 | 0.21729405 | 0.31234465 | 0.75477861 | NA | NA | NA | |
| ENSG000002 | 1.86612604 | -0.377671775 | 0.35581028 | -1.0614414 | 0.28848934 | NA | NA | NA | |
| ENSG000002 | 0.11934008 | 0.083564698 | 0.12760091 | 0.65489106 | 0.5125379 | NA | USP17L25 | | 728373 |
| ENSG000002 | 0.90344102 | -0.436658607 | 0.29584239 | -1.4759839 | 0.13994822 | NA | NA | NA | |
| ENSG000002 | 1.84328029 | -0.25725445 | 0.35933408 | -0.71592 | 0.47404075 | NA | NA | NA | |
| ENSG000002 | 0 | NA | NA | NA | NA | NA | GPM6BP1 | | 106480720 |
| ENSG000002 | 0.61656602 | -0.102985032 | 0.25681525 | -0.4010082 | 0.68841406 | NA | NA | NA | |
| ENSG000002 | 0.90027816 | -0.281064798 | 0.30047954 | -0.9353875 | 0.34958865 | NA | NA | NA | |
| ENSG000002 | 1.01045173 | -0.196178082 | 0.31280784 | -0.6271521 | 0.53055957 | NA | NA | NA | |
| ENSG000002 | 0 | NA | NA | NA | NA | NA | NA | NA | |

| | | | | | | | | | |
|---|---|---|---|---|---|---|---|---|---|
| ENSG000002 | 0.63091271 | -0.116973765 | 0.27178557 | -0.4303899 | 0.66691204 | NA | NA | NA | |
| ENSG000002 | 0.35273207 | -0.08884607 | 0.20605566 | -0.4311751 | 0.66634105 | NA | NA | NA | |
| ENSG000002 | 1.35697355 | -0.254033313 | 0.3368165 | -0.7542187 | 0.45071789 | NA | NA | NA | |
| ENSG000002 | 0.50630812 | -0.189615696 | 0.24175631 | -0.7843257 | 0.432849 | NA | NA | NA | |
| ENSG000002 | 1.02553564 | -0.190777554 | 0.31023961 | -0.6149362 | 0.53859688 | NA | NA | NA | |
| ENSG000002 | 1.18440403 | 0.259342666 | 0.32754006 | 0.79178914 | 0.42848363 | NA | NA | NA | |
| ENSG000002 | 1.09040483 | -0.673744073 | 0.30410349 | -2.2155092 | 0.02672513 | NA | NA | NA | |
| ENSG000002 | 1.74274346 | -0.198188279 | 0.35375604 | -0.56024 | 0.57531575 | NA | NA | NA | |
| ENSG000002 | 1.37846358 | -0.271959268 | 0.34172162 | -0.7958503 | 0.42611903 | NA | NA | NA | |
| ENSG000002 | 2.16156514 | -0.769642794 | 0.36301444 | -2.1201437 | 0.03399392 | NA | NA | NA | |
| ENSG000002 | 1.58227414 | 0.018651073 | 0.34959315 | 0.0533508 | 0.95745241 | NA | NA | NA | |
| ENSG000002 | 1.20058897 | -0.173112411 | 0.21872647 | -0.7914562 | 0.42867785 | NA | NA | NA | |
| ENSG000002 | 1.03282277 | -0.203095518 | 0.31365537 | -0.6475117 | 0.51730083 | NA | NA | NA | |
| ENSG000002 | 1.31601344 | 0.165829565 | 0.33360182 | 0.4970883 | 0.61912678 | NA | NA | NA | |
| ENSG000002 | 0.36351096 | 0.070290861 | 0.21590418 | 0.32556508 | 0.74475344 | NA | MBD3L2 | | 125997 |
| ENSG000002 | 1.85962776 | 0.126712546 | 0.35436033 | 0.35758107 | 0.72065685 | NA | NA | NA | |
| ENSG000002 | 1.87481434 | -0.532377536 | 0.35892309 | -1.4832636 | 0.1380044 | NA | NA | NA | |
| ENSG000002 | 0.7456715 | -0.195362308 | 0.2860302 | -0.6830129 | 0.49459871 | NA | NA | NA | |
| ENSG000002 | 1.63908652 | -0.396413512 | 0.34847008 | -1.1375826 | 0.2552948 | NA | NA | NA | |
| ENSG000002 | 0.76744227 | -0.18689009 | 0.2813993 | -0.6641455 | 0.50659717 | NA | NA | NA | |
| ENSG000002 | 1.62946863 | 0.003885623 | 0.34004538 | 0.01142678 | 0.99088295 | NA | NA | NA | |
| ENSG000002 | 1.3137724 | -0.492859173 | 0.32598826 | -1.5118924 | 0.13056124 | NA | USP17L1 | | 401447 |
| ENSG000002 | 0.36416385 | 0.068880976 | 0.20758164 | 0.33182596 | 0.74002068 | NA | NA | NA | |
| ENSG000002 | 2.0190624 | -0.450321023 | 0.36116952 | -1.2468412 | 0.21245574 | NA | NA | NA | |
| ENSG000002 | 1.91605623 | -0.802897711 | 0.35543123 | -2.2589397 | 0.02388714 | NA | NA | NA | |
| ENSG000002 | 0.11205239 | 0.082118251 | 0.12671962 | 0.64803108 | 0.51696485 | NA | NA | NA | |
| ENSG000002 | 2.10836849 | -0.239459647 | 0.36434592 | -0.6572316 | 0.51103202 | NA | NA | NA | |
| ENSG000002 | 0.11934008 | 0.083564698 | 0.12760091 | 0.65489106 | 0.5125379 | NA | NA | NA | |
| ENSG000002 | 1.67508589 | -0.039832107 | 0.34265889 | -0.1162442 | 0.90745901 | NA | NA | NA | |

| | | | | | | | | |
|---|---|---|---|---|---|---|---|---|
| ENSG000002 | 0.85638641 | -0.10060963 | 0.29104879 | -0.3456796 | 0.72958351 | NA | NA | NA |
| ENSG000002 | 0 | NA | NA | NA | NA | NA | CT47A4 | 728075 |
| ENSG000002 | 1.42949866 | -0.423742039 | 0.3379465 | -1.2538732 | 0.20988811 | NA | GPAA1P2 | 106481722 |
| ENSG000002 | 1.90805307 | -0.650726514 | 0.35593418 | -1.8282215 | 0.06751632 | NA | NA | NA |
| ENSG000002 | 0.37194025 | 0.067575365 | 0.21717998 | 0.31114914 | 0.75568724 | NA | NA | NA |
| ENSG000002 | 1.3803424 | -0.272958917 | 0.34050905 | -0.8016201 | 0.42277272 | NA | NA | NA |
| ENSG000002 | 1.64062958 | -0.69787911 | 0.34852804 | -2.0023615 | 0.04524587 | NA | TSEN15P2 | 106480781 |
| ENSG000002 | 0 | NA | NA | NA | NA | NA | NA | NA |
| ENSG000002 | 1.11558913 | -0.108321552 | 0.31618912 | -0.3425847 | 0.73191093 | NA | NA | NA |
| ENSG000002 | 1.11971733 | -0.27463679 | 0.32326001 | -0.8495848 | 0.39555597 | NA | NA | NA |
| ENSG000002 | 0.51911933 | -0.349228947 | 0.23754574 | -1.4701545 | 0.14151991 | NA | NA | NA |
| ENSG000002 | 0.85040642 | 0.053066201 | 0.29032665 | 0.18278102 | 0.85496984 | NA | NA | NA |
| ENSG000002 | 0.62496306 | -0.108686629 | 0.26501936 | -0.4101083 | 0.68172654 | NA | NA | NA |
| ENSG000002 | 1.01224622 | -0.206895503 | 0.31834465 | -0.6499104 | 0.51575009 | NA | NA | NA |
| ENSG000002 | 1.35688868 | -0.247306099 | 0.33185333 | -0.7452271 | 0.45613443 | NA | NA | NA |
| ENSG000002 | 1.73711284 | -0.320499854 | 0.35425958 | -0.9047034 | 0.36562252 | NA | NA | NA |
| ENSG000002 | 0 | NA | NA | NA | NA | NA | NA | NA |
| ENSG000002 | 1.16215289 | -0.59203245 | 0.32014781 | -1.8492472 | 0.06442212 | NA | NA | NA |
| ENSG000002 | 0.96015544 | 0.113786635 | 0.296576 | 0.38366771 | 0.70122475 | NA | KLF2P2 | 106480678 |
| ENSG000002 | 1.9417226 | -0.681702369 | 0.35884431 | -1.8997163 | 0.05747036 | NA | NA | NA |
| ENSG000002 | 1.48707683 | -0.3189847 | 0.34175641 | -0.9333686 | 0.35062969 | NA | NA | NA |
| ENSG000002 | 0 | NA | NA | NA | NA | NA | NA | NA |
| ENSG000002 | 0.87284005 | -0.120944163 | 0.3030077 | -0.3991455 | 0.68978599 | NA | NA | NA |
| ENSG000002 | 2.08673624 | -0.117964317 | 0.36556275 | -0.3226924 | 0.74692822 | NA | LOC105375266 | 105375266 |
| ENSG000002 | 1.41313326 | -0.729445817 | 0.33556176 | -2.173805 | 0.02971979 | NA | NA | NA |
| ENSG000002 | 0.51959753 | -0.202283781 | 0.25032396 | -0.808088 | 0.41903995 | NA | NA | NA |
| ENSG000002 | 1.119435 | -0.436060276 | 0.32298771 | -1.3500832 | 0.1769893 | NA | NA | NA |
| ENSG000002 | 0.38133351 | -0.102466351 | 0.2189713 | -0.4679442 | 0.63982449 | NA | NA | NA |
| ENSG000002 | 1.87937548 | -0.351893666 | 0.34606343 | -1.0168473 | 0.30922608 | NA | NA | NA |

| | | | | | | | | | |
|---|---|---|---|---|---|---|---|---|---|
| ENSG000002 | 1.65767662 | -0.61337965 | 0.33621774 | -1.8243524 | 0.06809881 | NA | | LOC101928628 | 101928628 |
| ENSG000002 | 1.1146275 | -0.408072454 | 0.31717208 | -1.2865964 | 0.198235 | NA | | NA | NA |
| ENSG000002 | 1.17835117 | -0.557572044 | 0.31343597 | -1.7789025 | 0.07525575 | NA | | NA | NA |
| ENSG000002 | 0.61816464 | -0.117186973 | 0.27074914 | -0.4328249 | 0.66514197 | NA | | LOC105376480 | 105376480 |
| ENSG000002 | 0 | NA | NA | NA | NA | NA | | NA | NA |
| ENSG000002 | 1.25439978 | -0.337683135 | 0.32662402 | -1.0338588 | 0.30120216 | NA | | NA | NA |
| ENSG000002 | 2.0424765 | -0.536203695 | 0.35400655 | -1.5146717 | 0.12985558 | NA | | NA | NA |
| ENSG000002 | 1.44576631 | 0.241448978 | 0.34500625 | 0.69983943 | 0.48402759 | NA | | NA | NA |
| ENSG000002 | 1.80909975 | -0.35207825 | 0.35091296 | -1.0033207 | 0.31570614 | NA | | NA | NA |
| ENSG000002 | 1.89595788 | -0.640359126 | 0.35474219 | -1.8051395 | 0.07105286 | NA | | NA | NA |
| ENSG000002 | 0.11934008 | 0.083564698 | 0.12760091 | 0.65489106 | 0.5125379 | NA | | NA | NA |
| ENSG000002 | 0.89236323 | -0.395354214 | 0.28664737 | -1.3792355 | 0.16782217 | NA | | NA | NA |
| ENSG000002 | 1.19456676 | 0.108532542 | 0.32746711 | 0.33143036 | 0.74031944 | NA | | NA | NA |
| ENSG000002 | 0.82226586 | -0.505896824 | 0.27464408 | -1.8420089 | 0.06547385 | NA | | ANKRD63 | 100131244 |
| ENSG000002 | 1.87529501 | -0.221131857 | 0.34802665 | -0.6353877 | 0.52517558 | NA | | NA | NA |
| ENSG000002 | 1.69254196 | -0.540105949 | 0.34714662 | -1.5558439 | 0.11974521 | NA | | NA | NA |
| ENSG000002 | 1.12236718 | -0.119233432 | 0.31642544 | -0.3768137 | 0.70631208 | NA | | NA | NA |
| ENSG000002 | 2.09735543 | -0.231560962 | 0.36026197 | -0.6427572 | 0.52038168 | NA | | NA | NA |
| ENSG000002 | 1.51510916 | -0.325340978 | 0.34084502 | -0.9545129 | 0.33982407 | NA | | NA | NA |
| ENSG000002 | 1.86909451 | 0.039550098 | 0.24201198 | 0.16342207 | 0.87018613 | NA | | NA | NA |
| ENSG000002 | 0.36351096 | 0.070290861 | 0.21590418 | 0.32556508 | 0.74475344 | NA | | NA | NA |
| ENSG000002 | 1.85237569 | -0.130348987 | 0.35801938 | -0.3640836 | 0.71579558 | NA | | NA | NA |
| ENSG000002 | 0 | NA | NA | NA | NA | NA | | USP9YP24 | 106478928 |
| ENSG000002 | 0 | NA | NA | NA | NA | NA | | ZNF736P5Y | 106480711 |
| ENSG000002 | 0.90047403 | -0.281185275 | 0.30049169 | -0.9357506 | 0.34940162 | NA | | NA | NA |
| ENSG000002 | 1.05292112 | -0.337947424 | 0.30412401 | -1.1112159 | 0.26647544 | NA | | NA | NA |
| ENSG000002 | 2.04170529 | 0.255272042 | 0.36533044 | 0.69874287 | 0.48471274 | NA | | NA | NA |
| ENSG000002 | 0 | NA | NA | NA | NA | NA | | NA | NA |
| ENSG000002 | 0.85399796 | -0.109832103 | 0.29091059 | -0.3775459 | 0.70576795 | NA | | NA | NA |

| | | | | | | | | |
|---|---|---|---|---|---|---|---|---|
| ENSG000002 | 0 | NA | NA | NA | NA | NA | NA | NA |
| ENSG000002 | 1.38817087 | -0.265266482 | 0.3352626 | -0.79122 | 0.42881561 | NA | NA | NA |
| ENSG000002 | 0.25370795 | -0.171331747 | 0.17411721 | -0.9840024 | 0.32511435 | NA | NA | NA |
| ENSG000002 | 1.20692591 | -0.176256011 | 0.31988068 | -0.5510055 | 0.58162992 | NA | NA | NA |
| ENSG000002 | 0 | NA | NA | NA | NA | NA | USP9YP20 | 106480717 |
| ENSG000002 | 0.62087103 | -0.279525375 | 0.26646004 | -1.049033 | 0.29416293 | NA | NA | NA |
| ENSG000002 | 2.11343659 | -0.259035969 | 0.36743824 | -0.7049783 | 0.48082377 | NA | NA | NA |
| ENSG000002 | 1.59218723 | -0.181593265 | 0.21005836 | -0.8644896 | 0.38731899 | NA | NA | NA |
| ENSG000002 | 2.02272361 | -0.431729024 | 0.35912688 | -1.2021629 | 0.2293004 | NA | NA | NA |
| ENSG000002 | 1.0053619 | -0.324036727 | 0.30386972 | -1.0663673 | 0.28625765 | NA | NA | NA |
| ENSG000002 | 0 | NA | NA | NA | NA | NA | NA | NA |
| ENSG000002 | 1.54129961 | -0.595821749 | 0.3356649 | -1.7750493 | 0.07588975 | NA | NA | NA |
| ENSG000002 | 1.61509883 | -0.257303039 | 0.34799405 | -0.7393892 | 0.4596707 | NA | NA | NA |
| ENSG000002 | 1.85186579 | -0.246279427 | 0.35736786 | -0.6891482 | 0.49072999 | NA | NA | NA |
| ENSG000002 | 0.9859487 | -0.166067641 | 0.3011363 | -0.55147 | 0.58131152 | NA | NA | NA |
| ENSG000002 | 1.12603755 | -0.433672183 | 0.32323244 | -1.3416728 | 0.1797021 | NA | NA | NA |
| ENSG000002 | 1.02639492 | -0.370661357 | 0.31595022 | -1.1731638 | 0.24073014 | NA | NA | NA |
| ENSG000002 | 0 | NA | NA | NA | NA | NA | NA | NA |
| ENSG000002 | 2.00191796 | 0.052728252 | 0.35544917 | 0.14834259 | 0.8820724 | NA | NA | NA |
| ENSG000002 | 1.70047206 | 0.079195837 | 0.35462347 | 0.22332373 | 0.82328356 | NA | NA | NA |
| ENSG000002 | 1.39511777 | -0.26134542 | 0.33059549 | -0.7905293 | 0.42921872 | NA | NA | NA |
| ENSG000002 | 1.99485491 | -0.200415589 | 0.36455131 | -0.5497596 | 0.58248426 | NA | NA | NA |
| ENSG000002 | 0.96380544 | -0.187512512 | 0.31093159 | -0.6030668 | 0.54646426 | NA | NA | NA |
| ENSG000002 | 0.70900933 | 0.127098089 | 0.27497685 | 0.46221378 | 0.64392803 | NA | NA | NA |
| ENSG000002 | 1.9629631 | -0.805426639 | 0.35520535 | -2.2674958 | 0.02335996 | NA | NA | NA |
| ENSG000002 | 1.96901439 | -0.182475304 | 0.36275406 | -0.5030276 | 0.61494487 | NA | NA | NA |
| ENSG000002 | 1.44894024 | -0.043712534 | 0.3430757 | -0.1274137 | 0.898613 | NA | NA | NA |
| ENSG000002 | 0 | NA | NA | NA | NA | NA | NA | NA |
| ENSG000002 | 0.49862008 | 0.145705772 | 0.2401745 | 0.6066663 | 0.54407239 | NA | NA | NA |

| | | | | | | | | |
|---|---|---|---|---|---|---|---|---|
| ENSG000002 | 0.51208364 | -0.022981898 | 0.24832117 | -0.0925491 | 0.92626179 | NA | NA | NA |
| ENSG000002 | 1.23059778 | -0.037795899 | 0.32853598 | -0.1150434 | 0.90841072 | NA | NA | NA |
| ENSG000002 | 1.61905289 | -0.271170812 | 0.35310407 | -0.7679629 | 0.44250926 | NA | NA | NA |
| ENSG000002 | 0.50684946 | -0.202346264 | 0.24895102 | -0.8127955 | 0.41633533 | NA | NA | NA |
| ENSG000002 | 0.59760982 | 0.066257965 | 0.26871197 | 0.24657616 | 0.80523626 | NA | NA | NA |
| ENSG000002 | 0.78247198 | -0.337783522 | 0.27652822 | -1.2215156 | 0.22189088 | NA | NA | NA |
| ENSG000002 | 1.12245652 | -0.401923574 | 0.31666974 | -1.26922 | 0.2043626 | NA | NA | NA |
| ENSG000002 | 1.97547204 | -0.192379642 | 0.36131279 | -0.5324463 | 0.59441696 | NA | NA | NA |
| ENSG000002 | 1.69340788 | -0.281299554 | 0.34141977 | -0.8239112 | 0.40999004 | NA | NA | NA |
| ENSG000002 | 1.71479255 | 0.181870804 | 0.34592856 | 0.52574672 | 0.5990642 | NA | NA | NA |
| ENSG000002 | 1.03831989 | -0.50780213 | 0.30797528 | -1.6488405 | 0.09918031 | NA | NA | NA |
| ENSG000002 | 0 | NA | NA | NA | NA | NA | NA | NA |
| ENSG000002 | 1.72208258 | -0.200017194 | 0.35828799 | -0.5582582 | 0.57666811 | NA | NA | NA |
| ENSG000002 | 2.03565689 | -0.733263514 | 0.36176583 | -2.026901 | 0.04267254 | NA | NA | NA |
| ENSG000002 | 0.9962727 | 0.116269009 | 0.30673009 | 0.37905968 | 0.70464355 | NA | NA | NA |
| ENSG000002 | 0 | NA | NA | NA | NA | NA | USP17L28 | 728400 |
| ENSG000002 | 0.22410479 | 0.147643983 | 0.16638082 | 0.88738586 | 0.37487119 | NA | NA | NA |
| ENSG000002 | 0.35870346 | 0.077368606 | 0.21515516 | 0.35959447 | 0.71915042 | NA | NA | NA |
| ENSG000002 | 1.67270454 | -0.389454022 | 0.34633816 | -1.1244907 | 0.26080493 | NA | NA | NA |
| ENSG000002 | 1.60211135 | -0.129173593 | 0.35251027 | -0.3664392 | 0.71403735 | NA | NA | NA |
| ENSG000002 | 0.66527186 | -0.43035014 | 0.25829332 | -1.6661296 | 0.09568761 | NA | NA | NA |
| ENSG000002 | 2.01169039 | -0.438651069 | 0.35902568 | -1.2217819 | 0.22179012 | NA | NA | NA |
| ENSG000002 | 1.27264104 | -0.361300261 | 0.33572482 | -1.0761798 | 0.28184686 | NA | NA | NA |
| ENSG000002 | 1.10865219 | -0.257460636 | 0.31695249 | -0.8123004 | 0.41661928 | NA | NA | NA |
| ENSG000002 | 1.72245315 | -0.32637547 | 0.35531622 | -0.9185493 | 0.35833136 | NA | NA | NA |
| ENSG000002 | 1.08602159 | -0.113284216 | 0.31514512 | -0.3594668 | 0.71924589 | NA | NA | NA |
| ENSG000002 | 1.3126691 | -0.626026282 | 0.32128955 | -1.9484801 | 0.05135755 | NA | NA | NA |
| ENSG000002 | 1.48591094 | -0.051486256 | 0.33899127 | -0.1518808 | 0.87928098 | NA | NA | NA |
| ENSG000002 | 0.34936227 | 0.241139221 | 0.20762307 | 1.16142789 | 0.24546793 | NA | MTHFD2P3 | 106481665 |

| | | | | | | | | |
|---|---|---|---|---|---|---|---|---|
| ENSG000002 | 1.23629545 | -0.32842872 | 0.32614123 | -1.0070138 | 0.31392814 | NA | NA | NA |
| ENSG000002 | 0.46956163 | 0.155192681 | 0.24412227 | 0.63571702 | 0.52496088 | NA | NA | NA |
| ENSG000002 | 1.5849588 | -0.241396032 | 0.34425933 | -0.701204 | 0.48317569 | NA | NA | NA |
| ENSG000002 | 1.1416647 | -0.256242155 | 0.31807371 | -0.8056062 | 0.42046994 | NA | HAR1B | 768097 |
| ENSG000002 | 0.85131777 | -0.102855071 | 0.29519776 | -0.3484277 | 0.72751902 | NA | NA | NA |
| ENSG000002 | 0.51316968 | -0.377777671 | 0.24556403 | -1.538408 | 0.12394889 | NA | NA | NA |
| ENSG000002 | 0.13280363 | -0.093394593 | 0.13115644 | -0.7120855 | 0.47641185 | NA | NA | NA |
| ENSG000002 | 0 | NA | NA | NA | NA | NA | NA | NA |
| ENSG000002 | 1.46042601 | 0.237536207 | 0.34335832 | 0.69180268 | 0.48906126 | NA | NA | NA |
| ENSG000002 | 1.81768266 | -0.441860191 | 0.34843997 | -1.2681099 | 0.2047587 | NA | NA | NA |
| ENSG000002 | 0.88223331 | -0.275481339 | 0.29958866 | -0.9195319 | 0.35781741 | NA | NA | NA |
| ENSG000002 | 1.53127268 | 0.282356421 | 0.34079174 | 0.82853072 | 0.40737001 | NA | NA | NA |
| ENSG000002 | 0 | NA | NA | NA | NA | NA | NA | NA |
| ENSG000002 | 1.08960563 | 0.035673201 | 0.3211968 | 0.11106338 | 0.91156609 | NA | NA | NA |
| ENSG000002 | 1.96481129 | -0.196469152 | 0.36218592 | -0.5424539 | 0.58750588 | NA | NA | NA |
| ENSG000002 | 2.01006143 | -0.327033162 | 0.36425975 | -0.8978021 | 0.36929107 | NA | NA | NA |
| ENSG000002 | 0.73729832 | 0.131121994 | 0.27696116 | 0.47343099 | 0.63590573 | NA | NA | NA |
| ENSG000002 | 1.33862231 | -0.113856913 | 0.33425572 | -0.3406282 | 0.73338352 | NA | NA | NA |
| ENSG000002 | 0.27408862 | -0.199826037 | 0.18669465 | -1.0703362 | 0.28446801 | NA | NA | NA |
| ENSG000002 | 1.43444059 | 0.227581366 | 0.33829252 | 0.67273543 | 0.50111562 | NA | NA | NA |
| ENSG000002 | 1.05940714 | 0.17785428 | 0.31416685 | 0.56611409 | 0.57131623 | NA | NA | NA |
| ENSG000002 | 0.14128499 | -0.097772014 | 0.13403235 | -0.7294658 | 0.4657168 | NA | NA | NA |
| ENSG000002 | 1.13388776 | -0.269928135 | 0.32225003 | -0.8376357 | 0.40223532 | NA | NA | NA |
| ENSG000002 | 1.68593941 | -0.557468807 | 0.34969686 | -1.5941487 | 0.11090273 | NA | NA | NA |
| ENSG000002 | 0.38036605 | -0.293832733 | 0.22119083 | -1.3284128 | 0.18404178 | NA | NA | NA |
| ENSG000002 | 0.23936338 | -0.005479057 | 0.17759977 | -0.0308506 | 0.9753887 | NA | NA | NA |
| ENSG000002 | 1.42309917 | -0.030609111 | 0.33064669 | -0.0925735 | 0.92624242 | NA | NA | NA |
| ENSG000002 | 1.59394848 | -0.243241277 | 0.34438853 | -0.706299 | 0.48000221 | NA | NA | NA |
| ENSG000002 | 0.87420842 | -0.278408432 | 0.29914995 | -0.9306652 | 0.3520268 | NA | KLF2P1 | 106480253 |

| | | | | | | | | |
|---|---|---|---|---|---|---|---|---|
| ENSG000002 | 0.64616918 | -0.423578009 | 0.25668375 | -1.6501941 | 0.09890325 | NA | NA | NA |
| ENSG000002 | 0.23936338 | -0.005479057 | 0.17759977 | -0.0308506 | 0.9753887 | NA | NA | NA |
| ENSG000002 | 0.38270188 | -0.272446727 | 0.2140009 | -1.2731102 | 0.20297894 | NA | NA | NA |
| ENSG000002 | 2.03875972 | -0.866098721 | 0.35860469 | -2.4151908 | 0.01572697 | NA | NA | NA |
| ENSG000002 | 0.11934008 | 0.083564698 | 0.12760091 | 0.65489106 | 0.5125379 | NA | NA | NA |
| ENSG000002 | 2.11419499 | -0.134861325 | 0.36763269 | -0.3668371 | 0.7137405 | NA | NA | NA |
| ENSG000002 | 1.87153098 | -0.117640039 | 0.35647656 | -0.3300078 | 0.74139408 | NA | NA | NA |
| ENSG000002 | 1.26393292 | -0.198687541 | 0.33139639 | -0.5995465 | 0.54880852 | NA | NA | NA |
| ENSG000002 | 1.713116 | 0.076069113 | 0.35098192 | 0.21673228 | 0.82841699 | NA | NA | NA |
| ENSG000002 | 1.99138598 | -0.169391343 | 0.35811927 | -0.4730026 | 0.63621133 | NA | NA | NA |
| ENSG000002 | 1.62556898 | -0.414799063 | 0.35313807 | -1.1746087 | 0.24015128 | NA | NA | NA |
| ENSG000002 | 2.05728332 | -0.719456352 | 0.35944386 | -2.001582 | 0.04532971 | NA | NA | NA |
| ENSG000002 | 0.49270076 | -0.021329091 | 0.24621941 | -0.0866264 | 0.93096851 | NA | NA | NA |
| ENSG000002 | 0 | NA | NA | NA | NA | NA | NA | NA |
| ENSG000002 | 0.8794426 | -0.118660638 | 0.30333647 | -0.3911849 | 0.69566059 | NA | NA | NA |
| ENSG000002 | 0.86683249 | 0.037236871 | 0.29129111 | 0.12783387 | 0.89828044 | NA | NA | NA |
| ENSG000002 | 1.90148086 | -0.808250349 | 0.3552033 | -2.2754584 | 0.02287844 | NA | OOEP-AS1 | 106478938 |
| ENSG000002 | 1.52359499 | -0.613719411 | 0.34062255 | -1.801758 | 0.07158349 | NA | NA | NA |
| ENSG000002 | 1.09749009 | -0.264509638 | 0.32106828 | -0.8238423 | 0.41002916 | NA | NA | NA |
| ENSG000002 | 1.39958092 | -0.277135818 | 0.33889798 | -0.8177559 | 0.41349659 | NA | NA | NA |
| ENSG000002 | 0.75025279 | -0.033352346 | 0.29080996 | -0.1146878 | 0.90869261 | NA | NA | NA |
| ENSG000002 | 0.90864082 | -0.400702233 | 0.2875379 | -1.3935632 | 0.16344955 | NA | NA | NA |
| ENSG000002 | 1.39144893 | -0.565373992 | 0.33648791 | -1.6802208 | 0.09291436 | NA | NA | NA |
| ENSG000002 | 0.62362749 | 0.019297443 | 0.15764368 | 0.12241178 | 0.90257291 | NA | NA | NA |
| ENSG000002 | 0.65445467 | -0.486865861 | 0.27154464 | -1.7929496 | 0.07298098 | NA | NA | NA |
| ENSG000002 | 0.37945415 | -0.107841678 | 0.21865587 | -0.4932027 | 0.62186934 | NA | NA | NA |
| ENSG000002 | 0 | NA | NA | NA | NA | NA | NA | NA |
| ENSG000002 | 1.16073713 | -0.148257522 | 0.20687098 | -0.7166666 | 0.47357984 | NA | NA | NA |
| ENSG000002 | 1.85489123 | -0.482967759 | 0.35303972 | -1.3680267 | 0.17130373 | NA | NA | NA |

| | | | | | | | | |
|---|---|---|---|---|---|---|---|---|
| ENSG000002 | 1.93184973 | -0.036673594 | 0.35020739 | -0.1047196 | 0.91659827 | NA | NA | NA |
| ENSG000002 | 0 | NA | NA | NA | NA | NA | NA | NA |
| ENSG000002 | 0.50681912 | -0.019233989 | 0.24032284 | -0.080034 | 0.93621025 | NA | NA | NA |
| ENSG000002 | 0 | NA | NA | NA | NA | NA | USP17L10 | 100287144 |
| ENSG000002 | 1.95849162 | -0.06433024 | 0.36243828 | -0.177493 | 0.8591212 | NA | NA | NA |
| ENSG000002 | 1.57866811 | -0.78075439 | 0.33890979 | -2.3037233 | 0.02123818 | NA | NA | NA |
| ENSG000002 | 1.20165802 | 0.102084619 | 0.32768603 | 0.3115318 | 0.75539637 | NA | NA | NA |
| ENSG000002 | 0 | NA | NA | NA | NA | NA | NA | NA |
| ENSG000002 | 2.11639284 | -0.251091618 | 0.36628816 | -0.685503 | 0.4930266 | NA | NA | NA |
| ENSG000002 | 1.74571099 | -0.326680355 | 0.35434604 | -0.9219247 | 0.35656787 | NA | NA | NA |
| ENSG000002 | 0 | NA | NA | NA | NA | NA | NA | NA |
| ENSG000002 | 0 | NA | NA | NA | NA | NA | NA | NA |
| ENSG000002 | 1.97721054 | 0.058191525 | 0.36336825 | 0.16014477 | 0.87276703 | NA | WASIR2 | 100132169 |
| ENSG000002 | 0.75964551 | -0.194136624 | 0.28087797 | -0.6911778 | 0.48945379 | NA | NA | NA |
| ENSG000002 | 1.08886842 | 0.03543426 | 0.3211724 | 0.11032785 | 0.91214937 | NA | NA | NA |
| ENSG000002 | 1.27515101 | -0.51710732 | 0.33236375 | -1.5558475 | 0.11974435 | NA | NA | NA |
| ENSG000002 | 1.14476753 | -0.438324541 | 0.3238073 | -1.3536586 | 0.17584532 | NA | NA | NA |
| ENSG000002 | 1.87401242 | -0.486255534 | 0.35154112 | -1.383211 | 0.16660018 | NA | NA | NA |
| ENSG000002 | 0.61172818 | 0.021749657 | 0.15747952 | 0.13811102 | 0.89015268 | NA | NA | NA |
| ENSG000002 | 1.73523347 | -0.320938501 | 0.35317731 | -0.9087178 | 0.36349911 | NA | NA | NA |
| ENSG000002 | 0.75441187 | -0.377475959 | 0.28823563 | -1.3096089 | 0.19032816 | NA | FCF1P9 | 106480444 |
| ENSG000002 | 0.37330862 | -0.105391168 | 0.21773449 | -0.4840352 | 0.62836086 | NA | NA | NA |
| ENSG000002 | 0.23276084 | -0.008011382 | 0.17603982 | -0.0455089 | 0.96370166 | NA | NA | NA |
| ENSG000002 | 0.75167514 | -0.031455075 | 0.28559977 | -0.1101369 | 0.9123008 | NA | NA | NA |
| ENSG000002 | 0 | NA | NA | NA | NA | NA | NA | NA |
| ENSG000002 | 0.37928001 | 0.070449581 | 0.21829793 | 0.32272217 | 0.74690566 | NA | MTND4LP1 | 106478909 |
| ENSG000002 | 0 | NA | NA | NA | NA | NA | NA | NA |
| ENSG000002 | 1.5311353 | -0.631853588 | 0.34256483 | -1.8444789 | 0.06511339 | NA | NA | NA |
| ENSG000002 | 0.26199343 | -0.194018184 | 0.18422801 | -1.0531416 | 0.29227609 | NA | LINC00278 | 100873962 |

| | | | | | | | | |
|---|---|---|---|---|---|---|---|---|
| ENSG000002 | 0.7164389 | -0.025867019 | 0.28328556 | -0.0913108 | 0.92724567 | NA | NA | NA |
| ENSG000002 | 0.72167308 | 0.133011302 | 0.2775391 | 0.47925248 | 0.63175902 | NA | NA | NA |
| ENSG000002 | 1.4913758 | -0.336306894 | 0.34674602 | -0.969894 | 0.33209931 | NA | DPP10-AS3 | 101927591 |
| ENSG000002 | 0 | NA | NA | NA | NA | NA | NA | NA |
| ENSG000002 | 0.97000709 | 0.121464882 | 0.3072071 | 0.39538436 | 0.69255925 | NA | NA | NA |
| ENSG000002 | 1.50668592 | -0.4784321 | 0.3439377 | -1.3910429 | 0.16421242 | NA | NA | NA |
| ENSG000002 | 2.11178535 | -0.258379625 | 0.36522468 | -0.7074539 | 0.47928445 | NA | NA | NA |
| ENSG000002 | 1.86515815 | -0.002143844 | 0.3595452 | -0.0059627 | 0.99524252 | NA | NA | NA |
| ENSG000002 | 1.51154741 | -0.34501085 | 0.34704344 | -0.9941431 | 0.32015319 | NA | NA | NA |
| ENSG000002 | 0.38270188 | -0.272446727 | 0.2140009 | -1.2731102 | 0.20297894 | NA | NA | NA |
| ENSG000002 | 0.602333 | 0.056422611 | 0.25511105 | 0.22116882 | 0.82496098 | NA | NA | NA |
| ENSG000002 | 0.11865494 | 0.083440866 | 0.12751478 | 0.65436232 | 0.5128784 | NA | NA | NA |
| ENSG000002 | 1.50146955 | -0.189827156 | 0.34505912 | -0.5501294 | 0.58223065 | NA | NA | NA |
| ENSG000002 | 0.88126585 | -0.455824927 | 0.30137169 | -1.5125008 | 0.1304065 | NA | NA | NA |
| ENSG000002 | 1.74736112 | -0.326172547 | 0.353703 | -0.9221651 | 0.35644249 | NA | NA | NA |
| ENSG000002 | 0.37422052 | -0.268731395 | 0.21273223 | -1.2632378 | 0.20650373 | NA | NA | NA |
| ENSG000002 | 0 | NA | NA | NA | NA | NA | NA | NA |
| ENSG000002 | 1.45351881 | -0.537848821 | 0.32995828 | -1.630051 | 0.10309073 | NA | NA | NA |
| ENSG000002 | 0.51299499 | -0.204672849 | 0.24959455 | -0.8200213 | 0.41220396 | NA | NA | NA |
| ENSG000002 | 1.75962887 | -0.554160684 | 0.34381948 | -1.611778 | 0.10701025 | NA | NA | NA |
| ENSG000002 | 0.93627854 | 0.237120892 | 0.28973418 | 0.81840843 | 0.41312401 | NA | NA | NA |
| ENSG000002 | 1.13711375 | -0.589529324 | 0.31927247 | -1.8464772 | 0.06482295 | NA | NA | NA |
| ENSG000002 | 2.05742112 | -0.32875026 | 0.35637517 | -0.9224836 | 0.35627637 | NA | NA | NA |
| ENSG000002 | 1.74516965 | -0.340540167 | 0.35799569 | -0.951241 | 0.34148207 | NA | NA | NA |
| ENSG000002 | 2.02608894 | -0.46441825 | 0.36352552 | -1.2775396 | 0.2014118 | NA | NA | NA |
| ENSG000002 | 1.11265832 | -0.271859025 | 0.32304286 | -0.8415571 | 0.40003588 | NA | NA | NA |
| ENSG000002 | 2.1019018 | -0.243299955 | 0.36138611 | -0.673241 | 0.50079396 | NA | NA | NA |
| ENSG000002 | 2.15991211 | -0.396565441 | 0.36726022 | -1.0797942 | 0.28023385 | NA | NA | NA |
| ENSG000002 | 1.76501188 | -0.721396549 | 0.34889432 | -2.0676649 | 0.03867154 | NA | NA | NA |

| | | | | | | | | |
|---|---|---|---|---|---|---|---|---|
| ENSG000002 | 1.94655396 | -0.35359427 | 0.34897435 | -1.0132386 | 0.31094622 | NA | LOC105373524 | 105373524 |
| ENSG000002 | 0.47687965 | -0.021216903 | 0.24445659 | -0.0867921 | 0.93083676 | NA | NA | NA |
| ENSG000002 | 1.03501659 | -0.697779481 | 0.30815271 | -2.2643951 | 0.02354983 | NA | AGBL5-AS1 | 100874031 |
| ENSG000002 | 0 | NA | NA | NA | NA | NA | USP17L29 | 728405 |
| ENSG000002 | 1.75594854 | -0.20411658 | 0.35861116 | -0.5691864 | 0.56922968 | NA | NA | NA |
| ENSG000002 | 1.74205886 | -0.193846913 | 0.35547927 | -0.5453114 | 0.58553934 | NA | DLG3-AS1 | 100873930 |
| ENSG000002 | 1.21882969 | -0.448703196 | 0.31853721 | -1.4086367 | 0.15894263 | NA | NA | NA |
| ENSG000002 | 0.12070844 | -0.088503695 | 0.12783931 | -0.6923042 | 0.48874631 | NA | NA | NA |
| ENSG000002 | 1.77027192 | -0.48752561 | 0.35821131 | -1.361 | 0.17351368 | NA | NA | NA |
| ENSG000002 | 1.94678819 | -0.061094204 | 0.36180158 | -0.1688611 | 0.86590592 | NA | NA | NA |
| ENSG000002 | 1.91148151 | 0.594931725 | 0.35808968 | 1.66140428 | 0.09663228 | NA | NA | NA |
| ENSG000002 | 1.937001 | -0.539807008 | 0.3591936 | -1.5028303 | 0.13288282 | NA | NA | NA |
| ENSG000002 | 1.61717407 | -0.2700383 | 0.35408399 | -0.7626391 | 0.44567865 | NA | NA | NA |
| ENSG000002 | 0.84648407 | 0.042140931 | 0.2962168 | 0.14226381 | 0.88687163 | NA | NA | NA |
| ENSG000002 | 0.95537965 | 0.121853116 | 0.31143989 | 0.39125725 | 0.69560709 | NA | NA | NA |
| ENSG000002 | 2.06815394 | -0.595260123 | 0.36189221 | -1.6448547 | 0.09999977 | NA | NA | NA |
| ENSG000002 | 1.76976339 | -0.444074667 | 0.35065164 | -1.2664269 | 0.20536028 | NA | NA | NA |
| ENSG000002 | 1.00766101 | -0.039578712 | 0.30716166 | -0.128853 | 0.89747394 | NA | NA | NA |
| ENSG000002 | 1.36093229 | -0.226897475 | 0.32065369 | -0.7076091 | 0.47918803 | NA | NA | NA |
| ENSG000002 | 0.84919046 | -0.109177322 | 0.296711 | -0.3679585 | 0.71290421 | NA | NA | NA |
| ENSG000002 | 1.88482569 | -0.259080107 | 0.35970488 | -0.7202574 | 0.47136652 | NA | NA | NA |
| ENSG000002 | 0.50516651 | -0.183853222 | 0.24168957 | -0.7606999 | 0.44683635 | NA | NA | NA |
| ENSG000002 | 0.82898056 | 0.04846601 | 0.29524784 | 0.16415365 | 0.86961019 | NA | NA | NA |
| ENSG000002 | 1.65437545 | -0.71407314 | 0.35044767 | -2.0376028 | 0.04158967 | NA | NA | NA |
| ENSG000002 | 1.23145706 | -0.183181855 | 0.326802 | -0.5605286 | 0.57511897 | NA | NA | NA |
| ENSG000002 | 0 | NA | NA | NA | NA | NA | NA | NA |
| ENSG000002 | 1.15444257 | -0.445335221 | 0.32404689 | -1.3742925 | 0.16935088 | NA | NA | NA |
| ENSG000002 | 1.21716456 | -0.038860888 | 0.32966871 | -0.1178786 | 0.90616385 | NA | NA | NA |
| ENSG000002 | 2.06815449 | -0.576451814 | 0.36040779 | -1.5994432 | 0.10972215 | NA | NA | NA |

| | | | | | | | | |
|---|---|---|---|---|---|---|---|---|
| ENSG000002 | 1.93671225 | 0.197864424 | 0.36080416 | 0.54839839 | 0.58341838 | NA | NA | NA |
| ENSG000002 | 0.96840792 | 0.117990867 | 0.31199888 | 0.37817722 | 0.70529895 | NA | NA | NA |
| ENSG000002 | 0.89720511 | -0.276906822 | 0.29874913 | -0.9268875 | 0.35398497 | NA | NA | NA |
| ENSG000002 | 0.87801812 | -0.27594325 | 0.29776051 | -0.9267288 | 0.35406734 | NA | NA | NA |
| ENSG000002 | 1.50397952 | -0.347958955 | 0.34806559 | -0.9996936 | 0.31745879 | NA | NA | NA |
| ENSG000002 | 2.03268991 | -0.573430607 | 0.36016458 | -1.5921349 | 0.11135439 | NA | NA | NA |
| ENSG000002 | 1.44235751 | -0.039885941 | 0.33818161 | -0.1179424 | 0.90611332 | NA | NA | NA |
| ENSG000002 | 1.6754665 | -0.245310295 | 0.34195866 | -0.7173683 | 0.47314689 | NA | NA | NA |
| ENSG000002 | 0.62821079 | -0.27685072 | 0.26713242 | -1.0363801 | 0.30002486 | NA | NA | NA |
| ENSG000002 | 1.22690218 | -0.19088789 | 0.32896214 | -0.5802731 | 0.56173045 | NA | NA | NA |
| ENSG000002 | 1.72436689 | -0.182187758 | 0.35355821 | -0.5152978 | 0.60634495 | NA | NA | NA |
| ENSG000002 | 0.60307021 | 0.056690296 | 0.25518062 | 0.22215753 | 0.82419125 | NA | NA | NA |
| ENSG000002 | 1.37695086 | -0.129027042 | 0.33989859 | -0.3796045 | 0.70423901 | NA | NA | NA |
| ENSG000002 | 1.37128408 | 0.02237469 | 0.3279987 | 0.06821579 | 0.94561386 | NA | NA | NA |
| ENSG000002 | 0.49512483 | -0.344267812 | 0.23617548 | -1.457678 | 0.14492931 | NA | NA | NA |
| ENSG000002 | 0.71900341 | -0.01905758 | 0.27706964 | -0.0687826 | 0.94516264 | NA | NA | NA |
| ENSG000002 | 0.40327843 | -0.281391716 | 0.21700406 | -1.2967118 | 0.19473038 | NA | NA | NA |
| ENSG000002 | 1.37411096 | -0.401268159 | 0.33686125 | -1.1911972 | 0.23357619 | NA | NA | NA |
| ENSG000002 | 1.14552805 | -0.11661481 | 0.31720969 | -0.3676269 | 0.71315146 | NA | UBE2V2P4 | 106480802 |
| ENSG000002 | 1.50150977 | -0.169400449 | 0.33414731 | -0.5069634 | 0.61218052 | NA | NA | NA |
| ENSG000002 | 1.37621599 | -0.525415668 | 0.32840439 | -1.5999045 | 0.10961976 | NA | NA | NA |
| ENSG000002 | 1.8390913 | -0.365626843 | 0.35515971 | -1.0294716 | 0.30325811 | NA | NA | NA |
| ENSG000002 | 1.61636768 | 0.000543672 | 0.34290018 | 0.00158551 | 0.99873495 | NA | NA | NA |
| ENSG000002 | 0 | NA | NA | NA | NA | NA | NA | NA |
| ENSG000002 | 1.63694869 | -0.245682904 | 0.34503751 | -0.7120469 | 0.47643571 | NA | NA | NA |
| ENSG000002 | 2.11489265 | -0.622512895 | 0.36127797 | -1.7230857 | 0.08487303 | NA | NA | NA |
| ENSG000002 | 1.09663081 | -0.108391837 | 0.32174573 | -0.3368866 | 0.73620236 | NA | F10-AS1 | 104413892 |
| ENSG000002 | 1.89763793 | -0.2678292 | 0.35887664 | -0.7462988 | 0.45548694 | NA | NA | NA |
| ENSG000002 | 1.03922732 | -0.354912528 | 0.31056143 | -1.1428094 | 0.25311773 | NA | NA | NA |

| | | | | | | | | |
|---|---|---|---|---|---|---|---|---|
| ENSG000002 | 0.24756242 | -0.187034346 | 0.18119817 | -1.0322088 | 0.30197431 | NA | NA | NA |
| ENSG000002 | 1.98814473 | -0.447436554 | 0.36095557 | -1.239589 | 0.21512745 | NA | NA | NA |
| ENSG000002 | 1.11169489 | -0.110683234 | 0.32072136 | -0.3451072 | 0.73001382 | NA | NA | NA |
| ENSG000002 | 2.04677573 | -0.11916507 | 0.35997037 | -0.3310413 | 0.74061326 | NA | NA | NA |
| ENSG000002 | 0.60914194 | -0.10149904 | 0.25613314 | -0.3962745 | 0.69190251 | NA | RARRES2P3 | 106481659 |
| ENSG000002 | 1.04330208 | -0.72334856 | 0.31208131 | -2.3178208 | 0.02045906 | NA | NA | NA |
| ENSG000002 | 0.81900821 | 0.351371426 | 0.2870767 | 1.22396357 | 0.22096597 | NA | NA | NA |
| ENSG000002 | 1.34900477 | 0.031469685 | 0.33906312 | 0.09281365 | 0.92605161 | NA | NEBL-AS1 | 100128511 |
| ENSG000002 | 1.01523282 | -0.347954946 | 0.30960905 | -1.1238526 | 0.26107555 | NA | NA | NA |
| ENSG000002 | 2.08604898 | -0.235869129 | 0.36231873 | -0.650999 | 0.51504714 | NA | NA | NA |
| ENSG000002 | 1.8764823 | -0.134840298 | 0.3602235 | -0.374324 | 0.70816328 | NA | NA | NA |
| ENSG000002 | 0 | NA | NA | NA | NA | NA | NA | NA |
| ENSG000002 | 1.96002203 | -0.310543747 | 0.3616578 | -0.8586674 | 0.39052406 | NA | NA | NA |
| ENSG000002 | 1.53119479 | -0.616074981 | 0.33925932 | -1.8159412 | 0.06937939 | NA | HS1BP3-IT1 | 100874343 |
| ENSG000002 | 1.51533942 | -0.184887375 | 0.33845498 | -0.5462687 | 0.58488123 | NA | NA | NA |
| ENSG000002 | 2.00625015 | -0.447268245 | 0.36257083 | -1.2336024 | 0.21735109 | NA | NA | NA |
| ENSG000002 | 2.02587988 | -0.421100305 | 0.35427468 | -1.1886266 | 0.23458664 | NA | NA | NA |
| ENSG000002 | 0.60264758 | 0.238025744 | 0.26986178 | 0.88202838 | 0.37776146 | NA | NA | NA |
| ENSG000002 | 0.11865494 | 0.083440866 | 0.12751478 | 0.65436232 | 0.5128784 | NA | NA | NA |
| ENSG000002 | 1.76500741 | -0.478920863 | 0.3568729 | -1.3419928 | 0.17959833 | NA | NA | NA |
| ENSG000002 | 0.3575613 | 0.072305152 | 0.21497462 | 0.33634273 | 0.73661243 | NA | NA | NA |
| ENSG000002 | 1.22466076 | -0.038989694 | 0.32399524 | -0.1203403 | 0.90421356 | NA | NA | NA |
| ENSG000002 | 1.83993414 | 0.250261294 | 0.35063027 | 0.71374697 | 0.47538366 | NA | NA | NA |
| ENSG000002 | 0 | NA | NA | NA | NA | NA | NA | NA |
| ENSG000002 | 1.52476748 | -0.790224644 | 0.34133162 | -2.3151229 | 0.02060621 | NA | NA | NA |
| ENSG000002 | 0.6313676 | -0.265579877 | 0.26046629 | -1.0196324 | 0.30790281 | NA | NA | NA |
| ENSG000002 | 0 | NA | NA | NA | NA | NA | NA | NA |
| ENSG000002 | 0 | NA | NA | NA | NA | NA | NA | NA |
| ENSG000002 | 0 | NA | NA | NA | NA | NA | NA | NA |

| | | | | | | | | |
|---|---|---|---|---|---|---|---|---|
| ENSG000002 | 0 | NA | NA | NA | NA | NA | NA | NA |
| ENSG000002 | 2.05316975 | -0.109941586 | 0.35771912 | -0.3073405 | 0.75858419 | NA | NA | NA |
| ENSG000002 | 0.97621968 | -0.304047064 | 0.29688408 | -1.0241272 | 0.30577519 | NA | NA | NA |
| ENSG000002 | 0.64144758 | -0.286694252 | 0.26813299 | -1.0692241 | 0.28496871 | NA | NA | NA |
| ENSG000002 | 0 | NA | NA | NA | NA | NA | DEFT1P2 | 100287083 |
| ENSG000002 | 1.36677121 | -0.403925432 | 0.33668423 | -1.1997159 | 0.23024968 | NA | NA | NA |
| ENSG000002 | 2.07874088 | -0.725611534 | 0.35990106 | -2.0161417 | 0.04378516 | NA | NA | NA |
| ENSG000002 | 0.11205239 | 0.082118251 | 0.12671962 | 0.64803108 | 0.51696485 | NA | NA | NA |
| ENSG000002 | 2.02351506 | 0.132919908 | 0.36280281 | 0.36636957 | 0.71408932 | NA | NA | NA |
| ENSG000002 | 0 | NA | NA | NA | NA | NA | USP9YP33 | 106478931 |
| ENSG000002 | 1.97003338 | -0.05985262 | 0.35881154 | -0.166808 | 0.86752116 | NA | NA | NA |
| ENSG000002 | 0.25260017 | -0.015978705 | 0.18049464 | -0.0885273 | 0.92945758 | NA | NA | NA |
| ENSG000002 | 0.9731987 | -0.328787898 | 0.30544532 | -1.0764215 | 0.28173881 | NA | NA | NA |
| ENSG000002 | 0 | NA | NA | NA | NA | NA | NA | NA |
| ENSG000002 | 2.06767132 | -0.24605012 | 0.36527516 | -0.6736021 | 0.50056431 | NA | NA | NA |
| ENSG000002 | 1.22690568 | 0.093435007 | 0.32840751 | 0.28450935 | 0.77602007 | NA | NA | NA |
| ENSG000002 | 0.61562241 | 0.056234209 | 0.25476066 | 0.22073349 | 0.82529995 | NA | NA | NA |
| ENSG000002 | 2.06545202 | -0.72985043 | 0.36104658 | -2.0214855 | 0.04322953 | NA | NA | NA |
| ENSG000002 | 1.96757222 | -0.1825724 | 0.36125348 | -0.5053859 | 0.61328786 | NA | NA | NA |
| ENSG000002 | 1.63550762 | -0.225067013 | 0.33872369 | -0.6644561 | 0.50639846 | NA | NA | NA |
| ENSG000002 | 2.12442634 | -0.515250962 | 0.36580036 | -1.4085578 | 0.15896595 | NA | NA | NA |
| ENSG000002 | 0.92882561 | -0.439371363 | 0.29719023 | -1.4784179 | 0.13929595 | NA | NA | NA |
| ENSG000002 | 0.38747904 | -0.104908247 | 0.21987514 | -0.4771265 | 0.63327207 | NA | NA | NA |
| ENSG000002 | 0.73636311 | -0.02696217 | 0.28461296 | -0.0947328 | 0.9245271 | NA | NA | NA |
| ENSG000002 | 2.09933996 | -0.126307224 | 0.36639987 | -0.3447251 | 0.73030106 | NA | NA | NA |
| ENSG000002 | 0.11205239 | 0.082118251 | 0.12671962 | 0.64803108 | 0.51696485 | NA | NA | NA |
| ENSG000002 | 0.36075696 | -0.085953824 | 0.20740622 | -0.4144226 | 0.67856464 | NA | NA | NA |
| ENSG000002 | 1.03000874 | -0.345463641 | 0.30781682 | -1.1223027 | 0.26173377 | NA | NA | NA |
| ENSG000002 | 0.35141577 | 0.074527565 | 0.21401461 | 0.34823588 | 0.72766304 | NA | NA | NA |

| | | | | | | | | | |
|---|---|---|---|---|---|---|---|---|---|
| ENSG000002 | 0.99969403 | -0.200271688 | 0.31387515 | -0.6380616 | 0.52343357 | NA | | C11orf98P1 | 106480289 |
| ENSG000002 | 1.97903434 | -0.193893436 | 0.36475394 | -0.5315732 | 0.5950216 | NA | | RAB9BP1 | 9366 |
| ENSG000002 | 0.36212533 | -0.235926492 | 0.20089643 | -1.1743688 | 0.24024735 | NA | | NA | NA |
| ENSG000002 | 0.73889482 | -0.029099289 | 0.29011813 | -0.1003015 | 0.92010495 | NA | | NA | NA |
| ENSG000002 | 0 | NA | NA | NA | NA | NA | | NA | NA |
| ENSG000002 | 0.91507974 | -0.461155367 | 0.30304372 | -1.5217454 | 0.1280729 | NA | | NA | NA |
| ENSG000002 | 1.23621304 | -0.173924502 | 0.32326694 | -0.5380213 | 0.59056234 | NA | | NA | NA |
| ENSG000002 | 1.00953983 | -0.046292298 | 0.31369112 | -0.1475729 | 0.88267987 | NA | | NA | NA |
| ENSG000002 | 0.78745417 | -0.533348008 | 0.2804936 | -1.9014623 | 0.05724148 | NA | | NA | NA |
| ENSG000002 | 1.8984039 | -0.503259015 | 0.35251802 | -1.4276122 | 0.15340352 | NA | | NA | NA |
| ENSG000002 | 1.70409036 | -0.169303169 | 0.3493244 | -0.4846589 | 0.62791835 | NA | | NA | NA |
| ENSG000002 | 0.73243662 | -0.016475534 | 0.26887492 | -0.0612758 | 0.95113954 | NA | | NA | NA |
| ENSG000002 | 0.59485136 | 0.227885061 | 0.26312451 | 0.86607312 | 0.38645007 | NA | | NA | NA |
| ENSG000002 | 0 | NA | NA | NA | NA | NA | | NA | NA |
| ENSG000002 | 0.12685397 | -0.090215768 | 0.12901919 | -0.699243 | 0.48440017 | NA | | NA | NA |
| ENSG000002 | 2.10583624 | -0.254631993 | 0.36669313 | -0.6944008 | 0.48743091 | NA | | NA | NA |
| ENSG000002 | 1.11037668 | -0.111093452 | 0.32067756 | -0.3464335 | 0.72901695 | NA | | NA | NA |
| ENSG000002 | 0 | NA | NA | NA | NA | NA | | NA | NA |
| ENSG000002 | 1.13312881 | -0.430296951 | 0.3208914 | -1.3409426 | 0.17993908 | NA | | NA | NA |
| ENSG000002 | 1.33397239 | 0.024028053 | 0.34167488 | 0.07032432 | 0.94393553 | NA | | NA | NA |
| ENSG000002 | 0.97803198 | 0.116610914 | 0.30004355 | 0.38864663 | 0.69753757 | NA | | NA | NA |
| ENSG000002 | 0.75096827 | -0.201604636 | 0.2863373 | -0.7040809 | 0.48138237 | NA | | NA | NA |
| ENSG000002 | 0 | NA | NA | NA | NA | NA | | NA | NA |
| ENSG000002 | 0 | NA | NA | NA | NA | NA | | NA | NA |
| ENSG000002 | 1.63290874 | -0.402496351 | 0.35101784 | -1.146655 | 0.25152425 | NA | | NA | NA |
| ENSG000002 | 1.23812377 | -0.188681269 | 0.32332512 | -0.5835651 | 0.55951292 | NA | | NA | NA |
| ENSG000002 | 0.84899458 | -0.103232959 | 0.29062518 | -0.35521 | 0.7224323 | NA | | NA | NA |
| ENSG000002 | 1.97006763 | -0.43191282 | 0.35921863 | -1.2023675 | 0.22922116 | NA | | NA | NA |
| ENSG000002 | 1.2246052 | -0.32665329 | 0.32764584 | -0.9969707 | 0.31877875 | NA | | NA | NA |

| | | | | | | | | |
|---|---|---|---|---|---|---|---|---|
| ENSG000002 | 0.25260017 | -0.015978705 | 0.18049464 | -0.0885273 | 0.92945758 | NA | NA | NA |
| ENSG000002 | 1.76834049 | -0.458524791 | 0.35176001 | -1.303516 | 0.19239867 | NA | NA | NA |
| ENSG000002 | 1.47756342 | 0.092453159 | 0.34020627 | 0.27175619 | 0.7858095 | NA | NA | NA |
| ENSG000002 | 1.82947433 | -0.771890815 | 0.35421943 | -2.1791317 | 0.02932188 | NA | KRTAP25-1 | 100131902 |
| ENSG000002 | 0 | NA | NA | NA | NA | NA | USP17L24 | 728369 |
| ENSG000002 | 1.97326095 | -0.188758328 | 0.36278492 | -0.5203037 | 0.60285193 | NA | NA | NA |
| ENSG000002 | 1.54917867 | 0.145381344 | 0.34486604 | 0.42155888 | 0.67334702 | NA | NA | NA |
| ENSG000002 | 1.73341065 | -0.557265577 | 0.34583618 | -1.6113571 | 0.10710192 | NA | NA | NA |
| ENSG000002 | 1.43581888 | -0.582434506 | 0.33866238 | -1.7198087 | 0.08546722 | NA | NA | NA |
| ENSG000002 | 1.92368205 | 0.072228847 | 0.35894987 | 0.20122266 | 0.84052447 | NA | NA | NA |
| ENSG000002 | 1.6095137 | -0.242336905 | 0.33943128 | -0.7139498 | 0.4752582 | NA | NA | NA |
| ENSG000002 | 1.62349374 | -0.416528747 | 0.35257363 | -1.1813951 | 0.23744582 | NA | NA | NA |
| ENSG000002 | 2.11705488 | -0.495718801 | 0.364138 | -1.3613487 | 0.17340352 | NA | NA | NA |
| ENSG000002 | 0.86344734 | 0.04862751 | 0.30222795 | 0.1608968 | 0.87217468 | NA | NA | NA |
| ENSG000002 | 1.76810164 | -0.30105573 | 0.34528558 | -0.8719036 | 0.38326099 | NA | NA | NA |
| ENSG000002 | 0 | NA | NA | NA | NA | NA | LINC00279 | 100873963 |
| ENSG000002 | 1.21205011 | 0.090635872 | 0.32570145 | 0.278279 | 0.7807982 | NA | NA | NA |
| ENSG000002 | 0.45549726 | 0.302229013 | 0.22838165 | 1.32335066 | 0.18571879 | NA | NA | NA |
| ENSG000002 | 1.33070347 | 0.023672364 | 0.33869437 | 0.069893 | 0.94427883 | NA | SEMA3B-AS1 | 101928931 |
| ENSG000002 | 0.74709385 | -0.19484779 | 0.28612562 | -0.6809869 | 0.4958798 | NA | NA | NA |
| ENSG000002 | 0.65561801 | -0.268676647 | 0.26257757 | -1.0232277 | 0.3062002 | NA | NA | NA |
| ENSG000002 | 1.52458887 | -0.327503744 | 0.34239607 | -0.9565056 | 0.33881688 | NA | NA | NA |
| ENSG000002 | 1.15632193 | -0.440729556 | 0.3241513 | -1.3596415 | 0.1739434 | NA | NA | NA |
| ENSG000002 | 0.77498842 | -0.384054853 | 0.28945545 | -1.3268185 | 0.18456873 | NA | NA | NA |
| ENSG000002 | 1.01298344 | -0.206632615 | 0.31837313 | -0.6490266 | 0.51632121 | NA | NA | NA |
| ENSG000002 | 1.80106876 | -0.776066742 | 0.35447762 | -2.1893251 | 0.02857322 | NA | NA | NA |
| ENSG000002 | 1.72117068 | -0.062936056 | 0.35384966 | -0.177861 | 0.85883214 | NA | NA | NA |
| ENSG000002 | 1.44315809 | -0.548671536 | 0.33129082 | -1.6561628 | 0.09768885 | NA | NA | NA |
| ENSG000002 | 2.12393866 | -0.382900839 | 0.36490784 | -1.0493083 | 0.29403623 | NA | NA | NA |

| | | | | | | | | |
|---|---|---|---|---|---|---|---|---|
| ENSG000002 | 0.26199343 | -0.194018184 | 0.18422801 | -1.0531416 | 0.29227609 | NA | NA | NA |
| ENSG000002 | 0.7557499 | -0.354842837 | 0.28293339 | -1.2541568 | 0.20978501 | NA | NA | NA |
| ENSG000002 | 1.14547441 | -0.241365636 | 0.31111854 | -0.7757996 | 0.43786732 | NA | NA | NA |
| ENSG000002 | 1.25935218 | -0.342579228 | 0.33221209 | -1.0312064 | 0.30244404 | NA | NA | NA |
| ENSG000002 | 0 | NA | NA | NA | NA | NA | USP17L13 | 100287238 |
| ENSG000002 | 0.76152487 | -0.199410494 | 0.28701953 | -0.6947628 | 0.48720396 | NA | LINC00112 | 54089 |
| ENSG000002 | 1.52273516 | -0.471655819 | 0.34218248 | -1.3783751 | 0.16808749 | NA | NA | NA |
| ENSG000002 | 1.35035085 | -0.262689591 | 0.33521442 | -0.7836465 | 0.43324759 | NA | NA | NA |
| ENSG000002 | 0.87420842 | -0.278408432 | 0.29914995 | -0.9306652 | 0.3520268 | NA | NA | NA |
| ENSG000002 | 0.11865494 | 0.083440866 | 0.12751478 | 0.65436232 | 0.5128784 | NA | NA | NA |
| ENSG000002 | 0 | NA | NA | NA | NA | NA | TTTY19 | 252952 |
| ENSG000002 | 0 | NA | NA | NA | NA | NA | USP9YP29 | 106480425 |
| ENSG000002 | 0.60370328 | 0.056896924 | 0.25523913 | 0.22291614 | 0.82360077 | NA | NA | NA |
| ENSG000002 | 0 | NA | NA | NA | NA | NA | NA | NA |
| ENSG000002 | 1.7931843 | -0.487327383 | 0.35753187 | -1.363032 | 0.17287242 | NA | LOC100289656 | 100289656 |
| ENSG000002 | 0.99992078 | -0.363103686 | 0.31497042 | -1.1528184 | 0.24898494 | NA | NA | NA |
| ENSG000002 | 0.24619405 | -0.007630671 | 0.17915639 | -0.0425922 | 0.96602658 | NA | NA | NA |
| ENSG000002 | 1.08863974 | 0.028312312 | 0.3149176 | 0.08990387 | 0.9283636 | NA | NA | NA |
| ENSG000002 | 2.02312004 | -0.459345395 | 0.36265011 | -1.2666352 | 0.20528576 | NA | NA | NA |
| ENSG000002 | 1.70636965 | 0.081119662 | 0.35866353 | 0.22617204 | 0.82106762 | NA | NA | NA |
| ENSG000002 | 2.07039306 | -0.11631542 | 0.36132878 | -0.3219102 | 0.74752073 | NA | NA | NA |
| ENSG000002 | 0 | NA | NA | NA | NA | NA | HSFY5P | 106478934 |
| ENSG000002 | 1.9869699 | 0.04287974 | 0.35627488 | 0.12035578 | 0.90420132 | NA | NA | NA |
| ENSG000002 | 1.41654973 | 0.084904719 | 0.33048743 | 0.25690756 | 0.79725013 | NA | NA | NA |
| ENSG000002 | 1.93046638 | 0.442608651 | 0.35417824 | 1.24967772 | 0.2114173 | NA | NA | NA |
| ENSG000002 | 1.97974279 | 0.065891665 | 0.36415168 | 0.18094566 | 0.85641023 | NA | NA | NA |
| ENSG000002 | 2.16381273 | -0.504143507 | 0.36437431 | -1.3835869 | 0.16648499 | NA | NA | NA |
| ENSG000002 | 1.53921969 | -0.612984237 | 0.33945057 | -1.805813 | 0.07094756 | NA | NA | NA |
| ENSG000002 | 1.6413163 | -0.547455998 | 0.34906357 | -1.5683562 | 0.11679804 | NA | NA | NA |

| | | | | | | | | |
|---|---|---|---|---|---|---|---|---|
| ENSG000002 | 0.40942396 | -0.284058841 | 0.21788566 | -1.303706 | 0.19233384 | NA | NA | NA |
| ENSG000002 | 0 | NA | NA | NA | NA | NA | NA | NA |
| ENSG000002 | 0.64830914 | -0.459484042 | 0.26530677 | -1.7318972 | 0.08329187 | NA | NA | NA |
| ENSG000002 | 0.14128499 | -0.097772014 | 0.13403235 | -0.7294658 | 0.4657168 | NA | NA | NA |
| ENSG000002 | 1.07357757 | 0.19487222 | 0.32462569 | 0.60029821 | 0.54830751 | NA | NA | NA |
| ENSG000002 | 2.08369274 | -0.719880774 | 0.35964833 | -2.0016241 | 0.04532517 | NA | LINC00945 | 101930746 |
| ENSG000002 | 1.1540403 | -0.24481916 | 0.31141981 | -0.7861387 | 0.43178624 | NA | NA | NA |
| ENSG000002 | 1.59842427 | -0.257909053 | 0.34985559 | -0.7371872 | 0.46100853 | NA | NA | NA |
| ENSG000002 | 0.14128499 | -0.097772014 | 0.13403235 | -0.7294658 | 0.4657168 | NA | NA | NA |
| ENSG000002 | 0.60213713 | 0.060404575 | 0.26266752 | 0.22996591 | 0.81811826 | NA | NA | NA |
| ENSG000002 | 1.58181713 | 0.015173312 | 0.35075901 | 0.04325851 | 0.96549546 | NA | KCND3-IT1 | 100874295 |
| ENSG000002 | 0.50405874 | -0.025925695 | 0.24745105 | -0.104771 | 0.91655752 | NA | CRB3P1 | 106480279 |
| ENSG000002 | 0.634074 | -0.448061355 | 0.26266228 | -1.7058458 | 0.08803679 | NA | NA | NA |
| ENSG000002 | 1.86237892 | -0.527225144 | 0.35824349 | -1.471695 | 0.14110327 | NA | NA | NA |
| ENSG000002 | 0 | NA | NA | NA | NA | NA | NA | NA |
| ENSG000002 | 0 | NA | NA | NA | NA | NA | NA | NA |
| ENSG000002 | 0 | NA | NA | NA | NA | NA | NA | NA |
| ENSG000002 | 1.15517978 | -0.429306277 | 0.32005601 | -1.3413474 | 0.1798077 | NA | NA | NA |
| ENSG000002 | 0.9842073 | -0.043104386 | 0.31268272 | -0.1378534 | 0.89035626 | NA | NA | NA |
| ENSG000002 | 0.50611225 | -0.202624884 | 0.24886748 | -0.8141879 | 0.41553732 | NA | NA | NA |
| ENSG000002 | 0.75799536 | -0.171531325 | 0.27367317 | -0.6267744 | 0.53080715 | NA | NA | NA |
| ENSG000002 | 0.99605901 | -0.318888985 | 0.30081979 | -1.0600665 | 0.28911435 | NA | NA | NA |
| ENSG000002 | 1.77824071 | -0.732550604 | 0.34895759 | -2.0992539 | 0.03579452 | NA | NA | NA |
| ENSG000002 | 0.62821079 | -0.27685072 | 0.26713242 | -1.0363801 | 0.30002486 | NA | NA | NA |
| ENSG000002 | 0 | NA | NA | NA | NA | NA | NA | NA |
| ENSG000002 | 0 | NA | NA | NA | NA | NA | NA | NA |
| ENSG000002 | 0 | NA | NA | NA | NA | NA | NA | NA |
| ENSG000002 | 1.18642971 | -0.404880396 | 0.31466053 | -1.2867213 | 0.19819146 | NA | NA | NA |
| ENSG000002 | 0 | NA | NA | NA | NA | NA | NA | NA |

| | | | | | | | | |
|---|---|---|---|---|---|---|---|---|
| ENSG000002 | 0 | NA | NA | NA | NA | NA | NA | NA |
| ENSG000002 | 1.22511565 | -0.191823281 | 0.33256442 | -0.5768004 | 0.56407431 | NA | NA | NA |
| ENSG000002 | 1.37821508 | -0.265546677 | 0.33582178 | -0.7907369 | 0.42909753 | NA | NA | NA |
| ENSG000002 | 1.13804237 | -0.257422017 | 0.312677 | -0.8232842 | 0.41034642 | NA | NA | NA |
| ENSG000002 | 0.11205239 | 0.082118251 | 0.12671962 | 0.64803108 | 0.51696485 | NA | NA | NA |
| ENSG000002 | 2.13373528 | -0.646206955 | 0.36543303 | -1.7683321 | 0.0770054 | NA | NA | NA |
| ENSG000002 | 1.97647622 | -0.316812944 | 0.36350577 | -0.8715486 | 0.38345469 | NA | NA | NA |
| ENSG000002 | 0.52742601 | -0.196769605 | 0.24405015 | -0.8062671 | 0.42008885 | NA | NA | NA |
| ENSG000002 | 1.71519537 | 0.064986311 | 0.35305484 | 0.1840686 | 0.85395964 | NA | NA | NA |
| ENSG000002 | 2.12801141 | -0.256246705 | 0.36580793 | -0.7004952 | 0.48361809 | NA | NA | NA |
| ENSG000002 | 1.44060268 | -0.037791893 | 0.33491308 | -0.1128409 | 0.91015669 | NA | NA | NA |
| ENSG000002 | 1.63863164 | -0.271928221 | 0.35334415 | -0.7695846 | 0.44154633 | NA | HSPE1P9 | 106480255 |
| ENSG000002 | 0 | NA | NA | NA | NA | NA | NA | NA |
| ENSG000002 | 2.00735951 | -0.448193018 | 0.36257705 | -1.2361318 | 0.21640958 | NA | NA | NA |
| ENSG000002 | 0 | NA | NA | NA | NA | NA | NA | NA |
| ENSG000002 | 1.85063539 | 0.001396344 | 0.35674268 | 0.00391415 | 0.99687697 | NA | NA | NA |
| ENSG000002 | 0.99372318 | -0.356597057 | 0.31328177 | -1.138263 | 0.25501067 | NA | NA | NA |
| ENSG000002 | 0.60950859 | 0.048840033 | 0.2633134 | 0.18548252 | 0.85285059 | NA | NA | NA |
| ENSG000002 | 1.96187966 | -0.43325399 | 0.35982523 | -1.2040678 | 0.22856336 | NA | NA | NA |
| ENSG000002 | 0.73448374 | -0.031861808 | 0.28448729 | -0.1119973 | 0.91082555 | NA | NA | NA |
| ENSG000002 | 1.58423727 | 0.018224352 | 0.35078113 | 0.05195363 | 0.95856564 | NA | NA | NA |
| ENSG000002 | 1.78925782 | -0.4536384 | 0.35201237 | -1.2887002 | 0.19750235 | NA | NA | NA |
| ENSG000002 | 1.72208313 | -0.1921382 | 0.35829253 | -0.5362607 | 0.59177839 | NA | NA | NA |
| ENSG000002 | 1.26412934 | -0.177084571 | 0.32404874 | -0.5464751 | 0.58473939 | NA | NA | NA |
| ENSG000002 | 0 | NA | NA | NA | NA | NA | NA | NA |
| ENSG000002 | 0.70163732 | 0.137651641 | 0.28236374 | 0.48749759 | 0.62590575 | NA | NA | NA |
| ENSG000002 | 1.51674259 | -0.793062058 | 0.34109994 | -2.3250137 | 0.02007123 | NA | ATG4AP1 | 106480786 |
| ENSG000002 | 0 | NA | NA | NA | NA | NA | USP9YP16 | 106478927 |
| ENSG000002 | 0.6265601 | -0.275936484 | 0.2670182 | -1.0333995 | 0.30141697 | NA | NA | NA |

| | | | | | | | | |
|---|---|---|---|---|---|---|---|---|
| ENSG000002 | 1.53756828 | 0.390728507 | 0.33757444 | 1.15745881 | 0.24708496 | NA | LOC105376111 | 105376111 |
| ENSG000002 | 2.02529617 | -0.730859848 | 0.36112062 | -2.0238663 | 0.04298391 | NA | NA | NA |
| ENSG000002 | 0.6275579 | -0.281336606 | 0.26701873 | -1.0536213 | 0.29205636 | NA | NA | NA |
| ENSG000002 | 0 | NA | NA | NA | NA | NA | NA | NA |
| ENSG000002 | 1.67057531 | -0.520377384 | 0.34317475 | -1.5163627 | 0.12942767 | NA | NA | NA |
| ENSG000002 | 0.50204195 | -0.181972198 | 0.24131623 | -0.7540819 | 0.45080006 | NA | NA | NA |
| ENSG000002 | 0.50611225 | -0.202624884 | 0.24886748 | -0.8141879 | 0.41553732 | NA | NA | NA |
| ENSG000002 | 2.00414403 | -0.325412836 | 0.36338349 | -0.895508 | 0.37051557 | NA | NA | NA |
| ENSG000002 | 1.50563213 | -0.201928769 | 0.34886506 | -0.5788163 | 0.56271316 | NA | NA | NA |
| ENSG000002 | 1.98820339 | -0.547156058 | 0.35704967 | -1.5324368 | 0.12541468 | NA | BMS1P22 | 106480334 |
| ENSG000002 | 0.14128499 | -0.097772014 | 0.13403235 | -0.7294658 | 0.4657168 | NA | NA | NA |
| ENSG000002 | 0.85524617 | 0.042708506 | 0.29061608 | 0.14695851 | 0.88316477 | NA | NA | NA |
| ENSG000002 | 1.72065865 | -0.321240449 | 0.35407705 | -0.9072614 | 0.36426857 | NA | NA | NA |
| ENSG000002 | 1.12352703 | -0.266790378 | 0.31963776 | -0.8346648 | 0.40390648 | NA | NA | NA |
| ENSG000002 | 0.14128499 | -0.097772014 | 0.13403235 | -0.7294658 | 0.4657168 | NA | NA | NA |
| ENSG000002 | 1.76962351 | -0.760787274 | 0.35316987 | -2.1541681 | 0.03122699 | NA | NA | NA |
| ENSG000002 | 0.13280363 | -0.093394593 | 0.13115644 | -0.7120855 | 0.47641185 | NA | NA | NA |
| ENSG000002 | 0.11865494 | 0.083440866 | 0.12751478 | 0.65436232 | 0.5128784 | NA | NA | NA |
| ENSG000002 | 1.88454336 | -0.382592186 | 0.35726468 | -1.0708928 | 0.28421762 | NA | NA | NA |
| ENSG000002 | 0 | NA | NA | NA | NA | NA | TTTY20 | 252951 |
| ENSG000002 | 0.38884741 | -0.29745092 | 0.22235466 | -1.3377319 | 0.18098386 | NA | NA | NA |
| ENSG000002 | 1.57695171 | 0.155180222 | 0.34887018 | 0.44480793 | 0.65645856 | NA | COL4A2-AS1 | 100874203 |
| ENSG000002 | 2.14846534 | -0.138530321 | 0.36508816 | -0.3794435 | 0.70435857 | NA | DUX4L50 | 106480350 |
| ENSG000002 | 0 | NA | NA | NA | NA | NA | NA | NA |
| ENSG000002 | 1.95577471 | 0.200420783 | 0.36432613 | 0.55011367 | 0.58224141 | NA | NA | NA |
| ENSG000002 | 2.11670997 | -0.473241451 | 0.36062034 | -1.3122983 | 0.18941952 | NA | NA | NA |
| ENSG000002 | 1.35634398 | 0.025462498 | 0.34050555 | 0.07477851 | 0.94039094 | NA | NA | NA |
| ENSG000002 | 1.98748132 | -0.308871938 | 0.35990357 | -0.8582075 | 0.3907779 | NA | NA | NA |
| ENSG000002 | 1.62212592 | -0.260927558 | 0.35195604 | -0.7413641 | 0.45847274 | NA | NA | NA |

| | | | | | | | | | |
|---|---|---|---|---|---|---|---|---|---|
| ENSG000002 | 1.26117254 | -0.200687202 | 0.32985872 | -0.6084035 | 0.54291989 | NA | NA | NA | |
| ENSG000002 | 0 | NA | NA | NA | NA | NA | NA | NA | |
| ENSG000002 | 1.96474713 | -0.182075048 | 0.35796163 | -0.508644 | 0.61100176 | NA | NA | NA | |
| ENSG000002 | 1.35131832 | -0.11487615 | 0.3287045 | -0.3494815 | 0.72672784 | NA | NA | NA | |
| ENSG000002 | 1.27028348 | -0.506206397 | 0.33016948 | -1.5331714 | 0.12523364 | NA | NA | NA | |
| ENSG000002 | 1.90216365 | -0.408481933 | 0.36118609 | -1.1309459 | 0.25807786 | NA | NA | NA | |
| ENSG000002 | 0.63982777 | -0.42118588 | 0.25611206 | -1.6445375 | 0.10006523 | NA | NA | NA | |
| ENSG000002 | 0.50204195 | -0.181972198 | 0.24131623 | -0.7540819 | 0.45080006 | NA | NA | NA | |
| ENSG000002 | 1.19810619 | -0.591329122 | 0.31849508 | -1.8566351 | 0.06336309 | NA | NA | NA | |
| ENSG000002 | 1.44579802 | 0.21013234 | 0.3362255 | 0.62497442 | 0.53198784 | NA | NA | NA | |
| ENSG000002 | 1.94641604 | -0.059249265 | 0.36083268 | -0.1642015 | 0.86957252 | NA | NA | NA | |
| ENSG000002 | 0.60987915 | -0.108304852 | 0.26372492 | -0.4106736 | 0.68131192 | NA | NA | NA | |
| ENSG000002 | 1.53130477 | -0.721113915 | 0.33199683 | -2.1720506 | 0.02985185 | NA | NA | NA | |
| ENSG000002 | 0 | NA | NA | NA | NA | NA | NA | NA | |
| ENSG000002 | 0 | NA | NA | NA | NA | NA | NA | NA | |
| ENSG000002 | 0 | NA | NA | NA | NA | NA | NA | NA | |
| ENSG000002 | 1.25110052 | -0.195549445 | 0.3310706 | -0.5906578 | 0.5547497 | NA | NA | NA | |
| ENSG000002 | 0 | NA | NA | NA | NA | NA | NA | NA | |
| ENSG000002 | 0.73906896 | -0.197744251 | 0.28558485 | -0.6924186 | 0.48867451 | NA | NA | NA | |
| ENSG000002 | 0.50135681 | -0.182211159 | 0.24123516 | -0.7553259 | 0.45005347 | NA | NA | NA | |
| ENSG000002 | 0.35733509 | 0.244646831 | 0.20894085 | 1.17089037 | 0.24164285 | NA | NA | NA | |
| ENSG000002 | 0 | NA | NA | NA | NA | NA | NA | NA | |
| ENSG000002 | 0 | NA | NA | NA | NA | NA | NA | NA | |
| ENSG000002 | 2.06012959 | -0.599127873 | 0.36306447 | -1.6501969 | 0.09890266 | NA | NA | NA | |
| ENSG000002 | 0.78731386 | -0.195162287 | 0.28101843 | -0.6944822 | 0.48737989 | NA | NA | NA | |
| ENSG000002 | 0.51322524 | -0.027081322 | 0.24101349 | -0.1123643 | 0.91053453 | NA | NA | NA | |
| ENSG000002 | 0 | NA | NA | NA | NA | NA | DEFB130 | | 245940 |
| ENSG000002 | 1.09688927 | -0.24574033 | 0.31115835 | -0.7897597 | 0.42966812 | NA | NA | NA | |
| ENSG000002 | 1.96332533 | -0.190077291 | 0.36466276 | -0.5212413 | 0.60219868 | NA | NA | NA | |

| | | | | | | | | |
|---|---|---|---|---|---|---|---|---|
| ENSG000002 | 1.09253878 | -0.252778792 | 0.3186311 | -0.7933274 | 0.42758708 | NA | NA | NA |
| ENSG000002 | 1.1377101 | -0.279768332 | 0.3238186 | -0.8639662 | 0.38760644 | NA | NA | NA |
| ENSG000002 | 1.34064199 | -0.372341102 | 0.32696759 | -1.1387707 | 0.25479882 | NA | NA | NA |
| ENSG000002 | 1.84308979 | 0.128605785 | 0.34706991 | 0.37054721 | 0.71097481 | NA | NA | NA |
| ENSG000002 | 1.30324201 | -0.682264372 | 0.33144118 | -2.058478 | 0.03954427 | NA | NA | NA |
| ENSG000002 | 0 | NA | NA | NA | NA | NA | NA | NA |
| ENSG000002 | 1.88334385 | -0.261285352 | 0.35925949 | -0.7272887 | 0.46704914 | NA | NA | NA |
| ENSG000002 | 1.70347075 | -0.059974447 | 0.35533438 | -0.1687831 | 0.86596723 | NA | NA | NA |
| ENSG000002 | 0.49150654 | -0.026274915 | 0.24607766 | -0.1067749 | 0.91496757 | NA | NA | NA |
| ENSG000002 | 1.48019197 | -0.480142374 | 0.34438093 | -1.3942188 | 0.16325154 | NA | NA | NA |
| ENSG000002 | 0.12070844 | -0.088503695 | 0.12783931 | -0.6923042 | 0.48874631 | NA | LINC01219 | 104355220 |
| ENSG000002 | 0.74521503 | -0.189818705 | 0.28603889 | -0.6636115 | 0.50693897 | NA | NA | NA |
| ENSG000002 | 1.84909414 | -0.047177932 | 0.24124604 | -0.1955594 | 0.84495504 | NA | MTATP6P7 | 106480798 |
| ENSG000002 | 1.76522583 | 0.139606992 | 0.3488622 | 0.40017804 | 0.68902539 | NA | NA | NA |
| ENSG000002 | 1.22328836 | -0.198664676 | 0.33383385 | -0.5951004 | 0.55177633 | NA | NA | NA |
| ENSG000002 | 1.08934312 | -0.106081292 | 0.31694868 | -0.3346955 | 0.73785481 | NA | VPS13A-AS1 | 100286938 |
| ENSG000002 | 1.13295412 | -0.268810023 | 0.31776627 | -0.8459363 | 0.39758828 | NA | NA | NA |
| ENSG000002 | 0.84785243 | -0.11359032 | 0.29663151 | -0.3829341 | 0.70176863 | NA | NA | NA |
| ENSG000002 | 0.73329399 | -0.327183738 | 0.2730203 | -1.1983861 | 0.23076673 | NA | NA | NA |
| ENSG000002 | 1.56824305 | 0.286968969 | 0.34907429 | 0.82208567 | 0.41102814 | NA | NA | NA |
| ENSG000002 | 0.50204195 | -0.181972198 | 0.24131623 | -0.7540819 | 0.45080006 | NA | MTCO1P5 | 106480318 |
| ENSG000002 | 1.73491553 | -0.190446559 | 0.35645099 | -0.5342854 | 0.5931441 | NA | NA | NA |
| ENSG000002 | 0.6054383 | 0.055876002 | 0.26295202 | 0.21249505 | 0.83172084 | NA | NA | NA |
| ENSG000002 | 0.48302518 | -0.023466322 | 0.24515318 | -0.0957211 | 0.92374212 | NA | NA | NA |
| ENSG000002 | 0.85166269 | -0.106217298 | 0.28902612 | -0.3675007 | 0.71324558 | NA | NA | NA |
| ENSG000002 | 0.14128499 | -0.097772014 | 0.13403235 | -0.7294658 | 0.4657168 | NA | LOC100133267 | 100133267 |
| ENSG000002 | 0.64218479 | -0.28641133 | 0.26819919 | -1.0679053 | 0.28556324 | NA | NA | NA |
| ENSG000002 | 2.0489914 | 0.133095689 | 0.36523272 | 0.36441338 | 0.71554935 | NA | NA | NA |
| ENSG000002 | 1.25171903 | -0.303078371 | 0.31785973 | -0.9534972 | 0.3403382 | NA | NA | NA |

| | | | | | | | | |
|---|---|---|---|---|---|---|---|---|
| ENSG000002 | 0 | NA | NA | NA | NA | NA | NA | NA |
| ENSG000002 | 0.98236074 | -0.042248734 | 0.31261001 | -0.1351484 | 0.89249456 | NA | NA | NA |
| ENSG000002 | 1.31149661 | -0.481780985 | 0.32466916 | -1.4839136 | 0.13783184 | NA | NA | NA |
| ENSG000002 | 1.46139155 | 0.221698647 | 0.3364943 | 0.65884816 | 0.50999328 | NA | FTH1P6 | 106481787 |
| ENSG000002 | 0.51316968 | -0.377777671 | 0.24556403 | -1.538408 | 0.12394889 | NA | NA | NA |
| ENSG000002 | 1.8801671 | -0.809246112 | 0.35549363 | -2.2764012 | 0.02282201 | NA | NA | NA |
| ENSG000002 | 0.89634975 | -0.435966242 | 0.29720041 | -1.46691 | 0.14240055 | NA | NA | NA |
| ENSG000002 | 1.65997965 | -0.684844784 | 0.34629253 | -1.9776481 | 0.04796841 | NA | NA | NA |
| ENSG000002 | 1.49734664 | -0.200002045 | 0.3487411 | -0.5734972 | 0.56630809 | NA | NA | NA |
| ENSG000002 | 0.62545232 | -0.114578523 | 0.27135287 | -0.4222492 | 0.67284316 | NA | NA | NA |
| ENSG000002 | 0 | NA | NA | NA | NA | NA | USP9YP15 | 106480713 |
| ENSG000002 | 1.16206657 | -0.110408346 | 0.30773295 | -0.3587797 | 0.71975987 | NA | NA | NA |
| ENSG000002 | 0 | NA | NA | NA | NA | NA | NA | NA |
| ENSG000002 | 1.6092914 | -0.384847491 | 0.34677697 | -1.1097839 | 0.26709217 | NA | NA | NA |
| ENSG000002 | 0.11131518 | 0.081951103 | 0.12663804 | 0.64712866 | 0.51754868 | NA | NA | NA |
| ENSG000002 | 0.75423719 | -0.191783781 | 0.2805177 | -0.683678 | 0.49417852 | NA | NA | NA |
| ENSG000002 | 0.3588776 | -0.099902505 | 0.21555794 | -0.4634601 | 0.6430346 | NA | NA | NA |
| ENSG000002 | 1.011563 | -0.039786304 | 0.30732304 | -0.1294609 | 0.896993 | NA | NA | NA |
| ENSG000002 | 0.24619405 | -0.007630671 | 0.17915639 | -0.0425922 | 0.96602658 | NA | USP17L11 | 100287178 |
| ENSG000002 | 1.79605547 | -0.097825575 | 0.34988184 | -0.279596 | 0.77978745 | NA | LOC728660 | 728660 |
| ENSG000002 | 1.94147786 | 0.046781329 | 0.35180255 | 0.1329761 | 0.89421228 | NA | NA | NA |
| ENSG000002 | 0 | NA | NA | NA | NA | NA | HSFY8P | 106480723 |
| ENSG000002 | 1.564261 | 0.018755705 | 0.34743327 | 0.05398362 | 0.95694822 | NA | NA | NA |
| ENSG000002 | 1.92481991 | -0.658827347 | 0.35674284 | -1.846785 | 0.0647783 | NA | NA | NA |
| ENSG000002 | 0.52165104 | -0.403921832 | 0.25252134 | -1.5995552 | 0.10969729 | NA | NA | NA |
| ENSG000002 | 0.74772501 | -0.357780902 | 0.28233051 | -1.2672414 | 0.205069 | NA | NA | NA |
| ENSG000002 | 0.64469531 | -0.422865637 | 0.25651704 | -1.6484894 | 0.09925228 | NA | NA | NA |
| ENSG000002 | 1.301451 | -0.33349633 | 0.32546053 | -1.0246906 | 0.30550922 | NA | NA | NA |
| ENSG000002 | 1.50370069 | -0.193021416 | 0.34197334 | -0.5644341 | 0.57245873 | NA | NA | NA |

| | | | | | | | | |
|---|---|---|---|---|---|---|---|---|
| ENSG000002 | 2.07734478 | -0.360769795 | 0.36231463 | -0.9957362 | 0.31937834 | NA | NA | NA |
| ENSG000002 | 0.96015681 | 0.284230535 | 0.312132 | 0.91061004 | 0.36250088 | NA | NA | NA |
| ENSG000002 | 0.35682409 | 0.072008558 | 0.21485623 | 0.33514763 | 0.73751372 | NA | NA | NA |
| ENSG000002 | 0.78039674 | -0.368519854 | 0.28442264 | -1.295677 | 0.1950868 | NA | RPS3AP1 | 101241875 |
| ENSG000002 | 1.40663835 | -0.423542215 | 0.33882934 | -1.2500163 | 0.21129358 | NA | NA | NA |
| ENSG000002 | 1.54616715 | -0.575342019 | 0.33190351 | -1.7334617 | 0.08301364 | NA | NA | NA |
| ENSG000002 | 0 | NA | NA | NA | NA | NA | NA | NA |
| ENSG000002 | 0.90147183 | -0.268817485 | 0.29320795 | -0.9168151 | 0.35923952 | NA | NA | NA |
| ENSG000002 | 2.10501177 | -0.483643234 | 0.36257207 | -1.333923 | 0.18222908 | NA | DEFA9P | 449492 |
| ENSG000002 | 0.96841839 | -0.214826771 | 0.19245321 | -1.1162546 | 0.26431318 | NA | NA | NA |
| ENSG000002 | 1.55642915 | 0.297272043 | 0.34675559 | 0.85729561 | 0.39128153 | NA | NA | NA |
| ENSG000002 | 1.1170134 | 0.034064043 | 0.31333691 | 0.10871379 | 0.9134295 | NA | CRIP1P1 | 106480793 |
| ENSG000002 | 1.37063647 | -0.105379934 | 0.32311993 | -0.3261326 | 0.74432406 | NA | NA | NA |
| ENSG000002 | 1.81074652 | -0.115319984 | 0.35657998 | -0.3234057 | 0.74638805 | NA | NA | NA |
| ENSG000002 | 1.03763475 | -0.508045393 | 0.30793986 | -1.6498202 | 0.09897972 | NA | NA | NA |
| ENSG000002 | 1.24719853 | -0.192800664 | 0.32950263 | -0.5851263 | 0.55846278 | NA | NA | NA |
| ENSG000002 | 2.08489305 | -0.373704372 | 0.3646985 | -1.024694 | 0.30550758 | NA | NA | NA |
| ENSG000002 | 0.69343828 | 0.300367975 | 0.27699246 | 1.08439043 | 0.27819173 | NA | NA | NA |
| ENSG000002 | 1.27355643 | -0.191482191 | 0.32869695 | -0.5825493 | 0.56019671 | NA | NA | NA |
| ENSG000002 | 0.24004852 | -0.005223787 | 0.17775844 | -0.029387 | 0.97655594 | NA | NA | NA |
| ENSG000002 | 1.89195175 | -0.64487746 | 0.35425068 | -1.8203987 | 0.06869832 | NA | NA | NA |
| ENSG000002 | 1.37492961 | 0.010411084 | 0.33480241 | 0.0310962 | 0.97519282 | NA | NA | NA |
| ENSG000002 | 1.13130398 | -0.273022612 | 0.32364448 | -0.843588 | 0.39889966 | NA | NA | NA |
| ENSG000002 | 1.37254618 | -0.273939765 | 0.34159844 | -0.801935 | 0.42259053 | NA | NA | NA |
| ENSG000002 | 0.85405407 | 0.203507704 | 0.29712682 | 0.68491867 | 0.49339524 | NA | NA | NA |
| ENSG000002 | 0.71254041 | 0.271406322 | 0.27001992 | 1.00513444 | 0.31483212 | NA | NA | NA |
| ENSG000002 | 1.73902412 | -0.33341962 | 0.35644265 | -0.9354089 | 0.34957762 | NA | NA | NA |
| ENSG000002 | 1.35907858 | -0.387486077 | 0.33034355 | -1.1729791 | 0.24080419 | NA | NA | NA |
| ENSG000002 | 0.65524745 | -0.109156678 | 0.2585811 | -0.4221371 | 0.67292494 | NA | NA | NA |

| | | | | | | | | | |
|---|---|---|---|---|---|---|---|---|---|
| ENSG000002 | 1.65702531 | -0.547152414 | 0.34858383 | -1.5696437 | 0.11649802 | NA | NA | NA | |
| ENSG000002 | 0.73975464 | -0.177073248 | 0.27958599 | -0.6333409 | 0.52651104 | NA | NA | NA | |
| ENSG000002 | 0 | NA | NA | NA | NA | NA | NA | NA | |
| ENSG000002 | 0.76445355 | -0.200993415 | 0.28720706 | -0.6998206 | 0.48403935 | NA | NA | NA | |
| ENSG000002 | 1.65235632 | -0.394042217 | 0.34810389 | -1.1319673 | 0.25764819 | NA | NA | NA | |
| ENSG000002 | 2.15586168 | 0.061876261 | 0.36601932 | 0.1690519 | 0.86575582 | NA | NA | NA | |
| ENSG000002 | 0.24004852 | -0.005223787 | 0.17775844 | -0.029387 | 0.97655594 | NA | NA | NA | |
| ENSG000002 | 1.51091838 | -0.052562884 | 0.34482662 | -0.1524328 | 0.8788456 | NA | NA | NA | |
| ENSG000002 | 2.0084113 | -0.31022095 | 0.36044418 | -0.8606629 | 0.38942371 | NA | LINC00028 | | 140875 |
| ENSG000002 | 1.84624064 | -0.747916917 | 0.34999162 | -2.1369566 | 0.03260152 | NA | NA | NA | |
| ENSG000002 | 1.81481873 | 0.006727863 | 0.35184576 | 0.01912163 | 0.98474408 | NA | NA | NA | |
| ENSG000002 | 0.4972498 | 0.131765009 | 0.23019643 | 0.5724025 | 0.56704932 | NA | NA | NA | |
| ENSG000002 | 0 | NA | NA | NA | NA | NA | NA | NA | |
| ENSG000002 | 1.74875122 | -0.322341267 | 0.35372304 | -0.9112815 | 0.36214706 | NA | NA | NA | |
| ENSG000002 | 1.8931866 | 0.471582499 | 0.36106613 | 1.30608345 | 0.19152418 | NA | NA | NA | |
| ENSG000002 | 0.24619405 | -0.007630671 | 0.17915639 | -0.0425922 | 0.96602658 | NA | NA | NA | |
| ENSG000002 | 0.25993993 | -0.013144386 | 0.18211683 | -0.0721756 | 0.94246219 | NA | NA | NA | |
| ENSG000002 | 1.97094474 | -0.177558072 | 0.35496085 | -0.5002187 | 0.61692106 | NA | LINC01046 | | 103689843 |
| ENSG000002 | 1.52045545 | -0.174664467 | 0.33699063 | -0.5183066 | 0.60424437 | NA | NA | NA | |
| ENSG000002 | 1.8125328 | -0.598113278 | 0.35161881 | -1.7010275 | 0.08893781 | NA | NA | NA | |
| ENSG000002 | 0.72398718 | -0.026990458 | 0.2774178 | -0.0972917 | 0.92249472 | NA | NA | NA | |
| ENSG000002 | 1.01381047 | -0.348485104 | 0.3095448 | -1.1257986 | 0.26025078 | NA | NA | NA | |
| ENSG000002 | 1.42294818 | -0.392729717 | 0.32886218 | -1.1942076 | 0.23239678 | NA | NA | NA | |
| ENSG000002 | 1.4836311 | -0.342527291 | 0.34659668 | -0.988259 | 0.32302582 | NA | HOTAIRM1 | | 100506311 |
| ENSG000002 | 0.83502003 | -0.102565745 | 0.28807063 | -0.3560437 | 0.72180781 | NA | NA | NA | |
| ENSG000002 | 1.5023809 | -0.329312153 | 0.34275623 | -0.9607766 | 0.33666453 | NA | LOC100130849 | | 100130849 |
| ENSG000002 | 0 | NA | NA | NA | NA | NA | NA | NA | |
| ENSG000002 | 1.12515845 | -0.271245119 | 0.32345284 | -0.8385925 | 0.40169803 | NA | NA | NA | |
| ENSG000002 | 1.43175522 | -0.140747054 | 0.32262289 | -0.4362587 | 0.66264902 | NA | NA | NA | |

| | | | | | | | | |
|---|---|---|---|---|---|---|---|---|
| ENSG000002 | 1.96719762 | -0.313323846 | 0.3601554 | -0.8699685 | 0.38431762 | NA | NA | NA |
| ENSG000002 | 0 | NA | NA | NA | NA | NA | NA | NA |
| ENSG000002 | 2.0105507 | -0.319673481 | 0.36254275 | -0.8817539 | 0.37790991 | NA | NA | NA |
| ENSG000002 | 0.59548251 | 0.058022206 | 0.26207208 | 0.22139789 | 0.82478264 | NA | NA | NA |
| ENSG000002 | 1.8309049 | -0.228193592 | 0.35082574 | -0.6504471 | 0.51540345 | NA | NA | NA |
| ENSG000002 | 1.23726291 | -0.197629439 | 0.33418696 | -0.5913739 | 0.55426994 | NA | NA | NA |
| ENSG000002 | 0.86916674 | -0.10847042 | 0.29614034 | -0.3662805 | 0.71415581 | NA | NA | NA |
| ENSG000002 | 1.43373372 | 0.090032932 | 0.34141101 | 0.26370834 | 0.79200467 | NA | NA | NA |
| ENSG000002 | 1.18699887 | 0.110083506 | 0.32724836 | 0.33639132 | 0.7365758 | NA | NA | NA |
| ENSG000002 | 1.48097678 | -0.186741118 | 0.33937881 | -0.5502439 | 0.5821521 | NA | NA | NA |
| ENSG000002 | 1.75233472 | -0.195271931 | 0.35496348 | -0.5501184 | 0.58223819 | NA | NA | NA |
| ENSG000002 | 1.02915338 | -0.534444235 | 0.31367491 | -1.7038157 | 0.08841552 | NA | NA | NA |
| ENSG000002 | 2.01874569 | -0.733176656 | 0.36105032 | -2.0306772 | 0.04228775 | NA | PAXBP1P1 | 106481704 |
| ENSG000002 | 0 | NA | NA | NA | NA | NA | NA | NA |
| ENSG000002 | 1.63749003 | -0.264191019 | 0.3521469 | -0.7502296 | 0.45311645 | NA | SYNJ2-IT1 | 100874300 |
| ENSG000002 | 1.08291876 | 0.0372422 | 0.3209687 | 0.11603063 | 0.90762827 | NA | NA | NA |
| ENSG000002 | 0 | NA | NA | NA | NA | NA | NA | NA |
| ENSG000002 | 1.13181251 | -0.287282343 | 0.32726178 | -0.8778365 | 0.38003244 | NA | NA | NA |
| ENSG000002 | 0.34948434 | 0.063159058 | 0.2051263 | 0.30790326 | 0.75815594 | NA | NA | NA |
| ENSG000002 | 1.80937544 | -0.610901226 | 0.35294102 | -1.7308876 | 0.08347182 | NA | NA | NA |
| ENSG000002 | 2.06955068 | -0.241114561 | 0.36495599 | -0.6606675 | 0.50882557 | NA | NA | NA |
| ENSG000002 | 1.13093341 | -0.122399657 | 0.32284929 | -0.3791232 | 0.70459637 | NA | NA | NA |
| ENSG000002 | 1.20478416 | 0.234807803 | 0.32404257 | 0.72462024 | 0.46868504 | NA | NA | NA |
| ENSG000002 | 0 | NA | NA | NA | NA | NA | PRYP5 | 106478933 |
| ENSG000002 | 0.60282172 | 0.050569991 | 0.26272347 | 0.19248372 | 0.84736332 | NA | NA | NA |
| ENSG000002 | 1.24432384 | -0.047223593 | 0.33260414 | -0.1419814 | 0.88709472 | NA | NA | NA |
| ENSG000002 | 1.15947198 | 0.563654806 | 0.32145923 | 1.75342548 | 0.079529 | NA | NA | NA |
| ENSG000002 | 1.90162521 | -0.777828177 | 0.35277617 | -2.2048773 | 0.0274627 | NA | NA | NA |
| ENSG000002 | 1.15461726 | -0.614542622 | 0.32342171 | -1.9001279 | 0.05741633 | NA | NA | NA |

| | | | | | | | | |
|---|---|---|---|---|---|---|---|---|
| ENSG000002 | 1.72066215 | -0.058528744 | 0.35383645 | -0.1654119 | 0.86861982 | NA | NA | NA |
| ENSG000002 | 1.14095783 | -0.421225193 | 0.31963334 | -1.3178387 | 0.18755764 | NA | NA | NA |
| ENSG000002 | 1.833997 | -0.235506402 | 0.35360213 | -0.6660209 | 0.50539776 | NA | NA | NA |
| ENSG000002 | 0.23202362 | -0.008302493 | 0.17586207 | -0.0472103 | 0.96234565 | NA | NA | NA |
| ENSG000002 | 0.75523498 | -0.185546569 | 0.27894532 | -0.6651718 | 0.5059406 | NA | NA | NA |
| ENSG000002 | 1.48345696 | -0.19332447 | 0.34718227 | -0.5568386 | 0.57763774 | NA | NA | NA |
| ENSG000002 | 1.7456484 | -0.179700705 | 0.35182248 | -0.510771 | 0.60951145 | NA | LINC00462 | 100129597 |
| ENSG000002 | 1.80018966 | -0.614715192 | 0.35439678 | -1.7345394 | 0.08282242 | NA | NA | NA |
| ENSG000002 | 1.42007319 | 0.209499352 | 0.33408157 | 0.62709042 | 0.53059997 | NA | NA | NA |
| ENSG000002 | 0 | NA | NA | NA | NA | NA | NA | NA |
| ENSG000002 | 2.10461032 | -0.256596013 | 0.36740085 | -0.6984089 | 0.48492153 | NA | NA | NA |
| ENSG000002 | 0 | NA | NA | NA | NA | NA | C4A-AS1 | 106478955 |
| ENSG000002 | 0.85200291 | -0.096788563 | 0.28906802 | -0.3348297 | 0.73775354 | NA | NA | NA |
| ENSG000002 | 1.5678233 | 0.019119616 | 0.34939241 | 0.05472247 | 0.95635957 | NA | NA | NA |
| ENSG000002 | 0 | NA | NA | NA | NA | NA | NA | NA |
| ENSG000002 | 1.97596871 | -0.17258211 | 0.3531425 | -0.4887039 | 0.62505136 | NA | NA | NA |
| ENSG000002 | 1.89149082 | -0.403841019 | 0.36078962 | -1.1193255 | 0.2630013 | NA | NA | NA |
| ENSG000002 | 1.51098488 | -0.492535031 | 0.34671306 | -1.420584 | 0.15543773 | NA | NA | NA |
| ENSG000002 | 1.39519797 | -0.24650209 | 0.3306073 | -0.7456039 | 0.45590673 | NA | NA | NA |
| ENSG000002 | 0 | NA | NA | NA | NA | NA | NA | NA |
| ENSG000002 | 0.27408862 | -0.199826037 | 0.18669465 | -1.0703362 | 0.28446801 | NA | NA | NA |
| ENSG000002 | 0 | NA | NA | NA | NA | NA | NA | NA |
| ENSG000002 | 1.36568325 | -0.244163851 | 0.33156049 | -0.7364082 | 0.46148234 | NA | NA | NA |
| ENSG000002 | 1.73295418 | -0.552392057 | 0.34585047 | -1.5971991 | 0.11022133 | NA | NA | NA |
| ENSG000002 | 0.4009426 | -0.302672773 | 0.2240145 | -1.3511303 | 0.17665371 | NA | NA | NA |
| ENSG000002 | 0.39499294 | -0.277908454 | 0.21584201 | -1.287555 | 0.19790091 | NA | NA | NA |
| ENSG000002 | 0.96087229 | 0.115852936 | 0.31168972 | 0.37169315 | 0.71012133 | NA | NA | NA |
| ENSG000002 | 1.39486165 | -0.572959208 | 0.33786735 | -1.6958111 | 0.08992165 | NA | NA | NA |
| ENSG000002 | 1.12654855 | -0.266995406 | 0.32351576 | -0.8252933 | 0.40920507 | NA | LOC105375342 | 105375342 |

| | | | | | | | | | |
|---|---|---|---|---|---|---|---|---|---|
| ENSG000002 | 1.29097213 | -0.489257293 | 0.32697388 | -1.4963192 | 0.13457049 | NA | NA | NA | |
| ENSG000002 | 0 | NA | NA | NA | NA | NA | TTTY18 | | 252950 |
| ENSG000002 | 0.53517016 | -0.075146746 | 0.15579947 | -0.4823299 | 0.6295716 | NA | NA | NA | |
| ENSG000002 | 1.17209098 | -0.396832703 | 0.31331288 | -1.26657 | 0.20530907 | NA | NA | NA | |
| ENSG000002 | 2.13854224 | -0.652703576 | 0.36471749 | -1.7896141 | 0.07351597 | NA | NA | NA | |
| ENSG000002 | 1.60917091 | -0.124941169 | 0.35362204 | -0.3533184 | 0.72384976 | NA | NA | NA | |
| ENSG000002 | 1.33118401 | -0.219052026 | 0.31987009 | -0.6848156 | 0.4934603 | NA | NA | NA | |
| ENSG000002 | 0.5226185 | -0.189580046 | 0.24357686 | -0.7783172 | 0.43638207 | NA | NA | NA | |
| ENSG000002 | 1.19714536 | -0.287932482 | 0.31231667 | -0.9219248 | 0.3565678 | NA | NA | NA | |
| ENSG000002 | 1.51985625 | 0.444661861 | 0.34759914 | 1.27923752 | 0.20081343 | NA | NA | NA | |
| ENSG000002 | 0.75327376 | -0.030668806 | 0.27945337 | -0.1097457 | 0.91261106 | NA | NA | NA | |
| ENSG000002 | 0.38651158 | -0.274258716 | 0.21461355 | -1.2779189 | 0.20127801 | NA | NA | NA | |
| ENSG000002 | 1.98971557 | -0.656424943 | 0.35290497 | -1.8600615 | 0.06287683 | NA | NA | NA | |
| ENSG000002 | 2.04975895 | 0.004440032 | 0.36441924 | 0.01218386 | 0.99027893 | NA | NA | NA | |
| ENSG000002 | 0.9982701 | -0.343188488 | 0.30892322 | -1.1109184 | 0.26660348 | NA | NA | NA | |
| ENSG000002 | 0 | NA | NA | NA | NA | NA | NA | NA | |
| ENSG000002 | 0.47826975 | -0.016435741 | 0.244632 | -0.0671856 | 0.94643397 | NA | NA | NA | |
| ENSG000002 | 2.04299567 | -0.179126767 | 0.35093467 | -0.5104277 | 0.60975187 | NA | NA | NA | |
| ENSG000002 | 1.37021481 | -0.502145937 | 0.32574554 | -1.5415282 | 0.12318828 | NA | NA | NA | |
| ENSG000002 | 1.97807484 | -0.310427933 | 0.36178388 | -0.858048 | 0.39086596 | NA | DSCAM-IT1 | | 100874326 |
| ENSG000002 | 0.36553413 | 0.069401489 | 0.20780452 | 0.33397488 | 0.73839851 | NA | NA | NA | |
| ENSG000002 | 0 | NA | NA | NA | NA | NA | NA | NA | |
| ENSG000002 | 0.87420842 | -0.278408432 | 0.29914995 | -0.9306652 | 0.3520268 | NA | NA | NA | |
| ENSG000002 | 2.08773529 | -0.479939067 | 0.36042699 | -1.3315847 | 0.1829967 | NA | NA | NA | |
| ENSG000002 | 1.55893912 | 0.160046037 | 0.35043569 | 0.45670588 | 0.64788247 | NA | NA | NA | |
| ENSG000002 | 0.63984896 | -0.285743295 | 0.26802755 | -1.0660967 | 0.28637991 | NA | NA | NA | |
| ENSG000002 | 0 | NA | NA | NA | NA | NA | TSPY4 | | 728395 |
| ENSG000002 | 2.02616275 | -0.318310511 | 0.36152854 | -0.8804575 | 0.37861153 | NA | NA | NA | |
| ENSG000002 | 1.24314998 | -0.037426317 | 0.32887045 | -0.1138026 | 0.90939427 | NA | NA | NA | |

| | | | | | | | | | |
|---|---|---|---|---|---|---|---|---|---|
| ENSG000002 | 1.7794905 | -0.48189743 | 0.35700977 | -1.3498158 | 0.17707507 | NA | NA | NA | |
| ENSG000002 | 0.79807549 | -0.536675498 | 0.28113813 | -1.9089389 | 0.05626998 | NA | NA | NA | |
| ENSG000002 | 0.87010195 | 0.048098246 | 0.29583509 | 0.16258466 | 0.87084547 | NA | NA | NA | |
| ENSG000002 | 1.30301525 | -0.509671631 | 0.33100847 | -1.539754 | 0.12362032 | NA | NA | NA | |
| ENSG000002 | 0.26062507 | -0.012888319 | 0.18226503 | -0.070712 | 0.94362699 | NA | NA | NA | |
| ENSG000002 | 0.64002365 | -0.456557672 | 0.26464873 | -1.7251459 | 0.08450119 | NA | NA | NA | |
| ENSG000002 | 0 | NA | NA | NA | NA | NA | NA | NA | |
| ENSG000002 | 1.48396941 | -0.428404958 | 0.33348295 | -1.2846383 | 0.19891872 | NA | NA | NA | |
| ENSG000002 | 0.37441639 | -0.268856313 | 0.21277471 | -1.2635727 | 0.20638344 | NA | NA | NA | |
| ENSG000002 | 1.08701079 | 0.188393495 | 0.321295 | 0.58635676 | 0.55763579 | NA | NA | NA | |
| ENSG000002 | 2.05522529 | 0.378485073 | 0.36243623 | 1.04428047 | 0.29635565 | NA | NA | NA | |
| ENSG000002 | 0.13280363 | -0.093394593 | 0.13115644 | -0.7120855 | 0.47641185 | NA | NA | NA | |
| ENSG000002 | 0 | NA | NA | NA | NA | NA | TTTY15 | | 64595 |
| ENSG000002 | 0.99260276 | -0.190905426 | 0.3121099 | -0.6116609 | 0.54076213 | NA | NA | NA | |
| ENSG000002 | 1.57803967 | 0.019806419 | 0.34952878 | 0.05666606 | 0.95481121 | NA | NA | NA | |
| ENSG000002 | 1.39719302 | -0.273663342 | 0.33763541 | -0.8105292 | 0.41763611 | NA | NA | NA | |
| ENSG000002 | 1.81023527 | -0.772303829 | 0.35397902 | -2.1817786 | 0.02912588 | NA | NA | NA | |
| ENSG000002 | 0.85400243 | -0.391827062 | 0.2851755 | -1.3739857 | 0.16944611 | NA | NA | NA | |
| ENSG000002 | 2.12828969 | -0.394056104 | 0.36697963 | -1.073782 | 0.28292042 | NA | NA | NA | |
| ENSG000002 | 1.38873786 | -0.38748962 | 0.33265318 | -1.1648457 | 0.24408147 | NA | NA | NA | |
| ENSG000002 | 1.52117092 | -0.342983509 | 0.34720467 | -0.9878424 | 0.32322982 | NA | NA | NA | |
| ENSG000002 | 0.12685397 | -0.090215768 | 0.12901919 | -0.699243 | 0.48440017 | NA | NA | NA | |
| ENSG000002 | 0.82103138 | 0.338029444 | 0.28129135 | 1.20170577 | 0.22947754 | NA | NA | NA | |
| ENSG000002 | 1.9116109 | -0.382455473 | 0.35620457 | -1.0736961 | 0.28295889 | NA | NA | NA | |
| ENSG000002 | 0.37330862 | -0.105391168 | 0.21773449 | -0.4840352 | 0.62836086 | NA | POTEB | | 100996331 |
| ENSG000002 | 0.26199343 | -0.194018184 | 0.18422801 | -1.0531416 | 0.29227609 | NA | NA | NA | |
| ENSG000002 | 0.87576928 | -0.100297505 | 0.2903629 | -0.3454212 | 0.72977773 | NA | NA | NA | |
| ENSG000002 | 0.77613058 | -0.361458377 | 0.28422771 | -1.2717211 | 0.20347223 | NA | NA | NA | |
| ENSG000002 | 1.38583504 | -0.278357211 | 0.34060417 | -0.8172455 | 0.41378817 | NA | NA | NA | |

| | | | | | | | | |
|---|---|---|---|---|---|---|---|---|
| ENSG000002 | 1.744733 | -0.305984969 | 0.35051895 | -0.8729484 | 0.38269119 | NA | NA | NA |
| ENSG000002 | 0.85063263 | -0.097257973 | 0.28899195 | -0.3365422 | 0.73646207 | NA | NA | NA |
| ENSG000002 | 0.89085264 | -0.117645689 | 0.29886997 | -0.393635 | 0.69385052 | NA | NA | NA |
| ENSG000002 | 0 | NA | NA | NA | NA | NA | NA | NA |
| ENSG000002 | 0.7731096 | -0.379060994 | 0.28942548 | -1.3097015 | 0.19029682 | NA | NA | NA |
| ENSG000002 | 1.38743421 | -0.258562509 | 0.33603156 | -0.769459 | 0.44162088 | NA | NA | NA |
| ENSG000002 | 1.51130393 | -0.603315541 | 0.33915994 | -1.7788526 | 0.07526393 | NA | NA | NA |
| ENSG000002 | 0.25370795 | -0.171331747 | 0.17411721 | -0.9840024 | 0.32511435 | NA | NA | NA |
| ENSG000002 | 1.24150789 | -0.33581852 | 0.32807869 | -1.0235914 | 0.30602832 | NA | NA | NA |
| ENSG000002 | 1.57692786 | -0.11824568 | 0.3447632 | -0.3429765 | 0.73161613 | NA | NA | NA |
| ENSG000002 | 0.97635956 | -0.027138674 | 0.29985599 | -0.0905057 | 0.92788537 | NA | NA | NA |
| ENSG000002 | 1.00037333 | -0.042011726 | 0.30685759 | -0.1369095 | 0.89110231 | NA | NA | NA |
| ENSG000002 | 1.74608155 | -0.482295806 | 0.3579921 | -1.3472247 | 0.17790787 | NA | NA | NA |
| ENSG000002 | 1.60135241 | -0.265349911 | 0.34907476 | -0.7601521 | 0.44716367 | NA | NA | NA |
| ENSG000002 | 0.8872155 | -0.45762078 | 0.30165688 | -1.5170242 | 0.12926059 | NA | NA | NA |
| ENSG000002 | 1.21691661 | -0.039714821 | 0.33190354 | -0.1196577 | 0.90475429 | NA | NA | NA |
| ENSG000002 | 0.24004852 | -0.005223787 | 0.17775844 | -0.029387 | 0.97655594 | NA | NA | NA |
| ENSG000002 | 1.91494049 | -0.633203781 | 0.35211753 | -1.798274 | 0.0721336 | NA | NA | NA |
| ENSG000002 | 1.72450665 | -0.455576465 | 0.35274234 | -1.2915276 | 0.19652079 | NA | NA | NA |
| ENSG000002 | 1.51718598 | -0.192267295 | 0.34220271 | -0.561852 | 0.57421683 | NA | NA | NA |
| ENSG000002 | 1.45106308 | 0.236155132 | 0.34512508 | 0.68425954 | 0.49381129 | NA | NA | NA |
| ENSG000002 | 0.80880231 | -0.346232408 | 0.27824793 | -1.2443306 | 0.21337792 | NA | NDUFA5P7 | 106481784 |
| ENSG000002 | 1.53093028 | -0.331587209 | 0.34324247 | -0.9660437 | 0.33402233 | NA | NA | NA |
| ENSG000002 | 0.24550891 | -0.007886114 | 0.17900103 | -0.0440563 | 0.96485956 | NA | NA | NA |
| ENSG000002 | 0.35346928 | -0.088570748 | 0.20618147 | -0.4295767 | 0.66750362 | NA | NA | NA |
| ENSG000002 | 1.24386491 | -0.20423389 | 0.33433595 | -0.6108643 | 0.54128944 | NA | NA | NA |
| ENSG000002 | 1.58762826 | -0.121626572 | 0.3492237 | -0.348277 | 0.72763219 | NA | FAM96AP1 | 106480777 |
| ENSG000002 | 1.20708807 | 0.253488137 | 0.32822455 | 0.77230096 | 0.4399362 | NA | NA | NA |
| ENSG000002 | 0 | NA | NA | NA | NA | NA | NA | NA |

| | | | | | | | | | |
|---|---|---|---|---|---|---|---|---|---|
| ENSG000002 | 1.86993629 | -0.376077947 | 0.35714671 | -1.0530069 | 0.29233783 | NA | NA | NA | |
| ENSG000002 | 0.77035114 | -0.189893091 | 0.27994117 | -0.6783321 | 0.49756113 | NA | NA | NA | |
| ENSG000002 | 0.73567988 | 0.119721393 | 0.26952818 | 0.44418878 | 0.65690611 | NA | NA | NA | |
| ENSG000002 | 0.85747595 | -0.11158464 | 0.29714457 | -0.3755231 | 0.7072715 | NA | NA | NA | |
| ENSG000002 | 0.87392664 | -0.422259382 | 0.29440295 | -1.4342906 | 0.15148935 | NA | NA | NA | |
| ENSG000002 | 0 | NA | NA | NA | NA | NA | NA | NA | |
| ENSG000002 | 1.10665714 | -0.230273625 | 0.30969732 | -0.7435441 | 0.45715231 | NA | NA | NA | |
| ENSG000002 | 0.46956163 | 0.155192681 | 0.24412227 | 0.63571702 | 0.52496088 | NA | NA | NA | |
| ENSG000002 | 1.48830221 | -0.323927221 | 0.34036864 | -0.9516953 | 0.34125156 | NA | NA | NA | |
| ENSG000002 | 0.77018092 | -0.359515013 | 0.28385235 | -1.2665564 | 0.20531394 | NA | NA | NA | |
| ENSG000002 | 1.61531836 | 0.013872231 | 0.348155 | 0.03984499 | 0.96821671 | NA | NA | NA | |
| ENSG000002 | 1.04312739 | -0.535440803 | 0.31265489 | -1.7125618 | 0.08679319 | NA | NA | NA | |
| ENSG000002 | 1.14846726 | -0.255399844 | 0.31035212 | -0.8229357 | 0.41054455 | NA | NA | NA | |
| ENSG000002 | 0.63874119 | -0.117901629 | 0.26613938 | -0.4430071 | 0.65776062 | NA | NA | NA | |
| ENSG000002 | 0 | NA | NA | NA | NA | NA | NA | NA | |
| ENSG000002 | 0.50544884 | -0.019721866 | 0.2401671 | -0.0821173 | 0.93455346 | NA | NA | NA | |
| ENSG000002 | 1.33693937 | -0.118619402 | 0.33569207 | -0.3533578 | 0.72382024 | NA | NA | NA | |
| ENSG000002 | 0.25351207 | -0.189965547 | 0.1824784 | -1.0410303 | 0.29786148 | NA | NA | NA | |
| ENSG000002 | 0 | NA | NA | NA | NA | NA | NA | NA | |
| ENSG000002 | 1.79088677 | -0.591130172 | 0.35023926 | -1.6877896 | 0.09145163 | NA | NA | NA | |
| ENSG000002 | 1.4818084 | -0.045469378 | 0.34558511 | -0.1315722 | 0.89532271 | NA | UGT1A13P | | 404204 |
| ENSG000002 | 2.1102771 | -0.389455918 | 0.36614302 | -1.0636716 | 0.28747751 | NA | NA | NA | |
| ENSG000002 | 0.96468646 | -0.179071102 | 0.30600965 | -0.5851812 | 0.55842586 | NA | NA | NA | |
| ENSG000002 | 0.75350052 | -0.19232973 | 0.28654979 | -0.6711913 | 0.50209866 | NA | NA | NA | |
| ENSG000002 | 1.35409247 | 0.021328581 | 0.3357919 | 0.06351726 | 0.94935462 | NA | NA | NA | |
| ENSG000002 | 2.05773122 | -0.000951476 | 0.36447159 | -0.0026106 | 0.99791707 | NA | NA | NA | |
| ENSG000002 | 1.42124405 | -0.582965417 | 0.33908582 | -1.7192267 | 0.08557309 | NA | DEFA11P | | 724068 |
| ENSG000002 | 0 | NA | NA | NA | NA | NA | NA | NA | |
| ENSG000002 | 0.47804353 | 0.152460376 | 0.23772494 | 0.64133101 | 0.52130765 | NA | NA | NA | |

| | | | | | | | | | |
|---|---|---|---|---|---|---|---|---|---|
| ENSG000002 | 1.58652103 | 0.018114431 | 0.35313074 | 0.05129667 | 0.95908912 | NA | NA | NA | |
| ENSG000002 | 2.06480125 | -0.557404556 | 0.35696194 | -1.5615238 | 0.11840021 | NA | NA | NA | |
| ENSG000002 | 2.07202585 | -0.001407105 | 0.36310071 | -0.0038752 | 0.99690801 | NA | LINC01191 | | 440900 |
| ENSG000002 | 0.89839934 | -0.272249175 | 0.29883702 | -0.911029 | 0.36228012 | NA | NA | NA | |
| ENSG000002 | 1.32729599 | -0.620254993 | 0.32007296 | -1.937855 | 0.05264091 | NA | NA | NA | |
| ENSG000002 | 1.57325979 | -0.794577221 | 0.3408123 | -2.3314218 | 0.01973113 | NA | NA | NA | |
| ENSG000002 | 0 | NA | NA | NA | NA | NA | LINC01505 | | 100996590 |
| ENSG000002 | 0.86660862 | -0.252242393 | 0.2931025 | -0.8605945 | 0.38946143 | NA | NA | NA | |
| ENSG000002 | 0.88647829 | -0.457898022 | 0.30161276 | -1.5181653 | 0.12897273 | NA | NA | NA | |
| ENSG000002 | 1.05834126 | 0.491531897 | 0.31046235 | 1.58322547 | 0.11337008 | NA | NA | NA | |
| ENSG000002 | 0.5953301 | 0.377753376 | 0.250069 | 1.51059659 | 0.13089126 | NA | NA | NA | |
| ENSG000002 | 0.76579158 | -0.196446233 | 0.28731812 | -0.6837238 | 0.49414961 | NA | NA | NA | |
| ENSG000002 | 0.67599868 | -0.271668661 | 0.26265684 | -1.0343102 | 0.30099116 | NA | NA | NA | |
| ENSG000002 | 1.1352905 | -0.256090704 | 0.31621303 | -0.8098676 | 0.41801625 | NA | NA | NA | |
| ENSG000002 | 1.12293025 | 0.030919025 | 0.32613242 | 0.09480513 | 0.92446962 | NA | NA | NA | |
| ENSG000002 | 1.55157546 | -0.632652325 | 0.34159827 | -1.8520361 | 0.06402064 | NA | NA | NA | |
| ENSG000002 | 1.68292483 | -0.697212 | 0.34843033 | -2.0010084 | 0.04539149 | NA | NA | NA | |
| ENSG000002 | 0.24619405 | -0.007630671 | 0.17915639 | -0.0425922 | 0.96602658 | NA | NA | NA | |
| ENSG000002 | 0 | NA | NA | NA | NA | NA | PRR20E | | 729250 |
| ENSG000002 | 1.08843122 | 0.035098336 | 0.32115863 | 0.1092866 | 0.91297517 | NA | NA | NA | |
| ENSG000002 | 0.86179473 | -0.108705225 | 0.2973844 | -0.3655377 | 0.71471003 | NA | NA | NA | |
| ENSG000002 | 1.83621677 | -0.44549188 | 0.34864781 | -1.2777705 | 0.20133037 | NA | NA | NA | |
| ENSG000002 | 1.40704329 | -0.431839374 | 0.34180188 | -1.2634201 | 0.20643824 | NA | NA | NA | |
| ENSG000002 | 0.2596576 | -0.193002035 | 0.1837913 | -1.0501152 | 0.29366517 | NA | NA | NA | |
| ENSG000002 | 1.3275966 | -0.108518268 | 0.32895384 | -0.329889 | 0.7414838 | NA | NA | NA | |
| ENSG000002 | 1.60732826 | -0.395910399 | 0.34954128 | -1.1326571 | 0.25735829 | NA | NA | NA | |
| ENSG000002 | 2.11308776 | -0.003973342 | 0.36527473 | -0.0108777 | 0.99132104 | NA | NA | NA | |
| ENSG000002 | 1.59109986 | 0.007668998 | 0.35089583 | 0.02185548 | 0.98256324 | NA | NA | NA | |
| ENSG000002 | 0.49241844 | -0.197108126 | 0.24744176 | -0.7965839 | 0.42569273 | NA | NA | NA | |

| | | | | | | | | | |
|---|---|---|---|---|---|---|---|---|---|
| ENSG000002 | 0.62023988 | -0.112292793 | 0.27094277 | -0.4144521 | 0.67854306 | NA | NA | NA | |
| ENSG000002 | 0.11205239 | 0.082118251 | 0.12671962 | 0.64803108 | 0.51696485 | NA | NA | NA | |
| ENSG000002 | 1.27125094 | -0.361134426 | 0.33440988 | -1.0799155 | 0.2801798 | NA | NA | NA | |
| ENSG000002 | 1.92197581 | -0.177290279 | 0.35980059 | -0.4927459 | 0.62219211 | NA | NA | NA | |
| ENSG000002 | 0.513452 | -0.199963756 | 0.2496885 | -0.8008529 | 0.42321682 | NA | NA | NA | |
| ENSG000002 | 0.12685397 | -0.090215768 | 0.12901919 | -0.699243 | 0.48440017 | NA | NA | NA | |
| ENSG000002 | 2.04111728 | -0.749351569 | 0.36336479 | -2.062257 | 0.03918328 | NA | NA | NA | |
| ENSG000002 | 1.71848657 | 0.213709217 | 0.35493764 | 0.60210356 | 0.54710521 | NA | NA | NA | |
| ENSG000002 | 0.85995209 | -0.423384572 | 0.29356981 | -1.4421938 | 0.1492477 | NA | NA | NA | |
| ENSG000002 | 0 | NA | NA | NA | NA | NA | NA | NA | |
| ENSG000002 | 0 | NA | NA | NA | NA | NA | NA | NA | |
| ENSG000002 | 2.15368013 | -0.520802971 | 0.36593542 | -1.4232101 | 0.15467527 | NA | NA | NA | |
| ENSG000002 | 1.76004695 | -0.174451705 | 0.34788029 | -0.5014705 | 0.61604003 | NA | NA | NA | |
| ENSG000002 | 0 | NA | NA | NA | NA | NA | NA | NA | |
| ENSG000002 | 0 | NA | NA | NA | NA | NA | NA | NA | |
| ENSG000002 | 0.94900386 | -0.030219346 | 0.30457487 | -0.0992181 | 0.92096509 | NA | NA | NA | |
| ENSG000002 | 0.89359129 | -0.291434556 | 0.3048858 | -0.955881 | 0.33913234 | NA | NA | NA | |
| ENSG000002 | 0 | NA | NA | NA | NA | NA | RBMY1A1 | | 5940 |
| ENSG000002 | 1.77795408 | 0.145647603 | 0.35606599 | 0.40904666 | 0.68250542 | NA | SLC25A5P4 | | 100131027 |
| ENSG000002 | 1.54799039 | -0.956985252 | 0.33818375 | -2.829779 | 0.00465802 | NA | NA | NA | |
| ENSG000002 | 0 | NA | NA | NA | NA | NA | NA | NA | |
| ENSG000002 | 0.88086495 | -0.113054583 | 0.29836514 | -0.3789135 | 0.70475209 | NA | NA | NA | |
| ENSG000002 | 0.4933859 | -0.021085811 | 0.24629532 | -0.0856119 | 0.93177494 | NA | KBTBD13 | | 390594 |
| ENSG000002 | 1.09526244 | 0.042338448 | 0.32137535 | 0.13174143 | 0.89518881 | NA | NA | NA | |
| ENSG000002 | 0.24411881 | -0.012946855 | 0.17864669 | -0.0724718 | 0.94222641 | NA | NA | NA | |
| ENSG000002 | 0 | NA | NA | NA | NA | NA | NA | NA | |
| ENSG000002 | 0 | NA | NA | NA | NA | NA | NA | NA | |
| ENSG000002 | 1.9802886 | -0.16581683 | 0.35572956 | -0.4661317 | 0.64112122 | NA | NA | NA | |
| ENSG000002 | 2.11867414 | -0.374689141 | 0.36347219 | -1.0308606 | 0.30260622 | NA | NA | NA | |

| | | | | | | | | |
|---|---|---|---|---|---|---|---|---|
| ENSG000002 | 0.60401594 | 0.058796518 | 0.26924067 | 0.21837904 | 0.8271338 | NA | NA | NA |
| ENSG000002 | 0.83349734 | 0.196566422 | 0.28582694 | 0.68771131 | 0.4916346 | NA | COX6CP12 | 106481682 |
| ENSG000002 | 0.12685397 | -0.090215768 | 0.12901919 | -0.699243 | 0.48440017 | NA | NA | NA |
| ENSG000002 | 1.70855895 | -0.492402094 | 0.33828145 | -1.4555989 | 0.14550354 | NA | NA | NA |
| ENSG000002 | 1.51235325 | -0.639714061 | 0.34406712 | -1.8592711 | 0.06298872 | NA | AP1B1P1 | 23782 |
| ENSG000002 | 1.58420694 | 0.151601688 | 0.34469798 | 0.43981021 | 0.66007457 | NA | NA | NA |
| ENSG000002 | 1.39033201 | -0.124733469 | 0.33683912 | -0.3703058 | 0.71115468 | NA | NA | NA |
| ENSG000002 | 2.01259156 | -0.443559282 | 0.36170456 | -1.2263027 | 0.22008478 | NA | NA | NA |
| ENSG000002 | 1.69053801 | 0.216998448 | 0.35287045 | 0.61495217 | 0.53858631 | NA | NA | NA |
| ENSG000002 | 1.04788337 | -0.532169807 | 0.31292209 | -1.7006463 | 0.08900942 | NA | NA | NA |
| ENSG000002 | 1.39514398 | -0.411293256 | 0.33726829 | -1.2194839 | 0.22266058 | NA | NA | NA |
| ENSG000002 | 0 | NA | NA | NA | NA | NA | NA | NA |
| ENSG000002 | 1.23333642 | -0.172492516 | 0.32318556 | -0.5337259 | 0.59353116 | NA | NA | NA |
| ENSG000002 | 1.979066 | -0.665606104 | 0.3561616 | -1.8688318 | 0.06164622 | NA | NA | NA |
| ENSG000002 | 0.85363187 | -0.268235387 | 0.2964963 | -0.9046837 | 0.36563295 | NA | NA | NA |
| ENSG000002 | 1.78407571 | -0.607710898 | 0.35311026 | -1.7210231 | 0.08524664 | NA | CHEK2P5 | 106479022 |
| ENSG000002 | 1.81242031 | 0.139872723 | 0.35908488 | 0.38952552 | 0.69688744 | NA | LINC01309 | 104326190 |
| ENSG000002 | 1.52114919 | -0.498506047 | 0.34683971 | -1.4372808 | 0.15063823 | NA | NA | NA |
| ENSG000002 | 1.24249654 | -0.05210282 | 0.3338948 | -0.1560456 | 0.87599706 | NA | NA | NA |
| ENSG000002 | 1.78906003 | -0.601453184 | 0.35252915 | -1.7061091 | 0.08798777 | NA | NA | NA |
| ENSG000002 | 1.13918452 | -0.279227964 | 0.32386779 | -0.8621665 | 0.3885959 | NA | NA | NA |
| ENSG000002 | 1.54558751 | -0.469353697 | 0.33991363 | -1.3808028 | 0.1673396 | NA | NA | NA |
| ENSG000002 | 1.64438487 | -0.273274458 | 0.35044561 | -0.7797914 | 0.4355137 | NA | NA | NA |
| ENSG000002 | 0 | NA | NA | NA | NA | NA | NA | NA |
| ENSG000002 | 1.00518722 | -0.196835355 | 0.31410736 | -0.6266499 | 0.53088873 | NA | NA | NA |
| ENSG000002 | 1.83046516 | 0.137954301 | 0.36107505 | 0.38206545 | 0.70241282 | NA | NA | NA |
| ENSG000002 | 1.84418113 | -0.250917282 | 0.35834645 | -0.7002086 | 0.48379702 | NA | NA | NA |
| ENSG000002 | 0.89020422 | -0.427620104 | 0.29524143 | -1.4483743 | 0.14751239 | NA | NA | NA |
| ENSG000002 | 1.34436694 | 0.16197479 | 0.33426969 | 0.48456319 | 0.62798623 | NA | NA | NA |

| | | | | | | | | |
|---|---|---|---|---|---|---|---|---|
| ENSG000002 | 0 | NA | NA | NA | NA | NA | NA | NA |
| ENSG000002 | 0 | NA | NA | NA | NA | NA | NA | NA |
| ENSG000002 | 0.38270188 | -0.272446727 | 0.2140009 | -1.2731102 | 0.20297894 | NA | NA | NA |
| ENSG000002 | 1.12082118 | -0.109217511 | 0.31637059 | -0.3452202 | 0.72992885 | NA | NA | NA |
| ENSG000002 | 1.58353487 | -0.374727065 | 0.34519982 | -1.0855367 | 0.27768403 | NA | NA | NA |
| ENSG000002 | 1.76258334 | -0.203302819 | 0.35775719 | -0.5682704 | 0.56985138 | NA | NA | NA |
| ENSG000002 | 1.44433112 | -0.702216627 | 0.3312797 | -2.1197092 | 0.03403058 | NA | NECAP1P2 | 106479026 |
| ENSG000002 | 1.6987454 | -0.688540313 | 0.34635805 | -1.9879437 | 0.04681791 | NA | NA | NA |
| ENSG000002 | 1.75819793 | -0.337542263 | 0.35662761 | -0.9464838 | 0.34390187 | NA | NA | NA |
| ENSG000002 | 0.62061044 | -0.252611122 | 0.25798806 | -0.9791582 | 0.32750182 | NA | NA | NA |
| ENSG000002 | 0.97388385 | -0.328542053 | 0.30547786 | -1.075502 | 0.28215002 | NA | NA | NA |
| ENSG000002 | 0 | NA | NA | NA | NA | NA | NA | NA |
| ENSG000002 | 1.82497039 | 0.005291002 | 0.35819318 | 0.01477136 | 0.98821458 | NA | NA | NA |
| ENSG000002 | 1.35119804 | -0.39151426 | 0.33291061 | -1.1760342 | 0.23958122 | NA | NA | NA |
| ENSG000002 | 1.9456487 | 0.069889241 | 0.3585413 | 0.19492661 | 0.8454504 | NA | NA | NA |
| ENSG000002 | 1.63772888 | -0.399201215 | 0.34788399 | -1.1475125 | 0.25116988 | NA | NA | NA |
| ENSG000002 | 1.62671059 | -0.3934156 | 0.34701224 | -1.1337226 | 0.25691095 | NA | NA | NA |
| ENSG000002 | 1.22639522 | -0.047895427 | 0.33215181 | -0.1441974 | 0.88534459 | NA | NA | NA |
| ENSG000002 | 0.48032325 | -0.179266136 | 0.23878452 | -0.7507444 | 0.45280651 | NA | NA | NA |
| ENSG000002 | 0.39388516 | -0.104610737 | 0.21253504 | -0.4922047 | 0.62257467 | NA | NA | NA |
| ENSG000002 | 0.61927241 | -0.278564577 | 0.26634985 | -1.0458597 | 0.29562581 | NA | NA | NA |
| ENSG000002 | 1.74106298 | -0.048699419 | 0.34299012 | -0.1419849 | 0.88709193 | NA | TMEM212-AS1 | 100874219 |
| ENSG000002 | 0 | NA | NA | NA | NA | NA | NA | NA |
| ENSG000002 | 0.85803847 | 0.039330813 | 0.29683333 | 0.13250134 | 0.89458777 | NA | RPEP5 | 106481798 |
| ENSG000002 | 0 | NA | NA | NA | NA | NA | USP9YP26 | 106478929 |
| ENSG000002 | 0.46341609 | 0.157328408 | 0.24338973 | 0.64640528 | 0.51801692 | NA | NA | NA |
| ENSG000002 | 0.36502313 | -0.093755175 | 0.2080862 | -0.4505593 | 0.6523072 | NA | NA | NA |
| ENSG000002 | 1.66758191 | -0.544710079 | 0.3487519 | -1.5618842 | 0.11831528 | NA | NA | NA |
| ENSG000002 | 1.15379023 | -0.424268504 | 0.32005247 | -1.3256217 | 0.18496503 | NA | NA | NA |

| | | | | | | | | |
|---|---|---|---|---|---|---|---|---|
| ENSG000002 | 1.88866572 | -0.387955462 | 0.35729024 | -1.0858272 | 0.27755545 | NA | NA | NA |
| ENSG000002 | 0.23868015 | 0.154625155 | 0.16997278 | 0.90970538 | 0.36297791 | NA | NA | NA |
| ENSG000002 | 1.62822799 | -0.508173708 | 0.34105279 | -1.4900148 | 0.13622034 | NA | NA | NA |
| ENSG000002 | 1.23826016 | -0.183159436 | 0.32247927 | -0.5679728 | 0.57005347 | NA | NA | NA |
| ENSG000002 | 1.35995835 | 0.030204184 | 0.32857538 | 0.09192467 | 0.92675789 | NA | NA | NA |
| ENSG000002 | 0.8650959 | -0.118179347 | 0.30263724 | -0.3904984 | 0.69616807 | NA | NA | NA |
| ENSG000002 | 1.26834387 | -0.722281176 | 0.31032746 | -2.3274807 | 0.01993969 | NA | NA | NA |
| ENSG000002 | 0 | NA | NA | NA | NA | NA | NA | NA |
| ENSG000002 | 0 | NA | NA | NA | NA | NA | NA | NA |
| ENSG000002 | 1.21893978 | -0.030488659 | 0.31957639 | -0.0954034 | 0.92399445 | NA | NA | NA |
| ENSG000002 | 0.71698024 | -0.023804784 | 0.27695399 | -0.0859521 | 0.93150447 | NA | NA | NA |
| ENSG000002 | 0 | NA | NA | NA | NA | NA | NA | NA |
| ENSG000002 | 1.63566486 | -0.114795325 | 0.34296661 | -0.3347128 | 0.73784172 | NA | NA | NA |
| ENSG000002 | 1.3416948 | -0.124403157 | 0.33579853 | -0.3704696 | 0.7110326 | NA | IFNA17 | 3451 |
| ENSG000002 | 0 | NA | NA | NA | NA | NA | NA | NA |
| ENSG000002 | 1.51952023 | -0.328334609 | 0.34305631 | -0.9570866 | 0.33852353 | NA | NA | NA |
| ENSG000002 | 0 | NA | NA | NA | NA | NA | NA | NA |
| ENSG000002 | 1.35679886 | -0.118636959 | 0.33949283 | -0.3494535 | 0.72674887 | NA | NA | NA |
| ENSG000002 | 2.16250389 | 0.058861565 | 0.35869172 | 0.16410071 | 0.86965187 | NA | LINC00676 | 101409253 |
| ENSG000002 | 1.36866912 | 0.636524772 | 0.32726521 | 1.9449815 | 0.05177721 | NA | NA | NA |
| ENSG000002 | 2.02545642 | -0.454907585 | 0.36321198 | -1.2524575 | 0.21040319 | NA | NA | NA |
| ENSG000002 | 1.11720736 | -0.118552941 | 0.32623891 | -0.363393 | 0.71631132 | NA | NA | NA |
| ENSG000002 | 0.38036605 | -0.293832733 | 0.22119083 | -1.3284128 | 0.18404178 | NA | NA | NA |
| ENSG000002 | 1.91881372 | -0.36543187 | 0.35143651 | -1.0398233 | 0.29842201 | NA | NA | NA |
| ENSG000002 | 1.71758953 | -0.539389865 | 0.34217244 | -1.5763685 | 0.1149409 | NA | NA | NA |
| ENSG000002 | 1.24907735 | -0.206105181 | 0.33446029 | -0.6162321 | 0.53774132 | NA | NA | NA |
| ENSG000002 | 1.06572736 | 0.036481253 | 0.31884695 | 0.11441619 | 0.90890788 | NA | LINC00163 | 727699 |
| ENSG000002 | 0 | NA | NA | NA | NA | NA | NA | NA |
| ENSG000002 | 0.63484558 | -0.278698901 | 0.26767877 | -1.0411692 | 0.29779701 | NA | NA | NA |

| | | | | | | | | |
|---|---|---|---|---|---|---|---|---|
| ENSG000002 | 1.40672659 | -0.714253651 | 0.33280933 | -2.1461347 | 0.03186224 | NA | NA | NA |
| ENSG000002 | 0 | NA | NA | NA | NA | NA | NA | NA |
| ENSG000002 | 0.49856397 | -0.186262775 | 0.24090455 | -0.7731808 | 0.43941538 | NA | NA | NA |
| ENSG000002 | 1.0225205 | -0.329020232 | 0.30463065 | -1.0800628 | 0.28011421 | NA | NA | NA |
| ENSG000002 | 1.58885729 | -0.485616847 | 0.33875727 | -1.4335245 | 0.15170801 | NA | NA | NA |
| ENSG000002 | 2.15314849 | -0.785771603 | 0.36425001 | -2.1572315 | 0.03098763 | NA | NA | NA |
| ENSG000002 | 0.49671898 | -0.027206884 | 0.23914704 | -0.1137663 | 0.90942302 | NA | NA | NA |
| ENSG000002 | 1.61234483 | -0.381412251 | 0.34562458 | -1.1035449 | 0.2697906 | NA | NA | NA |
| ENSG000002 | 1.09908871 | -0.254591417 | 0.31661755 | -0.8040976 | 0.4213406 | NA | NA | NA |
| ENSG000002 | 1.53238455 | -0.348251327 | 0.34549975 | -1.007964 | 0.31347173 | NA | NA | NA |
| ENSG000002 | 0 | NA | NA | NA | NA | NA | NA | NA |
| ENSG000002 | 1.0285005 | -0.53844435 | 0.31357068 | -1.7171387 | 0.08595383 | NA | LOC101929374 | 101929374 |
| ENSG000002 | 1.76557397 | -0.058433849 | 0.34946184 | -0.167211 | 0.86720406 | NA | NA | NA |
| ENSG000002 | 0 | NA | NA | NA | NA | NA | NA | NA |
| ENSG000002 | 1.14368149 | -0.122054925 | 0.32325304 | -0.3775832 | 0.70574023 | NA | NA | NA |
| ENSG000002 | 0.96494705 | -0.184243905 | 0.30600798 | -0.6020886 | 0.54711519 | NA | FABP5P13 | 106480712 |
| ENSG000002 | 1.33762506 | -0.111232745 | 0.33705645 | -0.3300122 | 0.74139074 | NA | LOC102724264 | 102724264 |
| ENSG000002 | 0.86455457 | -0.111874481 | 0.29589056 | -0.3780941 | 0.70536068 | NA | NA | NA |
| ENSG000002 | 1.73446009 | -0.064828898 | 0.35816092 | -0.1810049 | 0.85636369 | NA | NA | NA |
| ENSG000002 | 2.09730592 | -0.582003286 | 0.3578483 | -1.6263967 | 0.1038653 | NA | NA | NA |
| ENSG000002 | 1.33332397 | -0.237159467 | 0.32828582 | -0.7224176 | 0.47003775 | NA | NA | NA |
| ENSG000002 | 1.88520093 | -0.391378686 | 0.35811069 | -1.0928986 | 0.2744383 | NA | NA | NA |
| ENSG000002 | 0.63484558 | -0.278698901 | 0.26767877 | -1.0411692 | 0.29779701 | NA | NA | NA |
| ENSG000002 | 1.73663622 | -0.322878038 | 0.35359041 | -0.9131414 | 0.36116819 | NA | NA | NA |
| ENSG000002 | 0.75555403 | -0.349570212 | 0.28131632 | -1.2426233 | 0.21400668 | NA | NA | NA |
| ENSG000002 | 0 | NA | NA | NA | NA | NA | NA | NA |
| ENSG000002 | 1.48317622 | -0.203077551 | 0.34957911 | -0.5809202 | 0.56129427 | NA | NA | NA |
| ENSG000002 | 1.30055892 | -0.175103229 | 0.31389641 | -0.5578376 | 0.57695526 | NA | NA | NA |
| ENSG000002 | 0 | NA | NA | NA | NA | NA | NA | NA |

| | | | | | | | | |
|---|---|---|---|---|---|---|---|---|
| ENSG000002 | 0.54456342 | -0.359011254 | 0.24026449 | -1.4942335 | 0.1351146 | NA | NA | NA |
| ENSG000002 | 1.115587 | -0.258948766 | 0.31719322 | -0.8163755 | 0.41428542 | NA | NA | NA |
| ENSG000002 | 1.01557773 | -0.324951973 | 0.30170866 | -1.0770389 | 0.28146287 | NA | NA | NA |
| ENSG000002 | 0.26813897 | -0.197007041 | 0.18550319 | -1.0620143 | 0.2882292 | NA | NA | NA |
| ENSG000002 | 1.61410048 | -0.268666363 | 0.35071229 | -0.7660592 | 0.4436411 | NA | NA | NA |
| ENSG000002 | 2.10657558 | -0.122492431 | 0.36539598 | -0.335232 | 0.73745007 | NA | NA | NA |
| ENSG000002 | 2.10778695 | -0.239466144 | 0.36181903 | -0.6618395 | 0.50807406 | NA | NA | NA |
| ENSG000002 | 0.35961481 | -0.099625193 | 0.21567743 | -0.4619176 | 0.64414043 | NA | NA | NA |
| ENSG000002 | 0 | NA | NA | NA | NA | NA | LOC101929148 | 101929148 |
| ENSG000002 | 1.44435231 | -0.57075154 | 0.33637784 | -1.6967573 | 0.08974256 | NA | UBE2V1P7 | 106481746 |
| ENSG000002 | 0.23070733 | 0.167416683 | 0.17664244 | 0.94777156 | 0.34324576 | NA | NA | NA |
| ENSG000002 | 1.43041056 | -0.58068763 | 0.33856401 | -1.7151487 | 0.08631797 | NA | NA | NA |
| ENSG000002 | 1.48424782 | -0.049615676 | 0.3357842 | -0.1477606 | 0.8825317 | NA | NA | NA |
| ENSG000002 | 1.69332873 | 0.084984073 | 0.35452858 | 0.23971007 | 0.81055502 | NA | NA | NA |
| ENSG000002 | 0.97916955 | -0.186021397 | 0.30829886 | -0.6033801 | 0.54625586 | NA | NA | NA |
| ENSG000002 | 2.14226424 | -0.397388172 | 0.36788925 | -1.0801842 | 0.28006015 | NA | NA | NA |
| ENSG000002 | 0 | NA | NA | NA | NA | NA | NA | NA |
| ENSG000002 | 1.28656352 | -0.327126718 | 0.32590597 | -1.0037457 | 0.3155012 | NA | NA | NA |
| ENSG000002 | 1.50763165 | -0.473319851 | 0.341899 | -1.384385 | 0.1662406 | NA | NA | NA |
| ENSG000002 | 0.6009429 | 0.055606074 | 0.26256469 | 0.21178047 | 0.83227831 | NA | NA | NA |
| ENSG000002 | 2.10315359 | -0.370196523 | 0.36392996 | -1.0172191 | 0.30904919 | NA | NA | NA |
| ENSG000002 | 1.45912168 | -0.184521771 | 0.3404131 | -0.5420525 | 0.58778234 | NA | BSN-AS1 | 106478963 |
| ENSG000002 | 0.47804353 | 0.152460376 | 0.23772494 | 0.64133101 | 0.52130765 | NA | NA | NA |
| ENSG000002 | 1.82350294 | -0.932532844 | 0.35224761 | -2.6473787 | 0.00811184 | NA | NA | NA |
| ENSG000002 | 1.78717908 | -0.773036458 | 0.35434431 | -2.1815969 | 0.02913929 | NA | NA | NA |
| ENSG000002 | 1.33990031 | -0.117620203 | 0.33010931 | -0.3563068 | 0.72161079 | NA | NA | NA |
| ENSG000002 | 0.74521449 | -0.186736727 | 0.27823071 | -0.6711579 | 0.50211997 | NA | NA | NA |
| ENSG000002 | 1.54437657 | -0.978745771 | 0.34045755 | -2.8747953 | 0.0040429 | NA | NA | NA |
| ENSG000002 | 2.11454539 | -0.25541402 | 0.3638474 | -0.7019812 | 0.48269091 | NA | NA | NA |

| | | | | | | | | |
|---|---|---|---|---|---|---|---|---|
| ENSG000002 | 1.56420893 | 0.017383494 | 0.33652243 | 0.05165627 | 0.95880258 | NA | NA | NA |
| ENSG000002 | 0.37928001 | 0.070449581 | 0.21829793 | 0.32272217 | 0.74690566 | NA | NA | NA |
| ENSG000002 | 1.62263638 | -0.127689357 | 0.35378582 | -0.3609228 | 0.71815719 | NA | NA | NA |
| ENSG000002 | 0.88448035 | 0.031576389 | 0.29050026 | 0.1086966 | 0.91344314 | NA | NA | NA |
| ENSG000002 | 0.58150796 | 0.05262168 | 0.25154459 | 0.20919424 | 0.83429661 | NA | NA | NA |
| ENSG000002 | 0.22336757 | 0.163792188 | 0.17487321 | 0.93663397 | 0.34894687 | NA | NA | NA |
| ENSG000002 | 0.62357296 | -0.113393436 | 0.2648832 | -0.4280884 | 0.66858674 | NA | LOC105378614 | 105378614 |
| ENSG000002 | 1.78137323 | -0.752740668 | 0.35198312 | -2.1385704 | 0.03247048 | NA | NA | NA |
| ENSG000002 | 0 | NA | NA | NA | NA | NA | NA | NA |
| ENSG000002 | 0.796903 | -0.342445501 | 0.27749094 | -1.2340781 | 0.2171738 | NA | NA | NA |
| ENSG000002 | 1.34895312 | -0.48265618 | 0.32203902 | -1.4987506 | 0.13393834 | NA | NA | NA |
| ENSG000002 | 0.7278812 | -0.032565449 | 0.27770112 | -0.117268 | 0.9066477 | NA | NA | NA |
| ENSG000002 | 1.23712597 | -0.184015392 | 0.32329508 | -0.5691871 | 0.56922917 | NA | NA | NA |
| ENSG000002 | 1.97515992 | -0.182648712 | 0.36224546 | -0.5042126 | 0.61411201 | NA | NA | NA |
| ENSG000002 | 0.76559516 | -0.206287743 | 0.28724125 | -0.7181689 | 0.47265312 | NA | NA | NA |
| ENSG000002 | 0.24485602 | -0.012655878 | 0.17881722 | -0.0707755 | 0.94357643 | NA | NA | NA |
| ENSG000002 | 1.0667469 | 0.182727564 | 0.31609824 | 0.57807207 | 0.56321546 | NA | NA | NA |
| ENSG000002 | 1.4700232 | -0.190582096 | 0.34061599 | -0.5595219 | 0.57580561 | NA | NA | NA |
| ENSG000002 | 2.09612896 | -0.25736019 | 0.3680314 | -0.6992887 | 0.48437164 | NA | NA | NA |
| ENSG000002 | 1.83750532 | -0.381178077 | 0.35788958 | -1.0650717 | 0.28684346 | NA | NA | NA |
| ENSG000002 | 1.88403683 | -0.748870797 | 0.34890163 | -2.1463666 | 0.03184374 | NA | NA | NA |
| ENSG000002 | 1.25818829 | -0.512993105 | 0.33195581 | -1.5453657 | 0.12225784 | NA | NA | NA |
| ENSG000002 | 0 | NA | NA | NA | NA | NA | NA | NA |
| ENSG000002 | 0.36827086 | -0.266056975 | 0.21181163 | -1.2561018 | 0.20907905 | NA | NA | NA |
| ENSG000002 | 1.79931991 | 0.138634546 | 0.3552318 | 0.39026502 | 0.69634058 | NA | NA | NA |
| ENSG000002 | 0.73880837 | -0.163829537 | 0.26808589 | -0.6111084 | 0.54112783 | NA | NA | NA |
| ENSG000002 | 1.72039964 | -0.188633677 | 0.35481587 | -0.5316382 | 0.59497659 | NA | NA | NA |
| ENSG000002 | 1.62254801 | -0.395527367 | 0.34864926 | -1.1344563 | 0.25660317 | NA | NA | NA |
| ENSG000002 | 0.52779657 | -0.383508725 | 0.24707466 | -1.5521977 | 0.12061492 | NA | NA | NA |

| | | | | | | | | | |
|---|---|---|---|---|---|---|---|---|---|
| ENSG000002 | 0.41537361 | -0.286583533 | 0.21871501 | -1.3103057 | 0.19009243 | NA | NA | NA | |
| ENSG000002 | 0.83575724 | -0.109698178 | 0.29597774 | -0.3706298 | 0.71091327 | NA | NA | NA | |
| ENSG000002 | 1.30473979 | 0.170714185 | 0.33613724 | 0.50787049 | 0.61154417 | NA | NA | NA | |
| ENSG000002 | 2.12148212 | -0.133536529 | 0.36354117 | -0.3673216 | 0.71337913 | NA | NA | NA | |
| ENSG000002 | 2.09837545 | -0.012234499 | 0.36579863 | -0.033446 | 0.97331893 | NA | NA | NA | |
| ENSG000002 | 1.96360106 | 0.056052169 | 0.35990479 | 0.15574166 | 0.87623666 | NA | NA | NA | |
| ENSG000002 | 2.00348062 | -0.191571575 | 0.36329616 | -0.5273152 | 0.59797474 | NA | NA | NA | |
| ENSG000002 | 2.07353061 | -0.129280472 | 0.36556276 | -0.3536478 | 0.72360283 | NA | NA | NA | |
| ENSG000002 | 1.28911667 | 0.321468514 | 0.33490111 | 0.95989086 | 0.33711015 | NA | NA | NA | |
| ENSG000002 | 1.56346617 | -0.319648413 | 0.33898298 | -0.942963 | 0.34569982 | NA | NA | NA | |
| ENSG000002 | 1.03193963 | -0.341576984 | 0.30498076 | -1.1199952 | 0.26271581 | NA | NA | NA | |
| ENSG000002 | 0.61546271 | -0.255876753 | 0.25745011 | -0.9938887 | 0.32027706 | NA | NA | NA | |
| ENSG000002 | 1.1250392 | -0.148413522 | 0.20866282 | -0.71126 | 0.47692313 | NA | NA | NA | |
| ENSG000002 | 1.49752133 | -0.337857404 | 0.34684122 | -0.9740982 | 0.33000778 | NA | NA | NA | |
| ENSG000002 | 2.1620576 | -0.061529374 | 0.3635758 | -0.169234 | 0.86561262 | NA | NA | NA | |
| ENSG000002 | 1.8645618 | -0.265784875 | 0.36183814 | -0.7345408 | 0.46261921 | NA | NA | NA | |
| ENSG000002 | 0 | NA | NA | NA | NA | NA | NA | NA | |
| ENSG000002 | 1.73005541 | -0.191006944 | 0.35535576 | -0.5375091 | 0.59091602 | NA | NA | NA | |
| ENSG000002 | 1.02021501 | -0.529182659 | 0.31179354 | -1.6972214 | 0.08965482 | NA | LOC101929350 | | 101929350 |
| ENSG000002 | 1.35848572 | -0.373909609 | 0.32900779 | -1.1364765 | 0.25575721 | NA | NA | NA | |
| ENSG000002 | 1.51130049 | -0.184482704 | 0.33049739 | -0.5581972 | 0.57670976 | NA | NA | NA | |
| ENSG000002 | 0.51139849 | -0.021802213 | 0.24083338 | -0.0905282 | 0.92786748 | NA | NA | NA | |
| ENSG000002 | 0.95968308 | -0.159296336 | 0.29728489 | -0.5358373 | 0.592071 | NA | NA | NA | |
| ENSG000002 | 0.76693319 | -0.191420094 | 0.28134921 | -0.6803648 | 0.49627351 | NA | NA | NA | |
| ENSG000002 | 1.98588832 | 0.061936182 | 0.36159264 | 0.17128718 | 0.86399796 | NA | NA | NA | |
| ENSG000002 | 2.13543549 | -0.260506762 | 0.36683122 | -0.7101543 | 0.47760847 | NA | NA | NA | |
| ENSG000002 | 0 | NA | NA | NA | NA | NA | NA | NA | |
| ENSG000002 | 1.8730089 | 0.130524436 | 0.35696571 | 0.36564979 | 0.71462642 | NA | NA | NA | |
| ENSG000002 | 2.11796623 | -0.648909066 | 0.3655278 | -1.775266 | 0.07585398 | NA | NA | NA | |

| | | | | | | | | |
|---|---|---|---|---|---|---|---|---|
| ENSG000002 | 0 | NA | NA | NA | NA | NA | NA | NA |
| ENSG000002 | 1.87592169 | -0.127729202 | 0.3582146 | -0.3565717 | 0.72141244 | NA | NA | NA |
| ENSG000002 | 1.67845334 | 0.082956465 | 0.35652787 | 0.23267877 | 0.81601086 | NA | NA | NA |
| ENSG000002 | 1.70876338 | -0.04410642 | 0.34820032 | -0.1266697 | 0.89920185 | NA | NA | NA |
| ENSG000002 | 1.19166998 | 0.107653642 | 0.32738762 | 0.32882624 | 0.74228703 | NA | SUMO2P10 | 106481678 |
| ENSG000002 | 0.78141628 | -0.193205819 | 0.28064196 | -0.6884424 | 0.49117424 | NA | NA | NA |
| ENSG000002 | 1.63018508 | -0.688074544 | 0.34729216 | -1.9812556 | 0.04756262 | NA | NA | NA |
| ENSG000002 | 2.00275448 | -0.301466165 | 0.3581408 | -0.8417532 | 0.39992611 | NA | NA | NA |
| ENSG000002 | 1.94729918 | 0.069344562 | 0.36170595 | 0.19171529 | 0.84796523 | NA | NA | NA |
| ENSG000002 | 1.5964011 | -0.254568012 | 0.3469998 | -0.7336258 | 0.46317682 | NA | NA | NA |
| ENSG000002 | 0.95127263 | 0.445116954 | 0.30707994 | 1.44951493 | 0.14719383 | NA | NA | NA |
| ENSG000002 | 0 | NA | NA | NA | NA | NA | NA | NA |
| ENSG000002 | 0 | NA | NA | NA | NA | NA | NA | NA |
| ENSG000002 | 1.98276164 | -0.063602986 | 0.36008143 | -0.176635 | 0.8597951 | NA | NA | NA |
| ENSG000002 | 0 | NA | NA | NA | NA | NA | NA | NA |
| ENSG000002 | 0.86637994 | -0.271645032 | 0.29878654 | -0.9091609 | 0.36326522 | NA | NA | NA |
| ENSG000002 | 2.0647685 | -0.130395738 | 0.36468186 | -0.3575603 | 0.72067242 | NA | JARID2-AS1 | 100506681 |
| ENSG000002 | 1.40758408 | -0.405715559 | 0.33301861 | -1.218297 | 0.22311114 | NA | NA | NA |
| ENSG000002 | 0 | NA | NA | NA | NA | NA | NA | NA |
| ENSG000002 | 0 | NA | NA | NA | NA | NA | USP9YP27 | 106480716 |
| ENSG000002 | 1.66857579 | -0.253814557 | 0.34408407 | -0.7376527 | 0.4607255 | NA | NA | NA |
| ENSG000002 | 1.97186656 | -0.96528174 | 0.35367595 | -2.729283 | 0.00634722 | NA | NA | NA |
| ENSG000002 | 0.86091955 | -0.265574426 | 0.29690668 | -0.894471 | 0.37106993 | NA | NA | NA |
| ENSG000002 | 1.02829547 | -0.183417066 | 0.3030352 | -0.6052665 | 0.54500192 | NA | NA | NA |
| ENSG000002 | 0.63025982 | -0.115141786 | 0.26544572 | -0.4337677 | 0.66445712 | NA | NA | NA |
| ENSG000002 | 2.12948583 | -0.263307727 | 0.3679329 | -0.7156406 | 0.47421328 | NA | NA | NA |
| ENSG000002 | 0.97234079 | -0.032345511 | 0.30729297 | -0.1052595 | 0.91616988 | NA | NA | NA |
| ENSG000002 | 1.62579127 | -0.25449284 | 0.34607721 | -0.7353643 | 0.46211763 | NA | NA | NA |
| ENSG000002 | 1.97914197 | 0.062688381 | 0.36415504 | 0.1721475 | 0.86332157 | NA | NA | NA |

| | | | | | | | | |
|---|---|---|---|---|---|---|---|---|
| ENSG000002 | 1.50472023 | -0.058896045 | 0.34403552 | -0.1711918 | 0.86407299 | NA | NA | NA |
| ENSG000002 | 0.63296622 | -0.299126688 | 0.27328461 | -1.0945611 | 0.27370899 | NA | NA | NA |
| ENSG000002 | 1.58259706 | -0.61342477 | 0.33945101 | -1.8071084 | 0.07074538 | NA | NA | NA |
| ENSG000002 | 0 | NA | NA | NA | NA | NA | NA | NA |
| ENSG000002 | 0.3588776 | -0.099902505 | 0.21555794 | -0.4634601 | 0.6430346 | NA | NA | NA |
| ENSG000002 | 0.37422052 | -0.268731395 | 0.21273223 | -1.2632378 | 0.20650373 | NA | NA | NA |
| ENSG000002 | 1.0895213 | 0.039318981 | 0.32118777 | 0.12241743 | 0.90256844 | NA | NA | NA |
| ENSG000002 | 1.04335764 | -0.348999546 | 0.308338 | -1.1318733 | 0.25768771 | NA | NKX1-1 | 54729 |
| ENSG000002 | 0 | NA | NA | NA | NA | NA | NA | NA |
| ENSG000002 | 1.28949613 | -0.678631562 | 0.33112094 | -2.0494976 | 0.04041349 | NA | NA | NA |
| ENSG000002 | 0.75144647 | -0.037105481 | 0.28557336 | -0.1299333 | 0.89661922 | NA | NA | NA |
| ENSG000002 | 0.96605483 | -0.317561686 | 0.300273 | -1.0575766 | 0.29024854 | NA | NA | NA |
| ENSG000002 | 0.49196355 | -0.021608171 | 0.24613638 | -0.0877894 | 0.93004404 | NA | NA | NA |
| ENSG000002 | 0.48609822 | -0.022320526 | 0.23791393 | -0.0938177 | 0.92525401 | NA | NA | NA |
| ENSG000002 | 0 | NA | NA | NA | NA | NA | NA | NA |
| ENSG000002 | 1.23427577 | -0.05310361 | 0.33013745 | -0.160853 | 0.87220916 | NA | NA | NA |
| ENSG000002 | 1.31111612 | 0.333444614 | 0.33856756 | 0.98486877 | 0.32468854 | NA | NA | NA |
| ENSG000002 | 0.36576034 | -0.0934779 | 0.20820828 | -0.4489634 | 0.65345805 | NA | NA | NA |
| ENSG000002 | 0.73701545 | -0.03254472 | 0.28464749 | -0.1143334 | 0.90897349 | NA | NA | NA |
| ENSG000002 | 0.5008998 | -0.200150804 | 0.24832882 | -0.805991 | 0.42024801 | NA | NA | NA |
| ENSG000002 | 0.37945415 | -0.107841678 | 0.21865587 | -0.4932027 | 0.62186934 | NA | NA | NA |
| ENSG000002 | 0 | NA | NA | NA | NA | NA | NA | NA |
| ENSG000002 | 1.64331031 | -0.651608973 | 0.34172304 | -1.9068336 | 0.05654214 | NA | NA | NA |
| ENSG000002 | 0 | NA | NA | NA | NA | NA | NA | NA |
| ENSG000002 | 2.06359656 | 0.000484484 | 0.36725398 | 0.00131921 | 0.99894743 | NA | NA | NA |
| ENSG000002 | 1.93011025 | 0.19886927 | 0.35845824 | 0.55479063 | 0.57903788 | NA | NA | NA |
| ENSG000002 | 1.0978264 | 0.036185051 | 0.3152317 | 0.11478874 | 0.90861257 | NA | DPP10-AS2 | 106478969 |
| ENSG000002 | 0 | NA | NA | NA | NA | NA | NA | NA |
| ENSG000002 | 1.92525464 | -0.797531953 | 0.35364433 | -2.255181 | 0.02412197 | NA | NA | NA |

| | | | | | | | | | |
|---|---|---|---|---|---|---|---|---|---|
| ENSG000002 | 0.62087103 | -0.279525375 | 0.26646004 | -1.049033 | 0.29416293 | NA | NA | NA | |
| ENSG000002 | 1.45447635 | 0.232314644 | 0.3400851 | 0.68310738 | 0.49453899 | NA | NA | NA | |
| ENSG000002 | 1.26370671 | -0.044382136 | 0.32256985 | -0.1375892 | 0.89056507 | NA | NA | NA | |
| ENSG000002 | 0.23202362 | -0.008302493 | 0.17586207 | -0.0472103 | 0.96234565 | NA | NA | NA | |
| ENSG000002 | 1.02832581 | -0.356454241 | 0.31009328 | -1.1495065 | 0.25034718 | NA | NA | NA | |
| ENSG000002 | 1.49034183 | -0.047626511 | 0.34572716 | -0.1377575 | 0.89043207 | NA | NA | NA | |
| ENSG000002 | 0.77265259 | -0.365549345 | 0.28395139 | -1.2873659 | 0.19796678 | NA | NA | NA | |
| ENSG000002 | 0.83426872 | 0.335934032 | 0.28136048 | 1.19396312 | 0.23249239 | NA | NA | NA | |
| ENSG000002 | 0 | NA | NA | NA | NA | NA | USP17L27 | | 728393 |
| ENSG000002 | 1.87985264 | -0.39923971 | 0.36103649 | -1.1058154 | 0.26880643 | NA | NA | NA | |
| ENSG000002 | 0.53374623 | -0.385819457 | 0.24767828 | -1.5577444 | 0.11929385 | NA | NA | NA | |
| ENSG000002 | 1.55999291 | -0.106037007 | 0.34449508 | -0.3078041 | 0.75823138 | NA | NA | NA | |
| ENSG000002 | 0.26062507 | -0.012888319 | 0.18226503 | -0.070712 | 0.94362699 | NA | NA | NA | |
| ENSG000002 | 1.00634864 | -0.189808043 | 0.30944695 | -0.6133783 | 0.53962623 | NA | NA | NA | |
| ENSG000002 | 0.24004852 | -0.005223787 | 0.17775844 | -0.029387 | 0.97655594 | NA | NA | NA | |
| ENSG000002 | 0.35346928 | -0.088570748 | 0.20618147 | -0.4295767 | 0.66750362 | NA | NA | NA | |
| ENSG000002 | 1.72504352 | -0.194451754 | 0.35356283 | -0.5499779 | 0.58233451 | NA | NA | NA | |
| ENSG000002 | 1.50993267 | -0.058640793 | 0.3394253 | -0.1727649 | 0.86283619 | NA | NA | NA | |
| ENSG000002 | 1.97158102 | -0.038581591 | 0.34856807 | -0.110686 | 0.91186538 | NA | NA | NA | |
| ENSG000002 | 0 | NA | NA | NA | NA | NA | NA | NA | |
| ENSG000002 | 1.74468285 | -0.182828029 | 0.35377402 | -0.5167933 | 0.60530048 | NA | NA | NA | |
| ENSG000002 | 1.19400369 | -0.041196486 | 0.3290376 | -0.125203 | 0.90036286 | NA | NA | NA | |
| ENSG000002 | 1.51883455 | -0.33046353 | 0.34090248 | -0.9693785 | 0.33235639 | NA | NA | NA | |
| ENSG000002 | 1.79691687 | -0.096660775 | 0.3481258 | -0.2776605 | 0.781273 | NA | NA | NA | |
| ENSG000002 | 1.97403041 | -0.197599535 | 0.36519168 | -0.5410844 | 0.58844939 | NA | NA | NA | |
| ENSG000002 | 0.13280363 | -0.093394593 | 0.13115644 | -0.7120855 | 0.47641185 | NA | NA | NA | |
| ENSG000002 | 2.1195295 | -0.257631538 | 0.36440695 | -0.7069885 | 0.4795736 | NA | NA | NA | |
| ENSG000002 | 1.57300137 | -0.116759947 | 0.34813811 | -0.335384 | 0.73733545 | NA | NA | NA | |
| ENSG000002 | 0.50451363 | -0.188485218 | 0.24157334 | -0.7802401 | 0.43524954 | NA | NA | NA | |

| | | | | | | | | |
|---|---|---|---|---|---|---|---|---|
| ENSG000002 | 0 | NA | NA | NA | NA | NA | NA | NA |
| ENSG000002 | 0.14128499 | -0.097772014 | 0.13403235 | -0.7294658 | 0.4657168 | NA | NA | NA |
| ENSG000002 | 0 | NA | NA | NA | NA | NA | NA | NA |
| ENSG000002 | 1.4280763 | -0.411202489 | 0.33454863 | -1.2291262 | 0.21902449 | NA | APOA1-AS | 104326055 |
| ENSG000002 | 1.77875721 | -0.726730871 | 0.34814816 | -2.0874184 | 0.03685033 | NA | NA | NA |
| ENSG000002 | 1.37823524 | -0.25558768 | 0.33583709 | -0.7610466 | 0.44662923 | NA | NA | NA |
| ENSG000002 | 0.50613398 | -0.020909936 | 0.24769218 | -0.084419 | 0.93272327 | NA | NA | NA |
| ENSG000002 | 1.06390411 | 0.328405819 | 0.31292926 | 1.04945706 | 0.29396781 | NA | NA | NA |
| ENSG000002 | 0.11934008 | 0.083564698 | 0.12760091 | 0.65489106 | 0.5125379 | NA | NA | NA |
| ENSG000002 | 1.46063398 | 0.207980929 | 0.33577398 | 0.61940753 | 0.53564792 | NA | NA | NA |
| ENSG000002 | 1.99822629 | -0.305786737 | 0.35583608 | -0.8593472 | 0.390149 | NA | LCE6A | 448835 |
| ENSG000002 | 0.49270076 | -0.021329091 | 0.24621941 | -0.0866264 | 0.93096851 | NA | NA | NA |
| ENSG000002 | 1.92715344 | 0.192531266 | 0.36071082 | 0.53375517 | 0.59351093 | NA | NA | NA |
| ENSG000002 | 2.14995475 | -0.508686838 | 0.36480168 | -1.3944202 | 0.16319075 | NA | NA | NA |
| ENSG000002 | 0 | NA | NA | NA | NA | NA | NA | NA |
| ENSG000002 | 0.97848632 | -0.034332936 | 0.30756148 | -0.1116295 | 0.91111718 | NA | NA | NA |
| ENSG000002 | 0.36465257 | 0.064724075 | 0.21605106 | 0.29957767 | 0.76449932 | NA | NA | NA |
| ENSG000002 | 2.02187485 | -0.45119823 | 0.36212154 | -1.2459856 | 0.21276968 | NA | NA | NA |
| ENSG000002 | 0 | NA | NA | NA | NA | NA | CHEK2P4 | 106479021 |
| ENSG000002 | 0.77613058 | -0.361458377 | 0.28422771 | -1.2717211 | 0.20347223 | NA | NA | NA |
| ENSG000002 | 0 | NA | NA | NA | NA | NA | NA | NA |
| ENSG000002 | 1.43737085 | 0.373388061 | 0.34121004 | 1.09430562 | 0.27382096 | NA | NA | NA |
| ENSG000002 | 2.13556457 | 0.554312985 | 0.36443238 | 1.52103109 | 0.12825204 | NA | NA | NA |
| ENSG000002 | 1.35655092 | -0.118540236 | 0.33948808 | -0.3491735 | 0.72695907 | NA | NA | NA |
| ENSG000002 | 0.99963846 | -0.501648105 | 0.30628932 | -1.6378243 | 0.10145835 | NA | NA | NA |
| ENSG000002 | 0.14128499 | -0.097772014 | 0.13403235 | -0.7294658 | 0.4657168 | NA | NA | NA |
| ENSG000002 | 0.37304803 | -0.090842893 | 0.20939682 | -0.4338313 | 0.66441096 | NA | NA | NA |
| ENSG000002 | 1.49854032 | -0.196986069 | 0.34500799 | -0.5709609 | 0.56802615 | NA | NA | NA |
| ENSG000002 | 1.70557632 | -0.193958915 | 0.35509532 | -0.5462165 | 0.58491714 | NA | NA | NA |

| | | | | | | | | |
|---|---|---|---|---|---|---|---|---|
| ENSG000002 | 0.14128499 | -0.097772014 | 0.13403235 | -0.7294658 | 0.4657168 | NA | NA | NA |
| ENSG000002 | 0.96385943 | -0.029850629 | 0.30692108 | -0.0972583 | 0.92252126 | NA | NA | NA |
| ENSG000002 | 0.87392664 | -0.422259382 | 0.29440295 | -1.4342906 | 0.15148935 | NA | NA | NA |
| ENSG000002 | 0 | NA | NA | NA | NA | NA | NA | NA |
| ENSG000002 | 0.74268278 | -0.18879062 | 0.27975142 | -0.6748513 | 0.49977023 | NA | COX6CP18 | 106481684 |
| ENSG000002 | 0.73650691 | -0.025515431 | 0.27831123 | -0.0916795 | 0.92695269 | NA | NA | NA |
| ENSG000002 | 1.47703068 | -0.199271508 | 0.34842826 | -0.5719155 | 0.56737925 | NA | NA | NA |
| ENSG000002 | 1.34643971 | 0.016348827 | 0.32996227 | 0.04954757 | 0.96048293 | NA | NA | NA |
| ENSG000002 | 1.38370827 | -0.256465212 | 0.33357739 | -0.7688327 | 0.44199262 | NA | GSTM5P1 | 100505557 |
| ENSG000002 | 2.06453619 | -0.315793845 | 0.35880172 | -0.8801347 | 0.37878634 | NA | NA | NA |
| ENSG000002 | 2.01124307 | 0.113307812 | 0.34937901 | 0.32431202 | 0.74570182 | NA | NA | NA |
| ENSG000002 | 2.09764359 | -0.215181602 | 0.35700867 | -0.6027349 | 0.54668503 | NA | NA | NA |
| ENSG000002 | 1.93338777 | -0.056256463 | 0.36074208 | -0.1559465 | 0.87607519 | NA | NA | NA |
| ENSG000002 | 1.31345195 | 0.323456269 | 0.33547201 | 0.96418258 | 0.3349544 | NA | NA | NA |
| ENSG000002 | 1.54943768 | 0.014427779 | 0.34547476 | 0.04176218 | 0.96668829 | NA | ELFN1-AS1 | 101927125 |
| ENSG000002 | 0.97476197 | 0.104188468 | 0.30579771 | 0.34071042 | 0.73332159 | NA | NA | NA |
| ENSG000002 | 0 | NA | NA | NA | NA | NA | NA | NA |
| ENSG000002 | 0 | NA | NA | NA | NA | NA | NA | NA |
| ENSG000002 | 1.12984333 | -0.120430152 | 0.31668121 | -0.3802883 | 0.70373143 | NA | NA | NA |
| ENSG000002 | 1.50770737 | -0.184503415 | 0.34204295 | -0.5394159 | 0.58959989 | NA | NA | NA |
| ENSG000002 | 0.94363329 | -0.15212972 | 0.28750794 | -0.5291322 | 0.59671372 | NA | NA | NA |
| ENSG000002 | 1.34399735 | -0.245410528 | 0.33431611 | -0.7340673 | 0.46290771 | NA | NA | NA |
| ENSG000002 | 1.95417381 | -0.542389676 | 0.35586857 | -1.5241292 | 0.12747644 | NA | NA | NA |
| ENSG000002 | 1.75491174 | -0.743723805 | 0.35059808 | -2.1213003 | 0.03389654 | NA | NA | NA |
| ENSG000002 | 1.4658474 | -0.329268927 | 0.34489624 | -0.9546898 | 0.33973458 | NA | NA | NA |
| ENSG000002 | 1.90014742 | -0.502376925 | 0.35532042 | -1.4138701 | 0.15740004 | NA | NA | NA |
| ENSG000002 | 0 | NA | NA | NA | NA | NA | CT47A3 | 728082 |
| ENSG000002 | 0 | NA | NA | NA | NA | NA | NA | NA |
| ENSG000002 | 0.87252547 | -0.269732918 | 0.29750671 | -0.9066448 | 0.36459466 | NA | NA | NA |

| | | | | | | | | |
|---|---|---|---|---|---|---|---|---|
| ENSG000002 | 1.8206285 | -0.237409007 | 0.35221129 | -0.6740528 | 0.50027775 | NA | NA | NA |
| ENSG000002 | 1.03726418 | -0.330778546 | 0.30263006 | -1.0930129 | 0.27438815 | NA | NA | NA |
| ENSG000002 | 1.4658474 | -0.329268927 | 0.34489624 | -0.9546898 | 0.33973458 | NA | NA | NA |
| ENSG000002 | 1.8479186 | -0.23185045 | 0.3469745 | -0.668206 | 0.50400209 | NA | NA | NA |
| ENSG000002 | 0 | NA | NA | NA | NA | NA | CHCHD4P3 | 106480792 |
| ENSG000002 | 2.03572623 | -0.315042899 | 0.36012637 | -0.8748121 | 0.38167617 | NA | NA | NA |
| ENSG000002 | 0.83760627 | 0.05226442 | 0.28780949 | 0.1815938 | 0.85590151 | NA | NA | NA |
| ENSG000002 | 0.61126925 | -0.095064869 | 0.2547532 | -0.3731646 | 0.70902596 | NA | NA | NA |
| ENSG000002 | 2.15650276 | -0.641302364 | 0.36342465 | -1.7646089 | 0.07762952 | NA | NA | NA |
| ENSG000002 | 0.36576034 | -0.0934779 | 0.20820828 | -0.4489634 | 0.65345805 | NA | NA | NA |
| ENSG000002 | 0 | NA | NA | NA | NA | NA | NA | NA |
| ENSG000002 | 2.13534904 | -0.377599729 | 0.36444523 | -1.0360946 | 0.30015801 | NA | NA | NA |
| ENSG000002 | 0.24756242 | -0.187034346 | 0.18119817 | -1.0322088 | 0.30197431 | NA | NA | NA |
| ENSG000002 | 2.12538274 | -0.251777103 | 0.36478106 | -0.6902143 | 0.49005944 | NA | HNRNPDLP3 | 106481662 |
| ENSG000002 | 1.63729808 | -0.531027139 | 0.34711692 | -1.5298221 | 0.12606077 | NA | NA | NA |
| ENSG000002 | 2.04310429 | -0.449910371 | 0.36189736 | -1.2431988 | 0.21379458 | NA | NA | NA |
| ENSG000002 | 1.23262563 | -0.04206857 | 0.32628719 | -0.1289311 | 0.89741216 | NA | NA | NA |
| ENSG000002 | 2.16404086 | -0.493379268 | 0.36215554 | -1.3623408 | 0.17309035 | NA | NA | NA |
| ENSG000002 | 1.94793285 | -0.67991695 | 0.35793594 | -1.8995493 | 0.0574923 | NA | NA | NA |
| ENSG000002 | 1.46811404 | -0.051173118 | 0.347972 | -0.147061 | 0.88308386 | NA | NA | NA |
| ENSG000002 | 2.15187049 | -0.762260856 | 0.36192596 | -2.1061238 | 0.0351936 | NA | NA | NA |
| ENSG000002 | 1.90480359 | 0.323875026 | 0.35983741 | 0.90005935 | 0.36808867 | NA | NA | NA |
| ENSG000002 | 1.7746057 | -0.892127753 | 0.34851263 | -2.5598147 | 0.0104728 | NA | NA | NA |
| ENSG000002 | 1.09899889 | -0.106208641 | 0.31560672 | -0.3365221 | 0.73647717 | NA | NA | NA |
| ENSG000002 | 1.69644395 | -0.54450203 | 0.34799612 | -1.5646784 | 0.11765833 | NA | LOC101928381 | 101928381 |
| ENSG000002 | 0.11934008 | 0.083564698 | 0.12760091 | 0.65489106 | 0.5125379 | NA | NA | NA |
| ENSG000002 | 1.39898751 | -0.241051542 | 0.32306223 | -0.7461459 | 0.45557931 | NA | NA | NA |
| ENSG000002 | 1.89860829 | -0.483543904 | 0.3512269 | -1.376728 | 0.16859636 | NA | NA | NA |
| ENSG000002 | 0.87989748 | -0.286905324 | 0.30426178 | -0.9429555 | 0.34570366 | NA | NA | NA |

| | | | | | | | | |
|---|---|---|---|---|---|---|---|---|
| ENSG000002 | 0.23139247 | 0.167748926 | 0.17680352 | 0.94878726 | 0.34272882 | NA | NA | NA |
| ENSG000002 | 0.72788567 | -0.324716073 | 0.27260818 | -1.1911458 | 0.23359637 | NA | NA | NA |
| ENSG000002 | 0.62889593 | -0.276614073 | 0.26719437 | -1.0352541 | 0.30055024 | NA | NA | NA |
| ENSG000002 | 1.04087801 | -0.355695264 | 0.3106182 | -1.1451205 | 0.25215924 | NA | NA | NA |
| ENSG000002 | 2.10363784 | -0.133822085 | 0.36094628 | -0.3707535 | 0.71082116 | NA | NA | NA |
| ENSG000002 | 1.65682343 | -0.151402183 | 0.33822156 | -0.447642 | 0.65441157 | NA | NA | NA |
| ENSG000002 | 0.91049845 | -0.644235083 | 0.3003423 | -2.1450028 | 0.03195262 | NA | NA | NA |
| ENSG000002 | 2.01678502 | -0.567474257 | 0.35966515 | -1.5777849 | 0.11461502 | NA | NA | NA |
| ENSG000002 | 0.59943712 | 0.024381757 | 0.15730283 | 0.15499884 | 0.87682223 | NA | NA | NA |
| ENSG000002 | 1.88708422 | -0.686933325 | 0.35982283 | -1.9090876 | 0.05625079 | NA | NA | NA |
| ENSG000002 | 1.39150292 | -0.406713696 | 0.33644987 | -1.2088389 | 0.22672474 | NA | NA | NA |
| ENSG000002 | 1.35386379 | 0.016379895 | 0.33579817 | 0.04877899 | 0.96109543 | NA | NA | NA |
| ENSG000002 | 2.15177958 | -0.663942342 | 0.36625046 | -1.8128096 | 0.0698612 | NA | NA | NA |
| ENSG000002 | 0.84069466 | 0.360628166 | 0.29080316 | 1.2401109 | 0.21493438 | NA | NA | NA |
| ENSG000002 | 0.37692244 | -0.098341116 | 0.20997214 | -0.4683532 | 0.63953205 | NA | NA | NA |
| ENSG000002 | 0 | NA | NA | NA | NA | NA | CT47A1 | 728096 |
| ENSG000002 | 0 | NA | NA | NA | NA | NA | NA | NA |
| ENSG000002 | 0.88855353 | -0.453244102 | 0.30180501 | -1.5017779 | 0.13315447 | NA | NA | NA |
| ENSG000002 | 0.37304803 | -0.090842893 | 0.20939682 | -0.4338313 | 0.66441096 | NA | NA | NA |
| ENSG000002 | 1.53819623 | -0.481607645 | 0.3437391 | -1.4010849 | 0.1611887 | NA | NA | NA |
| ENSG000002 | 0.77248003 | -0.035615716 | 0.27170725 | -0.1310812 | 0.89571106 | NA | ITCH-AS1 | 106480737 |
| ENSG000002 | 0.61656602 | -0.102985032 | 0.25681525 | -0.4010082 | 0.68841406 | NA | NA | NA |
| ENSG000002 | 0.37899768 | -0.10176489 | 0.21865765 | -0.4654074 | 0.64163973 | NA | NA | NA |
| ENSG000002 | 2.03279324 | 0.621171104 | 0.35604855 | 1.74462471 | 0.08105022 | NA | TAS2R39 | 259285 |
| ENSG000002 | 0.99255069 | -0.182637364 | 0.30722096 | -0.5944821 | 0.5521897 | NA | NA | NA |
| ENSG000002 | 0.6974565 | 0.289376825 | 0.27577059 | 1.04933896 | 0.29402214 | NA | NA | NA |
| ENSG000002 | 0.88106997 | -0.429369843 | 0.29470046 | -1.4569704 | 0.14512455 | NA | NA | NA |
| ENSG000002 | 0 | NA | NA | NA | NA | NA | TSPY10 | 100289087 |
| ENSG000002 | 0.85912506 | -0.250823615 | 0.29092846 | -0.8621488 | 0.38860566 | NA | NA | NA |

| | | | | | | | | |
|---|---|---|---|---|---|---|---|---|
| ENSG000002 | 0 | NA | NA | NA | NA | NA | GPM6BP2 | 106480426 |
| ENSG000002 | 1.73560225 | -0.061126401 | 0.3581692 | -0.1706635 | 0.86448838 | NA | NA | NA |
| ENSG000002 | 0.63073857 | 0.056382307 | 0.26514218 | 0.21264933 | 0.83160049 | NA | NA | NA |
| ENSG000002 | 0 | NA | NA | NA | NA | NA | NA | NA |
| ENSG000002 | 1.59995233 | -0.12002124 | 0.3512179 | -0.3417287 | 0.73255506 | NA | NA | NA |
| ENSG000002 | 1.85041502 | 0.006762852 | 0.357803 | 0.01890105 | 0.98492004 | NA | NA | NA |
| ENSG000002 | 0.55665861 | -0.363599063 | 0.24151673 | -1.5054819 | 0.13220023 | NA | NA | NA |
| ENSG000002 | 1.857925 | 0.132731539 | 0.35788758 | 0.37087495 | 0.71073067 | NA | NA | NA |
| ENSG000002 | 0 | NA | NA | NA | NA | NA | NA | NA |
| ENSG000002 | 0.60264758 | 0.238025744 | 0.26986178 | 0.88202838 | 0.37776146 | NA | NA | NA |
| ENSG000002 | 1.63450343 | -0.122873475 | 0.35051497 | -0.3505513 | 0.72592501 | NA | NA | NA |
| ENSG000002 | 0.96630097 | 0.117434284 | 0.30291998 | 0.38767428 | 0.69825709 | NA | NA | NA |
| ENSG000002 | 0 | NA | NA | NA | NA | NA | NA | NA |
| ENSG000002 | 2.03297169 | -0.457564366 | 0.36217732 | -1.2633711 | 0.20645584 | NA | NA | NA |
| ENSG000002 | 2.10318797 | -0.254456465 | 0.36668504 | -0.6939374 | 0.48772146 | NA | LOC101928067 | 101928067 |
| ENSG000002 | 2.02503324 | -0.335071951 | 0.36377388 | -0.9210995 | 0.35699847 | NA | NA | NA |
| ENSG000002 | 0.90844494 | -0.459619446 | 0.30273017 | -1.5182479 | 0.12895191 | NA | NA | NA |
| ENSG000002 | 1.5420963 | -0.653482357 | 0.34571531 | -1.8902326 | 0.05872685 | NA | NA | NA |
| ENSG000002 | 1.81236475 | -0.118887896 | 0.35927888 | -0.330907 | 0.74071473 | NA | NA | NA |
| ENSG000002 | 1.59001616 | -0.122868465 | 0.3499316 | -0.3511214 | 0.72549729 | NA | NA | NA |
| ENSG000002 | 1.65339884 | -0.504774599 | 0.34164126 | -1.4774989 | 0.13954194 | NA | NA | NA |
| ENSG000002 | 0.26062507 | -0.012888319 | 0.18226503 | -0.070712 | 0.94362699 | NA | NA | NA |
| ENSG000002 | 2.10483261 | -0.131029424 | 0.36642798 | -0.3575858 | 0.72065334 | NA | NA | NA |
| ENSG000002 | 1.69113279 | 0.340696506 | 0.35076092 | 0.9713069 | 0.33139547 | NA | NA | NA |
| ENSG000002 | 1.59798762 | -0.246635287 | 0.34866714 | -0.707366 | 0.47933904 | NA | NA | NA |
| ENSG000002 | 1.34320752 | -0.256480577 | 0.33506417 | -0.765467 | 0.44399349 | NA | NA | NA |
| ENSG000002 | 1.78590109 | -0.750255448 | 0.35164877 | -2.1335364 | 0.03288075 | NA | NA | NA |
| ENSG000002 | 1.87646558 | -0.505414502 | 0.35571715 | -1.4208325 | 0.15536545 | NA | NA | NA |
| ENSG000002 | 1.49869327 | -0.490906417 | 0.34501183 | -1.4228684 | 0.15477431 | NA | NA | NA |

| | | | | | | | | |
|---|---|---|---|---|---|---|---|---|
| ENSG000002 | 1.60737587 | -0.129269249 | 0.35131807 | -0.367955 | 0.71290677 | NA | NA | NA |
| ENSG000002 | 0.37422052 | -0.268731395 | 0.21273223 | -1.2632378 | 0.20650373 | NA | NA | NA |
| ENSG000002 | 0.34931019 | 0.26109409 | 0.21538338 | 1.21222954 | 0.22542453 | NA | NA | NA |
| ENSG000002 | 0.39479706 | -0.300008631 | 0.22317059 | -1.3443018 | 0.17885081 | NA | NA | NA |
| ENSG000002 | 1.09204952 | -0.23168629 | 0.30913013 | -0.7494782 | 0.45356904 | NA | NA | NA |
| ENSG000002 | 0.65405377 | -0.105271043 | 0.2601228 | -0.4046975 | 0.6856999 | NA | NA | NA |
| ENSG000002 | 2.11830412 | -0.255716579 | 0.36630069 | -0.6981056 | 0.48511113 | NA | MTATP6P4 | 106480797 |
| ENSG000002 | 1.19964226 | 0.09505642 | 0.32069778 | 0.29640498 | 0.76692082 | NA | NA | NA |
| ENSG000002 | 0.49696748 | -0.016894442 | 0.23921338 | -0.070625 | 0.94369622 | NA | NA | NA |
| ENSG000002 | 1.9191248 | -0.524455889 | 0.35721865 | -1.4681649 | 0.14205943 | NA | NA | NA |
| ENSG000002 | 0 | NA | NA | NA | NA | NA | NA | NA |
| ENSG000002 | 0 | NA | NA | NA | NA | NA | NA | NA |
| ENSG000002 | 0.47872676 | -0.008834112 | 0.22824852 | -0.0387039 | 0.96912645 | NA | NA | NA |
| ENSG000002 | 2.03021578 | -0.734325989 | 0.3614618 | -2.0315452 | 0.04219972 | NA | NA | NA |
| ENSG000002 | 0.74661723 | -0.178670198 | 0.27282214 | -0.6548963 | 0.51253455 | NA | NA | NA |
| ENSG000002 | 0.70479626 | 0.296063265 | 0.27790032 | 1.06535775 | 0.28671407 | NA | NA | NA |
| ENSG000002 | 0 | NA | NA | NA | NA | NA | NA | NA |
| ENSG000002 | 0.76696407 | -0.358999402 | 0.28364792 | -1.2656514 | 0.2056379 | NA | NA | NA |
| ENSG000002 | 0 | NA | NA | NA | NA | NA | NA | NA |
| ENSG000002 | 1.79281178 | 0.265754159 | 0.35086635 | 0.75742277 | 0.44879662 | NA | NA | NA |
| ENSG000002 | 0 | NA | NA | NA | NA | NA | NA | NA |
| ENSG000002 | 1.9845851 | -0.315996189 | 0.36182408 | -0.8733421 | 0.38247666 | NA | LAMTOR5P1 | 106478910 |
| ENSG000002 | 0 | NA | NA | NA | NA | NA | NA | NA |
| ENSG000002 | 0.26560726 | -0.177019426 | 0.17672274 | -1.0016788 | 0.31649875 | NA | NA | NA |
| ENSG000002 | 0.26199343 | -0.194018184 | 0.18422801 | -1.0531416 | 0.29227609 | NA | FRGCA | 106481742 |
| ENSG000002 | 1.36673449 | -0.118867572 | 0.33633783 | -0.3534172 | 0.7237757 | NA | NA | NA |
| ENSG000002 | 0.12685397 | -0.090215768 | 0.12901919 | -0.699243 | 0.48440017 | NA | NA | NA |
| ENSG000002 | 1.62195124 | -0.12924716 | 0.3547583 | -0.3643246 | 0.71561566 | NA | NA | NA |
| ENSG000002 | 1.01663556 | -0.327349972 | 0.30258504 | -1.0818446 | 0.27932161 | NA | NA | NA |

| | | | | | | | | |
|---|---|---|---|---|---|---|---|---|
| ENSG000002 | 0 | NA | NA | NA | NA | NA | NA | NA |
| ENSG000002 | 1.44728955 | -0.041781897 | 0.34050399 | -0.122706 | 0.90233988 | NA | NA | NA |
| ENSG000002 | 2.12732627 | -0.259152278 | 0.36659163 | -0.7069236 | 0.47961396 | NA | NA | NA |
| ENSG000002 | 1.92553092 | -0.539210236 | 0.35875056 | -1.5030227 | 0.13283319 | NA | NA | NA |
| ENSG000002 | 0 | NA | NA | NA | NA | NA | NA | NA |
| ENSG000002 | 1.77556401 | -0.450290854 | 0.35186739 | -1.2797175 | 0.20064451 | NA | NA | NA |
| ENSG000002 | 1.24893993 | -0.319145424 | 0.32405324 | -0.9848549 | 0.32469535 | NA | NA | NA |
| ENSG000002 | 1.86817754 | -0.125515452 | 0.35814737 | -0.3504576 | 0.72599534 | NA | NA | NA |
| ENSG000002 | 0.99106886 | -0.189588774 | 0.30880259 | -0.6139481 | 0.5392496 | NA | NA | NA |
| ENSG000002 | 0 | NA | NA | NA | NA | NA | GAGE12B | 729428 |
| ENSG000002 | 0.74575582 | -0.19939939 | 0.28601262 | -0.6971699 | 0.48569647 | NA | NA | NA |
| ENSG000002 | 0.49856397 | -0.186262775 | 0.24090455 | -0.7731808 | 0.43941538 | NA | NA | NA |
| ENSG000002 | 1.64372146 | -0.132126386 | 0.34997365 | -0.3775324 | 0.705778 | NA | NA | NA |
| ENSG000002 | 0.63370343 | -0.298851213 | 0.27334848 | -1.0932975 | 0.2742632 | NA | NA | NA |
| ENSG000002 | 2.04438173 | -0.455259152 | 0.36132585 | -1.2599684 | 0.20768076 | NA | NA | NA |
| ENSG000002 | 0.99212614 | -0.190783629 | 0.30884287 | -0.6177369 | 0.53674881 | NA | NA | NA |
| ENSG000002 | 0.48962772 | -0.021045505 | 0.24589998 | -0.0855856 | 0.93179582 | NA | NA | NA |
| ENSG000002 | 0 | NA | NA | NA | NA | NA | NA | NA |
| ENSG000002 | 2.1689026 | 0.0561058 | 0.36758718 | 0.15263264 | 0.87868799 | NA | NA | NA |
| ENSG000002 | 2.11259849 | -0.004253135 | 0.36705178 | -0.0115873 | 0.99075489 | NA | LOC102724589 | 102724589 |
| ENSG000002 | 1.82124441 | -0.455616739 | 0.35123545 | -1.2971832 | 0.19456815 | NA | NA | NA |
| ENSG000002 | 1.356549 | -0.26734178 | 0.34004101 | -0.7862045 | 0.43174767 | NA | NA | NA |
| ENSG000002 | 0.12685397 | -0.090215768 | 0.12901919 | -0.699243 | 0.48440017 | NA | NA | NA |
| ENSG000002 | 1.71135184 | -0.059116984 | 0.35792836 | -0.1651643 | 0.86881467 | NA | NA | NA |
| ENSG000002 | 1.5694936 | 0.018099144 | 0.33909192 | 0.05337533 | 0.95743286 | NA | NA | NA |
| ENSG000002 | 1.58734752 | -0.11887256 | 0.34274493 | -0.3468251 | 0.72872269 | NA | NA | NA |
| ENSG000002 | 1.83934988 | 0.009266845 | 0.35557598 | 0.02606151 | 0.97920828 | NA | NA | NA |
| ENSG000002 | 0.88286 | -0.118727439 | 0.2925247 | -0.4058715 | 0.68483702 | NA | NA | NA |
| ENSG000002 | 1.01110407 | -0.204132122 | 0.31431391 | -0.649453 | 0.51604559 | NA | NA | NA |

| | | | | | | | | |
|---|---|---|---|---|---|---|---|---|
| ENSG000002 | 1.84649768 | -0.235772493 | 0.354526 | -0.6650358 | 0.50602758 | NA | NA | NA |
| ENSG000002 | 2.14844808 | -0.519661865 | 0.3658051 | -1.4205977 | 0.15543376 | NA | LINC00421 | 100287114 |
| ENSG000002 | 0.73701545 | -0.03254472 | 0.28464749 | -0.1143334 | 0.90897349 | NA | PCYT1B-AS1 | 100874088 |
| ENSG000002 | 0.38747904 | -0.104908247 | 0.21987514 | -0.4771265 | 0.63327207 | NA | NA | NA |
| ENSG000002 | 1.83732617 | -0.002123582 | 0.35831437 | -0.0059266 | 0.99527129 | NA | NA | NA |
| ENSG000002 | 1.38398902 | -0.241769191 | 0.32952078 | -0.7336994 | 0.46313198 | NA | RPL23AP95 | 106480338 |
| ENSG000002 | 2.15510057 | -0.639744468 | 0.36342463 | -1.7603223 | 0.07835318 | NA | NA | NA |
| ENSG000002 | 1.11811871 | -0.270222481 | 0.32173496 | -0.8398916 | 0.40096918 | NA | NA | NA |
| ENSG000002 | 1.1566925 | -0.602268935 | 0.32217505 | -1.8693841 | 0.06156939 | NA | NA | NA |
| ENSG000002 | 1.46971109 | -0.172089588 | 0.33762331 | -0.5097088 | 0.61025545 | NA | NA | NA |
| ENSG000002 | 1.6299544 | -0.272785719 | 0.35423406 | -0.770072 | 0.4412572 | NA | NA | NA |
| ENSG000002 | 0.96619526 | -0.027997535 | 0.29940688 | -0.09351 | 0.92549841 | NA | NA | NA |
| ENSG000002 | 0.50750235 | -0.197774682 | 0.24907427 | -0.794039 | 0.42717272 | NA | NA | NA |
| ENSG000002 | 2.028143 | -0.577012913 | 0.36073045 | -1.5995681 | 0.10969444 | NA | NA | NA |
| ENSG000002 | 1.35862424 | -0.263106077 | 0.34009514 | -0.7736249 | 0.43915263 | NA | NA | NA |
| ENSG000002 | 1.16764553 | -0.574676996 | 0.31593997 | -1.8189437 | 0.06892003 | NA | SETD6P1 | 106481727 |
| ENSG000002 | 1.46462284 | -0.168741843 | 0.33591311 | -0.5023378 | 0.61542995 | NA | NA | NA |
| ENSG000002 | 1.26375686 | -0.1922786 | 0.32993646 | -0.5827746 | 0.56004502 | NA | NA | NA |
| ENSG000002 | 1.01424417 | -0.667476388 | 0.30336331 | -2.2002542 | 0.02778887 | NA | NA | NA |
| ENSG000002 | 0.37404583 | -0.105111695 | 0.21784993 | -0.4824959 | 0.62945372 | NA | NA | NA |
| ENSG000002 | 1.9931875 | -0.577036047 | 0.36144158 | -1.596485 | 0.11038056 | NA | NA | NA |
| ENSG000002 | 0.74575582 | -0.19939939 | 0.28601262 | -0.6971699 | 0.48569647 | NA | NA | NA |
| ENSG000002 | 0.88052864 | -0.456101693 | 0.30132726 | -1.5136423 | 0.13011659 | NA | NA | NA |
| ENSG000002 | 0.8678023 | -0.271126522 | 0.29886542 | -0.907186 | 0.36430845 | NA | NA | NA |
| ENSG000002 | 1.40063662 | -0.42922408 | 0.34045438 | -1.2607389 | 0.20740293 | NA | NA | NA |
| ENSG000002 | 0.71575568 | 0.130993046 | 0.27711751 | 0.47269855 | 0.63642826 | NA | NA | NA |
| ENSG000002 | 2.0219864 | -0.431995646 | 0.3591215 | -1.2029234 | 0.22900598 | NA | NA | NA |
| ENSG000002 | 0.63250975 | -0.278024813 | 0.26750458 | -1.0393273 | 0.29865254 | NA | NA | NA |
| ENSG000002 | 1.22920768 | -0.045354498 | 0.33659976 | -0.1347431 | 0.89281499 | NA | NA | NA |

| | | | | | | | | |
|---|---|---|---|---|---|---|---|---|
| ENSG000002 | 1.41205114 | -0.694833161 | 0.33055349 | -2.1020294 | 0.0355507 | NA | NA | NA |
| ENSG000002 | 1.40584907 | -0.42385874 | 0.33881128 | -1.2510172 | 0.2109282 | NA | BAALC-AS2 | 157556 |
| ENSG000002 | 1.83772059 | 0.138229028 | 0.35830505 | 0.38578588 | 0.69965526 | NA | NA | NA |
| ENSG000002 | 0 | NA | NA | NA | NA | NA | LOC101929148 | 101929148 |
| ENSG000002 | 1.22237892 | 0.105958062 | 0.3298514 | 0.32122969 | 0.74803634 | NA | NA | NA |
| ENSG000002 | 0.99070221 | -0.346865624 | 0.31020477 | -1.1181827 | 0.26348895 | NA | NA | NA |
| ENSG000002 | 0.2596576 | -0.193002035 | 0.1837913 | -1.0501152 | 0.29366517 | NA | NA | NA |
| ENSG000002 | 1.96606397 | -0.307251552 | 0.36075704 | -0.8516855 | 0.39438865 | NA | NA | NA |
| ENSG000002 | 1.01201597 | -0.366610227 | 0.31542515 | -1.1622733 | 0.24512447 | NA | NA | NA |
| ENSG000002 | 1.03393055 | -0.34903534 | 0.31038672 | -1.1245176 | 0.26079349 | NA | NA | NA |
| ENSG000002 | 0.12685397 | -0.090215768 | 0.12901919 | -0.699243 | 0.48440017 | NA | NA | NA |
| ENSG000002 | 1.38532906 | -0.679925452 | 0.3286103 | -2.0690936 | 0.03853731 | NA | NA | NA |
| ENSG000002 | 0.75941929 | -0.032774607 | 0.27987107 | -0.1171061 | 0.90677598 | NA | NA | NA |
| ENSG000002 | 1.00359072 | -0.033603177 | 0.30863071 | -0.1088783 | 0.91329904 | NA | NA | NA |
| ENSG000002 | 0.86339122 | -0.275317148 | 0.2986111 | -0.9219924 | 0.35653256 | NA | NA | NA |
| ENSG000002 | 0 | NA | NA | NA | NA | NA | NA | NA |
| ENSG000002 | 0 | NA | NA | NA | NA | NA | NA | NA |
| ENSG000002 | 1.08609539 | 0.033698977 | 0.3147952 | 0.10705048 | 0.91474893 | NA | NA | NA |
| ENSG000002 | 2.0163474 | -0.066508681 | 0.36262248 | -0.1834103 | 0.85447613 | NA | USP17L8 | 392188 |
| ENSG000002 | 0.9554857 | 0.271521574 | 0.30572459 | 0.88812476 | 0.37447365 | NA | DPH3P2 | 106478926 |
| ENSG000002 | 0 | NA | NA | NA | NA | NA | TTTY12 | 83867 |
| ENSG000002 | 1.43845881 | 0.23331122 | 0.34368678 | 0.67884839 | 0.49723393 | NA | NA | NA |
| ENSG000002 | 1.47153592 | -0.335444897 | 0.34521383 | -0.9717018 | 0.33119892 | NA | NA | NA |
| ENSG000002 | 1.95730187 | -0.310070135 | 0.36069466 | -0.8596471 | 0.38998358 | NA | NA | NA |
| ENSG000002 | 1.95899018 | 0.071021565 | 0.35968897 | 0.19745272 | 0.84347328 | NA | NA | NA |
| ENSG000002 | 1.23740672 | -0.195332072 | 0.33287139 | -0.5868094 | 0.5573317 | NA | NA | NA |
| ENSG000002 | 0.59685088 | -0.09712515 | 0.25497858 | -0.3809149 | 0.70326637 | NA | NA | NA |
| ENSG000002 | 0.25145857 | -0.010118565 | 0.18031426 | -0.0561163 | 0.95524918 | NA | NA | NA |
| ENSG000002 | 1.61873831 | -0.406531264 | 0.35145983 | -1.1566934 | 0.24739765 | NA | NA | NA |

| | | | | | | | | | |
|---|---|---|---|---|---|---|---|---|---|
| ENSG000002 | 1.03445015 | -0.511543607 | 0.30773863 | -1.6622665 | 0.09645936 | NA | NA | NA | |
| ENSG000002 | 0.12070844 | -0.088503695 | 0.12783931 | -0.6923042 | 0.48874631 | NA | NA | NA | |
| ENSG000002 | 0 | NA | NA | NA | NA | NA | NA | NA | |
| ENSG000002 | 0.50818749 | -0.197539377 | 0.24915052 | -0.7928516 | 0.42786432 | NA | NA | NA | |
| ENSG000002 | 1.83384868 | -0.4337375 | 0.34687501 | -1.2504144 | 0.21114821 | NA | NA | NA | |
| ENSG000002 | 1.9643983 | 0.067199945 | 0.36027248 | 0.18652534 | 0.85203281 | NA | NA | NA | |
| ENSG000002 | 0.2596576 | -0.193002035 | 0.1837913 | -1.0501152 | 0.29366517 | NA | NA | NA | |
| ENSG000002 | 1.27423966 | -0.357749603 | 0.33450052 | -1.0695039 | 0.28484269 | NA | NA | NA | |
| ENSG000002 | 1.38176475 | -0.25494589 | 0.33353562 | -0.7643738 | 0.44464451 | NA | NA | NA | |
| ENSG000002 | 1.11037805 | 0.027858661 | 0.31736906 | 0.08778002 | 0.93005152 | NA | NA | NA | |
| ENSG000002 | 1.60888858 | -0.268211976 | 0.35398483 | -0.7576934 | 0.44863453 | NA | NA | NA | |
| ENSG000002 | 1.15290006 | -0.122560958 | 0.32203872 | -0.3805783 | 0.70351617 | NA | NA | NA | |
| ENSG000002 | 1.63592971 | -0.389398551 | 0.34788578 | -1.1193287 | 0.26299992 | NA | NA | NA | |
| ENSG000002 | 1.02523158 | -0.498388862 | 0.30490411 | -1.6345757 | 0.10213801 | NA | NA | NA | |
| ENSG000002 | 1.91566551 | -0.635930158 | 0.35175708 | -1.8078674 | 0.07062714 | NA | NA | NA | |
| ENSG000002 | 1.9263585 | -0.690802621 | 0.36014523 | -1.9181224 | 0.05509549 | NA | NA | NA | |
| ENSG000002 | 1.30970211 | -0.510858528 | 0.33117216 | -1.5425769 | 0.12293346 | NA | NA | NA | |
| ENSG000002 | 1.87207974 | -0.118515906 | 0.35461807 | -0.3342072 | 0.73822323 | NA | NA | NA | |
| ENSG000002 | 1.84134301 | -0.120101575 | 0.35114856 | -0.342025 | 0.7323321 | NA | NA | NA | |
| ENSG000002 | 1.24941366 | 0.091608253 | 0.32901705 | 0.27843011 | 0.78068221 | NA | NA | NA | |
| ENSG000002 | 0.53627793 | -0.360851627 | 0.24080453 | -1.4985251 | 0.13399688 | NA | NA | NA | |
| ENSG000002 | 0.63640983 | -0.455149466 | 0.26433333 | -1.7218769 | 0.08509182 | NA | NA | NA | |
| ENSG000002 | 0 | NA | NA | NA | NA | NA | NA | NA | |
| ENSG000002 | 0.34202251 | 0.257871122 | 0.21422266 | 1.20375278 | 0.22868515 | NA | KRTAP9-10P | | 106480422 |
| ENSG000002 | 1.91868973 | -0.37356457 | 0.35439814 | -1.0540816 | 0.29184554 | NA | NA | NA | |
| ENSG000002 | 1.40425492 | -0.726984337 | 0.33538482 | -2.1676125 | 0.03018818 | NA | NA | NA | |
| ENSG000002 | 1.24381284 | -0.204269206 | 0.33433463 | -0.6109723 | 0.54121791 | NA | NA | NA | |
| ENSG000002 | 0 | NA | NA | NA | NA | NA | NA | NA | |
| ENSG000002 | 0 | NA | NA | NA | NA | NA | NA | NA | |

| | | | | | | | | |
|---|---|---|---|---|---|---|---|---|
| ENSG000002 | 0.33394554 | 0.210095954 | 0.19490774 | 1.07792513 | 0.28106717 | NA | NA | NA |
| ENSG000002 | 1.57881114 | -0.376241677 | 0.34376593 | -1.0944705 | 0.27374867 | NA | NA | NA |
| ENSG000002 | 1.74562666 | -0.323374206 | 0.35435049 | -0.9125829 | 0.36146195 | NA | NA | NA |
| ENSG000002 | 1.10133417 | -0.109792096 | 0.3173723 | -0.345941 | 0.72938704 | NA | NA | NA |
| ENSG000002 | 0 | NA | NA | NA | NA | NA | NA | NA |
| ENSG000002 | 1.7184693 | -0.183012439 | 0.35460767 | -0.5160984 | 0.60578571 | NA | NA | NA |
| ENSG000002 | 1.60088734 | 0.017467333 | 0.34598189 | 0.05048627 | 0.95973489 | NA | NA | NA |
| ENSG000002 | 0.36827086 | -0.266056975 | 0.21181163 | -1.2561018 | 0.20907905 | NA | NA | NA |
| ENSG000002 | 1.38922712 | -0.390193442 | 0.33347541 | -1.1700816 | 0.24196814 | NA | NA | NA |
| ENSG000002 | 0.11131518 | 0.081951103 | 0.12663804 | 0.64712866 | 0.51754868 | NA | MBD3L5 | 284428 |
| ENSG000002 | 1.80682003 | -0.104888558 | 0.3527109 | -0.2973783 | 0.76617773 | NA | NA | NA |
| ENSG000002 | 1.61379697 | -0.485299164 | 0.33671325 | -1.4412833 | 0.14950468 | NA | NA | NA |
| ENSG000002 | 2.14340585 | -0.401293716 | 0.36788858 | -1.0908023 | 0.27535987 | NA | NA | NA |
| ENSG000002 | 1.24739441 | -0.185637638 | 0.32525832 | -0.5707391 | 0.56817651 | NA | NA | NA |
| ENSG000002 | 0 | NA | NA | NA | NA | NA | NA | NA |
| ENSG000002 | 0 | NA | NA | NA | NA | NA | NA | NA |
| ENSG000002 | 1.00501253 | -0.042485437 | 0.30871955 | -0.1376182 | 0.89054215 | NA | NA | NA |
| ENSG000002 | 0.61221498 | -0.115173106 | 0.27027142 | -0.4261387 | 0.6700068 | NA | NA | NA |
| ENSG000002 | 0 | NA | NA | NA | NA | NA | NA | NA |
| ENSG000002 | 1.82597368 | -0.116198025 | 0.35101236 | -0.3310369 | 0.74061665 | NA | NA | NA |
| ENSG000002 | 1.6977476 | -0.619550798 | 0.33653816 | -1.8409526 | 0.06562851 | NA | NA | NA |
| ENSG000002 | 1.54101026 | -0.341815556 | 0.34565856 | -0.9888821 | 0.32272084 | NA | NA | NA |
| ENSG000002 | 1.728112 | 0.312880092 | 0.34278253 | 0.91276558 | 0.36136584 | NA | NA | NA |
| ENSG000002 | 2.09974524 | -0.099662345 | 0.3540817 | -0.2814671 | 0.77835217 | NA | NA | NA |
| ENSG000002 | 1.44725675 | -0.048410432 | 0.34375798 | -0.1408271 | 0.88800655 | NA | RPL7L1P10 | 106480729 |
| ENSG000002 | 0.25260017 | -0.015978705 | 0.18049464 | -0.0885273 | 0.92945758 | NA | NA | NA |
| ENSG000002 | 2.15379691 | -0.490333799 | 0.36043705 | -1.3603868 | 0.17370756 | NA | NA | NA |
| ENSG000002 | 1.80769309 | -0.118010582 | 0.35436549 | -0.3330194 | 0.73911964 | NA | NA | NA |
| ENSG000002 | 1.91101558 | -0.536630242 | 0.35921888 | -1.4938809 | 0.13520674 | NA | SYP-AS1 | 100873921 |

| | | | | | | | | |
|---|---|---|---|---|---|---|---|---|
| ENSG000002 | 1.99455936 | -0.427816272 | 0.35849411 | -1.1933704 | 0.23272432 | NA | NA | NA |
| ENSG000002 | 1.88070843 | -0.788644485 | 0.35322685 | -2.2326856 | 0.02556969 | NA | NA | NA |
| ENSG000002 | 2.0382745 | -0.560548354 | 0.35674047 | -1.5713057 | 0.11611165 | NA | NA | NA |
| ENSG000002 | 0.72353016 | -0.033011912 | 0.2837491 | -0.1163419 | 0.90738157 | NA | NA | NA |
| ENSG000002 | 1.82560154 | -0.126045748 | 0.35939567 | -0.3507158 | 0.72580155 | NA | NA | NA |
| ENSG000002 | 1.88522713 | -0.510153742 | 0.35696336 | -1.4291487 | 0.15296149 | NA | NA | NA |
| ENSG000002 | 1.10047488 | 0.040290995 | 0.32155197 | 0.12530166 | 0.90028474 | NA | NA | NA |
| ENSG000002 | 0.3863157 | -0.274134926 | 0.21457207 | -1.277589 | 0.20139437 | NA | NA | NA |
| ENSG000002 | 1.5873784 | -0.262115472 | 0.35036288 | -0.7481257 | 0.45438434 | NA | NA | NA |
| ENSG000002 | 0.64218479 | -0.28641133 | 0.26819919 | -1.0679053 | 0.28556324 | NA | NA | NA |
| ENSG000002 | 0.24485602 | -0.012655878 | 0.17881722 | -0.0707755 | 0.94357643 | NA | NA | NA |
| ENSG000002 | 0.48624257 | -0.012072666 | 0.23796621 | -0.0507327 | 0.95953853 | NA | NA | NA |
| ENSG000002 | 1.93532023 | 0.069523363 | 0.36417315 | 0.19090744 | 0.84859811 | NA | BRWD1-IT1 | 106480743 |
| ENSG000002 | 1.08584744 | 0.036912291 | 0.32497036 | 0.11358664 | 0.90956548 | NA | NA | NA |
| ENSG000002 | 2.09040743 | -0.246164382 | 0.36425771 | -0.6757973 | 0.49916935 | NA | NA | NA |
| ENSG000002 | 2.05573426 | -0.108055489 | 0.3593451 | -0.3007012 | 0.76364238 | NA | NA | NA |
| ENSG000002 | 2.08673804 | -0.466576788 | 0.3589876 | -1.2997017 | 0.19370323 | NA | NA | NA |
| ENSG000002 | 1.74092412 | -0.180997604 | 0.34853823 | -0.5193049 | 0.60354815 | NA | NA | NA |
| ENSG000002 | 0.66156766 | -0.287991838 | 0.26979811 | -1.0674346 | 0.28577564 | NA | NA | NA |
| ENSG000002 | 0.87076481 | -0.120542517 | 0.29782815 | -0.4047385 | 0.68566975 | NA | NA | NA |
| ENSG000002 | 0.63324854 | -0.111307913 | 0.26569851 | -0.4189256 | 0.6752705 | NA | NA | NA |
| ENSG000002 | 1.52747833 | 0.595963878 | 0.34589735 | 1.7229501 | 0.08489755 | NA | NA | NA |
| ENSG000002 | 1.60365441 | -0.406902078 | 0.35125422 | -1.1584262 | 0.24669016 | NA | LOC101928932 | 101928932 |
| ENSG000002 | 2.16742208 | -0.255535605 | 0.36319579 | -0.7035754 | 0.48169726 | NA | NA | NA |
| ENSG000002 | 0 | NA | NA | NA | NA | NA | NA | NA |
| ENSG000002 | 1.3472142 | -0.245761065 | 0.33438871 | -0.7349562 | 0.46236615 | NA | NA | NA |
| ENSG000002 | 1.52407787 | -0.496440377 | 0.34623546 | -1.433823 | 0.15162277 | NA | NA | NA |
| ENSG000002 | 1.73335411 | -0.303093213 | 0.34631057 | -0.8752064 | 0.38146163 | NA | NA | NA |
| ENSG000002 | 2.03861498 | -0.106062976 | 0.3564883 | -0.2975216 | 0.76606831 | NA | NA | NA |

| | | | | | | | | |
|---|---|---|---|---|---|---|---|---|
| ENSG000002 | 2.13454979 | -0.233604998 | 0.36089235 | -0.6472983 | 0.51743891 | NA | NA | NA |
| ENSG000002 | 0.11205239 | 0.082118251 | 0.12671962 | 0.64803108 | 0.51696485 | NA | NA | NA |
| ENSG000002 | 1.12177083 | -0.42067392 | 0.31896151 | -1.3188861 | 0.18720718 | NA | NA | NA |
| ENSG000002 | 0.61881753 | -0.112804359 | 0.27082437 | -0.4165222 | 0.67702794 | NA | NA | NA |
| ENSG000002 | 0.73961029 | -0.187025619 | 0.27953219 | -0.6690665 | 0.50345308 | NA | NA | NA |
| ENSG000002 | 1.41119348 | 0.091159926 | 0.33585957 | 0.27142274 | 0.78606591 | NA | TUBB4BP3 | 106480746 |
| ENSG000002 | 1.12538466 | -0.443269451 | 0.32457208 | -1.3657042 | 0.17203183 | NA | NA | NA |
| ENSG000002 | 1.11060427 | -0.112386496 | 0.3086212 | -0.3641568 | 0.71574095 | NA | NA | NA |
| ENSG000002 | 0 | NA | NA | NA | NA | NA | USP9YP35 | 106480719 |
| ENSG000002 | 1.2452913 | 0.089941921 | 0.32035538 | 0.28075671 | 0.77889701 | NA | NA | NA |
| ENSG000002 | 0.36827086 | -0.266056975 | 0.21181163 | -1.2561018 | 0.20907905 | NA | NA | NA |
| ENSG000002 | 1.14666808 | -0.262494406 | 0.31825016 | -0.8248052 | 0.40948217 | NA | NA | NA |
| ENSG000002 | 1.50918984 | -0.477455164 | 0.34190438 | -1.3964582 | 0.16257657 | NA | NA | NA |
| ENSG000002 | 0.4009426 | -0.302672773 | 0.2240145 | -1.3511303 | 0.17665371 | NA | NA | NA |
| ENSG000002 | 1.14937448 | -0.390849179 | 0.31253631 | -1.2505721 | 0.21109064 | NA | NA | NA |
| ENSG000002 | 1.79179867 | -0.751444796 | 0.35171227 | -2.1365328 | 0.03263601 | NA | NA | NA |
| ENSG000002 | 0.98480654 | -0.179926523 | 0.306894 | -0.5862823 | 0.55768582 | NA | NA | NA |
| ENSG000002 | 0.6190462 | -0.100577229 | 0.26454309 | -0.3801922 | 0.70380272 | NA | NA | NA |
| ENSG000002 | 1.21935953 | 0.238929246 | 0.32445691 | 0.73639747 | 0.46148885 | NA | NA | NA |
| ENSG000002 | 1.37818125 | -0.424970497 | 0.34126631 | -1.2452753 | 0.21303058 | NA | NA | NA |
| ENSG000002 | 1.85553518 | -0.002859766 | 0.35616371 | -0.0080294 | 0.99359357 | NA | NA | NA |
| ENSG000002 | 0.89547065 | -0.261071432 | 0.29465241 | -0.8860319 | 0.37560034 | NA | NA | NA |
| ENSG000002 | 0 | NA | NA | NA | NA | NA | NA | NA |
| ENSG000002 | 0.11934008 | 0.083564698 | 0.12760091 | 0.65489106 | 0.5125379 | NA | NA | NA |
| ENSG000002 | 1.71507564 | -0.187408546 | 0.35345393 | -0.5302206 | 0.59595901 | NA | NA | NA |
| ENSG000002 | 2.04378943 | -0.444579534 | 0.36096969 | -1.2316257 | 0.21808895 | NA | NA | NA |
| ENSG000002 | 0.71348903 | -0.155897422 | 0.26621239 | -0.585613 | 0.55813564 | NA | NA | NA |
| ENSG000002 | 0 | NA | NA | NA | NA | NA | NA | NA |
| ENSG000002 | 0 | NA | NA | NA | NA | NA | NA | NA |

| | | | | | | | | |
|---|---|---|---|---|---|---|---|---|
| ENSG000002 | 0.37285215 | -0.099309896 | 0.2177345 | -0.4561055 | 0.64831414 | NA | NA | NA |
| ENSG000002 | 0.62087103 | -0.279525375 | 0.26646004 | -1.049033 | 0.29416293 | NA | NA | NA |
| ENSG000002 | 1.7771374 | -0.919795743 | 0.35135239 | -2.6178725 | 0.00884799 | NA | NA | NA |
| ENSG000002 | 0 | NA | NA | NA | NA | NA | NA | NA |
| ENSG000002 | 1.85249886 | -0.2529458 | 0.35737912 | -0.70778 | 0.47908188 | NA | NA | NA |
| ENSG000002 | 1.00971397 | -0.20046801 | 0.30956332 | -0.6475832 | 0.51725456 | NA | NA | NA |
| ENSG000002 | 0.51413714 | -0.199727707 | 0.24976394 | -0.7996659 | 0.42390439 | NA | NA | NA |
| ENSG000002 | 1.36710752 | -0.105686889 | 0.32986891 | -0.3203906 | 0.74867227 | NA | NA | NA |
| ENSG000002 | 0.77174124 | -0.198363094 | 0.28768104 | -0.6895244 | 0.49049333 | NA | NA | NA |
| ENSG000002 | 0.36827086 | -0.266056975 | 0.21181163 | -1.2561018 | 0.20907905 | NA | NA | NA |
| ENSG000002 | 1.68909819 | -0.158391379 | 0.3428159 | -0.4620304 | 0.64405949 | NA | NA | NA |
| ENSG000002 | 1.02192181 | -0.177492783 | 0.30275912 | -0.5862508 | 0.55770697 | NA | NA | NA |
| ENSG000002 | 0 | NA | NA | NA | NA | NA | NA | NA |
| ENSG000002 | 1.12740199 | -0.252735954 | 0.31510351 | -0.8020728 | 0.42251085 | NA | NA | NA |
| ENSG000002 | 0.8716979 | -0.114697509 | 0.29189776 | -0.3929373 | 0.69436581 | NA | NA | NA |
| ENSG000002 | 1.86075346 | -0.119667747 | 0.35449828 | -0.3375693 | 0.73568776 | NA | NA | NA |
| ENSG000002 | 0.50377641 | -0.188767022 | 0.24148558 | -0.7816906 | 0.4343964 | NA | NA | NA |
| ENSG000002 | 0.62911822 | -0.101139175 | 0.25635403 | -0.3945293 | 0.6931903 | NA | NA | NA |
| ENSG000002 | 0 | NA | NA | NA | NA | NA | USP9YP32 | 106480718 |
| ENSG000002 | 1.7124392 | -0.686575928 | 0.3458007 | -1.9854671 | 0.04709252 | NA | NA | NA |
| ENSG000002 | 1.36476977 | -0.264758132 | 0.34021671 | -0.7782044 | 0.43644854 | NA | NA | NA |
| ENSG000002 | 1.61648033 | 0.020029077 | 0.33981768 | 0.05894066 | 0.95299937 | NA | NA | NA |
| ENSG000002 | 1.37285828 | -0.251571808 | 0.33007607 | -0.7621631 | 0.44596265 | NA | NA | NA |
| ENSG000002 | 1.52180208 | -0.494830045 | 0.34687642 | -1.4265312 | 0.15371506 | NA | NA | NA |
| ENSG000002 | 0 | NA | NA | NA | NA | NA | DUX4L29 | 106481970 |
| ENSG000002 | 0 | NA | NA | NA | NA | NA | NA | NA |
| ENSG000002 | 1.36636839 | -0.257112959 | 0.33701504 | -0.7629124 | 0.44551562 | NA | NA | NA |
| ENSG000002 | 0.6190462 | -0.100577229 | 0.26454309 | -0.3801922 | 0.70380272 | NA | NA | NA |
| ENSG000002 | 0 | NA | NA | NA | NA | NA | NA | NA |

| | | | | | | | | |
|---|---|---|---|---|---|---|---|---|
| ENSG000002 | 1.39331915 | -0.255433824 | 0.33616037 | -0.759857 | 0.44734005 | NA | NA | NA |
| ENSG000002 | 1.98514954 | -0.061990507 | 0.3570795 | -0.1736042 | 0.86217652 | NA | NA | NA |
| ENSG000002 | 1.87945162 | -0.658066293 | 0.35676122 | -1.8445567 | 0.06510205 | NA | GRM7-AS2 | 105376946 |
| ENSG000002 | 1.90787254 | -0.351370114 | 0.34910878 | -1.0064775 | 0.31418595 | NA | NA | NA |
| ENSG000002 | 0.62775378 | -0.28146231 | 0.26704095 | -1.0540043 | 0.29188095 | NA | NA | NA |
| ENSG000002 | 1.72468134 | -0.559433729 | 0.34640791 | -1.6149566 | 0.1063201 | NA | NA | NA |
| ENSG000002 | 1.61496031 | -0.394144592 | 0.34756966 | -1.1340017 | 0.25679383 | NA | NA | NA |
| ENSG000002 | 1.66595104 | -0.501929724 | 0.3415304 | -1.4696487 | 0.14165691 | NA | NA | NA |
| ENSG000002 | 0 | NA | NA | NA | NA | NA | NA | NA |
| ENSG000002 | 1.64768466 | -0.397104218 | 0.34899081 | -1.1378644 | 0.25517711 | NA | NA | NA |
| ENSG000002 | 0 | NA | NA | NA | NA | NA | NA | NA |
| ENSG000002 | 1.22388918 | -0.185859517 | 0.32888627 | -0.5651179 | 0.57199358 | NA | NA | NA |
| ENSG000002 | 1.5104917 | -0.19425425 | 0.34208905 | -0.567847 | 0.57013884 | NA | NA | NA |
| ENSG000002 | 2.09296947 | -0.36246602 | 0.36270677 | -0.9993362 | 0.31763183 | NA | RNF216-IT1 | 100874342 |
| ENSG000002 | 1.46709609 | -0.051678647 | 0.33865686 | -0.1525989 | 0.87871463 | NA | NA | NA |
| ENSG000002 | 0.64332695 | -0.264269033 | 0.26152646 | -1.0104868 | 0.31226213 | NA | NA | NA |
| ENSG000002 | 1.76423348 | -0.199715391 | 0.35174367 | -0.5677867 | 0.57017978 | NA | LINC01143 | 104355141 |
| ENSG000002 | 1.49188625 | -0.19306624 | 0.34612868 | -0.5577875 | 0.57698953 | NA | NA | NA |
| ENSG000002 | 2.06511571 | -1.0479324 | 0.35955095 | -2.9145588 | 0.00356192 | NA | NA | NA |
| ENSG000002 | 0.36502313 | -0.093755175 | 0.2080862 | -0.4505593 | 0.6523072 | NA | AMY1A | 276 |
| ENSG000002 | 0.24141689 | -0.165257896 | 0.17127401 | -0.9648743 | 0.33460776 | NA | NA | NA |
| ENSG000002 | 0.47684931 | 0.147509143 | 0.23759598 | 0.62084023 | 0.53470475 | NA | NA | NA |
| ENSG000002 | 1.38529425 | -0.270070693 | 0.34061649 | -0.7928879 | 0.42784314 | NA | NA | NA |
| ENSG000002 | 0.12685397 | -0.090215768 | 0.12901919 | -0.699243 | 0.48440017 | NA | NA | NA |
| ENSG000002 | 1.32275642 | 0.475909376 | 0.33521483 | 1.41971455 | 0.1556908 | NA | NA | NA |
| ENSG000002 | 1.98603915 | -0.317699696 | 0.36126176 | -0.8794169 | 0.37917527 | NA | NA | NA |
| ENSG000002 | 0 | NA | NA | NA | NA | NA | NA | NA |
| ENSG000002 | 0 | NA | NA | NA | NA | NA | NA | NA |
| ENSG000002 | 0.48348219 | -0.018797026 | 0.24521266 | -0.076656 | 0.9388972 | NA | NA | NA |

| | | | | | | | | | |
|---|---|---|---|---|---|---|---|---|---|
| ENSG000002 | 0.50702415 | -0.349150675 | 0.23755647 | -1.4697586 | 0.14162714 | NA | NA | NA | |
| ENSG000002 | 1.72006133 | -0.063250708 | 0.35600688 | -0.1776671 | 0.85898443 | NA | NA | NA | |
| ENSG000002 | 1.05886635 | 0.186500081 | 0.31411261 | 0.59373637 | 0.55268846 | NA | NA | NA | |
| ENSG000002 | 0 | NA | NA | NA | NA | NA | NA | NA | |
| ENSG000002 | 1.22801345 | -0.046083825 | 0.32847408 | -0.1402967 | 0.88842558 | NA | NA | NA | |
| ENSG000002 | 1.73059537 | -0.294145385 | 0.34782268 | -0.8456763 | 0.39773337 | NA | NA | NA | |
| ENSG000002 | 1.92479371 | -0.539479704 | 0.35874322 | -1.5038046 | 0.13263168 | NA | NA | NA | |
| ENSG000002 | 1.39728393 | -0.379874022 | 0.32882606 | -1.155243 | 0.24799091 | NA | NA | NA | |
| ENSG000002 | 1.86968834 | -0.387699493 | 0.35971549 | -1.0777948 | 0.28112533 | NA | NA | NA | |
| ENSG000002 | 1.38378208 | -0.123478385 | 0.34130234 | -0.3617859 | 0.71751201 | NA | NA | NA | |
| ENSG000002 | 0 | NA | NA | NA | NA | NA | NA | NA | |
| ENSG000002 | 2.02823824 | 0.131638495 | 0.36338796 | 0.36225332 | 0.71716274 | NA | NA | NA | |
| ENSG000002 | 0 | NA | NA | NA | NA | NA | NA | NA | |
| ENSG000002 | 0.72008656 | 0.117012389 | 0.27576463 | 0.42431979 | 0.67133261 | NA | NA | NA | |
| ENSG000002 | 1.70913591 | -0.816699761 | 0.34124224 | -2.3933138 | 0.01669695 | NA | NA | NA | |
| ENSG000002 | 0.60717723 | -0.256800472 | 0.2582777 | -0.9942805 | 0.32008634 | NA | NA | NA | |
| ENSG000002 | 1.18967046 | 0.419416519 | 0.32845693 | 1.27693004 | 0.20162695 | NA | NA | NA | |
| ENSG000002 | 0.97697752 | -0.164936641 | 0.29502854 | -0.5590532 | 0.57612544 | NA | NA | NA | |
| ENSG000002 | 0.90064872 | -0.456924471 | 0.30237983 | -1.5110944 | 0.1307644 | NA | NA | NA | |
| ENSG000002 | 1.85529115 | -0.258652104 | 0.36000452 | -0.7184691 | 0.47246812 | NA | NA | NA | |
| ENSG000002 | 1.51297134 | -0.199300861 | 0.3445483 | -0.578441 | 0.56296642 | NA | KCNQ1DN | | 55539 |
| ENSG000002 | 2.05716653 | -0.723507499 | 0.36043496 | -2.0073178 | 0.04471584 | NA | NA | NA | |
| ENSG000002 | 1.80772795 | 0.041366854 | 0.25053766 | 0.16511232 | 0.86885558 | NA | NA | NA | |
| ENSG000002 | 2.15065971 | -0.51303684 | 0.36594054 | -1.4019678 | 0.16092487 | NA | NA | NA | |
| ENSG000002 | 0.24141689 | -0.165257896 | 0.17127401 | -0.9648743 | 0.33460776 | NA | NA | NA | |
| ENSG000002 | 1.98143825 | -0.43659261 | 0.35880614 | -1.2167925 | 0.22368318 | NA | NA | NA | |
| ENSG000002 | 1.99027608 | -0.185712891 | 0.36234466 | -0.5125311 | 0.60827939 | NA | NA | NA | |
| ENSG000002 | 0.75053511 | 0.123730904 | 0.27954741 | 0.44261152 | 0.65804675 | NA | NA | NA | |
| ENSG000002 | 0 | NA | NA | NA | NA | NA | CT47A5 | | 728072 |

| | | | | | | | | |
|---|---|---|---|---|---|---|---|---|
| ENSG000002 | 1.08889015 | 0.196991595 | 0.32375079 | 0.60846677 | 0.54287794 | NA | NA | NA |
| ENSG000002 | 0 | NA | NA | NA | NA | NA | NA | NA |
| ENSG000002 | 1.57351045 | -0.116531097 | 0.35086589 | -0.3321243 | 0.73979538 | NA | NA | NA |
| ENSG000002 | 1.91409182 | 0.165376574 | 0.35181037 | 0.47007305 | 0.63830283 | NA | NA | NA |
| ENSG000002 | 1.97940491 | -0.311812861 | 0.3602444 | -0.8655592 | 0.38673194 | NA | NA | NA |
| ENSG000002 | 0 | NA | NA | NA | NA | NA | NA | NA |
| ENSG000002 | 1.17746492 | -0.521763955 | 0.30622498 | -1.7038583 | 0.08840757 | NA | NA | NA |
| ENSG000002 | 0.25370795 | -0.171331747 | 0.17411721 | -0.9840024 | 0.32511435 | NA | NA | NA |
| ENSG000002 | 0.61241141 | -0.098777276 | 0.2639741 | -0.3741931 | 0.70826069 | NA | NA | NA |
| ENSG000002 | 2.04046494 | -0.716963054 | 0.36035234 | -1.9896168 | 0.04663317 | NA | NA | NA |
| ENSG000002 | 1.96025228 | -0.191146117 | 0.36510929 | -0.5235312 | 0.60060462 | NA | NA | NA |
| ENSG000002 | 1.99207058 | -0.180701739 | 0.3598299 | -0.5021866 | 0.6155363 | NA | NA | NA |
| ENSG000002 | 0.51550551 | -0.37860525 | 0.24578489 | -1.5403927 | 0.12346466 | NA | NA | NA |
| ENSG000002 | 1.59076994 | -0.380310648 | 0.34516894 | -1.1018102 | 0.27054421 | NA | OOSP1P1 | 106480251 |
| ENSG000002 | 0.62450659 | -0.10299827 | 0.26500891 | -0.3886597 | 0.69752793 | NA | NA | NA |
| ENSG000002 | 0.36556447 | -0.101943371 | 0.21659044 | -0.4706735 | 0.63787395 | NA | NA | NA |
| ENSG000002 | 1.22952225 | 0.098031846 | 0.32846601 | 0.29845355 | 0.76535702 | NA | NA | NA |
| ENSG000002 | 2.12277374 | -0.641021675 | 0.36405877 | -1.7607643 | 0.07827831 | NA | NA | NA |
| ENSG000002 | 0.75395699 | -0.187568606 | 0.28051187 | -0.6686655 | 0.50370888 | NA | NA | NA |
| ENSG000002 | 1.61398288 | -0.363732539 | 0.34203129 | -1.0634481 | 0.28757877 | NA | NA | NA |
| ENSG000002 | 2.15145465 | 0.311152076 | 0.3670568 | 0.84769462 | 0.39660806 | NA | NA | NA |
| ENSG000002 | 0 | NA | NA | NA | NA | NA | NA | NA |
| ENSG000002 | 0.96853054 | -0.032347758 | 0.30712832 | -0.1053233 | 0.9161193 | NA | NA | NA |
| ENSG000002 | 0 | NA | NA | NA | NA | NA | NA | NA |
| ENSG000002 | 0 | NA | NA | NA | NA | NA | NA | NA |
| ENSG000002 | 0.99423418 | -0.165628961 | 0.29972324 | -0.5526063 | 0.580533 | NA | NA | NA |
| ENSG000002 | 1.25229474 | -0.197677572 | 0.33455733 | -0.5908631 | 0.55461217 | NA | NA | NA |
| ENSG000002 | 1.15307475 | -0.274348016 | 0.32432702 | -0.8458993 | 0.3976089 | NA | NA | NA |
| ENSG000002 | 0 | NA | NA | NA | NA | NA | NA | NA |

| | | | | | | | | | |
|---|---|---|---|---|---|---|---|---|---|
| ENSG000002 | 1.10006994 | 0.034909074 | 0.31699605 | 0.11012463 | 0.91231053 | NA | NA | NA | |
| ENSG000002 | 2.14907318 | -0.521965117 | 0.36557783 | -1.4277811 | 0.15335487 | NA | NA | NA | |
| ENSG000002 | 0.72972831 | -0.025248506 | 0.28417722 | -0.0888477 | 0.92920291 | NA | NA | NA | |
| ENSG000002 | 1.70770555 | -0.053342324 | 0.35186091 | -0.1516006 | 0.87950196 | NA | NA | NA | |
| ENSG000002 | 1.01207154 | -0.046453419 | 0.31227181 | -0.1487596 | 0.88174336 | NA | NA | NA | |
| ENSG000002 | 0.88359721 | -0.118458047 | 0.2925649 | -0.4048949 | 0.68555474 | NA | NA | NA | |
| ENSG000002 | 0.11131518 | 0.081951103 | 0.12663804 | 0.64712866 | 0.51754868 | NA | NA | NA | |
| ENSG000002 | 1.23812377 | -0.188681269 | 0.32332512 | -0.5835651 | 0.55951292 | NA | NA | NA | |
| ENSG000002 | 1.96130407 | -0.061747447 | 0.36412931 | -0.1695756 | 0.86534391 | NA | NA | NA | |
| ENSG000002 | 0 | NA | NA | NA | NA | NA | NA | NA | |
| ENSG000002 | 0.73124048 | -0.17762673 | 0.27730728 | -0.6405412 | 0.52182083 | NA | NA | NA | |
| ENSG000002 | 0.5854929 | -0.091197421 | 0.25231745 | -0.3614392 | 0.71777113 | NA | NA | NA | |
| ENSG000002 | 1.56211767 | 0.291618756 | 0.34687011 | 0.84071457 | 0.40050786 | NA | NA | NA | |
| ENSG000002 | 1.3545514 | 0.153030227 | 0.32653492 | 0.46864889 | 0.63932063 | NA | NA | NA | |
| ENSG000002 | 0.61833933 | -0.261391558 | 0.25929785 | -1.0080745 | 0.31341869 | NA | NA | NA | |
| ENSG000002 | 1.79651487 | 0.1322309 | 0.35126661 | 0.37644028 | 0.70658961 | NA | NA | NA | |
| ENSG000002 | 0 | NA | NA | NA | NA | NA | NA | NA | |
| ENSG000002 | 1.77766724 | -0.20200568 | 0.35734339 | -0.5652985 | 0.57187075 | NA | NA | NA | |
| ENSG000002 | 0.84099142 | 0.050396409 | 0.3010945 | 0.16737738 | 0.86707312 | NA | NA | NA | |
| ENSG000002 | 1.43536241 | -0.563756537 | 0.33622193 | -1.6767394 | 0.09359346 | NA | LOC646938 | | 646938 |
| ENSG000002 | 1.02411116 | -0.351091141 | 0.3099654 | -1.1326785 | 0.25734929 | NA | NA | NA | |
| ENSG000002 | 1.82200596 | -0.229244588 | 0.35221484 | -0.6508658 | 0.51513313 | NA | NA | NA | |
| ENSG000002 | 0.87278449 | -0.433275285 | 0.29591838 | -1.4641716 | 0.14314707 | NA | NA | NA | |
| ENSG000002 | 1.20060623 | -0.040671408 | 0.33282946 | -0.1221989 | 0.90274146 | NA | NA | NA | |
| ENSG000002 | 0.54222759 | -0.38902668 | 0.24851205 | -1.5654238 | 0.11748356 | NA | NA | NA | |
| ENSG000002 | 0.75936373 | -0.356158746 | 0.28317817 | -1.2577196 | 0.20849317 | NA | NA | NA | |
| ENSG000002 | 1.80422374 | 0.395672117 | 0.35200417 | 1.12405519 | 0.26098962 | NA | NA | NA | |
| ENSG000002 | 0 | NA | NA | NA | NA | NA | NA | NA | |
| ENSG000002 | 1.62145577 | -0.667878769 | 0.34453476 | -1.9384946 | 0.05256291 | NA | NA | NA | |

| | | | | | | | | |
|---|---|---|---|---|---|---|---|---|
| ENSG000002 | 2.13742248 | -0.002780625 | 0.35684677 | -0.0077922 | 0.99378278 | NA | NA | NA |
| ENSG000002 | 0.25260017 | -0.015978705 | 0.18049464 | -0.0885273 | 0.92945758 | NA | NA | NA |
| ENSG000002 | 1.62881224 | -0.276616188 | 0.35421222 | -0.7809335 | 0.43484161 | NA | NA | NA |
| ENSG000002 | 0.4974562 | -0.028348769 | 0.24672499 | -0.1149003 | 0.90852417 | NA | NA | NA |
| ENSG000002 | 0.50479595 | -0.02564641 | 0.24753216 | -0.1036084 | 0.91748012 | NA | NA | NA |
| ENSG000002 | 0.23276084 | -0.008011382 | 0.17603982 | -0.0455089 | 0.96370166 | NA | NA | NA |
| ENSG000002 | 1.64493012 | -0.555071631 | 0.35028008 | -1.5846509 | 0.11304568 | NA | NA | NA |
| ENSG000002 | 0 | NA | NA | NA | NA | NA | NA | NA |
| ENSG000002 | 0.9854377 | -0.339886722 | 0.3083842 | -1.1021535 | 0.27039495 | NA | NA | NA |
| ENSG000002 | 1.45999933 | 0.087947657 | 0.34318002 | 0.25627266 | 0.79774031 | NA | NA | NA |
| ENSG000002 | 2.10307821 | -0.61105584 | 0.36063181 | -1.6944036 | 0.09018861 | NA | NA | NA |
| ENSG000002 | 1.21782851 | -0.167124347 | 0.32186807 | -0.5192325 | 0.60359865 | NA | NA | NA |
| ENSG000002 | 1.30850844 | -0.492384017 | 0.32826465 | -1.4999605 | 0.13362463 | NA | NA | NA |
| ENSG000002 | 0.36538978 | 0.059033792 | 0.20773944 | 0.28417229 | 0.77627835 | NA | NA | NA |
| ENSG000002 | 1.84031543 | 0.00870882 | 0.35932585 | 0.02423655 | 0.98066392 | NA | NA | NA |
| ENSG000002 | 0.93818799 | -0.409036498 | 0.28911747 | -1.4147761 | 0.15713415 | NA | NA | NA |
| ENSG000002 | 0.90827025 | -0.266474502 | 0.2935965 | -0.9076215 | 0.36407821 | NA | NA | NA |
| ENSG000002 | 0.52125014 | -0.010203417 | 0.15567753 | -0.065542 | 0.94774247 | NA | NA | NA |
| ENSG000002 | 0.50630812 | -0.189615696 | 0.24175631 | -0.7843257 | 0.432849 | NA | NA | NA |
| ENSG000002 | 0.48487421 | 0.137177479 | 0.22868862 | 0.59984392 | 0.54861026 | NA | NA | NA |
| ENSG000002 | 0.26560726 | -0.177019426 | 0.17672274 | -1.0016788 | 0.31649875 | NA | NA | NA |
| ENSG000002 | 0.37928001 | 0.070449581 | 0.21829793 | 0.32272217 | 0.74690566 | NA | SNORD121B | 101340252 |
| ENSG000002 | 0.38651158 | -0.274258716 | 0.21461355 | -1.2779189 | 0.20127801 | NA | RNU7-120P | 106480817 |
| ENSG000002 | 0.74435521 | -0.028370124 | 0.27885082 | -0.1017394 | 0.9189635 | NA | NA | NA |
| ENSG000002 | 1.50522719 | -0.198274157 | 0.34511609 | -0.5745144 | 0.56561977 | NA | NA | NA |
| ENSG000002 | 0.8650959 | -0.118179347 | 0.30263724 | -0.3904984 | 0.69616807 | NA | NA | NA |
| ENSG000002 | 0.72358278 | -0.021486528 | 0.27739644 | -0.0774578 | 0.93825933 | NA | NA | NA |
| ENSG000002 | 0.51327731 | -0.027046708 | 0.2410203 | -0.1122176 | 0.91065091 | NA | NA | NA |
| ENSG000002 | 0.48962772 | -0.021045505 | 0.24589998 | -0.0855856 | 0.93179582 | NA | RNU7-140P | 106479077 |

| | | | | | | | | |
|---|---|---|---|---|---|---|---|---|
| ENSG000002 | 0.39274356 | -0.107428268 | 0.22062504 | -0.4869269 | 0.62631013 | NA | NA | NA |
| ENSG000002 | 1.78728672 | -0.469327716 | 0.35431365 | -1.3246109 | 0.18530024 | NA | NA | NA |
| ENSG000002 | 0.76949578 | -0.359754798 | 0.28380236 | -1.2676244 | 0.20493211 | NA | NA | NA |
| ENSG000002 | 0.38651158 | -0.274258716 | 0.21461355 | -1.2779189 | 0.20127801 | NA | NA | NA |
| ENSG000002 | 0.12685397 | -0.090215768 | 0.12901919 | -0.699243 | 0.48440017 | NA | RNU7-180P | 106480830 |
| ENSG000002 | 0.23139247 | 0.167748926 | 0.17680352 | 0.94878726 | 0.34272882 | NA | NA | NA |
| ENSG000002 | 0.63825247 | -0.10803491 | 0.26614363 | -0.4059271 | 0.68479617 | NA | RNU7-165P | 106479087 |
| ENSG000002 | 0.12685397 | -0.090215768 | 0.12901919 | -0.699243 | 0.48440017 | NA | RNU6-1211P | 106480096 |
| ENSG000002 | 0.59009593 | 0.234584782 | 0.26263803 | 0.89318664 | 0.37175723 | NA | NA | NA |
| ENSG000002 | 0 | NA | NA | NA | NA | NA | NA | NA |
| ENSG000002 | 0.88695649 | -0.277948495 | 0.29981959 | -0.9270525 | 0.35389929 | NA | NA | NA |
| ENSG000002 | 1.72396195 | -0.196059071 | 0.35830941 | -0.5471781 | 0.58425637 | NA | MIR2053 | 100302225 |
| ENSG000002 | 0.26199343 | -0.194018184 | 0.18422801 | -1.0531416 | 0.29227609 | NA | RNA5SP311 | 106480358 |
| ENSG000002 | 0.12685397 | -0.090215768 | 0.12901919 | -0.699243 | 0.48440017 | NA | RNU7-171P | 106479089 |
| ENSG000002 | 0.87087479 | -0.249546264 | 0.28910243 | -0.863176 | 0.38804073 | NA | NA | NA |
| ENSG000002 | 0.11205239 | 0.082118251 | 0.12671962 | 0.64803108 | 0.51696485 | NA | RNU7-186P | 106480363 |
| ENSG000002 | 0 | NA | NA | NA | NA | NA | RNU6-437P | 106479759 |
| ENSG000002 | 1.50215469 | -0.192012845 | 0.34629373 | -0.5544797 | 0.57925059 | NA | SNORD42B | 26808 |
| ENSG000002 | 0.12685397 | -0.090215768 | 0.12901919 | -0.699243 | 0.48440017 | NA | NA | NA |
| ENSG000002 | 0.64099112 | -0.296274405 | 0.27396996 | -1.081412 | 0.27951389 | NA | RNU7-136P | 106479076 |
| ENSG000002 | 0.37304803 | -0.090842893 | 0.20939682 | -0.4338313 | 0.66441096 | NA | RNU7-157P | 106480826 |
| ENSG000002 | 0.12070844 | -0.088503695 | 0.12783931 | -0.6923042 | 0.48874631 | NA | NA | NA |
| ENSG000002 | 0.35961481 | -0.099625193 | 0.21567743 | -0.4619176 | 0.64414043 | NA | RNU7-130P | 106479074 |
| ENSG000002 | 0.39246123 | -0.276864021 | 0.21549122 | -1.2848042 | 0.19886073 | NA | NA | NA |
| ENSG000002 | 0.76810568 | -0.364419366 | 0.28364061 | -1.2847926 | 0.19886476 | NA | RNU6-893P | 106479961 |
| ENSG000002 | 1.13758803 | -0.120160334 | 0.32306217 | -0.3719418 | 0.70993617 | NA | NA | NA |
| ENSG000002 | 0.49512483 | -0.344267812 | 0.23617548 | -1.457678 | 0.14492931 | NA | RNU7-134P | 106480820 |
| ENSG000002 | 0.87050477 | -0.105667046 | 0.29784719 | -0.3547693 | 0.72276241 | NA | RNU7-144P | 106479079 |
| ENSG000002 | 0.99556582 | -0.037878896 | 0.31314051 | -0.1209645 | 0.90371912 | NA | RNU6-1014P | 106480013 |

| | | | | | | | | | |
|---|---|---|---|---|---|---|---|---|---|
| ENSG000002 | 0.28256998 | -0.18483211 | 0.18021686 | -1.0256094 | 0.30507573 | NA | | RNU7-169P | 106480827 |
| ENSG000002 | 0.26813897 | -0.197007041 | 0.18550319 | -1.0620143 | 0.2882292 | NA | | NA | NA |
| ENSG000002 | 0 | NA | NA | NA | NA | NA | | RNU6-498P | 106481338 |
| ENSG000002 | 0.4855357 | -0.181797071 | 0.23938927 | -0.7594203 | 0.44760118 | NA | | RNU6-1208P | 106480094 |
| ENSG000002 | 2.00277217 | -0.447582462 | 0.361997 | -1.2364259 | 0.2163003 | NA | | NA | NA |
| ENSG000002 | 0 | NA | NA | NA | NA | NA | | RNU6-749P | 106480606 |
| ENSG000002 | 0.89179679 | -0.274686446 | 0.29848248 | -0.9202766 | 0.35742822 | NA | | NA | NA |
| ENSG000002 | 0.63370343 | -0.298851213 | 0.27334848 | -1.0932975 | 0.2742632 | NA | | RNU6-221P | 106480561 |
| ENSG000002 | 0.85955119 | -0.106906951 | 0.29726276 | -0.3596379 | 0.71911795 | NA | | NA | NA |
| ENSG000002 | 0.36345889 | 0.070257596 | 0.21589421 | 0.32542602 | 0.74485866 | NA | | NA | NA |
| ENSG000002 | 0.5301324 | -0.3843196 | 0.24728844 | -1.5541349 | 0.12015224 | NA | | NA | NA |
| ENSG000002 | 0.37304803 | -0.090842893 | 0.20939682 | -0.4338313 | 0.66441096 | NA | | RNU6-104P | 106479612 |
| ENSG000002 | 2.078648 | 0.001433019 | 0.35504461 | 0.00403617 | 0.99677961 | NA | | NA | NA |
| ENSG000002 | 0.23868015 | 0.154625155 | 0.16997278 | 0.90970538 | 0.36297791 | NA | | NA | NA |
| ENSG000002 | 1.97898281 | -0.164511413 | 0.35498907 | -0.4634267 | 0.64305855 | NA | | RNU7-107P | 106479067 |
| ENSG000002 | 0.61955474 | -0.11252945 | 0.27088619 | -0.4154123 | 0.67784012 | NA | | RNU6-599P | 106481370 |
| ENSG000002 | 1.18359874 | 0.548108819 | 0.31782779 | 1.72454653 | 0.08460923 | NA | | NA | NA |
| ENSG000002 | 1.47966844 | -0.039907308 | 0.33479786 | -0.1191982 | 0.90511832 | NA | | RNU7-93P | 106480813 |
| ENSG000002 | 1.70779592 | -0.168235467 | 0.34813369 | -0.4832496 | 0.62891852 | NA | | RNU7-104P | 106480448 |
| ENSG000002 | 0 | NA | NA | NA | NA | NA | | RNU6-1193P | 106481556 |
| ENSG000002 | 1.25933044 | -0.50232137 | 0.33067667 | -1.5190711 | 0.12874461 | NA | | RNU1-88P | 106481619 |
| ENSG000002 | 1.08712992 | -0.233085789 | 0.30625434 | -0.7610857 | 0.44660589 | NA | | NA | NA |
| ENSG000002 | 0.12070844 | -0.088503695 | 0.12783931 | -0.6923042 | 0.48874631 | NA | | NA | NA |
| ENSG000002 | 0.36482725 | -0.102221582 | 0.21647267 | -0.4722147 | 0.63677353 | NA | | RNU6ATAC28P | 106479555 |
| ENSG000002 | 0.38651158 | -0.274258716 | 0.21461355 | -1.2779189 | 0.20127801 | NA | | RNU7-159P | 106481747 |
| ENSG000002 | 0.61491745 | 0.056458936 | 0.26379319 | 0.21402727 | 0.83052579 | NA | | NA | NA |
| ENSG000002 | 0.40942396 | -0.284058841 | 0.21788566 | -1.303706 | 0.19233384 | NA | | NA | NA |
| ENSG000002 | 1.62076148 | -0.38346237 | 0.34695102 | -1.1052349 | 0.26905783 | NA | | RNU7-167P | 106479088 |
| ENSG000002 | 1.30733403 | -0.473192856 | 0.3238236 | -1.4612674 | 0.14394208 | NA | | NA | NA |

| | | | | | | | | |
|---|---|---|---|---|---|---|---|---|
| ENSG000002 | 0.9674445 | 0.271109132 | 0.30627275 | 0.88518856 | 0.37605494 | NA | NA | NA |
| ENSG000002 | 0 | NA | NA | NA | NA | NA | SNORD4B | 26772 |
| ENSG000002 | 1.7469929 | -0.612038871 | 0.35379133 | -1.7299431 | 0.08364044 | NA | NA | NA |
| ENSG000002 | 0.77330548 | -0.35598868 | 0.28246534 | -1.2602915 | 0.20756421 | NA | RNU6-300P | 106479700 |
| ENSG000002 | 1.82389295 | -0.002887443 | 0.35397287 | -0.0081572 | 0.99349153 | NA | RNU6-775P | 106479909 |
| ENSG000002 | 0 | NA | NA | NA | NA | NA | SNORD97 | 692223 |
| ENSG000002 | 0.12070844 | -0.088503695 | 0.12783931 | -0.6923042 | 0.48874631 | NA | NA | NA |
| ENSG000002 | 0.71675348 | 0.133341687 | 0.28341107 | 0.47048864 | 0.63800595 | NA | NA | NA |
| ENSG000002 | 1.02886714 | -0.315044737 | 0.29925492 | -1.0527638 | 0.29244928 | NA | RNU7-126P | 106479073 |
| ENSG000002 | 0.48367807 | -0.017505926 | 0.23765649 | -0.0736606 | 0.94128043 | NA | NA | NA |
| ENSG000002 | 0 | NA | NA | NA | NA | NA | SNORD42A | 26809 |
| ENSG000002 | 0 | NA | NA | NA | NA | NA | SNORD54 | 26795 |
| ENSG000002 | 1.09933273 | 0.036042709 | 0.32152164 | 0.11210042 | 0.91074379 | NA | NA | NA |
| ENSG000002 | 0.88553256 | -0.426391846 | 0.29501377 | -1.4453286 | 0.1483656 | NA | RNU6-625P | 106479843 |
| ENSG000002 | 0.36484899 | 0.075158314 | 0.21612264 | 0.34775772 | 0.72802215 | NA | RNU6-333P | 106480571 |
| ENSG000002 | 1.38199096 | -0.379758313 | 0.32709868 | -1.1609901 | 0.24564594 | NA | RNU6-1037P | 106481508 |
| ENSG000002 | 0.11131518 | 0.081951103 | 0.12663804 | 0.64712866 | 0.51754868 | NA | NA | NA |
| ENSG000002 | 0.50107449 | -0.372905725 | 0.24426561 | -1.5266403 | 0.12685049 | NA | NA | NA |
| ENSG000002 | 0.70141111 | 0.318465679 | 0.28333314 | 1.12399728 | 0.26101418 | NA | RNU6-261P | 106481885 |
| ENSG000002 | 1.52325421 | -0.610368312 | 0.33833887 | -1.8040147 | 0.071229 | NA | RNU7-147P | 106480451 |
| ENSG000002 | 0 | NA | NA | NA | NA | NA | MIR1976 | 100302190 |
| ENSG000002 | 0.73041345 | -0.026515459 | 0.28960252 | -0.0915581 | 0.92704914 | NA | NA | NA |
| ENSG000002 | 0.24411881 | -0.012946855 | 0.17864669 | -0.0724718 | 0.94222641 | NA | NA | NA |
| ENSG000002 | 1.48231885 | 0.092073109 | 0.34632949 | 0.26585408 | 0.79035159 | NA | RNU7-96P | 106480361 |
| ENSG000002 | 1.25417411 | -0.174031465 | 0.32377511 | -0.5375072 | 0.59091729 | NA | RNU7-194P | 106480834 |
| ENSG000002 | 1.44042662 | -0.035634274 | 0.33813689 | -0.1053842 | 0.91607098 | NA | NA | NA |
| ENSG000002 | 1.72279861 | -0.32946057 | 0.3562811 | -0.9247209 | 0.35511111 | NA | NA | NA |
| ENSG000002 | 0.72996048 | 0.271217395 | 0.2689077 | 1.00858915 | 0.31317171 | NA | MIR1972-1 | 100302243 |
| ENSG000002 | 0.26813897 | -0.197007041 | 0.18550319 | -1.0620143 | 0.2882292 | NA | RNU7-164P | 106479086 |

| | | | | | | | | | |
|---|---|---|---|---|---|---|---|---|---|
| ENSG000002 | 0.61130363 | 0.061366006 | 0.26984585 | 0.22741134 | 0.82010391 | NA | | RNU7-113P | 106479068 |
| ENSG000002 | 0.9875388 | -0.181571486 | 0.30121583 | -0.6027953 | 0.54664487 | NA | | NA | NA |
| ENSG000002 | 0 | NA | NA | NA | NA | NA | | NA | NA |
| ENSG000002 | 1.32635298 | 0.031750353 | 0.33110907 | 0.09589092 | 0.92360721 | NA | | NA | NA |
| ENSG000002 | 0.75418566 | -0.181711922 | 0.28055616 | -0.6476847 | 0.51718893 | NA | | RNU7-155P | 106480825 |
| ENSG000002 | 1.39977926 | -0.113089787 | 0.32895198 | -0.3437881 | 0.73100563 | NA | | NA | NA |
| ENSG000002 | 0 | NA | NA | NA | NA | NA | | NA | NA |
| ENSG000002 | 0.87715884 | -0.112985904 | 0.29817966 | -0.3789189 | 0.7047481 | NA | | RNU7-141P | 106480822 |
| ENSG000002 | 0.8209774 | 0.219137605 | 0.30028134 | 0.7297743 | 0.46552815 | NA | | NA | NA |
| ENSG000002 | 0.86842953 | -0.108738482 | 0.29610105 | -0.3672344 | 0.71344421 | NA | | NA | NA |
| ENSG000002 | 0.28256998 | -0.18483211 | 0.18021686 | -1.0256094 | 0.30507573 | NA | | NA | NA |
| ENSG000002 | 1.11541635 | 0.181949205 | 0.32077866 | 0.56721107 | 0.57057079 | NA | | RNU7-92P | 106479061 |
| ENSG000002 | 2.09347992 | -0.240925408 | 0.36216831 | -0.6652305 | 0.50590307 | NA | | RNU7-182P | 106480831 |
| ENSG000002 | 0.38036605 | -0.293832733 | 0.22119083 | -1.3284128 | 0.18404178 | NA | | RNU7-152P | 106479081 |
| ENSG000002 | 0.45476004 | 0.3294872 | 0.23735363 | 1.38817005 | 0.16508527 | NA | | SNORD124 | 101340251 |
| ENSG000002 | 0.36416385 | 0.068880976 | 0.20758164 | 0.33182596 | 0.74002068 | NA | | SCARNA12 | 677777 |
| ENSG000002 | 1.19266753 | -0.391252932 | 0.31090412 | -1.258436 | 0.20823412 | NA | | NA | NA |
| ENSG000002 | 1.46314046 | -0.190522548 | 0.34125857 | -0.5582938 | 0.57664377 | NA | | NA | NA |
| ENSG000002 | 1.0908852 | 0.179690252 | 0.31995927 | 0.56160351 | 0.57438619 | NA | | RNU7-102P | 106479064 |
| ENSG000002 | 1.56964813 | -0.116029094 | 0.34465741 | -0.3366505 | 0.73638037 | NA | | RNU7-127P | 106480362 |
| ENSG000002 | 1.13406244 | -0.430764449 | 0.32352773 | -1.3314607 | 0.18303748 | NA | | NA | NA |
| ENSG000002 | 1.00587044 | -0.364802939 | 0.31519397 | -1.1573919 | 0.24711229 | NA | | NA | NA |
| ENSG000002 | 0.96103646 | 0.121982531 | 0.30514254 | 0.3997559 | 0.68933632 | NA | | NA | NA |
| ENSG000002 | 0.25993993 | -0.013144386 | 0.18211683 | -0.0721756 | 0.94246219 | NA | | NA | NA |
| ENSG000002 | 0.59852118 | -0.096713565 | 0.25514249 | -0.3790571 | 0.70464548 | NA | | RNU1-131P | 106480193 |
| ENSG000002 | 1.12521052 | -0.280538266 | 0.32709639 | -0.8576624 | 0.39107892 | NA | | NA | NA |
| ENSG000002 | 0.88667417 | -0.451945682 | 0.30012996 | -1.5058333 | 0.13210997 | NA | | RNU7-106P | 106479066 |
| ENSG000002 | 0.63893761 | -0.11408674 | 0.27243711 | -0.4187636 | 0.67538892 | NA | | NA | NA |
| ENSG000002 | 0.38494734 | -0.095409837 | 0.21124663 | -0.4516514 | 0.65152014 | NA | | NA | NA |

| | | | | | | | | |
|---|---|---|---|---|---|---|---|---|
| ENSG000002 | 0 | NA | NA | NA | NA | NA | NA | NA |
| ENSG000002 | 0.84534638 | -0.252050477 | 0.29012693 | -0.8687593 | 0.38497878 | NA | NA | NA |
| ENSG000002 | 0 | NA | NA | NA | NA | NA | RNU6-329P | 106479712 |
| ENSG000002 | 1.3497197 | -0.127076387 | 0.34062444 | -0.3730689 | 0.70909716 | NA | NA | NA |
| ENSG000002 | 2.15949882 | 0.31252367 | 0.36799136 | 0.84926905 | 0.3957316 | NA | SNORD121A | 100113379 |
| ENSG000002 | 0.7735878 | -0.177395638 | 0.27477067 | -0.6456134 | 0.51852974 | NA | RNU1-80P | 106480174 |
| ENSG000002 | 0.77495809 | -0.176906983 | 0.27486729 | -0.6436087 | 0.5198292 | NA | NA | NA |
| ENSG000002 | 0.63347722 | -0.105383092 | 0.26574722 | -0.3965539 | 0.69169647 | NA | NA | NA |
| ENSG000002 | 0.75597219 | -0.185264009 | 0.27899653 | -0.664037 | 0.50666665 | NA | NA | NA |
| ENSG000002 | 0.49031286 | -0.020802257 | 0.24597635 | -0.0845702 | 0.93260313 | NA | NA | NA |
| ENSG000002 | 1.48373929 | -0.050250217 | 0.34682779 | -0.1448852 | 0.8848015 | NA | RNU7-196P | 106479098 |
| ENSG000002 | 0 | NA | NA | NA | NA | NA | NA | NA |
| ENSG000002 | 0 | NA | NA | NA | NA | NA | RNU7-1 | 100147744 |
| ENSG000002 | 0.46273095 | 0.157073337 | 0.24331041 | 0.64556769 | 0.51855937 | NA | NA | NA |
| ENSG000002 | 0.53608206 | -0.386623811 | 0.24788931 | -1.5596631 | 0.11883952 | NA | NA | NA |
| ENSG000002 | 1.44569034 | -0.540008126 | 0.33142546 | -1.6293502 | 0.10323891 | NA | NA | NA |
| ENSG000002 | 0.85551123 | -0.243943637 | 0.28822537 | -0.8463642 | 0.39734961 | NA | NA | NA |
| ENSG000002 | 1.17350695 | -0.256970868 | 0.31123198 | -0.825657 | 0.40899868 | NA | NA | NA |
| ENSG000002 | 1.09045494 | 0.045114756 | 0.30931248 | 0.14585495 | 0.8840359 | NA | RNU6-953P | 106479988 |
| ENSG000002 | 0 | NA | NA | NA | NA | NA | NA | NA |
| ENSG000002 | 1.58973651 | -0.108744716 | 0.34803068 | -0.3124573 | 0.75469303 | NA | NA | NA |
| ENSG000002 | 0.14128499 | -0.097772014 | 0.13403235 | -0.7294658 | 0.4657168 | NA | NA | NA |
| ENSG000002 | 0.3588776 | -0.099902505 | 0.21555794 | -0.4634601 | 0.6430346 | NA | RNU7-173P | 106479090 |
| ENSG000002 | 1.27469455 | -0.518563938 | 0.33372084 | -1.5538854 | 0.12021176 | NA | RNU7-176P | 106481803 |
| ENSG000002 | 1.48365284 | -0.187334235 | 0.34406882 | -0.5444673 | 0.58611992 | NA | NA | NA |
| ENSG000002 | 0.4869258 | -0.176879301 | 0.23959723 | -0.738236 | 0.46037105 | NA | RNA5SP351 | 106480764 |
| ENSG000002 | 1.55299002 | 0.158923479 | 0.3428018 | 0.46360164 | 0.64293318 | NA | NA | NA |
| ENSG000002 | 1.06489173 | 0.510358507 | 0.31456425 | 1.6224301 | 0.10471129 | NA | RNU7-133P | 106479075 |
| ENSG000002 | 0.3934287 | -0.107186984 | 0.22072612 | -0.4856108 | 0.62724315 | NA | RNU7-187P | 106479095 |

| | | | | | | | | |
|---|---|---|---|---|---|---|---|---|
| ENSG000002 | 0.26199343 | -0.194018184 | 0.18422801 | -1.0531416 | 0.29227609 | NA | RNU6-420P | 106481769 |
| ENSG000002 | 0.63330253 | 0.047732352 | 0.25793088 | 0.1850587 | 0.853183 | NA | SCARNA10 | 692148 |
| ENSG000002 | 1.77977891 | 0.012618011 | 0.35387926 | 0.03565626 | 0.97155645 | NA | NA | NA |
| ENSG000002 | 1.77667004 | 0.247624222 | 0.34592219 | 0.71583793 | 0.47409143 | NA | NA | NA |
| ENSG000002 | 0.24411881 | -0.012946855 | 0.17864669 | -0.0724718 | 0.94222641 | NA | RNU7-95P | 106479062 |
| ENSG000002 | 1.11351973 | -0.258606322 | 0.31711803 | -0.8154892 | 0.41479235 | NA | NA | NA |
| ENSG000002 | 1.48958076 | -0.343974963 | 0.34668986 | -0.9921691 | 0.32111505 | NA | NA | NA |
| ENSG000002 | 0 | NA | NA | NA | NA | NA | SNORD108 | 338427 |
| ENSG000002 | 1.27306717 | -0.184389898 | 0.32429962 | -0.5685788 | 0.56964199 | NA | NA | NA |
| ENSG000002 | 0.74612639 | -0.351728697 | 0.28064337 | -1.2532942 | 0.21009867 | NA | RNU1-98P | 106481622 |
| ENSG000002 | 0.53374623 | -0.385819457 | 0.24767828 | -1.5577444 | 0.11929385 | NA | RNU6-956P | 106480404 |
| ENSG000002 | 1.96936268 | -0.446985458 | 0.36139648 | -1.2368285 | 0.21615077 | NA | NA | NA |
| ENSG000002 | 1.49672856 | -0.608871614 | 0.33880514 | -1.7971144 | 0.07231745 | NA | NA | NA |
| ENSG000002 | 1.31060512 | 0.177673151 | 0.33955993 | 0.52324535 | 0.60080353 | NA | NA | NA |
| ENSG000002 | 0.74365079 | -0.022786451 | 0.27880612 | -0.0817287 | 0.93486249 | NA | RNU7-138P | 106481801 |
| ENSG000002 | 0.51299499 | -0.204672849 | 0.24959455 | -0.8200213 | 0.41220396 | NA | MIR500B | 100422911 |
| ENSG000002 | 0.23868015 | 0.154625155 | 0.16997278 | 0.90970538 | 0.36297791 | NA | NA | NA |
| ENSG000002 | 0.61290067 | -0.091310841 | 0.25649069 | -0.3560006 | 0.72184011 | NA | RNU6-274P | 106479688 |
| ENSG000002 | 1.88624612 | -0.39208181 | 0.35926766 | -1.0913362 | 0.27512496 | NA | NA | NA |
| ENSG000002 | 0.90807438 | -0.279916171 | 0.29927035 | -0.9353288 | 0.34961889 | NA | NA | NA |
| ENSG000002 | 0.75232803 | -0.025593668 | 0.27939852 | -0.0916027 | 0.92701368 | NA | NA | NA |
| ENSG000002 | 0.8863036 | -0.294029201 | 0.30451173 | -0.965576 | 0.33425641 | NA | RNU7-170P | 106480453 |
| ENSG000002 | 1.92513666 | 0.067756322 | 0.35653667 | 0.19004026 | 0.84927758 | NA | NA | NA |
| ENSG000002 | 0.38884741 | -0.29745092 | 0.22235466 | -1.3377319 | 0.18098386 | NA | NA | NA |
| ENSG000002 | 1.90273065 | -0.533981154 | 0.35916096 | -1.4867461 | 0.13708188 | NA | NA | NA |
| ENSG000002 | 0.71903374 | -0.165977372 | 0.26905974 | -0.6168793 | 0.53731437 | NA | RNU7-185P | 106479094 |
| ENSG000002 | 0.50107449 | -0.372905725 | 0.24426561 | -1.5266403 | 0.12685049 | NA | NA | NA |
| ENSG000002 | 0.72848202 | -0.028325979 | 0.27774574 | -0.1019853 | 0.91876835 | NA | NA | NA |
| ENSG000002 | 0 | NA | NA | NA | NA | NA | RNU6-1132P | 106481937 |

| | | | | | | | | | |
|---|---|---|---|---|---|---|---|---|---|
| ENSG000002 | 0.58133382 | 0.231430989 | 0.26184763 | 0.88383838 | 0.37678347 | NA | | NA | NA |
| ENSG000002 | 0.48285104 | 0.156035981 | 0.24564443 | 0.63521075 | 0.52529097 | NA | | NA | NA |
| ENSG000002 | 1.2495561 | -0.045562567 | 0.33273362 | -0.1369341 | 0.89108291 | NA | | MIR1972-2 | 100422922 |
| ENSG000002 | 1.75406972 | -0.198121677 | 0.35767678 | -0.5539126 | 0.57963866 | NA | | RNU7-119P | 106479070 |
| ENSG000002 | 0.48416733 | -0.018553864 | 0.24528998 | -0.0756405 | 0.93970509 | NA | | RNU2-11P | 106481637 |
| ENSG000002 | 0 | NA | NA | NA | NA | NA | | SNORD125 | 100113380 |
| ENSG000002 | 0.95511906 | 0.126703664 | 0.31141718 | 0.40686151 | 0.6841097 | NA | | NA | NA |
| ENSG000002 | 0.38270188 | -0.272446727 | 0.2140009 | -1.2731102 | 0.20297894 | NA | | NA | NA |
| ENSG000002 | 1.10963947 | -0.111349078 | 0.32065306 | -0.3472572 | 0.72839811 | NA | | NA | NA |
| ENSG000002 | 0.38133351 | -0.102466351 | 0.2189713 | -0.4679442 | 0.63982449 | NA | | NA | NA |
| ENSG000002 | 0.49224375 | -0.024566489 | 0.23863564 | -0.1029456 | 0.91800615 | NA | | NA | NA |
| ENSG000002 | 1.26088187 | 0.435747621 | 0.32308882 | 1.34869297 | 0.1774356 | NA | | SNORA59A | 677885 |
| ENSG000002 | 1.65392057 | -0.553082479 | 0.35042133 | -1.5783357 | 0.11448852 | NA | | RNU7-195P | 106479097 |
| ENSG000002 | 0.9793139 | -0.164523313 | 0.30084235 | -0.5468755 | 0.58446426 | NA | | RNA5SP310 | 106480762 |
| ENSG000002 | 0.24411881 | -0.012946855 | 0.17864669 | -0.0724718 | 0.94222641 | NA | | NA | NA |
| ENSG000002 | 0.48032325 | -0.179266136 | 0.23878452 | -0.7507444 | 0.45280651 | NA | | RNU7-197P | 106480455 |
| ENSG000002 | 0.63159785 | -0.116736614 | 0.27184111 | -0.4294296 | 0.66761063 | NA | | SNORD109B | 338429 |
| ENSG000002 | 0.71763504 | 0.134026678 | 0.2755896 | 0.48632705 | 0.6267353 | NA | | RNU6-1218P | 106480099 |
| ENSG000002 | 0.66156766 | -0.287991838 | 0.26979811 | -1.0674346 | 0.28577564 | NA | | NA | NA |
| ENSG000002 | 1.58011491 | 0.023661747 | 0.34887317 | 0.06782335 | 0.94592626 | NA | | NA | NA |
| ENSG000002 | 1.20028966 | 0.248412721 | 0.32656762 | 0.76067775 | 0.44684957 | NA | | SNORA84 | 100124534 |
| ENSG000002 | 0.97109258 | -0.188894688 | 0.30794033 | -0.6134133 | 0.53960311 | NA | | NA | NA |
| ENSG000002 | 0.2596576 | -0.193002035 | 0.1837913 | -1.0501152 | 0.29366517 | NA | | RNU7-190P | 106479096 |
| ENSG000002 | 0.35210091 | 0.074791332 | 0.21412433 | 0.34928928 | 0.72687215 | NA | | RNU6-634P | 106479847 |
| ENSG000002 | 0.58958493 | 0.063416451 | 0.26802881 | 0.23660311 | 0.8129647 | NA | | RNU6-717P | 106479884 |
| ENSG000002 | 0 | NA | NA | NA | NA | NA | | RNU1-123P | 106481628 |
| ENSG000002 | 0 | NA | NA | NA | NA | NA | | SNORD5 | 692072 |
| ENSG000002 | 1.61054979 | -0.381219317 | 0.34346381 | -1.1099257 | 0.26703104 | NA | | NA | NA |
| ENSG000002 | 0 | NA | NA | NA | NA | NA | | NA | NA |

| | | | | | | | | |
|---|---|---|---|---|---|---|---|---|
| ENSG000002 | 1.07125221 | -0.498055016 | 0.30386257 | -1.6390799 | 0.10119663 | NA | NA | NA |
| ENSG000002 | 0.12685397 | -0.090215768 | 0.12901919 | -0.699243 | 0.48440017 | NA | NA | NA |
| ENSG000002 | 1.62733782 | -0.267750545 | 0.35088727 | -0.7630671 | 0.44542336 | NA | RN7SL499P | 106481059 |
| ENSG000002 | 0.35073255 | 0.241740392 | 0.2078501 | 1.16305162 | 0.24480856 | NA | NA | NA |
| ENSG000002 | 1.14507159 | -0.113208356 | 0.31886051 | -0.3550404 | 0.72255934 | NA | NA | NA |
| ENSG000002 | 2.0476335 | -0.311633744 | 0.35970508 | -0.866359 | 0.3862933 | NA | RPS26P55 | 100129650 |
| ENSG000002 | 1.49808386 | -0.194681002 | 0.34622677 | -0.5622933 | 0.57391619 | NA | RN7SL563P | 106479427 |
| ENSG000002 | 0.99166968 | -0.200962024 | 0.31760022 | -0.6327515 | 0.52689592 | NA | NA | NA |
| ENSG000002 | 0 | NA | NA | NA | NA | NA | NA | NA |
| ENSG000002 | 1.60524698 | -0.246485964 | 0.34316215 | -0.7182784 | 0.47258564 | NA | RN7SL442P | 106481043 |
| ENSG000002 | 1.62425481 | -0.114363931 | 0.34544483 | -0.3310628 | 0.74059706 | NA | NA | NA |
| ENSG000002 | 0.74686572 | -0.180243062 | 0.28006088 | -0.6435853 | 0.5198444 | NA | RN7SL546P | 106479420 |
| ENSG000002 | 0 | NA | NA | NA | NA | NA | NA | NA |
| ENSG000002 | 0.7523063 | -0.206231678 | 0.2915808 | -0.7072883 | 0.47938735 | NA | NA | NA |
| ENSG000002 | 0.52028267 | -0.202047067 | 0.25039858 | -0.8069018 | 0.41972305 | NA | NA | NA |
| ENSG000002 | 0.90761736 | -0.270783511 | 0.29354236 | -0.9224683 | 0.35628438 | NA | RN7SL578P | 106479433 |
| ENSG000002 | 0.94689828 | 0.124354263 | 0.31107421 | 0.39975755 | 0.6893351 | NA | NA | NA |
| ENSG000002 | 1.5549287 | -0.648161245 | 0.34474775 | -1.8801029 | 0.06009405 | NA | RN7SL757P | 106480530 |
| ENSG000002 | 0.7717195 | -0.360573024 | 0.28228954 | -1.2773163 | 0.20149061 | NA | RPS4XP23 | 106480153 |
| ENSG000002 | 1.76156827 | -0.606838076 | 0.35400444 | -1.7142104 | 0.0864901 | NA | NA | NA |
| ENSG000002 | 1.04369208 | 0.327645322 | 0.31213476 | 1.04969188 | 0.2938598 | NA | NA | NA |
| ENSG000002 | 0 | NA | NA | NA | NA | NA | NA | NA |
| ENSG000002 | 1.73239124 | -0.183897344 | 0.35167632 | -0.5229165 | 0.60103237 | NA | RN7SL184P | 106479288 |
| ENSG000002 | 1.53187601 | -0.34584511 | 0.3461793 | -0.9990346 | 0.31777791 | NA | NA | NA |
| ENSG000002 | 0.72244254 | 0.129271463 | 0.27760219 | 0.46567163 | 0.64145056 | NA | NA | NA |
| ENSG000002 | 0 | NA | NA | NA | NA | NA | NA | NA |
| ENSG000002 | 1.68613584 | -0.544828289 | 0.34901378 | -1.561051 | 0.11851173 | NA | NA | NA |
| ENSG000002 | 1.58771471 | 0.01410711 | 0.35414806 | 0.03983393 | 0.96822552 | NA | RN7SL11P | 106479229 |
| ENSG000002 | 0.73109859 | -0.024786187 | 0.2842681 | -0.087193 | 0.93051811 | NA | NA | NA |

| | | | | | | | | |
|---|---|---|---|---|---|---|---|---|
| ENSG000002 | 0.86518572 | -0.258761418 | 0.29124258 | -0.8884739 | 0.37428591 | NA | RN7SL854P | 106480383 |
| ENSG000002 | 0.47614243 | -0.020043108 | 0.23674202 | -0.0846622 | 0.93252992 | NA | NA | NA |
| ENSG000002 | 0.4009426 | -0.302672773 | 0.2240145 | -1.3511303 | 0.17665371 | NA | ATP6V1B1-AS1 | 101927750 |
| ENSG000002 | 1.80064613 | -0.619273413 | 0.35436897 | -1.7475385 | 0.08054398 | NA | NA | NA |
| ENSG000002 | 1.88756242 | -0.542678209 | 0.36080443 | -1.5040786 | 0.13256114 | NA | NA | NA |
| ENSG000002 | 1.59790117 | -0.368234707 | 0.34406416 | -1.0702501 | 0.28450674 | NA | RN7SL658P | 106480524 |
| ENSG000002 | 1.38961521 | 0.364606318 | 0.33432766 | 1.09056583 | 0.27546397 | NA | LLPH-AS1 | 103625681 |
| ENSG000002 | 0.25260017 | -0.015978705 | 0.18049464 | -0.0885273 | 0.92945758 | NA | NA | NA |
| ENSG000002 | 2.02585924 | -0.596163791 | 0.36336856 | -1.6406587 | 0.10086828 | NA | NA | NA |
| ENSG000002 | 0.55665861 | -0.363599063 | 0.24151673 | -1.5054819 | 0.13220023 | NA | RN7SL309P | 106481007 |
| ENSG000002 | 0.6567905 | -0.46248946 | 0.26597757 | -1.7388288 | 0.08206488 | NA | NA | NA |
| ENSG000002 | 1.01594437 | -0.180561054 | 0.29984811 | -0.6021751 | 0.54705761 | NA | RN7SL477P | 106479397 |
| ENSG000002 | 1.97124204 | -0.44500271 | 0.36178737 | -1.2300117 | 0.21869271 | NA | NA | NA |
| ENSG000002 | 0.7635422 | -0.029733906 | 0.28015406 | -0.1061341 | 0.91547594 | NA | NA | NA |
| ENSG000002 | 1.61783748 | -0.39132357 | 0.34688876 | -1.1280953 | 0.25927969 | NA | RN7SL87P | 106479255 |
| ENSG000002 | 1.67591947 | -0.566942013 | 0.3523632 | -1.6089706 | 0.10762278 | NA | NA | NA |
| ENSG000002 | 0 | NA | NA | NA | NA | NA | NA | NA |
| ENSG000002 | 1.03137711 | -0.522227534 | 0.30997739 | -1.6847278 | 0.0920411 | NA | NA | NA |
| ENSG000002 | 1.56275875 | -0.47158332 | 0.33873384 | -1.3921943 | 0.16386357 | NA | NA | NA |
| ENSG000002 | 1.0342145 | 0.501906792 | 0.31344364 | 1.60126648 | 0.10931791 | NA | NA | NA |
| ENSG000002 | 1.50227676 | -0.329387184 | 0.34275479 | -0.9609995 | 0.33655241 | NA | NA | NA |
| ENSG000002 | 0.89905168 | -0.281707301 | 0.30041599 | -0.9377241 | 0.34838623 | NA | NA | NA |
| ENSG000002 | 0.76394502 | -0.170709309 | 0.27234044 | -0.6268232 | 0.53077513 | NA | NA | NA |
| ENSG000002 | 0.99895681 | -0.192453284 | 0.30913013 | -0.622564 | 0.53357109 | NA | NA | NA |
| ENSG000002 | 1.01976012 | -0.369210986 | 0.31570354 | -1.1694864 | 0.24220773 | NA | NA | NA |
| ENSG000002 | 0 | NA | NA | NA | NA | NA | NA | NA |
| ENSG000002 | 0.63099704 | -0.107803684 | 0.25810428 | -0.4176749 | 0.67618481 | NA | RN7SL691P | 106481112 |
| ENSG000002 | 1.45652347 | 0.231143084 | 0.34401647 | 0.67189539 | 0.50165029 | NA | RN7SL569P | 106481077 |
| ENSG000002 | 0.12070844 | -0.088503695 | 0.12783931 | -0.6923042 | 0.48874631 | NA | RN7SL584P | 106480519 |

| | | | | | | | | |
|---|---|---|---|---|---|---|---|---|
| ENSG000002 | 0.7022164 | -0.022749028 | 0.26658526 | -0.0853349 | 0.93199514 | NA | RN7SL221P | 106480982 |
| ENSG000002 | 1.44156611 | -0.180766387 | 0.340862 | -0.5303213 | 0.59588917 | NA | NA | NA |
| ENSG000002 | 1.56553954 | 0.019723104 | 0.35052774 | 0.05626689 | 0.9551292 | NA | RN7SL333P | 106481014 |
| ENSG000002 | 2.0607841 | -0.003226549 | 0.36527114 | -0.0088333 | 0.99295214 | NA | NA | NA |
| ENSG000002 | 0.83284637 | 0.381014175 | 0.29569031 | 1.28855818 | 0.19755173 | NA | RN7SL583P | 106481082 |
| ENSG000002 | 1.55994276 | 0.023653895 | 0.33892441 | 0.06979106 | 0.94435996 | NA | NA | NA |
| ENSG000002 | 0.89820738 | -0.590908788 | 0.29086846 | -2.0315327 | 0.04220098 | NA | LOC102724728 | 102724728 |
| ENSG000002 | 1.2824387 | -0.514485804 | 0.33258611 | -1.5469251 | 0.1218813 | NA | NA | NA |
| ENSG000002 | 1.07591182 | 0.038345851 | 0.3144378 | 0.12195051 | 0.90293821 | NA | NA | NA |
| ENSG000002 | 0.26199343 | -0.194018184 | 0.18422801 | -1.0531416 | 0.29227609 | NA | NA | NA |
| ENSG000002 | 1.15967675 | -0.277549505 | 0.32304177 | -0.8591753 | 0.39024381 | NA | RPL30P5 | 100270978 |
| ENSG000002 | 1.71818698 | -0.324381421 | 0.35624355 | -0.9105608 | 0.36252682 | NA | NA | NA |
| ENSG000002 | 0 | NA | NA | NA | NA | NA | NA | NA |
| ENSG000002 | 1.27294672 | 0.641782045 | 0.33182152 | 1.93411819 | 0.0530986 | NA | RN7SL399P | 106479370 |
| ENSG000002 | 0.76215602 | -0.362469896 | 0.28326072 | -1.2796335 | 0.20067407 | NA | NA | NA |
| ENSG000002 | 2.01522078 | -0.399121622 | 0.35266201 | -1.1317398 | 0.25774386 | NA | RN7SL843P | 106481151 |
| ENSG000002 | 2.11978192 | -0.494505547 | 0.36263676 | -1.3636388 | 0.17268128 | NA | RN7SL12P | 106480926 |
| ENSG000002 | 2.09066857 | -0.2479615 | 0.36585151 | -0.6777654 | 0.49792045 | NA | RN7SL841P | 106481150 |
| ENSG000002 | 1.67207756 | -0.049420006 | 0.35262842 | -0.1401475 | 0.88854342 | NA | NA | NA |
| ENSG000002 | 0.737016 | -0.022718234 | 0.28466054 | -0.0798082 | 0.93638984 | NA | NA | NA |
| ENSG000002 | 0.95669594 | -0.03065973 | 0.29713163 | -0.1031857 | 0.9178156 | NA | NA | NA |
| ENSG000002 | 1.7341995 | -0.058315793 | 0.35445111 | -0.1645242 | 0.86931848 | NA | NA | NA |
| ENSG000002 | 1.45106308 | 0.236155132 | 0.34512508 | 0.68425954 | 0.49381129 | NA | RN7SL388P | 106479365 |
| ENSG000002 | 0 | NA | NA | NA | NA | NA | NA | NA |
| ENSG000002 | 1.62762249 | -0.554564227 | 0.35070199 | -1.5812976 | 0.11381 | NA | OR1J4 | 26219 |
| ENSG000002 | 1.96159031 | -0.178005385 | 0.36121097 | -0.4928017 | 0.6221527 | NA | NA | NA |
| ENSG000002 | 1.53440772 | -0.344085727 | 0.34554552 | -0.9957754 | 0.31935929 | NA | NA | NA |
| ENSG000002 | 1.25998278 | -0.497016518 | 0.32840001 | -1.5134485 | 0.13016577 | NA | RUVBL1-AS1 | 100874089 |
| ENSG000002 | 1.60627502 | -0.105498386 | 0.34300468 | -0.3075713 | 0.75840858 | NA | NA | NA |

| | | | | | | | | |
|---|---|---|---|---|---|---|---|---|
| ENSG000002 | 1.13983741 | -0.284518798 | 0.32751823 | -0.8687113 | 0.38500505 | NA | NA | NA |
| ENSG000002 | 0.27408862 | -0.199826037 | 0.18669465 | -1.0703362 | 0.28446801 | NA | NA | NA |
| ENSG000002 | 1.50045002 | -0.318184129 | 0.3405963 | -0.9341972 | 0.35020216 | NA | NA | NA |
| ENSG000002 | 1.24913581 | -0.326644859 | 0.32651273 | -1.0004047 | 0.31711471 | NA | NA | NA |
| ENSG000002 | 0.87535057 | -0.256239951 | 0.29181893 | -0.8780786 | 0.37990107 | NA | RN7SL241P | 106480988 |
| ENSG000002 | 1.49686707 | -0.410512681 | 0.3297232 | -1.2450221 | 0.21312362 | NA | NA | NA |
| ENSG000002 | 1.62770894 | -0.410641048 | 0.35216027 | -1.166063 | 0.24358899 | NA | NA | NA |
| ENSG000002 | 1.26672522 | -0.197627368 | 0.3277355 | -0.6030087 | 0.54650288 | NA | NA | NA |
| ENSG000002 | 1.84324591 | -0.386023894 | 0.35790552 | -1.0785637 | 0.28078228 | NA | NA | NA |
| ENSG000002 | 1.52829699 | -0.721594469 | 0.33139294 | -2.1774588 | 0.02944635 | NA | MEIKIN | 728637 |
| ENSG000002 | 0.35022155 | 0.06940789 | 0.21379411 | 0.32464827 | 0.74544729 | NA | NA | NA |
| ENSG000002 | 1.84978587 | -0.224600037 | 0.35097374 | -0.639934 | 0.5222155 | NA | NA | NA |
| ENSG000002 | 1.49005951 | -0.191223397 | 0.34728999 | -0.5506159 | 0.58189699 | NA | NA | NA |
| ENSG000002 | 1.49174582 | -0.463862824 | 0.34042303 | -1.362607 | 0.17300639 | NA | RN7SL193P | 106480493 |
| ENSG000002 | 1.96750325 | 0.058962297 | 0.36090932 | 0.1633715 | 0.87022594 | NA | NA | NA |
| ENSG000002 | 1.40886654 | -0.746617553 | 0.3374013 | -2.2128473 | 0.02690818 | NA | NA | NA |
| ENSG000002 | 1.06942738 | -0.316267841 | 0.29760586 | -1.062707 | 0.28791483 | NA | RN7SL668P | 106479466 |
| ENSG000002 | 1.78160621 | 0.016144971 | 0.34443563 | 0.0468737 | 0.96261389 | NA | NA | NA |
| ENSG000002 | 1.79956009 | -0.331726794 | 0.35385433 | -0.9374671 | 0.34851833 | NA | NA | NA |
| ENSG000002 | 1.73419559 | 0.174984219 | 0.34245584 | 0.51096871 | 0.60937296 | NA | RN7SL507P | 106479408 |
| ENSG000002 | 1.84669411 | -0.214119373 | 0.34917426 | -0.6132164 | 0.53973328 | NA | NA | NA |
| ENSG000002 | 2.15194567 | -0.467216851 | 0.35794541 | -1.305274 | 0.19179955 | NA | RN7SL692P | 106479476 |
| ENSG000002 | 1.74315287 | -0.442210313 | 0.350345 | -1.2622139 | 0.20687184 | NA | LOC105375115 | 105375115 |
| ENSG000002 | 0.97643282 | 0.115461059 | 0.30750612 | 0.37547565 | 0.70730676 | NA | IMP3P2 | 106480272 |
| ENSG000002 | 1.1284577 | -0.422016732 | 0.31918754 | -1.3221592 | 0.18611515 | NA | NA | NA |
| ENSG000002 | 1.33155223 | 0.025985345 | 0.33302238 | 0.07802883 | 0.93780512 | NA | TLR9 | 54106 |
| ENSG000002 | 0.9859487 | -0.166067641 | 0.3011363 | -0.55147 | 0.58131152 | NA | RN7SL63P | 106480939 |
| ENSG000002 | 0.76078766 | -0.199688475 | 0.28697083 | -0.6958494 | 0.48652316 | NA | RN7SL795P | 106479511 |
| ENSG000002 | 0 | NA | NA | NA | NA | NA | NA | NA |

| | | | | | | | | |
|---|---|---|---|---|---|---|---|---|
| ENSG000002 | 0.7585039 | -0.199073479 | 0.28684023 | -0.6940222 | 0.48766831 | NA | NA | NA |
| ENSG000002 | 0.84357575 | 0.054277865 | 0.28816962 | 0.18835388 | 0.85059925 | NA | RN7SL653P | 106481101 |
| ENSG000002 | 1.2077713 | 0.108521972 | 0.33289453 | 0.32599506 | 0.7444281 | NA | ITIH4-AS1 | 100873993 |
| ENSG000002 | 1.39997322 | -0.246729138 | 0.32988105 | -0.7479337 | 0.45450018 | NA | NA | NA |
| ENSG000002 | 1.35011254 | -0.098464479 | 0.33032421 | -0.2980844 | 0.76563878 | NA | NA | NA |
| ENSG000002 | 1.89345608 | -0.252590324 | 0.35669476 | -0.7081414 | 0.47885745 | NA | RN7SL255P | 106479315 |
| ENSG000002 | 1.08698905 | 0.031769778 | 0.32363216 | 0.09816632 | 0.92180023 | NA | NA | NA |
| ENSG000002 | 1.65861401 | 0.085577041 | 0.35235994 | 0.24286825 | 0.80810746 | NA | RN7SL213P | 106480979 |
| ENSG000002 | 0.51111617 | -0.186068642 | 0.2423498 | -0.7677689 | 0.44262448 | NA | RN7SL513P | 106481062 |
| ENSG000002 | 0.24004852 | -0.005223787 | 0.17775844 | -0.029387 | 0.97655594 | NA | NA | NA |
| ENSG000002 | 0.91741557 | -0.435441321 | 0.29663039 | -1.4679592 | 0.1421153 | NA | NA | NA |
| ENSG000002 | 1.32621501 | -0.094384581 | 0.32417667 | -0.2911517 | 0.77093532 | NA | NA | NA |
| ENSG000002 | 1.20546985 | 0.264892504 | 0.33301163 | 0.7954452 | 0.42635458 | NA | RN7SL225P | 106479304 |
| ENSG000002 | 0.35824644 | 0.072568381 | 0.21508272 | 0.33739754 | 0.73581724 | NA | NA | NA |
| ENSG000002 | 1.625828 | -0.543303186 | 0.34886327 | -1.5573528 | 0.11938675 | NA | RN7SL34P | 106481824 |
| ENSG000002 | 1.48103872 | -0.035095298 | 0.32393863 | -0.1083393 | 0.91372652 | NA | NA | NA |
| ENSG000002 | 1.54012924 | -0.332008438 | 0.34265238 | -0.9689366 | 0.33257681 | NA | NA | NA |
| ENSG000002 | 0 | NA | NA | NA | NA | NA | KRTAP9-2 | 83899 |
| ENSG000002 | 0 | NA | NA | NA | NA | NA | NA | NA |
| ENSG000002 | 0.11205239 | 0.082118251 | 0.12671962 | 0.64803108 | 0.51696485 | NA | NA | NA |
| ENSG000002 | 0.65382509 | -0.109680555 | 0.25845987 | -0.424362 | 0.67130181 | NA | NA | NA |
| ENSG000002 | 1.53275511 | -0.501635125 | 0.34700999 | -1.4455928 | 0.14829146 | NA | RN7SL864P | 106479536 |
| ENSG000002 | 1.27943946 | -0.323626087 | 0.32488075 | -0.9961381 | 0.31918307 | NA | RPL29P22 | 100271472 |
| ENSG000002 | 0.59490343 | 0.227912985 | 0.26313009 | 0.86616087 | 0.38640195 | NA | LOC101930094 | 101930094 |
| ENSG000002 | 2.13901886 | -0.624097677 | 0.36175428 | -1.7251978 | 0.08449184 | NA | NA | NA |
| ENSG000002 | 1.12577696 | -0.39284248 | 0.31389786 | -1.2514978 | 0.21075291 | NA | RN7SL394P | 106479367 |
| ENSG000002 | 0.99693611 | -0.035312292 | 0.30670838 | -0.1151331 | 0.90833961 | NA | NA | NA |
| ENSG000002 | 0 | NA | NA | NA | NA | NA | NA | NA |
| ENSG000002 | 1.5645742 | 0.022827458 | 0.34933815 | 0.06534487 | 0.94789941 | NA | RN7SL368P | 106479357 |

| | | | | | | | | |
|---|---|---|---|---|---|---|---|---|
| ENSG000002 | 0.12685397 | -0.090215768 | 0.12901919 | -0.699243 | 0.48440017 | NA | NA | NA |
| ENSG000002 | 0 | NA | NA | NA | NA | NA | RN7SL273P | 106481834 |
| ENSG000002 | 0.72409323 | 0.135832926 | 0.28390139 | 0.47845107 | 0.63232919 | NA | RN7SL748P | 106481125 |
| ENSG000002 | 0.22263036 | 0.146922267 | 0.16600367 | 0.88505432 | 0.37612734 | NA | NA | NA |
| ENSG000002 | 0.48555743 | -0.013775713 | 0.2454629 | -0.0561214 | 0.95524512 | NA | NA | NA |
| ENSG000002 | 1.25730059 | -0.034607525 | 0.31368213 | -0.1103267 | 0.91215026 | NA | NA | NA |
| ENSG000002 | 0.72235822 | 0.133238945 | 0.27758738 | 0.47998921 | 0.63123507 | NA | RN7SL420P | 106479377 |
| ENSG000002 | 0 | NA | NA | NA | NA | NA | NA | NA |
| ENSG000002 | 0.68448004 | -0.087655298 | 0.15827669 | -0.5538105 | 0.57970852 | NA | NA | NA |
| ENSG000002 | 1.21902219 | -0.179471509 | 0.32646092 | -0.5497488 | 0.58249165 | NA | NA | NA |
| ENSG000002 | 0 | NA | NA | NA | NA | NA | NA | NA |
| ENSG000002 | 2.12515815 | 0.432124663 | 0.36516269 | 1.18337571 | 0.23666028 | NA | NA | NA |
| ENSG000002 | 1.38893841 | -0.424120316 | 0.33954778 | -1.249074 | 0.211638 | NA | NA | NA |
| ENSG000002 | 2.07281513 | -0.00126673 | 0.36730159 | -0.0034487 | 0.9972483 | NA | NA | NA |
| ENSG000002 | 0.634074 | -0.448061355 | 0.26266228 | -1.7058458 | 0.08803679 | NA | NA | NA |
| ENSG000002 | 1.70341573 | -0.31349074 | 0.35167332 | -0.891426 | 0.37270067 | NA | NA | NA |
| ENSG000002 | 0.50681912 | -0.019233989 | 0.24032284 | -0.080034 | 0.93621025 | NA | NA | NA |
| ENSG000002 | 0 | NA | NA | NA | NA | NA | NA | NA |
| ENSG000002 | 0.39479706 | -0.300008631 | 0.22317059 | -1.3443018 | 0.17885081 | NA | NA | NA |
| ENSG000002 | 0.76403539 | -0.331533216 | 0.27524153 | -1.2045174 | 0.22838966 | NA | NA | NA |
| ENSG000002 | 1.78270951 | 0.137636754 | 0.3542814 | 0.38849557 | 0.69764933 | NA | NA | NA |
| ENSG000002 | 2.07197186 | -0.122424685 | 0.36290361 | -0.3373477 | 0.73585483 | NA | NA | NA |
| ENSG000002 | 0 | NA | NA | NA | NA | NA | NA | NA |
| ENSG000002 | 1.65360344 | -0.265847316 | 0.34704654 | -0.7660279 | 0.44365972 | NA | NA | NA |
| ENSG000002 | 0 | NA | NA | NA | NA | NA | NA | NA |
| ENSG000002 | 1.54248631 | 0.288272949 | 0.34749002 | 0.82958627 | 0.40677274 | NA | NA | NA |
| ENSG000002 | 1.97533407 | -0.320403459 | 0.36347807 | -0.8814932 | 0.37805092 | NA | NA | NA |
| ENSG000002 | 1.96485944 | 0.049879104 | 0.35369369 | 0.14102345 | 0.88785142 | NA | RN7SL146P | 106481827 |
| ENSG000002 | 0.9982701 | -0.343188488 | 0.30892322 | -1.1109184 | 0.26660348 | NA | NA | NA |

| | | | | | | | | |
|---|---|---|---|---|---|---|---|---|
| ENSG000002 | 1.7576354 | -0.475008493 | 0.35621854 | -1.3334749 | 0.182376 | NA | NA | NA |
| ENSG000002 | 1.35634398 | 0.025462498 | 0.34050555 | 0.07477851 | 0.94039094 | NA | NA | NA |
| ENSG000002 | 0.47821768 | -0.016471461 | 0.24462483 | -0.0673336 | 0.94631616 | NA | NA | NA |
| ENSG000002 | 1.65263652 | -0.413444779 | 0.35187923 | -1.1749622 | 0.24000985 | NA | RN7SL826P | 106479523 |
| ENSG000002 | 1.64151217 | -0.513463931 | 0.34272914 | -1.4981625 | 0.13409105 | NA | NA | NA |
| ENSG000002 | 0.70836193 | -0.024852369 | 0.26706475 | -0.0930575 | 0.92585791 | NA | RN7SL263P | 106480993 |
| ENSG000002 | 1.73349263 | -0.198885752 | 0.35643488 | -0.5579862 | 0.57685379 | NA | NA | NA |
| ENSG000002 | 1.38960237 | -0.493519355 | 0.32222451 | -1.5316009 | 0.12562094 | NA | NA | NA |
| ENSG000002 | 1.13685273 | 0.02586963 | 0.30831448 | 0.08390663 | 0.93313066 | NA | RN7SL771P | 106481132 |
| ENSG000002 | 1.96767973 | -0.424147289 | 0.35772095 | -1.1856932 | 0.23574349 | NA | MAGI1-AS1 | 100873983 |
| ENSG000002 | 0 | NA | NA | NA | NA | NA | NA | NA |
| ENSG000002 | 1.11485328 | 0.025028763 | 0.31584987 | 0.07924259 | 0.93683967 | NA | RN7SL297P | 106479330 |
| ENSG000002 | 0 | NA | NA | NA | NA | NA | RN7SL633P | 106479453 |
| ENSG000002 | 0 | NA | NA | NA | NA | NA | RN7SL621P | 106479449 |
| ENSG000002 | 1.37024068 | -0.419094537 | 0.33807547 | -1.2396479 | 0.21510566 | NA | NA | NA |
| ENSG000002 | 1.12612591 | -0.110015603 | 0.31653539 | -0.3475618 | 0.72816932 | NA | RN7SL520P | 106481065 |
| ENSG000002 | 0.853421 | 0.200392875 | 0.29549761 | 0.67815396 | 0.49767408 | NA | RN7SL223P | 106481754 |
| ENSG000002 | 1.88041909 | -0.520080748 | 0.35804457 | -1.4525587 | 0.14634633 | NA | RN7SL567P | 106481976 |
| ENSG000002 | 1.12518878 | -0.415709447 | 0.31749361 | -1.3093475 | 0.19041668 | NA | RN7SL865P | 106481154 |
| ENSG000002 | 1.20028966 | 0.248412721 | 0.32656762 | 0.76067775 | 0.44684957 | NA | NA | NA |
| ENSG000002 | 2.15937077 | -0.386895832 | 0.36533591 | -1.059014 | 0.28959342 | NA | RN7SL587P | 106479437 |
| ENSG000002 | 1.07830964 | -0.709323051 | 0.30979063 | -2.2896853 | 0.02203957 | NA | RN7SL794P | 106479510 |
| ENSG000002 | 1.41506819 | -0.416783209 | 0.33904439 | -1.229288 | 0.21896384 | NA | NA | NA |
| ENSG000002 | 0.11865494 | 0.083440866 | 0.12751478 | 0.65436232 | 0.5128784 | NA | NA | NA |
| ENSG000002 | 0.65084084 | -0.46034757 | 0.26550072 | -1.7338844 | 0.08293859 | NA | NA | NA |
| ENSG000002 | 2.09509499 | 0.000657136 | 0.36377309 | 0.00180644 | 0.99855867 | NA | NA | NA |
| ENSG000002 | 1.99545764 | -0.059863313 | 0.35475855 | -0.1687438 | 0.86599815 | NA | NA | NA |
| ENSG000002 | 0.14128499 | -0.097772014 | 0.13403235 | -0.7294658 | 0.4657168 | NA | NA | NA |
| ENSG000002 | 0.74003697 | -0.021372094 | 0.27854971 | -0.0767263 | 0.93884127 | NA | RN7SL187P | 106481829 |

| | | | | | | | | | |
|---|---|---|---|---|---|---|---|---|---|
| ENSG000002 | 1.18147977 | -0.254023892 | 0.31150519 | -0.8154724 | 0.41480195 | NA | | RN7SL717P | 106481118 |
| ENSG000002 | 1.87364048 | -0.501830633 | 0.35507323 | -1.4133159 | 0.15756287 | NA | | RN7SL100P | 106480948 |
| ENSG000002 | 1.66900235 | -0.71227597 | 0.35000084 | -2.0350693 | 0.0418439 | NA | | NA | NA |
| ENSG000002 | 1.47634554 | -0.194414508 | 0.34587384 | -0.5620966 | 0.57405021 | NA | | RN7SL764P | 106480531 |
| ENSG000002 | 0.38019136 | -0.10756138 | 0.21876967 | -0.491665 | 0.62295621 | NA | | NA | NA |
| ENSG000002 | 2.12059276 | 0.574851194 | 0.36554622 | 1.57258141 | 0.11581577 | NA | | RN7SL481P | 106481054 |
| ENSG000002 | 1.16636362 | -0.278924516 | 0.3232454 | -0.8628878 | 0.38819916 | NA | | NA | NA |
| ENSG000002 | 1.1486676 | -0.605126604 | 0.32183091 | -1.8802625 | 0.06007231 | NA | | NA | NA |
| ENSG000002 | 2.08436898 | -0.004630896 | 0.36279592 | -0.0127645 | 0.98981571 | NA | | NA | NA |
| ENSG000002 | 1.28895088 | -0.34715385 | 0.32929681 | -1.0542278 | 0.29177863 | NA | | NA | NA |
| ENSG000002 | 1.36403201 | -0.260050714 | 0.3347435 | -0.7768656 | 0.43723806 | NA | | RN7SL503P | 106481060 |
| ENSG000002 | 1.00339877 | -0.316062234 | 0.30116963 | -1.0494492 | 0.29397141 | NA | | NA | NA |
| ENSG000002 | 0.49076933 | -0.025124039 | 0.23846109 | -0.1053591 | 0.91609089 | NA | | NA | NA |
| ENSG000002 | 0 | NA | NA | NA | NA | NA | | LCE1F | 353137 |
| ENSG000002 | 1.27057027 | -0.661038152 | 0.32854268 | -2.0120313 | 0.04421664 | NA | | NA | NA |
| ENSG000002 | 0 | NA | NA | NA | NA | NA | | MTATP8P1 | 106480795 |
| ENSG000002 | 2.04145168 | -0.580218313 | 0.36081207 | -1.6080901 | 0.10781545 | NA | | NA | NA |
| ENSG000002 | 0.50630812 | -0.189615696 | 0.24175631 | -0.7843257 | 0.432849 | NA | | NA | NA |
| ENSG000002 | 0.88237128 | -0.115115451 | 0.29844458 | -0.385718 | 0.69970552 | NA | | NA | NA |
| ENSG000002 | 2.09151521 | -0.34242787 | 0.35842762 | -0.9553613 | 0.33939503 | NA | | NA | NA |
| ENSG000002 | 0.75441187 | -0.377475959 | 0.28823563 | -1.3096089 | 0.19032816 | NA | | NA | NA |
| ENSG000002 | 2.09539174 | -0.257583034 | 0.36802802 | -0.6999006 | 0.48398938 | NA | | KRTAP13-3 | 337960 |
| ENSG000002 | 0.34270765 | 0.258172734 | 0.21433185 | 1.20454674 | 0.22837833 | NA | | NA | NA |
| ENSG000002 | 0 | NA | NA | NA | NA | NA | | NA | NA |
| ENSG000002 | 0.93297772 | 0.27742623 | 0.30628238 | 0.9057858 | 0.36504924 | NA | | NA | NA |
| ENSG000002 | 0 | NA | NA | NA | NA | NA | | NA | NA |
| ENSG000002 | 0 | NA | NA | NA | NA | NA | | CSPG4P1Y | 114758 |
| ENSG000002 | 1.73913409 | -0.450424322 | 0.3529309 | -1.2762394 | 0.2018709 | NA | | NA | NA |
| ENSG000002 | 1.82064969 | -0.127145568 | 0.35835699 | -0.3548014 | 0.72273837 | NA | | NA | NA |

| | | | | | | | | |
|---|---|---|---|---|---|---|---|---|
| ENSG000002 | 0.60301814 | 0.060465317 | 0.26274989 | 0.230125 | 0.81799464 | NA | NA | NA |
| ENSG000002 | 1.83912121 | 0.005183108 | 0.36117207 | 0.0143508 | 0.98855011 | NA | RN7SL346P | 106479350 |
| ENSG000002 | 0.71894942 | -0.17346135 | 0.27644947 | -0.6274613 | 0.53035688 | NA | RN7SL38P | 106480932 |
| ENSG000002 | 0.88584714 | -0.288702375 | 0.30453821 | -0.9480005 | 0.3431292 | NA | NA | NA |
| ENSG000002 | 0.12685397 | -0.090215768 | 0.12901919 | -0.699243 | 0.48440017 | NA | NA | NA |
| ENSG000002 | 1.82577406 | -0.918509485 | 0.35134234 | -2.6142863 | 0.00894141 | NA | NA | NA |
| ENSG000002 | 1.41438304 | -0.417018261 | 0.33902866 | -1.2300384 | 0.21868274 | NA | NA | NA |
| ENSG000002 | 0.81226928 | 0.365915945 | 0.29070341 | 1.25872603 | 0.20812931 | NA | RN7SL837P | 106480536 |
| ENSG000002 | 0.86483744 | 0.050884827 | 0.29718268 | 0.17122407 | 0.86404758 | NA | MED28P2 | 106480445 |
| ENSG000002 | 1.03097621 | -0.184807282 | 0.30315411 | -0.609615 | 0.5421169 | NA | NA | NA |
| ENSG000002 | 1.38235918 | -0.127290928 | 0.33667319 | -0.3780845 | 0.70536779 | NA | NA | NA |
| ENSG000002 | 0.4009426 | -0.302672773 | 0.2240145 | -1.3511303 | 0.17665371 | NA | NA | NA |
| ENSG000002 | 1.01862043 | -0.175923532 | 0.30261218 | -0.5813498 | 0.56100472 | NA | RN7SL69P | 106480940 |
| ENSG000002 | 1.69101001 | -0.161263497 | 0.34636483 | -0.4655885 | 0.64151005 | NA | NA | NA |
| ENSG000002 | 0.58761574 | 0.20785012 | 0.25374465 | 0.81913104 | 0.41271166 | NA | KRTAP9-1 | 728318 |
| ENSG000002 | 0.51706583 | -0.070027487 | 0.15549871 | -0.4503413 | 0.65246439 | NA | RN7SL492P | 106479402 |
| ENSG000002 | 1.65260564 | -0.260121137 | 0.34647812 | -0.7507578 | 0.45279846 | NA | NA | NA |
| ENSG000002 | 1.28155768 | -0.49098552 | 0.32752079 | -1.4990973 | 0.1338484 | NA | NA | NA |
| ENSG000002 | 1.24235665 | -0.319018776 | 0.32386488 | -0.9850367 | 0.32460606 | NA | NA | NA |
| ENSG000002 | 0 | NA | NA | NA | NA | NA | NA | NA |
| ENSG000002 | 0.50704533 | -0.189333637 | 0.24184367 | -0.7828761 | 0.43369986 | NA | NA | NA |
| ENSG000002 | 1.20114702 | -0.046971503 | 0.32773889 | -0.1433199 | 0.88603755 | NA | NA | NA |
| ENSG000002 | 1.08580915 | 0.15326794 | 0.30380901 | 0.5044878 | 0.61391863 | NA | RN7SL445P | 106481044 |
| ENSG000002 | 1.20658755 | -0.553661937 | 0.31107561 | -1.7798308 | 0.07510366 | NA | NA | NA |
| ENSG000002 | 1.97331685 | 0.064090505 | 0.35867508 | 0.17868681 | 0.85818363 | NA | NA | NA |
| ENSG000002 | 0.8841642 | -0.254282913 | 0.29233359 | -0.8698382 | 0.38438884 | NA | RN7SL547P | 106479421 |
| ENSG000002 | 1.27690775 | -0.325612254 | 0.32566765 | -0.9998299 | 0.31739283 | NA | RN7SL98P | 106479259 |
| ENSG000002 | 1.60393865 | -0.123469648 | 0.35127246 | -0.3514925 | 0.72521887 | NA | NA | NA |
| ENSG000002 | 1.67576923 | 0.352930865 | 0.35049269 | 1.00695642 | 0.31395572 | NA | RN7SL403P | 106481034 |

| | | | | | | | | | |
|---|---|---|---|---|---|---|---|---|---|
| ENSG000002 | 1.89729072 | 0.539737577 | 0.35108001 | 1.53736345 | 0.12420433 | NA | | RN7SL150P | 106479276 |
| ENSG000002 | 0.49557738 | -0.021677348 | 0.2390401 | -0.090685 | 0.9277429 | NA | | NA | NA |
| ENSG000002 | 1.38817087 | -0.265266482 | 0.3352626 | -0.79122 | 0.42881561 | NA | | NA | NA |
| ENSG000002 | 1.10731863 | -0.543885597 | 0.31085259 | -1.7496576 | 0.08017741 | NA | | RN7SL808P | 106481861 |
| ENSG000002 | 0.33615718 | 0.211074506 | 0.19531107 | 1.08070938 | 0.27982641 | NA | | RN7SL666P | 106481854 |
| ENSG000002 | 2.09490483 | -0.573914862 | 0.35610543 | -1.6116431 | 0.10703962 | NA | | RN7SL580P | 106481081 |
| ENSG000002 | 1.93657837 | -0.370761683 | 0.3509641 | -1.0564091 | 0.29078133 | NA | | RN7SL305P | 106479334 |
| ENSG000002 | 1.30890934 | -0.853597831 | 0.32800414 | -2.6023996 | 0.00925739 | NA | | NA | NA |
| ENSG000002 | 1.68468374 | 0.080132771 | 0.34932435 | 0.22939361 | 0.81856301 | NA | | NA | NA |
| ENSG000002 | 0.61335714 | -0.10408451 | 0.2640241 | -0.3942235 | 0.69341602 | NA | | RN7SL310P | 106479336 |
| ENSG000002 | 0.74772108 | -0.029248394 | 0.27742891 | -0.1054266 | 0.91603729 | NA | | NA | NA |
| ENSG000002 | 0.23730987 | 0.1539805 | 0.16964526 | 0.90766167 | 0.364057 | NA | | NA | NA |
| ENSG000002 | 2.01851948 | -0.598677831 | 0.36330589 | -1.6478616 | 0.09938107 | NA | | NA | NA |
| ENSG000002 | 0.87921234 | -0.287138853 | 0.30422636 | -0.9438329 | 0.34525504 | NA | | NA | NA |
| ENSG000002 | 1.87525424 | -0.245101173 | 0.35400288 | -0.6923706 | 0.48870465 | NA | | NA | NA |
| ENSG000002 | 1.76711456 | -0.475622013 | 0.35528707 | -1.3386978 | 0.18066908 | NA | | NA | NA |
| ENSG000002 | 0.48367807 | -0.017505926 | 0.23765649 | -0.0736606 | 0.94128043 | NA | | RN7SL538P | 106480517 |
| ENSG000002 | 1.88944178 | -0.5273173 | 0.35907518 | -1.4685429 | 0.14195682 | NA | | NA | NA |
| ENSG000002 | 0 | NA | NA | NA | NA | NA | | NA | NA |
| ENSG000002 | 1.3526534 | -0.367978519 | 0.32884981 | -1.1189866 | 0.26314586 | NA | | NA | NA |
| ENSG000002 | 0 | NA | NA | NA | NA | NA | | NA | NA |
| ENSG000002 | 0 | NA | NA | NA | NA | NA | | NA | NA |
| ENSG000002 | 1.23310383 | 0.105175918 | 0.33355079 | 0.31532204 | 0.75251715 | NA | | NA | NA |
| ENSG000002 | 1.28951732 | -0.448440856 | 0.31759782 | -1.4119771 | 0.15795669 | NA | | NA | NA |
| ENSG000002 | 0 | NA | NA | NA | NA | NA | | NA | NA |
| ENSG000002 | 2.07461877 | -0.252761319 | 0.36577065 | -0.6910377 | 0.48954183 | NA | | RN7SL776P | 106479503 |
| ENSG000002 | 1.73574605 | -0.056448781 | 0.35004306 | -0.1612624 | 0.87188674 | NA | | NA | NA |
| ENSG000002 | 1.21293167 | 0.105176392 | 0.33170022 | 0.31708267 | 0.75118087 | NA | | NA | NA |
| ENSG000002 | 0 | NA | NA | NA | NA | NA | | NA | NA |

| | | | | | | | | |
|---|---|---|---|---|---|---|---|---|
| ENSG000002 | 1.55131488 | -0.642895689 | 0.34401509 | -1.8688009 | 0.06165053 | NA | NA | NA |
| ENSG000002 | 1.23615459 | -0.054776917 | 0.32265353 | -0.1697701 | 0.86519095 | NA | NA | NA |
| ENSG000002 | 0 | NA | NA | NA | NA | NA | NA | NA |
| ENSG000002 | 1.34829577 | -0.260597025 | 0.33661402 | -0.7741716 | 0.43882931 | NA | RN7SL231P | 106480984 |
| ENSG000002 | 2.05509129 | -0.750638522 | 0.36293724 | -2.0682323 | 0.03861819 | NA | NA | NA |
| ENSG000002 | 0 | NA | NA | NA | NA | NA | NA | NA |
| ENSG000002 | 1.718431 | -0.048371258 | 0.35111246 | -0.1377657 | 0.89042558 | NA | RN7SL23P | 106480482 |
| ENSG000002 | 0.35022155 | 0.06940789 | 0.21379411 | 0.32464827 | 0.74544729 | NA | NA | NA |
| ENSG000002 | 1.74013189 | -0.476069137 | 0.35703977 | -1.3333784 | 0.18240764 | NA | NA | NA |
| ENSG000002 | 0.97706452 | -0.025446604 | 0.30749601 | -0.0827543 | 0.93404694 | NA | RN7SL77P | 106479252 |
| ENSG000002 | 1.2952711 | -0.517255339 | 0.33289437 | -1.5538122 | 0.12022921 | NA | NA | NA |
| ENSG000002 | 1.20339695 | -0.183803417 | 0.32832469 | -0.5598221 | 0.57560079 | NA | NA | NA |
| ENSG000002 | 1.59728211 | -0.266742958 | 0.35162533 | -0.7585999 | 0.4480919 | NA | NA | NA |
| ENSG000002 | 2.15416105 | 0.189820477 | 0.36764794 | 0.51631046 | 0.60563759 | NA | NA | NA |
| ENSG000002 | 1.22994489 | -0.043758902 | 0.33357853 | -0.1311802 | 0.89563275 | NA | NA | NA |
| ENSG000002 | 0.40942396 | -0.284058841 | 0.21788566 | -1.303706 | 0.19233384 | NA | NA | NA |
| ENSG000002 | 1.58471374 | -0.554703162 | 0.32732847 | -1.6946377 | 0.09014417 | NA | NA | NA |
| ENSG000002 | 1.90884567 | 0.584193255 | 0.35640549 | 1.63912529 | 0.10118717 | NA | RN7SL521P | 106479413 |
| ENSG000002 | 1.12942461 | -0.263371317 | 0.31766218 | -0.8290924 | 0.4070521 | NA | NA | NA |
| ENSG000002 | 1.91092969 | -0.663925164 | 0.3565706 | -1.8619739 | 0.06260676 | NA | NA | NA |
| ENSG000002 | 1.72020568 | -0.053694414 | 0.35317016 | -0.1520355 | 0.87915891 | NA | NA | NA |
| ENSG000002 | 1.63385447 | -0.419137338 | 0.35380929 | -1.184642 | 0.23615903 | NA | NA | NA |
| ENSG000002 | 1.39912445 | -0.277403715 | 0.34086982 | -0.8138113 | 0.41575306 | NA | NA | NA |
| ENSG000002 | 0.96818171 | 0.282914018 | 0.31099265 | 0.90971286 | 0.36297396 | NA | RN7SL209P | 106480494 |
| ENSG000002 | 1.01549341 | -0.353204735 | 0.30957606 | -1.1409304 | 0.25389889 | NA | NA | NA |
| ENSG000002 | 0 | NA | NA | NA | NA | NA | NA | NA |
| ENSG000002 | 0.99309203 | -0.193195534 | 0.31363677 | -0.615985 | 0.53790445 | NA | RN7SL554P | 106479423 |
| ENSG000002 | 1.81075291 | -0.205239047 | 0.34483867 | -0.5951741 | 0.55172709 | NA | RN7SL591P | 106481084 |
| ENSG000002 | 1.96218709 | -0.401988161 | 0.35401549 | -1.1355101 | 0.25616166 | NA | NA | NA |

| | | | | | | | | |
|---|---|---|---|---|---|---|---|---|
| ENSG000002 | 0.48032325 | -0.179266136 | 0.23878452 | -0.7507444 | 0.45280651 | NA | NA | NA |
| ENSG000002 | 1.73850707 | -0.316925858 | 0.3492502 | -0.9074465 | 0.36417075 | NA | RN7SL415P | 106479375 |
| ENSG000002 | 1.33209995 | -0.087625155 | 0.32025913 | -0.273607 | 0.78438662 | NA | NA | NA |
| ENSG000002 | 1.66993816 | -0.042231509 | 0.34238646 | -0.1233446 | 0.90183425 | NA | NA | NA |
| ENSG000002 | 0.4833075 | 0.150471097 | 0.24570075 | 0.61241611 | 0.54026249 | NA | NA | NA |
| ENSG000002 | 0 | NA | NA | NA | NA | NA | NA | NA |
| ENSG000002 | 1.91305465 | -0.256794351 | 0.3568601 | -0.7195939 | 0.47177507 | NA | NA | NA |
| ENSG000002 | 0.37285215 | -0.099309896 | 0.2177345 | -0.4561055 | 0.64831414 | NA | NA | NA |
| ENSG000002 | 0.63352874 | -0.1085548 | 0.25832188 | -0.4202308 | 0.67431688 | NA | NA | NA |
| ENSG000002 | 0.86455457 | -0.111874481 | 0.29589056 | -0.3780941 | 0.70536068 | NA | RN7SL459P | 106479390 |
| ENSG000002 | 1.11123597 | -0.28165796 | 0.3266603 | -0.8622351 | 0.38855818 | NA | NA | NA |
| ENSG000002 | 1.22467846 | -0.178220993 | 0.32461535 | -0.5490221 | 0.5829903 | NA | RN7SL183P | 106480971 |
| ENSG000002 | 0.50107449 | -0.372905725 | 0.24426561 | -1.5266403 | 0.12685049 | NA | NA | NA |
| ENSG000002 | 0 | NA | NA | NA | NA | NA | NA | NA |
| ENSG000002 | 1.4942736 | -0.185102711 | 0.34104322 | -0.5427544 | 0.5872989 | NA | RN7SL709P | 106481116 |
| ENSG000002 | 1.13404071 | -0.601316557 | 0.32137655 | -1.8710654 | 0.06133601 | NA | NA | NA |
| ENSG000002 | 1.57010515 | -0.109262881 | 0.34199038 | -0.3194911 | 0.74935414 | NA | NA | NA |
| ENSG000002 | 2.14368605 | -0.382683064 | 0.36359464 | -1.0524992 | 0.29257058 | NA | NA | NA |
| ENSG000002 | 0.97794553 | -0.022432951 | 0.29810695 | -0.0752513 | 0.94001473 | NA | NA | NA |
| ENSG000002 | 1.35086185 | -0.117708124 | 0.33599767 | -0.3503242 | 0.72609539 | NA | NA | NA |
| ENSG000002 | 0.46227394 | 0.141974152 | 0.23581243 | 0.60206391 | 0.5471316 | NA | NA | NA |
| ENSG000002 | 0 | NA | NA | NA | NA | NA | NA | NA |
| ENSG000002 | 0.87577321 | -0.423224474 | 0.29448938 | -1.4371468 | 0.15067628 | NA | NA | NA |
| ENSG000002 | 1.71067117 | -0.307665784 | 0.35176413 | -0.8746366 | 0.38177167 | NA | RN7SL813P | 106480534 |
| ENSG000002 | 0.38036605 | -0.293832733 | 0.22119083 | -1.3284128 | 0.18404178 | NA | RN7SL259P | 106480992 |
| ENSG000002 | 1.91352555 | -0.652341081 | 0.35415563 | -1.8419616 | 0.06548076 | NA | NA | NA |
| ENSG000002 | 1.47720537 | -0.331046587 | 0.34463445 | -0.9605731 | 0.33676685 | NA | NA | NA |
| ENSG000002 | 1.99338283 | -0.57673388 | 0.36022755 | -1.6010266 | 0.10937103 | NA | NA | NA |
| ENSG000002 | 1.30287144 | -0.492259456 | 0.3272812 | -1.5040872 | 0.13255893 | NA | NA | NA |

| | | | | | | | | | |
|---|---|---|---|---|---|---|---|---|---|
| ENSG000002 | 1.90990623 | -0.529824287 | 0.35826476 | -1.4788624 | 0.13917709 | NA | | LOC284344 | 284344 |
| ENSG000002 | 1.1060196 | 0.032575485 | 0.31550625 | 0.1032483 | 0.91776591 | NA | | NA | NA |
| ENSG000002 | 1.35125415 | -0.10760767 | 0.33453465 | -0.3216638 | 0.74770744 | NA | | NA | NA |
| ENSG000002 | 0.86569672 | -0.108917689 | 0.29758624 | -0.3660038 | 0.71436225 | NA | | KRTAP10-5 | 386680 |
| ENSG000002 | 1.42121372 | -0.418340249 | 0.3391633 | -1.2334479 | 0.21740872 | NA | | NA | NA |
| ENSG000002 | 1.4721626 | -0.193553166 | 0.34141777 | -0.5669101 | 0.5707753 | NA | | LOC100507461 | 100507461 |
| ENSG000002 | 0.12070844 | -0.088503695 | 0.12783931 | -0.6923042 | 0.48874631 | NA | | NA | NA |
| ENSG000002 | 0 | NA | NA | NA | NA | NA | | NA | NA |
| ENSG000002 | 0.23730987 | 0.1539805 | 0.16964526 | 0.90766167 | 0.364057 | NA | | NA | NA |
| ENSG000002 | 0.89222292 | -0.117181559 | 0.29894016 | -0.39199 | 0.69506559 | NA | | RN7SL160P | 106480964 |
| ENSG000002 | 1.77320803 | -0.459417608 | 0.35181603 | -1.3058462 | 0.19160488 | NA | | RN7SL617P | 106481852 |
| ENSG000002 | 1.60188514 | 0.014449501 | 0.35332972 | 0.04089523 | 0.96737942 | NA | | NA | NA |
| ENSG000002 | 1.14206561 | -0.598469485 | 0.32172533 | -1.8601877 | 0.06285898 | NA | | RN7SL70P | 106479249 |
| ENSG000002 | 1.60708032 | -0.379782 | 0.34554948 | -1.0990669 | 0.27173888 | NA | | RN7SL570P | 106479430 |
| ENSG000002 | 1.75734916 | -0.335397835 | 0.35508463 | -0.9445575 | 0.34488486 | NA | | RN7SL494P | 106481057 |
| ENSG000002 | 0.12685397 | -0.090215768 | 0.12901919 | -0.699243 | 0.48440017 | NA | | NA | NA |
| ENSG000002 | 1.94562267 | -0.80777134 | 0.35512191 | -2.2746311 | 0.02292807 | NA | | NA | NA |
| ENSG000002 | 2.07312971 | 0.116760541 | 0.36312576 | 0.32154299 | 0.74779894 | NA | | RN7SL165P | 106479281 |
| ENSG000002 | 1.49174582 | -0.463862824 | 0.34042303 | -1.362607 | 0.17300639 | NA | | RN7SL191P | 106479292 |
| ENSG000002 | 0 | NA | NA | NA | NA | NA | | NA | NA |
| ENSG000002 | 1.20294206 | -0.04053012 | 0.33154045 | -0.1222479 | 0.9027027 | NA | | NA | NA |
| ENSG000002 | 0 | NA | NA | NA | NA | NA | | NA | NA |
| ENSG000002 | 0 | NA | NA | NA | NA | NA | | NA | NA |
| ENSG000002 | 1.40335891 | -0.232956775 | 0.32495156 | -0.7168969 | 0.4734377 | NA | | RN7SL39P | 106479239 |
| ENSG000002 | 0.84906839 | 0.05307851 | 0.30150448 | 0.17604551 | 0.86025819 | NA | | NA | NA |
| ENSG000002 | 0.62023988 | -0.112292793 | 0.27094277 | -0.4144521 | 0.67854306 | NA | | RN7SL836P | 106481149 |
| ENSG000002 | 1.58894319 | -0.368347662 | 0.34313446 | -1.0734791 | 0.28305623 | NA | | RN7SL553P | 106480518 |
| ENSG000002 | 1.59182227 | -0.234309587 | 0.34296262 | -0.6831928 | 0.49448503 | NA | | NA | NA |
| ENSG000002 | 1.68528185 | -0.506349751 | 0.34115718 | -1.4842125 | 0.13775256 | NA | | RN7SL443P | 106480510 |

| | | | | | | | | | |
|---|---|---|---|---|---|---|---|---|---|
| ENSG000002 | 1.41747944 | 0.384808308 | 0.3420775 | 1.12491557 | 0.26062481 | NA | | RN7SL801P | 106481139 |
| ENSG000002 | 1.04105269 | -0.509071869 | 0.30808165 | -1.6523927 | 0.09845452 | NA | | KRTAP5-8 | 57830 |
| ENSG000002 | 0 | NA | NA | NA | NA | NA | | NA | NA |
| ENSG000002 | 0.98346851 | -0.187046005 | 0.30848592 | -0.6063356 | 0.5442919 | NA | | RN7SL629P | 106479452 |
| ENSG000002 | 1.6617187 | -0.408984954 | 0.35137323 | -1.1639616 | 0.24443957 | NA | | NA | NA |
| ENSG000002 | 1.75076835 | -0.190443916 | 0.35116691 | -0.5423174 | 0.58759989 | NA | | RN7SL302P | 106479332 |
| ENSG000002 | 1.38006062 | -0.40823766 | 0.33835053 | -1.2065524 | 0.22760455 | NA | | NA | NA |
| ENSG000002 | 2.14599613 | -0.003826853 | 0.35243996 | -0.0108582 | 0.9913366 | NA | | NA | NA |
| ENSG000002 | 0.7182662 | -0.019331073 | 0.27704777 | -0.0697752 | 0.94437256 | NA | | NA | NA |
| ENSG000002 | 1.99355518 | -0.303784151 | 0.35577545 | -0.8538648 | 0.39317988 | NA | | LOC152225 | 152225 |
| ENSG000002 | 0.49101782 | -0.013272127 | 0.22978098 | -0.0577599 | 0.95393988 | NA | | NA | NA |
| ENSG000002 | 0.71299529 | 0.133310031 | 0.28316265 | 0.4707896 | 0.63779099 | NA | | NA | NA |
| ENSG000002 | 2.01398792 | -0.322927702 | 0.36200842 | -0.8920447 | 0.37236896 | NA | | RN7SL791P | 106481860 |
| ENSG000002 | 1.24200782 | -0.040995596 | 0.3273316 | -0.1252418 | 0.90033213 | NA | | NA | NA |
| ENSG000002 | 0.75982019 | -0.379788636 | 0.28855699 | -1.3161651 | 0.18811864 | NA | | NA | NA |
| ENSG000002 | 1.26205802 | -0.481972717 | 0.32618586 | -1.4776015 | 0.13951447 | NA | | NA | NA |
| ENSG000002 | 1.45525991 | -0.190570986 | 0.34428591 | -0.5535254 | 0.57990371 | NA | | NA | NA |
| ENSG000002 | 0 | NA | NA | NA | NA | NA | | NA | NA |
| ENSG000002 | 1.37305526 | -0.266740823 | 0.34037924 | -0.7836577 | 0.43324097 | NA | | NA | NA |
| ENSG000002 | 1.73363643 | -0.188833163 | 0.35140275 | -0.5373696 | 0.59101233 | NA | | NA | NA |
| ENSG000002 | 0.88007217 | -0.43061833 | 0.29637842 | -1.4529341 | 0.14624204 | NA | | NA | NA |
| ENSG000002 | 0.73411765 | -0.177781191 | 0.27747615 | -0.640708 | 0.52171241 | NA | | NA | NA |
| ENSG000002 | 1.09071553 | 0.041804406 | 0.31496857 | 0.13272564 | 0.89441036 | NA | | NA | NA |
| ENSG000002 | 0 | NA | NA | NA | NA | NA | | NA | NA |
| ENSG000002 | 0.27408862 | -0.199826037 | 0.18669465 | -1.0703362 | 0.28446801 | NA | | NA | NA |
| ENSG000002 | 1.97610565 | -0.193922676 | 0.36520443 | -0.5309976 | 0.59542044 | NA | | RN7SL234P | 106479307 |
| ENSG000002 | 0.82963345 | 0.054595074 | 0.3005004 | 0.18168054 | 0.85583344 | NA | | RN7SL205P | 106479297 |
| ENSG000002 | 1.11829689 | -0.114559094 | 0.31795914 | -0.360295 | 0.71862652 | NA | | RN7SL515P | 106481063 |
| ENSG000002 | 1.2321191 | -0.474711875 | 0.32535723 | -1.4590482 | 0.14455184 | NA | | NA | NA |

| | | | | | | | | |
|---|---|---|---|---|---|---|---|---|
| ENSG000002 | 1.60066569 | -0.373118948 | 0.34180128 | -1.0916254 | 0.2749978 | NA | RN7SL441P | 106479383 |
| ENSG000002 | 0 | NA | NA | NA | NA | NA | NA | NA |
| ENSG000002 | 1.34923043 | -0.122156988 | 0.33596234 | -0.3636032 | 0.71615432 | NA | NA | NA |
| ENSG000002 | 1.98576288 | -0.559707645 | 0.35772112 | -1.564648 | 0.11766546 | NA | NA | NA |
| ENSG000002 | 0.73684131 | 0.135920058 | 0.28475401 | 0.47732447 | 0.63313109 | NA | NA | NA |
| ENSG000002 | 1.50332664 | -0.347615721 | 0.34690172 | -1.0020582 | 0.31631547 | NA | NA | NA |
| ENSG000002 | 1.97596522 | -0.45435774 | 0.36315325 | -1.251146 | 0.21088121 | NA | NA | NA |
| ENSG000002 | 1.07988052 | -0.35043658 | 0.30032741 | -1.1668485 | 0.24327156 | NA | RN7SL182P | 106479287 |
| ENSG000002 | 0.48761094 | -0.176642206 | 0.23968055 | -0.7369902 | 0.46112833 | NA | NA | NA |
| ENSG000002 | 0.26062507 | -0.012888319 | 0.18226503 | -0.070712 | 0.94362699 | NA | NA | NA |
| ENSG000002 | 1.22339654 | 0.099643261 | 0.32394768 | 0.3075906 | 0.75839387 | NA | SSX2 | 6757 |
| ENSG000002 | 1.25301022 | -0.345213867 | 0.33059302 | -1.0442261 | 0.29638078 | NA | RN7SL586P | 106481083 |
| ENSG000002 | 0.61950267 | -0.112567707 | 0.27088102 | -0.4155614 | 0.67773094 | NA | NA | NA |
| ENSG000002 | 0.98463185 | -0.033962714 | 0.30023809 | -0.1131193 | 0.909936 | NA | NA | NA |
| ENSG000002 | 1.11018218 | 0.029474609 | 0.32188917 | 0.09156757 | 0.92704162 | NA | NA | NA |
| ENSG000002 | 1.38102808 | -0.252104713 | 0.33590409 | -0.7505259 | 0.45293804 | NA | RN7SL632P | 106481095 |
| ENSG000002 | 0.63982777 | -0.42118588 | 0.25611206 | -1.6445375 | 0.10006523 | NA | NA | NA |
| ENSG000002 | 0.37236288 | -0.09108266 | 0.20928647 | -0.4352057 | 0.66341314 | NA | NA | NA |
| ENSG000002 | 1.77450831 | -0.332666989 | 0.35571071 | -0.9352178 | 0.34967607 | NA | RN7SL369P | 106480694 |
| ENSG000002 | 1.22787412 | -0.326970115 | 0.32773501 | -0.9976661 | 0.31844128 | NA | NA | NA |
| ENSG000002 | 2.17118419 | -0.49103234 | 0.36090645 | -1.3605529 | 0.17365501 | NA | RN7SL58P | 106479245 |
| ENSG000002 | 0.73329399 | -0.327183738 | 0.2730203 | -1.1983861 | 0.23076673 | NA | NA | NA |
| ENSG000002 | 1.53275511 | -0.501635125 | 0.34700999 | -1.4455928 | 0.14829146 | NA | RN7SL338P | 106479347 |
| ENSG000002 | 0 | NA | NA | NA | NA | NA | NA | NA |
| ENSG000002 | 0.73926538 | -0.17468127 | 0.27788076 | -0.6286195 | 0.52959817 | NA | NA | NA |
| ENSG000002 | 0.11934008 | 0.083564698 | 0.12760091 | 0.65489106 | 0.5125379 | NA | RN7SL484P | 106479400 |
| ENSG000002 | 0.12070844 | -0.088503695 | 0.12783931 | -0.6923042 | 0.48874631 | NA | NA | NA |
| ENSG000002 | 1.33349866 | -0.371467015 | 0.32837315 | -1.1312344 | 0.25795644 | NA | KRTAP9-4 | 85280 |
| ENSG000002 | 2.06168102 | 0.246301147 | 0.36540154 | 0.67405612 | 0.50027564 | NA | NA | NA |

| | | | | | | | | |
|---|---|---|---|---|---|---|---|---|
| ENSG000002 | 1.01549396 | -0.343561364 | 0.30964813 | -1.1095219 | 0.26720512 | NA | NA | NA |
| ENSG000002 | 0.81892389 | 0.385469336 | 0.29640724 | 1.30047208 | 0.19343922 | NA | KRTAP5-4 | 387267 |
| ENSG000002 | 0.37441639 | -0.268856313 | 0.21277471 | -1.2635727 | 0.20638344 | NA | NA | NA |
| ENSG000002 | 1.1486676 | -0.605126604 | 0.32183091 | -1.8802625 | 0.06007231 | NA | RN7SL340P | 106481016 |
| ENSG000002 | 0 | NA | NA | NA | NA | NA | NA | NA |
| ENSG000002 | 1.25508546 | -0.340782337 | 0.33066942 | -1.0305832 | 0.30273634 | NA | NA | NA |
| ENSG000002 | 0.89478551 | -0.274894995 | 0.30023915 | -0.9155868 | 0.35988367 | NA | NA | NA |
| ENSG000002 | 1.6196537 | -0.253749824 | 0.34806131 | -0.7290377 | 0.46597858 | NA | RN7SL253P | 106479314 |
| ENSG000002 | 2.07911302 | -0.673070961 | 0.35309402 | -1.9062089 | 0.05662311 | NA | NA | NA |
| ENSG000002 | 0.61679627 | 0.054898003 | 0.27030252 | 0.20309838 | 0.83905814 | NA | RN7SL418P | 106480508 |
| ENSG000002 | 0.86897479 | -0.416382535 | 0.29162984 | -1.4277775 | 0.1533559 | NA | NA | NA |
| ENSG000002 | 1.57891877 | -0.118420858 | 0.35093838 | -0.3374406 | 0.73578478 | NA | NA | NA |
| ENSG000002 | 0 | NA | NA | NA | NA | NA | NA | NA |
| ENSG000002 | 1.63792722 | -0.248032832 | 0.34624867 | -0.7163431 | 0.47377955 | NA | RN7SL704P | 106481115 |
| ENSG000002 | 0.89225326 | -0.284002901 | 0.30005063 | -0.9465166 | 0.34388516 | NA | NA | NA |
| ENSG000002 | 0.61084662 | 0.056873999 | 0.26981345 | 0.21079009 | 0.83305108 | NA | RN7SL265P | 106479318 |
| ENSG000002 | 1.61712145 | -0.268869128 | 0.35075349 | -0.7665473 | 0.44335073 | NA | NA | NA |
| ENSG000002 | 0 | NA | NA | NA | NA | NA | NA | NA |
| ENSG000002 | 1.84119241 | -0.259843128 | 0.36026692 | -0.7212517 | 0.47075468 | NA | NA | NA |
| ENSG000002 | 1.48796253 | -0.323684291 | 0.34036184 | -0.9510005 | 0.34160411 | NA | NA | NA |
| ENSG000002 | 1.70643773 | -0.181369185 | 0.34814075 | -0.5209651 | 0.60239107 | NA | NA | NA |
| ENSG000002 | 1.0968914 | -0.111687087 | 0.32022735 | -0.3487744 | 0.72725872 | NA | RN7SL83P | 106479254 |
| ENSG000002 | 1.16448872 | -0.573771943 | 0.31672267 | -1.811591 | 0.07004942 | NA | RN7SL714P | 106480527 |
| ENSG000002 | 1.4998016 | -0.600250662 | 0.33877986 | -1.7718015 | 0.07642752 | NA | NA | NA |
| ENSG000002 | 1.3184038 | 0.329389692 | 0.3367683 | 0.97808997 | 0.32802983 | NA | RN7SL390P | 106479366 |
| ENSG000002 | 1.67820323 | -0.533607509 | 0.34651281 | -1.5399359 | 0.12357599 | NA | NA | NA |
| ENSG000002 | 1.684246 | 0.07255087 | 0.35039253 | 0.20705598 | 0.83596614 | NA | RN7SL504P | 106480515 |
| ENSG000002 | 0 | NA | NA | NA | NA | NA | NA | NA |
| ENSG000002 | 2.02050593 | -0.305946452 | 0.35827383 | -0.8539459 | 0.39313499 | NA | NA | NA |

| | | | | | | | | |
|---|---|---|---|---|---|---|---|---|
| ENSG000002 | 0.14128499 | -0.097772014 | 0.13403235 | -0.7294658 | 0.4657168 | NA | RN7SL319P | 106479339 |
| ENSG000002 | 1.54616659 | -0.633039583 | 0.34165853 | -1.8528429 | 0.06390488 | NA | NA | NA |
| ENSG000002 | 0 | NA | NA | NA | NA | NA | NA | NA |
| ENSG000002 | 1.60302675 | 0.009051827 | 0.34702078 | 0.02608439 | 0.97919002 | NA | NA | NA |
| ENSG000002 | 1.36682328 | -0.416597871 | 0.33983311 | -1.2258896 | 0.22024023 | NA | RN7SL170P | 106479283 |
| ENSG000002 | 1.29321759 | -0.351280895 | 0.33161099 | -1.0593162 | 0.2894558 | NA | RN7SL149P | 106480961 |
| ENSG000002 | 0.12685397 | -0.090215768 | 0.12901919 | -0.699243 | 0.48440017 | NA | NA | NA |
| ENSG000002 | 0.24550891 | -0.007886114 | 0.17900103 | -0.0440563 | 0.96485956 | NA | NA | NA |
| ENSG000002 | 0.9155362 | -0.460427183 | 0.30148657 | -1.5271897 | 0.12671384 | NA | RN7SL401P | 106480507 |
| ENSG000002 | 0.7381576 | -0.029362007 | 0.29007186 | -0.1012232 | 0.91937327 | NA | RN7SL363P | 106480504 |
| ENSG000002 | 0.85817878 | -0.255621562 | 0.29085906 | -0.8788503 | 0.37948247 | NA | NA | NA |
| ENSG000002 | 1.71932466 | -0.056233094 | 0.35706488 | -0.157487 | 0.87486101 | NA | NA | NA |
| ENSG000002 | 1.25829647 | -0.192907069 | 0.33126015 | -0.5823431 | 0.56033559 | NA | NA | NA |
| ENSG000002 | 1.44756783 | -0.182296863 | 0.34097252 | -0.534638 | 0.5929002 | NA | NA | NA |
| ENSG000002 | 0.52073914 | -0.194865995 | 0.24332694 | -0.8008402 | 0.42322415 | NA | NA | NA |
| ENSG000002 | 0.74222789 | -0.03475108 | 0.28498529 | -0.1219399 | 0.90294661 | NA | NA | NA |
| ENSG000002 | 1.49657114 | -0.05807026 | 0.33919224 | -0.1712016 | 0.86406524 | NA | NA | NA |
| ENSG000002 | 1.52390318 | -0.348549899 | 0.34604207 | -1.0072472 | 0.31381601 | NA | NA | NA |
| ENSG000002 | 0.61890185 | -0.110660695 | 0.2644872 | -0.4183972 | 0.67565676 | NA | RN7SL188P | 106479290 |
| ENSG000002 | 0.90563091 | -0.593331152 | 0.29128265 | -2.0369602 | 0.04165403 | NA | NA | NA |
| ENSG000002 | 1.09800109 | -0.101487914 | 0.31557046 | -0.3216014 | 0.74775466 | NA | NA | NA |
| ENSG000002 | 0.12070844 | -0.088503695 | 0.12783931 | -0.6923042 | 0.48874631 | NA | NA | NA |
| ENSG000002 | 1.77641691 | -0.450259823 | 0.34888328 | -1.2905744 | 0.19685131 | NA | RN7SL76P | 106479251 |
| ENSG000002 | 1.56775667 | -0.105964799 | 0.34337106 | -0.3086014 | 0.75762473 | NA | RN7SL655P | 106481102 |
| ENSG000002 | 0.60338479 | 0.225396395 | 0.26390915 | 0.85406814 | 0.39306724 | NA | RN7SL566P | 106481850 |
| ENSG000002 | 1.77738884 | -0.595026059 | 0.3512133 | -1.6942014 | 0.09022703 | NA | NA | NA |
| ENSG000002 | 1.28451002 | -0.182605852 | 0.32457998 | -0.5625912 | 0.57371326 | NA | RN7SL831P | 106481862 |
| ENSG000002 | 0.39246123 | -0.276864021 | 0.21549122 | -1.2848042 | 0.19886073 | NA | NA | NA |
| ENSG000002 | 1.85789274 | 0.135906165 | 0.35787748 | 0.37975612 | 0.70412646 | NA | RN7SL323P | 106479341 |

| | | | | | | | | |
|---|---|---|---|---|---|---|---|---|
| ENSG000002 | 0.46956163 | 0.155192681 | 0.24412227 | 0.63571702 | 0.52496088 | NA | NA | NA |
| ENSG000002 | 1.3542349 | -0.108972027 | 0.33459936 | -0.3256791 | 0.74466714 | NA | RN7SL26P | 106479234 |
| ENSG000002 | 1.51641102 | -0.057103185 | 0.34313273 | -0.1664172 | 0.86782864 | NA | NA | NA |
| ENSG000002 | 1.87611565 | -0.26457392 | 0.36054349 | -0.7338197 | 0.46305862 | NA | LOC255654 | 255654 |
| ENSG000002 | 1.00501253 | -0.042485437 | 0.30871955 | -0.1376182 | 0.89054215 | NA | NA | NA |
| ENSG000002 | 1.46846287 | -0.322961224 | 0.34140866 | -0.9459667 | 0.34416557 | NA | NA | NA |
| ENSG000002 | 0.98606438 | -0.168921331 | 0.29240132 | -0.5777037 | 0.56346416 | NA | RN7SL214P | 106479300 |
| ENSG000002 | 0.90280986 | -0.2778642 | 0.29901695 | -0.929259 | 0.35275487 | NA | NA | NA |
| ENSG000002 | 0 | NA | NA | NA | NA | NA | NA | NA |
| ENSG000002 | 0.51299499 | -0.204672849 | 0.24959455 | -0.8200213 | 0.41220396 | NA | NA | NA |
| ENSG000002 | 1.53563172 | -0.464426487 | 0.33893587 | -1.3702489 | 0.17060923 | NA | NA | NA |
| ENSG000002 | 0.11205239 | 0.082118251 | 0.12671962 | 0.64803108 | 0.51696485 | NA | RN7SL318P | 106481009 |
| ENSG000002 | 0.37899768 | -0.10176489 | 0.21865765 | -0.4654074 | 0.64163973 | NA | RN7SL416P | 106481037 |
| ENSG000002 | 0 | NA | NA | NA | NA | NA | NA | NA |
| ENSG000002 | 0 | NA | NA | NA | NA | NA | NA | NA |
| ENSG000002 | 1.34239818 | -0.245867132 | 0.33100757 | -0.742784 | 0.45761245 | NA | RN7SL267P | 106481833 |
| ENSG000002 | 0.71740691 | 0.131932464 | 0.26987097 | 0.48887238 | 0.62493205 | NA | NA | NA |
| ENSG000002 | 1.9277486 | -0.672384515 | 0.35837655 | -1.8761956 | 0.06062842 | NA | NA | NA |
| ENSG000002 | 1.84083232 | -0.691675513 | 0.34247767 | -2.0196222 | 0.04342259 | NA | DYNLL1P5 | 106480681 |
| ENSG000002 | 0 | NA | NA | NA | NA | NA | NA | NA |
| ENSG000002 | 0.14128499 | -0.097772014 | 0.13403235 | -0.7294658 | 0.4657168 | NA | NA | NA |
| ENSG000002 | 1.21967061 | 0.104974983 | 0.33321107 | 0.3150405 | 0.75273091 | NA | RN7SL361P | 106479355 |
| ENSG000002 | 1.89482444 | -0.390089318 | 0.35836853 | -1.0885144 | 0.27636807 | NA | NA | NA |
| ENSG000002 | 1.24290148 | -0.048225889 | 0.33390272 | -0.144431 | 0.88516017 | NA | RN7SL222P | 106481831 |
| ENSG000002 | 1.10389621 | -0.274942423 | 0.32274067 | -0.8518989 | 0.3942702 | NA | NA | NA |
| ENSG000002 | 2.11382889 | -0.239864922 | 0.36186985 | -0.6628486 | 0.50742753 | NA | RN7SL329P | 106479343 |
| ENSG000002 | 0.83325077 | 0.332656575 | 0.28215118 | 1.17900117 | 0.23839771 | NA | NA | NA |
| ENSG000002 | 0.71763504 | 0.134026678 | 0.2755896 | 0.48632705 | 0.6267353 | NA | NA | NA |
| ENSG000002 | 1.23655736 | -0.700007046 | 0.30737793 | -2.2773498 | 0.02276535 | NA | OR51B5 | 282763 |

| | | | | | | | | |
|---|---|---|---|---|---|---|---|---|
| ENSG000002 | 0.74677928 | -0.319192781 | 0.2716115 | -1.1751814 | 0.23992216 | NA | NA | NA |
| ENSG000002 | 1.17931863 | -0.399011436 | 0.31356755 | -1.2724896 | 0.20319922 | NA | NA | NA |
| ENSG000002 | 1.6557782 | -0.664964986 | 0.343494 | -1.9358853 | 0.05288175 | NA | NA | NA |
| ENSG000002 | 1.16016601 | -0.266619678 | 0.3186885 | -0.8366153 | 0.40280884 | NA | NA | NA |
| ENSG000002 | 0.89839879 | -0.285938669 | 0.30035378 | -0.9520062 | 0.34109383 | NA | NA | NA |
| ENSG000002 | 0.6009429 | 0.055606074 | 0.26256469 | 0.21178047 | 0.83227831 | NA | RN7SL215P | 106480980 |
| ENSG000002 | 1.10203858 | -0.110837856 | 0.31317725 | -0.3539141 | 0.72340323 | NA | NA | NA |
| ENSG000002 | 1.23436401 | -0.328230544 | 0.32444975 | -1.011653 | 0.31170402 | NA | NA | NA |
| ENSG000002 | 0.8388005 | 0.053439402 | 0.27876409 | 0.19170117 | 0.84797629 | NA | RN7SL97P | 106479258 |
| ENSG000002 | 0.90201709 | -0.64176087 | 0.29994787 | -2.1395747 | 0.03238916 | NA | NA | NA |
| ENSG000002 | 1.33452113 | 0.024638165 | 0.32797726 | 0.07512157 | 0.94011799 | NA | RN7SL594P | 106479439 |
| ENSG000002 | 1.55251127 | 0.022479466 | 0.34550665 | 0.06506233 | 0.94812438 | NA | RN7SL306P | 106481006 |
| ENSG000002 | 0.98159127 | -0.038717571 | 0.31257898 | -0.1238649 | 0.90142223 | NA | NA | NA |
| ENSG000002 | 0.51459361 | -0.192534203 | 0.24265108 | -0.7934611 | 0.42750921 | NA | NA | NA |
| ENSG000002 | 1.48118977 | -0.465126632 | 0.34021571 | -1.3671521 | 0.17157762 | NA | BGLAP | 632 |
| ENSG000002 | 0 | NA | NA | NA | NA | NA | NA | NA |
| ENSG000002 | 0.83238744 | 0.205712217 | 0.29592731 | 0.69514443 | 0.4869648 | NA | RN7SL456P | 106480511 |
| ENSG000002 | 0.9860346 | -0.036930222 | 0.30788545 | -0.1199479 | 0.9045244 | NA | NA | NA |
| ENSG000002 | 0 | NA | NA | NA | NA | NA | NA | NA |
| ENSG000002 | 0.74672329 | -0.032948972 | 0.27900439 | -0.1180948 | 0.90599253 | NA | RN7SL159P | 106479279 |
| ENSG000002 | 0 | NA | NA | NA | NA | NA | NA | NA |
| ENSG000002 | 0.86247932 | -0.11814524 | 0.29740301 | -0.3972564 | 0.69117842 | NA | NA | NA |
| ENSG000002 | 0.49492896 | -0.339454393 | 0.23477063 | -1.4458981 | 0.1482058 | NA | NA | NA |
| ENSG000002 | 0.51364788 | -0.186913858 | 0.24261662 | -0.7704083 | 0.44105773 | NA | NA | NA |
| ENSG000002 | 0.39462238 | -0.104329077 | 0.21264929 | -0.4906157 | 0.6236983 | NA | NA | NA |
| ENSG000002 | 0.89651997 | -0.277149519 | 0.29871254 | -0.9278135 | 0.35350433 | NA | NA | NA |
| ENSG000002 | 1.22280101 | -0.041426398 | 0.32138016 | -0.1289015 | 0.89743556 | NA | NA | NA |
| ENSG000002 | 0.46404725 | -0.01414962 | 0.22636165 | -0.0625089 | 0.95015757 | NA | RN7SL685P | 106479473 |
| ENSG000002 | 1.36958132 | 0.025612409 | 0.33949449 | 0.07544278 | 0.93986242 | NA | NA | NA |

| | | | | | | | | |
|---|---|---|---|---|---|---|---|---|
| ENSG000002 | 1.34550952 | -0.390586473 | 0.33404543 | -1.1692615 | 0.24229828 | NA | NA | NA |
| ENSG000002 | 1.74878348 | -0.344965495 | 0.35891464 | -0.9611352 | 0.33648419 | NA | RN7SL646P | 106481099 |
| ENSG000002 | 0.12070844 | -0.088503695 | 0.12783931 | -0.6923042 | 0.48874631 | NA | RN7SL561P | 106481075 |
| ENSG000002 | 1.69179619 | 0.206356438 | 0.35293023 | 0.58469471 | 0.55875301 | NA | NA | NA |
| ENSG000002 | 1.14120578 | -0.442826606 | 0.32509083 | -1.3621627 | 0.17314654 | NA | NA | NA |
| ENSG000002 | 0.6010317 | -0.250734806 | 0.2561147 | -0.9789942 | 0.32758284 | NA | RN7SL272P | 106480996 |
| ENSG000002 | 0 | NA | NA | NA | NA | NA | CT47A2 | 728090 |
| ENSG000002 | 1.47813846 | -0.342073523 | 0.34787654 | -0.9833188 | 0.32545058 | NA | NA | NA |
| ENSG000002 | 1.55518069 | -0.30987041 | 0.33622728 | -0.92161 | 0.35673206 | NA | NA | NA |
| ENSG000002 | 0 | NA | NA | NA | NA | NA | NA | NA |
| ENSG000002 | 0.63921993 | 0.0498278 | 0.25845554 | 0.19279061 | 0.84712296 | NA | RN7SL741P | 106481124 |
| ENSG000002 | 0 | NA | NA | NA | NA | NA | NA | NA |
| ENSG000002 | 2.0546692 | -0.570858736 | 0.35927905 | -1.5889007 | 0.11208281 | NA | NA | NA |
| ENSG000002 | 0 | NA | NA | NA | NA | NA | NA | NA |
| ENSG000002 | 1.18696662 | 0.106271494 | 0.31861976 | 0.33353705 | 0.73872893 | NA | NA | NA |
| ENSG000002 | 0 | NA | NA | NA | NA | NA | RN7SL800P | 106479512 |
| ENSG000002 | 1.47464278 | -0.18810567 | 0.34284914 | -0.5486543 | 0.58324268 | NA | NA | NA |
| ENSG000002 | 1.48730049 | -0.05837799 | 0.34399896 | -0.169704 | 0.86524295 | NA | NA | NA |
| ENSG000002 | 1.21870953 | -0.168425715 | 0.32276193 | -0.5218265 | 0.60179116 | NA | NA | NA |
| ENSG000002 | 0.99951988 | -0.038043584 | 0.31329561 | -0.1214303 | 0.90335021 | NA | NA | NA |
| ENSG000002 | 1.70585865 | -0.060657596 | 0.35585759 | -0.1704547 | 0.86465256 | NA | NA | NA |
| ENSG000002 | 0.89720511 | -0.276906822 | 0.29874913 | -0.9268875 | 0.35398497 | NA | NA | NA |
| ENSG000002 | 0.73055726 | -0.021363932 | 0.26874046 | -0.0794965 | 0.9366377 | NA | RN7SL789P | 106479508 |
| ENSG000002 | 1.68664629 | -0.398289277 | 0.34783842 | -1.1450411 | 0.25219215 | NA | GTF2A1L | 11036 |
| ENSG000002 | 1.83044125 | -0.617185834 | 0.35353078 | -1.7457768 | 0.08084974 | NA | NA | NA |
| ENSG000002 | 0 | NA | NA | NA | NA | NA | NA | NA |
| ENSG000002 | 1.46288199 | -0.048890071 | 0.345278 | -0.1415963 | 0.88739893 | NA | NA | NA |
| ENSG000002 | 1.70249131 | 0.353967075 | 0.35338075 | 1.00165918 | 0.31650823 | NA | RN7SL37P | 106479238 |
| ENSG000002 | 0.38036605 | -0.293832733 | 0.22119083 | -1.3284128 | 0.18404178 | NA | RN7SL156P | 106480963 |

| | | | | | | | | |
|---|---|---|---|---|---|---|---|---|
| ENSG000002 | 2.1218315 | -0.39338443 | 0.36780183 | -1.0695554 | 0.28481948 | NA | NA | NA |
| ENSG000002 | 0.76559516 | -0.206287743 | 0.28724125 | -0.7181689 | 0.47265312 | NA | MAGEA5 | 4104 |
| ENSG000002 | 1.60959546 | -0.113323767 | 0.34830353 | -0.3253592 | 0.74490921 | NA | NA | NA |
| ENSG000002 | 0 | NA | NA | NA | NA | NA | NA | NA |
| ENSG000002 | 0.26199343 | -0.194018184 | 0.18422801 | -1.0531416 | 0.29227609 | NA | NA | NA |
| ENSG000002 | 1.99784597 | -0.506722806 | 0.35073089 | -1.4447624 | 0.14852464 | NA | NA | NA |
| ENSG000002 | 1.72231126 | -0.188797729 | 0.35527339 | -0.5314153 | 0.59513099 | NA | NA | NA |
| ENSG000002 | 0.50750235 | -0.197774682 | 0.24907427 | -0.794039 | 0.42717272 | NA | NA | NA |
| ENSG000002 | 1.88943731 | -0.271188985 | 0.36201909 | -0.7491013 | 0.45379613 | NA | NA | NA |
| ENSG000002 | 1.85488676 | -0.25990449 | 0.36176791 | -0.7184288 | 0.47249292 | NA | RN7SL144P | 106480960 |
| ENSG000002 | 0.85792032 | -0.094687964 | 0.28939487 | -0.327193 | 0.74352195 | NA | RN7SL797P | 106481138 |
| ENSG000002 | 1.87493347 | -0.901197592 | 0.34799677 | -2.5896723 | 0.00960673 | NA | RN7SL372P | 106479358 |
| ENSG000002 | 0 | NA | NA | NA | NA | NA | NA | NA |
| ENSG000002 | 1.00174808 | -0.339205349 | 0.30909496 | -1.0974147 | 0.27246016 | NA | NA | NA |
| ENSG000002 | 0 | NA | NA | NA | NA | NA | NA | NA |
| ENSG000002 | 1.10368769 | -0.249842606 | 0.31428549 | -0.7949543 | 0.42664008 | NA | NA | NA |
| ENSG000002 | 1.11179849 | -0.115053625 | 0.31605856 | -0.3640263 | 0.71583838 | NA | NA | NA |
| ENSG000002 | 1.82251249 | 0.127483798 | 0.349517 | 0.36474277 | 0.71530343 | NA | NA | NA |
| ENSG000002 | 1.98458702 | -0.19379716 | 0.36449243 | -0.5316905 | 0.59494037 | NA | NA | NA |
| ENSG000002 | 1.24745287 | -0.314723622 | 0.32402291 | -0.9713005 | 0.33139865 | NA | NA | NA |
| ENSG000002 | 1.80299785 | -0.295755155 | 0.3433352 | -0.8614181 | 0.38900782 | NA | RN7SL164P | 106480965 |
| ENSG000002 | 0.77658704 | -0.341413701 | 0.27605383 | -1.236765 | 0.21617435 | NA | GNG10 | 2790 |
| ENSG000002 | 1.1491786 | -0.424387989 | 0.32149757 | -1.3200348 | 0.18682339 | NA | NA | NA |
| ENSG000002 | 0.46882441 | 0.14446209 | 0.23662046 | 0.6105224 | 0.54151581 | NA | RN7SL294P | 106479329 |
| ENSG000002 | 0.14128499 | -0.097772014 | 0.13403235 | -0.7294658 | 0.4657168 | NA | NA | NA |
| ENSG000002 | 0.62482117 | 0.05427805 | 0.26463902 | 0.20510222 | 0.83749226 | NA | RN7SL292P | 106481002 |
| ENSG000002 | 1.75686394 | -0.064635428 | 0.35588907 | -0.1816168 | 0.85588348 | NA | RN7SL862P | 106479535 |
| ENSG000002 | 2.06049569 | -0.448452953 | 0.35881634 | -1.249812 | 0.21136824 | NA | RN7SL132P | 106479270 |
| ENSG000002 | 1.49583447 | -0.053603955 | 0.34702273 | -0.1544681 | 0.87724063 | NA | NA | NA |

| | | | | | | | | | |
|---|---|---|---|---|---|---|---|---|---|
| ENSG000002 | 1.62881617 | -0.521092702 | 0.34379622 | -1.5157022 | 0.12959466 | NA | | RN7SL581P | 106479434 |
| ENSG000002 | 1.24267123 | -0.205275703 | 0.33726848 | -0.6086418 | 0.54276187 | NA | | NA | NA |
| ENSG000002 | 0.94646301 | 0.277536047 | 0.30691005 | 0.90429116 | 0.36584103 | NA | | NA | NA |
| ENSG000002 | 1.35651866 | -0.116573719 | 0.34076461 | -0.3420946 | 0.73227973 | NA | | NA | NA |
| ENSG000002 | 1.59819401 | -0.402474124 | 0.35054641 | -1.1481336 | 0.2509134 | NA | | NA | NA |
| ENSG000002 | 1.02832581 | -0.356454241 | 0.31009328 | -1.1495065 | 0.25034718 | NA | | RN7SL167P | 106480966 |
| ENSG000002 | 0.39462238 | -0.104329077 | 0.21264929 | -0.4906157 | 0.6236983 | NA | | NA | NA |
| ENSG000002 | 2.11494876 | -0.383660532 | 0.36617308 | -1.0477573 | 0.29475042 | NA | | RN7SL304P | 106479333 |
| ENSG000002 | 1.72136957 | 0.080692197 | 0.34811716 | 0.23179609 | 0.81669639 | NA | | NA | NA |
| ENSG000002 | 1.83362644 | -0.123699009 | 0.35946035 | -0.3441242 | 0.73075288 | NA | | RN7SL516P | 106479411 |
| ENSG000002 | 0.79827137 | -0.536769119 | 0.28115538 | -1.9091547 | 0.05624213 | NA | | NA | NA |
| ENSG000002 | 0.73750472 | -0.033552356 | 0.29002437 | -0.1156881 | 0.90789977 | NA | | NA | NA |
| ENSG000002 | 1.4781128 | -0.18040199 | 0.33806135 | -0.5336368 | 0.59359281 | NA | | NA | NA |
| ENSG000002 | 0 | NA | NA | NA | NA | NA | | NA | NA |
| ENSG000002 | 2.1663227 | -0.621564035 | 0.36140122 | -1.7198725 | 0.08545562 | NA | | NA | NA |
| ENSG000002 | 0 | NA | NA | NA | NA | NA | | NA | NA |
| ENSG000002 | 1.98697045 | 0.05397544 | 0.36046325 | 0.1497391 | 0.88097046 | NA | | RN7SL824P | 106481146 |
| ENSG000002 | 1.43977565 | 0.09052716 | 0.33396743 | 0.27106584 | 0.7863404 | NA | | NA | NA |
| ENSG000002 | 1.48479444 | -0.185918314 | 0.340878 | -0.5454101 | 0.58547147 | NA | | RN7SL432P | 106481041 |
| ENSG000002 | 1.10098534 | 0.193106283 | 0.32415947 | 0.59571384 | 0.55136641 | NA | | NA | NA |
| ENSG000002 | 2.11184244 | -0.497725768 | 0.36462591 | -1.3650313 | 0.17224323 | NA | | NA | NA |
| ENSG000002 | 0.61721891 | -0.105633141 | 0.26437344 | -0.3995603 | 0.68948037 | NA | | MRPL42P6 | 106480794 |
| ENSG000002 | 2.08659828 | -0.22618593 | 0.36054692 | -0.6273412 | 0.53043562 | NA | | RN7SL229P | 106479305 |
| ENSG000002 | 1.28616016 | -0.205070991 | 0.33199348 | -0.6176958 | 0.53677586 | NA | | NA | NA |
| ENSG000002 | 2.09374051 | -0.257078652 | 0.36595151 | -0.7024938 | 0.48237129 | NA | | RN7SL846P | 106479530 |
| ENSG000002 | 1.09683933 | -0.111720888 | 0.3202256 | -0.3488818 | 0.72717804 | NA | | NA | NA |
| ENSG000002 | 1.94406507 | -0.305933592 | 0.36131992 | -0.8467111 | 0.39715616 | NA | | NA | NA |
| ENSG000002 | 1.14440144 | -0.58687804 | 0.31959221 | -1.836334 | 0.06630828 | NA | | NA | NA |
| ENSG000002 | 1.83636082 | 0.000924276 | 0.35769333 | 0.00258399 | 0.99793828 | NA | | NA | NA |

| | | | | | | | | |
|---|---|---|---|---|---|---|---|---|
| ENSG000002 | 1.37826716 | -0.277192628 | 0.3404557 | -0.8141812 | 0.41554116 | NA | NA | NA |
| ENSG000002 | 2.15942748 | 0.302005727 | 0.36435068 | 0.82888752 | 0.40716806 | NA | RN7SL749P | 106479494 |
| ENSG000002 | 0 | NA | NA | NA | NA | NA | NA | NA |
| ENSG000002 | 2.03860285 | -0.317513423 | 0.35860152 | -0.8854213 | 0.37592945 | NA | RN7SL496P | 106481058 |
| ENSG000002 | 0 | NA | NA | NA | NA | NA | NA | NA |
| ENSG000002 | 2.09675347 | -0.717245673 | 0.3590659 | -1.9975321 | 0.04576741 | NA | NA | NA |
| ENSG000002 | 0.84223963 | 0.21011263 | 0.29483721 | 0.71263947 | 0.47606888 | NA | RN7SL180P | 106480970 |
| ENSG000002 | 1.50375276 | -0.2060895 | 0.34883083 | -0.5908007 | 0.55465397 | NA | RN7SL677P | 106479470 |
| ENSG000002 | 0 | NA | NA | NA | NA | NA | RN7SL244P | 106479311 |
| ENSG000002 | 0 | NA | NA | NA | NA | NA | RBMY1B | 378948 |
| ENSG000002 | 1.9233397 | -0.058706578 | 0.35539275 | -0.1651879 | 0.8687961 | NA | RN7SL812P | 106479518 |
| ENSG000002 | 0 | NA | NA | NA | NA | NA | NA | NA |
| ENSG000002 | 0 | NA | NA | NA | NA | NA | NA | NA |
| ENSG000002 | 1.19434 | 0.250311398 | 0.32638459 | 0.76692163 | 0.44312811 | NA | NA | NA |
| ENSG000002 | 1.84837011 | -0.116024409 | 0.35321784 | -0.3284783 | 0.74255002 | NA | NA | NA |
| ENSG000002 | 0.98121858 | -0.044597913 | 0.30604201 | -0.1457248 | 0.88413864 | NA | RN7SL384P | 106481028 |
| ENSG000002 | 1.60034263 | 0.279357199 | 0.34422686 | 0.81154969 | 0.41705007 | NA | NA | NA |
| ENSG000002 | 1.62125075 | -0.383141327 | 0.34696076 | -1.1042786 | 0.26947232 | NA | RN7SL868P | 106479537 |
| ENSG000002 | 0.72130252 | 0.311846629 | 0.28479648 | 1.09498063 | 0.27352512 | NA | NA | NA |
| ENSG000002 | 2.10221603 | -0.132436258 | 0.36486831 | -0.36297 | 0.71662728 | NA | NA | NA |
| ENSG000002 | 0.25370795 | -0.171331747 | 0.17411721 | -0.9840024 | 0.32511435 | NA | NA | NA |
| ENSG000002 | 1.4581886 | -0.186035297 | 0.34116604 | -0.5452925 | 0.58555233 | NA | NA | NA |
| ENSG000002 | 0.25351207 | -0.189965547 | 0.1824784 | -1.0410303 | 0.29786148 | NA | NA | NA |
| ENSG000002 | 0.12685397 | -0.090215768 | 0.12901919 | -0.699243 | 0.48440017 | NA | NA | NA |
| ENSG000002 | 1.1284577 | -0.422016732 | 0.31918754 | -1.3221592 | 0.18611515 | NA | LOC101926934 | 101926934 |
| ENSG000002 | 2.03402155 | -0.444607703 | 0.36031852 | -1.2339296 | 0.21722913 | NA | NA | NA |
| ENSG000002 | 0.38884741 | -0.29745092 | 0.22235466 | -1.3377319 | 0.18098386 | NA | RN7SL332P | 106479345 |
| ENSG000002 | 1.35762589 | -0.255921919 | 0.33557285 | -0.7626419 | 0.44567698 | NA | NA | NA |
| ENSG000002 | 0.25333738 | -0.015686291 | 0.1806605 | -0.0868275 | 0.93080867 | NA | RN7SL379P | 106480505 |

| | | | | | | | | |
|---|---|---|---|---|---|---|---|---|
| ENSG000002 | 2.09009297 | 0.123876222 | 0.36160241 | 0.34257576 | 0.73191764 | NA | NA | NA |
| ENSG000002 | 1.46632347 | -0.300835616 | 0.33457395 | -0.8991603 | 0.36856728 | NA | NA | NA |
| ENSG000002 | 0 | NA | NA | NA | NA | NA | RN7SL737P | 106481123 |
| ENSG000002 | 0.34270765 | 0.258172734 | 0.21433185 | 1.20454674 | 0.22837833 | NA | NA | NA |
| ENSG000002 | 0.60375535 | 0.060734453 | 0.26281561 | 0.23109149 | 0.81724372 | NA | NA | NA |
| ENSG000002 | 1.08704167 | 0.039971473 | 0.32110378 | 0.12448148 | 0.90093406 | NA | NA | NA |
| ENSG000002 | 0.84859411 | -0.386648446 | 0.28403842 | -1.3612541 | 0.17343342 | NA | NA | NA |
| ENSG000002 | 1.7501215 | -0.334098942 | 0.35707985 | -0.9356421 | 0.34945749 | NA | NA | NA |
| ENSG000002 | 1.9907003 | -0.184644669 | 0.36140808 | -0.5109036 | 0.60941859 | NA | RN7SL810P | 106481142 |
| ENSG000002 | 1.26236676 | -0.196422322 | 0.32989515 | -0.5954083 | 0.55157055 | NA | NA | NA |
| ENSG000002 | 0 | NA | NA | NA | NA | NA | RBMY2FP | 159162 |
| ENSG000002 | 0.34948434 | 0.063159058 | 0.2051263 | 0.30790326 | 0.75815594 | NA | NA | NA |
| ENSG000002 | 0.73218621 | -0.185627455 | 0.27901773 | -0.6652891 | 0.5058656 | NA | NA | NA |
| ENSG000002 | 1.3606772 | -0.39706911 | 0.3327926 | -1.1931428 | 0.23281344 | NA | RN7SL33P | 106479236 |
| ENSG000002 | 0.68351258 | -0.46544448 | 0.26656372 | -1.7460909 | 0.08079516 | NA | NA | NA |
| ENSG000002 | 1.48317229 | 0.081179148 | 0.3403246 | 0.23853447 | 0.81146658 | NA | NA | NA |
| ENSG000002 | 0.49059519 | 0.153206417 | 0.24651 | 0.62150184 | 0.53426948 | NA | NA | NA |
| ENSG000002 | 0.12685397 | -0.090215768 | 0.12901919 | -0.699243 | 0.48440017 | NA | RN7SL400P | 106481033 |
| ENSG000002 | 0.75387054 | -0.354739931 | 0.28113912 | -1.261795 | 0.20702257 | NA | NA | NA |
| ENSG000002 | 1.25619089 | -0.046159439 | 0.33070265 | -0.1395799 | 0.88899193 | NA | NA | NA |
| ENSG000002 | 0.14128499 | -0.097772014 | 0.13403235 | -0.7294658 | 0.4657168 | NA | NA | NA |
| ENSG000002 | 1.63892949 | -0.459617489 | 0.33365996 | -1.3775027 | 0.16835688 | NA | NA | NA |
| ENSG000002 | 0 | NA | NA | NA | NA | NA | NA | NA |
| ENSG000002 | 1.7757387 | -0.580687786 | 0.35042268 | -1.6571068 | 0.09749789 | NA | RN7SL643P | 106481098 |
| ENSG000002 | 0.84648854 | -0.247366387 | 0.2902103 | -0.8523694 | 0.39400908 | NA | RN7SL807P | 106481141 |
| ENSG000002 | 1.29413107 | -0.345270734 | 0.32943944 | -1.0480553 | 0.29461315 | NA | NA | NA |
| ENSG000002 | 1.6608484 | 0.333762338 | 0.34587644 | 0.96497564 | 0.33455701 | NA | UGT1A3 | 54659 |
| ENSG000002 | 1.24449661 | -0.339366582 | 0.33039402 | -1.0271572 | 0.30434646 | NA | RN7SL22P | 106480928 |
| ENSG000002 | 0.96585895 | -0.331548819 | 0.30509477 | -1.0867077 | 0.27716603 | NA | NA | NA |

| | | | | | | | | |
|---|---|---|---|---|---|---|---|---|
| ENSG000002 | 1.71656875 | -0.313035958 | 0.35298189 | -0.8868329 | 0.37516886 | NA | NA | NA |
| ENSG000002 | 0.24485602 | -0.012655878 | 0.17881722 | -0.0707755 | 0.94357643 | NA | NA | NA |
| ENSG000002 | 0.58747935 | 0.229296668 | 0.2624377 | 0.87371848 | 0.38227159 | NA | NA | NA |
| ENSG000002 | 0.61335714 | -0.10408451 | 0.2640241 | -0.3942235 | 0.69341602 | NA | NA | NA |
| ENSG000002 | 2.08863625 | -0.001270811 | 0.365152 | -0.0034802 | 0.99722319 | NA | RN7SL861P | 106479534 |
| ENSG000002 | 1.51023486 | 0.589363712 | 0.34437995 | 1.71137637 | 0.08701166 | NA | NA | NA |
| ENSG000002 | 0.90844494 | -0.459619446 | 0.30273017 | -1.5182479 | 0.12895191 | NA | TUBBP11 | 106480745 |
| ENSG000002 | 0 | NA | NA | NA | NA | NA | PPAN-P2RY11 | 692312 |
| ENSG000002 | 1.73556999 | -0.053286542 | 0.35060025 | -0.1519866 | 0.8791975 | NA | NA | NA |
| ENSG000002 | 0.72444206 | -0.19300978 | 0.28464216 | -0.6780787 | 0.49772181 | NA | RN7SL55P | 106479244 |
| ENSG000002 | 1.49055269 | -0.441980642 | 0.33681696 | -1.312228 | 0.18944322 | NA | NA | NA |
| ENSG000002 | 1.25394331 | -0.349615287 | 0.33203331 | -1.0529524 | 0.29236279 | NA | NA | NA |
| ENSG000002 | 1.5117221 | -0.492264752 | 0.34672723 | -1.4197465 | 0.15568151 | NA | RN7SL350P | 106481019 |
| ENSG000002 | 0.7265173 | -0.177755649 | 0.27865224 | -0.6379121 | 0.52353088 | NA | NA | NA |
| ENSG000002 | 0.62550439 | -0.114540246 | 0.27135801 | -0.4221001 | 0.67295194 | NA | NA | NA |
| ENSG000002 | 0.8868146 | -0.114915491 | 0.29866695 | -0.3847613 | 0.70041426 | NA | NA | NA |
| ENSG000002 | 1.35370856 | -0.525357833 | 0.32981516 | -1.5928856 | 0.11118585 | NA | NA | NA |
| ENSG000002 | 1.74715899 | -0.428815069 | 0.34913085 | -1.228236 | 0.2193584 | NA | RN7SL614P | 106481089 |
| ENSG000002 | 1.93828347 | -0.829905674 | 0.35801616 | -2.3180676 | 0.02044565 | NA | RN7SL571P | 106479431 |
| ENSG000002 | 0.90932596 | -0.400444177 | 0.28758106 | -1.3924567 | 0.16378414 | NA | NA | NA |
| ENSG000002 | 1.36315291 | -0.126523967 | 0.34089456 | -0.3711528 | 0.71052374 | NA | NA | NA |
| ENSG000002 | 1.72937027 | -0.197830041 | 0.35836046 | -0.5520421 | 0.58091954 | NA | NA | NA |
| ENSG000002 | 0.74752913 | -0.352552616 | 0.28071348 | -1.2559163 | 0.20914634 | NA | NA | NA |
| ENSG000002 | 2.11687965 | -0.398259563 | 0.36818347 | -1.0816878 | 0.27939128 | NA | NA | NA |
| ENSG000002 | 2.14211221 | -0.148503473 | 0.35040929 | -0.4238 | 0.67171164 | NA | NA | NA |
| ENSG000002 | 1.98518379 | -0.455763095 | 0.3632117 | -1.2548139 | 0.20954632 | NA | NA | NA |
| ENSG000002 | 0.74672329 | -0.032948972 | 0.27900439 | -0.1180948 | 0.90599253 | NA | RN7SL174P | 106479285 |
| ENSG000002 | 1.26123099 | -0.347053654 | 0.33223688 | -1.044597 | 0.29620925 | NA | RN7SL307P | 106480500 |
| ENSG000002 | 1.71528918 | -0.424841291 | 0.34852237 | -1.2189785 | 0.22285234 | NA | RN7SL738P | 106480695 |

| | | | | | | | | |
|---|---|---|---|---|---|---|---|---|
| ENSG000002 | 0.75281538 | -0.182194416 | 0.28046106 | -0.6496246 | 0.51593472 | NA | RN7SL601P | 106480520 |
| ENSG000002 | 1.13983741 | -0.284518798 | 0.32751823 | -0.8687113 | 0.38500505 | NA | NA | NA |
| ENSG000002 | 0.88089528 | -0.266461602 | 0.29382599 | -0.9068687 | 0.36447622 | NA | NA | NA |
| ENSG000002 | 1.85912314 | -0.117043986 | 0.35636786 | -0.3284359 | 0.74258213 | NA | RN7SL278P | 106479324 |
| ENSG000002 | 1.75774146 | -0.329044262 | 0.35554213 | -0.9254719 | 0.35472048 | NA | MCCC1-AS1 | 106480736 |
| ENSG000002 | 2.06889779 | -0.241750563 | 0.36413897 | -0.6638964 | 0.5067566 | NA | RN7SL173P | 106479284 |
| ENSG000002 | 1.32782932 | 0.154397852 | 0.3206153 | 0.48156732 | 0.63011335 | NA | RN7SL72P | 106480941 |
| ENSG000002 | 1.08974998 | 0.042005985 | 0.3123879 | 0.13446739 | 0.893033 | NA | NA | NA |
| ENSG000002 | 1.08519456 | 0.029889764 | 0.31477095 | 0.09495719 | 0.92434883 | NA | RN7SL89P | 106479256 |
| ENSG000002 | 1.5278073 | -0.181801415 | 0.3349155 | -0.5428277 | 0.58724843 | NA | NA | NA |
| ENSG000002 | 0.58298238 | 0.057700061 | 0.2609372 | 0.22112624 | 0.82499414 | NA | NA | NA |
| ENSG000002 | 0.35690842 | 0.061733055 | 0.20636749 | 0.29914137 | 0.76483218 | NA | NA | NA |
| ENSG000002 | 0.77154482 | -0.197957795 | 0.28162873 | -0.7029034 | 0.48211594 | NA | RN7SL107P | 106479262 |
| ENSG000002 | 0.11865494 | 0.083440866 | 0.12751478 | 0.65436232 | 0.5128784 | NA | RN7SL454P | 106479388 |
| ENSG000002 | 1.046515 | -0.336034665 | 0.3056447 | -1.0994291 | 0.27158096 | NA | NA | NA |
| ENSG000002 | 0.50750235 | -0.197774682 | 0.24907427 | -0.794039 | 0.42717272 | NA | RN7SL352P | 106479352 |
| ENSG000002 | 2.04239933 | -0.448315882 | 0.36095291 | -1.2420343 | 0.21422392 | NA | NA | NA |
| ENSG000002 | 2.02201418 | -0.194484779 | 0.3594942 | -0.5409956 | 0.58851061 | NA | NA | NA |
| ENSG000002 | 1.7223891 | 0.203076362 | 0.34653937 | 0.58601239 | 0.55786719 | NA | INGX | 27160 |
| ENSG000002 | 0.99807422 | -0.34307921 | 0.30891336 | -1.1106001 | 0.26674051 | NA | MNX1-AS1 | 645249 |
| ENSG000002 | 1.36887232 | -0.274177343 | 0.3402748 | -0.8057527 | 0.42038545 | NA | NA | NA |
| ENSG000002 | 1.14083521 | -0.265756092 | 0.31804191 | -0.8356009 | 0.40337948 | NA | LOC105376912 | 105376912 |
| ENSG000002 | 1.86364832 | -0.25940986 | 0.3600702 | -0.7204425 | 0.47125262 | NA | RN7SL337P | 106481015 |
| ENSG000002 | 2.04210594 | -0.435414954 | 0.35838029 | -1.2149523 | 0.22438429 | NA | KRTAP10-11 | 386678 |
| ENSG000002 | 0.77379475 | -0.378828349 | 0.28947278 | -1.3086838 | 0.19064148 | NA | NA | NA |
| ENSG000002 | 0.36465257 | 0.064724075 | 0.21605106 | 0.29957767 | 0.76449932 | NA | NA | NA |
| ENSG000002 | 0.26560726 | -0.177019426 | 0.17672274 | -1.0016788 | 0.31649875 | NA | NA | NA |
| ENSG000002 | 0.48555743 | -0.013775713 | 0.2454629 | -0.0561214 | 0.95524512 | NA | NA | NA |
| ENSG000002 | 0.63025982 | -0.115141786 | 0.26544572 | -0.4337677 | 0.66445712 | NA | NA | NA |

| | | | | | | | | |
|---|---|---|---|---|---|---|---|---|
| ENSG000002 | 0 | NA | NA | NA | NA | NA | TNFRSF6B | 8771 |
| ENSG000002 | 0.72995857 | 0.137808191 | 0.28428846 | 0.48474776 | 0.62785529 | NA | RN7SL111P | 106479263 |
| ENSG000002 | 1.20945424 | 0.09945328 | 0.32791193 | 0.30329265 | 0.76166684 | NA | NA | NA |
| ENSG000002 | 1.56084802 | -0.438759373 | 0.33454037 | -1.3115289 | 0.18967912 | NA | NA | NA |
| ENSG000002 | 0 | NA | NA | NA | NA | NA | NA | NA |
| ENSG000002 | 1.21842879 | -0.17650433 | 0.32443557 | -0.5440351 | 0.58641734 | NA | NA | NA |
| ENSG000002 | 0.97729265 | -0.028877505 | 0.30586432 | -0.0944128 | 0.92478125 | NA | NA | NA |
| ENSG000002 | 0.90298455 | -0.457507079 | 0.30248048 | -1.5125177 | 0.13040221 | NA | RN7SL649P | 106481100 |
| ENSG000002 | 0.88883586 | -0.273072391 | 0.29994637 | -0.9104041 | 0.36260945 | NA | RN7SL172P | 106480967 |
| ENSG000002 | 0.98090805 | 0.118158695 | 0.31251281 | 0.37809233 | 0.70536201 | NA | RN7SL108P | 106480950 |
| ENSG000002 | 1.12270158 | 0.022366281 | 0.31610236 | 0.07075645 | 0.94359159 | NA | RN7SL705P | 106481856 |
| ENSG000002 | 0.77330548 | -0.35598868 | 0.28246534 | -1.2602915 | 0.20756421 | NA | RN7SL838P | 106479527 |
| ENSG000002 | 1.09367592 | 0.02937719 | 0.32134385 | 0.0914198 | 0.92715903 | NA | NA | NA |
| ENSG000002 | 1.40044075 | -0.434142106 | 0.34165506 | -1.270703 | 0.20383433 | NA | NA | NA |
| ENSG000002 | 1.23672157 | -0.20137977 | 0.33594737 | -0.5994385 | 0.54888054 | NA | NA | NA |
| ENSG000002 | 1.43265602 | -0.396777836 | 0.3320354 | -1.1949865 | 0.23209229 | NA | NA | NA |
| ENSG000002 | 1.19569201 | 0.359686852 | 0.31102039 | 1.15647354 | 0.24748752 | NA | NA | NA |
| ENSG000002 | 0.61767784 | 0.056941233 | 0.25494708 | 0.22334531 | 0.82326677 | NA | NA | NA |
| ENSG000002 | 0.2596576 | -0.193002035 | 0.1837913 | -1.0501152 | 0.29366517 | NA | NA | NA |
| ENSG000002 | 0.99823784 | -0.315368334 | 0.30097328 | -1.0478283 | 0.29471771 | NA | RN7SL542P | 106479419 |
| ENSG000002 | 2.10614836 | -0.262791463 | 0.36570259 | -0.7185934 | 0.47239149 | NA | NA | NA |
| ENSG000002 | 1.76557151 | -0.198217155 | 0.35292968 | -0.5616336 | 0.57436571 | NA | NA | NA |
| ENSG000002 | 2.11070323 | -0.247521872 | 0.36185776 | -0.6840309 | 0.49395568 | NA | NA | NA |
| ENSG000002 | 1.83152533 | -0.215149465 | 0.34727182 | -0.619542 | 0.53555939 | NA | RN7SL526P | 106479415 |
| ENSG000002 | 0 | NA | NA | NA | NA | NA | NA | NA |
| ENSG000002 | 0.57336291 | 0.401178563 | 0.25679184 | 1.56227146 | 0.11822405 | NA | RN7SL834P | 106481148 |
| ENSG000002 | 1.84219021 | -0.258036133 | 0.35832656 | -0.7201144 | 0.47145454 | NA | NA | NA |
| ENSG000002 | 0.98384066 | -0.185725188 | 0.30683884 | -0.6052858 | 0.54498912 | NA | NA | NA |
| ENSG000002 | 0.88035395 | -0.280358811 | 0.29946114 | -0.936211 | 0.34916456 | NA | NA | NA |

| | | | | | | | | |
|---|---|---|---|---|---|---|---|---|
| ENSG000002 | 1.48063591 | 0.089602836 | 0.33881912 | 0.26445626 | 0.79142837 | NA | RN7SL761P | 106479499 |
| ENSG000002 | 1.00698171 | -0.189611479 | 0.30947391 | -0.6126897 | 0.54008153 | NA | NA | NA |
| ENSG000002 | 0 | NA | NA | NA | NA | NA | NA | NA |
| ENSG000002 | 0 | NA | NA | NA | NA | NA | NA | NA |
| ENSG000002 | 1.0143518 | -0.367132577 | 0.31550649 | -1.1636292 | 0.2445743 | NA | NA | NA |
| ENSG000002 | 1.2541656 | -0.183852882 | 0.3212534 | -0.5722986 | 0.56711967 | NA | RN7SL598P | 106481759 |
| ENSG000002 | 1.5120377 | -0.188181496 | 0.33301035 | -0.565092 | 0.57201117 | NA | RN7SL638P | 106479455 |
| ENSG000002 | 0.37199232 | 0.06760863 | 0.21718977 | 0.31128828 | 0.75558148 | NA | RN7SL105P | 106480487 |
| ENSG000002 | 1.6485445 | -0.520624788 | 0.34411595 | -1.5129342 | 0.13029637 | NA | NA | NA |
| ENSG000002 | 0 | NA | NA | NA | NA | NA | NA | NA |
| ENSG000002 | 1.47105929 | -0.323772535 | 0.34005361 | -0.952122 | 0.3410351 | NA | RN7SL393P | 106481756 |
| ENSG000002 | 0.58895377 | 0.229836184 | 0.2625711 | 0.87532932 | 0.38139475 | NA | NA | NA |
| ENSG000002 | 0.23816916 | -0.010712105 | 0.1772998 | -0.060418 | 0.9518227 | NA | NA | NA |
| ENSG000002 | 0 | NA | NA | NA | NA | NA | ZMYM6NB | 100506144 |
| ENSG000002 | 0.64007922 | -0.113235616 | 0.26626803 | -0.4252693 | 0.67064038 | NA | NA | NA |
| ENSG000002 | 0.62023988 | -0.112292793 | 0.27094277 | -0.4144521 | 0.67854306 | NA | NA | NA |
| ENSG000002 | 1.9886012 | -0.449559512 | 0.36052018 | -1.2469746 | 0.21240681 | NA | NA | NA |
| ENSG000002 | 1.25087027 | -0.355937019 | 0.33393285 | -1.065894 | 0.28647157 | NA | NA | NA |
| ENSG000002 | 0.75116469 | -0.191751385 | 0.28641359 | -0.6694912 | 0.50318218 | NA | NA | NA |
| ENSG000002 | 0 | NA | NA | NA | NA | NA | NA | NA |
| ENSG000002 | 1.25775514 | -0.199637515 | 0.33468688 | -0.5964904 | 0.55084766 | NA | RN7SL78P | 106481752 |
| ENSG000002 | 1.99108138 | -0.457410671 | 0.36331667 | -1.2589862 | 0.20803532 | NA | NA | NA |
| ENSG000002 | 0.52147635 | -0.19458195 | 0.24341208 | -0.7993932 | 0.42406247 | NA | SNRPCP10 | 106481701 |
| ENSG000002 | 1.24147701 | -0.19493898 | 0.32934838 | -0.5918929 | 0.55392229 | NA | NA | NA |
| ENSG000002 | 2.14038488 | -0.390145862 | 0.36580109 | -1.0665519 | 0.28617422 | NA | RN7SL189P | 106480973 |
| ENSG000002 | 1.24998925 | -0.356057169 | 0.33390887 | -1.0663304 | 0.28627433 | NA | NA | NA |
| ENSG000002 | 2.11995661 | -0.627894127 | 0.36294981 | -1.7299751 | 0.08363473 | NA | LOC100507210 | 100507210 |
| ENSG000002 | 1.52162739 | -0.34131812 | 0.34534673 | -0.9883346 | 0.32298881 | NA | RN7SL610P | 106479444 |
| ENSG000002 | 0.98415578 | -0.033874028 | 0.31268148 | -0.108334 | 0.91373077 | NA | NA | NA |

| | | | | | | | | |
|---|---|---|---|---|---|---|---|---|
| ENSG000002 | 1.00930958 | -0.192415952 | 0.30791607 | -0.6248974 | 0.53203839 | NA | RN7SL236P | 106480986 |
| ENSG000002 | 0.91411227 | -0.636940681 | 0.29906252 | -2.129791 | 0.03318887 | NA | RN7SL487P | 106481055 |
| ENSG000002 | 0.89887754 | -0.114830835 | 0.29928032 | -0.3836899 | 0.7012083 | NA | RN7SL419P | 106479376 |
| ENSG000002 | 0 | NA | NA | NA | NA | NA | NA | NA |
| ENSG000002 | 0.59032214 | 0.060325881 | 0.26160816 | 0.23059632 | 0.81762842 | NA | NA | NA |
| ENSG000002 | 1.42084261 | -0.267309516 | 0.33323168 | -0.8021732 | 0.42245278 | NA | RN7SL320P | 106481010 |
| ENSG000002 | 1.77670425 | -0.598696329 | 0.35308493 | -1.6956157 | 0.08995869 | NA | ELFN2 | 114794 |
| ENSG000002 | 1.68662902 | -0.863977128 | 0.34755657 | -2.4858605 | 0.01292386 | NA | NA | NA |
| ENSG000002 | 2.00072079 | -0.185747903 | 0.36240998 | -0.5125353 | 0.60827643 | NA | NA | NA |
| ENSG000002 | 1.76369269 | -0.203026023 | 0.3572087 | -0.5683681 | 0.56978506 | NA | NA | NA |
| ENSG000002 | 0.75984193 | -0.181352233 | 0.27925466 | -0.6494152 | 0.51607002 | NA | NA | NA |
| ENSG000002 | 0 | NA | NA | NA | NA | NA | NA | NA |
| ENSG000002 | 1.56403424 | 0.134396519 | 0.33673564 | 0.39911582 | 0.68980787 | NA | NA | NA |
| ENSG000002 | 1.22499695 | -0.305212122 | 0.32037967 | -0.9526576 | 0.34076359 | NA | NA | NA |
| ENSG000002 | 2.09308114 | 0.116326814 | 0.35964523 | 0.32344879 | 0.74635538 | NA | RN7SL308P | 106479335 |
| ENSG000002 | 0.84904665 | -0.109000923 | 0.29670143 | -0.3673758 | 0.71333873 | NA | RN7SL743P | 106479492 |
| ENSG000002 | 1.76314744 | 0.047150358 | 0.34154022 | 0.13805214 | 0.89019921 | NA | RN7SL647P | 106479458 |
| ENSG000002 | 1.60535461 | 0.004298224 | 0.34803832 | 0.01234986 | 0.99014649 | NA | RN7SL684P | 106479472 |
| ENSG000002 | 0.82366206 | -0.09834969 | 0.28736554 | -0.3422459 | 0.73216581 | NA | NA | NA |
| ENSG000002 | 0.60301814 | 0.060465317 | 0.26274989 | 0.230125 | 0.81799464 | NA | RN7SL402P | 106479371 |
| ENSG000002 | 0 | NA | NA | NA | NA | NA | RN7SL702P | 106481114 |
| ENSG000002 | 1.98653097 | -0.717995609 | 0.36020626 | -1.9932902 | 0.04622968 | NA | NA | NA |
| ENSG000002 | 1.81444403 | 0.138370972 | 0.34990749 | 0.39545016 | 0.6925107 | NA | NA | NA |
| ENSG000002 | 1.57199774 | 0.013366449 | 0.34878262 | 0.03832315 | 0.96943003 | NA | RN7SL447P | 106479386 |
| ENSG000002 | 1.84277838 | -0.222939195 | 0.34780245 | -0.6409937 | 0.52152675 | NA | LOC105372630 | 105372630 |
| ENSG000002 | 0 | NA | NA | NA | NA | NA | NA | NA |
| ENSG000002 | 0.98987518 | -0.181818895 | 0.3071081 | -0.5920355 | 0.55382682 | NA | NA | NA |
| ENSG000002 | 1.39380595 | -0.432924975 | 0.34152976 | -1.2676054 | 0.20493889 | NA | RN7SL799P | 106480382 |
| ENSG000002 | 1.09094174 | -0.10994409 | 0.32002774 | -0.3435455 | 0.73118812 | NA | NA | NA |

| | | | | | | | | |
|---|---|---|---|---|---|---|---|---|
| ENSG000002 | 1.48078831 | -0.187235175 | 0.34157148 | -0.5481581 | 0.58358334 | NA | MT1HL1 | 645745 |
| ENSG000002 | 1.91935505 | -0.39441336 | 0.35854235 | -1.1000468 | 0.27131174 | NA | NA | NA |
| ENSG000002 | 1.76794056 | -0.751452854 | 0.35254727 | -2.1314953 | 0.03304836 | NA | KRTAP19-3 | 337970 |
| ENSG000002 | 1.49253522 | 0.086087078 | 0.33903936 | 0.2539147 | 0.79956146 | NA | RN7SL424P | 106480376 |
| ENSG000002 | 1.78033926 | -0.459298747 | 0.35336536 | -1.2997843 | 0.19367491 | NA | NA | NA |
| ENSG000002 | 1.93874657 | -0.395449227 | 0.35256403 | -1.121638 | 0.26201638 | NA | RN7SL735P | 106480529 |
| ENSG000002 | 0.70007253 | 0.267174664 | 0.26906372 | 0.99297915 | 0.32072012 | NA | NA | NA |
| ENSG000002 | 0.49765262 | -0.018104365 | 0.24679454 | -0.073358 | 0.9415212 | NA | LCE3C | 353144 |
| ENSG000002 | 1.26014495 | -0.045127133 | 0.32781726 | -0.1376594 | 0.89050959 | NA | NA | NA |
| ENSG000002 | 0.91878393 | -0.601261151 | 0.29269512 | -2.0542234 | 0.03995408 | NA | NA | NA |
| ENSG000002 | 1.76065637 | -0.461254938 | 0.35356089 | -1.3045983 | 0.19202966 | NA | RN7SL148P | 106479275 |
| ENSG000002 | 0.90639143 | -0.265472269 | 0.29522373 | -0.899224 | 0.36853334 | NA | NA | NA |
| ENSG000002 | 1.81461021 | 0.004564983 | 0.35536575 | 0.01284587 | 0.98975076 | NA | RN7SL833P | 106479525 |
| ENSG000002 | 0.12070844 | -0.088503695 | 0.12783931 | -0.6923042 | 0.48874631 | NA | NA | NA |
| ENSG000002 | 1.37411096 | -0.401268159 | 0.33686125 | -1.1911972 | 0.23357619 | NA | NA | NA |
| ENSG000002 | 1.77122212 | -0.741788949 | 0.35077763 | -2.1146986 | 0.03445565 | NA | NA | NA |
| ENSG000002 | 0.58966925 | 0.051416892 | 0.25233406 | 0.20376516 | 0.83853702 | NA | RN7SL483P | 106479399 |
| ENSG000002 | 1.66770316 | -0.817697996 | 0.34234223 | -2.3885397 | 0.01691548 | NA | NA | NA |
| ENSG000002 | 1.72615789 | -0.447875817 | 0.35278967 | -1.2695264 | 0.20425336 | NA | NA | NA |
| ENSG000002 | 1.88508628 | -0.268829663 | 0.36148144 | -0.7436887 | 0.45706482 | NA | RN7SL659P | 106479462 |
| ENSG000002 | 0.96137925 | -0.03540024 | 0.29652198 | -0.1193849 | 0.90497044 | NA | RN7SL250P | 106479313 |
| ENSG000002 | 1.49234326 | -0.18689102 | 0.34491331 | -0.5418493 | 0.58792235 | NA | RN7SL508P | 106481061 |
| ENSG000002 | 0 | NA | NA | NA | NA | NA | DNAJC25-GNG1 | 552891 |
| ENSG000002 | 0.36690249 | -0.096999144 | 0.21683809 | -0.4473344 | 0.65463361 | NA | NA | NA |
| ENSG000002 | 0.64030597 | -0.296505743 | 0.27391229 | -1.0824843 | 0.27903741 | NA | NA | NA |
| ENSG000002 | 0.95423749 | 0.117516692 | 0.3114056 | 0.37737501 | 0.70589493 | NA | UBE2Q2P9 | 106480276 |
| ENSG000002 | 1.62243995 | -0.132699853 | 0.35151414 | -0.3775093 | 0.70579516 | NA | NA | NA |
| ENSG000002 | 0.7863464 | -0.370445737 | 0.28479156 | -1.3007609 | 0.1933403 | NA | ATP6V0CP2 | 106478961 |
| ENSG000002 | 1.7130387 | -0.309574838 | 0.35086541 | -0.8823179 | 0.37760491 | NA | NA | NA |

| | | | | | | | | | |
|---|---|---|---|---|---|---|---|---|---|
| ENSG000002 | 1.5152516 | -0.483825865 | 0.34405608 | -1.4062413 | 0.15965249 | NA | NA | NA | |
| ENSG000002 | 1.01742429 | -0.349661349 | 0.30969455 | -1.1290523 | 0.25887577 | NA | NA | NA | |
| ENSG000002 | 1.38483724 | -0.260265032 | 0.33519743 | -0.7764529 | 0.43748159 | NA | NA | NA | |
| ENSG000002 | 2.10400895 | -0.25678504 | 0.3662252 | -0.701167 | 0.48319877 | NA | NA | NA | |
| ENSG000002 | 0.85951681 | -0.245407694 | 0.28844915 | -0.8507832 | 0.39488978 | NA | RN7SL120P | | 106480953 |
| ENSG000002 | 1.15082929 | -0.41306503 | 0.3184191 | -1.297237 | 0.19454966 | NA | NA | NA | |
| ENSG000002 | 1.23950995 | 0.0950731 | 0.33022325 | 0.28790553 | 0.77341905 | NA | NA | NA | |
| ENSG000002 | 0 | NA | NA | NA | NA | NA | NA | NA | |
| ENSG000002 | 1.93786083 | -0.670183843 | 0.35744944 | -1.8749053 | 0.06080575 | NA | NA | NA | |
| ENSG000002 | 2.16882662 | -0.622036593 | 0.36136192 | -1.7213673 | 0.0851842 | NA | RN7SL766P | | 106479501 |
| ENSG000002 | 0 | NA | NA | NA | NA | NA | NA | NA | |
| ENSG000002 | 1.73970981 | -0.300109899 | 0.34927208 | -0.8592439 | 0.39020597 | NA | NA | NA | |
| ENSG000002 | 0 | NA | NA | NA | NA | NA | CSPG4P1Y | | 114758 |
| ENSG000002 | 1.55216581 | 0.021827088 | 0.34412013 | 0.06342869 | 0.94942514 | NA | RN7SL698P | | 106481113 |
| ENSG000002 | 2.00991887 | -0.676047072 | 0.35406112 | -1.9094078 | 0.0562095 | NA | NA | NA | |
| ENSG000002 | 0 | NA | NA | NA | NA | NA | NA | NA | |
| ENSG000002 | 0 | NA | NA | NA | NA | NA | NA | NA | |
| ENSG000002 | 0.96929809 | -0.187967297 | 0.30786581 | -0.6105494 | 0.54149791 | NA | NA | NA | |
| ENSG000002 | 0 | NA | NA | NA | NA | NA | NA | NA | |
| ENSG000002 | 1.57071973 | 0.021199108 | 0.35059506 | 0.06046608 | 0.95178443 | NA | NA | NA | |
| ENSG000002 | 2.02608545 | -0.730539885 | 0.36112885 | -2.0229341 | 0.04307994 | NA | NA | NA | |
| ENSG000002 | 1.95612299 | -0.068063026 | 0.36187128 | -0.1880863 | 0.85080901 | NA | NA | NA | |
| ENSG000002 | 1.06587117 | 0.032148086 | 0.30383018 | 0.10580939 | 0.91573359 | NA | NA | NA | |
| ENSG000002 | 0 | NA | NA | NA | NA | NA | NA | NA | |
| ENSG000002 | 2.05570993 | -0.314540742 | 0.35909023 | -0.8759379 | 0.38106379 | NA | NA | NA | |
| ENSG000002 | 1.57419505 | -0.123034932 | 0.34971556 | -0.3518143 | 0.72497754 | NA | NA | NA | |
| ENSG000002 | 0.87301474 | -0.285254425 | 0.30393182 | -0.9385474 | 0.34796316 | NA | NA | NA | |
| ENSG000002 | 1.09199745 | -0.244062525 | 0.31470623 | -0.7755249 | 0.43802955 | NA | NA | NA | |
| ENSG000002 | 0.61742743 | -0.111215928 | 0.26435729 | -0.4207031 | 0.67397191 | NA | RN7SL740P | | 106479491 |

| | | | | | | | | |
|---|---|---|---|---|---|---|---|---|
| ENSG000002 | 0 | NA | NA | NA | NA | NA | NA | NA |
| ENSG000002 | 1.12538521 | -0.40631212 | 0.31757681 | -1.2794137 | 0.20075142 | NA | RN7SL168P | 106480491 |
| ENSG000002 | 1.24967659 | -0.322621105 | 0.32490881 | -0.9929589 | 0.32072997 | NA | RN7SL395P | 106481031 |
| ENSG000002 | 1.52253929 | -0.494559475 | 0.34689046 | -1.4256935 | 0.15395683 | NA | NA | NA |
| ENSG000002 | 1.43366529 | 0.346437947 | 0.33072 | 1.04752643 | 0.29485685 | NA | RN7SL36P | 106479237 |
| ENSG000002 | 1.0310322 | -0.510514485 | 0.30759561 | -1.6596937 | 0.0969761 | NA | RN7SL814P | 106481143 |
| ENSG000002 | 1.57137208 | 0.015537357 | 0.34300485 | 0.04529778 | 0.96386996 | NA | RN7SL845P | 106481863 |
| ENSG000002 | 0.3588776 | -0.099902505 | 0.21555794 | -0.4634601 | 0.6430346 | NA | NA | NA |
| ENSG000002 | 0.37945415 | -0.107841678 | 0.21865587 | -0.4932027 | 0.62186934 | NA | NA | NA |
| ENSG000002 | 0.86592293 | -0.272159547 | 0.29714053 | -0.9159287 | 0.35970427 | NA | RN7SL687P | 106481110 |
| ENSG000002 | 0.9979551 | 0.096636333 | 0.30098085 | 0.32107137 | 0.74815631 | NA | LY6G6D | 58530 |
| ENSG000002 | 0.74547562 | -0.19522748 | 0.2860138 | -0.6825806 | 0.49487187 | NA | NA | NA |
| ENSG000002 | 1.59961739 | -0.247488265 | 0.34356358 | -0.7203565 | 0.47130551 | NA | KRTAP19-7 | 337974 |
| ENSG000002 | 0.36538978 | 0.059033792 | 0.20773944 | 0.28417229 | 0.77627835 | NA | NA | NA |
| ENSG000002 | 1.86342799 | -1.023809042 | 0.34322319 | -2.982925 | 0.00285508 | NA | RN7SL423P | 106479378 |
| ENSG000002 | 0.48142868 | 0.145033972 | 0.23815494 | 0.60898998 | 0.54253108 | NA | RN7SL636P | 106481097 |
| ENSG000002 | 1.51246089 | -0.345916564 | 0.34705516 | -0.9967193 | 0.3189008 | NA | NA | NA |
| ENSG000002 | 0.63853479 | 0.015623989 | 0.15783986 | 0.09898633 | 0.92114912 | NA | RN7SL359P | 106479354 |
| ENSG000002 | 0.74384667 | -0.022939696 | 0.27882226 | -0.0822735 | 0.93442919 | NA | RN7SL330P | 106481013 |
| ENSG000002 | 0.8863036 | -0.294029201 | 0.30451173 | -0.965576 | 0.33425641 | NA | RN7SL869P | 106481864 |
| ENSG000002 | 0 | NA | NA | NA | NA | NA | RBMY1D | 378949 |
| ENSG000002 | 2.09060798 | -0.249881681 | 0.36279782 | -0.6887629 | 0.49097249 | NA | RN7SL251P | 106480990 |
| ENSG000002 | 0.11934008 | 0.083564698 | 0.12760091 | 0.65489106 | 0.5125379 | NA | RN7SL314P | 106481836 |
| ENSG000002 | 1.16517386 | -0.573527426 | 0.31675334 | -1.8106436 | 0.07019603 | NA | RN7SL216P | 106480495 |
| ENSG000002 | 1.2591235 | -0.347788725 | 0.33217816 | -1.0469945 | 0.29510209 | NA | KRTAP5-7 | 440050 |
| ENSG000002 | 1.03254045 | -0.358491974 | 0.311878 | -1.1494622 | 0.25036544 | NA | NA | NA |
| ENSG000002 | 0.79212584 | -0.534732018 | 0.28076432 | -1.9045583 | 0.05683751 | NA | NA | NA |
| ENSG000002 | 1.01910723 | -0.373267329 | 0.31564045 | -1.1825713 | 0.2369791 | NA | RN7SL268P | 106479319 |
| ENSG000002 | 0.74677928 | -0.319192781 | 0.2716115 | -1.1751814 | 0.23992216 | NA | RPS3AP33 | 100271259 |

| | | | | | | | | |
|---|---|---|---|---|---|---|---|---|
| ENSG000002 | 0.61426904 | -0.250234224 | 0.25739856 | -0.9721664 | 0.33096779 | NA | NA | NA |
| ENSG000002 | 1.54007368 | -0.61819878 | 0.34089884 | -1.8134376 | 0.06976435 | NA | NA | NA |
| ENSG000002 | 1.97561639 | -0.193015303 | 0.3647349 | -0.5291934 | 0.59667129 | NA | NA | NA |
| ENSG000002 | 0.77288285 | -0.203654568 | 0.28771447 | -0.7078357 | 0.47904731 | NA | NA | NA |
| ENSG000002 | 0.38270188 | -0.272446727 | 0.2140009 | -1.2731102 | 0.20297894 | NA | NA | NA |
| ENSG000002 | 0 | NA | NA | NA | NA | NA | NA | NA |
| ENSG000002 | 1.99257774 | -0.274340858 | 0.35131975 | -0.7808865 | 0.43486924 | NA | NA | NA |
| ENSG000002 | 1.40607582 | -0.587042961 | 0.34003824 | -1.7264028 | 0.08427497 | NA | NA | NA |
| ENSG000002 | 1.93037727 | -0.652289508 | 0.35459815 | -1.8395175 | 0.0658391 | NA | UGT1A4 | 54657 |
| ENSG000002 | 1.57219574 | 0.148212195 | 0.34590769 | 0.42847326 | 0.66830659 | NA | NA | NA |
| ENSG000002 | 1.93500174 | 0.193754405 | 0.36226717 | 0.53483844 | 0.59276158 | NA | HDAC11-AS1 | 100874101 |
| ENSG000002 | 0 | NA | NA | NA | NA | NA | NA | NA |
| ENSG000002 | 2.03966519 | -0.110105632 | 0.36061038 | -0.3053313 | 0.76011385 | NA | RN7SL28P | 106480929 |
| ENSG000002 | 1.07767014 | 0.353806943 | 0.32011693 | 1.10524285 | 0.26905438 | NA | RN7SL125P | 106479267 |
| ENSG000002 | 0 | NA | NA | NA | NA | NA | NA | NA |
| ENSG000002 | 1.34192402 | -0.102841023 | 0.32846707 | -0.3130939 | 0.75420935 | NA | RN7SL380P | 106479361 |
| ENSG000002 | 1.98985098 | -0.201763309 | 0.36403926 | -0.554235 | 0.57941801 | NA | KRTAP4-2 | 85291 |
| ENSG000002 | 1.5076553 | -0.173764374 | 0.33589229 | -0.5173217 | 0.6049316 | NA | NA | NA |
| ENSG000002 | 1.66791822 | -0.237078351 | 0.33921785 | -0.698897 | 0.48461637 | NA | NA | NA |
| ENSG000002 | 1.23749316 | -0.046146052 | 0.33376755 | -0.1382581 | 0.89003647 | NA | RN7SL446P | 106479385 |
| ENSG000002 | 0 | NA | NA | NA | NA | NA | NA | NA |
| ENSG000002 | 0.74661723 | -0.178670198 | 0.27282214 | -0.6548963 | 0.51253455 | NA | NA | NA |
| ENSG000002 | 1.95160616 | -0.652655813 | 0.35358953 | -1.8458007 | 0.06492114 | NA | NA | NA |
| ENSG000002 | 0 | NA | NA | NA | NA | NA | NA | NA |
| ENSG000002 | 2.06965674 | -0.111392187 | 0.36229108 | -0.307466 | 0.7584887 | NA | NA | NA |
| ENSG000002 | 0.87867101 | -0.252341977 | 0.28952996 | -0.8715574 | 0.38344989 | NA | NA | NA |
| ENSG000002 | 0.60162804 | 0.055843814 | 0.26262531 | 0.21263683 | 0.83161024 | NA | NA | NA |
| ENSG000002 | 0.49512483 | -0.344267812 | 0.23617548 | -1.457678 | 0.14492931 | NA | NA | NA |
| ENSG000002 | 0 | NA | NA | NA | NA | NA | NA | NA |

| | | | | | | | | |
|---|---|---|---|---|---|---|---|---|
| ENSG000002 | 0.85587733 | -0.115937995 | 0.30218708 | -0.383663 | 0.70122826 | NA | RN7SL334P | 106480502 |
| ENSG000002 | 1.80251443 | -0.442550383 | 0.34903353 | -1.2679309 | 0.20482264 | NA | NA | NA |
| ENSG000002 | 1.21279425 | -0.025772442 | 0.31101055 | -0.0828668 | 0.93395747 | NA | NA | NA |
| ENSG000002 | 1.97246563 | -0.069156317 | 0.35949273 | -0.1923719 | 0.84745086 | NA | KRTAP20-1 | 337975 |
| ENSG000002 | 1.73579654 | -0.184445193 | 0.35367696 | -0.5215075 | 0.60201327 | NA | NA | NA |
| ENSG000002 | 0 | NA | NA | NA | NA | NA | NA | NA |
| ENSG000002 | 0.99741027 | -0.188865662 | 0.3074178 | -0.6143615 | 0.53897647 | NA | NA | NA |
| ENSG000002 | 0.85231557 | -0.109410774 | 0.29687299 | -0.3685441 | 0.7124676 | NA | NA | NA |
| ENSG000002 | 2.08952935 | -0.006316918 | 0.35399653 | -0.0178446 | 0.98576284 | NA | RN7SL280P | 106480498 |
| ENSG000002 | 1.85509528 | -0.244092256 | 0.35497122 | -0.6876396 | 0.49167979 | NA | RN7SL706P | 106479480 |
| ENSG000002 | 0.61258555 | -0.25554054 | 0.25718763 | -0.9935958 | 0.32041971 | NA | RN7SL724P | 106479487 |
| ENSG000002 | 2.03198303 | -0.743520029 | 0.3624674 | -2.0512742 | 0.04024025 | NA | NA | NA |
| ENSG000002 | 0.58275617 | 0.231943815 | 0.26197681 | 0.88536009 | 0.37596245 | NA | NA | NA |
| ENSG000002 | 0.49921686 | -0.164688411 | 0.23121313 | -0.7122797 | 0.47629162 | NA | NA | NA |
| ENSG000002 | 1.80641466 | 0.395645985 | 0.35433475 | 1.11658816 | 0.26417046 | NA | NA | NA |
| ENSG000002 | 2.0701991 | 0.002196043 | 0.36728234 | 0.00597917 | 0.99522934 | NA | NA | NA |
| ENSG000002 | 1.58483864 | 0.021821353 | 0.34093857 | 0.06400377 | 0.94896723 | NA | C4A | 720 |
| ENSG000002 | 0.50405874 | -0.025925695 | 0.24745105 | -0.104771 | 0.91655752 | NA | RN7SL870P | 106481155 |
| ENSG000002 | 0.49356059 | -0.178885509 | 0.24036594 | -0.7442215 | 0.45674246 | NA | NA | NA |
| ENSG000002 | 1.58702957 | 0.013921727 | 0.35413988 | 0.03931138 | 0.96864214 | NA | NA | NA |
| ENSG000002 | 2.0236128 | -0.195420361 | 0.35965253 | -0.5433588 | 0.5868828 | NA | NA | NA |
| ENSG000002 | 2.02710848 | -0.331274598 | 0.36379103 | -0.9106178 | 0.36249677 | NA | NA | NA |
| ENSG000002 | 0.84878552 | -0.106052095 | 0.28885264 | -0.3671495 | 0.71350752 | NA | NA | NA |
| ENSG000002 | 1.8246014 | 0.243059939 | 0.35246006 | 0.68960988 | 0.49043955 | NA | NA | NA |
| ENSG000002 | 1.34439102 | -0.087464311 | 0.31638498 | -0.276449 | 0.78220323 | NA | NA | NA |
| ENSG000002 | 1.02278109 | -0.350503169 | 0.30990351 | -1.1310074 | 0.25805198 | NA | NA | NA |
| ENSG000002 | 0.3863157 | -0.274134926 | 0.21457207 | -1.277589 | 0.20139437 | NA | NA | NA |
| ENSG000002 | 1.47389293 | -0.162878623 | 0.33053222 | -0.4927768 | 0.62217028 | NA | NA | NA |
| ENSG000002 | 0.50016259 | -0.200428691 | 0.24824437 | -0.8073846 | 0.41944491 | NA | NA | NA |

| | | | | | | | | |
|---|---|---|---|---|---|---|---|---|
| ENSG000002 | 0 | NA | NA | NA | NA | NA | NA | NA |
| ENSG000002 | 1.13451891 | -0.419945847 | 0.31942037 | -1.3147122 | 0.18860665 | NA | NA | NA |
| ENSG000002 | 1.67672084 | -0.538631688 | 0.34646118 | -1.5546668 | 0.12002544 | NA | NA | NA |
| ENSG000002 | 0 | NA | NA | NA | NA | NA | NA | NA |
| ENSG000002 | 0 | NA | NA | NA | NA | NA | NA | NA |
| ENSG000002 | 1.1461359 | -0.60452063 | 0.32175527 | -1.8788212 | 0.06026891 | NA | NA | NA |
| ENSG000002 | 0.87206901 | -0.264258287 | 0.29752107 | -0.8882002 | 0.37443306 | NA | NA | NA |
| ENSG000002 | 1.96806404 | -0.537237755 | 0.35600374 | -1.509079 | 0.1312786 | NA | NA | NA |
| ENSG000002 | 1.12717915 | -0.404639256 | 0.31427386 | -1.2875371 | 0.19790715 | NA | BTG4P1 | 106480330 |
| ENSG000002 | 0 | NA | NA | NA | NA | NA | TRIM39-RPP21 | 202658 |
| ENSG000002 | 1.30813132 | 0.029256405 | 0.32900949 | 0.08892268 | 0.92914336 | NA | NA | NA |
| ENSG000002 | 1.11060694 | 0.035289427 | 0.31568464 | 0.11178696 | 0.91099232 | NA | NA | NA |
| ENSG000002 | 1.64614608 | -0.398366726 | 0.35050266 | -1.1365584 | 0.25572296 | NA | NA | NA |
| ENSG000002 | 0.63370343 | -0.298851213 | 0.27334848 | -1.0932975 | 0.2742632 | NA | NA | NA |
| ENSG000002 | 1.35589744 | -0.602509263 | 0.31751843 | -1.8975568 | 0.05775449 | NA | NA | NA |
| ENSG000002 | 0.38747904 | -0.104908247 | 0.21987514 | -0.4771265 | 0.63327207 | NA | NA | NA |
| ENSG000002 | 1.13796051 | -0.100746853 | 0.30870372 | -0.3263545 | 0.74415615 | NA | NA | NA |
| ENSG000002 | 0.78588993 | -0.359597889 | 0.28326202 | -1.2694886 | 0.20426687 | NA | NA | NA |
| ENSG000002 | 1.62078981 | -0.11766912 | 0.34650239 | -0.3395911 | 0.7341645 | NA | NA | NA |
| ENSG000002 | 0 | NA | NA | NA | NA | NA | NA | NA |
| ENSG000002 | 1.66075124 | -0.567831844 | 0.35277847 | -1.6095989 | 0.10748545 | NA | NA | NA |
| ENSG000002 | 1.53842244 | -0.644916076 | 0.34375714 | -1.8760806 | 0.06064422 | NA | NA | NA |
| ENSG000002 | 1.36525849 | -0.276357526 | 0.34144562 | -0.8093749 | 0.41829952 | NA | NA | NA |
| ENSG000002 | 0.38019136 | -0.10756138 | 0.21876967 | -0.491665 | 0.62295621 | NA | NA | NA |
| ENSG000002 | 1.25610445 | -0.189972045 | 0.32974056 | -0.5761258 | 0.56453016 | NA | NA | NA |
| ENSG000002 | 1.35737795 | -0.258948028 | 0.33681651 | -0.7688104 | 0.44200588 | NA | NA | NA |
| ENSG000002 | 0.11865494 | 0.083440866 | 0.12751478 | 0.65436232 | 0.5128784 | NA | NA | NA |
| ENSG000002 | 0.82504769 | 0.203340619 | 0.29552703 | 0.68806097 | 0.49141439 | NA | NA | NA |
| ENSG000002 | 1.4135034 | -0.267501747 | 0.33579574 | -0.7966204 | 0.42567151 | NA | NA | NA |

| | | | | | | | | | |
|---|---|---|---|---|---|---|---|---|---|
| ENSG000002 | 0.65561801 | -0.268676647 | 0.26257757 | -1.0232277 | 0.3062002 | NA | NA | NA | |
| ENSG000002 | 0.50818749 | -0.197539377 | 0.24915052 | -0.7928516 | 0.42786432 | NA | NA | NA | |
| ENSG000002 | 1.47922216 | -0.187553887 | 0.34154431 | -0.5491349 | 0.58291291 | NA | NA | NA | |
| ENSG000002 | 1.18438204 | -0.164735679 | 0.21957314 | -0.7502542 | 0.45310159 | NA | NA | NA | |
| ENSG000002 | 0 | NA | NA | NA | NA | NA | NA | NA | |
| ENSG000002 | 0 | NA | NA | NA | NA | NA | NA | NA | |
| ENSG000002 | 1.58813735 | -0.123365421 | 0.35336011 | -0.349121 | 0.7269985 | NA | NA | NA | |
| ENSG000002 | 0.50565578 | -0.183478728 | 0.24174047 | -0.7589905 | 0.44785822 | NA | NA | NA | |
| ENSG000002 | 2.12972056 | -0.343171676 | 0.35870845 | -0.9566869 | 0.33872531 | NA | NA | NA | |
| ENSG000002 | 1.90131076 | -0.372326505 | 0.35612844 | -1.0454838 | 0.29579944 | NA | NA | NA | |
| ENSG000002 | 1.15074284 | -0.565431982 | 0.31544929 | -1.7924655 | 0.07305842 | NA | NA | NA | |
| ENSG000002 | 0.23868015 | 0.154625155 | 0.16997278 | 0.90970538 | 0.36297791 | NA | NA | NA | |
| ENSG000002 | 0.3588776 | -0.099902505 | 0.21555794 | -0.4634601 | 0.6430346 | NA | NA | NA | |
| ENSG000002 | 1.35335683 | 0.154836218 | 0.33449873 | 0.4628903 | 0.643443 | NA | NA | NA | |
| ENSG000002 | 1.84318332 | -0.237135623 | 0.35491433 | -0.6681489 | 0.50403854 | NA | NA | NA | |
| ENSG000002 | 0 | NA | NA | NA | NA | NA | NA | NA | |
| ENSG000002 | 1.2749248 | -0.346440767 | 0.33116321 | -1.046133 | 0.29549963 | NA | NA | NA | |
| ENSG000002 | 1.98086556 | -0.43577569 | 0.36054884 | -1.2086454 | 0.22679912 | NA | NA | NA | |
| ENSG000002 | 1.83521709 | -0.123384187 | 0.35575269 | -0.3468257 | 0.72872225 | NA | NA | NA | |
| ENSG000002 | 1.75390113 | 0.268932509 | 0.35359458 | 0.76056739 | 0.4469155 | NA | NA | NA | |
| ENSG000002 | 0 | NA | NA | NA | NA | NA | NA | NA | |
| ENSG000002 | 1.61936849 | -0.002428276 | 0.34024042 | -0.0071369 | 0.99430559 | NA | NA | NA | |
| ENSG000002 | 0.77925514 | -0.363111002 | 0.28443192 | -1.2766183 | 0.20173703 | NA | NA | NA | |
| ENSG000002 | 1.02340778 | -0.179728615 | 0.29417515 | -0.6109578 | 0.5412275 | NA | NA | NA | |
| ENSG000002 | 1.48737642 | 0.217983727 | 0.33931875 | 0.6424158 | 0.52060324 | NA | NA | NA | |
| ENSG000002 | 1.15386561 | -0.109241664 | 0.3074497 | -0.3553156 | 0.72235319 | NA | NA | NA | |
| ENSG000002 | 1.50215415 | -0.200434001 | 0.34628668 | -0.5788094 | 0.56271775 | NA | NA | NA | |
| ENSG000002 | 1.24261916 | -0.193315202 | 0.33085304 | -0.5842933 | 0.55902302 | NA | MTATP6P9 | | 106479044 |
| ENSG000002 | 0 | NA | NA | NA | NA | NA | NA | NA | |

| | | | | | | | | |
|---|---|---|---|---|---|---|---|---|
| ENSG000002 | 0.47551128 | 0.153262041 | 0.24481524 | 0.62603146 | 0.53129431 | NA | NA | NA |
| ENSG000002 | 0.53627793 | -0.360851627 | 0.24080453 | -1.4985251 | 0.13399688 | NA | RPL23AP94 | 106481971 |
| ENSG000002 | 0.97820612 | -0.032054347 | 0.31244016 | -0.1025936 | 0.91828556 | NA | NA | NA |
| ENSG000002 | 0.95443391 | 0.126494054 | 0.311389 | 0.40622519 | 0.68457715 | NA | NA | NA |
| ENSG000002 | 1.51784589 | -0.640420176 | 0.34342667 | -1.8647945 | 0.06221021 | NA | NENFP3 | 106481703 |
| ENSG000002 | 2.060834 | -0.590732997 | 0.36130714 | -1.6349884 | 0.10205147 | NA | NA | NA |
| ENSG000002 | 0.51433302 | -0.186673425 | 0.24269587 | -0.7691661 | 0.44179474 | NA | LOC102577426 | 102577426 |
| ENSG000002 | 1.19872687 | -0.042298905 | 0.3261394 | -0.1296958 | 0.89680712 | NA | NA | NA |
| ENSG000002 | 1.69239701 | 0.203825438 | 0.35025304 | 0.58193767 | 0.56060867 | NA | NA | NA |
| ENSG000002 | 0.88727107 | -0.118675059 | 0.29705507 | -0.3995053 | 0.68952095 | NA | NA | NA |
| ENSG000002 | 1.35333799 | -0.414806766 | 0.33884954 | -1.2241621 | 0.22089108 | NA | NA | NA |
| ENSG000002 | 1.35876805 | -0.249381023 | 0.33464223 | -0.7452168 | 0.45614065 | NA | NA | NA |
| ENSG000002 | 1.97832333 | -0.284271622 | 0.35678473 | -0.7967595 | 0.42559073 | NA | NA | NA |
| ENSG000002 | 0 | NA | NA | NA | NA | NA | NA | NA |
| ENSG000002 | 1.72828073 | -0.186930492 | 0.35533572 | -0.5260673 | 0.59884147 | NA | NA | NA |
| ENSG000002 | 1.19733677 | -0.044343054 | 0.32065918 | -0.1382872 | 0.89001346 | NA | NA | NA |
| ENSG000002 | 0.87875937 | 0.039011202 | 0.29019183 | 0.13443246 | 0.89306061 | NA | NA | NA |
| ENSG000002 | 0.59758809 | -0.103924321 | 0.26264093 | -0.3956897 | 0.69233393 | NA | NA | NA |
| ENSG000002 | 1.68250028 | -0.707119784 | 0.34905877 | -2.0257901 | 0.0427863 | NA | NA | NA |
| ENSG000002 | 0 | NA | NA | NA | NA | NA | PRYP6 | 106480721 |
| ENSG000002 | 1.55638351 | -0.617203046 | 0.3397454 | -1.8166634 | 0.06926866 | NA | NA | NA |
| ENSG000002 | 0.60338479 | 0.225396395 | 0.26390915 | 0.85406814 | 0.39306724 | NA | NA | NA |
| ENSG000002 | 1.71335327 | -0.189125014 | 0.35408457 | -0.5341239 | 0.59325584 | NA | TMEM110-MUS | 100526772 |
| ENSG000002 | 1.4832302 | -0.053389099 | 0.34561713 | -0.1544747 | 0.87723547 | NA | NA | NA |
| ENSG000002 | 0.59892612 | -0.099249839 | 0.26277623 | -0.3776972 | 0.70565557 | NA | NA | NA |
| ENSG000002 | 1.50896363 | -0.338145992 | 0.3451475 | -0.9797144 | 0.3272271 | NA | NA | NA |
| ENSG000002 | 1.35477973 | 0.17154319 | 0.33924722 | 0.50565835 | 0.61309651 | NA | NA | NA |
| ENSG000002 | 1.88264281 | -0.514164603 | 0.35626915 | -1.4431915 | 0.14896654 | NA | NA | NA |
| ENSG000002 | 0.77613058 | -0.361458377 | 0.28422771 | -1.2717211 | 0.20347223 | NA | ENPP7P3 | 106481658 |

| | | | | | | | | | |
|---|---|---|---|---|---|---|---|---|---|
| ENSG000002 | 0.12070844 | -0.088503695 | 0.12783931 | -0.6923042 | 0.48874631 | NA | NA | NA | |
| ENSG000002 | 1.17299841 | -0.253702555 | 0.31210121 | -0.8128855 | 0.41628369 | NA | NA | NA | |
| ENSG000002 | 0 | NA | NA | NA | NA | NA | NA | NA | |
| ENSG000002 | 0.61104304 | 0.059501982 | 0.25592296 | 0.23249958 | 0.81615001 | NA | NA | NA | |
| ENSG000002 | 1.11237812 | -0.273939681 | 0.32537936 | -0.8419086 | 0.39983912 | NA | NA | NA | |
| ENSG000002 | 1.69485005 | -0.303108875 | 0.35002829 | -0.8659554 | 0.38651464 | NA | NA | NA | |
| ENSG000002 | 1.98230483 | -0.066155734 | 0.36204233 | -0.1827293 | 0.85501044 | NA | GCSHP1 | | 106480441 |
| ENSG000002 | 2.05608074 | 0.001715409 | 0.36721733 | 0.00467137 | 0.9962728 | NA | NA | NA | |
| ENSG000002 | 0.85593289 | 0.188277766 | 0.29142981 | 0.6460484 | 0.51824801 | NA | NA | NA | |
| ENSG000002 | 0.3934287 | -0.107186984 | 0.22072612 | -0.4856108 | 0.62724315 | NA | NA | NA | |
| ENSG000002 | 1.40795464 | -0.591309903 | 0.34002702 | -1.7390086 | 0.08203325 | NA | NA | NA | |
| ENSG000002 | 0.49076933 | -0.025124039 | 0.23846109 | -0.1053591 | 0.91609089 | NA | NA | NA | |
| ENSG000002 | 0 | NA | NA | NA | NA | NA | NA | NA | |
| ENSG000002 | 1.91203703 | -0.247035924 | 0.35352705 | -0.6987752 | 0.48469255 | NA | NA | NA | |
| ENSG000002 | 2.03436553 | 0.250825422 | 0.36446225 | 0.68820686 | 0.49132252 | NA | NA | NA | |
| ENSG000002 | 1.88884041 | -0.528610079 | 0.35852451 | -1.4744043 | 0.14037277 | NA | NA | NA | |
| ENSG000002 | 0.49810908 | -0.023852705 | 0.24681213 | -0.0966432 | 0.92300978 | NA | NA | NA | |
| ENSG000002 | 0.76073209 | -0.532127553 | 0.28034426 | -1.8981218 | 0.05768004 | NA | NA | NA | |
| ENSG000002 | 1.16693278 | 0.249173764 | 0.32112384 | 0.77594291 | 0.4377827 | NA | NA | NA | |
| ENSG000002 | 1.59779366 | -0.103360734 | 0.3428762 | -0.3014521 | 0.76306981 | NA | NA | NA | |
| ENSG000002 | 0.38651158 | -0.274258716 | 0.21461355 | -1.2779189 | 0.20127801 | NA | NA | NA | |
| ENSG000002 | 1.62140832 | -0.225149894 | 0.3379833 | -0.6661569 | 0.50531084 | NA | NA | NA | |
| ENSG000002 | 0.37441639 | -0.268856313 | 0.21277471 | -1.2635727 | 0.20638344 | NA | NA | NA | |
| ENSG000002 | 2.12713274 | -0.648475037 | 0.36538403 | -1.7747766 | 0.07593478 | NA | NA | NA | |
| ENSG000002 | 1.01958598 | -0.197147843 | 0.31466692 | -0.6265287 | 0.53096824 | NA | NA | NA | |
| ENSG000002 | 1.36320498 | -0.129765453 | 0.34362016 | -0.3776421 | 0.70569646 | NA | NA | NA | |
| ENSG000002 | 1.50161336 | -0.182679522 | 0.34117103 | -0.5354485 | 0.59233975 | NA | NA | NA | |
| ENSG000002 | 1.85850004 | -0.264275859 | 0.36040206 | -0.7332807 | 0.46338727 | NA | NA | NA | |
| ENSG000002 | 1.90632274 | -0.679922527 | 0.35811711 | -1.8986039 | 0.05761658 | NA | NA | NA | |

| | | | | | | | | |
|---|---|---|---|---|---|---|---|---|
| ENSG000002 | 0 | NA | NA | NA | NA | NA | NA | NA |
| ENSG000002 | 2.15086768 | -0.522825462 | 0.36561274 | -1.4299979 | 0.15271762 | NA | NA | NA |
| ENSG000002 | 1.34900477 | 0.031469685 | 0.33906312 | 0.09281365 | 0.92605161 | NA | NA | NA |
| ENSG000002 | 1.36639617 | 0.012106915 | 0.33206415 | 0.03645957 | 0.97091592 | NA | NA | NA |
| ENSG000002 | 1.95381997 | -0.066009949 | 0.35642191 | -0.1852017 | 0.85307082 | NA | NA | NA |
| ENSG000002 | 1.01363632 | -0.187240698 | 0.3097588 | -0.6044726 | 0.54552949 | NA | NA | NA |
| ENSG000002 | 1.50554623 | -0.320994915 | 0.34283163 | -0.9363048 | 0.34911626 | NA | NA | NA |
| ENSG000002 | 0 | NA | NA | NA | NA | NA | NA | NA |
| ENSG000002 | 0.35733509 | 0.244646831 | 0.20894085 | 1.17089037 | 0.24164285 | NA | NA | NA |
| ENSG000002 | 0 | NA | NA | NA | NA | NA | NA | NA |
| ENSG000002 | 2.13087806 | -0.119534706 | 0.36262568 | -0.3296366 | 0.74167455 | NA | NA | NA |
| ENSG000002 | 1.21774364 | -0.184085733 | 0.32872053 | -0.5600068 | 0.57547479 | NA | NA | NA |
| ENSG000002 | 1.73463478 | -0.201485335 | 0.35929221 | -0.560784 | 0.57494477 | NA | NA | NA |
| ENSG000002 | 0 | NA | NA | NA | NA | NA | NA | NA |
| ENSG000002 | 0.49356059 | -0.178885509 | 0.24036594 | -0.7442215 | 0.45674246 | NA | NA | NA |
| ENSG000002 | 1.15535447 | -0.614270225 | 0.32345318 | -1.8991009 | 0.05755121 | NA | NA | NA |
| ENSG000002 | 1.35788895 | -0.10817513 | 0.33389561 | -0.3239789 | 0.74595402 | NA | NA | NA |
| ENSG000002 | 1.82584949 | -0.124017148 | 0.35791729 | -0.3464967 | 0.72896949 | NA | NA | NA |
| ENSG000002 | 0.52623233 | -0.191068033 | 0.24398404 | -0.7831169 | 0.43355844 | NA | NA | NA |
| ENSG000002 | 1.63775253 | -0.124241443 | 0.34868736 | -0.3563119 | 0.72160702 | NA | NA | NA |
| ENSG000002 | 1.2337717 | -0.328595146 | 0.32789279 | -1.002142 | 0.31627501 | NA | NA | NA |
| ENSG000002 | 2.03419625 | -0.586079032 | 0.36073654 | -1.6246733 | 0.10423219 | NA | NA | NA |
| ENSG000002 | 1.5085672 | -0.302709081 | 0.33698439 | -0.8982881 | 0.36903196 | NA | NA | NA |
| ENSG000002 | 0.98457787 | -0.19372119 | 0.31177452 | -0.6213503 | 0.53436917 | NA | NA | NA |
| ENSG000002 | 0.86486777 | -0.10897299 | 0.29754053 | -0.3662459 | 0.71418162 | NA | NA | NA |
| ENSG000002 | 1.08565549 | -0.249702686 | 0.3136269 | -0.7961775 | 0.42592886 | NA | NA | NA |
| ENSG000002 | 0 | NA | NA | NA | NA | NA | TNFSF12-TNFSI | 407977 |
| ENSG000002 | 1.75094516 | -0.192244429 | 0.3555729 | -0.5406611 | 0.58874121 | NA | NA | NA |
| ENSG000002 | 1.75375707 | -0.187345345 | 0.35122269 | -0.533409 | 0.59375049 | NA | NA | NA |

| | | | | | | | | |
|---|---|---|---|---|---|---|---|---|
| ENSG000002 | 1.45619355 | -0.181960282 | 0.34430666 | -0.5284832 | 0.59716401 | NA | FAM138E | 100124412 |
| ENSG000002 | 0.50474388 | -0.025682234 | 0.2475253 | -0.103756 | 0.91736299 | NA | NA | NA |
| ENSG000002 | 0.63250975 | -0.278024813 | 0.26750458 | -1.0393273 | 0.29865254 | NA | NA | NA |
| ENSG000002 | 1.02701343 | -0.481140796 | 0.30215201 | -1.5923799 | 0.11129937 | NA | NA | NA |
| ENSG000002 | 1.76899393 | -0.474063189 | 0.35581739 | -1.3323216 | 0.18275454 | NA | NA | NA |
| ENSG000002 | 0.22263036 | 0.146922267 | 0.16600367 | 0.88505432 | 0.37612734 | NA | NA | NA |
| ENSG000002 | 1.86758222 | -0.256909185 | 0.35853932 | -0.716544 | 0.47365554 | NA | ATP5J2-PTCD1 | 100526740 |
| ENSG000002 | 0.91661832 | -0.39754197 | 0.2849311 | -1.3952214 | 0.16294908 | NA | USP17L19 | 100287404 |
| ENSG000002 | 1.05206576 | -0.500849589 | 0.30607069 | -1.6363853 | 0.10175897 | NA | NA | NA |
| ENSG000002 | 1.94659014 | -0.6414905 | 0.35340339 | -1.8151793 | 0.06949637 | NA | NA | NA |
| ENSG000002 | 1.74787158 | -0.204545044 | 0.35761466 | -0.5719705 | 0.56734195 | NA | NA | NA |
| ENSG000002 | 0.60307021 | 0.056690296 | 0.25518062 | 0.22215753 | 0.82419125 | NA | USP17L22 | 100287513 |
| ENSG000002 | 0.38270188 | -0.272446727 | 0.2140009 | -1.2731102 | 0.20297894 | NA | NA | NA |
| ENSG000002 | 0 | NA | NA | NA | NA | NA | NA | NA |
| ENSG000002 | 0.5301324 | -0.3843196 | 0.24728844 | -1.5541349 | 0.12015224 | NA | NA | NA |
| ENSG000002 | 1.60732826 | -0.395910399 | 0.34954128 | -1.1326571 | 0.25735829 | NA | NA | NA |
| ENSG000002 | 1.70201311 | 0.204491869 | 0.34830539 | 0.5871051 | 0.55713313 | NA | NA | NA |
| ENSG000002 | 0.61750123 | 0.055806918 | 0.26401083 | 0.21138117 | 0.83258985 | NA | NA | NA |
| ENSG000002 | 2.05623344 | -0.736893769 | 0.36189302 | -2.0362199 | 0.04172828 | NA | NA | NA |
| ENSG000002 | 0.54319505 | -0.072630779 | 0.15595508 | -0.465716 | 0.64141878 | NA | NA | NA |
| ENSG000002 | 0.94498858 | 0.26020925 | 0.29960274 | 0.86851426 | 0.38511287 | NA | NA | NA |
| ENSG000002 | 1.97989809 | -0.574557914 | 0.36078789 | -1.5925089 | 0.11127041 | NA | NA | NA |
| ENSG000002 | 0.71294322 | 0.124158965 | 0.27526995 | 0.45104438 | 0.65195757 | NA | NA | NA |
| ENSG000002 | 1.94309314 | -0.185465964 | 0.3610882 | -0.5136306 | 0.60751025 | NA | NA | NA |
| ENSG000002 | 0 | NA | NA | NA | NA | NA | NA | NA |
| ENSG000002 | 0.58070266 | 0.427028859 | 0.26367036 | 1.61955579 | 0.10532773 | NA | NA | NA |
| ENSG000002 | 1.73091277 | -0.454415266 | 0.35132998 | -1.2934144 | 0.19586776 | NA | NA | NA |
| ENSG000002 | 0 | NA | NA | NA | NA | NA | NA | NA |
| ENSG000002 | 1.20803381 | 0.23956795 | 0.32411785 | 0.73913841 | 0.45982296 | NA | NA | NA |

| | | | | | | | | |
|---|---|---|---|---|---|---|---|---|
| ENSG000002 | 2.09555979 | -0.718714847 | 0.35974839 | -1.9978264 | 0.04573548 | NA | NA | NA |
| ENSG000002 | 0.6054383 | 0.055876002 | 0.26295202 | 0.21249505 | 0.83172084 | NA | NA | NA |
| ENSG000002 | 1.80472878 | 0.143370457 | 0.3544791 | 0.4044539 | 0.68587897 | NA | ZNF519P4 | 106481981 |
| ENSG000002 | 2.12280757 | -0.501036419 | 0.36231892 | -1.3828602 | 0.16670773 | NA | AARSP1 | 106480683 |
| ENSG000002 | 0.71034736 | 0.133412303 | 0.27671809 | 0.48212353 | 0.6297182 | NA | NA | NA |
| ENSG000002 | 1.09731595 | -0.108171341 | 0.32176864 | -0.3361774 | 0.7367371 | NA | NA | NA |
| ENSG000002 | 1.06082949 | 0.178283864 | 0.31421915 | 0.56738701 | 0.57045128 | NA | NA | NA |
| ENSG000002 | 1.08999793 | 0.042116199 | 0.31322934 | 0.13445803 | 0.8930404 | NA | FAM138D | 677784 |
| ENSG000002 | 1.81586234 | -0.099012525 | 0.34726301 | -0.2851226 | 0.77555023 | NA | TBCAP3 | 101290501 |
| ENSG000002 | 1.90931392 | -0.481562321 | 0.35133448 | -1.3706663 | 0.17047902 | NA | NA | NA |
| ENSG000002 | 1.41347306 | -0.114694236 | 0.32672634 | -0.3510407 | 0.72555783 | NA | NA | NA |
| ENSG000002 | 1.37272086 | -0.410866386 | 0.33817628 | -1.2149474 | 0.22438617 | NA | NA | NA |
| ENSG000002 | 0.67599868 | -0.271668661 | 0.26265684 | -1.0343102 | 0.30099116 | NA | NA | NA |
| ENSG000002 | 1.98561515 | -0.319962545 | 0.36271075 | -0.8821424 | 0.37769978 | NA | NA | NA |
| ENSG000002 | 1.13432303 | -0.435775362 | 0.32348252 | -1.3471373 | 0.17793603 | NA | C5orf66-AS1 | 101927953 |
| ENSG000002 | 1.70890838 | -0.306708796 | 0.34803948 | -0.8812471 | 0.37818408 | NA | NA | NA |
| ENSG000002 | 0.3863157 | -0.274134926 | 0.21457207 | -1.277589 | 0.20139437 | NA | NA | NA |
| ENSG000002 | 0.95743315 | -0.0327559 | 0.30495501 | -0.1074122 | 0.91446195 | NA | AIG1P1 | 106479028 |
| ENSG000002 | 1.95680711 | -0.176189604 | 0.35608999 | -0.4947895 | 0.62074869 | NA | NA | NA |
| ENSG000002 | 0.86523834 | -0.252727523 | 0.29302256 | -0.8624849 | 0.38842076 | NA | NA | NA |
| ENSG000002 | 0.12070844 | -0.088503695 | 0.12783931 | -0.6923042 | 0.48874631 | NA | USP17L17 | 100287327 |
| ENSG000002 | 1.6112784 | -0.121227729 | 0.35136646 | -0.3450179 | 0.73008089 | NA | NA | NA |
| ENSG000002 | 1.24156792 | -0.331006074 | 0.32809721 | -1.0088659 | 0.31303897 | NA | NA | NA |
| ENSG000002 | 0.38651158 | -0.274258716 | 0.21461355 | -1.2779189 | 0.20127801 | NA | NA | NA |
| ENSG000002 | 1.82605855 | -0.125343809 | 0.36123757 | -0.3469844 | 0.72860302 | NA | LOC101929696 | 101929696 |
| ENSG000002 | 1.76850713 | -0.306114203 | 0.35078762 | -0.8726483 | 0.38285483 | NA | EPPIN-WFDC6 | 100526773 |
| ENSG000002 | 0.12685397 | -0.090215768 | 0.12901919 | -0.699243 | 0.48440017 | NA | NA | NA |
| ENSG000002 | 1.91155746 | -0.50923827 | 0.35612684 | -1.4299351 | 0.15273565 | NA | NA | NA |
| ENSG000002 | 0.72833821 | -0.029759877 | 0.2840795 | -0.104759 | 0.91656707 | NA | NA | NA |

| | | | | | | | | | |
|---|---|---|---|---|---|---|---|---|---|
| ENSG000002 | 1.61629889 | -0.376336399 | 0.34436082 | -1.0928549 | 0.27445753 | NA | NA | NA | |
| ENSG000002 | 0 | NA | NA | NA | NA | NA | NA | NA | |
| ENSG000002 | 1.96981768 | -0.052858025 | 0.35337943 | -0.1495787 | 0.88109704 | NA | NA | NA | |
| ENSG000002 | 1.75492901 | -0.342946049 | 0.35808442 | -0.957724 | 0.33820194 | NA | NA | NA | |
| ENSG000002 | 1.27821959 | -0.186360163 | 0.3165237 | -0.5887716 | 0.55601451 | NA | NA | NA | |
| ENSG000002 | 1.49219891 | -0.195215168 | 0.34490594 | -0.5659954 | 0.57139692 | NA | UBE2V1P12 | | 106479048 |
| ENSG000002 | 0.49219223 | -0.014160136 | 0.23865874 | -0.0593321 | 0.95268755 | NA | NA | NA | |
| ENSG000002 | 0.471387 | -0.012999703 | 0.2361605 | -0.0550461 | 0.95610178 | NA | NA | NA | |
| ENSG000002 | 1.15558472 | -0.441010777 | 0.32412477 | -1.3606204 | 0.17363368 | NA | NA | NA | |
| ENSG000002 | 1.88436676 | -0.387271435 | 0.35725597 | -1.0840167 | 0.2783574 | NA | NA | NA | |
| ENSG000002 | 0 | NA | NA | NA | NA | NA | NA | NA | |
| ENSG000002 | 1.50067814 | -0.303231756 | 0.33440093 | -0.906791 | 0.36451733 | NA | NA | NA | |
| ENSG000002 | 0.11205239 | 0.082118251 | 0.12671962 | 0.64803108 | 0.51696485 | NA | NA | NA | |
| ENSG000002 | 1.73247309 | -0.325003809 | 0.35307834 | -0.9204864 | 0.35731864 | NA | NA | NA | |
| ENSG000002 | 1.78378112 | 0.269337883 | 0.35533034 | 0.75799293 | 0.44845522 | NA | NA | NA | |
| ENSG000002 | 0.64488726 | -0.109815036 | 0.26668753 | -0.4117742 | 0.68050496 | NA | NA | NA | |
| ENSG000002 | 2.10731279 | -0.124911801 | 0.36689137 | -0.3404599 | 0.73351021 | NA | NA | NA | |
| ENSG000002 | 0.98375084 | -0.039415757 | 0.31680944 | -0.1244147 | 0.90098692 | NA | NA | NA | |
| ENSG000002 | 1.98521762 | -0.312182382 | 0.35869365 | -0.8703315 | 0.38411928 | NA | NA | NA | |
| ENSG000002 | 1.2498363 | -0.048362168 | 0.33054017 | -0.1463125 | 0.88367468 | NA | NA | NA | |
| ENSG000002 | 0.38747904 | -0.104908247 | 0.21987514 | -0.4771265 | 0.63327207 | NA | NA | NA | |
| ENSG000002 | 2.14886925 | -0.227059621 | 0.35880053 | -0.6328297 | 0.52684488 | NA | NA | NA | |
| ENSG000002 | 0.71550581 | -0.024336771 | 0.27684741 | -0.0879068 | 0.92995075 | NA | NA | NA | |
| ENSG000002 | 2.09726609 | -0.011282349 | 0.36671825 | -0.0307657 | 0.97545638 | NA | NA | NA | |
| ENSG000002 | 1.1276317 | -0.121205536 | 0.31660604 | -0.3828276 | 0.70184758 | NA | NA | NA | |
| ENSG000002 | 0.50704533 | -0.189333637 | 0.24184367 | -0.7828761 | 0.43369986 | NA | NA | NA | |
| ENSG000002 | 0.86976701 | -0.115601083 | 0.29779035 | -0.3881962 | 0.69787084 | NA | NA | NA | |
| ENSG000002 | 1.08137625 | 0.346937677 | 0.31805049 | 1.09082579 | 0.27534955 | NA | NA | NA | |
| ENSG000002 | 1.86410725 | -0.115300774 | 0.35460202 | -0.3251554 | 0.74506344 | NA | NA | NA | |

| | | | | | | | | |
|---|---|---|---|---|---|---|---|---|
| ENSG000002 | 1.80368884 | -0.451191428 | 0.35102054 | -1.2853704 | 0.19866287 | NA | NA | NA |
| ENSG000002 | 2.00006735 | -0.203903521 | 0.36534196 | -0.5581169 | 0.57676456 | NA | NA | NA |
| ENSG000002 | 0 | NA | NA | NA | NA | NA | NA | NA |
| ENSG000002 | 0.40327843 | -0.281391716 | 0.21700406 | -1.2967118 | 0.19473038 | NA | NA | NA |
| ENSG000002 | 1.06256842 | -0.100315582 | 0.30858944 | -0.3250778 | 0.74512218 | NA | NA | NA |
| ENSG000002 | 0.6238537 | -0.107426552 | 0.26493731 | -0.4054791 | 0.68512535 | NA | NA | NA |
| ENSG000002 | 0.61332276 | -0.255253496 | 0.25725954 | -0.9922023 | 0.32109886 | NA | NA | NA |
| ENSG000002 | 1.86015073 | -0.266559179 | 0.36096312 | -0.7384665 | 0.46023101 | NA | NA | NA |
| ENSG000002 | 1.25799241 | -0.478796001 | 0.32450288 | -1.4754754 | 0.14008479 | NA | NA | NA |
| ENSG000002 | 1.19810619 | -0.591329122 | 0.31849508 | -1.8566351 | 0.06336309 | NA | NA | NA |
| ENSG000002 | 0.63731726 | -0.263777881 | 0.25938813 | -1.0169235 | 0.30918984 | NA | NA | NA |
| ENSG000002 | 1.11128995 | -0.116335392 | 0.32222196 | -0.3610412 | 0.71806867 | NA | NA | NA |
| ENSG000002 | 0.22410479 | 0.147643983 | 0.16638082 | 0.88738586 | 0.37487119 | NA | NA | NA |
| ENSG000002 | 2.08531781 | -0.343475285 | 0.35951392 | -0.955388 | 0.33938153 | NA | NA | NA |
| ENSG000002 | 1.63934711 | -0.419256313 | 0.35276792 | -1.1884763 | 0.23464582 | NA | NA | NA |
| ENSG000002 | 0.71282115 | 0.314277914 | 0.28418684 | 1.10588483 | 0.26877637 | NA | NA | NA |
| ENSG000002 | 1.64896658 | -0.641303271 | 0.33995451 | -1.8864385 | 0.05923589 | NA | NA | NA |
| ENSG000002 | 1.99537843 | -0.588825944 | 0.36177455 | -1.6276047 | 0.10360874 | NA | NA | NA |
| ENSG000002 | 0.63484558 | -0.278698901 | 0.26767877 | -1.0411692 | 0.29779701 | NA | NA | NA |
| ENSG000002 | 1.80687402 | 0.017734994 | 0.35127897 | 0.05048692 | 0.95973437 | NA | NA | NA |
| ENSG000002 | 1.21394908 | 0.098133254 | 0.32803809 | 0.29915201 | 0.76482406 | NA | NA | NA |
| ENSG000002 | 0.38036605 | -0.293832733 | 0.22119083 | -1.3284128 | 0.18404178 | NA | NA | NA |
| ENSG000002 | 0.61582881 | -0.11034135 | 0.26423602 | -0.4175863 | 0.67624959 | NA | NA | NA |
| ENSG000002 | 0.71294322 | 0.124158965 | 0.27526995 | 0.45104438 | 0.65195757 | NA | NA | NA |
| ENSG000002 | 0.99579395 | -0.04180985 | 0.3131507 | -0.1335135 | 0.89378728 | NA | NA | NA |
| ENSG000002 | 1.12640419 | -0.276028344 | 0.32347082 | -0.8533331 | 0.39347463 | NA | NA | NA |
| ENSG000002 | 1.43169013 | -0.402384605 | 0.33199597 | -1.2120165 | 0.22550605 | NA | NA | NA |
| ENSG000002 | 1.76845102 | -0.607852508 | 0.35407813 | -1.7167186 | 0.08603061 | NA | NA | NA |
| ENSG000002 | 1.228781 | -0.195375277 | 0.32900711 | -0.593833 | 0.55262381 | NA | MLLT10P2 | 106480724 |

| | | | | | | | | |
|---|---|---|---|---|---|---|---|---|
| ENSG000002 | 0.59032214 | 0.060325881 | 0.26160816 | 0.23059632 | 0.81762842 | NA | NA | NA |
| ENSG000002 | 1.68568704 | -0.048928874 | 0.35092665 | -0.1394276 | 0.88911223 | NA | NA | NA |
| ENSG000002 | 1.20294206 | -0.04053012 | 0.33154045 | -0.1222479 | 0.9027027 | NA | NA | NA |
| ENSG000002 | 1.02209595 | -0.350747276 | 0.30987289 | -1.1319069 | 0.25767356 | NA | NA | NA |
| ENSG000002 | 1.77081663 | -0.731979237 | 0.34885975 | -2.0982049 | 0.03588705 | NA | NA | NA |
| ENSG000002 | 1.95561151 | -0.291072269 | 0.35547193 | -0.8188333 | 0.41288151 | NA | NA | NA |
| ENSG000002 | 1.34029261 | -0.118991113 | 0.33915213 | -0.3508488 | 0.7257018 | NA | NA | NA |
| ENSG000002 | 1.23140307 | -0.320635076 | 0.32354003 | -0.9910214 | 0.32167515 | NA | NA | NA |
| ENSG000002 | 0 | NA | NA | NA | NA | NA | NA | NA |
| ENSG000002 | 0 | NA | NA | NA | NA | NA | NA | NA |
| ENSG000002 | 0.71755932 | -0.166538792 | 0.26894713 | -0.619225 | 0.53576813 | NA | NA | NA |
| ENSG000002 | 1.58495275 | -0.119416448 | 0.34796846 | -0.3431818 | 0.73146168 | NA | NA | NA |
| ENSG000002 | 2.16686294 | -0.639262944 | 0.36253723 | -1.7633029 | 0.07784942 | NA | NA | NA |
| ENSG000002 | 0.62413603 | 0.057496507 | 0.27089994 | 0.2122426 | 0.83191778 | NA | NA | NA |
| ENSG000002 | 1.70192867 | -0.300313555 | 0.3488284 | -0.8609206 | 0.38928178 | NA | NA | NA |
| ENSG000002 | 0.84699953 | -0.082240496 | 0.27963929 | -0.2940949 | 0.76868537 | NA | NA | NA |
| ENSG000002 | 0.11934008 | 0.083564698 | 0.12760091 | 0.65489106 | 0.5125379 | NA | NA | NA |
| ENSG000002 | 0.94300213 | -0.028401351 | 0.3043005 | -0.0933332 | 0.92563883 | NA | NA | NA |
| ENSG000002 | 1.22516772 | -0.19178798 | 0.33256576 | -0.5766919 | 0.56414761 | NA | NA | NA |
| ENSG000002 | 0 | NA | NA | NA | NA | NA | NA | NA |
| ENSG000002 | 1.2493105 | -0.473566726 | 0.32588276 | -1.4531813 | 0.14617343 | NA | NA | NA |
| ENSG000002 | 0 | NA | NA | NA | NA | NA | NA | NA |
| ENSG000002 | 2.15788698 | 0.189751605 | 0.36641089 | 0.51786563 | 0.60455202 | NA | NA | NA |
| ENSG000002 | 1.342999 | -0.258405154 | 0.33776945 | -0.7650341 | 0.44425123 | NA | NA | NA |
| ENSG000002 | 0.87830044 | -0.118101042 | 0.29822623 | -0.3960116 | 0.69209648 | NA | NA | NA |
| ENSG000002 | 0.5226185 | -0.189580046 | 0.24357686 | -0.7783172 | 0.43638207 | NA | NA | NA |
| ENSG000002 | 1.65309354 | -0.3937714 | 0.34811475 | -1.131154 | 0.25799026 | NA | NA | NA |
| ENSG000002 | 0 | NA | NA | NA | NA | NA | NA | NA |
| ENSG000002 | 1.49581666 | -0.458360632 | 0.3381922 | -1.3553259 | 0.17531376 | NA | NA | NA |

| | | | | | | | | | |
|---|---|---|---|---|---|---|---|---|---|
| ENSG000002 | 1.62992406 | -0.120073286 | 0.3497874 | -0.343275 | 0.73139156 | NA | NA | NA | |
| ENSG000002 | 0 | NA | NA | NA | NA | NA | NA | NA | |
| ENSG000002 | 1.56862251 | -0.647128954 | 0.3442784 | -1.8796676 | 0.06015339 | NA | NA | NA | |
| ENSG000002 | 0.60470109 | 0.059032226 | 0.26929751 | 0.21920821 | 0.82648786 | NA | KRT18P56 | | 106479027 |
| ENSG000002 | 1.64165069 | -0.392848254 | 0.34722933 | -1.1313798 | 0.25789527 | NA | NA | NA | |
| ENSG000002 | 2.04902523 | 0.25470419 | 0.36559944 | 0.69667555 | 0.48600586 | NA | NA | NA | |
| ENSG000002 | 0.668711 | -0.27443715 | 0.26201124 | -1.0474251 | 0.29490356 | NA | NA | NA | |
| ENSG000002 | 0.85704067 | 0.04415857 | 0.29677886 | 0.14879284 | 0.8817171 | NA | NA | NA | |
| ENSG000002 | 0 | NA | NA | NA | NA | NA | NA | NA | |
| ENSG000002 | 0.87033008 | 0.046821889 | 0.30257353 | 0.15474549 | 0.87702197 | NA | NA | NA | |
| ENSG000002 | 1.86962129 | 0.001037514 | 0.35797419 | 0.00289829 | 0.9976875 | NA | NA | NA | |
| ENSG000002 | 1.37271797 | 0.008145081 | 0.32805913 | 0.02482809 | 0.98019209 | NA | NA | NA | |
| ENSG000002 | 0.86689094 | -0.10433263 | 0.29765455 | -0.3505158 | 0.72595162 | NA | LOC100506406 | | 100506406 |
| ENSG000002 | 0.76018684 | -0.193725132 | 0.28091263 | -0.6896277 | 0.49042834 | NA | NA | NA | |
| ENSG000002 | 0.37171 | -0.104406568 | 0.21753023 | -0.4799635 | 0.63125336 | NA | NA | NA | |
| ENSG000002 | 1.71999525 | -0.171250873 | 0.34827953 | -0.4917052 | 0.62292773 | NA | NA | NA | |
| ENSG000002 | 0.64506141 | -0.288091388 | 0.2684363 | -1.0732207 | 0.28317211 | NA | NA | NA | |
| ENSG000002 | 1.98976563 | -0.310965998 | 0.36241366 | -0.8580416 | 0.39086949 | NA | NA | NA | |
| ENSG000002 | 0.37996515 | 0.064665424 | 0.21008146 | 0.3078112 | 0.758226 | NA | NA | NA | |
| ENSG000002 | 2.13145447 | -0.381260549 | 0.36442602 | -1.0461946 | 0.29547117 | NA | NA | NA | |
| ENSG000002 | 0.84975298 | 0.041735596 | 0.29639127 | 0.1408125 | 0.88801807 | NA | NA | NA | |
| ENSG000002 | 1.60743377 | -0.235262383 | 0.34164533 | -0.6886158 | 0.49106505 | NA | NA | NA | |
| ENSG000002 | 1.7235953 | -0.318120598 | 0.35411633 | -0.8983505 | 0.36899869 | NA | SAP18P1 | | 106480689 |
| ENSG000002 | 0.82945931 | 0.206551976 | 0.28980035 | 0.71273887 | 0.47600736 | NA | TRMT112P2 | | 106479033 |
| ENSG000002 | 1.13483349 | -0.274699772 | 0.32225896 | -0.8524194 | 0.39398138 | NA | NA | NA | |
| ENSG000002 | 1.10954911 | 0.027311429 | 0.3156612 | 0.08652134 | 0.93105199 | NA | NA | NA | |
| ENSG000002 | 0.84975353 | 0.05329801 | 0.30153883 | 0.17675339 | 0.8597021 | NA | NA | NA | |
| ENSG000002 | 1.61759862 | -0.255845394 | 0.35077483 | -0.7293722 | 0.46577404 | NA | NA | NA | |
| ENSG000002 | 2.16347386 | -0.383652933 | 0.36458343 | -1.0523049 | 0.29265968 | NA | NA | NA | |

| | | | | | | | | | |
|---|---|---|---|---|---|---|---|---|---|
| ENSG000002 | 0.63233507 | -0.116461163 | 0.27190168 | -0.4283209 | 0.66841754 | NA | | SNRPCP13 | 106481786 |
| ENSG000002 | 1.07002416 | -0.683165347 | 0.30569832 | -2.2347697 | 0.02543247 | NA | | LINC01377 | 102467074 |
| ENSG000002 | 0.12070844 | -0.088503695 | 0.12783931 | -0.6923042 | 0.48874631 | NA | | USP17L21 | 100287478 |
| ENSG000002 | 1.76563912 | -0.549740823 | 0.34392326 | -1.5984403 | 0.10994501 | NA | | NA | NA |
| ENSG000002 | 1.26888286 | -0.32856825 | 0.32543528 | -1.009627 | 0.31267402 | NA | | NA | NA |
| ENSG000002 | 2.11514072 | -0.137875539 | 0.3672086 | -0.3754693 | 0.70731151 | NA | | NA | NA |
| ENSG000002 | 1.21826771 | -0.241128973 | 0.22318837 | -1.0803832 | 0.27997157 | NA | | NA | NA |
| ENSG000002 | 1.51044121 | -0.060321155 | 0.33239638 | -0.1814736 | 0.85599589 | NA | | NA | NA |
| ENSG000002 | 2.0269126 | -0.32083724 | 0.36115919 | -0.8883541 | 0.37435032 | NA | | CCDC37-AS1 | 100506907 |
| ENSG000002 | 1.88238435 | -0.383955329 | 0.35764954 | -1.0735519 | 0.28302359 | NA | | NA | NA |
| ENSG000002 | 2.00180978 | -0.192007713 | 0.35990461 | -0.5334961 | 0.5936902 | NA | | NA | NA |
| ENSG000002 | 1.64865605 | -0.558959114 | 0.35160703 | -1.5897268 | 0.1118964 | NA | | NA | NA |
| ENSG000002 | 2.11965985 | -0.355740898 | 0.3610702 | -0.9852403 | 0.32450607 | NA | | NA | NA |
| ENSG000002 | 1.42181267 | -0.043324826 | 0.33116752 | -0.1308245 | 0.89591414 | NA | | MTRNR2L5 | 100463289 |
| ENSG000002 | 0.50611225 | -0.202624884 | 0.24886748 | -0.8141879 | 0.41553732 | NA | | NA | NA |
| ENSG000002 | 1.69760988 | -0.304519158 | 0.34924694 | -0.8719308 | 0.38324614 | NA | | LOC440117 | 440117 |
| ENSG000002 | 0.9705135 | -0.036451869 | 0.30553971 | -0.1193032 | 0.90503514 | NA | | NA | NA |
| ENSG000002 | 1.93872554 | 0.064763814 | 0.36255386 | 0.17863225 | 0.85822647 | NA | | RNF103-CHMP3 | 100526767 |
| ENSG000002 | 0.36827086 | -0.266056975 | 0.21181163 | -1.2561018 | 0.20907905 | NA | | NA | NA |
| ENSG000002 | 1.25638677 | -0.046299752 | 0.33070754 | -0.1400021 | 0.88865832 | NA | | NA | NA |
| ENSG000002 | 2.13717442 | -0.527044473 | 0.36762248 | -1.4336568 | 0.15167023 | NA | | NA | NA |
| ENSG000002 | 1.39059102 | -0.275355756 | 0.34070733 | -0.8081885 | 0.41898206 | NA | | KRT19P6 | 106480807 |
| ENSG000002 | 1.27013265 | -0.049395617 | 0.32808321 | -0.1505582 | 0.88032424 | NA | | LOC100288866 | 100288866 |
| ENSG000002 | 0.14128499 | -0.097772014 | 0.13403235 | -0.7294658 | 0.4657168 | NA | | NA | NA |
| ENSG000002 | 2.12004093 | -0.646910589 | 0.3648526 | -1.7730738 | 0.07621648 | NA | | NA | NA |
| ENSG000002 | 2.17166298 | 0.057703087 | 0.36470031 | 0.15822056 | 0.874283 | NA | | NA | NA |
| ENSG000002 | 1.99546275 | -0.569047865 | 0.35841455 | -1.5876807 | 0.11235855 | NA | | NA | NA |
| ENSG000002 | 1.68649997 | 0.221909951 | 0.35392117 | 0.62700388 | 0.53065669 | NA | | NA | NA |
| ENSG000002 | 0.70593786 | 0.286914504 | 0.27645988 | 1.03781607 | 0.2993557 | NA | | NA | NA |

| | | | | | | | | |
|---|---|---|---|---|---|---|---|---|
| ENSG000002 | 1.67856583 | -0.698158296 | 0.34793648 | -2.0065683 | 0.04479565 | NA | NA | NA |
| ENSG000002 | 1.44250574 | -0.576356875 | 0.33750957 | -1.7076756 | 0.08769655 | NA | NA | NA |
| ENSG000002 | 0.90027816 | -0.281064798 | 0.30047954 | -0.9353875 | 0.34958865 | NA | NA | NA |
| ENSG000002 | 1.46969147 | -0.17840941 | 0.34060814 | -0.5237967 | 0.60041997 | NA | NA | NA |
| ENSG000002 | 0 | NA | NA | NA | NA | NA | NA | NA |
| ENSG000002 | 0 | NA | NA | NA | NA | NA | NA | NA |
| ENSG000002 | 2.09171921 | -0.57160779 | 0.35853309 | -1.5942958 | 0.1108698 | NA | NA | NA |
| ENSG000002 | 0.36465257 | 0.064724075 | 0.21605106 | 0.29957767 | 0.76449932 | NA | NA | NA |
| ENSG000002 | 1.79372509 | -0.447622134 | 0.34945589 | -1.2809117 | 0.20022468 | NA | NA | NA |
| ENSG000002 | 2.16196562 | -0.520047773 | 0.36565221 | -1.422247 | 0.15495454 | NA | NA | NA |
| ENSG000002 | 0.76033119 | -0.194152683 | 0.28698065 | -0.6765358 | 0.49870053 | NA | NA | NA |
| ENSG000002 | 1.10460309 | -0.113252834 | 0.32048551 | -0.353379 | 0.72380436 | NA | NA | NA |
| ENSG000002 | 0.25351207 | -0.189965547 | 0.1824784 | -1.0410303 | 0.29786148 | NA | NA | NA |
| ENSG000002 | 2.13217049 | -0.520365624 | 0.36630558 | -1.4205779 | 0.1554395 | NA | NA | NA |
| ENSG000002 | 1.87290519 | -0.380740203 | 0.35817727 | -1.0629938 | 0.28778478 | NA | NA | NA |
| ENSG000002 | 0 | NA | NA | NA | NA | NA | C15orf38-AP3S2 | 100526783 |
| ENSG000002 | 1.39947775 | -0.888978406 | 0.33152088 | -2.6815156 | 0.00732895 | NA | NA | NA |
| ENSG000002 | 1.83069137 | -0.000397714 | 0.3611142 | -0.0011014 | 0.99912125 | NA | NA | NA |
| ENSG000002 | 0.2596576 | -0.193002035 | 0.1837913 | -1.0501152 | 0.29366517 | NA | NA | NA |
| ENSG000002 | 2.08782867 | -0.166594931 | 0.25506459 | -0.653148 | 0.51366086 | NA | NA | NA |
| ENSG000002 | 0.12685397 | -0.090215768 | 0.12901919 | -0.699243 | 0.48440017 | NA | NA | NA |
| ENSG000002 | 2.17147068 | -0.173099868 | 0.36411788 | -0.4753951 | 0.63450536 | NA | NA | NA |
| ENSG000002 | 0.98318777 | -0.200674067 | 0.3132137 | -0.6406938 | 0.52172166 | NA | NA | NA |
| ENSG000002 | 1.88192789 | -0.382165689 | 0.35825381 | -1.0667457 | 0.28608671 | NA | NA | NA |
| ENSG000002 | 1.95347156 | -0.285175917 | 0.35370427 | -0.8062552 | 0.4200957 | NA | NA | NA |
| ENSG000002 | 0.82907038 | -0.094230763 | 0.28039407 | -0.3360655 | 0.7368215 | NA | NA | NA |
| ENSG000002 | 1.41480705 | -0.392289816 | 0.33163215 | -1.1829065 | 0.23684622 | NA | NA | NA |
| ENSG000002 | 1.62648438 | -0.243134063 | 0.34338764 | -0.7080455 | 0.47891702 | NA | NA | NA |
| ENSG000002 | 0 | NA | NA | NA | NA | NA | NA | NA |

| | | | | | | | | |
|---|---|---|---|---|---|---|---|---|
| ENSG000002 | 0.14128499 | -0.097772014 | 0.13403235 | -0.7294658 | 0.4657168 | NA | NA | NA |
| ENSG000002 | 1.48884355 | -0.344233728 | 0.34667693 | -0.9929525 | 0.32073309 | NA | NA | NA |
| ENSG000002 | 1.3820774 | -0.269735082 | 0.34055326 | -0.7920496 | 0.42833173 | NA | NA | NA |
| ENSG000002 | 0.77333931 | -0.184645157 | 0.27296846 | -0.676434 | 0.49876512 | NA | NA | NA |
| ENSG000002 | 1.51428818 | -0.337606311 | 0.34592161 | -0.9759619 | 0.32908335 | NA | NA | NA |
| ENSG000002 | 1.70193059 | -0.170102026 | 0.34806599 | -0.4887063 | 0.62504968 | NA | NA | NA |
| ENSG000002 | 1.48064898 | -0.470953856 | 0.3433066 | -1.3718171 | 0.17012039 | NA | NA | NA |
| ENSG000002 | 1.1050601 | -0.110783624 | 0.32202153 | -0.3440255 | 0.7308271 | NA | NA | NA |
| ENSG000002 | 2.05952823 | -0.577164409 | 0.35823457 | -1.6111354 | 0.10715021 | NA | NA | NA |
| ENSG000002 | 1.90821686 | 0.327219881 | 0.35885342 | 0.9118483 | 0.36184858 | NA | HOXC-AS2 | 100874364 |
| ENSG000002 | 0 | NA | NA | NA | NA | NA | NA | NA |
| ENSG000002 | 1.01472237 | -0.5233652 | 0.3116574 | -1.6792966 | 0.09309426 | NA | NA | NA |
| ENSG000002 | 0.12685397 | -0.090215768 | 0.12901919 | -0.699243 | 0.48440017 | NA | ARPC4-TTLL3 | 100526693 |
| ENSG000002 | 1.25024069 | -0.052697936 | 0.32907245 | -0.1601408 | 0.87277014 | NA | TRMT112P5 | 106480784 |
| ENSG000002 | 0.61699269 | 0.06131205 | 0.26397339 | 0.23226602 | 0.8163314 | NA | NA | NA |
| ENSG000002 | 1.75501737 | -0.06033462 | 0.34950684 | -0.1726279 | 0.86294395 | NA | NA | NA |
| ENSG000002 | 1.36300051 | 0.158374217 | 0.32972302 | 0.48032503 | 0.6309963 | NA | NA | NA |
| ENSG000002 | 2.01911626 | -0.859696324 | 0.35819914 | -2.4000513 | 0.01639277 | NA | NA | NA |
| ENSG000002 | 1.55928578 | -0.797011497 | 0.34180144 | -2.3317968 | 0.01971138 | NA | NA | NA |
| ENSG000002 | 2.12166183 | -0.474357356 | 0.3613346 | -1.3127925 | 0.18925287 | NA | NA | NA |
| ENSG000002 | 1.72668369 | -0.056352834 | 0.3532454 | -0.1595289 | 0.87325223 | NA | NA | NA |
| ENSG000002 | 0.72335602 | 0.142054296 | 0.28912098 | 0.49133168 | 0.62319188 | NA | NA | NA |
| ENSG000002 | 0 | NA | NA | NA | NA | NA | NA | NA |
| ENSG000002 | 1.8233288 | -0.761937929 | 0.35312685 | -2.157689 | 0.03095202 | NA | YWHAEP4 | 106481785 |
| ENSG000002 | 0.75711827 | -0.523135555 | 0.27850213 | -1.8783898 | 0.06032786 | NA | NA | NA |
| ENSG000002 | 0.36576034 | -0.0934779 | 0.20820828 | -0.4489634 | 0.65345805 | NA | NA | NA |
| ENSG000002 | 1.20125308 | 0.093517087 | 0.32257669 | 0.28990653 | 0.77188775 | NA | NA | NA |
| ENSG000002 | 1.86502073 | -0.122469547 | 0.35919435 | -0.3409562 | 0.73313654 | NA | ALDH1L1-AS1 | 100874204 |
| ENSG000002 | 2.10427243 | -0.607163231 | 0.3606535 | -1.6835085 | 0.09227669 | NA | NA | NA |

| | | | | | | | | |
|---|---|---|---|---|---|---|---|---|
| ENSG000002 | 1.22940218 | -0.156024021 | 0.31501011 | -0.4952985 | 0.62038946 | NA | NA | NA |
| ENSG000002 | 1.77768898 | -0.05842832 | 0.34974954 | -0.1670576 | 0.86732472 | NA | NA | NA |
| ENSG000002 | 1.98364129 | -0.184527409 | 0.3623021 | -0.5093192 | 0.61052852 | NA | NA | NA |
| ENSG000002 | 2.03047883 | -0.559221863 | 0.35807629 | -1.5617394 | 0.11834939 | NA | NA | NA |
| ENSG000002 | 1.55907889 | -0.113538364 | 0.34448794 | -0.3295859 | 0.74171288 | NA | NA | NA |
| ENSG000002 | 1.85807838 | -0.619618227 | 0.35120213 | -1.7642781 | 0.07768517 | NA | NA | NA |
| ENSG000002 | 0 | NA | NA | NA | NA | NA | NA | NA |
| ENSG000002 | 0.12685397 | -0.090215768 | 0.12901919 | -0.699243 | 0.48440017 | NA | NA | NA |
| ENSG000002 | 0.58087681 | 0.226918272 | 0.26184027 | 0.86662863 | 0.38614552 | NA | NA | NA |
| ENSG000002 | 1.22534241 | -0.326376339 | 0.32766755 | -0.9960594 | 0.31922129 | NA | PSMC1P5 | 106480732 |
| ENSG000002 | 1.79505477 | 0.285422766 | 0.36082869 | 0.79102015 | 0.42893223 | NA | NA | NA |
| ENSG000002 | 1.82676406 | -0.102873127 | 0.35345758 | -0.291048 | 0.77101459 | NA | VWA8P1 | 106480787 |
| ENSG000002 | 0.74590018 | -0.179161319 | 0.28000754 | -0.6398446 | 0.52227362 | NA | NA | NA |
| ENSG000002 | 2.07569621 | -0.243638789 | 0.3653164 | -0.6669254 | 0.50481979 | NA | NA | NA |
| ENSG000002 | 0.90221296 | -0.620929665 | 0.29629503 | -2.0956466 | 0.03611355 | NA | NA | NA |
| ENSG000002 | 1.94547244 | 0.074938933 | 0.3639176 | 0.20592281 | 0.83685121 | NA | NA | NA |
| ENSG000002 | 1.93004771 | -0.359463669 | 0.35011899 | -1.02669 | 0.30456646 | NA | NA | NA |
| ENSG000002 | 1.10000061 | -0.401041023 | 0.31585546 | -1.2696979 | 0.20419224 | NA | NA | NA |
| ENSG000002 | 1.14325928 | -0.603209097 | 0.32167111 | -1.8752355 | 0.06076033 | NA | NA | NA |
| ENSG000002 | 2.06621067 | -0.126882356 | 0.36468539 | -0.3479228 | 0.72789817 | NA | NA | NA |
| ENSG000002 | 1.24061827 | -0.034391802 | 0.32274284 | -0.106561 | 0.91513725 | NA | NA | NA |
| ENSG000002 | 1.15965501 | -0.431490781 | 0.32019018 | -1.3476078 | 0.17778457 | NA | NA | NA |
| ENSG000002 | 0.26560726 | -0.177019426 | 0.17672274 | -1.0016788 | 0.31649875 | NA | NA | NA |
| ENSG000002 | 0.49170296 | -0.014565292 | 0.2386078 | -0.0610428 | 0.95132511 | NA | SNRPCP16 | 106480291 |
| ENSG000002 | 2.09724666 | -1.005418787 | 0.35588905 | -2.8250906 | 0.00472673 | NA | NA | NA |
| ENSG000002 | 1.10187997 | -0.366775624 | 0.30765232 | -1.1921757 | 0.23319236 | NA | NA | NA |
| ENSG000002 | 0 | NA | NA | NA | NA | NA | NA | NA |
| ENSG000002 | 1.0161791 | -0.327070024 | 0.30435441 | -1.0746354 | 0.28253799 | NA | NA | NA |
| ENSG000002 | 1.63495778 | -0.268805771 | 0.35098386 | -0.7658636 | 0.44375745 | NA | HSPE1P23 | 106481676 |

| | | | | | | | | |
|---|---|---|---|---|---|---|---|---|
| ENSG000002 | 1.55791742 | -0.640472441 | 0.34416341 | -1.8609545 | 0.0627506 | NA | NA | NA |
| ENSG000002 | 1.38097355 | -0.398489163 | 0.33326533 | -1.1957114 | 0.2318092 | NA | NA | NA |
| ENSG000002 | 1.66166259 | -0.7149154 | 0.34985767 | -2.0434464 | 0.04100826 | NA | NA | NA |
| ENSG000002 | 1.39545059 | -0.254137383 | 0.32975934 | -0.7706753 | 0.4408994 | NA | NA | NA |
| ENSG000002 | 0.25260017 | -0.015978705 | 0.18049464 | -0.0885273 | 0.92945758 | NA | NA | NA |
| ENSG000002 | 1.02255084 | -0.536784374 | 0.31334684 | -1.7130678 | 0.08670007 | NA | NA | NA |
| ENSG000002 | 0.60282172 | 0.050569991 | 0.26272347 | 0.19248372 | 0.84736332 | NA | NA | NA |
| ENSG000002 | 0.84423861 | -0.112252556 | 0.29643258 | -0.3786782 | 0.70492685 | NA | NA | NA |
| ENSG000002 | 1.97140174 | -0.176024643 | 0.35928035 | -0.4899367 | 0.62417867 | NA | NA | NA |
| ENSG000002 | 0.75304351 | -0.205958918 | 0.29162776 | -0.7062391 | 0.48003948 | NA | LOC101929034 | 101929034 |
| ENSG000002 | 0 | NA | NA | NA | NA | NA | NA | NA |
| ENSG000002 | 1.53163682 | -0.167063126 | 0.32710826 | -0.5107273 | 0.60954201 | NA | NA | NA |
| ENSG000002 | 0.94550601 | -0.633425358 | 0.29832934 | -2.1232419 | 0.03373358 | NA | NA | NA |
| ENSG000002 | 0.87787781 | 0.029373897 | 0.29013993 | 0.10124045 | 0.91935959 | NA | NA | NA |
| ENSG000002 | 0.82963345 | 0.054595074 | 0.3005004 | 0.18168054 | 0.85583344 | NA | NA | NA |
| ENSG000002 | 1.47525738 | -0.057081005 | 0.34808905 | -0.1639839 | 0.86974381 | NA | HOXC-AS1 | 100874363 |
| ENSG000002 | 1.58811561 | -0.26498561 | 0.35150561 | -0.7538588 | 0.45093399 | NA | NA | NA |
| ENSG000002 | 1.73481182 | -0.700783431 | 0.34594309 | -2.0257188 | 0.0427936 | NA | NA | NA |
| ENSG000002 | 0.12070844 | -0.088503695 | 0.12783931 | -0.6923042 | 0.48874631 | NA | NA | NA |
| ENSG000002 | 1.24981903 | -0.47868751 | 0.32585118 | -1.4690372 | 0.1418227 | NA | NA | NA |
| ENSG000002 | 2.01433007 | -0.182170297 | 0.35651943 | -0.5109688 | 0.60937292 | NA | NA | NA |
| ENSG000002 | 2.11414739 | -0.365200336 | 0.36345407 | -1.0048046 | 0.31499093 | NA | NA | NA |
| ENSG000002 | 2.15353632 | -0.518632028 | 0.36563875 | -1.4184274 | 0.15606601 | NA | NA | NA |
| ENSG000002 | 0 | NA | NA | NA | NA | NA | DUX4L51 | 106481740 |
| ENSG000002 | 1.32609184 | 0.029255275 | 0.33858658 | 0.08640412 | 0.93114517 | NA | NA | NA |
| ENSG000002 | 1.16155611 | -0.272927668 | 0.32311821 | -0.8446682 | 0.39829613 | NA | NA | NA |
| ENSG000002 | 0.65542213 | -0.28582049 | 0.26931472 | -1.061288 | 0.28855903 | NA | NA | NA |
| ENSG000002 | 1.12663445 | -0.119797104 | 0.32120224 | -0.3729647 | 0.7091747 | NA | CDK3 | 1018 |
| ENSG000002 | 1.26920136 | -0.491955005 | 0.32713564 | -1.5038258 | 0.13262623 | NA | NA | NA |

| | | | | | | | | |
|---|---|---|---|---|---|---|---|---|
| ENSG000002 | 0 | NA | NA | NA | NA | NA | NA | NA |
| ENSG000002 | 1.15188053 | -0.268324986 | 0.32004897 | -0.8383873 | 0.40181323 | NA | NA | NA |
| ENSG000002 | 0.62431017 | -0.119347761 | 0.27124333 | -0.4400026 | 0.65993525 | NA | NA | NA |
| ENSG000002 | 1.76620163 | -0.47506525 | 0.35690319 | -1.3310759 | 0.18316403 | NA | NA | NA |
| ENSG000002 | 1.37735422 | -0.267608881 | 0.34046407 | -0.7860121 | 0.43186039 | NA | NA | NA |
| ENSG000002 | 0.51987986 | -0.01050728 | 0.1556516 | -0.0675051 | 0.94617959 | NA | NA | NA |
| ENSG000002 | 1.74875314 | -0.191980903 | 0.35554752 | -0.5399585 | 0.58922566 | NA | NA | NA |
| ENSG000002 | 0.71103058 | -0.021840671 | 0.27652595 | -0.0789824 | 0.93704666 | NA | NA | NA |
| ENSG000002 | 1.9311075 | 0.192316466 | 0.36074227 | 0.5331132 | 0.59395522 | NA | NA | NA |
| ENSG000002 | 0 | NA | NA | NA | NA | NA | NA | NA |
| ENSG000002 | 2.13503839 | -0.754321769 | 0.36086127 | -2.0903373 | 0.03658751 | NA | NA | NA |
| ENSG000002 | 0.52125014 | -0.010203417 | 0.15567753 | -0.065542 | 0.94774247 | NA | NA | NA |
| ENSG000002 | 0.35067856 | 0.074230421 | 0.21389444 | 0.34704231 | 0.72855953 | NA | NA | NA |
| ENSG000002 | 1.59433899 | 0.156095043 | 0.34977897 | 0.44626767 | 0.65540391 | NA | NA | NA |
| ENSG000002 | 0.82963345 | 0.054595074 | 0.3005004 | 0.18168054 | 0.85583344 | NA | NA | NA |
| ENSG000002 | 0.23890637 | -0.010420674 | 0.17747379 | -0.0587167 | 0.95317776 | NA | NA | NA |
| ENSG000002 | 0.48257029 | 0.139654084 | 0.23829294 | 0.58606053 | 0.55783484 | NA | NA | NA |
| ENSG000002 | 2.04253964 | -0.734108478 | 0.36181104 | -2.0289831 | 0.04246001 | NA | NA | NA |
| ENSG000002 | 0.96726631 | 0.117798155 | 0.30709794 | 0.38358497 | 0.70128608 | NA | NA | NA |
| ENSG000002 | 0.99172175 | -0.193654422 | 0.31358032 | -0.6175592 | 0.53686592 | NA | NA | NA |
| ENSG000002 | 1.97755937 | -0.198011461 | 0.36313592 | -0.5452819 | 0.58555962 | NA | NA | NA |
| ENSG000002 | 1.35432135 | 0.028372984 | 0.3335299 | 0.08506879 | 0.9322067 | NA | NA | NA |
| ENSG000002 | 1.98994135 | -0.319835139 | 0.36241054 | -0.8825216 | 0.37749479 | NA | NA | NA |
| ENSG000002 | 1.60077736 | 0.146635182 | 0.34990191 | 0.41907512 | 0.67516123 | NA | NA | NA |
| ENSG000002 | 1.45087504 | -0.291369865 | 0.33338987 | -0.8739614 | 0.3821393 | NA | LOC100133461 | 100133461 |
| ENSG000002 | 1.26117254 | -0.200687202 | 0.32985872 | -0.6084035 | 0.54291989 | NA | NA | NA |
| ENSG000002 | 0.72967569 | -0.033529616 | 0.27782303 | -0.120687 | 0.90393899 | NA | NA | NA |
| ENSG000002 | 1.74722316 | -0.470974221 | 0.35508959 | -1.3263532 | 0.18472272 | NA | NA | NA |
| ENSG000002 | 0.9917791 | -0.123187141 | 0.20281927 | -0.607374 | 0.54360277 | NA | NA | NA |

| | | | | | | | | |
|---|---|---|---|---|---|---|---|---|
| ENSG000002 | 0.23202362 | -0.008302493 | 0.17586207 | -0.0472103 | 0.96234565 | NA | NA | NA |
| ENSG000002 | 1.32452705 | 0.162167183 | 0.33527931 | 0.48367788 | 0.6286145 | NA | NA | NA |
| ENSG000002 | 0.48228797 | -0.022292485 | 0.23747863 | -0.0938715 | 0.9252112 | NA | NA | NA |
| ENSG000002 | 0.66313191 | -0.429680649 | 0.25813351 | -1.6645675 | 0.09599908 | NA | NA | NA |
| ENSG000002 | 0.38056192 | -0.244105065 | 0.20391883 | -1.1970698 | 0.23127937 | NA | NA | NA |
| ENSG000002 | 2.02175222 | -0.316127905 | 0.3620603 | -0.8731361 | 0.38258892 | NA | NA | NA |
| ENSG000002 | 0 | NA | NA | NA | NA | NA | NA | NA |
| ENSG000002 | 0.84423861 | -0.112252556 | 0.29643258 | -0.3786782 | 0.70492685 | NA | NA | NA |
| ENSG000002 | 1.89057543 | -0.523088505 | 0.35696364 | -1.4653831 | 0.14281642 | NA | NA | NA |
| ENSG000002 | 2.15092422 | -0.78443297 | 0.36375421 | -2.1564918 | 0.03104528 | NA | NA | NA |
| ENSG000002 | 0.85380656 | -0.412184383 | 0.29077846 | -1.4175203 | 0.15633085 | NA | NA | NA |
| ENSG000002 | 0.602844 | 0.2204762 | 0.25672804 | 0.85879283 | 0.39045483 | NA | NA | NA |
| ENSG000002 | 0 | NA | NA | NA | NA | NA | CCDC169-SOHLI | 100526761 |
| ENSG000002 | 2.05915166 | -0.124517728 | 0.36460426 | -0.3415147 | 0.73271611 | NA | NA | NA |
| ENSG000002 | 1.39352363 | -0.588007859 | 0.3397524 | -1.7306952 | 0.08350614 | NA | NA | NA |
| ENSG000002 | 0.79670713 | -0.362882081 | 0.28391224 | -1.2781488 | 0.20119698 | NA | NA | NA |
| ENSG000002 | 1.86789679 | -0.1309401 | 0.35875329 | -0.3649865 | 0.7151215 | NA | LINC01017 | 102467075 |
| ENSG000002 | 0.98002938 | -0.337746284 | 0.30815809 | -1.0960163 | 0.27307164 | NA | NA | NA |
| ENSG000002 | 0.49150654 | -0.026274915 | 0.24607766 | -0.1067749 | 0.91496757 | NA | NA | NA |
| ENSG000002 | 1.37186103 | -0.281302253 | 0.34419406 | -0.8172781 | 0.41376954 | NA | NA | NA |
| ENSG000002 | 2.02896556 | -0.444957663 | 0.35978316 | -1.2367384 | 0.2161842 | NA | NA | NA |
| ENSG000002 | 0 | NA | NA | NA | NA | NA | NT5C1B-RDH14 | 100526794 |
| ENSG000002 | 0 | NA | NA | NA | NA | NA | USP17L20 | 100287441 |
| ENSG000002 | 1.16704876 | -0.268118945 | 0.31891204 | -0.8407301 | 0.40049917 | NA | NA | NA |
| ENSG000002 | 1.38011214 | -0.417019057 | 0.33830139 | -1.232685 | 0.2176933 | NA | NA | NA |
| ENSG000002 | 1.63311568 | -0.54068408 | 0.34898846 | -1.5492893 | 0.12131219 | NA | NA | NA |
| ENSG000002 | 0.55070895 | -0.361402396 | 0.24091882 | -1.5001003 | 0.13358842 | NA | NA | NA |
| ENSG000002 | 1.07949744 | 0.337913945 | 0.31345651 | 1.07802496 | 0.28102262 | NA | NA | NA |
| ENSG000002 | 0.52147635 | -0.19458195 | 0.24341208 | -0.7993932 | 0.42406247 | NA | NA | NA |

| | | | | | | | | |
|---|---|---|---|---|---|---|---|---|
| ENSG000002 | 0.47804353 | 0.152460376 | 0.23772494 | 0.64133101 | 0.52130765 | NA | NA | NA |
| ENSG000002 | 0.64235948 | -0.482579697 | 0.27062934 | -1.7831759 | 0.07455768 | NA | NA | NA |
| ENSG000002 | 1.94661247 | -0.048721619 | 0.35676842 | -0.1365637 | 0.89137566 | NA | NA | NA |
| ENSG000002 | 1.10445928 | -0.118876151 | 0.32584441 | -0.3648249 | 0.71524213 | NA | NA | NA |
| ENSG000002 | 2.04706639 | -0.738578923 | 0.36123003 | -2.0446222 | 0.04089212 | NA | NA | NA |
| ENSG000002 | 1.47857374 | -0.461029964 | 0.34018703 | -1.355225 | 0.17534588 | NA | NA | NA |
| ENSG000002 | 0.93497115 | -0.441253409 | 0.29749306 | -1.4832393 | 0.13801084 | NA | NA | NA |
| ENSG000002 | 2.12237224 | -0.822820604 | 0.35427436 | -2.322552 | 0.02020323 | NA | NA | NA |
| ENSG000002 | 1.73605659 | -0.198259283 | 0.35432913 | -0.5595342 | 0.57579717 | NA | NA | NA |
| ENSG000002 | 1.93823356 | -0.163708853 | 0.35282448 | -0.4639952 | 0.64265121 | NA | NA | NA |
| ENSG000002 | 1.89786523 | -0.389511713 | 0.35896895 | -1.0850847 | 0.27788414 | NA | NA | NA |
| ENSG000002 | 0.84512017 | -0.096634382 | 0.29040562 | -0.3327566 | 0.73931803 | NA | NA | NA |
| ENSG000002 | 1.49040779 | -0.479610099 | 0.34289783 | -1.3986968 | 0.16190392 | NA | NA | NA |
| ENSG000002 | 2.00235111 | -0.201223186 | 0.36411433 | -0.5526374 | 0.58051175 | NA | NA | NA |
| ENSG000002 | 0.47821768 | -0.016471461 | 0.24462483 | -0.0673336 | 0.94631616 | NA | NA | NA |
| ENSG000002 | 2.00576089 | -0.452559272 | 0.36342592 | -1.2452586 | 0.21303669 | NA | NA | NA |
| ENSG000002 | 1.72162557 | -0.193514853 | 0.35352713 | -0.5473833 | 0.58411542 | NA | NA | NA |
| ENSG000002 | 0 | NA | NA | NA | NA | NA | NA | NA |
| ENSG000002 | 1.98578124 | -0.182614062 | 0.36137751 | -0.5053277 | 0.6133287 | NA | NA | NA |
| ENSG000002 | 1.37900437 | -0.276934604 | 0.34047116 | -0.8133864 | 0.41599657 | NA | NA | NA |
| ENSG000002 | 0.11865494 | 0.083440866 | 0.12751478 | 0.65436232 | 0.5128784 | NA | NA | NA |
| ENSG000002 | 1.21073671 | -0.181310907 | 0.32853118 | -0.5518834 | 0.58102823 | NA | NA | NA |
| ENSG000002 | 1.24210816 | -0.330667888 | 0.32465358 | -1.0185253 | 0.30842837 | NA | NA | NA |
| ENSG000002 | 1.49183227 | -0.340989237 | 0.34673747 | -0.9834219 | 0.32539981 | NA | NA | NA |
| ENSG000002 | 0.37404583 | -0.105111695 | 0.21784993 | -0.4824959 | 0.62945372 | NA | NA | NA |
| ENSG000002 | 0.25351207 | -0.189965547 | 0.1824784 | -1.0410303 | 0.29786148 | NA | NA | NA |
| ENSG000002 | 0 | NA | NA | NA | NA | NA | NA | NA |
| ENSG000002 | 1.36287217 | -0.117834926 | 0.32813598 | -0.3591039 | 0.71951733 | NA | NA | NA |
| ENSG000002 | 0 | NA | NA | NA | NA | NA | NA | NA |

| | | | | | | | | |
|---|---|---|---|---|---|---|---|---|
| ENSG000002 | 1.12586341 | -0.26722627 | 0.32349262 | -0.826066 | 0.40876663 | NA | NA | NA |
| ENSG000002 | 1.34923098 | -0.122258384 | 0.34335912 | -0.3560656 | 0.72179141 | NA | NA | NA |
| ENSG000002 | 0.9235611 | -0.457490269 | 0.30194891 | -1.5151248 | 0.12974081 | NA | NA | NA |
| ENSG000002 | 1.31331343 | 0.162625071 | 0.32685802 | 0.4975404 | 0.61880802 | NA | NA | NA |
| ENSG000002 | 1.51370855 | -0.190703008 | 0.33996088 | -0.5609557 | 0.57482771 | NA | NA | NA |
| ENSG000002 | 1.10551499 | -0.250911746 | 0.31685714 | -0.7918766 | 0.42843265 | NA | NA | NA |
| ENSG000002 | 1.62317771 | -0.119275854 | 0.3475359 | -0.3432044 | 0.73144468 | NA | NA | NA |
| ENSG000002 | 0.62812434 | -0.408089682 | 0.2528562 | -1.61392 | 0.10654477 | NA | NA | NA |
| ENSG000002 | 0 | NA | NA | NA | NA | NA | NA | NA |
| ENSG000002 | 1.1040709 | -0.398262383 | 0.31343045 | -1.2706563 | 0.20385094 | NA | NA | NA |
| ENSG000002 | 0.71433332 | 0.130507939 | 0.27701556 | 0.47112132 | 0.6375541 | NA | NA | NA |
| ENSG000002 | 0 | NA | NA | NA | NA | NA | NA | NA |
| ENSG000002 | 0.49036493 | -0.020766486 | 0.24598337 | -0.0844223 | 0.93272066 | NA | NA | NA |
| ENSG000002 | 1.26233643 | -0.048019462 | 0.33086145 | -0.1451347 | 0.88460456 | NA | NA | NA |
| ENSG000002 | 0 | NA | NA | NA | NA | NA | WBP1LP4 | 106480779 |
| ENSG000002 | 1.01215978 | -0.340326264 | 0.30709292 | -1.1082192 | 0.26776717 | NA | NA | NA |
| ENSG000002 | 1.47318661 | -0.323818873 | 0.34149477 | -0.9482396 | 0.34300748 | NA | NA | NA |
| ENSG000002 | 0.51433302 | -0.186673425 | 0.24269587 | -0.7691661 | 0.44179474 | NA | NA | NA |
| ENSG000002 | 1.84810953 | -0.131143667 | 0.36224709 | -0.3620282 | 0.71733095 | NA | NA | NA |
| ENSG000002 | 0.98446631 | -0.179925643 | 0.30106788 | -0.5976248 | 0.55009028 | NA | NA | NA |
| ENSG000002 | 1.37275567 | -0.845086132 | 0.32632394 | -2.5897154 | 0.00960553 | NA | NA | NA |
| ENSG000002 | 1.84276537 | -0.125507433 | 0.3558245 | -0.3527229 | 0.72429622 | NA | NA | NA |
| ENSG000002 | 1.96782464 | -0.397924767 | 0.35412223 | -1.1236933 | 0.26114318 | NA | NA | NA |
| ENSG000002 | 1.96959348 | -0.30864874 | 0.36172682 | -0.8532647 | 0.39351251 | NA | NA | NA |
| ENSG000002 | 1.49501136 | -0.168207093 | 0.3364875 | -0.4998911 | 0.61715178 | NA | NA | NA |
| ENSG000002 | 1.83257575 | -0.226865849 | 0.35367102 | -0.6414601 | 0.52122379 | NA | NA | NA |
| ENSG000002 | 0.8706124 | 0.193012212 | 0.29224605 | 0.66044421 | 0.50896881 | NA | NA | NA |
| ENSG000002 | 1.81088424 | -0.462857384 | 0.3523343 | -1.3136881 | 0.18895118 | NA | NA | NA |
| ENSG000002 | 0.92492946 | -0.640001169 | 0.29955459 | -2.1365093 | 0.03263793 | NA | NA | NA |

| | | | | | | | | |
|---|---|---|---|---|---|---|---|---|
| ENSG000002 | 1.4090968 | -0.580153266 | 0.3388515 | -1.7121165 | 0.08687519 | NA | NA | NA |
| ENSG000002 | 2.00837245 | -0.188522922 | 0.35824575 | -0.5262391 | 0.59872209 | NA | NA | NA |
| ENSG000002 | 2.11807579 | -0.251190687 | 0.36525902 | -0.6877056 | 0.49163817 | NA | NA | NA |
| ENSG000002 | 0.48211382 | 0.155752635 | 0.24556107 | 0.63427249 | 0.525903 | NA | NA | NA |
| ENSG000002 | 1.29046359 | -0.494664122 | 0.329288 | -1.5022234 | 0.13303943 | NA | NA | NA |
| ENSG000002 | 0 | NA | NA | NA | NA | NA | NA | NA |
| ENSG000002 | 0.87457898 | -0.427920645 | 0.29435302 | -1.4537668 | 0.14601097 | NA | LOC101929573 | 101929573 |
| ENSG000002 | 2.13187318 | -0.253878795 | 0.36480731 | -0.6959257 | 0.48647534 | NA | NA | NA |
| ENSG000002 | 1.23171765 | -0.194591902 | 0.33056302 | -0.5886681 | 0.55608395 | NA | NA | NA |
| ENSG000002 | 1.89679363 | -0.495582395 | 0.35179784 | -1.4087136 | 0.15891987 | NA | NA | NA |
| ENSG000002 | 1.69005927 | 0.079279549 | 0.3566566 | 0.22228538 | 0.82409173 | NA | HULC | 728655 |
| ENSG000002 | 0.38133351 | -0.102466351 | 0.2189713 | -0.4679442 | 0.63982449 | NA | NA | NA |
| ENSG000002 | 0.49512483 | -0.344267812 | 0.23617548 | -1.457678 | 0.14492931 | NA | NA | NA |
| ENSG000002 | 0 | NA | NA | NA | NA | NA | NA | NA |
| ENSG000002 | 2.07674499 | -0.241538528 | 0.36049837 | -0.6700128 | 0.50284962 | NA | NA | NA |
| ENSG000002 | 2.13196197 | -0.516205532 | 0.36631525 | -1.4091838 | 0.15878081 | NA | NA | NA |
| ENSG000002 | 1.6038852 | -0.251359723 | 0.34993614 | -0.7183017 | 0.47257128 | NA | NA | NA |
| ENSG000002 | 2.05362421 | 0.250000493 | 0.36458239 | 0.68571741 | 0.49289133 | NA | NA | NA |
| ENSG000002 | 1.11703267 | 0.029979084 | 0.31759487 | 0.09439411 | 0.9247961 | NA | TMED7-TICAM2 | 100302736 |
| ENSG000002 | 1.49267958 | 0.081739118 | 0.33200505 | 0.24619841 | 0.80552864 | NA | NA | NA |
| ENSG000002 | 2.11268227 | -0.014956457 | 0.36239668 | -0.041271 | 0.96707989 | NA | NA | NA |
| ENSG000002 | 1.68429319 | -0.875351093 | 0.34880953 | -2.509539 | 0.01208889 | NA | NA | NA |
| ENSG000002 | 0.8678023 | -0.271126522 | 0.29886542 | -0.907186 | 0.36430845 | NA | B3GNTL1P1 | 106481968 |
| ENSG000002 | 1.99522705 | -0.183514183 | 0.35630105 | -0.5150537 | 0.60651548 | NA | LOC101927253 | 101927253 |
| ENSG000002 | 1.49943103 | -0.480277065 | 0.34471937 | -1.3932407 | 0.163547 | NA | NA | NA |
| ENSG000002 | 0 | NA | NA | NA | NA | NA | NA | NA |
| ENSG000002 | 1.33360574 | -0.11524404 | 0.33696956 | -0.3420013 | 0.73234988 | NA | NA | NA |
| ENSG000002 | 2.11730228 | -0.512374291 | 0.36543818 | -1.402082 | 0.16089077 | NA | NA | NA |
| ENSG000002 | 1.9913637 | -0.329464627 | 0.36486137 | -0.9029858 | 0.36653345 | NA | NA | NA |

| | | | | | | | | |
|---|---|---|---|---|---|---|---|---|
| ENSG000002 | 0.40191006 | -0.101653297 | 0.21376195 | -0.4755444 | 0.63439899 | NA | NA | NA |
| ENSG000002 | 1.54202792 | 0.149334685 | 0.34544098 | 0.43230159 | 0.66552223 | NA | NA | NA |
| ENSG000002 | 0.99900943 | -0.19112582 | 0.31388018 | -0.6089133 | 0.54258191 | NA | NA | NA |
| ENSG000002 | 1.32865635 | 0.03259023 | 0.33294158 | 0.09788573 | 0.92202303 | NA | NA | NA |
| ENSG000002 | 1.28431806 | -0.486187103 | 0.32764254 | -1.4838949 | 0.1378368 | NA | NA | NA |
| ENSG000002 | 1.73051242 | -0.187401968 | 0.35536173 | -0.5273555 | 0.59794673 | NA | NA | NA |
| ENSG000002 | 1.86521256 | -0.382320744 | 0.35709553 | -1.07064 | 0.28433135 | NA | NA | NA |
| ENSG000002 | 1.73122844 | -0.293953 | 0.3478309 | -0.8451032 | 0.39805323 | NA | NA | NA |
| ENSG000002 | 0 | NA | NA | NA | NA | NA | MSANTD3-TME | 100526694 |
| ENSG000002 | 1.92732596 | -0.541265725 | 0.36028105 | -1.502343 | 0.13300854 | NA | NA | NA |
| ENSG000002 | 0.63759958 | -0.112459832 | 0.26607281 | -0.4226656 | 0.67253924 | NA | NA | NA |
| ENSG000002 | 0 | NA | NA | NA | NA | NA | NA | NA |
| ENSG000002 | 1.34746226 | 0.295121875 | 0.32559864 | 0.90639776 | 0.36472536 | NA | NA | NA |
| ENSG000002 | 1.38504226 | -0.579005098 | 0.33833132 | -1.7113553 | 0.08701554 | NA | NA | NA |
| ENSG000002 | 1.12480962 | 0.033582263 | 0.32084407 | 0.10466848 | 0.91663887 | NA | NA | NA |
| ENSG000002 | 0.99487856 | -0.178928224 | 0.30155082 | -0.5933601 | 0.55294018 | NA | NA | NA |
| ENSG000002 | 0 | NA | NA | NA | NA | NA | DCANP1 | 140947 |
| ENSG000002 | 0.47781273 | -0.018045496 | 0.22812977 | -0.0791019 | 0.93695158 | NA | NA | NA |
| ENSG000002 | 0.59760982 | 0.066257965 | 0.26871197 | 0.24657616 | 0.80523626 | NA | NA | NA |
| ENSG000002 | 1.09313458 | 0.028487421 | 0.31980269 | 0.08907812 | 0.92901983 | NA | NA | NA |
| ENSG000002 | 1.96771411 | -0.31260139 | 0.36171036 | -0.8642312 | 0.38746087 | NA | NA | NA |
| ENSG000002 | 2.06635502 | -0.123501356 | 0.36696992 | -0.3365435 | 0.73646102 | NA | NA | NA |
| ENSG000002 | 0.98506768 | -0.166200817 | 0.30110626 | -0.5519673 | 0.58097074 | NA | NA | NA |
| ENSG000002 | 1.35572534 | -0.389302834 | 0.33186585 | -1.1730729 | 0.24076657 | NA | NA | NA |
| ENSG000002 | 0.87512032 | -0.447852863 | 0.29956856 | -1.4949929 | 0.1349163 | NA | NA | NA |
| ENSG000002 | 0.87898613 | -0.113387039 | 0.30333236 | -0.3738046 | 0.70854968 | NA | NA | NA |
| ENSG000002 | 0.97962848 | -0.029387251 | 0.305964 | -0.0960481 | 0.9234824 | NA | NA | NA |
| ENSG000002 | 0.88650003 | -0.284549442 | 0.30460367 | -0.9341629 | 0.35021988 | NA | NA | NA |
| ENSG000002 | 1.54174747 | -0.341551672 | 0.34567106 | -0.9880829 | 0.32311203 | NA | NA | NA |

| | | | | | | | | |
|---|---|---|---|---|---|---|---|---|
| ENSG000002 | 0.75070768 | -0.185851474 | 0.28031532 | -0.6630086 | 0.50732504 | NA | NA | NA |
| ENSG000002 | 0.74709385 | -0.19484779 | 0.28612562 | -0.6809869 | 0.4958798 | NA | NA | NA |
| ENSG000002 | 0.9882478 | 0.120464272 | 0.31280431 | 0.38511065 | 0.70015544 | NA | GTF2IP9 | 106481734 |
| ENSG000002 | 1.10964139 | 0.038843262 | 0.32434054 | 0.11976073 | 0.90467269 | NA | NA | NA |
| ENSG000002 | 1.82595767 | 0.138400887 | 0.35820322 | 0.38637533 | 0.69921872 | NA | NA | NA |
| ENSG000002 | 1.10277634 | -0.108641013 | 0.32042825 | -0.3390494 | 0.7345725 | NA | NA | NA |
| ENSG000002 | 2.00444672 | -0.170822669 | 0.3571006 | -0.4783601 | 0.63239395 | NA | NA | NA |
| ENSG000002 | 1.25362639 | -0.048535495 | 0.32479867 | -0.1494326 | 0.88121232 | NA | NA | NA |
| ENSG000002 | 1.93250403 | -0.68043289 | 0.35877025 | -1.89657 | 0.05788471 | NA | NA | NA |
| ENSG000002 | 1.255594 | -0.328431171 | 0.32505881 | -1.0103746 | 0.31231584 | NA | NA | NA |
| ENSG000002 | 1.25861442 | -0.347268399 | 0.33073335 | -1.0499951 | 0.29372036 | NA | NA | NA |
| ENSG000002 | 1.71639406 | -0.191591183 | 0.35674416 | -0.5370549 | 0.59122973 | NA | NA | NA |
| ENSG000002 | 0.40942396 | -0.284058841 | 0.21788566 | -1.303706 | 0.19233384 | NA | APITD1-CORT | 100526739 |
| ENSG000002 | 0.37125511 | 0.067313264 | 0.2170755 | 0.31009148 | 0.75649139 | NA | NA | NA |
| ENSG000002 | 0 | NA | NA | NA | NA | NA | NA | NA |
| ENSG000002 | 1.5817656 | 0.023402668 | 0.35202692 | 0.06647977 | 0.94699587 | NA | NA | NA |
| ENSG000002 | 0.77302316 | -0.486369413 | 0.27058578 | -1.7974685 | 0.07226128 | NA | LOC105374729 | 105374729 |
| ENSG000002 | 1.72775438 | -0.593892191 | 0.35207516 | -1.6868335 | 0.09163537 | NA | NA | NA |
| ENSG000002 | 0.61221498 | -0.115173106 | 0.27027142 | -0.4261387 | 0.6700068 | NA | NA | NA |
| ENSG000002 | 1.75882963 | -0.474273281 | 0.35682963 | -1.3291309 | 0.1838048 | NA | NA | NA |
| ENSG000002 | 1.04005435 | -0.513708114 | 0.3079622 | -1.6680882 | 0.09529821 | NA | NA | NA |
| ENSG000002 | 0.88374156 | -0.114651691 | 0.29851557 | -0.3840727 | 0.70092454 | NA | NA | NA |
| ENSG000002 | 1.96696226 | 0.059307945 | 0.35447087 | 0.16731402 | 0.86712298 | NA | NA | NA |
| ENSG000002 | 0 | NA | NA | NA | NA | NA | NA | NA |
| ENSG000002 | 0.49307133 | -0.179259067 | 0.24031383 | -0.7459374 | 0.45570524 | NA | NA | NA |
| ENSG000002 | 0 | NA | NA | NA | NA | NA | NA | NA |
| ENSG000002 | 1.93283254 | 0.307576859 | 0.35319244 | 0.8708478 | 0.38383726 | NA | NA | NA |
| ENSG000002 | 0 | NA | NA | NA | NA | NA | NA | NA |
| ENSG000002 | 1.62493783 | -0.259935194 | 0.35020605 | -0.742235 | 0.45794496 | NA | LOC100287592 | 100287592 |

| | | | | | | | | |
|---|---|---|---|---|---|---|---|---|
| ENSG000002 | 0.74501861 | -0.208803734 | 0.29111786 | -0.7172481 | 0.47322101 | NA | NA | NA |
| ENSG000002 | 1.41273236 | -0.416331642 | 0.33899902 | -1.2281205 | 0.21940174 | NA | RPL37P25 | 106481666 |
| ENSG000002 | 2.13525943 | -0.238464318 | 0.36089961 | -0.6607497 | 0.50877283 | NA | NA | NA |
| ENSG000002 | 1.78139497 | -0.565598064 | 0.34775043 | -1.6264482 | 0.10385435 | NA | NA | NA |
| ENSG000002 | 0 | NA | NA | NA | NA | NA | NA | NA |
| ENSG000002 | 0.90249529 | -0.431468428 | 0.2958793 | -1.4582583 | 0.14476937 | NA | NA | NA |
| ENSG000002 | 0.59618939 | 0.215036664 | 0.25454907 | 0.84477491 | 0.39823653 | NA | NA | NA |
| ENSG000002 | 1.75194634 | -0.458506471 | 0.35227185 | -1.30157 | 0.19306343 | NA | NA | NA |
| ENSG000002 | 1.83339776 | -0.130934567 | 0.36129811 | -0.3624004 | 0.71705287 | NA | NA | NA |
| ENSG000002 | 1.53189583 | -0.332481291 | 0.34325572 | -0.9686111 | 0.33273927 | NA | NA | NA |
| ENSG000002 | 2.05197619 | 0.37847517 | 0.36163067 | 1.04657929 | 0.29529366 | NA | NA | NA |
| ENSG000002 | 1.50654348 | -0.347922387 | 0.34695293 | -1.0027942 | 0.31596016 | NA | NA | NA |
| ENSG000002 | 1.93116655 | -0.655530941 | 0.3561283 | -1.8407157 | 0.06566324 | NA | PRELID3BP5 | 106480242 |
| ENSG000002 | 0 | NA | NA | NA | NA | NA | NA | NA |
| ENSG000002 | 0 | NA | NA | NA | NA | NA | NA | NA |
| ENSG000002 | 1.64779677 | -0.388678805 | 0.34805343 | -1.1167217 | 0.26411336 | NA | NA | NA |
| ENSG000002 | 0 | NA | NA | NA | NA | NA | USP17L9P | 391627 |
| ENSG000002 | 0.96894926 | 0.11678612 | 0.31050278 | 0.37611941 | 0.70682813 | NA | RN7SKP24 | 106480840 |
| ENSG000002 | 0.88526963 | 0.029214191 | 0.28333818 | 0.10310715 | 0.91787793 | NA | NA | NA |
| ENSG000002 | 1.2310865 | -0.046903466 | 0.33004793 | -0.1421111 | 0.88699227 | NA | NA | NA |
| ENSG000002 | 0.11205239 | 0.082118251 | 0.12671962 | 0.64803108 | 0.51696485 | NA | NA | NA |
| ENSG000002 | 0.85895037 | -0.111050564 | 0.29722404 | -0.3736258 | 0.70868275 | NA | NA | NA |
| ENSG000002 | 0.50565578 | -0.183478728 | 0.24174047 | -0.7589905 | 0.44785822 | NA | NA | NA |
| ENSG000002 | 0.8906785 | 0.039077521 | 0.29082116 | 0.13436959 | 0.89311033 | NA | RNU6-632P | 106479846 |
| ENSG000002 | 0.37194025 | 0.067575365 | 0.21717998 | 0.31114914 | 0.75568724 | NA | NA | NA |
| ENSG000002 | 1.59594409 | -0.260204762 | 0.34772192 | -0.7483128 | 0.45427151 | NA | RN7SKP67 | 106479125 |
| ENSG000002 | 0.13280363 | -0.093394593 | 0.13115644 | -0.7120855 | 0.47641185 | NA | NA | NA |
| ENSG000002 | 0.14128499 | -0.097772014 | 0.13403235 | -0.7294658 | 0.4657168 | NA | NA | NA |
| ENSG000002 | 0.23799501 | 0.170958857 | 0.17834799 | 0.95856902 | 0.33777591 | NA | RNU2-13P | 106480203 |

| | | | | | | | | |
|---|---|---|---|---|---|---|---|---|
| ENSG000002 | 0.48279897 | 0.145542372 | 0.23831717 | 0.6107087 | 0.54139244 | NA | RNU6-408P | 106479746 |
| ENSG000002 | 0.602844 | 0.2204762 | 0.25672804 | 0.85879283 | 0.39045483 | NA | RNU7-123P | 106479071 |
| ENSG000002 | 1.34037905 | 0.025732102 | 0.34018015 | 0.07564257 | 0.93970346 | NA | NA | NA |
| ENSG000002 | 0.70750265 | 0.13244101 | 0.27651219 | 0.47896988 | 0.63196006 | NA | NA | NA |
| ENSG000002 | 2.06906584 | -0.73386441 | 0.36143451 | -2.0304215 | 0.04231371 | NA | RNU6-488P | 106479782 |
| ENSG000002 | 1.74243347 | -0.052291499 | 0.35453119 | -0.1474948 | 0.8827415 | NA | NA | NA |
| ENSG000002 | 1.23985878 | -0.192714935 | 0.3278088 | -0.5878882 | 0.55660732 | NA | RN7SKP122 | 106479148 |
| ENSG000002 | 0.48436321 | -0.017262119 | 0.23773787 | -0.0726099 | 0.94211656 | NA | NA | NA |
| ENSG000002 | 0.36621735 | -0.097238239 | 0.21673053 | -0.4486596 | 0.65367721 | NA | RNU4ATAC13P | 106479543 |
| ENSG000002 | 0.6394784 | -0.110578951 | 0.25884399 | -0.4272031 | 0.66923141 | NA | RN7SKP13 | 106479100 |
| ENSG000002 | 0.79621786 | -0.342696846 | 0.27744022 | -1.2352097 | 0.21675247 | NA | NA | NA |
| ENSG000002 | 1.12886264 | -0.408853326 | 0.31684577 | -1.2903859 | 0.19691669 | NA | RNU7-124P | 106479072 |
| ENSG000002 | 1.23171765 | -0.194591902 | 0.33056302 | -0.5886681 | 0.55608395 | NA | NA | NA |
| ENSG000002 | 0.48009704 | -0.008352361 | 0.22842088 | -0.0365657 | 0.97083132 | NA | NA | NA |
| ENSG000002 | 1.85964164 | -0.264377138 | 0.3590626 | -0.7362982 | 0.46154926 | NA | RNU4ATAC11P | 106481158 |
| ENSG000002 | 0.84287024 | 0.04568409 | 0.30119185 | 0.15167771 | 0.87944114 | NA | NA | NA |
| ENSG000002 | 0.49128033 | 0.142942346 | 0.23932146 | 0.59728178 | 0.55031926 | NA | RN7SKP46 | 106480847 |
| ENSG000002 | 0.68546737 | 0.487906161 | 0.27704477 | 1.76110947 | 0.07821988 | NA | RNU4-29P | 106481181 |
| ENSG000002 | 0.76033119 | -0.194152683 | 0.28698065 | -0.6765358 | 0.49870053 | NA | RNU6-75P | 100873776 |
| ENSG000002 | 1.10475315 | 0.002822643 | 0.19797322 | 0.0142577 | 0.98862438 | NA | RNU6-999P | 106480008 |
| ENSG000002 | 1.38369721 | -0.117349721 | 0.33113115 | -0.3543905 | 0.72304628 | NA | NA | NA |
| ENSG000002 | 1.78502199 | -0.566712767 | 0.34721273 | -1.6321774 | 0.10264212 | NA | RNU6-848P | 106479941 |
| ENSG000002 | 0.75076379 | 0.127405659 | 0.27790738 | 0.45844647 | 0.64663171 | NA | RN7SKP298 | 106479228 |
| ENSG000002 | 0.25370795 | -0.171331747 | 0.17411721 | -0.9840024 | 0.32511435 | NA | NA | NA |
| ENSG000002 | 0.52028267 | -0.202047067 | 0.25039858 | -0.8069018 | 0.41972305 | NA | RNU6-138P | 106479627 |
| ENSG000002 | 0.23816916 | -0.010712105 | 0.1772998 | -0.060418 | 0.9518227 | NA | RNU6-470P | 106479773 |
| ENSG000002 | 1.13112929 | -0.122534329 | 0.32285603 | -0.3795324 | 0.70429254 | NA | NA | NA |
| ENSG000002 | 0 | NA | NA | NA | NA | NA | NA | NA |
| ENSG000002 | 1.1740522 | -0.607203744 | 0.32269269 | -1.881678 | 0.05987975 | NA | NA | NA |

| | | | | | | | | | |
|---|---|---|---|---|---|---|---|---|---|
| ENSG000002 | 0.98569829 | -0.345149663 | 0.30835238 | -1.1193352 | 0.26299718 | NA | | RNU6-1129P | 106481536 |
| ENSG000002 | 1.54408493 | 0.26949376 | 0.34107928 | 0.79012058 | 0.42945735 | NA | | NA | NA |
| ENSG000002 | 0.94461802 | 0.443161989 | 0.30679167 | 1.44450463 | 0.14859708 | NA | | RN7SKP242 | 106480908 |
| ENSG000002 | 0.23202362 | -0.008302493 | 0.17586207 | -0.0472103 | 0.96234565 | NA | | NA | NA |
| ENSG000002 | 0.67503122 | -0.469043005 | 0.26742254 | -1.7539397 | 0.07944084 | NA | | NA | NA |
| ENSG000002 | 0.61446491 | -0.254170746 | 0.25900567 | -0.9813328 | 0.32642867 | NA | | NA | NA |
| ENSG000002 | 0.41537361 | -0.286583533 | 0.21871501 | -1.3103057 | 0.19009243 | NA | | RNU6-1170P | 106480081 |
| ENSG000002 | 0.73980617 | -0.187156215 | 0.27954903 | -0.6694933 | 0.50318083 | NA | | NA | NA |
| ENSG000002 | 0 | NA | NA | NA | NA | NA | | NA | NA |
| ENSG000002 | 1.60908054 | 0.007628999 | 0.34808557 | 0.02191702 | 0.98251415 | NA | | RNU4-57P | 106479584 |
| ENSG000002 | 0.70305967 | 0.129084386 | 0.27453658 | 0.47019011 | 0.6382192 | NA | | NA | NA |
| ENSG000002 | 1.41861947 | 0.228075884 | 0.33797601 | 0.67482862 | 0.49978465 | NA | | RNU6-237P | 106479672 |
| ENSG000002 | 0 | NA | NA | NA | NA | NA | | NA | NA |
| ENSG000002 | 1.07788101 | -0.100547196 | 0.31484634 | -0.3193532 | 0.74945867 | NA | | RNU6-863P | 106481453 |
| ENSG000002 | 1.28581393 | 0.162392218 | 0.32616973 | 0.49787643 | 0.61857114 | NA | | NA | NA |
| ENSG000002 | 0.53471369 | -0.194084218 | 0.24487205 | -0.7925944 | 0.42801417 | NA | | NA | NA |
| ENSG000002 | 1.28161167 | -0.335264627 | 0.32738591 | -1.0240655 | 0.30580432 | NA | | NA | NA |
| ENSG000002 | 0.4869258 | -0.176879301 | 0.23959723 | -0.738236 | 0.46037105 | NA | | RNU6-1294P | 106480131 |
| ENSG000002 | 1.03763637 | 0.167683256 | 0.30502365 | 0.54973854 | 0.58249872 | NA | | NA | NA |
| ENSG000002 | 2.07136086 | 0.007007727 | 0.36013408 | 0.01945866 | 0.98447521 | NA | | RNU6-202P | 106479656 |
| ENSG000002 | 1.33955485 | -0.115577044 | 0.32758124 | -0.3528195 | 0.72422378 | NA | | RN7SKP132 | 106480874 |
| ENSG000002 | 0.40191006 | -0.101653297 | 0.21376195 | -0.4755444 | 0.63439899 | NA | | NA | NA |
| ENSG000002 | 1.54878475 | -0.47733489 | 0.34139541 | -1.3981878 | 0.16205669 | NA | | RNU6-983P | 106481490 |
| ENSG000002 | 1.85574604 | -0.39229374 | 0.35958773 | -1.0909542 | 0.27529304 | NA | | RN7SKP18 | 106480838 |
| ENSG000002 | 0 | NA | NA | NA | NA | NA | | SNORD116-26 | 100033438 |
| ENSG000002 | 0.53471369 | -0.194084218 | 0.24487205 | -0.7925944 | 0.42801417 | NA | | NA | NA |
| ENSG000002 | 0.49036493 | -0.020766486 | 0.24598337 | -0.0844223 | 0.93272066 | NA | | NA | NA |
| ENSG000002 | 1.02027057 | -0.195565578 | 0.30838824 | -0.6341538 | 0.52598044 | NA | | NA | NA |
| ENSG000002 | 0.40191006 | -0.101653297 | 0.21376195 | -0.4755444 | 0.63439899 | NA | | NA | NA |

| | | | | | | | | | |
|---|---|---|---|---|---|---|---|---|---|
| ENSG000002 | 1.98791852 | -0.315907168 | 0.36184154 | -0.8730539 | 0.3826337 | NA | | RN7SKP19 | 106479103 |
| ENSG000002 | 0.36553413 | 0.069401489 | 0.20780452 | 0.33397488 | 0.73839851 | NA | | NA | NA |
| ENSG000002 | 0.75341407 | -0.349034172 | 0.2811917 | -1.2412677 | 0.21450687 | NA | | NA | NA |
| ENSG000002 | 1.11513658 | -0.382209046 | 0.31128736 | -1.2278335 | 0.21950948 | NA | | NA | NA |
| ENSG000002 | 0.62496306 | -0.108686629 | 0.26501936 | -0.4101083 | 0.68172654 | NA | | RNU6-1114P | 106481531 |
| ENSG000002 | 0.11934008 | 0.083564698 | 0.12760091 | 0.65489106 | 0.5125379 | NA | | RNU6-319P | 106481280 |
| ENSG000002 | 0.46267888 | 0.146619746 | 0.23584798 | 0.62167056 | 0.53415852 | NA | | RNU6ATAC32P | 106479556 |
| ENSG000002 | 0.51959753 | -0.202283781 | 0.25032396 | -0.808088 | 0.41903995 | NA | | NA | NA |
| ENSG000002 | 0.52668879 | -0.197054386 | 0.24396589 | -0.8077129 | 0.4192559 | NA | | NA | NA |
| ENSG000002 | 1.37015636 | -0.415537848 | 0.33809401 | -1.2290601 | 0.21904929 | NA | | NA | NA |
| ENSG000002 | 1.75809029 | -0.611260928 | 0.35407271 | -1.7263712 | 0.08428065 | NA | | RN7SKP155 | 106479164 |
| ENSG000002 | 0 | NA | NA | NA | NA | NA | | RNU6-1334P | 106480148 |
| ENSG000002 | 1.83464989 | -0.224867272 | 0.35082797 | -0.6409616 | 0.52154761 | NA | | RNU6-732P | 106479890 |
| ENSG000002 | 0.64739724 | -0.288756313 | 0.26860627 | -1.075017 | 0.28236711 | NA | | RNU6-803P | 106481434 |
| ENSG000002 | 1.28198223 | -0.499509687 | 0.33054639 | -1.5111637 | 0.13074676 | NA | | NA | NA |
| ENSG000002 | 1.30375246 | -0.509384589 | 0.3310308 | -1.538783 | 0.12385727 | NA | | NA | NA |
| ENSG000002 | 0.37304803 | -0.090842893 | 0.20939682 | -0.4338313 | 0.66441096 | NA | | RNU6-772P | 106479908 |
| ENSG000002 | 0.51327731 | -0.027046708 | 0.2410203 | -0.1122176 | 0.91065091 | NA | | NA | NA |
| ENSG000002 | 0 | NA | NA | NA | NA | NA | | RNU6-507P | 106481341 |
| ENSG000002 | 0.53608206 | -0.386623811 | 0.24788931 | -1.5596631 | 0.11883952 | NA | | MIR449C | 100313923 |
| ENSG000002 | 1.47309781 | -0.056438252 | 0.34545403 | -0.1633741 | 0.87022387 | NA | | NA | NA |
| ENSG000002 | 0.26199343 | -0.194018184 | 0.18422801 | -1.0531416 | 0.29227609 | NA | | NA | NA |
| ENSG000002 | 0.76901758 | -0.527141184 | 0.27928891 | -1.8874404 | 0.05910111 | NA | | RNU6-1288P | 106480128 |
| ENSG000002 | 1.00060593 | -0.362869951 | 0.31500014 | -1.1519676 | 0.24933439 | NA | | NA | NA |
| ENSG000002 | 0.75523498 | -0.185546569 | 0.27894532 | -0.6651718 | 0.5059406 | NA | | MIR1343 | 100616437 |
| ENSG000002 | 0.33394554 | 0.210095954 | 0.19490774 | 1.07792513 | 0.28106717 | NA | | NA | NA |
| ENSG000002 | 1.25366311 | -0.352444185 | 0.33401964 | -1.0551601 | 0.29135212 | NA | | RNU6-518P | 106481344 |
| ENSG000002 | 1.38509837 | -0.273265904 | 0.34184847 | -0.7993773 | 0.42407168 | NA | | NA | NA |
| ENSG000002 | 1.57496664 | 0.012753414 | 0.34661237 | 0.03679446 | 0.97064889 | NA | | RNU2-69P | 106481656 |

| | | | | | | | | | |
|---|---|---|---|---|---|---|---|---|---|
| ENSG000002 | 2.15048714 | -0.25038313 | 0.3649016 | -0.6861662 | 0.49260833 | NA | | NA | NA |
| ENSG000002 | 1.1355707 | -0.26384213 | 0.31786779 | -0.8300373 | 0.40651769 | NA | | RNU6-615P | 106480595 |
| ENSG000002 | 0.37097279 | -0.10468565 | 0.21741418 | -0.4815033 | 0.63015882 | NA | | NA | NA |
| ENSG000002 | 0.46153673 | 0.141687071 | 0.23571996 | 0.60108219 | 0.54778525 | NA | | RNU2-45P | 106480217 |
| ENSG000002 | 0.47616417 | 0.157689149 | 0.24487722 | 0.64395189 | 0.51960663 | NA | | NA | NA |
| ENSG000002 | 1.38874254 | -0.401251321 | 0.33373002 | -1.2023231 | 0.22923836 | NA | | NA | NA |
| ENSG000002 | 1.85792308 | 0.003695152 | 0.35989544 | 0.01026729 | 0.99180803 | NA | | RNU6-475P | 106479776 |
| ENSG000002 | 0.11865494 | 0.083440866 | 0.12751478 | 0.65436232 | 0.5128784 | NA | | RNU6ATAC17P | 106481164 |
| ENSG000002 | 0.76380067 | -0.180699362 | 0.27231745 | -0.6635614 | 0.50697104 | NA | | NA | NA |
| ENSG000002 | 0.25351207 | -0.189965547 | 0.1824784 | -1.0410303 | 0.29786148 | NA | | RNU6-120P | 106479619 |
| ENSG000002 | 1.35537651 | -0.122555041 | 0.3423343 | -0.3579981 | 0.72034471 | NA | | RNU6-760P | 106481420 |
| ENSG000002 | 1.6087139 | -0.128555234 | 0.35361599 | -0.3635447 | 0.716198 | NA | | RN7SKP190 | 106480891 |
| ENSG000002 | 1.08114545 | 0.176555629 | 0.31500704 | 0.56048153 | 0.57515103 | NA | | RNU4-49P | 106481187 |
| ENSG000002 | 0.99106886 | -0.189588774 | 0.30880259 | -0.6139481 | 0.5392496 | NA | | NA | NA |
| ENSG000002 | 0.63159785 | -0.116736614 | 0.27184111 | -0.4294296 | 0.66761063 | NA | | NA | NA |
| ENSG000002 | 0.57683685 | 0.056087234 | 0.2526609 | 0.22198621 | 0.82432462 | NA | | NA | NA |
| ENSG000002 | 0 | NA | NA | NA | NA | NA | | SNORD116-27 | 100033439 |
| ENSG000002 | 0.6394784 | -0.110578951 | 0.25884399 | -0.4272031 | 0.66923141 | NA | | RNU6-916P | 106481468 |
| ENSG000002 | 0.13280363 | -0.093394593 | 0.13115644 | -0.7120855 | 0.47641185 | NA | | VTRNA2-2P | 106479538 |
| ENSG000002 | 0.98938783 | -0.030862894 | 0.30044856 | -0.1027227 | 0.91818304 | NA | | NA | NA |
| ENSG000002 | 0.75537934 | -0.187046077 | 0.28061071 | -0.6665679 | 0.50504821 | NA | | NA | NA |
| ENSG000002 | 0.12685397 | -0.090215768 | 0.12901919 | -0.699243 | 0.48440017 | NA | | RNU6-351P | 106479722 |
| ENSG000002 | 0.49819341 | -0.026649049 | 0.23931956 | -0.1113534 | 0.9113361 | NA | | RNU6-491P | 106479784 |
| ENSG000002 | 0 | NA | NA | NA | NA | NA | | RNU6-513P | 106481343 |
| ENSG000002 | 0.50479595 | -0.02564641 | 0.24753216 | -0.1036084 | 0.91748012 | NA | | RNU7-200P | 106480835 |
| ENSG000002 | 0 | NA | NA | NA | NA | NA | | NA | NA |
| ENSG000002 | 0.3854038 | -0.110128691 | 0.21952326 | -0.5016721 | 0.61589821 | NA | | RNU1-61P | 106481980 |
| ENSG000002 | 0 | NA | NA | NA | NA | NA | | RNU1-86P | 106480666 |
| ENSG000002 | 0.51570138 | -0.378721611 | 0.24581506 | -1.540677 | 0.12339542 | NA | | RNU7-105P | 106479065 |

| | | | | | | | | |
|---|---|---|---|---|---|---|---|---|
| ENSG000002 | 2.06448455 | -0.894219309 | 0.36151989 | -2.4734996 | 0.01337969 | NA | NA | NA |
| ENSG000002 | 0.26813897 | -0.197007041 | 0.18550319 | -1.0620143 | 0.2882292 | NA | NA | NA |
| ENSG000002 | 0.14128499 | -0.097772014 | 0.13403235 | -0.7294658 | 0.4657168 | NA | RNU6-276P | 106479689 |
| ENSG000002 | 0 | NA | NA | NA | NA | NA | NA | NA |
| ENSG000002 | 0 | NA | NA | NA | NA | NA | NA | NA |
| ENSG000002 | 0 | NA | NA | NA | NA | NA | RNU6-189P | 106479650 |
| ENSG000002 | 0.2596576 | -0.193002035 | 0.1837913 | -1.0501152 | 0.29366517 | NA | RNU6-871P | 106479950 |
| ENSG000002 | 0.87594342 | -0.270876514 | 0.29768468 | -0.9099444 | 0.36285183 | NA | RN7SKP179 | 106481815 |
| ENSG000002 | 0.34270765 | 0.258172734 | 0.21433185 | 1.20454674 | 0.22837833 | NA | NA | NA |
| ENSG000002 | 0.2596576 | -0.193002035 | 0.1837913 | -1.0501152 | 0.29366517 | NA | RNU7-129P | 106480450 |
| ENSG000002 | 0.49219223 | -0.014160136 | 0.23865874 | -0.0593321 | 0.95268755 | NA | RNU6-1278P | 106480124 |
| ENSG000002 | 0.49375647 | -0.17902118 | 0.24039617 | -0.7446923 | 0.45645774 | NA | NA | NA |
| ENSG000002 | 2.06101087 | -0.119390299 | 0.35779351 | -0.3336849 | 0.73861733 | NA | NA | NA |
| ENSG000002 | 0.85679081 | -0.111816085 | 0.29710779 | -0.3763485 | 0.7066578 | NA | RNU6-693P | 106481399 |
| ENSG000002 | 0.89606743 | -0.631542144 | 0.2981933 | -2.1178952 | 0.03418395 | NA | RNU6-1023P | 106480017 |
| ENSG000002 | 1.80976991 | 0.011822548 | 0.35530377 | 0.03327448 | 0.97345571 | NA | RN7SKP34 | 106479110 |
| ENSG000002 | 0.26560726 | -0.177019426 | 0.17672274 | -1.0016788 | 0.31649875 | NA | RNU6-1219P | 106480649 |
| ENSG000002 | 0 | NA | NA | NA | NA | NA | RNA5SP522 | 106479019 |
| ENSG000002 | 0.36621735 | -0.097238239 | 0.21673053 | -0.4486596 | 0.65367721 | NA | RNU6-496P | 106480584 |
| ENSG000002 | 0.8385697 | -0.098916793 | 0.29002443 | -0.3410637 | 0.73305566 | NA | RNU6-27P | 106478918 |
| ENSG000002 | 0.71689591 | -0.019796298 | 0.27694967 | -0.0714798 | 0.94301592 | NA | RNU6-617P | 106479839 |
| ENSG000002 | 0.64235948 | -0.482579697 | 0.27062934 | -1.7831759 | 0.07455768 | NA | RNU6-1095P | 106481525 |
| ENSG000002 | 0.65427998 | -0.290650429 | 0.26916107 | -1.0798383 | 0.28021419 | NA | RNU6-1102P | 106481527 |
| ENSG000002 | 0.50516651 | -0.183853222 | 0.24168957 | -0.7606999 | 0.44683635 | NA | RNU6-559P | 106481771 |
| ENSG000002 | 1.01119231 | -0.456160862 | 0.29536475 | -1.5443984 | 0.12249184 | NA | RNU6ATAC13P | 106479549 |
| ENSG000002 | 0.11865494 | 0.083440866 | 0.12751478 | 0.65436232 | 0.5128784 | NA | NA | NA |
| ENSG000002 | 1.71393616 | -0.048538376 | 0.35309638 | -0.137465 | 0.8906633 | NA | NA | NA |
| ENSG000002 | 0 | NA | NA | NA | NA | NA | NA | NA |
| ENSG000002 | 0.38747904 | -0.104908247 | 0.21987514 | -0.4771265 | 0.63327207 | NA | RNU6-1333P | 106481601 |

| | | | | | | | | | |
|---|---|---|---|---|---|---|---|---|---|
| ENSG000002 | 0.35824644 | 0.072568381 | 0.21508272 | 0.33739754 | 0.73581724 | NA | | RNU6-473P | 106479775 |
| ENSG000002 | 0.48761094 | -0.176642206 | 0.23968055 | -0.7369902 | 0.46112833 | NA | | NA | NA |
| ENSG000002 | 0.36556447 | -0.101943371 | 0.21659044 | -0.4706735 | 0.63787395 | NA | | RNU6-718P | 106481406 |
| ENSG000002 | 1.54787182 | -0.482811339 | 0.34462956 | -1.4009574 | 0.16122681 | NA | | RN7SKP141 | 106479157 |
| ENSG000002 | 1.64878547 | 0.345143193 | 0.34890479 | 0.98921885 | 0.32255608 | NA | | RNU6-991P | 106480004 |
| ENSG000002 | 0.48490454 | -0.018274975 | 0.24537428 | -0.074478 | 0.94063008 | NA | | NA | NA |
| ENSG000002 | 1.05259608 | 0.176836815 | 0.31141161 | 0.56785557 | 0.57013304 | NA | | RNU6-436P | 106481316 |
| ENSG000002 | 0.37236288 | -0.09108266 | 0.20928647 | -0.4352057 | 0.66341314 | NA | | NA | NA |
| ENSG000002 | 1.99036198 | -0.067689451 | 0.36298667 | -0.1864792 | 0.85206902 | NA | | RN7SKP157 | 106479165 |
| ENSG000002 | 0.62550439 | -0.114540246 | 0.27135801 | -0.4221001 | 0.67295194 | NA | | RNU6-1161P | 106480077 |
| ENSG000002 | 1.85989172 | -0.128624519 | 0.35764727 | -0.3596407 | 0.71911584 | NA | | NA | NA |
| ENSG000002 | 0.64602887 | -0.11525331 | 0.26675303 | -0.43206 | 0.66569782 | NA | | NA | NA |
| ENSG000002 | 0.11205239 | 0.082118251 | 0.12671962 | 0.64803108 | 0.51696485 | NA | | NA | NA |
| ENSG000002 | 0 | NA | NA | NA | NA | NA | | NA | NA |
| ENSG000002 | 1.3991915 | -0.682890278 | 0.32895271 | -2.0759528 | 0.03789832 | NA | | NA | NA |
| ENSG000002 | 1.29248038 | -0.351556288 | 0.33159098 | -1.0602107 | 0.28904878 | NA | | RNU2-27P | 106481642 |
| ENSG000002 | 0 | NA | NA | NA | NA | NA | | NA | NA |
| ENSG000002 | 0.4983681 | -0.186127524 | 0.2408746 | -0.7727154 | 0.4396908 | NA | | RNA5SP458 | 106479016 |
| ENSG000002 | 0.99510936 | -0.028734984 | 0.3007006 | -0.0955601 | 0.92386994 | NA | | NA | NA |
| ENSG000002 | 0.88377637 | -0.615319728 | 0.29536899 | -2.0832238 | 0.03723083 | NA | | NA | NA |
| ENSG000002 | 0.74268278 | -0.18879062 | 0.27975142 | -0.6748513 | 0.49977023 | NA | | NA | NA |
| ENSG000002 | 0.26560726 | -0.177019426 | 0.17672274 | -1.0016788 | 0.31649875 | NA | | RNU7-154P | 106481802 |
| ENSG000002 | 0.36007182 | -0.08619198 | 0.20729252 | -0.4157988 | 0.67755724 | NA | | RNU4ATAC8P | 106481157 |
| ENSG000002 | 0 | NA | NA | NA | NA | NA | | RNU6-927P | 106479976 |
| ENSG000002 | 0 | NA | NA | NA | NA | NA | | RNA5SP521 | 106479018 |
| ENSG000002 | 0.6265601 | -0.275936484 | 0.2670182 | -1.0333995 | 0.30141697 | NA | | RNU4ATAC14P | 106481159 |
| ENSG000002 | 0.28256998 | -0.18483211 | 0.18021686 | -1.0256094 | 0.30507573 | NA | | NA | NA |
| ENSG000002 | 0.49557738 | -0.021677348 | 0.2390401 | -0.090685 | 0.9277429 | NA | | RNU6-904P | 106479966 |
| ENSG000002 | 1.4062717 | -0.576073672 | 0.33808868 | -1.7039129 | 0.08839735 | NA | | NA | NA |

| | | | | | | | | | |
|---|---|---|---|---|---|---|---|---|---|
| ENSG000002 | 0.36391535 | 0.058449071 | 0.20749591 | 0.28168782 | 0.77818289 | NA | | RNU6-1194P | 106481557 |
| ENSG000002 | 0.34407602 | 0.071624546 | 0.21281613 | 0.33655599 | 0.73645163 | NA | | RNU2-30P | 106481643 |
| ENSG000002 | 0.66253513 | -0.036612812 | 0.15813333 | -0.2315313 | 0.81690208 | NA | | RNU6ATAC9P | 106479547 |
| ENSG000002 | 0.14128499 | -0.097772014 | 0.13403235 | -0.7294658 | 0.4657168 | NA | | NA | NA |
| ENSG000002 | 0.84397802 | -0.10543835 | 0.29479587 | -0.3576656 | 0.72059356 | NA | | NA | NA |
| ENSG000002 | 1.50059437 | -0.338024576 | 0.34689077 | -0.974441 | 0.32983764 | NA | | NA | NA |
| ENSG000002 | 1.94972576 | -0.835435172 | 0.35806072 | -2.3332221 | 0.01963649 | NA | | RNU6-581P | 106479823 |
| ENSG000002 | 0.33615718 | 0.211074506 | 0.19531107 | 1.08070938 | 0.27982641 | NA | | RNU6-982P | 106481979 |
| ENSG000002 | 0.25351207 | -0.189965547 | 0.1824784 | -1.0410303 | 0.29786148 | NA | | RNU6ATAC14P | 106481163 |
| ENSG000002 | 1.51900923 | -0.494935448 | 0.34616123 | -1.429783 | 0.1527793 | NA | | RN7SKP52 | 106479118 |
| ENSG000002 | 0 | NA | NA | NA | NA | NA | | RNA5SP253 | 106480760 |
| ENSG000002 | 0.46067745 | 0.331901404 | 0.23807331 | 1.39411428 | 0.1632831 | NA | | RNU6-1196P | 106480089 |
| ENSG000002 | 0.36621735 | -0.097238239 | 0.21673053 | -0.4486596 | 0.65367721 | NA | | RNU6-311P | 106479705 |
| ENSG000002 | 0.95088268 | -0.032588813 | 0.29685893 | -0.1097788 | 0.91258482 | NA | | RNU6-397P | 106479741 |
| ENSG000002 | 1.99068048 | -0.187848544 | 0.36141006 | -0.5197657 | 0.6032269 | NA | | RN7SKP288 | 106479223 |
| ENSG000002 | 0 | NA | NA | NA | NA | NA | | RNU6-162P | 106481226 |
| ENSG000002 | 0.3588776 | -0.099902505 | 0.21555794 | -0.4634601 | 0.6430346 | NA | | RNU6-287P | 106481269 |
| ENSG000002 | 0.98030723 | 0.114240896 | 0.31250032 | 0.36557049 | 0.7146856 | NA | | NA | NA |
| ENSG000002 | 1.93658151 | -0.291851802 | 0.35673299 | -0.818124 | 0.41328641 | NA | | NA | NA |
| ENSG000002 | 1.22140954 | -0.187849582 | 0.33029804 | -0.5687275 | 0.56954107 | NA | | NA | NA |
| ENSG000002 | 0.47029884 | 0.145033566 | 0.23680225 | 0.612467 | 0.54022882 | NA | | NA | NA |
| ENSG000002 | 0.8841642 | -0.254282913 | 0.29233359 | -0.8698382 | 0.38438884 | NA | | RNU6-150P | 106479632 |
| ENSG000002 | 0.13280363 | -0.093394593 | 0.13115644 | -0.7120855 | 0.47641185 | NA | | NA | NA |
| ENSG000002 | 0.74593051 | -0.356732965 | 0.28222667 | -1.2639945 | 0.206232 | NA | | NA | NA |
| ENSG000002 | 1.53918297 | -0.331835126 | 0.34125458 | -0.9723976 | 0.33085279 | NA | | RN7SKP276 | 106479218 |
| ENSG000002 | 2.00266612 | -0.566109286 | 0.35783843 | -1.5820248 | 0.1136439 | NA | | RNU6-1101P | 106480050 |
| ENSG000002 | 1.97712519 | -0.059644473 | 0.36161791 | -0.1649378 | 0.86899292 | NA | | NA | NA |
| ENSG000002 | 0.73807116 | -0.164114325 | 0.26803315 | -0.6122911 | 0.54034515 | NA | | RNU6-415P | 106481309 |
| ENSG000002 | 1.74278175 | -0.324355275 | 0.35365543 | -0.9171506 | 0.35906375 | NA | | RNU6-469P | 106481327 |

| | | | | | | | | |
|---|---|---|---|---|---|---|---|---|
| ENSG000002 | 0.14128499 | -0.097772014 | 0.13403235 | -0.7294658 | 0.4657168 | NA | RNU6-1041P | 106481509 |
| ENSG000002 | 1.4452703 | -0.301507468 | 0.3343969 | -0.9016455 | 0.3672452 | NA | NA | NA |
| ENSG000002 | 0.4923881 | -0.012789975 | 0.22994905 | -0.0556209 | 0.95564381 | NA | RNU6-864P | 106481776 |
| ENSG000002 | 0.956291 | -0.037707389 | 0.30658778 | -0.1229905 | 0.90211461 | NA | RNU2-53P | 106480220 |
| ENSG000002 | 0.54817724 | -0.365482324 | 0.24206593 | -1.5098462 | 0.13108267 | NA | RNU4-71P | 106481194 |
| ENSG000002 | 0.52125014 | -0.010203417 | 0.15567753 | -0.065542 | 0.94774247 | NA | RNU6-390P | 106479738 |
| ENSG000002 | 0.99240689 | -0.200699655 | 0.31762938 | -0.6318674 | 0.52747352 | NA | NA | NA |
| ENSG000002 | 1.08711493 | 0.167046079 | 0.30790354 | 0.5425273 | 0.5874553 | NA | NA | NA |
| ENSG000002 | 0.76015651 | -0.032508449 | 0.27992106 | -0.1161343 | 0.90754607 | NA | RNU6-947P | 106479985 |
| ENSG000002 | 0.25370795 | -0.171331747 | 0.17411721 | -0.9840024 | 0.32511435 | NA | RNU6-416P | 106480579 |
| ENSG000002 | 1.99064822 | -0.17995232 | 0.35927365 | -0.5008781 | 0.61645688 | NA | RN7SKP196 | 106479182 |
| ENSG000002 | 0 | NA | NA | NA | NA | NA | RNU6-327P | 106481887 |
| ENSG000002 | 0.14128499 | -0.097772014 | 0.13403235 | -0.7294658 | 0.4657168 | NA | RNU7-99P | 106480814 |
| ENSG000002 | 0.62889593 | -0.276614073 | 0.26719437 | -1.0352541 | 0.30055024 | NA | RNU6-277P | 106481266 |
| ENSG000002 | 0.37236288 | -0.09108266 | 0.20928647 | -0.4352057 | 0.66341314 | NA | RNU6-547P | 106481902 |
| ENSG000002 | 0.36345889 | 0.070257596 | 0.21589421 | 0.32542602 | 0.74485866 | NA | NA | NA |
| ENSG000002 | 1.82946991 | -0.229221458 | 0.35276006 | -0.6497943 | 0.51582513 | NA | RN7SKP12 | 106480836 |
| ENSG000002 | 0.99542786 | -0.173481066 | 0.30157371 | -0.5752526 | 0.56512046 | NA | NA | NA |
| ENSG000002 | 0.84434466 | 0.043418337 | 0.29446077 | 0.14745033 | 0.88277658 | NA | RNU4ATAC7P | 106480539 |
| ENSG000002 | 0.25214371 | -0.009862892 | 0.18046657 | -0.0546522 | 0.95641556 | NA | RNU6-1146P | 106480070 |
| ENSG000002 | 0 | NA | NA | NA | NA | NA | NA | NA |
| ENSG000002 | 0.60644002 | -0.257083821 | 0.25820471 | -0.9956589 | 0.31941589 | NA | NA | NA |
| ENSG000002 | 0.22410479 | 0.147643983 | 0.16638082 | 0.88738586 | 0.37487119 | NA | RNU6-346P | 106481768 |
| ENSG000002 | 1.50924541 | -0.200862267 | 0.34343908 | -0.5848556 | 0.55864482 | NA | NA | NA |
| ENSG000002 | 0.26813897 | -0.197007041 | 0.18550319 | -1.0620143 | 0.2882292 | NA | RNU6-917P | 106479971 |
| ENSG000002 | 0.49879422 | -0.023609326 | 0.24688725 | -0.095628 | 0.92381605 | NA | RNU6-191P | 106479651 |
| ENSG000002 | 0.23276084 | -0.008011382 | 0.17603982 | -0.0455089 | 0.96370166 | NA | RNU1-143P | 106481634 |
| ENSG000002 | 0 | NA | NA | NA | NA | NA | NA | NA |
| ENSG000002 | 0.24485602 | -0.012655878 | 0.17881722 | -0.0707755 | 0.94357643 | NA | NA | NA |

| | | | | | | | | | |
|---|---|---|---|---|---|---|---|---|---|
| ENSG000002 | 0.61907654 | -0.261106204 | 0.25936927 | -1.0066967 | 0.31408052 | NA | | RNU6-1232P | 106480105 |
| ENSG000002 | 1.76062054 | -0.033518918 | 0.18226913 | -0.1838979 | 0.85409352 | NA | | NA | NA |
| ENSG000002 | 2.09610722 | -0.384021234 | 0.36625907 | -1.0484962 | 0.29441007 | NA | | RNU6-523P | 106479798 |
| ENSG000002 | 0.24550891 | -0.007886114 | 0.17900103 | -0.0440563 | 0.96485956 | NA | | RNU4ATAC4P | 106479539 |
| ENSG000002 | 0.11205239 | 0.082118251 | 0.12671962 | 0.64803108 | 0.51696485 | NA | | NA | NA |
| ENSG000002 | 1.51721632 | -0.351388519 | 0.34711224 | -1.0123196 | 0.31138528 | NA | | RNU6ATAC39P | 106481172 |
| ENSG000002 | 0.11934008 | 0.083564698 | 0.12760091 | 0.65489106 | 0.5125379 | NA | | NA | NA |
| ENSG000002 | 1.23433423 | -0.179926905 | 0.32489222 | -0.5538049 | 0.57971238 | NA | | RNU6-970P | 106481486 |
| ENSG000002 | 0.11131518 | 0.081951103 | 0.12663804 | 0.64712866 | 0.51754868 | NA | | RNU6-1299P | 106480133 |
| ENSG000002 | 1.39537265 | -0.391776062 | 0.33360877 | -1.1743578 | 0.24025174 | NA | | RNU6-313P | 106480569 |
| ENSG000002 | 1.01503694 | -0.366898172 | 0.31553574 | -1.1627785 | 0.24491939 | NA | | NA | NA |
| ENSG000002 | 0.88126585 | -0.455824927 | 0.30137169 | -1.5125008 | 0.1304065 | NA | | RNU6-795P | 106480610 |
| ENSG000002 | 0 | NA | NA | NA | NA | NA | | NA | NA |
| ENSG000002 | 0.52147635 | -0.19458195 | 0.24341208 | -0.7993932 | 0.42406247 | NA | | NA | NA |
| ENSG000002 | 0.74410726 | -0.031277165 | 0.29043452 | -0.1076909 | 0.91424086 | NA | | RNU6-1338P | 106480150 |
| ENSG000002 | 0.5008998 | -0.200150804 | 0.24832882 | -0.805991 | 0.42024801 | NA | | NA | NA |
| ENSG000002 | 0 | NA | NA | NA | NA | NA | | NA | NA |
| ENSG000002 | 0.51550551 | -0.37860525 | 0.24578489 | -1.5403927 | 0.12346466 | NA | | RNU6-65P | 106480751 |
| ENSG000002 | 1.02434141 | -0.203341632 | 0.31483099 | -0.6458755 | 0.51835997 | NA | | NA | NA |
| ENSG000002 | 2.03114439 | -0.434810442 | 0.35694864 | -1.2181317 | 0.22317394 | NA | | RNU6-1225P | 106480650 |
| ENSG000002 | 0.35690842 | 0.061733055 | 0.20636749 | 0.29914137 | 0.76483218 | NA | | RNY3P16 | 106480686 |
| ENSG000002 | 1.14606939 | -0.120921155 | 0.3218241 | -0.3757368 | 0.70711258 | NA | | RNU6-1206P | 106481561 |
| ENSG000002 | 0 | NA | NA | NA | NA | NA | | RNA5SP315 | 106479005 |
| ENSG000002 | 0.11131518 | 0.081951103 | 0.12663804 | 0.64712866 | 0.51754868 | NA | | RNU6-1265P | 106481579 |
| ENSG000002 | 1.07719586 | -0.105604382 | 0.31956186 | -0.3304662 | 0.74104776 | NA | | MIR2278 | 100313780 |
| ENSG000002 | 0 | NA | NA | NA | NA | NA | | NA | NA |
| ENSG000002 | 1.00359017 | -0.045803433 | 0.31755408 | -0.1442382 | 0.88531237 | NA | | RNU6-155P | 106481224 |
| ENSG000002 | 2.12259365 | 0.425147895 | 0.36457963 | 1.16613177 | 0.24356117 | NA | | RNU6-479P | 106481331 |
| ENSG000002 | 0.740227 | 0.037763023 | 0.1591977 | 0.23720835 | 0.81249516 | NA | | NA | NA |

| | | | | | | | | |
|---|---|---|---|---|---|---|---|---|
| ENSG000002 | 1.98319746 | -0.178586406 | 0.35976905 | -0.4963918 | 0.61961801 | NA | RNU6-830P | 106480613 |
| ENSG000002 | 1.29319586 | -0.51524323 | 0.33145573 | -1.5544858 | 0.12006857 | NA | RN7SKP31 | 106480842 |
| ENSG000002 | 1.71588149 | -0.446494054 | 0.35116842 | -1.2714528 | 0.20356762 | NA | NA | NA |
| ENSG000002 | 0 | NA | NA | NA | NA | NA | RNU1-95P | 106481621 |
| ENSG000002 | 2.02773806 | -0.591430635 | 0.36222589 | -1.6327674 | 0.10251792 | NA | NA | NA |
| ENSG000002 | 0.87710676 | -0.118147501 | 0.30323109 | -0.3896286 | 0.69681121 | NA | NA | NA |
| ENSG000002 | 1.03419114 | -0.368499765 | 0.3147961 | -1.1705983 | 0.2417603 | NA | NA | NA |
| ENSG000002 | 0.98557813 | -0.03123057 | 0.30622151 | -0.1019869 | 0.91876711 | NA | NA | NA |
| ENSG000002 | 1.11536024 | -0.122613735 | 0.32235003 | -0.3803745 | 0.70366743 | NA | RNU6-720P | 106479885 |
| ENSG000002 | 0 | NA | NA | NA | NA | NA | RNU6-109P | 106481211 |
| ENSG000002 | 0.36212533 | -0.235926492 | 0.20089643 | -1.1743688 | 0.24024735 | NA | NA | NA |
| ENSG000002 | 0.98573592 | 0.043452226 | 0.19874519 | 0.21863284 | 0.82693607 | NA | NA | NA |
| ENSG000002 | 0 | NA | NA | NA | NA | NA | NA | NA |
| ENSG000002 | 1.7035451 | 0.084153852 | 0.35287333 | 0.23848176 | 0.81150746 | NA | RNU6-948P | 106481479 |
| ENSG000002 | 1.91653715 | -0.044768248 | 0.35613994 | -0.1257041 | 0.89996616 | NA | RNU2-10P | 106480201 |
| ENSG000002 | 0.40942396 | -0.284058841 | 0.21788566 | -1.303706 | 0.19233384 | NA | RNU6-752P | 106481417 |
| ENSG000002 | 2.10243419 | -0.004942436 | 0.35942776 | -0.0137508 | 0.98902876 | NA | RNU6-781P | 106479911 |
| ENSG000002 | 0.84827856 | 0.038186956 | 0.28846579 | 0.1323795 | 0.89468413 | NA | NA | NA |
| ENSG000002 | 1.13337922 | -0.250178375 | 0.31614289 | -0.7913459 | 0.42874219 | NA | NA | NA |
| ENSG000002 | 0.7523063 | -0.206231678 | 0.2915808 | -0.7072883 | 0.47938735 | NA | RNU6-751P | 106479898 |
| ENSG000002 | 0.13280363 | -0.093394593 | 0.13115644 | -0.7120855 | 0.47641185 | NA | NA | NA |
| ENSG000002 | 1.09823134 | 0.039512292 | 0.31269643 | 0.12635991 | 0.89944703 | NA | NA | NA |
| ENSG000002 | 0.48555743 | -0.013775713 | 0.2454629 | -0.0561214 | 0.95524512 | NA | NA | NA |
| ENSG000002 | 1.07011061 | -0.505926479 | 0.30679689 | -1.64906 | 0.09913534 | NA | RNU6-764P | 106479904 |
| ENSG000002 | 1.21961854 | 0.104947502 | 0.33320976 | 0.31495927 | 0.75279258 | NA | NA | NA |
| ENSG000002 | 0.85337128 | -0.249072601 | 0.29060646 | -0.8570787 | 0.3914014 | NA | NA | NA |
| ENSG000002 | 0 | NA | NA | NA | NA | NA | NA | NA |
| ENSG000002 | 0.9155362 | -0.460427183 | 0.30148657 | -1.5271897 | 0.12671384 | NA | RNA5SP473 | 106479017 |
| ENSG000002 | 1.67016569 | -0.521258123 | 0.34440535 | -1.5135018 | 0.13015224 | NA | NA | NA |

| | | | | | | | | | |
|---|---|---|---|---|---|---|---|---|---|
| ENSG000002 | 0.87875745 | -0.113785529 | 0.29825526 | -0.3815038 | 0.70282942 | NA | | NA | NA |
| ENSG000002 | 2.17286968 | -0.906608623 | 0.36109789 | -2.5107004 | 0.01204919 | NA | | MIR892C | 102466721 |
| ENSG000002 | 0.49356059 | -0.178885509 | 0.24036594 | -0.7442215 | 0.45674246 | NA | | RNU6-977P | 106480626 |
| ENSG000002 | 1.83990836 | -0.118259612 | 0.3518488 | -0.3361092 | 0.73678854 | NA | | NA | NA |
| ENSG000002 | 0.36502313 | -0.093755175 | 0.2080862 | -0.4505593 | 0.6523072 | NA | | RNU6-1049P | 106481779 |
| ENSG000002 | 0.76559516 | -0.206287743 | 0.28724125 | -0.7181689 | 0.47265312 | NA | | RNU4ATAC15P | 106479544 |
| ENSG000002 | 0 | NA | NA | NA | NA | NA | | NA | NA |
| ENSG000002 | 1.44384377 | -0.561509047 | 0.33562378 | -1.6730312 | 0.09432116 | NA | | NA | NA |
| ENSG000002 | 0.87608778 | -0.255953804 | 0.29186148 | -0.8769701 | 0.38050286 | NA | | NA | NA |
| ENSG000002 | 1.67797297 | -0.713643355 | 0.35011672 | -2.0383013 | 0.04151981 | NA | | NA | NA |
| ENSG000002 | 1.3482109 | -0.256967349 | 0.33291953 | -0.7718602 | 0.4401972 | NA | | NA | NA |
| ENSG000002 | 1.34531364 | -0.385790393 | 0.33279513 | -1.1592429 | 0.24635718 | NA | | RNU4-54P | 106479583 |
| ENSG000002 | 0.96931982 | -0.033717682 | 0.3120798 | -0.1080419 | 0.91396249 | NA | | NA | NA |
| ENSG000002 | 0 | NA | NA | NA | NA | NA | | RNU7-115P | 106479069 |
| ENSG000002 | 0.12070844 | -0.088503695 | 0.12783931 | -0.6923042 | 0.48874631 | NA | | RNU6-412P | 106481308 |
| ENSG000002 | 0.94717848 | 0.118876188 | 0.30957398 | 0.38399929 | 0.70097897 | NA | | NA | NA |
| ENSG000002 | 0.52193336 | -0.189821458 | 0.24349877 | -0.7795582 | 0.43565098 | NA | | RNU6-70P | 106478988 |
| ENSG000002 | 0.12685397 | -0.090215768 | 0.12901919 | -0.699243 | 0.48440017 | NA | | NA | NA |
| ENSG000002 | 1.9228541 | -0.76946389 | 0.35095114 | -2.19251 | 0.0283427 | NA | | RNU6-687P | 106479869 |
| ENSG000002 | 0.7731096 | -0.379060994 | 0.28942548 | -1.3097015 | 0.19029682 | NA | | NA | NA |
| ENSG000002 | 0.3513637 | 0.074494292 | 0.21400438 | 0.34809704 | 0.72776731 | NA | | NA | NA |
| ENSG000002 | 1.29320032 | -0.839186213 | 0.32625791 | -2.5721559 | 0.01010674 | NA | | NA | NA |
| ENSG000002 | 0.46748638 | 0.139734475 | 0.23646399 | 0.59093341 | 0.55456504 | NA | | NA | NA |
| ENSG000002 | 0.37441639 | -0.268856313 | 0.21277471 | -1.2635727 | 0.20638344 | NA | | NA | NA |
| ENSG000002 | 0.99101679 | -0.197748834 | 0.31353795 | -0.6307014 | 0.52823577 | NA | | RNU6-659P | 106481390 |
| ENSG000002 | 0.25993993 | -0.013144386 | 0.18211683 | -0.0721756 | 0.94246219 | NA | | RNU7-153P | 106479082 |
| ENSG000002 | 0.46153673 | 0.141687071 | 0.23571996 | 0.60108219 | 0.54778525 | NA | | NA | NA |
| ENSG000002 | 1.23220342 | -0.451222897 | 0.31895731 | -1.4146811 | 0.157162 | NA | | NA | NA |
| ENSG000002 | 0.11205239 | 0.082118251 | 0.12671962 | 0.64803108 | 0.51696485 | NA | | NA | NA |

| | | | | | | | | | |
|---|---|---|---|---|---|---|---|---|---|
| ENSG000002 | 0.72509495 | -0.178274683 | 0.2785491 | -0.6400117 | 0.52216499 | NA | | RNU4ATAC12P | 106481865 |
| ENSG000002 | 2.0549174 | -0.114295953 | 0.36000523 | -0.3174841 | 0.75087627 | NA | | RNU6-1110P | 106480638 |
| ENSG000002 | 0 | NA | NA | NA | NA | NA | | NA | NA |
| ENSG000002 | 0.12070844 | -0.088503695 | 0.12783931 | -0.6923042 | 0.48874631 | NA | | RNU6-426P | 106479754 |
| ENSG000002 | 0.26062507 | -0.012888319 | 0.18226503 | -0.070712 | 0.94362699 | NA | | RNU7-192P | 106480833 |
| ENSG000002 | 0.37831254 | -0.102004459 | 0.21855316 | -0.4667261 | 0.64069587 | NA | | RNU6-406P | 106481306 |
| ENSG000002 | 1.33397451 | 0.167459308 | 0.33547672 | 0.49916819 | 0.6176609 | NA | | NA | NA |
| ENSG000002 | 1.35145863 | -0.409428765 | 0.3363104 | -1.2174133 | 0.223447 | NA | | NA | NA |
| ENSG000002 | 0.12070844 | -0.088503695 | 0.12783931 | -0.6923042 | 0.48874631 | NA | | NA | NA |
| ENSG000002 | 1.25836298 | -0.629674495 | 0.32267023 | -1.951449 | 0.05100366 | NA | | RNU6-966P | 106480625 |
| ENSG000002 | 0 | NA | NA | NA | NA | NA | | NA | NA |
| ENSG000002 | 0.99428571 | -0.190008893 | 0.30893655 | -0.6150418 | 0.53852712 | NA | | NA | NA |
| ENSG000002 | 1.48821686 | -0.463200821 | 0.34159574 | -1.3559912 | 0.17510196 | NA | | RN7SKP238 | 106479201 |
| ENSG000002 | 0.37441639 | -0.268856313 | 0.21277471 | -1.2635727 | 0.20638344 | NA | | RNU6-589P | 106481367 |
| ENSG000002 | 0.76940933 | -0.516275266 | 0.27702168 | -1.8636638 | 0.06236894 | NA | | NA | NA |
| ENSG000002 | 2.15225832 | -0.520279318 | 0.3668723 | -1.4181483 | 0.15614748 | NA | | RNU6-875P | 106479953 |
| ENSG000002 | 1.6754302 | -0.535940125 | 0.34692304 | -1.5448387 | 0.12238529 | NA | | NA | NA |
| ENSG000002 | 0.74393257 | 0.128890331 | 0.285235 | 0.45187418 | 0.65135962 | NA | | NA | NA |
| ENSG000002 | 0.97658577 | -0.193587304 | 0.31296964 | -0.6185498 | 0.53621298 | NA | | NA | NA |
| ENSG000002 | 1.52835989 | 0.596599347 | 0.34461279 | 1.73121649 | 0.08341316 | NA | | RNU1-103P | 106481623 |
| ENSG000002 | 0.11934008 | 0.083564698 | 0.12760091 | 0.65489106 | 0.5125379 | NA | | NA | NA |
| ENSG000002 | 0 | NA | NA | NA | NA | NA | | RNA5SP520 | 106480771 |
| ENSG000002 | 1.96372972 | -0.186072224 | 0.36123463 | -0.5151007 | 0.60648263 | NA | | RNY4 | 6086 |
| ENSG000002 | 0.73516888 | -0.031630387 | 0.28453243 | -0.1111662 | 0.91148456 | NA | | NA | NA |
| ENSG000002 | 0.59604504 | 0.222764966 | 0.2632626 | 0.84617018 | 0.39745781 | NA | | NA | NA |
| ENSG000002 | 1.16887359 | -0.440583365 | 0.32232684 | -1.3668839 | 0.1716617 | NA | | RNU6-320P | 106480570 |
| ENSG000002 | 1.75934008 | -0.339420533 | 0.35813714 | -0.947739 | 0.34326235 | NA | | RN7SKP83 | 106479131 |
| ENSG000002 | 1.35509419 | -0.261618156 | 0.33801922 | -0.7739742 | 0.43894605 | NA | | RN7SKP244 | 106479203 |
| ENSG000002 | 0 | NA | NA | NA | NA | NA | | RNU6-184P | 106479648 |

| | | | | | | | | | |
|---|---|---|---|---|---|---|---|---|---|
| ENSG000002 | 0.11205239 | 0.082118251 | 0.12671962 | 0.64803108 | 0.51696485 | NA | | RNU6-1038P | 106480022 |
| ENSG000002 | 0 | NA | NA | NA | NA | NA | | NA | NA |
| ENSG000002 | 0.61039015 | 0.058968166 | 0.26339887 | 0.22387403 | 0.82285533 | NA | | NA | NA |
| ENSG000002 | 0.22410479 | 0.147643983 | 0.16638082 | 0.88738586 | 0.37487119 | NA | | RNU6-911P | 106479969 |
| ENSG000002 | 0.39499294 | -0.277908454 | 0.21584201 | -1.287555 | 0.19790091 | NA | | RNU6-964P | 106479992 |
| ENSG000002 | 2.0727551 | -0.0117576 | 0.35903351 | -0.0327479 | 0.97387561 | NA | | RNU6-1337P | 106481602 |
| ENSG000002 | 1.0053619 | -0.324036727 | 0.30386972 | -1.0663673 | 0.28625765 | NA | | RNU6-62P | 106478987 |
| ENSG000002 | 1.01412504 | -0.201798225 | 0.3129295 | -0.644868 | 0.51901272 | NA | | NA | NA |
| ENSG000002 | 1.07122 | 0.029177651 | 0.31428088 | 0.09283941 | 0.92603114 | NA | | RNU6-503P | 106479789 |
| ENSG000002 | 0.25145857 | -0.010118565 | 0.18031426 | -0.0561163 | 0.95524918 | NA | | RNU6-1061P | 106480033 |
| ENSG000002 | 0.51139849 | -0.021802213 | 0.24083338 | -0.0905282 | 0.92786748 | NA | | NA | NA |
| ENSG000002 | 0.52165104 | -0.403921832 | 0.25252134 | -1.5995552 | 0.10969729 | NA | | RNU6-1046P | 106480026 |
| ENSG000002 | 1.60399072 | -0.123438894 | 0.35127294 | -0.3514045 | 0.7252849 | NA | | RNU6-1256P | 106480114 |
| ENSG000002 | 1.94656448 | -0.540019716 | 0.35897196 | -1.5043507 | 0.13249109 | NA | | NA | NA |
| ENSG000002 | 1.96369589 | -0.312252291 | 0.36292143 | -0.8603854 | 0.38957664 | NA | | RPPH1-3P | 106479602 |
| ENSG000002 | 1.99361776 | -0.440306933 | 0.35846731 | -1.2283043 | 0.21933277 | NA | | RNU6ATAC12P | 106479548 |
| ENSG000002 | 1.08937537 | -0.108161702 | 0.31526431 | -0.3430826 | 0.73153631 | NA | | NA | NA |
| ENSG000002 | 0.71561133 | 0.119508664 | 0.27543591 | 0.43388919 | 0.66436891 | NA | | NA | NA |
| ENSG000002 | 0.50548109 | -0.025402968 | 0.24760631 | -0.1025942 | 0.91828506 | NA | | NA | NA |
| ENSG000002 | 1.58873212 | 0.009093771 | 0.34344913 | 0.02647778 | 0.97887626 | NA | | RNU6-506P | 106479791 |
| ENSG000002 | 0.11131518 | 0.081951103 | 0.12663804 | 0.64712866 | 0.51754868 | NA | | NA | NA |
| ENSG000002 | 1.12860205 | -0.412736221 | 0.31927353 | -1.2927355 | 0.19610255 | NA | | NA | NA |
| ENSG000002 | 0.88533669 | -0.426289624 | 0.29500043 | -1.4450475 | 0.14844456 | NA | | NA | NA |
| ENSG000002 | 1.54991588 | 0.14916522 | 0.34728367 | 0.42951982 | 0.66754497 | NA | | NA | NA |
| ENSG000002 | 0.25370795 | -0.171331747 | 0.17411721 | -0.9840024 | 0.32511435 | NA | | NA | NA |
| ENSG000002 | 0.48573157 | -0.181932918 | 0.23941992 | -0.7598905 | 0.44732006 | NA | | NA | NA |
| ENSG000002 | 2.15118727 | -0.606392733 | 0.36025279 | -1.6832423 | 0.0923282 | NA | | NA | NA |
| ENSG000002 | 0.55070895 | -0.361402396 | 0.24091882 | -1.5001003 | 0.13358842 | NA | | RNU6-725P | 106481408 |
| ENSG000002 | 0.97035145 | 0.102046536 | 0.29790912 | 0.34254251 | 0.73194266 | NA | | RNU6-358P | 106481291 |

| | | | | | | | | |
|---|---|---|---|---|---|---|---|---|
| ENSG000002 | 0.63296622 | -0.299126688 | 0.27328461 | -1.0945611 | 0.27370899 | NA | NA | NA |
| ENSG000002 | 0.76542102 | -0.036241538 | 0.28646241 | -0.1265141 | 0.89932498 | NA | RNU6-504P | 106481340 |
| ENSG000002 | 1.11732943 | -0.270532996 | 0.32170779 | -0.8409277 | 0.40038843 | NA | RNU6-314P | 106481278 |
| ENSG000002 | 0.11934008 | 0.083564698 | 0.12760091 | 0.65489106 | 0.5125379 | NA | RNU6-68P | 100873772 |
| ENSG000002 | 1.01449561 | -0.348241391 | 0.30957571 | -1.1248989 | 0.26063186 | NA | RNU6-1018P | 106481502 |
| ENSG000002 | 0.11131518 | 0.081951103 | 0.12663804 | 0.64712866 | 0.51754868 | NA | NA | NA |
| ENSG000002 | 1.21250947 | -0.319648435 | 0.3229796 | -0.9896862 | 0.32232754 | NA | RNU6-1004P | 106480009 |
| ENSG000002 | 0.6181343 | 0.056012223 | 0.26406445 | 0.21211573 | 0.83201675 | NA | RNU6-1007P | 106480629 |
| ENSG000002 | 0.74481413 | 0.129398785 | 0.27749152 | 0.46631618 | 0.6409892 | NA | NA | NA |
| ENSG000002 | 1.66009835 | -0.571388597 | 0.35274234 | -1.6198469 | 0.10526516 | NA | NA | NA |
| ENSG000002 | 0.13280363 | -0.093394593 | 0.13115644 | -0.7120855 | 0.47641185 | NA | RNU6-1291P | 106481588 |
| ENSG000002 | 1.26609461 | -0.046859047 | 0.32797502 | -0.1428738 | 0.88638983 | NA | RNU4ATAC6P | 106479540 |
| ENSG000002 | 1.68365527 | 0.212910182 | 0.35011006 | 0.60812358 | 0.54310551 | NA | NA | NA |
| ENSG000002 | 0.65493287 | -0.26892161 | 0.26251527 | -1.0244037 | 0.30564465 | NA | RNU7-183P | 106480454 |
| ENSG000002 | 0 | NA | NA | NA | NA | NA | NA | NA |
| ENSG000002 | 1.4847738 | -0.292542518 | 0.33492443 | -0.8734583 | 0.38241333 | NA | NA | NA |
| ENSG000002 | 1.72182199 | -0.188191978 | 0.3548309 | -0.5303709 | 0.59585481 | NA | RNU6-1087P | 106481522 |
| ENSG000002 | 0.50684946 | -0.202346264 | 0.24895102 | -0.8127955 | 0.41633533 | NA | NA | NA |
| ENSG000002 | 0.12685397 | -0.090215768 | 0.12901919 | -0.699243 | 0.48440017 | NA | RNU7-121P | 106480449 |
| ENSG000002 | 1.23589612 | 0.094974253 | 0.32864698 | 0.28898563 | 0.77259238 | NA | RNU6-100P | 106481208 |
| ENSG000002 | 0.5226185 | -0.189580046 | 0.24357686 | -0.7783172 | 0.43638207 | NA | RNU6-1012P | 106481500 |
| ENSG000002 | 0.23065526 | 0.167384503 | 0.17662711 | 0.94767164 | 0.34329664 | NA | RNU6-885P | 106481458 |
| ENSG000002 | 0.72102019 | 0.142484107 | 0.28898156 | 0.49305605 | 0.62197298 | NA | RNU7-179P | 106479092 |
| ENSG000002 | 0.37404583 | -0.105111695 | 0.21784993 | -0.4824959 | 0.62945372 | NA | NA | NA |
| ENSG000002 | 0.38494734 | -0.095409837 | 0.21124663 | -0.4516514 | 0.65152014 | NA | RNU6-1069P | 106480036 |
| ENSG000002 | 0.81036655 | -0.540716938 | 0.28190939 | -1.9180522 | 0.05510439 | NA | NA | NA |
| ENSG000002 | 0.25370795 | -0.171331747 | 0.17411721 | -0.9840024 | 0.32511435 | NA | RNU6-229P | 106481250 |
| ENSG000002 | 0.71422624 | -0.155614713 | 0.26626742 | -0.5844302 | 0.55893094 | NA | NA | NA |
| ENSG000002 | 0 | NA | NA | NA | NA | NA | RNU1-107P | 106480186 |

| | | | | | | | | |
|---|---|---|---|---|---|---|---|---|
| ENSG000002 | 0.71469997 | 0.309652873 | 0.28436427 | 1.08893033 | 0.27618461 | NA | NA | NA |
| ENSG000002 | 0.81084692 | 0.377874869 | 0.29444671 | 1.28333875 | 0.19937342 | NA | NA | NA |
| ENSG000002 | 1.20498304 | 0.390537823 | 0.31984522 | 1.22102127 | 0.22207797 | NA | NA | NA |
| ENSG000002 | 0.74313979 | -0.194647915 | 0.28587591 | -0.6808825 | 0.49594582 | NA | RNU6-1247P | 106481574 |
| ENSG000002 | 1.04312739 | -0.535440803 | 0.31265489 | -1.7125618 | 0.08679319 | NA | NA | NA |
| ENSG000002 | 0.91897981 | -0.610230583 | 0.29431969 | -2.0733597 | 0.03813882 | NA | NA | NA |
| ENSG000002 | 0.26199343 | -0.194018184 | 0.18422801 | -1.0531416 | 0.29227609 | NA | NA | NA |
| ENSG000002 | 0.47633831 | -0.018593199 | 0.22794174 | -0.08157 | 0.93498869 | NA | NA | NA |
| ENSG000002 | 0.99218013 | -0.040479937 | 0.31300325 | -0.1293275 | 0.89709849 | NA | NA | NA |
| ENSG000002 | 0.59032214 | 0.060325881 | 0.26160816 | 0.23059632 | 0.81762842 | NA | RNU6-344P | 106480572 |
| ENSG000002 | 1.06277085 | -0.508777778 | 0.30642404 | -1.6603716 | 0.09683972 | NA | RNU6-405P | 106479744 |
| ENSG000002 | 1.01207154 | -0.046453419 | 0.31227181 | -0.1487596 | 0.88174336 | NA | NA | NA |
| ENSG000002 | 0.67599868 | -0.271668661 | 0.26265684 | -1.0343102 | 0.30099116 | NA | RNU6-139P | 106481977 |
| ENSG000002 | 1.13483349 | -0.274699772 | 0.32225896 | -0.8524194 | 0.39398138 | NA | NA | NA |
| ENSG000002 | 1.13127118 | -0.246256639 | 0.30973008 | -0.7950685 | 0.42657364 | NA | NA | NA |
| ENSG000002 | 0.12070844 | -0.088503695 | 0.12783931 | -0.6923042 | 0.48874631 | NA | NA | NA |
| ENSG000002 | 1.72276827 | -0.189812517 | 0.35737379 | -0.5311316 | 0.59532758 | NA | NA | NA |
| ENSG000002 | 0.61016202 | 0.063256892 | 0.2633751 | 0.24017795 | 0.81019231 | NA | NA | NA |
| ENSG000002 | 0.96568426 | -0.18397265 | 0.30604131 | -0.6011367 | 0.54774897 | NA | NA | NA |
| ENSG000002 | 0.61927241 | -0.278564577 | 0.26634985 | -1.0458597 | 0.29562581 | NA | NA | NA |
| ENSG000002 | 0.76675905 | -0.030196205 | 0.28036461 | -0.1077033 | 0.91423102 | NA | RNU6-635P | 106481382 |
| ENSG000002 | 1.15230316 | -0.410693718 | 0.31767026 | -1.2928302 | 0.19606978 | NA | NA | NA |
| ENSG000002 | 0.36690249 | -0.096999144 | 0.21683809 | -0.4473344 | 0.65463361 | NA | RNU6-113P | 106479616 |
| ENSG000002 | 0.24004852 | -0.005223787 | 0.17775844 | -0.029387 | 0.97655594 | NA | MIR2115 | 100313840 |
| ENSG000002 | 0.25370795 | -0.171331747 | 0.17411721 | -0.9840024 | 0.32511435 | NA | RNU6-347P | 106479720 |
| ENSG000002 | 0 | NA | NA | NA | NA | NA | RNU2-57P | 106480222 |
| ENSG000002 | 0.49902098 | -0.194741582 | 0.24819886 | -0.7846192 | 0.43267688 | NA | RNU7-160P | 106479085 |
| ENSG000002 | 1.13477792 | -0.601040431 | 0.32140896 | -1.8700177 | 0.06148136 | NA | RNU6-551P | 106479811 |
| ENSG000002 | 0 | NA | NA | NA | NA | NA | RNU6-521P | 106481900 |

| | | | | | | | | |
|---|---|---|---|---|---|---|---|---|
| ENSG000002 | 0.75555403 | -0.349570212 | 0.28131632 | -1.2426233 | 0.21400668 | NA | RNU6-539P | 106481351 |
| ENSG000002 | 0 | NA | NA | NA | NA | NA | RNU6-1332P | 106480660 |
| ENSG000002 | 1.76579615 | -0.437315967 | 0.34658107 | -1.2617999 | 0.20702079 | NA | NA | NA |
| ENSG000002 | 1.33184954 | -0.251079119 | 0.33403178 | -0.7516624 | 0.45225412 | NA | RNU6-719P | 106481913 |
| ENSG000002 | 0.14128499 | -0.097772014 | 0.13403235 | -0.7294658 | 0.4657168 | NA | NA | NA |
| ENSG000002 | 2.02443688 | -0.596650306 | 0.36335645 | -1.6420523 | 0.10057916 | NA | RNU6-1178P | 106480083 |
| ENSG000002 | 1.72438862 | -0.045142008 | 0.35047983 | -0.1288006 | 0.89751544 | NA | RN7SKP49 | 106480848 |
| ENSG000002 | 0.62450659 | -0.10299827 | 0.26500891 | -0.3886597 | 0.69752793 | NA | NA | NA |
| ENSG000002 | 0.3863157 | -0.274134926 | 0.21457207 | -1.277589 | 0.20139437 | NA | NA | NA |
| ENSG000002 | 0.51570138 | -0.378721611 | 0.24581506 | -1.540677 | 0.12339542 | NA | NA | NA |
| ENSG000002 | 1.84068599 | -0.118263834 | 0.35951535 | -0.3289535 | 0.74219084 | NA | RN7SKP66 | 106480853 |
| ENSG000002 | 0.49902098 | -0.194741582 | 0.24819886 | -0.7846192 | 0.43267688 | NA | RNU1-142P | 106480197 |
| ENSG000002 | 1.38474954 | 0.020635631 | 0.33499352 | 0.06160009 | 0.9508813 | NA | RNU6-63P | 100873768 |
| ENSG000002 | 0.6567905 | -0.46248946 | 0.26597757 | -1.7388288 | 0.08206488 | NA | RNU4-77P | 106481196 |
| ENSG000002 | 0.96473853 | -0.169713303 | 0.3001574 | -0.5654144 | 0.57179196 | NA | RNU6-180P | 106481233 |
| ENSG000002 | 1.78191457 | -0.771676217 | 0.3542961 | -2.1780545 | 0.02940198 | NA | NA | NA |
| ENSG000002 | 0.39274356 | -0.107428268 | 0.22062504 | -0.4869269 | 0.62631013 | NA | RNU4ATAC3P | 106480753 |
| ENSG000002 | 0.86939644 | 0.035091061 | 0.29743942 | 0.11797717 | 0.90608575 | NA | NA | NA |
| ENSG000002 | 0.88223331 | -0.275481339 | 0.29958866 | -0.9195319 | 0.35781741 | NA | NA | NA |
| ENSG000002 | 0.49287545 | -0.179123382 | 0.24028356 | -0.7454666 | 0.45598967 | NA | RNA5SP206 | 106478999 |
| ENSG000002 | 0 | NA | NA | NA | NA | NA | RNU1-128P | 106480191 |
| ENSG000002 | 0.12685397 | -0.090215768 | 0.12901919 | -0.699243 | 0.48440017 | NA | NA | NA |
| ENSG000002 | 0.14128499 | -0.097772014 | 0.13403235 | -0.7294658 | 0.4657168 | NA | RNU6-1171P | 106481940 |
| ENSG000002 | 0.51911933 | -0.349228947 | 0.23754574 | -1.4701545 | 0.14151991 | NA | NA | NA |
| ENSG000002 | 0.25351207 | -0.189965547 | 0.1824784 | -1.0410303 | 0.29786148 | NA | NA | NA |
| ENSG000002 | 1.08365597 | 0.037481185 | 0.32099328 | 0.11676626 | 0.90704528 | NA | RNU6-783P | 106479912 |
| ENSG000002 | 1.61097647 | -0.249743101 | 0.34721057 | -0.7192843 | 0.47196579 | NA | NA | NA |
| ENSG000002 | 0.91049845 | -0.644235083 | 0.3003423 | -2.1450028 | 0.03195262 | NA | RNU6-267P | 106480565 |
| ENSG000002 | 0.37330862 | -0.105391168 | 0.21773449 | -0.4840352 | 0.62836086 | NA | NA | NA |

| | | | | | | | | |
|---|---|---|---|---|---|---|---|---|
| ENSG000002 | 0.12070844 | -0.088503695 | 0.12783931 | -0.6923042 | 0.48874631 | NA | NA | NA |
| ENSG000002 | 1.08030218 | 0.029874938 | 0.31206558 | 0.09573288 | 0.92373272 | NA | RNU6-965P | 106481484 |
| ENSG000002 | 0.14128499 | -0.097772014 | 0.13403235 | -0.7294658 | 0.4657168 | NA | RNU7-193P | 106481804 |
| ENSG000002 | 0.60947688 | 0.058372533 | 0.25419409 | 0.22963765 | 0.81837335 | NA | NA | NA |
| ENSG000002 | 0 | NA | NA | NA | NA | NA | RNA5SP254 | 106481793 |
| ENSG000002 | 0.60507165 | -0.101443751 | 0.26332021 | -0.3852486 | 0.70005322 | NA | NA | NA |
| ENSG000002 | 0.76624805 | -0.191662006 | 0.28130234 | -0.6813381 | 0.49565757 | NA | NA | NA |
| ENSG000002 | 0 | NA | NA | NA | NA | NA | SNORD36C | 26813 |
| ENSG000002 | 0.97548246 | -0.304345198 | 0.29684868 | -1.0252537 | 0.30524352 | NA | RNU6-1282P | 106481585 |
| ENSG000002 | 1.53377465 | -0.334386157 | 0.34253861 | -0.9761999 | 0.32896541 | NA | RNU7-149P | 106479080 |
| ENSG000002 | 0.53627793 | -0.360851627 | 0.24080453 | -1.4985251 | 0.13399688 | NA | NA | NA |
| ENSG000002 | 1.25440032 | -0.352175911 | 0.33403966 | -1.0542937 | 0.29174847 | NA | RNU6-307P | 106481276 |
| ENSG000002 | 1.25692756 | -0.055576391 | 0.33292159 | -0.1669354 | 0.8674209 | NA | RNU6-861P | 106479947 |
| ENSG000002 | 0.73957996 | -0.027414423 | 0.28481756 | -0.0962526 | 0.92331998 | NA | RNU6-567P | 106479818 |
| ENSG000002 | 1.34468194 | -0.267795944 | 0.33979385 | -0.788113 | 0.43063063 | NA | RNU6-256P | 106479680 |
| ENSG000002 | 0 | NA | NA | NA | NA | NA | NA | NA |
| ENSG000002 | 0.38270188 | -0.272446727 | 0.2140009 | -1.2731102 | 0.20297894 | NA | RNU4-18P | 106480545 |
| ENSG000002 | 0 | NA | NA | NA | NA | NA | NA | NA |
| ENSG000002 | 0.62178293 | -0.443481895 | 0.26161272 | -1.6951848 | 0.09004038 | NA | NA | NA |
| ENSG000002 | 0.58150796 | 0.05262168 | 0.25154459 | 0.20919424 | 0.83429661 | NA | RNU6-1153P | 106480643 |
| ENSG000002 | 0.24004852 | -0.005223787 | 0.17775844 | -0.029387 | 0.97655594 | NA | NA | NA |
| ENSG000002 | 0.64144758 | -0.286694252 | 0.26813299 | -1.0692241 | 0.28496871 | NA | RNU6-135P | 106479625 |
| ENSG000002 | 0.52028267 | -0.202047067 | 0.25039858 | -0.8069018 | 0.41972305 | NA | MIR514B | 100422847 |
| ENSG000002 | 0 | NA | NA | NA | NA | NA | NA | NA |
| ENSG000002 | 0.26199343 | -0.194018184 | 0.18422801 | -1.0531416 | 0.29227609 | NA | RNU7-143P | 106479078 |
| ENSG000002 | 0.71619095 | -0.024103923 | 0.27689662 | -0.0870503 | 0.93063156 | NA | RNA5SP468 | 106480769 |
| ENSG000002 | 1.66976292 | 0.076777114 | 0.35065149 | 0.21895562 | 0.82668462 | NA | RNU6-1224P | 106480101 |
| ENSG000002 | 1.49709466 | -0.484131846 | 0.34373039 | -1.4084639 | 0.15899375 | NA | RNU6-656P | 106479857 |
| ENSG000002 | 1.01660118 | -0.506439653 | 0.30700104 | -1.649635 | 0.09901762 | NA | RN7SKP82 | 106480462 |

| | | | | | | | | | |
|---|---|---|---|---|---|---|---|---|---|
| ENSG000002 | 0.59146429 | 0.065095808 | 0.26170762 | 0.24873486 | 0.80356588 | NA | | RNU7-137P | 106480821 |
| ENSG000002 | 0.9855236 | -0.193118322 | 0.30855488 | -0.62588 | 0.53139367 | NA | | NA | NA |
| ENSG000002 | 1.36896213 | -0.391444096 | 0.33056853 | -1.1841542 | 0.23635205 | NA | | RNU6-566P | 106481903 |
| ENSG000002 | 0.61172818 | 0.021749657 | 0.15747952 | 0.13811102 | 0.89015268 | NA | | NA | NA |
| ENSG000002 | 0.12070844 | -0.088503695 | 0.12783931 | -0.6923042 | 0.48874631 | NA | | RNU6-1150P | 106480072 |
| ENSG000002 | 0.62069689 | -0.101414475 | 0.26466902 | -0.3831747 | 0.70159023 | NA | | NA | NA |
| ENSG000002 | 0.51316968 | -0.377777671 | 0.24556403 | -1.538408 | 0.12394889 | NA | | RNU6-1191P | 106481555 |
| ENSG000002 | 0.23868015 | 0.154625155 | 0.16997278 | 0.90970538 | 0.36297791 | NA | | RNU6-1121P | 106481780 |
| ENSG000002 | 0.26813897 | -0.197007041 | 0.18550319 | -1.0620143 | 0.2882292 | NA | | NA | NA |
| ENSG000002 | 1.83866282 | -0.123484833 | 0.36045796 | -0.3425776 | 0.73191623 | NA | | RNU6-807P | 106479923 |
| ENSG000002 | 1.31758658 | -0.844883601 | 0.32688443 | -2.5846554 | 0.00974764 | NA | | NA | NA |
| ENSG000002 | 0.12685397 | -0.090215768 | 0.12901919 | -0.699243 | 0.48440017 | NA | | RNU6-1149P | 106481542 |
| ENSG000002 | 0.12685397 | -0.090215768 | 0.12901919 | -0.699243 | 0.48440017 | NA | | NA | NA |
| ENSG000002 | 1.89547679 | -0.397179688 | 0.35893188 | -1.1065601 | 0.26848418 | NA | | NA | NA |
| ENSG000002 | 0 | NA | NA | NA | NA | NA | | NA | NA |
| ENSG000002 | 0 | NA | NA | NA | NA | NA | | RNU1-40P | 106480158 |
| ENSG000002 | 1.55612642 | -0.322275913 | 0.33884761 | -0.9510939 | 0.34155669 | NA | | RNU6-1308P | 106480137 |
| ENSG000002 | 1.0280996 | -0.192572733 | 0.30871384 | -0.6237904 | 0.53276524 | NA | | RNU6-453P | 106479766 |
| ENSG000002 | 0 | NA | NA | NA | NA | NA | | RN7SKP282 | 106480919 |
| ENSG000002 | 0.13280363 | -0.093394593 | 0.13115644 | -0.7120855 | 0.47641185 | NA | | RNU2-24P | 106480207 |
| ENSG000002 | 0.39499294 | -0.277908454 | 0.21584201 | -1.287555 | 0.19790091 | NA | | NA | NA |
| ENSG000002 | 1.35651812 | -0.123628383 | 0.33948661 | -0.3641628 | 0.71573647 | NA | | RNU6-678P | 106479865 |
| ENSG000002 | 0.48624257 | -0.012072666 | 0.23796621 | -0.0507327 | 0.95953853 | NA | | NA | NA |
| ENSG000002 | 0.63435632 | -0.27906164 | 0.26763939 | -1.0426778 | 0.29709757 | NA | | RNU6-1136P | 106480066 |
| ENSG000002 | 0.13280363 | -0.093394593 | 0.13115644 | -0.7120855 | 0.47641185 | NA | | NA | NA |
| ENSG000002 | 0.99900888 | -0.200501448 | 0.31384716 | -0.6388506 | 0.52292012 | NA | | RNU7-111P | 106481800 |
| ENSG000002 | 0 | NA | NA | NA | NA | NA | | NA | NA |
| ENSG000002 | 0.37928001 | 0.070449581 | 0.21829793 | 0.32272217 | 0.74690566 | NA | | NA | NA |
| ENSG000002 | 0.7278812 | -0.032565449 | 0.27770112 | -0.117268 | 0.9066477 | NA | | RNU6-557P | 106479814 |

| | | | | | | | | | |
|---|---|---|---|---|---|---|---|---|---|
| ENSG000002 | 1.93324282 | -0.522652996 | 0.35627962 | -1.4669742 | 0.14238308 | NA | | NA | NA |
| ENSG000002 | 1.14749511 | -0.38473559 | 0.30932113 | -1.2438064 | 0.21357084 | NA | | NA | NA |
| ENSG000002 | 1.97058464 | -0.674781306 | 0.35750744 | -1.8874609 | 0.05909836 | NA | | NA | NA |
| ENSG000002 | 0 | NA | NA | NA | NA | NA | | RN7SKP88 | 106480860 |
| ENSG000002 | 0.12070844 | -0.088503695 | 0.12783931 | -0.6923042 | 0.48874631 | NA | | NA | NA |
| ENSG000002 | 2.08106836 | -6.17E-05 | 0.36593488 | -0.0001686 | 0.99986546 | NA | | RNU6-1088P | 106480044 |
| ENSG000002 | 0.6010317 | -0.250734806 | 0.2561147 | -0.9789942 | 0.32758284 | NA | | RNU6-782P | 106481917 |
| ENSG000002 | 0 | NA | NA | NA | NA | NA | | RNA5SP523 | 106480772 |
| ENSG000002 | 1.18805403 | -0.037079762 | 0.32291449 | -0.1148284 | 0.90858112 | NA | | NA | NA |
| ENSG000002 | 0 | NA | NA | NA | NA | NA | | NA | NA |
| ENSG000002 | 0.25351207 | -0.189965547 | 0.1824784 | -1.0410303 | 0.29786148 | NA | | NA | NA |
| ENSG000002 | 0.90344102 | -0.436658607 | 0.29584239 | -1.4759839 | 0.13994822 | NA | | NA | NA |
| ENSG000002 | 1.334095 | -0.114999414 | 0.33706443 | -0.3411793 | 0.73296859 | NA | | NA | NA |
| ENSG000002 | 0 | NA | NA | NA | NA | NA | | RNU1-97P | 106480182 |
| ENSG000002 | 1.63052029 | -0.378801816 | 0.34297559 | -1.1044571 | 0.26939492 | NA | | NA | NA |
| ENSG000002 | 0.11865494 | 0.083440866 | 0.12751478 | 0.65436232 | 0.5128784 | NA | | RNU6ATAC36P | 106480543 |
| ENSG000002 | 0.89404775 | -0.267563874 | 0.29281143 | -0.9137754 | 0.36083489 | NA | | RNU6-928P | 106479977 |
| ENSG000002 | 0.65101106 | -0.268990917 | 0.26053266 | -1.0324652 | 0.30185423 | NA | | RNU6-1234P | 106480106 |
| ENSG000002 | 0.96656582 | -0.164959882 | 0.30024482 | -0.5494179 | 0.58271869 | NA | | RNU6-579P | 106481364 |
| ENSG000002 | 0 | NA | NA | NA | NA | NA | | NA | NA |
| ENSG000002 | 0.48285104 | 0.156035981 | 0.24564443 | 0.63521075 | 0.52529097 | NA | | NA | NA |
| ENSG000002 | 0.49512483 | -0.344267812 | 0.23617548 | -1.457678 | 0.14492931 | NA | | NA | NA |
| ENSG000002 | 1.72188671 | -0.199902932 | 0.35828674 | -0.5579412 | 0.57688454 | NA | | MIR2276 | 100313842 |
| ENSG000002 | 1.11069463 | -0.252719333 | 0.31452977 | -0.803483 | 0.4216956 | NA | | NA | NA |
| ENSG000002 | 0.81748172 | 0.358474136 | 0.2894016 | 1.23867365 | 0.21546638 | NA | | RN7SKP209 | 106479188 |
| ENSG000002 | 0 | NA | NA | NA | NA | NA | | NA | NA |
| ENSG000002 | 0.95711411 | 0.109098762 | 0.30499913 | 0.35770188 | 0.72056643 | NA | | RNU7-110P | 106480815 |
| ENSG000002 | 1.45531198 | -0.1929097 | 0.34552376 | -0.558311 | 0.57663207 | NA | | RN7SKP277 | 106479219 |
| ENSG000002 | 0.25145857 | -0.010118565 | 0.18031426 | -0.0561163 | 0.95524918 | NA | | RNU6-175P | 106480558 |

| | | | | | | | | |
|---|---|---|---|---|---|---|---|---|
| ENSG000002 | 1.1832187 | -0.60909527 | 0.32297056 | -1.8859158 | 0.0593063 | NA | RNU6-295P | 106481272 |
| ENSG000002 | 1.31200574 | 0.024500742 | 0.33484789 | 0.07316977 | 0.94167102 | NA | RN7SKP116 | 106480869 |
| ENSG000002 | 0.9539769 | 0.115712872 | 0.30484181 | 0.37958334 | 0.70425473 | NA | SCARNA14 | 692149 |
| ENSG000002 | 0.3561712 | 0.067399341 | 0.21474131 | 0.31386296 | 0.75362513 | NA | RNU6-1198P | 106480090 |
| ENSG000002 | 0 | NA | NA | NA | NA | NA | NA | NA |
| ENSG000002 | 0.24619405 | -0.007630671 | 0.17915639 | -0.0425922 | 0.96602658 | NA | RN7SKP260 | 106479211 |
| ENSG000002 | 0.70141111 | 0.318465679 | 0.28333314 | 1.12399728 | 0.26101418 | NA | RNU6-612P | 106481374 |
| ENSG000002 | 0.6929292 | 0.29595641 | 0.27700004 | 1.06843454 | 0.28532454 | NA | NA | NA |
| ENSG000002 | 1.1610991 | -0.26645478 | 0.31872041 | -0.8360142 | 0.40314693 | NA | RNU6-1258P | 106480115 |
| ENSG000002 | 0.60219111 | 0.230513192 | 0.26377177 | 0.87391153 | 0.38216644 | NA | NA | NA |
| ENSG000002 | 0.5301324 | -0.3843196 | 0.24728844 | -1.5541349 | 0.12015224 | NA | RNU6-677P | 106481395 |
| ENSG000002 | 0 | NA | NA | NA | NA | NA | NA | NA |
| ENSG000002 | 1.12603755 | -0.433672183 | 0.32323244 | -1.3416728 | 0.1797021 | NA | RNU6-765P | 106479905 |
| ENSG000002 | 1.82697045 | -0.247084266 | 0.35819801 | -0.689798 | 0.49032123 | NA | NA | NA |
| ENSG000002 | 1.00783515 | -0.192961103 | 0.30785304 | -0.6267962 | 0.53079287 | NA | RNU6-971P | 106481926 |
| ENSG000002 | 0.89262382 | -0.459792846 | 0.30190855 | -1.522954 | 0.1277702 | NA | RNU6-980P | 106481489 |
| ENSG000002 | 0.74447728 | -0.175463996 | 0.27094248 | -0.6476061 | 0.51723977 | NA | RNU6-528P | 106479800 |
| ENSG000002 | 0.96226294 | 0.124254798 | 0.3068576 | 0.40492659 | 0.68553148 | NA | RNU7-151P | 106480824 |
| ENSG000002 | 0.95281706 | 0.266670486 | 0.30563367 | 0.87251672 | 0.38292655 | NA | NA | NA |
| ENSG000002 | 0.98306761 | 0.112072446 | 0.30614197 | 0.36607998 | 0.71430539 | NA | RNU6-850P | 106481919 |
| ENSG000002 | 0.27408862 | -0.199826037 | 0.18669465 | -1.0703362 | 0.28446801 | NA | NA | NA |
| ENSG000002 | 0.23202362 | -0.008302493 | 0.17586207 | -0.0472103 | 0.96234565 | NA | NA | NA |
| ENSG000002 | 0.82922851 | 0.045356974 | 0.28735996 | 0.15784027 | 0.87458266 | NA | NA | NA |
| ENSG000002 | 1.2549411 | -0.334872767 | 0.32844983 | -1.0195553 | 0.30793942 | NA | RNU7-116P | 106480816 |
| ENSG000002 | 0.60487577 | -0.101296586 | 0.26329793 | -0.3847223 | 0.70044317 | NA | NA | NA |
| ENSG000002 | 0.36538978 | 0.059033792 | 0.20773944 | 0.28417229 | 0.77627835 | NA | RNU6-143P | 106479629 |
| ENSG000002 | 1.42251508 | 0.358658409 | 0.33760477 | 1.0623618 | 0.28807147 | NA | RNU6-210P | 106479659 |
| ENSG000002 | 1.23823038 | -0.044428126 | 0.3302345 | -0.1345351 | 0.89297947 | NA | RNU6-703P | 106479876 |
| ENSG000002 | 1.35210759 | -0.114873994 | 0.33602078 | -0.3418657 | 0.73245193 | NA | RNU6-577P | 106479822 |

| | | | | | | | | | |
|---|---|---|---|---|---|---|---|---|---|
| ENSG000002 | 1.27350087 | -0.490478208 | 0.3288546 | -1.4914744 | 0.13583699 | NA | | RNU6-445P | 106479763 |
| ENSG000002 | 0.97453172 | -0.043521083 | 0.30739462 | -0.1415805 | 0.88741137 | NA | | NA | NA |
| ENSG000002 | 0.96663728 | 0.421022983 | 0.30038209 | 1.40162477 | 0.16102733 | NA | | NA | NA |
| ENSG000002 | 0.35067856 | 0.074230421 | 0.21389444 | 0.34704231 | 0.72855953 | NA | | RNU6-1092P | 106480046 |
| ENSG000002 | 0 | NA | NA | NA | NA | NA | | RNU6-255P | 106481258 |
| ENSG000002 | 0.61496952 | 0.052672128 | 0.25628412 | 0.2055224 | 0.837164 | NA | | RNU6-250P | 106479678 |
| ENSG000002 | 0.9854377 | -0.339886722 | 0.3083842 | -1.1021535 | 0.27039495 | NA | | RNU6-943P | 106480623 |
| ENSG000002 | 0.83157643 | 0.034789488 | 0.27753624 | 0.12535115 | 0.90024556 | NA | | NA | NA |
| ENSG000002 | 0.23730987 | 0.1539805 | 0.16964526 | 0.90766167 | 0.364057 | NA | | RNU6-714P | 106480603 |
| ENSG000002 | 1.87543297 | -0.120142994 | 0.35820176 | -0.3354059 | 0.7373189 | NA | | NA | NA |
| ENSG000002 | 2.16918275 | -0.363988707 | 0.3617273 | -1.0062517 | 0.31429451 | NA | | RNU4ATAC10P | 106479542 |
| ENSG000002 | 0.77427295 | -0.18871474 | 0.2818501 | -0.6695571 | 0.50314016 | NA | | NA | NA |
| ENSG000002 | 0.99967284 | -0.343913679 | 0.30897117 | -1.1130931 | 0.26566844 | NA | | NA | NA |
| ENSG000002 | 1.10056313 | -0.254040437 | 0.31667138 | -0.802221 | 0.42242512 | NA | | RNU6-182P | 106479647 |
| ENSG000002 | 0.38651158 | -0.274258716 | 0.21461355 | -1.2779189 | 0.20127801 | NA | | NA | NA |
| ENSG000002 | 0.13280363 | -0.093394593 | 0.13115644 | -0.7120855 | 0.47641185 | NA | | RNU6-835P | 106481445 |
| ENSG000002 | 1.27261931 | -0.516541857 | 0.33230424 | -1.5544245 | 0.12008319 | NA | | NA | NA |
| ENSG000002 | 0 | NA | NA | NA | NA | NA | | NA | NA |
| ENSG000002 | 0.37441639 | -0.268856313 | 0.21277471 | -1.2635727 | 0.20638344 | NA | | RN7SKP60 | 106480851 |
| ENSG000002 | 1.8647056 | -0.255687434 | 0.35851826 | -0.7131783 | 0.47573545 | NA | | RNU6-56P | 100873764 |
| ENSG000002 | 0.59704675 | -0.097265597 | 0.25500139 | -0.3814316 | 0.70288299 | NA | | NA | NA |
| ENSG000002 | 0.76764921 | -0.358759829 | 0.28369807 | -1.2645832 | 0.20602076 | NA | | NA | NA |
| ENSG000002 | 0.93708022 | -0.247252523 | 0.2826817 | -0.8746676 | 0.3817548 | NA | | NA | NA |
| ENSG000002 | 0.68584841 | -0.42838929 | 0.25774854 | -1.6620435 | 0.09650404 | NA | | NA | NA |
| ENSG000002 | 1.86251021 | 0.003705589 | 0.35993369 | 0.0102952 | 0.99178576 | NA | | RNU6-958P | 106481482 |
| ENSG000002 | 0.24550891 | -0.007886114 | 0.17900103 | -0.0440563 | 0.96485956 | NA | | NA | NA |
| ENSG000002 | 0.83577898 | 0.052536527 | 0.30082402 | 0.17464206 | 0.8613609 | NA | | RNU6-1006P | 106480010 |
| ENSG000002 | 0.50360227 | -0.018733935 | 0.23996924 | -0.0780681 | 0.93777391 | NA | | RNU6-345P | 106481288 |
| ENSG000002 | 1.31590134 | 0.143998319 | 0.32697711 | 0.44039266 | 0.65965274 | NA | | NA | NA |

| | | | | | | | | |
|---|---|---|---|---|---|---|---|---|
| ENSG000002 | 1.24634509 | -0.18966293 | 0.33095539 | -0.573077 | 0.56659254 | NA | RN7SKP233 | 106480475 |
| ENSG000002 | 1.95570772 | -0.522902444 | 0.35327893 | -1.4801405 | 0.13883576 | NA | RNU6-410P | 106481893 |
| ENSG000002 | 1.26584262 | -0.33574382 | 0.32695086 | -1.0268938 | 0.30447047 | NA | RNU6-388P | 106479737 |
| ENSG000002 | 0.59758809 | -0.103924321 | 0.26264093 | -0.3956897 | 0.69233393 | NA | NA | NA |
| ENSG000002 | 0.35682409 | 0.072008558 | 0.21485623 | 0.33514763 | 0.73751372 | NA | RNU6-148P | 106479631 |
| ENSG000002 | 0.37125511 | 0.067313264 | 0.2170755 | 0.31009148 | 0.75649139 | NA | NA | NA |
| ENSG000002 | 0.47684931 | 0.147509143 | 0.23759598 | 0.62084023 | 0.53470475 | NA | NA | NA |
| ENSG000002 | 0 | NA | NA | NA | NA | NA | NA | NA |
| ENSG000002 | 0.23202362 | -0.008302493 | 0.17586207 | -0.0472103 | 0.96234565 | NA | RNU6-1212P | 106481943 |
| ENSG000002 | 1.24794213 | -0.316844681 | 0.32486165 | -0.9753219 | 0.32940062 | NA | NA | NA |
| ENSG000002 | 0.73129447 | -0.021100375 | 0.26879465 | -0.0785 | 0.93743034 | NA | RNU6-419P | 106479751 |
| ENSG000002 | 2.11570579 | -0.371012239 | 0.36435183 | -1.0182801 | 0.30854484 | NA | NA | NA |
| ENSG000002 | 0.72833821 | -0.029759877 | 0.2840795 | -0.104759 | 0.91656707 | NA | NA | NA |
| ENSG000002 | 1.30420947 | -0.505365084 | 0.33108302 | -1.5263999 | 0.1269103 | NA | RNU2-46P | 106480218 |
| ENSG000002 | 0.75899316 | -0.198701114 | 0.28686906 | -0.6926544 | 0.48852648 | NA | NA | NA |
| ENSG000002 | 0.50016259 | -0.200428691 | 0.24824437 | -0.8073846 | 0.41944491 | NA | RNU1-117P | 106480189 |
| ENSG000002 | 0.74521503 | -0.189818705 | 0.28603889 | -0.6636115 | 0.50693897 | NA | NA | NA |
| ENSG000002 | 1.23373498 | -0.046143321 | 0.33367449 | -0.1382884 | 0.89001247 | NA | NA | NA |
| ENSG000002 | 0.12685397 | -0.090215768 | 0.12901919 | -0.699243 | 0.48440017 | NA | RNU6-1271P | 106480120 |
| ENSG000002 | 1.80297204 | 0.011156966 | 0.35053407 | 0.03182848 | 0.97460883 | NA | RNU6-375P | 106480575 |
| ENSG000002 | 1.25636504 | -0.203697155 | 0.33464405 | -0.608698 | 0.54272466 | NA | RNU6-570P | 106481361 |
| ENSG000002 | 1.3476984 | -0.436947566 | 0.3139726 | -1.3916742 | 0.16402108 | NA | RNU6-448P | 106479764 |
| ENSG000002 | 1.00151493 | -0.047469472 | 0.30854936 | -0.1538473 | 0.87773017 | NA | NA | NA |
| ENSG000002 | 0.11205239 | 0.082118251 | 0.12671962 | 0.64803108 | 0.51696485 | NA | RNA5SP243 | 106479001 |
| ENSG000002 | 1.04135621 | -0.183876889 | 0.30092995 | -0.6110289 | 0.54118047 | NA | RNU6-909P | 106481466 |
| ENSG000002 | 0.77476221 | -0.188343058 | 0.28188047 | -0.6681664 | 0.50402738 | NA | NA | NA |
| ENSG000002 | 1.08797476 | 0.04005403 | 0.32113533 | 0.12472632 | 0.90074022 | NA | RNU7-188P | 106480832 |
| ENSG000002 | 0 | NA | NA | NA | NA | NA | RNA5SP312 | 106480439 |
| ENSG000002 | 2.00064933 | -0.705421353 | 0.35907669 | -1.9645423 | 0.04946723 | NA | RN7SKP98 | 106480863 |

| | | | | | | | | | |
|---|---|---|---|---|---|---|---|---|---|
| ENSG000002 | 0 | NA | NA | NA | NA | NA | | RNU6-334P | 106479714 |
| ENSG000002 | 0.73599591 | -0.172781731 | 0.27032991 | -0.6391514 | 0.52272446 | NA | | RNU6-1137P | 106480067 |
| ENSG000002 | 0.12685397 | -0.090215768 | 0.12901919 | -0.699243 | 0.48440017 | NA | | RNU6-1213P | 106481563 |
| ENSG000002 | 1.5094201 | -0.349198913 | 0.3469948 | -1.006352 | 0.31424627 | NA | | RN7SKP197 | 106480894 |
| ENSG000002 | 0.96454265 | -0.178908235 | 0.3060021 | -0.5846634 | 0.55877405 | NA | | RNU6-430P | 106479756 |
| ENSG000002 | 0.11934008 | 0.083564698 | 0.12760091 | 0.65489106 | 0.5125379 | NA | | NA | NA |
| ENSG000002 | 0.8794426 | -0.118660638 | 0.30333647 | -0.3911849 | 0.69566059 | NA | | RNU6-1236P | 106481570 |
| ENSG000002 | 0.83037066 | 0.05283786 | 0.29532027 | 0.17891714 | 0.85800276 | NA | | RNU6-699P | 106479875 |
| ENSG000002 | 1.03698186 | -0.533697484 | 0.31242205 | -1.708258 | 0.08758847 | NA | | NA | NA |
| ENSG000002 | 1.17811607 | 0.415350982 | 0.32815682 | 1.26570884 | 0.20561734 | NA | | RNU6-548P | 106481354 |
| ENSG000002 | 0.78461194 | -0.338341767 | 0.27666213 | -1.2229421 | 0.22135156 | NA | | RNU6-685P | 106479868 |
| ENSG000002 | 0.12685397 | -0.090215768 | 0.12901919 | -0.699243 | 0.48440017 | NA | | RNU6-1096P | 106480700 |
| ENSG000002 | 0 | NA | NA | NA | NA | NA | | RNU6-303P | 106479702 |
| ENSG000002 | 0.36556447 | -0.101943371 | 0.21659044 | -0.4706735 | 0.63787395 | NA | | RNA5SP342 | 106479008 |
| ENSG000002 | 0.77498842 | -0.384054853 | 0.28945545 | -1.3268185 | 0.18456873 | NA | | RNU6-894P | 106481461 |
| ENSG000002 | 1.08380032 | 0.042247673 | 0.30634037 | 0.13791089 | 0.89031084 | NA | | NA | NA |
| ENSG000002 | 0.72853409 | -0.029918348 | 0.28409594 | -0.1053107 | 0.91612925 | NA | | NA | NA |
| ENSG000002 | 2.0410207 | 0.259579699 | 0.36442003 | 0.71230908 | 0.4762734 | NA | | RNU6-1003P | 106481778 |
| ENSG000002 | 0 | NA | NA | NA | NA | NA | | RNU6-201P | 106481240 |
| ENSG000002 | 0.85721536 | -0.106371576 | 0.29714523 | -0.3579784 | 0.72035947 | NA | | RNU6-158P | 106481225 |
| ENSG000002 | 0.2596576 | -0.193002035 | 0.1837913 | -1.0501152 | 0.29366517 | NA | | RNU6-789P | 106479915 |
| ENSG000002 | 1.16755729 | -0.267488923 | 0.31643592 | -0.8453178 | 0.39793341 | NA | | NA | NA |
| ENSG000002 | 0.24141689 | -0.165257896 | 0.17127401 | -0.9648743 | 0.33460776 | NA | | NA | NA |
| ENSG000002 | 0.58821656 | 0.242346449 | 0.26860002 | 0.90225774 | 0.36691997 | NA | | NA | NA |
| ENSG000002 | 1.92144957 | -0.351169025 | 0.34747014 | -1.0106452 | 0.31218628 | NA | | RNU6-365P | 106479728 |
| ENSG000002 | 1.48167043 | -0.162384961 | 0.33291012 | -0.4877742 | 0.6257098 | NA | | NA | NA |
| ENSG000002 | 1.8662968 | -0.258487211 | 0.36046951 | -0.7170848 | 0.47332177 | NA | | NA | NA |
| ENSG000002 | 1.21754113 | -0.568075434 | 0.31473407 | -1.8049378 | 0.07108441 | NA | | RNU6-64P | 100873769 |
| ENSG000002 | 0.23890637 | -0.010420674 | 0.17747379 | -0.0587167 | 0.95317776 | NA | | RNU6-218P | 106479663 |

| | | | | | | | | | |
|---|---|---|---|---|---|---|---|---|---|
| ENSG000002 | 0.48353618 | 0.145822039 | 0.2384055 | 0.61165553 | 0.54076569 | NA | | RNU6-1015P | 106481501 |
| ENSG000002 | 1.55167317 | 0.302892658 | 0.34940864 | 0.86687226 | 0.38601201 | NA | | RNU6-1223P | 106481566 |
| ENSG000002 | 1.06538045 | 0.481403452 | 0.31005682 | 1.55262979 | 0.1205116 | NA | | RNU7-172P | 106480828 |
| ENSG000002 | 1.35262251 | -0.249611865 | 0.33528559 | -0.7444754 | 0.45658893 | NA | | RNU6-1220P | 106481565 |
| ENSG000002 | 0.11865494 | 0.083440866 | 0.12751478 | 0.65436232 | 0.5128784 | NA | | NA | NA |
| ENSG000002 | 0 | NA | NA | NA | NA | NA | | NA | NA |
| ENSG000002 | 0.74846222 | -0.357497056 | 0.28238646 | -1.2659851 | 0.20551842 | NA | | NA | NA |
| ENSG000002 | 0.73978498 | -0.328755599 | 0.27348192 | -1.2021109 | 0.22932055 | NA | | RNU6-264P | 106481262 |
| ENSG000002 | 0.6313676 | -0.265579877 | 0.26046629 | -1.0196324 | 0.30790281 | NA | | RNU6-897P | 106479963 |
| ENSG000002 | 1.02536095 | -0.044289782 | 0.3078644 | -0.1438613 | 0.88560997 | NA | | NA | NA |
| ENSG000002 | 0 | NA | NA | NA | NA | NA | | RNU6-1314P | 106481595 |
| ENSG000002 | 1.75259475 | -0.184906904 | 0.34652911 | -0.533597 | 0.59362036 | NA | | NA | NA |
| ENSG000002 | 0.3593886 | 0.071604659 | 0.20679693 | 0.34625591 | 0.7291504 | NA | | NA | NA |
| ENSG000002 | 0.63026429 | -0.446551733 | 0.2623193 | -1.7023213 | 0.08869516 | NA | | RNU6-58P | 100873766 |
| ENSG000002 | 0.51276878 | -0.021314138 | 0.24098735 | -0.0884451 | 0.92952296 | NA | | RNU6-232P | 106480388 |
| ENSG000002 | 0.7164389 | -0.025867019 | 0.28328556 | -0.0913108 | 0.92724567 | NA | | RNU4ATAC9P | 106479541 |
| ENSG000002 | 0.76121029 | -0.351991847 | 0.28168635 | -1.2495879 | 0.21145014 | NA | | NA | NA |
| ENSG000002 | 1.81595842 | -0.122100928 | 0.35444732 | -0.3444826 | 0.73048339 | NA | | NA | NA |
| ENSG000002 | 0.8678023 | -0.271126522 | 0.29886542 | -0.907186 | 0.36430845 | NA | | NA | NA |
| ENSG000002 | 0.75984193 | -0.181352233 | 0.27925466 | -0.6494152 | 0.51607002 | NA | | RN7SKP258 | 106480912 |
| ENSG000002 | 0.37404583 | -0.105111695 | 0.21784993 | -0.4824959 | 0.62945372 | NA | | RNU6-993P | 106480005 |
| ENSG000002 | 1.32846494 | -0.054704825 | 0.18789689 | -0.2911428 | 0.77094211 | NA | | RN7SKP147 | 106480879 |
| ENSG000002 | 0.36827086 | -0.266056975 | 0.21181163 | -1.2561018 | 0.20907905 | NA | | NA | NA |
| ENSG000002 | 0.23799501 | 0.170958857 | 0.17834799 | 0.95856902 | 0.33777591 | NA | | NA | NA |
| ENSG000002 | 0.66654986 | -0.459691143 | 0.26528715 | -1.7328059 | 0.08313017 | NA | | RNU6-1057P | 106480030 |
| ENSG000002 | 0.50520035 | -0.028355362 | 0.23144993 | -0.1225119 | 0.90249365 | NA | | RNU6-1028P | 106480405 |
| ENSG000002 | 1.27309751 | -0.361931181 | 0.33444931 | -1.0821705 | 0.27917677 | NA | | NA | NA |
| ENSG000002 | 0.83074723 | -0.505049968 | 0.27443609 | -1.8403191 | 0.06572141 | NA | | RNA5SP70 | 106480436 |
| ENSG000002 | 1.47532601 | -0.327786713 | 0.34228073 | -0.9576546 | 0.33823696 | NA | | RN7SKP183 | 106480889 |

| | | | | | | | | |
|---|---|---|---|---|---|---|---|---|
| ENSG000002 | 0.37765965 | -0.098061994 | 0.21009083 | -0.46676 | 0.64067161 | NA | RNU6-165P | 106481227 |
| ENSG000002 | 1.00330785 | -0.201516375 | 0.31401882 | -0.6417334 | 0.52104628 | NA | RNU6-367P | 106481890 |
| ENSG000002 | 0.71552755 | 0.142356724 | 0.28330783 | 0.50248073 | 0.6153294 | NA | NA | NA |
| ENSG000002 | 0.26062507 | -0.012888319 | 0.18226503 | -0.070712 | 0.94362699 | NA | RNU6-111P | 106481212 |
| ENSG000002 | 0 | NA | NA | NA | NA | NA | NA | NA |
| ENSG000002 | 1.43763536 | 0.084896087 | 0.33395049 | 0.25421758 | 0.79932747 | NA | RNU1-54P | 106481611 |
| ENSG000002 | 0.24411881 | -0.012946855 | 0.17864669 | -0.0724718 | 0.94222641 | NA | RNU6-1073P | 106481518 |
| ENSG000002 | 1.37217561 | -0.123093098 | 0.33645815 | -0.3658497 | 0.71447727 | NA | NA | NA |
| ENSG000002 | 1.1295328 | 0.03166093 | 0.32251159 | 0.0981699 | 0.92179738 | NA | NA | NA |
| ENSG000002 | 1.95689802 | -0.276800114 | 0.35471569 | -0.7803436 | 0.43518866 | NA | RNU6-1231P | 106480104 |
| ENSG000002 | 0.59413396 | 0.214315572 | 0.25435676 | 0.84257864 | 0.39946414 | NA | RNU6-1315P | 106481950 |
| ENSG000002 | 1.38432287 | -0.124876923 | 0.33526157 | -0.3724761 | 0.7095384 | NA | NA | NA |
| ENSG000002 | 0.8662058 | -0.104563914 | 0.29761826 | -0.3513357 | 0.72533653 | NA | RNU1-45P | 106481610 |
| ENSG000002 | 1.00610069 | -0.197840257 | 0.31413529 | -0.6297932 | 0.52882992 | NA | RNU6-176P | 106479644 |
| ENSG000002 | 2.15546933 | -0.375331083 | 0.36454882 | -1.029577 | 0.30320861 | NA | RN7SKP283 | 106479221 |
| ENSG000002 | 0 | NA | NA | NA | NA | NA | NA | NA |
| ENSG000002 | 1.20769749 | -0.046413567 | 0.33166837 | -0.1399397 | 0.88870765 | NA | NA | NA |
| ENSG000002 | 0.47895489 | -0.016192723 | 0.24471014 | -0.066171 | 0.94724166 | NA | RN7SKP64 | 106480852 |
| ENSG000002 | 0 | NA | NA | NA | NA | NA | NA | NA |
| ENSG000002 | 1.64361383 | -0.421522054 | 0.3539203 | -1.1910084 | 0.23365028 | NA | RN7SKP40 | 106479113 |
| ENSG000002 | 0.51931521 | -0.354214731 | 0.23897171 | -1.4822455 | 0.13827499 | NA | RNU6-902P | 106480619 |
| ENSG000002 | 0.35067856 | 0.074230421 | 0.21389444 | 0.34704231 | 0.72855953 | NA | RNU6-731P | 106481411 |
| ENSG000002 | 0.37928001 | 0.070449581 | 0.21829793 | 0.32272217 | 0.74690566 | NA | RNU7-156P | 106479083 |
| ENSG000002 | 0.76015651 | -0.032508449 | 0.27992106 | -0.1161343 | 0.90754607 | NA | RNU6-1238P | 106480107 |
| ENSG000002 | 0.61950267 | -0.112567707 | 0.27088102 | -0.4155614 | 0.67773094 | NA | RNU6-464P | 106479771 |
| ENSG000002 | 0 | NA | NA | NA | NA | NA | NA | NA |
| ENSG000002 | 1.73089889 | -0.050404369 | 0.34922206 | -0.1443333 | 0.88523729 | NA | MIR2116 | 100313886 |
| ENSG000002 | 0.55665861 | -0.363599063 | 0.24151673 | -1.5054819 | 0.13220023 | NA | NA | NA |
| ENSG000002 | 0.74209047 | -0.163326534 | 0.27077347 | -0.6031851 | 0.54638553 | NA | RNU6-299P | 106480389 |

| | | | | | | | | |
|---|---|---|---|---|---|---|---|---|
| ENSG000002 | 1.12586341 | -0.26722627 | 0.32349262 | -0.826066 | 0.40876663 | NA | RNU4ATAC5P | 106481156 |
| ENSG000002 | 0.64878734 | -0.284003979 | 0.26878765 | -1.056611 | 0.29068916 | NA | RNU6-706P | 106479878 |
| ENSG000002 | 0.63324854 | -0.111307913 | 0.26569851 | -0.4189256 | 0.6752705 | NA | NA | NA |
| ENSG000002 | 1.45001964 | -0.741421222 | 0.33622168 | -2.2051559 | 0.02744315 | NA | NA | NA |
| ENSG000002 | 1.58213617 | -0.103660384 | 0.34558359 | -0.2999575 | 0.76420958 | NA | NA | NA |
| ENSG000002 | 1.41410072 | -0.584178567 | 0.34024575 | -1.7169313 | 0.08599173 | NA | NA | NA |
| ENSG000002 | 1.16385919 | 0.240581173 | 0.32107444 | 0.7493003 | 0.45367623 | NA | NA | NA |
| ENSG000002 | 0.26813897 | -0.197007041 | 0.18550319 | -1.0620143 | 0.2882292 | NA | NA | NA |
| ENSG000002 | 0 | NA | NA | NA | NA | NA | NA | NA |
| ENSG000002 | 0.36484899 | 0.075158314 | 0.21612264 | 0.34775772 | 0.72802215 | NA | RN7SKP289 | 106479224 |
| ENSG000002 | 1.4469388 | 0.095498156 | 0.34363939 | 0.27790224 | 0.7810874 | NA | NA | NA |
| ENSG000002 | 0 | NA | NA | NA | NA | NA | NA | NA |
| ENSG000002 | 0.49076933 | -0.025124039 | 0.23846109 | -0.1053591 | 0.91609089 | NA | RNU7-128P | 106480819 |
| ENSG000002 | 1.74262934 | -0.05171862 | 0.35340932 | -0.146342 | 0.88365143 | NA | RN7SKP57 | 106479120 |
| ENSG000002 | 1.643265 | -0.126510746 | 0.34910901 | -0.3623818 | 0.71706675 | NA | NA | NA |
| ENSG000002 | 1.61355577 | 0.006156923 | 0.35119336 | 0.01753143 | 0.98601266 | NA | NA | NA |
| ENSG000002 | 0.84312065 | 0.183806439 | 0.27924636 | 0.65822322 | 0.51039471 | NA | NA | NA |
| ENSG000002 | 0.37216701 | -0.099548723 | 0.21762847 | -0.4574251 | 0.64736554 | NA | RNU6-711P | 106479881 |
| ENSG000002 | 0.4009426 | -0.302672773 | 0.2240145 | -1.3511303 | 0.17665371 | NA | NA | NA |
| ENSG000002 | 0.38270188 | -0.272446727 | 0.2140009 | -1.2731102 | 0.20297894 | NA | RNU6-709P | 106479879 |
| ENSG000002 | 0.13280363 | -0.093394593 | 0.13115644 | -0.7120855 | 0.47641185 | NA | NA | NA |
| ENSG000002 | 0.64741897 | -0.110552013 | 0.26688537 | -0.4142303 | 0.67870543 | NA | RNU6-794P | 106480400 |
| ENSG000002 | 1.95997388 | -0.53152263 | 0.35382924 | -1.502201 | 0.13304521 | NA | RN7SKP295 | 106480924 |
| ENSG000002 | 1.80089408 | -0.622820023 | 0.35488913 | -1.7549707 | 0.07926432 | NA | RNU6-586P | 106480592 |
| ENSG000002 | 1.15532413 | -0.420043617 | 0.32014283 | -1.3120507 | 0.18950304 | NA | RN7SKP92 | 106479135 |
| ENSG000002 | 1.68429319 | -0.875351093 | 0.34880953 | -2.509539 | 0.01208889 | NA | NA | NA |
| ENSG000002 | 1.68565241 | -0.616996041 | 0.33633837 | -1.8344504 | 0.06658717 | NA | RNU6-1116P | 106480057 |
| ENSG000002 | 0.95699842 | 0.124732458 | 0.30495346 | 0.40902129 | 0.68252404 | NA | NA | NA |
| ENSG000002 | 1.36123972 | -0.248132127 | 0.33195021 | -0.747498 | 0.45476299 | NA | NA | NA |

| | | | | | | | | |
|---|---|---|---|---|---|---|---|---|
| ENSG000002 | 0.86962767 | -0.412029274 | 0.2917292 | -1.412369 | 0.15784135 | NA | RNA5SP106 | 106478994 |
| ENSG000002 | 0.11131518 | 0.081951103 | 0.12663804 | 0.64712866 | 0.51754868 | NA | RNU6-840P | 106479938 |
| ENSG000002 | 1.0214648 | -0.199205491 | 0.31322582 | -0.6359804 | 0.52478918 | NA | NA | NA |
| ENSG000002 | 0.83404005 | 0.381415969 | 0.29733374 | 1.28278739 | 0.19956657 | NA | RNU6-537P | 106479804 |
| ENSG000002 | 0.49076933 | -0.025124039 | 0.23846109 | -0.1053591 | 0.91609089 | NA | RNU6-1174P | 106481550 |
| ENSG000002 | 2.09602964 | 0.115349804 | 0.35758314 | 0.32258178 | 0.747012 | NA | NA | NA |
| ENSG000002 | 0.35022155 | 0.06940789 | 0.21379411 | 0.32464827 | 0.74544729 | NA | RNU1-104P | 106480184 |
| ENSG000002 | 0 | NA | NA | NA | NA | NA | NA | NA |
| ENSG000002 | 1.80836592 | 0.419437026 | 0.35800065 | 1.17160967 | 0.2413538 | NA | NA | NA |
| ENSG000002 | 1.12402076 | -0.549645385 | 0.31284511 | -1.756925 | 0.0789306 | NA | NA | NA |
| ENSG000002 | 0.26813897 | -0.197007041 | 0.18550319 | -1.0620143 | 0.2882292 | NA | NA | NA |
| ENSG000002 | 0.23070733 | 0.167416683 | 0.17664244 | 0.94777156 | 0.34324576 | NA | NA | NA |
| ENSG000002 | 1.2504997 | -0.20563256 | 0.33449627 | -0.6147529 | 0.5387179 | NA | RNU2-19P | 106481959 |
| ENSG000002 | 1.00262718 | -0.490562014 | 0.30482559 | -1.6093203 | 0.10754632 | NA | RNU7-161P | 106480452 |
| ENSG000002 | 2.05401189 | -0.116410927 | 0.36457272 | -0.3193078 | 0.74949309 | NA | RNU2-60P | 106481653 |
| ENSG000002 | 1.55919759 | 0.012569513 | 0.34424467 | 0.03651331 | 0.97087307 | NA | NA | NA |
| ENSG000002 | 1.4812974 | -0.194021419 | 0.34714695 | -0.5589029 | 0.57622802 | NA | NA | NA |
| ENSG000002 | 1.36400636 | -0.122858054 | 0.33253997 | -0.3694535 | 0.71178973 | NA | NA | NA |
| ENSG000002 | 0.3854038 | -0.110128691 | 0.21952326 | -0.5016721 | 0.61589821 | NA | OC90 | 729330 |
| ENSG000002 | 0.48367807 | -0.017505926 | 0.23765649 | -0.0736606 | 0.94128043 | NA | NA | NA |
| ENSG000002 | 0.5969266 | 0.215303566 | 0.25461873 | 0.84559202 | 0.39778038 | NA | NA | NA |
| ENSG000002 | 0.25260017 | -0.015978705 | 0.18049464 | -0.0885273 | 0.92945758 | NA | NA | NA |
| ENSG000002 | 0.49671898 | -0.027206884 | 0.23914704 | -0.1137663 | 0.90942302 | NA | NA | NA |
| ENSG000002 | 0.76605217 | -0.201845012 | 0.28729196 | -0.702578 | 0.48231878 | NA | NA | NA |
| ENSG000002 | 1.50489492 | -0.197131661 | 0.34633512 | -0.5691934 | 0.5692249 | NA | NA | NA |
| ENSG000002 | 1.2024856 | -0.035692698 | 0.33152587 | -0.1076619 | 0.91426391 | NA | NA | NA |
| ENSG000002 | 1.89712836 | -0.336722454 | 0.34718126 | -0.9698751 | 0.33210876 | NA | NA | NA |
| ENSG000002 | 2.11730228 | -0.512374291 | 0.36543818 | -1.402082 | 0.16089077 | NA | LOC102467222 | 102467222 |
| ENSG000002 | 1.35370566 | -0.121864955 | 0.33381266 | -0.3650699 | 0.71505923 | NA | NA | NA |

| | | | | | | | | | |
|---|---|---|---|---|---|---|---|---|---|
| ENSG000002 | 0.60259551 | 0.210653417 | 0.25675396 | 0.82044857 | 0.41196044 | NA | NA | NA | |
| ENSG000002 | 1.43914203 | 0.093195232 | 0.34738048 | 0.26827999 | 0.78848381 | NA | NA | NA | |
| ENSG000002 | 1.37106379 | -0.245791684 | 0.32784263 | -0.7497246 | 0.45342058 | NA | NA | NA | |
| ENSG000002 | 0.7381576 | -0.029362007 | 0.29007186 | -0.1012232 | 0.91937327 | NA | NA | NA | |
| ENSG000002 | 0.23936338 | -0.005479057 | 0.17759977 | -0.0308506 | 0.9753887 | NA | NA | NA | |
| ENSG000002 | 2.14840978 | -0.394969277 | 0.36724242 | -1.0755001 | 0.28215087 | NA | NA | NA | |
| ENSG000002 | 0.9700353 | -0.175695904 | 0.30041861 | -0.584837 | 0.55865735 | NA | NA | NA | |
| ENSG000002 | 1.12700501 | -0.272087911 | 0.32350711 | -0.841057 | 0.40031601 | NA | NA | NA | |
| ENSG000002 | 0.4923881 | -0.012789975 | 0.22994905 | -0.0556209 | 0.95564381 | NA | NA | NA | |
| ENSG000002 | 1.12956101 | -0.251078053 | 0.31054343 | -0.8085119 | 0.41879596 | NA | NA | NA | |
| ENSG000002 | 0.66313191 | -0.429680649 | 0.25813351 | -1.6645675 | 0.09599908 | NA | NA | NA | |
| ENSG000002 | 1.2456922 | -0.199865866 | 0.33439021 | -0.5977025 | 0.55003845 | NA | NA | NA | |
| ENSG000002 | 0.37441639 | -0.268856313 | 0.21277471 | -1.2635727 | 0.20638344 | NA | NA | NA | |
| ENSG000002 | 0.75825595 | -0.198978971 | 0.2868202 | -0.6937411 | 0.48784457 | NA | NA | NA | |
| ENSG000002 | 0.73069578 | 0.138063225 | 0.2843372 | 0.48556159 | 0.62727803 | NA | NA | NA | |
| ENSG000002 | 1.24369022 | -0.052540742 | 0.32286036 | -0.1627352 | 0.87072694 | NA | NA | NA | |
| ENSG000002 | 0.88744576 | -0.277587958 | 0.29984413 | -0.9257742 | 0.35456333 | NA | GUSBP3 | | 653188 |
| ENSG000002 | 0.7244638 | -0.023070588 | 0.28382879 | -0.0812835 | 0.93521652 | NA | NA | NA | |
| ENSG000002 | 0.85149191 | -0.248439926 | 0.28798297 | -0.8626896 | 0.38830813 | NA | NA | NA | |
| ENSG000002 | 1.81304754 | 0.266254347 | 0.35666521 | 0.74651056 | 0.45535905 | NA | NA | NA | |
| ENSG000002 | 0.51225778 | -0.204952203 | 0.24951191 | -0.8214125 | 0.41141135 | NA | NA | NA | |
| ENSG000002 | 1.66464684 | -0.402629917 | 0.34752535 | -1.1585627 | 0.24663447 | NA | NA | NA | |
| ENSG000002 | 1.80775913 | -0.46265829 | 0.35227551 | -1.3133422 | 0.18906767 | NA | NA | NA | |
| ENSG000002 | 1.01722842 | -0.363859965 | 0.3141749 | -1.1581446 | 0.24680503 | NA | COX6CP8 | | 106481680 |
| ENSG000002 | 0.75323938 | -0.18378281 | 0.27879518 | -0.6592037 | 0.50976498 | NA | NA | NA | |
| ENSG000002 | 1.09617325 | -0.105770848 | 0.30536495 | -0.3463752 | 0.72906075 | NA | NA | NA | |
| ENSG000002 | 1.68468315 | -0.382329964 | 0.34379296 | -1.1120936 | 0.26609789 | NA | NA | NA | |
| ENSG000002 | 1.77280418 | -0.461387804 | 0.35484412 | -1.3002549 | 0.19351362 | NA | NA | NA | |
| ENSG000002 | 1.83931708 | 0.004970726 | 0.3602897 | 0.01379647 | 0.98899236 | NA | NA | NA | |

| | | | | | | | | |
|---|---|---|---|---|---|---|---|---|
| ENSG000002 | 1.49274362 | -0.483453536 | 0.34365595 | -1.4067952 | 0.15948813 | NA | NA | NA |
| ENSG000002 | 0.22410479 | 0.147643983 | 0.16638082 | 0.88738586 | 0.37487119 | NA | NA | NA |
| ENSG000002 | 0.96724458 | -0.038194427 | 0.31199407 | -0.1224204 | 0.90256611 | NA | NA | NA |
| ENSG000002 | 0.35022155 | 0.06940789 | 0.21379411 | 0.32464827 | 0.74544729 | NA | NA | NA |
| ENSG000002 | 1.29689145 | -0.327626719 | 0.32535304 | -1.0069884 | 0.31394037 | NA | NA | NA |
| ENSG000002 | 0.74347611 | 0.134190709 | 0.2851946 | 0.47052332 | 0.63798117 | NA | NA | NA |
| ENSG000002 | 0.7859803 | -0.49392286 | 0.27219789 | -1.8145727 | 0.0695896 | NA | NA | NA |
| ENSG000002 | 2.06552799 | 0.004165841 | 0.36725819 | 0.01134308 | 0.99094972 | NA | NA | NA |
| ENSG000002 | 2.13424573 | -0.511185888 | 0.36556029 | -1.3983628 | 0.16200416 | NA | NA | NA |
| ENSG000002 | 0.59892612 | -0.099249839 | 0.26277623 | -0.3776972 | 0.70565557 | NA | NA | NA |
| ENSG000002 | 0.39499294 | -0.277908454 | 0.21584201 | -1.287555 | 0.19790091 | NA | NA | NA |
| ENSG000002 | 1.23055485 | -0.310371496 | 0.32185313 | -0.9643265 | 0.33488226 | NA | NA | NA |
| ENSG000002 | 1.65315362 | 0.080210039 | 0.34178728 | 0.23467824 | 0.81445848 | NA | NA | NA |
| ENSG000002 | 0.78941944 | -0.365615932 | 0.28339326 | -1.2901363 | 0.19700334 | NA | NA | NA |
| ENSG000002 | 1.81734826 | -0.59195635 | 0.35055719 | -1.6886156 | 0.09129312 | NA | NA | NA |
| ENSG000002 | 0.14128499 | -0.097772014 | 0.13403235 | -0.7294658 | 0.4657168 | NA | NA | NA |
| ENSG000002 | 1.53579782 | -0.325741392 | 0.34120826 | -0.9546703 | 0.33974445 | NA | NA | NA |
| ENSG000002 | 1.69686658 | -0.700716181 | 0.34864461 | -2.0098294 | 0.04444925 | NA | NA | NA |
| ENSG000002 | 1.39426488 | -0.259992325 | 0.3361724 | -0.7733899 | 0.43929169 | NA | NA | NA |
| ENSG000002 | 1.47657476 | -0.151311706 | 0.32871146 | -0.4603177 | 0.64528819 | NA | NA | NA |
| ENSG000002 | 0.63159785 | -0.116736614 | 0.27184111 | -0.4294296 | 0.66761063 | NA | NA | NA |
| ENSG000002 | 1.5474148 | -0.486658732 | 0.34459775 | -1.4122516 | 0.15787589 | NA | NA | NA |
| ENSG000002 | 1.54069356 | -0.596574656 | 0.33661146 | -1.7722946 | 0.07634567 | NA | NA | NA |
| ENSG000002 | 1.65732937 | -0.258929557 | 0.34856703 | -0.7428401 | 0.45757851 | NA | NA | NA |
| ENSG000002 | 1.4493108 | -0.173150206 | 0.34023366 | -0.5089156 | 0.6108114 | NA | NA | NA |
| ENSG000002 | 0.54456342 | -0.359011254 | 0.24026449 | -1.4942335 | 0.1351146 | NA | NA | NA |
| ENSG000002 | 1.48003509 | 0.089039087 | 0.34241999 | 0.26002888 | 0.7948415 | NA | NA | NA |
| ENSG000002 | 1.01910778 | -0.344738314 | 0.30979765 | -1.1127854 | 0.2658006 | NA | NA | NA |
| ENSG000002 | 1.74391986 | -0.604626907 | 0.35264337 | -1.7145563 | 0.08642661 | NA | LINC01298 | 619344 |

| | | | | | | | | | |
|---|---|---|---|---|---|---|---|---|---|
| ENSG000002 | 1.58841967 | 0.017054742 | 0.35084315 | 0.04861073 | 0.96122952 | NA | NA | NA | |
| ENSG000002 | 1.72757735 | -0.062821537 | 0.35715656 | -0.1758936 | 0.86037757 | NA | NA | NA | |
| ENSG000002 | 0.25351207 | -0.189965547 | 0.1824784 | -1.0410303 | 0.29786148 | NA | NA | NA | |
| ENSG000002 | 0.87235133 | -0.106572909 | 0.29793873 | -0.3577008 | 0.72056727 | NA | NA | NA | |
| ENSG000002 | 0.46062538 | 0.331874243 | 0.23806547 | 1.39404611 | 0.16330368 | NA | NA | NA | |
| ENSG000002 | 1.15264808 | -0.42869132 | 0.31997664 | -1.3397582 | 0.18032397 | NA | NA | NA | |
| ENSG000002 | 2.11389093 | -0.387841733 | 0.36567297 | -1.0606246 | 0.28886055 | NA | NA | NA | |
| ENSG000002 | 1.01458206 | -0.189427026 | 0.30816255 | -0.6146984 | 0.53875392 | NA | NA | NA | |
| ENSG000002 | 0.2596576 | -0.193002035 | 0.1837913 | -1.0501152 | 0.29366517 | NA | NA | NA | |
| ENSG000002 | 1.27304544 | -0.350976478 | 0.33109305 | -1.0600539 | 0.28912008 | NA | NA | NA | |
| ENSG000002 | 0.74621284 | -0.194950297 | 0.28606354 | -0.681493 | 0.49555961 | NA | NA | NA | |
| ENSG000002 | 1.95205462 | 0.063390019 | 0.36078903 | 0.1756983 | 0.86053097 | NA | NA | NA | |
| ENSG000002 | 2.07481754 | -0.551573827 | 0.35310331 | -1.5620749 | 0.11827033 | NA | NA | NA | |
| ENSG000002 | 1.49190799 | -0.044034518 | 0.3390866 | -0.1298622 | 0.89667548 | NA | NA | NA | |
| ENSG000002 | 0.95947167 | 0.284029678 | 0.31210451 | 0.9100467 | 0.36279788 | NA | NA | NA | |
| ENSG000002 | 1.28408939 | -0.48103971 | 0.32524329 | -1.479015 | 0.1391363 | NA | NA | NA | |
| ENSG000002 | 0.47872867 | 0.139368284 | 0.22792065 | 0.61147721 | 0.5408837 | NA | NA | NA | |
| ENSG000002 | 0.95968308 | -0.159296336 | 0.29728489 | -0.5358373 | 0.592071 | NA | NA | NA | |
| ENSG000002 | 1.21324275 | -0.041701149 | 0.32806816 | -0.1271112 | 0.89885236 | NA | NA | NA | |
| ENSG000002 | 1.40519672 | -0.396104325 | 0.33297421 | -1.1895946 | 0.23420577 | NA | NA | NA | |
| ENSG000002 | 1.79546457 | -0.760694324 | 0.35278991 | -2.1562247 | 0.03106612 | NA | NA | NA | |
| ENSG000002 | 1.60869271 | -0.265166257 | 0.35298385 | -0.7512136 | 0.45252413 | NA | NA | NA | |
| ENSG000002 | 1.99901748 | -0.183721855 | 0.35819573 | -0.5129091 | 0.60801489 | NA | NA | NA | |
| ENSG000002 | 1.09799917 | -0.249575148 | 0.31659137 | -0.7883195 | 0.43050985 | NA | HSPA8P13 | | 106480803 |
| ENSG000002 | 0.50681912 | -0.019233989 | 0.24032284 | -0.080034 | 0.93621025 | NA | NA | NA | |
| ENSG000002 | 2.08574999 | -0.129061342 | 0.3629875 | -0.3555531 | 0.72217525 | NA | NCRNA00250 | | 552853 |
| ENSG000002 | 0.63389931 | -0.262485172 | 0.25907614 | -1.0131584 | 0.3109845 | NA | NA | NA | |
| ENSG000002 | 1.12538466 | -0.443269451 | 0.32457208 | -1.3657042 | 0.17203183 | NA | NA | NA | |
| ENSG000002 | 2.14263426 | -0.515733698 | 0.36474449 | -1.4139588 | 0.15737399 | NA | NA | NA | |

| | | | | | | | | |
|---|---|---|---|---|---|---|---|---|
| ENSG000002 | 0.22263036 | 0.146922267 | 0.16600367 | 0.88505432 | 0.37612734 | NA | NA | NA |
| ENSG000002 | 0.60352914 | 0.217649947 | 0.25523451 | 0.85274497 | 0.39380075 | NA | NA | NA |
| ENSG000002 | 0.7523063 | -0.206231678 | 0.2915808 | -0.7072883 | 0.47938735 | NA | LOC101929217 | 101929217 |
| ENSG000002 | 0.35073255 | 0.241740392 | 0.2078501 | 1.16305162 | 0.24480856 | NA | NA | NA |
| ENSG000002 | 1.83868593 | 0.134150752 | 0.35771906 | 0.37501706 | 0.70764778 | NA | NA | NA |
| ENSG000002 | 0.60447083 | -0.105840125 | 0.26325248 | -0.402048 | 0.68764873 | NA | NA | NA |
| ENSG000002 | 0 | NA | NA | NA | NA | NA | NA | NA |
| ENSG000002 | 0.8794426 | -0.118660638 | 0.30333647 | -0.3911849 | 0.69566059 | NA | NA | NA |
| ENSG000002 | 0.25370795 | -0.171331747 | 0.17411721 | -0.9840024 | 0.32511435 | NA | NA | NA |
| ENSG000002 | 1.8493891 | -0.002050275 | 0.35941621 | -0.0057045 | 0.99544852 | NA | NA | NA |
| ENSG000002 | 1.90680148 | -0.540612913 | 0.36066657 | -1.4989271 | 0.13389254 | NA | NA | NA |
| ENSG000002 | 0.769666 | -0.190137122 | 0.27989473 | -0.6793165 | 0.49693731 | NA | NA | NA |
| ENSG000002 | 0.36345889 | 0.070257596 | 0.21589421 | 0.32542602 | 0.74485866 | NA | NA | NA |
| ENSG000002 | 0.59278059 | -0.088567558 | 0.25301851 | -0.3500438 | 0.72630584 | NA | NA | NA |
| ENSG000002 | 0.50955585 | -0.376243729 | 0.24515812 | -1.5346982 | 0.12485797 | NA | NA | NA |
| ENSG000002 | 1.95861695 | 0.056050195 | 0.35883507 | 0.15620044 | 0.87587503 | NA | NA | NA |
| ENSG000002 | 0 | NA | NA | NA | NA | NA | NA | NA |
| ENSG000002 | 0.23202362 | -0.008302493 | 0.17586207 | -0.0472103 | 0.96234565 | NA | NA | NA |
| ENSG000002 | 1.33303124 | 0.299781778 | 0.32520664 | 0.92181936 | 0.35662281 | NA | NA | NA |
| ENSG000002 | 0.73975464 | -0.177073248 | 0.27958599 | -0.6333409 | 0.52651104 | NA | NA | NA |
| ENSG000002 | 0.24619405 | -0.007630671 | 0.17915639 | -0.0425922 | 0.96602658 | NA | NA | NA |
| ENSG000002 | 0.8640981 | -0.100670617 | 0.28973712 | -0.347455 | 0.7282495 | NA | NA | NA |
| ENSG000002 | 1.92120162 | -0.39976339 | 0.35957996 | -1.111751 | 0.26624521 | NA | NA | NA |
| ENSG000002 | 1.71328273 | -0.060739881 | 0.35423517 | -0.1714677 | 0.86385605 | NA | NA | NA |
| ENSG000002 | 0.37236288 | -0.09108266 | 0.20928647 | -0.4352057 | 0.66341314 | NA | NA | NA |
| ENSG000002 | 2.04028971 | -0.606568134 | 0.36341429 | -1.6690817 | 0.09510119 | NA | NA | NA |
| ENSG000002 | 0.94644127 | 0.120225999 | 0.31106787 | 0.38649443 | 0.69913052 | NA | NA | NA |
| ENSG000002 | 1.86085952 | 0.003733622 | 0.35310906 | 0.01057357 | 0.99156367 | NA | NA | NA |
| ENSG000002 | 0.50497064 | -0.183718113 | 0.24165995 | -0.760234 | 0.44711472 | NA | NA | NA |

| | | | | | | | | |
|---|---|---|---|---|---|---|---|---|
| ENSG000002 | 0.27408862 | -0.199826037 | 0.18669465 | -1.0703362 | 0.28446801 | NA | PYDC2 | 152138 |
| ENSG000002 | 1.24309386 | -0.318736308 | 0.32388685 | -0.9840977 | 0.32506747 | NA | NA | NA |
| ENSG000002 | 1.12947668 | -0.273940106 | 0.32209379 | -0.8504979 | 0.3950483 | NA | NA | NA |
| ENSG000002 | 1.14556031 | -0.125036239 | 0.32180579 | -0.3885457 | 0.69761228 | NA | NA | NA |
| ENSG000002 | 1.09751182 | -0.099344728 | 0.31301128 | -0.3173839 | 0.75095235 | NA | NA | NA |
| ENSG000002 | 1.06515019 | 0.332237401 | 0.3137724 | 1.0588484 | 0.28966883 | NA | SUMO2P16 | 106480679 |
| ENSG000002 | 1.33377988 | -0.236843651 | 0.3252906 | -0.7280987 | 0.46655319 | NA | NA | NA |
| ENSG000002 | 0.51570138 | -0.378721611 | 0.24581506 | -1.540677 | 0.12339542 | NA | NA | NA |
| ENSG000002 | 0.5074159 | -0.309594122 | 0.22571446 | -1.3716185 | 0.17018225 | NA | NA | NA |
| ENSG000002 | 2.06627512 | -0.593980011 | 0.36225088 | -1.6396924 | 0.10106914 | NA | LOC101927822 | 101927822 |
| ENSG000002 | 0.2596576 | -0.193002035 | 0.1837913 | -1.0501152 | 0.29366517 | NA | NA | NA |
| ENSG000002 | 0.98159319 | 0.111594572 | 0.30607873 | 0.36459434 | 0.71541424 | NA | RN7SL260P | 106480497 |
| ENSG000002 | 1.34942686 | -0.115098062 | 0.33730626 | -0.3412272 | 0.73293257 | NA | NA | NA |
| ENSG000002 | 0.27408862 | -0.199826037 | 0.18669465 | -1.0703362 | 0.28446801 | NA | NA | NA |
| ENSG000002 | 2.05413576 | 0.362064714 | 0.35768001 | 1.01225873 | 0.31141436 | NA | NA | NA |
| ENSG000002 | 0.11205239 | 0.082118251 | 0.12671962 | 0.64803108 | 0.51696485 | NA | WBP1LP3 | 106479031 |
| ENSG000002 | 0.38651158 | -0.274258716 | 0.21461355 | -1.2779189 | 0.20127801 | NA | NA | NA |
| ENSG000002 | 0 | NA | NA | NA | NA | NA | NA | NA |
| ENSG000002 | 1.53710827 | -0.329782528 | 0.34197503 | -0.9643468 | 0.33487208 | NA | NA | NA |
| ENSG000002 | 0.87115711 | -0.109553848 | 0.29624207 | -0.3698119 | 0.71152263 | NA | NA | NA |
| ENSG000002 | 1.32518049 | 0.174381916 | 0.33524594 | 0.52016116 | 0.60295125 | NA | NA | NA |
| ENSG000002 | 1.74081257 | -0.199923959 | 0.35650882 | -0.5607826 | 0.57494572 | NA | NA | NA |
| ENSG000002 | 0.99491686 | -0.347261079 | 0.30873726 | -1.1247786 | 0.26068285 | NA | NA | NA |
| ENSG000002 | 1.75200032 | -0.334336234 | 0.35608635 | -0.938919 | 0.34777234 | NA | NA | NA |
| ENSG000002 | 0.77288285 | -0.203654568 | 0.28771447 | -0.7078357 | 0.47904731 | NA | NA | NA |
| ENSG000002 | 0.53608206 | -0.386623811 | 0.24788931 | -1.5596631 | 0.11883952 | NA | NA | NA |
| ENSG000002 | 1.81904247 | 0.128486873 | 0.34941283 | 0.36772225 | 0.71308035 | NA | NA | NA |
| ENSG000002 | 1.40526998 | -0.270869185 | 0.33780808 | -0.8018434 | 0.42264354 | NA | NA | NA |
| ENSG000002 | 1.82605855 | -0.125343809 | 0.36123757 | -0.3469844 | 0.72860302 | NA | NA | NA |

| | | | | | | | | |
|---|---|---|---|---|---|---|---|---|
| ENSG000002 | 2.15087215 | -0.768033019 | 0.36208936 | -2.1211146 | 0.03391216 | NA | NA | NA |
| ENSG000002 | 1.59594463 | -0.254160264 | 0.34863193 | -0.7290218 | 0.46598836 | NA | NA | NA |
| ENSG000002 | 1.11032653 | 0.037115625 | 0.3180473 | 0.11669844 | 0.90709902 | NA | NA | NA |
| ENSG000002 | 1.14234793 | -0.432863572 | 0.32377541 | -1.3369254 | 0.18124699 | NA | NA | NA |
| ENSG000002 | 1.81288622 | -0.538783431 | 0.34239871 | -1.5735557 | 0.11559021 | NA | NA | NA |
| ENSG000002 | 0.62023988 | -0.112292793 | 0.27094277 | -0.4144521 | 0.67854306 | NA | NA | NA |
| ENSG000002 | 1.03080599 | -0.331313494 | 0.30498096 | -1.0863416 | 0.27732791 | NA | NA | NA |
| ENSG000002 | 1.00720847 | -0.360508653 | 0.31528677 | -1.143431 | 0.2528597 | NA | NA | NA |
| ENSG000002 | 2.10928143 | -0.252437657 | 0.3657198 | -0.6902488 | 0.49003774 | NA | NA | NA |
| ENSG000002 | 0.59892612 | -0.099249839 | 0.26277623 | -0.3776972 | 0.70565557 | NA | NA | NA |
| ENSG000002 | 1.47170771 | -0.059367275 | 0.34348378 | -0.1728387 | 0.86277825 | NA | NA | NA |
| ENSG000002 | 0.24141689 | -0.165257896 | 0.17127401 | -0.9648743 | 0.33460776 | NA | NA | NA |
| ENSG000002 | 1.38165869 | -0.398246096 | 0.33328237 | -1.194921 | 0.23211791 | NA | NA | NA |
| ENSG000002 | 0.65903596 | -0.265924846 | 0.26125257 | -1.0178841 | 0.30873302 | NA | NA | NA |
| ENSG000002 | 1.00313371 | -0.039373136 | 0.31344169 | -0.1256155 | 0.90003629 | NA | NA | NA |
| ENSG000002 | 2.13950033 | 0.065631308 | 0.3578237 | 0.183418 | 0.85447005 | NA | NA | NA |
| ENSG000002 | 0.49782676 | -0.186543761 | 0.24081583 | -0.7746325 | 0.43855686 | NA | NA | NA |
| ENSG000002 | 0.63484558 | -0.278698901 | 0.26767877 | -1.0411692 | 0.29779701 | NA | NA | NA |
| ENSG000002 | 1.14900391 | -0.263004617 | 0.31832567 | -0.8262124 | 0.40868362 | NA | NA | NA |
| ENSG000002 | 1.21147337 | -0.167509099 | 0.31448826 | -0.5326402 | 0.59428265 | NA | NA | NA |
| ENSG000002 | 0.52779657 | -0.383508725 | 0.24707466 | -1.5521977 | 0.12061492 | NA | NA | NA |
| ENSG000002 | 0.85131777 | -0.102855071 | 0.29519776 | -0.3484277 | 0.72751902 | NA | NA | NA |
| ENSG000002 | 0.91371137 | -0.266813888 | 0.29560643 | -0.9025984 | 0.36673908 | NA | NA | NA |
| ENSG000002 | 1.3354851 | -0.113305101 | 0.3390529 | -0.3341812 | 0.73824284 | NA | NA | NA |
| ENSG000002 | 0.60401594 | 0.058796518 | 0.26924067 | 0.21837904 | 0.8271338 | NA | NA | NA |
| ENSG000002 | 0.89589274 | -0.433945216 | 0.29547227 | -1.4686495 | 0.14192787 | NA | NA | NA |
| ENSG000002 | 0.37859486 | 0.064147216 | 0.20986618 | 0.30565771 | 0.75986527 | NA | NA | NA |
| ENSG000002 | 1.49687002 | -0.197363176 | 0.34498101 | -0.5720987 | 0.56725514 | NA | NA | NA |
| ENSG000002 | 1.99881756 | -0.445416571 | 0.36006984 | -1.2370283 | 0.21607658 | NA | NA | NA |

| | | | | | | | | | |
|---|---|---|---|---|---|---|---|---|---|
| ENSG000002 | 1.10440721 | -0.118910289 | 0.32584268 | -0.3649316 | 0.71516247 | NA | | CARSP2 | 106481707 |
| ENSG000002 | 0.8648157 | -0.114173272 | 0.30263075 | -0.3772692 | 0.70597354 | NA | | NA | NA |
| ENSG000002 | 1.91799086 | -0.058439661 | 0.36198067 | -0.1614441 | 0.8717436 | NA | | NA | NA |
| ENSG000002 | 1.23111683 | -0.195852128 | 0.32906913 | -0.5951702 | 0.55172973 | NA | | NA | NA |
| ENSG000002 | 1.59221113 | 0.136408486 | 0.34487298 | 0.39553254 | 0.69244991 | NA | | NA | NA |
| ENSG000002 | 0.23065526 | 0.167384503 | 0.17662711 | 0.94767164 | 0.34329664 | NA | | NA | NA |
| ENSG000002 | 1.28129654 | -0.512277922 | 0.33116903 | -1.5468775 | 0.12189278 | NA | | NA | NA |
| ENSG000002 | 1.37015636 | -0.415537848 | 0.33809401 | -1.2290601 | 0.21904929 | NA | | NA | NA |
| ENSG000002 | 0.5226185 | -0.189580046 | 0.24357686 | -0.7783172 | 0.43638207 | NA | | NA | NA |
| ENSG000002 | 2.00726859 | -0.317330779 | 0.36102534 | -0.8789709 | 0.37941704 | NA | | NA | NA |
| ENSG000002 | 1.07597381 | -0.684802076 | 0.30593765 | -2.2383714 | 0.02519685 | NA | | NA | NA |
| ENSG000002 | 0.99850035 | -0.195348626 | 0.31384733 | -0.622432 | 0.53365784 | NA | | NA | NA |
| ENSG000002 | 1.99773654 | -0.426332662 | 0.35958953 | -1.1856092 | 0.23577666 | NA | | NA | NA |
| ENSG000002 | 2.0857787 | -0.867112838 | 0.3587175 | -2.4172583 | 0.01563792 | NA | | NA | NA |
| ENSG000002 | 1.28659386 | -0.4878471 | 0.32686521 | -1.4925023 | 0.1355675 | NA | | NA | NA |
| ENSG000002 | 0.50204195 | -0.181972198 | 0.24131623 | -0.7540819 | 0.45080006 | NA | | NA | NA |
| ENSG000002 | 0 | NA | NA | NA | NA | NA | | NA | NA |
| ENSG000002 | 0.26813897 | -0.197007041 | 0.18550319 | -1.0620143 | 0.2882292 | NA | | NA | NA |
| ENSG000002 | 1.44708048 | -0.044625451 | 0.33827384 | -0.1319211 | 0.89504671 | NA | | NA | NA |
| ENSG000002 | 1.37484371 | -0.119800231 | 0.33651052 | -0.3560074 | 0.72183503 | NA | | NA | NA |
| ENSG000002 | 0.11131518 | 0.081951103 | 0.12663804 | 0.64712866 | 0.51754868 | NA | | NA | NA |
| ENSG000002 | 0.48573157 | -0.181932918 | 0.23941992 | -0.7598905 | 0.44732006 | NA | | NA | NA |
| ENSG000002 | 0.39479706 | -0.300008631 | 0.22317059 | -1.3443018 | 0.17885081 | NA | | NA | NA |
| ENSG000002 | 1.76620109 | -0.483121488 | 0.35686654 | -1.3537876 | 0.17580415 | NA | | NA | NA |
| ENSG000002 | 0.76582541 | -0.037952969 | 0.27125181 | -0.1399178 | 0.8887249 | NA | | NA | NA |
| ENSG000002 | 0.48436321 | -0.017262119 | 0.23773787 | -0.0726099 | 0.94211656 | NA | | NA | NA |
| ENSG000002 | 0 | NA | NA | NA | NA | NA | | WBP1LP10 | 106480326 |
| ENSG000002 | 1.5438525 | -0.431844366 | 0.33015491 | -1.3080053 | 0.19087151 | NA | | NA | NA |
| ENSG000002 | 1.09236409 | -0.109466652 | 0.32007612 | -0.3420019 | 0.73234943 | NA | | NA | NA |

| | | | | | | | | |
|---|---|---|---|---|---|---|---|---|
| ENSG000002 | 2.16876404 | -0.519414782 | 0.36590784 | -1.4195235 | 0.15574645 | NA | NA | NA |
| ENSG000002 | 1.81197028 | -0.753867153 | 0.35124951 | -2.146244 | 0.03185352 | NA | NA | NA |
| ENSG000002 | 1.89498595 | -0.491628164 | 0.35039489 | -1.4030689 | 0.1605963 | NA | NA | NA |
| ENSG000002 | 0.11865494 | 0.083440866 | 0.12751478 | 0.65436232 | 0.5128784 | NA | NA | NA |
| ENSG000002 | 1.59548762 | -0.259699106 | 0.34929957 | -0.7434853 | 0.4571879 | NA | NA | NA |
| ENSG000002 | 1.74118368 | -0.31183838 | 0.35047853 | -0.8897503 | 0.37359997 | NA | NA | NA |
| ENSG000002 | 1.20657762 | 0.113032855 | 0.33285006 | 0.33959091 | 0.73416462 | NA | NA | NA |
| ENSG000002 | 1.39566358 | -0.551758429 | 0.33305499 | -1.6566586 | 0.09758851 | NA | NA | NA |
| ENSG000002 | 0 | NA | NA | NA | NA | NA | NA | NA |
| ENSG000002 | 1.48477326 | -0.33853051 | 0.34662953 | -0.9766349 | 0.32874992 | NA | NA | NA |
| ENSG000002 | 1.13932887 | -0.255730233 | 0.31799754 | -0.8041893 | 0.42128763 | NA | NA | NA |
| ENSG000002 | 1.11192111 | -0.28143253 | 0.32668269 | -0.8614859 | 0.38897049 | NA | NA | NA |
| ENSG000002 | 1.64819904 | -0.558988467 | 0.35095727 | -1.5927537 | 0.11121547 | NA | NA | NA |
| ENSG000002 | 0 | NA | NA | NA | NA | NA | NA | NA |
| ENSG000002 | 1.15037228 | -0.42327205 | 0.31994132 | -1.3229678 | 0.1858461 | NA | LOC154761 | 154761 |
| ENSG000002 | 0.67756292 | -0.43472079 | 0.25931557 | -1.6764161 | 0.09365672 | NA | NA | NA |
| ENSG000002 | 0.24619405 | -0.007630671 | 0.17915639 | -0.0425922 | 0.96602658 | NA | NA | NA |
| ENSG000002 | 0.91371137 | -0.266813888 | 0.29560643 | -0.9025984 | 0.36673908 | NA | NA | NA |
| ENSG000002 | 0.9752174 | -0.035570667 | 0.31232158 | -0.1138912 | 0.90932407 | NA | NA | NA |
| ENSG000002 | 0.23816916 | -0.010712105 | 0.1772998 | -0.060418 | 0.9518227 | NA | NA | NA |
| ENSG000002 | 0.79807549 | -0.536675498 | 0.28113813 | -1.9089389 | 0.05626998 | NA | NA | NA |
| ENSG000002 | 1.56387371 | -0.224462349 | 0.34094861 | -0.6583466 | 0.51031545 | NA | LOC105379715 | 105379715 |
| ENSG000002 | 0.24485602 | -0.012655878 | 0.17881722 | -0.0707755 | 0.94357643 | NA | NA | NA |
| ENSG000002 | 1.74259496 | -0.167055388 | 0.34560682 | -0.4833683 | 0.62883424 | NA | NA | NA |
| ENSG000002 | 1.10776202 | 0.033583502 | 0.32180346 | 0.10436029 | 0.91688344 | NA | NA | NA |
| ENSG000002 | 1.99750923 | -0.328824703 | 0.36442476 | -0.9023116 | 0.36689135 | NA | NA | NA |
| ENSG000002 | 0.45333769 | 0.328904543 | 0.23717926 | 1.38673401 | 0.16552289 | NA | NA | NA |
| ENSG000002 | 0.90452313 | -0.427819001 | 0.29348532 | -1.4577186 | 0.14491813 | NA | NA | NA |
| ENSG000002 | 2.1420077 | -0.12257609 | 0.36325686 | -0.3374364 | 0.73578794 | NA | NA | NA |

| | | | | | | | | |
|---|---|---|---|---|---|---|---|---|
| ENSG000002 | 2.00707718 | -0.589117616 | 0.36275006 | -1.6240318 | 0.10436903 | NA | NA | NA |
| ENSG000002 | 0 | NA | NA | NA | NA | NA | NA | NA |
| ENSG000002 | 0.63416044 | -0.278936162 | 0.26761738 | -1.0422946 | 0.2972751 | NA | NA | NA |
| ENSG000002 | 1.53417959 | -0.340496138 | 0.34555296 | -0.985366 | 0.32444432 | NA | NA | NA |
| ENSG000002 | 2.04628947 | 0.013040606 | 0.36293223 | 0.03593124 | 0.97133718 | NA | NA | NA |
| ENSG000002 | 1.74860879 | -0.206337742 | 0.35854024 | -0.575494 | 0.56495728 | NA | NA | NA |
| ENSG000002 | 1.01315812 | -0.362204918 | 0.31550826 | -1.1480045 | 0.25096669 | NA | NA | NA |
| ENSG000002 | 1.01454768 | -0.348198677 | 0.30957817 | -1.124752 | 0.26069412 | NA | NA | NA |
| ENSG000002 | 1.51496928 | -0.633419352 | 0.34246211 | -1.8496042 | 0.06437062 | NA | NA | NA |
| ENSG000002 | 0.24756242 | -0.187034346 | 0.18119817 | -1.0322088 | 0.30197431 | NA | NA | NA |
| ENSG000002 | 1.28266546 | -0.677345408 | 0.33094555 | -2.0466975 | 0.0406878 | NA | NA | NA |
| ENSG000002 | 1.15768583 | -0.259348877 | 0.31155972 | -0.8324211 | 0.40517131 | NA | NA | NA |
| ENSG000002 | 2.01419933 | -0.714278267 | 0.35988325 | -1.98475 | 0.0471723 | NA | NA | NA |
| ENSG000002 | 0.75665733 | -0.185020633 | 0.27904382 | -0.6630523 | 0.50729708 | NA | NA | NA |
| ENSG000002 | 1.90460947 | -0.490695898 | 0.35077578 | -1.3988876 | 0.16184668 | NA | NA | NA |
| ENSG000002 | 0.60680666 | -0.099379886 | 0.25590479 | -0.3883471 | 0.69775918 | NA | NA | NA |
| ENSG000002 | 1.13081471 | -0.264183285 | 0.3198881 | -0.8258616 | 0.40888263 | NA | NA | NA |
| ENSG000002 | 1.8504131 | -0.122851106 | 0.35860536 | -0.3425802 | 0.73191428 | NA | NA | NA |
| ENSG000002 | 2.08244508 | -0.834003243 | 0.35499916 | -2.3493104 | 0.01880822 | NA | NA | NA |
| ENSG000002 | 1.03598406 | -0.528870525 | 0.31247702 | -1.6925101 | 0.09054875 | NA | NA | NA |
| ENSG000002 | 2.09487862 | -0.492944675 | 0.36176794 | -1.3625991 | 0.17300889 | NA | NA | NA |
| ENSG000002 | 0.26560726 | -0.177019426 | 0.17672274 | -1.0016788 | 0.31649875 | NA | NA | NA |
| ENSG000002 | 1.76277921 | -0.198970664 | 0.35568908 | -0.5593949 | 0.57589223 | NA | NA | NA |
| ENSG000002 | 1.55373605 | -0.435615937 | 0.3303523 | -1.3186405 | 0.18728932 | NA | NA | NA |
| ENSG000002 | 0.35682409 | 0.072008558 | 0.21485623 | 0.33514763 | 0.73751372 | NA | NA | NA |
| ENSG000002 | 1.3477241 | -0.096158232 | 0.31923226 | -0.3012171 | 0.76324892 | NA | NA | NA |
| ENSG000002 | 1.29663946 | -0.684576067 | 0.33120234 | -2.0669421 | 0.03873961 | NA | NA | NA |
| ENSG000002 | 0.63435632 | -0.27906164 | 0.26763939 | -1.0426778 | 0.29709757 | NA | NA | NA |
| ENSG000002 | 0.6530863 | -0.285158107 | 0.26914708 | -1.059488 | 0.28937758 | NA | NA | NA |

| | | | | | | | | |
|---|---|---|---|---|---|---|---|---|
| ENSG000002 | 0.90434845 | -0.270382456 | 0.29337366 | -0.9216317 | 0.35672074 | NA | NA | NA |
| ENSG000002 | 0.11131518 | 0.081951103 | 0.12663804 | 0.64712866 | 0.51754868 | NA | NA | NA |
| ENSG000002 | 0.12070844 | -0.088503695 | 0.12783931 | -0.6923042 | 0.48874631 | NA | NA | NA |
| ENSG000002 | 1.1811652 | -0.427414722 | 0.32095046 | -1.3317156 | 0.18295367 | NA | NA | NA |
| ENSG000002 | 1.89368676 | -0.684650993 | 0.359902 | -1.9023262 | 0.05712853 | NA | NA | NA |
| ENSG000002 | 2.08087036 | -0.129722753 | 0.36704391 | -0.3534257 | 0.72376932 | NA | NA | NA |
| ENSG000002 | 1.47286948 | -0.053887558 | 0.34238377 | -0.1573893 | 0.87493801 | NA | LOC105375787 | 105375787 |
| ENSG000002 | 1.98145999 | -0.321632242 | 0.3632439 | -0.8854443 | 0.37591705 | NA | NA | NA |
| ENSG000002 | 1.01773887 | -0.205648565 | 0.31456741 | -0.6537504 | 0.51327264 | NA | PKMP4 | 106480443 |
| ENSG000002 | 2.09376726 | -0.311005612 | 0.35301493 | -0.8809985 | 0.37831864 | NA | NA | NA |
| ENSG000002 | 1.84669356 | -0.231642586 | 0.35292323 | -0.656354 | 0.51159636 | NA | NA | NA |
| ENSG000002 | 0.66184999 | -0.106729201 | 0.25913995 | -0.4118593 | 0.68044255 | NA | NA | NA |
| ENSG000002 | 0.11131518 | 0.081951103 | 0.12663804 | 0.64712866 | 0.51754868 | NA | NA | NA |
| ENSG000002 | 2.13580998 | -0.639052185 | 0.36414742 | -1.7549271 | 0.07927177 | NA | NA | NA |
| ENSG000002 | 1.10372261 | -0.093943638 | 0.31018134 | -0.3028668 | 0.76199135 | NA | NA | NA |
| ENSG000002 | 0.86860367 | -0.259912622 | 0.29143534 | -0.8918363 | 0.37248067 | NA | NA | NA |
| ENSG000002 | 2.01917237 | -0.595301253 | 0.36332748 | -1.6384702 | 0.10132364 | NA | NA | NA |
| ENSG000002 | 1.73876298 | -0.320571753 | 0.35171036 | -0.9114652 | 0.36205032 | NA | LOC101927392 | 101927392 |
| ENSG000002 | 2.08144552 | -0.003856616 | 0.36278002 | -0.0106307 | 0.99151807 | NA | NA | NA |
| ENSG000002 | 0.36212533 | -0.235926492 | 0.20089643 | -1.1743688 | 0.24024735 | NA | NA | NA |
| ENSG000002 | 1.85081592 | -0.262399194 | 0.36173477 | -0.7253911 | 0.46821212 | NA | NA | NA |
| ENSG000002 | 1.53998723 | -0.787561413 | 0.34078278 | -2.3110364 | 0.02083084 | NA | NA | NA |
| ENSG000002 | 0.83720133 | 0.04865582 | 0.28956773 | 0.16802915 | 0.86656035 | NA | NA | NA |
| ENSG000002 | 0.73796118 | -0.036030065 | 0.2783958 | -0.1294203 | 0.89702509 | NA | NA | NA |
| ENSG000002 | 1.4191041 | -0.552958101 | 0.33360646 | -1.6575162 | 0.09741516 | NA | NA | NA |
| ENSG000002 | 1.219166 | -0.183603128 | 0.32876002 | -0.5584716 | 0.5765224 | NA | NA | NA |
| ENSG000002 | 0.37171 | -0.104406568 | 0.21753023 | -0.4799635 | 0.63125336 | NA | NA | NA |
| ENSG000002 | 0.71469997 | 0.309652873 | 0.28436427 | 1.08893033 | 0.27618461 | NA | NA | NA |
| ENSG000002 | 0.34197044 | 0.237837536 | 0.20637034 | 1.15247924 | 0.2491242 | NA | NA | NA |

| | | | | | | | | |
|---|---|---|---|---|---|---|---|---|
| ENSG000002 | 0.66797379 | -0.274731044 | 0.26194525 | -1.048811 | 0.29426513 | NA | NA | NA |
| ENSG000002 | 0.83477726 | 0.358756326 | 0.29046689 | 1.23510229 | 0.21679244 | NA | NA | NA |
| ENSG000002 | 1.60309091 | 0.002079429 | 0.3382837 | 0.006147 | 0.99509544 | NA | NA | NA |
| ENSG000002 | 1.58495667 | -0.382353449 | 0.3450756 | -1.1080281 | 0.26784969 | NA | NA | NA |
| ENSG000002 | 0.74094832 | -0.192779168 | 0.28573214 | -0.6746849 | 0.49987596 | NA | NA | NA |
| ENSG000002 | 1.83757858 | -0.249982692 | 0.35724694 | -0.6997476 | 0.48408494 | NA | NA | NA |
| ENSG000002 | 0 | NA | NA | NA | NA | NA | NA | NA |
| ENSG000002 | 1.28758251 | -0.190542918 | 0.3238309 | -0.5884025 | 0.55626215 | NA | NA | NA |
| ENSG000002 | 2.06438933 | 0.244552217 | 0.36379679 | 0.67222204 | 0.50144235 | NA | NA | NA |
| ENSG000002 | 1.62434105 | 0.012009392 | 0.34828287 | 0.03448172 | 0.97249302 | NA | NA | NA |
| ENSG000002 | 0.73039619 | -0.459565217 | 0.26470411 | -1.7361469 | 0.08253787 | NA | NA | NA |
| ENSG000002 | 1.39995203 | -0.418959555 | 0.34006434 | -1.2320008 | 0.21794877 | NA | CTB-7E3.1 | 102557615 |
| ENSG000002 | 0.47092999 | -0.01913341 | 0.24378131 | -0.078486 | 0.9374415 | NA | NA | NA |
| ENSG000002 | 0.75207604 | -0.35881839 | 0.28263394 | -1.2695516 | 0.20424438 | NA | NA | NA |
| ENSG000002 | 1.60436128 | -0.254938379 | 0.34993711 | -0.7285263 | 0.46629149 | NA | NA | NA |
| ENSG000002 | 0 | NA | NA | NA | NA | NA | NA | NA |
| ENSG000002 | 0.83575724 | -0.109698178 | 0.29597774 | -0.3706298 | 0.71091327 | NA | NA | NA |
| ENSG000002 | 0.4009426 | -0.302672773 | 0.2240145 | -1.3511303 | 0.17665371 | NA | NA | NA |
| ENSG000002 | 0.74640871 | -0.195085107 | 0.28607993 | -0.6819252 | 0.49528626 | NA | NA | NA |
| ENSG000002 | 1.51238708 | -0.466894439 | 0.33931448 | -1.3759933 | 0.16882372 | NA | NA | NA |
| ENSG000002 | 0.50107449 | -0.372905725 | 0.24426561 | -1.5266403 | 0.12685049 | NA | NA | NA |
| ENSG000002 | 1.78026546 | -0.615192137 | 0.35416326 | -1.7370298 | 0.08238191 | NA | NA | NA |
| ENSG000002 | 0.50474388 | -0.025682234 | 0.2475253 | -0.103756 | 0.91736299 | NA | NA | NA |
| ENSG000002 | 0.87101522 | 0.044260014 | 0.29588867 | 0.14958333 | 0.88109336 | NA | NA | NA |
| ENSG000002 | 1.63783886 | -0.553112649 | 0.35085618 | -1.5764655 | 0.11491857 | NA | NA | NA |
| ENSG000002 | 0.37304803 | -0.090842893 | 0.20939682 | -0.4338313 | 0.66441096 | NA | NA | NA |
| ENSG000002 | 0 | NA | NA | NA | NA | NA | NA | NA |
| ENSG000002 | 0.71433332 | 0.130507939 | 0.27701556 | 0.47112132 | 0.6375541 | NA | NA | NA |
| ENSG000002 | 1.28627928 | -0.664965104 | 0.32894475 | -2.02151 | 0.043227 | NA | NA | NA |

| | | | | | | | | |
|---|---|---|---|---|---|---|---|---|
| ENSG000002 | 0.2596576 | -0.193002035 | 0.1837913 | -1.0501152 | 0.29366517 | NA | NA | NA |
| ENSG000002 | 1.68212775 | 0.082988992 | 0.34133241 | 0.24313247 | 0.80790278 | NA | NA | NA |
| ENSG000002 | 1.31818348 | -0.500910807 | 0.32881795 | -1.5233682 | 0.12766661 | NA | NA | NA |
| ENSG000002 | 1.02198385 | -0.332149583 | 0.30195792 | -1.0999863 | 0.27133808 | NA | NA | NA |
| ENSG000002 | 1.91160899 | -0.529955184 | 0.35820697 | -1.4794664 | 0.1390157 | NA | NA | NA |
| ENSG000002 | 1.38137244 | -0.259847714 | 0.33512312 | -0.7753799 | 0.43811524 | NA | NA | NA |
| ENSG000002 | 0.47616417 | 0.157689149 | 0.24487722 | 0.64395189 | 0.51960663 | NA | NA | NA |
| ENSG000002 | 0.48555743 | -0.013775713 | 0.2454629 | -0.0561214 | 0.95524512 | NA | NA | NA |
| ENSG000002 | 1.57858042 | -0.445199053 | 0.3352733 | -1.3278691 | 0.18422135 | NA | NA | NA |
| ENSG000002 | 1.69180262 | -0.67640394 | 0.34395248 | -1.9665622 | 0.0492337 | NA | NA | NA |
| ENSG000002 | 0.35022155 | 0.06940789 | 0.21379411 | 0.32464827 | 0.74544729 | NA | NA | NA |
| ENSG000002 | 1.1491786 | -0.424387989 | 0.32149757 | -1.3200348 | 0.18682339 | NA | NA | NA |
| ENSG000002 | 1.20111668 | 0.103094292 | 0.32916118 | 0.31320307 | 0.75412639 | NA | NA | NA |
| ENSG000002 | 1.53839585 | 0.271990586 | 0.34028385 | 0.799305 | 0.42411358 | NA | NA | NA |
| ENSG000002 | 0.62382145 | -0.09615089 | 0.25748875 | -0.3734178 | 0.70883749 | NA | NA | NA |
| ENSG000002 | 1.22895781 | -0.18029443 | 0.32473834 | -0.5551991 | 0.57875849 | NA | NA | NA |
| ENSG000002 | 0.12685397 | -0.090215768 | 0.12901919 | -0.699243 | 0.48440017 | NA | NA | NA |
| ENSG000002 | 0.81881783 | 0.196008695 | 0.28746236 | 0.68185863 | 0.49532834 | NA | NA | NA |
| ENSG000002 | 1.65377868 | -0.393539091 | 0.34812508 | -1.1304532 | 0.25828533 | NA | NA | NA |
| ENSG000002 | 2.1056621 | -0.122698439 | 0.36595838 | -0.3352798 | 0.73741406 | NA | NA | NA |
| ENSG000002 | 0.84338124 | 0.205156406 | 0.29492171 | 0.69563006 | 0.48666054 | NA | NA | NA |
| ENSG000002 | 1.88391379 | -0.122811032 | 0.3527457 | -0.3481574 | 0.72772196 | NA | NA | NA |
| ENSG000002 | 1.87990663 | -0.244150831 | 0.3551747 | -0.6874105 | 0.49182406 | NA | NA | NA |
| ENSG000002 | 1.98293229 | -0.424134557 | 0.35659789 | -1.1893917 | 0.23428558 | NA | NA | NA |
| ENSG000002 | 1.62051004 | -0.652811972 | 0.34300808 | -1.9031971 | 0.05701484 | NA | NA | NA |
| ENSG000002 | 2.14409546 | -0.64046275 | 0.36419075 | -1.7585915 | 0.07864692 | NA | NA | NA |
| ENSG000002 | 0.36576034 | -0.0934779 | 0.20820828 | -0.4489634 | 0.65345805 | NA | SNRPCP17 | 106481702 |
| ENSG000002 | 0.74438746 | -0.033988507 | 0.28512681 | -0.1192049 | 0.90511304 | NA | NA | NA |
| ENSG000002 | 0.98227641 | -0.039913906 | 0.31675261 | -0.1260097 | 0.89972423 | NA | NA | NA |

| | | | | | | | | | |
|---|---|---|---|---|---|---|---|---|---|
| ENSG000002 | 0.97772792 | -0.168848749 | 0.30077603 | -0.561377 | 0.57454055 | NA | NA | NA | |
| ENSG000002 | 2.03302534 | -0.310542574 | 0.35654923 | -0.8709669 | 0.38377224 | NA | NA | NA | |
| ENSG000002 | 1.37871344 | -0.107768106 | 0.32847956 | -0.3280816 | 0.74284995 | NA | NA | NA | |
| ENSG000002 | 0.48216589 | 0.145317976 | 0.23824352 | 0.60995564 | 0.54189119 | NA | NA | NA | |
| ENSG000002 | 0.24141689 | -0.165257896 | 0.17127401 | -0.9648743 | 0.33460776 | NA | NA | NA | |
| ENSG000002 | 0.25214371 | -0.009862892 | 0.18046657 | -0.0546522 | 0.95641556 | NA | NA | NA | |
| ENSG000002 | 2.09379504 | -0.118206944 | 0.36301013 | -0.3256299 | 0.74470441 | NA | NA | NA | |
| ENSG000002 | 0 | NA | NA | NA | NA | NA | NA | NA | |
| ENSG000002 | 1.90433144 | 0.069760681 | 0.35779046 | 0.19497636 | 0.84541146 | NA | NA | NA | |
| ENSG000002 | 1.61856609 | -0.11663296 | 0.3496376 | -0.3335824 | 0.73869468 | NA | MAFA-AS1 | | 104326051 |
| ENSG000002 | 0 | NA | NA | NA | NA | NA | SNORD87 | | 641648 |
| ENSG000002 | 0.59687261 | 0.065989748 | 0.26864914 | 0.24563543 | 0.80596446 | NA | NA | NA | |
| ENSG000002 | 0.11865494 | 0.083440866 | 0.12751478 | 0.65436232 | 0.5128784 | NA | NA | NA | |
| ENSG000002 | 0.37171 | -0.104406568 | 0.21753023 | -0.4799635 | 0.63125336 | NA | NA | NA | |
| ENSG000002 | 1.87159081 | -0.105694259 | 0.34851085 | -0.3032739 | 0.76168111 | NA | NA | NA | |
| ENSG000002 | 1.64256451 | -0.390543886 | 0.34797538 | -1.1223319 | 0.26172133 | NA | NA | NA | |
| ENSG000002 | 0.98868253 | -0.043402631 | 0.3063604 | -0.1416718 | 0.88733925 | NA | NA | NA | |
| ENSG000002 | 1.55342449 | -0.420198006 | 0.33034081 | -1.2720136 | 0.20336828 | NA | NA | NA | |
| ENSG000002 | 2.07149072 | -0.445964549 | 0.35462342 | -1.2575722 | 0.20854652 | NA | NA | NA | |
| ENSG000002 | 0.92315828 | -0.245138419 | 0.28498236 | -0.860188 | 0.3896854 | NA | NA | NA | |
| ENSG000002 | 1.37117644 | -0.250427004 | 0.33569274 | -0.7460007 | 0.45566702 | NA | NA | NA | |
| ENSG000002 | 1.75993932 | -0.413059767 | 0.34518933 | -1.196618 | 0.23145548 | NA | NA | NA | |
| ENSG000002 | 1.20703463 | 0.112249722 | 0.32780597 | 0.34242732 | 0.73202933 | NA | NA | NA | |
| ENSG000002 | 0.84836343 | 0.046896239 | 0.29631143 | 0.15826672 | 0.87424663 | NA | NA | NA | |
| ENSG000002 | 0.64833032 | -0.271392816 | 0.26190928 | -1.0362092 | 0.30010454 | NA | NA | NA | |
| ENSG000002 | 1.56179183 | -0.646004053 | 0.34416833 | -1.8770003 | 0.06051805 | NA | NA | NA | |
| ENSG000002 | 1.83701159 | -0.130668221 | 0.36044675 | -0.3625174 | 0.71696542 | NA | NA | NA | |
| ENSG000002 | 1.74943828 | -0.182840275 | 0.35114501 | -0.5206973 | 0.60257763 | NA | NA | NA | |
| ENSG000002 | 0.72258443 | -0.025922983 | 0.27568004 | -0.0940329 | 0.92508305 | NA | NA | NA | |

| | | | | | | | | | |
|---|---|---|---|---|---|---|---|---|---|
| ENSG000002 | 1.93115957 | 0.198761164 | 0.36338262 | 0.54697488 | 0.58439599 | NA | NA | NA | |
| ENSG000002 | 0.13280363 | -0.093394593 | 0.13115644 | -0.7120855 | 0.47641185 | NA | NA | NA | |
| ENSG000002 | 1.1393804 | -0.265120033 | 0.31799676 | -0.8337193 | 0.40443919 | NA | NA | NA | |
| ENSG000002 | 1.21696814 | -0.046967871 | 0.32817589 | -0.143118 | 0.88619701 | NA | NA | NA | |
| ENSG000002 | 1.87774357 | -0.521862277 | 0.35840595 | -1.4560648 | 0.14537471 | NA | NA | NA | |
| ENSG000002 | 0.6639035 | -0.267460023 | 0.26165968 | -1.0221675 | 0.30670162 | NA | NA | NA | |
| ENSG000002 | 1.15712331 | -0.40363464 | 0.31275461 | -1.2905794 | 0.19684958 | NA | NA | NA | |
| ENSG000002 | 0.11131518 | 0.081951103 | 0.12663804 | 0.64712866 | 0.51754868 | NA | NA | NA | |
| ENSG000002 | 1.19713229 | 0.24256158 | 0.32210783 | 0.75304465 | 0.45142309 | NA | NA | NA | |
| ENSG000002 | 0.63296622 | -0.299126688 | 0.27328461 | -1.0945611 | 0.27370899 | NA | NA | NA | |
| ENSG000002 | 0.88558813 | -0.11555507 | 0.29860532 | -0.3869826 | 0.69876907 | NA | NA | NA | |
| ENSG000002 | 0.89954094 | -0.281346535 | 0.30044017 | -0.9364478 | 0.34904268 | NA | NA | NA | |
| ENSG000002 | 0 | NA | NA | NA | NA | NA | TMX2-CTNND1 | | 100528016 |
| ENSG000002 | 0.71484432 | 0.299199048 | 0.27698049 | 1.08021704 | 0.28004554 | NA | NA | NA | |
| ENSG000002 | 0.96972421 | -0.038864349 | 0.31209406 | -0.1245277 | 0.90089749 | NA | NA | NA | |
| ENSG000002 | 1.09959567 | -0.389769313 | 0.31151695 | -1.2511978 | 0.21086233 | NA | NA | NA | |
| ENSG000002 | 0.63893761 | -0.11408674 | 0.27243711 | -0.4187636 | 0.67538892 | NA | NA | NA | |
| ENSG000002 | 1.5486522 | 0.297676444 | 0.34753002 | 0.85654887 | 0.39169425 | NA | NA | NA | |
| ENSG000002 | 1.00220454 | -0.34457114 | 0.30907251 | -1.1148554 | 0.26491241 | NA | NA | NA | |
| ENSG000002 | 0 | NA | NA | NA | NA | NA | NA | NA | |
| ENSG000002 | 0.74538918 | -0.35201633 | 0.2805873 | -1.2545697 | 0.209635 | NA | NA | NA | |
| ENSG000002 | 0.88163641 | -0.585934749 | 0.28999956 | -2.0204677 | 0.04333489 | NA | NA | NA | |
| ENSG000002 | 1.36796489 | -0.408585816 | 0.33668581 | -1.2135522 | 0.22491876 | NA | NA | NA | |
| ENSG000002 | 1.25092638 | -0.045145042 | 0.33276742 | -0.1356655 | 0.89208576 | NA | NA | NA | |
| ENSG000002 | 1.97715944 | -0.452580347 | 0.36350353 | -1.2450508 | 0.21311309 | NA | NA | NA | |
| ENSG000002 | 0 | NA | NA | NA | NA | NA | NA | NA | |
| ENSG000002 | 2.09570522 | -0.126224744 | 0.35916072 | -0.3514436 | 0.72525557 | NA | LOC105376633 | | 105376633 |
| ENSG000002 | 1.59671295 | -0.764873937 | 0.33662009 | -2.2722172 | 0.0230734 | NA | NA | NA | |
| ENSG000002 | 0.24756242 | -0.187034346 | 0.18119817 | -1.0322088 | 0.30197431 | NA | NA | NA | |

| | | | | | | | | | |
|---|---|---|---|---|---|---|---|---|---|
| ENSG000002 | 0.85066701 | 0.050294646 | 0.2964275 | 0.1696693 | 0.86527022 | NA | NA | NA | |
| ENSG000002 | 1.13972978 | -0.566869226 | 0.31589767 | -1.7944711 | 0.07273801 | NA | NA | NA | |
| ENSG000002 | 0.81036655 | -0.540716938 | 0.28190939 | -1.9180522 | 0.05510439 | NA | NA | NA | |
| ENSG000002 | 0.5681808 | 0.209349632 | 0.25180656 | 0.83139071 | 0.40575295 | NA | NA | NA | |
| ENSG000002 | 0.63188018 | 0.051086839 | 0.26523141 | 0.19261232 | 0.84726259 | NA | NA | NA | |
| ENSG000002 | 1.38992761 | -0.121500207 | 0.33824689 | -0.3592057 | 0.71944122 | NA | NA | NA | |
| ENSG000002 | 1.17934897 | -0.562521107 | 0.31338762 | -1.7949691 | 0.07265862 | NA | NA | NA | |
| ENSG000002 | 1.0999658 | 0.032769198 | 0.30969282 | 0.10581194 | 0.91573156 | NA | NA | NA | |
| ENSG000002 | 1.97982575 | -0.054108184 | 0.35657998 | -0.1517421 | 0.87939037 | NA | NA | NA | |
| ENSG000002 | 0.49833776 | -0.016406893 | 0.23937171 | -0.0685415 | 0.9453546 | NA | NA | NA | |
| ENSG000002 | 1.85368147 | -0.113753549 | 0.35046312 | -0.3245807 | 0.74549847 | NA | OMP | | 4975 |
| ENSG000002 | 1.99273207 | -0.462349603 | 0.36406952 | -1.2699487 | 0.20410292 | NA | NA | NA | |
| ENSG000002 | 0.34270765 | 0.258172734 | 0.21433185 | 1.20454674 | 0.22837833 | NA | NA | NA | |
| ENSG000002 | 1.65244402 | -0.695140292 | 0.34721417 | -2.0020505 | 0.04527931 | NA | NA | NA | |
| ENSG000002 | 1.28895088 | -0.34715385 | 0.32929681 | -1.0542278 | 0.29177863 | NA | NA | NA | |
| ENSG000002 | 1.74949895 | -0.463860613 | 0.3527404 | -1.3150198 | 0.18850325 | NA | NA | NA | |
| ENSG000002 | 0.49101782 | -0.013272127 | 0.22978098 | -0.0577599 | 0.95393988 | NA | NA | NA | |
| ENSG000002 | 1.99671701 | -0.573172447 | 0.3602703 | -1.5909511 | 0.11162057 | NA | NA | NA | |
| ENSG000002 | 1.48485916 | -0.191819348 | 0.34164686 | -0.561455 | 0.57448738 | NA | NA | NA | |
| ENSG000002 | 0.8747492 | -0.246588133 | 0.28492303 | -0.8654552 | 0.38678897 | NA | NA | NA | |
| ENSG000002 | 1.7481028 | -0.590305176 | 0.35165101 | -1.6786677 | 0.09321683 | NA | NA | NA | |
| ENSG000002 | 0.4855357 | -0.181797071 | 0.23938927 | -0.7594203 | 0.44760118 | NA | NA | NA | |
| ENSG000002 | 0.22997012 | 0.167051788 | 0.17646558 | 0.94665368 | 0.34381528 | NA | NA | NA | |
| ENSG000002 | 1.99406563 | -0.200668915 | 0.36454692 | -0.5504611 | 0.58200312 | NA | NA | NA | |
| ENSG000002 | 0.49765262 | -0.018104365 | 0.24679454 | -0.073358 | 0.9415212 | NA | NA | NA | |
| ENSG000002 | 0.76052707 | -0.169513888 | 0.27209919 | -0.6229856 | 0.53329396 | NA | NA | NA | |
| ENSG000002 | 2.15396458 | -0.237860689 | 0.35921976 | -0.6621593 | 0.50786916 | NA | NA | NA | |
| ENSG000002 | 1.47557642 | -0.176840367 | 0.34147835 | -0.517867 | 0.60455107 | NA | NA | NA | |
| ENSG000002 | 0 | NA | NA | NA | NA | NA | NA | NA | |

| | | | | | | | | |
|---|---|---|---|---|---|---|---|---|
| ENSG000002 | 1.65161699 | -0.566026567 | 0.35204623 | -1.6078188 | 0.10787486 | NA | NA | NA |
| ENSG000002 | 0.12685397 | -0.090215768 | 0.12901919 | -0.699243 | 0.48440017 | NA | NA | NA |
| ENSG000002 | 1.86285904 | -0.259680129 | 0.36006403 | -0.7212054 | 0.47078314 | NA | NA | NA |
| ENSG000002 | 1.72145198 | -0.051654144 | 0.35600615 | -0.1450934 | 0.88463713 | NA | NA | NA |
| ENSG000002 | 0 | NA | NA | NA | NA | NA | NA | NA |
| ENSG000002 | 1.40040637 | -0.560188263 | 0.33461142 | -1.6741457 | 0.09410197 | NA | NA | NA |
| ENSG000002 | 0 | NA | NA | NA | NA | NA | NA | NA |
| ENSG000002 | 0 | NA | NA | NA | NA | NA | NA | NA |
| ENSG000002 | 0.85551123 | -0.243943637 | 0.28822537 | -0.8463642 | 0.39734961 | NA | NA | NA |
| ENSG000002 | 0.48279897 | 0.145542372 | 0.23831717 | 0.6107087 | 0.54139244 | NA | INS | 3630 |
| ENSG000002 | 0.14128499 | -0.097772014 | 0.13403235 | -0.7294658 | 0.4657168 | NA | NA | NA |
| ENSG000002 | 0.60373362 | -0.112425234 | 0.26959077 | -0.4170218 | 0.67666245 | NA | NA | NA |
| ENSG000002 | 1.01491824 | -0.477517023 | 0.30159351 | -1.5833133 | 0.11335007 | NA | NA | NA |
| ENSG000002 | 1.3727426 | -0.250081818 | 0.33572739 | -0.7448955 | 0.4563349 | NA | NA | NA |
| ENSG000002 | 0 | NA | NA | NA | NA | NA | NA | NA |
| ENSG000002 | 1.768192 | -0.444008514 | 0.35063858 | -1.2662854 | 0.20541093 | NA | NA | NA |
| ENSG000002 | 1.45599767 | -0.18049182 | 0.34360175 | -0.5252937 | 0.59937906 | NA | NA | NA |
| ENSG000002 | 0.72540953 | -0.028108977 | 0.28387695 | -0.0990182 | 0.92112384 | NA | NA | NA |
| ENSG000002 | 1.40812933 | -0.746887747 | 0.33737747 | -2.2138045 | 0.02684223 | NA | NA | NA |
| ENSG000002 | 1.72661878 | -1.022466622 | 0.3438222 | -2.9738238 | 0.00294114 | NA | NA | NA |
| ENSG000002 | 0 | NA | NA | NA | NA | NA | NA | NA |
| ENSG000002 | 0 | NA | NA | NA | NA | NA | NA | NA |
| ENSG000002 | 0.60029448 | -0.251020599 | 0.2560415 | -0.9803903 | 0.3268935 | NA | NA | NA |
| ENSG000002 | 1.4970952 | -0.46580079 | 0.34174659 | -1.3630006 | 0.17288233 | NA | NA | NA |
| ENSG000002 | 0.11934008 | 0.083564698 | 0.12760091 | 0.65489106 | 0.5125379 | NA | NA | NA |
| ENSG000002 | 1.47400111 | 0.086711084 | 0.33300671 | 0.2603884 | 0.79456419 | NA | NA | NA |
| ENSG000002 | 0 | NA | NA | NA | NA | NA | NA | NA |
| ENSG000002 | 0.62160824 | -0.2792447 | 0.26652831 | -1.0477112 | 0.29477168 | NA | IGLL5 | 100423062 |
| ENSG000002 | 0 | NA | NA | NA | NA | NA | NA | NA |

| | | | | | | | | | |
|---|---|---|---|---|---|---|---|---|---|
| ENSG000002 | 1.63233532 | 0.658099484 | 0.35027367 | 1.87881517 | 0.06026973 | NA | NA | NA | |
| ENSG000002 | 1.55733996 | 0.153702177 | 0.35109057 | 0.43778498 | 0.66154216 | NA | NA | NA | |
| ENSG000002 | 2.06800359 | -0.24374219 | 0.36359611 | -0.6703652 | 0.50262499 | NA | NA | NA | |
| ENSG000002 | 0.11205239 | 0.082118251 | 0.12671962 | 0.64803108 | 0.51696485 | NA | NA | NA | |
| ENSG000002 | 0.24004852 | -0.005223787 | 0.17775844 | -0.029387 | 0.97655594 | NA | NA | NA | |
| ENSG000002 | 2.08864484 | -0.244632765 | 0.36333814 | -0.6732923 | 0.50076136 | NA | NA | NA | |
| ENSG000002 | 1.99383941 | -0.064358116 | 0.36115574 | -0.1782004 | 0.85856556 | NA | HMGB1P42 | | 100873906 |
| ENSG000002 | 1.77940405 | -0.599880557 | 0.35197379 | -1.704333 | 0.08831889 | NA | NA | NA | |
| ENSG000002 | 0 | NA | NA | NA | NA | NA | NA | NA | |
| ENSG000002 | 1.99461089 | -0.438059366 | 0.35891752 | -1.2205015 | 0.22227482 | NA | NA | NA | |
| ENSG000002 | 0.88419803 | -0.119996672 | 0.29852609 | -0.4019638 | 0.6877107 | NA | NA | NA | |
| ENSG000002 | 1.74818266 | -0.331524626 | 0.35436218 | -0.9355531 | 0.34950334 | NA | NA | NA | |
| ENSG000002 | 1.25143492 | -0.049944267 | 0.33278258 | -0.1500808 | 0.88070089 | NA | NA | NA | |
| ENSG000002 | 1.20495728 | -0.014378109 | 0.19611902 | -0.0733132 | 0.9415569 | NA | NA | NA | |
| ENSG000002 | 2.04922204 | -0.417104448 | 0.35348625 | -1.1799736 | 0.23801071 | NA | NA | NA | |
| ENSG000002 | 0.47761686 | -0.01948631 | 0.23692179 | -0.0822479 | 0.93444962 | NA | NA | NA | |
| ENSG000002 | 0 | NA | NA | NA | NA | NA | NA | NA | |
| ENSG000002 | 1.59659944 | -0.113601521 | 0.34812833 | -0.3263208 | 0.74418164 | NA | NA | NA | |
| ENSG000002 | 1.31033327 | -0.67503004 | 0.32944573 | -2.0489871 | 0.04046338 | NA | NA | NA | |
| ENSG000002 | 0 | NA | NA | NA | NA | NA | EEF1G | | 1937 |
| ENSG000002 | 0.47895489 | -0.016192723 | 0.24471014 | -0.066171 | 0.94724166 | NA | NA | NA | |
| ENSG000002 | 1.16360962 | -0.422169876 | 0.32040352 | -1.3176193 | 0.18763111 | NA | NA | NA | |
| ENSG000002 | 0 | NA | NA | NA | NA | NA | NA | NA | |
| ENSG000002 | 1.25150142 | -0.494905107 | 0.32817629 | -1.5080465 | 0.13154263 | NA | NA | NA | |
| ENSG000002 | 0 | NA | NA | NA | NA | NA | CKLF-CMTM1 | | 100529251 |
| ENSG000002 | 1.21094522 | -0.182249059 | 0.32623275 | -0.5586473 | 0.57640242 | NA | NA | NA | |
| ENSG000002 | 1.11195199 | -0.379564089 | 0.30936568 | -1.2269108 | 0.21985613 | NA | NA | NA | |
| ENSG000002 | 0 | NA | NA | NA | NA | NA | NA | NA | |
| ENSG000002 | 1.0128758 | -0.5006805 | 0.30693294 | -1.6312374 | 0.10284023 | NA | NA | NA | |

| | | | | | | | | |
|---|---|---|---|---|---|---|---|---|
| ENSG000002 | 1.61535316 | -0.381707575 | 0.34687676 | -1.1004127 | 0.27115236 | NA | NA | NA |
| ENSG000002 | 0.25260017 | -0.015978705 | 0.18049464 | -0.0885273 | 0.92945758 | NA | NA | NA |
| ENSG000002 | 0.14128499 | -0.097772014 | 0.13403235 | -0.7294658 | 0.4657168 | NA | NA | NA |
| ENSG000002 | 0.3854038 | -0.110128691 | 0.21952326 | -0.5016721 | 0.61589821 | NA | NA | NA |
| ENSG000002 | 0.12070844 | -0.088503695 | 0.12783931 | -0.6923042 | 0.48874631 | NA | NA | NA |
| ENSG000002 | 0.25214371 | -0.009862892 | 0.18046657 | -0.0546522 | 0.95641556 | NA | SYS1-DBNDD2 | 767557 |
| ENSG000002 | 0.49833776 | -0.016406893 | 0.23937171 | -0.0685415 | 0.9453546 | NA | NA | NA |
| ENSG000002 | 1.72782372 | -0.186922361 | 0.3535915 | -0.5286393 | 0.59705569 | NA | NA | NA |
| ENSG000002 | 0.78384035 | -0.532082522 | 0.28025057 | -1.8985957 | 0.05761765 | NA | NA | NA |
| ENSG000002 | 0.41537361 | -0.286583533 | 0.21871501 | -1.3103057 | 0.19009243 | NA | NA | NA |
| ENSG000002 | 0.27408862 | -0.199826037 | 0.18669465 | -1.0703362 | 0.28446801 | NA | NA | NA |
| ENSG000002 | 2.17255256 | -0.617931014 | 0.36035002 | -1.7148078 | 0.08638048 | NA | NA | NA |
| ENSG000002 | 2.16489042 | -0.048090472 | 0.36473661 | -0.1318499 | 0.89510305 | NA | NA | NA |
| ENSG000002 | 1.56457557 | 0.152696675 | 0.34937744 | 0.43705362 | 0.66207246 | NA | NA | NA |
| ENSG000002 | 1.56065023 | -0.622740106 | 0.34124714 | -1.8248947 | 0.06801692 | NA | NA | NA |
| ENSG000002 | 1.84026324 | -0.485564225 | 0.35071344 | -1.3845042 | 0.16620413 | NA | NA | NA |
| ENSG000002 | 1.3871175 | -0.570737883 | 0.33771825 | -1.6899824 | 0.09103133 | NA | NA | NA |
| ENSG000002 | 1.49846357 | -0.599749423 | 0.33793007 | -1.7747738 | 0.07593525 | NA | NA | NA |
| ENSG000002 | 0.12685397 | -0.090215768 | 0.12901919 | -0.699243 | 0.48440017 | NA | NDUFA3P2 | 106479051 |
| ENSG000002 | 2.05805184 | 0.005565278 | 0.36204635 | 0.01537173 | 0.98773562 | NA | NA | NA |
| ENSG000002 | 1.90014687 | -0.542890817 | 0.36060019 | -1.50552 | 0.13219043 | NA | NA | NA |
| ENSG000002 | 0 | NA | NA | NA | NA | NA | ATP6V1G2-DDX | 100532737 |
| ENSG000002 | 0 | NA | NA | NA | NA | NA | NA | NA |
| ENSG000002 | 1.65192454 | 0.001758843 | 0.33653528 | 0.00522633 | 0.99583001 | NA | NA | NA |
| ENSG000002 | 1.00627483 | -0.356078214 | 0.31079499 | -1.1457013 | 0.25191877 | NA | NA | NA |
| ENSG000002 | 1.61452112 | 0.001697766 | 0.34508291 | 0.00491988 | 0.99607452 | NA | NA | NA |
| ENSG000002 | 1.24221634 | -0.048433427 | 0.3338859 | -0.1450598 | 0.88466364 | NA | NA | NA |
| ENSG000002 | 0.5883907 | 0.051464553 | 0.2537826 | 0.20278992 | 0.83929924 | NA | NA | NA |
| ENSG000002 | 1.06648631 | 0.194823934 | 0.32055797 | 0.60776506 | 0.5433433 | NA | NA | NA |

| | | | | | | | | | |
|---|---|---|---|---|---|---|---|---|---|
| ENSG000002 | 0.11934008 | 0.083564698 | 0.12760091 | 0.65489106 | 0.5125379 | NA | NA | NA | |
| ENSG000002 | 1.24309578 | -0.162830752 | 0.31543908 | -0.5162035 | 0.60571229 | NA | NA | NA | |
| ENSG000002 | 0.90756584 | -0.264865892 | 0.29528719 | -0.8969773 | 0.36973105 | NA | NA | NA | |
| ENSG000002 | 1.69008155 | 0.218840819 | 0.35169813 | 0.62224049 | 0.53378374 | NA | NA | NA | |
| ENSG000002 | 1.36109783 | -0.102885772 | 0.32379468 | -0.31775 | 0.75067455 | NA | NA | NA | |
| ENSG000002 | 1.699401 | -0.044441655 | 0.3521556 | -0.1261989 | 0.89957447 | NA | NA | NA | |
| ENSG000002 | 1.89569265 | 0.595680439 | 0.35835464 | 1.66226519 | 0.09645961 | NA | NA | NA | |
| ENSG000002 | 1.50008584 | -0.300602671 | 0.33680253 | -0.892519 | 0.37211479 | NA | NA | NA | |
| ENSG000002 | 1.77679194 | -0.881267851 | 0.34727885 | -2.5376376 | 0.01116035 | NA | NA | NA | |
| ENSG000002 | 0.26062507 | -0.012888319 | 0.18226503 | -0.070712 | 0.94362699 | NA | NA | NA | |
| ENSG000002 | 1.39240725 | -0.112788704 | 0.32965251 | -0.3421442 | 0.73224235 | NA | NA | NA | |
| ENSG000002 | 1.59068012 | -0.255672263 | 0.34975155 | -0.7310111 | 0.46477237 | NA | NA | NA | |
| ENSG000002 | 1.88206373 | -0.383826282 | 0.35764084 | -1.0732172 | 0.28317369 | NA | NA | NA | |
| ENSG000002 | 1.22288746 | 0.104575244 | 0.33329427 | 0.3137625 | 0.75370143 | NA | NA | NA | |
| ENSG000002 | 1.41751344 | -0.51110642 | 0.3263863 | -1.5659555 | 0.11735904 | NA | NA | NA | |
| ENSG000002 | 1.67091554 | -0.55135363 | 0.3491007 | -1.5793541 | 0.11425486 | NA | NA | NA | |
| ENSG000002 | 1.7018766 | -0.309601586 | 0.3516571 | -0.8804076 | 0.37863854 | NA | NA | NA | |
| ENSG000002 | 1.86311963 | -0.261019356 | 0.35909653 | -0.726878 | 0.46730072 | NA | NA | NA | |
| ENSG000002 | 2.07709895 | -0.230878059 | 0.36109285 | -0.639387 | 0.52257123 | NA | NA | NA | |
| ENSG000002 | 1.10118982 | -0.123426475 | 0.3218848 | -0.3834492 | 0.70138672 | NA | NA | NA | |
| ENSG000002 | 1.78855829 | 0.415055423 | 0.35784011 | 1.15989073 | 0.2460933 | NA | NA | NA | |
| ENSG000002 | 1.87490024 | -0.358096796 | 0.3483122 | -1.0280914 | 0.30390682 | NA | NA | NA | |
| ENSG000002 | 0.47621624 | 0.147284149 | 0.23752141 | 0.62008787 | 0.53519994 | NA | NA | NA | |
| ENSG000002 | 1.19921613 | -0.04255523 | 0.32768066 | -0.129868 | 0.89667086 | NA | SLC2A13P1 | | 106480239 |
| ENSG000002 | 0.76061352 | -0.028091069 | 0.27995155 | -0.1003426 | 0.92007233 | NA | INMT-FAM188E | | 100526825 |
| ENSG000002 | 1.98672238 | -0.4529653 | 0.3618743 | -1.25172 | 0.21067191 | NA | NA | NA | |
| ENSG000002 | 1.25206449 | -0.356014866 | 0.33525225 | -1.0619313 | 0.28826687 | NA | NA | NA | |
| ENSG000002 | 2.14664498 | -0.245586961 | 0.36400414 | -0.6746818 | 0.49987797 | NA | NA | NA | |
| ENSG000002 | 1.46814787 | 0.07855674 | 0.34084651 | 0.23047541 | 0.81772237 | NA | NA | NA | |

| | | | | | | | | | |
|---|---|---|---|---|---|---|---|---|---|
| ENSG000002 | 1.22209413 | -0.193984853 | 0.32882754 | -0.5899288 | 0.55523835 | NA | NA | NA | |
| ENSG000002 | 1.78693114 | -0.750618755 | 0.35165863 | -2.1345097 | 0.03280108 | NA | NA | NA | |
| ENSG000002 | 1.72999984 | -0.445074135 | 0.35134477 | -1.2667732 | 0.2052364 | NA | NA | NA | |
| ENSG000002 | 2.13503446 | -0.501023197 | 0.36393191 | -1.3766949 | 0.1686066 | NA | NA | NA | |
| ENSG000002 | 1.85125065 | -0.382602692 | 0.3556663 | -1.075735 | 0.28204579 | NA | NA | NA | |
| ENSG000002 | 2.08469054 | -0.726667395 | 0.35993941 | -2.0188603 | 0.04350174 | NA | NA | NA | |
| ENSG000002 | 1.61401513 | -0.401833499 | 0.35076624 | -1.1455877 | 0.25196578 | NA | NA | NA | |
| ENSG000002 | 1.68767246 | 0.085697133 | 0.35269729 | 0.24297644 | 0.80802365 | NA | NA | NA | |
| ENSG000002 | 1.09469882 | -0.106289156 | 0.30531055 | -0.3481346 | 0.72773913 | NA | KRTAP5-9 | | 3846 |
| ENSG000002 | 0.24550891 | -0.007886114 | 0.17900103 | -0.0440563 | 0.96485956 | NA | NA | NA | |
| ENSG000002 | 1.75575714 | -0.453782978 | 0.35311134 | -1.2850989 | 0.19875774 | NA | NA | NA | |
| ENSG000002 | 1.15788618 | -0.576254973 | 0.31642482 | -1.8211434 | 0.06858506 | NA | NA | NA | |
| ENSG000002 | 1.64524524 | -0.370368528 | 0.34250714 | -1.0813454 | 0.2795435 | NA | NA | NA | |
| ENSG000002 | 1.58310559 | -0.635177541 | 0.3421539 | -1.8564089 | 0.0633953 | NA | NA | NA | |
| ENSG000002 | 0.36416385 | 0.068880976 | 0.20758164 | 0.33182596 | 0.74002068 | NA | NA | NA | |
| ENSG000002 | 0.38747904 | -0.104908247 | 0.21987514 | -0.4771265 | 0.63327207 | NA | NA | NA | |
| ENSG000002 | 0.26199343 | -0.194018184 | 0.18422801 | -1.0531416 | 0.29227609 | NA | NA | NA | |
| ENSG000002 | 0 | NA | NA | NA | NA | NA | NA | NA | |
| ENSG000002 | 2.149864 | 0.30500787 | 0.36429878 | 0.83724648 | 0.40245404 | NA | NA | NA | |
| ENSG000002 | 1.58250227 | 0.01536264 | 0.35076785 | 0.04379717 | 0.96506608 | NA | NA | NA | |
| ENSG000002 | 0.66527186 | -0.43035014 | 0.25829332 | -1.6661296 | 0.09568761 | NA | NA | NA | |
| ENSG000002 | 1.3218772 | 0.026872941 | 0.33850254 | 0.07938771 | 0.93672425 | NA | NA | NA | |
| ENSG000002 | 1.22186792 | -0.046303628 | 0.33337651 | -0.1388929 | 0.88953479 | NA | LOC100499484- | | 57653 |
| ENSG000002 | 0.11205239 | 0.082118251 | 0.12671962 | 0.64803108 | 0.51696485 | NA | NA | NA | |
| ENSG000002 | 1.50725036 | -0.179010952 | 0.33672297 | -0.5316268 | 0.59498451 | NA | NA | NA | |
| ENSG000002 | 0 | NA | NA | NA | NA | NA | NA | NA | |
| ENSG000002 | 0 | NA | NA | NA | NA | NA | MTND1P35 | | 106480433 |
| ENSG000002 | 1.64241569 | -0.126053881 | 0.33808729 | -0.3728442 | 0.70926441 | NA | NA | NA | |
| ENSG000002 | 0 | NA | NA | NA | NA | NA | SAA2-SAA4 | | 100528017 |

| | | | | | | | | |
|---|---|---|---|---|---|---|---|---|
| ENSG000002 | 0 | NA | NA | NA | NA | NA | PIGY | 84992 |
| ENSG000002 | 0 | NA | NA | NA | NA | NA | ZFP91-CNTF | 386607 |
| ENSG000002 | 1.71610727 | -0.062829593 | 0.35546941 | -0.1767511 | 0.85970394 | NA | NA | NA |
| ENSG000002 | 0.59072708 | 0.064826349 | 0.26164045 | 0.24776883 | 0.80431327 | NA | NA | NA |
| ENSG000002 | 0.34481323 | 0.065936758 | 0.20433332 | 0.32269215 | 0.7469284 | NA | NA | NA |
| ENSG000002 | 0.89566653 | -0.261188394 | 0.29466492 | -0.8863912 | 0.37540676 | NA | NA | NA |
| ENSG000002 | 2.05992874 | -0.120037769 | 0.36320876 | -0.3304925 | 0.74102787 | NA | NA | NA |
| ENSG000002 | 0.36351096 | 0.070290861 | 0.21590418 | 0.32556508 | 0.74475344 | NA | NA | NA |
| ENSG000002 | 0.86976701 | -0.115601083 | 0.29779035 | -0.3881962 | 0.69787084 | NA | NA | NA |
| ENSG000002 | 1.71318306 | -0.306498473 | 0.35025211 | -0.8750796 | 0.38153061 | NA | OR1D4 | 653166 |
| ENSG000002 | 1.83524488 | -0.119626342 | 0.35847437 | -0.3337096 | 0.7385987 | NA | NA | NA |
| ENSG000002 | 0 | NA | NA | NA | NA | NA | TBC1D26 | 353149 |
| ENSG000002 | 0.13280363 | -0.093394593 | 0.13115644 | -0.7120855 | 0.47641185 | NA | NA | NA |
| ENSG000002 | 1.49709466 | -0.484131846 | 0.34373039 | -1.4084639 | 0.15899375 | NA | NA | NA |
| ENSG000002 | 1.84581254 | -0.247128244 | 0.35846108 | -0.6894144 | 0.4905625 | NA | NA | NA |
| ENSG000002 | 1.21865107 | -0.045349274 | 0.33195206 | -0.1366139 | 0.89133597 | NA | NA | NA |
| ENSG000002 | 2.16031356 | -0.653702212 | 0.3658739 | -1.7866872 | 0.07398806 | NA | OVOL1-AS1 | 101927828 |
| ENSG000002 | 1.35449686 | 0.016914806 | 0.33724668 | 0.05015559 | 0.9599984 | NA | NA | NA |
| ENSG000002 | 0.98620929 | -0.180604596 | 0.30694877 | -0.5883868 | 0.55627272 | NA | NA | NA |
| ENSG000002 | 1.60504642 | -0.243795634 | 0.34578481 | -0.7050502 | 0.48077902 | NA | NA | NA |
| ENSG000002 | 1.33915258 | 0.024856283 | 0.33887024 | 0.07335045 | 0.94152725 | NA | NA | NA |
| ENSG000002 | 0.96175385 | 0.123457392 | 0.31018235 | 0.39801553 | 0.69061874 | NA | NA | NA |
| ENSG000002 | 1.86668236 | -0.75961029 | 0.35043573 | -2.1676165 | 0.03018787 | NA | NA | NA |
| ENSG000002 | 0 | NA | NA | NA | NA | NA | NA | NA |
| ENSG000002 | 1.24953436 | -0.188772213 | 0.32695743 | -0.5773602 | 0.56369615 | NA | NA | NA |
| ENSG000002 | 0.89364336 | -0.29139313 | 0.3048889 | -0.9557354 | 0.33920591 | NA | NA | NA |
| ENSG000002 | 2.15450379 | -0.379498205 | 0.36419301 | -1.042025 | 0.29740011 | NA | NA | NA |
| ENSG000002 | 2.11010296 | -0.262767389 | 0.36784741 | -0.714338 | 0.4750182 | NA | NA | NA |
| ENSG000002 | 1.78043623 | -0.460084367 | 0.35189804 | -1.3074366 | 0.19106448 | NA | NA | NA |

| | | | | | | | | |
|---|---|---|---|---|---|---|---|---|
| ENSG000002 | 1.26763712 | -0.360055083 | 0.33432432 | -1.0769635 | 0.28149658 | NA | NA | NA |
| ENSG000002 | 1.64071602 | -0.553158378 | 0.35089559 | -1.5764187 | 0.11492934 | NA | NA | NA |
| ENSG000002 | 0.47826975 | -0.016435741 | 0.244632 | -0.0671856 | 0.94643397 | NA | NA | NA |
| ENSG000002 | 0.57473127 | 0.229046573 | 0.26124356 | 0.8767549 | 0.38061978 | NA | NA | NA |
| ENSG000002 | 1.67785253 | 0.076289093 | 0.35260811 | 0.2163566 | 0.82870979 | NA | NA | NA |
| ENSG000002 | 2.03938282 | -0.708250863 | 0.35938015 | -1.9707568 | 0.0487517 | NA | NA | NA |
| ENSG000002 | 0.78039674 | -0.368519854 | 0.28442264 | -1.295677 | 0.1950868 | NA | LOC105369423 | 105369423 |
| ENSG000002 | 0.39274356 | -0.107428268 | 0.22062504 | -0.4869269 | 0.62631013 | NA | NA | NA |
| ENSG000002 | 1.38557657 | -0.118962565 | 0.33673859 | -0.3532787 | 0.72387953 | NA | NA | NA |
| ENSG000002 | 1.98079893 | -0.523986246 | 0.3534793 | -1.4823676 | 0.13824252 | NA | NA | NA |
| ENSG000002 | 1.47722711 | -0.185895229 | 0.34589495 | -0.5374326 | 0.59096881 | NA | GTF2IP11 | 106480346 |
| ENSG000002 | 0.61107338 | -0.109784645 | 0.27021411 | -0.4062876 | 0.68453129 | NA | MED28P5 | 106479053 |
| ENSG000002 | 1.70554653 | -0.046130348 | 0.35222655 | -0.1309678 | 0.89580075 | NA | LOC105375888 | 105375888 |
| ENSG000002 | 1.63194127 | -0.530970848 | 0.34713645 | -1.5295739 | 0.12612223 | NA | NA | NA |
| ENSG000002 | 0.74365025 | -0.035627946 | 0.29040344 | -0.1226843 | 0.90235709 | NA | NA | NA |
| ENSG000002 | 0.51322524 | -0.027081322 | 0.24101349 | -0.1123643 | 0.91053453 | NA | NA | NA |
| ENSG000002 | 1.2261988 | -0.05258316 | 0.32149802 | -0.1635567 | 0.87008012 | NA | NA | NA |
| ENSG000002 | 0.80188519 | -0.545785546 | 0.28296247 | -1.9288266 | 0.05375239 | NA | NA | NA |
| ENSG000002 | 0.50377641 | -0.188767022 | 0.24148558 | -0.7816906 | 0.4343964 | NA | NA | NA |
| ENSG000002 | 1.85602837 | -0.266656681 | 0.36258611 | -0.7354299 | 0.46207769 | NA | OR5M11 | 219487 |
| ENSG000002 | 2.11932502 | -0.005798683 | 0.35974275 | -0.016119 | 0.98713948 | NA | NA | NA |
| ENSG000002 | 0.61790405 | -0.105393408 | 0.26443323 | -0.3985634 | 0.69021494 | NA | NA | NA |
| ENSG000002 | 1.01549396 | -0.343561364 | 0.30964813 | -1.1095219 | 0.26720512 | NA | NA | NA |
| ENSG000002 | 1.96358592 | -0.183782436 | 0.36026676 | -0.5101288 | 0.60996126 | NA | NA | NA |
| ENSG000002 | 1.98521425 | -0.062905721 | 0.36205229 | -0.1737476 | 0.86206382 | NA | NA | NA |
| ENSG000002 | 1.83584825 | -0.244878553 | 0.35362533 | -0.6924802 | 0.4886358 | NA | NA | NA |
| ENSG000002 | 1.74988679 | -0.19039275 | 0.34653672 | -0.5494158 | 0.58272014 | NA | NA | NA |
| ENSG000002 | 1.45757918 | 0.090814441 | 0.34201602 | 0.26552686 | 0.79060362 | NA | NA | NA |
| ENSG000002 | 0 | NA | NA | NA | NA | NA | FXYD6-FXYD2 | 100533181 |

| | | | | | | | | |
|---|---|---|---|---|---|---|---|---|
| ENSG000002 | 1.6953585 | 0.190471316 | 0.34191531 | 0.55707162 | 0.5774785 | NA | PRR23D1 | 100131608 |
| ENSG000002 | 1.34806764 | -0.253330333 | 0.33517759 | -0.7558093 | 0.44976356 | NA | NA | NA |
| ENSG000002 | 1.11306326 | -0.264119337 | 0.32158908 | -0.8212945 | 0.41147855 | NA | NA | NA |
| ENSG000002 | 0.90875952 | -0.279672956 | 0.29930643 | -0.9344034 | 0.35009583 | NA | NA | NA |
| ENSG000002 | 0 | NA | NA | NA | NA | NA | NA | NA |
| ENSG000002 | 0.23816916 | -0.010712105 | 0.1772998 | -0.060418 | 0.9518227 | NA | NA | NA |
| ENSG000002 | 1.74164015 | -0.309936589 | 0.34843316 | -0.8895152 | 0.37372626 | NA | NA | NA |
| ENSG000002 | 2.12886252 | -0.557884598 | 0.35351023 | -1.5781286 | 0.11453607 | NA | NA | NA |
| ENSG000002 | 2.10783521 | -0.498638713 | 0.36223721 | -1.376553 | 0.16865049 | NA | NA | NA |
| ENSG000002 | 0.23868015 | 0.154625155 | 0.16997278 | 0.90970538 | 0.36297791 | NA | NA | NA |
| ENSG000002 | 1.33531096 | 0.035050503 | 0.33535166 | 0.10451865 | 0.91675777 | NA | NA | NA |
| ENSG000002 | 1.83311654 | -0.227450001 | 0.3508486 | -0.6482853 | 0.51680042 | NA | NA | NA |
| ENSG000002 | 0.4855357 | -0.181797071 | 0.23938927 | -0.7594203 | 0.44760118 | NA | NA | NA |
| ENSG000002 | 1.4789095 | -0.190264356 | 0.34154189 | -0.5570747 | 0.57747637 | NA | NA | NA |
| ENSG000002 | 1.23185562 | -0.046790222 | 0.32417903 | -0.1443345 | 0.88523633 | NA | NA | NA |
| ENSG000002 | 0.3561712 | 0.067399341 | 0.21474131 | 0.31386296 | 0.75362513 | NA | NA | NA |
| ENSG000002 | 1.64616727 | -0.235061281 | 0.3399778 | -0.6914018 | 0.48931305 | NA | NA | NA |
| ENSG000002 | 1.51334245 | -0.327663175 | 0.34429341 | -0.9516975 | 0.34125043 | NA | NA | NA |
| ENSG000002 | 0.84441329 | -0.252217771 | 0.29006899 | -0.8695096 | 0.38456846 | NA | NA | NA |
| ENSG000002 | 1.56528108 | 0.164282655 | 0.34934531 | 0.47025865 | 0.63817024 | NA | NA | NA |
| ENSG000002 | 0.95896122 | 0.117525033 | 0.30506085 | 0.38525112 | 0.70005137 | NA | NA | NA |
| ENSG000002 | 1.48092526 | -0.180691983 | 0.34080688 | -0.5301888 | 0.59598105 | NA | NA | NA |
| ENSG000002 | 0 | NA | NA | NA | NA | NA | NA | NA |
| ENSG000002 | 0.88744576 | -0.277587958 | 0.29984413 | -0.9257742 | 0.35456333 | NA | NA | NA |
| ENSG000002 | 1.47497718 | -0.054187923 | 0.34915756 | -0.1551962 | 0.87666665 | NA | NA | NA |
| ENSG000002 | 0.60717723 | -0.256800472 | 0.2582777 | -0.9942805 | 0.32008634 | NA | NA | NA |
| ENSG000002 | 2.01393819 | -0.743417264 | 0.36269193 | -2.049721 | 0.04039166 | NA | NA | NA |
| ENSG000002 | 0.96084004 | 0.11959348 | 0.31167742 | 0.38370915 | 0.70119403 | NA | NA | NA |
| ENSG000002 | 0.9932879 | -0.182365576 | 0.30725311 | -0.5935353 | 0.55282294 | NA | NA | NA |

| | | | | | | | | |
|---|---|---|---|---|---|---|---|---|
| ENSG000002 | 1.75772185 | -0.332296532 | 0.35614974 | -0.9330248 | 0.35080717 | NA | NA | NA |
| ENSG000002 | 1.01979292 | -0.328248738 | 0.30451396 | -1.0779431 | 0.28105913 | NA | NA | NA |
| ENSG000002 | 0.50818749 | -0.197539377 | 0.24915052 | -0.7928516 | 0.42786432 | NA | NA | NA |
| ENSG000002 | 0.65445467 | -0.486865861 | 0.27154464 | -1.7929496 | 0.07298098 | NA | NA | NA |
| ENSG000002 | 0 | NA | NA | NA | NA | NA | NA | NA |
| ENSG000002 | 0.74177143 | -0.029328749 | 0.28496727 | -0.1029197 | 0.91802669 | NA | NA | NA |
| ENSG000002 | 1.61218975 | -0.254999081 | 0.34795393 | -0.732853 | 0.4636481 | NA | NA | NA |
| ENSG000002 | 1.47273014 | -0.329060617 | 0.34456333 | -0.9550077 | 0.33957382 | NA | NA | NA |
| ENSG000002 | 0.12685397 | -0.090215768 | 0.12901919 | -0.699243 | 0.48440017 | NA | NA | NA |
| ENSG000002 | 1.74504703 | -0.190101415 | 0.35112888 | -0.5414007 | 0.58823144 | NA | NA | NA |
| ENSG000002 | 1.21275541 | 0.098784214 | 0.32363467 | 0.30523372 | 0.76018815 | NA | NA | NA |
| ENSG000002 | 1.78137144 | -0.32349994 | 0.35254008 | -0.917626 | 0.35881471 | NA | NA | NA |
| ENSG000002 | 0.48971204 | -0.022179634 | 0.22960609 | -0.0965986 | 0.92304514 | NA | NA | NA |
| ENSG000002 | 0.83628998 | 0.212075688 | 0.29449747 | 0.72012736 | 0.47144658 | NA | NA | NA |
| ENSG000002 | 0 | NA | NA | NA | NA | NA | NA | NA |
| ENSG000002 | 0.89262382 | -0.459792846 | 0.30190855 | -1.522954 | 0.1277702 | NA | NA | NA |
| ENSG000002 | 0.13280363 | -0.093394593 | 0.13115644 | -0.7120855 | 0.47641185 | NA | NA | NA |
| ENSG000002 | 1.42147376 | -0.413664687 | 0.33439552 | -1.2370521 | 0.21606774 | NA | NA | NA |
| ENSG000002 | 2.09635572 | -0.364494247 | 0.36389871 | -1.0016366 | 0.31651916 | NA | UBTFL10 | 106480293 |
| ENSG000002 | 0.14128499 | -0.097772014 | 0.13403235 | -0.7294658 | 0.4657168 | NA | NA | NA |
| ENSG000002 | 1.85463477 | -0.499852452 | 0.3537715 | -1.4129246 | 0.1576779 | NA | NA | NA |
| ENSG000002 | 1.5952846 | -0.353678583 | 0.33871887 | -1.0441656 | 0.29640879 | NA | NA | NA |
| ENSG000002 | 1.40609756 | -0.415102611 | 0.33886696 | -1.2249722 | 0.22058569 | NA | NA | NA |
| ENSG000002 | 0.14128499 | -0.097772014 | 0.13403235 | -0.7294658 | 0.4657168 | NA | NA | NA |
| ENSG000002 | 1.51203825 | -0.199432513 | 0.34453363 | -0.5788477 | 0.56269192 | NA | NA | NA |
| ENSG000002 | 1.06387863 | -0.719575284 | 0.31142394 | -2.3105972 | 0.02085511 | NA | NA | NA |
| ENSG000002 | 1.21560032 | 0.107641345 | 0.32954521 | 0.32663605 | 0.74394318 | NA | NA | NA |
| ENSG000002 | 2.13204842 | -0.379896227 | 0.36577619 | -1.038603 | 0.29898943 | NA | NA | NA |
| ENSG000002 | 1.87706138 | -0.268600027 | 0.36054951 | -0.744974 | 0.45628741 | NA | NA | NA |

| | | | | | | | | |
|---|---|---|---|---|---|---|---|---|
| ENSG000002 | 0 | NA | NA | NA | NA | NA | NA | NA |
| ENSG000002 | 0 | NA | NA | NA | NA | NA | NA | NA |
| ENSG000002 | 1.41868304 | -0.254620452 | 0.32728126 | -0.7779867 | 0.43657688 | NA | NA | NA |
| ENSG000002 | 0 | NA | NA | NA | NA | NA | NA | NA |
| ENSG000002 | 2.11233116 | -0.510635704 | 0.36541824 | -1.3974007 | 0.16229311 | NA | NA | NA |
| ENSG000002 | 0.76101442 | -0.375123817 | 0.28870464 | -1.2993342 | 0.19382926 | NA | NA | NA |
| ENSG000002 | 1.00720847 | -0.360508653 | 0.31528677 | -1.143431 | 0.2528597 | NA | NA | NA |
| ENSG000002 | 1.70816469 | 0.081698618 | 0.35091303 | 0.23281728 | 0.81590329 | NA | NA | NA |
| ENSG000002 | 1.54023867 | -0.464499874 | 0.33992122 | -1.3664927 | 0.17178436 | NA | NA | NA |
| ENSG000002 | 1.88636819 | -0.529386917 | 0.35805895 | -1.4784909 | 0.13927642 | NA | NA | NA |
| ENSG000002 | 1.52183921 | 0.399393753 | 0.33772846 | 1.18258837 | 0.23697232 | NA | NA | NA |
| ENSG000002 | 0.61335714 | -0.10408451 | 0.2640241 | -0.3942235 | 0.69341602 | NA | NA | NA |
| ENSG000002 | 1.21518105 | -0.040361122 | 0.32202593 | -0.125335 | 0.90025834 | NA | NA | NA |
| ENSG000002 | 1.60771664 | -0.099465804 | 0.33995837 | -0.2925823 | 0.76984144 | NA | NA | NA |
| ENSG000002 | 0 | NA | NA | NA | NA | NA | SEPT14P8 | 106480341 |
| ENSG000002 | 1.72746917 | -0.323025451 | 0.35157751 | -0.9187887 | 0.35820609 | NA | NA | NA |
| ENSG000002 | 1.11883823 | -0.118806031 | 0.3224635 | -0.3684325 | 0.71255078 | NA | NA | NA |
| ENSG000002 | 1.21048876 | -0.169659109 | 0.32164862 | -0.5274672 | 0.59786916 | NA | NA | NA |
| ENSG000002 | 1.47144767 | -0.04586973 | 0.34234718 | -0.133986 | 0.89341364 | NA | NA | NA |
| ENSG000002 | 1.4759734 | -0.182359634 | 0.34006405 | -0.5362509 | 0.59178519 | NA | LINC00681 | 101409254 |
| ENSG000002 | 1.12503583 | -0.124742832 | 0.32646715 | -0.3820992 | 0.70238779 | NA | NA | NA |
| ENSG000002 | 0.12070844 | -0.088503695 | 0.12783931 | -0.6923042 | 0.48874631 | NA | NA | NA |
| ENSG000002 | 0.76371634 | -0.201271588 | 0.28715856 | -0.7009075 | 0.48336075 | NA | NA | NA |
| ENSG000002 | 0.61175852 | -0.109548347 | 0.27027148 | -0.4053271 | 0.68523711 | NA | NA | NA |
| ENSG000002 | 0 | NA | NA | NA | NA | NA | NA | NA |
| ENSG000002 | 1.24916614 | -0.4827207 | 0.3257962 | -1.4816646 | 0.13842956 | NA | NA | NA |
| ENSG000002 | 0.49076933 | -0.025124039 | 0.23846109 | -0.1053591 | 0.91609089 | NA | SNRPCP5 | 106480290 |
| ENSG000002 | 1.60700976 | -0.24364707 | 0.34655855 | -0.7030473 | 0.48202625 | NA | NA | NA |
| ENSG000002 | 0 | NA | NA | NA | NA | NA | NEDD8-MDP1 | 100528064 |

| | | | | | | | | |
|---|---|---|---|---|---|---|---|---|
| ENSG000002 | 2.10384979 | -0.500208998 | 0.36405292 | -1.3740008 | 0.16944144 | NA | NA | NA |
| ENSG000002 | 1.82061798 | -0.118182172 | 0.35976564 | -0.3284977 | 0.74253541 | NA | NA | NA |
| ENSG000002 | 0.26062507 | -0.012888319 | 0.18226503 | -0.070712 | 0.94362699 | NA | NA | NA |
| ENSG000002 | 1.01352814 | -0.517235448 | 0.30927899 | -1.6723911 | 0.09444722 | NA | OR10V2P | 81343 |
| ENSG000002 | 1.87318751 | -0.241626772 | 0.3558165 | -0.6790769 | 0.49708913 | NA | NA | NA |
| ENSG000002 | 1.20891345 | 0.11260035 | 0.33291183 | 0.33822874 | 0.73519082 | NA | NA | NA |
| ENSG000002 | 0.58756367 | 0.225272418 | 0.26247383 | 0.85826621 | 0.39074548 | NA | NA | NA |
| ENSG000002 | 1.02004245 | -0.191554231 | 0.30838552 | -0.6211518 | 0.53449973 | NA | NA | NA |
| ENSG000002 | 1.47246912 | 0.228239214 | 0.34118014 | 0.66896982 | 0.50351473 | NA | NA | NA |
| ENSG000002 | 0.85630346 | 0.041530201 | 0.29067593 | 0.14287458 | 0.88638923 | NA | NA | NA |
| ENSG000002 | 1.21916654 | -0.160294895 | 0.32025137 | -0.5005284 | 0.61670309 | NA | NA | NA |
| ENSG000002 | 2.04953861 | -0.724731376 | 0.36008755 | -2.0126532 | 0.04415112 | NA | NA | NA |
| ENSG000002 | 1.48268941 | -0.043706189 | 0.34235645 | -0.1276628 | 0.89841582 | NA | NA | NA |
| ENSG000002 | 0.12685397 | -0.090215768 | 0.12901919 | -0.699243 | 0.48440017 | NA | NA | NA |
| ENSG000002 | 1.90270891 | -0.674588931 | 0.35749803 | -1.8869724 | 0.05916403 | NA | NA | NA |
| ENSG000002 | 1.01315758 | -0.371579399 | 0.31542085 | -1.1780432 | 0.23877937 | NA | NA | NA |
| ENSG000002 | 1.90267858 | -0.534020711 | 0.3591606 | -1.4868577 | 0.1370524 | NA | NA | NA |
| ENSG000002 | 0.85860546 | -0.094436395 | 0.28941414 | -0.3263019 | 0.74419592 | NA | NA | NA |
| ENSG000002 | 1.34505016 | 0.023121094 | 0.33899629 | 0.06820456 | 0.94562279 | NA | NA | NA |
| ENSG000002 | 1.47503116 | 0.091707547 | 0.34796104 | 0.26355694 | 0.79212134 | NA | NA | NA |
| ENSG000002 | 1.41244169 | 0.238144811 | 0.34002175 | 0.7003811 | 0.48368933 | NA | NA | NA |
| ENSG000002 | 0.97760868 | -0.292311475 | 0.29012275 | -1.0075442 | 0.31367335 | NA | NA | NA |
| ENSG000002 | 1.95902244 | 0.069485001 | 0.3617844 | 0.19206191 | 0.84769371 | NA | NA | NA |
| ENSG000002 | 1.21166799 | 0.245126167 | 0.32169559 | 0.76198175 | 0.4460709 | NA | NA | NA |
| ENSG000002 | 1.28861511 | -0.670124302 | 0.3297392 | -2.0322858 | 0.04212473 | NA | NA | NA |
| ENSG000002 | 1.25881029 | -0.304131235 | 0.31606804 | -0.9622334 | 0.33593236 | NA | NA | NA |
| ENSG000002 | 0 | NA | NA | NA | NA | NA | NA | NA |
| ENSG000002 | 0.13280363 | -0.093394593 | 0.13115644 | -0.7120855 | 0.47641185 | NA | NA | NA |
| ENSG000002 | 1.62588411 | -0.25771316 | 0.35088253 | -0.7344713 | 0.46266152 | NA | NA | NA |

| | | | | | | | | |
|---|---|---|---|---|---|---|---|---|
| ENSG000002 | 2.00684404 | -0.327510028 | 0.36251032 | -0.9034502 | 0.366287 | NA | NA | NA |
| ENSG000002 | 2.13854525 | -0.439676146 | 0.35459338 | -1.2399446 | 0.21499588 | NA | RERG-AS1 | 100873980 |
| ENSG000002 | 1.00176343 | -0.037709418 | 0.30691391 | -0.1228664 | 0.90221287 | NA | NA | NA |
| ENSG000002 | 2.15522318 | -0.805941274 | 0.36577361 | -2.2033882 | 0.0275674 | NA | NA | NA |
| ENSG000002 | 1.12472317 | -0.107711213 | 0.31479936 | -0.3421583 | 0.73223177 | NA | NA | NA |
| ENSG000002 | 1.12466864 | -0.248688732 | 0.30948572 | -0.8035548 | 0.42165414 | NA | NA | NA |
| ENSG000002 | 0.85726688 | -0.108645082 | 0.28935898 | -0.3754682 | 0.70731233 | NA | NA | NA |
| ENSG000002 | 0.3593886 | 0.071604659 | 0.20679693 | 0.34625591 | 0.7291504 | NA | NA | NA |
| ENSG000002 | 1.01085612 | -0.195998049 | 0.30962028 | -0.6330272 | 0.52671591 | NA | NA | NA |
| ENSG000002 | 0.25370795 | -0.171331747 | 0.17411721 | -0.9840024 | 0.32511435 | NA | NA | NA |
| ENSG000002 | 0.52028267 | -0.202047067 | 0.25039858 | -0.8069018 | 0.41972305 | NA | NA | NA |
| ENSG000002 | 0.99872711 | -0.343927824 | 0.31057383 | -1.1073948 | 0.26812329 | NA | NA | NA |
| ENSG000002 | 1.08943094 | 0.174177147 | 0.31532051 | 0.55238128 | 0.58068715 | NA | NA | NA |
| ENSG000002 | 1.60058583 | -0.630946203 | 0.33917481 | -1.860239 | 0.06285172 | NA | NA | NA |
| ENSG000002 | 0 | NA | NA | NA | NA | NA | NA | NA |
| ENSG000002 | 0.52742601 | -0.196769605 | 0.24405015 | -0.8062671 | 0.42008885 | NA | NA | NA |
| ENSG000002 | 1.30167776 | -0.491044392 | 0.32808669 | -1.496691 | 0.13447369 | NA | NA | NA |
| ENSG000002 | 1.49142979 | -0.178651993 | 0.34099374 | -0.5239157 | 0.60033714 | NA | NA | NA |
| ENSG000002 | 1.61654237 | -0.132748297 | 0.35269223 | -0.3763857 | 0.70663021 | NA | NA | NA |
| ENSG000002 | 2.12752449 | -0.622200099 | 0.3621947 | -1.7178608 | 0.085822 | NA | NA | NA |
| ENSG000002 | 1.967854 | -0.066249995 | 0.36250147 | -0.1827579 | 0.85498801 | NA | NA | NA |
| ENSG000002 | 1.96671686 | -0.3000914 | 0.35976681 | -0.8341275 | 0.40420913 | NA | KLRC4-KLRK1 | 100528032 |
| ENSG000002 | 1.87381654 | -0.522393809 | 0.35797487 | -1.459303 | 0.14448173 | NA | NA | NA |
| ENSG000002 | 0 | NA | NA | NA | NA | NA | NA | NA |
| ENSG000002 | 1.03538728 | -0.177797599 | 0.3015194 | -0.5896722 | 0.55541046 | NA | NA | NA |
| ENSG000002 | 1.72127729 | 0.078136913 | 0.35484759 | 0.22019852 | 0.82571655 | NA | NA | NA |
| ENSG000002 | 1.96878763 | -0.059465937 | 0.36367835 | -0.1635124 | 0.87011498 | NA | NA | NA |
| ENSG000002 | 1.35967705 | 0.018112271 | 0.33592088 | 0.05391826 | 0.95700029 | NA | NA | NA |
| ENSG000002 | 1.09686106 | 0.041632926 | 0.32142911 | 0.12952444 | 0.89694269 | NA | NA | NA |

| | | | | | | | | |
|---|---|---|---|---|---|---|---|---|
| ENSG000002 | 0 | NA | NA | NA | NA | NA | NA | NA |
| ENSG000002 | 0 | NA | NA | NA | NA | NA | NA | NA |
| ENSG000002 | 1.85584371 | 0.015495244 | 0.34840059 | 0.04447537 | 0.96452549 | NA | NA | NA |
| ENSG000002 | 1.72290815 | 0.071165111 | 0.341867 | 0.20816608 | 0.83509929 | NA | NA | NA |
| ENSG000002 | 1.27657549 | -0.322472478 | 0.32318142 | -0.9978064 | 0.31837326 | NA | NA | NA |
| ENSG000002 | 0.86432431 | -0.23665616 | 0.28166791 | -0.8401957 | 0.40079869 | NA | NA | NA |
| ENSG000002 | 0.97609651 | -0.191311697 | 0.31143602 | -0.6142889 | 0.53902443 | NA | NA | NA |
| ENSG000002 | 1.42967334 | -0.580972021 | 0.33854491 | -1.7160855 | 0.08614639 | NA | LOC101928416 | 101928416 |
| ENSG000002 | 0.49241844 | -0.197108126 | 0.24744176 | -0.7965839 | 0.42569273 | NA | NA | NA |
| ENSG000002 | 1.20536166 | -0.044428128 | 0.32785314 | -0.1355123 | 0.89220685 | NA | NA | NA |
| ENSG000002 | 0.24004852 | -0.005223787 | 0.17775844 | -0.029387 | 0.97655594 | NA | NA | NA |
| ENSG000002 | 1.61594951 | -0.121852525 | 0.35027455 | -0.3478772 | 0.72793242 | NA | NA | NA |
| ENSG000002 | 1.77067632 | -0.468046993 | 0.35364862 | -1.3234803 | 0.1856757 | NA | NA | NA |
| ENSG000002 | 1.53195144 | 0.428754307 | 0.34510894 | 1.24237381 | 0.21409867 | NA | NA | NA |
| ENSG000002 | 2.13909208 | -0.585320848 | 0.3575328 | -1.6371109 | 0.1016073 | NA | NA | NA |
| ENSG000002 | 0.23202362 | -0.008302493 | 0.17586207 | -0.0472103 | 0.96234565 | NA | NA | NA |
| ENSG000002 | 0.55665861 | -0.363599063 | 0.24151673 | -1.5054819 | 0.13220023 | NA | NA | NA |
| ENSG000002 | 1.85035307 | -0.124798911 | 0.35693514 | -0.3496403 | 0.72660866 | NA | NA | NA |
| ENSG000002 | 1.46902752 | -0.051273049 | 0.34659146 | -0.1479351 | 0.88239397 | NA | NA | NA |
| ENSG000002 | 0.96152572 | 0.124015489 | 0.3068254 | 0.40418913 | 0.68607365 | NA | TUBB4BP1 | 106478981 |
| ENSG000002 | 0.85723517 | -0.086923536 | 0.28025668 | -0.3101569 | 0.75644166 | NA | NA | NA |
| ENSG000002 | 1.87888518 | -0.546379549 | 0.36155605 | -1.511189 | 0.13074029 | NA | NA | NA |
| ENSG000002 | 1.55953994 | 0.155460642 | 0.34508079 | 0.45050506 | 0.65234631 | NA | NA | NA |
| ENSG000002 | 0.99121266 | -0.205080843 | 0.31756778 | -0.645786 | 0.51841792 | NA | NA | NA |
| ENSG000002 | 1.50054039 | -0.473507883 | 0.34311855 | -1.3800125 | 0.1675828 | NA | NA | NA |
| ENSG000002 | 0.46404725 | -0.01414962 | 0.22636165 | -0.0625089 | 0.95015757 | NA | NA | NA |
| ENSG000002 | 0.73696338 | -0.030982117 | 0.27833596 | -0.1113119 | 0.91136898 | NA | NA | NA |
| ENSG000002 | 1.5382866 | -0.628187816 | 0.3418731 | -1.8374883 | 0.06613784 | NA | NA | NA |
| ENSG000002 | 0 | NA | NA | NA | NA | NA | NA | NA |

| | | | | | | | | | |
|---|---|---|---|---|---|---|---|---|---|
| ENSG000002 | 0 | NA | NA | NA | NA | NA | NA | NA | |
| ENSG000002 | 2.09239251 | -0.120298083 | 0.36589293 | -0.3287795 | 0.74232238 | NA | NA | NA | |
| ENSG000002 | 0.61881753 | -0.112804359 | 0.27082437 | -0.4165222 | 0.67702794 | NA | NA | NA | |
| ENSG000002 | 0.84053441 | 0.044053894 | 0.29589149 | 0.1488853 | 0.88164414 | NA | NA | NA | |
| ENSG000002 | 1.20973656 | 0.246118768 | 0.32413044 | 0.75932013 | 0.44766108 | NA | NA | NA | |
| ENSG000002 | 1.93158221 | 0.07096901 | 0.36334174 | 0.19532303 | 0.84514007 | NA | MTRNR2L10 | | 100463488 |
| ENSG000002 | 1.6098555 | -0.128390163 | 0.35019902 | -0.3666206 | 0.71390207 | NA | NA | NA | |
| ENSG000002 | 1.99073255 | -0.195268243 | 0.36452812 | -0.535674 | 0.59218386 | NA | NA | NA | |
| ENSG000002 | 1.82599329 | -0.106562122 | 0.34326017 | -0.3104413 | 0.75622541 | NA | NA | NA | |
| ENSG000002 | 1.86291111 | -0.267872977 | 0.36263574 | -0.7386833 | 0.4600993 | NA | NA | NA | |
| ENSG000002 | 0 | NA | NA | NA | NA | NA | NA | NA | |
| ENSG000002 | 1.61966905 | 0.006586619 | 0.34514318 | 0.01908373 | 0.98477431 | NA | NA | NA | |
| ENSG000002 | 0.24485602 | -0.012655878 | 0.17881722 | -0.0707755 | 0.94357643 | NA | NA | NA | |
| ENSG000002 | 0.35346928 | -0.088570748 | 0.20618147 | -0.4295767 | 0.66750362 | NA | NA | NA | |
| ENSG000002 | 0.38747904 | -0.104908247 | 0.21987514 | -0.4771265 | 0.63327207 | NA | NA | NA | |
| ENSG000002 | 1.61982392 | -0.127389368 | 0.35032787 | -0.363629 | 0.71613508 | NA | NA | NA | |
| ENSG000002 | 0.7571138 | -0.203662327 | 0.28672441 | -0.7103069 | 0.47751384 | NA | NA | NA | |
| ENSG000002 | 1.07506855 | -0.103608381 | 0.31302785 | -0.3309877 | 0.74065374 | NA | NA | NA | |
| ENSG000002 | 1.74895762 | -0.467518338 | 0.35340327 | -1.3229033 | 0.18586755 | NA | NA | NA | |
| ENSG000002 | 2.12030207 | -0.650932749 | 0.36573121 | -1.7798119 | 0.07510675 | NA | NA | NA | |
| ENSG000002 | 1.84198573 | 0.008993401 | 0.35933746 | 0.02502773 | 0.98003284 | NA | NA | NA | |
| ENSG000002 | 1.60985605 | -0.126121805 | 0.35461449 | -0.3556589 | 0.72209602 | NA | NA | NA | |
| ENSG000002 | 0 | NA | NA | NA | NA | NA | NA | NA | |
| ENSG000002 | 0 | NA | NA | NA | NA | NA | NA | NA | |
| ENSG000002 | 0.88035449 | -0.282626859 | 0.30431514 | -0.9287309 | 0.35302858 | NA | NA | NA | |
| ENSG000002 | 0 | NA | NA | NA | NA | NA | NA | NA | |
| ENSG000002 | 1.48269802 | -0.338619972 | 0.34539346 | -0.9803891 | 0.32689411 | NA | NA | NA | |
| ENSG000002 | 1.60107063 | -0.387590367 | 0.34664862 | -1.1181073 | 0.26352115 | NA | NA | NA | |
| ENSG000002 | 2.13586162 | -0.133279979 | 0.36611114 | -0.3640424 | 0.71582635 | NA | NA | NA | |

| | | | | | | | | | |
|---|---|---|---|---|---|---|---|---|---|
| ENSG000002 | 0.52125014 | -0.010203417 | 0.15567753 | -0.065542 | 0.94774247 | NA | NA | NA | |
| ENSG000002 | 1.93358364 | -0.056813712 | 0.36132217 | -0.1572384 | 0.87505698 | NA | NA | NA | |
| ENSG000002 | 0.35273207 | -0.08884607 | 0.20605566 | -0.4311751 | 0.66634105 | NA | NA | NA | |
| ENSG000002 | 0.46136259 | 0.353033165 | 0.24467579 | 1.44286102 | 0.14905963 | NA | NA | NA | |
| ENSG000002 | 0.5974134 | 0.053016323 | 0.26223774 | 0.20216893 | 0.83978466 | NA | NA | NA | |
| ENSG000002 | 0.86091955 | -0.265574426 | 0.29690668 | -0.894471 | 0.37106993 | NA | NA | NA | |
| ENSG000002 | 1.03584375 | -0.18180248 | 0.30069578 | -0.604606 | 0.5454408 | NA | NA | NA | |
| ENSG000002 | 2.06356218 | -0.117724225 | 0.36163066 | -0.3255372 | 0.74477455 | NA | NA | NA | |
| ENSG000002 | 1.84236894 | -0.119716701 | 0.35793077 | -0.3344689 | 0.73802578 | NA | NA | NA | |
| ENSG000002 | 0.85058056 | -0.103122652 | 0.29515749 | -0.3493818 | 0.7268027 | NA | NA | NA | |
| ENSG000002 | 0 | NA | NA | NA | NA | NA | BBS1 | | 582 |
| ENSG000002 | 0.84702987 | -0.241400035 | 0.28775544 | -0.8389069 | 0.40152153 | NA | NA | NA | |
| ENSG000002 | 1.9058915 | -0.255647466 | 0.35720066 | -0.7156971 | 0.47417841 | NA | NA | NA | |
| ENSG000002 | 0.95978275 | 0.108273074 | 0.29920334 | 0.36187121 | 0.71744827 | NA | NA | NA | |
| ENSG000002 | 1.72279806 | -0.327451662 | 0.35342947 | -0.9264979 | 0.35418729 | NA | NA | NA | |
| ENSG000002 | 2.1718009 | -0.248947894 | 0.36320884 | -0.6854125 | 0.49308364 | NA | NA | NA | |
| ENSG000002 | 0 | NA | NA | NA | NA | NA | NA | NA | |
| ENSG000002 | 1.95416074 | -0.055925267 | 0.36358286 | -0.1538171 | 0.87775394 | NA | NA | NA | |
| ENSG000002 | 1.96136006 | -0.303265304 | 0.36072123 | -0.8407193 | 0.4005052 | NA | NA | NA | |
| ENSG000002 | 2.10552167 | -0.385685929 | 0.36633364 | -1.052827 | 0.29242031 | NA | NA | NA | |
| ENSG000002 | 1.85272899 | -0.6322097 | 0.35244179 | -1.7937989 | 0.07284527 | NA | NA | NA | |
| ENSG000002 | 1.74205831 | -0.198395254 | 0.35374867 | -0.5608367 | 0.57490883 | NA | NA | NA | |
| ENSG000002 | 2.14462398 | 0.305148412 | 0.36511328 | 0.83576365 | 0.40328788 | NA | NA | NA | |
| ENSG000002 | 1.98114341 | -0.067541129 | 0.36503153 | -0.1850282 | 0.85320691 | NA | NA | NA | |
| ENSG000002 | 0.76673731 | -0.201606703 | 0.28733642 | -0.7016399 | 0.48290375 | NA | NA | NA | |
| ENSG000002 | 0.85701948 | -0.100411347 | 0.29108473 | -0.3449557 | 0.73012765 | NA | NA | NA | |
| ENSG000002 | 0.84374989 | -0.097099741 | 0.29032625 | -0.3344504 | 0.73803969 | NA | NA | NA | |
| ENSG000002 | 0 | NA | NA | NA | NA | NA | NA | NA | |
| ENSG000002 | 1.73418224 | -0.455447971 | 0.35327875 | -1.2892028 | 0.19732758 | NA | NA | NA | |

| | | | | | | | | |
|---|---|---|---|---|---|---|---|---|
| ENSG000002 | 1.96078447 | 0.068367192 | 0.36024915 | 0.18977753 | 0.84948347 | NA | NA | NA |
| ENSG000002 | 0.49862008 | 0.145705772 | 0.2401745 | 0.6066663 | 0.54407239 | NA | NA | NA |
| ENSG000002 | 0 | NA | NA | NA | NA | NA | NA | NA |
| ENSG000002 | 0 | NA | NA | NA | NA | NA | NA | NA |
| ENSG000002 | 1.91110365 | 0.181279506 | 0.35730303 | 0.50735508 | 0.6119057 | NA | NA | NA |
| ENSG000002 | 1.84828367 | -0.264187418 | 0.35994251 | -0.7339712 | 0.46296632 | NA | NA | NA |
| ENSG000002 | 2.12999225 | -0.361310002 | 0.35936606 | -1.0054094 | 0.31469977 | NA | NA | NA |
| ENSG000002 | 0 | NA | NA | NA | NA | NA | NA | NA |
| ENSG000002 | 2.1319641 | -0.376818711 | 0.36577978 | -1.0301792 | 0.30292591 | NA | NA | NA |
| ENSG000002 | 1.98561593 | -0.606890764 | 0.34661833 | -1.7508905 | 0.07996476 | NA | LINC00273 | 649159 |
| ENSG000002 | 1.99978503 | -0.333266267 | 0.36443064 | -0.9144848 | 0.36046218 | NA | NA | NA |
| ENSG000002 | 1.51453613 | -0.327771442 | 0.34297273 | -0.9556779 | 0.33923501 | NA | NA | NA |
| ENSG000002 | 0 | NA | NA | NA | NA | NA | NA | NA |
| ENSG000002 | 0.84935071 | 0.20708171 | 0.29524145 | 0.70139781 | 0.48305479 | NA | NA | NA |
| ENSG000002 | 1.27674963 | -0.463675091 | 0.32030485 | -1.4476056 | 0.14772739 | NA | NA | NA |
| ENSG000002 | 1.49646508 | -0.203441151 | 0.34619367 | -0.5876513 | 0.55676639 | NA | NA | NA |
| ENSG000002 | 0.99883666 | 0.103468916 | 0.29836313 | 0.34678854 | 0.72875018 | NA | NA | NA |
| ENSG000002 | 2.1128099 | -0.367875824 | 0.36485083 | -1.008291 | 0.31331476 | NA | NA | NA |
| ENSG000002 | 1.59542291 | -0.23043303 | 0.33918703 | -0.6793686 | 0.49690431 | NA | NA | NA |
| ENSG000002 | 1.85519082 | 0.012536089 | 0.35782921 | 0.03503372 | 0.97205285 | NA | LOC100996671 | 100996671 |
| ENSG000002 | 1.40402232 | -0.419441453 | 0.33879845 | -1.2380265 | 0.21570624 | NA | NA | NA |
| ENSG000002 | 1.20621043 | -0.034310732 | 0.31030532 | -0.1105709 | 0.91195664 | NA | NA | NA |
| ENSG000002 | 0 | NA | NA | NA | NA | NA | STH | 246744 |
| ENSG000002 | 1.14350735 | 0.028754788 | 0.3112865 | 0.09237403 | 0.92640087 | NA | NA | NA |
| ENSG000002 | 1.10437687 | 0.037518038 | 0.31291646 | 0.11989794 | 0.904564 | NA | NA | NA |
| ENSG000002 | 2.10893907 | -0.377960046 | 0.36302794 | -1.0411321 | 0.29781424 | NA | NA | NA |
| ENSG000002 | 2.0917198 | -0.123938008 | 0.36428176 | -0.3402257 | 0.73368658 | NA | NA | NA |
| ENSG000002 | 1.18631565 | 0.260118898 | 0.32759183 | 0.79403354 | 0.4271759 | NA | NA | NA |
| ENSG000002 | 0.47821768 | -0.016471461 | 0.24462483 | -0.0673336 | 0.94631616 | NA | NA | NA |

| | | | | | | | | |
|---|---|---|---|---|---|---|---|---|
| ENSG000002 | 0 | NA | NA | NA | NA | NA | NA | NA |
| ENSG000002 | 1.53984692 | -0.498995773 | 0.34714443 | -1.4374299 | 0.15059588 | NA | NA | NA |
| ENSG000002 | 1.70469722 | -0.059087074 | 0.35477492 | -0.1665481 | 0.86772567 | NA | NA | NA |
| ENSG000002 | 1.14527853 | -0.253524287 | 0.31654035 | -0.8009225 | 0.42317651 | NA | NA | NA |
| ENSG000002 | 0 | NA | NA | NA | NA | NA | NA | NA |
| ENSG000002 | 0.48609822 | -0.022320526 | 0.23791393 | -0.0938177 | 0.92525401 | NA | NA | NA |
| ENSG000002 | 1.35695629 | -0.652420339 | 0.32469839 | -2.0093119 | 0.04450407 | NA | NA | NA |
| ENSG000002 | 2.00024149 | -0.329602045 | 0.36246634 | -0.9093315 | 0.36317519 | NA | NA | NA |
| ENSG000002 | 1.56358475 | -0.81293609 | 0.34376455 | -2.3648049 | 0.01803957 | NA | NA | NA |
| ENSG000002 | 0.92452856 | -0.244627232 | 0.28505648 | -0.8581711 | 0.390798 | NA | NA | NA |
| ENSG000002 | 1.52057402 | -0.625226184 | 0.34244016 | -1.8257969 | 0.06788086 | NA | NA | NA |
| ENSG000002 | 0.7164389 | -0.025867019 | 0.28328556 | -0.0913108 | 0.92724567 | NA | NA | NA |
| ENSG000002 | 2.1155163 | -0.469005762 | 0.36060872 | -1.3005946 | 0.19339725 | NA | NA | NA |
| ENSG000002 | 2.0405417 | -0.323623415 | 0.36219004 | -0.8935183 | 0.3715797 | NA | NA | NA |
| ENSG000002 | 1.53798929 | -0.334083 | 0.34335623 | -0.9729924 | 0.33055709 | NA | NA | NA |
| ENSG000002 | 0.89364336 | -0.29139313 | 0.3048889 | -0.9557354 | 0.33920591 | NA | NA | NA |
| ENSG000002 | 1.67514788 | -0.697414586 | 0.34789223 | -2.0046857 | 0.04499665 | NA | LOC105369593 | 105369593 |
| ENSG000002 | 1.30238409 | -0.333313798 | 0.32548668 | -1.0240474 | 0.30581287 | NA | NA | NA |
| ENSG000002 | 0.12685397 | -0.090215768 | 0.12901919 | -0.699243 | 0.48440017 | NA | NA | NA |
| ENSG000002 | 0.65493287 | -0.26892161 | 0.26251527 | -1.0244037 | 0.30564465 | NA | NA | NA |
| ENSG000002 | 0 | NA | NA | NA | NA | NA | NA | NA |
| ENSG000002 | 0.5983253 | -0.096573125 | 0.25511974 | -0.3785404 | 0.7050292 | NA | NA | NA |
| ENSG000002 | 0 | NA | NA | NA | NA | NA | NA | NA |
| ENSG000002 | 0.36345889 | 0.070257596 | 0.21589421 | 0.32542602 | 0.74485866 | NA | NA | NA |
| ENSG000002 | 0.81978317 | 0.195087942 | 0.28926628 | 0.67442337 | 0.50004219 | NA | KHDC1L | 100129128 |
| ENSG000002 | 0.97091844 | -0.035928459 | 0.31631496 | -0.1135844 | 0.90956722 | NA | NA | NA |
| ENSG000002 | 0.37148378 | 0.06737793 | 0.20875796 | 0.32275622 | 0.74687987 | NA | NA | NA |
| ENSG000002 | 0.11934008 | 0.083564698 | 0.12760091 | 0.65489106 | 0.5125379 | NA | NA | NA |
| ENSG000002 | 2.01055113 | -0.863812655 | 0.35912602 | -2.405319 | 0.01615835 | NA | NA | NA |

| | | | | | | | | |
|---|---|---|---|---|---|---|---|---|
| ENSG000002 | 1.6222097 | -0.269401897 | 0.35081785 | -0.7679253 | 0.44253158 | NA | NA | NA |
| ENSG000002 | 0.12685397 | -0.090215768 | 0.12901919 | -0.699243 | 0.48440017 | NA | NA | NA |
| ENSG000002 | 1.3633453 | -0.379609167 | 0.32908186 | -1.1535402 | 0.24868872 | NA | NA | NA |
| ENSG000002 | 0.74304943 | -0.034670049 | 0.26965489 | -0.1285719 | 0.89769638 | NA | NA | NA |
| ENSG000002 | 0.52807889 | -0.192145741 | 0.24416537 | -0.7869492 | 0.4313116 | NA | NA | NA |
| ENSG000002 | 0 | NA | NA | NA | NA | NA | NA | NA |
| ENSG000002 | 0.63777427 | -0.25910394 | 0.25947273 | -0.9985787 | 0.31799883 | NA | NA | NA |
| ENSG000002 | 1.29200218 | -0.50706363 | 0.3307463 | -1.5330894 | 0.12525384 | NA | NA | NA |
| ENSG000002 | 1.35199996 | -0.406130636 | 0.33556904 | -1.2102745 | 0.22617359 | NA | NA | NA |
| ENSG000002 | 1.1580564 | -0.430687338 | 0.32014615 | -1.3452835 | 0.17853369 | NA | NA | NA |
| ENSG000002 | 0.48376239 | -0.021735405 | 0.23765615 | -0.0914574 | 0.92712918 | NA | LRRC37A13P | 106479024 |
| ENSG000002 | 1.51843419 | -0.051357139 | 0.33955459 | -0.1512485 | 0.87977965 | NA | NA | NA |
| ENSG000002 | 0.3575613 | 0.072305152 | 0.21497462 | 0.33634273 | 0.73661243 | NA | OR11H12 | 440153 |
| ENSG000002 | 1.78117736 | -0.771950528 | 0.35428349 | -2.1789063 | 0.02933863 | NA | NA | NA |
| ENSG000002 | 0.64013129 | -0.113197991 | 0.26627307 | -0.4251199 | 0.67074925 | NA | NA | NA |
| ENSG000002 | 1.55093788 | 0.442729152 | 0.34700145 | 1.2758712 | 0.20200105 | NA | NA | NA |
| ENSG000002 | 1.10417696 | -0.259029211 | 0.31842451 | -0.8134713 | 0.41594787 | NA | NA | NA |
| ENSG000002 | 1.59750887 | -0.402698534 | 0.35053597 | -1.148808 | 0.25063515 | NA | NA | NA |
| ENSG000002 | 1.21902165 | -0.182430184 | 0.3227705 | -0.5652009 | 0.57193711 | NA | NA | NA |
| ENSG000002 | 1.24230067 | -0.051960162 | 0.3338897 | -0.1556207 | 0.87633198 | NA | HMGN2P6 | 643872 |
| ENSG000002 | 0 | NA | NA | NA | NA | NA | NA | NA |
| ENSG000002 | 0 | NA | NA | NA | NA | NA | NA | NA |
| ENSG000002 | 1.94840112 | 0.071129838 | 0.36074055 | 0.19717727 | 0.84368882 | NA | NA | NA |
| ENSG000002 | 0 | NA | NA | NA | NA | NA | NA | NA |
| ENSG000002 | 1.00789361 | -0.346076759 | 0.31094524 | -1.112983 | 0.26571574 | NA | NA | NA |
| ENSG000002 | 1.47747505 | -0.173614371 | 0.33931072 | -0.5116678 | 0.60888353 | NA | NA | NA |
| ENSG000002 | 0.38747904 | -0.104908247 | 0.21987514 | -0.4771265 | 0.63327207 | NA | NA | NA |
| ENSG000002 | 0.38036605 | -0.293832733 | 0.22119083 | -1.3284128 | 0.18404178 | NA | NA | NA |
| ENSG000002 | 0.87736735 | -0.118228369 | 0.29817545 | -0.3965061 | 0.69173174 | NA | NA | NA |

| | | | | | | | | |
|---|---|---|---|---|---|---|---|---|
| ENSG000002 | 1.52324587 | 0.148699819 | 0.34091662 | 0.4361765 | 0.66270868 | NA | NA | NA |
| ENSG000002 | 1.49183069 | -0.451958809 | 0.33898925 | -1.3332541 | 0.18244843 | NA | NA | NA |
| ENSG000002 | 1.61777276 | -0.402924208 | 0.35081251 | -1.1485457 | 0.25074335 | NA | NA | NA |
| ENSG000002 | 1.15788225 | -0.276667752 | 0.32299046 | -0.8565818 | 0.39167604 | NA | NA | NA |
| ENSG000002 | 1.26237315 | -0.320618557 | 0.32525868 | -0.9857341 | 0.32426363 | NA | NA | NA |
| ENSG000002 | 1.42539616 | 0.092331855 | 0.3414271 | 0.2704292 | 0.78683008 | NA | NA | NA |
| ENSG000002 | 1.28542137 | -0.336453547 | 0.3274863 | -1.027382 | 0.30424061 | NA | NA | NA |
| ENSG000002 | 1.06484609 | -0.526071742 | 0.31132635 | -1.6897758 | 0.09107085 | NA | NA | NA |
| ENSG000002 | 0.79007233 | -0.361217845 | 0.28349795 | -1.2741462 | 0.20261159 | NA | NA | NA |
| ENSG000002 | 2.14912972 | -0.786454697 | 0.36400977 | -2.1605318 | 0.03073152 | NA | NA | NA |
| ENSG000002 | 1.00953983 | -0.046292298 | 0.31369112 | -0.1475729 | 0.88267987 | NA | NA | NA |
| ENSG000002 | 1.4976445 | -0.400155862 | 0.32739771 | -1.2222317 | 0.22162002 | NA | NA | NA |
| ENSG000002 | 1.70446909 | -0.055013651 | 0.35365665 | -0.1555567 | 0.87638248 | NA | NA | NA |
| ENSG000002 | 1.75817674 | -0.4878537 | 0.35862403 | -1.3603486 | 0.17371964 | NA | NA | NA |
| ENSG000002 | 0.3867939 | -0.105148852 | 0.21977268 | -0.4784437 | 0.63233444 | NA | NA | NA |
| ENSG000002 | 0 | NA | NA | NA | NA | NA | NA | NA |
| ENSG000002 | 0.47207214 | -0.012756135 | 0.23624418 | -0.0539956 | 0.95693871 | NA | NA | NA |
| ENSG000002 | 2.01673103 | -0.727169247 | 0.36131231 | -2.0125781 | 0.04415903 | NA | NA | NA |
| ENSG000002 | 0.99863832 | -0.045375191 | 0.30675492 | -0.14792 | 0.88240589 | NA | NA | NA |
| ENSG000002 | 0 | NA | NA | NA | NA | NA | NA | NA |
| ENSG000002 | 1.73397138 | -0.055669106 | 0.35564581 | -0.1565296 | 0.87561557 | NA | NA | NA |
| ENSG000002 | 0 | NA | NA | NA | NA | NA | NA | NA |
| ENSG000002 | 0 | NA | NA | NA | NA | NA | MIR3179-2 | 100422886 |
| ENSG000002 | 1.46288199 | -0.048890071 | 0.345278 | -0.1415963 | 0.88739893 | NA | NA | NA |
| ENSG000002 | 2.08169952 | -0.122120634 | 0.36387863 | -0.3356082 | 0.73716638 | NA | NA | NA |
| ENSG000002 | 0.74883278 | -0.478142561 | 0.26884577 | -1.7785013 | 0.07532156 | NA | NA | NA |
| ENSG000002 | 1.47471446 | -0.16108471 | 0.32723169 | -0.492265 | 0.62253201 | NA | NA | NA |
| ENSG000002 | 0.23936338 | -0.005479057 | 0.17759977 | -0.0308506 | 0.9753887 | NA | NA | NA |
| ENSG000002 | 0.98147112 | 0.271592889 | 0.30689914 | 0.88495813 | 0.37617921 | NA | NA | NA |

| | | | | | | | | | |
|---|---|---|---|---|---|---|---|---|---|
| ENSG000002 | 0 | NA | NA | NA | NA | NA | NA | NA | |
| ENSG000002 | 0.2596576 | -0.193002035 | 0.1837913 | -1.0501152 | 0.29366517 | NA | NA | NA | |
| ENSG000002 | 1.62349374 | -0.416528747 | 0.35257363 | -1.1813951 | 0.23744582 | NA | NA | NA | |
| ENSG000002 | 2.01652818 | 0.129571769 | 0.35916537 | 0.36075797 | 0.71828039 | NA | NA | NA | |
| ENSG000002 | 1.63759766 | 0.006507776 | 0.34173721 | 0.01904322 | 0.98480663 | NA | NA | NA | |
| ENSG000002 | 1.70856559 | -0.17914245 | 0.35338303 | -0.5069356 | 0.61219998 | NA | NA | NA | |
| ENSG000002 | 1.62832937 | -0.361168113 | 0.34092743 | -1.0593695 | 0.28943154 | NA | NA | NA | |
| ENSG000002 | 2.02151429 | -0.199844995 | 0.35016891 | -0.5707103 | 0.56819606 | NA | NA | NA | |
| ENSG000002 | 1.15495357 | -0.275275876 | 0.32290101 | -0.8525086 | 0.39393188 | NA | NA | NA | |
| ENSG000002 | 1.50626328 | -0.340343774 | 0.34578128 | -0.9842747 | 0.32498046 | NA | NA | NA | |
| ENSG000002 | 1.60797668 | -0.128784231 | 0.35360701 | -0.3642016 | 0.71570749 | NA | LOC642426 | | 642426 |
| ENSG000002 | 0.38270188 | -0.272446727 | 0.2140009 | -1.2731102 | 0.20297894 | NA | NA | NA | |
| ENSG000002 | 1.63331156 | -0.530487249 | 0.34716036 | -1.5280755 | 0.12649379 | NA | NA | NA | |
| ENSG000002 | 2.15152057 | -0.51614298 | 0.36513548 | -1.4135657 | 0.15748947 | NA | NA | NA | |
| ENSG000002 | 0.12685397 | -0.090215768 | 0.12901919 | -0.699243 | 0.48440017 | NA | NA | NA | |
| ENSG000002 | 1.88898689 | -0.371650593 | 0.35536818 | -1.0458184 | 0.29564487 | NA | NA | NA | |
| ENSG000002 | 0 | NA | NA | NA | NA | NA | NA | NA | |
| ENSG000002 | 0.61750123 | 0.055806918 | 0.26401083 | 0.21138117 | 0.83258985 | NA | NA | NA | |
| ENSG000002 | 0 | NA | NA | NA | NA | NA | NA | NA | |
| ENSG000002 | 2.11637111 | -0.383210327 | 0.36618068 | -1.0465061 | 0.29532742 | NA | NA | NA | |
| ENSG000002 | 1.19810619 | -0.591329122 | 0.31849508 | -1.8566351 | 0.06336309 | NA | NA | NA | |
| ENSG000002 | 1.18094508 | 0.100575792 | 0.31368905 | 0.32062258 | 0.74849643 | NA | NA | NA | |
| ENSG000002 | 0.34344486 | 0.238503239 | 0.20662376 | 1.15428757 | 0.24838229 | NA | NA | NA | |
| ENSG000002 | 0.73311985 | -0.175350888 | 0.27912815 | -0.6282093 | 0.52986687 | NA | NA | NA | |
| ENSG000002 | 1.34131214 | 0.02550809 | 0.33891407 | 0.07526418 | 0.94000452 | NA | NA | NA | |
| ENSG000002 | 1.49456507 | -0.309855651 | 0.33671344 | -0.9202355 | 0.35744972 | NA | NA | NA | |
| ENSG000002 | 1.43376699 | 0.21047781 | 0.33707891 | 0.62441702 | 0.53235375 | NA | NA | NA | |
| ENSG000002 | 1.62605825 | -0.412157823 | 0.35262564 | -1.1688254 | 0.24247397 | NA | NA | NA | |
| ENSG000002 | 1.29453388 | -0.517535719 | 0.33287212 | -1.5547584 | 0.12000363 | NA | NA | NA | |

| | | | | | | | | |
|---|---|---|---|---|---|---|---|---|
| ENSG000002 | 0 | NA | NA | NA | NA | NA | NA | NA |
| ENSG000002 | 0.97834589 | -0.310389428 | 0.29697817 | -1.0451591 | 0.29594946 | NA | NA | NA |
| ENSG000002 | 2.16079338 | 0.058179923 | 0.36896302 | 0.15768497 | 0.87470504 | NA | KCCAT198 | 105369954 |
| ENSG000002 | 1.88466015 | -0.406707362 | 0.36157032 | -1.1248361 | 0.26065848 | NA | NA | NA |
| ENSG000002 | 0.84773228 | 0.181666502 | 0.2821682 | 0.64382344 | 0.51968993 | NA | NA | NA |
| ENSG000002 | 0.37199232 | 0.06760863 | 0.21718977 | 0.31128828 | 0.75558148 | NA | NA | NA |
| ENSG000002 | 1.00655716 | -0.18306274 | 0.30210179 | -0.6059638 | 0.54453881 | NA | NA | NA |
| ENSG000002 | 1.49219891 | -0.195215168 | 0.34490594 | -0.5659954 | 0.57139692 | NA | NA | NA |
| ENSG000002 | 1.42097861 | 0.494958104 | 0.33075578 | 1.49644582 | 0.13453752 | NA | NA | NA |
| ENSG000002 | 0.73564763 | 0.13375234 | 0.27852684 | 0.48021347 | 0.63107561 | NA | NA | NA |
| ENSG000002 | 0.89244914 | -0.284123598 | 0.30006291 | -0.9468801 | 0.34369988 | NA | NA | NA |
| ENSG000002 | 1.06267448 | 0.0355814 | 0.30553168 | 0.11645732 | 0.90729011 | NA | NA | NA |
| ENSG000002 | 0.34270765 | 0.258172734 | 0.21433185 | 1.20454674 | 0.22837833 | NA | NA | NA |
| ENSG000002 | 1.5057421 | -0.305858287 | 0.33801423 | -0.9048681 | 0.36553528 | NA | NA | NA |
| ENSG000002 | 2.09518268 | -0.253238357 | 0.36586973 | -0.6921544 | 0.48884035 | NA | NA | NA |
| ENSG000002 | 2.11070378 | -0.257058028 | 0.36716519 | -0.7001155 | 0.4838552 | NA | NA | NA |
| ENSG000002 | 1.83997857 | -0.258757048 | 0.35830775 | -0.7221642 | 0.47019351 | NA | NA | NA |
| ENSG000002 | 0 | NA | NA | NA | NA | NA | NA | NA |
| ENSG000002 | 0.36690249 | -0.096999144 | 0.21683809 | -0.4473344 | 0.65463361 | NA | NA | NA |
| ENSG000002 | 1.73836071 | -0.187256521 | 0.3543546 | -0.5284439 | 0.59719128 | NA | NA | NA |
| ENSG000002 | 0.83683077 | 0.192554645 | 0.28855806 | 0.66729949 | 0.50458085 | NA | NA | NA |
| ENSG000002 | 1.39220733 | -0.420060297 | 0.33854424 | -1.2407841 | 0.21468553 | NA | NA | NA |
| ENSG000002 | 1.79427481 | -1.078970133 | 0.3484265 | -3.0966936 | 0.00195692 | NA | NA | NA |
| ENSG000002 | 0 | NA | NA | NA | NA | NA | NA | NA |
| ENSG000002 | 1.57218093 | -0.089064213 | 0.33467943 | -0.266118 | 0.79014835 | NA | NA | NA |
| ENSG000002 | 1.89029872 | -0.261926988 | 0.3587274 | -0.7301561 | 0.46529479 | NA | NA | NA |
| ENSG000002 | 1.48693495 | -0.175081285 | 0.33948515 | -0.5157259 | 0.60604586 | NA | NA | NA |
| ENSG000002 | 1.23860094 | -0.19360319 | 0.33422907 | -0.579253 | 0.56241848 | NA | NA | NA |
| ENSG000002 | 1.52854451 | -0.188353488 | 0.33896106 | -0.5556788 | 0.57843043 | NA | NA | NA |

| | | | | | | | | |
|---|---|---|---|---|---|---|---|---|
| ENSG000002 | 0 | NA | NA | NA | NA | NA | NA | NA |
| ENSG000002 | 1.0502192 | -0.511171751 | 0.30845471 | -1.6572019 | 0.09747865 | NA | NA | NA |
| ENSG000002 | 0.86134219 | -0.388987513 | 0.28567108 | -1.3616622 | 0.17330452 | NA | NA | NA |
| ENSG000002 | 1.76457617 | -0.049497755 | 0.33998647 | -0.1455874 | 0.88424709 | NA | NA | NA |
| ENSG000002 | 1.47318268 | -0.051468704 | 0.34544979 | -0.1489904 | 0.8815612 | NA | NA | NA |
| ENSG000002 | 0.60299641 | -0.1063938 | 0.26311939 | -0.4043556 | 0.68595125 | NA | NA | NA |
| ENSG000002 | 0.34197044 | 0.237837536 | 0.20637034 | 1.15247924 | 0.2491242 | NA | NA | NA |
| ENSG000002 | 1.45203446 | 0.095127323 | 0.34050098 | 0.2793746 | 0.77995736 | NA | NA | NA |
| ENSG000002 | 1.16978549 | -0.610076399 | 0.3224802 | -1.8918259 | 0.05851418 | NA | NA | NA |
| ENSG000002 | 1.14206561 | -0.598469485 | 0.32172533 | -1.8601877 | 0.06285898 | NA | NA | NA |
| ENSG000002 | 0.76678939 | -0.201566736 | 0.28734031 | -0.7014913 | 0.48299644 | NA | NA | NA |
| ENSG000002 | 1.99463317 | -0.297834583 | 0.35875577 | -0.8301876 | 0.40643274 | NA | NA | NA |
| ENSG000002 | 2.012221 | -0.322858056 | 0.36343594 | -0.8883493 | 0.3743529 | NA | NA | NA |
| ENSG000002 | 0 | NA | NA | NA | NA | NA | NA | NA |
| ENSG000002 | 0.78227611 | -0.381453213 | 0.2899593 | -1.3155405 | 0.18832831 | NA | NA | NA |
| ENSG000002 | 1.85474632 | -0.485046983 | 0.35163575 | -1.3794018 | 0.1677709 | NA | NA | NA |
| ENSG000002 | 1.57971155 | 0.155889338 | 0.35184888 | 0.44305766 | 0.65772403 | NA | NA | NA |
| ENSG000002 | 2.02275587 | -0.438266383 | 0.35975209 | -1.2182456 | 0.22313067 | NA | NA | NA |
| ENSG000002 | 0.14128499 | -0.097772014 | 0.13403235 | -0.7294658 | 0.4657168 | NA | NA | NA |
| ENSG000002 | 0.37859486 | 0.064147216 | 0.20986618 | 0.30565771 | 0.75986527 | NA | NA | NA |
| ENSG000002 | 2.01621803 | -0.432439894 | 0.35864105 | -1.2057735 | 0.22790482 | NA | NA | NA |
| ENSG000002 | 1.12930254 | -0.11794969 | 0.32280241 | -0.3653928 | 0.71481818 | NA | NA | NA |
| ENSG000002 | 1.12447331 | -0.280802665 | 0.32707248 | -0.8585334 | 0.39059797 | NA | RPL3P13 | 106480199 |
| ENSG000002 | 1.73922246 | -0.170881525 | 0.34850814 | -0.4903229 | 0.62390542 | NA | NA | NA |
| ENSG000002 | 1.37240237 | -0.26222318 | 0.33713477 | -0.7777993 | 0.43668735 | NA | NA | NA |
| ENSG000002 | 0.75116414 | -0.201739256 | 0.28635357 | -0.7045111 | 0.48111457 | NA | NA | NA |
| ENSG000002 | 1.26530129 | -0.363844343 | 0.3355307 | -1.0843846 | 0.27819429 | NA | NA | NA |
| ENSG000002 | 0 | NA | NA | NA | NA | NA | NA | NA |
| ENSG000002 | 0 | NA | NA | NA | NA | NA | NA | NA |

| | | | | | | | | | |
|---|---|---|---|---|---|---|---|---|---|
| ENSG000002 | 0.60150597 | 0.230273195 | 0.26371223 | 0.87319879 | 0.38255474 | NA | NA | NA | |
| ENSG000002 | 1.80453127 | 0.107023397 | 0.34486911 | 0.31033048 | 0.75630965 | NA | NA | NA | |
| ENSG000002 | 0.80518849 | -0.337327945 | 0.27554191 | -1.2242346 | 0.22086373 | NA | NA | NA | |
| ENSG000002 | 0.62482117 | 0.05427805 | 0.26463902 | 0.20510222 | 0.83749226 | NA | NA | NA | |
| ENSG000002 | 0.9487771 | 0.114735664 | 0.306278 | 0.37461282 | 0.70794844 | NA | NA | NA | |
| ENSG000002 | 0.99872656 | -0.362859801 | 0.31343729 | -1.1576791 | 0.24699501 | NA | NA | NA | |
| ENSG000002 | 1.88116004 | 0.194089769 | 0.35521811 | 0.54639604 | 0.58479373 | NA | NA | NA | |
| ENSG000002 | 1.75182426 | -0.327060555 | 0.35504151 | -0.9211896 | 0.35695143 | NA | NA | NA | |
| ENSG000002 | 0.90435292 | -0.633941791 | 0.29858306 | -2.1231673 | 0.03373983 | NA | NA | NA | |
| ENSG000002 | 1.892949 | -0.653290597 | 0.35465843 | -1.8420275 | 0.06547112 | NA | NA | NA | |
| ENSG000002 | 0.95355427 | 0.27081259 | 0.3072828 | 0.88131387 | 0.37814796 | NA | NA | NA | |
| ENSG000002 | 0.52668879 | -0.197054386 | 0.24396589 | -0.8077129 | 0.4192559 | NA | NA | NA | |
| ENSG000002 | 0 | NA | NA | NA | NA | NA | LOC388242 | | 388242 |
| ENSG000002 | 0 | NA | NA | NA | NA | NA | NA | NA | |
| ENSG000002 | 0.85565111 | 0.046315327 | 0.2888886 | 0.16032244 | 0.87262708 | NA | NA | NA | |
| ENSG000002 | 1.14598294 | -0.273361555 | 0.32262649 | -0.8473004 | 0.3968277 | NA | NA | NA | |
| ENSG000002 | 1.47773756 | -0.048025903 | 0.3455223 | -0.1389951 | 0.88945403 | NA | NA | NA | |
| ENSG000002 | 0.90807438 | -0.279916171 | 0.29927035 | -0.9353288 | 0.34961889 | NA | NA | NA | |
| ENSG000002 | 0.25993993 | -0.013144386 | 0.18211683 | -0.0721756 | 0.94246219 | NA | NA | NA | |
| ENSG000002 | 0 | NA | NA | NA | NA | NA | NA | NA | |
| ENSG000002 | 1.58740013 | -0.123594321 | 0.35335093 | -0.3497778 | 0.72650544 | NA | NA | NA | |
| ENSG000002 | 2.04821993 | 0.511762756 | 0.36243896 | 1.41199708 | 0.15795082 | NA | NA | NA | |
| ENSG000002 | 1.12296059 | -0.125047233 | 0.32259195 | -0.3876328 | 0.69828776 | NA | NA | NA | |
| ENSG000002 | 0.79511008 | -0.182452322 | 0.27194202 | -0.6709236 | 0.50226919 | NA | NA | NA | |
| ENSG000002 | 1.77234662 | -0.476107951 | 0.35533987 | -1.3398664 | 0.18028879 | NA | NA | NA | |
| ENSG000002 | 0.47075585 | 0.160115757 | 0.24423976 | 0.65556794 | 0.51210216 | NA | NA | NA | |
| ENSG000002 | 1.89621646 | -0.236308296 | 0.35219 | -0.6709682 | 0.50224078 | NA | NA | NA | |
| ENSG000002 | 0.36075696 | -0.085953824 | 0.20740622 | -0.4144226 | 0.67856464 | NA | NA | NA | |
| ENSG000002 | 0.88732314 | -0.112890519 | 0.2910107 | -0.3879257 | 0.69807104 | NA | NA | NA | |

| | | | | | | | | |
|---|---|---|---|---|---|---|---|---|
| ENSG000002 | 0.37216701 | -0.099548723 | 0.21762847 | -0.4574251 | 0.64736554 | NA | NA | NA |
| ENSG000002 | 1.16744966 | -0.609564442 | 0.32241415 | -1.8906256 | 0.05867434 | NA | NA | NA |
| ENSG000002 | 1.19428601 | 0.108288433 | 0.33254341 | 0.32563698 | 0.74469903 | NA | NA | NA |
| ENSG000002 | 1.66692957 | -0.523473187 | 0.34581161 | -1.5137525 | 0.13008862 | NA | NA | NA |
| ENSG000002 | 0.97597848 | 0.260695916 | 0.29933222 | 0.87092502 | 0.38379509 | NA | NA | NA |
| ENSG000002 | 2.12117056 | -0.104359406 | 0.3566451 | -0.2926142 | 0.76981709 | NA | NA | NA |
| ENSG000002 | 0.87457898 | -0.427920645 | 0.29435302 | -1.4537668 | 0.14601097 | NA | NA | NA |
| ENSG000002 | 0.90090931 | -0.442334366 | 0.29735163 | -1.4875801 | 0.13686166 | NA | NA | NA |
| ENSG000002 | 1.749346 | -0.204079997 | 0.35762905 | -0.5706471 | 0.56823885 | NA | NA | NA |
| ENSG000002 | 1.89222803 | -0.389873882 | 0.35780416 | -1.0896293 | 0.27587649 | NA | NA | NA |
| ENSG000002 | 2.10815943 | -0.236371323 | 0.36224408 | -0.6525195 | 0.51406611 | NA | NA | NA |
| ENSG000002 | 0.59078107 | 0.217502869 | 0.25401457 | 0.85626139 | 0.39185321 | NA | NA | NA |
| ENSG000002 | 0.65561801 | -0.268676647 | 0.26257757 | -1.0232277 | 0.3062002 | NA | NA | NA |
| ENSG000002 | 1.86445362 | -0.52851389 | 0.35778667 | -1.477176 | 0.13962846 | NA | NA | NA |
| ENSG000002 | 0.64870089 | -0.415503931 | 0.25466679 | -1.6315591 | 0.1027724 | NA | NA | NA |
| ENSG000002 | 1.25775514 | -0.199637515 | 0.33468688 | -0.5964904 | 0.55084766 | NA | NA | NA |
| ENSG000002 | 0.14128499 | -0.097772014 | 0.13403235 | -0.7294658 | 0.4657168 | NA | NA | NA |
| ENSG000002 | 1.75387969 | -0.3264086 | 0.35506463 | -0.9192935 | 0.35794209 | NA | NA | NA |
| ENSG000002 | 1.59335171 | 0.015930255 | 0.34907026 | 0.04563624 | 0.96360018 | NA | DUX4L52 | 106480354 |
| ENSG000002 | 1.79078594 | 0.134625295 | 0.35626466 | 0.37788002 | 0.70551973 | NA | NA | NA |
| ENSG000002 | 1.9312282 | -0.057888285 | 0.36206904 | -0.1598819 | 0.87297411 | NA | NA | NA |
| ENSG000002 | 1.41912958 | 0.370220206 | 0.33870698 | 1.09303979 | 0.27437633 | NA | NA | NA |
| ENSG000002 | 0.74268725 | -0.479320364 | 0.26912835 | -1.7810103 | 0.07491077 | NA | NA | NA |
| ENSG000002 | 1.09529627 | 0.186091026 | 0.32158634 | 0.57866582 | 0.56281468 | NA | NA | NA |
| ENSG000002 | 0.75896283 | -0.026905877 | 0.27820585 | -0.0967121 | 0.92295502 | NA | NA | NA |
| ENSG000002 | 1.47872092 | 0.346289858 | 0.33337986 | 1.03872458 | 0.29893285 | NA | NA | NA |
| ENSG000002 | 1.61859642 | -0.263224321 | 0.35190845 | -0.7479909 | 0.45446564 | NA | NA | NA |
| ENSG000002 | 2.16737448 | -0.514028435 | 0.36572244 | -1.4055151 | 0.15986815 | NA | NA | NA |
| ENSG000002 | 1.51802058 | -0.748065573 | 0.33512991 | -2.232166 | 0.025604 | NA | NA | NA |

| | | | | | | | | |
|---|---|---|---|---|---|---|---|---|
| ENSG000002 | 1.63111828 | -0.112514548 | 0.34689971 | -0.3243432 | 0.74567825 | NA | NA | NA |
| ENSG000002 | 1.28881099 | -0.678857587 | 0.33109616 | -2.0503336 | 0.04033189 | NA | NA | NA |
| ENSG000002 | 2.08428951 | -1.003523544 | 0.35578788 | -2.820567 | 0.00479389 | NA | NA | NA |
| ENSG000002 | 0.23890637 | -0.010420674 | 0.17747379 | -0.0587167 | 0.95317776 | NA | NA | NA |
| ENSG000002 | 0.6567905 | -0.46248946 | 0.26597757 | -1.7388288 | 0.08206488 | NA | MEG8 | 79104 |
| ENSG000002 | 1.34624618 | -0.384563017 | 0.33001118 | -1.165303 | 0.24389637 | NA | NA | NA |
| ENSG000002 | 1.32114053 | 0.035163161 | 0.3384712 | 0.10388819 | 0.91725808 | NA | NA | NA |
| ENSG000002 | 0.4833075 | 0.150471097 | 0.24570075 | 0.61241611 | 0.54026249 | NA | NA | NA |
| ENSG000002 | 0.47001864 | 0.159831094 | 0.24415434 | 0.65463138 | 0.51270511 | NA | NA | NA |
| ENSG000002 | 1.62739436 | -0.536646854 | 0.34821659 | -1.5411295 | 0.12328527 | NA | NA | NA |
| ENSG000002 | 0 | NA | NA | NA | NA | NA | NA | NA |
| ENSG000002 | 2.06931754 | -0.005160944 | 0.36754837 | -0.0140415 | 0.98879684 | NA | NA | NA |
| ENSG000002 | 1.83912175 | 0.012555567 | 0.35407823 | 0.03545987 | 0.97171305 | NA | LOC105370586 | 105370586 |
| ENSG000002 | 0 | NA | NA | NA | NA | NA | NA | NA |
| ENSG000002 | 2.16201768 | -0.504497768 | 0.36435813 | -1.3846206 | 0.16616852 | NA | NA | NA |
| ENSG000002 | 0.84426089 | 0.055158544 | 0.2899695 | 0.19022188 | 0.84913526 | NA | RNASE12 | 493901 |
| ENSG000002 | 2.00732705 | -0.410099954 | 0.35535582 | -1.1540544 | 0.24847786 | NA | NA | NA |
| ENSG000002 | 2.04737262 | 0.129164772 | 0.36739101 | 0.35157303 | 0.72515849 | NA | NA | NA |
| ENSG000002 | 1.71887212 | -0.312219161 | 0.35294478 | -0.8846119 | 0.37636598 | NA | NA | NA |
| ENSG000002 | 1.9763536 | -0.186693228 | 0.36225636 | -0.5153622 | 0.60629996 | NA | NA | NA |
| ENSG000002 | 1.76116333 | -0.610457026 | 0.35397252 | -1.7245887 | 0.08460162 | NA | OR11H2 | 79334 |
| ENSG000002 | 1.03510304 | -0.536100349 | 0.31389681 | -1.7078872 | 0.08765727 | NA | NA | NA |
| ENSG000002 | 2.00166832 | -0.589115625 | 0.36182116 | -1.6281956 | 0.10348341 | NA | NA | NA |
| ENSG000002 | 1.06874833 | 0.034337083 | 0.3124672 | 0.1098902 | 0.91249645 | NA | NA | NA |
| ENSG000002 | 1.83937963 | -0.598478036 | 0.35107656 | -1.7046938 | 0.08825155 | NA | NA | NA |
| ENSG000002 | 1.98342368 | -0.325369924 | 0.3643433 | -0.8930312 | 0.37184048 | NA | NA | NA |
| ENSG000002 | 1.89720537 | 0.470496747 | 0.36054383 | 1.30496409 | 0.19190507 | NA | NA | NA |
| ENSG000002 | 1.78065384 | -0.330290654 | 0.35470758 | -0.9311632 | 0.35176914 | NA | NA | NA |
| ENSG000002 | 1.60408108 | -0.253298122 | 0.35059943 | -0.7224716 | 0.47000459 | NA | NA | NA |

| | | | | | | | | |
|---|---|---|---|---|---|---|---|---|
| ENSG000002 | 1.90171166 | -0.667057772 | 0.35817252 | -1.8623924 | 0.0625478 | NA | NA | NA |
| ENSG000002 | 1.06034078 | 0.196872923 | 0.32033316 | 0.61458802 | 0.53882683 | NA | NA | NA |
| ENSG000002 | 1.58238603 | 0.025560277 | 0.33239421 | 0.07689748 | 0.9387051 | NA | NA | NA |
| ENSG000002 | 1.93432086 | -0.055419354 | 0.3597353 | -0.1540559 | 0.87756564 | NA | NA | NA |
| ENSG000002 | 0.64759311 | -0.271681639 | 0.26184098 | -1.0375826 | 0.29946444 | NA | NA | NA |
| ENSG000002 | 0 | NA | NA | NA | NA | NA | NA | NA |
| ENSG000002 | 1.16859181 | -0.605351814 | 0.32253551 | -1.8768533 | 0.06053821 | NA | NA | NA |
| ENSG000002 | 0.61833933 | -0.261391558 | 0.25929785 | -1.0080745 | 0.31341869 | NA | NA | NA |
| ENSG000002 | 0.27408862 | -0.199826037 | 0.18669465 | -1.0703362 | 0.28446801 | NA | NA | NA |
| ENSG000002 | 0.98661019 | -0.487243331 | 0.30323516 | -1.6068167 | 0.10809457 | NA | NA | NA |
| ENSG000002 | 1.88448546 | -0.269486834 | 0.36060643 | -0.7473157 | 0.45487304 | NA | NA | NA |
| ENSG000002 | 0.86344734 | 0.04862751 | 0.30222795 | 0.1608968 | 0.87217468 | NA | NA | NA |
| ENSG000002 | 2.07725708 | -0.114307594 | 0.36292396 | -0.3149629 | 0.7527898 | NA | NA | NA |
| ENSG000002 | 1.62297077 | 0.011951637 | 0.35014735 | 0.03413316 | 0.97277096 | NA | NA | NA |
| ENSG000002 | 0.6346709 | -0.110789319 | 0.26582002 | -0.4167832 | 0.67683699 | NA | NA | NA |
| ENSG000002 | 1.65761871 | -0.540428908 | 0.34746116 | -1.555365 | 0.11985918 | NA | NA | NA |
| ENSG000002 | 0 | NA | NA | NA | NA | NA | NA | NA |
| ENSG000002 | 1.71890246 | -0.459280829 | 0.35377767 | -1.2982188 | 0.19421215 | NA | NA | NA |
| ENSG000002 | 1.26029791 | -0.330103414 | 0.32681551 | -1.0100604 | 0.31246634 | NA | NA | NA |
| ENSG000002 | 1.48747518 | -0.196741751 | 0.34605282 | -0.568531 | 0.56967447 | NA | NA | NA |
| ENSG000002 | 1.83015505 | -0.232336351 | 0.35403541 | -0.6562517 | 0.51166217 | NA | NA | NA |
| ENSG000002 | 0 | NA | NA | NA | NA | NA | NA | NA |
| ENSG000002 | 1.553873 | -0.483227676 | 0.34266892 | -1.4101882 | 0.15848411 | NA | NA | NA |
| ENSG000002 | 1.21503725 | -0.04601322 | 0.33625988 | -0.1368383 | 0.89115863 | NA | NA | NA |
| ENSG000002 | 1.10093135 | 0.036578177 | 0.32544536 | 0.11239422 | 0.91051084 | NA | SPECC1L-ADOR/ | 101730217 |
| ENSG000002 | 1.87577788 | -0.128429606 | 0.35881573 | -0.3579264 | 0.72039843 | NA | NA | NA |
| ENSG000002 | 1.43792445 | -0.747935355 | 0.33726791 | -2.2176297 | 0.02658009 | NA | NA | NA |
| ENSG000002 | 0.94552992 | 0.288810542 | 0.31146791 | 0.92725618 | 0.35379354 | NA | NA | NA |
| ENSG000002 | 0.38133351 | -0.102466351 | 0.2189713 | -0.4679442 | 0.63982449 | NA | NA | NA |

| | | | | | | | | |
|---|---|---|---|---|---|---|---|---|
| ENSG000002 | 0 | NA | NA | NA | NA | NA | NA | NA |
| ENSG000002 | 1.34465161 | -0.109742798 | 0.33438927 | -0.3281888 | 0.74276895 | NA | NA | NA |
| ENSG000002 | 1.00427532 | -0.044259671 | 0.31348448 | -0.1411862 | 0.88772288 | NA | NA | NA |
| ENSG000002 | 1.1063381 | -0.112945479 | 0.31586705 | -0.3575728 | 0.720663 | NA | NA | NA |
| ENSG000002 | 2.15775097 | -0.530901699 | 0.36770867 | -1.4438107 | 0.14879224 | NA | NA | NA |
| ENSG000002 | 1.56466687 | -0.754622406 | 0.33629023 | -2.2439617 | 0.02483486 | NA | NA | NA |
| ENSG000002 | 0.26199343 | -0.194018184 | 0.18422801 | -1.0531416 | 0.29227609 | NA | NA | NA |
| ENSG000002 | 0 | NA | NA | NA | NA | NA | TRIM6-TRIM34 | 445372 |
| ENSG000002 | 0.87987629 | -0.394300418 | 0.28674147 | -1.3751077 | 0.16909805 | NA | NA | NA |
| ENSG000002 | 1.81886069 | -0.233265741 | 0.35014937 | -0.6661892 | 0.50529015 | NA | NA | NA |
| ENSG000002 | 0.14128499 | -0.097772014 | 0.13403235 | -0.7294658 | 0.4657168 | NA | NA | NA |
| ENSG000002 | 0 | NA | NA | NA | NA | NA | NA | NA |
| ENSG000002 | 0.36345889 | 0.070257596 | 0.21589421 | 0.32542602 | 0.74485866 | NA | NA | NA |
| ENSG000002 | 0.63045625 | -0.105028713 | 0.26550735 | -0.3955774 | 0.69241679 | NA | NA | NA |
| ENSG000002 | 0.76881256 | -0.186406664 | 0.28149285 | -0.6622075 | 0.50783827 | NA | NA | NA |
| ENSG000002 | 0.72902335 | -0.029528511 | 0.28412507 | -0.1039279 | 0.9172266 | NA | NA | NA |
| ENSG000002 | 1.7804275 | -0.704677798 | 0.34639249 | -2.0343334 | 0.04191799 | NA | NA | NA |
| ENSG000002 | 1.61689621 | -0.662557645 | 0.3445149 | -1.9231611 | 0.05445983 | NA | NA | NA |
| ENSG000002 | 0.87916027 | -0.287180256 | 0.30422322 | -0.9439788 | 0.3451805 | NA | NA | NA |
| ENSG000002 | 2.11686004 | -0.343202406 | 0.35586113 | -0.9644279 | 0.33483143 | NA | NA | NA |
| ENSG000002 | 0.99138735 | -0.346626079 | 0.31023635 | -1.1172968 | 0.26386744 | NA | BCL2L2-PABPN1 | 100529063 |
| ENSG000002 | 0 | NA | NA | NA | NA | NA | SYNJ2BP-COX16 | 100529257 |
| ENSG000002 | 0.14128499 | -0.097772014 | 0.13403235 | -0.7294658 | 0.4657168 | NA | NA | NA |
| ENSG000002 | 1.9979899 | -0.050847832 | 0.35194895 | -0.144475 | 0.88512538 | NA | NA | NA |
| ENSG000002 | 0 | NA | NA | NA | NA | NA | NA | NA |
| ENSG000002 | 1.67980672 | 0.351835957 | 0.35312444 | 0.99635119 | 0.31907954 | NA | NA | NA |
| ENSG000002 | 1.27927584 | -0.347250326 | 0.33127351 | -1.0482285 | 0.29453336 | NA | NA | NA |
| ENSG000002 | 1.46047235 | -0.18261946 | 0.33900751 | -0.5386885 | 0.59010178 | NA | NA | NA |
| ENSG000002 | 1.12364573 | -0.124826171 | 0.32261416 | -0.3869209 | 0.69881474 | NA | COX6CP11 | 106480266 |

| | | | | | | | | |
|---|---|---|---|---|---|---|---|---|
| ENSG000002 | 1.22736366 | -0.435569425 | 0.31747286 | -1.3719895 | 0.17006671 | NA | NA | NA |
| ENSG000002 | 2.03896452 | 0.242314722 | 0.36216255 | 0.66907724 | 0.50344621 | NA | NA | NA |
| ENSG000002 | 0 | NA | NA | NA | NA | NA | NA | NA |
| ENSG000002 | 1.39468697 | -0.406894707 | 0.33508856 | -1.2142901 | 0.22463698 | NA | NA | NA |
| ENSG000002 | 0.74194557 | -0.177437562 | 0.27249639 | -0.6511556 | 0.51494603 | NA | NA | NA |
| ENSG000002 | 1.81435818 | -0.755955374 | 0.35246365 | -2.1447754 | 0.03197081 | NA | NA | NA |
| ENSG000002 | 0 | NA | NA | NA | NA | NA | NA | NA |
| ENSG000002 | 2.06405248 | -0.011179552 | 0.35760539 | -0.0312623 | 0.97506039 | NA | NA | NA |
| ENSG000002 | 0.99232044 | -0.334091058 | 0.30627138 | -1.0908334 | 0.27534619 | NA | NA | NA |
| ENSG000002 | 1.36013194 | -0.124803194 | 0.33955955 | -0.3675443 | 0.71321302 | NA | NA | NA |
| ENSG000002 | 1.12961712 | 0.026536278 | 0.31802993 | 0.08343956 | 0.93350203 | NA | NA | NA |
| ENSG000002 | 1.85386723 | -0.379154522 | 0.3559238 | -1.0652688 | 0.2867543 | NA | NA | NA |
| ENSG000002 | 0.13280363 | -0.093394593 | 0.13115644 | -0.7120855 | 0.47641185 | NA | NA | NA |
| ENSG000002 | 0.49879422 | -0.023609326 | 0.24688725 | -0.095628 | 0.92381605 | NA | LINC01193 | 348120 |
| ENSG000002 | 1.08686643 | 0.169473599 | 0.31524395 | 0.53759508 | 0.59085662 | NA | NA | NA |
| ENSG000002 | 0.23936338 | -0.005479057 | 0.17759977 | -0.0308506 | 0.9753887 | NA | C20orf141 | 128653 |
| ENSG000002 | 1.56219022 | -0.221254496 | 0.33754777 | -0.6554761 | 0.51216127 | NA | NA | NA |
| ENSG000002 | 0 | NA | NA | NA | NA | NA | NA | NA |
| ENSG000002 | 0 | NA | NA | NA | NA | NA | NA | NA |
| ENSG000002 | 0 | NA | NA | NA | NA | NA | NA | NA |
| ENSG000002 | 1.20524406 | -0.163997585 | 0.32148867 | -0.5101193 | 0.60996786 | NA | NA | NA |
| ENSG000002 | 2.11476197 | -0.231381078 | 0.3612895 | -0.6404312 | 0.52189228 | NA | NA | NA |
| ENSG000002 | 2.03475234 | 0.385871152 | 0.36414245 | 1.05967088 | 0.28929435 | NA | NA | NA |
| ENSG000002 | 1.42784851 | -0.393956117 | 0.33274354 | -1.1839632 | 0.23642762 | NA | NA | NA |
| ENSG000002 | 0.96505311 | -0.034239163 | 0.30529647 | -0.1121505 | 0.91070405 | NA | NA | NA |
| ENSG000002 | 0.86184872 | 0.047268168 | 0.29703329 | 0.15913424 | 0.87356311 | NA | NA | NA |
| ENSG000002 | 0 | NA | NA | NA | NA | NA | NA | NA |
| ENSG000002 | 0.56686451 | 0.359787491 | 0.24507414 | 1.46807612 | 0.14208355 | NA | NA | NA |
| ENSG000002 | 0.36690249 | -0.096999144 | 0.21683809 | -0.4473344 | 0.65463361 | NA | NA | NA |

| | | | | | | | | |
|---|---|---|---|---|---|---|---|---|
| ENSG000002 | 0 | NA | NA | NA | NA | NA | NA | NA |
| ENSG000002 | 0.25370795 | -0.171331747 | 0.17411721 | -0.9840024 | 0.32511435 | NA | NA | NA |
| ENSG000002 | 1.8275845 | -0.124793505 | 0.35569276 | -0.3508463 | 0.72570362 | NA | NA | NA |
| ENSG000002 | 1.49780366 | -0.193534955 | 0.34741115 | -0.5570776 | 0.57747444 | NA | KIAA0391 | 9692 |
| ENSG000002 | 0.73833175 | -0.187716843 | 0.27944393 | -0.6717514 | 0.50174199 | NA | NA | NA |
| ENSG000002 | 0 | NA | NA | NA | NA | NA | NA | NA |
| ENSG000002 | 0.11205239 | 0.082118251 | 0.12671962 | 0.64803108 | 0.51696485 | NA | NA | NA |
| ENSG000002 | 0.50613398 | -0.020909936 | 0.24769218 | -0.084419 | 0.93272327 | NA | NA | NA |
| ENSG000002 | 1.90674887 | -0.54508127 | 0.36009141 | -1.5137303 | 0.13009426 | NA | OR4C13 | 283092 |
| ENSG000002 | 1.71539626 | -0.184177624 | 0.3552017 | -0.5185156 | 0.60409858 | NA | NA | NA |
| ENSG000002 | 0 | NA | NA | NA | NA | NA | NA | NA |
| ENSG000002 | 0 | NA | NA | NA | NA | NA | NA | NA |
| ENSG000002 | 2.12109484 | -0.378513136 | 0.36515745 | -1.0365752 | 0.2999339 | NA | NA | NA |
| ENSG000002 | 1.44733715 | 0.20985347 | 0.33462366 | 0.6271328 | 0.53057219 | NA | LOC105370489 | 105370489 |
| ENSG000002 | 0 | NA | NA | NA | NA | NA | NA | NA |
| ENSG000002 | 0.83575724 | -0.109698178 | 0.29597774 | -0.3706298 | 0.71091327 | NA | NA | NA |
| ENSG000002 | 1.36608552 | -0.395910371 | 0.33209058 | -1.1921759 | 0.2331923 | NA | NA | NA |
| ENSG000002 | 1.58655584 | -0.386898247 | 0.34825588 | -1.1109597 | 0.26658569 | NA | MIR381HG | 378881 |
| ENSG000002 | 0.64947248 | -0.283765241 | 0.26884769 | -1.055487 | 0.29120265 | NA | NA | NA |
| ENSG000002 | 0.37945415 | -0.107841678 | 0.21865587 | -0.4932027 | 0.62186934 | NA | NA | NA |
| ENSG000002 | 2.16131327 | -0.471611698 | 0.35846092 | -1.3156572 | 0.18828911 | NA | NA | NA |
| ENSG000002 | 0.85177478 | -0.09428293 | 0.29078922 | -0.3242312 | 0.74576302 | NA | NA | NA |
| ENSG000002 | 0 | NA | NA | NA | NA | NA | NA | NA |
| ENSG000002 | 1.37063064 | 0.009900369 | 0.32800494 | 0.0301836 | 0.97592063 | NA | NA | NA |
| ENSG000002 | 1.70531978 | 0.079160504 | 0.35173752 | 0.22505561 | 0.82193601 | NA | NA | NA |
| ENSG000002 | 1.89080568 | -0.412073715 | 0.36242221 | -1.1369991 | 0.25553868 | NA | NA | NA |
| ENSG000002 | 1.54086825 | -0.762468499 | 0.33757825 | -2.2586423 | 0.02390565 | NA | NA | NA |
| ENSG000002 | 0.98851074 | -0.340153574 | 0.30851192 | -1.1025622 | 0.27021734 | NA | NA | NA |
| ENSG000002 | 0.76468381 | -0.03650702 | 0.28641547 | -0.1274618 | 0.89857494 | NA | NA | NA |

| | | | | | | | | |
|---|---|---|---|---|---|---|---|---|
| ENSG000002 | 1.8694272 | -0.383514947 | 0.35799153 | -1.0712962 | 0.28403629 | NA | NA | NA |
| ENSG000002 | 0 | NA | NA | NA | NA | NA | NA | NA |
| ENSG000002 | 1.37663683 | -0.240703883 | 0.33096392 | -0.7272813 | 0.46705363 | NA | NA | NA |
| ENSG000002 | 1.65002049 | -0.406847046 | 0.35122788 | -1.1583564 | 0.24671864 | NA | NA | NA |
| ENSG000002 | 2.11357376 | -0.514687852 | 0.34686363 | -1.4838334 | 0.13785311 | NA | NA | NA |
| ENSG000002 | 2.08810755 | -0.005527703 | 0.36223861 | -0.0152598 | 0.98782488 | NA | NA | NA |
| ENSG000002 | 2.12791913 | -0.262005517 | 0.36530693 | -0.7172202 | 0.47323823 | NA | NA | NA |
| ENSG000002 | 0 | NA | NA | NA | NA | NA | NA | NA |
| ENSG000002 | 2.12652043 | -0.011107453 | 0.36567319 | -0.0303754 | 0.9757677 | NA | NA | NA |
| ENSG000002 | 0 | NA | NA | NA | NA | NA | NA | NA |
| ENSG000002 | 2.14704588 | -0.515581895 | 0.36639453 | -1.4071768 | 0.15937496 | NA | NA | NA |
| ENSG000002 | 1.34510862 | -0.105832917 | 0.33439784 | -0.316488 | 0.75163211 | NA | NA | NA |
| ENSG000002 | 0.89244968 | -0.286343813 | 0.30487766 | -0.9392089 | 0.34762349 | NA | NA | NA |
| ENSG000002 | 0 | NA | NA | NA | NA | NA | NA | NA |
| ENSG000002 | 0.87278449 | -0.433275285 | 0.29591838 | -1.4641716 | 0.14314707 | NA | NA | NA |
| ENSG000002 | 0.8714177 | -0.116365569 | 0.29786968 | -0.3906593 | 0.69604906 | NA | NA | NA |
| ENSG000002 | 0.75964605 | -0.203618105 | 0.2920432 | -0.6972191 | 0.48566567 | NA | NA | NA |
| ENSG000002 | 0 | NA | NA | NA | NA | NA | NA | NA |
| ENSG000002 | 0 | NA | NA | NA | NA | NA | NA | NA |
| ENSG000002 | 0 | NA | NA | NA | NA | NA | NA | NA |
| ENSG000002 | 1.70554049 | 0.068476484 | 0.34626577 | 0.19775701 | 0.84323518 | NA | NA | NA |
| ENSG000002 | 1.41982362 | -0.417025197 | 0.33776472 | -1.2346618 | 0.21695639 | NA | NA | NA |
| ENSG000002 | 0.65445467 | -0.486865861 | 0.27154464 | -1.7929496 | 0.07298098 | NA | NF1P2 | 440225 |
| ENSG000002 | 0.36345889 | 0.070257596 | 0.21589421 | 0.32542602 | 0.74485866 | NA | NA | NA |
| ENSG000002 | 0.62362749 | 0.019297443 | 0.15764368 | 0.12241178 | 0.90257291 | NA | NA | NA |
| ENSG000002 | 1.79178531 | 0.270762489 | 0.35854415 | 0.75517196 | 0.45014581 | NA | NA | NA |
| ENSG000002 | 1.86582253 | -0.668774447 | 0.3578361 | -1.8689407 | 0.06163107 | NA | NA | NA |
| ENSG000002 | 0.3433388 | 0.065348438 | 0.20407693 | 0.32021472 | 0.74880557 | NA | NA | NA |
| ENSG000002 | 2.09132145 | -0.244682393 | 0.36585533 | -0.6687955 | 0.50362595 | NA | NA | NA |

| | | | | | | | | | |
|---|---|---|---|---|---|---|---|---|---|
| ENSG000002 | 1.49034128 | -0.055974128 | 0.34573636 | -0.1618983 | 0.87138594 | NA | NA | NA | |
| ENSG000002 | 0.88881412 | -0.432208381 | 0.29509244 | -1.4646542 | 0.14301529 | NA | NA | NA | |
| ENSG000002 | 2.08990047 | -0.118456374 | 0.36086217 | -0.3282593 | 0.74271559 | NA | NA | NA | |
| ENSG000002 | 1.77405346 | 0.263957598 | 0.35382429 | 0.74601323 | 0.45565942 | NA | NA | NA | |
| ENSG000002 | 0.7580257 | -0.35564649 | 0.28142375 | -1.2637401 | 0.20632332 | NA | NA | NA | |
| ENSG000002 | 1.28157941 | -0.344040162 | 0.33134684 | -1.0383083 | 0.29912656 | NA | LINC00524 | | 338002 |
| ENSG000002 | 0 | NA | NA | NA | NA | NA | TVP23C-CDRT4 | | 100533496 |
| ENSG000002 | 0.61721891 | -0.105633141 | 0.26437344 | -0.3995603 | 0.68948037 | NA | NA | NA | |
| ENSG000002 | 0 | NA | NA | NA | NA | NA | NA | NA | |
| ENSG000002 | 1.9259576 | -0.392194246 | 0.35860061 | -1.0936798 | 0.27409544 | NA | NA | NA | |
| ENSG000002 | 1.26275906 | -0.177543414 | 0.32401132 | -0.5479543 | 0.58372325 | NA | NA | NA | |
| ENSG000002 | 0.87257946 | -0.103222136 | 0.29019349 | -0.3557011 | 0.72206444 | NA | NA | NA | |
| ENSG000002 | 1.51479225 | -0.063244475 | 0.33712621 | -0.1875988 | 0.85119116 | NA | NA | NA | |
| ENSG000002 | 0 | NA | NA | NA | NA | NA | NA | NA | |
| ENSG000002 | 1.40425045 | -0.417736252 | 0.33743086 | -1.2379907 | 0.21571952 | NA | NA | NA | |
| ENSG000002 | 2.01483903 | -0.690316022 | 0.35699018 | -1.9337115 | 0.05314861 | NA | NA | NA | |
| ENSG000002 | 0.49201754 | 0.143225237 | 0.2394083 | 0.59824675 | 0.5496753 | NA | NA | NA | |
| ENSG000002 | 1.51497332 | -0.323041933 | 0.34161068 | -0.9456435 | 0.34433043 | NA | NA | NA | |
| ENSG000002 | 1.68223914 | -0.706123896 | 0.34835865 | -2.0270026 | 0.04266215 | NA | NA | NA | |
| ENSG000002 | 0.97040799 | -0.176839398 | 0.30626639 | -0.5774039 | 0.56366666 | NA | NA | NA | |
| ENSG000002 | 0 | NA | NA | NA | NA | NA | NA | NA | |
| ENSG000002 | 0.37330862 | -0.105391168 | 0.21773449 | -0.4840352 | 0.62836086 | NA | NA | NA | |
| ENSG000002 | 1.89416496 | -0.529098159 | 0.35910799 | -1.4733678 | 0.14065188 | NA | NA | NA | |
| ENSG000002 | 2.09655509 | -0.123459344 | 0.36535067 | -0.3379201 | 0.73542338 | NA | NA | NA | |
| ENSG000002 | 2.01768513 | 0.391624491 | 0.36528 | 1.07212137 | 0.28366552 | NA | NA | NA | |
| ENSG000002 | 0.71339969 | 0.108133429 | 0.2654293 | 0.4073907 | 0.68372106 | NA | NA | NA | |
| ENSG000002 | 1.63399827 | -0.378096921 | 0.34383293 | -1.099653 | 0.27148333 | NA | NA | NA | |
| ENSG000002 | 1.26419351 | -0.200936507 | 0.32993659 | -0.6090155 | 0.54251415 | NA | NA | NA | |
| ENSG000002 | 0.61924208 | -0.030634071 | 0.15758473 | -0.1943975 | 0.84586468 | NA | POC1B-GALNT4 | | 100528030 |

| | | | | | | | | |
|---|---|---|---|---|---|---|---|---|
| ENSG000002 | 1.7445385 | -0.200955104 | 0.35850545 | -0.5605357 | 0.5751141 | NA | NA | NA |
| ENSG000002 | 1.75524516 | -0.069474373 | 0.34937329 | -0.1988543 | 0.84237674 | NA | NA | NA |
| ENSG000002 | 0 | NA | NA | NA | NA | NA | NA | NA |
| ENSG000002 | 1.77904008 | -0.561918381 | 0.34715581 | -1.6186345 | 0.10552592 | NA | LOC100507277 | 100507277 |
| ENSG000002 | 0.51322524 | -0.027081322 | 0.24101349 | -0.1123643 | 0.91053453 | NA | NA | NA |
| ENSG000002 | 1.86304908 | -0.130340629 | 0.35642556 | -0.3656882 | 0.71459774 | NA | NA | NA |
| ENSG000002 | 0.13280363 | -0.093394593 | 0.13115644 | -0.7120855 | 0.47641185 | NA | NA | NA |
| ENSG000002 | 1.29179524 | -0.339378985 | 0.3276478 | -1.0358043 | 0.30029347 | NA | NA | NA |
| ENSG000002 | 1.37679115 | -0.416781345 | 0.33823161 | -1.2322365 | 0.21786074 | NA | GTF2IP22 | 106480347 |
| ENSG000002 | 0.3588776 | -0.099902505 | 0.21555794 | -0.4634601 | 0.6430346 | NA | NA | NA |
| ENSG000002 | 1.9759285 | -0.178336941 | 0.35567946 | -0.5013979 | 0.61609111 | NA | NA | NA |
| ENSG000002 | 1.44894024 | -0.043712534 | 0.3430757 | -0.1274137 | 0.898613 | NA | NA | NA |
| ENSG000002 | 1.80115521 | -0.608242364 | 0.35330775 | -1.7215653 | 0.08514829 | NA | LOC105370648 | 105370648 |
| ENSG000002 | 1.10270253 | -0.259576304 | 0.318371 | -0.8153265 | 0.41488545 | NA | NA | NA |
| ENSG000002 | 0 | NA | NA | NA | NA | NA | NDUFC2-KCTD1 | 100532726 |
| ENSG000002 | 0 | NA | NA | NA | NA | NA | NA | NA |
| ENSG000002 | 0.96052738 | 0.112067483 | 0.29922033 | 0.37453165 | 0.70800881 | NA | NA | NA |
| ENSG000002 | 1.73784814 | -0.452949593 | 0.35264014 | -1.2844527 | 0.19898359 | NA | SMIM6 | 100130933 |
| ENSG000002 | 1.86453849 | -0.507132834 | 0.35496552 | -1.4286819 | 0.15309567 | NA | NA | NA |
| ENSG000002 | 1.65395282 | -0.563835537 | 0.35164293 | -1.603432 | 0.10883931 | NA | NA | NA |
| ENSG000002 | 1.65118383 | -0.26783496 | 0.35231396 | -0.7602167 | 0.44712506 | NA | NA | NA |
| ENSG000002 | 0 | NA | NA | NA | NA | NA | NA | NA |
| ENSG000002 | 0.70553347 | 0.296317779 | 0.27795352 | 1.06606952 | 0.28639222 | NA | NA | NA |
| ENSG000002 | 1.4690254 | -0.189695972 | 0.34253587 | -0.5537989 | 0.57971651 | NA | NA | NA |
| ENSG000002 | 0.77453354 | -0.191441115 | 0.28019317 | -0.6832469 | 0.49445084 | NA | NA | NA |
| ENSG000002 | 1.35037259 | -0.123378075 | 0.34063885 | -0.3621961 | 0.71720546 | NA | LINC00644 | 101954204 |
| ENSG000002 | 0 | NA | NA | NA | NA | NA | NA | NA |
| ENSG000002 | 1.35879838 | -0.419418757 | 0.33964564 | -1.2348716 | 0.2168783 | NA | NA | NA |
| ENSG000002 | 0 | NA | NA | NA | NA | NA | NA | NA |

| | | | | | | | | |
|---|---|---|---|---|---|---|---|---|
| ENSG000002 | 2.07717664 | -0.574348544 | 0.35876086 | -1.6009231 | 0.10939396 | NA | NA | NA |
| ENSG000002 | 0.37945415 | -0.107841678 | 0.21865587 | -0.4932027 | 0.62186934 | NA | NA | NA |
| ENSG000002 | 0 | NA | NA | NA | NA | NA | NA | NA |
| ENSG000002 | 0.74094832 | -0.192779168 | 0.28573214 | -0.6746849 | 0.49987596 | NA | NA | NA |
| ENSG000002 | 2.117024 | -0.377375267 | 0.36570318 | -1.0319168 | 0.30211108 | NA | LOC104613533 | 104613533 |
| ENSG000002 | 0 | NA | NA | NA | NA | NA | NA | NA |
| ENSG000002 | 1.79870532 | 0.015810632 | 0.35733397 | 0.04424609 | 0.96470824 | NA | NA | NA |
| ENSG000002 | 0.89999741 | -0.26939482 | 0.29312597 | -0.9190411 | 0.35807406 | NA | NA | NA |
| ENSG000002 | 0.96909361 | 0.125770916 | 0.31048429 | 0.40507981 | 0.68541885 | NA | MED28P6 | 106480808 |
| ENSG000002 | 1.02360262 | -0.303371748 | 0.2955469 | -1.0264758 | 0.30466735 | NA | NA | NA |
| ENSG000002 | 1.74653856 | -0.457761877 | 0.35409892 | -1.2927514 | 0.19609705 | NA | NA | NA |
| ENSG000002 | 0.38614101 | -0.1098476 | 0.21963552 | -0.5001359 | 0.61697942 | NA | NA | NA |
| ENSG000002 | 1.73275385 | -0.299751361 | 0.34434149 | -0.8705061 | 0.38402389 | NA | NA | NA |
| ENSG000002 | 1.64842187 | -0.396835085 | 0.34900155 | -1.1370582 | 0.25551395 | NA | NA | NA |
| ENSG000002 | 1.56187365 | 0.020390715 | 0.34640192 | 0.05886433 | 0.95306017 | NA | NA | NA |
| ENSG000002 | 0.81864369 | 0.366817392 | 0.28936936 | 1.26764419 | 0.20492504 | NA | NA | NA |
| ENSG000002 | 0 | NA | NA | NA | NA | NA | NA | NA |
| ENSG000002 | 0.76989467 | -0.051525063 | 0.15948306 | -0.3230755 | 0.74663809 | NA | NA | NA |
| ENSG000002 | 1.67683082 | -0.699213878 | 0.34755387 | -2.0118144 | 0.0442395 | NA | NA | NA |
| ENSG000002 | 0.72353016 | -0.033011912 | 0.2837491 | -0.1163419 | 0.90738157 | NA | NA | NA |
| ENSG000002 | 1.35962204 | -0.244121083 | 0.33056224 | -0.7385026 | 0.46020907 | NA | NA | NA |
| ENSG000002 | 0.87852912 | -0.112522425 | 0.29825115 | -0.3772741 | 0.70596994 | NA | NA | NA |
| ENSG000002 | 0.11934008 | 0.083564698 | 0.12760091 | 0.65489106 | 0.5125379 | NA | NA | NA |
| ENSG000002 | 0.25333738 | -0.015686291 | 0.1806605 | -0.0868275 | 0.93080867 | NA | HMGN2P47 | 106480791 |
| ENSG000002 | 0 | NA | NA | NA | NA | NA | NA | NA |
| ENSG000002 | 1.633398 | -0.409678184 | 0.35273112 | -1.1614461 | 0.24546053 | NA | NA | NA |
| ENSG000002 | 2.06175254 | -0.356102795 | 0.36253585 | -0.9822554 | 0.32597405 | NA | NA | NA |
| ENSG000002 | 0 | NA | NA | NA | NA | NA | NA | NA |
| ENSG000002 | 1.34849568 | 0.026379016 | 0.33340344 | 0.07912041 | 0.93693685 | NA | NDUFA3P4 | 106480806 |

| | | | | | | | | |
|---|---|---|---|---|---|---|---|---|
| ENSG000002 | 1.89117624 | -0.548919525 | 0.3616431 | -1.5178487 | 0.12905253 | NA | NA | NA |
| ENSG000002 | 1.86399927 | -0.392491191 | 0.36003918 | -1.0901347 | 0.27565383 | NA | NA | NA |
| ENSG000002 | 1.48864767 | -0.323452158 | 0.34037489 | -0.9502821 | 0.34196894 | NA | NA | NA |
| ENSG000002 | 0.67141739 | -0.461190807 | 0.26562301 | -1.7362608 | 0.08251773 | NA | NA | NA |
| ENSG000002 | 2.04541625 | -0.721909875 | 0.36035358 | -2.0033376 | 0.04514106 | NA | NA | NA |
| ENSG000002 | 1.81355897 | -0.114272165 | 0.35869922 | -0.3185738 | 0.75004974 | NA | NA | NA |
| ENSG000002 | 1.37154446 | 0.015307295 | 0.32888008 | 0.0465437 | 0.96287691 | NA | NA | NA |
| ENSG000002 | 1.19372349 | -0.036218167 | 0.32599171 | -0.1111015 | 0.91153586 | NA | PAK6 | 56924 |
| ENSG000002 | 0.60558073 | -0.089745409 | 0.25581394 | -0.350823 | 0.72572115 | NA | NA | NA |
| ENSG000002 | 0.11131518 | 0.081951103 | 0.12663804 | 0.64712866 | 0.51754868 | NA | NA | NA |
| ENSG000002 | 0.12685397 | -0.090215768 | 0.12901919 | -0.699243 | 0.48440017 | NA | NA | NA |
| ENSG000002 | 0.73792735 | -0.192434872 | 0.28554696 | -0.6739167 | 0.50036426 | NA | ZHX1-C8orf76 | 100533106 |
| ENSG000002 | 1.09498169 | 0.037145301 | 0.32137443 | 0.11558263 | 0.90798333 | NA | NA | NA |
| ENSG000002 | 1.69077661 | -0.383859367 | 0.34388465 | -1.1162446 | 0.26431746 | NA | NA | NA |
| ENSG000002 | 1.94826124 | -0.167603262 | 0.35592839 | -0.4708904 | 0.637719 | NA | NA | NA |
| ENSG000002 | 0 | NA | NA | NA | NA | NA | NA | NA |
| ENSG000002 | 1.65952276 | -0.127351273 | 0.34329419 | -0.3709683 | 0.71066112 | NA | NA | NA |
| ENSG000002 | 0.81762258 | 0.046469806 | 0.2774599 | 0.16748296 | 0.86699005 | NA | IGHV1OR15-2 | 100287128 |
| ENSG000002 | 0.88379363 | -0.119722389 | 0.30355205 | -0.3944048 | 0.69328219 | NA | NA | NA |
| ENSG000002 | 1.33731048 | -0.24410306 | 0.33416604 | -0.7304843 | 0.46509418 | NA | NA | NA |
| ENSG000002 | 0.80655685 | -0.497227187 | 0.27284408 | -1.8223858 | 0.06839646 | NA | NA | NA |
| ENSG000002 | 2.113648 | -0.580763239 | 0.35736489 | -1.6251268 | 0.10413555 | NA | NA | NA |
| ENSG000002 | 0.42385498 | -0.262217565 | 0.21037102 | -1.2464529 | 0.21259817 | NA | NA | NA |
| ENSG000002 | 0.74504035 | -0.029746166 | 0.28517406 | -0.1043088 | 0.91692429 | NA | NA | NA |
| ENSG000002 | 1.0548264 | 0.04098023 | 0.30522441 | 0.13426262 | 0.89319491 | NA | NA | NA |
| ENSG000002 | 1.10163823 | 0.182687897 | 0.31572657 | 0.57862694 | 0.56284092 | NA | NA | NA |
| ENSG000002 | 1.63245173 | -0.396915032 | 0.34878294 | -1.1380001 | 0.25512043 | NA | NA | NA |
| ENSG000002 | 0 | NA | NA | NA | NA | NA | NA | NA |
| ENSG000002 | 0.5226185 | -0.189580046 | 0.24357686 | -0.7783172 | 0.43638207 | NA | NA | NA |

| | | | | | | | | |
|---|---|---|---|---|---|---|---|---|
| ENSG000002 | 2.00972312 | -0.191481634 | 0.35941432 | -0.5327602 | 0.5941996 | NA | NA | NA |
| ENSG000002 | 1.59724986 | -0.252899526 | 0.34774514 | -0.7272554 | 0.46706952 | NA | NA | NA |
| ENSG000002 | 0 | NA | NA | NA | NA | NA | TGIF2-C20orf24 | 100527943 |
| ENSG000002 | 2.13301534 | -0.258255947 | 0.36527603 | -0.7070159 | 0.47955662 | NA | NA | NA |
| ENSG000002 | 1.71186731 | -0.170026072 | 0.35115209 | -0.484195 | 0.62824752 | NA | NA | NA |
| ENSG000002 | 0.63398576 | -0.111029822 | 0.26576185 | -0.4177794 | 0.67610842 | NA | NA | NA |
| ENSG000002 | 0.64648534 | -0.040210356 | 0.15791988 | -0.254625 | 0.79901271 | NA | NA | NA |
| ENSG000002 | 0.38133351 | -0.102466351 | 0.2189713 | -0.4679442 | 0.63982449 | NA | NA | NA |
| ENSG000002 | 1.9952622 | -0.563274513 | 0.35888384 | -1.5695176 | 0.11652737 | NA | NA | NA |
| ENSG000002 | 0.37404583 | -0.105111695 | 0.21784993 | -0.4824959 | 0.62945372 | NA | OR4N3P | 390539 |
| ENSG000002 | 1.7725127 | -0.276771547 | 0.34073282 | -0.8122832 | 0.41662915 | NA | NA | NA |
| ENSG000002 | 0 | NA | NA | NA | NA | NA | NA | NA |
| ENSG000002 | 0.88567245 | -0.119461241 | 0.29860273 | -0.4000675 | 0.68910681 | NA | NA | NA |
| ENSG000002 | 1.37789501 | -0.254682927 | 0.33183591 | -0.7674966 | 0.44278633 | NA | NA | NA |
| ENSG000002 | 1.48821239 | -0.196496401 | 0.34606494 | -0.5678021 | 0.57016935 | NA | NA | NA |
| ENSG000002 | 0.87206846 | -0.254828244 | 0.28914816 | -0.8813068 | 0.37815178 | NA | NA | NA |
| ENSG000002 | 1.5550086 | -0.106140788 | 0.3430029 | -0.3094457 | 0.75698247 | NA | NA | NA |
| ENSG000002 | 0.75350052 | -0.19232973 | 0.28654979 | -0.6711913 | 0.50209866 | NA | NA | NA |
| ENSG000002 | 1.47290832 | -0.169414114 | 0.33525775 | -0.505325 | 0.6133306 | NA | NA | NA |
| ENSG000002 | 1.41973129 | 0.353702235 | 0.3303805 | 1.07059054 | 0.28435358 | NA | NA | NA |
| ENSG000002 | 1.08704358 | 0.179683414 | 0.31267802 | 0.57465956 | 0.56552156 | NA | NA | NA |
| ENSG000002 | 1.12387249 | -0.26114804 | 0.31747439 | -0.8225799 | 0.41074696 | NA | NA | NA |
| ENSG000002 | 1.87176304 | -0.393497867 | 0.36010104 | -1.0927429 | 0.27450669 | NA | NA | NA |
| ENSG000002 | 1.88997422 | 0.202196868 | 0.35822343 | 0.56444345 | 0.57245237 | NA | NA | NA |
| ENSG000002 | 1.25325817 | -0.349846863 | 0.33201408 | -1.0537109 | 0.29201529 | NA | NA | NA |
| ENSG000002 | 0 | NA | NA | NA | NA | NA | NA | NA |
| ENSG000002 | 1.37753284 | -0.703111727 | 0.33225231 | -2.1161981 | 0.03432796 | NA | NA | NA |
| ENSG000002 | 1.55549526 | -0.176805309 | 0.33126828 | -0.5337224 | 0.5935336 | NA | NA | NA |
| ENSG000002 | 0.98275304 | -0.031088941 | 0.30610247 | -0.1015638 | 0.91910289 | NA | NA | NA |

| | | | | | | | | |
|---|---|---|---|---|---|---|---|---|
| ENSG000002 | 0.49150654 | -0.026274915 | 0.24607766 | -0.1067749 | 0.91496757 | NA | NA | NA |
| ENSG000002 | 1.47677009 | -0.194810611 | 0.34842976 | -0.5591101 | 0.57608656 | NA | LOC102723335 | 102723335 |
| ENSG000002 | 0 | NA | NA | NA | NA | NA | NA | NA |
| ENSG000002 | 1.10147214 | 0.026740532 | 0.32160708 | 0.08314659 | 0.93373498 | NA | NA | NA |
| ENSG000002 | 0.24619405 | -0.007630671 | 0.17915639 | -0.0425922 | 0.96602658 | NA | NA | NA |
| ENSG000002 | 1.09635006 | -0.108985638 | 0.31719667 | -0.3435901 | 0.73115456 | NA | NA | NA |
| ENSG000002 | 1.6528324 | -0.413539469 | 0.35188113 | -1.1752249 | 0.23990475 | NA | HMGN1P26 | 100128015 |
| ENSG000002 | 1.96848412 | -0.287186361 | 0.35602452 | -0.8066477 | 0.41986948 | NA | NA | NA |
| ENSG000002 | 1.62697015 | -0.535391633 | 0.34821825 | -1.5375174 | 0.12416665 | NA | NA | NA |
| ENSG000002 | 1.90620416 | -0.261824466 | 0.35786421 | -0.7316308 | 0.46439396 | NA | BLID | 414899 |
| ENSG000002 | 0 | NA | NA | NA | NA | NA | NA | NA |
| ENSG000002 | 1.71977955 | -0.179292891 | 0.35080509 | -0.5110898 | 0.6092882 | NA | NA | NA |
| ENSG000002 | 0.96952834 | -0.038714098 | 0.31208463 | -0.12405 | 0.90127569 | NA | NA | NA |
| ENSG000002 | 1.1281427 | 0.026056294 | 0.31798091 | 0.08194295 | 0.93469208 | NA | NA | NA |
| ENSG000002 | 0.85038468 | -0.1098152 | 0.30192298 | -0.3637192 | 0.71606767 | NA | NA | NA |
| ENSG000002 | 0.6094788 | 0.215652144 | 0.25580422 | 0.84303591 | 0.39920836 | NA | NA | NA |
| ENSG000002 | 1.61615399 | -0.403695534 | 0.34883851 | -1.1572562 | 0.2471677 | NA | NA | NA |
| ENSG000002 | 0.64030597 | -0.296505743 | 0.27391229 | -1.0824843 | 0.27903741 | NA | NA | NA |
| ENSG000002 | 1.22971868 | 0.108141918 | 0.33003756 | 0.32766548 | 0.7431646 | NA | NA | NA |
| ENSG000002 | 0.37692244 | -0.098341116 | 0.20997214 | -0.4683532 | 0.63953205 | NA | NA | NA |
| ENSG000002 | 1.36980753 | -0.127713282 | 0.34374615 | -0.3715337 | 0.71024005 | NA | NA | NA |
| ENSG000002 | 0 | NA | NA | NA | NA | NA | NA | NA |
| ENSG000002 | 0 | NA | NA | NA | NA | NA | NA | NA |
| ENSG000002 | 0.98432992 | -0.196257029 | 0.31327313 | -0.6264726 | 0.53100501 | NA | NA | NA |
| ENSG000002 | 2.14707971 | -0.390537079 | 0.36650047 | -1.0655841 | 0.28661168 | NA | NA | NA |
| ENSG000002 | 0.84190278 | -0.111716136 | 0.29631247 | -0.3770214 | 0.70615772 | NA | NA | NA |
| ENSG000002 | 2.06574098 | -0.2236749 | 0.35954812 | -0.6221 | 0.53387609 | NA | NA | NA |
| ENSG000002 | 0.97957449 | -0.18738479 | 0.31158508 | -0.6013921 | 0.54757889 | NA | NA | NA |
| ENSG000002 | 1.34447746 | 0.032462084 | 0.33330198 | 0.09739541 | 0.92241239 | NA | NA | NA |

| | | | | | | | | |
|---|---|---|---|---|---|---|---|---|
| ENSG000002 | 1.43086059 | 0.38346503 | 0.34204623 | 1.12109124 | 0.26224903 | NA | NA | NA |
| ENSG000002 | 1.36287114 | -0.242407282 | 0.33149404 | -0.7312568 | 0.46462229 | NA | NA | NA |
| ENSG000002 | 0.7505118 | -0.182952521 | 0.27862903 | -0.6566169 | 0.51142729 | NA | NA | NA |
| ENSG000002 | 1.36232184 | -0.249522909 | 0.32899105 | -0.7584489 | 0.44818228 | NA | NA | NA |
| ENSG000002 | 1.82759659 | -0.124471205 | 0.35103915 | -0.3545793 | 0.72290481 | NA | NA | NA |
| ENSG000002 | 1.26456407 | -0.353388526 | 0.33229131 | -1.0634901 | 0.28755974 | NA | NA | NA |
| ENSG000002 | 0.87375195 | -0.2849797 | 0.30397068 | -0.9375236 | 0.34848927 | NA | NA | NA |
| ENSG000002 | 0 | NA | NA | NA | NA | NA | NA | NA |
| ENSG000002 | 0.69887885 | 0.294045978 | 0.27747344 | 1.05972656 | 0.28926901 | NA | NA | NA |
| ENSG000002 | 1.79434764 | -0.338075848 | 0.35603446 | -0.9495593 | 0.34233621 | NA | NA | NA |
| ENSG000002 | 1.5666817 | 0.023426087 | 0.34936625 | 0.06705309 | 0.94653944 | NA | NA | NA |
| ENSG000002 | 1.88257886 | -0.445480357 | 0.34449956 | -1.2931232 | 0.19596847 | NA | NA | NA |
| ENSG000002 | 1.39929859 | -0.42618076 | 0.33865945 | -1.2584346 | 0.20823464 | NA | NA | NA |
| ENSG000002 | 1.08089559 | 0.032937455 | 0.32090736 | 0.10263851 | 0.91824988 | NA | NA | NA |
| ENSG000002 | 1.28337179 | -0.480083604 | 0.32523365 | -1.4761191 | 0.13991192 | NA | LOC100131315 | 100131315 |
| ENSG000002 | 0.99739257 | -0.041004633 | 0.30837275 | -0.132971 | 0.89421632 | NA | NA | NA |
| ENSG000002 | 1.49134492 | -0.174427766 | 0.33559542 | -0.5197561 | 0.60323358 | NA | NA | NA |
| ENSG000002 | 0.48325543 | 0.139912547 | 0.23837401 | 0.58694547 | 0.55724033 | NA | NA | NA |
| ENSG000002 | 2.10996499 | -0.364162238 | 0.3634289 | -1.0020178 | 0.31633498 | NA | NA | NA |
| ENSG000002 | 0 | NA | NA | NA | NA | NA | NA | NA |
| ENSG000002 | 1.51860642 | -0.343407129 | 0.3459729 | -0.9925839 | 0.32091278 | NA | NA | NA |
| ENSG000002 | 0.50750235 | -0.197774682 | 0.24907427 | -0.794039 | 0.42717272 | NA | NA | NA |
| ENSG000002 | 0.5226185 | -0.189580046 | 0.24357686 | -0.7783172 | 0.43638207 | NA | NA | NA |
| ENSG000002 | 1.15574622 | -0.513082617 | 0.30367074 | -1.6896018 | 0.09110417 | NA | NA | NA |
| ENSG000002 | 0 | NA | NA | NA | NA | NA | NA | NA |
| ENSG000002 | 0.63182461 | -0.278261844 | 0.26744298 | -1.040453 | 0.29812951 | NA | NA | NA |
| ENSG000002 | 0.6265601 | -0.275936484 | 0.2670182 | -1.0333995 | 0.30141697 | NA | NA | NA |
| ENSG000002 | 0.71487411 | 0.121165254 | 0.27707044 | 0.43730848 | 0.66188765 | NA | NA | NA |
| ENSG000002 | 0 | NA | NA | NA | NA | NA | NA | NA |

| | | | | | | | | |
|---|---|---|---|---|---|---|---|---|
| ENSG000002 | 1.18560017 | 0.417535153 | 0.32612844 | 1.28027827 | 0.20044729 | NA | NA | NA |
| ENSG000002 | 1.52065992 | -0.498826159 | 0.34682955 | -1.4382458 | 0.15036432 | NA | NA | NA |
| ENSG000002 | 1.1400121 | -0.438041731 | 0.32510406 | -1.3473893 | 0.17785489 | NA | NA | NA |
| ENSG000002 | 0 | NA | NA | NA | NA | NA | NA | NA |
| ENSG000002 | 0 | NA | NA | NA | NA | NA | NA | NA |
| ENSG000002 | 0.78227611 | -0.381453213 | 0.2899593 | -1.3155405 | 0.18832831 | NA | NA | NA |
| ENSG000002 | 1.86173862 | -0.121468282 | 0.35598512 | -0.3412173 | 0.73294 | NA | AGGF1P6 | 106481737 |
| ENSG000002 | 1.88517507 | -0.485498184 | 0.35119186 | -1.3824301 | 0.16683969 | NA | LOC388456 | 388456 |
| ENSG000002 | 0.9472683 | -0.032708447 | 0.29669023 | -0.1102444 | 0.91221552 | NA | NA | NA |
| ENSG000002 | 1.23128292 | -0.036960747 | 0.32703796 | -0.1130167 | 0.91001732 | NA | NA | NA |
| ENSG000002 | 0.11934008 | 0.083564698 | 0.12760091 | 0.65489106 | 0.5125379 | NA | NA | NA |
| ENSG000002 | 1.73653533 | -0.059113995 | 0.35511135 | -0.1664661 | 0.86779017 | NA | NA | NA |
| ENSG000002 | 0.12685397 | -0.090215768 | 0.12901919 | -0.699243 | 0.48440017 | NA | NA | NA |
| ENSG000002 | 0.99667305 | -0.197371362 | 0.31225574 | -0.6320824 | 0.52733302 | NA | NA | NA |
| ENSG000002 | 2.12385467 | -0.358337543 | 0.36151851 | -0.9912011 | 0.3215874 | NA | NA | NA |
| ENSG000002 | 1.31704796 | -0.110280087 | 0.33297407 | -0.3311972 | 0.74049552 | NA | NA | NA |
| ENSG000002 | 1.69999578 | 0.078845735 | 0.35285128 | 0.22345316 | 0.82318283 | NA | NA | NA |
| ENSG000002 | 2.06689636 | -0.123177063 | 0.36697318 | -0.3356569 | 0.73712965 | NA | NA | NA |
| ENSG000002 | 1.95806026 | -0.16659548 | 0.35406765 | -0.4705188 | 0.63798441 | NA | NA | NA |
| ENSG000002 | 1.59150715 | -0.396564923 | 0.34931555 | -1.1352627 | 0.25626525 | NA | NA | NA |
| ENSG000002 | 0.94317018 | -0.617561148 | 0.29553776 | -2.0896185 | 0.03665209 | NA | NA | NA |
| ENSG000002 | 0 | NA | NA | NA | NA | NA | NA | NA |
| ENSG000002 | 0.23890637 | -0.010420674 | 0.17747379 | -0.0587167 | 0.95317776 | NA | NA | NA |
| ENSG000002 | 1.60153156 | -0.647543584 | 0.34102671 | -1.8988061 | 0.05758998 | NA | NA | NA |
| ENSG000002 | 0 | NA | NA | NA | NA | NA | NA | NA |
| ENSG000002 | 0 | NA | NA | NA | NA | NA | NA | NA |
| ENSG000002 | 1.20799997 | 0.108726669 | 0.32784248 | 0.33164302 | 0.74015883 | NA | NA | NA |
| ENSG000002 | 1.41224309 | -0.393793748 | 0.33314399 | -1.1820527 | 0.2371848 | NA | NA | NA |
| ENSG000002 | 0.12685397 | -0.090215768 | 0.12901919 | -0.699243 | 0.48440017 | NA | NA | NA |

| | | | | | | | | |
|---|---|---|---|---|---|---|---|---|
| ENSG000002 | 1.6565945 | -0.658690533 | 0.34335721 | -1.9183827 | 0.05506251 | NA | NA | NA |
| ENSG000002 | 0.35961481 | -0.099625193 | 0.21567743 | -0.4619176 | 0.64414043 | NA | NA | NA |
| ENSG000002 | 0.25214371 | -0.009862892 | 0.18046657 | -0.0546522 | 0.95641556 | NA | NA | NA |
| ENSG000002 | 1.62950185 | -0.535889679 | 0.34825305 | -1.5387939 | 0.12385461 | NA | NA | NA |
| ENSG000002 | 1.38396069 | -0.539882203 | 0.3328817 | -1.621844 | 0.10483675 | NA | NA | NA |
| ENSG000002 | 1.91839206 | -0.807101869 | 0.35586891 | -2.2679752 | 0.02333072 | NA | NA | NA |
| ENSG000002 | 0.61158383 | 0.053697626 | 0.26349596 | 0.20378918 | 0.83851825 | NA | NA | NA |
| ENSG000002 | 0 | NA | NA | NA | NA | NA | NA | NA |
| ENSG000002 | 0 | NA | NA | NA | NA | NA | NA | NA |
| ENSG000002 | 0.59966333 | -0.0989736 | 0.26284329 | -0.3765498 | 0.70650817 | NA | NA | NA |
| ENSG000002 | 1.10944497 | 0.029235822 | 0.32186539 | 0.09083245 | 0.92762572 | NA | NA | NA |
| ENSG000002 | 0 | NA | NA | NA | NA | NA | NA | NA |
| ENSG000002 | 1.4534549 | -0.041745268 | 0.34385584 | -0.1214034 | 0.90337153 | NA | NA | NA |
| ENSG000002 | 1.27227385 | -0.456968961 | 0.31878808 | -1.4334569 | 0.15172731 | NA | DUX4L46 | 106480349 |
| ENSG000002 | 1.39471785 | -0.583906516 | 0.33982626 | -1.7182501 | 0.085751 | NA | NA | NA |
| ENSG000002 | 0.60914194 | -0.10149904 | 0.25613314 | -0.3962745 | 0.69190251 | NA | NA | NA |
| ENSG000002 | 0.37422052 | -0.268731395 | 0.21273223 | -1.2632378 | 0.20650373 | NA | NA | NA |
| ENSG000002 | 0.58682646 | 0.225002094 | 0.26240694 | 0.85745481 | 0.39119358 | NA | NA | NA |
| ENSG000002 | 0.24485602 | -0.012655878 | 0.17881722 | -0.0707755 | 0.94357643 | NA | NA | NA |
| ENSG000002 | 0 | NA | NA | NA | NA | NA | NA | NA |
| ENSG000002 | 1.26649551 | -0.359657695 | 0.33558407 | -1.0717365 | 0.2838384 | NA | NA | NA |
| ENSG000002 | 0.88305642 | -0.119986999 | 0.30351552 | -0.3953241 | 0.69260372 | NA | LOC105447648 | 105447648 |
| ENSG000002 | 0.99121266 | -0.205080843 | 0.31756778 | -0.645786 | 0.51841792 | NA | NA | NA |
| ENSG000002 | 1.38429253 | 0.016764249 | 0.33499053 | 0.05004395 | 0.96008737 | NA | NA | NA |
| ENSG000002 | 0.75184983 | -0.181113777 | 0.28040834 | -0.645893 | 0.51834867 | NA | MRPS21P7 | 359771 |
| ENSG000002 | 1.12421217 | -0.266553834 | 0.31966139 | -0.8338631 | 0.40435814 | NA | LOC101928614 | 101928614 |
| ENSG000002 | 1.53349232 | -0.488143522 | 0.3443486 | -1.4175853 | 0.15631186 | NA | NA | NA |
| ENSG000002 | 0.68448004 | -0.087655298 | 0.15827669 | -0.5538105 | 0.57970852 | NA | NA | NA |
| ENSG000002 | 2.11807175 | -0.495027952 | 0.36230095 | -1.3663446 | 0.17183082 | NA | NA | NA |

| | | | | | | | | |
|---|---|---|---|---|---|---|---|---|
| ENSG000002 | 1.73426656 | -0.449607283 | 0.3513852 | -1.2795282 | 0.20071111 | NA | NA | NA |
| ENSG000002 | 0.12685397 | -0.090215768 | 0.12901919 | -0.699243 | 0.48440017 | NA | NA | NA |
| ENSG000002 | 0.81232135 | 0.378347255 | 0.29452843 | 1.28458651 | 0.19893681 | NA | NA | NA |
| ENSG000002 | 1.72297487 | -0.295481538 | 0.34571511 | -0.8546966 | 0.39271912 | NA | NA | NA |
| ENSG000002 | 0.2596576 | -0.193002035 | 0.1837913 | -1.0501152 | 0.29366517 | NA | NA | NA |
| ENSG000002 | 0.99140909 | -0.177582075 | 0.30718362 | -0.5780975 | 0.56319831 | NA | NA | NA |
| ENSG000002 | 1.80476049 | 0.13340901 | 0.35529691 | 0.37548599 | 0.70729907 | NA | NA | NA |
| ENSG000002 | 0.99377525 | -0.356554529 | 0.31328434 | -1.138118 | 0.25507123 | NA | NA | NA |
| ENSG000002 | 1.1018427 | -0.123614086 | 0.32575757 | -0.3794665 | 0.70434148 | NA | NA | NA |
| ENSG000002 | 2.09960639 | -0.244113829 | 0.36285128 | -0.6727655 | 0.50109648 | NA | LOC101929224 | 101929224 |
| ENSG000002 | 0 | NA | NA | NA | NA | NA | NA | NA |
| ENSG000002 | 1.02161775 | -0.508343841 | 0.30719954 | -1.6547676 | 0.09797165 | NA | NA | NA |
| ENSG000002 | 1.54499466 | -0.459370164 | 0.33783099 | -1.3597632 | 0.17390486 | NA | NA | NA |
| ENSG000002 | 0 | NA | NA | NA | NA | NA | NA | NA |
| ENSG000002 | 1.62297077 | 0.011951637 | 0.35014735 | 0.03413316 | 0.97277096 | NA | NA | NA |
| ENSG000002 | 1.51357058 | -0.320006764 | 0.34082526 | -0.9389174 | 0.34777315 | NA | NA | NA |
| ENSG000002 | 0.85152629 | -0.109713239 | 0.29682987 | -0.3696166 | 0.7116682 | NA | NA | NA |
| ENSG000002 | 0.12685397 | -0.090215768 | 0.12901919 | -0.699243 | 0.48440017 | NA | C1QL1P1 | 106480353 |
| ENSG000002 | 1.3397003 | -0.101311497 | 0.33198348 | -0.3051703 | 0.76023645 | NA | NA | NA |
| ENSG000002 | 0.52165104 | -0.403921832 | 0.25252134 | -1.5995552 | 0.10969729 | NA | NA | NA |
| ENSG000002 | 2.16119777 | 0.0522251 | 0.36657111 | 0.14246922 | 0.88670939 | NA | NA | NA |
| ENSG000002 | 1.36328931 | -0.122304576 | 0.3340225 | -0.3661567 | 0.71424815 | NA | NA | NA |
| ENSG000002 | 0.72441173 | -0.021440533 | 0.27745778 | -0.0772749 | 0.93840483 | NA | NA | NA |
| ENSG000002 | 1.7734526 | -0.187754049 | 0.34890216 | -0.5381281 | 0.59048863 | NA | NA | NA |
| ENSG000002 | 0.82554142 | -0.091292857 | 0.28500536 | -0.3203198 | 0.74872593 | NA | RARRES2P10 | 106481972 |
| ENSG000002 | 0.12685397 | -0.090215768 | 0.12901919 | -0.699243 | 0.48440017 | NA | NA | NA |
| ENSG000002 | 0.77191538 | -0.365835526 | 0.28389718 | -1.2886198 | 0.1975303 | NA | NA | NA |
| ENSG000002 | 0.12070844 | -0.088503695 | 0.12783931 | -0.6923042 | 0.48874631 | NA | NA | NA |
| ENSG000002 | 0.86976701 | -0.115601083 | 0.29779035 | -0.3881962 | 0.69787084 | NA | NA | NA |

| | | | | | | | | |
|---|---|---|---|---|---|---|---|---|
| ENSG000002 | 0 | NA | NA | NA | NA | NA | NA | NA |
| ENSG000002 | 0.39499294 | -0.277908454 | 0.21584201 | -1.287555 | 0.19790091 | NA | NA | NA |
| ENSG000002 | 0.97183034 | -0.184703938 | 0.31127565 | -0.5933774 | 0.55292861 | NA | NA | NA |
| ENSG000002 | 1.63797875 | -0.272102667 | 0.3521419 | -0.7727074 | 0.43969554 | NA | NA | NA |
| ENSG000002 | 0.91743893 | 0.428437087 | 0.30033686 | 1.42652185 | 0.15371776 | NA | NA | NA |
| ENSG000002 | 0 | NA | NA | NA | NA | NA | NA | NA |
| ENSG000002 | 0.76670294 | -0.358458639 | 0.28198122 | -1.2712146 | 0.20365232 | NA | DUX4L47 | 106481738 |
| ENSG000002 | 0.72026124 | -0.028762852 | 0.26797177 | -0.1073354 | 0.91452292 | NA | NA | NA |
| ENSG000002 | 0.12685397 | -0.090215768 | 0.12901919 | -0.699243 | 0.48440017 | NA | NA | NA |
| ENSG000002 | 1.42084315 | -0.264938788 | 0.33595227 | -0.7886203 | 0.43033394 | NA | NA | NA |
| ENSG000002 | 1.06484609 | -0.526071742 | 0.31132635 | -1.6897758 | 0.09107085 | NA | NA | NA |
| ENSG000002 | 1.63582207 | -0.66813564 | 0.34507532 | -1.9362024 | 0.05284292 | NA | NA | NA |
| ENSG000002 | 0 | NA | NA | NA | NA | NA | PMF1-BGLAP | 100527963 |
| ENSG000002 | 0.22997012 | 0.167051788 | 0.17646558 | 0.94665368 | 0.34381528 | NA | NA | NA |
| ENSG000002 | 0.99855242 | -0.202560922 | 0.31785587 | -0.6372729 | 0.52394712 | NA | NA | NA |
| ENSG000002 | 0.12070844 | -0.088503695 | 0.12783931 | -0.6923042 | 0.48874631 | NA | NA | NA |
| ENSG000002 | 2.07873488 | -0.33726626 | 0.35765246 | -0.943 | 0.34568092 | NA | NA | NA |
| ENSG000002 | 0 | NA | NA | NA | NA | NA | NA | NA |
| ENSG000002 | 0.97384712 | -0.036009546 | 0.31226566 | -0.115317 | 0.90819384 | NA | NA | NA |
| ENSG000002 | 1.75222954 | -0.283193835 | 0.34454928 | -0.8219255 | 0.41111931 | NA | NA | NA |
| ENSG000002 | 0.11865494 | 0.083440866 | 0.12751478 | 0.65436232 | 0.5128784 | NA | NA | NA |
| ENSG000002 | 0.71664742 | -0.027372205 | 0.26769121 | -0.1022529 | 0.91855593 | NA | NA | NA |
| ENSG000002 | 0.61753348 | 0.051696206 | 0.26401285 | 0.19580943 | 0.84475933 | NA | TPSP2 | 650474 |
| ENSG000002 | 1.93361802 | 0.065584925 | 0.35660971 | 0.18391234 | 0.85408223 | NA | C6orf229 | 101928603 |
| ENSG000002 | 1.68180816 | 0.219141928 | 0.35604184 | 0.61549488 | 0.53822795 | NA | TBC1D3G | 101060321 |
| ENSG000002 | 2.0164699 | -0.728405486 | 0.36102383 | -2.0176105 | 0.04363184 | NA | NA | NA |
| ENSG000002 | 1.52871865 | -0.329257078 | 0.33984177 | -0.9688541 | 0.33261801 | NA | FRG2GP | 106480321 |
| ENSG000002 | 0.65427998 | -0.290650429 | 0.26916107 | -1.0798383 | 0.28021419 | NA | NA | NA |
| ENSG000002 | 0.35682409 | 0.072008558 | 0.21485623 | 0.33514763 | 0.73751372 | NA | NA | NA |

| | | | | | | | | |
|---|---|---|---|---|---|---|---|---|
| ENSG000002 | 0 | NA | NA | NA | NA | NA | NA | NA |
| ENSG000002 | 0.23202362 | -0.008302493 | 0.17586207 | -0.0472103 | 0.96234565 | NA | NA | NA |
| ENSG000002 | 1.23194441 | -0.351295305 | 0.33347478 | -1.0534389 | 0.29213989 | NA | NA | NA |
| ENSG000002 | 1.45977649 | -0.041255879 | 0.33139421 | -0.1244919 | 0.90092585 | NA | NA | NA |
| ENSG000002 | 0.27408862 | -0.199826037 | 0.18669465 | -1.0703362 | 0.28446801 | NA | NA | NA |
| ENSG000002 | 1.48894961 | -0.188039367 | 0.34171463 | -0.5502819 | 0.58212605 | NA | NA | NA |
| ENSG000002 | 0.9731639 | 0.120899458 | 0.31218839 | 0.38726443 | 0.69856046 | NA | NA | NA |
| ENSG000002 | 0 | NA | NA | NA | NA | NA | NA | NA |
| ENSG000002 | 0.25993993 | -0.013144386 | 0.18211683 | -0.0721756 | 0.94246219 | NA | NA | NA |
| ENSG000002 | 0 | NA | NA | NA | NA | NA | NA | NA |
| ENSG000002 | 0.47070378 | 0.149674697 | 0.23683599 | 0.63197616 | 0.52740246 | NA | NA | NA |
| ENSG000002 | 1.31339988 | 0.317071934 | 0.33330606 | 0.95129363 | 0.34145534 | NA | NA | NA |
| ENSG000002 | 1.00703378 | -0.197702163 | 0.31417445 | -0.6292751 | 0.52916896 | NA | NA | NA |
| ENSG000002 | 0 | NA | NA | NA | NA | NA | NA | NA |
| ENSG000002 | 1.53429616 | -0.296147017 | 0.33118727 | -0.8941981 | 0.37121592 | NA | NA | NA |
| ENSG000002 | 0.74552961 | -0.029356539 | 0.28520244 | -0.1029323 | 0.91801671 | NA | NA | NA |
| ENSG000002 | 0 | NA | NA | NA | NA | NA | NA | NA |
| ENSG000002 | 0.97723811 | -0.188071196 | 0.30654628 | -0.6135165 | 0.53953489 | NA | NA | NA |
| ENSG000002 | 0 | NA | NA | NA | NA | NA | NA | NA |
| ENSG000002 | 0.49765262 | -0.018104365 | 0.24679454 | -0.073358 | 0.9415212 | NA | NA | NA |
| ENSG000002 | 1.44776829 | 0.099446088 | 0.33818833 | 0.29405535 | 0.76871561 | NA | NA | NA |
| ENSG000002 | 1.84837459 | -0.368615582 | 0.35453547 | -1.0397143 | 0.29847267 | NA | NA | NA |
| ENSG000002 | 0 | NA | NA | NA | NA | NA | NA | NA |
| ENSG000002 | 0 | NA | NA | NA | NA | NA | NA | NA |
| ENSG000002 | 0 | NA | NA | NA | NA | NA | NA | NA |
| ENSG000002 | 1.56389043 | 0.158157823 | 0.35280487 | 0.44828696 | 0.65394611 | NA | LOC101928880 | 101928880 |
| ENSG000002 | 1.35145917 | -0.397563524 | 0.33560016 | -1.1846345 | 0.23616199 | NA | NA | NA |
| ENSG000002 | 1.40083304 | -0.404961685 | 0.33666164 | -1.2028745 | 0.22902491 | NA | NA | NA |
| ENSG000002 | 0.50451363 | -0.188485218 | 0.24157334 | -0.7802401 | 0.43524954 | NA | NA | NA |

| | | | | | | | | | |
|---|---|---|---|---|---|---|---|---|---|
| ENSG000002 | 0.11865494 | 0.083440866 | 0.12751478 | 0.65436232 | 0.5128784 | NA | NA | NA | |
| ENSG000002 | 0.64713665 | -0.283099576 | 0.26867821 | -1.0536752 | 0.29203163 | NA | NA | NA | |
| ENSG000002 | 1.88064584 | -0.643000971 | 0.3553141 | -1.8096692 | 0.07034711 | NA | NA | NA | |
| ENSG000002 | 0.13280363 | -0.093394593 | 0.13115644 | -0.7120855 | 0.47641185 | NA | NA | NA | |
| ENSG000002 | 0.13280363 | -0.093394593 | 0.13115644 | -0.7120855 | 0.47641185 | NA | NA | NA | |
| ENSG000002 | 1.56534487 | -0.919467564 | 0.33407103 | -2.7523116 | 0.00591762 | NA | NA | NA | |
| ENSG000002 | 1.62170329 | -0.119748533 | 0.34751641 | -0.3445838 | 0.73040727 | NA | NA | NA | |
| ENSG000002 | 1.98411039 | -0.187265329 | 0.35978602 | -0.5204908 | 0.60272151 | NA | NA | NA | |
| ENSG000002 | 1.84486392 | 0.135777348 | 0.36032221 | 0.37682203 | 0.70630587 | NA | NA | NA | |
| ENSG000002 | 1.09231202 | -0.109500448 | 0.32007435 | -0.3421094 | 0.73226856 | NA | NA | NA | |
| ENSG000002 | 0.47821768 | -0.016471461 | 0.24462483 | -0.0673336 | 0.94631616 | NA | NA | NA | |
| ENSG000002 | 0 | NA | NA | NA | NA | NA | NA | NA | |
| ENSG000002 | 1.48916742 | -0.041732566 | 0.33196095 | -0.1257153 | 0.8999573 | NA | NA | NA | |
| ENSG000002 | 1.20711841 | 0.106664649 | 0.33471958 | 0.31866869 | 0.74997776 | NA | NA | NA | |
| ENSG000002 | 1.38418298 | -0.414109791 | 0.33840632 | -1.2237059 | 0.22106321 | NA | NA | NA | |
| ENSG000002 | 0.38884741 | -0.29745092 | 0.22235466 | -1.3377319 | 0.18098386 | NA | NA | NA | |
| ENSG000002 | 1.87190347 | -0.122276736 | 0.35817791 | -0.3413855 | 0.73281341 | NA | NA | NA | |
| ENSG000002 | 1.80157922 | -0.622585464 | 0.35489868 | -1.7542626 | 0.07938553 | NA | NA | NA | |
| ENSG000002 | 1.86946158 | -0.257629357 | 0.36011832 | -0.715402 | 0.47436069 | NA | NA | NA | |
| ENSG000002 | 0.61496952 | 0.052672128 | 0.25628412 | 0.2055224 | 0.837164 | NA | NA | NA | |
| ENSG000002 | 0.62001175 | -0.094588771 | 0.25713061 | -0.3678627 | 0.71297559 | NA | NA | NA | |
| ENSG000002 | 0.11865494 | 0.083440866 | 0.12751478 | 0.65436232 | 0.5128784 | NA | NA | NA | |
| ENSG000002 | 0.84104349 | 0.050427642 | 0.30109754 | 0.16747942 | 0.86699284 | NA | NA | NA | |
| ENSG000002 | 1.25890431 | -0.569902524 | 0.31195885 | -1.8268516 | 0.06772209 | NA | NA | NA | |
| ENSG000002 | 1.04904671 | -0.336692541 | 0.30574771 | -1.1012104 | 0.27080511 | NA | CYP3A52P | | 106478983 |
| ENSG000002 | 0.13280363 | -0.093394593 | 0.13115644 | -0.7120855 | 0.47641185 | NA | NA | NA | |
| ENSG000002 | 1.69462329 | -0.188293708 | 0.3555292 | -0.5296153 | 0.59637867 | NA | NA | NA | |
| ENSG000002 | 0 | NA | NA | NA | NA | NA | NA | NA | |
| ENSG000002 | 1.57371233 | -0.476550117 | 0.34046082 | -1.3997209 | 0.16159692 | NA | NA | NA | |

| | | | | | | | | |
|---|---|---|---|---|---|---|---|---|
| ENSG000002 | 0 | NA | NA | NA | NA | NA | NA | NA |
| ENSG000002 | 0 | NA | NA | NA | NA | NA | NA | NA |
| ENSG000002 | 0.28256998 | -0.18483211 | 0.18021686 | -1.0256094 | 0.30507573 | NA | NA | NA |
| ENSG000002 | 1.60544498 | -0.128253752 | 0.35357502 | -0.3627342 | 0.71680345 | NA | NA | NA |
| ENSG000002 | 0 | NA | NA | NA | NA | NA | NA | NA |
| ENSG000002 | 1.53686267 | -0.801862042 | 0.34266192 | -2.3400967 | 0.01927875 | NA | NA | NA |
| ENSG000002 | 1.67522735 | 0.347144769 | 0.35122805 | 0.98837428 | 0.32296938 | NA | NA | NA |
| ENSG000002 | 0 | NA | NA | NA | NA | NA | NA | NA |
| ENSG000002 | 1.91443157 | 0.075602079 | 0.35785216 | 0.21126624 | 0.83267953 | NA | NA | NA |
| ENSG000002 | 1.60693541 | -0.375915041 | 0.34190704 | -1.0994656 | 0.27156501 | NA | NA | NA |
| ENSG000002 | 1.08650033 | 0.039576325 | 0.32108644 | 0.12325754 | 0.90190316 | NA | NA | NA |
| ENSG000002 | 1.38450093 | -0.543171453 | 0.3315225 | -1.6384151 | 0.10133514 | NA | NA | NA |
| ENSG000002 | 0.24004852 | -0.005223787 | 0.17775844 | -0.029387 | 0.97655594 | NA | NA | NA |
| ENSG000002 | 0 | NA | NA | NA | NA | NA | NA | NA |
| ENSG000002 | 1.58327139 | 0.013159933 | 0.34961371 | 0.03764135 | 0.96997364 | NA | NA | NA |
| ENSG000002 | 0.87916027 | -0.287180256 | 0.30422322 | -0.9439788 | 0.3451805 | NA | NA | NA |
| ENSG000002 | 0.73402886 | 0.123685362 | 0.27843246 | 0.44422034 | 0.65688329 | NA | NA | NA |
| ENSG000002 | 1.86439545 | -0.87141181 | 0.34612039 | -2.5176552 | 0.01181389 | NA | NA | NA |
| ENSG000002 | 0.68678367 | 0.298085096 | 0.27650287 | 1.07805424 | 0.28100955 | NA | NA | NA |
| ENSG000002 | 2.16751425 | -0.794127724 | 0.3645316 | -2.178488 | 0.02936972 | NA | NA | NA |
| ENSG000002 | 0.50684946 | -0.202346264 | 0.24895102 | -0.8127955 | 0.41633533 | NA | NA | NA |
| ENSG000002 | 1.84574336 | 0.00748269 | 0.35604925 | 0.02101589 | 0.98323298 | NA | NA | NA |
| ENSG000002 | 1.96323888 | -0.314327074 | 0.36257015 | -0.8669414 | 0.38597411 | NA | NA | NA |
| ENSG000002 | 1.63664463 | -0.529571579 | 0.34591178 | -1.5309441 | 0.12578322 | NA | NA | NA |
| ENSG000002 | 0 | NA | NA | NA | NA | NA | NA | NA |
| ENSG000002 | 2.16346394 | 0.181745468 | 0.36638139 | 0.4960554 | 0.61985533 | NA | NA | NA |
| ENSG000002 | 0 | NA | NA | NA | NA | NA | NA | NA |
| ENSG000002 | 0.3863157 | -0.274134926 | 0.21457207 | -1.277589 | 0.20139437 | NA | NA | NA |
| ENSG000002 | 0.73218621 | -0.185627455 | 0.27901773 | -0.6652891 | 0.5058656 | NA | NA | NA |

| | | | | | | | | |
|---|---|---|---|---|---|---|---|---|
| ENSG000002 | 0.11131518 | 0.081951103 | 0.12663804 | 0.64712866 | 0.51754868 | NA | NA | NA |
| ENSG000002 | 1.20721069 | 0.102382424 | 0.3209191 | 0.31902876 | 0.7497047 | NA | NA | NA |
| ENSG000002 | 0.23730987 | 0.1539805 | 0.16964526 | 0.90766167 | 0.364057 | NA | NA | NA |
| ENSG000002 | 0.12070844 | -0.088503695 | 0.12783931 | -0.6923042 | 0.48874631 | NA | LOC100653233 | 100653233 |
| ENSG000002 | 1.1740522 | -0.607203744 | 0.32269269 | -1.881678 | 0.05987975 | NA | NA | NA |
| ENSG000002 | 1.41523343 | 0.226953182 | 0.33791962 | 0.67161884 | 0.50182638 | NA | NA | NA |
| ENSG000002 | 2.10019925 | -0.263888274 | 0.3684503 | -0.7162113 | 0.47386088 | NA | LOC105374896 | 105374896 |
| ENSG000002 | 0.9143946 | -0.461388428 | 0.3030046 | -1.522711 | 0.12783103 | NA | NA | NA |
| ENSG000002 | 2.09077619 | 0.004735598 | 0.36462863 | 0.01298746 | 0.9896378 | NA | NA | NA |
| ENSG000002 | 1.10902331 | -0.392361568 | 0.31327512 | -1.2524505 | 0.21040576 | NA | NA | NA |
| ENSG000002 | 0.24141689 | -0.165257896 | 0.17127401 | -0.9648743 | 0.33460776 | NA | NA | NA |
| ENSG000002 | 2.10162105 | -0.259405229 | 0.36517721 | -0.7103544 | 0.47748441 | NA | NA | NA |
| ENSG000002 | 0.73155506 | -0.031636845 | 0.28965816 | -0.1092213 | 0.91302696 | NA | NA | NA |
| ENSG000002 | 0 | NA | NA | NA | NA | NA | NA | NA |
| ENSG000002 | 1.62696726 | -0.131786303 | 0.34923566 | -0.3773564 | 0.70590878 | NA | NA | NA |
| ENSG000002 | 1.50283983 | -0.189399533 | 0.34508192 | -0.5488538 | 0.58310577 | NA | NA | NA |
| ENSG000002 | 0.96221032 | 0.120012748 | 0.31173331 | 0.38498532 | 0.70024829 | NA | NA | NA |
| ENSG000002 | 1.62675874 | -0.130239821 | 0.35156581 | -0.3704564 | 0.71104243 | NA | NA | NA |
| ENSG000002 | 1.44184843 | -0.047069336 | 0.34491978 | -0.1364646 | 0.89145402 | NA | NA | NA |
| ENSG000002 | 0.49970612 | -0.194507174 | 0.24827631 | -0.7834303 | 0.43337449 | NA | NA | NA |
| ENSG000002 | 0.93958026 | 0.290700617 | 0.31119409 | 0.9341457 | 0.35022874 | NA | NA | NA |
| ENSG000002 | 0 | NA | NA | NA | NA | NA | NA | NA |
| ENSG000002 | 0.38884741 | -0.29745092 | 0.22235466 | -1.3377319 | 0.18098386 | NA | NA | NA |
| ENSG000002 | 1.35017671 | -0.121748929 | 0.33935727 | -0.3587633 | 0.71977218 | NA | NA | NA |
| ENSG000002 | 0.90064872 | -0.456924471 | 0.30237983 | -1.5110944 | 0.1307644 | NA | NA | NA |
| ENSG000002 | 1.49643317 | -0.175013746 | 0.33568461 | -0.5213636 | 0.60211349 | NA | NA | NA |
| ENSG000002 | 0 | NA | NA | NA | NA | NA | NA | NA |
| ENSG000002 | 1.84965607 | -0.12486204 | 0.35756037 | -0.3492055 | 0.72693505 | NA | NA | NA |
| ENSG000002 | 1.11787076 | -0.2830715 | 0.32685819 | -0.8660376 | 0.38646952 | NA | NA | NA |

| | | | | | | | | |
|---|---|---|---|---|---|---|---|---|
| ENSG000002 | 1.63558948 | -0.403619216 | 0.35104903 | -1.1497517 | 0.25024615 | NA | NA | NA |
| ENSG000002 | 0 | NA | NA | NA | NA | NA | NA | NA |
| ENSG000002 | 1.07196169 | -0.233091677 | 0.30746181 | -0.7581159 | 0.44838162 | NA | NA | NA |
| ENSG000002 | 1.16465894 | -0.428215128 | 0.32039193 | -1.3365353 | 0.18137437 | NA | NA | NA |
| ENSG000002 | 0.61496952 | 0.052672128 | 0.25628412 | 0.2055224 | 0.837164 | NA | RARRES2P7 | 106481660 |
| ENSG000002 | 1.60550289 | -0.251893821 | 0.34712989 | -0.7256472 | 0.4680551 | NA | NA | NA |
| ENSG000002 | 0 | NA | NA | NA | NA | NA | NA | NA |
| ENSG000002 | 0 | NA | NA | NA | NA | NA | NA | NA |
| ENSG000002 | 1.13657242 | -0.571052974 | 0.31568984 | -1.8089052 | 0.07046574 | NA | NA | NA |
| ENSG000002 | 0 | NA | NA | NA | NA | NA | NA | NA |
| ENSG000002 | 0.87138578 | -0.090065353 | 0.2810864 | -0.3204188 | 0.74865091 | NA | NA | NA |
| ENSG000002 | 0 | NA | NA | NA | NA | NA | NA | NA |
| ENSG000002 | 1.86973986 | -0.397367747 | 0.36007336 | -1.1035744 | 0.26977779 | NA | NA | NA |
| ENSG000002 | 0 | NA | NA | NA | NA | NA | LOC100505948 | 100505948 |
| ENSG000002 | 0.49307133 | -0.179259067 | 0.24031383 | -0.7459374 | 0.45570524 | NA | LOC100130700 | 100130700 |
| ENSG000002 | 1.27876931 | -0.835839877 | 0.32588403 | -2.5648384 | 0.01032239 | NA | NA | NA |
| ENSG000002 | 1.70668814 | -0.051277511 | 0.35367458 | -0.144985 | 0.88472271 | NA | NA | NA |
| ENSG000002 | 0.73366064 | -0.185067619 | 0.27912381 | -0.6630306 | 0.50731097 | NA | CYP2C60P | 106480748 |
| ENSG000002 | 0.25351207 | -0.189965547 | 0.1824784 | -1.0410303 | 0.29786148 | NA | NA | NA |
| ENSG000002 | 1.83407026 | -0.108175144 | 0.34712954 | -0.3116276 | 0.75532356 | NA | NA | NA |
| ENSG000002 | 1.1615 | -0.608026814 | 0.32223433 | -1.8869088 | 0.05917259 | NA | NA | NA |
| ENSG000002 | 0 | NA | NA | NA | NA | NA | NA | NA |
| ENSG000002 | 0.59852118 | -0.096713565 | 0.25514249 | -0.3790571 | 0.70464548 | NA | NA | NA |
| ENSG000002 | 0.14128499 | -0.097772014 | 0.13403235 | -0.7294658 | 0.4657168 | NA | NA | NA |
| ENSG000002 | 0.60914194 | -0.10149904 | 0.25613314 | -0.3962745 | 0.69190251 | NA | NA | NA |
| ENSG000002 | 1.84575354 | -0.133236886 | 0.35797073 | -0.3722005 | 0.70974357 | NA | NA | NA |
| ENSG000002 | 1.21040389 | -0.17649383 | 0.32251355 | -0.5472447 | 0.58421063 | NA | NA | NA |
| ENSG000002 | 1.87121833 | -0.124787017 | 0.35976738 | -0.3468547 | 0.72870046 | NA | NA | NA |
| ENSG000002 | 1.94516996 | -0.062706706 | 0.36353602 | -0.172491 | 0.86305151 | NA | NA | NA |

| | | | | | | | | | |
|---|---|---|---|---|---|---|---|---|---|
| ENSG000002 | 0.38884741 | -0.29745092 | 0.22235466 | -1.3377319 | 0.18098386 | NA | NA | NA | |
| ENSG000002 | 0.26560726 | -0.177019426 | 0.17672274 | -1.0016788 | 0.31649875 | NA | NA | NA | |
| ENSG000002 | 0.25370795 | -0.171331747 | 0.17411721 | -0.9840024 | 0.32511435 | NA | NA | NA | |
| ENSG000002 | 1.7015909 | -0.172136655 | 0.34929585 | -0.4928105 | 0.62214651 | NA | NA | NA | |
| ENSG000002 | 0 | NA | NA | NA | NA | NA | NA | NA | |
| ENSG000002 | 1.38506345 | -0.389736396 | 0.32961011 | -1.1824164 | 0.23704052 | NA | NA | NA | |
| ENSG000002 | 1.07243542 | 0.173941236 | 0.31469486 | 0.55272983 | 0.58044842 | NA | NA | NA | |
| ENSG000002 | 2.08441034 | -0.731420015 | 0.36087812 | -2.0267785 | 0.04268507 | NA | NA | NA | |
| ENSG000002 | 0.86757417 | -0.249321606 | 0.2914133 | -0.8555601 | 0.39224112 | NA | NA | NA | |
| ENSG000002 | 1.22402769 | -0.038535276 | 0.32228373 | -0.1195694 | 0.90482425 | NA | NA | NA | |
| ENSG000002 | 0.48962772 | -0.021045505 | 0.24589998 | -0.0855856 | 0.93179582 | NA | FOXC2-AS1 | | 103752587 |
| ENSG000002 | 1.81127871 | 0.148312807 | 0.36172812 | 0.41001183 | 0.68179727 | NA | NA | NA | |
| ENSG000002 | 2.06433535 | 0.121315809 | 0.36534995 | 0.33205371 | 0.7398487 | NA | NA | NA | |
| ENSG000002 | 1.68546317 | -0.255368891 | 0.33580219 | -0.7604742 | 0.44697121 | NA | NA | NA | |
| ENSG000002 | 0.37441639 | -0.268856313 | 0.21277471 | -1.2635727 | 0.20638344 | NA | RARRES2P5 | | 106480231 |
| ENSG000002 | 2.01603985 | -0.577116484 | 0.36039536 | -1.6013427 | 0.10930103 | NA | NA | NA | |
| ENSG000002 | 0.14128499 | -0.097772014 | 0.13403235 | -0.7294658 | 0.4657168 | NA | NA | NA | |
| ENSG000002 | 2.10016891 | -0.129636974 | 0.36823254 | -0.3520519 | 0.72479932 | NA | NA | NA | |
| ENSG000002 | 0.37928001 | 0.070449581 | 0.21829793 | 0.32272217 | 0.74690566 | NA | NA | NA | |
| ENSG000002 | 0.38614101 | -0.1098476 | 0.21963552 | -0.5001359 | 0.61697942 | NA | NA | NA | |
| ENSG000002 | 1.0288454 | -0.447417526 | 0.29282381 | -1.5279411 | 0.12652717 | NA | NA | NA | |
| ENSG000002 | 1.15948033 | -0.270470398 | 0.31616038 | -0.8554848 | 0.39228282 | NA | NA | NA | |
| ENSG000002 | 0 | NA | NA | NA | NA | NA | NA | NA | |
| ENSG000002 | 0.98227641 | -0.039913906 | 0.31675261 | -0.1260097 | 0.89972423 | NA | NA | NA | |
| ENSG000002 | 0 | NA | NA | NA | NA | NA | NA | NA | |
| ENSG000002 | 0.63270563 | -0.25685676 | 0.259029 | -0.9916139 | 0.3213859 | NA | NA | NA | |
| ENSG000002 | 1.69263237 | -0.174368126 | 0.34425184 | -0.5065133 | 0.61249638 | NA | NA | NA | |
| ENSG000002 | 0.76198188 | -0.18457699 | 0.28103812 | -0.6567685 | 0.51132977 | NA | NA | NA | |
| ENSG000002 | 1.37114556 | -0.115102195 | 0.336432 | -0.3421262 | 0.73225594 | NA | NA | NA | |

| | | | | | | | | |
|---|---|---|---|---|---|---|---|---|
| ENSG000002 | 0.74590018 | -0.179161319 | 0.28000754 | -0.6398446 | 0.52227362 | NA | NA | NA |
| ENSG000002 | 1.01796912 | -0.045010794 | 0.30172252 | -0.1491794 | 0.88141205 | NA | NA | NA |
| ENSG000002 | 0 | NA | NA | NA | NA | NA | NA | NA |
| ENSG000002 | 1.86877644 | -0.260732598 | 0.36103599 | -0.722179 | 0.47018446 | NA | NA | NA |
| ENSG000002 | 0.74817755 | -0.035125537 | 0.27910954 | -0.1258486 | 0.89985179 | NA | NA | NA |
| ENSG000002 | 0.48281878 | 0.136430125 | 0.22843528 | 0.59723755 | 0.55034879 | NA | NAMPTP3 | 106480319 |
| ENSG000002 | 1.43100341 | -0.549123202 | 0.33258669 | -1.6510679 | 0.09872472 | NA | NA | NA |
| ENSG000002 | 1.70182061 | -0.057736957 | 0.35474516 | -0.1627562 | 0.87071043 | NA | NA | NA |
| ENSG000002 | 1.4678575 | 0.208663457 | 0.32869612 | 0.63482179 | 0.52554465 | NA | NA | NA |
| ENSG000002 | 0.58092888 | 0.226946341 | 0.26184598 | 0.86671692 | 0.38609713 | NA | NA | NA |
| ENSG000002 | 0 | NA | NA | NA | NA | NA | NA | NA |
| ENSG000002 | 0.63777427 | -0.25910394 | 0.25947273 | -0.9985787 | 0.31799883 | NA | NA | NA |
| ENSG000002 | 1.81060855 | -0.237016032 | 0.35461472 | -0.6683762 | 0.50389349 | NA | NA | NA |
| ENSG000002 | 0.13280363 | -0.093394593 | 0.13115644 | -0.7120855 | 0.47641185 | NA | NA | NA |
| ENSG000002 | 0.40942396 | -0.284058841 | 0.21788566 | -1.303706 | 0.19233384 | NA | NA | NA |
| ENSG000002 | 2.11821321 | -0.136278152 | 0.36546234 | -0.3728925 | 0.70922848 | NA | NA | NA |
| ENSG000002 | 2.01614748 | -0.322228954 | 0.36202258 | -0.8900797 | 0.37342309 | NA | NA | NA |
| ENSG000002 | 2.12399319 | -0.2587135 | 0.36791174 | -0.7031945 | 0.48193457 | NA | LMO7-AS1 | 101927155 |
| ENSG000002 | 0.13280363 | -0.093394593 | 0.13115644 | -0.7120855 | 0.47641185 | NA | NA | NA |
| ENSG000002 | 0 | NA | NA | NA | NA | NA | NA | NA |
| ENSG000002 | 1.11351973 | -0.258606322 | 0.31711803 | -0.8154892 | 0.41479235 | NA | NA | NA |
| ENSG000002 | 1.06487 | 0.354028983 | 0.31965489 | 1.10753502 | 0.26806268 | NA | NA | NA |
| ENSG000002 | 0.13280363 | -0.093394593 | 0.13115644 | -0.7120855 | 0.47641185 | NA | NA | NA |
| ENSG000002 | 0 | NA | NA | NA | NA | NA | NA | NA |
| ENSG000002 | 0.6009429 | 0.055606074 | 0.26256469 | 0.21178047 | 0.83227831 | NA | NA | NA |
| ENSG000002 | 1.53623309 | -0.489050181 | 0.3452922 | -1.4163372 | 0.15667681 | NA | NA | NA |
| ENSG000002 | 0 | NA | NA | NA | NA | NA | NA | NA |
| ENSG000002 | 0.11934008 | 0.083564698 | 0.12760091 | 0.65489106 | 0.5125379 | NA | NA | NA |
| ENSG000002 | 0.36391535 | 0.058449071 | 0.20749591 | 0.28168782 | 0.77818289 | NA | NA | NA |

| | | | | | | | | |
|---|---|---|---|---|---|---|---|---|
| ENSG000002 | 1.84022941 | -0.589201938 | 0.34681823 | -1.6988782 | 0.08934213 | NA | NA | NA |
| ENSG000002 | 0.83489988 | 0.207754519 | 0.29444023 | 0.70559147 | 0.48044223 | NA | NA | NA |
| ENSG000002 | 2.00900807 | -0.58654796 | 0.36188611 | -1.6208082 | 0.10505878 | NA | NA | NA |
| ENSG000002 | 0.25145857 | -0.010118565 | 0.18031426 | -0.0561163 | 0.95524918 | NA | NA | NA |
| ENSG000002 | 2.1096465 | -0.251080932 | 0.36571971 | -0.6865392 | 0.49237313 | NA | NA | NA |
| ENSG000002 | 0.87540264 | -0.25619911 | 0.29182199 | -0.8779294 | 0.37998202 | NA | NA | NA |
| ENSG000002 | 1.0229214 | -0.694426287 | 0.30767157 | -2.2570376 | 0.02400573 | NA | NA | NA |
| ENSG000002 | 2.06407765 | -0.198900849 | 0.35154151 | -0.5657962 | 0.57153233 | NA | NA | NA |
| ENSG000002 | 1.33110786 | 0.018438863 | 0.33666514 | 0.05476915 | 0.95632238 | NA | NA | NA |
| ENSG000002 | 0.12070844 | -0.088503695 | 0.12783931 | -0.6923042 | 0.48874631 | NA | NA | NA |
| ENSG000002 | 0.12070844 | -0.088503695 | 0.12783931 | -0.6923042 | 0.48874631 | NA | NA | NA |
| ENSG000002 | 1.13620185 | -0.418582057 | 0.31785913 | -1.3168791 | 0.18787914 | NA | NA | NA |
| ENSG000002 | 1.76858844 | -0.453126754 | 0.35061644 | -1.2923717 | 0.19622846 | NA | NA | NA |
| ENSG000002 | 1.7517339 | -0.18747841 | 0.34914056 | -0.5369712 | 0.59128753 | NA | SULT1C2P2 | 106480333 |
| ENSG000002 | 2.02473827 | -0.425581109 | 0.35617762 | -1.1948564 | 0.23214314 | NA | NA | NA |
| ENSG000002 | 0.34407602 | 0.071624546 | 0.21281613 | 0.33655599 | 0.73645163 | NA | NA | NA |
| ENSG000002 | 0.63984896 | -0.285743295 | 0.26802755 | -1.0660967 | 0.28637991 | NA | NA | NA |
| ENSG000002 | 0.25214371 | -0.009862892 | 0.18046657 | -0.0546522 | 0.95641556 | NA | NA | NA |
| ENSG000002 | 0.36690249 | -0.096999144 | 0.21683809 | -0.4473344 | 0.65463361 | NA | NA | NA |
| ENSG000002 | 0.90015946 | -0.421486757 | 0.29333848 | -1.4368614 | 0.15075737 | NA | NA | NA |
| ENSG000002 | 1.38253229 | -0.399548778 | 0.33485301 | -1.1932065 | 0.23278853 | NA | NA | NA |
| ENSG000002 | 1.70947886 | -0.183705213 | 0.35404193 | -0.5188798 | 0.60384456 | NA | NA | NA |
| ENSG000002 | 1.33625615 | 0.022367468 | 0.33684326 | 0.06640319 | 0.94705683 | NA | NA | NA |
| ENSG000002 | 2.05722073 | -0.565166967 | 0.35889818 | -1.5747279 | 0.11531927 | NA | NA | NA |
| ENSG000002 | 0.37899768 | -0.10176489 | 0.21865765 | -0.4654074 | 0.64163973 | NA | NA | NA |
| ENSG000002 | 0 | NA | NA | NA | NA | NA | NA | NA |
| ENSG000002 | 2.08482654 | 0.006223083 | 0.36459625 | 0.01706842 | 0.98638203 | NA | NA | NA |
| ENSG000002 | 0.14128499 | -0.097772014 | 0.13403235 | -0.7294658 | 0.4657168 | NA | NA | NA |
| ENSG000002 | 0.85473517 | -0.109563843 | 0.29095278 | -0.3765692 | 0.70649381 | NA | NA | NA |

| | | | | | | | | |
|---|---|---|---|---|---|---|---|---|
| ENSG000002 | 0.3513637 | 0.074494292 | 0.21400438 | 0.34809704 | 0.72776731 | NA | NA | NA |
| ENSG000002 | 1.71923979 | -0.058560874 | 0.35316853 | -0.1658157 | 0.86830202 | NA | NA | NA |
| ENSG000002 | 1.45787649 | -0.184735712 | 0.34363269 | -0.5375965 | 0.59085568 | NA | NA | NA |
| ENSG000002 | 1.70910829 | -0.055632207 | 0.35482031 | -0.1567898 | 0.87541051 | NA | NA | NA |
| ENSG000002 | 0.4889793 | -0.336999675 | 0.23406338 | -1.4397796 | 0.14992977 | NA | NA | NA |
| ENSG000002 | 1.35606165 | -0.122348055 | 0.34234773 | -0.3573795 | 0.72080774 | NA | CYP4A44P | 106481791 |
| ENSG000002 | 0.11131518 | 0.081951103 | 0.12663804 | 0.64712866 | 0.51754868 | NA | NA | NA |
| ENSG000002 | 0.96929809 | -0.187967297 | 0.30786581 | -0.6105494 | 0.54149791 | NA | NA | NA |
| ENSG000002 | 1.00513323 | -0.365079386 | 0.31516187 | -1.1583869 | 0.24670617 | NA | NA | NA |
| ENSG000002 | 0.971289 | -0.176731084 | 0.30630694 | -0.5769738 | 0.56395715 | NA | NA | NA |
| ENSG000002 | 0.27408862 | -0.199826037 | 0.18669465 | -1.0703362 | 0.28446801 | NA | NA | NA |
| ENSG000002 | 1.10668747 | -0.402397236 | 0.31609267 | -1.2730356 | 0.2030054 | NA | NA | NA |
| ENSG000002 | 1.49395958 | -0.322721792 | 0.34262522 | -0.941909 | 0.34623921 | NA | NA | NA |
| ENSG000002 | 1.22701228 | 0.236911839 | 0.32216551 | 0.73537306 | 0.46211232 | NA | NA | NA |
| ENSG000002 | 1.95367148 | -0.056229498 | 0.363579 | -0.1546555 | 0.8770929 | NA | NA | NA |
| ENSG000002 | 0 | NA | NA | NA | NA | NA | NA | NA |
| ENSG000002 | 0 | NA | NA | NA | NA | NA | NA | NA |
| ENSG000002 | 0.99852017 | -0.168361517 | 0.29991672 | -0.5613609 | 0.57455154 | NA | NA | NA |
| ENSG000002 | 0 | NA | NA | NA | NA | NA | NA | NA |
| ENSG000002 | 2.01950622 | -0.455497767 | 0.36209382 | -1.2579551 | 0.20840799 | NA | NA | NA |
| ENSG000002 | 1.35065737 | 0.158101903 | 0.32642966 | 0.48433681 | 0.62814685 | NA | NA | NA |
| ENSG000002 | 1.06387863 | -0.719575284 | 0.31142394 | -2.3105972 | 0.02085511 | NA | NA | NA |
| ENSG000002 | 0.11865494 | 0.083440866 | 0.12751478 | 0.65436232 | 0.5128784 | NA | NA | NA |
| ENSG000002 | 1.92308124 | 0.070112528 | 0.36057487 | 0.19444652 | 0.84582627 | NA | NA | NA |
| ENSG000002 | 1.38350368 | -0.543924518 | 0.33283348 | -1.6342242 | 0.10221179 | NA | NA | NA |
| ENSG000002 | 0.11865494 | 0.083440866 | 0.12751478 | 0.65436232 | 0.5128784 | NA | NA | NA |
| ENSG000002 | 1.60581554 | -0.245783102 | 0.34579186 | -0.7107834 | 0.47721849 | NA | NA | NA |
| ENSG000002 | 1.58694367 | -0.115978761 | 0.35104407 | -0.3303823 | 0.74111109 | NA | NA | NA |
| ENSG000002 | 2.00556447 | -0.471908058 | 0.36534472 | -1.2916789 | 0.19646838 | NA | NA | NA |

| | | | | | | | | |
|---|---|---|---|---|---|---|---|---|
| ENSG000002 | 1.4067827 | -0.4148678 | 0.33888281 | -1.224222 | 0.22086848 | NA | NA | NA |
| ENSG000002 | 1.2532061 | -0.36066144 | 0.33525109 | -1.075795 | 0.28201896 | NA | NA | NA |
| ENSG000002 | 0 | NA | NA | NA | NA | NA | NA | NA |
| ENSG000002 | 1.49518495 | -0.321384638 | 0.33924879 | -0.947342 | 0.34346453 | NA | NA | NA |
| ENSG000002 | 0.91467692 | -0.263883047 | 0.29394991 | -0.8977143 | 0.36933786 | NA | NA | NA |
| ENSG000002 | 2.14023802 | 0.176123937 | 0.36500509 | 0.4825246 | 0.62943333 | NA | NA | NA |
| ENSG000002 | 1.35005588 | -0.369636684 | 0.32550524 | -1.1355783 | 0.25613309 | NA | NA | NA |
| ENSG000002 | 0 | NA | NA | NA | NA | NA | NA | NA |
| ENSG000002 | 0 | NA | NA | NA | NA | NA | NA | NA |
| ENSG000002 | 1.35586578 | -0.123643012 | 0.34348517 | -0.3599661 | 0.71887249 | NA | NA | NA |
| ENSG000002 | 0.53471369 | -0.194084218 | 0.24487205 | -0.7925944 | 0.42801417 | NA | NA | NA |
| ENSG000002 | 1.66272968 | 0.454594845 | 0.34118213 | 1.33241106 | 0.18272515 | NA | NA | NA |
| ENSG000002 | 0.73206414 | -0.024147029 | 0.27799144 | -0.0868625 | 0.93078081 | NA | NA | NA |
| ENSG000002 | 0.76903877 | -0.364246514 | 0.2837119 | -1.2838606 | 0.19919075 | NA | NA | NA |
| ENSG000002 | 0 | NA | NA | NA | NA | NA | NA | NA |
| ENSG000002 | 0.64218479 | -0.28641133 | 0.26819919 | -1.0679053 | 0.28556324 | NA | NA | NA |
| ENSG000002 | 0 | NA | NA | NA | NA | NA | NA | NA |
| ENSG000002 | 0 | NA | NA | NA | NA | NA | NA | NA |
| ENSG000002 | 0.49036493 | -0.020766486 | 0.24598337 | -0.0844223 | 0.93272066 | NA | NA | NA |
| ENSG000002 | 1.05297929 | 0.032929523 | 0.30602908 | 0.1076026 | 0.91431094 | NA | NA | NA |
| ENSG000002 | 1.34738791 | 0.151833241 | 0.32776791 | 0.463234 | 0.64319665 | NA | NA | NA |
| ENSG000002 | 0.23816916 | -0.010712105 | 0.1772998 | -0.060418 | 0.9518227 | NA | NA | NA |
| ENSG000002 | 2.00949678 | -0.567008422 | 0.35897926 | -1.5795019 | 0.11422098 | NA | NA | NA |
| ENSG000002 | 1.35657265 | 0.029599531 | 0.33718584 | 0.08778403 | 0.93004833 | NA | NA | NA |
| ENSG000002 | 0 | NA | NA | NA | NA | NA | NA | NA |
| ENSG000002 | 2.10682352 | -0.125707864 | 0.36716019 | -0.3423788 | 0.73206585 | NA | NA | NA |
| ENSG000002 | 2.09156897 | 0.250364448 | 0.36472854 | 0.68644051 | 0.49243536 | NA | NA | NA |
| ENSG000002 | 1.34101212 | 0.024821509 | 0.33550035 | 0.07398356 | 0.94102347 | NA | NA | NA |
| ENSG000002 | 0 | NA | NA | NA | NA | NA | NA | NA |

| | | | | | | | | |
|---|---|---|---|---|---|---|---|---|
| ENSG000002 | 0 | NA | NA | NA | NA | NA | NA | NA |
| ENSG000002 | 0.71524468 | -0.026690489 | 0.26759971 | -0.0997404 | 0.92055046 | NA | NA | NA |
| ENSG000002 | 0 | NA | NA | NA | NA | NA | NA | NA |
| ENSG000002 | 1.58839794 | -0.124659972 | 0.35106994 | -0.3550859 | 0.72252527 | NA | NA | NA |
| ENSG000002 | 0 | NA | NA | NA | NA | NA | NA | NA |
| ENSG000002 | 0 | NA | NA | NA | NA | NA | NA | NA |
| ENSG000002 | 0.71758105 | -0.021228247 | 0.28336419 | -0.0749151 | 0.94028229 | NA | NA | NA |
| ENSG000002 | 1.33824548 | -0.088580102 | 0.31950729 | -0.2772397 | 0.78159607 | NA | NA | NA |
| ENSG000002 | 1.1949127 | 0.227884317 | 0.32126078 | 0.70934373 | 0.4781112 | NA | NA | NA |
| ENSG000002 | 0.83993806 | -0.234470211 | 0.28282127 | -0.8290402 | 0.40708165 | NA | TMEM249 | 340393 |
| ENSG000002 | 0.64947248 | -0.283765241 | 0.26884769 | -1.055487 | 0.29120265 | NA | NA | NA |
| ENSG000002 | 0.65240116 | -0.285397204 | 0.26908736 | -1.0606117 | 0.2888664 | NA | NA | NA |
| ENSG000002 | 1.34280258 | -0.263914262 | 0.3364866 | -0.7843232 | 0.43285047 | NA | NA | NA |
| ENSG000002 | 1.02983852 | -0.53421432 | 0.31370855 | -1.7029001 | 0.08858676 | NA | NA | NA |
| ENSG000002 | 2.08935227 | -0.120159174 | 0.36479644 | -0.3293869 | 0.74186325 | NA | NA | NA |
| ENSG000002 | 0.24485602 | -0.012655878 | 0.17881722 | -0.0707755 | 0.94357643 | NA | NA | NA |
| ENSG000002 | 0.87357726 | -0.11559895 | 0.29798362 | -0.3879373 | 0.69806246 | NA | NA | NA |
| ENSG000002 | 0.50565578 | -0.183478728 | 0.24174047 | -0.7589905 | 0.44785822 | NA | NA | NA |
| ENSG000002 | 2.00488179 | -0.311624178 | 0.36195781 | -0.8609406 | 0.38927075 | NA | NA | NA |
| ENSG000002 | 1.16332146 | -0.384547532 | 0.3098971 | -1.2408878 | 0.2146472 | NA | NA | NA |
| ENSG000002 | 1.34991612 | -0.122053174 | 0.34337244 | -0.3554542 | 0.72224935 | NA | NA | NA |
| ENSG000002 | 1.50398144 | -0.196243602 | 0.34632018 | -0.5666537 | 0.5709495 | NA | NA | NA |
| ENSG000002 | 1.85392121 | -0.233611056 | 0.35298452 | -0.6618167 | 0.50808869 | NA | NA | NA |
| ENSG000002 | 1.43137397 | -0.675464758 | 0.32734049 | -2.0634928 | 0.03906583 | NA | NA | NA |
| ENSG000002 | 0.48353618 | 0.145822039 | 0.2384055 | 0.61165553 | 0.54076569 | NA | NA | NA |
| ENSG000002 | 1.32939356 | 0.03197461 | 0.32864265 | 0.09729294 | 0.92249376 | NA | BEAN1-AS1 | 101927726 |
| ENSG000002 | 1.46930827 | -0.046467443 | 0.34659023 | -0.1340703 | 0.89334701 | NA | TTC39A-AS1 | 102724097 |
| ENSG000002 | 0.62450659 | -0.10299827 | 0.26500891 | -0.3886597 | 0.69752793 | NA | TUBAP4 | 106479041 |
| ENSG000002 | 2.15911879 | -0.652051249 | 0.36430198 | -1.7898647 | 0.07347567 | NA | NA | NA |

| | | | | | | | | |
|---|---|---|---|---|---|---|---|---|
| ENSG000002 | 0.7802226 | -0.200980091 | 0.2881869 | -0.697395 | 0.48555566 | NA | NA | NA |
| ENSG000002 | 0.23816916 | -0.010712105 | 0.1772998 | -0.060418 | 0.9518227 | NA | NA | NA |
| ENSG000002 | 2.06389657 | 0.000572058 | 0.36308026 | 0.00157557 | 0.99874288 | NA | NA | NA |
| ENSG000002 | 1.36309735 | -0.40756483 | 0.33658358 | -1.2108874 | 0.22593855 | NA | NA | NA |
| ENSG000002 | 0 | NA | NA | NA | NA | NA | NA | NA |
| ENSG000002 | 1.38213297 | 0.016472119 | 0.33640423 | 0.04896526 | 0.96094698 | NA | NA | NA |
| ENSG000002 | 0.48823762 | -0.022725119 | 0.22942258 | -0.0990535 | 0.92109576 | NA | NA | NA |
| ENSG000002 | 0.23276084 | -0.008011382 | 0.17603982 | -0.0455089 | 0.96370166 | NA | NA | NA |
| ENSG000002 | 0 | NA | NA | NA | NA | NA | NA | NA |
| ENSG000002 | 0 | NA | NA | NA | NA | NA | NA | NA |
| ENSG000002 | 0 | NA | NA | NA | NA | NA | NA | NA |
| ENSG000002 | 1.72256835 | -0.423794571 | 0.3460201 | -1.2247686 | 0.2206624 | NA | NA | NA |
| ENSG000002 | 0.5974134 | 0.053016323 | 0.26223774 | 0.20216893 | 0.83978466 | NA | NA | NA |
| ENSG000002 | 1.25338079 | -0.507294168 | 0.3318855 | -1.5285217 | 0.12638307 | NA | NA | NA |
| ENSG000002 | 1.00552498 | -0.313740355 | 0.2982185 | -1.0520486 | 0.29277725 | NA | NA | NA |
| ENSG000002 | 0.52779657 | -0.383508725 | 0.24707466 | -1.5521977 | 0.12061492 | NA | NA | NA |
| ENSG000002 | 1.56862251 | -0.647128954 | 0.3442784 | -1.8796676 | 0.06015339 | NA | NA | NA |
| ENSG000002 | 1.87381262 | -0.256726107 | 0.35961603 | -0.7138895 | 0.47529551 | NA | NA | NA |
| ENSG000002 | 1.13027001 | 0.031899322 | 0.32253476 | 0.09890197 | 0.92121611 | NA | LOC100507334 | 100507334 |
| ENSG000002 | 0 | NA | NA | NA | NA | NA | DYX1C1-CCPG1 | 100533483 |
| ENSG000002 | 1.72940115 | -0.327318985 | 0.35635447 | -0.9185208 | 0.35834629 | NA | NA | NA |
| ENSG000002 | 0.68584841 | -0.42838929 | 0.25774854 | -1.6620435 | 0.09650404 | NA | NA | NA |
| ENSG000002 | 0.51931521 | -0.354214731 | 0.23897171 | -1.4822455 | 0.13827499 | NA | NA | NA |
| ENSG000002 | 0 | NA | NA | NA | NA | NA | NA | NA |
| ENSG000002 | 0 | NA | NA | NA | NA | NA | NA | NA |
| ENSG000002 | 0 | NA | NA | NA | NA | NA | ISY1-RAB43 | 100534599 |
| ENSG000002 | 1.63251644 | -0.413369289 | 0.35161678 | -1.1756245 | 0.23974497 | NA | NA | NA |
| ENSG000002 | 1.11152075 | 0.039998422 | 0.31316442 | 0.12772339 | 0.89836788 | NA | NA | NA |
| ENSG000002 | 1.55165978 | -0.636385754 | 0.34289585 | -1.8559156 | 0.06346559 | NA | NA | NA |

| | | | | | | | | |
|---|---|---|---|---|---|---|---|---|
| ENSG000002 | 0 | NA | NA | NA | NA | NA | NA | NA |
| ENSG000002 | 1.16013164 | -0.426075742 | 0.32025662 | -1.3304198 | 0.18338001 | NA | CYP3A54P | 106480749 |
| ENSG000002 | 2.08977628 | -0.115536585 | 0.35954194 | -0.3213438 | 0.74794984 | NA | NA | NA |
| ENSG000002 | 2.12892723 | -0.624067104 | 0.3623704 | -1.7221801 | 0.08503689 | NA | NA | NA |
| ENSG000002 | 0.24141689 | -0.165257896 | 0.17127401 | -0.9648743 | 0.33460776 | NA | NA | NA |
| ENSG000002 | 1.17724339 | -0.392761515 | 0.31037203 | -1.265454 | 0.20570864 | NA | NA | NA |
| ENSG000002 | 1.74530996 | -0.608989086 | 0.35393803 | -1.7206094 | 0.08532173 | NA | DNM1P49 | 730058 |
| ENSG000002 | 1.56852826 | 0.022805797 | 0.35056486 | 0.06505443 | 0.94813066 | NA | NA | NA |
| ENSG000002 | 1.27859462 | -0.638984377 | 0.32407137 | -1.9717397 | 0.04863933 | NA | NA | NA |
| ENSG000002 | 0.37928001 | 0.070449581 | 0.21829793 | 0.32272217 | 0.74690566 | NA | CLN3 | 1201 |
| ENSG000002 | 0.48487229 | -0.012559814 | 0.237804 | -0.0528158 | 0.95787865 | NA | NA | NA |
| ENSG000002 | 0.38884741 | -0.29745092 | 0.22235466 | -1.3377319 | 0.18098386 | NA | NA | NA |
| ENSG000002 | 1.48909149 | -0.337952415 | 0.34481536 | -0.9800967 | 0.32703837 | NA | NA | NA |
| ENSG000002 | 2.08714009 | -0.130942531 | 0.36634749 | -0.3574271 | 0.72077207 | NA | NA | NA |
| ENSG000002 | 0.13280363 | -0.093394593 | 0.13115644 | -0.7120855 | 0.47641185 | NA | NA | NA |
| ENSG000002 | 0.52807889 | -0.192145741 | 0.24416537 | -0.7869492 | 0.4313116 | NA | NA | NA |
| ENSG000002 | 1.51943008 | -0.181859614 | 0.33611848 | -0.5410581 | 0.58846757 | NA | NA | NA |
| ENSG000002 | 0 | NA | NA | NA | NA | NA | NA | NA |
| ENSG000002 | 0 | NA | NA | NA | NA | NA | NA | NA |
| ENSG000002 | 0.6567905 | -0.46248946 | 0.26597757 | -1.7388288 | 0.08206488 | NA | LARP7P2 | 106481699 |
| ENSG000002 | 2.13956424 | -0.367984499 | 0.36320713 | -1.0131533 | 0.31098694 | NA | NA | NA |
| ENSG000002 | 1.85878394 | -0.005512786 | 0.3550627 | -0.0155262 | 0.98761236 | NA | NA | NA |
| ENSG000002 | 0.36007182 | -0.08619198 | 0.20729252 | -0.4157988 | 0.67755724 | NA | NA | NA |
| ENSG000002 | 0.13280363 | -0.093394593 | 0.13115644 | -0.7120855 | 0.47641185 | NA | NA | NA |
| ENSG000002 | 0.58019167 | 0.226675249 | 0.26177829 | 0.86590545 | 0.38654202 | NA | NA | NA |
| ENSG000002 | 1.7481028 | -0.590305176 | 0.35165101 | -1.6786677 | 0.09321683 | NA | NA | NA |
| ENSG000002 | 1.50392587 | -0.451173296 | 0.33838179 | -1.3333262 | 0.18242479 | NA | NA | NA |
| ENSG000002 | 2.15630045 | 0.177326439 | 0.36560678 | 0.48501956 | 0.62766247 | NA | NA | NA |
| ENSG000002 | 1.84870843 | -0.254506837 | 0.35897836 | -0.7089754 | 0.47833972 | NA | NA | NA |

| | | | | | | | | | |
|---|---|---|---|---|---|---|---|---|---|
| ENSG000002 | 0.89171192 | -0.270959061 | 0.29440303 | -0.9203678 | 0.3573806 | NA | NA | NA | |
| ENSG000002 | 0 | NA | NA | NA | NA | NA | NA | NA | |
| ENSG000002 | 1.36935106 | -0.119687654 | 0.34101958 | -0.35097 | 0.72561083 | NA | NA | NA | |
| ENSG000002 | 1.17294442 | -0.424432905 | 0.3191153 | -1.33003 | 0.1835084 | NA | NA | NA | |
| ENSG000002 | 1.72370136 | -0.181406647 | 0.35354709 | -0.5131046 | 0.60787813 | NA | NA | NA | |
| ENSG000002 | 0.50451363 | -0.188485218 | 0.24157334 | -0.7802401 | 0.43524954 | NA | NA | NA | |
| ENSG000002 | 1.71869105 | -0.063632581 | 0.35599307 | -0.1787467 | 0.85813661 | NA | NA | NA | |
| ENSG000002 | 1.33284734 | -0.228380831 | 0.32176135 | -0.7097833 | 0.47783853 | NA | NA | NA | |
| ENSG000002 | 1.63242999 | -0.552261937 | 0.34942688 | -1.5804793 | 0.11399715 | NA | NA | NA | |
| ENSG000002 | 1.83111613 | 0.005894095 | 0.35763853 | 0.01648059 | 0.98685098 | NA | NA | NA | |
| ENSG000002 | 1.5638687 | 0.011004998 | 0.34867394 | 0.03156243 | 0.974821 | NA | NA | NA | |
| ENSG000002 | 0 | NA | NA | NA | NA | NA | NA | NA | |
| ENSG000002 | 0.95947167 | 0.284029678 | 0.31210451 | 0.9100467 | 0.36279788 | NA | NA | NA | |
| ENSG000002 | 0 | NA | NA | NA | NA | NA | NA | NA | |
| ENSG000002 | 0.46079952 | 0.128217767 | 0.22562762 | 0.56827158 | 0.56985057 | NA | NA | NA | |
| ENSG000002 | 0.26813897 | -0.197007041 | 0.18550319 | -1.0620143 | 0.2882292 | NA | NA | NA | |
| ENSG000002 | 0.50127037 | -0.342056569 | 0.23551462 | -1.4523793 | 0.14639616 | NA | NA | NA | |
| ENSG000002 | 0.97177827 | -0.176371147 | 0.30632783 | -0.5757595 | 0.56477777 | NA | NA | NA | |
| ENSG000002 | 0 | NA | NA | NA | NA | NA | NA | NA | |
| ENSG000002 | 1.34518656 | 0.018507181 | 0.33413229 | 0.05538878 | 0.95582873 | NA | TRPV1 | | 7442 |
| ENSG000002 | 1.4090585 | -0.401410062 | 0.33418105 | -1.2011754 | 0.22968316 | NA | NA | NA | |
| ENSG000002 | 0.50611225 | -0.202624884 | 0.24886748 | -0.8141879 | 0.41553732 | NA | NA | NA | |
| ENSG000002 | 1.78376113 | -0.755098203 | 0.35225656 | -2.143603 | 0.03206471 | NA | NA | NA | |
| ENSG000002 | 0 | NA | NA | NA | NA | NA | NA | NA | |
| ENSG000002 | 2.07981465 | -0.00018131 | 0.36368836 | -0.0004985 | 0.99960223 | NA | NA | NA | |
| ENSG000002 | 0.76542102 | -0.036241538 | 0.28646241 | -0.1265141 | 0.89932498 | NA | NDUFA3P6 | | 106479052 |
| ENSG000002 | 1.89746029 | -0.388092881 | 0.35785163 | -1.0845078 | 0.27813971 | NA | NA | NA | |
| ENSG000002 | 1.01677587 | -0.697407278 | 0.30816713 | -2.2630813 | 0.02363068 | NA | NA | NA | |
| ENSG000002 | 1.48798426 | -0.198042911 | 0.34859967 | -0.5681099 | 0.56996037 | NA | NA | NA | |

| | | | | | | | | | |
|---|---|---|---|---|---|---|---|---|---|
| ENSG000002 | 0.84383421 | -0.106858859 | 0.29642339 | -0.360494 | 0.71847772 | NA | NA | NA | |
| ENSG000002 | 1.03950964 | -0.196451507 | 0.30916914 | -0.6354176 | 0.5251561 | NA | NA | NA | |
| ENSG000002 | 0 | NA | NA | NA | NA | NA | NA | NA | |
| ENSG000002 | 1.09785674 | -0.115298626 | 0.32025822 | -0.3600177 | 0.7188339 | NA | NA | NA | |
| ENSG000002 | 0.47001864 | 0.159831094 | 0.24415434 | 0.65463138 | 0.51270511 | NA | NA | NA | |
| ENSG000002 | 0.25351207 | -0.189965547 | 0.1824784 | -1.0410303 | 0.29786148 | NA | NA | NA | |
| ENSG000002 | 1.75114266 | 0.408177167 | 0.35541974 | 1.14843698 | 0.25078822 | NA | NA | NA | |
| ENSG000002 | 1.83650088 | -0.701652202 | 0.34525332 | -2.0322823 | 0.04212509 | NA | NA | NA | |
| ENSG000002 | 2.05605041 | 0.132574724 | 0.36645464 | 0.36177663 | 0.71751896 | NA | NA | NA | |
| ENSG000002 | 0 | NA | NA | NA | NA | NA | NA | NA | |
| ENSG000002 | 1.38666262 | -0.426962511 | 0.34142534 | -1.2505296 | 0.21110614 | NA | NA | NA | |
| ENSG000002 | 1.86766517 | -0.384428593 | 0.35812325 | -1.0734533 | 0.28306778 | NA | NA | NA | |
| ENSG000002 | 0.8444871 | -0.096832364 | 0.29036906 | -0.3334803 | 0.73877175 | NA | NA | NA | |
| ENSG000002 | 0 | NA | NA | NA | NA | NA | NA | NA | |
| ENSG000002 | 0.74296511 | -0.035856325 | 0.29036133 | -0.1234886 | 0.90172017 | NA | NA | NA | |
| ENSG000002 | 0 | NA | NA | NA | NA | NA | NA | NA | |
| ENSG000002 | 1.03959788 | -0.500829612 | 0.30639782 | -1.634573 | 0.10213858 | NA | NA | NA | |
| ENSG000002 | 0.35095876 | 0.063746151 | 0.20537869 | 0.31038348 | 0.75626936 | NA | NA | NA | |
| ENSG000002 | 1.61117077 | -0.36510523 | 0.34281536 | -1.06502 | 0.28686686 | NA | NA | NA | |
| ENSG000002 | 0 | NA | NA | NA | NA | NA | NA | NA | |
| ENSG000002 | 1.06560666 | 0.332160431 | 0.31544378 | 1.05299408 | 0.29234371 | NA | NA | NA | |
| ENSG000002 | 0 | NA | NA | NA | NA | NA | NA | NA | |
| ENSG000002 | 0 | NA | NA | NA | NA | NA | NA | NA | |
| ENSG000002 | 1.40815052 | -0.573272649 | 0.33678227 | -1.7022055 | 0.08871685 | NA | NA | NA | |
| ENSG000002 | 0 | NA | NA | NA | NA | NA | OVCA2 | | 124641 |
| ENSG000002 | 0.68473016 | 0.493998038 | 0.27846386 | 1.77401133 | 0.07606128 | NA | GLIS2-AS1 | | 101926896 |
| ENSG000002 | 1.47019789 | -0.329544498 | 0.34218361 | -0.9630634 | 0.33551573 | NA | NA | NA | |
| ENSG000002 | 2.04338665 | 0.128913439 | 0.36594051 | 0.35227977 | 0.72462846 | NA | NA | NA | |
| ENSG000002 | 0 | NA | NA | NA | NA | NA | NA | NA | |

| | | | | | | | | |
|---|---|---|---|---|---|---|---|---|
| ENSG000002 | 1.68977477 | -0.647136409 | 0.3402657 | -1.9018561 | 0.05718996 | NA | NA | NA |
| ENSG000002 | 0.48416733 | -0.018553864 | 0.24528998 | -0.0756405 | 0.93970509 | NA | NA | NA |
| ENSG000002 | 0.72944948 | 0.12611948 | 0.27810362 | 0.45349817 | 0.65019006 | NA | NA | NA |
| ENSG000002 | 1.91338524 | -0.369665564 | 0.35044615 | -1.0548427 | 0.29149727 | NA | NA | NA |
| ENSG000002 | 1.39104748 | -0.280145047 | 0.34070291 | -0.8222561 | 0.41093115 | NA | NA | NA |
| ENSG000002 | 0.49270076 | -0.021329091 | 0.24621941 | -0.0866264 | 0.93096851 | NA | NA | NA |
| ENSG000002 | 0 | NA | NA | NA | NA | NA | NA | NA |
| ENSG000002 | 0.11865494 | 0.083440866 | 0.12751478 | 0.65436232 | 0.5128784 | NA | NA | NA |
| ENSG000002 | 1.46990696 | -0.163157255 | 0.33216001 | -0.4912008 | 0.62328445 | NA | NA | NA |
| ENSG000002 | 1.67560489 | -0.69137683 | 0.34762621 | -1.9888512 | 0.04671762 | NA | NA | NA |
| ENSG000002 | 0 | NA | NA | NA | NA | NA | NA | NA |
| ENSG000002 | 2.04276797 | -0.730266695 | 0.3612629 | -2.0214273 | 0.04323555 | NA | NA | NA |
| ENSG000002 | 1.26219846 | -0.180348775 | 0.32399627 | -0.5566384 | 0.57777449 | NA | NA | NA |
| ENSG000002 | 2.04780415 | -0.750127794 | 0.36340643 | -2.0641566 | 0.03900287 | NA | NA | NA |
| ENSG000002 | 1.53958633 | -0.481006813 | 0.34449877 | -1.3962512 | 0.16263887 | NA | NA | NA |
| ENSG000002 | 1.36861385 | -0.119928995 | 0.34100495 | -0.3516928 | 0.72506864 | NA | NA | NA |
| ENSG000002 | 1.4400236 | 0.099093036 | 0.34475868 | 0.28742724 | 0.77378521 | NA | NA | NA |
| ENSG000002 | 0 | NA | NA | NA | NA | NA | NA | NA |
| ENSG000002 | 0 | NA | NA | NA | NA | NA | NA | NA |
| ENSG000002 | 2.00593503 | -0.601864523 | 0.36390569 | -1.6539025 | 0.09814733 | NA | NA | NA |
| ENSG000002 | 1.26484482 | -0.343545255 | 0.33091634 | -1.0381635 | 0.29919396 | NA | NA | NA |
| ENSG000002 | 0 | NA | NA | NA | NA | NA | NA | NA |
| ENSG000002 | 0 | NA | NA | NA | NA | NA | NA | NA |
| ENSG000002 | 0.62157207 | 0.018969676 | 0.15761573 | 0.12035395 | 0.90420277 | NA | NA | NA |
| ENSG000002 | 0.46896081 | 0.127080046 | 0.22668366 | 0.56060523 | 0.57506668 | NA | NA | NA |
| ENSG000002 | 0.36484899 | 0.075158314 | 0.21612264 | 0.34775772 | 0.72802215 | NA | NA | NA |
| ENSG000002 | 0.82826508 | 0.221452585 | 0.30065499 | 0.73656713 | 0.46138563 | NA | NA | NA |
| ENSG000002 | 0.61742743 | -0.111215928 | 0.26435729 | -0.4207031 | 0.67397191 | NA | NA | NA |
| ENSG000002 | 0.85635553 | 0.046060708 | 0.30187838 | 0.15258035 | 0.87872922 | NA | NA | NA |

| | | | | | | | | |
|---|---|---|---|---|---|---|---|---|
| ENSG000002 | 0.50016259 | -0.200428691 | 0.24824437 | -0.8073846 | 0.41944491 | NA | NA | NA |
| ENSG000002 | 1.72643754 | -0.426834153 | 0.34557704 | -1.2351346 | 0.21678042 | NA | NA | NA |
| ENSG000002 | 0 | NA | NA | NA | NA | NA | NA | NA |
| ENSG000002 | 0.87138578 | -0.090065353 | 0.2810864 | -0.3204188 | 0.74865091 | NA | NA | NA |
| ENSG000002 | 0 | NA | NA | NA | NA | NA | NA | NA |
| ENSG000002 | 1.00701205 | -0.369774986 | 0.31519008 | -1.1731809 | 0.24072326 | NA | NA | NA |
| ENSG000002 | 0 | NA | NA | NA | NA | NA | NA | NA |
| ENSG000002 | 1.95119576 | -0.286055501 | 0.35542128 | -0.804835 | 0.42091488 | NA | NA | NA |
| ENSG000002 | 1.7857843 | -0.685461487 | 0.34373113 | -1.9941793 | 0.04613247 | NA | NA | NA |
| ENSG000002 | 1.4947932 | -0.339341994 | 0.34490756 | -0.9838636 | 0.32518258 | NA | LOC105371079 | 105371079 |
| ENSG000002 | 1.5292256 | -0.46417942 | 0.34019914 | -1.3644344 | 0.17243088 | NA | NA | NA |
| ENSG000002 | 0.86592348 | -0.266194925 | 0.29880302 | -0.8908709 | 0.37299842 | NA | NA | NA |
| ENSG000002 | 1.87326737 | -0.003362528 | 0.35756971 | -0.0094038 | 0.99249693 | NA | NA | NA |
| ENSG000002 | 1.34331358 | -0.120743311 | 0.34049601 | -0.3546101 | 0.72288173 | NA | NA | NA |
| ENSG000002 | 0.24004852 | -0.005223787 | 0.17775844 | -0.029387 | 0.97655594 | NA | NA | NA |
| ENSG000002 | 0.74521503 | -0.189818705 | 0.28603889 | -0.6636115 | 0.50693897 | NA | MIR5011 | 100847002 |
| ENSG000002 | 1.93540902 | -0.183450568 | 0.36285716 | -0.5055724 | 0.61315686 | NA | NA | NA |
| ENSG000002 | 0.87067891 | -0.272676395 | 0.29901497 | -0.9119155 | 0.36181318 | NA | RN7SL134P | 106479271 |
| ENSG000002 | 1.40091949 | -0.2612501 | 0.33773762 | -0.7735298 | 0.43920893 | NA | MIR378H | 100616306 |
| ENSG000002 | 1.27489042 | -0.502315723 | 0.33033319 | -1.5206335 | 0.12835183 | NA | NA | NA |
| ENSG000002 | 1.48175229 | -0.3342258 | 0.34539111 | -0.9676734 | 0.33320749 | NA | MIR548AO | 100847068 |
| ENSG000002 | 1.26874489 | -0.510667863 | 0.33226442 | -1.5369321 | 0.12430994 | NA | MIR5584 | 100847089 |
| ENSG000002 | 1.19360333 | 0.257097352 | 0.32554177 | 0.78975228 | 0.42967246 | NA | NA | NA |
| ENSG000002 | 1.20680383 | -0.040319846 | 0.32788912 | -0.1229679 | 0.90213249 | NA | NA | NA |
| ENSG000002 | 1.90588958 | -0.409245826 | 0.36205078 | -1.1303548 | 0.25832676 | NA | NA | NA |
| ENSG000002 | 0 | NA | NA | NA | NA | NA | NA | NA |
| ENSG000002 | 0.64694077 | -0.282974687 | 0.26865662 | -1.0532951 | 0.29220579 | NA | NA | NA |
| ENSG000002 | 0.63828472 | -0.112219173 | 0.26613064 | -0.4216695 | 0.67326627 | NA | MIR3170 | 100422881 |
| ENSG000002 | 0.12685397 | -0.090215768 | 0.12901919 | -0.699243 | 0.48440017 | NA | MIR4673 | 100616242 |

| | | | | | | | | |
|---|---|---|---|---|---|---|---|---|
| ENSG000002 | 1.03831596 | -0.182155769 | 0.30347174 | -0.6002396 | 0.54834654 | NA | MIR4660 | 100616350 |
| ENSG000002 | 0.12685397 | -0.090215768 | 0.12901919 | -0.699243 | 0.48440017 | NA | MIR4747 | 100616337 |
| ENSG000002 | 0.85177478 | -0.09428293 | 0.29078922 | -0.3242312 | 0.74576302 | NA | RN7SL460P | 106481048 |
| ENSG000002 | 0 | NA | NA | NA | NA | NA | MIR4538 | 100616276 |
| ENSG000002 | 0.50377641 | -0.188767022 | 0.24148558 | -0.7816906 | 0.4343964 | NA | MIR3187 | 100422854 |
| ENSG000002 | 1.78723465 | -0.490970416 | 0.35836347 | -1.3700348 | 0.17067605 | NA | NA | NA |
| ENSG000002 | 0.5125401 | -0.027325673 | 0.24093659 | -0.1134144 | 0.90970204 | NA | NA | NA |
| ENSG000002 | 1.81353669 | -0.246133179 | 0.35456714 | -0.6941793 | 0.48756975 | NA | NA | NA |
| ENSG000002 | 0.74021111 | -0.193056035 | 0.28568202 | -0.6757724 | 0.49918514 | NA | NA | NA |
| ENSG000002 | 0.65445467 | -0.486865861 | 0.27154464 | -1.7929496 | 0.07298098 | NA | MIR4450 | 100616299 |
| ENSG000002 | 0.68351258 | -0.46544448 | 0.26656372 | -1.7460909 | 0.08079516 | NA | MIR4532 | 100616353 |
| ENSG000002 | 0.37236288 | -0.09108266 | 0.20928647 | -0.4352057 | 0.66341314 | NA | MIR5189 | 100847057 |
| ENSG000002 | 1.21136688 | 0.227555716 | 0.3217686 | 0.707203 | 0.47944034 | NA | MIR4455 | 100616111 |
| ENSG000002 | 1.75914666 | -0.069893169 | 0.2531133 | -0.2761339 | 0.7824452 | NA | NA | NA |
| ENSG000002 | 1.49088845 | -0.172511497 | 0.33729648 | -0.5114536 | 0.60903348 | NA | NA | NA |
| ENSG000002 | 2.04680738 | -0.557949506 | 0.3559602 | -1.5674491 | 0.11700975 | NA | NA | NA |
| ENSG000002 | 1.80500164 | -0.931289258 | 0.35274773 | -2.6400999 | 0.00828816 | NA | NA | NA |
| ENSG000002 | 0.86600938 | -0.114092409 | 0.29758829 | -0.3833901 | 0.70143054 | NA | MTATP6P3 | 106479043 |
| ENSG000002 | 0.95646569 | -0.181789565 | 0.30563476 | -0.5947935 | 0.55198153 | NA | NA | NA |
| ENSG000002 | 1.49943049 | -0.490632952 | 0.34502651 | -1.4220152 | 0.15502183 | NA | RN7SL155P | 106479278 |
| ENSG000002 | 1.22511565 | -0.191823281 | 0.33256442 | -0.5768004 | 0.56407431 | NA | NA | NA |
| ENSG000002 | 0 | NA | NA | NA | NA | NA | MIR4497 | 100616454 |
| ENSG000002 | 0 | NA | NA | NA | NA | NA | MIR5699 | 100847086 |
| ENSG000002 | 1.67620179 | -0.40287043 | 0.34935675 | -1.1531778 | 0.24883743 | NA | MIR4311 | 100422905 |
| ENSG000002 | 0.49401706 | -0.184896882 | 0.240355 | -0.7692658 | 0.44173553 | NA | MIR3668 | 100500879 |
| ENSG000002 | 0.12685397 | -0.090215768 | 0.12901919 | -0.699243 | 0.48440017 | NA | MIR548AN | 100616144 |
| ENSG000002 | 0.94846253 | -0.03116119 | 0.30623541 | -0.1017557 | 0.91895061 | NA | CRAT40 | 105827617 |
| ENSG000002 | 1.56882055 | 0.012893382 | 0.34438504 | 0.03743886 | 0.97013509 | NA | MIR378G | 100616321 |
| ENSG000002 | 0.37304803 | -0.090842893 | 0.20939682 | -0.4338313 | 0.66441096 | NA | MIR4526 | 100616130 |

| | | | | | | | | |
|---|---|---|---|---|---|---|---|---|
| ENSG000002 | 1.9502582 | -0.064383982 | 0.36484147 | -0.1764711 | 0.85992384 | NA | RN7SL122P | 106480954 |
| ENSG000002 | 0.85106983 | -0.104357458 | 0.29681775 | -0.3515877 | 0.72514752 | NA | MIR4463 | 100616389 |
| ENSG000002 | 1.34696625 | -0.225565435 | 0.32560059 | -0.6927673 | 0.48845561 | NA | RN7SL387P | 106481029 |
| ENSG000002 | 2.08212875 | -0.124498632 | 0.36295765 | -0.3430115 | 0.73158984 | NA | MIR5582 | 100847020 |
| ENSG000002 | 2.08992471 | -0.587035755 | 0.36088553 | -1.6266536 | 0.10381069 | NA | NA | NA |
| ENSG000002 | 0.95520338 | 0.116334841 | 0.30489442 | 0.38155779 | 0.7027894 | NA | RN7SL383P | 106479363 |
| ENSG000002 | 1.83815821 | -0.373111008 | 0.3568685 | -1.045514 | 0.29578547 | NA | RN7SL716P | 106479484 |
| ENSG000002 | 0.52125014 | -0.010203417 | 0.15567753 | -0.065542 | 0.94774247 | NA | MIR4704 | 100616205 |
| ENSG000002 | 1.51448406 | -0.327808957 | 0.34297202 | -0.9557892 | 0.33917874 | NA | MIR7853 | 102466866 |
| ENSG000002 | 0.81991957 | 0.167950405 | 0.27483774 | 0.61108931 | 0.54114046 | NA | MIR4270 | 100422868 |
| ENSG000002 | 0.23936338 | -0.005479057 | 0.17759977 | -0.0308506 | 0.9753887 | NA | MIR4665 | 100616288 |
| ENSG000002 | 0.34270765 | 0.258172734 | 0.21433185 | 1.20454674 | 0.22837833 | NA | MIR548X2 | 100616302 |
| ENSG000002 | 0.35824644 | 0.072568381 | 0.21508272 | 0.33739754 | 0.73581724 | NA | MIR4522 | 100616277 |
| ENSG000002 | 0.58765349 | 0.017801079 | 0.15713255 | 0.11328702 | 0.90980301 | NA | MIR4480 | 100616151 |
| ENSG000002 | 1.53921577 | -0.343325832 | 0.34630445 | -0.9913988 | 0.32149086 | NA | NA | NA |
| ENSG000002 | 1.12631775 | -0.412612603 | 0.31673218 | -1.3027177 | 0.19267117 | NA | NA | NA |
| ENSG000002 | 0.39462238 | -0.104329077 | 0.21264929 | -0.4906157 | 0.6236983 | NA | MIR4804 | 100616152 |
| ENSG000002 | 0.48009704 | -0.008352361 | 0.22842088 | -0.0365657 | 0.97083132 | NA | RN7SL823P | 106479522 |
| ENSG000002 | 0.97339458 | -0.314763209 | 0.30063658 | -1.0469891 | 0.29510463 | NA | MIR3915 | 100500915 |
| ENSG000002 | 0.50548109 | -0.025402968 | 0.24760631 | -0.1025942 | 0.91828506 | NA | MIR3185 | 100422978 |
| ENSG000002 | 0.22997012 | 0.167051788 | 0.17646558 | 0.94665368 | 0.34381528 | NA | NA | NA |
| ENSG000002 | 1.483717 | -0.195870546 | 0.3416256 | -0.5733486 | 0.56640872 | NA | RN7SL353P | 106481020 |
| ENSG000002 | 0.72578009 | -0.178034771 | 0.27859866 | -0.6390367 | 0.52279905 | NA | MIR4306 | 100422861 |
| ENSG000002 | 1.29356642 | -0.675780685 | 0.32977403 | -2.0492235 | 0.04044027 | NA | RN7SL178P | 106479286 |
| ENSG000002 | 0 | NA | NA | NA | NA | NA | NA | NA |
| ENSG000002 | 0.86961269 | 0.041702653 | 0.27978323 | 0.14905344 | 0.88151147 | NA | MIR3155A | 100422989 |
| ENSG000002 | 0.34197044 | 0.237837536 | 0.20637034 | 1.15247924 | 0.2491242 | NA | MIR378D1 | 100616201 |
| ENSG000002 | 0.47207214 | -0.012756135 | 0.23624418 | -0.0539956 | 0.95693871 | NA | MIR3919 | 100500803 |
| ENSG000002 | 0.51225778 | -0.204952203 | 0.24951191 | -0.8214125 | 0.41141135 | NA | NA | NA |

| | | | | | | | | |
|---|---|---|---|---|---|---|---|---|
| ENSG000002 | 0.35870346 | 0.077368606 | 0.21515516 | 0.35959447 | 0.71915042 | NA | MIR4777 | 100616317 |
| ENSG000002 | 1.53858619 | -0.061242766 | 0.33864499 | -0.1808465 | 0.85648805 | NA | MIR4802 | 100616274 |
| ENSG000002 | 0.12685397 | -0.090215768 | 0.12901919 | -0.699243 | 0.48440017 | NA | MIR4515 | 100616404 |
| ENSG000002 | 0 | NA | NA | NA | NA | NA | NA | NA |
| ENSG000002 | 1.1135718 | -0.269208314 | 0.32158437 | -0.8371312 | 0.40251882 | NA | MIR3135B | 100616218 |
| ENSG000002 | 0.62339882 | 0.057228974 | 0.27083985 | 0.2113019 | 0.8326517 | NA | MIR548AX | 100847063 |
| ENSG000002 | 1.99696453 | -0.066487292 | 0.36386039 | -0.1827275 | 0.85501185 | NA | NA | NA |
| ENSG000002 | 1.56804952 | -0.12136965 | 0.34963107 | -0.3471363 | 0.72848889 | NA | SNORA70D | 100379141 |
| ENSG000002 | 0.63025982 | -0.115141786 | 0.26544572 | -0.4337677 | 0.66445712 | NA | RN7SL470P | 106479394 |
| ENSG000002 | 0.49879422 | -0.023609326 | 0.24688725 | -0.095628 | 0.92381605 | NA | MIR4457 | 100616235 |
| ENSG000002 | 0.9860346 | -0.036930222 | 0.30788545 | -0.1199479 | 0.9045244 | NA | NA | NA |
| ENSG000002 | 2.1475257 | 0.180975154 | 0.36638926 | 0.4939423 | 0.62134693 | NA | MIR5581 | 100847010 |
| ENSG000002 | 0.40689225 | -0.283036689 | 0.21754773 | -1.3010326 | 0.19324729 | NA | MIR4632 | 100616438 |
| ENSG000002 | 0.97549973 | 0.124934136 | 0.31637382 | 0.39489404 | 0.69292109 | NA | NA | NA |
| ENSG000002 | 0.24550891 | -0.007886114 | 0.17900103 | -0.0440563 | 0.96485956 | NA | MIR3197 | 100423023 |
| ENSG000002 | 1.59551272 | -0.359267829 | 0.33926808 | -1.0589497 | 0.2896227 | NA | NA | NA |
| ENSG000002 | 0.7888781 | -0.345530318 | 0.27689228 | -1.2478871 | 0.21207239 | NA | MIR3161 | 100423000 |
| ENSG000002 | 0 | NA | NA | NA | NA | NA | MIR3621 | 100500811 |
| ENSG000002 | 1.66820915 | -0.397793585 | 0.34639678 | -1.1483755 | 0.25081357 | NA | NA | NA |
| ENSG000002 | 0.90395202 | -0.244245327 | 0.28659974 | -0.8522175 | 0.39409336 | NA | MIR5689 | 100846998 |
| ENSG000002 | 0.99073604 | -0.183192029 | 0.30134584 | -0.6079129 | 0.54324523 | NA | NA | NA |
| ENSG000002 | 0.74501916 | -0.179271543 | 0.27994274 | -0.6403865 | 0.52192138 | NA | MIR4639 | 100616269 |
| ENSG000002 | 1.6706227 | -0.513508966 | 0.3436982 | -1.4940694 | 0.13515748 | NA | NA | NA |
| ENSG000002 | 1.68465095 | 0.082069576 | 0.35753948 | 0.2295399 | 0.81844931 | NA | NA | NA |
| ENSG000002 | 1.36059087 | 0.021838238 | 0.32942957 | 0.06629107 | 0.94714609 | NA | NA | NA |
| ENSG000002 | 1.81441536 | 0.129890796 | 0.35861797 | 0.36219824 | 0.7172039 | NA | NA | NA |
| ENSG000002 | 0.62311649 | -0.107704099 | 0.26487294 | -0.4066255 | 0.68428304 | NA | NA | NA |
| ENSG000002 | 1.22920768 | -0.045354498 | 0.33659976 | -0.1347431 | 0.89281499 | NA | NA | NA |
| ENSG000002 | 0.62197881 | -0.414424795 | 0.25447679 | -1.6285367 | 0.10341113 | NA | NA | NA |

| | | | | | | | | | |
|---|---|---|---|---|---|---|---|---|---|
| ENSG000002 | 0.69333414 | 0.278653872 | 0.26842621 | 1.03810232 | 0.29922242 | NA | | MIR3937 | 100500822 |
| ENSG000002 | 0.99560255 | -0.313468749 | 0.30084953 | -1.0419453 | 0.29743705 | NA | | NA | NA |
| ENSG000002 | 0.24004852 | -0.005223787 | 0.17775844 | -0.029387 | 0.97655594 | NA | | MIR4643 | 100616174 |
| ENSG000002 | 0.91012788 | -0.438173655 | 0.29618858 | -1.4793739 | 0.13904041 | NA | | MIR4662B | 100616255 |
| ENSG000002 | 0.51322524 | -0.027081322 | 0.24101349 | -0.1123643 | 0.91053453 | NA | | MIR548AS | 100847092 |
| ENSG000002 | 1.34029261 | -0.118991113 | 0.33915213 | -0.3508488 | 0.7257018 | NA | | MIR3664 | 100500844 |
| ENSG000002 | 1.88044495 | -0.399376608 | 0.35920149 | -1.1118456 | 0.26620453 | NA | | MIR4277 | 100422966 |
| ENSG000002 | 2.13881578 | 0.676996707 | 0.3613835 | 1.87334702 | 0.06102048 | NA | | MIR548AV | 100847083 |
| ENSG000002 | 0.63026429 | -0.446551733 | 0.2623193 | -1.7023213 | 0.08869516 | NA | | MIR3133 | 100422942 |
| ENSG000002 | 1.95584121 | -0.184985603 | 0.36117318 | -0.5121798 | 0.60852519 | NA | | MIR4286 | 100422982 |
| ENSG000002 | 0.3842622 | -0.095651135 | 0.21113931 | -0.4530238 | 0.65053159 | NA | | SNORD43 | 26807 |
| ENSG000002 | 1.49284575 | -0.053005199 | 0.33911554 | -0.1563042 | 0.87579321 | NA | | NA | NA |
| ENSG000002 | 0 | NA | NA | NA | NA | NA | | SNORA4 | 619568 |
| ENSG000002 | 1.01938956 | -0.203779619 | 0.31312988 | -0.6507831 | 0.51518654 | NA | | NA | NA |
| ENSG000002 | 1.99943483 | -0.176354486 | 0.35639713 | -0.4948258 | 0.62072311 | NA | | MIR5590 | 100847069 |
| ENSG000002 | 1.24125122 | -0.60802898 | 0.31924221 | -1.9046008 | 0.05683198 | NA | | MIR3182 | 100422853 |
| ENSG000002 | 0.78754062 | -0.365675631 | 0.28493535 | -1.2833635 | 0.19936476 | NA | | NA | NA |
| ENSG000002 | 1.24241222 | -0.044837174 | 0.32447852 | -0.1381823 | 0.89009638 | NA | | MIR4473 | 100616229 |
| ENSG000002 | 0.40122492 | -0.101896709 | 0.21365861 | -0.4769136 | 0.63342361 | NA | | MIR3115 | 100422866 |
| ENSG000002 | 1.12213748 | -0.267307256 | 0.32188046 | -0.830455 | 0.40628155 | NA | | RN7SL457P | 106481047 |
| ENSG000002 | 0.60355893 | 0.050839769 | 0.26278931 | 0.19346209 | 0.84659709 | NA | | MIR5100 | 100847014 |
| ENSG000002 | 1.71340884 | 0.066853643 | 0.34896582 | 0.19157648 | 0.84807397 | NA | | NA | NA |
| ENSG000002 | 0 | NA | NA | NA | NA | NA | | NA | NA |
| ENSG000002 | 0.61836106 | -0.107182493 | 0.27082484 | -0.3957632 | 0.69227976 | NA | | MIR5684 | 100847071 |
| ENSG000002 | 1.88315189 | -0.526985157 | 0.35901937 | -1.467846 | 0.14214606 | NA | | MIR3675 | 100500876 |
| ENSG000002 | 1.1427661 | -0.262347508 | 0.31811969 | -0.8246818 | 0.40955228 | NA | | RN7SL426P | 106481842 |
| ENSG000002 | 0.60901987 | 0.058493723 | 0.26327949 | 0.22217349 | 0.82417883 | NA | | MIR548AW | 100846992 |
| ENSG000002 | 1.32967588 | 0.171132349 | 0.33311856 | 0.51372806 | 0.60744213 | NA | | NA | NA |
| ENSG000002 | 1.10946478 | 0.03244994 | 0.32034241 | 0.10129767 | 0.91931416 | NA | | MIR4439 | 100616207 |

| | | | | | | | | |
|---|---|---|---|---|---|---|---|---|
| ENSG000002 | 0.92648978 | -0.438774645 | 0.29708296 | -1.4769432 | 0.13969087 | NA | MIR378E | 100616498 |
| ENSG000002 | 0 | NA | NA | NA | NA | NA | MIR4635 | 100616479 |
| ENSG000002 | 1.74100844 | -0.191457571 | 0.35231184 | -0.5434321 | 0.58683232 | NA | MIR4698 | 100616486 |
| ENSG000002 | 1.97719382 | -0.310556375 | 0.36178215 | -0.8584071 | 0.39066769 | NA | RN7SL44P | 106480934 |
| ENSG000002 | 1.89946173 | -0.553727999 | 0.36218396 | -1.5288584 | 0.12629954 | NA | MIR4744 | 100616420 |
| ENSG000002 | 0.99329705 | -0.45577571 | 0.29583679 | -1.5406323 | 0.1234063 | NA | MIR4729 | 100616204 |
| ENSG000002 | 0.62638541 | -0.108168827 | 0.26514251 | -0.4079649 | 0.68329947 | NA | MIR4769 | 100616147 |
| ENSG000002 | 1.63555915 | -0.251419225 | 0.3462231 | -0.7261769 | 0.46773032 | NA | NA | NA |
| ENSG000002 | 0.71755932 | -0.166538792 | 0.26894713 | -0.619225 | 0.53576813 | NA | MIR3927 | 100500898 |
| ENSG000002 | 0.26199343 | -0.194018184 | 0.18422801 | -1.0531416 | 0.29227609 | NA | MIR4528 | 100616232 |
| ENSG000002 | 0 | NA | NA | NA | NA | NA | MIR4436B2 | 100847033 |
| ENSG000002 | 0.57945445 | 0.209016946 | 0.25293627 | 0.8263621 | 0.40859872 | NA | RN7SL774P | 106481761 |
| ENSG000002 | 1.22970141 | -0.281007102 | 0.31334497 | -0.8967979 | 0.36982678 | NA | MIR4303 | 100422924 |
| ENSG000002 | 0 | NA | NA | NA | NA | NA | NA | NA |
| ENSG000002 | 1.12213748 | -0.267307256 | 0.32188046 | -0.830455 | 0.40628155 | NA | MIR3925 | 100500885 |
| ENSG000002 | 0.47092999 | -0.01913341 | 0.24378131 | -0.078486 | 0.9374415 | NA | MIR4669 | 100616236 |
| ENSG000002 | 2.07349601 | 0.238830678 | 0.36332255 | 0.65735165 | 0.51095483 | NA | RN7SL555P | 106479424 |
| ENSG000002 | 0.97018123 | -0.034709227 | 0.31211164 | -0.1112077 | 0.91145162 | NA | MIR5000 | 100846995 |
| ENSG000002 | 1.47745332 | -0.333424677 | 0.34532097 | -0.96555 | 0.33426939 | NA | NA | NA |
| ENSG000002 | 0 | NA | NA | NA | NA | NA | NA | NA |
| ENSG000002 | 0.12685397 | -0.090215768 | 0.12901919 | -0.699243 | 0.48440017 | NA | MIR3670-1 | 100500910 |
| ENSG000002 | 1.75222474 | -0.061556294 | 0.35023559 | -0.1757568 | 0.86048499 | NA | MIR4462 | 100616413 |
| ENSG000002 | 1.11032249 | -0.257022845 | 0.31701311 | -0.8107641 | 0.41750117 | NA | MIR4448 | 100616127 |
| ENSG000002 | 1.67489993 | -0.718273656 | 0.35069948 | -2.0481173 | 0.04054851 | NA | TOMM20P3 | 106480301 |
| ENSG000002 | 0.52779657 | -0.383508725 | 0.24707466 | -1.5521977 | 0.12061492 | NA | RN7SL275P | 106480997 |
| ENSG000002 | 1.74465473 | -0.196282613 | 0.35377468 | -0.5548238 | 0.57901518 | NA | RN7SL32P | 106480930 |
| ENSG000002 | 2.03541383 | 0.125667073 | 0.36511006 | 0.34418957 | 0.73070373 | NA | RN7SL435P | 106479381 |
| ENSG000002 | 0.86253544 | 0.190406077 | 0.2917991 | 0.65252454 | 0.51406285 | NA | MIR548Y | 100500919 |
| ENSG000002 | 0.88286447 | -0.430377155 | 0.29478205 | -1.4599843 | 0.1442944 | NA | RN7SL850P | 106479532 |

| | | | | | | | | | |
|---|---|---|---|---|---|---|---|---|---|
| ENSG000002 | 0.11131518 | 0.081951103 | 0.12663804 | 0.64712866 | 0.51754868 | NA | | MIR4509-3 | 100616382 |
| ENSG000002 | 0.7731096 | -0.379060994 | 0.28942548 | -1.3097015 | 0.19029682 | NA | | MIR4290 | 100422963 |
| ENSG000002 | 1.25819276 | -0.810851032 | 0.32272528 | -2.5125117 | 0.01198751 | NA | | RN7SL678P | 106481108 |
| ENSG000002 | 0.49076933 | -0.025124039 | 0.23846109 | -0.1053591 | 0.91609089 | NA | | NA | NA |
| ENSG000002 | 0.25370795 | -0.171331747 | 0.17411721 | -0.9840024 | 0.32511435 | NA | | MIR1587 | 100616251 |
| ENSG000002 | 0.62178293 | -0.443481895 | 0.26161272 | -1.6951848 | 0.09004038 | NA | | MIR4760 | 100616148 |
| ENSG000002 | 0.60644002 | -0.257083821 | 0.25820471 | -0.9956589 | 0.31941589 | NA | | RN7SL121P | 106479265 |
| ENSG000002 | 0.37404583 | -0.105111695 | 0.21784993 | -0.4824959 | 0.62945372 | NA | | YWHAEP3 | 106480273 |
| ENSG000002 | 1.37679115 | -0.416781345 | 0.33823161 | -1.2322365 | 0.21786074 | NA | | MIR4499 | 100616304 |
| ENSG000002 | 0.3561712 | 0.067399341 | 0.21474131 | 0.31386296 | 0.75362513 | NA | | MIR4672 | 100616429 |
| ENSG000002 | 2.02540223 | -0.582936735 | 0.36108451 | -1.6144053 | 0.10643955 | NA | | MIR5705 | 100847027 |
| ENSG000002 | 0.11865494 | 0.083440866 | 0.12751478 | 0.65436232 | 0.5128784 | NA | | NA | NA |
| ENSG000002 | 1.01417711 | -0.193638607 | 0.30813876 | -0.6284137 | 0.52973299 | NA | | MIR5698 | 100847024 |
| ENSG000002 | 1.50995091 | -0.194302087 | 0.34641672 | -0.5608912 | 0.57487172 | NA | | RN7SL615P | 106479446 |
| ENSG000002 | 2.04666412 | -0.58162806 | 0.36084325 | -1.6118579 | 0.10699286 | NA | | NA | NA |
| ENSG000002 | 1.73605659 | -0.198259283 | 0.35432913 | -0.5595342 | 0.57579717 | NA | | RN7SL786P | 106479506 |
| ENSG000002 | 1.7758246 | -0.472622112 | 0.3553932 | -1.3298569 | 0.18356541 | NA | | NA | NA |
| ENSG000002 | 1.5430082 | -0.808703482 | 0.3434393 | -2.3547203 | 0.01853665 | NA | | RN7SL52P | 106480937 |
| ENSG000002 | 1.01224622 | -0.206895503 | 0.31834465 | -0.6499104 | 0.51575009 | NA | | MIR4717 | 100616241 |
| ENSG000002 | 0.85704121 | 0.053051046 | 0.29513872 | 0.17974953 | 0.85734921 | NA | | MIR4429 | 100616469 |
| ENSG000002 | 0.90958655 | -0.438566765 | 0.29615636 | -1.4808622 | 0.13864329 | NA | | NA | NA |
| ENSG000002 | 0.87801866 | -0.269992537 | 0.29941829 | -0.9017236 | 0.36720371 | NA | | MIR3938 | 100500875 |
| ENSG000002 | 0.27408862 | -0.199826037 | 0.18669465 | -1.0703362 | 0.28446801 | NA | | MIR4253 | 100422914 |
| ENSG000002 | 0 | NA | NA | NA | NA | NA | | NA | NA |
| ENSG000002 | 0.94063848 | -0.603768454 | 0.29306415 | -2.0601922 | 0.03938017 | NA | | RN7SL744P | 106479493 |
| ENSG000002 | 0.50548109 | -0.025402968 | 0.24760631 | -0.1025942 | 0.91828506 | NA | | RN7SL66P | 106480485 |
| ENSG000002 | 0.47826975 | -0.016435741 | 0.244632 | -0.0671856 | 0.94643397 | NA | | MIR4491 | 100616330 |
| ENSG000002 | 1.28266546 | -0.677345408 | 0.33094555 | -2.0466975 | 0.0406878 | NA | | RN7SL198P | 106479294 |
| ENSG000002 | 0.71831827 | -0.020963193 | 0.28341454 | -0.0739665 | 0.94103701 | NA | | MIR4472-2 | 100616309 |

| | | | | | | | | |
|---|---|---|---|---|---|---|---|---|
| ENSG000002 | 1.43159376 | 0.09032587 | 0.34136764 | 0.26459998 | 0.79131764 | NA | NA | NA |
| ENSG000002 | 1.46242552 | -0.045743876 | 0.34787871 | -0.1314938 | 0.89538473 | NA | RN7SL455P | 106479389 |
| ENSG000002 | 1.98894125 | 0.501347074 | 0.36059249 | 1.39034254 | 0.16442488 | NA | MIR4737 | 100616210 |
| ENSG000002 | 1.83578813 | 0.010665444 | 0.35970766 | 0.02965031 | 0.97634594 | NA | NA | NA |
| ENSG000002 | 1.62124883 | -0.528886788 | 0.34698757 | -1.524224 | 0.12745276 | NA | NA | NA |
| ENSG000002 | 1.12113968 | -0.263718236 | 0.319023 | -0.8266433 | 0.40843924 | NA | MIR5685 | 100847075 |
| ENSG000002 | 1.61329965 | -0.247588586 | 0.34797478 | -0.711513 | 0.47676639 | NA | NA | NA |
| ENSG000002 | 0.14128499 | -0.097772014 | 0.13403235 | -0.7294658 | 0.4657168 | NA | MIR4316 | 100422851 |
| ENSG000002 | 0 | NA | NA | NA | NA | NA | NA | NA |
| ENSG000002 | 1.20723243 | 0.226312084 | 0.3112845 | 0.72702652 | 0.46720971 | NA | NA | NA |
| ENSG000002 | 0.62499586 | -0.095529625 | 0.25759142 | -0.3708572 | 0.71074392 | NA | RN7SL531P | 106479416 |
| ENSG000002 | 0.75982019 | -0.379788636 | 0.28855699 | -1.3161651 | 0.18811864 | NA | NA | NA |
| ENSG000002 | 0.26199343 | -0.194018184 | 0.18422801 | -1.0531416 | 0.29227609 | NA | MIR4783 | 100616187 |
| ENSG000002 | 1.98974836 | -0.681152267 | 0.35682697 | -1.9089148 | 0.05627309 | NA | NA | NA |
| ENSG000002 | 1.49362127 | -0.17968766 | 0.33649792 | -0.5339934 | 0.59334612 | NA | NA | NA |
| ENSG000002 | 1.57342166 | 0.146423047 | 0.34531355 | 0.42402926 | 0.67154448 | NA | MIR5688 | 100847077 |
| ENSG000002 | 0.889521 | -0.259225314 | 0.29433702 | -0.8807092 | 0.37847525 | NA | MIR3681 | 100500884 |
| ENSG000002 | 1.66465131 | -0.670054411 | 0.3441006 | -1.9472631 | 0.05150319 | NA | MIR4666B | 100847047 |
| ENSG000002 | 0.95762957 | -0.023510508 | 0.30495995 | -0.0770938 | 0.93854896 | NA | RN7SL474P | 106481052 |
| ENSG000002 | 0.11865494 | 0.083440866 | 0.12751478 | 0.65436232 | 0.5128784 | NA | NA | NA |
| ENSG000002 | 0 | NA | NA | NA | NA | NA | MIR4689 | 100616421 |
| ENSG000002 | 0 | NA | NA | NA | NA | NA | MIR4688 | 100616368 |
| ENSG000002 | 0.12685397 | -0.090215768 | 0.12901919 | -0.699243 | 0.48440017 | NA | NA | NA |
| ENSG000002 | 1.84386655 | -0.348855426 | 0.3504885 | -0.9953406 | 0.31957064 | NA | NA | NA |
| ENSG000002 | 0.23070733 | 0.167416683 | 0.17664244 | 0.94777156 | 0.34324576 | NA | MIR4300 | 100422823 |
| ENSG000002 | 0.47872676 | -0.008834112 | 0.22824852 | -0.0387039 | 0.96912645 | NA | RN7SL784P | 106481134 |
| ENSG000002 | 0.11934008 | 0.083564698 | 0.12760091 | 0.65489106 | 0.5125379 | NA | MIR3180-4 | 100500852 |
| ENSG000002 | 0.83284445 | 0.207108582 | 0.29432888 | 0.70366381 | 0.48164216 | NA | MIR4795 | 100616161 |
| ENSG000002 | 0.96042029 | -0.159022485 | 0.2973194 | -0.534854 | 0.59275079 | NA | NA | NA |

| | | | | | | | | |
|---|---|---|---|---|---|---|---|---|
| ENSG000002 | 0 | NA | NA | NA | NA | NA | MIR3689D2 | 100616344 |
| ENSG000002 | 0.85058056 | -0.103122652 | 0.29515749 | -0.3493818 | 0.7268027 | NA | MIR3667 | 100500882 |
| ENSG000002 | 0.12070844 | -0.088503695 | 0.12783931 | -0.6923042 | 0.48874631 | NA | MIR3928 | 100500901 |
| ENSG000002 | 1.16497352 | -0.268776859 | 0.31715991 | -0.847449 | 0.39674488 | NA | NA | NA |
| ENSG000002 | 1.75022756 | -0.191508367 | 0.35556236 | -0.538607 | 0.59015807 | NA | NA | NA |
| ENSG000002 | 0.45476004 | 0.3294872 | 0.23735363 | 1.38817005 | 0.16508527 | NA | NA | NA |
| ENSG000002 | 1.91232195 | -0.051468351 | 0.36058564 | -0.1427354 | 0.88649912 | NA | MIR4670 | 100616351 |
| ENSG000002 | 0 | NA | NA | NA | NA | NA | MIR3689B | 100500906 |
| ENSG000002 | 1.44565813 | -0.047380275 | 0.34301925 | -0.1381272 | 0.89013992 | NA | NA | NA |
| ENSG000002 | 0.98503488 | -0.18396325 | 0.30855684 | -0.5962054 | 0.55103803 | NA | RN7SL665P | 106481104 |
| ENSG000002 | 0.73160713 | -0.030178129 | 0.28429061 | -0.1061524 | 0.91546144 | NA | MIR4305 | 100422940 |
| ENSG000002 | 0.72008656 | 0.117012389 | 0.27576463 | 0.42431979 | 0.67133261 | NA | MIR3189 | 100422943 |
| ENSG000002 | 0.51931521 | -0.354214731 | 0.23897171 | -1.4822455 | 0.13827499 | NA | NA | NA |
| ENSG000002 | 1.41640784 | 0.227467455 | 0.33793463 | 0.67311082 | 0.50087679 | NA | NA | NA |
| ENSG000002 | 0 | NA | NA | NA | NA | NA | NA | NA |
| ENSG000002 | 1.92436665 | 0.065161134 | 0.35961322 | 0.18119782 | 0.8562123 | NA | NA | NA |
| ENSG000002 | 1.00478385 | -0.047356386 | 0.30701583 | -0.1542474 | 0.87741469 | NA | NA | NA |
| ENSG000002 | 1.36429507 | -0.117134827 | 0.33628724 | -0.3483178 | 0.72760154 | NA | NA | NA |
| ENSG000002 | 1.94821108 | -0.053758153 | 0.36269993 | -0.1482166 | 0.88217183 | NA | MIR4693 | 100616457 |
| ENSG000002 | 0.11131518 | 0.081951103 | 0.12663804 | 0.64712866 | 0.51754868 | NA | MIR4701 | 100616262 |
| ENSG000002 | 0.24550891 | -0.007886114 | 0.17900103 | -0.0440563 | 0.96485956 | NA | NA | NA |
| ENSG000002 | 1.22897742 | -0.186516225 | 0.32902362 | -0.5668779 | 0.57079712 | NA | MIR4716 | 100616332 |
| ENSG000002 | 0.5810132 | 0.071210634 | 0.156906 | 0.45384265 | 0.64994208 | NA | MIR3168 | 100422878 |
| ENSG000002 | 0.11131518 | 0.081951103 | 0.12663804 | 0.64712866 | 0.51754868 | NA | MIR4456 | 100616381 |
| ENSG000002 | 0.87394783 | -0.26920868 | 0.29758521 | -0.904644 | 0.36565401 | NA | NA | NA |
| ENSG000002 | 0.73747438 | 0.126914074 | 0.27698383 | 0.4582003 | 0.64680855 | NA | NA | NA |
| ENSG000002 | 1.20863113 | -0.038158429 | 0.33303517 | -0.1145778 | 0.9087798 | NA | NA | NA |
| ENSG000002 | 1.10992159 | 0.032362872 | 0.31566451 | 0.102523 | 0.91834156 | NA | NA | NA |
| ENSG000002 | 2.06476633 | -0.7143351 | 0.35820103 | -1.9942296 | 0.04612697 | NA | RN7SL835P | 106479526 |

| | | | | | | | | |
|---|---|---|---|---|---|---|---|---|
| ENSG000002 | 0.11934008 | 0.083564698 | 0.12760091 | 0.65489106 | 0.5125379 | NA | NA | NA |
| ENSG000002 | 0.73727446 | -0.186424749 | 0.27938103 | -0.6672778 | 0.50459473 | NA | NA | NA |
| ENSG000002 | 1.95581088 | -0.063164641 | 0.36242132 | -0.1742851 | 0.8616414 | NA | NA | NA |
| ENSG000002 | 0.51570138 | -0.378721611 | 0.24581506 | -1.540677 | 0.12339542 | NA | MIR4322 | 100422925 |
| ENSG000002 | 0.70764096 | 0.275229799 | 0.26960301 | 1.02087063 | 0.30731573 | NA | MIR4767 | 100616467 |
| ENSG000002 | 0.60698135 | -0.252864344 | 0.25666105 | -0.9852073 | 0.32452224 | NA | RN7SL488P | 106479401 |
| ENSG000002 | 1.07728019 | -0.102834727 | 0.31310955 | -0.3284305 | 0.74258618 | NA | NA | NA |
| ENSG000002 | 0.90895539 | -0.266226439 | 0.29363402 | -0.9066607 | 0.36458623 | NA | RN7SL753P | 106479496 |
| ENSG000002 | 0.98164526 | 0.122915952 | 0.31661942 | 0.38821356 | 0.697858 | NA | MIR4786 | 100616417 |
| ENSG000002 | 0.97711604 | -0.034776569 | 0.30750244 | -0.1130936 | 0.90995632 | NA | NA | NA |
| ENSG000002 | 2.17294702 | -0.059874533 | 0.36841583 | -0.1625189 | 0.87089725 | NA | NA | NA |
| ENSG000002 | 1.08777833 | 0.032037272 | 0.32365735 | 0.09898515 | 0.92115006 | NA | MIR2467 | 100616360 |
| ENSG000002 | 0 | NA | NA | NA | NA | NA | NA | NA |
| ENSG000002 | 0 | NA | NA | NA | NA | NA | SNORD55 | 26811 |
| ENSG000002 | 1.77297777 | -0.617800329 | 0.35405859 | -1.7449099 | 0.08100056 | NA | MIR4433B | 102465833 |
| ENSG000002 | 1.31595587 | 0.274620962 | 0.32107521 | 0.85531661 | 0.39237589 | NA | NA | NA |
| ENSG000002 | 0.22263036 | 0.146922267 | 0.16600367 | 0.88505432 | 0.37612734 | NA | NA | NA |
| ENSG000002 | 1.45747155 | -0.180047101 | 0.33895069 | -0.5311897 | 0.59528736 | NA | MIR4694 | 100616426 |
| ENSG000002 | 0.80429645 | 0.358103509 | 0.2885739 | 1.24094214 | 0.21462712 | NA | MIR548AT | 100847030 |
| ENSG000002 | 0.3433388 | 0.065348438 | 0.20407693 | 0.32021472 | 0.74880557 | NA | NA | NA |
| ENSG000002 | 2.09885207 | -0.008100013 | 0.36469328 | -0.0222105 | 0.98228006 | NA | RN7SL154P | 106479277 |
| ENSG000002 | 0.36482725 | -0.102221582 | 0.21647267 | -0.4722147 | 0.63677353 | NA | MIR4801 | 100616435 |
| ENSG000002 | 0.85767182 | -0.111725312 | 0.29715703 | -0.3759807 | 0.70693124 | NA | RN7SL448P | 106481843 |
| ENSG000002 | 1.9915349 | -0.671063637 | 0.35488552 | -1.8909299 | 0.0586337 | NA | MIR5186 | 100847036 |
| ENSG000002 | 2.13017097 | -0.253902525 | 0.36531623 | -0.6950212 | 0.48704201 | NA | NA | NA |
| ENSG000002 | 1.64193088 | -0.379157963 | 0.34472914 | -1.0998721 | 0.27138785 | NA | NA | NA |
| ENSG000002 | 1.35623188 | 0.01146653 | 0.33209857 | 0.03452749 | 0.97245652 | NA | NA | NA |
| ENSG000002 | 0.26813897 | -0.197007041 | 0.18550319 | -1.0620143 | 0.2882292 | NA | MIR4417 | 100616489 |
| ENSG000002 | 2.01194271 | -0.172327727 | 0.35571751 | -0.4844511 | 0.62806579 | NA | MIR3660 | 100500825 |

| | | | | | | | | |
|---|---|---|---|---|---|---|---|---|
| ENSG000002 | 0 | NA | NA | NA | NA | NA | NA | NA |
| ENSG000002 | 0.12070844 | -0.088503695 | 0.12783931 | -0.6923042 | 0.48874631 | NA | RN7SL602P | 106479441 |
| ENSG000002 | 1.76332 | -0.194260514 | 0.34927484 | -0.5561824 | 0.57808618 | NA | MIR3134 | 100422990 |
| ENSG000002 | 0 | NA | NA | NA | NA | NA | MIR4648 | 100616116 |
| ENSG000002 | 0.92082245 | -0.265816145 | 0.2942672 | -0.9033156 | 0.36635844 | NA | MIR3122 | 100422947 |
| ENSG000002 | 1.11288857 | -0.117074117 | 0.32227187 | -0.3632775 | 0.7163976 | NA | NA | NA |
| ENSG000002 | 1.19904144 | 0.101070648 | 0.33134656 | 0.30503002 | 0.76034329 | NA | MIR4425 | 100616365 |
| ENSG000002 | 0.86091955 | -0.265574426 | 0.29690668 | -0.894471 | 0.37106993 | NA | NA | NA |
| ENSG000002 | 0.47144099 | 0.149959821 | 0.2369265 | 0.63293816 | 0.52677403 | NA | MIR4671 | 100616380 |
| ENSG000002 | 0.3593886 | 0.071604659 | 0.20679693 | 0.34625591 | 0.7291504 | NA | MIR4513 | 100616183 |
| ENSG000002 | 1.27726063 | -0.307477044 | 0.31803592 | -0.9667997 | 0.33364415 | NA | MIR5007 | 100846996 |
| ENSG000002 | 1.53408922 | -0.191208545 | 0.3375853 | -0.5664007 | 0.57112143 | NA | RN7SL208P | 106480978 |
| ENSG000002 | 0 | NA | NA | NA | NA | NA | MIR3180-5 | 100500916 |
| ENSG000002 | 0.11865494 | 0.083440866 | 0.12751478 | 0.65436232 | 0.5128784 | NA | MIR4736 | 100616220 |
| ENSG000002 | 0 | NA | NA | NA | NA | NA | RN7SL491P | 106480514 |
| ENSG000002 | 1.08465769 | -0.234371138 | 0.30882403 | -0.7589148 | 0.44790351 | NA | MIR548AL | 100616215 |
| ENSG000002 | 0.39274356 | -0.107428268 | 0.22062504 | -0.4869269 | 0.62631013 | NA | NA | NA |
| ENSG000002 | 1.29367406 | -0.349332307 | 0.32941062 | -1.0604768 | 0.28892775 | NA | MIR548AP | 100847084 |
| ENSG000002 | 0.11865494 | 0.083440866 | 0.12751478 | 0.65436232 | 0.5128784 | NA | MIR4470 | 100616484 |
| ENSG000002 | 0.96109045 | 0.265145083 | 0.30036775 | 0.88273485 | 0.37737955 | NA | MIR548AC | 100616384 |
| ENSG000002 | 1.25659371 | -0.187003675 | 0.32826121 | -0.5696795 | 0.56889512 | NA | NA | NA |
| ENSG000002 | 1.11034635 | 0.040157213 | 0.3156658 | 0.12721433 | 0.89877077 | NA | RN7SL718P | 106479485 |
| ENSG000002 | 2.01723314 | 0.132879989 | 0.35915966 | 0.3699747 | 0.71140134 | NA | MIR4653 | 100616117 |
| ENSG000002 | 0 | NA | NA | NA | NA | NA | MIR4283-1 | 100422917 |
| ENSG000002 | 0.91371137 | -0.266813888 | 0.29560643 | -0.9025984 | 0.36673908 | NA | MIR3183 | 100422835 |
| ENSG000002 | 1.34898303 | -0.113352555 | 0.33595423 | -0.3374048 | 0.7358118 | NA | NA | NA |
| ENSG000002 | 1.29831597 | 0.30667206 | 0.32894733 | 0.93228318 | 0.3511902 | NA | RN7SL560P | 106479426 |
| ENSG000002 | 1.57900659 | 0.148685141 | 0.34959497 | 0.42530686 | 0.67061299 | NA | NA | NA |
| ENSG000002 | 0.12685397 | -0.090215768 | 0.12901919 | -0.699243 | 0.48440017 | NA | NA | NA |

| | | | | | | | | |
|---|---|---|---|---|---|---|---|---|
| ENSG000002 | 2.06188438 | 0.129171243 | 0.35613477 | 0.36270326 | 0.71682657 | NA | NA | NA |
| ENSG000002 | 0 | NA | NA | NA | NA | NA | NA | NA |
| ENSG000002 | 0.76949578 | -0.359754798 | 0.28380236 | -1.2676244 | 0.20493211 | NA | MIR4520-1 | 100616401 |
| ENSG000002 | 0.99761933 | -0.195448883 | 0.31381005 | -0.6228254 | 0.53339924 | NA | MIR4794 | 100616338 |
| ENSG000002 | 0.25333738 | -0.015686291 | 0.1806605 | -0.0868275 | 0.93080867 | NA | MIR4656 | 100616465 |
| ENSG000002 | 1.4395126 | -0.046636385 | 0.34487761 | -0.1352259 | 0.89243327 | NA | MIR5682 | 100847043 |
| ENSG000002 | 2.09635921 | -0.125143556 | 0.36638793 | -0.3415603 | 0.73268185 | NA | MIR4714 | 100616432 |
| ENSG000002 | 0.37148378 | 0.06737793 | 0.20875796 | 0.32275622 | 0.74687987 | NA | MIR5008 | 100847072 |
| ENSG000002 | 0.38133351 | -0.102466351 | 0.2189713 | -0.4679442 | 0.63982449 | NA | RN7SL697P | 106480526 |
| ENSG000002 | 0.86592348 | -0.266194925 | 0.29880302 | -0.8908709 | 0.37299842 | NA | NA | NA |
| ENSG000002 | 1.60774643 | -0.260795783 | 0.34997509 | -0.7451839 | 0.45616058 | NA | MIR4298 | 100423021 |
| ENSG000002 | 1.32737039 | 0.029141272 | 0.33518644 | 0.08694048 | 0.93071882 | NA | MIR4650-2 | 100616331 |
| ENSG000002 | 1.15307475 | -0.274348016 | 0.32432702 | -0.8458993 | 0.3976089 | NA | NA | NA |
| ENSG000002 | 1.1480402 | 0.386522643 | 0.31797584 | 1.21557238 | 0.22414786 | NA | MIR3124 | 100422879 |
| ENSG000002 | 0 | NA | NA | NA | NA | NA | NA | NA |
| ENSG000002 | 1.57254648 | 0.01745119 | 0.35295437 | 0.04944319 | 0.96056611 | NA | NA | NA |
| ENSG000002 | 1.47321941 | -0.298654936 | 0.33492859 | -0.8916973 | 0.37255518 | NA | MIR3678 | 100500841 |
| ENSG000002 | 0.61950267 | -0.112567707 | 0.27088102 | -0.4155614 | 0.67773094 | NA | MIR5692A2 | 100847038 |
| ENSG000002 | 1.57684545 | 0.015330492 | 0.34526237 | 0.04440244 | 0.96458362 | NA | RN7SL596P | 106481085 |
| ENSG000002 | 0.35210091 | 0.074791332 | 0.21412433 | 0.34928928 | 0.72687215 | NA | MIR4778 | 100616464 |
| ENSG000002 | 1.61394187 | -0.486868866 | 0.33919373 | -1.435371 | 0.15118139 | NA | RN7SL25P | 106479233 |
| ENSG000002 | 0.37945415 | -0.107841678 | 0.21865587 | -0.4932027 | 0.62186934 | NA | NA | NA |
| ENSG000002 | 0.53590737 | -0.074911264 | 0.15581405 | -0.4807735 | 0.63067751 | NA | MIR548AR | 100847035 |
| ENSG000002 | 2.05961059 | 0.002854846 | 0.36302511 | 0.00786405 | 0.99372546 | NA | RN7SL405P | 106479373 |
| ENSG000002 | 0 | NA | NA | NA | NA | NA | NA | NA |
| ENSG000002 | 0 | NA | NA | NA | NA | NA | SNORD95 | 619570 |
| ENSG000002 | 0.12070844 | -0.088503695 | 0.12783931 | -0.6923042 | 0.48874631 | NA | MIR4257 | 100422997 |
| ENSG000002 | 0.25214371 | -0.009862892 | 0.18046657 | -0.0546522 | 0.95641556 | NA | NA | NA |
| ENSG000002 | 0.60487577 | -0.101296586 | 0.26329793 | -0.3847223 | 0.70044317 | NA | MIR4633 | 100616175 |

| | | | | | | | | |
|---|---|---|---|---|---|---|---|---|
| ENSG000002 | 0.61753348 | 0.051696206 | 0.26401285 | 0.19580943 | 0.84475933 | NA | MIR23C | 100500809 |
| ENSG000002 | 1.52638337 | -0.344681703 | 0.34728158 | -0.9925136 | 0.32094703 | NA | MIR4529 | 100616290 |
| ENSG000002 | 2.15746864 | -0.649862711 | 0.36505092 | -1.7801975 | 0.07504365 | NA | RN7SL15P | 106479230 |
| ENSG000002 | 1.12677241 | 0.02346003 | 0.31069857 | 0.07550736 | 0.93981104 | NA | MIR1254-2 | 100616217 |
| ENSG000002 | 1.76343373 | 0.388752651 | 0.35197186 | 1.10449925 | 0.26937663 | NA | NA | NA |
| ENSG000002 | 2.14087121 | 0.644096323 | 0.3572036 | 1.80316305 | 0.07136261 | NA | MIR5692B | 100847013 |
| ENSG000002 | 1.52954785 | 0.138449881 | 0.33815539 | 0.4094268 | 0.68222647 | NA | RN7SL326P | 106479342 |
| ENSG000002 | 0.50611225 | -0.202624884 | 0.24886748 | -0.8141879 | 0.41553732 | NA | MIR4487 | 100616222 |
| ENSG000002 | 0.63202049 | -0.257145926 | 0.25898277 | -0.9929075 | 0.32075505 | NA | RN7SL131P | 106480957 |
| ENSG000002 | 2.10359078 | -0.389596941 | 0.36772449 | -1.0594805 | 0.28938099 | NA | NA | NA |
| ENSG000002 | 1.99807176 | -0.189915927 | 0.35987756 | -0.5277237 | 0.5976911 | NA | MIR3159 | 100423016 |
| ENSG000002 | 1.02761033 | -0.192933176 | 0.30869441 | -0.6249973 | 0.53197282 | NA | MIR3173 | 100422981 |
| ENSG000002 | 0 | NA | NA | NA | NA | NA | MIR4685 | 100616482 |
| ENSG000002 | 1.62005461 | -0.539328085 | 0.34808935 | -1.5493955 | 0.12128667 | NA | RN7SL290P | 106481001 |
| ENSG000002 | 0.14128499 | -0.097772014 | 0.13403235 | -0.7294658 | 0.4657168 | NA | MIR5588 | 100847054 |
| ENSG000002 | 1.48941011 | 0.079269493 | 0.3390043 | 0.23383035 | 0.81511669 | NA | RN7SL592P | 106481851 |
| ENSG000002 | 1.6692326 | -0.559683748 | 0.35123993 | -1.5934514 | 0.11105897 | NA | NA | NA |
| ENSG000002 | 0.93036114 | 0.249957807 | 0.29629899 | 0.84359992 | 0.39889301 | NA | RN7SL732P | 106481122 |
| ENSG000002 | 0.63621395 | -0.448723671 | 0.2628158 | -1.7073694 | 0.08775341 | NA | MIR5591 | 100847065 |
| ENSG000002 | 1.66857971 | -0.569525961 | 0.35224664 | -1.6168386 | 0.10591313 | NA | NA | NA |
| ENSG000002 | 0.76693319 | -0.191420094 | 0.28134921 | -0.6803648 | 0.49627351 | NA | NA | NA |
| ENSG000002 | 0.37441639 | -0.268856313 | 0.21277471 | -1.2635727 | 0.20638344 | NA | MIR4271 | 100422952 |
| ENSG000002 | 0.48418907 | 0.150283598 | 0.23847389 | 0.63018889 | 0.52857101 | NA | MIR3152 | 100422869 |
| ENSG000002 | 1.74164014 | -0.328462156 | 0.35598827 | -0.9226769 | 0.35617561 | NA | NA | NA |
| ENSG000002 | 1.89951572 | -0.387426458 | 0.3578707 | -1.0825878 | 0.27899142 | NA | NA | NA |
| ENSG000002 | 0.75144647 | -0.037105481 | 0.28557336 | -0.1299333 | 0.89661922 | NA | MIR5194 | 100847051 |
| ENSG000002 | 0.14128499 | -0.097772014 | 0.13403235 | -0.7294658 | 0.4657168 | NA | MIR3171 | 100422830 |
| ENSG000002 | 1.09467764 | -0.24657427 | 0.31107311 | -0.792657 | 0.4279777 | NA | MIR4798 | 100616471 |
| ENSG000002 | 1.44395455 | -0.166398197 | 0.33742054 | -0.4931478 | 0.62190816 | NA | NA | NA |

| | | | | | | | | |
|---|---|---|---|---|---|---|---|---|
| ENSG000002 | 0.73727446 | -0.186424749 | 0.27938103 | -0.6672778 | 0.50459473 | NA | NA | NA |
| ENSG000002 | 0 | NA | NA | NA | NA | NA | NA | NA |
| ENSG000002 | 0.11934008 | 0.083564698 | 0.12760091 | 0.65489106 | 0.5125379 | NA | MIR4285 | 100422858 |
| ENSG000002 | 0.24619405 | -0.007630671 | 0.17915639 | -0.0425922 | 0.96602658 | NA | RN7SL204P | 106480977 |
| ENSG000002 | 0.37441639 | -0.268856313 | 0.21277471 | -1.2635727 | 0.20638344 | NA | NA | NA |
| ENSG000002 | 1.12540639 | -0.277199927 | 0.32577282 | -0.8508995 | 0.39482519 | NA | NA | NA |
| ENSG000002 | 2.1141969 | -0.007173588 | 0.35973057 | -0.0199416 | 0.98408999 | NA | NA | NA |
| ENSG000002 | 0.35141577 | 0.074527565 | 0.21401461 | 0.34823588 | 0.72766304 | NA | NA | NA |
| ENSG000002 | 1.12933637 | 0.022299862 | 0.32100953 | 0.06946791 | 0.94461717 | NA | MIR4255 | 100422898 |
| ENSG000002 | 0.64759311 | -0.271681639 | 0.26184098 | -1.0375826 | 0.29946444 | NA | MIR4752 | 100616171 |
| ENSG000002 | 2.00681659 | -0.546047617 | 0.3571966 | -1.5287033 | 0.12633802 | NA | NA | NA |
| ENSG000002 | 1.09478582 | 0.037301556 | 0.32136739 | 0.11607138 | 0.90759597 | NA | NA | NA |
| ENSG000002 | 1.42738959 | -0.584430898 | 0.33920311 | -1.7229526 | 0.08489709 | NA | NA | NA |
| ENSG000002 | 0 | NA | NA | NA | NA | NA | MIR3670-2 | 100846994 |
| ENSG000002 | 0 | NA | NA | NA | NA | NA | MIR3129 | 100422908 |
| ENSG000002 | 1.9751615 | -0.061081424 | 0.36254037 | -0.1684817 | 0.86620433 | NA | NA | NA |
| ENSG000002 | 0.86321866 | 0.038651795 | 0.2910859 | 0.13278484 | 0.89436354 | NA | MIR4638 | 100616342 |
| ENSG000002 | 0.47523053 | 0.12360509 | 0.22747273 | 0.54338422 | 0.58686529 | NA | MIR4718 | 100616195 |
| ENSG000002 | 0.7536964 | -0.192464368 | 0.28656601 | -0.6716232 | 0.50182363 | NA | NA | NA |
| ENSG000002 | 0.11205239 | 0.082118251 | 0.12671962 | 0.64803108 | 0.51696485 | NA | MIR4511 | 100616379 |
| ENSG000002 | 0 | NA | NA | NA | NA | NA | MIR4505 | 100616158 |
| ENSG000002 | 0.11865494 | 0.083440866 | 0.12751478 | 0.65436232 | 0.5128784 | NA | MIR3689C | 100616333 |
| ENSG000002 | 0.8899219 | -0.629664671 | 0.29788939 | -2.1137533 | 0.03453635 | NA | MIR3924 | 100500834 |
| ENSG000002 | 1.0768253 | 0.038783439 | 0.31922825 | 0.12149125 | 0.90330194 | NA | NA | NA |
| ENSG000002 | 1.14624353 | -0.287735931 | 0.32635368 | -0.881669 | 0.37795581 | NA | MIR3131 | 100422957 |
| ENSG000002 | 1.72823215 | 0.070733257 | 0.3491266 | 0.2026006 | 0.83944722 | NA | MIR3198-1 | 100423025 |
| ENSG000002 | 0.26813897 | -0.197007041 | 0.18550319 | -1.0620143 | 0.2882292 | NA | MIR3141 | 100422950 |
| ENSG000002 | 1.32558084 | -0.112237008 | 0.33166702 | -0.3384027 | 0.73505975 | NA | MIR4295 | 100422909 |
| ENSG000002 | 0.61767784 | 0.056941233 | 0.25494708 | 0.22334531 | 0.82326677 | NA | MIR4772 | 100616157 |

| | | | | | | | | |
|---|---|---|---|---|---|---|---|---|
| ENSG000002 | 1.80399315 | 0.265785235 | 0.35412875 | 0.75053278 | 0.45293389 | NA | NA | NA |
| ENSG000002 | 0 | NA | NA | NA | NA | NA | MIR4420 | 100616164 |
| ENSG000002 | 0 | NA | NA | NA | NA | NA | MIR4537 | 100616422 |
| ENSG000002 | 1.26140121 | -0.1915509 | 0.32760121 | -0.5847076 | 0.55874437 | NA | NA | NA |
| ENSG000002 | 1.27515101 | -0.51710732 | 0.33236375 | -1.5558475 | 0.11974435 | NA | MIR3148 | 100422876 |
| ENSG000002 | 1.28761676 | -0.669706094 | 0.32888846 | -2.0362712 | 0.04172313 | NA | NA | NA |
| ENSG000002 | 0.64608094 | -0.115215656 | 0.26675803 | -0.4319107 | 0.6658063 | NA | MIR4637 | 100616271 |
| ENSG000002 | 0.23202362 | -0.008302493 | 0.17586207 | -0.0472103 | 0.96234565 | NA | MIR4771-1 | 100616370 |
| ENSG000002 | 0.58623011 | -0.031196001 | 0.15711109 | -0.1985601 | 0.84260683 | NA | MIR4294 | 100422895 |
| ENSG000002 | 0.14128499 | -0.097772014 | 0.13403235 | -0.7294658 | 0.4657168 | NA | MIR378C | 100422867 |
| ENSG000002 | 1.59454538 | 0.011955234 | 0.35093312 | 0.03406699 | 0.97282373 | NA | NA | NA |
| ENSG000002 | 0.86435869 | -0.122981079 | 0.30693344 | -0.4006767 | 0.68865816 | NA | MIR4441 | 100616493 |
| ENSG000002 | 0.513452 | -0.199963756 | 0.2496885 | -0.8008529 | 0.42321682 | NA | NA | NA |
| ENSG000002 | 0.87394783 | -0.26920868 | 0.29758521 | -0.904644 | 0.36565401 | NA | MIR1273C | 100422821 |
| ENSG000002 | 0 | NA | NA | NA | NA | NA | NA | NA |
| ENSG000002 | 1.11746582 | -0.26364324 | 0.31724605 | -0.8310371 | 0.40595266 | NA | MIR5571 | 100847006 |
| ENSG000002 | 1.53574575 | -0.309525066 | 0.33588111 | -0.9215316 | 0.35677294 | NA | NA | NA |
| ENSG000002 | 0.75964605 | -0.203618105 | 0.2920432 | -0.6972191 | 0.48566567 | NA | NA | NA |
| ENSG000002 | 1.66368951 | 0.070253356 | 0.34023644 | 0.20648392 | 0.83641292 | NA | RN7SL468P | 106481050 |
| ENSG000002 | 0.63893761 | -0.11408674 | 0.27243711 | -0.4187636 | 0.67538892 | NA | MIR1273F | 100616156 |
| ENSG000002 | 0.36538978 | 0.059033792 | 0.20773944 | 0.28417229 | 0.77627835 | NA | RN7SL404P | 106479372 |
| ENSG000002 | 0.26560726 | -0.177019426 | 0.17672274 | -1.0016788 | 0.31649875 | NA | MIR4310 | 100423013 |
| ENSG000002 | 1.51567381 | -0.058770891 | 0.34614289 | -0.1697879 | 0.86517691 | NA | MIR5696 | 100847007 |
| ENSG000002 | 0.23868015 | 0.154625155 | 0.16997278 | 0.90970538 | 0.36297791 | NA | MIR3613 | 100500908 |
| ENSG000002 | 1.69510204 | -0.042285721 | 0.35011213 | -0.1207777 | 0.90386715 | NA | NA | NA |
| ENSG000002 | 1.62357861 | -0.399354304 | 0.34975376 | -1.1418156 | 0.25353067 | NA | RN7SL690P | 106479475 |
| ENSG000002 | 1.26123099 | -0.347053654 | 0.33223688 | -1.044597 | 0.29620925 | NA | MIR1273D | 100422970 |
| ENSG000002 | 0 | NA | NA | NA | NA | NA | NA | NA |
| ENSG000002 | 0.11131518 | 0.081951103 | 0.12663804 | 0.64712866 | 0.51754868 | NA | MIR3616 | 100500814 |

| | | | | | | | | |
|---|---|---|---|---|---|---|---|---|
| ENSG000002 | 0 | NA | NA | NA | NA | NA | NA | NA |
| ENSG000002 | 0 | NA | NA | NA | NA | NA | MIR4494 | 100616478 |
| ENSG000002 | 0.52623233 | -0.191068033 | 0.24398404 | -0.7831169 | 0.43355844 | NA | MIR4540 | 100616278 |
| ENSG000002 | 0.98663137 | -0.344974254 | 0.30839688 | -1.1186049 | 0.26330874 | NA | MIR378F | 100616492 |
| ENSG000002 | 1.95358715 | -0.047702047 | 0.3563767 | -0.1338529 | 0.89351891 | NA | NA | NA |
| ENSG000002 | 0.8841642 | -0.254282913 | 0.29233359 | -0.8698382 | 0.38438884 | NA | NA | NA |
| ENSG000002 | 1.28066751 | -0.19283298 | 0.32810378 | -0.5877195 | 0.5567206 | NA | RN7SL190P | 106479291 |
| ENSG000002 | 0.26813897 | -0.197007041 | 0.18550319 | -1.0620143 | 0.2882292 | NA | MIR4265 | 100422863 |
| ENSG000002 | 1.01611906 | -0.315839523 | 0.29868871 | -1.0574203 | 0.29031979 | NA | MIR4474 | 100616441 |
| ENSG000002 | 0.12685397 | -0.090215768 | 0.12901919 | -0.699243 | 0.48440017 | NA | MIR3181 | 100422972 |
| ENSG000002 | 0.34948434 | 0.063159058 | 0.2051263 | 0.30790326 | 0.75815594 | NA | MIR4730 | 100616359 |
| ENSG000002 | 0.26813897 | -0.197007041 | 0.18550319 | -1.0620143 | 0.2882292 | NA | RN7SL440P | 106481042 |
| ENSG000002 | 1.138622 | -0.436631135 | 0.3236206 | -1.3492068 | 0.17727054 | NA | MIR5094 | 100847059 |
| ENSG000002 | 1.52457781 | -0.18730661 | 0.34094561 | -0.5493739 | 0.5827489 | NA | NA | NA |
| ENSG000002 | 1.21045651 | -0.173549603 | 0.32622614 | -0.5319917 | 0.59473174 | NA | MIR4789 | 100616395 |
| ENSG000002 | 1.46549197 | 0.072807064 | 0.33149714 | 0.21963105 | 0.82615851 | NA | MIR4431 | 100616431 |
| ENSG000002 | 1.79875684 | 0.008121886 | 0.35940287 | 0.02259828 | 0.98197072 | NA | RN7SL630P | 106481094 |
| ENSG000002 | 0 | NA | NA | NA | NA | NA | MIR4436B1 | 100616123 |
| ENSG000002 | 0.23070733 | 0.167416683 | 0.17664244 | 0.94777156 | 0.34324576 | NA | NA | NA |
| ENSG000002 | 0.88360168 | -0.430087375 | 0.29482856 | -1.4587711 | 0.14462812 | NA | MIR5691 | 100847015 |
| ENSG000002 | 0.59621972 | 0.061674489 | 0.26859331 | 0.22962035 | 0.81838679 | NA | MIR5092 | 100847039 |
| ENSG000002 | 1.90942101 | -0.245260422 | 0.35155331 | -0.6976479 | 0.48539743 | NA | NA | NA |
| ENSG000002 | 0.50107449 | -0.372905725 | 0.24426561 | -1.5266403 | 0.12685049 | NA | MIR5089 | 100847067 |
| ENSG000002 | 1.36813957 | -0.559398039 | 0.33600892 | -1.6648309 | 0.09594649 | NA | MIR4780 | 100616447 |
| ENSG000002 | 0.73750472 | -0.033552356 | 0.29002437 | -0.1156881 | 0.90789977 | NA | MIR548W | 100422923 |
| ENSG000002 | 1.25365864 | -0.05381498 | 0.32916252 | -0.1634906 | 0.87013217 | NA | MIR4327 | 100422891 |
| ENSG000002 | 1.4516987 | -0.174999862 | 0.3410669 | -0.5130954 | 0.60788458 | NA | NA | NA |
| ENSG000002 | 0.75190136 | -0.191187334 | 0.28037014 | -0.6819105 | 0.49529557 | NA | NA | NA |
| ENSG000002 | 1.22946827 | -0.048659607 | 0.33356852 | -0.1458759 | 0.88401936 | NA | NA | NA |

| | | | | | | | | |
|---|---|---|---|---|---|---|---|---|
| ENSG000002 | 0.91680777 | 0.5912928 | 0.29592765 | 1.99809921 | 0.04570591 | NA | NA | NA |
| ENSG000002 | 1.05499636 | -0.338406447 | 0.30599274 | -1.1059297 | 0.26875696 | NA | MIR4792 | 100616448 |
| ENSG000002 | 0.60327873 | 0.055099477 | 0.26276039 | 0.20969476 | 0.83390592 | NA | MIR1273E | 100526648 |
| ENSG000002 | 1.17370499 | 0.391500887 | 0.32261844 | 1.21351055 | 0.22493468 | NA | RN7SL262P | 106479317 |
| ENSG000002 | 0.69549179 | 0.132602096 | 0.27562182 | 0.48110159 | 0.6304443 | NA | RN7SL622P | 106481092 |
| ENSG000002 | 0.63210694 | -0.105863447 | 0.26563058 | -0.3985364 | 0.69023486 | NA | MIR548S | 100422862 |
| ENSG000002 | 1.02536095 | -0.044289782 | 0.3078644 | -0.1438613 | 0.88560997 | NA | MIR4765 | 100616219 |
| ENSG000002 | 1.20220485 | -0.038574913 | 0.32623495 | -0.1182427 | 0.90587533 | NA | PPY2P | 23614 |
| ENSG000002 | 0.24756242 | -0.187034346 | 0.18119817 | -1.0322088 | 0.30197431 | NA | MIR4419B | 100616298 |
| ENSG000002 | 0.35870346 | 0.077368606 | 0.21515516 | 0.35959447 | 0.71915042 | NA | MIR4692 | 100616410 |
| ENSG000002 | 1.15003205 | -0.431577722 | 0.32180297 | -1.3411241 | 0.17988017 | NA | MIR3622B | 100500871 |
| ENSG000002 | 0.46415331 | 0.147192669 | 0.23603211 | 0.6236129 | 0.53288183 | NA | MIR3691 | 100500900 |
| ENSG000002 | 0.23139247 | 0.167748926 | 0.17680352 | 0.94878726 | 0.34272882 | NA | MIR4655 | 100616160 |
| ENSG000002 | 1.30414638 | 0.14919346 | 0.32145419 | 0.46412044 | 0.64256146 | NA | MIR4484 | 100616327 |
| ENSG000002 | 1.62104168 | -0.369279999 | 0.34231059 | -1.0787864 | 0.28068297 | NA | C1QTNF1-AS1 | 100507410 |
| ENSG000002 | 0.26062507 | -0.012888319 | 0.18226503 | -0.070712 | 0.94362699 | NA | NA | NA |
| ENSG000002 | 0.48555743 | -0.013775713 | 0.2454629 | -0.0561214 | 0.95524512 | NA | NA | NA |
| ENSG000002 | 1.21340829 | 0.105143959 | 0.32648814 | 0.32204527 | 0.7474184 | NA | NA | NA |
| ENSG000002 | 0.35961481 | -0.099625193 | 0.21567743 | -0.4619176 | 0.64414043 | NA | MIR4731 | 100616125 |
| ENSG000002 | 1.03393055 | -0.34903534 | 0.31038672 | -1.1245176 | 0.26079349 | NA | NA | NA |
| ENSG000002 | 0.90298455 | -0.457507079 | 0.30248048 | -1.5125177 | 0.13040221 | NA | NA | NA |
| ENSG000002 | 1.00926198 | -0.521357627 | 0.31145026 | -1.6739676 | 0.09413698 | NA | NA | NA |
| ENSG000002 | 0.36212533 | -0.235926492 | 0.20089643 | -1.1743688 | 0.24024735 | NA | NA | NA |
| ENSG000002 | 1.68938651 | -0.38788162 | 0.34384767 | -1.1280624 | 0.25929357 | NA | NA | NA |
| ENSG000002 | 0.78227611 | -0.381453213 | 0.2899593 | -1.3155405 | 0.18832831 | NA | NA | NA |
| ENSG000002 | 1.44771568 | 0.08779137 | 0.33416117 | 0.26272164 | 0.79276514 | NA | NA | NA |
| ENSG000002 | 1.96720421 | -0.430194623 | 0.35981254 | -1.1956077 | 0.2318497 | NA | MIR4301 | 100422855 |
| ENSG000002 | 0.73443167 | -0.030284643 | 0.27816347 | -0.1088735 | 0.91330279 | NA | MIR151B | 100616247 |
| ENSG000002 | 2.00556447 | -0.471908058 | 0.36534472 | -1.2916789 | 0.19646838 | NA | MIR5003 | 100847029 |

| | | | | | | | | | |
|---|---|---|---|---|---|---|---|---|---|
| ENSG000002 | 1.50942064 | -0.334327473 | 0.34516683 | -0.9685968 | 0.33274642 | NA | | MIR2681 | 100616110 |
| ENSG000002 | 1.44397127 | 0.229581444 | 0.34310267 | 0.66913336 | 0.50341041 | NA | | NA | NA |
| ENSG000002 | 0.80475538 | 0.532355785 | 0.28536748 | 1.86550967 | 0.06211 | NA | | RN7SL667P | 106479465 |
| ENSG000002 | 0.87457898 | -0.427920645 | 0.29435302 | -1.4537668 | 0.14601097 | NA | | MIR4262 | 100422996 |
| ENSG000002 | 1.13021602 | -0.11731561 | 0.32132097 | -0.3651041 | 0.71503369 | NA | | NA | NA |
| ENSG000002 | 0.12685397 | -0.090215768 | 0.12901919 | -0.699243 | 0.48440017 | NA | | MIR4674 | 100616301 |
| ENSG000002 | 0.4889793 | -0.336999675 | 0.23406338 | -1.4397796 | 0.14992977 | NA | | MIR3923 | 100500877 |
| ENSG000002 | 0.83922859 | -0.494704744 | 0.27219032 | -1.8174957 | 0.06914124 | NA | | MIR4486 | 100616118 |
| ENSG000002 | 2.0140419 | -0.199824544 | 0.36418084 | -0.5486959 | 0.58321414 | NA | | MIR3684 | 100500867 |
| ENSG000002 | 0 | NA | NA | NA | NA | NA | | MIR4283-2 | 100422848 |
| ENSG000002 | 0.66293603 | -0.489755159 | 0.27215878 | -1.7995199 | 0.07193647 | NA | | MIR4269 | 100423043 |
| ENSG000002 | 1.94818935 | -0.184053229 | 0.36207273 | -0.5083322 | 0.61122038 | NA | | MIR548AI | 100616347 |
| ENSG000002 | 1.59956913 | 0.016661946 | 0.34264071 | 0.04862804 | 0.96121572 | NA | | MIR4699 | 100616133 |
| ENSG000002 | 0.48436321 | -0.017262119 | 0.23773787 | -0.0726099 | 0.94211656 | NA | | NA | NA |
| ENSG000002 | 0 | NA | NA | NA | NA | NA | | SNORD84 | 692199 |
| ENSG000002 | 1.7091907 | -0.181142908 | 0.34864379 | -0.5195644 | 0.60336721 | NA | | NA | NA |
| ENSG000002 | 1.57763631 | 0.154084346 | 0.35299939 | 0.43650032 | 0.66247377 | NA | | NA | NA |
| ENSG000002 | 0.60029448 | -0.251020599 | 0.2560415 | -0.9803903 | 0.3268935 | NA | | MIR4472-1 | 100616268 |
| ENSG000002 | 1.3516208 | 0.026665674 | 0.33573061 | 0.07942581 | 0.93669394 | NA | | MIR4288 | 100422903 |
| ENSG000002 | 0.62178293 | -0.443481895 | 0.26161272 | -1.6951848 | 0.09004038 | NA | | MIR3132 | 100423039 |
| ENSG000002 | 0.6265601 | -0.275936484 | 0.2670182 | -1.0333995 | 0.30141697 | NA | | MIR4446 | 100616476 |
| ENSG000002 | 0.11865494 | 0.083440866 | 0.12751478 | 0.65436232 | 0.5128784 | NA | | NA | NA |
| ENSG000002 | 0.36502313 | -0.093755175 | 0.2080862 | -0.4505593 | 0.6523072 | NA | | MIR4686 | 100616126 |
| ENSG000002 | 1.24728733 | -0.495702954 | 0.32963509 | -1.503793 | 0.13263467 | NA | | NA | NA |
| ENSG000002 | 1.07926664 | 0.191104423 | 0.32101775 | 0.59530796 | 0.55163764 | NA | | NA | NA |
| ENSG000002 | 2.06157731 | -0.237731403 | 0.36049837 | -0.6594521 | 0.5096055 | NA | | RN7SL693P | 106479477 |
| ENSG000002 | 1.08837915 | 0.036305514 | 0.32505052 | 0.11169191 | 0.91106769 | NA | | NA | NA |
| ENSG000002 | 1.95364527 | 0.058575579 | 0.36080707 | 0.16234598 | 0.8710334 | NA | | MIR3935 | 100500891 |
| ENSG000002 | 1.86485409 | -0.259580257 | 0.35685906 | -0.7274027 | 0.46697927 | NA | | NA | NA |

| | | | | | | | | |
|---|---|---|---|---|---|---|---|---|
| ENSG000002 | 1.85395019 | 0.791954318 | 0.34075524 | 2.3241149 | 0.02011934 | NA | MIR4710 | 100616300 |
| ENSG000002 | 0.36621735 | -0.097238239 | 0.21673053 | -0.4486596 | 0.65367721 | NA | NA | NA |
| ENSG000002 | 0.12070844 | -0.088503695 | 0.12783931 | -0.6923042 | 0.48874631 | NA | MIR548AD | 100616475 |
| ENSG000002 | 0 | NA | NA | NA | NA | NA | NA | NA |
| ENSG000002 | 0.38747904 | -0.104908247 | 0.21987514 | -0.4771265 | 0.63327207 | NA | MIR3650 | 100500824 |
| ENSG000002 | 0 | NA | NA | NA | NA | NA | MIR3195 | 100422838 |
| ENSG000002 | 0 | NA | NA | NA | NA | NA | RN7SL199P | 106480693 |
| ENSG000002 | 0.36391535 | 0.058449071 | 0.20749591 | 0.28168782 | 0.77818289 | NA | NA | NA |
| ENSG000002 | 1.37072292 | 0.019921817 | 0.33469396 | 0.05952249 | 0.95253596 | NA | RN7SL411P | 106481035 |
| ENSG000002 | 1.3182972 | 0.164160307 | 0.3328777 | 0.49315502 | 0.62190305 | NA | MIR548AB | 100616336 |
| ENSG000002 | 1.26871264 | -0.459454802 | 0.32185801 | -1.4275077 | 0.15343359 | NA | MIR3137 | 100422926 |
| ENSG000002 | 0.25993993 | -0.013144386 | 0.18211683 | -0.0721756 | 0.94246219 | NA | MIR4291 | 100422927 |
| ENSG000002 | 1.25664578 | -0.19612168 | 0.33335069 | -0.5883344 | 0.55630786 | NA | MIR3136 | 100422859 |
| ENSG000002 | 0.69549179 | 0.132602096 | 0.27562182 | 0.48110159 | 0.6304443 | NA | MIR4493 | 100616319 |
| ENSG000002 | 0.49475427 | -0.18461639 | 0.24044438 | -0.7678133 | 0.44259811 | NA | NA | NA |
| ENSG000002 | 0.50548109 | -0.025402968 | 0.24760631 | -0.1025942 | 0.91828506 | NA | MIR4476 | 100616456 |
| ENSG000002 | 0.76989467 | -0.051525063 | 0.15948306 | -0.3230755 | 0.74663809 | NA | MIR4675 | 100616383 |
| ENSG000002 | 0 | NA | NA | NA | NA | NA | MIR3180-2 | 100422956 |
| ENSG000002 | 0.48353618 | 0.145822039 | 0.2384055 | 0.61165553 | 0.54076569 | NA | NA | NA |
| ENSG000002 | 1.0493073 | -0.350693879 | 0.30857123 | -1.1365087 | 0.25574373 | NA | NA | NA |
| ENSG000002 | 1.13776217 | -0.288912756 | 0.32743308 | -0.8823566 | 0.37758398 | NA | RN7SL391P | 106481030 |
| ENSG000002 | 0.71820774 | 0.113838472 | 0.26822874 | 0.42440818 | 0.67126816 | NA | MIR4999 | 100847049 |
| ENSG000002 | 0.12685397 | -0.090215768 | 0.12901919 | -0.699243 | 0.48440017 | NA | MIR4252 | 100422975 |
| ENSG000002 | 0.97709485 | -0.169048404 | 0.30074587 | -0.5620972 | 0.57404981 | NA | MIR3944 | 100500911 |
| ENSG000002 | 1.48254506 | -0.055488763 | 0.34927378 | -0.158869 | 0.87377211 | NA | MIR3128 | 100422824 |
| ENSG000002 | 0.51111617 | -0.186068642 | 0.2423498 | -0.7677689 | 0.44262448 | NA | MIR4324 | 100422979 |
| ENSG000002 | 0.11131518 | 0.081951103 | 0.12663804 | 0.64712866 | 0.51754868 | NA | NA | NA |
| ENSG000002 | 0 | NA | NA | NA | NA | NA | RN7SL656P | 106481760 |
| ENSG000002 | 1.88961592 | -0.675814517 | 0.35840742 | -1.8856042 | 0.05934832 | NA | NA | NA |

| | | | | | | | | | |
|---|---|---|---|---|---|---|---|---|---|
| ENSG000002 | 0.11865494 | 0.083440866 | 0.12751478 | 0.65436232 | 0.5128784 | NA | | MIR4261 | 100422929 |
| ENSG000002 | 1.73123237 | -0.54069076 | 0.34385215 | -1.5724513 | 0.11584592 | NA | | MIR4779 | 100616159 |
| ENSG000002 | 1.40458484 | -0.261817894 | 0.33401032 | -0.7838617 | 0.43312127 | NA | | MIR4459 | 100616233 |
| ENSG000002 | 1.17752964 | -0.600477683 | 0.32052695 | -1.8734078 | 0.0610121 | NA | | MIR4684 | 100616391 |
| ENSG000002 | 0.9841214 | -0.188498655 | 0.31177486 | -0.6045986 | 0.54544572 | NA | | NA | NA |
| ENSG000002 | 0.75281538 | -0.182194416 | 0.28046106 | -0.6496246 | 0.51593472 | NA | | NA | NA |
| ENSG000002 | 0.70158525 | 0.137622095 | 0.28235954 | 0.4874002 | 0.62597475 | NA | | MIR4308 | 100422984 |
| ENSG000002 | 0.70955616 | -0.022372494 | 0.27641835 | -0.0809371 | 0.93549199 | NA | | MIR4447 | 100616485 |
| ENSG000002 | 1.67336352 | -0.043301844 | 0.348518 | -0.1242456 | 0.90112078 | NA | | MIR3121 | 100423032 |
| ENSG000002 | 0.50377641 | -0.188767022 | 0.24148558 | -0.7816906 | 0.4343964 | NA | | RN7SL811P | 106479517 |
| ENSG000002 | 1.38651434 | -0.121275404 | 0.33032076 | -0.3671444 | 0.71351134 | NA | | MIR3941 | 100500866 |
| ENSG000002 | 1.37064562 | -0.4097894 | 0.33673971 | -1.2169322 | 0.22363 | NA | | NA | NA |
| ENSG000002 | 0.71507054 | 0.137980815 | 0.28329186 | 0.4870624 | 0.62621411 | NA | | MIR4733 | 100616266 |
| ENSG000002 | 0.97843234 | -0.194458003 | 0.31303937 | -0.6211934 | 0.53447236 | NA | | NA | NA |
| ENSG000002 | 1.40179467 | -0.110978901 | 0.32899174 | -0.3373304 | 0.73586788 | NA | | NA | NA |
| ENSG000002 | 0.98392902 | 0.10536175 | 0.30036947 | 0.35077383 | 0.72575803 | NA | | MIR3672 | 100500869 |
| ENSG000002 | 1.72142577 | 0.063678911 | 0.34782767 | 0.18307603 | 0.85473836 | NA | | MIR3680-1 | 100500917 |
| ENSG000002 | 1.83888043 | 0.012318341 | 0.35820808 | 0.03438879 | 0.97256713 | NA | | MIR4768 | 100616249 |
| ENSG000002 | 1.22067424 | -0.037804302 | 0.32387529 | -0.1167249 | 0.90707808 | NA | | NA | NA |
| ENSG000002 | 0.998356 | -0.204556518 | 0.31380715 | -0.6518542 | 0.51449521 | NA | | NA | NA |
| ENSG000002 | 1.67190342 | 0.083867128 | 0.34974617 | 0.23979427 | 0.81048975 | NA | | DTX2P1-UPK3BI | 441263 |
| ENSG000002 | 1.65266435 | 0.085187911 | 0.3495056 | 0.24373833 | 0.80743348 | NA | | MIR4443 | 100616407 |
| ENSG000002 | 0.60968328 | -0.108157951 | 0.26370283 | -0.4101509 | 0.68169526 | NA | | RN7SL518P | 106481847 |
| ENSG000002 | 0.84681846 | 0.201203398 | 0.29674539 | 0.67803377 | 0.49775028 | NA | | NA | NA |
| ENSG000002 | 1.97752162 | -0.068608753 | 0.36238126 | -0.1893275 | 0.84983611 | NA | | NA | NA |
| ENSG000002 | 1.53303743 | -0.344538344 | 0.34552183 | -0.9971536 | 0.31868995 | NA | | MIR3156-2 | 100422907 |
| ENSG000002 | 0.76576985 | -0.381700686 | 0.28891137 | -1.3211688 | 0.18644509 | NA | | MIR1269B | 100616494 |
| ENSG000002 | 0.25214371 | -0.009862892 | 0.18046657 | -0.0546522 | 0.95641556 | NA | | MIR4435-1 | 100616499 |
| ENSG000002 | 0.7731096 | -0.379060994 | 0.28942548 | -1.3097015 | 0.19029682 | NA | | MIR4436A | 100616399 |

| | | | | | | | | |
|---|---|---|---|---|---|---|---|---|
| ENSG000002 | 0.64332695 | -0.264269033 | 0.26152646 | -1.0104868 | 0.31226213 | NA | NA | NA |
| ENSG000002 | 0.49879422 | -0.023609326 | 0.24688725 | -0.095628 | 0.92381605 | NA | MIR548V | 100422850 |
| ENSG000002 | 1.61968246 | -0.538030546 | 0.34809174 | -1.5456573 | 0.12218734 | NA | MIR5697 | 100847055 |
| ENSG000002 | 1.37066736 | -0.269657602 | 0.34157405 | -0.7894558 | 0.42984567 | NA | MIR4304 | 100422931 |
| ENSG000002 | 2.09738969 | -0.61630598 | 0.36192845 | -1.7028393 | 0.08859815 | NA | RN7SL475P | 106481845 |
| ENSG000002 | 0 | NA | NA | NA | NA | NA | MIR3180-1 | 100422870 |
| ENSG000002 | 0.5226185 | -0.189580046 | 0.24357686 | -0.7783172 | 0.43638207 | NA | MIR4421 | 100616189 |
| ENSG000002 | 0.59027007 | 0.063652523 | 0.26808709 | 0.23743226 | 0.81232147 | NA | MIR3164 | 100422846 |
| ENSG000002 | 1.41643655 | -0.577476665 | 0.33904326 | -1.7032536 | 0.08852061 | NA | NA | NA |
| ENSG000002 | 1.96669066 | -0.193857699 | 0.36274477 | -0.534419 | 0.59305167 | NA | NA | NA |
| ENSG000002 | 1.36340141 | -0.116537575 | 0.33962753 | -0.3431335 | 0.73149805 | NA | NA | NA |
| ENSG000002 | 1.11562083 | -0.122963593 | 0.31786702 | -0.3868397 | 0.69887486 | NA | NA | NA |
| ENSG000002 | 0.71327762 | 0.293628257 | 0.27857918 | 1.05402081 | 0.29187339 | NA | MIR3918 | 100500851 |
| ENSG000002 | 1.79500079 | 0.136482188 | 0.35448045 | 0.38502036 | 0.70022233 | NA | RN7SL652P | 106479460 |
| ENSG000002 | 1.42388519 | -0.285194709 | 0.23615705 | -1.2076485 | 0.22718248 | NA | NA | NA |
| ENSG000002 | 1.35335526 | 0.022242081 | 0.34045005 | 0.06533141 | 0.94791013 | NA | PIGPP4 | 106480811 |
| ENSG000002 | 0.75301317 | -0.023177929 | 0.2703647 | -0.0857284 | 0.93168234 | NA | NA | NA |
| ENSG000002 | 0.48646878 | -0.181653651 | 0.2395107 | -0.7584365 | 0.44818973 | NA | MIR3978 | 100616491 |
| ENSG000002 | 0.4833075 | 0.150471097 | 0.24570075 | 0.61241611 | 0.54026249 | NA | MIR5708 | 100847056 |
| ENSG000002 | 0 | NA | NA | NA | NA | NA | NA | NA |
| ENSG000002 | 1.492365 | -0.039098491 | 0.33666475 | -0.1161348 | 0.90754572 | NA | MIR4756 | 100616225 |
| ENSG000002 | 1.82970719 | -0.490916732 | 0.35277586 | -1.3915826 | 0.16404884 | NA | MIR3659 | 100500801 |
| ENSG000002 | 0.11131518 | 0.081951103 | 0.12663804 | 0.64712866 | 0.51754868 | NA | MIR4471 | 100616451 |
| ENSG000002 | 0 | NA | NA | NA | NA | NA | MIR4695 | 100616120 |
| ENSG000002 | 0 | NA | NA | NA | NA | NA | MIR4539 | 100616374 |
| ENSG000002 | 1.76714736 | -0.453009625 | 0.35176583 | -1.2878159 | 0.19781006 | NA | NA | NA |
| ENSG000002 | 0.11205239 | 0.082118251 | 0.12671962 | 0.64803108 | 0.51696485 | NA | MIR548AG2 | 100616440 |
| ENSG000002 | 2.01085476 | -0.059486146 | 0.35965551 | -0.1653976 | 0.86863107 | NA | NA | NA |
| ENSG000002 | 1.45280413 | 0.092642493 | 0.33830276 | 0.27384492 | 0.7842038 | NA | MIR3139 | 100423017 |

| | | | | | | | | |
|---|---|---|---|---|---|---|---|---|
| ENSG000002 | 0.24756242 | -0.187034346 | 0.18119817 | -1.0322088 | 0.30197431 | NA | NA | NA |
| ENSG000002 | 1.33597245 | -0.248183435 | 0.33412978 | -0.7427756 | 0.45761757 | NA | NA | NA |
| ENSG000002 | 0.23070733 | 0.167416683 | 0.17664244 | 0.94777156 | 0.34324576 | NA | NA | NA |
| ENSG000002 | 1.98862752 | -0.054961686 | 0.36112394 | -0.1521962 | 0.8790322 | NA | MIR3612 | 100500817 |
| ENSG000002 | 0.99918357 | -0.363379837 | 0.31493811 | -1.1538135 | 0.24857664 | NA | NA | NA |
| ENSG000002 | 0.59146429 | 0.065095808 | 0.26170762 | 0.24873486 | 0.80356588 | NA | MIR3929 | 100500864 |
| ENSG000002 | 1.56209986 | -0.114662668 | 0.34530066 | -0.3320662 | 0.7398393 | NA | NA | NA |
| ENSG000002 | 1.56770069 | 0.151443087 | 0.34875491 | 0.4342393 | 0.66411468 | NA | NA | NA |
| ENSG000002 | 1.76332447 | -0.455787284 | 0.35318687 | -1.2904989 | 0.19687748 | NA | RN7SL279P | 106480998 |
| ENSG000002 | 1.01429973 | -0.367175034 | 0.31550396 | -1.1637731 | 0.24451596 | NA | MIR548AK | 100616488 |
| ENSG000002 | 0.36621735 | -0.097238239 | 0.21673053 | -0.4486596 | 0.65367721 | NA | MIR3151 | 100422992 |
| ENSG000002 | 0 | NA | NA | NA | NA | NA | MIR3690 | 100500894 |
| ENSG000002 | 0.66527186 | -0.43035014 | 0.25829332 | -1.6661296 | 0.09568761 | NA | MIR548AJ1 | 100616191 |
| ENSG000002 | 0.5854929 | -0.091197421 | 0.25231745 | -0.3614392 | 0.71777113 | NA | MIR4508 | 100616275 |
| ENSG000002 | 0.71578601 | -0.030116478 | 0.2832364 | -0.1063298 | 0.91532066 | NA | RN7SL708P | 106479481 |
| ENSG000002 | 0.24485602 | -0.012655878 | 0.17881722 | -0.0707755 | 0.94357643 | NA | MIR4267 | 100422994 |
| ENSG000002 | 1.76608251 | -0.063940428 | 0.35139981 | -0.1819592 | 0.85561475 | NA | NA | NA |
| ENSG000002 | 0 | NA | NA | NA | NA | NA | NA | NA |
| ENSG000002 | 1.35779666 | -0.123013911 | 0.33951292 | -0.3623247 | 0.71710941 | NA | MIR4460 | 100616325 |
| ENSG000002 | 0 | NA | NA | NA | NA | NA | NA | NA |
| ENSG000002 | 1.54373864 | 0.151110757 | 0.33885352 | 0.44594713 | 0.65563544 | NA | MIR4774 | 100616356 |
| ENSG000002 | 0.49013872 | 0.148314611 | 0.23918465 | 0.62008415 | 0.53520239 | NA | MIR548AE2 | 100616339 |
| ENSG000002 | 1.30993237 | -0.326900111 | 0.32273854 | -1.0128946 | 0.31111052 | NA | MIR4681 | 100616398 |
| ENSG000002 | 0.25333738 | -0.015686291 | 0.1806605 | -0.0868275 | 0.93080867 | NA | MIR4284 | 100422948 |
| ENSG000002 | 0.47781273 | -0.018045496 | 0.22812977 | -0.0791019 | 0.93695158 | NA | MIR3144 | 100422951 |
| ENSG000002 | 0.39462238 | -0.104329077 | 0.21264929 | -0.4906157 | 0.6236983 | NA | MIR4438 | 100616375 |
| ENSG000002 | 1.14410412 | -0.258048109 | 0.31817605 | -0.811023 | 0.41735244 | NA | NA | NA |
| ENSG000002 | 1.58358831 | -0.256777039 | 0.34913789 | -0.7354602 | 0.46205924 | NA | NA | NA |
| ENSG000002 | 1.76827845 | -0.315202425 | 0.35204918 | -0.8953363 | 0.37060732 | NA | MIR4427 | 100616390 |

| | | | | | | | | |
|---|---|---|---|---|---|---|---|---|
| ENSG000002 | 1.69284944 | -0.057761137 | 0.34293657 | -0.168431 | 0.86624425 | NA | RN7SL375P | 106479359 |
| ENSG000002 | 1.37800657 | -0.279862056 | 0.34319938 | -0.8154503 | 0.41481458 | NA | NA | NA |
| ENSG000002 | 0.52165104 | -0.403921832 | 0.25252134 | -1.5995552 | 0.10969729 | NA | MIR4506 | 100616140 |
| ENSG000002 | 0 | NA | NA | NA | NA | NA | MIR3670-3 | 103504726 |
| ENSG000002 | 0.95125324 | -0.170419074 | 0.29951539 | -0.5689827 | 0.56936789 | NA | RN7SL624P | 106480379 |
| ENSG000002 | 0.90001914 | -0.118344353 | 0.29770615 | -0.3975207 | 0.69098353 | NA | NA | NA |
| ENSG000002 | 1.57837598 | 0.290678276 | 0.34688856 | 0.83795866 | 0.40205392 | NA | NA | NA |
| ENSG000002 | 0 | NA | NA | NA | NA | NA | MIR3118-4 | 100422935 |
| ENSG000002 | 0 | NA | NA | NA | NA | NA | NA | NA |
| ENSG000002 | 2.09066801 | -0.232769323 | 0.36063397 | -0.6454448 | 0.51863897 | NA | NA | NA |
| ENSG000002 | 0.75350052 | -0.19232973 | 0.28654979 | -0.6711913 | 0.50209866 | NA | MIR3907 | 100500835 |
| ENSG000002 | 0.5226185 | -0.189580046 | 0.24357686 | -0.7783172 | 0.43638207 | NA | RN7SL376P | 106481025 |
| ENSG000002 | 0 | NA | NA | NA | NA | NA | NA | NA |
| ENSG000002 | 0.36075696 | -0.085953824 | 0.20740622 | -0.4144226 | 0.67856464 | NA | MIR3177 | 100423012 |
| ENSG000002 | 0.11865494 | 0.083440866 | 0.12751478 | 0.65436232 | 0.5128784 | NA | MIR4422 | 100616272 |
| ENSG000002 | 1.15828665 | -0.271306926 | 0.3186143 | -0.8515215 | 0.39447973 | NA | MIR548AG1 | 100616450 |
| ENSG000002 | 0.77126304 | -0.326709471 | 0.27331554 | -1.1953564 | 0.23194782 | NA | MIR3123 | 100422856 |
| ENSG000002 | 1.69687033 | 0.072565224 | 0.35174324 | 0.20630169 | 0.83655526 | NA | NA | NA |
| ENSG000002 | 1.88474796 | -0.116139055 | 0.3466521 | -0.3350306 | 0.73760202 | NA | CCND3P2 | 106480307 |
| ENSG000002 | 0.71809205 | 0.130302615 | 0.26820504 | 0.48583209 | 0.62708622 | NA | NA | NA |
| ENSG000002 | 0.23065526 | 0.167384503 | 0.17662711 | 0.94767164 | 0.34329664 | NA | MIR548X | 100422920 |
| ENSG000002 | 1.82223174 | 0.136892197 | 0.35917276 | 0.3811319 | 0.70310538 | NA | MIR4287 | 100422828 |
| ENSG000002 | 0 | NA | NA | NA | NA | NA | MIR3689D1 | 100616131 |
| ENSG000002 | 1.97252161 | -0.324501264 | 0.36347073 | -0.8927851 | 0.37197224 | NA | MIR4797 | 100616216 |
| ENSG000002 | 0.63270563 | -0.25685676 | 0.259029 | -0.9916139 | 0.3213859 | NA | NA | NA |
| ENSG000002 | 0.71225808 | 0.123930976 | 0.27522079 | 0.45029657 | 0.65249662 | NA | MIR5704 | 100847040 |
| ENSG000002 | 0.63159785 | -0.116736614 | 0.27184111 | -0.4294296 | 0.66761063 | NA | MIR4495 | 100616287 |
| ENSG000002 | 1.02504637 | -0.179071629 | 0.30288867 | -0.5912127 | 0.55437791 | NA | MIR4516 | 100616258 |
| ENSG000002 | 0 | NA | NA | NA | NA | NA | MIR3689A | 100500846 |

| | | | | | | | | |
|---|---|---|---|---|---|---|---|---|
| ENSG000002 | 0.74891476 | -0.03642447 | 0.28541666 | -0.1276186 | 0.89845082 | NA | MIR4489 | 100616284 |
| ENSG000002 | 1.14001155 | -0.441561817 | 0.32361681 | -1.3644588 | 0.1724232 | NA | NA | NA |
| ENSG000002 | 0.96075359 | -0.031985823 | 0.29649332 | -0.1078804 | 0.91409055 | NA | MIR4748 | 100616425 |
| ENSG000002 | 1.60132207 | -0.126335455 | 0.35008397 | -0.3608719 | 0.71819523 | NA | RN7SL73P | 106479250 |
| ENSG000002 | 0.11131518 | 0.081951103 | 0.12663804 | 0.64712866 | 0.51754868 | NA | NA | NA |
| ENSG000002 | 0 | NA | NA | NA | NA | NA | NA | NA |
| ENSG000002 | 1.22581614 | 0.103037727 | 0.33337321 | 0.3090762 | 0.75726356 | NA | RN7SL311P | 106481755 |
| ENSG000002 | 0.11205239 | 0.082118251 | 0.12671962 | 0.64803108 | 0.51696485 | NA | NA | NA |
| ENSG000002 | 2.04703256 | -0.87001969 | 0.35937933 | -2.4208952 | 0.01548234 | NA | NA | NA |
| ENSG000002 | 1.21403945 | -0.038037732 | 0.32808634 | -0.1159382 | 0.90770154 | NA | NA | NA |
| ENSG000002 | 0.14128499 | -0.097772014 | 0.13403235 | -0.7294658 | 0.4657168 | NA | MIR5692C1 | 100847082 |
| ENSG000002 | 0.3867939 | -0.105148852 | 0.21977268 | -0.4784437 | 0.63233444 | NA | MIR5195 | 100847062 |
| ENSG000002 | 1.73352701 | -0.063075644 | 0.35508672 | -0.1776345 | 0.85901004 | NA | MIR3146 | 100422967 |
| ENSG000002 | 1.84078959 | -0.124586934 | 0.3585237 | -0.3474999 | 0.72821582 | NA | NA | NA |
| ENSG000002 | 1.25548189 | -0.343625155 | 0.32842654 | -1.0462771 | 0.29543314 | NA | NA | NA |
| ENSG000002 | 1.89623791 | 0.311212776 | 0.35650311 | 0.87295951 | 0.38268515 | NA | NA | NA |
| ENSG000002 | 0.51931521 | -0.354214731 | 0.23897171 | -1.4822455 | 0.13827499 | NA | NA | NA |
| ENSG000002 | 1.16224125 | -0.258097503 | 0.31709387 | -0.8139467 | 0.41567549 | NA | NA | NA |
| ENSG000002 | 0 | NA | NA | NA | NA | NA | MIR4749 | 100616313 |
| ENSG000002 | 0.36502313 | -0.093755175 | 0.2080862 | -0.4505593 | 0.6523072 | NA | MIR4445 | 100616129 |
| ENSG000002 | 0.92122527 | -0.443181411 | 0.29845654 | -1.484911 | 0.13756739 | NA | MIR4319 | 100422829 |
| ENSG000002 | 0.36827086 | -0.266056975 | 0.21181163 | -1.2561018 | 0.20907905 | NA | NA | NA |
| ENSG000002 | 0.51550551 | -0.37860525 | 0.24578489 | -1.5403927 | 0.12346466 | NA | MIR4266 | 100423027 |
| ENSG000002 | 1.06623836 | 0.194978799 | 0.32054838 | 0.60826638 | 0.54301082 | NA | NA | NA |
| ENSG000002 | 0.38747904 | -0.104908247 | 0.21987514 | -0.4771265 | 0.63327207 | NA | MIR4725 | 100616449 |
| ENSG000002 | 0 | NA | NA | NA | NA | NA | NA | NA |
| ENSG000002 | 2.10384979 | -0.500208998 | 0.36405292 | -1.3740008 | 0.16944144 | NA | NA | NA |
| ENSG000002 | 0.4869258 | -0.176879301 | 0.23959723 | -0.738236 | 0.46037105 | NA | MIR4519 | 100616231 |
| ENSG000002 | 0.9868278 | -0.31141388 | 0.30042272 | -1.0365856 | 0.29992901 | NA | MIR5694 | 100847064 |

| | | | | | | | | |
|---|---|---|---|---|---|---|---|---|
| ENSG000002 | 2.02315677 | -0.729588439 | 0.36111451 | -2.0203797 | 0.04334401 | NA | NA | NA |
| ENSG000002 | 1.45503178 | -0.177156426 | 0.33890298 | -0.5227349 | 0.60115872 | NA | NA | NA |
| ENSG000002 | 1.74104325 | -0.599246202 | 0.35195429 | -1.7026251 | 0.08863826 | NA | NA | NA |
| ENSG000002 | 0.25370795 | -0.171331747 | 0.17411721 | -0.9840024 | 0.32511435 | NA | MIR4488 | 100616470 |
| ENSG000002 | 0 | NA | NA | NA | NA | NA | NA | NA |
| ENSG000002 | 0.94567181 | 0.111414102 | 0.29852007 | 0.37322148 | 0.70898361 | NA | MIR4764 | 100616295 |
| ENSG000002 | 0.61927241 | -0.278564577 | 0.26634985 | -1.0458597 | 0.29562581 | NA | LOC105371766 | 105371766 |
| ENSG000002 | 0.38019136 | -0.10756138 | 0.21876967 | -0.491665 | 0.62295621 | NA | MIR4309 | 100422954 |
| ENSG000002 | 1.11134148 | -0.042539648 | 0.20210323 | -0.2104848 | 0.83328935 | NA | NA | NA |
| ENSG000002 | 0.73058759 | -0.195089346 | 0.28504587 | -0.6844139 | 0.49371387 | NA | NA | NA |
| ENSG000002 | 1.12159669 | -0.266366458 | 0.32186057 | -0.8275834 | 0.40790649 | NA | NA | NA |
| ENSG000002 | 0 | NA | NA | NA | NA | NA | MIR4509-2 | 100616228 |
| ENSG000002 | 0 | NA | NA | NA | NA | NA | MIR4690 | 100616292 |
| ENSG000002 | 0.87321062 | -0.269491476 | 0.29754442 | -0.9057185 | 0.36508489 | NA | NA | NA |
| ENSG000002 | 0.84850532 | -0.109408574 | 0.2966738 | -0.3687841 | 0.71228867 | NA | MIR3649 | 100500816 |
| ENSG000002 | 0.64759311 | -0.271681639 | 0.26184098 | -1.0375826 | 0.29946444 | NA | NA | NA |
| ENSG000002 | 0.72353016 | -0.033011912 | 0.2837491 | -0.1163419 | 0.90738157 | NA | MIR4500 | 100616182 |
| ENSG000002 | 1.29572811 | -0.513295808 | 0.33294832 | -1.5416681 | 0.12315426 | NA | RN7SL140P | 106479272 |
| ENSG000002 | 1.29782671 | 0.323785094 | 0.33509513 | 0.96624827 | 0.33391996 | NA | MIR4771-2 | 100616165 |
| ENSG000002 | 0.51046328 | -0.173690131 | 0.23251175 | -0.7470166 | 0.45505355 | NA | MIR3617 | 100500897 |
| ENSG000002 | 1.62189916 | -0.124876346 | 0.35150325 | -0.3552637 | 0.72239205 | NA | MIR5693 | 100847003 |
| ENSG000002 | 0.73155506 | -0.031636845 | 0.28965816 | -0.1092213 | 0.91302696 | NA | RN7SL574P | 106481079 |
| ENSG000002 | 0.4794119 | -0.010064858 | 0.23715327 | -0.0424403 | 0.9661477 | NA | NA | NA |
| ENSG000002 | 1.65795894 | -0.527595581 | 0.34671329 | -1.5217057 | 0.12808285 | NA | MIR4432 | 100616473 |
| ENSG000002 | 0 | NA | NA | NA | NA | NA | NA | NA |
| ENSG000002 | 0.26062507 | -0.012888319 | 0.18226503 | -0.070712 | 0.94362699 | NA | MIR5700 | 100847031 |
| ENSG000002 | 0.13280363 | -0.093394593 | 0.13115644 | -0.7120855 | 0.47641185 | NA | MIR4326 | 100422945 |
| ENSG000002 | 0.62775378 | -0.28146231 | 0.26704095 | -1.0540043 | 0.29188095 | NA | NA | NA |
| ENSG000002 | 1.48770139 | -0.334435117 | 0.34248567 | -0.9764937 | 0.32881985 | NA | NA | NA |

| | | | | | | | | |
|---|---|---|---|---|---|---|---|---|
| ENSG000002 | 0.64739724 | -0.288756313 | 0.26860627 | -1.075017 | 0.28236711 | NA | MIR4525 | 100616196 |
| ENSG000002 | 0.13280363 | -0.093394593 | 0.13115644 | -0.7120855 | 0.47641185 | NA | MIR4440 | 100616397 |
| ENSG000002 | 0.36556447 | -0.101943371 | 0.21659044 | -0.4706735 | 0.63787395 | NA | MIR4423 | 100616481 |
| ENSG000002 | 1.97875359 | -0.199475394 | 0.36521843 | -0.5461811 | 0.58494148 | NA | NA | NA |
| ENSG000002 | 1.35881777 | 0.160003831 | 0.33382121 | 0.47930995 | 0.63171814 | NA | MIR5587 | 100847028 |
| ENSG000002 | 1.68575872 | -0.049079898 | 0.35278112 | -0.1391228 | 0.88935311 | NA | RN7SL366P | 106481023 |
| ENSG000002 | 1.23473917 | -0.160574055 | 0.31518934 | -0.5094527 | 0.61043496 | NA | NA | NA |
| ENSG000002 | 0.37404583 | -0.105111695 | 0.21784993 | -0.4824959 | 0.62945372 | NA | MIR4759 | 100616243 |
| ENSG000002 | 0.69298127 | 0.291824635 | 0.27540541 | 1.05961838 | 0.28931825 | NA | MIR4435-2 | 100616341 |
| ENSG000002 | 0.23936338 | -0.005479057 | 0.17759977 | -0.0308506 | 0.9753887 | NA | MIR4533 | 100616362 |
| ENSG000002 | 1.68447626 | 0.21031271 | 0.35394682 | 0.59419296 | 0.55238306 | NA | MIR4654 | 100616386 |
| ENSG000002 | 0.96198006 | -0.036154158 | 0.31177589 | -0.115962 | 0.90768265 | NA | MIR5190 | 100847080 |
| ENSG000002 | 0.65445467 | -0.486865861 | 0.27154464 | -1.7929496 | 0.07298098 | NA | NA | NA |
| ENSG000002 | 1.42001949 | -0.402994189 | 0.334136 | -1.2060783 | 0.22778729 | NA | MIR4711 | 100616409 |
| ENSG000002 | 0.95213246 | 0.27951741 | 0.30715626 | 0.91001698 | 0.36281355 | NA | MIR3646 | 100500813 |
| ENSG000002 | 1.61687001 | -0.531852553 | 0.34606381 | -1.5368627 | 0.12432694 | NA | NA | NA |
| ENSG000002 | 0.23816916 | -0.010712105 | 0.1772998 | -0.060418 | 0.9518227 | NA | RN7SL785P | 106480532 |
| ENSG000002 | 1.31291312 | -0.317409603 | 0.31906841 | -0.9948011 | 0.31983302 | NA | RN7SL612P | 106479445 |
| ENSG000002 | 0.69947967 | 0.276464724 | 0.26894493 | 1.02796035 | 0.30396848 | NA | YWHAEP2 | 106481688 |
| ENSG000002 | 0.89495965 | -0.460377077 | 0.30201036 | -1.5243751 | 0.12741503 | NA | RN7SL557P | 106479425 |
| ENSG000002 | 0 | NA | NA | NA | NA | NA | MIR3147 | 100422939 |
| ENSG000002 | 0.49763088 | -0.186408494 | 0.24078583 | -0.7741672 | 0.43883193 | NA | MIR4666A | 100616308 |
| ENSG000002 | 0.89273201 | -0.112833687 | 0.29897169 | -0.3774059 | 0.70587197 | NA | NA | NA |
| ENSG000002 | 1.5930892 | -0.115238794 | 0.34929276 | -0.3299204 | 0.74146014 | NA | RN7SL804P | 106481140 |
| ENSG000002 | 1.91383718 | -0.820456981 | 0.35682573 | -2.2993213 | 0.0214867 | NA | RN7SL480P | 106481757 |
| ENSG000002 | 0.52398687 | -0.355918066 | 0.23944695 | -1.4864172 | 0.1371688 | NA | MIR5095 | 100616458 |
| ENSG000002 | 0.26813897 | -0.197007041 | 0.18550319 | -1.0620143 | 0.2882292 | NA | MIR3186 | 100422944 |
| ENSG000002 | 0.99323392 | -0.342530258 | 0.30870366 | -1.1095763 | 0.26718166 | NA | NA | NA |
| ENSG000002 | 0.49902098 | -0.194741582 | 0.24819886 | -0.7846192 | 0.43267688 | NA | MIR1260B | 100422991 |

| | | | | | | | | | |
|---|---|---|---|---|---|---|---|---|---|
| ENSG000002 | 0.11205239 | 0.082118251 | 0.12671962 | 0.64803108 | 0.51696485 | NA | | MIR4661 | 100616245 |
| ENSG000002 | 1.62776292 | -0.251079657 | 0.34685626 | -0.7238724 | 0.4691441 | NA | | RN7SL163P | 106479280 |
| ENSG000002 | 1.94064065 | 0.300196268 | 0.3542331 | 0.84745402 | 0.39674211 | NA | | NA | NA |
| ENSG000002 | 0.59414895 | -0.083234893 | 0.15710218 | -0.5298137 | 0.59624109 | NA | | NA | NA |
| ENSG000002 | 0.11131518 | 0.081951103 | 0.12663804 | 0.64712866 | 0.51754868 | NA | | MIR5589 | 100847093 |
| ENSG000002 | 0.76884289 | -0.364133569 | 0.2836951 | -1.2835384 | 0.1993035 | NA | | MIR3926-1 | 100500870 |
| ENSG000002 | 0.75048147 | -0.022471992 | 0.27018513 | -0.0831726 | 0.93371432 | NA | | RN7SL371P | 106481839 |
| ENSG000002 | 0.37097279 | -0.10468565 | 0.21741418 | -0.4815033 | 0.63015882 | NA | | MIR3156-3 | 100423018 |
| ENSG000002 | 0.39499294 | -0.277908454 | 0.21584201 | -1.287555 | 0.19790091 | NA | | MIR3167 | 100422918 |
| ENSG000002 | 1.02181363 | -0.537060267 | 0.31330976 | -1.7141511 | 0.08650098 | NA | | MIR4323 | 100422980 |
| ENSG000002 | 1.36152204 | -0.118489928 | 0.33755921 | -0.3510197 | 0.72557358 | NA | | MIR3611 | 100500890 |
| ENSG000002 | 0.11131518 | 0.081951103 | 0.12663804 | 0.64712866 | 0.51754868 | NA | | MIR3178 | 100422974 |
| ENSG000002 | 2.16511408 | 0.182428624 | 0.36742458 | 0.49650631 | 0.61953724 | NA | | MIR5091 | 100847023 |
| ENSG000002 | 0.73374653 | -0.030517642 | 0.27811547 | -0.1097301 | 0.91262342 | NA | | NA | NA |
| ENSG000002 | 0.3867939 | -0.105148852 | 0.21977268 | -0.4784437 | 0.63233444 | NA | | MIR4279 | 100422874 |
| ENSG000002 | 1.57602042 | -0.257047148 | 0.34954738 | -0.7353714 | 0.46211333 | NA | | MIR4644 | 100616430 |
| ENSG000002 | 1.45508577 | -0.047933258 | 0.34776379 | -0.1378328 | 0.89037256 | NA | | NA | NA |
| ENSG000002 | 0.87073447 | 0.03701048 | 0.29150635 | 0.12696286 | 0.89896979 | NA | | MIR4475 | 100616289 |
| ENSG000002 | 1.13235385 | -0.267315938 | 0.31939562 | -0.836943 | 0.40262459 | NA | | NA | NA |
| ENSG000002 | 0.84223963 | 0.21011263 | 0.29483721 | 0.71263947 | 0.47606888 | NA | | MIR5096 | 100616427 |
| ENSG000002 | 0.73975464 | -0.177073248 | 0.27958599 | -0.6333409 | 0.52651104 | NA | | NA | NA |
| ENSG000002 | 0 | NA | NA | NA | NA | NA | | NA | NA |
| ENSG000002 | 0.50479595 | -0.02564641 | 0.24753216 | -0.1036084 | 0.91748012 | NA | | NA | NA |
| ENSG000002 | 1.60806155 | -0.12088724 | 0.35132345 | -0.344091 | 0.73077784 | NA | | NA | NA |
| ENSG000002 | 0.61175852 | -0.109548347 | 0.27027148 | -0.4053271 | 0.68523711 | NA | | MIR3683 | 100500886 |
| ENSG000002 | 0 | NA | NA | NA | NA | NA | | MIR3180-3 | 100422836 |
| ENSG000002 | 0.86367355 | -0.12320517 | 0.30690073 | -0.4014496 | 0.68808915 | NA | | MIR744 | 100126313 |
| ENSG000002 | 0 | NA | NA | NA | NA | NA | | MIR3118-1 | 100423008 |
| ENSG000002 | 1.99389094 | -0.072043691 | 0.36116791 | -0.1994742 | 0.8418918 | NA | | NA | NA |

| | | | | | | | | |
|---|---|---|---|---|---|---|---|---|
| ENSG000002 | 0.70616654 | 0.296520925 | 0.27799783 | 1.06663036 | 0.28613878 | NA | MIR5093 | 100847022 |
| ENSG000002 | 1.83239996 | -0.122097221 | 0.35797192 | -0.3410804 | 0.73304302 | NA | RN7SL510P | 106479410 |
| ENSG000002 | 0.47092999 | -0.01913341 | 0.24378131 | -0.078486 | 0.9374415 | NA | NA | NA |
| ENSG000002 | 0.72881641 | 0.133210785 | 0.28422864 | 0.46867475 | 0.63930214 | NA | MIR4668 | 100616114 |
| ENSG000002 | 1.75342076 | -0.475443632 | 0.3557527 | -1.3364442 | 0.18140413 | NA | RN7SL703P | 106479479 |
| ENSG000002 | 0.87118799 | -0.254593165 | 0.29334962 | -0.8678831 | 0.38545835 | NA | MIR5692A1 | 100847066 |
| ENSG000002 | 0 | NA | NA | NA | NA | NA | MIR3909 | 100500826 |
| ENSG000002 | 1.09094174 | -0.10994409 | 0.32002774 | -0.3435455 | 0.73118812 | NA | MIR4293 | 100422843 |
| ENSG000002 | 0.3641961 | 0.070436377 | 0.21601083 | 0.32607799 | 0.74436535 | NA | MIR4663 | 100616260 |
| ENSG000002 | 0 | NA | NA | NA | NA | NA | MIR3665 | 100500861 |
| ENSG000002 | 0.97026555 | -0.038461989 | 0.31211508 | -0.1232302 | 0.90192485 | NA | MIR4503 | 100616280 |
| ENSG000002 | 0 | NA | NA | NA | NA | NA | NA | NA |
| ENSG000002 | 0.54222759 | -0.38902668 | 0.24851205 | -1.5654238 | 0.11748356 | NA | MIR4281 | 100422962 |
| ENSG000002 | 0.58475569 | -0.031541521 | 0.15708832 | -0.2007885 | 0.84086399 | NA | NA | NA |
| ENSG000002 | 0 | NA | NA | NA | NA | NA | NA | NA |
| ENSG000002 | 1.86235251 | -0.612823687 | 0.35002846 | -1.7507824 | 0.07998339 | NA | MIR5002 | 100847021 |
| ENSG000002 | 1.90304118 | -0.639682492 | 0.35327854 | -1.8107029 | 0.07018685 | NA | NA | NA |
| ENSG000002 | 0.8942098 | -0.385640757 | 0.2837493 | -1.3590897 | 0.17411815 | NA | MIR4740 | 100616294 |
| ENSG000002 | 1.11792283 | -0.273754723 | 0.32320706 | -0.8469949 | 0.39699798 | NA | MIR4273 | 100422955 |
| ENSG000002 | 1.87307541 | -0.257699923 | 0.35755442 | -0.7207292 | 0.47107612 | NA | MIR3157 | 100422892 |
| ENSG000002 | 1.74878794 | -0.605481924 | 0.3540056 | -1.7103739 | 0.08719676 | NA | NA | NA |
| ENSG000002 | 0.36553413 | 0.069401489 | 0.20780452 | 0.33397488 | 0.73839851 | NA | MIR4636 | 100616326 |
| ENSG000002 | 0.5964156 | 0.058115253 | 0.26216088 | 0.22167783 | 0.82456469 | NA | NA | NA |
| ENSG000002 | 1.12809105 | -0.591823701 | 0.31973711 | -1.8509697 | 0.06417391 | NA | MIR4424 | 100616328 |
| ENSG000002 | 0.23816916 | -0.010712105 | 0.1772998 | -0.060418 | 0.9518227 | NA | NA | NA |
| ENSG000002 | 0.14128499 | -0.097772014 | 0.13403235 | -0.7294658 | 0.4657168 | NA | MIR4719 | 100616172 |
| ENSG000002 | 1.14234738 | -0.426269247 | 0.31963785 | -1.3336007 | 0.18233477 | NA | MIR5580 | 100847076 |
| ENSG000002 | 0.74416125 | 0.125178122 | 0.27745532 | 0.45116497 | 0.65187066 | NA | MIR4264 | 100422888 |
| ENSG000002 | 1.38418298 | -0.414109791 | 0.33840632 | -1.2237059 | 0.22106321 | NA | MIR4746 | 100616371 |

| | | | | | | | | |
|---|---|---|---|---|---|---|---|---|
| ENSG000002 | 1.25986263 | -0.189963562 | 0.32983839 | -0.5759292 | 0.56466303 | NA | RN7SL493P | 106479403 |
| ENSG000002 | 0.86455511 | -0.103797748 | 0.29753822 | -0.3488552 | 0.72719804 | NA | NA | NA |
| ENSG000002 | 0.62311649 | -0.107704099 | 0.26487294 | -0.4066255 | 0.68428304 | NA | NA | NA |
| ENSG000002 | 0.48646878 | -0.181653651 | 0.2395107 | -0.7584365 | 0.44818973 | NA | MIR3713 | 100500855 |
| ENSG000002 | 0 | NA | NA | NA | NA | NA | MIR3179-3 | 100423006 |
| ENSG000002 | 1.58854174 | -0.123335616 | 0.34991217 | -0.3524759 | 0.72448137 | NA | NA | NA |
| ENSG000002 | 1.21910809 | -0.038094184 | 0.32383109 | -0.117636 | 0.9063561 | NA | MIR2392 | 100616495 |
| ENSG000002 | 0 | NA | NA | NA | NA | NA | MIR3196 | 100423014 |
| ENSG000002 | 1.88084935 | -0.384831878 | 0.35474904 | -1.0848003 | 0.2780101 | NA | NA | NA |
| ENSG000002 | 1.43629708 | -0.387793915 | 0.32917866 | -1.1780652 | 0.23877063 | NA | RN7SL616P | 106481090 |
| ENSG000002 | 0.51396245 | -0.026802372 | 0.24109708 | -0.1111684 | 0.91148283 | NA | MIR5197 | 100846991 |
| ENSG000002 | 2.10643314 | -0.005750909 | 0.36472914 | -0.0157676 | 0.98741978 | NA | NA | NA |
| ENSG000002 | 0.12685397 | -0.090215768 | 0.12901919 | -0.699243 | 0.48440017 | NA | MIR4641 | 100616178 |
| ENSG000002 | 0.88396832 | -0.267921803 | 0.29812377 | -0.8986932 | 0.36881611 | NA | NA | NA |
| ENSG000002 | 0.40327843 | -0.281391716 | 0.21700406 | -1.2967118 | 0.19473038 | NA | MIR5681A | 100847058 |
| ENSG000002 | 0.35596481 | 0.219647334 | 0.19879223 | 1.10490904 | 0.26919901 | NA | NA | NA |
| ENSG000002 | 0.24004852 | -0.005223787 | 0.17775844 | -0.029387 | 0.97655594 | NA | MIR4479 | 100616480 |
| ENSG000002 | 1.00894293 | -0.325898326 | 0.30225001 | -1.0782409 | 0.28092626 | NA | MIR3201 | 100422916 |
| ENSG000002 | 0.51433302 | -0.186673425 | 0.24269587 | -0.7691661 | 0.44179474 | NA | MIR3934 | 100500873 |
| ENSG000002 | 0.11131518 | 0.081951103 | 0.12663804 | 0.64712866 | 0.51754868 | NA | MIR3689F | 100616212 |
| ENSG000002 | 1.47842325 | -0.034701816 | 0.33608332 | -0.1032536 | 0.91776169 | NA | MIR4452 | 100616463 |
| ENSG000002 | 0.67739041 | 0.460061681 | 0.27047584 | 1.7009345 | 0.08895529 | NA | MIR4715 | 100616474 |
| ENSG000002 | 0.86660862 | -0.252242393 | 0.2931025 | -0.8605945 | 0.38946143 | NA | MIR4268 | 100422959 |
| ENSG000002 | 1.23279977 | -0.188633248 | 0.32683417 | -0.5771528 | 0.56383626 | NA | NA | NA |
| ENSG000002 | 1.93674067 | -0.059777648 | 0.36228835 | -0.1650002 | 0.86894383 | NA | NA | NA |
| ENSG000002 | 1.40481564 | -0.113823378 | 0.33157149 | -0.3432846 | 0.73138438 | NA | MIR4650-1 | 100616310 |
| ENSG000002 | 1.89006689 | -0.512389901 | 0.35591373 | -1.4396464 | 0.14996748 | NA | NA | NA |
| ENSG000002 | 0.71037769 | -0.023915726 | 0.26721617 | -0.0894995 | 0.92868492 | NA | MIR4318 | 100422857 |
| ENSG000002 | 0.71543718 | 0.290127831 | 0.27713806 | 1.04687112 | 0.29515902 | NA | MIR4706 | 100616490 |

| | | | | | | | | | |
|---|---|---|---|---|---|---|---|---|---|
| ENSG000002 | 0.85363187 | -0.268235387 | 0.2964963 | -0.9046837 | 0.36563295 | NA | | MIR3619 | 100500828 |
| ENSG000002 | 1.62198604 | -0.528602896 | 0.34700038 | -1.5233496 | 0.12767125 | NA | | NA | NA |
| ENSG000002 | 0 | NA | NA | NA | NA | NA | | MIR5701-1 | 100847060 |
| ENSG000002 | 1.48735703 | -0.325014894 | 0.34250291 | -0.9489405 | 0.34265086 | NA | | NA | NA |
| ENSG000002 | 1.39486165 | -0.572959208 | 0.33786735 | -1.6958111 | 0.08992165 | NA | | RN7SL356P | 106479353 |
| ENSG000002 | 0.11131518 | 0.081951103 | 0.12663804 | 0.64712866 | 0.51754868 | NA | | MIR4530 | 100616163 |
| ENSG000002 | 1.34786453 | 0.161101144 | 0.33080005 | 0.48700459 | 0.62625508 | NA | | NA | NA |
| ENSG000002 | 1.52105047 | 0.441999362 | 0.34612555 | 1.27699145 | 0.20160527 | NA | | MIR5579 | 100847000 |
| ENSG000002 | 0.75825595 | -0.198978971 | 0.2868202 | -0.6937411 | 0.48784457 | NA | | NA | NA |
| ENSG000002 | 0.4680172 | 0.307231342 | 0.22996067 | 1.33601689 | 0.18154375 | NA | | MIR3166 | 100423040 |
| ENSG000002 | 0.86418455 | 0.048878028 | 0.30226458 | 0.16170611 | 0.8715373 | NA | | MIR4527 | 100616264 |
| ENSG000002 | 1.02391528 | -0.343605875 | 0.30757022 | -1.1171624 | 0.26392491 | NA | | MIR4478 | 100616312 |
| ENSG000002 | 0.72967624 | -0.02677813 | 0.28955569 | -0.0924801 | 0.92631662 | NA | | NA | NA |
| ENSG000002 | 1.32688516 | 0.326923281 | 0.33697764 | 0.97016314 | 0.33196518 | NA | | MIR4512 | 100616149 |
| ENSG000002 | 0.12070844 | -0.088503695 | 0.12783931 | -0.6923042 | 0.48874631 | NA | | NA | NA |
| ENSG000002 | 0 | NA | NA | NA | NA | NA | | RN7SL176P | 106480968 |
| ENSG000002 | 0.37899768 | -0.10176489 | 0.21865765 | -0.4654074 | 0.64163973 | NA | | MIR4703 | 100616423 |
| ENSG000002 | 0.3867939 | -0.105148852 | 0.21977268 | -0.4784437 | 0.63233444 | NA | | NA | NA |
| ENSG000002 | 0.37267746 | 0.067870636 | 0.21729405 | 0.31234465 | 0.75477861 | NA | | MIR4742 | 100616468 |
| ENSG000002 | 1.86681118 | -0.387373758 | 0.35811266 | -1.0817092 | 0.27938177 | NA | | MIR4642 | 100616352 |
| ENSG000002 | 2.11577029 | -0.383981611 | 0.36568825 | -1.0500245 | 0.29370686 | NA | | RN7SL202P | 106479295 |
| ENSG000002 | 0 | NA | NA | NA | NA | NA | | MIR3972 | 100616188 |
| ENSG000002 | 0.23890637 | -0.010420674 | 0.17747379 | -0.0587167 | 0.95317776 | NA | | MIR4754 | 100616168 |
| ENSG000002 | 0.35273207 | -0.08884607 | 0.20605566 | -0.4311751 | 0.66634105 | NA | | MIR3677 | 100500812 |
| ENSG000002 | 1.08274462 | 0.192373794 | 0.31962318 | 0.60187686 | 0.54725611 | NA | | NA | NA |
| ENSG000002 | 0.85473517 | -0.109563843 | 0.29095278 | -0.3765692 | 0.70649381 | NA | | MIR4299 | 100423026 |
| ENSG000002 | 1.1434295 | -0.434028495 | 0.32156178 | -1.3497515 | 0.1770957 | NA | | NA | NA |
| ENSG000002 | 0.62618953 | -0.114303343 | 0.27141404 | -0.4211401 | 0.67365276 | NA | | MIR3126 | 100423030 |
| ENSG000002 | 1.08658466 | 0.037148866 | 0.32499352 | 0.11430648 | 0.90899484 | NA | | NA | NA |

| | | | | | | | | |
|---|---|---|---|---|---|---|---|---|
| ENSG000002 | 0.63759958 | -0.112459832 | 0.26607281 | -0.4226656 | 0.67253924 | NA | NA | NA |
| ENSG000002 | 0.99007263 | -0.039612669 | 0.30806217 | -0.1285866 | 0.89768477 | NA | MIR3908 | 100500909 |
| ENSG000002 | 1.72874959 | -0.652461534 | 0.34125332 | -1.9119566 | 0.05588176 | NA | MIR3169 | 100422973 |
| ENSG000002 | 0.12070844 | -0.088503695 | 0.12783931 | -0.6923042 | 0.48874631 | NA | NA | NA |
| ENSG000002 | 0.11131518 | 0.081951103 | 0.12663804 | 0.64712866 | 0.51754868 | NA | MIR4739 | 100616170 |
| ENSG000002 | 0.7289911 | -0.025513702 | 0.28412774 | -0.0897966 | 0.92844886 | NA | MIR5692C2 | 100847017 |
| ENSG000002 | 0.48510042 | -0.016983889 | 0.23782629 | -0.071413 | 0.94306906 | NA | MIR4297 | 100422873 |
| ENSG000002 | 0.48991005 | 0.142435647 | 0.23916196 | 0.59556147 | 0.55146823 | NA | MIR4274 | 100422826 |
| ENSG000002 | 1.23931642 | -0.341209262 | 0.33024551 | -1.0331988 | 0.30151088 | NA | NA | NA |
| ENSG000002 | 0 | NA | NA | NA | NA | NA | NA | NA |
| ENSG000002 | 1.46899969 | -0.524106723 | 0.32757585 | -1.5999553 | 0.1096085 | NA | NA | NA |
| ENSG000002 | 2.01950868 | -0.31478859 | 0.3620498 | -0.8694621 | 0.38459442 | NA | NA | NA |
| ENSG000002 | 2.16805073 | 0.186297699 | 0.36789848 | 0.50638345 | 0.61258749 | NA | MIR3945 | 100500818 |
| ENSG000002 | 0.38133351 | -0.102466351 | 0.2189713 | -0.4679442 | 0.63982449 | NA | MIR4490 | 100616186 |
| ENSG000002 | 0.13280363 | -0.093394593 | 0.13115644 | -0.7120855 | 0.47641185 | NA | MIR4498 | 100616179 |
| ENSG000002 | 0.88123551 | -0.252774788 | 0.29217137 | -0.8651593 | 0.38695135 | NA | MIR4437 | 100616213 |
| ENSG000002 | 0.98292718 | -0.180673668 | 0.30434119 | -0.593655 | 0.55274289 | NA | RN7SL48P | 106481825 |
| ENSG000002 | 1.10941059 | -0.119301621 | 0.32064182 | -0.3720713 | 0.70983976 | NA | NA | NA |
| ENSG000002 | 1.80983654 | 0.135314371 | 0.3557567 | 0.38035649 | 0.70368081 | NA | NA | NA |
| ENSG000002 | 1.12287469 | -0.2670354 | 0.32190569 | -0.8295454 | 0.40679583 | NA | MIR4676 | 100616286 |
| ENSG000002 | 1.38441323 | -0.273485415 | 0.34183431 | -0.8000526 | 0.42368035 | NA | RN7SL14P | 106480481 |
| ENSG000002 | 0 | NA | NA | NA | NA | NA | MIR5695 | 100847016 |
| ENSG000002 | 0 | NA | NA | NA | NA | NA | NA | NA |
| ENSG000002 | 1.74192089 | -0.3044383 | 0.35050244 | -0.8685768 | 0.38507863 | NA | NA | NA |
| ENSG000002 | 1.59508672 | 0.011635934 | 0.34690222 | 0.0335424 | 0.97324205 | NA | MIR4757 | 100616307 |
| ENSG000002 | 0.63367309 | -0.098454409 | 0.25836134 | -0.3810725 | 0.70314943 | NA | MIR4708 | 100616176 |
| ENSG000002 | 0.62459249 | 0.048506127 | 0.26461252 | 0.18331002 | 0.85455477 | NA | MIR3135A | 100422901 |
| ENSG000002 | 0.72972831 | -0.025248506 | 0.28417722 | -0.0888477 | 0.92920291 | NA | NA | NA |
| ENSG000002 | 1.37726935 | -0.272772526 | 0.34044828 | -0.8012157 | 0.42300676 | NA | MIR4645 | 100616285 |

| | | | | | | | | | |
|---|---|---|---|---|---|---|---|---|---|
| ENSG000002 | 2.03234236 | 0.241755536 | 0.36270635 | 0.6665324 | 0.50507086 | NA | | MIR3661 | 100500905 |
| ENSG000002 | 0.38036605 | -0.293832733 | 0.22119083 | -1.3284128 | 0.18404178 | NA | | RN7SL524P | 106481066 |
| ENSG000002 | 1.29951607 | -0.672569239 | 0.32918457 | -2.0431372 | 0.04103886 | NA | | MIR5680 | 100847001 |
| ENSG000002 | 0.40327843 | -0.281391716 | 0.21700406 | -1.2967118 | 0.19473038 | NA | | MIR3680-2 | 100847041 |
| ENSG000002 | 0.12685397 | -0.090215768 | 0.12901919 | -0.699243 | 0.48440017 | NA | | MIR4464 | 100616109 |
| ENSG000002 | 1.63886946 | -0.49621599 | 0.34044791 | -1.4575386 | 0.14496775 | NA | | MIR3194 | 100422889 |
| ENSG000002 | 0.26199343 | -0.194018184 | 0.18422801 | -1.0531416 | 0.29227609 | NA | | NA | NA |
| ENSG000002 | 1.66972186 | -0.545039946 | 0.34878092 | -1.5627 | 0.11812317 | NA | | NA | NA |
| ENSG000002 | 1.26927336 | 0.082368649 | 0.31227105 | 0.26377293 | 0.7919549 | NA | | MIR4646 | 100616230 |
| ENSG000002 | 0.98403316 | 0.112873297 | 0.30784261 | 0.36665911 | 0.71387331 | NA | | MIR4444-2 | 100846999 |
| ENSG000002 | 0.83352905 | 0.20077793 | 0.29601317 | 0.67827363 | 0.49759822 | NA | | MIR3663 | 100500893 |
| ENSG000002 | 1.16401052 | -0.789014351 | 0.32059011 | -2.4611313 | 0.01384997 | NA | | NA | NA |
| ENSG000002 | 0.89666433 | -0.262300427 | 0.29296884 | -0.8953185 | 0.37061685 | NA | | MIR4419A | 100616177 |
| ENSG000002 | 1.22918594 | -0.200322978 | 0.333979 | -0.5998071 | 0.54863479 | NA | | NA | NA |
| ENSG000002 | 0.40122492 | -0.101896709 | 0.21365861 | -0.4769136 | 0.63342361 | NA | | MIR548H5 | 100616455 |
| ENSG000002 | 1.80664535 | 0.013350441 | 0.35696471 | 0.03739989 | 0.97016616 | NA | | MIR548AE1 | 100616305 |
| ENSG000002 | 1.53073833 | -0.642819973 | 0.3443703 | -1.8666533 | 0.06195001 | NA | | NA | NA |
| ENSG000002 | 0 | NA | NA | NA | NA | NA | | NA | NA |
| ENSG000002 | 0.22997012 | 0.167051788 | 0.17646558 | 0.94665368 | 0.34381528 | NA | | MIR3689E | 100616460 |
| ENSG000002 | 1.71729097 | 0.081198333 | 0.35437488 | 0.22913118 | 0.81876696 | NA | | RN7SL712P | 106481117 |
| ENSG000002 | 0.85803847 | 0.039330813 | 0.29683333 | 0.13250134 | 0.89458777 | NA | | NA | NA |
| ENSG000002 | 0.89887754 | -0.114830835 | 0.29928032 | -0.3836899 | 0.7012083 | NA | | MIR3910-1 | 100500821 |
| ENSG000002 | 0.98717271 | -0.344579472 | 0.30842128 | -1.1172364 | 0.26389327 | NA | | RN7SL123P | 106479266 |
| ENSG000002 | 1.56058769 | 0.017171635 | 0.35005106 | 0.04905466 | 0.96087574 | NA | | NA | NA |
| ENSG000002 | 0.72335602 | 0.142054296 | 0.28912098 | 0.49133168 | 0.62319188 | NA | | MIR4535 | 100616415 |
| ENSG000002 | 0.49201754 | 0.143225237 | 0.2394083 | 0.59824675 | 0.5496753 | NA | | MIR4634 | 100616202 |
| ENSG000002 | 1.4698168 | -0.046401635 | 0.33709447 | -0.1376517 | 0.89051567 | NA | | NA | NA |
| ENSG000002 | 2.00177829 | -0.665283715 | 0.35405861 | -1.8790214 | 0.06024157 | NA | | NA | NA |
| ENSG000002 | 0.75964605 | -0.203618105 | 0.2920432 | -0.6972191 | 0.48566567 | NA | | NA | NA |

| | | | | | | | | |
|---|---|---|---|---|---|---|---|---|
| ENSG000002 | 0.72558422 | -0.175184843 | 0.27691871 | -0.6326219 | 0.52698058 | NA | NA | NA |
| ENSG000002 | 2.12860874 | -0.515487365 | 0.36571397 | -1.409537 | 0.15867644 | NA | NA | NA |
| ENSG000002 | 1.23740672 | -0.195332072 | 0.33287139 | -0.5868094 | 0.5573317 | NA | NA | NA |
| ENSG000002 | 0.98352271 | -0.031897561 | 0.30613359 | -0.1041949 | 0.91701468 | NA | NA | NA |
| ENSG000002 | 0 | NA | NA | NA | NA | NA | NA | NA |
| ENSG000002 | 1.48545648 | -0.487522759 | 0.34563754 | -1.4105029 | 0.15839124 | NA | NA | NA |
| ENSG000002 | 0 | NA | NA | NA | NA | NA | NA | NA |
| ENSG000002 | 1.53779341 | -0.343813323 | 0.34628027 | -0.9928759 | 0.32077045 | NA | NA | NA |
| ENSG000002 | 0.22336757 | 0.163792188 | 0.17487321 | 0.93663397 | 0.34894687 | NA | NA | NA |
| ENSG000002 | 0 | NA | NA | NA | NA | NA | NA | NA |
| ENSG000002 | 0.7394356 | -0.033285101 | 0.2693877 | -0.1235584 | 0.90166497 | NA | NA | NA |
| ENSG000002 | 1.89740768 | -0.402893867 | 0.3593365 | -1.1212161 | 0.26219589 | NA | NA | NA |
| ENSG000002 | 1.53188048 | -0.615492 | 0.34076668 | -1.8061977 | 0.07088746 | NA | NA | NA |
| ENSG000002 | 1.47636728 | -0.048425652 | 0.34550009 | -0.140161 | 0.88853275 | NA | NA | NA |
| ENSG000002 | 0 | NA | NA | NA | NA | NA | NA | NA |
| ENSG000002 | 1.60302483 | -0.125036486 | 0.3485699 | -0.3587128 | 0.71980995 | NA | NA | NA |
| ENSG000002 | 2.03599129 | -0.594445783 | 0.36292543 | -1.6379282 | 0.10143668 | NA | NA | NA |
| ENSG000002 | 1.24998925 | -0.356057169 | 0.33390887 | -1.0663304 | 0.28627433 | NA | NA | NA |
| ENSG000002 | 0.74772501 | -0.357780902 | 0.28233051 | -1.2672414 | 0.205069 | NA | NA | NA |
| ENSG000002 | 1.44847058 | -0.037533779 | 0.33116572 | -0.1133384 | 0.90976231 | NA | NA | NA |
| ENSG000002 | 0.46887648 | 0.154938272 | 0.24404396 | 0.63487854 | 0.52550763 | NA | NA | NA |
| ENSG000002 | 0.22336757 | 0.163792188 | 0.17487321 | 0.93663397 | 0.34894687 | NA | NA | NA |
| ENSG000002 | 1.74457275 | -0.593936802 | 0.35158192 | -1.6893269 | 0.0911568 | NA | NA | NA |
| ENSG000002 | 1.55636198 | 0.130651716 | 0.32847523 | 0.39775211 | 0.69081292 | NA | NA | NA |
| ENSG000002 | 0.25351207 | -0.189965547 | 0.1824784 | -1.0410303 | 0.29786148 | NA | NA | NA |
| ENSG000002 | 0.84975353 | 0.05329801 | 0.30153883 | 0.17675339 | 0.8597021 | NA | NA | NA |
| ENSG000002 | 1.25963395 | -0.201596298 | 0.333412 | -0.6046462 | 0.5454141 | NA | NA | NA |
| ENSG000002 | 2.05815206 | 0.250941355 | 0.36318413 | 0.69094802 | 0.48959821 | NA | NA | NA |
| ENSG000002 | 1.23208821 | -0.320392633 | 0.32356093 | -0.990208 | 0.32207244 | NA | NA | NA |

| | | | | | | | | |
|---|---|---|---|---|---|---|---|---|
| ENSG000002 | 0 | NA | NA | NA | NA | NA | NA | NA |
| ENSG000002 | 0 | NA | NA | NA | NA | NA | NA | NA |
| ENSG000002 | 1.53461274 | -0.648329204 | 0.34437551 | -1.8826228 | 0.0597515 | NA | NA | NA |
| ENSG000002 | 2.03340304 | -0.31470972 | 0.35894785 | -0.8767561 | 0.38061913 | NA | NA | NA |
| ENSG000002 | 1.12993561 | -0.111392587 | 0.3166667 | -0.351766 | 0.72501374 | NA | NA | NA |
| ENSG000002 | 0.78039674 | -0.368519854 | 0.28442264 | -1.295677 | 0.1950868 | NA | NA | NA |
| ENSG000002 | 1.79543369 | -0.63205617 | 0.35638704 | -1.7735105 | 0.07614415 | NA | NA | NA |
| ENSG000002 | 0 | NA | NA | NA | NA | NA | NA | NA |
| ENSG000002 | 1.23439839 | -0.183286316 | 0.32321658 | -0.5670697 | 0.57066685 | NA | NA | NA |
| ENSG000002 | 1.13818672 | -0.260195218 | 0.31796998 | -0.8183012 | 0.41318522 | NA | NA | NA |
| ENSG000002 | 1.37618174 | -0.11419386 | 0.33507994 | -0.3407959 | 0.73325726 | NA | NA | NA |
| ENSG000002 | 0.87329706 | -0.111549593 | 0.29797379 | -0.3743604 | 0.70813619 | NA | NA | NA |
| ENSG000002 | 0 | NA | NA | NA | NA | NA | NA | NA |
| ENSG000002 | 0.11934008 | 0.083564698 | 0.12760091 | 0.65489106 | 0.5125379 | NA | NA | NA |
| ENSG000002 | 0 | NA | NA | NA | NA | NA | NA | NA |
| ENSG000002 | 0.64713665 | -0.283099576 | 0.26867821 | -1.0536752 | 0.29203163 | NA | NA | NA |
| ENSG000002 | 1.84169757 | 0.007443724 | 0.34953827 | 0.02129588 | 0.98300963 | NA | NA | NA |
| ENSG000002 | 1.64141843 | -0.121695091 | 0.34569204 | -0.3520332 | 0.72481333 | NA | LOC105372273 | 105372273 |
| ENSG000002 | 1.74971656 | -0.339255931 | 0.35749028 | -0.9489934 | 0.34262395 | NA | NA | NA |
| ENSG000002 | 0.968917 | 0.115248834 | 0.3055111 | 0.37723288 | 0.70600055 | NA | NA | NA |
| ENSG000002 | 0.53471369 | -0.194084218 | 0.24487205 | -0.7925944 | 0.42801417 | NA | NA | NA |
| ENSG000002 | 0.54817724 | -0.365482324 | 0.24206593 | -1.5098462 | 0.13108267 | NA | NA | NA |
| ENSG000002 | 1.10685301 | -0.085943616 | 0.20805917 | -0.413073 | 0.67955317 | NA | NA | NA |
| ENSG000002 | 1.29028891 | -0.330067658 | 0.32516344 | -1.0150823 | 0.3100666 | NA | NA | NA |
| ENSG000002 | 1.16401052 | -0.789014351 | 0.32059011 | -2.4611313 | 0.01384997 | NA | NA | NA |
| ENSG000002 | 0 | NA | NA | NA | NA | NA | NA | NA |
| ENSG000002 | 1.09157034 | 0.172271253 | 0.31540354 | 0.54619315 | 0.58493318 | NA | NA | NA |
| ENSG000002 | 0 | NA | NA | NA | NA | NA | NA | NA |
| ENSG000002 | 1.35305725 | -0.370720904 | 0.3254678 | -1.1390402 | 0.25468639 | NA | NA | NA |

| | | | | | | | | |
|---|---|---|---|---|---|---|---|---|
| ENSG000002 | 0.12685397 | -0.090215768 | 0.12901919 | -0.699243 | 0.48440017 | NA | NA | NA |
| ENSG000002 | 2.15809048 | 0.410417492 | 0.36395342 | 1.1276649 | 0.25946147 | NA | NA | NA |
| ENSG000002 | 0 | NA | NA | NA | NA | NA | NA | NA |
| ENSG000002 | 0.61887151 | 0.056280962 | 0.26412849 | 0.21308175 | 0.8312632 | NA | NA | NA |
| ENSG000002 | 0.60763424 | -0.248288277 | 0.25676438 | -0.9669888 | 0.33354964 | NA | NA | NA |
| ENSG000002 | 0.48302518 | -0.023466322 | 0.24515318 | -0.0957211 | 0.92374212 | NA | NA | NA |
| ENSG000002 | 1.85306138 | -0.126007381 | 0.36057149 | -0.3494657 | 0.72673969 | NA | NA | NA |
| ENSG000002 | 1.4664432 | -0.057533391 | 0.34339222 | -0.1675442 | 0.86694184 | NA | NA | NA |
| ENSG000002 | 0.98273577 | -0.447649295 | 0.29538788 | -1.5154626 | 0.12965529 | NA | NA | NA |
| ENSG000002 | 0 | NA | NA | NA | NA | NA | NA | NA |
| ENSG000002 | 0.54817724 | -0.365482324 | 0.24206593 | -1.5098462 | 0.13108267 | NA | NA | NA |
| ENSG000002 | 0.27408862 | -0.199826037 | 0.18669465 | -1.0703362 | 0.28446801 | NA | NA | NA |
| ENSG000002 | 1.21417584 | -0.0415988 | 0.32809337 | -0.1267895 | 0.89910699 | NA | NA | NA |
| ENSG000002 | 0.12070844 | -0.088503695 | 0.12783931 | -0.6923042 | 0.48874631 | NA | NA | NA |
| ENSG000002 | 1.60019408 | -0.622119847 | 0.33777232 | -1.8418319 | 0.06549974 | NA | NA | NA |
| ENSG000002 | 0 | NA | NA | NA | NA | NA | NA | NA |
| ENSG000002 | 0.45612841 | 0.144125125 | 0.23502944 | 0.61322157 | 0.53972984 | NA | NA | NA |
| ENSG000002 | 1.3666274 | -0.413667823 | 0.33914074 | -1.2197527 | 0.22255865 | NA | NA | NA |
| ENSG000002 | 0 | NA | NA | NA | NA | NA | NA | NA |
| ENSG000002 | 0 | NA | NA | NA | NA | NA | NA | NA |
| ENSG000002 | 0.47551128 | 0.153262041 | 0.24481524 | 0.62603146 | 0.53129431 | NA | NA | NA |
| ENSG000002 | 1.58487034 | 0.017560397 | 0.34461487 | 0.05095659 | 0.95936012 | NA | NA | NA |
| ENSG000002 | 1.10098342 | 0.036161261 | 0.3240723 | 0.11158393 | 0.91115332 | NA | NA | NA |
| ENSG000002 | 0.28256998 | -0.18483211 | 0.18021686 | -1.0256094 | 0.30507573 | NA | SNRPCP20 | 106480292 |
| ENSG000002 | 0 | NA | NA | NA | NA | NA | NA | NA |
| ENSG000002 | 1.11752139 | 0.020200712 | 0.31423506 | 0.06428535 | 0.94874302 | NA | NA | NA |
| ENSG000002 | 1.0792449 | 0.033612684 | 0.32085083 | 0.10476109 | 0.91656538 | NA | NA | NA |
| ENSG000002 | 0.24619405 | -0.007630671 | 0.17915639 | -0.0425922 | 0.96602658 | NA | MIR212 | 406994 |
| ENSG000002 | 0.12685397 | -0.090215768 | 0.12901919 | -0.699243 | 0.48440017 | NA | MIR132 | 406921 |

| | | | | | | | | | |
|---|---|---|---|---|---|---|---|---|---|
| ENSG000002 | 1.48895118 | -0.061053652 | 0.34830502 | -0.1752879 | 0.86085343 | NA | NA | NA | |
| ENSG000002 | 1.87056744 | -0.499255863 | 0.35460521 | -1.4079203 | 0.15915469 | NA | NA | NA | |
| ENSG000002 | 0.34948434 | 0.063159058 | 0.2051263 | 0.30790326 | 0.75815594 | NA | LCN6 | | 158062 |
| ENSG000002 | 1.02365627 | -0.195516125 | 0.31015284 | -0.6303864 | 0.52844182 | NA | NA | NA | |
| ENSG000002 | 1.21746132 | -0.318953807 | 0.32397646 | -0.9844969 | 0.32487128 | NA | NA | NA | |
| ENSG000002 | 0.26813897 | -0.197007041 | 0.18550319 | -1.0620143 | 0.2882292 | NA | NA | NA | |
| ENSG000002 | 0.72540953 | -0.028108977 | 0.28387695 | -0.0990182 | 0.92112384 | NA | NA | NA | |
| ENSG000002 | 0.36345889 | 0.070257596 | 0.21589421 | 0.32542602 | 0.74485866 | NA | NA | NA | |
| ENSG000002 | 0 | NA | NA | NA | NA | NA | NA | NA | |
| ENSG000002 | 0 | NA | NA | NA | NA | NA | NA | NA | |
| ENSG000002 | 0.75030486 | -0.031917739 | 0.28551165 | -0.1117914 | 0.91098883 | NA | NA | NA | |
| ENSG000002 | 1.72619069 | -0.395980536 | 0.34238238 | -1.1565447 | 0.24745841 | NA | NA | NA | |
| ENSG000002 | 1.75705823 | -0.173874989 | 0.34574287 | -0.5029026 | 0.61503275 | NA | LOC100288123 | | 100288123 |
| ENSG000002 | 0.22336757 | 0.163792188 | 0.17487321 | 0.93663397 | 0.34894687 | NA | NA | NA | |
| ENSG000002 | 0.61890185 | -0.110660695 | 0.2644872 | -0.4183972 | 0.67565676 | NA | NA | NA | |
| ENSG000002 | 1.71778204 | -0.316088628 | 0.35292634 | -0.8956221 | 0.37045463 | NA | NA | NA | |
| ENSG000002 | 0 | NA | NA | NA | NA | NA | NA | NA | |
| ENSG000002 | 0.42385498 | -0.262217565 | 0.21037102 | -1.2464529 | 0.21259817 | NA | NA | NA | |
| ENSG000002 | 1.1295328 | 0.03166093 | 0.32251159 | 0.0981699 | 0.92179738 | NA | NA | NA | |
| ENSG000002 | 1.81962406 | -0.603769278 | 0.35166791 | -1.7168734 | 0.08600231 | NA | NA | NA | |
| ENSG000002 | 1.04102181 | -0.355845985 | 0.31062511 | -1.1455802 | 0.25196888 | NA | NA | NA | |
| ENSG000002 | 1.34763448 | 0.030642251 | 0.33699628 | 0.09092757 | 0.92755015 | NA | NA | NA | |
| ENSG000002 | 0.99649891 | -0.037776854 | 0.31317903 | -0.1206238 | 0.90398899 | NA | NA | NA | |
| ENSG000002 | 0.12685397 | -0.090215768 | 0.12901919 | -0.699243 | 0.48440017 | NA | NA | NA | |
| ENSG000002 | 1.41756981 | 0.228577987 | 0.33482047 | 0.68268821 | 0.49480388 | NA | NA | NA | |
| ENSG000002 | 0 | NA | NA | NA | NA | NA | NA | NA | |
| ENSG000002 | 1.39565498 | -0.246273735 | 0.33223693 | -0.7412594 | 0.4585362 | NA | NA | NA | |
| ENSG000002 | 0 | NA | NA | NA | NA | NA | NA | NA | |
| ENSG000002 | 1.83220409 | -0.126900141 | 0.36128581 | -0.3512459 | 0.72540391 | NA | NA | NA | |

| | | | | | | | | | |
|---|---|---|---|---|---|---|---|---|---|
| ENSG000002 | 0 | NA | NA | NA | NA | NA | NA | NA | |
| ENSG000002 | 0 | NA | NA | NA | NA | NA | NA | NA | |
| ENSG000002 | 0.11865494 | 0.083440866 | 0.12751478 | 0.65436232 | 0.5128784 | NA | NA | NA | |
| ENSG000002 | 1.53389726 | -0.503293403 | 0.34815766 | -1.4455905 | 0.1482921 | NA | NA | NA | |
| ENSG000002 | 1.6954421 | -0.806922873 | 0.34021969 | -2.3717701 | 0.0177031 | NA | NA | NA | |
| ENSG000002 | 2.14596146 | 0.317478587 | 0.36899002 | 0.86039885 | 0.38956922 | NA | NA | NA | |
| ENSG000002 | 1.84808834 | -0.246687521 | 0.35671477 | -0.691554 | 0.48921746 | NA | NA | NA | |
| ENSG000002 | 1.52495889 | -0.49953834 | 0.34689671 | -1.4400204 | 0.14986163 | NA | NA | NA | |
| ENSG000002 | 0.87715884 | -0.112985904 | 0.29817966 | -0.3789189 | 0.7047481 | NA | NA | NA | |
| ENSG000002 | 1.51195235 | -0.34531695 | 0.34820182 | -0.991715 | 0.32133659 | NA | NA | NA | |
| ENSG000002 | 1.99821666 | -0.166995675 | 0.35635937 | -0.4686159 | 0.6393442 | NA | NA | NA | |
| ENSG000002 | 0 | NA | NA | NA | NA | NA | UPK3BL | | 100134938 |
| ENSG000002 | 2.11010296 | -0.262767389 | 0.36784741 | -0.714338 | 0.4750182 | NA | NA | NA | |
| ENSG000002 | 0.75213161 | -0.036873935 | 0.28561738 | -0.1291026 | 0.8972765 | NA | NA | NA | |
| ENSG000002 | 1.1098676 | -0.124685886 | 0.32928509 | -0.3786563 | 0.70494308 | NA | NA | NA | |
| ENSG000002 | 1.50139575 | -0.337079884 | 0.34502508 | -0.9769721 | 0.32858296 | NA | NA | NA | |
| ENSG000002 | 0.96932386 | 0.269984829 | 0.3039266 | 0.88832247 | 0.37436732 | NA | NA | NA | |
| ENSG000002 | 1.10872599 | -0.112000084 | 0.32214247 | -0.3476725 | 0.72808614 | NA | NA | NA | |
| ENSG000002 | 0.5834394 | 0.062235112 | 0.26097587 | 0.23847075 | 0.811516 | NA | NA | NA | |
| ENSG000002 | 0.6164817 | -0.105910369 | 0.26430836 | -0.4007076 | 0.68863541 | NA | NA | NA | |
| ENSG000002 | 2.01262135 | -0.579271845 | 0.35958503 | -1.6109454 | 0.10719163 | NA | NA | NA | |
| ENSG000002 | 1.81617603 | 0.006056106 | 0.35802683 | 0.01691523 | 0.98650424 | NA | NA | NA | |
| ENSG000002 | 1.34945911 | -0.120978957 | 0.33934193 | -0.3565105 | 0.7214583 | NA | NA | NA | |
| ENSG000002 | 0.50818749 | -0.197539377 | 0.24915052 | -0.7928516 | 0.42786432 | NA | NA | NA | |
| ENSG000002 | 1.05041507 | -0.532808254 | 0.31301235 | -1.7021956 | 0.0887187 | NA | NA | NA | |
| ENSG000002 | 1.26601731 | -0.475702359 | 0.32476489 | -1.4647592 | 0.14298663 | NA | NA | NA | |
| ENSG000002 | 1.90583402 | -0.672223342 | 0.3581713 | -1.8768208 | 0.06054266 | NA | NA | NA | |
| ENSG000002 | 0 | NA | NA | NA | NA | NA | NA | NA | |
| ENSG000002 | 1.35449741 | 0.026628571 | 0.34207475 | 0.0778443 | 0.9379519 | NA | NA | NA | |

| | | | | | | | | | |
|---|---|---|---|---|---|---|---|---|---|
| ENSG000002 | 2.02315834 | -0.566868906 | 0.35801081 | -1.5833849 | 0.11333376 | NA | NA | NA | |
| ENSG000002 | 0.96577016 | -0.036559874 | 0.30533124 | -0.1197384 | 0.90469038 | NA | NA | NA | |
| ENSG000002 | 0.85542244 | 0.045978532 | 0.30182892 | 0.15233309 | 0.87892423 | NA | NA | NA | |
| ENSG000002 | 0 | NA | NA | NA | NA | NA | NA | NA | |
| ENSG000002 | 0.25993993 | -0.013144386 | 0.18211683 | -0.0721756 | 0.94246219 | NA | NA | NA | |
| ENSG000002 | 1.72427405 | -0.192682301 | 0.35420041 | -0.5439923 | 0.58644676 | NA | NA | NA | |
| ENSG000002 | 0 | NA | NA | NA | NA | NA | NA | NA | |
| ENSG000002 | 1.12057716 | -0.431729854 | 0.32306664 | -1.3363492 | 0.18143515 | NA | NA | NA | |
| ENSG000002 | 0.62160824 | -0.2792447 | 0.26652831 | -1.0477112 | 0.29477168 | NA | NA | NA | |
| ENSG000002 | 0.22263036 | 0.146922267 | 0.16600367 | 0.88505432 | 0.37612734 | NA | NA | NA | |
| ENSG000002 | 2.01463868 | -0.454681072 | 0.36206017 | -1.2558163 | 0.20918257 | NA | NA | NA | |
| ENSG000002 | 1.48780958 | -0.057269665 | 0.34935487 | -0.1639298 | 0.86978643 | NA | NA | NA | |
| ENSG000002 | 1.00513323 | -0.365079386 | 0.31516187 | -1.1583869 | 0.24670617 | NA | NA | NA | |
| ENSG000002 | 0.50972607 | -0.173976743 | 0.2324237 | -0.7485327 | 0.4541389 | NA | APOC4 | | 346 |
| ENSG000002 | 0.14128499 | -0.097772014 | 0.13403235 | -0.7294658 | 0.4657168 | NA | NA | NA | |
| ENSG000002 | 0 | NA | NA | NA | NA | NA | NA | NA | |
| ENSG000002 | 1.16859181 | -0.605351814 | 0.32253551 | -1.8768533 | 0.06053821 | NA | NA | NA | |
| ENSG000002 | 1.00032518 | -0.338397872 | 0.30533777 | -1.1082739 | 0.26774355 | NA | NA | NA | |
| ENSG000002 | 0 | NA | NA | NA | NA | NA | NA | NA | |
| ENSG000002 | 0.73587384 | -0.025715051 | 0.2782674 | -0.0924113 | 0.92637126 | NA | NA | NA | |
| ENSG000002 | 1.8454935 | -0.127547697 | 0.362225 | -0.3521228 | 0.72474614 | NA | NA | NA | |
| ENSG000002 | 2.13497711 | -0.372522783 | 0.36409495 | -1.0231474 | 0.30623818 | NA | NA | NA | |
| ENSG000002 | 0.7102413 | -0.022139713 | 0.27646803 | -0.0800806 | 0.93617319 | NA | NA | NA | |
| ENSG000002 | 2.07200062 | -0.359665108 | 0.36120464 | -0.9957378 | 0.31937757 | NA | FAM215A | | 23591 |
| ENSG000002 | 1.89094948 | -0.397225026 | 0.359295 | -1.1055679 | 0.26891358 | NA | NA | NA | |
| ENSG000002 | 1.5728793 | 0.021494487 | 0.34787995 | 0.06178708 | 0.95073239 | NA | NA | NA | |
| ENSG000002 | 2.05769952 | 0.009549515 | 0.36497278 | 0.02616501 | 0.97912573 | NA | NA | NA | |
| ENSG000002 | 0 | NA | NA | NA | NA | NA | NA | NA | |
| ENSG000002 | 1.55001569 | -0.289963412 | 0.2370673 | -1.223127 | 0.22128173 | NA | NA | NA | |

| | | | | | | | | |
|---|---|---|---|---|---|---|---|---|
| ENSG000002 | 0.60447083 | -0.105840125 | 0.26325248 | -0.402048 | 0.68764873 | NA | NA | NA |
| ENSG000002 | 2.09088225 | 0.125855508 | 0.36281799 | 0.34688332 | 0.72867898 | NA | NA | NA |
| ENSG000002 | 1.11860797 | -0.273523937 | 0.32323041 | -0.8462197 | 0.3974302 | NA | NA | NA |
| ENSG000002 | 0.75030486 | -0.031917739 | 0.28551165 | -0.1117914 | 0.91098883 | NA | NA | NA |
| ENSG000002 | 0.88881412 | -0.432208381 | 0.29509244 | -1.4646542 | 0.14301529 | NA | NA | NA |
| ENSG000002 | 0.6275579 | -0.281336606 | 0.26701873 | -1.0536213 | 0.29205636 | NA | NA | NA |
| ENSG000002 | 1.58679986 | -0.109024318 | 0.34565784 | -0.3154111 | 0.75244954 | NA | NA | NA |
| ENSG000002 | 0.78210142 | -0.19296092 | 0.28068748 | -0.6874582 | 0.491794 | NA | NA | NA |
| ENSG000002 | 1.0143518 | -0.367132577 | 0.31550649 | -1.1636292 | 0.2445743 | NA | NA | NA |
| ENSG000002 | 1.49768104 | -0.051600495 | 0.34141791 | -0.1511359 | 0.87986853 | NA | LOC105372145 | 105372145 |
| ENSG000002 | 0.91146591 | -0.433639492 | 0.2963326 | -1.463354 | 0.14337054 | NA | NA | NA |
| ENSG000002 | 0.12070844 | -0.088503695 | 0.12783931 | -0.6923042 | 0.48874631 | NA | NA | NA |
| ENSG000002 | 1.97909437 | -0.181452634 | 0.36133342 | -0.5021751 | 0.61554438 | NA | NA | NA |
| ENSG000002 | 1.64375818 | -0.379379643 | 0.34398158 | -1.1029069 | 0.27006761 | NA | NA | NA |
| ENSG000002 | 1.62451519 | -0.127264679 | 0.35038999 | -0.3632087 | 0.71644902 | NA | NA | NA |
| ENSG000002 | 0.87645835 | -0.422982294 | 0.29453257 | -1.4361138 | 0.15096996 | NA | NDUFA3P1 | 100131388 |
| ENSG000002 | 1.03698186 | -0.533697484 | 0.31242205 | -1.708258 | 0.08758847 | NA | NA | NA |
| ENSG000002 | 1.18486939 | -0.597753117 | 0.32083796 | -1.8630998 | 0.06244823 | NA | NA | NA |
| ENSG000002 | 1.77025466 | -0.905837687 | 0.34993223 | -2.5886089 | 0.00963645 | NA | NA | NA |
| ENSG000002 | 0.59715281 | 0.054585582 | 0.25462173 | 0.21437912 | 0.83025142 | NA | NA | NA |
| ENSG000002 | 2.05101708 | -0.446270943 | 0.36101028 | -1.2361724 | 0.21639449 | NA | NA | NA |
| ENSG000002 | 1.23931195 | -0.05014886 | 0.32018778 | -0.1566233 | 0.87554175 | NA | NA | NA |
| ENSG000002 | 0 | NA | NA | NA | NA | NA | NA | NA |
| ENSG000002 | 0.63828472 | -0.112219173 | 0.26613064 | -0.4216695 | 0.67326627 | NA | NA | NA |
| ENSG000002 | 0.89364336 | -0.29139313 | 0.3048889 | -0.9557354 | 0.33920591 | NA | NA | NA |
| ENSG000002 | 1.55586391 | -0.47536156 | 0.34274802 | -1.3869126 | 0.1654684 | NA | CGB1 | 114335 |
| ENSG000002 | 0.99149499 | -0.042075156 | 0.31709968 | -0.1326875 | 0.89444055 | NA | NA | NA |
| ENSG000002 | 1.32022651 | 0.026467087 | 0.33354982 | 0.07934973 | 0.93675445 | NA | NA | NA |
| ENSG000002 | 1.72139799 | -0.190213933 | 0.35735974 | -0.5322758 | 0.59453501 | NA | NA | NA |

| | | | | | | | | |
|---|---|---|---|---|---|---|---|---|
| ENSG000002 | 0 | NA | NA | NA | NA | NA | POLR2J2 | 246721 |
| ENSG000002 | 1.36264246 | -0.272330572 | 0.34140546 | -0.7976749 | 0.42505917 | NA | NA | NA |
| ENSG000002 | 1.90303687 | 0.337841188 | 0.36130235 | 0.93506501 | 0.3497548 | NA | NA | NA |
| ENSG000002 | 1.39337067 | -0.249060324 | 0.32972868 | -0.7553493 | 0.45003944 | NA | LOC100996324 | 100996324 |
| ENSG000002 | 0.60264758 | 0.238025744 | 0.26986178 | 0.88202838 | 0.37776146 | NA | NA | NA |
| ENSG000002 | 1.37242356 | -0.12159583 | 0.33500524 | -0.362967 | 0.71662957 | NA | NA | NA |
| ENSG000002 | 1.75654778 | -0.316954171 | 0.35335336 | -0.8969893 | 0.36972465 | NA | NA | NA |
| ENSG000002 | 0.47761686 | -0.01948631 | 0.23692179 | -0.0822479 | 0.93444962 | NA | FDX1L | 112812 |
| ENSG000002 | 0.66751732 | -0.272810278 | 0.26356336 | -1.0350842 | 0.30062957 | NA | NA | NA |
| ENSG000002 | 0.37945415 | -0.107841678 | 0.21865587 | -0.4932027 | 0.62186934 | NA | NA | NA |
| ENSG000002 | 1.40658628 | -0.435684064 | 0.34176813 | -1.2747943 | 0.20238205 | NA | NA | NA |
| ENSG000002 | 1.05178344 | -0.716293747 | 0.31097042 | -2.3034144 | 0.02125554 | NA | NA | NA |
| ENSG000002 | 0.85960326 | -0.112061982 | 0.30237674 | -0.3706038 | 0.71093262 | NA | NA | NA |
| ENSG000002 | 0.83626824 | 0.050891293 | 0.29564717 | 0.17213523 | 0.86333122 | NA | NA | NA |
| ENSG000002 | 0.64813445 | -0.288472774 | 0.26867189 | -1.0736991 | 0.28295756 | NA | NA | NA |
| ENSG000002 | 0 | NA | NA | NA | NA | NA | LUZP6 | 767558 |
| ENSG000002 | 0 | NA | NA | NA | NA | NA | NA | NA |
| ENSG000002 | 1.96761978 | -0.354701456 | 0.34834199 | -1.0182564 | 0.30855613 | NA | NA | NA |
| ENSG000002 | 0.11865494 | 0.083440866 | 0.12751478 | 0.65436232 | 0.5128784 | NA | NA | NA |
| ENSG000002 | 0.99191762 | -0.182835207 | 0.30719334 | -0.5951796 | 0.55172343 | NA | NA | NA |
| ENSG000002 | 1.52476301 | -0.496208817 | 0.34624859 | -1.4330999 | 0.15182929 | NA | NA | NA |
| ENSG000002 | 1.57450825 | -0.113135538 | 0.34472456 | -0.3281911 | 0.74276716 | NA | NA | NA |
| ENSG000002 | 0.79229605 | -0.346664159 | 0.27711096 | -1.2509941 | 0.21093665 | NA | NA | NA |
| ENSG000002 | 0.47075585 | 0.160115757 | 0.24423976 | 0.65556794 | 0.51210216 | NA | NA | NA |
| ENSG000002 | 0.99021095 | 0.099515752 | 0.3006715 | 0.33097833 | 0.74066086 | NA | NA | NA |
| ENSG000002 | 0 | NA | NA | NA | NA | NA | NA | NA |
| ENSG000002 | 0.88741138 | -0.451663868 | 0.3001743 | -1.504672 | 0.13240842 | NA | NA | NA |
| ENSG000002 | 1.21252728 | 0.111258646 | 0.33300904 | 0.33410098 | 0.73830336 | NA | NA | NA |
| ENSG000002 | 1.46842862 | 0.086243982 | 0.34403482 | 0.25068388 | 0.80205853 | NA | NA | NA |

| | | | | | | | | | |
|---|---|---|---|---|---|---|---|---|---|
| ENSG000002 | 0 | NA | NA | NA | NA | NA | NA | NA | |
| ENSG000002 | 0.63484558 | -0.278698901 | 0.26767877 | -1.0411692 | 0.29779701 | NA | NA | NA | |
| ENSG000002 | 1.0109082 | -0.204003008 | 0.31430482 | -0.649061 | 0.51629894 | NA | NA | NA | |
| ENSG000002 | 0 | NA | NA | NA | NA | NA | NA | NA | |
| ENSG000002 | 1.84139232 | 0.008424072 | 0.35601359 | 0.02366222 | 0.98112204 | NA | EML2-AS1 | | 100287177 |
| ENSG000002 | 1.98546411 | 0.061774376 | 0.36003956 | 0.17157663 | 0.86377038 | NA | NA | NA | |
| ENSG000002 | 0.86047844 | 0.043802335 | 0.2891494 | 0.15148686 | 0.87959168 | NA | NA | NA | |
| ENSG000002 | 1.12671877 | -0.1084661 | 0.31400686 | -0.3454259 | 0.72977422 | NA | NA | NA | |
| ENSG000002 | 0.86014796 | -0.390905035 | 0.2847271 | -1.3729112 | 0.16977995 | NA | NA | NA | |
| ENSG000002 | 1.98792064 | -0.18755845 | 0.36287672 | -0.5168655 | 0.60525006 | NA | NA | NA | |
| ENSG000002 | 0.40122492 | -0.101896709 | 0.21365861 | -0.4769136 | 0.63342361 | NA | NA | NA | |
| ENSG000002 | 1.62989883 | -0.523238764 | 0.34584356 | -1.5129348 | 0.13029621 | NA | NA | NA | |
| ENSG000002 | 2.01309938 | 0.500011379 | 0.36134787 | 1.38373965 | 0.16643819 | NA | NA | NA | |
| ENSG000002 | 1.85443132 | -0.132413163 | 0.36146326 | -0.3663254 | 0.7141223 | NA | NA | NA | |
| ENSG000002 | 2.16577395 | -0.639086989 | 0.36405068 | -1.7554891 | 0.07917569 | NA | NA | NA | |
| ENSG000002 | 1.98521212 | -0.185951305 | 0.35938559 | -0.5174145 | 0.60486686 | NA | LOC100132174 | | 100132174 |
| ENSG000002 | 0.5226185 | -0.189580046 | 0.24357686 | -0.7783172 | 0.43638207 | NA | NA | NA | |
| ENSG000002 | 0 | NA | NA | NA | NA | NA | WBP1LP11 | | 106481969 |
| ENSG000002 | 0 | NA | NA | NA | NA | NA | NA | NA | |
| ENSG000002 | 0.26199343 | -0.194018184 | 0.18422801 | -1.0531416 | 0.29227609 | NA | NA | NA | |
| ENSG000002 | 1.85218429 | -0.397294627 | 0.36081697 | -1.1010974 | 0.27085426 | NA | NA | NA | |
| ENSG000002 | 0.12685397 | -0.090215768 | 0.12901919 | -0.699243 | 0.48440017 | NA | NA | NA | |
| ENSG000002 | 0 | NA | NA | NA | NA | NA | NA | NA | |
| ENSG000002 | 0.99030076 | -0.043579311 | 0.3080742 | -0.1414572 | 0.88750878 | NA | NA | NA | |
| ENSG000002 | 0.70295361 | -0.024684415 | 0.27593502 | -0.0894573 | 0.92871845 | NA | NA | NA | |
| ENSG000002 | 1.6280795 | -0.550857306 | 0.35073528 | -1.5705785 | 0.11628058 | NA | NA | NA | |
| ENSG000002 | 0 | NA | NA | NA | NA | NA | NA | NA | |
| ENSG000002 | 1.49114904 | -0.187602169 | 0.34313618 | -0.546728 | 0.58456559 | NA | NA | NA | |
| ENSG000002 | 1.29534865 | 0.476077007 | 0.3345312 | 1.42311689 | 0.15470227 | NA | NA | NA | |

| | | | | | | | | |
|---|---|---|---|---|---|---|---|---|
| ENSG000002 | 1.33354975 | 0.160125278 | 0.33549461 | 0.47728122 | 0.63316188 | NA | NA | NA |
| ENSG000002 | 0.96147311 | 0.113052294 | 0.30518434 | 0.37043936 | 0.71105515 | NA | NA | NA |
| ENSG000002 | 0.72514894 | -0.021174814 | 0.27751054 | -0.0763027 | 0.93917825 | NA | NA | NA |
| ENSG000002 | 0.37199232 | 0.06760863 | 0.21718977 | 0.31128828 | 0.75558148 | NA | MIR3188 | 100422833 |
| ENSG000002 | 1.90116641 | -0.394816167 | 0.35938662 | -1.0985834 | 0.27194983 | NA | NA | NA |
| ENSG000002 | 0.96094609 | 0.269157969 | 0.30599103 | 0.87962699 | 0.37906141 | NA | NA | NA |
| ENSG000002 | 1.00803158 | -0.182514305 | 0.3021677 | -0.6040166 | 0.54583261 | NA | NA | NA |
| ENSG000002 | 0.80998764 | 0.560099784 | 0.291126 | 1.92390848 | 0.05436606 | NA | NA | NA |
| ENSG000002 | 0.63754751 | -0.112497443 | 0.26606775 | -0.422815 | 0.67243022 | NA | NA | NA |
| ENSG000002 | 0.73309811 | -0.347757121 | 0.27972943 | -1.2431911 | 0.21379744 | NA | NA | NA |
| ENSG000002 | 0.34270765 | 0.258172734 | 0.21433185 | 1.20454674 | 0.22837833 | NA | SSX4 | 6759 |
| ENSG000002 | 0.8678023 | -0.271126522 | 0.29886542 | -0.907186 | 0.36430845 | NA | NA | NA |
| ENSG000002 | 0 | NA | NA | NA | NA | NA | NA | NA |
| ENSG000002 | 1.91101558 | -0.536630242 | 0.35921888 | -1.4938809 | 0.13520674 | NA | LOC374890 | 374890 |
| ENSG000002 | 0.58092888 | 0.226946341 | 0.26184598 | 0.86671692 | 0.38609713 | NA | NA | NA |
| ENSG000002 | 1.89880884 | -0.537288981 | 0.35911135 | -1.4961626 | 0.13461128 | NA | NA | NA |
| ENSG000002 | 0 | NA | NA | NA | NA | NA | NA | NA |
| ENSG000002 | 0.46870234 | 0.33514356 | 0.239033 | 1.4020807 | 0.16089115 | NA | BNIP3P30 | 106481694 |
| ENSG000002 | 0.40689225 | -0.283036689 | 0.21754773 | -1.3010326 | 0.19324729 | NA | NA | NA |
| ENSG000002 | 0 | NA | NA | NA | NA | NA | NA | NA |
| ENSG000002 | 2.16406017 | -0.062856395 | 0.3616215 | -0.1738182 | 0.86200835 | NA | NA | NA |
| ENSG000002 | 0 | NA | NA | NA | NA | NA | NA | NA |
| ENSG000002 | 1.39124391 | -0.274903686 | 0.34196402 | -0.8038965 | 0.42145673 | NA | NA | NA |
| ENSG000002 | 1.51769294 | -0.330247167 | 0.34227009 | -0.964873 | 0.33460842 | NA | NA | NA |
| ENSG000002 | 0.75992838 | -0.028323838 | 0.27990546 | -0.1011907 | 0.91939906 | NA | NA | NA |
| ENSG000002 | 0 | NA | NA | NA | NA | NA | NA | NA |
| ENSG000002 | 1.12245205 | -0.118465868 | 0.3210696 | -0.3689725 | 0.71214819 | NA | NA | NA |
| ENSG000002 | 1.1146275 | -0.408072454 | 0.31717208 | -1.2865964 | 0.198235 | NA | NA | NA |
| ENSG000002 | 0.24485602 | -0.012655878 | 0.17881722 | -0.0707755 | 0.94357643 | NA | NA | NA |

| | | | | | | | | | |
|---|---|---|---|---|---|---|---|---|---|
| ENSG000002 | 0.73580004 | -0.186985061 | 0.27927554 | -0.6695361 | 0.50315356 | NA | NA | NA | |
| ENSG000002 | 0 | NA | NA | NA | NA | NA | NA | NA | |
| ENSG000002 | 1.21600122 | -0.175821913 | 0.3243643 | -0.5420507 | 0.58778355 | NA | NA | NA | |
| ENSG000002 | 0 | NA | NA | NA | NA | NA | NA | NA | |
| ENSG000002 | 1.27149889 | -0.337771646 | 0.32710086 | -1.0326223 | 0.30178069 | NA | NA | NA | |
| ENSG000002 | 0.36212533 | -0.235926492 | 0.20089643 | -1.1743688 | 0.24024735 | NA | NA | NA | |
| ENSG000002 | 1.72382206 | -0.436865435 | 0.35057887 | -1.246126 | 0.21271815 | NA | NA | NA | |
| ENSG000002 | 0.12070844 | -0.088503695 | 0.12783931 | -0.6923042 | 0.48874631 | NA | OPN1MW | | 2652 |
| ENSG000002 | 1.95805285 | -0.175782213 | 0.35721297 | -0.4920936 | 0.62265318 | NA | NA | NA | |
| ENSG000002 | 0.2596576 | -0.193002035 | 0.1837913 | -1.0501152 | 0.29366517 | NA | NA | NA | |
| ENSG000002 | 0.83897464 | -0.092974912 | 0.28827292 | -0.3225239 | 0.74705582 | NA | NA | NA | |
| ENSG000002 | 2.14614608 | -0.370233358 | 0.36046936 | -1.0270869 | 0.30437954 | NA | NA | NA | |
| ENSG000002 | 1.68254537 | 0.223008734 | 0.35753376 | 0.62374175 | 0.5327972 | NA | NA | NA | |
| ENSG000002 | 1.70038753 | 0.069309723 | 0.33998574 | 0.20386068 | 0.83846237 | NA | NA | NA | |
| ENSG000002 | 1.1530742 | -0.279898271 | 0.3228241 | -0.8670303 | 0.38592542 | NA | NA | NA | |
| ENSG000002 | 1.82137246 | 0.271223214 | 0.35776318 | 0.75810825 | 0.44838618 | NA | NA | NA | |
| ENSG000002 | 1.70133135 | -0.058056265 | 0.35473909 | -0.1636591 | 0.86999955 | NA | NA | NA | |
| ENSG000002 | 1.23945885 | -0.451846523 | 0.32098982 | -1.4076662 | 0.15922994 | NA | NA | NA | |
| ENSG000002 | 2.05563974 | -0.216893233 | 0.3504696 | -0.6188646 | 0.53600555 | NA | NA | NA | |
| ENSG000002 | 1.2580136 | -0.355680482 | 0.332109 | -1.0709751 | 0.28418061 | NA | NA | NA | |
| ENSG000002 | 1.15071719 | -0.431334726 | 0.32182785 | -1.3402654 | 0.18015908 | NA | SCGB1C2 | | 653486 |
| ENSG000002 | 1.23708822 | -0.049480308 | 0.33242372 | -0.1488471 | 0.88167428 | NA | NA | NA | |
| ENSG000002 | 0 | NA | NA | NA | NA | NA | NA | NA | |
| ENSG000002 | 0.96821204 | 0.122680799 | 0.31609655 | 0.3881118 | 0.6979333 | NA | NA | NA | |
| ENSG000002 | 0.9827855 | -0.032151716 | 0.30610195 | -0.105036 | 0.91634727 | NA | NA | NA | |
| ENSG000002 | 0.85112135 | -0.11399948 | 0.29680331 | -0.384091 | 0.70091101 | NA | NA | NA | |
| ENSG000002 | 1.39189467 | -0.381849851 | 0.32978302 | -1.1578821 | 0.24691216 | NA | NA | NA | |
| ENSG000002 | 0.35022155 | 0.06940789 | 0.21379411 | 0.32464827 | 0.74544729 | NA | NA | NA | |
| ENSG000002 | 0.63984896 | -0.285743295 | 0.26802755 | -1.0660967 | 0.28637991 | NA | NA | NA | |

| | | | | | | | | |
|---|---|---|---|---|---|---|---|---|
| ENSG000002 | 1.50870517 | -0.178201353 | 0.33592909 | -0.5304731 | 0.59578394 | NA | NA | NA |
| ENSG000002 | 0 | NA | NA | NA | NA | NA | NA | NA |
| ENSG000002 | 0.41537361 | -0.286583533 | 0.21871501 | -1.3103057 | 0.19009243 | NA | NA | NA |
| ENSG000002 | 0.62408396 | 0.054009456 | 0.26457561 | 0.20413618 | 0.83824708 | NA | SSX2B | 727837 |
| ENSG000002 | 0.74291303 | -0.034519579 | 0.28502992 | -0.1211086 | 0.90360501 | NA | NA | NA |
| ENSG000002 | 0.72423759 | 0.136226018 | 0.27602641 | 0.4935253 | 0.62164147 | NA | NA | NA |
| ENSG000002 | 0 | NA | NA | NA | NA | NA | NA | NA |
| ENSG000002 | 0.74094832 | -0.192779168 | 0.28573214 | -0.6746849 | 0.49987596 | NA | NA | NA |
| ENSG000002 | 1.06117223 | -0.511487739 | 0.30715732 | -1.6652305 | 0.09586678 | NA | NA | NA |
| ENSG000002 | 1.39491373 | -0.572914614 | 0.33786881 | -1.6956718 | 0.08994805 | NA | NA | NA |
| ENSG000002 | 0 | NA | NA | NA | NA | NA | NA | NA |
| ENSG000002 | 0.34202251 | 0.257871122 | 0.21422266 | 1.20375278 | 0.22868515 | NA | NA | NA |
| ENSG000002 | 0.49264869 | -0.021364872 | 0.24621241 | -0.0867741 | 0.93085104 | NA | NA | NA |
| ENSG000002 | 1.66934432 | 0.454047908 | 0.343668 | 1.3211818 | 0.18644075 | NA | NA | NA |
| ENSG000002 | 0.23202362 | -0.008302493 | 0.17586207 | -0.0472103 | 0.96234565 | NA | NA | NA |
| ENSG000002 | 0.70725416 | 0.110312514 | 0.2649635 | 0.41633097 | 0.67716783 | NA | NA | NA |
| ENSG000002 | 1.06026088 | -0.33515468 | 0.30446995 | -1.1007808 | 0.27099206 | NA | NA | NA |
| ENSG000002 | 0.62522207 | -0.280659539 | 0.2668423 | -1.0517806 | 0.29290024 | NA | NA | NA |
| ENSG000002 | 0.26560726 | -0.177019426 | 0.17672274 | -1.0016788 | 0.31649875 | NA | NA | NA |
| ENSG000002 | 1.83510244 | 0.002703082 | 0.35929505 | 0.00752329 | 0.99399734 | NA | NA | NA |
| ENSG000002 | 1.32735312 | -0.358573398 | 0.3249457 | -1.1034871 | 0.26981568 | NA | NA | NA |
| ENSG000002 | 0.36391535 | 0.058449071 | 0.20749591 | 0.28168782 | 0.77818289 | NA | NA | NA |
| ENSG000002 | 0.23065526 | 0.167384503 | 0.17662711 | 0.94767164 | 0.34329664 | NA | NA | NA |
| ENSG000002 | 0.60287625 | 0.213311073 | 0.25519469 | 0.83587584 | 0.40322475 | NA | RAET1E-AS1 | 100652739 |
| ENSG000002 | 0 | NA | NA | NA | NA | NA | NA | NA |
| ENSG000002 | 1.63898047 | -0.549933347 | 0.34953821 | -1.5733139 | 0.11564614 | NA | NA | NA |
| ENSG000002 | 0.98893294 | 0.119051617 | 0.31132261 | 0.38240594 | 0.70216028 | NA | NA | NA |
| ENSG000002 | 0.5301324 | -0.3843196 | 0.24728844 | -1.5541349 | 0.12015224 | NA | NA | NA |
| ENSG000002 | 0 | NA | NA | NA | NA | NA | NA | NA |

| | | | | | | | | |
|---|---|---|---|---|---|---|---|---|
| ENSG000002 | 0.37148378 | 0.06737793 | 0.20875796 | 0.32275622 | 0.74687987 | NA | NA | NA |
| ENSG000002 | 0.35961481 | -0.099625193 | 0.21567743 | -0.4619176 | 0.64414043 | NA | NA | NA |
| ENSG000002 | 1.86339846 | -0.391506466 | 0.35907701 | -1.0903134 | 0.27557513 | NA | NA | NA |
| ENSG000002 | 0 | NA | NA | NA | NA | NA | NA | NA |
| ENSG000002 | 0 | NA | NA | NA | NA | NA | CTAG1A | 246100 |
| ENSG000002 | 0 | NA | NA | NA | NA | NA | WBP1LP6 | 106481797 |
| ENSG000002 | 1.13654676 | -0.397925026 | 0.31199719 | -1.2754122 | 0.20216338 | NA | NA | NA |
| ENSG000002 | 0.11205239 | 0.082118251 | 0.12671962 | 0.64803108 | 0.51696485 | NA | NA | NA |
| ENSG000002 | 0.86481516 | -0.118678381 | 0.29751956 | -0.3988927 | 0.68997227 | NA | COX6CP7 | 106480265 |
| ENSG000002 | 1.01884664 | -0.349130761 | 0.30975853 | -1.1271062 | 0.25969758 | NA | NA | NA |
| ENSG000002 | 1.41131001 | -0.368788171 | 0.32371478 | -1.139238 | 0.25460391 | NA | NA | NA |
| ENSG000002 | 0.24756242 | -0.187034346 | 0.18119817 | -1.0322088 | 0.30197431 | NA | NA | NA |
| ENSG000002 | 0 | NA | NA | NA | NA | NA | NA | NA |
| ENSG000002 | 0.61061849 | 0.057717611 | 0.2634075 | 0.21911908 | 0.82655728 | NA | BNIP3P33 | 106481695 |
| ENSG000002 | 0.61767784 | 0.056941233 | 0.25494708 | 0.22334531 | 0.82326677 | NA | NA | NA |
| ENSG000002 | 1.26937212 | -0.350081366 | 0.3323955 | -1.0532073 | 0.29224599 | NA | NA | NA |
| ENSG000002 | 0.12070844 | -0.088503695 | 0.12783931 | -0.6923042 | 0.48874631 | NA | HSFX2 | 100130086 |
| ENSG000002 | 0.94856858 | 0.118036465 | 0.30458884 | 0.38752722 | 0.69836594 | NA | NA | NA |
| ENSG000002 | 1.62810124 | -0.367956461 | 0.3437554 | -1.070402 | 0.28443841 | NA | LOC105372277 | 105372277 |
| ENSG000002 | 1.58748446 | -0.126855731 | 0.35335202 | -0.3590067 | 0.7195901 | NA | NA | NA |
| ENSG000002 | 1.58743396 | 0.008275249 | 0.34778787 | 0.02379396 | 0.98101696 | NA | NA | NA |
| ENSG000002 | 1.83487219 | -0.128367119 | 0.35988767 | -0.3566866 | 0.72132642 | NA | NA | NA |
| ENSG000002 | 0 | NA | NA | NA | NA | NA | NA | NA |
| ENSG000002 | 0.87719109 | -0.116929605 | 0.29817431 | -0.3921518 | 0.69494602 | NA | NA | NA |
| ENSG000002 | 1.00408391 | -0.29739137 | 0.29462096 | -1.0094033 | 0.31278125 | NA | NA | NA |
| ENSG000002 | 0 | NA | NA | NA | NA | NA | NA | NA |
| ENSG000002 | 1.85511489 | -0.254628275 | 0.35902946 | -0.7092128 | 0.47819245 | NA | NA | NA |
| ENSG000002 | 1.1930706 | -0.038295236 | 0.32136448 | -0.1191645 | 0.90514504 | NA | NA | NA |
| ENSG000002 | 0.35141577 | 0.074527565 | 0.21401461 | 0.34823588 | 0.72766304 | NA | NA | NA |

| | | | | | | | | | |
|---|---|---|---|---|---|---|---|---|---|
| ENSG000002 | 2.06516292 | 0.003184403 | 0.36161239 | 0.00880612 | 0.99297382 | NA | NA | NA | |
| ENSG000002 | 0 | NA | NA | NA | NA | NA | NA | NA | |
| ENSG000002 | 0.36690249 | -0.096999144 | 0.21683809 | -0.4473344 | 0.65463361 | NA | NA | NA | |
| ENSG000002 | 2.15229045 | -0.984270098 | 0.35309285 | -2.7875674 | 0.00531054 | NA | NA | NA | |
| ENSG000002 | 0.69887885 | 0.294045978 | 0.27747344 | 1.05972656 | 0.28926901 | NA | NA | NA | |
| ENSG000002 | 1.11018218 | 0.029474609 | 0.32188917 | 0.09156757 | 0.92704162 | NA | BNIP3P32 | | 106480283 |
| ENSG000002 | 1.57715185 | -0.089819248 | 0.33067622 | -0.2716229 | 0.78591195 | NA | NA | NA | |
| ENSG000002 | 0.70209433 | 0.134859374 | 0.27610772 | 0.48843028 | 0.6252451 | NA | NA | NA | |
| ENSG000002 | 1.50002738 | -0.192452605 | 0.34433845 | -0.5589054 | 0.57622632 | NA | NA | NA | |
| ENSG000002 | 1.74803183 | 0.06676494 | 0.35156243 | 0.1899092 | 0.84938028 | NA | NA | NA | |
| ENSG000002 | 0 | NA | NA | NA | NA | NA | ARHGEF18 | | 23370 |
| ENSG000002 | 0 | NA | NA | NA | NA | NA | NA | NA | |
| ENSG000002 | 1.00601425 | -0.360230169 | 0.31375627 | -1.148121 | 0.25091861 | NA | NA | NA | |
| ENSG000002 | 1.1032328 | -0.113696873 | 0.3204396 | -0.3548153 | 0.72272796 | NA | NA | NA | |
| ENSG000002 | 1.76567974 | 0.548162831 | 0.35516335 | 1.54341048 | 0.12273121 | NA | NA | NA | |
| ENSG000002 | 1.87100017 | -0.251882677 | 0.35650178 | -0.7065397 | 0.47985254 | NA | NA | NA | |
| ENSG000002 | 2.17209554 | -0.64566806 | 0.36432387 | -1.7722365 | 0.07635531 | NA | CSAG3 | | 389903 |
| ENSG000002 | 0.60810852 | 0.215175582 | 0.25567951 | 0.8415832 | 0.4000213 | NA | NA | NA | |
| ENSG000002 | 0.96853054 | -0.032347758 | 0.30712832 | -0.1053233 | 0.9161193 | NA | NA | NA | |
| ENSG000002 | 0.38747904 | -0.104908247 | 0.21987514 | -0.4771265 | 0.63327207 | NA | NA | NA | |
| ENSG000002 | 1.75390142 | -0.176716837 | 0.34988969 | -0.5050644 | 0.61351358 | NA | CT45A1 | | 541466 |
| ENSG000002 | 1.27515156 | -0.508192942 | 0.33246163 | -1.5285762 | 0.12636953 | NA | LOC79160 | | 79160 |
| ENSG000002 | 0.38494734 | -0.095409837 | 0.21124663 | -0.4516514 | 0.65152014 | NA | NA | NA | |
| ENSG000002 | 1.13477792 | -0.601040431 | 0.32140896 | -1.8700177 | 0.06148136 | NA | NA | NA | |
| ENSG000002 | 0.72992419 | -0.021564412 | 0.26869413 | -0.0802564 | 0.93603337 | NA | NA | NA | |
| ENSG000002 | 1.21184159 | 0.098929346 | 0.32946773 | 0.30027021 | 0.76397105 | NA | NA | NA | |
| ENSG000002 | 0 | NA | NA | NA | NA | NA | NA | NA | |
| ENSG000002 | 0.38884741 | -0.29745092 | 0.22235466 | -1.3377319 | 0.18098386 | NA | NA | NA | |
| ENSG000002 | 0.63621395 | -0.448723671 | 0.2628158 | -1.7073694 | 0.08775341 | NA | NA | NA | |

| | | | | | | | | |
|---|---|---|---|---|---|---|---|---|
| ENSG000002 | 0.11205239 | 0.082118251 | 0.12671962 | 0.64803108 | 0.51696485 | NA | NA | NA |
| ENSG000002 | 0 | NA | NA | NA | NA | NA | LOC100132304 | 100132304 |
| ENSG000002 | 0 | NA | NA | NA | NA | NA | NA | NA |
| ENSG000002 | 1.49348083 | -0.483176747 | 0.34367087 | -1.4059287 | 0.1597453 | NA | NA | NA |
| ENSG000002 | 1.48118977 | -0.465126632 | 0.34021571 | -1.3671521 | 0.17157762 | NA | NA | NA |
| ENSG000002 | 1.83521705 | -0.578417286 | 0.34855664 | -1.6594642 | 0.09702228 | NA | NA | NA |
| ENSG000002 | 1.11091939 | 0.027734448 | 0.31570842 | 0.0878483 | 0.92999725 | NA | NA | NA |
| ENSG000002 | 0 | NA | NA | NA | NA | NA | NA | NA |
| ENSG000002 | 0 | NA | NA | NA | NA | NA | NA | NA |
| ENSG000002 | 0 | NA | NA | NA | NA | NA | NA | NA |
| ENSG000002 | 1.36805133 | -0.252419869 | 0.33336878 | -0.7571791 | 0.44894258 | NA | NA | NA |
| ENSG000002 | 0.7731096 | -0.379060994 | 0.28942548 | -1.3097015 | 0.19029682 | NA | BNIP3P18 | 106481693 |
| ENSG000002 | 0.35961481 | -0.099625193 | 0.21567743 | -0.4619176 | 0.64414043 | NA | NA | NA |
| ENSG000002 | 1.39059156 | -0.239370327 | 0.32964325 | -0.7261496 | 0.46774704 | NA | NA | NA |
| ENSG000002 | 2.02445807 | -0.460239417 | 0.36345487 | -1.2662904 | 0.20540912 | NA | NA | NA |
| ENSG000002 | 0 | NA | NA | NA | NA | NA | NA | NA |
| ENSG000002 | 0.84512017 | -0.096634382 | 0.29040562 | -0.3327566 | 0.73931803 | NA | NA | NA |
| ENSG000002 | 1.97642169 | -0.444083401 | 0.36049816 | -1.2318604 | 0.21800125 | NA | NA | NA |
| ENSG000002 | 0.36556447 | -0.101943371 | 0.21659044 | -0.4706735 | 0.63787395 | NA | NA | NA |
| ENSG000002 | 1.33139308 | -0.246154413 | 0.33402786 | -0.7369278 | 0.46116626 | NA | NA | NA |
| ENSG000002 | 0 | NA | NA | NA | NA | NA | NA | NA |
| ENSG000002 | 0.54456342 | -0.359011254 | 0.24026449 | -1.4942335 | 0.1351146 | NA | NA | NA |
| ENSG000002 | 1.01700166 | -0.197891823 | 0.3098715 | -0.6386254 | 0.52306662 | NA | NA | NA |
| ENSG000002 | 1.64836091 | 0.354470302 | 0.35272684 | 1.00494282 | 0.31492439 | NA | NA | NA |
| ENSG000002 | 0.72985038 | -0.185024451 | 0.2788647 | -0.6634918 | 0.50701561 | NA | NA | NA |
| ENSG000002 | 1.31704536 | -0.053150148 | 0.18587552 | -0.2859449 | 0.77492036 | NA | NA | NA |
| ENSG000002 | 0 | NA | NA | NA | NA | NA | NA | NA |
| ENSG000002 | 1.28977845 | -0.511818077 | 0.33280871 | -1.5378747 | 0.12407927 | NA | NA | NA |
| ENSG000002 | 1.12586341 | -0.26722627 | 0.32349262 | -0.826066 | 0.40876663 | NA | NA | NA |

| | | | | | | | | | |
|---|---|---|---|---|---|---|---|---|---|
| ENSG000002 | 1.39241585 | -0.424892273 | 0.33852035 | -1.2551454 | 0.20942597 | NA | NA | NA | |
| ENSG000002 | 1.78971079 | -0.773474294 | 0.35437006 | -2.182674 | 0.02905983 | NA | NA | NA | |
| ENSG000002 | 2.064565 | -0.335329172 | 0.35872897 | -0.9347703 | 0.34990671 | NA | LINC00528 | | 200298 |
| ENSG000002 | 2.12211574 | -0.125182385 | 0.36316382 | -0.3446995 | 0.7303203 | NA | NA | NA | |
| ENSG000002 | 0 | NA | NA | NA | NA | NA | NA | NA | |
| ENSG000002 | 0 | NA | NA | NA | NA | NA | NA | NA | |
| ENSG000002 | 2.16784634 | 0.159751934 | 0.36115821 | 0.44233227 | 0.65824878 | NA | NA | NA | |
| ENSG000002 | 1.67537813 | -0.550991029 | 0.34959534 | -1.5760823 | 0.11500684 | NA | NA | NA | |
| ENSG000002 | 0.51225778 | -0.204952203 | 0.24951191 | -0.8214125 | 0.41141135 | NA | NA | NA | |
| ENSG000002 | 0 | NA | NA | NA | NA | NA | NA | NA | |
| ENSG000002 | 0.84264211 | 0.045261084 | 0.28988579 | 0.15613419 | 0.87592725 | NA | NA | NA | |
| ENSG000002 | 0.11865494 | 0.083440866 | 0.12751478 | 0.65436232 | 0.5128784 | NA | NA | NA | |
| ENSG000002 | 1.16907001 | -0.440005341 | 0.32456851 | -1.3556625 | 0.17520659 | NA | NA | NA | |
| ENSG000002 | 0.63387812 | -0.454271587 | 0.26413408 | -1.7198522 | 0.08545931 | NA | NA | NA | |
| ENSG000002 | 0 | NA | NA | NA | NA | NA | NA | NA | |
| ENSG000002 | 0.65427998 | -0.290650429 | 0.26916107 | -1.0798383 | 0.28021419 | NA | NA | NA | |
| ENSG000002 | 0.38019136 | -0.10756138 | 0.21876967 | -0.491665 | 0.62295621 | NA | NA | NA | |
| ENSG000002 | 0.70305967 | 0.129084386 | 0.27453658 | 0.47019011 | 0.6382192 | NA | NA | NA | |
| ENSG000002 | 1.20797824 | -0.040377975 | 0.3294209 | -0.1225726 | 0.90244556 | NA | NA | NA | |
| ENSG000002 | 1.50700049 | -0.348855324 | 0.34830803 | -1.0015713 | 0.3165507 | NA | NA | NA | |
| ENSG000002 | 0.49082195 | -0.014647497 | 0.23849846 | -0.0614155 | 0.95102832 | NA | NA | NA | |
| ENSG000002 | 1.01930366 | -0.364009021 | 0.31573733 | -1.1528856 | 0.24895734 | NA | LINC01233 | | 100128139 |
| ENSG000002 | 0.47895489 | -0.016192723 | 0.24471014 | -0.066171 | 0.94724166 | NA | NA | NA | |
| ENSG000002 | 0.78159097 | -0.381686531 | 0.28991249 | -1.3165577 | 0.18798691 | NA | NA | NA | |
| ENSG000002 | 1.38991035 | -0.567294384 | 0.33781713 | -1.6792943 | 0.0930947 | NA | NA | NA | |
| ENSG000002 | 1.1958246 | 0.092434516 | 0.32145728 | 0.28754837 | 0.77369247 | NA | NA | NA | |
| ENSG000002 | 1.60450372 | -0.346925203 | 0.33898747 | -1.023416 | 0.30611119 | NA | NA | NA | |
| ENSG000002 | 0.11865494 | 0.083440866 | 0.12751478 | 0.65436232 | 0.5128784 | NA | NA | NA | |
| ENSG000002 | 1.21577501 | -0.031644624 | 0.32372603 | -0.0977513 | 0.92212982 | NA | NXF2 | | 56001 |

| | | | | | | | | |
|---|---|---|---|---|---|---|---|---|
| ENSG000002 | 0.3854038 | -0.110128691 | 0.21952326 | -0.5016721 | 0.61589821 | NA | NA | NA |
| ENSG000002 | 1.12338514 | -0.115293234 | 0.31813306 | -0.3624057 | 0.71704888 | NA | NA | NA |
| ENSG000002 | 1.31974989 | 0.022741016 | 0.33846479 | 0.06718872 | 0.94643147 | NA | NA | NA |
| ENSG000002 | 0 | NA | NA | NA | NA | NA | OPN1MW3 | 101060233 |
| ENSG000002 | 0.2596576 | -0.193002035 | 0.1837913 | -1.0501152 | 0.29366517 | NA | NXF2B | 728343 |
| ENSG000002 | 2.03579487 | -0.58945328 | 0.3611372 | -1.6322142 | 0.10263437 | NA | NA | NA |
| ENSG000002 | 1.26618285 | -0.32180747 | 0.32536186 | -0.9890756 | 0.32262617 | NA | NA | NA |
| ENSG000002 | 0 | NA | NA | NA | NA | NA | NA | NA |
| ENSG000002 | 0.82877416 | 0.203407181 | 0.28803679 | 0.70618473 | 0.48007327 | NA | NA | NA |
| ENSG000002 | 0.92512534 | -0.60329841 | 0.29304014 | -2.0587569 | 0.03951753 | NA | NA | NA |
| ENSG000002 | 0 | NA | NA | NA | NA | NA | NA | NA |
| ENSG000002 | 1.91819721 | -0.595690969 | 0.34538047 | -1.7247384 | 0.08457462 | NA | NA | NA |
| ENSG000002 | 0 | NA | NA | NA | NA | NA | NA | NA |
| ENSG000002 | 0 | NA | NA | NA | NA | NA | NA | NA |
| ENSG000002 | 1.46379334 | -0.189329346 | 0.34265951 | -0.5525291 | 0.58058591 | NA | NA | NA |
| ENSG000002 | 2.07165179 | -0.128876954 | 0.36743475 | -0.3507479 | 0.72577751 | NA | NA | NA |
| ENSG000002 | 0 | NA | NA | NA | NA | NA | NA | NA |
| ENSG000002 | 0.97119864 | -0.036229316 | 0.30556973 | -0.1185632 | 0.90562144 | NA | NA | NA |
| ENSG000002 | 0.86336089 | -0.100933932 | 0.28969003 | -0.3484205 | 0.72752444 | NA | NA | NA |
| ENSG000002 | 0.91664398 | -0.637643136 | 0.29917608 | -2.1313306 | 0.03306191 | NA | NA | NA |
| ENSG000002 | 0.60914194 | -0.10149904 | 0.25613314 | -0.3962745 | 0.69190251 | NA | NA | NA |
| ENSG000002 | 0.51322524 | -0.027081322 | 0.24101349 | -0.1123643 | 0.91053453 | NA | NA | NA |
| ENSG000002 | 0.85066892 | 0.192756384 | 0.28685996 | 0.6719529 | 0.50161368 | NA | NA | NA |
| ENSG000002 | 1.83746043 | -0.31881254 | 0.34357227 | -0.9279344 | 0.35344157 | NA | NA | NA |
| ENSG000002 | 0.13280363 | -0.093394593 | 0.13115644 | -0.7120855 | 0.47641185 | NA | NA | NA |
| ENSG000002 | 1.85992397 | -0.137307278 | 0.36150637 | -0.3798198 | 0.70407918 | NA | NA | NA |
| ENSG000002 | 1.28881099 | -0.678857587 | 0.33109616 | -2.0503336 | 0.04033189 | NA | NA | NA |
| ENSG000002 | 0.52165104 | -0.403921832 | 0.25252134 | -1.5995552 | 0.10969729 | NA | NA | NA |
| ENSG000002 | 0 | NA | NA | NA | NA | NA | NA | NA |

| | | | | | | | | |
|---|---|---|---|---|---|---|---|---|
| ENSG000002 | 1.36022478 | -0.1024738 | 0.32805025 | -0.3123723 | 0.75475762 | NA | NA | NA |
| ENSG000002 | 2.04642744 | 0.127049563 | 0.36295297 | 0.35004415 | 0.72630557 | NA | NA | NA |
| ENSG000002 | 0 | NA | NA | NA | NA | NA | NA | NA |
| ENSG000002 | 0.50016259 | -0.200428691 | 0.24824437 | -0.8073846 | 0.41944491 | NA | NA | NA |
| ENSG000002 | 0.11131518 | 0.081951103 | 0.12663804 | 0.64712866 | 0.51754868 | NA | NA | NA |
| ENSG000002 | 2.02324267 | -0.570083438 | 0.35798665 | -1.5924712 | 0.11127887 | NA | NA | NA |
| ENSG000002 | 1.58694559 | 0.021224991 | 0.34193263 | 0.0620736 | 0.95050422 | NA | NA | NA |
| ENSG000002 | 1.73448885 | -0.312749103 | 0.34967034 | -0.8944113 | 0.37110186 | NA | NA | NA |
| ENSG000002 | 1.12589566 | -0.270975439 | 0.32347778 | -0.8376941 | 0.40220251 | NA | NA | NA |
| ENSG000002 | 1.13972978 | -0.566869226 | 0.31589767 | -1.7944711 | 0.07273801 | NA | NA | NA |
| ENSG000002 | 1.73800458 | -0.457572124 | 0.3526286 | -1.2976036 | 0.1944236 | NA | NA | NA |
| ENSG000002 | 0.81300649 | 0.360892625 | 0.28909106 | 1.24837006 | 0.21189557 | NA | NA | NA |
| ENSG000002 | 1.60413261 | -0.252332927 | 0.34710987 | -0.7269541 | 0.4672541 | NA | NA | NA |
| ENSG000002 | 1.84732079 | -0.115653458 | 0.35690273 | -0.3240476 | 0.74590203 | NA | NA | NA |
| ENSG000002 | 0 | NA | NA | NA | NA | NA | NA | NA |
| ENSG000002 | 0 | NA | NA | NA | NA | NA | WBP1LP7 | 106479030 |
| ENSG000002 | 1.03305145 | -0.180117028 | 0.30322729 | -0.5940001 | 0.55251207 | NA | NA | NA |
| ENSG000002 | 0.23868015 | 0.154625155 | 0.16997278 | 0.90970538 | 0.36297791 | NA | KLK9 | 284366 |
| ENSG000002 | 0.59072708 | 0.064826349 | 0.26164045 | 0.24776883 | 0.80431327 | NA | NA | NA |
| ENSG000002 | 0 | NA | NA | NA | NA | NA | NA | NA |
| ENSG000002 | 0.56744359 | 0.212023309 | 0.25327123 | 0.83713935 | 0.40251425 | NA | NA | NA |
| ENSG000002 | 0.75213216 | -0.027056401 | 0.28563056 | -0.0947252 | 0.92453313 | NA | NA | NA |
| ENSG000002 | 1.61071533 | -0.265956235 | 0.35180198 | -0.7559828 | 0.44965953 | NA | NA | NA |
| ENSG000002 | 0.99354849 | -0.198387564 | 0.31363452 | -0.6325438 | 0.52703161 | NA | NA | NA |
| ENSG000002 | 1.69509811 | 0.204311096 | 0.34558163 | 0.59120937 | 0.55438015 | NA | NA | NA |
| ENSG000002 | 1.37428511 | -0.560900371 | 0.33613803 | -1.6686608 | 0.09518461 | NA | NA | NA |
| ENSG000002 | 1.01429973 | -0.367175034 | 0.31550396 | -1.1637731 | 0.24451596 | NA | NA | NA |
| ENSG000002 | 0 | NA | NA | NA | NA | NA | MIR1470 | 100302127 |
| ENSG000002 | 0.78979 | -0.568364433 | 0.28747989 | -1.9770581 | 0.04803506 | NA | NA | NA |

| | | | | | | | | | |
|---|---|---|---|---|---|---|---|---|---|
| ENSG000002 | 1.23877508 | -0.352729432 | 0.333643 | -1.0572061 | 0.29041751 | NA | NA | NA | |
| ENSG000002 | 1.45141583 | -0.306949141 | 0.33584877 | -0.9139505 | 0.36074287 | NA | SSX4B | | 548313 |
| ENSG000002 | 1.1027546 | -0.270075398 | 0.32272512 | -0.8368589 | 0.40267187 | NA | NA | NA | |
| ENSG000002 | 0.12070844 | -0.088503695 | 0.12783931 | -0.6923042 | 0.48874631 | NA | NA | NA | |
| ENSG000002 | 1.1377839 | -0.120294899 | 0.32306886 | -0.3723506 | 0.70963179 | NA | NA | NA | |
| ENSG000002 | 1.27378265 | -0.36170194 | 0.33446752 | -1.0814262 | 0.27950758 | NA | NA | NA | |
| ENSG000002 | 0 | NA | NA | NA | NA | NA | NA | NA | |
| ENSG000002 | 1.34414115 | -0.247467045 | 0.33509881 | -0.7384898 | 0.46021688 | NA | NA | NA | |
| ENSG000002 | 1.50741866 | -0.196571959 | 0.34340825 | -0.5724148 | 0.567041 | NA | NA | NA | |
| ENSG000002 | 0.38133351 | -0.102466351 | 0.2189713 | -0.4679442 | 0.63982449 | NA | NA | NA | |
| ENSG000002 | 1.5368582 | -0.488003986 | 0.34418674 | -1.4178466 | 0.15623556 | NA | NA | NA | |
| ENSG000002 | 1.89464818 | -0.363456 | 0.35225469 | -1.0317989 | 0.30216634 | NA | NA | NA | |
| ENSG000002 | 0.62023988 | -0.112292793 | 0.27094277 | -0.4144521 | 0.67854306 | NA | NA | NA | |
| ENSG000002 | 0.59464496 | 0.377502009 | 0.25000244 | 1.50999329 | 0.13104514 | NA | NA | NA | |
| ENSG000002 | 0.85679081 | -0.111816085 | 0.29710779 | -0.3763485 | 0.7066578 | NA | NA | NA | |
| ENSG000002 | 1.19827198 | 0.094653604 | 0.32065671 | 0.29518673 | 0.76785124 | NA | NA | NA | |
| ENSG000002 | 1.27338175 | -0.050825052 | 0.32112615 | -0.1582713 | 0.87424302 | NA | NA | NA | |
| ENSG000002 | 1.64395014 | -0.124729669 | 0.34782168 | -0.3586023 | 0.7198926 | NA | NA | NA | |
| ENSG000002 | 1.16292448 | -0.438326075 | 0.32438726 | -1.3512432 | 0.17661754 | NA | NA | NA | |
| ENSG000002 | 0 | NA | NA | NA | NA | NA | COMMD3-BMI1 | | 100532731 |
| ENSG000002 | 1.67548631 | -0.238153003 | 0.34039676 | -0.6996336 | 0.48415617 | NA | NA | NA | |
| ENSG000002 | 1.15429179 | 0.573733503 | 0.32353296 | 1.77333868 | 0.0761726 | NA | NA | NA | |
| ENSG000002 | 0 | NA | NA | NA | NA | NA | RMRP | | 6023 |
| ENSG000002 | 1.88143415 | -0.131732774 | 0.35985023 | -0.3660767 | 0.71430787 | NA | NA | NA | |
| ENSG000002 | 1.39124391 | -0.274903686 | 0.34196402 | -0.8038965 | 0.42145673 | NA | NA | NA | |
| ENSG000002 | 0.11934008 | 0.083564698 | 0.12760091 | 0.65489106 | 0.5125379 | NA | NA | NA | |
| ENSG000002 | 0.62684242 | -0.109923117 | 0.27148862 | -0.4048903 | 0.68555813 | NA | NA | NA | |
| ENSG000002 | 1.53577663 | -0.4799736 | 0.34443932 | -1.3934925 | 0.16347092 | NA | NA | NA | |
| ENSG000002 | 0.74043787 | -0.345110653 | 0.28031763 | -1.2311414 | 0.21826997 | NA | NA | NA | |

| | | | | | | | | |
|---|---|---|---|---|---|---|---|---|
| ENSG000002 | 1.06974558 | 0.025566821 | 0.3026341 | 0.08448097 | 0.93267403 | NA | NA | NA |
| ENSG000002 | 1.2460266 | -0.044699362 | 0.32458299 | -0.1377132 | 0.89046709 | NA | NA | NA |
| ENSG000002 | 0 | NA | NA | NA | NA | NA | C7orf55-LUC7L2 | 100996928 |
| ENSG000002 | 0 | NA | NA | NA | NA | NA | NA | NA |
| ENSG000002 | 1.78525712 | 0.418380649 | 0.35774595 | 1.16949095 | 0.24220589 | NA | NA | NA |
| ENSG000002 | 1.82900898 | 0.00417155 | 0.35655444 | 0.01169962 | 0.99066527 | NA | NA | NA |
| ENSG000002 | 2.05129285 | 0.007875176 | 0.36259347 | 0.02171902 | 0.98267209 | NA | NA | NA |
| ENSG000002 | 0.84260965 | 0.045668382 | 0.28815332 | 0.1584864 | 0.87407354 | NA | NA | NA |
| ENSG000002 | 1.7688048 | 0.064585698 | 0.34622274 | 0.18654378 | 0.85201835 | NA | NA | NA |
| ENSG000002 | 0.12070844 | -0.088503695 | 0.12783931 | -0.6923042 | 0.48874631 | NA | NA | NA |
| ENSG000002 | 1.41592163 | -0.38914352 | 0.33034672 | -1.1779851 | 0.23880254 | NA | NA | NA |
| ENSG000002 | 1.72394021 | -0.32969541 | 0.35518575 | -0.9282338 | 0.35328632 | NA | NA | NA |
| ENSG000002 | 0 | NA | NA | NA | NA | NA | NA | NA |
| ENSG000002 | 1.20331263 | -0.169960277 | 0.32227968 | -0.5273689 | 0.59793744 | NA | NA | NA |
| ENSG000002 | 1.20773285 | -0.407489467 | 0.30970568 | -1.3157313 | 0.18826424 | NA | NA | NA |
| ENSG000002 | 0.82954308 | 0.186001077 | 0.28563285 | 0.65118938 | 0.51492424 | NA | C8orf44-SGK3 | 100533105 |
| ENSG000002 | 0.6044405 | 0.057164517 | 0.25530853 | 0.22390367 | 0.82283226 | NA | NA | NA |
| ENSG000002 | 1.13255185 | -0.106020367 | 0.30940887 | -0.3426546 | 0.73185833 | NA | NA | NA |
| ENSG000002 | 0 | NA | NA | NA | NA | NA | NA | NA |
| ENSG000002 | 1.89482444 | -0.390089318 | 0.35836853 | -1.0885144 | 0.27636807 | NA | NA | NA |
| ENSG000002 | 0.99881301 | -0.200372067 | 0.31383792 | -0.6384572 | 0.52317613 | NA | NA | NA |
| ENSG000002 | 0.62701656 | -0.28174368 | 0.26697332 | -1.0553252 | 0.29127661 | NA | NA | NA |
| ENSG000002 | 2.15619282 | -0.054099898 | 0.36163651 | -0.1495974 | 0.88108222 | NA | NA | NA |
| ENSG000002 | 0 | NA | NA | NA | NA | NA | SCARNA2 | 677766 |
| ENSG000002 | 1.37047148 | -0.254301541 | 0.33567148 | -0.7575906 | 0.44869613 | NA | NA | NA |
| ENSG000002 | 0 | NA | NA | NA | NA | NA | NA | NA |
| ENSG000002 | 0.85884431 | -0.256667227 | 0.290884 | -0.8823697 | 0.37757692 | NA | NA | NA |
| ENSG000002 | 1.70150603 | -0.193019225 | 0.35715742 | -0.5404318 | 0.58889929 | NA | NA | NA |
| ENSG000002 | 0.97399093 | -0.034019229 | 0.30569514 | -0.1112848 | 0.91139049 | NA | NA | NA |

| | | | | | | | | | |
|---|---|---|---|---|---|---|---|---|---|
| ENSG000002 | 1.60481775 | -0.259545969 | 0.34993878 | -0.7416896 | 0.45827539 | NA | NA | NA | |
| ENSG000002 | 1.58828115 | -0.115138367 | 0.34687579 | -0.3319297 | 0.73994236 | NA | NA | NA | |
| ENSG000002 | 0.74530094 | -0.035008937 | 0.28517589 | -0.1227626 | 0.90229508 | NA | NA | NA | |
| ENSG000002 | 0 | NA | NA | NA | NA | NA | NA | NA | |
| ENSG000002 | 0 | NA | NA | NA | NA | NA | RNU11 | | 26824 |
| ENSG000002 | 1.6176002 | -0.12055216 | 0.34876513 | -0.3456543 | 0.72960254 | NA | NA | NA | |
| ENSG000002 | 1.140098 | -0.276591795 | 0.32242116 | -0.8578587 | 0.3909705 | NA | NA | NA | |
| ENSG000002 | 1.41464151 | -0.567623775 | 0.33568601 | -1.6909366 | 0.09084891 | NA | NA | NA | |
| ENSG000002 | 0.72233649 | -0.027848463 | 0.28368037 | -0.0981685 | 0.92179853 | NA | NA | NA | |
| ENSG000002 | 0 | NA | NA | NA | NA | NA | NA | NA | |
| ENSG000002 | 0.75213216 | -0.027056401 | 0.28563056 | -0.0947252 | 0.92453313 | NA | NA | NA | |
| ENSG000002 | 0 | NA | NA | NA | NA | NA | NA | NA | |
| ENSG000002 | 1.71428691 | -0.180888391 | 0.35409225 | -0.510851 | 0.60945537 | NA | NA | NA | |
| ENSG000002 | 0 | NA | NA | NA | NA | NA | NA | NA | |
| ENSG000002 | 0.26560726 | -0.177019426 | 0.17672274 | -1.0016788 | 0.31649875 | NA | MINOS1-NBL1 | | 100532736 |
| ENSG000002 | 2.10427592 | -0.378163717 | 0.36563901 | -1.0342543 | 0.30101729 | NA | NA | NA | |
| ENSG000002 | 0.58417661 | 0.05870191 | 0.25337197 | 0.23168273 | 0.81678444 | NA | TERC | | 7012 |
| ENSG000002 | 1.75010886 | -0.288454681 | 0.34472592 | -0.8367653 | 0.40272451 | NA | NA | NA | |
| ENSG000002 | 2.0655919 | -0.412255881 | 0.35353828 | -1.1660856 | 0.24357985 | NA | NA | NA | |
| ENSG000002 | 1.74102206 | -0.740119654 | 0.35038342 | -2.1123136 | 0.03465956 | NA | NA | NA | |
| ENSG000002 | 0 | NA | NA | NA | NA | NA | BIVM-ERCC5 | | 100533467 |
| ENSG000002 | 2.03550394 | -0.445027926 | 0.36129034 | -1.2317737 | 0.21803364 | NA | NA | NA | |
| ENSG000002 | 0 | NA | NA | NA | NA | NA | NA | NA | |
| ENSG000002 | 0 | NA | NA | NA | NA | NA | NA | NA | |
| ENSG000002 | 1.23040135 | -0.048563667 | 0.33359206 | -0.145578 | 0.88425453 | NA | NA | NA | |
| ENSG000002 | 1.31237842 | 0.031572028 | 0.3333602 | 0.09470845 | 0.9245464 | NA | BNIP3P31 | | 106480282 |
| ENSG000002 | 0 | NA | NA | NA | NA | NA | NA | NA | |
| ENSG000002 | 0.86981908 | -0.120659197 | 0.30286715 | -0.3983898 | 0.69034284 | NA | NA | NA | |
| ENSG000002 | 1.22277982 | -0.187405241 | 0.3303356 | -0.5673177 | 0.57049834 | NA | NA | NA | |

| | | | | | | | | | |
|---|---|---|---|---|---|---|---|---|---|
| ENSG000002 | 0.36690249 | -0.096999144 | 0.21683809 | -0.4473344 | 0.65463361 | NA | NA | NA | |
| ENSG000002 | 1.64022868 | -0.395305296 | 0.34920247 | -1.1320232 | 0.25762468 | NA | NA | NA | |
| ENSG000002 | 1.5815348 | -0.123950171 | 0.35224687 | -0.3518844 | 0.72492498 | NA | NA | NA | |
| ENSG000002 | 2.02378345 | -0.595350998 | 0.3616042 | -1.6464162 | 0.09967811 | NA | NA | NA | |
| ENSG000002 | 0.12685397 | -0.090215768 | 0.12901919 | -0.699243 | 0.48440017 | NA | NA | NA | |
| ENSG000002 | 0.84960918 | 0.041275706 | 0.29474918 | 0.14003671 | 0.88863098 | NA | NA | NA | |
| ENSG000002 | 0 | NA | NA | NA | NA | NA | NA | NA | |
| ENSG000002 | 0 | NA | NA | NA | NA | NA | NA | NA | |
| ENSG000002 | 1.09000653 | -0.25629258 | 0.31628782 | -0.8103144 | 0.4177595 | NA | NA | NA | |
| ENSG000002 | 1.38532459 | -0.435904782 | 0.34254622 | -1.272543 | 0.20318027 | NA | NA | NA | |
| ENSG000002 | 1.6012 | 0.014441699 | 0.3543121 | 0.04075982 | 0.96748737 | NA | NA | NA | |
| ENSG000002 | 1.46990909 | -0.039874703 | 0.34091075 | -0.1169652 | 0.90688761 | NA | NA | NA | |
| ENSG000002 | 1.40945455 | 0.377969242 | 0.34061989 | 1.10965112 | 0.26714939 | NA | NA | NA | |
| ENSG000002 | 1.83891515 | 0.13184778 | 0.34701664 | 0.37994656 | 0.70398508 | NA | NA | NA | |
| ENSG000002 | 2.12546707 | -0.252056265 | 0.36390124 | -0.6926502 | 0.4885291 | NA | NA | NA | |
| ENSG000002 | 1.71007576 | 0.070222085 | 0.34546664 | 0.20326734 | 0.83892608 | NA | NA | NA | |
| ENSG000002 | 1.54875283 | -0.460781234 | 0.34001242 | -1.3551894 | 0.17535721 | NA | NA | NA | |
| ENSG000002 | 0.90551233 | -0.110719234 | 0.29373998 | -0.3769294 | 0.70622607 | NA | NA | NA | |
| ENSG000002 | 0.25260017 | -0.015978705 | 0.18049464 | -0.0885273 | 0.92945758 | NA | NA | NA | |
| ENSG000002 | 1.25301022 | -0.345213867 | 0.33059302 | -1.0442261 | 0.29638078 | NA | NA | NA | |
| ENSG000002 | 0.74410726 | -0.031277165 | 0.29043452 | -0.1076909 | 0.91424086 | NA | NA | NA | |
| ENSG000002 | 0.87989748 | -0.286905324 | 0.30426178 | -0.9429555 | 0.34570366 | NA | NA | NA | |
| ENSG000002 | 1.2543636 | -0.04995535 | 0.32917739 | -0.1517581 | 0.8793777 | NA | NA | NA | |
| ENSG000002 | 0 | NA | NA | NA | NA | NA | NA | NA | |
| ENSG000002 | 0.71598189 | -0.026426461 | 0.26765551 | -0.0987331 | 0.92135018 | NA | NA | NA | |
| ENSG000002 | 1.26809358 | -0.334315519 | 0.32452712 | -1.030162 | 0.30293395 | NA | NA | NA | |
| ENSG000002 | 0.60558265 | 0.061136384 | 0.2538335 | 0.24085231 | 0.80966959 | NA | NA | NA | |
| ENSG000002 | 1.19235458 | 0.096948725 | 0.32512227 | 0.29819159 | 0.76555694 | NA | NA | NA | |
| ENSG000002 | 0.82961171 | -0.100242807 | 0.28771855 | -0.3484058 | 0.72753546 | NA | BNIP3P9 | | 106480280 |

| | | | | | | | | |
|---|---|---|---|---|---|---|---|---|
| ENSG000002 | 1.69932371 | -0.399792482 | 0.34010414 | -1.1755002 | 0.23979466 | NA | NA | NA |
| ENSG000002 | 1.96529451 | -0.296344223 | 0.35859478 | -0.8264042 | 0.40857485 | NA | NA | NA |
| ENSG000002 | 1.84127885 | -0.12847556 | 0.36135937 | -0.3555341 | 0.72218954 | NA | NA | NA |
| ENSG000002 | 0.74342403 | 0.124923428 | 0.27740429 | 0.45032983 | 0.65247263 | NA | NA | NA |
| ENSG000002 | 1.92590203 | -0.663998138 | 0.35737953 | -1.8579635 | 0.06317419 | NA | NA | NA |
| ENSG000002 | 0.11131518 | 0.081951103 | 0.12663804 | 0.64712866 | 0.51754868 | NA | NA | NA |
| ENSG000002 | 1.86285712 | -0.398847061 | 0.36057028 | -1.1061562 | 0.2686589 | NA | NA | NA |
| ENSG000002 | 0 | NA | NA | NA | NA | NA | NA | NA |
| ENSG000002 | 0.80285266 | -0.364894835 | 0.28429052 | -1.283528 | 0.19930716 | NA | NA | NA |
| ENSG000002 | 0.23890637 | -0.010420674 | 0.17747379 | -0.0587167 | 0.95317776 | NA | NA | NA |
| ENSG000002 | 0.47821768 | -0.016471461 | 0.24462483 | -0.0673336 | 0.94631616 | NA | NA | NA |
| ENSG000002 | 1.90199398 | -0.519850222 | 0.35816547 | -1.4514247 | 0.14666162 | NA | NA | NA |
| ENSG000002 | 1.23245486 | -0.194334232 | 0.33058291 | -0.5878532 | 0.55663081 | NA | NA | NA |
| ENSG000002 | 2.12328577 | -0.390320648 | 0.36572025 | -1.0672656 | 0.28585191 | NA | NA | NA |
| ENSG000002 | 1.26374151 | -0.477320073 | 0.32629738 | -1.4628376 | 0.14351182 | NA | NA | NA |
| ENSG000002 | 1.7570907 | -0.200388452 | 0.35862498 | -0.5587688 | 0.57631952 | NA | NA | NA |
| ENSG000002 | 0.58958493 | 0.063416451 | 0.26802881 | 0.23660311 | 0.8129647 | NA | NA | NA |
| ENSG000002 | 1.81267933 | 0.008664178 | 0.35808369 | 0.02419596 | 0.9806963 | NA | NA | NA |
| ENSG000002 | 0 | NA | NA | NA | NA | NA | NA | NA |
| ENSG000002 | 2.05613473 | 0.132031705 | 0.36784388 | 0.35893408 | 0.71964441 | NA | NA | NA |
| ENSG000002 | 0.77860225 | -0.362322636 | 0.28273614 | -1.2814868 | 0.20002273 | NA | NA | NA |
| ENSG000002 | 1.90662871 | -0.24854402 | 0.35471144 | -0.7006935 | 0.48349428 | NA | NA | NA |
| ENSG000002 | 1.15284608 | -0.276016838 | 0.32283135 | -0.8549877 | 0.39255795 | NA | MREGP1 | 106480304 |
| ENSG000002 | 0 | NA | NA | NA | NA | NA | NA | NA |
| ENSG000002 | 1.23854887 | -0.184566991 | 0.32928293 | -0.5605119 | 0.57513035 | NA | NA | NA |
| ENSG000002 | 0 | NA | NA | NA | NA | NA | NA | NA |
| ENSG000002 | 1.79967224 | 0.133128368 | 0.35446818 | 0.37557213 | 0.70723501 | NA | NA | NA |
| ENSG000002 | 0.73425753 | 0.129277406 | 0.27843923 | 0.46429307 | 0.64243779 | NA | NA | NA |
| ENSG000002 | 0.73086992 | -0.031865141 | 0.2896153 | -0.1100258 | 0.91238895 | NA | NA | NA |

| | | | | | | | | |
|---|---|---|---|---|---|---|---|---|
| ENSG000002 | 0.50593756 | -0.029751655 | 0.24018594 | -0.1238693 | 0.90141879 | NA | NA | NA |
| ENSG000002 | 0.73309811 | -0.347757121 | 0.27972943 | -1.2431911 | 0.21379744 | NA | NA | NA |
| ENSG000002 | 2.12339574 | -0.516542782 | 0.36588678 | -1.4117558 | 0.15802188 | NA | DPPA3P3 | 106481963 |
| ENSG000002 | 0 | NA | NA | NA | NA | NA | NA | NA |
| ENSG000002 | 1.37442554 | -0.254383507 | 0.33575764 | -0.7576403 | 0.44866636 | NA | NA | NA |
| ENSG000002 | 0.95947167 | 0.284029678 | 0.31210451 | 0.9100467 | 0.36279788 | NA | NA | NA |
| ENSG000002 | 1.92291881 | -0.816674735 | 0.35580043 | -2.2953169 | 0.02171497 | NA | NA | NA |
| ENSG000002 | 1.71238364 | -1.043369817 | 0.34582616 | -3.0170356 | 0.0025526 | NA | NA | NA |
| ENSG000002 | 0.11205239 | 0.082118251 | 0.12671962 | 0.64803108 | 0.51696485 | NA | AZU1P1 | 106480244 |
| ENSG000002 | 1.14476753 | -0.438324541 | 0.3238073 | -1.3536586 | 0.17584532 | NA | NA | NA |
| ENSG000002 | 0.85473964 | -0.391529356 | 0.2852255 | -1.3727011 | 0.16984529 | NA | NA | NA |
| ENSG000002 | 0.12685397 | -0.090215768 | 0.12901919 | -0.699243 | 0.48440017 | NA | NA | NA |
| ENSG000002 | 0 | NA | NA | NA | NA | NA | NA | NA |
| ENSG000002 | 0.50127037 | -0.342056569 | 0.23551462 | -1.4523793 | 0.14639616 | NA | NA | NA |
| ENSG000002 | 0 | NA | NA | NA | NA | NA | NA | NA |
| ENSG000002 | 1.35939473 | -0.119638944 | 0.33471889 | -0.3574311 | 0.72076909 | NA | NA | NA |
| ENSG000002 | 0.87763084 | -0.569725011 | 0.28695805 | -1.9853947 | 0.04710057 | NA | NA | NA |
| ENSG000002 | 0 | NA | NA | NA | NA | NA | NA | NA |
| ENSG000002 | 0.50630812 | -0.189615696 | 0.24175631 | -0.7843257 | 0.432849 | NA | NA | NA |
| ENSG000002 | 1.78876855 | -0.439200213 | 0.34887394 | -1.258908 | 0.20806355 | NA | NA | NA |
| ENSG000002 | 0.75047955 | -0.181596022 | 0.28031293 | -0.6478332 | 0.51709284 | NA | NA | NA |
| ENSG000002 | 2.14981431 | -0.753112239 | 0.36048668 | -2.0891542 | 0.03669385 | NA | BNIP3P37 | 106480285 |
| ENSG000002 | 1.37695141 | -0.116581605 | 0.33655599 | -0.3463959 | 0.72904523 | NA | NA | NA |
| ENSG000002 | 1.45713928 | -0.184983197 | 0.3436199 | -0.5383367 | 0.59034463 | NA | NA | NA |
| ENSG000002 | 1.66591649 | 0.334670802 | 0.34918867 | 0.95842399 | 0.337849 | NA | NA | NA |
| ENSG000002 | 2.07293829 | -0.104390952 | 0.35901247 | -0.2907725 | 0.77122532 | NA | NA | NA |
| ENSG000002 | 0 | NA | NA | NA | NA | NA | NA | NA |
| ENSG000002 | 1.07573713 | 0.174934362 | 0.31229588 | 0.56015585 | 0.57537314 | NA | NA | NA |
| ENSG000002 | 1.12107965 | -0.096244815 | 0.20843335 | -0.4617534 | 0.64425815 | NA | NA | NA |

| | | | | | | | | |
|---|---|---|---|---|---|---|---|---|
| ENSG000002 | 0.52876404 | -0.191903645 | 0.2442426 | -0.7857092 | 0.4320379 | NA | NA | NA |
| ENSG000002 | 0 | NA | NA | NA | NA | NA | NA | NA |
| ENSG000002 | 0.87958291 | -0.415226017 | 0.29226182 | -1.420733 | 0.15539439 | NA | NA | NA |
| ENSG000002 | 1.45668281 | -0.171677581 | 0.33892903 | -0.5065296 | 0.61248493 | NA | NA | NA |
| ENSG000002 | 0 | NA | NA | NA | NA | NA | NA | NA |
| ENSG000002 | 1.49943453 | -0.177748882 | 0.33658942 | -0.5280882 | 0.59743815 | NA | NA | NA |
| ENSG000002 | 1.95706872 | -0.070701482 | 0.35994906 | -0.1964208 | 0.84428082 | NA | NA | NA |
| ENSG000002 | 1.91112985 | 0.07281046 | 0.35989223 | 0.20231184 | 0.83967294 | NA | SNRPGP19 | 106481686 |
| ENSG000002 | 0.96840847 | 0.127136949 | 0.31197202 | 0.40752677 | 0.68362114 | NA | NA | NA |
| ENSG000002 | 0 | NA | NA | NA | NA | NA | TCEB1P34 | 106481671 |
| ENSG000002 | 1.72515562 | -0.19136634 | 0.35421459 | -0.5402554 | 0.58902092 | NA | NA | NA |
| ENSG000002 | 1.21500691 | 0.105840007 | 0.32804571 | 0.32263798 | 0.74696942 | NA | NA | NA |
| ENSG000002 | 1.78705941 | -0.14581367 | 0.25842938 | -0.5642302 | 0.57259745 | NA | NA | NA |
| ENSG000002 | 0.52876404 | -0.191903645 | 0.2442426 | -0.7857092 | 0.4320379 | NA | NA | NA |
| ENSG000002 | 1.86965217 | -0.117906172 | 0.35457974 | -0.3325237 | 0.73949384 | NA | NA | NA |
| ENSG000002 | 0.14128499 | -0.097772014 | 0.13403235 | -0.7294658 | 0.4657168 | NA | NA | NA |
| ENSG000002 | 0.36690249 | -0.096999144 | 0.21683809 | -0.4473344 | 0.65463361 | NA | NA | NA |
| ENSG000002 | 1.62058095 | -0.122919643 | 0.34966654 | -0.3515339 | 0.72518784 | NA | NA | NA |
| ENSG000002 | 1.37800657 | -0.279862056 | 0.34319938 | -0.8154503 | 0.41481458 | NA | NA | NA |
| ENSG000002 | 1.19423394 | 0.103445962 | 0.32747311 | 0.31589147 | 0.75208489 | NA | NA | NA |
| ENSG000002 | 1.52037815 | -0.636815333 | 0.3442519 | -1.8498528 | 0.06433477 | NA | NA | NA |
| ENSG000002 | 0.48609822 | -0.022320526 | 0.23791393 | -0.0938177 | 0.92525401 | NA | NA | NA |
| ENSG000002 | 0.52184691 | -0.350291387 | 0.23784392 | -1.4727784 | 0.1408108 | NA | NA | NA |
| ENSG000002 | 0.99465627 | -0.342002294 | 0.30876944 | -1.10763 | 0.26802165 | NA | NA | NA |
| ENSG000002 | 0 | NA | NA | NA | NA | NA | URGCP-MRPS24 | 100534592 |
| ENSG000002 | 0.47958604 | -0.1795445 | 0.23869266 | -0.7521995 | 0.4519311 | NA | NA | NA |
| ENSG000002 | 0.6115323 | 0.059122672 | 0.25438264 | 0.2324163 | 0.81621469 | NA | NA | NA |
| ENSG000002 | 1.82874839 | 0.008304169 | 0.35519979 | 0.02337887 | 0.98134806 | NA | NA | NA |
| ENSG000002 | 1.01654519 | -0.189906583 | 0.30824946 | -0.6160808 | 0.53784117 | NA | NA | NA |

| | | | | | | | | |
|---|---|---|---|---|---|---|---|---|
| ENSG000002 | 1.5952826 | 0.011970736 | 0.34978003 | 0.03422361 | 0.97269884 | NA | NA | NA |
| ENSG000002 | 0 | NA | NA | NA | NA | NA | NA | NA |
| ENSG000002 | 1.11775206 | -0.395950926 | 0.3135858 | -1.2626558 | 0.2067129 | NA | CYP2C61P | 106480734 |
| ENSG000002 | 1.22260726 | 0.100882623 | 0.32557876 | 0.30985628 | 0.75667025 | NA | NA | NA |
| ENSG000002 | 0.49150654 | -0.026274915 | 0.24607766 | -0.1067749 | 0.91496757 | NA | NA | NA |
| ENSG000002 | 1.79290253 | -0.594591288 | 0.3521723 | -1.6883534 | 0.09134341 | NA | NA | NA |
| ENSG000002 | 1.24488444 | -0.048284428 | 0.32286194 | -0.1495513 | 0.88111862 | NA | NA | NA |
| ENSG000002 | 0 | NA | NA | NA | NA | NA | NA | NA |
| ENSG000002 | 1.32566517 | -0.113821351 | 0.32976148 | -0.3451627 | 0.72997208 | NA | NA | NA |
| ENSG000002 | 0.11865494 | 0.083440866 | 0.12751478 | 0.65436232 | 0.5128784 | NA | NA | NA |
| ENSG000002 | 1.1174143 | -0.254293808 | 0.31726797 | -0.8015111 | 0.42283581 | NA | NA | NA |
| ENSG000002 | 0.62775378 | -0.28146231 | 0.26704095 | -1.0540043 | 0.29188095 | NA | NA | NA |
| ENSG000002 | 0.24756242 | -0.187034346 | 0.18119817 | -1.0322088 | 0.30197431 | NA | NA | NA |
| ENSG000002 | 0.87187313 | -0.268035467 | 0.29910571 | -0.8961229 | 0.37018715 | NA | NA | NA |
| ENSG000002 | 1.00677787 | -0.069601965 | 0.19989884 | -0.3481859 | 0.72770055 | NA | NA | NA |
| ENSG000002 | 0.61290067 | -0.091310841 | 0.25649069 | -0.3560006 | 0.72184011 | NA | NA | NA |
| ENSG000002 | 1.63764298 | -0.553029585 | 0.35085396 | -1.5762387 | 0.1149708 | NA | NA | NA |
| ENSG000002 | 0.37945415 | -0.107841678 | 0.21865587 | -0.4932027 | 0.62186934 | NA | NA | NA |
| ENSG000002 | 1.69378574 | 0.087170925 | 0.35276108 | 0.24711038 | 0.8048228 | NA | NA | NA |
| ENSG000002 | 0 | NA | NA | NA | NA | NA | NA | NA |
| ENSG000002 | 0.72921923 | -0.029686978 | 0.28414149 | -0.1044796 | 0.91678879 | NA | NA | NA |
| ENSG000002 | 0.85300071 | -0.109179228 | 0.29690999 | -0.3677183 | 0.71308332 | NA | NA | NA |
| ENSG000002 | 2.03628358 | -0.58064392 | 0.35975717 | -1.6139884 | 0.10652993 | NA | NA | NA |
| ENSG000002 | 1.75928408 | -0.057526478 | 0.34877752 | -0.1649375 | 0.86899321 | NA | NA | NA |
| ENSG000002 | 1.39249369 | 0.016517394 | 0.32847761 | 0.05028469 | 0.95989552 | NA | BNIP3P15 | 106481692 |
| ENSG000002 | 0.36556447 | -0.101943371 | 0.21659044 | -0.4706735 | 0.63787395 | NA | NA | NA |
| ENSG000002 | 1.88158765 | -0.3786237 | 0.35724583 | -1.0598408 | 0.28921704 | NA | NA | NA |
| ENSG000002 | 1.32060362 | -0.464544037 | 0.32161794 | -1.4443972 | 0.14862729 | NA | RPL23AP92 | 106481731 |
| ENSG000002 | 0.77407652 | -0.195940532 | 0.28015013 | -0.6994126 | 0.48429421 | NA | NA | NA |

| | | | | | | | | |
|---|---|---|---|---|---|---|---|---|
| ENSG000002 | 1.92956041 | 0.172613342 | 0.35490458 | 0.48636549 | 0.62670805 | NA | NA | NA |
| ENSG000002 | 0 | NA | NA | NA | NA | NA | NA | NA |
| ENSG000002 | 0.49375647 | -0.17902118 | 0.24039617 | -0.7446923 | 0.45645774 | NA | NA | NA |
| ENSG000002 | 0.11934008 | 0.083564698 | 0.12760091 | 0.65489106 | 0.5125379 | NA | NA | NA |
| ENSG000002 | 0 | NA | NA | NA | NA | NA | NA | NA |
| ENSG000002 | 1.707792 | 0.079799601 | 0.35684066 | 0.22362811 | 0.82304669 | NA | DEAR | 102191832 |
| ENSG000002 | 1.14020798 | -0.42381935 | 0.32154805 | -1.3180591 | 0.18748385 | NA | NA | NA |
| ENSG000002 | 0.93183557 | 0.256167429 | 0.29896412 | 0.85685006 | 0.39152776 | NA | NA | NA |
| ENSG000002 | 0 | NA | NA | NA | NA | NA | HSPE1-MOB4 | 100529241 |
| ENSG000002 | 0 | NA | NA | NA | NA | NA | NA | NA |
| ENSG000002 | 0.25370795 | -0.171331747 | 0.17411721 | -0.9840024 | 0.32511435 | NA | NA | NA |
| ENSG000002 | 1.22713189 | -0.053406294 | 0.32408317 | -0.1647919 | 0.86910775 | NA | NA | NA |
| ENSG000002 | 1.41894009 | 0.224110326 | 0.33239722 | 0.67422443 | 0.50016864 | NA | NA | NA |
| ENSG000002 | 0 | NA | NA | NA | NA | NA | NA | NA |
| ENSG000002 | 1.8448946 | 0.004118639 | 0.35491561 | 0.01160456 | 0.99074111 | NA | NA | NA |
| ENSG000002 | 1.26834207 | -0.332587205 | 0.3270304 | -1.0169917 | 0.30915737 | NA | NA | NA |
| ENSG000002 | 1.10916422 | 0.019098902 | 0.30549398 | 0.0625181 | 0.95015025 | NA | NA | NA |
| ENSG000002 | 1.6868488 | -0.045723649 | 0.35344722 | -0.1293649 | 0.89706896 | NA | NA | NA |
| ENSG000002 | 0 | NA | NA | NA | NA | NA | NA | NA |
| ENSG000002 | 0 | NA | NA | NA | NA | NA | RPS10-NUDT3 | 100529239 |
| ENSG000002 | 1.95809115 | -0.316892634 | 0.36253604 | -0.8740997 | 0.38206398 | NA | NA | NA |
| ENSG000002 | 2.07176831 | -0.558518612 | 0.35765243 | -1.561624 | 0.1183766 | NA | NA | NA |
| ENSG000002 | 1.89835395 | -0.394573113 | 0.35836926 | -1.1010239 | 0.27088627 | NA | NA | NA |
| ENSG000002 | 1.59084974 | -0.628315443 | 0.34083129 | -1.8434793 | 0.06525905 | NA | NA | NA |
| ENSG000002 | 0.53627793 | -0.360851627 | 0.24080453 | -1.4985251 | 0.13399688 | NA | NA | NA |
| ENSG000002 | 0.85709274 | 0.044189217 | 0.29678189 | 0.14889458 | 0.88163681 | NA | NA | NA |
| ENSG000002 | 0 | NA | NA | NA | NA | NA | NA | NA |
| ENSG000002 | 0.88126585 | -0.455824927 | 0.30137169 | -1.5125008 | 0.1304065 | NA | NA | NA |
| ENSG000002 | 1.82090222 | 0.142894136 | 0.35754167 | 0.3996573 | 0.68940895 | NA | NA | NA |

| | | | | | | | | |
|---|---|---|---|---|---|---|---|---|
| ENSG000002 | 0.23276084 | -0.008011382 | 0.17603982 | -0.0455089 | 0.96370166 | NA | NA | NA |
| ENSG000002 | 0.11865494 | 0.083440866 | 0.12751478 | 0.65436232 | 0.5128784 | NA | NA | NA |
| ENSG000002 | 1.75293396 | -0.319935022 | 0.35443149 | -0.902671 | 0.36670054 | NA | NA | NA |
| ENSG000002 | 1.66126224 | -0.396258799 | 0.349488 | -1.1338266 | 0.25686728 | NA | NA | NA |
| ENSG000002 | 0.86825484 | 0.042287756 | 0.30247575 | 0.13980544 | 0.88881371 | NA | NA | NA |
| ENSG000002 | 1.28939062 | -0.256118772 | 0.20751792 | -1.2342007 | 0.21712811 | NA | NA | NA |
| ENSG000002 | 0.96770351 | 0.116484622 | 0.30545434 | 0.38134872 | 0.7029445 | NA | NA | NA |
| ENSG000002 | 0.76485795 | -0.196355366 | 0.28121884 | -0.6982298 | 0.48503351 | NA | NA | NA |
| ENSG000002 | 2.0343406 | -0.601530748 | 0.36391089 | -1.6529617 | 0.09833866 | NA | NA | NA |
| ENSG000002 | 1.19170007 | -0.599120801 | 0.32105333 | -1.8661099 | 0.06202599 | NA | NA | NA |
| ENSG000002 | 0 | NA | NA | NA | NA | NA | NA | NA |
| ENSG000002 | 0.88790222 | -0.250340812 | 0.28567466 | -0.8763144 | 0.38085914 | NA | NA | NA |
| ENSG000002 | 1.08268851 | -0.1130467 | 0.32127513 | -0.3518688 | 0.72493665 | NA | NA | NA |
| ENSG000002 | 0.12685397 | -0.090215768 | 0.12901919 | -0.699243 | 0.48440017 | NA | NA | NA |
| ENSG000002 | 2.05181365 | -0.966983843 | 0.35269359 | -2.7417108 | 0.00611201 | NA | NA | NA |
| ENSG000002 | 1.0360705 | -0.349467167 | 0.31046478 | -1.1256258 | 0.26032393 | NA | NA | NA |
| ENSG000002 | 0.38019136 | -0.10756138 | 0.21876967 | -0.491665 | 0.62295621 | NA | NA | NA |
| ENSG000002 | 1.48642395 | -0.341729736 | 0.34738227 | -0.9837282 | 0.32524917 | NA | NA | NA |
| ENSG000002 | 1.47480441 | 0.206332281 | 0.3352039 | 0.6155426 | 0.53819645 | NA | GPR75-ASB3 | 100302652 |
| ENSG000002 | 0.89606743 | -0.631542144 | 0.2981933 | -2.1178952 | 0.03418395 | NA | NA | NA |
| ENSG000002 | 1.22135747 | -0.187884833 | 0.33029671 | -0.5688365 | 0.56946709 | NA | NA | NA |
| ENSG000002 | 1.37686441 | -0.263013611 | 0.33502024 | -0.7850678 | 0.43241381 | NA | NA | NA |
| ENSG000002 | 0.70483298 | -0.019337048 | 0.27437201 | -0.0704775 | 0.94381362 | NA | NA | NA |
| ENSG000002 | 0.90711041 | -0.11066242 | 0.28229539 | -0.3920093 | 0.69505134 | NA | NA | NA |
| ENSG000002 | 1.38532705 | -0.237574352 | 0.32952432 | -0.7209615 | 0.47093319 | NA | NA | NA |
| ENSG000002 | 1.504037 | 0.069708517 | 0.33275332 | 0.20949008 | 0.83406568 | NA | FAM195DP | 106480278 |
| ENSG000002 | 0.36212533 | -0.235926492 | 0.20089643 | -1.1743688 | 0.24024735 | NA | NA | NA |
| ENSG000002 | 0.38056192 | -0.244105065 | 0.20391883 | -1.1970698 | 0.23127937 | NA | NA | NA |
| ENSG000002 | 1.22465919 | -0.170093555 | 0.32293524 | -0.526711 | 0.59839427 | NA | NA | NA |

| | | | | | | | | |
|---|---|---|---|---|---|---|---|---|
| ENSG000002 | 1.32819742 | -0.115612856 | 0.33889938 | -0.3411421 | 0.73299659 | NA | RAP2CP1 | 106480249 |
| ENSG000002 | 0.22263036 | 0.146922267 | 0.16600367 | 0.88505432 | 0.37612734 | NA | NA | NA |
| ENSG000002 | 1.19317666 | 0.100590244 | 0.32305124 | 0.31137551 | 0.75551517 | NA | NA | NA |
| ENSG000002 | 0.89316516 | -0.433131994 | 0.29533334 | -1.4665868 | 0.14248848 | NA | NA | NA |
| ENSG000002 | 0.50107449 | -0.372905725 | 0.24426561 | -1.5266403 | 0.12685049 | NA | NA | NA |
| ENSG000002 | 0 | NA | NA | NA | NA | NA | NA | NA |
| ENSG000002 | 0.36576034 | -0.0934779 | 0.20820828 | -0.4489634 | 0.65345805 | NA | NA | NA |
| ENSG000002 | 1.86240257 | -0.24753759 | 0.35746012 | -0.6924901 | 0.48862961 | NA | NA | NA |
| ENSG000002 | 0.3513637 | 0.074494292 | 0.21400438 | 0.34809704 | 0.72776731 | NA | NA | NA |
| ENSG000002 | 1.20046243 | -0.036546017 | 0.32328437 | -0.113046 | 0.90999406 | NA | NA | NA |
| ENSG000002 | 0.75070926 | -0.035821281 | 0.27927834 | -0.1282637 | 0.89794027 | NA | NA | NA |
| ENSG000002 | 1.08402654 | -0.102441396 | 0.3150702 | -0.3251383 | 0.74507639 | NA | NA | NA |
| ENSG000002 | 0.86253544 | 0.190406077 | 0.2917991 | 0.65252454 | 0.51406285 | NA | NA | NA |
| ENSG000002 | 0.96619526 | -0.027997535 | 0.29940688 | -0.09351 | 0.92549841 | NA | NA | NA |
| ENSG000002 | 1.76364351 | -0.565449477 | 0.34821528 | -1.6238503 | 0.10440777 | NA | NA | NA |
| ENSG000002 | 1.92695485 | -0.394536753 | 0.35631291 | -1.1072761 | 0.26817459 | NA | NA | NA |
| ENSG000002 | 0.75759255 | -0.029385622 | 0.28598041 | -0.102754 | 0.91815824 | NA | NA | NA |
| ENSG000002 | 0.26062507 | -0.012888319 | 0.18226503 | -0.070712 | 0.94362699 | NA | NA | NA |
| ENSG000002 | 0.12685397 | -0.090215768 | 0.12901919 | -0.699243 | 0.48440017 | NA | NA | NA |
| ENSG000002 | 0.49953144 | -0.023330163 | 0.24696918 | -0.0944659 | 0.92473908 | NA | CTHRC1P1 | 106480299 |
| ENSG000002 | 0.49782676 | -0.186543761 | 0.24081583 | -0.7746325 | 0.43855686 | NA | NA | NA |
| ENSG000002 | 0.33468275 | 0.234558662 | 0.20511328 | 1.14355667 | 0.25280754 | NA | NA | NA |
| ENSG000002 | 1.01087786 | -0.042074694 | 0.31374356 | -0.1341054 | 0.89331926 | NA | NA | NA |
| ENSG000002 | 0.9700353 | -0.175695904 | 0.30041861 | -0.584837 | 0.55865735 | NA | NA | NA |
| ENSG000002 | 2.1042481 | -0.858137929 | 0.35817339 | -2.3958729 | 0.01658084 | NA | HSPE1P19 | 106481675 |
| ENSG000002 | 0.64030597 | -0.296505743 | 0.27391229 | -1.0824843 | 0.27903741 | NA | NA | NA |
| ENSG000002 | 1.88608642 | -0.65256235 | 0.35596987 | -1.8331955 | 0.06677351 | NA | NA | NA |
| ENSG000002 | 1.2511314 | -0.336094849 | 0.32911873 | -1.0211964 | 0.30716141 | NA | NA | NA |
| ENSG000002 | 1.1084775 | -0.125106288 | 0.32596134 | -0.3838071 | 0.7011214 | NA | CLPTM1LP1 | 106480241 |

| | | | | | | | | |
|---|---|---|---|---|---|---|---|---|
| ENSG000002 | 0 | NA | NA | NA | NA | NA | NA | NA |
| ENSG000002 | 2.02660503 | 0.459090422 | 0.35205671 | 1.30402405 | 0.19222538 | NA | NA | NA |
| ENSG000002 | 0.38747904 | -0.104908247 | 0.21987514 | -0.4771265 | 0.63327207 | NA | NA | NA |
| ENSG000002 | 1.34222808 | 0.171160849 | 0.33340659 | 0.51336972 | 0.60769272 | NA | NA | NA |
| ENSG000002 | 0 | NA | NA | NA | NA | NA | SNRPGP17 | 106481685 |
| ENSG000002 | 1.89265279 | -0.392460306 | 0.35989573 | -1.0904834 | 0.27550027 | NA | NA | NA |
| ENSG000002 | 1.88503071 | -0.550925022 | 0.3620825 | -1.5215456 | 0.12812299 | NA | NA | NA |
| ENSG000002 | 0.96152572 | 0.124015489 | 0.3068254 | 0.40418913 | 0.68607365 | NA | NA | NA |
| ENSG000002 | 1.94395972 | 0.198596655 | 0.36346593 | 0.5463969 | 0.58479314 | NA | NA | NA |
| ENSG000002 | 0.50548109 | -0.025402968 | 0.24760631 | -0.1025942 | 0.91828506 | NA | NA | NA |
| ENSG000002 | 0.74072211 | -0.018990669 | 0.26947758 | -0.0704722 | 0.94381785 | NA | BNIP3P28 | 106481966 |
| ENSG000002 | 1.69689104 | 0.078096509 | 0.34528285 | 0.22618126 | 0.82106045 | NA | NA | NA |
| ENSG000002 | 1.65886796 | -0.259657335 | 0.34392446 | -0.7549836 | 0.45025881 | NA | NA | NA |
| ENSG000002 | 1.15834319 | -0.572105318 | 0.31651541 | -1.8075117 | 0.07068253 | NA | NA | NA |
| ENSG000002 | 1.50717464 | -0.4867564 | 0.34389465 | -1.415423 | 0.15694451 | NA | NA | NA |
| ENSG000002 | 0 | NA | NA | NA | NA | NA | NA | NA |
| ENSG000002 | 0 | NA | NA | NA | NA | NA | NA | NA |
| ENSG000002 | 0.76940933 | -0.516275266 | 0.27702168 | -1.8636638 | 0.06236894 | NA | NA | NA |
| ENSG000002 | 1.25251703 | -0.048242105 | 0.3268257 | -0.1476081 | 0.8826521 | NA | NA | NA |
| ENSG000002 | 1.7870666 | 0.014686191 | 0.35509518 | 0.04135846 | 0.96701013 | NA | NA | NA |
| ENSG000002 | 1.15054697 | -0.60063283 | 0.32198505 | -1.865406 | 0.06212452 | NA | NA | NA |
| ENSG000002 | 1.8463726 | -0.252348785 | 0.3583621 | -0.7041726 | 0.48132526 | NA | NA | NA |
| ENSG000002 | 1.27170741 | -0.355131065 | 0.33103791 | -1.0727806 | 0.28336955 | NA | NA | NA |
| ENSG000002 | 1.76030754 | -0.190533503 | 0.3539424 | -0.5383178 | 0.59035764 | NA | NA | NA |
| ENSG000002 | 1.34189122 | -0.124546688 | 0.343216 | -0.3628814 | 0.71669351 | NA | NA | NA |
| ENSG000002 | 0 | NA | NA | NA | NA | NA | PRR20FP | 106480416 |
| ENSG000002 | 0.24619405 | -0.007630671 | 0.17915639 | -0.0425922 | 0.96602658 | NA | NA | NA |
| ENSG000002 | 0.86435924 | -0.097823818 | 0.29150034 | -0.3355873 | 0.73718208 | NA | NA | NA |
| ENSG000002 | 0.51299499 | -0.204672849 | 0.24959455 | -0.8200213 | 0.41220396 | NA | NA | NA |

| | | | | | | | | |
|---|---|---|---|---|---|---|---|---|
| ENSG000002 | 0 | NA | NA | NA | NA | NA | NA | NA |
| ENSG000002 | 0 | NA | NA | NA | NA | NA | NA | NA |
| ENSG000002 | 1.42605952 | -0.539932205 | 0.33095432 | -1.6314403 | 0.10279745 | NA | NA | NA |
| ENSG000002 | 1.22792364 | 0.093195901 | 0.32239453 | 0.28907408 | 0.77252469 | NA | NA | NA |
| ENSG000002 | 1.11083507 | 0.034634703 | 0.32575494 | 0.10632134 | 0.9153274 | NA | NA | NA |
| ENSG000002 | 0.99582876 | -0.511618665 | 0.30857089 | -1.6580263 | 0.09731215 | NA | NA | NA |
| ENSG000002 | 1.02488667 | -0.501256812 | 0.30569507 | -1.6397282 | 0.1010617 | NA | NA | NA |
| ENSG000002 | 0.52125014 | -0.010203417 | 0.15567753 | -0.065542 | 0.94774247 | NA | NA | NA |
| ENSG000002 | 0 | NA | NA | NA | NA | NA | NA | NA |
| ENSG000002 | 0.88695649 | -0.277948495 | 0.29981959 | -0.9270525 | 0.35389929 | NA | NA | NA |
| ENSG000002 | 1.46715705 | -0.914635442 | 0.33423698 | -2.7364879 | 0.00620989 | NA | NA | NA |
| ENSG000002 | 0.87989748 | -0.286905324 | 0.30426178 | -0.9429555 | 0.34570366 | NA | NA | NA |
| ENSG000002 | 0.3433388 | 0.065348438 | 0.20407693 | 0.32021472 | 0.74880557 | NA | NA | NA |
| ENSG000002 | 1.591785 | 0.008336959 | 0.35321443 | 0.02360311 | 0.98116919 | NA | NA | NA |
| ENSG000002 | 2.08554393 | 0.003471593 | 0.36156567 | 0.00960156 | 0.99233918 | NA | NA | NA |
| ENSG000002 | 1.93628557 | 0.064066 | 0.36125639 | 0.17734219 | 0.85923961 | NA | NA | NA |
| ENSG000002 | 0.70084804 | 0.12831363 | 0.2743736 | 0.46766027 | 0.64002756 | NA | NA | NA |
| ENSG000002 | 1.14561238 | -0.125002439 | 0.32180743 | -0.3884386 | 0.69769146 | NA | NA | NA |
| ENSG000002 | 1.11445336 | -0.249054184 | 0.31717509 | -0.7852262 | 0.43232097 | NA | NA | NA |
| ENSG000002 | 1.79264297 | -0.477123894 | 0.35555182 | -1.341925 | 0.1796203 | NA | NA | NA |
| ENSG000002 | 1.56947241 | -0.100629725 | 0.33937243 | -0.2965171 | 0.76683522 | NA | NA | NA |
| ENSG000002 | 1.14257214 | -0.105031551 | 0.30704768 | -0.3420692 | 0.73229882 | NA | NA | NA |
| ENSG000002 | 1.70824689 | -0.052079366 | 0.35028057 | -0.148679 | 0.88180693 | NA | NA | NA |
| ENSG000002 | 0 | NA | NA | NA | NA | NA | NA | NA |
| ENSG000002 | 0.99853068 | -0.367441329 | 0.31487463 | -1.1669448 | 0.24323264 | NA | NA | NA |
| ENSG000002 | 1.49985716 | -0.338283614 | 0.34687798 | -0.9752237 | 0.32944933 | NA | NA | NA |
| ENSG000002 | 0.87306681 | -0.285213035 | 0.30393499 | -0.9384015 | 0.34803814 | NA | NA | NA |
| ENSG000002 | 0.59146429 | 0.065095808 | 0.26170762 | 0.24873486 | 0.80356588 | NA | NA | NA |
| ENSG000002 | 1.72948493 | -0.31801568 | 0.35033812 | -0.9077393 | 0.36401598 | NA | NA | NA |

| | | | | | | | | |
|---|---|---|---|---|---|---|---|---|
| ENSG000002 | 1.89146103 | -0.236172775 | 0.35412412 | -0.6669209 | 0.50482266 | NA | NA | NA |
| ENSG000002 | 0.88604301 | -0.272984982 | 0.29819841 | -0.9154475 | 0.35995677 | NA | NA | NA |
| ENSG000002 | 2.08712799 | -0.116764582 | 0.36473896 | -0.3201319 | 0.74886834 | NA | NA | NA |
| ENSG000002 | 1.47819245 | -0.180876279 | 0.34152603 | -0.529612 | 0.59638098 | NA | NA | NA |
| ENSG000002 | 0.39462238 | -0.104329077 | 0.21264929 | -0.4906157 | 0.6236983 | NA | NA | NA |
| ENSG000002 | 1.30005201 | 0.455607161 | 0.33142463 | 1.37469311 | 0.16922661 | NA | MZT1P1 | 106480302 |
| ENSG000002 | 0.11131518 | 0.081951103 | 0.12663804 | 0.64712866 | 0.51754868 | NA | UGT2A2 | 574537 |
| ENSG000002 | 0.75982019 | -0.379788636 | 0.28855699 | -1.3161651 | 0.18811864 | NA | NA | NA |
| ENSG000002 | 0.86472925 | -0.266964717 | 0.29711342 | -0.898528 | 0.36890414 | NA | NA | NA |
| ENSG000002 | 0.98238247 | 0.118633573 | 0.31257181 | 0.37954022 | 0.70428674 | NA | NA | NA |
| ENSG000002 | 2.11742435 | -0.625366758 | 0.36161511 | -1.7293712 | 0.08374268 | NA | NA | NA |
| ENSG000002 | 1.51400586 | -0.485483375 | 0.34531596 | -1.4059106 | 0.15975067 | NA | NA | NA |
| ENSG000002 | 0.14128499 | -0.097772014 | 0.13403235 | -0.7294658 | 0.4657168 | NA | NA | NA |
| ENSG000002 | 1.14606884 | -0.122967388 | 0.31470425 | -0.3907395 | 0.69598979 | NA | NA | NA |
| ENSG000002 | 1.00262718 | -0.490562014 | 0.30482559 | -1.6093203 | 0.10754632 | NA | NA | NA |
| ENSG000002 | 1.4688311 | -0.056791469 | 0.34091985 | -0.1665831 | 0.86769813 | NA | NA | NA |
| ENSG000002 | 1.81054095 | 0.133478437 | 0.35747574 | 0.3733916 | 0.70885701 | NA | SRXN1 | 140809 |
| ENSG000002 | 1.76639751 | -0.472009112 | 0.35632458 | -1.3246605 | 0.18528376 | NA | NA | NA |
| ENSG000002 | 2.00705098 | -0.455113149 | 0.36164039 | -1.2584688 | 0.20822226 | NA | NA | NA |
| ENSG000002 | 0.12685397 | -0.090215768 | 0.12901919 | -0.699243 | 0.48440017 | NA | NA | NA |
| ENSG000002 | 0.48646878 | -0.181653651 | 0.2395107 | -0.7584365 | 0.44818973 | NA | NA | NA |
| ENSG000002 | 0.77453354 | -0.191441115 | 0.28019317 | -0.6832469 | 0.49445084 | NA | NA | NA |
| ENSG000002 | 0.6256482 | -0.108446506 | 0.2650784 | -0.4091111 | 0.68245816 | NA | NA | NA |
| ENSG000002 | 0.11934008 | 0.083564698 | 0.12760091 | 0.65489106 | 0.5125379 | NA | NA | NA |
| ENSG000002 | 1.63431093 | -0.395459614 | 0.34711775 | -1.1392665 | 0.25459202 | NA | B3GNTL1P2 | 106480419 |
| ENSG000002 | 1.48955042 | -0.190244194 | 0.3448633 | -0.551651 | 0.58118747 | NA | NA | NA |
| ENSG000002 | 0.86569672 | -0.108917689 | 0.29758624 | -0.3660038 | 0.71436225 | NA | NA | NA |
| ENSG000002 | 0.11131518 | 0.081951103 | 0.12663804 | 0.64712866 | 0.51754868 | NA | NA | NA |
| ENSG000002 | 0 | NA | NA | NA | NA | NA | NA | NA |

| | | | | | | | | |
|---|---|---|---|---|---|---|---|---|
| ENSG000002 | 1.11380047 | -0.263847833 | 0.32161457 | -0.8203852 | 0.41199655 | NA | NA | NA |
| ENSG000002 | 0.13280363 | -0.093394593 | 0.13115644 | -0.7120855 | 0.47641185 | NA | NA | NA |
| ENSG000002 | 0 | NA | NA | NA | NA | NA | NA | NA |
| ENSG000002 | 1.52492855 | -0.311768802 | 0.33727314 | -0.9243808 | 0.35528811 | NA | NA | NA |
| ENSG000002 | 2.04359343 | -1.022118338 | 0.35794206 | -2.8555413 | 0.00429635 | NA | NA | NA |
| ENSG000002 | 0 | NA | NA | NA | NA | NA | NA | NA |
| ENSG000002 | 1.4271998 | -0.158350964 | 0.32622456 | -0.4854048 | 0.62738923 | NA | NA | NA |
| ENSG000002 | 1.25627914 | -0.328189132 | 0.32507914 | -1.0095669 | 0.31270284 | NA | NA | NA |
| ENSG000002 | 1.47585874 | -0.043848915 | 0.34548583 | -0.1269196 | 0.89900405 | NA | NA | NA |
| ENSG000002 | 0.63233507 | -0.116461163 | 0.27190168 | -0.4283209 | 0.66841754 | NA | NA | NA |
| ENSG000002 | 1.01595042 | -0.327600166 | 0.30255227 | -1.0827887 | 0.27890224 | NA | NA | NA |
| ENSG000002 | 0.23936338 | -0.005479057 | 0.17759977 | -0.0308506 | 0.9753887 | NA | NA | NA |
| ENSG000002 | 1.61760722 | -0.531566617 | 0.34607676 | -1.535979 | 0.12454354 | NA | NA | NA |
| ENSG000002 | 0 | NA | NA | NA | NA | NA | NA | NA |
| ENSG000002 | 0.74484447 | -0.03101441 | 0.29048039 | -0.1067694 | 0.91497194 | NA | NA | NA |
| ENSG000002 | 1.20182219 | 0.110447894 | 0.32328294 | 0.34164467 | 0.73261831 | NA | C8orf59P2 | 106480687 |
| ENSG000002 | 1.9478375 | -0.062077633 | 0.35565016 | -0.1745469 | 0.86143567 | NA | NA | NA |
| ENSG000002 | 0 | NA | NA | NA | NA | NA | NA | NA |
| ENSG000002 | 0.70616654 | 0.296520925 | 0.27799783 | 1.06663036 | 0.28613878 | NA | NA | NA |
| ENSG000002 | 1.61933755 | -0.499282237 | 0.34242364 | -1.4580834 | 0.14481755 | NA | SRGNP1 | 106480246 |
| ENSG000002 | 0.87078654 | 0.041646547 | 0.30259789 | 0.13763 | 0.89053285 | NA | NA | NA |
| ENSG000002 | 1.96367416 | -0.451760028 | 0.3630786 | -1.2442486 | 0.21340809 | NA | NA | NA |
| ENSG000002 | 0.23276084 | -0.008011382 | 0.17603982 | -0.0455089 | 0.96370166 | NA | NA | NA |
| ENSG000002 | 1.5391985 | -0.783334673 | 0.34032556 | -2.3017215 | 0.02135088 | NA | NA | NA |
| ENSG000002 | 0.49270076 | -0.021329091 | 0.24621941 | -0.0866264 | 0.93096851 | NA | NA | NA |
| ENSG000002 | 1.02067202 | -0.53213514 | 0.31335919 | -1.6981635 | 0.08947691 | NA | NA | NA |
| ENSG000002 | 2.08514045 | -0.343511195 | 0.3575538 | -0.9607259 | 0.33669001 | NA | NA | NA |
| ENSG000002 | 2.02767654 | 0.110956336 | 0.3562723 | 0.31143688 | 0.75546852 | NA | NA | NA |
| ENSG000002 | 1.47606671 | -0.051807616 | 0.33476648 | -0.1547575 | 0.87701251 | NA | RPL17P51 | 106480245 |

| | | | | | | | | | |
|---|---|---|---|---|---|---|---|---|---|
| ENSG000002 | 1.22304488 | -0.445795295 | 0.31903181 | -1.3973381 | 0.16231193 | NA | NA | NA | |
| ENSG000002 | 1.49383021 | -0.742702498 | 0.3346463 | -2.2193656 | 0.02646186 | NA | NA | NA | |
| ENSG000002 | 0.8970349 | -0.435726653 | 0.29724246 | -1.4658964 | 0.14267651 | NA | NA | NA | |
| ENSG000002 | 0.62547406 | 0.058623002 | 0.26469482 | 0.22147393 | 0.82472344 | NA | NA | NA | |
| ENSG000002 | 1.92576564 | -0.655315251 | 0.35568971 | -1.842379 | 0.06541973 | NA | NA | NA | |
| ENSG000002 | 0 | NA | NA | NA | NA | NA | NA | NA | |
| ENSG000002 | 0.36553413 | 0.069401489 | 0.20780452 | 0.33397488 | 0.73839851 | NA | NA | NA | |
| ENSG000002 | 1.68882784 | 0.206019445 | 0.34890701 | 0.59047092 | 0.55487498 | NA | NA | NA | |
| ENSG000002 | 0.62587496 | -0.276172896 | 0.26695604 | -1.0345257 | 0.30089045 | NA | NA | NA | |
| ENSG000002 | 1.50350132 | -0.485991284 | 0.34475344 | -1.4096778 | 0.15863483 | NA | NA | NA | |
| ENSG000002 | 1.96946289 | -0.165915077 | 0.35033783 | -0.473586 | 0.63579519 | NA | NA | NA | |
| ENSG000002 | 0.50955585 | -0.376243729 | 0.24515812 | -1.5346982 | 0.12485797 | NA | NA | NA | |
| ENSG000002 | 0.49699973 | -0.021154807 | 0.23920592 | -0.0884376 | 0.92952885 | NA | NA | NA | |
| ENSG000002 | 0.48399264 | 0.150724011 | 0.24577677 | 0.61325572 | 0.53970727 | NA | MEMO1P5 | | 106481668 |
| ENSG000002 | 0.36621735 | -0.097238239 | 0.21673053 | -0.4486596 | 0.65367721 | NA | NA | NA | |
| ENSG000002 | 0 | NA | NA | NA | NA | NA | NA | NA | |
| ENSG000002 | 0.27408862 | -0.199826037 | 0.18669465 | -1.0703362 | 0.28446801 | NA | NA | NA | |
| ENSG000002 | 1.64156087 | -0.245573439 | 0.34629732 | -0.7091405 | 0.47823732 | NA | NA | NA | |
| ENSG000002 | 0.49375647 | -0.17902118 | 0.24039617 | -0.7446923 | 0.45645774 | NA | NA | NA | |
| ENSG000002 | 1.39083897 | -0.26720986 | 0.33751508 | -0.7916975 | 0.42853705 | NA | NA | NA | |
| ENSG000002 | 0.52028267 | -0.202047067 | 0.25039858 | -0.8069018 | 0.41972305 | NA | NA | NA | |
| ENSG000002 | 1.57802241 | -0.348604125 | 0.33843473 | -1.0300483 | 0.30298732 | NA | NA | NA | |
| ENSG000002 | 0.49150654 | -0.026274915 | 0.24607766 | -0.1067749 | 0.91496757 | NA | WBP1LP12 | | 106480327 |
| ENSG000002 | 1.76440817 | -0.339306792 | 0.35559494 | -0.9541947 | 0.33998511 | NA | NA | NA | |
| ENSG000002 | 0 | NA | NA | NA | NA | NA | NA | NA | |
| ENSG000002 | 0.63370343 | -0.298851213 | 0.27334848 | -1.0932975 | 0.2742632 | NA | NA | NA | |
| ENSG000002 | 1.50138473 | 0.248016459 | 0.3292363 | 0.75330837 | 0.45126463 | NA | NA | NA | |
| ENSG000002 | 1.40791847 | -0.249710685 | 0.33003278 | -0.7566239 | 0.44927523 | NA | NA | NA | |
| ENSG000002 | 0.87118991 | -0.097962296 | 0.29011903 | -0.3376624 | 0.7356176 | NA | NA | NA | |

| | | | | | | | | | |
|---|---|---|---|---|---|---|---|---|---|
| ENSG000002 | 1.11399635 | -0.253234712 | 0.31714974 | -0.7984705 | 0.42459751 | NA | NA | NA | |
| ENSG000002 | 0.60470109 | 0.059032226 | 0.26929751 | 0.21920821 | 0.82648786 | NA | NA | NA | |
| ENSG000002 | 0.26199343 | -0.194018184 | 0.18422801 | -1.0531416 | 0.29227609 | NA | PPIAL4E | | 730262 |
| ENSG000002 | 1.19841531 | -0.02443865 | 0.31899191 | -0.0766121 | 0.93893211 | NA | NA | NA | |
| ENSG000002 | 0.39499294 | -0.277908454 | 0.21584201 | -1.287555 | 0.19790091 | NA | NA | NA | |
| ENSG000002 | 2.0254038 | -0.473479942 | 0.36500429 | -1.2971901 | 0.1945658 | NA | NA | NA | |
| ENSG000002 | 1.38469343 | -0.27706962 | 0.34182927 | -0.8105497 | 0.4176243 | NA | NA | NA | |
| ENSG000002 | 2.08838426 | -0.257780816 | 0.3666109 | -0.7031455 | 0.48196506 | NA | NA | NA | |
| ENSG000002 | 0.79303327 | -0.346366603 | 0.27716593 | -1.2496723 | 0.21141926 | NA | NA | NA | |
| ENSG000002 | 0.41537361 | -0.286583533 | 0.21871501 | -1.3103057 | 0.19009243 | NA | NA | NA | |
| ENSG000002 | 1.00808365 | -0.16095894 | 0.2934225 | -0.5485569 | 0.58330956 | NA | NA | NA | |
| ENSG000002 | 1.12128212 | -0.387004649 | 0.31237234 | -1.238921 | 0.21537476 | NA | LARP7P1 | | 106480287 |
| ENSG000002 | 1.04193371 | -0.537615225 | 0.31415985 | -1.7112792 | 0.08702958 | NA | NA | NA | |
| ENSG000002 | 0 | NA | NA | NA | NA | NA | TCEB1P35 | | 106480250 |
| ENSG000002 | 1.46233908 | -0.182274016 | 0.34370933 | -0.5303144 | 0.59589395 | NA | NA | NA | |
| ENSG000002 | 0.11205239 | 0.082118251 | 0.12671962 | 0.64803108 | 0.51696485 | NA | NA | NA | |
| ENSG000002 | 1.60342819 | -0.266468027 | 0.35392088 | -0.7529028 | 0.45150832 | NA | NA | NA | |
| ENSG000002 | 0.37304803 | -0.090842893 | 0.20939682 | -0.4338313 | 0.66441096 | NA | NA | NA | |
| ENSG000002 | 0.62160824 | -0.2792447 | 0.26652831 | -1.0477112 | 0.29477168 | NA | NA | NA | |
| ENSG000002 | 0.70679769 | 0.126328142 | 0.27481541 | 0.45968362 | 0.64574333 | NA | NA | NA | |
| ENSG000002 | 1.05559314 | -0.674291479 | 0.30432019 | -2.2157303 | 0.02670998 | NA | NA | NA | |
| ENSG000002 | 1.74404923 | -0.191137824 | 0.35376894 | -0.54029 | 0.58899705 | NA | HSPE1P14 | | 106481674 |
| ENSG000002 | 1.12370172 | -0.408147645 | 0.31667423 | -1.2888565 | 0.19744797 | NA | NA | NA | |
| ENSG000002 | 0.77288285 | -0.203654568 | 0.28771447 | -0.7078357 | 0.47904731 | NA | NA | NA | |
| ENSG000002 | 1.12452729 | -0.114036451 | 0.32113983 | -0.3550991 | 0.72251538 | NA | NA | NA | |
| ENSG000002 | 0.51533082 | -0.192250929 | 0.24273717 | -0.7920127 | 0.42835324 | NA | NA | NA | |
| ENSG000002 | 0.72423567 | -0.015070868 | 0.26826514 | -0.056179 | 0.95519921 | NA | NA | NA | |
| ENSG000002 | 0.38651158 | -0.274258716 | 0.21461355 | -1.2779189 | 0.20127801 | NA | NA | NA | |
| ENSG000002 | 0.60487577 | -0.101296586 | 0.26329793 | -0.3847223 | 0.70044317 | NA | NA | NA | |

| | | | | | | | | |
|---|---|---|---|---|---|---|---|---|
| ENSG000002 | 1.21386531 | 0.110470538 | 0.32799987 | 0.33680056 | 0.73626725 | NA | NA | NA |
| ENSG000002 | 1.53696818 | -0.181310467 | 0.21383689 | -0.8478914 | 0.39649845 | NA | NA | NA |
| ENSG000002 | 1.929375 | -0.378995624 | 0.35335696 | -1.0725574 | 0.28346974 | NA | BNIP3P19 | 106480281 |
| ENSG000002 | 1.72236491 | -0.060415539 | 0.35602972 | -0.1696924 | 0.86525205 | NA | NA | NA |
| ENSG000002 | 1.75455044 | 0.495204871 | 0.3435291 | 1.44152236 | 0.14943717 | NA | NA | NA |
| ENSG000002 | 2.05517076 | 0.252917145 | 0.36563728 | 0.69171596 | 0.48911572 | NA | NA | NA |
| ENSG000002 | 0.12685397 | -0.090215768 | 0.12901919 | -0.699243 | 0.48440017 | NA | NA | NA |
| ENSG000002 | 1.99785469 | -0.318177615 | 0.36190987 | -0.8791626 | 0.37931313 | NA | NA | NA |
| ENSG000002 | 0.72170533 | 0.13625606 | 0.28374667 | 0.48020321 | 0.6310829 | NA | BNIP3P35 | 106480284 |
| ENSG000002 | 1.95187506 | -0.164297955 | 0.35370597 | -0.4645043 | 0.64228646 | NA | NA | NA |
| ENSG000002 | 0.7546942 | -0.187287667 | 0.28056327 | -0.6675416 | 0.50442622 | NA | NA | NA |
| ENSG000002 | 1.94286638 | -0.064106012 | 0.36177793 | -0.1771971 | 0.85935355 | NA | BNIP3P36 | 106481696 |
| ENSG000002 | 1.4244848 | 0.243991456 | 0.34461746 | 0.70800666 | 0.47894112 | NA | NA | NA |
| ENSG000002 | 0.89020422 | -0.427620104 | 0.29524143 | -1.4483743 | 0.14751239 | NA | NA | NA |
| ENSG000002 | 1.6729368 | -0.663317925 | 0.34297445 | -1.9340156 | 0.05311121 | NA | NA | NA |
| ENSG000002 | 0 | NA | NA | NA | NA | NA | NA | NA |
| ENSG000002 | 1.35649693 | -0.267378179 | 0.34003998 | -0.786314 | 0.43168358 | NA | LSM1P2 | 106480288 |
| ENSG000002 | 1.26811586 | -0.181051647 | 0.32580723 | -0.5557017 | 0.57841477 | NA | NA | NA |
| ENSG000002 | 1.92539103 | -0.822972705 | 0.35691622 | -2.3057868 | 0.02112255 | NA | NA | NA |
| ENSG000002 | 0.50593756 | -0.029751655 | 0.24018594 | -0.1238693 | 0.90141879 | NA | NA | NA |
| ENSG000002 | 0.36690249 | -0.096999144 | 0.21683809 | -0.4473344 | 0.65463361 | NA | NA | NA |
| ENSG000002 | 0.12070844 | -0.088503695 | 0.12783931 | -0.6923042 | 0.48874631 | NA | NA | NA |
| ENSG000002 | 1.63801355 | -0.673849099 | 0.34454437 | -1.9557687 | 0.0504924 | NA | NA | NA |
| ENSG000002 | 0.73443167 | -0.030284643 | 0.27816347 | -0.1088735 | 0.91330279 | NA | NA | NA |
| ENSG000002 | 1.62511017 | 0.009816246 | 0.34732162 | 0.0282627 | 0.97745263 | NA | NA | NA |
| ENSG000002 | 0.47753253 | -0.016714525 | 0.24454657 | -0.068349 | 0.94550778 | NA | NA | NA |
| ENSG000002 | 1.50076852 | -0.493899365 | 0.34653022 | -1.4252707 | 0.15407897 | NA | NA | NA |
| ENSG000002 | 1.92276832 | -0.338709616 | 0.3464499 | -0.9776583 | 0.32824336 | NA | NA | NA |
| ENSG000002 | 1.52575167 | -0.196474102 | 0.34158956 | -0.575176 | 0.5651723 | NA | NA | NA |

| | | | | | | | | |
|---|---|---|---|---|---|---|---|---|
| ENSG000002 | 1.76001938 | -0.196196334 | 0.35619741 | -0.5508079 | 0.5817654 | NA | NA | NA |
| ENSG000002 | 1.33528868 | -0.123277646 | 0.34033079 | -0.3622289 | 0.71718099 | NA | NA | NA |
| ENSG000002 | 1.61782887 | -0.116867367 | 0.34962813 | -0.3342619 | 0.73818193 | NA | NA | NA |
| ENSG000002 | 0.76101442 | -0.375123817 | 0.28870464 | -1.2993342 | 0.19382926 | NA | NA | NA |
| ENSG000002 | 1.60151457 | 0.14490715 | 0.34872613 | 0.41553281 | 0.6777519 | NA | NA | NA |
| ENSG000002 | 1.12717915 | -0.404639256 | 0.31427386 | -1.2875371 | 0.19790715 | NA | NA | NA |
| ENSG000002 | 1.62469043 | -0.237047909 | 0.34468625 | -0.6877208 | 0.49162861 | NA | NA | NA |
| ENSG000002 | 0.7009541 | 0.298692198 | 0.27758308 | 1.07604614 | 0.28190662 | NA | RNU1-29P | 106480154 |
| ENSG000002 | 0 | NA | NA | NA | NA | NA | NA | NA |
| ENSG000002 | 0.23868015 | 0.154625155 | 0.16997278 | 0.90970538 | 0.36297791 | NA | NA | NA |
| ENSG000002 | 1.95755036 | -0.309458748 | 0.36253886 | -0.8535878 | 0.39333341 | NA | NA | NA |
| ENSG000002 | 0 | NA | NA | NA | NA | NA | NA | NA |
| ENSG000002 | 0.85135215 | 0.050516499 | 0.29646393 | 0.17039678 | 0.8646981 | NA | NA | NA |
| ENSG000002 | 0 | NA | NA | NA | NA | NA | NA | NA |
| ENSG000002 | 1.56306383 | -0.224476294 | 0.33757899 | -0.6649593 | 0.50607651 | NA | NA | NA |
| ENSG000002 | 1.52299183 | -0.199450958 | 0.34540132 | -0.577447 | 0.56363752 | NA | NA | NA |
| ENSG000002 | 0.49375647 | -0.17902118 | 0.24039617 | -0.7446923 | 0.45645774 | NA | NA | NA |
| ENSG000002 | 1.12267826 | -0.276164768 | 0.32186163 | -0.8580233 | 0.39087962 | NA | NA | NA |
| ENSG000002 | 0.48211382 | 0.155752635 | 0.24556107 | 0.63427249 | 0.525903 | NA | NA | NA |
| ENSG000002 | 0 | NA | NA | NA | NA | NA | NA | NA |
| ENSG000002 | 0.37148378 | 0.06737793 | 0.20875796 | 0.32275622 | 0.74687987 | NA | SNORA51 | 677831 |
| ENSG000002 | 0 | NA | NA | NA | NA | NA | NA | NA |
| ENSG000002 | 1.51533592 | -0.458366313 | 0.33949102 | -1.3501574 | 0.1769655 | NA | RN7SKP290 | 106480480 |
| ENSG000002 | 2.09075391 | -0.1332164 | 0.36709146 | -0.362897 | 0.71668181 | NA | NA | NA |
| ENSG000002 | 2.08921184 | -0.380547344 | 0.36653712 | -1.0382232 | 0.29916615 | NA | NA | NA |
| ENSG000002 | 1.11141065 | -0.413696525 | 0.31865285 | -1.2982671 | 0.19419555 | NA | RNU6-94P | 106481206 |
| ENSG000002 | 1.87734213 | -0.271033617 | 0.36274029 | -0.7471837 | 0.45495271 | NA | NA | NA |
| ENSG000002 | 1.43893352 | 0.089604248 | 0.33804374 | 0.26506702 | 0.79095783 | NA | LOC389332 | 389332 |
| ENSG000002 | 1.33583449 | -0.360658207 | 0.32515845 | -1.1091768 | 0.26735392 | NA | NA | NA |

| | | | | | | | | |
|---|---|---|---|---|---|---|---|---|
| ENSG000002 | 0.2596576 | -0.193002035 | 0.1837913 | -1.0501152 | 0.29366517 | NA | NA | NA |
| ENSG000002 | 0 | NA | NA | NA | NA | NA | NA | NA |
| ENSG000002 | 1.73890149 | -0.188127921 | 0.3543555 | -0.5309017 | 0.59548693 | NA | NA | NA |
| ENSG000002 | 1.51360304 | -0.337830777 | 0.34590961 | -0.9766447 | 0.32874509 | NA | NA | NA |
| ENSG000002 | 1.00962807 | -0.3256485 | 0.30228304 | -1.0772966 | 0.28134777 | NA | NA | NA |
| ENSG000002 | 0.99850035 | -0.195348626 | 0.31384733 | -0.622432 | 0.53365784 | NA | NA | NA |
| ENSG000002 | 0 | NA | NA | NA | NA | NA | NA | NA |
| ENSG000002 | 2.1152209 | -0.101468936 | 0.35545916 | -0.2854588 | 0.77529268 | NA | NA | NA |
| ENSG000002 | 0.87728091 | -0.262583863 | 0.29191055 | -0.8995354 | 0.36836756 | NA | MIR4791 | 100616291 |
| ENSG000002 | 0.12070844 | -0.088503695 | 0.12783931 | -0.6923042 | 0.48874631 | NA | MIR4466 | 100616154 |
| ENSG000002 | 0.3867939 | -0.105148852 | 0.21977268 | -0.4784437 | 0.63233444 | NA | NA | NA |
| ENSG000002 | 1.21986703 | 0.108680849 | 0.32817014 | 0.33117226 | 0.74051437 | NA | NA | NA |
| ENSG000002 | 2.07755613 | 0.346464974 | 0.35865314 | 0.96601684 | 0.33403576 | NA | NA | NA |
| ENSG000002 | 0.59760982 | 0.066257965 | 0.26871197 | 0.24657616 | 0.80523626 | NA | NA | NA |
| ENSG000002 | 0.39499294 | -0.277908454 | 0.21584201 | -1.287555 | 0.19790091 | NA | RNU6-86P | 106479604 |
| ENSG000002 | 0.28256998 | -0.18483211 | 0.18021686 | -1.0256094 | 0.30507573 | NA | RNU6-85P | 106481203 |
| ENSG000002 | 1.49390751 | -0.322759299 | 0.34262449 | -0.9420205 | 0.34618214 | NA | NA | NA |
| ENSG000002 | 2.12094994 | -0.392429193 | 0.36619049 | -1.0716532 | 0.28387585 | NA | NA | NA |
| ENSG000002 | 1.99216797 | -0.670852548 | 0.35489329 | -1.8902937 | 0.05871869 | NA | NA | NA |
| ENSG000002 | 0 | NA | NA | NA | NA | NA | NA | NA |
| ENSG000002 | 1.40222085 | 0.523231425 | 0.33818539 | 1.54717334 | 0.12182145 | NA | LOC101929733 | 101929733 |
| ENSG000002 | 0.50479595 | -0.02564641 | 0.24753216 | -0.1036084 | 0.91748012 | NA | NA | NA |
| ENSG000002 | 1.92740586 | -0.052380843 | 0.36166004 | -0.1448345 | 0.88484156 | NA | NA | NA |
| ENSG000002 | 0.12070844 | -0.088503695 | 0.12783931 | -0.6923042 | 0.48874631 | NA | NA | NA |
| ENSG000002 | 0 | NA | NA | NA | NA | NA | NA | NA |
| ENSG000002 | 1.64598349 | -0.256960288 | 0.34376065 | -0.7474977 | 0.4547632 | NA | NA | NA |
| ENSG000002 | 0 | NA | NA | NA | NA | NA | SNORD58B | 26790 |
| ENSG000002 | 2.05683397 | -0.228112867 | 0.35679247 | -0.6393433 | 0.52259964 | NA | NA | NA |
| ENSG000002 | 0.86156797 | 0.043923168 | 0.3021415 | 0.14537284 | 0.88441651 | NA | NA | NA |

| | | | | | | | | | |
|---|---|---|---|---|---|---|---|---|---|
| ENSG000002 | 1.9193556 | -0.368470123 | 0.35440639 | -1.0396825 | 0.29848743 | NA | | RN7SL827P | 106481762 |
| ENSG000002 | 0 | NA | NA | NA | NA | NA | | NA | NA |
| ENSG000002 | 0.99098645 | -0.03510399 | 0.3114328 | -0.1127177 | 0.91025437 | NA | | NA | NA |
| ENSG000002 | 0.62522207 | -0.280659539 | 0.2668423 | -1.0517806 | 0.29290024 | NA | | NA | NA |
| ENSG000002 | 0.61175852 | -0.109548347 | 0.27027148 | -0.4053271 | 0.68523711 | NA | | RNU1-30P | 106480662 |
| ENSG000002 | 1.36443538 | -0.386083084 | 0.33208625 | -1.1625988 | 0.24499231 | NA | | NA | NA |
| ENSG000002 | 0.49150709 | -0.015856992 | 0.24611816 | -0.0644284 | 0.94862914 | NA | | RNU6-87P | 106481204 |
| ENSG000002 | 1.15119539 | -0.278942585 | 0.32424946 | -0.8602716 | 0.38963937 | NA | | MIR139 | 406931 |
| ENSG000002 | 0.13280363 | -0.093394593 | 0.13115644 | -0.7120855 | 0.47641185 | NA | | NA | NA |
| ENSG000002 | 1.39881337 | -0.12015792 | 0.3306301 | -0.363421 | 0.71629042 | NA | | NA | NA |
| ENSG000002 | 0.93702465 | -0.643555212 | 0.30011803 | -2.1443404 | 0.03200562 | NA | | NA | NA |
| ENSG000002 | 0.74752913 | -0.352552616 | 0.28071348 | -1.2559163 | 0.20914634 | NA | | NA | NA |
| ENSG000002 | 0.62684242 | -0.109923117 | 0.27148862 | -0.4048903 | 0.68555813 | NA | | NA | NA |
| ENSG000002 | 1.52478475 | -0.343998406 | 0.34725831 | -0.9906124 | 0.32187486 | NA | | NA | NA |
| ENSG000002 | 0.71148951 | 0.1280784 | 0.26770406 | 0.47843279 | 0.6323422 | NA | | RN7SL828P | 106481147 |
| ENSG000002 | 0.2596576 | -0.193002035 | 0.1837913 | -1.0501152 | 0.29366517 | NA | | MIR502 | 574504 |
| ENSG000002 | 1.55652732 | -0.652191112 | 0.34527435 | -1.888907 | 0.0589043 | NA | | NA | NA |
| ENSG000002 | 1.76644778 | -0.048582966 | 0.3424601 | -0.1418646 | 0.88718695 | NA | | NA | NA |
| ENSG000002 | 0 | NA | NA | NA | NA | NA | | NA | NA |
| ENSG000002 | 1.54078001 | -0.498814846 | 0.34716111 | -1.4368397 | 0.15076356 | NA | | NA | NA |
| ENSG000002 | 0 | NA | NA | NA | NA | NA | | NA | NA |
| ENSG000002 | 0.71489639 | 0.303414967 | 0.2785771 | 1.08915976 | 0.27608344 | NA | | NA | NA |
| ENSG000002 | 1.38689129 | -0.389719961 | 0.33342324 | -1.1688446 | 0.24246623 | NA | | NA | NA |
| ENSG000002 | 1.05431122 | -0.354787196 | 0.31117408 | -1.1401566 | 0.25422105 | NA | | NA | NA |
| ENSG000002 | 0 | NA | NA | NA | NA | NA | | NA | NA |
| ENSG000002 | 1.93275661 | 0.068126346 | 0.35902384 | 0.18975438 | 0.8495016 | NA | | NA | NA |
| ENSG000002 | 1.43020578 | 0.23756372 | 0.34475316 | 0.68908351 | 0.49077072 | NA | | NA | NA |
| ENSG000002 | 2.12075853 | -0.642859428 | 0.36428601 | -1.7647107 | 0.0776124 | NA | | NA | NA |
| ENSG000002 | 0.71217273 | -0.016761464 | 0.27491005 | -0.0609707 | 0.95138253 | NA | | NA | NA |

| | | | | | | | | |
|---|---|---|---|---|---|---|---|---|
| ENSG000002 | 2.14935708 | -0.24732679 | 0.35854883 | -0.6897995 | 0.49032031 | NA | NA | NA |
| ENSG000002 | 1.7595451 | -0.625250393 | 0.3560428 | -1.7561102 | 0.07906959 | NA | NA | NA |
| ENSG000002 | 0 | NA | NA | NA | NA | NA | NA | NA |
| ENSG000002 | 0.62618953 | -0.114303343 | 0.27141404 | -0.4211401 | 0.67365276 | NA | NA | NA |
| ENSG000002 | 0.59100728 | 0.060563842 | 0.26166992 | 0.23145129 | 0.81696422 | NA | NA | NA |
| ENSG000002 | 1.80755435 | 0.270215202 | 0.35869058 | 0.75333787 | 0.45124691 | NA | NA | NA |
| ENSG000002 | 1.41182208 | 0.347640587 | 0.3335292 | 1.0423093 | 0.2972683 | NA | NA | NA |
| ENSG000002 | 1.95001473 | -0.307241875 | 0.36136022 | -0.8502371 | 0.39519326 | NA | RN7SL803P | 106479514 |
| ENSG000002 | 1.96955229 | 0.168033873 | 0.35711632 | 0.47052981 | 0.63797654 | NA | NA | NA |
| ENSG000002 | 0 | NA | NA | NA | NA | NA | NA | NA |
| ENSG000002 | 0.23936338 | -0.005479057 | 0.17759977 | -0.0308506 | 0.9753887 | NA | MIR3666 | 100500896 |
| ENSG000002 | 1.45531253 | -0.180705924 | 0.34358968 | -0.5259352 | 0.5989332 | NA | RN7SL726P | 106479488 |
| ENSG000002 | 0.23936338 | -0.005479057 | 0.17759977 | -0.0308506 | 0.9753887 | NA | NA | NA |
| ENSG000002 | 2.0612677 | -0.890247014 | 0.36114884 | -2.4650419 | 0.01369972 | NA | NA | NA |
| ENSG000002 | 0.60925012 | 0.227481777 | 0.26441857 | 0.8603094 | 0.38961851 | NA | NA | NA |
| ENSG000002 | 0.72304144 | -0.021905802 | 0.27736023 | -0.0789796 | 0.93704884 | NA | NA | NA |
| ENSG000002 | 0.98506713 | -0.203217809 | 0.31733922 | -0.6403804 | 0.52192534 | NA | RNU6-89P | 106480550 |
| ENSG000002 | 0.94398495 | 0.423666649 | 0.30180104 | 1.40379455 | 0.16038004 | NA | NA | NA |
| ENSG000002 | 0 | NA | NA | NA | NA | NA | SNORD96A | 619571 |
| ENSG000002 | 0 | NA | NA | NA | NA | NA | NA | NA |
| ENSG000002 | 0.24619405 | -0.007630671 | 0.17915639 | -0.0425922 | 0.96602658 | NA | NA | NA |
| ENSG000002 | 0.23730987 | 0.1539805 | 0.16964526 | 0.90766167 | 0.364057 | NA | NA | NA |
| ENSG000002 | 1.41131001 | -0.368788171 | 0.32371478 | -1.139238 | 0.25460391 | NA | NA | NA |
| ENSG000002 | 0.89905168 | -0.281707301 | 0.30041599 | -0.9377241 | 0.34838623 | NA | NA | NA |
| ENSG000002 | 1.54199759 | 0.304886177 | 0.35039071 | 0.87013202 | 0.38422826 | NA | RNU6-90P | 106481205 |
| ENSG000002 | 1.0020304 | -0.171114558 | 0.30187219 | -0.5668444 | 0.57081991 | NA | NA | NA |
| ENSG000002 | 0.88012774 | -0.113321964 | 0.2983266 | -0.3798587 | 0.70405029 | NA | SNORD114-21 | 767599 |
| ENSG000002 | 0 | NA | NA | NA | NA | NA | NA | NA |
| ENSG000002 | 1.15976107 | -0.270751624 | 0.3186645 | -0.8496448 | 0.39552262 | NA | NA | NA |

| | | | | | | | | |
|---|---|---|---|---|---|---|---|---|
| ENSG000002 | 1.38830726 | -0.261210757 | 0.33527012 | -0.7791054 | 0.43591764 | NA | NA | NA |
| ENSG000002 | 0.37194025 | 0.067575365 | 0.21717998 | 0.31114914 | 0.75568724 | NA | RNU4-89P | 106481200 |
| ENSG000002 | 1.54319127 | 0.295285893 | 0.34865906 | 0.84691875 | 0.39704041 | NA | NA | NA |
| ENSG000002 | 0.11865494 | 0.083440866 | 0.12751478 | 0.65436232 | 0.5128784 | NA | NA | NA |
| ENSG000002 | 1.84100002 | 0.001020968 | 0.35834275 | 0.00284914 | 0.99772672 | NA | NA | NA |
| ENSG000002 | 1.32230442 | 0.163230412 | 0.32006174 | 0.50999664 | 0.61005382 | NA | NA | NA |
| ENSG000002 | 0.50544884 | -0.019721866 | 0.2401671 | -0.0821173 | 0.93455346 | NA | NA | NA |
| ENSG000002 | 1.77009778 | -0.328138374 | 0.3556718 | -0.9225875 | 0.35622219 | NA | NA | NA |
| ENSG000002 | 0.4009426 | -0.302672773 | 0.2240145 | -1.3511303 | 0.17665371 | NA | RNU6-92P | 106479607 |
| ENSG000002 | 2.1581736 | -0.646462462 | 0.36507143 | -1.7707835 | 0.07659671 | NA | IFNL4 | 101180976 |
| ENSG000002 | 1.05710352 | -0.328100859 | 0.30041929 | -1.0921431 | 0.2747702 | NA | NA | NA |
| ENSG000002 | 2.10821308 | -0.120579908 | 0.36153911 | -0.3335183 | 0.73874308 | NA | NA | NA |
| ENSG000002 | 0 | NA | NA | NA | NA | NA | NA | NA |
| ENSG000002 | 0.48399264 | 0.150724011 | 0.24577677 | 0.61325572 | 0.53970727 | NA | NA | NA |
| ENSG000002 | 0.87870381 | -0.256084916 | 0.29376523 | -0.8717332 | 0.38335392 | NA | RN7SL778P | 106479504 |
| ENSG000002 | 1.64239841 | -0.537656935 | 0.34725306 | -1.5483145 | 0.12154661 | NA | FAM47E-STBD1 | 100631383 |
| ENSG000002 | 2.10355652 | 0.001896714 | 0.36231278 | 0.00523502 | 0.99582308 | NA | NA | NA |
| ENSG000002 | 1.50531755 | -0.340053488 | 0.34696033 | -0.9800933 | 0.32704008 | NA | NA | NA |
| ENSG000002 | 1.79875684 | 0.008121886 | 0.35940287 | 0.02259828 | 0.98197072 | NA | NA | NA |
| ENSG000002 | 0.42385498 | -0.262217565 | 0.21037102 | -1.2464529 | 0.21259817 | NA | NA | NA |
| ENSG000002 | 1.75439069 | -0.171644103 | 0.34731634 | -0.4942011 | 0.6211642 | NA | NA | NA |
| ENSG000002 | 0.86501158 | -0.114317285 | 0.30264298 | -0.3777298 | 0.7056313 | NA | NA | NA |
| ENSG000002 | 0.11934008 | 0.083564698 | 0.12760091 | 0.65489106 | 0.5125379 | NA | RNU6-91P | 106479606 |
| ENSG000002 | 1.49193832 | -0.195393233 | 0.34731618 | -0.5625803 | 0.57372073 | NA | NA | NA |
| ENSG000002 | 1.5090157 | -0.338108593 | 0.34514827 | -0.9796039 | 0.32728168 | NA | NA | NA |
| ENSG000002 | 0.38651158 | -0.274258716 | 0.21461355 | -1.2779189 | 0.20127801 | NA | MIR432 | 574451 |
| ENSG000002 | 0.4009426 | -0.302672773 | 0.2240145 | -1.3511303 | 0.17665371 | NA | SNORD115-40 | 100033814 |
| ENSG000002 | 0.73292343 | -0.195671801 | 0.28518655 | -0.6861186 | 0.49263832 | NA | SNORD113-5 | 767565 |
| ENSG000002 | 0 | NA | NA | NA | NA | NA | NA | NA |

| | | | | | | | | |
|---|---|---|---|---|---|---|---|---|
| ENSG000002 | 1.60168926 | 0.014192027 | 0.35102341 | 0.04043043 | 0.96774997 | NA | NA | NA |
| ENSG000002 | 2.08661362 | 0.002641956 | 0.36461501 | 0.00724588 | 0.99421868 | NA | NA | NA |
| ENSG000002 | 0.64002365 | -0.456557672 | 0.26464873 | -1.7251459 | 0.08450119 | NA | NA | NA |
| ENSG000002 | 1.3007103 | -0.668394863 | 0.32930215 | -2.0297312 | 0.04238387 | NA | RNU6-88P | 106479605 |
| ENSG000002 | 2.13949978 | 0.060099898 | 0.36142878 | 0.16628421 | 0.86793329 | NA | NA | NA |
| ENSG000002 | 1.74648148 | -0.198111462 | 0.35264942 | -0.5617802 | 0.57426578 | NA | NA | NA |
| ENSG000002 | 1.14801526 | -0.592052842 | 0.32046828 | -1.8474616 | 0.06468027 | NA | NA | NA |
| ENSG000002 | 1.00749271 | -0.030678216 | 0.29858593 | -0.102745 | 0.91816535 | NA | RN7SL840P | 106479528 |
| ENSG000002 | 1.79653912 | -0.326479903 | 0.35234542 | -0.9265904 | 0.35413922 | NA | NA | NA |
| ENSG000002 | 1.88904893 | -0.523675528 | 0.35694799 | -1.467092 | 0.14235103 | NA | NA | NA |
| ENSG000002 | 0.11865494 | 0.083440866 | 0.12751478 | 0.65436232 | 0.5128784 | NA | MIR4787 | 100616138 |
| ENSG000002 | 1.96649261 | -0.813967951 | 0.35562501 | -2.2888378 | 0.02208878 | NA | NA | NA |
| ENSG000002 | 0.72390073 | -0.168754891 | 0.26943439 | -0.6263302 | 0.53109839 | NA | NA | NA |
| ENSG000002 | 1.32454687 | 0.157197944 | 0.32967738 | 0.47682357 | 0.63348776 | NA | NA | NA |
| ENSG000002 | 1.63390599 | -0.368725177 | 0.3410098 | -1.0812744 | 0.27957506 | NA | MUSTN1 | 389125 |
| ENSG000002 | 0 | NA | NA | NA | NA | NA | NA | NA |
| ENSG000002 | 0 | NA | NA | NA | NA | NA | ZSWIM8-AS1 | 100507331 |
| ENSG000002 | 0 | NA | NA | NA | NA | NA | NA | NA |
| ENSG000002 | 0 | NA | NA | NA | NA | NA | NA | NA |
| ENSG000002 | 1.72051779 | -0.066410618 | 0.35384739 | -0.1876815 | 0.85112631 | NA | NA | NA |
| ENSG000002 | 1.79378163 | -0.752018676 | 0.35224437 | -2.1349345 | 0.03276636 | NA | NA | NA |
| ENSG000002 | 0.38036605 | -0.293832733 | 0.22119083 | -1.3284128 | 0.18404178 | NA | NA | NA |
| ENSG000002 | 1.36272836 | -0.124564244 | 0.33479715 | -0.3720588 | 0.70984904 | NA | NA | NA |
| ENSG000002 | 0 | NA | NA | NA | NA | NA | NA | NA |
| ENSG000002 | 1.25641711 | -0.20366184 | 0.33464535 | -0.6085901 | 0.54279619 | NA | NA | NA |
| ENSG000002 | 0 | NA | NA | NA | NA | NA | NA | NA |
| ENSG000002 | 1.10679511 | -0.10882473 | 0.31588207 | -0.3445106 | 0.73046231 | NA | NA | NA |
| ENSG000002 | 1.23974118 | -0.308544999 | 0.32214724 | -0.9577763 | 0.33817556 | NA | NA | NA |
| ENSG000002 | 1.12057716 | -0.431729854 | 0.32306664 | -1.3363492 | 0.18143515 | NA | LOC105373098 | 105373098 |

| | | | | | | | | |
|---|---|---|---|---|---|---|---|---|
| ENSG000002 | 1.72208313 | -0.1921382 | 0.35829253 | -0.5362607 | 0.59177839 | NA | NA | NA |
| ENSG000002 | 0 | NA | NA | NA | NA | NA | NA | NA |
| ENSG000002 | 0 | NA | NA | NA | NA | NA | NA | NA |
| ENSG000002 | 1.02869637 | -0.531493232 | 0.31211448 | -1.7028791 | 0.0885907 | NA | NA | NA |
| ENSG000002 | 1.37946138 | -0.261299418 | 0.3358556 | -0.7780112 | 0.43656241 | NA | NA | NA |
| ENSG000002 | 1.62420062 | -0.26683825 | 0.35257669 | -0.7568233 | 0.44915574 | NA | NA | NA |
| ENSG000002 | 2.14481714 | -0.239035907 | 0.35763325 | -0.6683828 | 0.50388928 | NA | NA | NA |
| ENSG000002 | 1.74101057 | -0.061097187 | 0.35516016 | -0.1720271 | 0.86341619 | NA | NA | NA |
| ENSG000002 | 0 | NA | NA | NA | NA | NA | NA | NA |
| ENSG000002 | 0.23936338 | -0.005479057 | 0.17759977 | -0.0308506 | 0.9753887 | NA | STAG3L5P-PVRI | 101752399 |
| ENSG000002 | 0.3561712 | 0.067399341 | 0.21474131 | 0.31386296 | 0.75362513 | NA | NA | NA |
| ENSG000002 | 0 | NA | NA | NA | NA | NA | NA | NA |
| ENSG000002 | 0 | NA | NA | NA | NA | NA | NA | NA |
| ENSG000002 | 1.23985878 | -0.192714935 | 0.3278088 | -0.5878882 | 0.55660732 | NA | NA | NA |
| ENSG000002 | 0 | NA | NA | NA | NA | NA | NA | NA |
| ENSG000002 | 0.11205239 | 0.082118251 | 0.12671962 | 0.64803108 | 0.51696485 | NA | NA | NA |
| ENSG000002 | 1.10243803 | 0.030209146 | 0.31540552 | 0.09577875 | 0.9236963 | NA | NA | NA |
| ENSG000002 | 1.43048598 | 0.235981764 | 0.34557405 | 0.68286888 | 0.4946897 | NA | NA | NA |
| ENSG000002 | 2.10759457 | -0.006547356 | 0.36703559 | -0.0178385 | 0.98576771 | NA | NA | NA |
| ENSG000002 | 0.11865494 | 0.083440866 | 0.12751478 | 0.65436232 | 0.5128784 | NA | NA | NA |
| ENSG000002 | 1.13246732 | -0.104946942 | 0.30940357 | -0.3391911 | 0.73446576 | NA | KRTAP10-7 | 386675 |
| ENSG000002 | 0.7367675 | -0.032420367 | 0.28462732 | -0.1139046 | 0.9093134 | NA | NA | NA |
| ENSG000002 | 0.24619405 | -0.007630671 | 0.17915639 | -0.0425922 | 0.96602658 | NA | NA | NA |
| ENSG000002 | 1.55168544 | -0.786023188 | 0.34053914 | -2.3081728 | 0.02098953 | NA | NA | NA |
| ENSG000002 | 1.22938182 | -0.198012034 | 0.33266234 | -0.5952343 | 0.55168687 | NA | NA | NA |
| ENSG000002 | 1.48403604 | -0.329166394 | 0.34378436 | -0.9574793 | 0.33832541 | NA | NA | NA |
| ENSG000002 | 1.12352703 | -0.266790378 | 0.31963776 | -0.8346648 | 0.40390648 | NA | NA | NA |
| ENSG000002 | 0.73730024 | 0.273640027 | 0.26944423 | 1.01557206 | 0.30983322 | NA | NA | NA |
| ENSG000002 | 1.15558472 | -0.441010777 | 0.32412477 | -1.3606204 | 0.17363368 | NA | NA | NA |

| | | | | | | | | | |
|---|---|---|---|---|---|---|---|---|---|
| ENSG000002 | 1.7785081 | 0.219575254 | 0.3395198 | 0.64672297 | 0.51781125 | NA | NA | NA | |
| ENSG000002 | 1.94058666 | 0.193438982 | 0.36121015 | 0.53553031 | 0.59228321 | NA | NA | NA | |
| ENSG000002 | 1.47725744 | -0.331009196 | 0.34463524 | -0.9604624 | 0.33682254 | NA | NA | NA | |
| ENSG000002 | 1.28075184 | -0.196477303 | 0.32810234 | -0.5988293 | 0.54928671 | NA | NA | NA | |
| ENSG000002 | 1.26760486 | -0.332867286 | 0.32700917 | -1.0179142 | 0.30871871 | NA | NA | NA | |
| ENSG000002 | 1.89959969 | -0.37725044 | 0.35499105 | -1.0627041 | 0.28791616 | NA | NA | NA | |
| ENSG000002 | 0.38614101 | -0.1098476 | 0.21963552 | -0.5001359 | 0.61697942 | NA | NA | NA | |
| ENSG000002 | 0.36553413 | 0.069401489 | 0.20780452 | 0.33397488 | 0.73839851 | NA | NA | NA | |
| ENSG000002 | 0 | NA | NA | NA | NA | NA | NA | NA | |
| ENSG000002 | 0 | NA | NA | NA | NA | NA | NA | NA | |
| ENSG000002 | 0.14128499 | -0.097772014 | 0.13403235 | -0.7294658 | 0.4657168 | NA | NA | NA | |
| ENSG000002 | 1.67339168 | -0.855093669 | 0.34614539 | -2.470331 | 0.01349881 | NA | NA | NA | |
| ENSG000002 | 0.82491027 | 0.049523814 | 0.27973895 | 0.17703582 | 0.85948025 | NA | NDST2 | | 8509 |
| ENSG000002 | 0.3934287 | -0.107186984 | 0.22072612 | -0.4856108 | 0.62724315 | NA | NA | NA | |
| ENSG000002 | 1.13999036 | -0.594959047 | 0.32011792 | -1.8585621 | 0.06308923 | NA | NA | NA | |
| ENSG000002 | 0.72765946 | -0.158679159 | 0.26971653 | -0.5883183 | 0.5563187 | NA | NA | NA | |
| ENSG000002 | 0 | NA | NA | NA | NA | NA | NA | NA | |
| ENSG000002 | 0 | NA | NA | NA | NA | NA | NA | NA | |
| ENSG000002 | 0 | NA | NA | NA | NA | NA | NA | NA | |
| ENSG000002 | 2.08499664 | -0.35597078 | 0.36061136 | -0.9871314 | 0.32357824 | NA | NA | NA | |
| ENSG000002 | 0 | NA | NA | NA | NA | NA | NA | NA | |
| ENSG000002 | 0.75304351 | -0.205958918 | 0.29162776 | -0.7062391 | 0.48003948 | NA | NA | NA | |
| ENSG000002 | 0 | NA | NA | NA | NA | NA | RBAK-RBAKDN | | 100533952 |
| ENSG000002 | 1.44204196 | 0.363245068 | 0.33922525 | 1.07080789 | 0.28425582 | NA | NA | NA | |
| ENSG000002 | 0.37404583 | -0.105111695 | 0.21784993 | -0.4824959 | 0.62945372 | NA | NA | NA | |
| ENSG000002 | 0.66202413 | -0.272682834 | 0.26145371 | -1.0429488 | 0.296972 | NA | NA | NA | |
| ENSG000002 | 0 | NA | NA | NA | NA | NA | NA | NA | |
| ENSG000002 | 0 | NA | NA | NA | NA | NA | NA | NA | |
| ENSG000002 | 1.87045938 | -0.267167191 | 0.36188187 | -0.7382718 | 0.46034928 | NA | NA | NA | |

| | | | | | | | | |
|---|---|---|---|---|---|---|---|---|
| ENSG000002 | 1.48177402 | -0.177077913 | 0.34082074 | -0.5195632 | 0.60336806 | NA | NA | NA |
| ENSG000002 | 0 | NA | NA | NA | NA | NA | NA | NA |
| ENSG000002 | 0 | NA | NA | NA | NA | NA | FAM95C | 100289137 |
| ENSG000002 | 0.23816916 | -0.010712105 | 0.1772998 | -0.060418 | 0.9518227 | NA | NA | NA |
| ENSG000002 | 0 | NA | NA | NA | NA | NA | NA | NA |
| ENSG000002 | 0 | NA | NA | NA | NA | NA | NA | NA |
| ENSG000002 | 0 | NA | NA | NA | NA | NA | NA | NA |
| ENSG000002 | 1.73084077 | 0.07052463 | 0.34914585 | 0.20199189 | 0.83992306 | NA | NA | NA |
| ENSG000002 | 1.44503745 | -0.570503452 | 0.33639559 | -1.6959303 | 0.08989909 | NA | NA | NA |
| ENSG000002 | 2.10776871 | -0.136898387 | 0.36717466 | -0.3728427 | 0.70926553 | NA | NA | NA |
| ENSG000002 | 0.49902098 | -0.194741582 | 0.24819886 | -0.7846192 | 0.43267688 | NA | NA | NA |
| ENSG000002 | 0 | NA | NA | NA | NA | NA | NA | NA |
| ENSG000002 | 1.58624318 | -0.346165988 | 0.33869313 | -1.0220638 | 0.3067507 | NA | NA | NA |
| ENSG000002 | 1.06697557 | 0.195204077 | 0.32057318 | 0.60892205 | 0.54257611 | NA | NA | NA |
| ENSG000002 | 1.91181096 | -0.171035465 | 0.355645 | -0.4809163 | 0.63057601 | NA | NA | NA |
| ENSG000002 | 2.07483191 | 0.119480767 | 0.36589691 | 0.32654216 | 0.7440142 | NA | NA | NA |
| ENSG000002 | 0 | NA | NA | NA | NA | NA | NA | NA |
| ENSG000002 | 1.83095034 | -0.597823064 | 0.35115412 | -1.7024521 | 0.08867065 | NA | NA | NA |
| ENSG000002 | 0.96614127 | -0.175719289 | 0.30359707 | -0.5787911 | 0.56273013 | NA | NA | NA |
| ENSG000002 | 1.11764263 | -0.255560168 | 0.3172709 | -0.8054951 | 0.42053401 | NA | NA | NA |
| ENSG000002 | 0 | NA | NA | NA | NA | NA | NA | NA |
| ENSG000002 | 0.9691234 | -0.04234686 | 0.31053688 | -0.1363666 | 0.89153147 | NA | NA | NA |
| ENSG000002 | 1.15025183 | 0.397095352 | 0.32094708 | 1.23726116 | 0.21599014 | NA | NA | NA |
| ENSG000002 | 0.73957996 | -0.027414423 | 0.28481756 | -0.0962526 | 0.92331998 | NA | NA | NA |
| ENSG000002 | 1.54478831 | 0.157777591 | 0.34840031 | 0.45286295 | 0.65064743 | NA | NA | NA |
| ENSG000002 | 1.22511565 | -0.191823281 | 0.33256442 | -0.5768004 | 0.56407431 | NA | NA | NA |
| ENSG000002 | 0.98048192 | -0.037096279 | 0.3110075 | -0.1192778 | 0.9050553 | NA | NA | NA |
| ENSG000002 | 0.71489639 | 0.303414967 | 0.2785771 | 1.08915976 | 0.27608344 | NA | NA | NA |
| ENSG000002 | 0.8403082 | 0.19488034 | 0.28875592 | 0.67489643 | 0.49974157 | NA | NA | NA |

| | | | | | | | | |
|---|---|---|---|---|---|---|---|---|
| ENSG000002 | 0 | NA | NA | NA | NA | NA | NA | NA |
| ENSG000002 | 1.23623231 | -0.173079689 | 0.31939678 | -0.5418955 | 0.58789048 | NA | NA | NA |
| ENSG000002 | 0.23139247 | 0.167748926 | 0.17680352 | 0.94878726 | 0.34272882 | NA | NA | NA |
| ENSG000002 | 0.3867939 | -0.105148852 | 0.21977268 | -0.4784437 | 0.63233444 | NA | NA | NA |
| ENSG000002 | 0 | NA | NA | NA | NA | NA | NA | NA |
| ENSG000002 | 1.60749848 | -0.271967322 | 0.35396023 | -0.7683556 | 0.44227596 | NA | NA | NA |
| ENSG000002 | 0.8417069 | -0.10578696 | 0.2902031 | -0.3645273 | 0.71546427 | NA | NA | NA |
| ENSG000002 | 0.62550439 | -0.114540246 | 0.27135801 | -0.4221001 | 0.67295194 | NA | NA | NA |
| ENSG000002 | 0 | NA | NA | NA | NA | NA | NA | NA |
| ENSG000002 | 0.79807549 | -0.536675498 | 0.28113813 | -1.9089389 | 0.05626998 | NA | NA | NA |
| ENSG000002 | 0.25351207 | -0.189965547 | 0.1824784 | -1.0410303 | 0.29786148 | NA | NA | NA |
| ENSG000002 | 0.35141577 | 0.074527565 | 0.21401461 | 0.34823588 | 0.72766304 | NA | NA | NA |
| ENSG000002 | 0.47781273 | -0.018045496 | 0.22812977 | -0.0791019 | 0.93695158 | NA | NA | NA |
| ENSG000002 | 0 | NA | NA | NA | NA | NA | NA | NA |
| ENSG000002 | 0 | NA | NA | NA | NA | NA | NA | NA |
| ENSG000002 | 0.62408396 | 0.054009456 | 0.26457561 | 0.20413618 | 0.83824708 | NA | NA | NA |
| ENSG000002 | 1.61919532 | -0.383840683 | 0.34692875 | -1.1063963 | 0.26855503 | NA | NA | NA |
| ENSG000002 | 0.61907654 | -0.261106204 | 0.25936927 | -1.0066967 | 0.31408052 | NA | NA | NA |
| ENSG000002 | 0.77453354 | -0.191441115 | 0.28019317 | -0.6832469 | 0.49445084 | NA | NA | NA |
| ENSG000002 | 0 | NA | NA | NA | NA | NA | NA | NA |
| ENSG000002 | 1.23489674 | -0.036433174 | 0.32258638 | -0.1129408 | 0.91007747 | NA | NA | NA |
| ENSG000002 | 1.27975404 | -0.195822174 | 0.3303429 | -0.5927846 | 0.55332533 | NA | NA | NA |
| ENSG000002 | 0.84929652 | 0.044636708 | 0.29027382 | 0.15377449 | 0.87778755 | NA | NA | NA |
| ENSG000002 | 1.47039835 | -0.040159569 | 0.34214274 | -0.1173766 | 0.90656159 | NA | NA | NA |
| ENSG000002 | 1.20246386 | -0.172422066 | 0.32140615 | -0.5364616 | 0.59163955 | NA | NA | NA |
| ENSG000002 | 1.09313458 | 0.028487421 | 0.31980269 | 0.08907812 | 0.92901983 | NA | NA | NA |
| ENSG000002 | 1.40191909 | -0.718185475 | 0.33415131 | -2.1492823 | 0.03161203 | NA | NA | NA |
| ENSG000002 | 1.76070844 | -0.483419283 | 0.35777428 | -1.3511851 | 0.17663615 | NA | NA | NA |
| ENSG000002 | 0.12070844 | -0.088503695 | 0.12783931 | -0.6923042 | 0.48874631 | NA | NA | NA |

| | | | | | | | | |
|---|---|---|---|---|---|---|---|---|
| ENSG000002 | 1.30324363 | 0.020456195 | 0.32804752 | 0.06235741 | 0.95027821 | NA | NA | NA |
| ENSG000002 | 0 | NA | NA | NA | NA | NA | NA | NA |
| ENSG000002 | 2.14163701 | -0.507458873 | 0.36425819 | -1.3931296 | 0.1635806 | NA | NA | NA |
| ENSG000002 | 0.49031286 | -0.020802257 | 0.24597635 | -0.0845702 | 0.93260313 | NA | NA | NA |
| ENSG000002 | 1.60954147 | -0.25002593 | 0.34882196 | -0.7167723 | 0.47351459 | NA | NA | NA |
| ENSG000002 | 0.24485602 | -0.012655878 | 0.17881722 | -0.0707755 | 0.94357643 | NA | NA | NA |
| ENSG000002 | 0.50107449 | -0.372905725 | 0.24426561 | -1.5266403 | 0.12685049 | NA | NA | NA |
| ENSG000002 | 0.37899768 | -0.10176489 | 0.21865765 | -0.4654074 | 0.64163973 | NA | NA | NA |
| ENSG000002 | 0 | NA | NA | NA | NA | NA | NA | NA |
| ENSG000002 | 0 | NA | NA | NA | NA | NA | NA | NA |
| ENSG000002 | 0.72423759 | 0.136226018 | 0.27602641 | 0.4935253 | 0.62164147 | NA | NA | NA |
| ENSG000002 | 1.80749854 | -0.433851818 | 0.34706935 | -1.2500436 | 0.21128363 | NA | NA | NA |
| ENSG000002 | 1.37431501 | -0.118781198 | 0.33088499 | -0.3589803 | 0.71960981 | NA | NA | NA |
| ENSG000002 | 1.77741083 | 0.016130546 | 0.34894485 | 0.04622664 | 0.96312961 | NA | NA | NA |
| ENSG000002 | 0.49921686 | -0.164688411 | 0.23121313 | -0.7122797 | 0.47629162 | NA | NA | NA |
| ENSG000002 | 0 | NA | NA | NA | NA | NA | NA | NA |
| ENSG000002 | 1.05067566 | -0.505361849 | 0.30593435 | -1.6518637 | 0.09856234 | NA | LINC01396 | 103695434 |
| ENSG000002 | 1.93054094 | -0.18037625 | 0.35837212 | -0.5033211 | 0.61473853 | NA | NA | NA |
| ENSG000002 | 1.85830361 | -0.256735458 | 0.35498518 | -0.7232287 | 0.46953942 | NA | NA | NA |
| ENSG000002 | 2.10503241 | -0.360620328 | 0.36098842 | -0.9989803 | 0.31780422 | NA | NA | NA |
| ENSG000002 | 1.99944067 | -0.32039257 | 0.36335995 | -0.8817498 | 0.37791211 | NA | NA | NA |
| ENSG000002 | 0 | NA | NA | NA | NA | NA | NA | NA |
| ENSG000002 | 1.13932696 | -0.422633486 | 0.32117748 | -1.3158877 | 0.18821175 | NA | NA | NA |
| ENSG000002 | 0 | NA | NA | NA | NA | NA | NA | NA |
| ENSG000002 | 2.1372498 | -0.253878962 | 0.3656199 | -0.6943795 | 0.48744425 | NA | NA | NA |
| ENSG000002 | 0 | NA | NA | NA | NA | NA | NA | NA |
| ENSG000002 | 1.67147806 | -0.398209217 | 0.34835427 | -1.1431156 | 0.25299058 | NA | NA | NA |
| ENSG000002 | 2.01259156 | -0.443559282 | 0.36170456 | -1.2263027 | 0.22008478 | NA | NA | NA |
| ENSG000002 | 0.53471369 | -0.194084218 | 0.24487205 | -0.7925944 | 0.42801417 | NA | NA | NA |

| | | | | | | | | |
|---|---|---|---|---|---|---|---|---|
| ENSG000002 | 2.0606665 | -0.115910612 | 0.3654145 | -0.3172031 | 0.75108949 | NA | NA | NA |
| ENSG000002 | 1.74857496 | -0.32352841 | 0.35437725 | -0.9129492 | 0.36126928 | NA | NA | NA |
| ENSG000002 | 2.10366459 | -0.244970024 | 0.36518619 | -0.6708086 | 0.50234249 | NA | NA | NA |
| ENSG000002 | 0 | NA | NA | NA | NA | NA | NA | NA |
| ENSG000002 | 0.82720725 | 0.186483195 | 0.28549554 | 0.65319127 | 0.51363298 | NA | NA | NA |
| ENSG000002 | 1.11262607 | -0.253709345 | 0.31710029 | -0.8000918 | 0.42365761 | NA | NA | NA |
| ENSG000002 | 1.55000233 | 0.284426842 | 0.34599541 | 0.8220538 | 0.41104628 | NA | NA | NA |
| ENSG000002 | 1.00396074 | -0.194820594 | 0.31255594 | -0.6233143 | 0.53307798 | NA | NA | NA |
| ENSG000002 | 0.73998086 | -0.354752955 | 0.28183273 | -1.2587358 | 0.20812577 | NA | NA | NA |
| ENSG000002 | 0 | NA | NA | NA | NA | NA | NA | NA |
| ENSG000002 | 1.59198781 | -0.095431775 | 0.33672461 | -0.2834119 | 0.77686107 | NA | MIR6129 | 102465137 |
| ENSG000002 | 1.40604494 | -0.395593903 | 0.3314182 | -1.1936397 | 0.23261896 | NA | NA | NA |
| ENSG000002 | 0.38133351 | -0.102466351 | 0.2189713 | -0.4679442 | 0.63982449 | NA | NA | NA |
| ENSG000002 | 1.52174651 | -0.789238544 | 0.34117451 | -2.3132987 | 0.02070622 | NA | NA | NA |
| ENSG000002 | 2.06926836 | -0.346190744 | 0.36013626 | -0.9612771 | 0.33641285 | NA | FOXD4L6 | 653404 |
| ENSG000002 | 1.65785344 | -0.689563313 | 0.34771471 | -1.9831295 | 0.04735296 | NA | NA | NA |
| ENSG000002 | 1.43692665 | -0.734330147 | 0.33605041 | -2.1851785 | 0.02887576 | NA | NA | NA |
| ENSG000002 | 0 | NA | NA | NA | NA | NA | NA | NA |
| ENSG000002 | 1.61059271 | -0.126682837 | 0.34901474 | -0.3629727 | 0.71662525 | NA | NA | NA |
| ENSG000002 | 1.60808649 | 0.803444912 | 0.34632546 | 2.3199129 | 0.02034559 | NA | NA | NA |
| ENSG000002 | 0 | NA | NA | NA | NA | NA | NA | NA |
| ENSG000002 | 1.48317676 | -0.187428871 | 0.34599132 | -0.5417155 | 0.58801449 | NA | NA | NA |
| ENSG000002 | 0 | NA | NA | NA | NA | NA | NA | NA |
| ENSG000002 | 0 | NA | NA | NA | NA | NA | NA | NA |
| ENSG000002 | 0 | NA | NA | NA | NA | NA | SNORA44 | 677825 |
| ENSG000002 | 1.12584167 | -0.433570142 | 0.32322501 | -1.341388 | 0.17979452 | NA | NA | NA |
| ENSG000002 | 0.45476004 | 0.3294872 | 0.23735363 | 1.38817005 | 0.16508527 | NA | NA | NA |
| ENSG000002 | 1.98453091 | -0.454640841 | 0.362405 | -1.2545104 | 0.20965655 | NA | NA | NA |
| ENSG000002 | 0.22997012 | 0.167051788 | 0.17646558 | 0.94665368 | 0.34381528 | NA | MIR6812 | 102465487 |

| | | | | | | | | | |
|---|---|---|---|---|---|---|---|---|---|
| ENSG000002 | 1.15162909 | -0.550605944 | 0.3125357 | -1.7617377 | 0.07811362 | NA | NA | NA | |
| ENSG000002 | 1.73893767 | -0.469292518 | 0.35500398 | -1.3219359 | 0.18618948 | NA | NA | NA | |
| ENSG000002 | 0.22336757 | 0.163792188 | 0.17487321 | 0.93663397 | 0.34894687 | NA | NA | NA | |
| ENSG000002 | 0.12070844 | -0.088503695 | 0.12783931 | -0.6923042 | 0.48874631 | NA | NA | NA | |
| ENSG000002 | 0 | NA | NA | NA | NA | NA | NA | NA | |
| ENSG000002 | 1.35490278 | -0.5184575 | 0.32938367 | -1.5740231 | 0.11548211 | NA | NA | NA | |
| ENSG000002 | 1.14006766 | -0.11922429 | 0.32163471 | -0.3706823 | 0.71087418 | NA | NA | NA | |
| ENSG000002 | 1.48787664 | -0.425543431 | 0.33074358 | -1.2866264 | 0.19822453 | NA | NA | NA | |
| ENSG000002 | 2.04406324 | -0.301185583 | 0.35316945 | -0.8528076 | 0.39376603 | NA | NA | NA | |
| ENSG000002 | 0 | NA | NA | NA | NA | NA | SNORA78 | | 677844 |
| ENSG000002 | 0 | NA | NA | NA | NA | NA | NA | NA | |
| ENSG000002 | 0.37216701 | -0.099548723 | 0.21762847 | -0.4574251 | 0.64736554 | NA | NA | NA | |
| ENSG000002 | 0.11934008 | 0.083564698 | 0.12760091 | 0.65489106 | 0.5125379 | NA | NA | NA | |
| ENSG000002 | 0.51775097 | -0.069813375 | 0.15551247 | -0.4489246 | 0.65348604 | NA | MIR7160 | | 102465695 |
| ENSG000002 | 1.99558248 | -0.29731926 | 0.35188653 | -0.8449294 | 0.39815028 | NA | NA | NA | |
| ENSG000002 | 0 | NA | NA | NA | NA | NA | NA | NA | |
| ENSG000002 | 0.49810908 | -0.023852705 | 0.24681213 | -0.0966432 | 0.92300978 | NA | NA | NA | |
| ENSG000002 | 1.41480705 | -0.392289816 | 0.33163215 | -1.1829065 | 0.23684622 | NA | NA | NA | |
| ENSG000002 | 1.15558472 | -0.441010777 | 0.32412477 | -1.3606204 | 0.17363368 | NA | MIR6776 | | 102465465 |
| ENSG000002 | 0.11865494 | 0.083440866 | 0.12751478 | 0.65436232 | 0.5128784 | NA | NA | NA | |
| ENSG000002 | 0.34344486 | 0.238503239 | 0.20662376 | 1.15428757 | 0.24838229 | NA | RN7SL386P | | 106479364 |
| ENSG000002 | 2.15638653 | -0.642737827 | 0.36425007 | -1.764551 | 0.07763925 | NA | NA | NA | |
| ENSG000002 | 0 | NA | NA | NA | NA | NA | MIR6511A4 | | 102465684 |
| ENSG000002 | 0 | NA | NA | NA | NA | NA | NA | NA | |
| ENSG000002 | 0.49475427 | -0.18461639 | 0.24044438 | -0.7678133 | 0.44259811 | NA | NA | NA | |
| ENSG000002 | 0 | NA | NA | NA | NA | NA | MIR4726 | | 100616153 |
| ENSG000002 | 1.91291668 | -0.375332966 | 0.35622486 | -1.0536406 | 0.29204751 | NA | NA | NA | |
| ENSG000002 | 0 | NA | NA | NA | NA | NA | NA | NA | |
| ENSG000002 | 0.47029884 | 0.145033566 | 0.23680225 | 0.612467 | 0.54022882 | NA | NA | NA | |

| | | | | | | | | |
|---|---|---|---|---|---|---|---|---|
| ENSG000002 | 0 | NA | NA | NA | NA | NA | WBP1LP8 | 106480325 |
| ENSG000002 | 0 | NA | NA | NA | NA | NA | NA | NA |
| ENSG000002 | 0.91673043 | -0.435686733 | 0.29658919 | -1.4689906 | 0.14183534 | NA | MIR8065 | 102465867 |
| ENSG000002 | 0 | NA | NA | NA | NA | NA | MIR6792 | 102465475 |
| ENSG000002 | 0 | NA | NA | NA | NA | NA | NA | NA |
| ENSG000002 | 0.48279897 | 0.145542372 | 0.23831717 | 0.6107087 | 0.54139244 | NA | NA | NA |
| ENSG000002 | 1.08695871 | 0.166972473 | 0.30789757 | 0.54229877 | 0.5876127 | NA | NA | NA |
| ENSG000002 | 1.03254045 | -0.358491974 | 0.311878 | -1.1494622 | 0.25036544 | NA | NA | NA |
| ENSG000002 | 1.00976659 | -0.188211986 | 0.30796956 | -0.6111383 | 0.54110804 | NA | NA | NA |
| ENSG000002 | 0 | NA | NA | NA | NA | NA | NA | NA |
| ENSG000002 | 0.22997012 | 0.167051788 | 0.17646558 | 0.94665368 | 0.34381528 | NA | NA | NA |
| ENSG000002 | 0.36007182 | -0.08619198 | 0.20729252 | -0.4157988 | 0.67755724 | NA | NA | NA |
| ENSG000002 | 0 | NA | NA | NA | NA | NA | NA | NA |
| ENSG000002 | 2.0728672 | -0.001231289 | 0.36686219 | -0.0033563 | 0.99732209 | NA | NA | NA |
| ENSG000002 | 0 | NA | NA | NA | NA | NA | MIR6760 | 102465455 |
| ENSG000002 | 0 | NA | NA | NA | NA | NA | NA | NA |
| ENSG000002 | 1.39816048 | -0.128308196 | 0.33477497 | -0.383267 | 0.70152181 | NA | NA | NA |
| ENSG000002 | 0.71512261 | 0.122662368 | 0.26971035 | 0.45479296 | 0.6492582 | NA | NA | NA |
| ENSG000002 | 0.81133402 | -0.339316464 | 0.27594092 | -1.2296707 | 0.21882045 | NA | NA | NA |
| ENSG000002 | 0.63393368 | -0.111067416 | 0.26575676 | -0.4179288 | 0.67599914 | NA | NA | NA |
| ENSG000002 | 0 | NA | NA | NA | NA | NA | RN7SL343P | 106481018 |
| ENSG000002 | 1.97008757 | 0.070058529 | 0.35755986 | 0.19593511 | 0.84466096 | NA | NA | NA |
| ENSG000002 | 0.14128499 | -0.097772014 | 0.13403235 | -0.7294658 | 0.4657168 | NA | NA | NA |
| ENSG000002 | 1.70668622 | -0.176814936 | 0.35010151 | -0.5050391 | 0.61353141 | NA | NA | NA |
| ENSG000002 | 0.56533801 | 0.392832502 | 0.25455097 | 1.5432371 | 0.12277326 | NA | NA | NA |
| ENSG000002 | 0.85021 | 0.04998197 | 0.30595275 | 0.163365 | 0.87023106 | NA | NA | NA |
| ENSG000002 | 0.11131518 | 0.081951103 | 0.12663804 | 0.64712866 | 0.51754868 | NA | RNA5-8S5 | 100008587 |
| ENSG000002 | 0 | NA | NA | NA | NA | NA | NA | NA |
| ENSG000002 | 0.51139849 | -0.021802213 | 0.24083338 | -0.0905282 | 0.92786748 | NA | NA | NA |

| | | | | | | | | |
|---|---|---|---|---|---|---|---|---|
| ENSG000002 | 1.95075194 | -0.306998422 | 0.36136539 | -0.8495513 | 0.39557462 | NA | NA | NA |
| ENSG000002 | 0 | NA | NA | NA | NA | NA | NA | NA |
| ENSG000002 | 1.88912721 | -0.644601107 | 0.35539127 | -1.8137787 | 0.06971181 | NA | NA | NA |
| ENSG000002 | 0.11205239 | 0.082118251 | 0.12671962 | 0.64803108 | 0.51696485 | NA | NA | NA |
| ENSG000002 | 1.69748334 | 0.077566048 | 0.35616438 | 0.2177816 | 0.82759928 | NA | MIR6075 | 102466103 |
| ENSG000002 | 0.23070733 | 0.167416683 | 0.17664244 | 0.94777156 | 0.34324576 | NA | NA | NA |
| ENSG000002 | 2.12527511 | -0.519521672 | 0.36686395 | -1.4161153 | 0.15674173 | NA | NA | NA |
| ENSG000002 | 0 | NA | NA | NA | NA | NA | NA | NA |
| ENSG000002 | 1.56591666 | -0.44918888 | 0.33670789 | -1.3340611 | 0.18218385 | NA | NA | NA |
| ENSG000002 | 0 | NA | NA | NA | NA | NA | MIR6790 | 102465473 |
| ENSG000002 | 0.76747453 | -0.201328414 | 0.28738472 | -0.7005536 | 0.48358162 | NA | NA | NA |
| ENSG000002 | 1.18956283 | 0.100490079 | 0.32124916 | 0.3128104 | 0.75442471 | NA | NA | NA |
| ENSG000002 | 0.35022155 | 0.06940789 | 0.21379411 | 0.32464827 | 0.74544729 | NA | NA | NA |
| ENSG000002 | 0 | NA | NA | NA | NA | NA | NA | NA |
| ENSG000002 | 0.51459361 | -0.192534203 | 0.24265108 | -0.7934611 | 0.42750921 | NA | MIR6126 | 102465134 |
| ENSG000002 | 0 | NA | NA | NA | NA | NA | MIR6765 | 102465458 |
| ENSG000002 | 0 | NA | NA | NA | NA | NA | NA | NA |
| ENSG000002 | 0.23868015 | 0.154625155 | 0.16997278 | 0.90970538 | 0.36297791 | NA | NA | NA |
| ENSG000002 | 1.28568643 | -0.655539254 | 0.32753507 | -2.0014323 | 0.04534583 | NA | NA | NA |
| ENSG000002 | 1.85306296 | 0.001301262 | 0.36128744 | 0.00360174 | 0.99712624 | NA | NA | NA |
| ENSG000002 | 0.39462238 | -0.104329077 | 0.21264929 | -0.4906157 | 0.6236983 | NA | NA | NA |
| ENSG000002 | 0 | NA | NA | NA | NA | NA | NA | NA |
| ENSG000002 | 0.72779688 | -0.028558914 | 0.27769729 | -0.1028419 | 0.91808846 | NA | NA | NA |
| ENSG000002 | 0.99463061 | -0.184102047 | 0.30484099 | -0.6039281 | 0.54589143 | NA | NA | NA |
| ENSG000002 | 0 | NA | NA | NA | NA | NA | NA | NA |
| ENSG000002 | 0.63114139 | -0.097702348 | 0.25814354 | -0.3784807 | 0.70507353 | NA | NA | NA |
| ENSG000002 | 0.3934287 | -0.107186984 | 0.22072612 | -0.4856108 | 0.62724315 | NA | RN7SL673P | 106480525 |
| ENSG000002 | 0.99625097 | -0.037659479 | 0.31316771 | -0.1202534 | 0.90428243 | NA | NA | NA |
| ENSG000002 | 1.31926559 | -0.455966196 | 0.31845966 | -1.4317864 | 0.15220497 | NA | NA | NA |

| | | | | | | | | | |
|---|---|---|---|---|---|---|---|---|---|
| ENSG000002 | 0.12070844 | -0.088503695 | 0.12783931 | -0.6923042 | 0.48874631 | NA | | MIR7843 | 102465832 |
| ENSG000002 | 0 | NA | NA | NA | NA | NA | | SNORD115-13 | 100033450 |
| ENSG000002 | 0.61927241 | -0.278564577 | 0.26634985 | -1.0458597 | 0.29562581 | NA | | MIR6842 | 102465507 |
| ENSG000002 | 0 | NA | NA | NA | NA | NA | | NA | NA |
| ENSG000002 | 1.78759632 | 0.131548658 | 0.34908825 | 0.37683497 | 0.70629625 | NA | | NA | NA |
| ENSG000002 | 0.87838868 | -0.38610365 | 0.28281027 | -1.3652391 | 0.1721779 | NA | | NA | NA |
| ENSG000002 | 0 | NA | NA | NA | NA | NA | | NA | NA |
| ENSG000002 | 0.70961015 | 0.140347015 | 0.28290789 | 0.49608732 | 0.6198328 | NA | | NA | NA |
| ENSG000002 | 1.35588751 | 0.029841394 | 0.33920725 | 0.08797393 | 0.9298974 | NA | | NA | NA |
| ENSG000002 | 0 | NA | NA | NA | NA | NA | | NA | NA |
| ENSG000002 | 1.96600999 | -0.445959311 | 0.36200977 | -1.2318985 | 0.21798699 | NA | | NA | NA |
| ENSG000002 | 0 | NA | NA | NA | NA | NA | | NA | NA |
| ENSG000002 | 0.62675597 | -0.08300713 | 0.15756539 | -0.5268107 | 0.59832506 | NA | | NA | NA |
| ENSG000002 | 2.01785553 | -0.462417682 | 0.36340646 | -1.2724531 | 0.20321218 | NA | | NA | NA |
| ENSG000002 | 0.38036605 | -0.293832733 | 0.22119083 | -1.3284128 | 0.18404178 | NA | | MIR6859-2 | 102465909 |
| ENSG000002 | 0.11934008 | 0.083564698 | 0.12760091 | 0.65489106 | 0.5125379 | NA | | NA | NA |
| ENSG000002 | 0 | NA | NA | NA | NA | NA | | NA | NA |
| ENSG000002 | 0.14128499 | -0.097772014 | 0.13403235 | -0.7294658 | 0.4657168 | NA | | NA | NA |
| ENSG000002 | 1.41475306 | -0.570815196 | 0.33695745 | -1.6940275 | 0.09026006 | NA | | NA | NA |
| ENSG000002 | 1.55567413 | -0.109382958 | 0.34512851 | -0.316934 | 0.75129368 | NA | | NA | NA |
| ENSG000002 | 0.39274356 | -0.107428268 | 0.22062504 | -0.4869269 | 0.62631013 | NA | | NA | NA |
| ENSG000002 | 0 | NA | NA | NA | NA | NA | | MIR6798 | 102466738 |
| ENSG000002 | 0.37236288 | -0.09108266 | 0.20928647 | -0.4352057 | 0.66341314 | NA | | NA | NA |
| ENSG000002 | 0 | NA | NA | NA | NA | NA | | NA | NA |
| ENSG000002 | 0.12070844 | -0.088503695 | 0.12783931 | -0.6923042 | 0.48874631 | NA | | NA | NA |
| ENSG000002 | 0 | NA | NA | NA | NA | NA | | NA | NA |
| ENSG000002 | 0.58765349 | 0.017801079 | 0.15713255 | 0.11328702 | 0.90980301 | NA | | MIR8068 | 102466876 |
| ENSG000002 | 1.11969614 | -0.406844482 | 0.31653256 | -1.2853164 | 0.19868175 | NA | | NA | NA |
| ENSG000002 | 2.05850924 | -0.738246018 | 0.36154534 | -2.0419182 | 0.04115964 | NA | | NA | NA |

| | | | | | | | | |
|---|---|---|---|---|---|---|---|---|
| ENSG000002 | 1.7116637 | -0.480288119 | 0.3328984 | -1.4427469 | 0.14909179 | NA | NA | NA |
| ENSG000002 | 1.09435054 | 0.190580096 | 0.32153387 | 0.59272168 | 0.55336743 | NA | NA | NA |
| ENSG000002 | 0 | NA | NA | NA | NA | NA | NA | NA |
| ENSG000002 | 0 | NA | NA | NA | NA | NA | MIR6877 | 102465528 |
| ENSG000002 | 0 | NA | NA | NA | NA | NA | NA | NA |
| ENSG000002 | 0 | NA | NA | NA | NA | NA | RN7SL722P | 106479486 |
| ENSG000002 | 0 | NA | NA | NA | NA | NA | NA | NA |
| ENSG000002 | 0 | NA | NA | NA | NA | NA | MIR6853 | 102466201 |
| ENSG000002 | 1.47964726 | -0.180036679 | 0.34078803 | -0.5282952 | 0.59729447 | NA | RN7SL733P | 106479489 |
| ENSG000002 | 0.53969588 | -0.357270382 | 0.23978341 | -1.4899712 | 0.13623181 | NA | NA | NA |
| ENSG000002 | 0.62636422 | -0.258424308 | 0.2600681 | -0.9936794 | 0.32037899 | NA | NA | NA |
| ENSG000002 | 1.12194105 | -0.27643682 | 0.32183639 | -0.8589359 | 0.3903759 | NA | NA | NA |
| ENSG000002 | 0 | NA | NA | NA | NA | NA | NA | NA |
| ENSG000002 | 0 | NA | NA | NA | NA | NA | NA | NA |
| ENSG000002 | 1.27651992 | -0.662487327 | 0.32869433 | -2.0155119 | 0.04385104 | NA | NA | NA |
| ENSG000002 | 0.81017068 | -0.540624039 | 0.28189242 | -1.9178381 | 0.05513154 | NA | NA | NA |
| ENSG000002 | 0 | NA | NA | NA | NA | NA | NA | NA |
| ENSG000002 | 1.71679399 | 0.062407713 | 0.35079921 | 0.17790152 | 0.85880032 | NA | NA | NA |
| ENSG000002 | 0.75992838 | -0.028323838 | 0.27990546 | -0.1011907 | 0.91939906 | NA | NA | NA |
| ENSG000002 | 0.86987519 | 0.192770645 | 0.29220536 | 0.65970949 | 0.50944028 | NA | NA | NA |
| ENSG000002 | 0.11131518 | 0.081951103 | 0.12663804 | 0.64712866 | 0.51754868 | NA | RN7SL106P | 106479261 |
| ENSG000002 | 0.74435521 | -0.028370124 | 0.27885082 | -0.1017394 | 0.9189635 | NA | NA | NA |
| ENSG000002 | 0.59687261 | 0.065989748 | 0.26864914 | 0.24563543 | 0.80596446 | NA | NA | NA |
| ENSG000002 | 1.81167961 | -0.119081425 | 0.35927314 | -0.331451 | 0.74030388 | NA | NA | NA |
| ENSG000002 | 0.35596481 | 0.219647334 | 0.19879223 | 1.10490904 | 0.26919901 | NA | NA | NA |
| ENSG000002 | 1.84121873 | 0.138622411 | 0.35016948 | 0.39587234 | 0.69219921 | NA | NA | NA |
| ENSG000002 | 1.0113864 | -0.045143544 | 0.30728843 | -0.1469094 | 0.88320357 | NA | NA | NA |
| ENSG000002 | 1.72706827 | -0.064149221 | 0.35326294 | -0.1815906 | 0.85590404 | NA | NA | NA |
| ENSG000002 | 0 | NA | NA | NA | NA | NA | MIR3670-4 | 103504740 |

| | | | | | | | | |
|---|---|---|---|---|---|---|---|---|
| ENSG000002 | 0.38133351 | -0.102466351 | 0.2189713 | -0.4679442 | 0.63982449 | NA | NA | NA |
| ENSG000002 | 1.37914872 | -0.271739879 | 0.34173587 | -0.7951752 | 0.42651161 | NA | NA | NA |
| ENSG000002 | 0 | NA | NA | NA | NA | NA | MIR6813 | 102466741 |
| ENSG000002 | 0.11131518 | 0.081951103 | 0.12663804 | 0.64712866 | 0.51754868 | NA | NA | NA |
| ENSG000002 | 0 | NA | NA | NA | NA | NA | NA | NA |
| ENSG000002 | 0.35596481 | 0.219647334 | 0.19879223 | 1.10490904 | 0.26919901 | NA | NA | NA |
| ENSG000002 | 0.86894445 | -0.248832029 | 0.29149308 | -0.8536464 | 0.39330092 | NA | NA | NA |
| ENSG000002 | 2.0559766 | 0.001879898 | 0.36263355 | 0.00518401 | 0.99586377 | NA | MIR4727 | 100616416 |
| ENSG000002 | 0 | NA | NA | NA | NA | NA | NA | NA |
| ENSG000002 | 1.08772681 | 0.040180652 | 0.32112645 | 0.12512408 | 0.90042532 | NA | MIR6507 | 102465253 |
| ENSG000002 | 0 | NA | NA | NA | NA | NA | MIR6724-2 | 103504727 |
| ENSG000002 | 0 | NA | NA | NA | NA | NA | NA | NA |
| ENSG000002 | 1.22878547 | -0.450307556 | 0.31886017 | -1.4122415 | 0.15787887 | NA | NA | NA |
| ENSG000002 | 0.23890637 | -0.010420674 | 0.17747379 | -0.0587167 | 0.95317776 | NA | MIR6514 | 102465257 |
| ENSG000002 | 0 | NA | NA | NA | NA | NA | NA | NA |
| ENSG000002 | 0 | NA | NA | NA | NA | NA | NA | NA |
| ENSG000002 | 0 | NA | NA | NA | NA | NA | NA | NA |
| ENSG000002 | 0.56744359 | 0.212023309 | 0.25327123 | 0.83713935 | 0.40251425 | NA | NA | NA |
| ENSG000002 | 0 | NA | NA | NA | NA | NA | RN7SL539P | 106479418 |
| ENSG000002 | 0 | NA | NA | NA | NA | NA | NA | NA |
| ENSG000002 | 0 | NA | NA | NA | NA | NA | SNORD1C | 677850 |
| ENSG000002 | 0.38056192 | -0.244105065 | 0.20391883 | -1.1970698 | 0.23127937 | NA | NA | NA |
| ENSG000002 | 0.87926845 | 0.044006275 | 0.29196642 | 0.15072375 | 0.88019364 | NA | NA | NA |
| ENSG000002 | 1.54280692 | 0.279608508 | 0.34191268 | 0.81777753 | 0.41348423 | NA | RN7SL787P | 106479507 |
| ENSG000002 | 0.60076876 | 0.230005242 | 0.26364708 | 0.87239822 | 0.38299117 | NA | FAM27AP1 | 106481705 |
| ENSG000002 | 0.26813897 | -0.197007041 | 0.18550319 | -1.0620143 | 0.2882292 | NA | OR4M2 | 390538 |
| ENSG000002 | 0 | NA | NA | NA | NA | NA | MIR6777 | 102465466 |
| ENSG000002 | 0.85809446 | -0.24599379 | 0.28838825 | -0.8529952 | 0.39366197 | NA | MIR6081 | 102466518 |
| ENSG000002 | 0.86031874 | -0.256095975 | 0.29097097 | -0.8801427 | 0.378782 | NA | NA | NA |

| | | | | | | | | | |
|---|---|---|---|---|---|---|---|---|---|
| ENSG000002 | 0.38036605 | -0.293832733 | 0.22119083 | -1.3284128 | 0.18404178 | NA | NA | NA | |
| ENSG000002 | 1.09840357 | -0.254828384 | 0.31659239 | -0.80491 | 0.42087161 | NA | NA | NA | |
| ENSG000002 | 1.39873607 | -0.572455657 | 0.33675573 | -1.6999136 | 0.08914718 | NA | NA | NA | |
| ENSG000002 | 0.67141739 | -0.461190807 | 0.26562301 | -1.7362608 | 0.08251773 | NA | NA | NA | |
| ENSG000002 | 0.52165104 | -0.403921832 | 0.25252134 | -1.5995552 | 0.10969729 | NA | MIR6774 | | 102466732 |
| ENSG000002 | 0.38270188 | -0.272446727 | 0.2140009 | -1.2731102 | 0.20297894 | NA | NA | NA | |
| ENSG000002 | 1.35154508 | -0.256593355 | 0.33447105 | -0.7671616 | 0.44298542 | NA | NA | NA | |
| ENSG000002 | 0.87050422 | -0.120430766 | 0.30290137 | -0.3975907 | 0.69093192 | NA | NA | NA | |
| ENSG000002 | 0.81363764 | 0.207625688 | 0.29485537 | 0.70416113 | 0.48133244 | NA | NA | NA | |
| ENSG000002 | 0.13280363 | -0.093394593 | 0.13115644 | -0.7120855 | 0.47641185 | NA | NA | NA | |
| ENSG000002 | 0 | NA | NA | NA | NA | NA | NA | NA | |
| ENSG000002 | 2.1573804 | -0.393969893 | 0.36684707 | -1.073935 | 0.28285183 | NA | RNA5SP439 | | 106479014 |
| ENSG000002 | 0.85817878 | -0.255621562 | 0.29085906 | -0.8788503 | 0.37948247 | NA | RN7SL585P | | 106479436 |
| ENSG000002 | 0.99849451 | -0.013680769 | 0.19890195 | -0.0687815 | 0.94516357 | NA | NA | NA | |
| ENSG000002 | 0.13280363 | -0.093394593 | 0.13115644 | -0.7120855 | 0.47641185 | NA | MIR8071-1 | | 102465871 |
| ENSG000002 | 0 | NA | NA | NA | NA | NA | NA | NA | |
| ENSG000002 | 1.01226796 | -0.034445764 | 0.303293 | -0.1135726 | 0.90957664 | NA | NA | NA | |
| ENSG000002 | 0.87280567 | -0.239192171 | 0.28216319 | -0.8477086 | 0.39660025 | NA | NA | NA | |
| ENSG000002 | 1.27475011 | -0.194872149 | 0.32794824 | -0.5942162 | 0.55236754 | NA | NA | NA | |
| ENSG000002 | 0 | NA | NA | NA | NA | NA | H2AFB1 | | 474382 |
| ENSG000002 | 0.26062507 | -0.012888319 | 0.18226503 | -0.070712 | 0.94362699 | NA | RN7SL248P | | 106479312 |
| ENSG000002 | 0 | NA | NA | NA | NA | NA | NA | NA | |
| ENSG000002 | 0.59829496 | 0.063152461 | 0.26232549 | 0.24074084 | 0.80975598 | NA | NA | NA | |
| ENSG000002 | 1.39858312 | -0.274480122 | 0.33959349 | -0.8082608 | 0.41894044 | NA | NA | NA | |
| ENSG000002 | 0.92247506 | -0.105203374 | 0.28310368 | -0.3716072 | 0.7101853 | NA | NA | NA | |
| ENSG000002 | 0.40942396 | -0.284058841 | 0.21788566 | -1.303706 | 0.19233384 | NA | NA | NA | |
| ENSG000002 | 1.59955978 | -0.60473962 | 0.33768046 | -1.7908636 | 0.07331519 | NA | NA | NA | |
| ENSG000002 | 0.87289212 | -0.115830741 | 0.29794765 | -0.3887621 | 0.69745217 | NA | NA | NA | |
| ENSG000002 | 1.05226381 | 0.191973105 | 0.31854925 | 0.60264811 | 0.54674281 | NA | NA | NA | |

| | | | | | | | | |
|---|---|---|---|---|---|---|---|---|
| ENSG000002 | 1.72669922 | -0.440858892 | 0.35143928 | -1.2544383 | 0.20968274 | NA | NA | NA |
| ENSG000002 | 0.61767592 | -0.10105625 | 0.26442364 | -0.3821755 | 0.70233116 | NA | NA | NA |
| ENSG000002 | 1.39283903 | -0.554511425 | 0.33532997 | -1.6536292 | 0.09820288 | NA | NA | NA |
| ENSG000002 | 0.26199343 | -0.194018184 | 0.18422801 | -1.0531416 | 0.29227609 | NA | TBC1D3H | 729877 |
| ENSG000002 | 0 | NA | NA | NA | NA | NA | NA | NA |
| ENSG000002 | 0 | NA | NA | NA | NA | NA | NA | NA |
| ENSG000002 | 0 | NA | NA | NA | NA | NA | NA | NA |
| ENSG000002 | 0 | NA | NA | NA | NA | NA | RN7SL818P | 106479520 |
| ENSG000002 | 0.75944963 | -0.20423668 | 0.28685914 | -0.7119755 | 0.47647995 | NA | NA | NA |
| ENSG000002 | 1.70816119 | -0.165240853 | 0.3486098 | -0.4739994 | 0.6355003 | NA | NA | NA |
| ENSG000002 | 0.12070844 | -0.088503695 | 0.12783931 | -0.6923042 | 0.48874631 | NA | GGT2 | 728441 |
| ENSG000002 | 2.09538727 | -0.007582174 | 0.36697677 | -0.0206612 | 0.98351593 | NA | NA | NA |
| ENSG000002 | 0.72477837 | 0.128795829 | 0.27776445 | 0.46368724 | 0.64287184 | NA | MIR6728 | 102465436 |
| ENSG000002 | 2.00838956 | -0.431092333 | 0.35795529 | -1.2043189 | 0.22846633 | NA | MIR7702 | 102465800 |
| ENSG000002 | 0 | NA | NA | NA | NA | NA | SNORA73A | 6080 |
| ENSG000002 | 0.48418907 | 0.150283598 | 0.23847389 | 0.63018889 | 0.52857101 | NA | NA | NA |
| ENSG000002 | 1.28519716 | -0.636511691 | 0.32433393 | -1.9625196 | 0.04970203 | NA | NA | NA |
| ENSG000002 | 0.23936338 | -0.005479057 | 0.17759977 | -0.0308506 | 0.9753887 | NA | NA | NA |
| ENSG000002 | 1.47085324 | -0.163866128 | 0.33520565 | -0.4888525 | 0.6249461 | NA | MIR6069 | 102464824 |
| ENSG000002 | 0 | NA | NA | NA | NA | NA | NA | NA |
| ENSG000002 | 0 | NA | NA | NA | NA | NA | NA | NA |
| ENSG000002 | 0.39246123 | -0.276864021 | 0.21549122 | -1.2848042 | 0.19886073 | NA | NA | NA |
| ENSG000002 | 1.45622738 | -0.048705913 | 0.34068075 | -0.1429664 | 0.88631669 | NA | NA | NA |
| ENSG000002 | 1.55372668 | 0.160151764 | 0.34919983 | 0.45862498 | 0.6465035 | NA | NA | NA |
| ENSG000002 | 0 | NA | NA | NA | NA | NA | MIR6864 | 102465521 |
| ENSG000002 | 0.11934008 | 0.083564698 | 0.12760091 | 0.65489106 | 0.5125379 | NA | MIR5088 | 100847074 |
| ENSG000002 | 0 | NA | NA | NA | NA | NA | SNORA71E | 677821 |
| ENSG000002 | 0 | NA | NA | NA | NA | NA | NA | NA |
| ENSG000002 | 0 | NA | NA | NA | NA | NA | MIR6749 | 102466727 |

| | | | | | | | | |
|---|---|---|---|---|---|---|---|---|
| ENSG000002 | 2.03061467 | -0.594611954 | 0.36252826 | -1.640181 | 0.10096754 | NA | NA | NA |
| ENSG000002 | 1.18350265 | 0.241897219 | 0.32532449 | 0.74355675 | 0.45714468 | NA | NA | NA |
| ENSG000002 | 1.2403354 | -0.187582817 | 0.32782703 | -0.5722006 | 0.5671861 | NA | NA | NA |
| ENSG000002 | 0.71712404 | -0.025635761 | 0.28333197 | -0.0904796 | 0.92790611 | NA | NA | NA |
| ENSG000002 | 0 | NA | NA | NA | NA | NA | NA | NA |
| ENSG000002 | 0 | NA | NA | NA | NA | NA | NA | NA |
| ENSG000002 | 2.06880628 | -0.552708062 | 0.35470326 | -1.5582266 | 0.11917953 | NA | NA | NA |
| ENSG000002 | 0.38270188 | -0.272446727 | 0.2140009 | -1.2731102 | 0.20297894 | NA | NA | NA |
| ENSG000002 | 0 | NA | NA | NA | NA | NA | RN7SL853P | 106481153 |
| ENSG000002 | 0.50611225 | -0.202624884 | 0.24886748 | -0.8141879 | 0.41553732 | NA | NA | NA |
| ENSG000002 | 1.87265085 | -0.258390736 | 0.35858152 | -0.7205913 | 0.47116099 | NA | NA | NA |
| ENSG000002 | 0.47821768 | -0.016471461 | 0.24462483 | -0.0673336 | 0.94631616 | NA | NA | NA |
| ENSG000002 | 1.56497647 | -0.117635739 | 0.34670541 | -0.3392959 | 0.7343868 | NA | NA | NA |
| ENSG000002 | 0 | NA | NA | NA | NA | NA | NA | NA |
| ENSG000002 | 1.55065364 | 0.163921399 | 0.35142488 | 0.46644791 | 0.64089493 | NA | NA | NA |
| ENSG000002 | 0.74231222 | -0.037174701 | 0.27870433 | -0.133384 | 0.89388969 | NA | NA | NA |
| ENSG000002 | 0 | NA | NA | NA | NA | NA | NA | NA |
| ENSG000002 | 0.60968775 | -0.401217092 | 0.25115369 | -1.5974963 | 0.11015512 | NA | MIR6801 | 102466984 |
| ENSG000002 | 1.22465727 | -0.313628678 | 0.32335806 | -0.9699114 | 0.33209065 | NA | NA | NA |
| ENSG000002 | 0.35273207 | -0.08884607 | 0.20605566 | -0.4311751 | 0.66634105 | NA | NA | NA |
| ENSG000002 | 1.86154432 | -0.001201837 | 0.35791213 | -0.0033579 | 0.99732078 | NA | NA | NA |
| ENSG000002 | 0.23936338 | -0.005479057 | 0.17759977 | -0.0308506 | 0.9753887 | NA | MIR6885 | 102465533 |
| ENSG000002 | 0 | NA | NA | NA | NA | NA | NA | NA |
| ENSG000002 | 1.54718613 | -0.501625296 | 0.34658118 | -1.447353 | 0.14779809 | NA | NA | NA |
| ENSG000002 | 0.51180131 | -0.185828522 | 0.2424294 | -0.7665263 | 0.44336319 | NA | NA | NA |
| ENSG000002 | 0 | NA | NA | NA | NA | NA | MIR6515 | 102466659 |
| ENSG000002 | 0.11865494 | 0.083440866 | 0.12751478 | 0.65436232 | 0.5128784 | NA | NA | NA |
| ENSG000002 | 0 | NA | NA | NA | NA | NA | RN7SL196P | 106479293 |
| ENSG000002 | 0 | NA | NA | NA | NA | NA | NA | NA |

| | | | | | | | | |
|---|---|---|---|---|---|---|---|---|
| ENSG000002 | 0.11205239 | 0.082118251 | 0.12671962 | 0.64803108 | 0.51696485 | NA | NA | NA |
| ENSG000002 | 0 | NA | NA | NA | NA | NA | NA | NA |
| ENSG000002 | 1.02644699 | -0.370618883 | 0.31595273 | -1.1730201 | 0.24078776 | NA | NA | NA |
| ENSG000002 | 0.25214371 | -0.009862892 | 0.18046657 | -0.0546522 | 0.95641556 | NA | NA | NA |
| ENSG000002 | 1.95586879 | -0.163816387 | 0.35333558 | -0.4636283 | 0.64291404 | NA | NA | NA |
| ENSG000002 | 0.37404583 | -0.105111695 | 0.21784993 | -0.4824959 | 0.62945372 | NA | NA | NA |
| ENSG000002 | 0.49970612 | -0.194507174 | 0.24827631 | -0.7834303 | 0.43337449 | NA | NA | NA |
| ENSG000002 | 0.47895489 | -0.016192723 | 0.24471014 | -0.066171 | 0.94724166 | NA | NA | NA |
| ENSG000002 | 0.3854038 | -0.110128691 | 0.21952326 | -0.5016721 | 0.61589821 | NA | MIR1273H | 102466247 |
| ENSG000002 | 0 | NA | NA | NA | NA | NA | NA | NA |
| ENSG000002 | 0.9818194 | -0.042650985 | 0.31258931 | -0.1364442 | 0.89147017 | NA | NA | NA |
| ENSG000002 | 0.84975353 | 0.05329801 | 0.30153883 | 0.17675339 | 0.8597021 | NA | NA | NA |
| ENSG000002 | 1.969645 | -0.326664791 | 0.36425514 | -0.8968022 | 0.3698245 | NA | NA | NA |
| ENSG000002 | 1.01270111 | -0.3663759 | 0.31545448 | -1.1614224 | 0.24547016 | NA | NA | NA |
| ENSG000002 | 0 | NA | NA | NA | NA | NA | NA | NA |
| ENSG000002 | 0.12070844 | -0.088503695 | 0.12783931 | -0.6923042 | 0.48874631 | NA | NA | NA |
| ENSG000002 | 0.74279096 | 0.133965577 | 0.28515073 | 0.46980618 | 0.6384935 | NA | NA | NA |
| ENSG000002 | 0.97630502 | -0.188217358 | 0.30650371 | -0.6140786 | 0.53916342 | NA | MIR6832 | 102466745 |
| ENSG000002 | 0.61946829 | -0.085161233 | 0.1574564 | -0.540856 | 0.58860686 | NA | NA | NA |
| ENSG000002 | 2.09928735 | -0.137015167 | 0.36823357 | -0.3720877 | 0.70982759 | NA | NA | NA |
| ENSG000002 | 0.23936338 | -0.005479057 | 0.17759977 | -0.0308506 | 0.9753887 | NA | NA | NA |
| ENSG000002 | 0 | NA | NA | NA | NA | NA | NA | NA |
| ENSG000002 | 1.37712555 | -0.275905675 | 0.34168334 | -0.8074894 | 0.41938458 | NA | NA | NA |
| ENSG000002 | 1.19157962 | 0.257848452 | 0.33136629 | 0.77813725 | 0.4364881 | NA | NA | NA |
| ENSG000002 | 2.15155336 | -0.503944995 | 0.36400024 | -1.3844634 | 0.16621662 | NA | NA | NA |
| ENSG000002 | 1.28821366 | -0.358772376 | 0.33281566 | -1.0779913 | 0.28103765 | NA | NA | NA |
| ENSG000002 | 1.11852365 | -0.279165329 | 0.32689835 | -0.8539821 | 0.39311492 | NA | FAM74A6 | 653123 |
| ENSG000002 | 0.50341032 | -0.342798738 | 0.23572799 | -1.4542131 | 0.14588723 | NA | NA | NA |
| ENSG000002 | 0.23816916 | -0.010712105 | 0.1772998 | -0.060418 | 0.9518227 | NA | SEBOX | 645832 |

| | | | | | | | | |
|---|---|---|---|---|---|---|---|---|
| ENSG000002 | 0.14128499 | -0.097772014 | 0.13403235 | -0.7294658 | 0.4657168 | NA | NA | NA |
| ENSG000002 | 0.9779238 | -0.181055242 | 0.30825798 | -0.5873497 | 0.55696885 | NA | NA | NA |
| ENSG000002 | 0 | NA | NA | NA | NA | NA | SNORD28 | 9300 |
| ENSG000002 | 0 | NA | NA | NA | NA | NA | NA | NA |
| ENSG000002 | 1.47711501 | -0.189449798 | 0.34150875 | -0.5547436 | 0.57907004 | NA | NA | NA |
| ENSG000002 | 0.26199343 | -0.194018184 | 0.18422801 | -1.0531416 | 0.29227609 | NA | MIR6889 | 102466758 |
| ENSG000002 | 0 | NA | NA | NA | NA | NA | NA | NA |
| ENSG000002 | 1.38780422 | -0.399836427 | 0.33341859 | -1.1992026 | 0.2304492 | NA | NA | NA |
| ENSG000002 | 1.16896557 | 0.084988484 | 0.30745821 | 0.27642288 | 0.78222328 | NA | NA | NA |
| ENSG000002 | 0 | NA | NA | NA | NA | NA | RN7SL545P | 106481848 |
| ENSG000002 | 1.55985048 | 0.017202414 | 0.3517514 | 0.04890503 | 0.96099498 | NA | NA | NA |
| ENSG000002 | 0.50722002 | -0.375409304 | 0.24493426 | -1.5326941 | 0.12535123 | NA | ZYXP1 | 106480342 |
| ENSG000002 | 0.28256998 | -0.18483211 | 0.18021686 | -1.0256094 | 0.30507573 | NA | NA | NA |
| ENSG000002 | 2.16935961 | 0.058514724 | 0.3664255 | 0.15969065 | 0.87312477 | NA | NA | NA |
| ENSG000002 | 0 | NA | NA | NA | NA | NA | SNORA16A | 692073 |
| ENSG000002 | 1.58193974 | -0.121759283 | 0.35328266 | -0.3446512 | 0.73035663 | NA | NA | NA |
| ENSG000002 | 0 | NA | NA | NA | NA | NA | RNU2-1 | 6066 |
| ENSG000002 | 1.35358551 | 0.167537872 | 0.33923804 | 0.49386523 | 0.62140136 | NA | NA | NA |
| ENSG000002 | 0.73869839 | -0.035763555 | 0.2784475 | -0.1284391 | 0.89780147 | NA | NA | NA |
| ENSG000002 | 0 | NA | NA | NA | NA | NA | NA | NA |
| ENSG000002 | 1.4163501 | -0.739047635 | 0.33693248 | -2.1934592 | 0.02827431 | NA | RIMBP3B | 440804 |
| ENSG000002 | 0.65427998 | -0.290650429 | 0.26916107 | -1.0798383 | 0.28021419 | NA | NA | NA |
| ENSG000002 | 1.23091235 | 0.102041544 | 0.32849205 | 0.31063627 | 0.75607715 | NA | NA | NA |
| ENSG000002 | 0 | NA | NA | NA | NA | NA | NA | NA |
| ENSG000002 | 0.48624257 | -0.012072666 | 0.23796621 | -0.0507327 | 0.95953853 | NA | RNU6-866P | 106481454 |
| ENSG000002 | 0.27408862 | -0.199826037 | 0.18669465 | -1.0703362 | 0.28446801 | NA | TBC1D3 | 729873 |
| ENSG000002 | 1.83572755 | -0.001986967 | 0.35334881 | -0.0056232 | 0.99551332 | NA | NA | NA |
| ENSG000002 | 2.03402548 | -0.734260037 | 0.3613899 | -2.0317669 | 0.04217726 | NA | NA | NA |
| ENSG000002 | 0.98708839 | -0.340680839 | 0.30844566 | -1.1045085 | 0.26937263 | NA | NA | NA |

| | | | | | | | | | |
|---|---|---|---|---|---|---|---|---|---|
| ENSG000002 | 0 | NA | NA | NA | NA | NA | | MIR6867 | 102465523 |
| ENSG000002 | 0 | NA | NA | NA | NA | NA | | NA | NA |
| ENSG000002 | 0.9928639 | -0.178222784 | 0.30724023 | -0.5800763 | 0.56186314 | NA | | NA | NA |
| ENSG000002 | 0.50722002 | -0.375409304 | 0.24493426 | -1.5326941 | 0.12535123 | NA | | RN7SL719P | 106481119 |
| ENSG000002 | 0.78247198 | -0.337783522 | 0.27652822 | -1.2215156 | 0.22189088 | NA | | NA | NA |
| ENSG000002 | 0 | NA | NA | NA | NA | NA | | SNORD109A | 338428 |
| ENSG000002 | 0.23276084 | -0.008011382 | 0.17603982 | -0.0455089 | 0.96370166 | NA | | NA | NA |
| ENSG000002 | 0.78130864 | -0.565696235 | 0.28699154 | -1.9711251 | 0.04870956 | NA | | NA | NA |
| ENSG000002 | 0 | NA | NA | NA | NA | NA | | NA | NA |
| ENSG000002 | 0.14128499 | -0.097772014 | 0.13403235 | -0.7294658 | 0.4657168 | NA | | RN7SKP236 | 106480906 |
| ENSG000002 | 0 | NA | NA | NA | NA | NA | | NA | NA |
| ENSG000002 | 1.35009026 | -0.269708286 | 0.33990144 | -0.7934897 | 0.42749258 | NA | | NA | NA |
| ENSG000002 | 0.73906896 | -0.197744251 | 0.28558485 | -0.6924186 | 0.48867451 | NA | | NA | NA |
| ENSG000002 | 1.25758099 | -0.04149421 | 0.3292537 | -0.126025 | 0.89971211 | NA | | NA | NA |
| ENSG000002 | 1.45766508 | 0.229729776 | 0.34527202 | 0.6653588 | 0.50582103 | NA | | NA | NA |
| ENSG000002 | 1.05831952 | 0.352178843 | 0.3194342 | 1.10250826 | 0.27024077 | NA | | NA | NA |
| ENSG000002 | 1.21417797 | 0.109330819 | 0.33172076 | 0.32958691 | 0.74171212 | NA | | NA | NA |
| ENSG000002 | 0.50681912 | -0.019233989 | 0.24032284 | -0.080034 | 0.93621025 | NA | | NA | NA |
| ENSG000002 | 0.11131518 | 0.081951103 | 0.12663804 | 0.64712866 | 0.51754868 | NA | | MIR6761 | 102465456 |
| ENSG000002 | 1.13272441 | -0.440635094 | 0.32484009 | -1.3564677 | 0.17495039 | NA | | NA | NA |
| ENSG000002 | 1.72454103 | -0.313507061 | 0.35155303 | -0.8917774 | 0.37251223 | NA | | NA | NA |
| ENSG000002 | 0.83486763 | 0.214468614 | 0.29601929 | 0.72450892 | 0.46875335 | NA | | NA | NA |
| ENSG000002 | 0 | NA | NA | NA | NA | NA | | NA | NA |
| ENSG000002 | 0 | NA | NA | NA | NA | NA | | NA | NA |
| ENSG000002 | 0.84745153 | 0.201391656 | 0.2967784 | 0.6785927 | 0.49739597 | NA | | MIR7974 | 102465856 |
| ENSG000002 | 0 | NA | NA | NA | NA | NA | | NA | NA |
| ENSG000002 | 1.26225913 | -0.470078123 | 0.32335897 | -1.4537346 | 0.1460199 | NA | | NA | NA |
| ENSG000002 | 0.39499294 | -0.277908454 | 0.21584201 | -1.287555 | 0.19790091 | NA | | NA | NA |
| ENSG000002 | 0.61881753 | -0.112804359 | 0.27082437 | -0.4165222 | 0.67702794 | NA | | NA | NA |

| | | | | | | | | |
|---|---|---|---|---|---|---|---|---|
| ENSG000002 | 0.23202362 | -0.008302493 | 0.17586207 | -0.0472103 | 0.96234565 | NA | NA | NA |
| ENSG000002 | 0 | NA | NA | NA | NA | NA | NA | NA |
| ENSG000002 | 0.12685397 | -0.090215768 | 0.12901919 | -0.699243 | 0.48440017 | NA | NA | NA |
| ENSG000002 | 0 | NA | NA | NA | NA | NA | TCEB3CL2 | 100506888 |
| ENSG000002 | 0.11865494 | 0.083440866 | 0.12751478 | 0.65436232 | 0.5128784 | NA | RN7SL627P | 106481093 |
| ENSG000002 | 0.74268278 | -0.18879062 | 0.27975142 | -0.6748513 | 0.49977023 | NA | NA | NA |
| ENSG000002 | 1.35495388 | 0.022324065 | 0.34323608 | 0.06503997 | 0.94814217 | NA | NA | NA |
| ENSG000002 | 1.97781996 | -0.193806837 | 0.35934753 | -0.5393298 | 0.5896593 | NA | NA | NA |
| ENSG000002 | 1.52216818 | -0.315463624 | 0.33428737 | -0.9436899 | 0.34532812 | NA | NA | NA |
| ENSG000002 | 1.8569413 | -0.247823058 | 0.35303589 | -0.701977 | 0.48269353 | NA | NA | NA |
| ENSG000002 | 0 | NA | NA | NA | NA | NA | NA | NA |
| ENSG000002 | 2.00497106 | -0.434556391 | 0.35855854 | -1.2119538 | 0.22553006 | NA | NA | NA |
| ENSG000002 | 0.62684242 | -0.109923117 | 0.27148862 | -0.4048903 | 0.68555813 | NA | NA | NA |
| ENSG000002 | 0.49264869 | -0.021364872 | 0.24621241 | -0.0867741 | 0.93085104 | NA | MIR6128 | 102465135 |
| ENSG000002 | 0.25370795 | -0.171331747 | 0.17411721 | -0.9840024 | 0.32511435 | NA | NA | NA |
| ENSG000002 | 0 | NA | NA | NA | NA | NA | NA | NA |
| ENSG000002 | 0 | NA | NA | NA | NA | NA | NA | NA |
| ENSG000002 | 0 | NA | NA | NA | NA | NA | F8A2 | 474383 |
| ENSG000002 | 0.61816464 | -0.117186973 | 0.27074914 | -0.4328249 | 0.66514197 | NA | NA | NA |
| ENSG000002 | 1.69458848 | 0.208067111 | 0.35295161 | 0.58950605 | 0.55552186 | NA | NA | NA |
| ENSG000002 | 2.02628537 | -0.451774831 | 0.3626911 | -1.2456187 | 0.2129044 | NA | NA | NA |
| ENSG000002 | 1.11672861 | -0.278122138 | 0.32315071 | -0.8606577 | 0.38942661 | NA | MIR6768 | 102465460 |
| ENSG000002 | 0.86390222 | -0.107975723 | 0.29749683 | -0.3629475 | 0.71664412 | NA | MIR6078 | 102464829 |
| ENSG000002 | 1.08701133 | 0.194779844 | 0.31975983 | 0.6091442 | 0.54242886 | NA | NA | NA |
| ENSG000002 | 0.4009426 | -0.302672773 | 0.2240145 | -1.3511303 | 0.17665371 | NA | NA | NA |
| ENSG000002 | 0 | NA | NA | NA | NA | NA | MIR6772 | 102465463 |
| ENSG000002 | 1.59879483 | -0.382665378 | 0.34662871 | -1.1039633 | 0.26960905 | NA | NA | NA |
| ENSG000002 | 0 | NA | NA | NA | NA | NA | NA | NA |
| ENSG000002 | 0.7418235 | -0.027672983 | 0.27868031 | -0.0993001 | 0.9209 | NA | NA | NA |

| | | | | | | | | |
|---|---|---|---|---|---|---|---|---|
| ENSG000002 | 0 | NA | NA | NA | NA | NA | MIR6762 | 102465457 |
| ENSG000002 | 1.12521052 | -0.280538266 | 0.32709639 | -0.8576624 | 0.39107892 | NA | NA | NA |
| ENSG000002 | 0 | NA | NA | NA | NA | NA | MIR7152 | 102465689 |
| ENSG000002 | 1.57376244 | 0.161986098 | 0.34947409 | 0.4635139 | 0.64299605 | NA | NA | NA |
| ENSG000002 | 1.08769593 | 0.178511065 | 0.31523925 | 0.56627169 | 0.5712091 | NA | NA | NA |
| ENSG000002 | 0.75982019 | -0.379788636 | 0.28855699 | -1.3161651 | 0.18811864 | NA | NA | NA |
| ENSG000002 | 0.35690842 | 0.061733055 | 0.20636749 | 0.29914137 | 0.76483218 | NA | NA | NA |
| ENSG000002 | 0.24756242 | -0.187034346 | 0.18119817 | -1.0322088 | 0.30197431 | NA | NA | NA |
| ENSG000002 | 0 | NA | NA | NA | NA | NA | NA | NA |
| ENSG000002 | 0 | NA | NA | NA | NA | NA | MIR6788 | 102466735 |
| ENSG000002 | 1.50580627 | -0.3481815 | 0.3469402 | -1.0035778 | 0.31558214 | NA | NA | NA |
| ENSG000002 | 0.60282172 | 0.050569991 | 0.26272347 | 0.19248372 | 0.84736332 | NA | NA | NA |
| ENSG000002 | 0 | NA | NA | NA | NA | NA | NA | NA |
| ENSG000002 | 0 | NA | NA | NA | NA | NA | NA | NA |
| ENSG000002 | 2.05357912 | -0.601368194 | 0.36300938 | -1.6566189 | 0.09759655 | NA | NA | NA |
| ENSG000002 | 0.13280363 | -0.093394593 | 0.13115644 | -0.7120855 | 0.47641185 | NA | RN7SL422P | 106481038 |
| ENSG000002 | 0.83705698 | 0.037685274 | 0.28530005 | 0.13208997 | 0.89491313 | NA | NA | NA |
| ENSG000002 | 0 | NA | NA | NA | NA | NA | NA | NA |
| ENSG000002 | 0 | NA | NA | NA | NA | NA | NA | NA |
| ENSG000002 | 0 | NA | NA | NA | NA | NA | NA | NA |
| ENSG000002 | 1.45577091 | -0.045643567 | 0.3439022 | -0.1327225 | 0.89441283 | NA | NA | NA |
| ENSG000002 | 0 | NA | NA | NA | NA | NA | NA | NA |
| ENSG000002 | 0 | NA | NA | NA | NA | NA | NA | NA |
| ENSG000002 | 0.25351207 | -0.189965547 | 0.1824784 | -1.0410303 | 0.29786148 | NA | NA | NA |
| ENSG000002 | 0.77339385 | -0.029713334 | 0.27178516 | -0.1093265 | 0.91294349 | NA | NA | NA |
| ENSG000002 | 0.85177478 | -0.09428293 | 0.29078922 | -0.3242312 | 0.74576302 | NA | NA | NA |
| ENSG000002 | 0 | NA | NA | NA | NA | NA | NA | NA |
| ENSG000002 | 2.10445061 | -0.482837652 | 0.36184358 | -1.3343823 | 0.18207861 | NA | SEPT14P17 | 106480344 |
| ENSG000002 | 0.34999534 | 0.241411141 | 0.20772605 | 1.16216113 | 0.24517002 | NA | NA | NA |

| | | | | | | | | |
|---|---|---|---|---|---|---|---|---|
| ENSG000002 | 0 | NA | NA | NA | NA | NA | NA | NA |
| ENSG000002 | 1.62830818 | -0.507524421 | 0.34386609 | -1.4759362 | 0.13996101 | NA | NA | NA |
| ENSG000002 | 1.89351207 | -0.552546357 | 0.36214262 | -1.52577 | 0.12706715 | NA | NA | NA |
| ENSG000002 | 0.2596576 | -0.193002035 | 0.1837913 | -1.0501152 | 0.29366517 | NA | NA | NA |
| ENSG000002 | 0 | NA | NA | NA | NA | NA | RNA5-8S5 | 100008587 |
| ENSG000002 | 1.87654935 | -0.540593626 | 0.36072128 | -1.4986464 | 0.1339654 | NA | NA | NA |
| ENSG000002 | 1.01861852 | -0.350116737 | 0.31136779 | -1.1244475 | 0.26082322 | NA | NA | NA |
| ENSG000002 | 1.69042074 | -0.074378515 | 0.24858607 | -0.2992063 | 0.76478265 | NA | NA | NA |
| ENSG000002 | 1.11809752 | -0.409040804 | 0.31729248 | -1.2891601 | 0.19734244 | NA | NA | NA |
| ENSG000002 | 0 | NA | NA | NA | NA | NA | MIR7150 | 102465688 |
| ENSG000002 | 0.5008998 | -0.200150804 | 0.24832882 | -0.805991 | 0.42024801 | NA | TBC1D3I | 102724862 |
| ENSG000002 | 0.88881412 | -0.432208381 | 0.29509244 | -1.4646542 | 0.14301529 | NA | MIR8083 | 102466879 |
| ENSG000002 | 0.11934008 | 0.083564698 | 0.12760091 | 0.65489106 | 0.5125379 | NA | NA | NA |
| ENSG000002 | 1.03419114 | -0.368499765 | 0.3147961 | -1.1705983 | 0.2417603 | NA | NA | NA |
| ENSG000002 | 0.27408862 | -0.199826037 | 0.18669465 | -1.0703362 | 0.28446801 | NA | NA | NA |
| ENSG000002 | 0 | NA | NA | NA | NA | NA | NA | NA |
| ENSG000002 | 0.25351207 | -0.189965547 | 0.1824784 | -1.0410303 | 0.29786148 | NA | NA | NA |
| ENSG000002 | 0 | NA | NA | NA | NA | NA | MIR6771 | 102465462 |
| ENSG000002 | 0.14128499 | -0.097772014 | 0.13403235 | -0.7294658 | 0.4657168 | NA | NA | NA |
| ENSG000002 | 0 | NA | NA | NA | NA | NA | MIR6735 | 102465440 |
| ENSG000002 | 0.86316501 | -0.108242685 | 0.2974574 | -0.3638931 | 0.71593786 | NA | NA | NA |
| ENSG000002 | 0.11934008 | 0.083564698 | 0.12760091 | 0.65489106 | 0.5125379 | NA | RNU11 | 26824 |
| ENSG000002 | 0 | NA | NA | NA | NA | NA | NA | NA |
| ENSG000002 | 0.73719014 | -0.192711607 | 0.28549663 | -0.6750048 | 0.4996727 | NA | NA | NA |
| ENSG000002 | 0.12685397 | -0.090215768 | 0.12901919 | -0.699243 | 0.48440017 | NA | MIR6750 | 102466192 |
| ENSG000002 | 1.53255923 | -0.501544651 | 0.34700709 | -1.4453441 | 0.14836126 | NA | NA | NA |
| ENSG000002 | 0.11934008 | 0.083564698 | 0.12760091 | 0.65489106 | 0.5125379 | NA | MIR6876 | 102465527 |
| ENSG000002 | 1.18236551 | -0.032029269 | 0.32487177 | -0.0985905 | 0.92146342 | NA | MIR8056 | 102465862 |
| ENSG000002 | 1.70005777 | -0.524144109 | 0.34411158 | -1.5231807 | 0.12771351 | NA | NA | NA |

| | | | | | | | | |
|---|---|---|---|---|---|---|---|---|
| ENSG000002 | 0 | NA | NA | NA | NA | NA | SNORA17 | 677804 |
| ENSG000002 | 1.38093525 | -0.26204044 | 0.33588484 | -0.7801496 | 0.4353028 | NA | MIR8052 | 102466873 |
| ENSG000002 | 1.91001441 | -0.25479363 | 0.35785635 | -0.7119997 | 0.47646494 | NA | NA | NA |
| ENSG000002 | 2.0116241 | -0.572307256 | 0.36038238 | -1.5880556 | 0.11227377 | NA | NA | NA |
| ENSG000002 | 0.13280363 | -0.093394593 | 0.13115644 | -0.7120855 | 0.47641185 | NA | NA | NA |
| ENSG000002 | 0.35664995 | 0.244349217 | 0.20882949 | 1.17008962 | 0.2419649 | NA | NA | NA |
| ENSG000002 | 1.39932437 | 0.01429156 | 0.32609395 | 0.04382651 | 0.96504269 | NA | NA | NA |
| ENSG000002 | 0.23202362 | -0.008302493 | 0.17586207 | -0.0472103 | 0.96234565 | NA | MIR6753 | 102465451 |
| ENSG000002 | 1.16998136 | -0.602258787 | 0.32106012 | -1.8758443 | 0.06067666 | NA | NA | NA |
| ENSG000002 | 1.53400494 | 0.28220105 | 0.34439488 | 0.81941128 | 0.4125518 | NA | NA | NA |
| ENSG000002 | 0 | NA | NA | NA | NA | NA | RNU6-538P | 106479805 |
| ENSG000002 | 0 | NA | NA | NA | NA | NA | NA | NA |
| ENSG000002 | 0 | NA | NA | NA | NA | NA | LOC102723655 | 102723655 |
| ENSG000002 | 0.85021 | 0.04998197 | 0.30595275 | 0.163365 | 0.87023106 | NA | MIR8086 | 102465880 |
| ENSG000002 | 1.12099979 | -0.549336909 | 0.31273209 | -1.7565735 | 0.07899053 | NA | NA | NA |
| ENSG000002 | 1.36104831 | -0.52248844 | 0.33002792 | -1.5831643 | 0.11338401 | NA | NA | NA |
| ENSG000002 | 0 | NA | NA | NA | NA | NA | SNORD25 | 9303 |
| ENSG000002 | 0.840795 | 0.038310401 | 0.29427135 | 0.13018733 | 0.89641822 | NA | NA | NA |
| ENSG000002 | 0.51413714 | -0.199727707 | 0.24976394 | -0.7996659 | 0.42390439 | NA | BSNDP1 | 106480329 |
| ENSG000002 | 0 | NA | NA | NA | NA | NA | NA | NA |
| ENSG000002 | 0 | NA | NA | NA | NA | NA | NA | NA |
| ENSG000002 | 1.23447165 | -0.053244199 | 0.3301425 | -0.1612764 | 0.8718757 | NA | NA | NA |
| ENSG000002 | 0.38133351 | -0.102466351 | 0.2189713 | -0.4679442 | 0.63982449 | NA | NA | NA |
| ENSG000002 | 0.11131518 | 0.081951103 | 0.12663804 | 0.64712866 | 0.51754868 | NA | MIR6825 | 102466199 |
| ENSG000002 | 0.13280363 | -0.093394593 | 0.13115644 | -0.7120855 | 0.47641185 | NA | NA | NA |
| ENSG000002 | 0.46478446 | -0.015417091 | 0.23534337 | -0.0655089 | 0.9477688 | NA | NA | NA |
| ENSG000002 | 0.96221032 | 0.120012748 | 0.31173331 | 0.38498532 | 0.70024829 | NA | NA | NA |
| ENSG000002 | 0 | NA | NA | NA | NA | NA | SNORD50B | 692088 |
| ENSG000002 | 0 | NA | NA | NA | NA | NA | NA | NA |

| | | | | | | | | |
|---|---|---|---|---|---|---|---|---|
| ENSG000002 | 0.78979 | -0.568364433 | 0.28747989 | -1.9770581 | 0.04803506 | NA | NA | NA |
| ENSG000002 | 1.71456732 | -0.176899626 | 0.35162137 | -0.5030969 | 0.61489613 | NA | NA | NA |
| ENSG000002 | 0.7102413 | -0.022139713 | 0.27646803 | -0.0800806 | 0.93617319 | NA | MIR6766 | 102466983 |
| ENSG000002 | 0 | NA | NA | NA | NA | NA | MIR6782 | 102465469 |
| ENSG000002 | 1.02009643 | -0.042232876 | 0.30764618 | -0.1372774 | 0.89081151 | NA | NA | NA |
| ENSG000002 | 0.61272598 | 0.058460947 | 0.26359129 | 0.22178634 | 0.82448022 | NA | MIR6504 | 102465251 |
| ENSG000002 | 0.75944963 | -0.20423668 | 0.28685914 | -0.7119755 | 0.47647995 | NA | MIR6087 | 102464835 |
| ENSG000002 | 1.10974498 | 0.025527862 | 0.31007394 | 0.0823283 | 0.93438565 | NA | NA | NA |
| ENSG000002 | 0 | NA | NA | NA | NA | NA | NA | NA |
| ENSG000002 | 0.34936227 | 0.241139221 | 0.20762307 | 1.16142789 | 0.24546793 | NA | SNORD116-22 | 100033433 |
| ENSG000002 | 0 | NA | NA | NA | NA | NA | RN7SL203P | 106479296 |
| ENSG000002 | 0.12070844 | -0.088503695 | 0.12783931 | -0.6923042 | 0.48874631 | NA | NA | NA |
| ENSG000002 | 0.36502313 | -0.093755175 | 0.2080862 | -0.4505593 | 0.6523072 | NA | NA | NA |
| ENSG000002 | 0.88493524 | -0.124764995 | 0.30358944 | -0.4109662 | 0.68109733 | NA | RN7SL663P | 106481103 |
| ENSG000002 | 0 | NA | NA | NA | NA | NA | NA | NA |
| ENSG000002 | 1.02869637 | -0.531493232 | 0.31211448 | -1.7028791 | 0.0885907 | NA | NA | NA |
| ENSG000002 | 0 | NA | NA | NA | NA | NA | MIR6888 | 102465535 |
| ENSG000002 | 1.63177923 | -0.413631025 | 0.35160613 | -1.1764045 | 0.23943328 | NA | NA | NA |
| ENSG000002 | 0 | NA | NA | NA | NA | NA | SCARNA16 | 677781 |
| ENSG000002 | 1.52180208 | -0.494830045 | 0.34687642 | -1.4265312 | 0.15371506 | NA | NA | NA |
| ENSG000002 | 0 | NA | NA | NA | NA | NA | NA | NA |
| ENSG000002 | 1.01976012 | -0.369210986 | 0.31570354 | -1.1694864 | 0.24220773 | NA | NA | NA |
| ENSG000002 | 0.25370795 | -0.171331747 | 0.17411721 | -0.9840024 | 0.32511435 | NA | NA | NA |
| ENSG000002 | 1.7580257 | -0.053016335 | 0.343208 | -0.1544729 | 0.87723688 | NA | NA | NA |
| ENSG000002 | 0.58087681 | 0.226918272 | 0.26184027 | 0.86662863 | 0.38614552 | NA | NA | NA |
| ENSG000002 | 1.73633946 | -0.060909861 | 0.35817601 | -0.1700557 | 0.86496636 | NA | NA | NA |
| ENSG000002 | 0.75899316 | -0.198701114 | 0.28686906 | -0.6926544 | 0.48852648 | NA | KCNMB2 | 10242 |
| ENSG000002 | 0.12685397 | -0.090215768 | 0.12901919 | -0.699243 | 0.48440017 | NA | MIR6814 | 102465488 |
| ENSG000002 | 0.64081643 | -0.112957221 | 0.26633067 | -0.424124 | 0.6714754 | NA | MIR6815 | 102465489 |

| | | | | | | | | |
|---|---|---|---|---|---|---|---|---|
| ENSG000002 | 0.11934008 | 0.083564698 | 0.12760091 | 0.65489106 | 0.5125379 | NA | NA | NA |
| ENSG000002 | 1.62748218 | -0.25414236 | 0.34610885 | -0.7342845 | 0.46277537 | NA | NA | NA |
| ENSG000002 | 0.76169956 | -0.374892198 | 0.28875268 | -1.2983159 | 0.19417881 | NA | NA | NA |
| ENSG000002 | 0.49261432 | -0.167130763 | 0.23040378 | -0.725382 | 0.46821769 | NA | NA | NA |
| ENSG000002 | 1.95275821 | -0.05102728 | 0.36093071 | -0.1413769 | 0.88757217 | NA | NA | NA |
| ENSG000002 | 1.71650349 | -0.308348496 | 0.35042623 | -0.8799241 | 0.37890041 | NA | NA | NA |
| ENSG000002 | 0.11205239 | 0.082118251 | 0.12671962 | 0.64803108 | 0.51696485 | NA | NA | NA |
| ENSG000002 | 2.08759552 | -0.250545658 | 0.36660924 | -0.6834134 | 0.49434567 | NA | MIR6740 | 102465443 |
| ENSG000002 | 0.48627291 | -0.18151782 | 0.23948009 | -0.7579662 | 0.44847119 | NA | MIR6745 | 102466726 |
| ENSG000002 | 1.76679799 | -0.189888795 | 0.3488242 | -0.5443682 | 0.58618814 | NA | NA | NA |
| ENSG000002 | 1.94360806 | -0.306719679 | 0.36059815 | -0.8505858 | 0.39499946 | NA | NA | NA |
| ENSG000002 | 0 | NA | NA | NA | NA | NA | NA | NA |
| ENSG000002 | 0.62023988 | -0.112292793 | 0.27094277 | -0.4144521 | 0.67854306 | NA | NA | NA |
| ENSG000002 | 1.12029194 | -0.115847267 | 0.30810116 | -0.376004 | 0.70691392 | NA | NA | NA |
| ENSG000002 | 0.23202362 | -0.008302493 | 0.17586207 | -0.0472103 | 0.96234565 | NA | RN7SL544P | 106481070 |
| ENSG000002 | 0 | NA | NA | NA | NA | NA | NA | NA |
| ENSG000002 | 1.35911296 | -0.271436398 | 0.34008239 | -0.7981489 | 0.42478406 | NA | NA | NA |
| ENSG000002 | 0 | NA | NA | NA | NA | NA | MIR4734 | 100616203 |
| ENSG000002 | 1.31157313 | 0.323574016 | 0.33399366 | 0.96880286 | 0.33264356 | NA | NA | NA |
| ENSG000002 | 0.98218997 | -0.171595247 | 0.29833824 | -0.5751701 | 0.56517624 | NA | NA | NA |
| ENSG000002 | 1.96992924 | -0.059906394 | 0.35881146 | -0.1669579 | 0.86740321 | NA | NA | NA |
| ENSG000002 | 1.3358895 | -0.108143767 | 0.3300016 | -0.3277068 | 0.74313336 | NA | NA | NA |
| ENSG000002 | 0.73961029 | -0.187025619 | 0.27953219 | -0.6690665 | 0.50345308 | NA | NA | NA |
| ENSG000002 | 0 | NA | NA | NA | NA | NA | MIR6511A1 | 102466268 |
| ENSG000002 | 0 | NA | NA | NA | NA | NA | NA | NA |
| ENSG000002 | 0.61084662 | 0.056873999 | 0.26981345 | 0.21079009 | 0.83305108 | NA | NA | NA |
| ENSG000002 | 0.24004852 | -0.005223787 | 0.17775844 | -0.029387 | 0.97655594 | NA | MIR6780A | 102466195 |
| ENSG000002 | 2.11874849 | -0.230874641 | 0.36080416 | -0.6398891 | 0.52224472 | NA | NA | NA |
| ENSG000002 | 0 | NA | NA | NA | NA | NA | NA | NA |

| | | | | | | | | |
|---|---|---|---|---|---|---|---|---|
| ENSG000002 | 0.75537934 | -0.187046077 | 0.28061071 | -0.6665679 | 0.50504821 | NA | NA | NA |
| ENSG000002 | 1.56357586 | 0.018563757 | 0.34742377 | 0.05343261 | 0.95738723 | NA | NA | NA |
| ENSG000002 | 1.6500784 | -0.544273459 | 0.34921182 | -1.5585768 | 0.11909657 | NA | NA | NA |
| ENSG000002 | 0.6558786 | -0.270452548 | 0.26091264 | -1.0365636 | 0.29993928 | NA | NA | NA |
| ENSG000002 | 0.36553413 | 0.069401489 | 0.20780452 | 0.33397488 | 0.73839851 | NA | NA | NA |
| ENSG000002 | 0.14128499 | -0.097772014 | 0.13403235 | -0.7294658 | 0.4657168 | NA | NA | NA |
| ENSG000002 | 0 | NA | NA | NA | NA | NA | NA | NA |
| ENSG000002 | 0 | NA | NA | NA | NA | NA | NA | NA |
| ENSG000002 | 0.26813897 | -0.197007041 | 0.18550319 | -1.0620143 | 0.2882292 | NA | NA | NA |
| ENSG000002 | 1.79895272 | 0.007772129 | 0.35563069 | 0.0218545 | 0.98256402 | NA | NA | NA |
| ENSG000002 | 0.86341296 | -0.106740575 | 0.29584124 | -0.3608036 | 0.71824629 | NA | NA | NA |
| ENSG000002 | 0 | NA | NA | NA | NA | NA | NA | NA |
| ENSG000002 | 0.23276084 | -0.008011382 | 0.17603982 | -0.0455089 | 0.96370166 | NA | NA | NA |
| ENSG000002 | 0.12685397 | -0.090215768 | 0.12901919 | -0.699243 | 0.48440017 | NA | MIR5787 | 102464817 |
| ENSG000002 | 0.77339193 | -0.188827122 | 0.28178772 | -0.6701042 | 0.50279139 | NA | MIR8088 | 102466880 |
| ENSG000002 | 1.48437236 | -0.048449776 | 0.34368555 | -0.1409712 | 0.88789267 | NA | RN7SL680P | 106480380 |
| ENSG000002 | 1.71018382 | -0.178809142 | 0.35339981 | -0.5059684 | 0.61287883 | NA | NA | NA |
| ENSG000002 | 2.04418586 | -0.469573979 | 0.36388576 | -1.2904434 | 0.19689675 | NA | NA | NA |
| ENSG000002 | 1.2490779 | -0.188309517 | 0.32955773 | -0.5714007 | 0.56772805 | NA | MIR6503 | 102465250 |
| ENSG000002 | 1.50555538 | -0.541147972 | 0.3280405 | -1.6496377 | 0.09901707 | NA | NA | NA |
| ENSG000002 | 1.67916218 | -0.38098994 | 0.34015632 | -1.1200437 | 0.26269514 | NA | NA | NA |
| ENSG000002 | 0 | NA | NA | NA | NA | NA | NA | NA |
| ENSG000002 | 0.75005637 | -0.040089581 | 0.27923102 | -0.1435714 | 0.88583895 | NA | NA | NA |
| ENSG000002 | 1.6164903 | -0.131170226 | 0.35143764 | -0.373239 | 0.70897058 | NA | NA | NA |
| ENSG000002 | 0.49264869 | -0.021364872 | 0.24621241 | -0.0867741 | 0.93085104 | NA | NA | NA |
| ENSG000002 | 0 | NA | NA | NA | NA | NA | MIR6868 | 102466753 |
| ENSG000002 | 0.35067856 | 0.074230421 | 0.21389444 | 0.34704231 | 0.72855953 | NA | NA | NA |
| ENSG000002 | 0 | NA | NA | NA | NA | NA | NA | NA |
| ENSG000002 | 0 | NA | NA | NA | NA | NA | NA | NA |

| | | | | | | | | | |
|---|---|---|---|---|---|---|---|---|---|
| ENSG000002 | 1.71886765 | -0.058431223 | 0.35492313 | -0.1646306 | 0.86923471 | NA | | MIR6124 | 102466906 |
| ENSG000002 | 1.48157972 | -0.050275115 | 0.34558757 | -0.1454772 | 0.88433411 | NA | | NA | NA |
| ENSG000002 | 1.15677895 | -0.414679213 | 0.31861263 | -1.3015153 | 0.19308213 | NA | | NA | NA |
| ENSG000002 | 0 | NA | NA | NA | NA | NA | | MIR6770-3 | 102466259 |
| ENSG000002 | 1.81580836 | -0.239223591 | 0.35642427 | -0.6711765 | 0.50210811 | NA | | NA | NA |
| ENSG000002 | 2.01586674 | -0.330334126 | 0.36292286 | -0.9102048 | 0.36271452 | NA | | NA | NA |
| ENSG000002 | 0 | NA | NA | NA | NA | NA | | NA | NA |
| ENSG000002 | 0.6089678 | 0.061849836 | 0.269667 | 0.22935634 | 0.81859197 | NA | | MIR6882 | 102465531 |
| ENSG000002 | 1.36568517 | -0.113122812 | 0.33631545 | -0.3363593 | 0.73659997 | NA | | NA | NA |
| ENSG000002 | 1.92231369 | 0.198661613 | 0.35965758 | 0.5523632 | 0.58069954 | NA | | NA | NA |
| ENSG000002 | 1.78075123 | -0.903597683 | 0.34966752 | -2.5841625 | 0.00976158 | NA | | NA | NA |
| ENSG000002 | 0 | NA | NA | NA | NA | NA | | MIR6862-1 | 102465520 |
| ENSG000002 | 1.10240769 | 0.175236718 | 0.31327322 | 0.55937344 | 0.57590688 | NA | | NA | NA |
| ENSG000002 | 0.76812741 | -0.197020246 | 0.28745118 | -0.6854042 | 0.49308889 | NA | | NA | NA |
| ENSG000002 | 0.47826975 | -0.016435741 | 0.244632 | -0.0671856 | 0.94643397 | NA | | NA | NA |
| ENSG000002 | 0.63416044 | -0.278936162 | 0.26761738 | -1.0422946 | 0.2972751 | NA | | NA | NA |
| ENSG000002 | 0 | NA | NA | NA | NA | NA | | NA | NA |
| ENSG000002 | 1.60635633 | -0.266441455 | 0.35061146 | -0.7599337 | 0.44729423 | NA | | MIR6085 | 102464834 |
| ENSG000002 | 1.41807889 | 0.210783445 | 0.3232807 | 0.6520137 | 0.51439233 | NA | | MIR7852 | 102465839 |
| ENSG000002 | 0.11205239 | 0.082118251 | 0.12671962 | 0.64803108 | 0.51696485 | NA | | MIR6809 | 102465485 |
| ENSG000002 | 1.11695886 | -0.127417906 | 0.3262173 | -0.3905921 | 0.69609875 | NA | | NA | NA |
| ENSG000002 | 1.13753596 | -0.120194247 | 0.32306052 | -0.3720487 | 0.70985659 | NA | | NA | NA |
| ENSG000002 | 1.13181306 | -0.268887597 | 0.32367719 | -0.8307277 | 0.40612749 | NA | | MIR6130 | 102466967 |
| ENSG000002 | 0 | NA | NA | NA | NA | NA | | NA | NA |
| ENSG000002 | 1.78654951 | -0.481018565 | 0.35649633 | -1.3492946 | 0.17724237 | NA | | NA | NA |
| ENSG000002 | 1.79773918 | -0.429386147 | 0.34645746 | -1.2393618 | 0.21521153 | NA | | NA | NA |
| ENSG000002 | 1.24560438 | -0.457258208 | 0.32205621 | -1.4198087 | 0.15566339 | NA | | C17orf98 | 388381 |
| ENSG000002 | 1.46922394 | -0.04007797 | 0.34089874 | -0.1175656 | 0.90641185 | NA | | NA | NA |
| ENSG000002 | 1.49188434 | -0.340951916 | 0.34673826 | -0.9833121 | 0.32545388 | NA | | NA | NA |

| | | | | | | | | | |
|---|---|---|---|---|---|---|---|---|---|
| ENSG000002 | 0.13280363 | -0.093394593 | 0.13115644 | -0.7120855 | 0.47641185 | NA | NA | NA | |
| ENSG000002 | 0.76810568 | -0.364419366 | 0.28364061 | -1.2847926 | 0.19886476 | NA | NA | NA | |
| ENSG000002 | 0.72509495 | -0.178274683 | 0.2785491 | -0.6400117 | 0.52216499 | NA | NA | NA | |
| ENSG000002 | 0.49833776 | -0.016406893 | 0.23937171 | -0.0685415 | 0.9453546 | NA | NA | NA | |
| ENSG000002 | 1.78503925 | -0.188529402 | 0.3502487 | -0.5382729 | 0.59038863 | NA | NA | NA | |
| ENSG000002 | 1.30449184 | 0.162759991 | 0.3308162 | 0.49199522 | 0.62272272 | NA | NA | NA | |
| ENSG000002 | 1.35196963 | -0.265359328 | 0.33995197 | -0.7805789 | 0.43505021 | NA | NA | NA | |
| ENSG000002 | 2.02714128 | -0.292430137 | 0.35427965 | -0.8254218 | 0.40913218 | NA | MIR6718 | | 102465430 |
| ENSG000002 | 0 | NA | NA | NA | NA | NA | MIR6804 | | 102465482 |
| ENSG000002 | 0.23799501 | 0.170958857 | 0.17834799 | 0.95856902 | 0.33777591 | NA | NA | NA | |
| ENSG000002 | 1.40461709 | -0.257942958 | 0.33079934 | -0.7797566 | 0.43553417 | NA | MIR6508 | | 102466972 |
| ENSG000002 | 0 | NA | NA | NA | NA | NA | SNORD115-26 | | 100033802 |
| ENSG000002 | 0 | NA | NA | NA | NA | NA | SNORD116-4 | | 100033416 |
| ENSG000002 | 1.22557212 | -0.190226416 | 0.32892623 | -0.5783255 | 0.56304441 | NA | NA | NA | |
| ENSG000002 | 1.08205735 | 0.036356747 | 0.31466476 | 0.11554121 | 0.90801616 | NA | NA | NA | |
| ENSG000002 | 1.99638124 | -0.801541692 | 0.35324211 | -2.2691 | 0.02326224 | NA | NA | NA | |
| ENSG000002 | 1.7978004 | -0.72247297 | 0.34794237 | -2.0764156 | 0.03785553 | NA | NA | NA | |
| ENSG000002 | 0 | NA | NA | NA | NA | NA | TCEB3CL | | 728929 |
| ENSG000002 | 1.80310946 | 0.131780796 | 0.35741571 | 0.36870454 | 0.71234797 | NA | NA | NA | |
| ENSG000002 | 0 | NA | NA | NA | NA | NA | RN7SKP175 | | 106479173 |
| ENSG000002 | 0.22336757 | 0.163792188 | 0.17487321 | 0.93663397 | 0.34894687 | NA | NA | NA | |
| ENSG000002 | 1.25915733 | -0.1777897 | 0.31595633 | -0.5627034 | 0.57363687 | NA | NA | NA | |
| ENSG000002 | 1.45166198 | 0.092612761 | 0.34302322 | 0.26998977 | 0.78716812 | NA | NA | NA | |
| ENSG000002 | 0 | NA | NA | NA | NA | NA | NA | NA | |
| ENSG000002 | 1.35728971 | 0.019959594 | 0.33586883 | 0.05942675 | 0.95261221 | NA | NA | NA | |
| ENSG000002 | 1.24904701 | -0.048634427 | 0.33052005 | -0.1471452 | 0.88301744 | NA | RN7SL341P | | 106481017 |
| ENSG000002 | 0.62138203 | -0.094107809 | 0.25725582 | -0.3658141 | 0.71450379 | NA | NA | NA | |
| ENSG000002 | 2.04105023 | -0.316121665 | 0.35900385 | -0.8805523 | 0.37856018 | NA | NA | NA | |
| ENSG000002 | 0.87508998 | -0.254813262 | 0.29356359 | -0.8680002 | 0.38539421 | NA | NA | NA | |

| | | | | | | | | |
|---|---|---|---|---|---|---|---|---|
| ENSG000002 | 0 | NA | NA | NA | NA | NA | NA | NA |
| ENSG000002 | 0.69282314 | 0.115007774 | 0.26385302 | 0.43587818 | 0.66292512 | NA | NA | NA |
| ENSG000002 | 0 | NA | NA | NA | NA | NA | NA | NA |
| ENSG000002 | 0.13280363 | -0.093394593 | 0.13115644 | -0.7120855 | 0.47641185 | NA | NA | NA |
| ENSG000002 | 0.88584714 | -0.288702375 | 0.30453821 | -0.9480005 | 0.3431292 | NA | NA | NA |
| ENSG000002 | 0.24141689 | -0.165257896 | 0.17127401 | -0.9648743 | 0.33460776 | NA | NA | NA |
| ENSG000002 | 1.12282262 | -0.256369936 | 0.31745204 | -0.8075864 | 0.41932875 | NA | NA | NA |
| ENSG000002 | 0 | NA | NA | NA | NA | NA | NA | NA |
| ENSG000002 | 0.12685397 | -0.090215768 | 0.12901919 | -0.699243 | 0.48440017 | NA | NA | NA |
| ENSG000002 | 0 | NA | NA | NA | NA | NA | MIR6793 | 102466737 |
| ENSG000002 | 2.13196155 | -0.008622193 | 0.36148367 | -0.0238522 | 0.98097048 | NA | NA | NA |
| ENSG000002 | 0 | NA | NA | NA | NA | NA | NA | NA |
| ENSG000002 | 0 | NA | NA | NA | NA | NA | NA | NA |
| ENSG000002 | 0.76633237 | -0.195742052 | 0.28129046 | -0.6958716 | 0.48650922 | NA | NA | NA |
| ENSG000002 | 0.12685397 | -0.090215768 | 0.12901919 | -0.699243 | 0.48440017 | NA | MIR6851 | 102465512 |
| ENSG000002 | 0.50684946 | -0.202346264 | 0.24895102 | -0.8127955 | 0.41633533 | NA | MIR6796 | 102465477 |
| ENSG000002 | 0.12070844 | -0.088503695 | 0.12783931 | -0.6923042 | 0.48874631 | NA | NA | NA |
| ENSG000002 | 0.27408862 | -0.199826037 | 0.18669465 | -1.0703362 | 0.28446801 | NA | NA | NA |
| ENSG000002 | 0 | NA | NA | NA | NA | NA | NA | NA |
| ENSG000002 | 0 | NA | NA | NA | NA | NA | SNORD112 | 692215 |
| ENSG000002 | 1.01110407 | -0.204132122 | 0.31431391 | -0.649453 | 0.51604559 | NA | HIST1H4G | 8369 |
| ENSG000002 | 0 | NA | NA | NA | NA | NA | NA | NA |
| ENSG000002 | 0.59348108 | 0.227402675 | 0.26300367 | 0.86463689 | 0.38723812 | NA | NA | NA |
| ENSG000002 | 0.86911467 | -0.102688647 | 0.29002244 | -0.3540714 | 0.72328538 | NA | MIR6839 | 102465505 |
| ENSG000002 | 0 | NA | NA | NA | NA | NA | MIR6744 | 102466725 |
| ENSG000002 | 0.25145857 | -0.010118565 | 0.18031426 | -0.0561163 | 0.95524918 | NA | MIR8076 | 102466253 |
| ENSG000002 | 1.11060481 | -0.120723725 | 0.32603316 | -0.3702805 | 0.71117349 | NA | NA | NA |
| ENSG000002 | 0.73998086 | -0.354752955 | 0.28183273 | -1.2587358 | 0.20812577 | NA | NA | NA |
| ENSG000002 | 0 | NA | NA | NA | NA | NA | NA | NA |

| | | | | | | | | |
|---|---|---|---|---|---|---|---|---|
| ENSG000002 | 1.57214154 | 0.013707765 | 0.3519177 | 0.03895162 | 0.96892896 | NA | NA | NA |
| ENSG000002 | 0.26199343 | -0.194018184 | 0.18422801 | -1.0531416 | 0.29227609 | NA | MIR6724-4 | 103504733 |
| ENSG000002 | 1.73451463 | 0.077925152 | 0.3543554 | 0.21990677 | 0.82594376 | NA | NA | NA |
| ENSG000002 | 0.50818749 | -0.197539377 | 0.24915052 | -0.7928516 | 0.42786432 | NA | NA | NA |
| ENSG000002 | 1.88166091 | -0.257451017 | 0.35924538 | -0.7166439 | 0.47359383 | NA | NA | NA |
| ENSG000002 | 0 | NA | NA | NA | NA | NA | MIR6795 | 102465476 |
| ENSG000002 | 2.10116671 | -0.133346178 | 0.36824095 | -0.3621166 | 0.71726486 | NA | NA | NA |
| ENSG000002 | 1.78470294 | -0.485597772 | 0.35745446 | -1.3584885 | 0.17430873 | NA | NA | NA |
| ENSG000002 | 1.67901237 | -0.045909277 | 0.34377167 | -0.1335458 | 0.89376171 | NA | NA | NA |
| ENSG000002 | 0.74672329 | -0.032948972 | 0.27900439 | -0.1180948 | 0.90599253 | NA | MIR6088 | 102464836 |
| ENSG000002 | 1.22832454 | -0.180338481 | 0.32304069 | -0.5582531 | 0.57667153 | NA | NA | NA |
| ENSG000002 | 0.36484899 | 0.075158314 | 0.21612264 | 0.34775772 | 0.72802215 | NA | RBM27 | 54439 |
| ENSG000002 | 0 | NA | NA | NA | NA | NA | NA | NA |
| ENSG000002 | 1.16699477 | -0.444508723 | 0.32445779 | -1.3700048 | 0.17068541 | NA | NA | NA |
| ENSG000002 | 0.12070844 | -0.088503695 | 0.12783931 | -0.6923042 | 0.48874631 | NA | NA | NA |
| ENSG000002 | 0.86372754 | 0.042498317 | 0.29713559 | 0.14302668 | 0.88626911 | NA | MIR6505 | 102466657 |
| ENSG000002 | 1.21182679 | -0.162810359 | 0.32001501 | -0.5087585 | 0.6109215 | NA | NA | NA |
| ENSG000002 | 1.99817422 | -0.28825422 | 0.35012271 | -0.8232948 | 0.41034035 | NA | NA | NA |
| ENSG000002 | 0.24004852 | -0.005223787 | 0.17775844 | -0.029387 | 0.97655594 | NA | RN7SL185P | 106479289 |
| ENSG000002 | 0 | NA | NA | NA | NA | NA | NA | NA |
| ENSG000002 | 0.37285215 | -0.099309896 | 0.2177345 | -0.4561055 | 0.64831414 | NA | NA | NA |
| ENSG000002 | 1.97733358 | -0.578841802 | 0.36074873 | -1.6045567 | 0.10859139 | NA | NA | NA |
| ENSG000002 | 1.24637543 | -0.354972115 | 0.33382112 | -1.0633603 | 0.2876186 | NA | NA | NA |
| ENSG000002 | 1.14729924 | -0.438926243 | 0.32388124 | -1.3552074 | 0.1753515 | NA | MIR8064 | 102465866 |
| ENSG000002 | 1.77487833 | -0.444276437 | 0.34869266 | -1.2741204 | 0.20262075 | NA | NA | NA |
| ENSG000002 | 0 | NA | NA | NA | NA | NA | NA | NA |
| ENSG000002 | 1.69535603 | 0.073400544 | 0.35574294 | 0.20633029 | 0.83653292 | NA | NA | NA |
| ENSG000002 | 0.48302518 | -0.023466322 | 0.24515318 | -0.0957211 | 0.92374212 | NA | NA | NA |
| ENSG000002 | 1.71025045 | -0.047964475 | 0.34823124 | -0.1377374 | 0.89044794 | NA | NA | NA |

| | | | | | | | | |
|---|---|---|---|---|---|---|---|---|
| ENSG000002 | 0.48416733 | -0.018553864 | 0.24528998 | -0.0756405 | 0.93970509 | NA | NA | NA |
| ENSG000002 | 0.24004852 | -0.005223787 | 0.17775844 | -0.029387 | 0.97655594 | NA | MIR6500 | 102466656 |
| ENSG000002 | 0.46278494 | 0.309182332 | 0.23065454 | 1.34045629 | 0.18009705 | NA | MIR6818 | 102466742 |
| ENSG000002 | 0 | NA | NA | NA | NA | NA | NA | NA |
| ENSG000002 | 1.34250974 | -0.226422784 | 0.32380855 | -0.6992489 | 0.48439648 | NA | NA | NA |
| ENSG000002 | 0.28256998 | -0.18483211 | 0.18021686 | -1.0256094 | 0.30507573 | NA | NA | NA |
| ENSG000002 | 1.71818409 | 0.07008176 | 0.35540379 | 0.19718912 | 0.84367955 | NA | NA | NA |
| ENSG000002 | 0.8129525 | 0.207408747 | 0.29481735 | 0.70351608 | 0.48173419 | NA | NA | NA |
| ENSG000002 | 0.47735839 | 0.152206097 | 0.23764319 | 0.64048164 | 0.52185952 | NA | MIR6068 | 102464823 |
| ENSG000002 | 1.93835889 | -0.060444892 | 0.36348726 | -0.1662916 | 0.86792745 | NA | NA | NA |
| ENSG000002 | 0.62450659 | -0.10299827 | 0.26500891 | -0.3886597 | 0.69752793 | NA | NA | NA |
| ENSG000002 | 0.24756242 | -0.187034346 | 0.18119817 | -1.0322088 | 0.30197431 | NA | NA | NA |
| ENSG000002 | 1.0904332 | -0.104919227 | 0.3200143 | -0.3278579 | 0.74301908 | NA | NA | NA |
| ENSG000002 | 0 | NA | NA | NA | NA | NA | MIR6720 | 102466720 |
| ENSG000002 | 0 | NA | NA | NA | NA | NA | NA | NA |
| ENSG000002 | 0 | NA | NA | NA | NA | NA | NA | NA |
| ENSG000002 | 0 | NA | NA | NA | NA | NA | NA | NA |
| ENSG000002 | 1.03080599 | -0.331313494 | 0.30498096 | -1.0863416 | 0.27732791 | NA | MIR8054 | 102465860 |
| ENSG000002 | 0.12685397 | -0.090215768 | 0.12901919 | -0.699243 | 0.48440017 | NA | NA | NA |
| ENSG000002 | 0.73961029 | -0.187025619 | 0.27953219 | -0.6690665 | 0.50345308 | NA | NA | NA |
| ENSG000002 | 2.14386679 | -0.616104642 | 0.36047612 | -1.7091414 | 0.08742477 | NA | RN7SL43P | 106479241 |
| ENSG000002 | 1.03877852 | 0.169424175 | 0.30326417 | 0.55866862 | 0.5763879 | NA | NA | NA |
| ENSG000002 | 0 | NA | NA | NA | NA | NA | NA | NA |
| ENSG000002 | 1.09043266 | -0.115661355 | 0.32153261 | -0.3597189 | 0.71905736 | NA | NA | NA |
| ENSG000002 | 0 | NA | NA | NA | NA | NA | NA | NA |
| ENSG000002 | 1.38617335 | -0.381686926 | 0.32966611 | -1.1577985 | 0.24694628 | NA | NA | NA |
| ENSG000002 | 1.23308155 | -0.055137891 | 0.32422604 | -0.17006 | 0.86496292 | NA | NA | NA |
| ENSG000002 | 0 | NA | NA | NA | NA | NA | NA | NA |
| ENSG000002 | 1.82061935 | 0.006062894 | 0.35581332 | 0.01703954 | 0.98640507 | NA | NA | NA |

| | | | | | | | | |
|---|---|---|---|---|---|---|---|---|
| ENSG000002 | 0.65465054 | -0.455468986 | 0.2643416 | -1.7230318 | 0.08488278 | NA | NA | NA |
| ENSG000002 | 0.23070733 | 0.167416683 | 0.17664244 | 0.94777156 | 0.34324576 | NA | MIR6807 | 102465976 |
| ENSG000002 | 1.54948115 | 0.279795156 | 0.34112538 | 0.82021207 | 0.41209522 | NA | NA | NA |
| ENSG000002 | 0 | NA | NA | NA | NA | NA | NA | NA |
| ENSG000002 | 0.11934008 | 0.083564698 | 0.12760091 | 0.65489106 | 0.5125379 | NA | NA | NA |
| ENSG000002 | 1.58576726 | 0.279151486 | 0.3452308 | 0.8085938 | 0.41874883 | NA | NA | NA |
| ENSG000002 | 0.63891587 | -0.285900823 | 0.2679393 | -1.0670358 | 0.28595567 | NA | NA | NA |
| ENSG000002 | 0.66156766 | -0.287991838 | 0.26979811 | -1.0674346 | 0.28577564 | NA | NA | NA |
| ENSG000002 | 0 | NA | NA | NA | NA | NA | NA | NA |
| ENSG000002 | 0 | NA | NA | NA | NA | NA | NA | NA |
| ENSG000002 | 0 | NA | NA | NA | NA | NA | MIR6724-1 | 102465433 |
| ENSG000002 | 1.74701676 | -0.306501628 | 0.35056681 | -0.874303 | 0.38195324 | NA | NA | NA |
| ENSG000002 | 1.3362904 | -0.360592604 | 0.32323588 | -1.1155711 | 0.26460577 | NA | TBC1D3F | 84218 |
| ENSG000002 | 0.73564763 | 0.13375234 | 0.27852684 | 0.48021347 | 0.63107561 | NA | NA | NA |
| ENSG000002 | 1.39537265 | -0.391776062 | 0.33360877 | -1.1743578 | 0.24025174 | NA | RN7SL210P | 106479298 |
| ENSG000002 | 1.49692401 | -0.05682441 | 0.3446045 | -0.1648975 | 0.86902468 | NA | NA | NA |
| ENSG000002 | 0.48945358 | 0.14806159 | 0.23910487 | 0.61923286 | 0.53576297 | NA | NA | NA |
| ENSG000002 | 0.36465257 | 0.064724075 | 0.21605106 | 0.29957767 | 0.76449932 | NA | MIR6880 | 102466204 |
| ENSG000002 | 1.4011457 | -0.418152711 | 0.33874423 | -1.2344202 | 0.21704637 | NA | NA | NA |
| ENSG000002 | 0 | NA | NA | NA | NA | NA | NA | NA |
| ENSG000002 | 1.14920894 | -0.604762374 | 0.32185513 | -1.8789894 | 0.06024594 | NA | NA | NA |
| ENSG000002 | 0 | NA | NA | NA | NA | NA | NA | NA |
| ENSG000002 | 0.60438651 | -0.101682894 | 0.2632591 | -0.3862465 | 0.69931415 | NA | NA | NA |
| ENSG000002 | 0.97470641 | -0.190151385 | 0.30809904 | -0.6171762 | 0.53711853 | NA | NA | NA |
| ENSG000002 | 0 | NA | NA | NA | NA | NA | NA | NA |
| ENSG000002 | 0.8723296 | -0.285487742 | 0.3038961 | -0.9394255 | 0.34751233 | NA | NA | NA |
| ENSG000002 | 0 | NA | NA | NA | NA | NA | RN7SL327P | 106481012 |
| ENSG000002 | 0 | NA | NA | NA | NA | NA | SNORA24 | 677809 |
| ENSG000002 | 1.22306215 | -0.043529992 | 0.33645225 | -0.1293794 | 0.89705744 | NA | NA | NA |

| | | | | | | | | |
|---|---|---|---|---|---|---|---|---|
| ENSG000002 | 0 | NA | NA | NA | NA | NA | SNORD27 | 9301 |
| ENSG000002 | 1.59208557 | 0.01498229 | 0.34475309 | 0.04345803 | 0.96533642 | NA | NA | NA |
| ENSG000002 | 0.86435869 | -0.122981079 | 0.30693344 | -0.4006767 | 0.68865816 | NA | NA | NA |
| ENSG000002 | 0 | NA | NA | NA | NA | NA | NA | NA |
| ENSG000002 | 0.60355893 | 0.050839769 | 0.26278931 | 0.19346209 | 0.84659709 | NA | NA | NA |
| ENSG000002 | 1.22804379 | -0.198209482 | 0.33046028 | -0.5997982 | 0.54864073 | NA | RN7SL301P | 106480499 |
| ENSG000002 | 1.93697326 | -0.299719452 | 0.35894385 | -0.8350037 | 0.40371561 | NA | NA | NA |
| ENSG000002 | 0.48536101 | -0.024029524 | 0.24539296 | -0.0979226 | 0.92199373 | NA | LOC102724728 | 102724728 |
| ENSG000002 | 0 | NA | NA | NA | NA | NA | NA | NA |
| ENSG000002 | 0 | NA | NA | NA | NA | NA | RNU12 | 267010 |
| ENSG000002 | 0.89452438 | -0.261834816 | 0.29286276 | -0.8940529 | 0.37129356 | NA | MIR4477A | 100616184 |
| ENSG000002 | 1.689178 | -0.388967976 | 0.34538377 | -1.1261907 | 0.26008482 | NA | RN7SL197P | 106480975 |
| ENSG000002 | 0.62701656 | -0.28174368 | 0.26697332 | -1.0553252 | 0.29127661 | NA | NA | NA |
| ENSG000002 | 1.01586397 | -0.506728927 | 0.30696171 | -1.6507887 | 0.09878172 | NA | NA | NA |
| ENSG000002 | 0.5008998 | -0.200150804 | 0.24832882 | -0.805991 | 0.42024801 | NA | NA | NA |
| ENSG000002 | 1.7121268 | -0.188361943 | 0.35342223 | -0.5329657 | 0.59405729 | NA | SPATA31A7 | 26165 |
| ENSG000002 | 0 | NA | NA | NA | NA | NA | NA | NA |
| ENSG000002 | 0.11934008 | 0.083564698 | 0.12760091 | 0.65489106 | 0.5125379 | NA | NA | NA |
| ENSG000002 | 0 | NA | NA | NA | NA | NA | NA | NA |
| ENSG000002 | 1.12652681 | -0.433333667 | 0.32325023 | -1.3405518 | 0.18006603 | NA | NA | NA |
| ENSG000002 | 0 | NA | NA | NA | NA | NA | NA | NA |
| ENSG000002 | 0 | NA | NA | NA | NA | NA | NA | NA |
| ENSG000002 | 0.26560726 | -0.177019426 | 0.17672274 | -1.0016788 | 0.31649875 | NA | NA | NA |
| ENSG000002 | 0 | NA | NA | NA | NA | NA | NA | NA |
| ENSG000002 | 1.67733484 | 0.320748167 | 0.34332843 | 0.93423129 | 0.3501846 | NA | NA | NA |
| ENSG000002 | 0.35961481 | -0.099625193 | 0.21567743 | -0.4619176 | 0.64414043 | NA | NA | NA |
| ENSG000002 | 0.26813897 | -0.197007041 | 0.18550319 | -1.0620143 | 0.2882292 | NA | MIR7156 | 102466995 |
| ENSG000002 | 1.10640226 | -0.117524395 | 0.31416674 | -0.3740829 | 0.70834266 | NA | MIR7976 | 102465857 |
| ENSG000002 | 0.74549928 | 0.118820701 | 0.26777766 | 0.44372895 | 0.65723856 | NA | RN7SL620P | 106479448 |

| | | | | | | | | |
|---|---|---|---|---|---|---|---|---|
| ENSG000002 | 0 | NA | NA | NA | NA | NA | NA | NA |
| ENSG000002 | 1.58014972 | -0.357753002 | 0.34225929 | -1.0452689 | 0.29589869 | NA | NA | NA |
| ENSG000002 | 0.38064837 | -0.102706251 | 0.21886737 | -0.4692625 | 0.63888201 | NA | NA | NA |
| ENSG000002 | 1.7243865 | -0.171921967 | 0.34880737 | -0.4928851 | 0.62209375 | NA | NA | NA |
| ENSG000002 | 1.91727785 | 0.192538323 | 0.35349022 | 0.54467793 | 0.58597505 | NA | NA | NA |
| ENSG000002 | 0 | NA | NA | NA | NA | NA | NA | NA |
| ENSG000002 | 1.57177489 | -0.113999338 | 0.34536581 | -0.3300829 | 0.74133734 | NA | MIR6747 | 102465447 |
| ENSG000002 | 1.84003119 | -0.247736536 | 0.3572683 | -0.6934187 | 0.4880468 | NA | NA | NA |
| ENSG000002 | 0.85200291 | -0.096788563 | 0.28906802 | -0.3348297 | 0.73775354 | NA | NA | NA |
| ENSG000002 | 0 | NA | NA | NA | NA | NA | NA | NA |
| ENSG000002 | 1.75564448 | -0.470939545 | 0.35395562 | -1.3305045 | 0.1833521 | NA | NA | NA |
| ENSG000002 | 1.25338079 | -0.507294168 | 0.3318855 | -1.5285217 | 0.12638307 | NA | NA | NA |
| ENSG000002 | 0.11131518 | 0.081951103 | 0.12663804 | 0.64712866 | 0.51754868 | NA | MIR6855 | 102466750 |
| ENSG000002 | 1.85989652 | -0.349597445 | 0.35142325 | -0.9948045 | 0.31983134 | NA | MIR6133 | 102465139 |
| ENSG000002 | 0.23890637 | -0.010420674 | 0.17747379 | -0.0587167 | 0.95317776 | NA | NA | NA |
| ENSG000002 | 0 | NA | NA | NA | NA | NA | NA | NA |
| ENSG000002 | 1.10958528 | -0.267975363 | 0.32146244 | -0.8336133 | 0.40449893 | NA | MIR548AY | 102465247 |
| ENSG000002 | 0 | NA | NA | NA | NA | NA | NA | NA |
| ENSG000002 | 0 | NA | NA | NA | NA | NA | NA | NA |
| ENSG000002 | 0.36556447 | -0.101943371 | 0.21659044 | -0.4706735 | 0.63787395 | NA | NA | NA |
| ENSG000002 | 1.18449027 | 0.389577602 | 0.32053136 | 1.21541182 | 0.22420906 | NA | NA | NA |
| ENSG000002 | 0 | NA | NA | NA | NA | NA | NA | NA |
| ENSG000002 | 0 | NA | NA | NA | NA | NA | SNORA43 | 677824 |
| ENSG000002 | 0 | NA | NA | NA | NA | NA | NA | NA |
| ENSG000002 | 0 | NA | NA | NA | NA | NA | RN7SL1 | 6029 |
| ENSG000002 | 0 | NA | NA | NA | NA | NA | MIR6871 | 102466912 |
| ENSG000002 | 0 | NA | NA | NA | NA | NA | MIR6090 | 102466104 |
| ENSG000002 | 0 | NA | NA | NA | NA | NA | NA | NA |
| ENSG000002 | 1.14149002 | -0.112339957 | 0.31536579 | -0.3562211 | 0.72167497 | NA | NA | NA |

| | | | | | | | | |
|---|---|---|---|---|---|---|---|---|
| ENSG000002 | 0 | NA | NA | NA | NA | NA | MIR6794 | 102466196 |
| ENSG000002 | 0.35802023 | 0.220517756 | 0.19914056 | 1.1073473 | 0.2681438 | NA | NA | NA |
| ENSG000002 | 0 | NA | NA | NA | NA | NA | NA | NA |
| ENSG000002 | 2.10785695 | -0.386965882 | 0.3663112 | -1.0563856 | 0.29079207 | NA | NA | NA |
| ENSG000002 | 0.67141739 | -0.461190807 | 0.26562301 | -1.7362608 | 0.08251773 | NA | NA | NA |
| ENSG000002 | 0.25260017 | -0.015978705 | 0.18049464 | -0.0885273 | 0.92945758 | NA | NA | NA |
| ENSG000002 | 0.13280363 | -0.093394593 | 0.13115644 | -0.7120855 | 0.47641185 | NA | NA | NA |
| ENSG000002 | 1.8770598 | -0.400820081 | 0.36068047 | -1.1112886 | 0.26644415 | NA | RN7SL820P | 106480535 |
| ENSG000002 | 0.12070844 | -0.088503695 | 0.12783931 | -0.6923042 | 0.48874631 | NA | NA | NA |
| ENSG000002 | 1.61850763 | 0.008647223 | 0.35124957 | 0.02461846 | 0.9803593 | NA | NA | NA |
| ENSG000002 | 1.83824253 | -0.380922223 | 0.35789653 | -1.0643362 | 0.28717644 | NA | NA | NA |
| ENSG000002 | 0.69280713 | 0.465428716 | 0.27165989 | 1.7132773 | 0.08666154 | NA | NA | NA |
| ENSG000002 | 1.82884263 | -0.584710494 | 0.34846625 | -1.6779545 | 0.09335599 | NA | NA | NA |
| ENSG000002 | 1.1477557 | -0.404006993 | 0.31326092 | -1.2896821 | 0.19716107 | NA | NA | NA |
| ENSG000002 | 0.65354277 | -0.290934597 | 0.26909603 | -1.0811553 | 0.27962804 | NA | NA | NA |
| ENSG000002 | 0.885162 | -0.288936305 | 0.30450304 | -0.9488782 | 0.34268255 | NA | NA | NA |
| ENSG000002 | 0.49196355 | -0.021608171 | 0.24613638 | -0.0877894 | 0.93004404 | NA | MIR8057 | 102466874 |
| ENSG000002 | 0 | NA | NA | NA | NA | NA | NA | NA |
| ENSG000002 | 0.36827086 | -0.266056975 | 0.21181163 | -1.2561018 | 0.20907905 | NA | NA | NA |
| ENSG000002 | 0 | NA | NA | NA | NA | NA | NA | NA |
| ENSG000002 | 0 | NA | NA | NA | NA | NA | NA | NA |
| ENSG000002 | 1.82298552 | -0.120369461 | 0.35289965 | -0.3410869 | 0.73303814 | NA | NA | NA |
| ENSG000002 | 1.33002471 | -0.105723789 | 0.3340578 | -0.3164835 | 0.75163553 | NA | NA | NA |
| ENSG000002 | 0 | NA | NA | NA | NA | NA | NA | NA |
| ENSG000002 | 0.98868253 | -0.043402631 | 0.3063604 | -0.1416718 | 0.88733925 | NA | NA | NA |
| ENSG000002 | 0.12685397 | -0.090215768 | 0.12901919 | -0.699243 | 0.48440017 | NA | MIR6799 | 102465479 |
| ENSG000002 | 0.85363187 | -0.268235387 | 0.2964963 | -0.9046837 | 0.36563295 | NA | NA | NA |
| ENSG000002 | 2.11798508 | -0.135596457 | 0.36694674 | -0.3695263 | 0.7117355 | NA | RN7SL126P | 106479268 |
| ENSG000002 | 1.5411844 | -0.480800029 | 0.34246059 | -1.4039572 | 0.16033158 | NA | LOC102723475 | 102723475 |

| | | | | | | | | |
|---|---|---|---|---|---|---|---|---|
| ENSG000002 | 1.45144357 | -0.561836942 | 0.33365562 | -1.6838828 | 0.09220433 | NA | NA | NA |
| ENSG000002 | 0.23139247 | 0.167748926 | 0.17680352 | 0.94878726 | 0.34272882 | NA | NA | NA |
| ENSG000002 | 0 | NA | NA | NA | NA | NA | NA | NA |
| ENSG000002 | 0 | NA | NA | NA | NA | NA | MIR6511A3 | 102465683 |
| ENSG000002 | 0 | NA | NA | NA | NA | NA | NA | NA |
| ENSG000002 | 0 | NA | NA | NA | NA | NA | SNORD107 | 91380 |
| ENSG000002 | 1.25317385 | -0.329098508 | 0.32661895 | -1.0075916 | 0.31365057 | NA | NA | NA |
| ENSG000002 | 2.05162207 | -0.225521006 | 0.35934436 | -0.6275902 | 0.53027242 | NA | MIR8079 | 102465876 |
| ENSG000002 | 1.30637567 | -0.022576701 | 0.19197004 | -0.1176053 | 0.90638038 | NA | MIR2909 | 100422969 |
| ENSG000002 | 1.89817769 | -0.383564836 | 0.35608867 | -1.077161 | 0.28140833 | NA | NA | NA |
| ENSG000002 | 0 | NA | NA | NA | NA | NA | MIR6858 | 102465517 |
| ENSG000002 | 0 | NA | NA | NA | NA | NA | NA | NA |
| ENSG000002 | 0.11131518 | 0.081951103 | 0.12663804 | 0.64712866 | 0.51754868 | NA | NA | NA |
| ENSG000002 | 2.08893066 | 0.113762343 | 0.36284476 | 0.31352897 | 0.75387882 | NA | NA | NA |
| ENSG000002 | 1.05755841 | -0.52878758 | 0.31097085 | -1.700441 | 0.089048 | NA | NA | NA |
| ENSG000002 | 1.40636006 | -0.263197598 | 0.33641956 | -0.7823493 | 0.43400933 | NA | NA | NA |
| ENSG000002 | 0.39479706 | -0.300008631 | 0.22317059 | -1.3443018 | 0.17885081 | NA | MIR145 | 406937 |
| ENSG000002 | 0.88647829 | -0.457898022 | 0.30161276 | -1.5181653 | 0.12897273 | NA | NA | NA |
| ENSG000002 | 1.57965811 | 0.022292425 | 0.34764855 | 0.06412345 | 0.94887193 | NA | NA | NA |
| ENSG000002 | 2.12235165 | -0.475915603 | 0.36018893 | -1.3212944 | 0.18640323 | NA | NA | NA |
| ENSG000002 | 0.36482725 | -0.102221582 | 0.21647267 | -0.4722147 | 0.63677353 | NA | NA | NA |
| ENSG000002 | 1.45938921 | -0.286714713 | 0.33305783 | -0.8608556 | 0.38931755 | NA | NA | NA |
| ENSG000002 | 0.24756242 | -0.187034346 | 0.18119817 | -1.0322088 | 0.30197431 | NA | NA | NA |
| ENSG000002 | 0 | NA | NA | NA | NA | NA | NA | NA |
| ENSG000002 | 0.25351207 | -0.189965547 | 0.1824784 | -1.0410303 | 0.29786148 | NA | NA | NA |
| ENSG000002 | 1.26319571 | -0.191866549 | 0.32732869 | -0.5861587 | 0.55776889 | NA | NA | NA |
| ENSG000002 | 1.03968825 | -0.674629002 | 0.30443993 | -2.2159675 | 0.02669373 | NA | NA | NA |
| ENSG000002 | 0 | NA | NA | NA | NA | NA | NA | NA |
| ENSG000002 | 0.99832566 | -0.040897058 | 0.30841276 | -0.1326049 | 0.89450582 | NA | NA | NA |

| | | | | | | | | |
|---|---|---|---|---|---|---|---|---|
| ENSG000002 | 0 | NA | NA | NA | NA | NA | NA | NA |
| ENSG000002 | 0.66116485 | -0.036912824 | 0.15811556 | -0.2334547 | 0.81540833 | NA | NA | NA |
| ENSG000002 | 0.76903877 | -0.364246514 | 0.2837119 | -1.2838606 | 0.19919075 | NA | NA | NA |
| ENSG000002 | 0.47821768 | -0.016471461 | 0.24462483 | -0.0673336 | 0.94631616 | NA | FAM25C | 644054 |
| ENSG000002 | 1.47153592 | -0.335444897 | 0.34521383 | -0.9717018 | 0.33119892 | NA | NA | NA |
| ENSG000002 | 0.69486063 | 0.296704413 | 0.27549718 | 1.07697805 | 0.28149007 | NA | NA | NA |
| ENSG000002 | 0.26560726 | -0.177019426 | 0.17672274 | -1.0016788 | 0.31649875 | NA | NA | NA |
| ENSG000002 | 0 | NA | NA | NA | NA | NA | NA | NA |
| ENSG000002 | 1.45596679 | -0.04414747 | 0.34066642 | -0.1295915 | 0.89688963 | NA | NA | NA |
| ENSG000002 | 0.25370795 | -0.171331747 | 0.17411721 | -0.9840024 | 0.32511435 | NA | NA | NA |
| ENSG000002 | 0 | NA | NA | NA | NA | NA | NA | NA |
| ENSG000002 | 1.98395395 | -0.187792606 | 0.3619346 | -0.5188578 | 0.60385988 | NA | NA | NA |
| ENSG000002 | 1.32705931 | 0.175438565 | 0.3386452 | 0.51806009 | 0.60441634 | NA | NA | NA |
| ENSG000002 | 1.14303307 | -0.416611372 | 0.31974486 | -1.3029494 | 0.19259204 | NA | NA | NA |
| ENSG000002 | 1.33739426 | -0.258453728 | 0.3349294 | -0.7716663 | 0.4403121 | NA | MIR548AZ | 102466162 |
| ENSG000002 | 0.24141689 | -0.165257896 | 0.17127401 | -0.9648743 | 0.33460776 | NA | NA | NA |
| ENSG000002 | 1.14570062 | -0.41319844 | 0.31741795 | -1.3017488 | 0.19300227 | NA | NA | NA |
| ENSG000002 | 0 | NA | NA | NA | NA | NA | MIR6829 | 102465977 |
| ENSG000002 | 0 | NA | NA | NA | NA | NA | NA | NA |
| ENSG000002 | 1.1293348 | -0.115361668 | 0.31666339 | -0.3643038 | 0.71563118 | NA | NA | NA |
| ENSG000002 | 0.86523834 | -0.252727523 | 0.29302256 | -0.8624849 | 0.38842076 | NA | NA | NA |
| ENSG000002 | 0 | NA | NA | NA | NA | NA | NA | NA |
| ENSG000002 | 0.25214371 | -0.009862892 | 0.18046657 | -0.0546522 | 0.95641556 | NA | NA | NA |
| ENSG000002 | 2.09217627 | -0.131168201 | 0.36614517 | -0.358241 | 0.72016299 | NA | NA | NA |
| ENSG000002 | 0.64759311 | -0.271681639 | 0.26184098 | -1.0375826 | 0.29946444 | NA | NA | NA |
| ENSG000002 | 0.72495306 | -0.021021293 | 0.27749391 | -0.0757541 | 0.93961476 | NA | NA | NA |
| ENSG000002 | 0.63347722 | -0.105383092 | 0.26574722 | -0.3965539 | 0.69169647 | NA | NA | NA |
| ENSG000002 | 1.13841293 | -0.397597343 | 0.31209391 | -1.273967 | 0.20267511 | NA | NA | NA |
| ENSG000002 | 1.61003019 | -0.272478546 | 0.35399098 | -0.769733 | 0.44145828 | NA | NA | NA |

| | | | | | | | | |
|---|---|---|---|---|---|---|---|---|
| ENSG000002 | 0.39499294 | -0.277908454 | 0.21584201 | -1.287555 | 0.19790091 | NA | RN7SL462P | 106479391 |
| ENSG000002 | 0.87141612 | -0.272396215 | 0.29905577 | -0.9108542 | 0.36237218 | NA | NA | NA |
| ENSG000002 | 0 | NA | NA | NA | NA | NA | NA | NA |
| ENSG000002 | 0 | NA | NA | NA | NA | NA | NA | NA |
| ENSG000002 | 1.36455663 | -0.669484424 | 0.32724571 | -2.0458158 | 0.0407745 | NA | NA | NA |
| ENSG000002 | 0.74248691 | -0.185910538 | 0.27806603 | -0.6685841 | 0.50376079 | NA | NA | NA |
| ENSG000002 | 1.15808188 | 0.553609168 | 0.32024623 | 1.72869847 | 0.08386308 | NA | NA | NA |
| ENSG000002 | 0.7513944 | -0.035588106 | 0.27932511 | -0.1274075 | 0.8986179 | NA | NA | NA |
| ENSG000002 | 0 | NA | NA | NA | NA | NA | NA | NA |
| ENSG000002 | 0.12685397 | -0.090215768 | 0.12901919 | -0.699243 | 0.48440017 | NA | RN7SL453P | 106481046 |
| ENSG000002 | 0 | NA | NA | NA | NA | NA | NA | NA |
| ENSG000002 | 0.64795976 | -0.039872043 | 0.15793999 | -0.2524506 | 0.8006928 | NA | NA | NA |
| ENSG000002 | 1.21438883 | -0.319590162 | 0.32460293 | -0.9845572 | 0.3248416 | NA | NA | NA |
| ENSG000002 | 0.58953286 | 0.060024281 | 0.26153531 | 0.22950737 | 0.8184746 | NA | RN7SL565P | 106481076 |
| ENSG000002 | 1.52936996 | -0.482444624 | 0.34431278 | -1.4011813 | 0.16115987 | NA | NA | NA |
| ENSG000002 | 0.47826975 | -0.016435741 | 0.244632 | -0.0671856 | 0.94643397 | NA | NA | NA |
| ENSG000002 | 0.63296622 | -0.299126688 | 0.27328461 | -1.0945611 | 0.27370899 | NA | NA | NA |
| ENSG000002 | 0.4553912 | 0.143837295 | 0.23493577 | 0.61224092 | 0.54037837 | NA | NA | NA |
| ENSG000002 | 1.78660295 | -0.319030526 | 0.34847132 | -0.9155144 | 0.35992163 | NA | MIR6895 | 102465539 |
| ENSG000002 | 1.55646587 | 0.029406816 | 0.33499793 | 0.08778208 | 0.93004988 | NA | NA | NA |
| ENSG000002 | 0 | NA | NA | NA | NA | NA | RNU6-467P | 106479772 |
| ENSG000002 | 0.70232246 | 0.137878926 | 0.28241054 | 0.4882216 | 0.62539289 | NA | MIR8055 | 102465861 |
| ENSG000002 | 0.2596576 | -0.193002035 | 0.1837913 | -1.0501152 | 0.29366517 | NA | SPATA31A5 | 727905 |
| ENSG000002 | 0.11205239 | 0.082118251 | 0.12671962 | 0.64803108 | 0.51696485 | NA | MIR6737 | 102465441 |
| ENSG000002 | 0.50704533 | -0.189333637 | 0.24184367 | -0.7828761 | 0.43369986 | NA | NA | NA |
| ENSG000002 | 1.96711462 | -0.606079421 | 0.34765871 | -1.7433173 | 0.08127821 | NA | RN7SL322P | 106479340 |
| ENSG000002 | 0.11205239 | 0.082118251 | 0.12671962 | 0.64803108 | 0.51696485 | NA | PHBP6 | 106478940 |
| ENSG000002 | 2.12899657 | -0.262043412 | 0.36750683 | -0.7130301 | 0.47582716 | NA | NA | NA |
| ENSG000002 | 0 | NA | NA | NA | NA | NA | NA | NA |

| | | | | | | | | |
|---|---|---|---|---|---|---|---|---|
| ENSG000002 | 1.80066732 | -0.450367046 | 0.34899394 | -1.2904724 | 0.19688669 | NA | NA | NA |
| ENSG000002 | 0 | NA | NA | NA | NA | NA | MIR6893 | 102466271 |
| ENSG000002 | 0 | NA | NA | NA | NA | NA | NA | NA |
| ENSG000002 | 1.35430546 | -0.252021854 | 0.33453959 | -0.7533394 | 0.45124598 | NA | NA | NA |
| ENSG000002 | 0 | NA | NA | NA | NA | NA | NA | NA |
| ENSG000002 | 0 | NA | NA | NA | NA | NA | SNORD64 | 347686 |
| ENSG000002 | 0 | NA | NA | NA | NA | NA | NA | NA |
| ENSG000002 | 1.56156507 | -0.479926515 | 0.34160686 | -1.4049089 | 0.16004837 | NA | NA | NA |
| ENSG000002 | 0.14128499 | -0.097772014 | 0.13403235 | -0.7294658 | 0.4657168 | NA | MIR6499 | 102465246 |
| ENSG000002 | 0 | NA | NA | NA | NA | NA | NA | NA |
| ENSG000002 | 1.14459284 | -0.281267894 | 0.32403159 | -0.8680262 | 0.38538001 | NA | NA | NA |
| ENSG000002 | 1.03464603 | -0.540193964 | 0.31380249 | -1.7214458 | 0.08516996 | NA | NA | NA |
| ENSG000002 | 0.58486175 | 0.058939796 | 0.25343823 | 0.23256079 | 0.81610248 | NA | MIR7154 | 102465691 |
| ENSG000002 | 0.60373362 | -0.112425234 | 0.26959077 | -0.4170218 | 0.67666245 | NA | MIR6769A | 102466731 |
| ENSG000002 | 0 | NA | NA | NA | NA | NA | NA | NA |
| ENSG000002 | 0 | NA | NA | NA | NA | NA | NA | NA |
| ENSG000002 | 0.26560726 | -0.177019426 | 0.17672274 | -1.0016788 | 0.31649875 | NA | NA | NA |
| ENSG000002 | 0.8876937 | -0.277667413 | 0.29985958 | -0.9259915 | 0.35445041 | NA | NA | NA |
| ENSG000002 | 0.98232848 | -0.03988105 | 0.31675486 | -0.1259051 | 0.89980705 | NA | RN7SL856P | 106479533 |
| ENSG000002 | 1.05773309 | -0.717898212 | 0.31119291 | -2.3069234 | 0.02105909 | NA | MIR6083 | 102464832 |
| ENSG000002 | 1.55062581 | -0.329911017 | 0.34145304 | -0.9661973 | 0.33394545 | NA | NA | NA |
| ENSG000002 | 0 | NA | NA | NA | NA | NA | NA | NA |
| ENSG000002 | 0.25370795 | -0.171331747 | 0.17411721 | -0.9840024 | 0.32511435 | NA | NA | NA |
| ENSG000002 | 0.72540953 | -0.028108977 | 0.28387695 | -0.0990182 | 0.92112384 | NA | NA | NA |
| ENSG000002 | 1.33562554 | 0.169296667 | 0.33326239 | 0.50799811 | 0.61145467 | NA | NA | NA |
| ENSG000002 | 0.23799501 | 0.170958857 | 0.17834799 | 0.95856902 | 0.33777591 | NA | NA | NA |
| ENSG000002 | 1.01796563 | -0.363579607 | 0.31420679 | -1.1571348 | 0.24721729 | NA | NA | NA |
| ENSG000002 | 0 | NA | NA | NA | NA | NA | NA | NA |
| ENSG000002 | 0 | NA | NA | NA | NA | NA | NA | NA |

| | | | | | | | | |
|---|---|---|---|---|---|---|---|---|
| ENSG000002 | 0 | NA | NA | NA | NA | NA | NA | NA |
| ENSG000002 | 1.58762826 | -0.121626572 | 0.3492237 | -0.348277 | 0.72763219 | NA | NA | NA |
| ENSG000002 | 0.27408862 | -0.199826037 | 0.18669465 | -1.0703362 | 0.28446801 | NA | NA | NA |
| ENSG000002 | 1.37852617 | -0.401576066 | 0.33319609 | -1.2052244 | 0.22811667 | NA | NA | NA |
| ENSG000002 | 0 | NA | NA | NA | NA | NA | NA | NA |
| ENSG000002 | 0 | NA | NA | NA | NA | NA | NA | NA |
| ENSG000002 | 0.73824193 | -0.03188941 | 0.28472511 | -0.1120007 | 0.91082286 | NA | NA | NA |
| ENSG000002 | 1.47733262 | -0.049544019 | 0.34105754 | -0.1452659 | 0.88450096 | NA | NA | NA |
| ENSG000002 | 0.96129492 | -0.036373653 | 0.31174745 | -0.1166767 | 0.90711628 | NA | NA | NA |
| ENSG000002 | 0.11131518 | 0.081951103 | 0.12663804 | 0.64712866 | 0.51754868 | NA | RNA5-8S5 | 100008587 |
| ENSG000002 | 0.11865494 | 0.083440866 | 0.12751478 | 0.65436232 | 0.5128784 | NA | NA | NA |
| ENSG000002 | 0.25214371 | -0.009862892 | 0.18046657 | -0.0546522 | 0.95641556 | NA | NA | NA |
| ENSG000002 | 0.51396245 | -0.026802372 | 0.24109708 | -0.1111684 | 0.91148283 | NA | NA | NA |
| ENSG000002 | 1.23643925 | -0.334106796 | 0.32794682 | -1.0187835 | 0.30830577 | NA | NA | NA |
| ENSG000002 | 1.85088008 | -0.252713026 | 0.35673811 | -0.7083993 | 0.47869733 | NA | NA | NA |
| ENSG000002 | 0.11865494 | 0.083440866 | 0.12751478 | 0.65436232 | 0.5128784 | NA | NA | NA |
| ENSG000002 | 1.35725588 | -0.116332488 | 0.34077944 | -0.3413718 | 0.73282371 | NA | NA | NA |
| ENSG000002 | 1.15037228 | -0.42327205 | 0.31994132 | -1.3229678 | 0.1858461 | NA | NA | NA |
| ENSG000002 | 1.25761133 | -0.183192392 | 0.32553071 | -0.5627499 | 0.57360517 | NA | NA | NA |
| ENSG000002 | 0.75878622 | -0.032974418 | 0.27982875 | -0.1178378 | 0.90619614 | NA | NA | NA |
| ENSG000002 | 0.69064756 | 0.464680151 | 0.27149568 | 1.71155635 | 0.08697846 | NA | NA | NA |
| ENSG000002 | 0.26813897 | -0.197007041 | 0.18550319 | -1.0620143 | 0.2882292 | NA | MIR7158 | 102465693 |
| ENSG000002 | 0.91627341 | -0.460143215 | 0.30152955 | -1.5260303 | 0.12700233 | NA | NA | NA |
| ENSG000002 | 1.06788693 | 0.037171252 | 0.31892133 | 0.11655304 | 0.90721425 | NA | NA | NA |
| ENSG000002 | 0 | NA | NA | NA | NA | NA | MIR6821 | 102465493 |
| ENSG000002 | 0.47075585 | 0.160115757 | 0.24423976 | 0.65556794 | 0.51210216 | NA | NA | NA |
| ENSG000002 | 0.24619405 | -0.007630671 | 0.17915639 | -0.0425922 | 0.96602658 | NA | NA | NA |
| ENSG000002 | 1.87686393 | -0.393741216 | 0.3591855 | -1.0962058 | 0.27298874 | NA | NA | NA |
| ENSG000002 | 0.35664995 | 0.244349217 | 0.20882949 | 1.17008962 | 0.2419649 | NA | NA | NA |

| | | | | | | | | |
|---|---|---|---|---|---|---|---|---|
| ENSG000002 | 1.56503042 | -0.448851562 | 0.33668994 | -1.3331303 | 0.18248905 | NA | NA | NA |
| ENSG000002 | 0.4923881 | -0.012789975 | 0.22994905 | -0.0556209 | 0.95564381 | NA | NA | NA |
| ENSG000002 | 1.53944198 | -0.489518483 | 0.3444428 | -1.4211895 | 0.15526168 | NA | MIR8073 | 102465872 |
| ENSG000002 | 0 | NA | NA | NA | NA | NA | MIR6752 | 102465450 |
| ENSG000002 | 0.37285215 | -0.099309896 | 0.2177345 | -0.4561055 | 0.64831414 | NA | MIR6780B | 102466746 |
| ENSG000002 | 0 | NA | NA | NA | NA | NA | NA | NA |
| ENSG000002 | 0.24619405 | -0.007630671 | 0.17915639 | -0.0425922 | 0.96602658 | NA | NA | NA |
| ENSG000002 | 2.16076087 | -0.366659142 | 0.36182649 | -1.0133563 | 0.31089001 | NA | NA | NA |
| ENSG000002 | 0 | NA | NA | NA | NA | NA | NA | NA |
| ENSG000002 | 0 | NA | NA | NA | NA | NA | SNORD26 | 9302 |
| ENSG000002 | 1.00797951 | -0.191736877 | 0.30785515 | -0.6228152 | 0.53340594 | NA | NA | NA |
| ENSG000002 | 1.98960087 | -0.790052656 | 0.35268126 | -2.2401322 | 0.02508234 | NA | NA | NA |
| ENSG000002 | 0.48348219 | -0.018797026 | 0.24521266 | -0.076656 | 0.9388972 | NA | NA | NA |
| ENSG000002 | 0.14128499 | -0.097772014 | 0.13403235 | -0.7294658 | 0.4657168 | NA | NA | NA |
| ENSG000002 | 1.23182762 | -0.282512936 | 0.3117001 | -0.9063614 | 0.3647446 | NA | NA | NA |
| ENSG000002 | 1.41392603 | -0.433110204 | 0.34193037 | -1.266662 | 0.20527617 | NA | MIR7159 | 102466816 |
| ENSG000002 | 0 | NA | NA | NA | NA | NA | NA | NA |
| ENSG000002 | 0.53590737 | -0.074911264 | 0.15581405 | -0.4807735 | 0.63067751 | NA | MIR6730 | 102466722 |
| ENSG000002 | 1.66611292 | 0.342498026 | 0.3491452 | 0.98096158 | 0.32661169 | NA | NA | NA |
| ENSG000002 | 0.23868015 | 0.154625155 | 0.16997278 | 0.90970538 | 0.36297791 | NA | NA | NA |
| ENSG000002 | 0.36391535 | 0.058449071 | 0.20749591 | 0.28168782 | 0.77818289 | NA | NA | NA |
| ENSG000002 | 1.93922527 | -0.508803916 | 0.35456287 | -1.4350175 | 0.15128212 | NA | NA | NA |
| ENSG000002 | 0 | NA | NA | NA | NA | NA | SNORD115-46 | 100873857 |
| ENSG000002 | 0.51931521 | -0.354214731 | 0.23897171 | -1.4822455 | 0.13827499 | NA | NA | NA |
| ENSG000002 | 0.22336757 | 0.163792188 | 0.17487321 | 0.93663397 | 0.34894687 | NA | NA | NA |
| ENSG000002 | 0.3854038 | -0.110128691 | 0.21952326 | -0.5016721 | 0.61589821 | NA | NA | NA |
| ENSG000002 | 0.37194025 | 0.067575365 | 0.21717998 | 0.31114914 | 0.75568724 | NA | NA | NA |
| ENSG000002 | 0.37945415 | -0.107841678 | 0.21865587 | -0.4932027 | 0.62186934 | NA | NA | NA |
| ENSG000002 | 2.13185391 | -0.224461156 | 0.35608412 | -0.6303599 | 0.52845913 | NA | MIR7151 | 102466814 |

| | | | | | | | | |
|---|---|---|---|---|---|---|---|---|
| ENSG000002 | 1.93444756 | -0.639276878 | 0.35195708 | -1.816349 | 0.06931686 | NA | NA | NA |
| ENSG000002 | 0 | NA | NA | NA | NA | NA | MIR6729 | 102466982 |
| ENSG000002 | 0 | NA | NA | NA | NA | NA | NA | NA |
| ENSG000002 | 0 | NA | NA | NA | NA | NA | NA | NA |
| ENSG000002 | 0 | NA | NA | NA | NA | NA | NA | NA |
| ENSG000002 | 0.64099112 | -0.296274405 | 0.27396996 | -1.081412 | 0.27951389 | NA | NA | NA |
| ENSG000002 | 1.35795409 | -0.672093525 | 0.3270152 | -2.0552364 | 0.03985618 | NA | MIR6074 | 102464827 |
| ENSG000002 | 0.60259551 | 0.210653417 | 0.25675396 | 0.82044857 | 0.41196044 | NA | NA | NA |
| ENSG000002 | 1.09011808 | -0.234601567 | 0.30817899 | -0.761251 | 0.44650716 | NA | LOC101929572 | 101929572 |
| ENSG000002 | 2.14383095 | -0.371543117 | 0.36449384 | -1.0193399 | 0.30804162 | NA | NA | NA |
| ENSG000002 | 0.13280363 | -0.093394593 | 0.13115644 | -0.7120855 | 0.47641185 | NA | RN7SL238P | 106480987 |
| ENSG000002 | 0.25370795 | -0.171331747 | 0.17411721 | -0.9840024 | 0.32511435 | NA | NA | NA |
| ENSG000002 | 2.14295331 | -0.646987243 | 0.36450335 | -1.774983 | 0.07590069 | NA | NA | NA |
| ENSG000002 | 1.3635761 | -0.242319402 | 0.33313008 | -0.7274017 | 0.46697989 | NA | NA | NA |
| ENSG000002 | 0.75941929 | -0.032774607 | 0.27987107 | -0.1171061 | 0.90677598 | NA | NA | NA |
| ENSG000002 | 0.53590737 | -0.074911264 | 0.15581405 | -0.4807735 | 0.63067751 | NA | NA | NA |
| ENSG000002 | 0 | NA | NA | NA | NA | NA | NA | NA |
| ENSG000002 | 0 | NA | NA | NA | NA | NA | MIR7106 | 102466222 |
| ENSG000002 | 0 | NA | NA | NA | NA | NA | NA | NA |
| ENSG000002 | 0.91049845 | -0.644235083 | 0.3003423 | -2.1450028 | 0.03195262 | NA | NA | NA |
| ENSG000002 | 1.23675437 | -0.17354407 | 0.32328255 | -0.5368186 | 0.59139296 | NA | MIR8080 | 102465877 |
| ENSG000002 | 1.03236631 | -0.192432806 | 0.31051435 | -0.6197228 | 0.53544033 | NA | NA | NA |
| ENSG000002 | 0 | NA | NA | NA | NA | NA | NA | NA |
| ENSG000002 | 1.14492724 | -0.120564886 | 0.31718999 | -0.3801031 | 0.70386892 | NA | NA | NA |
| ENSG000002 | 0.50955585 | -0.376243729 | 0.24515812 | -1.5346982 | 0.12485797 | NA | NA | NA |
| ENSG000002 | 1.67026965 | 0.455590913 | 0.34448968 | 1.32250962 | 0.18599851 | NA | MIR6797 | 102465478 |
| ENSG000002 | 0.49970612 | -0.194507174 | 0.24827631 | -0.7834303 | 0.43337449 | NA | NA | NA |
| ENSG000002 | 0.8842501 | -0.124993706 | 0.30355585 | -0.4117651 | 0.6805116 | NA | NA | NA |
| ENSG000002 | 1.27051373 | -0.350395479 | 0.33103085 | -1.0584979 | 0.28982849 | NA | NA | NA |

| | | | | | | | | |
|---|---|---|---|---|---|---|---|---|
| ENSG000002 | 0 | NA | NA | NA | NA | NA | MIR4509-1 | 100616223 |
| ENSG000002 | 0.99852208 | -0.028779217 | 0.2990393 | -0.0962389 | 0.92333083 | NA | NA | NA |
| ENSG000002 | 0.12685397 | -0.090215768 | 0.12901919 | -0.699243 | 0.48440017 | NA | NA | NA |
| ENSG000002 | 1.41949857 | 0.091158225 | 0.33767287 | 0.26996017 | 0.7871909 | NA | NA | NA |
| ENSG000002 | 0 | NA | NA | NA | NA | NA | NA | NA |
| ENSG000002 | 1.61176465 | -0.252397141 | 0.34452069 | -0.7326037 | 0.46380016 | NA | RN7SL769P | 106479502 |
| ENSG000002 | 0.48416733 | -0.018553864 | 0.24528998 | -0.0756405 | 0.93970509 | NA | NA | NA |
| ENSG000002 | 1.65414869 | -0.570119657 | 0.35267309 | -1.6165669 | 0.10597179 | NA | NA | NA |
| ENSG000002 | 0.36007182 | -0.08619198 | 0.20729252 | -0.4157988 | 0.67755724 | NA | MIR7848 | 102466865 |
| ENSG000002 | 0 | NA | NA | NA | NA | NA | NA | NA |
| ENSG000002 | 0 | NA | NA | NA | NA | NA | NA | NA |
| ENSG000002 | 1.6439844 | -0.497197663 | 0.33865443 | -1.4681564 | 0.14206175 | NA | NA | NA |
| ENSG000002 | 1.32250835 | -0.114741739 | 0.33536751 | -0.3421373 | 0.73224756 | NA | NA | NA |
| ENSG000002 | 0.90001914 | -0.118344353 | 0.29770615 | -0.3975207 | 0.69098353 | NA | NA | NA |
| ENSG000002 | 0 | NA | NA | NA | NA | NA | NA | NA |
| ENSG000002 | 1.64382234 | -0.400712001 | 0.34796549 | -1.1515855 | 0.24949145 | NA | NA | NA |
| ENSG000002 | 0.24756242 | -0.187034346 | 0.18119817 | -1.0322088 | 0.30197431 | NA | NA | NA |
| ENSG000002 | 0.47477407 | 0.142502628 | 0.23735343 | 0.60038158 | 0.54825196 | NA | NA | NA |
| ENSG000002 | 0.12070844 | -0.088503695 | 0.12783931 | -0.6923042 | 0.48874631 | NA | NA | NA |
| ENSG000002 | 0 | NA | NA | NA | NA | NA | NA | NA |
| ENSG000002 | 0.24756242 | -0.187034346 | 0.18119817 | -1.0322088 | 0.30197431 | NA | NA | NA |
| ENSG000002 | 0 | NA | NA | NA | NA | NA | NA | NA |
| ENSG000002 | 0 | NA | NA | NA | NA | NA | MIR3179-4 | 103504729 |
| ENSG000002 | 0.89387361 | -0.116349482 | 0.29739608 | -0.3912274 | 0.69562919 | NA | NA | NA |
| ENSG000002 | 0 | NA | NA | NA | NA | NA | RMRP | 6023 |
| ENSG000002 | 0 | NA | NA | NA | NA | NA | RNU6-1192P | 106480088 |
| ENSG000002 | 0 | NA | NA | NA | NA | NA | MIR8071-2 | 102466889 |
| ENSG000002 | 0 | NA | NA | NA | NA | NA | NA | NA |
| ENSG000002 | 0 | NA | NA | NA | NA | NA | NA | NA |

| | | | | | | | | |
|---|---|---|---|---|---|---|---|---|
| ENSG000002 | 0.55070895 | -0.361402396 | 0.24091882 | -1.5001003 | 0.13358842 | NA | NA | NA |
| ENSG000002 | 0.3863157 | -0.274134926 | 0.21457207 | -1.277589 | 0.20139437 | NA | NA | NA |
| ENSG000002 | 0.12070844 | -0.088503695 | 0.12783931 | -0.6923042 | 0.48874631 | NA | NA | NA |
| ENSG000002 | 0.74430313 | -0.030011485 | 0.2851257 | -0.105257 | 0.91617186 | NA | NA | NA |
| ENSG000002 | 0.22410479 | 0.147643983 | 0.16638082 | 0.88738586 | 0.37487119 | NA | MIR6763 | 102465975 |
| ENSG000002 | 0.25145857 | -0.010118565 | 0.18031426 | -0.0561163 | 0.95524918 | NA | NA | NA |
| ENSG000002 | 0 | NA | NA | NA | NA | NA | MIR6779 | 102465467 |
| ENSG000002 | 1.03355999 | -0.202825642 | 0.31368466 | -0.6465909 | 0.51789675 | NA | HNRNPCL3 | 649330 |
| ENSG000002 | 1.23854248 | -0.049964253 | 0.32875967 | -0.151978 | 0.87920425 | NA | RN7SL88P | 106480945 |
| ENSG000002 | 0.49287545 | -0.179123382 | 0.24028356 | -0.7454666 | 0.45598967 | NA | MIR8084 | 102467005 |
| ENSG000002 | 1.45397745 | 0.072984679 | 0.33433297 | 0.21829938 | 0.82719586 | NA | NA | NA |
| ENSG000002 | 0.46882441 | 0.14446209 | 0.23662046 | 0.6105224 | 0.54151581 | NA | MIR6131 | 102465138 |
| ENSG000002 | 0.47758652 | 0.147793567 | 0.23768536 | 0.6218034 | 0.53407115 | NA | NA | NA |
| ENSG000002 | 0 | NA | NA | NA | NA | NA | NA | NA |
| ENSG000002 | 0 | NA | NA | NA | NA | NA | NA | NA |
| ENSG000002 | 1.26123099 | -0.347053654 | 0.33223688 | -1.044597 | 0.29620925 | NA | NA | NA |
| ENSG000002 | 0.47092999 | -0.01913341 | 0.24378131 | -0.078486 | 0.9374415 | NA | NA | NA |
| ENSG000002 | 1.92945544 | 0.0739297 | 0.3621293 | 0.20415277 | 0.83823412 | NA | NA | NA |
| ENSG000002 | 0 | NA | NA | NA | NA | NA | NA | NA |
| ENSG000002 | 0.53374623 | -0.385819457 | 0.24767828 | -1.5577444 | 0.11929385 | NA | NA | NA |
| ENSG000002 | 0 | NA | NA | NA | NA | NA | NA | NA |
| ENSG000002 | 0 | NA | NA | NA | NA | NA | MIR6089 | 102464837 |
| ENSG000002 | 0.22410479 | 0.147643983 | 0.16638082 | 0.88738586 | 0.37487119 | NA | NA | NA |
| ENSG000002 | 0.59278059 | -0.088567558 | 0.25301851 | -0.3500438 | 0.72630584 | NA | NA | NA |
| ENSG000002 | 0.61890185 | -0.110660695 | 0.2644872 | -0.4183972 | 0.67565676 | NA | NA | NA |
| ENSG000002 | 0.61816464 | -0.117186973 | 0.27074914 | -0.4328249 | 0.66514197 | NA | NA | NA |
| ENSG000002 | 0.11865494 | 0.083440866 | 0.12751478 | 0.65436232 | 0.5128784 | NA | NA | NA |
| ENSG000002 | 1.11838176 | -0.107372848 | 0.31626856 | -0.339499 | 0.73423388 | NA | NA | NA |
| ENSG000002 | 0.3593886 | 0.071604659 | 0.20679693 | 0.34625591 | 0.7291504 | NA | MIR6509 | 102465254 |

| | | | | | | | | |
|---|---|---|---|---|---|---|---|---|
| ENSG000002 | 2.01048015 | -0.19265104 | 0.3615401 | -0.5328622 | 0.594129 | NA | NA | NA |
| ENSG000002 | 1.59728266 | -0.260243232 | 0.35224588 | -0.7388113 | 0.46002159 | NA | NA | NA |
| ENSG000002 | 0 | NA | NA | NA | NA | NA | NA | NA |
| ENSG000002 | 0.85724569 | -0.273352963 | 0.29829209 | -0.9163936 | 0.35946048 | NA | F8A3 | 474384 |
| ENSG000002 | 0 | NA | NA | NA | NA | NA | NA | NA |
| ENSG000002 | 1.76363904 | -0.328839599 | 0.35516013 | -0.9258911 | 0.35450258 | NA | NA | NA |
| ENSG000002 | 0 | NA | NA | NA | NA | NA | SNORA9 | 677798 |
| ENSG000002 | 1.11971733 | -0.27463679 | 0.32326001 | -0.8495848 | 0.39555597 | NA | NA | NA |
| ENSG000002 | 1.04290118 | -0.351036066 | 0.31073619 | -1.1296916 | 0.25860619 | NA | NA | NA |
| ENSG000002 | 0.39499294 | -0.277908454 | 0.21584201 | -1.287555 | 0.19790091 | NA | NA | NA |
| ENSG000002 | 0 | NA | NA | NA | NA | NA | SNORD22 | 9304 |
| ENSG000002 | 0.67004903 | -0.269626612 | 0.26217122 | -1.0284371 | 0.30374426 | NA | RN7SL270P | 106480995 |
| ENSG000002 | 0.12685397 | -0.090215768 | 0.12901919 | -0.699243 | 0.48440017 | NA | NA | NA |
| ENSG000002 | 0.12070844 | -0.088503695 | 0.12783931 | -0.6923042 | 0.48874631 | NA | MIR8063 | 102466875 |
| ENSG000002 | 1.25611359 | -0.440864227 | 0.31658662 | -1.3925548 | 0.16375445 | NA | NA | NA |
| ENSG000002 | 0.12685397 | -0.090215768 | 0.12901919 | -0.699243 | 0.48440017 | NA | NA | NA |
| ENSG000002 | 0 | NA | NA | NA | NA | NA | RPPH1 | 85495 |
| ENSG000002 | 0.11131518 | 0.081951103 | 0.12663804 | 0.64712866 | 0.51754868 | NA | NA | NA |
| ENSG000002 | 0.11934008 | 0.083564698 | 0.12760091 | 0.65489106 | 0.5125379 | NA | NA | NA |
| ENSG000002 | 2.05841946 | -0.33020136 | 0.35775566 | -0.9229801 | 0.35601757 | NA | NA | NA |
| ENSG000002 | 0 | NA | NA | NA | NA | NA | NA | NA |
| ENSG000002 | 0.3588776 | -0.099902505 | 0.21555794 | -0.4634601 | 0.6430346 | NA | NA | NA |
| ENSG000002 | 1.77106807 | -0.420030044 | 0.34298931 | -1.2246155 | 0.22072011 | NA | NA | NA |
| ENSG000002 | 0 | NA | NA | NA | NA | NA | MIR3118-3 | 100422844 |
| ENSG000002 | 0.49168123 | -0.197384925 | 0.247356 | -0.7979791 | 0.4248826 | NA | NA | NA |
| ENSG000002 | 0.23816916 | -0.010712105 | 0.1772998 | -0.060418 | 0.9518227 | NA | MIR6784 | 102465470 |
| ENSG000002 | 0.47781273 | -0.018045496 | 0.22812977 | -0.0791019 | 0.93695158 | NA | NA | NA |
| ENSG000002 | 0.46938748 | 0.335416007 | 0.2391134 | 1.40274868 | 0.16069179 | NA | MIR7854 | 102465840 |
| ENSG000002 | 1.64347002 | -0.398160519 | 0.34852632 | -1.1424116 | 0.25328296 | NA | NA | NA |

| | | | | | | | | |
|---|---|---|---|---|---|---|---|---|
| ENSG000002 | 0 | NA | NA | NA | NA | NA | NA | NA |
| ENSG000002 | 0.84585738 | -0.096367014 | 0.29044834 | -0.3317871 | 0.74005002 | NA | RN7SL556P | 106481073 |
| ENSG000002 | 0 | NA | NA | NA | NA | NA | NA | NA |
| ENSG000002 | 0 | NA | NA | NA | NA | NA | NA | NA |
| ENSG000002 | 0.48416733 | -0.018553864 | 0.24528998 | -0.0756405 | 0.93970509 | NA | NA | NA |
| ENSG000002 | 0 | NA | NA | NA | NA | NA | NA | NA |
| ENSG000002 | 2.06547592 | 0.00417593 | 0.36681796 | 0.0113842 | 0.99091692 | NA | NA | NA |
| ENSG000002 | 0.12685397 | -0.090215768 | 0.12901919 | -0.699243 | 0.48440017 | NA | RN7SL489P | 106481056 |
| ENSG000002 | 2.17329296 | 0.049701766 | 0.36737185 | 0.13529008 | 0.89238253 | NA | NA | NA |
| ENSG000002 | 0.98255661 | -0.042398807 | 0.31261927 | -0.1356244 | 0.8921182 | NA | NA | NA |
| ENSG000002 | 0.11934008 | 0.083564698 | 0.12760091 | 0.65489106 | 0.5125379 | NA | NA | NA |
| ENSG000002 | 2.05990326 | -1.03560266 | 0.35862663 | -2.8876903 | 0.00388082 | NA | NA | NA |
| ENSG000002 | 0 | NA | NA | NA | NA | NA | NA | NA |
| ENSG000002 | 0.78039674 | -0.368519854 | 0.28442264 | -1.295677 | 0.1950868 | NA | NA | NA |
| ENSG000002 | 0 | NA | NA | NA | NA | NA | RNU6-489P | 106481334 |
| ENSG000002 | 0.49375647 | -0.17902118 | 0.24039617 | -0.7446923 | 0.45645774 | NA | RNU1-6P | 106480152 |
| ENSG000002 | 1.3291109 | -0.112616888 | 0.33551486 | -0.3356539 | 0.73713184 | NA | NA | NA |
| ENSG000002 | 1.2594037 | -0.346954158 | 0.33075547 | -1.0489748 | 0.29418971 | NA | NA | NA |
| ENSG000002 | 1.62284532 | -0.690793076 | 0.34714128 | -1.9899479 | 0.04659667 | NA | NA | NA |
| ENSG000002 | 0.48624257 | -0.012072666 | 0.23796621 | -0.0507327 | 0.95953853 | NA | NA | NA |
| ENSG000002 | 1.16078452 | -0.413015931 | 0.31797011 | -1.2989143 | 0.19397333 | NA | NA | NA |
| ENSG000002 | 1.77399338 | -0.198543541 | 0.35408106 | -0.5607291 | 0.57498225 | NA | NA | NA |
| ENSG000002 | 0 | NA | NA | NA | NA | NA | SNORD49A | 26800 |
| ENSG000002 | 1.28127536 | -0.67275127 | 0.32946654 | -2.0419411 | 0.04115737 | NA | NA | NA |
| ENSG000002 | 1.65265771 | -0.25663169 | 0.3450911 | -0.7436636 | 0.45708002 | NA | NA | NA |
| ENSG000002 | 0.37171 | -0.104406568 | 0.21753023 | -0.4799635 | 0.63125336 | NA | NA | NA |
| ENSG000002 | 0 | NA | NA | NA | NA | NA | MIR6724-3 | 103504739 |
| ENSG000002 | 0.87903765 | -0.1121152 | 0.29232114 | -0.3835344 | 0.70132361 | NA | NA | NA |
| ENSG000002 | 1.62898693 | -0.41565237 | 0.35317787 | -1.1768925 | 0.23923843 | NA | NA | NA |

| | | | | | | | | |
|---|---|---|---|---|---|---|---|---|
| ENSG000002 | 1.07719586 | -0.105604382 | 0.31956186 | -0.3304662 | 0.74104776 | NA | NA | NA |
| ENSG000002 | 0.36621735 | -0.097238239 | 0.21673053 | -0.4486596 | 0.65367721 | NA | MIR8058 | 102465863 |
| ENSG000002 | 0.60215886 | 0.217171948 | 0.25510838 | 0.8512929 | 0.39460667 | NA | MIR8053 | 102467004 |
| ENSG000002 | 0.3701135 | 0.066858436 | 0.20853829 | 0.32060509 | 0.74850968 | NA | MIR6881 | 102465530 |
| ENSG000002 | 0 | NA | NA | NA | NA | NA | NA | NA |
| ENSG000002 | 0 | NA | NA | NA | NA | NA | NA | NA |
| ENSG000002 | 1.26242467 | -0.334755824 | 0.32685874 | -1.0241606 | 0.30575945 | NA | NA | NA |
| ENSG000002 | 0.23202362 | -0.008302493 | 0.17586207 | -0.0472103 | 0.96234565 | NA | NA | NA |
| ENSG000002 | 0.49031286 | -0.020802257 | 0.24597635 | -0.0845702 | 0.93260313 | NA | NA | NA |
| ENSG000002 | 0 | NA | NA | NA | NA | NA | REXO1L2P | 100288527 |
| ENSG000002 | 2.07617688 | 0.01392685 | 0.35316254 | 0.03943468 | 0.96854383 | NA | NA | NA |
| ENSG000002 | 1.48843915 | -0.323078211 | 0.34176994 | -0.945309 | 0.34450112 | NA | NA | NA |
| ENSG000002 | 1.89625127 | -0.631931287 | 0.35362781 | -1.7869955 | 0.07393822 | NA | NA | NA |
| ENSG000002 | 0 | NA | NA | NA | NA | NA | NA | NA |
| ENSG000002 | 0.23868015 | 0.154625155 | 0.16997278 | 0.90970538 | 0.36297791 | NA | MIR3687-1 | 100500815 |
| ENSG000002 | 0 | NA | NA | NA | NA | NA | NA | NA |
| ENSG000002 | 0 | NA | NA | NA | NA | NA | NA | NA |
| ENSG000002 | 0 | NA | NA | NA | NA | NA | NA | NA |
| ENSG000002 | 0.26062507 | -0.012888319 | 0.18226503 | -0.070712 | 0.94362699 | NA | NA | NA |
| ENSG000002 | 1.47196259 | -0.173138847 | 0.3399893 | -0.5092479 | 0.61057845 | NA | NA | NA |
| ENSG000002 | 0.24619405 | -0.007630671 | 0.17915639 | -0.0425922 | 0.96602658 | NA | NA | NA |
| ENSG000002 | 0 | NA | NA | NA | NA | NA | NA | NA |
| ENSG000002 | 0 | NA | NA | NA | NA | NA | RN7SL286P | 106479327 |
| ENSG000002 | 1.46165994 | -0.548913898 | 0.33173564 | -1.6546727 | 0.09799091 | NA | NA | NA |
| ENSG000002 | 0.25333738 | -0.015686291 | 0.1806605 | -0.0868275 | 0.93080867 | NA | NA | NA |
| ENSG000002 | 0 | NA | NA | NA | NA | NA | RN7SL628P | 106479451 |
| ENSG000002 | 0 | NA | NA | NA | NA | NA | NA | NA |
| ENSG000002 | 0 | NA | NA | NA | NA | NA | NA | NA |
| ENSG000002 | 1.48491068 | -0.190529729 | 0.34022433 | -0.5600121 | 0.57547121 | NA | NA | NA |

| | | | | | | | | |
|---|---|---|---|---|---|---|---|---|
| ENSG000002 | 0 | NA | NA | NA | NA | NA | MIR6894 | 102466759 |
| ENSG000002 | 0.90112692 | -0.259399981 | 0.29324956 | -0.8845708 | 0.37638819 | NA | MIR6165 | 102465141 |
| ENSG000002 | 0.63939408 | -0.113476381 | 0.26621037 | -0.4262658 | 0.66991416 | NA | NA | NA |
| ENSG000002 | 1.74348671 | -0.326933603 | 0.35539771 | -0.9199091 | 0.35762024 | NA | NA | NA |
| ENSG000002 | 0.26062507 | -0.012888319 | 0.18226503 | -0.070712 | 0.94362699 | NA | RN7SL321P | 106480501 |
| ENSG000002 | 0 | NA | NA | NA | NA | NA | NA | NA |
| ENSG000002 | 0.87531619 | -0.427631553 | 0.29439996 | -1.452553 | 0.1463479 | NA | NA | NA |
| ENSG000002 | 0 | NA | NA | NA | NA | NA | NA | NA |
| ENSG000002 | 0.6265601 | -0.275936484 | 0.2670182 | -1.0333995 | 0.30141697 | NA | NA | NA |
| ENSG000002 | 0 | NA | NA | NA | NA | NA | NA | NA |
| ENSG000002 | 0.51570138 | -0.378721611 | 0.24581506 | -1.540677 | 0.12339542 | NA | NA | NA |
| ENSG000002 | 1.26091642 | -0.494721778 | 0.33002771 | -1.4990311 | 0.13386558 | NA | RN7SL802P | 106479513 |
| ENSG000002 | 0 | NA | NA | NA | NA | NA | BSNDP3 | 106481725 |
| ENSG000002 | 1.02163894 | -0.355029704 | 0.30982402 | -1.1459076 | 0.25183338 | NA | NA | NA |
| ENSG000002 | 1.3686461 | -0.120391855 | 0.33842704 | -0.3557395 | 0.72203569 | NA | NA | NA |
| ENSG000002 | 0 | NA | NA | NA | NA | NA | SNORD65 | 692106 |
| ENSG000002 | 1.85833037 | -0.382097 | 0.35905324 | -1.0641792 | 0.28724752 | NA | NA | NA |
| ENSG000002 | 0.90047403 | -0.281185275 | 0.30049169 | -0.9357506 | 0.34940162 | NA | RN7SL495P | 106479404 |
| ENSG000002 | 0.71749069 | 0.124407623 | 0.27556104 | 0.45147029 | 0.65165063 | NA | NA | NA |
| ENSG000002 | 0 | NA | NA | NA | NA | NA | NA | NA |
| ENSG000002 | 0 | NA | NA | NA | NA | NA | NA | NA |
| ENSG000002 | 1.10272372 | -0.110611827 | 0.31320149 | -0.3531651 | 0.72396469 | NA | NF1P10 | 106480348 |
| ENSG000002 | 1.26444358 | -0.04787209 | 0.31534043 | -0.1518108 | 0.87933615 | NA | NA | NA |
| ENSG000002 | 0.41537361 | -0.286583533 | 0.21871501 | -1.3103057 | 0.19009243 | NA | MIR8077 | 102465875 |
| ENSG000002 | 0.12685397 | -0.090215768 | 0.12901919 | -0.699243 | 0.48440017 | NA | NA | NA |
| ENSG000002 | 0.72037177 | -0.172939881 | 0.27655343 | -0.6253398 | 0.53174804 | NA | NA | NA |
| ENSG000002 | 0.37831254 | -0.102004459 | 0.21855316 | -0.4667261 | 0.64069587 | NA | NA | NA |
| ENSG000002 | 0 | NA | NA | NA | NA | NA | NA | NA |
| ENSG000002 | 1.97224388 | -0.17018628 | 0.3549626 | -0.4794485 | 0.63161961 | NA | NA | NA |

| | | | | | | | | |
|---|---|---|---|---|---|---|---|---|
| ENSG000002 | 0.49101782 | -0.013272127 | 0.22978098 | -0.0577599 | 0.95393988 | NA | NA | NA |
| ENSG000002 | 0 | NA | NA | NA | NA | NA | NA | NA |
| ENSG000002 | 0.36351096 | 0.070290861 | 0.21590418 | 0.32556508 | 0.74475344 | NA | NA | NA |
| ENSG000002 | 0.36212533 | -0.235926492 | 0.20089643 | -1.1743688 | 0.24024735 | NA | MIR6134 | 102465140 |
| ENSG000002 | 0.12685397 | -0.090215768 | 0.12901919 | -0.699243 | 0.48440017 | NA | NA | NA |
| ENSG000002 | 1.74951017 | -0.181108072 | 0.35200332 | -0.5145067 | 0.60689777 | NA | NA | NA |
| ENSG000002 | 0 | NA | NA | NA | NA | NA | NA | NA |
| ENSG000002 | 0 | NA | NA | NA | NA | NA | NA | NA |
| ENSG000002 | 0 | NA | NA | NA | NA | NA | RN7SL428P | 106479379 |
| ENSG000002 | 0 | NA | NA | NA | NA | NA | NA | NA |
| ENSG000002 | 1.08470976 | -0.246686027 | 0.31443436 | -0.784539 | 0.43272388 | NA | NA | NA |
| ENSG000002 | 1.41779794 | 0.235922072 | 0.34013493 | 0.69361319 | 0.48792482 | NA | MIR6806 | 102465484 |
| ENSG000002 | 0.969498 | 0.11559493 | 0.30553578 | 0.37833516 | 0.70518163 | NA | MIR7845 | 102465835 |
| ENSG000002 | 0.99607628 | 0.114016091 | 0.31313774 | 0.36410843 | 0.71577704 | NA | RN7SL575P | 106479432 |
| ENSG000002 | 1.91705011 | -0.519645401 | 0.35709678 | -1.4551949 | 0.14561533 | NA | NA | NA |
| ENSG000002 | 0.52165104 | -0.403921832 | 0.25252134 | -1.5995552 | 0.10969729 | NA | NA | NA |
| ENSG000002 | 0 | NA | NA | NA | NA | NA | NA | NA |
| ENSG000002 | 0 | NA | NA | NA | NA | NA | NA | NA |
| ENSG000002 | 0.37079864 | 0.073139042 | 0.21703538 | 0.33699134 | 0.73612343 | NA | NA | NA |
| ENSG000002 | 2.06306786 | -0.003587211 | 0.35803694 | -0.0100191 | 0.99200604 | NA | NA | NA |
| ENSG000002 | 0 | NA | NA | NA | NA | NA | NA | NA |
| ENSG000002 | 0.11205239 | 0.082118251 | 0.12671962 | 0.64803108 | 0.51696485 | NA | NA | NA |
| ENSG000002 | 0.89495965 | -0.460377077 | 0.30201036 | -1.5243751 | 0.12741503 | NA | NA | NA |
| ENSG000002 | 0.27408862 | -0.199826037 | 0.18669465 | -1.0703362 | 0.28446801 | NA | NA | NA |
| ENSG000002 | 0.99294768 | -0.182417453 | 0.30147696 | -0.6050792 | 0.54512634 | NA | NA | NA |
| ENSG000002 | 0.8560732 | -0.109326917 | 0.29544606 | -0.3700402 | 0.71135254 | NA | NA | NA |
| ENSG000002 | 0.46748638 | 0.139734475 | 0.23646399 | 0.59093341 | 0.55456504 | NA | NA | NA |
| ENSG000002 | 2.02388951 | -0.437843005 | 0.35799064 | -1.2230571 | 0.22130813 | NA | NA | NA |
| ENSG000002 | 1.30235401 | 0.331140002 | 0.33838791 | 0.97858106 | 0.32778702 | NA | NA | NA |

| | | | | | | | | |
|---|---|---|---|---|---|---|---|---|
| ENSG000002 | 0 | NA | NA | NA | NA | NA | RN7SL469P | 106480512 |
| ENSG000002 | 1.00718673 | -0.525923654 | 0.31127369 | -1.689586 | 0.0911072 | NA | NA | NA |
| ENSG000002 | 0.53121452 | -0.133405847 | 0.15544486 | -0.8582198 | 0.39077113 | NA | NA | NA |
| ENSG000002 | 1.48889774 | -0.17899992 | 0.33799392 | -0.5295951 | 0.5963927 | NA | NA | NA |
| ENSG000002 | 1.28960376 | -0.343195717 | 0.32933085 | -1.0421001 | 0.29736527 | NA | NA | NA |
| ENSG000002 | 0.89498139 | -0.275015684 | 0.30025139 | -0.9159514 | 0.35969239 | NA | NA | NA |
| ENSG000002 | 1.98825628 | -0.438922408 | 0.35994446 | -1.219417 | 0.22268595 | NA | NA | NA |
| ENSG000002 | 0 | NA | NA | NA | NA | NA | NA | NA |
| ENSG000002 | 0.26062507 | -0.012888319 | 0.18226503 | -0.070712 | 0.94362699 | NA | NA | NA |
| ENSG000002 | 0 | NA | NA | NA | NA | NA | NA | NA |
| ENSG000002 | 0.86065896 | -0.231025277 | 0.28410078 | -0.8131807 | 0.41611446 | NA | NA | NA |
| ENSG000002 | 0.83760436 | -0.093441283 | 0.28819276 | -0.3242319 | 0.74576248 | NA | NA | NA |
| ENSG000002 | 0.12070844 | -0.088503695 | 0.12783931 | -0.6923042 | 0.48874631 | NA | NA | NA |
| ENSG000002 | 1.01159334 | -0.203772921 | 0.31433242 | -0.648272 | 0.51680901 | NA | MIR6739 | 102466724 |
| ENSG000002 | 0 | NA | NA | NA | NA | NA | NA | NA |
| ENSG000002 | 0.24141689 | -0.165257896 | 0.17127401 | -0.9648743 | 0.33460776 | NA | NA | NA |
| ENSG000002 | 0 | NA | NA | NA | NA | NA | NA | NA |
| ENSG000002 | 0 | NA | NA | NA | NA | NA | MIR6770-2 | 102465908 |
| ENSG000002 | 1.69379571 | -0.059406488 | 0.35354727 | -0.1680298 | 0.86655981 | NA | NA | NA |
| ENSG000002 | 0.71621269 | 0.14258294 | 0.28335356 | 0.50319798 | 0.61482509 | NA | NA | NA |
| ENSG000002 | 0.64878734 | -0.284003979 | 0.26878765 | -1.056611 | 0.29068916 | NA | NA | NA |
| ENSG000002 | 1.98972891 | -0.058886987 | 0.35950888 | -0.1637984 | 0.86988983 | NA | NA | NA |
| ENSG000002 | 0 | NA | NA | NA | NA | NA | NA | NA |
| ENSG000002 | 0.12070844 | -0.088503695 | 0.12783931 | -0.6923042 | 0.48874631 | NA | MIR6856 | 102465515 |
| ENSG000002 | 1.14550632 | -0.278636311 | 0.32258745 | -0.8637544 | 0.38772285 | NA | NA | NA |
| ENSG000002 | 0.99232044 | -0.334091058 | 0.30627138 | -1.0908334 | 0.27534619 | NA | NA | NA |
| ENSG000002 | 0.36690249 | -0.096999144 | 0.21683809 | -0.4473344 | 0.65463361 | NA | MIR6827 | 102466744 |
| ENSG000002 | 0 | NA | NA | NA | NA | NA | NA | NA |
| ENSG000002 | 0 | NA | NA | NA | NA | NA | NA | NA |

| | | | | | | | | | |
|---|---|---|---|---|---|---|---|---|---|
| ENSG000002 | 0 | NA | NA | NA | NA | NA | | RNA5-8S5 | 100008587 |
| ENSG000002 | 2.02412635 | -0.415910591 | 0.35579623 | -1.1689573 | 0.24242085 | NA | | NA | NA |
| ENSG000002 | 1.08863974 | 0.028312312 | 0.3149176 | 0.08990387 | 0.9283636 | NA | | NA | NA |
| ENSG000002 | 0 | NA | NA | NA | NA | NA | | H2AFB3 | 83740 |
| ENSG000002 | 1.5127972 | -0.054912483 | 0.34091179 | -0.1610753 | 0.87203406 | NA | | NA | NA |
| ENSG000002 | 0.88991743 | -0.27958823 | 0.29835725 | -0.9370921 | 0.34871117 | NA | | NA | NA |
| ENSG000002 | 0.80783485 | -0.539945266 | 0.28176122 | -1.9163222 | 0.0553241 | NA | | NA | NA |
| ENSG000002 | 1.93452328 | -0.377751332 | 0.35409031 | -1.066822 | 0.28605223 | NA | | NA | NA |
| ENSG000002 | 0.72332164 | -0.023862463 | 0.26820519 | -0.0889709 | 0.92910502 | NA | | NA | NA |
| ENSG000002 | 0.49998845 | -0.018666443 | 0.24702698 | -0.0755644 | 0.93976567 | NA | | MIR6719 | 102465974 |
| ENSG000002 | 0.86065896 | -0.231025277 | 0.28410078 | -0.8131807 | 0.41611446 | NA | | NA | NA |
| ENSG000002 | 0.73654172 | -0.473859347 | 0.26792886 | -1.7686013 | 0.07696043 | NA | | RN7SL261P | 106479316 |
| ENSG000002 | 0.94560372 | 0.432893913 | 0.30174255 | 1.43464655 | 0.15138784 | NA | | NA | NA |
| ENSG000002 | 1.08749601 | -0.112768333 | 0.31519801 | -0.3577698 | 0.72051558 | NA | | NA | NA |
| ENSG000002 | 0.72377866 | -0.02164005 | 0.2774131 | -0.0780066 | 0.93782281 | NA | | NA | NA |
| ENSG000002 | 0.13280363 | -0.093394593 | 0.13115644 | -0.7120855 | 0.47641185 | NA | | NA | NA |
| ENSG000002 | 0.52668879 | -0.197054386 | 0.24396589 | -0.8077129 | 0.4192559 | NA | | NA | NA |
| ENSG000002 | 0 | NA | NA | NA | NA | NA | | RN7SL763P | 106481130 |
| ENSG000002 | 0 | NA | NA | NA | NA | NA | | NA | NA |
| ENSG000002 | 0 | NA | NA | NA | NA | NA | | MIR6786 | 102465471 |
| ENSG000002 | 0 | NA | NA | NA | NA | NA | | SNORD116-21 | 100033432 |
| ENSG000002 | 0.96280709 | -0.170529837 | 0.29744015 | -0.5733249 | 0.56642474 | NA | | NA | NA |
| ENSG000002 | 0 | NA | NA | NA | NA | NA | | NA | NA |
| ENSG000002 | 1.14479841 | -0.513560224 | 0.30498702 | -1.6838756 | 0.0922057 | NA | | MIR6738 | 102465442 |
| ENSG000002 | 0.47092999 | -0.01913341 | 0.24378131 | -0.078486 | 0.9374415 | NA | | NA | NA |
| ENSG000002 | 0.49879422 | -0.023609326 | 0.24688725 | -0.095628 | 0.92381605 | NA | | NA | NA |
| ENSG000002 | 0.37996515 | 0.064665424 | 0.21008146 | 0.3078112 | 0.758226 | NA | | NA | NA |
| ENSG000002 | 0.48761094 | -0.176642206 | 0.23968055 | -0.7369902 | 0.46112833 | NA | | NA | NA |
| ENSG000002 | 0 | NA | NA | NA | NA | NA | | NA | NA |

| | | | | | | | | |
|---|---|---|---|---|---|---|---|---|
| ENSG000002 | 0.51316968 | -0.377777671 | 0.24556403 | -1.538408 | 0.12394889 | NA | NA | NA |
| ENSG000002 | 0 | NA | NA | NA | NA | NA | MIR5701-2 | 100847005 |
| ENSG000002 | 0 | NA | NA | NA | NA | NA | MIR6811 | 102465486 |
| ENSG000002 | 0.60489751 | 0.065499153 | 0.26291219 | 0.24912938 | 0.8032607 | NA | SNORD30 | 9299 |
| ENSG000002 | 0.62601485 | 0.045160216 | 0.25727761 | 0.17553107 | 0.86066236 | NA | NA | NA |
| ENSG000002 | 0 | NA | NA | NA | NA | NA | H2AFB2 | 474381 |
| ENSG000002 | 1.23589454 | -0.045458417 | 0.33372795 | -0.136214 | 0.89165211 | NA | NA | NA |
| ENSG000002 | 0 | NA | NA | NA | NA | NA | NA | NA |
| ENSG000002 | 0 | NA | NA | NA | NA | NA | SCARNA15 | 677778 |
| ENSG000002 | 0.47614243 | -0.020043108 | 0.23674202 | -0.0846622 | 0.93252992 | NA | NA | NA |
| ENSG000002 | 0.61121718 | -0.095100843 | 0.25474802 | -0.3733134 | 0.70891523 | NA | NA | NA |
| ENSG000002 | 0.11205239 | 0.082118251 | 0.12671962 | 0.64803108 | 0.51696485 | NA | MIR6817 | 102466198 |
| ENSG000002 | 0 | NA | NA | NA | NA | NA | NA | NA |
| ENSG000002 | 1.87071137 | 0.004343927 | 0.35365836 | 0.01228283 | 0.99019996 | NA | NA | NA |
| ENSG000002 | 1.29092006 | -0.489302153 | 0.3269723 | -1.4964636 | 0.13453288 | NA | NA | NA |
| ENSG000002 | 0 | NA | NA | NA | NA | NA | SNORA76C | 677842 |
| ENSG000002 | 0 | NA | NA | NA | NA | NA | MIR6846 | 102465509 |
| ENSG000002 | 1.21947866 | -0.176971665 | 0.32191945 | -0.549739 | 0.58249843 | NA | MIR8081 | 102465995 |
| ENSG000002 | 0 | NA | NA | NA | NA | NA | NA | NA |
| ENSG000002 | 0.46346816 | 0.146936161 | 0.23594779 | 0.62274862 | 0.53344973 | NA | MIR6746 | 102465446 |
| ENSG000002 | 1.94237549 | -0.65484049 | 0.35585346 | -1.8401971 | 0.06573931 | NA | NA | NA |
| ENSG000002 | 0 | NA | NA | NA | NA | NA | NA | NA |
| ENSG000002 | 0.84717078 | 0.166456873 | 0.27950973 | 0.59553159 | 0.55148819 | NA | MIR7850 | 102465838 |
| ENSG000002 | 1.12905852 | -0.417975625 | 0.31924322 | -1.3092702 | 0.19044284 | NA | NA | NA |
| ENSG000002 | 1.08863816 | -0.114785919 | 0.32147416 | -0.3570611 | 0.72104606 | NA | MIR8089 | 102465882 |
| ENSG000002 | 0.95623947 | -0.025982677 | 0.29893 | -0.0869189 | 0.93073595 | NA | NA | NA |
| ENSG000002 | 0 | NA | NA | NA | NA | NA | NA | NA |
| ENSG000002 | 0.62522207 | -0.280659539 | 0.2668423 | -1.0517806 | 0.29290024 | NA | MIR8067 | 102465869 |
| ENSG000002 | 0 | NA | NA | NA | NA | NA | NA | NA |

| | | | | | | | | | |
|---|---|---|---|---|---|---|---|---|---|
| ENSG000002 | 1.93352966 | -0.187328234 | 0.36285296 | -0.5162649 | 0.60566944 | NA | NA | NA | |
| ENSG000002 | 0 | NA | NA | NA | NA | NA | TERC | | 7012 |
| ENSG000002 | 0 | NA | NA | NA | NA | NA | NA | NA | |
| ENSG000002 | 0 | NA | NA | NA | NA | NA | NA | NA | |
| ENSG000002 | 1.43487924 | -0.166034263 | 0.33284024 | -0.4988407 | 0.6178916 | NA | NA | NA | |
| ENSG000002 | 0 | NA | NA | NA | NA | NA | NA | NA | |
| ENSG000002 | 0.25333738 | -0.015686291 | 0.1806605 | -0.0868275 | 0.93080867 | NA | RN7SL478P | | 106481053 |
| ENSG000002 | 0 | NA | NA | NA | NA | NA | SNORD3D | | 780854 |
| ENSG000002 | 0.98352408 | 0.108681701 | 0.30783243 | 0.35305474 | 0.7240474 | NA | NA | NA | |
| ENSG000002 | 0.48536101 | -0.024029524 | 0.24539296 | -0.0979226 | 0.92199373 | NA | NA | NA | |
| ENSG000002 | 1.87778187 | -0.639175434 | 0.35426254 | -1.8042422 | 0.07119334 | NA | NA | NA | |
| ENSG000002 | 0.83540841 | 0.214732553 | 0.30106421 | 0.71324504 | 0.47569414 | NA | NA | NA | |
| ENSG000002 | 0 | NA | NA | NA | NA | NA | NA | NA | |
| ENSG000002 | 0 | NA | NA | NA | NA | NA | NA | NA | |
| ENSG000002 | 0.50611225 | -0.202624884 | 0.24886748 | -0.8141879 | 0.41553732 | NA | NA | NA | |
| ENSG000002 | 0 | NA | NA | NA | NA | NA | NA | NA | |
| ENSG000002 | 1.46507134 | -0.187460129 | 0.34128954 | -0.5492701 | 0.58282013 | NA | NA | NA | |
| ENSG000002 | 1.4988897 | -0.489708237 | 0.34652817 | -1.4131845 | 0.15760149 | NA | NA | NA | |
| ENSG000002 | 0 | NA | NA | NA | NA | NA | SNORA67 | | 26781 |
| ENSG000002 | 0 | NA | NA | NA | NA | NA | NA | NA | |
| ENSG000002 | 0.37441639 | -0.268856313 | 0.21277471 | -1.2635727 | 0.20638344 | NA | NA | NA | |
| ENSG000002 | 0.81904596 | 0.191911473 | 0.28749143 | 0.66753806 | 0.5044285 | NA | NA | NA | |
| ENSG000002 | 0.48991005 | 0.142435647 | 0.23916196 | 0.59556147 | 0.55146823 | NA | NA | NA | |
| ENSG000002 | 0.8602884 | -0.106640097 | 0.29730239 | -0.3586924 | 0.71982524 | NA | NA | NA | |
| ENSG000002 | 0 | NA | NA | NA | NA | NA | NA | NA | |
| ENSG000002 | 1.15441223 | -0.248970101 | 0.31144772 | -0.7993961 | 0.42406076 | NA | NA | NA | |
| ENSG000002 | 0 | NA | NA | NA | NA | NA | NA | NA | |
| ENSG000002 | 0 | NA | NA | NA | NA | NA | NA | NA | |
| ENSG000002 | 0 | NA | NA | NA | NA | NA | NA | NA | |

| | | | | | | | | |
|---|---|---|---|---|---|---|---|---|
| ENSG000002 | 1.63729808 | -0.531027139 | 0.34711692 | -1.5298221 | 0.12606077 | NA | NA | NA |
| ENSG000002 | 0.49810908 | -0.023852705 | 0.24681213 | -0.0966432 | 0.92300978 | NA | NA | NA |
| ENSG000002 | 0.36212533 | -0.235926492 | 0.20089643 | -1.1743688 | 0.24024735 | NA | MIR6731 | 102465437 |
| ENSG000002 | 1.4511857 | 0.091650059 | 0.34049657 | 0.26916588 | 0.78780204 | NA | NA | NA |
| ENSG000002 | 0.25214371 | -0.009862892 | 0.18046657 | -0.0546522 | 0.95641556 | NA | NA | NA |
| ENSG000002 | 0.8067549 | 0.196250619 | 0.28671901 | 0.68447021 | 0.4936783 | NA | NA | NA |
| ENSG000002 | 0 | NA | NA | NA | NA | NA | NA | NA |
| ENSG000002 | 0 | NA | NA | NA | NA | NA | NA | NA |
| ENSG000002 | 0 | NA | NA | NA | NA | NA | NA | NA |
| ENSG000002 | 1.02685138 | -0.357028872 | 0.31002754 | -1.1516037 | 0.24948395 | NA | TEX28 | 1527 |
| ENSG000002 | 0.11205239 | 0.082118251 | 0.12671962 | 0.64803108 | 0.51696485 | NA | NA | NA |
| ENSG000002 | 1.26501951 | -0.457740716 | 0.32175758 | -1.422626 | 0.15484459 | NA | NA | NA |
| ENSG000002 | 1.28912556 | -0.515686166 | 0.33274927 | -1.549774 | 0.12119577 | NA | MIR6072 | 102465944 |
| ENSG000002 | 0.99172366 | -0.034851242 | 0.31146264 | -0.1118954 | 0.91090633 | NA | NA | NA |
| ENSG000002 | 0 | NA | NA | NA | NA | NA | NA | NA |
| ENSG000002 | 0 | NA | NA | NA | NA | NA | MIR6726 | 102465434 |
| ENSG000002 | 0.83281412 | 0.367184123 | 0.29024071 | 1.26510206 | 0.20583474 | NA | NA | NA |
| ENSG000002 | 0 | NA | NA | NA | NA | NA | NA | NA |
| ENSG000002 | 0.12685397 | -0.090215768 | 0.12901919 | -0.699243 | 0.48440017 | NA | NA | NA |
| ENSG000002 | 1.09204505 | 0.032365052 | 0.31671742 | 0.10218905 | 0.91860662 | NA | NA | NA |
| ENSG000002 | 0 | NA | NA | NA | NA | NA | NA | NA |
| ENSG000002 | 0 | NA | NA | NA | NA | NA | SNORD115-7 | 100033444 |
| ENSG000002 | 0.7513944 | -0.035588106 | 0.27932511 | -0.1274075 | 0.8986179 | NA | NA | NA |
| ENSG000002 | 0.85684479 | 0.046448936 | 0.3019 | 0.15385537 | 0.87772377 | NA | NA | NA |
| ENSG000002 | 0.36351096 | 0.070290861 | 0.21590418 | 0.32556508 | 0.74475344 | NA | NA | NA |
| ENSG000002 | 0.7265173 | -0.177755649 | 0.27865224 | -0.6379121 | 0.52353088 | NA | NA | NA |
| ENSG000002 | 1.13437702 | -0.273246191 | 0.32373902 | -0.8440323 | 0.39865136 | NA | MIR8060 | 102465864 |
| ENSG000002 | 0.93064401 | 0.432413493 | 0.30097419 | 1.43671289 | 0.15079959 | NA | NA | NA |
| ENSG000002 | 0.34407602 | 0.071624546 | 0.21281613 | 0.33655599 | 0.73645163 | NA | MIR6757 | 102466193 |

| | | | | | | | | |
|---|---|---|---|---|---|---|---|---|
| ENSG000002 | 0.62812434 | -0.408089682 | 0.2528562 | -1.61392 | 0.10654477 | NA | NA | NA |
| ENSG000002 | 0.24756242 | -0.187034346 | 0.18119817 | -1.0322088 | 0.30197431 | NA | NA | NA |
| ENSG000002 | 0 | NA | NA | NA | NA | NA | NA | NA |
| ENSG000002 | 0 | NA | NA | NA | NA | NA | NA | NA |
| ENSG000002 | 0 | NA | NA | NA | NA | NA | SNORD116-28 | 100033820 |
| ENSG000002 | 0.88286447 | -0.430377155 | 0.29478205 | -1.4599843 | 0.1442944 | NA | RN7SL186P | 106480972 |
| ENSG000002 | 0.12070844 | -0.088503695 | 0.12783931 | -0.6923042 | 0.48874631 | NA | NA | NA |
| ENSG000002 | 1.60516658 | -0.526845398 | 0.34543717 | -1.5251555 | 0.12722032 | NA | NA | NA |
| ENSG000002 | 0 | NA | NA | NA | NA | NA | PIK3R3 | 8503 |
| ENSG000002 | 1.84028105 | -0.125901454 | 0.3621848 | -0.3476166 | 0.72812812 | NA | NA | NA |
| ENSG000002 | 0 | NA | NA | NA | NA | NA | NA | NA |
| ENSG000002 | 0.98371858 | -0.032043013 | 0.30614283 | -0.1046669 | 0.91664015 | NA | NA | NA |
| ENSG000002 | 0 | NA | NA | NA | NA | NA | NA | NA |
| ENSG000002 | 1.23877116 | -0.043943195 | 0.3264539 | -0.1346077 | 0.89292209 | NA | NA | NA |
| ENSG000002 | 0 | NA | NA | NA | NA | NA | NA | NA |
| ENSG000002 | 1.74876621 | -0.731159165 | 0.34907356 | -2.0945704 | 0.0362092 | NA | MIR8062 | 102465865 |
| ENSG000002 | 0 | NA | NA | NA | NA | NA | NA | NA |
| ENSG000002 | 0.97544574 | -0.035137544 | 0.30742933 | -0.1142947 | 0.90900418 | NA | NA | NA |
| ENSG000002 | 1.74385281 | -0.191547272 | 0.34979553 | -0.5475978 | 0.58396808 | NA | NA | NA |
| ENSG000002 | 2.08785394 | -0.872747812 | 0.35970397 | -2.4262946 | 0.01525388 | NA | NA | NA |
| ENSG000002 | 0.63393368 | -0.111067416 | 0.26575676 | -0.4179288 | 0.67599914 | NA | NA | NA |
| ENSG000002 | 0 | NA | NA | NA | NA | NA | NA | NA |
| ENSG000002 | 1.02070585 | -0.339157921 | 0.30449216 | -1.1138478 | 0.26534449 | NA | NF1P11 | 106481736 |
| ENSG000002 | 1.02278109 | -0.350503169 | 0.30990351 | -1.1310074 | 0.25805198 | NA | NA | NA |
| ENSG000002 | 0.11205239 | 0.082118251 | 0.12671962 | 0.64803108 | 0.51696485 | NA | NA | NA |
| ENSG000002 | 0 | NA | NA | NA | NA | NA | NA | NA |
| ENSG000002 | 0.82032451 | 0.203072529 | 0.29363926 | 0.69157145 | 0.4892065 | NA | NA | NA |
| ENSG000002 | 0 | NA | NA | NA | NA | NA | NA | NA |
| ENSG000002 | 0 | NA | NA | NA | NA | NA | MIR6857 | 102465516 |

| | | | | | | | | |
|---|---|---|---|---|---|---|---|---|
| ENSG000002 | 0 | NA | NA | NA | NA | NA | NA | NA |
| ENSG000002 | 2.1272707 | -0.50545043 | 0.36468849 | -1.3859786 | 0.16575345 | NA | NA | NA |
| ENSG000002 | 0.48418907 | 0.150283598 | 0.23847389 | 0.63018889 | 0.52857101 | NA | NA | NA |
| ENSG000002 | 0 | NA | NA | NA | NA | NA | NA | NA |
| ENSG000002 | 0.24485602 | -0.012655878 | 0.17881722 | -0.0707755 | 0.94357643 | NA | NA | NA |
| ENSG000002 | 0 | NA | NA | NA | NA | NA | MIR6511A2 | 102466812 |
| ENSG000002 | 0.24485602 | -0.012655878 | 0.17881722 | -0.0707755 | 0.94357643 | NA | RN7SL536P | 106481068 |
| ENSG000002 | 0.25351207 | -0.189965547 | 0.1824784 | -1.0410303 | 0.29786148 | NA | MIR6783 | 102466734 |
| ENSG000002 | 0.96838673 | -0.033819419 | 0.31203989 | -0.1083817 | 0.9136929 | NA | NA | NA |
| ENSG000002 | 0.88086495 | -0.113054583 | 0.29836514 | -0.3789135 | 0.70475209 | NA | NA | NA |
| ENSG000002 | 2.06566753 | 0.246977624 | 0.36409971 | 0.67832413 | 0.4975662 | NA | NA | NA |
| ENSG000002 | 0 | NA | NA | NA | NA | NA | RNA5-8S5 | 100008587 |
| ENSG000002 | 1.49789135 | -0.452352609 | 0.33762042 | -1.339826 | 0.18030193 | NA | NA | NA |
| ENSG000002 | 1.85216255 | -0.51980234 | 0.35664993 | -1.4574581 | 0.14498996 | NA | NA | NA |
| ENSG000002 | 0 | NA | NA | NA | NA | NA | RN7SL725P | 106481120 |
| ENSG000002 | 0 | NA | NA | NA | NA | NA | SCARNA2 | 677766 |
| ENSG000002 | 0.63091271 | -0.116973765 | 0.27178557 | -0.4303899 | 0.66691204 | NA | NA | NA |
| ENSG000002 | 0.99761878 | -0.204823835 | 0.31377682 | -0.6527692 | 0.5139051 | NA | NA | NA |
| ENSG000002 | 0 | NA | NA | NA | NA | NA | SNORD1A | 677848 |
| ENSG000002 | 0.14128499 | -0.097772014 | 0.13403235 | -0.7294658 | 0.4657168 | NA | LOC642131 | 642131 |
| ENSG000002 | 0 | NA | NA | NA | NA | NA | MIR6803 | 102466739 |
| ENSG000002 | 0 | NA | NA | NA | NA | NA | MIR6770-1 | 102465461 |
| ENSG000002 | 0 | NA | NA | NA | NA | NA | SNORA61 | 677838 |
| ENSG000002 | 1.92150208 | -0.484729967 | 0.34638625 | -1.3993915 | 0.16169562 | NA | NA | NA |
| ENSG000002 | 0.40327843 | -0.281391716 | 0.21700406 | -1.2967118 | 0.19473038 | NA | NA | NA |
| ENSG000002 | 0 | NA | NA | NA | NA | NA | MIR6816 | 102465490 |
| ENSG000002 | 0.50016259 | -0.200428691 | 0.24824437 | -0.8073846 | 0.41944491 | NA | MIR7856 | 102465842 |
| ENSG000002 | 0.52398687 | -0.355918066 | 0.23944695 | -1.4864172 | 0.1371688 | NA | NA | NA |
| ENSG000002 | 0 | NA | NA | NA | NA | NA | NA | NA |

| | | | | | | | | |
|---|---|---|---|---|---|---|---|---|
| ENSG000002 | 0 | NA | NA | NA | NA | NA | NA | NA |
| ENSG000002 | 0.27408862 | -0.199826037 | 0.18669465 | -1.0703362 | 0.28446801 | NA | NA | NA |
| ENSG000002 | 0.46748638 | 0.139734475 | 0.23646399 | 0.59093341 | 0.55456504 | NA | TBC1D3C | 414060 |
| ENSG000002 | 1.80140508 | -0.474076003 | 0.35565596 | -1.3329624 | 0.18254416 | NA | NA | NA |
| ENSG000002 | 0.84150242 | 0.209871342 | 0.29479735 | 0.71191733 | 0.47651597 | NA | NA | NA |
| ENSG000002 | 0 | NA | NA | NA | NA | NA | NA | NA |
| ENSG000002 | 0 | NA | NA | NA | NA | NA | MIR6802 | 102465481 |
| ENSG000002 | 0 | NA | NA | NA | NA | NA | RNU6-156P | 106479635 |
| ENSG000002 | 2.1302464 | 0.431070789 | 0.36518924 | 1.18040385 | 0.23783963 | NA | NA | NA |
| ENSG000002 | 0.27408862 | -0.199826037 | 0.18669465 | -1.0703362 | 0.28446801 | NA | NA | NA |
| ENSG000002 | 0 | NA | NA | NA | NA | NA | NA | NA |
| ENSG000002 | 0.12685397 | -0.090215768 | 0.12901919 | -0.699243 | 0.48440017 | NA | NA | NA |
| ENSG000002 | 0 | NA | NA | NA | NA | NA | MIR6862-2 | 102465907 |
| ENSG000002 | 1.24930706 | -0.055619659 | 0.31217822 | -0.1781664 | 0.85859232 | NA | NA | NA |
| ENSG000002 | 2.07392558 | 0.013167267 | 0.361532 | 0.03642075 | 0.97094687 | NA | NA | NA |
| ENSG000002 | 2.11607244 | -0.24303252 | 0.36248865 | -0.6704555 | 0.50256745 | NA | NA | NA |
| ENSG000002 | 0.11205239 | 0.082118251 | 0.12671962 | 0.64803108 | 0.51696485 | NA | MIR6754 | 102466728 |
| ENSG000002 | 0.71171572 | -0.021607924 | 0.27657551 | -0.0781267 | 0.93772729 | NA | NA | NA |
| ENSG000002 | 0.12685397 | -0.090215768 | 0.12901919 | -0.699243 | 0.48440017 | NA | NA | NA |
| ENSG000002 | 0.48302518 | -0.023466322 | 0.24515318 | -0.0957211 | 0.92374212 | NA | MIR7155 | 102466815 |
| ENSG000002 | 1.47667973 | -0.052854427 | 0.34027978 | -0.1553264 | 0.87656402 | NA | NA | NA |
| ENSG000002 | 1.62930598 | -0.529432581 | 0.34710688 | -1.5252725 | 0.12719114 | NA | NA | NA |
| ENSG000002 | 0.97709677 | -0.02688482 | 0.29988949 | -0.0896491 | 0.92856607 | NA | NA | NA |
| ENSG000002 | 0.12070844 | -0.088503695 | 0.12783931 | -0.6923042 | 0.48874631 | NA | NA | NA |
| ENSG000002 | 1.30170559 | 0.035484257 | 0.33232235 | 0.10677662 | 0.9149662 | NA | NA | NA |
| ENSG000002 | 1.209681 | -0.038034844 | 0.32186044 | -0.1181719 | 0.90593149 | NA | NA | NA |
| ENSG000002 | 0.91741557 | -0.435441321 | 0.29663039 | -1.4679592 | 0.1421153 | NA | NA | NA |
| ENSG000002 | 0 | NA | NA | NA | NA | NA | NA | NA |
| ENSG000002 | 0 | NA | NA | NA | NA | NA | NA | NA |

| | | | | | | | | |
|---|---|---|---|---|---|---|---|---|
| ENSG000002 | 0.11934008 | 0.083564698 | 0.12760091 | 0.65489106 | 0.5125379 | NA | NA | NA |
| ENSG000002 | 0 | NA | NA | NA | NA | NA | NA | NA |
| ENSG000002 | 1.66743811 | -0.56883085 | 0.35285888 | -1.6120633 | 0.10694816 | NA | NA | NA |
| ENSG000002 | 1.16859181 | -0.605351814 | 0.32253551 | -1.8768533 | 0.06053821 | NA | NA | NA |
| ENSG000002 | 0.26560726 | -0.177019426 | 0.17672274 | -1.0016788 | 0.31649875 | NA | MIR6854 | 102465514 |
| ENSG000002 | 0.14128499 | -0.097772014 | 0.13403235 | -0.7294658 | 0.4657168 | NA | MIR6512 | 102465255 |
| ENSG000002 | 0 | NA | NA | NA | NA | NA | NA | NA |
| ENSG000002 | 0.47280936 | -0.012478229 | 0.23633508 | -0.0527989 | 0.95789215 | NA | MIR6819 | 102465491 |
| ENSG000002 | 1.3520217 | -0.265322928 | 0.339953 | -0.7804694 | 0.43511461 | NA | NA | NA |
| ENSG000002 | 0 | NA | NA | NA | NA | NA | RN7SL331P | 106479344 |
| ENSG000002 | 0.50818749 | -0.197539377 | 0.24915052 | -0.7928516 | 0.42786432 | NA | NA | NA |
| ENSG000002 | 1.8300435 | -0.25038501 | 0.35927504 | -0.6969173 | 0.48585452 | NA | NA | NA |
| ENSG000002 | 1.6819823 | 0.073966752 | 0.35294141 | 0.20957233 | 0.83400148 | NA | MIR6070 | 102464825 |
| ENSG000002 | 0.89597919 | -0.262547092 | 0.2929304 | -0.8962781 | 0.37010427 | NA | NA | NA |
| ENSG000002 | 1.58779243 | -0.093985027 | 0.33493364 | -0.2806079 | 0.77901115 | NA | NA | NA |
| ENSG000002 | 0 | NA | NA | NA | NA | NA | MIR6781 | 102465468 |
| ENSG000002 | 0 | NA | NA | NA | NA | NA | MIR6892 | 102465538 |
| ENSG000002 | 0 | NA | NA | NA | NA | NA | NA | NA |
| ENSG000002 | 1.49737944 | -0.177633615 | 0.33966529 | -0.5229666 | 0.60099747 | NA | NA | NA |
| ENSG000002 | 0.49810908 | -0.023852705 | 0.24681213 | -0.0966432 | 0.92300978 | NA | NA | NA |
| ENSG000002 | 0.73787528 | -0.192474738 | 0.28554293 | -0.6740658 | 0.50026945 | NA | NA | NA |
| ENSG000002 | 0 | NA | NA | NA | NA | NA | MIR6769B | 102466202 |
| ENSG000002 | 1.70707652 | 0.216006713 | 0.35417707 | 0.6098834 | 0.54193905 | NA | NA | NA |
| ENSG000002 | 0.72170533 | 0.13625606 | 0.28374667 | 0.48020321 | 0.6310829 | NA | NA | NA |
| ENSG000002 | 0 | NA | NA | NA | NA | NA | NA | NA |
| ENSG000002 | 0 | NA | NA | NA | NA | NA | NA | NA |
| ENSG000002 | 1.12154462 | -0.255713863 | 0.31740557 | -0.8056376 | 0.42045183 | NA | MIR8087 | 102465881 |
| ENSG000002 | 0.11934008 | 0.083564698 | 0.12760091 | 0.65489106 | 0.5125379 | NA | NA | NA |
| ENSG000002 | 1.53734087 | -0.640429689 | 0.34452023 | -1.858903 | 0.06304089 | NA | NA | NA |

| | | | | | | | | |
|---|---|---|---|---|---|---|---|---|
| ENSG000002 | 0.96198006 | -0.036154158 | 0.31177589 | -0.115962 | 0.90768265 | NA | NA | NA |
| ENSG000002 | 1.68064482 | 0.07810023 | 0.35263203 | 0.22147798 | 0.82472028 | NA | NA | NA |
| ENSG000002 | 0.14128499 | -0.097772014 | 0.13403235 | -0.7294658 | 0.4657168 | NA | RN7SL759P | 106479498 |
| ENSG000002 | 1.69278986 | 0.197308019 | 0.33857459 | 0.58276086 | 0.5600543 | NA | NA | NA |
| ENSG000002 | 1.51379679 | -0.468102903 | 0.34081159 | -1.3734947 | 0.16959861 | NA | NA | NA |
| ENSG000002 | 0.86228792 | -0.414834899 | 0.2912593 | -1.4242803 | 0.15436534 | NA | HOXD4 | 3233 |
| ENSG000002 | 0.59032214 | 0.060325881 | 0.26160816 | 0.23059632 | 0.81762842 | NA | MIR6082 | 102466953 |
| ENSG000002 | 0 | NA | NA | NA | NA | NA | NA | NA |
| ENSG000002 | 0.35682409 | 0.072008558 | 0.21485623 | 0.33514763 | 0.73751372 | NA | MIR6865 | 102465522 |
| ENSG000002 | 1.25870087 | -0.195291381 | 0.32980107 | -0.592149 | 0.5537508 | NA | NA | NA |
| ENSG000002 | 0.634074 | -0.448061355 | 0.26266228 | -1.7058458 | 0.08803679 | NA | NA | NA |
| ENSG000002 | 0 | NA | NA | NA | NA | NA | MIR6810 | 102466197 |
| ENSG000002 | 1.38492369 | -0.129859802 | 0.34289402 | -0.378717 | 0.70489801 | NA | RN7SL607P | 106481088 |
| ENSG000002 | 0 | NA | NA | NA | NA | NA | NA | NA |
| ENSG000002 | 0 | NA | NA | NA | NA | NA | NA | NA |
| ENSG000002 | 0 | NA | NA | NA | NA | NA | NA | NA |
| ENSG000002 | 0.74250864 | -0.029063449 | 0.2850158 | -0.1019714 | 0.91877941 | NA | MIR8070 | 102465870 |
| ENSG000002 | 0 | NA | NA | NA | NA | NA | NA | NA |
| ENSG000002 | 0.24756242 | -0.187034346 | 0.18119817 | -1.0322088 | 0.30197431 | NA | NA | NA |
| ENSG000002 | 1.66190879 | 0.192431212 | 0.34581783 | 0.55645255 | 0.57790152 | NA | NA | NA |
| ENSG000002 | 0.13280363 | -0.093394593 | 0.13115644 | -0.7120855 | 0.47641185 | NA | MIR6736 | 102466191 |
| ENSG000002 | 0 | NA | NA | NA | NA | NA | RNU6-368P | 106479729 |
| ENSG000002 | 0.11205239 | 0.082118251 | 0.12671962 | 0.64803108 | 0.51696485 | NA | NA | NA |
| ENSG000002 | 0.14128499 | -0.097772014 | 0.13403235 | -0.7294658 | 0.4657168 | NA | NA | NA |
| ENSG000002 | 1.49125318 | -0.195603369 | 0.34730499 | -0.5632034 | 0.57329636 | NA | NA | NA |
| ENSG000002 | 0.35022155 | 0.06940789 | 0.21379411 | 0.32464827 | 0.74544729 | NA | NA | NA |
| ENSG000002 | 0.3934287 | -0.107186984 | 0.22072612 | -0.4856108 | 0.62724315 | NA | NA | NA |
| ENSG000002 | 0 | NA | NA | NA | NA | NA | NA | NA |
| ENSG000002 | 0.12070844 | -0.088503695 | 0.12783931 | -0.6923042 | 0.48874631 | NA | MIR6080 | 102464831 |

| | | | | | | | | |
|---|---|---|---|---|---|---|---|---|
| ENSG000002 | 1.26205802 | -0.481972717 | 0.32618586 | -1.4776015 | 0.13951447 | NA | NA | NA |
| ENSG000002 | 0.11131518 | 0.081951103 | 0.12663804 | 0.64712866 | 0.51754868 | NA | NA | NA |
| ENSG000002 | 0.72142459 | 0.121470654 | 0.27585246 | 0.44034645 | 0.6596862 | NA | NA | NA |
| ENSG000002 | 1.44472287 | -0.72190573 | 0.33414399 | -2.160463 | 0.03073685 | NA | NA | NA |
| ENSG000002 | 1.85810361 | -0.257422559 | 0.36094013 | -0.7132002 | 0.4757219 | NA | RN7SL113P | 106480951 |
| ENSG000002 | 0 | NA | NA | NA | NA | NA | NA | NA |
| ENSG000002 | 0 | NA | NA | NA | NA | NA | NA | NA |
| ENSG000002 | 0.61195439 | -0.094823693 | 0.25481619 | -0.3721259 | 0.70979915 | NA | NA | NA |
| ENSG000002 | 1.75484077 | -0.067795806 | 0.35468183 | -0.1911454 | 0.84841167 | NA | NA | NA |
| ENSG000002 | 0.89404775 | -0.267563874 | 0.29281143 | -0.9137754 | 0.36083489 | NA | MIR6077 | 102466225 |
| ENSG000002 | 0.7169499 | 0.142838539 | 0.28340334 | 0.50401148 | 0.61425331 | NA | MIR6775 | 102465464 |
| ENSG000002 | 1.48380579 | -0.470530976 | 0.34222785 | -1.3749056 | 0.16916072 | NA | TBC1D3E | 102723859 |
| ENSG000002 | 0 | NA | NA | NA | NA | NA | NA | NA |
| ENSG000002 | 0 | NA | NA | NA | NA | NA | NA | NA |
| ENSG000002 | 0.67503122 | -0.469043005 | 0.26742254 | -1.7539397 | 0.07944084 | NA | RN7SL429P | 106479380 |
| ENSG000002 | 1.00376432 | -0.198698202 | 0.309319 | -0.6423731 | 0.52063097 | NA | NA | NA |
| ENSG000002 | 0.46204773 | 0.332451567 | 0.23823694 | 1.39546606 | 0.16287534 | NA | C11orf98 | 102288414 |
| ENSG000002 | 2.15305289 | -0.588342306 | 0.35764221 | -1.6450584 | 0.09995778 | NA | NA | NA |
| ENSG000002 | 1.15919855 | -0.44209514 | 0.32423497 | -1.3635023 | 0.17272425 | NA | MIR8074 | 102465873 |
| ENSG000002 | 0.11205239 | 0.082118251 | 0.12671962 | 0.64803108 | 0.51696485 | NA | NA | NA |
| ENSG000002 | 0.48761094 | -0.176642206 | 0.23968055 | -0.7369902 | 0.46112833 | NA | NA | NA |
| ENSG000002 | 0.52147635 | -0.19458195 | 0.24341208 | -0.7993932 | 0.42406247 | NA | NA | NA |
| ENSG000002 | 0.72173567 | -0.030452891 | 0.27726234 | -0.1098342 | 0.91254086 | NA | LOC102724151 | 102724151 |
| ENSG000002 | 1.27216284 | -0.501653305 | 0.33026616 | -1.5189365 | 0.12877849 | NA | NA | NA |
| ENSG000002 | 1.70563478 | -0.30178422 | 0.34934547 | -0.8638561 | 0.38766694 | NA | NA | NA |
| ENSG000002 | 0 | NA | NA | NA | NA | NA | NA | NA |
| ENSG000002 | 0.75030486 | -0.031917739 | 0.28551165 | -0.1117914 | 0.91098883 | NA | NA | NA |
| ENSG000002 | 0.8333232 | 0.335105949 | 0.27829048 | 1.20415888 | 0.22852818 | NA | NA | NA |
| ENSG000002 | 0.24004852 | -0.005223787 | 0.17775844 | -0.029387 | 0.97655594 | NA | NA | NA |

| | | | | | | | | |
|---|---|---|---|---|---|---|---|---|
| ENSG000002 | 0 | NA | NA | NA | NA | NA | NA | NA |
| ENSG000002 | 1.03257133 | -0.495560084 | 0.30529011 | -1.6232432 | 0.10453743 | NA | NA | NA |
| ENSG000002 | 1.12837338 | -0.418218994 | 0.31921684 | -1.3101408 | 0.19014822 | NA | NA | NA |
| ENSG000002 | 1.70782571 | -0.310126585 | 0.34888794 | -0.8889003 | 0.37405667 | NA | NA | NA |
| ENSG000002 | 0.51316968 | -0.377777671 | 0.24556403 | -1.538408 | 0.12394889 | NA | MIR6826 | 102465496 |
| ENSG000002 | 0 | NA | NA | NA | NA | NA | NA | NA |
| ENSG000002 | 1.60356796 | -0.500898474 | 0.34047672 | -1.471168 | 0.14124568 | NA | NA | NA |
| ENSG000002 | 1.09199298 | 0.033727334 | 0.32128099 | 0.10497768 | 0.91639352 | NA | NA | NA |
| ENSG000002 | 1.52002877 | -0.342920528 | 0.34599755 | -0.9911068 | 0.32163342 | NA | NA | NA |
| ENSG000002 | 0.68946223 | -0.438924761 | 0.26029181 | -1.6862796 | 0.09174197 | NA | MIR8059 | 102466918 |
| ENSG000002 | 0.74686572 | -0.180243062 | 0.28006088 | -0.6435853 | 0.5198444 | NA | NA | NA |
| ENSG000002 | 0.73787528 | -0.192474738 | 0.28554293 | -0.6740658 | 0.50026945 | NA | NA | NA |
| ENSG000002 | 1.58016856 | 0.15766925 | 0.35073598 | 0.44953828 | 0.6530434 | NA | HIST1H2AM | 8336 |
| ENSG000002 | 0.47895489 | -0.016192723 | 0.24471014 | -0.066171 | 0.94724166 | NA | NA | NA |
| ENSG000002 | 1.37231805 | -0.266998709 | 0.3403637 | -0.7844512 | 0.43277541 | NA | NA | NA |
| ENSG000002 | 0.25370795 | -0.171331747 | 0.17411721 | -0.9840024 | 0.32511435 | NA | NA | NA |
| ENSG000002 | 0.62197881 | -0.414424795 | 0.25447679 | -1.6285367 | 0.10341113 | NA | NA | NA |
| ENSG000002 | 1.22787412 | -0.326970115 | 0.32773501 | -0.9976661 | 0.31844128 | NA | NA | NA |
| ENSG000002 | 0 | NA | NA | NA | NA | NA | RN7SL82P | 106480944 |
| ENSG000002 | 1.24628898 | -0.500111357 | 0.32956494 | -1.5174896 | 0.12914312 | NA | NA | NA |
| ENSG000002 | 0.12070844 | -0.088503695 | 0.12783931 | -0.6923042 | 0.48874631 | NA | NA | NA |
| ENSG000002 | 0 | NA | NA | NA | NA | NA | NA | NA |
| ENSG000002 | 1.21375768 | -0.17004178 | 0.32176435 | -0.5284668 | 0.59717538 | NA | NA | NA |
| ENSG000002 | 0 | NA | NA | NA | NA | NA | SNORD116-20 | 100033431 |
| ENSG000002 | 1.3637285 | -0.527714073 | 0.32896441 | -1.6041677 | 0.1086771 | NA | NA | NA |
| ENSG000002 | 0.62775378 | -0.28146231 | 0.26704095 | -1.0540043 | 0.29188095 | NA | NA | NA |
| ENSG000002 | 0.76940933 | -0.516275266 | 0.27702168 | -1.8636638 | 0.06236894 | NA | NA | NA |
| ENSG000002 | 0.97592182 | -0.038695461 | 0.30578089 | -0.1265464 | 0.89929945 | NA | NA | NA |
| ENSG000002 | 0.25333738 | -0.015686291 | 0.1806605 | -0.0868275 | 0.93080867 | NA | NA | NA |

| | | | | | | | | |
|---|---|---|---|---|---|---|---|---|
| ENSG000002 | 0.79075747 | -0.360972421 | 0.28354666 | -1.2730618 | 0.20299612 | NA | NA | NA |
| ENSG000002 | 0.72902782 | -0.31270445 | 0.27032771 | -1.1567606 | 0.24737018 | NA | NA | NA |
| ENSG000002 | 1.24862481 | -0.486697553 | 0.32658191 | -1.4902772 | 0.13615138 | NA | NA | NA |
| ENSG000002 | 0.35664995 | 0.244349217 | 0.20882949 | 1.17008962 | 0.2419649 | NA | MIR6859-4 | 103504738 |
| ENSG000002 | 1.11189992 | -0.379607586 | 0.30936377 | -1.227059 | 0.21980043 | NA | NA | NA |
| ENSG000002 | 1.77112069 | -0.463269582 | 0.35434835 | -1.3073846 | 0.19108212 | NA | NA | NA |
| ENSG000002 | 0.4716672 | -0.017393026 | 0.23619084 | -0.0736397 | 0.94129707 | NA | RN7SL660P | 106479463 |
| ENSG000002 | 0.98432992 | -0.196257029 | 0.31327313 | -0.6264726 | 0.53100501 | NA | NA | NA |
| ENSG000002 | 0.98508887 | -0.033772222 | 0.31272058 | -0.1079949 | 0.91399975 | NA | NA | NA |
| ENSG000002 | 0.91049845 | -0.644235083 | 0.3003423 | -2.1450028 | 0.03195262 | NA | MIR6722 | 102465431 |
| ENSG000002 | 1.20977328 | -0.031093283 | 0.32643902 | -0.0952499 | 0.92411634 | NA | NA | NA |
| ENSG000002 | 0 | NA | NA | NA | NA | NA | NA | NA |
| ENSG000002 | 1.70231471 | 0.341348713 | 0.35124261 | 0.97183173 | 0.33113427 | NA | NA | NA |
| ENSG000002 | 1.01082229 | -0.342469623 | 0.30946802 | -1.1066398 | 0.2684497 | NA | NA | NA |
| ENSG000002 | 0.38064837 | -0.102706251 | 0.21886737 | -0.4692625 | 0.63888201 | NA | NA | NA |
| ENSG000002 | 1.19744283 | 0.094183373 | 0.32319366 | 0.29141467 | 0.7707342 | NA | NA | NA |
| ENSG000002 | 0 | NA | NA | NA | NA | NA | NA | NA |
| ENSG000002 | 0 | NA | NA | NA | NA | NA | NA | NA |
| ENSG000002 | 0.39499294 | -0.277908454 | 0.21584201 | -1.287555 | 0.19790091 | NA | NA | NA |
| ENSG000002 | 0.23070733 | 0.167416683 | 0.17664244 | 0.94777156 | 0.34324576 | NA | MIR6071 | 102466516 |
| ENSG000002 | 0 | NA | NA | NA | NA | NA | NA | NA |
| ENSG000002 | 0 | NA | NA | NA | NA | NA | NA | NA |
| ENSG000002 | 0.50630812 | -0.189615696 | 0.24175631 | -0.7843257 | 0.432849 | NA | NA | NA |
| ENSG000002 | 1.96474521 | -0.296879144 | 0.35743851 | -0.830574 | 0.40621432 | NA | NA | NA |
| ENSG000002 | 0.61332276 | -0.255253496 | 0.25725954 | -0.9922023 | 0.32109886 | NA | NA | NA |
| ENSG000002 | 0.11934008 | 0.083564698 | 0.12760091 | 0.65489106 | 0.5125379 | NA | MIR6823 | 102465494 |
| ENSG000002 | 0 | NA | NA | NA | NA | NA | NA | NA |
| ENSG000002 | 0 | NA | NA | NA | NA | NA | NA | NA |
| ENSG000002 | 0.23202362 | -0.008302493 | 0.17586207 | -0.0472103 | 0.96234565 | NA | NA | NA |

| | | | | | | | | |
|---|---|---|---|---|---|---|---|---|
| ENSG000002 | 0 | NA | NA | NA | NA | NA | VTRNA2-1 | 100126299 |
| ENSG000002 | 0 | NA | NA | NA | NA | NA | NA | NA |
| ENSG000002 | 0 | NA | NA | NA | NA | NA | RN7SL640P | 106479456 |
| ENSG000002 | 1.12561491 | -0.27633688 | 0.32344409 | -0.8543575 | 0.39290694 | NA | NA | NA |
| ENSG000002 | 0.11934008 | 0.083564698 | 0.12760091 | 0.65489106 | 0.5125379 | NA | NA | NA |
| ENSG000002 | 0 | NA | NA | NA | NA | NA | NA | NA |
| ENSG000002 | 0.11205239 | 0.082118251 | 0.12671962 | 0.64803108 | 0.51696485 | NA | LOC102723655 | 102723655 |
| ENSG000002 | 0 | NA | NA | NA | NA | NA | MIR6879 | 102466756 |
| ENSG000002 | 0 | NA | NA | NA | NA | NA | NA | NA |
| ENSG000002 | 0 | NA | NA | NA | NA | NA | NA | NA |
| ENSG000002 | 0 | NA | NA | NA | NA | NA | NA | NA |
| ENSG000002 | 1.14812235 | -0.26675062 | 0.31578363 | -0.8447259 | 0.39826387 | NA | NA | NA |
| ENSG000002 | 1.21545988 | -0.183571371 | 0.3286588 | -0.5585469 | 0.57647096 | NA | NA | NA |
| ENSG000002 | 1.72849117 | -0.056350991 | 0.35439123 | -0.1590079 | 0.87366268 | NA | NA | NA |
| ENSG000002 | 1.46677602 | -0.054930233 | 0.34795377 | -0.1578665 | 0.87456202 | NA | NA | NA |
| ENSG000002 | 1.97006818 | -0.410258617 | 0.35649983 | -1.1507961 | 0.24981612 | NA | NA | NA |
| ENSG000002 | 1.98942877 | -0.585091029 | 0.36137576 | -1.6190655 | 0.10543317 | NA | NA | NA |
| ENSG000002 | 1.35567382 | -0.392356685 | 0.33300986 | -1.1782134 | 0.23871154 | NA | NA | NA |
| ENSG000002 | 0 | NA | NA | NA | NA | NA | NA | NA |
| ENSG000002 | 2.07722675 | 0.009810244 | 0.35997087 | 0.02725288 | 0.97825804 | NA | NA | NA |
| ENSG000002 | 1.39385994 | -0.267440722 | 0.33757841 | -0.7922329 | 0.42822489 | NA | NA | NA |
| ENSG000002 | 0.25351207 | -0.189965547 | 0.1824784 | -1.0410303 | 0.29786148 | NA | NA | NA |
| ENSG000002 | 0.11131518 | 0.081951103 | 0.12663804 | 0.64712866 | 0.51754868 | NA | NA | NA |
| ENSG000002 | 2.06123149 | 0.130543477 | 0.3598105 | 0.36281175 | 0.71674552 | NA | NA | NA |
| ENSG000002 | 1.51239617 | -0.295597146 | 0.33240344 | -0.8892722 | 0.3738568 | NA | NA | NA |
| ENSG000002 | 0.71322555 | 0.289371075 | 0.27698025 | 1.04473541 | 0.29614527 | NA | NA | NA |
| ENSG000002 | 1.90062507 | -0.403706283 | 0.36118823 | -1.1177172 | 0.26368781 | NA | NA | NA |
| ENSG000002 | 0.63530205 | -0.267258942 | 0.26077599 | -1.0248602 | 0.30542915 | NA | NA | NA |
| ENSG000002 | 1.08450942 | 0.029677929 | 0.31474659 | 0.0942915 | 0.9248776 | NA | NA | NA |

| | | | | | | | | |
|---|---|---|---|---|---|---|---|---|
| ENSG000002 | 0 | NA | NA | NA | NA | NA | NA | NA |
| ENSG000002 | 1.33813359 | -0.107539089 | 0.32917882 | -0.326689 | 0.74390315 | NA | NA | NA |
| ENSG000002 | 1.60895564 | -0.657376496 | 0.34255791 | -1.919023 | 0.05498143 | NA | NA | NA |
| ENSG000002 | 1.37235477 | -0.491856638 | 0.32352825 | -1.5202896 | 0.1284382 | NA | NA | NA |
| ENSG000002 | 0 | NA | NA | NA | NA | NA | NA | NA |
| ENSG000002 | 1.25325817 | -0.349846863 | 0.33201408 | -1.0537109 | 0.29201529 | NA | NA | NA |
| ENSG000002 | 0.12685397 | -0.090215768 | 0.12901919 | -0.699243 | 0.48440017 | NA | NA | NA |
| ENSG000002 | 1.59633404 | 0.146499616 | 0.34983884 | 0.41876315 | 0.67538924 | NA | NA | NA |
| ENSG000002 | 0 | NA | NA | NA | NA | NA | NA | NA |
| ENSG000002 | 0.73824193 | -0.03188941 | 0.28472511 | -0.1120007 | 0.91082286 | NA | NA | NA |
| ENSG000002 | 0 | NA | NA | NA | NA | NA | NA | NA |
| ENSG000002 | 1.00610015 | -0.196494509 | 0.30776743 | -0.6384513 | 0.52317996 | NA | NA | NA |
| ENSG000002 | 1.36049914 | 0.023462067 | 0.33592726 | 0.0698427 | 0.94431886 | NA | NA | NA |
| ENSG000002 | 0.99511328 | -0.337711937 | 0.3088181 | -1.0935626 | 0.27414684 | NA | NA | NA |
| ENSG000002 | 1.40522238 | -0.527533335 | 0.33004523 | -1.5983668 | 0.10996137 | NA | NA | NA |
| ENSG000002 | 1.70696642 | -0.188913946 | 0.35448071 | -0.5329315 | 0.59408098 | NA | NA | NA |
| ENSG000002 | 0.66908156 | -0.466940971 | 0.26696073 | -1.7490999 | 0.08027376 | NA | NA | NA |
| ENSG000002 | 1.37368036 | -0.250883402 | 0.33222767 | -0.7551551 | 0.45015595 | NA | NA | NA |
| ENSG000002 | 1.44516887 | -0.039473119 | 0.32414564 | -0.1217759 | 0.90307652 | NA | NA | NA |
| ENSG000002 | 0.23730987 | 0.1539805 | 0.16964526 | 0.90766167 | 0.364057 | NA | NA | NA |
| ENSG000002 | 0.76491057 | -0.186165194 | 0.2812379 | -0.6619492 | 0.5080038 | NA | NA | NA |
| ENSG000002 | 1.24022989 | -0.335209323 | 0.32804282 | -1.0218463 | 0.30685367 | NA | NA | NA |
| ENSG000002 | 2.02995284 | -0.322473627 | 0.36211061 | -0.890539 | 0.37317652 | NA | NA | NA |
| ENSG000002 | 1.97665036 | -0.444545691 | 0.36145383 | -1.2298824 | 0.21874116 | NA | NA | NA |
| ENSG000002 | 0.75349997 | -0.202315365 | 0.28648959 | -0.7061875 | 0.48007155 | NA | NA | NA |
| ENSG000002 | 1.33594211 | -0.105461736 | 0.33566323 | -0.3141891 | 0.7533774 | NA | NA | NA |
| ENSG000002 | 0 | NA | NA | NA | NA | NA | NA | NA |
| ENSG000002 | 0.36827086 | -0.266056975 | 0.21181163 | -1.2561018 | 0.20907905 | NA | NA | NA |
| ENSG000002 | 2.01205546 | -0.431646754 | 0.35792574 | -1.2059673 | 0.22783009 | NA | NA | NA |

| | | | | | | | | | |
|---|---|---|---|---|---|---|---|---|---|
| ENSG000002 | 0.73580004 | -0.186985061 | 0.27927554 | -0.6695361 | 0.50315356 | NA | NA | NA | |
| ENSG000002 | 1.44785207 | 0.092135371 | 0.34366377 | 0.26809742 | 0.78862433 | NA | NA | NA | |
| ENSG000002 | 1.66408274 | -0.68206046 | 0.34588492 | -1.9719289 | 0.04861772 | NA | NA | NA | |
| ENSG000002 | 0.11934008 | 0.083564698 | 0.12760091 | 0.65489106 | 0.5125379 | NA | NA | NA | |
| ENSG000002 | 0 | NA | NA | NA | NA | NA | NA | NA | |
| ENSG000002 | 0.23276084 | -0.008011382 | 0.17603982 | -0.0455089 | 0.96370166 | NA | NA | NA | |
| ENSG000002 | 0 | NA | NA | NA | NA | NA | NA | NA | |
| ENSG000002 | 0.74572549 | -0.027904447 | 0.27894518 | -0.1000356 | 0.92031607 | NA | NA | NA | |
| ENSG000002 | 0.85300071 | -0.109179228 | 0.29690999 | -0.3677183 | 0.71308332 | NA | NA | NA | |
| ENSG000002 | 0.86321654 | -0.117877844 | 0.2974425 | -0.3963046 | 0.6918803 | NA | NA | NA | |
| ENSG000002 | 0.37996515 | 0.064665424 | 0.21008146 | 0.3078112 | 0.758226 | NA | NA | NA | |
| ENSG000002 | 0 | NA | NA | NA | NA | NA | NA | NA | |
| ENSG000002 | 1.37405442 | -0.114029445 | 0.33087109 | -0.3446341 | 0.7303695 | NA | NA | NA | |
| ENSG000002 | 2.10016891 | -0.129636974 | 0.36823254 | -0.3520519 | 0.72479932 | NA | NA | NA | |
| ENSG000002 | 0.87018964 | -0.249794079 | 0.28906262 | -0.8641521 | 0.38750432 | NA | NA | NA | |
| ENSG000002 | 0 | NA | NA | NA | NA | NA | NA | NA | |
| ENSG000002 | 2.17249001 | -0.060554533 | 0.36567569 | -0.1655963 | 0.86847468 | NA | OR8D2 | | 283160 |
| ENSG000002 | 1.21486256 | 0.102211625 | 0.33309619 | 0.30685318 | 0.75895513 | NA | NA | NA | |
| ENSG000002 | 0.5974134 | 0.053016323 | 0.26223774 | 0.20216893 | 0.83978466 | NA | NA | NA | |
| ENSG000002 | 1.48226328 | -0.178148279 | 0.34159672 | -0.5215164 | 0.60200711 | NA | NA | NA | |
| ENSG000002 | 0.11934008 | 0.083564698 | 0.12760091 | 0.65489106 | 0.5125379 | NA | NA | NA | |
| ENSG000002 | 1.23191407 | -0.183141071 | 0.32910664 | -0.5564794 | 0.57788316 | NA | NA | NA | |
| ENSG000002 | 1.57603769 | 0.150830465 | 0.35070032 | 0.43008362 | 0.66713481 | NA | NA | NA | |
| ENSG000002 | 0.80450335 | -0.337587828 | 0.275493 | -1.2253953 | 0.22042632 | NA | NA | NA | |
| ENSG000002 | 1.37638676 | -0.403435194 | 0.33613345 | -1.2002233 | 0.23005264 | NA | NA | NA | |
| ENSG000002 | 1.8473162 | -0.381544386 | 0.35562561 | -1.0728822 | 0.28332397 | NA | NA | NA | |
| ENSG000002 | 0.53969588 | -0.357270382 | 0.23978341 | -1.4899712 | 0.13623181 | NA | NA | NA | |
| ENSG000002 | 1.60758728 | -0.528972704 | 0.34677669 | -1.5253987 | 0.12715969 | NA | NA | NA | |
| ENSG000002 | 1.1705478 | -0.236006741 | 0.20041441 | -1.1775937 | 0.23895863 | NA | NA | NA | |

| | | | | | | | | |
|---|---|---|---|---|---|---|---|---|
| ENSG000002 | 0.84975353 | 0.05329801 | 0.30153883 | 0.17675339 | 0.8597021 | NA | NA | NA |
| ENSG000002 | 0.60373362 | -0.112425234 | 0.26959077 | -0.4170218 | 0.67666245 | NA | NA | NA |
| ENSG000002 | 1.6859369 | 0.082070205 | 0.35334084 | 0.23226923 | 0.81632891 | NA | NA | NA |
| ENSG000002 | 0 | NA | NA | NA | NA | NA | NA | NA |
| ENSG000002 | 0 | NA | NA | NA | NA | NA | NA | NA |
| ENSG000002 | 1.5009432 | -0.632359692 | 0.34319911 | -1.8425447 | 0.06539552 | NA | NA | NA |
| ENSG000002 | 1.25508546 | -0.340782337 | 0.33066942 | -1.0305832 | 0.30273634 | NA | NA | NA |
| ENSG000002 | 0 | NA | NA | NA | NA | NA | NA | NA |
| ENSG000002 | 0 | NA | NA | NA | NA | NA | NA | NA |
| ENSG000002 | 1.97066253 | 0.184518963 | 0.3590467 | 0.51391354 | 0.60731244 | NA | NA | NA |
| ENSG000002 | 0.12685397 | -0.090215768 | 0.12901919 | -0.699243 | 0.48440017 | NA | NA | NA |
| ENSG000002 | 0 | NA | NA | NA | NA | NA | NA | NA |
| ENSG000002 | 1.13218308 | -0.4267924 | 0.32125178 | -1.3285293 | 0.18400331 | NA | NA | NA |
| ENSG000002 | 0 | NA | NA | NA | NA | NA | NA | NA |
| ENSG000002 | 1.54211804 | -0.471840969 | 0.34252711 | -1.3775288 | 0.1683488 | NA | NA | NA |
| ENSG000002 | 0 | NA | NA | NA | NA | NA | NA | NA |
| ENSG000002 | 1.08920069 | 0.030457512 | 0.31734001 | 0.09597754 | 0.92353841 | NA | NA | NA |
| ENSG000002 | 1.61580358 | -0.25263435 | 0.34668194 | -0.7287208 | 0.46617246 | NA | NA | NA |
| ENSG000002 | 1.1811652 | -0.427414722 | 0.32095046 | -1.3317156 | 0.18295367 | NA | NA | NA |
| ENSG000002 | 0.11934008 | 0.083564698 | 0.12760091 | 0.65489106 | 0.5125379 | NA | NA | NA |
| ENSG000002 | 2.04356267 | -0.297263158 | 0.35468025 | -0.8381159 | 0.40196561 | NA | NA | NA |
| ENSG000002 | 1.48260117 | 0.220371158 | 0.33922463 | 0.64963195 | 0.51592999 | NA | NA | NA |
| ENSG000002 | 2.07988967 | -0.348372369 | 0.36024462 | -0.9670439 | 0.33352208 | NA | NA | NA |
| ENSG000002 | 1.11736031 | -0.409329296 | 0.31726361 | -1.2901867 | 0.19698583 | NA | NA | NA |
| ENSG000002 | 1.96758937 | -0.301663103 | 0.35677855 | -0.8455192 | 0.39782104 | NA | NA | NA |
| ENSG000002 | 1.23422153 | -0.721407117 | 0.31063344 | -2.3223743 | 0.02021279 | NA | NA | NA |
| ENSG000002 | 1.80800787 | 0.011587924 | 0.35249827 | 0.0328737 | 0.9737753 | NA | NA | NA |
| ENSG000002 | 0 | NA | NA | NA | NA | NA | NA | NA |
| ENSG000002 | 2.091263 | -0.129384064 | 0.36682021 | -0.3527179 | 0.72429992 | NA | NA | NA |

| | | | | | | | | |
|---|---|---|---|---|---|---|---|---|
| ENSG000002 | 0 | NA | NA | NA | NA | NA | NA | NA |
| ENSG000002 | 0.64579862 | -0.287808079 | 0.26850213 | -1.0719024 | 0.28376388 | NA | NA | NA |
| ENSG000002 | 1.38683676 | -0.557841476 | 0.33430983 | -1.6686362 | 0.0951895 | NA | NA | NA |
| ENSG000002 | 0.39479706 | -0.300008631 | 0.22317059 | -1.3443018 | 0.17885081 | NA | NA | NA |
| ENSG000002 | 0.72238856 | -0.029280768 | 0.28909343 | -0.1012848 | 0.91932439 | NA | NA | NA |
| ENSG000002 | 1.3344294 | 0.028072596 | 0.34282922 | 0.08188507 | 0.9347381 | NA | NA | NA |
| ENSG000002 | 0.51316968 | -0.377777671 | 0.24556403 | -1.538408 | 0.12394889 | NA | NA | NA |
| ENSG000002 | 1.22926379 | 0.240074781 | 0.322237 | 0.7450255 | 0.45625631 | NA | NA | NA |
| ENSG000002 | 1.60004113 | -0.3854436 | 0.348364 | -1.1064393 | 0.26853644 | NA | NA | NA |
| ENSG000002 | 1.36969543 | -0.122774524 | 0.32829842 | -0.3739723 | 0.70842491 | NA | NA | NA |
| ENSG000002 | 0.78745417 | -0.533348008 | 0.2804936 | -1.9014623 | 0.05724148 | NA | NA | NA |
| ENSG000002 | 0.34931019 | 0.26109409 | 0.21538338 | 1.21222954 | 0.22542453 | NA | NA | NA |
| ENSG000002 | 2.17274843 | -0.637384387 | 0.363759 | -1.7522161 | 0.07973665 | NA | NA | NA |
| ENSG000002 | 1.02155304 | -0.514261121 | 0.30968768 | -1.6605798 | 0.09679786 | NA | OR5L1 | 219437 |
| ENSG000002 | 0.85740732 | 0.173972714 | 0.28277537 | 0.6152329 | 0.53840092 | NA | NA | NA |
| ENSG000002 | 1.12062476 | -0.12299574 | 0.32100767 | -0.3831551 | 0.70160474 | NA | NA | NA |
| ENSG000002 | 0 | NA | NA | NA | NA | NA | LOC105369274 | 105369274 |
| ENSG000002 | 0.25333738 | -0.015686291 | 0.1806605 | -0.0868275 | 0.93080867 | NA | NA | NA |
| ENSG000002 | 1.61722751 | -0.129491573 | 0.3502962 | -0.3696631 | 0.71163354 | NA | LINC00244 | 64433 |
| ENSG000002 | 0.23890637 | -0.010420674 | 0.17747379 | -0.0587167 | 0.95317776 | NA | NA | NA |
| ENSG000002 | 0.61153039 | -0.091790886 | 0.25636364 | -0.3580495 | 0.72030624 | NA | NA | NA |
| ENSG000002 | 0.25214371 | -0.009862892 | 0.18046657 | -0.0546522 | 0.95641556 | NA | NA | NA |
| ENSG000002 | 0.12685397 | -0.090215768 | 0.12901919 | -0.699243 | 0.48440017 | NA | NA | NA |
| ENSG000002 | 1.21773917 | 0.099290079 | 0.33184246 | 0.29920848 | 0.76478098 | NA | NA | NA |
| ENSG000002 | 1.62807449 | -0.256897507 | 0.34342292 | -0.74805 | 0.45443002 | NA | NA | NA |
| ENSG000002 | 0 | NA | NA | NA | NA | NA | OR4N3P | 390539 |
| ENSG000002 | 1.30073149 | -0.495479308 | 0.32802182 | -1.5105071 | 0.13091407 | NA | NA | NA |
| ENSG000002 | 1.08240831 | -0.107607168 | 0.31973967 | -0.3365462 | 0.73645903 | NA | NA | NA |
| ENSG000002 | 0.36075696 | -0.085953824 | 0.20740622 | -0.4144226 | 0.67856464 | NA | NA | NA |

| | | | | | | | | |
|---|---|---|---|---|---|---|---|---|
| ENSG000002 | 1.7030669 | -0.050063184 | 0.34814866 | -0.1437983 | 0.88565975 | NA | NA | NA |
| ENSG000002 | 0 | NA | NA | NA | NA | NA | NA | NA |
| ENSG000002 | 0.49082195 | -0.014647497 | 0.23849846 | -0.0614155 | 0.95102832 | NA | NA | NA |
| ENSG000002 | 0.36465257 | 0.064724075 | 0.21605106 | 0.29957767 | 0.76449932 | NA | NA | NA |
| ENSG000002 | 1.42258208 | -0.586085133 | 0.34040368 | -1.7217356 | 0.08511743 | NA | NA | NA |
| ENSG000002 | 1.20020129 | -0.041812868 | 0.32618054 | -0.1281893 | 0.89799914 | NA | NA | NA |
| ENSG000002 | 0 | NA | NA | NA | NA | NA | NA | NA |
| ENSG000002 | 1.47896157 | -0.187701101 | 0.3401217 | -0.5518645 | 0.58104117 | NA | NA | NA |
| ENSG000002 | 0.96687848 | -0.166120343 | 0.2976221 | -0.5581586 | 0.57673606 | NA | NA | NA |
| ENSG000002 | 0.23139247 | 0.167748926 | 0.17680352 | 0.94878726 | 0.34272882 | NA | NA | NA |
| ENSG000002 | 1.03510304 | -0.536100349 | 0.31389681 | -1.7078872 | 0.08765727 | NA | NA | NA |
| ENSG000002 | 1.19658801 | -0.030470349 | 0.32316077 | -0.0942885 | 0.92487997 | NA | NA | NA |
| ENSG000002 | 1.04096625 | -0.704125123 | 0.30912422 | -2.2778064 | 0.02273812 | NA | NA | NA |
| ENSG000002 | 0 | NA | NA | NA | NA | NA | NA | NA |
| ENSG000002 | 1.03092064 | -0.48177817 | 0.3023184 | -1.5936118 | 0.11102301 | NA | NA | NA |
| ENSG000002 | 0.63099704 | -0.107803684 | 0.25810428 | -0.4176749 | 0.67618481 | NA | NA | NA |
| ENSG000002 | 2.04540845 | 0.013023044 | 0.36255398 | 0.03592029 | 0.97134592 | NA | NA | NA |
| ENSG000002 | 1.25024069 | -0.052697936 | 0.32907245 | -0.1601408 | 0.87277014 | NA | NA | NA |
| ENSG000002 | 0.38133351 | -0.102466351 | 0.2189713 | -0.4679442 | 0.63982449 | NA | NA | NA |
| ENSG000002 | 0.49241844 | -0.197108126 | 0.24744176 | -0.7965839 | 0.42569273 | NA | NA | NA |
| ENSG000002 | 0.72967569 | -0.033529616 | 0.27782303 | -0.120687 | 0.90393899 | NA | NA | NA |
| ENSG000002 | 1.71908389 | -0.043232707 | 0.348314 | -0.1241199 | 0.90122032 | NA | NA | NA |
| ENSG000002 | 1.99796625 | -0.320873497 | 0.36335048 | -0.8830964 | 0.37718419 | NA | NA | NA |
| ENSG000002 | 1.92348134 | -0.663157137 | 0.35561251 | -1.8648307 | 0.06220513 | NA | NA | NA |
| ENSG000002 | 1.8946503 | -0.241795157 | 0.35416223 | -0.6827243 | 0.49478108 | NA | NA | NA |
| ENSG000002 | 0 | NA | NA | NA | NA | NA | NA | NA |
| ENSG000002 | 1.78237478 | 0.006458977 | 0.35060201 | 0.01842253 | 0.98530178 | NA | NA | NA |
| ENSG000002 | 2.00243991 | -0.44102509 | 0.3601112 | -1.2246914 | 0.22069152 | NA | NA | NA |
| ENSG000002 | 0.96563274 | -0.161087644 | 0.2975649 | -0.541353 | 0.58826431 | NA | NA | NA |

| | | | | | | | | |
|---|---|---|---|---|---|---|---|---|
| ENSG000002 | 1.02553509 | -0.20565425 | 0.3133652 | -0.6562766 | 0.51164616 | NA | NA | NA |
| ENSG000002 | 0 | NA | NA | NA | NA | NA | NA | NA |
| ENSG000002 | 0.83165023 | 0.205469485 | 0.29588711 | 0.6944185 | 0.4874198 | NA | NA | NA |
| ENSG000002 | 0 | NA | NA | NA | NA | NA | DUX4L28 | 106481728 |
| ENSG000002 | 1.56218631 | 0.015705296 | 0.34428974 | 0.04561651 | 0.96361591 | NA | NA | NA |
| ENSG000002 | 0 | NA | NA | NA | NA | NA | NA | NA |
| ENSG000002 | 2.13038879 | -0.115097347 | 0.35934241 | -0.3202999 | 0.74874098 | NA | NA | NA |
| ENSG000002 | 1.5702971 | 0.156282621 | 0.35060475 | 0.44575157 | 0.65577671 | NA | NA | NA |
| ENSG000002 | 0 | NA | NA | NA | NA | NA | NA | NA |
| ENSG000002 | 0.64081643 | -0.112957221 | 0.26633067 | -0.424124 | 0.6714754 | NA | NA | NA |
| ENSG000002 | 1.32609184 | 0.029255275 | 0.33858658 | 0.08640412 | 0.93114517 | NA | NA | NA |
| ENSG000002 | 0 | NA | NA | NA | NA | NA | NA | NA |
| ENSG000002 | 1.44910829 | -0.573886873 | 0.33767976 | -1.6995003 | 0.08922495 | NA | NA | NA |
| ENSG000002 | 1.79352773 | 0.241199941 | 0.34840342 | 0.69230073 | 0.48874849 | NA | NA | NA |
| ENSG000002 | 1.92896618 | 0.07430968 | 0.36301973 | 0.20469874 | 0.83780751 | NA | NA | NA |
| ENSG000002 | 1.42696695 | -0.414798717 | 0.3357521 | -1.2354315 | 0.21666996 | NA | NA | NA |
| ENSG000002 | 0 | NA | NA | NA | NA | NA | ECSCR | 641700 |
| ENSG000002 | 0 | NA | NA | NA | NA | NA | NA | NA |
| ENSG000002 | 0.88650003 | -0.284549442 | 0.30460367 | -0.9341629 | 0.35021988 | NA | NA | NA |
| ENSG000002 | 1.92650286 | -0.655026762 | 0.35569876 | -1.8415211 | 0.06554524 | NA | NA | NA |
| ENSG000002 | 2.08397978 | -0.360980286 | 0.36266369 | -0.9953582 | 0.31956207 | NA | NA | NA |
| ENSG000002 | 0.49879422 | -0.023609326 | 0.24688725 | -0.095628 | 0.92381605 | NA | NA | NA |
| ENSG000002 | 0.37171 | -0.104406568 | 0.21753023 | -0.4799635 | 0.63125336 | NA | NA | NA |
| ENSG000002 | 0.98637078 | -0.307257407 | 0.29736015 | -1.0332837 | 0.30147114 | NA | NA | NA |
| ENSG000002 | 1.77925539 | 0.422812619 | 0.35824448 | 1.18023483 | 0.23790683 | NA | NA | NA |
| ENSG000002 | 1.71692233 | 0.34028986 | 0.35108229 | 0.96925955 | 0.33241571 | NA | NA | NA |
| ENSG000002 | 0.60831938 | -0.248045635 | 0.25683119 | -0.9657925 | 0.33414803 | NA | NA | NA |
| ENSG000002 | 0 | NA | NA | NA | NA | NA | NA | NA |
| ENSG000002 | 1.11536471 | -0.407782747 | 0.31720104 | -1.2855656 | 0.19859471 | NA | NA | NA |

| | | | | | | | | |
|---|---|---|---|---|---|---|---|---|
| ENSG000002 | 0.73618842 | 0.131739793 | 0.28472287 | 0.46269481 | 0.64358314 | NA | NA | NA |
| ENSG000002 | 0.65445467 | -0.486865861 | 0.27154464 | -1.7929496 | 0.07298098 | NA | NA | NA |
| ENSG000002 | 0.87833324 | -0.112382167 | 0.29823906 | -0.3768191 | 0.70630806 | NA | NA | NA |
| ENSG000002 | 1.33277871 | 0.028059221 | 0.34001823 | 0.0825227 | 0.93423107 | NA | LOC100128002 | 100128002 |
| ENSG000002 | 0 | NA | NA | NA | NA | NA | NA | NA |
| ENSG000002 | 0 | NA | NA | NA | NA | NA | NA | NA |
| ENSG000002 | 1.05567959 | -0.534683329 | 0.31319853 | -1.7071706 | 0.08779035 | NA | NA | NA |
| ENSG000002 | 0.7456715 | -0.195362308 | 0.2860302 | -0.6830129 | 0.49459871 | NA | NA | NA |
| ENSG000002 | 0.62752756 | -0.103354502 | 0.26525132 | -0.3896475 | 0.69679725 | NA | OR5D13 | 390142 |
| ENSG000002 | 0 | NA | NA | NA | NA | NA | NA | NA |
| ENSG000002 | 0 | NA | NA | NA | NA | NA | LOC441956 | 441956 |
| ENSG000002 | 1.34945911 | -0.120978957 | 0.33934193 | -0.3565105 | 0.7214583 | NA | NA | NA |
| ENSG000002 | 0.25370795 | -0.171331747 | 0.17411721 | -0.9840024 | 0.32511435 | NA | NA | NA |
| ENSG000002 | 0.23936338 | -0.005479057 | 0.17759977 | -0.0308506 | 0.9753887 | NA | PPIAL4F | 728945 |
| ENSG000002 | 0.50016259 | -0.200428691 | 0.24824437 | -0.8073846 | 0.41944491 | NA | NA | NA |
| ENSG000002 | 0 | NA | NA | NA | NA | NA | NA | NA |
| ENSG000002 | 0.89450319 | -0.455035162 | 0.30208731 | -1.5063035 | 0.13198929 | NA | NA | NA |
| ENSG000002 | 1.3976162 | -0.407614919 | 0.33734213 | -1.2083131 | 0.22692684 | NA | NA | NA |
| ENSG000002 | 0.36277375 | 0.069994797 | 0.21578753 | 0.32436906 | 0.74565864 | NA | NA | NA |
| ENSG000002 | 0.49264869 | -0.021364872 | 0.24621241 | -0.0867741 | 0.93085104 | NA | LOC101926951 | 101926951 |
| ENSG000002 | 1.52994108 | -0.628172037 | 0.34124077 | -1.840847 | 0.06564399 | NA | NA | NA |
| ENSG000002 | 0 | NA | NA | NA | NA | NA | NA | NA |
| ENSG000002 | 1.37092795 | -0.262747859 | 0.33710238 | -0.7794304 | 0.4357262 | NA | NA | NA |
| ENSG000002 | 0.36277375 | 0.069994797 | 0.21578753 | 0.32436906 | 0.74565864 | NA | NA | NA |
| ENSG000002 | 0.73755679 | -0.032120828 | 0.28468017 | -0.1128313 | 0.91016432 | NA | NA | NA |
| ENSG000002 | 1.74259496 | -0.167055388 | 0.34560682 | -0.4833683 | 0.62883424 | NA | NA | NA |
| ENSG000002 | 1.19585272 | 0.106470748 | 0.32057198 | 0.33212743 | 0.73979304 | NA | NA | NA |
| ENSG000002 | 1.77602048 | -0.451631303 | 0.35116697 | -1.2860871 | 0.19841267 | NA | NA | NA |
| ENSG000002 | 0.75099 | -0.033089548 | 0.29085543 | -0.1137663 | 0.90942305 | NA | NA | NA |

| | | | | | | | | |
|---|---|---|---|---|---|---|---|---|
| ENSG000002 | 1.1218209 | 0.026552797 | 0.3086592 | 0.08602626 | 0.93144554 | NA | NA | NA |
| ENSG000002 | 1.00871672 | -0.182276483 | 0.30219843 | -0.6031682 | 0.54639681 | NA | NA | NA |
| ENSG000002 | 1.83616729 | -0.380396169 | 0.35683782 | -1.0660198 | 0.28641471 | NA | NA | NA |
| ENSG000002 | 0.37404583 | -0.105111695 | 0.21784993 | -0.4824959 | 0.62945372 | NA | NA | NA |
| ENSG000002 | 0.63435632 | -0.27906164 | 0.26763939 | -1.0426778 | 0.29709757 | NA | NA | NA |
| ENSG000002 | 0.71373251 | 0.133566383 | 0.28321275 | 0.47161148 | 0.63720413 | NA | NA | NA |
| ENSG000002 | 0 | NA | NA | NA | NA | NA | NA | NA |
| ENSG000002 | 1.72648761 | -0.053707371 | 0.34418069 | -0.1560441 | 0.87599825 | NA | NA | NA |
| ENSG000002 | 1.38854143 | -0.413680213 | 0.33711226 | -1.227129 | 0.21977412 | NA | NA | NA |
| ENSG000002 | 0.97862876 | -0.164762438 | 0.30081039 | -0.5477286 | 0.58387831 | NA | NA | NA |
| ENSG000002 | 0 | NA | NA | NA | NA | NA | NA | NA |
| ENSG000002 | 1.91452232 | -0.820222852 | 0.35683609 | -2.2985984 | 0.02152776 | NA | NA | NA |
| ENSG000002 | 1.28314366 | -0.466907545 | 0.32314738 | -1.4448749 | 0.14849302 | NA | NA | NA |
| ENSG000002 | 0.3854038 | -0.110128691 | 0.21952326 | -0.5016721 | 0.61589821 | NA | NA | NA |
| ENSG000002 | 0.49953144 | -0.023330163 | 0.24696918 | -0.0944659 | 0.92473908 | NA | NA | NA |
| ENSG000002 | 1.77840446 | -0.197245063 | 0.35522669 | -0.5552653 | 0.57871319 | NA | NA | NA |
| ENSG000002 | 0.48573157 | -0.181932918 | 0.23941992 | -0.7598905 | 0.44732006 | NA | NA | NA |
| ENSG000002 | 0 | NA | NA | NA | NA | NA | NA | NA |
| ENSG000002 | 1.21728663 | -0.188097604 | 0.32870337 | -0.5722412 | 0.56715859 | NA | NA | NA |
| ENSG000002 | 0.85584699 | 0.046156439 | 0.28890047 | 0.15976588 | 0.8730655 | NA | NA | NA |
| ENSG000002 | 2.06659225 | -0.852534422 | 0.35790124 | -2.3820382 | 0.01721711 | NA | NA | NA |
| ENSG000002 | 1.87133691 | -0.527791505 | 0.35891071 | -1.4705371 | 0.14141636 | NA | NA | NA |
| ENSG000002 | 1.82730834 | 0.132639487 | 0.35475161 | 0.37389396 | 0.70848321 | NA | NA | NA |
| ENSG000002 | 0 | NA | NA | NA | NA | NA | NA | NA |
| ENSG000002 | 0.58953286 | 0.060024281 | 0.26153531 | 0.22950737 | 0.8184746 | NA | NA | NA |
| ENSG000002 | 0 | NA | NA | NA | NA | NA | NA | NA |
| ENSG000002 | 1.91500402 | -0.379222848 | 0.35512899 | -1.0678454 | 0.28559027 | NA | NA | NA |
| ENSG000002 | 0.73292343 | -0.195671801 | 0.28518655 | -0.6861186 | 0.49263832 | NA | NA | NA |
| ENSG000002 | 0 | NA | NA | NA | NA | NA | NA | NA |

| | | | | | | | | |
|---|---|---|---|---|---|---|---|---|
| ENSG000002 | 0 | NA | NA | NA | NA | NA | NA | NA |
| ENSG000002 | 2.0069884 | -0.320814383 | 0.36102906 | -0.888611 | 0.3742122 | NA | NA | NA |
| ENSG000002 | 1.01654519 | -0.189906583 | 0.30824946 | -0.6160808 | 0.53784117 | NA | NA | NA |
| ENSG000002 | 1.72987723 | -0.322868351 | 0.35417673 | -0.9116024 | 0.36197806 | NA | NA | NA |
| ENSG000002 | 1.26347803 | -0.052108982 | 0.32941685 | -0.1581855 | 0.8743106 | NA | NA | NA |
| ENSG000002 | 1.77854422 | -0.43397569 | 0.34621891 | -1.2534719 | 0.210034 | NA | NA | NA |
| ENSG000002 | 0 | NA | NA | NA | NA | NA | NA | NA |
| ENSG000002 | 0.51396245 | -0.026802372 | 0.24109708 | -0.1111684 | 0.91148283 | NA | NA | NA |
| ENSG000002 | 1.12686312 | -0.109757665 | 0.31656055 | -0.3467193 | 0.72880218 | NA | NA | NA |
| ENSG000002 | 0.75711827 | -0.523135555 | 0.27850213 | -1.8783898 | 0.06032786 | NA | LOC100506596 | 100506596 |
| ENSG000002 | 1.13337922 | -0.250178375 | 0.31614289 | -0.7913459 | 0.42874219 | NA | NA | NA |
| ENSG000002 | 2.02241955 | -0.744872014 | 0.36274405 | -2.0534369 | 0.04003023 | NA | NA | NA |
| ENSG000002 | 0.88800828 | -0.112655767 | 0.29104807 | -0.3870693 | 0.69870492 | NA | NA | NA |
| ENSG000002 | 2.15982566 | -0.511008763 | 0.36433281 | -1.4025878 | 0.16073978 | NA | NA | NA |
| ENSG000002 | 1.61996637 | -0.238931568 | 0.34010828 | -0.7025162 | 0.48235731 | NA | NA | NA |
| ENSG000002 | 0.24411881 | -0.012946855 | 0.17864669 | -0.0724718 | 0.94222641 | NA | NA | NA |
| ENSG000002 | 1.50509836 | -0.584798244 | 0.33528818 | -1.744166 | 0.08113015 | NA | NA | NA |
| ENSG000002 | 0.75327376 | -0.030668806 | 0.27945337 | -0.1097457 | 0.91261106 | NA | NA | NA |
| ENSG000002 | 0.26062507 | -0.012888319 | 0.18226503 | -0.070712 | 0.94362699 | NA | NA | NA |
| ENSG000002 | 1.84651303 | 0.00726527 | 0.36206686 | 0.0200661 | 0.98399064 | NA | NA | NA |
| ENSG000002 | 0.25260017 | -0.015978705 | 0.18049464 | -0.0885273 | 0.92945758 | NA | NA | NA |
| ENSG000002 | 0 | NA | NA | NA | NA | NA | NA | NA |
| ENSG000002 | 0.6265601 | -0.275936484 | 0.2670182 | -1.0333995 | 0.30141697 | NA | NA | NA |
| ENSG000002 | 0.14128499 | -0.097772014 | 0.13403235 | -0.7294658 | 0.4657168 | NA | NA | NA |
| ENSG000002 | 1.49219836 | -0.188133737 | 0.34035392 | -0.5527591 | 0.58042836 | NA | NA | NA |
| ENSG000002 | 1.00432739 | -0.044227129 | 0.31348668 | -0.1410814 | 0.88780566 | NA | NA | NA |
| ENSG000002 | 1.00771892 | -0.189340575 | 0.30950532 | -0.6117522 | 0.54070169 | NA | NA | NA |
| ENSG000002 | 1.14254381 | -0.416954641 | 0.31972632 | -1.3040986 | 0.19219997 | NA | NA | NA |
| ENSG000002 | 1.93136635 | -0.969646782 | 0.35471816 | -2.7335696 | 0.00626519 | NA | LOC388572 | 388572 |

| | | | | | | | | | |
|---|---|---|---|---|---|---|---|---|---|
| ENSG000002 | 0 | NA | NA | NA | NA | NA | NA | NA | |
| ENSG000002 | 1.23858078 | -0.173466357 | 0.31623255 | -0.5485405 | 0.58332084 | NA | NA | NA | |
| ENSG000002 | 1.00143296 | -0.491564337 | 0.30388723 | -1.617588 | 0.10575142 | NA | NA | NA | |
| ENSG000002 | 0.24141689 | -0.165257896 | 0.17127401 | -0.9648743 | 0.33460776 | NA | NA | NA | |
| ENSG000002 | 0.64081643 | -0.112957221 | 0.26633067 | -0.424124 | 0.6714754 | NA | NA | NA | |
| ENSG000002 | 1.87683359 | -0.253828983 | 0.35861781 | -0.707798 | 0.47907069 | NA | NA | NA | |
| ENSG000002 | 1.12635212 | -0.285304152 | 0.32710432 | -0.8722115 | 0.38309301 | NA | NA | NA | |
| ENSG000002 | 1.51493894 | -0.475711221 | 0.34336455 | -1.3854407 | 0.16591775 | NA | PCDH20 | | 64881 |
| ENSG000002 | 1.23563608 | 0.107020853 | 0.32709793 | 0.32718291 | 0.74352955 | NA | NA | NA | |
| ENSG000002 | 0.12070844 | -0.088503695 | 0.12783931 | -0.6923042 | 0.48874631 | NA | NA | NA | |
| ENSG000002 | 0.38019136 | -0.10756138 | 0.21876967 | -0.491665 | 0.62295621 | NA | NA | NA | |
| ENSG000002 | 1.59691547 | -0.381805743 | 0.34525976 | -1.1058507 | 0.26879115 | NA | NA | NA | |
| ENSG000002 | 0.12685397 | -0.090215768 | 0.12901919 | -0.699243 | 0.48440017 | NA | NA | NA | |
| ENSG000002 | 1.53087629 | -0.484141869 | 0.34432839 | -1.4060469 | 0.15971019 | NA | NA | NA | |
| ENSG000002 | 1.54397567 | -0.631342607 | 0.34306121 | -1.8403206 | 0.06572118 | NA | C18orf12 | | 84322 |
| ENSG000002 | 0.98315743 | -0.034471163 | 0.30017172 | -0.1148381 | 0.90857342 | NA | NA | NA | |
| ENSG000002 | 0.12685397 | -0.090215768 | 0.12901919 | -0.699243 | 0.48440017 | NA | NA | NA | |
| ENSG000002 | 1.58679932 | -0.124008457 | 0.35104639 | -0.3532538 | 0.72389821 | NA | NA | NA | |
| ENSG000002 | 0.35870346 | 0.077368606 | 0.21515516 | 0.35959447 | 0.71915042 | NA | NA | NA | |
| ENSG000002 | 0 | NA | NA | NA | NA | NA | NA | NA | |
| ENSG000002 | 1.09615206 | -0.261892015 | 0.31813153 | -0.8232193 | 0.4103833 | NA | NA | NA | |
| ENSG000002 | 1.72518595 | -0.314251337 | 0.35155459 | -0.8938906 | 0.37138043 | NA | NA | NA | |
| ENSG000002 | 0.36482725 | -0.102221582 | 0.21647267 | -0.4722147 | 0.63677353 | NA | NA | NA | |
| ENSG000002 | 0.62638541 | -0.108168827 | 0.26514251 | -0.4079649 | 0.68329947 | NA | NA | NA | |
| ENSG000002 | 0.70964048 | -0.026383209 | 0.2764226 | -0.0954452 | 0.92396122 | NA | NA | NA | |
| ENSG000002 | 0.86097162 | -0.265533134 | 0.29690985 | -0.8943224 | 0.3711494 | NA | NA | NA | |
| ENSG000002 | 0 | NA | NA | NA | NA | NA | ACTR3BP2 | | 440888 |
| ENSG000002 | 1.22673196 | -0.32084456 | 0.32424527 | -0.9895119 | 0.32241273 | NA | NA | NA | |
| ENSG000002 | 0.87394783 | -0.26920868 | 0.29758521 | -0.904644 | 0.36565401 | NA | NA | NA | |

| | | | | | | | | |
|---|---|---|---|---|---|---|---|---|
| ENSG000002 | 1.63607977 | -0.247805442 | 0.34035875 | -0.7280713 | 0.46656993 | NA | NA | NA |
| ENSG000002 | 2.08260482 | -0.131088144 | 0.3650614 | -0.3590852 | 0.71953136 | NA | NA | NA |
| ENSG000002 | 0.12685397 | -0.090215768 | 0.12901919 | -0.699243 | 0.48440017 | NA | NA | NA |
| ENSG000002 | 1.49228268 | -0.192775328 | 0.34100862 | -0.5653093 | 0.57186344 | NA | NA | NA |
| ENSG000002 | 1.24292322 | 0.097776166 | 0.32448007 | 0.30133181 | 0.76316149 | NA | NA | NA |
| ENSG000002 | 0.77749894 | -0.529922771 | 0.27983286 | -1.8937117 | 0.05826328 | NA | NA | NA |
| ENSG000002 | 1.13479911 | -0.396496401 | 0.31196121 | -1.2709798 | 0.20373583 | NA | NA | NA |
| ENSG000002 | 0.8250799 | -0.098366843 | 0.15985981 | -0.6153319 | 0.53833555 | NA | NA | NA |
| ENSG000002 | 0 | NA | NA | NA | NA | NA | NA | NA |
| ENSG000002 | 0 | NA | NA | NA | NA | NA | NA | NA |
| ENSG000002 | 0.87875533 | -0.262010202 | 0.29199538 | -0.8973094 | 0.36955383 | NA | NA | NA |
| ENSG000002 | 1.15649662 | -0.602182096 | 0.32216736 | -1.8691592 | 0.06160066 | NA | NA | NA |
| ENSG000002 | 1.11508004 | -0.122935201 | 0.32616764 | -0.376908 | 0.70624196 | NA | NA | NA |
| ENSG000002 | 0 | NA | NA | NA | NA | NA | NA | NA |
| ENSG000002 | 1.68847167 | -0.545237057 | 0.34904433 | -1.5620854 | 0.11826787 | NA | NA | NA |
| ENSG000002 | 1.38803098 | -0.575669893 | 0.33843478 | -1.7009773 | 0.08894724 | NA | NA | NA |
| ENSG000002 | 1.30103771 | 0.484398986 | 0.33595818 | 1.44184309 | 0.14934664 | NA | NA | NA |
| ENSG000002 | 0 | NA | NA | NA | NA | NA | NA | NA |
| ENSG000002 | 0.49059519 | 0.153206417 | 0.24651 | 0.62150184 | 0.53426948 | NA | NA | NA |
| ENSG000002 | 0.69356035 | 0.125831047 | 0.27383501 | 0.4595141 | 0.64586503 | NA | NA | NA |
| ENSG000002 | 1.73910844 | -0.30909306 | 0.35055131 | -0.8817341 | 0.37792059 | NA | NA | NA |
| ENSG000002 | 1.36334934 | -0.115024594 | 0.33831839 | -0.3399892 | 0.73386468 | NA | NA | NA |
| ENSG000002 | 1.04563791 | -0.705264854 | 0.30928943 | -2.2802747 | 0.0225914 | NA | NA | NA |
| ENSG000002 | 1.74150272 | -0.408871455 | 0.34495944 | -1.185274 | 0.23590913 | NA | NA | NA |
| ENSG000002 | 0.49224375 | -0.024566489 | 0.23863564 | -0.1029456 | 0.91800615 | NA | NA | NA |
| ENSG000002 | 1.50784534 | -0.051470843 | 0.34298047 | -0.1500693 | 0.88070993 | NA | NA | NA |
| ENSG000002 | 1.51123241 | 0.063069963 | 0.33236993 | 0.18975833 | 0.84949851 | NA | NA | NA |
| ENSG000002 | 0.78822576 | -0.365434278 | 0.2849842 | -1.2822966 | 0.19973861 | NA | NA | NA |
| ENSG000002 | 1.74382302 | -0.056334783 | 0.35408017 | -0.1591018 | 0.87358869 | NA | NA | NA |

| | | | | | | | | | |
|---|---|---|---|---|---|---|---|---|---|
| ENSG000002 | 0.40327843 | -0.281391716 | 0.21700406 | -1.2967118 | 0.19473038 | NA | NA | NA | |
| ENSG000002 | 1.84407306 | 0.007231391 | 0.35667287 | 0.02027458 | 0.98382434 | NA | NA | NA | |
| ENSG000002 | 1.54035792 | -0.313412425 | 0.3383681 | -0.926247 | 0.35431765 | NA | NA | NA | |
| ENSG000002 | 2.06493709 | -0.603423804 | 0.36300211 | -1.6623149 | 0.09644965 | NA | NA | NA | |
| ENSG000002 | 1.70149531 | -0.008042655 | 0.18245574 | -0.04408 | 0.96484061 | NA | NA | NA | |
| ENSG000002 | 0.72714846 | -0.325005647 | 0.27254781 | -1.1924721 | 0.2330762 | NA | NA | NA | |
| ENSG000002 | 1.02887106 | -0.710128537 | 0.31010642 | -2.2899511 | 0.02202415 | NA | NA | NA | |
| ENSG000002 | 1.58625853 | -0.114646943 | 0.34987393 | -0.3276807 | 0.74315308 | NA | NA | NA | |
| ENSG000002 | 0.98626136 | -0.188884989 | 0.31185466 | -0.6056828 | 0.54472544 | NA | LINC01101 | | 84931 |
| ENSG000002 | 0.51987986 | -0.01050728 | 0.1556516 | -0.0675051 | 0.94617959 | NA | NA | NA | |
| ENSG000002 | 1.03624465 | -0.533979215 | 0.31238521 | -1.7093614 | 0.08738403 | NA | NA | NA | |
| ENSG000002 | 0.86046309 | -0.246260481 | 0.29102415 | -0.8461857 | 0.39744914 | NA | NA | NA | |
| ENSG000002 | 1.73913409 | -0.450424322 | 0.3529309 | -1.2762394 | 0.2018709 | NA | NA | NA | |
| ENSG000002 | 1.08123582 | 0.03980283 | 0.31460953 | 0.12651502 | 0.89932426 | NA | NA | NA | |
| ENSG000002 | 0.98508887 | -0.033772222 | 0.31272058 | -0.1079949 | 0.91399975 | NA | NA | NA | |
| ENSG000002 | 0.60148871 | -0.081011337 | 0.15721804 | -0.5152802 | 0.60635726 | NA | NA | NA | |
| ENSG000002 | 0.25214371 | -0.009862892 | 0.18046657 | -0.0546522 | 0.95641556 | NA | NA | NA | |
| ENSG000002 | 2.17053287 | -0.384097046 | 0.36426733 | -1.0544373 | 0.29168277 | NA | NA | NA | |
| ENSG000002 | 0.65465054 | -0.455468986 | 0.2643416 | -1.7230318 | 0.08488278 | NA | NA | NA | |
| ENSG000002 | 0 | NA | NA | NA | NA | NA | NA | NA | |
| ENSG000002 | 0 | NA | NA | NA | NA | NA | NA | NA | |
| ENSG000002 | 2.01025731 | -0.314286529 | 0.36104928 | -0.870481 | 0.38403761 | NA | NA | NA | |
| ENSG000002 | 0 | NA | NA | NA | NA | NA | NA | NA | |
| ENSG000002 | 0 | NA | NA | NA | NA | NA | SNORA43 | | 677824 |
| ENSG000002 | 0 | NA | NA | NA | NA | NA | SNORA16A | | 692073 |
| ENSG000002 | 0.39479706 | -0.300008631 | 0.22317059 | -1.3443018 | 0.17885081 | NA | NA | NA | |
| ENSG000002 | 1.35862369 | -0.263526682 | 0.33683477 | -0.7823619 | 0.43400193 | NA | NA | NA | |
| ENSG000002 | 0 | NA | NA | NA | NA | NA | NA | NA | |
| ENSG000002 | 0.95918935 | 0.12025443 | 0.31160872 | 0.38591485 | 0.69955973 | NA | NA | NA | |

| | | | | | | | | |
|---|---|---|---|---|---|---|---|---|
| ENSG000002 | 0 | NA | NA | NA | NA | NA | NA | NA |
| ENSG000002 | 0 | NA | NA | NA | NA | NA | NA | NA |
| ENSG000002 | 0 | NA | NA | NA | NA | NA | NA | NA |
| ENSG000002 | 1.51690366 | -0.319323274 | 0.33867181 | -0.9428694 | 0.34574773 | NA | NA | NA |
| ENSG000002 | 0 | NA | NA | NA | NA | NA | NA | NA |
| ENSG000002 | 0 | NA | NA | NA | NA | NA | NA | NA |
| ENSG000002 | 0 | NA | NA | NA | NA | NA | NA | NA |
| ENSG000002 | 1.22072789 | 0.09899665 | 0.32822358 | 0.30161346 | 0.76294675 | NA | NA | NA |
| ENSG000002 | 0 | NA | NA | NA | NA | NA | NA | NA |
| ENSG000002 | 0 | NA | NA | NA | NA | NA | NA | NA |
| ENSG000002 | 0.80032095 | -0.34356993 | 0.27770516 | -1.2371752 | 0.21602205 | NA | LOC727924 | 727924 |
| ENSG000002 | 0 | NA | NA | NA | NA | NA | NA | NA |
| ENSG000002 | 0 | NA | NA | NA | NA | NA | NA | NA |
| ENSG000002 | 1.19941201 | -0.04208655 | 0.32615861 | -0.1290371 | 0.89732832 | NA | NA | NA |
| ENSG000002 | 1.25141318 | -0.197758596 | 0.32960691 | -0.5999832 | 0.54851746 | NA | NA | NA |
| ENSG000002 | 0 | NA | NA | NA | NA | NA | NA | NA |
| ENSG000002 | 0 | NA | NA | NA | NA | NA | NA | NA |
| ENSG000002 | 0.11865494 | 0.083440866 | 0.12751478 | 0.65436232 | 0.5128784 | NA | NA | NA |
| ENSG000002 | 1.16056691 | -0.608217967 | 0.32219504 | -1.8877322 | 0.05906191 | NA | NA | NA |
| ENSG000002 | 1.04105269 | -0.509071869 | 0.30808165 | -1.6523927 | 0.09845452 | NA | NA | NA |
| ENSG000002 | 0.36553413 | 0.069401489 | 0.20780452 | 0.33397488 | 0.73839851 | NA | NA | NA |
| ENSG000002 | 0 | NA | NA | NA | NA | NA | NA | NA |
| ENSG000002 | 0 | NA | NA | NA | NA | NA | NA | NA |
| ENSG000002 | 0.34344486 | 0.238503239 | 0.20662376 | 1.15428757 | 0.24838229 | NA | NA | NA |
| ENSG000002 | 0.23868015 | 0.154625155 | 0.16997278 | 0.90970538 | 0.36297791 | NA | NA | NA |
| ENSG000002 | 1.37886056 | -0.235492673 | 0.3263588 | -0.721576 | 0.4705552 | NA | NA | NA |
| ENSG000002 | 0 | NA | NA | NA | NA | NA | NA | NA |
| ENSG000002 | 0.11865494 | 0.083440866 | 0.12751478 | 0.65436232 | 0.5128784 | NA | MIR3653 | 100500833 |
| ENSG000002 | 0 | NA | NA | NA | NA | NA | RPS4Y2 | 140032 |

| | | | | | | | | | |
|---|---|---|---|---|---|---|---|---|---|
| ENSG000002 | 1.81834607 | -0.614718712 | 0.35341171 | -1.7393841 | 0.08196723 | NA | NA | NA | |
| ENSG000002 | 0 | NA | NA | NA | NA | NA | SNORD3D | | 780854 |
| ENSG000002 | 0 | NA | NA | NA | NA | NA | SNORD60 | | 26788 |
| ENSG000002 | 0 | NA | NA | NA | NA | NA | PI4K2B | | 55300 |
| ENSG000002 | 0 | NA | NA | NA | NA | NA | NA | NA | |
| ENSG000002 | 0.13280363 | -0.093394593 | 0.13115644 | -0.7120855 | 0.47641185 | NA | NA | NA | |
| ENSG000002 | 0.23276084 | -0.008011382 | 0.17603982 | -0.0455089 | 0.96370166 | NA | NA | NA | |
| ENSG000002 | 0 | NA | NA | NA | NA | NA | NA | NA | |
| ENSG000002 | 0.86096715 | 0.037689904 | 0.29699504 | 0.12690415 | 0.89901626 | NA | NA | NA | |
| ENSG000002 | 0.11131518 | 0.081951103 | 0.12663804 | 0.64712866 | 0.51754868 | NA | NA | NA | |
| ENSG000002 | 1.3819091 | -0.225557013 | 0.3244294 | -0.6952422 | 0.48690353 | NA | NA | NA | |
| ENSG000002 | 0.72969798 | 0.127609973 | 0.27082704 | 0.47118624 | 0.63750774 | NA | NA | NA | |
| ENSG000002 | 0.12685397 | -0.090215768 | 0.12901919 | -0.699243 | 0.48440017 | NA | MIR3651 | | 100500918 |
| ENSG000002 | 0 | NA | NA | NA | NA | NA | NA | NA | |
| ENSG000002 | 2.1552793 | -0.50829327 | 0.3651778 | -1.3919063 | 0.16395078 | NA | NA | NA | |
| ENSG000002 | 0 | NA | NA | NA | NA | NA | NA | NA | |
| ENSG000002 | 1.63517076 | -0.542202409 | 0.34829675 | -1.5567254 | 0.11953569 | NA | LINC00254 | | 64735 |
| ENSG000002 | 0.51931521 | -0.354214731 | 0.23897171 | -1.4822455 | 0.13827499 | NA | NA | NA | |
| ENSG000002 | 0.98776906 | -0.044467267 | 0.31282942 | -0.1421454 | 0.88696515 | NA | NA | NA | |
| ENSG000002 | 0 | NA | NA | NA | NA | NA | NA | NA | |
| ENSG000002 | 0 | NA | NA | NA | NA | NA | SNORA76C | | 677842 |
| ENSG000002 | 0.26560726 | -0.177019426 | 0.17672274 | -1.0016788 | 0.31649875 | NA | NF1P2 | | 440225 |
| ENSG000002 | 0 | NA | NA | NA | NA | NA | NA | NA | |
| ENSG000002 | 1.43965946 | -0.727040857 | 0.33492476 | -2.1707588 | 0.02994941 | NA | NA | NA | |
| ENSG000002 | 1.84116207 | -0.119501858 | 0.35804391 | -0.3337631 | 0.73855831 | NA | NA | NA | |
| ENSG000002 | 0.24485602 | -0.012655878 | 0.17881722 | -0.0707755 | 0.94357643 | NA | NA | NA | |
| ENSG000002 | 1.99754361 | -0.187865587 | 0.36047272 | -0.5211645 | 0.60225218 | NA | NA | NA | |
| ENSG000002 | 0 | NA | NA | NA | NA | NA | SCARNA4 | | 677771 |
| ENSG000002 | 0.34270765 | 0.258172734 | 0.21433185 | 1.20454674 | 0.22837833 | NA | NA | NA | |

| | | | | | | | | |
|---|---|---|---|---|---|---|---|---|
| ENSG000002 | 0 | NA | NA | NA | NA | NA | TGFB2-OT1 | 103611157 |
| ENSG000002 | 0.61582881 | -0.11034135 | 0.26423602 | -0.4175863 | 0.67624959 | NA | NA | NA |
| ENSG000002 | 2.17210001 | -0.905855788 | 0.36110026 | -2.5085991 | 0.0121211 | NA | PISRT1 | 140464 |
| ENSG000002 | 1.73593843 | -0.317249515 | 0.35316968 | -0.8982921 | 0.36902987 | NA | LOC105374847 | 105374847 |
| ENSG000002 | 0 | NA | NA | NA | NA | NA | NA | NA |
| ENSG000002 | 1.95001418 | -0.312396657 | 0.36064201 | -0.8662237 | 0.3863675 | NA | NA | NA |
| ENSG000002 | 0 | NA | NA | NA | NA | NA | NA | NA |
| ENSG000002 | 1.89488393 | -0.362733663 | 0.35072615 | -1.0342361 | 0.30102578 | NA | DBET | 100419743 |
| ENSG000002 | 1.96011432 | -0.294027627 | 0.35741065 | -0.8226605 | 0.41070108 | NA | NA | NA |
| ENSG000002 | 1.91739246 | -0.346319943 | 0.3487073 | -0.9931537 | 0.32063506 | NA | NA | NA |
| ENSG000002 | 1.85151641 | 0.001612662 | 0.35842386 | 0.00449932 | 0.99641008 | NA | NA | NA |
| ENSG000002 | 0 | NA | NA | NA | NA | NA | NA | NA |
| ENSG000002 | 0 | NA | NA | NA | NA | NA | NA | NA |
| ENSG000002 | 0.23890637 | -0.010420674 | 0.17747379 | -0.0587167 | 0.95317776 | NA | MIR664A | 100302234 |
| ENSG000002 | 0 | NA | NA | NA | NA | NA | NA | NA |
| ENSG000002 | 0.37194025 | 0.067575365 | 0.21717998 | 0.31114914 | 0.75568724 | NA | NA | NA |
| ENSG000002 | 0.66683218 | -0.273056421 | 0.26350214 | -1.0362588 | 0.30008141 | NA | LOC100996720 | 100996720 |
| ENSG000002 | 1.36360938 | -0.124475121 | 0.33481579 | -0.371772 | 0.71006264 | NA | NA | NA |
| ENSG000002 | 0.3641961 | 0.070436377 | 0.21601083 | 0.32607799 | 0.74436535 | NA | NA | NA |
| ENSG000002 | 0 | NA | NA | NA | NA | NA | SNORA17 | 677804 |
| ENSG000002 | 1.54494121 | -0.605969974 | 0.33881538 | -1.788496 | 0.07369601 | NA | NA | NA |
| ENSG000002 | 1.88235009 | 0.005600757 | 0.35595332 | 0.01573453 | 0.98744618 | NA | LINC00602 | 441177 |
| ENSG000002 | 0.11205239 | 0.082118251 | 0.12671962 | 0.64803108 | 0.51696485 | NA | NA | NA |
| ENSG000002 | 1.52444451 | -0.289447762 | 0.32909772 | -0.8795192 | 0.37911983 | NA | NA | NA |
| ENSG000002 | 0.71192962 | 0.137101895 | 0.15853758 | 0.86479112 | 0.38715344 | NA | NA | NA |
| ENSG000002 | 0 | NA | NA | NA | NA | NA | SNORD38B | 94163 |
| ENSG000002 | 0 | NA | NA | NA | NA | NA | NA | NA |
| ENSG000002 | 0.78273257 | -0.363992205 | 0.28297036 | -1.2863263 | 0.19832922 | NA | NA | NA |
| ENSG000002 | 0 | NA | NA | NA | NA | NA | GIMAP1-GIMAF | 100527949 |

| | | | | | | | | | |
|---|---|---|---|---|---|---|---|---|---|
| ENSG000002 | 1.05852587 | -0.327535763 | 0.30048082 | -1.0900388 | 0.27569604 | NA | | C1QTNF9B-AS1 | 542767 |
| ENSG000002 | 0 | NA | NA | NA | NA | NA | | SNORA76A | 677830 |
| ENSG000002 | 0.40327843 | -0.281391716 | 0.21700406 | -1.2967118 | 0.19473038 | NA | | NA | NA |
| ENSG000002 | 0.13280363 | -0.093394593 | 0.13115644 | -0.7120855 | 0.47641185 | NA | | NA | NA |
| ENSG000002 | 0.86543421 | -0.227847958 | 0.28173184 | -0.8087405 | 0.4186644 | NA | | NA | NA |
| ENSG000002 | 0.66293603 | -0.489755159 | 0.27215878 | -1.7995199 | 0.07193647 | NA | | NA | NA |
| ENSG000002 | 0 | NA | NA | NA | NA | NA | | TMEM265 | 100862671 |
| ENSG000002 | 0.82171461 | 0.207400687 | 0.29369875 | 0.70616811 | 0.4800836 | NA | | LOC100506563 | 100506563 |
| ENSG000002 | 0.49307133 | -0.179259067 | 0.24031383 | -0.7459374 | 0.45570524 | NA | | NA | NA |
| ENSG000002 | 0 | NA | NA | NA | NA | NA | | NA | NA |
| ENSG000002 | 0 | NA | NA | NA | NA | NA | | NA | NA |
| ENSG000002 | 0.4716672 | -0.017393026 | 0.23619084 | -0.0736397 | 0.94129707 | NA | | NA | NA |
| ENSG000002 | 1.24449852 | -0.186214924 | 0.3294393 | -0.5652481 | 0.57190506 | NA | | NA | NA |
| ENSG000002 | 0.76715995 | -0.377154177 | 0.28907496 | -1.3046933 | 0.19199729 | NA | | NA | NA |
| ENSG000002 | 0.88351289 | -0.120228751 | 0.29849063 | -0.402789 | 0.68710343 | NA | | NA | NA |
| ENSG000002 | 1.75648933 | -0.198300087 | 0.35208864 | -0.5632107 | 0.57329143 | NA | | NA | NA |
| ENSG000002 | 0.12685397 | -0.090215768 | 0.12901919 | -0.699243 | 0.48440017 | NA | | NA | NA |
| ENSG000002 | 0 | NA | NA | NA | NA | NA | | NA | NA |
| ENSG000002 | 0 | NA | NA | NA | NA | NA | | NA | NA |
| ENSG000002 | 0 | NA | NA | NA | NA | NA | | NA | NA |
| ENSG000002 | 0.52668879 | -0.197054386 | 0.24396589 | -0.8077129 | 0.4192559 | NA | | NA | NA |
| ENSG000002 | 0 | NA | NA | NA | NA | NA | | NA | NA |
| ENSG000002 | 0.24756242 | -0.187034346 | 0.18119817 | -1.0322088 | 0.30197431 | NA | | CYP3A7-CYP3A5 | 100861540 |
| ENSG000002 | 0.11934008 | 0.083564698 | 0.12760091 | 0.65489106 | 0.5125379 | NA | | NA | NA |
| ENSG000002 | 0 | NA | NA | NA | NA | NA | | NA | NA |
| ENSG000002 | 0.99593584 | -0.189409988 | 0.30735381 | -0.6162604 | 0.53772265 | NA | | NA | NA |
| ENSG000002 | 0.62775378 | -0.28146231 | 0.26704095 | -1.0540043 | 0.29188095 | NA | | NA | NA |
| ENSG000002 | 0 | NA | NA | NA | NA | NA | | NA | NA |
| ENSG000002 | 0.7459517 | -0.174898099 | 0.27105011 | -0.6452611 | 0.51875798 | NA | | NA | NA |

| | | | | | | | | | |
|---|---|---|---|---|---|---|---|---|---|
| ENSG000002 | 2.09994504 | -0.327113849 | 0.35617922 | -0.9183968 | 0.35841117 | NA | LOC401442 | | 401442 |
| ENSG000002 | 0 | NA | NA | NA | NA | NA | NA | NA | |
| ENSG000002 | 0.24485602 | -0.012655878 | 0.17881722 | -0.0707755 | 0.94357643 | NA | NA | NA | |
| ENSG000002 | 1.57047571 | -0.234527413 | 0.34263415 | -0.6844835 | 0.49366991 | NA | NA | NA | |
| ENSG000002 | 0.47684931 | 0.147509143 | 0.23759598 | 0.62084023 | 0.53470475 | NA | NA | NA | |
| ENSG000002 | 0 | NA | NA | NA | NA | NA | NA | NA | |
| ENSG000002 | 0 | NA | NA | NA | NA | NA | NA | NA | |
| ENSG000002 | 0 | NA | NA | NA | NA | NA | NA | NA | |
| ENSG000002 | 0.52876404 | -0.191903645 | 0.2442426 | -0.7857092 | 0.4320379 | NA | NA | NA | |
| ENSG000002 | 1.10514442 | -0.114466106 | 0.32202152 | -0.355461 | 0.72224422 | NA | NA | NA | |
| ENSG000002 | 2.00425167 | -0.069541602 | 0.36026112 | -0.1930311 | 0.84693461 | NA | NA | NA | |
| ENSG000002 | 0 | NA | NA | NA | NA | NA | FAM138F | | 641702 |
| ENSG000002 | 0.49105007 | -0.020523257 | 0.24605963 | -0.0834077 | 0.9335274 | NA | NA | NA | |
| ENSG000002 | 1.34649233 | 0.026593111 | 0.33697902 | 0.07891622 | 0.93709926 | NA | NA | NA | |
| ENSG000002 | 1.90166659 | 0.328127886 | 0.35880347 | 0.91450589 | 0.36045109 | NA | NA | NA | |
| ENSG000002 | 0.47821768 | -0.016471461 | 0.24462483 | -0.0673336 | 0.94631616 | NA | NA | NA | |
| ENSG000002 | 0 | NA | NA | NA | NA | NA | NA | NA | |
| ENSG000002 | 0 | NA | NA | NA | NA | NA | NA | NA | |
| ENSG000002 | 1.14189092 | -0.442592986 | 0.32511543 | -1.3613411 | 0.17340593 | NA | NA | NA | |
| ENSG000002 | 0.49356059 | -0.178885509 | 0.24036594 | -0.7442215 | 0.45674246 | NA | NA | NA | |
| ENSG000002 | 1.00422133 | -0.202520951 | 0.31404679 | -0.6448751 | 0.51900814 | NA | NA | NA | |
| ENSG000002 | 1.63131416 | -0.110996465 | 0.34553568 | -0.3212301 | 0.74803602 | NA | NA | NA | |
| ENSG000002 | 0.25145857 | -0.010118565 | 0.18031426 | -0.0561163 | 0.95524918 | NA | NA | NA | |
| ENSG000002 | 0 | NA | NA | NA | NA | NA | NA | NA | |
| ENSG000002 | 0.91507974 | -0.461155367 | 0.30304372 | -1.5217454 | 0.1280729 | NA | NA | NA | |
| ENSG000002 | 0.38884741 | -0.29745092 | 0.22235466 | -1.3377319 | 0.18098386 | NA | NA | NA | |
| ENSG000002 | 0 | NA | NA | NA | NA | NA | NA | NA | |
| ENSG000002 | 0 | NA | NA | NA | NA | NA | NA | NA | |
| ENSG000002 | 0.3588776 | -0.099902505 | 0.21555794 | -0.4634601 | 0.6430346 | NA | NA | NA | |

| | | | | | | | | |
|---|---|---|---|---|---|---|---|---|
| ENSG000002 | 1.76050205 | -0.290539011 | 0.34599216 | -0.8397271 | 0.40106139 | NA | NA | NA |
| ENSG000002 | 1.49583605 | 0.076322459 | 0.33911636 | 0.22506275 | 0.82193046 | NA | NA | NA |
| ENSG000002 | 1.26299983 | -0.18893732 | 0.32566252 | -0.5801629 | 0.56180476 | NA | NA | NA |
| ENSG000002 | 1.73673176 | -0.0477403 | 0.34927831 | -0.1366827 | 0.89128161 | NA | NA | NA |
| ENSG000002 | 0 | NA | NA | NA | NA | NA | NA | NA |
| ENSG000002 | 1.05828919 | 0.508552028 | 0.31432706 | 1.61790724 | 0.10568259 | NA | NA | NA |
| ENSG000002 | 1.01354988 | -0.31059533 | 0.2985882 | -1.040213 | 0.29824096 | NA | NA | NA |
| ENSG000002 | 0.12685397 | -0.090215768 | 0.12901919 | -0.699243 | 0.48440017 | NA | NA | NA |
| ENSG000002 | 0 | NA | NA | NA | NA | NA | NA | NA |
| ENSG000002 | 0.12070844 | -0.088503695 | 0.12783931 | -0.6923042 | 0.48874631 | NA | NA | NA |
| ENSG000002 | 0.3588776 | -0.099902505 | 0.21555794 | -0.4634601 | 0.6430346 | NA | NA | NA |
| ENSG000002 | 1.78957048 | -0.4764796 | 0.35563998 | -1.3397807 | 0.18031664 | NA | NA | NA |
| ENSG000002 | 0.22997012 | 0.167051788 | 0.17646558 | 0.94665368 | 0.34381528 | NA | NA | NA |
| ENSG000002 | 0 | NA | NA | NA | NA | NA | NA | NA |
| ENSG000002 | 0.73513663 | -0.025980961 | 0.27821552 | -0.0933843 | 0.92559827 | NA | NA | NA |
| ENSG000002 | 0.49856397 | -0.186262775 | 0.24090455 | -0.7731808 | 0.43941538 | NA | NA | NA |
| ENSG000002 | 1.50101646 | -0.449496802 | 0.33699745 | -1.3338285 | 0.18226006 | NA | NA | NA |
| ENSG000002 | 0.48302518 | -0.023466322 | 0.24515318 | -0.0957211 | 0.92374212 | NA | NA | NA |
| ENSG000002 | 1.67588913 | -0.386215687 | 0.34638869 | -1.1149778 | 0.26485995 | NA | NA | NA |
| ENSG000002 | 2.15884806 | 0.436401904 | 0.36661349 | 1.19035963 | 0.23390507 | NA | NA | NA |
| ENSG000002 | 0 | NA | NA | NA | NA | NA | NA | NA |
| ENSG000002 | 1.46809231 | -0.198220508 | 0.34692863 | -0.5713582 | 0.56775688 | NA | NA | NA |
| ENSG000002 | 1.36932933 | -0.277688424 | 0.34304044 | -0.8094918 | 0.41823231 | NA | NA | NA |
| ENSG000002 | 0.73567797 | -0.025561802 | 0.27825106 | -0.091866 | 0.92680454 | NA | NA | NA |
| ENSG000002 | 0.13280363 | -0.093394593 | 0.13115644 | -0.7120855 | 0.47641185 | NA | NA | NA |
| ENSG000002 | 1.87760272 | -0.262497908 | 0.35862447 | -0.7319576 | 0.46419444 | NA | NA | NA |
| ENSG000002 | 1.7596761 | 0.407580718 | 0.35592703 | 1.14512438 | 0.25215764 | NA | NA | NA |
| ENSG000002 | 0.85036295 | -0.238800028 | 0.28345998 | -0.8424471 | 0.39953775 | NA | NA | NA |
| ENSG000002 | 0 | NA | NA | NA | NA | NA | NA | NA |

| | | | | | | | | |
|---|---|---|---|---|---|---|---|---|
| ENSG000002 | 1.03561349 | -0.367980092 | 0.31485625 | -1.1687241 | 0.2425148 | NA | NA | NA |
| ENSG000002 | 0.11131518 | 0.081951103 | 0.12663804 | 0.64712866 | 0.51754868 | NA | NA | NA |
| ENSG000002 | 1.50861872 | -0.318685671 | 0.33934278 | -0.9391261 | 0.34766599 | NA | NA | NA |
| ENSG000002 | 0.36465257 | 0.064724075 | 0.21605106 | 0.29957767 | 0.76449932 | NA | NA | NA |
| ENSG000002 | 0.51225778 | -0.204952203 | 0.24951191 | -0.8214125 | 0.41141135 | NA | NA | NA |
| ENSG000002 | 1.56206314 | 0.142070138 | 0.34515842 | 0.41160849 | 0.68062641 | NA | NA | NA |
| ENSG000002 | 0 | NA | NA | NA | NA | NA | NA | NA |
| ENSG000002 | 1.71400616 | -0.187906786 | 0.35518682 | -0.5290365 | 0.59678014 | NA | NA | NA |
| ENSG000002 | 0 | NA | NA | NA | NA | NA | NA | NA |
| ENSG000002 | 0 | NA | NA | NA | NA | NA | NA | NA |
| ENSG000002 | 0 | NA | NA | NA | NA | NA | NA | NA |
| ENSG000002 | 0.49082386 | 0.135175125 | 0.22941993 | 0.58920394 | 0.55572448 | NA | NA | NA |
| ENSG000002 | 1.13369188 | -0.282768575 | 0.3273404 | -0.8638365 | 0.38767774 | NA | NA | NA |
| ENSG000002 | 0.37285215 | -0.099309896 | 0.2177345 | -0.4561055 | 0.64831414 | NA | NA | NA |
| ENSG000002 | 0.53608206 | -0.386623811 | 0.24788931 | -1.5596631 | 0.11883952 | NA | NA | NA |
| ENSG000002 | 0 | NA | NA | NA | NA | NA | NA | NA |
| ENSG000002 | 0.54222759 | -0.38902668 | 0.24851205 | -1.5654238 | 0.11748356 | NA | NA | NA |
| ENSG000002 | 0.26560726 | -0.177019426 | 0.17672274 | -1.0016788 | 0.31649875 | NA | NA | NA |
| ENSG000002 | 0 | NA | NA | NA | NA | NA | NA | NA |
| ENSG000002 | 0.40191006 | -0.101653297 | 0.21376195 | -0.4755444 | 0.63439899 | NA | NA | NA |
| ENSG000002 | 0.37899768 | -0.10176489 | 0.21865765 | -0.4654074 | 0.64163973 | NA | NA | NA |
| ENSG000002 | 0 | NA | NA | NA | NA | NA | NA | NA |
| ENSG000002 | 2.17338013 | -0.780534474 | 0.36356897 | -2.1468677 | 0.03180382 | NA | NA | NA |
| ENSG000002 | 0 | NA | NA | NA | NA | NA | NA | NA |
| ENSG000002 | 2.08335054 | -0.12066096 | 0.36585346 | -0.3298068 | 0.74154595 | NA | NA | NA |
| ENSG000002 | 0.634074 | -0.448061355 | 0.26266228 | -1.7058458 | 0.08803679 | NA | NA | NA |
| ENSG000002 | 1.11209579 | -0.413453992 | 0.31867971 | -1.2973967 | 0.19449474 | NA | NA | NA |
| ENSG000002 | 0.36465257 | 0.064724075 | 0.21605106 | 0.29957767 | 0.76449932 | NA | NA | NA |
| ENSG000002 | 1.89713037 | -0.813536444 | 0.35673438 | -2.2805104 | 0.02257744 | NA | NA | NA |

| | | | | | | | | |
|---|---|---|---|---|---|---|---|---|
| ENSG000002 | 1.28454035 | -0.342030838 | 0.32920541 | -1.0389587 | 0.29882393 | NA | NA | NA |
| ENSG000002 | 0.11865494 | 0.083440866 | 0.12751478 | 0.65436232 | 0.5128784 | NA | NA | NA |
| ENSG000002 | 0.26062507 | -0.012888319 | 0.18226503 | -0.070712 | 0.94362699 | NA | NA | NA |
| ENSG000002 | 0.98964842 | -0.039843885 | 0.31290384 | -0.1273359 | 0.89867456 | NA | NA | NA |
| ENSG000002 | 0 | NA | NA | NA | NA | NA | NA | NA |
| ENSG000002 | 1.48887443 | -0.430503153 | 0.33281253 | -1.2935305 | 0.19582764 | NA | NA | NA |
| ENSG000002 | 1.5080562 | -0.477960846 | 0.34396532 | -1.3895611 | 0.16466218 | NA | NA | NA |
| ENSG000002 | 0.12685397 | -0.090215768 | 0.12901919 | -0.699243 | 0.48440017 | NA | NA | NA |
| ENSG000002 | 0.66654986 | -0.459691143 | 0.26528715 | -1.7328059 | 0.08313017 | NA | NA | NA |
| ENSG000002 | 2.04701542 | -0.665772516 | 0.35341742 | -1.8838135 | 0.05959021 | NA | NA | NA |
| ENSG000002 | 0.47029884 | 0.145033566 | 0.23680225 | 0.612467 | 0.54022882 | NA | NA | NA |
| ENSG000002 | 0.23890637 | -0.010420674 | 0.17747379 | -0.0587167 | 0.95317776 | NA | NA | NA |
| ENSG000002 | 2.09929373 | -0.233633353 | 0.36070421 | -0.6477145 | 0.51716962 | NA | NA | NA |
| ENSG000002 | 0.71347404 | 0.29874652 | 0.27688408 | 1.07895881 | 0.28060609 | NA | NA | NA |
| ENSG000002 | 0.75982019 | -0.379788636 | 0.28855699 | -1.3161651 | 0.18811864 | NA | NA | NA |
| ENSG000002 | 1.11097538 | -0.258164454 | 0.31924678 | -0.8086674 | 0.41870651 | NA | NA | NA |
| ENSG000002 | 0 | NA | NA | NA | NA | NA | NA | NA |
| ENSG000002 | 0.25370795 | -0.171331747 | 0.17411721 | -0.9840024 | 0.32511435 | NA | NA | NA |
| ENSG000002 | 1.02214994 | -0.19080937 | 0.30847374 | -0.6185595 | 0.53620657 | NA | NA | NA |
| ENSG000002 | 0.60606945 | -0.106717248 | 0.26337649 | -0.405189 | 0.68533862 | NA | NA | NA |
| ENSG000002 | 0 | NA | NA | NA | NA | NA | NA | NA |
| ENSG000002 | 0.12685397 | -0.090215768 | 0.12901919 | -0.699243 | 0.48440017 | NA | NA | NA |
| ENSG000002 | 0.78461194 | -0.338341767 | 0.27666213 | -1.2229421 | 0.22135156 | NA | NA | NA |
| ENSG000002 | 0 | NA | NA | NA | NA | NA | NA | NA |
| ENSG000002 | 0.62041457 | -0.27371262 | 0.26650407 | -1.0270486 | 0.3043976 | NA | NA | NA |
| ENSG000002 | 0.7228673 | 0.135772152 | 0.27592955 | 0.49205369 | 0.62268139 | NA | NA | NA |
| ENSG000002 | 0.76747453 | -0.201328414 | 0.28738472 | -0.7005536 | 0.48358162 | NA | NA | NA |
| ENSG000002 | 0.97516342 | -0.185907671 | 0.30812709 | -0.6033474 | 0.54627762 | NA | NA | NA |
| ENSG000002 | 0.62411876 | -0.406499733 | 0.2524666 | -1.6101129 | 0.10737321 | NA | NA | NA |

| | | | | | | | | |
|---|---|---|---|---|---|---|---|---|
| ENSG000002 | 0.12685397 | -0.090215768 | 0.12901919 | -0.699243 | 0.48440017 | NA | NA | NA |
| ENSG000002 | 1.68571712 | -0.699472994 | 0.34802911 | -2.0098118 | 0.04445112 | NA | NA | NA |
| ENSG000002 | 1.74539833 | -0.328395192 | 0.35602917 | -0.9223828 | 0.35632893 | NA | NA | NA |
| ENSG000002 | 0.14128499 | -0.097772014 | 0.13403235 | -0.7294658 | 0.4657168 | NA | NA | NA |
| ENSG000002 | 1.62662077 | -0.25846459 | 0.34815348 | -0.7423869 | 0.45785295 | NA | NA | NA |
| ENSG000002 | 0.48078026 | -0.157692509 | 0.2289571 | -0.6887426 | 0.49098527 | NA | NA | NA |
| ENSG000002 | 0.61377977 | -0.254411229 | 0.25893928 | -0.9825131 | 0.32584714 | NA | NA | NA |
| ENSG000002 | 0.94626713 | 0.289037131 | 0.31149838 | 0.92789289 | 0.35346313 | NA | NA | NA |
| ENSG000002 | 0.73499474 | 0.129533122 | 0.27849067 | 0.46512554 | 0.64184156 | NA | NA | NA |
| ENSG000002 | 0.72995644 | -0.027793814 | 0.27785056 | -0.1000315 | 0.9203193 | NA | NA | NA |
| ENSG000002 | 0.11865494 | 0.083440866 | 0.12751478 | 0.65436232 | 0.5128784 | NA | NA | NA |
| ENSG000002 | 0 | NA | NA | NA | NA | NA | NA | NA |
| ENSG000002 | 1.08787061 | 0.038182591 | 0.31487164 | 0.12126399 | 0.90348194 | NA | NA | NA |
| ENSG000002 | 0.12070844 | -0.088503695 | 0.12783931 | -0.6923042 | 0.48874631 | NA | NA | NA |
| ENSG000002 | 1.57140488 | 0.021523662 | 0.35173115 | 0.0611935 | 0.9512051 | NA | NA | NA |
| ENSG000002 | 0.37441639 | -0.268856313 | 0.21277471 | -1.2635727 | 0.20638344 | NA | NA | NA |
| ENSG000002 | 1.95157395 | -0.190529077 | 0.35899695 | -0.5307262 | 0.59560855 | NA | NA | NA |
| ENSG000002 | 0 | NA | NA | NA | NA | NA | NA | NA |
| ENSG000002 | 0 | NA | NA | NA | NA | NA | NA | NA |
| ENSG000002 | 0 | NA | NA | NA | NA | NA | NA | NA |
| ENSG000002 | 0.23890637 | -0.010420674 | 0.17747379 | -0.0587167 | 0.95317776 | NA | NA | NA |
| ENSG000002 | 0.95702876 | -0.027534525 | 0.30493537 | -0.0902963 | 0.92805179 | NA | NA | NA |
| ENSG000002 | 0.64195858 | -0.101091878 | 0.25907734 | -0.3901996 | 0.69638895 | NA | NA | NA |
| ENSG000002 | 1.38896015 | -0.259266138 | 0.3360614 | -0.7714844 | 0.44041984 | NA | NA | NA |
| ENSG000002 | 0.26813897 | -0.197007041 | 0.18550319 | -1.0620143 | 0.2882292 | NA | NA | NA |
| ENSG000002 | 0 | NA | NA | NA | NA | NA | NA | NA |
| ENSG000002 | 2.01606954 | -0.420236588 | 0.35795381 | -1.1739967 | 0.24039636 | NA | NA | NA |
| ENSG000002 | 1.70307683 | -0.703406555 | 0.3482457 | -2.0198571 | 0.04339821 | NA | NA | NA |
| ENSG000002 | 0.94317018 | -0.617561148 | 0.29553776 | -2.0896185 | 0.03665209 | NA | NA | NA |

| | | | | | | | | |
|---|---|---|---|---|---|---|---|---|
| ENSG000002 | 0.74772501 | -0.357780902 | 0.28233051 | -1.2672414 | 0.205069 | NA | NA | NA |
| ENSG000002 | 0.59829496 | 0.063152461 | 0.26232549 | 0.24074084 | 0.80975598 | NA | NA | NA |
| ENSG000002 | 0 | NA | NA | NA | NA | NA | NA | NA |
| ENSG000002 | 1.66670089 | -0.569099 | 0.35284722 | -1.6128765 | 0.10677134 | NA | NA | NA |
| ENSG000002 | 0.23202362 | -0.008302493 | 0.17586207 | -0.0472103 | 0.96234565 | NA | NA | NA |
| ENSG000002 | 0 | NA | NA | NA | NA | NA | NA | NA |
| ENSG000002 | 1.17509509 | 0.415540235 | 0.32806253 | 1.26664947 | 0.20528065 | NA | NA | NA |
| ENSG000002 | 0.26062507 | -0.012888319 | 0.18226503 | -0.070712 | 0.94362699 | NA | NA | NA |
| ENSG000002 | 0.12685397 | -0.090215768 | 0.12901919 | -0.699243 | 0.48440017 | NA | NA | NA |
| ENSG000002 | 0 | NA | NA | NA | NA | NA | NA | NA |
| ENSG000002 | 0.25260017 | -0.015978705 | 0.18049464 | -0.0885273 | 0.92945758 | NA | NA | NA |
| ENSG000002 | 0 | NA | NA | NA | NA | NA | NA | NA |
| ENSG000002 | 0.86703071 | -0.38206962 | 0.28215912 | -1.3540927 | 0.17570679 | NA | NA | NA |
| ENSG000002 | 0 | NA | NA | NA | NA | NA | NA | NA |
| ENSG000002 | 0.23799501 | 0.170958857 | 0.17834799 | 0.95856902 | 0.33777591 | NA | NA | NA |
| ENSG000002 | 1.52439692 | -0.60214654 | 0.33792549 | -1.7818914 | 0.07476694 | NA | NA | NA |
| ENSG000002 | 1.24449852 | -0.186214924 | 0.3294393 | -0.5652481 | 0.57190506 | NA | NA | NA |
| ENSG000002 | 2.10771561 | -0.246553586 | 0.36547509 | -0.6746112 | 0.49992282 | NA | NA | NA |
| ENSG000002 | 0.76650706 | -0.381422914 | 0.28896362 | -1.3199686 | 0.1868455 | NA | NA | NA |
| ENSG000002 | 1.32202155 | 0.034283893 | 0.3335762 | 0.10277679 | 0.91814012 | NA | NA | NA |
| ENSG000002 | 0 | NA | NA | NA | NA | NA | NA | NA |
| ENSG000002 | 0.52876404 | -0.191903645 | 0.2442426 | -0.7857092 | 0.4320379 | NA | NA | NA |
| ENSG000002 | 0 | NA | NA | NA | NA | NA | NA | NA |
| ENSG000002 | 0 | NA | NA | NA | NA | NA | NA | NA |
| ENSG000002 | 0 | NA | NA | NA | NA | NA | NA | NA |
| ENSG000002 | 1.21705403 | 0.095468158 | 0.32813128 | 0.29094501 | 0.77109338 | NA | NA | NA |
| ENSG000002 | 1.707281 | -0.061515199 | 0.35789164 | -0.1718822 | 0.86353015 | NA | NA | NA |
| ENSG000002 | 0.62069689 | -0.101414475 | 0.26466902 | -0.3831747 | 0.70159023 | NA | NA | NA |
| ENSG000002 | 0.36465257 | 0.064724075 | 0.21605106 | 0.29957767 | 0.76449932 | NA | NA | NA |

| | | | | | | | | |
|---|---|---|---|---|---|---|---|---|
| ENSG000002 | 0.2596576 | -0.193002035 | 0.1837913 | -1.0501152 | 0.29366517 | NA | NA | NA |
| ENSG000002 | 0.37899768 | -0.10176489 | 0.21865765 | -0.4654074 | 0.64163973 | NA | NA | NA |
| ENSG000002 | 1.9904846 | -0.187742184 | 0.36140994 | -0.5194716 | 0.60343194 | NA | NA | NA |
| ENSG000002 | 0 | NA | NA | NA | NA | NA | NA | NA |
| ENSG000002 | 1.57716203 | -0.242161803 | 0.34182512 | -0.7084377 | 0.47867348 | NA | NA | NA |
| ENSG000002 | 0.37285215 | -0.099309896 | 0.2177345 | -0.4561055 | 0.64831414 | NA | NA | NA |
| ENSG000002 | 0 | NA | NA | NA | NA | NA | NA | NA |
| ENSG000002 | 0.38494734 | -0.095409837 | 0.21124663 | -0.4516514 | 0.65152014 | NA | NA | NA |
| ENSG000002 | 1.90134055 | -0.543539827 | 0.36005342 | -1.5096088 | 0.13114326 | NA | NA | NA |
| ENSG000002 | 0 | NA | NA | NA | NA | NA | NA | NA |
| ENSG000002 | 0.76576985 | -0.381700686 | 0.28891137 | -1.3211688 | 0.18644509 | NA | NA | NA |
| ENSG000002 | 0.58486175 | 0.058939796 | 0.25343823 | 0.23256079 | 0.81610248 | NA | NA | NA |
| ENSG000002 | 0.49763088 | -0.186408494 | 0.24078583 | -0.7741672 | 0.43883193 | NA | NA | NA |
| ENSG000002 | 0.37831254 | -0.102004459 | 0.21855316 | -0.4667261 | 0.64069587 | NA | NA | NA |
| ENSG000002 | 0.48032325 | -0.179266136 | 0.23878452 | -0.7507444 | 0.45280651 | NA | NA | NA |
| ENSG000002 | 0.6176233 | -0.104294753 | 0.25690244 | -0.4059703 | 0.68476445 | NA | NA | NA |
| ENSG000002 | 0.48536101 | -0.024029524 | 0.24539296 | -0.0979226 | 0.92199373 | NA | NA | NA |
| ENSG000002 | 0.12685397 | -0.090215768 | 0.12901919 | -0.699243 | 0.48440017 | NA | NA | NA |
| ENSG000002 | 2.14451363 | -0.532115693 | 0.36764108 | -1.4473782 | 0.14779103 | NA | NA | NA |
| ENSG000002 | 0.9860346 | -0.036930222 | 0.30788545 | -0.1199479 | 0.9045244 | NA | NA | NA |
| ENSG000002 | 0 | NA | NA | NA | NA | NA | NA | NA |
| ENSG000002 | 0.843097 | -0.112320797 | 0.30154593 | -0.3724832 | 0.70953311 | NA | NA | NA |
| ENSG000002 | 0.12685397 | -0.090215768 | 0.12901919 | -0.699243 | 0.48440017 | NA | NA | NA |
| ENSG000002 | 0.11131518 | 0.081951103 | 0.12663804 | 0.64712866 | 0.51754868 | NA | NA | NA |
| ENSG000002 | 0 | NA | NA | NA | NA | NA | NA | NA |
| ENSG000002 | 0 | NA | NA | NA | NA | NA | NA | NA |
| ENSG000002 | 0 | NA | NA | NA | NA | NA | NA | NA |
| ENSG000002 | 0.34862505 | 0.240809381 | 0.20749856 | 1.16053522 | 0.24583096 | NA | NA | NA |
| ENSG000002 | 1.22536125 | 0.246634366 | 0.32877712 | 0.75015673 | 0.45316032 | NA | NA | NA |

| | | | | | | | | |
|---|---|---|---|---|---|---|---|---|
| ENSG000002 | 0.51931521 | -0.354214731 | 0.23897171 | -1.4822455 | 0.13827499 | NA | NA | NA |
| ENSG000002 | 1.91055911 | -0.528642836 | 0.35866609 | -1.4739136 | 0.14050485 | NA | NA | NA |
| ENSG000002 | 1.5729165 | -0.112195534 | 0.33979128 | -0.3301896 | 0.74125673 | NA | NA | NA |
| ENSG000002 | 1.20592631 | 0.248522032 | 0.32822199 | 0.75717668 | 0.44894402 | NA | NA | NA |
| ENSG000002 | 0 | NA | NA | NA | NA | NA | NA | NA |
| ENSG000002 | 0.73798346 | 0.133274977 | 0.27868582 | 0.47822663 | 0.63248891 | NA | NA | NA |
| ENSG000002 | 0.23816916 | -0.010712105 | 0.1772998 | -0.060418 | 0.9518227 | NA | NA | NA |
| ENSG000002 | 0.9350833 | 0.117782235 | 0.303968 | 0.38748235 | 0.69839915 | NA | NA | NA |
| ENSG000002 | 1.05811313 | 0.491830643 | 0.30956612 | 1.58877415 | 0.11211139 | NA | NA | NA |
| ENSG000002 | 0.63298795 | -0.105769843 | 0.26571024 | -0.3980646 | 0.69058256 | NA | NA | NA |
| ENSG000002 | 1.53614272 | -0.330894781 | 0.34258525 | -0.9658757 | 0.33410637 | NA | NA | NA |
| ENSG000002 | 0.12070844 | -0.088503695 | 0.12783931 | -0.6923042 | 0.48874631 | NA | NA | NA |
| ENSG000002 | 0.12070844 | -0.088503695 | 0.12783931 | -0.6923042 | 0.48874631 | NA | NA | NA |
| ENSG000002 | 1.15444257 | -0.445335221 | 0.32404689 | -1.3742925 | 0.16935088 | NA | NA | NA |
| ENSG000002 | 0 | NA | NA | NA | NA | NA | NA | NA |
| ENSG000002 | 0 | NA | NA | NA | NA | NA | NA | NA |
| ENSG000002 | 0.6439198 | -0.261315757 | 0.26001163 | -1.0050157 | 0.31488931 | NA | NA | NA |
| ENSG000002 | 0.12685397 | -0.090215768 | 0.12901919 | -0.699243 | 0.48440017 | NA | NA | NA |
| ENSG000002 | 0.59141222 | 0.061254288 | 0.25406951 | 0.24109263 | 0.80948332 | NA | NA | NA |
| ENSG000002 | 0.86384824 | -0.270954282 | 0.29866119 | -0.9072296 | 0.36428538 | NA | NA | NA |
| ENSG000002 | 0.26199343 | -0.194018184 | 0.18422801 | -1.0531416 | 0.29227609 | NA | NA | NA |
| ENSG000002 | 0.37194025 | 0.067575365 | 0.21717998 | 0.31114914 | 0.75568724 | NA | NA | NA |
| ENSG000002 | 0.95884106 | 0.426815259 | 0.30074743 | 1.41918175 | 0.15584603 | NA | NA | NA |
| ENSG000002 | 0.38270188 | -0.272446727 | 0.2140009 | -1.2731102 | 0.20297894 | NA | NA | NA |
| ENSG000002 | 0.3863157 | -0.274134926 | 0.21457207 | -1.277589 | 0.20139437 | NA | NA | NA |
| ENSG000002 | 0 | NA | NA | NA | NA | NA | NA | NA |
| ENSG000002 | 0.46136259 | 0.353033165 | 0.24467579 | 1.44286102 | 0.14905963 | NA | NA | NA |
| ENSG000002 | 0 | NA | NA | NA | NA | NA | NA | NA |
| ENSG000002 | 0.12070844 | -0.088503695 | 0.12783931 | -0.6923042 | 0.48874631 | NA | NA | NA |

| | | | | | | | | |
|---|---|---|---|---|---|---|---|---|
| ENSG000002 | 0.11131518 | 0.081951103 | 0.12663804 | 0.64712866 | 0.51754868 | NA | NA | NA |
| ENSG000002 | 0.60419063 | -0.101535721 | 0.26323679 | -0.3857201 | 0.69970398 | NA | NA | NA |
| ENSG000002 | 0.36212533 | -0.235926492 | 0.20089643 | -1.1743688 | 0.24024735 | NA | NA | NA |
| ENSG000002 | 0.37996515 | 0.064665424 | 0.21008146 | 0.3078112 | 0.758226 | NA | NA | NA |
| ENSG000002 | 0.58486175 | 0.058939796 | 0.25343823 | 0.23256079 | 0.81610248 | NA | NA | NA |
| ENSG000002 | 1.22531207 | -0.166149882 | 0.32295293 | -0.5144709 | 0.60692281 | NA | NA | NA |
| ENSG000002 | 0 | NA | NA | NA | NA | NA | NA | NA |
| ENSG000002 | 0.25260017 | -0.015978705 | 0.18049464 | -0.0885273 | 0.92945758 | NA | NA | NA |
| ENSG000002 | 0.12070844 | -0.088503695 | 0.12783931 | -0.6923042 | 0.48874631 | NA | NA | NA |
| ENSG000002 | 0.61972943 | -0.273948374 | 0.26644131 | -1.0281753 | 0.30386737 | NA | NA | NA |
| ENSG000002 | 1.93970135 | -0.503462811 | 0.35293791 | -1.4264912 | 0.15372662 | NA | NA | NA |
| ENSG000002 | 0.73884274 | -0.027679667 | 0.28476881 | -0.0972005 | 0.92256718 | NA | NA | NA |
| ENSG000002 | 0 | NA | NA | NA | NA | NA | NA | NA |
| ENSG000002 | 2.04304817 | -0.733478072 | 0.36123924 | -2.0304496 | 0.04231086 | NA | NA | NA |
| ENSG000002 | 0.34948434 | 0.063159058 | 0.2051263 | 0.30790326 | 0.75815594 | NA | NA | NA |
| ENSG000002 | 1.97692664 | -0.167042771 | 0.3503982 | -0.4767227 | 0.63355961 | NA | NA | NA |
| ENSG000002 | 0.61175852 | -0.109548347 | 0.27027148 | -0.4053271 | 0.68523711 | NA | NA | NA |
| ENSG000002 | 1.07002416 | -0.683165347 | 0.30569832 | -2.2347697 | 0.02543247 | NA | NA | NA |
| ENSG000002 | 1.14524573 | -0.273634142 | 0.3226019 | -0.8482099 | 0.39632106 | NA | NA | NA |
| ENSG000002 | 0.76445355 | -0.200993415 | 0.28720706 | -0.6998206 | 0.48403935 | NA | NA | NA |
| ENSG000002 | 0.65084084 | -0.46034757 | 0.26550072 | -1.7338844 | 0.08293859 | NA | NA | NA |
| ENSG000002 | 1.52652368 | -0.63376727 | 0.34366754 | -1.844129 | 0.06516435 | NA | NA | NA |
| ENSG000002 | 0.79602198 | -0.363127991 | 0.28386389 | -1.2792328 | 0.2008151 | NA | NA | NA |
| ENSG000002 | 1.88522659 | -0.542225803 | 0.36077223 | -1.5029588 | 0.13284967 | NA | NA | NA |
| ENSG000002 | 0.39246123 | -0.276864021 | 0.21549122 | -1.2848042 | 0.19886073 | NA | NA | NA |
| ENSG000002 | 0.48302518 | -0.023466322 | 0.24515318 | -0.0957211 | 0.92374212 | NA | NA | NA |
| ENSG000002 | 0.35210091 | 0.074791332 | 0.21412433 | 0.34928928 | 0.72687215 | NA | NA | NA |
| ENSG000002 | 0.27408862 | -0.199826037 | 0.18669465 | -1.0703362 | 0.28446801 | NA | NA | NA |
| ENSG000002 | 1.38413091 | -0.42642298 | 0.34137831 | -1.2491215 | 0.21162064 | NA | NA | NA |

| | | | | | | | | |
|---|---|---|---|---|---|---|---|---|
| ENSG000002 | 0 | NA | NA | NA | NA | NA | NA | NA |
| ENSG000002 | 0 | NA | NA | NA | NA | NA | NA | NA |
| ENSG000002 | 0 | NA | NA | NA | NA | NA | NA | NA |
| ENSG000002 | 0.74194557 | -0.177437562 | 0.27249639 | -0.6511556 | 0.51494603 | NA | NA | NA |
| ENSG000002 | 0.9903553 | 0.111893664 | 0.30398842 | 0.36808528 | 0.71280965 | NA | NA | NA |
| ENSG000002 | 1.55080442 | -0.810642258 | 0.34355497 | -2.3595708 | 0.01829609 | NA | NA | NA |
| ENSG000002 | 0.23276084 | -0.008011382 | 0.17603982 | -0.0455089 | 0.96370166 | NA | NA | NA |
| ENSG000002 | 0.37267746 | 0.067870636 | 0.21729405 | 0.31234465 | 0.75477861 | NA | NA | NA |
| ENSG000002 | 1.71955088 | -0.199511941 | 0.35826258 | -0.5568875 | 0.57760431 | NA | NA | NA |
| ENSG000002 | 0.14128499 | -0.097772014 | 0.13403235 | -0.7294658 | 0.4657168 | NA | NA | NA |
| ENSG000002 | 0.38019136 | -0.10756138 | 0.21876967 | -0.491665 | 0.62295621 | NA | NA | NA |
| ENSG000002 | 0 | NA | NA | NA | NA | NA | NA | NA |
| ENSG000002 | 0.72135459 | 0.274397577 | 0.27068711 | 1.01370759 | 0.3107223 | NA | NA | NA |
| ENSG000002 | 0.85767182 | -0.111725312 | 0.29715703 | -0.3759807 | 0.70693124 | NA | NA | NA |
| ENSG000002 | 0.75759255 | -0.029385622 | 0.28598041 | -0.102754 | 0.91815824 | NA | NA | NA |
| ENSG000002 | 0.34270765 | 0.258172734 | 0.21433185 | 1.20454674 | 0.22837833 | NA | NA | NA |
| ENSG000002 | 1.222377 | -0.043741271 | 0.33643593 | -0.1300137 | 0.89655561 | NA | NA | NA |
| ENSG000002 | 0.38747904 | -0.104908247 | 0.21987514 | -0.4771265 | 0.63327207 | NA | NA | NA |
| ENSG000002 | 1.24992612 | -0.173710215 | 0.31750666 | -0.5471073 | 0.58430501 | NA | NA | NA |
| ENSG000002 | 0.24004852 | -0.005223787 | 0.17775844 | -0.029387 | 0.97655594 | NA | NA | NA |
| ENSG000002 | 1.12840618 | -0.384999221 | 0.31177849 | -1.2348486 | 0.21688687 | NA | NA | NA |
| ENSG000002 | 0.25214371 | -0.009862892 | 0.18046657 | -0.0546522 | 0.95641556 | NA | NA | NA |
| ENSG000002 | 0.12685397 | -0.090215768 | 0.12901919 | -0.699243 | 0.48440017 | NA | NA | NA |
| ENSG000002 | 0.25145857 | -0.010118565 | 0.18031426 | -0.0561163 | 0.95524918 | NA | NA | NA |
| ENSG000002 | 0.36827086 | -0.266056975 | 0.21181163 | -1.2561018 | 0.20907905 | NA | NA | NA |
| ENSG000002 | 0.98392553 | -0.191024511 | 0.31327738 | -0.6097616 | 0.54201976 | NA | NA | NA |
| ENSG000002 | 1.856647 | 0.127057242 | 0.35360747 | 0.35931719 | 0.71935781 | NA | NA | NA |
| ENSG000002 | 1.22223849 | -0.173647622 | 0.32201569 | -0.5392521 | 0.58971295 | NA | NA | NA |
| ENSG000002 | 0 | NA | NA | NA | NA | NA | NA | NA |

| | | | | | | | | |
|---|---|---|---|---|---|---|---|---|
| ENSG000002 | 1.13351719 | -0.121407813 | 0.32142371 | -0.3777189 | 0.70563942 | NA | NA | NA |
| ENSG000002 | 0 | NA | NA | NA | NA | NA | NA | NA |
| ENSG000002 | 1.24381284 | -0.204269206 | 0.33433463 | -0.6109723 | 0.54121791 | NA | NA | NA |
| ENSG000002 | 1.12290852 | -0.125081176 | 0.32259025 | -0.3877401 | 0.69820838 | NA | NA | NA |
| ENSG000002 | 0.87995168 | -0.095346645 | 0.28343201 | -0.3364004 | 0.73656894 | NA | NA | NA |
| ENSG000002 | 0 | NA | NA | NA | NA | NA | NA | NA |
| ENSG000002 | 0 | NA | NA | NA | NA | NA | NA | NA |
| ENSG000002 | 0 | NA | NA | NA | NA | NA | NA | NA |
| ENSG000002 | 0 | NA | NA | NA | NA | NA | NA | NA |
| ENSG000002 | 0 | NA | NA | NA | NA | NA | NA | NA |
| ENSG000002 | 0 | NA | NA | NA | NA | NA | NA | NA |
| ENSG000002 | 0 | NA | NA | NA | NA | NA | NA | NA |
| ENSG000002 | 0 | NA | NA | NA | NA | NA | NA | NA |
| ENSG000002 | 0 | NA | NA | NA | NA | NA | NA | NA |
| ENSG000002 | 0 | NA | NA | NA | NA | NA | NA | NA |
| ENSG000002 | 0 | NA | NA | NA | NA | NA | NA | NA |
| ENSG000002 | 0 | NA | NA | NA | NA | NA | NA | NA |
| ENSG000002 | 0 | NA | NA | NA | NA | NA | NA | NA |
| ENSG000002 | 0 | NA | NA | NA | NA | NA | NA | NA |
| ENSG000002 | 0 | NA | NA | NA | NA | NA | NA | NA |
| ENSG000002 | 0 | NA | NA | NA | NA | NA | NA | NA |
| ENSG000002 | 0 | NA | NA | NA | NA | NA | NA | NA |
| ENSG000002 | 0.49064726 | 0.142718762 | 0.23924906 | 0.596528 | 0.55082255 | NA | NA | NA |
| ENSG000002 | 0 | NA | NA | NA | NA | NA | NA | NA |
| ENSG000002 | 2.14181328 | -0.491022019 | 0.36144248 | -1.3585067 | 0.17430297 | NA | NA | NA |
| ENSG000002 | 0 | NA | NA | NA | NA | NA | NA | NA |
| ENSG000002 | 0 | NA | NA | NA | NA | NA | NA | NA |
| ENSG000002 | 0 | NA | NA | NA | NA | NA | NA | NA |
| ENSG000002 | 0.12685397 | -0.090215768 | 0.12901919 | -0.699243 | 0.48440017 | NA | NA | NA |

| | | | | | | | | |
|---|---|---|---|---|---|---|---|---|
| ENSG000002 | 0 | NA | NA | NA | NA | NA | NA | NA |
| ENSG000002 | 0 | NA | NA | NA | NA | NA | NA | NA |
| ENSG000002 | 0 | NA | NA | NA | NA | NA | NA | NA |
| ENSG000002 | 0 | NA | NA | NA | NA | NA | NA | NA |
| ENSG000002 | 0 | NA | NA | NA | NA | NA | NA | NA |
| ENSG000002 | 0 | NA | NA | NA | NA | NA | NA | NA |
| ENSG000002 | 1.76787351 | -0.318876435 | 0.35204034 | -0.9057952 | 0.36504428 | NA | NA | NA |
| ENSG000002 | 1.32770635 | 0.295378931 | 0.32675783 | 0.90396895 | 0.36601187 | NA | NA | NA |
| ENSG000002 | 0 | NA | NA | NA | NA | NA | NA | NA |
| ENSG000002 | 0 | NA | NA | NA | NA | NA | NA | NA |
| ENSG000002 | 0.49031286 | -0.020802257 | 0.24597635 | -0.0845702 | 0.93260313 | NA | NA | NA |
| ENSG000002 | 0.51433302 | -0.186673425 | 0.24269587 | -0.7691661 | 0.44179474 | NA | NA | NA |
| ENSG000002 | 0 | NA | NA | NA | NA | NA | NA | NA |
| ENSG000002 | 0 | NA | NA | NA | NA | NA | NA | NA |
| ENSG000002 | 0.14128499 | -0.097772014 | 0.13403235 | -0.7294658 | 0.4657168 | NA | NA | NA |
| ENSG000002 | 0 | NA | NA | NA | NA | NA | NA | NA |
| ENSG000002 | 0 | NA | NA | NA | NA | NA | NA | NA |
| ENSG000002 | 0 | NA | NA | NA | NA | NA | NA | NA |
| ENSG000002 | 0 | NA | NA | NA | NA | NA | NA | NA |
| ENSG000002 | 0 | NA | NA | NA | NA | NA | NA | NA |
| ENSG000002 | 0 | NA | NA | NA | NA | NA | NA | NA |
| ENSG000002 | 0 | NA | NA | NA | NA | NA | NA | NA |
| ENSG000002 | 0 | NA | NA | NA | NA | NA | NA | NA |
| ENSG000002 | 0 | NA | NA | NA | NA | NA | NA | NA |
| ENSG000002 | 0 | NA | NA | NA | NA | NA | NA | NA |
| ENSG000002 | 0 | NA | NA | NA | NA | NA | NA | NA |
| ENSG000002 | 0 | NA | NA | NA | NA | NA | NA | NA |
| ENSG000002 | 0 | NA | NA | NA | NA | NA | NA | NA |
| ENSG000002 | 0 | NA | NA | NA | NA | NA | NA | NA |

| | | | | | | | | |
|---|---|---|---|---|---|---|---|---|
| ENSG000002 | 0 | NA | NA | NA | NA | NA | NA | NA |
| ENSG000002 | 0 | NA | NA | NA | NA | NA | NA | NA |
| ENSG000002 | 0 | NA | NA | NA | NA | NA | NA | NA |
| ENSG000002 | 0 | NA | NA | NA | NA | NA | NA | NA |
| ENSG000002 | 0 | NA | NA | NA | NA | NA | NA | NA |
| ENSG000002 | 0 | NA | NA | NA | NA | NA | NA | NA |
| ENSG000002 | 0 | NA | NA | NA | NA | NA | NA | NA |
| ENSG000002 | 0 | NA | NA | NA | NA | NA | NA | NA |
| ENSG000002 | 0 | NA | NA | NA | NA | NA | NA | NA |
| ENSG000002 | 1.94645685 | -0.820167656 | 0.35644286 | -2.3009794 | 0.0213928 | NA | NA | NA |
| ENSG000002 | 0 | NA | NA | NA | NA | NA | NA | NA |
| ENSG000002 | 0 | NA | NA | NA | NA | NA | NA | NA |
| ENSG000002 | 0 | NA | NA | NA | NA | NA | NA | NA |
| ENSG000002 | 0 | NA | NA | NA | NA | NA | NA | NA |
| ENSG000002 | 0 | NA | NA | NA | NA | NA | NA | NA |
| ENSG000002 | 0.55070895 | -0.361402396 | 0.24091882 | -1.5001003 | 0.13358842 | NA | NA | NA |
| ENSG000002 | 0.92950186 | 0.447250693 | 0.30599053 | 1.46164881 | 0.14383747 | NA | NA | NA |
| ENSG000002 | 0.65427998 | -0.290650429 | 0.26916107 | -1.0798383 | 0.28021419 | NA | NA | NA |
| ENSG000002 | 0.6346709 | -0.110789319 | 0.26582002 | -0.4167832 | 0.67683699 | NA | NA | NA |
| ENSG000002 | 0 | NA | NA | NA | NA | NA | NA | NA |
| ENSG000002 | 0 | NA | NA | NA | NA | NA | NA | NA |
| ENSG000002 | 0 | NA | NA | NA | NA | NA | NA | NA |
| ENSG000002 | 0 | NA | NA | NA | NA | NA | NA | NA |
| ENSG000002 | 0 | NA | NA | NA | NA | NA | NA | NA |
| ENSG000002 | 1.09615653 | -0.504731812 | 0.30299024 | -1.6658352 | 0.09574623 | NA | NA | NA |
| ENSG000002 | 0 | NA | NA | NA | NA | NA | NA | NA |
| ENSG000002 | 0.3867939 | -0.105148852 | 0.21977268 | -0.4784437 | 0.63233444 | NA | NA | NA |
| ENSG000002 | 0 | NA | NA | NA | NA | NA | NA | NA |
| ENSG000002 | 0 | NA | NA | NA | NA | NA | NA | NA |

| | | | | | | | | |
|---|---|---|---|---|---|---|---|---|
| ENSG000002 | 0 | NA | NA | NA | NA | NA | NA | NA |
| ENSG000002 | 1.3949098 | -0.258918702 | 0.33541491 | -0.7719356 | 0.44015259 | NA | NA | NA |
| ENSG000002 | 0 | NA | NA | NA | NA | NA | NA | NA |
| ENSG000002 | 0 | NA | NA | NA | NA | NA | NA | NA |
| ENSG000002 | 0 | NA | NA | NA | NA | NA | NA | NA |
| ENSG000002 | 0 | NA | NA | NA | NA | NA | NA | NA |
| ENSG000002 | 0 | NA | NA | NA | NA | NA | NA | NA |
| ENSG000002 | 0 | NA | NA | NA | NA | NA | NA | NA |
| ENSG000002 | 0.49105007 | -0.020523257 | 0.24605963 | -0.0834077 | 0.9335274 | NA | NA | NA |
| ENSG000002 | 0 | NA | NA | NA | NA | NA | NA | NA |
| ENSG000002 | 0 | NA | NA | NA | NA | NA | NA | NA |
| ENSG000002 | 0 | NA | NA | NA | NA | NA | NA | NA |
| ENSG000002 | 0 | NA | NA | NA | NA | NA | NA | NA |
| ENSG000002 | 0 | NA | NA | NA | NA | NA | NA | NA |
| ENSG000002 | 0 | NA | NA | NA | NA | NA | NA | NA |
| ENSG000002 | 0.35067856 | 0.074230421 | 0.21389444 | 0.34704231 | 0.72855953 | NA | NA | NA |
| ENSG000002 | 0 | NA | NA | NA | NA | NA | NA | NA |
| ENSG000002 | 0 | NA | NA | NA | NA | NA | NA | NA |
| ENSG000002 | 0.22263036 | 0.146922267 | 0.16600367 | 0.88505432 | 0.37612734 | NA | NA | NA |
| ENSG000002 | 0 | NA | NA | NA | NA | NA | NA | NA |
| ENSG000002 | 0.49170296 | -0.014565292 | 0.2386078 | -0.0610428 | 0.95132511 | NA | NA | NA |
| ENSG000002 | 0 | NA | NA | NA | NA | NA | NA | NA |
| ENSG000002 | 0 | NA | NA | NA | NA | NA | NA | NA |
| ENSG000002 | 0 | NA | NA | NA | NA | NA | NA | NA |
| ENSG000002 | 0 | NA | NA | NA | NA | NA | NA | NA |
| ENSG000002 | 0.37928001 | 0.070449581 | 0.21829793 | 0.32272217 | 0.74690566 | NA | NA | NA |
| ENSG000002 | 0 | NA | NA | NA | NA | NA | NA | NA |
| ENSG000002 | 0 | NA | NA | NA | NA | NA | NA | NA |
| ENSG000002 | 0 | NA | NA | NA | NA | NA | NA | NA |

| | | | | | | | | |
|---|---|---|---|---|---|---|---|---|
| ENSG000002 | 0.75441187 | -0.377475959 | 0.28823563 | -1.3096089 | 0.19032816 | NA | NA | NA |
| ENSG000002 | 1.09483981 | 0.195139949 | 0.32395196 | 0.60237311 | 0.5469258 | NA | NA | NA |
| ENSG000002 | 0 | NA | NA | NA | NA | NA | NA | NA |
| ENSG000002 | 0.9235611 | -0.457490269 | 0.30194891 | -1.5151248 | 0.12974081 | NA | NA | NA |
| ENSG000002 | 0.24619405 | -0.007630671 | 0.17915639 | -0.0425922 | 0.96602658 | NA | NA | NA |
| ENSG000002 | 0 | NA | NA | NA | NA | NA | NA | NA |
| ENSG000002 | 0.8212576 | 0.176872863 | 0.27789754 | 0.63646789 | 0.5244715 | NA | NA | NA |
| ENSG000002 | 0 | NA | NA | NA | NA | NA | NA | NA |
| ENSG000002 | 0.37330862 | -0.105391168 | 0.21773449 | -0.4840352 | 0.62836086 | NA | NA | NA |
| ENSG000002 | 0 | NA | NA | NA | NA | NA | NA | NA |
| ENSG000002 | 0 | NA | NA | NA | NA | NA | NA | NA |
| ENSG000002 | 0 | NA | NA | NA | NA | NA | NA | NA |
| ENSG000002 | 0 | NA | NA | NA | NA | NA | NA | NA |
| ENSG000002 | 0 | NA | NA | NA | NA | NA | NA | NA |
| ENSG000002 | 0 | NA | NA | NA | NA | NA | NA | NA |
| ENSG000002 | 1.01930311 | -0.354487838 | 0.309735 | -1.1444875 | 0.25242152 | NA | NA | NA |
| ENSG000002 | 0 | NA | NA | NA | NA | NA | NA | NA |
| ENSG000002 | 0 | NA | NA | NA | NA | NA | NA | NA |
| ENSG000002 | 0 | NA | NA | NA | NA | NA | NA | NA |
| ENSG000002 | 0 | NA | NA | NA | NA | NA | NA | NA |
| ENSG000002 | 0.11205239 | 0.082118251 | 0.12671962 | 0.64803108 | 0.51696485 | NA | NA | NA |
| ENSG000002 | 0 | NA | NA | NA | NA | NA | NA | NA |
| ENSG000002 | 0.13280363 | -0.093394593 | 0.13115644 | -0.7120855 | 0.47641185 | NA | NA | NA |
| ENSG000002 | 2.04752341 | -0.739975963 | 0.36180246 | -2.0452486 | 0.04083036 | NA | NA | NA |
| ENSG000002 | 0 | NA | NA | NA | NA | NA | NA | NA |
| ENSG000002 | 0 | NA | NA | NA | NA | NA | NA | NA |
| ENSG000002 | 0 | NA | NA | NA | NA | NA | NA | NA |
| ENSG000002 | 0.12685397 | -0.090215768 | 0.12901919 | -0.699243 | 0.48440017 | NA | NA | NA |
| ENSG000002 | 0 | NA | NA | NA | NA | NA | NA | NA |

| | | | | | | | | |
|---|---|---|---|---|---|---|---|---|
| ENSG000002 | 0 | NA | NA | NA | NA | NA | NA | NA |
| ENSG000002 | 0 | NA | NA | NA | NA | NA | NA | NA |
| ENSG000002 | 0 | NA | NA | NA | NA | NA | NA | NA |
| ENSG000002 | 0 | NA | NA | NA | NA | NA | NA | NA |
| ENSG000002 | 0 | NA | NA | NA | NA | NA | NA | NA |
| ENSG000002 | 0 | NA | NA | NA | NA | NA | NA | NA |
| ENSG000002 | 0.11865494 | 0.083440866 | 0.12751478 | 0.65436232 | 0.5128784 | NA | NA | NA |
| ENSG000002 | 0 | NA | NA | NA | NA | NA | NA | NA |
| ENSG000002 | 0 | NA | NA | NA | NA | NA | NA | NA |
| ENSG000002 | 0 | NA | NA | NA | NA | NA | NA | NA |
| ENSG000002 | 0 | NA | NA | NA | NA | NA | NA | NA |
| ENSG000002 | 0 | NA | NA | NA | NA | NA | NA | NA |
| ENSG000002 | 0 | NA | NA | NA | NA | NA | NA | NA |
| ENSG000002 | 0 | NA | NA | NA | NA | NA | NA | NA |
| ENSG000002 | 0 | NA | NA | NA | NA | NA | NA | NA |
| ENSG000002 | 0 | NA | NA | NA | NA | NA | NA | NA |
| ENSG000002 | 0 | NA | NA | NA | NA | NA | NA | NA |
| ENSG000002 | 0 | NA | NA | NA | NA | NA | NA | NA |
| ENSG000002 | 0 | NA | NA | NA | NA | NA | NA | NA |
| ENSG000002 | 0 | NA | NA | NA | NA | NA | NA | NA |
| ENSG000002 | 0 | NA | NA | NA | NA | NA | NA | NA |
| ENSG000002 | 0 | NA | NA | NA | NA | NA | NA | NA |
| ENSG000002 | 0 | NA | NA | NA | NA | NA | NA | NA |
| ENSG000002 | 0 | NA | NA | NA | NA | NA | NA | NA |
| ENSG000002 | 0 | NA | NA | NA | NA | NA | NA | NA |
| ENSG000002 | 0 | NA | NA | NA | NA | NA | NA | NA |
| ENSG000002 | 0 | NA | NA | NA | NA | NA | NA | NA |
| ENSG000002 | 0.13280363 | -0.093394593 | 0.13115644 | -0.7120855 | 0.47641185 | NA | NA | NA |
| ENSG000002 | 0 | NA | NA | NA | NA | NA | NA | NA |

| | | | | | | | | |
|---|---|---|---|---|---|---|---|---|
| ENSG000002 | 0 | NA | NA | NA | NA | NA | NA | NA |
| ENSG000002 | 0 | NA | NA | NA | NA | NA | NA | NA |
| ENSG000002 | 0 | NA | NA | NA | NA | NA | NA | NA |
| ENSG000002 | 0 | NA | NA | NA | NA | NA | NA | NA |
| ENSG000002 | 1.9151964 | -0.681069485 | 0.35918216 | -1.8961673 | 0.05793792 | NA | NA | NA |
| ENSG000002 | 0 | NA | NA | NA | NA | NA | NA | NA |
| ENSG000002 | 0.85818091 | -0.107369534 | 0.29718939 | -0.3612832 | 0.71788775 | NA | NA | NA |
| ENSG000002 | 0 | NA | NA | NA | NA | NA | NA | NA |
| ENSG000002 | 0 | NA | NA | NA | NA | NA | NA | NA |
| ENSG000002 | 0 | NA | NA | NA | NA | NA | NA | NA |
| ENSG000002 | 0 | NA | NA | NA | NA | NA | NA | NA |
| ENSG000002 | 0 | NA | NA | NA | NA | NA | NA | NA |
| ENSG000002 | 0 | NA | NA | NA | NA | NA | NA | NA |
| ENSG000002 | 0 | NA | NA | NA | NA | NA | NA | NA |
| ENSG000002 | 0 | NA | NA | NA | NA | NA | NA | NA |
| ENSG000002 | 0 | NA | NA | NA | NA | NA | NA | NA |
| ENSG000002 | 0 | NA | NA | NA | NA | NA | NA | NA |
| ENSG000002 | 0 | NA | NA | NA | NA | NA | NA | NA |
| ENSG000002 | 0 | NA | NA | NA | NA | NA | NA | NA |
| ENSG000002 | 0.13280363 | -0.093394593 | 0.13115644 | -0.7120855 | 0.47641185 | NA | NA | NA |
| ENSG000002 | 0 | NA | NA | NA | NA | NA | NA | NA |
| ENSG000002 | 0 | NA | NA | NA | NA | NA | NA | NA |
| ENSG000002 | 0.26813897 | -0.197007041 | 0.18550319 | -1.0620143 | 0.2882292 | NA | NA | NA |
| ENSG000002 | 0 | NA | NA | NA | NA | NA | NA | NA |
| ENSG000002 | 0 | NA | NA | NA | NA | NA | NA | NA |
| ENSG000002 | 0 | NA | NA | NA | NA | NA | NA | NA |
| ENSG000002 | 0 | NA | NA | NA | NA | NA | NA | NA |
| ENSG000002 | 0.88033276 | -0.429659423 | 0.29465382 | -1.4581838 | 0.14478989 | NA | NA | NA |
| ENSG000002 | 0.11131518 | 0.081951103 | 0.12663804 | 0.64712866 | 0.51754868 | NA | NA | NA |

| | | | | | | | | |
|---|---|---|---|---|---|---|---|---|
| ENSG000002 | 0 | NA | NA | NA | NA | NA | NA | NA |
| ENSG000002 | 0 | NA | NA | NA | NA | NA | NA | NA |
| ENSG000002 | 0 | NA | NA | NA | NA | NA | NA | NA |
| ENSG000002 | 0 | NA | NA | NA | NA | NA | NA | NA |
| ENSG000002 | 0 | NA | NA | NA | NA | NA | NA | NA |
| ENSG000002 | 0.11934008 | 0.083564698 | 0.12760091 | 0.65489106 | 0.5125379 | NA | NA | NA |
| ENSG000002 | 0 | NA | NA | NA | NA | NA | NA | NA |
| ENSG000002 | 0 | NA | NA | NA | NA | NA | NA | NA |
| ENSG000002 | 0 | NA | NA | NA | NA | NA | NA | NA |
| ENSG000002 | 0 | NA | NA | NA | NA | NA | NA | NA |
| ENSG000002 | 0.63504146 | -0.278824355 | 0.26770076 | -1.0415524 | 0.29761927 | NA | NA | NA |
| ENSG000002 | 0 | NA | NA | NA | NA | NA | NA | NA |
| ENSG000002 | 0 | NA | NA | NA | NA | NA | NA | NA |
| ENSG000002 | 0 | NA | NA | NA | NA | NA | NA | NA |
| ENSG000002 | 0 | NA | NA | NA | NA | NA | NA | NA |
| ENSG000002 | 0 | NA | NA | NA | NA | NA | NA | NA |
| ENSG000002 | 0 | NA | NA | NA | NA | NA | NA | NA |
| ENSG000002 | 0 | NA | NA | NA | NA | NA | NA | NA |
| ENSG000002 | 0 | NA | NA | NA | NA | NA | NA | NA |
| ENSG000002 | 0 | NA | NA | NA | NA | NA | NA | NA |
| ENSG000002 | 0 | NA | NA | NA | NA | NA | NA | NA |
| ENSG000002 | 0 | NA | NA | NA | NA | NA | NA | NA |
| ENSG000002 | 0 | NA | NA | NA | NA | NA | NA | NA |
| ENSG000002 | 0 | NA | NA | NA | NA | NA | NA | NA |
| ENSG000002 | 0.11131518 | 0.081951103 | 0.12663804 | 0.64712866 | 0.51754868 | NA | NA | NA |
| ENSG000002 | 0 | NA | NA | NA | NA | NA | NA | NA |
| ENSG000002 | 0 | NA | NA | NA | NA | NA | NA | NA |
| ENSG000002 | 0 | NA | NA | NA | NA | NA | NA | NA |
| ENSG000002 | 0.63387812 | -0.454271587 | 0.26413408 | -1.7198522 | 0.08545931 | NA | NA | NA |

| | | | | | | | | |
|---|---|---|---|---|---|---|---|---|
| ENSG000002 | 0 | NA | NA | NA | NA | NA | NA | NA |
| ENSG000002 | 1.09431154 | -0.375382938 | 0.30731864 | -1.2214779 | 0.22190512 | NA | NA | NA |
| ENSG000002 | 0 | NA | NA | NA | NA | NA | NA | NA |
| ENSG000002 | 1.42598902 | 0.077883783 | 0.33348119 | 0.23354775 | 0.81533609 | NA | NA | NA |
| ENSG000002 | 0 | NA | NA | NA | NA | NA | NA | NA |
| ENSG000002 | 0 | NA | NA | NA | NA | NA | NA | NA |
| ENSG000002 | 0 | NA | NA | NA | NA | NA | NA | NA |
| ENSG000002 | 0.23799501 | 0.170958857 | 0.17834799 | 0.95856902 | 0.33777591 | NA | NA | NA |
| ENSG000002 | 0 | NA | NA | NA | NA | NA | NA | NA |
| ENSG000002 | 0 | NA | NA | NA | NA | NA | NA | NA |
| ENSG000002 | 0.23868015 | 0.154625155 | 0.16997278 | 0.90970538 | 0.36297791 | NA | NA | NA |
| ENSG000002 | 0 | NA | NA | NA | NA | NA | NA | NA |
| ENSG000002 | 0 | NA | NA | NA | NA | NA | NA | NA |
| ENSG000002 | 0 | NA | NA | NA | NA | NA | NA | NA |
| ENSG000002 | 1.12014188 | -0.249241995 | 0.3148547 | -0.7916096 | 0.42858837 | NA | NA | NA |
| ENSG000002 | 0 | NA | NA | NA | NA | NA | NA | NA |
| ENSG000002 | 0 | NA | NA | NA | NA | NA | NA | NA |
| ENSG000002 | 0 | NA | NA | NA | NA | NA | NA | NA |
| ENSG000002 | 0 | NA | NA | NA | NA | NA | NA | NA |
| ENSG000002 | 0 | NA | NA | NA | NA | NA | NA | NA |
| ENSG000002 | 0 | NA | NA | NA | NA | NA | NA | NA |
| ENSG000002 | 0.39388516 | -0.104610737 | 0.21253504 | -0.4922047 | 0.62257467 | NA | NA | NA |
| ENSG000002 | 0 | NA | NA | NA | NA | NA | NA | NA |
| ENSG000002 | 0.65451023 | -0.109436795 | 0.25851814 | -0.4233235 | 0.67205928 | NA | NA | NA |
| ENSG000002 | 0 | NA | NA | NA | NA | NA | NA | NA |
| ENSG000002 | 0 | NA | NA | NA | NA | NA | NA | NA |
| ENSG000002 | 0 | NA | NA | NA | NA | NA | NA | NA |
| ENSG000002 | 0 | NA | NA | NA | NA | NA | NA | NA |
| ENSG000002 | 0.25333738 | -0.015686291 | 0.1806605 | -0.0868275 | 0.93080867 | NA | NA | NA |

| | | | | | | | | |
|---|---|---|---|---|---|---|---|---|
| ENSG000002 | 0 | NA | NA | NA | NA | NA | NA | NA |
| ENSG000002 | 0 | NA | NA | NA | NA | NA | NA | NA |
| ENSG000002 | 0 | NA | NA | NA | NA | NA | NA | NA |
| ENSG000002 | 0.75830802 | -0.198939032 | 0.28682413 | -0.6935924 | 0.48793787 | NA | NA | NA |
| ENSG000002 | 0 | NA | NA | NA | NA | NA | NA | NA |
| ENSG000002 | 0 | NA | NA | NA | NA | NA | NA | NA |
| ENSG000002 | 0 | NA | NA | NA | NA | NA | NA | NA |
| ENSG000002 | 0 | NA | NA | NA | NA | NA | NA | NA |
| ENSG000002 | 0.52193336 | -0.189821458 | 0.24349877 | -0.7795582 | 0.43565098 | NA | NA | NA |
| ENSG000002 | 0 | NA | NA | NA | NA | NA | NA | NA |
| ENSG000002 | 1.90660506 | -0.539170999 | 0.3591709 | -1.5011545 | 0.13331562 | NA | NA | NA |
| ENSG000002 | 0.24756242 | -0.187034346 | 0.18119817 | -1.0322088 | 0.30197431 | NA | NA | NA |
| ENSG000002 | 0.38133351 | -0.102466351 | 0.2189713 | -0.4679442 | 0.63982449 | NA | NA | NA |
| ENSG000002 | 0 | NA | NA | NA | NA | NA | NA | NA |
| ENSG000002 | 0 | NA | NA | NA | NA | NA | NA | NA |
| ENSG000002 | 0 | NA | NA | NA | NA | NA | NA | NA |
| ENSG000002 | 0 | NA | NA | NA | NA | NA | NA | NA |
| ENSG000002 | 0.12070844 | -0.088503695 | 0.12783931 | -0.6923042 | 0.48874631 | NA | NA | NA |
| ENSG000002 | 0 | NA | NA | NA | NA | NA | NA | NA |
| ENSG000002 | 0.39499294 | -0.277908454 | 0.21584201 | -1.287555 | 0.19790091 | NA | NA | NA |
| ENSG000002 | 0 | NA | NA | NA | NA | NA | NA | NA |
| ENSG000002 | 0 | NA | NA | NA | NA | NA | NA | NA |
| ENSG000002 | 0 | NA | NA | NA | NA | NA | NA | NA |
| ENSG000002 | 0 | NA | NA | NA | NA | NA | NA | NA |
| ENSG000002 | 0 | NA | NA | NA | NA | NA | NA | NA |
| ENSG000002 | 0 | NA | NA | NA | NA | NA | NA | NA |
| ENSG000002 | 0 | NA | NA | NA | NA | NA | NA | NA |
| ENSG000002 | 0 | NA | NA | NA | NA | NA | NA | NA |
| ENSG000002 | 0 | NA | NA | NA | NA | NA | NA | NA |

| | | | | | | | | |
|---|---|---|---|---|---|---|---|---|
| ENSG000002 | 0 | NA | NA | NA | NA | NA | NA | NA |
| ENSG000002 | 0 | NA | NA | NA | NA | NA | NA | NA |
| ENSG000002 | 0 | NA | NA | NA | NA | NA | NA | NA |
| ENSG000002 | 0 | NA | NA | NA | NA | NA | NA | NA |
| ENSG000002 | 2.11254587 | -0.011795326 | 0.36476888 | -0.0323364 | 0.97420375 | NA | NA | NA |
| ENSG000002 | 0 | NA | NA | NA | NA | NA | NA | NA |
| ENSG000002 | 0 | NA | NA | NA | NA | NA | NA | NA |
| ENSG000002 | 0 | NA | NA | NA | NA | NA | NA | NA |
| ENSG000002 | 0 | NA | NA | NA | NA | NA | NA | NA |
| ENSG000002 | 0 | NA | NA | NA | NA | NA | NA | NA |
| ENSG000002 | 0 | NA | NA | NA | NA | NA | NA | NA |
| ENSG000002 | 0.23139247 | 0.167748926 | 0.17680352 | 0.94878726 | 0.34272882 | NA | NA | NA |
| ENSG000002 | 0.39499294 | -0.277908454 | 0.21584201 | -1.287555 | 0.19790091 | NA | NA | NA |
| ENSG000002 | 0 | NA | NA | NA | NA | NA | NA | NA |
| ENSG000002 | 1.99054004 | -0.424877042 | 0.35708212 | -1.1898581 | 0.23410218 | NA | NA | NA |
| ENSG000002 | 0 | NA | NA | NA | NA | NA | NA | NA |
| ENSG000002 | 0 | NA | NA | NA | NA | NA | NA | NA |
| ENSG000002 | 0 | NA | NA | NA | NA | NA | NA | NA |
| ENSG000002 | 0 | NA | NA | NA | NA | NA | NA | NA |
| ENSG000002 | 0 | NA | NA | NA | NA | NA | NA | NA |
| ENSG000002 | 0 | NA | NA | NA | NA | NA | NA | NA |
| ENSG000002 | 1.92014379 | -0.36311226 | 0.3487513 | -1.0411782 | 0.29779284 | NA | NA | NA |
| ENSG000002 | 0 | NA | NA | NA | NA | NA | NA | NA |
| ENSG000002 | 0 | NA | NA | NA | NA | NA | NA | NA |
| ENSG000002 | 0 | NA | NA | NA | NA | NA | NA | NA |
| ENSG000002 | 0 | NA | NA | NA | NA | NA | NA | NA |
| ENSG000002 | 0 | NA | NA | NA | NA | NA | NA | NA |
| ENSG000002 | 0 | NA | NA | NA | NA | NA | NA | NA |
| ENSG000002 | 0 | NA | NA | NA | NA | NA | NA | NA |

| | | | | | | | | |
|---|---|---|---|---|---|---|---|---|
| ENSG000002 | 0 | NA | NA | NA | NA | NA | NA | NA |
| ENSG000002 | 0 | NA | NA | NA | NA | NA | NA | NA |
| ENSG000002 | 0 | NA | NA | NA | NA | NA | NA | NA |
| ENSG000002 | 0.63389931 | -0.262485172 | 0.25907614 | -1.0131584 | 0.3109845 | NA | NA | NA |
| ENSG000002 | 1.18831517 | -0.033774112 | 0.32504355 | -0.1039064 | 0.91724361 | NA | NA | NA |
| ENSG000002 | 0 | NA | NA | NA | NA | NA | NA | NA |
| ENSG000002 | 0 | NA | NA | NA | NA | NA | NA | NA |
| ENSG000002 | 0 | NA | NA | NA | NA | NA | NA | NA |
| ENSG000002 | 0 | NA | NA | NA | NA | NA | NA | NA |
| ENSG000002 | 0 | NA | NA | NA | NA | NA | NA | NA |
| ENSG000002 | 0 | NA | NA | NA | NA | NA | NA | NA |
| ENSG000002 | 0 | NA | NA | NA | NA | NA | NA | NA |
| ENSG000002 | 0.50016259 | -0.200428691 | 0.24824437 | -0.8073846 | 0.41944491 | NA | NA | NA |
| ENSG000002 | 0 | NA | NA | NA | NA | NA | NA | NA |
| ENSG000002 | 0 | NA | NA | NA | NA | NA | NA | NA |
| ENSG000002 | 0 | NA | NA | NA | NA | NA | NA | NA |
| ENSG000002 | 1.3764944 | -0.125774528 | 0.34115888 | -0.3686685 | 0.71237484 | NA | NA | NA |
| ENSG000002 | 0.12070844 | -0.088503695 | 0.12783931 | -0.6923042 | 0.48874631 | NA | NA | NA |
| ENSG000002 | 0 | NA | NA | NA | NA | NA | NA | NA |
| ENSG000002 | 0.14128499 | -0.097772014 | 0.13403235 | -0.7294658 | 0.4657168 | NA | NA | NA |
| ENSG000002 | 0 | NA | NA | NA | NA | NA | NA | NA |
| ENSG000002 | 0 | NA | NA | NA | NA | NA | NA | NA |
| ENSG000002 | 0.11205239 | 0.082118251 | 0.12671962 | 0.64803108 | 0.51696485 | NA | NA | NA |
| ENSG000002 | 0 | NA | NA | NA | NA | NA | NA | NA |
| ENSG000002 | 0 | NA | NA | NA | NA | NA | NA | NA |
| ENSG000002 | 0 | NA | NA | NA | NA | NA | NA | NA |
| ENSG000002 | 1.31129026 | 0.177859054 | 0.33957385 | 0.52377135 | 0.60043758 | NA | NA | NA |
| ENSG000002 | 1.23262955 | -0.351067317 | 0.33349376 | -1.0526953 | 0.29248066 | NA | NA | NA |
| ENSG000002 | 0 | NA | NA | NA | NA | NA | NA | NA |

| | | | | | | | | |
|---|---|---|---|---|---|---|---|---|
| ENSG000002 | 0 | NA | NA | NA | NA | NA | NA | NA |
| ENSG000002 | 0 | NA | NA | NA | NA | NA | NA | NA |
| ENSG000002 | 0 | NA | NA | NA | NA | NA | NA | NA |
| ENSG000002 | 0 | NA | NA | NA | NA | NA | NA | NA |
| ENSG000002 | 0 | NA | NA | NA | NA | NA | NA | NA |
| ENSG000002 | 0.23070733 | 0.167416683 | 0.17664244 | 0.94777156 | 0.34324576 | NA | NA | NA |
| ENSG000002 | 0.26062507 | -0.012888319 | 0.18226503 | -0.070712 | 0.94362699 | NA | NA | NA |
| ENSG000002 | 0 | NA | NA | NA | NA | NA | NA | NA |
| ENSG000002 | 0 | NA | NA | NA | NA | NA | NA | NA |
| ENSG000002 | 0 | NA | NA | NA | NA | NA | NA | NA |
| ENSG000002 | 0 | NA | NA | NA | NA | NA | NA | NA |
| ENSG000002 | 0 | NA | NA | NA | NA | NA | NA | NA |
| ENSG000002 | 0 | NA | NA | NA | NA | NA | NA | NA |
| ENSG000002 | 0.96838673 | -0.033819419 | 0.31203989 | -0.1083817 | 0.9136929 | NA | NA | NA |
| ENSG000002 | 0 | NA | NA | NA | NA | NA | NA | NA |
| ENSG000002 | 0 | NA | NA | NA | NA | NA | NA | NA |
| ENSG000002 | 0 | NA | NA | NA | NA | NA | NA | NA |
| ENSG000002 | 0.92970663 | -0.439225344 | 0.29724343 | -1.4776621 | 0.13949825 | NA | NA | NA |
| ENSG000002 | 0 | NA | NA | NA | NA | NA | NA | NA |
| ENSG000002 | 0 | NA | NA | NA | NA | NA | NA | NA |
| ENSG000002 | 0 | NA | NA | NA | NA | NA | NA | NA |
| ENSG000002 | 0.61158383 | 0.053697626 | 0.26349596 | 0.20378918 | 0.83851825 | NA | NA | NA |
| ENSG000002 | 0 | NA | NA | NA | NA | NA | NA | NA |
| ENSG000002 | 0 | NA | NA | NA | NA | NA | NA | NA |
| ENSG000002 | 1.72390851 | -0.306614679 | 0.35191799 | -0.8712674 | 0.38360815 | NA | NA | NA |
| ENSG000002 | 0 | NA | NA | NA | NA | NA | NA | NA |
| ENSG000002 | 0 | NA | NA | NA | NA | NA | NA | NA |
| ENSG000002 | 0 | NA | NA | NA | NA | NA | NA | NA |
| ENSG000002 | 0 | NA | NA | NA | NA | NA | NA | NA |

| | | | | | | | | |
|---|---|---|---|---|---|---|---|---|
| ENSG000002 | 0 | NA | NA | NA | NA | NA | NA | NA |
| ENSG000002 | 0 | NA | NA | NA | NA | NA | NA | NA |
| ENSG000002 | 0 | NA | NA | NA | NA | NA | NA | NA |
| ENSG000002 | 0 | NA | NA | NA | NA | NA | NA | NA |
| ENSG000002 | 0 | NA | NA | NA | NA | NA | NA | NA |
| ENSG000002 | 0 | NA | NA | NA | NA | NA | NA | NA |
| ENSG000002 | 0 | NA | NA | NA | NA | NA | NA | NA |
| ENSG000002 | 0 | NA | NA | NA | NA | NA | NA | NA |
| ENSG000002 | 0 | NA | NA | NA | NA | NA | NA | NA |
| ENSG000002 | 0.11205239 | 0.082118251 | 0.12671962 | 0.64803108 | 0.51696485 | NA | NA | NA |
| ENSG000002 | 0 | NA | NA | NA | NA | NA | NA | NA |
| ENSG000002 | 0 | NA | NA | NA | NA | NA | NA | NA |
| ENSG000002 | 0 | NA | NA | NA | NA | NA | NA | NA |
| ENSG000002 | 0.11205239 | 0.082118251 | 0.12671962 | 0.64803108 | 0.51696485 | NA | NA | NA |
| ENSG000002 | 0 | NA | NA | NA | NA | NA | NA | NA |
| ENSG000002 | 0 | NA | NA | NA | NA | NA | NA | NA |
| ENSG000002 | 0 | NA | NA | NA | NA | NA | NA | NA |
| ENSG000002 | 0 | NA | NA | NA | NA | NA | NA | NA |
| ENSG000002 | 0 | NA | NA | NA | NA | NA | NA | NA |
| ENSG000002 | 0 | NA | NA | NA | NA | NA | NA | NA |
| ENSG000002 | 0 | NA | NA | NA | NA | NA | NA | NA |
| ENSG000002 | 0.62775378 | -0.28146231 | 0.26704095 | -1.0540043 | 0.29188095 | NA | NA | NA |
| ENSG000002 | 1.62079139 | 0.007211665 | 0.3459427 | 0.02084642 | 0.98336817 | NA | NA | NA |
| ENSG000002 | 0.23276084 | -0.008011382 | 0.17603982 | -0.0455089 | 0.96370166 | NA | NA | NA |
| ENSG000002 | 1.81908992 | -0.23187151 | 0.35198552 | -0.658753 | 0.5100544 | NA | NA | NA |
| ENSG000002 | 0.36345889 | 0.070257596 | 0.21589421 | 0.32542602 | 0.74485866 | NA | NA | NA |
| ENSG000002 | 0.96326411 | -0.170256409 | 0.30008628 | -0.5673582 | 0.57047085 | NA | NA | NA |
| ENSG000002 | 0 | NA | NA | NA | NA | NA | NA | NA |
| ENSG000002 | 0 | NA | NA | NA | NA | NA | NA | NA |

| | | | | | | | | |
|---|---|---|---|---|---|---|---|---|
| ENSG000002 | 0 | NA | NA | NA | NA | NA | NA | NA |
| ENSG000002 | 1.18805403 | -0.037079762 | 0.32291449 | -0.1148284 | 0.90858112 | NA | NA | NA |
| ENSG000002 | 1.44430024 | -0.570796634 | 0.33637647 | -1.6968982 | 0.0897159 | NA | NA | NA |
| ENSG000002 | 0.48441528 | -0.017227682 | 0.23774503 | -0.0724628 | 0.94223357 | NA | NA | NA |
| ENSG000002 | 0 | NA | NA | NA | NA | NA | NA | NA |
| ENSG000002 | 1.99210304 | -0.176661921 | 0.35747097 | -0.4941994 | 0.6211654 | NA | NA | NA |
| ENSG000002 | 0.75982019 | -0.379788636 | 0.28855699 | -1.3161651 | 0.18811864 | NA | NA | NA |
| ENSG000002 | 0.76696407 | -0.358999402 | 0.28364792 | -1.2656514 | 0.2056379 | NA | NA | NA |
| ENSG000002 | 0.4620998 | 0.332478694 | 0.23824476 | 1.39553412 | 0.16285483 | NA | NA | NA |
| ENSG000002 | 0.84488757 | 0.195433437 | 0.29075995 | 0.67214704 | 0.50149009 | NA | NA | NA |
| ENSG000002 | 0 | NA | NA | NA | NA | NA | NA | NA |
| ENSG000002 | 0 | NA | NA | NA | NA | NA | NA | NA |
| ENSG000002 | 0 | NA | NA | NA | NA | NA | NA | NA |
| ENSG000002 | 0 | NA | NA | NA | NA | NA | NA | NA |
| ENSG000002 | 1.01773887 | -0.205648565 | 0.31456741 | -0.6537504 | 0.51327264 | NA | NA | NA |
| ENSG000002 | 0.75506476 | -0.355081327 | 0.28288244 | -1.2552257 | 0.20939682 | NA | NA | NA |
| ENSG000002 | 0.73833175 | -0.187716843 | 0.27944393 | -0.6717514 | 0.50174199 | NA | NA | NA |
| ENSG000002 | 0.60355948 | 0.064278365 | 0.2692118 | 0.23876504 | 0.81128778 | NA | NA | NA |
| ENSG000002 | 1.36072884 | 0.155763424 | 0.33611541 | 0.46342245 | 0.64306159 | NA | NA | NA |
| ENSG000002 | 0 | NA | NA | NA | NA | NA | NA | NA |
| ENSG000002 | 0.13280363 | -0.093394593 | 0.13115644 | -0.7120855 | 0.47641185 | NA | NA | NA |
| ENSG000002 | 0 | NA | NA | NA | NA | NA | NA | NA |
| ENSG000002 | 0.35801832 | 0.071082974 | 0.20657063 | 0.34410977 | 0.73076373 | NA | NA | NA |
| ENSG000002 | 0 | NA | NA | NA | NA | NA | NA | NA |
| ENSG000002 | 0 | NA | NA | NA | NA | NA | NA | NA |
| ENSG000002 | 0.40327843 | -0.281391716 | 0.21700406 | -1.2967118 | 0.19473038 | NA | NA | NA |
| ENSG000002 | 0 | NA | NA | NA | NA | NA | NA | NA |
| ENSG000002 | 1.45425819 | 0.084221553 | 0.34198209 | 0.24627474 | 0.80546956 | NA | NA | NA |
| ENSG000002 | 1.97422629 | -0.199603395 | 0.36591594 | -0.5454898 | 0.58541673 | NA | NA | NA |

| | | | | | | | | |
|---|---|---|---|---|---|---|---|---|
| ENSG000002 | 0.35870346 | 0.077368606 | 0.21515516 | 0.35959447 | 0.71915042 | NA | NA | NA |
| ENSG000002 | 1.24540851 | -0.446148995 | 0.31944238 | -1.3966494 | 0.16251904 | NA | NA | NA |
| ENSG000002 | 0.64081643 | -0.112957221 | 0.26633067 | -0.424124 | 0.6714754 | NA | NA | NA |
| ENSG000002 | 0 | NA | NA | NA | NA | NA | NA | NA |
| ENSG000002 | 0.50684946 | -0.202346264 | 0.24895102 | -0.8127955 | 0.41633533 | NA | NA | NA |
| ENSG000002 | 2.04043406 | -0.579430364 | 0.36080686 | -1.6059295 | 0.10828941 | NA | NA | NA |
| ENSG000002 | 0 | NA | NA | NA | NA | NA | NA | NA |
| ENSG000002 | 1.96702505 | -0.066286572 | 0.36249724 | -0.1828609 | 0.85490716 | NA | NA | NA |
| ENSG000002 | 0.5226185 | -0.189580046 | 0.24357686 | -0.7783172 | 0.43638207 | NA | NA | NA |
| ENSG000002 | 0 | NA | NA | NA | NA | NA | NA | NA |
| ENSG000002 | 1.63539752 | -0.697931504 | 0.34850792 | -2.0026274 | 0.0452173 | NA | NA | NA |
| ENSG000002 | 0.61221553 | -0.098630193 | 0.26395208 | -0.373667 | 0.70865205 | NA | NA | NA |
| ENSG000002 | 1.80897768 | -0.236771892 | 0.35594858 | -0.6651857 | 0.50593175 | NA | NA | NA |
| ENSG000002 | 0 | NA | NA | NA | NA | NA | NA | NA |
